# Functional exploration of *in vivo* and *in vitro* lignocellulose-fed rumen bacterial microbiomes reveals novel enzymes involved in polysaccharide breakdown

**DOI:** 10.1101/2024.03.15.585145

**Authors:** L. Ufarté, E. Laville, A. Lazuka, S. Lajus, E. Bouhajja, D. Cecchini, A. Rizzo, E. Amblard, E. Drula, V. Lombard, N. Terrapon, B. Henrissat, M. Cleret, D.P. Morgavi, C. Dumon, P. Robe, C. Klopp, S. Bozonnet, G. Hernandez-Raquet, G. Potocki-Veronese

**Affiliations:** TBI, Université de Toulouse, CNRS, INRAE, INSA, Toulouse, France; CNRS, UMR 7257, Aix-Marseille Université, F-13288 Marseille, France; USC 1408 AFMB, INRAE, F-13288 Marseille, France; Université Clermont Auvergne, INRAE, VetAgro Sup, UMR Herbivores, F-63122 Saint-Genes-Champanelle, France; Givaudan SA, Toulouse, France; Plateforme Bio-informatique Toulouse Genopole, UBIA INRAE, BP 52627, Castanet-Tolosan, France

**Keywords:** activity-based metagenomics, bovine rumen microbiome, plant cell wall polysaccharides, glycoside hydrolases

## Abstract

**Background:** Plant cell walls are the main carbon sources for ruminal bacteria, which have evolved to produce sophisticated multi-functional enzyme cocktails in response to the structural diversity of lignocelulloses. Since a large proportion of ruminal bacteria are not yet cultured, we developed a high-throughput activity-based metagenomic approach to gain insight into this enzymatic diversity.

**Results:** A multi-step screening methodology was implemented to identify metagenomic clones acting on polysaccharides and polyaromatic compounds. This approach was used to explore the functional potential of two different microbial consortia derived from *in vivo* and *in vitro* enrichments of the bovine rumen microbiome on wheat straw. One hundred and sixty-eight fosmid clones were isolated from libraries. Five to seven times more β-mannanase and β-glucanase clones, and seven times less xylanase clones were obtained from the *in vitro* enrichment compared to the *in vivo* one.

The sequencing of 51 fosmids, covering in total 1.4 Gb of metagenomic DNA, enabled the identification of various novel glycoside-hydrolases, esterases and oxidoreductases mostly encoded by unknown bacterial genera. Functional analysis showed that most of the identified xylanases belonged to Firmicutes members that were not enriched in the fermenter, while most cellulases and mannanases originate from Bacteroidetes.

**Conclusion:** These enzymes, that, for most of them, had not been previously identified by in depth-metagenome sequencing, present a high potential for biotechnological applications, as they could be used alone or in cocktails to break down plant cell walls. The relationships established between enzyme function and taxonomy highlight the complementary roles played by ruminal Firmicutes and Bacteroidetes in plant cell wall degradation.

## 1. Background

The bovine ruminal microbiota is a highly complex ecosystem mainly composed of bacteria, which represent the largest microbial biomass with 10^10^-10^11^ cells per gram of rumen content, and by bacteriophages, archaea, protozoa and fungi (1). Due to the paucity of genes coding for polysaccharide-degrading enzymes (the so-called Carbohydrate Active enZymes, or CAZymes (2)) in the genome of cattle, ruminants, like other mammals and some insects, depend on these symbiotic microorganisms colonizing their digestive tract to break down plant cell walls (PCW), their main dietary resource. This complex polymeric PCW consists of a cellulose scaffold cross-linked with hemicelluloses, pectins, lignins and some proteins. The composition and structural complexity of these biopolymers, their physical state and their distribution, which vary for each plant type, tissue and growth stage, make the complete degradation of PCW a complex issue (3).

Ruminal bacteria are key players in PCW degradation and fermentation, producing volatile fatty acids which are directly absorbed through the stomach wall to supply their host with 60 to 80% of their energy requirements (4). Ruminal bacteria have evolved sophisticated mechanisms for PCW and energy storage polysaccharides degradation. Indeed, those encoded by the Polysaccharide Utilization Loci (PULs, (5)) first described in Bacteroidetes (6,7) and further also in Gram positive bacteria (8,9), or the cellulosome-like structures described in cultivated Firmicutes (10,11) orchestrate the binding, transport and degradation of polysaccharides. These features of the rumen microbiome could be exploited for lignocellulose bioconversion to produce synthons and biofuels, which remains a scientific and economic challenge in the context of depleted fossil resources and fighting global warming (12).

Since only a minor fraction of the ruminal bacterial species is cultured (13), sequence-based metagenomics has been used extensively in the past decade to study their plant biomass degradation potential (14–17). These studies revealed that the ruminal bacteriome is one of the richest and most diverse in terms of CAZy-encoding genes. This specificity has been exploited many times to mine for novel PCW polysaccharide-degrading enzymes, using activity-based metagenomics. This approach indeed provides a direct link between DNA and encoded protein functions, allowing the discovery of novel enzyme families and/or novel functions. In total, more than one hundred novel glycoside-hydrolases were identified by applying this approach to the ruminal bacteriome (1). The efficiency of bovine ruminal bacteria in degrading PCW has also been exploited by engineering mixed cultures derived from rumen content. This *in vitro* microbiome enrichment strategy, using lignocellulosic material like wheat straw as the sole carbon source in controlled bioreactors, was recently used to obtain a stable hyper-hydrolytic microbial consortium (18). Microbial diversity was significantly affected during enrichment, with an increase in xylanase and cellulase activities and a decrease in β-glucosidase activities. Nevertheless, the enzymes that are ultimately responsible for lignocellulose degradation in this enriched microbiota have not been fully identified. Metatranscriptomics and metaproteomics could provide additional information about expressed functional genes and the proteins produced, yet none of these methods would provide experimental proof of their function.

In this study, we thus used activity-based metagenomics in order to identify key enzymes involved in lignocellulose degradation by two hyper-lignocellulolytic bacterial consortia derived from bovine rumen. These microbiomes were obtained after *in vitro* and *in vivo* enrichments on wheat straw, an agricultural residue which is a valuable feedstock for European biorefineries to produce second-generation biofuels (12). A multi-step high-throughput screening strategy was first implemented, to access not only glycoside-hydrolases, but also esterases and oxidoreductases, all of which are necessary to completely break down PCW. Taxonomic and functional profiles of both enriched microbiomes were then compared, in the light of the biochemical and metagenomic data that are now available on these semi-natural and artificially enriched ruminal ecosystems.

## 2. Results

### 2.1 Diversity analysis of samples used for metagenomic libraries

The ‘*in vivo* enriched’ (IVVE) rumen sample was obtained from a mix of the ruminal contents of two cows fed with a diet enriched in wheat straw. The ‘*in vitro* enriched’ (IVTE) rumen sample was obtained from the ruminal contents of the same cows before dietary enrichment, but was enriched *in vitro* in a bioreactor, following the method described by (18). The microbial diversity of the IVVE- and IVTE-enriched rumen samples was analyzed by 16S rRNA amplicon sequencing. For the purposes of comparison, a representative sample of the original rumen microbiota (before enrichment) was also included. Sequencing data showed significant changes in microbial community composition, particularly in the IVTE samples (Figure 1A). Compared to the original rumen which displays a Shannon diversity index of 5.91 and an observed richness of 919 Operational taxonomic units (OTUs), diversity decreased strongly in IVTE (Shannon of 2.87, and 255 OTUs). In contrast, a lesser impact on diversity was observed in IVVE despite the wheat straw-fed diet imposed to the cows (Shannon 5.85 and 876 OTUs). This undoubtedly resulted from the different environmental conditions imposed in the bioreactor. Microbial community composition was strongly modified in the *in-vitro* and, to a lesser extent, *in-vivo* enrichments, with an increase in both cases in Bacteroidetes-related members at the expense of Firmicutes (Figure 1B and C), while the latter phylum was the more abundant in the original rumen microbiome.

**Figure 1.**
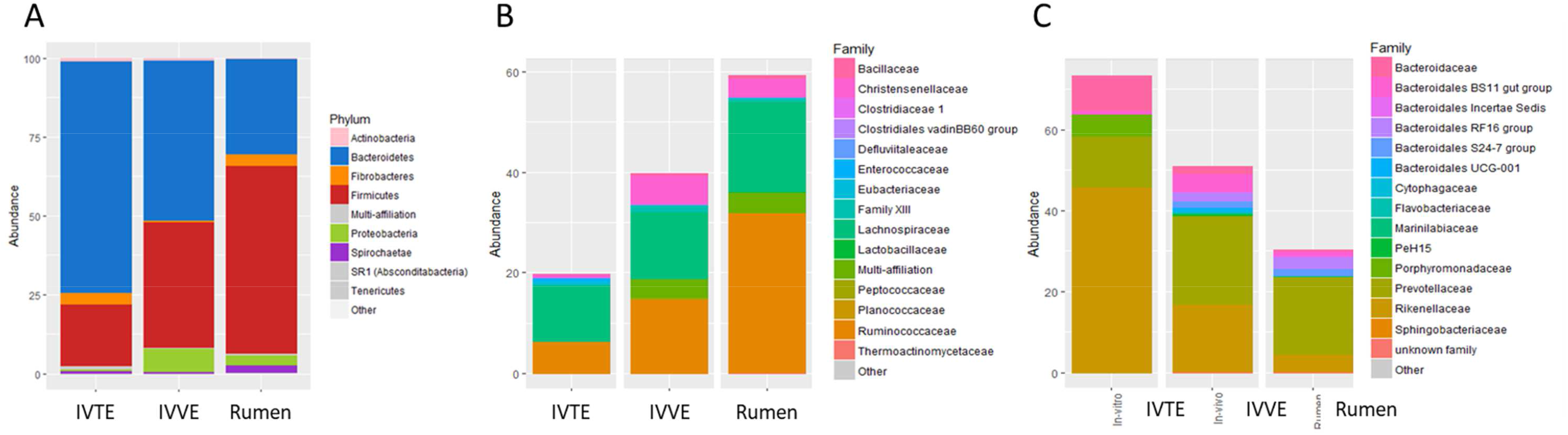
Taxonomic composition in the *in vitro* enriched (IVTE), *in vivo* enriched (IVVE) and initial rumen (Rumen) microbial communities. Relative abundance distribution of A) all phyla, B) families within Firmicutes phylum, and C) families within the Bacteroidetes phylum.

### 2.2 Primary high-throughput screening of the metagenomic libraries

The IVVE metagenomic library consisted of 19,968 *Escherichia coli* fosmid clones, each comprising a 30-40 kb DNA insert. The IVTE library consisted of 20,352 *Escherichia coli* fosmid clones. Each library thus covers around 700 Mbp of metagenomic DNA, the equivalent of more than one hundred bacterial genomes. Both libraries were screened for the degradation of PCW polysaccharide-degrading enzymes, esterases, and oxidoreductases. After a primary screening, a total of 168 clones were isolated from both libraries, corresponding to 207 hits, some of them being active on more than one substrate (Table 1). One hundred and twenty-nine hits were obtained from the IVTE library, while only 78 were obtained from the IVVE library. In addition, the distribution of enzymes’ activities retrieved from the two libraries was different, too.

**Table 1:** Number of hits obtained from the primary screening of the IVVE and IVTE metagenomic libraries.

| Activity | Substrate | Hit number<br>IVVE | Hit number<br>IVTE |
| --- | --- | --- | --- |
| esterase | Tween 20 | 26 (0.13%) | 3 (0.01%) |
| endo-1,4-β-D-xylanase | AZCL-xylan<br>(birchwood) | 30 (0.15%) | 4 (0.02%) |
| endo-1,4-β-D-mannanase | AZO-Carob<br>galactomannan | 9 (0.05%) | 47 (0.20%) |
| endo-(1,4)-(1,3)-β-D-glucanase | AZCL-Barley<br>β-glucan | 10 (0.05%) | 26 (0.12%) |
| endo-1,4-β-D-glucanase<br>(cellulase) | AZO-CM-cellulose | 0 (0%) | 44 (0.22%) |
| oxidoreductase | Lignin alkali | 3 (0.02%) | 3 (0.01%) |
|  | ABTS | 0 (0%) | 2 (0.01%) |
| Total hit number |  | 78 | 129 |

Hemicellulases were screened using AZCL-xylan and AZO-Carob Galactomannan chromogenic substrates. From the IVVE library, 30 hits (yield 0.15%) were active on the first, while 9 hits (yield 0.05%) were found on the latter. Conversely, for the IVTE library, 4 hits (yield 0.02%) and 47 hits (yield 0.20%), respectively, were found on these substrates. β-glucanase activities, including (1,4)-β-D-glucanase (cellulase) and 1,4-1,3-β-D-glucanase (lichenase) activities, were screened using the AZO-Carboxy Methyl-Cellulose (Azo-CMC) and AZCL-Barley β-glucan substrates. Here again, the primary screening results differed between the two libraries. From the IVVE library, 10 hits (yield 0.05%) were active on AZCL-Barley β-glucan, while none was found on Azo-CMC. For the IVTE library, 26 hits (yield 0.12%) and 44 hits (yield 0.22%), respectively, were found on these substrates. Eight of the hit clones were positive on both substrates.

In addition, the rumen libraries were screened for esterase activities, since PCW components harbor acetyl substituents that impede glycosidase access to the polymer backbone. Ester linkages are also involved in covalent binding between lignin and carbohydrates (19–21). Esterases are therefore key enzymes for breaking down PCW components and they were screened using Tween 20 (polyethylene glycol sorbitan monolaurate, an ester with C12 chain length), a substrate known to be easily degraded by esterases, including carbohydrate-esterases (22). Following this primary screening, 26 hits (yield 0.13%) were isolated from the IVVE library, while only 3 (yield 0.01%) were found for the IVTE library.

Finally, screening for bacterial oxidoreductases, which may be involved in ruminal lignin degradation, was also performed on both libraries using alkaline lignin (AL), a degradation product of lignin, and ABTS, a chromogenic substrate usually used for characterizing oxidases acting on polyaromatics, especially laccases. From the IVVE library, only 3 clones (0.02%) were found to be active on AL, and none reacted on ABTS. For the IVTE bioreactor enrichment, 3 hits (0.01%) were found on AL, and 2 (0.01%) on ABTS. None of these hit clones was found to be active with both substrates.

### 2.3 Discrimination screening

To refine the specificity profiling of the CAZymes responsible for the activities identified by the primary screening and to assess more largely the functional potential of each positive hits, the 168 hit clones were further tested on other chromogenic hemicellulosic (AZCL-arabinoxylan, AZCL-arabinan, AZCL-xyloglucan) and cellulosic substrates (AZCL-hydroxy-ethyl-cellulose (AZCL-HEC), AZO-α cellulose and AZO-Avicel).

Of the 168 hit clones from the primary screening, 76 were active on more than one polysaccharidic substrate (Table 2 and Figure 2). In particular, 32 of the 34 clones active on AZCL-xylan were also active on AZCL-arabinoxylan. In contrast, one clone active on AZO-Carob galactomannan and one active on AZCL-Barley β-glucan were found to be active on AZCL-arabinan.

**Table 2:** Number of hits obtained from secondary screening of the hit clones.

|  | Activity | Substrate | Hit<br>number<br>IVVE | Hit<br>number<br>IVTE |
| --- | --- | --- | --- | --- |
| esterases | esterase | *para*-nitrophenyl<br>acetate | 16 | 3 |
|  | esterase | *para*-nitrophenyl<br>butyrate | 20 | 3 |
|  | lipase | *para*-nitrophenyl<br>palmytate | 3 | 3 |
| polysaccharidases | endo-1,4-β-D-arabino-xylanase | AZCL-arabinoxylan<br>(wheat) | 30 | 16 |
| | endo-1,5-α-L-arabinanase | AZCL-debranched<br>arabinan | 1 | 1 |
|  | endo-cellulase | AZCL-xyloglucan<br>(tamarind) | 2 | 14 |
|  | endo-cellulase | AZCL-HE Cellulose | 3 | 13 |
|  | endo-cellulase | AZO-Avicel | 0 | 0 |
| | endo-cellulase | AZO-α cellulose | 0 | 0 |
| oxidoreductases | oxidoreductase | ABTS | 3 | 5 |

**Figure 2.**
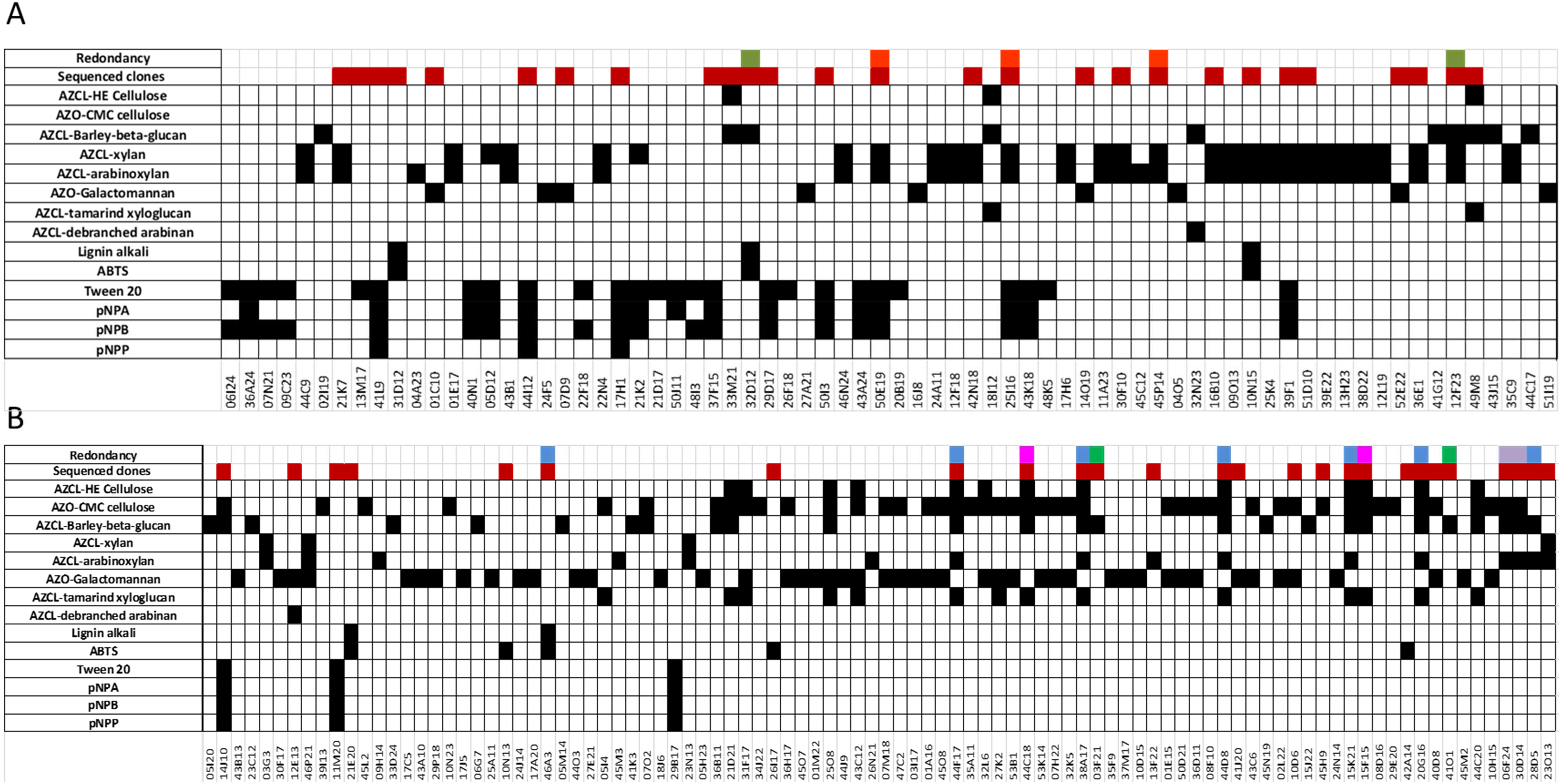
Hit clone activities on substrates of the primary and secondary screenings in the IVVE (A) and IVTE (B) libraries. The sequenced clones are indicated in red and the redundant clones share the same colors.

Regarding β-glucanase activities, most of those selected on Azo-CMC were also positive on AZCL-HEC, another chemically modified amorphous, soluble cellulose. Unfortunately, no active clone was found on the chromogenic substrates representing the most resistant fractions of cellulose, namely AZO-α cellulose (the insoluble amorphous alkali-resistant cellulose fraction, with a very high degree of polymerization) and AZO-Avicel (a highly crystalline cellulose fraction). The amorphous parts of cellulose are indeed the easiest to degrade, and this fraction was probably the first to be degraded by the IVTE consortium. In contrast, α-crystalline and crystalline parts of cellulose are known to be highly resistant to most cellulases (23–25).

In addition, 14 of the 44 clones obtained from the IVTE library on CMC were also active on AZCL-xyloglucan, a hemicellulosic component. At this stage, one might think that most of the cellulases retrieved here thus present great flexibility to amorphous substrates (CMC, HEC, xyloglucan, β-(1,3)-(1,4)-glucans). The dual specificity for xyloglucanases/cellulases is a trait that has already been observed for many enzymes, like those belonging to GH5, GH8 and GH44 families (26–28). Nevertheless, the ability of these clones to degrade both cellulosic substrates and xyloglucan might also result from the expression of various GH-encoding genes present on the same metagenomic DNA insert. This is further analyzed in the section below presenting the results of the sequence analysis.

All the primary screening hits found to be positive on Tween 20 were also tested on *para*-nitrophenyl acetate (*p*NPA), *para*-nitrophenyl butyrate (*p*NPB), and *para*-nitrophenyl palmitate (*p*NPP). Activity levels were low (between 0.4 and 2x10^−4^ U/mL of extract), since in fosmids, gene expression is not controlled by strong promoters, but rather, by sequences randomly scattered across the metagenomic DNA insert, which are recognized as promoters by *E. coli* (29). Although activity levels were low, they allowed us to distinguish clones depending on their substrate specificity. Twenty-four clones were active only on *p*NPA and/or *p*NPB, confirming the presence of esterases. Only six clones were active on *p*NPP, a lipase-specific substrate acting on long fatty acid chains, concurrently or otherwise with *p*NPA and/or *p*NPB. These enzymes may be involved in microbial lipolysis mechanisms in the cow rumen, which are still not well understood.

Regarding clones mined by the oxidoreductase screening, concentrated enzymatic extracts were tested on the chromogenic polycyclic compound, ABTS, to further discriminate activities. Unsurprisingly, measured activities were again low (between 0.1 and 0.8 U/L of culture). However, the activity displayed by these eight clones was at least three times greater than for the negative control (an *E. coli* clone containing an empty fosmid), which is itself able to break down ABTS slightly, thanks to native *E. coli* oxidoreductase activities.

### 2.4 Sequencing and data analysis

Based on their multiple or complementary activities, we selected 27 IVVE and 26 IVTE clones for sequencing of their metagenomic DNA inserts (Figure 2, Table S1). As in the original population of hits, xylanase activity was predominant for the sequenced clones from the IVVE library, whereas glucanase and mannanase activities predominated for the clones from the IVTE library (Figure 2).

A total of 2 Mb of metagenomic DNA was sequenced, with an average coverage of 40 times. No sequence redundancy was observed between IVVE and IVTE clones, while several cases of partial sequence redundancy were observed between clones from the same library. In the IVTE library, four loci, between 9 and 35 kb, were indeed found to be redundant in 13 clones (Figure 2, Table S1): one locus in the two clones 15F15 and 44C18; one locus in the seven clones 15K21, 20G16, 28D5, 38A17, 44D8, 44F17 and 46A3; one locus in the two clones 3F21 and 41O1; and one locus in the two clones 6F24 and 30D14. In the IVVE library, two clones (12F23 and 32D12) share highly similar sequences (> 98% identity and > 70% contig coverage), and three other clones (50E19, 45P14 and 25I16) are partially redundant. The very high redundancy of the clones in the IVTE library might be due to lower microbial diversity in the IVTE sample than in the IVVE sample as shown previously by the Shannon index.

#### 2.4.1 Taxonomic annotation

Firstly, the BLASTN analysis against the non-redundant database of the NCBI revealed that two IVVE clones (50E19 and 45P14) present high sequence identity (> 80 % identity, > 50% coverage) with metagenomic contigs obtained from ruminal samples derived from Chinese Holstein cattle (30), and five more (16B10, 25I16, 42N18, 45P14 and 50E19) from Jersey cows (16). These sequences, which belong to xylanase-active clones, were assigned to the Firmicutes phylum.

In addition, very high sequence identity (> 99% identity, > 90% coverage) was also found between IVTE contigs and genomic loci of the human gut bacterium *Enterococcus faecium* T110 (clone 45H9), or human gut metagenomic sequences (clone 10D6, corresponding to the sequence GU942953.1, of a metagenomic clone active on AZCL-β-glucan and yeast β-glucan) assigned to the *Bacteroides* genus (31). These results indicate that our activity-based functional metagenomics strategy allowed us to discover major fibrolytic pathways that are common to different gut microbiomes.

Furthermore, we compared our sequences to MAGs (metagenome-assembled genomes) of the bovine rumen obtained from cows from the same herd than rumen content donors (32). In the IVVE library, six clones presented similarities to three MAGs (clones 25I16, 45P14 and 50E19 to MGS205, clones 7D9 and 14O19 to MGS147, and 52E22 to MGS052). In the IVTE library, because of the high sequence redundancy among the hit clones, 11 clones (23O13, 10N13, 15K21, 20G16, 28D5, 38A17, 40D8, 44D8, 44F17, 46A3 and 21E20) match a single MAG (MGS049). The three MAGs matching clones from the IVVE library are annotated as uncultured Clostridiales. According to Li *et al*. (2020) (32), MGS147 and MGS052 are similar to MAGs from the Scottish cattle dataset of Stewart *et al*. (2019) (33), while MGS205 is specific to the dataset of Li *et al*. (2020) (32). MGS049, to which the 11 clones from the IVTE library match, is similar to the genome of a Bacteroidales bacterium from the Hungate 1000 genomes collection (34).

Finally, MEGAN sequence analysis revealed that the most probable common ancestors of the sequences in the IVVE library are Firmicutes (16 of the 25 clones in the IVVE library, Table S1). Sequences were too distantly related to those of sequenced genomes to annotate them below the phylum level, except for two clones (41L9 and 44I12) which were assigned to Clostridiales and a further two (1C10 and 31D12) to *Faecalibacterium prausnitzii*, a species which is fairly well represented in the ruminal microbiota (32,35). Regarding the other clones in this library, the assignments matched bacteria from environmental samples, with no further taxonomic information, apart from their similarity to genome sequences from the bovine ruminal metagenome (16). The assignment of the 26 IVTE clones showed that most of the clones (18 out of 26) were assigned to the Bacteroidetes phylum which was, indeed, the most abundant phylum in the *in vitro* enriched microbiota. In this population, it was possible to assign the sequences more precisely. Some clones were indeed assigned to the *Prevotella* (6F24 and 30D14) or *Bacteroides* genus (15K21 and 12E13) and even at the species rank to *Bacteroides graminisolvens* (13F22 and 41J20) and *Bacteroides uniformis* (10D6). Only five clones (11M20,14J10,15F15, 44C18 and 45H9) were assigned to the Firmicutes phylum, of which two were assigned to the Lachnospiraceae family. Clone 21E20 with an oxidase activity was assigned at species level to *Paenirhodobacter enshiensis* from the Proteobacteria phylum.

#### 2.4.2 Functional annotation

All clones harbor at least one ORF coding for an enzyme that is likely responsible for the detected activities, i.e., glycoside hydrolases (GHs), carbohydrate esterases (CEs) or oxidoreductases (Table S1). The genes coding for GHs are mainly organized into functional clusters, surrounded by other genes having complementary and accessory functions for plant cell wall harvesting, degradation and metabolization: complementary GHs, CEs, carbohydrate transporters, isomerases, kinases, and epimerases. In addition, these contigs also contain several sortases and signal peptidases, related to secretion and addressing of membrane-bound proteins involved in the degradation of extracellular substrates like polysaccharides.

In total, we found 153 non-redundant CAZyme-encoding modules (several proteins being multimodular), including 111 GHs, 26 carbohydrate-binding modules (CBM), 9 carbohydrate esterases (CE) and 8 glycosyltransferases (GT) (Table S1). These sequences were not reported before among biochemically characterized CAZymes, except the GH11 identified in clone 23O13 (ORF31, from nucleotides 33261 to 34256), which is 100 % identical to the xylanase C from *Fibrobacter succinogenes subsp. succinogenes* S85 (accession number ACX76028.1). However, the 23O13 sequence was not assigned to any particular phylum or genus, since 48 % of its nucleotide sequence (from nucleotides 17326 to 34323) shares more that 90 % identity in seven different stretches with a part of the genome of *Fibrobacter succinogenes subsp. succinogenes* S85, while 27 % of the sequence (from nucleotides 3 to 9778) shares 98% identity with a part of the genome of Bacteroidales bacterium WCE2008 : Ga0070252_15. The 23O13 sequence is thus probably issued from an unknown bacterium which exchanged genes with both Fibrobacter and Bacteroidales by horizontal gene transfers. Among the 147 non-redundant CAZyme-encoding sequences, 52 present less than 50% identity with characterized enzymes (Table S1). Most of the GHs belong to families known to contain enzymes active on the substrates of the primary screening. They are classified in the GH5 (subfamilies 2, 4, 7 and 37), GH10, GH11, GH16, GH26, and GH55 families. The remaining GH modules correspond to unscreened functions which in several cases are complementary to the screened functions for PCW degradation.

We found 68 GH modules (including 2 redundant sequences) in the IVVE library sequences, of which GH10 is the most prominent (15 contigs). The GH10 family containing many xylanases, we hypothesize that the GH10 sequences that we identified may be involved in the predominant 1,4-β-D-xylanase activity in the IVVE microbiome screening (18). The GH10 modules were often linked to a CBM22, reported to be associated with xylanases (36). Families GH5_2/ GH5_4 (6 contigs) and GH26 (4 contigs) are less frequent in the IVVE dataset. Although these families are multifunctional, they contain β-glucanases and 1,4-β-D-mannanases, respectively. The GH5_2/ GH5_4 and GH26 sequences of the IVVE clones may thus be responsible for the screened β-glucanase and 1,4-β-D-mannanase activities.

In the IVTE library, we found 72 GH modules (including 27 redundant sequences); those belonging to the GH26 (11 contigs) and GH5_2/GH5_4/GH5_7/GH5_25/GH5_37 (22 contigs) families were the most frequent. As for the IVVE clones, these GH encoding sequences are probably involved in the prominent 1,4-β-D-mannanase and the β-1,3/1,4-glucanase activities in the IVTE microbiome screening (18). In a few clones from the library, β-1,3/1,4-glucanase activity may also be attributed to the GH16 family (clones 3F21, 40D8 and 41O1). In clone 23O13, the β-xylanase activity may be attributed to a protein annotated as GH11 and/or GH5_4.

Finally, one clone displaying esterase activity (50I3 from IVVE) possesses two genes encoding CEs, including a sequence classified in the CE1 family, which contains acetylxylan-esterases. This gene belongs to a multigenic cluster likely dedicated to xylan degradation. The remaining clones with esterase activity harbor putative esterases, not necessarily targeting carbohydrate esters. The eight oxidase clones contain diverse genes coding for oxidoreductases, dehydrogenases, methyltransferases, and acyltransferases known to be involved in aromatics degradation.

#### 2.4.3 Sequence prevalence and abundance in the bovine rumen microbiome

After identifying the presence of our metagenomic sequences in the bovine rumen gene catalog, we looked for their prevalence and abundance in the independent dataset of 77 cattle from two different genetic stocks (32) (Table S1). Two contigs from the IVVE library (13M17 and 31D12) and five from the IVTE library (42A4, 3F21, 10D6, 13F22 and 41O1) have no homologous genes in the rumen gene catalog. No more than 41% of the protein sequences in the IVVE library and 34% of those in the IVTE library have a homolog in the rumen gene catalog. Furthermore, the number of occurrences was reduced to 14% for the IVVE library and 21% for the IVTE in the datasets obtained from 77 animals.

Of the GH genes likely involved in the screened activities (GH5, GH10, GH11, GH16, and GH26), 67.8% were found in the rumen gene catalog and only 39% in the 77-cattle database, the most prevalent gene being a GH10 from an unknown bacterium present in 49 metagenomic samples out of 77 (table S1). A GH26 assigned to *Faecalibacterium prauznitzii* was found in 41 samples, and the GH5_7/4 and GH16 assigned to Bacteroides were found in 41, 35 and 40 samples, respectively. The remaining target genes (including the GH11 identified in clone 23O13, which is identical to the xylanase C from *Fibrobacter succinogenes subsp. succinogenes* S85, a common rumen species) are mostly present in the catalog but absent or poorly represented in the 77-cattle database. Curiously, the genes we assigned to well characterized species known to thrive in plant glycan-enriched environments like *Bacteroides uniformis, Bacteroides graminisolvens* and *Prevotella* (37,38) are absent in the catalog. This is in keeping with the absence of *B. uniformis* and *B. graminisolvens* in the list of the most abundant strains in the rumen reported by Li *et al*. (32), while the absence in the catalog of the GH5_4 that we assigned to *Prevotella*, the most abundant genus in the rumen (39.2 %), is unexpected.

## 3. Discussion

We constructed two libraries of metagenomic clones from two bovine ruminal microbiomes obtained after *in vivo* and *in vitro* enrichment on wheat straw. Libraries were screened for esterase, oxidase and glycoside-hydrolase activities in order to identify enzymes involved in PCW degradation.

Regarding oxidase activity screening, ABTS-based assays were previously set up in liquid medium, allowing the detection of activity by either extracellular fungal oxidases (39) or intracellular recombinant enzymes expressed in *E. coli* (40). However, to our knowledge, no screen has previously been described for oxidases on a solid medium using this specific substrate. Here, we developed a high-throughput method to screen oxidase activity on solid plates. We transferred the recombinant clones from the culture medium to the screening medium, which is lethal to *E. coli* cells because of the presence of high amounts of Cu and Mn, using a PVDF membrane. Like other solid medium assays, this method means enzyme libraries can be screened at a throughput of 400,000 assays per week, using a simple colony picker system. The low number of oxidase hits is in accordance with the few data available in the literature regarding bacterial rumen oxidases acting on cyclic compounds, which are likely not abundant in this anaerobic ecosystem (41). It is difficult to know what role is played by these oxidoreductases in plant cell wall degradation by the rumen microbiota. Nevertheless, it is possible that the presence, abundance and functional importance of bacterial ligninases in the rumen may be overestimated in studies like those of Ausec *et al*. (42) and Strachan *et al*. (43), as suggested by the absence of lignin degradation by the rumen-derived consortium enriched on wheat straw by Lazuka *et al*. (18). In addition, the anaerobic conditions of plant biomass degradation by the IVVE and IVTE consortia do not favor oxidative reactions, although not all oxidoreductases need oxygen (41), even if some digestive reactions may partially occur in aerobic conditions when ruminal content is regurgitated during rumination.

Regarding glycoside-hydrolase hits, the proportions obtained in the present study are in the same range as those obtained from the screening of the human-colon-metagenomic library for endo-β-glucanase (0.06%) and xylanase (0.02 %) activities, except for the yield of xylanase-producing clones, which was 7.5 times higher in the IVVE library than in the human colon-derived library (31). By comparing the results obtained from the IVTE library with those from the IVVE library, a much lower yield of hit clones active on AZCL-xylan was observed, along with a higher hit yield on AZCL-galactomannan, AZCL-β-glucan and Azo-CMC. These findings raise a number of questions regarding the low number of cellulase hits from the IVVE library and the low number of esterase and xylanases hits from the IVTE library. To examine the enrichment efficiency, we compared our results to data from Hess *et al*. (15), who counted all CAZyme-encoding genes in 268 Gb of rumen microbiota DNA sequences. We expressed our results and those of Hess et al. as the number of genes of given CAZy families (clustered by families containing cellulases, xylanases and mannanases, as referenced in Hess *et al*. (44)) per 100 Mb (Table 3). Of course, we analyzed only one sample of each enrichment conditions, and the number of genes identified in the present study is highly biased by metagenomic library construction, heterologous gene expression in *E. coli*, screening stringency and hit clone selection for sequencing. However, this comparison shows that *in vivo* enrichment on wheat straw followed by library construction and functional screening did not increase the yield in cellulase-encoding genes, while the number of genes encoding xylanases and mannanases increased by a factor of 8. On the contrary, *in vitro* enrichment and library construction increased the yield of cellulase-encoding genes by a factor of 9, and mannanase-encoding genes by a factor of 54.

**Table 3:** Comparison of the occurrence of genes encoding CAZymes involved in PCW degradation in rumen metagenomes.

| Functions of CAZymes, according to the clustering of Hess *et al.*, 2011 (doi.org/10.1126/science.1200387) | CAZy number/ 100 MBp in Hess *et al.*, 2011 (doi.org/10.1126/science.1200387) | Hit number/ 100 MBp IVVE in this study | Hit number/ 100 MBp IVTE in this study |
| --- | --- | --- | --- |
| Cellulases<br>GH5/6/7/9/44/45/48 | 1 | 1.14 | 9 |
| Xylanases<br>GH8/10/11 | 0.57 | 4.35 | 0.57 |
| Mannanases<br>GH26 | 0.14 | 1.13 | 7.5 |

The duration and the rate of substrate degradation *in vivo* and *in vitro* might explain these differences. In the bovine rumen, the transit time is indeed between 20 and 48h, while each step of the *in vitro* enrichment on wheat straw lasted 7 days, repeated for 10 cycles needed to stabilize the ecosystem (18). Previous studies on the diversity of carbohydrate degradation machinery in the rumen state that hydrolysis of PCW appears to be a dynamic process, involving a succession of specialized microbial communities responsible for a sequential attack on polysaccharides (45,46,47). In the rumen, the PCW components that are degraded first are probably the readily accessible hemicellulose side chains, principally xylan, thus freeing up access to the polymer backbone of the most soluble hemicelluloses. The crystalline parts of cellulose are then degraded, if fermentation and hydrolysis duration allow, which would be the case in the fermenter enrichment process but not in the rumen enrichment.

Our results partially corroborate the results of Lazuka *et al*. (18), who observed that around 60% of the initial cellulose and hemicelluloses were metabolized at the end of *in vitro* enrichment on wheat straw. The IVTE consortium is thus particularly efficient at breaking down hemicelluloses and cellulose; this is consistent with the high rate of hit clones we obtained from the IVTE library on Azo-CMC, AZCL-Barley β-glucan and AZO-Carob Galactomannan. Regarding xylan degradation, Lazuka *et al*. (18) observed an increase of xylanase and/or xylosidase activity during enrichment, which however did not reach the level of activity in the *in vivo* enriched microbiome. In the present study, the xylanase hit rate was 7.5 times lower in the IVTE library than in the IVVE one. The Firmicutes-producing xylanases evidenced in the IVVE library were thus probably not selected during enrichment in the fermenter. During *in vitro* enrichment, Bacteroidetes indeed significantly increased at the expense of the Firmicutes phylum (Figure 1). In accordance with changes in microbial diversity, the majority of the hit clones in the IVTE library come from Bacteroidetes, suggesting that these are the best adapted for hydrolysis of cellulose and mannans under our experimental conditions. However, we cannot exclude the inability of certain microorganisms to survive in bioreactors, as mentioned by Lazuka *et al*. (18), who showed a sharp decrease in microbial diversity during transition between the natural environment of the rumen and the artificial conditions of the bioreactor. Regarding the *in vivo* enrichment, the absence of Bacteroidetes in our selection from the IVVE clone hits was unexpected since i) Bacteroidetes account for around 50% of the IVVE microbiota, ii) they are known for their involvement in plant cell wall polysaccharide degradation, especially hemicelluloses, and iii) their genes are well expressed in *E. coli*, as shown by the high number of Bacteroidetes clones in the IVTE library, and in other functional metagenomics studies (31). We thus assume that we missed Bacteroidetes hemicellulase sequences in our study, since we randomly sequenced only 11 clones among the 30 ones that were active on xylan.

## 4. Conclusions

In the present work, activity-based metagenomics was used to explore the functional potential of bacterial consortia derived from the bovine ruminal ecosystem enriched *in vivo* and *in vitro* on wheat straw. A multi-step high-throughput screening strategy was developed to identify metagenomic clones that are able to break down polysaccharidic and polyaromatic PCW components. This generic approach, which is applicable to any natural or artificial microbiome, allowed us to assign metagenomic sequences to particular taxons highlighting novel enzymes involved in PCW degradation derived from uncultivated ruminal bacteria of low prevalence. Most of these sequences have not been previously identified by sequencing cultured rumen bacteria or bovine rumen metagenomes, despite the latter being sequenced in great depth. The structure-function relationships of the most original glycoside-hydrolases, carbohydrate esterases and oxidases encoded by these metagenomic loci will be further investigated in order to unravel their specific role in PCW catabolism.

## 5. Methods

### 5.1 Chemicals

Tween 20, Pefabloc, *para*-nitrophenyl acetate (*p*NPA), *para*-nitrophenyl butyrate (*p*NPB), *para*-nitrophenyl palmytate (*p*NPP), *para*-nitrophenol (*p*NP), 2,2′-Azino-bis(3-ethylbenzothiazoline-6-sulfonic acid) diammonium salt (ABTS), and alkaline lignin were purchased from Sigma Aldrich (France). AZCL-xylan (birchwood), AZCL-xyloglucan (tamarind), AZCL-HE Cellulose, AZCL-arabinoxylan (wheat), AZCL-debranched arabinan, AZCL-Barley β-glucan, AZO-CM Cellulose, AZO-Avicel, AZO-α cellulose, AZO-Carob Galactomannan were purchased from Megazyme (Ireland).

### 5.2 Metagenomic DNA sampling and library construction

Two non-producing Holstein dairy cows were used in this study. Prior to enrichment, the cows were fed a standard dairy cow ration composed of corn silage (64% DM), hay (6% DM) and concentrate (30% DM). They were fed *ad libitum* once a day in the morning. For seven weeks, both cows received a diet of 80% wheat straw, 12% concentrate and 8% beetroot molasses before sampling of their rumen content. The rumen content was collected from the cows via a rumen cannula. Samples were taken from various parts of the rumen and manually homogenized. Subsamples (30g) were immediately frozen in liquid nitrogen prior to storage at −80 °C. To construct the IVTE library, the two cows were sampled before being subjected to the wheat straw-rich diet. This sample was used to inoculate the bioreactors fed only with wheat straw, as described by Lazuka *et al*. (18). For IVVE, the same two cows were fed for seven weeks with wheat straw-enriched food prior to sampling and library construction.

Two metagenomic libraries were constructed in the *Escherichia coli* strain from rumen bacterial DNAs extracted from the bioreactor (IVTE library) or directly from the rumen bacterial DNA of the cows, after the diet (IVVE library). Briefly, total metagenomic DNA was extracted from homogenized samples containing both solid and liquid fractions, and fragmented. Fragments sized between 30 and 40 kb were isolated as previously described (Tasse et al., 2010) and cloned into pEPIFOS-5 fosmids (Epicentre Technologies). EPI100 *E. coli* cells were then transfected to obtain a library of 20,352 and 19,968 clones from the rumen samples from the bioreactor and rumen enrichment respectively. Recombinant clones were transferred to 53 or 52 384-well microtiter plates containing Luria Bertani (LB) medium, supplemented with 12.5 mg/L chloramphenicol and 8% glycerol. After 22 hours of growth at 37°C without shaking, the plates were stored for a long duration at -80°C.

### 5.3 Microbial diversity analysis

The microbial diversity of rumen, IVVE and IVTE samples was analysed by genomic DNA extraction followed by sequencing of a 460bp fragment of the V3-V4 variable region of the 16S rRNA gene, using the procedures described by Auer *et al*. (47). Sequencing was performed by the GenoToul Genomics and Transcriptomics facility (Auzeville, France) using MiSeq Illumina V.3 reagents according to the manufacturer’s instructions. Joined-pair reads (76,000 reads on average per sample, minimum 36,794) were pre-processed and filtered for chimera removal using the Frogs pipeline (48). Operational Taxonomic Units (OTUs) were defined on a random subsample of 35,000 high-quality sequences using Swarm implemented in the Frogs pipeline; OTUs displaying fewer than 10 sequences across the dataset were removed. Taxonomic classification was based on Silva 128.

### 5.4 Primary high-throughput screening of the metagenomic libraries

The metagenomic libraries were gridded on 22 cm x 22 cm trays containing solid agar medium, using an automated microplate gridder (K2, KBiosystem, Basildon, UK). Three major activities were screened.

Esterase/lipase screening medium was formed of solid LB medium supplemented with 12.5 mg/L chloramphenicol and 1% (w/v) final concentration of Tween 20. The assay plates were incubated for 3 days at 37°C until a powdery halo appeared, indicating the presence of a positive clone.

The screening media for polysaccharide degrading activity were LB media supplemented with 12.5 mg/L chloramphenicol and with chromogenic substrates (AZCL-xylan, AZCL-Barley β-glucan, AZO-CM Cellulose, AZO-Carob Galactomannan) at a final concentration of 0.2% (w/v). After incubation of the plates at 37°C, for plates containing AZCL-substrates, positive clones were visually detected after 1 to 10 days of incubation by the presence of a blue halo resulting from the production of colored oligosaccharides from which the color diffused around the bacterial colonies. For AZO-polysaccharides, clear halos were observed around the positive clones.

Oxidoreductase activity screening took place on two different substrates. The first medium was a minimal medium supplemented with 12.5 mg/L chloramphenicol, 15 g/L agar and 1% (w/v) filter-sterilized alkaline lignin as the sole carbon source (other than the agar itself). The minimal medium was composed of a final concentration of salts (3.6 g/L Na_2_HPO_4_, H_2_O; 0.62 g/L KH_2_PO_4_; 0.11 g/L NaCl; 0.42 g/L NH_4_Cl), 2 mM MgSO_4_, 0.03 mM CaCl2, other salts (15 mg/L Na_2_EDTA, 2H_2_O; 4.5 mg/L ZnSO_4_, 7H_2_O; 0.3 mg/L CoCl_2_, 6H_2_O; 1 mg/L MnCl_2_, 4H_2_O; 1 mg/L H_3_BO_3_; 0.4 mg/L Na_2_MoO_4_, 2H_2_O; 3 mg/L FeSO_4_, 7H_2_O; 0.3 mg/L CuSO_4_, 5H_2_O), 0.04 g/L leucine and 0.1 g/L thiamine hypochloride. The assay plates were incubated at 37°C until the growth of hit clones was visualized.

The other screening method used ABTS as the substrate. An initial layer was composed of solid LB medium with 12.5 mg/L chloramphenicol, on which was deposited a 0.22 µM Durapore PVDF membrane (Millipore) that had previously been autoclaved. The clones were gridded and grown on the membrane. After growth, the membrane was moved to another plate, containing 15 g/L agar, 12.5 mg/L chloramphenicol, 0.1M sodium citrate pH 4.5, 1 mM CuSO_4_ 5H_2_O, 1 mM MnCl_2_ 4H_2_O and 1mM ABTS. Positive hits turned green or grey, sometimes with the presence of halos.

### 5.4 Discrimination screening

The hit clones obtained after primary screening for esterase activity were grown in microplates containing 200µL of LB medium for pre-culture, overnight at 37°C. These pre-cultures were used to inoculate deep well microplates containing 1.6 mL of LB liquid medium supplemented with 12.5 mg/L chloramphenicol, which were subsequently incubated overnight at 37°C. Cells were pelleted by centrifugation for 30 minutes at 4°C and 1,760 x g. Each pellet was re-suspended in 250 µL of lysis buffer (100 mM HEPES pH 7.5, 1 mM Pefabloc and 666.7 µl/L Lysonase) and incubated for one hour at 37°C and 600 rpm (shaking throw 25 mm) in a Multitron Pro shaker (INFORS HT) followed by a freeze/unfreeze cycle at -80°C in order to perform cell lysis. Cellular extracts were isolated by centrifugation for 30 minutes at 4°C and 1,760 x g, and stored at 4°C for less than 24 h before use.

Esterase/lipase activity of the cytoplasmic extracts was estimated by monitoring the hydrolysis of *p*NP esters into the corresponding acid and *p*-nitrophenol. Assays were performed in 96-well microtiter plates on *p*NPA, *p*NPB and *p*NPP at 1 mM final concentration. *p*NP was used for the standard curve. For *p*NPA and *p*NPB, 195 µL of cellular extract was mixed with 5µL of substrate dissolved at 40 mM in 2-methyl-2-butanol (2M2B). For *p*NPP, 175µL of cellular extract was mixed with 25µL of substrate dissolved at 8 mM in isopropanol. Enzymatic activity was determined by following the absorbance increase at 405 nm for 30 min at 30°C, in a microplate spectrophotometer (BioTek™ Eon™ Microplate Spectrophotometers, Colmar, France).

The hit clones obtained after primary screening for carbohydrate-degrading activity were further characterized using chromogenic substrates in solid medium (AZO-Avicel, AZO HE cellulose, AZO-α cellulose, AZCL arabinoxylan, AZCL-xyloglucan, AZCL debranched arabinan) at a final concentration of 0.2% (w/v). After incubation of plates at 37°C, for plates containing AZCL-substrates, positive clones were visually detected after 1 to 10 days of incubation by the presence of a blue halo resulting from the production of colored oligosaccharides from which the color diffused around the bacterial colonies. For AZO-polysaccharides, clear halos were observed around the positive clones.

The hit clones obtained after primary screening for oxidoreductase activity were grown at 37°C in 20 mL LB medium, with orbital shaking at 120 rpm. After 16 h, cells were harvested by centrifuging for 5 min at 5,000 rpm, and re-suspended in activity buffer to obtain a final OD_600nm_ of 80. Cell lysis was performed using sonication. Cell debris was centrifuged at 13,000 rpm for 10 min and the cytoplasmic extracts were filtered with a 0.20 μm Minisart RC4 syringe filter. Enzymatic reactions were carried out at 30°C, adding 5mM ABTS as a colored substrate, in 96-well microtiter plate assays. Each well contained 20 µl of concentrated cytoplasmic extract, 5 mM ABTS, 0 or 10 mM CuSO_4_, 0 or 3% H_2_O_2_ and 100 mM sodium citrate buffer pH 4.5. Enzymatic activity was determined by following the absorbance increase at 420 nm for 30 min, measured using a microplate spectrophotometer (BioTek™ Eon™ Microplate Spectrophotometers, Colmar, France). ABTS-oxidizing activity was expressed as µmol/min/L_culture_, using an ABTS extinction coefficient value of ε_ABTS, 420nm_ = 36,000 M^-1^.cm^-1^.

### 5.5 Sequencing and data analysis

Fosmid DNA of the hit clones was extracted with the NucleoBond Xtra Midi kit from Macherey-Nagel (France) following supplier recommendations. Fosmids were then sequenced with the MiSeq technology, on the Genotoul platform (http://get.genotoul.fr/). Read assembly was performed using Masurca (http://www.genome.umd.edu/masurca.html). The contigs were cleaned from the vector pCC1FOS sequence using Crossmatch (http://bozeman.mbt.washington.edu/phredphrapconsed.html). Detection of ORFs and functional annotation was performed using the RAST software (49–51).

Sequences of contigs were compared to the Metagenome Assembled Genomes (MAGs) obtained from the work of Li et al. (32) by BLASTN analysis (E-value=0; identity ≥90%).

CAZyme-encoding genes were identified by BLAST analysis of the predicted ORFs against the sequences included in the CAZy database (http://www.cazy.org) using a cut-off E-value of 7.10^−6^ followed by visual inspection and alignment with known CAZy families. CAZyme functions were predicted by BLASTP (version 2.3.0+) (52) comparison against a library of experimentally characterized sequences from the CAZy database (release 11/09/2023). The best hit was retained.

The homolog sequences of the genes identified in the contigs sequences were searched for using BLASTP, E-value=0, identity ≥90%, in a translated catalog of 13 million reference genes resulting from the sequencing of the rumen content of 5 Holstein cows and 5 Charolais bulls (32). The gene richness in bovine rumen was assessed by recovering the occurrence frequency data for homologous sequences in the catalog from the gene frequency dataset of 77 cattle from two independent genetic stocks (32).

The most likely taxonomic origin of the contigs was retrieved using MEGAN v3.2.1 (53), based on ORF BLASTX analysis against the non-redundant NCBI protein database (E-value ≤10^−8^, identity ≥90%, query length coverage ≥50%). Contigs were assigned to a phylum, order, class, genus or species only if at least 50% of their ORFs were assigned to the same level.

## Supporting information

Sequences

## Ethics approval

The cows involved in this study were reared according to the national standards set by the legislation on animal care (Certificate of Authorization to Experiment on Living Animals, No. 004495, Ministry of Agriculture, France).

## Availability of data and materials

The sequence data generated and analysed during the current study are available in the ‘European Nucleotide Archive’ (http://www.ebi.ac.uk/ena/data/view/OZ022632-OZ022684).

## Authors’ contributions

G.P.V., G.H.R., E.L., C.D., S.B. and D.M. conceptualized and supervised the study. L.U. set up the functional screening methodology, performed all screening experiments and functional annotation of sequences. C.K. and E.L. analyzed the sequences. A.L. performed the IVTE enrichment experiments and 16S sequence analyses. D.C., A.R., E. A., M.C. were involved in primary and validation screening campaigns. S.L. and E.B. performed the discrimination screening. E.D., V.L., N.T. and B.H. annotated the CAZy encoding genes. L.U. and E.L. wrote the original draft; G.P.V., G.H.R., D.M. and B.H. reviewed and edited the manuscript; G.P.V., S.B., G.H.R., C.D., C.K. and D.M. obtained the funding.

## Table legends

Table 3: Enrichment in genes encoding CAZymes involved in PCW degradation by functional metagenomics, in comparison to gene abundance in the bovine rumen metagenome, according to Hess et al. (15).

## Supplementary data

Table S1: Functional and taxonomic annotation of metagenomic sequences.

