## Supplemental Table 1 for "Functional exploration of *in vivo* and *in vitro* lignocellulose-fed rumen bacterial microbiomes reveals novel enzymes involved in polysaccharide breakdown"

| clone ID | Accession | Contig size | Taxonomic assignation MAG ID and number of base pairs of coverage | Activities | ORFs number Final | functional annotation (RAST) ORFs in orange are the ones assumed to be responsible for the activity | Start | Stop | Strand | Presence in the bovine rumen metagenome (Li et all, 2020) | Prevalence (number of individuals harbouring the gene in the 77 cattle database (Li et al., 2020)) | CAZy annotation | Redundancy of CAZy sequences | Most similar biochemically characterized proteins listen in the Cazy DB | | | |
|---|---|---|---|---|---|---|---|---|---|---|---|---|---|---|---|---|---|
| | | | | | | | | | | | | | | NCBI Accession | %ID | evalue | function description (EC) |
| IVVE | | | | | | | | | | | | | | | | | |
| | | | | | 1 | hypothetical protein | 53 | 1762 | - | | | | | | | | |
| | | | | | 2 | N-Acetyl-D-glucosamine ABC transport system, permease protein 2 | 1822 | 2826 | - | x | | | | | | | |
| | | | | | 3 | binding-protein-dependent transport systems inner membrane component | 2847 | 3773 | - | x | 3 | | | | | | |
| | | | | | 4 | hypothetical protein | 3770 | 6361 | - | x | 2 | GH129 | 14O19(5):100%cov, 99%id | BAL14929.1 | 32.31 | 0.19 | a-N-acetylgalactosaminidase (3.2.1.49) |
| | | | | | 5 | FIG01031734: hypothetical protein | 6376 | 7008 | - | x | | | | | | | |
| | | | | | 6 | NHL repeat containing protein | 7005 | 8486 | - | | | | | | | | |
| | | | | | 7 | binding-protein-dependent transport systems inner membrane component | 8498 | 9487 | - | x | | | | | | | |
| | | | | | 8 | binding-protein-dependent transport systems inner membrane component | 9484 | 10416 | - | x | 60 | | | | | | |
| | | | | | 9 | N-Acetyl-D-glucosamine ABC transport system, sugar-binding protein | 10418 | 13402 | - | x | | | | | | | |
| | | | | | 10 | Mannan endo-1,4-beta-mannosidase precursor (EC 3.2.1.78) | 13439 | 14911 | - | x | 41 | CBM35-GH26 | 14O19 (11): 100% cov, 99% id | QIZ03395.1 | 57.81 | 2E-126 | endo-b-1,4-mannanase (3.2.1.78) |
| | | | | | 11 | Polygalacturonase | 15231 | 16637 | - | x | | GH28 | 14O19 (12): 100% cov, 98% id | AAO79251.1 | 29.27 | 2E-38 | rhamnogalacturonan a-1,2-galacturonohydrolase (3.2.1.173) |
| | | | | | 12 | Putative isomerase | 16653 | 18083 | - | x | | | | | | | |
| | | | | | 13 | ABC transporter, permease protein | 18323 | 19276 | + | x | 34 | | | | | | |
| | | | | | 14 | ABC transporter, permease protein | 19293 | 20225 | + | | | | | | | | |
| | | | | | 15 | Transcriptional regulator, AraC family | 20240 | 21145 | + | x | | | | | | | |
| | | | | | 16 | ABC transporter, substrate-binding protein | 21207 | 22712 | + | x | 26 | | | | | | |
| | | | | | 17 | ABC transporter, substrate-binding protein | 22829 | 24376 | + | x | | | | | | | |
| | | | | | 18 | Transcriptional regulator, AraC family | 24380 | 26647 | + | x | | | | | | | |
| | | | | | 19 | putative glycosyl hydrolase, family 43 | 26657 | 27499 | + | x | 3 | GH43_17 | 14O19 (20): 100% cov, 99% id | ADQ05609.1 | 49.12 | 2E-87 | a-L-arabinofuranosidase (3.2.1.55) |
| | | | | | 20 | Rhamnulokinase (EC 2.7.1.5) | 27546 | 28826 | + | x | 33 | | | | | | |
| | | | | | 21 | L-rhamnose isomerase (EC 5.3.1.14) | 28823 | 30067 | + | x | 27 | | | | | | |
| | | | | | 22 | Rhamnulose-1-phosphate aldolase (EC 4.1.2.19) | 30075 | 30911 | + | x | 26 | | | | | | |
| | | | | | 23 | Alpha-galactosidase (EC 3.2.1.22) | 30908 | 32305 | + | x | | GH4_3 | 1C10 (24): 100% cov, 100% id | CAB12532.1 | 52.37 | 1E-166 | exo-polygalacturonase (3.2.1.67) |
| 1C10-IVVE | | 32,580 bp | 1 **Firmicutes** *Feacalibacterium prausnitzii* | AZO-galacto-mannan | 24 | hypothetical protein | 32406 | 32570 | - | x | | | | | | | |
| | | | | | 1 | Rubredoxin | 64 | 216 | + | | | | | | | | |
| | | | | | 2 | Homoserine kinase (EC 2.7.1.39) | 232 | 831 | + | | | | | | | | |
| | | | | | 3 | ABC transporter substrate-binding protein - sugar transport | 912 | 2279 | - | x | 31 | | | | | | |
| | | | | | 4 | hypothetical protein | 2312 | 4021 | - | x | 6 | | | | | | |
| | | | | | 5 | sugar ABC transporter permease | 4081 | 5085 | - | x | 38 | | | | | | |
| | | | | | 6 | Sugar ABC transporter permease | 5104 | 6030 | - | x | 2 | | | | | | |
| | | | | | 7 | hypothetical protein | 6027 | 8618 | - | x | 19 | GH129 | | BAL14929.1 | 23.32 | 5,00E-06 | a-N-acetylgalactosaminidase (3.2.1.49) |
| | | | | | 8 | hypothetical protein | 8633 | 9265 | - | | | | | | | | |
| | | | | | 9 | hypothetical protein | 9262 | 10743 | - | x | | | | | | | |
| | | | | | 10 | binding-protein-dependent transport systems inner membrane component | 10755 | 11735 | - | x | 4 | | | | | | |
| | | | | | 11 | binding-protein-dependent transport systems inner membrane component | 11741 | 12673 | - | x | 13 | | | | | | |
| | | | | | 12 | N-Acetyl-D-glucosamine ABC transport system, sugar-binding protein | 12675 | 15644 | - | x | | | | | | | |
| | | | | | 13 | Mannan endo-1,4-beta-mannosidase precursor (EC 3.2.1.78) | 15699 | 17072 | - | x | | CBM35-GH26 | | QIZ03395.1 | 56.56 | 2E-122 | endo-b-1,4-mannanase (3.2.1.78) |
| | | | | | 14 | hypothetical protein | 17425 | 17544 | - | | | | | | | | |
| | | | | | 15 | lipoprotein, putative | 17519 | 18970 | - | x | | GH37 | | AAC76544.1 | 23.61 | 0.037 | a,a-trehalase (3.2.1.28) |
| | | | | | 16 | ABC transporter, permease protein | 19300 | 20166 | + | x | | | | | | | |
| | | | | | 17 | ABC transporter, permease protein | 20183 | 21121 | + | x | 1 | | | | | | |
| | | | | | 18 | hypothetical protein | 21135 | 22040 | + | x | 11 | | | | | | |
| | | | | | 19 | ABC transporter, substrate-binding protein | 22092 | 23597 | + | x | | | | | | | |
| | | | | | 20 | hypothetical protein | 23714 | 25261 | + | x | 38 | | | | | | |
| | | | | | 21 | Transcriptional regulator, AraC family | 25266 | 27512 | + | x | | | | | | | |
| | | | | | 22 | putative glycosyl hydrolase, family 43 | 27537 | 28379 | + | x | 11 | GH43_17 | | ADQ05609.1 | 46.48 | 2E-81 | a-L-arabinofuranosidase (3.2.1.55) |
| | | | | | 23 | Alpha-galactosidase (EC 3.2.1.22) | 28476 | 29870 | + | x | | GH4 | | CAB12532.1 | 51.92 | 1E-164 | exo-polygalacturonase (3.2.1.67) |
| | | | | | 24 | hypothetical protein | 30093 | 30305 | + | | | | | | | | |
| | | | | | 25 | DNA mismatch repair protein [MutS] | 30364 | 32088 | - | x | | | | | | | |
| | | | | | 26 | Cys-tRNA(Pro) deacylase YbaK | 32225 | 32698 | + | | | | | | | | |
| | | | | | 27 | Exopolysaccharide biosynthesis protein related to N-acetylglucosamine-1-phosphodiester alpha-N-acetylglucosaminidase | 32790 | 33797 | + | x | | | | | | | |
| | | | | | 28 | Prolipoprotein diacylglyceryl transferase (EC 2.4.99.-) | 33846 | 34937 | + | x | | | | | | | |
| | | | | | 29 | Ferredoxin | 34940 | 36094 | + | x | 16 | | | | | | |
| | | | | | 30 | hypothetical protein | 36326 | 36781 | + | | | | | | | | |
| | | | | | 31 | Thioredoxin reductase (EC 1.8.1.9) | 36955 | 37881 | - | x | 5 | | | | | | |
| 7D9-IVVE | | 38,453 bp | 1 **Firmicutes RumenMGS147_scaffold_9 4,786 bp** | AZO-galacto-mannan | 32 | Ribonucleotide reductase of class III (anaerobic), large subunit (EC 1.17.4.2) | 38002 | 38418 | - | | | | | | | | |
| | | | | | 1 | Phenylalanyl-tRNA synthetase beta chain (EC 6.1.1.20) | 44 | 934 | + | x | | | | | | | |
| | | | | | 2 | N-Acetyl-D-glucosamine ABC transport system, sugar-binding protein | 1134 | 4019 | + | x | 24 | | | | | | |
| | | | | | 3 | binding-protein-dependent transport systems inner membrane component | 4041 | 4985 | + | x | 6 | | | | | | |
| | | | | | 4 | binding-protein-dependent transport systems inner membrane component | 4987 | 6225 | + | x | | | | | | | |
| | | | | | 5 | hypothetical protein | 6238 | 7785 | + | x | 6 | | | | | | |
| | | | | | 6 | hypothetical protein | 7751 | 8446 | + | | | | | | | | |
| | | | | | 7 | hypothetical protein | 8456 | 11038 | + | x | 17 | GH129 | | BAL14929.1 | 21.99 | 0.27 | a-N-acetylgalactosaminidase (3.2.1.49) |
| | | | | | 8 | hypothetical protein | 11035 | 12063 | + | x | 8 | | | | | | |
| | | | | | 9 | putative sugar uptake ABC transporter permease protein | 12056 | 13066 | + | x | | | | | | | |
| | | | | | 10 | hypothetical protein | 13221 | 14678 | + | x | 64 | | | | | | |
| | | | | | 11 | Endo-1,4-beta-xylanase A precursor (EC 3.2.1.8) | 14824 | 16377 | + | x | | CBM22-GH10 | | AHF24999.1 | 84.78 | 0.0 | endo-b-1,4-xylanase (3.2.1.8) |
| | | | | | 12 | Chitobiose phosphorylase (EC 2.4.1.-)//cellobiose phosphorylase | 16555 | 18942 | + | x | 25 | GH94 | | QCO92991.1 | 66.08 | 0.0 | cellodextrin phosphorylase (2.4.1.49) |
| | | | | | 13 | COG1720: Uncharacterized conserved protein | 18944 | 19612 | + | | | | | | | | |
| | | | | | 14 | Phosphoserine aminotransferase (EC 2.6.1.52) | 19773 | 20849 | + | x | 18 | | | | | | |
| | | | | | 15 | D-3-phosphoglycerate dehydrogenase (EC 1.1.1.95) | 20987 | 22162 | + | x | 1 | | | | | | |
| | | | | | 16 | DNA-directed RNA polymerase beta' subunit (EC 2.7.7.6) | 22220 | 25789 | - | x | 28 | | | | | | |
| | | | | | 17 | DNA-directed RNA polymerase beta subunit (EC 2.7.7.6) | 25810 | 29688 | - | x | 60 | | | | | | |
| | | | | | 18 | DNA internalization-related competence protein ComEC/Rec2 | 29922 | 30923 | + | x | 24 | | | | | | |
| | | | | | 19 | Firmicutes ribosomal L7Ae family protein | 31017 | 31331 | + | | | | | | | | |
| | | | | | 20 | SSU ribosomal protein S12p (S23e) | 31396 | 31770 | + | | | | | | | | |
| | | | | | 21 | SSU ribosomal protein S7p (S5e) | 31976 | 32446 | + | | | | | | | | |
| | | | | | 22 | Translation elongation factor G | 32546 | 34612 | + | x | 39 | | | | | | |
| | | | | | 23 | Translation elongation factor Tu | 34654 | 35868 | + | x | | | | | | | |
| | | | | | 24 | LSU ribosomal protein L33p | 35890 | 36051 | + | | | | | | | | |
| | | | | | 25 | Preprotein translocase subunit SecE (TC 3.A.5.1.1) | 36089 | 36370 | + | | | | | | | | |
| | | | | | 26 | Transcription antitermination protein NusG | 36386 | 36922 | + | | | | | | | | |
| | | | | | 27 | LSU ribosomal protein L11p (L12e) | 37068 | 37493 | + | | | | | | | | |
| 10N15- IVVE | LT674548 | 39,312 bp | 1 **Bacterium** | AZCL-xylan, AZCL-arabinoxylan, lignin alkali | 28 | LSU ribosomal protein L1p (L10Ae) | 37563 | 38252 | + | | | GH3 | | ABG59531.1 | 23.93 | 0.46 | b-glucosidase (3.2.1.21) |
| | | | | | 1 | Cytochrome c-type biogenesis protein Ccs1/ResB | 729 | 1757 | - | x | 6 | | | | | | |
| | | | | | 2 | Fumarate reductase flavoprotein subunit (EC 1.3.99.1) | 1800 | 4724 | - | x | | | | | | | |
| | | | | | 3 | Electron transport complex protein RnfG | 4762 | 5940 | - | x | | | | | | | |
| | | | | | 4 | Electron transport complex protein RnfB | 5953 | 6747 | - | x | 57 | | | | | | |
| | | | | | 5 | Electron transport complex protein RnfA | 6762 | 7340 | - | | | | | | | | |
| | | | | | 6 | Electron transport complex protein RnfE | 7337 | 8095 | - | | | | | | | | |
| | | | | | 7 | Electron transport complex protein RnfD | 8097 | 9638 | - | x | | | | | | | |
| | | | | | 8 | Electron transport complex protein RnfC | 9635 | 10984 | - | x | | | | | | | |
| | | | | | 9 | Sensory box/GGDEF family protein | 11020 | 12921 | - | x | | | | | | | |
| | | | | | 10 | hypothetical protein | 13193 | 14566 | + | x | | | | | | | |
| | | | | | 11 | Lysophospholipase (EC 3.1.1.5); Monoglyceride lipase (EC 3.1.1.23); putative | 14617 | 15501 | - | x | | | | | | | |
| | | | | | 12 | Malate permease | 15679 | 16608 | + | x | 14 | | | | | | |
| | | | | | 13 | conserved hypothetical protein 374 | 16611 | 17753 | - | x | | | | | | | |
| | | | | | 14 | Neopullulanase (EC 3.2.1.135) | 17814 | 19889 | - | x | | GH13_21 | 32D12 (33): 100% cov, 99% id; 38D22 (27): 100% cov, 99% id; 51D10 (3): 100% cov, 95% id | BAA02471.1 | 34.47 | 7E-80 | a-amylase (3.2.1.1) |
| | | | | | 15 | Beta-1,3-glucosyltransferase | 19925 | 20746 | - | | | GT2 | 32D12 (14): 100% cov, 99% id; 38D22 (28): 100% cov, 99% id | | | | |

| | | | | | | | | | | | | | | | | | |
|---|---|---|---|---|---|---|---|---|---|---|---|---|---|---|---|---|---|
| | | | | | 16 | Endo-1,4-beta-xylanase A precursor (EC 3.2.1.8) | 20895 | 22409 | - | x | | CBM22-GH10 | 38D22 (29): 100% cov, 99% id; 32D12 (35): 100% cov, 99% id | AHF26235.1 | 67.63 | 6E-162 | endo-b-1,4-xylanase (3.2.1.8) |
| | | | | | 17 | hypothetical protein | 22395 | 22508 | + | | | | | | | | |
| 12F23 IVVE | | 25,530 bp | 1 Bacterium | AZCL-xylan, AZCL-arabinoxylan, AZCL-Barley beta glucan | 18 | Alcohol dehydrogenase (EC 1.1.1.1) | 22612 | 25479 | - | x | | | | | | | |
| | | | | | 1 | hypothetical protein | 41 | 742 | + | | | GH50 | | QZN05586.1 | 32.35 | 0.82 | b-agarase (3.2.1.81) |
| | | | | | 2 | ATPase | 739 | 2247 | + | | | | | | | | |
| | | | | | 3 | Beta-glucosidase (EC 3.2.1.21) | 2332 | 3642 | + | | | GH1 | 41L9 (9): 100% cov, 97% id | CBL17177.1 | 46.31 | 3E-131 | b-glucosidase (3.2.1.21) |
| | | | | | 4 | hypothetical protein | 3657 | 4520 | + | | | | | | | | |
| | | | | | 5 | benzyl alcohol dehydrogenase | 4623 | 5717 | - | | | | | | | | |
| | | | | | 6 | hypothetical protein | 5959 | 6864 | - | | | | | | | | |
| | | | | | 7 | hypothetical protein | 6885 | 7265 | - | | | | | | | | |
| | | | | | 8 | hypothetical protein | 7262 | 7705 | - | | | | | | | | |
| | | | | | 9 | cobalamin synthesis protein, P47K | 7686 | 8750 | - | | | | | | | | |
| | | | | | 10 | Aromatic hydrocarbon utilization transcriptional regulator CatR (LysR family) | 9122 | 10009 | + | | | | | | | | |
| | | | | | 11 | Xanthine dehydrogenase iron-sulfur subunit (EC 1.17.1.4) / Xanthine dehydrogenase, molybdenum binding subunit (EC 1.17.1.4) | 10124 | 12949 | + | | | | | | | | |
| | | | | | 12 | Glutamate synthase [NADPH] small chain (EC 1.4.1.13) | 13025 | 15127 | + | | | | | | | | |
| | | | | | 13 | hypothetical protein | 15148 | 15324 | + | | | | | | | | |
| | | | | | 14 | 2-Hydroxy-6-Oxo-6-Phenylhexa-2,4-Dienoate hydrolase | 15396 | 16169 | - | | | | | | | | |
| | | | | | 15 | Esterase/lipase | 16191 | 17096 | - | | | | | | | | |
| | | | | | 16 | hypothetical protein | 17241 | 18578 | - | | | | | | | | |
| | | | | | 17 | hypothetical protein | 18863 | 22315 | + | | | | | | | | |
| | | | | | 18 | transcriptional regulator, LysR family | 22353 | 23258 | - | | | | | | | | |
| | | | | | 19 | hypothetical protein | 23433 | 24071 | + | | | | | | | | |
| | | | | | 20 | Deblocking aminopeptidase (EC 3.4.11.-) | 24523 | 25221 | + | | | | | | | | |
| | | | | | 21 | amidohydrolase | 25218 | 26507 | + | | | | | | | | |
| | | | | | 22 | Oligopeptide ABC transporter, periplasmic oligopeptide-binding protein OppA (TC 3.A.1.5.1) | 26461 | 28059 | + | | | | | | | | |
| | | | | | 23 | Dipeptide transport system permease protein DppB (TC 3.A.1.5.2) | 28062 | 29012 | + | | | | | | | | |
| | | | | | 24 | Dipeptide transport system permease protein DppC (TC 3.A.1.5.2) | 29005 | 29868 | + | | | | | | | | |
| 13M17 - IVVE | LT674547 | 30,963 bp | 1 Firmicutes | Tween 20 | 25 | Oligopeptide transport ATP-binding protein OppD (TC 3.A.1.5.1) | 29879 | 30898 | + | | | | | | | | |
| | | | | | 1 | hypothetical protein | 527 | 1000 | + | | | | | | | | |
| | | | | | 2 | hypothetical protein | 967 | 2676 | - | x | | | | | | | |
| | | | | | 3 | N-Acetyl-D-glucosamine ABC transport system, permease protein 2 | 2736 | 3740 | - | x | 3 | | | | | | |
| | | | | | 4 | binding-protein-dependent transport systems inner membrane component | 3761 | 4687 | - | x | 2 | | | | | | |
| | | | | | 5 | hypothetical protein | 4684 | 7275 | - | x | | GH129 | 1C10(4):100%cov, 99%id | BAL14929.1 | 21.61 | 0.22 | a-N-acetylgalactosaminidase (3.2.1.49) |
| | | | | | 6 | hypothetical protein | 7290 | 7922 | - | | | | | | | | |
| | | | | | 7 | hypothetical protein | 7919 | 9400 | - | x | 1 | | | | | | |
| | | | | | 8 | binding-protein-dependent transport systems inner membrane component | 9412 | 10401 | - | x | 60 | | | | | | |
| | | | | | 9 | binding-protein-dependent transport systems inner membrane component | 10398 | 11330 | - | x | | | | | | | |
| | | | | | 10 | N-Acetyl-D-glucosamine ABC transport system, sugar-binding protein | 11332 | 14316 | - | x | 41 | | | | | | |
| | | | | | 11 | Mannan endo-1,4-beta-mannosidase( EC:3.2.1.78 ) | 14353 | 15825 | - | x | | CBM35-GH26 | 1C10 (10): 100% cov, 99% id | QIZ03395.1 | 57.50 | 9E-126 | endo-b-1,4-mannanase (3.2.1.78) |
| | | | | | 12 | Polygalacturonase | 16145 | 17551 | - | x | | GH28 | 1C10 (11): 100% cov, 98% id | AAO79251.1 | 29.06 | 6E-38 | rhamnogalacturonan a-1,2-galacturonohydrolase (3.2.1.173) |
| | | | | | 13 | Long-chain-fatty-acid--CoA ligase (EC 6.2.1.3) | 17567 | 18997 | - | x | 34 | | | | | | |
| | | | | | 14 | Predicted beta-xyloside ABC transporter, permease component | 19235 | 20188 | + | x | 49 | | | | | | |
| | | | | | 15 | ABC transporter, permease protein | 20205 | 21143 | + | x | 33 | | | | | | |
| | | | | | 16 | Transcriptional regulator, AraC family | 21186 | 22091 | + | x | 26 | | | | | | |
| | | | | | 17 | ABC transporter, substrate-binding protein | 22153 | 23658 | + | x | | | | | | | |
| | | | | | 18 | hypothetical protein | 23775 | 25322 | + | x | | | | | | | |
| | | | | | 19 | Transcriptional regulator, AraC family | 25326 | 27593 | + | x | 3 | | | | | | |
| | | | | | 20 | putative glycosyl hydrolase, family 43 | 27603 | 28445 | + | x | 33 | GH43_17 | 1C10 (19): 100% cov, 99% id | ADQ05609.1 | 49.12 | 6E-88 | a-L-arabinofuranosidase (3.2.1.55) |
| | | | | | 21 | Rhamnulokinase (EC 2.7.1.5) | 28492 | 29772 | + | x | 27 | | | | | | |
| | | | | | 22 | L-rhamnose isomerase (EC 5.3.1.14) | 29769 | 31025 | + | x | 26 | | | | | | |
| | | | | | 23 | Rhamnulose-1-phosphate aldolase (EC 4.1.2.19) | 31022 | 31858 | + | x | | | | | | | |
| | | | | | 24 | Alpha-galactosidase (EC 3.2.1.22) | 31855 | 33252 | + | x | | GH4 | 1C10 (23): 100% cov, 99% id | CAB12532.1 | 52.37 | 8E-167 | exo-polygalacturonase (3.2.1.67) |
| | | | | | 25 | FIG00615029: hypothetical protein | 33353 | 34918 | - | x | | | | | | | |
| 14O19- IVVE | | 35,658 bp | 1 Firmicutes | AZO-galacto-mannan | 26 | Lipolytic enzyme, G-D-S-L precursor | 34952 | 35581 | + | | | | | | | | |
| | | | | | 1 | binding-protein-dependent transport systems inner membrane component | 32 | 898 | + | x | 2 | | | | | | |
| | | | | | 2 | hypothetical protein | 913 | 2445 | + | x | 3 | | | | | | |
| | | | | | 3 | hypothetical protein | 2414 | 3106 | + | | | | | | | | |
| | | | | | 4 | hypothetical protein | 3122 | 5695 | + | x | 7 | GH68_2 | | BAB72022.1 | 24.57 | 0.029 | b-fructofuranosidase / invertase (3.2.1.26) |
| | | | | | 5 | binding-protein-dependent transport systems inner membrane component | 5692 | 6672 | + | x | | | | | | | |
| | | | | | 6 | Alpha-glucoside transport system permease protein AglG | 6665 | 7675 | + | x | 12 | | | | | | |
| | | | | | 7 | hypothetical protein | 7684 | 7803 | + | | | | | | | | |
| | | | | | 8 | ABC transporter substrate-binding protein - sugar transport | 7831 | 9273 | + | x | 11 | | | | | | |
| | | | | | 9 | Endo-1,4-beta-xylanase A precursor (EC 3.2.1.8) | 9412 | 10968 | + | x | 49 | CBM22-GH10 | | AHF24999.1 | 81.90 | 0.0 | endo-b-1,4-xylanase (3.2.1.8) |
| | | | | | 10 | Endo-1,4-beta-xylanase A precursor (EC 3.2.1.8) | 11167 | 12330 | + | x | 19 | GH5_4 | | AHF24998.1 | 75.00 | 0.0 | endo-b-1,4-glucanase (3.2.1.4) |
| | | | | | 11 | Chitobiose phosphorylase (EC 2.4.1.-) | 12435 | 14822 | + | x | | GH94 | | QCO92991.1 | 66.83 | 0.0 | cellodextrin phosphorylase (2.4.1.49) |
| | | | | | 12 | COG1720: Uncharacterized conserved protein | 14948 | 15610 | + | | | | | | | | |
| | | | | | 13 | Phosphoserine aminotransferase (EC 2.6.1.52) | 15812 | 16888 | + | x | 11 | | | | | | |
| | | | | | 14 | D-3-phosphoglycerate dehydrogenase (EC 1.1.1.95) | 17068 | 18243 | + | x | 29 | | | | | | |
| | | | | | 15 | DNA-directed RNA polymerase beta' subunit (EC 2.7.7.6) | 18368 | 21937 | - | x | 28 | | | | | | |
| | | | | | 16 | DNA-directed RNA polymerase beta subunit (EC 2.7.7.6) | 21958 | 25824 | - | x | 60 | | | | | | |
| | | | | | 17 | DNA internalization-related competence protein ComEC/Rec2 | 26073 | 27059 | + | x | 2 | | | | | | |
| | | | | | 18 | Firmicutes ribosomal L7Ae family protein | 27231 | 27476 | + | | | | | | | | |
| | | | | | 19 | SSU ribosomal protein S12p (S23e) | 27541 | 27915 | + | | | | | | | | |
| | | | | | 20 | SSU ribosomal protein S7p (S5e) | 28126 | 28596 | + | | | | | | | | |
| | | | | | 21 | Translation elongation factor G | 28696 | 30762 | + | x | | | | | | | |
| | | | | | 22 | Translation elongation factor Tu | 30832 | 32046 | + | x | | | | | | | |
| | | | | | 23 | LSU ribosomal protein L33p | 32068 | 32229 | + | | | | | | | | |
| | | | | | 24 | hypothetical protein | 32257 | 32547 | + | | | | | | | | |
| | | | | | 25 | Transcription antitermination protein NusG | 32565 | 33101 | + | | | | | | | | |
| 16B10-IVVE | | 34,263 bp | 1 Bacterium | AZCL-xylan, AZCL-arabinoxylan | 26 | LSU ribosomal protein L11p (L12e) | 33248 | 33673 | + | | | | | | | | |
| | | | | | 1 | hypothetical protein | 530 | 694 | + | | | | | | | | |
| | | | | | 2 | hypothetical protein | 691 | 849 | + | | | | | | | | |
| | | | | | 3 | tRNA proofreading protein STM4549 | 868 | 1347 | - | | | | | | | | |
| | | | | | 4 | hypothetical protein | 1589 | 1816 | + | | | | | | | | |
| | | | | | 5 | ABC transporter, ATP-binding protein | 1861 | 2598 | + | x | | | | | | | |
| | | | | | 6 | ABC transporter permease protein | 2591 | 4222 | + | | | | | | | | |
| | | | | | 7 | Lipopolysaccharide biosynthesis protein-like protein | 4282 | 5937 | - | x | | | | | | | |
| | | | | | 8 | Carbamoylphosphate synthase large subunit (split gene in MJ) | 5985 | 7169 | - | | | | | | | | |
| | | | | | 9 | hypothetical protein | 7174 | 7998 | - | | | | | | | | |
| | | | | | 10 | hypothetical protein | 8143 | 8625 | - | x | | | | | | | |
| | | | | | 11 | Cystathionine beta-lyase, Bsu PatB (EC 4.4.1.8) | 8693 | 9874 | - | | | | | | | | |
| | | | | | 12 | hypothetical protein | 9920 | 10684 | - | | | | | | | | |
| | | | | | 13 | Branched-chain amino acid transport protein AzlC | 10842 | 11534 | + | | | | | | | | |
| | | | | | 14 | Branched-chain amino acid transport protein azlD | 11531 | 11854 | + | x | | | | | | | |
| | | | | | 15 | Dehydrogenase | 12102 | 13253 | - | | | | | | | | |
| | | | | | 16 | Transcriptional regulator, MerR family | 13464 | 14195 | - | | | | | | | | |
| | | | | | 17 | Uridine kinase family protein YggC homolog | 14283 | 15101 | - | | | | | | | | |
| | | | | | 18 | spermidine acetyltransferase | 15098 | 15580 | - | | | | | | | | |
| | | | | | 19 | FIG01116832: hypothetical protein | 15630 | 15950 | - | | | | | | | | |
| | | | | | 20 | Putative Nudix hydrolase YfcD (EC 3.6.-.-) | 15980 | 16516 | - | x | 8 | | | | | | |
| | | | | | 21 | Mg-protoporphyrin O-methyltransferase (EC 2.1.1.11) | 16589 | 17518 | - | | | | | | | | |
| | | | | | 22 | hypothetical protein | 17825 | 18226 | + | x | | | | | | | |
| | | | | | 23 | response regulator | 18361 | 21465 | - | | | | | | | | |
| | | | | | 24 | Cell surface protein | 21620 | 22837 | - | | | | | | | | |
| | | | | | 25 | hypothetical protein | 22834 | 23241 | - | | | | | | | | |
| | | | | | 26 | hypothetical protein | 23330 | 23842 | + | x | | | | | | | |
| | | | | | 27 | Manganese-dependent inorganic pyrophosphatase (EC 3.6.1.1) | 23864 | 25024 | - | | | | | | | | |
| | | | | | 28 | acetyltransferase, GNAT family | 25034 | 26077 | - | | | | | | | | |
| | | | | | 29 | Aminoglycoside phosphotransferase | 26108 | 26905 | - | | | | | | | | |
| | | | | | 30 | hypothetical protein | 26954 | 27118 | - | | | | | | | | |
| | | | | | 31 | hypothetical protein | 27382 | 27861 | - | | | | | | | | |
| | | | | | 32 | hypothetical protein | 27868 | 28662 | - | | | | | | | | |
| | | | | | 33 | probable methyltransferase | 29050 | 29622 | + | | | | | | | | |
| | | | | | 34 | Metal transporter, ZIP family | 29718 | 30497 | + | x | | | | | | | |
| | | | | | 35 | hypothetical protein | 30559 | 30729 | - | | | | | | | | |

| | | | | | | | | | | | | | | | | | |
|---|---|---|---|---|---|---|---|---|---|---|---|---|---|---|---|---|---|
| | | | | | 36 | Sensory box histidine kinase/response regulator | 30847 | 34113 | - | x | | | | | | | |
| | | | | | 37 | hypothetical protein | 34199 | 34396 | - | x | | | | | | | |
| | | | | | 38 | hypothetical protein | 34448 | 34588 | - | x | 17 | | | | | | |
| | | | | | 39 | DNA-binding response regulator, AraC family | 34578 | 35531 | - | x | | | | | | | |
| | | | | | 40 | hypothetical protein | 35528 | 35746 | - | | | | | | | | |
| | | | | | 41 | Xylose isomerase (EC 5.3.1.5) | 35705 | 36742 | + | | | | | | | | |
| | | | | | 42 | Ribokinase (EC 2.7.1.15) | 36929 | 38032 | + | | | | | | | | |
| | | | | | 43 | Fructose-bisphosphate aldolase class II (EC 4.1.2.13) | 38143 | 38988 | + | | | | | | | | |
| 17H1- IVVE | | 39,405 bp | 1 **Firmicutes** | Tween 20, pNPA, pNPB, pNPP | 44 | hypothetical protein | 39064 | 39270 | + | | | | | | | | |
| | | | | | 1 | Sodium:alanine symporter | 266 | 1906 | - | x | 2 | | | | | | |
| | | | | | 2 | hypothetical protein | 1997 | 2116 | - | | | | | | | | |
| | | | | | 3 | Probable transcriptional regulator | 2123 | 3076 | + | x | | | | | | | |
| | | | | | 4 | hypothetical protein | 3191 | 4477 | + | x | | | | | | | |
| | | | | | 5 | FIG01032461: hypothetical protein | 4499 | 5236 | + | | | | | | | | |
| | | | | | 6 | FIG01032596: hypothetical protein | 5278 | 5949 | + | | | | | | | | |
| | | | | | 7 | hypothetical protein | 5974 | 7236 | + | | | | | | | | |
| | | | | | 8 | hypothetical protein | 7229 | 7423 | + | | | | | | | | |
| | | | | | 9 | hypothetical protein | 7447 | 7581 | - | | | | | | | | |
| | | | | | 10 | hypothetical protein | 7614 | 8531 | - | x | | GH18 | | BAO37115.1 | 29.58 | 0.75 | chitinase (3.2.1.14) |
| | | | | | 11 | Alpha-galactosidase (EC 3.2.1.22) | 8582 | 9889 | - | x | | GH4 | | BAB05947.1 | 68.42 | 0.0 | a-galactosidase (3.2.1.22) |
| | | | | | 12 | Endo-1,4-beta-xylanase A precursor (EC 3.2.1.8) | 10181 | 11734 | + | | | CBM22-GH10 | | AHF25590.1 | 89.64 | 0.0 | endo-b-1,4-xylanase (3.2.1.8) |
| 21K7-IVVE | | 13,661 bp | 1 **Firmicutes** | AZCL-xylan, AZCL-arabinoxylan | 13 | hypothetical protein | 11704 | 13425 | - | | | | | | | | |
| | | | | | 1 | Na+ driven multidrug efflux pump | 13 | 1116 | + | x | | | | | | | |
| | | | | | 2 | Transposase | 1591 | 2886 | + | x | | | | | | | |
| | | | | | 3 | no significant homology | 3054 | 3266 | - | | | | | | | | |
| | | | | | 4 | Homoserine kinase (EC 2.7.1.39) | 3337 | 3993 | - | | | | | | | | |
| | | | | | 5 | Rubredoxin | 4058 | 4210 | - | | | | | | | | |
| | | | | | 6 | Uncharacterized protein MJ1614 | 4371 | 5207 | + | x | | | | | | | |
| | | | | | 7 | hypothetical protein | 5209 | 6930 | + | x | | | | | | | |
| | | | | | 8 | ABC-type polysaccharide transport system, permease component | 7185 | 8159 | + | x | | | | | | | |
| | | | | | 9 | Predicted beta-xyloside ABC transporter, permease component | 8175 | 9047 | + | x | | | | | | | |
| | | | | | 10 | hypothetical protein | 9106 | 10716 | + | x | | | | | | | |
| | | | | | 11 | Transcriptional regulator, AraC family | 10834 | 11760 | - | x | | | | | | | |
| | | | | | 12 | Oxidoreductase | 11931 | 13091 | + | x | | | | | | | |
| | | | | | 13 | oxidoreductase domain protein | 13093 | 14172 | + | x | | | | | | | |
| | | | | | 14 | Gluconokinase (EC 2.7.1.12) / oxidoreductase domain | 14176 | 15357 | + | x | | | | | | | |
| | | | | | 15 | Phage repressor | 15504 | 16256 | + | x | | | | | | | |
| | | | | | 16 | hypothetical protein | 16269 | 17660 | + | x | | | | | | | |
| | | | | | 17 | hypothetical protein | 17678 | 17944 | + | | | | | | | | |
| | | | | | 18 | putative esterase | 17950 | 18735 | - | x | | | | | | | |
| | | | | | 19 | Endo-1,4-beta-xylanase A precursor (EC 3.2.1.8) | 18777 | 19832 | - | x | | GH10 | 45P14 (37), 50E19 (15): 100% cov, 100 % id | AEW47914.1 | 82.81 | 0.0 | endo-b-1,4-xylanase (3.2.1.8) |
| | | | | | 20 | Acetyl-coenzyme A synthetase (EC 6.2.1.1) | 19914 | 22187 | - | x | | | | | | | |
| | | | | | 21 | DNA mismatch repair protein [MutS] | 22641 | 24356 | + | x | | | | | | | |
| | | | | | 22 | hypothetical protein | 24483 | 26360 | + | x | | | | | | | |
| | | | | | 23 | Ribosome protection-type tetracycline resistance related proteins, group 2 | 26389 | 29022 | - | x | | | | | | | |
| | | | | | 24 | hypothetical protein | 29192 | 30460 | + | x | | | | | | | |
| | | | | | 25 | hypothetical protein | 30465 | 31160 | + | | | | | | | | |
| | | | | | 26 | hypothetical protein | 31258 | 33780 | + | x | | | | | | | |
| | | | | | 27 | ABC transporter, ATP-binding protein | 33767 | 34705 | + | x | | | | | | | |
| | | | | | 28 | hypothetical protein | 34698 | 35615 | + | x | | | | | | | |
| 25I16-IVVE | LT674546 | 37,436 bp | 1 **Bacterium RumenMGS205_scaffold_35 11,466bp** | AZCL-xylan, AZCL-arabinoxylan, Tween 20, pNPA, pNPB | 29 | hypothetical protein | 35636 | 37135 | + | x | | | | | | | |
| | | | | | 1 | Pyrophosphate-energized proton pump (EC 3.6.1.1) | 1220 | 3379 | - | x | | | | | | | |
| | | | | | 2 | Sugar phosphate isomerases/epimerases | 3424 | 4314 | - | x | | | | | | | |
| | | | | | 3 | Gluconokinase (EC 2.7.1.12) / oxidoreductase domain | 4346 | 5557 | - | x | | | | | | | |
| | | | | | 4 | Transcriptional regulator of rhamnose utilization, AraC family | 5696 | 6550 | + | | | | | | | | |
| | | | | | 5 | Beta-hexosaminidase (EC 3.2.1.52) | 6690 | 8738 | + | | | GH20 | | AGU10374.1 | 35.70 | 2E-76 | b-N-acetylhexosaminidase (3.2.1.52) |
| | | | | | 6 | MATE efflux family protein | 8784 | 10196 | + | | | | | | | | |
| | | | | | 7 | hypothetical protein | 10230 | 11345 | - | | | | | | | | |
| | | | | | 8 | Esterase/lipase | 11382 | 12242 | - | | | | | | | | |
| | | | | | 9 | LacX protein, plasmid | 12290 | 13207 | - | | | | | | | | |
| | | | | | 10 | Lactoylglutathione lyase and related lyase | 13535 | 13999 | - | | | | | | | | |
| | | | | | 11 | hypothetical protein | 14400 | 15200 | + | | | | | | | | |
| | | | | | 12 | hypothetical protein | 15260 | 16681 | + | | | | | | | | |
| | | | | | 13 | Amino acid ABC transporter, permease protein | 16674 | 17348 | + | | | | | | | | |
| | | | | | 14 | putative amino acid ABC transporter, ATP-binding protein | 17345 | 18091 | + | | | | | | | | |
| | | | | | 15 | Phosphohydrolase | 18134 | 18880 | - | | | | | | | | |
| | | | | | 16 | hypothetical protein | 19040 | 19180 | + | | | | | | | | |
| | | | | | 17 | Ribosomal subunit interface protein | 19266 | 19814 | - | | | | | | | | |
| | | | | | 18 | 8-amino-7-oxononanoate synthase (EC 2.3.1.47) | 20224 | 21468 | + | x | | | | | | | |
| | | | | | 19 | hypothetical protein | 21443 | 21769 | - | | | | | | | | |
| | | | | | 20 | Potassium voltage-gated channel subfamily KQT; possible potassium channel, VIC family | 21821 | 22840 | + | | | | | | | | |
| | | | | | 21 | heat shock protein, Hsp20 family | 23007 | 23435 | + | | | | | | | | |
| | | | | | 22 | Adenine phosphoribosyltransferase (EC 2.4.2.7) | 23505 | 24059 | - | | | | | | | | |
| | | | | | 23 | Nucleoside:H+ symporter:Major facilitator superfamily | 24200 | 25360 | - | | | | | | | | |
| | | | | | 24 | COG family: phosphoserine phosphatase | 25539 | 26135 | - | | | | | | | | |
| | | | | | 25 | Exodeoxyribonuclease III (EC 3.1.11.2) | 26132 | 26959 | - | | | | | | | | |
| | | | | | 26 | LysM-repeat proteins and domains | 27007 | 27573 | - | | | CBM50-CBM50-CBM50 | | | | | |
| | | | | | 27 | Usp domain | 27681 | 28064 | - | | | | | | | | |
| | | | | | 28 | hypothetical protein | 28061 | 28576 | - | | | | | | | | |
| 29D17-IVVE | LT674545 | 29,355 bp | 1 **Firmicutes** | Tween 20, pNPA, pNPB | 29 | Tryptophanyl-tRNA synthetase (EC 6.1.1.2) | 28722 | 29327 | - | | | | | | | | |
| | | | | | 1 | hypothetical protein | 865 | 1383 | - | | | | | | | | |
| | | | | | 2 | hypothetical protein | 1538 | 3238 | - | | | | | | | | |
| | | | | | 3 | Signal peptidase I (EC 3.4.21.89) | 3818 | 4414 | - | | | | | | | | |
| | | | | | 4 | hypothetical protein | 4490 | 4894 | - | | | | | | | | |
| | | | | | 5 | Solute-binding protein family 5 domain | 4917 | 7001 | - | | 4 | | | | | | |
| | | | | | 6 | hypothetical protein | 7039 | 7578 | - | | | | | | | | |
| | | | | | 7 | Sortase A, LPXTG specific | 7562 | 8608 | - | | | | | | | | |
| | | | | | 8 | Peptidase S26A, signal peptidase I | 8605 | 9516 | - | x | 9 | | | | | | |
| | | | | | 9 | Endo-1,4-beta-xylanase A precursor (EC 3.2.1.8) | 9720 | 11276 | - | x | 6 | CBM22-GH10 | | AHF24999.1 | 74.55 | 0.0 | endo-b-1,4-xylanase (3.2.1.8) |
| | | | | | 10 | hypothetical protein | 11408 | 12901 | - | x | | | | | | | |
| | | | | | 11 | ABC transporter type 1, transmembrane domain MetI-like | 13077 | 14087 | - | x | | | | | | | |
| | | | | | 12 | ABC transporter type 1, transmembrane domain MetI-like | 14080 | 15090 | - | x | | | | | | | |
| | | | | | 13 | hypothetical protein | 15087 | 17666 | - | x | | | | | | | |
| | | | | | 14 | Yip1 domain | 17682 | 18377 | - | | | | | | | | |
| | | | | | 15 | FIG01032054: hypothetical protein | 18352 | 19884 | - | x | | | | | | | |
| | | | | | 16 | binding-protein-dependent transport systems inner membrane component | 19904 | 21130 | - | x | | | | | | | |
| | | | | | 17 | binding-protein-dependent transport systems inner membrane component | 21132 | 22076 | - | x | | | | | | | |
| | | | | | 18 | N-Acetyl-D-glucosamine ABC transport system, sugar-binding protein | 22098 | 24983 | - | x | 35 | | | | | | |
| | | | | | 19 | Phenylalanyl-tRNA synthetase beta chain (EC 6.1.1.20) | 25185 | 27593 | - | x | 11 | | | | | | |
| | | | | | 20 | Phenylalanyl-tRNA synthetase alpha chain (EC 6.1.1.20) | 27963 | 29051 | - | x | | | | | | | |
| 30F10-IVVE | | 30,169 bp | 1 **Bacterium** | AZCL-xylan, AZCL-arabinoxylan | 21 | Chromosomal replication initiator protein DnaA | 29120 | 30139 | - | x | | | | | | | |
| | | | | | 1 | Hypothetical lipoprotein | 1398 | 2540 | - | | | | | | | | |
| | | | | | 2 | hypothetical protein | 2610 | 2744 | + | | | | | | | | |
| | | | | | 3 | hypothetical protein / (cupin [Ruminococcaceae bacterium AB4001]) | 2896 | 3228 | + | | | | | | | | |
| | | | | | 4 | MG(2+) CHELATASE FAMILY PROTEIN / ComM-related protein | 3260 | 4792 | + | | | | | | | | |
| | | | | | 5 | hypothetical protein | 4812 | 5969 | + | | | | | | | | |
| | | | | | 6 | hypothetical protein | 6042 | 6458 | - | | | | | | | | |
| | | | | | 7 | hypothetical protein | 6738 | 7637 | - | | | | | | | | |
| | | | | | 8 | Exopolysaccharide biosynthesis protein related to N-acetylglucosamine-1-phosphodiester alpha-N-acetylglucosaminidase | 7673 | 8581 | - | | | | | | | | |
| | | | | | 9 | Ribosomal RNA small subunit methyltransferase C (EC 2.1.1.52) | 8748 | 9398 | - | | | | | | | | |
| | | | | | 10 | Histone acetyltransferase HPA2 and related acetyltransferases | 9391 | 9903 | - | | | | | | | | |
| | | | | | 11 | Beta-galactosidase (EC 3.2.1.23) | 9940 | 12879 | - | | | GH2 | | AJP16550.1 | 44.88 | 2E-65 | b-galactosidase (3.2.1.23) |
| | | | | | 12 | Ribulose-phosphate 3-epimerase (EC 5.1.3.1) | 12902 | 13549 | - | | | | | | | | |
| | | | | | 13 | Lipid A export ATP-binding/permease protein MsbA | 13747 | 15549 | + | | | | | | | | |
| | | | | | 14 | Lipid A export ATP-binding/permease protein MsbA | 15542 | 17443 | + | | | | | | | | |
| | | | | | 15 | Methionine synthase activation domain (EC 2.1.1.13) | 17443 | 18105 | + | | | | | | | | |

| | | | | | | | | | | | | | | | | | |
|---|---|---|---|---|---|---|---|---|---|---|---|---|---|---|---|---|---|
| | | | | | 16 | 5-methyltetrahydrofolate--homocysteine methyltransferase (EC 2.1.1.13) | 18092 | 20509 | + | | | | | | | | |
| | | | | | 17 | Metallo-dependent hydrolases, subgroup C | 20585 | 21739 | + | | | | | | | | |
| 31D12-IVVE | LT674549 | 22,580 bp | 1 **Firmicutes** *Feacalibacterium prausnitzii* | Lignin alkali | 18 | Putative Holliday junction resolvase YggF | 21736 | 22533 | + | | | | | | | | |
| | | | | | 1 | Endoglucanase E precursor (EC 3.2.1.4) (EgE) (Endo-1,4-beta-glucanase E) (Cellulase E) | 6 | 830 | + | x | | CE2 | 38D22 (1): 100% cov, 100% id | | | | |
| | | | | | 2 | Substrate-specific component RibU of riboflavin ECF transporter | 1108 | 1707 | + | | | | | | | | |
| | | | | | 3 | Fibronectin/fibrinogen-binding protein | 1727 | 3466 | + | | | | | | | | |
| | | | | | 4 | hypothetical protein | 3516 | 4067 | - | | | | | | | | |
| | | | | | 5 | Phosphoglycolate phosphatase (EC 3.1.3.18) | 4113 | 4763 | - | | | | | | | | |
| | | | | | 6 | 2-oxoglutarate oxidoreductase, delta subunit, putative (EC 1.2.7.3) | 4935 | 5138 | + | | | | | | | | |
| | | | | | 7 | 2-oxoglutarate oxidoreductase, alpha subunit (EC 1.2.7.3) | 5154 | 6230 | + | x | | | | | | | |
| | | | | | 8 | 2-oxoglutarate oxidoreductase, beta subunit (EC 1.2.7.3) | 6233 | 6970 | + | | | | | | | | |
| | | | | | 9 | 2-oxoglutarate oxidoreductase, gamma subunit (EC 1.2.7.3) | 6974 | 7510 | + | | | | | | | | |
| | | | | | 10 | putative xylanase | 7591 | 8808 | - | | | CE17 | 38D22 (10): 100% cov, 100% id; | | | | |
| | | | | | 11 | Cytochrome c-type biogenesis protein CcsA/ResC | 8825 | 9640 | - | | | | | | | | |
| | | | | | 12 | Cytochrome c-type biogenesis protein Ccs1/ResB | 9649 | 10677 | - | | | | | | | | |
| | | | | | 13 | Fumarate reductase flavoprotein subunit (EC 1.3.99.1) | 10720 | 13644 | - | x | | GH29 | | AIC77302.1 | 27.66 | 0.80 | a-L-fucosidase (3.2.1.51) |
| | | | | | 14 | hypothetical protein | 13682 | 14839 | - | x | | | | | | | |
| | | | | | 15 | Electron transport complex protein RnfB | 14852 | 15646 | - | x | 15 | | | | | | |
| | | | | | 16 | Electron transport complex protein RnfA | 15661 | 16239 | - | | | | | | | | |
| | | | | | 17 | Electron transport complex protein RnfE | 16236 | 16994 | - | | | | | | | | |
| | | | | | 18 | Electron transport complex protein RnfD | 16996 | 18537 | - | | | | | | | | |
| | | | | | 19 | Electron transport complex protein RnfC | 18534 | 19859 | - | | | | | | | | |
| | | | | | 20 | Sensory box/GGDEF family protein | 19919 | 21820 | - | | | | | | | | |
| | | | | | 21 | hypothetical protein | 22092 | 23465 | + | | | | | | | | |
| | | | | | 22 | Lysophospholipase (EC 3.1.1.5); Monoglyceride lipase (EC 3.1.1.23); putative | 23516 | 24439 | - | | | | | | | | |
| | | | | | 23 | hypothetical protein | 24419 | 24556 | + | | | | | | | | |
| | | | | | 24 | Malate permease | 24579 | 25508 | + | | | | | | | | |
| | | | | | 25 | hypothetical protein | 25511 | 26416 | - | | | | | | | | |
| | | | | | 26 | hypothetical protein | 26413 | 26652 | - | x | | | | | | | |
| | | | | | 27 | Neopullulanase (EC 3.2.1.135) | 26713 | 28788 | - | | | GH13_20 | 38D22 (27): 100% cov, 100% id; 12F23 (16): 100% cov, 99% id; 51D10 (3): 100% cov, 95% id | BAA02473.1 | 33.92 | 3E-80 | neopullulanase (3.2.1.135), cyclomaltodextrinase(3.2.1.54) |
| | | | | | 28 | Beta-1,3-glucosyltransferase | 28824 | 29645 | - | x | | GT2 | 38D22 (27): 100% cov, 100% id; 12F23 (17): 100% cov, 99% id | | | | |
| | | | | | 29 | Endo-1,4-beta-xylanase A precursor (EC 3.2.1.8) | 29793 | 31307 | - | x | | CBM22-GH10 | 38D22 (29): 100% cov, 99% id; 12F23 (18): 100% cov, 99% id | AHF26235.1 | 66.76 | 1E-159 | endo-b-1,4-xylanase(3.2.1.8) |
| | | | | | 30 | hypothetical protein | 31293 | 31406 | + | | | | | | | | |
| 32D12-IVVE | LT674550 | 33,736 bp | 1 **Bacterium** | Lignin alkali, AZCL-barley beta glucan | 31 | Collagen-binding A protein-like | 31510 | 33693 | - | | | | | | | | |
| | | | | | 1 | Adenosylcobinamide-phosphate guanylyltransferase (EC 2.7.7.62) | 81 | 617 | + | | | | | | | | |
| | | | | | 2 | Cobalamin synthase | 614 | 1366 | + | | | | | | | | |
| | | | | | 3 | hypothetical protein | 1363 | 1731 | + | | | | | | | | |
| | | | | | 4 | hypothetical protein | 1728 | 2309 | + | | | | | | | | |
| | | | | | 5 | Cob(I)alamin adenosyltransferase (EC 2.5.1.17) | 2311 | 2832 | + | | | | | | | | |
| | | | | | 6 | hypothetical protein | 2863 | 3234 | + | | | | | | | | |
| | | | | | 7 | FIG022979: MoxR-like ATPases | 3262 | 4206 | + | | | | | | | | |
| | | | | | 8 | hypothetical protein | 4203 | 5270 | + | | | | | | | | |
| | | | | | 9 | hypothetical protein | 5286 | 6506 | + | | | | | | | | |
| | | | | | 10 | exo-b-1,3-glucanase (3.2.1.58) | 6752 | 7450 | - | | | GH3 | | ACE85841.1 | 35.00 | 0.31 | exo-b-1,3-glucanase (3.2.1.58) |
| | | | | | 11 | Endo-1,4-beta-xylanase A precursor (EC 3.2.1.8) | 7472 | 8401 | - | | | GH5_2 | | ABA42185.1 | 54.25 | 5E-109 | endo-b-1,4-glucanase (3.2.1.4) |
| | | | | | 12 | hypothetical protein | 8426 | 9832 | - | x | 18 | | | | | | |
| | | | | | 13 | Beta-galactosidase (EC 3.2.1.23) | 9895 | 11832 | - | | | GH42 | | AAN05452.1 | 47.66 | 0.0 | b-galactosidase(3.2.1.23),a-L-arabinopyranosidase (3.2.1.-) |
| | | | | | 14 | hypothetical protein | 11991 | 13547 | - | | | | | | | | |
| | | | | | 15 | hypothetical protein | 13653 | 14348 | - | | | | | | | | |
| | | | | | 16 | hypothetical protein | 14368 | 14733 | - | | | | | | | | |
| | | | | | 17 | hypothetical protein | 14891 | 15124 | + | | | | | | | | |
| | | | | | 18 | hypothetical protein | 15127 | 15333 | + | | | | | | | | |
| | | | | | 19 | hypothetical protein | 15330 | 15710 | + | | | | | | | | |
| | | | | | 20 | hypothetical protein | 15715 | 16011 | + | | | | | | | | |
| | | | | | 21 | hypothetical protein | 16061 | 16192 | + | | | | | | | | |
| | | | | | 22 | hypothetical protein | 16255 | 16437 | + | | | | | | | | |
| | | | | | 23 | hypothetical protein | 17045 | 17299 | + | | | | | | | | |
| | | | | | 24 | hypothetical protein | 17292 | 17651 | + | | | | | | | | |
| | | | | | 25 | Protein export cytoplasm protein SecA ATPase RNA helicase (TC 3.A.5.1.1) | 17664 | 18479 | + | | | | | | | | |
| | | | | | 26 | hypothetical protein | 18472 | 19017 | + | | | | | | | | |
| | | | | | 27 | hypothetical protein | 19087 | 19560 | + | | | | | | | | |
| | | | | | 28 | hypothetical protein | 19572 | 20216 | + | | | | | | | | |
| | | | | | 29 | hypothetical protein | 20254 | 21120 | + | | | | | | | | |
| | | | | | 30 | Single-stranded DNA-binding protein | 21135 | 21536 | + | | | | | | | | |
| | | | | | 31 | hypothetical protein | 21551 | 21937 | + | | | | | | | | |
| | | | | | 32 | hypothetical protein | 21934 | 22338 | + | | | | | | | | |
| | | | | | 33 | hypothetical protein | 22340 | 22600 | + | | | | | | | | |
| | | | | | 34 | hypothetical protein | 22593 | 23132 | + | | | | | | | | |
| | | | | | 35 | hypothetical protein | 23117 | 23332 | + | | | | | | | | |
| | | | | | 36 | hypothetical protein | 23311 | 23517 | + | | | | | | | | |
| | | | | | 37 | phage Gp37Gp68 family protein | 23486 | 24337 | - | | | | | | | | |
| | | | | | 38 | hypothetical protein | 24342 | 24551 | + | | | | | | | | |
| | | | | | 39 | hypothetical protein | 24548 | 24823 | + | | | | | | | | |
| | | | | | 40 | hypothetical protein | 25045 | 25233 | + | | | | | | | | |
| | | | | | 41 | hypothetical protein | 25223 | 25417 | + | | | | | | | | |
| | | | | | 42 | hypothetical protein | 25513 | 26031 | + | | | | | | | | |
| | | | | | 43 | hypothetical protein | 26167 | 26781 | + | | | | | | | | |
| | | | | | 44 | hypothetical protein | 26778 | 27296 | + | | | | | | | | |
| | | | | | 45 | Phage terminase, large subunit | 27299 | 29158 | + | x | | | | | | | |
| | | | | | 46 | hypothetical protein | 29181 | 29423 | + | | | | | | | | |
| | | | | | 47 | hypothetical protein | 29440 | 31092 | + | | | | | | | | |
| | | | | | 48 | Prophage Clp protease-like protein | 31085 | 32218 | + | | | | | | | | |
| | | | | | 49 | hypothetical protein | 32245 | 32901 | + | | | | | | | | |
| | | | | | 50 | Putative capsid protein of prophage | 32937 | 33974 | + | | | | | | | | |
| | | | | | 51 | hypothetical protein | 33995 | 34345 | + | | | | | | | | |
| | | | | | 52 | hypothetical protein | 34342 | 34500 | + | | | | | | | | |
| 33M21-IVVE | | 35,197 bp | 1 **Firmicutes** | AZCL-Barley beta glucan, AZCL-HE cellulose | 53 | hypothetical protein | 34500 | 34844 | + | | | | | | | | |
| | | | | | 1 | alpha-galactosidase (3.2.1.22) | 149 | 2038 | + | | | GH27 | | BAB83765.1 | 23.08 | 0.50 | a-galactosidase (3.2.1.22) |
| | | | | | 2 | hypothetical protein | 2264 | 4207 | + | | | | | | | | |
| | | | | | 3 | hypothetical protein / (metallophosphoesterase [Subdoligranulum sp. 4_3_54A2FAA] ) | 4204 | 4659 | + | | | | | | | | |
| | | | | | 4 | hypothetical protein / (metallophosphoesterase [Faecalibacterium prausnitzii]) | 4659 | 4844 | + | | | | | | | | |
| | | | | | 5 | N-acetylglucosamine-1-phosphate uridyltransferase (EC 2.7.7.23) / Glucosamine-1-phosphate N-acetyltransferase (EC 2.3.1.157) | 4937 | 5680 | - | | | | | | | | |
| | | | | | 6 | conserved hypothetical protein | 5705 | 6349 | - | | | | | | | | |
| | | | | | 7 | pH adaptation potassium efflux system protein D 2 | 6416 | 8647 | - | | | | | | | | |
| | | | | | 8 | hypothetical protein | 8641 | 8856 | - | | | | | | | | |
| | | | | | 9 | NADH-ubiquinone oxidoreductase chain L (EC 1.6.5.3) | 8868 | 10094 | - | | | | | | | | |
| | | | | | 10 | NADH-ubiquinone oxidoreductase chain L (EC 1.6.5.3) | 10105 | 11559 | - | | | | | | | | |
| | | | | | 11 | PH adaptation potassium efflux system protein D 1 | 11578 | 13062 | - | | | | | | | | |
| | | | | | 12 | putative Na(+) H(+) antiporter subunit C | 13066 | 13440 | - | | | | | | | | |
| | | | | | 13 | putative Na(+) H(+) antiporter subunit B | 13440 | 14396 | - | | | | | | | | |
| | | | | | 14 | NADH dehydrogenase subunit 5 | 14380 | 14652 | - | | | | | | | | |
| | | | | | 15 | hypothetical protein | 14640 | 14966 | - | | | | | | | | |
| | | | | | 16 | Na(+) H(+) antiporter subunit F | 14968 | 15246 | - | | | | | | | | |
| | | | | | 17 | Na(+) H(+) antiporter subunit E | 15257 | 15721 | - | | | | | | | | |
| | | | | | 18 | Hypothetical similar to thiamin biosynthesis lipoprotein ApbE | 16044 | 17135 | + | | | | | | | | |
| | | | | | 19 | hypothetical protein | 17143 | 17664 | + | | | | | | | | |
| | | | | | 20 | hypothetical protein | 17792 | 21991 | + | | | | | | | | |
| | | | | | 21 | Electron transport complex protein RnfC | 22034 | 23383 | + | x | 1 | | | | | | |
| | | | | | 22 | Electron transport complex protein RnfD | 23380 | 24324 | + | | | | | | | | |
| | | | | | 23 | Electron transport complex protein RnfE | 24321 | 24959 | + | | | | | | | | |
| | | | | | 24 | Electron transport complex protein RnfA | 24956 | 25540 | + | | | | | | | | |

| | | | | | | | | | | | | | | | | | |
|---|---|---|---|---|---|---|---|---|---|---|---|---|---|---|---|---|---|
| | | | | | 25 | Electron transport complex protein RnfB | 25556 | 26347 | + | | | | | | | | |
| | | | | | 26 | Endo-1,4-beta-xylanase A precursor (EC 3.2.1.8) | 26455 | 28059 | - | | | CBM22-GH10 | | AHF24999.1 | 73.59 | 1E-180 | endo-b-1,4-xylanase (3.2.1.8) |
| | | | | | 27 | hypothetical protein | 28172 | 29623 | - | | | | | | | | |
| | | | | | 28 | binding-protein-dependent transport systems inner membrane component | 29781 | 30791 | - | x | 42 | | | | | | |
| | | | | | 29 | binding-protein-dependent transport systems inner membrane component | 30784 | 32190 | - | | | | | | | | |
| | | | | | 30 | a-N-acetylgalactosaminidase (3.2.1.49) | 32190 | 34763 | - | | | GH129 | | BAL14929.1 | 23.48 | 0.017 | a-N-acetylgalactosaminidase (3.2.1.49) |
| | | | | | 31 | hypothetical protein | 34779 | 35474 | - | | | | | | | | |
| 36E1-IVVE | | 36,794 bp | 1 **Firmicutes** | AZCL-xylan, AZCL-arabinoxylan | 32 | NHL repeat containing protein | 35443 | 36765 | - | | | | | | | | |
| | | | | | 1 | hypothetical protein | 673 | 1515 | + | | | | | | | | |
| | | | | | 2 | hypothetical protein | 1654 | 2151 | + | | | | | | | | |
| | | | | | 3 | Polygalacturonase (EC 3.2.1.15) | 2148 | 3704 | + | | | GH28 | | AAO76125.1 | 30.09 | 2E-51 | exo-polygalacturonase (3.2.1.67) |
| | | | | | 4 | 4-deoxy-L-threo-5-hexosulose-uronate ketol-isomerase (EC 5.3.1.17) | 3719 | 4561 | + | x | 2 | | | | | | |
| | | | | | 5 | 2-deoxy-D-gluconate 3-dehydrogenase (EC 1.1.1.125) | 4576 | 5373 | + | x | | | | | | | |
| | | | | | 6 | Beta-lactamase class C and other penicillin binding proteins | 5431 | 6570 | + | | | | | | | | |
| | | | | | 7 | putative alpha-dextrin endo-1, 6-alpha-glucosidase | 6575 | 7342 | - | | | | | | | | |
| | | | | | 8 | hypothetical protein | 7374 | 8384 | - | | | | | | | | |
| | | | | | 9 | NAD-dependent protein deacetylase of SIR2 family | 8391 | 9122 | - | | | | | | | | |
| | | | | | 10 | Alanyl-tRNA synthetase (EC 6.1.1.7) | 9254 | 11224 | - | x | 3 | | | | | | |
| | | | | | 11 | hypothetical protein | 11270 | 11386 | + | | | | | | | | |
| | | | | | 12 | Deblocking aminopeptidase (EC 3.4.11.-) | 11585 | 12577 | - | | | | | | | | |
| | | | | | 13 | Deblocking aminopeptidase (EC 3.4.11.-) | 12594 | 13607 | - | | | | | | | | |
| | | | | | 14 | Deblocking aminopeptidase (EC 3.4.11.-) | 13598 | 14635 | - | | | | | | | | |
| | | | | | 15 | Esterase/lipase-like protein | 14675 | 15550 | - | | | | | | | | |
| | | | | | 16 | Fibronectin/fibrinogen-binding protein | 15578 | 17317 | - | | | | | | | | |
| | | | | | 17 | Rubredoxin | 17433 | 17585 | - | | | | | | | | |
| | | | | | 18 | Conserved membrane protein | 17774 | 18556 | + | | | | | | | | |
| | | | | | 19 | hypothetical protein | 18907 | 19479 | + | | | | | | | | |
| | | | | | 20 | hypothetical protein | 19479 | 20552 | + | x | | | | | | | |
| | | | | | 21 | ABC transporter, membrane spanning protein [sugar] | 20565 | 21455 | + | x | 7 | | | | | | |
| | | | | | 22 | hypothetical protein | 21480 | 23543 | + | x | 12 | | | | | | |
| | | | | | 23 | hypothetical protein | 23669 | 24262 | + | | | | | | | | |
| | | | | | 24 | hypothetical protein | 24259 | 24726 | + | | | | | | | | |
| | | | | | 25 | Maltose/maltodextrin transport ATP-binding protein MalK (EC 3.6.3.19) | 24739 | 25812 | + | | | | | | | | |
| | | | | | 26 | hypothetical protein | 25815 | 26369 | + | | | | | | | | |
| | | | | | 27 | hypothetical protein | 26379 | 27032 | + | | | | | | | | |
| | | | | | 28 | Rhamnogalacturonides degradation protein RhiN | 27055 | 28188 | + | | | GH105 | | AAO76118.1 | 41.53 | 9E-99 | d-4,5-unsaturated a-galacturonidase (3.2.1.-) |
| | | | | | 29 | Transcriptional regulator, AraC family | 28163 | 28963 | - | | | | | | | | |
| | | | | | 30 | Multiple sugar ABC transporter, ATP-binding protein | 29109 | 30185 | + | | | | | | | | |
| | | | | | 31 | hypothetical protein | 30372 | 32816 | - | | | | | | | | |
| | | | | | 32 | Ribosomal subunit interface protein | 32898 | 33428 | - | | | | | | | | |
| | | | | | 33 | hypothetical protein | 33901 | 34746 | + | | | | | | | | |
| | | | | | 34 | hypothetical protein | 34960 | 35127 | - | | | | | | | | |
| 37F15-IVVE | LT674544 | 35,496 bp | 1 **Firmicutes** | Tween 20, pNPA, pNPB | 35 | hypothetical protein | 35323 | 35442 | + | | | | | | | | |
| | | | | | 1 | Phosphopentomutase (EC 5.4.2.7) | 376 | 2358 | - | | | | | | | | |
| | | | | | 2 | Segregation and condensation protein B | 2444 | 2995 | - | | | | | | | | |
| | | | | | 3 | Segregation and condensation protein A | 2982 | 3737 | - | | | | | | | | |
| | | | | | 4 | FIG004556: membrane metalloprotease | 3742 | 4497 | - | | | | | | | | |
| | | | | | 5 | hypothetical protein | 4565 | 4702 | + | | | | | | | | |
| | | | | | 6 | Endo-1,4-beta-xylanase A precursor (EC 3.2.1.8) | 4728 | 6275 | - | x | | CBM22-GH10 | | AHF24999.1 | 70.62 | 1E-176 | endo-b-1,4-xylanase (3.2.1.8) |
| | | | | | 7 | ABC transporter substrate-binding protein - sugar transport | 6468 | 7937 | - | x | | | | | | | |
| | | | | | 8 | binding-protein-dependent transport systems inner membrane component | 8098 | 9108 | - | | | | | | | | |
| | | | | | 9 | binding-protein-dependent transport systems inner membrane component | 9101 | 10087 | - | x | | | | | | | |
| | | | | | 10 | hypothetical protein | 10084 | 12660 | - | x | | GH129 | | BAL14929.1 | 24.04 | 0.004 | a-N-acetylgalactosaminidase (3.2.1.49) |
| | | | | | 11 | hypothetical protein | 12675 | 13379 | - | | | | | | | | |
| | | | | | 12 | FIG01032054: hypothetical protein | 13348 | 14895 | - | | | | | | | | |
| | | | | | 13 | binding-protein-dependent transport systems inner membrane component | 14945 | 16171 | - | x | | | | | | | |
| | | | | | 14 | binding-protein-dependent transport systems inner membrane component | 16173 | 17120 | - | | | | | | | | |
| | | | | | 15 | N-Acetyl-D-glucosamine ABC transport system, sugar-binding protein | 17143 | 20034 | - | x | | | | | | | |
| | | | | | 16 | Pyrroline-5-carboxylate reductase (EC 1.5.1.2) | 20443 | 21246 | - | | | | | | | | |
| | | | | | 17 | hypothetical protein | 21417 | 22265 | + | | | | | | | | |
| | | | | | 18 | hypothetical protein | 22452 | 22622 | - | | | | | | | | |
| | | | | | 19 | COG0613, Predicted metal-dependent phosphoesterases (PHP family) | 22711 | 23421 | - | | | | | | | | |
| 39F1- IVVE | LT674543 | 24,784 bp | 1 **Bacterium** | AZCL-xylan, AZCL-arabinoxylan, Tween 20, pNPA, pNPB | 20 | FIG176548: DRTGG domain-containing protein | 23423 | 23770 | - | | | | | | | | |
| | | | | | 1 | Formate dehydrogenase-O, major subunit (EC 1.2.1.2) | 1589 | 3988 | - | | | | | | | | |
| | | | | | 2 | Xanthine dehydrogenase, molybdenum binding subunit (EC 1.17.1.4) | 4006 | 6312 | - | | | | | | | | |
| | | | | | 3 | Xanthine dehydrogenase iron-sulfur subunit (EC 1.17.1.4) | 6314 | 6829 | - | x | 5 | | | | | | |
| | | | | | 4 | hypothetical protein | 6858 | 7397 | - | | | | | | | | |
| | | | | | 5 | Methyltransferase | 7592 | 8473 | + | | | | | | | | |
| | | | | | 6 | Cell surface protein | 8542 | 9768 | + | | | | | | | | |
| | | | | | 7 | FIG00262634: hypothetical protein | 9765 | 11237 | + | | | | | | | | |
| | | | | | 8 | FIG01077095: hypothetical protein | 11234 | 12742 | + | | | | | | | | |
| | | | | | 9 | Beta-glucosidase (EC 3.2.1.21) | 12827 | 14137 | + | | | GH1 | 13M17 (3): 100% cov, 97% id | CBL17177.1 | 46.09 | 5E-130 | b-glucosidase (3.2.1.21) |
| | | | | | 10 | Uncharacterized conserved protein | 14152 | 15015 | + | | | | | | | | |
| | | | | | 11 | benzyl alcohol dehydrogenase | 15102 | 16196 | - | | | | | | | | |
| | | | | | 12 | CTP:molybdopterin cytidylyltransferase | 16427 | 17356 | - | | | | | | | | |
| | | | | | 13 | hypothetical protein | 17353 | 17733 | - | | | | | | | | |
| | | | | | 14 | hypothetical protein | 17730 | 18143 | - | | | | | | | | |
| | | | | | 15 | cobalamin synthesis protein, P47K | 18154 | 19230 | - | | | | | | | | |
| | | | | | 16 | Aromatic hydrocarbon utilization transcriptional regulator CatR (LysR family) | 19589 | 20476 | - | | | | | | | | |
| | | | | | 17 | Xanthine dehydrogenase iron-sulfur subunit (EC 1.17.1.4) / Xanthine dehydrogenase, molybdenum binding subunit (EC 1.17.1.4) | 20591 | 23416 | + | | | | | | | | |
| | | | | | 18 | Glutamate synthase [NADPH] small chain (EC 1.4.1.13) | 23423 | 25594 | + | | | | | | | | |
| | | | | | 19 | hypothetical protein | 25626 | 25802 | - | | | | | | | | |
| | | | | | 20 | 2-Hydroxy-6-Oxo-6-Phenylhexa-2,4-Dienoate hydrolase | 25892 | 26662 | - | | | | | | | | |
| | | | | | 21 | Esterase/lipase | 26684 | 27589 | - | | | | | | | | |
| | | | | | 22 | Phenylacetate-coenzyme A ligase (EC 6.2.1.30) | 27734 | 29071 | - | | | | | | | | |
| | | | | | 23 | hypothetical protein | 29392 | 32844 | + | | | | | | | | |
| | | | | | 24 | transcriptional regulator, LysR family | 32883 | 33788 | - | | | | | | | | |
| | | | | | 25 | hypothetical protein | 33963 | 34403 | + | | | | | | | | |
| | | | | | 26 | HNH endonuclease | 34743 | 36230 | + | | | | | | | | |
| | | | | | 27 | hypothetical protein | 36507 | 37688 | - | | | | | | | | |
| | | | | | 28 | Mobile element protein | 38041 | 39429 | + | | | | | | | | |
| | | | | | 29 | hypothetical protein | 39401 | 39532 | - | | | | | | | | |
| | | | | | 30 | hypothetical protein | 39668 | 40918 | + | | | | | | | | |
| | | | | | 31 | Mobile element protein | 41085 | 42311 | - | | | | | | | | |
| | | | | | 32 | Mobile element protein | 42521 | 42760 | + | | | | | | | | |
| 41L9- IVVE | | 44,123 bp | 1 **Firmicutes** Clostridiales | Tween 20, pNPA, pNPB, pNPP | 33 | hypothetical protein | 42781 | 43176 | + | | | | | | | | |
| | | | | | 1 | hypothetical protein | 402 | 797 | - | | | | | | | | |
| | | | | | 2 | hypothetical protein | 831 | 2867 | - | | | | | | | | |
| | | | | | 3 | Proline-rich protein | 2899 | 3663 | - | | | | | | | | |
| | | | | | 4 | hypothetical protein | 3660 | 4625 | - | | | | | | | | |
| | | | | | 5 | hypothetical protein | 4622 | 5545 | - | x | | | | | | | |
| | | | | | 6 | Endo-1,4-beta-xylanase A precursor (EC 3.2.1.8) | 5714 | 7270 | - | x | | CBM22-GH10 | | AHF24999.1 | 87.76 | 0.0 | endo-b-1,4-xylanase (3.2.1.8) |
| | | | | | 7 | ABC transporter substrate-binding protein - sugar transport | 7427 | 8854 | - | x | 6 | | | | | | |
| | | | | | 8 | Alpha-glucoside transport system permease protein AglG | 9011 | 10021 | - | | | | | | | | |
| | | | | | 9 | binding-protein-dependent transport systems inner membrane component | 10014 | 10994 | - | x | 2 | | | | | | |
| | | | | | 10 | hypothetical protein | 10991 | 13588 | - | x | | | | | | | |
| | | | | | 11 | hypothetical protein | 13604 | 14299 | - | x | | | | | | | |
| | | | | | 12 | NHL repeat containing protein | 14265 | 15800 | - | x | 3 | | | | | | |
| | | | | | 13 | binding-protein-dependent transport systems inner membrane component | 15815 | 17053 | - | x | | | | | | | |
| | | | | | 14 | binding-protein-dependent transport systems inner membrane component | 17055 | 17999 | - | x | 5 | | | | | | |
| | | | | | 15 | N-Acetyl-D-glucosamine ABC transport system, sugar-binding protein | 18023 | 20908 | - | x | | | | | | | |
| | | | | | 16 | Phenylalanyl-tRNA synthetase beta chain (EC 6.1.1.20) | 21103 | 23508 | - | x | 4 | | | | | | |
| | | | | | 17 | Phenylalanyl-tRNA synthetase alpha chain (EC 6.1.1.20) | 23601 | 24623 | - | x | | | | | | | |
| | | | | | 18 | hypothetical protein | 24645 | 24776 | + | x | | | | | | | |
| | | | | | 19 | Chromosomal replication initiator protein DnaA | 24830 | 26158 | - | | | | | | | | |
| | | | | | 20 | hypothetical protein / (patatin [Cyanothece sp. PCC 7822]) | 26577 | 27083 | + | | | | | | | | |
| | | | | | 21 | Zinc ABC transporter, periplasmic-binding protein ZnuA | 27143 | 28036 | + | | | | | | | | |

| | | | | | | | | | | | | | | | | | |
|---|---|---|---|---|---|---|---|---|---|---|---|---|---|---|---|---|---|
| | | | | | 22 | hypothetical protein | 28094 | 28876 | + | | | | | | | | |
| | | | | | 23 | Cell division protein FtsW | 28873 | 30006 | + | x | 2 | | | | | | |
| | | | | | 24 | hypothetical protein | 30032 | 30331 | + | | | | | | | | |
| | | | | | 25 | hypothetical protein | 30359 | 31135 | + | | | | | | | | |
| | | | | | 26 | hypothetical protein | 31139 | 31771 | + | | | | | | | | |
| | | | | | 27 | hypothetical protein | 33847 | 33984 | - | | | | | | | | |
| 42N18-IVVE | | 37,103 bp | 1 **Bacterium** | AZCL-xylan, AZCL-arabinoxylan | 28 | hypothetical protein | 34020 | 34145 | - | | | | | | | | |
| | | | | | 1 | RNA polymerase sigma-28 factor | 524 | 1249 | + | | | | | | | | |
| | | | | | 2 | hypothetical protein | 1389 | 2348 | + | | | | | | | | |
| | | | | | 3 | IS481 family transposase ISAnsp2 | 2629 | 3813 | + | | | | | | | | |
| | | | | | 4 | 4-hydroxy-tetrahydrodipicolinate synthase | 4238 | 5125 | - | | | | | | | | |
| | | | | | 5 | hypothetical protein | 5207 | 6139 | - | | | | | | | | |
| | | | | | 6 | Sensor histidine kinase RcsC | 6449 | 9496 | + | | | | | | | | |
| | | | | | 7 | hypothetical protein probable lipase/esterase | 9894 | 10781 | + | | | | | | | | |
| | | | | | 8 | hypothetical protein | 10774 | 11562 | + | | | | | | | | |
| | | | | | 9 | Alpha-galactosidase A | 11879 | 13189 | + | | | GH27_2 | | 4NX0 | 58.40 | 3E-135 | a-L-fucosidase (3.2.1.51) |
| | | | | | 10 | Alpha-xylosidase | 13340 | 15562 | + | x | | GH31_3 | | AAC76680.1 | 68.06 | 7E-159 | a-xylosidase (3.2.1.177) |
| | | | | | 11 | Intracellular exo-alpha-(1->5)-L-arabinofuranosidase | 15593 | 17203 | + | | | GH51_1 | | ALK02878.1 | 40.45 | 2E-102 | a-L-arabinofuranosidase (3.2.1.55) |
| | | | | | 12 | hypothetical protein | 17499 | 19073 | + | x | | | | | | | |
| | | | | | 13 | putative multiple-sugar transport system permease YteP | 19160 | 20119 | + | | | | | | | | |
| | | | | | 14 | L-arabinose transport system permease protein AraQ | 20136 | 21035 | + | x | | | | | | | |
| | | | | | 15 | Intracellular exo-alpha-L-arabinofuranosidase 2 | 21328 | 22818 | + | | | GH51_1 | | ACE73681.1 | 55.78 | 1E-161 | a-L-arabinofuranosidase (3.2.1.55) |
| | | | | | 16 | hypothetical protein | 22938 | 25133 | + | | | | | | | | |
| | | | | | 17 | hypothetical protein | 25182 | 26657 | + | | | | | | | | |
| | | | | | 18 | Aminopeptidase PepS | 26849 | 28084 | + | | | | | | | | |
| | | | | | 19 | Lactaldehyde reductase | 28281 | 29426 | + | x | 8 | | | | | | |
| | | | | | 20 | hypothetical protein | 29829 | 31673 | - | | | | | | | | |
| | | | | | 21 | Putative peroxiredoxin bcp | 31907 | 32359 | - | | | | | | | | |
| | | | | | 22 | hypothetical protein | 32731 | 33891 | + | | | | | | | | |
| | | | | | 23 | hypothetical protein | 33878 | 34663 | + | | | | | | | | |
| | | | | | 24 | hypothetical protei | 34670 | 35065 | - | | | | | | | | |
| | | | | | 25 | Osmoregulated proline transporter OpuE | 35264 | 36703 | + | | | | | | | | |
| | | | | | 26 | Cyclic beta-(1,2)-glucan synthase NdvB | 36831 | 39149 | + | | | GT84 | | | | | |
| 44I12-IVVE | LT674542 | 40,627 bp | **Firmicute** Clostridiales | Tween 20, pNPA, pNPB, pNPP | 27 | hypothetical protein | 39151 | 39813 | - | | | GH29 | | ADV49759.1 | 47.49 | 8E-121 | a-L-fucosidase (3.2.1.51) |
| | | | | | 1 | hypothetical protein | 200 | 709 | - | | | | | | | | |
| | | | | | 2 | hypothetical protein | 743 | 1189 | - | | | | | | | | |
| | | | | | 3 | Rubrerythrin | 1662 | 2204 | - | | | | | | | | |
| | | | | | 4 | ABC transporter, periplasmic spermidine putrescine-binding protein PotD (TC 3.A.1.11.1) | 2336 | 3835 | - | | | | | | | | |
| | | | | | 5 | Spermidine Putrescine ABC transporter permease component potC (TC_3.A.1.11.1) | 3832 | 4635 | - | | | | | | | | |
| | | | | | 6 | Spermidine Putrescine ABC transporter permease component PotB (TC 3.A.1.11.1) | 4632 | 5420 | - | x | 3 | | | | | | |
| | | | | | 7 | Putrescine transport ATP-binding protein PotA (TC 3.A.1.11.1) | 5431 | 6876 | - | | | | | | | | |
| | | | | | 8 | Transcriptional regulator, MerR family, near polyamine transporter | 6928 | 7500 | - | | | | | | | | |
| | | | | | 9 | hypothetical protein | 7730 | 10381 | - | | | | | | | | |
| | | | | | 10 | hypothetical protein | 10378 | 11946 | - | | | | | | | | |
| | | | | | 11 | hypothetical protein | 11957 | 12805 | - | | | | | | | | |
| | | | | | 12 | MoxR-like ATPase in aerotolerance operon | 12795 | 13772 | - | x | | | | | | | |
| | | | | | 13 | hypothetical protein | 13789 | 15282 | - | x | | | | | | | |
| | | | | | 14 | hypothetical protein | 15309 | 16226 | - | x | | | | | | | |
| | | | | | 15 | ABC transporter, ATP-binding protein | 16219 | 17157 | - | x | | | | | | | |
| | | | | | 16 | hypothetical protein | 17144 | 19666 | - | x | | | | | | | |
| | | | | | 17 | hypothetical protein | 19764 | 20459 | - | | | | | | | | |
| | | | | | 18 | hypothetical protein | 20464 | 21732 | - | x | | | | | | | |
| | | | | | 19 | Ribosome protection-type tetracycline resistance related proteins, group 2 | 21902 | 24535 | + | x | | | | | | | |
| | | | | | 20 | hypothetical protein | 24564 | 26441 | - | x | | | | | | | |
| | | | | | 21 | DNA mismatch repair protein [MutS] | 26568 | 28283 | - | x | | | | | | | |
| | | | | | 22 | Acetyl-coenzyme A synthetase (EC 6.2.1.1) | 28737 | 31010 | + | x | | | | | | | |
| | | | | | 23 | Endo-1,4-beta-xylanase A precursor (EC 3.2.1.8) | 31092 | 32147 | + | x | | GH10 | 25I16 (19), 50E19 (15): 100% cov, 100 % id | AEW47914.1 | 82.81 | 0.0 | endo-b-1,4-xylanase (3.2.1.8) |
| | | | | | 24 | putative esterase | 32189 | 32974 | + | x | | | | | | | |
| | | | | | 25 | hypothetical protein | 32980 | 33246 | - | x | | | | | | | |
| | | | | | 26 | hypothetical protein | 33264 | 34655 | - | x | | | | | | | |
| | | | | | 27 | Phage repressor | 34668 | 35420 | - | x | 15 | | | | | | |
| | | | | | 28 | Gluconokinase (EC 2.7.1.12) / oxidoreductase domain | 35567 | 36748 | - | x | | | | | | | |
| | | | | | 29 | oxidoreductase domain protein | 36752 | 37831 | - | x | | | | | | | |
| | | | | | 30 | Oxidoreductase | 37833 | 38993 | - | x | | | | | | | |
| | | | | | 31 | Transcriptional regulator, AraC family | 39164 | 40090 | - | x | | | | | | | |
| | | | | | 32 | hypothetical protein | 40208 | 41818 | - | x | | | | | | | |
| | | | | | 33 | Predicted beta-xyloside ABC transporter, permease component | 41877 | 42749 | - | x | | | | | | | |
| 45P14-IVVE | | 43,328 bp | 1 **Bacterium RumenMGS205_scaffold_27 13,128bp** | AZCL-xylan, AZCL-arabinoxylan | 34 | ABC-type polysaccharide transport system, permease component | 42765 | 43286 | - | | | | | | | | |
| | | | | | 1 | hypothetical protein | 406 | 678 | + | | | | | | | | |
| | | | | | 2 | hypothetical protein | 657 | 2150 | - | | | | | | | | |
| | | | | | 3 | hypothetical protein | 2286 | 2708 | - | | | | | | | | |
| | | | | | 4 | N-carbamoylputrescine amidase (3.5.1.53) | 3192 | 4073 | - | | | | | | | | |
| | | | | | 5 | acetyltransferase, GNAT family | 4089 | 4523 | - | | | | | | | | |
| | | | | | 6 | hypothetical protein | 4534 | 6117 | - | | | | | | | | |
| | | | | | 7 | Endo-1,4-beta-xylanase A precursor (EC 3.2.1.8) | 6198 | 8033 | - | | | DOC-GH5_4 | | AEV59736.1 | 43.06 | 3E-102 | endo-b-1,4-glucanase (3.2.1.4) |
| | | | | | 8 | hypothetical protein | 8297 | 9454 | + | | | | | | | | |
| | | | | | 9 | tRNA-specific adenosine-34 deaminase (EC 3.5.4.-) | 9464 | 9961 | + | | | | | | | | |
| | | | | | 10 | UDP-glucose 4-epimerase (EC 5.1.3.2) | 10191 | 11138 | + | | | | | | | | |
| | | | | | 11 | conserved hypothetical protein | 11243 | 12583 | + | x | | | | | | | |
| | | | | | 12 | N-acetylglucosamine-6-phosphate deacetylase (EC 3.5.1.25) | 12608 | 14794 | + | | | CE9 | | | | | |
| | | | | | 13 | D-alanine--D-alanine ligase (EC 6.3.2.4) | 14866 | 15918 | + | | | | | | | | |
| | | | | | 14 | UDP-N-acetylmuramoylalanyl-D-glutamyl-2,6-diaminopimelate--D-alanyl-D-alanine ligase (EC 6.3.2.10) | 15921 | 17342 | + | | | | | | | | |
| | | | | | 15 | hypothetical protein | 17740 | 19311 | - | | | | | | | | |
| | | | | | 16 | Beta-xylosidase (EC 3.2.1.37) | 19534 | 21090 | + | | | CBM22-CE1 | | | | | |
| | | | | | 17 | Membrane-associated phospholipid phosphatase | 21461 | 22141 | - | | | | | | | | |
| | | | | | 18 | Rhomboid family protein | 22190 | 22804 | - | | | | | | | | |
| | | | | | 19 | hypothetical protein | 22801 | 25215 | - | | | | | | | | |
| | | | | | 20 | FIG002343: hypothetical protein | 25256 | 26551 | - | | | | | | | | |
| | | | | | 21 | FIG022979: MoxR-like ATPases | 26548 | 27489 | - | | | | | | | | |
| | | | | | 22 | hypothetical protein | 27520 | 27975 | - | | | | | | | | |
| | | | | | 23 | Superoxide reductase (EC 1.15.1.2) | 27972 | 28325 | - | | | | | | | | |
| | | | | | 24 | hypothetical protein | 28388 | 29137 | - | | | | | | | | |
| | | | | | 25 | hypothetical protein | 29125 | 29307 | + | | | | | | | | |
| | | | | | 26 | Glutamyl-tRNA synthetase (EC 6.1.1.17) | 29343 | 30986 | - | | | | | | | | |
| | | | | | 27 | hypothetical protein | 31349 | 32278 | - | | | | | | | | |
| 49M8-IVVE | | 33,187 bp | 1 **Firmicutes** | AZCL-Barley beta glucan, AZCL-HE cellulose, AZCL-tamarind xyloglucan | 28 | hypothetical protein | 32275 | 33135 | - | | | | | | | | |
| | | | | | 1 | Rubredoxin | 173 | 325 | - | | | | | | | | |
| | | | | | 2 | Uncharacterized protein MJ1614 | 486 | 1322 | + | x | | | | | | | |
| | | | | | 3 | hypothetical protein | 1324 | 3045 | + | x | | | | | | | |
| | | | | | 4 | ABC-type polysaccharide transport system, permease component | 3300 | 4274 | + | x | | | | | | | |
| | | | | | 5 | Predicted beta-xyloside ABC transporter, permease component | 4290 | 5162 | + | x | | | | | | | |
| | | | | | 6 | hypothetical protein | 5221 | 6831 | + | x | | | | | | | |
| | | | | | 7 | Transcriptional regulator, AraC family | 6949 | 7917 | - | x | | | | | | | |
| | | | | | 8 | Oxidoreductase | 8046 | 9206 | + | x | | | | | | | |
| | | | | | 9 | oxidoreductase domain protein | 9208 | 10287 | + | x | | | | | | | |
| | | | | | 10 | Gluconokinase (EC 2.7.1.12) / oxidoreductase domain | 10291 | 11472 | + | x | | | | | | | |
| | | | | | 11 | Phage repressor | 11619 | 12371 | + | x | 15 | | | | | | |
| | | | | | 12 | hypothetical protein | 12384 | 13775 | + | x | | | | | | | |
| | | | | | 13 | hypothetical protein | 13793 | 14059 | + | | | | | | | | |
| | | | | | 14 | putative esterase | 14065 | 14850 | - | x | | | | | | | |
| | | | | | 15 | Endo-1,4-beta-xylanase A precursor (EC 3.2.1.8) | 14892 | 15947 | - | x | | GH10 | 45P14 (37), 25I16 (19): 100% cov, 100 % id | AEW47914.1 | 82.81 | 0.0 | endo-b-1,4-xylanase (3.2.1.8) |
| | | | | | 16 | Acetyl-coenzyme A synthetase (EC 6.2.1.1) | 16029 | 18302 | - | x | | | | | | | |
| | | | | | 17 | DNA mismatch repair protein [MutS] | 18756 | 20471 | + | x | | | | | | | |

| | | | | | | | | | | | | | | | | | |
|---|---|---|---|---|---|---|---|---|---|---|---|---|---|---|---|---|---|
| | | | | | 18 | hypothetical protein | 20598 | 22475 | + | x | | | | | | | |
| | | | | | 19 | Ribosome protection-type tetracycline resistance related proteins, group 2 | 22504 | 25137 | - | x | | | | | | | |
| | | | | | 20 | hypothetical protein | 25307 | 26575 | + | x | | | | | | | |
| | | | | | 21 | hypothetical protein | 26580 | 27275 | + | | | | | | | | |
| | | | | | 22 | hypothetical protein | 27373 | 29895 | + | x | | | | | | | |
| | | | | | 23 | ABC transporter, ATP-binding protein | 29882 | 30820 | + | x | | | | | | | |
| | | | | | 24 | hypothetical protein | 30813 | 31730 | + | x | | | | | | | |
| | | | | | 25 | hypothetical protein | 31751 | 33250 | + | x | | | | | | | |
| | | | | | 26 | MoxR-like ATPase in aerotolerance operon | 33267 | 34244 | + | x | | | | | | | |
| | | | | | 27 | hypothetical protein | 34234 | 35082 | + | x | | | | | | | |
| 50E19-IVVE | LT674541 | 36,732 bp | 1 **Bacterium RumenMGS205_scaffold_35 11,466bp** | AZCL-xylan, AZCL-arabinoxylan, Tween 20, pNPA, pNPB | 28 | hypothetical protein | 35093 | 36661 | + | x | | | | | | | |
| | | | | | 1 | Pili retraction protein pilT | 30 | 596 | + | | | | | | | | |
| | | | | | 2 | N-acetylmuramoyl-L-alanine amidase (EC 3.5.1.28) | 694 | 2115 | + | | | | | | | | |
| | | | | | 3 | Mobile element protein | 2289 | 3752 | + | | | | | | | | |
| | | | | | 4 | tmRNA-binding protein SmpB | 4299 | 4772 | - | | | | | | | | |
| | | | | | 5 | Gamma-glutamyl phosphate reductase (EC 1.2.1.41) | 5008 | 6282 | + | | | | | | | | |
| | | | | | 6 | hypothetical protein | 6414 | 6620 | - | | | | | | | | |
| | | | | | 7 | hypothetical protein | 6624 | 6911 | - | | | | | | | | |
| | | | | | 8 | hypothetical protein | 7199 | 7354 | - | | | | | | | | |
| | | | | | 9 | hypothetical protein | 7528 | 7779 | + | | | | | | | | |
| | | | | | 10 | putative rRNA methylase | 8278 | 9120 | + | | | | | | | | |
| | | | | | 11 | hypothetical protein | 9194 | 11386 | - | | | | | | | | |
| | | | | | 12 | hypothetical protein | 11922 | 12767 | - | x | | | | | | | |
| | | | | | 13 | hypothetical protein | 12786 | 13688 | - | x | | | | | | | |
| | | | | | 14 | Cys-tRNA(Pro) deacylase YbaK | 13772 | 14257 | - | | | | | | | | |
| | | | | | 15 | hypothetical protein | 14351 | 15724 | - | x | | | | | | | |
| | | | | | 16 | N-acetylglucosamine-6-phosphate deacetylase (EC 3.5.1.25) | 15853 | 17001 | - | x | | CE9 | | | | | |
| | | | | | 17 | glycoside hydrolase, family 43 | 17218 | 18744 | - | | | GH43_10 | | ACY69973.1 | 62.08 | 4E-118 | a-L-arabinofuranosidase (3.2.1.55) |
| | | | | | 18 | Endo-1,4-beta-xylanase A precursor (EC 3.2.1.8) | 18707 | 21388 | - | | | GH10-GH43_10-CBM6 | | | | | |
| | | | | | 19 | Endo-1,4-beta-xylanase A precursor (EC 3.2.1.8) | 21508 | 22581 | - | | | CE1 | | | | | |
| | | | | | 20 | putative esterase | 22756 | 23589 | - | x | | | | | | | |
| | | | | | 21 | oxidoreductase domain protein | 23773 | 24873 | - | x | | | | | | | |
| | | | | | 22 | Predicted rhamnose oligosaccharide ABC transport system, permease component | 25077 | 25919 | - | x | 2 | | | | | | |
| | | | | | 23 | Predicted rhamnose oligosaccharide ABC transport system, permease component 2 | 25937 | 26863 | - | x | | | | | | | |
| | | | | | 24 | Predicted rhamnose oligosaccharide ABC transport system, substrate-binding component | 26967 | 28301 | - | x | | | | | | | |
| | | | | | 25 | Oxidoreductase | 28565 | 29719 | - | x | | GH179 | | QFQ13829.1 | 23.15 | 0.30 | b-N-acetylhexosaminidase (3.2.1.52) |
| | | | | | 26 | lipoprotein | 29814 | 31406 | - | x | | | | | | | |
| | | | | | 27 | binding-protein-dependent transport systems inner membrane component | 31614 | 32528 | - | x | | | | | | | |
| | | | | | 28 | Xylose ABC transporter, permease component | 32542 | 33486 | - | x | | | | | | | |
| 50I3-IVVE | LT674540 | 34,579 bp | 1 **Firmicutes** | Tween 20, pNPA, pNPB | 29 | Neuraminidase NanP | 33599 | 34561 | - | | | GH123 | | ABR39859.1 | 24.79 | 0,0002 | b-N-acetylgalactosaminidase (3.2.1.53) |
| | | | | | 1 | hypothetical protein | 709 | 1638 | + | x | | | | | | | |
| | | | | | 2 | Integral membrane protein | 1641 | 2786 | - | x | | | | | | | |
| | | | | | 3 | Neopullulanase (EC 3.2.1.135) | 2849 | 4924 | - | x | | GH13_21 | 32D12 (33): 100% cov, 95% id; 12F23 (16): 100% cov, 95% id; 38D22 (27): 100% cov, 95% id | BAA02471.1 | 34.85 | 4E-81 | a-amylase(3.2.1.1) |
| | | | | | 4 | Beta-1,3-glucosyltransferase | 4961 | 5782 | - | x | | GT2 | | | | | |
| | | | | | 5 | Endo-1,4-beta-xylanase A precursor (EC 3.2.1.8) | 5933 | 7459 | - | x | | CBM22-GH10 | | AHF25644.1 | 66.86 | 2E-161 | endo-b-1,4-xylanase (3.2.1.8) |
| | | 9,817 bp | 1 **Bacterium** | | 6 | hypothetical protein | 7650 | 9800 | - | | | | | | | | |
| | | | | | 1 | Glycosyltransferase | 270 | 1331 | + | x | | | | | | | |
| | | | | | 2 | hypothetical protein | 1522 | 3423 | + | x | | | | | | | |
| 51D10- IVVE | | 6,846 bp | 2 **Bacterium** | AZCL-xylan, AZCL-arabinoxylan | 3 | hypothetical protein | 3459 | 5852 | + | x | | | | | | | |
| | | | | | 1 | hypothetical protein | 417 | 1334 | - | | | | | | | | |
| | | | | | 2 | hypothetical protein | 1336 | 1623 | - | | | | | | | | |
| | | | | | 3 | Ribosyl nicotinamide transporter, PnuC-like | 1620 | 2312 | - | | | | | | | | |
| | | | | | 4 | Nicotinamide-nucleotide adenylyltransferase, NadR family (EC 2.7.7.1) / Ribosylnicotinamide kinase (EC 2.7.1.22) | 2305 | 3324 | - | | | | | | | | |
| | | | | | 5 | COG2152 predicted glycoside hydrolase | 3533 | 4558 | - | | | GH130_2 | | VCV21229.1 | 75.90 | 9E-166 | b-1,4-mannooligosaccharide phosphorylase (2.4.1.319) |
| | | | | | 6 | ABC transporter substrate-binding protein - sugar transport | 4732 | 6102 | - | x | 13 | | | | | | |
| | | | | | 7 | hypothetical protein | 6189 | 7922 | - | x | 37 | | | | | | |
| | | | | | 8 | N-Acetyl-D-glucosamine ABC transport system, permease protein 2 | 7985 | 8956 | - | x | 20 | | | | | | |
| | | | | | 9 | binding-protein-dependent transport systems inner membrane component | 8971 | 9891 | - | x | | | | | | | |
| | | | | | 10 | hypothetical protein | 9888 | 12485 | - | x | 72 | GH129 | | | | | |
| | | | | | 11 | FIG01031734: hypothetical protein | 12502 | 13137 | - | x | 6 | | | | | | |
| | | | | | 12 | NHL repeat containing protein / (gluconolactonase [Lachnoclostridium phytofermentans]) | 13134 | 14609 | - | | | | | | | | |
| | | | | | 13 | binding-protein-dependent transport systems inner membrane component | 14622 | 15551 | - | x | 1 | | | | | | |
| | | | | | 14 | binding-protein-dependent transport systems inner membrane component | 15553 | 16485 | - | x | 15 | | | | | | |
| | | | | | 15 | N-Acetyl-D-glucosamine ABC transport system, sugar-binding protein | 16487 | 19489 | - | x | 21 | | | | | | |
| | | | | | 16 | Mannan endo-1,4-beta-mannosidase precursor (EC 3.2.1.78) | 19515 | 20996 | - | x | 22 | CBM35-GH26 | | QIZ03395.1 | 55.97 | 1E-127 | endo-b-1,4-mannanase (3.2.1.78) |
| | | | | | 17 | hypothetical protein | 21391 | 21594 | + | x | 1 | | | | | | |
| | | | | | 18 | hypothetical protein / (gDSL-like Lipase/Acylhydrolase [Eubacterium eligens CAG:72]) | 21877 | 22686 | - | | | | | | | | |
| | | | | | 19 | hypothetical protein | 22727 | 23929 | - | | | | | | | | |
| | | | | | 20 | hypothetical protein | 23993 | 24151 | + | | | | | | | | |
| | | | | | 21 | Phage protein | 24205 | 25533 | - | | | | | | | | |
| | | | | | 22 | hypothetical protein | 25785 | 25931 | + | | | | | | | | |
| | | | | | 23 | hypothetical protein | 26230 | 26760 | - | x | 50 | | | | | | |
| | | | | | 24 | Glycogen synthase, ADP-glucose transglucosylase (EC 2.4.1.21) | 26870 | 28345 | - | | | GT5 | | | | | |
| | | | | | 25 | hypothetical protein | 28356 | 28538 | - | | | | | | | | |
| | | | | | 26 | Uridine kinase family protein | 28628 | 29410 | + | | | | | | | | |
| | | | | | 27 | hypothetical protein | 29559 | 30704 | + | | | | | | | | |
| | | | | | 28 | hypothetical protein | 30717 | 31187 | + | | | | | | | | |
| | | | | | 29 | hypothetical protein | 31184 | 31819 | + | | | | | | | | |
| | | | | | 30 | hypothetical protein | 31861 | 32052 | + | x | 21 | | | | | | |
| | | | | | 31 | hypothetical protein | 32077 | 32844 | + | | | | | | | | |
| | | | | | 32 | hypothetical protein | 32875 | 33657 | + | | | | | | | | |
| | | | | | 33 | hypothetical protein | 33694 | 33984 | + | | | | | | | | |
| | | | | | 34 | GCN5-related N-acetyltransferase | 34001 | 34447 | + | | | | | | | | |
| | | | | | 35 | hypothetical protein | 34515 | 34706 | + | | | GH3 | | AEF56854.1 | 45.00 | 0.011 | xylan b-1,4-xylosidase (3.2.1.37) |
| 52E22-IVVE | | 35,389 bp | 1 **Firmicutes RumenMGS052_scaffold_132 8,277 bp** | AZO-galacto-mannan | 36 | hypothetical protein | 34810 | 35031 | + | | | | | | | | |
| | | | | | 1 | Acyltransferase | 3083 | 4321 | + | | | | | | | | |
| | | | | | 2 | FIG053235: Diacylglucosamine hydrolase like | 4468 | 4971 | + | | | | | | | | |
| | | | | | 3 | Beta-glucosidase (EC 3.2.1.21) | 5109 | 7262 | - | | | GH3 | 41O1 (9): 100% cov, 100% id; | ZP_02070067.1 | 72.40 | 0.0 | exo-b-1,3-glucanase (3.2.1.58) |
| | | | | | 4 | Beta-glucanase precursor (EC 3.2.1.73) | 7275 | 8324 | - | | | GH16_3 | 41O1 (10): 100% cov, 100% id; | WP_072066679.1 | 80.95 | 2E-138 | endo-b-1,3-glucanase / laminarinase (3.2.1.39) |
| | | | | | 5 | hypothetical protein | 8308 | 8442 | - | | | | | | | | |
| | | | | | 6 | FIG00412289: hypothetical protein | 8446 | 9810 | - | | | | | | | | |
| | | | | | 7 | RagB/SusD domain protein | 9831 | 11300 | - | | | | | | | | |
| | | | | | 8 | hypothetical protein | 11254 | 11373 | + | | | | | | | | |
| | | | | | 9 | TonB family protein / TonB-dependent receptor | 11374 | 14385 | - | | | | | | | | |
| | | | | | 10 | FIG00417103: hypothetical protein | 14689 | 17538 | - | | | | | | | | |
| | | | | | 11 | Fe-S OXIDOREDUCTASE (1.8.-.-) | 17691 | 18398 | - | | | | | | | | |
| | | | | | 12 | FIG00898434: hypothetical protein | 18528 | 20591 | - | | | | | | | | |
| | | | | | 13 | hypothetical protein | 20602 | 21285 | - | | | | | | | | |
| | | | | | 14 | FIG00896812: hypothetical protein | 21518 | 22123 | + | | | | | | | | |
| | | | | | 15 | 23S rRNA methyltransferase (EC 2.1.1.-) | 22115 | 22645 | - | | | | | | | | |
| | | | | | 16 | Quinolinate synthetase (EC 2.5.1.72) | 23167 | 24165 | + | | | | | | | | |
| | | | | | 17 | Immunoreactive 84kD antigen PG93 | 24169 | 26457 | - | | | | | | | | |
| | | | | | 18 | Nucleoside 5-triphosphatase RdgB (dHAPTP, dITP, XTP-specific) (EC 3.6.1.15) | 26514 | 27089 | - | | | | | | | | |
| | | | | | 19 | putative transporter | 27168 | 28043 | - | | | | | | | | |
| 3F21-IVTE | | 30,213 bp | 1 **Bacteroidetes** Bacteroidales | AZCL-Barley beta glucan | 20 | Leucyl-tRNA synthetase (EC 6.1.1.4) | 28043 | 30211 | - | | | | | | | | |
| | | | | | 1 | hypothetical protein | 1027 | 2538 | - | x | 52 | | | | | | |
| | | | | | 2 | hypothetical protein | 2568 | 3572 | - | x | 3 | | | | | | |
| | | | | | 3 | hypothetical protein | 3586 | 3792 | - | | | | | | | | |
| | | | | | 4 | hypothetical protein | 3841 | 4011 | - | | | | | | | | |

| | | | | | | | | | | | | | | | | | |
|---|---|---|---|---|---|---|---|---|---|---|---|---|---|---|---|---|---|
| | | | | | 5 | transposase | 4169 | 5239 | + | x | | | | | | | |
| | | | | | 6 | FIG00937925: hypothetical protein | 5405 | 6115 | - | | | | | | | | |
| | | | | | 7 | Ynd | 6108 | 6869 | - | | | | | | | | |
| | | | | | 8 | hypothetical protein | 7408 | 7665 | - | | | | | | | | |
| | | | | | 9 | hypothetical protein | 7640 | 7756 | - | | | | | | | | |
| | | | | | 10 | Hypothetical Nudix-like regulator | 7910 | 8665 | + | | | | | | | | |
| | | | | | 11 | hydrolase, putative | 8668 | 9465 | + | | | | | | | | |
| | | | | | 12 | Mobile element protein | 9875 | 11110 | - | | | | | | | | |
| | | | | | 13 | Beta-glucosidase (EC 3.2.1.21) | 11571 | 13883 | - | | | GH3 | 30D14 (6): 100%cov, 100%id | ALJ47761.1 | 61.88 | 1,00E-163 | b-glucosidase (3.2.1.21) |
| | | | | | 14 | hypothetical protein | 13883 | 14623 | - | | | | | | | | |
| | | | | | 15 | FIG00937602: hypothetical protein | 14762 | 16504 | - | | | | | | | | |
| | | | | | 16 | FIG01289191: hypothetical protein | 16616 | 19795 | - | | | | | | | | |
| | | | | | 17 | DNA mismatch repair protein precursor (EC 3.2.1.4):B-glucanase | 19849 | 21096 | - | | | GH5_4 | 30D14 (2): 100%cov, 100%id | ALJ47761.1 | 47.27 | 4,00E-116 | lichenase / endo-b-1,3-1,4-glucanase (3.2.1.73) |
| 6F24 IVTE | | 21,494 bp | 1<br>**Bacteroidetes** Prevotella | AZO-CM cellulose, AZCL-Barley beta glucan | 18 | FIG00938042: hypothetical protein | 21306 | 21458 | - | | | | | | | | |
| | | | | | 1 | hypothetical protein | 104 | 265 | + | | | | | | | | |
| | | | | | 2 | Putative ion-channel protein | 289 | 1293 | - | | | | | | | | |
| | | | | | 3 | Cellobiose phosphorylase (EC 2.4.1.-) | 1312 | 3798 | - | | | GH94 | | ABD80580.1 | 76.64 | 0.0 | cellobiose phosphorylase (2.4.1.20) |
| | | | | | 4 | Beta-galactosidase (EC 3.2.1.23) | 4022 | 6466 | + | | | GH2 | | AAY50033.1 | 45.85 | 6E-67 | b-galactosidase (3.2.1.23) |
| | | | | | 5 | Transcriptional regulator of various polyols utilization, AraC family | 6792 | 7658 | + | | | | | | | | |
| | | | | | 6 | FIG00937556: hypothetical protein | 7779 | 9014 | + | | | CE7 | | | | | |
| | | | | | 7 | Endo-1,4-beta-mannosidase | 9106 | 10398 | + | | | GH5_7 | | AAS19695.1 | 43.93 | 4E-109 | b-mannosidase (3.2.1.25) |
| | | | | | 8 | Endoglucanase 5A (EC 3.2.1.4) (Endo-1,4-beta-glucanase 5A) (Alkaline cellulase) | 10556 | 11482 | + | | | GH5_2 | | AGL50932.1 | 63.41 | 1E-136 | endo-b-1,4-glucanase (3.2.1.4) |
| | | | | | 9 | hypothetical protein | 11546 | 11659 | + | | | | | | | | |
| | | | | | 10 | TonB family protein / TonB-dependent receptor | 11717 | 14983 | + | | | | | | | | |
| | | | | | 11 | putative outer membrane protein, probably involved in nutrient binding | 15004 | 16725 | + | | | | | | | | |
| | | | | | 12 | FIG00417257: hypothetical protein | 16730 | 18043 | + | | | | | | | | |
| | | | | | 13 | FIG00410329: hypothetical protein | 18065 | 20050 | + | | | | | | | | |
| | | 21,977 bp | 1<br>**Bacteroidetes** *Bacteroides uniformis* | | 14 | Mannan endo-1,4-beta-mannosidase B precursor (EC 3.2.1.78) | 20118 | 21815 | + | | | GH26 | 12E13 (7): 100% cov, 100% id | ARK08557.1 | 53.96 | 2,00E-69 | endo-b-1,4-mannanase (3.2.1.78) |
| | | | | | 1 | hypothetical protein | 403 | 1125 | - | | | | | | | | |
| | | | | | 2 | Di-/tripeptide transporter | 1125 | 2687 | - | | | | | | | | |
| | | | | | 3 | Lipid A biosynthesis lauroyl acyltransferase (EC 2.3.1.-) | 2786 | 3700 | + | | | | | | | | |
| 10D6 IVTE | | 5,173 bp | 2<br>**Bacteroidales** (Bacteroidetes) | AZO-CM cellulose, AZO Galactomannan | 4 | hypothetical protein | 3712 | 4722 | + | | | | | | | | |
| | | | | | 1 | hypothetical protein | 38 | 664 | + | | | | | | | | |
| | | | | | 2 | hypothetical protein | 690 | 1625 | + | | | | | | | | |
| | | | | | 3 | TonB family protein / TonB-dependent receptor | 2265 | 5618 | + | x | | | | | | | |
| | | | | | 4 | FIG00937915: hypothetical protein | 5627 | 7282 | + | x | 2 | | | | | | |
| | | | | | 5 | FIG00938925: hypothetical protein | 7301 | 8206 | + | x | | | | | | | |
| | | | | | 6 | hypothetical protein | 8218 | 9012 | + | x | 6 | | | | | | |
| | | | | | 7 | hypothetical protein | 9019 | 10353 | + | | | | | | | | |
| | | | | | 8 | hypothetical protein | 10710 | 12101 | + | | | | | | | | |
| | | | | | 9 | metallo-beta-lactamase family protein | 12083 | 12892 | + | x | 43 | | | | | | |
| | | | | | 10 | Cystathionine beta-lyase, Bsu PatB (EC 4.4.1.8) | 12889 | 13116 | - | | | | | | | | |
| | | | | | 11 | HIT family protein | 13321 | 13722 | - | | | | | | | | |
| | | | | | 12 | Transcription elongation factor GreA | 13728 | 14192 | - | | | | | | | | |
| | | | | | 13 | Aspartate aminotransferase (EC 2.6.1.1) | 14409 | 15716 | - | | | | | | | | |
| | | | | | 14 | Indolepyruvate oxidoreductase subunit IorB (EC 1.2.7.8) | 15697 | 16275 | - | | | | | | | | |
| | | | | | 15 | Indolepyruvate oxidoreductase subunit IorA (EC 1.2.7.8) | 16279 | 17931 | - | | | | | | | | |
| | | | | | 16 | Aliphatic amidase AmiE (EC 3.5.1.4) | 17910 | 18692 | - | | | | | | | | |
| | | | | | 17 | Aspartate-semialdehyde dehydrogenase (EC 1.2.1.11) | 18733 | 19743 | - | x | | | | | | | |
| | | | | | 18 | Probable exoglucanase a (1,4-beta-cellobiosidase) protein (EC 3.2.1.91) | 20198 | 25573 | + | | | | | | | | |
| | | | | | 19 | hypothetical protein | 25580 | 26773 | + | | | | | | | | |
| | | | | | 20 | hypothetical protein | 26791 | 28212 | + | | | | | | | | |
| | | | | | 21 | hypothetical protein | 28260 | 28559 | + | | | | | | | | |
| | | | | | 22 | Na+/H+ antiporter | 29313 | 31316 | + | | | | | | | | |
| | | | | | 23 | Lipoprotein releasing system ATP-binding protein LolD | 31345 | 32043 | + | | | | | | | | |
| | | | | | 24 | FIG00938655: hypothetical protein | 32070 | 32936 | + | | | | | | | | |
| | | | | | 25 | Octaprenyl diphosphate synthase (EC 2.5.1.90) / Dimethylallyltransferase (EC 2.5.1.1 | 33028 | 33924 | + | | | | | | | | |
| | | | | | 26 | Deoxyribose-phosphate aldolase (EC 4.1.2.4) | 33986 | 34927 | - | | | | | | | | |
| | | | | | 27 | D-tyrosyl-tRNA(Tyr) deacylase (EC 3.6.1.n1) | 34945 | 35397 | - | | | | | | | | |
| | | | | | 28 | Excinuclease ABC subunit C | 35394 | 37256 | - | | | | | | | | |
| | | | | | 29 | tRNA uridine 5-carboxymethylaminomethyl modification enzyme GidA | 37299 | 39194 | - | | | | | | | | |
| | | | | | 30 | nitroreductase | 39303 | 39917 | - | | | | | | | | |
| | | | | | 31 | Fumarate hydratase class I, anaerobic (EC 4.2.1.2) | 40220 | 41872 | + | x | | | | | | | |
| | | | | | 32 | Ribosome recycling factor | 41902 | 42465 | + | | | | | | | | |
| | | | | | 33 | Ribosome small subunit-stimulated GTPase EngC | 42473 | 43402 | + | | | | | | | | |
| | | | | | 34 | hypothetical protein | 43788 | 44300 | - | | | | | | | | |
| 10N13-IVTE | | 46,551 bp | 1<br>**Bacteroidetes** Bacteroidales RumenMGS049_scaffold_103 2,009 bp | ABTS | 35 | Glutamine synthetase type III, GlnN (EC 6.3.1.2) | 44544 | 46541 | - | x | | | | | | | |
| | | | | | 1 | hypothetical protein | 3 | 1190 | - | x | | | | | | | |
| | | | | | 2 | D-3-phosphoglycerate dehydrogenase (EC 1.1.1.95) | 1231 | 2178 | - | | | | | | | | |
| | | | | | 3 | Methyltransferase corrinoid activation protein | 2231 | 2665 | - | | | | | | | | |
| | | | | | 4 | Probable poly(beta-D-mannuronate) O-acetylase (EC 2.3.1.-) | 2846 | 4186 | + | | | | | | | | |
| | | | | | 5 | N-Acetylneuraminate cytidylyltransferase (EC 2.7.7.43) / (GDSL family lipase [[Clostri | 4196 | 5122 | + | | | | | | | | |
| | | | | | 6 | Flavoredoxin | 5190 | 5765 | - | | | | | | | | |
| | | | | | 7 | hypothetical protein | 5838 | 6158 | - | | | | | | | | |
| | | | | | 8 | Aldose 1-epimerase (EC 5.1.3.3) | 6169 | 7125 | - | | | | | | | | |
| | | | | | 9 | 2,4-dienoyl-CoA reductase [NADPH] (EC 1.3.1.34) | 7277 | 9229 | + | x | | | | | | | |
| | | | | | 10 | hypothetical protein | 9312 | 10736 | - | | | | | | | | |
| | | | | | 11 | Zn-dependent hydrolases of the metallo-beta-lactamase superfamily | 10813 | 11448 | - | | | | | | | | |
| | | | | | 12 | hypothetical protein / (thioesterase family protein [Firmicutes bacterium CAG:466]) | 11497 | 11970 | - | | | | | | | | |
| | | | | | 13 | hypothetical protein | 12023 | 12280 | - | | | | | | | | |
| | | | | | 14 | hypothetical protein | 12309 | 12821 | - | | | | | | | | |
| | | | | | 15 | Na(+)-linked D-alanine glycine permease | 13048 | 14448 | - | x | | | | | | | |
| | | | | | 16 | Phosphohydrolase / (metallophosphoesterase [Lachnospiraceae bacterium FE2018]) | 14859 | 15605 | - | | | | | | | | |
| | | | | | 17 | L-fuco-beta-pyranose dehydrogenase (EC 1.1.1.122) | 15700 | 16689 | - | x | | | | | | | |
| | | 17,653 bp | 1<br>**Firmicutes** | | 18 | Hydrolase (HAD superfamily) | 16705 | 17280 | - | | | | | | | | |
| | | | | | 1 | ABC transporter ATP-binding protein | 25 | 777 | + | | | | | | | | |
| | | | | | 2 | putative cell wall-binding protease | 1058 | 2212 | - | x | | | | | | | |
| | | | | | 3 | S-layer protein | 2228 | 3262 | - | | | | | | | | |
| | | | | | 4 | Alpha/beta superfamily hydrolase / (feruloyl esterase [Butyrivibrio sp. AE2015]) | 3549 | 4289 | + | | | | | | | | |
| | | | | | 5 | putative rRNA methylase | 4321 | 5163 | + | | | | | | | | |
| | | | | | 6 | Phenylacetic acid degradation protein PaaD, thioesterase | 5294 | 5746 | + | | | | | | | | |
| | | | | | 7 | hypothetical protein | 5773 | 6462 | + | | | | | | | | |
| | | | | | 8 | hypothetical protein | 6592 | 6882 | + | | | | | | | | |
| | | | | | 9 | hypothetical protein | 6896 | 8857 | + | x | | | | | | | |
| | | | | | 10 | hypothetical protein | 9327 | 9485 | - | | | | | | | | |
| | | | | | 11 | hypothetical protein | 9570 | 9866 | - | | | | | | | | |
| | | | | | 12 | Late competence protein ComEC, DNA transport | 9880 | 11007 | + | x | | | | | | | |
| | | | | | 13 | hypothetical protein | 11033 | 11581 | + | | | | | | | | |
| | | | | | 14 | Potassium voltage-gated channel subfamily KQT; possible potassium channel, VIC far | 11782 | 12519 | + | | | | | | | | |
| | | | | | 15 | hypothetical protein | 12628 | 12843 | + | | | | | | | | |
| | | | | | 16 | 2-dehydropantoate 2-reductase (EC 1.1.1.169) | 12951 | 13853 | + | x | | | | | | | |
| | | | | | 17 | Permease of the drug/metabolite transporter (DMT) superfamily | 13862 | 14749 | + | | | | | | | | |
| 11M20-IVTE | | 15,380 bp | 2<br>**Firmicutes** Lachnospiraceae | Tween 20, pNPA, pNPB, pNPP | 18 | hypothetical protein | 14868 | 15080 | + | | | | | | | | |
| | | | | | 1 | Alpha-N-arabinofuranosidase 2 (EC 3.2.1.55) | 15 | 929 | + | | | GH51 | | AAO75455.1 | 86.10 | 2E-118 | a-L-arabinofuranosidase (3.2.1.55) |
| | | | | | 2 | Transketolase (EC 2.2.1.1) | 999 | 3011 | + | | | | | | | | |
| | | | | | 3 | Ribose 5-phosphate isomerase B (EC 5.3.1.6) | 3113 | 3544 | + | | | | | | | | |
| | | | | | 4 | Beta-xylosidase (EC 3.2.1.37) | 3663 | 5552 | - | | | GH43_4 | | AAO75467.1 | 68.47 | 0.0 | endo-a-1,5-L-arabinanase (3.2.1.99) |
| | | | | | 5 | Putative outer membrane protein, probably involved in nutrient binding | 5622 | 7421 | - | | | | | | | | |
| | | | | | 6 | TonB-dependent receptor | 7446 | 10580 | - | | | | | | | | |
| | | | | | 7 | putative outer membrane protein, probably involved in nutrient binding | 10674 | 12440 | - | | | | | | | | |
| | | 13,214 bp | 1<br>**Bacteroidetes** *Bacteroides graminisolvens* | | 8 | TonB-dependent receptor | 12446 | 13165 | - | | | | | | | | |
| | | | | | 1 | putative DNA-binding protein | 318 | 1505 | + | | | | | | | | |
| | | | | | 2 | hypothetical protein | 1548 | 1661 | + | | | | | | | | |
| | | | | | 3 | hypothetical protein | 1828 | 2472 | - | | | | | | | | |
| | | | | | 4 | FIG00416167: hypothetical protein | 2621 | 3490 | - | | | | | | | | |
| | | | | | 5 | putative outer membrane protein, probably involved in nutrient binding | 3505 | 5274 | - | | | | | | | | |
| | | | | | 6 | SusC, outer membrane protein involved in starch binding | 5287 | 8535 | - | | | | | | | | |
| 12E13-IVTE | | 10,713 bp | 2<br>**Bacteroidetes** *Bacteroides uniformis* | AZO-galacto-mannan, AZCL-debranched arabinan | 7 | Mannan endo-1,4-beta-mannosidase B precursor (EC 3.2.1.78) | 8941 | 10638 | - | | | GH26 | 10D6 (14): 100% cov, 100% id | ARK08557.1 | 53.96 | 2E-69 | endo-b-1,4-mannanase (3.2.1.78) |
| | | | | | 1 | putative outer membrane protein, probably involved in nutrient binding | 4 | 1998 | + | | | | | | | | |

| | | | | | | | | | | | | | | | | | |
|---|---|---|---|---|---|---|---|---|---|---|---|---|---|---|---|---|---|
| | | | | | 2 | FIG00938343: hypothetical protein | 2041 | 3837 | + | | | | | | | | |
| | | | | | 3 | FIG00939126: hypothetical protein | 3868 | 4902 | + | | | | | | | | |
| | | | | | 4 | Endo-1,4-beta-xylanase A precursor (EC 3.2.1.8) | 4965 | 7139 | + | | | GH10-CBM4 | | EDV05072.1 | 86.76 | 2E-170 | endo-b-1,4-xylanase (3.2.1.8) |
| | | | | | 5 | hypothetical protein | 7146 | 7274 | - | | | | | | | | |
| | | | | | 6 | Endo-1,4-beta-xylanase A precursor (EC 3.2.1.8) | 7228 | 9450 | + | | | GH10-CBM48 | | ADN44261.1 | 68.15 | 3E-168 | endo-b-1,4-xylanase (3.2.1.8) |
| | | | | | 7 | DNA-binding response regulator, AraC family | 9595 | 13485 | - | | | | | | | | |
| | | | | | 8 | Endoglucanase D precursor (EC 3.2.1.4) | 13889 | 16414 | + | | | GH9 | | ACL75133.1 | 21.15 | 0,00003 | endo-b-1,4-glucanase (3.2.1.4) |
| | | | | | 9 | Endo-1,4-beta-xylanase A precursor (EC 3.2.1.8) | 16459 | 17583 | + | | | CBM48-CE1 | | | | | |
| | | | | | 10 | Endo-1,4-beta-xylanase A precursor (EC 3.2.1.8) | 17638 | 20157 | + | | | GH43_2-CBM6-GH8 | | EDV05843.1 | 85.57 | 0.0 | oligosaccharide reducing-end xylanase (3.2.1.156) |
| | | | | | 11 | Hypothetical glycoside hydrolase, family 43, similar to arabinosidase | 20221 | 21183 | - | | | GH43_29 | | AAO75476.1 | 53.45 | 1E-102 | a-1,2-L-arabinofuranosidase (3.2.1.-),a-L-arabinofuranosidase (3.2.1.55) |
| | | | | | 12 | Pyridoxamine 5'-phosphate oxidase (EC 1.4.3.5) | 21674 | 22165 | + | | | | | | | | |
| | | | | | 13 | hypothetical protein | 22187 | 22603 | - | | | | | | | | |
| | | 24,657 bp | 1 **Bacteroidetes** *Bacteroides graminisolvens* | | 14 | transcriptional regulator | 22852 | 23058 | + | | | | | | | | |
| | | | | | 1 | SusD, outer membrane protein | 1282 | 3024 | - | | | | | | | | |
| 13F22-IVTE | | 4,084 bp | 2 **Bacteroidetes** | AZCL-arabinoxylan | 2 | putative outer membrane protein, probably involved in nutrient binding | 3043 | 3864 | - | | | | | | | | |
| | | | | | 1 | Amino acid ABC transporter, ATP-binding protein | 32 | 625 | + | | | | | | | | |
| | | | | | 2 | putative transmembrane protein | 644 | 1156 | + | | | | | | | | |
| | | | | | 3 | Possible glyoxylase family protein (Lactoylglutathione lyase) (EC 4.4.1.5) | 1179 | 1565 | + | | | | | | | | |
| | | | | | 4 | O-acetylhomoserine sulfhydrylase (EC 2.5.1.49) / O-succinylhomoserine sulfhydrylase | 1754 | 3049 | + | x | | | | | | | |
| | | | | | 5 | Cysteine synthase (EC 2.5.1.47) | 3190 | 4122 | + | x | | | | | | | |
| | | | | | 6 | hypothetical protein | 4174 | 5907 | - | x | | | | | | | |
| | | | | | 7 | Endoglucanase (EC 3.2.1.4) | 6107 | 7135 | + | x | | GH5_37 | | ADX05739.1 | 59.13 | 5E-153 | endo-b-1,4-glucanase (3.2.1.4) |
| | | | | | 8 | Anthranilate synthase, aminase component (EC 4.1.3.27) | 7463 | 8911 | + | x | | | | | | | |
| | | | | | 9 | Anthranilate synthase, amidotransferase component (EC 4.1.3.27) | 8913 | 9482 | + | | | | | | | | |
| | | | | | 10 | Glutamate synthase [NADPH] large chain (EC 1.4.1.13) | 9707 | 14236 | + | | | | | | | | |
| | | | | | 11 | Glutamate synthase [NADPH] small chain (EC 1.4.1.13) | 14253 | 15716 | + | | | | | | | | |
| | | | | | 12 | Glutamine synthetase type I (EC 6.3.1.2) | 15803 | 17020 | + | | | | | | | | |
| | | | | | 13 | Response regulator receiver and ANTAR domain protein | 17041 | 17622 | + | | | | | | | | |
| | | | | | 14 | putative esterase | 17661 | 18488 | - | | | | | | | | |
| | | | | | 15 | Carotenoid cis-trans isomerase (EC 5.2.-.-) | 18687 | 20168 | + | | | | | | | | |
| | | | | | 16 | hypothetical protein | 20178 | 20741 | + | | | | | | | | |
| | | | | | 17 | Glutamine synthetase type III, GlnN (EC 6.3.1.2) | 20871 | 22973 | + | | | | | | | | |
| | | | | | 18 | Asparagine synthetase [glutamine-hydrolyzing] (EC 6.3.5.4) | 23031 | 24617 | + | x | 10 | | | | | | |
| | | | | | 19 | CTP synthase (EC 6.3.4.2) | 24829 | 25860 | + | | | | | | | | |
| | | 27,156 bp | 1 **Firmicutes** Lachnospiraceae | | 20 | CTP synthase (EC 6.3.4.2) | 25844 | 26437 | + | | | | | | | | |
| | | | | | 1 | Fructose-bisphosphate aldolase class II (EC 4.1.2.13) | 127 | 270 | + | | | | | | | | |
| | | | | | 2 | hypothetical protein | 425 | 994 | + | | | | | | | | |
| | | | | | 3 | hypothetical protein | 1370 | 2146 | - | | | | | | | | |
| | | | | | 4 | hypothetical protein | 2146 | 2676 | - | | | | | | | | |
| 14J10-IVTE | | 2,993 bp | 2 **Bacteroidetes** **Bacteroides** | AZCL-Barley beta glucan, AZO-CM cellulose,Tween 20 | 5 | hypothetical protein | 2705 | 2968 | - | | | | | | | | |
| | | | | | 1 | hypothetical protein | 1140 | 3074 | + | | | | | | | | |
| | | | | | 2 | Alanine racemase (EC 5.1.1.1) | 3046 | 4215 | - | | | | | | | | |
| | | | | | 3 | hypothetical protein | 4248 | 4871 | - | | | | | | | | |
| | | | | | 4 | Cellulase( EC:3.2.1.4 ) | 4881 | 6245 | - | x | | GH5_4 | 44C18 (4): 100% cov, 100% id; 44C20 (4) 100% cov, 100% id | ABX42426.1 | 44.47 | 7E-110 | endo-b-1,4-glucanase (3.2.1.4) |
| | | | | | 5 | Ribosome small subunit-stimulated GTPase EngC | 6299 | 7171 | - | | | | | | | | |
| | | | | | 6 | Serine/threonine protein kinase PrkC, regulator of stationary phase | 7183 | 9264 | - | x | | | | | | | |
| | | | | | 7 | Protein serine/threonine phosphatase PrpC, regulation of stationary phase | 9292 | 10020 | - | | | | | | | | |
| | | | | | 8 | Ribosomal RNA large subunit methyltransferase N (EC 2.1.1.-) | 10066 | 11103 | - | x | | | | | | | |
| | | (11,640 bp) | 1 **Firmicutes** | | 9 | Ribosomal RNA small subunit methyltransferase B (EC 2.1.1.-) | 11103 | 11627 | - | | | | | | | | |
| | | | | | 1 | Transcription termination factor Rho | 30 | 266 | + | | | | | | | | |
| | | | | | 2 | Choline-sulfatase (EC 3.1.6.6) | 421 | 1866 | + | | | | | | | | |
| | | | | | 3 | Signal recognition particle, subunit Ffh SRP54 (TC 3.A.5.1.1) | 2060 | 3382 | + | | | | | | | | |
| | | | | | 4 | Methylenetetrahydrofolate dehydrogenase (NADP+) (EC 1.5.1.5) / Methenyltetrahyd | 3394 | 4275 | + | | | | | | | | |
| | | | | | 5 | Capsule biosynthesis protein capA | 4283 | 5428 | + | | | | | | | | |
| | | | | | 6 | Fe-S oxidoreductase | 6053 | 7747 | + | | | | | | | | |
| | | | | | 7 | Glutamine synthetase type I (EC 6.3.1.2) | 7809 | 9311 | - | | | | | | | | |
| | | | | | 8 | hypothetical protein | 9340 | 9486 | + | | | | | | | | |
| | | | | | 9 | 3-oxoacyl-[acyl-carrier-protein] synthase, KASIII (EC 2.3.1.180) | 9527 | 10453 | - | | | | | | | | |
| | | | | | 10 | hypothetical protein | 10622 | 10792 | - | | | | | | | | |
| | | | | | 11 | hypothetical protein | 11135 | 11305 | + | | | | | | | | |
| | | | | | 12 | hypothetical protein | 11558 | 11692 | - | | | | | | | | |
| | | | | | 13 | Arsenical resistance operon repressor | 11912 | 12238 | + | | | | | | | | |
| | | | | | 14 | Arsenical pump-driving ATPase (EC 3.6.3.16) | 12309 | 12548 | + | | | | | | | | |
| | | | | | 15 | hypothetical protein | 12558 | 13001 | + | | | | | | | | |
| | | | | | 16 | Cytochrome c biogenesis protein | 13003 | 13710 | + | | | | | | | | |
| | | | | | 17 | Arsenical resistance operon trans-acting repressor ArsD | 13745 | 14071 | + | | | | | | | | |
| | | | | | 18 | Arsenical pump-driving ATPase (EC 3.6.3.16) | 14097 | 15809 | + | | | | | | | | |
| 15F15-IVTE | | 16,634 bp | 2 **Bacteroidetes** **Bacteroides** | AZCL-Barley beta glucan, AZO-CM cellulose, AZCL Tamarind xyloglucan, AZCL-HE cellulose | 19 | Arsenate reductase (EC 1.20.4.1) | 15814 | 16242 | + | | | | | | | | |
| | | | | | 1 | SSU ribosomal protein S21p | 490 | 684 | - | | | | | | | | |
| | | | | | 2 | hypothetical protein | 992 | 2002 | + | x | 69 | | | | | | |
| | | | | | 3 | NAD-dependent protein deacetylase of SIR2 family | 1999 | 2706 | + | | | | | | | | |
| | | | | | 4 | LSU ribosomal protein L21p | 2830 | 3141 | + | | | | | | | | |
| | | | | | 5 | LSU ribosomal protein L27p | 3161 | 3421 | + | | | | | | | | |
| | | | | | 6 | Seryl-tRNA synthetase (EC 6.1.1.11) | 3505 | 4776 | + | x | | | | | | | |
| | | | | | 7 | Crossover junction endodeoxyribonuclease RuvC (EC 3.1.22.4) | 4783 | 5346 | + | | | | | | | | |
| | | | | | 8 | ThiF domain protein | 5339 | 6085 | - | x | 14 | | | | | | |
| | | | | | 9 | Apolipoprotein N-acyltransferase (EC 2.3.1.-) | 6082 | 7587 | - | x | 2 | | | | | | |
| | | | | | 10 | Putative threonine efflux protein | 7855 | 8421 | - | | | | | | | | |
| | | | | | 11 | hypothetical protein | 8423 | 9205 | - | | | | | | | | |
| | | | | | 12 | Ribosomal large subunit pseudouridine synthase D (EC 4.2.1.70) | 9207 | 10238 | + | x | | | | | | | |
| | | | | | 13 | Cytoplasmic axial filament protein CafA and Ribonuclease G (EC 3.1.4.-) | 10256 | 11827 | - | x | | | | | | | |
| | | | | | 14 | DNA-binding protein HU | 12056 | 12328 | - | | | | | | | | |
| | | | | | 15 | ATP-dependent protease La (EC 3.4.21.53) Type I | 12572 | 14956 | + | x | | | | | | | |
| | | | | | 16 | Uracil-DNA glycosylase, family 1 | 14956 | 15624 | + | | | | | | | | |
| | | | | | 17 | Phosphopantetheine adenylyltransferase (EC 2.7.7.3) | 15621 | 16094 | + | | | | | | | | |
| | | | | | 18 | hypothetical protein | 16096 | 16290 | - | | | | | | | | |
| | | | | | 19 | hypothetical protein | 16333 | 17565 | + | x | 4 | | | | | | |
| | | | | | 20 | DJ-1/YajL/PfpI superfamily, includes chaperone protein YajL (former ThiJ), parkinsoni | 17554 | 18126 | - | | | | | | | | |
| | | | | | 21 | UDP-N-acetylmuramoylalanyl-D-glutamyl-2,6-diaminopimelate--D-alanyl-D-alanine li | 18236 | 19348 | - | x | | | | | | | |
| | | | | | 22 | probable metal-dependent peptidase | 19707 | 20381 | + | | | | | | | | |
| | | | | | 23 | hypothetical protein | 20378 | 20671 | - | | | | | | | | |
| | | | | | 24 | tRNA-specific adenosine-34 deaminase (EC 3.5.4.-) | 20688 | 21122 | - | | | | | | | | |
| | | | | | 25 | hypothetical protein | 21243 | 21473 | - | | | | | | | | |
| | | | | | 26 | Transcriptional regulator, AraC family | 21558 | 22130 | + | | | | | | | | |
| | | | | | 27 | Endo-1,4-beta-mannosidase | 22237 | 24699 | + | x | 41 | GH5_7-CE7 | 20G16 (18): 97% cov, 100% id; 38A17 (18): 100% cov, 100% id; 44D8 (13): 97%cov, 100% id; 44F17 (19): 97% cov, 100% id | AAS19695.1 | 41.77 | 1E-99 | b-mannosidase (3.2.1.25) |
| | | | | | 28 | TonB family protein / TonB-dependent receptor | 24787 | 27936 | + | x | 9 | | | | | | |
| | | | | | 29 | putative outer membrane protein, probably involved in nutrient binding | 27949 | 29697 | + | x | 24 | | | | | | |
| | | | | | 30 | FIG00417257: hypothetical protein | 29710 | 31014 | + | x | 7 | | | | | | |
| | | | | | 31 | hypothetical protein | 31021 | 33447 | + | x | | | | | | | |
| | | | | | 32 | Cellulase( EC:3.2.1.4 ) | 33542 | 35068 | + | x | 35 | GH5_4 | 20G16 (23): 100% cov, 100% id; 46A3 (18): 100% cov, 100% id; 44D8 (18): 100%cov, 100% id; 44F17 (19): 100% cov, 100% id | CAJ19139.1 | 67.53 | 0.0 | lichenase / endo-b-1,3-1,4-glucanase (3.2.1.73) |
| | | | | | 33 | Mannan endo-1,4-beta-mannosidase | 35185 | 36213 | + | x | 3 | GH26 | 20G16 (25):100% cov, 100% id; 28D5 (3): 100% cov, 100% id; 38A17 (24): 100% cov, 100% id; 44D8 (19): 100% cov, 100% id; 44F17 (13): 99% cov, 100% id; 46A3 (4): 99% cov, 100% id | AIT97142.1 | 49.85 | 1E-106 | exo-b-1,4-mannobiohydrolase / mannobiose-producing exo-b-mannanase (3.2.1.100) |

| | | | | | | | | | | | | | | | | | |
|---|---|---|---|---|---|---|---|---|---|---|---|---|---|---|---|---|---|
| | | | | | 34 | COG2152 predicted glycoside hydrolase | 36291 | 37253 | + | x | 10 | GH130 | 20G16 (25): 100% cov, 100% id; 28D5 (4): 100% cov, 100% id; 38A17 (25): 100% cov, 100% id; 44F17 (12): 100% cov, 100% id; 46A3 (5): 100% cov, 100% id; 44D8 (20): 100% cov, 100% id | AAS19693.1 | 70.82 | 5E-109 | |
| | | | | | 35 | COG2152 predicted glycoside hydrolase | 37214 | 37465 | + | | | GH130 | | AAS19693.1 | 68.63 | 7E-18 | |
| 15K21-IVTE | | 39,574 bp | 1 **Bacteroidetes** Bacteroides RumenMGS049_scaffold_83 18,169 bp | AZO-CM cellulose, AZCL-HE cellulose, AZCL-Barley beta glucan, AZCL-Tamarind xyloglucan, AZCL-arabinoxylan | 36 | Xyloside transporter XynT | 37486 | 38913 | + | x | | | | | | | |
| | | | | | 1 | putative glycosyltransferase | 10 | 870 | - | | | GT14 | | | | | |
| | | | | | 2 | Succinoglycan biosynthesis protein | 871 | 1950 | - | x | 11 | | | | | | |
| | | | | | 3 | Ribosomal large subunit pseudouridine synthase D (EC 4.2.1.70) | 1930 | 2634 | - | | | | | | | | |
| | | | | | 4 | Cytoplasmic axial filament protein CafA and Ribonuclease G (EC 3.1.4.-) | 2652 | 4223 | - | x | | | | | | | |
| | | | | | 5 | DNA-binding protein HU | 4452 | 4724 | - | | | | | | | | |
| | | | | | 6 | ATP-dependent protease La (EC 3.4.21.53) Type I | 5010 | 7352 | + | x | | | | | | | |
| | | | | | 7 | Uracil-DNA glycosylase, family 1 | 7352 | 8020 | + | | | | | | | | |
| | | | | | 8 | Phosphopantetheine adenylyltransferase (EC 2.7.7.3) | 8017 | 8490 | + | | | | | | | | |
| | | | | | 9 | hypothetical protein | 8492 | 8710 | - | | | | | | | | |
| | | | | | 10 | Thiol:disulfide interchange protein tlpA | 8714 | 9961 | + | x | 4 | | | | | | |
| | | | | | 11 | DJ-1/YajL/PfpI superfamily, includes chaperone protein YajL (former ThiJ), parkinsoni | 9950 | 10522 | - | | | | | | | | |
| | | | | | 12 | UDP-N-acetylmuramoylalanyl-D-glutamyl-2,6-diaminopimelate--D-alanyl-D-alanine li | 10632 | 11744 | - | x | 4 | | | | | | |
| | | | | | 13 | probable metal-dependent peptidase | 12103 | 12777 | + | | | | | | | | |
| | | | | | 14 | hypothetical protein | 12774 | 13067 | - | | | | | | | | |
| | | | | | 15 | tRNA-specific adenosine-34 deaminase (EC 3.5.4.-) | 13084 | 13518 | - | | | | | | | | |
| | | | | | 16 | hypothetical protein | 13639 | 13869 | - | | | | | | | | |
| | | | | | 17 | Transcriptional regulator, AraC family | 13954 | 14526 | + | | | | | | | | |
| | | | | | 18 | Endo-1,4-beta-mannosidase | 14696 | 17095 | + | x | 41 | GH5_7-CE7 | 15K21 (27): 100% cov, 100% id; 38A17 (18): 100% cov, 100% id; 44D8 (13): 100%cov, 100% id; 44F17 (19): 99% cov, 100% id | AAS19695.1 | 41.77 | 6E-100 | b-mannosidase (3.2.1.25) |
| | | | | | 19 | TonB family protein / TonB-dependent receptor | 17153 | 20332 | + | x | 9 | | | | | | |
| | | | | | 20 | putative outer membrane protein, probably involved in nutrient binding | 20345 | 22093 | + | x | 24 | | | | | | |
| | | | | | 21 | FIG00417257: hypothetical protein | 22106 | 23410 | + | x | 7 | | | | | | |
| | | | | | 22 | hypothetical protein | 23417 | 25843 | + | x | | | | | | | |
| | | | | | 23 | Cellulase( EC:3.2.1.4 ) | 25938 | 27464 | + | x | 35 | GH5_4 | 15K21 (32): 100% cov, 100% id; 46A3(3): 100% cov, 100% id; 44D8 (18): 100%cov, 100% id; 44F17 (14): 99% cov, 100% id | CAJ19139.1 | 67.53 | 0.0 | lichenase / endo-b-1,3-1,4-glucanase (3.2.1.73) |
| | | | | | 24 | Mannan endo-1,4-beta-mannosidase | 27578 | 28609 | + | x | 3 | GH26 | 15K21 (33): 100% cov, 100% id; 46A3(4): 100% cov, 100% id; 44D8 (19): 100%cov, 100% id; 44F17 (13): 99% cov, 100% id | AIT97142.1 | 49.85 | 7E-107 | exo-b-1,4-mannobiohydrolase / mannobiose-producing exo-b-mannanase (3.2.1.100) |
| | | | | | 25 | COG2152 predicted glycoside hydrolase | 28667 | 29842 | + | x | 10 | GH130_1 | 28D5 (4): 100% cov, 100% id; 38A17 (25): 100% cov, 100% id; 44F17 (12): 99% cov, 100% id; 46A3 (5): 99% cov, 100% id; 44D8 (20): 100% cov, 100% id | AAS19693.1 | 69.49 | 4E-139 | b-1,4-mannosylglucose phosphorylase (2.4.1.281) |
| | | | | | 26 | Xyloside transporter XynT | 29863 | 31290 | + | x | 16 | | | | | | |
| | | | | | 27 | DNA polymerase I (EC 2.7.7.7) | 31846 | 34494 | + | x | 27 | | | | | | |
| 20G16-IVTE | | 35,118 bp | 1 **Bacteroidetes** Bacteroidales RumenMGS049_scaffold_83 12,691 bp | AZO-CM cellulose, AZCL-HE cellulose, AZCL-Barley beta glucan, AZCL-Tamarind xyloglucan, AZCL-arabinoxylan | 28 | RNA polymerase sigma factor RpoE | 34466 | 35071 | + | | | | | | | | |
| | | | | | 1 | OmpA family outer membrane protein | 2606 | 3292 | - | | | | | | | | |
| | | | | | 2 | Glucosamine-6-phosphate deaminase (EC 3.5.99.6) | 3374 | 5296 | - | x | | | | | | | |
| | | | | | 3 | hypothetical protein | 5535 | 6131 | - | | | | | | | | |
| | | | | | 4 | putative outer membrane protein, probably involved in nutrient binding | 6214 | 7995 | - | x | 19 | | | | | | |
| | | | | | 5 | SusC, outer membrane protein involved in starch binding | 8002 | 11127 | - | x | 14 | | | | | | |
| | | | | | 6 | Hypothetical sugar kinase, ROK family | 11212 | 12045 | - | x | | | | | | | |
| | | | | | 7 | Xylose operon regulatory protein | 12133 | 13290 | - | | | | | | | | |
| | | | | | 8 | Beta-hexosaminidase (EC 3.2.1.52) | 13394 | 15634 | - | x | 12 | GH20 | | AEW20137.1 | 56.57 | 2E-179 | b-N-acetylhexosaminidase (3.2.1.52) |
| | | | | | 9 | Alkaline phosphatase (EC 3.1.3.1) | 15667 | 18771 | - | x | 70 | GH2 | | AAX62629.2 | 36.70 | 5E-57 | exo-b-1,4-glucosaminidase (3.2.1.165) |
| | | | | | 10 | Small protein containing transglutaminase-like domain | 18870 | 19316 | - | | | | | | | | |
| | | | | | 11 | Lipid-A-disaccharide synthase (EC 2.4.1.182) | 19343 | 20497 | - | | | GT19 | | | | | |
| | | | | | 12 | Protein of unknown function DUF1009 clustered with KDO2-Lipid A biosynthesis gen | 20494 | 21312 | - | | | | | | | | |
| | | | | | 13 | Acyl-[acyl-carrier-protein]--UDP-N-acetylglucosamine O-acyltransferase (EC 2.3.1.129 | 21309 | 22103 | - | | | | | | | | |
| | | | | | 14 | 3-hydroxyacyl-[acyl-carrier-protein] dehydratase, FabZ form (EC 4.2.1.59) | 22135 | 22602 | - | | | | | | | | |
| | | | | | 15 | Outer membrane protein H precursor | 22745 | 23458 | - | | | | | | | | |
| | | | | | 16 | Outer membrane protein assembly factor YaeT precursor | 23467 | 25848 | - | | | | | | | | |
| | | | | | 17 | Membrane-associated zinc metalloprotease | 26009 | 27379 | - | | | | | | | | |
| | | | | | 18 | 1-deoxy-D-xylulose 5-phosphate reductoisomerase (EC 1.1.1.267) | 27383 | 28606 | - | | | | | | | | |
| | | | | | 19 | Phosphatidate cytidylyltransferase (EC 2.7.7.41) | 28608 | 29420 | - | | | | | | | | |
| | | | | | 20 | Undecaprenyl diphosphate synthase (EC 2.5.1.31) | 29417 | 30169 | - | | | | | | | | |
| | | | | | 21 | Ribosome recycling factor | 30185 | 30748 | - | | | | | | | | |
| | | | | | 22 | Uridine monophosphate kinase (EC 2.7.4.22) | 30868 | 31599 | - | | | | | | | | |
| | | | | | 23 | tRNA dimethylallyltransferase (EC 2.5.1.75) | 31681 | 32580 | + | | | | | | | | |
| | | | | | 24 | Transcriptional regulator, AraC family | 32774 | 33535 | + | | | | | | | | |
| | | | | | 25 | hypothetical protein | 33545 | 33913 | - | | | | | | | | |
| | | | | | 26 | Aldehyde dehydrogenase (EC 1.2.1.3) | 34122 | 35651 | - | | | | | | | | |
| | | | | | 27 | hypothetical protein | 35706 | 35867 | + | | | | | | | | |
| 21E20-IVTE | | 37,465 bp | 1 **Proteobacteria** *Paenirhodobacter enshiensis* **RumenMGS049_scaffold_2 16,318 bp** | lignin alkali | 28 | Succinate-semialdehyde dehydrogenase [NAD] (EC 1.2.1.24); Succinate-semialdehyd | 35988 | 37379 | - | | | | | | | | |
| | | | | | 1 | hypothetical protein | 347 | 652 | + | | | | | | | | |
| | | | | | 2 | hypothetical protein | 652 | 981 | + | | | | | | | | |
| | | | | | 3 | hypothetical protein | 1210 | 1545 | - | | | | | | | | |
| | | | | | 4 | FIG00523589: hypothetical protein | 1528 | 1725 | + | | | | | | | | |
| | | | | | 5 | hypothetical protein | 2236 | 2790 | + | | | | | | | | |
| | | | | | 6 | Transcriptional regulator, PadR family | 2899 | 3219 | + | | | | | | | | |
| | | | | | 7 | hypothetical protein | 3225 | 3794 | + | | | | | | | | |
| | | | | | 8 | hypothetical protein | 3914 | 4777 | - | | 10 | | | | | | |
| | | | | | 9 | hypothetical protein | 4886 | 5515 | + | | | | | | | | |
| | | | | | 10 | Endo-1,4-beta-xylanase A precursor (EC 3.2.1.8) | 5584 | 7191 | + | x | | GH5_4-CBM72 | | ACA61140.1 | 43.10 | 4E-91 | endo-b-1,4-glucanase (3.2.1.4) |
| | | | | | 11 | Potassium uptake protein TrkH | 7262 | 8719 | - | x | 37 | | | | | | |
| | | | | | 12 | Putative outer membrane protein, probably involved in nutrient binding | 8750 | 10426 | - | x | 5 | | | | | | |
| | | | | | 13 | TonB-dependent receptor | 10440 | 13223 | - | x | 5 | | | | | | |
| | | | | | 14 | FIG00405972: hypothetical protein | 13245 | 15311 | - | x | 5 | | | | | | |
| | | | | | 15 | TonB-dependent receptor | 15319 | 15717 | - | | | | | | | | |
| | | | | | 16 | Ferric uptake regulation protein FUR | 15733 | 16059 | + | | | | | | | | |
| | | | | | 17 | Putative metal chaperone, involved in Zn homeostasis, GTPase of COG0523 family | 16096 | 17253 | + | x | | | | | | | |
| | | | | | 18 | aminoglycoside phosphotransferase | 17347 | 18252 | - | | | | | | | | |
| | | | | | 19 | CRISPR-associated protein, Csn1 family | 18567 | 21662 | + | x | | | | | | | |
| | | | | | 20 | CRISPR-associated protein Cas1 | 21659 | 22579 | + | x | | | | | | | |
| | | | | | 21 | CRISPR-associated protein Cas2 | 22612 | 22968 | + | | | | | | | | |
| | | | | | 22 | Arabinogalactan endo-1,4-beta-galactosidase | 23531 | 24682 | - | x | | GH53 | | ADM99400.1 | 32.36 | 5E-40 | endo-b-1,4-galactanase (3.2.1.89) |
| | | | | | 23 | hypothetical protein | 25119 | 26456 | + | x | | | | | | | |
| | | | | | 24 | Uracil-DNA glycosylase, family 1 | 26473 | 27150 | + | | | | | | | | |
| | | | | | 25 | Cobalt-zinc-cadmium resistance protein | 27271 | 28407 | + | x | | | | | | | |
| | | | | | 26 | hypothetical protein | 28458 | 28784 | + | | | | | | | | |
| | | | | | 27 | Aspartyl-tRNA(Asn) amidotransferase subunit A (EC 6.3.5.6) @ Glutamyl-tRNA(Gln) a | 28784 | 30202 | + | x | 23 | | | | | | |
| | | | | | 28 | hypothetical protein | 30268 | 30957 | + | | | | | | | | |
| | | | | | 29 | hypothetical protein | 30989 | 31192 | + | | | | | | | | |
| | | | | | 30 | Inosine-5'-monophosphate dehydrogenase (EC 1.1.1.205) | 31532 | 32989 | - | x | | | | | | | |
| | | | | | 31 | Endo-1,4-beta-xylanase A precursor (EC 3.2.1.8) (1,4-beta-D-xylan xylanohydrolase) ( | 33261 | 34256 | - | x | | GH11 | | ACX76028.1 | 100.00 | 2E-174 | endo-b-1,4-xylanase (3.2.1.8) |
| 23O13-IVTE | | 35,397 bp | 1 **Bacteria** **RumenMGS049_scaffold_62 8,109 bp** | AZCL-xylan, AZCL-arabinoxylan | 32 | hypothetical protein | 34327 | 35394 | - | | | | | | | | |
| | | | | | 1 | Chromate transporter | 33 | 506 | + | | | | | | | | |
| | | | | | 2 | probable chromate transport protein, putative | 503 | 1078 | + | | | | | | | | |
| | | | | | 3 | putative integral membrane transport protein | 1307 | 2515 | + | | | | | | | | |
| | | | | | 4 | Sugar phosphate isomerases/epimerases | 2560 | 3453 | + | | | | | | | | |
| | | | | | 5 | hypothetical protein | 3802 | 3933 | + | | | | | | | | |
| | | | | | 6 | Putative phosphate ABC transporter, phosphate-binding component | 4003 | 4836 | - | | | | | | | | |
| | | | | | 7 | Ferric siderophore transport system, periplasmic binding protein TonB | 4839 | 5654 | - | | | | | | | | |
| | | | | | 8 | Biopolymer transport protein ExbD/TolR | 5676 | 6332 | - | | | | | | | | |
| | | | | | 9 | Biopolymer transport protein ExbD/TolR | 6347 | 6952 | - | | | | | | | | |
| | | | | | 10 | MotA/TolQ/ExbB proton channel family protein | 6978 | 7790 | - | | | | | | | | |

| | | | | | | | | | | | | | | | | | |
|---|---|---|---|---|---|---|---|---|---|---|---|---|---|---|---|---|---|
| | | | | | 11 | hypothetical protein | 7910 | 8311 | - | | | | | | | | |
| | | | | | 12 | hypothetical protein | 11678 | 11797 | + | | | | | | | | |
| | | | | | 13 | hypothetical protein | 14289 | 14420 | + | | | | | | | | |
| | | | | | 14 | Probable poly(beta-D-mannuronate) O-acetylase (EC 2.3.1.-)//Aminobenzoate degradation :<br>Ethylbenzene degradation | 14832 | 16247 | - | | | | | | | | |
| | | | | | 15 | putative periplasmic protein | 16288 | 17133 | - | | | | | | | | |
| | | | | | 16 | putative periplasmic protein | 17180 | 18439 | - | | | | | | | | |
| | | | | | 17 | FIG00936253: hypothetical protein | 18618 | 18911 | - | | | | | | | | |
| | | | | | 18 | Cobalt-zinc-cadmium resistance protein CzcA; Cation efflux system protein CusA | 18908 | 22066 | - | | | | | | | | |
| | | | | | 19 | Membrane fusion protein of RND family multidrug efflux pump | 22079 | 23095 | - | | | | | | | | |
| | | | | | 20 | Outer membrane efflux protein precursor | 23130 | 24467 | - | | | | | | | | |
| | | | | | 21 | Transcriptional regulator, TetR family | 24502 | 25134 | - | | | | | | | | |
| | | | | | 22 | FIG00405371: hypothetical protein | 25263 | 25895 | - | | | | | | | | |
| | | | | | 23 | LSU ribosomal protein L17p | 26423 | 26914 | - | | | | | | | | |
| | | | | | 24 | DNA-directed RNA polymerase alpha subunit (EC 2.7.7.6) | 26918 | 27910 | - | x | | | | | | | |
| | | | | | 25 | SSU ribosomal protein S4p (S9e) | 27922 | 28527 | - | | | | | | | | |
| | | | | | 26 | SSU ribosomal protein S11p (S14e) | 28634 | 29023 | - | | | | | | | | |
| | | | | | 27 | SSU ribosomal protein S13p (S18e) | 29035 | 29415 | - | | | | | | | | |
| | | | | | 28 | LSU ribosomal protein L36p | 29449 | 29565 | - | | | | | | | | |
| | | | | | 29 | Translation initiation factor 1 | 29574 | 29792 | - | | | | | | | | |
| | | | | | 30 | Methionine aminopeptidase (EC 3.4.11.18) | 29807 | 30598 | - | | | | | | | | |
| | | | | | 31 | Preprotein translocase secY subunit (TC 3.A.5.1.1) | 30607 | 31956 | - | | | | | | | | |
| | | | | | 32 | LSU ribosomal protein L15p (L27Ae) | 31961 | 32407 | - | | | | | | | | |
| | | | | | 33 | LSU ribosomal protein L30p (L7e) | 32435 | 32611 | - | | | | | | | | |
| | | | | | 34 | SSU ribosomal protein S5p (S2e) | 32623 | 33144 | - | | | | | | | | |
| | | | | | 35 | LSU ribosomal protein L18p (L5e) | 33144 | 33488 | - | | | | | | | | |
| | | | | | 36 | LSU ribosomal protein L6p (L9e) | 33511 | 34065 | - | | | | | | | | |
| | | | | | 37 | SSU ribosomal protein S8p (S15Ae) | 34083 | 34478 | - | | | | | | | | |
| | | | | | 38 | SSU ribosomal protein S14p (S29e) @ SSU ribosomal protein S14p (S29e), zinc-indepe | 34531 | 34800 | - | | | | | | | | |
| | | | | | 39 | LSU ribosomal protein L5p (L11e) | 34805 | 35362 | - | | | | | | | | |
| | | | | | 40 | LSU ribosomal protein L24p (L26e) | 35365 | 35682 | - | | | | | | | | |
| | | | | | 41 | LSU ribosomal protein L14p (L23e) | 35702 | 36067 | - | | | | | | | | |
| | | | | | 42 | SSU ribosomal protein S17p (S11e) | 36070 | 36327 | - | | | | | | | | |
| | | | | | 43 | LSU ribosomal protein L29p (L35e) | 36341 | 36535 | - | | | | | | | | |
| | | | | | 44 | LSU ribosomal protein L16p (L10e) | 36541 | 36975 | - | | | | | | | | |
| | | | | | 45 | SSU ribosomal protein S3p (S3e) | 36998 | 37723 | - | | | | | | | | |
| | | | | | 46 | LSU ribosomal protein L22p (L17e) | 37735 | 38139 | - | | | | | | | | |
| | | | | | 47 | SSU ribosomal protein S19p (S15e) | 38166 | 38435 | - | | | | | | | | |
| | | | | | 48 | LSU ribosomal protein L2p (L8e) | 38455 | 39279 | - | | | | | | | | |
| | | | | | 49 | LSU ribosomal protein L23p (L23Ae) | 39281 | 39571 | - | | | | | | | | |
| | | | | | 50 | LSU ribosomal protein L4p (L1e) | 39584 | 40210 | - | | | | | | | | |
| | | | 1 | | 51 | LSU ribosomal protein L3p (L3e) | 40210 | 40827 | - | | | | | | | | |
| 26I17-IVTE | | 41,253 bp | **Bacteroidetes** Bacteroidaceae | ABTS | 52 | SSU ribosomal protein S10p (S20e) | 40848 | 41153 | - | | | | | | | | |
| | | | | | 1 | hypothetical protein | 167 | 2233 | + | x | | | | | | | |
| | | | | | 2 | Cellulase( EC:3.2.1.4 ) | 2328 | 3854 | + | x | 35 | GH5_4 | 38A17 (23): 100% cov, 100% id | CAJ19139.1 | 67.53 | 0.0 | lichenase / endo-b-1,3-1,4-glucanase (3.2.1.73) |
| | | | | | 3 | Mannan endo-1,4-beta-mannosidase | 3968 | 4999 | + | x | 3 | GH26 | 15K21 (33): 99% cov, 100% id;<br>38A17 (24): 100% cov, 100% id;<br>44D8 (19): 100% cov, 100% id;<br>44F17 (13): 98% cov, 100% id;<br>46A3 (4): 98% cov, 100% id | AIT97142.1 | 49.85 | 1E-106 | exo-b-1,4-mannobiohydrolase / mannobiose-producing exo-b-mannanase (3.2.1.100) |
| | | | | | 4 | COG2152 predicted glycoside hydrolase | 5077 | 6252 | + | x | 10 | GH130 | 20G16 (25): 100% cov, 100% id;<br>38A17 (25): 100% cov, 100% id;<br>44F17 (12): 99% cov, 100% id;<br>46A3 (5): 99% cov, 100% id; 44D8 (20): 100% cov, 100% id | AAS19693.1 | 69.49 | 4E-139 | b-1,4-mannosylglucose phosphorylase (2.4.1.281) |
| | | | | | 5 | Xyloside transporter XynT | 6273 | 7700 | + | x | 16 | | | | | | |
| | | | | | 6 | DNA polymerase I (EC 2.7.7.7) | 8185 | 9375 | + | x | | | | | | | |
| | | | | | 7 | DNA polymerase I (EC 2.7.7.7) | 9372 | 10916 | + | x | 27 | | | | | | |
| | | | | | 8 | RNA polymerase sigma factor RpoE | 10888 | 11493 | + | | | | | | | | |
| | | | | | 9 | rRNA small subunit 7-methylguanosine (m7G) methyltransferase GidB | 11490 | 12152 | + | | | | | | | | |
| | | | 1 | | 10 | ZINC PROTEASE (EC 3.4.99.-) | 12631 | 15408 | + | | 37 | | | | | | |
| | | | **Bacteroidetes** Bacteroidales | | 11 | iron (III) ABC transporter ATP-binding | 15422 | 16081 | - | | | | | | | | |
| | | | RumenMGS049_scaffold_83 | | 12 | Vitamin B12 ABC transporter, permease component BtuC | 16078 | 17079 | - | | 19 | | | | | | |
| | | 17,825 bp | 1,103 bp | | 13 | hypothetical protein | 17088 | 17759 | - | | | | | | | | |
| | | | | | 1 | hypothetical protein | 422 | 1354 | + | | 9 | | | | | | |
| | | | | | 2 | putative secretory protein | 1869 | 2651 | - | | | | | | | | |
| | | | | | 3 | Alpha-1,2-mannosidase | 2661 | 4937 | - | | 34 | GH92 | | AAO79068.1 | 66.93 | 0.0 | a-mannosidase (3.2.1.24) |
| | | | 2 | | 4 | putative outer membrane protein, probably involved in nutrient binding | 4990 | 6924 | - | | 27 | | | | | | |
| | | | **Bacteroidetes** Bacteroidales | AZCL-Barley beta | 5 | TonB-dependent receptor | 6972 | 10127 | - | | 10 | | | | | | |
| | | | RumenMGS049_scaffold_63 | glucan, AZCL- | 6 | DNA-binding response regulator, AraC family | 10312 | 14082 | - | | 12 | | | | | | |
| 28D5-IVTE | | 14,524 bp | 11,862 bp | arabinoxylan | 7 | RNA methyltransferase, TrmA family | 14192 | 14503 | + | | | | | | | | |
| | | | | | 1 | FIG00938042: hypothetical protein | 13 | 1584 | + | | | | | | | | |
| | | | | | 2 | lichenase / endo-b-1,3-1,4-glucanase (3.2.1.73) | 1791 | 3041 | + | | | GH5_4 | 6F24 (17): 100% cov, 100% id | CAJ19146.1 | 47.27 | 4E-116 | lichenase / endo-b-1,3-1,4-glucanase (3.2.1.73) |
| | | | | | 3 | FIG01289191: hypothetical protein | 3095 | 6274 | + | | | | | | | | |
| | | | | | 4 | FIG00937602: hypothetical protein | 6386 | 8128 | + | | | | | | | | |
| | | | | | 5 | hypothetical protein | 8267 | 9007 | + | | | | | | | | |
| | | | | | 6 | Beta-glucosidase (EC 3.2.1.21) | 9007 | 11319 | + | | | GH3 | 6F24 (13): 100% cov, 100% id | ALJ47761.1 | 61.88 | 1E-163 | b-glucosidase (3.2.1.21) |
| | | | | | 7 | Mobile element protein | 11780 | 13015 | + | | | | | | | | |
| | | | | | 8 | hydrolase, putative | 13425 | 14222 | - | | | | | | | | |
| | | | | | 9 | Hypothetical Nudix-like regulator | 14225 | 14980 | - | | | | | | | | |
| | | | | | 10 | hypothetical protein | 15134 | 15250 | - | | | | | | | | |
| | | | | | 11 | hypothetical protein | 15210 | 15482 | + | | | | | | | | |
| | | | | | 12 | Ynd | 16102 | 16782 | + | | | | | | | | |
| | | | | | 13 | FIG00937925: hypothetical protein | 16775 | 17485 | + | | | | | | | | |
| | | | | | 14 | transposase | 17651 | 18721 | - | x | | | | | | | |
| | | | | AZO-CM cellulose, | 15 | hypothetical protein | 19098 | 19304 | + | | | | | | | | |
| | | | | AZCL-Barley beta | 16 | hypothetical protein | 19318 | 20322 | + | x | 3 | | | | | | |
| | | | 1 | glucan, AZCL- | 17 | hypothetical protein | 20322 | 21863 | + | x | 52 | | | | | | |
| 30D14-IVTE | | 23,779 bp | **Bacteroidetes** Prevotella | arabinoxylan | 18 | hypothetical protein | 22629 | 23006 | + | | | | | | | | |
| | | | | | 1 | Putative threonine efflux protein | 267 | 833 | + | | | | | | | | |
| | | | | | 2 | hypothetical protein | 835 | 1617 | + | x | 5 | | | | | | |
| | | | | | 3 | Ribosomal large subunit pseudouridine synthase D (EC 4.2.1.70) | 1619 | 2650 | + | x | 2 | | | | | | |
| | | | | | 4 | Cytoplasmic axial filament protein CafA and Ribonuclease G (EC 3.1.4.-) | 2668 | 4239 | - | x | 18 | | | | | | |
| | | | | | 5 | DNA-binding protein HU | 4468 | 4740 | - | | | | | | | | |
| | | | | | 6 | ATP-dependent protease La (EC 3.4.21.53) Type I | 4963 | 7368 | + | x | 22 | | | | | | |
| | | | | | 7 | Uracil-DNA glycosylase, family 1 | 7368 | 8036 | + | | | | | | | | |
| | | | | | 8 | Phosphopantetheine adenylyltransferase (EC 2.7.7.3) | 8033 | 8506 | + | | | | | | | | |
| | | | | | 9 | hypothetical protein | 8508 | 8726 | - | | | | | | | | |
| | | | | | 10 | Thiol:disulfide interchange protein tlpA | 8730 | 9977 | + | x | 4 | | | | | | |
| | | | | | 11 | DJ-1/YajL/PfpI superfamily, includes chaperone protein YajL (former ThiJ), parkinsoni | 9966 | 10538 | - | | | | | | | | |
| | | | | | 12 | UDP-N-acetylmuramoylalanyl-D-glutamyl-2,6-diaminopimelate--D-alanyl-D-alanine li | 10648 | 11760 | - | x | 4 | | | | | | |
| | | | | | 13 | probable metal-dependent peptidase | 12119 | 12793 | + | | | | | | | | |
| | | | | | 14 | hypothetical protein | 12790 | 13083 | - | | | | | | | | |
| | | | | | 15 | tRNA-specific adenosine-34 deaminase (EC 3.5.4.-) | 13100 | 13534 | - | | | | | | | | |
| | | | | | 16 | hypothetical protein | 13655 | 13885 | - | | | | | | | | |
| | | | | | 17 | Transcriptional regulator, AraC family | 14003 | 14542 | + | | | | | | | | |
| | | | | | 18 | Endo-1,4-beta-mannosidase | 14649 | 17111 | + | x | 41 | GH5_7-CE7 | 20G16 (18): 97% cov, 100% id;<br>15K21 (27): 100% cov, 100% id;<br>44D8 (13): 97%cov, 100% id; 44F17 (19): 97% cov, 100% id | AAS19695.1 | 41.77 | 1E-99 | b-mannosidase (3.2.1.25) |
| | | | | | 19 | TonB family protein / TonB-dependent receptor | 17169 | 20348 | + | x | 9 | | | | | | |
| | | | | | 20 | putative outer membrane protein, probably involved in nutrient binding | 20361 | 22109 | + | x | 24 | | | | | | |
| | | | | | 21 | FIG00417257: hypothetical protein | 22122 | 23426 | + | x | 7 | | | | | | |
| | | | | | 22 | hypothetical protein | 23433 | 25859 | + | x | | | | | | | |
| | | | | | 23 | Cellulase( EC:3.2.1.4 ) | 25954 | 27480 | + | x | 35 | GH5_4 | 28D5 (2): 100% cov, 100% id | CAJ19139.1 | 67.53 | 0.0 | lichenase / endo-b-1,3-1,4-glucanase (3.2.1.73) |

| | | | | | | | | | | | | | | | | | |
|---|---|---|---|---|---|---|---|---|---|---|---|---|---|---|---|---|---|
| | | | | | 24 | Mannan endo-1,4-beta-mannosidase | 27594 | 28625 | + | x | 3 | GH26 | 15K21 (33): 99% cov, 100% id; 28D5 (3): 100% cov, 100% id; 44D8 (19): 100% cov, 100% id; 44F17 (13): 98% cov, 100% id; 46A3 (4): 98% cov, 100% id | AIT97142.1 | 49.85 | 1E-106 | exo-b-1,4-mannobiohydrolase / mannobiose-producing exo-b-mannanase (3.2.1.100) |
| | | | | | 25 | COG2152 predicted glycoside hydrolase | 28703 | 29878 | + | x | 10 | GH130 | 20G16 (25): 100% cov, 100% id; 28D5 (4): 100% cov, 100% id; 44F17 (12): 99% cov, 100% id; 46A3 (5): 99% cov, 100% id; 44D8 (20): 100% cov, 100% id | AAS19693.1 | 69.49 | 4E-139 | b-1,4-mannosylglucose phosphorylase (2.4.1.281) |
| | | | | | 26 | Xyloside transporter XynT | 29899 | 31326 | + | x | 16 | | | | | | |
| | | | | | 27 | DNA polymerase I (EC 2.7.7.7) | 31895 | 34543 | + | x | 27 | | | | | | |
| | | | | | 28 | RNA polymerase sigma factor RpoE | 34530 | 35120 | + | | | | | | | | |
| | | | | | 29 | rRNA small subunit 7-methylguanosine (m7G) methyltransferase GidB | 35117 | 35779 | + | | | | | | | | |
| 38A17-IVTE | | 37,749 bp | 1 **Bacteroidetes** Bacteroidales **RumenMGS049_scaffold_83 12,691 bp** | AZO-CM cellulose, AZCL Tamarind xyloglucan, AZCL-HE cellulose, AZCL-Barley beta glucan, AZCL-arabinoxylan | 30 | hypothetical protein | 36417 | 36683 | - | | | | | | | | |
| | | | | | 1 | SusC, outer membrane protein involved in starch binding | 103 | 846 | + | | | | | | | | |
| | | | | | 2 | SusD, outer membrane protein | 865 | 2451 | + | x | 36 | | | | | | |
| | | | | | 3 | hypothetical protein | 2461 | 4842 | + | x | 26 | | | | | | |
| | | | | | 4 | Alpha-amylase (EC 3.2.1.1) | 5108 | 7348 | + | x | 6 | GH13_36-CBM58 | | AAO78803.1 | 48.47 | 3E-102 | a-amylase (3.2.1.1), neopullulanase (3.2.1.135) |
| | | | | | 5 | Neopullulanase (EC 3.2.1.135) | 7362 | 10502 | + | x | 18 | GH13_46 | | ADF53136.1 | 46.69 | 5E-169 | a-amylase (3.2.1.1) |
| | | | | | 6 | Alpha-glucosidase SusB (EC 3.2.1.20) | 10559 | 12838 | + | x | 50 | GH97 | | AEW20125.1 | 57.72 | 0.0 | a-glucosidase (3.2.1.20) |
| | | | | | 7 | hypothetical protein | 12851 | 13171 | + | | | | | | | | |
| | | | | | 8 | ABC transporter ATP-binding protein | 13182 | 13823 | + | | | | | | | | |
| | | | | | 9 | hypothetical protein | 13816 | 14001 | - | | | | | | | | |
| | | | | | 10 | Uridine kinase (EC 2.7.1.48) | 14036 | 15754 | - | x | 45 | | | | | | |
| | | | | | 11 | Excinuclease ABC subunit A | 15751 | 18540 | - | x | | | | | | | |
| | | | | | 12 | Hypothetical metal-binding enzyme, YcbL homolog | 18540 | 19247 | - | | | | | | | | |
| | | | | | 13 | Deoxyadenosine kinase (EC 2.7.1.76) / Deoxyguanosine kinase (EC 2.7.1.113) | 19317 | 19964 | + | | | | | | | | |
| | | | | | 14 | Deoxyadenosine kinase (EC 2.7.1.76) / Deoxyguanosine kinase (EC 2.7.1.113) | 19955 | 20575 | + | | | | | | | | |
| | | | | | 15 | Metal transporter, ZIP family | 20582 | 21364 | + | | | | | | | | |
| | | | | | 16 | ABC-type multidrug transport system, permease component | 21586 | 22800 | - | x | | | | | | | |
| | | | | | 17 | ABC-type multidrug transport system, permease component | 22802 | 23971 | - | x | 31 | | | | | | |
| | | | | | 18 | FIG00937622: hypothetical protein | 23976 | 24965 | - | x | 1 | | | | | | |
| | | | | | 19 | Type I secretion system, outer membrane component LapE | 24979 | 26457 | - | x | 6 | | | | | | |
| | | | | | 20 | Transcriptional regulator, AraC family | 26745 | 27662 | + | x | 17 | | | | | | |
| | | | | | 21 | Transcriptional regulator, AraC family | 27665 | 28435 | + | x | 66 | | | | | | |
| | | | | | 22 | TonB-dependent receptor | 28541 | 30790 | + | x | | | | | | | |
| | | | | | 23 | hypothetical protein | 30808 | 31260 | + | | | | | | | | |
| | | | | | 24 | Thiol:disulfide interchange protein tlpA | 31580 | 32608 | - | x | | | | | | | |
| | | | | | 25 | Glucokinase (EC 2.7.1.2) | 32785 | 33744 | - | x | 4 | | | | | | |
| | | | | | 26 | Predicted glucose transporter in maltodextrin utilization gene cluster | 33771 | 35051 | - | x | 23 | | | | | | |
| | | | | | 27 | Periplasmic beta-glucosidase (EC 3.2.1.21) | 35158 | 37410 | - | x | 49 | GH3 | | ZP_02070067.1 | 68.47 | 2E-166 | exo-b-1,3-glucanase (3.2.1.58) |
| 40D8-IVTE | | 38,624 bp | 1 **Bacteroidetes** Bacteroidales RumenMGS049_scaffold_128 15,191 bp | AZCL-Barley beta glucan | 28 | Beta-glucanase precursor (EC 3.2.1.73) | 37422 | 38405 | - | x | 40 | GH16_3 | | WP_072066679.1 | 52.36 | 4E-77 | endo-b-1,3-glucanase / laminarinase (3.2.1.39) |
| | | | | | 1 | FIG00410329: hypothetical protein | 25 | 2022 | + | | | | | | | | |
| | | | | | 2 | Mannan endo-1,4-beta-mannosidase B precursor (EC 3.2.1.78) | 2244 | 3524 | + | | | GH26 | | ARK08557.1 | 62.97 | 2E-146 | endo-b-1,4-mannanase (3.2.1.78) |
| | | | | | 3 | Beta-glucosidase (EC 3.2.1.21) | 3635 | 5917 | + | | | GH3 | | | | | |
| | | | | | 4 | Mannan endo-1,4-beta-mannosidase | 6010 | 7044 | + | | | GH26 | | CAH06517.1 | 74.40 | 0.0 | exo-b-1,4-mannobiohydrolase / mannobiose-producing exo-b-mannanase (3.2.1.100) |
| | | | | | 5 | COG2152 predicted glycoside hydrolase | 7077 | 8267 | + | | | GH130 | | AAS19693.1 | 71.86 | 2E-146 | b-1,4-mannosylglucose phosphorylase (2.4.1.281) |
| | | | | | 6 | Cation symporter | 8287 | 9720 | + | | | | | | | | |
| | | | | | 7 | N-acylglucosamine 2-epimerase (EC 5.1.3.8) | 9717 | 10907 | + | | | | | | | | |
| | | | | | 8 | hypothetical protein | 10914 | 11399 | - | | | | | | | | |
| | | | | | 9 | Ferredoxin | 11507 | 12526 | - | | | | | | | | |
| | | | | | 10 | Transcriptional regulator, AraC family | 12538 | 13347 | - | | | | | | | | |
| | | | | | 11 | Isochorismatase (EC 3.3.2.1) | 13351 | 13908 | - | | | | | | | | |
| | | | | | 12 | Predicted nucleotide-binding protein | 14921 | 16189 | - | | | | | | | | |
| | | | | | 13 | hypothetical protein | 16203 | 16391 | - | | | | | | | | |
| | | | | | 14 | FIG00896557: hypothetical protein | 16741 | 17868 | - | | | | | | | | |
| | | | | | 15 | FIG00403577: hypothetical protein | 17980 | 20352 | - | | | | | | | | |
| | | | | | 16 | TonB-dependent receptor | 20404 | 23046 | - | | | | | | | | |
| | | 23,867 bp | 1 **Bacteroidetes** *Bacteroides graminisolvens* | | 17 | NADH-ubiquinone oxidoreductase chain N (EC 1.6.5.3) | 23311 | 23688 | - | | | | | | | | |
| | | | | | 1 | Transcriptional regulator, PadR family | 86 | 424 | + | | | | | | | | |
| | | | | | 2 | hypothetical protein | 435 | 1415 | + | | | | | | | | |
| | | | | | 3 | hypothetical protein | 1425 | 2180 | + | | | | | | | | |
| 41J20-IVTE | | 4,322 bp | 2 ***Lachnospiraceae*** ( Firmicutes; Clostridia; Clostridiales) | AZO-galactomannan | 4 | two-component response regulator | 2233 | 4257 | - | | | | | | | | |
| | | | | | 1 | Osmosensitive K+ channel histidine kinase KdpD (EC 2.7.3.-) | 347 | 1810 | + | | | | | | | | |
| | | | | | 2 | Two-component system response regulator | 1807 | 2499 | + | | | | | | | | |
| | | | | | 3 | CRISPR-associated protein, Csn1 family | 2724 | 7178 | + | | | | | | | | |
| | | | | | 4 | CRISPR-associated protein Cas1 | 7252 | 8193 | + | | | | | | | | |
| | | | | | 5 | CRISPR-associated protein Cas2 | 8235 | 8540 | + | | | | | | | | |
| | | | | | 6 | Acyltransferase | 12475 | 12966 | + | | | | | | | | |
| | | | | | 7 | Acyltransferase | 12957 | 13712 | + | | | | | | | | |
| | | | | | 8 | FIG053235: Diacylglucosamine hydrolase like | 13859 | 14362 | + | | | | | | | | |
| | | | | | 9 | Beta-glucosidase (EC 3.2.1.21) | 14403 | 16652 | - | | | GH3 | 3F21 (4): 100% cov, 100% id; | ZP_02070067.1 | 72.40 | 0.0 | exo-b-1,3-glucanase (3.2.1.58) |
| | | | | | 10 | Beta-glucanase precursor (EC 3.2.1.73) | 16665 | 17762 | - | | | GH16_3 | 3F21 (3): 100% cov, 100% id; | WP_072066679.1 | 80.95 | 3E-138 | endo-b-1,3-glucanase / laminarinase (3.2.1.39) |
| | | | | | 11 | FIG00412289: hypothetical protein | 17836 | 19200 | - | | | | | | | | |
| | | | | | 12 | RagB/SusD domain protein | 19221 | 20645 | - | | | | | | | | |
| | | | | | 13 | hypothetical protein | 20644 | 20763 | + | | | | | | | | |
| | | 22,594 bp | 1 **Bacteroidetes** Bacteroidales | | 14 | TonB family protein / TonB-dependent receptor | 20764 | 22581 | - | | | | | | | | |
| | | | | | 1 | L-rhamnose-proton symporter | 17 | 808 | + | | | | | | | | |
| | | | | | 2 | Rhamnulose-1-phosphate aldolase (EC 4.1.2.19) | 845 | 1639 | + | | | | | | | | |
| | | | | | 3 | putative glycosyslhydrolase | 1650 | 4871 | + | | | GH106 | | BAD12237.1 | 31.34 | 3E-76 | a-L-rhamnosidase (3.2.1.40) |
| | | | | | 4 | Alfa-L-rhamnosidase (EC 3.2.1.40) | 4916 | 6715 | + | | | GH78 | | AAO76108.1 | 26.44 | 3E-30 | a-L-rhamnosidase (3.2.1.40) |
| 41O1-IVTE | | 10,243 bp | 2 **Bacteroidetes; Bacteroidales** | AZCL-Barley beta glucan | 5 | Alfa-L-rhamnosidase (EC 3.2.1.40) | 6785 | 9319 | + | | | GH78 | | ACI19983.1 | 32.87 | 1E-42 | a-L-rhamnosidase (3.2.1.40) |
| | | | | | 1 | hypothetical protein | 122 | 448 | - | | | | | | | | |
| | | | | | 2 | hypothetical protein | 448 | 645 | - | | | | | | | | |
| | | | | | 3 | Prophage LambdaBa04, Gp54 | 642 | 1004 | - | | | | | | | | |
| | | | | | 4 | hypothetical protein | 1410 | 1589 | - | | | | | | | | |
| | | | | | 5 | hypothetical protein | 1586 | 1915 | - | | | | | | | | |
| | | | | | 6 | hypothetical protein | 1905 | 2117 | - | | | | | | | | |
| | | | | | 7 | CDP-glycerol: N-acetyl-beta-D-mannosaminyl-1,4-N-acetyl-D-glucosaminyldiphospho | 2110 | 3081 | - | | | | | | | | |
| | | | | | 8 | hypothetical protein | 3081 | 3674 | - | | | | | | | | |
| | | | | | 9 | hypothetical protein | 3667 | 4386 | - | | | | | | | | |
| | | | | | 10 | hypothetical protein | 4383 | 4781 | - | | | GT8 | | | | | |
| | | | | | 11 | hypothetical protein | 4782 | 4979 | - | | | | | | | | |
| | | | | | 12 | hypothetical protein / (oxidoreductase [Sphingobium sp. Ant17]) | 4988 | 5194 | - | | | | | | | | |
| | | | | | 13 | Phage terminase, large subunit | 5203 | 6918 | - | | | | | | | | |
| | | | | | 14 | hypothetical protein | 6905 | 7063 | - | | | | | | | | |
| | | | | | 15 | hypothetical protein | 7072 | 7341 | - | | | | | | | | |
| | | | | | 16 | Phage portal protein | 7554 | 8876 | + | | | | | | | | |
| | | | | | 17 | Phage head maturation protease | 8892 | 9581 | + | | | | | | | | |
| | | | | | 18 | Phage major capsid protein #Fam0024 | 9602 | 10906 | + | | | | | | | | |
| | | | | | 19 | hypothetical protein | 10918 | 11148 | + | | | | | | | | |
| | | | | | 20 | hypothetical protein | 11217 | 11549 | + | | | | | | | | |
| | | | | | 21 | hypothetical protein | 11550 | 11918 | + | | | | | | | | |
| | | | | | 22 | hypothetical protein | 11915 | 12319 | + | | | | | | | | |
| | | | | | 23 | hypothetical protein | 12319 | 12651 | + | | | | | | | | |
| | | | | | 24 | Phage major tail protein | 12644 | 13249 | + | | | | | | | | |
| | | | | | 25 | hypothetical protein | 13262 | 13624 | + | | | | | | | | |
| | | | | | 26 | hypothetical protein | 13668 | 13835 | + | | | | | | | | |
| | | | | | 27 | Phage tail length tape-measure protein | 13847 | 18889 | + | | | | | | | | |
| | | 22,822 bp | 1 **Bacteria** | | 28 | hypothetical protein | 18901 | 22011 | + | | | | | | | | |
| | | | | | 1 | Spore maturation protein A | 477 | 1052 | - | | | | | | | | |
| | | | | | 2 | hypothetical protein | 1184 | 2125 | + | | | | | | | | |
| | | | | | 3 | LSU m3Psi1915 methyltransferase RlmH | 2199 | 2675 | - | | | | | | | | |
| | | | | | 4 | Zn-dependent hydrolase (beta-lactamase superfamily) | 2687 | 3505 | - | | | | | | | | |
| | | | | | 5 | Malate permease | 3634 | 4569 | + | | | | | | | | |
| | | | | | 6 | UDP-N-acetylglucosamine 1-carboxyvinyltransferase (EC 2.5.1.7) | 4615 | 5880 | - | | | | | | | | |
| 42A14-IVTE | | 6,910 bp | 2 **Firmicute** | ABTS | 7 | Ribonuclease P protein component (EC 3.1.26.5) | 6396 | 6755 | + | | | | | | | | |
| | | | | | 1 | hypothetical protein | 18 | 665 | + | | | | | | | | |

| | | | | | | | | | | | | | | | | | |
|---|---|---|---|---|---|---|---|---|---|---|---|---|---|---|---|---|---|
| | | | | | 2 | Alanine racemase (EC 5.1.1.1) | 637 | 1821 | - | | | | | | | | |
| | | | | | 3 | hypothetical protein | 1839 | 2462 | - | | | | | | | | |
| | | | | | 4 | Endo-1,4-beta-xylanase A precursor (EC 3.2.1.8) | 2472 | 3836 | - | x | | GH5_4 | 15F15 (4): 100% cov, 100% id;<br>44C20 (4) 100% cov, 100% id | ABX42426.1 | 44.47 | 7E-110 | endo-b-1,4-glucanase (3.2.1.4) |
| | | | | | 5 | Ribosome small subunit-stimulated GTPase EngC | 3890 | 4762 | - | x | | | | | | | |
| | | | | | 6 | Serine/threonine protein kinase PrkC, regulator of stationary phase | 4774 | 6855 | - | x | | | | | | | |
| | | | | | 7 | Protein serine/threonine phosphatase PrpC, regulation of stationary phase | 6883 | 7611 | - | | | | | | | | |
| | | | | | 8 | Ribosomal RNA large subunit methyltransferase N (EC 2.1.1.-) | 7657 | 8694 | - | | | | | | | | |
| | | | | | 9 | 16S rRNA (cytosine(967)-C(5))-methyltransferase (EC 2.1.1.176) | 8694 | 9026 | - | | | | | | | | |
| | | | | | 10 | Ribosomal RNA small subunit methyltransferase B (EC 2.1.1.-) | 9026 | 10042 | - | x | | | | | | | |
| | | | | | 11 | Methionyl-tRNA formyltransferase (EC 2.1.2.9) | 10039 | 10983 | - | x | | | | | | | |
| | | | | | 12 | Peptide deformylase (EC 3.5.1.88) | 10983 | 11468 | - | | | | | | | | |
| | | | | | 13 | Helicase PriA essential for oriC/DnaA-independent DNA replication | 11487 | 13709 | - | x | | | | | | | |
| | | | | | 14 | Single-stranded-DNA-specific exonuclease RecJ (EC 3.1.-.-) | 13755 | 15530 | - | x | | | | | | | |
| | | 17,351 bp | 1<br>**Firmicutes** | | 15 | Protein-export membrane protein SecD (TC 3.A.5.1.1) / Protein-export membrane pr | 15534 | 17303 | - | x | | | | | | | |
| | | | | | 1 | putative outer membrane protein, probably involved in nutrient binding | 25 | 2934 | + | x | 38 | | | | | | |
| | | | | | 2 | putative outer membrane protein, probably involved in nutrient binding | 2971 | 4782 | + | x | 6 | | | | | | |
| 44C18-IVTE | | 6,528 bp | 2<br>Bacteroidetes; Bacteroidia;<br>Bacteroidales | AZCL-Barley beta glucan, AZO-CM cellulose, AZCL Tamarind xyloglucan, AZCL-HE cellulose | 3 | hypothetical protein | 4779 | 4913 | + | | | | | | | | |
| | | | | | 1 | Cytoplasmic axial filament protein CafA and Ribonuclease G (EC 3.1.4.-) | 373 | 1944 | - | x | 18 | | | | | | |
| | | | | | 2 | DNA-binding protein HU | 2173 | 2445 | - | | | | | | | | |
| | | | | | 3 | ATP-dependent protease La (EC 3.4.21.53) Type I | 2731 | 5073 | + | x | 22 | | | | | | |
| | | | | | 4 | Uracil-DNA glycosylase, family 1 | 5073 | 5741 | + | | | | | | | | |
| | | | | | 5 | Phosphopantetheine adenylyltransferase (EC 2.7.7.3) | 5738 | 6211 | + | | | | | | | | |
| | | | | | 6 | Thiol:disulfide interchange protein tlpA | 6204 | 7682 | + | x | 4 | | | | | | |
| | | | | | 7 | DJ-1/YajL/PfpI superfamily, includes chaperone protein YajL (former ThiJ), parkinsoni | 7671 | 8243 | - | | | | | | | | |
| | | | | | 8 | UDP-N-acetylmuramoylalanyl-D-glutamyl-2,6-diaminopimelate--D-alanyl-D-alanine li | 8353 | 9465 | - | x | 4 | | | | | | |
| | | | | | 9 | probable metal-dependent peptidase | 9824 | 10498 | + | | | | | | | | |
| | | | | | 10 | hypothetical protein | 10495 | 10788 | - | | | | | | | | |
| | | | | | 11 | tRNA-specific adenosine-34 deaminase (EC 3.5.4.-) | 10805 | 11239 | - | | | | | | | | |
| | | | | | 12 | Transcriptional regulator of various polyols utilization, AraC family | 11369 | 12247 | + | x | 12 | | | | | | |
| | | | | | 13 | Endo-1,4-beta-mannosidase | 12417 | 14816 | + | x | 41 | GH5_7-CE7 | 15K21 (27): 100% cov, 100% id;<br>38A17 (18): 100% cov, 100% id;<br>20G16 (18): 100%cov, 100% id;<br>44F17 (19): 99% cov, 100% id | AAS19695.1 | 41.77 | 6E-100 | b-mannosidase (3.2.1.25) |
| | | | | | 14 | TonB family protein / TonB-dependent receptor | 14904 | 18053 | + | x | 9 | | | | | | |
| | | | | | 15 | putative outer membrane protein, probably involved in nutrient binding | 18081 | 19814 | + | x | 24 | | | | | | |
| | | | | | 16 | FIG00417257: hypothetical protein | 19875 | 21131 | + | x | 7 | | | | | | |
| | | | | | 17 | hypothetical protein | 21138 | 23564 | + | x | | | | | | | |
| | | | | | 18 | Cellulase( EC:3.2.1.4 ) | 23626 | 25185 | + | x | 35 | GH5_4 | 20G16 (18): 97% cov, 100% id;<br>46A3 (3): 100% cov, 100% id;<br>15K21 (32): 97%cov, 100% id;<br>44F17 (19): 97% cov, 100% id | CAJ19139.1 | 67.53 | 0.0 | lichenase / endo-b-1,3-1,4-glucanase (3.2.1.73) |
| | | | | | 19 | Mannan endo-1,4-beta-mannosidase | 25188 | 26330 | + | x | 3 | GH26 | 15K21 (33): 100% cov, 100% id;<br>28D5 (3): 100% cov, 100% id;<br>44F17(13): 100% cov, 100% id;<br>38A17 (24): 100% cov, 100% id;<br>46A3 (4): 100% cov, 100% id | AIT97142.1 | 49.85 | 9E-107 | exo-b-1,4-mannobiohydrolase / mannobiose-producing exo-b-mannanase (3.2.1.100) |
| | | | | | 20 | COG2152 predicted glycoside hydrolase | 26408 | 27583 | + | x | 10 | GH130_1 | 20G16 (25): 100% cov, 100% id;<br>28D5 (4): 100% cov, 100% id;<br>44F17 (12): 99% cov, 100% id;<br>46A3 (5): 99% cov, 100% id; 38A17<br>(25): 100% cov, 100% id | AAS19693.1 | 69.49 | 4E-139 | b-1,4-mannosylglucose phosphorylase (2.4.1.281) |
| | | | | | 21 | Xyloside transporter XynT | 27604 | 29031 | + | x | 16 | | | | | | |
| | | | | | 22 | DNA polymerase I (EC 2.7.7.7) | 29516 | 32248 | + | x | 27 | | | | | | |
| | | | | | 23 | RNA polymerase sigma factor RpoE | 32235 | 32825 | + | | | | | | | | |
| | | | | | 24 | rRNA small subunit 7-methylguanosine (m7G) methyltransferase GidB | 32822 | 33484 | + | | | | | | | | |
| | | | | | 25 | ZINC PROTEASE (EC 3.4.99.-) | 33963 | 35471 | + | x | 37 | | | | | | |
| | | | | | 26 | Voltage-gated chloride channel family protein | 35530 | 37128 | - | | | | | | | | |
| | | | | | 27 | FIG00405231: hypothetical protein | 37201 | 37353 | - | | | | | | | | |
| | | | | | 28 | putative outer membrane protein, probably involved in nutrient binding | 37497 | 39041 | - | | | | | | | | |
| | | | | | 29 | putative anti-sigma factor | 39047 | 39907 | - | | | | | | | | |
| | | | | | 30 | RNA polymerase ECF-type sigma factor | 39904 | 40455 | - | | | | | | | | |
| | | | | | 31 | Ketol-acid reductoisomerase (EC 1.1.1.86) | 40507 | 41700 | - | | | | | | | | |
| | | | | | 32 | Vitamin B12 ABC transporter, permease component BtuC | 42003 | 43031 | - | | | | | | | | |
| 44D8-IVTE | | 43,948 bp | 1<br>**Bacteroidetes Bacteroidales**<br>**RumenMGS049_scaffold_83**<br>**12,691 bp** | AZO-CM cellulose, AZCL-Barley beta glucan, AZCL Tamarind xyloglucan, AZCL-arabinoxylan, AZCL-HE cellulose | 33 | Vitamin B12 ABC transporter, B12-binding component BtuF | 43113 | 43931 | - | | | | | | | | |
| | | | | | 1 | Citrate lyase alpha chain (EC 4.1.3.6) | 171 | 1694 | - | x | 26 | | | | | | |
| | | | | | 2 | Citrate lyase beta chain (EC 4.1.3.6) | 1691 | 2581 | - | x | 13 | | | | | | |
| | | | | | 3 | Citrate lyase gamma chain, acyl carrier protein (EC 4.1.3.6) | 2588 | 2866 | - | | | | | | | | |
| | | | | | 4 | DNA recombination protein RmuC | 2979 | 4298 | - | x | 5 | | | | | | |
| | | | | | 5 | Vitamin B12 ABC transporter, permease component BtuC | 4597 | 5367 | + | | | | | | | | |
| | | | | | 6 | iron (III) ABC transporter ATP-binding | 5364 | 6023 | + | | | | | | | | |
| | | | | | 7 | ZINC PROTEASE (EC 3.4.99.-) | 6037 | 8814 | - | x | 37 | | | | | | |
| | | | | | 8 | rRNA small subunit 7-methylguanosine (m7G) methyltransferase GidB | 9293 | 9955 | - | | | | | | | | |
| | | | | | 9 | RNA polymerase sigma factor RpoE | 9952 | 10542 | - | | | | | | | | |
| | | | | | 10 | DNA polymerase I (EC 2.7.7.7) | 10529 | 13261 | - | x | 27 | | | | | | |
| | | | | | 11 | Xyloside transporter XynT | 13746 | 15173 | - | x | 16 | | | | | | |
| | | | | | 12 | COG2152 predicted glycoside hydrolase | 15194 | 16360 | - | x | 10 | GH130_1 | 20G16 (25): 100% cov, 100% id;<br>28D5 (4): 100% cov, 100% id; 44D8<br>(20): 100% cov, 100% id; 46A3 (5):<br>100% cov, 100% id; 38A17 (25):<br>100% cov, 100% id | AAS19693.1 | 69.49 | 3E-139 | b-1,4-mannosylglucose phosphorylase (2.4.1.281) |
| | | | | | 13 | Mannan endo-1,4-beta-mannosidase | 16447 | 17466 | - | x | 3 | GH26 | 15K21 (33): 100% cov, 100% id;<br>28D5 (3): 100% cov, 100% id; 44D8<br>(19): 100% cov, 100% id; 38A17<br>(24): 100% cov, 100% id; 46A3 (4):<br>100% cov, 100% id | AIT97142.1 | 49.85 | 1E-106 | exo-b-1,4-mannobiohydrolase / mannobiose-producing exo-b-mannanase (3.2.1.100) |
| | | | | | 14 | Cellulase( EC:3.2.1.4 ) | 17592 | 19118 | - | x | 35 | GH5_4 | 20G16 (23): 100% cov, 100% id;<br>46A3 (3): 100% cov, 100% id; 44D8<br>(18): 100%cov, 100% id; 15K21<br>(32): 100% cov, 100% id | CAJ19139.1 | 67.53 | 0.0 | lichenase / endo-b-1,3-1,4-glucanase (3.2.1.73) |
| | | | | | 15 | hypothetical protein | 19213 | 21639 | - | x | | | | | | | |
| | | | | | 16 | FIG00417257: hypothetical protein | 21646 | 22902 | - | x | 7 | | | | | | |
| | | | | | 17 | putative outer membrane protein, probably involved in nutrient binding | 22963 | 24711 | - | x | 24 | | | | | | |
| | | | | | 18 | TonB family protein / TonB-dependent receptor | 24724 | 27873 | - | x | 9 | | | | | | |
| | | | | | 19 | Endo-1,4-beta-mannosidase | 27961 | 30354 | - | x | 41 | GH5_7-CE7 | 20G16 (18): 100% cov, 100% id;<br>38A17 (18): 100% cov, 100% id;<br>44D8 (13): 100%cov, 100% id;<br>15K21 (27): 100% cov, 100% id | AAS19695.1 | 41.77 | 6E-100 | b-mannosidase (3.2.1.25) |
| | | | | | 20 | Transcriptional regulator of various polyols utilization, AraC family | 30530 | 31414 | - | x | 12 | | | | | | |
| | | | | | 21 | tRNA-specific adenosine-34 deaminase (EC 3.5.4.-) | 31538 | 31972 | + | | | | | | | | |
| | | | | | 22 | hypothetical protein | 31989 | 32282 | + | | | | | | | | |
| | | | | | 23 | probable metal-dependent peptidase | 32279 | 32947 | + | | | | | | | | |
| | | | | | 24 | UDP-N-acetylmuramoylalanyl-D-glutamyl-2,6-diaminopimelate--D-alanyl-D-alanine li | 33312 | 34424 | + | x | 4 | | | | | | |
| | | | | | 25 | DJ-1/YajL/PfpI superfamily, includes chaperone protein YajL (former ThiJ), parkinsoni | 34534 | 35106 | + | | | | | | | | |
| 44F17-IVTE | | 35,340 bp | 1<br>**Bacteroidetes Bacteroidales**<br>**RumenMGS049_scaffold_83**<br>**12,691 bp** | AZO-CM cellulose, AZCL-Barley beta glucan, AZCL Tamarind xyloglucan, AZCL-arabinoxylan, AZCL-HE cellulose | 26 | hypothetical protein | 35095 | 35331 | - | | | | | | | | |
| | | | | | 1 | widely conserved MoxR-like protein in magnesium chelatase family | 286 | 1284 | - | x | | | | | | | |
| | | | | | 2 | FIG028593: membrane protein | 1397 | 2362 | - | | | | | | | | |
| | | | | | 3 | Guanine deaminase (EC 3.5.4.3) | 2377 | 2835 | - | | | | | | | | |
| | | | | | 4 | hypothetical protein | 2850 | 3833 | - | x | | | | | | | |
| | | | | | 5 | Alanyl-tRNA synthetase (EC 6.1.1.7) | 3850 | 5085 | - | x | | | | | | | |
| | | | | | 6 | hypothetical protein | 5381 | 5557 | - | | | | | | | | |
| | | | | | 7 | Inosine-5'-monophosphate dehydrogenase (EC 1.1.1.205) | 5801 | 7261 | - | | 39 | | | | | | |
| | | | | | 8 | hypothetical protein | 7312 | 7827 | - | | | | | | | | |
| | | | | | 9 | Endo-1,4-beta-xylanase A precursor (EC 3.2.1.8) | 7860 | 9128 | - | | | GH5_4 | | ADD61809.1 | 52.17 | 4E-131 | endo-b-1,3(4)-glucanase / lichenase-laminarinase(3.2.1.6), |
| | | | | | 10 | Metallo-beta-lactamase family protein, RNA-specific | 9137 | 10744 | - | x | | | | | | | |
| | | | | | 11 | Transcription regulator [contains diacylglycerol kinase catalytic domain] | 10981 | 11877 | - | | 17 | | | | | | |
| | | | | | 12 | hypothetical protein | 11890 | 12042 | - | | | | | | | | |

| | | | | | | | | | | | | | | | | | |
|---|---|---|---|---|---|---|---|---|---|---|---|---|---|---|---|---|---|
| | | | 1 | | 13 | Pyruvate:ferredoxin oxidoreductase, beta subunit (EC 1.2.7.1) | 12098 | 13036 | - | x | | | | | | | |
| | | 14,385 bp | **Firmicutes** Clostridiales | | 14 | Pyruvate:ferredoxin oxidoreductase, alpha subunit (EC 1.2.7.1) | 13065 | 14246 | - | x | | | | | | | |
| | | | | | 1 | Cof protein:HAD-superfamily hydrolase, subfamily IIB | 1552 | 2385 | + | | | | | | | | |
| | | | | | 2 | Cell division protein FtsI [Peptidoglycan synthetase] (EC 2.4.1.129) | 2688 | 4841 | - | | | | | | | | |
| | | | | | 3 | hypothetical protein | 4874 | 4990 | - | | | | | | | | |
| | | | 2 | | 4 | Transcriptional regulator, RpiR family | 5147 | 6007 | - | | | | | | | | |
| | | | ***Enterococcus facecium*** | | 5 | Threonyl-tRNA synthetase (EC 6.1.1.3) | 6217 | 8160 | - | | | | | | | | |
| | | | (Firmicutes; Bacilli; | | 6 | hypothetical protein | 8157 | 8270 | - | | | | | | | | |
| | | | Lactobacillales; | | 7 | 4-oxalocrotonate tautomerase (EC 5.3.2.-); Xylose transport system permease protei | 8496 | 8711 | - | | | | | | | | |
| 45H9-IVTE | | 1,161 bp | Enterococcaceae; Enterococcus) | AZO-CM cellulose | 8 | cadmium-translocating P-type ATPase | 8722 | 10545 | - | | | | | | | | |
| | | | | | 1 | hypothetical protein | 56 | 379 | + | | | | | | | | |
| | | | | | 2 | hypothetical protein | 398 | 2812 | + | x | | | | | | | |
| | | | | | 3 | Cellulase( EC:3.2.1.4 ) | 2907 | 4433 | + | x | 35 | GH5_4 | 20G16 (23): 100% cov, 100% id;<br>44F17 (14): 100% cov, 100% id;<br>44D8 (18): 100%cov, 100% id;<br>15K21 (32): 100% cov, 100% id | CAJ19139.1 | 67.53 | 0.0 | lichenase / endo-b-1,3-1,4-glucanase (3.2.1.73) |
| | | | | | 4 | Mannan endo-1,4-beta-mannosidase | 4559 | 5578 | + | x | 3 | GH26 | 15K21 (33): 100% cov, 100% id;<br>28D5 (3): 100% cov, 100% id; 44D8<br>(19): 100% cov, 100% id; 38A17<br>(24): 100% cov, 100% id; 44F17<br>(13): 100% cov, 100% id | AIT97142.1 | 49.85 | 1E-106 | exo-b-1,4-mannobiohydrolase / mannobiose-producing exo-b-mannanase (3.2.1.100) |
| | | | | | 5 | COG2152 predicted glycoside hydrolase | 5665 | 6831 | + | x | 10 | GH130 | 20G16 (25): 100% cov, 100% id;<br>28D5 (4): 100% cov, 100% id; 44D8<br>(20): 100% cov, 100% id; 44F17<br>(12): 100% cov, 100% id; 38A17<br>(25): 100% cov, 100% id | AAS19693.1 | 69.49 | 3E-139 | b-1,4-mannosylglucose phosphorylase (2.4.1.281) |
| | | | | | 6 | Predicted beta-glucoside transporter, GPH family | 7057 | 8280 | + | x | 16 | | | | | | |
| | | | | | 7 | DNA polymerase I (EC 2.7.7.7) | 8765 | 11497 | + | x | 27 | | | | | | |
| | | | | | 8 | RNA polymerase sigma factor RpoE | 11484 | 12074 | + | | | | | | | | |
| | | | | | 9 | rRNA small subunit 7-methylguanosine (m7G) methyltransferase GidB | 12071 | 12733 | + | | 37 | | | | | | |
| | | | | | 10 | ZINC PROTEASE (EC 3.4.99.-) | 13212 | 15989 | + | x | | | | | | | |
| | | | | | 11 | iron (III) ABC transporter ATP-binding | 16003 | 16662 | - | | | | | | | | |
| | | | | | 12 | Vitamin B12 ABC transporter, permease component BtuC | 16659 | 17660 | - | x | 19 | | | | | | |
| | | | | | 13 | DNA recombination protein RmuC | 17729 | 19048 | + | x | 5 | | | | | | |
| | | | | | 14 | Citrate lyase gamma chain, acyl carrier protein (EC 4.1.3.6) | 19161 | 19439 | + | | | | | | | | |
| | | | | | 15 | Citrate lyase beta chain (EC 4.1.3.6) | 19467 | 20336 | + | x | 13 | | | | | | |
| | | | | | 16 | Citrate lyase alpha chain (EC 4.1.3.6) | 20333 | 21856 | + | x | 26 | | | | | | |
| | | | | | 17 | Apo-citrate lyase phosphoribosyl-dephospho-CoA transferase (EC 2.7.7.61) | 21856 | 22359 | + | | | | | | | | |
| | | | | | 18 | 2-(5''-triphosphoribosyl)-3'-dephosphocoenzyme-A synthase (EC 2.7.8.25) | 22349 | 24079 | + | x | 16 | | | | | | |
| | | | | | 19 | Putative enzyme | 24113 | 26350 | + | x | 15 | | | | | | |
| | | | | | 20 | hypothetical protein | 26414 | 26530 | + | | | | | | | | |
| | | | | | 21 | Isocitrate dehydrogenase [NADP] (EC 1.1.1.42) | 26544 | 27755 | + | | 29 | | | | | | |
| | | | | | 22 | ABC transporter ATP-binding protein | 27755 | 29431 | + | x | 33 | | | | | | |
| | | | | | 23 | Low-specificity L-threonine aldolase (EC 4.1.2.48) | 29440 | 30465 | - | x | 47 | | | | | | |
| | | | | | 24 | hypothetical protein | 30470 | 30598 | - | | | | | | | | |
| | | | 1 | | 25 | Adenylate kinase (EC 2.7.4.3) | 30575 | 31717 | + | x | 2 | | | | | | |
| | | | **Bacteroidetes** Bacteroidales | | 26 | hypothetical protein | 31948 | 33498 | + | x | | | | | | | |
| | | | RumenMGS049_scaffold_83 | | 27 | Membrane-bound lytic murein transglycosylase D precursor (EC 3.2.1.-) | 33500 | 34918 | - | x | 14 | GH23-CBM50 | | AAD08609.1 | 40.85 | 8E-27 | peptidoglycan lytic transglycosylase / peptidoglycan lyase (4.2.2.-) |
| 46A3-IVTE | | 35,712 bp | 1,682 bp | lignin alkali, ABTS | 28 | hypothetical protein | 34966 | 35712 | - | x | 1 | | | | | | |
