## Supplementary material for "Functional exploration of *in vivo* and *in vitro* lignocellulose-fed rumen bacterial microbiomes reveals novel enzymes involved in polysaccharide breakdown": Sequences

```
ID   XXX; XXX; linear; XXX; XXX; XXX;  XXX.
XX
AC   PRJEB67889
XX
DE   Uncultured bacterium DNA fragment from in vivo enriched cow rumen, clone
rum-1c10
XX
FH   Key             Location/Qualifiers
FH
FT   source          1..32580
FT                   /organism="uncultured bacterium"
FT                   /environmental_sample
FT                   /mol_type="genomic DNA"
FT                   /isolation_source="Cow rumen"
FT                   /clone="rum-1c10"
FT                   /note="AZO-galacto-mannan active clone"
FT   CDS             complement(53..1762)
FT                   /db_xref="SEED:fig|6666666.127271.peg.1"
FT                   /translation="MKKLISWLLVAVMAVGMCSWASADPVNALDFEDGVFAFLGVSAAK
FT                   PNADASTALEVVDYNGSKALRVAAQGIPYVALNLEGLAGEKLADVAAVTFDIGVDKAAD
FT                   GKFYAVSGVVYSYTGENADENKADWSVYLEKKNPRNVKIQFKAPFVAGAGNYIMISRED
FT                   QAGGEPATFYLDNIQFLDAEGNAIALDPTAEYVSEASEKDLSNLVALTNAVEFPDFHKS
FT                   AGAWAQDGLEMPQEIIDALVPGSVVEVEYASADGSMWIVMPWATAGWMRVGQGTAAINN
FT                   SKTIAQIPYEMIEALCGEDKSTWGAMLQCESASDWEVFAVRVGQRANRIVLKNAVEFPG
FT                   FTKSADAWAQDGLEMPQEIIDALVPGSVVEITYSSEDGDIWLVMPWAEAGWMRVSQGTA
FT                   AKMGGKAYITYEEIAALCGEDKSTWGAMMQCEGSSPWEVYGIRVGQKAEFFGLTNLVEF
FT                   PGFTKSADAWAQDGLEMPQEIIDALVPGSVVTISYESEDGNMWLVMPWAAAGWMRVGND
FT                   GADVADGKIAQVTYEQIEALCGEDKSTWGAMMQCESSSPWNVYAVAVGQAIK"
FT                   /product="hypothetical protein"
FT   CDS             complement(1822..2826)
FT                   /db_xref="SEED:fig|6666666.127271.peg.2"
FT                   /db_xref="GO:0005215"
FT                   /db_xref="GO:0006810"
FT                   /db_xref="GO:0016020"
FT                   /translation="MQTIHPAQNGKITWQGFRNSPAYANQRDKTLNGMKNLVIGLFRLI
FT                   VIVGISYVILAPVINIISNSFFSRQDSINPMVFTIPINPTLERYSMAVKYLNYLPILGN
FT                   TILYVFGVTLIQLLICSMVGYGFARFRFPLKSVLFAFVLVMIVIPLNTIQFPLYTTFRF
FT                   FNPLGIVGLFNGGKTINLLQTAWPTFIMSVFGCGLRSGLYIYIFNQFFRGLPKEIEEAA
FT                   LVDGAGTARTFFTIMMPNAAPAVITVAVFSIVWQYNDSFFANTFNISDSILVTRKLESL
FT                   INVIANAEKVLTIREQQLYFDAGVVLVLVPVVVIYLVLQRRFIEGVERSGIVG"
FT                   /product="N-Acetyl-D-glucosamine ABC transport system,
FT                   permease protein 2"
FT   CDS             complement(2847..3773)
FT                   /db_xref="SEED:fig|6666666.127271.peg.3"
FT                   /translation="MTGIRRKNRIFHLSYEQRNAVRGYWFILPWLIGFLVFYAGCLVQL
FT                   GEFSLSRIGLDAATGMTKEYIGFGNYIEAFTSHATFKQTLTSSVMDMLIDLPMILFFSL
FT                   FVAMLLNRKFKGRALVRAIFFLPIILSADAVGSAITRATELVNAGVPSANLETAQAGTV
FT                   SVAYYMQLFGDLAIPESVLGYIVDAVNRISSIIKGSGVQIILFIAALQSIPGSLYEVAR
FT                   IEGATGYESFWKVTFPMVMPHMITNTIYTVVDRFSSSEVIRLATRTYKELYNYGLSSAF
FT                   SVVSTLITILILGLIVYLLNRRTFYYN"
FT                   /product="binding-protein-dependent transport systems
FT                   inner membrane component"
FT   CDS             complement(3770..6361)
FT                   /db_xref="SEED:fig|6666666.127271.peg.4"
```

```
FT                   /translation="MNEKNRLTLPTAFRRVLWIIAGLAAAAAVLFLARYVIHYRAYDEY
FT                   RQYIVQPAAKTEAEALKPLKDPENAVPGFVLAAQNDRLGLYLKEKTAEIALYDRKSGLT
FT                   VYSNPQDAADDPVARSGLNQANLKSQFILNYLDTNSREGTPWSSYAKCVENGQIEYLQL
FT                   ENGFRAVYTLSNEKLMLVPQQMTAEWYGILSASGKKQAAKSYVLDEETGLYVLKSQGVT
FT                   ARHRQQIDADARKAGFTMEDYAEMQALAEEEESEAAESTSFVVALDYTLTPDGLTVTIP
FT                   HAELAETGGAKIRTIQLLPFFGAAGTAEKGVIVVPDGSGALIRFNNGKNTAPQYNQNIY
FT                   DLDLIDSDITATQNMQSARLALFGICREDCGLLVSCGRGAALASVTADVAGRNNSYNYA
FT                   YFTFGLRRTDTLMVAGEEAIVAEQDSYPVDCAVSYRILDGEYTGYNGLARAVRESLLAD
FT                   GSLTLKTEKSGDIPFYCDILCGVRETAHWMGVQYLRVMPMTTFAQAEEILSGLRQEQVG
FT                   NLRVNLQGWMNGGYYHDPVNHVSVQGGLGGERGLQSLRTAAAETGGKIYPDAALQQVTE
FT                   IAKGFLRSEEASRYYAKGYVAEFGTVSPVSMRRTATMGFTERAYMLLSPRFLPRYAACL
FT                   ASAAERLGLDALSLRDLGNEVHADKRRTNVISREADLDLVRHAFGVIGAGERELMVSGG
FT                   NDYSFPYVRHVINAPVEATMFAIVDEQIPLWEMILHGSADYCGSPLNLMQSEDRRAALL
FT                   HLIEYGASTHYTFTWQDAADMKYTGLNNNYATTFSSWKEEAAESYRFVNGALAGVSGAE
FT                   MLRHDRLSDTLARVTYSNGTVLYVNSGEHEAEADGYRIPAMDYLAVGGENE"
FT                   /product="GH129"
FT   CDS             complement(6376..7008)
FT                   /db_xref="SEED:fig|6666666.127271.peg.5"
FT                   /translation="MNRMGIKDKWQHFFHTVSHPADGYYWIRHQEKGSLWIAFLMVLLY
FT                   GIVFSMNRISSSFIVNDIEPRTVNLPAELAGVILLYLILCVGNWSVTCLMDGEGRFKDI
FT                   LIAAGYALFPMMAATAAATAVSHLVAENEEAFYTLLIGLGTAYTVIMLVIGIMQVHNYT
FT                   FGKTLITLFLTVVAMLIIIFLGLLVINFITQVYSFFRGIYMELIFRV"
FT                   /product="FIG01031734: hypothetical protein"
FT   CDS             complement(7005..8486)
FT                   /db_xref="SEED:fig|6666666.127271.peg.6"
FT                   /translation="MKQVKRILCLMLLACLAAGSACAADSLPYDCYNYDHWNNILYTPA
FT                   PYVPDGLVSGATLRLEGTPVGAFRNPQDLCVSADGDVYIADTGNNRIVVLAPDLKTVRR
FT                   IITGFEAEGGTQTFNAPTGVALSEKHRLYIADSLNRRIVVLEPDGTFVKYVRDPQSEVL
FT                   DEGYVFTPLRVSVDYADRVYVIAQNMFQGIMVFSGEGDFTGFFGTISVKISLWEKFWRK
FT                   LATKEERSKQQLFIPTEFTGIDVDDEGFVYASSKDTDGLQAVRRLNPKGEDVIRKGPRG
FT                   NLGGDRVFGSVGNYAGPSQIVDVVYRGHGAYSLLDGRRGRIFTYDHEGNLLYIFGGIGT
FT                   QAGTFQTPAAIEKLGSSLLVLDSQQGNLTVFAETEYGALINRAVSLRFDGDESQAVPLW
FT                   QEVLRLDEINELANTGIGKAYLSAGDNANAMLYLRRGMNREYYSVAFKRWRNERLRNNI
FT                   SLILSGVVAAAALAVLWVKAVRPAIRKKGKRRDPS"
FT                   /product="NHL repeat containing protein"
FT   CDS             complement(8498..9487)
FT                   /db_xref="SEED:fig|6666666.127271.peg.7"
FT                   /translation="MTGMEKAKISRLFSRMPHRKPAELSPEEETALRQKRLAKIYRKMK
FT                   RQSGKKKLNRSTAGNAVLFILMGICGVFMVLPLVMILNNAVKPLDELYQYPPRIFVRNP
FT                   TLNNFTDLYVLLNESWVPFSRYILNTIIITVGGMTGHVIIASMAAYPLAKNHFPGKKIL
FT                   FSVVVLSMMFSWTVTQIPQYLIISWLGINNTYAALVLPAWSFGMGLYLMKQFMEQLPDS
FT                   LMESARLQGANEWQIFWRIVMPNVKPAWLTLAIFQFQQMWANTGSMFLRSEELKPLQYA
FT                   LQQITSGGVSRAGAGAAVTFLIAAVPIIFFLICQSSIMETMTTSGMKD"
FT                   /product="binding-protein-dependent transport systems
FT                   inner membrane component"
FT   CDS             complement(9484..10416)
FT                   /db_xref="SEED:fig|6666666.127271.peg.8"
FT                   /translation="MSMAIRNGRSMRKKDTLWHQVRINRASYLLMAPYFTLFFLFTVLP
FT                   VLMSVYLSFNYFNMLEMPRWVGWANYIKLLLEDDIFIISLRNTLLFAVVTGPVSYLLCL
FT                   LFAWIINEFRPKVRAFLTLVFYAPSICGNAYMIWNLILTGDRYGYLNGILLRTGILNEP
FT                   ILWMQTEKFVMPVLIVVQLWLSLGTGFLSFIAGLQTVDKGLYEAAAIDGVKNRWQELWF
FT                   VTLPAMKPQLMFGAVMQITQSFAVADISIALAGNPSVNYSGATIVTHLLDYGSTRLDMG
FT                   YASAIATILFLLMVGTNKLVQRILRRVGE"
FT                   /product="binding-protein-dependent transport systems
```

```
FT                   inner membrane component"
FT   CDS             complement(10418..13402)
FT                   /db_xref="SEED:fig|6666666.127271.peg.9"
FT                   /db_xref="GO:0005215"
FT                   /db_xref="GO:0006810"
FT                   /translation="MRKKTGIVLAMLLALAAGTAMAATDLGIREYEDQVSTYSINPDIP
FT                   SYAEYCAAHGTAARPEKEIVLEAADTVRYAEEGGENAPLLKAQAEGRDGVSVLTGEEAV
FT                   IEWDFEAEEAGWYDLEVEYYPWPGRNAEIQRAFFLDGALPYRELALVNFSRVWRNDLHP
FT                   EIVRSTAEDGTEEIRWLRDNQGNELKPSPAEAPEWRTSLLHDSNGYISESLKLYLDRGA
FT                   HTLTVLSLREPMLIRRMCFRAGTDVKSYAPPEAGGAANRVIRIEAEAASGTSSQMLYPV
FT                   QDQSSRSVSPSSPKYLLNNSIGGNAWKSAGQWIEWRFSVPEEGNYAISLYDKQNFIRGT
FT                   DVYRKIYIDGEVPFAEFQAAAFPYTQDWRLETLSGEPGEPYLIHLAAGEHTLRMEVVLG
FT                   EMAAVIRQVQDCVLQLNGIYRQVLYITGVSPDPYRDYQIERSLPGLEAQLQMVKKDLRS
FT                   AIEALEKTAGHRSDKLTVLKTMDDQLTELIRDQERFTEVLSSYKTNVRACGNWITQVLG
FT                   QPLQIDRIFVHTADVRPEISGTDFLSGTGFELSRLYSSFTINYNQIGNVAEKTEDSRVI
FT                   TLWVGTGRDQANVIKALIDEHFTPKTGISVNVQLVDMNTLLRATLSGQGPDVAIQVANT
FT                   TGIAGAVMNTGNDTPVNYGLRNAVMDLTQFDDFAEVAERFAPSALVPFSFNGAAYALPD
FT                   TQTFPMMFYRKDILAEIGLEVPETWDDVKVAMSVLSKNQMEFGMLPGEQTFAMLLFQAG
FT                   GEYYTENGDASALDSDIAVNVFRRYCDFYTDYKLDMATSVEERFRTGECPIIITDYTTY
FT                   NNLQVSAPDIKGLWDFTHVPGTLREDGTLDTTTGCSGLADIIMSATKEPEACWEFLKWW
FT                   TGAEIQTLYGREMESLMGASARVATANLEALAGLNWPIRDYRELAKQFSHVRGIPQVPG
FT                   GYYTWRNVNNAFYAVTTGSASTGRTEFGNTPREELMDKIYYINAEITYKRTEFGLPVAE
FT                   TGEGK"
FT                   /product="N-Acetyl-D-glucosamine ABC transport system,
FT                   sugar-binding protein"
FT   CDS             complement(13439..14911)
FT                   /db_xref="SEED:fig|6666666.127271.peg.10"
FT                   /translation="MKSSMKSICGIAVAVMTLLFIILSGTAEETGETMLVEAETGELKG
FT                   HTAVAGNGGNQWVEGFKTAGEDSVSVEVPVSREGFYDIAVIQASQGGHKENPVLLDGQN
FT                   IGSAVTDETTFGKSVLEHIWLSAGMHTLGIGTSWGHIRLDGFELKPSAALPEDLYTVPE
FT                   KMCVGEASPEARKLYDWLRECYGKKILSGQQCAGGMYGMENQAIWRATGGDYPAVLGLD
FT                   MIDYSPSRVEHGGESDKAVPYAVEYWNQGGIVTFCWHWNAPSPYLKEPWYSGFYTEHTS
FT                   FNLGRVMDGEDEEGYRLLNRDIDAVAAQLLRLKEAGVPVLWRPLHEASGGWFWWGASGK
FT                   EAYLKLYHLLFDRLTHEYGLNNLIWIWNGQDPEWYPGDDEVDMIGIDIYPGEHVYDSQS
FT                   ASFIPLTRIAGERKMIVLSENGCIPDPEKVFRDGTVWGYWCTWGGEFVIRNTKFNKPSE
FT                   QYTETDMVRKAYSDTRVVTRKDLPDFRAADGD"
FT                   /product="CBM35-GH26"
FT                   /EC_number="3.2.1.78"
FT   CDS             complement(15231..16637)
FT                   /db_xref="SEED:fig|6666666.127271.peg.11"
FT                   /translation="METIREGKEPSWCRDLKTNMARMAQELIKVARERTDGGPVFTPEE
FT                   FGYQPGETATEAIQGAIEAAAEQGGIVRLGNGDYVSGTLVMRSGVCLEVRAGSRLLGST
FT                   DLRDYPEHHAARLTVQDTSMGMHQSLLFAEGCENICLRGGGVIDGRGSPAHFPGEETAQ
FT                   GTPGRPFLIRMIDCRNVHIVDLTMKDAACWMQNYLNCENLLLEGLTVRNHANYNNDGMD
FT                   IDGCRNVVIRNCRVSSGDDALCFKGAGQRELSRVLVEDCDFYSACNAVKVGTDTQGDFR
FT                   QVYIRRCRIGGLAEDPSGLKHACADSGISLEMVDGGTLENFLIEDIHMIRAWSPFFLRL
FT                   ENRGRVKPGDPAPGPGTLRRVLFSDIRGRETGPRGSYLLGIPEKPIEDIAFRRVLLRQE
FT                   ASRRPLRKEKDFDEMRGVYPDAHMIDPVGDAPAYALWARHVRGLSMDGYEVIPEGKERR
FT                   PAYIFTDTEF"
FT                   /product="GH28"
FT   CDS             complement(16653..18083)
FT                   /db_xref="SEED:fig|6666666.127271.peg.12"
FT                   /translation="MLPETSVSFRTEDEKLQRVYDAAEKKCLLNLKDFSGDTVLVEGGG
FT                   YEKIWLETQPMGGEMYALRNLEAAKNNILLFMKHQREDGRMPGSIQHINGKTEPQFNKY
```

```
FT                   QGFCFPDPALNLYYLMGKDGEYLEQLKEALIRFDACLWRTRDVSGDGLLSSFCVYDTGE
FT                   DLAVRYGDAPCWWEEDEPPEGYGVVPMASMDVTSWSFAARRAAAEICRIQGDPAFTDWE
FT                   KKAGDVADALRRRLWDARRGALYDRDRTGKTINILCHNTLRCMYWGSVSRDMADRFVKE
FT                   HLLNASEFWTPFPLPSVAADDPAFRNAPENNWSGQPEGLTYQRAITALERYGYARIVTA
FT                   LGKKLINAVDANGCRFTQQYDPFTGKASLVDEADHRPVSGDSSEKVQDAYGPTMLACLE
FT                   YIAHRFGIHPHMGQVWFSLDRGKPYEYEAAFYGHRYGIRSDGKKAEIRADGKDIGRFSC
FT                   GRRIVTNENGDFLRTELI"
FT                   /product="Putative isomerase"
FT   CDS             18323..19276
FT                   /db_xref="SEED:fig|6666666.127271.peg.13"
FT                   /translation="MSLSSSPLAYRGGIRHKMAMTLRRDWQLWVLMLPCLAFFIIFCYF
FT                   PMYGVQIAFRDYKFNLGITGSRWVGLQHFQKFFNSYYCGRMLRNTFLLNVYGLLFGFPV
FT                   PIILAILLNQLKSKRFKSFAQTAIYVPHFISIVVMIGMIYLFFSPTNGIVNHLITAAGG
FT                   KSIYFIMEPGWFRPLFIGSAIWQDAGWNTILYIATLTSVDPQLYEAATIDGATRWQKIK
FT                   SIDIPHLIPITVMMLILSCGSLLSSDTQKALLLQTSGNIATSDVLGVYIFNVGIAASKA
FT                   QHSYAAAIGLCLNAVSFIIVMGVNAIARKMSDISLF"
FT                   /product="ABC transporter, permease protein"
FT   CDS             19293..20225
FT                   /db_xref="SEED:fig|6666666.127271.peg.14"
FT                   /translation="MTTLAAKRRNRIRETRRDRIFNTFNLIFWIIILLIVLYPLWLILI
FT                   CSVSDPDAVLQAKVLIWPVDFSLMGYEAVLQNNELLGSYLNSIFYTVAGSALSVIVSVA
FT                   AAYSLSRKFAGKKLVNLAITFTMFFSGGLIPIFLNVRDLGLYDTRAVMILMNLVSVWNL
FT                   MVARTYIQTSIPNDLYEAAVIDGADHFTYFFKCVLPLSGTIIAVLSVYYGVARWNDYFT
FT                   AQVMLRSRSLFPLQRILKEILTSLTSTGSSDTFFAAYADNLGGLQAALRKAEVAKYCCI
FT                   VVATVPAVLLYLFMQKYFVKGVMIGSLKG"
FT                   /product="ABC transporter, permease protein"
FT   CDS             20240..21145
FT                   /db_xref="SEED:fig|6666666.127271.peg.15"
FT                   /translation="MSQMKGVEVSARAARHIDGYRVLEGKREYVTYQGDSSIRIWYSDI
FT                   PWRYETHDHSAVEIVLVLEGTVTYSVEGTSYPVHKGEVLIVPSNTSHSLEMGEGSSRYL
FT                   FLFETDTIMTMRDIKSMVLYLHKPFHLRDESDAHKQIRELLLRAKDFYEKQELMWNTAC
FT                   YSCILQVYATLGRHYLIGIRSRTGDAARSMDTEVINAVMTYINDHYREDLRLDDVAGFS
FT                   GFSRFYFSRSFKQQTGYSFKDYLCQKRLQAAMDLLTSTNLSMRGIAVESGFGSTASFNR
FT                   AFRACKGCTPTQYRAVYGML"
FT                   /product="Transcriptional regulator, AraC family"
FT   CDS             21207..22712
FT                   /db_xref="SEED:fig|6666666.127271.peg.16"
FT                   /translation="MKKLVSLLLACCMLLSLSAAFAEEKTELTAFQYALENQNTDFNGL
FT                   WFFDELEKKTNTHVTFQMVKDADWATNSNLMFAVPDQMPDMILRHAVDVEEYGVSQGLL
FT                   LPLDEYLTEEIMPNYVSRLNINNAGDSIPASDGKTYYIGFLMAQNVNHTGTWYVNQKEL
FT                   DKLGKTAPATIDEVTDLLRAMKADGIKWPFSASYTQFGPETIWNQFASFGVPENTYYYF
FT                   IDADNKVQLTFAQDGWRACVEWLHQLYEEELMDPESLTQDSNLWANKVNNGEVGYFCYL
FT                   RLINTAIKAENVENFKSILPPAADGFKACVSQILEVPEAGAYLTKNCKDPVAALKWIDA
FT                   QLETETMMVSLNGKVGEQIVLNDEGKYEVINIPEDNGLYQFVPVTMGQFFAPGDYYTSI
FT                   YQMAPHRVERYEDSKWYAEAGVLEPKSFQYLRDLSKLSNDDATEAATIYTDLQKLVEES
FT                   FANFVRNGVTDESWNDFQAQAKSIGSDRYIELYQNAYDAYLAK"
FT                   /product="ABC transporter, substrate-binding protein"
FT   CDS             22829..24376
FT                   /db_xref="SEED:fig|6666666.127271.peg.17"
FT                   /translation="MNRTVKITCLLAAALFLLTGCAVSRPGEGSAPPGASLTALQYEID
FT                   NIAVDFSSMWYYRQIEQQTGVHVDFSEVKDSEWSSSVGLAFAKGKMPDMILRGSLDVEE
FT                   YGVTRHLLVPLDEYIDKGFLPNYAARMDQAGLREQLTASDGHIYQLGFLISQGVNTNGH
FT                   FFINKTWLDKLNLQVPRTVDQLTEVLRRFRSDDPNGNGLQDEIPLEFTFDDNTTGIYNL
FT                   FSFFGLPLNEDYVYVDNTGNVRFAPEEDAFFETLTWLHDLYLEGLLDIDFISQGSSIWA
```

```
FT                   AKVNEGNTGMFSYWRLQNTALSPGTAANYQVMVPVHVEGRNACLPRNMDIIEFGAALTT
FT                   DNRDIEASLRWLDAQFETENMLVSQNGPVGDTLIRREDGRYEVSYVPPDNELYQSVPVI
FT                   CGQFFAPAEYYASVYVPAAHRLEKSAYCDLYEQAGVLEPVSGKLLTAVAPKTTAQSAET
FT                   TRLKAKLKSLVDAEIVAMVTRGVTDESREAFRSKLKENGSVEYRQVYQQIYDRWKEK"
FT                   /product="ABC transporter, substrate-binding protein"
FT   CDS             24380..26647
FT                   /db_xref="SEED:fig|6666666.127271.peg.18"
FT                   /translation="MIRKSRTSVFIRYALSYIAVVLLLFCSITGYLYVRLSGEAREEII
FT                   DNQINRLSRIASQHESYISAMLNTAEEIGLSPDIEFFRYNEEPWKAYDLQLQMVPYTTT
FT                   ASTFCDQVYLYFFGDDMIYSSSSSMTLSLFARMMRYEFIPMENLVSRIASTDRMTILPA
FT                   QQVSSTLVDSSRVVTFLVPLGSAPGSGKGILLFLIKDSVYQSLFADAIDDRVNTYIYQG
FT                   DDLLSSSEKLSLPRDRVEASQDTPSRLFRWENQDWTQVTFSGRNWGLTYTAVLRNADIN
FT                   SSIMQQLFSILAILPVFVFLSLLLALWMARRHAKPIRAITGMLSRETDETPRDELQQIS
FT                   SGISRLTTRNQELSSRLERALPLQRHDFVFRFVKGRFETRQEAVAAGQAVGLEIDRPVY
FT                   AVILCSGSAGNDHPFDLSLPPFSRLSGYSGAGVELVALKANLYLVFSDDPKVLTGMAET
FT                   LRCECAAGGERCITSISALHTDFTEAPSAYLEAAAAYENRFVMGSQKVLCYGDISPDLE
FT                   EILPQARKITTSISQALTLGSREVLDVRISDLLHFLKNTSMSPFAFRMIYSEVIRSLVH
FT                   THTAALSGQGSVQETYNIFTLTSCQSIDDLDELLRRLCDRLITGESASGEKVPDEEDEI
FT                   SQVVRYMEEHFSDPEISITALADSLELSTTRFSLSFKEKAGMSPLEYLTLLRVEHAKEL
FT                   LESSDLTIRDISTQVGYYDSGSFIRRFKQVTGETPLQYRRSRNPKKENGIPDRDASVPT
FT                   EE"
FT                   /product="Transcriptional regulator, AraC family"
FT   CDS             26657..27499
FT                   /db_xref="SEED:fig|6666666.127271.peg.19"
FT                   /translation="MIHTDQINIRDPYVLVHNGTYYLYGTRGPDCWGPADGFDVYTGRD
FT                   LVSWEGPRVCFRNDGSFWADRNYWAPEVHPWQGMFYMFASFKSDARRRGTAILRSPSPL
FT                   GPFEPWSDGPVTPADWECLDGTFYASADGTPWIVFCHEWVQAGDGEVLAMRLTDDLKAS
FT                   AGTPVLLFRGSDAPWSHAVRHSSGITGFVTDGPFLWRGKDGSLLCLWSGFSEKGYAQGL
FT                   AVSDNGEIDGHFSQADPLFLEDGGHGMLFRSKEGRTLLALHSPNTHLLERPRFVPVDL"
FT                   /product="GH43_17"
FT   CDS             27546..28826
FT                   /db_xref="SEED:fig|6666666.127271.peg.20"
FT                   /db_xref="GO:0008993"
FT                   /translation="MHYVLAIDIGASSGRHIVGWREGSGIRTEEVYRFPNGITEQDGSL
FT                   TWDTDALLFHVKAGIDKALEKYPSIQSLSIDTWAVDYVLMNGGSPVLPCYAYRDSRTEA
FT                   VIGEVHEKIPFPELYRRTGIQYQPFNTIYQLYADKVSGRLDQADGFLMIPEYLMYRLCD
FT                   AESHEYTNATTTGLVSAETGEYDPGIISALGLPARFFGPLRQPGTVIGEYRGIRVMLCA
FT                   THDTGSAVEGIPMEGCQPYISSGTWSLLGVKTERPVITEAGRLANWSNEGGAGYNRYQK
FT                   NIMGMWLMNRLRAELCPDRSWDEIMAEAAEKHFDHLVDVNDPVFLAPESMKAAFDSRLS
FT                   HPPRCIAGYFRCACRSLADGYRRAVEEMEQNTGRRYDRLYIVGGGAKNTYLNRLTEEST
FT                   GKQVTALPIEATALGNIRIQLKGGTDT"
FT                   /product="Rhamnulokinase (EC 2.7.1.5)"
FT                   /EC_number="2.7.1.5"
FT   CDS             28823..30067
FT                   /db_xref="SEED:fig|6666666.127271.peg.21"
FT                   /db_xref="GO:0008740"
FT                   /translation="MSYEESRKRYAALGVDTDAVIRLLRTVPVSLHCWQGDDVRGFDTD
FT                   PSKPLTGGIQTTGNYPGRARTPEELMADLDMVLSLIPGTPKMNLHASYAIFENGEWADR
FT                   NRLQPKHFRKWAAFCKERNLGCDFNPTFFSHPKCDPLTLSSPNAETRKFWIEHGKACIR
FT                   ISSYLAEELGFPCVMNIWTGDGFKDIPGDRIGPRIRYRDAIDEILREPYDFSRVKPCIE
FT                   SKVFGIGVEAYTAGSAEFALCYAAMNRDKCIPLMDNGHYHPTEVVSDKIPALLTFFREI
FT                   ALHITRPIRWDSDHVVLFDDETREIAREIVRCGGLDGRVHIALDYFDASINRISAWTTG
FT                   YRNLQKALLYALLQPHEDMKKLQDKGQFSRLMVLQEEIKTMPFGDIWAEYCRSAESPPT
FT                   ANGSRWWKNMKRKCW"
```

```
FT                   /product="L-rhamnose isomerase (EC 5.3.1.14)"
FT                   /EC_number="5.3.1.14"
FT   CDS             30075..30911
FT                   /db_xref="SEED:fig|6666666.127271.peg.22"
FT                   /db_xref="GO:0008994"
FT                   /translation="MKDILAAPFMTEMIRTATAMYDHGWDERNGGNISLMLDEKEAGEY
FT                   LDMHSVLRTIPTGFSMPELDGKTFLVTGTGKYFRNIQYEPEVNLGIIRLRDCGQQADLL
FT                   WGYSDGGQFTSEFPAHMMSHSARLKVNPANRVIMHCHPANLLAMTYVHTLDEREFTRTL
FT                   WQMCTECVVVFPDGVNVLPWMLCGTNEIGEATAEKMKTARLVVWSQHGIYGAGKDLDET
FT                   FGLIETAEKAAEIYMKIAHLPRVNTITDEQMHQLEKRFGIKAREGYLEEPDRGTDK"
FT                   /product="Rhamnulose-1-phosphate aldolase (EC 4.1.2.19)"
FT                   /EC_number="4.1.2.19"
FT   CDS             30908..32305
FT                   /db_xref="SEED:fig|6666666.127271.peg.23"
FT                   /db_xref="GO:0004557"
FT                   /translation="MKQHAGKNIRELKVAYIGGGSRGWAWGFMKDLALDGEICGTVRLY
FT                   DIDREAADRNRIIGEKISAHPDAVSRWAYTVSDSLQEALTGADFVVISILPGTFDEMES
FT                   DVHLPERLGVWQPVGDTVGAGGFMRSMRTIPMFVTIGEAIRDYAPEAWVINYTNPMSLC
FT                   VRTLYEVFPSIRAFGCCHEVFGTQKLLCSMLKTGEGIDGVKRQDLYTTVTGINHFTWLT
FT                   SASWKGTDLYPLYARFAEKYAESGYNEGGDDNWMNSHFSCAQRVKFDLFRRYGVIAAAG
FT                   DRHLAEFVAPWYTRDPETVREWKFSLTPVSWRKQDLKEKLQMSDDLVSGKKEITLNGSG
FT                   EEGHLLLKALLGLGDMVSNVNIPNRGQIPNLPLGAVVETNALFGLNRIDPVYAGPVPAS
FT                   ILPLITRHVLNQENTLTAALRCDRKLGLTTFLNDPQLAAVTPKDGEKLFNDMLENQRIF
FT                   LPEKWFE"
FT                   /product="GH4_3"
FT                   /EC_number="3.2.1.22"
FT   CDS             complement(32406..32570)
FT                   /db_xref="SEED:fig|6666666.127271.peg.24"
FT                   /translation="MTGDAEALELPLRKHPVTGETIRIPPLYRRNGETVFEGRAMPGKY
FT                   ELCRIIKRK"
FT                   /product="hypothetical protein"
XX
SQ   Sequence 32580 BP; 6799 A; 10036 C; 8728 G; 7017 T; 0 other;
     attgctgtta ttttttttcat aagtcagttt ctcctttcca gaaatcaggt catcatttga        60
     tcgcctggcc cacggccaca gcataaacat tccaggggga ggagctttcg cactgcatca       120
     tggcgcccca ggtggacttg tcctcgccgc agagggcttc gatctgttca taagtcacct       180
     gcgcgatttt gccgtccgcc acgtccgcgc cgtcgttgcc aacgcgcatc cagccggcgg       240
     ctgcccaggg catgaccagc cacatattgc cgtcttcgga ttcataggag atggtcacaa       300
     cgctgccggg aaccaaggcg tcgatgatct cctggggcat ttccaggccg tcctgcgccc       360
     aggcgtcagc ggacttggtg aagccgggga attcaaccag gttggtcagg ccgaagaact       420
     ctgccttctg gcccacgcgg atgccgtaaa cctcccaggg agaggagcct tcgcactgca       480
     tcatggcgcc ccaggtggac ttgtcctcac cgcacagcgc agcgatctct tcataggtga       540
     tataggcctt gccgcccatc ttggcagcgg ttccctggga cacgcgcatc cagcctgctt       600
     ccgcccaggg catgaccagc cagatatcgc cgtcctcgga gctgtaggtg atctccacca       660
     cagagccggg caccagcgcg tcaatgatct cctgcggcat ttccaggccg tcctgcgccc       720
     aggcgtcagc ggacttggta aagccgggga attcaaccgc gttcttcagc acgatccggt       780
     ttgcccgctg acccacgcgc acggcaaaga cttcccagtc ggaagcgctt tcgcactgca       840
     gcatggcacc ccaggtggat ttgtcttcac cgcagagggc ttcgatcatc tcatagggaa       900
     tctgggcaat ggtcttgctg ttgttgatgg cagcagttcc ctggccgaca cgcatccagc       960
     cggcggttgc ccagggcatg acgatccaca tggagccatc cgcactcgcg tattccactt      1020
     ccaccacgga gccgggcacc agcgcatcaa taatctcctg cggcatttcc aggccgtcct      1080
     gtgcccaggc accggcagac ttgtggaaat ccgggaattc aacggcgttc gtcagggcaa      1140
     caaggttgct cagatccttt tcggaagcct ccgacacgta ttccgcagtc ggatccaggg      1200
     cgatcgcgtt tccttccgcg tccaggaact gaatgttgtc caggtagaag gtcgcgggtt      1260
```

ctccgccggc ctggtcttcc ctggagatca tgatgtagtt gcccgcgccg gcaacaaacg 1320
gcgccttgaa ctgaatcttt acattgcgcg gatttttctt ttccagatag acagaccagt 1380
cggccttgtt ctcatccgcg ttctcgccgg tgtagctgta taccacgccg gagaccgcgt 1440
agaacttgcc gtccgcggcc ttgtcaacac cgatgtcaaa cgtgacagcg gcaacgtcag 1500
ccagtttttc gccggccagg ccctcaagat tcagggccac ataggggatc ccctgcgcgg 1560
caacgcgcag cgccttggaa ccgttgtagt caacaacctc cagggccgtt gaggcatctg 1620
cgttcggttt cgcggcggaa acaccgagga aggcaaaaac gccgtcttcg aagtcaagcg 1680
cgttcaccgg atcggcagac gcccaggagc acataccgac tgccatgacg gcaaccagca 1740
accaggaaat gagtttcttc acaggacttt cctcctttta ttttttaccc gatcccgggt 1800
aattaacaat tgaaagcggg ctcagcccac aataccggaa cgttccacac cctcgataaa 1860
ccgccgctgc agcaccaggt agatcaccac caccggcacc agaaccagaa ccacgccggc 1920
gtcaaaatac agctgctgct cgcggatggt cagcaccttt tccgcattgg cgatcacgtt 1980
gatcagcgat tccagtttcc gggtcacgag gatggaatcg gaaatgttga aggtgttggc 2040
gaaaaaggaa tcgttatact gccagacgat ggagaacaca gcaaccgtaa tgaccgccgg 2100
agccgcgttg ggcatcatga tggtgaaaaa cgtccgggct gtgccggcgc cgtccaccag 2160
ggccgcctcc tcaatctcct tcggcaggcc gcggaaaaac tggttgaaga tatagatgta 2220
caagccggac cgcaggccgc agccgaaaac ggacatgatg aacgtcggcc aggcggtctg 2280
cagcaggttg atcgtcttcc cgccgttgaa aagtcctaca attcccaggg gattaaagaa 2340
gcggaaggtc gtataaagcg ggaactggat tgtattcagc ggaatgacga tcatcacaag 2400
cacaaaagca aaaagcacgc tcttcagcgg aaagcggaaa cgggcaaaac cgtaaccgac 2460
catggagcag atcagcagct ggatcagggt caccccgaac acatacagaa tcgtattgcc 2520
caggatgggc aggtaattca ggtatttgac tgccatgcta tagcgttcca gcgtcgggtt 2580
gatcgggatc gtgaatacca tggggttaat ggagtcctgc cgggaaaaga aggagttcga 2640
gatgatattg atcaccgggg cgaggatgac atagctgatc ccgacgatca cgatcagccg 2700
gaaaagaccg atcaccaggt tcttcatccc gttcagggtt ttatcccgct gattggcgta 2760
ggcgggagaa ttccggaatc cctgccaggt gatttttccg ttctgtgccg gatggattgt 2820
ctgcatgttc tttctccctc ctcccgtcag ttgtaataga aggtccgccg gttcagcagg 2880
taaacgatca gccccaggat caggatggtg atcagtgtcg agaccacaga gaacgctgac 2940
gacaggccgt agttgtacag ttccttgtag gtgcgcgttg ccagtctgat cacctcactg 3000
gaggaaaacc ggtccacgac cgtgtaaatc gtgtttgtga tcatgtgagg cataaccatg 3060
ggaaaagtca ccttccagaa ggattcatac cctgtcgctc cttcaatgcg ggccacctca 3120
tacagggaac ccggaatgga ctgcagcgcc gcgatgaaca ggatgatctg cacgcccgat 3180
cccttgatga tggacgagat ccggttcacc gcatccacaa tgtagccaag cacggattcc 3240
ggaatggcca ggtcgccgaa aagctgcatg taataggcta cggagacggt ccccgcctgc 3300
gcggtttcca ggtttgcgga gggcacgccg gcattcacca gttccgtcgc ccgggtgatg 3360
gcagagccca ccgcgtccgc gctcaggata atcggcagga agaagatcgc gcgtaccagc 3420
gccctgcctt tgaacttccg gttcaggagc atcgccacaa aaaggctgaa gaacaggatc 3480
atcggcaggt cgatcagcat atccatcaca gaggatgtca gtgtctgctt gaaggtggcg 3540
tggcttgtga atgcctcgat ataattgccg aagccgatat actcctttgt catacccgtg 3600
gcggcgtcca ggccgatccg ggacaggctg aattctccca gctgcaccag gcacccggca 3660
taaaacacca ggaagccgat cagccacgga aggatgaacc agtatccgcg cacggcgttc 3720
ctctgctcat aggaaagatg aaaaatacgg tttttctgc ggatgccagt cattcgtttt 3780
ctcctccgac cgccagataa tccattgccg ggatccggta tccgtccgcc tcagcttcat 3840
gctccccgct gttcacatac agcaccgtgc cgttggaata cgtcacgcgg gccagcgtat 3900
cgctgagccg gtcatgccgg agcatctccg cgccgctcac ccggctagc gctccgttca 3960
caaagcggta gctttccgct gcctcctcct tccatgaaga gaaggtcgtg gcgtaattgt 4020
tgttcagccc ggtgtacttc atatccgccg cgtcctgcca ggtaaatgtg tagtgcgttg 4080
acgcgccgta ttcgatcagg tgcagcaggg ctgcccgccg gtcctcgctc tgcatcaggt 4140
tcagcggcga gccgcagtaa tcagctgaac cgtggaggat catctcccac agcggaatct 4200
gctcatccac aatagcgaac atcgtcgcct ccaccggcgc gttgatcacg tgccggacat 4260
aggggaagct gtaatcgttg ccgccggaaa ccatcagttc ccgctctccg gcgccgataa 4320
cgccgaaggc atgcctgacc aggtccaggt ccgcttcgcg gctgatgacg ttcgtccgcc 4380
gcttgtccgc atggacctcg tttcccaggt cccggagtga aagcgcgtcc aggcccagcc 4440
gctccgccgc cgatgccaga caggcggcat accggggcag gaaacggggg gacagaagca 4500

tatatgcccg ctccgtaaag cccatggtgg ccgtacgcct catggaaacc ggactgacgg 4560
taccgaattc cgccacatag cccttggcgt aataccggct ggcttcctcg ctgcgcagga 4620
atcctttcgc gatctcggtc acctgctgca gggctgcgtc cggataaatc tttcctccgg 4680
tttcagcagc tgccgtgcgc aggctttgca atcctctttc cccgccgagt ccgccctgca 4740
cagagacgtg gttcaccgga tcgtggtaat atccgccgtt catccagccc tgcaggttca 4800
cccgcaggtt cccgacctgc tcctgacgga gtccggacag aatctcctcc gcctgggcga 4860
aggtcgtcat cggcataacc cggagatact gaacacccat ccagtgcgcc gtttcccgaa 4920
cgccgcagag aatgtcgcag tagaagggga tatcgccgga cttttccgtt ttcagtgtca 4980
ggctgccgtc cgccaggagg gattcccgca cagccctggc cagtccgtta tagccggtat 5040
attccccgtc caggatccgg taggatacgg cgcagtccac cgggtaggaa tcctgttccg 5100
ccacgatcgc ttcctctccg gccaccatca gggtatcggt ccgtctcagg ccgaaggtaa 5160
agtaggcgta gttatagctg ttgttccttc ccgccacgtc cgcggtgaca gacgccagtg 5220
ccgcgcccct gccgcaggag accagcaggc cgcagtcctc ccggcagatg ccgaaaagag 5280
ccagcctggc ggactgcata ttctgcgtgg cggtaatatc gctgtcgatc agatccaggt 5340
cataaatgtt ctggttatac tggggcgcgg tattcttgcc gttgttgaaa cggatcaggg 5400
cgccgctgcc gtcgggaacc acgatcacac ccttctccgc cgtaccggcc gcgccgaaga 5460
acggcagcag ctggatggtc cggatctttg ccccgcctgt ctccgcgagt tccgcgtggg 5520
ggatggtgac cgtcagccca tccggcgtca gcgtgtaatc cagggcaacc acaaaggaag 5580
tgctttccgc cgcttcggat tcctcttcct ctgccagcgc ctgcatctcc gcgtaatcct 5640
ccatggtgaa accggccttg cgcgcgtccg cgtcgatctg ctgacggtgg cgggccgtga 5700
cgccctggga cttcagcaca tacagaccgg tttcctcatc cagaacatag gatttggccg 5760
cctgcttctt tccggaagcc gacaggatgc cgtaccattc cgccgtcatc tgctgcggta 5820
caagcatcag cttttcattg gacagtgtat aaaccgcccg gaaaccgttt tccagctgaa 5880
ggtattcaat ctgcccgttc tccacgcact tggcgtagga gctccagggt gtaccttccc 5940
gggagtttgt atccaggtag ttcaggataa actggctctt caggtttgcc tggttcagtc 6000
cagaccgggc caccgggtca tccgccgcgt cctgcggatt ggaatagacg gtcagcccgc 6060
tcttcctgtc atacagcgcg atctccgcgg tttttttcctt caggtagagt cccagccggt 6120
cattctgtgc cgcgagaacg aatcccggca ctgcgttctc cggatccttc aggggcttca 6180
gggcttccgc ttccgttttc gccgcgggct gaacaatata ctgacggtac tcgtcatacg 6240
cccggtagtg gatcacgtac cgggcaagga acagcaccgc ggcagccgca gccaggcccg 6300
cgatgatcca cagtacacgc cggaaggcgg tcgggagggt cagcctgttt ttctcattca 6360
ttcagcattc cctccttaca cacggaaaat cagttccata tagatccccc ggaaaaacga 6420
atacacctgc gtaatgaaat tgatcaccag cagtcccagg aaaataatga tcagcatcgc 6480
caccaccgtc agaaacagtg tgatcagcgt tttgccgaag gtatagttat gcacctgcat 6540
gatcccgatc accagcatga tcacggtgta ggccgtaccc agtccgatca gcagggtgta 6600
gaatgcctcc tcattctccg ccaccaggtg ggaaacagcc gtcgctgccg cggtggccgc 6660
catcatggga aacaaggcgt aacccgctgc gatcaggatg tctttaaacc ttccctctcc 6720
gtccatcagg caggtcacgg accagttgcc gacacagagg atgagataca gcagaatgac 6780
gcctgccagc tccgccggca ggttcacagt gcggggctca atatcgttga caataaagct 6840
ggaggagatc cggttcatgg agaacacgat tccgtacagg agcaccatca ggaaggcgat 6900
ccacaggctg ccctttttcct gatgccggat ccagtaatac ccgtctgccg gatggctgac 6960
cgtatggaaa aaatgctgcc atttgtcctt gattcccatc cggttcatga cggatccctc 7020
ctcttccctt ttttgcggat cgccggccgg accgccttca cccagagcac ggccaacgca 7080
gccgcggcga caacgccgga aaggatcagg ctgatattgt tccgcagccg ttcattcctc 7140
caccgcttga aggcgacgga gtaatactcc cggttcattc cgcgccgcag gtacagcatg 7200
gcgttcgcgt tgtctcccgc ggacaggtag gccttgccga tcccggtatt cgccagttcg 7260
ttgatctcgt ccagccgcag cacttcctgc cacagcggaa ccgcctggct ttcatccccg 7320
tcaaaccgca ggctgaccgc ccggttgatc agggcgccgt attccgtctc tgcgaatacg 7380
gtcaggttcc cctgctggga atccaggacg agcaggctgc ttcccagctt ctcaatggcc 7440
gcgggcgtct ggaaggtccc ggcctgcgtg ccgatgccgc caaagatgta cagcaggttc 7500
ccttcatggt cataggtaaa gatcctgccc cgccggccgt ccagcaggga ataggctccg 7560
tggccgcggt acaccacgtc cacgatctgg ctcgggccgg catagttgcc gaccgaaccg 7620
aacacccggt cgccgcccag atttccgcgg gggcccttcc ggatcacgtc ttcacccttc 7680
ggattcagcc ggcggacagc ctgcaggccg tccgtatcct tgctggacgc gtaaacaaag 7740

ccctcgtcat ccacatcgat accggtaaat tcagtgggaa taaacagctg ctgtttgctc 7800
cgttcctcct tggtcgccag cttccgccag aacttttccc aaagggagat cttcacgctg 7860
atcgttccga aaaagccggt gaaatctcct tccccgctga aaaccatgat gccctggaac 7920
atgttctgtg cgatcacata gacccggtcc gcgtaatcca cagacacccg caggggagtg 7980
aacacatagc cctcgtccag cacctcgctc tgcgggtccc gcacatactt gacaaaggtt 8040
ccgtccggtt caagcacaac gatccggcgg ttcaggctgt ccgcgatata cagcctgtgt 8100
ttttccgaca gtgcgacgcc ggtgggcgcg ttgaatgtct gcgtgccgcc ctccgcctca 8160
aagccggtga tgatccgcct gaccgtcttc aggtccggcg ccagcaccac gatccggttg 8220
ttcccggtat ccgcgatata cacatccccg tccgccgaca cgcacaggtc ctggggattc 8280
cggaatgcgc caacgggtgt tccttccagc ctgagcgtgg caccgctgac cagtccgtcc 8340
ggaacgtagg gcgccggggt atacaggata ttgttccagt ggtcatagtt gtaacagtca 8400
tacggcaggg aatccgcggc gcacgcgctt cccgcggcca ggcaggccag aagcatcagg 8460
cacaggatcc gtttcacttg cttcatgctc cgcctcctca gtccttcatg ccggaggtcg 8520
tcatggtctc catgatcgag ctctggcaga tcaggaaaaa gataatcggg accgccgcga 8580
tcaggaaggt caccgccgct ccggcgcccg cccggctcac accgccggag gtaatctgct 8640
gcagggcgta ctgcaggggc ttcagttcct cggaacgcag gaacatgctg cccgtattgg 8700
cccacatctg ctggaactgg aagatcgcca gcgtgagcca ggccggcttc acgttcggca 8760
tcacgatccg ccagaagatc tgccattcat tggcaccctg cagccgtgcg ctttccatca 8820
gggaatccgg cagctgctcc atgaactgct tcatcaggta caggcccatg ccgaaagacc 8880
atgccggcag caccagcgcg gcataggtgt tgttgatccc gagccaggag atgatcaggt 8940
actgcggaat ctgggtcacc gtccaggaga acatcatgga cagtacaacc acggaaaaca 9000
ggatcttttt gccgggaaaa tgattcttgg ccaggggata ggccgccata gacgcgatga 9060
tcacatgacc ggtcattccc ccaaccgtaa tgatgatggt attcaggata taccgggaga 9120
acggaaccca gctttcattc agcagcacat acaggtctgt gaagttattc agcgtgggat 9180
tccgcacaaa gatcctcggc ggatactggt acagttcgtc cagcggcttc accgcgttgt 9240
taaggatcat gaccagcggc aggaccataa acacgccgca gatccccatc aggataaaca 9300
ggaccgcgtt gccggccgtg gaccggttca gtttttttctt tccggactgc cgtttcattt 9360
tccggtagat cttcgccagg cgcttctgcc gcagggctgt ttcctcctcg ggctcagct 9420
ccgcaggttt ccggtgcggc atgcgggaaa acagccggct gatctttgcc ttttccatac 9480
cggtcattca cccacccttc tgagaatccg ctgcaccagc ttgtttgttc ccaccatcag 9540
caggaacagg attgtcgcga tcgccgaggc ataccccata tccagtcggg tggaaccgta 9600
atccagcaga tgggtcacaa tggtcgcgcc ggaataattg accgacggat tgcccgccaa 9660
cgcaatggag atatccgcca cggcaaaaga ctgcgtgatc tgcatcaccg cgccgaacat 9720
cagctgcggc ttcatcgcgg gcagggtcac aaaccacagc tcctgccacc ggttcttcac 9780
gccgtcgatc gccgccgctt catagaggcc cttgtccacc gtctgcaggc ccgcgataaa 9840
ggacaggaag ccggttccca gggacagcca cagctgtaca acgatcagca ccggcatgac 9900
aaacttttcg gtctgcatcc agaggatcgg ctcgttcagg atacccgtcc gcagcagaat 9960
gccgttcagg tatccgtacc ggtcgcctgt caggatcagg ttccagatca tatacgcgtt 10020
gccgcagatg gacggcgcgt agaaaacaag cgtcagaaaa gcccgcactt tcggcctgaa 10080
ctcattgatg atccatgcaa acagcaggca cagcagatag ctgaccggtc cggtcaccac 10140
ggcgaatagc agggtattcc gcagggagat gatgaaaata tcatcctcca gcagcagctt 10200
gatataattg gcccagccga cccagcgggg catctccagc atgttgaaat agttgaacga 10260
caggtaaaca gacatcagca cgggcagcac tgtaaagagg aaaaacaggg taaaatacgg 10320
cgccatcagc aggtaggatg ccctgttgat ccggacctga tgccacaggg tatccttttt 10380
ccgcatgctc cgtccgttcc ggattgccat gctcatctca tttcccctcc cctgtttctg 10440
cgacaggcag tccgaattct gtccgtttat aggtaatctc cgcgttaatg taataaatct 10500
tgtccatcag ttcctcgcgg ggcgtattgc cgaactcggt gcgtcccgtg ctggcgctgc 10560
ccgtagtcac tgcgtagaag gcgttgttca cattgcgcca ggtgtaataa ccgccgggca 10620
cctgggggat ccccctcaca tggctgaact gctttgccag ttcccggtag tcccggatgg 10680
gccagttcag tcccgccagg gcttcaaggt tcgcggtggc cacccgtgcg gaggcgccca 10740
tcagtgactc catttcccgt ccgtacagcg tctggatctc ggcgccggtc caccacttca 10800
ggaattccca gcaggcttcc ggttcctttg tcgcgctcat gatgatatcc gcaagaccgg 10860
agcagccggt tgttgtgtcc agcgttccgt cttcccggag ggtaccgggc acatgggtga 10920
aatcccacag tcccttgata tccggcgcgg atacctgcag gttgttgtag gttgtataat 10980

ccgtaatgat gatcgggcat tcacccgtac ggaagcgttc ttccaccgag gtggccatgt 11040
ccagcttgta atctgtataa aaatcgcagt accggcggaa cacattgacc gcgatatcgc 11100
tgtccagtgc ggacgcgtct ccgttctccg tgtagtactc gcctccggcc tggaacagca 11160
gcatggcaaa agtctgctcc ccgggcagca tgccgaattc catctggttc ttgctcagca 11220
cgctcatggc caccttgacg tcgtcccagg tttccggcac ttccaggccg atctcagcca 11280
ggatgtcctt ccggtagaac atcatgggga aagtctgcgt gtccggcagc gcgtaagccg 11340
cgccgttaaa ggagaaggga accagggcgc tcggagcgaa ccgctccgcc acctcggcaa 11400
aatcatcaaa ctgcgtgaga tccataacgg cgttgcgcag gccgtagttc accggcgtat 11460
cgttcccggt attcatcacc gcgcccgcga tccccgtggt attcgcgacc tgaatagcca 11520
cgtccggccc ctggccggac agcgtggcgc gcagcagcgt gttcatatcc accagctgga 11580
cgttcacgct gatcccggtc ttcggcgtaa aatgctcgtc gatcagggcc ttgatcacat 11640
tagcctggtc ccgtcccgtg ccgacccaca gcgtgatcac ccgggagtcc tccgttttct 11700
ccgccacatt gccgatctgg ttgtaattga tagtgaagga gctgtataaa cgggataatt 11760
cgaaacccgt accggaaagg aaatccgtcc cgctgatctc cggacggaca tcggcagtat 11820
ggacaaagat cctgtcaatc tgcagcggct gacccagcac ctgggtgatc cagttgccgc 11880
aggcacgtac gttggtctta taggaagaaa gcacctcggt gaagcgttcc tgatcccgga 11940
tcagctctgt cagctggtca tccatcgtct tcagcacagt cagcttgtcg ctccggtgcc 12000
ctgctgtttt ttccagcgcc tcaatggccg accgaaggtc cttcttaacc atttgcagct 12060
gtgcttccag tcccggcagg cttctttcga tctggtaatc ccggtacgga tccggcgaaa 12120
caccggtgat ataaagcacc tgccggtaaa tcccgttcag ctgcagcaca cagtcctgca 12180
cctgccggat gacagctgcc atttcgccca gtacaacctc catccgcagg gtatgttccc 12240
ctgccgccag atggatcagg tacggttctc ccggctctcc ggacagcgtt tccaggcgcc 12300
agtcctgggt atagggaaac gcggccgcct ggaattcggc aaacggcact tctccgtcaa 12360
tatagatctt gcgatacacg tctgttccgc ggataaaatt ctgcttatcg taaagtgaga 12420
tcgcgtagtt tccttcctcc ggcacggaaa aacgccactc gatccactgt cccgcgcttt 12480
tccacgcgtt gccgccgatg ctgttgttca gcagatactt cgggctggac ggggaaacag 12540
atcgggagga ctgatcctgc acagggtaca gcatctggga ggaggtgccg gacgccgctt 12600
ccgcctcaat ccggatcacc cggttcgcgg ctccgcccgc ttccggcggg gcatacgact 12660
tcacgtccgt acccgcccgg aaacacatac gccggatcag catcggttcc cgcagggaca 12720
gcaccgtcag cgtatgcgcg ccccggtcca ggtaaagctt caggctttcg ctgatatatc 12780
cgtttgagtc atggaggaga gaagtgcgcc actccggcgc ttccgccgga gaaggtttca 12840
gttcgttccc ctggttatcc ctgagccagc ggatctcttc cgttccgtcc tcagccgtgc 12900
tgcggacgat ctccggatga aggtcgttcc gccatacccg ggaaaagttc accagcgcaa 12960
gctccctgta cggaagagca ccgtccagga aaaaggcccg ctggatctca gcgttccgtc 13020
ccggccaggg atagtattcc acctccaggt cataccaccc ggcctcttca gcctcaaaat 13080
cccactcgat gactgcttcc tcacccgtca gcaccgagac tccatcccgt ccttcagcct 13140
gtgccttcag caggggagcg ttctctccgc cttcctccgc gtaccggacg gtatccgccg 13200
cctccaggac gatctccttt tcaggccggg ccgccgttcc atgggccgcg cagtattccg 13260
cataggaggg aatgtccggg tttatggaat aagtgctcac ctggtcctca tactcccgga 13320
tacccagatc ggtggcagcc atcgccgtgc cggctgccag cgccagcagc atcgccagca 13380
caattcccgt tttctttctc acttgttctt tcctccggtc cattcgcacg cggcagactc 13440
agtctccgtc cgccgcgcgg aaatcgggca gatccttccg ggtcacgacc cgcgtgtccg 13500
aatacgcctt ccggaccata tccgtttctg tatactgctc tgaaggcttg ttgaacttgg 13560
tattccggat cacaaattcg ccgccccagg tgcaccagta tccccagacc gtcccgtccc 13620
ggaacacctt ctccggatcc ggaatgcagc cgttctcgga cagcacgatc atcttccgtt 13680
ctcccgcgat ccgcgtcagc ggtatgaagg acgcgctctg ggagtcataa acatgttccc 13740
cgggatagat gtcgatgccg atcatatcca cctcgtcatc ccccggatac cactccggat 13800
cctgtccgtt ccagatccag atcaggttgt tgaggccgta ttcatgggtc agccggtcaa 13860
aaagcaggtg atacagtttc aggtatgctt cctttccgga ggctccccac cagaaccacc 13920
cgccgctcgc ttcatgcagc ggccgccaca gcaccggcac cccggcctcc ttcagccgca 13980
gcagctgcgc cgccaccgcg tcgatatccc tgttcaggag ccggtatccc tcttcatcct 14040
ctccgtccat aaccctgccc aggttgaagc tggtatgctc ggtataaaag cccgaatacc 14100
agggttcctt cagatagggc gatggtgcgt tccagtgcca gcaaaaggtc acgatcccgc 14160
cctggttcca gtattcaacc gcgtacggga ccgccttgtc gctctcgccg ccatgctcca 14220

cgcggctggg ggaataatcg atcatgtcca gtccgagaac cgcgggataa tcaccgcccg 14280
ttgcccgcca gattgcctgg ttttccattc cgtacattcc gccggcacac tgctggccgg 14340
aaaggatctt tttcccgtag cattcccgca gccagtcata cagcttccgt gcttccggtg 14400
aagcttcgcc gacgcacatt ttttccggaa cggtatacag gtcctccggc agcgcagcgg 14460
aaggcttcag ctcaaacccg tccagccgga tatgtcccca gctggtaccg atacccaggg 14520
tatgcatacc tgcggacagc cagatatgtt ccagcacgct tttcccgaac gtcgtttcat 14580
ccgtaaccgc gcttccgata ttctgcccgt ccaggagcac cgggttttcc ttgtgcccgc 14640
cctggcttgc ctggataacc gcgatatcat aaaaaccttc ccgggagacc ggcacctcta 14700
ccgaaacaga atcttccccg gcagttttaa agccttccac ccactgattc ccgccgtttc 14760
cggcaacggc agtatgtccc ttcagttcac ctgtttccgc ttccactaac atcgtttctc 14820
ccgtctcttc tgctgtcccg gagagaatga tgaaaagcag ggtcatgacc gctactgcga 14880
ttccgcatat gcttttcata ctgcttttca tttccgttct ctccctgttt tgttagtttg 14940
ttaacctatt tgtaaacctt ttgttaacta attttagcga aagaaaaatc gcctgtcaag 15000
acaatttttg cacatatatt gggctttgct ggtcatttat tgttatcttc cttccggaaa 15060
agacatgtgt tttcttgttt cgcgggttat attggttaac ttattagtta acaaattaat 15120
acatgatttt cttctgtata ggccgcccct ccggatccgc cgggtgtcgc agacctgcag 15180
aacgcaaaag ggagagccgc tatggctctc ccttttaccg cactatccga ttaaaattct 15240
gtgtccgtaa agatatacgc cggtcgtcgt tccttcccct ccgggatcac ttcatacccg 15300
tccatggaca aaccccgcac atggcgcgcc caaagggcat aagccggggc gtctcccacc 15360
ggatcgatca tatgggcgtc cggatacacg ccgcgcattt catcaaaatc cttttctttc 15420
cgcaaaggcc ggcgggaagc ctcctgccgc aggaggaccc ggcggaaagc gatgtcctct 15480
atcggctttt cagggatccc cagcagatag gaaccgcggg ggccagtctc ccgtccgcgg 15540
atatccgaaa acagaacccg gcgcagggtc cccgggccgg gtgccggatc ccccggtttg 15600
acccttccgc ggttttccag ccgcaggaag aatgggctcc aggcgcggat catatggata 15660
tcctcgatca gaaaattctc cagcgttccg ccgtccacca tttcaaggga gatcccgctg 15720
tccgcgcagg catgcttcag cccggacgga tcctccgcca ggccgccgat ccggcagcgt 15780
cggatataca cctgccggaa atccccctgg gtatctgttc ccaccttcac cgcgttgcac 15840
gcggaataaa aatcgcagtc ttccaccaaa accctgctca gttcccgctg gcctgcgcct 15900
ttaaagcaga gcgcgtcatc cccgctgctg acccggcagt tccggatcac cacattccgg 15960
catccgtcta tatccatccc gtcattgtta tagttggcat ggttgcggac cgtaagccct 16020
tccagcagca ggttttcaca gttcaggtag ttctgcatcc agcaggcggc gtctttcatg 16080
gtcaggtcca cgatatgaac gttccggcag tcgatcatcc ggatcaggaa cggccggccc 16140
ggcgtaccct gggccgtctc ctccccggga aaatgcgccg ggcttccccg cccgtcgatg 16200
acgcctccgc cccgcaggca gatattctca catccttcgg caaagagcag cgactggtgc 16260
atacccatac tggtatcctg gaccgtcagc cgcgcggcgt gatgctccgg atagtcgcga 16320
agatccgtgc tgcccagcag ccgggagcct gcccggacct ccaggcatac gccggagcgc 16380
attaccagcg ttccgctgac gtaatctccg ttcccgagcc gcacgatccc gccctgctcc 16440
gctgccgctt cgatcgctcc ctggatggct tcggtcgccg tttccccggg ctggtacccg 16500
aattcctccg gcgtaaagac cggccccccg tctgtccgtt cccgggcaac ctttatcagt 16560
tcctgcgcca tccgcgccat gtttgtcttg agatcccggc accaggaagg ttcctttccc 16620
tcccgaatgg tctccatact gtttcctccg ttctaaatca gctctgtccg caggaagtcc 16680
ccgttttcat tcgttacgat gcgtctgccg cagctgaacc tgccgatatc ctttccgtca 16740
gcccggatct ccgctttctt tccatcgctc cggatcccgt agcgatggcc gtaaaaggcc 16800
gcttcatatt catatggttt tccgcggtca aggctgaacc acacctgccc catatgcgga 16860
tggattccga agcggtgggc aatatactcc aggcaggcga gcattgtcgg gccgtaggca 16920
tcctggacct tttcgctgct gtctccgctg accggccgat ggtccgcctc gtccaccagc 16980
gatgccttcc cggtaaacgg gtcatattgc tgggtaaacc ggcagccgtt cgcgtcaaca 17040
gcattgatca gtttcttccc cagagctgtc acgatccggg catacccgta ccgctcaagg 17100
gcggtgatcg cccgctggta ggtcaggccc tccggctgtc cgctccagtt attctccggc 17160
gcgttccgga aagccgggtc atccgcggca acgctcggca gggggaacgg tgtccagaat 17220
tccgaagcgt tcagcaggtg ttccttcaca aaccggtctg ccatatcccg ggaaacgctt 17280
ccccagtaca tgcaccgcag cgtgttatgg cacaggatat tgatcgtttt tcccgtccgg 17340
tcccggtcat acagcgcgcc ccgcctggca tcccacaggc gccttcgcag ggcatccgcc 17400
acgtctcccg ccttcttttc ccagtccgta aaagccggat ccccctgaat gcggcagatc 17460

tctgccgccg cgcggcgtgc cgcgaagctc cagctcgtca catccatgct ggccatggga 17520
accaccccgt atccttccgg cggttcatcc tcctcccacc agcagggcgc gtccccgtag 17580
cggaccgcca gatcctctcc cgtatcatac acacagaagg aagaaagcag cccgtccccc 17640
gaaacatccc tcgtcctcca taagcaggcg tcaaagcgga tcagtgcttc cttcagctgt 17700
tccaggtatt ccccatcttt tcccatcagg taatacaggt tcagggccgg atcagggaag 17760
cagaatccct ggtatttatt gaactgcggt tctgtttttc cattgatatg ctggatggag 17820
ccgggcatcc gcccgtcctc ccgctgatgt ttcataaaaa ggagaatatt gttcttcgcg 17880
gcctcaaggt tccgcagggc atacatttcc ccgcccatcg gctgcgtttc cagccagatc 17940
ttttcatacc cgcctccttc caccagcacg gtgtcgccgc tgaaatcctt caggttcagg 18000
aggcactttt tttctgccgc gtcataaacc cgctgcagtt tctcatcttc cgttcggaag 18060
ctcacgcttg tctccggcag catttttttgt ctttcccttt cagtgtgttt tgtcccttat 18120
ataattaggt agattttatc atttcttgcg aaatcagaac agttacaaaa aaactgcata 18180
ttttttttctt ttcatgaaat caaggaaaat caacggtatg caaaattttt gcttctttct 18240
tttccggtgc actttcgttc ataatcggga tgacagagca atcgttgaca aattcactca 18300
acccccctgaa ggaggatatc ccatgagtct ttcctcatcc ccgctggctt atcgcggagg 18360
gatccggcac aaaatggcta tgaccctgcg cagggactgg cagctgtggg tactgatgct 18420
gccctgcctg gctttcttta ttatcttctg ttatttcccg atgtacggcg tgcagatcgc 18480
tttccgtgac tataagttta acctgggaat caccggcagt cgatgggttg ggctgcagca 18540
ttttcaaaag ttctttaaca gctactactg cggaagaatg cttcgcaaca ccttcctgct 18600
gaatgtctac ggtctccttt tcggtttccc ggtcccgatc atcctggcaa tcctgctgaa 18660
ccagctgaaa agcaagcgct ttaaaagttt tgcccagacg gcaatctatg tgccccattt 18720
tatttccata gtggtcatga tcggtatgat ctatcttttt ttctccccga caaacggcat 18780
cgtgaaccac ctgatcaccg ccgccggggg aaagagtatc tatttcatta tggaacccgg 18840
atggttccgg ccgctgttta tcggatcagc catctggcag gatgcaggct ggaacacgat 18900
cctgtatatc gcgaccctga ccagtgttga tccccagctc tatgaagccg cgaccatcga 18960
cggggccacc cgctggcaga agatcaaaag catcgatatc ccccacctga tccccatcac 19020
ggtgatgatg ctgatcctca gctgcggctc cctgctttca tccgacacgc agaaagccct 19080
gctgcttcag accagcggca acatagccac ttccgatgtc ctgggcgtct atatcttcaa 19140
cgttggtatc gccgcctcaa aagcgcagca ctcatatgcc gccgctatcg gcctgtgcct 19200
caacgccgtc agtttcatca tcgtcatggg agtcaacgcg atcgcccgta agatgagtga 19260
tatcagcctg ttctgagaaa gaggggaaaa acatgacaac ccttgcagcc aaaagacgga 19320
accgcatccg tgaaacccgg agagaccgga tcttcaatac gtttaacctt attttctgga 19380
tcatcatcct gctcatcgtt ctttacccccc tctggctcat cctcatctgc tcagtgtctg 19440
acccggatgc cgttctgcag gcaaaggtcc tgatctggcc ggttgatttc tccctcatgg 19500
gatatgaggc agtcctgcag aacaacgaac tgctgggcag ttacctgaac tccatcttct 19560
ataccgttgc cggctccgcc ctgagcgtga tcgtttcggt agcggccgcc tactccctct 19620
cccggaagtt tgccgggaaa aagcttgtca atctggccat taccttcacc atgttcttct 19680
ccggcggcct gatcccgatc ttcctgaacg tcagggatct gggcctttat gacacccggg 19740
ccgtcatgat cctgatgaac ctggtctctg tctggaacct gatggtggct cgcacctaca 19800
ttcagaccag catccccaac gatctttatg aggcggctgt gatcgacggc gcagaccatt 19860
tcacctattt tttcaaatgc gtcctgcccc tttccggcac gatcatagcg gtgctctccg 19920
tttattacgg cgttgcccgc tggaatgatt atttcactgc ccaggtcatg ctgcgcagcc 19980
gcagcctgtt ccccctgcag cgtatcctca aggagattct gacgtccctg acctccacag 20040
gatcctccga taccttcttc gcggcttacg cggacaatct gggcggtctg caggccgcgc 20100
tgcgcaaggc cgaagtcgcc aaatactgct gcatcgtggt ggccaccgtg cctgctgtgc 20160
tgctctatct cttcatgcag aaatacttcg tcaaaggcgt catgatcgga tccctgaaag 20220
gataaggagg tacgtcccga tgtctcagat gaaaggggtg gaggtgtccg cccgggccgc 20280
ccgccatata gacggttatc gggtgctgga gggcaaacgt gaatatgtga cctaccaggg 20340
tgattcctcc atccgcatct ggtattcaga cattccatgg cgttacgaaa cccatgacca 20400
ttcggctgtt gagatcgtcc tggtcctgga gggaacggtc acctactccg tagagggaac 20460
gagttatccc gtacacaagg gggaagtgct gatcgtgccg tccaacacgt ctcattccct 20520
ggaaatgggc gagggcagca gccggtacct tttcctgttt gaaaccgaca ccatcatgac 20580
catgcgggat atcaaatcca tggtcctgta tcttcataaa cctttccacc tccgggatga 20640
atcagatgcc cataagcaga tcagggaact gcttcttcgg gcgaaggatt tttacgaaaa 20700

gcaggagctg atgtggaata cggcgtgtta cagctgcatt ctccaggtat atgccaccct 20760
gggccggcac tacctgatcg gaatcaggtc ccggaccggt gatgcggcac ggagcatgga 20820
cacagaggtg atcaacgctg tgatgaccta tatcaatgat cattaccgcg aggatctgcg 20880
cctggatgat gttgccgggt tttccggatt cagccgtttc tatttctccc ggtctttcaa 20940
gcagcagaca ggatactctt ttaaggatta tctgtgccag aaacggctgc aggccgccat 21000
ggacctgctc accagcacca atctgtccat gcgggggata gccgtcgaaa gcggctttgg 21060
cagcacagca tccttcaacc gcgcctttcg cgcctgcaaa ggctgtaccc ccacccaata 21120
ccgggccgtt tacggaatgc tgtaaagtgc cttgcctgat cggtatctag ggctgtatgc 21180
cctaaaataa aaaggaggaa aaacccatga agaaactcgt ttccctgctg ctggcatgct 21240
gtatgctgct cagcctgagt gctgccttcg ctgaagagaa gacggaactc accgctttcc 21300
agtacgcgct ggaaaaccag aacaccgatt tcaacggcct atggttcttt gacgaactgg 21360
aaaagaagac caatacccac gtgaccttcc agatggtgaa ggacgcggac tgggcgacca 21420
actccaacct gatgttcgct gtgccggacc agatgcccga catgatcctc cgtcacgcgg 21480
tggacgtgga agaatacggc gtctcccagg gcctgctgct tcccctcgat gaatacctga 21540
ccgaggagat catgcccaac tatgtcagcc gcctgaacat taacaacgcc ggcgactcca 21600
tccccgccag cgacggcaag acctattaca tcggcttcct catggcccag aacgtcaacc 21660
ataccggtac ctggtatgtg aaccagaagg aactggacaa gctgggaaag accgctcccg 21720
ccaccattga tgaagtgact gatctgctcc gcgccatgaa ggctgacggg atcaagtggc 21780
ccttctccgc gtcctacacc cagttcggac cggaaaccat ctggaaccag tttgcttcct 21840
tcggcgttcc ggaaaatacc tattactact tcatcgacgc tgacaacaag gtccagctca 21900
cctttgccca ggacggctgg cgtgcctgtg ttgaatggct gcaccagctg tatgaagagg 21960
aactgatgga tcccgaatcc ctgacccagg actccaacct gtgggccaac aaggtgaaca 22020
acggcgaagt cggttacttc tgctacctgc gcctgatcaa caccgccatc aaggctgaga 22080
acgtggagaa cttcaagtcc atcctcccgc ccgctgcgga cggattcaag gcctgcgttt 22140
cccagatcct ggaggttccg gaagccggcg cctacctgac aaagaactgc aaggatcccg 22200
ttgctgccct gaagtggatc gatgcccagc tggaaaccga aaccatgatg gtctccctga 22260
acggcaaggt cggcgagcag atcgtgctca acgatgaagg caagtatgaa gtcatcaaca 22320
tccccgaaga caacggcctg tatcagttcg ttccggttac catgggtcag ttcttcgccc 22380
ccggtgacta ttacacctcc atctaccaga tggctcccca ccgcgtggaa cgctatgaag 22440
acagcaagtg gtacgccgaa gccggcgtgc tggagcccaa gagcttccag tatctccgcg 22500
atctgagcaa gctgagcaac gatgacgcca ccgaagccgc caccatctac accgacctgc 22560
agaagctggt tgaggaatcc ttcgccaact tcgtccgcaa tggtgtaacg gatgagagct 22620
ggaacgattt ccaggctcag gccaaatcca tcggttccga tcgttacatc gagctgtacc 22680
agaatgctta cgatgcctat ctggcaaagt aaatccttga acgccgcggc ctgagatgtt 22740
aaaatgcaag catcagaaaa gccgccgggc atcgccgcag gaattgattc cgggggcgat 22800
gccctttgc cgtatagagg gagtacatat gaaccgtacc gtaaagatca cctgcctgct 22860
ggctgccgcc ctgtttctgc tgacagggtg tgctgtttca cggccgggag agggctctgc 22920
ccctcccggc gcatccctta ccgcgctgca gtatgagatc gacaatatcg ccgtggactt 22980
cagcagtatg tggtattacc ggcagatcga gcagcagaca ggcgttcatg ttgacttcag 23040
tgaagtcaag gattccgaat ggagcagctc cgtcggtctg gcctttgcca agggaaagat 23100
gccggacatg atcctgcggg gatccctgga tgtggaggaa tacggcgtga cccgccatct 23160
tctggtcccg ctggatgaat atatcgacaa gggcttcctg cccaactacg cggcccgtat 23220
ggaccaggcc ggtctgcgtg agcagcttac cgcttctgac ggccatattt accagctcgg 23280
attcctcatc tcccagggtg tcaacaccaa cgggcatttc ttcatcaaca aaacatggct 23340
ggacaagctg aatctgcagg tcccccggac cgtggaccag ctgacggaag tgctcaggcg 23400
tttccggtcg gatgatccca acggaaacgg tcttcaggat gagatccccc tggaattcac 23460
ctttgatgac aataccaccg gcatatacaa tctcttctcc tttttcggcc tgcccctgaa 23520
cgaggattat gtctatgtgg acaataccgg aaacgtacgt tttgcgccgg aggaggacgc 23580
cttttttgag acgctcacct ggcttcacga tctgtatctg gaaggtctcc tggacattga 23640
ttttatcagc cagggcagca gcatctgggc tgccaaggtc aacgaaggaa ataccggcat 23700
gttcagctac tggcggctcc agaacaccgc cctcagcccc ggcactgccg ccaattacca 23760
ggtgatggtt ccggtgcacg ttgagggaag gaatgcctgc cttccgagaa acatggacat 23820
cattgagttc ggcgccgccc tgaccacgga caacagggat atcgaggcca gtctccgctg 23880
gctggatgcc cagtttgaaa cggaaaacat gctcgtctcc cagaacggcc ccgtcggcga 23940

cacgctcatc cggcgggagg acggtcggta tgaggtcagc tatgtcccgc cggacaacga 24000
gctgtatcag tccgttccgg tcatctgcgg ccagtttttt gccccggctg aatattacgc 24060
ctctgtctat gtcccggccg cccaccggct ggaaaagtcc gcctactgcg atctgtatga 24120
gcaggccggc gtgctggaac ccgtctccgg caagctgctg acagccgtgg cgccgaaaac 24180
aacagcccag tccgccgaaa ccacccgcct gaaggcaaag ctgaagtccc tggtggacgc 24240
ggagatcgtt gccatggtca cccggggtgt aacggatgaa agccgcgaag ccttcaggtc 24300
aaagctgaaa gaaaacggaa gcgtggaata ccgtcaggtt taccagcaga tctatgaccg 24360
ctggaaggag aaataaaccg tgatcagaaa gagccggacc tctgtcttca tccggtatgc 24420
gctgtcgtat atcgctgttg tgctgctgct gttctgcagc atcacgggat acctgtatgt 24480
ccgtctcagc ggagaagccc gggaagagat cattgataac cagatcaacc ggctttcccg 24540
gatcgccagc cagcatgaaa gctatatctc cgccatgctg aataccgccg aggagatcgg 24600
tctcagtccg gatatcgagt tcttccgcta caatgaagag ccctggaaag cctatgacct 24660
ccagctccag atggtcccct acaccacaac ggccagcacc ttctgtgatc aggtttatct 24720
gtatttcttc ggggatgaca tgatctattc ctcatcatcc tccatgaccc tttccctttt 24780
tgcgcgcatg atgcgctatg agtttatccc gatggaaaat cttgtctccc ggattgcgag 24840
cactgaccgg atgaccattc tccccgccca gcaggtttcc tccacccttg tggacagttc 24900
ccgggttgtc accttcctgg tgccgctggg gtcggctccc ggatccggaa agggcatcct 24960
gctcttcctg atcaaggaca gcgtctacca gtcgcttttt gccgacgcca ttgacgaccg 25020
tgtcaacacc tatatctacc agggggatga cctgctgtcc tcctcggaga agctttccct 25080
tccccgggac cgggtggaag cctcccagga cactccttcc cgcctgttcc gatgggaaaa 25140
ccaggactgg acccaggtca ctttctccgg cagaaactgg ggactgacct atactgcggt 25200
cctccgaaac gccgatatca actcctccat catgcagcag ctcttcagca tcctggccat 25260
cctgccggta tttgtctttc tgagccttct gcttgctctc tggatggccc ggcgccatgc 25320
taagcccatc cgggcgatca ccggtatgct ttcgcgggaa acagatgaaa caccccgcga 25380
tgagctccag cagatctcct ccggcatcag ccggctcacc acccggaacc aggagctgtc 25440
ctcccggctg gaacgcgccc tgcccctcca gcggcatgat tttgtcttcc gctttgtgaa 25500
aggacgcttt gagaccaggc aggaagccgt ggctgcaggg caggctgtag ggctggagat 25560
cgaccggcct gtatacgcgg tcatcctttg cagcggctct gccggaaatg atcatccctt 25620
tgatctcagc ctgccgccct tcagccgcct ctccggctat tccggtgccg gcgtggaact 25680
ggttgccctg aaggccaacc tttaccttgt tttctccgat gacccgaagg tgcttaccgg 25740
catggcggaa acgctgcgct gcgaatgcgc tgccggcgga gagcgctgta tcacctccat 25800
ttccgccctg cataccgatt tcacggaagc cccttccgcc tatctggagg cagcagccgc 25860
ctatgaaaac cggtttgtca tgggcagcca gaaggtactc tgctacggcg atatctcccc 25920
tgacctggag gagatccttc cccaggcccg caagatcaca acctccatca gtcaggcgct 25980
gaccctggga agccgggaag ttctggacgt ccggatcagc gacctgctgc atttcctgaa 26040
gaacaccagc atgtctccct tcgcgttccg gatgatctac agcgaggtga tccgttccct 26100
ggtgcacacg catacggcgg cgctgtccgg acagggctcc gttcaggaaa cctacaacat 26160
cttcactctc acctcctgcc agtccattga tgacctggat gaactcctgc gcaggctgtg 26220
cgaccgcctg atcaccggag aatcagcttc cggggagaaa gttccggacg aggaggatga 26280
gatcagccag gttgtccgct acatggaaga gcacttcagc gatccggaga tctccatcac 26340
cgccctggcg gactccctgg agctttccac cacccggttc agcctttcct tcaaggaaaa 26400
ggccgggatg tctccgctgg aatacctgac gctcctgcgg gtggaacacg ccaaggaact 26460
gctggaatcc agcgacctga ctatccggga catcagcacg caggtcggct attacgactc 26520
cggctccttt atccgccgct tcaagcaggt aacaggcgaa acgcccctgc agtaccggcg 26580
cagcagaaac ccgaaaaaag agaacggcat tccggaccgt gatgcctccg tcccgacaga 26640
ggagtaaaac catcctatga ttcatactga tcagatcaat attcgggatc cctatgtcct 26700
tgtgcacaac ggcacctatt atctttacgg cacccgcggc cccgactgct ggggacccgc 26760
ggacggtttc gacgtatata ccggccggga tcttgtgtcc tgggaaggcc cccgcgtctg 26820
tttccggaat gacggctcct tctgggccga ccggaattac tgggctccgg aagtgcatcc 26880
ctggcaggga atgttttata tgttcgccag cttcaaaagt gacgcccgcc ggcgcggcac 26940
ggccatcctg cggtccccct ctcccctggg cccctttgag ccctggtccg acggtccggt 27000
tacccccgcg gactgggaat gtcttgacgg aacattctac gcttccgcgg acggcactcc 27060
ctggatcgtg ttctgccatg aatgggtcca ggccggcgac ggagaagttc tcgccatgcg 27120
cctgacggat gacctgaagg cgtcggccgg tacaccggtt ctgctcttcc ggggttccga 27180

cgctccctgg agccatgccg tacggcattc ctccggcata accggtttcg tcacggacgg 27240
tcctttttg tggcgcggaa aagacggctc cctgctctgc ctctggtccg gcttttcaga 27300
aaaaggatat gcccaggggc tggctgtttc tgacaacggg gagatcgacg ggcatttttc 27360
acaggctgat ccgcttttcc tggaggacgg cggccacggt atgcttttcc gttcgaagga 27420
aggcagaacc ctcctggctc tccacagtcc gaacacccat ctgctggagc gtccccgctt 27480
tgtaccggtg gatctctgat cccgcaaagg atctcctgaa atctcccttt atgaaaggat 27540
aaaacatgca ttacgttctg gcgatcgaca tcggcgcgtc cagcggccgg cacattgtgg 27600
gctggcggga aggaagcggg atccgaacag aggaggtcta tcgctttccc aacggcatca 27660
cggaacagga cggcagcctg acctgggata ccgatgccct gcttttccat gtcaaagcag 27720
gcattgacaa agccctggaa aaatacccct ccatccaaag cctttccatt gatacctggg 27780
ccgtggatta tgtgctgatg aacggcggca gccctgtcct tccctgctat gcctaccgtg 27840
acagccgtac ggaagcagtg atcggagagg ttcacgaaaa gattcccttt ccggagctgt 27900
accggcgcac cggcatccag taccagccct tcaacaccat ctatcagttg tatgccgaca 27960
aggtttccgg gcggctggat caggccgatg gtttcctgat gatcccggaa tacctgatgt 28020
accggctgtg cgatgcggaa agccatgaat acaccaatgc cacgaccacc ggcctggtca 28080
gcgcggaaac cggggaatac gatcccggaa tcatcagcgc cctgggcctg cccgcgcgtt 28140
ttttcgggcc tctccggcag cccggcaccg tgatcgggga atacaggggg atcagggtta 28200
tgctctgcgc cacccatgat accggaagcg cggtggaagg cattcccatg gaaggctgcc 28260
agccttatat ctccagcggc acctggagcc tgctgggggt gaagaccgaa cggccggtca 28320
tcactgaagc cggccggctg gccaactgga gcaatgaagg cggcgccgga tataaccgtt 28380
accagaagaa catcatgggt atgtggctca tgaaccggct ccgggcggag ctgtgtccgg 28440
acagatcctg ggatgagatc atggccgaag cggccgaaaa gcattttgat cacctggtgg 28500
atgtgaatga cccggtcttt ctggcaccgg aaagcatgaa ggccgccttt gattccaggc 28560
tttcccatcc gccccggtgt attgccggct acttccggtg cgcctgcagg tccctcgccg 28620
acggataccg ccgggctgtt gaggaaatgg agcagaacac cgggcggcgc tatgacaggc 28680
tgtacatcgt cggcggcgga gcaaagaaca catacctgaa ccgcctgact gaagaatcga 28740
ccggaaagca ggtgaccgcc ctgccgatcg aagctaccgc tctgggcaat atacgcattc 28800
agctgaaagg tggaactgat acatgagtta tgaagaatcc aggaaacggt atgcggccct 28860
gggggtggat accgatgccg tcatccgtct cctcaggaca gtgcccgtca gccttcactg 28920
ctggcagggt gatgatgtcc gcggttttga taccgatccc tccaaacccc tgaccggcgg 28980
catccagacc accggcaatt atcccggccg ggccaggacg ccggaagagc tgatggccga 29040
cctggatatg gttctgagcc tgatccccgg cacgccgaag atgaacctgc atgccagcta 29100
tgccatcttt gaaaacggtg aatgggcaga ccggaacagg ctgcagccga aacatttccg 29160
gaaatgggca gccttctgca aagaacggaa ccttggctgt gattttaatc ccaccttctt 29220
ctcccatccg aaatgcgatc cgctgaccct ttccagtccg aatgccgaaa cccggaaatt 29280
ctggattgag cacggcaagg cctgtatccg catcagcagc tacctggcgg aggaactggg 29340
cttcccctgc gtcatgaaca tctggaccgg cgatgggttc aaggacatcc ccggcgaccg 29400
tatcggcccc cggatccgtt accgcgacgc cattgacgag atcctgcggg agccctatga 29460
cttcagccgg gtgaagcctt gcatcgaaag caaagtgttc ggcatcggcg tggaggccta 29520
tacagcaggc agcgcagagt tcgccctgtg ctatgccgcc atgaaccggg acaagtgcat 29580
ccccctgatg gacaacggtc actaccatcc cactgaggta gtcagcgata agatccctgc 29640
cctgctgact ttcttccggg agatcgccct gcacataacc cggccgatcc gctgggacag 29700
tgaccatgtg gtcctgttcg acgatgaaac ccgggaaatt gcccgggaga tcgttcgctg 29760
cggcggtctg gacggccgtg tacatatcgc cctggattac tttgatgcca gcatcaaccg 29820
catatccgcc tggacaaccg gataccggaa cctgcagaag gcattgctct atgccctgct 29880
gcagccccat gaggacatga aaaaactgca ggataagggg cagttcagcc ggctgatggt 29940
cctgcaggaa gaaataaaaa ccatgccctt cggggacatc tgggcggaat actgccgctc 30000
tgcggaaagc ccgccgacgg cgaatggttc ccgctggtgg aagaatatga aaagaaagtg 30060
ctggtgaacc gcaaatgaaa gatatcctgg ccgccccctt tatgacggaa atgatacgca 30120
cagcgacagc catgtatgac catggctggg atgaacgcaa cggcggcaat atttccctga 30180
tgctggatga aaaggaagcc ggggaatacc tggatatgca cagtgtcctg cgaaccatcc 30240
cgacaggctt ttccatgccg gaactggacg gaaaaacctt cctggtcacc ggcaccggca 30300
aatacttcag aaatatacag tatgaaccgg aggtcaatct ggggatcatc cgtctgcggg 30360
actgcggcca gcaggcggac ctgctctggg gatactcgga cggcggacag tttaccagtg 30420

```
        agttccctgc ccatatgatg agccattcag cgcggctgaa ggtgaacccg gcaaaccggg     30480
        tgatcatgca ctgccatccg gcaaacctgc tggccatgac ctatgtgcac acactggatg     30540
        aacgggagtt tacccgaacc ctctggcaga tgtgcactga atgcgtggtt gttttcccgg     30600
        acggtgtaaa cgtgcttccc tggatgctgt gcggcaccaa tgagatcggt gaagccacag     30660
        cagagaaaat gaagaccgcg cgcctggtag tctggagcca gcacggtatc tacggagcag     30720
        gcaaagacct ggacgaaacc ttcggcctga tcgaaacggc ggaaaaggct gccgagatct     30780
        atatgaagat cgcgcatctg ccccgggtca acacgatcac ggatgaacag atgcaccagt     30840
        tggaaaagcg gttcggcatc aaagcccggg agggatatct ggaggaaccg gatcggggga     30900
        cggacaaatg aaacagcatg caggaaaaaa catccgggag ctgaaagttg cctatatcgg     30960
        cggcggttcc cgcggctggg cctggggctt tatgaaggat cttgccctgg acggcgaaat     31020
        ctgcggtacc gtacgcctgt atgatattga ccgggaagcc gcagaccgga acaggatcat     31080
        cggcgaaaag atcagcgccc atccggatgc tgtttcccgc tgggcttata cagtctctga     31140
        ttccctgcag gaagccctga ccggagcgga ctttgtggtc atctccatcc tgcccggcac     31200
        ctttgatgaa atggaatcag acgtccatct gccggagcgg ctgggggtct ggcagccggt     31260
        gggtgatacg gtcggcgccg gcggattcat gcgttccatg cgcaccatcc ccatgtttgt     31320
        caccatcggc gaggccatcc gggattacgc gccggaagcc tgggtgatca actataccaa     31380
        tcccatgtcc ctttgtgtgc gcaccctcta tgaggttttt ccatcgatcc gggccttcgg     31440
        ctgctgccat gaggtcttcg gcacccagaa gctgctctgc tccatgctga aaaccgggga     31500
        agggatcgac ggggtaaaac ggcaggacct gtatacaaca gtcaccggca ttaaccactt     31560
        cacctggctg accagcgcct cctggaaggg gacggacctc tatcccctct atgcccggtt     31620
        tgcggaaaag tacgcagaaa gcggctacaa cgaaggcggg gatgacaact ggatgaattc     31680
        tcacttcagc tgtgcccagc gggtaaagtt tgatctgttc cggcgttacg ggtcattgc      31740
        cgctgccggg gaccggcacc tggcagagtt cgtggcgccc tggtatacac gagatcccga     31800
        aacagtccgg gaatggaaat tcagcctgac cccgtttcc  tggcggaagc aggacctgaa     31860
        ggaaaaactg cagatgtcgg atgacctggt cagcggaaag aaggaaatca ccctgaacgg     31920
        ttccggggag gaagggcacc tgctgctgaa agccctgctc ggtctcggcg atatggtatc     31980
        caatgtcaac attcccaacc ggggccagat tcccaatctg cccctgggcg ctgtggtgga     32040
        aaccaacgcc ctcttcgggc tgaaccgtat tgatcccgta tacgcggggc cggttcccgc     32100
        ttccatcctg ccgctcatca cccggcatgt gctgaatcag gaaaacaccc tgaccgctgc     32160
        cctccgctgt gaccggaagc tgggcctgac caccttcctc aatgacccgc agctggccgc     32220
        agttaccccg aaggacgggg aaaagctctt taacgacatg ctggaaaacc agcggatctt     32280
        cctgccggaa aaatggtttg aataactccg gacatccgaa agaatcatat aatcaggaaa     32340
        cccgcttctg atttttatca gaagcgggtt ttttccgcca ttcttcattt ttcattcttc     32400
        atttttcatt tccgctttat aatcctgcac agttcatatt tcccgggcat ggcccgtcct     32460
        tcaaacaccg tctctccgtt ccgccggtac agcggcggga tcctgatggt ttccccggtc     32520
        accggatgct tccgaagcgg cagctcaagc gcttccgcat cgccggtcac atggatcagg     32580
//
ID   XXX; XXX; linear; XXX; XXX; XXX;  XXX.
XX
AC   PRJEB67889
XX
DE   Uncultured bacterium DNA fragment from in vivo enriched cow rumen, clone
rum-7d9
XX
FH   Key             Location/Qualifiers
FH
FT   source          1..38453
FT                   /organism="uncultured bacterium"
FT                   /environmental_sample
FT                   /mol_type="genomic DNA"
FT                   /isolation_source="Cow rumen"
FT                   /clone="rum-7d9"
FT                   /note="AZO-galacto-mannan active clone"
FT   CDS             64..216
```

```
FT                   /translation="MKYECPCGYIYDPEVGDPEGGIAAGTAWEDVPEDWVCPVCGLGKD
FT                   AFTPV"
FT                   /product="Rubredoxin"
FT   CDS             232..831
FT                   /db_xref="GO:0004413"
FT                   /translation="MNIVCLDLEGVLVPEIWIAFSQASGIPELKRTTRDEPDYDKLMKW
FT                   RLGILREHGLGLKDIQDTIATIDPMPGAKEFLDTLRSETQAIILSDTFTQFASPLMKKL
FT                   GWPTIFCNELEVGEDGMITGYRIRVEKSKLTTVRALQSIGFETIATGDSYNDLGMIQAS
FT                   KAGILFRSTEQIKKDYPELPACETYDELLAQIRKAL"
FT                   /product="Homoserine kinase (EC 2.7.1.39)"
FT                   /EC_number="2.7.1.39"
FT   CDS             complement(912..2279)
FT                   /translation="MKKLTAVLLALIMVLSMAVTALAEDPRNITIALWWDIYYDSNDES
FT                   WEDNEAATGKETDLMRFDNVKNIEDTYGVTYEFQNLTYAGVQDSVNNSILAGEPDCDVY
FT                   MVELGWGVPAVMNGYAMDLRDVLDPDDPLLTHNDTVMNYVNLESGAVSLLTVNGAEDQV
FT                   AATYPLAFNLQMIEEANLEDPRELAARGEWTWDKFREYCIALTKDNDGDGVTDVYGYGA
FT                   WITDCLPYWYMSNGTTLASTPKENLSSVEMGETLKFLQDLWLTDKAAYPLPEENGWDVC
FT                   RWLYRDKKVAFTTTAAWIMANYDDYNWDGKAETTLDFDMVFVDYPIGPHGNAETNATKI
FT                   AAGSFYFIPVGIEDPKLVYDVFRAYQNWYHDDISLRDDPEELEWWYTTTSDKLDLQEWN
FT                   FEIMKKMGEKTVVDFAGTVLGEMPLVEFINGDLTPAQLQEEYKQVVQDTLDQIFGA"
FT                   /product="ABC transporter substrate-binding protein -
FT                   sugar transport"
FT   CDS             complement(2312..4021)
FT                   /translation="MKKIISWLLVAVLAIGMCSWASADPVNVLDFEDGVFAFLGVSAVK
FT                   PNADAAASLEVVDYNGSKALRVAAQGIPYVALNLEGLAGEKLADVAGVSFDIGVDKAAD
FT                   GKFYAVSGVVYSYTGENADENKADWSVYMEKKNPRNVKIAFKAPFVAGAGNYVMISRED
FT                   QAGGEPATFYLDNIRFLDAEGNAIALDPAAVYVAEASERDLSNLVALTNAVEFPDFHKS
FT                   AGAWAQDGLEMPQEIIDALVPGSVVEVEYASADGSMWIVMPWATAGWMRVGQGTAAINN
FT                   SKNIAQIPYEMIEALCGEDKSTWGAMLQCESASDWEVFSVRVGQRANRIVLKNAVEFPG
FT                   FTKSADAWAQDGLEMPQEIIDALVPGSVVEITYSSEDGDIWLVMPWAEAGWMRVSQGTA
FT                   TRIGGKAYITYEEIAALCGEDKATWGAMMQCEGSSPWEVYSVRVGQKAEFFGLTNLVEF
FT                   PGFTKSADAWAQDGLEMPQEIIDALVPGSVVTISYESEDGNMWLVMPWATAGWMRVGND
FT                   GADVADGKIAQVTYEQIEALCGEDKSTWGAMMQCESSSAWNVYAVAVGQAIK"
FT                   /product="hypothetical protein"
FT   CDS             complement(4081..5085)
FT                   /translation="MHTGNAAQSGKITWQGFRNSPAYANRRDKTLTGMKNLVLGIFRLI
FT                   VIVGISYVILAPVINIIANSFFSRQDSINPMVFTIPISPTLERYTMAIKYLDYLPILGN
FT                   TILYVFGITLIQLLICSMVGYGFARYRFPLKGLLFGFVLVMIVIPLNTIQFPLYTTFRF
FT                   FNPLGLVGLFNGGEGINLLQTAWPTVVMSVFGCGLRSGLYIYIFNQFFRGLPKEIEEAA
FT                   LVDGAGTARTYFTIMMPNAAPAVITVAVFSIVWQYNDSFFANTFNISDSILVTRKLESL
FT                   INVIANAEKVLTIREQQLYFDAGVVLVLVPVVVIYLVLQRRFIEGVERSGIVG"
FT                   /product="sugar ABC transporter permease"
FT   CDS             complement(5104..6030)
FT                   /translation="MTGIRRKNRIFHLSYEQKNAVRGLWFILPWLIGFLVFYAGCLVQL
FT                   GEFSLSRIGLDAAAGMTKEYIGFGNYIEAFTSHATFKQTLASSVIDMLIDLPMIIFFSL
FT                   FVAMLLNRRFKGRALVRAIFFLPIILSADAVGSAITRATELVNAGVSSASLETAQAGTV
FT                   SVAYYMDLFGDLAIPQSVLGYIVDAVNRISSIIKGSGVQIILFIAALQSIPGSLYEVAR
FT                   IEGATGYESFWKVTFPMVMPHIITNTIYTVVDRFSSGEVIKLATRTYKELYNYGLSSAF
FT                   SVVSTLITILILGLIVYLLNRRTFYYN"
FT                   /product="Sugar ABC transporter permease"
FT   CDS             complement(6027..8618)
FT                   /translation="MSEKNRLPLPTAVRRALWILAALVVIGAGVFLGWYLIHYRGYDEY
FT                   RLYVQPPAEKTEAAALKTRKDPENSVPGFALAAQNDNQALYLNEKTAEVALRDRKSGRV
FT                   VYSNPQDAAEDPVARSGLNQGNLRSQFILNYLDTNSREGTPWSSYAKCVENGQIEYLQL
```

```
FT                   ENGFRAVYTLSNEKLMLVPRQMTAEWFEVLSNAGRKQAAKSYVLDEESGLYVLKSQGVT
FT                   ARNRQQIDADARTAGFTMEDYAEMEALAEEEAAEAAESTSFVIALDYTLTPEGLTVTIP
FT                   HAEIMEAGGGKIRSIQLLPFFGAAGTAETGGIVVPDGSGALINFNNGKNTAPQYNQAIY
FT                   DLDLIDSDFTATQNMQSARLALFGICREDYSLLASCGRGATLASVTADVAGRNNSYNYA
FT                   YFTFSLRRTDTLIVAGEEAIVAEQDAYPVDCAVTYRILDGDYTGYNGLAKAVRESLLSS
FT                   GTLKLKAETSGDIPFYCDILGGVRETAHWMGVQYLRVMPMTTFAQAEEMMSSLRGEQVG
FT                   NLRVNLQGWMNGGYYHDPVSHVSVLNELGGEKGLKRLCLAAAENGDRIYPDAALQLVTE
FT                   IAKGFMRSEEASRYYAKGYAAELGVVNPVSMRRMATLGFAERGYMLLSPRFLPRYAALL
FT                   AASADRLGLDALSLRDLGNEVHADKRRTNVISREADLDLVKQAFGVIGAGERELMVSGG
FT                   NDYSFPYVRHVINAPVEATMFAIVDEQIPLWELILHGSADYCGSPLNLMQSENRRATLL
FT                   HLIEYGASTHYTFTWKDAADMKYTGLNNNYATTFSSWKEEAAESYRFVNGALSRVSGAE
FT                   MLRHDRLSDTLARVTYSNGTVLYVNSGEREAAADGYRIPAMDYLAVGGDKE"
FT                   /product="GH129"
FT   CDS             complement(8633..9265)
FT                   /translation="MNRGTMKAKWQHFFHTVSHPADGYYWIRHQEKGSLWIAFLMVILY
FT                   GVAFSMNRISSSFIVNDIEPRTVNLPAELAGVILLYFILCVGNWSVTCLMEGEGRFRDI
FT                   LIAVGYALFPMLVTTAAATAVSHLVAENEEAFYTLLLGLGTAYTVVMLLIGIMQVHNYT
FT                   FGKTLVTLFLTVISMLIIIFLGLLVINFITQVYSFFRGIYMELVFRV"
FT                   /product="hypothetical protein"
FT   CDS             complement(9262..10743)
FT                   /translation="MKQLKSILCLVLLACLAFGNALAADSLPYDCYNYDHWNNILYTPA
FT                   PYVPDGLVSGATLRFEGAPIGAFRNPQDLCVSPDGDVYIADTGNNRIVVLADDLKTVRR
FT                   IIIGFEAEGAAQAFNAPSGVAISEKHQLYIADSLNRRIVVLDPDGTFVKYVQNPRSEVL
FT                   DEGYVFTPLRVSVDYADRVYVIAQNMFQGIMVFSEAGDFTGFFGTISVKISLWEKFWRK
FT                   LATKEERGKQQLFIPTEFTGIDVDDEGFVYASSKDTDGLQAVRRLNPKGEDVIRKGPHG
FT                   NLGGDRVFGSVGNYAGPSQIVDVVYRGHGTYSLLDSRRGRIFTYDHEGNLLYIFGGIGS
FT                   QAGTFLTPTAIEQLGSRLLVLDSQQGNLTVFAQTEYGSLINRAVALRFDGDETQAVPLW
FT                   QEGLRLDEINELANTGIGKAYLSAEDNANAMTYLRRGMNREYYSVAFKRWRNERLRNNI
FT                   SFILSGVVAAAVLAVLWVKAVRPAIRKKAKRRNLS"
FT                   /product="hypothetical protein"
FT   CDS             complement(10755..11735)
FT                   /translation="MEKAKNSRLLPAWLHRNPVELTAEQEEILRQKRLAKIHRKMKRKS
FT                   GKKKLNRSTAGNAALFILMGICGVFMVLPLVMILNNAVKPLDELYQYPPRIFVRNPTLN
FT                   NFTDLYVLLNESWVPFSRYVLNTIIITLGGMIGHVIIASMAAYPLAKNHFPGKKILFSV
FT                   VVLSMMFSWTVTQIPQYLIISWLGINNTYAALVLPAWSFGMGLYLMKQFMEQLPDSLME
FT                   SARLQGANEWQIFWRIVMPNVKPAWLTLAIFQFQQMWANTGSMFLRSEELKPLQYALQQ
FT                   ITSGGVSRAGAGAAVTFLIAAVPIIFFLICQSSIMETMTTSGMKD"
FT                   /product="binding-protein-dependent transport systems
FT                   inner membrane component"
FT   CDS             complement(11741..12673)
FT                   /translation="MSMAIRKERIVPRKDTLWHQVRINKASYLLMAPYFILFFLFTVLP
FT                   VLMSVYLSFNYFNMLEPPRWVGWANYIKLLLEDDIFIISLRNTLLFAVITGPVSYLLCL
FT                   LFAWIINEFKPRVRAFLTLVFYAPSICGNAYMIWNLILTGDRYGYLNGVLLRMGFLNEP
FT                   ILWMQTEKFVMPVLIVVQLWLSLGTGFLSFIAGLQTVDKGLYEAAAIDGVKNRWQELWF
FT                   VTLPAMKPQLMFGAVMQITQSFAVADISIALAGNPSVNYSGATIVPHLLDYGSTRLDMG
FT                   YASAIATILFLLMVGTNKLVQRILRRVGE"
FT                   /product="binding-protein-dependent transport systems
FT                   inner membrane component"
FT   CDS             complement(12675..15644)
FT                   /db_xref="GO:0005215"
FT                   /db_xref="GO:0006810"
FT                   /translation="MVLALLLALAAGAAQAATGFGISEYEEQVSTYSINPAVPSYADYC
FT                   AEHEMTAGPEKEIVLEASDAVRYEEEDGKNAPVFMEQAEGREGVSVLTGEEAVIEWAFE
FT                   AEESGWYDLEVEYYPWPGRNAEIQRAFFVDGALPYRELALVNFSRVWRNDLHPDVSRFT
```

```
FT                   AEDGAEEIRWLRDNQGNELKPSPAEAPEWMTSLLHDSNGYISESLKVFLPRGPHTLSVL
FT                   SLREPMLIRRLCFRAGADAQPYRKPEGESGAAGQIIRIEAEAASRTSSQMLYPVQDQSS
FT                   CSVSPSSPRYLLNNSIGGNAWKSAGQWIEWRFTVAEEGNYAISLYDKQNFIRGTDVYRK
FT                   IYIDGEVPFVEFQAATFPYTQDWRLETLSGEQDEPYLIHLSAGEHTLRMEVVLGGMAAV
FT                   IRQVQDCVLQLNGIYRQVLYITGVSPDPYRDYQIERSLPGLEAQLRKVKADLQTAIGAL
FT                   EETAGHRSDKLTVLKTMDDQLTELIRDQERFTEVLSSYKTNVRACGNWITQVLGQPLQI
FT                   DRIFVHTADVRPEISGRDFLSGTGFEASRLYSSFTINYNQIGNVAEKSDDSRVITLWVG
FT                   TGRDQANVIKALIDEHFTPQTGINVNVQLVDMNTLLRATLSGQGPDVAIQVANTTGIAG
FT                   AVMNTGNDTPVNYGLRNAVMDLTQFEDFEEVAKRFAPSALVPFSFNGASYALPDTQTFP
FT                   MMFYRKDILAEIGLAVPETWDDVKVAMSVLSKNQMEFGMLPGEQTFAMLLFQAGGEYYT
FT                   ENGDASALDSDIAVNVFRRYCDFYTDYKLDMATSAEERFRTGECPIIITDYTTYNNLQV
FT                   SAPDIKGLWDFTHVPGTLREDGTLDTTTGCSGLADIIMSATKEPEACWSFLKWWTSAEI
FT                   QTLYGREMESLMGASARVATANLEALAGLSWPIRDYRELARQFSHVKGIPQVPGGYYTW
FT                   RNVNNAFYAVTTGSASSGRSENGNTPREELMDKIFYINAEITYKRTEFGLPVANAGEGK
FT                   "
FT                   /product="N-Acetyl-D-glucosamine ABC transport system,
FT                   sugar-binding protein"
FT   CDS             complement(15699..17072)
FT                   /translation="MIVEAETGKPEGHTVIAGDGENQWVEGFHTAGEDGIITEVTVSRE
FT                   GFYDIAVIQASQGGHKENPVLLDGRNIGSAVTDDTAFGKSVLEHIYLSAGTHTLGIGTS
FT                   WGYIRVDRFELTPSAALPADLYTIPEKMSVGEATPEARRLYDWLRENYGKKILSGQQCD
FT                   GGMFGMENQAIWRATGGDYPAVLGLDMMEYSHSRVEHGGESDRAVPYAIEYWNKGGIVT
FT                   FCWHWNAPSPYLKEPWYSGFYTKYTSFNLARVMNGEDEEGYRLLNQDIDAIAAQLLRLK
FT                   AAGVPVLWRPLHEASGGWVWGGACGKEAYLKLYHLLFDRLTNEYGLNNLIWIWNAQDPD
FT                   WYPGDDEVDLIGTDIYPGERVYDSQSASFLPLTRIAGERKMIVLSENGCIPDPEQVFRD
FT                   GTVWGYWCTWGGEFVLRNTKFNKPSEQYTETAMIRKAYSDERVVTRKDLPDFRTRDEK"
FT                   /product="CBM35_GH26"
FT                   /EC_number="3.2.1.78"
FT   CDS             complement(17425..17544)
FT                   /translation="MKARERKSEPEKSEYQQLYAGGREKPPAFLRAEKHMAFL"
FT                   /product="hypothetical protein"
FT   CDS             complement(17519..18970)
FT                   /translation="MLPETSVFFRTEDEKLQRVYDAAEEKCRFNLKDFGGDTVLVEGGG
FT                   YEKIWLETQPMGGEMYALRSLEAARNNITLFMRHQRADGRLPGSIQCTGGKVEPQFNKY
FT                   QGFCFPFPALNLYYLIGKDRDYLAELKESLIRFDRCLWNTRHVSGDGLLSSFCVYDTGE
FT                   DLALRYGDAPCWWEGDEPPEGYQVVPMASMDVTSWSFAARRTVAEICRLQGDRSADAWE
FT                   RKARMVADALRRRLWDRKRGALYDRDRTGKTVDILCHNTLRCMYWGSISRGMAARFVKE
FT                   HLLNPAEFRTPFPLPSVAVNDPAFRNAPENNWSGQPEGLTYQRAILALERYGYNRLVTG
FT                   LGRKLIDAVYANGCRFTQQYDPFTGKASLVRMSDHQPAEEGSGEAVQDAYGPTMLACLE
FT                   YIAHHYGIHPHLGQVWFSLGSGAPCAYEAAFCDRHYAIRSDGQRAEIRVDGKALGSWKC
FT                   GRRVITDENGRFLRTELIEGAGAEI"
FT                   /product="lipoprotein, putative"
FT   CDS             19300..20166
FT                   /translation="MLLLPCLAFFIVFCYFPMYGVQIAFRDYKAAFGITGSQWVGLKYF
FT                   EKFFNSYYCGRMFANTFLLNLYGLLFGFPIPILLAIMLNQLNSKRFKGFAQTAIYVPHF
FT                   ISTVVLAGMIYLFFSPTNGIINHLITAAGGKSIYFIMEPQWFRPLFIGSDIWQNAGWNT
FT                   ILYIATLTSIDPQLYEAATIDGATRWEKIRHIDIPHLIPIAVMMLILNCGSLLSSNTDK
FT                   ALLLQTSGNMATSDIFGVYVYSVGIAAQKAQYSYASAIGLCLNLINFAIIMGVNSISRR
FT                   MSDISLF"
FT                   /product="ABC transporter, permease protein"
FT   CDS             20183..21121
FT                   /translation="MTVTAVKAKKQNRIRETRKDRIFNAINTLFWVLVLFLVLYPLWLI
FT                   LIASVSDPDAVLQAKVLIWPVDFSLMGYDAVLQNSELLNSYVNSVFYTVVGSALSVIVS
FT                   LAAAYALSRGFAGKKFVNLAITFTMFFSGGLIPIFLNIRDLGLYNTRTVMILMNLVSVW
```

```
FT                   NLMVARTYIQTSIPNDLYEAAVIDGADHFTYFFKCVLPLSGTIIAVLSVYYGVARWNDY
FT                   FTGLVMLRDRSLYPLQLVLREILASLTSTGSSDTFFAAYADNLGGLQAALRKAEVAKYC
FT                   CIVVATVPAVVLYVFMQKYFVKGVMIGSLKG"
FT                   /product="ABC transporter, permease protein"
FT   CDS             21135..22040
FT                   /translation="MSQIRSTNDSARAARYEDGFRVLQGRREYVVTYQADSSIRIWYSDI
FT                   PWRYEAHNHSAVEILLTLEGTVTYTVEDKLYQVRKGEILIVPPDTLHSLSMGEGSSRYL
FT                   FLFEPDAIMSMRDIKYMASYFNKPFHLRDGSDAHVRIRELLLRANDAYEKREMLWNTLC
FT                   YSCILRIYAILGQHYLGSVRPHTGDSTGNMDAEVVNAVMTYVNNHYREDLTLEEVAQFA
FT                   GFSRYYFSRSFKKQTGYSFKEYLCQKRVQVAMDLLIRTNRSMRDVALESGFGSVATFNR
FT                   VFREKKGCTPSQYRTIYGIP"
FT                   /product="hypothetical protein"
FT   CDS             22092..23597
FT                   /translation="MKKLVSLLLACCMLLSLSAAFAEEKVELTAFQYALENQNTDFNGL
FT                   WFFDELEKKTNTHVTFQMVKDADWATNVNLMFAVPDQMPDMIIRHAVDVEEYGVSQGLL
FT                   LPLDEYLTEEIMPNYVSRLNINNAGDSIPASDGKTYYIGFLMAQNVNHTGTWYVNQKEL
FT                   EKLGKTAPTTIAETTELLRAMKADGIKWPFSASYTQFGPETIWNQFASFGVPENTYYYF
FT                   IDADNKVQFTCAQDGWRACVEWLHQLYEEELMDPESLTQDSNLWANKVNDGEVGYFCYL
FT                   RLINTAIKAENVENFKSILPPAADGFKACVSQILEVPEAGAYLTKNCKDPVAALKWIDA
FT                   QLETETMMVSLNGKVGEQIVLNDEGKYEVIDIPADNGLYQFVPVTMGQFFAPGDYYTSI
FT                   YQMAPHRVERYEDSKWYAESGVLEPKSFQYLRDLSKLSNEDATEAADIYTELQKLVEES
FT                   FANFVRNGVTDESWNNFQEQAKAIGSERYIELYQNAYDAYLAK"
FT                   /product="ABC transporter, substrate-binding protein"
FT   CDS             23714..25261
FT                   /translation="MKRTVKIICLLAAALLLMTGCSAPPRDEGSPAPAISLSALQYEIE
FT                   NIAVDFSSMWYFRQIEQQTGVHVDFNEVKDSEWSSSVGLAFAQGKMPDMILRGSLDVEE
FT                   YGVKRHLLVPLDEYIEKGFLPNYAARMNQAGLREQLTASDGHIYQLGFLISQDVNTNGH
FT                   FFINKTWLDRLKLPVPETVEQLTEVLRHFREDDPNGNGIQDEIPLEFTLDDNTTGIYNL
FT                   FSFFGLPLNEDYVYADKAGRVHFAPEEDAFFDTLSWLHQLYQEKLLDIDFISQGGSIWA
FT                   AKVNEGSTGMFSYWRLQNTALNPGVAGNYEIMLPVHADGKAACLPRNMDIIEFGAALTK
FT                   DNRNIEASLRWLDAQFETENMLVSQNGPVGDTLIRREDGRYEVSYVPPDNELYQAVPVI
FT                   CGQFFAPADYYASVYVPAAHRLEKAAYCDLYERSGVLEPISAKLLTTVAPKTTDQSAET
FT                   TRLKAKLKSLVDAEIVSMVTQGVTEEGRAAFRESLKAGGSADYRQVYQQIYDRWKEK"
FT                   /product="hypothetical protein"
FT   CDS             25266..27512
FT                   /translation="MIRKSRTSVFIRYALSYIAVVLLLFLSITGYLYIRLSSKTREEII
FT                   DNQINRLSRIAAQHESYISAMLNTAEEIGLSPDIEFFRYEEEPWKAYDLQLKMVPYTTT
FT                   SSTFCDQVYLYFFGDDRVYSSSSSMSLSLFARMMRYEFIPMESLVSRIANTDRLTILPA
FT                   QQVSSTLVDSSRVVTFLVPLGSAPGSGKGVLLFLIKDSIYQSLFADAINDHINTYIHQG
FT                   DELLSSSEELSLPRDQVKVAQETASSLFRWDNQDWTQVTFSGRNWGLSYTAVLRNADIN
FT                   SSIMQQLFSILAILPVFVFLSLLLALWMARRHAQPIRAITGMLSRETDETPRDELQQIS
FT                   TGISRLTSRNQELTSRLDRALPVQRHDFVFKFVKDRFPTREEAVVAGKAVGLEIDRPAY
FT                   AVILCSGSGGNDHPFDLGAPPFDRFPGCSGAGVELVALKANLYLVFSDDPETLPDLAET
FT                   LRSEGAAGGERCITSISALHSDFTEAPSAYLEAAAAYENRFVMGSQKVLCYGDISSNLE
FT                   EILPQARKITNSISQALTLGSRDVLDVRISDLLHFLKNTNMSPFAFRMIYSEVIRSLVH
FT                   THTAALSGEGSVQEIYNIFTLTSCQSIDDLDELLRRLCDRLLLEEPDSCEIAPEEEDEI
FT                   SQVVHYMEEHFSDPEMSITALADSLELSTTRFSLSFKEKTGMSPLEYLTLLRVEHAKEL
FT                   LEMTDLTIRDISARAGYYDSGSFIRRFKQVTGETPAQYRRSRNLKKETPEGGKN"
FT                   /product="Transcriptional regulator, AraC family"
FT   CDS             27537..28379
FT                   /translation="MIHTEEINVRDPFVLVHEGQYYLYGTRGATCWGPADGFDVYTGRD
FT                   LHTWEGPAVCFHNDGTFWADRNYWAPEVHPWQGMFYMFASFKSDARRRGTAILRSASPL
FT                   GPFEPWSDGPVTPADWECLDGTFHVSSDGVPYIVFCHEWVQAGDGEILAMRLTGDLKAA
FT                   AENPFLLFRGSDAPWARPVHHSSGITGYVTDGPFLWRRKDGTLLCLWSGFSEKGYAQGL
```

```
FT                   AVSDNGEIDGHFSQIDPLFLEDGGHGMLFRDLEGGVYLTLHSPNTHLLERPRFIPVSL"
FT                   /product="GH43_17"
FT   CDS             28476..29870
FT                   /db_xref="GO:0004557"
FT                   /translation="MNQTTKNVRELTVAYIGGGSRGWAWGFMKDLALDGEICGTVRLYD
FT                   IDREAAERNRIIGEKISAHPDTASRWAYKVSNSLQEALTGADFVVISILPGTFDEMESD
FT                   VHLPERLGIWQPVGDTVGAGGFMRAMRTIPMFVTIGEAIRDYAPEAWVINYTNPMSLCV
FT                   RTLYEVFPEIRAFGCCHEVFGTQKLLCSMLKTEEGIDGVKRQDLYTTVTGINHFTWLTS
FT                   ATWQGRDLFPLYARFVEKYAESGYDEGGDDNWMNSHFNCAQRVKFDLFRRYGVIAAAGD
FT                   RHLAEFVAPWYTRNPETVRKWKFSLTPVSWRKQDLKERLQMSEDLVSGKKEITLNGSGE
FT                   EGHLLLKALLGLGDMVSNVNIPNRGQIPNLPLGAVVETNALFGLNRIDPVYAGPVPASI
FT                   LPLITRHVLNQENTLTAALRCDRKLGLTTFLNDPQLSGVTPADGEKLFNEMLENQRAYL
FT                   PVKWFE"
FT                   /product="GH4"
FT   CDS             30093..30305
FT                   /translation="MAEQYVTGETLVGEVVTKYPEAIEILLSIGMHCLGCPASRNESLQ
FT                   DACAVHGIPAEQVIEAINEKIAENK"
FT                   /product="hypothetical protein"
FT   CDS             complement(30364..32088)
FT                   /translation="MKQTKLSILFPRQDEVTYREIPEESWHDLGMDALTEKVAAQPQEE
FT                   PLLRRVMMSLTADPAVAAFRSDVFDDILRHPEIRERLMKLLDKVKMFYDYGVVNRHEGD
FT                   ETGIWDLMHRLEEYHDYILTVEAIRECLSDKDLVSEGLTTLRDTVEQIYQEKGFAALRK
FT                   DVEEMRVAASEIRSLTVGINLNDRFEAISMGLVSVNAKPFVKSSILKNFLASMSPRDEI
FT                   RKEADWSGSYNYYPANTEVGLLQSVGQFVESSVILRNPLAAMSLSRIPAADGTANVPRQ
FT                   MDGAASMLTSRIARKLRDTLGQYLNVSVKEIADLIPELVFYTRWAEYIEKKQKAGWKFC
FT                   KPQARLNRQGAACMKAAGLYNLKLIATEKPENVVPNDLVFDPEKRVYVLTGANRGGKTT
FT                   VTQAVGQLFILAQSGIPVPAESFEYDPADRVLTHFPADEDKTLDLGRLGEECRRFRELF
FT                   VSCTPESLLLLNETFSTTSFEEGYFIAADAVRAILGRGTRTIYNTHMHKLAQDLDTVIN
FT                   TGEAEGKAVSLVAETREGKHSFRVVIAPPEGQSFARDIAEKYGVTYESLTASVPSEN"
FT                   /product="DNA mismatch repair protein [MutS]"
FT   CDS             32225..32698
FT                   /translation="MGKQDKTNCMRVLDSKKIAYSAHLYEADPTLTGEDIAHLLNEEPS
FT                   QVFKTLVTVGKPQKYYVFVIPVNAELNLKKAAAAAGEKSVSMIPQRDLLPLTGYVHGGC
FT                   SPIGMKKHFPSFIDESASGLSRIFVSAGRVGCQVELAPADLIQIASLTPAALI"
FT                   /product="Cys-tRNA(Pro) deacylase YbaK"
FT   CDS             32790..33797
FT                   /translation="MTKKRTLLWLIPLIVLILLIPFLVPSALPYAGAEEMPVFPSVELS
FT                   NPEPEPVVLEPNPKYKNSMVSRYAPHKDAFTSEPLGYQDGTICVKIEKRTVVNGNGGKT
FT                   AVYFTWVKIADPSQLRTSTNQPYPSKQLVAATSLARRERSVLAINGDYFCDRTEGIVYR
FT                   NGELLRNSVFNTGYGYDALIIDLNGNFHILLKPDPSDFEPYEGCIAHSFIFGPALVVDG
FT                   KMQVLDGNNYASSPGMGLHKYLQRQAICQMDNLTYLMITAEGPEQSKDGGLTGQEMAQL
FT                   AYDCGAVQAYNLDGGSSSWLVLGTDRINATRGKKRDIQDIIYFVTAEAEPVPAE"
FT                   /product="Exopolysaccharide biosynthesis protein related
FT                   to N-acetylglucosamine-1-phosphodiester alpha-N-
FT                   acetylglucosaminidase"
FT   CDS             33846..34937
FT                   /translation="MNTILSIGPLNLTPYGLAVLAGVLAGALLCLLNKKVFQLLPLTIL
FT                   GAVVLGHAFWTFLRLDGYMDEISSFAFMFQPWLGGYTLYGALIGGTLGALAGAKLSGIR
FT                   PLEALDTLAPAACAVLFFCRLGEYYTGEGVGAEALESLSFFPIASEGLNSGFWLYAVWF
FT                   WEAIAALVILILLLCRRKNAREGELTVLFVTLLGTSQILLEQFRKDNFVRLNSFVRFTQ
FT                   IAAVLTLLALMIVLVIRRRPDRKKTILSFAVLVAAAGAVVMAEFVFDKPQMETLLYIAL
FT                   AATAVLFGLMLIAFRGKAGKLPAGLFGLFTGTLILLYLLATVDPDSSGFLYTILLDLMM
FT                   LSSLTAIGVTLMTLYHSESALEK"
FT                   /product="Prolipoprotein diacylglyceryl transferase (EC
```

```
FT                   2.4.99.-)"
FT   CDS             34940..36094
FT                   /db_xref="GO:0005506"
FT                   /db_xref="GO:0006118"
FT                   /db_xref="GO:0009055"
FT                   /translation="MTLTKEETLMSQKSSPVYFTDFRTVADVSQGVKLQKLCRRAGIDK
FT                   IDFAGKFAAIKMHFGELGNFAYLRPNYVKAVADLIKELGGKPFLTDCNTLYPGSRKNAV
FT                   DHLYNAEVNGFNSVTTGCYIIIGDGLKGTDDITVPVPGGEYCKEAYIGRALYDADIIIS
FT                   LNHFKGHEMAGFGGAVKNLGMGGGSRAGKMQQHSDGKPVVDQALCRSCHKCARECGSDA
FT                   ITYKSGKAYIMQDICKGCGRCIGACAFDAIHPSFDTAADMLGRKMAEYTAAICAGKPSF
FT                   HITLIMDVSPNCDCHGENDAPILPNIGMLASFDPVALDQACADLCLAAEPLPNSQLSDN
FT                   LAKPGWKHHHDHFKDSNPNVSWRETLAHGEKIGLGTRTYELIKI"
FT                   /product="Ferredoxin"
FT   CDS             36326..36781
FT                   /translation="MSTFLPAVFGENMMDLFDDFDRDFFRGFGRPERALYGKNAARMMK
FT                   TDVKETDEGYELAVDLPGMKKDEIHLDLQNGYLTISTQKNLENKEEKNGKLLRQERYTG
FT                   TMQRSFYVGDNLTEEDVQAKYEDGVLTVKLPKKEAKKDPEKKQILIA"
FT                   /product="hypothetical protein"
FT   CDS             complement(36955..37881)
FT                   /db_xref="GO:0004791"
FT                   /translation="MGEKMTDIIVIGGGPAGMTAALYAQRNGKNALVLEKHGFGGQITY
FT                   SPKVENWPGTAQMSGNEYAEALLDQIMNQGAEVDLAEVVRIEDHGDYKRVFTDDGLERD
FT                   AKAVIVAAGVKHRMLGLPGENELVGEGISFCAVCDGDFYTGKTVCVAGGGNSALQEAIL
FT                   LSGKCEKVIMLQDLPEFTGEKKLQEILFSRPNVEKYTGVKITGLNTEEGILRSVTVKPQ
FT                   DGGEERQVACDGLFVAIGLIPENDAFAHLADLNKWGYFDSDERCETKTPGVFVAGDCRS
FT                   KTVRQLTTAAADGATAALAACRYIDSL"
FT                   /product="Thioredoxin reductase (EC 1.8.1.9)"
FT                   /EC_number="1.8.1.9"
FT   CDS             complement(38002..38418)
FT                   /db_xref="GO:0008998"
FT                   /translation="MQNWNDGKAQEFKDRKVYDIGHSRLTHTGPVMATAPAAEPVKEEP
FT                   AAQPEKENKGCNCARAILFVSATCPNCKLAISMLEKAGFLFKKVLATDNPELTNQYGVK
FT                   QAPTLVVLGQNPDEYTKYKGVSEIKGMLTAKQVG"
FT                   /product="Ribonucleotide reductase of class III
FT                   (anaerobic), large subunit (EC 1.17.4.2)"
FT                   /EC_number="1.17.4.2"
XX
SQ   Sequence 38453 BP; 7868 A; 11877 C; 10630 G; 8078 T; 0 other;
     gagccagaaa agaaacggcg tttcgccgct tccatctgag ataaaaagaa aggatgaaaa        60
     tcgatgaagt acgaatgtcc ctgcggctac atctatgatc ctgaagttgg cgatcccgaa       120
     ggcggcattg ctgccggcac tgcctgggaa gacgtcccgg aggattgggt ctgccccgtc       180
     tgcggtctcg gaaaggatgc cttcactccc gtctgaggct gattattgtc tatgaatatt       240
     gtatgtcttg accttgaagg ggtgctggtg cctgagatct ggattgcctt ctcccaggcc       300
     agcggcatcc cggaactgaa acgcaccacc cgggacgagc cggattatga caagctgatg       360
     aagtggcggc tgggcatcct gcgggagcac ggcctgggcc tgaaggatat tcaggacacc       420
     attgccacca tcgatcccat gcccggcgcg aaagaattcc tggataccct ccggagtgaa       480
     acccaggcga tcatcctgtc cgataccttt acccagttcg cttctcccct gatgaagaag       540
     ctgggctggc ccaccatctt ctgcaatgag ctggaggtcg gtgaggacgg catgatcacc       600
     ggttaccgca tccgggtgga aaagagcaag ctgaccaccg tcagggcgct gcagagcatc       660
     ggctttgaaa ccatcgccac cggtgacagc tacaatgacc tgggcatgat ccaggcaagc       720
     aaggccggta tcctgttccg ctccaccgag cagattaaaa aggattatcc ggagctgccc       780
     gcctgcgaaa cctatgatga gctgctggcg cagatccgca aggcgctgta aaacaaaacc       840
     aaagaaccaa aagagggaga accgcagaat tgcggctctc ccttttcatt ttggaaaaca       900
     ggtttccgga atcaggctcc gaagatctgg tccagggtat cctgaaccac ctgcttgtat       960
```

tcctcctgca gctgggcggg ggtcaggtcg ccgttgatga attccaccag cggcatctct 1020
cccagcacgg tgccggcaaa gtcaaccacc gtcttttcgc ccatcttctt catgatttca 1080
aagttccact cctgcaggtc cagcttgtcg gaggtagtgg tgtaccacca ctccagttct 1140
tccggatcat cccggaggct gatgtcatca tgataccagt tctggtaggc acggaagaca 1200
tcgtaaacca gtttgggatc ctcgatgccg acggggatga agtagaagct gccggcagcg 1260
atcttggtgg cgttcgtttc cgcattgccg tggggtccga tggggtaatc cacaaagacc 1320
atgtcgaagt ccagggtggt ttcggccttg ccgtcccagt tatagtcatc atagttggcc 1380
atgatccagg ccgcggtggt ggtgaaggct accttcttgt cgcggtacag ccagcggcag 1440
acatcccagc cgttctcctc gggcagcgga taagccgcct tgtccgtgag ccacagatcc 1500
tggaggaact tcagggtttc gcccatctcc acagaggaca ggttttcctt gggggtggag 1560
gccagcgttg tgccgttgga catataccag tagggcaggc agtccgtgat ccaggcgccg 1620
tagccgtaaa cgtccgtgac gccgtcgccg tcgttatcct tggtcagggc gatgcagtac 1680
tcccggaact tgtcccaggt ccactcgccg cgggcagcca gttcccgggg atcctccagg 1740
ttggcttcct cgatcatctg caggttaaag gccagcggat aagtcgcggc cacctggtct 1800
tccgcaccgt tgacagtgag cagggagaca gcgccgcttt ccaggttcac atagttcatg 1860
acggtgtcgt tatgggtgag cagcggatca tcgggatcca gcacgtcccg caggtccatg 1920
gcatagccgt tcatcacagc aggaacaccc cagcccagct ccaccatgta cacatcgcag 1980
tccggttcac cggccaggat ggagttgttc acggaatcct gcacgccggc ataggtcagg 2040
ttctggaatt cgtaggttac gccgtaggta tcctcaatgt tcttcacgtt gtcgaagcgc 2100
atgaggtctg tttccttacc ggtggcggct tcattgtctt cccaggattc gtcgttggag 2160
tcatagtaga tatcccacca gagcgcgatg gtgatattcc ggggatcttc cgccagcgct 2220
gttacagcca tggaaagtac catgatcagt gccagcagta ccgctgtcag tttttcata 2280
agtcaatttc tcctttccag aaatctggtt attacttgat tgcctggccc acggccacag 2340
cataaacatt ccaggcagag gagctttcgc actgcatcat agcgccccag gtggacttgt 2400
cctcaccgca gagcgcctcg atctgctcat aggtcacctg cgcgattttg ccgtccgcca 2460
catccgcgcc gtcgttgccc acgcgcatcc agccggcggt tgcccagggc atgaccagcc 2520
acatattgcc gtcctcggat tcataggaga tggttaccac ggaaccggga accagggcgt 2580
cgatgatttc ctgcggcatt tccaggccgt cctgcgccca ggcgtcagca gacttggtga 2640
agccggggaa ttcaaccaga ttggtcaggc cgaagaactc cgccttctgg cccacacgga 2700
cgctgtaaac ctcccaggga gaggagccct cgcactgcat catggcaccc caggtggcct 2760
tgtcctcgcc gcacagcgca gcgatctctt cataggtgat ataggccttg ccgccgatcc 2820
tggtggcagt accctgggat acgcgcatcc agccggcttc cgcccagggc atgaccagcc 2880
agatatcgcc gtcttcggag ctgtaggtga tctccaccac agaaccgggc accagggcgt 2940
caatgatttc ctgcggcatt tccaggccgt cctgcgccca ggcgtcggcg gacttggtga 3000
agccggggaa ttcaaccgcg ttcttcagca caatgcggtt ggcccgctgg cccacgcgca 3060
cggagaagac ttcccagtca gaagcgcttt cgcactgcag catggcaccc caggtggact 3120
tgtcctcgcc gcagagcgcc tcgatcatct catagggaat ctgggcgatg ttcttgctgt 3180
tgttgatggc agcggttccc tggcccacgc gcatccagcc ggcggttgcc cagggcatga 3240
caatccacat ggaaccgtcc gcgctggcat attccacttc cacaacggag ccaggcacca 3300
gggcatcaat gatttcctgc ggcatttcca ggccgtcctg cgcccaggcg ccggcggact 3360
tgtggaaatc cgggaactca acagcgttcg tcagggctac gaggttgctc agatcccgct 3420
cggaagcttc cgccacatag accgcggcgg gatccagcgc gatggcgttt ccttccgcat 3480
ccaggaaccg gatgttgtcc aggtagaagg tggcgggttc gccgccggcc tggtcttcac 3540
gggagatcat gacataattg cccgcaccgg cgacaaacgg cgccttgaag gcaatcttta 3600
cattgcgcgg attcttcttc tccatataga cagaccagtc ggccttgttc tcatccgcgt 3660
tttcaccggt gtagctgtaa accacgccgg aaaccgcgta gaacttgccg tccgcggcct 3720
tgtcaacacc gatatcaaag gagacaccgg caacatccgc cagcttttcg ccggcgaggc 3780
cctccagatt cagggccaca taggggatcc cctgagcggc gacgcgcagc gccttggaac 3840
cgttgtaatc cacaacttcc agggacgcgg cggcatccgc gttcggtttg acggcggaga 3900
cgccgaggaa ggcaaacacg ccgtcctcaa agtcaagcac gttcaccgga tcggcggacg 3960
cccaggagca cataccgatt gccaagacgg caaccagcaa ccaggaaatg attttcttca 4020
cgggactttc ctccttttat tttttacccg atcccgggta attaacaaat cagcgcgggc 4080
tcagcccaca atgccggagc gttccacgcc ctcgataaac cgccgctgga gcaccaggta 4140
gatgaccaca acgggaacca gcaccagcac cacgccggcg tcaaaataga gctgctgctc 4200

ccggatggtg agcaccttct ccgcgttggc aatgacgttg atgagggact ccagcttccg 4260
ggtgacgagg atggaatcgg aaatgttgaa ggtgttggca aaaaaggaat cgttatactg 4320
ccagacgatg gagaacaccg cgaccgtgat gaccgccggc gccgcgttcg gcatcatgat 4380
ggtgaaatag gtgcgcgcgg tgcccgcgcc gtccaccagg gcagcctcct caatctcctt 4440
gggcaggccg cggaaaaact gattgaaaat atagatatac aggccggacc ggaggccgca 4500
gccgaagacg gacatgacca ccgttggcca ggcggtctgg agcaggttga ttccctcgcc 4560
gccgttgaag agccccacaa ggcccagggg attgaagaaa cggaaggttg tataaagagg 4620
gaactggatc gtgttcagcg ggatcacgat catgaccagc acaaagccga acagcagccc 4680
cttcagcggg aaccggtagc gggcgaagcc gtaaccgacc atggagcaga tgagcagctg 4740
gatgagggtg atgccgaaca catacaggat ggtgttgccg aggatcggca ggtaatccag 4800
gtatttgatc gccatggtat agcgctccag cgtggggctg atgggaatgg tgaagaccat 4860
gggattgatg gaatcctggc gggaaaagaa ggagttggcg atgatattga tcaccggcgc 4920
caggatgaca tagctgatgc ccacaatgac gatcagccgg aagatgccca gcaccaggtt 4980
cttcatgccg gtgagggttt tatcccgccg gttggcatag gcgggagaat tccggaagcc 5040
ctgccaggtg atttttccgc tttgcgcggc gttgccggtg tgcatgttct ttctccctcc 5100
ccgtcagttg taatagaagg tgcgccggtt gagcaggtag acgataagcc ccaggatgag 5160
gatggtgatc agcgtggaga ccacggagaa cgcggaagac aggccgtagt tatacagctc 5220
cttataggtg cgcgttgcca gtttgatcac ttccccggag gaaaaccggt ccaccacggt 5280
atagatcgtg ttggtgataa tgtgcggcat gaccatgggg aaggtgacct tccagaagga 5340
ctcatagccc gtggccccct cgatccgggc cacctcgtag agggagccgg gaatggactg 5400
gagcgccgcg atgaacagga tgatctgcac gcccgaaccc ttgatgatgg acgaaatccg 5460
gttgaccgcg tccacaatgt agccgaggac ggactgcgga atggccaggt cgccgaaaag 5520
atccatgtaa taggccacgg agacggtacc cgcctgggcg gtttccaggc ttgcggagga 5580
aacgccggca ttcaccagct ccgttgcccg ggtgatggcg gaacccaccg cgtccgcgct 5640
gaggatgatg ggcaggaaga agatggcgcg taccagcgcc cggcctttga accgccggtt 5700
caggagcatg gccacaaaca ggctgaagaa aatgatcatc ggcaggtcga tcagcatatc 5760
gatcacggag gaagccagcg tctgcttgaa ggtggcgtgg ctggtgaagg cctcaatata 5820
attgccgaaa ccgatgtatt ccttcgtcat tcccgcggcg gcgtccaggc cgatgcggga 5880
caggctgaac tctcccagct gcaccaggca tccggcgtag aacaccagga agccgatcag 5940
ccaggggaga atgaaccaga gcccgcgcac ggcgttcttc tgctcatagg aaagatgaaa 6000
aatacgattc tttctgcgga tgccagtcat tctttatctc ctccgaccgc caggtaatcc 6060
attgccggga tccggtatcc gtccgctgcg gcctcacgct ccccgctgtt cacatacagc 6120
acggtgccgt tggaataggt tacgcgggcc agcgtatcgc tgagccggtc atggcggagc 6180
atctccgcgc cgctcacccg ggacagggcg ccgttcacaa agcggtagct ttccgccgct 6240
tcctccttcc aggaggagaa ggtggtcgcg taattgttgt tgagcccggt gtacttcata 6300
tctgccgcgt ccttccaggt gaacgtgtaa tgcgtggacg cgccgtattc aatgaggtga 6360
agcagggttg cccgccggtt ctcgctctgc atcaggttca ggggggatcc gcagtaatcc 6420
gccgatccgt ggaggatcag ctcccacagc gggatctgtt catccacaat ggcgaacatt 6480
gtcgcttcca ccggcgcgtt gattacatgc cggacatagg gaaaactgta atcattgccg 6540
ccggacacca tgagctcccg ctccccggcg ccgatgacac cgaaggcttg cttcaccagg 6600
tccaggtcag cctcccggct gatgacgttc gtgcggcgct tgtccgcgtg gacctcgttg 6660
cccagatccc gcagggagag tgcgtccagg cccagccggt ccgccgaggc cgccagcagt 6720
gccgcatacc gtggcaggaa ccggggagac aggagcatat agccccgctc tgcaaagccc 6780
agggtggcca tgcgccgcat ggacacggga ttgaccacgc ccagctcggc cgcgtagccc 6840
ttggcgtaat accggctggc ctcctcactg cgcataaagc ccttcgcgat ctccgtgacc 6900
agctgcagcg ccgcgtccgg atagatccgg tccccgttct ccgccgccgc caggcacagg 6960
cgcttgaggc ccttttcccc gcccagctcg ttcagcacgg acacatggct gaccgggtcg 7020
tgataatagc cgccgttcat ccagccctgc aggttgaccc gcaggttgcc gacctgctcc 7080
ccgcgcaggc tggacatcat ttcctccgcc tgggcaaagg tggtcatggg catcacccgc 7140
aggtactgca cacccatcca gtgcgcggtt tcccgcacgc cgccgaggat atcgcagtaa 7200
aagggaatat cgccggatgt ttcggccttc agcttcaggg ttccggacga gagcagagat 7260
tcccgcacgg cctttgccag gccgttatag ccggtataat ccccgtccag gatcctgtag 7320
gtcacggcgc agtccaccgg ataggcgtcc tgttccgcga cgatggcctc ctcgccggcc 7380
acgatgaggg tatccgtcct tctcaggctg aaagtgaaat aggcatagtt gtaactgttg 7440

ttccggcccg ccacatccgc cgtgacggaa gccagggtcg caccccctgcc gcaggaggcc 7500
aggaggctgt aatcctcccg gcagatgccg aagagtgcca gccgggccga ctgcatattc 7560
tgggttgcgg tgaaatcact gtcgatcaga tccaggtcat agattgcctg gttatactgg 7620
ggcgcggtgt tcttgccatt gttgaaattg atcagggcgc cgcttccgtc cggcaccacg 7680
atgccgccgg tctccgccgt tccggccgcg ccgaagaagg gcagcagctg gatggagcgg 7740
atctttccgc cgccggcttc catgatctcc gcatgaggga tcgtgacggt gagtccctcc 7800
ggcgtgagcg tgtaatccag ggcgatgaca aaggaagtgc tctccgcggc ttccgcggcc 7860
tcttcctccg ccagcgcctc catctccgca taatcctcca tggtgaagcc ggccgtccgc 7920
gcgtccgcgt caatctgctg gcggttgcgc gccgtgacac cctgggactt gagcacatac 7980
aggccgctct cctcatccag gacataggat ttggccgcct gcttcctgcc ggcgttcgac 8040
agaacctcaa accattccgc ggtcatctgg cgcggcacca gcatcagctt ctcattagac 8100
agggtataga ccgcccggaa gccgttctcc agctgaaggt attcaatctg cccgttctcc 8160
acgcactttg cgtaggagct ccagggcgta ccctcgcggg agtttgtatc caggtaattg 8220
aggataaact ggctccgcag gttgccttgg ttcagcccgg agcgggccac cggatcctcc 8280
gccgcgtcct gcgggttgga atagaccaca cgtccgcttt tccggtcccg cagggcgacc 8340
tccgcggtct tttcattcag gtacagggcc tggttgtcat tctgcgccgc cagggcgaag 8400
ccgggcacgc tgttttccgg atccttccgg gtcttcaggg ccgcggcttc cgttttctcc 8460
gcgggcggct gcacatagag ccggtattcg tcataccccc ggtaatgaat caggtaccag 8520
cccaggaaca cccccgcgcc aatgaccacc agcgccgcga ggatccacag cgcgcgccgg 8580
accgcggtcg ggagggggcag cctgtttttc tcactcattc tgctttccct ccttacacac 8640
ggaaaaccag ttccatatag ataccccgga agaacgaata cacctgggtg ataaagttga 8700
tcaccaggag cccgaggaaa atgatgatca gcatggagat cacggtgagg aacagcgtga 8760
ccagcgtttt gccgaaggtg tagttatgca cctgcatgat tccgatgagc agcatgacca 8820
ccgtgtaggc ggtacccagc cccaggagga gggtatagaa cgcctcctca ttctccgcca 8880
ccaggtggga aaccgcagtc gccgccgcgg tagtgaccag catgggaaac agcgcgtagc 8940
ccaccgcgat gaggatgtcc ctgaaccttc cctcaccctc catgaggcag gtgacggacc 9000
agttgcccac gcagagaatg aaatacagca ggatgacccc ggccagttcc gccgggaggt 9060
tcaccgtgcg gggttcaata tcgttgacaa taaagctgga ggaaatccgg ttcatggaaa 9120
aggcgacgcc gtacaggatc accatgagga aggcgatcca caggctgccc ttttcctggt 9180
gccggatcca gtaatatccg tccgccggat ggctgactgt atggaagaaa tgctgccatt 9240
tggccttcat tgtcccccgg ttcatgacag gttcctcctc tttgcctttt tgcggattgc 9300
cggccgcacc gctttcaccc agagcaccgc cagcacggcc gccgcgacca cgccggaaag 9360
aatgaagctg atattgttcc gcagccgctc attcctccac cgcttgaaag cgacggaata 9420
atactcccgg ttcatgcccc gccgcaggta ggtcatggcg tttgcgttat cctccgcgga 9480
caggtaagcc ttgccgatgc cggtgttggc cagctcgttg atttcatcca gccgcagccc 9540
ctcctgccac agcggaaccg cctgggtttc atccccgtca aaccgcagag ccaccgcccg 9600
gttgatcagc gagccgtatt ccgtctgcgc gaaaacggtc aggttcccct gctgggaatc 9660
cagcacaagc aggcggctgc ccagctgctc aatggccgtg ggcgtgagga aggtgcccgc 9720
ctgggagccg atgccgccga agatatacag caggttgcct tcatggtcat aggtaaagat 9780
ccggccccgg cggctgtcca gcagggagta ggtgccgtgg ccgcggtaca ccacgtccac 9840
aatctggctc gggccggcat aattgccgac ggaaccgaag acccggtccc cgcccaggtt 9900
gccgtggggt cctttgcgga tcacgtcctc cccctttcggg ttcagccggc ggacagcctg 9960
caggccgtcc gtatccttgc tggaggcgta gacaaagccc tcgtcatcca catcgatgcc 10020
ggtgaactcc gtggggataa acagctgctg tttgccccgc tcttccttgg tggccagctt 10080
ccgccagaac ttttcccaga gggagatctt cacgctgatg gtgccgaaaa agccggtgaa 10140
atcgcctgcc tcgctgaaaa ccatgatgcc ctggaacatg ttctgggcga tcacatagac 10200
gcgatccgca tagtccacgg acacccgcag gggcgtgaac acatacccct cgtccagcac 10260
ctcgctccgg gggttctgca catatttgac aaaggttccg tccggatcca gcaccacgat 10320
gcggcggttc aggctgtccg caatgtacag ctgatgcttt tcggagatcg ccacgccgct 10380
gggggcgttg aacgcctggg cggcgccctc cgcctcaaag ccgatgatga tccgcctgac 10440
cgtcttcagg tcgtccgcca ggaccacaat ccggttgttg ccggtatccg cgatatacac 10500
atccccgtcc ggcgaaacgc acaggtcctg gggattgcgg aaggcgccga tcggcgcgcc 10560
ttcaaaccgc agcgtggcgc cgctgaccag cccgtccggc acatagggggg ccggggtgta 10620
aaggatgttg ttccagtggt catagttgta gcagtcatag ggcagggaat ccgccgccaa 10680

agcgttcccg aaggccaggc aggccagaag caccaggcac aggatggatt tcagttgctt 10740
catgctccgc ctcctcagtc cttcatgccg gaagtcgtca tggtctccat gatcgagctc 10800
tggcagatca ggaaaaagat gatcgggacc gccgcgatca ggaaagtgac cgccgccccg 10860
gcgcccgcac ggctcacccc gccggaggta atctgctgca gggcatactg caggggcttc 10920
agctcctcgg agcggaggaa catgctgccc gtgttcgccc acatctgctg gaactggaaa 10980
atggccagcg tcagccaggc cggtttcaca ttcggcatga cgatgcgcca gaaaatctgc 11040
cattcattgg cgccctgcag acgagcgctt tccatcaggg aatccggcag ctgctccata 11100
aactgcttca tcaggtacag gcccatgccg aaggaccagg cgggcagcac cagcgcggca 11160
taggtgttgt tgatgccgag ccaggaaatg atcaggtact gcgggatctg cgtcacggtc 11220
caggaaaaca tcatggacag cacaaccacg gaaaacagga tctttttgcc ggggaaatga 11280
ttcttggcca ggggataagc cgccatggag gcaatgatca catggccaat catgccgccc 11340
agggtgatga tgatggtatt cagcacatac cgggagaagg gcacccagct ttcattcagc 11400
aggacgtaaa gatctgtgaa attgttcagc gtggggttcc gcacaaagat ccgcggcgga 11460
tactggtaca gttcgtccag ggttttcacc gcgttgttga ggatcatgac cagcggcagc 11520
accataaaca cgccgcagat gcccatgaga ataaacagtg ccgcgttgcc ggcggtggac 11580
cggttcagct ttttctttcc ggatttccgc ttcattttcc ggtggatttt tgccagccgc 11640
ttctgccgca ggatctcctc ctgctccgcg gtcagttcca caggattcct gtgcagccag 11700
gcaggcagca accggctgtt tttcgccttt tccataccgg ttattcacct acccttctga 11760
gaatccgctg caccagtttg ttggttccga ccatgagcag gaacaggatg gtcgcgatgg 11820
ccgaggcata gcccatgtcc aggcgggtgg aaccgtaatc cagcaggtgc gggacgatgg 11880
ttgcgccgga ataattgacc gagggattgc cggccagggc aatggaaata tccgccaccg 11940
cgaaggactg ggtgatctgc atcaccgcgc cgaacatcag ctggggtttc atggccggca 12000
gggtcacaaa ccacagctcc tgccaccggt ttttaacgcc gtcgattgcc gccgcctcat 12060
agaggccctt gtccaccgtc tgcaggcccg cgataaagga caggaagcca gtgcccaggg 12120
acagccacag ctggacgacg atcagcaccg gcatgacaaa cttttcggtc tgcatccaga 12180
ggatcggctc attcaggaag cccatccgca gcaggacgcc gttcaggtat ccgtaacggt 12240
cgccggtgag gatcaggttc cagatcatat acgcgttgcc gcagatggac ggggcgtaga 12300
aaaccagcgt caggaacgcc cgcacccggg gcttgaactc attgatgatc caggcgaaga 12360
gcaggcacag gaggtagctg accggtccgg tgatcaccgc gaagagcagg gtattgcgca 12420
gggagatgat gaaaatatca tcctccagca gcagcttgat atagttggcc cagccgaccc 12480
agcggggcgg ctccagcatg ttgaaataat tgaacgaaag gtaaacagac atcaacacgg 12540
gcagcaccgt gaagaggaaa aacaggatga aatacggcgc catcagcagg taggatgcct 12600
tgttgatccg gacctggtgc cacagggtat ccttccgcgg cacgatccgt tctttccgga 12660
ttgccatgct catctcactt tccctcccct gcgttcgcga caggcagtcc gaattctgtc 12720
cgtttatagg taatctccgc attgatgtaa aaaatcttgt ccatcagttc ctcacggggc 12780
gtgttgccgt tctcgctgcg gccgctgctg gcgctgcccg tggtgaccgc gtagaacgcg 12840
ttgttcacat tgcgccaggt gtaatagccg ccgggtacct gggggatgcc cttcacgtgg 12900
ctgaactgcc tggccagctc ccggtaatcc cggatgggcc agctcagccc cgccagggct 12960
tccaggttcg cggttgccac ccgcgcggag gcgcccatca gcgattccat ttcccggccg 13020
tacagggtct ggatctccgc gctggtccac cacttgagga agctccagca ggcttccggc 13080
tctttggtgg cgctcatgat gatatccgcc aggccggagc agccggttgt cgtatccagc 13140
gttccgtcct cccgcagggt gccgggcaca tgggtaaaat cccacaggcc cttgatatcc 13200
ggggcggata cctgcaggtt gttgtaggtt gtgtaatccg tgatgataat cgggcactcg 13260
cccgtgcgga agcgctcctc cgccgaggtg gccatatcca gcttgtaatc cgtatagaaa 13320
tcgcagtagc ggcggaaaac gttgaccgcg atatcgctgt ccagcgcgga ggcgtcgccg 13380
ttctccgtgt agtattcccc gccggcctgg aacagcagca tggcgaaggt ctgctccccg 13440
ggcagcatgc cgaactccat ctggttcttg ctcagcacgc tcatggccac cttcacgtcg 13500
tcccaggttt ccgggacggc caggccgatt tcagccagga tatccttgcg gtagaacatc 13560
atggggaagg tctgcgtatc cggcagcgcg taggaggcac cgttgaagga gaagggcacc 13620
agggcgctgg gggcgaaccg cttcgccacc tcctcaaagt cctcaaactg cgtcaggtcc 13680
atcacggcgt tccgcaggcc gtagttcacc ggggtatcgt tgccggtgtt catcaccgcg 13740
ccggcaatac ccgtggtgtt cgccacctgg atggccacgt ccggcccctg gccggagagg 13800
gtggcgcgga gcagcgtgtt catatccacc agctggacgt tcacgttgat cccggtctgc 13860
ggggtgaaat gctcgtcgat cagggccttg atcacgttgg cctggtcccg tcccgtgccg 13920

acccagaggg tgatgacccg ggaatcatcg gacttctcgg ccacgttgcc gatctggtta 13980
tagttgatgg tgaaggagct gtagagccgg gacgcctcaa agccggtgcc ggaaaggaaa 14040
tccctgccgc tgatctccgg ccggacgtcc gcggtgtgga caaaaatccg gtcgatctgc 14100
agcggctggc ccagcacctg ggtgatccag ttgccgcagg cgcgcacgtt cgtcttgtag 14160
gaagaaagca cctccgtaaa gcgctcctgg tcccggatca gctccgtgag ctggtcgtcc 14220
atggttttca ggacggtcag cttgtcgctg cggtggcccg ccgtctcttc cagcgcgccg 14280
atcgcggtct gcaggtccgc cttgaccttc cgcagctgcg cctccagccc gggcaggctt 14340
ctttcaatct ggtaatcccg gtacgggtcc ggcgaaacgc cggtgatata cagcacctgc 14400
cggtaaatgc cgttgagctg cagcacgcag tcctgcacct ggcggatgac ggccgccatg 14460
ccgcccagca ccacctccat ccgcagggta tgctccccg cggaaaggtg gatcaggtaa 14520
ggctcatcct gttccccgga gagggtctcc aggcgccagt cctgggtata gggaaaggtc 14580
gccgcctgga actcgacaaa cggaacctcg ccgtcaatat agattttgcg gtacacgtcc 14640
gtgccgcgga taaaattctg cttatcgtac agggaaatgg cgtagttccc ttcctccgcc 14700
accgtgaagc gccactcaat ccactggccg gcgctcttcc acgcgttgcc gccgatgctg 14760
ttgttcagca ggtagcgtgg actggaggga gaaacggaac aggaggactg gtcctgcacc 14820
ggatacagca tctgggagga ggtcctggag gccgcctccg cctcgatccg gatgatctgc 14880
ccggcagcgc cggattcccc ttccggtttc cggtagggct gcgcgtccgc tcccgcccgg 14940
aagcagaggc gccggatcag catcggctcc cgcagggaca ggaccgagag cgtatgcggg 15000
ccgcggggca ggaaaacctt caggctttcg ctgatgtagc cgtttgaatc atgcaggagg 15060
gaagtcatcc actccggcgc ttcggcggga gagggcttca gctcattgcc ctgattgtcc 15120
cggagccacc ggatttcctc cgccccgtcc tccgccgtga agcggctgac gtccggatgc 15180
aggtcattcc gccagacccg ggaaaagttc accagcgcca gctcccggta agggagcgcg 15240
ccgtccacaa aaaaggcccg ctggatttcc gcattccggc ccggccacgg atagtattcc 15300
acctccaggt cgtaccagcc ggattcctcc gcttcaaagg cccactcgat gaccgcttcc 15360
tcgcccgtaa ggaccgaaac gccctcccgg ccttccgcct gctccatgaa aacaggcgcg 15420
ttcttgccgt cttcctcctc gtagcggacg gcgtcagacg cctccagcac gatctccttt 15480
tccggcccgg ccgtcatttc atgttccgcg cagtaatccg cataggaagg aacagccgga 15540
ttgatggaat aagtgctcac ctgttcttca tattcactta tgccaaagcc ggtggcagcc 15600
tgcgcggcgc ccgccgccag cgccagcagc agggccagca ccaatccagt ccactttctc 15660
acttgtttat tcctccggtc catccgcgta cagcggcgtc acttttcatc cctggtacgg 15720
aaatcgggca ggtcctttcg ggtgacgacc cgttcatccg aataggcttt ccggatcatg 15780
gccgtttccg tatactgctc cgacggcttg ttgaatttcg tgttccgcag cacaaactcg 15840
ccgccccagg tgcaccagta tccccagacg gttccgtccc ggaacacctg ttccggatcc 15900
ggaatgcacc cgttctccga aagcacaatc atcttccgct ctcccgcgat ccgcgtcagc 15960
ggcaggaagg atgcgctctg ggagtcatag acccgttccc cgggatatat atccgttccg 16020
atgagatcca cctcatcgtc ccccggatac cagtccggat cctgggcgtt ccagatccag 16080
atcaggttgt tcagtccgta ttcattggtc agccggtcaa acagcaggtg gtacagcttc 16140
aggtacgctt ctttcccgca tgcccccccc cagacccagc cgccgctggc ctcatgcagc 16200
ggccgccaca gcaccggcac gccggccgcc ttcagccgga gcagctgcgc cgcgatcgcg 16260
tcgatgtcct gattcaacag ccggtatcct tcttcgtcct cgccgttcat caccctggcc 16320
aggttgaagc ttgtgtattt ggtataaaaa ccggaatacc agggttcctt caggtaaggg 16380
gacggagcgt tccagtgcca gcagaaagtg acgatgccgc ccttgttcca gtattcaatg 16440
gcatagggca ccgcccggtc gctttcgccg ccgtgctcca cacggctgtg ggaatactcc 16500
atcatgtcca ggccgagcac agcgggataa tcgccgccgg ttgcccgcca gattgcctgg 16560
ttttccatcc cgaacatgcc gccgtcgcac tgctggccgg agagaatctt tttgccatag 16620
ttttcccgca gccagtcata cagcctccgg gcctccgggg ttgcctcccc gacgctcatc 16680
ttttcgggaa tggtataaag atctgccgga agggccgcgg agggtgtcag ctcaaaccgg 16740
tccacccgga tatatcccca gcttgtgccg attcccagcg tatgggtgcc ggcggacagg 16800
taaatatgct ccagcacgct cttcccgaag gcggtatcat ccgttaccgc gcttccgata 16860
ttccggccgt ccaggagcac cgggttttcc ttgtgcccgc cctggcttgc ctggatcacc 16920
gcgatatcat aaaagccttc ccggctcacc gtgacttccg ttatgatccc gtcttcaccg 16980
gctgtgtgga aaccttccac ccattggttt tcgccatccc cagcgatcac ggtatggcct 17040
tccggctttc ctgtttccgc ctccacgatc atcgtctccc ctgttccttt cacggataca 17100
gggagcagta cgcaaagcag catcataccc attgcggcca ttccgcaaaa gctgcttttc 17160

acgcttttca tgccggttac tctccctgtt taattaatct tgttaactta tttgttaacc 17220
tgttgtaaac taaattttat cggaacaaaa aacgcctgtc aagacaattt ttgcacacaa 17280
ataacgctat gcttgtcgtt tattgcgttt ttgctcccgt gaaaaacgca tggcttcttg 17340
tttttgaact tatttagatt aacaaattag ttaacaaata aatccctgct tttcttctgt 17400
tccttctccg gtcacagggg ctgtttacag gaaagccatg tgcttttccg cacgtaaaaa 17460
agccggcggt ttttcccggc cgccggcata caactgctga tattcacttt tttccggttc 17520
agatttccgc tcccgcgcct tcaatcagct ccgtccgcag gaaccgtccg ttttcatccg 17580
tgatgacgcg cctgccgcat ttccagctgc ccagtgcttt cccgtccaca cggatttccg 17640
ccctctgccc gtcggaccgg atggcgtaat gacggtcaca gaaggcagcc tcatacgcac 17700
agggcgcgcc gctccccagg ctgaaccaca cctgtcccag gtgcggatgg atgccgtaat 17760
ggtgtgcaat atattcaagg caggccagca tggtgggccc gtaggcgtcc tgcaccgctt 17820
ccccgctgcc ttcctccgcc ggctgatgat ccgacatgcg caccagagag gctttgccgg 17880
tgaagggatc atattgctgg gtaaaccggc agccgttcgc gtaaaccgcg tcaatcagtt 17940
tccggcccag ccccgtcacc agccggttat acccgtagcg ttccagggca aggatcgccc 18000
gctggtaggt gaggccctcc ggctggccgc tccagttgtt ctccggcgcg ttgcggaaag 18060
ccggatcatt gaccgccacg ctgggaagcg ggaaaggcgt ccggaattcc gcgggattca 18120
gcaggtgctc ctttacaaag cgggccgcca ttccccggga aatgctgccc cagtacatgc 18180
accgtagcgt attgtggcac aggatgtcca ccgttttccc cgtccggtcc cggtcataca 18240
gggcgccgcg cttcctgtcc cacagccggc ggcggagcgc atccgccacc atacgggctt 18300
tccgctccca ggcgtcagcg gaccgatccc cctgcaggcg gcagatttca gccaccgtcc 18360
ggcgggccgc aaagctccag cttgtcacgt ccatgctggc catgggaacc acctgatatc 18420
cttccggcgg ttcatcccct tcccaccagc agggcgcgtc cccgtaccgc agggccaggt 18480
cctcccccgt gtcatagacg cagaaggagg aaagcagtcc gtcgccggaa acatggcgcg 18540
tattccacag gcaccggtca aagcggatca ggctctcctt cagctctgcc aggtaatccc 18600
tgtcttttcc gattaagtaa tacaggttca gggccgggaa gggaaaacag aacccctggt 18660
atttgttgaa ctggggttcc actttgccgc cggtgcactg gatggagccc ggcaggcgcc 18720
catctgcccg ctggtggcgc atgaacaggg tgatgttgtt ccgggccgcc tccaggctgc 18780
gcagggcata catctcgccg cccatgggct gggtttccag ccagattttt tcatatccgc 18840
cgccttccac cagcaccgta tcgccgccga aatccttcag gttaaagcgg catttctctt 18900
ccgccgcgtc ataaacacgc tgaagctttt catcctccgt gcggaaaaaa acgcttgttt 18960
caggcagcat ttttcagcct tcccttccgg tttttattcg gaatttccac tggttgaatt 19020
ctatcattta ttgcgatttt tggacagatg cgaaaaaaat gcgcatcccc cgaaatccga 19080
aaaaaccttt atttttcaac aatgtgcaaa attttagctt cttccctttc agaagatttg 19140
ccgtcataat gagagcgaaa attgagcaac gaaaaatcat atcgccccga atatcccgaa 19200
ggaggattcc tgatgagtgt ttccgcatcc acccccgcgc atcgcggcgg actcaggtac 19260
agatttggat tagccctgcg ccgtgactgg cagttgtggg tgctgctgct gccctgcctg 19320
gcattcttca ttgtattctg ttatttcccg atgtacggcg tacagattgc cttccgggac 19380
tataaagccg ccttcggcat taccggcagc cagtgggtcg gactgaaata tttcgaaaaa 19440
ttcttcaaca gctattactg cgggcggatg tttgcgaaca ccttcctgct gaacctctac 19500
ggcctgctgt tcggtttccc gatcccgatc ctgctggcga tcatgctgaa ccagctgaac 19560
agcaaacggt tcaagggctt tgcccagacc gccatctatg tgccgcactt tatttccacc 19620
gttgtgctgg ctggcatgat ctaccttttc ttttcgccca ccaacggcat catcaatcac 19680
ctgatcaccg ctgccggagg gaaaagcatc tatttcatta tggaacccca atggttccgg 19740
ccgctgttca tcggatcgga catctggcag aacgccggat ggaacaccat cctctatatc 19800
gccacactga ccagtatcga tccccagctt tacgaggcag ccaccattga cggcgccacc 19860
cgctgggaaa agatccggca tattgatatt ccccacctga tccccatcgc ggtgatgatg 19920
ctgatcctca actgcggatc gctcctctcc tccaacacgg acaaggccct gctgctgcag 19980
accagcggca acatggccac ctcggacatc ttcggcgtct acgtgtacag cgtcggcatc 20040
gctgcccaga aggcccagta ttcctacgcc tccgccattg gcctgtgcct gaacctgatc 20100
aacttcgcca tcatcatggg agtcaacagc atttcccgca ggatgagcga tatcagcctg 20160
ttctgagaaa ggaggaagaa ggatgactgt tacggccgtg aaagcaaaaa agcaaaaccg 20220
gatccgcgaa acccggaagg accggatctt taacgcaatc aacaccctgt tctgggtcct 20280
ggtcctgttc ctcgtcctgt atccactgtg gctgatcctc atcgcttccg tctcagaccc 20340
ggacgcggtc ctgcaggcaa aggtcctgat ctggccggtc gatttttccc tgatgggata 20400

tgacgcggtg ctccagaaca gcgaactgct gaacagctat gtgaactccg tcttctacac 20460
ggtggtcggt tccgccctga gcgtaatcgt ttccctggct gccgcctacg cgctctcccg 20520
cgggtttgca ggcaaaaagt ttgtcaacct ggccatcacg tttaccatgt tcttttccgg 20580
cggcctgatc cctatcttcc tgaacatccg ggatctcggg ctgtacaaca cccggacagt 20640
catgatcctg atgaacctgg tttccgtctg gaacctgatg gttgcaagga cctatatcca 20700
gaccagcatt ccaaacgacc tgtatgaagc cgccgtcatc gacggggcgg atcactttac 20760
ctatttcttc aagtgcgtgc tgccgctgtc cggcaccatc atcgcggtgc tttcggtcta 20820
ctacggcgtt gcccgctgga acgattattt caccggcctg gtcatgctgc gggaccgcag 20880
cctgtatccc ctgcagctgg tgctgcggga aatcctggcc tccctcacct ccacaggctc 20940
ctccgatacc ttcttcgcgg cttacgcgga taacctgggc ggcctgcagg cggcgctgcg 21000
caaggccgag gtggccaaat actgctgcat cgtggtggcc actgtgcccg cggttgtgct 21060
gtatgtcttc atgcagaaat actttgtcaa aggcgtcatg atcgggtctt tgaagggata 21120
aagggggtaa accggtgtca cagatcagga gcacgaacga ttccgcccgg gccgcgcgct 21180
atgaggacgg cttccgggtt ctgcagggcc ggcgggaata cgtgacctac caggcggact 21240
catccatccg gatctggtat tcggatattc cctggcgcta cgaagcccat aaccactccg 21300
cggtggaaat cctcctgacg ctggagggca cggtgaccta taccgttgag gacaaactct 21360
accaggtccg caagggggaa atcctgatcg tccccccgga caccctgcat tccctgtcca 21420
tgggtgaagg cagcagccgg tatcttttcc tgtttgagcc tgacgccatc atgtccatgc 21480
gggatatcaa gtacatggct tcgtatttca acaaaccttt ccatctccgg gatggatcgg 21540
atgcccatgt gcgcatccgg gaactgctgc tccgggccaa tgacgcctat gagaagcggg 21600
aaatgctttg gaacaccctg tgctacagct gcattctccg catctacgcc atcctgggac 21660
agcattacct gggcagcgtc cggccccata ccggggacag cacgggaaac atggacgccg 21720
aggttgtcaa cgcagtcatg acctatgtca acaaccatta ccgggaggat cttaccctgg 21780
aggaagtagc gcagtttgcc ggcttcagcc gctactattt ctcccgttcc ttcaaaaagc 21840
aaacgggata ttcctttaaa gagtacctgt gccagaagcg ggtacaggta gccatggacc 21900
tgctcatccg gaccaaccgt tccatgcggg acgttgcgct ggagagcggc ttcggcagcg 21960
tggccacctt taaccgggta ttccgggaaa agaaaggctg cacgccctcc cagtacagga 22020
cgatttacgg aatcccgtga gattggtaat caagggcaca gtgccctgaa ataaaaaagg 22080
aggaaagacc catgaagaaa ctcgtttccc tgctgttggc atgctgtatg ctgctcagcc 22140
tgagcgccgc cttcgctgaa gagaaggtag agctcaccgc cttccagtac gcgctggaaa 22200
accagaacac cgatttcaac ggcctctggt tcttcgacga actggaaaag aaaaccaaca 22260
cccacgtcac cttccagatg gtgaaggacg ctgactgggc aaccaacgtc aacctgatgt 22320
tcgccgtgcc ggatcagatg cccgacatga tcatccgcca cgcggtggat gtggaagaat 22380
acggcgtctc ccaaggcctg ctgcttcccc tcgatgaata cctgaccgag gagatcatgc 22440
ccaactacgt cagccgcctg aacatcaaca acgccggcga ctccatcccc gccagcgacg 22500
gcaaaaccta ctacatcggc ttcctgatgg cccagaacgt gaaccatacc ggcacctggt 22560
atgtgaacca gaaggaactg gagaagctgg gcaagaccgc tcccaccacc attgccgaaa 22620
ccaccgagct gctccgcgcc atgaaggctg acggaatcaa gtggcccttc tccgcgtcct 22680
acacccagtt cggaccggaa accatctgga accagttcgc ctccttcggc gtaccggaaa 22740
acacctacta ctacttcatc gacgctgaca acaaggtcca gttcacctgc gcccaggacg 22800
gctggcgtgc ctgcgtggaa tggctgcatc agctgtatga agaggaactg atggatccgg 22860
aatccctgac ccaggactcc aacctctggg ccaacaaggt caacgacggc gaagtcggct 22920
acttctgcta cctgcgcctg atcaacaccg ccatcaaggc tgagaacgtg gagaacttca 22980
agtccatcct gcctcccgcc gctgacggat tcaaggcctg tgtgtcccag atcctggaag 23040
tgccggaagc cggcgcctac ctgaccaaga actgcaagga tcccgtggct gccctgaagt 23100
ggatcgacgc ccagctggaa accgaaacca tgatggtttc cctgaacggc aaggtcggcg 23160
agcagatcgt gctcaacgac gaaggcaagt atgaagtcat cgatatcccc gcggacaacg 23220
gcctgtatca gttcgttccg gttactatgg gccagttctt cgcccccggc gactactaca 23280
cctccattta ccagatggca cctcaccgcg tggaacgcta tgaagacagc aagtggtacg 23340
ctgaatccgg cgtgctcgag cccaagagct tccagtacct gcgcgacctg agcaagctga 23400
gcaacgaaga cgctactgag gctgctgaca tctacaccga gctgcagaag ctggtggaag 23460
aatccttcgc caacttcgtg cgcaacggcg tcacggatga aagctggaat aatttccagg 23520
agcaggccaa ggcgatcggt tccgagcgct acatcgaact gtaccagaac gcctatgacg 23580
cctacctggc caagtaagaa tcttgaacac cacggctggt gatgttagaa tagaagcacc 23640

gaaaagccgc cgggcatcgc cgcaggattt gtttccgggg gcgatgccct tttgccgtat 23700
agagggagta cctatgaagc gtaccgttaa aatcatctgc ctgctggccg cggcattgct 23760
cctgatgaca ggctgctccg cgcctccgcg ggacgagggt tcccccgccc ccgccatatc 23820
cctgtctgcc cttcagtacg aaattgaaaa catcgccgtg gatttcagca gcatgtggta 23880
tttccggcag attgaacagc agaccggcgt ccatgtggat ttcaacgagg tcaaggattc 23940
cgaatggagc agctccgtag gcctggcgtt tgcccagggc aaaatgccgg atatgattct 24000
ccggggttcc ctggacgtgg aagaatacgg tgtgaaacgg catctcctgg tgccgctgga 24060
tgaatatatt gaaaaaggct tcctgcccaa ctatgcagcc cgcatgaacc aggccggcct 24120
gcgggaacag ctcaccgctt cggacggaca catctaccag ctgggcttcc tgatctccca 24180
ggacgtcaac accaacggac atttcttcat caacaaaacc tggctggacc ggctgaagct 24240
gcccgttccc gaaacggtgg agcagctgac ggaggtgctc cggcacttcc gggaggatga 24300
tcccaacggc aacggcatcc aggacgagat tcccctggag tttaccctgg atgacaacac 24360
caccggcata tacaacctgt tttccttctt cggcctgccc ctgaacgagg actacgttta 24420
cgcggacaag gccggccggg tgcattttgc cccggaagag gacgcctttt ttgacacgct 24480
atcctggctc caccagctgt atcaggaaaa actcctggac attgatttca tcagccaggg 24540
aggcagtatc tgggctgcca aagtcaacga gggcagcacc ggcatgttca gctactggcg 24600
gctgcagaac accgccctga atcccggcgt cgccggaaac tatgagatca tgctcccggt 24660
ccatgctgat ggaaaggccg cctgcctgcc ccggaacatg gacatcattg aattcggcgc 24720
cgccctcacc aaagacaacc ggaacattga agcaagcctg cgctggctgg acgcccagtt 24780
tgaaacggaa aacatgctgg tttcccagaa cggccccgtc ggggacacgc tcatccggcg 24840
ggaagacggc cgctatgagg tcagctatgt gcccccggac aatgaactgt accaggctgt 24900
tccggtcatc tgcggccagt tttcgcgcc cgcggactat tatgcttccg tttatgtgcc 24960
tgccgcccac cggctggaaa aagcagccta ctgcgacctg tatgagcgct ccggtgtgct 25020
ggagcccatc tccgccaagc tgctgaccac cgtggcaccc aagaccaccg accagtccgc 25080
cgaaaccacc cgcctgaagg cgaagctgaa atccctggtg gatgcggaaa tcgtgtccat 25140
ggtcacgcag ggagtcacgg aagaaggccg cgcggcattc cgtgagagcc tgaaggcagg 25200
cggaagcgct gactaccgcc aggtatacca gcagatctat gaccgctgga aggagaagta 25260
aaaccgtgat cagaaagagc cgcacctccg tctttatccg gtatgcgctg tcgtatatcg 25320
ctgttgtgct gctgctgttc ctgagcatca ccggctatct ctatatccgg ctcagcagca 25380
aaacccggga ggaaatcatt gacaaccaga tcaaccggct gtcccggatt gccgcccagc 25440
acgaaagcta tatctccgcc atgctgaata ctgccgagga aatcggtctc agtcctgata 25500
tcgagttttt ccgttatgag gaagagccct ggaaagccta tgacctgcag ctgaagatgg 25560
tgccctatac caccaccagt agcaccttct gcgaccaggt ataccttttac ttcttcgggg 25620
atgaccgggt ttattcttcc tcttcttcca tgagcctttc ccttttgcg cggatgatgc 25680
gctatgaatt tatcccgatg gaaagcctgg tttcccggat tgcaaacacc gaccggctga 25740
ccattctgcc ggcccagcag gtatcctcca ccctggtgga cagttcccgg gtggttacct 25800
tcctggtgcc gctgggctcc gcccccggct ccggaaaagg cgtcctgctg ttcctgatca 25860
aagacagcat ttaccagtcg ctctttgccg atgccatcaa cgaccatatc aacacctata 25920
tccaccaggg cgatgaactc ctctcctcct ctgaggagct gtcccttccc cgggatcagg 25980
tgaaggttgc ccaggaaacc gcctccagcc tcttccggtg ggacaaccag gactggaccc 26040
aggttacttt ctccggccgg aactgggggc tgtcctatac ggcggttctc cgcaacgccg 26100
atatcaattc ctccatcatg cagcagcttt tcagcatcct ggccatcctg ccggtctttg 26160
ttttcctgag cctgctgctt gccctgtgga tggcgcggcg tcatgcccag cccatccggg 26220
cgatcaccgg catgctctca cgggaaacgg acgaaacgcc ccgggatgaa ctccagcaga 26280
tttcaaccgg catcagccgg ctcacctccc gcaaccagga gctgacctcc cgcctggacc 26340
gggccctgcc cgtccagcgg cacgattttg ttttcaaatt tgtcaaggac cgcttcccca 26400
cccgggaaga agcagtggtg gccggaaagg ccgtgggcct ggagatcgac cgcccggcct 26460
acgcggtgat cctgtgcagc ggttccgggg gaaacgacca tcccttcgac ctgggcgccc 26520
cgccctttga tcgcttcccc ggctgttccg gcgcaggggt ggaactggtc gccctgaaag 26580
ccaatcttta cctggtcttc tccgatgatc cggagaccct gccggacctg gcagaaacgc 26640
tccgcagcga aggcgccgcc ggcggggagc ggtgtatcac ctccatctcc gccctgcatt 26700
ccgatttcac ggaagctcct tccgcctacc tggaagcagc tgccgcctat gaaaaccggt 26760
ttgtcatggg cagccagaaa gtgctctgct acggcgatat ttcctccaac ctggaggaga 26820
tccttcccca ggcccggaag atcaccaact ccatcagcca ggcgctgacc ctgggaagcc 26880

gggatgtgct ggatgtccgg atcagcgacc tgctgcattt cctgaagaac accaacatgt 26940
cgcccttcgc tttccggatg atctacagtg aagtgatccg ttccctggtc cacacccata 27000
ccgccgccct ttccggggaa ggctctgtgc aggaaatcta caatatcttc acgctgactt 27060
cctgccagtc catcgacgac ctggacgagc tgctgcgccg gctttgcgac cgcctgctcc 27120
tggaggagcc ggattcctgc gaaatcgcgc ctgaggagga ggacgagatc agccaggtgg 27180
tccactatat ggaggagcac ttcagcgatc cggaaatgtc catcaccgcc ctggcagact 27240
ccctggagct ttccaccacc cgcttcagtc tttcctttaa ggaaaagacc ggcatgtccc 27300
ccctggaata cctgacgctc ctgcgggtgg agcacgcgaa ggagctgctg gaaatgaccg 27360
acctgaccat ccgggatatc agtgcccggg caggctatta tgattccggc tccttcatcc 27420
gccgcttcaa gcaggtcacc ggcgaaacgc ccgcccagta ccgccgcagc cggaacctga 27480
aaaaggaaac gccggaaggc ggaaaaaact gattatcatc catcaaggag ttttccatga 27540
ttcacacaga ggaaatcaac gtccgggatc cgtttgtgct ggtccacgaa ggacagtatt 27600
acctgtacgg cacccgcggg gctacctgct ggggacctgc ggacggcttt gacgtctata 27660
ccggccggga cctgcacacc tgggaaggtc cggccgtctg tttccataat gacggcacct 27720
tctgggccga ccggaattac tgggcgcccg aggtgcatcc ctggcagggc atgttctata 27780
tgttcgccag cttcaaaagc gacgcgcgca ggcgcggaac cgccatcctg cgttccgcct 27840
ctcccctcgg tccctttgaa ccctggtccg acggccctgt caccccggca gactgggaat 27900
gcctcgacgg caccttccac gtctcctccg acggcgtacc ctatatcgtg ttctgccacg 27960
aatgggtgca ggcaggagac ggtgaaatcc tggccatgcg cctgaccggg gacctgaagg 28020
ctgccgcgga aaacccottc ctgctcttcc gcggctctga cgctccctgg gcccggccgg 28080
tccatcactc ctccggcatc accggctatg tgacggacgg acctttcctg tggcgccgga 28140
aggacggaac cctgctctgc ctgtggtcag gattctcgga aaaaggctat gcccagggcc 28200
tggccgtttc ggacaacggc gaaatagacg ggcacttttc ccagatcgat cccctctttc 28260
tggaagacgg cggccacggc atgctcttcc gggacctgga aggcggcgtg tacctgaccc 28320
ttcacagtcc gaacacccac ctgctggagc gtccccggtt tatcccggtc agcctgtaaa 28380
attcttattc cgttgaaagg atatccggag ggaaaggcga acaataatca gtgcctttcc 28440
tccggttttc atcaatcact ctcaggaggc agcatatgaa tcagacaacc aagaatgttc 28500
gcgaactcac ggtagcctat atcggcggcg gttcccgggg ctgggcctgg ggctttatga 28560
aggaccttgc cctggacggc gaaatctgcg gtaccgtacg cctgtatgat attgaccggg 28620
aagccgcgga gcggaaccgg atcatcggtg aaaaaatcag cgcccacccg gacacggcgt 28680
cccgctgggc ctacaaggtc agcaattccc tgcaggaagc gctgaccggg gcggattttg 28740
tggtcatctc catcctgccc ggcacctttg acgaaatgga atcagacgtc cacctgccgg 28800
agcgcctcgg catctggcag ccggtgggcg acacggtcgg cgccggcgga ttcatgcgcg 28860
ccatgcgtac catccccatg tttgtcacca tcggcgaagc catccgggat tacgctccgg 28920
aagcctgggt tatcaactat actaacccca tgtccctttg cgtgcgcacc ctctacgagg 28980
tctttccgga gatccgggcc tttggctgct gccatgaggt tttcggcacc cagaagctgc 29040
tctgctccat gctgaaaacc gaagagggca tcgacggcgt gaagcgccag gatctgtata 29100
ccaccgtcac cggtatcaac cacttcacct ggctgacctc tgccacctgg cagggacgcg 29160
acctgttccc cctctatgcc cgcttcgtgg agaaatacgc ggaaagtggc tatgatgaag 29220
gcggggacga caactggatg aactcccact tcaactgcgc ccagcgcgtg aagtttgacc 29280
tgttccgccg ctacggcgtg atcgccgcag ccggagaccg gcacctggcc gagtttgtgg 29340
ctccctggta tacccggaat ccggaaaccg tccggaaatg gaaattcagc ctgacccccg 29400
tctcctggcg gaagcaggac ctgaaggaac ggctgcagat gtcggaagac ctggtcagcg 29460
gaaagaagga aatcaccctg aacggttccg gggaggaagg ccacctgctg ctgaaagccc 29520
tcctcggcct cggggatatg gtgtccaatg tcaacatccc caaccgcggc cagattccca 29580
acctgcccct gggcgccgtt gtggaaacca acgccctctt cgggctgaac cgcattgatc 29640
ccgtatacgc ggggccggtg cccgcctcca tccttccgct catcacccgg catgtgctga 29700
accaggaaaa caccctgacc gccgccctcc gctgcgaccg gaagctgggt ctcaccacct 29760
tcctgaatga tccgcagctg tccggcgtca ccccggcgga cggagaaaag ctcttcaacg 29820
aaatgctgga aaaccagcgc gcctacctgc cggtaaaatg gtttgaataa ccctgaatat 29880
caggatcaac aaaacagctt ctgagttttc tcagaagctg ttttttccgt cattcttcat 29940
tcttcatttg gagggtttcc tccatcgccc ttccctccag cacccgtctc cccgtttctt 30000
tcaccgattg atttataagg gatatgactt gtttttttatt caaggatagt gtatgatgag 30060
ccagtaactc agtatgtagg agggatttta ccatggctga acagtatgtt accggtgaaa 30120

cactggttgg cgaagttgtt accaagtacc ccgaagcgat cgagatcctg ctgtccatcg 30180
gcatgcactg cctgggctgc cccgcttccc gcaatgaatc cctgcaggat gcctgcgcgg 30240
ttcacggcat tcccgcggag caggtcatcg aagcaatcaa cgaaaagatc gctgagaaca 30300
agtaagttct gaccgacaat aaggccggag ggattttccc tccggccttt gttatgccct 30360
gattcagttt tctgacggca cggaagccgt caggctttcg taggtcacac cgtatttttc 30420
cgcgatatcc ctggcgaaag actgtccttc cggcggagcg atcacaaccc ggaaggaatg 30480
ctttccttcc ctggtttccg ccaccagaga aaccgccttc ccttcggctt cgcccgtatt 30540
gatcaccgtg tccaggtcct gggccagctt gtgcatgtgc gtattgtaga ttgtgcgggt 30600
accccggccg aggatggccc ggacggcatc cgccgcaata aagtatccct cttcaaagga 30660
tgtggtggag aaggtctcgt tcaggagcag caggctttcc ggtgtgcagc tgacaaacag 30720
ttcccggaac cgccggcatt cctcgcccag ccgtcccagg tccagcgtct tgtcctcatc 30780
cgcggggaaa tgggtcagca cccggtccgc cgggtcatac tcaaagcttt ccgccggtac 30840
cggaataccg ctctgggcca ggataaacag ctggccgacc gcctgagtca ccgtggtttt 30900
gccgccccgg ttggcgccag tcaggacgta tacccgtttt tccggatcaa agaccaggtc 30960
attgggtacc acgttctccg gcttttcggt ggcaatcagc ttcaggttgt acagtcccgc 31020
tgccttcatg caggccgccc cctgccggtt cagcctggcc tgcggcttgc agaacttcca 31080
gcctgctttc tgtttcttct caatgtattc cgcccaccgg gtataaaaca ccagttccgg 31140
tatcaggtcg gcgatttcct tgacgctcac attcaggtac tgtcccagtg tgtcccgcag 31200
cttccgggca atccgggaag tcagcatgga cgccgcgccg tccatctgcc gcggcacatt 31260
ggcagtgccg tccgccgcgg ggatccggga cagggacatg gccgccagcg gattgcggag 31320
aatcaccgag ctctccacaa actgaccgac agactgaagc agccccacct ccgtattggc 31380
gggataatag ttatagcttc cgctccagtc cgcctctttc cggatctcat ccctggggga 31440
catggacgcc aggaagttct tcaggatgct ggacttcaca aaaggctttg cgttgacgga 31500
aaccagtccc atactgatcg cctcaaagcg gtcattcagg ttgatgccta ccgtcaggct 31560
ccggatttcg gaagcggcaa cccgcatttc ctccacgtcc ttgcgcagtg cggcaaaccc 31620
tttttcctgg tagatctgct ccacggtgtc ccgcagcgtg gtcagacctt ccgaaaccag 31680
atccttatcc gacaggcact cccggatggc ttccacggtc aggatatagt cgtgatactc 31740
ctccagccgg tgcatcaggt cccagatgcc ggtctcatcg ccctcatgac ggttgaccac 31800
cccgtagtca tagaacatct tgaccttatc cagcagcttc atcagccgct cccggatttc 31860
cggatggcgc aggatgtcgt caaaaacgtc gctccggaaa gccgccactg ccgggtccgc 31920
cgtcaggctc atcatcaccc ggcggagcag cggttcttcc tggggctgcg ccgccacttt 31980
ttccgtcagc gcgtccatac ccaggtcgtg ccagctttct tccgggattt cccggtaggt 32040
cacttcatcc tgccggggaa acaatatact cagcttcgtc tgcttcattt cattatcctc 32100
atttgaacgg ttcttgcgga tctgattata tctgatccgc ccgctcccgt aaagtccgcg 32160
cggcggatga cagcctgtac gcaatatgat acaataatcc ccggccggaa aggaggcctt 32220
tgccatggga aaacaggaca aaaccaactg tatgcgggta ctggacagca aaaagattgc 32280
ctacagcgct catttatatg aagcggatcc gaccctgacc ggggaagata tcgcccacct 32340
gctgaatgag gagccctccc aggttttcaa aaccctggtt accgtcggca agcctcagaa 32400
gtattatgtg tttgtgatcc cggtcaatgc agaactgaac ctgaaaaagg ccgccgctgc 32460
cgccggtgaa aaatccgtga gcatgattcc ccagcgggat ctgcttccgc tgaccggcta 32520
tgtgcacggg ggctgctcac ctatcggaat gaagaagcat ttcccttcgt ttattgatga 32580
gagtgcttcc ggtctctccc ggatctttgt cagtgcgggc cgcgtcggct gccaggtgga 32640
actggctccc gcggatctga ttcagattgc ttccctgacg cccgcagccc tgatttgatt 32700
gactttccgc aaagcggagt gataaactgt aatgtaatgt tttttgcatt tttaccgcca 32760
ttcccggcgt taactgatgg aggttcctca tgacgaaaaa gcgtaccctg ctgtggctga 32820
tcccgctgat tgtgctgatt ctgctgatcc cgttcctggt tccttccgcc ctgccctatg 32880
cgggtgcaga agagatgcct gttttccctt ccgtggagct gtccaacccg gaaccggaac 32940
cggttgtgct ggagccgaat cccaagtaca agaattccat ggtctcccgg tatgccccgc 33000
acaaggacgc cttcacgtcc gaacccctgg gataccagga cggtaccatc tgcgtaaaga 33060
tcgagaagag aacggtcgtc aacggcaacg gcggcaagac cgcggtgtat ttcacctggg 33120
taaagattgc cgatccttcc cagctccgca ccagcaccaa ccagccttat ccttccaagc 33180
agctcgtcgc cgcaacctcc ctcgcccgtc gtgaacgctc cgtcctggcc atcaacggcg 33240
actatttctg cgaccggacg gaaggcatcg tctaccggaa cggtgaactg ctgcggaaca 33300
gcgttttcaa taccggttac ggatatgatg cattgattat cgacctgaac ggcaatttcc 33360

atatcctgct gaaacccgat cccagcgact tcgaacctta cgaaggctgt atcgcccaca 33420
gcttcatctt cgggcccgcg cttgtcgtcg acggcaagat gcaggttctg gacggcaaca 33480
attatgcaag ttcccccggc atggggctcc ataaatacct gcagcgccag gccatctgcc 33540
agatggataa cctgacctac ctgatgatca ctgcggaagg tcctgaacag tccaaggacg 33600
gcggcctcac cggccaggaa atggcccagc tggcctatga ctgcggtgcg gtccaggctt 33660
ataacctgga cggcggttcc tccagctggc tggtgctcgg taccgaccgg atcaacgcca 33720
cccgcggcaa gaagcgggat attcaggata tcatttactt tgtcactgct gaagcggaac 33780
cggttcccgc ggaataaccc aggagctgtc ccacagccca ccgtcaacaa aggaagcgtc 33840
agcctatgaa taccatcctg tccatcggac ccctgaacct gaccccctac ggactggcag 33900
tgctggccgg cgtactggcc ggtgcattgc tgtgcctgct gaacaaaaag gtctttcagc 33960
tcctgccgct gacaatcctg ggagcggtcg tcctggggca tgccttctgg actttccttc 34020
gtctcgacgg ctatatggat gaaatcagtt ccttcgcctt catgttccag ccctggctgg 34080
gcggctatac gctttacggt gccctcatcg gcggcaccct tggtgcgctg gccggtgcga 34140
agctgagcgg catccgtccc ctggaagcac tggacaccct ggctcccgca gcctgtgcag 34200
tcctcttttt ctgccgtctt ggagaatact ataccggcga gggtgtcggc gctgaagcgc 34260
tggaatccct ctccttcttc cccattgcca gcgaagggct gaattccggt ttctggctgt 34320
acgcggtctg gttctgggaa gccatcgccg cgttggtcat cctgatcctg ctgctgtgcc 34380
gccggaagaa cgcccgcgaa ggagagctta ccgtcctgtt cgttacgctg ctcggtacaa 34440
gccagatcct gctggaacag ttccggaaag acaacttcgt ccgtctgaat tccttcgttc 34500
gtttcaccca gatcgccgcg gtcctgaccc tgctggccct gatgatcgtc ctggtcatcc 34560
gccgtaggcc cgaccggaaa aagaccatcc tgagctttgc ggtgctcgtt gccgctgccg 34620
gcgcggtcgt aatggctgag ttcgtgtttg acaagcccca gatggaaacc ctgctgtata 34680
tcgcgctggc cgccacagcc gtgctgttcg gcctgatgct catcgccttc cgggggaaag 34740
ccggaaagct gcccgccggc ctgttcggcc tgttcaccgg cacgctgatc ctgctttacc 34800
tgctggccac ggtcgatccg gattcctccg gattcctgta caccattctg ctggacctga 34860
tgatgctgtc ctccctgacc gccatcggcg tcaccttaat gaccttgtac cattccgaat 34920
ctgcactgga aaagtaaata tgaccctgac aaaggaggaa actcttatgt cccagaagtc 34980
ttcccccgtc tatttcaccg atttccggac cgtagctgac gtcagccagg gcgtcaagct 35040
ccagaagctc tgccgccggg ccggaataga caagatcgat tttgccggga aattcgctgc 35100
cattaaaatg cacttcggtg agttgggcaa ctttgcctat ctccggccga actatgtcaa 35160
ggcagtggca gacctgatca aggaactggg cggaaagccc ttccttaccg actgcaacac 35220
cctctatccc ggcagccgga aaaacgccgt agaccatctg tacaacgcag aagtcaacgg 35280
ttttaacagc gtgaccaccg gctgctacat catcatcggt gacgggctga aggggacgga 35340
tgatatcacc gtaccggtcc ccggcggtga atactgcaag gaagcctata tcggccgtgc 35400
cctctacgac gcggacatca ttatttccct gaatcatttc aagggtcatg aaatggccgg 35460
cttcggcggt gcggtcaaga atctcggtat gggcggcggc agccgggccg ggaaaatgca 35520
gcagcacagc gacggcaagc cggttgttga ccaggccctg tgccgttcct gccacaagtg 35580
cgcccgggag tgcggatcgg acgccatcac ctataaaagc ggcaaggcct atatcatgca 35640
ggatatctgc aagggctgcg gccggtgcat cggcgcctgt gcctttgatg ccatccatcc 35700
gtcttttgac accgcggcgg atatgctggg ccgcaagatg gcagaataca ccgccgcgat 35760
ctgtgccggg aagccttcct tccatatcac cctgatcatg gatgtttccc ccaactgcga 35820
ctgccacggg gaaaacgatg cgcccatcct gccgaatatc ggcatgctgg cctcctttga 35880
tccggtagcc ctggaccagg cctgtgcgga tctgtgcctg gccgcggaac cgctgcccaa 35940
cagccagctg tctgacaacc tggcaaaacc cggctggaag catcatcacg atcacttcaa 36000
ggacagcaac ccgaacgtca gctggcggga aaccctcgcc cacggggaaa agatcggcct 36060
gggtacccga acctatgaac tgatcaaaat ctgatcagcg cgaaataatc agaaaaattc 36120
agcattttga agtatttaag agctttctgc ccgttttcgg cagaaagctc ttcttattga 36180
ctatttatgt ctttgactat ctttgacttt taagtataat ataatcagca caagaaagag 36240
agtgcggtgg ccagtgacca cggaaagtaa ttccccggaa ccgcaataag caacggaacc 36300
ggatcaaaag aatggagggt tttatatgag tactttcctg cctgctgttt tcggtgaaaa 36360
tatgatggat ctgtttgatg acttcgaccg tgatttcttc cgcggcttcg gccgtccgga 36420
aagagcgctc tacggtaaga acgccgcccg gatgatgaaa accgacgtga aggaaaccga 36480
cgaaggctat gagctggcgg tggatctgcc cggtatgaag aaggatgaaa ttcatctgga 36540
tcttcagaac ggctacctga ccatctccac ccagaagaac ctggagaaca aggaagaaaa 36600

```
     gaacggcaag ctgctccgcc aggagcggta taccggcacc atgcagcgca gcttctacgt     36660
     gggcgacaac ctgacagagg aggatgtgca ggccaaatac gaagacggtg tcctgaccgt     36720
     caagcttccc aagaaggaag ccaagaagga tcccgaaaag aaacaaatcc tcattgcata     36780
     atctccaacc aacaaaggag cggctctctg agccgctcct tttatttttc agtttttctc     36840
     acatccggcg tcagccggat atttcatatc gcgaagcgat atttcacatt gagccgcaag     36900
     cggctcaata tttcatactg ccgaaggcag tatttcattt tcaaccggac ttcttcacag     36960
     actatcaata tacctgcagg ctgccagcgc tgctgtcgca ccgtccgccg ccgcggtagt     37020
     cagctgacgt accgtcttgc tgcggcagtc cccgccacaa aagacgccgg gggtctttgt     37080
     ctcgcagcgc tcatcgctgt caaagtatcc ccacttgttc agatccgcca aatgcgcgaa     37140
     agcatcgttt tccggaatca ggccgatggc aacaaacagg ccgtcacagg caacctgccg     37200
     ttcttccccg ccgtcctgcg gcttcacggt cacgctccgg agaatgccct cctccgtgtt     37260
     caggcctgtg atcttcacgc cggtgtattt ttccacattc ggacggctga acaggatctc     37320
     ctgcagcttc ttttcaccgg tgaattccgg caggtcctgc agcataatca ccttttcgca     37380
     cttaccggac aacaggatcg cctcctgcag ggcggagttt ccgccgccgg ccacgcatac     37440
     ggtttttccg gtatagaaat ccccgtcgca cacggcgcag aagctgatgc cctcgcccac     37500
     cagctcgttt tcccccggca ggcccagcat ccggtgcttc acgcccgccg ccacaataac     37560
     agcctttgcg tcccgctcca ggccgtcgtc tgtaaatacc cgcttatagt ccccgtgatc     37620
     ttcgatccgg acaacctccg ccaggtccac ttccgcgccc tggttcatga tctggtccag     37680
     cagcgcctcg gcgtattcat tgccgctcat ctgggcggta cccggccagt tttccacctt     37740
     ggggctgtag gtaatctgtc cgccgaaacc gtgcttttcc agcaccagtg cgttctttcc     37800
     gttccgctgc gcgtacagcg cggccgtcat gcccgcgggg cctccgccga tcacgatgat     37860
     atccgtcatt ttttcaccca tcctttgaaa aagggacgcg ccgcgtccct ttttctcatt     37920
     ttgtaatcct atcggagtgg aacgaaaacc ataatccggt ttccgacttc ctacctccta     37980
     cttcctacct catacctctt attatccgac ctgttttgcc gtcagcatac ccttgatttc     38040
     ggatacaccc ttgtacttgg tgtattcatc gggattctgg ccgagcacca ccagcgtggg     38100
     cgcctgcttc acgccgtact ggttggtcag ttcgggattg tccgtcgcca gcaccttctt     38160
     gaacaggaag ccggccttct ccagcatact gattgccagc ttgcagttgg ggcaggtggc     38220
     actgacaaac aggatcgcgc gggcacagtt gcagcccttg ttttccttct caggctgggc     38280
     agcgggctct tccttcacgg gttcggctgc cggtgcggtg gccatcacag ggccggtatg     38340
     ggtcaggcgg ctgtgaccga tgtcataaac cttgcggtct ttgaattcct gggccttgcc     38400
     gtcgttccag ttctgcacag gacgatagta accggtgatc cggctgtaaa cct             38453
//
ID   XXX; XXX; linear; XXX; XXX; XXX;  XXX.
XX
AC   PRJEB32780
XX
DE   Uncultured bacterium DNA fragment from in vivo enriched cow rumen, clone
rum-12f23
XX
FH   Key             Location/Qualifiers
FH
FT   source          1..25530
FT                   /organism="uncultured bacterium"
FT                   /environmental_sample
FT                   /mol_type="genomic DNA"
FT                   /isolation_source="Cow rumen"
FT                   /clone="rum-12f23"
FT                   /note="AZCL-xylan, AZCL-arabinoxylan, AZCL-Barley beta
glucan active clone"
FT   CDS             complement(729..1757)
FT                   /translation="MTILRKIRKILGSMPLAITLLVMLAAACAVCSAVPQGQSQEWYAS
FT                   QFGERTAAVIIALHADDAYHSWWFLTLSGFLCLSLLMCNLIRVKPLIRRTRKEPGNIAG
FT                   IWGAWVCHLGILLLIIGFALGQMTQEEYAVSGLPGDTLPLAETGLQVTVDDFTIDWRED
FT                   GAAGQYTAELTVRGADGTEEHGTASVNHPAELCGYEFFQNSAGWAADVTVEKNGEILQK
```

```
FT                   KTLCVQEYMPMADDPDIVVMFYDFYPNYDGETGTVSPMASTRPEKPGYLYMVFYQGEIR
FT                   GMNVLEADDEIIIDQEYTVRFSNPSNYTLLMVKRDRFSWMAMLGAAIVMAGLVMAFYLR
FT                   KR"
FT                   /product="Cytochrome c-type biogenesis protein Ccs1/ResB"
FT   CDS             complement(1800..4724)
FT                   /db_xref="GO:0000104"
FT                   /translation="MKKLICLLLAMTVMLFAASAWAEPAPAETLFTPGTYQAEAQGMLS
FT                   TVKVWVTVDEQEIIAVLIDATDETETLGGVAAGMLSAAIRDAQTPNVDTISGATVTSKA
FT                   VIQAVTEALVAAGADIDVLDANRRDAGGDSGAKPEKTIDTEIVIIGAGGAGMTAAIMVK
FT                   QAGKDFVLLEKMPYAGGNTTKATGGMNAAETHYQKEQGIEDSVALFVQDTITGGHDLND
FT                   PALVEVMAKESSNAIDWLDSIGAELPKISFSGGASVNRIHAPADGSGVGEFLVERFSAK
FT                   LDELGVEIMYNTAATELLTDAEGRICGVKAEGKDYAYTFNCKAVILATGGFGANEEMYT
FT                   QYRPDLKGTVTTNAPGATGDGIVMAQALGADLVDIEQIQLHPTVEQSTSMLITESVRGD
FT                   GAILVNQSGKRFTNELLTRDAVSAAELEQEGHYAYIIFDQNLRDHLKAVEKYVKTGITV
FT                   QADTIEGLAELISVDPAVLAKTLADWNECVKNQEDPEFGRTTGMNADLTTAPYYAIKIA
FT                   PGIHHTMGGVKIDTSAHVINTEGNVIPGLYAAGEVTGGVHGGNRLGGNAVADIVIFGRI
FT                   AAETALSEMDGTAVPETAEEPAAEPEPAKEETAQVPPSVISRVAVTMGQNSENNIVTSY
FT                   LEQDPYLTNIYEGYGFAKDYGSARGHTYTLEDVSKTLRPHAKANCITCKTDDFARLVNT
FT                   EGIEVYSRAFDDVFPQMKNPISCYTCHGDGMGMDGQLTVSHSYVKTALGDAAAEIPAAI
FT                   LACGQCHIEYYFTPSDSETMMPYHSKAEMTPEAILAYYDAMGFYDWEQPGTGTKMLKAQ
FT                   HPEMETVLAGKHAANFGMTCADCHMPRETTDDGIEYSSHYFVSPLTNETLLNSCTKCHG
FT                   TAENLISLVQNVQVKVTARETEVGNKLSALKDKLTEAAAGGKLSEEQLEAVRKLHREAQ
FT                   WYFDFCYVENSEGAHNSSLSLTCLETSEAKIDEAMALIGEEQGV"
FT                   /product="Fumarate reductase flavoprotein subunit (EC
FT                   1.3.99.1)"
FT                   /EC_number="1.3.99.1"
FT   CDS             complement(4762..5940)
FT                   /translation="MNNFEKKQWLRLILCALVLIAASLFLTGIIGGGHMTYTLTEQTDA
FT                   PAVSGTAVEETEQGFGGTVTVHATLDGKTVQALTIDTPDETPGLGQRASEEAFTEQFIG
FT                   KEGPFTNGENGVDALSGATVTSNAVLKALNRAITGEDAAEPVQEAAEEPAPVAEEEPAA
FT                   EEPAAEEPATEEPAAAESETAGTAVEETEQGFGGNVTVKATLDGKTVKALTIDTPNETP
FT                   GLGQRASEEAFISQFIGKEGPFTYGENGVDALTGATVTSNAVLKALNRAITGEDAAEEP
FT                   KAEEAAAPAEEAKAEEAGRVIYGSYMAKRETNFSTIRVYADSKGGKITDCRITSEAKME
FT                   NSDFLTEEIREAWAKAIVENGTADVDVITGATLKFSSEAVVDAVNEILGRVQ"
FT                   /product="Electron transport complex protein RnfG"
FT   CDS             complement(5953..6747)
FT                   /translation="MNIILVTTLVIAVIGVCVGAGLVFTGKKFAVEVDEREVAVRSKLP
FT                   GNNCGACGYAGCDALAAAIAKGEAPVNACPVGGAASAEAIGDIMGITAGALERKVAFVA
FT                   CKGPCDVTRNQGNYIGIKDCRAAVLAGINVTDCNYGCLGFGSCVEVCPEKAISIRNGVA
FT                   VVNRNRCVSCGLCAKTCPRGLITLIPESRIVRVQCSNRDKGPQVKKVCTAGCIGCGLCV
FT                   KQCEFEAIEFDGTLARINPDKCTLCGKCAAKCPVKAITTTAV"
FT                   /product="Electron transport complex protein RnfB"
FT   CDS             complement(6762..7340)
FT                   /translation="MNGLIMIMVGSALISNVVLNQFLGICSFLGVSKQMKASASLGVAV
FT                   IFVITIASAVASLLYDYVLKPLGMDFMKTIIFILVIAALVQIVEMFLKKKSPGVYQALG
FT                   ISLPLITTNCAVLGVALNNVQDGYNFIECVTSGFGTALGYTIAIILLAGIRSRINEKDL
FT                   PAPLRGAPIVMIAAALMSIAFMGFGGLPH"
FT                   /product="Electron transport complex protein RnfA"
FT   CDS             complement(7337..8095)
FT                   /translation="MKKYFERLWNGLVKENPVLVLMLGMCPALAVSTQATNGIGMGLST
FT                   LAVLVLSNLVISLLRKVIPDQVRLPAYIVIVASLVTVTELLIKAYIPGIYEALGIYIPL
FT                   IVVNCIILGRAEAYANKHTPLLSVMDGIGMGLGFTIALTIAGAIRELLGAGKIFGFQIL
FT                   PEGFGMTIFIQPPGAFLVFALIIAVMNAIGIKTRQRQLVESGCDGNCALCSTVCSDRKE
FT                   ETAGAEEQQKALQEAPKAIEAAVKEGGEKK"
```

```
FT                   /product="Electron transport complex protein RnfE"
FT   CDS             complement(8097..9638)
FT                   /translation="MSEALNISSSPHVRDRWTTPYIMRVVLLSLMPATCVGIIAYGWQA
FT                   FAIIALAVVSAVLTEWVFNKIVHKPNTIWDGSAAVTGLLLALSLGPRTPLYLPVIGSVF
FT                   AIAVVKGCFGGLGKNFINPALAARCFLLISFANAMSIKPILDATSSATPMGALKMYETV
FT                   DITRMFLGTADGVIGSSILALLIGGLVLWSMDIIHGQICFSVLGGFTLFIALFGGQGFD
FT                   IKFLLAHLCGGGVVMGAFFMATDYVTSPVSRLGQFIYGCLIGVLGGVFRLYGNTADSFS
FT                   YAIIIGNVCTPLIDTYIVQKPLAYRKIGKTRPVKREPFRIPKPVIALTLIAAIAGVALS
FT                   GVYSMTKGTIELQELQKKTAAYKEVCPDAEIFDVVDKAEEMIAAQDEKASPKINEFYVA
FT                   KSADGQVAGYAVSVTAKGFGGDVTMALGLTPDGKINKIAFTELNETAGLGMRADEDSFK
FT                   GQFAGKGGELSLVKGTAGGGQEISALTGATVTSTAVVNGVNAGLAFCEEAAKGGN"
FT                   /product="Electron transport complex protein RnfD"
FT   CDS             complement(9635..10984)
FT                   /translation="MRNTFRGGVHPEERKELSREARLRLFNPKGEMVFPLQQHIGKPAK
FT                   PLVQRGDEVLTGQKIAEADGFVSAHIYSSCSGKVKAIEKRRVLGGGMIDCIVIDNDGQF
FT                   TPFADFSQKESANGLSNDEIIGRVQEAGIVGLGGAGFPTHVKLKPKDPDAIQYIIANGA
FT                   ECEPYLTCNDQLMRTQSTAIAEGLELVLRLFPNAEGVICIEENKPEAIHAMQIAVSGKP
FT                   KIRVQEMITKYPQGGERSLIQAVTGVDFPISKLPADVGCIVQNVGTLYAIQRAVLYREP
FT                   LFSQCFTVSGDAVKEPGNFFVRVGTSYAELLEECGGIKEECKAAKALAGGPMMGIAIGS
FT                   LDIPIQKQNNGLTVMAEDPVEEAEKLMTACLHCGRCTTVCPVGLMPQLMADAAIAGDLA
FT                   RYEKKLYGLECIQCGSCTFICPAKRPLMQVFKQAKAEIAAAKKREGGNQK"
FT                   /product="Electron transport complex protein RnfC"
FT   CDS             complement(11020..12921)
FT                   /translation="MGSGKQQYRIAFFTVDWNYELVESTLHGLKQYVDDHEDVYIYIFD
FT                   CLGKDLDNAKDKSEYVIFDLPDLRKFDGVLVQGNQLVLKRVRDDIERRIVEAGVPAVSI
FT                   GAPMKGCALIRFDNRAAQKAIANHVIREHHASRLVYLAGNTENGSPEAGERIDGFVDAC
FT                   RENNIPEENVRIVPATWRTSDGADLAERWMQEGLELPDAFICANDEMAIGLMTALKRHE
FT                   VRIPGDVIVTGFDNLAGSELSSPRLSTVGGDNRNQNYRAMDMLISLIDGKDCGQEVITD
FT                   FHTICSESCGCPETARPGTIRELYFQQTRFLKDFYNLQGQMAEDLFECNDLQELAETVG
FT                   KNRRIFGCQDIYLCFNDYYFDHFDKSEWPEEEKPFGEEMILTVRKRGCRSGDGNEKYDF
FT                   IRFRTEELLPEPLLRRERFMMFYPLHYNTYSIGYIALDGISEAAKMNLHESIFNFLEIA
FT                   IENVRKKELLRQFNETLDELYVRDSLTGLYNRFGLKRFGQETFDWLMERDHGAQVLFTD
FT                   MDDMKLVNDLYGHNTGDEALKASAWILSECCDPEDVIIRYGGDEFVIIARWDEKDLKER
FT                   ILATADEYNRNSGMPFQLGFSIGTCSAEASEGKTMDDCIKVADARMYEVKTERKAGR"
FT                   /product="Sensory box/GGDEF family protein"
FT   CDS             13193..14566
FT                   /translation="MCPSAGAARSGSSTLVSGDYEYPLQDNGTAVITRFTGNTDVVVIP
FT                   ARLDGIPVSALGEKSFAWCPSLLDVTIPEGITSIGDYAFLYCTGLAAVFMPDSLTDIGR
FT                   NPFTACSTLADIIVSDDHPALLFEDGVLFSRQDYRLVTFTAARPDGPYSVPDGTRTIGA
FT                   SAFYSCRNLTALTMPESVSAIGQKAFGLCVSLQSVSLPDSVDFIGASAFFGCRGLREMS
FT                   VPASVTSIADRTFEDCTNLSELHLPKTVRSIGNRAFMNCSSLEAVDLPPRIREISRSAF
FT                   AGCESLSRITLPGRVTRIGDGAFLNCMNLESIDIPAGVRSIGRDAFSGCRCLREISLPP
FT                   ECGEIGTGAFRSCFALVSFRVPDSAETIPEEIFAGCTALAQVYIPSGVRSIGSHAFLAC
FT                   EQLTEIIIPDSVRSIGSQAFFGCLNLARVSLPDPPADIHPDAFEGCERLAGTIPDLVP"
FT                   /product="hypothetical protein"
FT   CDS             complement(14617..15501)
FT                   /db_xref="GO:0004622"
FT                   /db_xref="GO:0047372"
FT                   /translation="MCEISAHIYTPDDGFFETVMVIHHGMAEHQKRYEEFIRFLCDHGI
FT                   AVYMHDMANHGESCQDPELKGWFGKKGGWNALIEDYREMVMRARQEHPACRLIVMGHSM
FT                   GSFIVRMYVSRYPKDGFAGAVFMGTGGPNPAAGAGKAVSAMIGAVRGKKHRSSLMNTMA
FT                   FGTYGNKFEGRTPFDWLTRDKAIVDRYIADPYCGFMFPVQGMNDLVRLNAASNTAAWAR
FT                   DIPAELPILLVSGGLDPVGDYGAGVQKVGELLAETGHTRVTTKIYPECRHEVLNEINRD
FT                   EVMNDITEWIRGI"
```

```
FT                   /product="Lysophospholipase (EC 3.1.1.5); Monoglyceride
FT                   lipase (EC 3.1.1.23); putative"
FT                   /EC_number="3.1.1.5"
FT                   /EC_number="3.1.1.23"
FT   CDS             15679..16608
FT                   /translation="MSTFSVTLGNVLLTLLYMSAGFLLSKAKKVVPDHMSSISSILLYI
FT                   CGPCMFLDTIVNMDYSPSQTVSMLLFLVITLAVMGLVLLLIFLLLGKRRKDFAFRMMAI
FT                   ASVMGNVGFFGLPIVKAVFPAAPEAAAYSCIFCASMNIVAWTLCVFILTDDRKYMSVKA
FT                   SLLNPTVLPVALAMILYALGAKNRMPQLVLGGIHSVGSISTPLCMFILGIRLSTMTMKS
FT                   FFTSPVPWLASFGKLLVFPLISYGIVLLLPLSDVFRASILLLSATPCASIILNLSEIHH
FT                   HGQDIAARCALLSTLLSVLTIPLLSLLL"
FT                   /product="Malate permease"
FT   CDS             complement(16611..17753)
FT                   /translation="MSHTIKRVLSALVIVISITAVIIIAFSNPEMKNAWKALASMDLKW
FT                   VLGLFLCWAAYAGFEGCGTWSYLRSKGYRINLGRVIGVALIGFYYSNITPSAAGGQPMQ
FT                   VNSLRKAGIPVAYGTTAVTIRFVSNQLTICLMTLGLMIANRTFAYNQLGGLIWAARIGW
FT                   AINFIAVPLVLLAAFKRNLIQAVTERLIALLAKMHLVRDKNATIAKISEILDSYHAAMI
FT                   ALIHSPGQILLQVLFSALSMLGMTGSIVFTYHAFELHGTPWYQILTMSCLLFVSASYTP
FT                   LPGASGAQEGGFMAYFAGIFTDGTIGVGLLIWRFFTYYMFLIVGVGTVLNDKRLIRLEK
FT                   KRRQAEERLSGTAEDHAGETGDASPETPEPAGGPEGNGKA"
FT                   /product="conserved hypothetical protein 374"
FT   CDS             complement(17814..19889)
FT                   /db_xref="GO:0031216"
FT                   /translation="MVVSQNVLSAVFSDETPDYRFPAEPDPGDSVTIRIRVAKNSAKRV
FT                   IVIFDSMSVGTQMENTRSDDYFDYYETSLVCNDQEVMYRFLIECHDDGRIAYDKSGARL
FT                   EEGGTPDFNSVYAFRFIPGFHVPEWAKGAVQYQIFTDRFCNGDPSNDVTDNEYYYTIGH
FT                   AKHISEWDALPTDTDIRSFYGGDLQGIMDKLDYLQDIGIEVLYLNPIFVSPSSHKYDTQ
FT                   DYEHVDPHFGVITDDVEHEMQLWEKHNGFAPRYIRRVTSTENLEKSDELFARLCQEMHR
FT                   RGMRIILDGVFNHCGSFNKWMDHEGIYLGKAGFLPGAFQSVHSPYRHYFHFNESGNGRS
FT                   PAYEGWWGYSTLPKLNYEESPELREQILKIAEKWMSPPYSIDGWRLDVAADLGHTRECN
FT                   HSFWQDFRKRVKAANPDALIIAEHYGDPTEWLNGREWDTVMNYDAFMEPVSWFLTGMEK
FT                   HSDSYRDDLYQNGSAFFGIMREKMAHLKYPSLITAMNELSNHDHSRFLTRTNRMVGRMT
FT                   TVGSGAAAAGINKGIFREAVTIQMTWPGAPTVYYADEAGQGGWTDPDNRRTYPWGHEDQ
FT                   DLISLHRDLIALRKDLPVLACGSLKQLLADYGRIAYARFNGNCRCIVAVNNTGGWTDFR
FT                   LFVRDAGAKDGETYFRRILTTQESHETGCEPLGAATDGYLTFDMPPYSAAVISNRE"
FT                   /product="GH13"
FT                   /EC_number="3.2.1.135"
FT   CDS             complement(19925..20746)
FT                   /translation="MEEQPLVSIVIPAYNAEKTLKRMMDSVTGQTWQNTQIILVDDGSR
FT                   DNTLQIAREAAERDARIQVHTQENKGVSQARNTGMRFCTGKYIRFLDADDTLPPESTEI
FT                   LVNKAEKEESDLVIGGYKECLGSREFRFNLENRDDTMTTLEMLPMLCRKANTYFYGVLW
FT                   NKLFRRDIIEDMGLVFQDGLTYGEDFAFHVTYLKKAERISFVKEYLYDYHRSMGSMTFR
FT                   QAFSCVTHPLRNIQTKVRLYHFLKDMYICRNEYGKYRKKLWLYLFKVGLDK"
FT                   /product="GT2"
FT   CDS             complement(20895..22409)
FT                   /db_xref="GO:0031176"
FT                   /translation="MTAAHAGQETLVYASAFGAGTDLWYARGAQKVYHTTEATLRTEGR
FT                   TSDWHSPGRNFELAAGTRYDMSVEVLQNEAETATLMVSVQVTKEDGSPDWKNLFKGEAK
FT                   KGEWTTLSGSYTPDPSEDYILYVETVGAPELSYEIRNFRVVAPDGVPDVQPTATPAPEF
FT                   EAVEEMPSLKDLYAGKFDFGTALPRNAFNDMKLLMLVKEQFNILTPENEMKPDAILDVY
FT                   ASRKLATEDETAVAVKFDACKTLLRFAQANGLKVHGHTLLWHNQTPEALFHEGYDTSKP
FT                   LASREVMLGRMENYIRQVLTWTEENYPGVIVSWDVVNEAIDDGTNKLRTRANWYKTVGP
FT                   DYLARAFEYARKYAAEGVLLYYNDYNTAYSGKLYGIFDLLKSLMAEGNIDGYGFQMHHS
FT                   LGEPSMDMITRAVEKIASLGLKLRVSELDINAGKPTEQNFQSQKNKYKQVMKLMLRFKD
```

```
FT                   QTEAVQVWGVIDTMSWRRDGYPLLFDKNMNPKPAFFGVAEAAQEDQ"
FT                   /product="CBM22-GH10"
FT   CDS             22395..22508
FT                   /translation="MGGRHSCHHQHHGCQYDPDSSHRSILSPAHPASVVPI"
FT                   /product="hypothetical protein"
FT   CDS             complement(22612..25479)
FT                   /db_xref="GO:0004022"
FT                   /translation="MTNSHKTEETEVKVKKIWDDADNQDGYRTDDVTVNLLADGEVIDT
FT                   VKLSEENNWSYGWTKLDKKAAGKDIDYTVNEDEVPEYTTKIDKAEDGTFTYTVTNSHET
FT                   EKTQVEVEKIWDDNDNAEGFRPESVTVKLLANGEEAGTATLNEDNEWKYTWTGLDKNSG
FT                   GAEIEYTVTEDQLENYKVPVIEKVSETGWAYTVTNSRDFEETEVNVTKVWEDAENQDGC
FT                   RPEVITVNLLADGETIDTVELREDDNWSFSWSQLQKYRNGEAIDYTVTEEPVENYETQI
FT                   TKAEDGTFTYTVTNSHETEKTQVEVEKIWEDKDNVKGFRPANVTVKLLANGEEAGEATL
FT                   DADNEWKYTWTDLDKYSEGEEIEYTVEEEQAAGYLKPVIMKTAEDAWVFTVTNTQTEVS
FT                   VSKVDIANGDELPGAHIEILREATEADAEDENTKYADETKTLVIVEEWISEKEPHVIEG
FT                   LLTDTEYILRETVAPNGYDIATDTKFTIDEEGNVTSTGSVTEDGILLVEDSMLFVTAAV
FT                   KKVWDDGNNQDGVRPLTLRVNLLANGKKIDSVVLSMANGWMASIGNLPMVDADQKDIEY
FT                   TWEEEAPGNRYTLTSQEKKGTLTVITNSRPPEETGVSVQKIWDDSENAYRTRPESIRVQ
FT                   LYADGEACGEEVILNAANNWKAEWTGLDKFANADGQSGHAAEVIYTVEELNMPEGYKAT
FT                   VTGSAATGFVIRNVLERGKLIIEKTFEFGEVEPNEPDDTPMDIPVIKTWNDNGNKDGNR
FT                   PANVTVRLLADGTEIASAELNAAGGWRTVFTGLPRYNGEKKITYTITEDPVAWYEAEIN
FT                   GFNIRNNYVPEVTSATVRKVWDDNNNAEKMRPSSIHVTLSNGTTVLLSEENNWTATINN
FT                   LPTRVNGKPVTYTWTEQTVIAYKLSKTERQGNTTIFTNVFTQVPKIPEGPKKPTTPEGT
FT                   WQKFEEYTTPLGVQTIINHVGDCFD"
FT                   /product="Alcohol dehydrogenase (EC 1.1.1.1)"
FT                   /EC_number="1.1.1.1"
XX
SQ   Sequence 25530 BP; 4635 A; 8025 C; 6669 G; 6201 T; 0 other;
     cttccctttc catccgcgca taatccgcag atgcaggtat accgcgtaaa tgatccaggt        60
     caccagggac caggtttcct tcggatccca gctccagtag cttccccagc atttcttggc       120
     ccagaaagcc ccggtgataa tcgtaattgt caggaacagc aggcccaggc acacactgcg       180
     gtattcgatc acgtccagct tgtcccggtc cggaatatgc tggtccatga atccgtcctt       240
     ccggatcttt tcccgtacca ggaaaatgaa ccccagcaca aagctgacgc cgaaggccca       300
     gtaactcaga atggccgtgg agacgtggaa taccagccag ttgctctgca gtgccggcat       360
     cagcgcatga atttcacggc tctgcatggc ggcatatccc ataatcagga atactaccgg       420
     agcggcaaac acgccaagga aactgaaccg gtacttcgcg ataaagatca ggctcaccag       480
     gcacagtgcc cacgcaaagc aggttgcaaa ctcatactga ttggtcatgg gaacccgccc       540
     cgccgcgact ccccgcacaa tcagggcggc agtatgggca gccaggccgc ctgtctgaat       600
     cagggaggcc gcctttgccg cggcgttttt cttcagtccg gcaaacacaa agtacagcag       660
     catcgccagg aagtatccgg ccatggcaat cgtgaacagg atattctccg cctgttccat       720
     tcagatcgtc acctctttcg cagatagaag gccattacca gcccggccat cacgatggcg       780
     gcgcccagca tggccatcca ggaaaaccgg tcgcgtttta ccatcagcag cgtgtagttg       840
     gaagggttgc tgaaccgcac ggtatattcc tggtcgatta taatttcgtc atccgcttcc       900
     agcacattca ttccccggat ctccccctgg tagaaaacca tgtacaggta tccgggtttt       960
     tccggccggg tggatgccat cggggataca gttcccgttt ccccgtcata gttcggatag      1020
     aagtcgtaga acattaccac gatatcggga tcatccgcca tcggcatata ttcctgcacg      1080
     cacagggtct tcttctgcag gatctccccg tttttctcca cggtcacgtc tgccgcccat      1140
     cctgcggaat tctggaagaa ttcataccgg cagagttccg cgggatggtt cacgctggca      1200
     gtcccgtgct cctctgttcc gtcagcaccg cgcaccgtca gttcagcggt atactgcccg      1260
     gctgccccgt cttcccgcca gtcgattgta aagtcatcga ctgtaacctg caggccggtt      1320
     tccgccagcg gaagggtatc tcccggcagt ccgctgaccg cgtattcctc ctgggtcatc      1380
     tgccccagtg caaatccgat aatcagcagc aggatcccca ggtggcatac ccaggcaccc      1440
     cagattcccg cgatatttcc cggttctttc cgtgtgcggc ggatcagcgg tttcacccgg      1500
     atcaggttgc acatcagcag gctcaggcag aggaaaccgc tcagcgtcag gaaccaccag      1560
```

ctgtggtagg cgtcatcggc atgaagggcg ataatcaccg ccgccgtccg ctcaccgaac 1620
tggcttgcat accactcctg gctctgtccc tgcggaacag ctgaacaaac ggcgcatgcc 1680
gcggcaagca tcaccaggag ggtgatggcc agcggcatgg agccaaggat ctttctgatt 1740
tttctgagta tcgtcatatt accccgtgtt cagtccggcg gaggctttca cctccggttt 1800
tatacaccct gttcttcgcc gatcagcgcc atggcctcat cgatctttgc ctcggatgtc 1860
tccaggcagg tcagggacag gctggaattg tgcgcgccct ctgagttctc cacgtagcag 1920
aagtcaaagt accactgcgc ttcccgatgc agtttgcgca cagcctccag ctgttcttcg 1980
ctgagctttc cgcccgccgc tgcctcggtc agcttgtcct tcaaagcact cagcttgttg 2040
ccaacctccg tctcgcgcgc ggtcaccttc acctgaacat tctgcaccag ggaaatcaag 2100
ttttccgcgg tcccatggca ctttgtgcag gaattcagga gcgtctcatt cgtcaggggg 2160
ctcacaaagt agtggctgct gtattcaatg ccgtcatccg tggtttccct cggcatatgg 2220
cagtccgcgc atgtcatgcc gaaatttgcc gcgtgttttc cggccagcac cgtctccatc 2280
tccggatgct gggctttcag catcttcgtc ccggtgcccg gctgctccca atcgtagaat 2340
cccatggcat catagtaagc caggattgcc tccggcgtca tctctgcctt gctgtggtac 2400
ggcatcattg tctcgctgtc ggagggggtg aagtaatact caatatggca ctgtccgcag 2460
gccaggatcg cggcgggaat ttctgccgca gcgtctccca gcgctgtctt cacatagctg 2520
tggctgaccg tcagctgccc gtccattccc attccgtccc cgtggcatgt atagcaggaa 2580
atggggttct tcatctgcgg gaaaacatcg tcaaatgcgc gggaatatac ctcaatcccc 2640
tcggtattca ccagccgtgc aaaatcatcc gtcttgcagg taatgcagtt tgcttttgcg 2700
tggggccgca gcgtcttgga tacatcctcc agcgtatagg tgtggccgcg ggcgctgccg 2760
tagtccttcg caaacccgta tccttcatag atatttgtca ggtacggatc ctgttccagg 2820
taactggtca cgatattgtt ttcggaattc tgccccatgg tcaccgcaac cctggaaatc 2880
acagacggag gaacctgggc agtttcttcc ttcgccggtt ccggttccgc agcaggttct 2940
tccgctgttt ccggaactgc cgtaccgtcc atttcgctga gtgctgtttc cgccgcgatc 3000
cggccgaaga ttacaatatc cgccaccgcg ttaccgccca ggcggttccc gccgtgcacg 3060
ccgccggtca cctcgcctgc tgcatacagt ccgggaatca cattgccttc ggtgttgatt 3120
acatgcgcgg aagtgtcgat cttcacacca cccatggtgt ggtggatacc gggagcgatc 3180
ttgatcgcat agtacggcgc agtcgtcagg tccgcgttca taccggtggt ccggccgaat 3240
tccggatcct cctggttctt cacgcactcg ttccagtccg ccagtgtctt ggccagcacg 3300
gcaggatcaa cggaaatcag ttccgccaga ccttcgatgg tgtccgcctg gacggtaatg 3360
cccgtcttga cgtacttctc gaccgccttc aggtgatccc gcaggttctg gtcaaagatg 3420
atgtaggcgt aatggccttc ctgctccagc tcggcggcgg ataccgcgtc gcgggtcagt 3480
agttcattgg tgaagcgttt gccgctctgg tttaccagga tggcgccgtc gccccgcaca 3540
gattcggtaa tcagcatgga ggtcgactgc tccacggtgg gatgcagctg gatctgttcg 3600
atatccacga ggtccgcgcc cagcgcctgt gccatcacga tgccgtcacc cgttgcgccg 3660
ggagcgttcg tggtcacagt gcccttcagg tccggccggt actgggtata catttcctca 3720
ttggcgccga agccgcctgt tgccaggata accgccttgc agttgaaggt gtaggcataa 3780
tccttgcctt ccgccttcac gccgcagatc cggccctccg cgtcggtcag cagttccgtc 3840
gcggctgtgt tgtacataat ctctacgccc agctcgtcca gcttggcgct gaagcgttcc 3900
accaggaatt cacccacgcc ggagccgtcc gccggggcat ggatccggtt cacggatgcg 3960
ccgccggaga aggaaatctt cggcagctcc gcaccgatgg agtccagcca gtcaatggca 4020
ttggaggatt ccttcgccat cacttccacc agggccgggt cgttcaggtc atggccgccc 4080
gtgatggtgt cctggacaaa cagtgcaacc gagtcttcaa ttccctgttc cttctggtaa 4140
tgggtttcgg ccgcgttcat accgccggtc gctttggtgg tgtttccgcc cgcataaggc 4200
attttctcca gcaggacgaa atccttcccc gcctgcttca ccataatcgc ggcggtcatt 4260
ccggcgccgc cggcgccgat gatgacgatt tcggtatcaa tggtcttttc gggttttgcc 4320
ccggagtcgc cgccggcatc gcggcggttg gcatccagca cgtcaatgtc cgcgccggcc 4380
gcaaccagtg cctccgtaac cgcctggatc acagccttac tggttaccgt ggcgccgctg 4440
atcgtatcca cgttcggggt ctgggcgtcc cggatggcag cggacagcat gccggctgcc 4500
acaccgccca gtgtttcggt ttcatcggtc gcatcaatca gtacagcgat gatttcctgt 4560
tcatccaccg taacccatac ctttacggta ctgagcattc cctgtgcctc cgcctggtag 4620
gtaccgggtg taaacagggt ttccgcaggc gccggttccg cccaggcgga tgcagcgaac 4680
agcatgaccg tcatggccag cagcaggcag atcagtttct tcatgatggt cctccttctc 4740
gtctgttcag gtcaacaaca cttactggac tcttcccagg atctcattta cagcgtcaac 4800

aaccgcttcc gaggagaact tcagggtcgc gccggtgatc acatcgacat cggccgttcc 4860
gttctccaca atggccttgg cccatgcttc cctgatctct tccgtcagga agtcgctgtt 4920
ttccattttg gcttcagagg taatccggca gtccgtgatc ttgccgccct tggaatcggc 4980
ataaacccgg atggtgctga agttggtttc ccgctttgcc atatagctgc cgtaaatcac 5040
ccgtccggct tcttccgcct ttgcctcttc agccggcgca gcagcttctt ccgccttcgg 5100
ttcttccgcg gcatcctcgc cggtgatcgc ccggttcagg gctttcagca cggcgttcga 5160
ggtcacggtc gcgcccgtca gggcatccac gccgttttca ccataggtga acgggccttc 5220
cttgccgata aactgactga taaatgcttc ttcggaagcc cgctggccca gtccgggcgt 5280
ttcattcggc gtatcaatcg tcagcgcctt tacggtcttt ccgtccagcg tagccttgac 5340
cgttacattc ccaccgaatc cctgttcggt ttcttccacg gccgttccgg cagtctccga 5400
ttcagccgcc gcaggttcct cagtcgcggg ttcttccgct gccggttcct cagcagctgg 5460
ttcttcttcc gcaacaggag caggctcttc tgctgcttcc tgcaccggtt ccgcggcatc 5520
ctcgccggtg atcgcccggt tcagggcttt cagcacggcg ttcgatgtca cggtcgcgcc 5580
ggacagcgcg tcaactccgt tctccccatt ggtgaacggg ccttccttgc cgataaactg 5640
ttccgtaaac gcttcttcgg aagccctctg gcccaggccg gggtttcat ccggcgtgtc 5700
aatcgtcagc gcctgcacgg tttttccgtc cagcgttgcg tgtacggtca ccgtaccgcc 5760
gaatccctgt tcagtctctt ccacggcggt tccgctcaca gccggggcgt ctgtttgctc 5820
cgtcagcgta tacgtcatgt gtcccccgcc gatgatcccc gtcaggaaca gggaggcggc 5880
aatcaggacc agggcacata gaatcaaccg gagccactgt ttcttttcaa agttgttcat 5940
ggccatctct ccttatacag cggtcgtggt aattgctttc accggacatt tcgccgcgca 6000
ctttccgcac agcgtgcact tgtccggatt gatccgggcc agcgttccgt caaactcaat 6060
cgcttcgaac tcacactgct tcacgcacag gccgcatccg atacagccag ccgtgcatac 6120
cttctttacc tggggaccct tgtccctgtt cgaacactgc acccggacaa tccggctttc 6180
cggaatcagg gtgatcagtc cccgcgggca tgtcttggcg cacaggccgc agctgacgca 6240
ccggttccgg ttcaccaccg cgacgccgtt ccggatggaa attgccttct ccgggcagac 6300
ttccacgcag gagccgaatc ccaggcagcc gtagttgcag tcggttacgt tgatgccggc 6360
cagcacggcg gcacggcagt ccttgattcc gatatagttg ccctggttgc gggtcacgtc 6420
gcaggggccc ttgcacgcca caaacgccac cttgcgttcc agtgccccgg cggtgattcc 6480
catgatatcg ccgatggctt ccgctgaagc ggcgccgccc accgggcagg cattcaccgg 6540
cgcttctccc ttggcgatcg ccgcagccag cgcgtcgcat cccgcatatc cgcaggcgcc 6600
gcagttattg cccggcagct tggagcggac cgccacttcg cgttcatcca cctcaacggc 6660
gaatttcttg cccgtaaaga ccaggccggc gccgacgcac acgccgatga ccgcaatgac 6720
cagtgtggtt accagaatga tattcattcg tacacacctc ctcagtgcgg cagcccgccg 6780
aagcccataa acgcgatgga catcagcgcg gcggcaatca tgacgatcgg ggcgccccgc 6840
agcggcgccg ggagatcctt ctcgttgatc cgggaacgga ttccggccag caggataatg 6900
gcgatggtgt agcccagcgc cgtcccgaaa ccgctggtaa cgcattcgat gaagttatat 6960
ccgtcctgca cgttgttcag ggccacgccc agcaccgcgc agttggtggt gatcagcggc 7020
agggagatgc ccagcgcctg gtacaccccg ggggactttt tcttcaggaa catttccaca 7080
atctgaacca gcgcggcaat caccaggatg aagatgatgg tcttcatgaa gtccatgcca 7140
agcggcttca gcacatagtc atacagcagg ctggccacgg cgctggcaat ggtgataacg 7200
aagatgacgg ccacgcccag gctggcggat gccttcatct gtttggaaac accgaggaag 7260
gaacagattc ccaggaactg gttcagcacg acgttgctga tgagggcgct gcccaccata 7320
atcatgatca gtccgttcac ttcttctcgc ctccttcctt cacggcagcc tcaatggctt 7380
tgggcgcctc ctgcagcgct ttctgctgct cttccgcgcc ggcggtttct tccttccggt 7440
cgctgcacac ggtggagcac agcgcgcagt tgccgtcgca tccgctttcc accagctggc 7500
gctggcgggt cttgatgccg atcgcgttca ttacggcgat gatcagcgcg aataccagga 7560
aagctcccgg aggctggatg aaaatcgtca ttccgaagcc ttccggcagg atctggaatc 7620
cgaagatctt tccggcgccc aacagctcgc ggatcgcgcc ggcgatcgtc agggcaatcg 7680
tgaaaccgag gcccatgccg ataccgtcca tcacggaaag cagcggcgta tgcttgttgg 7740
cataggcttc cgcccggccg aggatgatgc agttcaccac gatcagcgga atgtagatac 7800
ccagcgcttc ataaataccg ggaatatagg ccttgatcag cagctccgtc accgtcacca 7860
ggctggcgac gatcacgata tacgcgggaa ggcgtacctg gtccggaatc accttccgca 7920
gcaggctgat caccaggttg ctcagtacca gcaccgccag cgtgctcagg cccatgccga 7980
taccgttggt ggcctgagtg gaaaccgcca gggccgggca catgcccagc atcaggacca 8040

gtaccgggtt ttccttcacc aggccgttcc agaggcgctc gaaatacttc ttcatctcaa 8100
ttgcctccct tcgctgcttc ttcgcagaat gcaagtccgg cgttcacgcc gtttaccaca 8160
gcagtggagg tgaccgtcgc gccggtcagc gcgctgatct cctgtccgcc gcctgcagta 8220
cccttgacca ggctcagctc tccgcccttg ccggcgaact ggcccttgaa cgaatcctca 8280
tccgccctca taccgaggcc ggccgtctca ttcagttctg taaaggcaat cttattgatc 8340
tttccgtccg gcgtcaggcc gagcgccatg gtcacgtccc cgccgaatcc cttggcggtt 8400
acgctcacgg catagccggc cacctggccg tctgcggatt tggcgacgta gaattcgttg 8460
atcttcgggg aggctttttc atcctgcgcc gcgatcattt cttccgcctt gtccaccacg 8520
tcgaagattt ccgcgtccgg gcatacttcc ttataggctg cggtcttctt ctgcagctcc 8580
tgcagttcaa ttgttccctt ggtcatggag tagacaccgc tcagcgccac gcccgcgatg 8640
gccgcgatca gcgtcagggc gatcaccggc ttcgggatcc ggaacggctc ccgctttacc 8700
gggcgggtct tgccgatctt ccggtaggcc agcggcttct ggacgatgta ggtatcaatc 8760
aggggcgtgc atacgttgcc aatgatgatg gcgtagctga aggagtcggc cgtgttgccg 8820
tacaggcgga acacgccgcc cagcacgccg atcaggcatc cgtagatgaa ctgtcccagc 8880
cggctgaccg gactggtcac atagtccgtc gccatgaaga acgcgcccat gaccacgccg 8940
ccgccgcaca ggtgcgccag caggaatttg atgtcgaagc cctgtccgcc gaacagggcg 9000
ataaacaggg tgaatccgcc cagcacggag aagcagatct ggccgtggat gatatccatg 9060
ctccacagca ccaggccgcc gatcagcagc gccaggatgg agcttccgat cacgccgtcc 9120
gccgtgccga ggaacatccg ggtaatatcc accgtctcat acatcttcag ggctcccatg 9180
ggggtcgcgg aagatgttgc gtccaggatg ggcttgatgc tcatcgcgtt cgcgaaggag 9240
atcagcagga agcaccgggc cgccagcgca gggttgatga agttcttgcc cagtccgccg 9300
aagcagccct tgacaaccgc gatcgcgaat acggaaccga tcaccggcag gtacagcggc 9360
gtccgcggtc ccagcgacag ggccagcagc aggccggtca ccgccgcgct tccgtcccag 9420
atggtgttcg gcttatggac gattttgttg aatacccatt ccgtcagcac cgcggaaacc 9480
accgccagcg cgatgatcgc gaatgcctgc cagccgtatg cgatgattcc gacgcaggtc 9540
gccggcatca ggctcagcaa caccacccgc atgatatacg gggtcgtcca tctgtccctt 9600
acatgggggg atgaggaaat attcagcgcc tcgctcattt ctggttcccc ccttcccgtt 9660
tcttcgcggc ggcaatttcc gccttggctt gtttgaacac ctgcatcagg ggccgtttcg 9720
ccgggcagat gaaggtgcag ctgccgcact ggatgcattc cagaccgtac agcttcttct 9780
cataccgggc caggtccccg gcgatcgccg cgtcggccat cagctggggc atcagtccca 9840
ccgggcagac cgtggtgcag cgtccgcagt gcaggcatgc ggtcatcagc ttctccgctt 9900
cctccaccgg atcctccgcc atcacggtca gtccgttatt ctgcttctgg atcggaatgt 9960
ccaggctgcc gattgcgata cccatcatcg ggccgccagc cagcgctttg gccgccttgc 10020
attcttcctt gattccgccg cattcttcca gcagctccgc gtagctcgtg cccacccgca 10080
caaagaagtt gccgggttcc tttaccgcgt cgccgctcac cgtgaagcac tggctgaaca 10140
gcggttcacg gtacaggacc gcgcgctgga tcgcatacag cgtacctacg ttctgcacaa 10200
tgcatcccac gtccgccggc agcttgctga tcgggaaatc gacgcccgtc accgcctgga 10260
tcaggctgcg ctcaccgccc tggggatact tcgtgatcat ttcctgcacg cggatcttcg 10320
gctttccgga aacggcgatc tgcatcgcat gaatcgcttc cggcttgttc tcctcaatgc 10380
agatcacgcc ttccgcgttc gggaacagcc ggagcaccag ctccagtcct tccgcgatgg 10440
ccgtggactg cgtgcgcatc agctggtcgt tgcaggtcag gtacggctcg cattccgcgc 10500
cgttggcaat gatgtactga attgcgtccg gatctttggg cttcagcttc acatgcgtcg 10560
ggaatccggc accgcccagg cccacaatgc cggcctcctg cacgcggccg atgatttcgt 10620
cgttggaaag accgttcgcg ctctccttct gcgaaaaatc agcgaaggga gtaaactgcc 10680
cgtcattgtc gatcacgata cagtcaatca tgccgccgcc cagcacccgg cgtttttcaa 10740
tggctttcac cttgccggag cagctggaat agatatgggc ggaaacaaat ccgtccgcct 10800
ccgctatttt ttgtcctgtc agaacctcgt ctccccgttg gaccagcggt tttgcgggct 10860
tcccgatgtg ctgctggagc gggaatacca tttccccttt cgggttgaac aggcgcagcc 10920
gggcctcacg gctcagctcc ttccgctcct ccgggtgcac cccgccgcga aacgtgtttc 10980
tcactgtgat actggctcct ttccgtttcg attattgatt tatcttccgg ctttccgctc 11040
ggttttgacc tcgtacatcc gggcgtcggc gaccttgata cagtcgtcca tcgtctttcc 11100
ttccgaagcc tccgcactgc atgtgccgat ggagaatcca agctggaacg gcatccccga 11160
attccggttg tactcgtctg cggttgccag gatccgttcc ttcaggtctt tctcgtccca 11220
ccgggcaatg atcacgaatt catcgccgcc gtaccggatg atgacgtctt ccggatcgca 11280

gcattccgac aggatccatg ccgatgcctt cagcgcctca tcacccgtgt tatggccgta 11340
cagatcattg accagcttca tgtcgtccat atccgtaaag aggacctggg cgccgtggtc 11400
cctttccatc agccagtcaa acgtttcctg cccgaagcgc ttcagtccga accggttgta 11460
cagtccggtc aggctgtccc gcacatacag ctcgtccagc gtttcgttga actggcggag 11520
cagttccttc ttccgcacgt tttcaatggc gatttccagg aagttgaata tactctcatg 11580
caggttcatc ttcgccgctt cgctgattcc gtccagcgcg atatacccga tggaatatgt 11640
gttgtaatgc agcgggtaga acatcataaa ccgttcccgc ctgagcaggg gctccggcag 11700
gagctcctcc gtacggaacc ggatgaaatc gtatttctcg ttcccgtccc cgctccggca 11760
tccccgtttc cggacggtca ggatcatctc ctcgccgaag ggtttttcct cctccggcca 11820
ttcactcttg tcaaaatggt cgaaataata gtcgttgaaa cacaggtaga tatcctggca 11880
gccgaaaatc cgccggtttt tgccgacggt ttccgccagt tcctgcaggt cgttgcattc 11940
gaacaggtct tccgccatct ggccctgcag gttgtaaaaa tccttcagga accgggtctg 12000
ctggaaatag agttcccgga ttgtacccgg ccgggcggtc tcggggcatc cgcaggattc 12060
ggagcatatc gtatggaagt ccgtaatgac ttcctgcccg cagtccttcc cgtcgatcag 12120
gctgatcagc atgtccatcg cccggtaatt ctggttccgg ttgtctccgc ccaccgtgga 12180
aagacgcggg ctggaaagct cggaaccggc caggttgtca aacccggtta cgatcacgtc 12240
cccggggatc cggacttcat ggcgcttcag cgctgtcatc agtccgatcg ccatttcatc 12300
attcgcgcag atgaatgcgt cgggcagttc aagcccttcc tgcatccatc tttccgcaag 12360
atccgcgccg tcgcttgtac gccaggtggc gggaacaatc cggacgttct cctccggaat 12420
attgttttcc cggcatgcgt ccacaaaccc gtctatccgc tccccggcct ccgggcttcc 12480
gttctccgtg ttgccggcca ggtagaccag ccgcgatgca tggtgttccc ggatcacatg 12540
gttcgcgatg gccttctggg ctgcgcggtt gtcaaagcgg atcagggcgc accctttcat 12600
cggagccccg atgctgactg ccggaacccc ggcttccaca atccggcgtt cgatatcgtc 12660
gcgcacccgt ttcagcacga gctggtttcc ctgcaccagc acaccgtcaa actttctcag 12720
gtccggcagg tcaaaaatca catattcact tttgtctttt gcattgtcca ggtccttgcc 12780
caggcagtca aaaatataaa tatagacgtc ctcatggtca tccacatact gcttcaggcc 12840
atgcagcgta ctttcaacca attcataatt ccagtcaacc gtaaaaaagg cgatccggta 12900
ctgttgcttg cctgaaccca tgcttcagtc tccttcttga tctcctgtgt tttcggtatc 12960
cctgttcccg gctgaaatgg tgatgcccgg ggcatctttt acggtccttt tattgtaaga 13020
tgattcgccc gcaatgtcaa tatttgatta ggcttccggc ctgtttccgg aagcgaaaat 13080
ccatttgacg gaacgaagga aaaccgatat cataactata cattattgtg aggtgaagac 13140
catgcgtaaa ctcccggcat tattgctggc ctgtatgctg tgtcttcttt tcgtgtgccc 13200
gtcagccggg gctgcccgtt caggcagcag caccctcgtc agcggggatt acgaatatcc 13260
cctccaggat aacggcaccg ccgtgatcac acgctttacg ggaacacgg acgttgtggt 13320
gattcccgcc cggcttgacg gcattcccgt ctccgccctg gggaaaagt cctttgcctg 13380
gtgtccgtcg ctgctggacg tcaccattcc cgagggaatc acatccatcg gggattatgc 13440
cttcctctac tgtaccggcc tggctgccgt ctttatgccg gacagcctga ccgatatcgg 13500
ccggaatccg ttcaccgcat gcagtacact ggcagatatt atcgtatcgg atgatcatcc 13560
ggcgctcctg ttcgaagacg gcgtgctttt cagccggcag gattaccggc tggttacctt 13620
taccgccgca cgcccggacg gaccgtactc cgtcccggac ggaacccgga ccatcggtgc 13680
ctctgcgttt tactcctgcc gcaatctgac agcactgacg atgccggaat ccgtatccgc 13740
catcgggcag aaagcattcg gcctctgtgt cagcctgcag tctgtttcct tgccggacag 13800
cgttgatttc attggcgctt cagcgttctt cggatgccgc ggcctgcgcg aaatgagcgt 13860
tccggcttcc gtcacttcca tcgccgacag gacgtttgaa gactgcacaa acctcagcga 13920
actgcatctt ccgaagacgg tccggtccat cgggaaccgt gctttcatga actgttcgtc 13980
gctggaggcc gttgatcttc cccgcggat ccgggaaatc agccggagcg cctttgccgg 14040
atgcgaaagc ctgtcccgca ttaccctgcc gggacgggtc acccggatcg gcgacggggc 14100
tttcctgaac tgtatgaacc tggaaagcat tgatattccg gcaggcgtcc gatccatcgg 14160
cagggacgcc ttttccggct gccgttgtct ccgggaaatc agcctgccgc ccgaatgcgg 14220
tgaaatcggt acaggcgctt tcaggagctg ctttgctttg gtttccttcc gtgttccgga 14280
ttccgccgaa accatcccgg aggaaatatt cgccggctgt actgcccttg ctcaggtata 14340
cattccgtcc ggtgtccgtt ccatcggaag ccatgcgttc ctggcctgtg aacagcttac 14400
ggaaatcatc attccggaca gcgtccgttc catcggcagc caggcattct tcggatgcct 14460
gaacctggct cgcgtctctc ttccggatcc gcccgccgat attcaccccg acgcatttga 14520

ggggtgtgag cggcttgccg gcaccatccc ggaccttgtt ccctgaaaac ctgcatcagc 14580
agtcaggcgc ccgccggaac ggcaggcgcc tggtttttat attccccgga tccattcagt 14640
gatatcgttc atcacttcat cccggttgat ctcattcagc acttcatggc ggcattccgg 14700
ataaattttc gtggtgacgc gtgtatgccc ggtttccgcc agcagttccc cgaccttctg 14760
cacccctgcg ccgtaatcgc ctaccgggtc caggccgccg ctcaccagca ggatgggaag 14820
ttccgccgga atatcccggg cccatgcggc cgtattggat gcagcattga gccggaccag 14880
gtcgttcatg ccctggaccg gaaacataaa tccgcagtac ggatccgcta tataccggtc 14940
cacaatcgcc ttatcgcggg tcagccagtc aaatggcgtc ctcccctcaa acttattccc 15000
gtacgtgccg aacgccatgg tattcatcag gctgctcctg tgcttctttc cgcgcactgc 15060
gccgatcatc gcggagactg cctttccggc gcctgccgcc ggattggggc cgccggttcc 15120
cataaagacc gcccctgcga atccatcctt cggataacgg gatacataca tccgcacaat 15180
aaacgatccc atggaatggc ccatcacaat cagccggcat gccgggtgct cctgtcgtgc 15240
gcgcatcacc atttcccggt aatcctcaat cagggcgttc catccgccct tttttccgaa 15300
ccagcccttc agttccggat cctggcagga ttcgccgtgg ttggccatat catgcatgta 15360
tacggcaatc ccgtgatcac acaggaagcg gataaactct tcatatcgtt tctggtgttc 15420
cgccattccg tggtggatca ccatcaccgt ctcgaaaaaa ccgtcgtccg gagtataaat 15480
atgggcgctg atctcacaca cacctgtttt ggacggaaaa tgaaacggta tattcttcac 15540
gactgtccct ccccgttctt ttctttggtt gtgttctcct gccccatttg taccacattt 15600
ttccggcccg cgcaaccgtt tctgttgctt atccggcagg aggcagtgta tactgatact 15660
gttgtgggag gcgaagctgt gagtactttt tctgtaaccc tgggcaatgt gctgctgacg 15720
ctgctgtata tgtcagcggg tttcctgctg tccaaagcga aaaaggtcgt accggatcat 15780
atgagcagta tcagttcgat cctgctgtac atctgcggcc cctgcatgtt tctggacacc 15840
attgtcaaca tggattattc tccgtcccag acggtcagca tgctgctgtt cctggtaatc 15900
acactcgctg tcatgggact ggttctcctg ctgatctttc tgctgctcgg caagcggcgg 15960
aaggattttg ccttccgcat gatggccatc gcatccgtca tgggaaatgt cggcttcttc 16020
ggcctgccga ttgtcaaagc tgtattcccc gccgcgccgg aagctgccgc ctattcctgt 16080
atcttctgtg catccatgaa tatcgtggcc tggacgctgt gcgtcttcat cctcacggat 16140
gaccggaaat acatgtccgt caaagcatcc ctgctgaacc cgaccgtgct gcccgttgcg 16200
cttgcaatga tcctgtatgc cctgggcgca aaaaaccgga tgccccagct tgtgctcggg 16260
ggcatccatt ctgtcggcag tatttccacg ccgctgtgta tgtttattct cggaatccgt 16320
ctttccacca tgacgatgaa atcattcttc acatcaccgg tcccgtggct ggcatccttc 16380
gggaagctgc tcgtttttcc gctgatcagc tatggaatcg tgctgctgct tccgctttca 16440
gatgtgttcc gtgccagcat cctgctgctt tccgccacgc cgtgcgccag tattatcctg 16500
aatctttcgg aaatccatca ccacggccag gacatcgctg caaggtgtgc gctcctgtcc 16560
acactgctca gcgtgctgac gattccgctc ctgagcctgc tcctctgatc tcaggctttc 16620
ccgtttccct caggcccgcc ggcaggttcc ggcgtttcgg gagatgcgtc ccctgtttca 16680
ccggcatgat cttccgctgt tcccgacagg cgttcctccg cctgtcttct tttttctcc 16740
agccggatca gccgcttgtc gttcagtacg gttcccacgc ccacaatcag gaacatgtaa 16800
taggtgaaga accgccagat cagcaggccg acgccgatgg tgccgtccgt gaatatcccg 16860
gcgaaatagg ccatgaatcc gccctcctgg gcaccgctcg cgccgggaag gggggtgtag 16920
ctggcggata caaacagcag gcagctcatc gtcaggatct ggtaccacgg ggttccgtgc 16980
agctcgaacg catgataggt gaatacgatg gagcctgtca ttcccagcat actcagcgcg 17040
ctgaacagaa cctgcagcag gatctggccg ggggaatgga tcagagcgat cattgcggca 17100
tgataggaat ccagaatctc ggaaatcttc gcgatcgtgg cgttcttgtc ccgcaccagg 17160
tgcattttcg ccagcagcgc gatcagtctt tcggtcactg cctggatcag gttccgtttg 17220
aacgcagcaa gcagtaccag cggaactgcg ataaagttga tcgcccaccc gatgcgcgca 17280
gcccagatca agccgcccag ctgattatag gcgaacgtcc ggttggcaat cataagaccc 17340
agcgtcatca ggcagatcgt cagctggttg gatacgaagc ggatcgtcac cgccgtggtt 17400
ccgtatgcca ccgggatccc cgccttccgc aggctgttca cctgcatcgg ctgtccccg 17460
gctgcgctgg gagtgatgtt gctgtaatag aatccgatca gggccacgcc gatcacgcgg 17520
cccaggttaa tcctgtatcc tttggaccga aggtagctcc atgtgccgca gccttcaaac 17580
ccggcatatg ccgcccagca gaggaacagg cccagtaccc acttgaggtc cattgaagca 17640
agcgccttcc aggcgttctt catctcaggg ttactgaaag cgataataat cactgcagtg 17700
atggaaatta caatcaccag tgcactgagt acccgcttga tcgtgtggct catgtccgac 17760

tccttgctgt catgataatc cgtgttccgt atagtgtttt gctgtattcc gccttattcc 17820
cggttggaga tcactgctgc ggaatacggc ggcatatcaa atgtcaggta cccgtccgtt 17880
gcggcgccca gcggctcgca tccggtttca tgggattcct gcgttgtcag gatccggcgg 17940
aaataagttt ccccgtcctt tgcgccggcg tcccgcacaa acaggcggaa atccgtccat 18000
ccgccggtgt tgttcaccgc gacaatgcag cggcagttcc cgttaaaccg tgcgtaggcg 18060
atccggccgt aatccgccag cagctgtttc aggctcccgc acgccagcac gggcaggtcc 18120
ttccgcaggg cgatcaggtc ccggtgcagg ctgatcaggt cctgatcctc atggccccag 18180
ggatatgtcc gccggttgtc cggatccgtc cagcccccct gtcccgcctc gtccgcataa 18240
tatacggtcg gtgcgcccgg ccaggtcatc tggatggtaa cggcttcccg gaaaatgcct 18300
ttgttgattc ctgcagcggc ggcgccggaa ccgactgtcg tcatccggcc gaccatccgg 18360
ttggtccgtg tcaggaaccg ggaatggtcg tggttgctca gttcgttcat tgcggtaatc 18420
aggctcgggt atttcaggtg ggccattttt tcccgcatga tgccgaagaa tgcgctgccg 18480
ttctggtaca gatcatcccg gtaggaatcg gagtgttttt ccatccccgt cagaaaccag 18540
gataccggct ccataaacgc atcgtaattc atcacggtat cccattcccg gccgttcagc 18600
cattcggtgg gatcaccgta atgctctgcg ataatcagtg cgtcgggatt ggcggctttc 18660
acccgcttcc ggaaatcctg ccagaagctg tggttgcatt ccctggtgtg tccgaggtcc 18720
gccgcaacgt ccagccgcca tccgtcaata ctgtacggcg gcgacatcca tttctcggca 18780
attttcagga tctgctcccg cagctccggc gattcctcat agttcagctt cggaagggtg 18840
gaataacccc accatccttc ataggcggga gaccggccat tgccggattc gttgaaatgg 18900
aagtagtgcc tgtacgggct gtgaacgctc tggaaggcac ccggcaggaa gccggctttc 18960
cccaggtaaa tcccctcatg gtccatccat ttattgaagg aaccgcagtg gttgaacaca 19020
ccgtccagga taatccgcat tccgcgcctg tgcatctcct ggcacagcct ggcgaacagc 19080
tcgtcgcttt tctccaggtt ctccgtggag gtgacccgcc ggatataccg gggtgcaaat 19140
ccgttgtgtt tctcccacag ctgcatttca tgctcaacgt cgtcggtaat cacgccgaag 19200
tggggatcca catgctcata atcctgcgtg tcgtacttat ggctcgacgg ggaaacaaag 19260
atcggattca ggtacagcac ctcaatcccg atatcctgca ggtaatccag cttgtccata 19320
attccctgca ggtccccacc gtaaaaggac cggatatccg tatccgtcgg cagggcatcc 19380
cattcactga tatgctttgc gtgcccgatc gtataataat actcgttgtc tgtcacgtcg 19440
ttggacggat cgccgttgca gaagcggtct gtgaagatct ggtactgcac ggcgcctttg 19500
gcccattccg gcacatggaa tccgggaatg aacctgaatg catagaccga attaaaatcc 19560
ggcgtaccgc cctcttccag gcgcgcgccg ctcttgtcat aggcaatccg gccatcgtca 19620
tggcattcaa tcaggaaacg gtacatcacc tcctggtcgt tgcataccag gctggtttca 19680
taataatcaa aataatcgtc gctgcgggtg ttctccatct gcgtgcctac gctcatggaa 19740
tcaaaaatga caataacccg tttcgcgctg ttcttcgcca cccggatccg gatggtaacg 19800
gaatcaccgg gatccggctc agccggaaaa cggtaatccg gcgtttcatc gctgaacaca 19860
gcggataaaa cgttctgact tacaaccatg aatgcctgct cctcttatca agtctgtccg 19920
gccatcactt gtccagtccg accttaaaca gatacagcca aagttttttt ctgtatttcc 19980
catactcatt ccggcagata tacatgtcct tcaggaaatg gtacaggcgg actttggtct 20040
ggatgttccg cagcggatgc gtcacgcagc tgaatgcctg ccggaacgtc atacttccca 20100
tggaccggtg gtaatcatac agatattcct ttacaaacga gatccgttcc gcttttttca 20160
ggtaggtaac gtggaaagca aagtcttccc cgtatgtgag cccgtcctgg aagaccagtc 20220
ccatatcctc tatgatatcc cgccggaaca gcttgttcca caggacgccg taaaaatacg 20280
tattggcctt ccggcacagc atcggcagca tttccaaagt ggtcatcgta tcgtcccggt 20340
tctccaggtt gaaccggaat tcccggctcc ccaggcactc cttgtacccg ccgatcacca 20400
ggtctgactc ttctttttcc gccttgttca ccaggatttc ggtggattcc ggaggcagcg 20460
tatcgtctgc gtccaggaac cggatatatt tccccgtgca gaaccgcatt ccggtatttc 20520
tcgcctggga tacgcctttg ttttcctggg tatgcacctg gatccgtgca tcccgttctg 20580
cggcttcccg tgcaatctgc agggtattat ccctgctgcc gtcgtccacc aggataatct 20640
gggtgttctg ccaggtctgt cccgtcacgc tgtccatcat gcgtttcagc gtcttctccg 20700
cgttgtacgc cggaatcacg atgctgacaa ggggctgttc ttccaactct gcatacctct 20760
ctcccaatgg atcatatgca aaagcataca ggttattgta ccatggtttt cccggaaaag 20820
gaagtgtaaa taaagaacgg ggcctgatcg ctcagtcccc gccgatatgc aatggccggt 20880
cagaatggtt cctgttactg gtcttcctgt gccgcctcgg ccacgccgaa gaacgcaggc 20940
ttcgggttca tgttcttgtc aaacagcagc ggatagccgt ccctgcgcca gctcatggta 21000

tcaataacgc cccagacctg cacggcctcc gtctggtctt taaaccggag catcagcttc 21060
ataacctgct tatacttgtt cttctgcgac tggaaattct gctctgtggg tttcccggca 21120
ttgatatcca gttcgctgac ccgcagtttg aggcccagtg atgcgatctt ttccacagcg 21180
cgggtaatca tatccatgga gggttctccc aggctgtgat gcatctggaa tccgtatccg 21240
tcaatattgc cttcagccat caggctcttc agcaggtcga agatcccgta cagctttccg 21300
ctgtatgcgg tgttgtaatc gttatagtac agcagcacgc cttccgccgc gtacttccgg 21360
gcatactcaa acgcccgcgc cagatagtcc ggcccgacgg tcttgtacca gtttgctctg 21420
gtccgcagct tgttcgtccc gtcgtcaatc gcttcgttga ccacgtccca ggaaacgata 21480
acgcccggat agttttcctc ggtccaggtc agcacctggc ggatgtagtt ctccatcctg 21540
cccagcatca cttcgcggct ggccagcggt ttcgaagtgt cgtatccctc atggaaaagc 21600
gcttccggcg tctggttgtg ccagagcagc gtatggccgt gcaccttgag cccgttcgcc 21660
tgcgcaaacc gaagcagcgt cttgcacgca tcgaacttca cagctaccgc cgtttcatcc 21720
tccgttgcca gcttccggct ggcatatacg tccaggatgg catccggctt catctcgttt 21780
tccggcgtca ggatattgaa ctgctccttc accagcatca gcagcttcat atcgttgaac 21840
gcattccggg gaagcgccgt gccaaagtcg aattttcccg catacaggtc tttcagggaa 21900
ggcatctctt caacagcttc aaactccgga gccggcgtcg cagtcggctg gacgtccggt 21960
actccgtccg gcgcaactac ccggaaattc cggatttcat agctcagttc cggagcaccg 22020
acagtctcca catacaggat atagtcttcg gagggatccg gcgtgtagct gcccgacagg 22080
gtcgtccact cgcctttctt tgcttcgcct ttgaacaggt tcttccagtc cggggatccg 22140
tcctctttcg ttacctgcac ggataccatc agcgtggccg tctccgcttc attctggagc 22200
acttcaacgc tcatgtcata tctggttccc gccgccagtt cgaaatttcg tcccggggaa 22260
tgccagtcgc tggttcttcc ttccgtccgg agggttgcct ccgttgtgtg atataccttc 22320
tgggcgcccc gcgcatacca gagatctgtt ccggctccga aagcggaagc gtataccagt 22380
gtttcctgcc cggcatgggc ggccgtcatt cctgtcatca tcaacatcat ggctgtcagt 22440
atgatcccga ttcttctcat cggtctatcc tttccccggc acatccggcg tctgttgtgc 22500
ctatatgata cacggaatcc ccctttgaat caatatttgt tcgtacaaat tctgtggaaa 22560
aacaaaacaa aatagggccc tttccttttg cggaaagggc ccgggattct gttaatcgaa 22620
gcagtcaccc acgtggttga taatggtctg aacacccaac ggtgttgtgt attcctcaaa 22680
cttctgccat gttccttccg gtgtggtcgg cttcttcggt ccttccggaa tcttcggcac 22740
ctgggtgaat acgttggtga agatggtcgt gttgccttgc cgttccgtct tactcagctt 22800
gtacgcgatc acagtctgct ctgtccatgt gtaggtaacc ggcttgccgt ttacccgggt 22860
gggcaggttg ttgatcgtcg cggtccagtt gttctcctcg ctcaggagta cggtggtccc 22920
gttgctcagc gtcacatgga tgctgctcgg ccgcatcttc tccgcgttgt tgttgtcatc 22980
ccataccttc cggaccgttg cgctggtcac ttccggcaca tagttgttcc ggatgttgaa 23040
tccgttgatc tctgcctcgt accaggccac aggatcctcg gtaatggtgt aggtgatctt 23100
cttctcgccg ttgtagcgcg gcagtccggt aaataccgtt ctccagccgc cggctgcatt 23160
cagctcggca ctggcgatct ccgttccgtc tgccagcagg cggaccgtaa cgttcgccgg 23220
ccggttcccg tccttgttcc cgttgtcatt ccaggtcttg atcacgggaa tgtccatcgg 23280
ggtgtcatcc ggttcattgg gctcgacctc accgaactcg aaggtctttt cgataatcag 23340
ttttccgcgc tccaggacgt tccggatcac gaatccggtc gcggcgctgc cggtcacggt 23400
tgccttgtag ccttccggca tgttcagctc ttccaccgta tagatgactt ccgctgcatg 23460
gccgctctgt ccgtccgcat tggcgaactt gtccagtccg gtccattctg ccttccagtt 23520
attcgccgcg ttcagtatta cctcttcgcc gcaggcttct ccgtcagcat acagctgcac 23580
acggatcgat tcaggcctgg tcctgtaagc gttttcgctg tcatcccaga tcttctgtac 23640
gcttactccg gtttcttccg gcggacggct gttggtgatg acggtcagtg ttcccttctt 23700
ctcctggctg gtcagcgtgt atctgttgcc gggcgcttcc tcttcccagg tatactcgat 23760
atccttctga tccgcgtcca ccatcggcag gttgccgatc gatgccatcc agccgtttgc 23820
catgctcagt accacgctgt cgatcttctt gccgttcgcc agcaggttca ccctcagggt 23880
cagcggccgt acgccgtcct ggttgttccc gtcgtcccag accttcttca ccgcggccgt 23940
tacgaagagc atgctgtctt ccaccagcag gatgccgtct tcggtcaccg agcctgtgga 24000
tgtcacattg ccttcttcgt caatcgtaaa cttcgtatcc gttgcgatgt cgtatccgtt 24060
cggagccacg gtttcgcgca ggatgtactc cgtatccgtc agcaggcctt cgatgacgtg 24120
cggttctttc tcagagatcc actcttccac aatgacgagt gtctttgttt catccgcata 24180
cttcgtgttc tcatcctcag cgtccgcttc tgttgcttca cgcaggatct cgatatgcgc 24240

```
        acccggcagt tcgtcgccgt ttgcgatatc caccttgctg acgctcacct cggtctgggt     24300
        gtttgtaacc gtgaataccc atgcatcttc tgcggtcttc atgatgacag gcttcagata     24360
        accggctgcc tgttcttctt caaccgtgta ctcgatttct tcgccttcgg agtacttatc     24420
        caggtcggtc caggtgtact tccattcgtt gtccgcatcc agcgtcgctt cgccggcttc     24480
        ctcaccgttc gccagcagct tcacggtcac gttcgccgga cggaagccct tcacgttgtc     24540
        cttgtcttcc cagatctttt cgacttcaac ctgtgtcttc tccgtctcat gactgttcgt     24600
        cacggtatac gtgaaggtac cgtcttcagc ctttgtgatc tgcgtctcgt agttctcgac     24660
        cggttcttcg gtgaccgtgt agtcaatcgc ttcaccgttc cgatacttct gcagctgact     24720
        ccagctgaag ctccagttgt cgtcttctct cagctcgact gtatcgatag tctcgccgtc     24780
        agccagcagg ttcactgtga tgacttccgg acggcagcca tcctggttct ccgcgtcttc     24840
        ccagaccttc gtgacgttca cttcggtctc ttcgaaatca cggctgttgg taacagtgta     24900
        ggcccatccg gtctcggaca ctttctcaat gaccggtacc ttgtagttct ccagctggtc     24960
        ttccgtaacc gtgtactcga tctcagcgcc accggagttc ttgtccaggc ctgtccaggt     25020
        atacttccat tcgttgtctt cattcagcgt cgccgtgccg gcttcctcac cgttcgccag     25080
        cagcttcacg gttacgcttt caggacggaa gccttcagca ttgtcattgt catcccagat     25140
        cttttcgact tcgacctgcg tcttttccgt ctcatggctg ttggttaccg tgtaggtgaa     25200
        ggtgccgtcc tcggccttat cgatcttcgt ggtgtactca ggcacttcgt cctcgttcac     25260
        cgtgtaatcg atatccttgc ccgcagcctt cttgtccagc ttcgtccagc cgtagctcca     25320
        gttgttctcc tcgctcagct tcacggtgtc gatcacttca ccgtccgcca gcaggttcac     25380
        tgtcacgtcg tccgtccggt agccgtcctg gttatccgcg tcgtcccaga tcttcttaac     25440
        ctttacctcg gtttcttcgg tcttgtggct gttcgtcacc gtgtaggtga aggtgccgtc     25500
        ctcggcctta tcgatcttcg tggtgtactc                                      25530
//
ID   XXX; XXX; linear; XXX; XXX; XXX;  XXX.
XX
AC   PRJEB67889
XX
DE   Uncultured bacterium DNA fragment from in vivo enriched cow rumen, clone
rum-14o19
XX
FH   Key             Location/Qualifiers
FH
FT   source          1..35658
FT                   /organism="uncultured bacterium"
FT                   /environmental_sample
FT                   /mol_type="genomic DNA"
FT                   /isolation_source="Cow rumen"
FT                   /clone="rum-14o19"
FT                   /note="AZO-galacto-mannan active clone"
FT   CDS             527..1000
FT                   /translation="MGEQRIVGIQNIPQVHGIAVHDGGNAPAQLHHVHVTVRFTGQDGV
FT                   VHGILHASIGKVLELIGHAVGVLNVLHIIETHEIGILAGCGFIVFPGFIVGVIVDIPPE
FT                   TDGNIPGIFRQRRYSQGEHHGQSQQHCCYFFHKSVSPFQKSGHHLIAWPTATA"
FT                   /product="hypothetical protein"
FT   CDS             complement(967..2676)
FT                   /translation="MKKLISWLLVAVMAVGMCSWASADPVNALDFEDGVFAFLGVSAAK
FT                   PNADASTALEVVDYNGSKALRVAAQGIPYVALNLEGLAGEKLADVAAVTFDIGVDKAAD
FT                   GKFYAVSGVVYSYTGENADENKADWSVYLEKKNPRNVKIQFKAPFVAGAGNYIMISRED
FT                   QAGGEPATFYLDNIQFLDAEGNAIALDPTAEYVSEASEKDLSNLVALTNAVEFPDFHKS
FT                   AGAWAQDGLEMPQEIIDALVPGSVVEVEYASADGSMWIVMPWATAGWMRVGQGTAAINN
FT                   SKTIAQIPYEMIEALCGEDKSTWGAMLQCESASDWEVFAVRVGQRANRIVLKNAVEFPG
FT                   FTKSADAWAQDGLEMPQEIIDALVPGSVVEITYSSEDGDIWLVMPWAEAGWMRVSQGTA
FT                   AKMGGKAYITYEEIAALCGEDKATWGAMMQCEGSSPWEVYGIRVGQKAEFFGLTNLVEF
FT                   PGFTKSADAWAQDGLEMPQEIIDALVPGSVVTISYESEDGNMWLVMPWAAAGWMRVGND
```

```
FT                   GADVADGKIAQVTYEQIEALCGEDKSTWGAMMQCESSSPWNVYAVAVGQAIK"
FT                   /product="hypothetical protein"
FT   CDS             complement(2736..3740)
FT                   /db_xref="GO:0005215"
FT                   /db_xref="GO:0006810"
FT                   /db_xref="GO:0016020"
FT                   /translation="MQTIHPAQNGKITWQGFRNSPVYANQRDKTLNGMKNLVIGLFRLI
FT                   VIVGISYVILAPVINIISNSFFSRQDSINPMVFTIPINPTLERYSMAVKYLNYLPILGN
FT                   TILYVFGVTLIQLLICSMVGYGFARFRFPLKSVLFAFVLVMIVIPLNTIQFPLYTTFRF
FT                   FNPLGIVGLFNGGKTINLLQTAWPTFIMSVFGCGLRSGLYIYIFNQFFRGLPKEIEEAA
FT                   LVDGAGTARTFFTIMMPNAAPAVITVAVFSIVWQYNDSFFANTFNISDSILVTRKLESL
FT                   INVIANAEKVLTIREQQLYFDAGVVLVLVPVVVIYLVLQRRFIEGVERSGIVG"
FT                   /product="N-Acetyl-D-glucosamine ABC transport system,
FT                   permease protein 2"
FT   CDS             complement(3761..4687)
FT                   /translation="MTGIRRKNRIFHLSYEQRNAVRGYWFILPWLIGFLVFYAGCLVQL
FT                   GEFSLSRIGLDAATGMTKEYIGFGNYIEAFTSHATFKQTLTSSVMDMLIDLPMILFFSL
FT                   FVAMLLNRKFKGRALVRAIFFLPIILSADAVGSAITRATELVNAGVSSANLETAQAGTV
FT                   SVAYYMQLFGDLAIPESVLGYIVDAVNRISSIIKGSGVQIILFIAALQSIPGSLYEVAR
FT                   IEGATGYESFWKVTFPMVMPHMITNTIYTVVDRFSSSEVIRLATRTYKELYNYGLSSAF
FT                   SVVSTLITILILGLIVYLLNRRTFYYN"
FT                   /product="binding-protein-dependent transport systems
FT                   inner membrane component"
FT   CDS             complement(4684..7275)
FT                   /translation="MNEKNRLTLPTAFRRVLWIIAGLAAAAAVLFLARYVIHYRAYDEY
FT                   RQYIVQPAAKTEAEALKPLKDPENAVPGFVLAAQNDRLGLYLKEKTAEIALYDRKSGLT
FT                   VYSNPQDAADDPVARSGLNQANLKSQFILNYLDTNSREGTPWSSYAKCVENGQIEYLQL
FT                   ENGFRAVYTLSNEKLMLVPQQMTAEWYGILSASGKKQAAKSYVLDEETGLYVLKSQGVT
FT                   ARHRQQIDADAHKAGFTMEDYAEMQALAEEEESEAAESTSFVVALDYTLTPDGLTVTIP
FT                   HAELAETGGAKIRTIQLLPFFGAAGTAEKGVIVVPDGSGALIRFNNGKNTAPQYNQNIY
FT                   DLDLIDSDITATQNMQSARLALFGICREDCGLLVSCGRGAALASVTADVAGRNNSYNYA
FT                   YFTFGLRRTDTLMVAGEEAIVAEQDSYPVDCAVSYRILDGEYTGYNGLARAVRESLLAD
FT                   GSLTLKTEKSGDIPFYCDILCGVRETAHWMGVQYLRVMPMTTFAQAEEILSGLRQEQVG
FT                   NLRVNLQGWMNGGYYHDPVNHVSVQDGLGGERGLQSLRAAAAETGGKIYPDAALQQVTE
FT                   IAKGFLRSEEASRYYAKGYVAEFGTVSPVSMRRTATMGFTERAYMLLSPRFLPRYAASL
FT                   ASAAERLGLDALSLRDLGNEVHADKRRTNVISREADLDLVRHAFGVIGAGERELMVSGG
FT                   NDYSFPYVRHVINAPVEATMFAIVDEQIPLWEMILHGSADYCGSPLNLMQSEDRRAALL
FT                   HLIEYGASTHYTFTWQDAADMKYTGLNNNYATTFSSWKEEAAESYRFVNGALAGVSGAE
FT                   MLRHDRLSDTLARVTYSNGTVLYVNSGEHEAEADGYRIPAMDYLAVGGENE"
FT                   /product="GH129"
FT   CDS             complement(7290..7922)
FT                   /translation="MNRMGIKDKWQHFFHTVSHPADGYYWIRHQEKGSLWIAFLMVLLY
FT                   GIVFSMNRISSSFIVNDIEPRTVNLPAELAGVILLYLILCVGNWSVTCLMDGEGRFKDI
FT                   LIAAGYALFPMMAATAAATAVSHLVAENEEAFYTLLIGLGTAYTVIMLVIGIMQVHNYT
FT                   FGKTLITLFLTVVAMLIIIFLGLLVINFITQVYSFFRGIYMELIFRV"
FT                   /product="hypothetical protein"
FT   CDS             complement(7919..9400)
FT                   /translation="MKQVKRILCLMLLACLAAGSACAADSLPYDCYNYDHWNNILYTPA
FT                   PYVPDGLVSGATLRLEGEPVGAFRNPQDLCVSADGDVYIADTGNNRIVVLAPDLKTVRR
FT                   IITGFEAEGGTQTFNAPTGVALSEKHRLYIADSLNRRIVVLEPDGTFVKYVRDPQSEVL
FT                   DEGYVFTPLRVSVDYADRVYVIAQNMFQGIMVFSGEGDFTGFFGTISVKISLWEKFWRK
FT                   LATKEERSKQQLFIPTEFTGIDVDDEGFVYASSKDTDGLQAVRRLNPKGEDVIRKGPRG
FT                   NLGGDRVFGSVGNYAGPSQIVDVVYRGHGAYSLLDGRRGRIFTYDHEGNLLYIFGGIGT
FT                   QAGTFQTPAAIEKLGSSLLVLDSQQGNLTVFAETEYGALINRAVSLRFDGDESQAVPLW
```

```
FT                   QGVLRLDEINELANTGIGKAYLSAGDNANAMLYLRRGMNREYYSVAFKRWRNERLRNNI
FT                   SLILSGVVAAAALAVLWVKAVRPAIRKKGKRRDPS"
FT                   /product="hypothetical protein"
FT   CDS             complement(9412..10401)
FT                   /translation="MTGMEKAKISRLLPRMPHRKPAELSPEEETALRQKRLAKIYRKMK
FT                   RQSGKKKLNRSTAGNAVLFILMGICGVFMVLPLVMILNNAVKPLDELYQYPPRIFVRNP
FT                   TLNNFTDLYVLLNESWVPFSRYILNTIIITVGGMTGHVIIASMAAYPLAKNHFPGKKIL
FT                   FSVVVLSMMFSWTVTQIPQYLIISWLGINNTYAALVLPAWSFGMGLYLMKQFMEQLPDS
FT                   LMESARLQGANEWQIFWRIVMPNVKPAWLTLAIFQFQQMWANTGSMFLRSEELKPLQYA
FT                   LQQITSGGVSRAGAGAAVTFLIAAVPIIFFLICQSSIMETMTTSGMKD"
FT                   /product="binding-protein-dependent transport systems
FT                   inner membrane component"
FT   CDS             complement(10398..11330)
FT                   /translation="MSMAIRNERSMRKKDTLWHQVRINRASYLLMAPYFTLFFLFTVLP
FT                   VLMSVYLSFNYFNMLEMPRWVGWANYIKLLLEDDIFIISLRNTLLFAVVTGPVSYLLCL
FT                   LFAWIINEFRPKVRAFLTLVFYAPSICGNAYMIWNLILTGDRYGYLNGILLRTGILNEP
FT                   ILWMQTEKFVMPVLIVVQLWLSLGTGFLSFIAGLQTVDKGLYEAAAIDGVKNRWQELWF
FT                   VTLPAMKPQLMFGAVMQITQSFAVADISIALAGNPSVNYSGATIVTHLLDYGSTRLDMG
FT                   YASAIATILFLLMVGTNKLVQRILRRVGE"
FT                   /product="binding-protein-dependent transport systems
FT                   inner membrane component"
FT   CDS             complement(11332..14316)
FT                   /db_xref="GO:0005215"
FT                   /db_xref="GO:0006810"
FT                   /translation="MRKKTGIVLAMLLALAAGTAMAATDLGIREYEDQVSTYSINPDIP
FT                   SYAEYCAAHGTAARPEKEIVLEAADTVRYAEEGGENAPLLKAQAEGRDGVSVLTGEEAV
FT                   IEWDFEAEEAGWYDLEVEYYPWPGRNAEIQRAFFLDGALPYRELALVNFSRVWRNDLHP
FT                   EIVRSTAEDGTEEIRWLRDNQGNELKPSPAEAPEWRTSLLHDSNGYISESLKLYLDRGA
FT                   HTLTVLSLREPMLIRRMCFRAGTDVKPYAPPEAGGAANRVIRIEAEAASGTSSQMLYPV
FT                   QDQSSRSVSPSSPKYLLNNSIGGNAWKSAGQWIEWRFSVPEEGNYAISLYDKQNFIRGT
FT                   DVYRKIYIDGEVPFAEFQAAAFPYTQDWRLETLSGEQGEPYLIHLAAGEHTLRMEVVLG
FT                   EMAAVIRQVQDCVLQLNGIYRQVLYITGVSPDPYRDYQIERSLPGLEAQLQTVKKDLRS
FT                   AIEALEKTAGHRSDKLTVLKTMDDQLTELIRDQERFTEVLSSYKTNVRACGNWITQVLG
FT                   QPLQIDRIFVHTADVRPEISGTDFLSGTGFELSRLYSSFTINYNQIGNVAEKTEDSRVI
FT                   TLWVGTGRDQANVIKALIDEHFTPKTGISVNVQLVDMNTLLRATLSGQGPDVAIQVANT
FT                   TGIAGAVMNTGNDTPVNYGLRNAVMDLTQFDDFAEVAERFAPSALVPFSFNGAAYALPD
FT                   TQTFPMMFYRKDILAEIGLEVPETWDDVKVAMSVLSKNQMEFGMLPGEQTFAMLLFQAG
FT                   GEYYTENGDASALDSDIAVNVFRRYCDFYTDYKLDMATSVEERFRTGECPIIITDYTTY
FT                   NNLQVSAPDIKGLWDFTHVPGTLREDGTLDTTTGCSGLADIIMSATKEPEACWEFLKWW
FT                   TGAEIQTLYGREMESLMGASARVATANLEALAGLNWPIRDYRELAKQFSHVRGIPQVPG
FT                   GYYTWRNVNNAFYAVTTGSASTGRTEFGNTPREELMDKIYYINAEITYKRTEFGLPVAE
FT                   TGEGK"
FT                   /product="N-Acetyl-D-glucosamine ABC transport system,
FT                   sugar-binding protein"
FT   CDS             complement(14353..15825)
FT                   /translation="MKSSMKSICGIAAAVMTLLFIILSGTAEETGETMLVEAETGELKG
FT                   HTAVAGNGGNQWGEGFKTAGEDSVSVEVLVSREGFYDIAVIQASQGGHKENPVLLDGQN
FT                   IGSAVTDETTFGKSVLEHIWLSAGMHTLGVGTSWGHIRLDGFELKPSAALPEDLYTVPE
FT                   KMCVGEASPEARKLYDWLRECYGKKILSGQQCAGGMYGMENQAIWRATGGDYPAVLGLD
FT                   MIDYSPSRVEHGGESDKAVPYAVEYWNQGGIVTFCWHWNAPSPYLKEPWYSGFYTEHTS
FT                   FNLGRVMNGEDEEGYRLLNRDIDAVAAQLLRLKEAGVPVLWRPLHEASGGWFWWGASGK
FT                   EAYLKLYHLLFDRLTHEYGLNNLIWIWNGQDPEWYPGDDEVDMIGIDIYPGEHVYDSQS
FT                   ASFIPLTRIAGERKMIVLSENGCIPDPEKVFRDGTVWGYWCTWGGEFVIRNTKFNKPSE
FT                   QYTETDMVRKAYSDTRVVTRKDLPDFRAADGD"
```

```
FT                   /product="CBM35-GH26"
FT   CDS             complement(16145..17551)
FT                   /translation="METIREGKEPSWCRELKTNMARMAQELIKVVRERTDGGPVFTPEE
FT                   FGYQPGETATEAIQGAIEAAAEQSGIVRLGNGDYVSGTLVMRSGVCLEVRSGSRLLGST
FT                   DLRDYPEHHAARLTVQDTSMGMHQSLLFAEGCENICLRGGGVIDGRGSPAHFPGEETAQ
FT                   GTPGRPFLIRMIDCRNVHIADLTMKDAACWMQNYLNCENLLLEGLTVRNHANYNNDGMD
FT                   IDGCRNVVIRNCRVSSGDDALCFKGAGQRELSRVLVEDCDFYSACNAVKVGTDTQGDFR
FT                   QVYIRRCRIGGLAEDPSGLKHACADSGISLEMVDGGTLENFLIEDIHMIRAWSPFFLRL
FT                   ENRGRVKPGDPAPGPGTLRRVLFSDIRGRETGPRGSYLLGIPEKPIEDIAFRRVLLRRE
FT                   ASRRPVRKEKDFDEMRGVYPDAHMIDPVGDAPAYALWARHVRGLSLDGYEVIPEGKERR
FT                   PAYIFTDTEF"
FT                   /product="GH28"
FT   CDS             complement(17567..18997)
FT                   /db_xref="GO:0004467"
FT                   /translation="MLPETSVSFRTEDEKLQRVYDAAEKKCLLNLKDFSGDTVLVEGGG
FT                   YEKIWLETQPMGGEMYALRNLEAAKNNILLFMKHQREDGRMPGSIQHINGKTEPQFNKY
FT                   QGFCFPDPALNLYYLMGKDREYLEQLKEALIRFDACLWRTRDVSGDGLLSSFCVYDTGE
FT                   DLAVRYGDAPCWWEEDEPPEGYGVVPMASMDVTSWSFAARRAAAEICRIQGDPAFTDWE
FT                   KKAGDVADALRRRLWDARRGALYDRDRTGKTIDILCHNTLRCMYWGSVSRDMADRFVKE
FT                   HLLNASEFWTPFPLPSVAADDPAFRNAPENNWSGQPEGLTYQRAITALERYGYARIVTA
FT                   LGKKLIDAVDANGCRFTQQYDPFTGKASLVDEADHRPVSGDSGEKVQDAYGPTMLACLE
FT                   YIAHRFGIHPHMGQVWFSLDRGKPYEYEAAFYGHRYGIRSDGKKAEIRADGKDIGRFSC
FT                   GRRIVTNENGDFLRTELI"
FT                   /product="Long-chain-fatty-acid--CoA ligase (EC 6.2.1.3)"
FT                   /EC_number="6.2.1.3"
FT   CDS             19235..20188
FT                   /translation="MSLSSSPLAYRGGIRHRIAMTLRRDWQLWVLLLPCLVFFIVFCYF
FT                   PMYGIQIAFKDYKAVLKIEGSPWVGLKHFEKFFGSYYCGRMFLNTFLLNFYGLLFGFPV
FT                   PIILAILLNQLNSKRFKGFAQTAVYVPHFISTVVLAGMIYLFFSPTNGIINHLITAAGG
FT                   KSIYFIMEPGWFRPLFIGSDIWQNAGWNTILYIATLTGIDPQLYEAATIDGATRWEKIK
FT                   HIDIPHLIPIAVMMLILNSGSLLQSNTDKALLLQTSGNMATSDIFGVYVYNVGIAAPKA
FT                   QYSYASAIGLCLNIINFVIIMGVNAVSRKMSDISLF"
FT                   /product="Predicted beta-xyloside ABC transporter,
FT                   permease component"
FT   CDS             20205..21143
FT                   /translation="MTVLSLKHRKRNRIRETRKDRIFNTFNLLFWILVLFIVLYPLWLI
FT                   LIASVSDPDAVLQAKVLLWPVDFSLMGYETVLQNSELLRSYVNSIFYTVVGSALSVIIT
FT                   LAAAYALSRKFAGKKLVNLAITFTMFFSGGLIPIFLNVLNLGLWNTRAIMILMNLVSVW
FT                   NLMVARTYIQTTIPEDLYEAAVIDGSNHFTYFFKCVLPLSGTIVAVLSVYYGVARWNDY
FT                   FTAQVMLRDRALFPLQRVLREILASLTSTGSSDTFFAAYADNLGGLQAALRKAEVAKYC
FT                   CIVVATVPAVFLYIFMQKYFVKGVMIGSVKG"
FT                   /product="ABC transporter, permease protein"
FT   CDS             21186..22091
FT                   /translation="MSQKKGVEVSARAARHIDGYRVLEGKREYVTYQGDSSIRIWYSDI
FT                   PWRYETHDHSAVEIVLVLEGTVTYSVEGTSYPVRKGEVLIVPSNTPHSLEMGEGSSRYL
FT                   FLFETDTIMTMRDIKSMVLYLHKPFHLRDESDAHKQIRELLLRAKDFYEKQELMWNTAC
FT                   YSCILQVYATLGRHYLIGIRSRTGDTVRSMDTEVINAVMTYINDHYREDLRLDDVAGFS
FT                   GFSRFYFSRSFKQQTGYSFKDYLCQKRLQAAMDLLTSTNLSMRGIAVESGFGSTASFNR
FT                   AFRACKGCTPTQYRAVYGML"
FT                   /product="Transcriptional regulator, AraC family"
FT   CDS             22153..23658
FT                   /translation="MKKLVSLLLACCMLLSLSAAFAEEKTELTAFQYALENQNTDFNGL
FT                   WFFDELENKTNTHVTFQMVKDADWATSVNLMFAVPDQMPDMILRHAVDVEEYGVSQGLL
FT                   LPLDEYLTEEIMPNYVSRLNINNAGDSIPASDGKTYYIGFLMAQNVNHTGTWYVNQKEL
```

```
FT                   DKLGKTAPATIDEVTELLRAMKADGIKWPFSASYTQFGPETIWNQFASFGVPENTYYYF
FT                   IDADNKVQLTFAQDGWRACVEWLHQLYEEELMDPESLTQDSNLWANKVNDGEVGYFCYL
FT                   RLINTAIKAENVENFKSILPPAADGFKACVSQILEVPEAGAYLTKNCKDPVAALKWIDA
FT                   QLETETMMVSLNGKVGEQIVLNDEGKYEVINIPEDNGLYQFVPVTMGQFFAPGDYYTSI
FT                   YQMAPHRVERYEDSKWYAEAGVLEPKSFQYLRDLSKMSNDDATEAATLYTDLQKLVEES
FT                   FANFVRNGVTDESWNEFQAQAKAIGSDRYIELYQNAYNAYLAK"
FT                   /product="ABC transporter, substrate-binding protein"
FT   CDS             23775..25322
FT                   /translation="MNRTVKITCLLAAALFLLTGCAVSRPGEGSAPPGASLTALQYEID
FT                   NIAVDFSSMWYYRQIEQQTGVHVDFSEVKDSEWSSSVGLAFAKGKMPDMILRGSLDVEE
FT                   YGVTRHLLVPLDEYIDKGFLPNYAARMDQAGLREQLTASDGHIYQLGFLISQGVNTNGH
FT                   FFINKTWLDKLNLQVPRTVDQLTEVLRRFRSDDPNGNGLQDEIPLEFTFDDNTTGIYNL
FT                   FSFFGLPLNEDYVYVDNTGNVRFAPEEDAFFETLTWLHDLYLEGLLDIDFISQGSSIWA
FT                   AKVNEGNTGMFSYWRLQNTALSPGTAANYQVMVPVHVEGRNACLPRNMDIIEFGAALTT
FT                   DNRDIEASLRWLDAQFETENMLVSQNGPVGDTLIRREDGRYEVSYVPPDNELYQSVPVI
FT                   CGQFFAPAEYYASVYVPAAHRLEKSAYCDLYEQAGVLEPVSGKLLTAVAPKTTAQSAET
FT                   TRLKAKLKSLVDAEIVSMVTRGVTDESREAFRSKLKENGSVEYRQVYQQIYDRWKEK"
FT                   /product="hypothetical protein"
FT   CDS             25326..27593
FT                   /translation="MIRKSRTSVFIRYALSYIAVVLLLFCSITGYLYVRLSGEAREEII
FT                   DNQINRLSRIASQHESYISAMLNTAEEIGLSPDIEFFRYNEEPWKAYDLQLQMVPYTTT
FT                   ASTFCDQVYLYFFGDDMIYSSSSSMTLSLFARMMRYEFIPMENLVSRIASTDRMTILPA
FT                   QQVSSTLVDSSRVVTFLVPLGSAPGSGKGILLFLIKDSVYQSLFADAIDDRVNTYIYQG
FT                   DDLLSSSEKLSLPRDRVEASQDAPSRLFRWENQDWTQVTFSGRNWGLTYTAVLRNADIN
FT                   SSIMQQLFSILAILPVFVFLSLLLALWMARRHAKPIRAITGMLSRETDETPRDELQQIS
FT                   SGISRLTTRNQELSSRLERALPLQRHDFVFRFVKGRFETRQEAVAAGQAVGLEIDRPVY
FT                   AVILCSGSAGNDHPFDLSLPPFSRLSGYSGAGVELVALKANLYLVFSDDPKVLTGMAET
FT                   LRCECAAGGERCITSISALHTDFTEAPSAYLEAAAAYENRFVMGSQKVLCYGDISPDLE
FT                   EILPQARKITTSISQALTLGSREVLDVRISDLLHFLKNTSMSPFAFRMIYSEVIRSLVH
FT                   THTAALSGQGSVQETYNIFTLTSCQSIDDLDELLRRLCDRLITGESASGEKVPDEEDEI
FT                   SQVVRYMEEHFSDPEISITALADSLELSTTRFSLSFKEKAGMSPLEYLTLLRVEHAKEL
FT                   LESSDLTIRDISTQVGYYDSGSFIRRFKQVTGETPLQYRRGRNPKKENGIPDRDASVPT
FT                   EE"
FT                   /product="Transcriptional regulator, AraC family"
FT   CDS             27603..28445
FT                   /translation="MIHTDQINIRDPYVLVHNGTYYLYGTRGPDCWGPADGFDVYTGQD
FT                   LVSWEGPRVCFRNDGSFWADRNYWAPEVHPWQGMFYMFASFKSDARRRGTAILRSPSPL
FT                   GPFEPWSDGPVTPADWECLDGTFYVSADGTPWIVFCHEWVQAGDGEVLAMRLTDDLKAS
FT                   AGTPVLLFRGSDAPWSHAVRHSSGITGFVTDGPFLWRGKDGSLLCLWSGFSEKGYAQGL
FT                   AVSDNGEIDGHFSQADPLFLEDGGHGMLFRSKEGRTLLALHSPNTHLLERPRFVPVDL"
FT                   /product="GH43_17"
FT   CDS             28492..29772
FT                   /db_xref="GO:0008993"
FT                   /translation="MHYVLAIDIGASSGRHIVGWREGSGIRTEEVYRFPNGITEQDGSL
FT                   TWDTDALLFHVKAGIDKALEKYPSIQSLSIDTWAVDYVLMNGGSPVLPCYAYRDSRTEA
FT                   VIGEVHEKMPFPELYRRTGIQYQPFNTLYQLYADKVSGRLDQADGFLMIPEYLMYRLCG
FT                   AESHEYTNATTTGLVSAETGEYDPGIISALGLPARFFGPLRQPGTVIGEYRGIRVMLCA
FT                   THDTGSAVEGIPMEGCQPYISSGTWSLLGVKTERPVITEAGRLANWSNEGGAGYNRYQK
FT                   NIMGMWLMNRLRAELCPDRPWDEIMAEAAEKHFDHLVDVNDPVFLAPESMKAAFDSRLP
FT                   HPPRCIAGYFRCACRSLADGYRRAVEEMEQNTGRRYDRLYIVGGGAKNTYLNRLTEEST
FT                   GKQVTALPIEATALGNIRIQLKGGTDT"
FT                   /product="Rhamnulokinase (EC 2.7.1.5)"
FT                   /EC_number="2.7.1.5"
FT   CDS             29769..31025
```

```
FT                   /db_xref="GO:0008740"
FT                   /translation="MSYEESRKRYAALGVDTDAVIRLLRAVPVSLHCWQGDDVRGFDTD
FT                   PSKPLTGGIQTTGNYPGRARTPEELMADLDMVLSLIPGTPKMNLHASYAIFENGEWADR
FT                   NRLQPKHFRKWAAFCKERNLGCDFNPTFFSHPKCDPLTLSSPNAETRKFWIEHGKACIR
FT                   ISSYLAEELDFPCVMNIWTGDGFKDIPGDRIGPRIRYRDAIDEILREPYDFSRVKPCIE
FT                   SKVFGIGVEAYTTGSAEFALCYAAMNRDKCIPLMDNGHYHPTEVVSDKIPALLTFFREI
FT                   ALHITRPIRWDSDHVVLYDDETREIAREIVRCGGLDGRVHIALDYFDASINRISAWATG
FT                   YRNLQKALLYALLQPHEEMKKMQDKGQFSRLMVLQEEIKTMPFGEIWAEYCRFCGKPAD
FT                   GEWFPLVEEYEKKVLVNRK"
FT                   /product="L-rhamnose isomerase (EC 5.3.1.14)"
FT                   /EC_number="5.3.1.14"
FT   CDS             31022..31858
FT                   /db_xref="GO:0008994"
FT                   /translation="MKDILAAPFMTEMIRTATAMYDHGWDERNGGNISLMLDEKEAGEY
FT                   LDMRNVLRTIPTGFSMPELDGKIFLVTGTGKYFRNIQYEPEVNLGIIRLRDCGQQADLL
FT                   WGYSDGGQFTSEFPAHMMSHSARLKMNPANRVIMHCHPANLLAMTYVHALDEREFTRTL
FT                   WQMCTECVVVFPDGVNVLPWMLCGTNEIGEATAEKMKTTRLVVWSQHGIYGAGKDLDET
FT                   FGLIETAEKAAEIYMKIAHLPRVNTITDEQMHQLEKRFGIKAREGYLEEPDRGTDK"
FT                   /product="Rhamnulose-1-phosphate aldolase (EC 4.1.2.19)"
FT                   /EC_number="4.1.2.19"
FT   CDS             31855..33252
FT                   /db_xref="GO:0004557"
FT                   /translation="MKQHAGKNIRELKVAYIGGGSRGWAWGFMKDLALDGEICGTVRLY
FT                   DIDREAADRNRIIGEKISAHPDAVSRWAYTVSDSLQEALTGADFVVISILPGTFDEMES
FT                   DVHLPERLGVWQPVGDTVGAGGFMRSMRTIPMFVTIGEAIRDYAPEAWVINYTNPMSLC
FT                   VRTLYEVFPSIRAFGCCHEVFGTQKLLCSMLKTGEGIDRVKRQDLYTTVTGINHFTWLT
FT                   SASWKGTDLYPLYARFAEKYAESGYSEGGDDNWMNSHFSCAQRVKFDLFRRYGVIAAAG
FT                   DRHLAEFVAPWYTRDPETVREWKFSLTPVSWRKQDLKEKLQMSDDLVSGKKEITLNGSG
FT                   EEGHLLLKALLGLGDMVSNVNIPNRGQIPNLPLGAVVETNALFGLNRIDPVYAGPVPAS
FT                   ILPLITRHVLNQENTLTAALCCDRKLGLTTFLNDPQLAAVTPKDGEKLFNDMLENQRTF
FT                   LPEKWFE"
FT                   /product="GH4"
FT                   /EC_number="3.2.1.22"
FT   CDS             complement(33353..34918)
FT                   /translation="MAGGLTTTIRTPEGDEPKQRLFDTFCYNDEKMLAELRKVCSFIGG
FT                   HFDEFIIDDFFFTNCTCEACRRERDEYNRERGITDGSWKKYRLHLMEKVSREDVIAPAK
FT                   AANPKCRITIKYPNWAESYQETGYNPAAQRKLFDRIYTGTEARDTVTTDQHLPRYLSYS
FT                   LMTYFENMWPEHNGGGWFDPFDLHITEQYLEQAYLTAFSKPKEIMLFCFQMLTDNPFIP
FT                   PLGWHLDKLDEVLDYCGNPVGIQCYLPDNCQGEDNIQDFLGMCGLPVMLTPYWPEKAEY
FT                   ILLTRSSACDPDVVEKLEKYVAAGGKALVTSGFVEDTMDRGIQRMTSIRFTGRRIRGRN
FT                   YRVENAGVWPSTEFPRGTEEIGIPVCEFRNNATWAVVKAAESEESFAILLRDTYGKGQM
FT                   LTLAVPDNFPDLYKLPGKVLTRIRREFPAGGVYLESKAGISLFPYDNDTFVLYPYVASG
FT                   AQPATILIHVTGDAEALELPLRKHPVTGEAIRMQPLYRRNGETVFEGRAMPGKYELCRI
FT                   IKRK"
FT                   /product="FIG00615029: hypothetical protein"
FT   CDS             34952..35581
FT                   /translation="MQHAYDQARDFGQLCRYRAANAALTERPRVVFAGDSITDNWINLD
FT                   PSFFTHGLVDRGISGQTTPQLLVRFRQDVIDLHPRAVHIMIGTNDIAGNTGAAAMETVE
FT                   GNIASMADLARAHGIRVILASVPPAGAFPWAKDKQPVPQIAALNRWIAAYARSNGFTYV
FT                   DYHPVLATPDGAMKPGLASDGVHPTAAGYAAMRPLAEAAIEKALGR"
FT                   /product="Lipolytic enzyme, G-D-S-L precursor"
XX
SQ   Sequence 35658 BP; 7394 A; 10994 C; 9610 G; 7660 T; 0 other;
     ccttgccgtc ccagttatag tcgtcatagt tggccatgat ccaggcagcg gtggtggtga        60
```

aggctacctt cttgtcacgg tacagccagc ggcagacatc ccagccgttc tcctcgggca 120
gcggataggc cgccttgtcc gtaagccaca gatcctgcag gaacttcagg gtttcaccca 180
tctccacaga ggagaggttc tccttgggag aggaggccag cgttgtgccg ttggacatat 240
accagtaagg caggcagtcc gtgatccagg cgccgtagcc gtaaacatcc gtgacgccgt 300
cgccgtcgtt atccttggtc agggcaatgc agtactcccg gaacttgtcc caggtccact 360
cgccgcgggc agccaggtcc cgggggtcct ccaggttggc ttcctcgatc atctgcaggt 420
tgaaagccag gggataggtg gcggccacct ggtcctccgc gccgttgacg gtcagcaggg 480
agacagcgcc gctttccagc gccacataat tcatgacggt gtcgttatgg gtgagcagcg 540
gatcgtcggg atccagaaca tcccgcaggt ccatggcata gccgttcatg acggcgggaa 600
cgccccagcc cagctccacc atgtacacgt cacagtccgg ttcaccggcc aggatggagt 660
tgttcacgga atcctgcacg ccagcatagg taaggttctg gaactcatag gtcacgccgt 720
aggtgttctc aatgttcttc acattatcga aacgcatgag atcggaatcc ttgccggttg 780
cggcttcatt gtcttcccag gattcatcgt tggagtcata gtagatatcc caccagagac 840
cgatggtaat attccgggga tcttccgcca gcgccgttac agccagggag agcaccatgg 900
ccagagccag cagcattgct gttatttttt tcataagtca gtttctcctt tccagaaatc 960
aggtcatcat ttgatcgcct ggcccacggc cacagcataa acattccagg gggaggagct 1020
ttcgcactgc atcatggcgc cccaggtgga cttgtcctcg ccgcagaggg cttcgatctg 1080
ctcataggtc acctgcgcga ttttgccgtc cgccacgtcc gcgccatcgt tgccaacgcg 1140
catccagccg gctgctgccc agggcatgac cagccacata ttgccgtcct cggactcata 1200
ggagatggtt accacggagc cgggaaccag agcgtcgatg atttcctggg gcatttccag 1260
gccgtcctgc gcccatgcgt cggcggactt ggtgaagccg gggaattcaa ccaggttggt 1320
caggccgaag aactctgcct tctggcccac gcggatgccg taaacctccc agggagagga 1380
gccttcgcac tgcatcatgg cgccccaggt ggctttgtcc tcgccgcaaa gtgcagcgat 1440
ctcttcatag gtgatatagg ccttgccacc catcttggca gcggttccct gggacacgcg 1500
catccagccg gcttccgccc agggcatgac cagccagata tcgccgtcct cggagctgta 1560
ggtgatctcc accacagagc cgggcaccag cgcgtcaatg atctcctgcg gcatttccag 1620
gccgtcctgc gcccaggcgt ccgcggactt ggtaaagccg gggaattcaa ccgcgttctt 1680
cagcacgatc cggtttgccc gctgacccac gcgcacggca aagacttccc agtcggaagc 1740
gctttcgcac tgcagcatgg caccccaggt ggatttgtct tcaccgcaga gggcttcgat 1800
catctcatag ggaatctggg caatggtctt gctgttgttg atggcagcag ttccctggcc 1860
gacacgcatc cagccagcgg ttgcccaggg catgacgatc cacatggagc cgtccgcact 1920
cgcgtattcc acttccacca cggagccggg caccagcgca tcaataatct cctgcggcat 1980
ttccaggccg tcctgtgccc aggcaccggc agacttgtgg aaatccggga attcaacggc 2040
gttcgtcagg gcaacgaggt tgctcagatc cttttcggaa gcctccgaca cgtattccgc 2100
agtcggatcc agggcgatcg cgtttccttc cgcgtccagg aactgaatgt tgtccaggta 2160
gaaggtcgcg ggttctccgc cggcctggtc ttccctggag atcatgatgt agttgcccgc 2220
gccggcaaca aacggcgcct tgaactgaat ctttacattg cgcggatttt tcttttccag 2280
atagacagac cagtcggcct tgttctcatc cgcgttctcg ccggtgtagc tgtataccac 2340
gccggagacc gcgtagaact tgccgtccgc ggccttgtca acgccgatgt caaacgtgac 2400
agcggcaacg tcagccagtt tttcgccggc caggccctca agattcaggg ccacataggg 2460
gatcccctgc gcggcaacgc gcagtgcctt ggaaccgttg tagtcaacaa cctccagggc 2520
cgttgaggca tctgcgttcg gtttcgcagc ggaaacaccg aggaaggcaa aaacgccatc 2580
ttcgaagtca agcgcgttca ccggatcggc agacgcccag gagcacatac cgactgccat 2640
aacggcaacc agcaaccagg aaatgagttt cttcacagga ctttcctcct tttatttttt 2700
acccgatccc gggtaattaa caattgaaag cgggctcagc ccacaatacc ggaacgttcc 2760
acgccctcga taaaccgccg ctgcagcacc aggtagatca ccaccaccgg caccaggacc 2820
agaaccacgc cggcgtcaaa atacagctgc tgttcgcgga tggtcagcac cttttccgca 2880
ttggcgatca cgttgatcag cgattccagc ttcctggtca cgaggatgga atcggaaatg 2940
ttgaaggtgt tggcaaaaaa ggaatcgttg tactgccaga cgatggagaa cacggcgacc 3000
gtaatgaccg ccggagccgc attgggcatc atgatggtga aaaacgtccg ggctgtgccg 3060
gcgccgtcca ccagggccgc ctcctcaatc tccttcggca ggccgcggaa aaactggttg 3120
aagatataga tgtacaggcc ggaccgcagg ccgcagccga aaacggacat gatgaacgtc 3180
ggccaggcag tctgcagcag gttgatcgtc ttcccgccat tgaaaagtcc tacaattccc 3240
aggggattaa agaagcggaa ggtcgtataa agcgggaact ggatcgtatt cagcggaatg 3300

acgatcatca caagcacaaa agcaaaaagc acgctcttca gcggaaagcg gaaacgggca 3360
aaaccgtaac cgaccatgga gcagatcagc agctggatca gggtcacccc gaacacatac 3420
agaatcgtat tgcccaggat gggcaggtaa ttcaggtatt tgactgccat gctatagcgt 3480
tccagcgtcg ggttgatcgg gatcgtgaat accatggggt taatggagtc ctgccgggaa 3540
aagaaggagt tcgagatgat attgatcacc ggagcgagaa tgacatagct gatcccgacg 3600
atcacgatca gccggaaaag accgatcacc aggttcttca tcccgttcag ggttttatcc 3660
cgctgattgg cgtagacggg agaattccgg aatccctgcc aggtgatttt tccgttctgt 3720
gccggatgga ttgtctgcat gttctttctc cctcctcccg tcagttgtaa tagaaggtcc 3780
gccggttcag caggtaaacg atcagcccca ggatcaggat ggtgatcagt gtcgagacca 3840
cagagaacgc tgacgacagg ccgtagttgt acagttcctt gtaggtgcgc gttgccagtc 3900
tgatcacctc actggaggaa aaccggtcca cgaccgtgta gatcgtgttt gtgatcatgt 3960
gcggcataac catgggaaaa gtcaccttcc agaaggattc ataccctgtc gctccttcaa 4020
tgcgggccac ctcatacagg gaacccggaa tggactgcag cgccgcgatg aacaggatga 4080
tctgcacgcc cgatcccttg atgatggacg agatccggtt caccgcatcc acaatgtagc 4140
caagcacgga ttccggaatg gccaggtcgc cgaaaagctg catgtaatag gctacggaga 4200
cggtccccgc ctgcgcggtt tccaggtttg cggaggacac gccggcattc accagttccg 4260
tcgcccgggt gatggcggag cccaccgcgt ccgcgctcag gataatcggc aggaagaaga 4320
tcgcgcgtac cagcgccctg cctttgaact tccggttcag gagcatcgcc acaaaaaggc 4380
tgaagaacag gatcatcggc aggtcgatca gcatatccat cacagaggat gtcagtgtct 4440
gcttgaaggt ggcgtggctt gtgaatgcct cgatataatt gccgaagccg atatactcct 4500
ttgtcatacc cgtggcggcg tccaggccga tccgggacag gctgaattct cccagctgca 4560
ccaggcaccc ggcataaaac accaggaagc cgatcagcca cggaaggatg aaccagtatc 4620
cgcgcacggc gttcctctgc tcataggaaa gatgaaaaat acggtttttt ctgcggatgc 4680
cagtcattcg ttttctcctc cgaccgccag ataatccatt gccgggatcc ggtatccgtc 4740
cgcctcagct tcatgctccc cgctgttcac atacagcacc gtgccgttgg aatacgtcac 4800
gcgggccagc gtatcgctga gccggtcatg ccggagcatc tccgcgccgc tcaccccagc 4860
cagcgctccg ttcacaaagc ggtagctttc cgctgcctcc tccttccatg aagagaaggt 4920
cgtggcgtaa ttgttgttca gcccggtgta cttcatatcc gccgcgtcct gccaggtaaa 4980
tgtgtagtgc gttgacgcgc cgtattcgat caggtgcagc agggctgccc gccggtcctc 5040
gctctgcatc aggttcagcg gcgagccgca gtaatcagct gaaccgtgga ggatcatctc 5100
ccacagcgga atctgctcat ccacaatagc gaacatcgtc gcctccaccg gcgcgttgat 5160
cacgtgccgg acatagggga agctgtaatc gttgccgccg gaaaccatca gttcccgctc 5220
tccggcgccg ataacgccga aggcatgcct gaccaggtcc aggtccgctt cgcggctgat 5280
gacgttcgtc cgccgcttgt ccgcatggac ctcgtttccc aggtcccgga gtgaaagcgc 5340
gtccaggccc agccgctccg ccgccgatgc cagactggcg gcataccggg gcaggaaacg 5400
gggggacagg agcatatacg cccgctccgt aaatcccatg gtggccgtac gcctcatgga 5460
aaccggactg acggtaccga attccgccac atagcccttg gcgtaatacc ggctggcttc 5520
ctcgctgcgc aggaatcctt tcgcgatctc ggtcacctgc tgcagggctg cgtccggata 5580
aatctttcct ccggtttcag cggctgccgc gcgcaggctt tgcaatcctc tttccccgcc 5640
gagtccgtcc tgcacagaga cgtggttcac cggatcgtgg taatatccgc cgttcatcca 5700
gccctgcagg ttcacccgca ggttccctac ctgctcctga cggagcccgg acaaaatctc 5760
ctccgcctgg gcgaaggtcg tcatcggcat aacccggaga tactgaacac ccatccagtg 5820
cgccgtttcc cgaacgccgc agagaatgtc gcagtagaag gggatatcgc cggacttttc 5880
cgttttcagt gtcaggctgc cgtccgcgag cagggattcc cgcacagccc ttgccagtcc 5940
gttatagccg gtatattccc cgtccaggat ccggtaggat acggcgcagt ccaccgggta 6000
ggaatcctgt tccgccacga tcgcttcctc tccggccacc atcagggtat cggtccgtct 6060
caggccgaag gtaaagtagg cgtagttata gctgttgttc cttcccgcca catccgcggt 6120
gacagacgcc agtgccgcgc ctctgccgca ggagaccagc aggccgcagt cctcccggca 6180
gatgccgaaa agcgccagcc tggcggactg catattctgc gtggcggtaa tatcgctgtc 6240
gatcagatcc aggtcataaa tgttctggtt atactggggc gcggtattct tgccgttgtt 6300
gaaacggatc agggcgccgc tgccgtcggg aaccacgatc acacccttct ccgccgtacc 6360
ggccgcgccg aagaacggca gcagctggat ggtccggatc tttgctccgc ccgtctccgc 6420
gagttccgcg tgggggatgg tgaccgtcag cccatccggc gtcagcgtgt aatccagggc 6480
aaccacaaag gaagtgcttt ccgccgcttc ggattcctct tcctctgcca gcgcctgcat 6540

ctccgcgtaa tcctccatgg tgaaaccggc cttgtgcgcg tccgcgtcga tctgctgacg 6600
gtggcgggcc gtgacgccct gggacttcag cacatacaga ccggtttcct catccagaac 6660
ataggatttg gccgcctgct tctttccgga agccgacagg atgccgtacc attccgccgt 6720
catctgctgc ggtacaagca tcagcttttc attggacagt gtataaaccg cccggaaacc 6780
gttttccagc tgaaggtatt caatctgccc gttctccacg cacttggcgt aggagctcca 6840
gggtgtacct tcccgggagt ttgtatccag gtagttcagg ataaactggc tcttcaggtt 6900
tgcctggttc agtccggacc gggccaccgg gtcatctgcc gcgtcctgcg gattggaata 6960
gacggtcagc ccgctcttcc tgtcatacag cgcaatctcc gcggtttttt ccttcaggta 7020
gagtcccagc cggtcattct gtgccgcgag aacgaatccc ggcactgcgt tctccggatc 7080
cttcaggggc ttcagggctt ccgcttccgt tttcgccgcg ggctgaacaa tatactgacg 7140
gtactcgtca tacgcccggt agtggatcac gtaccgggca aggaacagca ccgcggcagc 7200
cgcagccagg cccgcgatga tccacagtac acgccggaag gcggtcggga gggtcagcct 7260
gtttttctca ttcattcagc attccctcct tacacacgga aaatcagttc catatagatc 7320
ccccggaaaa acgaatacac ctgcgtaata aaattgatca ccagcagtcc caggaaaata 7380
atgatcagca tcgccaccac cgtcagaaac agtgtgatca gcgttttacc gaaggtatag 7440
ttatgcacct gcatgatccc gatcaccagc atgatcacgg tgtaggctgt acccaggccg 7500
atcagcaggg tgtaaaacgc ctcctcattc tccgccacca ggtgggaaac agccgtcgcc 7560
gccgcggtgg ccgccatcat gggaaacaag gcgtaacccg ctgcgatcag gatgtcttta 7620
aaccttccct ctccgtccat caggcaggtc acggaccagt tgccgacaca gaggatgaga 7680
tacagcagaa tgacgcctgc cagctccgcc ggcaggttca cggtgcgggg ctcaatatcg 7740
ttgacaataa agctggaaga gatccggttc atggagaaca cgattccgta caggagcacc 7800
atcaggaagg cgatccacag gctgcccttt tcctgatgcc ggatccagta atacccgtct 7860
gccggatggc tgaccgtatg gaaaaaatgc tgccatttgt ccttgattcc catccggttc 7920
atgacggatc cctcctcttc ccttttttgc ggatcgccgg ccggaccgcc ttcacccaga 7980
gcacggccag tgcagccgca gcgacaacgc cggaaaggat caggctgata ttgttccgca 8040
gccgttcatt cctccaccgc ttaaaggcga cggagtaata ctcccggttc attccgcgcc 8100
gcaggtacag catggcgttc gcgttatctc ccgcggacag gtaggccttg ccgatcccgg 8160
tattcgccaa ttcgttgatc tcgtccagcc gcagcactcc ctgccacagc ggaaccgcct 8220
ggctttcatc cccgtcaaac cgcaggctga ccgcccggtt gatcagggcg ccgtattccg 8280
tctctgcgaa tacagtcagg ttcccctgct gggaatccag gacgagcagg ctgcttccca 8340
gcttctcaat ggccgcgggc gtctggaagg tcccggcctg cgtgccgatg ccgccaaaga 8400
tgtacagcag gtttccttca tggtcatagg taaagatcct gcccctccgg ccgtccagca 8460
gggaataggc tccgtggccg cggtacacca cgtccacgat ctggctcggg ccggcatagt 8520
tgccgaccga accgaacacc cggtcaccgc ccagatttcc gcgggggccc ttccggatca 8580
cgtcttcacc cttcggattc agccggcgga cagcctgcag gccgtccgta tccttgctgg 8640
acgcgtaaac aaagccctcg tcatccacat cgataccggt aaattcagtg ggaataaaca 8700
gctgctgttt gctccgctcc tccttggtcg ccagcttgcg ccagaacttt tcccacaggg 8760
agatcttcac gctgatcgtt ccgaaaaagc cggtgaaatc tccttccccg ctgaaaacca 8820
tgatgccctg gaacatgttc tgtgcgatca catagacccg gtccgcgtaa tccacagaca 8880
cccgcagggg agtgaacaca tagccctcgt ccagcacctc gctctgcggg tcccgcacat 8940
acttgacaaa ggttccgtcc ggttcaagca caacgatccg gcggttcagg ctgtccgcga 9000
tatacagcct gtgtttttcc gacagtgcga cgccggtggg cgcgttgaat gtctgcgtgc 9060
cgccctccgc ctcaaagccg gtgatgatcc gcctgaccgt cttcaggtcc ggcgccagca 9120
ccacgatccg gttgttcccg gtatccgcga tatacacatc cccgtccgcc gacacgcaca 9180
ggtcctgggg attccggaag gcgccgacag gctccccttc cagcctgagc gtggcaccgc 9240
tgaccagtcc gtccggaacg tagggcgccg ggtatacag gatattgttc cagtggtcat 9300
agttgtaaca gtcatacggc agggaatccg cggcgcacgc gcttcccgcg gccaggcagg 9360
ccagaagcat caggcacagg atccgtttca cttgcttcat gctccgcctc ctcagtcctt 9420
catgccggag gtcgtcatgg tctccatgat cgagctctgg cagatcagga aaaagatgat 9480
cgggaccgcc gcgatcagga aggtcaccgc cgccccggcg cccgcccggc tcacaccgcc 9540
ggaggtaatc tgctgcaggg cgtactgcag gggcttcagc tcctcggaac gcaggaacat 9600
gctgcccgta ttggcccaca tctgctggaa ctggaagatc gccagcgtga gccaggccgg 9660
cttcacgttc ggcatcacga tccgccagaa gatctgccat tcattggcgc cctgcagccg 9720
cgcgctttcc atcagggaat ccggcagctg ctccatgaac tgcttcatca ggtacaggcc 9780

catgccgaaa gaccatgccg gcagcaccag cgcggcatag gtgttgttga tcccgagcca 9840
ggagatgatc aggtactgcg gaatctgggt caccgtccag gagaacatca tggacagtac 9900
aaccacggaa aacaggatct ttttgccggg aaaatgattc ttggccaggg gataggccgc 9960
catggacgcg atgatcacat gaccggtcat tccccccacc gtaatgatga tggtattcag 10020
gatataccgg gagaacggaa cccagctttc attcagcagc acatacaggt ctgtaaagtt 10080
attcagcgtg ggattccgca caaagatcct cggcggatac tggtacagtt cgtccagcgg 10140
cttcaccgcg ttgttgagga tcatgaccag cggcaggacc ataaacacgc cgcagatccc 10200
catcaggata aacaggaccg cgttgccggc cgtggaccgg ttcagttttt tctttccgga 10260
ctgccgtttc attttccggt agatcttcgc caggcgcttc tgccgcaggg ctgtttcctc 10320
ctcggggctc agctccgcag gtttccggtg cggcatgcgg ggaagcagcc ggctgatctt 10380
tgccttttcc ataccggtca ttcacccacc cttctgagaa tccgctgcac cagcttgttt 10440
gttcccacca tcagcaggaa caggattgtc gcgatcgccg aggcataccc catatccagc 10500
cgggtggaac cgtaatccag cagatgggtc acaatggtcg cgccggaata attgaccgac 10560
ggattgcccg ccagcgcaat ggagatatcc gccacggcaa aagactgcgt gatctgcatc 10620
accgcgccga acatcagctg cggcttcatc gcgggcaggg tcacaaacca cagctcctgc 10680
caccggttct tcacgccgtc gatcgccgcc gcttcataga ggcccttgtc caccgtctgc 10740
aggcccgcga taaaggacag gaagccggtt cccagggaca gccacagctg tacaacgatc 10800
agcaccggca tgacaaactt ttcggtctgc atccagagga tcggctcgtt caggataccc 10860
gtccgcagca gaatgccgtt caggtatccg taccggtcgc ctgtcaggat caggttccag 10920
atcatatacg cgttgccgca gatggacggc gcgtagaaaa caagcgtcag aaaagcccgc 10980
actttcggcc tgaactcatt gatgatccat gcaaacagca ggcacagcag atagctgacc 11040
ggtccggtca ccacggcgaa tagcagggta ttccgcaggg agatgatgaa aatatcatcc 11100
tccagcagaa gcttgatata attggcccag ccgacccagc ggggcatctc cagcatgttg 11160
aaataattga acgacaggta aacagacatc agcacgggca gcactgtaaa gaggaaaaac 11220
agggtaaaat acggcgccat cagcaggtag gatgccctgt tgatccggac ctgatgccac 11280
agggtatcct tttccgcat gctccgttcg ttccggattg ccatgctcat ctcatttccc 11340
ctcccctgtt tctgcgacag gcagtccgaa ttctgtccgt ttataggtaa tctccgcgtt 11400
aatgtaataa atcttgtcca tcagttcctc acggggcgta ttgccgaact cggtgcgtcc 11460
cgtgctggcg ctgcccgtag tcactgcgta gaaggcgttg ttcacatttc gccaggtgta 11520
ataaccgccg ggcacctggg ggattcccct cacatggctg aactgctttg ccagttcccg 11580
gtagtcccgg atgggccagt tcagtcccgc cagggcttca aggttcgcgg tggccacccg 11640
tgcggaggcg cccatcagtg actccatttc ccgtccgtac agcgtctgga tctccgcgcc 11700
ggtccaccac ttcaggaatt cccagcaggc ttccggttcc tttgtcgcgc tcatgatgat 11760
atccgcaaga ccggagcagc cggttgttgt gtccagcgtt ccgtcttccc ggagggtacc 11820
gggcacatgg gtgaaatccc acagtccctt gatatccggc gcggatacct gcaggttgtt 11880
gtaggttgta taatccgtaa tgatgatcgg gcattcaccc gtacggaagc gttcttccac 11940
cgaggtggcc atgtccagct tgtaatctgt ataaaaatcg cagtaccggc ggaacacatt 12000
gaccgcgata tcgctgtcca gtgcggacgc gtctccgttc tccgtgtagt actcgcctcc 12060
ggcctggaac agcagcatgg caaaagtctg ctccccgggc agcatgccga attccatctg 12120
gttcttgctc agcacgctca tggccacctt gacgtcgtcc caggtttccg gcacttccag 12180
gccgatctca gccaggatgt ccttccggta gaacatcatg gggaaagtct gcgtgtccgg 12240
cagcgcgtaa gccgcgccgt taaaggagaa gggaaccagg gcgctcggag cgaaccgctc 12300
cgccacctcg gcaaaatcat caaactgcgt gagatccata acggcgttgc gcaggccgta 12360
gttcaccggc gtatcgttcc cggtattcat caccgcgccc gcgatccccg tggtattcgc 12420
gacctgaatg gccacgtccg gcccctggcc ggacagcgtg gcgcgcagca gcgtgttcat 12480
atccaccagc tggacgttca cgctgatccc ggtcttcggc gtaaaatgct cgtcgatcag 12540
ggctttgatc acattggcct ggtcccgtcc cgtgccgacc cacagcgtga tcacccggga 12600
gtcctccgtt ttctccgcca cattgccgat ctggttgtaa ttgatagtga aggagctgta 12660
taaacgggat aattcgaaac ccgtaccgga aaggaaatcc gtcccgctga tctccggacg 12720
gacgtcggcg gtatggacaa agatcctgtc aatctgcagc ggctgaccca gcacctgggt 12780
gatccagttg ccgcaggcac gtacgttggt cttataggaa gaaagcacct cggtgaaacg 12840
ttcctggtcc cggatcagct ctgtcagctg gtcatccatc gtcttcagca cagtcagctt 12900
gtcgctccgg tgtcctgctg ttttttccag cgcctcaatg gccgaccgca ggtccttctt 12960
aaccgtttgc agctgtgctt ccagtcccgg caggcttctt tcgatctggt aatcccggta 13020

cggatccggc gaaacaccgg tgatataaag cacctgccgg taaatcccgt tcagctgcag 13080
cacacagtcc tgcacctgcc ggatgacagc tgccatttcg cccagtacaa cctccatccg 13140
cagggtatgt tcccctgccg ccagatggat caggtacggt tctccctgct ccccggacag 13200
cgtttccagg cgccagtcct gggtataggg aaacgcggcc gcctggaatt cggcaaacgg 13260
cacttctccg tcaatataga tcttgcggta cacgtctgtt ccgcggataa aattctgctt 13320
atcgtaaagt gagatcgcgt agtttccttc ctccggcacg gaaaaacgcc actcgatcca 13380
ctgtcccgcg cttttccacg cgttgccgcc gatgctgttg ttcagcagat acttcgggct 13440
ggacggggaa acagatcggg aggactgatc ctgcacaggg tacagcatct gggaggaggt 13500
gccggacgcc gcttccgcct caatccggat cacccggttc gcggctccgc ccgcttccgg 13560
cggggcatac ggcttcacgt ccgtacccgc ccggaaacac atacgccgga tcagcatcgg 13620
ttcccgcagg gacagtaccg tcagcgtatg cgcgccccgg tccaggtaaa gcttcaggct 13680
ttcgctgata tatccgtttg agtcatggag gagagaagtg cgccactccg gcgcttccgc 13740
cggagaaggt ttcagttcgt ttccctggtt atccctgagc cagcggatct cttccgttcc 13800
gtcctcagcc gtgctgcgga cgatctccgg atgaaggtca ttccgccata cccgggaaaa 13860
gttcaccagc gcaagctccc tgtacggaag ggcaccgtcc aggaaaaagg ccctctggat 13920
ctcagcgttc cgtcccggcc agggatagta ttccacctcc aggtcatacc acccggcctc 13980
ttcagcctca aaatcccact cgatgactgc ttcctcaccc gtcagcaccg agactccatc 14040
ccgtccttca gcctgtgcct tcagcagggg agcgttctct ccgccttcct ccgcgtaccg 14100
gacggtatcc gccgcctcca ggacgatctc cttttcaggc cgggccgccg ttccatgggc 14160
cgcgcagtat tccgcatagg agggaatgtc cgggtttatg gaataagtgc tcacctggtc 14220
ctcatactcc cggataccca gatcggtggc agccatcgcc gtgccggctg ccagcgccag 14280
cagcatcgcc agcacaattc ccgttttctt tctcacttgt tctttcctcc ggtccattcg 14340
cacgcggcag actcagtctc cgtccgccgc gcggaaatca ggcagatcct tccgggtcac 14400
gacccgcgtg tccgaatacg ccttccggac catatccgtt tctgtatact gctctgaagg 14460
cttgttgaac ttggtattcc ggatcacaaa ttcgccgccc caggtgcacc agtatcccca 14520
gaccgtcccg tcccggaaca ccttctccgg atccggaatg cagccgttct cggacagcac 14580
gatcatcttc cgttccccg cgatccgcgt cagcggtatg aaggacgcgc tctgggagtc 14640
ataaacgtgt tccccgggat agatgtcgat gccgatcata tccacctcgt catcccccgg 14700
ataccactcc ggatcctgtc cgttccagat ccagatcagg ttgttgaggc cgtattcatg 14760
ggtcagccgg tcaaaaagca ggtgatacag tttcaggtat gcttcctttc cggaggctcc 14820
ccaccagaac cacccgccgc tcgcttcatg cagcggccgc cacagcaccg gcaccccggc 14880
ctccttcagc cgcagcagct gcgccgccac cgcgtcgata tccctgttca ggagccggta 14940
tccctcttca tcctctccgt tcataaccct gcccaggttg aagctggtat gctcggtata 15000
aaagcccgaa taccagggtt ccttcagata gggcgatggt gcgttccagt gccagcaaaa 15060
ggtcacgatc ccgccctggt tccagtattc aaccgcgtac gggaccgcct tgtcgctctc 15120
accgccatgc tccacgcggc tgggggaata atcgatcatg tccagtccga gaaccgcggg 15180
ataatcaccg cccgttgccc gccagattgc ctggttttcc attccgtaca ttccgccggc 15240
acactgctgg ccggaaagga tctttttccc gtagcattcc cgcagccagt catacagctt 15300
tcgcgcttcc ggtgaggctt caccgacgca catttttttcc ggaacggtat acaggtcctc 15360
cggcagcgca gcggaaggct tcagctcaaa cccgtccagc cggatatgtc cccagctggt 15420
accgacaccc agggtatgca tacctgcgga cagccagata tgttccagca cgcttttccc 15480
gaacgtcgtt tcatccgtaa ccgcgctgcc gatattctgc ccgtccagga gcaccgggtt 15540
ttccttgtgc ccgccctggc ttgcctggat aaccgcgata tcataaaaac cttcccggga 15600
gaccagcacc tctaccgaaa cagaatcctc cccggcagtt ttaaagcctt ccccccactg 15660
attcccgccg tttccggcaa cggcagtatg tcccttcagt tcacctgttt ccgcttccac 15720
taacatcgtt tctcccgtct cttctgctgt cccggagaga atgatgaaaa gcagggtcat 15780
gaccgctgct gcgattccgc atatgctttt catactgctt ttcatttccg ttctctccct 15840
gttttgttag tttgttaacc tatttgtaaa ccaattgtta actaatttta gcgaaagaaa 15900
aatcgcctgt caagacaatt tttgcacata tattgggctt tgctggtcat ttattgttat 15960
cttgtttccg gaaaagacat gtgttttctt gtttcgcggg ttatattggt taacttatta 16020
gttaacaaat taatacatga ttttcttctg tataggccgt ccctccggat ccgccgggtg 16080
tcgcaggtct gcagaacgca aacgggagag ccgccatggc tctcccgttt accgcactat 16140
ccggttaaaa ttctgtgtcc gtaaagatat acgccggtcg tcgttccttc ccctccggga 16200
tcacttcata gccgtccagg gacaaacccc gtacatggcg cgcccaaagg gcataggccg 16260

gggcgtctcc caccggatcg atcatatggg cgtccggata cacgccgcgc atttcatcaa 16320
aatccttttc tttccgcaca ggccggcggg aagcctcccg ccgcaggagg acccggcgga 16380
aagcgatgtc ctctatcggc ttttcaggga tccccagcag ataggaaccg cgggggcctg 16440
tctcccgtcc gcggatatcc gaaaacagaa cccggcgcag ggtccccggg ccgggtgccg 16500
gatcccccgg tttgaccctt cctcggtttt ccagccgcag gaagaacggg ctccaggcgc 16560
ggatcatatg gatatcctcg atcagaaaat tctccagcgt tccgccgtcc accatttcaa 16620
gggagatccc gctgtccgcg caggcatgct tcagcccgga cggatcctcc gccaggccgc 16680
cgatccggca gcgacggata tacacctgcc ggaaatcccc ctgggtatct gttcccacct 16740
tcaccgcgtt gcacgcggaa taaaaatcgc agtcttctac cagaaccctg ctcagttccc 16800
gctggcctgc gcctttaaag cagagcgcgt catccccgct gctgacccgg cagttccgga 16860
tcaccacatt ccggcatccg tctatatcca tcccgtcatt gttatagttg gcatggttgc 16920
ggaccgtaag cccttccagc agcaggtttt cacagttcag gtagttctgc atccagcagg 16980
cggcgtcttt catggtcagg tccgcgatat gaacgttccg gcagtcgatc atccggatca 17040
ggaacggccg gcccggcgta ccctgggccg tctcctcccc gggaaaatgc gccgggcttc 17100
cccgcccgtc aatgacgcct ccgccccgca ggcagatatt ctcacatcct tcggcaaaga 17160
gcagcgactg gtgcataccc atactggtat cctggaccgt cagccgcgcg gcgtgatgct 17220
ccggatagtc gcgaagatcc gtgctgccca gcagccggga gcctgaccgg acctccaggc 17280
atacgccgga gcgcattacc agcgttccgc tgacgtaatc tccgttcccg agccgcacga 17340
tcccgctctg ctccgctgcc gcttcgatcg ctccctggat ggcttcggtc gccgtttccc 17400
cgggctggta cccgaattcc tccggcgtaa agaccggccc cccgtctgtc cgttcccgga 17460
caacctttat cagttcctgc gccatccgcg ccatgtttgt cttcagttcc cggcaccagg 17520
aaggttcctt tccctcccga atggtctcca tactgtttcc tccgttctaa atcagctctg 17580
tccgcaggaa gtccccgttt tcattcgtta cgatgcgtct gccgcagctg aacctgccga 17640
tatcctttcc atcagcccgg atctctgctt tctttccatc gctccggatc ccgtagcgat 17700
ggccgtaaaa ggccgcttca tattcatatg gttttccgcg gtcaaggctg aaccacacct 17760
gccccatatg cggatggatt ccgaagcggt gggcaatata ctccaggcag gcgagcattg 17820
tcgggccgta ggcatcctgg accttttcgc cgctgtctcc gctgaccggc cgatggtccg 17880
cctcgtccac cagcgaagcc ttcccggtaa acgggtcata ttgctgggta aaccggcagc 17940
cgttcgcgtc aacagcatcg atcagtttct tccccagggc tgtcacgatc cgggcatacc 18000
cgtaccgctc aagggcagtg atcgcccgtt ggtaggtaag gccttccggc tgtccgctcc 18060
agttattctc cggcgcgttc cggaaagccg ggtcatccgc ggcaacgctc ggcaggggga 18120
acggtgtcca gaattccgaa gcgttcagca ggtgttcctt cacaaaccgg tctgccatat 18180
cccgggaaac gcttccccaa tacatgcagc gtagcgtgtt atggcacagg atatcgatcg 18240
tttttcccgt ccggtcccgg tcatacagcg cgccccgcct ggcatcccac aggcgcctgc 18300
gcagcgcatc cgccacgtct cccgccttct tttcccagtc cgtaaaagcc ggatccccct 18360
gaatgcggca gatctccgcc gccgcgcggc gcgccgcgaa gctccagctc gtcacatcca 18420
tgctggccat gggaaccacc ccgtatcctt ccggcggttc atcctcctcc caccagcagg 18480
gcgcatcccc gtagcggacc gccaggtcct ctcccgtatc ataaacacag aaggaggaaa 18540
gcagcccgtc ccccgaaacg tccctcgtcc tccataagca ggcgtcaaag cggatcagcg 18600
cttccttcag ctgttccagg tattccctgt cttttcccat caggtaatac aggttcagcg 18660
ccggatcggg gaagcagaac ccctggtatt tattgaactg cggttctgtt tttccgttga 18720
tatgctggat ggagccgggc atccgcccgt cctcccgctg atgtttcata aaaaggagaa 18780
tattgttctt cgcggcctca agattccgca gggcatacat ttccccgccc atcggctgcg 18840
tttccagcca gatcttttca tacccgcctc cttccaccag cacggtgtcg ccgctgaaat 18900
ccttcaggtt caggaggcac tttttttctg ccgcgtcata aacccgctgc agtttctcat 18960
cttccgttcg gaagctcacg cttgtctcag gcagcatttt ttgtctttcc cttcagtgtg 19020
ttttgtccct tatataatta ggtagatttt atcatttctt gcgaaatcag aacagccaca 19080
aaaaaactgc acattttttt cttttcgcga aatcaaggaa aatcaacggt atgcaaaatt 19140
ttcgcttctt tcttttctgg tgcgttttcg tccataatct ggataagaaa gcaatcgttg 19200
acaaatccac ttaactcccg aaggaggata ccccatgagt ctttcctcat ccccgctggc 19260
ttatcgcgga gggatccggc acagaatagc tatgaccctg cgcagggact ggcagctgtg 19320
ggtactgctg ctcccctgtc tggttttctt tattgtcttc tgttacttcc ccatgtacgg 19380
tatccagatc gcattcaagg attataaggc agtactcaaa attgagggaa gcccctgggt 19440
aggcctgaaa cattttgaaa aattctttgg cagctattac tgcggaagaa tgttcctgaa 19500

cacttttctg ctgaacttct acggcctgct gttcggtttc cccgtcccga tcatactggc 19560
catcctgctg aatcagctga acagcaagcg cttcaaggga tttgcccaga ccgcggtcta 19620
tgtgcctcac ttcatttcca ccgtggtgct ggccggaatg atttacctct ttttttcccc 19680
aaccaacggc atcatcaacc acctgattac tgccgccggc ggaaaaagca tctactttat 19740
tatggaaccc ggctggttcc ggccgctctt catcggttcg gatatctggc agaacgcggg 19800
atggaacacc attctttata tcgcgaccct caccggtatt gatccccagc tgtacgaggc 19860
cgcaactatc gacggcgcca cccgctggga aaagatcaag catattgata tcccccatct 19920
gattcccatt gcggtcatga tgctgatcct gaacagcggt tccctgctgc aatccaacac 19980
ggacaaggcg ctgctcctgc agaccagcgg caacatggcc acctcggata tctttggtgt 20040
ctatgtctac aacgtcggca ttgccgcccc caaggcccag tattcctatg cttccgccat 20100
cggcctttgt ctgaacatca tcaacttcgt cattattatg ggcgtgaacg ctgtctcccg 20160
taaaatgagc gacatcagcc tgttctgaga aaggagggaa aaggatgact gtactgtccc 20220
ttaaacatag gaagcgcaac cggatccgtg aaacccggaa ggacaggatc tttaacactt 20280
tcaatctcct cttctggatc ctggtcctgt ttattgtcct ctatcctctc tggctgatcc 20340
tgattgcctc cgtctcggat ccggatgcgg tgctgcaggc aaaagtcctg ctgtggccgg 20400
ttgacttttc cctgatggga tacgagaccg ttttacagaa cagcgagctg ctgcgcagct 20460
atgtgaactc cattttctac accgttgtcg gttcagccct gagcgttatc atcaccctgg 20520
ccgccgccta tgcgctgtcc cgtaaatttg ccggcaaaaa gctggtaaac ctggccatta 20580
cttttaccat gttcttctcc ggcggcctga tcccgatctt cctgaacgtc ctgaacctgg 20640
ggctgtggaa cacccgggca atcatgatcc tgatgaacct ggtctccgtc tggaacctga 20700
tggtagcccg aacctatatc cagaccacca ttccggaaga tctttacgaa gctgctgtca 20760
tcgacggatc caaccatttc acctacttct tcaaatgcgt cctgcccctg tccggcacga 20820
ttgtcgcggt gctttcagtg tattacgggg ttgcccggtg gaatgactat ttcaccgccc 20880
aggtcatgct gcgcgaccgt gcgctgttcc ccctgcagag ggttctccgg gaaatcctgg 20940
cctccctcac ttccaccggt tcttccgata ccttcttcgc agcctacgcg gataacctgg 21000
gcggcctgca ggcagctctc cgcaaggcgg aagtcgcgaa gtactgctgt attgtggtag 21060
ctactgttcc ggctgttttc ctgtacatct tcatgcagaa gtactttgtc aaaggcgtca 21120
tgatcgggtc tgtcaaagga taatgcgttt taatcagaag gtttccggaa aggaggtgcg 21180
gcccgatgtc tcagaagaaa ggggtggagg tgtccgcccg ggccgcccgc catatagacg 21240
gttatcgggt actggagggc aaacgtgaat atgtgaccta ccagggtgat tcctccatcc 21300
gcatctggta ttcagacatt ccatggcgtt atgaaaccca tgaccattcg gctgttgaga 21360
tcgtcctggt cctggaggga acagtcacct actctgtaga gggaacgagt tatcccgtac 21420
gcaagggaga agtgctgatc gtaccgtcca acacgcccca ttccctggaa atgggcgaag 21480
gcagcagccg ctaccttttc ctttttgaaa ccgacacaat catgaccatg cgggatatca 21540
aatccatggt cctgtatctt cataaacctt tccacctccg ggatgaatca gatgcccata 21600
agcaaatcag ggaactgctt cttcgggcga aggatttcta cgaaaagcag gaactgatgt 21660
ggaatacggc atgctacagc tgcattctcc aggtatatgc cacgctgggc cggcactacc 21720
tgatcggaat caggtcccgg accggcgata cggttcggag catggacaca gaggtgatca 21780
acgctgtgat gacctatatc aatgatcatt accgcgaaga cctgcgtctg gatgatgttg 21840
ccggtttttc cggattcagc cgtttctatt tttcccggtc tttcaagcag cagaccggat 21900
actcctttaa ggattatctg tgccagaaac ggctgcaggc tgccatggat ctgctcacca 21960
gcaccaatct gtccatgcgg gggatagccg tcgaaagcgg ctttggcagc acggcatcct 22020
tcaaccgcgc cttccgcgcc tgcaaaggct gtacccccac ccaataccgg gccgtttacg 22080
gaatgctgta aagcgccttg tctgatcggt atcaagggct gtatgccctg aaataaaaag 22140
gaggaaaaac ccatgaagaa actcgtttcc ctgctgctgg catgctgtat gctgctcagc 22200
ctgagcgccg ccttcgctga agagaagacg gaactcaccg ccttccagta cgcgctggaa 22260
aatcagaaca ccgatttcaa cggtctctgg ttcttcgacg agctggaaaa caagaccaac 22320
acccacgtga ccttccagat ggtgaaggac gcggactggg ccaccagcgt gaacctgatg 22380
ttcgccgtgc cggaccagat gcctgacatg atcctccgtc acgcggtgga tgtggaagaa 22440
tacggtgtct cccagggtct gctgcttccc ctcgatgaat acctgaccga ggagatcatg 22500
cccaactacg tcagccgcct gaacatcaac aacgccggcg actccatccc cgccagcgac 22560
ggcaagacct attacatcgg cttcctcatg gcccagaacg tcaaccatac cggtacctgg 22620
tatgtgaacc agaaggaact ggacaagctg ggaaagaccg ctcccgccac cattgatgaa 22680
gtgaccgaac tgctccgcgc catgaaggct gacgggatca agtggccctt ctccgcatcc 22740

tacacccagt tcggaccgga aaccatctgg aaccagtttg cttccttcgg cgttccggaa 22800
aatacctatt actacttcat cgacgctgac aacaaggtcc agctcacctt tgcccaggac 22860
ggctggcgtg cctgtgttga atggctgcac cagctgtatg aagaggaact gatggatccg 22920
gaatccctga cccaggactc caacctgtgg gccaacaaag tcaacgacgg cgaagtcggc 22980
tacttctgct acctgcgcct gatcaacacc gccatcaagg ctgaaaacgt ggaaaacttc 23040
aagtccatcc tcccgcctgc cgcggacgga ttcaaggcct gcgtttccca gatcctggag 23100
gttccggaag ccggcgccta cctgacaaag aactgcaagg atcccgttgc cgccctgaag 23160
tggatcgatg cccagctgga aaccgaaact atgatggtct ccctcaacgg caaggtcggc 23220
gagcagatcg tgctcaacga tgaaggcaag tatgaagtca tcaacattcc cgaagacaac 23280
ggcctgtatc agttcgttcc ggttaccatg ggccagttct tcgctcccgg tgactattac 23340
acctccatct accagatggc tcctcaccgt gtggaacgct atgaagacag caagtggtac 23400
gccgaagccg gcgtgctgga acccaagagc ttccagtatc tccgcgacct gagcaagatg 23460
agcaacgatg acgccaccga agccgccacc ctctacaccg acctgcagaa gctggttgag 23520
gaatccttcg ccaacttcgt ccgcaatggt gtaacggatg agagctggaa cgagttccag 23580
gctcaggcca aggccatcgg ttccgaccgt tacatcgagc tgtaccagaa tgcttacaat 23640
gcctatctgg caaagtaaat ccttgaacgt tgcggcctga gatgttaaaa tgcaagcatc 23700
agaaaagccg ccgggcatcg ccgcaggaat tgattccggg ggcgatgccc ttttgccgta 23760
tagagggagt acatatgaac cgtaccgtaa agatcacctg cctgttggct gccgccctgt 23820
ttctgctgac agggtgtgct gtttcacggc cgggagaggg ctctgcccct cccggcgcat 23880
cccttaccgc gctgcagtat gagatcgaca atatcgccgt ggacttcagc agtatgtggt 23940
attaccggca gatcgaacag cagacaggcg ttcatgttga cttcagtgaa gtaaaggatt 24000
ccgaatggag cagctccgtc ggtctggcct ttgccaaggg aaagatgccg gacatgatcc 24060
tgcggggatc cctggatgtg gaggaatacg gcgtaacccg ccatcttctg gtcccgctgg 24120
atgaatatat cgacaagggc ttcctgccca actacgcggc ccgtatggac caggccggtc 24180
tgcgggagca gcttaccgct tctgacggcc atatttacca gctcggattc ctcatctccc 24240
agggtgtcaa caccaacggg catttcttta tcaacaaaac atggctggac aagctgaacc 24300
tgcaggtccc ccggaccgtg gaccagctga cggaagtgct caggcgtttc cggtcggatg 24360
atcccaacgg taacggcctt caggatgaga ttcccctgga attcaccttt gatgacaata 24420
ccaccggcat atacaatctc ttctcctttt tcggcctgcc cctgaacgag gattatgtct 24480
atgtggacaa taccggaaac gtacgttttg cgccggagga ggacgccttt tttgagacgc 24540
tcacctggct tcacgatctg tatctggaag gtctcctgga cattgatttt atcagccagg 24600
gcagcagcat ctgggccgcc aaggtcaacg aaggaaatac cggcatgttc agctactggc 24660
ggctccagaa caccgccctc agccccggca ctgccgccaa ttaccaggtg atggttccgg 24720
tgcacgttga gggaaggaat gcctgcctgc cgagaaacat ggacatcatt gagttcggcg 24780
ccgccctgac cacggacaac agggatatcg aggccagtct ccgctggctg gatgcccagt 24840
ttgaaacgga aaacatgctc gtctcccaga acggccccgt cggcgacacg ctcatccggc 24900
gggaggacgg ccggtatgag gtcagctatg ttccgccgga caacgagctg tatcagtccg 24960
ttccggtcat ctgcggccag tttttgccc cggctgaata ttacgcctct gtctatgtcc 25020
cggccgccca ccggctggaa aagtccgcct actgcgatct gtatgagcag gccggcgtgc 25080
tggaacccgt ctccggcaag ctgctgacag ccgtggcgcc gaaaacaaca gcccagtccg 25140
ccgaaaccac ccgcctgaag gcaaagctga agtccctggt ggacgcggag atcgtttcca 25200
tggtcacccg gggtgtaacg gatgaaagcc gcgaagcctt caggtcaaag ctgaaagaaa 25260
acggaagcgt ggaataccgt caggtttacc agcagatcta tgaccgctgg aaggagaaat 25320
aaaccgtgat cagaaagagc cggacctctg tcttcatccg gtatgcgctg tcgtatatcg 25380
ctgttgtgct gctgctgttc tgcagcatca cgggatacct gtatgtccgt ctcagcggag 25440
aagcccggga agagatcatt gataaccaga tcaaccggct ttcccggatc gccagccagc 25500
atgaaagcta tatctccgcc atgctgaata ccgccgagga gatcggtctc agtccggata 25560
tcgagttctt ccgctacaat gaagagccct ggaaagccta tgacctccag ctccagatgg 25620
tcccctacac cacaacggcc agcaccttct gtgatcaggt ttatctgtat ttcttcgggg 25680
atgacatgat ctattcctca tcatcctcca tgaccctttc ccttttgcg cgcatgatgc 25740
gctatgagtt tatcccgatg gaaaatcttg tctcccggat tgcgagcact gaccggatga 25800
ccattctccc cgcccagcag gtttcctcca cccttgtgga cagttcccgg gttgtcacct 25860
tcctggtgcc gctggggtcg gctcccggat ccggaaaggg catcctgctg ttcctgatca 25920
aggacagcgt ctaccagtcg ctttttgccg acgccattga cgaccgtgtc aacacctata 25980

tctaccaggg ggatgacctg ctgtcctcct cggagaagct ttcccttccc cgagaccggg 26040
tggaagcctc ccaggacgct ccttcccgcc tgttccgatg ggaaaaccag gactggaccc 26100
aggtcacttt ctccggcaga aactgggggc tgacctatac tgcggtcctc cgcaacgccg 26160
atatcaactc gtccatcatg cagcagctct tcagcatcct ggccatcctg ccggtatttg 26220
tcttcctgag ccttctgctt gctctctgga tggcccggcg ccatgctaag ccaatccggg 26280
cgatcaccgg tatgctttca cgggaaacag atgaaacacc ccgcgatgag ctccagcaga 26340
tctcctccgg catcagccgg ctcaccaccc ggaaccagga gctgtcctcc cggctggaac 26400
gcgccttgcc cctccaacgg catgattttg tcttccgctt tgtgaaagga cgctttgaga 26460
ccaggcagga agccgtggct gcagggcagg ctgtggggct ggagatcgac cggcctgtat 26520
acgcggtcat cctttgcagc ggctctgccg gaaatgatca tccctttgat ctcagcctgc 26580
cgcccttcag ccgcctctcc ggctattccg gtgccggcgt ggaactggtt gccctgaagg 26640
ccaaccttta ccttgttttc tccgatgacc cgaaggtgct taccggcatg gcggaaacgc 26700
tgcgctgcga atgcgctgcc ggcggagagc gctgtatcac ctccatttcc gccctgcata 26760
ccgatttcac ggaagcccct tccgcctatc tggaggcagc agccgcctat gaaaaccggt 26820
ttgtcatggg cagccagaag gtactctgct acggcgatat ctcccctgac ctggaggaga 26880
tccttcccca ggcccgcaag atcacaacct ccatcagtca ggcgttgacc ctgggaagcc 26940
gggaagttct ggacgtccgg atcagcgacc tgctgcattt cctgaagaac accagcatgt 27000
ctcccttcgc attccggatg atctacagcg aggtgatccg ttccctggtg cacacgcata 27060
cggcggcgct gtccggacag ggctccgttc aggaaaccta caacatcttc actctcacct 27120
cctgccagtc cattgatgac ctggatgaac tcctgcgcag gctgtgcgac cgcctgatca 27180
ccggagaatc agcttccgga gagaaagttc cggacgagga ggatgagatc agccaggttg 27240
tccgctacat ggaagagcac ttcagcgatc cggagatctc catcaccgcc ctggcggact 27300
ccctggagct ttcaaccacc cggttcagcc tttccttcaa ggaaaaggcc gggatgtctc 27360
cgctggaata cctgacgctc ctgcgggtgg aacacgccaa ggaactgctg gaatccagcg 27420
acctgactat ccgggacatc agcacgcagg tcggctatta cgactccggc tcctttatcc 27480
gccgcttcaa gcaggtaaca ggcgaaacgc ccctgcagta ccggcgcgga agaaatccga 27540
aaaaagagaa cggcattccg gaccgggatg cctccgtccc gacagaggag taaaaccatc 27600
ctatgattca tactgatcag atcaatattc gggatcccta tgtccttgtg cacaacggca 27660
cctattatct ttacggcacc cgcggccccg actgttgggg acccgcggac ggtttcgacg 27720
tgtataccgg acaggatctt gtgtcctggg aaggcccccg cgtctgtttc cggaatgacg 27780
gctccttctg ggccgaccgg aattactggg ctccggaagt gcatccctgg cagggaatgt 27840
tttatatgtt cgccagcttc aaaagtgacg cccgccggcg cggcacggcc atcctgcggt 27900
ccccctctcc cctgggcccc tttgaaccct ggtccgacgg tccggttacc cccgcggact 27960
gggaatgtct tgacggaaca ttctacgttt ccgcggacgg taccccctgg atcgtattct 28020
gccatgaatg ggtccaagcc ggcgacggag aagttctcgc catgcgcctg acggatgacc 28080
tgaaggcgtc ggccggtaca ccggttctgc tcttccgggg ttccgacgct ccctggagcc 28140
atgccgtacg gcattcctcc ggcataaccg gtttcgtcac ggacggtcct tttttgtggc 28200
gcggaaaaga cggctccctg ctctgcctct ggtccggctt ttcagaaaaa ggatatgccc 28260
aggggctggc tgtttccgac aacggggaga tcgacgggca tttttcacag gctgatccgc 28320
ttttcctgga ggacggcggc cacggtatgc ttttccgttc gaaggaaggc agaaccctcc 28380
tggctctcca cagtccgaac acccatctgc tggagcgtcc ccgctttgta ccggtggatc 28440
tctgatcccg caaaggatct cctgaaatct ccctttatga aaggataaaa catgcattac 28500
gttctggcga tcgacatcgg cgcgtccagc ggccggcaca ttgtgggctg gcgggaagga 28560
agcgggatcc gtacagagga ggtctatcgc tttcccaacg gcatcacgga acaggacggc 28620
agcctgacct gggataccga tgccctgctt ttccatgtca aagcaggcat tgacaaagcc 28680
ctggaaaaat acccctccat ccaaagcctt tccattgata cctgggccgt ggattatgtg 28740
ctgatgaatg gcggcagccc tgtccttccc tgctatgcct accgtgacag ccgtacggaa 28800
gcagtgatcg gagaggttca cgaaaagatg ccttttccgg agctgtaccg gcgcaccggc 28860
atccagtacc agcccttcaa caccctctac cagctgtatg ccgacaaggt ttccgggcgg 28920
ctggatcagg ccgatggttt cctgatgatc ccggaatacc tgatgtaccg gctgtgcggt 28980
gcggaaagcc atgaatacac caatgccacg accaccggcc tggtcagtgc ggaaaccggg 29040
gaatacgatc ccggaatcat cagcgccctg ggcctgcccg cgcgtttttt cgggcctctc 29100
cggcagcccg gcaccgtgat cggggaatac agggggatca gggttatgct ctgcgccacc 29160
catgataccg gaagcgcggt ggaaggcatt cccatggaag gctgccagcc ttatatctcc 29220

agcggcacct ggagcctgct gggggtgaag accgaacggc cggtcatcac tgaagccggc 29280
cggctggcca actggagcaa tgaaggcggt gccggatata accgttacca gaagaacatc 29340
atgggtatgt ggctcatgaa ccggctccgg gcggagctgt gtccggacag accctgggat 29400
gagatcatgg ccgaagcggc cgaaaagcat tttgatcacc tggtggatgt gaatgacccg 29460
gtcttcctgg caccggaaag catgaaggcc gcctttgatt ccaggcttcc ccatccgccc 29520
cggtgtattg ccggctactt ccggtgcgcc tgcaggtccc tcgccgacgg ataccgccgg 29580
gctgttgagg aaatggagca gaacaccggg cggcgctatg acaggctgta catcgtcggc 29640
ggcggagcga agaacacata cctgaaccgc ctgaccgaag aatcgaccgg aaagcaggtg 29700
accgccctgc cgatcgaagc taccgctctg ggaaatatac gcattcagct gaaaggtgga 29760
actgatacat gagttatgaa gaatccagga aacggtatgc agccctgggt gtggataccg 29820
atgccgtcat ccgtctcctc agggcagtgc ctgtcagcct tcactgctgg cagggtgatg 29880
atgtccgcgg ttttgatacc gatccctcca agcccctgac cggcggcatc cagaccaccg 29940
gcaattaccc cggccgggcc aggacgccgg aagagctgat ggctgacctg gatatggtcc 30000
tgagcctgat ccccggcacg ccgaagatga acctgcatgc cagctatgcc atctttgaaa 30060
acggtgaatg ggcagaccgg aacaggctgc agccgaaaca tttccggaaa tgggcagctt 30120
tctgcaaaga acggaacctt ggctgtgatt ttaaccccac cttcttctcc catccgaaat 30180
gcgatccgct gaccctttcc agtccgaatg ccgaaacccg gaaattctgg attgagcacg 30240
gcaaggcctg tatccgcatt agcagctacc tggcggagga actggacttc ccctgcgtca 30300
tgaacatctg gaccggcgac gggttcaagg acatccccgg cgaccgtata ggcccccgga 30360
tccgttaccg tgacgccatt gacgagatcc tgcgggaacc ctatgacttc agccgggtga 30420
agccctgcat cgaaagcaaa gtgttcggca tcggcgtgga ggcctataca acaggcagcg 30480
cagagttcgc cctgtgctat gccgccatga accgggacaa gtgcatcccc ctgatggaca 30540
acggtcacta ccatcccacc gaggtagtca gcgataagat ccccgccctg ctgactttct 30600
tccgggaaat cgccctgcat ataacccggc cgatccgctg ggacagtgac catgtggtcc 30660
tgtatgacga tgaaacccgg gaaattgccc gggagatcgt tcgctgcggc ggtctggacg 30720
gccgtgtgca tatcgccctg gattactttg atgccagcat caaccgcata tccgcctggg 30780
caaccggata ccggaacctg cagaaagcac tgctctatgc tctgctgcag ccccatgagg 30840
aaatgaaaaa gatgcaggat aaggggcagt tcagccggct gatggtcctg caggaagaaa 30900
taaaaaccat gcccttcggg gaaatctggg cggaatactg ccgcttctgc ggaaagcccg 30960
ccgacggcga atggttcccg ctggtggaag aatatgaaaa gaaagtgctg gtgaaccgca 31020
aatgaaagat atcctggccg ccccctttat gacggaaatg atacgcaccg cgacagccat 31080
gtatgaccat ggctgggatg aacgcaacgg cggcaatatt tccctgatgc tggatgaaaa 31140
ggaagccggg gaatacctgg atatgcgcaa tgtcctgcga accatcccga caggcttttc 31200
tatgccggaa ctggacggga aaatctttct ggttaccggc acaggcaaat acttcagaaa 31260
tatacagtat gaaccggagg tcaatctggg gatcatccgt ctgcgggact gcgggcagca 31320
ggcagacctg ctctggggat actcggacgg cggacagttt accagtgagt tccctgccca 31380
tatgatgagc cattcagcgc ggctgaagat gaacccggca aaccgggtga tcatgcactg 31440
ccatccggcg aacctgctgg ccatgaccta tgtgcacgca ctggatgaac gggagtttac 31500
ccgaaccctc tggcagatgt gcactgaatg cgtggttgtt ttcccggacg gtgtaaacgt 31560
gcttccctgg atgctgtgcg gcaccaatga gatcggcgaa gccacagcag agaagatgaa 31620
gaccacgcgc ctggtggtct ggagccagca cggcatctac ggagcaggca aagacctgga 31680
cgaaaccttc ggcctgatcg aaacggcgga aaaggctgcc gagatctata tgaagatcgc 31740
gcatctgccc cgggtcaaca cgatcacgga tgaacagatg caccagttgg aaaagcggtt 31800
cggcatcaaa gcccgggagg gatatctgga ggaaccggat cgggggacgg acaaatgaaa 31860
cagcatgcag gaaaaaacat ccgggagctg aaagttgcct atatcggcgg cggttcccgc 31920
ggctgggcct ggggctttat gaaggatctt gccctggacg gcgaaatctg cggtaccgta 31980
cgcctgtatg atattgaccg ggaagccgca gaccggaaca ggatcatcgg cgaaaagatc 32040
agcgcccatc cggatgctgt ttcccgctgg gcttatacag tctctgattc cctgcaggaa 32100
gccctgaccg gagcggactt tgtggtcatc tccatcctgc ccggcacctt tgatgaaatg 32160
gaatcagacg tccatctgcc ggagcggctg ggggtctggc agccggtggg cgatacggtc 32220
ggcgccggcg gattcatgcg ttccatgcgc accatcccca tgtttgtcac catcggcgag 32280
gccatccggg attacgcgcc ggaagcctgg gtgatcaact ataccaatcc catgtccctt 32340
tgtgtgcgca ccctctatga ggtttttccg tcgatccggg ccttcggctg ctgccatgag 32400
gtcttcggca cccagaagct gctctgctcc atgctgaaaa ccggggaagg gatcgacagg 32460

gtaaaacggc aggacctgta tacaacagtc accggcatta accacttcac ctggctgacc 32520
agcgcctcct ggaaggggac ggacctctat cccctctatg cccggtttgc ggaaaagtac 32580
gcggaaagcg gctacagcga aggcggggat gacaactgga tgaattctca cttcagctgt 32640
gcccagcggg taaagtttga tctgttccgg cgttacgggg tcattgccgc tgccggggac 32700
cggcacctgg cagagttcgt ggcgccctgg tatacacggg atcccgaaac agtccgggaa 32760
tggaaattca gcctgacccc cgtttcctgg cggaagcagg acctgaagga aaaactgcag 32820
atgtcggatg acctggtcag cggaaagaag gaaatcaccc tgaacggttc cggggaggaa 32880
gggcacctgc tgctgaaagc cctgctcggt ctcggcgata tggtgtccaa tgtcaacatt 32940
cccaaccggg gccagattcc caatctgccc ctgggcgccg tggtggaaac caacgccctc 33000
ttcgggctga accgtattga tcccgtatac gcggggccgg ttcccgcctc catcctgccg 33060
ctcatcaccc ggcatgtgct gaatcaggaa aataccctga ccgctgccct ctgctgtgac 33120
cgcaaactgg gcctgaccac cttcctcaat gacccgcagc tggccgcagt taccccgaag 33180
gacggggaaa agctctttaa cgacatgctg gaaaaccagc ggaccttcct gcccgaaaaa 33240
tggtttgaat aactccggac atccgaaaga atcatataat caggaaaccc gcttctgatt 33300
tttatcagaa gcgggttttt tccgccattc ttcatttttc attcttcatt tttcatttcc 33360
gctttataat cctgcacagt tcatatttcc cgggcatggc ccgtccttca aataccgttt 33420
ctccattccg ccggtacagc ggctgcatcc ggatggcttc tccggtcaca ggatgcttcc 33480
gcagcggcag ctccagcgct tccgcatcgc cagtcacatg gatcaggatc gttgccggct 33540
gcgctcccga agccacatac ggatagagga caaaagtatc attatcgtaa ggaaagaggc 33600
ttatacccgc cttgctttcc aggtagacgc cgcccgcggg aaactccctg cggatccggg 33660
tcaggacttt tcccggcagc ttgtacagat ccgggaagtt gtccggtacc gccagggtca 33720
gcatctgtcc ctttccgtaa gtatcccgca gcaggatggc gaagctttcc tcggattcag 33780
cagctttcac caccgcccag gtggcattat tccggaattc gcacaccgga ataccgatct 33840
cttccgtacc ccggggaaat tctgtcgagg gccatactcc cgcgttctcc acccggtaat 33900
tccggccccg gatcctgcgg ccggtgaaac ggatggaggt catccgctgg attccccggt 33960
ccatggtatc ttccacgaag ccgctggtca ccagcgcttt gccgccagcc gccacatact 34020
tctccagctt ttccaccaca tccggatcgc aggcggagga acgggtcagc aggatatact 34080
ccgccttttc cggccagtag ggtgtcagca tcaccggcag gccgcacata cccaggaaat 34140
cctggatgtt gtcctcaccc tggcagttgt ccggcaggta acactgaatg cccaccggat 34200
tgccgcagta atccagcacc tcatccagtt tgtccaggtg ccagcccagc ggcggaatga 34260
atggattgtc cgtcagcatc tggaagcaga acagcatgat ctccttcggt ttggagaagg 34320
cggtcaggta cgcctgctcc aggtactgtt ccgtgatatg caggtcaaag ggatcgaacc 34380
agcctccgcc gttgtgttcc ggccacatgt tctcaaaata agtcatcagg gaatagctca 34440
ggtagcgggg caggtgctgg tctgttgtta cggtatcccg ggcttccgtt ccggtgtaga 34500
tccggtcgaa cagcttccgc tgcgccgcag gattgtatcc ggtttcctgg tagctctccg 34560
cccagttggg atacttgatg gtgatccggc atttcggatt ggctgccttc gccggggcga 34620
tcacgtcctc ccggctcacc ttttccatca ggtgaaggcg gtatttcttc cagctgccgt 34680
ccgtaatccc gcgctcccgg ttatattcat cccgttcccg gcggcaggcc tcgcaggtac 34740
agttggtaaa gaagaagtcg tcaataatga actcgtcaaa atgcccgccg ataaagctgc 34800
agaccttccg gagttccgcc agcatctttt catcgttgta gcagaaggtg tcaaacaggc 34860
gctgcttcgg ttcatcccct tccggggttc ggatcgtggt ggtcaggccg ccggccaccg 34920
ccacccttcg catggcaagg tcgattgggc ggtgcagcac gcctatgatc aggcacgcga 34980
tttcggccag ctttgccggt accgtgccgc caatgccgcg ctgaccgagc ggccgcgcgt 35040
cgtcttcgcg ggcgattcga tcaccgacaa ctggatcaac ctcgacccct cgttcttcac 35100
ccatggactc gtcgatcgcg gcatcagcgg acagacgacg ccgcagctgc tcgtccgctt 35160
ccgccaggac gtgatcgacc tgcacccgcg cgcggtccac atcatgatcg gcaccaacga 35220
catcgccgga aacaccggtg ccgcggcgat ggagacggtg gaaggcaata tcgcgagcat 35280
ggcggacctc gcccgagcgc atggcatccg ggtgatcctg gcctccgttc cgcccgccgg 35340
agcctttccc tgggccaagg acaagcagcc cgtaccgcag atcgccgcgc tcaaccgctg 35400
gatcgccgcc tatgcgagga gcaacggctt cacctatgtc gactatcatc ccgtgctcgc 35460
gacaccggac ggcgcgatga agcccgggct ggccagcgat ggcgtccatc cgaccgcggc 35520
gggctatgcg gcgatgcgcc cccttgccga ggcggcaatc gaaaaggcgc tcggtcgatg 35580
aaccgcttca tcatgacgct cgccctgctg ctcgccggca ccgcgccggt cgcggcgcag 35640
cggatggcga cgctcgat 35658

```
//
ID   XXX; XXX; linear; XXX; XXX; XXX;  XXX.
XX
AC   PRJEB67889
XX
DE   Uncultured bacterium DNA fragment from in vivo enriched cow rumen, clone
rum-16b10
XX
FH   Key             Location/Qualifiers
FH
FT   source          1..34263
FT                   /organism="uncultured bacterium"
FT                   /environmental_sample
FT                   /mol_type="genomic DNA"
FT                   /isolation_source="Cow rumen"
FT                   /clone="rum-16b10"
FT                   /note="AZCL-xylan, AZCL-arabinoxylan active clone"
FT   CDS             32..898
FT                   /translation="MPRKHLITRRHPNRSIAGDIIIYLLLLLVAIAMVFPLVFLIGSSL
FT                   KPLDELFRFPPPVWPSHPTMDNFSDLFVTMGQSWVPFSRYLVNTIIITVVGTFGHLVVA
FT                   GMAAFVLSKYEFPGGKLFFAVATTCLMFTGVVTDIPNYLILSRLGLVDTYWALILPACA
FT                   APIGLFLMKQFMEGLPTALIEAATIDGASRFRVFWSIVMPNVKPAWLTMIIFSVQALWN
FT                   NRAATIIYSEAKKPLVYALQQIQAGGVARTGQMAAANVIIVAVPILIFVFSQSRILETM
FT                   ASSGIKD"
FT                   /product="binding-protein-dependent transport systems
FT                   inner membrane component"
FT   CDS             913..2445
FT                   /translation="MMKKTISFLLVLLMLLSIAGALADDSFNMNKDGYSTSYTYGYDYW
FT                   GDVQESPDAYRVETVLDSVSLGLDVRMSKPQSLYVRNNDLYIVDTLNNRIIQVEYKDHA
FT                   YKLVRIIDHMTGGNPETFNNPYDVFVDENDSIYVADYNNNRVVMMDKDLNWIKEFTKPT
FT                   DATFDQSIDFLPKKIVVDVAGRVYALVTNVNKGIVKYEADTTFTGFIGATPVSVSMVDY
FT                   IWKKYFMTQAQRDASPSFVPTEYENLYMDKDGFIYATITNFSEYDLKSDKAKPIRRLNG
FT                   LGNDILVKNDRYPPIGDLWWMEGDSTYFGPSKMTDVTVFDNDIYVVLDRVRGRLFGYDN
FT                   QGVMLWAFGTRGNIEGAFTSAVSIEHMGTDLFVLDQLENSVTVFTPTEYGNKIYDAILK
FT                   YQEGEYDESAAVWEDVLKQNANYPMAFRGIGRTVLRQNKYEEAMEYFKMAHDRENYGRV
FT                   FKLYRKEWIEKNILWIFLGLAVLLIVPLVIGRVKRMKWEVIMHEQSKVRKNG"
FT                   /product="hypothetical protein"
FT   CDS             2414..3106
FT                   /translation="MNRAKSVKMDKAASATARKAWWSKYKDSLRYALHVITHPFDGFWD
FT                   LTHEHRGTLAAANTFLILFLITRVLKLLCTSFQFISTPVQHINVFEEMGSLLLPFLILC
FT                   LANWAMTTLFDGKGRFIDIYIGMCYALVPYVLIQLPMILVSNMLTFEEGSLYSVMLSIS
FT                   IVWCVFLVFVGLMEIHDYGPGKTFIFIIVTIVGAAVIIFLILVFFSLLSDAVGFFVSLY
FT                   REASYRLN"
FT                   /product="hypothetical protein"
FT   CDS             3122..5695
FT                   /translation="MRKKPILLRLLPWIIVLVLIAGAYIVLDKIYSSPGHSFARDTKII
FT                   SYEGDGKPLTMENDQLLFEMDGSTTQFSVTNKETGKVWYSNPEDRANDTTATGKTTKEY
FT                   LSCTLNAHYYNDISETEVNNYTKSIMNQNYEIQQEADGSIRVNYTLGEKVFMVPNAITK
FT                   ENMDALLANLKKKDQGKLKNHYSLVETKKLDKQKKKDEIIALYPSVLEQDLYILKSDRT
FT                   TEQKKQAEQLLASAGYTEEDFARDQELQAGEKSNSGPVFNISVIYRLEGNDLVVSVPYS
FT                   ELTCSSDYPLVYIDMLPMFGAGGKKQDGFILVPEGGGAVINYNNGKLSQNPYYANMYGW
FT                   DYGTKRTSVTGETENAFPVFGMSQEDGSFICIMEGADSYGGVCADISGRFNDYNTVYGR
FT                   YNVLHFDSVEVDEKKSARLFLMYENEIPDDTIVQRYRFLNDNGYVNMAEAYGEYLRAKP
FT                   EMRGECASEEMPVNVELIGAINKTEIKFGLPVKSVVATTTFYQAESIMNDLLNAGVRDL
```

```
FT                   NLRYTGWCNGGVQQRVLTSIHVEGGLGGENGMKKLIASAAEKNVDLYFDGITCFAYDSG
FT                   LFNGFNSLADAARATTRTVIKLYAYDIVTYRLSTWMDNPYFLVKPSYAKNCVNNLINGL
FT                   KNRGVAGIAFRDVGNLLSADYQDSGTVTREQAKAMDIQTLEDSVATGMKVMIKEGNAYS
FT                   IPYADLITDMNLSGNAYALLDYGIPFYQIAIHGLKDYTGEAINLAGDTQTLLLESAEYG
FT                   AGLNFSFMQKDTKVLQDSMYSCYASAYYTPWKDEAIAMITRYQKEMSGLNRQAITGHER
FT                   LNEDVAVTTYADGTQVYVNYGDREFTIGSVTVPGRDYLVKRGGGQ"
FT                   /product="hypothetical protein"
FT   CDS             5692..6672
FT                   /translation="MSKKNRKYREHWYSGITEAFQMFVPGNKLYRTIPAKRARTGFLFI
FT                   LPFIIGFIFFMLKPLYVSMQMSLSRYVINTGEMTWNNFYNYNYAFNTDPDYNRMLVDEI
FT                   SRMAINVIATLVMSFVIAVILNQKFKGRILCRIIFFLPVILSSGVLPGIETSSSSLNLI
FT                   NSMSQAMGDSSGVNISAALENLLSVSGVASQVFDVIFQMIDAIYDIVMASGIQIIVFLS
FT                   GLQSISPSLYEAADVEGCSAWESFWKITFPMVSPLLLVNCIYTIIDFFMKNNNQVMEHI
FT                   NNTYMNGVRMDLGIAAAESWIYFGVAIVFIGISSFIISRAVKSYE"
FT                   /product="binding-protein-dependent transport systems
FT                   inner membrane component"
FT   CDS             6665..7675
FT                   /translation="MSKDNTIFIGEHSSFRERNRRSGGYLMKRRIKEISVSVIRGLLMF
FT                   GLCFMIISPMISRISMSLMDEKDLYDTTIVLIPRNVTMDNFTAVARATAMPTSMINTLW
FT                   ISILSSILQIISCTLVAYGFARYEFPLKKFWFACVIALIIIPPQTIQSALYRTFTRFDF
FT                   LGIITLINGEPLNIRKSIWPYLLMCLTCMGLKDGLYIYMLRQYFKNVPESLEEAAYVDG
FT                   CGTMHTFVKIMLPDALPIIASCFLFSFVWQWTDLFYSRNFLSGYTLYAPQLSSMLDRIA
FT                   DELSKMAGTPVTPSPGRQLQLISVGVLFCCIPLIILYCFTQRTFVESLAMTGSKE"
FT                   /product="Alpha-glucoside transport system permease
FT                   protein AglG"
FT   CDS             7684..7803
FT                   /translation="MRKMYEPILEKMDKPVYNKPEQWGFRINNVPIQISAGNL"
FT                   /product="hypothetical protein"
FT   CDS             7831..9273
FT                   /translation="MKKIMALVLAALMLLGCCSFAAAEEVPAEYPEIIEGLDFGGATVY
FT                   INDWYSTGERAEEPTEEQQAQYDYWDWLEKTYNVKLVETKLGDWDGMVAELQNIVSNKD
FT                   NSELRIIGVSGGFCGPVLANGLFAEWTYGLENFNEATVKFMTLGGKCYGVCGGKFFEPR
FT                   QMVFFNKKVLEDAGIDWNEIYDAQADGSWTWEKMESYMDAVKADKDNDGELDVFALTGN
FT                   FDDGVVGLVVSNFGDFYEFDENGKLYYSADSDATLAALNRIQEWNRNYYRPYVNWDDYK
FT                   QFWPEGNVAFLIGQSYEGFNGNDVVNQVEEWGGVCLPKGPNADAYTHAADNNIFGIPAV
FT                   YDDETSLKLQQIYTLYRKNPVIDADTDAWATRLYELTDERAVDETYAYLRENGTIMNFN
FT                   YIGDRNSTIGPNLTWGIMGGAAAEQVEAARLAMEDMAAVFNGDKTQEQVDAEKAAREAA
FT                   AAEEAAAEEGTEEAPADEGAAE"
FT                   /product="ABC transporter substrate-binding protein -
FT                   sugar transport"
FT   CDS             9412..10968
FT                   /db_xref="GO:0031176"
FT                   /translation="MGKKILAFLMILSLLGGCVSALADESKTIYTSDFSKDEDGWYGRG
FT                   AKSFRTADGTLKTVDRQSSWNSPGRDFDLIEGGKYQLSAEVKQDDMDSASFMISIAHSA
FT                   EGVETYENLAFGTAKKGEWVTLKGEYIAGAFDRSVLYVETTGSDTLDFEIRNFTVIAPE
FT                   GEPEPKPTEPPMVIEEAENIPSLKEIYAGKFDFGSAAPQMVFTNPKWMNLITEQFNILT
FT                   PENEMKPDSVLDVQASKKLVEETGDETSAAVHFDAAKALLRFAQSNGIKVHGHVLIWHS
FT                   QTPEAFFHEGYDPKKPLVTREVMLGRMENYIKGVFEYLDENYPGVVVSWDVLNEAIDDS
FT                   TNWLRNSNWLKIIGDDYPNKAYEFARKYAPEGTLLYYNDYNTAMPMKLKGIVKLLNKLI
FT                   PEGNIDGYGFQMHHGVTFPTIKQIQTAVETIAALNIRLRVSELDVTVSNNSEASFAKQA
FT                   KYYADVMKILIAHSDQFEAVQVWGLTDMMSWRSRQYPLLFDGRGNPKPAFWAVADPESV
FT                   Q"
FT                   /product="CBM22-GH10"
FT   CDS             11167..12330
```

```
FT                   /db_xref="GO:0031176"
FT                   /translation="MSSGLADPPVQTEGITVPVIEYKGEFKIPDTEAMAMLRDMKCGWN
FT                   LGNTFDAYNGYTGHFSGTGMESSWVGVKTSKKLIEAIKEAGFNTIRIPVSWHNHVDAND
FT                   VIDQEWMDRVKEVAGWALDLGMYVIVNVHHDNDVHYVYPDAQHYERSAAYLISVWKQMA
FT                   EAFKDCDDHLILESMNEPRLVGTQFEWSWNNYSPDCKAAAKYINQLNQLFVDTVRATGG
FT                   NNATRYLAVPAYCAAPANAADYAFQLPEDTADHRIIVTAHAYTPYNVALNLNSSDRTFD
FT                   LDKDSGKKSDIDGVIQKVYSRFIKYGTPVIMDEFGALDKDGNTQDRVNLTAYYVAAAAA
FT                   RGIPCVWWDNHAFGGSGERFGLINRSTLEWVYPDIVLAIQANCLKDK"
FT                   /product="GH5_4"
FT   CDS             12435..14822
FT                   /translation="MKYGYFDEKAREYVITNPNTPAPWANYLGSPEYGAIVTQNAGGYS
FT                   FVKSGAAGRILRYTFNQFDEPGRYIYLRDEETGDFWSASWRPVCKPLDEYKTECHHGTS
FT                   YTEIVSEYNGIRSRALYYVPMGATHEVWRISLENLTDRERKISVFGYAELTTESLYTQD
FT                   LVNMQYTQFITKTEFHDSFILEQINEFNYPNPDGTTGRERFFGLAGVPVASYTGRREQF
FT                   LGRNHFDKPQAVLDGKCDNSLNYNANPCGALQFSAALKPGEKKTAAFLLGQKTVFEAKA
FT                   IMERYEKTEETVDKELKELIAYWHGELENLKISTPDKNFDAMINTWNAFQCFTTFVWSR
FT                   AASLVYCGERNGYGYRDTVQDIQGIMHLDPELARKQLTFMLSAQVHHGAGLPLVKFTHN
FT                   AGHEDTPEQESYVKATGHPSYRADDALWLFPTVYKYIAETGNTAYLDEEIPFADKDTGT
FT                   VREHLKRAINFSMTHLGPHGMPAGLHADWNDCLVLGDQGESTFVAFQLYYALNIMKEFC
FT                   ENEPEYVKYLDETAAKLLKILNDLCFEGDRFIRGFRDTGDIIGSKKDNEAAMWLNPQSW
FT                   AVISRAATPEQAEAILNTANEKLNTPYGLELMQPSYRFEYFEGARMRLFNPGTKENGGI
FT                   FCQPQGWAILAEALCGHGNRAFKYYEESSPASFNDDADRRVIEPYVHGQFIEGHESPFA
FT                   GRSHVHWLTGTASTVMVGCVEGILGLRPTTKGILISPAIPGEWDGFTMEKVFRGKKLHI
FT                   KVDNAAHKEGNPEKIILNGVERPAGLIPEGDLQTENEITVIM"
FT                   /product="GH94"
FT   CDS             14948..15610
FT                   /translation="MKTIARIRNAYDGKFGVPRQSGLVEEVVSTIVFEPEFRVAEALRG
FT                   IEEFSHLWLIWAFDRMEREGWSPTVRPPRLGGNQRIGVFATRSPYRPNAIGLSCVRLLG
FT                   IEKGKDGTVLKVAGADLMNGTPIYDIKPYLPYADCRPEATGGFTDRTEKRTVTVEIPEE
FT                   TTKKMNPEDRKALEAVLREDPRPAYQDDPERIYAFEFGGRHVEFRVADQVLSVIRIV"
FT                   /product="COG1720: Uncharacterized conserved protein"
FT   CDS             15812..16888
FT                   /db_xref="GO:0004648"
FT                   /translation="MERVYNFSPGPSQLALPVLEKAQKDLVCYGDTGMSVMEMSHRSKM
FT                   YTDIYDKTVADLRDLMNIPESYDVVFLQGGATQQFSAVPLNLMVNKKADYIDSGNFAHL
FT                   AAEEGKRYGEVNVVASSREDVYTYIPDLDAIKFDDDADYVHITQNNTIYGTRFIELPKC
FT                   KAPIVCDASSMILSEEMDVTKYGAIYAGAQKNIGPSGLCVLIVRHDLVGKAMDICPKLL
FT                   NWQIQVEKASMYNTPNTWGIYLAGLTFEWLKDIGGVKAVEAVNKEKAAMLYDFLDNSKL
FT                   FKATAQPKYRSRMNVTFVTGDADKDAAFVKAAAAEGLVNLKGHRSVGGMRASIYNAMPI
FT                   EGVQKLVAFMKKFEAENA"
FT                   /product="Phosphoserine aminotransferase (EC 2.6.1.52)"
FT                   /EC_number="2.6.1.52"
FT   CDS             17068..18243
FT                   /db_xref="GO:0004617"
FT                   /translation="MYKIQTLNAISDIIHTQLSADKYTVSKDEPVPDAILVRSAAMHEM
FT                   EFGKELLAIGRAGAGVNNIPIDRCSKEGIVVFNTPGANANAVAELVIAGLMMSGRKIAD
FT                   GIDWVKSLKGKGDEVAKLVEKGKSQFVGAELRGKTLGVIGLGAIGSIVANAASRGLGMN
FT                   VVGFDPFISVESAWSLSTTIHRAASEDEVIAQADYITIHMPLNDKTRGSFNAEKIAKMK
FT                   DGACLLNFARGELVNTADLLAALASGKLERYVTDFPNDELIGAEKVVCIPHLGASTPES
FT                   EENCARMAAAEIRDYLENGSIHNSVNYPEVLLGDPEAVRVLVLHENIPNMISNITAAAA
FT                   KEGINIENMVNKSKKDMSVTVMEMEELPSKHALDTLSELPGIIRIRTFTKE"
FT                   /product="D-3-phosphoglycerate dehydrogenase (EC
FT                   1.1.1.95)"
FT                   /EC_number="1.1.1.95"
```

```
FT   CDS             complement(18368..21937)
FT                   /db_xref="GO:0003899"
FT                   /translation="MFELSNLDSIQIGMASPEQILAWSHGEVTKPETINYRTLKPERDG
FT                   LYCERIFGPTKDWECNCGKYKRIKFKDKDKICDKCGVQVTRAKVRRERMGHIQLAAPVS
FT                   HIWYFKGIPSRMGMLLDISPRNLEKVLYFANFIVLDPGNTEETGLKRLELINDARYREV
FT                   EAKCGKGSFVAKMGAEAVQDLLMELDLDKLSADLKAEIAQLTEKERDRETEGQKRARAV
FT                   KRLEVVESFRTSHNKPEWMILTVVPVIPPDLRPLIQLDGGRFATSDLNDLYRRVINRNN
FT                   RLKRLLELGAPDIIVRNEKRMLQESVDALIDNGRRGRPVTGPGNRALKSLSDLLRGKQG
FT                   RFRQNLLGKRVDYSGRSVIVVGPELKLYQCGLPKEMALELFKPFVMERLVTLKLSHNVK
FT                   SAKRAVEKIRPEVWDILEEVIRDHPVLLNRAPTLHRLGIQAFEPILVEGKAIKLHPLAC
FT                   TAFNADFDGDQMAVHVPLSMEAQAEARFLMLAHNNILKPQDGRPVISPSQDMVIGCYYL
FT                   TIVHDGAKGSGRVFSSEDEAIMAYSLGQITLQSPIKVRLTRSFGDETGSKLIDTTLGRL
FT                   IFNDALPQDLGFVKRECLDDMFKLEVDELVGKKQLSRIVDQCFHTQGEHRTAQVLDKIK
FT                   ALGFKYSTIGAITVSVADIKVPEVKKTMLADADEAVRKVNNMYHRGLLSEDERYNRVIN
FT                   IWTECTANLRNQILPNLDTFNPIRMMTDSGARGSSAQVSQLAGMRGLMASPSGKTIELP
FT                   IRANFREGLTVLEFFLSSHGSRKSLADTALRTADSGYLTRRLVDVSQEVIVREDDCFEA
FT                   RGEKVRGITVAELMSGKQSIESLEDRLRGRTAAEDIIDPATGEVLVRLNEPIDYKMARE
FT                   IVARGITKATVRSVLTCRTENGVCARCYGENMAHGGKVDVGEAVGIIAAQAIGEPGTQL
FT                   TMRTFHTGGIAGADDITQGLPRVEELFEARKPKRDAILAEIAGTVSIQETKKRRELVIT
FT                   SDDDPKDTKSYQITYGSRLKVQVGDHVEAGDELTEGSVNPHDLLRILGVKAVQEYLLKE
FT                   VLSVYRLQGVGVSDKHIEIIIRQMLRKVRVEDSGDTDLLTGSLVDIFRFEEANRKAIME
FT                   GGTPAIAKRILLGITKASLATESFLSAASFQETTRVLTEAAIRGKIDPLIGLKENVILG
FT                   KLIPAGTGMKRYKDIEIKDSYSF"
FT                   /product="DNA-directed RNA polymerase beta' subunit (EC
FT                   2.7.7.6)"
FT                   /EC_number="2.7.7.6"
FT   CDS             complement(21958..25824)
FT                   /db_xref="GO:0003899"
FT                   /translation="MGKLRERMSFSRIDEVLEMPNLIEVQKNSYQRFLDTDLREVLADV
FT                   SPITDYNGNLVLEFVDYSLDEPKNSIARCRERDITYAAPLKVFVRLKNKETGEIKESDV
FT                   FMGDFPLMTDHGTFIINGAERVIVSQLVRSPGVYFDVAYDKAGKKLYSSQIIPNRGAWL
FT                   EYETDSNDVLWVRIDRARKLPITVIIRALGYGSDAEIIELMGEDERLMATIQKGDNAKT
FT                   REEGLIEIYKRQKPGEPASLESATNLFHSLFFDPKRYDLMRVGRYKYNKKLSIATRIHN
FT                   HVAAEDIINPQTGEVMIKKGDSIPLDVAREIQNAGVNSVLLDCDGQIRKVVGNNFVDAS
FT                   AYLPFNPKDAGILEMVHLPTLKQIMASVEPDQLFDAVKENVAELVPKHIIPDDIISSVN
FT                   YMLGLPYGIGSTDDIDHLGNRRLRSVGELLQNQIRIGLSRLERVVRERMSIQDSTDIKP
FT                   GDLINIRPVSAAIKEFFGSSQLSQFMDQPNPLAELTHKRRLSALGPGGLNRDRASFEVR
FT                   DVHYSHYARICPIETPEGPNIGLIGSLATYARINEYGFIEAPYRRVDKEHGKVLDEIDY
FT                   MTADEEDLYKVATANEPLNADGTFVQDRITVRDKAEIVEVPVSQVDFMDVSPRQVVSVA
FT                   TAMVPFLENDDATRALMGSNMQRQAVPLLVPEAPLVATGMEYRAAHDSGVVILAKEDGV
FT                   VEKVDATQIIIRDNFGERHTYNLTKFARSNQSTCINQHPVVSAGQKIEAGDLLADGPST
FT                   EKGEIGLGRNALIGFMTWEGYNYEDAVLLSEKLVKEDIYTSIHIEEFESEARETKLGPE
FT                   EITRDIPNISDDAVKDLDEGGIVRIGAEVHAGDILVGKVTPKGETELTAEERLLRAIFG
FT                   EKAREVRDTSLRVPHGEGGIVVDVKVFTRENKDELSPGVNQMVRIYIAQKRKISVGDKM
FT                   AGRHGNKGVVSRILREEDMPFLPDGTPLQIVLNPLGVPSRMNLGQVLEVHLGLACRALG
FT                   WHIATPVFDGATYEEILDHLELAEKKMAAPEDENDPHVNEYHFHNGKPLRLDLDEEGKA
FT                   LFRQGKIRVRDGRTGDYFENPVTVGIMYFLKLHHLVDDKMHARSTGPYSLVTQQPLGGK
FT                   AQFGGQRFGEMEVWALEAYGAANTLQEILTVKSDDTVGRVKTYEAIIKGQNIPNPGVPE
FT                   SFKVLLKEMQALSLDIRVLDAARNEIEIPELDPEDMPQPETMRANLLGKTEADDEPVAA
FT                   EESEELEQAADEAFSMSEDDLSFGADDESTADSDDFSVSDENAEDLEDFSVEDLSDLDL
FT                   DDDI"
FT                   /product="DNA-directed RNA polymerase beta subunit (EC
FT                   2.7.7.6)"
FT                   /EC_number="2.7.7.6"
```

```
FT   CDS             26073..27059
FT                   /translation="MRLRGLFRTVILILTLALTLCAGAAEEAIPPSPEPPAGNPAGRPT
FT                   PAPTAVPTETIVDHTHQPGLYPDFSFMPDRKLLEIWFPNITDADEAVLIYDGDVWMIDC
FT                   GDEKAGARGARLLQQLGIKKIDRLFTSHLHHDHINGLALTDDTAKIGEVLICFSPDLTP
FT                   SGLKLIQVAEEHHIPVKEYGDGDRFTMGDGAVELLFIKNNEQDLDMNNQSAVTRITYGQ
FT                   RTILFMADMEQPGQIAMFKKGDPELLKCDIVKYPHHAKTDMYTPFFNAMGAKLAIVTST
FT                   EGRKDGGQTGLANRKLPAVFTFVGGQFTHLVTDGEYWLCERVEITVK"
FT                   /product="DNA internalization-related competence protein
FT                   ComEC/Rec2"
FT   CDS             27231..27476
FT                   /translation="MEMLKAAGSRKVVGTKQVLRALKAGTVARVYVCSDADTFIYQQVV
FT                   RSAEEAGIPCVRAASMKELGMICGVDVPTAAAGLLR"
FT                   /product="Firmicutes ribosomal L7Ae family protein"
FT   CDS             27541..27915
FT                   /db_xref="GO:0003735"
FT                   /db_xref="GO:0005622"
FT                   /db_xref="GO:0005840"
FT                   /db_xref="GO:0006412"
FT                   /translation="MPTINQLVRKGRERATKKPTAPILQGCPQKRGVCLSVKTQTPKKP
FT                   NSALRKIARIRLTNSIEGTCYIPGIGHNLQEHSVVLIRGGRVRDLPGVRYHIIRGALDT
FT                   AGVAKRMQARSLYGAKRPKK"
FT                   /product="SSU ribosomal protein S12p (S23e)"
FT   CDS             28126..28596
FT                   /db_xref="GO:0003735"
FT                   /db_xref="GO:0005622"
FT                   /db_xref="GO:0005840"
FT                   /db_xref="GO:0006412"
FT                   /translation="MPRRGFVPKREVLPDPVYGTTVVTKLINQIMLDGKRGVAQNVCYQ
FT                   AFETVAEKTGKAATDVFQEALNNVAPQLEVKARRVGGATYQVPIEIRPERRQTLALRWI
FT                   VDFARKRGEKTMAERLAGELLDASNNTGAAVKRKEEMHRMAEANKAFAHYRW"
FT                   /product="SSU ribosomal protein S7p (S5e)"
FT   CDS             28696..30762
FT                   /db_xref="GO:0005525"
FT                   /db_xref="GO:0006412"
FT                   /translation="MAHIDAGKTTTTERILFYTGKNHRIGETHEGTATMDWMAQEQERG
FT                   ITITSAATTCFWHDPHDPANTNKQYRINIIDTPGHVDFTVEVERSLRVLDGAVSVFCAK
FT                   GGVEPQSETVWKQAETYRVPRMAYVNKMDITGADFFRVVDMMKDRLGANAVPIQLPIGK
FT                   ENTFRGIIDLVRMRAEIYYDDLGQDVRDEEIPADMQEIADEYRANLIEKVAETDDTLME
FT                   KFLEGEELTEEEIRSAIRKCTIDCSMNPVLCGTSYRNKGVQPLLDAVIDYMPSPLDIPA
FT                   IEGTDPDNPEKKMERHPSDDEPFSALAFKIMVDPFVGKLAFFRVYSGKLSAGNYVMNST
FT                   KQKRERISRIMLMHANHREEVENIYTGEIAGAVGLKDTTTGDTLCDENHQIILESMVFP
FT                   DPVIEVAIEPKTKAGQEKMTYALQRLAEEDPTFKTYTNQETGQTIIAGMGELHLEIIVD
FT                   RLLREFKVEATVGKPQVTYKETIRKPAKAEGRYVRQTGGHGQYGHCWIEIEPQEPGKGY
FT                   EFESRIVGGVIPKEFIGPIDQGIQEAAKSGAIAGYEVVNFKAAVTDGSYHEVDSSEMAY
FT                   KIAASMAFKEAVRKADPCLMEPMMKVEIIVPDTYLGDVMGNVSSRRGRVEGTEVRGQDQ
FT                   IIHAYVPLSEMFGYTTDLRSRTQGRGMFTMQFDHYEEVPKSVAEKIIGKKGAD"
FT                   /product="Translation elongation factor G"
FT   CDS             30832..32046
FT                   /db_xref="GO:0005525"
FT                   /db_xref="GO:0006412"
FT                   /translation="MAKGRYERTKPHVNIGTIGHVDHGKTTLTAAITMTLAVKGEAQAM
FT                   RYDEIDKAPEEKARGITINTAHVEYETAKRHYAHVDCPGHADYVKNMITGAAQMDGAIL
FT                   VVSAPDGPMPQTREHILLARQVGVPYIVVFMNKTDMMDDEELLELVEMEIRELLSSYDF
FT                   PGDEIPIIKGSALKTLEYLQNGGTDVDNAPECKCIWELMDAVDSYIPEPERATDQPFLM
```

```
FT                   PVEDVFSISGRGTVATGRVERGTVKVSDPVEIVGLMDKPRNTVVTGVEMFHKLLDQAEA
FT                   GDNIGALLRGIQRNEVERGQVLAKPGSIHPHTHCIGQVYVLTKEEGGRHTPFFNGYRPQ
FT                   FYFRTTDVTGNIKLPDGVEMVMPGDNIDMEITLITPIAMEQGLRFAIREGGRTVGSGVV
FT                   AKVIE"
FT                   /product="Translation elongation factor Tu"
FT   CDS             32068..32229
FT                   /db_xref="GO:0003735"
FT                   /db_xref="GO:0005622"
FT                   /db_xref="GO:0005840"
FT                   /db_xref="GO:0006412"
FT                   /translation="MANAARNKVCLACTECKQRNYNNMKNKKNTPDRIELMKYCPFCKK
FT                   HTTHRETK"
FT                   /product="LSU ribosomal protein L33p"
FT   CDS             32257..32547
FT                   /translation="MKEVSDVKKTAENTEKAVKKSGESKLSKIISWFKNLPKRISKPFK
FT                   NMYYELKKVTWPSKEKLIAYSVIVLVFMLFMGIVIGLLDMGASALVRMLIA"
FT                   /product="hypothetical protein"
FT   CDS             32565..33101
FT                   /db_xref="GO:0003711"
FT                   /translation="MAEAKKEPCWYVIHTYSGYENKVKDTLEKSVENNGMQDLILEVRV
FT                   PMEEVVEIRNGKRVKSTRKVYPGYVLVHMIETSESWYVVRNTRGVTGFVGPDSKPVPLT
FT                   QEEVDMMLNTEQSSVDFGFAIGENVRILSGPLENFSGVVEDVDTVRGKLTVKVQMFLGR
FT                   EMPVEVDLDQVEKED"
FT                   /product="Transcription antitermination protein NusG"
FT   CDS             33248..33673
FT                   /db_xref="GO:0003735"
FT                   /db_xref="GO:0005622"
FT                   /db_xref="GO:0005840"
FT                   /db_xref="GO:0006412"
FT                   /translation="MAKKVQAYIKLQVPAGKATPAPPIGPALGQHGVNIPGFCKEFNDR
FT                   TAKENGMIIPVVITVYTDRTFTFITKTPPAPVLIKKELNLQKASGRPNKEKVGKLTKEQ
FT                   VKKIAEIKMPDLNAGSIEAAMSMVAGTARSMGVTVEE"
FT                   /product="LSU ribosomal protein L11p (L12e)"
XX
SQ   Sequence 34263 BP; 8287 A; 9127 C; 9872 G; 6977 T; 0 other;
     acaggcggaa cgcctgggcg agaagattga ggtgccgagg aagcatctga tcacccgcag        60
     acatccgaac cgttccatcg ccggcgatat catcatctat ctgctgcttc ttcttgtagc       120
     cattgcaatg gtgttcccgc tggtgttcct gatcggttcc tccctgaagc ccctggatga       180
     actgttccgg ttcccgccgc ccgtatggcc cagccatccg acgatggaca acttctccga       240
     cctgtttgtg acgatgggcc agagctgggt gcctttcagc cggtacctgg tgaacaccat       300
     catcattacc gtggtcggca ccttcggcca cctggtggtc gccggcatgg ccgcattcgt       360
     gctttccaag tatgaattcc ccggcggaaa gctgttcttc gctgttgcca ccacatgtct       420
     gatgttcacc ggcgtcgtta cggatattcc gaactacctg atcctgagcc gcctgggcct       480
     ggtggacacc tactgggcgc tgatcctgcc ggcctgcgcc gcgccgatcg gactgttcct       540
     gatgaagcag ttcatggaag gcctgccgac cgcactgatc gaagcggcga caattgacgg       600
     cgccagccgg ttccgggttt tctggagtat tgtgatgccc aacgtaaaac ccgcctggct       660
     gacgatgatc atcttcagcg tgcaggcgct gtggaacaac cgggccgcga cgatcatcta       720
     ttccgaagcg aagaaacccc tggtctacgc actgcagcag atccaggccg gcggcgtggc       780
     ccgtaccggt cagatggctg cagccaacgt gatcattgtt gccgttccga tcctgatctt       840
     cgtgttcagc cagagccgga tcctggagac catggcctcc tccggcatca aggactaaga       900
     ggggagggca taatgatgaa gaaaacaatc tccttcctgc ttgttctgct gatgctgctg       960
     tccatagccg gcgcactggc ggacgacagc ttcaacatga acaaggacgg ctacagcact      1020
     tcctatacct acggttacga ctactggggc gatgtgcagg aatccccgga cgcataccgc      1080
     gtcgagaccg tgctggacag cgtcagcctg ggacttgacg ttagaatgag caagccccag      1140
```

agcctgtacg tccggaacaa cgacctgtat attgtcgata ccctgaacaa ccggatcatt 1200
caggtcgagt acaaggatca tgcgtacaag ctggtccgga tcattgacca tatgaccggc 1260
ggaaatccgg aaacattcaa caatccttac gatgtgtttg tggacgagaa cgacagcatc 1320
tatgttgccg actacaacaa caaccgcgtc gtgatgatgg acaaggacct gaactggatc 1380
aaggaattta cgaaaccgac ggacgccacc tttgaccaga gtatcgactt tcttcccaaa 1440
aagatcgtag tggatgttgc aggacgcgta tacgcgctgg ttaccaacgt gaacaagggt 1500
attgtcaagt atgaggcgga taccaccttc actgggttta tcggcgccac gccggtgagc 1560
gtttcgatgg tggactatat ctggaaaaag tacttcatga cccaggccca gcgggacgcg 1620
tctccctcct ttgtcccgac ggaatatgaa aatctctata tggacaagga cggattcatc 1680
tacgcgacca ttaccaattt cagcgaatac gacctgaaat cagacaaagc gaaaccgatc 1740
cgcaggctga atggcctggg caacgatatc ctggtcaaga acgaccgtta tcctcccatc 1800
ggcgacctgt ggtggatgga aggcgactcc acctatttcg gaccttcgaa gatgacggac 1860
gtcaccgtgt ttgacaacga tatttacgtg gtcctggaca gggtacgcgg ccgcctgttc 1920
ggctatgaca accagggcgt catgctgtgg gctttcggta cccgaggcaa tatcgaaggg 1980
gcatttacca gcgctgtcag cattgaacac atgggaaccg acctgtttgt gctggatcag 2040
ctggagaaca gcgtcacggt gtttacgccc acggaatacg gcaacaagat ctatgacgcc 2100
atcctgaaat accaggaagg cgaatatgac gaaagcgcgg cggtctggga agacgtgctg 2160
aaacagaacg ccaactaccc gatggcattc cgcggaatcg gccggacggt actccgccag 2220
aacaagtacg aagaggccat ggaatacttc aagatggccc acgaccggga aaactacgga 2280
cgggtattca aactctaccg gaaagaatgg attgagaaga acatcctgtg gatcttcctg 2340
ggcctggctg tgctgctgat tgtgccgctg gttatcggaa gggtcaagag aatgaaatgg 2400
gaggtgatca tgcatgaaca gagcaaagtc cgtaaaaatg gataaagccg cctcggctac 2460
agcacgtaaa gcctggtgga gcaagtacaa ggactccctg cgttacgcgc tgcacgtgat 2520
cacccacccg ttcgacggtt tctgggatct gacccacgaa cacagaggca cgctggcagc 2580
cgccaacacg ttcctgatcc tgttcctgat tacgcgggtg ttaaaactgc tgtgcaccag 2640
cttccagttt atcagcacgc ctgttcagca catcaacgtg tttgaggaaa tgggttcgct 2700
gctgctgccg ttcctgattc tgtgcctggc caactgggct atgactaccc tgttcgacgg 2760
taaaggacgg tttatcgaca tctatatcgg catgtgctac gcactggttc cgtatgttct 2820
gatccagctg ccgatgatcc tggtcagcaa catgctgaca tttgaggagg ggagccttta 2880
cagcgttatg ctgagcatca gcatcgtctg gtgcgtgttc ctggtgtttg tcggcttgat 2940
ggagatccat gattacggtc ccggcaaaac cttcatcttc atcattgtca ccattgtcgg 3000
cgcggctgtg atcatattcc tgattctcgt attcttcagc ttgctgagcg acgcggttgg 3060
cttcttcgta agtttgtacc gtgaagccag ttaccgactg aattaaggag ggatggcaaa 3120
aatgagaaaa aaacccatcc ttttacggct gcttccgtgg atcatcgtgc ttgtactgat 3180
cgccggtgcc tatattgtgc ttgataagat ctattcatct cccggccaca gctttgcccg 3240
ggatacgaaa atcatcagct atgaaggcga cgggaagccg ctgaccatgg aaaacgatca 3300
gttgctcttt gaaatggacg gatcgaccac tcagttctcg gttaccaata aggaaaccgg 3360
gaaggtatgg tattccaatc cggaagaccg cgcaaacgat acgacagcca caggtaagac 3420
aaccaaggaa tacctgtcct gcacgctgaa tgcgcattat tacaatgaca tcagcgaaac 3480
agaagtgaac aactacacaa aatccatcat gaaccagaac tatgaaatcc agcaggaagc 3540
ggacggctcg atccgggtca actacacgct gggtgagaaa gtgttcatgg tgcccaacgc 3600
aataaccaag gaaaacatgg acgctctcct ggccaacctg aaaaagaagg accagggcaa 3660
actgaagaac cattactccc tggttgaaac aaaaaaactc gataaacaga agaagaagga 3720
cgagattatc gccctgtatc ccagtgtgct ggagcaggac ctgtacatcc tgaaatcgga 3780
ccggaccacc gagcaaaaga aacaggcaga acagctgctg gccagcgcgg gatacactga 3840
ggaagacttt gcgagggacc aggagctaca ggccggcgag aaatcaaaca gcggaccggt 3900
atttaacatt agcgtgattt accgcctgga agggaacgac ctggtggtat ccgttccgta 3960
cagcgaactg acctgctctt cggattatcc gctggtatat atcgacatgc tgccgatgtt 4020
cggcgcgggc gggaagaaac aggatggatt catcctggtg ccggaaggcg gcggcgccgt 4080
catcaactac aataacggca agctgtccca gaatccctat tatgccaaca tgtacggctg 4140
ggactacgga acgaagcgga cgagcgttac cggggaaacg gagaacgctt tccccgtgtt 4200
cggcatgagc caggaagacg gttctttcat ctgcattatg gaaggcgctg attcctacgg 4260
cggcgtatgc gccgatatct ccggcagatt caacgattac aatacggtat acggccggta 4320
caatgttctg cacttcgaca gcgtggaagt ggacgagaaa aatccgcac gcctgttcct 4380

gatgtacgaa aacgaaatcc cggacgatac tattgttcag cgttaccgtt tcctgaacga 4440
caacggctat gtgaacatgg ctgaagccta tggagagtat ctccgtgcga agcccgaaat 4500
gagaggcgaa tgtgcatccg aagagatgcc ggtgaacgtg gaactgatcg gcgcgatcaa 4560
caagacagaa atcaaattcg gtctgccggt caagtccgtg gtagccacga ccacctttta 4620
tcaagcggaa agcattatga acgacctgct gaacgcaggc gtcagggacc tgaatctccg 4680
ctataccgga tggtgcaacg gcggcgttca gcagcgggtg cttaccagca tccatgttga 4740
aggcggcctg ggcggtgaaa acggcatgaa gaaactgatt gccagcgccg cggaaaagaa 4800
cgtggacctg tattttgacg gtatcacatg cttcgcttat gacagcggtc tcttcaacgg 4860
attcaactca ctggccgacg cagcccgggc tacgacccgg accgtgatca agctgtatgc 4920
atacgatatt gttacatacc ggttgtcgac ctggatggat aatccatatt ttctggtgaa 4980
accgtcctac gccaagaact gtgtaaacaa cctgatcaac ggcctgaaga accgtggcgt 5040
ggcaggtatc gcgttccgcg atgtcggcaa cctgctgagc gctgattatc aggacagcgg 5100
caccgtgacc cgcgaacagg ccaaggcaat ggatattcag acccttgagg attcagtggc 5160
gacgggcatg aaggtgatga tcaaggaagg taacgcgtac tccattccgt atgccgatct 5220
gatcacagac atgaaccttt ccggcaacgc ctacgctttg ctggattacg gtattccttt 5280
ctaccagatt gcgatccatg gcctgaagga ctataccggt gaggcgatca acctggcagg 5340
agatacacag acgcttctgc tggaatcagc cgaatacggc gcgggcctga acttctcctt 5400
tatgcagaag gataccaagg tgctgcagga ttccatgtac agctgctacg cctcagccta 5460
ttacacgccc tggaaagacg aagcgatcgc catgatcacc cggtatcaga aggaaatgtc 5520
cggcctgaac cgccaggcga ttaccgggca tgaaagacta aatgaagacg ttgccgttac 5580
cacttacgca gacggaaccc aggtgtatgt gaattacgga gacagggagt tcacgatagg 5640
ttcagttacc gtgcccggac gcgattatct tgtgaaaaga ggtggcggac aatgagtaag 5700
aagaaccgga aataccggga acactggtat agcggcatta cggaagcttt ccagatgttt 5760
gtcccgggaa acaaactcta cagaaccatt ccggccaaaa gggcccggac cggtttcctg 5820
tttatcctgc cgtttatcat cggatttatc ttcttcatgc tgaagccact ctatgtttcc 5880
atgcagatga gcctcagccg gtatgtgatc aataccggtg agatgacctg gaacaatttc 5940
tacaactaca actacgcgtt caacacggat ccggattaca accggatgct ggtggacgag 6000
atcagccgga tggcaatcaa cgtgatcgcc acgctggtta tgagctttgt gattgcggtt 6060
attctgaacc agaagtttaa gggccggatt ctttgccgga ttatcttctt cctgcccgta 6120
atcctgtcat ccggcgtgct gcccggcatt gaaacctcca gttcgtccct gaacctgatc 6180
aacagcatgt cacaggccat gggggattcg tccggcgtca acatctccgc cgcgctggaa 6240
aacctgctgt cagtatccgg cgtggccagc caggtttttg acgtcatatt ccagatgatt 6300
gacgcaatct atgacatcgt gatggccagc ggaatccaga tcatcgtatt cctttccgga 6360
ctgcagtcga tatcgccctc cctgtatgag gccgcggacg tggaaggctg ctccgcatgg 6420
gaatccttct ggaagatcac cttcccgatg gtcagccccc ttctgctggt caactgcatc 6480
tataccatca tcgacttctt catgaagaac aacaaccagg tcatggagca tatcaataac 6540
acctatatga acggcgtccg gatggatctg ggcatcgcag ccgccgaaag ctggatctat 6600
ttcggcgtgg cgattgtttt catcggcatc agctccttca ttatctcaag ggcggtgaaa 6660
tcctatgagt aaagacaata cgatctttat cggcgagcac agcagcttcc gggaaagaaa 6720
ccgccgcagc ggcggttacc tgatgaaacg caggatcaag gagatcagtg tgtccgtcat 6780
ccgcggactg ctgatgttcg gcctttgctt catgatcatc tcgccgatga tcagccggat 6840
ctccatgagc ctgatggatg aaaaggatct gtacgatact acgatcgttc tcatccccccg 6900
gaacgtgaca atggataact ttacggccgt cgcgcgcgcc accgctatgc cgacatccat 6960
gattaatacg ctgtggattt ccatcctgag cagtattctt cagattattt cctgcacact 7020
ggtggcttac ggattcgcca gatatgagtt cccgctgaag aaattctggt ttgcctgcgt 7080
catcgcgctg attattattc cgccccagac aatccagagc gccctttacc gcacatttac 7140
acgctttgat ttcctgggca tcatcacgct gatcaacggg gaaccgctga atatcaggaa 7200
atccatctgg ccctatctgc tgatgtgcct gacctgcatg ggcctgaagg acggactgta 7260
tatctacatg ctgcggcagt atttcaagaa tgttcctgaa agcctggaag aggcggctta 7320
tgtggacggc tgcggcacaa tgcacacctt tgtgaaaatc atgctgccgg acgccctgcc 7380
gataatcgcc agctgcttcc tgttcagttt tgtatggcag tggacggatt tgttctattc 7440
ccgcaacttt ctgtcaggct atacactgta cgcaccgcaa ctcagttcga tgctcgaccg 7500
gatcgcagat gaactctcca aaatggccgg cacaccggtt accccatctc ccggcaggca 7560
gctgcagctg atctctgttg gcgtgctgtt ctgctgcatt ccgctgatca tcctgtactg 7620

cttcacacag cggacattcg tggaaagcct tgccatgacc gggtcgaagg aataagacac 7680
aaaatgcgaa aaatgtacga acctatactt gaaaaaatgg acaaaccggt ctataataaa 7740
cctgaacagt gggggttccg catcaacaat gtacctatac agatttccgc gggaaatctg 7800
taaggaaata aaaaaggagg ttttctcaat atgaagaaaa tcatggctct ggtactggct 7860
gctctgatgc tgctgggctg ctgcagcttc gctgccgcgg aagaagttcc ggctgaatat 7920
ccggaaatca tcgaaggtct tgattttggc ggcgcaaccg tctacatcaa tgactggtat 7980
tccaccggcg aacgcgctga agaacccact gaagaacagc aggctcagta tgactactgg 8040
gattggctgg aaaaaaccta caacgttaag ctggttgaga ccaagctggg cgactgggac 8100
ggcatggtgg ccgaactcca gaacatcgtt tccaacaagg acaacagcga actgcgtatc 8160
atcggcgtgt ccggcggctt ctgcggcccg gtgctcgcta acggactgtt cgccgagtgg 8220
acctacggtc tggagaactt caacgaagcg accgttaagt tcatgaccct gggcggcaag 8280
tgctacggtg tctgcggcgg caagttcttc gagccccgcc agatggtatt cttcaacaag 8340
aaggttctgg aagatgccgg catcgactgg aacgagatct atgacgctca ggccgacgga 8400
agctggacct gggaaaagat ggaaagctac atggacgctg taaaggctga caaagacaat 8460
gacggtgaac tcgacgtatt tgccctgacc ggcaacttcg atgatggcgt tgttggcctg 8520
gttgtcagca acttcggtga tttctatgaa ttcgacgaga acggcaaact gtactactct 8580
gctgacagcg atgccactct ggctgccctg aaccgcatcc aggagtggaa ccggaactac 8640
tatcgtccct atgtcaattg ggatgactat aagcagttct ggccggaagg aaacgttgcg 8700
ttcctcatcg gacagtccta tgaaggtttc aacggcaacg acgtcgtgaa ccaggttgaa 8760
gagtggggcg gcgtatgcct gcccaaggga cccaacgcgg atgcttacac ccatgcggcg 8820
gacaacaaca tcttcggcat cccggctgtg tatgacgatg agacttccct gaagctccag 8880
cagatctaca ccctgtatcg caagaatccc gtcattgacg ctgatactga tgcttgggcg 8940
acccgtctgt atgagctgac cgacgaacgc gctgttgacg aaacctacgc ttatctgcgc 9000
gagaacggca ccatcatgaa cttcaactac atcggcgacc gcaacagcac catcggcccg 9060
aacctcacct ggggtatcat gggcggcgct gctgctgagc aggttgaagc tgcacgtctg 9120
gccatggaag acatggcggc tgtgttcaac ggcgacaaga ctcaggagca ggttgatgct 9180
gaaaaggccg cgcgcgaagc cgctgctgcc gaagaagctg ctgctgaaga aggcactgaa 9240
gaagcccccg ctgatgaagg cgctgctgaa taagcaacaa ttcctttggt ttcccggacc 9300
gtccgaaagg acggtccttt ctttttcaga aaagcttgaa gaaaactccg ggagaggata 9360
caatatgtac gaacaagtcg gattcagatc cggaggaaag gatggaacag catgggcaaa 9420
aagatactgg cttttctgat gattctctcg ctgcttggcg gatgcgtatc agcactggca 9480
gacgaatcaa aaacgattta tacttccgat ttcagcaaag atgaagacgg atggtacggc 9540
cggggagcga agtctttccg taccgcggac ggcacgctga aaactgttga caggcaaagc 9600
agctggaatt ctccgggacg ggactttgac ctgatcgagg gcgggaaata ccagctgagc 9660
gcagaggtca aacaggacga catggacagc gccagcttca tgatttccat tgcacattca 9720
gccgagggag tggaaaccta tgagaacctg gcgttcggga cggcgaaaaa aggcgaatgg 9780
gttacactga aaggtgaata tattgccgga gcgttcgatc gcagcgtgct gtatgtggag 9840
accaccggtt cggacacgct ggattttgaa atccgcaatt ttaccgtcat cgccccggaa 9900
ggagagcctg aacccaagcc cacagagcct ccgatggtga ttgaggaagc ggaaaacatc 9960
cccagcctga aggaaatcta cgcaggtaag tttgatttcg gttctgccgc tccgcagatg 10020
gtgtttacca atccgaagtg gatgaacctg attacggaac agttcaatat cctgacgccg 10080
gaaaacgaaa tgaaaccgga ttccgtactg gacgttcagg cgagcaaaaa actggtggaa 10140
gaaaccgggg atgagacatc cgctgcagtt cattttgacg cggccaaagc gctgctgagg 10200
tttgcccaga gcaacgggat caaggtgcac ggtcatgtgc tgatctggca cagccagacg 10260
ccggaagctt tcttccatga aggctatgat ccgaagaagc cgctggttac ccgtgaggtt 10320
atgctcgggc ggatggaaaa ctacatcaag ggcgtgtttg aatacettga tgaaaactat 10380
ccgggcgtgg tcgtttcctg ggacgtgctg aatgaagcga tcgacgacag taccaactgg 10440
ctgcggaaca gcaactggct gaagatcatc ggcgacgatt atccgaacaa agcctatgaa 10500
tttgcccgta agtacgcgcc ggaaggcacg ctgctttatt acaacgacta caacaccgct 10560
atgcccatga agctgaaagg aatcgtgaag cttctgaaca agctgattcc ggaaggcaat 10620
atcgacggct acgggttcca gatgcaccac ggggtcactt tcccgaccat caagcagatc 10680
cagaccgcgg tcgagacgat cgccgcgctg aacatccgcc tgcgggtcag cgagctggac 10740
gttacggttt ccaacaattc ggaagcgtct tttgccaagc aggcgaagta ttacgcggat 10800
gttatgaaga tcctgatcgc ccacagcgac cagtttgaag ccgtccaggt atggggactg 10860

accgatatga tgagctggag aagccggcaa tacccgcttc tctttgacgg aagaggtaac 10920
ccgaaacccg cattctgggc ggttgccgac ccggaatccg ttcagtaaaa aactgatcaa 10980
agagccacga aagccatttc aacacgggag agcaaagacg ctctcccgtg ttattccata 11040
caggcggatt gaagaaacaa cccggttcgt atataataac agacggaaga accgaaaaag 11100
ggaggaaaac aaatgcggag caaaagacta tatgctgtta tgctcgcttt aggaatcatg 11160
atctgcgtgt cttccggact ggcggatccg ccggtccaga cggagggaat caccgtgccc 11220
gtgattgaat acaagggcga attcaagatt ccggataccg aagccatggc catgctgcgg 11280
gacatgaagt gcggctggaa cctggggaac acgtttgacg catataacgg atacaccggc 11340
catttttccg gaaccggtat ggaaagcagc tgggtcggcg tcaaaaccag caaaaagctg 11400
atcgaggcca tcaaggaagc gggattcaac acaatccgga ttccggtgag ctggcataac 11460
catgtggacg cgaacgacgt gatcgatcag gaatggatgg accgggtgaa agaagtggcc 11520
ggatgggcgc ttgatctggg catgtatgtg atcgtgaacg ttcaccatga caatgacgtc 11580
cattatgtgt atccggacgc acagcattat gaacgttcgg cggcttatct gatttcggtg 11640
tggaaacaga tggcggaagc gtttaaagac tgcgacgatc atctgatcct ggaatcgatg 11700
aacgaaccga gactggtggg gacgcagttt gagtggagct ggaacaacta cagtcctgac 11760
tgcaaggcag ctgccaaata tatcaatcag cttaaccagc tgtttgtgga tacagtccgt 11820
gccaccggcg gcaataacgc aacgcgctat ctggcggtac cggcttactg cgcggccccg 11880
gcaaatgcgg cggattacgc tttccagctg ccagaagaca cggcggatca ccggatcatt 11940
gtgaccgccc acgcttatac gccgtataac gtcgccttaa acctgaacag ctctgacaga 12000
acatttgacc tggacaaaga cagcggcaag aaaagcgata ttgacggcgt gatccagaag 12060
gtttacagcc gttttatcaa atacggcacg ccggtgatta tggatgagtt cggcgcgctg 12120
gacaaggacg gaaacacgca ggaccgggtg aatcttaccg catactacgt tgccgcggca 12180
gccgccaggg gaattccctg cgtctggtgg gacaaccacg ctttcggcgg aagcggcgaa 12240
cgcttcggcc tgatcaaccg gtcgaccctg gaatgggtct atcctgatat tgtactggct 12300
atccaggcta actgcctgaa ggacaaatga gggcagaaaa gggcggttca ggaagaactg 12360
ccctgctttt tacggttgcg tgtaaacgtt tactatgata agatgacaac agaagataca 12420
aggaggaaca acaaatgaaa tacggctatt ttgatgaaaa agccagggaa tacgtcatta 12480
cgaatcccaa tacgccggcg ccctgggcca actacctcgg ctcgccggaa tacggcgcaa 12540
tcgtaacgca gaacgcaggc ggttacagct ttgtgaagag cggagcggcc ggccggattc 12600
ttcgctatac gttcaaccag tttgatgagc ccggacggta tatctacctg cgggacgaag 12660
agacgggcga tttctggtcc gccagctggc ggccggtgtg caaaccgctg gatgaataca 12720
agacggagtg ccatcatggc acaagctata ccgaaatcgt cagcgaatac aacggcatcc 12780
gcagtcgcgc actgtactat gttccgatgg gcgcgaccca tgaagtatgg cggatcagcc 12840
tggagaacct gacggacagg gaacggaaga tcagcgtgtt cggttacgct gaactgacga 12900
cggaaagcct gtatacgcag gacctggtga atatgcagta tacgcagttc atcacaaaga 12960
ctgagttcca tgacagcttt atcctggaac agatcaatga gttcaattat ccgaatcctg 13020
acggaactac cggcagggaa cgtttcttcg gcctggccgg ggtgcctgtt gccagctata 13080
ccggacggcg cgaacagttc ctggggcgga accactttga caaaccgcag gcggtgctgg 13140
acggaaaatg cgacaacagc ctgaactata acgcgaatcc ctgcggcgcg ctgcagttca 13200
gcgccgcact gaagccgggg gaaaagaaga ccgcggcatt cctgcttggc cagaagacgg 13260
tttttgaggc gaaagccatc atggaacgtt atgaaaagac cgaagaaacc gtggacaaag 13320
agctgaagga actgatcgcc tactggcacg gcgaactgga aaacctgaag atcagcacac 13380
ccgacaagaa cttcgacgcg atgatcaaca cctggaacgc cttccagtgc ttcaccacct 13440
ttgtgtggag ccgcgcggcc agcctggtat actgcggtga gcggaacgga tacggctacc 13500
gtgataccgt gcaggacatc cagggcatca tgcacctgga tccggaactg gcgcggaaac 13560
agctgacgtt catgctgagc gcccaggtac atcacggcgc gggcctgccg ctggtaaaat 13620
tcacgcacaa cgccggccat gaagacacac cggaacagga aagctacgta aaagcgacgg 13680
gccatccgag ctaccgggcg gacgacgcgc tctggctgtt cccgacggtc tacaagtata 13740
tcgccgaaac cggaaacacc gcttacctgg atgaggagat tccgttcgcg gataaggata 13800
ccggcacagt gcgggagcac ctgaagcgcg caatcaactt cagcatgacc catctgggac 13860
cgcacggtat gcccgcaggc ctgcacgccg actggaacga ctgcctggtc ctcggcgatc 13920
agggcgagag tacgtttgtc gcgttccagc tgtattacgc gctgaacatc atgaaggagt 13980
tctgcgagaa cgaaccggag tatgtgaaat acctggatga aaccgccgcg aagctgctga 14040
agatcctgaa cgatctgtgc tttgagggtg accgttttat ccgcggtttc cgggatacgg 14100

gcgatatcat cggcagcaaa aaggacaacg aagccgctat gtggctgaat ccccagagct 14160
gggcggtcat cagccgcgcg gcaacgccgg aacaggcgga agctatcctg aatacggcga 14220
acgagaagct gaatactcct tacgggcttg aactgatgca gcccagctac cggtttgagt 14280
actttgaagg cgcccgcatg agactgttca atccgggaac gaaagaaaac ggcggtatct 14340
tctgccagcc ccagggatgg gcgatcctgg ctgaggcact gtgcggacac ggtaaccgcg 14400
cattcaagta ctatgaggag agcagtcccg catccttcaa cgacgacgca gaccgccggg 14460
tgatcgaacc ctatgttcac ggacagttca ttgagggaca tgagagcccc tttgccggca 14520
gaagccatgt gcactggctg accggtaccg ccagcactgt catggtaggc tgcgtggaag 14580
gtatactggg cctgaggccc acgacaaagg gaatcctgat ttctcccgcg attcccggcg 14640
aatgggacgg gttcacgatg gaaaaggtct tccgcgggaa gaaactgcat atcaaggttg 14700
acaatgccgc gcacaaagaa gggaatcctg aaaagattat cctgaacggc gttgagcggc 14760
ctgccgggct gattccggag ggggatctgc agacggaaaa cgaaatcacg gtgattatgt 14820
aatactggat tctgattatt ccggaaagct catgctttcc ggaatttatt aggggattct 14880
ttgaatttgt ccaggataac ctgtcggata tccaaacagt ttttgaacag gatgaaaacg 14940
gagaaatatg aagactattg ccagaatcag aaacgcatat gacgggaagt tcggtgttcc 15000
ccggcagagc gggctggtgg aggaagtcgt ctccacgatt gtgttcgagc cggaattccg 15060
ggttgccgaa gcgctgcggg gcattgagga gttttcccat ctctggctga tctgggcttt 15120
tgaccggatg gaaagggaag gatggtctcc cacggtccgg ccgccccggc tcggaggaaa 15180
ccaacgaatc ggcgtgttcg cgacacggtc accataccgg cccaacgcga tcggactcag 15240
ctgcgtcagg ctgctgggca ttgaaaaagg gaaggacggc acagtgctga aggtggcagg 15300
ggccgatctg atgaacggta cgccgatcta tgacattaag ccgtatctgc cctacgcgga 15360
ctgcagaccg gaggcaacag gcggctttac cgaccggacg gaaaaaagaa cggtgacagt 15420
ggagatcccg gaggaaacaa cgaaaaagat gaatccggaa gacaggaaag ccctggaagc 15480
agtactgaga gaagatccgc ggcccgcata ccaggacgac ccggaacgga tctacgcttt 15540
tgaattcgga ggaaggcatg tggagttcag ggtggctgat caggtgctga gcgttatcag 15600
gattgtctga cagcaagtga acccggcccg cacggccgcg acatggccga ctgtaacagc 15660
aaagagaagc tggagtcatc atacggcttc tttttttgtt tttgaactgc atttggggtt 15720
gctttttatt gggaaatatg gcatactagt cgcgtggtta aaaaataacc ctgtgggaca 15780
aaaacgtccc agttgatgag gaggatgaac tatggaacgt gtttacaact tttccccggg 15840
accttcccag ctggctctgc ctgtgctcga aaaagcccag aaggacctgg tttgctatgg 15900
cgataccggc atgtccgtga tggaaatgag ccaccgcagc aaaatgtata cggatatcta 15960
tgacaagact gtcgctgatc tccgcgatct gatgaatatt cctgaaagtt atgacgtcgt 16020
gttcctgcag ggcggcgcga cccagcagtt ctctgctgtt ccgctgaacc tgatggtaaa 16080
taaaaaggct gattatatcg acagcggcaa ctttgcccac ctggccgccg aagagggaaa 16140
acgctacggt gaagtgaatg ttgtggcttc cagccgtgag gacgtatata cctatattcc 16200
ggatctggac gccatcaagt ttgacgacga tgcggactac gtccatatta cccagaataa 16260
cacgatttac ggcacccggt tcatcgaact gcccaagtgc aaggcaccta tcgtgtgcga 16320
cgcgagcagc atgatcctct ctgaagagat ggacgtaacg aaatacggcg ccatctatgc 16380
cggcgcacag aagaacatcg gaccgagcgg actgtgtgta ctgattgtgc ggcacgacct 16440
ggtgggcaag gcaatggata tctgcccgaa gctgctgaac tggcagatcc aggttgaaaa 16500
ggccagcatg tacaacacac cgaatacctg gggcatctac ctggccggcc tgaccttcga 16560
atggctgaag gacatcggcg gcgtaaaggc tgttgaagct gtgaacaagg aaaaggccgc 16620
catgctgtat gatttcctgg acaacagcaa gctgttcaaa gcgacggctc agcccaagta 16680
ccgcagccgg atgaacgtca cctttgtcac cggagacgcc gacaaggatg ccgcgtttgt 16740
caaggcggcc gcggctgaag gtctcgtgaa cctgaagggc catcgctctg tcggcggtat 16800
gcgcgccagc atttacaacg cgatgccgat cgaaggcgtt cagaaattgg ttgccttcat 16860
gaagaagttt gaagctgaaa acgcataaat aaggaataat ccttttcccc ctgcccccat 16920
ccccatgctc taaaagttcc gtgatgcaaa ttgatttgca tggggacggg gattatttag 16980
ttggcagggg ctccacccct gtaaccctga ctgtaatgag agggaaagaa taggaacaga 17040
atcatttatt gaggaggaaa acaggaaatg tataaaattc agacgctgaa tgcgatttcg 17100
gacattatcc atacccagct gagcgccgat aagtataccg tgagcaagga cgagccggtg 17160
ccggacgcga tcctggtccg gagcgccgcg atgcatgaga tggaattcgg caaggaactg 17220
ctggccatcg gccgtgccgg cgcgggcgtg aacaacattc cgatcgaccg ctgctccaag 17280
gaaggcatcg tggtattcaa cacccccgga gccaacgcga acgccgtggc cgagctggta 17340

atcgccggac tcatgatgag cggacggaag atcgccgacg gcattgactg ggtgaaatcc 17400
ctgaagggca agggcgacga agttgccaag ctcgttgaaa aaggaaagag ccagtttgtc 17460
ggcgccgagc tgcgggggaa gaccctgggc gtgatcggcc tgggtgccat cggttccatt 17520
gtggccaacg cggccagccg cggccttggc atgaacgtcg tcggctttga ccctttcatc 17580
tccgttgaga gcgcctggag cctgagcaca accatccatc gcgccgcctc ggaagacgag 17640
gtgatcgcgc aagctgatta catcaccatt catatgccgc tgaacgacaa gacgcgcggc 17700
tccttcaacg cagagaagat cgctaagatg aaggacggcg cctgcctgct gaattttgcc 17760
cgcggcgagc tggtgaacac ggctgacctg ctggcggctc tcgcaagcgg caaactggag 17820
cgctatgtga cagatttccc gaacgatgaa ctgatcggcg cggagaaggt ggtctgcatt 17880
ccgcacctgg gagccagcac gccggaaagc gaagagaact gcgccaggat ggcagcggcg 17940
gaaatccgcg attacctgga gaacggcagc attcacaaca gcgtgaacta ccctgaagtt 18000
ctgctgggcg acccggaggc ggtgcgcgta ctggtgctgc atgagaatat cccgaatatg 18060
atttccaata ttaccgccgc ggcggccaaa gagggaatca acatcgaaaa catggtcaac 18120
aagagtaaga aggatatgtc ggtgaccgtg atggagatgg aggaactgcc gagcaagcat 18180
gcgctggata cgctgagcga gctgcccggg attatcagaa tccggacgtt tacgaaagag 18240
taagggaaaa gattactcca tataaaaagg gaatgcggaa atgcattcct tttttaatac 18300
gaaaatgact gttacagata aacatatgaa aaagaggaaa gcggctgctt tcctcttttg 18360
atcatactca gaagctgtag gaatccttga tttcgatatc cttgtagcgc ttcatgccgg 18420
taccggcggg gatcagctta ccaaggatga cgttctcctt caggccgatc agcggatcga 18480
tcttaccgcg gatggcggct tccgtcagga cgcgcgtggt ctcctggaag gaggcggcgg 18540
acaggaagct ttcggtggcc agggaagcct tggtgatacc gagcaggata cgcttggcga 18600
tagccggcgt tccgccctcc ataatggctt tccggttggc ttcctcaaag cggaagatat 18660
ccacgaggga accggtcagc agatcggtgt ctccgctgtc ctccacgcga accttgcgca 18720
gcatctggcg aatgatgatc tcgatatgct tgtcgctcac gccgacgccc tggaggcgat 18780
agacggagag gacttccttg agcaggtatt cctgcacagc tttcacaccg aggatccgga 18840
gcaggtcatg ggattgacg gaaccttcag tcagttcgtc gccagcttcc acatgatcgc 18900
cgacctgaac cttcaggcgg gaaccatagg tgatctgata ggacttggtg tccttgggat 18960
catcgtcgga cgtgatcacc agttcacggc gcttcttggt ttcctggatg gaaaccgtgc 19020
cggcgatctc cgcaaggatg gcgtcacgct tcggcttgcg ggcttcgaac agttcctcga 19080
cgcgggggaag accctgggtg atgtcgtctg cacccgcgat accgccggta tggaaggtac 19140
gcattgtcag ctgggtgccg ggttcaccga ttgcctgggc ggcaatgatg ccgaccgctt 19200
caccgacgtc caccttgccg ccgtgggcca tgttctcgcc atagcagcgg gcgcagacac 19260
cgttttcagt gcggcaggta aggaccgaac ggaccgtagc tttggtgatg cccctggcaa 19320
cgatctcacg ggccatctta taatcgatcg gctcgttcag ccggaccagc acttcgccgg 19380
tggccggatc gatgatgtct tctgcagcgg tacggccgcg gagacggtct tccagggatt 19440
caatggactg cttgccgctc atgagctcgg caacggtgat gccgcggacc ttttcgccgc 19500
gggcttcgaa acaatcgtcc tcccggacga tgacttcctg ggaaacgtcc accagacggc 19560
gggtcagata acccgagtca gcggtacgca gggcggtatc ggccagggac ttccgggaac 19620
cgtgagagga gaggaagaac tccagcaccg tcaggccctc acggaagttc gcgcgaatag 19680
gcagctcgat ggtcttaccg gaaggggagg ccatcaggcc gcgcatgccg gccagctggg 19740
atacctgagc ggaagaaccg cgggcgccgg agtcggtcat catgcggatc gggttgaagg 19800
tatccagatt cggcagaatc tgattcctca ggttagcggt gcactcagtc cagatattga 19860
taacacggtt gtaacgctca tcctcggaca gaagaccgcg gtgatacata ttgttcacct 19920
tgcggacagc ctcgtcggca tcggccagca ttgtcttctt cacttcaggc accttgatgt 19980
cggcaacgga tacggtgatg gcgccgatgg tggagtactt gaagccgagg gccttgatct 20040
tgtccagtac ttgggcggtc ctgtgttcac cctgggtgtg gaagcactgg tcaacgatcc 20100
tggagagctg ctttttgccg accagctcat ccacctcaag cttgaacata tcgtccaggc 20160
attcacgttt gacaaagcca agatcctgcg gcagcgcgtc gttgaagatc agacggccca 20220
gcgtggtatc gatcagttta gaaccggttt cgtcaccgaa agagcgggtc aggcgtacct 20280
tgatgggaga ctgcagggta atctgaccga gggagtaagc catgatggct tcatcttcag 20340
aggagaagac ccggccggat cccttggcac cgtcatgaac gatggtcagg tagtagcagc 20400
cgataaccat gtcctgagac ggagagatga ccggacggcc gtcctgaggc ttcaggatgt 20460
tgttatgggc cagcatcagg aagcgggctt ccgcctgggc ttccatggaa agcggtacgt 20520
gtacggccat ctggtcgccg tcaaagtcgg cgttgaaagc ggtacaggcc agggggtgca 20580

gcttgatggc cttgccttcg accaggatcg gctcgaaagc ctggataccc agacggtgca 20640
gggtaggcgc acggttcagc agcacaggat gatcccggat gacttcttcc agaatgtccc 20700
agacctcggg gcggatcttt tcgaccgcac gcttggcgga tttgacatta tgggagagtt 20760
tcagggtgac caggcgctcc atgacgaagg gcttgaacag ctcaagcgcc atttccttcg 20820
gcaggccgca ctggtaaagc ttgagttcgg ggccgacgac gataacgctg cggccggaat 20880
agtcaacgcg cttgcccagc aggttctgac ggaaacggcc ctgcttaccg cggagcagat 20940
ccgacaggga ctttagggca cggtttccgg gaccggtgac cggacggccg cggcggccgt 21000
tgtcgatcag ggcgtccaca gattcctgca gcatacgctt ttcgttgcgg acgatgatgt 21060
ccgggggcgcc cagctccagc aatctcttca ggcggttgtt acggttgatg acccggcggt 21120
acagatcgtt caggtcggag gtggcgaaac ggccgccgtc gagctggatc aggggacgga 21180
gatccggggg gatgaccggt acaacagtaa ggatcatcca ctcgggctta ttgtggctgg 21240
tacggaagga ttcgacgacc tcgagccgtt tgacggcacg ggcacgcttc tggccttcgg 21300
tttcacggtc gcgctctttt tcggtcagct gggcaatctc agccttcaga tcggcggaga 21360
gcttatccag gtcgagctcc atgagcagat cctgaaccgc ttcagcgccc atcttggcga 21420
cgaaagatcc tttgccgcac ttcgcttcga cttcacggta gcgggcgtca ttgatcaact 21480
caagccgctt cagacccgtt tcctcggtgt tgccgggatc gagcacgatg aaattggcga 21540
aatagagcac tttttccagg ttgcgggggg aaatatctag cagcatgccc atccggctgg 21600
gaatgccctt gaaataccag atgtggctga caggagccgc cagctggata tggcccatgc 21660
gctcacggcg gactttggcc cgggtgacct gcacgccgca cttgtcacag atcttgtcct 21720
tgtccttgaa cttgatccgt ttatatttac cgcagttaca ttcccaatcc ttggtgggtc 21780
cgaagatgcg ttcgcagtac agaccgtccc gttcgggctt cagcgtgcgg tagttgatgg 21840
tttcgggttt ggttacttca ccgtgggacc acgcgaggat ctgctccgga gaggccatac 21900
cgatctgaat ggagtcaaga ttagaaagtt cgaacaaaag gggcacactc ccttcgatta 21960
aatatcatcg tcaaggtcaa ggtctgacag atcttctacg ctgaagtctt caagatcttc 22020
cgcattctcg tcggatacag agaagtcatc gctgtccgcc gtgctttcat catcagcgcc 22080
gaaggacaga tcgtcctcgg acatgctgaa ggcttcgtca gcggcctgct caagctcttc 22140
actctcttcg gcagcaaccg gttcgtcatc cgcttcggtc ttgccgagga gattcgcgcg 22200
catggtttca ggctgcggca tatcctcggg atcgagctcg gggatttcga tctcgttccg 22260
ggcagcgtcg agcacgcgga tatcgaggct cagggcctgc atttccttca gcaggacttt 22320
gaagctctcc ggtacgccgg ggttcggaat gttctgtccc ttaatgatcg cttcataggt 22380
cttgacgcgg ccgacggtat cgtcggactt cacggtcagg atctcctgca gggtgttggc 22440
agcgccgtag gcttccagcg cccagacttc catttcaccg aagcgctggc cgccgaactg 22500
ggctttaccg cccagaggct gctgggtaac aaggctgtag ggaccggtag accgggcatg 22560
catcttgtcg tcaaccaggt ggtgcagttt caggaagtac atgataccga cggtaaccgg 22620
gttctcgaag tagtcgccgg tgcggccgtc gcgcacacgg attttgccct ggcggaagag 22680
cgccttgcct tcttcgtcca gatccaggcg caggggcttg ccgttatgga aatgatactc 22740
atttacatgg ggatcgttct catcttcggg cgcggccatc ttcttttcgg ccagttcgag 22800
gtggtccagg atttcctcat aggtggcgcc gtcgaagacc ggggtggcaa tgtgccagcc 22860
cagcgcccgg caggccaggc caaggtgcac ttcaagcacc tgaccgaggt tcatacggga 22920
gggaacgccc agcgggttca gcacgatctg caggggagtg ccgtccggaa ggaagggcat 22980
atcctcttca cgcaggatgc gggaaacaac gcccttgttg ccgtgacggc cggccatctt 23040
atcgccgacg gagattttcc gcttctgggc gatatagatc cggaccatct ggttgacgcc 23100
gggggagagc tcgtctttat tctcgcgggt gaagaccttg acgtcgacga cgatgccgcc 23160
ttcaccgtga ggtacgcgca gggaagtatc ccggacctcg cgggcctttt caccgaagat 23220
ggcgcgcagc aggcgctctt cagcggtcag ctcggtttcg cccttcgggg tgaccttgcc 23280
gaccaggatg tcgccggcat ggacttccgc accgatgcgg acgataccgc cctcatccag 23340
atccttgacg gcgtcatcgg agatgttcgg gatgtcgcgg gtaatctctt ccggcccaag 23400
cttggtttca cgggcttcgg attcgaactc ctcaatatgg atcgaggtgt agatatcttc 23460
cttaaccagc ttctcgctca gcaggacagc gtcttcatag ttatagcctt cccaggtcat 23520
gaaaccgatc agggcgttgc ggcccagacc gatctcaccc ttctccgtgg agggaccgtc 23580
cgccagtaga tcgccggctt cgatcttctg tcccgcggag accaccgggt gctgattgat 23640
gcaggtggac tggttggaac gggcaaactt ggtcaggttg taggtgtggc gttcgccgaa 23700
gttgtcccgg atgatgatct gggtggcgtc cactttttcc acaacgccgt cttccttcgc 23760
caggataaca acgccggagt cgtgggcggc gcggtattcc ataccggtgg caacgagggg 23820

cgcttcgggt accagcagcg gcacagcctg gcgctgcatg ttggaaccca tcagggcgcg 23880
ggtggcgtcg tcgttctcca ggaagggaac catggcggtg gccacggaca caacctggcg 23940
cgggctgacg tccatgaagt caacctggga cacagggact tcgacgattt cggccttgtc 24000
acggacagtg atccggtcct gaacaaaagt tccgtcggcg ttcaggggtt cgttggcggt 24060
ggcgaccttg tagaggtctt cctcgtcagc ggtcatgtaa tcgatctcgt cgaggacctt 24120
tccgtgttcc ttatccacac ggcggtaggg ggcttcgatg aaaccgtatt cgttgatgcg 24180
ggcgtaggta gccagagaac cgatcagacc gatgttcggg ccttcaggcg tctcgatcgg 24240
gcagatgcgg gcatagtggc tgtaatgcac gtcgcgaacc tcaaaggagg cacggtcacg 24300
gttcagaccg ccggggccca gggcgctcag acggcgcttg tgggtcagct cagccagcgg 24360
gttgggctgg tccatgaact gggacagctg ggaagaaccg aagaactctt taatcgcggc 24420
gcttaccgga cggatgttga tcagatcgcc gggcttgata tcggtggaat cctgaatgga 24480
catccgctca cggaccacgc gctccaggcg gctcaggcca atacggatct gattctgaag 24540
gagttcgccg acgctgcgca ggcggcggtt gccaagatgg tcgatgtcat ccgtggaacc 24600
gatgccgtag ggcagaccca gcatatagtt aacagaggaa atgatgtcgt cgggaataat 24660
gtgcttcggc acgagttcgg cgacgttttc ctttacggcg tcgaaaagct gatccggttc 24720
cacggaagcc atgatctgct tcagggtggg cagatggacc atttccagga tgccggcatc 24780
cttcggattg aagggcagat aggcggaggc gtcgacaaag ttgttgccga ctaccttacg 24840
gatctggccg tcacaatcga gcaggacaga attcacaccg gcgttctgga tctcacgcgc 24900
cacatccagc gggatggaat cgcccttttt gatcatgact tcgccggtct ggggattgat 24960
gatatcttcc gccgcaacat ggttgtgaat ccgcgtcgcg atcgaaagct tcttattata 25020
cttatagcgg ccgacacgca tcaggtcgta gcgcttggga tcgaagaaga gggaatggaa 25080
gagattggta gcgctttcaa ggctggcggg ttcgccgggc ttctggcgct tatagatctc 25140
aatgaggcct tcttctctcg tcttggcgtt gtcgcccttc tgaatggtgg ccatcaggcg 25200
ctcgtcttcg cccatgagct cgatgatttc ggcatcggaa ccgtatccga gcgcacggat 25260
aatcacggtg atgggcagtt tgcgggcacg gtcaatacgg acccagagca catcgttgga 25320
gtcggtctca tattcgagcc aggcaccgcg gttcgggatg atctgggaag agtaaagctt 25380
tttgccggct ttgtcgtagg cgacgtcaaa atagacgcca ggcgaacgga ccagctggct 25440
gacgattacg cgttcggcac cgttgatgat aaaggtgcca tggtcggtca tgaggggaaa 25500
gtcgcccata aaaacgtcgg attccttgat ttcaccggtc tccttattct tcaggcgaac 25560
aaaaaccttc agcggcgccg cataggtaat atcccgttcg cggcagcggg cgatggagtt 25620
tttcggctcg tccagagaat agtccacgaa ttccaggacc aggttcccat tgtagtcggt 25680
aatgggagaa acatcggcca ggacttcgcg cagatcggta tcgagaaagc gctgatagga 25740
gttcttctgt acctcaatca ggttgggcat ctcaagaact tcatcaatgc gtgagaagct 25800
catacgttcc cggagcttcc ccatgtggac ctcgtgtacc atgttatgaa atcacccctt 25860
atattgtccg accagatacg ccctgacggc cggtaaaaac ggacttgcgt ttcacacaga 25920
acgtgtgtaa aataaggaag tccgcagtct taaccgggca tcaagacgat actgatgcaa 25980
ttataaattc taacatagta tcagatgaaa gtcaacgata ttttatccaa tattcaatat 26040
attctatata tttaaatagg aaggaataag aagtgagact cagaggactg ttccggactg 26100
tgatcctgat tctgaccctg gccctgacgc tgtgcgcggg cgccgcggaa gaagcgattc 26160
cgccttctcc ggaaccgccg gcagggaacc cagccggccg accgaccccg gcgccgacgg 26220
cagtgcccac ggagacgatc gttgaccaca cccatcagcc cggtctctac cctgactttt 26280
cctttatgcc ggacagaaaa ctgctcgaaa tctggtttcc gaatataaca gacgctgatg 26340
aagcggtcct gatttacgac ggagacgtct ggatgatcga ctgcggagac gaaaaagccg 26400
gggcacgggg cgcacgcctg cttcagcagc tcggaatcaa aaaaatcgac cggctgttca 26460
cctcccatct gcatcatgac catatcaacg gcctggccct gacggacgac acggccaaga 26520
tcggcgaggt cctgatctgc ttttccccgg accttacgcc cagcggactg aaactgattc 26580
aggtcgcaga ggaacatcat atccccgtga aggaatacgg agatggagac cgtttcacca 26640
tgggggacgg cgcggtcgaa cttcttttca tcaaaaacaa cgaacaggac ctggacatga 26700
acaaccagag cgctgtgacc cggattacct acggccagcg gacgattctc ttcatggcgg 26760
atatggaaca gccggggcag attgccatgt tcaaaaaggg tgatcccgaa ctgctgaagt 26820
gcgatatcgt caagtatccc catcatgcca aaacagatat gtacactcct ttctttaacg 26880
ccatgggcgc gaaactggcg atcgtgacat cgacagaagg cagaaaggac ggcggccaga 26940
ccggcctggc caaccggaaa ctgccggctg tgttcacttt tgtcggcgga cagttcaccc 27000
acctggttac ggacggggaa tactggctgt gcgagcgcgt ggagataaca gtgaaatgaa 27060

acagttcgcc cggtgaccgg aaaacaggct gccgcatgca aatttacaga taataaaacg 27120
ttgacaagcc tttacaacaa tgatatgata gtcgagttgc agtatgctcc gggtgaagtg 27180
cgcccttttt tgagacgccg gaacccgatg cggaaggagg tctgaaacgc atggagatgc 27240
tgaaagcggc cggcagccgg aaggtcgtcg gaacgaaaca ggtgctccgc gcgctgaagg 27300
ccggaaccgt cgccagggtg tatgtctgca gcgacgcgga tacctttatt tatcagcagg 27360
tggtccgaag cgccgaagag gccggcattc cctgcgtcag ggccgcgtcc atgaaagagc 27420
ttgggatgat ctgcggagtg gacgtaccca ccgctgccgc gggtctgctc aggtaatcaa 27480
aacgcctttt ggatcctgag ggttgcggga tcttaaggat attattcagg aggtgctcta 27540
atgcccacaa ttaatcagct tgtccgtaag ggccgtgaac gggcgaccaa aaagcccacc 27600
gctcctattc ttcagggatg cccccagaag cgcggtgtgt gcctgagcgt caaaacgcag 27660
acgcccaaga agcctaactc tgctctgcgg aaaatcgcga gaatccgcct gacgaattcc 27720
atcgaaggta cctgctacat ccccggtatc ggccacaacc tgcaggaaca cagcgttgtg 27780
ctgatccgcg gcggccgtgt gcgggacctg cccggcgttc gttatcacat catccgcggt 27840
gctctggata ccgccggtgt ggccaagcgg atgcaggcgc gttccctgta tggcgcaaag 27900
cgcccgaaga agtaagccat tacgctgccg acgcagcctg aaacggtgat gatccgcgcc 27960
tgacagtccg ttgttcttcg gaacaggaat gacggaaaga caggaagcag gcggaggaag 28020
ccggccggtc tgttacgtga ccggggccgt gtgatgctga cacagggccg gaaggagagg 28080
gagcagaaca cttacatcag gattatcaat tgaggaggga aacctatgcc ccgtcgcggt 28140
tttgtaccga aacgtgaagt gctgccggat cccgtttacg gaaccacggt agttacaaaa 28200
cttattaacc agattatgct ggacggaaag cgcggagtcg cgcagaacgt ctgctaccag 28260
gcttttgaga ccgttgccga gaagaccggc aaggctgcga ctgacgtttt ccaggaagcc 28320
ctgaacaatg tcgctccgca gctggaagtc aaagcccgcc gtgttggcgg cgccacctac 28380
caggtaccta ttgaaatccg tcccgagcgc cgtcagaccc tcgccctgcg ctggatcgtc 28440
gactttgccc ggaaacgcgg tgaaaagacc atggctgaac gcctggccgg cgaactgctg 28500
gacgcttcca ataataccgg cgccgcagtg aagcgcaagg aagaaatgca ccgtatggcc 28560
gaagcaaaca aggcattcgc tcattaccgc tggtaatcgg catttcacct gcatcattca 28620
agcagctgtt gaaaggagca attgcccagt atgcccagaa gccatccact agacagagtc 28680
cggaacatcg gtattatggc tcatatcgat gccggaaaaa cgacaaccac cgagcggatc 28740
ctgttctaca caggcaagaa ccaccgcatc ggtgagacgc atgaaggtac ggccaccatg 28800
gactggatgg cccaggaaca ggaacgcggc attacgatca catccgctgc caccacctgc 28860
ttctggcatg atccgcatga cccggccaat accaacaagc agtatcggat caacatcatt 28920
gacacgcccg gccacgtgga ctttacggtc gaggtggagc gcagccttcg cgtgctggac 28980
ggcgcagtca gcgtcttctg tgccaagggc ggcgtggagc cccagagcga gactgtctgg 29040
aaacaggcgg aaacctaccg ggttccccgt atggcctatg tcaataagat ggatatcacc 29100
ggcgcggatt tcttccgcgt ggttgatatg atgaaggacc gcctcggcgc caacgccgtt 29160
ccgattcagc tgccgatcgg caaggaaaac accttccgcg gcattatcga cctggttcgt 29220
atgcgcgcag agatctatta cgacgatctc ggacaggacg tacgcgacga agaaattccc 29280
gctgacatgc aggaaatcgc cgacgaatac cgcgccaacc tgatcgaaaa agtggctgag 29340
acagacgaca ccctgatgga aaaattcctg gaagggggaag aactgaccga ggaagagatc 29400
cgcagcgcca tccgtaagtg caccatcgac tgctcgatga accctgtgct gtgcggtacc 29460
agttaccgta acaagggcgt gcagccgctg ctggacgcgg tgatcgatta catgccgagc 29520
ccgctggata ttcccgcgat tgaaggcacc gatcccgaca atccggagaa gaaaatggaa 29580
cgccatcctt ccgatgacga accgttctcc gcactggcgt tcaagatcat ggttgatccg 29640
tttgtcggca aactggcgtt cttccgtgtt tacagcggaa agctgagcgc cgggaactat 29700
gtgatgaatt ccacgaagca gaagcgtgag cgcatcagcc gtattatgct gatgcacgcc 29760
aaccaccggg aagaggttga gaacatctat accggtgaaa ttgccggcgc cgtgggcctg 29820
aaagatacca cgacaggcga taccctttgc gacgagaatc atcagattat cctggaaagc 29880
atggtcttcc cggatcccgt tatcgaagtg gccatcgagc ccaagaccaa agccggtcag 29940
gagaagatga cctatgcgct tcagcgcctt gctgaagagg atccgacatt caagacctac 30000
accaaccagg aaacaggcca gaccatcatc gccggcatgg gcgaactcca cctggagatc 30060
atcgtggacc gcctgctgcg cgaattcaag gtcgaggcaa ccgtgggcaa gccccaggtt 30120
acctacaagg aaaccatccg caagcccgcc aaggctgaag gacgctatgt ccggcagacc 30180
ggcggacacg gccagtacgg ccactgctgg atcgagatcg aaccccagga acccggcaag 30240
ggctacgagt tcgagagccg gatcgtcggc ggcgtgattc cgaaggaatt catcggaccg 30300

atcgaccagg gcatccagga agccgcaaag agcggcgcga tcgccggcta cgaggttgtg 30360
aacttcaaag ccgcagttac ggacggttcc tatcacgaag tggactcctc cgaaatggcg 30420
tacaagatcg ccgcgagcat ggctttcaag gaagccgtcc gcaaggcaga tccgtgcctg 30480
atggagccca tgatgaaagt ggagatcatc gttccggata cgtacctcgg cgacgtgatg 30540
ggcaatgtgt ccagccgccg cggacgggtc gaaggcacgg aagtgcgcgg ccaggatcag 30600
atcatccacg cctatgtccc gctgagcgag atgttcggat acaccacgga tcttcgttcc 30660
cgtacccagg gccgcggcat gttcaccatg cagtttgacc attacgagga agtgccgaag 30720
tccgtcgcgg aaaagatcat cggcaaaaag ggcgcggact gagaagaacg caccatgcac 30780
aataaatcat gacaattatc tacggactga tttacaagga ggaaaaatcc aatggcaaaa 30840
ggacgttatg aacggacaaa gccccatgta aacatcggca ccatcggtca cgtcgaccat 30900
ggcaagacca ccctgaccgc ggcgatcacg atgacgctgg cagtgaaggg cgaagcgcag 30960
gccatgcgct atgacgaaat tgacaaggct ccggaagaga aagcgcgcgg aatcacgatc 31020
aacaccgccc acgttgagta cgagacggcg aagcgccact acgctcacgt tgactgcccc 31080
ggccacgctg actatgttaa gaacatgatc accggcgcgg ctcagatgga cggcgcgatc 31140
ctggtcgtgt ctgctcccga cggcccgatg ccccagaccc gtgagcacat cctgctcgcc 31200
cgtcaggtcg gcgtgcctta catcgttgtg ttcatgaaca agacggacat gatggacgac 31260
gaagaactgc tcgagctggt cgaaatggaa atccgtgagc tgctgagcag ctacgacttc 31320
ccgggcgacg aaatcccgat catcaaggga tccgctctga agaccctgga gtacctgcag 31380
aacggcggca ccgacgtgga caacgcgccc gagtgcaaat gcatctggga actgatggac 31440
gcggtcgaca gctacattcc ggaaccggaa cgcgcgaccg accagccctt cctgatgccc 31500
gttgaagacg tgttctccat ttccggccgc ggcaccgtgg ctacgggccg tgtcgagcgc 31560
ggcaccgtga aggtgtctga cccggtggaa atcgtcggcc tgatggacaa gccccgcaac 31620
acggttgtta ccggtgttga aatgttccac aagctgctgg accaggcgga agccggcgac 31680
aacatcggcg ccctgctgcg tggtatccag aggaacgaag tcgagcgcgg ccaggttctg 31740
gcgaagcccg gcagcattca tccccacacc cactgcatcg gccaggtgta cgtgctgacc 31800
aaggaagaag gcggacgtca tactcccttc ttcaacggct atcgtcccca gttctacttc 31860
cggacgactg acgttaccgg caacatcaag ctgcccgacg gcgtggaaat ggttatgccc 31920
ggcgacaaca tcgacatgga aatcaccctg atcaccccca tcgctatgga gcagggactg 31980
cgcttcgcta tccgcgaagg cggccgtacc gttggttccg gcgttgtggc aaaggtcatt 32040
gagtaatcaa ttcaggaggg atcccaaatg gcaaacgccg cacgtaacaa ggtttgcctg 32100
gcttgcactg aatgcaaaca gcgcaactac aacaacatga agaacaagaa gaacactccg 32160
gatcgcattg agctcatgaa gtattgccca ttctgcaaga agcacacaac tcatcgcgag 32220
accaagtgat ccaacaccta tcaacgctgt gaggatgtga aagaagtgtc tgatgttaag 32280
aagacggctg aaaatactga aaaggccgta aagaaatccg gagagagcaa gctgagcaag 32340
attatcagct ggttcaagaa cctgcccaag cggatctcta agcctttcaa gaacatgtat 32400
tatgagctga agaaggttac gtggcccagc aaagagaagc tgatcgccta ttccgttatc 32460
gtgctcgttt tcatgctctt catgggcatc gtgatcggcc tgctggacat gggcgcgtcc 32520
gcgttggtca gaatgctgat cgcgtaacat aggagagaag aaacatggct gaagcaaaga 32580
aggagccttg ctggtacgtt attcacactt attccggcta tgagaataag gtgaaagaca 32640
cgctggaaaa atccgtcgag aacaacggca tgcaggacct gatccttgag gttcgggtac 32700
cgatggaaga agttgtcgag atccggaacg gaaagcgcgt caagagcacc cgtaaggttt 32760
atcccggata tgtgctggtg cacatgatcg agaccagtga gagctggtat gtcgtgagga 32820
atacccgcgg cgtgaccggc tttgtgggcc cggattccaa accggttccg ctgacgcagg 32880
aagaagtcga catgatgctc aataccgagc agtccagcgt ggacttcgga tttgcgatcg 32940
gagagaacgt acggatcctt tccggcccgc tggaaaattt cagcggtgtt gtggaagacg 33000
tcgatactgt ccgcggcaag ctgacggtca aggttcagat gttcctcggc cgcgagatgc 33060
cggtggaagt agatctcgac caggtcgaga aggaagactg atttccactc ctgtgaattc 33120
tgttcgctgc aacgggaaca gtccggacag actgttccct gcatcgccct gtcttatggc 33180
aaggcgcgac gtgggaggaa gctgacagct tccgctctac cacaatattt atggaggtgc 33240
cactcaaatg gcaaagaaag tacaggctta catcaagctg caggttcctg ccggcaaggc 33300
aacacccgca ccgcctatcg gtcccgctct gggccagcat ggcgtgaaca ttcccggatt 33360
ctgcaaggaa ttcaacgacc gtaccgccaa ggaaaacggc atgatcatcc cggtggtcat 33420
taccgtttac accgaccgta ccttcacctt catcaccaag acgcccccgg cgcccgtgct 33480
gatcaagaag gaactgaacc tgcagaaggc cagcggccgc cccaacaagg aaaaggttgg 33540

```
     caagctgacg aaggaacagg tcaagaagat cgctgagatc aagatgcccg acctgaacgc     33600
     cggcagcatc gaagctgcca tgagcatggt cgccggaacc gcccgcagca tgggcgtgac     33660
     cgtcgaagag tgacgtaccg gggcaatgcc cctgatcatg tgggaggaca tagcgccgtt     33720
     aagaccacat aggaggaata aaaaatgaaa cacggtaaga aatataccga cagcgtaaag     33780
     ctgatcgacc acctgaagca gtacgacccc gaagaggccg ttgacctggt gctccagacc     33840
     ggcaaagcca agtttgacga gaccgtcgag atctccgtcc gactgggcgt tgaccccgc      33900
     cacgctgacc agcaggtccg cggcgctgtg gttctgccgc acggaaccgg ccgcaccatc     33960
     cgcgtgctgg ttatcgccaa gggcgataag gccaaggaag ccgaagaagc cggcgcggat     34020
     tacgtcggcg ctgaagagat gatccagaag atccagcagg agaactggtt cgagtttgac     34080
     gtctgcgtag ccacgcctga tatgatgggt atggtcggcc gtatcggccg cgtgctcggc     34140
     ccgaagggcc tgatgcccaa cccgaagtcc ggcaccgtaa cgatggatat cgcccgcgcg     34200
     gtgaacgaca tcaaagccgg taaagtggaa taccgcctgg acaagaccgc gatcatccac     34260
     tgc                                                                   34263
//
ID   XXX; XXX; linear; XXX; XXX; XXX;  XXX.
XX
AC   PRJEB67889
XX
DE   Uncultured bacterium DNA fragment from in vivo enriched cow rumen, clone
rum-17h1
XX
FH   Key             Location/Qualifiers
FH
FT   source          1..39405
FT                   /organism="uncultured bacterium"
FT                   /environmental_sample
FT                   /mol_type="genomic DNA"
FT                   /isolation_source="Cow rumen"
FT                   /clone="rum-17h1"
FT                   /note="Tween 20, pNPA, pNPB, pNPP active clone"
FT   CDS             530..694
FT                   /db_xref="SEED:fig|6666666.127272.peg.1"
FT                   /translation="MLDTIRTGIELTEPVNRQAPEQAVPKAAEMFPCFAVKMVLKGSEY
FT                   LLEPDDRLF"
FT                   /product="hypothetical protein"
FT   CDS             691..849
FT                   /db_xref="SEED:fig|6666666.127272.peg.2"
FT                   /translation="MISPEGRFAALLKEFGIPFEEYEPERFGICGFKVPENESSSDKTR
FT                   SACFGGR"
FT                   /product="hypothetical protein"
FT   CDS             complement(868..1347)
FT                   /db_xref="SEED:fig|6666666.127272.peg.3"
FT                   /translation="MAFEKAKAYLEGFGLGEKIIVTEHSSATVAEAAEAIGCEESMIAK
FT                   TLSFLQDEKPVLILADGTARIDNRKYKDRFGCKARMIPAGLVEQLVGHDIGGVCPFGVN
FT                   PGVSVYLDESLRRHETVYPAVGTDHSGVKLTIPELERCSGFLDWVDVCKQAEQPS"
FT                   /product="tRNA proofreading protein STM4549"
FT   CDS             1589..1816
FT                   /db_xref="SEED:fig|6666666.127272.peg.4"
FT                   /translation="MKKLENNPLLYGIVNCLLSAIGVVIVCFIISLVKKDESFSDVISG
FT                   TYMLIILIGGSIFAGISAYLKAKNKDKKEM"
FT                   /product="hypothetical protein"
FT   CDS             1861..2598
FT                   /db_xref="SEED:fig|6666666.127272.peg.5"
FT                   /translation="MLQINHYTKQYSVNGPKAVDDLTLHIRPGEIYGFLGHNGAGKTTT
```

```
FT                   IKACCGILDFTEGEILINGMSIKKQPLECKKIMAYIPDNPDLYEFMSGIRYLNFIGDVY
FT                   GIPTDVREKRIAEYAEDMGITDALAEKVASYSHGMKQKLAIVAALMHEPKLLIMDEPFV
FT                   GLDPKAAFILKKFMRRFCDNGGAIFFSTHVLDVAEKLCDKIAIIKGGKLLADGDIETVR
FT                   GNESLEEVFLELVDDMKVEEENA"
FT                   /product="ABC transporter, ATP-binding protein"
FT   CDS             2591..4222
FT                   /db_xref="SEED:fig|6666666.127272.peg.6"
FT                   /db_xref="GO:0005215"
FT                   /db_xref="GO:0006810"
FT                   /db_xref="GO:0016020"
FT                   /translation="MLKILLKSRLLAIADQFSGQSKGKKAATITSHITLIVLGLLLMVG
FT                   AAVGVNALLGGTVAALAPQGLTWGFYAAAGGLSLIASLFLTMFYAQGVVFEAKDNEMLL
FT                   SMPIPPSAILASRIGSVYLMNLIAPVVIMFATGYTVAVRASGISVLSVVFLVLCTLLLT
FT                   LLTATITCLLAWLVSIITRRTRKKTLVQLLLSLAAMGLLYFVTSNVNRDLLASIKDNAV
FT                   SIADFFRSKLYPLYVMGQAIAESSWLQLLIFAACCIVPFVLVYFLLSKSFIRIVTARST
FT                   AAKLKYEATSLKSSSLVWAMTKKDLTRFFNSSPYMMNAGIGLLYTLGWTIFNVFSGNSV
FT                   IKGLFEQGTTVPDAGPYMILVSAYTLSFFAGLTTISGCSVSVEGKNLWILKSLPVRAKG
FT                   ILIGKFLSHLVLAAPVSIICSILYLLSMPSPDVIGIAAIFLMPLAANVLCALTGVISNL
FT                   FFGKLNYPSIAKAAKSSASSLIPMLVTTVLVVGTGAVYLFFLRETVSVRIFTVAVILFL
FT                   LAVDAGLYFFLGSSAVQKRWEALGNS"
FT                   /product="ABC transporter permease protein"
FT   CDS             complement(4282..5937)
FT                   /db_xref="SEED:fig|6666666.127272.peg.7"
FT                   /translation="MRKSVAIIFISMATVLAVVMGVLYVSNNADKAKMIETLTTSVKDK
FT                   TEKIGSLNSDSAEKDEEITRLKSEIADRDERIDSLNADAEDKAAQIENLNAENADKAAQ
FT                   IDTLKTAVSDRDKTIKALNAENADKAAQIDTLKAAVSERDKEIEALNADVTDMGSQIET
FT                   MYAENADKSEQIDTLKAAVSDRDKANETLDADVTDKGSQIEVLNADVNDKTAQIDALKA
FT                   DAADKSEQIGALKAAIAERDRKIEALNADITDKGSQIEALNADVTDRDAQIEDLNTENA
FT                   DKAAQIDTLKAAAAERDKEIETLKTDAADKTARIEAMNADAAEKANQIDTLKAAVSDRD
FT                   KQIEALNAAVSDRDKQIEALNADVTDKNAQIEDLNTENADKAAQINTITAGGIAASGVF
FT                   ETQKAAVGLSESGKEPDEPHEVKTENGTYRLTVAEVRKLDGDLWQMLAKDKGDHYMIIA
FT                   VHGEIENVDFTSRTGDTLPASLVTEEMEVKDQAGTPLAFFHISGGSDGEYAVNPDVRPG
FT                   TTQKVSYTYFADKDLTEVSIRMANGETLTVPVAE"
FT                   /product="Lipopolysaccharide biosynthesis protein-like
FT                   protein"
FT   CDS             complement(5985..7169)
FT                   /db_xref="SEED:fig|6666666.127272.peg.8"
FT                   /translation="MNYVFISPHFPHTYWQFCDRLHRNGINVLGIGDAPYDGLEEPLKA
FT                   ALTEYYRVDSLEDYDQVYRAVAFFAFKYGKIDWVESMNEYWLEQDARLRTDFNIITGIR
FT                   SDRISFIKEKSLMKKLYLEAGIPTARQHVVSDREAGKSFLREVGYPVIVKPDVGVGATR
FT                   TWKLENDADLEAFYDNLPAVPYVMEEFIQGEICSYDAILDSRCEPLFESMTVWPPVMDI
FT                   VNKDLDLMYYTCPEVPEKLRALGRKTVKAFGVDRRFVHLEFFRLTKARKNLGKTGDFVA
FT                   LEVNMRPAGGYTPDMMDFAHSTDVYQIYADMVAFDTRKLPESTEHFYCVYASRKDGHTY
FT                   ARTHEEIMARYGGRMMMQEEMPPMNWPQMGRYMYTARLQTFEEAQEFIRFVQGA"
FT                   /product="Carbamoylphosphate synthase large subunit (split
FT                   gene in MJ)"
FT   CDS             complement(7174..7998)
FT                   /db_xref="SEED:fig|6666666.127272.peg.9"
FT                   /translation="MYNKKASSRADGRRSGKEQKKAGRCSMNREELTHYSQSLDREMHI
FT                   MVYGQGGVPFLCFPTQDSMCHNYEDFGMIHHLADFIEDNKIQLFVVDTVDRESWSARGW
FT                   DNEFRAGRQEQYFRYITGEVIPLIRSRNPEVPAVTGFSMGADHAVITFLRRPDLFRGVI
FT                   ALSGVYDADYFFDGWMNPTLYDNSPERFLPNMPGDHPWISLYNERKMILCCGQGAWEED
FT                   GVRTLRHLERIFHEKGIEAWCDFWGCDVNHDWPWWFKQMRYFVPFMINEQRN"
FT                   /product="hypothetical protein"
```

```
FT   CDS             complement(8143..8625)
FT                   /db_xref="SEED:fig|6666666.127272.peg.10"
FT                   /translation="MILAAALLMVSAAYAESAVDVIMATGTAQAFTDEAVPAGDRPAAP
FT                   PAGAGAKASLGDSPAAIIIYKDGASGLPDASFDCGLAAQNMVIAASAKGYGVKIVSSPA
FT                   RTLNGENHDALCEKLGVDPSMQAVAVLLIGKPDTGVDGASGATTRDPYETKTSIIE"
FT                   /product="hypothetical protein"
FT   CDS             complement(8693..9874)
FT                   /db_xref="SEED:fig|6666666.127272.peg.11"
FT                   /translation="MAERNLDFDRIISRKNTRSVKYDLAARKGYPEDILPLWVADMDFR
FT                   TSSYVEDALGELTRHNIYGYSNTLPGDGFFEAVSGWMKRHHGWDVDPEWHVKTPGVCFA
FT                   VSMAIRAFTEPGDAVIIQEPVYYPFSSLIRQNGRKVVSSDLVPDAEGNYSMDPEDFERK
FT                   VAEHHVKLFILCSPHNPVGRVWTGDELRTIGRICASHGVIVFSDEIHSDFVWKGEHSVF
FT                   QEADPSFREFTVTATAPSKTFNLAGLQQSNIFIPNPGLRRGFAEEISRTGFDEPTVFGI
FT                   AAAQAAYENGDTWYAAMKTYVRDNIRFADEFVRERLPGVRMNQPEGTYLIWLDFRGTGM
FT                   DAAELDDVILRKARLWLDSGSMFGKAGEGFQRINTACPRSILREAMERLESVL"
FT                   /product="Cystathionine beta-lyase, Bsu PatB (EC 4.4.1.8)"
FT                   /EC_number="4.4.1.8"
FT   CDS             complement(9920..10684)
FT                   /db_xref="SEED:fig|6666666.127272.peg.12"
FT                   /translation="MIFLKTGGKMIIRDDEIRLSAVLEEPEGSGRHPLAILLHGFGSAK
FT                   DRTHNILTAAAMREKGCATLRLDLYGHGESGGEFRKHTLYKWISNTMAVIDYARELGYT
FT                   EIYLSGHSQGGLTAALVAGMEADRISGLILRAPAFMIPQAARDGCMLGYRFDPDHVPES
FT                   VTVAEDVVLDGNYIRVAQTIHAEDAADRYRGPVLILHGDEDDVVPPEDSARMAKRYSNC
FT                   EMHMIPGETHHFDRHPEQMQEIIRNWMEQQRS"
FT                   /product="hypothetical protein"
FT   CDS             10842..11534
FT                   /db_xref="SEED:fig|6666666.127272.peg.13"
FT                   /translation="MKTVKAAFVRSLPVMAGYIILGIGFGILAHKAGYGLPWALAMSLF
FT                   IFAGSMQFVGVGLLAGGASVLTVILTTVMVNARHLFYSISMLRHYRNAGPWKPYLVFSL
FT                   TDETYSLLCDGKTPEGTDPDRYRFLVSVFNHSYWVLGSVIGSVVGGILPFSAAGIEFSM
FT                   TALFIASFTEQWITAKNHIPAAAGLLGTLLCLVVFGPGQFLIPAMLLITLLLTLLKSRL
FT                   DSGKKVDA"
FT                   /product="Branched-chain amino acid transport protein
FT                   AzlC"
FT   CDS             11531..11854
FT                   /db_xref="SEED:fig|6666666.127272.peg.14"
FT                   /translation="MNSSAYSALLVAVMSAVTILLRAAPFLVFKKKAPAYIMYLGQVLP
FT                   AAIIGMLVIYCLKDTNLTAAPFGAPELIAGLLVAVLHAWKRNILLSILAGTVAYMLLIQ
FT                   LVF"
FT                   /product="Branched-chain amino acid transport protein
FT                   azlD"
FT   CDS             complement(12102..13253)
FT                   /db_xref="SEED:fig|6666666.127272.peg.15"
FT                   /translation="MRDFQIGILGCGVISRTYASDIKAFYPRLHIAACADVRPESARKL
FT                   AEEFGIPKACTPEELLGDPEIEIIIDLTPPQFHVELNRQIIQAGKHLFSEKPFAQTTRE
FT                   AEELVALAAEKGVKIGSAPDTFLASGLQSMRYYLDTGLIGKPFFVTANMTTFGAETWHP
FT                   APAPFYRKGSGPLYDMAPYYLSALVALFGPAESVGAFSAKPSETRHVYSGPQAGTEIPA
FT                   EIPTHYSAILRMRSGVIANLNVSFDIYRSSLPMFEVYGDGGTLAYPDPNFGGGTPKVYR
FT                   KEQFTDTVWQHTPESEARRKTFRDLPELFTRAKDYSRGIGVLDLAAAIERNGRNRANGE
FT                   MIIHITEIIEGLIRSAESGSFVQLRTTCGRPEPLKPGCAPDEI"
FT                   /product="Dehydrogenase"
FT   CDS             complement(13464..14195)
FT                   /db_xref="SEED:fig|6666666.127272.peg.16"
FT                   /translation="MSQYTTGEMAKECGVTVRTVQFYDQKGILAPSALTEGGRRLYSEG
```

```
FT                   DLKKMKIICFLRDTGLSLDTIGQLLKEEDPGSVISIFLEQQEQVLRSEIEERRGKMDRL
FT                   AELKKGLKDTPAFSLETIGDIAAVMENKKKLSAVHRKMVLSAIPLGILQWASILLWILK
FT                   GIWWPFVVWAVVAVPYAIWAVRYYYRNVAYICPQCHTVFRPAMKEFFFANHTPKTRKLT
FT                   CVKCGSKRFCVETWGGELKQE"
FT                   /product="Transcriptional regulator, MerR family"
FT   CDS             complement(14283..15101)
FT                   /db_xref="SEED:fig|6666666.127272.peg.17"
FT                   /translation="MKAGGHQDRFGKTGANIPWTENSMIWHTVTINGIAVEAGYSERAV
FT                   NEIFAPLLEKLTEKQRAAGRRILVMLAAPPGAGKSTLVSFLHRLAEERRDAGRIQAIGM
FT                   DGFHRRQEYLLTHDTERDGKKIRMVEIKGAPETFDLEALTKRVKSLAAGEETKWPVYDR
FT                   LLHNPVEDAVRIDGDIVLLEGNYLLLDRDGWRDLPSFADYTISVRADEDLLRKRLVDRR
FT                   IKTGVGEEAAVRFVDFSDMPNARLCLSCSMRADLQLEIDEQGDYRMIRGK"
FT                   /product="Uridine kinase family protein YggC homolog"
FT   CDS             complement(15098..15580)
FT                   /db_xref="SEED:fig|6666666.127272.peg.18"
FT                   /translation="MTGQEAERAQAVRLIPVTEDNWMEIASLDVKEHQKSYVAPAVGIL
FT                   ARGYVVYRGCNAKIFAFEGDGTIVGMALVREFTDEPSGYDLQQFMVDRRYQNRGYGSRAL
FT                   RLILDELRKEGRYDHVELCVKKADEEAIRLYEKHGFADSGYVDEDLPDSLNMICRL"
FT                   /product="spermidine acetyltransferase"
FT   CDS             complement(15630..15950)
FT                   /db_xref="SEED:fig|6666666.127272.peg.19"
FT                   /translation="MASGREYLDFILEQLSGLDGITYRQMMGEYILYMNGRIIGGIYDD
FT                   RFLVKPTRSALAMMPDADRECPYEGAKEMLLADNVDNREFLQELLTAMYDELPAPKQQK
FT                   KK"
FT                   /product="FIG01116832: hypothetical protein"
FT   CDS             complement(15980..16516)
FT                   /db_xref="SEED:fig|6666666.127272.peg.20"
FT                   /translation="MELFDLYTANHEKTGRTMVRGESTPEGFYRMVVHVCIFDPEGRML
FT                   IQQRQPFKRGWSNLWDISVGGCAVSGDSSQSAAERETREELGLAIDLKNARPTLTVHFE
FT                   DGFDDIYVLTMPEDLSSLKLQYEEVQAVRWATKDEILRMIDDGRFIPYEKSLIELLFFR
FT                   RDHRSSHTKGDPTND"
FT                   /product="Putative Nudix hydrolase YfcD (EC 3.6.-.-)"
FT   CDS             complement(16589..17518)
FT                   /db_xref="SEED:fig|6666666.127272.peg.21"
FT                   /db_xref="GO:0046406"
FT                   /translation="MDYSQKISERQWNLPDKGELESRREATYMDDIIQKDHIQRKLLEN
FT                   LDGIGTVFDGGAGSGRFSILLAKQGCRVTHFDISRPMIEKARELAAREGVLDRITFVKG
FT                   ALEDLSAYRDRAFDLVISFDAPVSYTYPNQNKVIGELVRIADKRIMISVSSRLGSLPYL
FT                   ANPIQKNQFILDEKADDPFVQWCVANREQMISDFSFNKQNAMKLLADGLMGGEEEIALY
FT                   EQGGAPWCITYTFMPDELEKILSGYGVKNIRLSGPGAYGRTVPREILTRIMKDPKQKED
FT                   FLDFCYRYDSSPYVCGMGKDNLLAIGER"
FT                   /product="Mg-protoporphyrin O-methyltransferase (EC
FT                   2.1.1.11)"
FT                   /EC_number="2.1.1.11"
FT   CDS             17825..18226
FT                   /db_xref="SEED:fig|6666666.127272.peg.22"
FT                   /translation="MSVLQMNFVRGEDSIRRINDALAGMCLSDVRIRDSRGCLATRKDF
FT                   DENGQVKAVTYSYLFDSIALETTGDETMGTRTWRIETAVFLKGSPNAPQNTDAFFDSWS
FT                   AGHPDRSVRVIQWITFDKADVFIVLSRGR"
FT                   /product="hypothetical protein"
FT   CDS             complement(18361..21465)
FT                   /db_xref="SEED:fig|6666666.127272.peg.23"
FT                   /translation="MKKSVPHILLTFLIAFFAYFAMGQTARASYMSNKTEIQVNLEVSG
FT                   AEALCQTEDGYVWIAQYSGLTRYDSKEYVTYKSFEYEGQEYNIINVTALAAEGNVLYVA
```

```
FT                   TSDHLYVVYKDYHFELLATADGVITDIVLDPASGVLYICTRENGGIVFNLETGTKSTIPG
FT                   TEGKAIRDFAVGYEKGKYYYQTNEGVYDQDGKEILQNSKIMELYAYDTILYIGEDSGII
FT                   YRYDMKNEAFLDELSVPDQINKLLYSEADQILFVACEKDGIYCIDFSSGAPVISLAGNL
FT                   DNKSQLMDLMIDYQGNLWVASHFIGTSGVSIITRNALTELLYDDQIWQSLAEPPAYDRN
FT                   VYAVERYGNILYIVCSNRVFMYDLTRKEILPDNNLMKAIDEYAERRIEESQAKGDSNSS
FT                   FAYSPKDVEVFRNKLYFAFVNIGLVEYNPDTEQVVIYDYDYISNHLGTLVNDPDIALAD
FT                   KIRSMRSFDDALILGTSRGIMRFDGETFSIINVGANVLYINKTKDGKILFDQTKGLFVV
FT                   DDDFTTVTEIPTVKEITGNRLKFLVDGDDLYYTLNSRLFRTNPGKGDGVSEEIAIPYVK
FT                   GSIVELAKIRFTGADGKPEYKYVIGSQTQLYVTDSLAGTSLTEYESYDAINGLQPIIAN
FT                   TSGYYDESEMKYYLQSTNGIYVYEPNVGRESDLPVKVAASSVDLDGEPHYGDSISLEKD
FT                   VSRVTINLSVFGFRPNNGFTSYYKLDGVDSEYTSYTDGNRSITYTNLPGGSYDFHYYVT
FT                   DENNKESNHILIHLEKEKRFVEQWWFWAILSVLAGAIVFLISSLVIRHKTKQSLKRQLE
FT                   YKNITLESLQAIARTIDAKDEYTNGHSMRVSSYSRLIAQNMGMDQDEVENVYYIALLHD
FT                   IGKIAIPDSILNKPGRLTDEEFKIMKSHTTRGAEILRGISTIPQIIEGAKSHHEKYDGS
FT                   GYPEGLSGENIPLVARIICCADCFDAMASKRVYKAPFSQEVIIGEFERCSGTQFDPEIA
FT                   KIVIGMIASGELEPNAADNIYLGSDGKTHRIKEGEKREDTENSAS"
FT                   /product="response regulator"
FT   CDS             complement(21620..22837)
FT                   /db_xref="SEED:fig|6666666.127272.peg.24"
FT                   /translation="MKRIAVFLAAALYLALCMMIPASAEDGTIETTMETDGFRYRLLED
FT                   GSAEILKYVGKELQLVIPETLDGHPVTSIGEEAFDNYRYSMEDVTIPASVTHIGSGAFQ
FT                   LTDLAEIEIPAGVVSMDENPFGYCTELKTITVAEGNPVFYVEDGVLFCREDHSLVCFPN
FT                   LAADPGEDELATYTVPAGTEIIRADAFGGVETLSDILLPDSLRVIGPQAFGFCWGLDRI
FT                   VIPDSVTEIGDAAFDTCKYLGRVVLPKGLKKIPPMLFYECDSLYEAEIPETVTEIGDSA
FT                   FDFCGSLEHARIPDGVTVIKTGVFEGCSSLKSIEIPDSVTEIGAFAFCDCSALGSIQIP
FT                   AGVTKIGDRAFAYCSSLKELVIPDSVQAIGENAFDNCPEDLVILCGPDSCAKAYCEQNG
FT                   IAFRVE"
FT                   /product="Cell surface protein"
FT   CDS             complement(22834..23241)
FT                   /db_xref="SEED:fig|6666666.127272.peg.25"
FT                   /translation="MNDGFFTGIIYQNKRIRNRIQAEAGNAPYNRYEYAFVPHAEEQDI
FT                   ALPTENYHMISRRKNRKWLPAGISEPASRVNAPMARACFMNPEKVEKKSHDYTGVIVKK
FT                   IDTGNTDRGNKTPGSIVLPAEHEPEARRIPT"
FT                   /product="hypothetical protein"
FT   CDS             23330..23842
FT                   /db_xref="SEED:fig|6666666.127272.peg.26"
FT                   /translation="MVELDNERMKQILHDETAETEDLVTILRGIYTRYMNLYERYFADI
FT                   EALNDDRITEFAKYHEETKSLIRYYYMDIPQDICAGISEFEEKVSGRLLGREWKMNLFD
FT                   VYDEFREKNRVRNKSEDYYKAEFAKLALKEFYKAMENIFRDGFGTYSQTVKDVLGGITG
FT                   LLFGKKD"
FT                   /product="hypothetical protein"
FT   CDS             complement(23864..25024)
FT                   /db_xref="SEED:fig|6666666.127272.peg.27"
FT                   /db_xref="GO:0004427"
FT                   /translation="MNNHCEQYPGQRRYCEPGTNKKEKGDPMIKKKYIIPLLVLFLAFS
FT                   GMAARADGTGFADVLRSVDYGDQVTYVVGHKSPDADTIGTAIAYAHLLRQLGINAKAAA
FT                   TAKVNQETKYALDFYGLEQPEILDNAAGRQFVLVDHSEYSQALDGMKNARIVGIVDHHG
FT                   IGDIRNTERIAVISLPAGATASIVYKLYLDCNVGIPRDIARVMLMSILSDTKGMTSNVT
FT                   ALDRDAYNYLLSVADIDDIDALYAGMKAAKASYEGMTASGIFYNDYKEYLAGEYTFGIG
FT                   DVYTDSEENATALAQKIEAAFPSIYASNNLDMLFCMVSTDDMTRLIWYGEDAEKAVYDS
FT                   FPDYDGSGYLVFTPKASRKAKIVPPLTAALEKWTESGSRSRPAADD"
FT                   /product="Manganese-dependent inorganic pyrophosphatase
FT                   (EC 3.6.1.1)"
FT                   /EC_number="3.6.1.1"
```

```
FT   CDS             complement(25034..26077)
FT                   /db_xref="SEED:fig|6666666.127272.peg.28"
FT                   /translation="MIYYSDGILVIRTMEEADAQIFTDEEAEQGWHSDISKYLTRLRDQ
FT                   AEGKCVSLTAVYEGHPAGYINVYKTGLGGAFSGKGWPEIVDFGVLERFQRNGIGSKLMD
FT                   IAEQIAGQYADTVWLGVGLHNGYGSAQRMYVKRGYIPDGTGVWYRGRPCEQYETEIEND
FT                   DSLQLFFSKRLCTARKEDGLKAIVGLRRMTDEEYRRFKEYSVSDYAEDLTRGKGFTREQ
FT                   ALEKAEGEFRGMLPDGQDTVNQFLMTVEDKKAGKEVGWIWFSYEEGYDGSPQVFLNDFL
FT                   IFKPERRKGYAKAALHEMEQMAKADGRTESVLYVWAHNVPGFNLYQKCGYETVTQGDSG
FT                   THMKKEL"
FT                   /product="acetyltransferase, GNAT family"
FT   CDS             complement(26108..26905)
FT                   /db_xref="SEED:fig|6666666.127272.peg.29"
FT                   /translation="MKDIMNQAPASIAAVLGNKAGKTDRIGKSGAQVLLFDDYVLKIRP
FT                   ADGWDTADVKVLQWLAGKAPVPQVAAHEVLNGQDWLLMTRIRGKELCDPDMMSHPALLL
FT                   DCMAEALHTLWTIPVAGCPFERTVADNLSHAEAAILSGRFDPSGCEPETFGPGGFESPE
FT                   KLLDWLKTHLPPQDRVVTHGDFCLPNIFTDGKRFAGLIDVGNAGAGDRWMDLALGWRSL
FT                   KHNSDGHYGTVCPNIDPDDLFRAAGVPKDEEKLRYYILLDELN"
FT                   /product="Aminoglycoside phosphotransferase"
FT   CDS             complement(26954..27118)
FT                   /db_xref="SEED:fig|6666666.127272.peg.30"
FT                   /translation="MKLRPELVPGMDSQTFREYYYLKEELADFCRANHLPVSGGKQELT
FT                   ERVFPSMYP"
FT                   /product="hypothetical protein"
FT   CDS             complement(27382..27861)
FT                   /db_xref="SEED:fig|6666666.127272.peg.31"
FT                   /translation="MMKQQIINDTFALQIPDGFEILTEDDLRKMYRNAGDPFRWGVRDA
FT                   ENHVIILALWKKYPALLSWTLDLKAIVKKNEQLTGKAHAAHGYRFLEFFSMQAGDEKAE
FT                   GYRFSYDKEGMTQVCSNCLVRDGRTIYAFMRVGREENLETDREAFRRMMETLAYA"
FT                   /product="hypothetical protein"
FT   CDS             complement(27868..28662)
FT                   /db_xref="SEED:fig|6666666.127272.peg.32"
FT                   /translation="MKKMISILLVIMMLVAIACTASADEAPQPEAGKKFESDWAIAGGL
FT                   AQIVYEEEGYRVTLDITKDDGTGSIWEYSCYYHEDTDSLSSVSSSRTDYRINPDTGDKV
FT                   IGEKVYEGLDTEKGTEFTIGTDGFLTWKDGHDDAGAGLKFANIGRFEGVWKNEAEETDA
FT                   EFMWNGTTADEMFYTVYILRGKTDGDTYSSFLMNGTYDPATGKLTANGTCTVFRKNAEG
FT                   EYTPEEDGETYDAVFSKTADGKVLFETDNGIELEYDMLGSQG"
FT                   /product="hypothetical protein"
FT   CDS             29050..29622
FT                   /db_xref="SEED:fig|6666666.127272.peg.33"
FT                   /translation="MESIRPGKPGSRREDSGGRGFSLLPAIAPEKAVDLGCGGGRNAGE
FT                   LLKKYPKALVTAIDYSGLSVEKAKAFNQAAIDKGRCIVRQGDVSDLRLPAVSFDLATAF
FT                   ETVYFWPGLAECFGQVAKVLKPGGYFLICNESDGLDDAGKKFEKIIDGMRCHTAEEIEK
FT                   ALRAAGFTEVTRYRYPSKPWIAVLAKK"
FT                   /product="probable methyltransferase"
FT   CDS             29718..30497
FT                   /db_xref="SEED:fig|6666666.127272.peg.34"
FT                   /translation="MNPDYKSWILIPFVGTSLGAAMVFFLKKQISAGLQKVLTGFAAGV
FT                   MVAASFWSLLQPSLESSASMGRLAFLPAAVGFLIGMGFLLALDILTPHMHMDRQHEGPQ
FT                   SRLRRTTKLILAVTLHNIPEGMAVGVVCAGFLAGDTGISAAGALALAIGIAIQNFPEGA
FT                   IVSMPLLAEGMPKGKTFLMGVLSGAVEPVAAVITLLAAGLVTPVMPYFLSFAAGAMMYV
FT                   VVEELIPEMSEGKHSNTGTVAFALGFVVMMILDVALG"
FT                   /product="Metal transporter, ZIP family"
FT   CDS             complement(30559..30729)
FT                   /db_xref="SEED:fig|6666666.127272.peg.35"
```

```
FT                   /translation="MYRLRYARAGRQPGRLCQKAQGSKLPLRAVRFCDHQPVQEGFVMD
FT                   STQGRLTVLFE"
FT                   /product="hypothetical protein"
FT   CDS             complement(30847..34113)
FT                   /db_xref="SEED:fig|6666666.127272.peg.36"
FT                   /translation="MSYSERRNIRRICAMIITVAALLSVMTLPVSVTAETKTVRVGYYE
FT                   NEVFQEGAREGSVKTGYAYEYYRKLSEYTGWDYQYVYGEFGELYQMLLDGEIDLLAGLA
FT                   WREERAAIIGYPKAVMGRESYYLIKHDTDAGITAEPSTLNGYRIGVLDSAMVSVLNQYL
FT                   DEHHVTAEVIAYPDYKQLFGAFDSRDVDVLAAESDGARGREHSEVLTVFGTSDYYLCVN
FT                   IHRPDLLEELNSAQTLLATEEPNYLNSLSAKYYSVSVTARAFSRAERDWMDNHNALQVG
FT                   YLENYLPYSDTDRQGSATGIVKDIIPRILSSLGMDSISVTYKGYRSYDDMIAGINSGEI
FT                   DVAFPVSGGLYYSEESGIYQSNPVSSSPAELVYKGEFSDETTAHFAVNENNRMQFYFVR
FT                   TNYPDAEITFYPSSEDCLAAVLSGKAGCATLNGLRANDILRNRKYEDLSLRQTSYNDAR
FT                   CFGVGIGNEGLLKLLNRGINILGSDYAQNISYRYTGGLYSYDFIDAIRDNQAIVGTAIL
FT                   AVFAIILFLLVRDRKHKKAEIGEKETARLELEKKNKELAESQQALSDALIAAEHANRAK
FT                   TTFLNNMSHDIRTPMNAIIGFTALAASHIDNREQVQDYLGKISVSSQHLLSLINDVLDM
FT                   SRIESGKVKIEEAEVHLPDIIHDLRTIIQSNVLSKQMELFIDTQDVRHEDIITDKLRLN
FT                   QVLLNILSNAIKFTPAGGIISFRVIEKPSPAADFANFEFRIKDNGIGMSEEFQKTIFDA
FT                   FTREKTSTVSGIQGTGLGMAITKSIVDMMGGVITVNSEVGKGSEFIVQLPCRISGASAK
FT                   FEPLTELNSLRALVADDDTDTCLSICSMLREIGMRPDWTNYGKEAVVRAKEAMDQADEF
FT                   RVYIIDWLMPDLNGIETVRRIRKVIGDNTPIIILTAYDWTDIEEEAKKAGVTAFCAKPL
FT                   FMSELRNILAKSFTASRDSQEKKEETPVLPHFSGKRILLAEDNEMNQMIAEAILTEAGL
FT                   TVDVVGDGTDAVEKMRDAPAGTYDVILMDIQMPKMNGYEAAKQIRALEDPRKATIPIVA
FT                   VTANAFEEDRKTAMEAGMNGHLAKPYDIPAMMETLRTLLE"
FT                   /product="Sensory box histidine kinase/response regulator"
FT   CDS             complement(34199..34396)
FT                   /db_xref="SEED:fig|6666666.127272.peg.37"
FT                   /translation="MEYKIRTAVQADEGTIRELFPEMLRAIYHTDDVKGCGDGDPDRFR
FT                   CGNGDRIYAAEDGEFDWIIL"
FT                   /product="hypothetical protein"
FT   CDS             complement(34448..34588)
FT                   /db_xref="SEED:fig|6666666.127272.peg.38"
FT                   /translation="MLNNRFSAVCRTHYEADPCNHGFRSRQRSVCDVLFGEYDQKESRR
FT                   E"
FT                   /product="hypothetical protein"
FT   CDS             complement(34578..35531)
FT                   /db_xref="SEED:fig|6666666.127272.peg.39"
FT                   /translation="MKKEFEVVSQETFKYLHVFLVNLASRTPHIHKEIELGLVLKGKLT
FT                   VREGKNAWSLDRDEMYLINSMEAHEFVADDPDTLILAIQISPKILMPFLSDAALLRFMT
FT                   EPPVYGFFREKQNVWQNLRNDCMELARYYHARPDGFEFDCFHLISHIVLLLRRHIPIAS
FT                   LSQKDYNTVRRRMDRIASITDYIDQNYQRKLLLEEIAEMNNLTLTHVSHLFKDTLGITF
FT                   QDYLKEKRFEHALSLFENTDGTILEITVESGFSDERYLTKLFHERMGCSPKEYRKKQSQ
FT                   AQNRPRNAVINTQKIYNDEEAIAFLGCPAEPVLVAE"
FT                   /product="DNA-binding response regulator, AraC family"
FT   CDS             complement(35528..35746)
FT                   /db_xref="SEED:fig|6666666.127272.peg.40"
FT                   /translation="MNIPGADRKLKLRHCCILLLLMEAFRFICLNHSRIFEQKPVLVCE
FT                   KTPINLRFLCLTFFSPDLMISGGRIEK"
FT                   /product="hypothetical protein"
FT   CDS             35705..36742
FT                   /db_xref="SEED:fig|6666666.127272.peg.41"
FT                   /db_xref="GO:0009045"
FT                   /translation="MSQFKFSVGPWNVHTGADSYGPETRNAIDLETKFAAFKKLGFDAV
FT                   QFHDDDAVPNMNNLSEEEIKTEARKVKALLDKYDLKAEFVAPRLWMDPHTTDGGFTSTS
```

```
FT                   KEDREFAMWRAYRSIDIARELGCDLIVLWLAREGTLCAEAKSPVTATKQLVEAIDNMLR
FT                   YDATIRVCIEPKPNEPIDRSFAGTIGHVMGISAATIDPKRVGANLESAHAILAGLDPAN
FT                   EIGFALAFGKLFTVHLNDQNGIRYDQDKSFGVENLRSAFNQVKVLVENNYGSRGEYVGL
FT                   DVKAMRTTSDEDSYEHLKNSLAIFKAMEEKVKKYDYDYAGKLIAEKKFEQLEMYTVGLI
FT                   MGEAL"
FT                   /product="Xylose isomerase (EC 5.3.1.5)"
FT                   /EC_number="5.3.1.5"
FT   CDS             36929..38032
FT                   /db_xref="SEED:fig|6666666.127272.peg.42"
FT                   /db_xref="GO:0004747"
FT                   /translation="MGKVISAGHICLDITPVFPSGHRYDRIGDLLVPGKLIRTDAASVH
FT                   TGGSVANTGLALKLLGNDVTLLGKVGCDAFGGMVQSILAGYGASGLIADEQSSTSYSVV
FT                   LAVPGMDRIFLHNPGANDTFSSSDIPEETLADAVLFHFGYPTLMKRMYEAGGAELKNLF
FT                   MRVKSKGIATSLDLAAVDPQSDAGKADWPAILGSVLPYVDFFVPSFEELCYMLDRSCYD
FT                   ALAAKGGDMTEQEGILQTADILAQKCLDLGCGAVLIKCGLSGMVYRTGCRERIAQVGAR
FT                   LELYADEWTDRSGAQPCYRAEVVRSATGAGDVSIAAYLTALMSGETPEVCAKIAAAEGA
FT                   ASVSSYDALGGVLPLDELKKRIAEYEG"
FT                   /product="Ribokinase (EC 2.7.1.15)"
FT                   /EC_number="2.7.1.15"
FT   CDS             38143..38988
FT                   /db_xref="SEED:fig|6666666.127272.peg.43"
FT                   /db_xref="GO:0004332"
FT                   /translation="MLVNMKEMLAKAQKEQYGIGFFNAVNVEMARAVIETAEELHAPVI
FT                   VGTAEVLLPAMSLERVAEYLIPMAQKASVPVAVHYDHGLTFEKCMEALKLGFSSVMYDC
FT                   STASYEVNLDRVAEMVRICHATGVTVEGELGHVGDNEGAGKLENPSDYYTDPAVAADFV
FT                   QKTGVDALAVAVGNAHGDYRFPPKLDFERITAIREATGTPLVLHGGSGLSDDDFREAVR
FT                   RGVCKINIFTDLDKAGKAGIEEGLKKGAASMMGLIPYEIEAMKEVVRNKIGLFGSIGHA
FT                   "
FT                   /product="Fructose-bisphosphate aldolase class II (EC
FT                   4.1.2.13)"
FT                   /EC_number="4.1.2.13"
FT   CDS             39064..39270
FT                   /db_xref="SEED:fig|6666666.127272.peg.44"
FT                   /translation="MIQVQGDVNFPEFALPGQEVQFDGSAGSADRQPKESSGHRKVPIL
FT                   SGSLSGLAQAIRGDPACRPYPFI"
FT                   /product="hypothetical protein"
XX
SQ   Sequence 39405 BP; 8013 A; 11506 C; 10371 G; 9515 T; 0 other;
     agtgtggcac acccggcggg attcgattcc gttcgctcaa tggtcgcctg aagcccttgg        60
     agcgggcttc attctccctt tcgcgctaac aaactcgagt cgcttccgct atgcggacag       120
     cccaccgggc tgacgcttct ctcgtttgcc acggccgccg tggattctga tctgcagggc       180
     agatcaatct aaaaccggcc atccataaag gatgaccggc ttcagatggc acacccggcg       240
     ggattcgaac ccacggcctg ccgcttagga ggcggccgct ctatcctact gagctacggg       300
     tgcatagtat agaagcaagc gaaattatat gctttacagc atcttttgtc aatccgaacc       360
     tgtagacata gaatgttcta tagcatttca ggccttgatt cttgtgcagt ctcctgtggc       420
     tgaagtggat ttttctgcct gtccatgata taaaaaacaa acattaacaa gtgagggccg       480
     taccttgagc cgaaaagtga acaaactgac gtgcactgcg ggacccggga tgctggacac       540
     catccgtacc gggattgaac tgacggagcc ggtgaaccgg caggcgccgg agcaggccgt       600
     cccaaaggcc gcggaaatgt tcccctgttt cgccgtaaag atggtgctga agggatcgga       660
     gtatctgctt gagccggacg accggctgtt ttgatttcac cggaagggcg gttcgccgcg       720
     ctgctgaagg aattcggaat cccgtttgag gaatatgaac cggaacggtt tggcatatgc       780
     ggctttaagg ttccggaaaa tgaatcatcc tcggataaga cccggagtgc atgcttcggc       840
     ggccggtaat cgtatcatgc gattctctca ggatggttgt tccgcctgtt tgcagacatc       900
     cacccagtcc agaaagccgc tgcagcgttc cagttccggt atggtcagtt tcactccgct       960
```

gtggtcggtg cccacggcgg gataaacggt ctcatgcctc cggaggcttt cgtccaggta 1020
tacgctgacc cctggattga caccgaaggg acagacgccc ccgatatcat gcccgaccag 1080
ctgttccacc aggcccgcag gaatcatcct ggccttgcag ccaaaccggt ctttgtattt 1140
ccggttgtcg atacgggccg tgccgtccgc aaggatcaga accggctttt catcctgcag 1200
gaacgaaagc gtctttgcaa tcatcgactc ctcgcagccg atggcttccg ccgcttcggc 1260
aaccgtagca ctgctgtgct cagtgacgat aatcttctct ccgagtccga agccttccag 1320
gtacgctttt gctttttcga atgccattgc cgtcatcctc ccctgccccg cggtccgctg 1380
tccgccatcg ggcaccgtaa aagaatagca ggtcggagcc cggctgtaaa ccgcggatta 1440
cccggagcgg cgccgcgtgt atgctgcagg ggctattttt gcaggaaaaa atgcgtgaca 1500
acgcccggaa ctatatattt ccgaaggcaa atatgctaca atacaacatc attattccga 1560
aacggaatgc attgaaggag gttattttat gaaaaaactg gagaacaatc cgctcctcta 1620
cggcatcgtg aactgcctgc tcagcgcgat aggcgtcgtc atcgtttgtt ttatcatcag 1680
cctggtcaag aaagacgagt ctttttccga tgttatcagc ggaacctata tgctgatcat 1740
cctgatcggc ggcagcattt ttgccggcat ttccgcgtac cttaaggcga agaacaaaga 1800
caagaaggaa atgtaatctt ttcctccggg cgtaaatcga ataaaggaga aaacaagttc 1860
atgcttcaga tcaatcatta cacaaaacag tacagcgtga acggccccaa ggccgtggat 1920
gacctgacgc tccatatccg gcccggcgag atttatggtt tcctgggcca caacggtgcg 1980
ggcaagacca ccacgatcaa agcctgctgc ggcatcctgg attttaccga aggggagatc 2040
ctcatcaacg gcatgtccat caaaaaacag ccgctggaat gcaagaaaat catggcgtac 2100
atcccggata acccagacct gtacgagttc atgagcggta tccgctacct gaactttatc 2160
ggggacgttt acggcattcc cacagacgtg cgggaaaagc gcatcgccga gtacgccgaa 2220
gacatgggaa tcaccgacgc gctggcggaa aaagtcgctt cctactccca cggtatgaag 2280
cagaaactgg ccattgtggc tgcactgatg catgagccga aactcctgat catggacgag 2340
ccctttgtcg gcctggatcc caaggccgcc tttatcctga agaagttcat gcgcaggttc 2400
tgtgacaacg gcggggcaat cttcttctcc acccatgtgc tggatgtcgc ggaaaaactg 2460
tgcgacaaaa tcgcgatcat caaaggcgga aagctgcttg ctgacggaga tatcgaaacc 2520
gtcaggggca acgaaagcct ggaagaagtc ttcctcgaac tggtggatga tatgaaggtg 2580
gaggaagaaa atgcttaaaa tactgctgaa aagccggttg cttgcaatcg cggaccagtt 2640
ttccggccag tcgaagggca aaaaagcagc cacaatcacc agccatatca ctctcatcgt 2700
cttgggcctc ctgctcatgg tcggcgccgc cgtgggagtc aacgcgctgc tcggtggaac 2760
ggttgccgcc ctcgctcccc aggggctgac ctggggcttc tatgccgccg ccggcggact 2820
cagcctcatt gcctccctgt ttctgaccat gttctatgcg cagggcgtcg tctttgaggc 2880
aaaggacaat gaaatgctcc tgtccatgcc catcccgccc tccgccatcc tcgcaagccg 2940
catcggatcg gtatatctca tgaacctgat cgcccctgtg gtgattatgt tcgcgaccgg 3000
atacacggtt gcggtgagag ccagcggcat ctccgtgctc agcgtggtct tcctggtact 3060
gtgcacgctg ctgctgacgc tccttacagc cacgataacc tgcctgctcg cctggctggt 3120
ttcgatcatc acaaggcgca cgcggaaaaa gacgctggta cagctgctcc tgtccctggc 3180
cgccatgggt ctgctctatt ttgtgacgag caacgtcaac cgcgacctgc tggcgtccat 3240
caaggacaac gccgtctcca tcgcggactt tttccggagt aaactgtatc ccctgtacgt 3300
catgggacag gccattgcgg aaagcagctg gctgcagctg ctgatcttcg cggcctgctg 3360
tatcgttcct ttcgtgctgg tatacttcct gctgagcaag agctttatcc ggatcgtcac 3420
agcccggagc accgcggcaa aactgaagta tgaagcgacg tccctgaaga gttcctccct 3480
ggtctgggcc atgacaaaga aagacctgac ccggtttttc aacagttccc cctatatgat 3540
gaacgccggc attggcctgc tgtataccct gggctggacc atttttaacg tttttttccgg 3600
caactcggtg atcaagggat tgtttgagca ggggacaaca gtacctgatg ccggccccta 3660
tatgatcctg gtctcggcct atacgctgag tttcttcgcc ggcctgacca ccatctccgg 3720
ctgttccgtt tccgtggagg gcaagaacct ctggatcctg aagagcctgc cggtgcgcgc 3780
aaagggaatc ctgatcggaa aattcctttc ccatctggtg ctggcggcgc ccgtgagtat 3840
catctgctca atcctgtacc tgctgtccat gccgtctccc gatgtgatcg gcatcgccgc 3900
catcttcctg atgccgctgg ccgccaacgt cctttgcgca ctgaccggcg tgatcagcaa 3960
cctgtttttc ggcaagctga attatccgag cattgccaaa gcggcgaaga gcagcgccag 4020
ctccctcatc cccatgctgg tcaccaccgt gctcgtcgtg ggcacaggtg cggtgtactt 4080
attcttcctg cgggaaacgg tatccgtcag gatattcacc gtcgctgtca tcctcttcct 4140
gctcgcggtg gatgccggtc tgtatttctt cctcggatcc tccgccgtgc agaaacgctg 4200

ggaagcgctg ggcaattcct gatctgttta tggccatata aaagcgcggg ctgcgcatct 4260
gcagcccgcg ctttttttcgg tttattccgc aaccggaacc gtcagtgttt ccccgtttgc 4320
catcctgatg gacacctccg tcaggtcctt gtccgcgaag taggtatagg agaccttctg 4380
ggttgttccc ggcctgacgt ccggattcac agcatactcc ccgtcgcttc cgccggaaat 4440
gtggaagaaa gccagcggcg ttccggcctg gtccttcact tccatctctt cggtcacgag 4500
ggatgccggc aatgtatccc cggtcctgga tgtgaagtcc acgttctcaa tttccccatg 4560
cacggcgatg atcatataat gatcgccctt gtctttggcc agcatctgcc acaggtctcc 4620
gtcaagcttc ctgacctccg cgacggtcag ccggtaggtg ccgttctccg ttttgacctc 4680
atgcggttcg tccggttcct tcccggattc cgacagtccc acagctgcct tctgcgtttc 4740
aaaaacaccg gatgccgcga tcccgccggc tgttatcgta ttgatctgcg ccgctttgtc 4800
cgcgttttcc gtgttcaggt cctcaatctg agcatttttg tccgtgacat ccgcgttcag 4860
cgcctcaatc tgcttatccc ggtccgatac cgccgcgttc agcgcctcaa tctgcttatc 4920
ccggtccgat actgccgcct tcagcgtgtc aatctggttt gccttttcag cggcatccgc 4980
gttcatggct tcaatccgcg cggtcttgtc tgccgcgtcc gttttcaggg tctctatttc 5040
cttgtcacgc tccgctgctg ccgccttcag cgtgtcgatc tgcgccgctt tgtccgcgtt 5100
ttccgtgttc aggtcctcta tctgtgcgtc cctgtccgtc acgtccgcgt tcagcgcctc 5160
aatctgtgag cccttgtccg tgatgtccgc gttcagcgcc tcaattttcc tgtcccgctc 5220
cgcaatcgcc gccttcagcg cgcctatctg tttccgatttg tccgccgcgt ccgctttcag 5280
ggcatcaatc tgcgcggtct tgtcattcac gtccgcgttc aggacttcaa tctgcgaccc 5340
cttgtccgtc acgtccgcgt ccagggtctc gtttgccttg tcccggtccg ataccgccgc 5400
cttcagcgta tcgatctgtt ccgatttgtc cgcgttttcc gcgtacatgg tttcaatctg 5460
cgaacccatg tccgtcacgt ccgcgttcag ggcttcaatc tccttgtccc gctccgatac 5520
tgccgccttc agcgtgtcga tctgcgccgc tttgtccgcg ttttccgcgt tcagggcttt 5580
gattgttttg tcccgatccg ataccgccgt cttcagcgtg tcgatctgcg ccgctttgtc 5640
cgcattttcc gcgttcaggt tttcaatctg cgcggccttg tcctccgcgt ccgcgttcag 5700
cgagtcaatc cgctcatccc tgtcagcgat ttccgatttc agccgcgtga tctcctcgtc 5760
cttttccgcg gaatccgaat tcagcgaccc gatttttttcc gtcttgtcct tcacggacgt 5820
ggtcagtgtt tcgatcatct tcgccttgtc cgcgttgttg ctgacataca gcacgcccat 5880
cacaaccgcc agaactgtcg ccatcgaaat gaagatgatt gccacgctct ttctcatgtt 5940
tccgttcctt ctttcctgtc gattcgttcc ggcccggttt gtcttcaggc gccctggaca 6000
aaccggataa actcctgcgc ttcctcaaat gtctgcaggc gcgcggtata catataccgt 6060
cccatctgcg gccagttcat cggcggcatt tcctcctgca tcatcatccg gccgccgtaa 6120
cgggccatga tctcctcatg cgtccgggca taggtatgcc cgtccttccg gctggcatac 6180
acgcagtaga aatgctccgt ggattccggc agcttccggg tgtcaaaagc aaccatgtcc 6240
gcgtagatct ggtaaacgtc cgtggaatgg gcaaagtcca tcatgtccgg cgtatagcct 6300
cccgcggggc gcatatttac ttccagcgcc acaaaatccc cggtcttccc gaggtttttg 6360
cgggccttcg tcaggcggaa aaactccagg tgcacaaacc ggcggtccac accgaaagcc 6420
ttcaccgtct tccggcccag cgcccgcagc ttttccggga cttccgggca ggtgtagtac 6480
atcaggtcca ggtccttgtt cacaatgtcc attaccggcg gccaaaccgt catgctttca 6540
aacagcggtt cgcaccggga atccaggatc gcgtcatagg agcagatttc gccctggata 6600
aactcctcca tcacataggg aaccgccggc aggttatcat aaaaagcttc caggtccgcg 6660
tcgttttcca gcttccaggt ccgcgttgcc ccgacgccca catccggctt gacgatcacc 6720
ggatacccga cttcccggag aaagcttttc cccgcttccc tgtcggatac cacatgctgc 6780
cgggccgtcg ggattcccgc ctccaggtac agctttttca tcaggctctt ttccttgatg 6840
aaactgatcc ggtccgaccg gatcccggtg atgatattga agtccgtccg caggcgggcg 6900
tcctgctcca gccagtattc gttcatggac tcaacccagt cgatctttcc gtacttaaag 6960
gcgaaaaaag caaccgcccg gtatacctga tcgtaatcct ccaggctgtc cacccggtaa 7020
tactccgtca atgcggcctt cagcggttct tccaggccgt catacggcgc gtctccgatg 7080
cccagtacat tgatgccgtt ccggtggagc cggtcgcaga attgccagta tgtgtgggga 7140
aaatggggag aaataaagac atagttcatt gtttcagttc ctttgttcgt taatcataaa 7200
ggggacgaaa taccgcatct gcttaaacca ccagggccag tcgtggttga cgtcgcagcc 7260
ccagaaatca caccaggcct caatcccttt ctcatggaaa atccgctcca ggtgccgcag 7320
cgtgcgtacg ccgtcctctt cccaggcgcc ctggccgcag caaaggatca tcttcctttc 7380
gttgtacagg ctgatccacg gatggtcccc gggcatgttg ggcaggaagc gttccggaga 7440

attgtcatac agcgtcgggt tcatccaccc gtcaaagaaa taatccgcgt catatacgcc 7500
cgacagggcg atcactcccc tgaagaggtc aggccggcgg agaaacgtaa tcaccgcgtg 7560
gtcggcgccc atggagaaac cggttacggc cggaacctcc ggattgcggg aacggatcag 7620
cgggatgact tccccggtaa tgtaccggaa atactgctcc tggcgcccgg cacggaactc 7680
gttgtcccat cccctggctg accagctttc cctgtccacg gtatccacca caaaaagctg 7740
tattttattg tcttcaataa aatccgccag gtgatggatc atgccgaaat cctcatagtt 7800
atggcacatg gaatcctggg tcggaaagca caggaagggc acgccgccct gcccgtaaac 7860
catgatatgc atttcccggt ccagggattg gctgtaatga gtcaactctt ctctgttcat 7920
actgcacctt ccagcctttt tctgttcctt tccggagcgg cggccgtccg cgcggctgct 7980
tgccttttta ttatacacat tcttgggaca gatggaagtc cctgcccgat ttccgcatgc 8040
gcgaaaggcc ttcgaccctt ctgtgccgcg aaaaaaagca gcccccgtaa gggggctgcc 8100
ccggaagaag aaagtgtttc cgaaaaccgc tgtttcagcc ggtcattcaa tgatgctggt 8160
cttcgtctca tagggatccc gggtcgtcgc gccgctggcc ccgtccacac cggtatccgg 8220
cttgccgatc agcaggacgg ccaccgcctg catggaagga tccacaccca gcttttcgca 8280
cagggcgtcg tggttttctc cgttcagcgt cctggccggg gaagaaacga tcttcacccc 8340
gtaccccttg gcggaagccg caatcaccat gttctgcgcg gcaagtccgc agtcaaagct 8400
ggcgtccggt aatccggaag cgccgtcttt gtaaataatg atcgcagcgg gcgagtctcc 8460
cagtgaagcc ttcgcgccgg cgcctgccgg aggcgcggcg ggcctgtccc ctgcgggaac 8520
cgcctcatcc gtgaaggcct gcgcggttcc ggtggccata atgacgtcaa ccgcggactc 8580
ggcataagcg gcggaaacca tcagcagcgc agctgccagg atcatagtcc ccaaagctaa 8640
attgaaactg aaacccgttc ccggcgggga gggtataccg cctcgggccc tttcacagta 8700
cgctttccag cctttccatg gcttcccgca ggatactccg cgggcaggcc gtgttaatcc 8760
tctggaagcc ttccccggct ttcccgaaca tcgatccgct gtccagccac aggcgtgcct 8820
tgcggaggat cacgtcgtcc agttcggccg cgtccatccc ggtgcccctg aaatccagcc 8880
agatcaggta ggttccttcc ggctggttca tacgcacgcc tggcagccgc tccctgacaa 8940
attcatccgc gaaacggata ttgtcccgga cgtatgtttt catggccgca taccatgtat 9000
cgccgttttc ataggccgcc tgcgccgccg caatcccgaa gaccgtcggt tcgtcgaaac 9060
cggtccggct gatctcctcc gcaaagccgc gtctcaggcc gggattcgga ataaaaatat 9120
tggactgctg gagtccggcc aggttaaacg tcttgctggg ggccgtcgcc gtcacggtaa 9180
actcccggaa ggaaggatcc gcctcctgga atacgctgtg ctccccttc cagacaaagt 9240
ccgaatggat ctcgtcgctg aacacgatta ccccgtgcga cgcgcagatc cggccgattg 9300
tccgcagttc gtccccggtc cagaccctgc cgaccggatt gtgcgggctg cagaggatga 9360
acagcttcac atggtgttcc gccaccttcc gctcaaagtc ttccggatcc atggaatagt 9420
ttccttccgc gtccgggacc aggtcggagc tgaccacttt ccggccgttc tgccggatca 9480
ggctgctgaa aggataatag accggctcct gaatgatgac ggcgtccccg ggctcggtaa 9540
aggcccggat ggccatcgag actgcaaagc acacgcccgg ggtcttgacg tgccattccg 9600
ggtccacgtc ccagccgtga tgccgtttca tccagccgct gaccgcctca aaaaagccgt 9660
ctccgggcaa tgtattgctg tatccgtaga tattgtggcg cgtcagttcc ccgagcgcgt 9720
cctccacata ggaggatgtc ctgaagtcca tatccgccac ccacaggggc aggatatcct 9780
ccgggtatcc tttccgggcg gcaagatcat acttcacgct ccgggtgttc ttccggctga 9840
tgatccggtc aaaatcaagg tttctctctg ccatatccgc atgcctcccc attttccctg 9900
ccggtaaaca ccggtgctgt tatgatcttt gctgttccat ccagttccgg atgatctcct 9960
gcatctgttc cggatgccgg tcaaagtgat gggtttcccc gggaatcata tgcatttcgc 10020
agttgctgta gcgtttcgcc atccgcgcgg aatcctccgg gggaacaacg tcgtcctcat 10080
ccccgtgaag gatcaggacc ggccccctgt agcggtccgc cgcgtcctcc gcgtggattg 10140
tctgcgccac gcggatatag tttccgtcca gcaccacgtc ctcggccacc gtcacgctct 10200
ccggaacatg gtccgggtca aaccggtatc ccagcatgca gccgtccctg gcagcctgcg 10260
gaatcatgaa cgccggggcg cggaggatca gaccgctgat ccggtccgct tccatacctg 10320
ccaccagcgc ggcggtcagg ccgccctggg aatggccgga aaggtagatc tccgtatacc 10380
cgagctcccg ggcatagtcg attaccgcca tggtattgga gatccatttg tacagggtat 10440
gcttccggaa ctctccgccg ctttcgccat ggccgtacag gtcaaggcgg agggtggcgc 10500
accctttctc ccgcatggcg gccgcagtaa ggatgttgtg cgtcctgtcc ttcgcgctgc 10560
caaagccatg gagcaggatt gccagcggat gcctgccgct gccctcaggc tcttccagca 10620
ccgcgctcag ccggatttca tcatctcgga taatcatttt ccctccggtt ttcaggaata 10680

tcattcatac acgcgccggt tcttcttccg tcagcatctt atccccatgt gtacctgaag 10740
tcaacatgtg caccgtaaac gcttgactta gaataggctc aaggctgtat actggagtgg 10800
cggcgttgca tcgccgtaaa tcaccgttcg gagaagaaac aatgaaaact gtaaaggctg 10860
cttttgtcag gtcccttcct gtcatggccg gatatattat cctcggaatc ggcttcggga 10920
tcctcgcgca caaagccggc tacggcctgc cgtgggccct ggcgatgagc ctgtttatct 10980
ttgctggttc catgcagttt gtgggcgtgg ggttgctggc cggcggcgcc tccgtgctga 11040
cagtcatcct caccaccgtc atggtcaacg cccggcacct gttctacagc atttccatgc 11100
ttcgccatta ccggaacgcg ggtccctgga aaccctacct ggtcttttcc ctgacagatg 11160
aaacctattc cctgctgtgc gacggaaaaa caccggaggg gacggatccg gaccgctacc 11220
gtttcctggt gtccgtcttc aaccattcct actgggtgct cggaagcgtc atcggaagcg 11280
ttgtcggcgg catcctgcct ttttccgccg caggcatcga gttctccatg accgccctgt 11340
tcatcgcatc cttcacagag cagtggataa cggcaaagaa ccatattccc gccgccgcgg 11400
gcctgttggg aacactgctg tgcctggtcg ttttcgggcc cgggcagttc ctgattcccg 11460
ccatgctgct gattacgctc cttttaaccc tgctcaagag ccggcttgac tccggaaaga 11520
aggtggacgc gtgaacagtt ccgcttattc cgccctgctg gtggctgtca tgtccgcggt 11580
aacgatcctg ctccgggccg cgccgttcct ggttttcaaa aagaaagccc cggcgtacat 11640
tatgtacctg gggcaggtcc ttcccgcagc aatcatcggt atgctcgtga tttactgcct 11700
gaaggataca aatctgactg cagcgccctt cggggcgccg gaactgatcg caggccttct 11760
ggtcgccgta ctgcacgcat ggaaaaggaa tatcctgctg agcatcctgg cgggcaccgt 11820
tgcgtatatg ctgctgatcc agctggtctt ctgagggaag cggcggccgt atcggatccg 11880
ggcattcccg gcccgatacc ggttccgcgc tcatccggcc ggacttcccc gagctgaaac 11940
aatggccttt cctgtggcag cagacggaat atgtgctggg cggcggttcc tcccgcttca 12000
tgttcctggt cctggccgtg cgcgaactca tgcggggctg accggcaaac tgaaaacgga 12060
gagcggcggg aatcctccct ccgctctttc gtatgctcct gtcagatttc atcaggggca 12120
cagcccggtt tcaggggttc cggcctgccg caggtggtgc gcagctgcac gaagctgccg 12180
ctttccgcgg accggatcag cccttcgata atctctgtga tatggataat catttccccg 12240
ttggcccggt tcctgccgtt ccgctcaatg gcggccgcca ggtcaagcac accgatgccc 12300
cgggagtaat cctttgcccg ggtaaacagt tccggcaggt cccggaatgt tttccgccgc 12360
gcttcgcttt ccggcgtgtg ctgccagacc gtgtcggtaa actgttcttt ccggtacacc 12420
ttcggcgtgc cgccgccgaa attcggatcc ggataggcca gcgtgccgcc gtctccgtac 12480
acttcaaaca tcggcaggct ggagcggtaa atgtcaaaac tgacgttcag gttggcgatc 12540
accccgctgc gcatccgcaa aatggccgaa taatgcgtcg ggatttccgc gggaatctcc 12600
gtgcctgcct gggggccgga ataaacatgg cgtgtttccg acggtttcgc gctgaacgcg 12660
ccgacgcttt ccgccgggcc gaacagcgca accagcgccg acaggtaata cggcgccatg 12720
tcatacagcg gaccgcttcc tttccggtag aaaggcgcgg gcgccggatg ccaggtctcc 12780
gcgccgaacg tcgtcatgtt ggccgtcacg aaaaacggtt tcccgatcag tccggtatcc 12840
aggtaatacc gcatgctctg cagtccggaa gccaggaacg tatccggcgc ggaaccgatc 12900
ttcacgcctt tttccgcggc aagcgccaca agctcctccg cttcgcgggt cgtctgcgcg 12960
aacggtttct cgctgaacag gtgcttgcct gcctggataa tctgccggtt cagctccaca 13020
tggaactgcg gcggcgtcag gtcgatgata atctcaatct ccggatcccc cagcagctct 13080
tccggtgtgc aggccttcgg aatgccgaat tcctccgcca gtttccgcgc ggactcaggc 13140
cggacatccg cgcaggcggc gatatgcagc ctgggataaa aggccttgat atccgaagca 13200
taggtcctgc tgatgacgcc gcatccgaga ataccgatct gaaaatcccg catctgtatt 13260
cctccctccg gttctgtgta aagccatttt acagattgaa gccgactcta agtcaagcgt 13320
ttcctgccgg gcgctgcctc tttccgccgg gtaggttcca tggtgcttcc tttccgcaaa 13380
aaaagaacag tgctcccgcg cgcagcggaa cgcactgttc gcccggcccg gggaaagccg 13440
tcaggctgtc cccggctgcc ggatcattcc tgtttgagct ctcctcccca ggtttccacg 13500
cagaatctct tgctgccgca ttttacgcag gtcagtttgc gggttttggg ggtatggtta 13560
gcgaagaaaa attccttcat tgccgggcgg aataccgtat ggcactgggg gcagatgtag 13620
gccacattcc gataataata ccggacggcc cagattgcgt aaggaaccgc cacaacggcc 13680
cagacaacaa acggccacca gatccctttc aggatccaca gcaggatgga agcccattgc 13740
aggataccca gcggtattgc ggacaggacc attttcctgt gcaccgcgct cagtttcttc 13800
ttgttttcca taacagccgc tatgtcacct atggtttcca gggagaacgc gggcgtatcc 13860
ttcaggcctt ttttcagttc cgccagtctg tccatctttc cccggcgttc ttcgatctca 13920

ctgcgcagca cctgttcctg ctgttccagg aagatcgaaa tgacgctgcc gggatcctct 13980
tccttcagca gctgcccgat ggtatccagc gacaggcctg tgtcccgcag gaagcagatg 14040
attttcattt tcttcaggtc gccttcggaa tacagccttc ttccgccttc ggtcagcgcg 14100
ctgggagcca ggattccttt ctggtcataa aactgaactg tccggactgt gacgccgcac 14160
tctttcgcca tttctcccgt ggtgtactgt gacatagcca tgacctcctt tcggcgccat 14220
cctaccgcat tacgtaacgt tacaagcaac aggtgacgca gggggatttt tatttttttgg 14280
aatcatttcc cccgaatcat gcggtaatcg ccctgttcgt caatttccag ctgcaggtct 14340
gcccgcatgg aacagctgag gcacagccgg gcattgggca tgtcgctgaa atccacaaac 14400
cggactgccg cctcttcccc gaccccggtt ttgatccgcc tgtcaaccag gcgtttccgc 14460
agcaggtcct catccgccct cacggagatc gtataatccg cgaaggaggg caggtcccgc 14520
catccgtccc ggtccagcag caggtagttt ccttccagca gcacaatgtc cccgtcaatc 14580
cgaaccgcgt cttccaccgg attatgcagc agcctgtcgt aaaccggcca cttcgtttcc 14640
tctccggccg ccaggctttt cacgcgcttt gtcagcgcct ccaggtcaaa ggtctccggc 14700
gcgcctttga tttccaccat ccggatcttt ttcccgtccc gctctgtgtc atgggtcagc 14760
aggtattcct gcctcctgtg gaagccgtcc atgccgatgg cctggatccg tccggcatcc 14820
cggcgttctt ccgccaggcg gtgcaaaaag ctcaccagtg tgcttttccc cgccccgggc 14880
ggcgcggcca gcatcaccag gattctcctg cctgccgccc gctgcttttc cgtcagtttc 14940
tccagcagcg gcgcgaagat ttcgttcacc gcgcgctcac tgtatccggc ctcaaccgct 15000
attccgttaa tcgtcactgt atgccagatc atgctgttct ccgtccaggg tatattcgcg 15060
cctgttttcc caaaccggtc ctgatgcccg cctgctttca caggcggcag atcatgttca 15120
gtgaatccgg caggtcttcg tcaacatatc cggaatccgc aaaaccgtgc ttttcataca 15180
ggcggatagc ttcttcgtcc gcctttttca cgcagagctc cacatgatca taacggcctt 15240
ctttccgcag ctcatccagg atgagccgca aggcccggga gccgtacccc ctgttctggt 15300
accgcctgtc caccataaac tgctgcaggt catagccgga cggttcatcc gtgaattccc 15360
tgaccagcgc catgccgaca atggtcccgt ccccctcaaa cgcgaagatc ttcgcgttgc 15420
agccccggta aacgtagccc cgcgccagga tgccgaccgc gggggccaca tagctcttct 15480
ggtgctcctt cacgtccagt gaggcaattt ccatccagtt gtcttcagtc acgggaatca 15540
gcctgaccgc ctgcgcccgt tcagcttcct gccctgtcat ccgcgttcat ccttttcctt 15600
tcaaaagccc gcgtgtttcg gttccgttgt tatttcttct tctgctgctt cggagcgggc 15660
agttcatcgt acatcgcagt cagcagctcc tgcaggaatt cccggttgtc cacgttgtcg 15720
gcgagcagca tttccttcgc gccttcatag gggcattccc ggtccgcgtc cggcatcatg 15780
gccagcgccg accgggtggg tttgaccagg aaccggtcgt catagatgcc tccgataatc 15840
ctgccgttca tatacaggat gtattctccc atcatctgcc tgtaagtaat cccgtccagg 15900
cccgaaagct gttccaggat aaaatccagg tattccctgc ctgatgccat tccgtatccc 15960
tcctgcgccg tcctgtgctt taatcgtttg tcggatcgcc tttggtatgc gagctgcgat 16020
ggtccctgcg gaagaaaagc agctcaatca ggcttttttc atagggaatg aaacgcccgt 16080
cgtcaatcat ccggaggatc tcatccttcg tggcccagcg gaccgcctgt acttcctcgt 16140
actgaagctt cagggaagaa aggtcctccg gcatcgtcag cacataaata tcgtcaaagc 16200
cgtcctcaaa atgaactgtc agggtaggcc gggcgttttt caggtcaatg gccaggccca 16260
gttcttcccg ggtctcccgc tcggcggctg actggctgct gtcccctgac accgcgcatc 16320
cgccgacact gatatcccag agattcgacc agccccgttt gaacggctgg cgctgctgga 16380
tcagcatcct gccttcagga tcaaagatac agacatggac caccatcctg tagaagcctt 16440
ccggcgtgga ttcgccgcgc accatggtac ggccggtttt ctcatggttt gcggtataca 16500
ggtcaaacag ttccatacag ctttcctcct ccttcgttcc cggccggcct tcctccgcgc 16560
ccggctggtc atggactgtt tcccggtgtc atctttcacc gattgccagc aggttgtctt 16620
tccccatccc gcagacgtac ggactggaat cataccggta gcagaaatcc aggaagtctt 16680
ccttctgctt cgggtctttc atgatccggg tcaggatttc ccggggaacg gtcctcccat 16740
acgcgcccgg gcctgacagg cgaatgttct ttacgccgta tccggacagg atttttcca 16800
gttcatccgg catgaaggta taggtaatgc accagggcgc gcctccctgt tcatacaggg 16860
cgatctcttc ctcgccgccc atcaggccgt ccgccagcag cttcattgcg ttctgtttgt 16920
tgaatgaaaa gtcgctgatc atctgttcac ggttggcgac gcaccattgc acaaaaggat 16980
catcggcctt ttcgtccagg ataaactggt ttttctggat cggattcgcc aggtacggca 17040
gggaacccag cctgctggac acgctgatca taatccgctt gtccgcaatc cgcaccagtt 17100
cgccgatcac tttgttctgg ttgggatatg tataggaaac cggcgcgtca aaagagatca 17160

ccaggtcgaa cgcccggtcg cgataggcgc tcaggtcctc cagcgcgccc ttcacaaagg 17220
ttatcctgtc aagaacgcct tcccgcgccg ccagctcgcg cgctttttcg atcatcggcc 17280
gggaaatatc aaaatgggtc acgcggcagc cctgcttcgc cagcaggatc gagaaccgtc 17340
cgcttccggc gccgccgtcg aacaccgtgc cgattccgtc caggttctcc agcagttttc 17400
gctggatatg gtctttctgg atgatgtcgt ccatataggt tgcttcccgg cggctttcca 17460
gttctccctt gtccggaagg ttccactgcc gttctgaaat tttctggctg taatccatat 17520
ccctttccct tcatgatgtt gcccggggcc tgttgctttt gccgtccggg ccgtcttttc 17580
cggccatcat tcacattaaa tgcgatccgt cccgtttgtc aagacgggcg gccgtttctt 17640
ttttccccct tcggccgccg tccgcgaagc cggcataccg gccgtgcgtt cccgggcaga 17700
ttccggaaac agggccggcg gcgggcagga aacgatctgc aattttcctc cggggcgtct 17760
gataacatgc cgtatgtaat aaaaccggcc ctccgatggg ccgggagaaa ggaagatgtg 17820
aaaaatgtca gtgctgcaga tgaactttgt tcgcggagaa gacagtatcc ggaggatcaa 17880
tgatgccctg gccggaatgt gcctgagcga cgtgcgcatc cgggacagcc gcggatgcct 17940
cgcgaccagg aaggactttg acgaaaacgg gcaggtgaaa gccgtaacct attcctacct 18000
gtttgacagc atcgcgctgg aaaccacggg tgacgaaacc atgggaaccc gcacctggcg 18060
gatcgaaacc gctgttttcc tcaagggatc tcccaatgcg ccccaaaata cggacgcttt 18120
tttcgacagc tggagcgcgg ggcatccgga caggtccgtc cgcgtcatcc agtggattac 18180
ctttgacaag gcggacgtgt ttatcgtcct cagccgggga cggtaaaaag aggatccggt 18240
acgttcagat cccgctgctg cgcagcacct catagctgat tcctgccaga tgcagttcat 18300
ccgggaagca cctgaaggcg cgttttgccg gccatattct ttcatttctg tatgcgcggg 18360
tcaggaagcg ctgttttctg tgtcttcccg cttctcgccc tctttgatcc ggtgcgtttt 18420
gccgtcgctt ccgagataga tattatccgc ggcattcggt tccagctctc cgcttgcgat 18480
cataccgatg acaattttgg cgatttccgg atcaaactgg gtgccggagc agcgttcaaa 18540
ctctccgata atgacttcct gcgagaacgg cgccttgtag acgcgtttgg aggccatcgc 18600
gtcaaagcag tccgcgcagc atatgatcct ggcaaccagc gggatgtttt caccgctgag 18660
tccttcggga tagccggatc cgtcatactt ctcatgatgg gatttcgcgc cttcaataat 18720
ctggggaatg gtcgagattc ccctcaggat ttcggcgccg cgggtcgtat ggctcttcat 18780
gatcttgaat tcttcgtcgg tcagcctgcc cggcttgttc agaatgctgt cgggaatcgc 18840
aatttttccg atatcatgaa gcagggcgat ataataaacg ttttcgacct cgtcctgatc 18900
catgcccatg ttttgcgcaa tcagcctgga atacgaactg acgcgcatgg aatggccgtt 18960
tgtatattcg tccttggcgt cgatggtccg ggcaattgcc tggagagatt caagggtgat 19020
gttcttgtat tccagctggc gcttcaggga ctgcttggtc ttgtggcgga tgaccaggga 19080
cgaaatcagg aacacgatcg cgcccgcgag cacagagagg atggcccaga accaccactg 19140
ttcaacaaag cgcttttcct tctccaggtg gatcaggatg tggttggatt ccttgttgtt 19200
ttcatccgtc acataataat ggaaatcgta gcttcctccg gggaggttgg tataagtgat 19260
gcttcggttc ccgtccgtat agctcgtgta ttcgctgtcc acaccgtcca gcttgtaata 19320
ggaggtgaag ccgttgttcg gccggaatcc gaaaacggaa aggttgatgg tcacacggga 19380
tacgtctttc tccaggctga tgctgtcccc ataatggggc tccccgtcga ggtcaacaga 19440
agaagcggcg actttgaccg gaaggtcgct ttctcttccc acgttcggct cgtagacata 19500
gatgccgttg gtggactgga gatagtattt catttccgat tcgtcgtaat agccggaggt 19560
gttggcaatg atcggctgca ggccgttgat tgcgtcatag gattcatatt ctgtcaggct 19620
ggtgcctgcc agcgaatcgg taacgtacag ctgggtctgg ctcccgatta catatttgta 19680
ttcgggtttg ccgtccgcgc ccgtgaacct gatctttgcc agttcaacga tggagccttt 19740
gacatacgga atcgcgattt cttcggaaac cccgtcaccc tttccgggat tcgtcctgaa 19800
gagcctgctg ttcagcgtgt agtagaggtc atccccgtcc accaggaatt tcagccggtt 19860
gcccgtgatt tcttttacag tgggaatctc cgtcacggtg gtgaaatcat cgtctacgac 19920
gaacaggccc ttggtctggt caaacaggat ctttccatcc ttcgtcttgt tgatatacag 19980
aacattggcg cctacgttga tgatggagaa agtttcaccg tcaaagcgca tgatacccct 20040
ggaagtaccg aggatcagcg cgtcgtcaaa gctccgcatc gaacggattt tatccgccag 20100
cgcaatatcc gggtcattga cgagcgtgcc caggtggttg ctgatataat cataatcgta 20160
gatcacgacc tgttccgtat ccgggttata ttccaccagg ccgatattga caaaagcaaa 20220
atagagcttg ttcctgaaaa cctcaacatc cttcggcgaa taggcgaaag agctgttgct 20280
gtctcccttg gcctggcttt cctctatcct gcgctcggca tattcgtcga tggccttcat 20340
gagattgttg tccggcagga tttccttgcg tgtcaggtca tacatgaaaa ccctgttgga 20400

gcagacaata tacaggatat ttccgtaccg ttccaccgcg tagacgttcc ggtcatatgc 20460
gggaggctcg gccagggact gccagatctg atcgtcatac agcagttctg tcagggcatt 20520
cctggtaata atggaaacgc cggacgtgcc gataaagtgg gaggctaccc aaaggttccc 20580
ctggtaatca atcatcaggt ccatcagctg gctcttgttg tccaggtttc ccgccaggga 20640
aataaccggt gcgccggagc tgaaatcaat acagtagatg ccgtccttct cacaggccac 20700
aaacaggatc tgatccgctt cggagtacag aagcttgttg atctggtccg gaacggaaag 20760
ctcgtccagg aaggcctcgt ttttcatatc gtaccggtaa atgattccgc tgtcttcgcc 20820
gatatacagg atggtgtcat aggcatacag ctccatgatc tttgaattct gcagaatctc 20880
ttttccgtcc tggtcataga cgccttcgtt cgtctggtag taatacttcc ctttctcata 20940
cccgaccgcg aaatcccgga tggctttccc ctctgtcccg ggaatagtgc tttttgtgcc 21000
ggtctcaagg ttgaaaacaa tgccgccgtt ttcccgggta cagatgtaca gcacgccgct 21060
cgccggatcc aggacgatgt ccgtgatgac gccgtccgcc gtggccagca gttcaaaatg 21120
ataatccttg tatacgtaca ggtggtcgga ggttgcgacg tacagcacat tgccttcggc 21180
cgccagggca gtgacattga tgatgttgta ttcctggccc tcatattcaa agctcttgta 21240
ggtcacgtat tctttggaat cgtaccgggt cagtccggaa tactgggcaa tccagacata 21300
cccgtcctcc gtctggcaca gagcctcggc gccggagacc tcaagattga cctggatctc 21360
cgtcttgttg ctcatatatg acgcccgggc tgtctgtccc atggcaaaat aagcgaagaa 21420
cgcaatcaga aaggtcagca gtatatgcgg aacgcttttt ttcatacttt cctcctgcct 21480
gtcagggttc tgctctttgc gctacagcaa ctccggcagg ttccggatcg gcagtctgct 21540
cacagatggc ctgttgccat aatacaatag attggcagaa aagaaaatat cctttttttc 21600
gccagggtct ccgtgggcgt cattccaccc ggaaggcgat gccgttctgc tcgcagtacg 21660
ccttcgcgca gctgtccgga ccgcagagga tcaccaggtc ctccgggcag ttatcaaaag 21720
cgttttcacc gatagcctgc acgctgtccg gaatcaccag ttccttcagt gaactgcagt 21780
atgcgaaggc cctgtccccg atcttcgtaa ccccggcggg gatctgaatg ctgcccaggg 21840
cggagcagtc gcagaaggca aaggcgccga tttcagtcac actgtccgga atctcaatgg 21900
atttgaggga ggagcagcct tcaaagacac cggttttgat tactgtgaca ccatccggga 21960
ttcgggcatg ctccaggctg ccgcagaaat cgaacgcgga gtccccgatc tccgtaactg 22020
tttccggaat ctctgcctca tacaggctgt cgcattcata gaacagcatg gggggaatct 22080
tcttcaggcc cttcggaagc accaccctgc caaggtattt gcaggtgtca aaggcggcgt 22140
ccccgatctc cgtgacgctg tccggaatga caatgcggtc cagtccccag cagaagccga 22200
aggcctgcgg cccgatgacc cggagggaat ccggaaggag gatatcggac agggtttcca 22260
cacctccgaa agcatcggcg cggataatct ccgttcctgc cggaaccgta tatgtcgcaa 22320
gctcgtcttc ccccggatca gctgccaggt tggggaagca caccagggaa tgatcctccc 22380
ggcagaacag gacgccgtcc tccacataaa acacgggatt cccctcggca accgtaatgg 22440
ttttcagttc cgtgcagtag ccgaacggat tctcatccat gctgaccacg cccgccggaa 22500
tttcgatttc cgccaggtcg gtcagctgga aggcgccgga cccgatatgc gtaaccgaag 22560
ccggaatggt gacatcctcc atgctgtatc tgtaattatc aaaggcttcc tcgccgattg 22620
atgtgacggg atggccgtcc agggtctccg gaatcaccag ctgaagttcc ttgccaacgt 22680
atttcaggat ctcagccgat ccgtcctcca gcaggcggta acggaaaccg tcggtttcca 22740
tggtggtttc tatcgtgccg tcctcggccg acgcggggat catcatgcac agcgcgaggt 22800
acagcgccgc tgccaggaat acagcgatcc ttttcatgtc ggaatcctcc ttgcttcagg 22860
ttcatgttct gcaggtaaga cgatggatcc gggggttttg ttccccctgt cagtatttcc 22920
cgtatcgatc tttttcacaa tcacgcccgt gtaatcatgg ctcttttttt caaccttctc 22980
cgggttcatg aaacacgccc gggccatcgg ggcgtttacc cggctggccg gttccgaaat 23040
ccctgctggc agccatttcc ggttcttcct tctgctgatc atgtgatagt tctccgtagg 23100
cagcgcgata tcctgttctt ccgcgtgcgg gacaaatgca tattcgtacc ggttatacgg 23160
cgcattcccg gcttctgcct gaatgcggtt gcggatgcgt ttgttctggt atataatacc 23220
tgtaaagaaa ccatcgttca ttgcttataa tcggataggc ggctgtcagg cgggaactgc 23280
cgttcagctg cgggaaccgg acaattgctg acgatgaagg aggcccacca tggtagaact 23340
ggacaatgaa agaatgaagc agatcctgca cgatgagacg gcggaaacgg aagacctggt 23400
aacgattctt cgcggtatct ataccaggta tatgaacctg tacgagcgtt attttgcaga 23460
cattgaagcg ctgaacgatg acaggattac tgagttcgca aaatatcatg aagaaaccaa 23520
aagcctgatc cggtattatt acatggatat tccccaggat atctgcgcgg gaatcagtga 23580
attcgaggaa aaggtcagcg gcagactgct ggggcgcgaa tggaagatga atcttttcga 23640

cgtctatgac gaattcaggg aaaaaaaccg ggtccgaaac aagagtgagg actattataa 23700
ggcagagttt gcaaaactgg ccctgaaaga attctataag gccatggaaa acattttcag 23760
ggacggtttc ggtacttaca gccagaccgt caaggatgtg cttggcggga tcacaggact 23820
gctgttcggc aaaaaagact gatgagtcat ggggacggtt ctgttaatca tctgcggccg 23880
gccggctccg gctgcctgat tcagtccatt tttcaagcgc agcggtcagc gggggcacga 23940
tcttcgcctt ccgggaagcc tttggggtga agaccaggta tccgctgccg tcatagtccg 24000
gaaaactgtc gtatacggct ttctccgcgt cttcgccgta ccagatgagc cgtgtcatgt 24060
catccgtgga aaccatgcaa aacagcatat ccagattgtt ggacgcataa atggacggaa 24120
atgcggcctc tattttctgc gccagcgctg tggcgttctc ctctgagtct gtatacacgt 24180
cgccgatgcc aaaggtgtac tcgccggcaa ggtattcctt gtaatcattg tagaaaatcc 24240
cggacgcggt catcccttca taggacgcct ttgctgcttt cattcccgcg tacagtgcgt 24300
caatatcatc gatatctgca acagaaagga gataattata tgcatcccgg tcaagcgccg 24360
tgacgttgct ggtcattccc ttggtatcgg acaaaatgct catgagcatc acccgggcaa 24420
tatcccttgg aatgcccaca ttgcagtcca ggtacagttt gtaaacgatg gacgctgttg 24480
cgcccgccgg aagggagatg acagcaatgc gctccgtgtt ccggatatcg ccgattccgt 24540
gatggtctac aattcctacg atccgggcat ttttcatgcc gtccagtgcc tgtgaatact 24600
cactgtggtc caccaggaca aactgtctcc ctgcggcgtt gtccaggatc tcaggctgtt 24660
ccaggccata gaaatccagt gcgtatttcg tttcctggtt gacttttgcg gtagcggctg 24720
cttttgcgtt gatcccgagc tgccgcagca aatgcgcgta tgcgatcgcg gttccgattg 24780
tatctgcgtc cgggctctta tggccgacga cgtaagtcac ctgatccccg taatccaccg 24840
agcgcaggac atccgcaaaa cccgttccat ccgcgcgggc tgccatgccg gaaaaggcaa 24900
ggaagagtac aagcagcgga atgatatatt ttttctttat catcggatca cctttctctt 24960
ttttgtttgt cccgggttcg caatatctcc gctgtccggg atactgttca caatgattat 25020
tcataaaaca gactcacagc tctttttttca tgtgtgtccc gctgtcgccc tgcgtaacgg 25080
tttcatatcc gcatttctga tacaggttaa atccggggac attgtgcgcc cagacataaa 25140
ggacactctc cgtacggccg tcggctttcg ccatctgttc catttcatgg agcgcggcct 25200
tcgcgtatcc ctttcttctc tccggtttaa aaatcaaaaa gtcgttcaga aaaacctgcg 25260
ggctgccgtc atagccttcc tcataagaaa accagatcca gccaacttct ttgcctgcct 25320
ttttatcttc cacggtcatc aggaactgat tcacagtgtc ctggccgtca gggagcatcc 25380
cgcggaattc cccttcggcc ttctccaaag cctgctccct tgtgaatcct ttccctctgg 25440
tcaaatcctc tgcatagtcg gaaacgctgt attctttaaa acgccggtat tcctcatccg 25500
tcattctgcg cagacccacg atggctttca ggccatcttc tttcctggca gtgcagagcc 25560
ttttggagaa gaaaagctgg agactgtcat cgttttctat ttcggtttca tattgttcac 25620
atggcctgcc ccgataccat acgcctgtgc cgtccgggat atatccccgt ttgacgtaca 25680
tccgctgggc gcttccataa ccgttgtgaa ggccgacgcc gagccagacc gtatccgcgt 25740
attgcccggc gatctgttcg gcaatatcca tgagcttact gccgattcca tttctctgga 25800
acctttccag cacgccaaag tcaacaatct caggccagcc ttttccgctg aacgcaccgc 25860
cgaggcctgt tttgtaaaca ttgatgtatc cggccggatg tccttcatat acagccgtca 25920
gtgagacaca tttcccttcc gcctggtccc tgagtcttgt caggtatttg gagatgtctg 25980
aatgccagcc ctgttccgcc tcttcatctg taaagatttg agcgtctgct tcttccatgg 26040
tgcggatcac cagtatgccg tcactgtaat aaatcatgtt tcctcctgaa aaaagctgtt 26100
tccgtcctta attcaattcg tccagcagga tgtagtaccg gagtttctct tcatctttcg 26160
gaaccccggc agcccggaac agatcatccg gatcaatatt cggacagact gttccatagt 26220
gtccgtcgct gttatgtttg aggcttctcc agcccagggc cagatccatc cagcggtcac 26280
cggctccggc attgcccacg tcaatcagtc ccgcaaaacg cttgccgtct gtgaagatgt 26340
tcggcaggca aaagtccccg tgcgtgacca cgcggtcctg cggaggcaga tgggttttca 26400
gccagtccag gagtttttcc gggctttcaa agccgcccgg gccgaaggtt tccggttcgc 26460
agccggaagg gtcaaagcgt ccggacagga ttgccgcttc agcatgggaa aggttatccg 26520
ccacagttct ttcaaacggg cagcctgcca cgggaatcgt ccacaaggta tgcaaggctt 26580
cagccataca atccagcagc agtgcgggat ggctcatcat atctgggtca cacagctctt 26640
ttccccggat gcgcgtcata agcagccagt cctgtccgtt cagcacctca tgggccgcga 26700
cctgcggaac aggcgccttt ccggcaagcc actgcagaac cttcacgtct gccgtatccc 26760
agccatccgc cggacggatt ttcagcacgt agtcatcaaa aagcaatacc tgtgccccgg 26820
attttccaat cctgtccgtt ttccctgcct tgttgcccag gacagcagcg atcgaagccg 26880

gtgcctgatt catgatgtct ttcatatccg ttcctgattc tccagcactg ccagatctga 26940
ccgctcatac cggttatggg tacattgaag gaaaaacccg ctccgtcagc tcctgttttc 27000
ctccccgaaac cggaagatgg ttggcccggc agaaatccgc cagttcttcc ttcaggtaat 27060
aatactcccg gaaggtttgg ctgtccatgc cgggaaccag ttccggccgt agcttcatgt 27120
tgtcctcctg atgcatttta tccagtgcgc ggctgtccgg aatgcggcaa atacattcgt 27180
tcccatcgga ttcactcttc tctcacatac cgggaatact gactggatca tacctgcagt 27240
atcggaacac tgtcaagttt acattccccg gacataatca gtgaaggcat caaacagcgc 27300
ctgtcccgga ggatcctttc ggatcgttcc ggggcaggcg ttttgttact gtttattcca 27360
ttcttcctgc agccttccgg atcaggcgta tgccagcgtt tccatcatcc ggcggaatgc 27420
ttccctgtcc gtttccaggt tttcttcccg gccgacgcgc atgaaagcgt aaatggtcct 27480
cccgtccctg accaggcagt tgctgcagac ctgcgtcatc ccttccttgt cataggaaaa 27540
gcgatagcct tccgctttct catcgccggc ctgcatggag aagaattcca ggaaccggta 27600
accatgcgcc gcatgggcct taccggtcag ctgctcgttc ttcttcacga tcgctttcag 27660
gtccagcgtc caggagagga gcgcgggata ctttttccac agggccagaa taatcacatg 27720
gttttccgca tcccgtacgc cccacctgaa cggatccccg gcgttcctgt acattttgcg 27780
caggtcgtct tccgtcagga tttcaaagcc gtcaggaatc tgcagtgcaa aagtatcatt 27840
gattatctgt tgtttcatca tgatgcatca gccctgcgat cccagcatat cgtactccag 27900
ctcgatgccg ttgtccgtct caaacagcac tttgccatct gccgttttgg agaagaccgc 27960
gtcgtacgtc tctccgtctt cctccggggt atattcacct tccgcgttct tcctgaacac 28020
tgtgcaggta ccgttcgcgg tcagcttgcc ggtcgccgga tcataggtgc cgttcatcag 28080
gaaagaagag tacgtatcgc cgtccgtttt tccgcggagg atatatacgg tgtagaacat 28140
ctcatcagcc gttgtgccat tccacatgaa ctctgcgtcc gtctcttccg cttcgttctt 28200
ccacacgccc tcgaaccggc cgatattggc aaacttcagg cccgcgcccg cgtcatcgtg 28260
gccgtctttc catgtcagga agccgtccgt gccgatcgtg aactcggtac ccttctctgt 28320
atcaagtcct tcgtacacct tttcaccgat aaccttgtcg ccggtgtccg gattaatccg 28380
gtaatccgtc cgcgaggagg aaacggaaga caggctgtcc gtatcctcat gatagtagca 28440
ggagtattcc cagatggagc cggttccgtc atcctttgta atatccagcg tcacccggta 28500
accttcttct tcatagacga tctgcgcaag accgccggcg atcgcccagt cgctctcgaa 28560
tttctttccg gcttcgggct ggggagcctc gtccgccgat gcggtgcagg caatcgcgac 28620
cagcatcatg attaccagca ggatactgat catttttttc atttttgttg cctcctttgt 28680
gtcctgacag tatttggctg cttttcagtt ctccatgatc ttgatacgta tttcccgcag 28740
gagatggcag tcaattctgt agaaatttca ggccccgcat catctgacag gttaaatgcg 28800
gccgatgagt tttcttcat gtgccgaaag gctgcatttt gctcttgaaa tccataaaaa 28860
atgttttata atagatcgtg tgtttgatat ggctaactca agcgcacaag gcgaaagagg 28920
tccagtcgac tgcaaatctt cttttttttgg tgaaaggtta gttaaggcta attgccaaac 28980
gtttccgttt gcactgaaaa gaagcggcgg catgatactg gcacaggtcc gtttcctcgt 29040
gatcccttga tggaaagcat ccgtccggga aagcccggca gccggcggga agattccggc 29100
ggccggggct tttcccttct ccccgccatc gcgcctgaaa aggctgtgga cctgggctgc 29160
ggcggcgggc gcaacgccgg agaactgctg aagaaatacc cgaaggcgct tgtgacggcc 29220
atcgattatt ccggtctgtc cgtcgaaaaa gcgaaagctt ttaatcaggc ggccatcgac 29280
aaaggccgct gcatcgtgcg gcagggcgat gtgtccgacc tccgcctgcc cgccgtctcc 29340
ttcgacctgg ccacagcctt cgagacggtt tacttctggc cggggctggc ggaatgcttc 29400
gggcaggtgg caaaggtgct gaagccgggc ggatatttcc tgatctgcaa tgagtcggac 29460
ggcctggacg atgccggcaa aaagttcgag aaaatcatcg acggcatgcg gtgccacaca 29520
gcggaggaaa tcgaaaaagc gctccgggca gccggcttca ccgaggtgac gcgctatcgt 29580
tatccgtcaa agccctggat tgccgtgctg gcgaagaagt gagcagaaaa gcgtatcatg 29640
aaaagaagat cctcggtctg tccggcagtc tgtactgaca cggcgaaatc atccaaaaat 29700
gaacggaggc tgataacatg aatccggatt acaaaagctg gatcctgatt cccttcgtcg 29760
gcaccagcct cggggcggca atggtgttct tcctgaagaa gcagatctct gccggcttgc 29820
agaaggttct gaccggcttc gcggcaggtg tcatggtggc ggcctccttc tggagcctgc 29880
tgcagccctc gctggaaagc agtgcttcca tgggccgtct ggccttcctg cccgcggcgg 29940
tcggcttcct gatcggcatg ggcttcctgc tggcgctgga catactgact ccccacatgc 30000
acatggacag gcagcacgag ggcccgcaga gcaggctcag gcgcaccacc aaactgatcc 30060
tggcggtcac gctgcacaac attcccgagg gcatggcggt gggcgtggtg tgcgcgggtt 30120

tccttgccgg agataccggc atcagcgccg ccggcgccct ggccctggcc atcggcatcg 30180
ccatccagaa cttcccggag ggcgctatcg tctccatgcc gctgctggcg gaaggcatgc 30240
caaagggcaa aaccttcctg atgggcgtgc tgtccggcgc ggtggagccg gtggccgcgg 30300
tcatcacact gctggccgcg ggactggtga ccccggtcat gccctatttc ctgtccttcg 30360
ctgccggggc gatgatgtat gtggtggtgg aggaactgat ccccgaaatg tcggagggca 30420
aacactccaa taccggcacc gttgcttttg ccctgggctt tgtggtgatg atgatcctgg 30480
atgtggcact gggttgaaaa ccggcatatt cgccccgctg atgatactgc cgggtgacat 30540
gcttcgccgg cgcggctgct attcaaaaag caccgtcagc cttccctgag tactgtccat 30600
aacgaaaccc tcctgtaccg gttgatgatc acagaaacgg acagcccgga ggggcagttt 30660
acttccctgt gccttctggc acagacggcc gggctgcctg ccggccctgg cataccgcag 30720
ccggtacata ttgcgttttt attctgtgat gatatacggc gcgtttccgc gcgttatccc 30780
tgcaggactg tatccttcga ttggtcacct cctgtcaggt gtttttttcgg tccctgccgc 30840
aattgtttat tccagcaggg ttctcagggt ttccatcatg gcgggaatat cgtagggttt 30900
tgccagatgg ccgttcatgc cggcctccat ggcagtcttc cggtcctcct caaaggcatt 30960
ggccgtgacg gccacgatag ggattgttgc tttccgggga tcttccagcg cacggatctg 31020
tttggccgct tcataaccat tcattttcgg catctggata tccatcagga taacatcata 31080
tgtccccgcg ggagcgtctc tcattttctc cacggcgtcc gttccatcac cgacaacgtc 31140
aactgttaat cccgcttctg tcaggattgc ttccgcgatc atctggttca tttcattgtc 31200
ttccgccagc agtatgcgct tcccgctgaa atgtggcagc acaggtgttt cttccttctt 31260
ttcctgtgaa tcccgtgatg ctgtaaacga ctttgcgaga atattcctca gttcagacat 31320
aaacaacggt ttggcgcaga aggctgtcac cccggctttt ttcgcttctt cctcaatatc 31380
cgtccagtca tatgcagtca ggatgataat cggcgtattg tcaccgatta ccttccggat 31440
cctccggact gtttcaatgc cgttcaggtc cggcatcagc cagtcaatga tatagacccg 31500
gaattcgtcc gcctgatcca tcgcttcttt cgcccggacc acggcttcct tcccgtaatt 31560
cgtccagtcc gggcgcatcc cgatttccct cagcattgag cagatggaca ggcatgtgtc 31620
cgtatcgtca tcagccacca gcgcacggag gctgttcaat tccgtcagcg gttcgaactt 31680
tgcggatgcg ccgctgattc tgcaaggcag ctggacaatg aattcacttc cctttccgac 31740
ttcagaattc actgtgatca cgccgcccat catatctaca atgcttttgg tgatcgccat 31800
accaaggcct gtgccctgta tgccgctgac tgtgcttgtt ttttcgcggg tgaaagcatc 31860
aaagattgtt ttctggaatt cctcgctcat accgattccg ttatccttga tccggaattc 31920
aaagttggcg aagtcggccg ccggcgaagg cttttcgatc acccggaagc tgatgatacc 31980
acctgcggga gtaaacttaa tcgcgttcga aaggatattc agcagcacct ggttcagccg 32040
gagcttgtca gtaatgatat cctcatgcct gacatcctgc gtatcgataa acagttccat 32100
ctgtttggac aggacattgg actggatgat cgtgcggagg tcatggataa tatccggtaa 32160
atgcacctcc gcttcttcaa tcttgacctt tccgctctca atccggctca tatccagcac 32220
atcgttgatc agcgacagga gatgctggct ggaaacagaa atcttcccga gataatcctg 32280
cacctgttcc ctgttgtcaa tatgggacgc ggccagcgcg gtgaagccta tgatggcgtt 32340
cataggcgta cggatatcat gggacatgtt gttgaggaat gtcgtcttgg cgcggttggc 32400
atgctccgcc gcaatcagtg cgtctgacag cgcctgctgg ctctcagcga gttctttatt 32460
cttcttctcc agttcaagcc tggctgtttc cttttccccg atctccgctt tcttgtgttt 32520
acggtcccgg accagcagga agagaatgat cgcgaatacg gccaggattg ccgtgccgac 32580
aatcgcctga ttgtctctga tggcatctat aaagtcgtaa gagtacaggc cgccggtata 32640
acggtatgaa atattctgcg cgtagtcgct gccaaggatg ttgattcccc ggttcagcag 32700
tttcagcagc ccttcgttgc cgattcccac accaaagcac cgggcgtcat tgtaactggt 32760
ctggcggagc gaaagatctt cgtatttccg gttacgcaga atatcattgg ctctcaggcc 32820
gttcagggtc gcgcatcccg cttttccgga cagcacggct gccaggcagt cctcgctcga 32880
cggataaaaa gtgatttccg cgtccgggta attcgttcgc acaaaataga actgcatgcg 32940
gttattttca ttcactgcga aatgcgctgt tgtctcatcg ctgaattcgc ctttgtaaac 33000
cagttcggcc ggcgaggaag ataccggatt ggattgatag ataccgctct cttcggaata 33060
ataaaggccg ccgctgaccg ggaaagccac gtcaatttca ccgctgttga tccctgcaat 33120
catatcatcg tagcttctgt agcctttgta tgtgacggaa atgctgtcca ttcctaagga 33180
agacaggatt ctgggaatga tatccttcac gataccggtc gcgcttccct gccggtccgt 33240
gtcactgtat ggcaggtaat tttccagata accgacctga agcgcgttat gattatccat 33300
ccagtcccgt tccgcccggg aaaacgctct ggccgtaacg ctgactgaat aatacttcgc 33360

gctcagggag ttcaggtaat tcggctcttc cgtcgcaagc agggtttgtg ccgagttcag 33420
ttcttccagc aggtcgggcc ggtgaatatt cacgcagaga taataatcgg atgtgccgaa 33480
aactgtcagc acctctgagt gttcccttcc gcgcgcgccg tcgctttccg ccgccaggac 33540
gtccacatca cgggaatcaa aggcgccgaa caattgtttg taatcgggat aagcgatcac 33600
ttccgccgtt acatgatgct catccaggta ctgattcagc acgctgacca tggcgctgtc 33660
cagcactccg atcctgtatc cgttcagcgt tgagggctca gcggtaatgc cggcatcggt 33720
atcatgcttt ataagataat aagattctct gcccataaca gccttcggat atccgataat 33780
cgccgcccgc tcttcacgcc acgccaggcc ggcgaggaga tcgatttccc cgtccaggag 33840
catctggtac agttcgccga attctccgta aacatattga taatcccagc ccgtatactc 33900
agaaagcttg cggtaatact cataggcgta gcccgtcttg accgagcctt cccgggcgcc 33960
ttcctggaac acctcattct catagtaacc tacacggacc gtttttgttt ctgccgtgac 34020
ggaaaccggt aaagtcatca ctgaaagcaa agcagcaacc gtaataatca ttgcacagat 34080
tctgcggata ttgcgtcttt ctgaatagct catgttcggt ctccttatat ggctgttgcc 34140
cgtttgttcc ggaaaagccg gcactcatgc tgtatcctga gttgattata taaaattatt 34200
acaaaataat ccagtcaaat tccccgtctt ccgccgcata gatccggtcg ccgtttccgc 34260
accggaaccg gtccggatcc ccgtctccgc agcccttcac atcatccgtg tgatatatcg 34320
cccgcagcat ttccgggaac agttcacgga ttgttccttc gtccgcctga acggcggttc 34380
tgattttata ctccatcgtt attcctcatt ccctccttca atccccggcg tattcatatc 34440
agcctgttta ttccctgcgt gattctttct gatcgtattc accgaacaga acgtcacaaa 34500
cggagcgctg cctgctgcgg aaaccgtgat tgcagggatc tgcttcgtaa tgtgtccggc 34560
aaaccgccga aaacctgtta ttcagcaaca agaaccggct cagccgggca gccgaggaat 34620
gcgatcgctt cctcatcatt atatattttc tgggtgttaa taacagcatt cctgggccgg 34680
ttctgcgcct gggactgttt ttttctgtac tctttgggag aacagcccat tctttcatgg 34740
aacagctttg tcaggtaacg ttcatcggaa aaaccgcttt ccaccgtgat ttcaagaatt 34800
gttccatccg tattctcaaa aaggctgagc gcgtgttcaa accgtttttc tttcagataa 34860
tcctgaaaag taatcccgag cgtatcctta aacaggtggg atacatgggt cagcgtcaga 34920
ttgttcattt cggcaatctc ttccagcaga agctttcgct ggtagttctg atcaatatag 34980
tctgtaatgg aagcaatccg gtccattcgt ctgcggacgg tattatagtc cttctggctg 35040
agggaagcga tggggatatg gcgtctcagc aacaatacaa tatgggaaat gagatggaag 35100
cagtcaaact caaaaccgtc cggtctggca tgatagtatc tcgccagttc catgcagtca 35160
ttgcgcagat tctgccagac attctgtttc tcacgaaaaa agccataaac aggcggttct 35220
gtcataaaac gcaggagagc cgcgtcagaa agaaacggca tcagtatttt gggagagatc 35280
tggattgcca gaatcagggt gtccggatca tccgccacaa attcatgggc ttccatggag 35340
ttgatcaggt acatctcatc ccggtccagg ctccaggcgt tttttccttc ccgaaccgtc 35400
agcttccctt ttaataccag gccgagttcg atttctttgt gtatatgcgg tgttcttgac 35460
gccagattca ccaaaaatac gtgcaggtat ttaaaagtct cctgggatac cacttcaaat 35520
tcctttttca tttttcgatc ctcccacctg agatcataag atccggcgaa aaaaacgtca 35580
agcataaaaa tcgcaaatta atcggtgttt tttcgcaaac taatacaggt ttttgttcaa 35640
agattctgct atgattcaga cagatgaagc gaaaagcttc catcaagaga aggaggatgc 35700
agcaatgtct cagtttaagt tttctgtcgg cccctggaat gttcataccg gtgctgattc 35760
ctatggtccc gaaacccgta acgccatcga tctggaaacc aagttcgccg ctttcaaaaa 35820
gctcggcttt gacgccgtcc agttccatga tgacgatgct gtcccgaaca tgaacaatct 35880
ctctgaagaa gagatcaaga ccgaagcgcg gaaggtcaaa gctctgctgg acaagtatga 35940
cctgaaggcc gagttcgtcg cgcccccgcct gtggatggat ccccacacca ccgacggcgg 36000
attcacctcc accagcaagg aagaccgcga gttcgccatg tggcgtgcct accgctccat 36060
cgacatcgcc cgggagcttg gctgcgacct gatcgtgctg tggcttgccc gtgaaggaac 36120
cctgtgcgcg gaagcaaaga gccctgtgac cgccacgaag cagctggtcg aagcgattga 36180
caatatgctg cgctatgacg caaccatccg cgtgtgcatc gagcccaagc ccaacgaacc 36240
catcgatcgt tccttcgccg gaaccatcgg ccatgtgatg ggcatctccg ccgccacgat 36300
tgatccgaag cgggtcggcg cgaacctgga aagcgcccat gcgattctgg ccggactgga 36360
tcccgccaac gagatcggct tcgccctggc gttcggcaag ctgttcaccg ttcatctgaa 36420
cgatcagaac ggtatccgtt atgaccagga caagtccttc ggcgtggaaa acctgcgctc 36480
cgccttcaat caggtgaaag tcctggtaga gaacaactac ggctccaggg gtgagtatgt 36540
cggcctggat gtcaaggcaa tgcgcaccac ctccgacgag gacagctacg agcacctgaa 36600

gaactccctg gcgatcttca aggccatgga agaaaaggtg aagaagtatg actatgatta 36660
tgccggcaaa ctgatcgcgg agaaaaagtt cgagcagctg gaaatgtata ctgtcggcct 36720
gatcatgggc gaagccctgt aaaccgccag tggcggaagt ctttgcaggg ctgaggtcat 36780
ccgaaacagg cagaggcgga tgtaacgcct ctgcgccgat tgaggatgcg gagatcgact 36840
gataaacgct tcagcctgct ggctgcggat gttccaaagg agcatgcagg cggcaggctg 36900
tttttgaaag aaagcagagg tcaggaaaat gggcaaagtc atctcagccg ggcatatttg 36960
tctggatatt acgccggttt ttccctccgg gcacagatat gaccggatcg gcgacctgct 37020
ggtaccgggc aagctgatcc gcacggatgc cgccagtgtt catacaggcg gcagcgtggc 37080
aaataccggc cttgcgctga aactgcttgg aaacgacgta acgctgctgg ggaaagtcgg 37140
ctgcgacgcc tttggcggaa tggttcagag tatcctggcc ggatacggag ccagcggact 37200
gatcgcggat gagcagagct ccacttccta ttccgttgtg ctggctgttc cgggaatgga 37260
ccgtattttc ctgcataatc ccggcgcaaa cgacacattc tcctcttcgg atattccgga 37320
ggaaacgctt gcggacgcgg tcctgttcca tttcggttat cccacgctga tgaagcggat 37380
gtatgaggca ggcggcgcgg aactgaagaa cctgttcatg cgggtaaagt caaaaggaat 37440
cgccaccagc ctggaccttg cagctgttga tccgcagagc gacgcaggga aggcagactg 37500
gccggcgatt ctgggcagtg tgctgccgta tgtggatttc tttgttccca gtttcgagga 37560
gctctgctac atgctggatc gttcctgcta tgacgcgctg gccgcaaagg gcggcgatat 37620
gaccgagcag gaaggcattc tgcaaacagc ggatatactg gcacagaaat gcctggatct 37680
gggatgcggc gccgtactca tcaaatgcgg cctgtccggc atggtttacc gtaccggctg 37740
cagggagcgg atagcacagg tcggcgccag gctggaactg tatgcggatg aatggacaga 37800
ccggtcaggc gcacagccct gttatcgggc ggaagttgtt cgcagtgcca ccggcgcggg 37860
cgatgtgagc atcgcggctt acctgactgc cctgatgagc ggcgaaacac cggaggtctg 37920
cgcgaagatt gccgccgcgg aaggtgcggc atccgtcagc agctacgacg ccctcggcgg 37980
agtcctgccg ctggatgaac tgaagaaaag aatcgcggaa tacgaaggat gaagatccga 38040
tgactgccgg tggcaggaat ctcatcgaat ctgaattcat tcgaaacgag cgggctgccc 38100
ggtgggcagt cgcgaagatt aagaatgcgg aggaaagaaa tcatgctggt aaacatgaaa 38160
gaaatgctgg ccaaagcgca gaaggaacag tacggcatcg gtttttttcaa tgcggtgaat 38220
gtggagatgg cccgtgcggt cattgaaaca gcggaagaac tgcacgcgcc ggtgatcgtc 38280
ggcacagcgg aagttcttct tcccgcgatg tccctggaac gggttgcgga atatctgatt 38340
cccatggcac agaaagccag tgtgccggtt gcggtgcatt acgaccacgg gctcaccttt 38400
gagaagtgca tggaagccct gaagctggga ttttcctcag tcatgtatga ctgctccacc 38460
gcttcctatg aggttaatct tgatcgcgtt gccgagatgg tccggatctg ccacgccacg 38520
ggcgtaaccg tggaaggtga actgggtcat gtgggcgata acgaaggtgc ggggaaactg 38580
gagaacccct ccgattacta taccgatccg gcagtggcgg cggattttgt acagaaaacc 38640
ggcgtggacg cactggcggt ggctgtcgga aacgctcacg gcgattacag gttcccgccg 38700
aaactggact ttgaacggat caccgcgatc cgggaagcaa ccggaacacc gcttgtactg 38760
cacgggggca gcggcctttc cgacgacgat ttccgggaag cggtcaggcg cggcgtctgc 38820
aagatcaaca tctttactga tctggacaaa gccggtaaag ccgggatcga ggaaggcctg 38880
aaaaaaggag cggccagcat gatgggactg atcccatatg agatcgaagc gatgaaagaa 38940
gtggtccgga ataagatcgg gctcttcggc tccatcgggc atgcctgaaa cagaaattgc 39000
ttttctgaca aggaacaact gatcacacat ggaggcagaa gcaatcgcgg catagtattc 39060
cggatgatcc aggttcaggg cgatgtcaat ttcccggaat ttgccctccc gggacaggaa 39120
gtgcagtttg acggcagtgc tggttctgct gaccggcagc caaaggaatc cagcggtcac 39180
agaaaggttc cgatcctttc cggatcgtta tccgggctgg cgcaagcgat acggggcgat 39240
cctgcgtgcc ggccgtatcc gtttatctga gcattgaatt caccgcgtca atataaactt 39300
cgctgtgttt ggtatggatg acatgatcgt ttgtatccgt ttcaaccagg cggcagtttt 39360
ccccgatata ggataccgcg cgttcagcct gctcgcgggt tgcgg 39405

//

ID XXX; XXX; linear; XXX; XXX; XXX; XXX.

XX

AC PRJEB67889

XX

DE Uncultured bacterium DNA fragment from in vivo enriched cow rumen, clone rum-21k7

```
XX
FH   Key             Location/Qualifiers
FH
FT   source          1..13661
FT                   /organism="uncultured bacterium"
FT                   /environmental_sample
FT                   /mol_type="genomic DNA"
FT                   /isolation_source="Cow rumen"
FT                   /clone="rum-21k7"
FT                   /note="AZCL-xylan, AZCL-arabinoxylan active clone"
FT   CDS             complement(266..1906)
FT                   /translation="MKKLPVFLIPALLTGVASAEAVLDETAAAGEAAVGFFDKLLNWIT
FT                   VGNDAVNGVVWGVPALVLLAFVGILMTVLTKVFQVSHFGHWFKKTIGAVFKDKHVTSHT
FT                   KDKSISQFQSLCTALAATVGTGNIVGVSGAIMVGGPGAVFWMWVIAFFGMMTNYSENVL
FT                   GIFYRRKNSKGEWSGGAMYYLRDGLGSRKGCKTIGKVLAWLFSLFCLIAAFGIGNMTQV
FT                   NSISENMLAVFKVPTWVSGLVIMILVGLVVVGGLKRIASVTEKIVPFMVILYMIGTIVI
FT                   FILHINQIGAVFSSIFRSAFAAQSAAGGIVGYGMKLAIEQGMKRGVFSNEAGLGSSVMV
FT                   HSNSNVKEPVKQGMWGIFEVFADTIIVCTLTAFAVLSSGLVNLETGVTAATFTNHAGEV
FT                   IELSKANLVSSVFNQQFGFIGSAFIAVSIMLFAFSTVLGWSHYGTKACEYLLGEKRTIP
FT                   YRVLFVLLVFGGAVMGNNLAWDLADTFNGLMMIPNLIGVIALSGLVAKITKNYVNRNRR
FT                   GIKEPPMYSAFPDVQAEQEQAPDSPEDMA"
FT                   /product="Sodium:alanine symporter"
FT   CDS             complement(1997..2116)
FT                   /translation="MSFQLYATGYRRNPRVSAFLVFFQPYSIPERLTGNGSAK"
FT                   /product="hypothetical protein"
FT   CDS             2123..3076
FT                   /translation="MAVVKRSFQIRQGGNGLKIYFAPMEGVTDGILRQTHHRIFGGVDV
FT                   YCLPFHKLTQSLSLLTREIRDISPDENEGLNVLPQALTRDPEQLSAWLYYVSECGYTTA
FT                   DLNIGCPSATVTKRGRGSGMLRDTAYLRSFFDRVFSNTLPVKLSVKTRIGWESPEEWPG
FT                   ILDILASYPFAHVTIHTRTTREQYTGRTHPEAFELAVRKGIPHPVYNGDLRTTDDVKQL
FT                   TERCPETETVMIGRGLLADPALARRLRGGAEAGTDELEAWYNTLYWGWKDRFGETIALG
FT                   RIKKLMEWPVGEDIRRKRLLRRAGDIENCVRAAIGE"
FT                   /product="Probable transcriptional regulator"
FT   CDS             3191..4477
FT                   /translation="MKMKRTFTRMAAAVTAVFLFAAMILTGGSGIMTSASAENGTPVYS
FT                   SSDLFTDRDLKQTADLDKAVTLTAEDGKEIRITEAGTYVLTGTASNATVYVEAGDDDKV
FT                   QLVLQDLHITNSDFPCIYVYNADKVFITTAADSSLTVNGTFRTDGSINTDGAVFSRTDL
FT                   VMNGTAELTIQSMDNGVVCKDDLKITGGSYTITAGSKAFEANDSIRIKDGTFSLRAGTD
FT                   GFHAENSDDDSKGYIYIGGGTMTIACGDDGIHATSVVQIDDGTLTITAREGLEGSFIQV
FT                   NGGTINITASDDGINAARKSTAVEVAVEFNGGTVNVTMGGGDTDGVDSNGSLYVNGGVI
FT                   SVNGNSTFDYDGMVSFNGGTVYINGQQASSIPNQMMGGGGRGGWGDGRNGGWNMPQDNG
FT                   GSTPPADGWNGSTGDNGGRGGRGGGRRGW"
FT                   /product="hypothetical protein"
FT   CDS             4499..5236
FT                   /translation="MGTETGSAIQVMKRYELKYLLDRGQTAFLVSCLRGHMEPDRYGLT
FT                   SIASLYYDTPDFRLVRASLDKPDFKEKLRLRSYGIATEESPVFLELKRKASGIVYKRRV
FT                   QTTLPDVRRFFAGGNGLFDDRQISGELEWFRDSYGLLRPACLIIYDRVAYFEPGGDLRL
FT                   TIDFNPRYRTDAMNLTSSMDGFPLLPDGDTILEIKVQGSMPLWLTAILDAGQIYKTSFS
FT                   KYGEAYSRQSVGPAVLPGFRIAG"
FT                   /product="FIG01032461: hypothetical protein"
FT   CDS             5278..5949
FT                   /translation="MFNSIYSASVTPVQLLLTAAAALCSGVLYSWLISFRIRANRRFFI
FT                   VTSVLPLVVGLVLTFINGNIGAGIAFGGAFALTRFRSAQGTSDEITAVLIAMAAGCAFG
FT                   MGYAAYGVLILVCMGLILLALSGVKLFDHSSMHSEKLLNITIPESLDYTRAFEEPFARY
```

```
FT                   LEQAENVGVKTTGMGSMYRLSFKIRMRNPAEEKKLIDELRTRNSNLEIAVLPYAEPQLQ
FT                   L"
FT                   /product="FIG01032596: hypothetical protein"
FT   CDS             5974..7236
FT                   /translation="MKNKHRIIRNLLLDAAAAGLVLVVFAAFHHVLPRAQQGLNVQIVK
FT                   PDSTAETAVAEIREAETAAEIPEAKKAEEQPAEMKAEAETEPDTATSATQVEEEKTGFR
FT                   PTVVTSASSGTADSGRRGPSQRPGSRKSGRGGGNMMNSKGGSGLDGEDGTAEDSAGTAD
FT                   STPLAEKFADKFTESVTVTGNSYTSPDISITVDEVNDGNLTYYIADIYVRDITCFQTAL
FT                   AQDTYGSGFRDSIEDMAVLNNALLAVNGDYYGNTSEGVVIRNGMIYRANPTDCDICVLY
FT                   YDGTMKVMSGDSFDLEDAIEDGAWQAWTFGPALLDIGGQPIASFSSTNRIISANPRTAI
FT                   GYYEPGHYCLVVVDGRGESAGISLPRLSQLFYDLGCTAAYNLDGGNSSIMVWGNEVINN
FT                   PSGGGRESSDALLIAEVRNYE"
FT                   /product="hypothetical protein"
FT   CDS             7229..7423
FT                   /translation="MSKVQGLFNHLTVMFGMVFLVFLVLDQFNPMMNFVDNSISRWMLA
FT                   AFCICGISRSILCWRKNDS"
FT                   /product="hypothetical protein"
FT   CDS             complement(7447..7581)
FT                   /translation="MNISRRFVYGIYPEWKVYVIPRNTLIWRSAFSFIPFVIAVFSSK"
FT                   /product="hypothetical protein"
FT   CDS             complement(7614..8531)
FT                   /translation="MKRRISVQTDGIFQPCGVDEGMRLISRAGFDCVDLDLIYISSGSI
FT                   RKGIIEGEFTLPEEQFMEEIIRPIRDAGRKYGVGFGQAHAPFPSWIPLDEEMNAHLMQA
FT                   FRMCLDACRFLDCPYLVIHPAFTDYRNRLDAAAEHEINIRLYSSLIPDAKRTGVKILLE
FT                   NMFVSWKAKRVEAVCADMGEACAYIDELNAIAGEEIFGFCYDCGHAALLGKDQGKAIRT
FT                   LGHRLLALHLHDNDGVSDSHRFPYYGIIDWETVLRALHEIGYSGTLNFETFNEMDSHDP
FT                   ALIPKLLELLGETGRLFASKMDGE"
FT                   /product="hypothetical protein"
FT   CDS             complement(8582..9889)
FT                   /db_xref="GO:0004557"
FT                   /translation="MPKITFMGAGSTVFARNVLGDSMLTPALEDSVISLYDIDPQRLKE
FT                   SEMMLTAINNNLGGKAKIEAWCGVENRKAALRGADYVVNAIQVGFYKPCTVTDFEVPKK
FT                   YGLRQTIGDTLGIGGIFRALRTIPVMFDFARDMEEVCPNAWFLNYTNPMAMLSGAMQRF
FT                   TGVKTVGLCHSVQHCVPKLMHLAGMEYDSRVQWKIAGINHQAWLLEVTKDGEDLYPEIK
FT                   RRCAAITEKHDDMVRLEIMKRFGYYVTESSEHSSEYMPYFIKDKYPELIDRFNIPLDEY
FT                   PRRCENQIARWEKQRDELIANPNLTHTRSNEYASRIMEAMEINEPYKFGGNVLNTGLIT
FT                   NLPKDACVEVPCFADASGVTPAFVGDLPEQCAAMNRTNINVQLLTILAAVTLKKDYIYQ
FT                   AAMMDPHCQSELSIDDIVAMCDDLIEAHRGWLPEYH"
FT                   /product="GH4"
FT                   /EC_number="3.2.1.22"
FT   CDS             10181..11734
FT                   /db_xref="GO:0031176"
FT                   /translation="MKKLAIILLAVIMMAACISPAMAARYYTSRFAAGTDGWYARGAQS
FT                   VYRTTEGTLRTEGRTSDWHSPGRDFVLVEGCLYKLSVEVYQAEADSAEFMISVAHSKDG
FT                   NETYENPARGTVKKGEWTKIEGTYTAGNYDRSVLYVETLGAPQLSFEIRNFIVDAPNGI
FT                   PKPQPAEPPMEIEAVAIEDMPSIREAYEGKFDFGAAVPQQAFMNKQLRALMAKQFSILT
FT                   PENELKPDSVLDVQASKSLVHNTGDQTAVAVHFDAAKGVLSFAKANGLKVHGHVLVWHS
FT                   QTPEEFFHEGYDRSKPLLSREVMLGRLENYIREVLTQTEEMYPGVIVSWDVVNEAIDDG
FT                   TNWLRKTSPWTRTIGEDHVARAFEFARKYAAEGVLLYYNDYNTPVPAKLAGITKLLKQL
FT                   MAEGNIDGYGFQMHYSNGDPSIAQISNAVSQIASLGLKLRVSELDIGAGTSESALMQQK
FT                   ARYKEIMQLMLQYADQTEAVQVWGLTDGMSWRDGQYPLLFNDQLYPKPAFYGVLEAAEE
FT                   "
FT                   /product="CBM22-GH10"
FT   CDS             complement(11704..13425)
```

```
FT                   /translation="MRLAHPGQLRAFRDLHVVAPEQLQVLPEQVAEDIGVHAAALFDDP
FT                   IHLTLVPCFPQVRHEVVVIGIAHDVGEHPAAESLIRAGLGQVVGQQQFVHRGRRFHQVG
FT                   GKPGVIVGLGSVGVHALPGMPQLMGQGEYVRHPVVPVQQDEGVLPVLAGTEAARALADV
FT                   PGEIHPAFPVGPVYHFHIILTEDRQGIPDVFMRFRDRHLRVEPGIHRDLQVGKGHFVQP
FT                   VDLFQEPGVPAHMRRQALGDQVDLAVVHFPRYLLVKQQAVKHVVKVPELGHDLVLGHVG
FT                   VVRGGRGVPVFRDRVGEGLESRPAHRGVRFQLVVVLEEGIAQRIFLSVHGEGIKGDIRQ
FT                   LHRIVYGVGHIQRFAEGRQHLLGFRGQHMRLSGNQVLQVDPASLCLLHRGLQHVVRRQQ
FT                   DFTVHEGRVPLQADHLVAHLFIQLAQLVVRVVRFHRQPRIAPDFRRSNPDLLHLVQAID
FT                   NILRGVQFTGICIDNFFHRCLRFFQGIPHLLFVFKNQGKVPCFLHGYVFPGTQFHIFPL
FT                   SRPSGRILFLLPAYLSYRRCGISSIFFPPPGYRKSAASAALSCKRGAYSSAASSTP"
FT                   /product="hypothetical protein"
XX
SQ   Sequence 13661 BP; 3089 A; 3720 C; 4255 G; 2597 T; 0 other;
     cccgccgtga ttctgaaccc actgtgggtc cgaatatccg caatctcaat cttcgcactg        60
     ctcgcttgga gcgcttcgca ttgctcgttt cgcaaggaat cgcgcgcgca ccgcgggcgg       120
     aattcccgcc tcgcgcaccg cacacggcag gactcgctca ggtcgctttt gtcttcgaca       180
     aatgccccac aggggcaccg cttccttcgc tccccgccgg gcaataaaaa acggctgccg       240
     tttccggcaa ccgggaatga acggattaag ccatatcttc cggggagtcg gggggcctgtt       300
     cctgttcggc ctggacatcc gggaaggcgg agtacatcgg cggctccttg atgccgcggc       360
     ggttccggtt gacgtagttc tttgtgatct tggccaccag gccggacagg gcgatgacgc       420
     cgatcaggtt ggggatcatc atcaggccgt tgaaggtgtc ggccaggtcc caggcgaggt       480
     tgttgcccat aaccgcaccg ccgaagacca gcagcacgaa caggacacgg taggggatgg       540
     tccgcttttc accgagcagg tattcgcagg ccttggtgcc gtaatggctc cagccgagca       600
     cggtggagaa ggcgaacagc atgatcgaga ccgcgatgaa cgcggaaccg atgaagccga       660
     actgctggtt gaagaccgag gaaacgaggt tcgccttgct gagctcgatg acctcgccgg       720
     catggttggt aaaggttgcc gcggtcacac cggtttccag gttgaccagg ccggaagaca       780
     ggacggcgaa ggcggtcagc gtgcatacga taatcgtatc cgcaaagacc tcgaagatgc       840
     cccacatacc ctgcttgacg ggctccttga cattggagtt ggagtggacc atgaccgagg       900
     aaccgaggcc ggcttcattg gagaagacgc cccgtttcat accttgctcg atggccagct       960
     tcatgccgta gccgacaata ccgccggcag cggactgggc ggcgaaagcg ctccggaaga      1020
     tggaggagaa cactgcgccg atctggttaa tatgcaggat gaagatcacg atggtgccga      1080
     tcatatagag gatgaccatg aagggcacga tcttttcggt cacggaagcg atccgcttca      1140
     ggccgccgac caccaccagg ccgaccagga tcatgatgac caggccggac acccaggtcg      1200
     ggaccttgaa cacagccagc atgttctcgg agatggagtt gacctgcgtc atgttgccga      1260
     taccgaaggc agcgatcagg cagaacaggg agaagagcca ggccagcact ttgccgatgg      1320
     ttttgcagcc cttgcgggag ccgagaccgt cgcgcaggta gtacatggca ccgccggacc      1380
     attcaccctt ggagttcttg cggcggtaga agatacccag cacgttttcg gaatagttgg      1440
     tcatcatgcc gaagaacgcg atgacccaca tccagaatac cgcaccgggg ccgccgacca      1500
     tgatggcgcc ggaaacgccc acgatattgc cggtgccgac ggtggccgcc agcgctgtgc      1560
     acagggactg gaactgcgaa atgctcttgt ccttggtatg gctggtaacg tgcttatcct      1620
     tgaacacggc accgatggtc tttttgaacc agtggccgaa atgggacacc tggaacacct      1680
     tcgtcaggac cgtcatcagg atgccgacga acgccagcag caccagggcg gggacgcccc      1740
     agaccacacc gttgaccgca tcattgccga cggtgatcca gttcaggagt ttgtcgaaaa      1800
     agccgacggc cgcttcaccg gcggcggcag tttcgtccag gaccgcttcc gcggaagcaa      1860
     cgcccgtcag cagcgcggga atgaggaata cagggagctt cttcatcttc tccagcctcg      1920
     cttttgtgaa ttgttgaaat aaaaacgtct tcgaggattg cctgttctcg aagacgccct      1980
     tgccttccgg aaaagtttac tttgcggaac cgttgcctgt caatcgttcc gggatgctgt      2040
     acggctgaaa aaacacaagg aatgcagaaa cacggggatt ccggcggtat ccggttgcat      2100
     acagctgaaa ggacatgata aaatggccgt cgtgaaacgg tcctttcaga ttcgacaggg      2160
     aggaaacggc ttgaagatct atttcgcgcc gatggaaggc gttacggacg gaatcctgcg      2220
     gcagacccac caccggatat tcgggggcgt ggacgtatac tgcctgccgt ttcacaagct      2280
     gacgcagtcc ctgtccctgc tgacgaggga gatcagagat atttcaccgg atgaaaatga      2340
     aggactgaac gtgctgccac aggcgctgac gcgggacccg gaacagctgt ccgcctggct      2400
     gtactatgtg agcgaatgcg gatatacgac tgcagacctg aatatcggat gtccgtccgc      2460
```

gacggtgaca aaacggggac ggggcagcgg gatgctgcgg gataccgcat acctgcgatc 2520
ctttttcgac cgggtgtttt ccaatacact gcccgtaaag ctttccgtga agacccggat 2580
cggatgggaa agcccggagg aatggccggg aatcctggat attttagctt catacccgtt 2640
tgcgcatgtg accatccaca cgcggaccac ccgggaacag tataccggga ggacgcatcc 2700
ggaggcgttt gaactggcgg tgcggaaggg aattccgcac ccggtttaca acggagacct 2760
gcgtacaacg gacgacgtga agcagctgac cgaacgctgc ccggaaacag agacggtgat 2820
gatcggccgg gggctgctgg cggatccggc tctggcccgc cggctgcgcg gcggcgcgga 2880
agccgggacg gatgaactgg aggcctggta caatacgctg tactggggct ggaaagaccg 2940
gttcggcgag accattgccc tgggaagaat caagaagctg atggagtggc cggttgggga 3000
ggacatccgg cggaaaaggc tgctgcggag ggccggcgat attgaaaact gcgtgcgcgc 3060
agcaatcgga gaataaaggg attccggccg gtctgatccg gccggaaatt ttcttttttt 3120
gagtttaggt tcattttagg aacctctttt acaatggcac catcaacgaa aggaaaggcg 3180
gtgctccaat atgaaaatga agcgcacatt taccaggatg gcggcagctg ttacagccgt 3240
attcctgttt gcagcaatga tccttaccgg aggttccgga ataatgacct ctgcctcggc 3300
ggagaacgga acgccggtat acagcagcag cgacctgttt accgaccggg acctgaaaca 3360
gacggcggac ctggacaagg cggttaccct gaccgcggaa gacgggaagg aaatccggat 3420
tacggaagcc gggacctacg tgctgacggg tacagcctcc aacgccacag tctatgtgga 3480
agcgggggac gatgacaagg tgcagctggt gctgcaggac ctgcacatta ccaacagcga 3540
tttcccctgt atctacgtgt acaacgcgga caaggtgttc atcacgactg cggcagacag 3600
ctccctgacg gtgaacggga cattccggac ggacgggagc atcaacacag acggcgcggt 3660
tttcagcagg acggacctgg tgatgaacgg gacagctgaa ctgacgatcc agtccatgga 3720
caacggcgtg gtatgcaagg atgacctgaa gatcaccggc ggaagctata ccatcacggc 3780
gggctccaag gcatttgaag ccaacgactc catccggatc aaggacggaa cgttcagcct 3840
ccgggccgga acggacgggt tccacgccga gaacagcgac gatgacagca agggatatat 3900
ctacatcggc ggaggcacca tgaccattgc ctgcggggat gacggcatac acgcgacatc 3960
cgtggtgcag attgatgacg gaacgctgac aatcacagcg agggaaggcc ttgagggatc 4020
ctttatccag gtgaacggag gcacgatcaa tatcacggct tcggatgacg gcatcaacgc 4080
ggcccggaag tccaccgcgg ttgaggtggc agttgaattc aacgggggaa ccgtgaacgt 4140
caccatgggc ggcggggaca ccgacggagt ggattccaac ggcagcctgt acgtgaacgg 4200
cggcgtaatc agcgtgaacg gaaacagcac cttcgactat gacggaatgg tttcgttcaa 4260
cgggggaacg gtttatatca acggccagca ggcgtcctcc atcccgaacc agatgatggg 4320
cggcggaggc cgcggcggct ggggagacgg ccggaacggc ggatggaata tgccccagga 4380
caacggaggc agcactccgc ctgcggacgg atggaacggc agcacaggcg ataacggcgg 4440
ccgcggcgga cgcgggggcg gccggagagg ctggtaaaaa aggaaaggag aatccaccat 4500
gggtacagag acaggaagtg ccattcaggt gatgaaacgc tatgaactga aatacctgct 4560
ggaccggggg cagacggctt tcctggtcag ctgcctcagg ggacatatgg aaccggaccg 4620
gtacgggctg acatcgattg catcgctgta ctatgacaca ccggatttcc ggctggtccg 4680
ggcatcgctg gacaaacccg atttcaagga aaagctccgg cttcgatcct atggaatcgc 4740
aacggaagaa tcacctgtat tcctggaact gaaacggaag gcttccggga ttgtctacaa 4800
gcggcgggtc cagacgaccc tgccggatgt gcggcggttc tttgccgggg gaaacggact 4860
gtttgatgac cggcagatca gcggggaact ggaatggttc cgcgacagct acggactgct 4920
ccggccggca tgcctgatca tctatgaccg ggtggcgtat tttgaacccg gcggggacct 4980
gcggctgacg attgatttca atccccggta ccggacggat gccatgaacc tgaccagctc 5040
catggacggg tttccgctcc tgccggacgg agacaccatc ctggaaatca aggtgcaggg 5100
ctcaatgccc ctgtggctga ccgcgatcct ggatgccggg cagatttaca agaccagctt 5160
ctccaaatac ggagaggcat acagccggca gtctgtagga ccggctgtgc ttccgggatt 5220
ccggattgcc ggatgagcaa tccgcaaata cgaaacgcaa gtgaggagaa aaaggctatg 5280
ttcaattcga tatattccgc gtcggtgaca ccggtgcagc tgctgctgac ggctgctgcc 5340
gcactgtgct ccggggttct gtattcctgg ctgatcagtt tccggatccg ggcaaaccgc 5400
cggttcttta ttgtgacatc ggtactgccg ctggtggtcg gcctggtgct gacgtttatc 5460
aacggcaaca tcggcgccgg aattgctttc ggcggggcat tcgcactgac ccggttccgt 5520
tccgcccagg gaacatcgga cgagattacg gcagtcctga ttgccatggc agccggatgc 5580
gccttcggaa tgggttatgc ggcctacggt gtgctgatcc tggtatgcat gggcctgatt 5640
ctcctggccc tttccggcgt aaagctgttt gaccacagca gcatgcacag cgagaaactg 5700

ctgaacatca cgattccgga atccctggac tatacgcggg catttgaaga gcctttcgcc 5760
cgctacctgg agcaggcgga gaacgtgggc gtgaaaacca ccggaatggg cagcatgtac 5820
cggctgtcat tcaaaatccg gatgaggaat ccggcggagg aaaagaagct gattgatgag 5880
ctgcggaccc ggaacagcaa cctggagatt gcagtcctgc cttatgcgga accgcagctg 5940
cagctgtaat ccctcccgaa aggagagaaa cgtatgaaga acaaacaccg gattatccgg 6000
aacctgctgc tggatgcggc agcggcgggg ctggtgctgg tggtctttgc ggcattccac 6060
catgtactgc ccgcgcgca gcagggcttg aacgtacaga ttgtgaaacc ggattcaacg 6120
gcagaaacgg ctgtcgctga gatcagggaa gcggaaaccg cggcggagat cccggaagcg 6180
aaaaaggccg aagaacagcc tgcagaaatg aaagcggagg ctgaaacgga acccgatacg 6240
gccacctcgg ccacacaggt ggaagaagag aaaaccggat tccggccgac cgtcgtgaca 6300
agcgcttcct ctggaaccgc tgattcgggc cggcgcggcc cttcccagag accgggaagc 6360
cggaaaagcg gaagaggcgg cggaaacatg atgaattcca agggcggaag cggcctggac 6420
ggggaggacg gaacggcgga agacagcgcc gggactgcgg acagtacccc gctggccgaa 6480
aagtttgcgg acaagtttac ggaatccgtt acggtgaccg gaaacagcta taccagcccg 6540
gatatttcaa tcacggtgga tgaagtgaac gacgggaacc tgacctacta catcgcggat 6600
atctacgtgc gggatatcac ctgcttccaa actgcgctgg cgcaggacac ctacggctcc 6660
ggcttccggg acagcattga ggatatggcc gtcctgaaca acgccctgct ggcggtgaac 6720
ggggattact acggcaatac cagcgagggt gtggtgatcc ggaacgggat gatctaccgg 6780
gcgaatccga cggactgtga catatgcgtg ctgtactatg acgggaccat gaaggtgatg 6840
agcggagact ccttcgacct ggaggacgcc attgaggacg gtgcgtggca ggcctggacc 6900
ttcgggccgg cgctgctgga tatcggcggg cagcccattg cttccttctc cagcacaaac 6960
cggatcattt ccgccaatcc acggaccgcc atcggatact acgaaccggg ccattactgc 7020
ctggtggtgg tggacggacg gggagaatcc gcgggaatca gcctcccccg gctgagccag 7080
ctgttctatg acctgggatg taccgcggcg tataacctgg acgggggcaa ttcctccatc 7140
atggtatggg ggaacgaagt gatcaacaat ccgtccggcg gcggacggga aagcagcgat 7200
gcgctgctga ttgcggaggt gcggaattat gagtaaagtg caggggctgt tcaatcacct 7260
gaccgtgatg ttcgggatgg tattcctggt gttcctggtg ctggaccagt tcaacccgat 7320
gatgaacttt gtggacaact cgatttcccg gtggatgctt gctgcgttct gcatctgcgg 7380
aatcagccgg agcatcctgt gctggcggaa aaacgattcc tgaaaatagg gggaatgaaa 7440
gcacagctac tttgagctga acaccgcgat gacaaacgga atgaaggaaa acgcgctgcg 7500
ccatatcagg gtgttccgcg ggatcacgta aaccttccat tccgggtata ttccatatac 7560
gaagcggcgg gaaatgttca tttcccgccg cttctgtatt atccggctga agattattct 7620
ccatccatct tcgacgcgaa caggcgtccg gtttctccca gcagttccag cagtttcgga 7680
atgagggctg gatcgtggga gtccatttcg ttgaaggtct caaaattgag ggtgccgctg 7740
tagccgatct catgaagggc gcggagcacc gtttcccagt cgatgattcc gtagtagggg 7800
aaacggtggc tgtcgctgac gccgtcgttg tcgtgcaggt gcagggcaag gagccggtga 7860
cccagggtgc ggatagcctt cccctggtct tttcccagca gggcggcgtg gccgcagtca 7920
tagcagaagc cgaagatttc ctccccggca atggcattca gctcatcaat gtacgcgcag 7980
gcctcgccca tgtcggcgca gacggcctcg acgcgctttg ccttccagga gacaaacatg 8040
ttctccagca ggattttgac acccgtgcgc ttcgcgtcgg ggatcaggga tgaatagagc 8100
cggatattga tctcatgttc cgcggcggcg tccagcctgt ttctgtaatc cgtgaatgcg 8160
ggatggatca ccaggtaagg acagtccagg aaacggcatg catccaggca catccggaag 8220
gcctgcatca ggtgggcgtt catctcctca tccaggggga tccaggaggg gaagggggca 8280
tgcgcctgtc cgaaaccgac gccgtatttc cggcccgcat cgcggatggg ccggatgatc 8340
tcctccataa actgttcctc cggcagggtg aattctccct cgatgattcc cttgcggatg 8400
ctgccggacg aaatatagat caggtccagg tccacgcagt caaagccggc ccggctgatg 8460
agccgcatgc cttcatcgac cccgcagggc tggaagattc cgtctgtctg tacgctgatc 8520
cgcctcttca tgaacaggcc tcccgtttcc ttttttccgg atatggcgcc gcggccaacc 8580
gtcagtggta ttccggcagc cagccccggt gggcttcaat caggtcgtcg cacatggcga 8640
cgatatcgtc gatggacagc tcgctctggc agtggggatc catcatggca gcctggtaga 8700
tgtaatcctt tttcagtgta acggcggcca ggatggtgag cagctggacg ttgatgttgg 8760
tgcggttcat ggcggcgcac tgctcgggca ggtctcccac aaaggcggga gtcacgccgg 8820
aagcgtcggc aaaacagggc acttccacgc aggcgtcctt gggcaggttg gtgatcaggc 8880
cggtattgag cacattgccg ccgaacttgt agggctcgtt gatctccatc gcttccatga 8940

tccggctggc gtactcgttg ctccgggtat gggtcaggtt cggattggcg atgagctcgt 9000
cccgctgctt ttcccaccgg gcgatctggt tttcgcagcg ccggggatat tcgtccaggg 9060
ggatgttgaa ccggtcaatc agttccggat acttgtcctt gatgaaatag ggcatgtatt 9120
ccgagctgtg ctcgctggat tcggtgacat aatagccgaa gcgcttcatg atttcgagac 9180
gcaccatgtc gtcgtgcttt tccgtgatgg ccgcgcagcg gcgtttgatc tctggataca 9240
ggtcctctcc gtctttggta acctccagca gccaggcctg gtggttgatg cctgcgatct 9300
tccactgcac ccggctgtca tattccattc cggccaggtg catcagcttc gggacacagt 9360
gctgcacgct gtggcacagg cccaccgttt tgacgccggt aaaccgctgc atcgcaccgg 9420
acagcatggc catggggttg gtgtagttga ggaaccaggc gttgggacac acttcttcca 9480
tgtccctggc aaagtcaaac atcacaggga tcgtgcgcag ggcccggaag atgcccccga 9540
tgcccagggt gtcgccgatg gtctggcgca ggccgtactt tttgggcacc tcgaaatccg 9600
tcacggtaca gggcttatag aaccccacct ggatggcgtt gaccacgtaa tcggccccgc 9660
gcagggcagc cttccggttc tccacgccgc accaggcttc aattttcgcc ttgccgccca 9720
gattgttgtt gatggccgtc agcatcattt cggattcctt caggcgctgg ggatcgatat 9780
cgtacaggga gattacgctg tcctccagcg ccggggtgag catgctgtcc cccagcacgt 9840
tccttgcgaa cacggtggaa cccgcgccca taaatgtgat tttcggcata tccagatcct 9900
cctttgagac gaatcaaaat cctccgggaa gggaacgatg aacaatcatt cagctccgat 9960
catatggcat tctcgctttg caggcagtat atcatagaaa atgacttttt accagagaag 10020
tacaggctcc tgttcgtgga tcggcttacc ttgttttgct gcccggcaca cccgggaacc 10080
tgtacagcgg cgcatggaac ggaacaggtt gaaatcaggc ggcattccgg gtaaactaat 10140
ggcagtacga accgggacag cccggggaaa ggacaaacga atgaagaagc tagcgattat 10200
tctgcttgcg gtaatcatga tggcagcctg catcagcccg gcaatggcgg caagatacta 10260
tacatcccgg tttgcggcag ggacggacgg atggtatgcc cgcggtgcac agtccgtgta 10320
ccgcacgacg gagggcaccc tgcggaccga ggggcggacc agcgactggc attcccccgg 10380
acgggatttt gtcctggtgg aaggatgcct gtacaagctg agcgtggaag tataccaggc 10440
tgaggcggac agcgcggagt tcatgatttc tgtcgcacac tcaaaggacg gaaatgagac 10500
atatgagaac ccggcccgcg gcaccgtaaa gaaaggtgaa tggacgaaga ttgaaggcac 10560
ctatacggcc gggaactacg accgcagcgt gctgtatgtt gaaacgcttg gagcgccgca 10620
gctgagcttt gaaatcagga atttcattgt tgatgctccg aacggcattc cgaagccgca 10680
gcctgctgaa cccccgatgg agatcgaagc ggtggccatc gaggacatgc ccagcatcag 10740
ggaagcctat gaaggaaagt tcgatttcgg cgcggcggtt ccgcagcagg cttttatgaa 10800
taagcagctg cgggcactga tggccaaaca gttcagtatc ctgacaccgg aaaatgagct 10860
gaagccggat tccgtgctgg atgtgcaggc cagcaagagc ctggtacata atacgggaga 10920
ccagacagcg gttgccgtcc actttgacgc ggcgaagggt gtgctgagct ttgccaaggc 10980
aaacggcctg aaggtgcacg gccatgtgct ggtatggcac agccagacgc cggaggaatt 11040
cttccacgag ggctatgacc ggagcaagcc gctgctcagc cgcgaggtca tgctgggccg 11100
gctggaaaac tacatcaggg aagtcctgac gcagaccgaa gagatgtatc cgggcgtgat 11160
tgtcagctgg gacgtggtga acgaagcgat tgacgacggg acgaactggc tgcggaaaac 11220
ctctccctgg acccggacca tcggcgagga ccatgtggcg cgcgccttcg agtttgcccg 11280
gaagtacgcc gcggaaggtg tgctgctgta ctacaacgac tacaatacgc ccgttcccgc 11340
caagctggct ggaatcacaa agctgctgaa gcagctgatg gcagaaggca acatcgacgg 11400
ctacggattc cagatgcact acagcaatgg cgatccgtcc atcgcgcaga tttccaacgc 11460
cgtgagccag atcgcctccc tcgggctgaa gctgcgcgtg agcgagctgg acatcggtgc 11520
cggcacgtcg gaatcggcct tgatgcagca gaaggcaagg tacaaggaaa tcatgcagct 11580
gatgctgcag tatgcggacc agacggaagc cgtacaggtg tggggcctga cggatggcat 11640
gagctggcgc gacggtcagt atccgctgct gttcaacgat cagctgtatc cgaagcccgc 11700
cttttacggc gtgctggaag cggcggaaga ataagcacct cttttacagg aaagcgctgc 11760
agatgcagcg cttttcctgt atcccggcgg cggaaaaaag attgaagaaa tcccgcatcg 11820
gcggtaagat agataagccg gaagcaaaaa caaaatacgc ccggaagggc gggaaagagg 11880
aaatatatgg aattgtgttc ccggaaagac gtacccgtgg aggaaacatg ggacctttcc 11940
ctgatttttg aaaacgaaca gcagatgtgg gatgccctgg aagaaacgaa ggcagcggtg 12000
aaaaaagttg tcgatacata tgccggtaaa ctgaacaccg cggagaatat tgtcaattgc 12060
ctggacgaga tggagcagat ccggattgct tcgtcgaaaa tctggagcta tacggggctg 12120
gcggtggaaa cggactacac ggacaacaag ctgcgcgagc tggatgaaaa ggtgggcgac 12180

gagatgatcc gcctggagcg ggacacgtcc ttcgtggaca gtgaaatcct gctggcgccg 12240
gacgacgtgc tgcaggccgc ggtggagaag gcaaagggat gccggatcta cctgaaggac 12300
ctgattgcca gaaaggcgca tatgctgtcc ccggaaaccg agaaggtgct ggcggccctc 12360
ggcaaaacgc tgaatgtgcc ctacaccgta tacaatacga tgaagctggc ggatatctcc 12420
tttgattcct tcaccgtgga cggaaaggaa tatccgctgg gctattccct cttcgaggac 12480
aactacgagc tggaagcgga caccgcggtg cgccgggcgg ctttccaggc cttctccgac 12540
acgatcgcga aatacaggaa caccacggcc accgcgtaca acgccaacgt ggcccaggac 12600
aagatcatgt ccgagctccg gcactttgac aacgtgtttg acagcctgtt gtttgaccag 12660
aaggtaacgc gggaaatgta cgaccgccag atcgacctga tcaccgagcg cctggcgccg 12720
catatgcgcc ggtacgcccg gctcctgaaa aagatctaca ggctggacaa aatgaccttt 12780
gccgacctga agatcgcggt ggatcccggg ttcgacccga aggtgacgat cgcggaatcg 12840
cataaataca tccgggatgc cctggcgatc ctcggtgagg attatgtgga aatggtagac 12900
cgggcctacc gggaacgctg ggtggatttc gcccggaacg tcggcaagag cacgggcggc 12960
ttctgttccg gccagtaccg gaagaactcc ttcatcctgc tgaactggaa cgaccggatg 13020
gcggacgtat tcaccctggc ccatgagctg gggcatgccg ggcagggcat gtacaccgac 13080
gctgcccagt cctactatga cacctggctt tccacctacc tggtggaagc gccttccacg 13140
atgaacgaac tgctgctggc ccaccacctg tccaagacca gcacggataa gcgattccgc 13200
cgctgggtgc tctccaacgt cgtgagcaat acctattacc acaacttcgt gacgcacctg 13260
cgggaagcat ggtaccagcg tgaggtgtat cggatcatcg aagagggcgg cagcgtgaac 13320
gccgatatcc tcagcgacct gttccggaag aacctggagc tgttctgggg cgacgacgtg 13380
gagatcccgg aaggcgcgga gctgacctgg atgcgccagc cgcactacta tatgggactg 13440
tactcctata cctacagcgc cggcctgacc gtggcgaccc aggcgctgct tcggatcgaa 13500
aaggaaggcg aacctgccgt gcaggactgg aagaaattcc tggctgccgg ctgcacccag 13560
gacccggtgg gactggcgaa gatcgccgga tttgatatca ccacggacgc gccgctgaac 13620
gcgaccatcg actatatcgg atccctgatc gacgagatct g 13661
//
ID XXX; XXX; linear; XXX; XXX; XXX; XXX.
XX
AC PRJEB67889
XX
DE Uncultured bacterium DNA fragment from in vivo enriched cow rumen, clone
rum-30f10
XX
FH Key Location/Qualifiers
FH
FT source 1..30169
FT /organism="uncultured bacterium"
FT /environmental_sample
FT /mol_type="genomic DNA"
FT /isolation_source="Cow rumen"
FT /clone="rum-30f10"
FT /note="AZCL-xylan, AZCL-arabinoxylan active clone"
FT CDS complement(865..1383)
FT /translation="MKKILALVMMLCLLCGAAMAEEVTWADIEPQLAENGLSGDFVVFD
FT QLGLKVWLPTGMNAAAVSEEDAASGRLAAFAADDGSAYFVVDAVNMGEMTIDQLLANVS
FT NNETVTDAEIVVNNGLSVVSYKNTASDCLTAALVDTNSNVIMFTMGPASAEGAAMVFYF
FT IMSSLQPAQ"
FT /product="hypothetical protein"
FT CDS complement(1538..3238)
FT /translation="MKKLISLVLALAMIMMVGAAFAADPVNNRTADTTISITGLDEGDK
FT VALYKIIEWDGFWKPATGYSGLTQTEVDDIVKQNHETGKEIANAAGWTDAYINKVITST
FT LPNAVTLTDAQATIGANGTFSLEVEPGMYYAVVTPKDGTKVYKPMFISADYSATPATHT
FT VGAKYETITINKTATDQTNNHDAGANYDEANTDANSYNTVNIGDTVDFEITTTIPKFPA
FT SFTSPVFKVSDTLNGLTLVTGEGKIKVYAGETLLTAGTEYTLNPTPTSTTFTVDFKSDY

```
FT                   IKSLTAAQAIKVVYQGIVDNTATYEINMKKNTAEVRFSNNPSDNKGANKLVDETRHYTF
FT                   DINGIISGEANRPGTDLIKVGIDDKGNEILETIQLPNQHWAGALQGAKFTLWGEEGKTN
FT                   KIRENLQSDEKGQIVITGLDAGTYWLEETEAPAGFIKSNKLAKIEIIPVFEKKTVAAHE
FT                   ETVEGIKVTVPTYSYAVLKSYTIKINGDITSTYTVKNEGTVDAIDDDQIAAGSESGKPT
FT                   NDALGKIQNPKGVELPSTGGIGTTIFYIVGGILLIGAAVILVARRKAQD"
FT                   /product="hypothetical protein"
FT   CDS             complement(3818..4414)
FT                   /db_xref="GO:0006508"
FT                   /db_xref="GO:0008236"
FT                   /translation="MAKQPKNPALPELNEIESEMSQVRSKSRFRQALKGTFGTFLVVAA
FT                   VAVIIAFIFLPVLRITNGHNMEPGLQPGDIVILYKTPDVKKGEIAAFYFNNKLLLRRII
FT                   ATAGDEVSIDEKGYVKVNGEILEEDAYISEHALGQCDLEFPFRVPTAQIFVLGDNRDYA
FT                   LDSRATNFGCIDQDEIYGKPLARIYPFNSLTWYGF"
FT                   /product="Signal peptidase I (EC 3.4.21.89)"
FT                   /EC_number="3.4.21.89"
FT   CDS             complement(4490..4894)
FT                   /translation="MAKNMLKSYGRFELLELIYSMRKENLELKERCDAAEKEVVEVRRQ
FT                   TEKQLDNARRDYENRIEELRMQGAAKDLQIRMKKIEEQLRMIQRLTSLDIDTRDMPTEE
FT                   EIEALAEQEEKEREEKEREEKEREGKQAKA"
FT                   /product="hypothetical protein"
FT   CDS             complement(4917..7001)
FT                   /translation="MIKRLIAVLLAGMLLMSIVPVQAEEEATEEPAEVTATPEPLPPYP
FT                   DRNYEELVVGNPTPMDGKFFTGMWGNSTTDIDVRTLVNSYYLTTWAYDTGIFHANPVVV
FT                   SGAVVVVNDDDGSRRYVFRLMDDLFYSDGTKINAWDYAFSALFQASPLIAELGGEPLDL
FT                   SYLVGYEDYYSGRTPYFAGVKVTDDRTIEFDVKAEYLPYFFELYRLGFLPYPIHEIAPS
FT                   CKVYDDGEGAYIANADPEVQETIFTKELLENTVMDPDSGYLSHPTVGSGPYVMTSWDGE
FT                   FATFEINKYFKGNELGYLPTIPKLRFTLAKNEDMIEKLEADEFQLLNKVTRRDSIDKGM
FT                   ELTGKGNHTVTNYPRIGLSFIVFTPDMPALQEQNVRKAIAYCMDKDKLREEYTLNYGIA
FT                   MDGLIGIGQWMYGMVNGTEDFPVELPENPTAAEEQAYADEMAEWEALSLDGVLHYELDV
FT                   KKAVSLLEKNGWTLNDQGDTFRPGEDEVRCKYIQGELVKLDLTCAYPVTNVMAEYMDTF
FT                   FVPYLAEAGIKLTLLPLDMKELFRSYNEKDIEEIDMFYLGDDFNIEFDPQLFFLAGDPE
FT                   APHEDTLAWAHTQMAEYARLMCETEPSDALGFVKKWITFQEKLSDLLPLIPIYSNVYYD
FT                   FYTTDLTNYDIRYYITWGDAIVPSEYHNVFQELLDQGINPYAPAEEEDDLDDDLEDWDW
FT                   "
FT                   /product="hypothetical protein"
FT   CDS             complement(7039..7578)
FT                   /translation="MAAENNGQIDITIGSVKSLFDRNGIHLKAYLVATGEYGRWETLST
FT                   FADIQLFSSQDDGSAWVNKSLAQLHTTIQNQGLQATQEGISQNGKISFKSLRHGIYYIE
FT                   FARGPENLDVKPLLLAVPDRTGRISYNTNAKLEFNPPVPPIDVPHRRNEHLVPIDDYET
FT                   ALGLGNIQMHVGVCFE"
FT                   /product="hypothetical protein"
FT   CDS             complement(7562..8608)
FT                   /translation="MRKLIIGLLVFLLAAALAAYLLYPTVSDQLAGLRDGNLMRDYRAK
FT                   AAALDEEAVQEHFQQAREYNDAIESFHPEDVFTSGNIRTDLDYQSRLNIHSGIIGELII
FT                   PAIRVSLPVYHQSSETPATAKLVHLDGSSLPADGTKENIVLAGPGVLKAEGILGQLNLT
FT                   DDRMLEDLDSLTPGDVMILTVMDRTMVYRVSEVQMLSAAGLKELDLTPGEEDRQLTVIS
FT                   RHQDRRLLVRSSGIPIAEARVLLEGEDKASYPDNWKNVLLLGSPVLLLGLLIIWIIERF
FT                   KMRSYKLPDEGRQTAARERKRKAKLENLTTDTERVEDENEKDKPENPVPGDPADAGNDQ
FT                   PAGDGSGE"
FT                   /product="Sortase A, LPXTG specific"
FT   CDS             complement(8605..9516)
FT                   /translation="MEIYHPRDEELLREMRRVRRADKRKRLAWGLSILLILSAAAGLFV
FT                   FHRYYQLAVMHGPAMGDSLPEGSVVLVRRTESGVTYPAGSILLYEKQFAEPVDLTILGP
FT                   KGKIRDYCRYKLYRDVGTTRQYYSTQEGTFRWTESESEADEFETSPEGILHLETDHLPN
```

```
FT                   GDYALKEVYASWGQNLMPDPVPFTVFNPVRTMLKRVLAAPGDRLILSPYQETQVNSRDI
FT                   SHLNTSGRTADAELVTRRVIVGNDEYFVQGDQLSLSRDSRDPDYDNVRLDEIQGRAEFA
FT                   LWPLRIFGDLTGASGGREGAGP"
FT                   /product="hypothetical protein"
FT   CDS             complement(9720..11276)
FT                   /db_xref="GO:0031176"
FT                   /translation="MLKRILLCLLTLCLLASCLAAAAEPVSVYTSDFSKDEDGWYGRGA
FT                   QSVRTKDNTLKTTGRSSDWNSPGRDFDLVEGGVYDLSVEVKQDELDSASFMISIAHTAE
FT                   GMETYENLAFGNAKKGEWATLKGSYTAGAFDRSVLYVETTGAADLSYEIRNFTVTAPQG
FT                   VPEAKPTEPPMEITEAESIPSLKEVYAGKFDFGTAAPRNVFMDPRWTNLMKEQFSILTP
FT                   ENEMKPDSVLDVDGSRTLARETGDPASVKVRFNAAKPLLDFASRNGLKVHGHTLLWHSQ
FT                   TPQAFFHEDYDSRKPLVTREVMLGRMENYIKGVFAYLDENYPGVVVSWDVLNEAIDDGT
FT                   NWLRKSSWYKVVGEDYASYAFKFARQYAPADVILCYNDYNTAYPAKLKGIVRLLKQLQE
FT                   DGNIDGYGFQMHHRVGSPSIGQIRNAVKTIADLGLKLRVSELDVGTTGRTETAFNKQAK
FT                   YYADVMRILIDHADQFEAVEVWGLTDLMSWRSKDFPLLFDGAGNPKPAFFAVADPDGWQ
FT                   K"
FT                   /product="CBM22-GH10"
FT   CDS             complement(11408..12901)
FT                   /translation="MKKITALVMAAIMMLVCFSFAAAEELPEGYPAIIEGLDMGGQTVY
FT                   IYDWWTQDDEEHSTRVADPDDDTQKLYDYRDWLEKTYNVKIVNCAWDKGGWQESPQELA
FT                   NMVNDGKADQLCLIIIDPNFAGGPLANGLLEEWTIDLTGPQWLSSVSEFMTKDGKVYGV
FT                   LAADEEPREVIFFNKKVLEDAGIDYNEIYDLQKSGEWTWDKFEEYMEKVQQDKDNDGIN
FT                   DIWALTGNGDRLLRGCIYGNGATIFGDKDGKLSITADSDACLKGIAKRQEWDRFMPQKA
FT                   DFAPDGEWNWFEGFWKQGTTAFFVGQNYEGIGNPDNLMRSCDFDWGYVAFPKGPDADDY
FT                   LYAAVDNICVIPAKCYDADTVKKLEQIYALYTYPTPGVDQETAWIGDKYEYADDRAVDE
FT                   TMAMLRQPQHAVGNKTYLLGDENTVLGSTLMWNLGSGTPAEIVEGARDVWQFRCDVFNG
FT                   DKTADDLQTFLDEKAAEAEAAAAAAAEEAAEGEGEEKAE"
FT                   /product="hypothetical protein"
FT   CDS             complement(13077..14087)
FT                   /translation="MSKNNTIFIGENKSFWERNRRSNGYLLKRRAGQVTISVIRALLMF
FT                   GLCFMIIQPLITRFSVSLMAAEDIYDSTIRLLPRNVTLDNYRIVAELTSMPNSMINTAW
FT                   ISLVMSICQVISTTLVAYGFARYEFPLKKFWFACVVLLIIIPPQTIQTALFMSFTHFDI
FT                   FGIFKATTGSDINLRSSLIPYTIMCLTCMGLKDGLYIYMLRQYFKSVPESLEEAAYVDG
FT                   CGTMRTFVKIMLPDAVPTIASCFLFSFVWQWTDLFYTRNFISGYKIYSRELSNITSIMS
FT                   KYFARNATENVTVPLGRQMQLIGIGVFVCCVPLIILYAFTQRTFVQSLAMTGSKE"
FT                   /product="hypothetical protein"
FT   CDS             complement(14080..15090)
FT                   /translation="MKKTKKYNHWYEGITEAFKMYIPGNKTYHTIQARRARTGTLFILP
FT                   LIIGFLAFMVRPLILSLQMSFSKVVVVEGTMSPFNIVLNLFGSNVTVPFGNYNYAFVSD
FT                   PDFNGLLVNEIGRMAINTIATLVMSFVIAVILNQNFKGRTLCRIIFFLPVILSSGVLPG
FT                   IESQNQYFNLMTSISESVNESAGVNLSQALQNLLSVSGVAREVFDVIFQMIDAIYDIVM
FT                   ASGIQIIIFLSGLQSISPSLYEAADVEGCSAWESFWKITFPMVSPLLLVNCIYTIIDFF
FT                   MKNDNTVMEHINTVMYGTQMDFGAASAESWIYFGVALLFIGISTFIITRAVKSYE"
FT                   /product="hypothetical protein"
FT   CDS             complement(15087..17666)
FT                   /translation="MDRNKKKIITRLIIWLVCLAALAALIIFVFVPIYSEREVSSGRDP
FT                   HLIFYEGDGAPVTIENEHLLLEMDGATTQFKVTNKDTGKIWYSNPQTRDKDPIAHGVNS
FT                   DFLSSTLTLTYTVSGNETELNNYRYSMENQNFNIIRENDTTIRLEYAIGKIERTYHIPL
FT                   AIPVERYKELTGQMSGGDKKKVSSFYSLYDQKKLDKADNLDELKAMYPEIESGNLYILK
FT                   SGTNTAAKEKIEGYFQKIGYDQEQYEADLQYVAQKEENHGPFFNATMILRLEGNQMIVE
FT                   VPYNEIMYEAAYPLTYVSVLPMFGAAGTDEEGFIFIPEGSGGIIKYNNGKLAQSAYYAN
FT                   LYGWDYGMERTEAVSETENTFAVFGMGQPDGSFICALEGASSYGGIYADIAGRFNSFNT
FT                   VYCKYNLIHYGQFKLSGRTNEQVLMYENTIPDDTVIQKYTFLDTDNYVGMAEAYGDYLR
FT                   GKEEFKKEQADPEVPIHVELIGAINKTIPKMGMPVDSTIAVTKFSEAQDIMNELLDQGV
```

```
FT                   KHLNLRMTGWSNGGVRQRVLTSIHTAGELGGDSALKTLAAAAKDKGVALYLDGISCFAY
FT                   DSGLFEGFIPTSHAARFTTRELVKLYAYDIVTYQQNTDGENYYLVKPKIAADFANRLID
FT                   GVKDRNAAGVAFRDIGNLLSADYYKNDTVTREQVKALNIKSLQKAQELGLNISIKKGND
FT                   YAAAYADLITDMNLTGRKYSIIDQAVPFYQIALHGLRNYTGEAINLSADFQNALLESAE
FT                   YGAGLNFSFMKEDTNILQDSAYSCYTAASYDRWKDQVLPMILRYQKEMSGLNSQRITGH
FT                   EFLTPEVHVTTYEDGTKVYVNYGRKDYRKGAVQIPAREYLVERRGKQ"
FT                   /product="hypothetical protein"
FT   CDS             complement(17682..18377)
FT                   /translation="MSTVKSVSDQAVKAGSGKTKAWFSRYGQSLHYALHVITHPFDGFW
FT                   DLVHEKKGSIAAANTFLILFLVTRVLKLIATNFQFISAPIQHINIFEEMASLLLPFIVL
FT                   CVANWAMTTLFEGKGRFKDIYMGMCYALVPYIIIQLPMILVSNMLTYEEGSFYSVLLSI
FT                   SVIWCVFLVFVGLMEIHDYGPGKTFIFLIVTVVGAGVILFLMLVFFSLLSDAVSFFVSF
FT                   YREIVFRLN"
FT                   /product="hypothetical protein"
FT   CDS             complement(18352..19884)
FT                   /translation="MKKLISFLMVLALALSAAGALADDNFDMKDDGFNTSYTYGYDYWG
FT                   DVLMSPDPYRVELVIDSTTLGLENLNNKTINKPQSIFVRDKDVYVVDSGNNRILQIRYT
FT                   EEGASLIRIIEEIKGAEPSSFSNPTDVFADEKGNLYVADQNNNRVVMMDKDLNFLKLFD
FT                   KPTDATFNQSSSFLPSKIAVDVAGRVYVLATNVNKGFMKYESDTTFTGYIGANPVKVNM
FT                   ADYIWKRYFMSKEQRAASENFVPTEYENMYMDEKGFIYATTLVFSEYELLSDVAKPIRR
FT                   LNGIGSDILIKNDRYPPIGDLDWKAESLQYGPSKMADITALKNEMYVALDRTRGRLFGY
FT                   DSQGVMLWAFGTKGSMDGTFGRAIALDHMGYDLLVVDEMKNNITIFTPTEYGQLIYDAT
FT                   DTYLNGDYDRSAELWEQVMKMNSNYSLAFRGIGRSLLRQEDYQGAMDYFQRAHAREEYG
FT                   RAFKQYRKIWVEKNVWWIVLILAVILIVPLVLGRIKRTKWEVVMHEHSKVRK"
FT                   /product="FIG01032054: hypothetical protein"
FT   CDS             complement(19904..21130)
FT                   /translation="MSEMNSVRSNGTLQLINGKKFRGYLQADKEGAAFYAKGKAEPVVQ
FT                   WHYARMNRMDNIRRGGTTSQITVILKDDSRYVMELAQGLKCYSHMDKNWTDKPVVKKTR
FT                   GDLLHEQAERLGQKIEVPKKHLITRRHPNRSIGGDIGIYLLLILFAVAMVFPIVFLIGS
FT                   SLKPLSDLFRFPPPVWPSQPTMDNFSDLFVTMGQSWVPFSRYLVNTLLITIVGTFGHLV
FT                   VAGMAAFVLAKYEFPGGRFFFGVVTTCLMFSGYVTGIPNYLIMSRLGMVDTYWALILPA
FT                   FAAPIGLFLMKQTMETLPTALIEAATIDGASRFRIFWSIVMPNVKPAWLTMIIFSVQSL
FT                   WNNAAATVIYSEAKKTLVYALQQIQAGGVARTGQMAAANVIIFIVPVIIFILSESQILE
FT                   TMASSGIKE"
FT                   /product="binding-protein-dependent transport systems
FT                   inner membrane component"
FT   CDS             complement(21132..22076)
FT                   /translation="MQSQSFTQRYLTMRKEKMQHGWTLAKRNKGCYLFLAPYALLFFTF
FT                   FILPIGTSIFYSFTYYNILEPPRFIGFQNYINLVLQDEVFLTAVKNTFVCAVIIGPVGY
FT                   MLSFLFAWFINELPKWIRSVAVVIFYAPSLGGAAFTVFSIIFRGDAYGWFNSILMEFGI
FT                   IEAPRLWLTDPRYMMTIVIIVSLWMSMGTGFLSFVAGFQGIDRSMYEAGYVDGVRNRWQ
FT                   ELYHITLPSMKPMLLFGAVMSITSSFNISDVPRILCGYPSTDYAARTVVTHLFDYGFSR
FT                   FEMGYASAIATFLFLVMILCKKAITALLGRVGT"
FT                   /product="binding-protein-dependent transport systems
FT                   inner membrane component"
FT   CDS             complement(22098..24983)
FT                   /db_xref="GO:0005215"
FT                   /db_xref="GO:0006810"
FT                   /translation="MSRKWRNILIGVLAVALIVVLITLLSGGTENYRAKYEGTDLSADV
FT                   TGIGRSNTYGAYVEQYAALPAVTEPVKVDIAAFEGTGGQTVPEGVLTSDESELTWKVEV
FT                   PKEGLYNIKLDYLTTESRGIDIERELAINGEVPFSGASTLCFSRLWTDAGEVRKDNQGN
FT                   DIRPTQVERFEQQSAYCRDDMGYQIEPYAFYFKEGENELTIRAVNEPLILCGINLLPIA
FT                   KFPAYTEYAAGQKEASMTDEGKNFTLTVQGEHAKRRSQPSLYARYDRSSSLTDPYSVTN
FT                   TILNYIGGDPWTHPGEWIEWDFEVPEDGFYNISIKARQIYQRGALSARTVYIDGVIPFE
```

```
FT                   DLEAVPFGYNTSWEMRTLGDKEGNPFRFYLTKGAHTIRMEVTLGEMGPILKDVEDSIFR
FT                   LNQIYRKILVLTGVNPDRFRDYNLNQVYPEVIQAMDLESKRLYKIVDDIVATTGEKSDR
FT                   AASAQTLANQLETFVESNERITESFVNFKDNITALGTAMQNMSESKLDVDLIMITGENA
FT                   KVPEVHENIFQSIGHEIRSCASSFFVDYNSLGDKYDDTDDVLEIWITTGRDQSTVLKTM
FT                   VDDTFTARTGIKVNVKLVQVDAILPAVVAGNGPDIVLSVSGWFAVNYAMRHAVEDLTQF
FT                   EDFEEVVKPFYKSILDPLVYNDSKHKGIYGLPETQNFPLLFYRTDVMEELGLPIPQTWD
FT                   ELIAELPTIQGNSLTVAVNFPDIAVADISILGSMIYQEGGYIYDEDAKRTLIDSEAGVA
FT                   AFKQYTSLYNDYGLPVVFDFVSRFRTGEMPLGVASYATFNTLMVSAPEIRGLWDFTLFP
FT                   GTKQADGTIDRTAQTDGLCCMMIATQDEKVKHNAWEFMKWWTGAEAQTRFGREMEAVLG
FT                   SSARYQTANRDAIKQLAWSNKQLKVLEEQMAHTRGFPEIAGGYSTTRHITNAIRRVINT
FT                   KEDQRETLLNYARTINEEIKIKRREFNLPVD"
FT                   /product="N-Acetyl-D-glucosamine ABC transport system,
FT                   sugar-binding protein"
FT   CDS             complement(25185..27593)
FT                   /db_xref="GO:0004826"
FT                   /translation="MKVPYSWLKEYVDIDITAEELEEKLFGCGFEVEELIDLGKEISRV
FT                   VVGEVLSAEPVEGTHLFICQVNCGEYGNPIQIVTGAPNVYAGMHTPAALDNSTLPGGIT
FT                   IKAKPLKGIPSDGMLCSGDELGLNEDLYPGAEVYGLLDLPKETVPGTDIREVVGLNDYI
FT                   FDISVTANRPDCQSVLGIAREVAAALGKELKMPKTDYTVSDYVFPGLDITVEAPDLCPR
FT                   YLGHGVRNITVGESPRWMKRHLALCGLRSISNVVDITNYVLLEIGQPMHAFDMDQLQER
FT                   RIIVRRAKEGEKIQTLDEKEFTLTPENLVICDGNRPVALAGIMGGLNSEITENTTQLLF
FT                   EAAKFARDNVRRTSRALGQSSDSSARFEKGISEYITELGMARALHLIQELDCGEITSSA
FT                   FDCSAGAPREGKRFKASLRRINDILGIEVPAGDVLRILKNLNFEVVRDGDTLDVTAPRY
FT                   REDIEVGEPDLAEEVIREYGYDHVVPTFLKAATVTSGGKTPAQQRRERTCRILCGQGYH
FT                   EVCTLSFYADADLDFLRIPEDAPERNVLRIRNPISANLAIMRPLLAPSLLNIVVENLKK
FT                   GNNEGRIFEISNIYVPREGGELPDERPHLGFAAFGDREDFFTVKGTVEALGKALGLTFD
FT                   VERAADVPWLHPGIAAYILCEGERIGCFGKLANEVTGELKLHKDAKSNHKIFLGEIDYT
FT                   AMMAHVAASFRFRPLSAFPPVVRDLALTVAEETPCGDLMHEIARACPRVSDVELFDIYR
FT                   GEQIGEGRKSMAFKITFEAEDKPLLPEELDKYIKKILGNLKFKLGAEIR"
FT                   /product="Phenylalanyl-tRNA synthetase beta chain (EC
FT                   6.1.1.20)"
FT                   /EC_number="6.1.1.20"
FT   CDS             complement(27963..29051)
FT                   /db_xref="GO:0004826"
FT                   /translation="MQKYNAAPFHRSRFLFVNMEEKMEQRIKALREQVEQRIGQVDSKE
FT                   KLAAFWQEFLGKKGSVADLMKGLGAVAKEDRPAMGKVINDFKVAVEKKYQALSARMEER
FT                   ELAARNCREQVDITLPGTEMPLGGLHPLTLVKNEMINVFAGMGFEIFEGPEIEDDDHNF
FT                   TRLNVPKDHPSRDMQDTFYLDNDLLLRTQTSGGQIRVMDMEKPPIKVLVPGRVFRSDSD
FT                   ATHSPMFHQMEGLVVDKGITLCDLQGMLDRFVQQIFDGDVRTRLRPSYFPFTEPSVEVD
FT                   VSCFECGGKGCSLCKHTGWIEVLGGGVVHPHVLENCGIDPNVYSGIAFGIGIERITMLK
FT                   YGINNIGLLFENDLRFLKQFKE"
FT                   /product="Phenylalanyl-tRNA synthetase alpha chain (EC
FT                   6.1.1.20)"
FT                   /EC_number="6.1.1.20"
FT   CDS             complement(29120..30139)
FT                   /translation="MNNRYTFESFVVGNANRFAQAAALAVAESPGEAYNPLFIYGGVGL
FT                   GKTHLMQAIGNFIHQADPKKKILYMTSESFTNELISAIQMKKTYEFREKIRRVDVLMVD
FT                   DVQFIAGRESTQQEFFNTFNELHNENKQIILTSDKPPKDIQRLEERLCSRFEWGLVADI
FT                   QRPDVETRVAILREKTTQDNIRVPDEVLQLIAGKIDSNIRELEGCLTRLVAYSTLVKEP
FT                   ITLDLCERALKEIFDNRRHKQITAELIMQTVTDYYGLSFEDMISPTRKREITVPRQIAM
FT                   YLTRELTGMSLPQIGSVFGGRDHTTVLHSCKTVEGNMASSPDVKAVVEDIKTLVRTAQ"
FT                   /product="Chromosomal replication initiator protein DnaA"
XX
SQ   Sequence 30169 BP; 6023 A; 8427 C; 8639 G; 7080 T; 0 other;
```

```
atccgggatc aggcctgccg ccatgccgtg gaaaacggag gccgtcatca tcagggcgga        60
ggtgccgtcc acaaagcttt cgtcttcatc caggagatca tgaaagcgcc catcctcggt       120
ttggtatggg atcatagcct gcagcaggga aaggaacatt tctgttgctt cctgctgcag       180
cgctgcctcc ccctggacct tcagcgtgtc cgccacgcgg atcaggccca gcagcgccca       240
tccgttcccc gtggcccatc tcttccgccg cacaaaccgg ccctgttccg tgtccacaat       300
gtggaaaaac aaaccggttt ccgcatcccg cagatgccgg atatagcccc ggatctgttc       360
catagcttcc tttgtcctgc ccatcaccgc cagaaaaggc ggcaccatat acaggccgtc       420
aatccacagc tggcggtcgg aatatccctt ttcaaatgag acattgttat ggcagatgat       480
gccgtcctcc gtccgcggtg cctggtccat caggtaggcc agcatgcgtt ccgcgccctc       540
caggaaaagc ctgtctccgg tcttttcata agcccgcagg cagacttccc cattggctgc       600
cggatccgac accgccggtt ccccgccgat catagccagc ctgccatccg ctccctgccg       660
ttgaatggcg tcataggcca tgggaatcca gagttcttcc cggccggctt catacagagc       720
ctgggcgcag acgccctgct cccagggata ccgctgcata gccagcatgg ccaggattac       780
caattccgta tgcgtttcca tatttccgct ccctaaaaaa acccgaccct ccgtgaagag       840
gatcgggttg tctgtttgga ttccttactg cgcaggctgc agagaagaca tgatgaagta       900
gaaaaccatg gcagcgcctt ccgcggaagc aggtcccatg gtgaacatga tgacgttgct       960
gttggtatcc accagagcag cggtcaggca gtctgaagcg gtgttcttgt agctcacaac      1020
gctcagtccg ttgttgacca cgatctcagc atcggtcacc gtttcattat tgctgacatt      1080
cgcaagcagc tgatcaatcg tcatttcgcc catattgacc gcgtccacaa caaaatatgc      1140
agacccgtca tctgcggcaa aggcagccag gcggccgctg gcagcatcct cttcagatac      1200
cgcggcagcg ttcatgccgg tgggcagcca taccttcagg cccagctgat cgaagacgac      1260
gaaatcgccg gacaggccat tctcagccaa ctgcggctcg atatcggccc aggtcacttc      1320
ttcagccatc gcggcgccgc agagcaggca cagcatcatc acaagagcaa gaatcttttt      1380
catatggggt aatctttttc atatggggta acctccttca taaacatcgg ttattattgt      1440
atggtttttc cggaggatgt caatagaaaa atcccgatcc cctgtaaagg ggaccgggat      1500
gaggtttacc tcagttcatt cgtaatccgt agaaggatta gtcctgagcc ttacggcggg      1560
ccaccaggat caccgcagcg ccaatcagca ggatgccgcc gacgatgtag aagatcgtgg      1620
taccgatgcc accggtgctg ggaagctcaa cgcccttcgg gttctgaatc ttaccaagag      1680
catcatttgt gggtttgcca gactctgaac cagcagcgat ctgatcgtca tcgatagcat      1740
cgacagtgcc ttcattcttc acagtgtaag tagaagtaat gtctccatta atcttgatgg      1800
tgtaggactt cagaactgca tagctgtatg tgggaacagt gactttgata ccttcaacag      1860
tttcttcatg agcagctaca gttttctttt caaacacggg gatgatttca atcttggcaa      1920
gcttattgct cttaatgaat ccagcaggag cttcagtctc ttcaagccag taggtgcctg      1980
catcaagacc agtgatcaca atctgaccct tttcatcgga ttgcagattc tctctaatct      2040
tgttcgtctt gccctcttca ccccagaggg taaacttggc accctgcaga gcaccggccc      2100
aatgctggtt gggaagctga atggtttcaa ggatttcatt acctttgtcg tcaataccga      2160
ccttaatcag gtcagtacca ggacggttag cttcgccgct aataattccg ttaatatcaa      2220
aggtataatg acgggtttca tcgaccagtt tgttagcgcc cttattatca gaaggattgt      2280
tggagaagcg aacttcagct gtgtttttct tcatgttgat ctcataggtc gctgtattgt      2340
caacaatacc ctgatacaca accttgatag cctgcgcagc agtcagactc ttgatgtaat      2400
cagatttaaa atcaacagtg aacgtggtgc tggtgggggt aggattcagt gtatactctg      2460
ttcctgcagt aaggagtgtt tcaccggcat aaaccttaat tttgccttca ccagttacaa      2520
gagtcagacc attcagagta tcagatacct tgaacacagg agatgtgaag ctggccggga      2580
acttcggaat tgtggtggtg atttcaaaat caacagtatc accgatattc accgtattgt      2640
aagaattagc gtcagtattc gcttcatcat aattggctcc cgcatcatgg ttgttcgtct      2700
gatcagtcgc tgtcttatta atggtgatgg tctcatactt ggcaccaaca gtatgagtgg      2760
caggagttgc actataatcg gcggaaataa acatcggctt ataaaccttg gtaccatcct      2820
taggggtaac tacagcatag tacatgccag gctccacttc aaggctgaag gttccgtttg      2880
ctccgattgt agcttgtgca tccgtaaggg taactgcgtt cggaagtgtg gaagtgatga      2940
ctttattgat atacgcatca gtccagccgg ctgcatttgc aatttcctta ccggtctcat      3000
gattctgttt aacaatgtca tcgacttctg tctgggttaa tccggagtat ccggttgcag      3060
gtttccaaaa gccatcccac tcgatgatct tatacagtgc aactttatca ccttcatcca      3120
gaccagtaat ggaaatggtt gtgtcagccg tacgattgtt gacagggtca gcagcaaacg      3180
cagcgccaac catcatgatc atcgccaggg ccagtaccag ggaaataagt ttcttcatgg      3240
```

ggggttcctc actttctttt ttattcgttc cttgagttaa cgtgtttatc acctggggta 3300
tttgtcatct caaatggatg actttgggtt tccggggttt ggttttgaat caaactggtt 3360
cggttgcccc tcctttcgga ttctcccagg ttctgaagca ggaggtttga cgtcctgcgg 3420
gaaccggttt tgtccggctt ctttgtgccg ggtggtcacg gagtacagat cactccattg 3480
cacgatgtaa ttatatccgc ggaaagggct tttgtcaatg cattgcaagg ctttgcaagg 3540
ggttgaaaag ttacttcgag catcccggaa aatcggttac atgagggttt ttgagcgatt 3600
gggtgtaact gtctttccgg aatgtaactg ggtcgtgtga atgcaccctc tgcttcactc 3660
ggatgaagtt ttggctattt tcaaggccta tagacggcaa agcccctcca tgtaactctt 3720
atgttaccat acatggaggg gctttttgcc tgtattgagg ggacaaaatt gtcttgttaa 3780
gaatttgtta aatatttatt aaatatacga cttattttca aaaaccatac caagtgaggc 3840
tgttaaaggg gtaaatacgg gccaaaggct tcccatatat ttcatcctgg tcaatacagc 3900
caaaattggt ggcccggctg tccagggcgt aatcccggtt gtcgccaagg acaaaaatct 3960
gcgcggtggg aacccggaag ggaaactcca ggtcgcactg gcccagggca tgttccgaga 4020
tgtaagcgtc ctcttccagg atttccccgt tcaccttgac atagcctttt tcgtcaattg 4080
atacttcatc tcccgctgtg gcgatgatcc ggcgcagaag cagtttgttg ttaaagtaga 4140
aagcagcgat ttctcctttt tttacatccg gtgttttgta caggatgaca atgtccccgg 4200
gctgaaggcc gggttccatg ttatggccgt tggtgatgcg cagcaccggc aggaaaatga 4260
aggcgatgat cacggcgaca gcagcaacca caagaaatgt tccaaaggtg cccttcaggg 4320
cttgcctgaa tcggcttttg ctccgaacct gactcatttc agattcgatt tcattcaact 4380
ccggtaaggc agggttcttt ggctgtttcg ccatgcgttt ttcaccagct ttgtcttatg 4440
tgatagaggg atttctgttc ggcctgatgg ccaggatccg gatgggttat caggccttgg 4500
cctgttttcc ttcccgttcc ttttcttctc tttcctttc ttctctttcc ttttcttcct 4560
gttccgccag ggcttcgatc tcttcctcgg tgggcatgtc ccgggtatcg atatccaggg 4620
aggtcaggcg ctggatcatg cgcagctgtt cctcgatctt cttcatgcgg atctgcaggt 4680
ccttggccgc accctgcata cgcagttcct cgatgcggtt ttcatagtcc cgccgggcgt 4740
tgtccagctg tttttccgtc tggcggcgaa cttccacaac ttccttctcc gccgcgtcgc 4800
agcgctcctt cagttccagg ttttccttgc gcatggagta aatcagctcc agcagctcga 4860
agcgcccgta gcttttcagc atgttcttgg ccatgcggtt tttcctccgt taaaccttac 4920
cagtcccagt cttccaaatc gtcgtccagg tcatcctctt cttcggcggg agcgtaggga 4980
ttgatgccct gatccagcag ttcctggaac acgttatgat attctgaggg aacgatggca 5040
tcgccccagg tgatataata acggatatcg tagttggtca ggtctgtggt atagaagtca 5100
taatagacat tggaataaat cggaatcagg ggcagcagat cgctgagctt ctcctggaag 5160
gtgatccact tcttgacgaa gccgagggcg tcgctgggct cggtttcgca catcaggcgg 5220
gcatattccg ccatctgggt atgcgcccag gccagggtgt cctcgtgcgg ggcctccggg 5280
tcgcccgcca ggaagaagag ctggggatcg aactcgatat tgaagtcatc cccgagatag 5340
aacatgtcga tctcctcgat gtccttctcg ttataggaac ggaagagctc cttcatgtcc 5400
aggggcagca gagtcagctt aatgccggcc tccgccaggt agggcacaaa gaaggtgtcc 5460
atgtattccg ccatcacgtt ggtgaccgga taggcgcagg tcaggtccag cttcaccagt 5520
tcgccctgga tatatttgca gcgaacctca tcctcgccgg ggcggaaggt gtcaccctgg 5580
tcgttcaggg tccagccgtt cttttccagc aggctgacgg ccttttttac atccagctca 5640
taatggagca caccgtccag ggacagtgct tcccattccg ccatttcgtc tgcataggcc 5700
tgttcttccg ccgcggtggg gttctccggc agttccacgg ggaaatcttc cgttccgttg 5760
accatgccgt acatccactg gccgataccg atgaggccat ccatggcgat accgtaattg 5820
agggtgtatt cctcccggag tttgtccttg tccatgcagt aggcaatggc cttgcggacg 5880
ttctgctcct gaagggcggg catatccggg gtaaagacga taaaggaaag gccgatgcgg 5940
ggatagttgg tcacggtatg gttgcccttg ccggtcagct ccatgccttt gtcgatggaa 6000
tcccgccggg taaccttgtt cagcagctgg aattcatccg cctccagctt ttcgatcata 6060
tcctcgttct ttgccagggt aaagcgcagc ttcgggatgg tgggaagata gcccaattca 6120
ttgcccttga aatacttgtt gatttcaaag gtggcaaact ccccgtccca ggatgtcatg 6180
acgtagggac cgctgcctac ggtaggatgg gacaggtaac cgctgtccgg atccatgact 6240
gtgttttcca gcagttcttt ggtaaagatt gtttcctgga cctccggatc cgcgttggca 6300
atataggccc cttccccgtc gtcatagact ttgcaggaag gcgcgatttc gtggatggga 6360
taaggcagga aacccaggcg gtacagttcg aagaaatagg gcaggtattc cgccttgaca 6420
tcgaattcga tggtcctgtc atccgttacc ttcacgccgg caaaataggg cgtgcggccg 6480

gaataataat cctcatagcc caccagatag gaaaggtcca agggctcgcc gcccagctcc 6540
gcgatcaggg gtgaagcctg gaacagggcg gaaaaggcgt agtcccaggc gttgatcttc 6600
gtgccgtcgg aatagaaaag atcatccatc aggcggaaaa catagcggcg ggaaccgtca 6660
tcgtcgttca cgaccaccac ggcgccgctg accaccaccg ggttggcatg gaaaatgccg 6720
gtatcatagg cccaggtagt caggtaatag gaattcacaa gggtacggac atcgatatcc 6780
gttgtggaat tgccccacat gccggtaaag aacttgccgt ccatgggagt ggggttgccc 6840
accaccagct cctcatagtt ccggtccgga taaggcggaa ggggttcggg ggtggcggtg 6900
acctccgccg gttcctcagt ggcttcttcc tccgcctgga ccgggacgat gctcatcagg 6960
agcatcccgg ccagcagtac cgcaatcagt cgtttaatca cgtgatttct ccccttgtca 7020
ttcccgtaaa ggatgtattt actcgaagca gacgcctacg tgcatctgga tattgcccag 7080
gcccaacgcg gtttcatagt catcgatggg caccaggtgc tcatttctgc gatggggcac 7140
atcgatgggc ggaacggggg ggttaaactc aagctttgca ttggtgttgt agctgatccg 7200
gccggttctg tccggaacag ccagcagcaa aggcttcaca tccaggtttt caggcccccg 7260
ggcaaactcg atataataga tgccatggcg cagggacttg aaggagattt tgccgttctg 7320
gctgatgccc tcctgggttg cctgcaggcc ctggttctgg atcgtggtat gcagctgggc 7380
cagggatttg ttaacccagg cggaaccgtc gtcctgggaa gagaaaagct ggatgtcagc 7440
aaacgtactg agggtttccc agcggccgta ttctcctgta gccaccaggt acgctttgag 7500
atggataccg ttccgatcga acaggctctt gacgctgccg atggtgatat cgatttggcc 7560
gttattctcc gctgccatcg ccggcaggct ggtcattacc agcatcagca ggatcgccag 7620
ggacaggatt ctcaggcttg tctttttcat tttcgtcctc caccctttca gtgtcggttg 7680
tcaggttttc cagtttagct ttgcgttttc tttcccgggc ggcggtctgc cggccctcgt 7740
cgggaagttt gtaggagcgc attttgaagc gttctatgat ccagatgatg agcagtccca 7800
gaaggagtac cgggctgccc agcaagagga cattcttcca gttatccgga taggaggctt 7860
tgtcctcccc ttccagcagc accctggctt ccgcgatggg aatgccggaa gagcggacca 7920
ggagcctccg gtcctggtgg cgggagatta ccgtcagctg ccggtcttcc tctccggggg 7980
tcaggtccag ttccttaagt cccgcggcgg agagcatctg cacctcgctg acccggtaga 8040
ccatggtccg gtccatgacc gtcaggatca tcacatcgcc gggggtcagg ctgtccaggt 8100
cctccagcat ccggtcatcc gtgaggttca gctgtcccag aatgccttcc gccttcagga 8160
ccccgggacc ggccaggacg atgttctcct tcgttccgtc cgccggcagg gagctgccgt 8220
ccaggtggac cagcttggcg gtggccgggg tctcgctgct ctggtggtaa accggcaggg 8280
aaacccggat ggcgggaatg atcagctctc cgatgatgcc ggaatggatg ttcagccggc 8340
tctggtagtc caggtccgta cggatgttcc cgctggtaaa cacatcctcc ggatggaagg 8400
actcgatggc gtcgttgtac tcccgggcct gttggaagtg ttcctgcacc gcttcctcat 8460
ccagggcggc ggccttggcc cggtaatccc tcatcaggtt gccgtcccgc aggccggcca 8520
gctggtcgga aaccgtggga taaagcaggt aagccgccag ggctgccgcc agcaggaaaa 8580
ccagcaggcc gatgatcagc tttctcatgg cccggctccc tcccttccgc cggaggcgcc 8640
ggtcaggtca ccgaaaatcc gcaagggcca gagggcgaat tccgcccgtc cctggatttc 8700
gtccagccgg acgttgtcat agtccggatc ccggctgtcc cgggaaaggc tcagctggtc 8760
gccctggaca aaatactcat cgtttccgac gatgaccctt cgggtgacca gttccgcgtc 8820
cgcggttcgt ccggaggtgt tcaggtggct gatgtcccgg ctgttcacct gggtttcctg 8880
gtaggggctc aggatcagcc ggtcccccgg cgccgccagg acccgcttga gcatggtgcg 8940
cacgggattg aacacggtaa agggaacggg atccggcatc aggttctggc cccaggaggc 9000
gtaaacctct ttcagggcgt agtccccgtt gggcaggtgg tccgtttcca gatggaggat 9060
gccctccggg ctggtttcga actcatccgc ctcgctctcc gattcggtcc agcggaaggt 9120
cccctcctgc gtggaataat actggcgggt ggtgcccaca tcccggtaga gcttgtaccg 9180
gcagtagtcc cggatcttgc cctttggccc gaggatcgtc agatccaccg gttcggcgaa 9240
ctgcttttca tacaggagaa tgctgcccgc cggataggtt acaccgcttt ccgtcctccg 9300
caccaggact acgcttccct ccggaaggga gtcccccatg gcggggccgt gcatcacggc 9360
cagctggtag taacggtgga agacgaacag gcctgcggca gcgctgagga tcagcagaat 9420
tgaaagaccc caggccagac gcttgcgctt gtctgcccgg cgtacccgcc gcatctcccg 9480
aagcagctcc tcatcccggg gatgatagat ctccatccgt tacactccgt attcctgttt 9540
ccggcaatca gcggccgtct ggatggttac cggaaataaa atgggcattt cgatacaaat 9600
aaccaatgtt agtatataac agctgttaca tcttttcaag cacaattcct tgcggaaaaa 9660
gggaaaaaaa cagagggaaa gcgtccttcc gggtgctttc cctcctgtat aaagctgttt 9720

tacttctgcc agccatccgg atctgccacg gcaaagaagg ccggtttcgg gttgcccgcg 9780
ccgtcaaaga gcagcgggaa atccttggaa cgccaggaca tcaggtccgt gagtccccag 9840
acttccacgg cttcaaactg gtccgcgtgg tcgatcagga tgcgcatgac atccgcgtaa 9900
tacttggcct gtttgttaaa ggcggtttcc gtccggccgg tggtgcccac gtccagctcg 9960
ctgacacgga gcttcaggcc caggtccgcg atggttttta cggcgttccg gatctggccg 10020
atgctggggc tgcctacccg gtgatgcatc tggaagccgt aaccgtcgat gttgccgtcc 10080
tcctgcagct gtttgagcag gcggacgatg cccttcagct tggccggata ggcggtattg 10140
taatcgttgt agcagaggat cacatccgcc ggggcgtact ggcgggcaaa tttgaaggca 10200
tagctggcat agtcctcgcc caccaccttg taccagctgc ttttgcgcag ccagttggtc 10260
ccgtcgtcaa tggcctcgtt cagcacatcc caggaaacca caacgccggg atagttctcg 10320
tccaggtagg caaagacgcc cttgatatag ttttccatcc ggcccagcat cacttcccgg 10380
gtcaccaggg gcttccggga atcatagtcc tcgtggaaga aggcctgcgg cgtctggctg 10440
tgccagagca gcgtatggcc gtggaccttg aggccgttgc gggaggcaaa atccagcagg 10500
ggcttcgcgg cgttgaagcg caccttcacg gaggcgggat cccccgtttc ccgggccagc 10560
gtgcggctgc cgtccacatc cagcacggag tcgggtttca tctcgttttc cggggtcagg 10620
atactgaact gctccttcat caagttggtc caccggggat ccataaagac gttccggggc 10680
gccgcagtgc cgaagtcgaa cttgccggca taaacttcct tcaggctggg aatgctttcc 10740
gcctcggtaa tctccatggg cggttcggta ggcttggctt ccggaacacc ctggggtgcg 10800
gtaacggtga aattgcggat ttcataggag aggtccgccg cgccggtggt ttccacatac 10860
agcacggacc ggtcgaaggc cccggcggta tagcttccct tcagcgtcgc ccactcgccc 10920
tttttcgcgt tgccgaaggc caggttctca taggtttcca tgccttcggc ggtatgggcg 10980
atggagatca tgaaggaagc gctgtccagc tcatcctgtt taacctctac ggagagatca 11040
tacacgccgc cctccacgag gtcaaaatcc cgtccgggag aattccagtc actggagcgg 11100
ccggtagttt tcagggtgtt gtccttcgtc cggacagact gggctccgcg gccgtaccag 11160
ccgtcctcgt ccttgctgaa gtccgaggtg taaacggaaa caggttccgc cgcggccgca 11220
aggcaggagg ccagcaggca gagggttaac aggcaaagca ggatacgttt caacatgttt 11280
cagttacatc ctttccgggc agcgcccagg ttgttcgtac acattttaca tggtttcgtg 11340
ccggacttca agggtcaaaa aaggaccgct ctttcgagcg gtccggggta aaccgaatga 11400
ttgctgatta ttcagccttc tcttcgcctt cgccttcagc agcttcttca gcagcggcag 11460
cggcagcagc ttcagcttcg gcagccttct catccaggaa ggtctgcaga tcgtcagcag 11520
tcttgtctcc gttgaacacg tcgcagcgga actgccacac gtcacgggcg ccttcgacga 11580
tctcggcggg ggtgccgctt cccaggttcc acatcagggt gctgcccagg acggtgttct 11640
cgtcgcccag cagataggtc ttgttgccaa cagcgtgctg ggctgacgc agcatagcca 11700
tggtctcgtc cacagcgcgg tcatcagcat actcgtactt gtcgccgatc caggccgtct 11760
cctgatccac accgggggtg ggataggtgt acagagcgta gatctgctcc agcttcttga 11820
cggtgtctgc atcatagcac ttggcgggga tcacgcagat gttgtcaaca gcggcataca 11880
gataatcgtc agcatcgggt cccttcggga aggccacata gccccagtcg aagtcgcagc 11940
tccgcatgag gttgtcgggg ttgccgatac cttcgtagtt ctggcctacg aagaaagcgg 12000
tcgtgccctg cttccagaag ccttcgaacc agttccactc accatcggga gcgaagtcag 12060
ccttctgagg catgaaccga tcccactcct gacgctttgc aataccсttc aggcaagcat 12120
cgctgtcagc ggtgatggac agtttgccgt ccttgtcgcc gaagatggtg gcgccgttgc 12180
cgtaaatgca gccgcgcagc aggcggtcgc cgttgccggt cagggcccag atgtcgttga 12240
taccgtcgtt gtccttatcc tgctggacct tttccatgta ctcttcgaac ttgtcccagg 12300
tccattcgcc gctcttctgg agatcataga tctcgttgta gtcgatgccg gcatcttcca 12360
gaaccttctt gttgaagaag attacttcgc ggggttcttc atcagcggcc aggacgccgt 12420
acactttgcc gtccttggtc atgaattcgg aaacggagct cagccactgc gggccggtca 12480
ggtcgatggt ccactcttcc aacaggccgt tagccagggg gccgccggcg aagttggggt 12540
cgatgatgat caggcacagc tgatcggctt taccatcgtt aaccatgttg gccagttcct 12600
gggggctctc ctgccagccg cccttgtccc atgcgcagtt gacgatcttg acgttgtagg 12660
tcttttcgag ccagtcacga tagtcataca gcttctgggt gtcgtcatcg ggatcggcga 12720
cgcgggtgga gtgctcttcg tcgtcctggg tccaccagtc atagatgtaa acagtctgtc 12780
cgcccatgtc caggccttcg atgatggccg gatagccttc gggcagttct tctgcggcag 12840
cgaagctgaa gcatacgagc atcatgatcg cagccataac cagggcagtg attttcttca 12900
tactgagaaa acctcctttt ttatttcctt acaggactcc aacagagcct gtataggtac 12960

attggtagaac ggggcacccc gttgataggg taaattatag accattcttt ccattttttc 13020
aagagtcgga tcgcacaatt tctgcatttt gttcgaacaa tttctgcatt ttgtgcttat 13080
tccttggaac cggtcatggc caggctctga acgaaagtcc gctgggtaaa cgcgtacagg 13140
atgatcagcg gcacgcagca gacgaacacg ccgataccga tcagctgcat ctgccggccc 13200
agcggaaccg tcacgttttc cgtggcgttc cgggcgaagt atttgctcat gatggaggtg 13260
atattgctca gctctctgga gtaaatcttg tagccggaga taaagttccg cgtatagaac 13320
aggtcggtcc actgccacac aaagctgaac aggaagcagc tggcgatggt gggcaccgcg 13380
tccggaagca tgatcttcac gaaggtgcgc atggtaccgc agccgtccac ataggccgcc 13440
tcttccaggc tttccggcac gctcttgaaa tactgccgga gcatgtagat gtacagtccg 13500
tccttcaggc ccatacaggt caggcacatg atggtataag gaatcaggga ggaccgcagg 13560
ttgatgtcgc tgccggtggt tgccttgaag attccgaaga tatcaaagtg cgtaaaggac 13620
atgaacagcg ctgtctggat ggtctggggc gggatgatga tcagcagcac cacgcaggca 13680
aaccagaact ttttcagcgg gaattcatac cgggcaaatc cgtaggccac cagtgtggtg 13740
gaaatgacct ggcagatgga catcaccagg gaaatccacg ccgtgttgat catggagttc 13800
ggcatggagg tcagctccgc cacgatccgg tagttgtcca gggtgacgtt gcgtggcagc 13860
aggcggatgg tggaatcgta gatatcctcc gcagccatca ggctgacgga gaaacgggtg 13920
atcaggggct ggatgatcat gaagcaaagg ccgaacatca gcagcgcgcg gatgacggag 13980
atcgttacct gccctgccct gcgcttgagc agatagccgt tgctgcggcg gttccgctcc 14040
cagaagcttt tgttttcgcc gatgaagatc gtattgtttt tactcataag acttcaccgc 14100
ccttgtgatg atgaatgtgc tgataccgat aaacagcagc gccacgccga agtaaatcca 14160
gctttcagcg gaggcggcgc cgaagtccat ctgggtaccg tacatcaccg tgttgatatg 14220
ctccatgacc gtattatcgt tcttcatgaa gaagtcgata atcgtataga tgcagttcac 14280
cagcagcagg gggctgacca tggggaaagt gatcttccag aaggattccc aggcggagca 14340
gccttccacg tccgccgcct cgtacaggga gggagagatg gactgcaggc cgctgaggaa 14400
gatgatgatc tggatgccgg atgccatgac gatgtcataa atggcatcga tcatctggaa 14460
gatgacgtcg aagacctccc gggctacgcc tgatacggag agcaggttct gcaaagcctg 14520
cgagaggttc acacccgccg attcgttcac cgactcggaa atgctggtca tcaggttgaa 14580
gtactggttc tggctttcaa tgccgggcag cacgccggag gacaggatca caggcaggaa 14640
gaaaatgatc cggcacaggg ttcttccctt gaagttctgg ttcaggatca ccgcgattac 14700
gaaactcatc accagggtgg cgatggtgtt gatcgccata cggccgatct cgttcaccag 14760
cagcccgttg aagtcagggt cggaaacaaa ggcgtagttg taattgccga agggaaccgt 14820
tacattgctt ccgaacaggt tcaggacgat gttgaagggg ctcatggttc cttcaaccac 14880
cacgaccttg ctgaagctca tctgaaggga caggatgagc gggcgcacca tgaaggcaag 14940
gaaaccaatg atcagcggca ggatgaagag cgttccggtc cgggcgcgcc tggcctggat 15000
ggtgtgatag gttttattgc cgggaatgta cattttgaac gcttccgtga tgccttcgta 15060
ccagtggttg tacttctttg ttttcttcac tgcttgcccc tcctttcaac cagatattca 15120
cgggccggaa tttgaaccgc gcccttccgg taatccttgc ggccgtagtt gacatacacc 15180
ttggtgccgt cctcataggt ggtcacgtgc acctccgggg taaggaactc atggcctgtg 15240
atccgctggc tgttcaggcc ggacatttcc ttctggtaac ggaggatcat cggcaggacc 15300
tggtccttcc accggtcata ggacgcggcg gtatagcagc tgtaagcgga atcctgcagg 15360
atattggtat cttccttcat gaaggagaag ttcaggcctg cgccgtattc cgcgctctcc 15420
agcaaagcgt tctggaagtc cgcggagagg ttgatggctt cgccggtata attccgcagt 15480
ccgtgaagcg cgatctggta gaagggtacg gcctggtcga taatggagta tttgcggccg 15540
gtcaggttca tgtcggtgat caggtccgca taggccgccg cgtagtcgtt gcccttcttg 15600
atgctgatgt tcaggccgag ctcctgggct ttctggaggg acttgatatt cagcgccttg 15660
acctgttccc gggtgacggt gtcgttcttg taatagtccg cgctgagcag gttgccgata 15720
tcgcggaatg ccacgccggc agcgttgcgg tccttcacgc cgtcgatcag ccggttggcg 15780
aaatcggcag cgatcttggg cttcaccagg tagtagttct cgccgtccgt gttctgctgg 15840
taggtgacga tgtcatacgc gtaaagcttg acaagctcgc gggtggtgaa gcgggccgcg 15900
tgggaggtcg gaataaagcc ttcaaacagg ccgctgtcat aggcaaagca actgatgccg 15960
tccaggtaca gggccacgcc cttatccttt gcggcggcgg ccagggtctt cagggcggaa 16020
tccccgccga gctcgccggc ggtatggatg ctggtcagca cccgctggcg gacgccgccg 16080
ttgctccagc cggtcatgcg caggttcagg tgcttgacac cctggtccag cagctcgttc 16140
atgatatcct gggcttcgct gaacttcgtc acggcaatgg tggaatccac cggcatgccc 16200

atcttcggga tggtcttgtt gatggcgccg atcagttcca catggatcgg gacttccgga 16260
tccgcctgtt ccttcttgaa ctcttcctta ccgcggagat agtcgccgta ggcttccgcc 16320
atgccgacat agttatcggt atctaggaag gtatacttct ggatgacggt atcgtccgga 16380
atggtgttct cgtacatgag cacctgctca ttggtgcggc cggaaagctt gaactggccg 16440
taatggatca ggttgtactt gcagtacacc gtgttgaagc tgttgaaccg tccggcgata 16500
tcagcgtaga tgccgccgta ggagctggcg ccttccaggg cgcagatgaa ggatccgtcg 16560
ggctggccca tgccgaagac ggcaaaggtg ttctcggtct cgctgacggc ttccgttctc 16620
tccatgccgt aatcccatcc gtacaggttg gcatagtaag cgctctgggc cagtttgccg 16680
ttgttgtact tgatgatgcc gccgctgcct tccggaatga agatgaaacc ttcctcatcc 16740
gttccggctg cgccgaacat cggcagaacc gatacatagg tcagcggata cgccgcttca 16800
tacatgatct cgttgtaggg aacttcgacg atcatctggt tcccttccag gcggagaatc 16860
atggtggcgt tgaagaacgg tccgtggttt tcctctttct gggcgacata ctggaggtca 16920
gcctcgtact gttcctggtc atagccgatt ttctggaagt atccttcgat cttttccttg 16980
gcagccgtat tggtgccgga cttcaggatg tacaggtttc cgctttcgat ctccgggtac 17040
atcgctttca gttcatccag gttgtccgcc ttgtccagtt tcttctggtc atagagggag 17100
tagaaggagg aaaccttctt cttgtctccg ccggacattt ggccggtcag ttccttgtag 17160
cgctccaccg gaatcgccag gggatgtgg taagtccgct cgatcttgcc gatggcgtac 17220
tccaggcgga tggtggtatc gttttcccgg atgatgttga agttctggtt ctccatggaa 17280
tagcgatagt tgttcagttc cgtctcgttg ccggagaccg tgtaggtcag ggtcagcgtg 17340
gaggaaagga aatcgctgtt cacgccatgg gcaatcgggt ccttatcccg ggtctgggga 17400
ttggagtacc agatcttgcc ggtatccttg ttggtcactt tgaactgggt ggtggccccg 17460
tccatttcca gcagcaggtg ttcattttca atggtcaccg gcgccccgtc tccttcatag 17520
aagatcaggt gcggatcccg gccggaggag acttccctct ccgaatagat cggaacgaag 17580
acgaagatga tcagggcggc cagggcagcc aggcaaacca gccagatgat caggcgggtg 17640
atgatctttt tcttgtttct gtccatcaat cttcccctc cttaattcaa tctgaagacg 17700
atctcgcgat agaagctgac gaagaaggat accgcatcac tgagcaggct gaagaacacc 17760
agcatcagga agaggatcac gcccgctccc accacggtca cgatcaggaa gatgaaggtt 17820
ttgccggggc cgtaatcgtg gatctccatc aggccgacga agaccaggaa tacgcaccag 17880
atcaccgaga tgctcagaag cacggaatag aaacttccct cttcataggt cagcatgttg 17940
ctgacgagga tcatcggcag ctggataata atgtagggca ccagggcgta gcacatgccc 18000
atgtagatgt ccttgaaacg gcccttgcct tcgaacaggg tggtcatggc ccagttcgcg 18060
acgcacagga cgatgaaggg cagcagcagg gaggccattt cctcaaagat gttgatatgc 18120
tggatgggtg cggaaatgaa ctggaagttg gtggcaatca gtttcagcac ccgggtgacc 18180
aggaacagga tcaggaaggt gtttgccgcg gcaatgcttc ccttcttttc atgaaccagg 18240
tcccagaatc cgtcaaacgg atgggtgatc acatgcagcg cgtaatgcag ggactggccg 18300
tagcggctga accaggcttt ggttttgccg ctgcccgcct tgaccgcctg atcacttacg 18360
gactttactg tgctcatgca ttaccacctc ccatttcgtt ctcttgatgc gtcccagaac 18420
caggggaacg atcagtatca ccgccaggat cagtacaatc caccagacgt tcttctccac 18480
ccagatcttc cggtattgct tgaacgcccg gccgtattcc tcccgcgcgt gagcgcgctg 18540
gaagtaatcc atggccccct ggtaatcctc ctgccggagc agggaccgtc cgatgccgcg 18600
gaaggccagg gagtagttgg agttcatctt catgacctgc tcccagagtt ccgcgctccg 18660
gtcatagtca ccgttgaggt aggtatcggt ggcgtcatag atcagctggc cgtattccgt 18720
cggggtgaag atggtgatgt tgttcttcat ctcatccacc accagcaggt catagcccat 18780
gtgatccagc gcgatggccc ggccgaaagt gccgtccatg ctgcccttgg taccgaaggc 18840
ccacagcatg acgccctggc tgtcatagcc gaacaggcgt ccgcgggtcc ggtccagcgc 18900
cacatacatt tcgtttttca gcgccgtgat atccgccatc ttggaggggc cgtactggag 18960
gctttccgcc ttccagtcca ggtcgccgat ggggggataa cggtcgttct tgatcaggat 19020
gtcactgccg atgccattga ggcggcggat cggcttcgcc acgtccgaca gaagctcgta 19080
ctcgctgaac acgagggtgg tagcatagat gaatcctttt tcatccatat acatgttctc 19140
gtattcggta ggcacgaagt tttcggaagc ggcccgctgt tccttggaca tgaagtaccg 19200
tttccagata tagtcggcca tgttgacctt aacggggttg gcgccgatat agcctgtgaa 19260
ggtggtatcg ctttcatatt tcatgaagcc cttgttgacg ttggtcgcca gcacgtaaac 19320
ccggcctgcc acgtccaccg cgatcttgct gggcaggaag gaggaggact ggttaaaggt 19380
ggcgtcggtg ggcttgtcaa acagcttcag gaagttgagg tctttgtcca tcatcacaac 19440

ccggttgttg ttctggtcgg caacgtaaag gttgcccttt tcgtccgcga aaacatccgt 19500
cgggttgctg aaggaggaag gctcagcgcc cttgatctct tcgatgatcc ggatcaggct 19560
ggcgccctct tcggtgtagc ggatctggag gatgcggttg ttgccggaat cgaccacata 19620
cacatccttg tcccggacga agatgctctg gggcttgttg atggttttgt tgttgaggtt 19680
ttccaggccc agggtggtgc tgtcgatgac cagctccaca cggtacggat ccggggacat 19740
gagcacatct ccccagtagt cgtatccgta ggtataggat gtgttgaagc cgtcatcctt 19800
catgtcaaag ttgtcatccg ccagcgcgcc ggcagcggag agcgccagcg ccaggaccat 19860
cagaaaggaa atcagttttt tcatttttgc ttcctccctc cccttactcc ttgataccgg 19920
aggacgccat ggtctccagg atctggctct cgctgaggat gaagatgatg accggcacga 19980
tgaagatgat cacgttggcc gcggccatct ggcctgtacg ggctacgccg cctgcctgga 20040
tctgctgcag cgcgtagacc agggttttct tcgcctcgga gtaaatcacg gtggccgcgg 20100
cgttgttcca cagggactgt acgctgaaga tgatcatggt cagccaggcg ggcttcacgt 20160
tgggcatcac gatggaccag aagatccgga agcggctggc gccgtcgatg gtggcagcct 20220
cgatcagcgc ggtgggcagg gtttccatgg tctgcttcat caggaacagg ccgatgggcg 20280
ccgcgaacgc cggcaggatc agggcccagt aggtatccac catgcccagg cgggacatga 20340
tcagatagtt cgggataccg gtcacataac cggagaacat caggcacgtg gtaaccacgc 20400
cgaagaagaa gcggccgccg ggaaactcat acttggccag cacgaaggcg gccatgccgg 20460
ccaccaccag gtggccgaag gtacccacaa tcgtgatcag cagggtgttc accaggtagc 20520
ggctgaaggg cacccagctc tggcccatgg tgacaaacag atcggagaag ttgtccatgg 20580
tcggctggct gggccacacg ggcggcggga accggaacag gtcgctcagg ggcttcaggg 20640
aggaaccgat caggaacaca atcgggaaaa ccatggccac ggcgaagagg atcagcagca 20700
gatagatgcc gatatcgccg ccaatggaac ggttgggatg gcgccgggtg atcagatgct 20760
tcttcggcac ctcaatcttc tggcccaggc gctccgcctg ttcgtggagg agatcgccgc 20820
gggtcttttt gaccacaggc ttgtccgtcc agtttttgtc catgtggctg tagcacttca 20880
gtccctgggc cagctccatc acatagcggg agtcgtcctt gaggataacg gtgatctggc 20940
tggtggtgcc gccccggcgg atattgtcca tccggttcat gcgcgcgtaa tgccactgca 21000
cgacaggttc agccttgccc tttgcataga aggcggcgcc ttccttgtca gcctgcaggt 21060
aaccgcggaa ctttttgccg ttgatcagct gaagggtacc gttggatctg acggaattca 21120
tttcgctcat atcaggtgcc caccсttccc agcaatgcgg taattgcttt tttgcaaaga 21180
atcatgacca ggaacaggaa ggtggcgatg gcgctggcat agcccatctc gaaccggctg 21240
aaaccgtagt caaacaggtg ggtaaccacc gtgcgggccg cgtagtccgt ggaaggataa 21300
ccgcacagga ttctcggaac gtcactgata ttgaaggaag aagtaatgct catcaccgcg 21360
ccgaacagca gcatgggctt catgctgggc agggtgatgt gatacagttc ctgccagcgg 21420
ttgcggacgc cgtccacata gcccgcttca tacatgctgc ggtcgatgcc ctggaaaccg 21480
gccacaaagc tcaggaagcc ggtgcccatg ctcatccaca ggctgacgat gatcacgatg 21540
gtcatcatgt agcggggatc ggtgagccac aggcggggcg cttcaatgat gccgaactcc 21600
atcaggatgg agttgaacca gccgtaggcg tcgccgcgga agatgatgga gaacacggtg 21660
aaagccgcgc cgcccaggga gggggcatag aagatgacca ccgccacgga gcggatccac 21720
ttgggcagct cgttgatgaa ccaggcgaac aggaaggaca gcatatatcc taccggtccg 21780
atgatcacgg cgcagacgaa ggtgttcttt accgcggtca ggaagacctc atcctgcagc 21840
accaggttga tatagttctg gaagccgatg aaccgcgggg gttccaggat gttatagtag 21900
gtaaagctgt aaaagatgga agtaccgatg ggcaggatga agaaggtaaa gaacaggagc 21960
gcgtagggcg ccaggaacag gtagcaaccc ttgttccgct tggccagcgt ccagccgtgc 22020
tgcatctttt ccttccgcat ggtcaggtat ctttgggtaa aggactgtga ctgcatgctt 22080
gcttcctcct tcctttctta atccaccggc aggttgaatt cccgccgctt gatcttgatc 22140
tcttcgttaa tggtccgcgc gtagttcagc agggtctcac gctggtcttc cttggtgttg 22200
atgacgcggc ggatagcgtt tgtgatgtgg cgcgtggtgg aatagccgcc cgcgatttcc 22260
gggaagcccc gggtgtgggc catctgttcc tccaggacct tcagctgttt gttggaccag 22320
gccagctgct tgatggcatc ccggttcgcc gtctggtaac gggcggaaga accgaggaca 22380
gcttccatct cgcggccgaa gcgggtctgg gcttcagcgc cggtccacca cttcatgaac 22440
tcccaggcgt tgtgcttcac cttttcatcc tgggtggcga tcatcatgca gcacaggccg 22500
tccgtctggg cagtgcggtc gatggtgccg tccgcctgct tcgtgccggg gaagagggtg 22560
aaatcccaca ggccccggat ttccggtgcg gaaaccatca gggtgttgaa ggtggcatag 22620
gaagctacgc ccagcggcat ttcgccggta cggaagcggc tgacaaagtc gaacaccacc 22680

ggcaggccgt agtcgttata caggctggta tactgtttga aggcggctac gccggcttcg 22740
ctgtcgatca gcgtgcgctt ggcgtcctcg tcatagatat atccgccttc ctggtagatc 22800
atggatccca ggatggagat atctgcaacg gcaatgtccg ggaagttgac agccacggtc 22860
aggctgttgc cctggatggt gggcagctcg gcaatgagct cgtcccaggt ctggggaatc 22920
ggcagtccca gttcttccat aacgtcggtg cggtagaaca gcagcgggaa gttctgggtt 22980
tccggcaggc cgtagatgcc cttgtgtttg ctgtcgttgt aaaccagcgg atccaggatg 23040
ctcttgtaga agggcttgac aacctcttca aagtcctcga actgggtcag gtcttccacg 23100
gcatggcgca tggcgtagtt gacggcgaac cagccgctga cggaaagcac gatgtccggt 23160
ccgttgccgg ccacgacggc gggcaggatg gcgtccacct ggaccagttt gacgttgacc 23220
ttaatgccgg tccgggcggt gaaggtatca tccaccatgg tcttcagcac tgtggactgg 23280
tcgcggccgg tggtgatcca gatttccagc acgtcgtcgg tatcgtcgta tttgtcgccc 23340
agggagttgt aatccacaaa gaaggaactc gcgcaggagc ggatctcgtg accgatgctc 23400
tggaagatgt tctcgtgcac ctcgggcacc ttggcgttct ctccggtgat catgatcagg 23460
tccacgtcca gtttgctttc ggacatgttc tgcatggcgg tgcccagtgc ggtgatgttg 23520
tccttgaagt tgacaaagga ctccgtgatg cgctcgttgc tctccacgaa ggtttccagc 23580
tggttggcca gcgtctgggc ggaagcggcg cggtcgctct tttcaccggt ggtggcgacg 23640
atgtcgtcca cgatcttgta cagccgcttg ctctccaggt ccatggcctg gataacctcg 23700
ggatagacct ggttcaggtt gtaatcgcgg aaccggtccg gattgacgcc cgtgagcacc 23760
aggatcttgc ggtagatctg gttgaggcgg aagatggagt cttccacatc cttcaggatg 23820
ggacccatct cacccagggt tacctccatg cggatggtat gggcgccctt ggtcagatag 23880
aagcggaagg gattgccctc cttgtcgccc agggtgcgca tttcccagct ggtgttgtag 23940
ccaaagggaa cggcctccag gtcttcgaag ggaatcacgc cgtcgatata cacggtacgt 24000
gcggacaggg cgccgcgctg atagatctgc cgcgccttga tggagatatt gtagaagccg 24060
tcctccggca cttcaaaatc ccattcgatc cattcgccgg gatgggtcca cggatcgccg 24120
ccgatatagt tcaggatggt gttggtcaca gaatagggat ccgtcaagga ggaggagcgg 24180
tcataccggg catagaggct cggctggctg cgccgcttgg catgttcgcc ctgtacggtc 24240
agagtgaagt tctttccctc atcggtcatg gaagcttcct tctggcccgc ggcatactcc 24300
gtataggcgg ggaacttcgc aatgggcagc aggttgatgc cgcagaggat cagcggctcg 24360
ttcaccgcgc ggatggtcag ttcgttctcc ccttctttaa aatagaaggc atagggttca 24420
atctggtatc ccatgtcatc gcggcagtac gcgctctgct gttcaaagcg ttccacctgg 24480
gtgggacgga tgtcgttccc ctggttgtcc ttgcggacct ctcccgcgtc ggtccaaagg 24540
cgcgagaaac acagggtgct ggcgccggaa aaaggtactt ctccgttgat ggccagttcc 24600
cgctcaatat caataccccg gctctctgtg gtcagatagt cgagcttgat attgtacagt 24660
ccctccttgg gcacctccac cttccaggtg agttcggact catctgaggt aaggacgcct 24720
tccggaaccg tttgtccgcc ggtcccttcg aaagcggcaa tatccacttt cacaggttcc 24780
gtcaccgcgg gaagggcagc gtactgctcc acataggctc cgtaggtgtt gctccgcccg 24840
atgccggtga catcggcgga gaggtctgtc ccttcgtatt tcgccctgta gttctcagtc 24900
ccgccgctca gcagcgtgat gagcacaaca atcagggcga ccgcaaggac accgatgaga 24960
atgttccgcc attttcttga catgagttca tctcccattc acgacgtttt gtttttgcgc 25020
cgcaaaaatc gtgttgatga ggtttttgaa attggaaaga ctgcataaaa taatgcccaa 25080
atgattatat cggggtttgt gcaaagtgtc aacggatcaa aaaaaagctg cgcagttctt 25140
gcgcagcctc ccgaaattgt atggacaact tgagaatgca atgcttagcg gatctcggcc 25200
cccagtttga acttcaggtt gccgaggatt ttttttatgt atttatccag ttcttccggc 25260
agcaggggtt tgtcctccgc ctcaaaggtg atcttgaagg ccatggactt gcggccttcg 25320
ccgatctgct cgccgcggta gatgtcaaac agctccacgt cgctcacccg cggacaggcc 25380
cgggcaatct cgtgcatcag gtcgccgcag gggtttcct ccgccacggt cagcgccagg 25440
tcgcgcacca ccggcgggaa ggcggacagg gggcggaagc ggaagctggc cgccacgtgg 25500
gccatcatgg ccgtatagtc aatctcgccc aggaagatct tgtggttgga tttggcgtcc 25560
ttgtgcagtt tcagctcgcc ggtcacttca ttggccagct tgccgaagca gccgatccgt 25620
tcgccttcgc acaggatgta tgccgcaatg cccgggtgca gccacggcac atccgcggcc 25680
cgctccacgt caaaggtgag gcccagggcc ttccccaggg cttccacggt gcccttgacg 25740
gtgaagaagt cctcccggtc gccgaaggcg gcaaagccca gatggggccg ttcgtccgga 25800
agctctcccc cctcccgggg cacatagatg ttgctgattt cgaaaatccg gccctcgttg 25860
ttgcccttct tcaggttttc caccacaatg ttcagcaggc tcggggccag cagcggccgc 25920

ataatggcca ggttggcgga aatggggttc cggatgcgca gcacgttccg ctccggggcg 25980
tcctccggaa tccgcaggaa gtccaggtcg gcatccgcgt agaaggacag ggtgcatact 26040
tcatggtacc cctggccgca gaggatgcgg caggtccgtt cgcggcgctg ctgggcgggt 26100
gtctttccgc cgctggtgac ggtggccgcc ttcaggaagg tgggcacaac atggtcatag 26160
ccgtattccc ggatcacttc ctccgccagg tccggctcgc ccacttcgat gtcctcccgg 26220
taacgaggcg cggtcacatc cagggtgtcg ccgtcccgga ccacttcaaa gttcaggttc 26280
ttcaggatcc ggagcacatc cccggcgggc acttcaatgc ccaggatgtc gttgatccgc 26340
cgcaggctgg ccttgaagcg cttcccttcc cggggtgcgc cggcgctgca gtcaaacgcg 26400
ctggaggtaa tctcgccgca gtccagctcc tggatcaggt gcagggcccg ggccatgccc 26460
agctccgtaa tatattcgct gatgcccttt tcaaaccggg cggaagaatc gctgctctgg 26520
cccagggcgc gggaagtccg ccgtacattg tcccgggcga atttggctgc ctcaaacagc 26580
agctgggtgg tgttctccgt aatctcgctg ttcaggccgc ccataatgcc cgccagggcc 26640
acaggacggt tgccgtcgca gatcaccagg ttttccggtg taagggtgaa ttccttctcg 26700
tccagggtct ggatcttctc cccttccttc gcccggcgga caatgatgcg ccgctcctgc 26760
agctggtcca tgtcaaaggc gtgcatgggc tggccgatct ccagcagcac atagttggtg 26820
atgtccacca cgttggaaat gctccgcagg ccgcacaggg ccaggtggcg cttcatccac 26880
cgggggctct cccccacggt gatgttccgg accccgtggc ccaggtaacg gggcacagg 26940
tccggcgctt ccacggtgat atccagtccg gggaacacat agtcggaaac ggtgtagtcc 27000
gttttcggca ttttcagctc cttgccaagt gcggccgcca cttcccgggc aatgcccagg 27060
acgctctggc agtccggccg gttggctgta acggaaatat caaagatata gtcgttgagg 27120
cccaccactt cccggatgtc cgtgccggga acggtttcct ttggcagatc cagcaggccg 27180
tatacctccg cgccgggata caggtcctcg ttgaggccca gttcatcccc ggagcagagc 27240
atgccgtcgg agggaatgcc cttcaggggc ttcgccttga tggtgatgcc gccgggtagc 27300
gtggagttgt ccagggccgc cggggtatgc atgccggcat agacgttggg ggcgccggtc 27360
acaatctgga tcggattgcc gtattccccg cagttcacct ggcagataaa caggtgggtt 27420
ccttccacgg gctccgcgga aagcacttcg cccaccacca cgcggctgat ctccttgccc 27480
aggtcgatca gctcttccac ttcaaaaccg cagccgaaga gtttctcttc cagttcttcc 27540
gcggtaatgt caatatccac atattccttt aaccagctgt atggaacctt catgatgccg 27600
atactccttt atctcatcct taccaacggt atcatccaaa acggtgtctt ccctcaacaa 27660
agggtcatcc ctcaacaaaa ggtgtcatcc tgagcgcagc gaaggatctc aggcgctgca 27720
attcccgatc tcatgaaccg caggaacatt cccccggtgt catcctgagc gcagcgaagg 27780
atctcaaacg ccgcaaaccc cgatctcttc agccgcagga acactccccc ggtgtcatcc 27840
tgagcgcagc gaaggatctc aggcgccgca attcccgatc tcatgaaccg caggaacact 27900
cccccggtgt catcctgagc gcagcgaagg atctcatttg aaacatgttc cttcttcttt 27960
tctcattcct tgaactgctt caggaacctg agatcgttct caaacagcag cccgatgttg 28020
ttaatcccgt acttgagcat ggtaatccgc tcgatgccga taccaaaggc gatgccggag 28080
tacacgttcg ggtcaatgcc gcagttctcc agcacatggg ggtgaaccac gccgccgccc 28140
agcacttcaa tccatccggt atgcttgcac aggctgcagc ccttgccgcc gcattcaaag 28200
cagctcacgt ccacttccac gctgggttcg gtgaagggga aatagctggg ccgcaggcgg 28260
gtgcggacat cgccgtcaaa gatctgctgc acaaaccggt ccagcatgcc ctgcaggtcg 28320
cacagggtaa tgcccttgtc caccaccagg ccttccatct ggtggaacat gggggaatgg 28380
gtggcgtcgc tgtcggagcg gaaaacccgt cccggcacca gtaccttgat gggggggcttt 28440
tccatatcca tcacccggat ctggccgccg ctggtctggg tgcgcagcag caggtcgttg 28500
tccaggtaga aggtgtcctg catatcccgg gagggatggt ccttgggcac gttcaggcgg 28560
gtgaagttat ggtcgtcgtc ctcaatttcc gggccttcga agatttcaaa gcccatgccg 28620
gcgaaaacgt tgatcatctc gtttttcacc agggtcaggg gatgcaggcc gcccaggggc 28680
atttccgtgc ccggcagggt gatgtccacc tgttcccgac agttgcgggc cgccagctcc 28740
ctctcttcca ttctggcgct cagggcctgg tactttttct ccacggcgac cttgaaatcg 28800
ttgatcacct tgcccatggc gggacggtct tccttggcca ccgcgcccag gcccttcatc 28860
aggtccgcaa cggaaccctt tttgcccagg aattcctgcc agaaggcagc cagtttctcc 28920
ttggaatcca cctggccgat gcgctgttca acctgttccc tcagggcttt gattctctgc 28980
tccattttct cctccatgtt tacaaaaaga aagcggctcc gatggaacgg agccgcgttg 29040
tatttctgca cccagtaccg ctttccatcg ttgcgatatc aagcgtcact atatcacgcg 29100
gaaggcccga ttgtcaatct tattgggcgg tccggaccag ggttttgatg tcctccacca 29160

```
     cggccttcac atccgggctg gaagccatgt ttccctccac ggtcttgcag ctgtgcaata     29220
     cggtggtatg gtccctgccc ccgaaaacgc tgccgatctg gggcaggctc atgcctgtca     29280
     gttcccgggt caggtacatg gcgatctgcc ggggcacggt aatctcccgt ttccgggtgg     29340
     gactgatcat atcctcaaag ctcagcccgt agtaatccgt cacggtctgc atgatcagct     29400
     ccgccgtgat ctgcttgtgg cgccggttgt caaagatctc cttcagcgcc cgttcgcaca     29460
     ggtccagggt gatgggttcc tttaccaggg tggaataggc caccaggcgt gtcaggcagc     29520
     cctccagctc ccggatatta ctgtcgatct ttccggcaat cagctgaagc acttcgtccg     29580
     gcacccggat attgtcctgg gtggtttttt cccgcaggat cgccacccgg gtttccacgt     29640
     ccggccgctg gatatctgcc accaggcccc actcaaaccg gctgcagagg cgttcctcca     29700
     gccgctggat atccttgggg ggcttgtcgc tggtcaggat tatttgcttg ttctcgttat     29760
     gcagctcgtt aaaggtgttg aaaaactcct gctgggtgct ttcccgtccg gcaatgaact     29820
     ggacatcgtc caccatgagc acgtccaccc ggcggatttt ctcccggaac tcgtaggttt     29880
     tcttcatctg gatggcgctg atcagctcgt tggtaaagct ttcgctggtc atatacagga     29940
     tctttttctt cggatccgcc tggtggatga agttgccgat ggcctgcatc aggtgggttt     30000
     tgcccaggcc cacgccgcca tagatgaaca ggggattata ggcctctccg gggctttccg     30060
     ccaccgccag ggccgccgcc tgggcaaagc ggttggcgtt gcccaccaca aagctttcaa     30120
     aagtgtaccg gttgttcaaa tggggatcgt tgtcctccgg ctcggcggg                 30169
//
ID   XXX; XXX; linear; XXX; XXX; XXX;  XXX.
XX
AC   PRJEB67889
XX
DE   Uncultured bacterium DNA fragment from in vivo enriched cow rumen, clone
rum-33m21
XX
FH   Key             Location/Qualifiers
FH
FT   source          1..35197
FT                   /organism="uncultured bacterium"
FT                   /environmental_sample
FT                   /mol_type="genomic DNA"
FT                   /isolation_source="Cow rumen"
FT                   /clone="rum-33m21"
FT                   /note="AZCL-Barley beta glucan, AZCL-HE cellulose active
clone"
FT   CDS             81..617
FT                   /translation="MSAYISGGSKSGKSMLAQRIARAMAGDGPLYYVATMLPGDDEDAA
FT                   RIARHRREREGWGFVMLEQARDLPDCLDKADPRGAFLVDSVTALLANEMFGPDGIHAEA
FT                   PAKVADDLAAFLRRAPKAVFVSDFIFSDAALYDSLTEAYRKGLAGVDRRMAALCDTVVE
FT                   AVAGRFILHKGELSI"
FT                   /product="Adenosylcobinamide-phosphate guanylyltransferase
FT                   (EC 2.7.7.62)"
FT                   /EC_number="2.7.7.62"
FT   CDS             614..1366
FT                   /db_xref="GO:0008818"
FT                   /db_xref="GO:0009236"
FT                   /translation="MKRFILALCMCFSNFTAIPCPYRPWDEDARELMIACLPIVGAVIG
FT                   FLWYLLALLAGRWLPAAHAALIVALPWLLTGFMHLDGFMDACDALLSWRPLDQRLRILK
FT                   DPHAGSFAVIAMALLALFSWGAASRMGPKDDLRVLLFIPVVSRCCSAFCVLALKPLSAS
FT                   QYAGREDVPVQRLAIFAMWMFCVVASALWLKARAVVLLVETAAYAAAMAWAYRTLKGVS
FT                   GDLAGFALSIGECAALVALANQIARAAL"
FT                   /product="Cobalamin synthase"
FT   CDS             1363..1731
FT                   /translation="MIFIYGGSWQGKLDFAKAQYGLEDGEIQICDEATEALDAGARCLA
```

FT                   YVEKWALNRVRAGVDPLEAFRVRHDAIVIATDISCGVVPVDATLRAWREASGRLSAQIA
FT                   ARSEAVWRLFCGLPERLK"
FT                   /product="hypothetical protein"
FT   CDS             1728..2309
FT                   /translation="MRTLYVVRHSLTEGNERRLYCGSTDLPLTGAGRALAKARAKDRPL
FT                   PPCDLRVTSGMARTDETLFLLTGRTPDIVIADLAEMRFGAFEMRGYEELREDADYRRWI
FT                   GDCMGAGLVRCPGGESQSEYRARVLRGGAALLALEWQSALAVIHGGSIANLMDAWFPGE
FT                   GKGFYDWQPGPCEGYAVRFDGDAPREYEVL"
FT                   /product="hypothetical protein"
FT   CDS             2311..2832
FT                   /db_xref="GO:0008817"
FT                   /translation="MEIGQFQVYTGDGKGKTTAALGLALRAAGAGLRVYIGQFMKAGET
FT                   SEHASLRRLGITVEQFGSGKGLLIRREADADDRACACEGLMKVRRAMTGGAYDLVIADE
FT                   INVAMACALLEEADLLALADARPEGVELVFTGRGATEAIIDRADLVTEMRAVRHYFRDK
FT                   GLPARTGIEK"
FT                   /product="Cob(I)alamin adenosyltransferase (EC 2.5.1.17)"
FT                   /EC_number="2.5.1.17"
FT   CDS             2863..3234
FT                   /translation="MRFKAILAEAARKRYPLSILLMVIGVLAIPFPYLFVHFGLELTTV
FT                   WWTILSIGLGVILGCVIIGLGSLVEDVHDISMHTVGFDLELGELDEEDYEDEEDAPEAE
FT                   NEQPEAEPEQPEEAPADAE"
FT                   /product="hypothetical protein"
FT   CDS             3262..4206
FT                   /translation="MDTQRFIDYRDRVVGAVSKAIVGKDDAIGKVLICYVCGGHVLLED
FT                   VPGTGKTMLLRAFARAIGGSFSRVQFTPDLQPSDLTGINFYSQKEAEFRFRPGPLFHDV
FT                   ILADEVNRATPRTQAALLEAMEERQITVDGDTYPLGGRFMVMATQNPLESYGTFPLPEA
FT                   QIDRFFMRLSLGYMKRDEELSVLSRRDTVDILPEIGQVVTDEDTAFIRANYRGVRVSDE
FT                   VAGYLMDLIEATRGGEFSCGVSTRGAIALYKAVQASAALSGRDYALPEDIRRMAPCVLA
FT                   HRVAAGGGLRRADAEAKLNHLIDRIPVPVEGAV"
FT                   /product="FIG022979: MoxR-like ATPases"
FT   CDS             4203..5270
FT                   /translation="MIAFLLMLAAVIAGAELLSLRRGLDGVEYDASPSKPLAGPDEPLE
FT                   LVTVVTNRRRRFLPFIRLNEDVPDALVVSGDLRVEDVKNQRSALRSTVYMMPRQRLTRR
FT                   TPFSIRQRGRYLFLGATLEGGDFMGLSTASRRVELERELVILPALADCGALREVLGGWM
FT                   GERSVNRFIMEDPVLTLGYRDYTGREPMKQIAWRQTARMGRMMVRCQDHTVERTVTVML
FT                   NAHTFAFGSYGRKMLERALSLCRDVCQQLEERRIPYAFISNLRVIGAAEGFGEVGDGLG
FT                   QAHLATVLEGLGRADYGSRDSLEGLFDRALSRANLGRGHIYITPVDRDIHHDLVDRLRA
FT                   ATGEAPLCIAADEYE"
FT                   /product="hypothetical protein"
FT   CDS             5286..6506
FT                   /translation="MSFAYALTFFCDALLYYGAIGCMGLLGACPANLYWAPVILLAACW
FT                   LSGRWTGRGKPWLRWLPTAAAIPAMLVISNWPGRLATLPMAAYLPLYIYNNRRAPDYDY
FT                   AADRFRHSLIGMGVALFLGALFRAASWKRGLPYLFLYFTLNMTLLRLLRHDDRVARSRR
FT                   FRVLNLAGVALVCAAGFALSQPAVIAAVRVAWNWFLENIVVNLLTALAFVLQYLLYALA
FT                   WLITRIFGEGALGIGGLPDLSPTEGYQSTLPRTPAEVRMLPVWVQWAVKGIGIALLAVL
FT                   AFVILRALSRRVSRTSTDSGNDERESLDAEAPREPRRLLRRRDPQQGVRRWYRRALALI
FT                   RARNGRVAPTMNTLQIQQENAETVDYDAMDALRELYLPVRYGGHEATREDVARAKEAYE
FT                   RLRKARD"
FT                   /product="hypothetical protein"
FT   CDS             complement(6752..7450)
FT                   /translation="MRIIFVRHGEPDYANDCLTGTGHEQAEACAERLAGEGIAEIYASP
FT                   CGRAAETAEHTARRLHLPVTTLDYMHEISWGGEGIPSEGHPWTLGDALIETEDFDFYAS
FT                   DWREHPFFNGNRATDYDRMIAGKIDEFLLGYGYKRAGTRYFSANGSDQTIAIFSHGGSG
FT                   ACALSRILSLPFPYVCSVMPYDFTSIIIVDLPAILAGYIHPRLELFNDCAHIHRDSQGP

```
FT                   LIQQKPDAAM"
FT                   /product="GH3"
FT   CDS             complement(7472..8401)
FT                   /db_xref="GO:0031176"
FT                   /translation="MATYFEQHGALSVSGTRLVDAAGQPVQLRGVSTLGLAWYPEFVNE
FT                   AGFATLRDDWGANTVRLAMYTCEEGGYMTGGDKAALEALIEKGVAICAKLGMYAIVDWH
FT                   ILSDGDPHIHMAAAEDFFRRMSARLADQPHVLYEICNEPNGAGVTWPVVKDYAERIIPI
FT                   IRENAPKAVILCGTPTWSQDVDHAAKDPLDGGNIMYSLHFYAATHKDGLRAKAEEALAA
FT                   GLPIFIDEFSICDASGDGVIDYDSAAAWVDLIRKHGLSYIAWSLSNRDESAALVRADCP
FT                   KTDSWAESDLTETGRWLRCLLRRDANLG"
FT                   /product="GH5_2"
FT                   /EC_number="3.2.1.8"
FT   CDS             complement(8426..9832)
FT                   /translation="MKKLHMGPAHCALDIGDGSERAEYVNQDYLLHKLGRPHRAVNIMY
FT                   TYYPHDAEWPARISEAWKDKSVNFAWDYPYDDYFPYGANGQPFEQMKDIRRHGQDVLLT
FT                   LTLDCGLSDDELRAVARQLRPFGRMMIRVNHECCGDWFQHNKRYTYPEVGAFFVRFAGI
FT                   IHQEAPQVRVIFCGGFCGEDGHVEKEEAFRDCYAAADLWSWDSYPALHYGWPYDIAEVG
FT                   GGRYKADPIGMIVESFARTHARATELAGESKPMITAEFNTDGDVTGPRMQGESVVRFAH
FT                   YWRDNRGDWFKSISLYQFRDRGRLGLECEDPNNSAVGVEQPMLAEYRDQVLNDPWFMPG
FT                   MAEGEAASFPMSMRWGGSEDADGLAMPIEFEGNPEFCEMTVEQEIGLMVELNGRWFYKA
FT                   PGTKTIDLMSAFFSKPLDGPQTLTLRVFAPPAEGVNPDDGSDDWMANYRATLDEPPKMR
FT                   IRYEVPGIVR"
FT                   /product="hypothetical protein"
FT   CDS             complement(9895..11832)
FT                   /db_xref="GO:0004565"
FT                   /translation="MRIGVDYYPEHWDRALWREDADLMAATGVKVVRLGEFAWSRLEPR
FT                   EGEYDFGWLDEVIALMAGRGIDVVLGTPTSCPPLWLYEAHPEIAQQHYTGLREPTGIRG
FT                   HRCYESPVFRDYAAKIVTRLAERYGRLKHVIAWQVDNEVEANFCCCEVCVGRFRDWLKA
FT                   RYGTLEKLNAAFHNDVWSGDWSSWTQITPPVKNHPMVWYNPALVLDYQRFAAGSTAGYV
FT                   AFQRDLIRSIIPGARITTNFWFCEHMPELYGMYRDLDFASYDNYPSTRLNQASHAFHLD
FT                   FTRGLRRQNFWIMEQLSGLPGCWMNMQRAPQPGMIKGYALQAFARGADAVVHFRWRSAV
FT                   GGAEMFWHGLLDHDNVPGRRFREFGSLCEAVEKLKPAEGTSVMSQVAILFGMDNEFAFK
FT                   SQLQTEGMDYFEQLTAWHRAFAALGVNVDVIDENADLTGYRVVVAPTMYVRAPEAVAAL
FT                   HAFAQRGGTVLLTCRSGVKDINNNCVMAPLPGDYADMVGARVVEYDSLENDTASIDMGG
FT                   RTWSARRWADVLALDTAEPLATYADQYYAGSPAATVNRWGGGRVYYVGTVPEDAACRHL
FT                   AERMLQEARVPFLPDLPVGVEVSTRAGGDRRFTFIFNNNPEPVTLEFMGRELALGPYAV
FT                   RVPELSEEAL"
FT                   /product="GH42"
FT                   /EC_number="3.2.1.23"
FT   CDS             complement(11991..13547)
FT                   /translation="MKAVIYARYSSHSQTEQSIEGQLRDNYAWAQQQGISVVGEYIDRH
FT                   LSGTKDQRPDFQRMIADAAKGQFELVIVWKLDRFARNRYDSAIYKAKLKKYGVRVVSVK
FT                   ENITDQPEGIILEGLLESMAEYYSANLSQNIRRGQRESIAKGTYLGGTTPYGYKALNGK
FT                   LVIDDKTAPMIRYLFEHYAKGEARKDIIDELNRRGYCNRYGKPLSPNSFSHVLSNTAYI
FT                   GEYRYNGEVVPGLSEPMIDPEIFARVQDRLKQAARAPASKKAKVSYLLQGKAYCGYCGT
FT                   AMVGESGRSHTGAKYHYYACHNKKKRRGCHKKNERKDFIEWYVVEQTVQYILTPGRLEQ
FT                   VAQAVVEQYKKDFSDSQVEELEKALAQIDREMDKLVDALIEAPKAAHKKIYDRMEVLDA
FT                   QKSDIEIDLAKLRIAQGIRLTEDEVRAWIKKFTNGDPLEEEFRRRIIDVFINCIYLYDD
FT                   RVVIFYNIRGGKQVSYIDLINSSDLPVEDESSDLKSDAVPAKSPENTAFSGLFFCVYPR
FT                   R"
FT                   /product="hypothetical protein"
FT   CDS             complement(13653..14348)
FT                   /translation="MRRNDDMILTTSSSIEGYKITAQCGLVFGETVFKHGFLSRLGASV
FT                   SNTIDTLALGSREMSGSMGLIEEARAFAYDKMIRQAKDLGANAIIAIDSDNTIGGDIMY
```

```
FT                   ISLYGTAVRAVSEKDYEKQIAAEREAAMQREAEMERQAQERKQRLDELKERRSAGDFAA
FT                   EEKLLSEIASYDSVMQIWQSWSALGLDVKYPDITSRIKNSKETERMYGKMAGEANAIKK
FT                   MINEAIFEE"
FT                   /product="hypothetical protein"
FT   CDS             complement(14368..14733)
FT                   /translation="MSIGTRIKERREELGITQVELAARLGVSKGAIGNYETDANSPKAS
FT                   IMYKIFDILQCDANYLYQDEVGTLESTMPPADERMLVSLYRKLNDEGKAKLVDYADDLV
FT                   KSEKYVRFHPSGMGKEA"
FT                   /product="hypothetical protein"
FT   CDS             14891..15124
FT                   /translation="MAKDPIAAGIQYIMAERGLIQRAVAERAGYTQQQFSDMMNDRKTI
FT                   KATDIVPISKALGVSIQEVYDAGARFMNREAV"
FT                   /product="hypothetical protein"
FT   CDS             15127..15333
FT                   /translation="MADKYIQVGVTALRNPATGDFLPAVPLYIKAEGDAPEAEEKLIED
FT                   IGNLLAHRMKAYIDGCKKAGVDL"
FT                   /product="hypothetical protein"
FT   CDS             15330..15710
FT                   /translation="MKIIAVDFDGTLCKADWPEIGAPNLRLIDWLKERRRTGDKLILWT
FT                   CRCGEMLEKAVMWCSVWGLTFDAVNANLPENVKQYGNDCRKVYADIYLDDKNGYIGRIS
FT                   PFPIYIRPEREDMKAPCWTARG"
FT                   /product="hypothetical protein"
FT   CDS             15715..16011
FT                   /translation="MSAKRFHVARPDELQTLTKYAKTNAPTVARTARIWKSAKAKKGKT
FT                   RWDNGVYVGFSGDRQRRAALFRVTGDDELVEFATLKYNLSLLWCEQHLRRVYE"
FT                   /product="hypothetical protein"
FT   CDS             16061..16192
FT                   /translation="MGAAEALAEALRKGLPPGLTAQRPCRNAFGALMGRAVPWGNDQ"
FT                   /product="hypothetical protein"
FT   CDS             16255..16437
FT                   /translation="MAWIMYTMRCFTLAFARSADADWAEAIPPAHRKKPTFSAIPPAGR
FT                   CIASGERDHTAATGE"
FT                   /product="hypothetical protein"
FT   CDS             17045..17299
FT                   /translation="MDSIEEIKIAAPGPRTCRICATAHRPEQPHDRDSLYYQNWFFKRY
FT                   RHFPSWADAMAHCTEAVQAEFTATLAKRGIAIEKGGDDG"
FT                   /product="hypothetical protein"
FT   CDS             17292..17651
FT                   /translation="MDSDWIEMSRSPIARECPDAPIIVWHVYSGAMVERLERARKNRFM
FT                   THWREIDNGAWIDARERKPTQEDADAYDCVISQDVWGEVAMAGWHRFDREPRLARWQTP
FT                   PPAPPNICELRNRIR"
FT                   /product="hypothetical protein"
FT   CDS             17664..18479
FT                   /db_xref="GO:0003676"
FT                   /db_xref="GO:0004386"
FT                   /db_xref="GO:0005515"
FT                   /db_xref="GO:0005524"
FT                   /db_xref="GO:0016020"
FT                   /db_xref="GO:0017038"
FT                   /translation="MNALANTETHLPSRLDELARQAQLFAQNACMNLLQLGRVLAEARP
FT                   MIPYGEFDGWCRQNAKMSKRTAEQYMQAYAEFGLNTKIAELGTSKVIKLLPMSEDEREQ
FT                   LLAENDVGSMSTRQLDEAIRKQRGQLLKEARAEAKEEIERHAETAKALLKEKLRLEREA
FT                   AEHDEDMAALQADYNAVRDELLAYKSADARGEDEHKATDTLTPDAFTSAVNAFIGKCCR
FT                   LPQMGRTFSAMPPEEKEIYEQSLCAIEKWAESARIAMNSVSFEEAIFVD"
```

```
FT                   /product="Protein export cytoplasm protein SecA ATPase RNA
FT                   helicase (TC 3.A.5.1.1)"
FT   CDS             18472..19017
FT                   /translation="MTDIIKHDSATPALPQETVEIISVVVRDMMAPIMQGVAKLLENNT
FT                   QAMEQIAAAQQATSERIAALEKRVRLQTPLSKSQENCINGAIRAKAKELLDAHGFSDDK
FT                   KAVTKLCGIIRKSVLARYGVGSLREAPAYDYDTAMKQVAMWNDRLTIRDVEREARNRAE
FT                   AMEEAVMELAKRSARLDG"
FT                   /product="hypothetical protein"
FT   CDS             19087..19560
FT                   /translation="MDKIRLSASDRESLMRLNVALEILETEGDQLARRLRVIKDGKRDY
FT                   GLIKSRINKLMECMIETIPEDQIMTYVRSLKMTSYTIGIKKPGPMGRNEKDYGMWLSYE
FT                   VINALLTGCHDHCMMCSEDKAERRACKLRKALATIPNDAPERDDNDCPFYTLM"
FT                   /product="hypothetical protein"
FT   CDS             19572..20216
FT                   /translation="MLISELAKQVHENAVAHGWWEENREAPEINALIHSEWSEALEEAR
FT                   AGRPLVWYDIYDRGDDCGGGGRCSAMVNEGVCSGRNGKCERNHKPEGVAVELIDGCIRI
FT                   LDYMGELGVTVQDCDSEADSEIEGLYTDEIMAEAEQPTAHVVAWLHMLTSEALSDAREN
FT                   ALLGDMNPMKLLSAMALALSWVKKQGLDPLAILLEKHEYNKTRPYKHGKKF"
FT                   /product="hypothetical protein"
FT   CDS             20254..21120
FT                   /translation="MDVNYLIEINGLDDWEFGNPLSATAYKVMRKLLMIANRQRFPERI
FT                   PISNTRLCFLVGCTEKSLIAARQQLIQRGLIDYKGKKKQTPVYTIHYFSHDPRYNRKNY
FT                   SYNGSYDGSMNGSYDGRMDGGYTGGRSINRSYGELEDCEEEDLGNDNPTTGTLTPVTAL
FT                   LHNPPPSLPSVDVLQGNRYPPRNRPLAGAETWDYAAAMSFFQNPDIQRMYGGGAGTIER
FT                   ILNSDAFPIKLVGYAMDKTYQRNRMYPEPLGNPAAYTLRLLNDWHEQGFHTPEDVQESK
FT                   DDWMHFG"
FT                   /product="hypothetical protein"
FT   CDS             21135..21536
FT                   /db_xref="GO:0003697"
FT                   /translation="MNKLTIIGNLTEHPQAREVSTSNGPATVCNFNVATNRYVRGQKRT
FT                   TYFRVSCWGKQAENAMKYLLKGRKVAVVGPVEASAYTAQDGSVRARMEVYADEIEYLSS
FT                   RQDAGEQAPPPQAPGYGEFTEVDDDELPF"
FT                   /product="Single-stranded DNA-binding protein"
FT   CDS             21551..21937
FT                   /translation="MADLYERRMVTARRPHKCHLCGLTIPEGHDYIREKWRDDGFREVK
FT                   RHIHCDAMLDEFLNSDSFVGEYYDDDVWEWIREKCLDLCGYEGREDCIDNPFACEQMID
FT                   TISNLNSRKAALDSLNLFRMRTGQ"
FT                   /product="hypothetical protein"
FT   CDS             21934..22338
FT                   /translation="MKSDATKWVEDVLAVLKQAASNHKDAMEAEKGCDDKTQDLLHELE
FT                   LVPHSYHERSGIAKELAAVRRKRRQAKDAHELLGPLVKWIGAQKTAIDALTRALGEMRK
FT                   IDEKRDNRLYYKRADGNHEIIRKQEDQEAK"
FT                   /product="hypothetical protein"
FT   CDS             22340..22600
FT                   /translation="MNEQEMRKKVLDGLTCCDALHNGGVVTCGTECPYGIADGIDRGCR
FT                   GKLIPDALALINEMDITIRALMGEFGDSCDVDSHCGGGKDG"
FT                   /product="hypothetical protein"
FT   CDS             22593..23132
FT                   /translation="MADRENVIKGLKCCRWGICFACPYNDGVHGNTGCKKKLVDDALEL
FT                   LNALDVTPEELERLKKCRHECKIDCLLEHYDAIKAERDALLEKQEPRLMVAKDFEDNPN
FT                   VDGDGYLPCWIECNEIEVERAIRFGAIQEGETLDGWTEVSKAELLPTDLSYNPNVRHWT
FT                   ARPSEEQRRKTPWPDA"
FT                   /product="hypothetical protein"
FT   CDS             23117..23332
```

```
FT                   /translation="MAGCLRIRDSVDIDIYVLPDTARCRCTGISPLDMDACPMRRFDYF
FT                   GMDCWPGECDEYTEDKEGEPWNDYSD"
FT                   /product="hypothetical protein"
FT   CDS             23311..23517
FT                   /translation="MERLLRLKTPSERRQDIQRELAFLERRKQYARSVYVKQLQEIAEQ
FT                   ENRLMKRLERIGIDGRKDENRLV"
FT                   /product="hypothetical protein"
FT   CDS             23486..24337
FT                   /translation="MGAKTKIDWCDASWNPVTGCLHGCDYCYARGIANRFGGHLRGEIK
FT                   DIMTPDGEIARVELVAVELTEPMHRNEAMGSGLMRVGPIIPYPYDFLPTFHRYKLDEPR
FT                   HWKKPRSIFVCSMADLFGEWVPDRWIRDVFDACAEAPQHRYLFLTKNPKRYIEAHPWYL
FT                   QRIQRPSKAFEAWFGYSYDGFGNSEIRKALKRQYSLHQFISIEPMRGAIWEDEINAITA
FT                   TCDWVIVGAETGNRKGKVVPDKRWIMKVAEQCEDSGIPVFMKESLRGIMGEDFRHETPW
FT                   ED"
FT                   /product="phage Gp37Gp68 family protein"
FT   CDS             24342..24551
FT                   /translation="MNKKPTPQELIAAIHAKCLECSGGSRKEVHTCKLTDCPLWLYRRG
FT                   EPRERLAQAKGQIDVFDIIQKEGA"
FT                   /product="hypothetical protein"
FT   CDS             24548..24823
FT                   /translation="MMTNRERMIQDFVEMSSEKLAETIRELSEDIIGENLREMVCDDCH
FT                   AEHGGRCVFSDNEGNEEADPDDVYDGCPTSMAEWLDKECKREHLIQ"
FT                   /product="hypothetical protein"
FT   CDS             25045..25233
FT                   /translation="MKVRVTVMTENNAPAEKAGKTKEERNATAKLAWNVIISLLQTVFL
FT                   AYDETIELESVEVLDDE"
FT                   /product="hypothetical protein"
FT   CDS             25223..25417
FT                   /translation="MTNKMNIRCCDTCRYGLGGGYDNCRINLEAECAAGEFEAWEPREP
FT                   DRKIIHYEDEISEWEPKDK"
FT                   /product="hypothetical protein"
FT   CDS             25513..26031
FT                   /translation="MARSGNVKINIGCEFVEKEIKPVVIHNKDIPLLADVLPIMQEICQ
FT                   IERRRDWQRDRMLNITQHLTGMPGGGRTPKGLDEAFALLSELDEEHEKRCKEYVRQLRT
FT                   AQKILNAIESRSMRTFVIMKYVMNVSDSEIRQELNMTRTGFNRARRCVESAERMADVVW
FT                   RERYILAKC"
FT                   /product="hypothetical protein"
FT   CDS             26167..26781
FT                   /translation="MAQLFTMDIDASDAQETIMRMKNAMTEVQFERAMYGIFQRTGRHV
FT                   SKILREDLPRKYEVLPGDISAAVGNPSLTTGFGGVGCTIPIRAARGTIGSKYRASGGAH
FT                   GWNSLKRRYKVKARIIKGVQSTLPDKWHSGYAEFRNLGSKLGGLTFARSTRTRGPIMKI
FT                   TGIAIPQMPLNRSEADVQNDIKEYLEKQIAERFMAMMRIGK"
FT                   /product="hypothetical protein"
FT   CDS             26778..27296
FT                   /translation="MSVEMTKKDLANIAGYSYHQLYNIDRDLPADKKLFVMGEGGKYDL
FT                   AFFVQRWVEYNVHRETDDTEDLDTVKARHEAVKIEKTELEVAKMRGALIDVQDVKRLWG
FT                   DVANTVMQNMIHLASKLAPMLLMMDNVEVISNLIDEEVRKTLADIAQCPLPSYAADDET
FT                   DESEEDEEV"
FT                   /product="hypothetical protein"
FT   CDS             27299..29158
FT                   /translation="MSALSELARYTFSMFRPPAKQTVSEWADENRVLVSESSAEPGAWR
FT                   TDRAPYQREIMDAFTQPGIWQIVIMASAQVGKSEIELNMMGCAIDNDPGPMLYIQPTDK
FT                   VAEDYSKRRIAPMINACPTLRDKVFKARSRDAMNTITMKTFPGGSLAIIGANSPADLSS
FT                   KPVRYIFMDETDRFPASAGTEGDPQELAERRTETFRHNRKIVKTSTPTIKGASRIETDY
```

```
FT                   MNGTQEEWHTECPHCHTYSYIRFADIHFEKEDFTNQNGDEDYHVKTVTWRCPICKRDIG
FT                   EYECKRLPAKWVSKNPKAKENGIRSFRLNAFMSPWSDWKNIVWKFLKARKDATKLQTFY
FT                   NTILGESWEIHHHDGLDEKLYQRREHYNAEVPTGVLLLTMGMDTQDNRLEYEVVGWDRH
FT                   GQSWGINRGVIPGRPDSQGVWDEVDNLLDREWKSTNGIKFRILATFIDSGGHYTTTIYK
FT                   QCARRAAKKIWPIKGEKGEGKDECRPMKRGQGDGAKFMLGVDAGKAGIMSDAQIEEPGP
FT                   NYMHFPIDYRAGYNMEYFKGLISERLEVHRRGGQGVAVWEQFYERNEPLDCRNYARAAY
FT                   RYFNWRFDELERYINGEEEPQITTRQQETKRKQRRVVSRGIQV"
FT                   /product="Phage terminase, large subunit"
FT   CDS             29181..29423
FT                   /translation="MAAISAYSLEEAREMLGLWKNAEKALASGQVTSYRVGTRECTLLD
FT                   LEDIRAAITYFGNLVEALSGTVRTKRVTRVVPRDL"
FT                   /product="hypothetical protein"
FT   CDS             29440..31092
FT                   /translation="MNMKPNLRERALFLFNPKRGNDAYNKRLREEQEQEQARGQRTPGS
FT                   PIMSYGSYGASTTKNSMLGWIVEAGNAEDNIDLYSSTLRQRARDLYAGGGLARSGPNTL
FT                   TTSVCGWGIMPKPKIDGDYLGMTDEAREETERNILREWKLWAENKLCDAERQQNFYDLQ
FT                   QLAFLSMLMSGDVFVLFGMKPNNRTPYQTTIRLLEADRICTPDSSGDSESVETEGGGRI
FT                   IDGVEIDKEGVVIRYYVASRSPIAGNDSSELTWQPIEAFGADTGYENILHIMTHERPEQ
FT                   RRGVPFVAAEIEALKQFSRYMNAELAGKLVSAMLTVFITSKDDDGKAGMEDAVNEDEKV
FT                   TDDQLKLELAPGAIYDLPPGKGIETVDPKRSDTQFEVFVNTCIMVIASSMGIPKEVLVK
FT                   KYESNYTAARAALLDFWRTVKVYRRRFNNAFNQPIYEQWLTEAVAAGRIEAPGFFDDPA
FT                   IRQAWCGCEWMGASMGHVDPLKEVNAAERRIANNITSQEQEASEYNGNDWAAIVRQRKR
FT                   EVEATRDMTGDTATQEPEAPTPDKDEEEGQEDE"
FT                   /product="hypothetical protein"
FT   CDS             31085..32218
FT                   /translation="MNKERYRIRFDMKAEGDEAEVMVYSEIANEKWWGDETTTGDFDKA
FT                   LKEARKSGATKLNLRINSGGGEVYAAVAMRSMIINAGFESVRVMIEGLCASAATLFATI
FT                   PGARVVIAEGSEFMIHNPMTIAWGNAEEIGKTVDHLHKMEEQFHGMYAAKTGATEDTIK
FT                   AWMDAETWFTAKEACDYGFCDELLTSEPVAACVTAQEMGVMKALYKAVPDGIAIKDPSP
FT                   TPTVSNGNPVAGLPTEIKNHEEEDSHMDMKDITLDQLRADNPALFTQVQQEAVAAERQR
FT                   IEDIDALTMEGYEAMAEEAKRNGTSALDFQKQVVAAQKQKPKDFIANRQKETAPAANVP
FT                   GGAPQNGKTDEQEIMDNAKEIAAFASEFSGNNNESMF"
FT                   /product="Prophage Clp protease-like protein"
FT   CDS             32245..32901
FT                   /translation="MSNLYETIGTSTPDQILAKVEADPIAVNLLPGQGEIKRGTLLYKN
FT                   AAGFYLPAGSGQISTSYDLVVLNETVDTGNDGTAVAEVAAAYREGTFIDGKVKYDNSGT
FT                   PTPVTEAHKVVLRLFGIKFNQSVESSGTFENGPATITYKANNDVTPAEADVAIEAQRGS
FT                   SYTILNNSDSKLSFTAPATKSFSKWNTKADGSGTDYAAAASYTASADLTLYAIWA"
FT                   /product="hypothetical protein"
FT   CDS             32937..33974
FT                   /translation="MDLYDVRSQLKAIELMPRECSVLADFFCKEEGVQNQKKAIYDFRK
FT                   GVKAMAPTVHPGTGGVIMERTGYETREIGFCTIAPERLIEDQNLEGRMFGEDVLGAMTP
FT                   EQREKKMLAKDLTEMRQAIQRRREWMVRQVLLTGKLSIFEYTNEGRNIEASMIADYHFT
FT                   NNFTPTTPWNQSGADIVSDMEAIYDLVYEGLGDVEKIVMAPNVWAALRHNDAFMKKLDM
FT                   RNVDMGDIKTKYRGSGLRFLGHNSDGVEMYSLSGTFIDDDGLKKPVMPAGTLIAGGSDI
FT                   LRMPYGPVTQVEETGPNARHKTYIKKEVPLRHGSIDGNSIMNRLTSCPTVIPFNVDAWA
FT                   VAKVI"
FT                   /product="Putative capsid protein of prophage"
FT   CDS             33995..34345
FT                   /translation="MYIATTYIGGEYTPGETLPDDFPAEKLAWLIRAGAVRKAAPAPSL
FT                   PDCEPEAMEEAEAPQEEPEANLDEIDEEAEAPEIDVMAGIVKDADEPEGKTKTASRTRA
FT                   ASKKVPKGGKTK"
FT                   /product="hypothetical protein"
FT   CDS             34342..34500
```

```
FT                   /translation="MKVLELKTKKQIEVNESYGARLIEHGMAILPPVISRVPVQKSEKA
FT                   AVKKGDA"
FT                   /product="hypothetical protein"
FT   CDS             34500..34844
FT                   /translation="MALKDRIAEDINRIFLQMDHFAETHYWNGEEITCVPDEEEALKRK
FT                   NNNVNDISWDSNTRKILIHVNESEFPGHEPPEPNTHVFFDRKPMRVLDVNINMGVLDIL
FT                   LESRDPREVM"
FT                   /product="hypothetical protein"
XX
SQ   Sequence 35197 BP; 7634 A; 10158 C; 11249 G; 6156 T; 0 other;
     atgggcacct ttgacgaggc cgggatcgac gacagctatc tttcggacat ccgcggcaac        60
     cgccgccaca tgggggaagc atgagcgcgt acatttccgg cgggagcaag agcggcaagt       120
     ccatgctggc gcagcggatc gcccgcgcca tggccggcga cgggccgctg tactacgtgg       180
     ccacgatgct gccgggcgac gatgaggacg cggcacggat cgcacggcac cgccgggagc       240
     gggagggctg gggctttgtc atgctggagc aggcccggga ccttccggat tgcctggata       300
     aggccgatcc ccggggcgcg ttcctggtgg acagcgtgac ggcgctgctg gccaacgaga       360
     tgttcggccc ggacggtatt catgcggaag ctccggccaa ggtggcggac gatctggcgg       420
     cgtttctcag gcgcgcgccg aaggcggtgt tcgtctcgga cttcatcttc tccgacgcgg       480
     ccctctacga cagcctcacc gaggcctatc gcaaaggact ggccggcgtc gatcgaagga       540
     tggcggcgct ctgcgacacc gtggtcgagg ccgtcgccgg gcggttcata ctgcacaagg       600
     gggaattgtc gatatgaagc ggttcatact ggcgctctgc atgtgctttt ccaacttcac       660
     cgccataccc tgcccctatc gcccctggga cgaggacgcg cgggagctga tgatcgcctg       720
     cctgcccatc gtcggcgcgg tgatcggctt tctctggtac ctgctggcgc tgctggcggg       780
     gcgctggctg cccgcggccc acgccgcgct gatcgtggcg ctgccctggc tgctgaccgg       840
     cttcatgcac ctggacggct tcatggacgc ctgcgacgcg ctgctctcct ggcggccgct       900
     ggatcagcgg ctcaggatcc tgaaggaccc ccacgccggc tccttcgccg tgatcgccat       960
     ggcgctgctg gccctgttca gctggggcgc ggcgagccgc atgggcccga aggacgacct      1020
     gcgggtgctg ctgttcatac ccgtcgtctc ccgttgctgc tccgccttct gtgtgctggc      1080
     gctgaagccc ctgagcgcaa gccagtacgc cgggcgcgag gacgtgcccg tccagcggct      1140
     ggcgatattc gccatgtgga tgttctgcgt ggtggcctct gcgctgtggc tgaaggcgcg      1200
     cgcggtggtg ctgctcgtgg agaccgccgc ctacgccgcc gcgatggcct gggcctatcg      1260
     cacgctgaag ggcgtatccg gcgacctggc gggctttgcg ctgtccatcg ggagtgcgc      1320
     cgcgctggtg gcgctggcca accagatcgc gagggccgcg ctatgatctt catctacggg      1380
     ggaagctggc agggaaagct ggacttcgcg aaggcgcaat acggccttga ggatggcgag      1440
     atccaaatct gcgacgaggc caccgaggcg ctggatgccg gcgcgcggtg cctggcctat      1500
     gtggagaagt gggcgctgaa ccgcgtccgc gcgggcgtgg acccgctcga ggcgttcagg      1560
     gttcggcacg acgccatcgt catcgccacg gacatctcct gcggcgtggt gcctgtggac      1620
     gcgacgctca gggcctggcg cgaggccagc gggcggctga gcgcgcagat cgccgcgcgg      1680
     agcgaggccg tctggcggct gttctgcggg cttccggagc gtttgaaatg aggacgctgt      1740
     acgtcgttcg ccacagcctg accgagggca acgagcgccg cctctactgc ggcagcaccg      1800
     acctgccgct cacgggcgcg ggccgggcgc tggcgaaggc gcgcgcgaag gatcgaccgc      1860
     tgccgccatg cgatctccgc gtcacaagcg gcatggcccg aacggacgag acgctcttcc      1920
     tgctgacggg gcggacgccg gacatcgtga tcgcagatct cgccgagatg cgcttcgggg      1980
     cctttgagat gcgcggctac gaggaactcc gggaggacgc ggactaccgg cgctggatcg      2040
     gcgactgcat gggggcgggt cttgtccgct gccccggcgg ggaatcccag tccgaatatc      2100
     gcgccagggt actgcggggc ggcgcggcgc tgctggccct tgaatggcag tctgcgctgg      2160
     cggtgatcca cggcggatcc atcgccaacc tgatggatgc ctggttcccc ggcgagggca      2220
     agggctttta cgactggcag cctgggccct gcgagggcta cgcggtacgg ttcgacggcg      2280
     acgcgccgcg ggaatacgag gtgctgtgac atggaaatcg gtcagtttca ggtgtacacc      2340
     ggtgacggca agggcaagac cacggcggcg ctgggcctgg cgctgcgggc cgccggcgcg      2400
     ggcctcaggg tgtacatcgg gcagttcatg aaggccgggg agaccagcga gcacgcctcg      2460
     ctgcgcaggc tcggcatcac ggtcgagcag ttcggctccg gcaaggggct gctgatccgc      2520
     cgggaggccg atgcggacga ccgcgcctgt gcctgcgaag ggctgatgaa ggtgcgccgg      2580
     gccatgaccg gcggcgccta cgacctggtg atcgccgacg agatcaacgt cgccatggcc      2640
```

tgcgccctgc tcgaggaggc ggacctgctg gccctggcgg acgcgaggcc ggagggcgtg 2700
gagctggtgt tcaccggccg cggtgcgacg gaggcaatca tcgaccgcgc cgacctggtc 2760
accgagatgc gggccgtccg gcactatttc cgggacaagg gcctgccagc ccgaacgggc 2820
atagaaaaat agaacataat ctctatcaag gaggtcatac ccatgagatt caaggcaatc 2880
ctcgcggagg cggcgcgcaa gcgctatccc ctctccatcc tgctgatggt gatcggcgtg 2940
ctggcaatcc ccttccccta cctgttcgtc cacttcggcc tggagctgac gaccgtatgg 3000
tggaccatcc tgtccatcgg tctgggcgtg atcctgggct gcgtgatcat cggcctcggc 3060
tcgctggtgg aggacgtgca cgacatctcc atgcacaccg tgggcttcga cctggagctt 3120
ggcgagctgg acgaggagga ctacgaggac gaggaggacg cgccggaggc ggagaacgag 3180
cagcccgagg cggagccgga gcagcccgaa gaagcccccg ccgacgccga ataagcgcaa 3240
aggcgcaccg gaggatcccc catggatacc cagagattca tcgattacag agaccgcgtg 3300
gtcggcgccg tctcgaaggc catcgtcggc aaggacgacg ccatcgggaa ggtgctgatc 3360
tgctatgtct gcggcggcca cgtgctgctg gaggacgtgc cgggcacggg caagacgatg 3420
ctgctgcggg cctttgcgcg ggccatcggc ggcagcttca gccgcgtgca gttcacgccg 3480
gacctgcagc cctccgacct gacgggcatc aacttttaca gccagaagga ggcggagttc 3540
cgcttccgtc cggggccgct gtttcacgac gtgatcctgg ccgacgaggt gaaccgcgcc 3600
acgccgcgca cccaggcggc cctgctggag gccatggagg agcggcagat caccgtcgac 3660
ggggacacct atcccctggg cgggcgcttt atggtgatgg cgacccagaa cccgctggaa 3720
tcctatggca cgtttccgct gccggaggcg cagatcgacc gattcttcat gcgcctctcg 3780
ctgggctaca tgaagcggga cgaggagctc tccgtgctct cccgccggga cacggtggac 3840
atcctcccgg agatcggcca ggtcgtcacc gacgaggaca ccgccttcat ccgcgccaac 3900
tatcgcggcg tgcgcgtttc ggacgaagtg gccggctacc tgatggacct gatcgaggcc 3960
acgcgcggcg gcgagttcag ctgcggcgtg tccacccgcg gggccatcgc gctctacaag 4020
gcggtgcagg cctcggcggc gctctccggg cgggactacg ccctgccgga ggacatccgc 4080
cggatggcgc cctgcgtgct ggcccaccgc gtcgccgcgg gcggcgggct gcgccgggcg 4140
gacgcggagg cgaagctcaa ccacctgatc gaccgcatcc cggtgccggt ggaaggcgct 4200
gtatgatcgc gtttctgctg atgctggccg cggtgatcgc cggggcggag ctcctgtccc 4260
tgcggcgggg gctggacggc gtggaatacg acgcctcccc cagcaaaccg ctggccgggc 4320
cggacgagcc gctggagctg gtgacggtgg tcacgaaccg caggcggcgc ttcctgccct 4380
tcatccggct gaacgaggac gtgccggacg cgctggtcgt gtccggcgat ttgcgcgtcg 4440
aggacgtgaa gaaccagcgc agcgcactgc gcagcacggt gtacatgatg ccgcgccagc 4500
gcctgacccg gcgcacgccc ttctcgatcc gccaacgcgg gcgctatctg ttcctcggcg 4560
cgacgctgga gggcggcgac ttcatgggcc tgagcacggc ctcccgccgc gtggagctgg 4620
agcgcgagct tgtgatcctt cccgccctcg ccgattgcgg cgcgctgcgg gaggtgctgg 4680
gcggctggat gggcgagcgc tcggtgaacc gcttcatcat ggaggacccg gtgctgacgc 4740
tgggctatcg cgactacacg ggccgcgagc cgatgaagca gatcgcctgg aggcagacgg 4800
cgcgcatggg gcgcatgatg gtgcgctgcc aggaccacac ggtggagcgc accgtgaccg 4860
tgatgctgaa cgcccacacc tttgccttcg gaagctatgg caggaagatg ctggaaaggg 4920
cgctttcgct gtgccgcgac gtgtgccagc agctggagga gcgccgcatc ccctatgcct 4980
tcatctccaa cctccgcgtc atcggcgcgg cggagggctt cggcgaggtg ggcgacggcc 5040
tgggccaggc ccacctggcc accgtgctgg aggggttggg acgcgccgat tatggcagtc 5100
gggattcgct ggagggcctc ttcgaccggg cgctctcgcg ggcgaacctg ggccgcggcc 5160
acatctatat cacccccgtc gaccgggaca tccaccacga cctggtcgat cggctgcggg 5220
ccgccaccgg cgaggcgccg ctgtgcatcg ccgcggacga atacgaatga cgaaaggagc 5280
cacccatgag cttcgcctac gccctcacgt tcttctgcga cgcgctgttg tactatggcg 5340
cgatcggctg catggggctg ctgggcgcat gccccgcgaa cctgtactgg gcgccggtga 5400
tcctgctggc ggcctgctgg ctcagcgggc ggtggacggg gcgcggcaaa ccgtggctca 5460
ggtggctgcc gacggccgcc gccatccccg cgatgctcgt gatctccaac tggccgggtc 5520
ggctcgccac gctgccgatg gcagcctacc tgcccctcta tatctacaac aaccgccgcg 5580
cgccggacta cgactacgcg gcggatcgct ttcgccacag cctgatcggc atgggcgtgg 5640
cgctgttcct gggagcgctg ttccgggcgg ccagctggaa gcggggcctg ccctacctgt 5700
tcctgtactt cacgctgaac atgacgctgc tgcgcctgct gcgccacgac gatcgcgtgg 5760
cccgctcccg ccggttccgg gtgctgaacc tggcaggcgt ggcgctggtc tgcgcggcgg 5820
gcttcgcgct gtcccagccc gccgtcattg cggcggtccg ggtggcctgg aactggttcc 5880

tggagaacat cgtggtgaac cttctgacgg cactggcatt cgtgcttcaa tacctgctct 5940
atgccctcgc ctggctgatc acgcgcatct ttggcgaagg ggcgctgggc atcggcggcc 6000
tgccagacct gtcgcccacg gagggatacc agtcgacgct tccccgcacc cccgcggagg 6060
tgcgcatgct gcccgtctgg gtgcagtggg ccgtgaaggg catcgggatc gcgctgctgg 6120
ccgtcctggc cttcgtcatc ctgcgggcgc tctcccgccg ggtcagccgg acgtcgaccg 6180
actccggcaa tgacgagcgg gaatcgctgg acgcggaggc gccccgggag ccccggcgcc 6240
tgctgcgacg ccgcgacccc cagcagggag tgcggcgctg gtaccggcgg gcgctcgcgc 6300
tgatccgcgc ccggaacggc cgtgtcgcgc cgacgatgaa cacgctgcag atccagcagg 6360
agaacgccga aaccgtggat tacgacgcca tggacgccct gcgcgagttg tacctgcccg 6420
tgcgctatgg cgggcacgag gcgacccggg aggacgtcgc ccgcgcgaaa gaagcctatg 6480
agcggctgcg aaaggccagg gattgaatcg cgcatcccca aaccgggccc gcaccggaaa 6540
taccagatcg acaaatcccc cgcttcgccc ggatggcaga gcatctgtca tcccgggtaa 6600
gcgggggatt tgttgggttt gtcagttttt gtgggatggt gatgaacacc cgcaggcagg 6660
gagcccacgc cgtgaaaggc gtccctgccg ccatcatcgg gaggcggcag ccgctctcaa 6720
ttcccaattt ctcgtttctg attgtaattc atcacatcgc cgcgtccggc ttttgctgga 6780
tcagcgggcc ctgggaatcc ctgtgaatgt gggcgcaatc gttgaacagc tccagccggg 6840
gatggatata gcccgccagg atggccggga ggtccacgat gatgatggac gtgaagtcat 6900
agggcatcac ggagcagacg tacgggaacg gcagcgacag gatgcgggac agcgcgcagg 6960
cgccggagcc gccatggctg aagatggcga tggtctgatc ggagccgttc gcactgaaat 7020
agcgggtgcc ggcgcgtttg tagccgtagc ccagcaggaa ttcgtcgatc ttccccgcga 7080
tcatgcggtc gtagtccgtg gccctgttgc cgttgaagaa gggatgctcg cgccagtccg 7140
aggcatagaa gtcaaaatcc tccgtctcga tcagcgcgtc gcccagcgtc cagggatgcc 7200
cttccgacgg gatcccctcg ccgccccagc tgatctcgtg catgtagtcc agcgtcgtga 7260
ccggcaggtg gagccgccgc gccgtgtgtt cggcggtctc ggcggcccgt ccgcagggcg 7320
aagcgtagat ctcggcaatg ccctcgcccg ccaggcgctc ggcgcaggcc tccgcctgct 7380
cgtggcctgt cccggtcagg cagtcgttcg cgtagtcggg ttccccgtgc ctcacaaata 7440
tgatgcgcat gtgtcctccg tcgtccggcg atcagccgag gttcgcgtcc cggcgcagca 7500
agcagcgcag ccagcgaccc gtttcggtca ggtccgattc ggcccagctg tcggtcttcg 7560
ggcagtccgc gcgcaccagc gcagcgctct cgtcgcggtt ggacaggctc catgcgatgt 7620
aggagaggcc gtgcttccgg atcaggtcga cccacgccgc ggcggaatcg tagtcgatca 7680
cgccgtcgcc ggaggcgtcg cagatgctga attcgtcgat gaatatcgga aggcccgcgg 7740
ccagcgcctc ctcagccttg gccctcaacc cgtccttgtg ggtggcggcg tagaagtgca 7800
gcgaatacat gatgttcccg ccgtccagcg ggtcctttgc tgcgtggtcc acatcctggg 7860
accaggtggg cgtgccgcac aggatcacgg ccttcggggc gttctcccgg atgatgggga 7920
tgatgcgctc ggcatagtcc ttcaccaccg gccaggtgac gcccgcgccg ttgggctcgt 7980
tgcagatctc atagagcacg tggggctggt ccgccaggcg cgcgctcatg cggcggaaga 8040
aatcctcagc ggcggccatg tggatgtgcg gatcgccgtc gctcaggatg tgccagtcca 8100
cgatggcgta catgcccagc ttcgcgcaga tggcgacgcc cttctcgatc agcgcctcca 8160
gcgcggcttt gtcgccgccg gtcatgtagc cgccctcctc gcaggtgtac atcgccaggc 8220
gcacggtgtt cgcgccccag tcgtcccgca gggtcgcaaa acctgcctcg ttcacaaact 8280
ccggatacca ggccaggccc agggtgctca cgccccgcag ctgcaccggc tgtcccgccg 8340
cgtccacaag cctggttccc gaaacgctca gcgcgccgtg ctgttcaaag taagttgcca 8400
tgcgattcct ctcctttgat gttcgttatc tcacgatccc cggcacctcg tagcggatgc 8460
gcatcttcgg gggctcgtcg agcgtcgcgc ggtagttcgc catccagtcg tcgctgccgt 8520
cgtccgggtt gacgccctcg gcgggcggcg cgaacacccg cagggtcagc gtctgcggcc 8580
cgtcgagggg cttgctgaag aaggcgctca tcaggtcgat ggtcttcgtg ccgggggcct 8640
tgtagaacca gcggccgttc agctcgacca tcaggccgat ctcctgctcc acggtcatct 8700
cgcagaactc cgggttgccc tcgaattcga tgggcatggc gagaccgtcg gcatcctccg 8760
agccgcccca gcgcatcgac atcgggaagg acgccgcctc gccctcggcc atgccgggca 8820
tgaaccaggg atcgttcagc acctggtcgc ggtattcggc cagcatcggc tgctccacgc 8880
ccacggcgct gttgttcggg tcctcgcact ccaggcccag gcggccccgg tcgcggaact 8940
gatagaggct gatggacttg aaccagtccc cccggttgtc ccgccagtag tgggcaaagc 9000
gcaccacgga ttcgccctgc atcctcgggc cggtcacgtc gccgtcggtg ttgaactcgg 9060
cggtgatcat cggcttgctc tcccccgcca gctccgtggc ccgggcgtgg gtccgggcaa 9120

agctctcgac gatcatgccg atcgggtcgg ccttgtagcg gccgccgccc acctcggcga 9180
tgtcataggg ccagccatag tgcagcgccg gatagctgtc ccaactccac agatccgcgg 9240
cggcatagca gtcccggaac gcctcttcct tctcaacatg gccgtcttcc ccgcagaagc 9300
cgccgcagaa gatgacgcgc acctgcggcg cttcctggtg gatgatgccg gcgaagcgca 9360
cgaagaaggc cccgacctcc ggataggtat agcgcttgtt gtgctggaac caatcgccgc 9420
agcactcgtg gttcacgcgg atcatcatcc gcccgaaggg ccgaagctgc cgggcaacgg 9480
cccgaagctc atcgtcggag aggccgcagt ccagcgtcag cgtcagcagc acgtcctggc 9540
cgtggcggcg gatgtccttc atctgctcaa agggctggcc gttcgcgcca tagggaaagt 9600
aatcgtcata gggatagtcc caggcgaagt tcacgctctt gtccttccag gcctcggaga 9660
tgcgggcggg ccactccgcg tcgtggggat agtaggtata catgatgttc acggcgcggt 9720
gggggcgccc cagcttgtgc agcaggtaat cctggttgac gtactcagcg cgctcgctgc 9780
cgtcgccgat gtccagcgcg cagtgcgcgg ggcccatgtg cagcttcttc atcgtctcac 9840
ctcatatggt tgatgctaca attataatcg aatcccgggg gcgtttcaag ggttcagag 9900
cgcttcttcg gaaagctccg gcacccggac ggcatagggc ccgagggcga gctcgcggcc 9960
catgaattcc agcgtcacgg gctcgggatt gttgttgaag atgaacgtga agcggcggtc 10020
gccgccggcg cgcgtcgaga cctcgacgcc cacgggcagg tccggcagaa agggcacgcg 10080
cgcctcctga agcatccgtt cggccagatg cctgcaggcc gcgtcctccg gcaccgtgcc 10140
gacgtaatac acccggccgc cgccccagcg gttcaccgtc gcggcggggc tcccggcata 10200
gtactggtcg gcgtaggtcg ccagcggttc ggcggtatcc agcgccagta cgtccgccca 10260
gcgccgcgcc gaccaggtcc tgccgcccat atcgatgctg gccgtgtcgt tctccaggga 10320
atcgtactcg accacgcgcg cgccgaccat gtccgcgtag tcgccgggaa ggggcgccat 10380
cacgcagttg ttgttgatgt ccttcacgcc gctgcggcag gtgagcagca ccgtgccgcc 10440
ccgctgcgcg aaggcgtgga gcgccgccac ggcctcggga gcgcggacgt acatcgtcgg 10500
ggcgacgacc acccgatagc ccgtcaggtc ggcgttttcg tcgatcacgt ccacgttcac 10560
gcccagcgcg gcaaacgccc ggtgccaggc ggtcagctgc tcgaagtaat ccatgccctc 10620
ggtctgcagc tggctcttga aggcgaattc attgtccatg ccaaacagga tcgccacctg 10680
agacatcaca gacgtgccct ccgccggctt cagcttctcc accgcctcgc agaggctccc 10740
gaattcccgg aagcgccggc cgggcacatt gtcgtggtcg agcaagccgt gccagaacat 10800
ctccgcgccg cccaccgcgc tgcgccagcg gaagtggacc acggcgtccg cgccgcgggc 10860
gaaagcctgc agcgcgtatc ccttgatcat ccccggctgc ggcgcgcgct gcatgttcat 10920
ccagcagccg ggcaggccgc tgagctgctc catgatccag aagttctgcc gcctgaggcc 10980
gcgggtgaag tccaggtgga aggcgtggga tgcctggttc agccgggtag agggatagtt 11040
gtcataggac gcgaagtcca ggtcccggta catgccatag agctccggca tgtgctcgca 11100
gaaccagaag tttgtggtga tcctggcgcc gggaatgatc gagcggatca aatcccgctg 11160
gaaggccacg tacccggcgg tgctcccggc ggcaaagcgc tggtagtcca gcaccagcgc 11220
cgggttgtac cagaccatcg ggtggttctt caccggtgga gtaatttggg tccagctgga 11280
ccagtcaccg ctccacacgt cgttgtggaa ggccgcattc agcttctcca gcgtgccgta 11340
tcgcgccttc agccagtcgc ggaagcggcc cacgcacacc tcgcagcagc agaagttcgc 11400
ctccacctcg ttgtccacct gccaggcgat cacgtgcttc aaccgcccat agcgctccgc 11460
gaggcgggtg acgatcttcg cggcatagtc gcggaacacc ggcgattcat agcagcgatg 11520
gccgcggatg cccgtcggct cgcgcaggcc ggtgtagtgc tgctgggcaa tctccggatg 11580
cgcctcgtag agccagagcg gcgggcagct ggtgggcgtg cccagcacca cgtcgatgcc 11640
gcgcccggcc atcagggcga tcacttcgtc cagccagccg aagtcgtatt ccccttcccg 11700
gggttccagg cggctccagg cgaattcccc caggcgcacg accttcacgc cggtcgccgc 11760
catgagatct gcgtcctcgc gccagagcgc cctgtcccag tgttccggat agtaatccac 11820
gccgatgcgc atgggcaagt ctcccccctgc attggaatgt tttgacagta ttataccatg 11880
attttcaccg taatcccttc gggcttccgg cgaaccgcgc aaccgccgca cttcatgcgg 11940
ccggttcgct aatccttatt ataccgtgtt tcaggattgc gcgcaaggga tcagcgtcgg 12000
ggataaacac aaaagaaaag ccccgaaaac gctgtgtttt cggggctttt tgctggtaca 12060
gcatccgatt tcaaatccga actttcgtcc tctactggaa ggtcggagga attgatgagg 12120
tcgatgtagg agacttgttt tccaccgcgg atattataga atatcacaac gcggtcgtca 12180
taaaggtata tgcagtttat gaacacgtcg atgatgcggc ggcggaattc ctcttccagg 12240
ggatcgccat tggtgaactt ctttatccag gcgcgcactt catcttccgt gaggcgtatg 12300
ccctgggcaa tgcgcagctt ggcaagatcg atctcaatat ccgacttctg ggcatcgaga 12360

acttccattc gatcatagat tttcttatgg gcagcctttg gcgcttcgat cagcgcgtcc 12420
accagcttgt ccatctcccg gtcgatctgc gcaagtgcct tttcgagttc ttcaacctgg 12480
ctatccgaaa aatccttctt atactgctct acgacggcct gggcgacctg ctccaggcgg 12540
cccggcgtga ggatgtattg gaccgtctgc tcgacgacat accattcgat gaaatccttg 12600
cgctcattct tcttgtggca gccgcgacgc tttttcttgt tatggcatgc gtaataatga 12660
tatttcgcgc cggtatggga ccttccgctc tcccccacca ttgccgtgcc gcaatagccg 12720
caataggctt tcccctggag cagataggac acctttgcct ttttcgacgc cggagcgcgg 12780
gcggcctgct ttaggcgatc ctggacgcgg gcaaagattt cgggatcaat catcggctcc 12840
gaaaggccgg gaaccacttc cccattatac ctgtattcgc cgatataggc ggtattggac 12900
agaacatggg agaagctgtt tggagacagg ggcttgccgt agcgattgca atagccgcgc 12960
cggttcagct cgtcgatgat gtccttccgc gcctcgccct tggcgtaatg ctcgaagaga 13020
tagcggatca tgggcgcggt tttatcgtct atgacgagtt tcccattaag cgccttatag 13080
ccatagggcg tcgtgccgcc cagataggtg cccttggcga tgctctcgcg ctggccgcgc 13140
cggatattct gggaaaggtt tgccgaatag tattccgcca tggactccag cagcccttcg 13200
aggatgatgc cctccggctg atcggtgatg ttctccttga cagagacgac gcggacgccg 13260
tatttttttca gcttcgcctt atagatggca ctgtcatagc gatttctggc gaagcgatcc 13320
agcttccaca caatgaccaa ctcaaactgg ccctttgcgg cgtctgcgat catgcgctga 13380
aaatccgggc gctgatcctt cgtgccggac aggtggcgat caatatactc gcccacgacg 13440
gatatgccct gctgctgcgc ccaggcataa ttatcccgca actggccttc gatggattgt 13500
tcagtttggc tgtgagacga atacctggca taaataacgg ctttcatgaa attctccttg 13560
ctttttgccg aaggctgtgc tataataacc ttgcgattgg acaattggtg ttggcttttc 13620
gcatcccgtt ccgggcaacc ggggcgggtt ttttattcct cgaaaattgc ttcgttaatc 13680
atcttcttta tcgcatttgc ttcccctgcc atcttgccgt acatccgctc cgtttccttg 13740
gaattcttga ttcgactggt aatatcggga tactttacgt ccaggcctaa cgcggaccag 13800
gattgccaaa tctgcatgac ggaatcataa cttgcgatct ctgacaacag cttctcttcg 13860
gcggcaaaat ctccagccga ccgccgttcc ttcaattcat cgaggcgctg cttcctctcc 13920
tgggcctgcc gctccatttc cgcctctcgc tgcatggcag cttcccgctc ggcggcaatc 13980
tgtttctcat agtccttttc cgaaaccgcg cggaccgctg tgccatatag ggagatgtac 14040
ataatatctc cgccaatggt attatcggaa tcaatggcaa tgatggcgtt tgcgcccaaa 14100
tccttcgcct ggcggatcat cttatcatag gcaaaggccc gcgcttcctc gatcaggccc 14160
attgagccgg acatttcccg agagcccaag gcaagcgtgt caatggtatt ggacacactc 14220
gcgccaagcc tggagaggaa gccgtgcttg aaaaccgttt ccccgaagac gagaccgcat 14280
tgtgcggtga tcttatatcc ttcaatcgag gaggaagttg tcaggatcat atcatcattt 14340
ctgcgcatgg attatgcccc ctatcgccta tgcttccttg cccattccag acggatgaaa 14400
cctcacatat ttttctgact ttaccaggtc atccgcgtag tccaccagct ttgccttacc 14460
ctcatcgttc agcttgcgat agagcgacac aagcattctt tcatccgcag gcggcatcgt 14520
ggactccagc gttccaacct catcctgata aagataattc gcgtcgcatt gaagaatatc 14580
aaatatctta tacattatgc tggccttcgg agaattggca tcggtttcat agtttccgat 14640
agcccctttg ctcaccccta gccgagccgc caactccacc tgggtaatgc ccaactcctc 14700
gcgcctttcc ttaatacggg ttccgatgct cataatcaac acctcttcac caagcatttt 14760
atgatattta tattgaaatg tcaatataaa tgtcaagatt tgctgagatt taagtattga 14820
ctgttaagtt attctgtgct atcatatgcc tacactcaga tttactgaga tttggcagga 14880
gggtgataag atggcaaaag acccaatcgc ggctggcatc caatacatca tggccgaacg 14940
cggcctgatt cagcgcgccg ttgccgagcg cgccggttat acccagcagc aattcagcga 15000
catgatgaac gaccgcaaaa ccatcaaagc gactgacatc gtgcccattt caaaagcctt 15060
gggcgtgagc atccaggaag tatacgacgc cggagcgaga ttcatgaacc gagaggcggt 15120
gtaatcatgg ctgataaata catacaggtc ggcgtcacgg ccctgcgcaa tcctgccact 15180
ggcgattttc tgcccgctgt gcccctctac atcaaggcag agggtgacgc gccggaggct 15240
gaggagaagc tgatcgagga catcggcaat ctgctggccc accgcatgaa ggcttacatc 15300
gacggctgca aaaaggcagg ggtcgatctg tgaagatcat cgctgtcgat tttgacggga 15360
cgctgtgcaa ggcggactgg ccggagatcg gtgcgcccaa cctgcggttg atcgactggc 15420
taaaagagcg caggcggacc ggggacaagc tgatcctttg gacatgccga tgcggcgaga 15480
tgctggagaa agccgtgatg tggtgcagcg tgtgggggtt gacgtttgac gccgtgaacg 15540
cgaatttgcc ggagaacgtg aagcaatacg ggaacgactg ccggaaggtg tatgcggaca 15600

tctacctgga cgacaagaac ggttacatcg ggcggatttc gcccttcccc atctatatcc 15660
gcccggagcg cgaggacatg aaagcgccct gctggacggc gagaggataa tgctatgagc 15720
gcaaagagat tccatgtggc gaggcccgat gagcttcaaa cgcttacgaa atatgccaag 15780
acaaacgctc caacggtggc ccggaccgcc cggatatgga aaagcgcgaa ggcaaagaag 15840
ggcaaaacgc gctgggacaa tggcgtgtat gtgggcttct ccggggaccg gcaacgcagg 15900
gcagcgctct tccgggtgac gggcgacgac gagctcgtgg agtttgcgac gctgaaatac 15960
aacctgtcgc tgctgtggtg cgagcagcac ctgcgaaggg tgtacgaata ggattcaatg 16020
cccgatgaca gggctggcgg cccggaaagg cgggcattac atgggggcag cggaggcgct 16080
tgcggaagca ctacgcaagg gcctgccgcc cgggctcacg gcccaacgcc cctgtcgcaa 16140
cgccttcggc gcgctgatgg gccgcgccgt gccatggggc aacgatcaat aagaccgagc 16200
gcgagacgcc ggattgagca caattcatcc gcccatcggg aaggagggcg ctgcttggcc 16260
tggatcatgt acaccatgcg ttgtttcacc cttgccttcg ccaggtcggc ggacgcggat 16320
tgggccgagg cgattccacc cgcgcatcgc aagaagccca cgtttagcgc aatcccgccc 16380
gcaggccggt gtatagcatc cggcgaacgg gatcacacgg cagcgacggg cgagtagatg 16440
cagcccggcg cgggcaaagg attttggcta tgggaaagat cggcggggcc gagtggttag 16500
ccccgccgag gtatgggaat gtagcgcagt ttggaagcgc ggttgggatg aaggacgggc 16560
cattgattga tcctctattg gcaccaccac ccagcaggtc gccggttcga ttccggccat 16620
tcccaccaag cggagccatg gttacggttt acccatggcc gagcaaaaag ccggaggagc 16680
gcggcggccc gtgtggtttg ccggacgccg cgccctctaa aaaaccgaca ggatatggga 16740
atgtggcgta atttggtaac gcacgttggc ggtgcaccta accagcggtt gtcggttcga 16800
ttccggccat tctcaccagt gccggtggcg gttatcggct gtcgccacgg cggtacaacc 16860
gaaacgacgc ggacagagcg tctccgtgaa agacggatgg cggcccggaa agacgggcat 16920
cctgctggga tggctcaatg gcagagcaga tgatctgtaa tcataaggtt cggggttcaa 16980
atccccgtcc cagctccaaa gcgcccgcga aagcgggcat tcttcaccag gagagcaatg 17040
aaacatggac agcatagagg aaatcaagat cgcggcacct gggccgagga catgccggat 17100
atgcgccacg gcacacaggc cggagcagcc ccacgacagg gacagcctat actatcaaaa 17160
ctggttcttc aagcgctaca gacacttccc atcctgggca gacgccatgg cgcattgcac 17220
cgaggccgtg caggccgagt ttacggcgac gctggcgaag cgcggcatcg ccattgagaa 17280
gggaggcgac gatggatagc gattggatcg aaatgagcag aagtccaatc gcaagggaat 17340
gccctgacgc gccaataatc gtatggcacg tctacagcgg cgcgatggtg gaacgactgg 17400
aacgggcgag gaagaacagg ttcatgaccc actggcggga aatcgacaac ggggcatgga 17460
tcgacgcacg agaacgcaag cccacccagg aggacgccga cgcctatgat tgcgtaatat 17520
cgcaggacgt gtggggcgaa gtggctatgg ccggatggca caggttcgac cgcgagccgc 17580
gcctggcgcg ctggcagacg ccaccccctg ccccaccgaa catctgcgaa ttgcgaaaca 17640
gaatccgata ggaggtatcg acattgaacg cattagccaa caccgaaacc catctgccgt 17700
ccaggttgga cgagctggca aggcaagcgc agttgtttgc ccagaacgcc tgtatgaacc 17760
ttttgcaact tggccgggtg ctcgccgagg ccaggcccat gatcccctat ggcgagtttg 17820
acggatggtg cagacagaac gcgaagatga gcaaacggac ggcggagcag tacatgcagg 17880
cctatgccga atttggcctg aacaccaaga tcgcagagct tgggacgagc aaggtcatca 17940
agctgctgcc catgagcgag gacgagcgcg aacagctatt ggctgagaac gacgtgggtt 18000
ccatgtccac ccggcagctc gacgaggcaa tccgcaaaca gcgcgggcag cttctcaaag 18060
aagcacgggc cgaggcgaag gaggagatcg agcgccacgc cgagaccgcc aaggcccttt 18120
tgaaggagaa gctgcgccta gagcgcgagg ccgccgaaca cgacgaggac atggccgcgc 18180
tgcaagcgga ttacaacgcc gtgagggatg aactgctggc ctataagagc gcagacgcgc 18240
gtggcgaaga tgaacacaag gccacggaca cgctgacccc tgacgccttc accagcgccg 18300
tgaacgcctt catcgggaaa tgctgccgcc tgccgcagat gggcaggact ttcagcgcca 18360
tgccccccgga ggagaaggaa atctacgagc aatccctatg cgccattgag aagtgggccg 18420
agagcgcccg catcgccatg aacagcgttt cattcgagga ggcgattttc gttgactgat 18480
attatcaagc acgacagcgc gacacctgcc cttccccagg aaacggtgga gatcatctca 18540
gtggtcgtgc gcgacatgat ggcccccatc atgcagggcg tcgccaaact gctggaaaac 18600
aacactcagg ccatggagca gatcgccgcc gcgcagcagg ccacgagcga acggatcgcc 18660
gcgctggaaa agcgcgtgcg cctacaaacc cctttgagca aatcccagga gaattgcatc 18720
aacggagcca tccgggccaa ggccaaggag cttctggacg ctcacggctt ttcggatgac 18780
aaaaaggctg tgacaaagct gtgcggcatc atccgcaaga gcgtgctggc ccgctatggc 18840

gtgggcagcc ttcgcgaggc cccagcctat gactatgaca ccgctatgaa gcaggtcgcc 18900
atgtggaacg accggcttac cattcgggac gttgagaggg aggcgcgaaa ccgtgcggaa 18960
gccatggaag aagcggttat ggagctggct aaacgatctg cgcgtttgga tggttaaccg 19020
gcggatgcgg gcaagcctga ttaaccgtgt aagcgcggtg tgtgtgaggc tacggaggga 19080
ctggtgatgg acaagatcag attgtccgcc agcgaccgtg agagcctgat gcggctgaac 19140
gtagcattag agatactgga gaccgagggg gaccagctcg cccgtcggct tcgagttatc 19200
aaggacggca agcgcgatta cgggcttatc aaatccagaa tcaacaagct gatggagtgc 19260
atgatcgaga cgatccccga agaccagatc atgacctatg ttcgctcatt gaagatgacg 19320
agctatacca tcgggatcaa aaagccaggc ccaatgggcc ggaatgagaa ggactacggc 19380
atgtggttga gctatgaggt tatcaacgcc ctgctgaccg gctgccacga ccattgcatg 19440
atgtgttcgg aggacaaggc cgagcgccgc gcctgcaagc tgcgaaaggc gctggcaacc 19500
atacccaatg acgcgccgga aagagacgac aacgattgcc cattttacac gctgatgtga 19560
ggaggatcac catgcttatt tctgaattgg caaagcaggt acacgagaac gccgtggcac 19620
acggctggtg ggaggagaac cgggaagcgc cagagatcaa cgcactgata cactctgaat 19680
ggagcgaagc gctggaggag gcgcgggcgg gcaggccgct tgtctggtat gacatctatg 19740
accggggcga cgattgcggc ggcggcggac gctgcagcgc gatggtcaac gaaggtgtat 19800
gctcggggcg gaacggaaaa tgcgagcgca accacaagcc ggagggcgtg gccgtggagc 19860
tgatcgacgg ctgcatccgc atcctggact acatgggcga gctgggagtg accgtgcagg 19920
attgcgacag cgaggcggac agcgagatcg aagggctcta caccgacgag atcatggccg 19980
aagccgaaca gcccacggcc catgtggtgg cgtggcttca catgctgact tccgaagcgc 20040
tttccgacgc gagagagaac gcgctattgg gcgacatgaa ccccatgaag ctattgagcg 20100
cgatggcgct ggcgctatcc tgggtgaaaa aacaggggct tgatccgctt gcgatactgc 20160
tggagaagca cgaatacaac aagacccgcc cgtacaagca cggcaagaaa ttctagcaaa 20220
acggagccaa caccaaaaag gcggtacatt gcaatggatg tgaattatct gatcgagata 20280
aacggattgg acgattggga gtttggcaat ccactctccg caactgctta caaggttatg 20340
cgaaaactgt tgatgatcgc aaaccgccag cggtttcctg agaggattcc gatatccaac 20400
accagactgt gtttcctcgt cggatgcacg gaaaaaagcc tgatcgccgc ccgccaacag 20460
cttattcaac gtggactgat cgactacaag gggaagaaaa agcagacccc agtctatacc 20520
attcactatt tttcccatga tccgcgctat aacaggaaaa attacagtta taacgggagt 20580
tatgacggga gtatgaatgg gagttatgac gggcgtatgg acgggggtta tacgggggga 20640
aggtcaataa atagatcata cggcgaactg gaggattgtg aggaggaaga tttgggcaat 20700
gacaacccca caacaggaac gctaacgcct gttactgctc tccttcataa cccacccccct 20760
tccctacctt ccgtggatgt actacagggg aacaggtatc cgccgaggaa tcgcccccctg 20820
gcaggcgcgg agacatggga ctatgccgcc gcaatgtcat tttttcaaaa ccctgacatc 20880
cagcggatgt acggcggcgg ggctgggacc atcgagagaa tcctgaacag cgacgccttc 20940
ccgatcaagc tggtgggcta cgccatggac aagacctatc agcgcaacag gatgtatccc 21000
gaaccgcttg gaaaccctgc tgcatacacc ctgcggcttt tgaacgactg gcacgagcag 21060
ggatttcata ccccggagga tgtgcaggag agcaaggacg actggatgca ctttggctga 21120
tggagggatg aggcatgaac aagctgacga tcatcgggaa tctgacggag cacccgcagg 21180
cgcgggaagt gagcaccagc aacgggcccg cgacggtgtg caatttcaac gtggcgacga 21240
accgctatgt gcgcgggcaa aagcggacga cttatttccg cgtgtcctgc tgggggaagc 21300
aggcggaaaa cgccatgaag tatctgctga aaggcaggaa agtggccgtg gtcggccccg 21360
tggaggccag cgcctacacc gcccaggacg gatcggtgcg ggcccggatg gaagtgtacg 21420
cagacgagat cgagtatttg agcagcaggc aggatgccgg agagcaggca ccgccgccgc 21480
aagcccccgg ctacggcgag tttaccgaag tggacgatga cgaattgcct ttctgacagg 21540
aggacgcacg atggctgatt tatacgagcg gcgcatggtg acggccagga ggccgcacaa 21600
atgccacctt tgcgggctga cgatcccgga agggcatgac tacatccggg agaaatggcg 21660
cgatgatggc ttccgcgagg tcaaacggca catccactgc gacgcgatgc tggacgagtt 21720
tctgaacagc gacagctttg tcggcgaata ctacgatgac gacgtgtggg agtggatcag 21780
ggagaaatgc ctggacctgt gcggctatga agggcgggag gattgcatcg acaatccctt 21840
tgcttgcgag cagatgatcg acacgatctc gaacctcaac agccggaagg cggcacttga 21900
ttccctgaat ctattccgaa tgaggaccgg gcaatgaaga gcgacgcgac aaagtgggtg 21960
gaggacgtgc tggcggtgct taaacaagcc gcgagcaacc acaaggatgc catggaggcg 22020
gaaaaggggt gcgacgacaa gacgcaggac ttgcttcacg aactggagct ggtgccccac 22080

agctaccacg agcgcagcgg gatcgccaag gagctggcgg cagtgcggcg gaagaggcgg 22140
caggccaagg acgcgcacga gctgctgggg ccgctggtga agtggatcgg ggcgcagaag 22200
accgccatcg acgcgctgac gcgggcgttg ggcgagatgc gcaagatcga cgagaagcgg 22260
gacaaccggc tttactacaa gcgagcggac gggaaccacg agatcatccg caagcaagag 22320
gatcaggagg ctaaatagca tgaacgagca ggagatgcgg aagaaggtcc tagatgggtt 22380
gacgtgctgc gacgcgctgc acaatggcgg cgtagtgact tgcgggacgg agtgccccta 22440
tgggatcgcc gacgggattg accggggatg ccggggcaag ctgattcccg acgccctggc 22500
gcttatcaac gagatggaca tcaccatccg ggcgctgatg ggcgaattcg gggacagctg 22560
cgacgtggac agccattgcg gaggtgggaa ggatggctga tcgggagaat gttatcaagg 22620
gtctgaaatg ctgtcgatgg ggaatttgct tcgcatgccc atacaatgac ggcgtgcacg 22680
ggaacacagg ctgcaagaaa aagctggtag acgatgccct tgaactgctc aatgccttgg 22740
acgtgacgcc ggaggagctg gagcggctga aaaagtgcag gcacgagtgc aagatcgact 22800
gtctgctgga gcattacgac gctatcaagg ctgagcgcga cgcgctgctg gagaagcagg 22860
agccgaggct tatggtcgcc aaggactttg aggacaaccc aaatgtggac ggggacggct 22920
atttgccctg ctggatcgag tgcaacgaga ttgaagtgga gcgggctata cgctttggtg 22980
ccattcagga aggcgaaacg ctggacggct ggacagaggt ttccaaggct gaactgttgc 23040
ccaccgacct ttcatacaac ccgaacgtgc gccactggac cgccaggccg agcgaggagc 23100
agaggaggaa aacgccgtgg ccggatgcct gagaatcaga gacagcgttg atattgatat 23160
ctacgttttg ccggacacgg caagatgcag gtgcaccggc atatctccgc tggatatgga 23220
cgcctgcccc atgcgccggt ttgattattt tgggatggat tgctggccgg gcgagtgtga 23280
cgagtacacc gaggacaagg agggcgaacc gtggaacgat tactcagact gaaaacgccg 23340
agcgagaggc ggcaggacat acaaagagag ctggccttcc tggaacgcag gaagcaatac 23400
gcccggagcg tctatgtgaa gcagcttcaa gagatcgcag agcaggaaaa ccgactgatg 23460
aagagacttg agaggattgg aatagatggg cgcaaagacg aaaatcgact ggtgtgacgc 23520
cagctggaac cccgtgacgg gctgcctgca tgggtgcgat tactgctatg ctcgggggat 23580
cgccaatcgg ttcggcggtc acctgcgcgg agagatcaag gacatcatga cgccggatgg 23640
ggagattgcc cgcgtggaac tggtggcggt ggaattgaca gagccaatgc accgaaacga 23700
ggctatggga agcgggctga tgcgcgtggg gccgatcatt ccatatccat acgacttttt 23760
accgaccttt catcgctata agttggacga accccggcac tggaagaagc cgaggagcat 23820
attcgtgtgc agcatggcgg acctgttcgg ggagtgggtg ccggacaggt ggatcaggga 23880
cgtgttcgac gcctgcgcgg aagcgccgca gcatcggtat ttgttcctaa cgaagaaccc 23940
gaagcgatac atagaggccc atccctggta tctacagcgc atacagcggc ccagcaaggc 24000
ctttgaagca tggtttggct acagctacga cggcttcgga aacagcgaga ttcgcaaggc 24060
gctcaagcgg caatactcgc tgcatcaatt catatccatc gagcccatgc ggggcgcgat 24120
atgggaggac gagatcaacg cgatcactgc gacctgcgat tgggtcatcg tcggggccga 24180
gacgggaaac cgcaaaggca aggtggtgcc ggacaaacgc tggatcatga aggtcgccga 24240
gcaatgcgag gattccggga tccccgtgtt catgaaggag agcctgcgcg ggatcatggg 24300
cgaggacttc cggcatgaaa cgccatggga ggactgatga cgtgaacaag aagccaacac 24360
cacaagagtt gatcgcggcc atccatgcga aatgcctgga gtgcagcggg gggagccgga 24420
aggaagtgca cacctgcaag ctgacggatt gcccactatg gctctaccgg cgaggagagc 24480
cgagggagag gctggcacag gccaagggac agatcgacgt attcgacatc atacagaagg 24540
agggcgcatg atgacgaacc gggagaggat gattcaggac tttgtggaga tgagcagcga 24600
aaagctggcc gagacgatcc gggaattgag cgaggacatc atcggcgaaa atcttcggga 24660
aatggtatgt gacgattgtc acgccgagca tggcgggcga tgcgtgttct ctgacaatga 24720
ggggaatgag gaggcagacc cggacgatgt gtatgacggt tgccccacca gcatggctga 24780
atggctggac aaggagtgca aacgggagca cctgatccaa taacgaggca cgagcatgaa 24840
ggtgagcata tgtcggaagt gccgtcactt ccgggagagg cgctggagcc atcgctatga 24900
accgaggggc tatcacccca tcggcatgac gcacgtctat ggctactgtt tgttgcggaa 24960
ggtgcgatgt atggaagtaa agaaagatga ttgcacaccc ggtcagacaa ctatattcga 25020
cgtagcccag gggagggac aaacatgaaa gtcagagtta ccgtaatgac agagaataac 25080
gcccctgctg aaaaagcggg aaaaaccaaa gaagaacgaa atgcgaccgc gaaattagct 25140
tggaatgtga taatttccct cttacaaact gtgtttcttg cttacgatga aacaatagaa 25200
ctggaatctg tggaggtact cgatgacgaa taaaatgaat attcggtgct gtgatacgtg 25260
ccgttacggc cttggcggcg gctatgacaa ctgccggatc aacctggagg cggagtgtgc 25320

cgcaggtgaa tttgaggcat gggaaccaag ggagccggat agaaagatca tccattatga 25380
ggatgaaatc tctgaatggg agccgaaaga taaataaggc gcgaacgtag tcatgcatcg 25440
atgaaatgcg cgaacgttcg cgcatttgaa tgaagcccca ggcgattatg cgcaacaatg 25500
gagggtttgg aaatggcaag gagcggcaac gtaaagatca atattggttg cgagttcgtg 25560
gagaaggaga tcaagccggt ggtgatccac aacaaggaca ttcccctgct ggccgacgtt 25620
cttcccataa tgcaggagat atgccagatc gagcggcgac gcgactggca gcgggacagg 25680
atgctcaaca tcacccagca tttgaccggg atgcccggcg gaggccggac gccgaaagga 25740
ctagatgaag cctttgccct gctttcggag ctggacgaag agcatgagaa gcggtgcaag 25800
gaatatgtgc gccagcttcg gaccgcgcaa aagatactga acgccatcga gagtcggagc 25860
atgcggactt tcgtgataat gaaatacgtg atgaacgtgt cagacagcga gattcgccag 25920
gagcttaaca tgacgcggac gggcttcaac cgggcgaggc gctgcgtgga gagcgcagag 25980
cgcatggcgg acgtggtatg gcgcgagcga tatatactcg caaaatgtta ataagttatg 26040
agtttgttaa aaagtgttga aatgacaagg gctttatgct aaaatgctat cgtcggtaga 26100
agtgtaagga agccgggaaa cgcaagttcc cggctttccc catttttggc gcatggaggc 26160
gataccgtgg cgcagctatt cacaatggac atcgacgctt ctgacgcgca agagacaatc 26220
atgcgcatga aaaacgccat gaccgaagtg cagtttgagc gggccatgta cggcattttc 26280
cagcggaccg gtcgccatgt gtccaagata ttgagagaag accttccaag gaagtatgag 26340
gtactgcccg gcgacatcag cgctgcggtg ggaaatccat cactgacaac gggcttcggc 26400
ggcgttggct gcacgatccc tattcgcgcg gcccgtggaa ccatcggcag caagtacagg 26460
gccagcggcg gtgcccatgg ttggaacagc cttaaacgga gatacaaggt caaggcgagg 26520
atcatcaagg gtgttcagag cactttgcct gacaaatggc attccggtta cgcggaattc 26580
aggaaccttg gatcgaagct gggaggcctt acgtttgcca ggagcacgcg gacacgcggc 26640
cccatcatga agatcactgg cattgccatt ccacagatgc cgctcaaccg ttcggaagcg 26700
gacgtgcaaa acgacatcaa agaatacctg gagaagcaga tcgcggagcg attcatggcg 26760
atgatgagga ttggaaaatg agcgtcgaaa tgaccaaaaa ggacctggca aatatcgcag 26820
gttactccta ccaccaatta tacaacatcg accgtgatct tccggcagat aaaaaattgt 26880
tcgtcatggg cgagggcggg aaatacgacc tcgctttttt tgtgcaacgt tgggtggaat 26940
acaacgtgca cagggagact gatgacacgg aggacctgga cacagtgaag gctcgccacg 27000
aggccgtgaa gatcgaaaaa acggagctgg aagtggctaa gatgcgcggt gctttgatcg 27060
atgtacagga cgtgaagcgg ctatggggcg atgtggccaa tacggtgatg caaaacatga 27120
tccacctggc gagcaagctg gctccgatgc tgctgatgat ggacaacgtt gaggttattt 27180
caaacctgat cgatgaggaa gtgcgcaaaa ccctggcgga tattgcgcaa tgcccgctgc 27240
ccagctacgc ggcagacgac gagaccgatg aaagcgagga ggatgaggag gtgtaaaggt 27300
gagcgcgtta tctgaactcg cccggtacac cttttcaatg ttcaggccac cggccaaaca 27360
gactgtttcg gagtgggccg atgaaaaccg cgtgctggtt tcggagagca gcgcggagcc 27420
tggcgcatgg cgcacagacc gtgccccata ccagcgcgag atcatggacg ccttcaccca 27480
gcccggcatt tggcagatcg tcatcatggc aagcgcccag gtgggcaaga gcgaaatcga 27540
gctcaacatg atgggatgcg ccatcgacaa cgatcccggc ccgatgctgt acatccagcc 27600
gacggacaag gtggccgagg attattccaa aaggcgcatc gcaccgatga ttaacgcctg 27660
ccctaccctg cgtgacaagg tattcaaggc gcgaagccga gacgcgatga acaccatcac 27720
catgaagacc tttcccggcg ggagcctggc gatcatcgga gcaaacagcc ccgccgacct 27780
gtccagcaag ccggtgagat acatattcat ggacgagacc gaccgcttcc cggccagcgc 27840
cggaaccgag ggcgatccgc aggaattggc cgagagacga accgaaacct tccggcacaa 27900
tcgaaagatc gtcaagactt ccacgccgac cataaagggg gcgtccagga tcgagactga 27960
ctacatgaac ggcacgcagg aggagtggca cactgaatgc ccgcattgcc acacctacag 28020
ttacatccgc tttgccgata ttcacttcga gaaggaagac ttcaccaacc agaacggcga 28080
cgaggattat catgtaaaga ctgtcacctg gcgctgcccg atatgcaagc gggatattgg 28140
cgaatacgag tgcaagcgcc tgcccgcgaa atgggtaagc aagaatccaa aggccaagga 28200
aaacggcatc cgcagctttc gcctgaatgc cttcatgtcg ccctggagcg actggaagaa 28260
catcgtgtgg aagttcttga aagcccgcaa ggacgcgaca aagctccaaa ctttctacaa 28320
cacgatcctt ggcgaaagct gggaaatcca ccatcacgac ggcttggacg aaaagctgta 28380
tcagcgacgc gagcattaca acgcggaagt gccgacaggc gtgctgctgc tgaccatggg 28440
catggacacg caggacaacc gcctggaata cgaggttgtg ggctgggacc ggcacgggca 28500
aagctggggc atcaaccgag gcgtaatccc aggaaggcca gattcgcaag gcgtatggga 28560

tgaagtggac aacctgctgg atcgagaatg gaaatcgacg aacgggatca agttccgcat 28620
actggcgacg ttcatcgact ccggcggcca ttacactacg acgatctaca agcaatgcgc 28680
ccggagggcc gccaagaaaa tatggcccat caagggagaa aaaggagaag ggaaagacga 28740
atgcaggccc atgaagcgcg gccagggcga cggagcgaaa ttcatgctgg gcgtggacgc 28800
gggcaaggca ggcatcatgt ctgacgcgca gatcgaggaa cccggaccga actacatgca 28860
cttccccatc gactaccgcg ccggttacaa catggaatac ttcaagggat tgatctcaga 28920
gcgattggaa gtgcaccggc gcggcggaca gggcgtggcc gtgtgggaac agttctacga 28980
gagaaatgaa ccactggatt gcaggaacta cgcccgcgcc gcctatcgct acttcaactg 29040
gcgattcgac gagctggagc gatacatcaa cggcgaggaa gaaccgcaga tcacaaccag 29100
gcaacaggaa accaagcgga agcagcggcg cgttgtgagc cgtggaatac aggtatagag 29160
catgagagga gcgtgaggac atggcggcca tatccgcata cagcctggag gaagccaggg 29220
aaatgttggg actatggaaa aatgcggaaa aagcgctggc gagcggacaa gtgaccagct 29280
atcgcgtggg aacgcgggag tgtacgcttt tggacctgga ggacattcgg gccgcgatca 29340
cctacttcgg caaccttgtg gaagcccttt ccggcacggt gagaaccaag cgcgtgacgc 29400
gcgtggtgcc gcgcgatttg taaaggagcg tgaatcctga tgaatatgaa gccaaatttg 29460
agggagcgcg ccctattcct gttcaaccca aagcgcggga acgacgccta caacaagcgc 29520
ctgcgggaag aacaggagca ggaacaggcc agaggccaga ggacgccagg cagtccaatc 29580
atgagctacg gcagctatgg cgcgagtacg acgaagaaca gcatgctcgg ctggatcgtg 29640
gaggccggta acgcggagga caacatcgac ctttactcct ccacccttcg gcagcgcgca 29700
cgcgacctgt acgcaggcgg cgggcttgcc cgcagcgggc cgaacaccct gacaacctcc 29760
gtttgcggct ggggcatcat gcccaagcct aagatcgacg gcgactacct gggcatgacg 29820
gacgaggcgc gggaggaaac cgagcgcaac atactgcgcg aatggaagct gtgggcggaa 29880
aacaaactgt gcgacgccga acgccagcag aacttctacg acctgcagca gctcgccttc 29940
ctgagcatgc tgatgagcgg cgacgtgttc gtgctgttcg ggatgaagcc caacaatcgg 30000
acgccctacc agaccacgat ccggctgctg gaggcggacc gcatctgcac cccggattcc 30060
agcggcgaca gtgagagcgt ggagacggag ggcggagggc gcatcatcga cggcgtggag 30120
atcgacaagg aaggcgtggt gatccgctat tacgtcgcca gcagaagccc gattgcggga 30180
aacgactcca gcgaattgac gtggcagccc atcgaagcct ttggcgcgga cacgggatat 30240
gaaaacatac tgcacatcat gacgcacgaa aggccggagc agcggcgcgg cgttccgttc 30300
gtggccgccg agatcgaagc cctgaaacaa ttttcccgct atatgaacgc ggaattggct 30360
ggcaagctgg tgagcgctat gctgaccgtg ttcatcacga gcaaggacga cgacggcaag 30420
gcgggcatgg aggacgccgt gaacgaggac gaaaaggtta cagacgatca actgaaactg 30480
gaactcgccc ccggcgcgat ctacgacctg ccgcccggaa agggcatcga aacggtagac 30540
ccgaagcgca gcgacacgca gtttgaggta ttcgtgaaca cctgcatcat ggtgattgcc 30600
tccagcatgg gcatcccgaa ggaagtgctg gtgaagaaat acgagagcaa ctacaccgcc 30660
gcccgtgccg cgctgctgga cttctggcgg acggtgaagg tgtaccgcag gcgattcaac 30720
aacgctttca accaacccat ttacgaacaa tggctgaccg aggccgtggc cgccggacgc 30780
atcgaagcgc ccggcttttt tgatgacccg gccattcgcc aggcatggtg cggctgcgaa 30840
tggatgggcg cgagcatggg ccacgtcgat ccgctgaaag aggtgaacgc ggcagagcga 30900
cgcattgcca acaacatcac cagccaggag caggaggcca gcgaatacaa cggcaacgat 30960
tgggcggcca tcgtgcgcca gcgcaagcgt gaagtggaag ccacgcggga catgacaggc 31020
gacaccgcca cccaggagcc ggaggcaccg acgcctgaca aggacgagga ggagggacaa 31080
gaggatgaat aaggagcgat accgtatccg cttcgacatg aaggccgagg gcgatgaggc 31140
cgaggttatg gtgtacagcg aaatcgccaa cgaaaagtgg tggggcgacg agaccaccac 31200
gggcgacttc gacaaggccc tgaaagaggc ccggaaaagc ggcgcgacca agctgaacct 31260
gcgcatcaac agcggcggcg gcgaggtgta cgccgccgtt gccatgcgga gcatgattat 31320
caacgcaggc tttgagagcg tgcgcgtgat gatcgaggga ctttgcgcca gcgccgcgac 31380
gctgtttgcc acgattcccg gcgctcgggt ggtgatcgcc gagggcagcg agttcatgat 31440
ccacaacccg atgacgattg cctggggcaa cgccgaggaa atcgggaaaa ccgtggatca 31500
ccttcacaag atggaggagc agtttcacgg catgtacgcc gccaagaccg gggccaccga 31560
ggacacgatc aaggcctgga tggacgcgga gacctggttc acggccaagg aagcctgcga 31620
ttatggattc tgtgacgagc tgctgacgag cgagccagtg gccgcctgcg tgacggcgca 31680
ggagatgggc gtgatgaagg ccctctacaa ggccgtgccg gatgggatcg ccataaaaga 31740
cccatccccc acccccactg tcagtaacgg aaaccctgtt gccgggttgc cgactgaaat 31800

aaaaaatcac gaggaggaag attcgcacat ggacatgaag gacatcacgc tggatcagct 31860
tcgtgcggat aacccggccc tgtttacgca ggtacagcag gaagccgtgg ccgccgagcg 31920
ccagcgcatc gaggacatcg acgcgctgac gatggagggc tatgaagcga tggccgagga 31980
ggccaagcgc aatggcacgt ctgccctgga cttccagaag caggtcgtgg cagcccagaa 32040
gcagaagccc aaggacttca tcgccaaccg ccagaaggaa accgcacccg cagcgaatgt 32100
gccgggcggc gctccccaga atggcaagac cgacgagcag gagatcatgg acaacgccaa 32160
ggaaatcgcg gcatttgcca gtgagttctc cggcaacaac aacgagagca tgttctgata 32220
catgcggaaa ggagcaaaaa agccatgagc aatctgtatg agaccatcgg caccagcacc 32280
ccggaccaga ttctggcgaa ggtggaagcc gatcccattg ccgtaaacct tctgcccggc 32340
cagggtgaga tcaagcgcgg cacgctgctg tacaagaacg ccgccggatt ctatctcccg 32400
gcaggcagcg gccagatcag caccagctat gacctggttg tgctgaacga gaccgtggac 32460
actggcaacg acggaaccgc cgtggccgag gtggccgccg cgtaccgcga gggcaccttc 32520
atcgacggca aggtaaagta tgacaacagc ggcacgccca cccccgtgac cgaggcgcac 32580
aaggtcgtgc tgcggctgtt cggcatcaag ttcaaccaga gtgttgagag ttccggcacg 32640
tttgaaaacg ggcccgcgac gatcacctac aaggccaaca acgacgtgac ccctgccgag 32700
gccgacgtgg cgattgaggc acagcgcggc agcagctaca ccatcctgaa caacagcgac 32760
tcgaagctga gtttcaccgc ccccgccacc aagagtttca gcaagtggaa taccaaggct 32820
gacggcagcg gcacggatta tgcggctgcg gccagctaca cggcgagcgc cgacctgacg 32880
ctttacgcga tctgggcctg atgagcgaaa atcaacagtg aaaggagaat acgaatatgg 32940
atctgtatga tgttcggagc cagctcaaag ccattgagct gatgccccgt gagtgcagcg 33000
ttctggccga cttcttctgc aaggaggagg gcgttcagaa ccagaaaaag gccatctacg 33060
acttccgcaa gggcgtgaag gcgatggccc ccaccgtgca tcccggcacc ggcggcgtca 33120
tcatggagcg caccggctac gagacccgcg aaatcggctt ctgcaccatc gccccggagc 33180
gcctgatcga ggatcagaac cttgaaggcc ggatgttcgg cgaggatgtg ctgggcgcta 33240
tgacgccgga gcagcgcgag aagaagatgc tggccaaaga cctgaccgag atgcgccagg 33300
ccattcagcg ccgcagggag tggatggtgc gccaggtgct cctgaccggc aagctgagca 33360
tcttcgagta caccaacgag ggccgcaata tcgaggcctc catgatcgcc gactatcact 33420
tcacgaacaa cttcaccccc accactccct ggaaccagtc tggcgctgac atcgtgagcg 33480
acatggaagc gatctatgac ctggtgtacg agggcctggg tgacgtggag aagatcgtga 33540
tggccccgaa cgtgtgggcc gcgctgcggc ataacgacgc cttcatgaag aagctggaca 33600
tgcgcaacgt ggacatgggc gacatcaaga cgaagtatcg cggcagcgga cttcgcttcc 33660
tgggccacaa ttccgacggc gtggagatgt acagcctgag cggcacgttc atcgacgacg 33720
acggcctaaa gaagcccgtc atgcccgccg gtacgctgat cgcgggcggc agcgacattc 33780
tgcggatgcc ctacggcccc gtgacccagg tggaggagac cggccccaac gcgcgtcaca 33840
agacctacat caagaaggaa gtgccgctgc gccacggcag catcgacggc aacagcatca 33900
tgaaccggct gacgagctgc ccgacggtga tcccgttcaa cgtggacgcc tgggccgttg 33960
cgaaggtgat ctgacgcgga aaggatgagg cgtaatgtat atcgctacca cctacatcgg 34020
cggcgagtat actccgggcg aaacccttcc cgatgacttt cccgcagaaa agctggcatg 34080
gctgattcgg gcgggcgcgg tacgcaaggc cgcgcctgcc ccttccctgc ccgattgcga 34140
gccggaagcg atggaggagg cggaagcgcc ccaggaggaa ccggaagcca acctggacga 34200
gatcgacgag gaggcggaag ccccggagat cgacgtgatg gcggggatcg tgaaggatgc 34260
tgacgagccg gaaggcaaga ccaagacggc atccaggacc cgcgccgcga gcaagaaagt 34320
gccgaaagga ggtaaaacca agtgaaggtg ctggagctta aaaccaagaa gcagatcgag 34380
gtgaacgagt cctacggcgc acgactgatc gaacacggca tggcgatcct gccgcccgtc 34440
atctctcgcg ttcccgttca gaagagcgag aaggccgcag tgaaaaaggg tgatgcctaa 34500
tggccctaaa agaccggatt gcggaggaca tcaaccgcat cttcctgcaa atggaccact 34560
ttgcagaaac ccactactgg aacggcgagg agattacatg cgttcccgat gaagaagaag 34620
cgctgaagcg gaaaaacaac aacgtgaacg acatctcctg ggattccaac acccggaaga 34680
tattgattca cgtcaatgaa tcggagtttc cgggccatga acccccggag ccgaacaccc 34740
acgttttttt cgacaggaag cccatgcgcg tattggatgt caatatcaat atgggcgtgc 34800
tggacatact tcttgaatcg cgcgatccga gggaggttat gtaatatgcg gttttcggaa 34860
aggttcattg gcctgaaaaa ctggacatac ctggagctgt gcaaggatcg ggagatgaaa 34920
gctcccgccg agaagatgga catcggcgtt attcatacgc aggagccaac ggttcacctt 34980
gcatgggcac ctgcccgcgt ggacaaggcc ggaaacctgc aagaaatccc actgagcacc 35040

```
     gttcctggaa tcctcgtgat gcccaatcaa gcctacgcga agtatatgga ggagaagcgg     35100
     tttgaccgct acaacaacat acacaggccg aaggagatgg ggcaacacct ggcaataacc     35160
     atactgttcg gcgtctatga acctggaatc cggctgc                              35197
//
ID   XXX; XXX; linear; XXX; XXX; XXX;  XXX.
XX
AC   PRJEB67889
XX
DE   Uncultured bacterium DNA fragment from in vivo enriched cow rumen, clone
rum-36e1
XX
FH   Key             Location/Qualifiers
FH
FT   source          1..36794
FT                   /organism="uncultured bacterium"
FT                   /environmental_sample
FT                   /mol_type="genomic DNA"
FT                   /isolation_source="Cow rumen"
FT                   /clone="rum-36e1"
FT                   /note="AZCL-xylan, AZCL-arabinoxylan active clone"
FT   CDS             149..2038
FT                   /translation="MHFRLRKPYGPIPHVFIVVKAHFFENLYQRHDLYFTQTGVFFFLL
FT                   NIQQAQIHIQVGIGIIIHIHLSDKCFASFKIQLFHLIGLPLMHIDGIRVQQHRGGKTVH
FT                   FPYNPGMGGIGDIINQKVFRHGGAQADFTGGKIFRHPVIGIPHPVQNIAFFQKAKHFFP
FT                   ARCISKAFPFFKRKLVGGAFEMTQQNIQIVRIQKRIFSRLRKKVLRMRDDELVQCAGGC
FT                   HKEGGAHAVATSGTAGLLPGAGNGSGISAQNAGFQGADVYAQFQRVGTHYGLDIPGPQP
FT                   LLNLPPFRWQVSAPISPYLAAVANGVGYQLFELPGHHLDLKPAFGKDNALDSVAHQHGS
FT                   DIPRFRKHAPADTELDIHHRRIVENIVFFTGRSAVIVNQHHAAARQRFRQLLRVLDGGA
FT                   AKDEPGMASVKITEPDQAAQDIGQIAAENAAIGVDFVHHDVFQVFKQLDPFGMLRQDTA
FT                   VKHVRIGDHDMPGLPDRPSGRGRGIAVVCIGFDIRPELLDQFVKLAHLIGGERFGGKQI
FT                   QCPGVRVFQDFSQNRQIVTKGFSGSGGCYHHIILPFLGSGKAGSLMGVQLMHAVLFQSR
FT                   RKPRVNAGRPGSIFRSGGRNCFPGGDIVHKGRIVFQFFQKLRQIHGLSSFGYI"
FT                   /product="GH27"
FT   CDS             2264..4207
FT                   /translation="MSRNDAIEQYNAALRMGRSYYKACTAKGEDPFPKVLEHLIEKQET
FT                   GNSVKIGLVDIPASRIVGTWMEGRKAAFAGNFMPLLDPDSEFGGKWISLCESHLDEGGI
FT                   RDPITCIEYLGYFYVQEGNKRVSVLRSFDSPSIPGIVTRLLPLPGDDPRIQLYFEFLQF
FT                   YKVSKTYLVMFSKPGSYLKLLAALGFEPDQEWSEDARRSFSSEFRRFEIIFDHLNSEKL
FT                   SLTSADALLAFLQVHSFQALRKMTNDELKTALSALWPDIRLLSEGEPVSVNPVPEEKEK
FT                   GLLSRILGGPRLHVGFFYDCDPQKSVWASAHAQGQHHLEQKFGSTVRVSSFVCTDEETD
FT                   ETMADAIQNKGVNVIFTTSPALMDNCRRIASKYRNAAIFNCSLSMPYAGVRSYYCRLYE
FT                   GKYIAGAIAGAMAKNDRIGYVANYPIMGVIAAVNAFALGARLTNPRARISLKWSCLPGN
FT                   PILEFKNEGISVISNRDESGGNGHLSWDLGTYMVNPDGSLQALASSRWNWGSYYEKTVQ
FT                   SLLNGGIDALKDSSHAVNDWWGMNSGVVSVEMGEGLPDGLRNLGRILKNGIVKGDIDPF
FT                   LCPIRNQKGVLISDGTHPFSLEELMRMDWLCDIVDGSIPPFEEILPRSQNLVRLLGLYR
FT                   EHIPPKTEEAAL"
FT                   /product="hypothetical protein"
FT   CDS             4204..4659
FT                   /translation="MKILLIADEESKYLWDYYQPGRLSGYNCILSCGDLKQEYLEFLVT
FT                   MAGIPLYYVHGNHDKGYVDFPPEGCDCLDDDLVIVDGIRVLGLGGCIRYNPGPYQYTER
FT                   EMAQRIRRAERKIRKAGGVDMSLPMLPLAVMVTPMITLTADLKRFCR"
FT                   /product="hypothetical protein"
FT   CDS             4659..4844
FT                   /translation="MEKYRPRYLIHGHVHQSYGHNLPRSIQYGDTTIFNAVGWHTLDIE
```

```
FT                   PCASVPAKPIFSFFRH"
FT                   /product="hypothetical protein"
FT   CDS             complement(4937..5680)
FT                   /db_xref="GO:0019134"
FT                   /db_xref="GO:0003977"
FT                   /translation="MERTKAVILAAGKGSRLTSEKDEIPKALRRVGDKTLIEHVLDGIN
FT                   FVQPEDITIVIGVMGEKIRQYLGEKYHYAWQKEQKGTAHAMLCAEETASGFDGPVLCLY
FT                   CDMPLLSRETYARMVREHLETGAANTLLAAKIHPIPAFGRLIRNEKGELCDIIEESACT
FT                   DEQKRIDEVNVGIQVMQGSIMWDALKKVDNLNPKHEYYLTGVVRVIYAMGLRQHAVITA
FT                   DSPEYWGVNTMEELLKVENYLKNRK"
FT                   /product="N-acetylglucosamine-1-phosphate
FT                   uridyltransferase (EC 2.7.7.23) / Glucosamine-1-phosphate
FT                   N-acetyltransferase (EC 2.3.1.157)"
FT                   /EC_number="2.7.7.23"
FT                   /EC_number="2.3.1.157"
FT   CDS             complement(5705..6349)
FT                   /translation="MRWLDKLERKLGRHYIRNLMQYLCIAMFGVFILEYLPLPRSAWAL
FT                   LYFNRDLILQGEVWRLITFIFLPPSGELLWILLSLYFYYFLGTGLENHWGSARFNIYYA
FT                   IGVLGNIVAGFLTGFATNSYLNLTLLLAFAVLYPDMEIMLFFILPVKLRWIGGAWGLYL
FT                   IYQLIVSPWAYRVSLLFSLLPFILFFGKQAWLQAKYDSRKFVRWVQNLRGH"
FT                   /product="conserved hypothetical protein"
FT   CDS             complement(6416..8647)
FT                   /translation="MLSVMLFSPVLGALVCPWAGRKNGAVRELLLRLFTIAETLFSLFF
FT                   FLKVREGQAYYLEAPGLVGLGLSFRVDAFRAMYAFLACFMWAMSCQFSREYFSGHGRHM
FT                   GRYCVFTLLTLCGVVGVFYSDDLYTAFVFFEIMSIASYPWVAHEQTPGAMRAAATYLGV
FT                   SVACGMITLMGMFLCWRETGSLAFSALRNAGENPSLYLPASLMLVGYCAKAGMFPLHIW
FT                   LPKAHPVAPAPASALLSGMLTKTGLFGVIAIGLNLMGENSLFGHILLALGLITMTLGAV
FT                   LAVFSVNLKRTLACSSLSQIGYITVGLSCTVLLGHHGALAAAGSLGHMLNHSLLKLCLF
FT                   LCAGAVYMRTHTLDLTRLQGYGRGKIVLHAVFLLGALGLMGVPMLNGYTSKTMIHEALV
FT                   ELAEETAGFSVYRLCEWIFLFAAGLTTAYMLKLYICLFWQKNRDEKTQAGYDALNGKEM
FT                   SLRSRLILILTVLPLLAIGVFPDHTLLPLAEKCAPFFNRPPLAHGLAFFSLTNLAGGAV
FT                   SLAIGGIVYWFFVRKLLYDPKTGYVDRWPRRLDLEELFYRPVFVRFLPRAACGAAAFLD
FT                   HIPECRLIRLWIPKAICSVFTFLGSIVENRLLLRWIPKGIASAAAALDRLPESRFFSRI
FT                   LPGMVTALGRMMDELTDHMLLFIRQMFLTDRQELLQEKEDGMISRLASACAEKLHGIGL
FT                   LPARYLPRREEDDPRYGTYLTNAISFGLLLCAAGLVAAILYGFIRIGAG"
FT                   /product="pH adaptation potassium efflux system protein D
FT                   2"
FT   CDS             complement(8641..8856)
FT                   /translation="MLACVYIFTVLFPAFFMKPLIAAGEPEIHDPGPCMRVSLAVLCAI
FT                   LLTLMVFSDRLIAFLQMLAGEVAGSC"
FT                   /product="hypothetical protein"
FT   CDS             complement(8868..10094)
FT                   /db_xref="GO:0008137"
FT                   /translation="MDKMLCALFPLIWGGALSRIPAAGRKTREVLAVFGSCVASALSWG
FT                   LVLGGSDSRLELFQLTETLSWVLHLDGTGKIFLAIAAVLWPLSTLYAVEYMRHEKREGH
FT                   FFSWYLMAYGCVILLAAADNLFSLYIAYEFLTLMTVPLVWHKRDEESNRAALQYMLYLL
FT                   GGGALSFVSLMGLSAYGTGQFLPGGNMPAMGNAELMLCLAGFLGFGVKAAVLPLCRWLP
FT                   RASAAPTPVTALLHAAAVVNAGVFSVARFLYFAVGTKVLAGTWVQTVFLGLSGCTAVYG
FT                   AAMAVRDRHLKRRLAWSTVSNLSYMLFGISLLTEQGLTAGLAHMVFHSCMKIILFFCAG
FT                   AVLTQTGRTQVNQMQGLGLRMPLVFGAFSLAGAALAGVPPMPGFASKYLLITAPFSREA
FT                   PERCGVDYA"
FT                   /product="NADH-ubiquinone oxidoreductase chain L (EC
FT                   1.6.5.3)"
FT                   /EC_number="1.6.5.3"
```

```
FT   CDS             complement(10105..11559)
FT                   /db_xref="GO:0008137"
FT                   /translation="MKELLLILTVFCPVALGCLFRMMNWSGKKLRMASVASVVLGAVLA
FT                   AVSAQFSGSQVCLHWTDMLAFSLRVDDTGRVWLLLIAVIWPGVALYATEYLEHDQNPQR
FT                   FFLFYTVTQGMLHGLALSGNLVTFYMFYEAMTLLTVPLVLHNMDREAVAAAIKYLIYSV
FT                   IGASAALAGIFFIASVGGTGDFAAGPLSDAQIQGREGMALGIFLAMLAGFSVKAGMFPL
FT                   HGWLPTAHPVAPAPASAVLSGVITKMGVLGVIRVIFCIAGVERLRGTWVQMTLLALSVL
FT                   TILMGSLLALKEKKLKKRLAYSSVSQVSYVLFGLFTMTEYGFVGAVLHVIFHSLMKNTL
FT                   FMAAGSVIHKTELTQVDDMDGLGRRMPWTYAFFTVASLGLVGIPPTGGFISKWNLAQGA
FT                   LATGLNYSWIGPAVLLISAVLTAGYLFTVVIRGCFPGADVPEGPACEAGLSMLLPMGVY
FT                   SALIVLLGIFSGPIVTYLDQLAHQLL"
FT                   /product="NADH-ubiquinone oxidoreductase chain L (EC
FT                   1.6.5.3)"
FT                   /EC_number="1.6.5.3"
FT   CDS             complement(11578..13062)
FT                   /translation="MLAWQNNVPLFMIISPLIGAALCSAFRERAARAVVAAIAALECTG
FT                   MTVLLLFTLREGSFVYPMGSIGAPFGNELRIGPVESATGLVLAAVMSLSLLGGRQMLKR
FT                   DVPPNRQSLYAAVICMLTAALPAMLFTNDLFTGYVFIEITTIAACALISANGRGNTLFA
FT                   AVRYMIMNLLGSGMFLLGLSVLYSLTGHLLFPQLGESVRRLTARGAYPVPLYLSFLLMT
FT                   VGIAIKSALYPFHTWVPNAYSSATAASGAILSSLISKMYIFLLLKLLLRAAGREVFAYW
FT                   INHILMLYAVSGIIMGSVHAIRQHHLNRMVAYSSVAQIGYIFLGISLGSSSGYAAAVFH
FT                   MLSHSAAKSLLFLSSNRLKTAAGGSDLLSDLYGSARQEPIAALAFTVGACSLVGVPLLG
FT                   GFASKVYLTQAGVQAGGWRTPVVLAALAVSTLLNVAYMLRSLLTLYSEKEAGSPGAGRD
FT                   RCFSAAAVLLLGINVVLGVCGGQIMTWVEKGLQVFL"
FT                   /product="PH adaptation potassium efflux system protein D
FT                   1"
FT   CDS             complement(13066..13440)
FT                   /translation="MLAVIYEHLDEFTAMALFGIGFTQLLLDKNLIKKIIGFSMMDTGI
FT                   YLFLASYGYIAGRTAPIYADGSMDTARYVNPLPAGLVLTGIVVGVSVTAVMLGLSVRLY
FT                   QLYHTLDVDEIAERIRGEGK"
FT                   /product="putative Na(+) H(+) antiporter subunit C"
FT   CDS             complement(13440..14396)
FT                   /translation="MAKENKKEASFPFREWLENGGQSLDKVMQDALKAEAGTLPGGTGR
FT                   KRKRKKQSAGMKGKQRVHPTAYAETWWYRLSAVLICLALTGLLFFAVQRMPRFGAADTL
FT                   VDSEVTRFYVEHTLEDTGAVNMVTGIILDYRGFDTLGETNVLFIAVSTVLLLLSLRGEA
FT                   NEAERLREGEYDRHFEPHNDVVLQSAARLITPLSIVLGFYIVLNGHLSPGGGFSGGAVL
FT                   GAGLILYQNAFGYQKTELFFNYRVFRWVSVCALLFYSLSKGYHFFTGANHLDSGLSVGI
FT                   AGRILSGGLLLPLNMAVSCVVACTMYALYTMFRKGEF"
FT                   /product="putative Na(+) H(+) antiporter subunit B"
FT   CDS             complement(14380..14652)
FT                   /translation="MEALNVILLLFVIVCALAVSLTKNLLSAVIIFMGQGLAMSAIWIL
FT                   LESPDLAVTEAAVGTGITSILLFIALRKIGAIQEREEKSKDGQGE"
FT                   /product="NADH dehydrogenase subunit 5"
FT   CDS             complement(14640..14966)
FT                   /translation="MLENVRLILFLALLLAGLLIQAASIVGVNRFRYSANRLHAAGMGD
FT                   TLGLFLLALAGAVYLGVDALTLKILCVVALFWITSPVCTHLIAKLIRETDDVQFRTEAK
FT                   EWKP"
FT                   /product="hypothetical protein"
FT   CDS             complement(14968..15246)
FT                   /db_xref="GO:0006811"
FT                   /db_xref="GO:0015075"
FT                   /db_xref="GO:0016021"
FT                   /translation="MAMQASLAVLGLFLFVCLYRAARGPSTSDRILAINMTSTVTMGII
FT                   LILALLKKEGYLMDIALIYAMLSFLAVVLLCRIYIGVHRAKTGKEGE"
```

```
FT                   /product="Na(+) H(+) antiporter subunit F"
FT   CDS             complement(15257..15721)
FT                   /db_xref="GO:0006812"
FT                   /db_xref="GO:0008324"
FT                   /db_xref="GO:0016021"
FT                   /translation="MVLLVFLFWIFLNGRVTTETVLLGMGIALLVLMFAFHACAWTWRK
FT                   ETAFYRSLPWILAFLFTLLREIVKANLRMLPLIFGGKPDPVVCTVRTGLRTRMGRMLLS
FT                   NAITLTPGTITLDCGQEELKVHCLTPALASELEDNAFENKLRRIEVVFLG"
FT                   /product="Na(+) H(+) antiporter subunit E"
FT   CDS             16044..17135
FT                   /translation="MQKHLFRAVFPLALALVLAFSCTVPRAAAEEEYFKFSRLIPDVFD
FT                   TKTTLIGFAESQESFDRAADEVTAQLTAYDRIFDAYTPYEGLNNLYYVNRHAGEEPVSV
FT                   PTDLFELLSWCKARWEEGLRVTNVAMGAVLSIWHEYREAGLNDPEHAELPPEDMLLAAN
FT                   AHVSLDDVVLDSEKQTVFFSDPGLTLDIGAVAKGFAADKVNGFLSHEMPSYLLSLGGNV
FT                   YAGAPPLDGRQNWAVGVQDPLTDELSLALGKTEIMDIMDVNRLTVVTSGDYWRYYLVDG
FT                   VKYHHIIDPVTLMPARQMISVTVICQSSTLADFLSTTFFALSYEDGLQLADRLDGVDVM
FT                   WVLPDGSVHCTDGMIRYARSLKQ"
FT                   /product="Hypothetical similar to thiamin biosynthesis
FT                   lipoprotein ApbE"
FT   CDS             17143..17664
FT                   /translation="MRNVSKTKRAAANMAVIGIFVLIVLLLLFVLPHRLNPDLEQTQED
FT                   IQRQMASLASSPKPGEETPPVPSQNPDAEAYLTITLNGSLWRIEPLGEEREIRVEQDDG
FT                   AVNVIHLLPNGYYMASSTCDNQLCVQEGTVTVENYQRRILGPAVYCLPHHLALELVVPF
FT                   HTPDPAAPDI"
FT                   /product="hypothetical protein"
FT   CDS             17792..21991
FT                   /translation="MKKKLPGWAVLLIITLAAGLALGGTYALTKDPIDRQTLAAAENSR
FT                   RAALPEADSFEEVSLAEGADVDWAYAGQKNGETIGYVAQKTVKGFGGKVEVIAGVDTRK
FT                   SPDSFTLGGIAVGGASFSETAGLGARAKEAAFSDQFIGKTAPVSYIKAGGEATESTVDA
FT                   ITSATITTTAVVNGVNDIVKYVKAGILGIAGIEMPPKPADGVYSASEKGFRGPVYVEAA
FT                   FDNAGKVTYIKVGDDSFAEDIGIGVIEPEFMIQFIGKSMPLDTSDIDMIAGATISSTAA
FT                   LKALNTAYAVAGGAEIVVPESPTLPEKPAEGVYEASAQGFGGPVQVEAAFDADGRMTYI
FT                   AIGDDAFAETAGLGAKALEPEFQAQFIGKQMPLELTDIDAIAGATITSTAVVSALNSAY
FT                   NASQGIVEEAPAAPVLPEKPAEGVYEASAQGFGGPVKVEAAFGADGQIAYIAIGDDAFA
FT                   ETAGLGAKALEPEFQAQFIGKQMPLELSDIDAIAGATITSTAVISALNDAYNASQGVVE
FT                   EAPAAPVLPEKPAEGVYEASAQGFGGPVKVEAAFDGDGQIAFIAIGDDAFAETAGLGAK
FT                   ALEPEFQAQFIGKQMPLELSDIDAIAGATITSTAVVNALNDAYNASQGVVEEAPPVPVL
FT                   PEKPAEGVYEASAQGFAGPVKVEAAFDGDGQIAYIAIGDDAFAETAGLGAKALEPEFQA
FT                   QFIGKQMPLELSDIDAIAGATITSTAVVSALNDAFIASQRTAEDISNAISSLPEKPAEG
FT                   VYEASAQGFGGPVKVEAAFDAEGKITYISIGDDAFAETAGLGAKALEPEFQAQFIGKQI
FT                   PLTLDDIDAITGATITSTAVVTALNEAYTASQGVVEEAPAAPVLPEKPAEGVYEASAQG
FT                   FGGPVKVEAAFDAEGKITYITIGDDAFAETAGLGAKALEPEFQAQFIGKQMPLELTDID
FT                   AIAGATITSTAVVSALNDAFIASQGTAEEISNGISSLPEKPAEGVYEASAQGFGGPVKV
FT                   EAAFDGDGQISFIAIGDDAFAETAGLGAKALEPEFQAQFIGKQMPLELSDIDAIAGATI
FT                   TSTAVISALNDAYNASQGIVEEAPAAPVLPEKPAEGVYEASAQGFAGPVKVEAAFDADG
FT                   KIAYIAIGDDSFAETAGLGAKALEPEFQAQFIGKQMPLELTDIDAIAGATITSTAVISA
FT                   LNDAFSASQGTAEDISNAISSLPEKPAEGVYEASAQGFGEPVKVEAAFDAEGKITYISI
FT                   GDDSFAETAGFGAKALEPEFQAQFIGKQMPLTLDDIDAITGATITSTALVTALNDAYTA
FT                   SQGTAEEISNEISSLPEKPAEGVYEASAQGFGGPVKVEAAFDAEGKITYISIGDESFAE
FT                   TAGFGAKALEPEFQAQFIGKQIPLTLDDIDAITGATITSTAVVTALNDAYQQTVLD"
FT                   /product="hypothetical protein"
FT   CDS             22034..23383
FT                   /translation="MLTKTFKRGIHPLHRMHEGKGLTRDLPVREYVSDTVVISMAMNIG
FT                   APAKPCVKKGDHVDIGQVIGEAQGFMSVPVHASVSGEVIAVDAIPSLGAGNVQAVTIKN
```

```
FT                   DFAETWVELHPLGSVEQVNPKDVIPAIQAAGITGMGGASFPTHVKLTFREGAFCDTVIV
FT                   NGAECETHLTADYRLMLEYAQRIVDGLRAVMLALGVKKGVIGIEDNKPEAIIAMEQASK
FT                   GRAGVSVQPLKTKYPQGGEKQLIEAITGRQVPSGGLPIDAHVVVLNVGTCAAIADAIID
FT                   GKPLISRICTVTGCVRQPSNLLLRIGTITEDIIGACGGYSEPAGKVVFGGMMTGICIPN
FT                   DSMPILKSTNGIVVLNDKDSRSKDESPCIHCGRCVAACPIGLNPYQIKVAADKSDFAAA
FT                   KKLHVMDCILCGSCAYECPSRRWLTPSFKYCKDRITAEAKAAAAKGGAKK"
FT                   /product="Electron transport complex protein RnfC"
FT   CDS             23380..24324
FT                   /translation="MSLRISTAPHMHNPLTTQRLMQYVFIALVPTAAAGVYTFGLKALL
FT                   VLAVATCTAVLSEYLWQKIAGQPVRIGDFSAAVTGLILGLNLPSTAPWWLAVIGSAFAV
FT                   IIVKQLFGGLGDNFLNPAMAARAVLLASWPVHMTAAAYAAPSFFSGVDALASATPLGAQ
FT                   SSFTMTQLFLGQIPGTIGEVCKAAILLGLLFLLVMRVISWRIPVIMAGSVFLCSMLIGG
FT                   MTPGAALAAVLSGGVLFGAVFMATDYVTSPITPLGQMLYAFMIGLIIVLIRTFGSYPEG
FT                   VTYGILLMNVATPLIDRFLPRAIYGYQKEGKKA"
FT                   /product="Electron transport complex protein RnfD"
FT   CDS             24321..24959
FT                   /translation="MSEKKSLGKVLFNGIVPENPTFRLVLGTCPTLAVTTSALNGLGMG
FT                   AAATFVLVCSNIAISLLRKFIPDQVRIPSFIVVICTFVTMVQLLMKAFLPALNESLGIF
FT                   IPLIVVNCIILARAEAFASKNTPIASAADGLGMGIGFTLALTLMGCIRELIGSGSVFGV
FT                   NVLGASYEPMLLIVLASGGFLTFGLLLGVFNLIVKKIERKRAVKEAQTA"
FT                   /product="Electron transport complex protein RnfE"
FT   CDS             24956..25540
FT                   /translation="MSIVEILLFMVSCILTHNFIFNRFLGCCPFLGVSGKVETSVSMGL
FT                   AVTFVMTIASLFCWLIYNLLLIPLGLEYMNTISFILVIAALVQFVEMFLKKSSPTLYSA
FT                   LGIYLPLITTNCAVLGVATLNMNEGYSLLQSVLNGTFSALGFLLAIVLMAGVRERLESS
FT                   KIPACMKGFPITLVSAGLMAVAFMGFSGLVP"
FT                   /product="Electron transport complex protein RnfA"
FT   CDS             25556..26347
FT                   /translation="MDMTILYAGILLGALGLIFGGVLSFASKKFHVETDERIARVRECL
FT                   GGANCGACGYAGCDAFAEAVVRGEAPINGCAPASAKGVQAIGEIMGQQADSGERMVARV
FT                   LCQGENGIAKERYQYDGYQSCATAAGLAGGPKDCRFACLGLGDCMDKCAFGAISMQNGI
FT                   AHVDPDKCTACGACVKTCPRSVLKLLPVSSNVIVRCRNSDIARVARSVCMNACIGCGRC
FT                   KKECQYEAITVENGFAHIDPSKCIRCGACATVCPCKCITIN"
FT                   /product="Electron transport complex protein RnfB"
FT   CDS             complement(26455..28059)
FT                   /db_xref="GO:0031176"
FT                   /translation="MKKKMNSGIIKQKGERMNNMKKILSAVLLLALVFCCLPGRAETEA
FT                   VYASDYSGGEDGWYGRGAVCYALDSILYTEGRQSDWNSPGHDFDLIAGASYCISAQVLQ
FT                   NELDSASFMISVAHSKDGAESYENLAHGTAKKGEWTTLTGSYTAGDFDRFVLYVETTGS
FT                   PSLSFAMKDFVMTAPEGLPEPKPTEAPMVIEAAGEMPSLKEIYADLFDFGAAAPQQAFR
FT                   DIRLMRLMKEQFSILTPENELKPDAVLDVYSSKKLAETGDETAAAVRFDAAKALLNFAQ
FT                   RNGLKVHGHVLVWHSQTPEAFFHEHYDTKQPLVSREVMLGRLENYIRGVMEYLDANYPG
FT                   VVVSWDVVNEAIDDGTGWLRKNSSWYKVVGEDFAARAFEYARKYAPEGTLLYYNDYNTA
FT                   MAGKRKGIVRLLESLIPEGNIDGYGFQMHHSAGSPSMQQIADSVKDIAALGLRLRVSEL
FT                   DVGVDNRTETSLKKQAQKYSWIMKLLRRYSDQFEAVQVWGLTDNMSWRSTSYPLLFDAS
FT                   LNPKPAFWAVADPDSVE"
FT                   /product="CBM22-GH10"
FT                   /EC_number="3.2.1.8"
FT   CDS             complement(28172..29623)
FT                   /translation="MKKIVSLILVAMMLLACCSFSALAEVPEGYPEIIEGLDFGGQTIW
FT                   IYDWWSSDDENHSARTSEPDEETALLYAYRDWLEETYNCKIVESALSDWAGNPTELANM
FT                   VMNGDSSKLCLVAVAADFAGTPLANDLYMPWQIDLSGEKWNKATSEFCTKNGQVYGVHV
FT                   GKTEPRSCIFFNKRVLSEAGIDPESIYDAQADGTWTWDMMLSIMEQVQRDTDNDGIFDI
FT                   YGMTGSGDDLATGLIFSNNASFFDYNEEGKLAITANSQAALDALQMRKTMWENYAAPQP
```

```
FT                   EGSNWDWFKDFWEQGTTAFYAGQTWQGFNDNAEMSDMEDEWGAVMYPKGPNADTYLAMA
FT                   CDNVYGIPNVYDDETSLKIQQIYDLYTNDTPGVDTEDAWIGNKYLFTDERAVDETYAMM
FT                   REDEHSVATKTSLLGSTNDVLGSALLWGPIGSMTPAEAVESATPVWQDMLDVFNGDKTQ
FT                   EQIDAEHAAAEAAAAEAAAEEVPAE"
FT                   /product="hypothetical protein"
FT   CDS             complement(29781..30791)
FT                   /translation="MSKDNTVFIGKNKSFKERNRRSGGYLMKRQIKRGCISVARALLMF
FT                   GLCFMIIQPMLTRFSMSFMAEKDLYDSTIILLPRNVTLENYKIVADLTDFPGTMINTLW
FT                   ISILVSICQVVAATLVSYGFARFEFPFKKFWFGCVVLLIIIPPQTIQTSLYMTFANFDL
FT                   FGIIQGLTGKTMNLRSSLTPYLLMSLTCMGLKNGLFIYMLRQYFKAVPESLEEAAYVDG
FT                   CSTLHTFFKIMLPDAVPTIASCFLFSFVWQWTDLFYARNFLTGYPTFAIRMSTMVSRMG
FT                   RYFSKDATQSVIVPNGRQLQLISIAVLFCCIPLIILYCFTQRTFVESLAMSGSKE"
FT                   /product="binding-protein-dependent transport systems
FT                   inner membrane component"
FT   CDS             complement(30784..32190)
FT                   /translation="MGGFVYRFLQQPTGIVLLGFAACLAVTAAFVFFLWKVFEKAGEKG
FT                   WKALIPLYNLYILWKTGWNGVRFFHALCVMLAGAYILDILTSFGPSVRLVGEIIFWLCA
FT                   AFAVAYSVMMGIHMGHRFGKSTVFGAVALGVLPIVGLGILAFDRSDYQYNRDTAGEPAA
FT                   QNSNYRITTIHERRAMTGYFFILPFIIGFLLFMVKPLVFSMQMSLNEVKLKAGGGFEMT
FT                   WNNFNNYHYALQTDPNFNQYLVDEMGRMAINTIATLVMSFVIAVILNQKFKGRVLCRVI
FT                   FFLPVILSSGVLPGIESSNEFYNMMSDISNSVQNSSGINISLALQNLLSVSGVAGEIFD
FT                   VIFQMIDAIYDIVMASGIQIIVFLSGLQSISPSLYEAADVEGCSAWESFWKITFPMVSP
FT                   LLLVNCIYTIIDFFMKNDNRVIERINTIMYGVQLNFGVASAMSWIYFGVALGFIGIGTF
FT                   IITRAVKSYE"
FT                   /product="binding-protein-dependent transport systems
FT                   inner membrane component"
FT   CDS             complement(32190..34763)
FT                   /translation="MKQKKSVLRWLIPLILVLAALAALVVFVFVPIYSQKEDTIGRPPS
FT                   VLNYAGDGKPLTMENDALYFEMDAGTTQFKITDKKTGKTWYSNPTDRESDPVARGVNKE
FT                   TLSSTLNVPYTATGGEVELNNYTYSMMNQTYDILQQEDGSIRVDYAIGKIEKTYLMPTA
FT                   ITAERYNEITGRMSKATKKKVSSSYSLYEPDKLDKKENKDEIIALYPSVVEQALYILKS
FT                   DTTTANKAKIEGYFAEAEYTQEDYEIDQQLVAGTRSNNGPVFNASVIYRLDGGDLLVEV
FT                   PYESLCCESSSPLTYISVLPMFGAAGTAQEGFMLIPEGGGALIHYNNGKLSQNAYYANL
FT                   YGWDYATERKEAVSETRNAFPVFGMGQPDGSFICIVEGASSYCGINADIAGRFNSYNFV
FT                   YGKYNVLHYDRFNVSNRTAQLLYMYESAIPDDVLIQRYCFLEGDSYVDMANLYGEYLRR
FT                   TPEMKNDTASIDMPVNVELIGAINKTVPRFGIPVDSVIPTTTFAQAGDIMNELVSSGVR
FT                   DLNLRMTGWANGGVRQKVLTGVHVLRQLGGENAMKKLISDAKARDVDLYFDGINCFAYS
FT                   SGVFDGFLPFSHAARFTTREQVKLYAYDIVTYTQADWMDTYYLVRPEYAHTNAGRLLDG
FT                   LKNRSAVGVAFRDIGNLLSADYYNRSTVTRQQVMDMDVQTLKDAVAKELKISIKEGNDY
FT                   AVPYADLITDMNLTGNAYAIIDRRIPFYQIALHGMKNYTGGAINLAGDYVTALLECAEY
FT                   GAGLEFSFMAEDTTVIQDSAYSCYTASGYERWKEQVIPAILRYQREMEGLNRQTIVNHE
FT                   YLTDEVVATTYADGTKVFVNYSVTDYETGGNTVPARDYMVERGVK"
FT                   /product="GH129"
FT   CDS             complement(34779..35474)
FT                   /translation="MSTIKSARAAEAAERAENRKAWWNRYKESFRYAFHVITHPFDGFW
FT                   DLAHEGKGNLAVANTFLFLFLMVRVLKLMFTSFQFISAPTQYINIFEEAASLALPFLVL
FT                   CLANWAMTTLFDGKGRFKDIYIAMCYALVPYILIQSIMVLVSNIITYEEGGFYSVFLGI
FT                   SLLWCAFLVFVGLMEVHDYGPGKTFIFLIVTVVGACIILFLVLVFFSLLSDAASYFISL
FT                   YREIIYRLY"
FT                   /product="hypothetical protein"
FT   CDS             complement(35443..36765)
FT                   /translation="MRISRPQSLFVRNSDLYIVDTNNNRILQVHWDGAEYTLTRVIDRI
FT                   NGCEVTSLNKPYDVFVDQDDSIYVADFGNYRVVMMDKDLNYIKVFVKPDDPTFDQKLDF
FT                   QPKKIAVDVAGRVYALVNNVNKGFVKYENDGTFTGYIGANTVSVNMADYIWKRYFQTKE
```

```
FT                   QRAASESFVPTEYENMYIDQYGFIYATNTVFSEYDLLWDNAKPIRRLNGVGSDILIKND
FT                   RYPPIGDLKWIEGGAASGPSKMTDITVLDNDIYIAVDRVRGRIFGYDSQGVMLWAFGTK
FT                   GNISGAFTSAISIEHMGYDLFVLDQLENSVTVFTPTEYGKTIYDAIETYLRGEYDASAD
FT                   LWRQVMKLNANYPLAFRGIGRALLRQDDYKGAMAYFEKAHDRENYGRAFKLFRKEWVEQ
FT                   NIWWLALIIAAVLVIPLALGRVKKMKWEVIAHEHNKVRKGG"
FT                   /product="NHL repeat containing protein"
XX
SQ   Sequence 36794 BP; 8525 A; 10372 C; 10043 G; 7854 T; 0 other;
     atacggcaat gtcccgcgga gagcacatga gatacagggg cgagaggatc ctgagcagat        60
     tggcaattcc cagcatgccg ctttgaagat cgtccttttc atattgtgcg gaaatgctgt       120
     ctttcaccgt aagccagcag gcagtggtat gcatttccgt ctccggaagc cgtacgggcc       180
     catacccac gttttcattg ttgtcaaagc gcattttttt gaaaaccttt accagcgcca       240
     cgacctttac ttcacccaga ccggcgtttt cttcttcctt ctcaatatcc agcaggctca       300
     gattcacatt caggtcggca tcggtataat aatccacatt cacctttcgg ataaatgctt       360
     tgcaagcttc aaaatccagc ttttccacct gataggtctg cccctcatgc atatagatgg       420
     cattcgggtg cagcagcatc ggggcggtaa aacggtccat ttcccctaca atcctgggat       480
     gggcgggatc ggtgatatca ttaatcagaa agttttccgt catggcggtg cgcaggctga       540
     tttcactggc gggaaaatct tccgccatcc agtgataggt atcccccacc ctgtgcagaa       600
     tattgctttc ttccagaaag ccaagcattt cttccccgcc cgctgcattt ccaaggcttt       660
     ccccttcttt aaaaggaagc tcgtaggcgg cgcatttgaa atgactcagc agaatataca       720
     gattgtccgg attcaaaagc gcattttcag caggctgcgt aaaaaagtac tcaggatgcg       780
     tgacgatgaa ctggtccaat gcgccggagg atgccacaag gaaggtggcg ctcacgccgt       840
     ggcgacgtcc ggcacggccg gactgctgcc aggtgctggc aatggttccg ggatatccgc       900
     acaaaacgca ggcttccagg gcgccgatgt ctatgcccag ttccagcgcg ttggtactca       960
     ctacggcctg gatattcccg gcccgcagcc cctgctcaat ctcccgccgt tccgatggca      1020
     ggtatccgcc ccgatatccc cgtaccttgc cgctgttgcc aacggggtcg gataccagct      1080
     ctttgagctg ccgggtcatc acctcgacct gaagcctgct tttggcaaag acaatgctct      1140
     ggactccgtt gcacaccagc atggaagcga tattcctcgt ttcagaaagc acgctccggc      1200
     ggataccgag ctggatattc accaccggcg gattgtagaa aatatagtgt ttttcaccgg      1260
     aaggagcgcc gttattgtca atcagcatca cgctgcggcc cgtcagcgtt tccgccagct      1320
     cctgcgggtt ttggatggtg gcgctgcaaa ggatgaaccg gggatggctt ccgtaaaaat      1380
     cacagagccg gatcaggcgg cgcaggacat tggccagatt gctgccgaaa acgccgcgat      1440
     aggtgtggat ttcgtccacc acgatgtatt tcaggttttc aaacagcttg acccatttgg      1500
     tatgctgagg caggataccg ctgtgaagca tgtccggatt ggtgaccacg atatgcccgg      1560
     cctgccggac cgcccgtcgg gccgcggccg gggtatcgcc gtcgtatgta taggttttga      1620
     tatccgcccg gagctgctcg atcagttcgt aaagctcgct cacctgatcg gcggagagcg      1680
     ctttggtggg aaacagatac agtgccctgg cgtccgggtt tttcaggatt tcagccagaa      1740
     caggcagatt gtaacaaagg gtttttccgg aagcggtggg tgttaccacc acataatcct      1800
     tcccttcctg ggaagcggca aagcaggaag cctgatgggt gtacagctga tgcacgccgt      1860
     gcttttccag agccgccgga agccgcgggt caatgccggc cggccaggga gcatattccg      1920
     cagcggcgga cggaactgtt ttccaggcgg tgacattgtc cataaaggca ggatcgtttt      1980
     tcagtttttc cagaagctgc ggcaaattca cggcctgtcc tccttcggat atatctgaaa      2040
     ctgcgctcgc aggaacaagt gttcgtttcg atgtccttcc ctattatatc ctttctttgt      2100
     gcgctttgca agctgttcct gctttcgagc cgtatttcag cttgcaacgc cagtctattc      2160
     ctgatacaat aaagacgttc tttccggtgc ggtacggctg tgcgcggaaa gaaaagcaga      2220
     cagaaagacg gatgaatcat ccggcaagat ataggaggca acgatgagca gaaacgatgc      2280
     gattgaacaa tacaatgccg cgctgcgcat gggaaggagc tattacaaag cctgtaccgc      2340
     gaaaggcgaa gacccctttc ccaaggtgct ggaacacctg attgaaaaac aggaaacggg      2400
     caactctgta aaaatcggtc tggtggatat ccccgccagc agaatcgtcg gcacctggat      2460
     ggagggacgg aaggcagcct ttgccggcaa ttttatgccg ctgctggacc cggacagcga      2520
     gtttggcgga aaatggatca gtctgtgtga gtcccatctg gacgagggag gcatccggga      2580
     tcccatcacc tgcatcgagt atctgggcta tttctatgtg caggagggca ataaacgggt      2640
     cagtgtgctc aggagcttcg attctcccag catcccgggc attgtgacgc gtttgctgcc      2700
     tctccctggc gacgatcccc ggattcagct gtattttgag tttcttcagt tttataaagt      2760
```

cagtaaaacc tatctggtta tgttcagcaa accgggcagt tatctgaagc tgctggccgc 2820
tttgggcttt gaaccggacc aggaatggtc ggaagatgcc aggcggagct tcagcagcga 2880
attccgccgg tttgaaatca tattcgatca tctgaacagt gaaaagcttt cgctcacctc 2940
ggcggatgct ctgctggcat tcctgcaggt gcattcgttc caggccctca ggaaaatgac 3000
aaacgatgaa ctgaagactg ccctttccgc cctgtggccc gatatacggc tgctgtcgga 3060
aggagaaccc gtgtctgtca atccggtgcc tgaggaaaag gaaaaaggcc tgctgagcag 3120
gatactgggc ggtccccgcc tgcatgtcgg ctttttttat gactgtgacc cgcagaaaag 3180
cgtctgggcc tccgcgcatg cccagggaca gcatcatctg gaacagaagt tcggctctac 3240
ggtgcgtgtt tcatccttcg tctgcaccga cgaggagaca gacgaaacca tggcagacgc 3300
cattcagaac aagggagtca atgttatatt caccacaagc ccgctctga tggacaattg 3360
ccggcgcatc gcgtccaaat accggaatgc cgccatcttc aactgctcgc tttccatgcc 3420
ttacgcgggc gtacgcagct attactgccg tctgtatgaa ggaaaataca tcgcaggcgc 3480
catcgccgga gcaatggcca aaaatgaccg tatcggctat gtggccaatt atccgatcat 3540
gggcgtgatt gccgcggtga acgcttttgc tctgggcgcc aggctcacca atcccagagc 3600
ccgcatcagc ctgaaatgga gctgcctccc cggaaatccc attctggaat tcaagaacga 3660
aggcatcagc gtcatttcaa atcgggatga aagcggcgga aacggccatc tttcctggga 3720
ccttggcacc tatatggtca acccggacgg cagcctgcag gctttggctt cttcccgctg 3780
gaactggggc agctattacg aaaaaacggt ccagtccctg ctgaacggcg gcattgacgc 3840
actgaaggac agcagccatg cggtcaatga ctggtggggg atgaactccg gcgtggtcag 3900
cgtggagatg ggcgagggcc tcccggatgg tctccggaat ctgggccgga ttctgaaaaa 3960
cggcatcgtc aaaggcgata ttgatccttt cctctgcccc atacggaatc agaagggcgt 4020
gctgatcagt gacgggaccc atcccttcag cctggaagaa ctgatgcgga tggactggct 4080
gtgcgatatt gtggacggct ccattcctcc gtttgaggaa atcctgccgc gttcccagaa 4140
cctggtacgt ctgctggggc tttaccggga acacattccg cccaaaacgg aggaagccgc 4200
gctatgaaaa tcctgctgat tgccgatgaa gaaagcaaat atctgtggga ttattatcag 4260
cccggccggc tgagcggata caactgcatt ctgtcctgcg gggacctgaa gcaggaatac 4320
ctcgaatttc tggtcaccat ggctggcatt cccctgtatt acgtgcatgg gaatcatgac 4380
aaaggttatg tggacttccc gccggaaggc tgcgactgtc tggacgatga cctggtcatt 4440
gtggacggaa tacgcgtcct ggggctgggc ggatgcatcc ggtacaatcc ggggccttat 4500
cagtacacag aaagagagat ggcgcagcga atccgccgcg ccgaaagaaa aatccggaaa 4560
gcgggagggg tggatatgtc attacccatg ctcccccctcg cggttatggt gacgccgatg 4620
ataacgctca ccgcggattt gaagcgtttc tgccgctgat ggaaaagtac cgtccccgat 4680
atctgataca cggtcatgtg catcagagtt acggacacaa tctgccccgc agcatccagt 4740
acggcgatac cacgattttc aacgccgttg gctggcacac gctggacatt gagccctgcg 4800
cctccgtccc ggcaaagccc atttttttctt ttttccgtca ttgaggccat gggaaatgcg 4860
gatgccataa taatgccgcc gcgcccgtat ccggttcctg ccggaaaaca cgcgcggcgg 4920
cggctttttt ggtttttttac tttctgtttt tcagataatt ttccaccttg agcagttctt 4980
ccatggtgtt cacgccccaa tactccgggg aatccgctgt tatcacagcg tgctgcctga 5040
gccccatggc gtaaatgacc cggacgacgc ccgtcaggta atactcatgc ttgggattca 5100
ggttgtccac tttttttagg gcatcccaca tgatgctgcc ctgcatcacc tgaatacccca 5160
cattaacttc gtcaatccgt ttctgttcgt cggtgcaggc gctttcttct atgatatcgc 5220
aaagctctcc tttttcattc cgaatcagcc ggccaaaggc gggaatggga tgaatcttcg 5280
cggccaggag cgtattggcc gctccggttt ccagatgctc cctcaccatg cgggcatagg 5340
tttccctgga aagcagcggc atatcgcagt aaaggcaaag caccggaccg tcaaatccgg 5400
aggctgtctc ctccgcgcac agcatggcgt gagccgtccc cttttgctcc ttctgccagg 5460
cataatgata cttttctccc agatactgac ggattttttc ccccatgacg ccgatcacga 5520
tggtaatgtc ttccggctgc acaaaattga tgccgtcaag cacatgttct atcagggttt 5580
tatcccccac cctgcgcaac gccttgggaa tctcgtcttt ttcacttgtc aggcgggaac 5640
cttttcccgc ggcgagaatg accgcttttg ttctttccat attccggctc tcctttgcat 5700
atactcaatg gccccgcagg ttctgcaccc atcgaacgaa ttttcgggaa tcatactttg 5760
cctgcagcca cgcctgcttc ccgaaaaaca gaatgaaggg aagcagcgaa aacagaaggg 5820
ataccctata ggcccaaggg ctgacgatca gctggtatat cagatacagg ccccaggcgc 5880
cgccgatcca gcgcagcttg acaggcagaa tgaaaaacag catgatttcc atgtcgggat 5940
agaggaccgc aaaagccaaa agtagcgtca ggttcagata gctgttggta gcgaagcccg 6000

tcaaaaagcc ggccacgata tttcccagaa caccgatggc ataataaatg ttgaaccttg 6060
cgctgcccca gtggttttcc agccctgtgc cgaggaagta ataaaaatac aggctcagca 6120
ggatccagag cagctcgccc gagggcggca gaaaaatgaa ggtgatcagg cgccagacct 6180
cgccctgcag aatcagatcc ctgttgaaat acagaagtgc ccacgcgctg cggggtagcg 6240
gcagatactc cagaatgaat acgccaaaca tggcgatgca caaatactgc atcagattcc 6300
ggatataatg ccgtcccagt ttcctttcca gtttatccag ccatcgcatt ttccccgcct 6360
ccaactgata taaatattat atcgcctttt ccccggttcg tcaagacggg cctgttcagc 6420
cggcgccgat acggataaaa ccgtacagga tcgccgccac cagaccggcg gcgcacagca 6480
gcagaccaaa ggaaatggcg ttcgtcagat aggtaccgta cctgggatca tcctcctccc 6540
tgcgcggaag atatctggcg ggcagaagcc cgattccatg cagcttttcc gcgcacgcgc 6600
tggccagacg ggaaatcatt ccgtcttctt tttcctgcag caattcctga cggtcagtca 6660
gaaacatctg tcggataaaa agcagcatat gatccgtcag ctcgtccatc atccttccca 6720
gcgcagttac cataccggga agaatgcggg aaaagaatct ggattccggc agtctgtcca 6780
aagcggccgc agcggaagct attcctttcg gaatccagcg cagcagcaaa cggttttcca 6840
cgatactccc cagaaaagtg aacaccgagc agatcgcttt gggaatccag agcctgatca 6900
ggcggcactc cggaatatga tccaggaaag ctgccgctcc acaggccgcc cggggcagaa 6960
accgtacaaa aacaggccgg taaaacagtt cctccagatc cagacggcgg ggccagcggt 7020
ccacgtatcc ggttttcgga tcataaagca gcttccggac gaaaaaccaa taaacgattc 7080
caccgatcgc caggctgacg gcgcctcctg ccagatttgt cagcgaaaag aaagcgaggc 7140
catgggccag aggcggacgg ttaaaaaacg gtgcgcactt ttcagccagc ggcaaaagag 7200
tatgatcggg aaatacgccg atcgccagca gcggcaggac ggtcaggatc aggatgagcc 7260
gggaacggag agacatctct tttccgttca gcgcgtcata tccggcctgc gtttttttcat 7320
ccctgttctt ttgccagaac agacagatat acagcttgag catataggcg gttgtcagcc 7380
cggctgcaaa cagaaaaatc cattcacaca gccggtagac agaaaagccg gccgtttctt 7440
ccgccagctc caccagcgcc tcgtgaatca tggttttact ggtatatccg ttcagcatcg 7500
gcacgcccat caagcccagg gcgcccagca ggaaaacagc gtgcagcacg atttttccac 7560
ggccatatcc ctgaagccgg gtgagatcca gcgtatgcgt gcgcatataa acggcgccgg 7620
cgcataaaaa gaggcacagc ttgagcagcg aatgattcag catgtgtccc aatgagccgg 7680
cggcagccag cgcaccgtga tgccccagca gcaccgtgca ggaaagcccg accgtaatat 7740
agccgatctg ggacagagaa gagcaggcaa gcgttctctt caggttaacg gaaaatacag 7800
ccagcactgc gcccagcgtc atggtgatca atcccagcgc cagcagaatg tgcccgaaca 7860
gactgttttc tcccatcaga ttgagcccga tcgcgatgac cccgaagagc cctgtttttg 7920
tgagcatgcc cgacagcaga gcgctggccg gagccggagc gacgggatgc gcctttggca 7980
gccagatatg caggggaaac atgcctgcct tggcacagta acccaccagc atgagagaag 8040
ccggaagata cagggaaggg ttttctcccg cgttcctcaa agccgaaaaa gccaggctgc 8100
ccgtttcccg ccagcataaa aacatgccca tcagcgtgat catgccgcag gctaccgata 8160
cccccagata agtggcggca gcccgcatcg cgcccggcgt ctgttcatgt gctacccagg 8220
gataggacgc gatggacatg atttcaaaaa agacaaatgc ggtatacaga tcgtctgaat 8280
aaaacacacc caccacgccg caaagcgtca gcagtgtaaa aacacagtat cgtcccatgt 8340
gtctgccgtg gccggaaaaa tattccctgg aaaactgaca ggacatggcc cacatgaagc 8400
aggcgagaaa cgcatacatc gcgcggaaag cgtccacgcg gaaggacagc cccaatccaa 8460
ccagtccggg cgcttccaga taatacgcct gtccttcccg gaccttcagg aagaaaaaga 8520
gagaaaacag ggtttccgct atcgtaaaca gccgcagcag cagttcccgg accgctccgt 8580
tttttcttcc tgcccaggga cataccagag cgcccagaac gggagaaaac agcataacgc 8640
tcagcatgat cccgccacct ccccggccag catctgaaga aaggcaatga gcctgtctga 8700
aaaaaccatc aatgtcagca ggatggcgca aaggacagcc agagaaacac gcatgcaggg 8760
tccagggtca tggatttccg gttctccggc cgctataagc ggtttcatga aaaaagcggg 8820
aaacaggacg gtaaaaatat aaacgcaggc cagcaccgcg gccagaatca ggcataatcc 8880
accccacagc gctccggcgc ttccctgctg aaaggcgcgg tgatcagcag atatttgctc 8940
gcaaagcccg gcataggcgg cactcccgcc agggctgctc cggccaggga aaacgctccg 9000
aataccagcg gcatgcgcag gccaaggccc tgcatctggt ttacctgtgt ccggcccgtc 9060
tgggtgagga cggcgcctgc gcagaaaaag agaatgattt tcatgcagct gtggaaaacc 9120
atgtgggcaa gcccggctgt cagtccctgt tcggtgagga gagaaatgcc aaagagcata 9180
tagctcagat tgctcacggt gctccaggcg agcctccgct tcagatgcct gtcccgtacg 9240

gccatggcgg ccccataaac cgctgtgcat ccgctcagcc ccagaaaaac agtctgtacc 9300
caggtgccgg ccagtacttt tgttcccaca gcaaaataca gaaatcttgc cacggaaaac 9360
acgcctgcgt tgacgaccgc ggcagcatgc agcagcgccg tcaccggggt gggggcagcc 9420
gatgcgcgcg gcagccagcg gcacagcggc aaaacggccg cttttacacc gaagcccaga 9480
aaaccggcca ggcacagcat caattcagca ttccccatcg ccggcatgtt tccgcccggc 9540
agaaactgtc ccgttccata ggcgctcagc cccatcaggg aaacgaaaga aagcgcgccg 9600
ccgccaagca ggtacagcat atactgcagc gcggcacggt tggattcttc atcccgtttg 9660
tgccagacaa gcggcacggt catcagtgtc aggaattcat aggctatata cagggaaaac 9720
agattgtctg ccgctgccag caggatgacg cagccataag ccatgaggta ccaggaaaag 9780
aaatgacctt cccttttttc atgacgcatg tattccaccg catacagcgt tgacagaggc 9840
caaagcacgg cagcaatggc aagaaatatt tttcccgttc catccagatg cagaacccat 9900
gaaagcgttt ccgtcagctg aaaaagctcc agcctgctgt cggatcctcc caggaccagc 9960
ccccatgaca gcgcggaagc gacacaggaa ccgaatacag cgagcacttc tctggttttc 10020
cgtccggcag ccggtattcg ggacagcgcg cctccccaga tcagcgggaa cagggcgcaa 10080
agcattttat ccatggctgt ttcctcacag cagctgatga gccagctgat ccaggtatgt 10140
tacgatcggg ccggagaaaa tgcccagcag cacaatgagc gccgaataaa cgcccatggg 10200
cagcagcatg ctgagcccgg cctcgcaggc agggccttcc ggcacatcgg cgccaggaaa 10260
acatccccgg atcactaccg taaacaggta ccccgccgtg agcaccgcgc tgatcaaaag 10320
tactgccggg ccgatccaac tatagttcaa tcccgtggcc agcgctccct gggccagatt 10380
ccatttgctg ataaaacccc cggtaggcgg aatgcccacc agtcccagag aagccacagt 10440
aaagaaagca taggtccacg gcatgcggcg gcccaacccg tccatatcat ccacctgggt 10500
cagttccgtt ttatgaatca cagagccggc agccatgaag agcgtgtttt tcatcagcga 10560
atggaagatc acatgaagca ctgccccgac aaagccgtat tccgtcatgg taaacaggcc 10620
gaaaagaaca taggagacct gggaaacaga ggaatatgcc agacgctttt tcagcttttt 10680
ttccttgagc gccagcagcg agcccatcag aatcgtcagg accgaaaggg ccagcagcgt 10740
catctgcacc caggttccgc gcagcctctc tacgccggca atgcagaaaa tcacccggat 10800
gacgcccaac acgcccattt tggtgatcac accggaaagt acggcgctgg ccggggccgg 10860
ggccaccgga tgcgccgtag gcagccagcc gtgcagcggg aacatgccgg ctttcacgct 10920
gaagcctgcc agcatggcca ggaaaatgcc caaggccatt ccttccctgc cctggatctg 10980
cgcgtctgag agagggcctg cggcgaaatc tcccgttcct cccacggatg cgataaagaa 11040
aatgccggcc agtgccgccg aagcaccgat gacagaataa atcaggtatt tgatagcagc 11100
cgccacagct tccctgtcca tattgtgaag cacaaggggc acagtcagca gcgtcatggc 11160
ttcgtagaac atgtaaaacg taaccagatt gccgctcagc gccagcccgt gcagcatgcc 11220
ctgtgtgacc gtatagaaca aaaagaatcg ctgcgggttc tggtcgtgct ccaggtattc 11280
cgtagcatac agggccacac ccggccagat gacggcgatc agaagcagcc agaccctgcc 11340
tgtatcgtct acccgcaggg aaaaggcaag catatccgtc cagtgcaggc acacctggct 11400
gccggaaaac tgcgcactga cagcagccag tacggcgccc agaaccacgg aagccacgga 11460
ggccattctg agtttttttc ctgaccagtt catcatccgg aacagacagc ccagcgcaac 11520
cggacaaaac acagtcagaa tcagcagaag ctccttcatt ttcgtttcct cccgcaatta 11580
tagaaatact tgcagtcctt tttcaaccca tgtcatgatc tgcccgccgc aaacgccaag 11640
caccacgttg atccccagaa gcagcacagc ggcggcggaa aagcaccgat cccgtccggc 11700
gcccgggctg ccggcctctt tttcgctgta cagcgtcagc aggctccgca gcatataggc 11760
aacgttcagc agcgtactga cggccagcgc ggccagcacc acaggcgttc tccagccgcc 11820
agcctgtacg cctgcctggg tcaggtacac tttgctggcg aagcccccca gcagcggcac 11880
tcccaccaga gagcacgcgc caacggtaaa cgccagcgcg gcgatcggct cctgcctggc 11940
cgaaccgtac agatcagaga gcaggtcgct ccctccggcg gccgtcttca ggcggttgga 12000
ggacaggaac agcagggact tggccgctga atgagacagc atatgaaaaa cagcggcggc 12060
atagccggaa gaagagccca gggatattcc cagaaaaatg tatccgatct gggcgacaga 12120
ggaataagca accattctgt tcagatgatg ctggcggatg gcgtgaactg accccatgat 12180
gatgcctgag actgcataga gcattaggat gtgattgatc caataggcga acacttccct 12240
tccggccgcc cgcaggagca gcttgagcag caggaaaata tacatcttgg aaatcagaga 12300
agataaaatg gccccggagg cggccgttgc gctgctgtaa gcattcggca cccacgtgtg 12360
aaagggatac aaagcgcttt ttatggcaat tccgacggtc atcagcagga aagacagata 12420
aagcggcacg ggataagcgc cgcgggcggt cagccggcga acgctttccc ccagctgagg 12480

aaagagcaga tggcccgtca ggctgtaaag cacagacagc cccagcaaaa acatgccgct 12540
ccccagcaga ttcatgatca tataccgtac cgcagcgaac agggtgtttc ccctgccgtt 12600
cgcgctgatc agggcgcagg ccgcgatcgt ggtgatttca ataaacacat agcccgtgaa 12660
caggtcgttg gtaaagagca tggcgggcag ggccgcggtc agcatacaga tgacggctgc 12720
atacaggctc tgacggttgg gcggcacatc tctcttcagc atctgccgcc cgcccagcag 12780
cgacagcgac atcaccgccg ccaggaccag tcctgtcgca gattccacgg ggccaatccg 12840
cagctcattg ccgaatggtg ctccgatgct tcccatcgga tagacaaagc tgccctccct 12900
gagcgtaaac agcagaagga cagtcatgcc ggtgcattcc agagccgcta tggctgccac 12960
gaccgcccgc gcggctcttt ccctgaaagc agagcacagc gctgcgccga tcagcggact 13020
gatgatcatg aacaaaggga cattgttctg ccacgccagc atcctttatt tcccctctcc 13080
gcggatgcgt tccgctattt catccacatc gagcgtgtga tacagctgat aaagccttac 13140
gctcagcccc agcatcaccg ctgtgaccga aacgcccacg acgatgcccg tcagcaccag 13200
cccggcggga agcgggttca cataacgggc tgtgtccatg ctgccgtccg cataaatcgg 13260
ggcggtccgc ccagcgatat agccatagga tgccaggaaa agatatatgc cggtatccat 13320
catggaaaag ccgatgattt tcttgatcag attcttatcc agtaaaagct gggtgaaccc 13380
gatgccgaac agcgccatgg ctgtgaattc atccagatgt tcataaatga ctgccagcat 13440
cagaactctc ctttccggaa catggtataa agggcataca tcgtgcaggc caccacacag 13500
ctgaccgcca tgttcaacgg cagcagcagg cctcccgaca ggattctgcc ggcaatgccc 13560
accgaaagac cgctgtccag atgattggcg ccggtaaaaa aatggtaacc tttggataag 13620
gaataaaaca gcagcgcgca caccgatacc cagcgaaaaa ccctatagtt gaaaaacagc 13680
tctgtttttt gatatccgaa ggcgttctga tacagaatca gcccggcccc caggacagcg 13740
cctcccgaaa aaccgccgcc cggggaaaga tggccgttca gaacgatata aaagcccaga 13800
acaatcgaca gcggcgtgat cagcctggct gcagactgaa gaaccacatc gttatgcggc 13860
tcaaagtgcc tgtcatattc tccttcgcga aggcgctccg cttcattcgc ctccccgcga 13920
agactcagca gcagcaggac cgtgctgaca gctatgaaca gcacattggt ctcccccagg 13980
gtgtcaaagc cccgataatc cagaatgata ccggtgacca tattgacggc gcccgtgtcc 14040
tccagagtgt gttccacata aaagcgggtc acttcgctgt ccaccagcgt atcggcagcc 14100
ccaaagcgcg gcatccgctg caccgcaaag aacaggagac ctgtgagggc cagacagatc 14160
agcacagcgc tgagacggta ccaccaggtt tccgcatacg ccgtcggatg cacacgctgt 14220
tttcctttca ttccggctga ctgttttttt cttttccttt ttctcccagt cccgccaggc 14280
agggttccgg cttctgcctt cagcgcatcc tgcatgacct tatccaggct ttgtccgccg 14340
ttttccagcc actcccggaa aggaaaagac gcttcttttt tattctcctt ggccatcttt 14400
gctttttTcc tctctttcct ggatcgcgcc gattttcctg agcgcaatga acaggagaat 14460
cgaagtaatg cctgtgccga cggctgcttc ggtaacggcc agatcgggcg attccagcag 14520
gatccagatc gcagacatgg ccagcccctg tcccataaag atgatgaccg cagacagaag 14580
gttcttcgtc aggctgacgg ccagcgcaca gacgatgaca aataacagca gaatgacatt 14640
caaggcttcc attcttttgc ctccgtgcga aattgtacat catccgtttc ccggatcagt 14700
ttcgcgatca gatgggtgca gactggactg gtaatccaga aaagcgcgac cacacacagg 14760
attttcaggg tcagagcatc aacgccgaga taaaccgctc ccgccagggc aagcagaaaa 14820
agccccagcg tatcgcccat tcccgctgca tgcagacgat tcgccgaata ccggaatcga 14880
ttcaccccca caatggaagc cgcctggatc agcagtcctg ccagcagaag cgccaggaaa 14940
agaatcagac gaacgttttc aagcattcta ttccccttcc ttcccggtct ttgcccgatg 15000
aacgccgata tagatacggc atagcagcac aacagccaga aagctgagca tggcgtaaat 15060
caaagcaata tccatcaaat atccttcttt tttcagcagg gcaagaatca ggatgatccc 15120
catggtgact gtgctggtca tgttgatggc cagaatccgg tccgatgtgg acggaccgcg 15180
cgctgccctg tacaggcaca caaacagaaa cagccccagc accgccagcg acgcctgcat 15240
agccatcccg tatactctat ccaaggaaca ccacctcgat tctgcgcagt ttgttttcaa 15300
acgcattgtc ctccagctcc gatgccagag ccggtgtcag gcagtgcacc ttcagttcct 15360
cctgtccgca gtccagcgta atggtgcccg gggtcagcgt gatggcgttg gacaggagca 15420
tccgacccat cctggtgcgc aggccggttc tgactgtgca tacgacgggg tcaggctttc 15480
ctccgaatat caggggcagc atccggagat tggctttgac gatctcccgg agcagcgtaa 15540
aaagaaatgc gaggatccac ggaagagaac ggtagaaggc tgtttctttc cgccaggtcc 15600
atgcgcaggc atgaaaagcg aacatcagga ccaacagcgc tatgcccatg cccagcaaaa 15660
cagtttccgt ggtcacgcgg ccgttcagga agatccagaa cagaaacacc agcagaacca 15720

tgccgctcct cttttccggc ggacacatcc gcctgttctt tggtttccca ttgaaatccg 15780
tctgttttcc gccaacagta tacacgaaag agcatcgttt ctcaagcccc gccgtcctga 15840
aatcttcggt tctgattatt gcagatgaat ccggatgaca gccgccgcat gaccccatga 15900
atggaaaaaa tacgatgaat gcatggaaaa acagcttgta acctcttctc atctttgcta 15960
taatactgga agaaatattg ccttcattcc atcagaaggc aatcaatttc atgcatcatt 16020
tatcccgcag tgaggttacc ataatgcaaa aacacttgtt ccgggctgtt tttccccttg 16080
ccctcgcgtt ggtactggct ttctcatgca ctgttccccg cgcggccgcc gaagaagaat 16140
acttcaaatt ttcaaggctg attcccgatg tttttgacac aaaaaccacg ctgatcggct 16200
ttgcggaaag ccaggaaagc tttgacagag ccgccgacga agtgaccgcg cagctgaccg 16260
cttatgaccg tatttttgat gcctataccc cttatgaggg cctgaataat ctttattatg 16320
tgaaccggca tgccggcgag gagccggtat ctgttccgac ggacctgttt gagctgcttt 16380
cctggtgtaa agcgcgctgg gaggaaggac tgcgtgtcac caatgtagcc atgggagccg 16440
ttctttcgat ctggcatgaa tacagggaag ccggactcaa tgatccggag catgctgaat 16500
tgcctcccga ggatatgctg ctggccgcca atgcgcacgt cagcctcgat gacgttgttc 16560
tggattcgga aaaacagacc gttttcttct ccgatcccgg cctgacgttg gacatcggcg 16620
ccgtggccaa aggctttgca gccgacaagg tcaacgggtt tctctcccat gaaatgccgt 16680
cttatctgct gagccttggc ggcaatgtct atgccggcgc tccccctctg gacggacggc 16740
agaactgggc agtgggcgta caggatccgc tgacggatga gctttccctg gcgcttggca 16800
agacagaaat catggatatt atggatgtca accgcctgac cgtagttacc agcggcgatt 16860
actggcggta ttatcttgta gacggcgtga agtatcatca catcattgac ccggtcactc 16920
tgatgcccgc aaggcagatg atttccgtga ccgtcatttg ccaaagctcc acgctggctg 16980
attttctcag caccactttt ttcgcgcttt cctatgaaga tggcctgcag ctggccgacc 17040
ggctggacgg agtggacgta atgtgggtgc ttccggacgg ctctgttcat tgcacggacg 17100
gaatgatccg ctacgcccgg tccctgaagc agtgaggcca ccatgcggaa tgtcagcaaa 17160
accaaacggg cagcggcaaa tatggcggtc atcggtattt tcgtgctcat tgtactgctg 17220
cttctttttg tccttcccca ccgcctgaac cccgatctgg agcaaacgca ggaagacatc 17280
caaaggcaga tggcttctct cgcatccagt ccgaagccag gagaggaaac cccgcctgtt 17340
ccttctcaga acccggacgc tgaagcgtac ctcaccatca cgctcaacgg cagcctttgg 17400
cgtattgagc ctctcggaga agaacgggag atcagggtgg aacaggatga cggtgcggtg 17460
aatgtgatcc atcttctgcc gaacggctac tatatggctt catcaacctg cgacaatcag 17520
ttatgcgttc aggagggcac cgtaacggtg gaaaactatc agcgccgcat tctgggacct 17580
gcggtttatt gtcttccgca tcatctggcg ctggagcttg tcgtgccctt ccatacgccg 17640
gatcccgccg cgccggatat ctgacacggg taaaacaagc atctgtattc ccatgggtca 17700
gaaaactgac cttttcccgc tgtgaaggga ataaataaca tcgaatctcc ccctgcttca 17760
gggagagatg gaacaaggag gacggaacca cgtgaagaaa aagcttcccg gctgggctgt 17820
gctgctcatc atcaccctgg cagcaggcct tgccctgggc gggacctacg ccctgaccaa 17880
ggaccccatt gacaggcaga ctctcgctgc ggctgaaaac agccgcaggg ctgcgttgcc 17940
ggaagcagat tcctttgagg aggtttcgct ggcagagggg gcagatgtgg attgggccta 18000
cgccggacag aagaatggcg aaaccatcgg atacgtcgcc cagaagactg taaaaggctt 18060
cggcggcaag gtggaggtta tagcgggcgt ggatacccgg aaatccccgg attccttcac 18120
ccttggaggc attgccgtgg gcggggccag cttttccgaa acggccggtc ttggcgcgcg 18180
tgccaaggaa gctgcttttt ctgatcagtt tatcggaaaa accgctcctg tcagctatat 18240
caaggctggc ggtgaagcca ccgaaagcac agtggatgcc atcacctccg ccaccatcac 18300
aacaacggcg gttgtgaacg gtgtcaacga catcgtcaaa tatgtcaagg ccgggatcct 18360
gggcattgcc ggtatagaaa tgcccccaa accggcggac ggtgtgtaca gcgcttctga 18420
aaaaggattc cgcggtcctg tgtatgtgga agccgccttt gacaatgccg gaaaagtcac 18480
ctatatcaag gtcggtgacg acagctttgc tgaggatatc ggcatcggtg tcatcgagcc 18540
tgaattcatg attcagttca tcggtaaatc catgccgctg gatacgtccg acatcgatat 18600
gattgccggc gcaaccattt cctccacagc agcattgaaa gcgctcaata cggcttacgc 18660
ggtggcaggc ggtgcggaga tcgtggttcc agagtccccc acactgccgg aaaagccggc 18720
ggaaggcgtt tatgaagcgt ctgcgcaggg cttcggcggg cccgtccagg tggaagccgc 18780
ctttgacgcg gacggacgaa tgacctatat cgccatcggt gacgatgcct ttgccgagac 18840
ggcagggctg ggagcaaagg ccctggagcc cgaattccag gcccaattca tcggcaagca 18900
gatgccgctg gagctgaccg atatcgacgc gatcgccggg gctacaatta ccagcaccgc 18960

tgtcgtaagc gcgctgaaca gcgcgtataa cgccagccag ggcattgtgg aagaagcgcc 19020
cgctgctccg gttctgcctg aaaagccggc agaaggcgtt tatgaagcgt ccgcgcaggg 19080
ctttggcggg ccggtcaagg tggaagcagc ctttggcgcg gacggacaga ttgcctatat 19140
agccatcggc gacgatgcct ttgccgagac cgccgggctg ggagcaaagg ccctggagcc 19200
cgagttccag gcccagttca tcggtaagca gatgccgctg gaactgagtg atatcgacgc 19260
gatcgccgga gcgacaatta ccagcaccgc tgtcatcagc gcgctgaacg atgcctataa 19320
cgccagtcag ggcgttgtgg aagaagcgcc cgctgctccg gttctgcctg aaaagccggc 19380
agaaggcgtt tatgaagcgt ccgcgcaggg ctttggcggg ccggtcaagg tggaagctgc 19440
ctttgacggg gacggacaga ttgcctttat agccatcggc gatgatgcct ttgccgagac 19500
ggcagggctg ggagcaaagg ccctggagcc cgagttccag gcccagttca tcggtaagca 19560
gatgccgctg gaactgagcg atatcgacgc gatcgccgga gcgacaatta ccagcacagc 19620
tgtggtcaat gcattgaacg atgcctataa cgccagccag ggcgttgtgg aagaagcgcc 19680
ccccgttccg gtactgcctg aaaaaccagc ggaaggcgtt tatgaagcgt ccgcccaggg 19740
cttcgccgga ccggtcaagg tggaagctgc ctttgacggg gacggacaga ttgcctatat 19800
cgccatcggt gatgatgcct ttgccgagac cgccggactg ggagctaagg ccctggaacc 19860
cgagttccag gcccagttca tcggtaagca gatgccgctg gaactgagcg atatcgacgc 19920
gatcgccgga gcgacaatta ccagtaccgc tgtcgtcagc gcgctgaacg atgcctttat 19980
cgccagtcag agaactgcgg aagatatatc caatgcaata agcagcctgc ctgaaaagcc 20040
agcagaaggc gtttatgaag cgtccgcgca gggcttcggc gggccggtca aggtggaagc 20100
cgcgtttgat gcggaaggaa aaatcaccta tatcagcatc ggcgacgatg cctttgccga 20160
gacggctggg ctgggagcaa aggccctgga gcccgagttc caggcccagt tcatcggcaa 20220
acagataccg ctgactctgg acgatatcga tgcgatcacg ggcgcgacga ttaccagcac 20280
tgcagtggtt accgccctga acgaagccta caccgccagt cagggcgttg tggaagaagc 20340
gcccgctgct ccggttctgc ctgaaaagcc ggcagaaggc gtttatgaag cgtccgcgca 20400
gggctttggc gggccggtca aggtggaagc tgcgtttgat gcggaaggaa aaatcaccta 20460
tatcaccatc ggcgacgatg cctttgccga gacggcaggg ctgggagcaa aggccctgga 20520
gcccgagttc caggcccagt tcatcggtaa acagatgccg ctggagctta ccgatatcga 20580
cgcgattgcc ggagcaacaa ttaccagcac cgctgtcgtc agcgcgctga acgatgcatt 20640
tatcgccagc cagggaactg cagaagaaat atccaatgga ataagcagcc tgcctgagaa 20700
gccggcggaa ggcgtttatg aggcgtccgc gcagggcttt ggtggcccgg tgaaggtgga 20760
agccgccttt gacggggacg gacagatttc ctttattgcc atcggcgacg atgcctttgc 20820
cgagacggca gggctgggag caaaggccct ggagcccgag ttccaggcgc agttcatcgg 20880
taagcagatg ccgctggaac tgagcgatat cgacgcgatc gccggagcga caattaccag 20940
caccgctgtc atcagcgcgc tgaacgatgc ctataatgcc agccagggca ttgtggaaga 21000
agcgcccgct gctccggttc tgcctgaaaa gccggcagaa ggcgtttatg aagcgtccgc 21060
ccagggcttc gccggaccgg tcaaggtgga agccgcgttt gatgcggacg gaaagattgc 21120
ctatattgcc atcggcgatg acagctttgc cgagaccgca gggctgggag cgaaggccct 21180
ggagcccgag ttccaggcgc agttcatcgg taagcagatg ccgctggagc tgactgatat 21240
cgacgcgatt gccggagcga caattaccag caccgctgtc atcagcgcgc tgaacgatgc 21300
ctttagcgcc agtcagggaa ctgcggaaga tatatccaat gcaataagca gcctgcctga 21360
aaaaccagca gaaggcgttt atgaagcgtc cgcgcagggc ttcggcgagc cggtcaaggt 21420
ggaagccgcg tttgatgcgg aagggaaaat cacctatatc agcatcggcg atgactcctt 21480
tgccgagacg gcgggattcg gcgcaaaggc gctggaaccc gagttccagg cgcagttcat 21540
cggcaaacag atgccgctga ctctggacga tatcgatgcg atcacgggcg cgacgattac 21600
cagcactgca ttggttaccg ccctgaacga cgcctacacc gccagtcagg gaactgcgga 21660
agaaatatcc aatgaaataa gtagcctgcc tgaaaagccg gcggaaggcg tttatgaagc 21720
gtctgcgcag ggctttggcg gcccggtgaa ggtggaagct gcgtttgatg cggaaggaaa 21780
aatcacctat atcagcatcg gtgacgaatc ctttgccgag acggcgggat tcggcgcaaa 21840
ggcgttggag cccgagttcc aggcgcagtt catcggcaaa cagataccgc tgactctgga 21900
tgatatcgat gcgatcacgg gtgcgacgat taccagcacc gcagtggtta ccgctctgaa 21960
cgacgcttat cagcagacgg tcctcgacta actgatcgag ccgggcgaga cagaaaagaa 22020
ggaggcacgg acattgctta cgaaaacatt caaacggggt attcatcccc tgcaccggat 22080
gcatgagggc aaaggcctca cgcgcgatct tccggtaagg gagtatgtat ctgatactgt 22140
agtgatttcc atggccatga atatcggtgc gcctgccaag ccatgcgtca aaaaaggaga 22200

ccatgtggac atcggacagg tcatcgggga agcgcagggc ttcatgagtg taccggttca 22260
cgccagtgta tccggagaag tgattgcagt ggatgccatt ccttcccttg gcgccggaaa 22320
cgtgcaggct gtcaccatca aaaatgattt tgccgaaacc tgggttgaat tgcatcctct 22380
gggcagtgtg gaacaggtga atccgaagga tgtgattccc gccattcagg cggcaggtat 22440
taccggcatg ggcggagcct cttttcccac ccacgtcaag ctcaccttcc gtgaaggcgc 22500
attctgcgac actgttattg tcaacggcgc ggaatgcgaa acccatctga cggcagatta 22560
ccggctgatg ctggagtatg cccagcgcat tgtagatggg ctgcgggccg tcatgctggc 22620
gcttggcgtg aaaaaaggcg tgatcggcat tgaggacaac aaaccggaag ccataattgc 22680
tatggaacag gcgtcgaagg ggcgtgcagg cgtaagcgtt cagccgctga aaaccaagta 22740
tccgcagggg ggcgaaaagc agctgattga agccattacc ggccgtcagg taccctccgg 22800
tggtctgccc atcgacgctc atgtggtggt gctgaatgtc ggcacctgtg cggccattgc 22860
ggacgccatc atagacggca agccgcttat cagccgcatc tgcaccgtaa ccggctgtgt 22920
acggcagccc tccaatctgc ttttgcgcat aggcaccatt acagaagata tcatcggcgc 22980
ctgcggcggc tactctgaac cggctggaaa agtggtcttt ggcggcatga tgacgggcat 23040
ctgcattccc aatgattcca tgcccattct caaatcaacc aacggcattg tggtgctcaa 23100
tgacaaagac tccaggagca aggacgaaag cccctgcata cactgcgggc gctgcgtcgc 23160
agcctgcccg atcggcctga atccctatca gatcaaggtg gcagcggata aaagcgattt 23220
tgccgctgcc aaaaagcttc atgtgatgga ttgcattctt tgcggctcat gcgcttatga 23280
atgcccgtca cgcagatggc tgactccgtc cttcaaatac tgtaaggacc gcattaccgc 23340
agaagcgaaa gctgctgccg cgaaaggagg agcaaagaaa tgagcctgag aatctctact 23400
gccccccata tgcacaatcc gctgaccacc cagcggctga tgcagtatgt attcatcgct 23460
ctggtgccga ctgccgccgc aggtgtttac acattcggcc tgaaagccct gctggtgctg 23520
gccgtcgcca cctgcacggc tgtcctgtcc gaatacctct ggcagaaaat agccggtcag 23580
cccgttcgga tcggtgattt ttctgccgct gtcaccggtc tgatcctggg cctgaatctc 23640
ccctctacag ctccgtggtg gctggctgtg attggcagcg cctttgcggt catcatcgtg 23700
aaacagcttt ttggcggcct gggagacaat tttctgaatc cagccatggc tgcccgcgcg 23760
gtgctgcttg cttcctggcc tgtacacatg acagctgccg cttatgccgc gccgtccttc 23820
ttttccggcg ttgatgccct ggcttccgcc acaccgctgg gcgcacagag ttctttcacc 23880
atgacccagc tgtttctcgg gcagattccc ggcacgatcg gcgaagtatg caaggcagct 23940
attctgctgg ggcttctttt ccttctggtc atgcgcgtta tttcctggcg catcccggtc 24000
atcatggcag gcagtgtatt cctttgctcc atgctgatcg gcggcatgac gcccggtgcc 24060
gcgctggcag ctgttctttc cggcggcgtg cttttcgggg cggtctttat ggctacggac 24120
tatgtcacct cccccattac gccgttgggt cagatgctgt atgctttcat gatcggcctg 24180
attattgtcc tgatccggac atttggcagc tatccggaag gcgttaccta tggcatcctg 24240
ctgatgaatg tagccacgcc tctcattgac cgattcctgc cccgcgccat ctatggctat 24300
caaaaggagg gtaaaaaagc atgagcgaaa aaaagagcct gggcaaggtg ctctttaacg 24360
gcatcgttcc tgagaacccc acattccgcc tggtgcttgg tacgtgtccc accctggcgg 24420
tgacaaccag cgccctgaac ggtctgggta tgggggcagc cgctacattt gtgctggtat 24480
gctccaatat cgctatttct cttctccgaa agttcattcc ggaccaggtg cgcatcccct 24540
ccttcatcgt cgtgatctgt acctttgtca ccatggttca gctgctgatg aaagcctttt 24600
tgcccgcttt gaatgaaagc cttggcattt tcatcccgct gattgtggtc aactgcatta 24660
tactggccag ggcagaagca tttgccagca aaaatactcc gatcgcatca gctgcggacg 24720
gcctggggat gggcatcggc ttcacgctgg cccttacgct gatgggctgc atccgggaac 24780
tgatcggcag cggctcagtc tttggcgtga atgtgcttgg cgcttcctat gagcccatgc 24840
tgctgattgt gctggcatcc ggcggcttcc tgacattcgg cctgcttttg ggcgtgttta 24900
atctgattgt caagaaaatt gagcgcaaac gcgcagtgaa ggaggcgcag acggcatgag 24960
tatcgttgag atcctgctgt tcatggtcag ctgcatcctg acccataact tcatttttaa 25020
ccgttttctg ggctgctgtc ccttcctggg agtatccggc aaggttgaaa cctccgtcag 25080
catgggcctg gctgttacat ttgtcatgac cattgcttcc ctgttctgct ggctgattta 25140
taacctgctc ctgatccctt tggggctgga gtacatgaac acaatttcct ttatcctggt 25200
cattgctgcg ctcgtacagt ttgtggaaat gttcctgaaa aaaagcagcc cgaccctgta 25260
cagtgccctg ggaatctatc tgcccctgat taccaccaac tgcgccgtgc tcggcgtggc 25320
cacgctgaat atgaacgaag gctacagcct gctgcagtcc gtgctgaacg gaacgttttc 25380
cgcgctgggc tttctgctgg ccatcgtgct gatggcgggc gtgcgcgagc gcctggagag 25440

cagcaaaatt ccggcctgta tgaaaggctt tcctattacc ctggtatccg ccggattgat 25500
ggctgttgcc ttcatgggct tcagcggtct ggtgccataa ggaggacatg agaaaatgga 25560
tatgacaatc ctgtatgccg gcattctgct tggcgcgctg ggccttattt tcggcggcgt 25620
tctgtccttc gccagcaaaa aattccatgt ggaaacggat gagcgcatcg cccgggtgcg 25680
tgaatgtctg ggcggggcaa actgcggcgc ctgcggatac gccggatgtg atgcttttgc 25740
cgaagcagtc gtacggggag aagcgccgat caacggctgc gctcccgcca gcgccaaggg 25800
cgtccaggcc atcggtgaaa tcatgggaca gcaggcggac tccggcgagc gcatggtggc 25860
gcgcgtcctg tgccagggcg aaaacggcat cgccaaggag cgctatcagt atgacggata 25920
tcagagctgt gccacagcgg ccggtctggc cggcggcccc aaggactgcc gcttcgcctg 25980
cctgggattg ggagactgca tggataagtg tgctttcggc gccatttcca tgcagaacgg 26040
cattgcccat gttgatccgg ataaatgcac agcctgcggc gcatgcgtga aaacatgccc 26100
ccgcagcgtc ctgaagcttc ttcctgtttc ttccaatgta atcgttcgct gccgtaacag 26160
cgatattgcc cgtgttgccc gttccgtctg catgaatgcc tgcatcggct gcgggcgctg 26220
caagaaagag tgccagtacg aagccattac ggttgagaac gggtttgccc atatcgatcc 26280
gtccaaatgc attcgctgcg gcgcctgcgc tactgtatgc ccctgcaaat gcattaccat 26340
taactgacct ttatcgtttg gaacaaattg aaagcggatg gcacgaaagt ccatccgctt 26400
tcagtttgtc cctgtttctc ctgtccttgc agaaaccaat ccggatatat ctgcttattc 26460
cacggaatcc gggtcagcta cagcccagaa tgcgggcttt gggttgaggg atgcgtcaaa 26520
aagaagcgga tagctggtgg agcgccagct catgttatcc gtcagccccc aaacctggac 26580
agcctcaaat tgatcgctgt atcggcgcag cagcttcatg atccaggaat acttctgcgc 26640
ctgttttttc aggctggttt ctgtcctgtt atccaccccc acatccagct cgctcacccg 26700
caggcgcagc cccagcgcgg caatatcctt cacagaatcc gcaatctgct gcatggaggg 26760
actgcctgcg ctgtgatgca tctggaagcc atacccatcg atattgccct ccggaatcag 26820
ggattccagc aggcggacaa tgcccttccg cttaccggcc atggctgtat tataatcgtt 26880
atagtacagc aaggttccct ccggcgcata tttccgggcg tattcgaagg cgcgggcggc 26940
aaagtcctct ccgaccacct tataccagga ggagtttttt cgcagccagc ctgtgccgtc 27000
atcgatcgct tcattcacca catcccagga aaccacaacg ccgggataat tggcatccag 27060
atactccatg acacccctga tatagttttc aaggcggccc agcataacct ccctgctgac 27120
cagcggctgc ttcgtatcat aatgctcatg gaagaaagct tccggggtct ggctgtgcca 27180
taccagcaca tgaccatgca ctttgaggcc gttcctctgg gcaaagttca gcagggcttt 27240
ggcagcgtca aaacgaacgg ctgcggccgt ttcatctcct gtctccgcca gtttttttgct 27300
gctgtacaca tccagcactg cgtcgggctt caattcattt tccggagtca gtatgctgaa 27360
ctgttctttc atcagccgca tcagccggat atcccggaaa gcctgctgag gcgcggcggc 27420
gccgaaatcg aacaaatcgg catagatttc tttcagggac ggcatttccc ccgctgcttc 27480
aatcaccatc ggtgcctcgg tgggcttggg ttcaggcaat ccttccggcg ccgtcatgac 27540
aaaatccttc atggcgaagg aaagggacgg cgaaccggtg gtttccacgt acagaacgaa 27600
ccgatcgaaa tcgcctgcgg tgtaactgcc agtcagcgtg gtccattctc cctttttttgc 27660
ggtgccgtgc gccagatttt catagctttc cgccccgtct ttcgaatgag caacagaaat 27720
catgaagctg gcgctgtcca gttcattctg gagcacctgc gcgctgatac aatagctggc 27780
ccctgcaatc agatcaaagt catgaccggg agaattccaa tcgctctgcc gcccctcggt 27840
atacagaata ctgtccagcg catagcagac ggcgcctctg ccgtaccagc cgtcttcgcc 27900
cccggaataa tccgaagcgt aaacagcttc tgtttctgcc ctgcccggaa gacagcagaa 27960
aaccagtgcc aacagcagga cagccgacag gattttttttc atattgttca tcctttcgcc 28020
tttttgcttt attatacctg aattcatttt ctttttcaat tgtttcttcc ctgctgacaa 28080
aaccattgca ccaagaggaa aaggccgctg tgctttcgca cagcggcccg gaaagcggat 28140
tcataatccg cagatctgct tatccggcag attattccgc aggaacctct tccgcagcag 28200
cttcagcggc agcggcttcg gcagcagcgt gctcagcatc aatctgctcc tgggtcttat 28260
cgccgttgaa cacatccagc atatcctgcc agacgggggt agcagattcg accgcttcgg 28320
cgggagtcat gctgccaatg gggccccaca gcagggcaga accaagcacg tcgttggtgg 28380
agcccagcag ggaggtcttg gtggcgacac tgtgttcatc ctcgcgcatc atggcatagg 28440
tttcatccac ggcgcgctcg tcagtgaaca gatacttgtt gccgatccag gcatcctcgg 28500
tgtcgacacc gggagtatcg ttggtgtaga ggtcatagat ctgctgaatc ttcaggctgg 28560
tttcgtcatc gtacacgttg ggaatgccgt acacgttgtc gcaggccatg gccagataag 28620
tatcggcatt gggtcccttg ggatacatga cagcgcccca ctcgtcttcc atgtcagaca 28680

tttcagcgtt gtcgttgaag ccctgccagg tctggccggc atagaaagca gtggtgccct 28740
gctcccagaa atccttgaac cagtcccagt tggagccttc aggctgagga gcggcatagt 28800
tttcccacat ggtcttgcgc atctgaagag catccagagc agcctggctg ttggcggtga 28860
tggccagctt gccttcttcg ttgtagtcaa agaaggaagc gttgttggag aagatgaggc 28920
cggtggccag atcgtcgccg gaaccggtca tgccgtagat atcgaagatg ccgtcgttgt 28980
cggtgtcacg ctgcacctgc tccatgatgc tcagcatcat atcccaggtc caggtgccgt 29040
cagcctgcgc atcgtagatg gattcgggat cgatgcccgc ttcggacagt acgcgcttgt 29100
tgaagaaaat gcaggaacgg ggctcagtct tgccgacatg cacgccgtag acctggccat 29160
tcttggtgca gaactcggag gtagctttgt tccacttttc gccgctcagg tcaatctgcc 29220
agggcatata cagatcgttg gccagagggg tgccggcaaa gtcagcagcc acggccacca 29280
ggcagagctt ggagctgtcg ccgttcatga ccatgttggc cagctcggtg gggttgcctg 29340
cccaatcgct cagagcggat tcaacaatct tgcagttgta ggtctcttcc agccagtcac 29400
gataagcgta aagcagcgcg gtttcttcat cgggttcaga ggtacgggcg ctgtggttct 29460
catcatcact gctccaccag tcgtaaatcc agatggtctg cccgccgaaa tccaggcctt 29520
caatgatttc gggatagcct tcgggaacct cggccagagc gctgaagctg cagcaagcca 29580
gcagcatcat ggcaaccaga atcagggata cgatttttctt catactggga aaacctcctt 29640
ttttatttcc atacgaacat cccgcggacg tccgtacagt gacggaatga aagatagccg 29700
gcttgtctgt tcgcggaata ttataaagca gattccacaa aaaatcaagc cattattctg 29760
ggaaaaacca tcatcattct tcattctttg gagccggaca tggccaggct ttccacaaag 29820
gtacgctgcg tgaagcagta caggataatc agcggaatgc agcagaacag cacggcaatg 29880
gaaatcagct gcagctggcg gccgttgggg acgatgacgg actgtgtggc atccttggag 29940
aaataacggc ccatcctgct caccatggtg ctcatgcgga tggcgaaggt gggatagccg 30000
gtcaggaagt tgcgggcgta gaacaggtcc gtccactgcc agacaaagct gaacaggaag 30060
cagctggcaa tggtgggaac agcgtcaggc agcatgattt tgaaaaaggt atgcagagtg 30120
ctgcagccat ccacatacgc cgcttcctcc aggctttcgg gcacagcttt gaaatactgg 30180
cgcagcatat agataaacag gccgtttttc aaccccatgc aggtcaggct catcagcaga 30240
taaggcgtca gggaggagcg caggttcatg gtctttcctg tgagcccctg gatgatcccg 30300
aagagatcga aattggcaaa cgtcatgtac agggaggtct gaatggtctg aggcgggatg 30360
atgatcagca gcacgacgca gccgaaccag aactttttaa acgggaactc aaaccgcgcg 30420
aaaccatagg acaccagcgt agcggcgacc acctggcaga tggataccag gatggaaatc 30480
cagagcgtat tgatcatggt gccgggaaag tccgtcagat ccgccacgat tttatagttt 30540
tccagcgtca cattccgcgg tagcagaatg atggtggaat cgtacaggtc cttttccgcc 30600
ataaagctca tggaaaaacg ggtgagcatc ggctggatga tcataaagca aaggccgaac 30660
atcagcaggg cgcgggcaac ggaaatgcag ccccgcttga tttgccgctt catcaggtaa 30720
ccgccgctgc ggcggttgcg ctccttaaag cttttgtttt ttccaatgaa aaccgtattg 30780
tccttactca taggatttca ccgccccttgt gatgataaag gttccgatgc cgatgaaccc 30840
gagggcaacc ccgaagtaga tccagctcat ggcggaagcg acgccgaagt tcagctgcac 30900
accgtacata atggtgttga tgcgctctat gacccggtta tcgttcttca tgaagaagtc 30960
gatgatggta tagatgcagt tgaccagaag aagcgggctg accatgggga acgtgatctt 31020
ccagaaggat tcccaggcag aacagccttc cacgtccgcc gcttcataca gagaaggaga 31080
aatggactgg aggccgctga gaaatacaat gatctgaatg ccgctggcca tgacgatgtc 31140
ataaatcgca tcgatcatct ggaaaatcac gtcgaatatc tcgccggcga caccggatac 31200
cgaaagcaga ttctgcagcg ccaaagaaat gttgatgccc gacgaattct gaacgctgtt 31260
ggaaatatcg ctcatcatgt tgtagaattc gttggagctt tcaatgccgg gcagcacgcc 31320
ggaggacaga atcacgggca ggaagaaaat gacgcggcac agcacccgtc ccttgaactt 31380
ctgattcaga atgaccgcga tcacaaagct catgaccagg gtggcaatgg tgttaatggc 31440
catacggccc atttcatcca ccaggtactg gttgaagttt ggatctgtct gcagggcgta 31500
gtgatagttg ttgaagttat tccatgtcat ttcaaagccg ccgccagcct tgagctttac 31560
ttcattcagg ctcatctgca tggagaacac caggggcttg accatgaaca gcagaaagcc 31620
gatgatgaag ggcagaatga agaagtagcc ggtcatcgcc cgtctctcgt ggatggtggt 31680
gatcctgtaa ttggaattct gggcagcagg ctcgcccgcc gtgtcgcggt tatactggta 31740
atcggaccgg tcaaaggcca gtatccccag cccgacgatg ggcaggacac caagggccac 31800
agcgccgaac accgtggatt ttccaaagcg atggcccatg tgtatgccca tcatgacgct 31860
gtaagccact gcaaacgccg cacaaagcca gaagatgatc tcaccaacca gccgaacgga 31920

agggccgaag gaagtaagaa tatccagaat atacgcgccg gccagcataa cgcataaagc 31980
gtggaaaaac cggacgccat tccagcctgt cttccacagg atatacaggt tgtacagggg 32040
gatcagcgct ttccagcctt tttctccggc tttttcgaat accttccaga ggaagaaaac 32100
gaaggccgct gtcacagcca ggcaggcagc aaagccgagc agaacgatgc ccgtgggctg 32160
ctgaagaaaa cgatacacaa agccgcccat tattttaccc ccctttcaac catatagtca 32220
cgggcaggca ccgtatttcc accggtttcg tagtccgtga cagaatagtt gacgaagacc 32280
ttcgtaccat ccgcataggt ggtcgccacc acctcatccg tcaggtattc gtggttgaca 32340
atcgtctggc ggtttaggcc ttccatttca cgctgatacc gaaggatcgc gggaataacc 32400
tgttctttcc agcgctcata gccggaagcg gtatagcagc tgtaggcgga gtcctgtatg 32460
accgtggtat cctccgccat aaaggagaac tccaggccgg ccccgtattc ggcgcactcc 32520
agcagagcgg tcacgtaatc gcccgccagg ttgatggcac cgccggtata gttcttcata 32580
ccatgaaggg caatctgata gaaggggatg cgcctgtcga tgatggcata cgcgttgccc 32640
gtcagattca tatcggtgat caggtccgca tagggaaccg cgtagtcatt gccttccttg 32700
atggatatct tcagttcttt cgccacggca tccttcaggg tctgcacgtc catgtccatc 32760
acctgctggc gggtgacggt gctgcggttg tagtaatcgg cgctgagcag attgccgatg 32820
tcccggaagg cgacgccaac cgcagagcgg tttttcaggc cgtcaagcag ccggcctgcg 32880
ttcgtgtggg cgtactcggg acgcaccaga taataggtat ccatccagtc agcctgggta 32940
taggtgacga tatcgtaggc ataaagcttc acctgttccc gggtggtaaa gcgcgcggca 33000
tgggagaagg gcaggaagcc gtcaaaaacg ccgctgctgt aggcaaagca gttgatcccg 33060
tcaaaataca ggtccacatc cctggctttt gcgtccgaaa tcagcttttt catggcgttt 33120
tctccgccaa gctgcctgag cacatgaacg ccggtgagca ccttctgccg cacaccgccg 33180
ttggcccagc cggtcatgcg aaggttcaga tcccttactc cgctgctgac cagctcgttc 33240
ataatatccc ccgcctgggc gaaagtggtg gtgggaatca cggaatcgac gggaatgcca 33300
aagcgcggaa cggttttatt gatcgcgccg atcaattcca cattgacggg catatcgatg 33360
ctggcggtat catttttcat ttcgggggtt ctgcgcagat actcgccgta aagattggcc 33420
atgtcaacgt aactgtcccc ctccaggaaa cagtaccgct gaatcagcac atcatccgga 33480
atggcgcttt catacatata cagcagctga gcggtgcggt tggacacgtt gaagcggtcg 33540
taatgcagca cattgtactt gccatatacg aaattgtagc tgttgaagcg tccggcaata 33600
tcagcgttaa tgccgcagta ggaactggcg ccttccacga tgcagatgaa ggagccgtcg 33660
ggctggccca tgccaaacac cgggaaagcg ttcctggttt cgctgacagc ttctttccgc 33720
tccgtggcat aatcccagcc gtaaaggttg gcgtaatagg cgttctgcga cagcttgcca 33780
ttattgtaat gaatcagcgc ccccccctcct tcgggaatca gcatgaaccc ttcctgcgct 33840
gttccggcgg cgccgaacat cggaagtacg ctgatatagg tcagcgggct ggagctttcg 33900
cagcaaaggc tctcgtaggg cacctcaacc agcagatcac cgccgtccag ccgatagatc 33960
acgctggcat tgaagacagg cccgttattg ctgcgggttc ccgccaccag ctgctgatca 34020
atctcataat cctcctgcgt atactccgct tccgcgaaat agccttcaat tttggcctta 34080
ttggcggtcg tggtatcgga tttcaggata tacagcgcct gttccaccac ggaaggatac 34140
aaagcgatga tctcgtcctt gttttctttt ttatccagct tatccggttc atagagagaa 34200
tagctggagc ttaccttttt cttggtggcc ttggacattc tgccggtaat ctcgttgtat 34260
ctctccgccg tgatggctgt gggcatcaga tacgtctttt caattttgcc gatcgcgtaa 34320
tccacccgga tggaaccgtc ctcctgctgc agaatgtcat aggtctggtt catcatggaa 34380
tacgtgtaat tattcagttc cacttcgccg cctgtggcgg tataggggac gttcagggtg 34440
gatgaaaggg tttccttgtt gacaccgcgg gccacgggat cgctttcccg atccgtcggg 34500
ttggaatacc aggttttgcc tgttttttttg tccgtgattt tgaattgggt ggtaccggca 34560
tccatttcaa aatacagcgc gtcgttttcc atggtcagcg gcttgccgtc accggcataa 34620
ttcagcacgg aaggcggcct gccgatggta tcctctttct gggaatagat ggggacaaac 34680
acgaagacca ccagggcggc cagggcggcc agaacaagaa tcagcggaat cagccagcgg 34740
aggacggact ttttctgttt cattttcatc attcctcctt aatacagcct gtagatgatt 34800
tcccggtaca gactgatgaa atagcttgcc gcatcgctca gaaggctgaa gaacaccagc 34860
accaggaaca gtatgataca ggcgcccacc accgtcacaa tcaggaaaat gaaggttttt 34920
ccagggccgt aatcgtgtac ctccatcagc cccacaaaaa ccaggaaggc gcaccagagc 34980
aggctgatcc ccaggaaaac ggaatagaag ccgccctctt cataggtgat gatattgctg 35040
accagcacca tgatggactg gatcaggata tagggcacca gcgcatagca catggcaata 35100
tagatatcct tgaacctgcc cttgccgtca aacagcgtgg tcatggccca gtttgccagg 35160

```
        cacagcacca ggaacggcag cgccagagac gccgcttctt caaaaatatt gatatactgg     35220
        gtgggggcgg aaatgaactg gaagctcgtg aacatcagct tgagaacacg aaccatcagg     35280
        aaaagaaaca ggaaagtgtt ggctacggcc aggtttccct ttccttcatg ggccaggtcc     35340
        caaaagccgt caaacggatg ggtgattaca tgaaacgcgt agcggaagga ttccttatac     35400
        cggttccacc atgcttttct gttttcagcc ctctcggccg cttcagccgc ccttgcggac     35460
        tttattgtgc tcatgtgcga tcacctccca cttcattttt ttcaccctgc cgagcgccag     35520
        cggaatgacc agcaccgccg cgatgatcag cgccagccac cagatgttct gttccaccca     35580
        ttccttccgg aagagcttaa aagcccggcc gtagttttcc cggtcatgcg ctttttcgaa     35640
        ataggccatg gcccctttgt aatcatcctg acggagcaaa gcccgtccga tgccgcggaa     35700
        agcaagggga taattggcat tgagcttcat gacctgacgc cacagatccg cgctggcgtc     35760
        atattcgccg cgcagatagg tttcaatcgc gtcgtagatc gttttgccgt attccgtggg     35820
        agtaaacacg gtcacgctgt tttccagctg atccagcacg aagagatcat accccatgtg     35880
        ctctatgctg atggcgctgg tgaatgcccc gctgatgttg cctttggtgc cgaaggccca     35940
        cagcatcacg ccctggctgt catagccgaa aatgcgtccg cggaccctgt caacggcaat     36000
        gtagatgtca ttgtccagca cggtgatatc cgtcatcttg ctggggcctg atgccgcgcc     36060
        gccttcgatc cacttcaggt cgccgatggg aggatatcgg tcgtttttta tgaggatatc     36120
        gctgcccacc ccgttcaggc ggcggatggg tttggcattg tcccataaaa ggtcatattc     36180
        gctgaagaca gtattggtgg catagatgaa gccgtactgg tcgatataca tgttttcgta     36240
        ttccgtggga acgaagcttt cagaagcggc ccgctgctcc ttggtctgga aatagcgctt     36300
        ccagatgtag tccgccatat tgacggaaac agtgttcgca ccgatgtagc cggtaaacgt     36360
        tccgtcattt tcgtatttca caaagccctt gttgacgttg ttcaccaggg catatacgcg     36420
        tccggccaca tccacggcga ttttcttcgg ctgaaaatcc agcttctgat caaacgtggg     36480
        atcgtcaggc ttgacaaata ccttgatata attcaggtcc ttgtccatca tgacgacgcg     36540
        atagttgcca aagtcggcca catatatgct gtcatcctgg tccacaaaaa catcgtaagg     36600
        cttattgagg ctggtcacct cgcacccgtt gatgcggtca atcacccggg tgagagtgta     36660
        ctcagcgccg tcccagtgca cctgaaggat gcggttgttg ttggtatcca cgatatacag     36720
        atcgctgttg cggacgaaca ggctctgggg acggctgatc ctcacgccgc ccaggccgtc     36780
        aaggcccagg gata                                                       36794
//
ID   XXX; XXX; linear; XXX; XXX; XXX;  XXX.
XX
AC   PRJEB67889
XX
DE   Uncultured bacterium DNA fragment from in vivo enriched cow rumen, clone
rum-41l9
XX
FH   Key             Location/Qualifiers
FH
FT   source          1..44123
FT                   /organism="uncultured bacterium"
FT                   /environmental_sample
FT                   /mol_type="genomic DNA"
FT                   /isolation_source="Cow rumen"
FT                   /clone="rum-41l9"
FT                   /note="Tween 20, pNPA, pNPB, pNPP active clone"
FT   CDS             complement(1589..3988)
FT                   /db_xref="SEED:fig|6666666.127274.peg.1"
FT                   /db_xref="GO:0008863"
FT                   /translation="MKTFGYIRPGSYEEASGIIRQGEGRIDPIAAGTDVLGSYKDRILP
FT                   TYPEAVVSLKSIPHTSYIREEDGQLAIGAGTTLKTLSESPLVKEKALALSEAAHSVASP
FT                   IIRSTATIGGNVCQDVRCWYYRYPDNVGGCLTCRRKGGDTCYAIAGENRYHSIFGGMST
FT                   GCGSACQQACPAGTDIPAYMAKVRAGDFDGAAEIFFRYNPMPMMTSRVCPHVCMDSCNQ
FT                   RVYGESVNIPAVERALGDYILSHKEKFYAAPKTSTGKKVAIIGAGPGGLTAAFYLRQAG
FT                   NEVTIYDAHEKPGGVLMYGIPHYRLPKSIVEEFCLALRDIMGIRFVMNTRVGEDLTVEA
```

```
FT                   IKAENDSVYFGTGAWRQPVLGLDGENLTSFGLNFLEEVNTYLEKAIGEEVLVCGGGNVA
FT                   MDVALTSKRLGAKTVKLVCLEQRDEMPADESEIRMAEEEGVQVMNGWGLGRVLQENGKV
FT                   CGLEAKRCLSVRDENGRFAPTYDEEDTMVVRSDFIILATGQGVDVSFLPESLSAQLKTP
FT                   RGLIGADIESGRTREAGYYAGGDAVTGPNLAIRAIRAGRSAARSMIRDLGEALPEKGEQ
FT                   ERFTHYDVKGVEETHAHRQKELPVEERTLTGEDASSLTAQEAMEEAKRCLHCACYAVNA
FT                   SDLTPVLVMLGATLVTTEREIGAKELFTSKLTVQQVLHLGELVKEIRIPVPTGVTHYDK
FT                   RRVRDAIDFAIVSLAERFVVEEGTITEASLVMGGVAPVPYELSGVEAFLKGKKITKELA
FT                   KEAGELAMAEATPMRENAYKVFMAKDVLYEAICRAGGFEKEEIPVL"
FT                   /product="Formate dehydrogenase-O, major subunit (EC
FT                   1.2.1.2)"
FT                   /EC_number="1.2.1.2"
FT   CDS             complement(4006..6312)
FT                   /db_xref="SEED:fig|6666666.127274.peg.2"
FT                   /translation="MFEEMKEKVKQSEHEYRYIGKHANRAEADELVTGKRQFLDDLFMP
FT                   GMLTVRVLRSPYPNCLIKSIDTEKARQVPGVVGVCTYKEAPDWKCGLPNHRRVLDSHMR
FT                   FTGDSVALVAAETLEAADEALEKIEVEYEVLPFVLDCEEALKPDAPQLYEEYPGNEIPK
FT                   TTPFSEEPFNQIFVGNVEEAFKKSKYVQEGYMRYENMAHPLPPESPGCIASWTGEKAVT
FT                   VWYNGGAPHLQKFNTEASIPGVAVRVIACQSGGSYGSKQLIPHMILQAILMAKITGKPC
FT                   KFIMDRNEHMLSYELRQGSRIRGRIGMTEDGIINAVEGMWYTDSGMSSTYAQAMTAVGL
FT                   GECQAFLGKCKDWALDTKLVATNHSSQAPIRGFGGQECKSVLVPMVCTAIRAAGMDPLE
FT                   VFRNNFVKDGDRYIWRDNLWWEVHSVDYTRAFDETAKAFGWKEKWKGWETPTWVSEDGR
FT                   YAIGVGVGVHGNADIGEDPTEGEVKLHPFGSVTVEICIGETGGAQRLAVQKMVAEILKV
FT                   PLSGVQVVDPDTHSTRYDAGMIGSRGTITMGTTAVNAAKDARRKLLEMAALKLNANVED
FT                   LDTEDFLVFNKNMPQMRLPWIAITGTPEMTINGTYSYKQNFDKPNFALYLAEVCVNLET
FT                   GEAKLIRAIEGSDVGQIIDPVNIRMQAEGSFGSAGGDTGLFEEMVMDRKLGRFMNGNMI
FT                   DYKWRTFNNFPRFDTVLLESEWDTEAFHAVGFGEITPSPAPAAILMAVSNAIGVEMTEY
FT                   PCTPTKILRALGKIE"
FT                   /product="Xanthine dehydrogenase, molybdenum binding
FT                   subunit (EC 1.17.1.4)"
FT                   /EC_number="1.17.1.4"
FT   CDS             complement(6314..6829)
FT                   /db_xref="SEED:fig|6666666.127274.peg.3"
FT                   /translation="MEQKMEMVNLTVNGKAYRFLVGDFEGSIRPSETLLDTLRERLHLT
FT                   AAKRSCDHGACGCCTVIVDGEAVASCMLLTADCDGKSVVTLEGLADPLTGELSPLQRAF
FT                   ADNSAFQCGFCTPGIIMSTQALLWKNPSPTEEELTEALSGNYCRCISHYQVIRTVMAFV
FT                   KQINEEGA"
FT                   /product="Xanthine dehydrogenase iron-sulfur subunit (EC
FT                   1.17.1.4)"
FT                   /EC_number="1.17.1.4"
FT   CDS             complement(6858..7397)
FT                   /db_xref="SEED:fig|6666666.127274.peg.4"
FT                   /translation="MNFYKNHFGMIVSSVIALCVSLIMSTSAIFVDHLTFTVPLWIRNW
FT                   GTAFLVITICGMIFPLTNWSFALGKKLGLKPETLPHVLLENTVATLVFNLAATLVLAGV
FT                   NIFSNPEISAAIEGGFLPFSSPMDMYVKTVLRDLPIMFVISFIAAFFITKLAIAIAKKS
FT                   LGLDTLATEESPQRKG"
FT                   /product="hypothetical protein"
FT   CDS             7592..8473
FT                   /db_xref="SEED:fig|6666666.127274.peg.5"
FT                   /translation="MKTVIIIGENNLMAAAADLLNTKEMKLVGFGDAREEAWNVFEKDG
FT                   TLKEEITSLPVMPIEMVSSFEPDLCLVATLDPERNEAFKYLLYRSSFFGEVLFLYDLAS
FT                   EFSIRTASLRRLCRRLDMISVEGCVAEVGCGRGDTSWQLNALLPERRLYLFDTFAGFDE
FT                   RDIAKEAELGTSQAKAGDMAYKEENRLLSRMARPEQVVLKKGYFPETASVAESEHFALV
FT                   LLDACLYAPTLAGLTFFFPRMSPGGVILLPGYEDDAFRGVCRAVADFEEAYGRLLLCPL
FT                   GDPLGTVCIVHP"
```

```
FT                   /product="Methyltransferase"
FT   CDS             8542..9768
FT                   /db_xref="SEED:fig|6666666.127274.peg.6"
FT                   /translation="MKEKDIFTLEKKGGDLVLVSYQGHLPRVEIPPGVTLIGPYAFRRC
FT                   FSLEELLIPESVIRIEAEAFNGCHGLREIFFSEGLREIGHRAFWCCSALRSVTLPESLT
FT                   RVGSRAFECCSALRRVRFRSEALQVDEYAFSETPYFEEKLRRAQEIREQARGGLSFYNA
FT                   HGDASVKAGAYEQLSLPEGVTHIDNWAYAGGIFESVTLPGSLRTLGMCAFKDCKRLKRV
FT                   SLSPDTKCNYRVALGPSDGIFSGCSALEEVILRGPLRDFTWYDADRPEILHGFDRERTF
FT                   LHCTSLKRLIAWEIPLEQIPAPWRQLALNGFVSDPEREEHYLPDIARDYHQALLGFRDQ
FT                   LIRRYRLQGGYALAQYLTSQGAIREEEVDELIRLASAKGDPEAAALLFAYKRDHFRERS
FT                   FARSLFDAL"
FT                   /product="Cell surface protein"
FT   CDS             9765..11237
FT                   /db_xref="SEED:fig|6666666.127274.peg.7"
FT                   /translation="MNPNEYTAEELAREILTLARNQLLIRLRFLEPALFALDLCPDPEG
FT                   TLSTDGFFLYYGFLPVIRRYQAKEEAIPFDYLHLVLHCLFHHPFVGPAIDRVCWDTATD
FT                   IAVCAAILELDLPYVQSGREERQRELLTALKKDVPLLTAEKLYHFFRDLGTEPERLLTL
FT                   REAFVSDDHERWYTHARTESESEERLPDPEGKDGDESGQEARPEAKEETKEEGEKKPGN
FT                   QGGAAPRGREEAEEANALSPTAEDPSETWRELSGRVQTDLETLSLEYGEKAGTLLRSLK
FT                   EVTRERVDYGAFLRAFCVPREVVRTSLEEFDYIFYTYGLRLYGNVPLVEPLEYSDQKRI
FT                   RDLAIVIDTSASVAGDLVEAFLRKTFTLLQARDTFLDTFNVHLIQCDAEVQRDDKATTL
FT                   AELETLLESLELKGFGGTDFRPAFTYLEELSRQGEFTELTGVLYFTDGDGLYPKDPPPW
FT                   QTAFVFLEDSYEDRDVPVWAIKLILEKEEVLS"
FT                   /product="FIG00262634: hypothetical protein"
FT   CDS             11234..12742
FT                   /db_xref="SEED:fig|6666666.127274.peg.8"
FT                   /translation="MNIKEAKEQIKIAMSAYLARDEWGNYEIPRSGQRPVFLLGAPGIG
FT                   KTQIIEQIAAELDVPLVSYSMTHHTRQSALGLPFIASRTYQGQPMQVSEYTMSEILSSV
FT                   YDIMEATGREEGILFLDEINCVSETLTPSMLQFLQFKTFGRHALPEGWLVVTAGNPPEY
FT                   NRTVREFDIVTLDRLKKIEVEPDFEAWKEYAYRADIHPAILSYLQVKKDHFYHVETRVS
FT                   GKSFVTARGWEDLSRMIRLCEKKELPVDRKLIGQYLQDEKISRDFSAYYDLFRKYRSDY
FT                   QVEAILAGTWSEEIEERARAARFDEKIALLGLLLHSLSDRCQAIASEEDSLLLILEVLQ
FT                   KVKERGGEGEGAGAGKREGEEKEELRKGTESLQEKIAQKEKKGTLALAEKRTIQRAHAF
FT                   LSSLAREAEGFEDCRRAYKKETDSLQEKAQETRQSLGQAFAFLEKVYPQGQEMLLFVTE
FT                   LTVDEATARFLAHYGCESYDRHSRKLLFQERRKDLSERAKRFLA"
FT                   /product="FIG01077095: hypothetical protein"
FT   CDS             12827..14137
FT                   /db_xref="SEED:fig|6666666.127274.peg.9"
FT                   /db_xref="GO:0008422"
FT                   /translation="MLFPETFLWGAASAATQVEGAWDEDGKCPSIWDVAGDHIKNGDTT
FT                   QVACDHYHRYKEDVALMKSLGLRSYRFSVNWCRIQPQPGVVNEKGIGFYKDLVAELKKA
FT                   GIKPLLTLYHWDLPLWAQEEGGWTNPKIVDWFLDYCQVVVDALSEEVDCWMPFNEPLCF
FT                   LDLGYAMGMHAPFLTDESLFRKAARHMLLAHGRVVSMIRERAKTTPSIGIAMAASTLIP
FT                   EDESPEAIKAAAKESFEAPFGERGNSLYMDPICRGKASPTMKDLLSEEDLALISAPIDF
FT                   IGLNVYQPANVWQGEERYDKSAFPQTMMGWVIDPRCLYWTIRHYWDRYQLPLMVTENGM
FT                   AAEDSVSPDGAVHDPVRVQFLDGFLAQVGRALQENIPVLGYQHWSVMDNFEWCEGYGPR
FT                   FGLIYVDYKTQRRFLKDSALHYAHLIRTRGADLKTGR"
FT                   /product="Beta-glucosidase (EC 3.2.1.21)"
FT                   /EC_number="3.2.1.21"
FT   CDS             14152..15015
FT                   /db_xref="SEED:fig|6666666.127274.peg.10"
FT                   /translation="MKAKALAFLLLPLTLCATFAFASEPAPAREAQELIEELVVDIGQY
FT                   KEEAKSHVEGLLEELSLADPDLGSRWERIADLWLSLDADLTLHEGILPDGLPDSEELCL
FT                   VALGFQLEPDGTMKEELLGRLGVVLESAKKYPKAYVLCTGGGTAAENPDATEAGEMAKW
```

```
FT                   LLEHGVEEDRLIVEDQSLTTAQNAIFSYDILTTRYPQVKQLAIVSSDYHIATGHLLFEA
FT                   EAILRCDTPGEEALTVVSNAAWQAPSGELSSMFQAGALIELSGDTETAFEIYYDTYDIH
FT                   ALPLLK"
FT                   /product="Uncharacterized conserved protein"
FT   CDS             complement(15102..16196)
FT                   /db_xref="SEED:fig|6666666.127274.peg.11"
FT                   /translation="MIKIRAGITREPQAPYQIVDVDLAEPKAGEVLVKNVASGLCHTDE
FT                   FGRTIGLPMPLVLGHEGAGIVQQVGPDVHGIEVGDHVAFSYATCGYCKNCISGKPFYCL
FT                   NFNAINFGGVSANGDTKLFENGKPISMFFGQSSLASHSVVNARNVVKVDKDVDLALIAP
FT                   FGCGIQTGSGAVLNVLQPEMDETIAIFGCGAVGMSAIMAAAVAGCEKIIAVGGNPDSLK
FT                   LAMELGATHTVNRKEVEDVPATIREITGGGVNYAIDTSGNGKMILNAIRSTAYNGTIIP
FT                   LAPTGVIENFDIGSEVLMQMRTVKGIDEGESNPKFFIPKLIRLYKEGRFPVDKIITKFK
FT                   FEDLEAARQASLSGKVIKAVVTMD"
FT                   /product="benzyl alcohol dehydrogenase"
FT   CDS             complement(16427..17356)
FT                   /db_xref="SEED:fig|6666666.127274.peg.12"
FT                   /translation="MTDRRTGAVIVAAGVPEENGGEPLHPLGSISVAERIVATLQQAGV
FT                   GRIVFVTGYHAERLEHQLSRKGVVFLRNERFETTQMIESARIGLSYLQASCDWILFTPV
FT                   DIPLFTADTVRSLMKTGARLACPVYEGTPGHPILIGSGLLPVLLGDTGEGGLQGAIERC
FT                   GEAMVRVEVPDPGILLHTGRGEEVSSLLEEHNRQLTRPVVTVSLAREKPFFDEQMALLF
FT                   HQIEETGSVRLACQRLQISYSSGWNMIRNLESQTDGQLVSRSQGGADPKRSGLTPEGKE
FT                   FLERFDAYARQVRENAQLLFGWFFEDLL"
FT                   /product="CTP:molybdopterin cytidylyltransferase"
FT   CDS             complement(17353..17733)
FT                   /db_xref="SEED:fig|6666666.127274.peg.13"
FT                   /translation="MIKIGCMLEEKGRKIAARFEQTRQVLLVDAESGETLGEVNAEGWS
FT                   EEELARWVLQEDCEAILTGPMQRKPFEIVAEEGLITRYDARGLEPLEAIRRMNAYRLPI
FT                   IPDYEGGTGCHRHSHGHGPDMP"
FT                   /product="hypothetical protein"
FT   CDS             complement(17730..18143)
FT                   /db_xref="SEED:fig|6666666.127274.peg.14"
FT                   /translation="MDAKKAVKRHLMRGSCCSEALIRFGLELLQEDEDERLTSAASALC
FT                   NGMFSGGVCGGLTGGCLLLGMVDRSAAPELCQEFCEWFDVTVASRYASVYCRDILGEDP
FT                   WNKIERCQPMIMEIADKCQELLVEHGLIEEGEG"
FT                   /product="hypothetical protein"
FT   CDS             complement(18154..19230)
FT                   /db_xref="SEED:fig|6666666.127274.peg.15"
FT                   /translation="MRRFLVCSGFLGAGKTTTMMALEKECRQRGIGASIISNDLGSRGL
FT                   ADHAYLSALGCSCSQMTGDCICDQRENLVSRLRRLYEEGSQIVFSDIPGFGVGALEHVY
FT                   HRLAKDYPGEFTLAPFTVVVERASLEKIMGEESASTLPDDLAYILDAQLKEADLILLNK
FT                   CDLLSDEEREDCLAFLARTWPDTPVLCLSAATGEGASAVMEYLLTHGSQMKKPDLGSGG
FT                   VAFRKALGKMSESYLQYYARVCCEQMDADAYLCDLAREIRDGLRKPLWDIPHLKLFGRA
FT                   EDGSFSKVDLLGIGKEPVLTHPMGKPCVDLAVQLNASAACPWGEFSRIVNGAIDRVSAA
FT                   HQLSVVRFFEESFGIMGS"
FT                   /product="cobalamin synthesis protein, P47K"
FT   CDS             19589..20476
FT                   /db_xref="SEED:fig|6666666.127274.peg.16"
FT                   /translation="METKTLEYFRVVYQEQNMHRAADKLYISQQGLSRTIQTLEKELEC
FT                   KLFERDSLGMHPTTAGDRLYESVRKIQSELDGLHRDIEELTTGKKILHVPAAYGVLHLL
FT                   YPAVRQFSEAQPKIEVRWAEHIDQECEQLLSQQQVELILTVLGEDNPAFDIHPLMTRRI
FT                   MLLVFEGHPLYEKEAVSVQDLKNEKLLTVGHGYQMFATLQKACLKAGFYPYIEGEISEI
FT                   NLCHKLVHLGEGLGLTVDFIADMVDQKNVRALHFVEDCFTWRVGIVRNKKTPLSPAAKA
FT                   FWKHIINWPCLDNK"
FT                   /product="Aromatic hydrocarbon utilization transcriptional
```

```
FT                   regulator CatR (LysR family)"
FT   CDS             20591..23416
FT                   /db_xref="SEED:fig|6666666.127274.peg.17"
FT                   /translation="MGLKRKYLNINGVERIVIYDPATDTLAETLRRHGLLGVKVGCGKG
FT                   MCGACNVILNGTLVRSCNRKMDKVEEFSTIETIEGLGLASNLHPLQLAWIFHGGAQCGF
FT                   CTPGFIVSAKALLDQNPSPTREEVRDWFQANKNVCRCTGYKPLVDAVMDAAAVMRGEKS
FT                   VKDIIYDPASDPEVYHSSCPRPTAVAKVMGQFEYGDDVRHHMPEDTLHLAIVQPRKYSH
FT                   AKILSIDVSEAEAMPGVVKVVTAKDVKGTNRILEPLPHPTTTIVGNENPIFCDDIIYRY
FT                   GDTVGVVVADSTEHARAAAEKVKVELEPLTEYMSYVEAVAPGAQSIYPNSPSNIYMHQP
FT                   LIKGEDTRDVIDESYCSVEGSFYAQKEPHLSVEGDVLQSYYDDEGRVVVQCKTQALGWN
FT                   TGAIAEGLGLPMDKLRLIMNPAGAAFGWGTFAGDSAIVAAAMMAVGRPVTLTMTYEEFM
FT                   HFSGKRTSNFVNARMSCDEKGKLTGLEFDVGCDHGAYFSNALPELEALVRFLGYPYVFP
FT                   NVRGMGRMGISNHSFGTAYRGLGSPQCYTPCESLMDDLARKMGIDPFEFRYMNAARPGD
FT                   LTVNSYPYVEPTIIQTLDKMRPIYEAAKKEYSGKETEDKAYGIGLACGGFMCNLGIIDR
FT                   SEVALELNKDGTITHYNSWEDVGQGGDIGTLVHTVKALEPLGIKPEQVRMIMNDLGRTP
FT                   DSGIAAGSRSHFMNGNATIDAANKLMDAMRKPDGTYRTYDEMVAENIPTKYFGVYETIG
FT                   KGSIPLDPKTGVGSPNNCMNYGCFLAVVEVDKKSGATKVIKYVAVGDVGVIGNKLAIDG
FT                   QAYGGISHTIGFALSEEYGDLKKGGSMAGAGVPYIKDIPDDIELYYVETPREHGPHGSS
FT                   GASELYQSAGHMAVLNGIRDAVGVRLYEMPARPERVKAAMEAKARNEEIKPEKYYLGPD
FT                   MFDLIDAMQEN"
FT                   /product="Xanthine dehydrogenase iron-sulfur subunit (EC
FT                   1.17.1.4) / Xanthine dehydrogenase, molybdenum binding
FT                   subunit (EC 1.17.1.4)"
FT                   /EC_number="1.17.1.4"
FT   CDS             23423..25594
FT                   /db_xref="SEED:fig|6666666.127274.peg.18"
FT                   /db_xref="GO:0004355"
FT                   /translation="MEISEYKSFAEKCFRDTPPPCAAVCPLSLPVRDFVRLMQKGSYRS
FT                   AYRLYRDQVIFPSLIARICPSPCSGVCIRGKHPGDTSVDLKALELACVKRMEGKKPERY
FT                   ALPKKQETIAVIGGGLSGLACAYRLSSFGYRVTLYERSSLLGGSVLAELDPAWVQEELR
FT                   REFSAVDCDLRTEKELTSLEGIEADAFYVATGAGGEAFEDRPDVFRGGALLGADHLTAL
FT                   SQGLWAATAIETYLKVGRREEGPPQPTLKPDERYDRLHYDYTVPEPKDGKGEAEAFRCL
FT                   RCQCSECYQVCPLMEKEKYFPQKMCVEVILTLKPNMSKRTSVRMLMGCTDCKRCRDVCP
FT                   EHIDMGQCLEQARTDFYESGAMAPAFHDYWLQDMDFSLSEEASLLYLPDEKAPCDLLFF
FT                   PGCQLGASDPEYVQRAYEALQRESQNPALSLGCCGVPARWAGMPQKALAIAEEIRQTWE
FT                   RLGKPRLLTGCPTCLSNLKQAIPEMEVEDLYTWFLAHPKALPPLKAQGERLTVLDPCSS
FT                   AEAPQMQRAVRTLLEQAGYQVEDPEPDIGCCGFGGHIYSSDPELHHAFATRRTKGKEGI
FT                   FAAYCANCRDVFAAEGSDARHVLGLLLGMEEENRFPPELDLRRENRRRLKRCFTGEEEP
FT                   PVKDMLQIMIPAELIAKMDRELLLREQVAETITHAEETSEKILDEADSLWYAHRRFGTV
FT                   TLWTAYRREGDNVQVEKVYAHRVEIREEG"
FT                   /product="Glutamate synthase [NADPH] small chain (EC
FT                   1.4.1.13)"
FT                   /EC_number="1.4.1.13"
FT   CDS             25626..25802
FT                   /db_xref="SEED:fig|6666666.127274.peg.19"
FT                   /translation="MICCKCQVPMEEKKTTFLYLGHEVTYPVLRCPVCGQAFIPEELVD
FT                   GKIREVEALLEEK"
FT                   /product="hypothetical protein"
FT   CDS             complement(25892..26662)
FT                   /db_xref="SEED:fig|6666666.127274.peg.20"
FT                   /translation="MRFDIKGNEAGKTLMLLPGTACDYQTNFGTVLERLGEKYHLICVN
FT                   YDGFDGSDLIFPDMITVTEKIEAYIQDHYGGRIDGAIGSSLGSSFVGQLVQREKIHIDH
FT                   AIFGSPDLDQSGKCSAWIQSKLVVPLLTSFTKNEKKRAKTREKLKSFFEMSDETADRFM
FT                   GCFAKFSPQSIKNEYYSDLLTWLREDIHVEGTKVHFIYANKMGEKYLKRYKKYFRDPDI
```

```
FT                   REFDMQHEQWLFGGESYAKPVLEAIDAWMEETEE"
FT                   /product="2-Hydroxy-6-Oxo-6-Phenylhexa-2,4-Dienoate
FT                   hydrolase"
FT   CDS             complement(26684..27589)
FT                   /db_xref="SEED:fig|6666666.127274.peg.21"
FT                   /translation="MSMKYELLKRLVRAIGIKKQWVGKSTEELLEACRKKNARNPIPDL
FT                   EDPDFEISRMEVMGFPVLKLIHKQPSGRANLFLIGGGMVRAPRPGSIKKALRFAKETGL
FT                   DVFVPYYPLCTDYPLTKAYEMIHETYKLMLREYEADQISVLGTSSGGNLALGMVPYIND
FT                   GHGDVPMPGYILAISPGTCVDTEEEWQRMLELDKKDVAIPAQYMKTAVEVMRHGDESVP
FT                   SYMLWLQRGDFSNSPRVTLLYGSDETLYACAPSFEAAMKRYGVDYRLIVGEGMFHCYPV
FT                   FPVCKEAKEGWDLMVRLMKE"
FT                   /product="Esterase/lipase"
FT   CDS             complement(27734..29071)
FT                   /db_xref="SEED:fig|6666666.127274.peg.22"
FT                   /db_xref="GO:0047475"
FT                   /translation="MAEGKKESAQGKKESGLGVAVDELVAQRMGIEVRDLTQERVEAYQ
FT                   LEAFRRTLAYAREKGDFYRAHLGGIDLANIQSLQDLAQVPTTCEEDLSGEEWRFACIDA
FT                   SSVQRVVTVPTTGTRGKRKRLSFSEEDQRRAIDFINRGYLTMDCTEGERMLIYMSGNAP
FT                   GSIGDLVKQAMEPLHMDIRVVGPVTDIAKAYEELMDFAPSVVEAIPWHMSGLARYGSRF
FT                   GNPERAFIRSVNLSADVVPDAIVERLERLWGCSVHRHYGSTEMCIFGGVECARHGGYHL
FT                   RPCDILLEIPETDEQGRGEILMTTLDREAMPLIRYRTGDIGRMTGERCGCGCQIKRLEK
FT                   VYGRESSRITLGSASFFLSDLAEALYGRDEVIDFEVRLTGERLLEVTVLTLPGEGMELS
FT                   AVEERLSSLAGLRKALEEGACVLKLTQRATDSYPEGRNLKKTVVRI"
FT                   /product="Phenylacetate-coenzyme A ligase (EC 6.2.1.30)"
FT                   /EC_number="6.2.1.30"
FT   CDS             29392..32844
FT                   /db_xref="SEED:fig|6666666.127274.peg.23"
FT                   /translation="MADDLGRVIPAIGSFFTELSSRMDVRAAVLEYRDVLEDGPASTWL
FT                   HQVGGSPWSSSPVEVLSSLFSIVPSGGGDAPESLPAALTVLTNGELSFREQAQKLVFVF
FT                   TDDTYQSTNGPSLDELAGYLQQSGIRVSVFADPSTFDASWEAPGRAIESQAMAAPQEED
FT                   EDLLSFALSGFESESFASEDLLGEDMASEEEMDLFFTDDLLAAEAPEPAETAAGISYRS
FT                   LAESTGGILADLSGDVEGSLFAYASTLFSEEEGILGEGTIAGTEAEASPSSKKAIYILP
FT                   GYLASRLYDEQGQEIWLDLDALMADILQYTQEGVTPSSILKLSPDGTGTRVHVDMARDL
FT                   YGVLNNTQPLVEMLKATCSGVFDVVFFPYNWTADLNEAARSLEEDIEAKGYEDVVFVTH
FT                   STGGLLAATYLARSQENRDKVEKAILIAAPLYGTLSALLPAETGTRPDMTSTIAMLTGL
FT                   PSSVLNAWVRDVTGNSPTIYQLFPSWEYLVNTGTLYRDEKDHSLLKPVASPEQMYALLN
FT                   HSRNLNPNLTDGNERSHAAHRESVLGGDILKILSQVDTLLIGSSIGVPTPAVAIYRDTK
FT                   GFSSLDEVIYQKKGDGTILGQSAFLCDAQGVPTLPWLDFPGLGHTALVGDTRVLTEVVR
FT                   QIMAPYGGVADLPAPAEEPYPPEVLQELSSMESMVKFIFESNEEVTPVIVDPNGYTISL
FT                   TDASLVYTVLSTTPGDYRSQLYLPASGYQVRLQTQAELSALKGKISVSTVDKDGYRSAT
FT                   SVYRLTEVSPSGGISLPTFRLPTEALTAEQLSALGSSQGTLAATEPATGWQIPETLSLD
FT                   ELGVATVLPVSGADIESGKLRPETLFFESQNESLVSVDPSGKIVGLAPGTAVVTATEPL
FT                   SGYTLSCQVNVAGYPETVSLADLRMKTGETVSLTPSFAPEGALKASVTYEVEDPSLARI
FT                   EQDVLTALAPGKTQIRAIALGGAETRFTLVVDRPEVNLSKPKKPTYTVTRSGVKLTWKP
FT                   VANADLYQVVYRGLPGKKVTRLTTTSDLSFTDTTAEPGVLYEYSVRAKSYKTREASYVSG
FT                   GKSKPTKAVWLPVPQSLALQGTSLTWETVPAGDGYEILFSDQKDLTEGTLVKVKGAGKS
FT                   KRTLAKLGLTKGTWYVKIRSYKTVPGLGKVRSSWSKKLKVKRK"
FT                   /product="hypothetical protein"
FT   CDS             complement(32883..33788)
FT                   /db_xref="SEED:fig|6666666.127274.peg.24"
FT                   /translation="MNLTHLRYFYVLAKYEHYGKAAEELHITQPSLSKAIRCLEGELGT
FT                   YLFEKDGRGVRLTRQGKRYLKFVSEGLTSLDRGLEQLQREQSLSEGYLDIGLISSISYL
FT                   EFPQWVHGFRQTEGKNLYLSLHTDTSTALVQDLRQGKYDLVFCTETEDQDVAFAHVLEQ
FT                   PLACVVPQGHPFALWPSVDIHELDGLSFVAHTPDSRMSKITSRLCQEAGIQLRSVSEAR
```

```
FT                   EDRTILGLVRQGLGLAIMTLSPEVHGDGFVAIPLTGTSYHRYVSMGWLKTVARSEMAEC
FT                   FRRYILSSTASVRQETDQER"
FT                   /product="transcriptional regulator, LysR family"
FT   CDS             33963..34403
FT                   /db_xref="SEED:fig|6666666.127274.peg.25"
FT                   /translation="MKHSMISRLLVWSMAALALCGAPAQAEALADGQVSMTLEEFLEKS
FT                   EEEWFLTGKKEYRVQAMMVSKETRFHNELEGTDYTVTDDGVTVLLKGTLGERWASQLSK
FT                   VILTYTKTDGSELTEEDFAQKDTYVNYPAPKGTGLDPRSFTP"
FT                   /product="hypothetical protein"
FT   CDS             34743..36230
FT                   /db_xref="SEED:fig|6666666.127274.peg.26"
FT                   /translation="MRRLLDNGKAKVVQTIPFTIQLLYEPDTHYVQPIVLGIDPGREHI
FT                   GLTATTKKGRAVFKARFDTHNGEVTKHMSKRRMYRQASRHGERLRRKRRAKKYGTLMKC
FT                   GEVDRKLPGYENGTVKVKGILNTEARFNNRKRKPGWMTPTARHLLECHENIVMFVERII
FT                   PIREISIEVARFDFVAMSYPNARYQMYENGPMKGYDSQLEALRDIQKGKCLLCQKGPVE
FT                   QKHHMRPRSLGGSDTLGNLVGLCNSCHDKIHQKQGKALERLSKKKEGFNKKYAAASVLN
FT                   QIMGRFIDWCTDRYPGKVTLTNGKATKDYRDAHGLVKDHHVDAQCIAMCACGYGAEWDG
FT                   SNLHCYVIRKYRRHDRAKVKAQVSRAYKIGKTIVAKNRKPAMVASVVRDGKMAYTKQKE
FT                   PALSDWYEEQVQKHGEQEAMRMRSQLVVAPSYRRYNDVKRTLPGTIFYHEGKRYVKSGQ
FT                   IRPDDSHLYYQAEGVDSRNFPAKDCRIFYNKGIVVMS"
FT                   /product="HNH endonuclease"
FT   CDS             complement(36507..37688)
FT                   /db_xref="SEED:fig|6666666.127274.peg.27"
FT                   /translation="MTDLVEDEGTAYYQYQISWADSFLEANGYDLQTLLPLEGTSIAYG
FT                   VLDENVFAVDGEEQPLTMEERHAMIDERAKRLNVKKSSEEQIEVGSDRTYEMEPDPSKG
FT                   VKISLDGVGAKRQKDSRPKMSGTESDPTFVRDENDGPEDYTRAPESKKRPKVETAVGHV
FT                   EHDGKKYVFAGKNMFDTCKMILAFLISHNLLENNLLVFFTDGGKDIRYCIDNIFRIYHP
FT                   IEILDWFHLRKHCTEALSMSLKGGKANRKMQYEVKRRLFRILWSGNVQNAICYLQSLDG
FT                   NYLKPGDKINELVGYLRKNEGRIPCYALRKSMGLQVSSNRVEKANDLIVSSRQKGDSMS
FT                   WSREGSWGLANVTVKYLNHEADVYRETGVVSFKMYEGYGKVFDLRSKEMAKIG"
FT                   /product="hypothetical protein"
FT   CDS             38041..39429
FT                   /db_xref="SEED:fig|6666666.127274.peg.28"
FT                   /translation="MARKASPIILSNADVNSLSILSTLSQDPDLARRAKVILECGKGLS
FT                   NKEVSAVIGMNEADVGRWRKVFLADGIDGLKGKQRGGDHRSDAPDETLYTRLNELLGQE
FT                   GRDWTVQELVQELNSTANIVSAALRKQGINLQRQRQWYIPTQDELIPKTLDVVGLYITK
FT                   AEQAIIVRCTQEALLECHGEFITRNRELSEDYLHYDGDITLAEAINTAVHHVKDTSRMV
FT                   PVSLSEFLNRTISAFPVKDGYEYHILICSPKNSGYQGARLKDIYQSWADDSRRWLHLVD
FT                   SKINELADRNQLHPAEELQKALELYLDKCIETTAPLIWRKKPDQPATAEHQAAHGQEKE
FT                   ENPIILKNGLESSPFLDELRELLKRYLASGLESPEIVKCGFISFAYDQDEVRVQLEDDN
FT                   GTLPGASDVDFGSLEAWKDGLTAVEKALINTRNRAGVDAAAMVSDLLKKNRCCRAKRGE
FT                   GSRE"
FT                   /product="Mobile element protein"
FT   CDS             complement(39401..39532)
FT                   /db_xref="SEED:fig|6666666.127274.peg.29"
FT                   /translation="MCAYPFNQPTTFDAFIFTQMPCYADREPDGKTSELTLYFPRHA"
FT                   /product="hypothetical protein"
FT   CDS             39668..40918
FT                   /db_xref="SEED:fig|6666666.127274.peg.30"
FT                   /translation="MREHQKQVLTSFGFGEDGLWPKDKEFPDELRNLDMEYIYFDPSMI
FT                   MEQAGYFPLPVIEFKEGQRSEPETGLHIRKMKRRDERLLVPDDQREQWLKGYILYINTQ
FT                   ADRSPFATIRHPARVERHAESVVNIFIDGDLVGEQAEHRVPGGKLVLRDEKPTIKHIDI
FT                   RIEIENGGRYNITSVDLDDAYQRLLAVLLETGLITRYFHFFVDGEKTLTKAIEKYFANW
FT                   NYSIFLDFFHIEEKIKTQTSLGIKGRRVASPWEEPEYYTRGPKKGEIKKQPMTSLSRTY
```

```
FT                   ASVLKTMVFYGNAKEAKEYIEHIPEEDIQDKAALQVLVNYLEERESMLACYTLRKHAGL
FT                   KNSSNASELCNELAVSSRQKVDDRMHWREDGSAALAAITVLYMNRADKRWFEQHNVEFE
FT                   LYYVEPTKTRKIKTQAA"
FT                   /product="hypothetical protein"
FT   CDS             complement(41085..42311)
FT                   /db_xref="SEED:fig|6666666.127274.peg.31"
FT                   /translation="MPKRVSPIVINETDVIMLRRIDDGTVVCGQDIVKRARALLLMSDG
FT                   RQIKDVAAEVGMRENTITDIRRRFLEAGIDCLYDKPKSGRPITYDPDQLEASIDLRIQQ
FT                   CVAQNVPVPSAKELATELKAPQNVIRDILQKKDIVAERGPHYWRFTMSDGTVPKLADMC
FT                   GLYLSPTEQVLVVKAWTIEKNMVIDLDLITDTSAVVETRTKAIASKIEALIDVSGKVDM
FT                   ADALTVFAESGKTVRLGKKDSAISFVKGILPNGIKASNIEYHIFTYGVSLVENNRSLLP
FT                   DTYLHVFKTQEQWMSQFEIIATNMYSGIQGKEKAAKVRDSIYRYLRTAKPNTEPFEWVK
FT                   VTKNEEGDQKPMALTEPKAQDDFSRSAPGSIRFTAQFVDDEHNVITVRVHIKTLLHRLN
FT                   SKPEAAGNI"
FT                   /product="Mobile element protein"
FT   CDS             42521..42760
FT                   /db_xref="SEED:fig|6666666.127274.peg.32"
FT                   /translation="METDKDHIHILLDYKVTDCVADIAKILKMVSTNTIWKNYPRMMKR
FT                   NFWREHTLWSDGYFACSIGEASKETIEQYIQNQG"
FT                   /product="Mobile element protein"
FT   CDS             42781..43176
FT                   /db_xref="SEED:fig|6666666.127274.peg.33"
FT                   /translation="MPILFIAVQAQEKAICFFSCAWNLFSILSKAHSSQHLKVLGFLGR
FT                   WIINLLTKSQPDSYYARRVPVEISVTVETAWGDVLHTNLPSTNHGEGDYLVCRVDENDG
FT                   PDLSDVWVVHGLLFTETYDLPDEPDPS"
FT                   /product="hypothetical protein"
XX
SQ   Sequence 44123 BP; 9542 A; 13382 C; 12276 G; 8923 T; 0 other;
     tccaaccagg ccctctcccc cacgagggag gcaaaggcct cgatctgtcc gtttgacagg        60
     tccagaggat gggccagggt ccagcctccc ccatccgggt tccctgggac ctgggccttc       120
     cttcccctct catccaccca gcgcccttcc agacgggcca ccagacttcc atccgcccgg       180
     gcaaagacca gcccctccag aagggccgag agggccgggc tttggagcca ctccaaaaga       240
     gtctcctccg aaaagctcct cttctccagg aaggcccggt acaaccttct ttggacttcc       300
     ctttccagga ccttttgctc ccttcgcccc tccctttcca gaagggcgaa aagcttccag       360
     aaaagctcct ccgaatcgat gccctgcacc ctggcatagg ctgccagctc cccattcacg       420
     gccgcgaaca cggcggcctc cggatccttg cttagcagga tggccttctg taggacctgc       480
     aggcagatcc tctcctgctc cccagacgaa tcagaaagct ctgcatgcaa gccgatccag       540
     ccggagagct cctcgcctcc ggccatccga aggagaggca ccagcctccg ggggtaggaa       600
     aggaccttcc ttctcccctc cttgggatct cccagttcct gcagggctct ttgcaggctt       660
     tccttcgaaa tcaaagaggc cagctcatcc gccaccggat cgggcaggag gtctttgtcc       720
     ggccgggaga ggtctgcctg gagaagatat cgatagagga aaaagcgcag cagctcgggg       780
     ggagccattc ttccttcctt ttcttccttc tctccctctt ctatcctcct gtccccccttc       840
     ccggcatagc ggatgggcgg cagcgccccg atccccgcct gggagaggac cttttcgatg       900
     tccgccaggg aagcctcccc ccggatccac cgtaggagcc gcttatcctc cgaggaagcc       960
     tccctccggt ccgggaagcg ttccgcctcg aggacatcga gaagcttttt ttcctccgcc      1020
     gaaacccgct ggctccgggc ataggaaaag agggccgctg ccttctcctc ctccaaagcc      1080
     gaaaggattc cccgacgctg cagggcttcg atggcctcca ggatgcagcc cttccggcag      1140
     agcagagccg gatccagcag gagctcctgc cggatatacg aaaggatctc ctccctctgc      1200
     ccggagtcat agagcccttc ccactccggg tgcaaaagaa agcccaggca gagcacctcc      1260
     tgctcttcct tctccttccc ccggaaacgg acaaaggggg cacagagggc cttcagctcc      1320
     ctggcccccg gcaagatccg catcagggga gtgtggcagg gcaccctcgt cagccagacc      1380
     acctcggccc ccggggcgat tccccagaag gccgaggggc ccaccgaccg gacactctgc      1440
     ccgacgaaca ccttgcgaag gctcgagcac ccgaggaagg ccccctcccc gacgctctct      1500
     accccctcct ccagccaaac ctcctgaagg cccaagtccc cccggaaggc ataggcaggg      1560
```

atctgttttt cttgaagatg ccggctgctc acaggaccgg gatctcttcc ttctcaaagc 1620
cacccgcccg gcagatggcc tcgtagagca catcctttgc catgaagacc ttgtaggcgt 1680
tctccctcat gggcgtcgcc tccgccatgg ccagctctcc ggcttccttg gccagctcct 1740
tcgtgatctt cttgcccttt aaaaaggcct cgacaccgga gagctcatag ggcaccggtg 1800
ccactccgcc catgaccaga gaggcttccg tgatggtacc ctcctcgacc acgaagcgtt 1860
cggccaggga gacgatggca aagtcgatgg catccctcac ccgcctcttg tcatagtggg 1920
tcacgccagt cggcaccgga atccggatct cctttacgag ctctcccagg tgcaggacct 1980
gctggacggt cagcttcgag gtgaagagct ccttcgcccc gatctcccgc tccgtcgtga 2040
ccaaggtcgc ccccagcatc accagcaccg gggtgaggtc cgaagcgttc acggcatagc 2100
aggcacagtg cagacacctt ttggcctcct ccatggcctc ctgggccgtc agggaggagg 2160
catcctcccc ggtcagggtc ctctcctcga cggggagctc cttctgccgg tgcgcatggg 2220
tctcttccac tccctttacg tcatagtggg tgaaccgctc ctgctcgccc ttctccggca 2280
acgcctcgcc caggtcccgg atcatggacc gggccgccga acggccggcc cggatcgccc 2340
ggattgccag gttcggcccg gtgaccgcat cgccgccggc ataatagccc gcctcccggg 2400
tccggccgga ctcgatgtcc gccccgatca ggcctcgggg ggtctttaac tgtgccgaaa 2460
ggctttcggg caggaaggag acatccaccc cctggccggt ggccaggatg atgaaatcgg 2520
agcgcaccac catggtgtcc tcttcatcgt aggtcggagc aaagcggcca ttctcatccc 2580
tcacggagag gcatctcttg gcctcgaggc cacagacctt cccgttctcc tgcaggaccc 2640
ggcccaggcc ccagccattc atgacctgca cgccctcttc ctcggccatc cggatctcgc 2700
tctcatccgc cggcatctca tccctctgct cgaggcagac gagctttacg gtctttgccc 2760
ccagccgctt cgaggtcagg gccacatcca tggccacgtt tccgccgccg cagaccagga 2820
cctcctcgcc gatggccttc tccaggtagg tgttgacctc ctcaaggaag tttaagccaa 2880
agctggtcag gttctccccg tccaggccca ggaccggctg cctccaggca ccggtgccaa 2940
agtagacgct gtcgttctcg gccttgatgg cctccaccgt gaggtcctct ccgacccggg 3000
tgttcatgac aaagcggatt cccatgatgt cccgaagggc caggcagaac tcctccacga 3060
tggacttcgg caggcgatag tgggggatgc catacatcaa gaccccaccc ggcttttcgt 3120
gggcatcgta gatggtcacc tcattgcccg cctggcgcag atagaaggcc gccgtcaggc 3180
cgccagggcc ggcgccgatg atcgccacct tcttgccggt ggaggtcttc ggggccgcat 3240
agaacttctc tttatgggaa aggatgtagt cgccgagggc ccgctccacc gcagggatgt 3300
tcacggactc gccgtagacc ctctggttgc acgaatccat gcagacatgg gggcagaccc 3360
ggctggtcat catgggcatc gggttatacc gaaagaagat ctcggccgcc ccgtcaaagt 3420
cgccggcgcg gaccttggcc atgtaggccg ggatgtcggt ccccgccgga caggcctgct 3480
ggcaggcgga cccgcagccg gtgctcatgc caccgaagat ggaatgatac cggttctctc 3540
cggcgatggc atagcaggta tccccgccct tgcggcggca agtgaggcac ccacccacat 3600
tgtctggata gcgataatac cagcagcgca catcctgaca gacgttcccg ccgatcgtgg 3660
cggtgctgcg gatgatcggg gaggccaccg aatgggccgc ctccgaaaga gccagggcct 3720
tttccttcac caggggggctc tcgctcaggg tcttcagcgt cgtcccggcc ccgatggcca 3780
gctggccatc ctcctcgcgg atgtaggagg tgtgggggat cgacttcagg ctcaccaccg 3840
cctcggggta ggtcggaagg atccggtcct tatagcttcc cagcacatcc gtcccggctg 3900
cgatcgggtc gatgcggccc tctccctgcc ggatgatgcc ggaggcctcc tcgtagctgc 3960
ccggtcggat atatccaaat gtcttcatcg ctccttgcct ccttcctact cgatcttccc 4020
cagggcccgc agaatcttcg tgggcgtaca gggatactcg gtcatctcca cgccgatggc 4080
gttgctgacc gccatcagga tggccgccgg agccggcgac ggggtgatct cgccaaagcc 4140
caccgcgtgg aaggcctcgg tatcccactc gctctcgagc agcacggtgt caaagcgggg 4200
gaagttgttg aaggtcctcc acttatagtc gatcatgttt ccgttcatga accggcccag 4260
cttcctgtcc atgaccatct cctcgaacag gcccgtatcc ccgccggcgg agccgaagga 4320
cccctcggcc tgcatacgga tgtttacagg gtcgatgatc tggcccacat ccgaaccctc 4380
gatggctcgg atcagcttcg cctcgccggt ctcgaggttc acgcagacct cggccaggta 4440
gagggcaaag ttgggcttgt caaagttctg cttgtagctg tacgtcccgt tgatggtcat 4500
ctccggggtg ccggtgatgg cgatccaggg aaggcgcatc tgaggcatgt tcttgttgaa 4560
gaccaggaaa tcctcggtat cgaggtcctc gacgttggca ttgagcttca gggccgccat 4620
ctcgagaagc tttctcctgg catccttggc cgcgttcacg gccgtcgttc ccatggtgat 4680
ggtcccccctc gagccgatca tgccggcatc gtagcgggtg gaatgggtgt cgggatcgac 4740
cacctgcacg ccgctaagcg gcaccttcaa gatctcggcc accatcttct ggaccgccag 4800

cctctgggcc ccgccggtct cgccgatgca gatctccacg gtcaccgagc cgaagggatg 4860
gagcttgacc tcaccctcgg tgggatcctc gccgatgtcc gcattgccat ggacgcccac 4920
gcccacgccg atggcgtagc ggccgtcctc agagacccag gtcggggttt cccagccctt 4980
ccatttttcc ttccagccaa aggccttggc cgtctcatca aaggcccggg tgtaatccac 5040
cgagtggacc tcccaccaca ggttgtcgcg ccagatgtac cggtccccgt ccttgacaaa 5100
gttgttgcgg aagacctcca aaggatccat gcccgccgcc cggatggcgg tgcagaccat 5160
gggcaccagg accgacttgc actcctgccc gccaaagcca cggatcgggg cctgggagga 5220
atggttggtg gccaccagct tggtatccag cgcccagtcc ttgcacttgc caaggaaggc 5280
ctggcattcc ccgaggccga cagcggtcat ggcctgggcg taggtggagc tcatgccgga 5340
gtcggtgtac cacatgccct ccacggcatt gatgatcccg tcctcggtca tgccgatccg 5400
tccccggatc cggctgccct ggcgaagctc atagctgagc atgtgctcgt tgcggtccat 5460
gatgaacttg cagggcttcc cggtgatctt cgccatcagg atggcctgca agatcatgtg 5520
ggggatcagc tgcttggagc cataggaccc gccggactgg caggcgatga cccgcaccgc 5580
gaccccgggg atggaagcct cggtattgaa cttctgcagg tggggagctc cgccgttgta 5640
ccagacggtc acggcctttt ccccggtcca gctggcgatg cagcccgggc tttccggcgg 5700
gagcggatgg gccatgttct catagcgcat gtacccctcc tggacatact tgctcttctt 5760
gaaggcctcc tctacgttcc ccacgaagat ctggttgaag ggctcctccg agaagggagt 5820
cgtcttgggg atctcattgc ccgggtattc ctcgtacagc tggggagcgt ccggcttcag 5880
ggcctcctcg cagtccagca cgaagggaag gacctcgtac tccacctcga tcttctcgag 5940
ggcctcatcg gccgcctcca gggtctcggc ggccaccagg gcgacactgt ccccggtgaa 6000
gcgcatatgg gagtccagca ccctccggtg attcggaagg ccgcacttcc agtccggggc 6060
ttctttatac gtacagaccc ccaccacccc cggcacctgc cgggcctttt cggtatcgat 6120
gctcttgatc agacagttgg gatagggaga gcgcagcacc ctcacggtca gcatccccgg 6180
catgaaaagg tcgtcgagga actgcctctt acccgtgacc agctcgtcgg cctcggcccg 6240
gttggcatgc ttgccgatgt agcggtactc gtgctcgctc tgcttcacct tctccttcat 6300
ttcttcaaac atatcatgca ccttcctcgt tgatctgctt tacaaacgcc atgaccgtcc 6360
ggatgacctg gtaatggctg atgcaccggc agtagttgcc cgacagggct tcggtgagct 6420
cctcctcggt gggcgagggg tttttccata agagagcctg ggtgctcatg atgatccccg 6480
gcgtacaaaa gccgcactgg aaggcagagt tgtcggcaaa ggccctctgc agggggggaaa 6540
gctccccagt gagaggatcg gccagcccct ctaaggtgac gacgctcttt ccatcgcagt 6600
cggcggtcag cagcatgcag gaggccaccg cctccccgtc cacgatcacg gtgcagcagc 6660
cgcaggcccc gtggtcgcag cttctcttgg ccgcggtcag atgcagcctt tccctcagcg 6720
tatccagaag ggtctcggaa gggcggatgc tcccttcaaa atcccccacc aggaagcgat 6780
aggccttccc gttcacggtg aggttcacca tttccatctt ctgttccatc ttcacttgct 6840
cctttctgct tccgtcttca gcccttcctc tgcggggact cctcggtggc cagggtatcc 6900
aggcccaggc tcttcttcgc gatggcgatg gccagcttgg tgatgaagaa ggcagcgatg 6960
aagctgatga cgaacatgat cggcaggtcc cgaaggacgg tcttcacata catgtccatg 7020
ggcgaggaaa agggaaggaa gcctccctcg atggccgcag agatctccgg gttggagaaa 7080
atgttgacgc cggccagcac cagagtggcg gccagattaa agaccagggt ggccacggtg 7140
ttctcgagaa gaacatgggg cagggtctcg ggcttcaggc ccagcttttt gcccagggcg 7200
aaggaccagt tggtcagggg aaagatcatg ccacagatgg tgatgaccag gaaggcggtg 7260
ccccagttcc tgatccagag cgggaccgtg aaggtcagat ggtcgacgaa gatcgccgag 7320
gtggacatga tcagcgagac gcaaagggcg atgaccgagg aaacgatcat gccaaagtgg 7380
ttcttgtaaa agttcatgga agcctccttt ggtgtggatg aaatgaaaag caacatatgg 7440
atatgtacag atgttttcat catctccctt tccctgcgga tttcccagta gcccaggcta 7500
cagctgatgc acgggataca ggtatgtaag ggcgtccggg tgtagactaa agggaaagaa 7560
aacaggatag aaagaaagga cggaccgatc catgaagact gtgatcatca tcggagaaaa 7620
caacctgatg gccgccgcgg cggacctgct gaacacgaag gaaatgaagc ttgtgggctt 7680
tggagatgcc cgggaggagg cctggaacgt ctttgagaag gacggcaccc tcaaagaaga 7740
gatcacctcc ctgcccgtca tgcccatcga gatggtctcc tccttcgagc cggacctctg 7800
cctggtggcc accctggacc cggagaggaa cgaggccttc aagtacctgc tctaccgctc 7860
cagctttttt ggcgaggtgc tcttcctcta cgacctggcc tcggaattct ccatccggac 7920
cgcttccctt cgccgcctct gccgccggct ggatatgatc tcagtggagg gctgcgtggc 7980
cgaggtgggc tgcgggcggg gggatacctc ctggcagctg aacgccctgc tgccagagag 8040

gaggctttac ctcttcgaca cctttgccgg ctttgatgaa agggacatcg ccaaagaagc 8100
ggagcttggc acctcccagg ccaaggccgg ggacatggcc tacaaggaag aaaaccgcct 8160
cctctcccgc atggcccggc ccgagcaggt ggtcctgaag aagggatatt tcccggagac 8220
cgcctcggtc gcagaatcgg agcattttgc cctggtcctc ctggacgcct gcctctatgc 8280
ccccaccctg gcgggcctca ccttttttctt cccccggatg agtccaggtg gcgtgatcct 8340
cttgccaggc tatgaggacg atgccttccg ggcgtgtgc cgggcggtgg cggactttga 8400
agaggcatac ggccgcctgc tcctttgccc cctgggcgac cccttgggca cggtctgcat 8460
cgtccatccc tgaaagaggg aagagagaga agaagagaag gaaaagagga ggggagaaag 8520
aagagggagg gaagaaagga tatgaaagaa aaagacatct tcaccctgga aaagaaggga 8580
ggagacctcg tcctggtctc ctatcagggc catcttcccc gggtcgagat ccccccgggg 8640
gtgaccctca tcggccccta cgccttccgg cgctgctttt ccctggagga gctcctgatc 8700
ccggagagcg tgatccgcat cgaggcggag gccttcaacg gctgccatgg gctaagggag 8760
atcttctttt ccgaaggact gcgggagatc gggcaccggg ccttctggtg ctgctcagcc 8820
cttcgaagcg tgaccttgcc cgagagcctg acgcgggttg gctcccgggc ctttgagtgc 8880
tgctcggccc ttcgaagggt ccggttccga agcgaagccc ttcaggtgga tgagtatgcc 8940
ttcagcgaga ccccctactt tgaagagaag ctccgccgag cccaggagat ccgagagcag 9000
gcccggggag gcctttcctt ctacaacgcg cacggggacg cctcggtgaa ggccggagca 9060
tacgagcagc tctcccttcc ggaggggggtg acccacatcg acaactgggc ctacgccggc 9120
gggatctttg agagcgtgac cctgcccggt tccctgcgca ccctgggcat gtgcgccttc 9180
aaggactgca aacgcctgaa gagggtctcc ctttccccgg ataccaaatg caattatcgg 9240
gtggcactgg gaccctccga cgggatcttc tccggctgct ccgctctgga ggaggtcatc 9300
ctgcgtggcc ccctgcggga ctttacctgg tatgatgccg accggcccga gatcctgcac 9360
ggcttcgacc gggagcggac cttcctgcat tgcacctccc tgaagcgcct gatagcctgg 9420
gagatccccc tggagcagat ccccgccccc tggcggcagc tggccctaaa cggctttgtc 9480
tcggacccgg agcgggagga gcattacctg ccggatatcg ccagggacta ccatcaggcc 9540
ctgctcggct ttcgggacca gctgatccgc cgctaccgcc ttcagggagg ctatgccctg 9600
gcccagtacc tcaccagcca gggagcgatc cgcgaggagg aggtcgacga gctgatccgc 9660
ctggcctctg ccaaaggaga cccggaggct gccgcccttc ttttcgctta caaacgagac 9720
catttccggg aacggtcctt tgcccgttcc ctttttgatg ccttatgaat ccgaacgagt 9780
ataccgcaga agaactggcc agagaaatcc tgacccttgc aaggaaccag ctgctgatcc 9840
ggctgcgttt cctcgagccg gccctctttg cactggacct ttgccctgac cccgagggga 9900
ccctctcgac ggatgggttc ttcctctact atggcttcct tccggtcatc cggcgctacc 9960
aggccaaaga agaggccatc cccttcgact acctgcacct ggtcctccac tgccttttcc 10020
accatccctt cgtaggtccg gccatcgatc gggtctgctg ggacaccgcg accgacatcg 10080
cggtctgcgc ggccatcctg gagctggacc ttccctatgt gcaaagcggc cgggaggaga 10140
ggcagaggga gctcctgacg gccctcaaaa aggatgtgcc cctccttacc gctgagaagc 10200
tctaccactt cttccgggac ctgggaaccg agccggagag gctcctcacc ctgcgggagg 10260
cctttgtcag cgacgaccac gaaagatggt acacccatgc ccgcacggaa tccgagtccg 10320
aggaaaggct tccggacccg gaagggaagg acggggacga gtccggacag gaggcaaggc 10380
ccgaggctaa agaggagacc aaagaggagg gagaaaaaaa gcccggaaac cagggagggg 10440
ctgccccccg gggaagggaa gaggccgagg aggccaacgc cctctccccc accgccgaag 10500
accccctccga aacctggagg gagctctccg gaagggtgca gacggacctg gaaactcttt 10560
ccctcgagta cggggaaaag gccggcaccc tccttcgctc cctcaaggag gtgacccggg 10620
agagggtcga ctatggcgcc ttccttcgcg ccttttgcgt cccccgggag gtggtgcgca 10680
cctccctgga ggaattcgac tacatctttt acacctacgg cctgcgcctc tatgggaacg 10740
taccccctggt ggagcccttg gaatactcgg accagaagcg gatccgggac ctggccatcg 10800
tgatcgacac ctcggcctcc gtggccgggg acctggtgga ggccttcctt cgaaagacct 10860
tcaccctcct gcaggcccgg gataccttcc tggacacctt caacgtccac ctcatccagt 10920
gcgacgccga ggtccagaga gatgacaagg ccaccaccct ggccgagctc gaaaccctcc 10980
tcgaaagcct ggagctcaag ggctttggcg gcacggactt tcgcccggcc ttcacctacc 11040
tcgaggagct ctcccgccag gggggaattca ccgagctgac cggcgtcctc tatttcaccg 11100
atggggacgg cctctatccc aaggaccccc cgccctggca gacggccttc gtcttcctgg 11160
aggactccta tgaagaccgg gatgtcccgg tctgggccat caagctgata ctggaaaagg 11220
aagaggtact ctcatgaaca tcaaagaagc gaaggaacag atcaagatcg cgatgagcgc 11280

ctacctggcc cgggacgaat ggggcaacta tgagatcccc cgctccgggc aaaggccggt 11340
gttcctcctg ggcgcccccg ggatcggcaa gacccagatc atcgagcaga tcgcagcgga 11400
gctggatgtg cccctggtgt cctattcaat gacccaccac acccgccaga gcgccctggg 11460
ccttcctttc atcgccagcc ggacctacca gggacagccg atgcaggtct cggaatatac 11520
catgagcgag attctctcca gtgtctacga catcatggaa gccaccggcc gggaggaggg 11580
gatcttgttc ctggatgaga tcaactgcgt ctccgagacc ctgacccccct ccatgctcca 11640
gttcctgcag ttcaagacct ttggccgcca cgcccttccg gagggctggc tggtggtcac 11700
cgccggcaac ccgccggaat acaaccgcac ggtgcgggag ttcgacatcg tgaccctcga 11760
ccggctcaag aagatcgagg tggagccgga ctttgaggcc tggaaggaat atgcctaccg 11820
ggcggatatc catccggcca tcctctccta cctccaggtc aaaaaggacc acttctacca 11880
cgtggagacc agggtctccg gcaagagctt cgtcacggcc cggggctggg aggacctctc 11940
ccgaatgatc cgcctctgcg aaaaaaaaga gctcccggtg gaccggaagc tgatcggcca 12000
gtatctgcag gacgaaaaga tcagccggga cttttccgcc tactacgacc tcttccgcaa 12060
gtaccggtcg gattaccagg tggaagccat cctggccggg acctggtctg aggagatcga 12120
agagcgggcc agggcggcca ggtttgatga gaagatcgcc cttctgggcc ttcttcttca 12180
cagcctctcc gaccgctgcc aggccatcgc ctccgaggag gactccctcc ttttgatcct 12240
ggaggtgctc caaaaagtga aagaaagagg aggagaggga gagggagcag gagcaggaaa 12300
aagggaagga gaagaaaagg aagagctccg gaaagggacc gagagcctgc aggagaagat 12360
cgcccagaag gaaaagaagg gcaccctggc cctcgccgaa aagaggacca tccagcgcgc 12420
ccacgccttc ctctcctccc tcgcccggga ggccgagggc ttcgaggact gccggagggc 12480
ctataaaaaa gagaccgatt ccttacagga gaaagcccag gagaccaggc aaagcctcgg 12540
gcaggccttt gccttcctcg aaaaggtcta tccccagggg caggagatgc tccttttcgt 12600
cacggagctg acggtggacg aggccaccgc ccgcttcctt gcccactatg gctgcgagtc 12660
ctatgaccgg cacagccgga agctcctctt ccaggaaagg agaaaggacc tctctgagag 12720
ggccaaacgc ttcctggcct gagcttgacg aagccccctt cccccggtat aacgatggta 12780
ccgaacatca accgctcaca cgcaaagagg aacaggagga accgacatgc tttttcccga 12840
aacatttctc tggggagctg ccagcgccgc cacgcaggtg gagggagcct gggacgagga 12900
tggcaagtgc ccgagcatct gggatgtggc aggagaccac atcaaaaacg gggacaccac 12960
ccaggtggcc tgcgaccact atcatcgcta taaagaagac gtggcactga tgaaaagcct 13020
ggggcttcgc tcctaccgct tttccgtcaa ctggtgccgc atccagccac agcccggcgt 13080
cgtgaacgag aaggggatcg gcttttataa ggacctggtg gccgagctga agaaggcagg 13140
aatcaagccc ctgctcaccc tctatcactg ggacctgccc ctctgggccc aagaggaagg 13200
cggttggacg aatccgaaga tcgtagactg gttcctggac tactgccagg tggtggtcga 13260
tgccctctcc gaggaggtgg actgctggat gcccttcaac gagcccttat gcttcctgga 13320
cctgggctat gccatgggga tgcacgctcc cttccttacg gatgagagcc tctttcgaaa 13380
ggcggcccgg cacatgctcc tggcccatgg cagggtcgtc tccatgatcc gggagagagc 13440
caaaaccact ccctccatcg ggatcgccat ggcggcctcc accctcatcc cggaagacga 13500
aagccccgaa gccataaagg ccgcagcaaa ggaaagcttt gaagcccccct ttggcgagcg 13560
gggcaacagc ctctacatgg accccatctg ccggggaaag gccagcccga cgatgaagga 13620
cctcttaagc gaggaggacc ttgccctcat ctcggccccg atcgacttca tcggcctgaa 13680
cgtctaccag cccgccaatg tctggcaggg cgaggagcgc tatgacaagt ccgccttccc 13740
ccagacgatg atgggctggg tcatcgatcc cagatgtctg tactggacga tccgccacta 13800
ctgggaccgc tatcagctgc ccctcatggt caccgagaac gggatggcgg ccgaggacag 13860
cgtctccccg gacggagccg tgcacgaccc ggtgcgggtg cagttcctgg atgggttcct 13920
cgcccaggta ggccgggccc ttcaggagaa catcccggtg ctgggctacc agcactggag 13980
cgtcatggac aactttgagt ggtgcgaagg ctacggtccc cgcttcggcc tcatctatgt 14040
ggattacaag acccagaggc gctttttgaa ggactccgcc ctccactatg cccacctcat 14100
ccggacccgg ggcgcggacc tgaagaccgg aagataagga ggaagcctgt catgaaagcc 14160
aaagctctcg cttttctcct cctgcccctc accctgtgcg caacctttgc cttcgcctcc 14220
gagccggccc cggcccggga ggctcaggag ctcatcgagg agctggtggt ggatatcggc 14280
cagtacaaag aggaagccaa gtcccatgtc gaaggccttc tggaggagct ctccctggcc 14340
gaccctgatc tgggaagccg ctgggaacgg atcgcagatc tgtggctttc gctggatgcg 14400
gacctgaccc ttcacgaagg catcctgccc gatggccttc ccgattccga ggagctctgc 14460
ctcgtcgccc tgggctttca gctcgagcca gacgggacga tgaaggagga gctgcttggg 14520

cggcttgggg tcgtcctgga aagcgcgaag aagtatccga aggcctatgt cctttgcacc 14580
ggaggcggga ccgcggcgga aaaccctgac gccaccgaag ccggcgagat ggccaaatgg 14640
ctcctggagc acggcgtaga ggaagaccgg ctgatcgtgg aggaccagtc gctcaccacg 14700
gcccagaatg ccatcttttc ctatgacatc ctgaccaccc gctatcccca ggtgaagcag 14760
ctggcgatcg tatccagcga ttatcacatc gcgaccggcc acctgctttt tgaggcagag 14820
gccatcctgc gctgtgacac gccgggagag gaagccctca cggtcgtctc gaacgccgcc 14880
tggcaggcac cctcgggaga gctctcgtcc atgtttcagg cgggagccct gatcgagctt 14940
tccggggaca ccgaaacggc ctttgaaatc tactatgaca cctacgacat ccatgctttg 15000
ccactcctga aatgacaaaa agccaggtcc cctgccaaaa ggcaggggac ctggctatct 15060
ttttacgttt tcgtttttc acttcgattt cggacgtacc ctcagtccat ggtcacgacg 15120
gccttgatca ccttcccgga gagggaagcc tgacgggcag cctcgaggtc ctcaaatttg 15180
aacttggtga tgatcttgtc caccgggaag cggccttcct tgtacagacg gatcagcttc 15240
gggatgaaga acttgggatt ggactcgccc tcgtcgatgc ccttcacggt cctcatctgc 15300
atcaggacct ccgaaccgat atcgaagttc tcgatcacgc ccgtgggggc cagcgggatg 15360
atcgtcccat tgtaggccgt ggaacggatg gcgttcagga tcatcttccc gtttccggag 15420
gtatcgatgg catagttcac gccgccaccc gtgatctcac ggatcgtggc cggcacatcc 15480
tccacttcct tgcggttgac ggtatgggtc gcgccgagct ccatcgcgag cttcaggctg 15540
tccggattgc cgccgacggc gatgatcttc tcgcatccgg ccaccgcggc cgccatgatc 15600
gcgctcatgc ccacagcccc gcaaccaaag atggcaatcg tctcatccat ttccggctga 15660
aggacgttca gcacggctcc ggatccggtc tggatcccgc acccaaaggg agcgatcaga 15720
gccaggtcca catccttgtc caccttcacg acgttccgtg cgttcacgac agaatggctc 15780
gccagggagg actgcccaaa gaacatggaa atcggcttgc cgttttcaaa cagcttcgta 15840
tccccgttcg ccgaaacccc accgaaattg atggcgttga agttcaggca atagaatggc 15900
ttgccggaga tacagttctt acagtagccg caggtcgcat aggaaaaggc cacatggtcc 15960
cctacctcga tgccatggac atccgggccc acctgctgca cgatgccggc gccctcatgg 16020
ccgagcacaa gaggcatcgg caggccgatc gtccggccaa actcatccgt atggcaaagg 16080
ccgctggcga cattcttcac gagcacctcg cccgccttcg gctcagccag gtccacatcc 16140
acgatctgat acggggcctg cggttcacgt gtgattccag ctcgaatctt gatcataaga 16200
aatcctcctc atagtatatt cgaccatttg tacgcactgg ttacctctga cgatgcggcc 16260
cttcgaaagc ggtaggacgc gtcagttggg aaggattgcc aacctccata cattttatga 16320
aaatggcctc acggaagcct tttccttatc atacgaaaac cagacaggaa agggcaaaca 16380
acaaatgctt gtcatggggg accagatcgt tgtcaccaca aagatcctac aggaggtcct 16440
caaagaacca gccaaaaaga agctgggcgt tttcccggac ctgcctcgca taggcgtcaa 16500
agcgctcgag gaattccttc ccctccgggg tcagcccgct gcgcttgggg tccgcgcctc 16560
cctgggagcg gctgaccagc tgtccgtcgg tctgggactc caggttccgg atcatgttcc 16620
acccactgga gtaggagatc tgaagccgct ggcaggccag ccggaccgac ccggtctcct 16680
cgatctggtg gaacaaaagg gccatctgct catcaaaaaa gggcttttcc ctggccaagg 16740
acaccgtcac caccggccgg gtcagctgtc ggttgtgctc ttccaaaagg gaagagacct 16800
cctccccccct ccccgtatgc agcaggatgc cggggtccgg cacctccacc cggaccatgg 16860
cctccccaca gcgctcgatc gccccctgca ggccaccctc gccggtgtcc ccaaggagca 16920
ccggaagcag cccggaaccg atcaggatgg gatggccagg ggtcccctca tagaccggac 16980
aggccagcct ggccccggtc ttcataagcg agcgcaccgt atccgccgtg aacaaaggga 17040
tgtccaccgg cgtgaacaag atccagtcgc aggaagcctg caggtaggaa aggccgatgc 17100
gggccgactc gatcatctgc gtcgtctcaa agcgttcgtt tctcagaaag acgactccct 17160
tcctggaaag ctggtgctcg agcctctcgg catgatagcc ggtgacgaac acgatccgcc 17220
cgaccccggc ctgctgaagg gtcgccacga tccgctcggc gacggagatg ctccccaggg 17280
ggtgcagcgg ctctccccg ttctcttccg ggacccctgc ggcgacgatc accgccccgg 17340
tccttcggtc tgtcatggca tgtccggacc gtgtccatgg gagtggcgat ggcagcccgt 17400
gcctccttca tagtccggga tgatcgggag ccggtaggca ttcatccggc ggatcgcctc 17460
caggggctct agccccctcg catcataccg ggtgatcagc ccctcctctg ccacgatctc 17520
aaacggcttt ctttgcatcg gcccggtgag gatggcctca cagtcctcct ggagcaccca 17580
gcgcgccagc tcctcctcgc tccacccttc ggcgttcacc tccccgaggg tctccccgct 17640
ctcggcatcg accaggagga cctggcgcgt ctgctcgaac cgggctgcaa tcttgcgtcc 17700
cttttcctcc aacatgcatc cgatcttgat catccttctc cctcctcgat cagcccatgc 17760

tccaccagca gctcctgaca cttgtcggcg atttccatga tcatgggctg gcagcgctcg 17820
atcttattcc aggggtcctc ccccaggata tccctacagt agacagaagc ataccgggag 17880
gcaacggtga catcgaacca ctcacaaaac tcctgacaaa gctccggcgc agcggaccgg 17940
tccaccatgc caaggagcag gcagcccccg gtcaggcctc cgcagacacc cccgctgaac 18000
atcccattgc acagggcact ggcggcagag gtcagacgct cgtcctcgtc ctcctgaagg 18060
agctcgagcc caaagcggat cagagcctcg ctgcagcagc tccccctcat cagatgccgt 18120
tttacggctt tcttcgcatc cataccctcc ccctcatgat cccatgattc caaagctttc 18180
ctcaaaaaac ctcacgaccg acagctgatg agcggcagat accctgtcga tggccccgtt 18240
gacgatgcgg gagaattccc cccaggggca ggccgcagag gcattgagct gcacggccag 18300
atccacgcag ggctttccca tggggtgggt caggaccggc tcctttccga tccccagaag 18360
gtcgaccttg gaaaagcttc catcctccgc ccgcccaaag agcttcaggt gagggatatc 18420
ccagagcggc ttcctcaggc catcccggat ttccctggcc aggtcgcaca gataggcgtc 18480
cgcgtccatc tgctcgcagc agaccctggc gtaatactga agatacgatt cgctcatctt 18540
ccccagggcc ttccgaaaag ccaccccgcc ggagcccaga tccggctttt tcatctgcga 18600
gccgtgggtc aggaggtact ccatcacggc cgatgccccc tctcctgtgg cagccgaaag 18660
gcagaggacg ggcgtatccg gccaggtcct ggcaagaaag gccaggcaat cctctctttc 18720
ctcatcggaa agaaggtcac atttattgag gaggatcaga tccgcctcct tcagctgggc 18780
atccagtata taggccaggt catccggcag ggtggaggca ctttcctctc ccatgatctt 18840
ttccagcgag gccctctcca ccaccaccgt aaagggggcc agggtaaact cccccggata 18900
gtccttcgcc agccgatgat agacatgctc cagagccccg accccaaaac cgggtatatc 18960
ggaaaagacg atctgggacc cctcctcgta cagccggcgc agcctcgata ccaggttttc 19020
tctctgatca cagatacagt cccctgtcat ctgcgagcag gagcagccca gggcgcttaa 19080
gtaggcatgg tccgccaggc ccctgctccc caggtcgtta gagatgatcg aagccccgat 19140
ccccctttgc ctgcactcct tttccaaagc catcatggtg gtcgtcttgc cggcccccag 19200
gaaaccgcta cacaccaaga atctgcgcat atcctccctc ccttcagatc cattcgttcc 19260
taagaatttt agcacaaagc aagaagaggc aacctccacg gatggtggat acccaacaac 19320
aaattgttgt catctttccc caacttgtat ccatttataa gttgataaag ggcagaagga 19380
ccttgtatac tgacagaaag agaaaggtct caaagggccc tcgtgcacga caggcatgga 19440
ctcctgccct tgacttccct catccagaga tcgatcaaat ccactgaaaa caagttcacg 19500
gcatctatca tccacctttt gctcaaagtc catgtgtttt tatccggcac gatgccagac 19560
aaggttgaac gcccccagga gaagaaaagt ggaaacaaaa acattggaat atttccgggt 19620
cgtctatcag gaacagaaca tgcaccgggc cgcggataag ctatacattt cgcagcaggg 19680
cctcagccgg acgatccaga ccctggagaa ggagctcgag tgcaagctct ttgaaagaga 19740
cagcctgggc atgcatccca ccaccgccgg cgaccggctg tatgagtccg tccggaagat 19800
ccagagcgag ctggacggtc tgcatcggga tatcgaagag ctgacgaccg gaaaaaagat 19860
cctccatgtg cccgctgcct atggagtcct gcatcttctc tatccggccg tccggcagtt 19920
cagcgaagcc cagcccaaga tcgaggtccg ctgggcggag catatcgacc aggaatgcga 19980
acagctgctc agccagcagc aggtggagct gatcctcacc gtcctggggg aggataaccc 20040
cgccttcgac atccatcccc tgatgacccg gaggatcatg ctcctggtct tcgaaggcca 20100
tccgctctat gagaaggagg ccgtcagcgt ccaggacctg aagaacgaaa agctgctcac 20160
ggtgggccat ggataccaga tgttcgccac tctgcagaag gcctgcctga aggccggctt 20220
ctatccctat atcgaagggg agatcagcga gatcaacctc tgccacaagc tggtccatct 20280
gggagagggg ctgggcctga cggtggattt catcgccgac atggtcgacc agaaaaacgt 20340
gagggctctt cactttgtcg aagactgctt tacctggagg gtcggcatcg tccggaacaa 20400
aaaaacaccc ctgtcccctg cggccaaggc cttctggaag catatcataa actggccctg 20460
cctggataac aagtaggggc aggcgcaaaa gccatcatcc gaaccatgtc ttcccgtttc 20520
ccggcgggat atggatatat aggaaccaaa gaggcagctg gaaaacaaca gaaaaaagga 20580
gaaaatgact atgggtctta agagaaagta cctgaacatc aacggtgtag agcgcatcgt 20640
catctacgat cccgcgaccg ataccctggc cgaaaccctg cgccgccatg gcctgctggg 20700
cgtgaaggtg ggctgtggaa agggcatgtg tggggcctgc aacgtgatct tgaacggcac 20760
cctggtccgc tcctgcaacc ggaagatgga taaggtggag gaattcagca ccatcgagac 20820
catcgagggc ctgggcctcg cctccaacct gcatccgctc cagctggcct ggatcttcca 20880
cggcggcgcc cagtgcggtt tctgcacccc gggcttcatc gtctccgcca aggctcttct 20940
ggatcagaac cccagcccca cccgggagga ggtccgcgac tggttccagg ccaacaagaa 21000

cgtctgtcgc tgtacgggct acaagcccct ggtcgatgct gtcatggatg ccgccgcggt 21060
catgcggggc gagaagagcg tgaaggacat catctatgat cctgccagcg atcccgaggt 21120
gtatcacagc agctgtccgc gtcccacggc cgtagccaag gtcatgggcc agttcgaata 21180
tggagacgat gtccgccatc acatgcccga ggataccctg catctggcca tcgtccagcc 21240
caggaagtat tcccatgcga agatcctttc catcgatgtc tccgaggcag aggccatgcc 21300
cggcgtcgtg aaggtcgtca ccgccaagga tgtcaagggc accaaccgga tcttagagcc 21360
ccttccccat ccgaccacga ccatcgtggg caatgagaac ccgatcttct gcgatgacat 21420
catctatcgc tatggcgata ccgtcggcgt cgtcgtggcc gacagcaccg agcatgcccg 21480
ggccgccgcc gagaaggtca aggtggagct cgagccgctg accgaataca tgagctacgt 21540
ggaggccgtg gcccccggtg cccagagcat ctatcccaac tccccctcca atatctacat 21600
gcatcagccg ttgatcaagg gagaggatac ccgggacgtc atcgacgagt cctactgctc 21660
ggtcgaagga agcttctacg cccagaagga gccgcatctg tccgtcgaag gcgatgtgct 21720
ccagagctac tatgacgacg agggccgggt ggtcgtgcag tgcaagaccc aggctctggg 21780
ctggaacacc ggtgccatcg ccgaaggcct tggcctgccg atggacaagc tccgcctgat 21840
catgaacccc gccggtgccg ccttcggatg gggcaccttc gccggtgaca gcgccatcgt 21900
tgccgccgcc atgatggccg tgggccgtcc ggtcaccctg acgatgacct atgaggagtt 21960
catgcacttc agtggcaagc gtacctccaa ctttgtcaat gcccggatgt cctgcgatga 22020
gaagggcaag ctcaccggtc tggaattcga tgtgggctgc gaccatggcg cctacttctc 22080
caatgccctg ccggagctgg aagccctggt gcgtttcctt gggtaccctt atgtcttccc 22140
gaatgtccgt ggcatgggcc gcatgggcat ctccaaccac tccttcggca ccgcgtaccg 22200
cggccttggc tcgccccagt gctacactcc ctgcgagagc ctgatggacg acttggcccg 22260
caagatgggc atcgatccct tcgagttcag gtatatgaac gccgcccgtc ccggggacct 22320
gacggtcaac agctatcctt acgtggagcc gaccatcatc cagaccctgg acaagatgcg 22380
ccccatctat gaggcggcca agaaggaata cagcggcaag gagaccgagg acaaggccta 22440
tggcatcggc ctcgcctgcg gcggcttcat gtgcaacctg ggcatcatcg accgctccga 22500
ggtagccctg gagctgaaca aggatggcac catcacccat tacaattcct gggaggatgt 22560
gggccagggt ggcgacatcg gcacgctggt gcacacggtc aaggctctcg agcctctggg 22620
catcaagccg gagcaggtcc gcatgatcat gaacgacctc gggcgtaccc ccgacagcgg 22680
catcgcagcc ggaagccgca gccacttcat gaacggcaat gccaccatcg atgcggccaa 22740
caagctcatg gatgccatga gaaagcccga tgggacctat cgcacctatg acgagatggt 22800
cgccgaaaac atccccacca agtacttcgg tgtctacgag accatcggca agggctccat 22860
cccgctcgat cccaagacgg gtgtcggctc tcccaacaac tgcatgaact atggctgctt 22920
cctggccgtg gtcgaggtgg ataagaagtc cggcgccacc aaggtcatca agtacgtggc 22980
cgtcggcgac gtgggcgtga tcggcaacaa gctggccatc gatgggcagg cctacggcgg 23040
catctcccat accatcggct tcgctctgtc ggaggagtat ggagatctca agaagggcgg 23100
ctccatggcc ggagcgggcg tcccgtacat caaggacatc cccgatgaca tcgagctgta 23160
ctatgtcgag actccgcgcg agcatggacc gcacggatcg agtggcgcct ccgagctcta 23220
tcagtccgcc gggcacatgg ccgtgctcaa cggcatccgc gatgccgtgg gcgtccgtct 23280
ctatgagatg cccgccaggc cggagagggt caaggctgcc atggaggcca aggccaggaa 23340
cgaggagatc aagcccgaga agtattacct ggggccggat atgttcgacc tcatcgatgc 23400
catgcaggag aactgatccc gaatggagat aagtgaatac aaaagcttcg ccgagaagtg 23460
tttccgggac acaccgccgc cttgtgcggc ggtgtgtcct ctctcgctcc ccgtccgcga 23520
ctttgtccgg ctgatgcaaa aaggatcgta tcgcagtgcc tatcgcctgt accgcgacca 23580
ggtgatcttt ccctccctca tcgcccggat ctgcccctct ccctgctccg gggtctgcat 23640
cagggggaag catccgggag atacgagcgt ggacctaaaa gccctggaac ttgcctgcgt 23700
gaagcggatg gagggaaaaa agccggagcg ttatgccctt cccaaaaagc aggagacgat 23760
cgccgtcatc ggcggggggc tctctggcct ggcctgtgcc tatcggctct cttccttcgg 23820
ataccgcgtg accctgtatg agcgctcctc ccttctggga ggatccgtgc tcgccgagct 23880
cgatcccgcc tgggtacagg aggagctgcg ccgggagttt tcggcggtgg actgcgacct 23940
gcgaaccgag aaggagctca cgagcctgga agggatcgag gctgatgcct tctatgtggc 24000
caccggagcg ggaggggaag ccttcgagga tcggccggat gtctttcgcg ggggcgccct 24060
gctgggggca gaccacctga cggccctctc ccaggggctt tgggcagcca cggccatcga 24120
gacctacctg aaggtcggcc ggagggagga gggacccccc cagcccacct taaagcccga 24180
cgagcgctat gaccggctcc actatgacta cacggtcccg gagccgaagg atgggaaggg 24240

ggaagcagag gcctttcgct gcctgcgctg ccagtgcagc gaatgctacc aggtctgccc 24300
cctgatggaa aaggaaaaat actttcctca gaaaatgtgc gtcgaggtca tcctgaccct 24360
caagcccaac atgtccaagc gcaccagcgt gcggatgctc atgggctgca ccgactgcaa 24420
aaggtgccgg gatgtctgcc cggagcacat cgacatggga cagtgcctgg agcaggcccg 24480
cacggacttt tatgagagtg gggccatggc cccggccttc catgactact ggctgcagga 24540
tatggacttt tctctttcgg aggaagcctc cctgctctat ctgccagatg aaaaggcccc 24600
ctgcgacctg ctgtttttcc cgggctgcca gctgggcgcc tccgatccgg aatacgtcca 24660
aagagcctat gaggccctcc aaagggaaag ccagaacccg gccctttccc tggggtgctg 24720
cggcgtcccg gcccgctggg ccggcatgcc ccaaaaggcc ctggccatcg cagaggagat 24780
ccgccagacc tgggaaaggc ttggaaagcc caggctcctg accggttgcc ccacctgcct 24840
ttcaaacctg aagcaggcga ttccggagat ggaggtggag gatctctata cctggtttct 24900
ggcccatccc aaggctctgc cgcccttaaa ggcacagggg gagcggctca ctgtgctcga 24960
tccctgctcc tcggccgagg ccccgcagat gcaaagggcc gtgcgcaccc ttctggagca 25020
ggccggctac caggtcgaag accccgagcc agatatcggc tgctgtggat tcggagggca 25080
tatctactcc tccgaccccg agcttcacca tgcctttgcc acccgccgga ccaaggggaa 25140
ggaaggaatc ttcgccgcct actgcgcgaa ctgccgggat gtgttcgccg cagaggggag 25200
cgacgccagg catgtgctcg gcctgctcct gggcatggaa gaagaaaacc gtttcccacc 25260
ggagctggac cttcgaaggg aaaaccgaag gagactgaaa agatgtttca ccggggagga 25320
ggaaccgccc gtgaaggata tgctgcagat catgatcccg gctgagctga tcgccaagat 25380
ggaccgggag ctccttttgc gcgagcaggt ggcagaaacc attacccatg cagaggagac 25440
ctcggaaaag atcctggatg aggccgattc cctttggtac gcccatcggc gctttggcac 25500
cgtcactctc tggacggcct accgccggga gggggataat gtccaggtgg aaaaggtcta 25560
tgcccaccgg gtcgagatcc gggaggaagg atgaagaagg ataaggaaag ggaggagagc 25620
atcggatgat ctgttgcaag tgccaggtcc ccatggagga aaagaagacc accttcctct 25680
acctgggcca tgaggtcacc taccccgtgc tgcgctgtcc cgtctgcggg caggctttca 25740
tccccgagga gctcgtcgat ggaaagatcc gggaggtgga ggctcttctg gaggagaagt 25800
gataagagta aagaggaggc tggccaaaag gccagcctcc tctgctgatg tactggaacg 25860
tattcctccc acaaaacaga cgccgaaagg cttactcttc tgtctcttcc atccacgcat 25920
cgattgcttc cagtaccggc ttggcatagg actccccgcc aaagagccac tgttcgtgct 25980
gcatgtcaaa ctcccggatg tccgggtccc ggaaatattt cttatatcgt ttcagatatt 26040
tttcacccat tttgttggca tagatgaaat ggacctttgt gccctccaca tggatgtcct 26100
ccctcagcca ggtgagcagg tccgagtagt actcgttttt gatggactga gggctgaact 26160
tcgcaaagca gcccatgaac cggtccgccg tctcatcgct catctcgaaa aagctcttga 26220
gcttttctcg ggtctttgcc ctcttcttct cgttcttcgt aaagctggtc aaaagaggga 26280
ccaccagctt ggactggatc catgcgctgc attttccgct ctgatccaga tccgggctgc 26340
caaagatcgc gtgatcgata tggatctttt cccgctggac cagctggccg acgaagctgc 26400
tgcccaggga agagccgatc gccccgtcga tcctcccgcc atagtgatcc tggatgtatg 26460
cttctatctt ctccgtgacc gtgatcatgt cgggaaagat taggtcgctg ccgtcgaacc 26520
cgtcatagtt gacgcagatc aggtgatatt tctctcccag cctttcgagc accgtgccga 26580
agttcgtctg atagtcgcag gccgtccccg gcagcagcat caacgtcttt cccgcttcgt 26640
ttccttttat gtcaaatcgc acagctgcac ctcctcacac ctctcactcc ttcataaggc 26700
ggaccatcag atcccagccc tctttggcct ctttgcagac cgggaatacc ggataacagt 26760
ggaacatgcc ctccccgacg atcagccgat aatccacccc gtacctcttc atcgccgctt 26820
caaaggacgg agcacaggcg tatagggttt catcgctgcc atagagtaag gtgacccttg 26880
ggctgttcga gaagtcgccc ctctgcagcc aaagcatata ggaggggacg ctctcatctc 26940
catgccgcat gacctccacc gctgttttca tatactgggc cgggatcgcc acatcctttt 27000
tgtccaactc cagcatcctc tgccactcct cctcggtatc cacacaggta ccgggagaga 27060
tcgccaggat ataccccggc atcgggacat ccccatgccc atcatttatg taaggcacca 27120
tcccaagagc gagattccca ccggaggagg tccccaggac agagatctga tccgcttcat 27180
actctctcag catgagcttg taggtctcat ggatcatctc ataggccttt gtcagcgggt 27240
agtctgtgca aagcggataa taggggacaa agacatccag accggtctct ttggcaaagc 27300
gcagcgcctt tttgatcgag cctggtctcg gcgcacggac catgccgcct ccgatgagaa 27360
agaggttcgc cctcccggag ggctgcttat ggatcagctt caggacagga aagcccatga 27420
cctccatccg gctgatctca aatcaggat cctccaggtc cgggatcgga ttcctggcgt 27480

ttttctttct gcaggcttcc agcagctcct ctgtgctttt gcccacccat tgctttttga 27540
tcccaatcgc cctgaccaga cgtttcaata gctcatactt catcgacatg atgtccgctc 27600
ccttcagtta gtattatcta actttaggtc attatagtcc actcccggat gaaaagcaag 27660
aaagggaaac ggcacgatcc ctttcgtacc gtgcctccct cctttgtctt cattccccgg 27720
gagcttggcc gcttcagatc cggaccacgg tcttcttgag attccttccc tccggatagg 27780
aatcggtagc cctctgggta agcttcagca cgcaggcccc ctcctccagg gcctttcgaa 27840
ggcccgcaag ggaggagagc ctctcctcca ccgctgaaag ctccatcccc tcccctggca 27900
gggtcaggac cgtcacctcc aggagccttt cccctgtcag gcgcacctcg aagtcgatca 27960
cctcgtcccg gccatagagg gcctcggcca ggtccgagag gaagaaggaa gcactcccca 28020
gggtgatccg gctgctctcc cgcccgtaca ccttttccag ccttttgatc tggcagccac 28080
agccgcatcg ttcccccgtc atccggccga tgtccccggt tctgtagcgg atgaggggca 28140
tggcctcccg gtccagggtc gtcatcagga tctcccctct tccctgttca tccgtctcgg 28200
ggatctccag gaggatgtcg caggggcgaa gatgatatcc cccgtgtctg gcgcattcca 28260
ctcccccgaa gatgcacatc tccgtggagc cgtaatgccg gtgcaccgag cagccccaca 28320
gcctctcgag ccgctctacg atggcatccg ggaccacgtc cgcgctcagg ttcacgctgc 28380
ggatgaaggc cctctccgga ttcccgaacc ggctcccata ccgggccagc ccggacatgt 28440
gccaggggat cgcctccacc accgaggggg caaagtccat cagctcttca taggcctttg 28500
cgatgtccgt gaccggcccc acgacccgga tgtccatgtg caggggctcc atcgcctgtt 28560
tgaccaggtc cccgatgctc ccgggggcat tcccggacat gtagatgagc atcctttccc 28620
cctcggtgca atccatcgtg agataccccc gattgatgaa gtcgatggcg cgcctttggt 28680
cctcctcgga aaaagaaagc cttttcctct tcccccgggt cccggtggtg gggacggtca 28740
cgacgcgctg cacggaagag gcatcgatgc acgcaaagcg ccattcctct ccggaaaggt 28800
cctcctcgca ggtggtcggc acctgggcca aatcctgcag gctctggata ttcgccaagt 28860
cgatgcctcc cagatgggcc cggtaaaaat ccccctttc cctggcatag gcgagcgtcc 28920
ttcgaaaggc ctccagctgg taggcctcca ccctctcctg ggtcaggtcc cggacctcga 28980
ttcccatgcg ctgggccacg agctcatcca cggccacccc aaggccgctc tcttttttcc 29040
cttgcgcgct ttcttttttc ccttccgcca tctctttttt ccctcctttt tctgctatac 29100
tctttcctga agcatgtcct gtccgttact gggacaggat gcccaccctg actcatctta 29160
ccatggaaag gagcggtcac catcaaatcc aaagcatttt tacgccccat tttctgcctt 29220
tttctgaccc tgtccctgat cctgggaagc ctgagcgcgg catctgccgc tgacctaggc 29280
tttaccgacg gatcggaggg attcctcctc gaaacggcag cccctgccga gacgcaggtc 29340
cttcccgcgc cccaggccga catcctcttc tgcctcgatg ccacggcctc gatggccgat 29400
gacctcggtc gggtgatccc ggccatcggc tcgttcttta cggagctctc ctcccggatg 29460
gacgtacggg ccgccgtcct ggaataccgg gatgtcctcg aggacgggcc agcatccacc 29520
tggctgcacc aggtgggggg ctccccctgg tcatcgagcc ccgtagaggt cctctcttcc 29580
ctcttctcca tcgtcccctc tgggggcggg gatgccccgg aatccctgcc ggcagccctc 29640
actgtgctga caaacgggga gctttccttc cgtgagcagg cacagaagct ggtctttgtg 29700
ttcacggacg atacctacca aagcactaac ggcccctctc tggacgagct ggccgggtat 29760
ctgcagcaaa gcggcatccg cgtatccgtc tttgccgacc cctccacctt cgacgcgagc 29820
tgggaagccc ccggccgggc catcgaaagc caggccatgg ctgccccaca ggaggaggat 29880
gaggacctgc tctcctttgc cctgtcgggc tttgaaagcg agagctttgc atcggaggac 29940
ctccttggcg aagacatggc ttcggaggag gagatggatc tcttcttcac cgacgacctt 30000
ctggccgccg aggccccaga accggccgag acagcggccg ggatatccta ccgctccctt 30060
gccgaatcca ccggcgggat cctggccgac ctttccgggg atgtggaagg cagccttttc 30120
gcttacgcct ccaccctctt ctctgaggag gaagggatac tgggagaagg gacgatagcg 30180
ggtacggagg cagaggccag cccctcctcc aaaaaagcca tctacatcct tccgggctac 30240
ctggcttccc gcctctatga cgagcagggc caggagatct ggctcgatct ggacgccctg 30300
atggcggaca tccttcagta tacccaggaa ggggtgacgc cctcctccat cctgaagctc 30360
tcccccgacg ggaccgggac gagggtccat gtcgacatgg cccgcgacct ctatggggtg 30420
ctgaacaaca cccagcccct ggtcgagatg ctaaaggcca cctgctccgg tgtcttcgac 30480
gtcgtcttct tcccctacaa ctggacagcc gacttaaacg aagccgcccg gagcctggag 30540
gaggacatcg aagcaaaggg ctatgaggac gtcgtctttg tcacccacag caccggcggc 30600
ctgctggctg ccacctacct ggccaggagc caggaaaacc gggacaaggt cgaaaaggcc 30660
attttgatcg cggctcccct gtatggcacc ctgtccgccc tgctgcccgc cgagaccggg 30720

acccggccgg atatgaccag caccatcgcc atgctgaccg gccttccctc ctcggtcctg 30780
aacgcctggg tgagggatgt gacggggaac tcccccacga tctaccagct cttcccctcc 30840
tgggagtatc tggtgaacac gggaacgctc taccgggatg aaaaagacca ctccctcctg 30900
aagccggtgg cttcccccga gcagatgtat gccctgctaa accacagccg caacctgaac 30960
ccgaacctga cagacggaaa cgaaagaagc cacgcggccc atcgggaaag cgtcctgggc 31020
ggagacatcc tgaagatcct cagccaggtg gacaccctgc tgatcggcag ctccatcggc 31080
gtccccaccc cggccgtcgc catctaccgg gataccaaag ggttcagctc cctggacgag 31140
gtcatctacc agaaaaaggg cgatggcacc atcctgggcc agagcgcttt cctctgcgat 31200
gcccagggcg tgcccaccct tccctggctg gatttcccgg gccttggcca caccgctctg 31260
gtgggagata cccgggtcct gacggaggtc gtaaggcaga tcatggcacc ctatgggggc 31320
gtagcggacc tgcccgctcc cgcagaagag ccctatccgc cggaggtgct ccaggagctt 31380
tcctccatgg agtccatggt caaatttatc ttcgagagca acgaagaggt cacgcccgtc 31440
atcgtggacc cgaacgggta taccatttcc ctgacggatg cctccctggt ctacaccgtg 31500
ctttccacca ccccggggga ctaccgaagc cagctgtacc tgccggcctc gggctatcag 31560
gtgcgcctgc agacccaggc agagctttct gccctgaagg gaaagatctc ggtctccacc 31620
gtcgacaagg acggatatcg ctcggccacc tccgtctacc ggctgacgga ggtttcccca 31680
tccggaggca tctcccttcc gaccttccgt ctgcccacag aggccctgac cgccgaacag 31740
ctctcggctc ttggatcctc ccaggggacc ttggcggcca cagagcccgc caccggctgg 31800
cagatccccg agaccctttc cctggatgag cttggagttg cgaccgtgct gcctgtgagc 31860
ggagccgaca tcgaaagcgg aaagctccgc cccgagaccc tcttcttcga gtcccagaat 31920
gagagcctcg tttcggtcga tccttccggg aaaatcgtcg gcctcgcccc cgggacggcg 31980
gtggtgactg ccaccgagcc cctgagcggc tatactcttt cctgccaggt gaacgtcgcc 32040
ggctacccgg aaacggtaag cctggcggat cttcggatga agaccggcga aacggtctcc 32100
ctgacgccct cctttgcacc ggagggtgcc ctgaaggcct ccgtcaccta tgaggtggaa 32160
gacccgagcc tggcccgcat cgagcaggat gtcctgaccg ccctcgcccc cggaaagacc 32220
cagatccggg ccatcgccct cggcggggcc gagacacgct tcaccctcgt cgtcgaccgg 32280
ccggaggtca acctgtctaa gcccaaaaag ccgacgtata cggtgacaag gagcggcgtg 32340
aagctcacct ggaagccggt cgcgaatgca gacctctatc aggtctaccg cggcctaccc 32400
ggcaaaaagg tgacccggct gaccaccacc tcggaccttt cctttaccga tacgaccgcc 32460
gagcccggcg tgctctacga gtattctgta agggccaagt cctataagac cagagaggcc 32520
agctatgtct ccggcggcaa gtccaaaccc actaaggctg tgtggctgcc cgttccccag 32580
agcctcgctt tgcagggcac gtccctgacc tgggagacgg tcccggccgg tgacggctat 32640
gagatcctct tctccgacca gaaggacctg accgaaggga ccctcgtcaa ggtcaaggga 32700
gccggcaaga gcaagcggac gcttgccaag ctgggcctga ccaaggggac ctggtatgtg 32760
aagataagaa gctacaaaac cgttcccggc ctgggcaagg tccgaagcag ctggagcaaa 32820
aagctgaagg tgaaacgcaa ataacacgca agaaagggcc tcgcatctgc gaagcccttt 32880
ctttatctct cttggtctgt ttcctgtcgg accgaagcgg tcgaggagag gatatatcgg 32940
cggaagcatt ccgccatttc cgaccgagcc acggtcttga gccagcccat cgaaacatag 33000
cgatggtatg aggtgccggt cagagggatg gccacaaagc catccccgtg gacctcgggg 33060
ctcagggtca tgatcgcaag gcccaggccc tggcgcacca gtcccaggat cgtccggtcc 33120
tccctggcct ccgataccga acggagctgg attccggcct cctggcacag gcgggaggtg 33180
atcttgctca tacgggaatc cggggtatga gcgacaaagg aaagcccgtc cagctcgtgg 33240
atgtccacag aaggccacaa ggcaaaggga tggccctggg gcactacgca ggccagaggc 33300
tgttccagga cgtgggcgaa ggccacgtcc tgatcctccg tctctgtgca aaagaccagg 33360
tcatactttc cctggcgcag gtcctggacc agggccgtgc tggtgtccgt atgcagggaa 33420
agatacagat tcttcccctc cgtctggcga aagccatgga cccactgggg gaattccagg 33480
taagagatcg agctgatcag cccgatatcc agataccctt ccgagaggga ctgttccctt 33540
tgcagctgct ccaggccccg gtccaggctg gtaagcccct ccgagacaaa tttcagatat 33600
cgcttgccct gccttgtcag ccttacgccc cggccatcct tttcaaacag ataggtgcca 33660
agctccccct ccaggcaacg gattgccttg ctgaggctcg gctgggtgat gtggagctct 33720
tcggcggctt tgccatagtg ttcatacttg gccagcacat agaaatatcg caggtgggtg 33780
aggttcatgg aatccctcct tcgtgacagt ttttctcagc cagtcggcca aacgcaaaac 33840
gatcacctcc tagcttacca tgttttctgc ccagcggtaa gggattttgt gccaaaggct 33900
gacaacggcc caggacggga gtatgataga cgagagcttg tccacaaaaa gaggaaacga 33960

agatgaaaca cagcatgatt tcccggctcc ttgtatggag catggcggcg ctcgcccttt 34020
gtggcgcccc cgcacaggca gaagccctgg cagatggcca ggtgtccatg accctggagg 34080
aattcctcga gaagagcgag gaggaatggt tcctgaccgg aaaaaaggaa taccgcgtcc 34140
aggcgatgat ggtgtccaag gagacccgct tccacaatga gctggagggg accgattaca 34200
ccgtgaccga tgacggcgtc accgtcctcc tgaagggcac cctcggggaa cggtgggcgt 34260
cacagctttc aaaggtgatc ctcacctaca ccaagacaga cggaagcgag cttaccgaag 34320
aagacttcgc ccagaaggac acctatgtca actacccggc acctaaaggt accgggcttg 34380
atccaaggag tttcactcct tgacaagcct gtactttcaa ggaccgtgta gttgagcagt 34440
ccaatgatcc ctcgtgggga atccaccttc gggcggagga accaacgtaa gccacttgcc 34500
cttcggggta ggtatgcgga aacagatcac tgtcatgtga tgccaagcat ggcacgctga 34560
ggtaggggga ccaatggtat ggcgcatcga ctcaatcctc cagcatgggc taccatagac 34620
atgttggagg tgagagaatt aactcttcaa gggagttttg taacatgaaa ccagagatct 34680
gtcttgttta cgtgcttgac gcatccggta agccactcat gcctacgaag aggttcggca 34740
aggtgcgcag gcttctggac aacggaaagg caaaggtcgt ccagaccatt cctttcacta 34800
tccagcttct ttacgagcca gacacgcatt atgtccagcc tatcgttttg ggaatcgatc 34860
cgggaaggga acatatcgga ctcacggcga caacaaaaaa aggcagggcg gtctttaagg 34920
ctcgattcga tacccataat ggggaagtta cgaaacatat gtcaaagcgg agaatgtatc 34980
gtcaggcttc caggcatgga gaaaggcttc gccgcaaacg cagggcaaag aagtacggaa 35040
cgctcatgaa atgcggggag gtggacagaa agcttccggg ttacgaaaat ggcactgtca 35100
aggtaaaagg gatccttaac actgaggcca gattcaataa caggaaaagg aagcccggct 35160
ggatgactcc tacagccagg catctgctgg aatgccatga aaacatcgtc atgttcgtgg 35220
aacggattat cccgatcagg gaaatttcaa tcgaggtggc gaggtttgat ttcgtggcaa 35280
tgagttaccc gaatgcaaga taccagatgt atgagaacgg gccgatgaaa ggttatgatt 35340
cccagcttga agccctgagg gacattcaaa aagggaagtg tcttctctgc cagaaggggc 35400
ccgtggagca aaagcaccac atgaggccga gatccctcgg cgggtcggat acgctcggaa 35460
atcttgttgg cctctgcaac tcatgccacg acaagattca tcagaaacag gggaaggcat 35520
tagaaaggct ctcgaaaaag aaggagggct tcaacaaaaa gtatgcggcc gcatcggtct 35580
tgaaccagat catgggaagg ttcatcgact ggtgtacaga tagatatcct ggaaaggtga 35640
cgttgacgaa tggcaaagcc accaaggatt acagggatgc ccatgggctt gtaaaggacc 35700
atcatgttga tgcccagtgc atcgccatgt gtgcctgtgg ctacggtgct gaatgggatg 35760
gttccaacct gcactgctat gtgatccgca aatatcggag gcatgatagg gccaaggtca 35820
aagcccaggt ctctcgggcc tataagattg gcaaaacgat tgtggcaaag aacaggaagc 35880
ctgccatggt ggcatcagtg gtcagagacg gaaagatggc ttacacgaaa cagaaggagc 35940
ctgccttgtc cgattggtat gaggaacagg tccagaaaca cggagaacag gaagccatgc 36000
gtatgcgcag ccagcttgtc gtggcacctt cctatcgaag atataatgac gtgaagcgga 36060
ctcttcctgg aaccatcttc taccatgaag ggaagcgata tgtcaaaagc ggacagatca 36120
ggcctgatga ctcgcatcta tactatcagg ctgaaggagt ggatagtcga aactttcctg 36180
cgaaagactg caggattttt tacaacaagg gcatcgtggt gatgtcatga atgatggaaa 36240
ataccgtagg gggtaacata acaagtacag gcttatggtg catttaatct ttgttgtgaa 36300
gtaccgcaag aggctattcg tcactgagaa tgtcgtggat gaccttaagc agagccttca 36360
cgatatctcc gaagcgaaag gttggaaaat ccttcgacct gtgcggtcgt aaaagtaacc 36420
gatgttataa cccctggggt aatctcggcc tttagccgga ccccccaggg cgttgttatg 36480
gttcatcatc attcgaatag tatgcatcaa cctatttttg ccatttcttt cgaccttaaa 36540
tcgaaaacct tgccatatcc ttcgtacatc ttaaaagata cgactccagt ctcacggtaa 36600
acatcagctt cgtgattcag gtacttcact gtcacattgg caagacccca actaccttca 36660
cgagaccagc tcatggaatc gcccttctgt cgggaagaaa cgataaggtc attcgccttt 36720
tccactctgt tgctggaaac ctgaagcccc atgctcttgc gaagagcata gcaaggaata 36780
cgcccttcat ttttccgaag atacccgacc agttcattga ttttatcacc gggtttgaga 36840
tagtttccat ctaacgactg tagatagcaa atagcattct gtacattccc tgaccacagt 36900
atccggaata gccttcgctt tacctcatat tgcatcttcc gattggcctt accacctttc 36960
aggctcatgc ttaaagcttc ggtacaatgc ttacgcagat ggaaccaatc caggatttct 37020
atgggatgat aaatcctaaa gatattgtct atgcaatatc gaatgtcttt tcccccatcc 37080
gtaaagaaaa ccagaaggtt attctccaga aggttatgag agattaaaaa tgccagtatc 37140
attttacagg tatcgaacat gttcttgcct gcaaacacat atttcttgcc atcatgttca 37200

acatgtccca cagctgtttc tactttgggc cgttttttgg actcgggtgc tcgggtatag 37260
tcttcaggac cgtcgttttc atcccggacg aacgtaggat cgctttcagt gccgctcatc 37320
tttggccggg aatctttctg acgttttgca cccactccat ccagggaaat cttcaccccc 37380
ttcgaaggat caggttccat ctcatatgtt ctgtccgaac cgacttcaat ctgttcctcc 37440
gaagacttct tcacattgag ccgtttggct ctttcgtcaa tcatcgcatg ccgttcttcc 37500
atagtcaaag gttgctcttc tccatccacg gcaaacacat tctcatcaag gaccccataa 37560
gcgatgctgg ttccttcgag cggaaggagc gtttgaaggt catagccatt agcctccaag 37620
aagctgtcgg cccatgaaat ctgatactga taatatgctg tgccctcatc ctcgactaaa 37680
tctgtcatag tccgtaactt gatgtcatcc agcccttcct cgcggtggag cagtgcgtta 37740
aggagacggt tagtaactcg tccactattt tcaatcatcc cttgttccat atagcccaac 37800
aaggaattgc tccatagata acctgattcc tttgaccctt tatgtatttc gctccatata 37860
ggggcaacca ggataccatc ttcgctttga atcttggcta tacggagccc gtcatcactt 37920
gtttttttc gagctgtacg tccagtcaac tatattgacc aacacctaag ttattgactt 37980
ggaatcaaaa atgggataga atgcagggag aaaaaacata taaggaggtg taaacaccgg 38040
atggctcgca aggcatcacc aattatattg agcaatgccg atgttaactc cctttctatc 38100
ctatcaacat tatcacagga ccctgacttg gcccgtcggg caaaagtgat tctggaatgc 38160
ggtaagggcc tctctaacaa agaggtatct gcagtcattg gaatgaatga ggccgatgta 38220
gggcgatggc ggaaggtttt tctagcagat gggatagatg ggctaaaagg gaagcagcgc 38280
ggaggtgatc atcgctctga cgctccagat gagaccttat acactcggct caatgagctt 38340
ctggggcaag aaggaagaga ctggactgtt caggaacttg tccaggaatt gaattcaacc 38400
gcgaacattg tctctgctgc tttacgtaag cagggaatca atctccagcg ccaaaggcaa 38460
tggtatatac caacccagga tgagctcatt cccaaaacat tggatgtggt tggattgtat 38520
atcacaaagg cagaacaagc catcattgtc cgctgtacac aggaggcact tttagagtgc 38580
catggagagt tcatcaccag gaacagggag ctgtcagagg actacctcca ttacgatgga 38640
gacattacgc tggctgaagc aataaataca gccgtccatc atgtgaagga tacttcccgc 38700
atggtccctg tttcattatc ggagtttttg aacaggacca tctccgcatt tccggtaaag 38760
gacggatatg agtatcatat tcttatctgc tctccaaaga acagcggata tcaaggagca 38820
cgcctgaaag atatatatca gtcatgggca gacgattcca ggcggtggct ccatctggtg 38880
gacagtaaga tcaatgaact ggcagaccgg aatcagttgc atccggcaga ggagctacag 38940
aaggcactgg aactgtacct tgacaaatgc atagagacaa cagctccact tatctggcgg 39000
aagaaaccag atcagccggc aacggctgag catcaagctg cgcatgggca agagaaagag 39060
gaaaatccga ttattttaaa aaacggactg gaaagctctc cctttcttga cgaattgagg 39120
gaactcctga aacgttatct tgcctcagga cttgagtcgc ctgagattgt taaatgtggc 39180
ttcatttcct ttgcctatga tcaagatgaa gtgcgcgtcc agttggagga cgacaacggc 39240
acccttccag gagcgtctga tgtagatttc ggttccttag aagcatggaa ggatggattg 39300
accgccgtgg aaaaggctct gataaacact cgaaacagag ctggtgtgga tgctgcagca 39360
atggtgtctg atttattaaa aaaaaacagg tgctgtcggg ctaagcgtgg cgagggaagt 39420
agagagtgag ctcggacgtt tttccgtctg gctccctgtc ggcatagcac ggcatctgcg 39480
taaagatgaa cgcatcaaat gtggtgggtt gattgaacgg atatgcgcac acgcttctga 39540
tacctcctac cgaacagcat gtaagaattt caacctcgat tatcatcgag aagaagcagg 39600
atgcgagatt ccgctcatga cattctcaga taccgttgtt catgagggac agcggttaat 39660
ctcgaagatg cgtgaacatc agaaacaggt ccttaccagt tttggttttg gagaggatgg 39720
tctatggccg aaagataagg agtttcccga tgagctccgc aacctggaca tggagtatat 39780
ctattttgac ccgtccatga ttatggaaca ggccggatac ttcccccttc cagtgattga 39840
gtttaaagaa gggcagcgct ccgaacctga aaccggatta cacattcgga aaatgaagag 39900
gagggacgaa agattacttg tccctgatga tcagcgagag caatggctca aaggctatat 39960
tctttatatc aatacgcagg cggataggag tccttttgca accatcaggc atccggcccg 40020
cgttgagaga catgccgaat cggtggtcaa catctttatc gatggtgacc ttgtaggcga 40080
gcaggcggaa catcgtgtcc caggcggtaa gctagtcctt cgtgatgaaa aacctacaat 40140
aaaacatata gatatccgga tcgaaataga aaacggagga agatataata tcacttctgt 40200
tgatttggac gatgcatacc aaagattact ggctgtactg ctagaaacag gattgataac 40260
gcgatatttt catttctttg tcgatggaga aaagacgtta actaaagcaa tcgaaaaata 40320
ttttgcaaat tggaactata gtatattttt ggatttcttt catatcgagg aaaagataaa 40380
gacacaaaca agccttggta ttaagggcag gcgggttgca agcccttggg aagaacctga 40440

atactacacc agaggtccca aaaagggaga gattaaaaaa cagccaatga catcactgag 40500
taggacgtat gcctcagtat tgaagaccat ggttttctat ggcaacgcaa aagaagccaa 40560
ggaatacata gaacatatcc ccgaagagga tatacaggac aaagcagcct tgcaggtatt 40620
ggtcaactat ctggaagaga gggagtccat gctggcctgc tataccctcc ggaagcacgc 40680
aggcttaaag aattccagta atgcctcgga gctttgcaat gagctagcag tcagttcccg 40740
ccagaaggtt gatgacagga tgcactggag ggaggatggc agtgcagcat tagcggctat 40800
cactgtcctt tatatgaacc gggctgacaa gcgatggttt gaacagcata atgtggaatt 40860
tgagctgtac tatgtagagc cgaccaagac aaggaaaatc aagactcagg ctgcgtaagc 40920
gagagatggg ggcctaaatg aataatttag gttcccatca atttagtgga ctcgccgtac 40980
agctcgtttt tttttagcga tttcatggag gtacgtctcg ctaagggctc tggatgcttc 41040
tcgcgaagca ctgatcaccg cctgttctac cttatcaagt gaattcaaat attcccggcg 41100
gcttccggtt tcgaattcag cctgtgtaat agggttttta tatgaacacg aacagtgatg 41160
acattatgtt catcgtctac gaactgtgca gtgaatctga tagaaccagg tgctgatcgg 41220
gaaaagtcat cctgagcttt aggttctgtc aaagccatgg gtttctggtc gccttcctcg 41280
ttctttgtca ctttcaccca ttcgaacggt tcggtgttcg gtttagcggt acgcaggtaa 41340
cggtatatac tatctctgac cttggctgct ttctcctttc cctgaatgcc agagtacata 41400
ttagttgcta ttatctcaaa ttgagacatc cattgttcct gtgtcttgaa gacgtgcagg 41460
taggtatccg gaagaaggga acgattgttt tctaccaaag aaacaccata tgtgaatata 41520
tggtattcaa tgttgctcgc ttttatcccg tttgggagga ttcctttcac gaatgaaata 41580
gccgaatcct ttttcccgag ccgaacagtc tttcctgact ccgcaaagac agtcagtgca 41640
tctgccatat caaccttgcc tgagacatca ataagggctt caatcttaga agcaatcgcc 41700
ttggttcttg tctctactac tgcagaggta tccgtaatca ggtcgaggtc gattaccata 41760
ttcttttcta tcgtccaagc cttgaccaca agaacctgtt cggtgggcga cagatataag 41820
ccgcacatat ctgccaattt aggtacggta ccgtcagaca ttgtaaatcg ccagtagtgc 41880
ggacctcgtt cagcaactat gtctttcttc tgcaagatgt cacggataac attctgtggt 41940
gccttaagtt ccgtagcaag ttccttagca gatggcacag gaacgttctg ggcaacacac 42000
tgctggattc gaaggtcgat tgaagcttct aactgatctg gatcatatgt gataggccgt 42060
ccagatttcg gcttgtcata tagacagtca atccccgcct ccaggaaacg ccgacgtata 42120
tccgtgatgg tgttctccct cataccaacc tcagcagcca cgtcctttat ttgcctgcca 42180
tcggacataa gcagtagtgc ccgggctctc tttacgatat cctggccaca gacaacggtt 42240
ccatcatcga ttcttctcag cattataaca tcggtttcat taatgacaat tggcgaaact 42300
cgttttggca tatcccattg cccctttcat aatggatatg ccatattata cagagtgaga 42360
atggcttcat caacaataat cgcatgaaag ttcctgattt gacagcaaga gaagcctcgg 42420
aaccatacca gcgccatggg gagtctggct aaagggcgag attagcccag gggttataac 42480
atcggttact tttacgaccg cacaggtcga aatccttgag atggagaccg ataaggatca 42540
cattcacatc ctgttggatt ataaggtgac ggattgtgtg gccgacattg caaaaatcct 42600
taagatggtc tccacgaaca cgatatggaa gaactatcca cgtatgatga aaaggaattt 42660
ctggagggag catacccttt ggtcggatgg atactttgct tgtagcatcg gcgaggcttc 42720
caaggaaacc atagaacaat acattcagaa ccaaggctga ggtttatagc aaaaggaagc 42780
atgcccatcc tcttcattgc cgtccaagca caggaaaaag ccatttgctt tttttcctgt 42840
gcttggaatc tattctctat tctctcaaag gcccattcat cccagcacct aaaggtactg 42900
ggttttctgg gccgatggat tataaacctg ctcacaaagt cccagccgga ttcctattat 42960
gcgagaaggg tcccggtgga gatctcggtg acggtagaga cagcctgggg cgatgtgctg 43020
cacacgaacc tgccaagcac caaccatggg gagggagact atctcgtctg ccgggtggat 43080
gaaaacgacg ggccggacct ttcggatgtc tgggtcgtcc atggcctcct ctttacggaa 43140
acctatgacc tcccggacga gccggacccg tcctgaccat ggaatgctga ttgcgcttgg 43200
ggctggtgaa agggtataat aaggccactg tgcgcgacag gcctatggac cagtcgccca 43260
ccaaagcaca aatgtagact acataagagg aaacctatgc gaaacgaaaa gctgctggag 43320
gccctgaccc agatttgggg agtctccggc tatgagaccg aggtaagaaa gtttataaaa 43380
gaagaggtcc gctcttacgc cgatgagatg tggaccgatg ccgtggggaa cctgattgtc 43440
ctcaaaaaag ggaccggggg acccacctcg agaaagatca tgtacgcggc ccacatggac 43500
cagatcgggt tcatggtcaa gaccatcgag gacaatggcc tgatccgcat ctgcaatatg 43560
ggttggaact gggccggctc ggcgtacaac gagcggatga tcttccgaaa cggcaccgtc 43620
ggggtcgtgg gctgctatgg ccccatcgag gaggccaaga acaagatcga gaacctgttc 43680

```
     gtggacatcg gggcctccgg caaggacgag accacaaagc tcgtgaaaat cggggactgc     43740
     tgcacctatg agggcagcta tgtccacctg gcgaccggaa agctctgctc cgtggccctg     43800
     gatgaccgcc tgggctgcta tcagctgatc gaggccctga aggaaaaccc cggcacctat     43860
     cccaacgact gttactactg ctttgtcgtc caggaggagc tcggctgccg tggctcccag     43920
     gtggccgccc agcagatcct gccggatatc gggatcgcgg tggacatcac cccggcccac     43980
     gattatccca atgacctgac gggcagcaac aagattggca aggggatcgg catcaagatc     44040
     tgcgacccct ccgtggtcag cgatgagacg gtggtcgccg tcatggaaca gctctgcgag     44100
     gagcatcaga tcccgaacca gag                                             44123
//
ID   XXX; XXX; linear; XXX; XXX; XXX;  XXX.
XX
AC   PRJEB67889
XX
DE   Uncultured bacterium DNA fragment from in vivo enriched cow rumen, clone
rum-42n18
XX
FH   Key             Location/Qualifiers
FH
FT   source          1..37103
FT                   /organism="uncultured bacterium"
FT                   /environmental_sample
FT                   /mol_type="genomic DNA"
FT                   /isolation_source="Cow rumen"
FT                   /clone="rum-42n18"
FT                   /note="AZCL-xylan, AZCL-arabinoxylan active clone"
FT   CDS             complement(402..797)
FT                   /translation="MAKNMLKNYGRFELLELIYNMRKENLELKERCDAAERQVVEIRLQ
FT                   TEKQLDNARRDYENRIEELRMQGAAKDLQIRMKKIEEQLRLLQKLTMVDIDLPEMPTEE
FT                   EIEQLAEQEAREREELYTEPGDNLKSE"
FT                   /product="hypothetical protein"
FT   CDS             complement(831..2867)
FT                   /translation="MNMKRMISILLTCILLFSVFVVHAEEQVEATNEPVLSYPDRNYEE
FT                   LTVGNATPMDGKFFTGMWGNATTDIDVRTLVNSYYLTIWGYDTGFFKANPVTVAGLSVV
FT                   DNTKRYRKYRFQLADDLFYSDGTQITAWDYAFSVLFQIAPEISELGGLPMDLSYLEGYE
FT                   EYVSGQVPYLAGVRVFDDLNIEFSVKPEALPYFFELYRLGFLPYPIHEIAPGCKVFDEG
FT                   NGVYIGNEDPDVIEKIFSTELLKATVMDPDNGYLSHPTVGSGPYVLTSWDGTTCTFEIN
FT                   PHFKGNQEGFKPSIPKLRFTLAENDTMIEKLEAGEFQLLNKVVRKDVIEKGSALVSAGK
FT                   GFALTNYPRVGLSFIVFTPDSPALQEQNVRQAINWCMDKDAMRDDYTHLYGMTMDGMIG
FT                   IGQWMYGMVMGTEDYPEILPDNPTEYEQEEYDERITAWKNLSLDGVKHYELDVKKACEL
FT                   LDANGWTMNENGERFTAGRDEVRCKKIHGEMVTLDLTCAYPAENIMAAAMEEYFIPYLK
FT                   EAGIRLNLVPMEMKELLRAYNDRDIEDIDMFYLADDFNIEFDPQLFFLAGDPNVVKEDT
FT                   LAWAHARMARYARRMCETVPSDALGFVEKWITFQEKLSELLPLIPVYSNVYYDFYTSDL
FT                   MNYDIIKYITWGDAIVPANYYNVYQSMVDMGIDPTVTEEEEIP"
FT                   /product="hypothetical protein"
FT   CDS             complement(2899..3663)
FT                   /translation="MMRRRVLSILMILILLFSCMSALAETQNENDGWLSITVGEDRSGF
FT                   DPEGIRMTIYLLATGDYGNWKMVEEFSDITVFTRSDGSASVDITLGQIRNRIADRKIKP
FT                   VEEAVSDKSGKLEFKNLAHGIYYMEMTAGPDKLKMSPMLLSVPNKNGSIQVRAVAKYEY
FT                   ETPTPSPTPSPRPTMTPFIPEETPTTDETPAPPTKEPVATPSDSPDPEKTPVPQHVPTL
FT                   VPKPDEYPVPLEDYEAALGLWNIQTHVGVCYE"
FT                   /product="Proline-rich protein"
FT   CDS             complement(3660..4625)
FT                   /translation="MKKLFAWILALLLLVVVIGHFLYPVLSDQLARRRDAEMMWEYREK
FT                   TSAMGKEQREKLFAEAAEWNKGLEEIRMEDIFTTGITRTTRDYQNHMNVHSGIIAELVI
```

```
FT                   PDIGVSLPVYHLSTETPATYKLVHVNTSSLPADGSRENIILAGPGVLKAEGILGDVGLT
FT                   DDRILEDLDKLIPGSLVILNVLDRTMVYRVKGVQMLSPVGLKELDLTPDEGEEKLTLVS
FT                   QRKDQRLLVQAERIPIREARTLLAENDQATFPANWQNVLLLGCPVLLAGLLVLLIIEKI
FT                   RGRAYRLPGEGRMADQELRERNARETINQIETEISEDDET"
FT                   /product="hypothetical protein"
FT   CDS             complement(4622..5545)
FT                   /translation="MEIYHPRDEELLREMKRVRRSDRRRRLCRGLIILLTLSIAVGLFV
FT                   FLRYYQLAVAHGTGMGDTLPEGSLLLIRKPEAGQTYQAGDVILYEKRLAKPVELTILNS
FT                   KGKTRRYCRYVLYRDVGTTRQYYATTEGGAAWLSTVSGADIFESTEEGTVSIGTENLKN
FT                   GEYWLKEVEASYGQALLTDPIPFAVVNPVKTQLKRVLAGPGERVVLSPYAETRVNGLLI
FT                   STDYTSGRTEDAAPEGRRLIMARDRYFVQGDQLSLSVDSRETDYSTVSDGEILGRAEFA
FT                   LWPLQCFGNLTGRQTVVAGTETEAAE"
FT                   /product="hypothetical protein"
FT   CDS             complement(5714..7270)
FT                   /db_xref="GO:0031176"
FT                   /translation="MGKKILVFLMILFLLGSCVSALADDAKTIYTSDFSKDADGWYGRG
FT                   AQSARTGDNTLKTIGRQSSWNSPGRDFDLVEGGKYNLSVEVKQDDLDSANFMISIAHSV
FT                   DGMETYENLAFGKAKKGEWTTLTGSYTAGHFDRSVLYVETTGADTLDFEIRAFTVTAPE
FT                   GVPEPKPTELPMVIEEADSVPSLKEIYAEKFDFGSAAPQMVFRDPKWMNLMQEQFSILT
FT                   PENEMKPDAVLDVNASKALLKETGDETAVAVHFDAAKALLRFAQNNGIKVHGHVLIWHS
FT                   QTPEVFFHEGYDAKKPLVSREVMLGRMENYIKGVFEYLDANYPGVVVSWDVLNEAIDDG
FT                   SNWLRNSNWKKIIGEDYPNFAYAYARKYAPEGTKLYYNDYNTAVGGKLRGIVKLLNTLI
FT                   PEGNIDGYGFQMHHNVSFPSIQQIRTAVETIGALGLRLRVSELDVTVSNNSEASFKKQA
FT                   KYYADVMKILIDHADQFEAVQVWGLTDMMSWRGSQFPLLFDSRGNPKPAFWAVADPENY
FT                   Q"
FT                   /product="CBM22-GH10"
FT                   /EC_number="3.2.1.8"
FT   CDS             complement(7427..8854)
FT                   /translation="MKKIMALVLAALMVLGCCSFAAAEDIPEEYPEIIEGLDFGGATVY
FT                   INDWYSSGERAEEPTEEQQAQYDYWDWLEATYNVKLVEIKLGDWDGMVAELQNIVSNKD
FT                   NSELRIVGVSGGFCGPVLANGLFSEWTYGLENFNKATVDFMTIGGKCYGVCGGKFFEPR
FT                   QCVFFNKKVLEDAGIKWEEIYDAQADGSWTWEKMESYMDAVKADKDNDGELDVFALTGN
FT                   FDDGVVGLVVSNNGDFYEKTEEGKLVYSADSDNTLAALNRIQEWNRNYYRPYVNWDDYK
FT                   QFWPEGNVAFLIGQSYEGFNGNDVVNNVEEWGCVCLPKGPAADAYTSAADNNVFGIPAV
FT                   YDEETSLKLQEIYTLYRKNPVIDADEDAWATRLYELTDERAVEETYAYLRENGTIMNFN
FT                   YIGDRNSTIGPNLTWGIMGGAASEQVEAARLAMEDMAAVFNGDKTQEQVDAEKAEREAA
FT                   AAAEAAAAEAAEEPAAE"
FT                   /product="ABC transporter substrate-binding protein -
FT                   sugar transport"
FT   CDS             complement(9011..10021)
FT                   /translation="MSKDNTIFIGKNKSFKERNRRSGGYLMKRRIKEVSISVFRALLMF
FT                   GLCFMIISPMISRLSMSLMDEKDLYDTTIVLLPRNITLDNFTAVARATSLPTSMLNTLW
FT                   ISILSSICQVVACTLVAYGFARFEFPLKKFWFACVIALIIIPPQTIQSALYRTFTRFDL
FT                   FGLITLFNGQPLNIRKSLWPYLMMCLTCMGLKDGLYIYMLRQYFKNVPESLEEAAYVDG
FT                   CGTMHTFLKIMLPDAVPIIASCFLFSFVWQWTDLFYSRNFLSGYKLYAPELSSITDRIA
FT                   DELSKMAGTPVTPSAGRSLQLISIGVLFCCIPLIIILYCFTQRTFVESLAMTGSKE"
FT                   /product="Alpha-glucoside transport system permease
FT                   protein AglG"
FT   CDS             complement(10014..10994)
FT                   /translation="MSKKNRKYREHWYSGITEAFQIFFPGSRFYHTIPARRARTGVLFI
FT                   LPFIIGFIAFMVKPLAVSLQMSFSRYIINTGEMTYNGLYNYNYAFNTDPDFNRMLLDEI
FT                   SRMAINVIATLVMSFVIAVILNQKFKGRILCRIIFFLPVILSSGVLPGIETNSSSLNLL
FT                   NKMSEAMGDSSGVNIAASIEDLLKVSGVGQGVFDVVFKMINSIYDIVMASGIQIIVFLS
FT                   GLQSISPSLYEAADVEGCSAWESFWKITFPMVSPLLLVNCIYTIIDFFMKNNNQVMEHI
```

```
FT                   NNIYMNGVRMDLGIAAAESWIYFGVAIVFIGISAFIISRAVKSYE"
FT                   /product="binding-protein-dependent transport systems
FT                   inner membrane component"
FT   CDS             complement(10991..13588)
FT                   /translation="MRKKSILRRLIPWIIAIVLIAGAAFVLNKIYTSPGKSFARETKII
FT                   YFEGDGKPLTMENEELLFEMDGATSQFKVTNKDTGKVWFSNPENRENDKIARGTTLEYL
FT                   SSTLNVSYYNDITKTEMNNFTGSIKNQSFEILQEEDGSIRVNYSLGKNAGKRYIIPNAM
FT                   TMERYSEIMEKMGGEKSKDAKKFKNHYSLYKTKDLNGKNKDKKDELLALYPSLEEQDLY
FT                   ILKSDRKDEQKKQAEDLLKKAGYTEEDYQKDQELMAGEKDSTGPVFNISVLYRLEGKDL
FT                   VVSVPYSDIACSSDYPLVFVDVLPMFGAAGTDQEGFTLVPEGGGAIIRYNNGRLSQSSY
FT                   YANMYGWDYGTKRETATSETENAFPVFGMSQEDGSFICIMESADSYGGVCADISGRLNE
FT                   YNTVFGRYNILHSDNVEVDENKSARLFLMYEKQIPDDTLVQRYRFLDGNGYVSMAQTYG
FT                   EYLRAKPEMRGECASEEMPVNVELVGAINKVEVKLGIPVEATVATTTFEQADSIMNDFL
FT                   EGGVKDLNLRMTGWCNGGAMQRVLTSIHVEGCLGGESGMKKLIASAAEKGVDLYFDGIS
FT                   CFAYRSGILNGFNTLSDSARATTRDLIKLYPYYIVTYEMSTWMDKYPYYLVKPSYAQRC
FT                   ASNLINGLKDRGAAGIAFRDIGNLLSADYQDSNTVTREQAKAQDTQTLEDAVATGMKVI
FT                   IKEGNAYAIPYADLITDMNLSGNAYALLDYSVPFYQIAIHGLKDYTGEALNMAGDPQTL
FT                   LLESAEYGAGLNFTFMEKDTRVLKDTLFSCYTSSHYAPWKDEVIAMITRYQKEMSGLNR
FT                   QAITGHERLNEDVAVTTYADGTQVYVNYSTKDYRAGSVTVPGRDYLVKRGGGQ"
FT                   /product="hypothetical protein"
FT   CDS             complement(13604..14299)
FT                   /translation="MSKAKSAETLINAEPVSGGKTSWAEYKRSLKYALYVITHPFDGFW
FT                   DLNREHKGSLAAANTFMALFLITRVLKILCTSFQFFDAAVQHINVFEQMASLLLPFLIL
FT                   CLANWAMTTLFEGKGRFKDIYIGMCYALVPYILIQLPMILVSNMLTYEEGSLYSVMLSI
FT                   SVVWCVFLVFVGLMEIHDYGPGKTFIFIIVTIVGACVIIFLALVCLSLLSDAIGFFVSL
FT                   YREAAYRLN"
FT                   /product="hypothetical protein"
FT   CDS             complement(14265..15800)
FT                   /translation="MMKKAVSFLMILMLLFTACTALADDSFNVNKDGYSTSYTYGYDYW
FT                   SDVQESPDAYRVETVIDSVDLGLDVRMNKPQSLFARNNDLYVADTFNNRILQLEYRNGQ
FT                   AKLKRIIDHVTGGTPETFNTPYDVYVDADGNIYVADYGNLRVVMMDKNLNWIKEFTKPA
FT                   DATFDQSLDFLPKKITVDVAGRLYALVTNINKGIVKYEADTTFTGFIGATPVQFSTFEY
FT                   IWKKYFMTQEQRDASTSFVPTEYENLYMDKDGFIYTTLTNFEEGALDAGQVKPIRRLNG
FT                   LGNDILIANDRYAPIGDLWWRAGESAYFGPARMTDITVFDNDIYIVLDRIRGRLFGYDN
FT                   QGIMLWAFGTRGNIEGAFTSPISLEHIGTDLFVLDQLENSITVFTPTEYGLKIYDAINK
FT                   YQDGDYDESAEVWQDVLKQNANYPMAFRGIGRTILRQNKYEEAMEYFKMAHDRENYGRA
FT                   FKLYRKEWIENNALWIFLGVVLLLIVPLILGRIKRMKWEVIMHEQSKIRRNAD"
FT                   /product="NHL repeat containing protein"
FT   CDS             complement(15815..17053)
FT                   /translation="MSEMNTVKSNGTLQLIGGKKYRGELVAGEKGAAFYAKGQKEPIAE
FT                   WHYARMNRMDNIRRGGTTSQITVILKDDTRYVFELAQGLKLYSHMDKNWADKPVKVKTR
FT                   GDILHEQAERLGEKIQVPKKHLMARRHPNRSIAGDVIIYVILALVAVAMLFPLIFLLGS
FT                   SLKPLDELFRFPPPVWPTRPTMDNFSDLYVTMGQSWVPFSRYLVNTVFITVVGTFGHLV
FT                   IAGMAAFVLAKYDFPGGRLFFVLATTFLMFASGVVTDIPNYLIMRRLCLVDTYWALILP
FT                   AFAAPIGLFLMKQFMEGLPTALIEAATIDGAGRFRVFWSIVMPNVKPAWLTMIIFSVQS
FT                   LWNNPAGTIIYSEAKKPLVVYALQQIQGGVMVNIARTGQVAAANVIIVAVPVLIFIFSQS
FT                   RILETMASSGIKD"
FT                   /product="binding-protein-dependent transport systems
FT                   inner membrane component"
FT   CDS             complement(17055..17999)
FT                   /translation="MQSQSFARRYLTMRREKIAHGWTLAKRNKGCYLFLAPYAILFITF
FT                   FILPICTSIFYSFTYYNILEPARFIGLQNYTNLFLQDEIFLTSIKNTFVCAVIIGPVGY
FT                   ILSFLFAWFINELPKWLRAVAVIVFYIPSIGGAAFEVFNTIFRGDAYGWFNALLMEYGM
FT                   INAPVLWLTDPRYMLTIVIIVSLWMSMGTGFLSFVAGFQGIDRSMYEAGYVDGVRNRWQ
```

```
FT                   ELYHITLPSMKPMLLFGAVMSITSSFNVGAVPRALCGFPSTDYAARTIVTHLFDYGYTR
FT                   FEMGYASAIAVILFVAMILCKKAITALLGRVGT"
FT                   /product="binding-protein-dependent transport systems
FT                   inner membrane component"
FT   CDS             complement(18023..20908)
FT                   /db_xref="GO:0005215"
FT                   /db_xref="GO:0006810"
FT                   /translation="MSRKCRNILIGVLVAVVVIGGIWLLSRNSENFRSKYEGADLSTDV
FT                   SGIGRSNTYTSYLEKYAELPAVSEAVEVDLASFEGNGGEVCADGLLTADNSELSWKVDV
FT                   PKAGLYNIRLEYLTTESRGVDIEREIAINGEVPFSGASTLCFSRLWTDATEVRKDNQGN
FT                   DIRPTQEERFEKQSAYCKDDMGYQTEPYAFYFNEGENELTIRAVNEPVILCSISLTPIL
FT                   KSPTYEEYTAAQPEAAMSEEGLNYCQTIQGEAATLRSTPSLYGRYDRSSSQTVPYSVSN
FT                   SILNYIGGEPWTHPGEWIQWDFEVPEDGYYNISVKARQMYQRGALSARSVYIDGEIPFT
FT                   DLEAVTFSYSTGWDMRTLGDKDGNPYRFWLTKGSHTIRMEVTMGKMGPVLKSVEDSIFR
FT                   LNQIYRKILVLTGASPDKYRDYNLKRVYPEVIEAMDLESKRLYKIVDDTVAITGEKSDR
FT                   VASAQTLAVQLETFVKKNERITESFQNFRDNITSLGTAMQNMAESKLDVDLIMVTGENV
FT                   KVPEVHENIFQSIGHEIRSCLSSYTVDYNSLGDKYEDSDDVVEIWITTGRDQSTVLKNL
FT                   VDNTFTAKTGIKVNIKLVLQDAILPAVVAGNGPDVVLSLSGWFAVNYAMRNAVEDLTQF
FT                   EDFEEVVKPFTKSILEPLTYNDSKKIGIYGLPETQEFAVLFYRKDVMEELGLAIPQTWD
FT                   ELIAELPTIQGNSLTVGINFPDIRVVDNSVLNSMIYQNGGEIYDKEAKHTLVDSEAGVA
FT                   GFKQYTSLYNDYGLPVVFDFPSRFRSGEMPLGIQNYNVYNTLMVSAPEIRGLWDFTLFP
FT                   GTVKEDGTIDHTAQTSGLCCMMIASDNETKKKNAWEFMKWWVGADAQVRFGREMESILG
FT                   ASARYQTANQDALKQLAWSDKQLKVLQTQVAQTKGFPEIAGGYSTQRHLINAVRKVINT
FT                   KEDQRETLLTYARTINEEIKIKRREFNLPVD"
FT                   /product="N-Acetyl-D-glucosamine ABC transport system,
FT                   sugar-binding protein"
FT   CDS             complement(21103..23508)
FT                   /db_xref="GO:0004826"
FT                   /translation="MKVPYSWLKEYVDIDITAEELQEKLFGCGFEVEELIDLGKEISRV
FT                   VVGEVLSCEPVEGTHLQICQVNCGSYGNPIQIVTGAPNVYAGMHTPAALDNSTLPGGIT
FT                   IKAKPLKGIPSDGMLCSGEELGLNDDLYPGAEVYGLMDLPKDTVPGTDIREVVGMNDYI
FT                   FDVTITANRPDCQSILGIAREVAAALGKELKMPATDYTVSDYVFPSLDITVEAPDLCPR
FT                   YLGHGVRNITVGESPRWMKRNLALCGLRSISNVVDITNYVLLEIGQPMHAFDMDQLQDR
FT                   RIIVRRAKDGEKIQTLDEKEFTLTPDNLVICDGNRPVALAGIMGGLNSEITENTTQLLF
FT                   EAAKFARDNVRKTSRALGQSSDSSARFEKGISEYITELGMARALHLIQELGCGEITSST
FT                   FDCSAGAPREGKRFTASLKRINAILGIEVPAEEVLRILKNLNFEVKRDGDTLDVTAPRY
FT                   REDIEVGEPDLAEEVIREYGYDHVIPTFLKAATVTSGGKTPAQQRRDRVCRILCGQGYH
FT                   EAYTMSFYSDSDLDFLRIPEGAPERNVLRIKNPISVNLSIMRPLLAPSLVNVVVENLKK
FT                   GNNEGRIFEISNIYIPRDGELPDERPHLGFAAFGDAEDFFTVKGTVEALGEALGLSFDV
FT                   ERASDVPWLHPGIAAYILCEGERIGCFGKLANEVTGELKLHKDAKSNHKIFLGEIDYNA
FT                   MMNHMAASFRYRPLSVFPSVVRDLALTVAEETACGDLMKEIAKACPRVSDVELFDIYRG
FT                   EQIGEGRKSMAFKITFEADDKPLTPEELDKYIKKILGNLKFKLGAEIR"
FT                   /product="Phenylalanyl-tRNA synthetase beta chain (EC
FT                   6.1.1.20)"
FT                   /EC_number="6.1.1.20"
FT   CDS             complement(23601..24623)
FT                   /db_xref="GO:0004826"
FT                   /translation="MEQKIKALQEQMEQQISQIDTKEKLAEFWQEFLGKKGAIADLMKG
FT                   LGAVAKEDRPAMGKVINEFKVLVEEKYQALSAKMDELELAARNRREQVDITLPGKGLPL
FT                   GGLHPLTLVKNEMINVFASMGFDIFEGPEIEDDDHNFTRLNVPKDHPSRDMQDTFYLEN
FT                   ELLLRTQTSGGQIRVMDMQKPPIKVLVPGRVFRSDSDATHSPMFHQMEGLVVDKGITLC
FT                   DLQGMLDKFVQQIFDADVRTRLRPSYFPFTEPSVEVDVSCFECGGKGCSLCKHTGWIEV
FT                   LGGGVVHPKVLENCGIDPNVYSGIAFGIGIERITMLKYGINNIGLLFENDMRFLKQFKE
FT                   "
```

```
FT                   /product="Phenylalanyl-tRNA synthetase alpha chain (EC
FT                   6.1.1.20)"
FT                   /EC_number="6.1.1.20"
FT   CDS             24645..24776
FT                   /translation="MICLIKTITGDPVITNYRNYELWIIQHKMSILSWFLSVFAPHI"
FT                   /product="hypothetical protein"
FT   CDS             complement(24830..26158)
FT                   /translation="MNMESLWEQTCALLSREMNYVSYSTWVDGNMVPGMLEDDTLFVVV
FT                   KMERMISMIQNKYADLIERKLTEVAGKAIRIVLLTKDEMKARQETDEDSGKPDDTDPHL
FT                   NPKYTFESFVVGNGNRFAHAAALAVAESPAEAYNPLFIYGGVGLGKTHLMQAIGHFIHQ
FT                   SDPKKKILYMTSENFTNELISAIQMKKTYEFREKIRKVDVLMVDDIQFIAGRESTQQEF
FT                   FNTFNELHNDNKQIILTSDKPPKDIQRLEERLCSRFEWGLVADIQRPDVDTRVAILREK
FT                   TVQDNIEVPDDVLQLIAGKIDSNIRELEGCLTRLVAYSTLVKEPITVDLCERALKEIFD
FT                   SKRHKQITSELIMQTVSDYYGLSMDDMTGPTRKREITVPRQIAMYLTREMTGMSLPQIG
FT                   TVFGGRDHTTVLHSCKTVEANMTANTDIRAVVEDIKTLVKNAQ"
FT                   /product="Chromosomal replication initiator protein DnaA"
FT   CDS             26577..27083
FT                   /translation="MIEAWHVREALRFRFGSALVDRPGIWEKAAALLRNFAPYRDTFSD
FT                   LAASLEDVLFNTVYEQLGPSMGAQMDNGTVRRIRSTEFRDASDDVMGVLFDHLKVYSVT
FT                   YDSLHQYCMATGSFAAMRVLYTRYADFMPASERKIIARIIRDSRPRDVWEPWMDPEDVP
FT                   PAPSR"
FT                   /product="hypothetical protein"
FT   CDS             27143..28036
FT                   /db_xref="GO:0005488"
FT                   /translation="MKKILALLLVLALFLPAVATAETIVTSFYPIWIMTLNLTEGIETV
FT                   KVVNLAEPTTGCLHDYTLQNSDMVTLSQADALLVNGAGMESFLPVVTGAYPNLPVIDAT
FT                   VGLPFLHESDTVEIGEAEEGEEVNSHLWLDPQRAAGMAANLAEGLISLMPDREAEILEN
FT                   LHAYQERLQALDETIREETKDVDRKVIIFHEAFPYFAEACGLTAAAIVNKEPEDILPTA
FT                   QLVRVTELIFNTGIPMPLILKSAEEDPSVNTLVNETGIPVCELDPMTSGPENPPLDYYE
FT                   SIMMQNVKTLLNAIAQ"
FT                   /product="Zinc ABC transporter, periplasmic-binding
FT                   protein ZnuA"
FT   CDS             28094..28876
FT                   /translation="MFSVSPEIYSVLSQASRWIFILLALMLLFLAFSWHHTDRKERRDR
FT                   FKNLPGAGTVGELVVLSGSDELPVDTWFPVPREGVLGSLRSCDLVIPCPGVSSHHLDFS
FT                   WQDGTGLLIRPRVGCEVLIDGVPVTRRGMDMPLVHGSVLQIGSAVLRLQLFAALSHTNR
FT                   TFIPPQPAVGSFSDPMPQPVPVPVQYPQEPVNAQPVFCPPPYQEPVPLQIPVEPPMPVQ
FT                   PVPQPPVPEQLPVQEGSPLAPAAPRRKRADRWKEDWSE"
FT                   /product="hypothetical protein"
FT   CDS             28873..30006
FT                   /translation="MTKQKSSARFRFSGVMLFLALFFFLVPALQAGDYRLYMLAAGVPC
FT                   AMLLCCTILARAFSLDRLILALSLFFCSVGIASLAMTAPDTALVQAVSCGIGIIVLIVG
FT                   AILIRSLSGSLLTAISAAFLGLLLLAAKLIAPTFTLPATQPALTLLLIAFAVLLSREGP
FT                   ISAAGLGIAAAVLLMLRGETADALIWGLTILLLLFAADGRLLVVLPSLAAFALLFYGMY
FT                   SMQTQTLVYPEALSADTLKAVGALGADLLPEGLAVPEKGSLFSRLLGHYGLVFTGLTVL
FT                   LYLPLILRGTTVASYARMRFHAILAMGICLLLGLSTLASVLSFFGFLPFTGLEMPLLTT
FT                   SLPSLCAHLFLAGILCGISGRNDSDLAEDAHLAMLAK"
FT                   /product="Cell division protein FtsW"
FT   CDS             30032..30331
FT                   /translation="MKQMRSRFRLLTLLIVCAFLLTIVLCTGHVLKTTGISLSSLSSIS
FT                   LPVPGQSASPDPSALPDPSPAPESTPDPWAGFSVENTPPGTDNPPEEEYNVVGL"
FT                   /product="hypothetical protein"
FT   CDS             30359..31135
FT                   /translation="MIQFKRALALMLVLILCCSSSLAAKKKTTEYTAREITTEIVQEIP
```

```
FT                   AEIQNMLDIAYEQLIETDGKNLKEKNKFTKWRNNYTFEWCGGFVTWCMLEAGIPMEEKN
FT                   KIQDGEVEGLFHVKEAGVGKLYEGYAKLNRITRVPQKGYIAVYGNEGSGGSTPYYHVGL
FT                   VYDVEKLSDGKYRITTIEGNVKGHTVKMYVRDYDLNLASDKKQKPKDMSLVPEEERNRE
FT                   ETNIFSYGYAYTKKHMYITIFLMPWIPEESQESTEE"
FT                   /product="hypothetical protein"
FT   CDS             31139..31771
FT                   /translation="MLWMYAVCALSIFVCLPFFLYYKKALHMKLAAAFKSLGTLCAASL
FT                   ALTAAIRLDPQYWVCFAALFLHAVADWILEFNMYIGTGFFIAGHICYIGFFTHLCPVSG
FT                   IHLIAALCLLGILVFLLWRWRKPIGKTIPLIAVYGTILSLASACAIACFSLNSLQGQFI
FT                   AVGGALFFISDVIILARLLFTAGRSVDWAIMITYYAAQLLFGLSCLL"
FT                   /product="hypothetical protein"
FT   CDS             complement(33847..33984)
FT                   /translation="MSSRWREKLLVYLKVPVPETDAGGRGENPETNGRTLVKELGKMTP
FT                   "
FT                   /product="hypothetical protein"
FT   CDS             complement(34020..34145)
FT                   /translation="MYIPALPKNEAVTQKDSLSGVMVNSVQALRLERSEVRVPLS"
FT                   /product="hypothetical protein"
XX
SQ   Sequence 37103 BP; 7784 A; 10742 C; 9085 G; 9492 T; 0 other;
     ttcttccaga aattctccgt ttaccttcac atatcccttc tcatcaattt ccaccttatc        60
     cccttcccgc gcgataatgc ggcgaagaag cagcttgttg ttgaaataga aggcacagac       120
     gtcgccggtc ttaacatcat ctgtttttgt aaggataact atatccccgg gctgaagtcc       180
     cggttccata ttatgtccgt tcgtaatccg tagtaccggc aggaagatga acgcgataat       240
     caccgccagg gccgccacca ccacaaaggt tccgaaggta cctttcagcg cctgtctgaa       300
     tttccccttg ctgcgaacct ggctcatctc ggtttcaatc tccttcagat ccggcagcgt       360
     atcggctttc cttgttttca cggccatggc agttccctca tttactcaga tttcaggttg       420
     tctcccggtt cggtgtacag ttcttcccgt tccctggctt cctgctccgc cagctgttct       480
     atttcttcct cggtaggcat ctccggcaag tcaatatcca ccatcgtcag tttctgcagc       540
     agccgcaact gttcctcgat cttcttcatg cggatctgca ggtccttagc cgcgccctgc       600
     atccgaagct cttcaatccg gttctcgtag tccctacggg cattgtccag ctgtttctct       660
     gtctgcaggc ggatttccac aacctgccgc tccgccgcat cacagcgttc cttgagctcc       720
     aggttttcct tccgcatgtt gtagattagt tccaacagtt caaaacgccc gtagtttttc       780
     agcatgtttt ttgccatgcg tcggttcctc cgttatctgt cctgatgctt ttacggtatt       840
     tcctcttcct cggtcaccgt gggatcgata cccatatcaa ccatggactg atagacatta       900
     tagtagtttg cagggacaat cgcgtcgccc caggtaatgt acttgatgat atcgtagttc       960
     atcagatcgc tggtgtaaaa atcataataa acattggaat aaaccggaat cagcggcagc      1020
     agttcgctaa gtttttcctg gaaagtgatc catttttcca caaagccaag tgcgtcactc      1080
     ggtactgtct cacacatccg acgtgcgtat ctcgccatcc gggcatgagc ccatgccaac      1140
     gtatcttcct ttactacatt cggatcaccg gccagaaaaa agagctgggg atcaaactca      1200
     atattgaagt catccgccag atagaacatg tctatatctt cgatatcccg atcattgtat      1260
     gcacggagca attccttcat ctccatgggt accaggttca gccgaatacc agcttctttc      1320
     agataaggaa taaagtattc ttccatggcc gctgccatga tgttttctgc cggataagcg      1380
     caggtcagat ccagcgtaac catttccccg tggatcttct tgcagcgcac ctcatcccgt      1440
     cctgcagtga agcgttcacc gttctcgttc atcgtccagc cattcgcatc cagcagctcg      1500
     caggcttttt tcacatccag ttcataatgc tttacgccat ccaatgagag attcttccag      1560
     gctgtgattc tctcatcata ctcttcctgc tcatattctg tagggttatc gggcagaatt      1620
     tccggataat cctctgtccc cattaccatg ccatacatcc actgtccgat gccgatcata      1680
     ccatccatag tcatgccgta cagatgagta tagtcatccc gcatggcatc tttatccata      1740
     caccagttta tagcctggcg cacattctgc tcctgcaggg cggggctgtc cggtgtgaaa      1800
     actataaagg aaaggcctac acgtggatag ttggtcagtg cgaatccttt acccgccgaa      1860
     accaaagcag aacccttctc aatcacgtcc ttgcgtacta ccttgttcaa cagctgaaac      1920
     tcacccgctt ccagcttttc aatcatggta tcgttctctg ccaatgtgaa acgaagtttc      1980
     ggaatggaag gcttaaaccc ctcctggttt cccttgaaat ggggattgat ctcaaacgta      2040
```

caggttgttc cgtcccagga tgtgagtaca taggggccgc ttcccactgt gggatgggat 2100
agataaccgt tatccggatc catcacagtt gccttcagga gttccgtact gaaaatcttc 2160
tcaatcacgt ccggatcttc atttccgata tagactccat tcccctcgtc aaaaactttg 2220
cagccgggtg cgatctcatg aataggatat ggcagaaaac caagacgata cagttcaaaa 2280
aaatacggaa gtgcttccgg cttaaccgaa aactcaatgt tcagatcatc gaacactctg 2340
acgccggcca gataaggcac ctgtccggaa acatattcct catacccttc caggtaggac 2400
aaatccatgg gcagcccgcc caattcactg atctccggtg caatctggaa cagtacagag 2460
aacgcatagt cccaggcagt aatctgggtt ccgtcggaat agaacaggtc atctgccagc 2520
tggaaacggt attttctata ccttttcgta ttatccacta cgctcagtcc ggcaactgtt 2580
acaggattag ctttaaaaaa gcctgtatca tagccccaga tagtcagata ataggagttg 2640
accagtgtgc ggacgtcaat gtcagttgta gcgtttcccc acatgcccgt aaagaatttg 2700
ccgtccatcg gtgtggcatt tccaactgtc agttcctcat aattcctgtc cggatacgaa 2760
agaacaggtt cgtttgtcgc ttccacttgt tcctccgcgt gcacaacaaa cacactgaag 2820
agcaggatgc atgtcagcaa tatagagatc attcgtttca tgttcatctg tctcccgtgt 2880
ttttctatct gccctgattt attcgtagca aacgcctaca tgtgtctgaa tattccacag 2940
tcccagtgcc gcttcataat cttccagtgg cacagggtat tcgtccggtt tgggtaccaa 3000
tgtgggaaca tgctgcggaa ccggcgtttt ttccggatcc ggtgagtcgc tcggagtcgc 3060
caccggctcc ttggtcggag gtgccggtgt ctcgtccgtt gtaggggttt cttccggaat 3120
aaacggtgtc atagtcggtc ttggtgacgg agttgggctt ggtgtcggcg tttcatattc 3180
atatttagcc actgcacgga cctgaatact gccattcttg ttcggtacag acaaaagcat 3240
cgggctcatt ttcagcttat ccggtcccgc agtcatttcc atgtaataaa tcccatgcgc 3300
aaggttttta aactcaagct tgccgctctt atcactgact gcctcttcca caggttttat 3360
ttttcggtca gcaatccggt ttcgaatctg accgagcgtt atatctacag atgctgaacc 3420
gtcacttcgt gtaaaaacgg tgatatcgct gaattcctcc accatcttcc agttaccgta 3480
atctccggtt gccagcagat aaatagtcat gcggattcct tccggatcga atccgctccg 3540
gtcctcaccc actgtaatgc tcagccagcc gtcgttctca ttctgcgtct cggccagagc 3600
tgacatgcag gaaaacagca gtatgagaat catcaggata ctcagcactc tccgtctcat 3660
catgtctcat catcctcact gatttcggtt tcaatctggt ttatcgtttc ccgggcgttt 3720
ctttcacgta gttcctgatc tgccattctt ccctcaccgg gaagccgata ggcgcgtccc 3780
ctgatctttt ctataatcaa tagcaccagc aagcccgcca gcagcaccgg acagcccagc 3840
agtaatacat tctgccagtt tgccgggaag gttgcctggt cattttccgc cagcagcgtc 3900
cgtgcctccc ggatcgggat ccgctccgcc tgaaccagca gccgctgatc cttccgctgg 3960
gaaaccagcg tcagcttctc ctccccttca tccggtgtca gatccagctc tttcagtccg 4020
acgggagaca gcatctgaac gcctttaacc cggtagacca tcgtccggtc cagcacattc 4080
agaatcacca ggctgccggg aatcagcttg tccaggtcct ccagtatccg gtcatccgtc 4140
agtccgacat cccccaggat gccttcagcc ttcagcacgc caggtcctgc gaggatgatg 4200
ttttcccggc ttccgtccgc cggcagggag cttgtgttta cgtgtaccag tttatatgtt 4260
gccggcgttt cagtgctcag gtggtatacc ggcagcgaaa ccccgatatc cggtatcacc 4320
agctccgcaa tgatgccgga atggacattc atgtggttct ggtaatcccg ggttgttctt 4380
gtaatcccgg ttgtgaagat atcctccatc cggatttctt ccagcccctt gttccattcc 4440
gccgcttccg caaacagttt ttcacgctgc tctttgccca tggccgaggt cttttcccgg 4500
tattcccaca tcatctccgc gtcacggcgc cgggccagct gatcggataa cacaggatac 4560
aggaaatgcc ctatgacgac gaccagcagc aaaagggcca gaatccaggc aaacagtttt 4620
ttcattccgc cgcctccgtt tccgtaccgg cgacaaccgt ctgtctgcct gtcaggtttc 4680
cgaagcactg cagcggccag agcgcaaatt ctgcccgccc caggatctct ccgtcagaaa 4740
ccgtactgta gtctgtttcc cggctgtcca cagaaagact cagctgatct ccctggacaa 4800
aataccggtc cctggccatg atcagccggc gcccttccgg cgccgcgtcc tctgtccggc 4860
cggaggtata atctgtgctg attagcagcc cgttcacccg cgtctcggcg taggggctca 4920
gtaccacccg ttccccgggg ccggccagca ctcgcttgag ctgtgttttg accggattga 4980
caaccgcaaa cggaatcgga tccgtcagca gcgcctgtcc gtaggaggct tctacctcct 5040
tcagccagta ttcaccgttc ttcaggttct ccgtaccgat actgaccgtt ccctcttcgg 5100
tgctttcaaa gatgtctgcg ccggacactg tggataacca ggccgcgccc ccttctgtgg 5160
ttgcgtaata ctgtctggtg gttcccacgt cacggtaaag cacgtaccgg cagtatcgcc 5220
gggtttttcc cttggaattc aggatcgtca gctcaaccgg cttagctagt ctcttttcat 5280

acaggataac atccccagcc tgatatgtct gaccagcctc cggtttgcgt atgagcagca 5340
ggctcccctc cggcagcgta tcgcccatgc ctgttccatg tgccacagcc agctggtaat 5400
accggaggaa tacaaaaaga ccgacggcaa tgctcagggt cagcaaaatg atcaggcccc 5460
ggcagaggcg tctccgtctg tcggaacgtc tcactcgctt catttcccga agcagttctt 5520
catctctggg atgatagatt tccatccgtt ctctccctgt ccggtattcc ggaataatcc 5580
ggtcaaacag ttaccggaaa aaaacagatt attttatata taaccatttt gagtataacg 5640
cgattaactt tccttttcaa gtttttttcca actcgtacca aacgaaaaca ggagggtttc 5700
ccctcctgtt tcattactgg tagttttcag gatctgcaac cgcccagaat gcgggtttcg 5760
gatttcccct gctgtcaaac agcagcggga actggcttcc ccgccagctc atcatatccg 5820
tcagtcccca tacctggacc gcctcaaact gatccgcgtg atcaatgagt attttcatga 5880
cgtccgcata atactttgcc tgtttcttaa aggaagcttc actgttattg cttaccgtca 5940
cgtccagctc acttacccga agccgcagtc ccagcgcgcc gatagtttcc accgccgtcc 6000
tgatctgctg gatagagggg aaagaaacat tgtgatgcat ctggaatcca taaccgtcaa 6060
tgtttccttc cggaatcagg gtgttcagca gcttcacaat gccccggagc ttgcccccga 6120
cagcggtatt gtaatcgtta tagtagagtt tcgttccttc cggagcatat ttccgggcgt 6180
atgcataggc gaagttcgga taatcctcac cgatgatctt tttccagttg ctgttccgca 6240
gccagttgct gccgtcgtca atcgcctcgt tcagcacgtc ccaggaaacg accacgcccg 6300
gatagttcgc gtcaaggtat tcaaaaacgc ccttgatata gttctccatc cgtcccagca 6360
tcacttcccg gcttaccagc ggctttttgg cgtcgtagcc ctcatggaag aatacttccg 6420
gcgtctggct gtgccagatc agcacatggc cgtggacttt gatcccgttg ttctgcgcaa 6480
accgcagcag ggctttggcc gcgtcaaaat gaaccgcgac ggctgtttcg tccccggtct 6540
ccttcagcag cgccttgctt gcattcacat ccagcacggc atccggcttc atttcgtttt 6600
caggcgtcag gatactgaac tgttcctgca tcaggttcat ccatttcgga tcacggaata 6660
ccatctgggg agcggcggag ccgaagtcaa acttttccgc gtaaatctcc ttcaggctcg 6720
gcacgctgtc cgcctcttca atcaccatgg gcagctcggt cggcttgggt tccggaacgc 6780
cttccggtgc ggtaaccgta aaggcacgga tttcaaaatc aagcgtatcg gcacctgtgg 6840
tctctacata cagcacactg cggtcaaaat gaccggcggt atagctgcct gtcagggtgg 6900
tccattcgcc cttcttcgct ttgccgaaag caaggttttc ataggtttcc attccatcca 6960
cgctgtgggc aatggaaatc atgaaattgg cgctgtccag gtcgtcctgc ttcacttcaa 7020
cgctcaggtt atatttgccg ccttccacca ggtcaaagtc ccgtcccggg gaattccagc 7080
tgctctgccg tccgatggtt ttcagcgtat tgtcacccgt cctggcagac tgtgcgccgc 7140
ggccgtacca tccgtccgca tccttgctga agtcagaggt gtagatcgtc tttgcgtcat 7200
ccgccagcgc ggaaacacag cttcccagca ggaaaagaat catcaggaaa accagaatct 7260
ttttacccat ggtattaggt cctttctgac tcgaatttgt tcgtacatat tctacctgct 7320
caggccgcat tattcaaggt gtaatgcatc ataaaaacca taattctgaa aatgaaaaaa 7380
aggaccgccc ttgcgggcgg tccgggaaac cgaaagattg tccgacttat tcagcagcgg 7440
gttcttcagc ggcttcagcg gcagcagctt cagcggcagc agcggcttcg cgctcggcct 7500
tttcagcgtc aacctgttcc tgagtcttat cgccgttgaa cacagcagcc atatcttcca 7560
tggccagacg tgcagcttca acctgctcgg aagcagcgcc gcccatgatg ccccaggtca 7620
ggttcgggcc gatggtgctg ttgcggtctc cgatgtagtt gaagttcatg atggtgccgt 7680
tctcacgcag ataagcgtag gtctcttcga cagcgcgctc atcggtcagt tcgtacaggc 7740
gggtggccca tgcatcttca tccgcgtcga tgacggggtt cttccgatac agggtgtaga 7800
tctcctgcag cttcagagag gtctcctcgt cgtacacggc gggaataccg aacacgttgt 7860
tgtcagcagc gctggtgtag gcatcagcag cgggtccctt gggcaagcag acgcagcccc 7920
actcttcgac gttgttcaca acgtcgttgc cgttgaagcc ttcataggac tgaccaatca 7980
ggaaagcaac gttgccttcc ggccagaact gcttatagtc atcccagttg acatagggac 8040
gatagtagtt gcggttccac tcctggatgc gattcagagc cgccagggtg ttgtcgctgt 8100
cagcggagta aaccagtttg ccttcttcgg tcttctcata gaagtcgccg ttgttgctga 8160
caaccaggcc cacaacgccg tcgtcgaagt taccggtcag ggcaaagacg tcgagttcac 8220
cgtcgttatc cttgtcggcc ttcacagcgt ccatgtagct ttccatcttt tcccaggtcc 8280
agcttccgtc ggcctgagcg tcataaatct cttcccactt gatgccggcg tcttccagaa 8340
ccttcttgtt gaagaacacg cactgacggg gctcgaagaa cttgccgccg caaacgccgt 8400
agcacttgcc gccgatggtc atgaaatcaa cagtcgcctt attgaagttc tccaggccat 8460
aggtccactc agagaacagg ccgttcgcga gcacggggcc acagaagccg ccggacacgc 8520

cgacgatgcg cagttcgctg ttatccttgt ttgaaacgat attctgcagt tcagcaacca 8580
tgccgtccca gtcgcccagc ttgatttcaa cgagcttcac attgtaggtt gcttccagcc 8640
aatcccagta gtcatactga gcctgctgtt cttcagtggg ttcttcagcg cgttcgccgg 8700
aagaatacca atcattgata tacacagtag cgccaccgaa gtcgaggcct tcgatgattt 8760
ccggatattc ttccggaatg tcttccgcag cagcgaagct gcagcagccc agaaccatca 8820
gagcggccaa taccagagcc atgattttct tcataatgag aaaacctcct tttttatttc 8880
cttacagatt tcccgcggaa atctgtatag gtgcaattgt aaatcgcgga tcccccgctg 8940
ttcaggacaa ttatagaccg tattttccat ttaatcaagt acaagtatgt acatttttcg 9000
cattttgtgc ttattctttt gatcccgtca tcgccaggct ctccacgaac gtccgctgcg 9060
tgaagcagta caggatgatc agcggaatac agcagaacag cacgccgatg gagatcagct 9120
gaagggaacg tcccgcggac ggtgtcacag gcgtacccgc cattttggac agttcatcgg 9180
caatccggtc cgtaatggaa ctcagttcag gagcatagag cttatatcct gacaggaagt 9240
tccgggaata gaacagatcc gtccactgcc acacaaagct gaacaggaaa cagctggcga 9300
taatcggtac cgcatcgggc agcatgatct tcaggaaagt atgcatggtc ccgcagccgt 9360
ccacatacgc agcttcttcc aggctttccg gcacgttctt gaaatactgc ctcagcatgt 9420
agatatacag gccgtctttc agtcccatac aggtcagaca catcatcaga tatggccaca 9480
gggatttgcg gatgttcagc ggctgtccgt tgaacagcgt aatgaggccg aaaaggtcaa 9540
accgggtaaa cgtgcggtaa agggcgctct ggatcgtctg cggcgggatg atgatcaagg 9600
caatgacgca ggcaaaccag aactttttca gcggaaactc aaaccgggcg aacccgtagg 9660
ccaccagtgt gcaggcaacc acctggcaga tactgctcag gattgagatc cacagggtgt 9720
tcagcatgga tgtaggcagg gatgttgccc gggcaaccgc tgtaaagtta tccagcgtta 9780
tattccgggg cagcagcacg atagtggtat cgtacaggtc cttttcatcc atcaggctca 9840
tggaaagacg gctgatcatc ggggaaataa tcataaagca caggccgaac atcagcaggg 9900
cccggaacac ggagatactt acctccttga tcctgcgctt catcaggtat ccgccgctgc 9960
ggcggttccg ttccttgaag ctcttgttct ttccgataaa gatcgtatta tctttactca 10020
taggatttca ccgcccttga gatgatgaaa gcgctgatgc cgataaagac gatcgcaacg 10080
ccgaaataga tccagctttc agccgccgca atgcccaggt ccatccggac gccgttcata 10140
tagatattgt tgatatgctc catgacctgg ttgttgttct tcatgaagaa gtcgatgatc 10200
gtatagatgc agttcaccag gagcagcgga ctgaccatcg ggaaggtgat cttccagaag 10260
gattcccatg cggagcagcc ttccacgtcc gcagcctcat aaagcgacgg ggaaatggac 10320
tgcagtccgg aaaggaatac gatgatctga ataccgctgg ccatcacgat gtcgtaaatc 10380
gagttgatca tcttgaaaac aacatcaaac acaccctggc cgacgccgga taccttcagc 10440
agatcctcaa tggatgccgc aatgttgacc cccgatgaat cccccatcgc ttccgacatt 10500
ttgttgagca ggttcaggga actgctgttg gtttcaatgc cgggcagcac gccggaagac 10560
aggatcaccg gcaggaagaa gataatccgg cagagaatcc ggcccttgaa tttctggttc 10620
aggatcacgg cgatcacgaa actcatgact agcgtcgcga tgacgttgat agccatgcgg 10680
ctgatctcgt ccagcagcat cctgttgaaa tccgggtctg tattgaacgc gtagttgtaa 10740
ttatacaggc cgttgtacgt catctcacct gtattgatga tataccggct gaagctcatc 10800
tgcagtgaaa cagccagcgg cttcaccatg aaggcgatga aaccgatgat gaacggaagg 10860
ataaacagga cgccggtgcg ggcacggcgg gcagggatcg tatgataaaa cctgctgccc 10920
gggaagaaaa tctggaaggc ttccgtaatg ccgctgtacc agtgttcccg gtatttccgg 10980
ttcttcttgc tcattgtccg ccacctcttt tcaccaggta atcccgtccg ggtactgtta 11040
cgcttccggc cctgtaatcc tttgtgctgt agtttacata aacctgtgtg ccgtccgcgt 11100
atgtggtcac cgcaacatcc tcattcagtc tctcatgccc ggtaattgcc tggcggttca 11160
gcccggacat ttccttctgg taccgggtga tcatcgcgat gacttcatct ttccagggtg 11220
cgtaatggga tgatgtatag cagctgaaca gagtatcctt cagcaccctc gtatccttct 11280
ccataaaggt gaagttcagt ccggcgccgt attctgcgga ttccagcagc agcgtctggg 11340
ggtcaccggc catgttcagc gcctctcccg tataatcctt caggccgtgt atggcaatct 11400
ggtagaaggg aacgctgtaa tccagcaggg cataggcatt gccgctcagg ttcatgtccg 11460
tgatcaggtc cgcgtaagga atcgcgtatg cgtttccttc cttgatgatg accttcatgc 11520
ccgttgccac agcgtcttcc agtgtctgag tatcctgggc cttggcctgt tcgcgcgtca 11580
ccgtattgct gtcctggtag tctgcactga gcaggttccc gatatcacgg aacgcaatgc 11640
cggctgcccc tcggtccttc aggccgttga tcaggttgga cgcgcaccgc tgggcatagg 11700
atggcttcac caggtagtac gggtacttgt ccatccaggt ggacatttca taagtgacaa 11760

tataataggg atacagtttg atcaaatccc gtgtcgttgc ccgtgctgaa tcagacagtg 11820
tgttgaatcc gttcaggatg ccgctgcgat aggcaaaaca ggaaatgccg tcaaagtaca 11880
ggtccacgcc cttttccgcg gcgctggcaa tcagcttctt catgccgctc tcaccgccca 11940
ggcagccttc cacatgaatg ctggtgagca ccctctgcat ggcgccgccg ttgcaccagc 12000
ccgtcatccg gaggttcagg tccttgacgc cgccctcgag gaagtcgttc atgatgctgt 12060
cggcctgctc aaaagtcgtg gtggctacag tggcctccac cggtatgccc agtttgacct 12120
caaccttgtt tattgcgccg accagttcca cgtttaccgg catttcttca gacgcgcatt 12180
caccccgcat ttcaggtttg gcgcgcaggt attcgccgta ggtctgcgcc atgctgacgt 12240
atccgttgcc gtccaggaag cggtaacgct ggaccagcgt atcatccggg atctgctttt 12300
cgtacatcag gaacaggcgc gcagacttat tctcatccac ttccacattg tcggaatgca 12360
ggatattgta tcttccgaaa accgtgttgt actcgttcag ccgtccggaa atatcggcgc 12420
acacgccgcc gtacgagtct gcgctttcca taatgcagat gaaggaaccg tcctcctggc 12480
tcatgccgaa gacagggaat gcgttttcgg tttcactggt agcggtttcc cgcttcgtac 12540
catagtccca gccgtacata ttggcatagt aggaactctg ggaaagcctt ccgttgttat 12600
agcggataat ggctccgccg ccttccggca ccagggtaaa tccttcctgg tccgttccag 12660
cggcgccgaa catcggcagc acgtccacaa aaaccagcgg atagtccgag gaacaggcga 12720
tatcgctgta cggcacggat accaccaggt ccttgccttc cagccggtac aggacgctga 12780
tgttgaacac cgggccggtg ctgtcctttt cgccggccat cagttcctgg tccttctggt 12840
aatcttcctc ggtatatccg gctttttca ggaggtcctc cgcctgtttt ttctgctcgt 12900
ctttccggtc tgatttcagg atgtacagat cctgctcctc aaggctggga tacaaggcaa 12960
gcagttcatc ctttttatcc ttgtttttcc cgttcaggtc ctttgtctta tacagcgaat 13020
agtggttctt gaatttttc gcgtccttgg atttctcccc gcccatcttc tccatgatct 13080
cgctgtatct ttccatggtc atggcattgg gaataatata ccgcttaccg gcatttttac 13140
ccagggagta gttaacccgt atcgaaccgt cctcttcctg caggatctca aagctctggt 13200
ttttgatgga tcctgtaaag ttgttcattt ccgtcttggt gatatcattg taataagaca 13260
cattcagtgt gctggacagg tattccagcg tggtgccccg cgcgatcttg tcattctcgc 13320
ggttctccgg gttggaaaac catacсttgc ccgtatcctt gttggtcacc ttgaactggc 13380
tggtcgcccc gtccatttca aagagcagtt cctcattttc catggtcagc ggcttcccgt 13440
cgccttcaaa atagataatc tttgtttccc tggcaaagct ctttcccgga ctggtataaa 13500
ttttgttaag cacaaacgcc gccccggcga tcagtacaat tgcaatgatc cacggaatca 13560
gccgcctcag gatggatttc tttctcatcg tcttcatccc tccttaattc agccggtaag 13620
cagcttcacg gtacaggctc acaaagaaac cgatcgcgtc gctcagcagg ctcaggcaaa 13680
caagggccag gaaaatgatt acacaggcgc ctacgattgt gacaataatg aagataaagg 13740
tttttcccgg gccgtagtca tgaatctcca tcaaaccgac aaacacgagg aaaacgcacc 13800
agaccacact gatcgaaagc atgacgctat agaggctccc ctcttcatac gtcagcatgt 13860
tgctgaccag gatcatcgga agctgaatca ggatataggg caccagcgca tagcacatac 13920
cgatgtagat atccttgaaa cggccttttc cttcaaacag agtggtcatc gcccagttgg 13980
ccaggcaaag gatcaggaac gggagcagca gtgaagccat ctgctcaaaa acattgatat 14040
gctgtactgc cgcgtcaaag aactggaaac tggtgcacag gattttcagc acacgggtga 14100
tcaggaaaag agccatgaac gtattggcgg cagccaggct tcccttatgc tcacggttca 14160
ggtcccagaa tccgtcaaag ggtgagtaa tcacataaag cgcgtatttc agggaacgct 14220
tatattctgc ccaggatgtt ttcccgccgc tgacgggttc ggcattaatc agcgtttctg 14280
cggattttgc tttgctcatg catgatcacc tcccacttca ttctcttgat tcttccgagg 14340
atcagcggca ctatcagcag cagcaccact ccgaggaaaa tccacagagc gttgttttca 14400
atccattcct tccggtacag tttgaatgcc ctgccgtagt tttcccggtc atgcgccatc 14460
ttgaagtatt ccatggcttc ttcgtacttg ttctgtctca gtatcgtacg gccgatgccg 14520
cggaaagcca tgggataatt ggcgttttgc ttcagcacat cctgccagac ttccgcactt 14580
tcgtcgtaat cgccgtcctg gtatttattg atggcgtcgt aaattttcag gccgtattcc 14640
gtcggtgtaa agactgtaat gctgttttcc agctggtcca gcacgaacag gtctgttccg 14700
atgtgctcca ggctgatagg gctggtaaaa gcgccctcaa tattgccgcg ggtaccgaag 14760
gcccacagca taatgccctg gttgtcataa ccaaacaggc gtccgcggat ccggtccagg 14820
acgatatata tatcattgtc aaaaacggta atgtcagtca tccttgccgg gccgaagtag 14880
gcagattcac ccgctctcca ccacaggtcg ccgatgggag catagcggtc gtttgcgatc 14940
aggatatcat ttcccaggcc gttcagcctg cggatcggtt tgacctgccc cgcgtccagg 15000

gcgccttctt cgaagtttgt cagcgtggta tagatgaatc cgtccttatc catgtaaagg 15060
ttttcatatt ccgtcggaac aaaggatgtg gaagcgtccc gctgttcctg ggtcatgaaa 15120
tactttttcc agatgtattc aaaggtcgag aactgtacag gcgtagcgcc gatgaatccg 15180
gtaaacgtgg tgtctgcctc atacttcaca atacctttgt ttatgtttgt aaccagggca 15240
tacaagcgtc ccgcaacgtc caccgtaatc ttctttggca ggaaatccag gctctggtca 15300
aaggttgcgt ccgcaggttt cgtgaattcc ttgatccagt tcaggttttt atccatcata 15360
accacgcgca ggtttccgta atccgcaacg tagatgtttc cgtccgcgtc cacatatacg 15420
tcgtacggag tattgaacgt ctcaggcgtc ccccctgtca catgatcaat gatccttttg 15480
agtttcgcct ggccgttcct gtattcaagc tgaaggatcc ggttgttgaa cgtgtctgca 15540
acgtacaggt cgttgttccg cgcaaacagg ctctggggct tgttcatacg cacgtccagg 15600
ccgaggtcaa cgctgtcgat gaccgtctca acccggtagg cgtccgggga ttcctggacg 15660
tcagaccagt aatcataccc gtaggtatag gatgtgctgt agccatcctt gtttacgttg 15720
aagctgtcgt ccgccagcgc agtgcatgcc gtgaacagca gcatcaggat catcagaaag 15780
gaaaccgctt ttttcatcat catttcctcc cctcttagtc cttgatgccg gaggaggcca 15840
tggtctccag gatccggctc tggctgaaaa tgaagatgag gaccggcacc gcgacgataa 15900
tcacattggc tgcggccacc tggccagtac gggcaatgtt gaccatgaca ccgccctgga 15960
tctgctgcag cgcgtacacc agcggcttct tcgcctcgga ataaatgatg gttcctgccg 16020
ggttgttcca cagggactgc acgctgaaga tgatcatcgt cagccaggca ggcttcacgt 16080
tgggcatcac gatgctccag aatactcgga aacggccggc gccgtcgatg gtcgccgcct 16140
cgatcagggc ggtgggcagg ccttccataa actgcttcat caggaacagg ccgatcggtg 16200
ctgcaaaggc cggcaggatc agcgcccagt aggtatccac caggcacagt cgtctcataa 16260
tcaggtagtt agggatgtcc gtaaccacgc cggaagcgaa catcaggaag gtggtagcca 16320
gtacgaagaa cagcctgccg cccgggaaat catacttggc cagcacgaaa gctgccatgc 16380
cggcgatcac cagatggccg aaggtgccga ctacggtaat gaatactgtg ttcaccaggt 16440
accggctgaa gggaacccag ctctggccca tcgttacata cagatcagag aaattgtcca 16500
ttgtcggacg ggtgggccat acgggcggcg ggaaccggaa cagctcgtcc aggggtttca 16560
ggctggaacc cagcaggaaa atcaggggga acagcatggc gaccgcaacc agcgccagga 16620
tgacgtaaat gataacatct cccgcaatgg accggttggg atgcctccgg gccatcagat 16680
gtttcttcgg cacctggatc ttctcaccca gccgttctgc ctgctcatgc aggatatcac 16740
cccgggtctt gaccttcaca ggtttgtccg cccagttttt gtccatgtgg ctgtaaagct 16800
tcaggccctg ggccagttca aatacatagc gggtgtcatc cttcagaata acggtgatct 16860
ggctggtggt accgccccgc cggatgttgt ccatccggtt catccgggca taatgccact 16920
cagcgatggg ctccttctgt ccttttgcgt agaaagccgc tcctttttca ccggcgacca 16980
gttcaccgcg gtactttta ccgccgatca gctgcagggt accgttggat tttacggtgt 17040
tcatctcact catatcaggt acccaccctt cccagcaatg cggtgattgc tttcttgcaa 17100
agaatcatgg ccacgaacag gattaccgca atggcgctgg catagcccat ctcgaagcgt 17160
gtatagccat agtcgaacag gtgggtaaca atcgtgcggg ccgcatagtc cgtggatggg 17220
aatccgcaca gagcccgcgg cacggcgccg acgttgaagg aggaagtaat gctcataacc 17280
gcgccgaaca gcagcatggg cttcatgctc ggcagggtga tatgatacag ttcctgccag 17340
cggttccgga cgccgtccac gtatccggct tcatacatgc tccggtcgat gccctggaaa 17400
ccggccacaa aactcaggaa gccggtaccc atgctcatcc acaggctgac gatgatgacg 17460
atcgtcagca tataccgggg atctgtcagc cacaggaccg gggcattaat catgccgtat 17520
tccatcagca gggcgttaaa ccaaccgtaa gcgtcgccgc ggaaaatcgt gttgaaaacc 17580
tcgaaggcgg cgccgccgat ggaagggatg taaaagacga taaccgccac tgcccgcagc 17640
catttcggca gctcgttgat aaaccaggca aacaggaagg aaaggatata tccaaccggc 17700
ccgataatca cggcgcacac gaatgtattc ttaatggatg tcaggaaaat ctcatcctgc 17760
aggaacaggt ttgtatagtt ctgcaggccg ataaaacggg caggctccag gatattataa 17820
taagtaaagc tgtagaaaat cgatgtgcag atcggcagaa tgaagaaggt aatgaaaagg 17880
atcgcgtagg gcgccaggaa caggtagcat cccttgttcc gctttgccag cgtccatccg 17940
tgtgcaatct tttccctccg cattgtcagg tatctccggg caaaagactg tgactgcatg 18000
tcagcttccc tccttccctt tctcagtcaa ccggcaggtt gaattcccgc cgtttgatct 18060
tgatttcttc gttgatggtg cgtgcatagg tcagcagtgt ctcacgctgg tcttccttgg 18120
tgttaatcac cttacggacc gcgttgatca gatggcgctg cgttgagtat ccgccggcaa 18180
tctccgggaa gcccttggtc tgtgccacct gtgtctgcag caccttcagc tgcttgtcag 18240

accatgccag ctgtttcagt gcatcctggt tggctgtctg gtaccgggcg gaagcaccaa 18300
ggatgctttc catttcacgg ccgaagcgga cctgtgcgtc cgcgccgacc caccacttca 18360
tgaattccca tgcgtttttc tttttcgttt cgttgtctga cgcaatcatc atacagcaca 18420
ggccgcttgt ctgggccgta tgatcaatcg ttccgtcctc tttcaccgtg cccgggaaca 18480
gggtaaagtc ccacaggccg cggatttccg gagcggaaac catgagggta ttatagacat 18540
tgtagttctg gatacccagc ggcatctcgc cgctgcggaa gcggctcggg aaatcgaata 18600
ccacgggcag gccgtaatcg ttatacaggc tcgtatactg cttgaacccc gccacgccgg 18660
cctcgctgtc caccagggtg tgcttggctt ccttatcgta gatttctccg ccgttctgat 18720
aaatcatcga gttcagcacg gagttgtcca caaccctgat gtccgggaaa ttgataccga 18780
ctgtcaggct gttgccctgg atcgtcggca gctccgcgat cagttcgtcc caggtttgcg 18840
gaattgccag tcccagttct tccatgacgt ccttgcggta gaacagcacg gcgaattcct 18900
gcgtctcagg caggccgtaa atgccgatct tcttgctgtc gttataagtc agcggttcaa 18960
gaatactttt tgtgaaaggt ttgaccactt cttcaaaatc ttcaaactgg gtcaggtcct 19020
ccaccgcgtt acgcatggca tagttcacgg caaaccatcc gctcaggctc agcacgacgt 19080
ccggtccgtt tccggccaca acagccggca ggatggcgtc ctgcaggacc agcttgatgt 19140
ttaccttgat tccggtcttc gccgtaaagg tgttgtcaac caggttcttc agcaccgtac 19200
tctggtcgcg gcctgtggtg atccagattt ccaccacgtc gtctgaatct tcatatttgt 19260
cgcccaggct gttatagtcc actgtatagg aactcaggca gctccggatc tcatgtccaa 19320
tactctggaa gatgttctca tgcacttcgg gcaccttgac gttctcgcct gtcaccatga 19380
tcaggtccac gtccagcttg ctctcagcca tattctgcat tgccgtgccg agggaggtaa 19440
tgttatcccg gaaattctgg aaagattcag taatccgttc attcttttta acgaaagtct 19500
caagctgtac tgccagcgtc tgcgcggaag caacccggtc gctcttttcc ccggtgattg 19560
ccaccgtatc atccacaatc ttgtacaggc gcttgctttc caggtccatc gcctcgataa 19620
cctcaggata aacccgcttc agattataat cgcggtactt gtccggactg gcgcctgtca 19680
gcaccaggat cttacggtag atctgattca ggcggaagat gctgtcttcc acggatttca 19740
gcacaggtcc catcttaccc atggtcactt ccatgcggat ggtatggctg ccttttgtga 19800
gccagaaacg gtaaggattg ccgtccttgt cacccagggt acgcatatcc catccggtgc 19860
tgtaactgaa ggtgacggct tccagatcgg taaagggaat ctccccgtca atatatacgg 19920
atcgtgcgga cagggcgccg cgctggtaca tctgccgggc tttgacggag atgttgtagt 19980
atccgtcttc cggcacttca aaatcccact ggatccattc acccggatga gtccatggct 20040
cgccgccgat ataattcagg attgaatttg atacagaata aggcacagtc tgggaggaac 20100
tgcggtcata gcgtccgtaa aggctcggcg tgctgcgcag cgtggcggcc tctccctgga 20160
tcgtctggca gtagttcagg ccttcttcac tcatggcggc ttcaggctgc gccgccgtgt 20220
attcctcgta agtcggagac ttcaggatgg gtgtcagtga aatggaacac agaattacag 20280
gctcgttaac agcccggata gtcagctcat tctcaccctc attgaaatag aaagcatagg 20340
gttccgtctg gtatcccata tcatccttgc agtacgcgct ctgtttttcg aagcgctcct 20400
cctgcgtggg acggatatca tttccctggt tgtccttgcg cacttcggtc gcgtcggtcc 20460
acaggcggga gaaacaaagc gtgcttgcgc cggagaaagg cacttccccg ttgatcgcga 20520
tttcacgctc aatgtccaca ccgcggcttt ccgtcgtcag gtactccagg cggatattgt 20580
acaggccggc tttaggtacg tccaccttcc agctcagttc ggaattgtca gcagtcagca 20640
gaccgtcagc acaaacctcc ccgccgtttc cctcaaagga agcgagatca acctcaacgg 20700
cctcactgac tgcgggcaac tccgcgtact tttccagata ggatgtgtaa gtattgctcc 20760
gtccgatgcc gctgacatct gtggacagat cagcgccctc gtacttgctc ctgaagtttt 20820
cagagttccg gctgagaagc cagattccgc caatcacaac aacagccacg aggacgccga 20880
tgagaatgtt ccgacatttc cttgacatga gttcatctcc cacttgcgac gtttgcacgt 20940
cagaaaaatc agatccctga tactgtgaaa ttggtaagaa ttacactaaa aaaccgtaaa 21000
taattatatc ggttatatca tattgtgtca attgtacgta caaaaaagcg acgcccgctt 21060
gcgtcgcttt ccggtaatgt tttcagtgtt gaaaaatctg gtttatcgaa tttcagctcc 21120
gagtttgaac ttcagattcc caagaatttt ttttatgtat ttatcaagtt cttcaggcgt 21180
caggggcttg tcatccgcct caaaggtaat tttgaaagcc atactcttac gtccctcacc 21240
gatctgctcg cccctgtaga tgtcgaacag ctccacatcg ctgacccgcg ggcaggcctt 21300
ggcaatttcc ttcatcaggt ctccgcaggc tgtttcctcc gccactgtca gggccaggtc 21360
gcgcaccacg gacgggaaca cggacagcgg acggtaccgg aagctggcag ccatgtggtt 21420
catcatggca ttgtaatcaa tttcgcccag gaagatcttg tgattggact tggcgtcctt 21480

gtgcagcttc agctcaccgg tcacttcatt ggccagcttg ccgaaacagc cgatcctttc 21540
tccttcacaa aggatatagg cggcgattcc gggatgcagc cacggcacgt cggaagcccg 21600
ttccacatca aaggacagtc cgagggcttc gcccagcgct tccactgttc ctttcacagt 21660
gaagaaatct tctgcgtcgc cgaaggctgc gaaaccaaga tgcggccttt catccggcag 21720
ctctccgtcc cgcggaatat agatattgct gatctcgaaa atccggccct cgttgttgcc 21780
cttcttcagg ttctccacca caacgttcac caggctgggt gccagcaggg gcctcatgat 21840
ggacaggttg acggaaatcg ggtttttgat gcgcagcaca ttccgctccg gcgctccttc 21900
cggaatccgc aggaaatcca ggtcgctatc ggaatagaag gacatcgtgt aggcttcatg 21960
gtatccctgt ccgcacagga tccggcaaac ccggtccctg cgctgctgcg caggagtctt 22020
accgccgctg gtcacggtcg ccgccttcag gaaggtgggg atcacgtgat cataaccgta 22080
ctcgcggatc acttcctccg ccagatccgg ttcgccgact tcaatatcct cgcggtaacg 22140
cggagcagtc acgtccagcg tatcgccgtc ccgctttact tcaaagttca ggttcttcag 22200
gatccgcagc acttcctccg ccggcacctc aataccgagg atcgcgttga tccgcttcag 22260
gctggcggta aaccgcttcc cttcccgcgg tgccccggcg ctgcagtcaa aggtgctgga 22320
ggtgatttca ccgcagccca gttcctggat aaggtgcagg gcgcgggcca tacccagctc 22380
cgtgatatat tcgctgatgc ctttctcaaa ccgggcggaa gagtcactgc tctgtcccag 22440
tgcacgggag gttttccgca cgttgtcccg ggcaaactta gccgcttcga aaagcagctg 22500
ggttgtgttt tccgtgattt cgctgttcag gccgcccatg atacccgcca gggccacggg 22560
gcggtttccg tcgcagatta ccagattgtc gggtgtcagg gtgaattcct tctcgtccag 22620
cgtttggatc ttctcgccgt cctttgcgcg gcggacaata atccggcggt cctgcagctg 22680
gtccatgtcg aaggcatgca tcggctggcc gatctccagc agcacatagt tcgtgatatc 22740
aaccacgttg gaaatgctgc gcagtccgca cagggccagg ttgcgcttca tccaccgggg 22800
gctctccccc acagtgatgt tccgtacgcc gtgtcccaga tagcgggggc acaggtccgg 22860
cgcttccacg gtaatatcca ggctggggaa aacataatcg ctgaccgtgt agtccgtcgc 22920
gggcattttc agttccttgc caagggcggc agcaacctca cgggcaatgc ccagaatgct 22980
ctggcagtcc ggacggtttg ctgtaatggt aacatcgaag atataatcat tcattccgac 23040
gacttcacgg atatccgttc cgggaaccgt atccttcggc agatccataa ggccgtagac 23100
ctccgcgccg ggatacaggt catcgttcag acccagttct tcaccggagc agagcatacc 23160
gtcggaagga ataccсttca ggggttttgc cttgatggtg attccgcctg gcagcgtgga 23220
attgtccagt gcggcaggcg tatgcatacc ggcgtataca ttgggagcgc cggtcacaat 23280
ctggatgggg ttgccgtagg aaccgcagtt cacctggcag atctgcaggt gtgtgccttc 23340
caccggttcg caggagagca cttcgcccac gaccacccgg ctgatctcct tccccaggtc 23400
gatcagctct tccacttcaa atccgcagcc gaacagcttc tcctggagct cttccgctgt 23460
gatatcaata tccacatatt ccttcagcca gctgtaaggt actttcatcg tttatctctc 23520
ctgctctgaa acatcaatca tgatcattat tcattcttaa gttctcagta ttcattattc 23580
attgcaattc aaagaatgtc ttattctttg aactgcttta agaatctcat atcgttctca 23640
aacagcagtc cgatattatt gatgccgtac ttcagcatgg tgatccgctc aataccgatg 23700
ccgaacgcga taccggagta tacgttcgga tcaatgccgc agttctccag caccttggga 23760
tgaaccacgc cgccgccgag cacctcgatc cagccggtat gcttgcacag gctgcatccc 23820
ttgccgccgc actcaaagca gctcacgtcc acttcaacgc tcggttcagt gaaggggaaa 23880
tagctggggc ggagccgggt acgcacgtct gcgtcaaaaa tctgctgcac aaacttgtcc 23940
agcataccct gcaggtcgca cagggttatg cccttgtcca ccaccaggcc ttccatctga 24000
tggaacatgg gagaatgggt cgcgtcgctg tcggaacgga aaacccgtcc gggaaccagc 24060
accttgatcg gcggcttctg catgtccatc acgcggatct ggccgccgct ggtctgcgta 24120
cgcagcagca gttcattttc caggtagaaa gtatcctgca tatcacggga aggatgatcc 24180
ttcggcacgt tcaggcgggt aaagttgtgg tcgtcatcct cgatttccgg tccttcgaaa 24240
atatcgaagc ccatgctggc aaaaacgttg atcatttcat tcttcaccag cgtcagggga 24300
tgcagtccgc ccagcggcag tccctttccg ggcagggtaa tatccacctg ttcccggcgg 24360
ttgcgggcag ccagctcaag ctcgtccatc tttgcgctca gggcctggta cttttcttcc 24420
accagcacct tgaactcatt gatgactttg cccatagccg gccggtcttc cttggccacg 24480
gcacccaggc ccttcatcag atcggcgatg gcacccttct tgcccagaaa ctcctgccag 24540
aattcagcca gcttttcctt tgtgtcgatc tgactgatct gctgttccat ctgttcctgc 24600
agtgccttga tcttctgctc catgtgctcc actcccgtgc aatcatgatt tgtctcatca 24660
aaacgatcac cggtgatcct gttatcacca attaccgtaa ttatgaatta tggattatac 24720

agcataaaat gtcaattctt tcttggtttt tatcagtatt tgcgcctcat atataacaga 24780
aggcggagct gttcagcccc gccctctgtt cattgtgcat tatgcattat tactgtgcgt 24840
tcttcaccag tgttttgatg tcctccacca ctgcccggat atcggtgttc gctgtcatgt 24900
tcgcttccac cgtcttacag ctgtgaagca ccgtcgtatg atcccggccg ccgaatacgg 24960
ttccgatctg gggcaggctc atgcctgtca tctcccgggt caggtacatg gcgatctgcc 25020
ggggcaccgt gatttcccgt ttccgggtag gaccggtcat gtcatccatg gacagtccgt 25080
aatagtcgct gacggtctgc atgatcaact cggaagtgat ctgcttatgc cgcttggaat 25140
caaaaatctc tttcagggcc cgctcgcaca ggtccacggt gatcggttcc tttaccagtg 25200
tggaataagc caccagccgg gtcaggcagc cttccagttc acggatgttg ctgtcaatct 25260
tgccggcaat cagctgcagt acgtcgtccg gcacttcgat attgtcctgt acggtcttct 25320
cgcgcaggat agccacgcgg gtatccacgt ccggacgctg gatatccgcc acaagacccc 25380
actcgaagcg gctgcacagc cgttcctcca ggcgttggat atccttcggc ggtttatcgc 25440
tggtcaggat aatctgcttg ttatcgttgt gcagttcatt gaaggtattg aagaactcct 25500
gctgggtact ctcacgtccg gcgatgaact ggatatcgtc caccatcagc acgtccacct 25560
tccggatctt ttcccggaat tcataggtct tcttcatctg gatcgcgctg atcagctcgt 25620
tggtaaagtt ctcactggtc atgtacagga ttttcttctt cggatcggac tgatggatga 25680
aatgtccgat cgcctgcatc aggtgtgtct tacccagacc cacgccgccg taaatgaaca 25740
gcgggttata ggcttcagcg gggctttccg ccactgccag ggccgcggca tgggcaaagc 25800
ggttgccgtt gccgaccacg aagctctcaa aggtatactt agggttgaga tggggatccg 25860
tatcatccgg ctttcccgaa tcttcatccg tttcctggcg ggccttcatc tcatccttgg 25920
tcagcagcac aatacggatg gccttgccgg ccacctcggt cagtttccgc tcgatcagat 25980
ccgcatactt gttctggatc atggagatca tgcgttccat tttcaccacg acaaacagag 26040
tatcgtcctc cagcattccg ggcaccatat tgccgtcaac ccacgtgctg taggacacgt 26100
agttcatttc ccgcgacagc agggcacacg tctgctccca gagcgactcc atgttcatca 26160
atacattctc cattcatctg gttattaaaa cgcaaaagcg atgatttgct gcatatgctg 26220
caaaaggcgg cgaaaaaagc gcacgaggga tcgtgagctg tggataaccg ggctgtggaa 26280
aactggcctt ttgcaaaccg gctcgactat catagcaaaa acattgagta atttcaaggt 26340
tcttaccctg tggataaaac gattaaaatg ttagaattct ttcatttttt ctgtcaaggt 26400
tttgtatttg ttccggatga ctccactaga tttatggtat aggccgtttt tttgcatcgt 26460
ttttttcctg ttttttgtat tcgacgacaa aatcaccgtg aaaagttaac aattacttca 26520
ttttatgata tcatacagta tacattatat aaggaaacta ttacggaggg atgggcatga 26580
tcgaagcctg gcatgtgagg gaagccctgc gtttccgctt cgggtctgcc cttgtggatc 26640
gtcccgggat ctgggagaag gctgccgcgc ttttgcggaa cttcgcgcct taccgggata 26700
ccttctccga tcttgccgcc agccttgagg atgtactctt caacacggtc tatgagcagc 26760
tcggcccctc catgggggca cagatggata acggtactgt acgccggatc cgttcaaccg 26820
agttcaggga cgcctccgat gatgtgatgg gcgttctttt tgatcatctg aaggtctatt 26880
cggtcaccta tgactccctg catcagtact gcatggccac cggctccttc gccgcgatgc 26940
gcgtgctcta tacccgctat gccgacttta tgcccgcctc ggagcggaaa atcatcgcca 27000
ggatcatccg cgacagccgc ccccgtgacg tatgggaacc ctggatggat ccggaggatg 27060
ttccgccggc accgtcacgt taatgtacaa tgtacaatgc ataaagcata gtgatgaatt 27120
ctgaattgaa atgaggttct ccatgaaaaa aatccttgcc cttctgcttg tccttgcgct 27180
tttccttccg gccgttgcca ctgccgagac gattgtcacc agcttttatc cgatctggat 27240
tatgaccctg aacctgactg aaggaatcga aaccgtcaaa gtcgtgaacc tggcggagcc 27300
gacaacagga tgcctgcatg attataccct gcagaactcc gacatggtca ccctttcaca 27360
ggcggatgcc ctgctggtca acggcgccgg aatggaatcc ttccttccgg tcgttaccgg 27420
agcctatccc aacctgccgg tcattgacgc gaccgtcggc cttcctttcc tccatgaatc 27480
agacaccgtg gaaatcggcg aagccgaaga aggtgaagaa gtgaattccc atctctggct 27540
ggatccccag cgggccgccg gcatggccgc caacctggcg gaagggctga tcagcctgat 27600
gccggacagg gaggcggaaa tcctggaaaa cctgcacgcg tatcaggaac gccttcaggc 27660
cctggatgaa accatccggg aagaaaccaa agacgtggac aggaaagtca ttattttcca 27720
tgaggctttc ccctactttg cggaagcctg cggtctgact gccgcggcta ttgtcaataa 27780
ggaaccggaa gacatcctgc cgaccgcgca gcttgtccgg gtaacggaac tcatcttcaa 27840
caccggcatc ccgatgccgc tgatcctgaa atcagctgaa gaggacccct ccgtcaatac 27900
cctggtcaat gagaccggca ttcccgtctg cgaactggat cccatgacca gcggtcctga 27960

aaacccgccc ctggattact atgaatccat catgatgcag aacgtcaaaa ccctgttaaa 28020
cgctatagct cagtaaaact ctttataatt ctgaattctt aattctgaat tctgacttaa 28080
gaaaggatca tctatgttca gcgtaagtcc ggaaatctac tcggttcttt cccaggcttc 28140
acgctggatt tttattctcc ttgcgctcat gctcctgttc ctggcctttt cctggcatca 28200
tacagacagg aaggagcgcc gggaccggtt caaaaatctt ccgggcgccg gaaccgtcgg 28260
tgaactggtt gttctctccg gcagtgatga actgcctgtg gatacctggt tccccgttcc 28320
ccgggaaggc gtgctcggtt cactgcgctc ctgcgatctg gttatcccct gccccggcgt 28380
cagctcccac catctggatt tctcctggca ggacgggaca ggtctcctga tccgtccccg 28440
ggttggctgc gaagtcctga tcgatggagt acccgtcacg cgcaggggca tggatatgcc 28500
gctggttcat ggatccgtgc tgcagatcgg cagtgccgta cttcggctgc agctttttgc 28560
tgccctttct cataccaacc ggacctttat tccccccgcag cctgccgtcg gctccttctc 28620
tgatccgatg ccgcagcctg ttccggttcc tgtgcagtat ccgcaggagc ccgtgaatgc 28680
gcagccagtt ttctgcccgc ctccttatca ggagccggtt ccgcttcaga ttcccgtcga 28740
gccgcccatg ccggttcaac ccgttccgca gccgcctgtc ccggaacagc tgcccgtaca 28800
ggaaggatcg cccctggcgc ctgctgcccc gcgtcgtaaa cgcgctgatc gctggaagga 28860
ggactggagc gaatgacaaa acagaaatcc tctgcccgtt tccgcttcag cggcgttatg 28920
ctgttccttg ccctttttt cttcctggtc ccggcgcttc aggcaggaga ttaccggctc 28980
tatatgctgg ccgcaggcgt tccctgcgcc atgctgttgt gctgcaccat cctggcccgg 29040
gcgttctccc tggacaggct gatcctggct ttatccctct tcttctgttc tgtggggatt 29100
gcgtccctgg ccatgactgc gccggatact gctctggttc aggcggtcag ctgcggcatc 29160
gggattattg tcctgatcgt cggagccatc ctgatccgtt ccctctccgg ttccctgctc 29220
accgcgatct ccgctgcttt cctcggcctg cttctgttgg ccgcaaagct gattgcgccc 29280
actttcaccc tgccggccac gcagcctgcg ctgaccctgc tgctgatcgc cttcgccgtt 29340
ctcctttcac gggaaggtcc catctccgcg gcggggcttg gaattgccgc agccgtcctg 29400
ctgatgcttc ggggtgaaac agcggatgcc ctgatctggg gtcttaccat cctgctgctg 29460
ctcttcgccg ctgacgggcg cctgctggtc gtcctgcctt ccctggcggc ctttgcgctg 29520
ctgttctacg gcatgtactc catgcagacg caaacgcttg tttatccgga agcgttatcc 29580
gcggataccc tgaaagctgt cggcgccctc ggcgcggatc tgcttccgga gggactggcc 29640
gttccggaga aaggttccct gttctcccgc ctgcttggcc attacgggct ggttttcacc 29700
ggattgacgg tgctgcttta tcttccgctc atcctgcggg gaaccacggt ggcttcctac 29760
gcccgcatgc gtttccatgc gattctcgct atgggaatct gcctgctttt gggattgtct 29820
acgctggcct ccgtattgtc cttcttcggt ttcctgcctt ttaccgggct ggaaatgccg 29880
cttttgacaa cgagcctgcc ctcgctctgc gcgcacctct tcctcgccgg cattctctgc 29940
ggcatatccg gccgcaatga ttccgacctg gcagaggatg cccatctggc catgctggcc 30000
aagtaacctc ctgtactgaa aggattatcg tatgaaacag atgcgttccc gtttccgcct 30060
gcttacgctg ctgattgtct gcgccttttt gctgacgatt gtgctgtgta cgggacacgt 30120
tctgaaaacg accggaatct ccctttcttc cctctcctcc atctcacttc cggttcccgg 30180
tcagtctgcg tccccggatc cttccgcatt gccggatcct tctcccgctc cggaaagcac 30240
acccgatccc tgggcaggtt tttccgtcga aaacacgccc ccggggactg acaatccccc 30300
tgaagaagag tataatgtag tgggcctctg attccccgtg aaaggaatga attaaccaat 30360
gattcagttt aaacgcgccc tggcgcttat gcttgttttg atcctctgct gctcctcttc 30420
cctggctgcg aagaagaaga ccacggaata cacagcccgg gagatcacca cggaaattgt 30480
ccaggagatc cctgctgaaa tccagaatat gctggacatc gcctatgagc agctgattga 30540
aaccgatggc aaaaacctga aggaaaagaa taaattcacc aaatggcgga acaactacac 30600
ctttgagtgg tgcggcggtt tcgtgacctg gtgcatgctg gaagccggca tccccatgga 30660
ggaaaagaac aaaatccagg acggcgaggt tgaaggcctg ttccatgtaa aggaagccgg 30720
cgtcggcaag ctctatgaag gctatgccaa actgaaccgg attacccgcg ttccccagaa 30780
gggctatatt gccgtctacg gaaatgaagg ttccggcggc tccacgcctt attatcatgt 30840
aggcctcgtt tacgatgtgg agaagctttc cgacggcaaa taccggatca ccaccatcga 30900
aggcaacgta aaaggacata ccgtcaaaat gtatgtccgc gattatgatc tgaatctggc 30960
ttctgacaaa aagcagaagc ccaaggatat gtcccttgtt cccgaggaag aacgcaaccg 31020
ggaggaaacc aacattttct cctacggcta tgcttacacc aagaagcata tgtacatcac 31080
catcttcctc atgccctgga tcccggaaga atctcaggag tcaacggagg aataatccat 31140
gctgtggatg tacgctgtat gcgctctgtc catctttgtc tgccttccct tctttctgta 31200

ttacaaaaaa gcgctgcaca tgaagcttgc agctgctttt aaatcgcttg gaaccctctg 31260
tgcggcaagt ctggcattga ctgccgccat tcgtctggat ccacagtact gggtctgttt 31320
tgcggccctg ttccttcatg ccgttgcaga ctggattctg gaattcaaca tgtatatcgg 31380
gaccggattc tttattgccg gtcatatctg ttatatcggt ttctttaccc atctctgccc 31440
ggtcagcggg attcatctga ttgcagccct ctgcctgctg ggaatcttgg ttttcctcct 31500
ctggcgctgg cggaagccca tcggcaaaac tataccgctc atcgcggttt acggcacgat 31560
cctgagcctt gcctctgcct gtgccatcgc ctgtttctcc ctcaacagcc tccagggcca 31620
gttcatcgcc gtcggcggcg ccctcttctt catcagcgac gtgatcattc tcgcccgcct 31680
gctcttcact gccggccgtt ccgtcgactg ggccatcatg atcacctact acgccgctca 31740
gctcctcttc ggcctctcct gccttcttta atgcaaataa tcacaatata ttatatcgga 31800
acggcctctt gatttttgaa tcatgatccg tatttctgtc aagccttcgc tatcgccatt 31860
tcatgggctt gacagcaaat cccggatcat gagtgacggt agacaagagg ccgttccctt 31920
tttatgttcc gtttatcaat gtacttgtca tcggtataag gtattaaagt atccttgctc 31980
tgatgcctat gggcatcaga gctctgtgtt cggtatggaa caggggagcg aaggaagcgg 32040
tgacccagtg ggtcatttgc cgtaaggcaa aagcgccctg agcgagtcaa ccgaaacgag 32100
cagagggctg aagcgccccc tcgatgccct ctgcgacgat tgaggttgcg gggggacctc 32160
ttcttctccc ctctcaccgc ctgtttttct aaatccattt ggctacaaag tattttagca 32220
gttgcgtaag caactgccta tttggaatcc ggggagtgat ccgagtggtg ccccagtggg 32280
gcattcgcca caggcgaaaa cgaggtgagc gagtcaatcg aaacgagcag aggaccgaag 32340
cgtccccgcg aggttctctg cgacgattga gattgcgggg gtgcgccgcc gagattccac 32400
agaaaaaacc gcctgtatta aacaagcggt ttttctctgt ggaatccggc ggcgacctac 32460
tctcccgggc cgttaccagc caagtaccat cggcactgaa gagcttaact tctgtgttcg 32520
gtatgggaac aggtgggacc tcttcgtcat tgccaccgaa aatggttcag tgtccttgcg 32580
cacactgaca actgcacagc tacaatattc aggaattatc tgacctgtta gtttatctgt 32640
taccttaatt ggtctctgga ccaatcatga aatcaagccc tcgacctatt agtatcatca 32700
agctccatgc gttaccgcac ttccaccgat gacctatcta cctggtagtc ctccaggggt 32760
cttacttgct tgcgcaatgg gagtcttaat cttgaggtgg gcttcacgct tagatgcctt 32820
cagcgtttat cccgtccgca tttcgcttcc ctgctatgcc gttggcacga caacagttgc 32880
accagtgatg cgtccattcc ggtcctctcg tactaggaac agctcctctc atgactccta 32940
cgcccacgat ggatagggac cgaactgtct cacgacgttc tgaacccagc tcgcgtgccg 33000
ctttaattgg cgaacagcca aacccttggg acctgcttca gccccaggat gcgacgagcc 33060
gacatcgagg tgccaaacct ccccgtcgat gtggactctt gggggagatc agcctgttat 33120
ccccgaggta gcttttatcc gttaagcgac ggctttttcca ctcaataccg ccggatcact 33180
aagcccgact ttcgtctctg ctcgaattgt cactctcgca gtcaggcacc cttctgcctt 33240
tgcactcttc gaatgatttc catccattct gagggtacct ttgggcgcct ccgttactct 33300
ttcggaggcg accgccccag tcaaactgtc cacctgacac tgtccattac ccggtttacg 33360
ggtccatgtt agaattccag taacagaaga gtggtatccc aacagcggct ccgctaagac 33420
cgaagtccta acatcctagc ctcccaccta tcctgtgcat ccgttaccga aatccaatat 33480
caggctacag taaagctcta cggggtcttt ccgtccagtc gcgggtaatg ggcatcttca 33540
cccatacttc aatttcaccg ggcccccctgt tgagacagcg ctcaaatcgt tacaccattc 33600
gtgcgggtcg gaacttaccc gacaaggaat ttcgctacct taggaccgtt atagttacgg 33660
ccgccgttta ctggggcttc ggttcgatgc ttcgattgct ctaacatgtt cccttaacct 33720
tccagcaccg ggcaggtgtc agcacctata cgtcagctct cgctttcgca gatacctgtg 33780
tttttgctaa acagtcgctt gagcctattc tctgcggcct ctctcgaggc accccttatc 33840
ccgaagttac ggggtcattt tgccgagttc cttaacaagg gttctcccgt tcgtctcagg 33900
attctctcct cgcccacctg cgtcggtttc cggtacgggt accttcagat atactagcag 33960
cttttctcgc cagcgtgaac tcatctgctt ccctacttga tttcggtcca cgtcacacct 34020
cagcttaaag gaacgcgtac ttcactacgt tccaacctca gtgcttgtac ggagttgacc 34080
atcaccccgc tcagactatc cttctgtgtc actgcttcat tcttcggtag tgcaggaata 34140
tacacctgct gtccatcgcc tacgacttcc gtcctcggct taggccccga cttaccctgg 34200
gtggatgaac cttccccagg aaaccttggg ctttcgacgg cgaagtttct cgcttcgctc 34260
tcgctactca taccggcatt ctctcttctg tgcggtccac cgtgcctctc gatacggctt 34320
cagcccccac agaacgctcc tctaccacgc gcttaaagcg catccatcgc ttcggtgtca 34380
cgttttagcc ccggacattt tcggcgcaca accactcgac cagtgagcta ttacgcactc 34440

```
        tttaaatgtg tggctgcttc tgagccaaca tcctggttgt ctgtgcaatt ccacatcctt     34500
        ctccacttaa cgtgtacttt gggaccttag cggttgatct ggggtgttcc ccttttgccc     34560
        gcggaactta tctctcgcgg actgactccc atggtttgca ttgtatggta ttcgcagttt     34620
        gatagagttt ggtaggattt gagtcccccg cgcccattca gtgctctacc cccatcaatt     34680
        atccatgagg ctagccctaa agctatttcg aggagaacca gctatctccg ggttcgattg     34740
        gaatttctcc cctatccaca gctcatcccc gccttttca acagacgtct ggttcggtcc     34800
        tccatgggat tttactccca cttcaacctg gccatggata ggtcacccgg tttcgggtct     34860
        acgtcatgca actatcgccc tattcaaact cgctttcgct tcggctccgg accttcagtc     34920
        cttaacctcg ctgcataacg taactcgccg gtccgttcta caaaaagtac gagatcgtac     34980
        atatcgtgta ctttctctgc ttgtaaacac agggtttcag gttctctttc actcccctcc     35040
        cggggttctt ttcacctttc cctcacggta ctatgcgcta tcggtcacca tatcgtattt     35100
        aggcttggat ggtggtccac cctgcttccc gccgggtttc acgtgccctg cggtactctg     35160
        gtatcagtta gctggttgac tctttcgctt acggggctgt taccctcttt ggctgagctt     35220
        tccatctctc ttcggctaaa gtcttccata cacgttactg acccacaacc ccggggagca     35280
        tgccccccgg tttggcctct ttcccgttcg ctcgccgcta cttaggaaat ctttaaatta     35340
        atttctcttc ctgcgggtac ttagatgttt cagttccccg cgtgccctcc tgtaagacta     35400
        tggattcatc ttacggtgac aggttcttca cctgccgggt ttccccattc ggatgtctac     35460
        ggatcgatgc ctgcttgcgg ctccccgtag cttatcgcag cttgccacgt ccttcgtcgg     35520
        cgtatggtgc cagggcatcc accctgtgct ctttgtagct tgatttctcg tcgttcttga     35580
        tcctgagacc aattctagta aaattgtttt acctttaact ggtctttcct gaatcttgtt     35640
        tctttgctgt gcagttgtca aggtgcgcgt gcctcttctc aaagtgagaa agagtgtggg     35700
        tcgcagtgga ctcgaaccac cgacctcccg cttatcaggc gggtgctcta accgactgag     35760
        ctagcgaccc atgtgagtgg agattagcgg gatcgaaccg ctgacctcct gcttgcaagg     35820
        caggcgctct cccagctgag ctaaaccccc acataatcgg cactattcat ttaagctcgg     35880
        agcttcgcga gccctgaaga cgatacagag ttttgcgcgc tgtaaggaac agctcaattc     35940
        tttttcgaac gttcgttgac gttccgtttt gcttctttga agctttttaa tgacgaccgg     36000
        taatccggtc gatcgacctc ggtttcggcg tccacctttc ggcttctgct ttctccctag     36060
        aaaggaggtg atccagccgc accttccgat acggctacct tgttacgact tcaccccagt     36120
        cattggtctt gccttaggcg gctggctcct tgcggttacc tcaccgactt tgggcactcc     36180
        caactcccat ggtgtgacgg gcggtgtgta caaggcccgg gaacgtattc accgcggcat     36240
        gctgatccgc gattactagc aactccgact tcatgtgggc gggttgcagc ccacaatccg     36300
        aactgggacc actttttga gatccgctct cccttacggg ttcgctcctc tctgtggtgg     36360
        ccattgtagc acgtgtgtag cccaggtcat aaggggcatg atgatttgac gtcgtcccca     36420
        ccttcctccg agttgtcccc ggcagtctct tcagagtcct cggcattacc cgttagtaac     36480
        tgaaaataag ggttgcgctc gttgcgggac ttaacccaac atctcacgac acgagctgac     36540
        gacaaccatg caccacctgt ctcagtatga gcaagctcac gatgtatctc tacaagtttt     36600
        actggatgtc aagacctggt aaggttcttc gcgttgcgtc gaattaaacc acatgctccg     36660
        ctgcttgtgc gggcccccgt caattccttt gagtttcaac cttgcggccg tactccccag     36720
        gcggaatgct tattgtgtta actgcggcac agaaggggtc gatacccccct acacctagca     36780
        ttcatcgttt acagtgtgga ctaccagggt atctaatcct gtttgctccc cacactttcg     36840
        cgcctcagcg tcagttacag tccagtaagt cgccttcgcc actggtgttc ctcccaatct     36900
        ctacgcattt caccgctaca ctgggaattc cacttacctc tcctgcactc aagcccgaca     36960
        gtatccaaag caattcccac cttaaaagca ggactttcac tccagactta ccgagccgcc     37020
        tacgcgccct ttacgcccaa tcattccgga caacgctcgc cccctacgta ttaccgcggc     37080
        tgctggcacg tagttagccg ggg                                             37103
//
ID   XXX; XXX; linear; XXX; XXX; XXX;  XXX.
XX
AC   PRJEB67889
XX
DE   Uncultured bacterium DNA fragment from in vivo enriched cow rumen, clone
rum-45p14
XX
FH   Key             Location/Qualifiers
```

```
FH
FT   source          1..43328
FT                   /organism="uncultured bacterium"
FT                   /environmental_sample
FT                   /mol_type="genomic DNA"
FT                   /isolation_source="Cow rumen"
FT                   /clone="rum-45p14"
FT                   /note="AZCL-xylan, AZCL-arabinoxylan active clone"
FT   CDS             complement(200..709)
FT                   /translation="MEMINTLKVNYTNHTIVMDRTFAKLAQDTRSDEYAHLQAVRKDYP
FT                   SFKVVQRTIRKNSDKKTYQGLTYEYMEGYIMRHGTEEQRLANLRKYAELREIAECQGKR
FT                   YRYPVIKSWFLEQYPEIVKFGIEEDASVASPVTANAVVKPQTNELKVVENDTEDTAKEQ
FT                   TEAIGA"
FT                   /product="hypothetical protein"
FT   CDS             complement(743..1189)
FT                   /translation="MTSITINAKNRTIELSNTFAKAASLFGSEEYRQLQEARRDYPSYR
FT                   VVTLKQKGTAKAEFAKLSFKFMDKYVKANATTDDLKAEYRSLRGLKEDWTKDENGIPVD
FT                   FQIVKDWFLNAFSEFEQFHNERKALLEKIQREKAARLAARKVAA"
FT                   /product="hypothetical protein"
FT   CDS             complement(1662..2204)
FT                   /translation="MKWVCPVCGYVHEGPEPPEKCPVCKVPGEKFIKQEGEMTWAAEHV
FT                   IGIAKDVPEEIKAGLRSNFEGECCEVGMYLAMGRAAAREGYPEIAEYWKTAAFEEAEHA
FT                   AKFAELLGEVVSPSTKENLRVRVEAENGATQGKTDLAKKAKELGLDAIHDTVHEMARDE
FT                   ARHGKAFKGLLERYFGK"
FT                   /product="Rubrerythrin"
FT   CDS             complement(2336..3835)
FT                   /translation="MMNTKRMLSAALLLCLLGTLLCPAFSLAAKEETITLRICNWEEYI
FT                   DQGDWGEDEVIDLESGDIFGENSLVDDFETWYEETYGKKVHVEYSTFGTNEDLYNMLTI
FT                   GDVYDLVCPSEYMIMKLMAEDKVQPLSEDFFDGENENNYYIRGVSPYIRNIFETHDIGG
FT                   VTWDKCAAGYMWGVTGFLYNPEVLEEENAATWNLLENRDYNRQITMKDNVRDAYFAAVG
FT                   ALKSEQLTSEDFTGRGDYVLALQDEMNDVSQETVDAAQEYLQKIRDNLYSFETDSGKAD
FT                   MITGKVVANYQWSGDAVYAMDTAEEDDYILNFAVPQECTNIYFDGWIMLKSGINGNAEK
FT                   QQAAEAFINFISRPDNVIRNMYYIGYTSAISGGEDERIFEYLDWNYGAEEDEEEVVDYP
FT                   VGYFFAGEEDNEDYVLSVPEEQLHRQLSAQYPSQEVIRRASIMVYFNEEQSERINRMWI
FT                   RIRCFNIQSVPVWVWIAIGAVIILGVLFGYQRKLSKQRQYS"
FT                   /product="ABC transporter, periplasmic spermidine
FT                   putrescine-binding protein PotD (TC 3.A.1.11.1)"
FT   CDS             complement(3832..4635)
FT                   /db_xref="GO:0005215"
FT                   /db_xref="GO:0006810"
FT                   /db_xref="GO:0016020"
FT                   /translation="MTRKIVRIAFPILVLLFLYVPILVLAFYSITDATQIGSFRNFSFQ
FT                   NYVTLFNTEELRNMIIGTILLALGAALIATILGTLGAIGAFYSGKRAGSAIATVNNIPV
FT                   INADVVTGFSLCVLLVVVFGIAKDTYIPLVIGHVTLCAPFVYLSVIPKLRQMDSSLYEA
FT                   ALDLGAKPSKALLKVVIPQIFSGMISGFALAVTLSLDDYFIATYTKPATFDTISTYVVN
FT                   ATKGSQTQIKTALWALSTLIFLLVIVFVVVMNIRSWRATKKGEKA"
FT                   /product="Spermidine Putrescine ABC transporter permease
FT                   component potC (TC_3.A.1.11.1)"
FT   CDS             complement(4632..5420)
FT                   /translation="MRFSRKTLGIPYAVFLLLFVVAPLLVLFYYAFTNGQGVFTLKNLT
FT                   DFFTDPNTLGTLAYSFAVSLATTVLCLLIAYPVAYILATSKLKAKSVIIMIFILPMWIN
FT                   FSLRVTALKEILTIIEGNLAFYPFLNSVLGMTYDFLPFMILPIYNSIVKLDISLNEAAR
FT                   DLGANGRMAFLRVTLPLSLPGIISGVSMVFLPAMTNYVVLDMLYNSTYIMGSLIGSYFN
FT                   AYNWHGGSMIALVLLGIICVFSLITNGKEDEEAKTGGALL"
```

```
FT                   /product="Spermidine Putrescine ABC transporter permease
FT                   component PotB (TC 3.A.1.11.1)"
FT   CDS             complement(5431..6876)
FT                   /translation="MEKQENIIELKHISRTFSDGFKAVSDFNLEVKRGEFVTFLGPSGC
FT                   GKTTTLRMIAGFDMPTEGDILLNGRDISKLPPNERPINTVFQRYALFPHMNVYENIAFG
FT                   LKLKKMPKDEIKKKVKKVLELVDLEGFEKRSVPTLSGGQQQRVAIARALVNEPEILLLD
FT                   EPLGALDLKMRKEMQIELKAMHDELGITFIYVTHDQEEALTMSDKIVVMSEGKIQQIGT
FT                   PEDIYNEPQNAFVADFIGDSNIFNGIMTGNMTARFAGGQFTCVDDVPEGTQITAVLRPE
FT                   DIELTAPENGQIRGKVISVIFKGIHYEIIVQSGRYEVVIQSTKSAEVGSEVGMRVEPDG
FT                   IHIMIAEDHTNVFPVDMNRNHRLEFNEQELDVSLTQLIKGSRRNENHELLDENGEVIDP
FT                   EKIHIIAAIEPQDIRMTDDVDDEQVLIEGTISNLIYKGDHYSYVINTDVGQDFIVDDEY
FT                   LWNMGDQVGLILPLDKMRFTVKK"
FT                   /product="Putrescine transport ATP-binding protein PotA
FT                   (TC 3.A.1.11.1)"
FT   CDS             complement(6928..7500)
FT                   /translation="MAGKTMDAERVMTQVGTKLRDLRKQNGLTQQELADRAELTKGFIS
FT                   QLEQGQVTPSLVTLLDLIECLGTTPSDFFKTSGAEKVVFGDRDYFEKIDDAGNSKVWVV
FT                   PSAQSNQMEPILVTLQPHGSLPVDDPHNGEEFGYVLSGKIALYRQETRYDVKAGESFYY
FT                   EADKTHRIENPGSRPARFIWVSTPPSF"
FT                   /product="Transcriptional regulator, MerR family, near
FT                   polyamine transporter"
FT   CDS             complement(7730..10381)
FT                   /translation="MSFEIAHPLRLLVIPVCMALVLWMAHLQRSRSLKERVSHILRHVL
FT                   IILTALAFAGLSILTASPDRTAWLVLDASASVREEEVVSLARQALQEAGNDRKTGIIVF
FT                   GRHAAVEKSIGGNTEWTGLHVSADRSASNLGEALSLASALLPTDSNGGIAVISDGAVTG
FT                   AAEWADSRTGVPVNTLQVARKTGPDAQVTEISVPDTLYQGQKYTTLVTVYTNTAGEATL
FT                   LLSENHETPRTQRVTLRKGENTFAFESVATRSGVIPCEAQILLDGDTVAPNNLNGTYTV
FT                   VSGEMNVLLAEGKSGEGEELKKMLESAGMKVRVIPVSMLSEQAADLWAYHAIALVNVDA
FT                   EQLSEAQAEALDEAVRELGCGLAVFGGDSSYALGGYRGSLIEKMLPVSIDVKNKMDLPS
FT                   TALVIAIDKSGSMSDSSWGLTRLQLAREAACSALEILNERDSAGVIAFDDTGKWVVPLS
FT                   PVTDVAAMQEKVRTIRLGGGTAFYSPLKMAKDALENVNAQYKHVIFLTDGEAGDTGYED
FT                   VVKEMAEKGITVTTVAVGDGSDTAGMRKLAELGKGRMYYAGPFDSLPKIFTKETLMISG
FT                   SYVQNRTFTPIVTDPSMTSFDGFPVLDGYLATTEKPLATVSLSSNRKEPVLAWWQYGAG
FT                   KVAAWTSDVQGGWTGSFLNWEDAPAFFSGILSYILPVNTRSGDAELEDGKLRFTADAED
FT                   ELMSRAAKAEARIIRPDGTQETVELNQVNARTFEGEADTSQAGAYAVHVSVRDRLGNEL
FT                   LATDSGAVLSWSREYDLRTTDDGILAQLSASSGGIATENTSELLRFPDTSARKRRDLTW
FT                   ILALLAGLIFLFDVAQRRLDWLHEPEKKEEPEAEKPVKPEKKKKPEKKQPEKPQEKAAD
FT                   VLWENLQKKKRL"
FT                   /product="hypothetical protein"
FT   CDS             complement(10378..11946)
FT                   /translation="MLTFAAPQFAWFLLLPAGIVVLYLLRRKYLPKQVPSTFLWRTAIR
FT                   DHAANKPLQRLRKNILLPVQFLAALVLALALMQPNLTGREASRTVMIFDLSASMQTVSA
FT                   GKSRLDEAKDRAEELLQGLPPGEEITILAAGTEVRQTLTGSTDREDVRQAIRSLSCGRE
FT                   GADLEKAVSLAEAIRREETDRGTRIVVFSDSYTPKAGIAGIHAGRGEENRGVYTLTAEE
FT                   GSAYARICNFGSDCRITVACEADGKLCEAKETEIPAGETAGVLFTIPENAAKVEVYIRE
FT                   PDALMADNRAEAPVERSTVRTVALTGDNLFLESALKVREDLRVVRTSEEAPAETEADLY
FT                   IYGTSPLLFSRDPSKTAFTWSEEEKEPAGSLTAEESPMTEGLTLRDVALRACKPLTGGK
FT                   AAIRAGSDVLAAYTDSEAAIGFLLQDSNLPLKYDFPVLVQNILNMLLPRSAAEVPETEA
FT                   PVAREESDVRTVARDTEGADAAAAGSRGQDLTGILMGVFLALLMIEFILAREPWSRRKG
FT                   KGAAA"
FT                   /product="hypothetical protein"
FT   CDS             complement(11957..12805)
FT                   /translation="MLSDAFLSRLDTLQLAMKGRASGGAGGTRRSRQTGSSAEFSDYRE
FT                   YIPGDDIRRIDWNAMARFDKIFLKLFMEEQESAVTVLLDGSGSMGEKWETARQAAEAVS
```

```
FT                   YLALTGGDRLRVICLQEGGNRMSPQLSGRQAYARLEQFLDTCAPGGAGRLTETVMRTEG
FT                   LRRGLCFLISDCYEEGGIGTALDSLRYRGQECAVIQPLSAFELDPALDGAVKLTDSETG
FT                   EAINLLADREALENYRQTLENFLREVRETCFRRETPYALLDAGKNFEEEWIPLISQSGM
FT                   I"
FT                   /product="hypothetical protein"
FT   CDS             complement(12795..13772)
FT                   /translation="MNTNQSIEAFAEKYAKIRTEIAKEMIGQEDVVEHLLLAVIAGGNV
FT                   LLEGVPGLGKTHLVRILSKTLDLPFSRIQFTPDLMPADITGTNILIRTEEGSSFRFQQG
FT                   PLFASIVLADEINRATPKTQAALLEAMQEHSVTVAGETMKLPEPYFVLATQNPVEQEGT
FT                   YPLPEAQLDRFLFKVLVPFPNLKELGGIMNLTIGSAAQEAGKAADAAELLDMRETARQV
FT                   PVADSVQESALRLVLATHPETEDAPEITKKYLRFGASPRAAQAILSTARVRALEQGRYN
FT                   VAYEDIRYVAPASLRHRLALNFAAVTEGKDMESIVAELVENYVK"
FT                   /product="MoxR-like ATPase in aerotolerance operon"
FT   CDS             complement(13789..15282)
FT                   /translation="MKGYDGMNELLKALRPVKNRIRLGRLWCGAATGLAAGAAAALVLL
FT                   AVTTFVPLGGRGGIAGAAVAGGALLGAVINALRPVDSREAARAADRCGLRERTVTALEM
FT                   ADRPPADGKAAELVVAQRQDACEHLQALDARKIRLQAPKKRLLAAATLLVLCAGTLLIP
FT                   GGGDRIVSERKILQEKTARMAEEIDEAEAAEEAGKTEKEKAELRKLTADLKRELEDSRD
FT                   DVDALVAVDKAESRLEEIRQKTAGDAVSELADALRDAGMDAAAQALENGDAAEMSATLS
FT                   EINTDELKEAAKGLSGEAQEMAEQMAQAAQQGEMSDAQMQAMQSGSSQASVMQQALAQM
FT                   KAALGAAQGASSQQQQSSNSNSSGSGSGGSGNQGQNAAGAGSGTTNEEEQGGGSGNKSS
FT                   ASRGNRPPEYKEGEYETIYDPEKVNTASRDVMTEQQQLGKDSVQIETGPGKGSLEGNVP
FT                   YRQVIGEYAETEAQAAESAHLTREQKEWVDEYFRQLTDE"
FT                   /product="hypothetical protein"
FT   CDS             complement(15309..16226)
FT                   /translation="MLNPILSFSATRRMRSFKTLLIAAAYVVVMLAVTLLIMGRIFGNE
FT                   ATFYSIRNSVTCYQAQMVTQFILILLIAPAMTSGAIAGERERQTLELLLVTNTRSFRIV
FT                   IGKAMESFAMLALLIVCGFPVMCLTMVAGAVTLPQILIGELFLLLIAFASVSVGVFSST
FT                   LARSTVASGVISYLILLAIGGLTLIPLIMDFPVEVTSVVYDTSSLAALAPSDALKMISP
FT                   VLLCNPGYALIALLHGQTGMLTSTMEYREIGRLLCEFRLMDKAGGETLALISCGIILVL
FT                   AVLLLLAAGFLLRKNSLKPAKRTK"
FT                   /product="hypothetical protein"
FT   CDS             complement(16219..17157)
FT                   /translation="MLSLEKLTKKYGDFTALNGLSLEIADGQLHGFVGPNGAGKTTTMR
FT                   ILATLLKPTSGTATIDGVDVVRSGTEARRLIGYMPDFFGVYDSLKCWEYLDFYARCYRI
FT                   PSADRKKMTERLLDLVQLSEKKDAFVDSLSRGMKQRLCLARSLIHDPKLLILDEPASGM
FT                   DPRARAEMKTILRTLREMGKTVLISSHILPELSEMCDSLTILDHGQLVFSGSVEALSEK
FT                   MNGNAPLEIRLQEGCGEAAVDMAVACLKELPFVKEILQEGTCVLKVRTEGGEAEHMQML
FT                   REMISRNIPVCDFHRAPMNLEKVFMEVTGDA"
FT                   /product="ABC transporter, ATP-binding protein"
FT   CDS             complement(17144..19666)
FT                   /translation="MRKKAVRWFAFGLVICLLAWAGAAAADGLHTEVENPQLEMEVRLG
FT                   YDGQVTYGKAIPVRVTVRNNGGDLEAMLAVNGYVNTAKYDRFETALSVPAGGERTVVLP
FT                   VVAQTRQNTFTVEIVKDGKVISAVNSTPSGVINPNAMMIGVLSTRPRNLANLDINQEND
FT                   VLYRYEFWQTVALTPETLPENPKLLDSFGMIVLDDTDPALLTEKQQQALKDWIVSGHVL
FT                   IAGGGSTAPRNLAFLGDLTKLQASDFTVSDGVLSEVESLAGQKHSGRKTEIALAKLEGA
FT                   NPLISDSEGNGLVWLEGAGSGRIITLAWEAGNAALNAESAMHIFFQQLLIKMDPNMYSK
FT                   LLYPGEESSALYSAGDDTKIQVRNSLPAAVAVIAGAVVICGVMWIILKKRSKTQWMWAV
FT                   IPAAALAAAAVGTVMAGSSMLNRPGTATTVNLVQNAEGRATRYVFVTAAAPRPGLHRYS
FT                   MNGEDLEAQVYDDGYYYWAEEEDTAELKEPSQLRVVQITGENGSSALNANTPWEVVRLS
FT                   SVRTSEQEGQVEAQIWMESDGLHGTVRNGLPYSLKEGVVLSVYGFVRVPALAPGESREF
FT                   AMVAKDAPDPSNPVFEDGVMLRNVTSGVYMVLSGMRENSDTNDYGGRESILSGMMNSAY
FT                   DQLTRDNSAGTRENVIFVYCAEPEGDFASPLMVDGKEVEGGTVLPMYTAKIEYLKIGKT
FT                   GVVFHAPGMDHAVRCEIDSSGMPAGDMAEDPNGGKLYYNYFPLSEHPTFRFTLENLDRI
```

```
FT                   EIEKLSVGVEDWYANDVACFVLNVELKTWVEVQMNTPLKQPGKYLDEQGNLYCQFRPKV
FT                   SDSYMDIPEPSLTLEGKIKEGGETHAEP"
FT                   /product="hypothetical protein"
FT   CDS             complement(19764..20459)
FT                   /translation="MYCVYCGVRLQDGVTECPLCHTEVQVAPVKRPAEEAPYSRKYPQP
FT                   EKQSGKYLVLGLVTIIFAAVALSCLIFCLKSRGEVGWSGIAALGMALGWVWLVLPLLFS
FT                   RWKPMIFLPIDFACLAGYLLYICVNTGGHWFLSFAFPVTMIAGILTLAGVAMMRYIVQG
FT                   RLRLMSLLVIAIGLSFMLVEFFQHITFRTPMFVWSLYCMIAFCMIGLFLFVASFIPSLR
FT                   AALRRSFFF"
FT                   /product="hypothetical protein"
FT   CDS             complement(20464..21732)
FT                   /translation="MKKQWYRLDNAALIFPAVMRKNWNNVFRVSVTLKNRVDPEILGRA
FT                   AEDLLPRFPTIYVRLKTGFFWYYLETVDEAPKAEEDYAWPLTPMTRRKLNKCCIRILYY
FT                   QNRIAVEFFHSVTDGTGGMAFLQNLAARYLTLKEGAVIPPEGNIVDPAEPPKEEEARDC
FT                   FLQCAAKGKVSRAEANAYHTRGNPEEGPFRHIVTGIIPTEALLAKAHAEGVTVTAWLAA
FT                   VMAEAVLARMKADPLAGKRSHIKITVPVNLRKTFGLNTLRNFTLAVNIGFDPRLGEYTR
FT                   SEICRLMHHQLAAETIPQRMAGRVATNVDLQRCLPLRLVPLPLKTLAMRLVYTLSGESK
FT                   GSLNISNMGVVKLPEAMAPYVERFDFIIGVQYSYPNNCSVVSWQGKTYISMIRGIRDSE
FT                   LERLFFSSLVEMGIPVDIESNRR"
FT                   /product="hypothetical protein"
FT   CDS             21902..24535
FT                   /translation="MATGPVIGLCAHVDAGKTTLSESMLFLSGAVRRQGRVDHGDAFLD
FT                   TDRMEKDRGITIFSKEARLSWNHTDLTLMDTPGHTDFSGETERALNVLDAAVLVISAAD
FT                   GVQPHTRTLWRLLEQRGLPVILFLNKTDLPHDAEALFASLSRELSGQIVAFPSPDPEKL
FT                   ALCDETCLDAYLREGEIPERMIHAMVAGRRIFPLFSGSALRNEGVEPLLDFLARFDPRP
FT                   APPAAFGARVYKVARDPQGARLAFLRITGGTLKARDLLTLKSPEGETLWAEKAAEIRLY
FT                   SGARYTAVQEVSAGQVCSVVGLSKALPGDGLGTDPGRKEQELRPCYACRVVLPPGADLH
FT                   YVLNCLETLQEEEPLIQVEYVEPRREIRVHSMGDVYLEVLRAQLADRFGLDVSFAETTV
FT                   LYRETIEAPVEGAGHYEPLRHYAEVHLLIAPLPRGSGLVCDSSVSTDDLALNWQRLIVT
FT                   HLREKVHVGVLTGSPVTDLHITLIAGKAHLKHTEGGDFRQATYRALRQGLMKARSILLE
FT                   PWMTLDITVPEDCVGRVMSDLSLMGGRFDAPEDAGETLRRLSAAVPASACADYGRQLAV
FT                   FTKGRGSLTAAFLDWEPCADPEKVIREKGYDPRRDVWNTPDSVFCSHGAGVTVPWDEAD
FT                   AHMHLPLLKDLSRREVPAPAAGGAASAYHGTREEDLALERIFERTYGPVKARQLVAAPS
FT                   AAAEKQREAAPAPASDGEILLIDGYNVLHAWDEWKPFLPDRLEDARDALRDLLCDYAGA
FT                   TGKPVILVFDAYAVPDNPGKAEKYKNIYVIYTREAQTADAFIEQTTYYGRNTARIRVVT
FT                   SDRPEQLIASGNAALRTSAREFHAEVNGVRDGIAAFIARNRLSRPVRTLEAAYKEAWKK
FT                   SREPQK"
FT                   /product="Ribosome protection-type tetracycline resistance
FT                   related proteins, group 2"
FT   CDS             complement(24564..26441)
FT                   /translation="MYAGDLVVLGVCSIMLILLFFSFKVRSRAFRMFLSLVGILMAAAA
FT                   THVTFHTLLKYRPNLPTGLFYGFQSLYHGLLLFMLYQFAAYICQVTGLTKEERKVYHTI
FT                   GGMLILVVLAAELFVNFSGSGIQLGDGAASAGGNRIFMIGYGLFIVMLSMLMIQVRHHL
FT                   YRRMMNGFYMTMVFAVAMLAIAKLNGQDSFTTATFLFPVVAIMYLLHSNPYDAQLGSND
FT                   LRGLEGLIRYSDDKGKTYLYLSLYMRELDETARQMPGDLQAIVRRFTAQYFRSSVMFRV
FT                   GNGHLLMMVRKERNPDYEARTEAILRAFDQEYQNFKLDYKIVIGQTTAELNRNNEYVSF
FT                   IRSIHTRIPENTIYRVQPDDIDKFRKYEAILHQLADIAGQKNLDDPRVLVYCQPVLNIR
FT                   TGKYDTAEALMRLQLPEEGLVYPDRFIALAEENGFIHVLTEIILHKVCKAVKALNEEGY
FT                   CIDRVSVNVSAMELRNQHFCMDISSIIDRSGIDGKQIAIEMTESRNERDFLLMKEKIAE
FT                   LKQKGITFYLDDFGTGYSNMERILELPFDIIKFDRSMVVAGAQDKRSEQVLGSLADLFA
FT                   KLDYAVLFEGVETEQDEEMCRECCASYLQGYKYSRPVPMEQLKEFLTKK"
FT                   /product="hypothetical protein"
FT   CDS             complement(26568..28283)
FT                   /translation="MKANKLSLLFPDQDRITYGMIPEESWHDLGLDAVAEKVAAQPAEL
```

```
FT                   MMIQQVMRSLTPDPEVTAFRCGVFEDILRHPEIRERMLKLLDRVKTFYDYGVVRKHQGD
FT                   ESGLWDLMHRLEEYHDYIVTVEEIRECLAEKDLQSDGMIRLREEVDSIYRDNGFAALRK
FT                   DVEEMRITASEVKSLTVGINLNDRFEAINMGLITVNSKPFTRSGLLKNFLTAVTPKDEI
FT                   HPEADWNKSYSYYLANAGVGLADGVGQFVERGLVMRNPIAAMSMARIPDADGSASVPRQ
FT                   MDSAATMLTSRITRRLREMLGKYLNVSVKEISELIPELVFYTRWAEYIEKCRAAGWKFC
FT                   TPEIRKNPEGSACAEAEGFYNLKLISAEKPENVVTNNLAFDREKRAYILTGANRGGKTT
FT                   ITQAVGQLFLLAQSGISVPADRFVFDPADQVLTHFPADEDKTLDLGRLGEECRRFRELY
FT                   GKSTGKSLILLNETFSTTSFEEGYFIAVDAVKAILRRDTRTIYNTHMHKLAKELDSEIN
FT                   TEGLPGKAVSLVAETRDGKNSFRVKIAPPEGKSFAREIAVKYGVTYEALTQEGQ"
FT                   /product="DNA mismatch repair protein [MutS]"
FT   CDS             28737..31010
FT                   /db_xref="GO:0003987"
FT                   /translation="MMQTQTDLAQLKAIASRLREMREIMGWTAEEMAGKTEVTPEEYLL
FT                   YEAGEADMPFSFIYKAAQAFNMELTELLEGHNAFLSTYTVTRRGKGETTAKEEGIAISN
FT                   LAPKFRDKAAEPYWVRYEYSEKQQNEPIHTTTHHGQEFDLVLSGSLKVQIGDHTEILHE
FT                   GDSIYYNSSLPHGMIAVDSKDCVFLAMILPGEGPQVETAAVTAPKAARYEGHLVAEKFI
FT                   DAKEDDQGRLQAISFPNKETFNFGFDVVDEIARQYPDKLAMLYVDKHHEERRFTFKDVK
FT                   DASNQCANYFTSLGIKRGDRVMLVLKRHYQFWFCMVALHKLGAIAIPATNQLKEHDFEY
FT                   RFNAAGVKALICTADGDTAEIAERAAARCPSVETLIIVNGTREGWRSLNDEYPLFTRRY
FT                   HRPEDASCGDDPALMFFTSGTTGNPKMAQHKHTYALGHYVTAKYWHCCERDGLHFTISD
FT                   TGWGKSLWGKLYGQWLCEGAVFVYDFDRFDAADILPMFKKYNITTFCAPPTMLRMFIKV
FT                   DLANYDLSSIHHMTTAGEALNPEVFRQFEKATGLQIMEGFGQTETTLTIGNLFGGVPKI
FT                   GSMGKANPQYDVDIVDPDGNPVATGEVGEIVIHTDKSAPCGLFREYYLDEEKTKDAWHD
FT                   GLYHTGDTAWRDEDGYFWYVSRIDDVIKSSGYRIGPFEIESVIMELPYVLECGVSAEPD
FT                   EVRGQIVKASIVLTRGTEGTEELKKEIQQYVKEHTAPYKYPRKIVFRDELPKTISGKIQ
FT                   RNLL"
FT                   /product="Acetyl-coenzyme A synthetase (EC 6.2.1.1)"
FT                   /EC_number="6.2.1.1"
FT   CDS             31092..32147
FT                   /db_xref="GO:0031176"
FT                   /translation="MISATQAPSLCKAYEPYFRVGAALSSWLVKDPELVEIVCRHFNSI
FT                   TADNQMKPDSVLNLEGTLAQGDPCHAATDFTRVDALLSFARDHQLSVRYHVLAWHNQTP
FT                   VWFFKKDWENDREAPFAPKEVILARLENYIRDVMQHVNTCFPGVVYTWDVVNEAIEPDQ
FT                   DGPGLYRTRSPWFTAAGQDFLPAAFRAARRYAAPGQTLCYNDYNAFEPVKRDAILDVLK
FT                   MLKAENLVDTMGMQGHYLLPHLDIAACETAARAYAALGLKLQVTELDIHCNSDDEAHVA
FT                   ALAEAYRSWFAMIKKLVKEGIPVEAVTFWGVTDADSWLPGFRKEPSYPLLITADRKTKP
FT                   AFDAVLREAEL"
FT                   /product="GH10"
FT                   /EC_number="3.2.1.8"
FT   CDS             32189..32974
FT                   /db_xref="GO:0008800"
FT                   /db_xref="GO:0030655"
FT                   /db_xref="GO:0046677"
FT                   /translation="MARLQIEYFSNALRRSTTFEMIIPNDFPAGVPWEAGKAHSRPMST
FT                   LFLLHGYTGKAGNWVPDGLPEKYNFAIVMPTAENSFYLNGEATGTAYQTMVGEELVNYV
FT                   RKTFHLASGPENTCIAGLSMGGFGALHTGLAYPETFGKIGALSSALIVHEVATMKPGGG
FT                   NAVANYAYYRACFGDLETVLERDANPEVLARKLKASGAKIPEIYMCCGTEDFLLQNNRQ
FT                   MHSFLEAEGIPHEYHESKGTHDMVFWREYIAKIVEWMFA"
FT                   /product="putative esterase"
FT   CDS             complement(32980..33246)
FT                   /translation="MRKAESNWDLRRIDRSFPYLRVLMRGRPARREDDPKHPPMDRGHR
FT                   AKIFAPFDALDGFSDLIGQKNRLREETALEKTEENICMPWEDP"
FT                   /product="hypothetical protein"
FT   CDS             complement(33264..34655)
```

```
FT                   /translation="MQYLCIDLKSFYASVECVARNLNPLEANLLVADESRTDKTICLAV
FT                   SPSLKAKGVSSRPRLFEAKQAIRLYEAAHKTKVEYIVAPPRMAEYLRVSARAYEIYRKF
FT                   LSEKDIHVYSIDECFIDLTPYLHLCRAEAERAGVSPAHFLAMRLIRAVLAETGITATAG
FT                   IGTNLYLAKVGMDIVAKKAKPDRDGVRIAELGEEDYRLLLWEHRPLTDFWQVGPGTARR
FT                   LMTHGMQTMGDVAAMAVANPEGLYRLFGVNAEILMDHAWGVEPVGMSDIKSYRPSARSF
FT                   STGQVLPRPYRFEESLLVFREMAEQAGIKLVEKRMRIPAAMWWISFDPASLEENPGYTG
FT                   PVCLDFYGRLHPVHVGGTVRFRPETNSRTVILEALDRAFREKVDPGLLVRRLGLALTDP
FT                   REEGCALQMDMFSDPDAQDREFRMQEAMLEVRGRFGSNLLVRGMNMLDGATAMERNKQI
FT                   GGHRA"
FT                   /product="hypothetical protein"
FT   CDS             complement(34668..35420)
FT                   /translation="MRYDAEKARERNEIGARLTQARRSLRLTQGELAEALALRGVRVQT
FT                   AAVSKWEKGDSVPSAYQLAAVCDALRIPLFGEEEKLNEEGRKMLAGYRGYLESQSAYRS
FT                   RPMHREPVVAMKVSTLPASAGFGEALDNNLFESQTFPASSVPAGASFAVRVSGDSMEPV
FT                   LHDGQYAWVEECSRLLPGEVGLFVVDGEGYIKVYAEQEPEGDEVENFTDSEGIRHLQPV
FT                   LVSCNTAYDPKVITSGTDFRIVGRVLNA"
FT                   /product="Phage repressor"
FT   CDS             complement(35567..36748)
FT                   /db_xref="GO:0046316"
FT                   /translation="MIQGTKVRLGIIGIGNMGSEHCRLILGGACPETELCAVADLRQER
FT                   REWARGNLPENVSIYTDGRELIASGRCDAVLIAVPHFQHEPLAVAALEAGLHVLCEKPA
FT                   AVQADRAEKMIRAAEKAGKTLTFMFNQRTNCMYRAVKELLDSGEMGAMKRMNWIITDWY
FT                   RTQRYYDSGSWRATWAGEGGGVLLNQCPHQLDLLIWLCGMPCAVTARCLEGKWHHIEVE
FT                   DDVTAFLEFPEGATGVFVASTGDLPGTNRLEIDCERGKIVCENGQVKVWRLGANEREIC
FT                   FSSDDPWYREEIRESLLETDGENPQHAGVINAFAAHLLRGEKPVAGAEDGLRALQLSNA
FT                   IHLSGWTGGRVGIPVSPEAFNRELERRIAASGTHKTEDITYDTDHSRSGRRMH"
FT                   /product="Gluconokinase (EC 2.7.1.12) / oxidoreductase
FT                   domain"
FT                   /EC_number="2.7.1.12"
FT   CDS             complement(36752..37831)
FT                   /translation="MNVSDGMTYAPKGKPQPVVKGREFVFSAVRLDHGHIYGMCNGLTE
FT                   AGGTLKYVYDPDKKKVAAFLKAFPQARAANSEEEALNDPETQMIAGAAVTSERCALGLR
FT                   AMDAGKDYFTDKAPFTTLEQLASAREAVKRTGKKYMVYYAERLHDEGSILAGYIAHSGE
FT                   IGKVISVTGFGPHRLGAPGRPAWFFEREKYGGILCDIGSHQFEQFLYYAQEEDATVTAS
FT                   RIANFAHPEYPELEDFGDAQLTGKNGAAGYFRLDWFTPDGLRTWGDGRMFILGTKGFVE
FT                   IRKYVNLASEKDGGGHVYLVNDQGEKYLNAAGATGYPFFGQLILDCLNRTENAMTQEHA
FT                   FKAAELCLLAQKNAIRIGE"
FT                   /product="oxidoreductase domain protein"
FT   CDS             complement(37833..38993)
FT                   /translation="MIRVAIVGTGGISHAHIKAYMQFPERCKIVALVDIIPGKAQQVKE
FT                   QYGLDAEVFLDHHEILEKDIDLVDVCTPPFVHAEISINALRSGKNVVCEKPMAASLEEC
FT                   DAMLKARDESGKKLSIIAQNRFRLPVRNLKALLDNGMAGKVRHVTVNSLWFRGHSYYDL
FT                   WWRGTWEKEGGGCTLNHAVHHIDMLGWMMGTPEKVTSVLANTGHDNAEVEDLSVSIAEY
FT                   PGALAVVTASVVDHGEDQALIFQCEKAKIAFPYSVFASQPQPNGFPLAEPDESFAKQAE
FT                   EFLASLPPVPYEMHTGQLENVLHALETDSAPAITGEDGRRTIELITAIYKAGSSHQAVT
FT                   LPIPPEDPFYTVEGIRGAVPHFYEKTTSVESQDGSLSFGNFKKGEK"
FT                   /product="Oxidoreductase"
FT   CDS             39164..40090
FT                   /translation="MQLIQAEMTRAPFLPMGQQGALGYEAHHNLDKTFAVTQYHCHDYY
FT                   EFYIHIRGGEYMGVDNQLFLLKPNQVFILPPFSMHGLSCTREMHDYERAYLNLSPEVMR
FT                   VLGCGQIDLDQFFRSQASGGVCTYQLTDADAEQFVAWIRKIRDNQQDESFENDVYHRFQ
FT                   NYSLMMSLMSLLCRTTRRTDPVEGDAFGSSVIQEILEYINNHYTQPIRIENLARSFGVS
FT                   VSWLSHEFSRFTNRSVYDYVLSRRVMLARQQMLGSASLNDIAYQCGFNDYSNFLRSFGK
FT                   IVGMSPSQYRKRLKAYREKEIDARIEF"
```

```
FT                   /product="Transcriptional regulator, AraC family"
FT   CDS             complement(40208..41818)
FT                   /translation="MKRILALVLALALFATLGITSAFAADDGFVDGKFTETRTITVRLF
FT                   QRGDNVPETSPWAEYIKNGMKEKYNVNVEFSVCGRWDDETDLGNLLAAQNAPDVIYTYA
FT                   YPTVLNYADMGGIIDLAPYIEEYKDLLPNLFNLLGEENVYWDKDPESGKLWALETRLVN
FT                   NARINTFIRKDWLDKLGMALPTNKAEFEACLVAFRDNAELLLGADAAKMVPYSTSQDIG
FT                   WRNNILTVSFVPDAITDEELYIRGFDDRQQFLPGAKEAMRLLNKWYNEGLVWKDFALYG
FT                   DGDTTEDDNLKAGFVGAFQHNYDYPYRNGEDSIEANIKRAAGEDAEFVAVDCFENDAGI
FT                   YRKYLGSTVDRKVCFPTTNKEVLASLLYLDFISQPDTILFLQTGVEGKNHERAENGAYM
FT                   MKAVDASDPYTQASLNNIDFTMTCNGQYLGEYTEASFAYGYAPVNPEVVQTAYAVAMNG
FT                   GRVPGHFSLPAIEAETGVGTTLSDERNSFLNKAVTAPVDQFDAVYDAGVENYMAIGGEA
FT                   IMNERAEKLEAATGYKFEK"
FT                   /product="hypothetical protein"
FT   CDS             complement(41877..42749)
FT                   /translation="MIKSKSTKFGDWVIVFICILLIFICLVPMLNVLARSLSDTKYLIR
FT                   NEVLLWPKGLNLDAYRTVLGNERYIHSLWYTALLTVGCTVLSLFMTVICAYPLTYDQLK
FT                   GRKFLNGFILFTMYFSAGMIPHYVMLSNMGMLETVWVLVIPSCLSVYNMIIMRSSFYSV
FT                   PESLREAAEIDGAGPIRILVQVYLPLSKPVLATLALFYAVGRWNGFSDALLYIKKNREL
FT                   YPIQLYLYNILQNASSGEAAAQEGFAMPGLSETLKMASVMFATVPILLVYPWLQRYFVS
FT                   GVTIGAVKG"
FT                   /product="Predicted beta-xyloside ABC transporter,
FT                   permease component"
FT   CDS             complement(42765..43286)
FT                   /translation="MADNDGLVNILNKQMTGNPDAVIPFLSNPSWWVGSYIGLGIWKEA
FT                   GWGTIIYLAALTSISPELYEAAAVDGAGRFRRIWHVTLPGLRPTIVVLLIMRLGYILGS
FT                   EFDRPLALSNKLVTDVSTVLSIYVYQKGINGGGQFSLTTAVGLFQSVVGVLFLFGANFL
FT                   AKRFGERGIM"
FT                   /product="ABC-type polysaccharide transport system,
FT                   permease component"
XX
SQ   Sequence 43328 BP; 8377 A; 13921 C; 11763 G; 9267 T; 0 other;
     acatagataa acaaaacagc acatgccttt cggcactgct gaagtgaatc tcttatcctc        60
     atcttctata attattatac cagaaattac attccgattc aaatacttga ctgctcctaa       120
     catttgattt cccaaaaatg aaaaacccct tctcgcttta tgagaaaggt ctttcacgca       180
     ttttcatgga gcggggcaat catgcaccga tagcttcggt ctgttccttt gcagtatcct       240
     cagtgtcgtt ctcaacaacc ttgagttcat tagtctgcgg tttcactaca gcatttgctg       300
     taacagggct tgctacagaa gcatcctctt cgatcccgaa tttgacaatt tcaggatact       360
     gttcgaggaa ccaactcttg atgacaggat accgatagcg tttgccctga cattctgcga       420
     tttcacgcag ttctgcgtat ttccgcagat tggcaagccg ctgctcttcc gtgccgtgac       480
     gcatgatata accttccatg tactcgtaag tcaaaccctg ataggttttc ttatccgagt       540
     ttttgcggat tgtccgctgt acaaccttaa agctgggata atccttcctg acagcctgaa       600
     ggtgggcata ctcatcgctt ctagtatcct gtgccagttt tgcaaacgtc cgatccataa       660
     cgatggtgtg gttggtgtag ttcactttca gtgtgttaat catttccatg tgctcctttc       720
     aatctcatac cttaggaagg ggtcaggcag ccactttgcg ggctgccaga cgagctgcct       780
     tttccctctg gatcttttcc agcaaagcct tccgctcgtt gtggaactgt tcgaactcgg       840
     agaaggcatt caagaaccag tccttgacga tctggaaatc aacaggaatt ccattttcat       900
     ccttcgtcca atcctctttc aaaccacgca gactgcgata ctcagccttg agatcatcag       960
     tcgtagcatt agccttgaca tacttgtcca tgaacttgaa ggacagtttg gcgaactcgg      1020
     ccttggcagt gcccttctgc ttcagggtga caacccgata gctcggatag tcgcggcggg      1080
     cttcctgcaa ctgcctatac tcttcagatc caaagaggga ggcggccttg gcgaaagtgt      1140
     tgctcagttc gatggtgcgg tttttggcgt tgatggtgat gctagtcatt ttagactcct      1200
     ttcaattggc gatatattag gctcgcctgc ccttattatc tttggcttcg ggtctctttc      1260
     gttcccttca ccttacatac atagtataac cgatactcat gtaaaacatg gcgttataat      1320
     ggtacatgtc aagataactg aaaaagaatt tccccgttgc ttttgcggga attgtgatct      1380
```

gcggaatctt gttatgaaga tgtagtggag atgctgagtt ctgcctttgg atttcttcta 1440
aacatgtaca ggacatgtga ccgagatgct attgttttca aggctttcag gtgataaaat 1500
gggctcgcga aagggtacaa cattttggaa gtacaacttt tgggaaaatg ctgtttttga 1560
aggaggggcg gtacaactaa agtacaactt tttgcaccga agatgatcac gaaaaaagga 1620
ttccctcatt gcgagagaat cccctgaatc ccttgaaatt attacttccc gaagtaccgc 1680
tccagcaggc ccttgaaggc tttgccgtgc cgggcttcgt cgcgggccat ttcatgcacg 1740
gtgtcatgga tggcatccag gccgagttcc ttggctttct tggccaggtc ggtctttccc 1800
tgggtggcgc cgttttcggc ttccacgcgg acgcgcaggt tttccttggt gctgggggaa 1860
acaacttcgc ccagcagctc ggcaaacttg gcagcgtgct ccgcttcctc gaacgcggcg 1920
gtcttccagt attccgcgat ctccggatag ccttcacggg cagcagcgcg gcccatggcc 1980
aggtacatac cgacttcaca gcactcccct tcaaagttgg agcgcaggcc ggccttgatc 2040
tcttccggca catccttggc gatgccgatc acatgttcgg cggcccaggt catttcgcct 2100
tcctgcttga tgaacttttc accgggcact ttgcagaccg ggcacttctc cggcggttcg 2160
ggtccttcat gtacgtagcc gcatacaggg catacccact tcataatttt ttacctccaa 2220
atattttatt tttcttttat tcggcacttg ccgtgaatgg attatatcat ccggcagaga 2280
aaaaagcaaa caggcagcgc aaaaatctgc gctgcctgtt ttttacggta ttctgtcagc 2340
tgtactgccg ctgtttggac aatttccgct ggtatccgaa gaggacaccg aggataatca 2400
ccgcgccgat ggcaatccag acccataccg gaaccgactg gatattgaag caccggatcc 2460
ggatccacat gcggttgatc cgctcactct gctcctcatt gaaatatacc atgatggagg 2520
cccggcggat gacttcctgc gacggatact gggcggacag ctgccggtgc agctgttcct 2580
ccggaacgga caggacatag tcctcattgt cctcttcccc ggcgaagaaa tatcccaccg 2640
gataatcgac gacttcctcc tcgtcttcct ccgcgccgta gttccagtcc aggtattcga 2700
agatccgctc atcctcaccg ccggagatcg cggaggtgta tccgatatag tacatattgc 2760
ggatgacatt gtccgggcgg gagataaagt tgataaacgc ctccgccgcc tgctgtttct 2820
ccgcgtttcc gttgattccg ctcttcagca tgatccatcc gtcaaagtag atgttcgtgc 2880
attcctgcgg aacagcgaag ttcaggatat aatcatcctc ttccgccgta tccatggcat 2940
acaccgcgtc gccggaccac tggtagttgg ccacgacctt gccggtgatc atatccgcct 3000
tgccggagtc ggtttcgaag gagtacaggt tatcgcggat cttctgcagg tattcctgcg 3060
ccgcgtccac cgtctcctgg gaaacatcgt tcatttcatc ctgcagggcc agcacataat 3120
ctcccctgcc ggtaaagtcc tccgacgtca gctgttcgga tttcagcgcg ccgaccgccg 3180
cgaaataggc gtcccgcacg ttgtccttca tggtgatctg gcggttgtaa tccctgtttt 3240
ccagcaggtt ccaggtagcg gcgttctctt cctcgagcac ttccggatta tacaggaatc 3300
cggtgacacc ccacatgtag cccgccgcgc acttgtccca ggtcacgccg ccgatatcat 3360
gggtttcgaa gatattgcgg atataggggg atacgccccg gatatagtag ttattctcat 3420
tctccccgtc gaagaagtcc tccgacagcg gctgcacctt gtcctccgcc atgagtttca 3480
tgatcatgta ctcggagggg cagaccagat cgtacacatc gccgatggtc agcatgttgt 3540
acaggtcttc attggtgccg aaggtggaat actccacatg gaccttcttc ccataggttt 3600
cctcatacca ggtttcgaaa tcatcgacca gggaattctc gccgaagata tcgccgcttt 3660
ccaggtcgat cacttcatcc tcgccccagt cgccctggtc gatatattcc tcccagttgc 3720
agatccgcag ggtgatggtt tcttccttcg ccgcaaggga gaaggccggg cacagcaggg 3780
tgccgagcag gcacagcagc agggcggcgc tcagcatgcg tttcgtgttc atcatgcttt 3840
ctccccccttc ttcgtcgccc gccaggagcg gatattcatc accactacga agacgatcac 3900
cagcaggaag atcagggtgg acagggccca cagggcggtc ttgatctggg tctgggagcc 3960
ttttgtggcg ttcaccacat aggtgctgat cgtatcgaac gtagccggct tggtataggt 4020
cgcgatgaaa tagtcatcca gggaaagggt gaccgccagg gcgaaaccgg agatcatacc 4080
ggagaagatc tgggggatca caaccttgag cagggccttg ctgggcttcg cgcccaggtc 4140
cagcgccgcc tcatacaggc tggaatccat ctgccggagc ttcggtatca cggacaggta 4200
cacaaacggc gcgcacagcg tcacatgccc gatcaccagc ggaatgtagg tatccttggc 4260
gatgccgaac acgaccacca gcagcacgca cagggagaag cctgtcacaa cgtccgcgtt 4320
gataaccgga atattgttga cggtggcgat ggcggaaccg gcccgcttcc cggagtagaa 4380
ggcgccgatg gcacccagcg taccgaggat cgtggcgatc agggccgcgc ccagcgccag 4440
caggattgtt ccgataatca tgttccgcag ttcctcggtg ttgaacagcg tcacataatt 4500
ctggaaggag aaattccgga aggaaccgat ctgcgtcgcg tccgtaatgc tgtagaaggc 4560
cagcaccagg atcgggacat acaggaagag cagcaccagg atcgggaacg caatgcgtac 4620

aatctttctt gtcacagcag cgcacctccc gtcttcgctt cttcatcttc cttaccgttg 4680
gtgatcaggg agaagacaca gatgataccc agcagcacca gcgcgatcat ggatccgccg 4740
tgccagttgt aggcattgaa gtagctgccg atcaggctgc ccatgatata ggtgctgttg 4800
tacagcatat ccagcaccac gtagttcgtc atagccggca ggaaaaccat ggacacgccg 4860
ctgatgattc ccggcaggga gagcggcagc gtcacccgca ggaaggccat gcgcccgttt 4920
gcgcccaggt cccgggccgc ctcattcagc gagatatcca gcttcacgat cgagttgtaa 4980
atcggcagga tcatgaaggg caggaaatcg taggtcatac ccagcacgga attcaggaag 5040
ggatagaagg ccaggttgcc ctcgatgatg gtcaggatct ccttcagggc ggtaacgcgc 5100
agggagaagt tgatccacat gggcagtatg aagatcatga taatgaccga tttggccttc 5160
agcttgctgg tggccaggat ataggccacc ggataagcga tcagcaggca cagcaccgtg 5220
gtcgccaggg aaacggcgaa gctgtaagcc agggttccca gggtattcgg atccgtaaag 5280
aaatccgtca ggtttttcag ggtgaatact ccctgcccgt tcgtaaacgc ataatagaac 5340
agtaccagca gcggcgccac cacgaatagc agcaggaaca cagcatacgg tatgcccagc 5400
gtttttctgg aaaatctcat atgctcctcc ttacttcttc accgtaaagc gcatcttgtc 5460
cagcggcagg atcaggccga cctggtcccc catgttccac aggtattcat cgtccacgat 5520
gaagtcctgt cccacatcgg tattgatcac gtagctgtag tggtcgccct tgtagatcag 5580
gttgctgatg gtgccttcga tcaggacctg ctcatcgtcc acatcgtccg tcatgcggat 5640
atcctggggt tcgatggccg cgatgatatg gatcttttca ggatcgatca cttcgccgtt 5700
ctcatccagc agctcgtgat tctcgttccg gcggctgccc ttgatcagct gcgtcaggct 5760
gacatccagc tcctgctcgt tgaattccag gcggtggttc ctgttcatat ccacggggaa 5820
cacattggtg tggtcctcgg cgatcatgat atggatgccg tccggctcca cccgcatgcc 5880
gacttcgctg ccgacctccg cggatttcgt gctctggatc accacttcat agcggccgga 5940
ctgcacgatg atttcatagt ggattccctt aaagatcacg gaaatgacct taccgcggat 6000
ctgtccgttt tccggcgcgg tcagttcgat atcctccgga cggagaaccg cggtgatctg 6060
cgtgccttcc ggcacatcat ccacgcaggt gaactggccg ccggcaaacc gcgccgtcat 6120
gttgccggtc atgatgccgt tgaagatgtt gctgtcgccg ataaagtccg cgacaaaggc 6180
gttctgcggc tcattgtaga tatcctcagg cgtgccgatc tgctggattt tgccttcgga 6240
catgaccacg atcttgtcgc tcatggtcag ggcttcctcc tggtcgtgcg tcacgtaaat 6300
gaaggtaatg cccagctcat cgtgcatcgc cttcagttcg atctgcattt ccttgcgcat 6360
cttcaggtcc agggcgccga ggggctcatc cagcaggagg atttccggct cattcaccag 6420
ggcgcgggcg attgccacac gctgctgctg gccgccggac agcgtgggaa cggagcgctt 6480
ctcgaagcct tccaggtcca ccagctccag caccttcttg acctttttct tgatttcatc 6540
cttcggcatc tttttcagct tcaggccaaa ggcgatattt tcatacacat tcatatgcgg 6600
gaacagcgcg tagcgctgga acaccgtatt gatcggccgc tcgttgggcg gcagcttgga 6660
gatatcccgt ccgttcagca ggatatcgcc ttccgtgggc atatcaaaac cggcgatcat 6720
acgaagggtc gtggtttttc cgcaaccgga cggcccgagg aacgtgacga attcgccgcg 6780
ctttacttcc aggttgaaat cagagacagc cttgaacccg tcgctgaagg tccgtgaaat 6840
gtgcttcagt tcaataatgt tttcctgctt ttccatgtga gaaatatctc cttcccctgt 6900
ctgaaaacta aaaagctgat acccgattta aaaactcgga ggtgtggaca cccagatgaa 6960
ccgcgccggc cggctgcccg gattctcgat ccggtgcgtc ttgtccgctt cataatagaa 7020
gctctccccc gccttgacgt catagcgggt ttcctgccgg taaagcgcga tcttcccgct 7080
gagcacgtag ccgaactcct ccccgttgtg gggatcgtcc acaggaagtg aaccgtgggg 7140
ctggagcgtc accaggatcg gttccatctg gttgctctgg gcggaaggga ccacccagac 7200
cttgctgttt ccggcgtcgt caatcttctc gaaataatcc cggtccccga agaccacttt 7260
ttccgctccc gacgtcttga agaaatcgga aggggtcgtt cccaggcact cgatcagatc 7320
cagcagcgtc accagcgacg gcgttacctg gccctgttcc agctgggaaa taaagccctt 7380
ggtcagctcc gcgcggtccg ccagctcctg ctgggtcagc ccgttctgtt tgcgcagatc 7440
ccggagtttt gtgcctacct gcgtcataac tctctccgcg tccattgttt ttcccgccat 7500
ccgcctttct gtcttaagtc caagtaattc taaacacgga atccattttt gctaaacctc 7560
gcaaacaata gcactcctaa actaaaatgt caatattact aaactttttt cttttgtatt 7620
tttttgttct ttgtgtacag tccatggaca gaagctcccg gaacccccga aaaacaaaga 7680
aaatccgggg ctcccatatg gggagccccg gtgttttccg ggtggttttt tacagccgct 7740
tctttttctg cagattctcc cacagcacgt ccgccgcttt ctcctgcggc ttctccggct 7800
gcttcttttc cggtttcttc ttcttttccg gcttcaccgg tttctccgct tccggctctt 7860

ccttcttctc cggctcatgc agccagtcga gccggcgctg cgccacgtca aacaggaaga 7920
tgaggccggc cagcagcgcc aggatccacg tcaggtccct gcgcttgcgc gcgctcgtat 7980
ccgggaaacg caggagctcc gatgtgttct ccgtcgcgat gcctccgctt gatgcgctca 8040
gctgcgccag gatcccgtca tcggtggtgc gcaggtcata ttccctgctc caggaaagca 8100
ccgcaccgct gtccgtagcc aggagctcat tccccagccg gtcccgcaca gacacgtgca 8160
ccgcataggc gccggcctgg gaggtatccg cttcgccctc aaaggtccgg gcattcacct 8220
ggttcagttc cacggtttcc tgggtgccgt ccggccggat aatccgggct tccgccttcg 8280
ccgcgcggct catcagctcg tcctccgcgt ccgcggtgaa gcgcagcttc ccgtcctcca 8340
gttccgcatc gccgctgcgg gtattcaccg gcaggatata ggacaggatt ccgctgaaga 8400
aagccggcgc gtcctcccag ttcaggaagc ttccggtcca gccgccctgc acgtcgctgg 8460
tccaggcggc caccttgcct gcgccatact gccaccaggc cagcaccggt tctttccggt 8520
tgctggagag ggagaccgtt gccagcggtt tttctgttgt cgccaggtat ccgtccagca 8580
ccgggaatcc gtcaaagctt gtcatggaag gatccgtgac aatgggcgtg aaggtccggt 8640
tctgcacata ggatccggag atcatcaggg tttccttggt gaaaatcttc ggcaggctgt 8700
cgaaggggcc ggcgtaatac atccgtccct tgccgagctc cgccagcttc cgcatgcccg 8760
ccgtatccga tccgtcgccg accgccacgg tcgttacggt gatgcccttc tccgccattt 8820
ctttcacgac gtcctcatat ccggtatccc cggcttcgcc gtccgtcagg aaaatcacat 8880
gcttgtactg cgcgttcaca ttttccagcg cgtccttcgc catcttcagc ggagaataga 8940
aggccgtacc gccgcccagg cggatggtcc ggaccttttc ctgcatggcc gccacatccg 9000
tcaccgggct caggggtacc acccatttcc cggtatcgtc aaaggcgatg acaccggcgc 9060
tgtcccgctc gttcaggatc tccagcgcgc tgcaggctgc ctcccgggcc agctgcagcc 9120
gggtcagtcc ccaggagctg tccgacatgg agccggattt gtcgatcgcg atcaccagcg 9180
ccgtggaagg caggtccatc ttgttcttga catcaatgga taccggcaac atcttttcaa 9240
tcaggctgcc ccggtatccg ccgagggcat agctggaatc accgccgaac accgccaggc 9300
cgcagcccag ctcccgcacg gcctcgtcca gcgcttccgc ctgtgcttcc gacagctgtt 9360
ccgcatccac gttcaccagg gcgatcgcgt gatacgccca caagtccgcg gcctgctcgc 9420
tgagcatgct gaccgggatc acgcggacct tcatgcccgc gctctccagc atctttttca 9480
gctcttcgcc ttccccgctc ttcccttccg ccaggaggac attcatctcg ccggagacga 9540
cggtataggt gccgttcagg ttattcggcg ccaccgtatc cccgtccagc aggatctgcg 9600
cctcgcaggg aatgacgccg ctgcgcgtgg caacggactc aaaggcaaag gtattctctc 9660
ccttccgcag ggtcacccgc tgcgtccggg gcgtttcatg gttttcactc agcagaagtg 9720
tcgcttcccc tgccgtgttt gtgtaaaccg tcaccagggt cgtatacttc tggccctggt 9780
acagcgtatc cggtacgctg atttccgtca cctgggcatc cggtcccgtt ttccgggcca 9840
cctgcagggt gttcaccggc acgccggtcc ggctgtccgc ccattccgcg gcgccggtta 9900
ccgcgccatc gctgatgacg gcaatgccgc cgttggaatc cgtgggcagc agcgccgagg 9960
ccagggacag cgcctctccc aggttgctgg cgctgcggtc ggccgagaca tgcagcccgg 10020
tccattccgt attcccgccg atggattttt ccaccgccgc atgccggccg aaaacgataa 10080
taccggtctt ccggtcattc cccgcttcct gcagggcctg ccgggccagg gaaaccactt 10140
cctcctcccg cacggaggcg gaagcgtcca gcaccagcca ggcggtccgg tccgggctcg 10200
ccgtcaggat actcagtccc gcgaaggcca gtgccgtcag gatgatcagc acatggcgca 10260
ggatatggga aaccctttcc ttcaggctcc tgctcctctg cagatgtgcc atccacagca 10320
ccagcgccat gcagaccggg atgaccagca gccggagcgg atgcgcaatc tcaaagctca 10380
tgcggcagcc cccttccctt tcctgcggct ccacggttca cgggccagga taaactcaat 10440
catcagcagc gcgaggaaca cacccatcag gattccggtc aggtcctggc cgcggcttcc 10500
ggcggctgcg gcgtccgcgc cttccgtatc ccgcgccacc gtacgcacat cgctctcctc 10560
ccgggccacc ggcgcttccg tctccggaac ctccgccgcg ctccggggca ggagcatatt 10620
cagaatgttc tgcaccagga caggaaaatc atatttcagg ggcaggttgg aatcctgcag 10680
aaggaacccg atggctgcct cgctgtccgt ataggccgcc agcacgtcgc ttccggcccg 10740
gatcgcggct tttccgccgg tgagcggctt gcaggcgcgc aacgccacat cccgcagggt 10800
caggccctcc gtcatcgggg attcctccgc cgtcaggctg ccggccggct ccttctcctc 10860
ttcgctccag gtgaaagccg tctttgaggg atcccgggag aacagcaggg gcgaggtccc 10920
gtaaatatac aggtcggctt ccgtctccgc cggggcttcc tcgctcgtcc gcacaacccg 10980
caggtcttcc cggaccttca gcgcgctttc caggaacagg ttatccccgg tcagcgccac 11040
agtccgcacc gtgctgcgct ccaccggcgc ttccgcccgg ttatccgcca tgagcgcgtc 11100

cggctcccgg atatagactt ccacctttgc ggcattttcc ggaatggtga acagcacccc 11160
cgccgtttca cccgcgggaa tctccgtttc ctttgcttca cacagtttgc cgtccgcctc 11220
gcaggccacc gtgatcctgc agtcgctgcc gaaattgcag atccgcgcgt aggcggatcc 11280
ctcctccgcg gtcagcgtat acacgccgcg gttctcttca ccccgcccgg catggatccc 11340
ggcgatgccc gccttcggtg tatagctgtc cgagaaaaca acaatccgcg ttccccggtc 11400
cgtttcctcc cggcggatcg cttccgccag ggaaaccgct ttttccaggt cggcgccttc 11460
ccgtccgcag gacaggctcc ggatggcctg gcgcacatcc tcccggtccg tgcttccggt 11520
cagggtctgg cggacttccg ttccggccgc aaggatcgtg atttcctctc ccggcggcag 11580
gccctgcagc agttcctccg cccggtcctt cgcctcgtcc agccggctct ttcccgcgct 11640
gaccgtctgc atactcgcgg acaggtcaaa aatcatcacg gtccgggaag cctccctgcc 11700
ggtcaggttc ggctgcatca gcgccagcgc cagcaccagc gccgccagga actgcaccgg 11760
aagcaggatg tttttccgca ggcgctgcag cggcttgttg gcggcatggt cccggatggc 11820
tgtccgccac aggaaggtgc tcggaacctg cttcggcaga tatttccgcc gcagcagata 11880
cagcaccaca atcccggccg ggagcagcag aaaccacgca aactgcgggg cagcaaatgt 11940
cagcatgata acctctctag atcattccgc tctgtgaaat cagcggaatc cattcctctt 12000
caaaattctt tccggcatcc agcagcgcat agggcgtttc acgccggaag caggtttcgc 12060
ggacctcccg caggaaattc tccagcgtct gccggtagtt ttccaaggct tcccggtccg 12120
ccaggagatt gatggcctcc ccggtctcgc tgtccgtcag cttcaccgca ccgtccaggg 12180
ccggatccag ctcaaaggcg gagagcggct ggatgaccgc gcattcctgc ccgcggtaac 12240
gcaggctgtc cagcgccgtg ccgattccgc cttcctcata gcagtcactg atcaggaagc 12300
agagtccccg ccgcaggcct tcggtccgca tcacggtctc cgtcagcctg ccggcgccgc 12360
cgggggcgca ggtatccagg aactgctcca gccgggcata ggcctggcgc ccggacagct 12420
ggggcgacat gcggtttccg ccttcctgca ggcagatcac ccgcagccgg tccccgccgg 12480
tgagcgccag gtagctgacg gcctccgccg cctgccgcgc cgtttcccat ttttcgccca 12540
tgcttccgct gccgtccagc aggaccgtca ccgccgattc ctgctcttcc atgaacagct 12600
tcaggaagat tttgtcgaac cgggccatgg cgttccagtc aatgcggcgg atatcgtccc 12660
cgggaatata ctcccggtaa tcggagaatt ccgccgagga accggtctgc cgggaccgcc 12720
gggttccgcc ggcgccgccg gaagcgcggc ccttcatcgc cagctgcagc gtgtccagcc 12780
gggacagaaa cgcgtcactt aacatagttt tccaccagtt ccgccacgat cgattccata 12840
tccttgccct cggtcacggc ggcaaaattc agggccaggc ggtgccgcag gcttgccggc 12900
gccacatagc ggatatcctc ataggccacg ttgtagcggc cctgctccag cgcccgcaca 12960
cgggccgtgg acaggatcgc ctgcgccgcc cgggggctgg cgccgaagcg caggtacttc 13020
ttcgtaattt ccggcgcatc ctccgtttcc ggatgggtag ccagcaccag ccggagcgcg 13080
ctttcctgca cgctgtccgc cacaggcacc tgccgcgcgg tctcccgcat atccagcagt 13140
tccgccgcat ccgccgcttt cccggcttcc tgcgccgcgc tgccgatggt caggttcatg 13200
atgccgccca gctccttcag gttcgggaag ggcaccagca ccttgaacag gaaacggtcc 13260
aactgcgcct cgggcagggg gtaggtacct tcctgctcca ccgggttctg cgtcgccagc 13320
acgaagtagg gttccggcag cttcatggtt tccccggcca cggtaacgct gtgctcctgc 13380
atagcttcca gcagggccgc ctgggtcttg ggcgtcgcgc ggttgatttc atccgccagc 13440
acgatggaag caaacagcgg tccctgctgg aagcggaagc tgcttccctc ctccgtacgg 13500
atcaggatgt tcgtacccgt gatatccgcg ggcatcaggt ccggcgtaaa ctggatccgg 13560
gagaaaggca gatccaacgt cttggaaagg atccggacca gatgggtttt tcccaggccg 13620
ggaacgcctt ccagcagcac gttgccgccg gcgatgaccg ccagcagcag gtgttccacc 13680
acgtcctcct ggccgatcat ttcctttgcg atttccgtgc ggatcttcgc atatttttcc 13740
gcgaacgctt cgatagactg attcgtgttc atgtttcgtt ccttcctctc attcatcggt 13800
gagctggcgg aaatactcat ccacccattc cttctgttcc cgggtcagat gcgccgattc 13860
cgccgcctgc gcttccgttt ccgcgtattc cccgatcacc tgccggtagg gcacattccc 13920
ttccaggctt cctttgccgg gaccggtttc gatctgcacg gagtcttttc ccagctgctg 13980
ctgttccgtc attacgtccc gggaagcggt attcaccttt tccgggtcat aaatcgtttc 14040
gtattcgcct tccttgtatt ccggcggccg gtttcccctg gacgcagagg acttgtttcc 14100
gctgcctccg ccctgttcct cctcattggt ggtgccgctg ccggcgccgg ccgcattctg 14160
tccctgattg ccgctgccgc ccgagccgct gccggaggag ttgctgttgc tgctctgctg 14220
ttgctgcgag gaagcgccct gtgcggcgcc gagcgcggct ttcatctgcg ccagggcctg 14280
ctgcatcacg ctggcctggc tgcttccgga ctgcatggcc tgcatctgcg catcggacat 14340

ttcgccctgc tgcgcagcct gcgccatctg ttccgccatt tcctgcgctt ccccgctcag 14400
gcctttcgcc gcctctttga gttcatcggt attgatttcc gacagtgtcg cggacatttc 14460
cgccgcgtcc ccgttctcca gggcctgggc cgccgcgtcc attccggcat cccgcagcgc 14520
gtccgccagc tcgctgacgg catccccggc cgtcttctgc cggatttcct ccagccggct 14580
ttccgctttg tcgaccgcca ccagcgcgtc cacatcgtcc cggctgtcct ccagctcccg 14640
tttcaggtcc gccgtcagtt tccgcagctc cgccttttcc ttctcggtct ttccggcttc 14700
ctccgcggct tcggcttcat cgatttcctc cgccatgcgg gcggtctttt cctgcaggat 14760
tttgcgttcc gacacgatcc ggtccccgcc gccgggaatc agcagcgtcc cggcgcaaag 14820
caccagcagc gtcgcggcgg ccagcagccg tttcttcggc gcctgcagcc ggatcttccg 14880
ggcgtccagc gcctgcaggt gctcgcaggc atcctgccgc tgggccacga caagctccgc 14940
agccttcccg tccgccggtg gccggtccgc catttccagc gcggtcacgg tccgctcccg 15000
caggccgcag cgatccgcgg cgcgggcagc ttcccggctg tccaccggac gcagggcgtt 15060
gattacggcc cccagcagcg cgcctccggc cacggcggcc ccggcaatcc ccccccggcc 15120
ccccagcggc acaaaggttg tcaccgccag cagcaccagt gccgctgccg cgccggccgc 15180
aagcccggtc gcagcgccgc accaaagccg cccgagccgg atccggttct tcaccggacg 15240
cagggctttc agaagctcat tcattccgtc gtaccctttc atcagcattc atctgtcatt 15300
acaatttttt acttggttct ttttgccggt ttcagggagt tcttccggag caggaatccc 15360
gccgcaagca gcagcaggac cgccagcacc aggatgattc cgcaggaaat caacgccagg 15420
gtttcgccgc ccgccttgtc catcagccgg aactcgcaca gcagccggcc tatttcacgg 15480
tattccatcg tgctggtgag catccccgtc tgtccgtgca ggagggcgat cagcgcatat 15540
cccggattgc acagcagcac cggggaaatc atcttcagcg catcactggg ggcaagcgcc 15600
gccaggctgc tcgtgtcata gacaacggag gtaacctcca cgggaaaatc cattatcagg 15660
gggatcagcg tcaggccgcc gatggccagg agaatcaggt agctgatgac gccgctggcc 15720
accgtggagc gcgccagcgt ggaagaaaac accccgacgc tcaccgacgc aaaagcgatc 15780
agcagcagga agagttcccc gatcaggatc tgcggcagcg tgactgcgcc ggccaccatc 15840
gtcaggcaca tcaccgggaa gccgcagaca atcagcagcg ccagcatggc gaagctttcc 15900
atggctttgc cgatcacgat ccggaaggac cgggtgtttg tcaccagcag caattccagc 15960
gtctgccgtt cccgctcccc ggcaatggcg ccggaggtca tcgccggcgc gatcaggagg 16020
atcaggataa actgggtgac catctgcgcc tggtagcagg tgacgctgtt gcggatggaa 16080
tagaaggtag cctcgttccc aaagatccgc cccataatca gcagcgtgac cgccagcatc 16140
accaccacat aggcagctgc gatcagcagc gttttgaaag accgcatccg gcgggttgcg 16200
gagaaggaca gaatcggatt aagcatctcc tgtcacctcc atgaatactt tttcgagatt 16260
catcggcgcg cggtggaaat cgcacaccgg gatattccgg gaaatcattt cccgcagcat 16320
ctgcatatgc tctgcttcgc cgccttccgt gcggaccttc agcacgcagg ttccctcctg 16380
caggatctcc ttcacaaagg gaagctcctt caggcaggcc accgccatat ccaccgccgc 16440
ctcaccgcag ccttcctgca ggcggatttc cagcggggca ttcccgttca ttttctccga 16500
aagcgcttcc acgctgccgg agaacaccag ctgaccgtgg tccaggatcg tcagggaatc 16560
gcacatctcg ctcagttccg gcaggatgtg ggaagagatc aggacggttt tccccatttc 16620
ccgcagggtc cgcaggatcg ttttcatttc cgcccgggcc cggggatcca tgccggaagc 16680
cggttcgtcc aggatcagca gcttcggatc atggatcagg ctccgcgcca ggcaaagccg 16740
ctgcttcatg ccccgggaca gggaatccac aaacgcgtcc ttcttttcgg acagctgcac 16800
caggtccagc aggcgttccg tcattttctt ccggtccgcc gaaggaatcc ggtagcaccg 16860
ggcatagaaa tccagatact cccagcactt caggctgtca tagacgccga agaagtccgg 16920
catatagccg atcagccgcc gtgcctcggt tccgctgcgg accacatcca ccccgtcgat 16980
cgtcgccgtt ccggaagtgg gcttgagcag cgttgccagg atccgcatgg tcgtggtctt 17040
tcccgcgccg ttgggcccca caaaaccgtg gagctgcccg tccgcaatct ccagggacag 17100
tccgttcagc gcggtgaaat caccgtattt cttcgtcagt ttctcaaggc tcagcatgcg 17160
tctcccctcc ttcctttatt tttccctcca gggtcagtga cggctccgga atatccatgt 17220
agctgtccga caccttgggg cggaactggc agtacaggtt cccctgttca tccaggtatt 17280
tcccgggctg cttgagcggg gtgttcatct gcacctccac ccaggttttc agttccacat 17340
tcagcacaaa gcacgccacg tcgttggcgt accagtcctc cacgccgacg gacagctttt 17400
caatctcaat ccggtccagg ttctccaggg taaagcggaa agtgggatgt tctgacagcg 17460
ggaaataatt gtaatacagt tttccgccgt tcggatcctc agccatatcg cccgccggca 17520
tgcccgacga atcgatctcg cagcgcaccg catgatccat tccgggagca tggaatacca 17580

cgcctgtttt cccgatcttc aggtactcta ttttcgccgt gtacatcggc agcaccgtgc 17640
cgccttccac ttccttcccg tccaccatga gcggcgaagc gaaatctcct tccggttccg 17700
cacagtaaac gaaaatcacg ttttcgcggg tccccgcact gttgtcacgc gtcagctggt 17760
cataggcgct gttcatcatc ccgctcagga tcgattcccg gccgccgtaa tcattcgtat 17820
ccgaattctc ccgcatgccg gagagcacca tataaacccc ggaggtcaca ttccgcagca 17880
tcacgccgtc ctcaaacacc ggattggagg gatcaggcgc atccttggcg accatggcaa 17940
attccctgct ttccccgggt gccagtgccg gaacacggac aaagccataa acgctcagca 18000
ccacgccttc cttcagggaa tagggcaggc cgttccggac ggtaccgtgc aggccgtcgc 18060
tctccatcca gatctgggcc tccacctgac cttcctgctc cgacgtccgt acggaggaca 18120
ggcggacgac ctcccagggc gtattcgcat tcagcgccga agagccgttc tccccggtga 18180
tctgaaccac ccgcagctgg gacggttcct tcagctccgc ggtgtcttcc tcctccgccc 18240
agtagtaata gccgtcgtca tagacctgcg cttccaggtc ctccccgttc atgctgtaac 18300
ggtgcaggcc cggccgcggc gccgccgcgg tcacaaacac atagcgggtg gctcttcctt 18360
ccgcattctg caccaggtta acggttgtcg cggtccccgg ccggttcagc atcgagcttc 18420
cagccatcac ggtccccacc gccgctgccg ccagcgccgc cgccggaatc accgcccaca 18480
tccactgggt tttactgcgc ttcttcagga tgatccacat cacgccgcag atcaccacag 18540
ccccggcaat gaccgccacc gcggccggca ggctgttccg gacctgtatt tttgtatcat 18600
cgcccgcgga atacagggcg ctggactctt ccccgggata cagcagcttg ctgtacatat 18660
tgggatccat tttgatcagc agctgctgga agaaaatatg catggcgctt tccgcgttca 18720
gcgccgcatt gccggcctcc caggcaaggg taatgatccg cccgctgccg gcgccttcca 18780
gccagaccag gccgttgccc tcgctgtctg aaatcagggg attcgcgcct tcgagctttg 18840
ccagggcgat ctccgttttc cttccggaat gtttctggcc cgcaaggctc tccacctcgg 18900
aaagcacgcc gtccgatacc gtgaaatcag aagcctgcag cttcgtcagg tcgcccagga 18960
aagccaggtt gcgcggcgcg gtgctcccgc cgccggcaat cagcacatgc ccggaaacga 19020
tccagtcctt cagcgcctgc tgctgttttt ccgtcagcag ggcaggatcc gtatcatcca 19080
gcacaatcat tccgaaggaa tccagcagtt tcggattctc cgggagggtc tccggcgtca 19140
gcgccacggt ctgccagaat tcatagcggt agagcacatc gttctcctgg ttgatatcca 19200
gattcgccag gttccgcggc cgggtggaca gtacgccgat catcatcgcg ttgggattga 19260
tcacgccgga aggtgtgctg ttcacagcgg agatgacttt tccgtcctta acgatctcca 19320
ccgtaaaggt attctgccgg gtctgggcca ccacgggcag caccaccgta cgctcgccgc 19380
ccgccggaac ggacagcgcc gtttcaaagc ggtcgtattt cgcggtgttc acatatccgt 19440
tcaccgccag catggcttcc aggtccccgc cgttattgcg gacggtcacc cgcaccggga 19500
tggctttccc gtaggtcacc tgtccgtcat acccgagccg gacttccatc tccagctgcg 19560
ggttttccac ttccgtatgc agcccgtctg ccgcggccgc tccggcccac gccagcaggc 19620
aaatcaccag gccgaaggca aaccacctta cagctttttt gcgcatacca atacctcctc 19680
gtgagcattg ctcactcatt tatacgctga aacaaccgga tttgttccat gaaggcggaa 19740
aattcatctt ccgttcattt tcctcagaag aagaaactcc tccgcagtgc cgcccgcagc 19800
gagggaataa acgacgcgac aaacaggaac agcccgatca tgcagaacgc gatcatacag 19860
tacagcgacc agacgaacat gggcgtccgg aacgtgatat gctggaagaa ctccaccagc 19920
atgaagctca gcccgatggc aatcaccagc aggctcatca gccgcagccg tccctgcacg 19980
atatagcgca tcattgccac gcctgccagc gtcaggatac cggcaatcat ggtcaccggg 20040
aaggcgaagg acaggaacca gtgcccgccc gtattcacgc agatatacag cagataccct 20100
gccaggcagg caaaatcaat cggcaggaaa atcatcggtt tccaccggga aaacagcagc 20160
ggcagcacca gccagaccca gcccagcgcc atacccagcg ccgcgattcc gctccagccg 20220
acttcccccc tgctcttcag gcagaaaatc aggcaggaca gcgcgaccgc cgcaaaaata 20280
atggtcacca ggcccagcac caggtatttc ccgctctgct tctccggctg cggatacttc 20340
ctggaatagg gagcttcctc cgccggccgc ttcaccggcg ccacctgtac ctccgtatga 20400
cacagggggc actccgtcac gccatcctgc agccgcaccc cgcaatatac acaatacatt 20460
cactcaccgc ctgttgcttt caatatccac aggtatgccc atctccacca gcgaggagaa 20520
aaacagccgt tccagctccg aatcccggat cccgcggatc atgctgatat aggtctttcc 20580
ctgccaggaa accacggagc agttattcgg ataggaatac tgcaccccga taataaaatc 20640
aaaccgttcc acatatgggg ccattgcctc cggcagcttc accacaccca tgttggaaat 20700
attcaggctt cccttgcttt ccccgctcag ggtgtaaacc agccgcatgg ccagggtctt 20760
cagcggcagg ggcaccagcc gcagcggcag gcagcgctgg agatccacgt tggtcgcgac 20820

ccggccggcc atccgctgtg gaattgtttc cgccgccagc tgatggtgca tcaggcggca 20880
gatttcgctg cgggtatatt cccccagccg cgggtcaaac ccgatgttga ccgccagggt 20940
aaagttccgc agggtattca ggccgaaggt tttccgcagg ttcaccggca cggtgatctt 21000
gatatggctc cgtttcccgg ccaggggatc cgccttcatc cgggccagca ccgcctccgc 21060
catcaccgcg gccagccatg ccgtcactgt gacgccttcc gcgtgtgcct tcgccagcag 21120
cgcctccgtc ggaatgattc cggttacgat atgccggaag gggccttctt ccggattgcc 21180
ccgcgtatgg taggcattcg cctccgcacg ggaaaccttt cctttcgccg cgcactgcag 21240
gaagcagtcc cgggcttcct cctccttcgg cggctccgcc ggatcgacaa tatttccctc 21300
cggcgggatc accgcgcctt ccttcagcgt caggtatctt gcagccaggt tctgcagaaa 21360
ggccatgcct cccgttccgt ctgtcacgga atggaagaac tccacagcga tgcggttttg 21420
ataatacaga atgcgaatgc agcatttgtt cagcttccgc ctggtcatcg gtgtcagcgg 21480
ccaggcgtaa tcttcctccg ccttcggtgc ttcgtccaca gtctccaggt aataccagaa 21540
aaaaccggtt ttcagccgca catagatcgt gggaaaccgg ggcagcagat cctccgcagc 21600
gcggcccagg atctccggat ccaccctgtt cttcagcgta actgaaaccc ggaaaacatt 21660
attccagttc tttcgcatca cggcggggaa aatcagcgcc gcgttatcca gccggtacca 21720
ttgcttcttc atggttctct ctcgtttcat tcgtcgggat accacccgtc attataccat 21780
atttatacat tccgctcaac cggaagcgcc cggattgtcc gtacaaaaca gcttttgttc 21840
tttcctttcc cggcctgtta tgatacgatg cacatacaaa agcggaaggg aggaaacagc 21900
tatggcaacg ggaccggtga tcggactgtg cgcccatgtg gacgcgggca aaaccacgct 21960
ctctgaatcc atgcttttcc tctccggcgc cgtccgccgc cagggccgcg tggatcatgg 22020
cgacgccttc ctcgataccg accggatgga gaaggaccgc ggcattacga tcttttccaa 22080
ggaagcccgt ctctcctgga accatacgga cctgacgctg atggacacgc cggggcacac 22140
cgacttttcc ggcgagaccg aacgcgccct gaatgtgctg gacgcggccg tgctcgtcat 22200
cagcgccgcc gacggcgtcc agccgcacac ccgtacgctc tggcggctgc tggagcagcg 22260
cgggctgccc gtgattctct tcctgaacaa aaccgatctg ccccatgacg cggaagcgct 22320
tttcgcttcc ctcagccggg agctttccgg ccagatcgtt gcatttccct ccccggatcc 22380
ggagaagctg gccctgtgcg acgaaacctg cctggacgcc tatctccggg agggtgaaat 22440
cccggaacgg atgatccatg ccatggtagc cggccgcagg atcttcccgc tgttctcggg 22500
ctccgccctg cggaacgagg gtgtggaacc cctgctggac ttcctggccc gcttcgatcc 22560
gcggccggct ccgcccgccg cgttcggcgc ccgggtttac aaagtggcca gggatccgca 22620
gggcgcacgc cttgccttcc tgcgcatcac cggcggaacg ctgaaggcgc gggacctgct 22680
gacgctgaaa agcccggagg gtgaaacgct ctgggcagaa aaggctgccg agatccggct 22740
gtattccggc gcgcgctata cggccgtgca ggaagtttcc gccggacagg tctgcagtgt 22800
cgtcgggctc agcaaagcgc tgcccgggga cgggctgggc accgatcccg gccggaagga 22860
gcaggagctc cgcccctgct atgcctgccg cgtggtgctg ccgcccggcg ccgatctgca 22920
ttatgtgctg aactgcctgg aaaccctgca ggaggaagag ccgctgatcc aggtggaata 22980
tgtggaaccc cgaagggaaa tccgggtcca ttccatgggc gatgtatacc tggaagtcct 23040
gcgcgcccag ctggcggacc gcttcgggct ggatgtctcc tttgccgaaa ccacagtgct 23100
ctaccgggag acgattgaag ccccggtgga gggcgccggc cactacgagc cgctgcgcca 23160
ctacgcggag gtccacctgc tgattgcgcc gctgccccgc ggcagcggac tggtctgcga 23220
ttcctccgtg tccacggacg acctggccct caactggcag cggctcatcg tgacccacct 23280
ccgggagaag gtgcacgtgg gcgtactcac cggttccccc gtcaccgatc tgcatatcac 23340
gctgattgcc ggaaaagcgc atctcaagca taccgaaggc ggagatttcc gccaggccac 23400
ctaccgcgcc ctgcggcagg ggctgatgaa ggcccgcagc atcctgctgg agccgtggat 23460
gacgctggac atcaccgtac cggaggactg tgtcggccgg gtgatgagcg acctgagcct 23520
catgggcggc cgcttcgacg cgccggagga cgccggggaa accctgcgcc gcctgtccgc 23580
ggcggttccc gcctccgcct gcgcggacta cgggcggcag ctggccgtct tcaccaaagg 23640
ccggggcagc ctgaccgccg ccttcctgga ctgggaaccc tgcgccgatc cggaaaaagt 23700
gatccgcgaa aaaggatatg atccccgccg ggatgtctgg aacaccccgg attccgtctt 23760
ctgctcccac ggcgccggcg tcaccgtccc ctgggacgag gcggacgccc acatgcacct 23820
gccgctgctg aaggatcttt cccgccggga ggttcccgcc cccgccgcgg gcggcgccgc 23880
ttccgcctac cacggcaccc gggaggaaga cctggccctg gaaagaatct ttgagcgcac 23940
ctacgggccg gtgaaggccc gccagctggt ggccgctccc tcggctgcgg cggaaaagca 24000
gcgggaggcg gcacccgctc cggcctccga cggtgaaatc ctgctgattg acggatacaa 24060

cgtcctgcat gcctgggacg aatggaagcc gttcctgccc gaccggctgg aggatgcccg 24120
ggatgccctg cgcgacctgc tgtgcgatta cgccggcgcc accggcaaac cggtaatcct 24180
ggtgttcgac gcctatgccg tgccggacaa tcccgggaag gccgagaaat acaagaacat 24240
ctatgtcatc tacacccggg aggcccagac ggcggacgcc ttcattgaac agacgaccta 24300
ttacggccgg aacaccgccc gcatccgggt ggtcacctcc gaccggccgg agcagctcat 24360
cgcctccggc aacgcggccc tgcgcacctc ggcgcgggaa ttccacgcgg aagtcaacgg 24420
cgtccgggac gggatcgccg cgttcatcgc ccggaaccgg ctttcccggc cggtacgcac 24480
cctggaagcc gcctacaagg aagcctggaa aaagagcagg gagcctcaga agtgaggctc 24540
cctgttcttt atacatgcag gatctatttt tttgtcagga attctttcag ctgttccatc 24600
ggcaccggcc gtgaatattt atatccctgc aggtaggaag cgcagcattc cctgcacatc 24660
tcctcatcct gctccgtttc cacgccttca aacagcaccg cgtaatccag cttggcaaac 24720
aggtctgcca ggctgcccag cacctgctcc gaacgcttgt cctgcgcgcc ggccaccacc 24780
atgctccggt caaacttgat gatatcgaag ggcagctcca ggatccgctc catattgctg 24840
tagccggtcc cgaaatcgtc caggtagaag gtgatcccct tctgcttgag ttcggcgatc 24900
ttttccttca tgaggaggaa gtcccgctca ttccgggatt ccgtcatttc aatggcaatc 24960
tgtttgccgt cgatcccgct ccggtcgata atgctgctga tatccatgca gaaatgctgg 25020
ttccgcagct ccatcgcgga aacattgacg ctgacgcggt caatgcagta gccctcctcg 25080
ttcagcgcct ttaccgcctt gcagacctta tgcaggataa tttcggtcag cacgtggata 25140
aacccgttct cttccgccag cgcgataaag cggtccggat aaacgaggcc ttcctccggc 25200
agctgcagcc gcatcagcgc ctccgcggta tcgtacttcc cggtgcggat attcagcacc 25260
ggctggcagt aaaccagcac ccgcggatca tccaggttct tttgcccggc aatatccgcc 25320
agctgatgca ggatcgcttc atacttccgg aatttgtcga tgtcgtccgg ctgcacccgg 25380
tagatcgtgt tttccggaat ccgggtatgg atgctccgga taaagctgac atattcgttg 25440
ttccggttca gttccgctgt cgtctggccg atcacaattt tgtaatccag cttgaagttc 25500
tgatattcct ggtcgaacgc ccgcaggatc gcttcggtcc gcgcctcata gtccgggttc 25560
cgctccttcc ggaccatcat cagcaggtgg ccgtttccga cccggaacat aacggaagag 25620
cggaaatact gcgccgtaaa acggcggacg atcgcctgca ggtcgccggg catctggcgt 25680
gcggtctcat ccagctcccg catgtagagg ctcaggtaca gatatgtctt gcctttatcg 25740
tcgctgtagc ggatcaggcc ttccagtccg cgcaggtcat tggatcccag ctgcgcgtca 25800
tagggattgg aatgcagcag gtacataatg gcaaccaccg ggaagaggaa ggtggcggtc 25860
gtaaacgaat cctgtccgtt cagcttggca atggcgagca tcgccacggc gaagaccatg 25920
gtcatgtaga aaccgttcat catccgacgg tacagatggt ggcgcacctg gatcatcagc 25980
atggacagca tcacgatgaa cagcccgtat ccgatcatga atatccggtt cccgccggcg 26040
gaagccgcgc cgtctcccag ctgtatcccg ctgccggaga aattgacaaa gagctccgca 26100
gccagcacca caaggatcag catcccgccg atggtatggt ataccttccg ttcctccttc 26160
gtcaggccgg tgacctggca gatgtaggca gcgaactgat acagcatgaa cagcagcagc 26220
ccgtgataca ggctctgaaa cccgtagaac agtccggtcg gcaggtttgg ccggtatttc 26280
agcagggtat ggaaggttac gtgggttgcc gccgcggcca tcaggatgcc gaccagggaa 26340
aggaacatcc ggaacgccct ggagcgcacc ttgaaggaga agaacagcaa aatcagcata 26400
atggaacaga cccccagaac caccaggtcg cctgcataca tatactgctc cagctgaaga 26460
gactgtggca ttccccttcc cctttccgcg tatattactc agttgttgtt ctgtaattgt 26520
atcatacaaa ccgacatctg aaaagaccgg ggattcttcc cttgcggtta ctgcccttcc 26580
tgcgtcagcg cttcataggt tacgccgtat ttgacggcga tttcccgggc aaaggatttc 26640
ccttccggcg gcgcgatctt cacccggaag gagttctttc cgtcccgggt ttccgccacc 26700
agggagaccg ctttcccggg caggccctcc gtgttgatct cgctgtccag ttccttcgcc 26760
agcttatgca tatgcgtatt gtaaatcgtg cgggtatccc tgcgcaggat ggctttcacc 26820
gcgtccaccg caatgaagta tccttcctca aaggaggtgg tggagaaagt ctcgttcagc 26880
aggatcaggc tctttccggt gctcttcccg tacagttccc ggaaccggcg gcattcctcg 26940
cccagccgcc ccaggtccag cgtcttgtcc tcatccgccg ggaaatgggt gagcacctgg 27000
tccgccggat caaacacgaa ccggtccgcc ggaacggaga taccgctctg cgccagcagg 27060
aaaagctgtc ccaccgcctg tgtgatcgtg gtttttccgc cgcggttcgc gccggtcagg 27120
atgtatgccc gcttctcccg gtcaaaggcc aggttgttgg taaccacatt ctccggtttt 27180
tccgcggaaa tcagcttcag gttgtagaat ccctccgcct ccgcgcaggc ggagccttcc 27240
gggtttttcc ggatctccgg cgtgcagaac ttccatccgg ccgcccggca cttctcaata 27300

tattccgccc agcgggtata gaacaccagc tccggaatca gttcggaaat ctccttcacg 27360
ctgacgttca ggtatttccc cagcatctcc cgcagcctgc gcgtgatccg ggaggtcagc 27420
atcgtcgcgg cgctgtccat ctggcgggga acggaagcgg atccgtccgc atccgggatc 27480
cgggccatgg acatcgccgc gatgggattg cgcatcacca ggccccgctc cacgaactgt 27540
cccacgccgt ccgcgagtcc cacccccgcg ttcgccagat aatagctgta gcttttgttc 27600
cagtccgcct ccggatggat ctcatccttc ggggtcacgg cggtcaggaa gttcttcagc 27660
aggccggagc gggtgaaggg cttcgaattc accgtgatca ggcccatatt gatggcttca 27720
aagcggtcgt tcaggttgat gccgacggtc aggcttttca cttccgacgc ggtgatccgc 27780
atctcctcca catccttgcg cagcgcggca aaaccgttat cccggtaaat actgtcaacc 27840
tcttcccgca ggcggatcat tccgtccgac tgcaggtcct tctccgccag gcactcgcgg 27900
atctcctcca ccgtgacgat atagtcgtgg tattcctcca ggcgatgcat caggtcccac 27960
aggccggact cgtcccccctg gtgcttccgt accacgccgt aatcatagaa ggtcttcacc 28020
cggtccagca gcttcagcat ccgctcccgg atttccggat ggcgcaggat atcctcaaac 28080
acgccgcacc ggaacgcggt gacttccgga tccggcgtca ggctccgcat cacctgctgg 28140
atcatcatca gctccgccgg ctgcgccgcc accttttccg ccacggcgtc cagtcccagg 28200
tcgtgccagc tttcctccgg gatcatccca taggtaatcc ggtcctggtc cggaaacaaa 28260
agactcagtt tattcgcttt catgcctgcc tccgctgtgt tgttccttta caaagagcgt 28320
atcacttccc gccgcggttt tcaagcatct gcaagcgcct ccgccggctg tacatccgaa 28380
aaacaaaaga atttccgaaa cgcccttgac agtcaccggg aggcggtgtt acaatgcgtg 28440
caatttcata caggcgatga cgaagacgga acagcgggag gccccgcaga gagccgggat 28500
tggtgcgaac cggcgggctg aagcttcatc ccatcccttc cgagccggag atccgaacgg 28560
gctttcctca gtaggcgtct cccgtgactg ctgcgtcaat gcataaggca tgacagtgcc 28620
tgaagaggtg tggaaacaca caatttgagt ggtaccgcga gtcagtatga cccgtctcaa 28680
agctgagacg ggtcatttta tttcaccaac gatttgtttt tcaaaggagg aaaacaatga 28740
tgcagaccca gacagatctt gcccagctga aagccatcgc ttcccgactg cgtgaaatgc 28800
gggagatcat gggatggacc gcggaggaaa tggccggaaa aaccgaggtc accccggagg 28860
aatacctgct gtacgaagca ggcgaggcgg acatgccctt ctccttcatc tacaaggccg 28920
cccaggcctt caacatggag ctgacggagc tgctggaagg acacaacgcc ttcctgtcca 28980
cctatacggt gacccgccgc ggcaagggcg aaaccaccgc gaaggaggaa ggcatcgcca 29040
tctcgaacct cgccccgaag ttccgggaca aggcggcgga gccttactgg gtgcgctatg 29100
aatattccga aaagcagcag aacgagccga tccacacgac gacgcaccac ggccaggaat 29160
ttgacctggt cctctccggc agcctgaagg tgcagatcgg cgaccatacc gaaatcctgc 29220
atgaaggcga cagcatctac tacaattcct ccctgcccca cggcatgatc gccgtggaca 29280
gcaaggactg cgtcttcctg gcgatgatcc tgcccggcga gggcccgcag gtggaaactg 29340
ccgccgtgac cgcgccgaaa gccgcgcggt acgagggaca cctcgtcgcc gagaaattca 29400
tcgacgcgaa ggaagacgac cagggacggc tccaggccat ctccttcccg aacaaggaaa 29460
ccttcaactt cggctttgac gttgtggacg aaatcgcccg ccagtatccg gacaagctgg 29520
ccatgctcta tgtggacaag catcacgagg agcgccgctt caccttcaag gatgtcaagg 29580
acgcttccaa ccagtgcgcc aactacttca cttccctggg catcaaacgg ggcgaccggg 29640
tgatgctggt gctcaagcgg cattaccagt tctggttctg catggtggcc ctgcacaagc 29700
tgggcgccat cgccatcccc gccaccaacc agctgaagga acacgacttt gaataccgct 29760
tcaacgccgc cggcgtgaag gcgctgatct gcaccgcgga cggggatacc gccgagatcg 29820
ccgagcgcgc ggccgccagg tgtccctccg tggaaacgct gatcatcgtc aacggcaccc 29880
gggagggctg gcgcagcctg aacgacgaat acccgctgtt cacccgccgc taccaccgtc 29940
cggaggatgc ttcctgcggc gacgatccgg ccctgatgtt cttcacctcc ggcaccaccg 30000
gaaacccgaa gatggcgcag cataagcaca cctacgccct gggccattat gtcaccgcca 30060
agtactggca ctgctgcgag cgcgacggac tgcacttcac catctccgat accggctggg 30120
gcaagtccct gtggggcaaa ctctacggcc agtggctgtg cgaaggcgcg gtcttcgtct 30180
acgactttga caggtttgac gcggcggaca tcctgccgat gttcaaaaag tacaatatca 30240
cgaccttctg cgcgccgccc accatgctcc ggatgttcat caaggtggac ctggccaact 30300
acgacctcag ttccatccat catatgacga ccgccggcga agcgctgaac ccggaagtct 30360
tccgccagtt cgaaaaggcc accggcctcc agatcatgga aggcttcggc cagacggaaa 30420
ccaccctgac catcggcaac ctgttcggcg gcgtgccgaa aatcggctcc atgggcaagg 30480
ccaacccgca atacgatgtg gacatcgtgg atccggacgg caaccctgtt gccaccggtg 30540

aagtcggcga gatcgtgatt cacacggaca agtccgcccc ctgcggcctg ttccgggaat 30600
attacctgga cgaagaaaaa acgaaggacg cctggcacga cggcctgtac cacacgggtg 30660
acaccgcctg gcgggatgaa gacggctatt tctggtatgt cagccgcatc gacgacgtga 30720
tcaagtcctc cggctaccgg atcgggccgt tcgaaatcga aagcgtcatc atggaactcc 30780
cctacgtgct ggaatgcggc gtctccgccg agccggatga agtccgcggc cagatcgtca 30840
aggcatccat tgtcctgacc aggggaacgg aaggcaccga ggagctgaaa aaggaaatcc 30900
agcagtatgt caaggagcac accgctccct ataaatatcc gcgcaagatc gttttccggg 30960
acgagcttcc gaagaccatt tccggcaaaa tccagcgcaa tctcctctga tccgcttttc 31020
tgcgcggaac gcttgcggcg ttccgcgttc ttttgatata ctgactgtat gattcaggag 31080
gtgaaccccc tatgatttct gcaacacagg ccccttccct ctgcaaagca tatgaaccgt 31140
atttccgcgt cggtgccgcc ctcagcagct ggctggtgaa agatcccgag ctcgtggaaa 31200
tcgtatgccg ccatttcaat tccatcacag cggacaacca gatgaagccg gacagcgtgc 31260
tgaacctgga aggaacgctg gcgcagggcg atccctgcca tgcggccacg gacttcaccc 31320
gggtggacgc gctgctttcc ttcgcccggg accatcagct gtccgtgcgc tatcatgtcc 31380
tggcctggca caaccagacg cctgtctggt tctttaaaaa ggactgggag aacgaccggg 31440
aggcgccttt cgcgccgaag gaagtgatcc ttgcccggct ggaaaactat atccgggatg 31500
tcatgcagca tgtgaacacc tgcttccccg gcgtggtcta cacctgggat gtggtgaacg 31560
aagcgattga gccggaccag gacggtcccg gcctgtaccg gacccgcagc ccctggttta 31620
ctgctgccgg acaggatttc ctgcccgccg ccttccgggc tgcccgcaga tacgctgccc 31680
cggggcagac gctgtgctac aacgactata acgcctttga gcccgtcaag cgggacgcca 31740
tcctcgacgt gctgaaaatg ctgaaggcgg aaaacctggt ggacacgatg ggaatgcaag 31800
gccattatct cctgccccac ctggatatcg ccgcctgcga gaccgcggcc cgggcctatg 31860
ctgccctggg gctgaagctg caggtcaccg agctggacat ccactgcaac agcgatgatg 31920
aagcgcatgt cgccgcgctg gccgaagcct accgttcctg gttcgccatg atcaaaaagc 31980
tggtgaagga aggcatcccg gtggaagccg tcaccttctg ggcgtaacg gacgcggaca 32040
gctggctgcc cggtttccgc aaggagccga gctatcccct cctcatcacc gcggaccgga 32100
aaaccaaacc cgcttttgac gctgtgctca gagaagcgga actgtaaaac atatcaacct 32160
gtcatagatt gctgaaagga gtttccccat ggccagactg cagatcgaat acttctccaa 32220
tgcccttcgc cgttcaacca cctttgaaat gatcatcccc aacgattttc ccgcgggcgt 32280
tccgtgggaa gccggcaagg cgcattcccg ccccatgtcc accctgttcc tcctgcatgg 32340
ctacaccgga aaagccggca actgggtgcc ggacggcctg ccggaaaagt ataacttcgc 32400
tatcgtcatg cccacggcgg aaaactcctt ctatctgaac ggggaagcca ccggcaccgc 32460
ctaccagacc atggtggggg aagagctggt gaattatgtc cgcaagacct tccacctggc 32520
cagcggcccg gaaaacacct gcatcgccgg cctgtccatg ggcggtttcg gcgcgctgca 32580
taccggcctc gcctatccgg agaccttcgg caaaatcggc gccctctcct ccgcgctcat 32640
cgtgcatgaa gtcgcgacca tgaagcccgg cggcggcaac gccgtcgcga actacgccta 32700
ctaccgggcc tgcttcgggg atctggaaac cgtcctggag cgggatgcga accccgaagt 32760
cctcgcccgc aaactgaaag cctccggcgc gaagattccg gagatctata tgtgctgcgg 32820
cacggaggac ttcctcctgc agaacaaccg ccagatgcac agcttcctcg aggcggaagg 32880
cattccgcac gaatatcacg aaagcaaggg cacccatgat atggtcttct ggcgcgaata 32940
tatcgcgaag atcgtggaat ggatgttcgc ctgaggtcct tacggatctt cccagggcat 33000
acagatattc tcctctgttt tttccagagc cgtctcctca cggaggcggt ttttctgtcc 33060
gatcaggtcc gaaaaaccgt ccagcgcgtc aaagggcgca aaaatcttgg cccggtgtcc 33120
tcggtccatg ggcggatgct tcggatcgtc ctcccggcgt gccggccgtc cccgcatcag 33180
tacccgcagg tacggaaaag agcggtcgat ccgccgcaga tcccagttgg attcagcctt 33240
ccgcaccgtc ccgcctcctt acgctatgcc ctgtgtccgc ctatttgttt atttctttcc 33300
atcgctgtgg cgccgtccag catattcatc ccgcgcacca gcagattgct tccaaaccgg 33360
ccccgcactt ccagcatcgc ttcctgcatg cggaattccc ggtcctgggc atcaggatcc 33420
gagaacatat ccatctgcag ggcgcagcct tcctcccggg gatccgtcag cgccagtccg 33480
agccggcgca ccagcagtcc cggatccacc ttttcccgga aggcccggtc cagtgcctcc 33540
aggatcaccg tgcggctgtt cgtctccggc cggaagcgca ccgttccgcc cacgtgcacc 33600
ggatgcagcc gcccgtaaaa atccaggcat accggccccg tatatcccgg gttctcctcc 33660
aggctggccg gatcaaagct gatccaccac attgccgcgg gaatccgcat ccgcttctcc 33720
accagtttga tccccgcctg ctccgccatt tcccggaaca ccagcagcga ttcctcaaac 33780

cggtacggcc ggggcagcac ctgcccggtg gaaaagctgc gcgcggacgg ccggtacgac 33840
ttgatatcgc tcatccccac gggttccacg ccccaggcgt gatccatcag gatctccgcg 33900
ttcaccccga acaggcggta cagcccttcc gggtttgcca cggccatggc cgcaacatcc 33960
cccatagtct gcatgccatg ggtcatcagg cggcgtgccg tccccggccc tacctgccag 34020
aaatccgtca gcggccggtg ttcccacaga aggagccggt aatcctcctc ccccagttcc 34080
gcgatgcgca caccatcccg gtccggcttc gcctttttgg ccacaatatc catgcccacc 34140
ttggccaggt acaggttcgt cccgatgccg gccgtcgcgg tgattcccgt ctccgccagc 34200
acggcgcgga tcagccgcat cgccagaaaa tgcgccggag aaactccggc gcgttccgcc 34260
tccgcccggc acaggtgaag gtaaggcgtc agatcaatga aacattcatc aatggaataa 34320
acgtggatat ccttttccga caggaatttc cggtagattt cgtaggctct cgcggatacc 34380
cgcaggtatt ccgccatccg cggcggtgcc acaatgtatt ccaccttcgt tttgtgcgcc 34440
gcttcataca gccggatggc ctgtttcgcc tcaaacaggc ggggacggga tgagaccccc 34500
tttgccttga gcgacgggga aactgccagg cagatcgtct tgtctgtccg gctttcatcc 34560
gccaccagca ggtttgcctc cagcgggttc aggttccggg ccacgcattc caccgaggca 34620
tagaaggact tcaggtcgat gcaaagatac tgcacgggat tgccccctta tgcattcagt 34680
accctgccga cgatccggaa gtccgtcccg gaggtgatca ccttcggatc atatgcggtg 34740
ttgcaggaaa ccagtaccgg ctggagatga cggatccctt cactgtccgt aaaattctcc 34800
acctcgtccc cctccggctc ctgctcggcg tacaccttga tatatccttc cccgtccact 34860
acaaacaggc ctacctcccc ggggagcagc cggctgcatt cctccaccca ggcatactgc 34920
ccgtcgtgca ggaccggctc catactgtcc ccgctgacgc ggacggcaaa ggaagctccc 34980
gccggcaccg aggaagccgg gaaagtctgg ctttcaaaca ggttgttgtc cagcgcctcg 35040
ccgaatccgg ccgaggccgg gagcgtgctg accttcatcg ccaccacagg ctccctgtgc 35100
atcgggcggc tgcggtacgc cgactggctt tccaggtagc cccggtagcc tgccagcatc 35160
ttccgccctt cctcgttcag cttttcctcc tcgccgaaga gcggaatccg cagcgcgtcg 35220
cacaccgccg ccagctgata ggcgctggga acggaatctc ccttttccca tttgctgacc 35280
gccgccgtct gtacccggac gccgcggagc gcgagcgctt ccgccagctc accctgcgtc 35340
agccgcaggg acctgcgcgc ctgcgtcagc cgcgcgccga tctcgttccg ctcccgtgcc 35400
ttttccgcgt cgtaccgcat ccctgtccct ccgtatatcc gctttctttg tctggagtat 35460
aatccccgga aaacggattt gtcaataaaa aattatcctg taaggataat tttttatcaa 35520
ttttatctta ttcggatatt ttcagaagga aagggttatc ttccgttcag tgcattcgcc 35580
ggccgctgcg gctgtggtct gtatcgtagg tgatgtcttc ggttttatgt gtcccggaag 35640
cggcaatccg gcgctccagc tcccggttga acgcctcagg ggacacggga atcccgaccc 35700
gcccgccggt ccatccggac aggtgaatgg cattggacag ctgcagcgcg cgaagcccgt 35760
cctccgcccc ggccaccggt ttttccccgc gcagcaggtg cgccgcgaaa gcgttgatca 35820
cgcccgcgtg ctgcggattc tccccgtcgg tttccagcag ggattcccgg atctcttccc 35880
ggtaccacgg atcatcgctg ctgaaacaga tctcccgctc attcgccccc aggcgccaga 35940
ccttcacctg accgttctcg cagacgatct ttccgcgttc gcagtcaatt tccagccggt 36000
tggttccggg caggtccccg gtggacgcca caaaaacgcc tgtcgccccc tccgggaact 36060
ccaggaaggc cgttacgtcg tcctccacct caatatggtg ccattttccc tccaggcacc 36120
gggccgtcac cgcgcagggc atcccgcaca gccagatcag caggtccagc tgatgggggc 36180
actggttcag cagcacgccg ccgccttccc ctgcccaggt ggcccgccag cttccggaat 36240
cgtagtagcg ctgcgtccgg taccagtcgg taatgatcca gttcatccgc ttcatggcgc 36300
ccatttcgcc gctgtccagc agttctttca ccgcccggta catacagttg gtgcgctggt 36360
tgaacatgaa ggtcagggtt ttcccggcct tctccgccgc gcggatcatc ttttccgccc 36420
ggtccgcctg caccgccgcc ggcttttcgc acagcacatg caggcccgcc tccagcgcag 36480
ccaccgccag cggttcatgc tgaaaatgcg gcacggcaat cagcaccgcg tcgcagcggc 36540
cggaagcgat cagctcccgg ccatctgtat aaatgcttac attttcaggc aggttcccgc 36600
gggcccactc ccggcgctcc tgccgcaggt ccgccacggc gcacagttct gtctccgggc 36660
atgctccgcc caggatcagc cggcagtgtt cgctgcccat gttgccgata ccgatgatgc 36720
ccagacggac ctttgttccc tgtatcatgg tttattctcc gatccggatg gcgttcttct 36780
gcgccagcag gcacagttcc gccgccttga aggcatgctc ctgggtcatg gcgttttccg 36840
tgcggttcag gcagtccaga atcagctgcc cgaagaaggg atatcctgtt gcgcctgccg 36900
cgttcaggta tttttctccc tggtcattca ccaggtatac atgtccgccg ccgtcttttt 36960
cggacgcaag attgacatac ttgcggattt ccacaaaacc tttggtcccc aggataaaca 37020

tccggccgtc accccaggtc cgcaggccgt ccggagtaaa ccagtccagc cggaaatagc 37080
cggctgcccc gttcttcccg gtcagctgcg cgtcaccgaa atcttccagc tccggatatt 37140
ccggatgcgc gaaattggca atccggctgg ccgtcacggt ggcgtcttcc tcctgcgcat 37200
aatacaagaa ctgttcgaac tgatggctgc cgatatcgca gaggatcccg ccgtattttt 37260
cccgctcaaa gaaccaggca ggccgtcccg gcgcacccag ccggtgcgga ccgaagccgg 37320
tgacggaaat caccttgccg atctccccgg aatgcgcgat gtatcccgcc aggatggagc 37380
cttcgtcatg caggcgctcc gcgtaataca ccatatattt ctttccggtg cgtttcaccg 37440
cttcccgggc agatgccaac tgctccagcg tcgtgaaagg cgccttgtcc gtaaagtaat 37500
cctttccggc gtccatggcc cgcagtccga gggcgcaccg ttcgctggtg accgccgcgc 37560
cggcgatcat ctgcgtttcc ggatcattca gtgcttcctc ctcgctgttg gctgcccggg 37620
cctgggggaa agctttcagg aacgccgcca ccttttttctt gtccggatca taaacatatt 37680
tcagggtgcc gccggcctct gtcagcccgt tgcacattcc gtaaatatgt ccgtggtcca 37740
gccgtacggc ggaaaagaca aattccctgc ccttgaccac cggctgcggt ttccctttcg 37800
gcgcatacgt catgccgtcc gaaacgttca ttttattttt ccccccttctt gaaattgcca 37860
aagctcaggc tgccgtcctg gctctccacc gacgtggttt tctcatagaa atgcggaacc 37920
gcaccgcgga ttccctccac ggtatagaaa ggatcctccg gagggatcgg cagcgtcacg 37980
gcctgatgcg aagatcccgc cttgtagatc gcggtaatca gctcaatcgt cctgcggccg 38040
tcctcgccgg tgatcgccgg agcggaatcc gtttccagcg cgtgcagcac attctccagc 38100
tggccggtat gcatctcata cggcaccggc ggcagcgacg caaggaattc ctccgcctgt 38160
ttcgcaaagg attcatccgg ctccgccagc gggaagccgt tgggctgcgg ctgcgaagca 38220
aacacgctgt acgggaacgc aatctttgcc ttctcgcact ggaagatcag cgcctggtcc 38280
tccccgtggt ccacgacgga agccgtcacc accgccaggg cgcccggata ttccgcgatg 38340
gaaaccgaca ggtcttccac ctcggcattg tcatggccgg tgtttgccag cacactggtc 38400
accttctccg gcgtacccat catccagccc agcatgtcga tatggtgcac cgcgtgattc 38460
agggtgcatc cgccgccttc cttctcccag gtgccccgcc accacaggtc atagtagctg 38520
tggccccgga accacaggga attcaccgtc acatggcgca ccttgccggc catgccgtta 38580
tccagcagcg ctttcaggtt gcgcaccggc aggcggaaac ggttctgcgc gatgatggac 38640
agtttcttcc cgctctcatc ccgggccttc agcatggcgt cacattcctc cagggaggct 38700
gccatgggtt tttcgcatac aacattcttc ccgctccgca gggcgttgat gctgatctcc 38760
gcatgcacaa acggcggggt gcagacatcc accaggtcga tgtccttctc caggatctcg 38820
tggtgatcca ggaacacttc cgcgtccagc ccgtactgtt cctttacctg ctgcgctttt 38880
ccgggaataa tgtccaccag ggcgacgatt ttgcaccgct ccggaaattg catatatgct 38940
ttgatatgtg cgtgggaaat tccgccggtt ccgacgattg cgacccgaat catacactgc 39000
acctctctga atgaaagctt tcgttgattt taaagcgttt gttccgatat aatcaggttc 39060
ttattatctg gatacagtat agcttgtatg aggttgacaa tactatataa atcatgtttc 39120
aatgtatgca gatcctgcac aatctgaaac cggaggaacc gccatgcagc tgatacaggc 39180
cgaaatgacc cgggctcctt tccttcccat gggacagcag gggcgctgg gatatgaagc 39240
ccatcataat ctcgacaaaa cctttgcggt tacgcaatac cactgccatg attattatga 39300
attctacatt catatccgcg gcggggaata tatgggtgtg gataaccagc tgttcctgct 39360
gaaacccaac caggttttca ttcttccgcc tttttccatg cacggtctgt cctgtacccg 39420
ggaaatgcat gattacgagc gcgcttatct gaacctttct ccggaggtca tgcgcgttct 39480
cggctgcggg cagatcgacc tggatcagtt cttccgttcc caggcatccg gcggcgtctg 39540
tacctatcag ctgacagacg cggacgcgga acagtttgtc gcatggatcc ggaagatccg 39600
ggacaaccag caggacgaat cctttgagaa cgacgtttat caccgtttcc agaactactc 39660
cctgatgatg agcctgatgt ccctgctctg ccggaccaca cgcaggaccg accctgtgga 39720
aggcgacgct ttcggcagca gcgtgattca ggaaatcctg gaatatatca acaaccacta 39780
tacccagccg atccggattg aaaatctggc ccgcagcttc ggtgtcagcg tttcctggct 39840
ttcccatgaa ttttcccgct ttaccaaccg cagcgtctat gactatgtcc tctcccgccg 39900
ggtcatgctg gcccgccagc agatgctggg ctccgcatcg ctcaatgata tcgcctacca 39960
gtgcggattc aatgattact ccaatttcct acgttctttc ggaaaaatag tcggcatgtc 40020
tcccagtcag taccggaagc ggctgaaagc ctatcgggaa aaggaaatcg acgcccggat 40080
agagttctga taaaaaactc ccttcacgcg gcggtttttc gcctcttatg aagggagtca 40140
tttggatgct tgtatatccc ggaccgggtt catctcctga acccgatccg ggattttatg 40200
tcaggaatta tttctcgaac ttgtaaccgg tcgctgcttc cagcttctcg gcgcgttcgt 40260

tcatgatggc ttcgccgccg atcgccatgt agttttccac accagcgtca taaacagcgt 40320
caaactgatc cacaggagca gtcacagcct tgttcaggaa ggagttgcgt tcgtcgctca 40380
gcgtggtgcc aacgccggtt tccgcctcaa tcgccggcag gctgaagtgt ccgggtacgc 40440
ggccgccgtt catggcgaca gcataagcgg tctggacaac ctcgggattc acgggagcat 40500
agccgtaagc gaagctggct tcggtgtatt cacccagata ctggccgttg caggtcatgg 40560
tgaagtcaat gttgttcagg ctggcctggg tgtagggatc ggaagcgtcc acggccttca 40620
tcatgtaggc accgttctcg gcacgctcat ggttcttgcc ttccacaccg gtctgcagga 40680
acaggatggt atcgggctgg gagatgaagt ccaggtacag caggctcgcc agcacttcct 40740
tgttcgtggt ggggaagcaa accttgcggt ccacggtgct gcccaggtac ttccggtaga 40800
taccggcgtc attttcgaag cagtcgaccg cgacaaactc agcgtcttcg ccggcagcgc 40860
gcttgatgtt cgcttcgatg gaatcttcgc cgttacggta aggatagtca tagttgtgct 40920
ggaaagcacc cacaaagccg gccttcaggt tgtcgtcttc ggtggtgtcg ccgtcgccgt 40980
acagagcgaa gtctttccag accaggcctt cattgtacca tttgttcagc aggcgcatgg 41040
cttctttggc gccgggcagg aactgctgcc tgtcatcgaa accgcggatg tagagttctt 41100
catcggtgat agcgtccgga acgaaggata cagtcaggat gttgttgcgc cagccgatat 41160
cctggctggt gctgtagggc accatcttgg cagcatcagc gcccagcagg agctcagcgt 41220
tgtcgcggaa ggcaaccagg caggcttcga attcagcctt gttggtcggc agggccatgc 41280
ccagcttgtc cagccagtcc ttacggatga aggtattgat acgggcattg ttcaccagac 41340
gggtttccag cgcccacagt tttccgctct cgggatcctt gtcccagtaa acgttttctt 41400
cgcccagcag gttgaagagg ttggggagca gatccttgta ttcttcaatg tagggagcca 41460
ggtcgatgat gccgcccatg tccgcatagt tcaggaccgt gggataggcg taggtgtaga 41520
tgacgtcggg agcgttctgg gcggccagca ggttgcccag gtccgtctca tcatcccagc 41580
ggccgcagac ggagaattca acgttcacgt tgtacttttc cttcatgccg ttcttgatat 41640
attcggccca gggagaagtc tcgggaacat tgtcaccgcg ctggaaaaga cgcacggtaa 41700
tggtgcgggt ttcggtgaac tttccgtcca cgaagccgtc gtccgctgca aatgcggaag 41760
taatgcccag cgtggcgaac agtgccagcg ccagaaccag agccaggata cgtttcatgt 41820
tttcttcctc cttttatttt atttcaaggg tttcgtaccc ttaaaatacg attggattat 41880
cccttgaccg cgccgatggt cacgccgctg acgaaataac gctgcagcca tggatatacc 41940
agcaggatcg gcaccgtcgc aaacatgacg gaagccatct tcagcgtttc gctcaggcct 42000
ggcatggcga atccttcctg ggccgccgcc tcgccgctgc tggcattctg caggatattg 42060
tacaggtaca gctgaatggg atacagttct ctgttcttct tgatgtaaag cagggcgtca 42120
gagaatccgt tccaccgtcc gaccgcatag aatagtgcca gggtggccag cacgggtttg 42180
ctcagcggca gataaacctg gaccagaatc cggatcggtc ccgcgccgtc gatctcggct 42240
gcctcccgca gggattccgg tacgctgtag aagcttgacc gcataataat catgttgtac 42300
acgctcaggc agctcggaat caccagaacc cagaccgttt ccagcatgcc catgttgctg 42360
agcattacgt agtggggaat catgccggcg ctgaagtaca tcgtgaaaag gataaatccg 42420
ttcaggaatt tccggccttt cagctgatca taagtcagcg gataggcgca gatgaccgtc 42480
ataaacagac tcagcaccgt acatccgacc gtcagcaggg ctgtatacca cagagagtgt 42540
atgtaccgtt cgtttcccag aaccgtccgg taggcgtcca ggttcagccc tttaggccaa 42600
agcagcactt cattgcggat caggtacttt gtatccgaca gggaacgggc cagcacgttc 42660
agcatcggca ctaagcagat aaagatcaga aggatacaga tgaacacgat tacccaatcg 42720
ccgaatttcg tggatttcga tttaatcatg tttctccccc tcccttacat gatgccgcgc 42780
tcgccgaagc gcttggccag gaaattggct ccgaagagga agaggacgcc gactacggac 42840
tggaacaggc cgacggcggt cgtcaggctg aactgcccgc cgccgtttat gcctttctgg 42900
tacacataga tggaaagcac cgtcgaaacg tcggtgacca gtttattgct taaggccagc 42960
ggccggtcaa actcgcttcc caaaatgtag cccaatcgca taatcagcag tacaacgatc 43020
gttggacgca ggcctggcag cgtcacatgc cagatgcggc ggaaacgccc ggcaccgtcc 43080
acggcggcag cttcataaag ctcggggctg atgctggtca gcgcagccag atagatgatc 43140
gtgccccagc cggcctcctt ccagatgccg aggccgatat agcttcccac ccaccaactg 43200
ggattgctca ggaatggaat caccgcgtcc ggattgcccg tcatctgttt gttcaggatg 43260
ttgaccaggc cgtcattatc cgccaacagg cgcagcgcca gggaggaaat gatgatccag 43320
ctcaggaa 43328

//

ID XXX; XXX; linear; XXX; XXX; XXX; XXX.

```
XX
AC   PRJEB67889
XX
DE   Uncultured bacterium DNA fragment from in vivo enriched cow rumen, clone
rum-49m8
XX
FH   Key             Location/Qualifiers
FH
FT   source          1..33187
FT                   /organism="uncultured bacterium"
FT                   /environmental_sample
FT                   /mol_type="genomic DNA"
FT                   /isolation_source="Cow rumen"
FT                   /clone="rum-49m8"
FT                   /note="AZCL-Barley beta glucan, AZCL-HE cellulose,
AZCL-tamarind xyloglucan active clone"
FT   CDS             406..678
FT                   /translation="MTAAGDFSMKTAIGSPYRPRCESAVPIAFAFRCSAENVSCILSSA
FT                   KAILSGTRGAVCSRYSKTFVCMMIPFYQAVRFQKPAFPDYDGVGG"
FT                   /product="hypothetical protein"
FT   CDS             complement(657..2150)
FT                   /translation="MKRSSRLSSGSIFRTAYALGGLIVLAALILVLISLIAGFGADPAA
FT                   DRLPAGPDYAVSQLSVWDGEGVPDPSAFLNDNTKSFVSTARYLLAPEESIGDQDIAILM
FT                   QLGDGTTRTENAVLTIREPVIRYEIGSAETAAEVLGKGYENAKFSQPLSSFTEIGTYPI
FT                   DVDIDGKTMHFTLVAQDTTPPVVQLKENLAFYVNQKLTVDDFIESCQDISEVEFHFSEV
FT                   PVTAAEGAFSIQIIATDAAGNSVTTDAVYYIGGDGLPPEFTGVGVMQTIAGVPVDYLRG
FT                   VQAIDANDGETAVQVTEPANFSIRTPGDYEILYTSMDKAGNRAEATGTLRVLASLDEVE
FT                   KLTSEDVARIGDYIVGTLKDGTEKTDKRAYLRRIYYYVQNHMFYKDNHDILDWEHAAVV
FT                   AITLGYGDCRNYYSFARLLLTCAGFENMMVEHEKPNEWSNAHYWNLVKVDGGWYHFDTT
FT                   PRVNRSEFFLWTDEQMDNYSAMNNNCFERDKSIYPPTPS"
FT                   /product="hypothetical protein"
FT   CDS             complement(2286..2708)
FT                   /translation="MKIIGTLFVCAALLTSCGGQNPDSEAPIQITAAPDSTVTTALTET
FT                   EAAPAAAETTAKAKKKETEISSETAKTETTARSTEAKTAKTETEPQKTETAAPKQKEST
FT                   ASPAQTKTTAAPETQKTEPVQTEEDDGLIYLPVIPL"
FT                   /product="hypothetical protein"
FT   CDS             complement(3192..4073)
FT                   /translation="MSRKITCAAVQMACSRDPKESMAKAERLIRLAAEKGAQIILLPEL
FT                   FERQYFCQERRYEYYAFAQRTEENPAVRRCAELAAELGVVLPVSFYERDGNQLFNSLAV
FT                   IDADGSLLGVYRKTHIPDDHYYQEKFYFSPGNTGFKVWKTKYAVIGAGICWDQWFPETA
FT                   RSLAVLGAELMFFPTAIGAEPILGTESGLHWQLTMQGHAAANMIPVIAANRIGLEKVTP
FT                   CPENGGQESALSFYGMSFMTNGIGQIAESADTAEETVLLHSYDLDALAQARLEWGLFRD
FT                   RRPECYGAITKP"
FT                   /product="N-carbamoylputrescine amidase (3.5.1.53)"
FT   CDS             complement(4089..4523)
FT                   /translation="MVRELEPQDLDALLQLYTHLHGNPIPEKSAALSALWQSILQDPNH
FT                   HIIGAFEDGRLAASCVCVIVPNLTHEQRPYALVENVVTDAAYRRRGLASACLNFAKELA
FT                   LRENCYKLMLMTGAKDEGTLHFYEQAGYNRTDKTAFIQWI"
FT                   /product="acetyltransferase, GNAT family"
FT   CDS             complement(4534..6117)
FT                   /translation="MELTRGTRVKLPEHLSTAAEFTVELKTAGGAVYDCSCFGVDKEGK
FT                   LSDDRYMVFYNQPQSPEGALKWTEIPGGSRWQVSLDRLPQTIDKLVFTVSIDGAGTMGQ
FT                   IASHSAVLLQGGQPRLTLSLSGADFKNERAIISVEIYRKDVWRLAATARGFNGGLGDLL
FT                   RFFGGEEAEAAPQPAPVSASAPRPAAPPRPAPMPSPQPMTPPRPAPMPSPQPMTPPRPA
```

```
FT                   PMPSPQPMTPPRPAPMPSPQQQNSHPVFGSPAFNAANPARQTPVFGAPPAQPQTPFPAQ
FT                   QPFPAQQPAQAPVSSGKVSLEKKLERGAPALVSLAKPIMNELKRCGLENEIARVALVLD
FT                   ISGSMSGRFANGTVQEIVNKVLPLAVQFDDDGALDFWYYGSRCRRMPEVTMANYTGAVP
FT                   DNWKKLMSSLGYGNHEPAVMKEVIDEYKNSRLPAYVIFVTDGGVGSENAIKKLLVESSR
FT                   YPIFWQFVGVGGSGYGVLERLDTMDGRFIDNANFFALDDFRTVSNEELYRRLLNEFPGW
FT                   LREIRRAGMI"
FT                   /product="hypothetical protein"
FT   CDS             complement(6198..8033)
FT                   /db_xref="GO:0031176"
FT                   /translation="MKNAKLIRLAGATAAALLAFAPLSAFPVNAAGSMTSKELSVCNYY
FT                   KELDVEVKRGEPYDTSLAALGVNLSEKVTQIWIKFEMDIESGLPAMPAIGYTSPGHIDP
FT                   TTGEQNDWYGDGLWLQPPASEGTIVIDIPEEDPLQNEFSVQMWGEKDKTIDKYQLKGIG
FT                   VVTSGIADIGIRTRTGDLNDDKTVNLADAEALAKWLTEGTEPAAPANGDLDRDNKLTAV
FT                   DLTLLKRGILDGSFNQTIDSGETAMEFVKHLRLGWNLGNTLDAQIRGAYQNPKQAETAW
FT                   GNPETTRELIDGVKAAGFNTVRLPVSWGEKMNSNYEIDGAWMDRVQEIVDYVIDNGMYC
FT                   ILNIHHDNGYEINQQFVEPSCPYFYPTNAHWEHSEKFVTAVWTQVSERFKSYGDHLIFE
FT                   TLNEPRMIGHRNQWWIDPNSADCKEAMDCVNKLNAAGLKAIRNSGGNNAKRFVMLPSYA
FT                   ASADQANLDGVVLPEDDHIIASAHAYNPQPFALQVTGTNKWSESNSSDTSAILTAFANL
FT                   KARYIDKGIPVIMGEFGAMNKDNDDDRAAWAKYYVSNADKYGIVCVWWDNNYFQAPVDD
FT                   NSGESFGLMNRNSNTVQFPKVMEALLEATAGRAEN"
FT                   /product="DOC-GH5_4"
FT                   /EC_number="3.2.1.8"
FT   CDS             8297..9454
FT                   /translation="MSEPHVRRFRRKKPVPKRIIDVVADFVCSSVNPTYNQAYRQWRAN
FT                   SKAPCMYRFRENTEERVFVSGKAVEPPSAARAETQALTRSSERIGAALLLFLAVGIGSE
FT                   LLMTAIFRAFGSDARFDLFSLSPSGSQPVVTLILCIASLLKYGVPILFLCRTCKLPPRV
FT                   LFPVSVGGFPELIAAFGFGMITASVYALTSADESVRTAQRLFEYSDSADGLAFALYGAF
FT                   EMLAVSLLAELFLRGAVLPVLRQFGDSFAVACTAAAAFFFPNPLHGRICELLLGLFCGY
FT                   LLLQSGSLLKCVLVRMVYVALEFAMLILVYAAGRMPLWQYALLLLTVGLVLLSFYARIR
FT                   RDRLLLQNRHTQLSDARKTACVLQAPTVLPWAAISLLFTVLQLFR"
FT                   /product="hypothetical protein"
FT   CDS             9464..9961
FT                   /db_xref="GO:0008270"
FT                   /db_xref="GO:0016787"
FT                   /translation="MFLDDFAYMREAIALAEEAGRAGEVPVGAVIVDRNTGAVIGRGRN
FT                   RVEADRSPTAHAEMLAIAEAAKLRGSRLTDCSLYVTLEPCPMCSGAIVMARITRVIFGA
FT                   YDERQGGVCSAAKLLSLPQSHRPRVTAGLLEEPCAALLQQFFRTIRGEESDKPQETEQF
FT                   SE"
FT                   /product="tRNA-specific adenosine-34 deaminase (EC 3.5.4.-
FT                   )"
FT   CDS             10191..11138
FT                   /db_xref="GO:0003978"
FT                   /translation="MATIMVTGGCGMIGDHVCSGLLKKGNTVIAVDRKESDYNKGKENF
FT                   YFRQAGPADQAKYGEIFDEFEIEFVVHIACTADNDIGPVIEKPQMDECAAMDRFIYKFA
FT                   LGKKIRKFILVSTTQVYKTPESREPVREDDTVKPVSNYGKMKAASEAAFAADTRPSKEM
FT                   LRCVIRVPQVYTLNFTANLMSRITDPKDHTIFVFRTGEYGFHMCCVHNISDFVIGYLRQ
FT                   ADAVSYSGVYNVADSSLIMASEIITFMKEHHRLGPVLQKANLSRSLKDRLFGSAEEKNN
FT                   YRYLDTATIDKNFMFDTTKASRICPFRWNITNTK"
FT                   /product="UDP-glucose 4-epimerase (EC 5.1.3.2)"
FT                   /EC_number="5.1.3.2"
FT   CDS             11243..12583
FT                   /translation="MIQPILDHDFVPAVYFNRDFLAKANRPFSIAVEREDGLVYRRDTR
FT                   LGDDFAENCRYAERLIKTILWCAGGWKVMLAGDHAVFEYIRDAYTKDGLRAFDVDFWST
FT                   TYERPFEVEERTFANLPQSCARPLSIGRNLAGCRVGFDAGGSDIKVAAVVDGETVWAEE
```

```
FT                   IIWLPKLAEDISYHKEHIKAAILKAAEHMPRFDAVGVSTAGVLVANRAMVSHLFLKVPK
FT                   DTQGEACKNVYVDVLNELGVPYAVANDGDVAALAASMALDVNGVLGIAMGTSQAGGYVD
FT                   ANGSVTGWHNELAFVPVDYNPDSPVDEWSGDYGCGVKYFSQDAVIRLAPKAGIELDESL
FT                   TLAEKLKAVQNVLAEGREGAKDIFRTIGTYFGYAIANYADFYDISYLQISGRVTSGLGG
FT                   DLIIEKAKEVLNKEFPELAAKIQFILPDEMDRRVGQAVAAASLPAIG"
FT                   /product="conserved hypothetical protein"
FT   CDS             12608..14794
FT                   /db_xref="GO:0008448"
FT                   /translation="MILKNLLLNGELTDITVENGKIAAIAPASPKTGGIDCSGLSVIPG
FT                   LVDMHTHGCVGYDTMDAHFEEMSVFLAQNGTTSWLPTTMTMDFNSIRRVTEAPRNVTGA
FT                   QILGFHMEGPYINEKRKGAQNGKFIKAPDLAEFRKLPDMRVVTIAPELPGSEEFIRECG
FT                   ALVCIGHTDADYETTLKAIAAGANCLTHTFNAMPGIHHREPGPIGAALKENIYVQAITD
FT                   GLHLHPAIVLMLYKLFGPERMVIISDAMRATGLGDGDYEFGGQTITVKDGVARTQDGAI
FT                   AGSTSTLWQCVKRAAEFGVPFEDAVRMATRTPADMIGAPEKGRIEVGADADLVLLDRNL
FT                   EIERVMIAGQFIEERDEERRRYLEEKEHVKKICSLFHTEGSFLAYETLHQGNINATYRV
FT                   DYRLESGQLKSYVLQKVNTYVFRNPKEIMENIRKVTEHLRGKVRRLNKPALHFHRTDSG
FT                   ENYYIEDDATFWRLENWIEGIGYDTCDSLDILRNAGEAFGEFQNQLSDFDASSLFTTIP
FT                   DFHNTAKRLETFFAAVEADPCGRAAQAQDEIAFLRENRALACELTEMLERGEVPLRVTH
FT                   NDTKINNVLFSRETGAPLTVIDLDTVMPGLAMHDFGDAVRFAASTAREDEPDVSKIAFD
FT                   LQKYRAFAEGFIGKTADALTENEIRTMALGAITITIELASRFLADYLTGDQYFKTAYPE
FT                   HNLVRARAQIALAKDMLAKRTRMESIVREIVAGV"
FT                   /product="CE9"
FT                   /EC_number="3.5.1.25"
FT   CDS             14866..15918
FT                   /db_xref="GO:0008716"
FT                   /translation="MQTVLTLFGGVSPEHDVSLVSVTAVLKNMRKENRRVLPVGITKEG
FT                   KWLLFTGENYDDIAANRWQEQSTPAFLSPERGTGLCILRDGALETVPVDVVFPVLHGEN
FT                   GEDGSLQGLLQIAGIPYVGCGICASASSMDKSVTKLIAATTGVRQAKWLTLTRRAFAED
FT                   AQAMCRTVLETLGLPLFVKPCSTGSSVGVSKVKTEAELLPALETAFRFDKKILAEEFIN
FT                   GREIETAVLGNDAPKAAAAGEIVAGAEFYDYETKYITSTSEAIIPAPIPADKMQEVRER
FT                   AVQIYSALGCAGLSRVDFFLTRDTNELVFNEINTMPGFTPISMYPKMWAADGLPFDELI
FT                   DRVIALALED"
FT                   /product="D-alanine--D-alanine ligase (EC 6.3.2.4)"
FT                   /EC_number="6.3.2.4"
FT   CDS             15921..17342
FT                   /translation="MAQLLAPADAEMTAAELASVLGTKLRNAENPDIAVKGFSTDTRTV
FT                   AAGNCFLALKGENFNGNLYAEQAVEKGAVLCILSEEPKEPIPFFLVSDTVQAYADIARH
FT                   FMQKRKAAGLRVVGVTGSSGKTTVKDMTAHVLSHHFRTICTTGNFNNHIGVPRTVLGIP
FT                   ADADAAVVEMGMNHTGEIAHLTRIAQPDAAIITNIGTAHIGNLGSQENIFRAKMEITEG
FT                   MGEPKTMIFPAEDAFLSTAAERFGAHLDLHFSSRTGLQNAELKAEIITENADDTCFTVR
FT                   FGEEAAEVRLPMTGLHNVTDALLAVHAGLLLGMPLAECAEALADFTPGAMRSERLSAGK
FT                   FTVIRDCYNANPEAMEAALKSLGTAAAGALRIAVLGNMNELGAFAGERHRELGAFCRTH
FT                   TDAAFFCGENYRDFAEGYGSSGTAFAAQEEMIPAVLAFLRQQTQPAFVLIKASRGLHLE
FT                   KSCEAVLAEFGSTVL"
FT                   /product="UDP-N-acetylmuramoylalanyl-D-glutamyl-2,6-
FT                   diaminopimelate--D-alanyl-D-alanine ligase (EC 6.3.2.10)"
FT                   /EC_number="6.3.2.10"
FT   CDS             complement(17740..19311)
FT                   /translation="MQKIKRVVSAAAAAALSLTASGVLALPAGAAGDDAPSLGLGVIAG
FT                   GKSVGIQLVVTNGIGSVLTLDGEEIEITESPQTAPLLVPAKEFTNQYSYKLTKDGETLK
FT                   SDSFSVAEYLNVLMNDAQYSKKYGALARAMLIYGGMAQRYFNPDIAKADCADSGIELTD
FT                   SDLAAVPDDPFDKAAFQEKFADAPVDYYGMNLSLVGQTIFRLFFQKKDSATDAEADAYM
FT                   KDVSLSMGASAPNGAEGFYEVSIPVPASDLSKDLSLIYDGQTENFRLTQYITAALQEEE
FT                   PLANVCKALYAYGNAVNAFHEEEPEEETWYTGYATVYTASAEGGSANLDQFAEDRGYIT
```

```
FT                   CAVNDEMYAGGKMAGAYLEVHNENDDTVVNVLVTNKDGNNNMQLGDLDFLEKDYSAIGT
FT                   PTGQKTAIKWRVIPLDTTDPVIFRFKSGSSIWWTEVQIQNHRYPIVKAEIKKNGEYVEM
FT                   ERQGDYNMFLAADSDHGLPKLDDSGPYEFRITDINGEVIEESVDIDLSSAISSEAFVEG
FT                   ENQFME"
FT                   /product="hypothetical protein"
FT   CDS             19534..21090
FT                   /db_xref="GO:0009044"
FT                   /translation="MKRKWTISAAGALAACCVCPAVSAAETEVMHCGFEDGIAGWQARG
FT                   SGTSVDVSDAASAEGASSAAVTGRGAAWHGIQYAAELDAGAKIRVSAQVMQKSGESVHF
FT                   KMTMQYGSGDSAVYDTFAEGDAASGEWLTLSAENYTVGSGTSPVLYFETESGSCDFWLD
FT                   EVILTQTGVSIKPSTKYRRGDADHSGKVEQADAEALCRFLQDGETEVFADTVDLDGDEA
FT                   LTAADLTILKRMLLYPEPEPEADPEYMSKIRDQVTANVPDSVLRGAGGTTEHITYFSST
FT                   ANREKGANVWLPPGYSQAETYPVFYVNHGYGGDEYAMMDGNGIREIATNLIQSGDAVPM
FT                   IIVFTDQYSDPAHPKQTGNGQADVPGYDLFVEDLPNSLMPYIESHYPVKTGRENTAVAG
FT                   FSMGGRESLYIGMKCCGKVGYVGGAAPAPGIFPTKDQFMDHPGVMSKEDMRIDPPYQPY
FT                   LLLIAGGTNDGMVNDFPLQYSNLFTEHGTENIYLSVPGGNHDKSTVIPLMYNFIKLIFK
FT                   A"
FT                   /product="CBM22-CE1"
FT                   /EC_number="3.2.1.37"
FT   CDS             complement(21461..22141)
FT                   /translation="MYFDWEFQFLNYLQDHIRNAFMDWFMPFITMFGDGGIFWIIVTLL
FT                   LFIPKRTRKYAHVSAIALIFCVVGGNLILKNVIARSRPFWIDNGHPLARIAYTIQDGIT
FT                   QYAAGHKTMTTTAPEGHTLIALLVKAPSDYCFPSGHTQASFAAATSICMWNRKAGIPAL
FT                   ILACLVAFSRMYLYVHYPTDVLFGMFCGIVYAFLALWICSKLFKNKDWDGCKGAKPAAK
FT                   APRH"
FT                   /product="Membrane-associated phospholipid phosphatase"
FT   CDS             complement(22190..22804)
FT                   /translation="MKNFLSKFQYNSPVILTFALLSALSLLLGWLTKGVSTDFLFCVYR
FT                   TRLLDPLMYVRMILYVFGHANYEHFIGNFTIILLVGPMLEEKYGSKVLLKMMLVTTVVT
FT                   GIVQVIFFPGTALLGASGIAFMLILLSSFANFQRGRIPITLILVVIIFLLGEIVDGLFK
FT                   SDNVSQMGHLIGGLCGTLFGWAITRSKEPSGDVSVGLDKLP"
FT                   /product="Rhomboid family protein"
FT   CDS             complement(22801..25215)
FT                   /translation="MFSALREHPVVKYLTEPRRIPMYYAVTVVAGIFYHYAPGLLWLWI
FT                   PLTLLLTAGLFRLFDFVREHHFLGGLAFALVSFLFLAASTYFMKIGFSEPIFSPADGSK
FT                   QIDFFIWFMTPQSVLETLYLGYTLALYLLFVFFIAFIAYYFTFVQYRVLMSFMVMVFPL
FT                   MIYAKEDENAPVPMIIILLAGWFAVMIYCRQAHAEDKTVVQRYAPKTQSKLSMPKKKSQ
FT                   FSRTKPEMLDGAFLGSAGIYIAAASILILVIPKPTVQADRSVLDTTLDMSTLSDYLMNA
FT                   ISGFTDSSDGGNYSPMNFRRTLYYARCKEPLNLRMRTFTNYSYENDEWQAGEYDRKPEV
FT                   NEKWIMKGGLRTLQVSPDPRELAAVIQSAAQRDAAFAERWNLTAFAQTELETDYTFDLE
FT                   VEAEISGNFVYPVPNYVKQATLTARLATTTDELDPFFSSAGIMFRFNGPRPFQENYSTV
FT                   YVSPQFAYTKAVQTLVNSLDEDEYFYMLLELIDMTEGEEQEIAQAAYDDCVEALLYSTT
FT                   VTSETPDSVAELAASLTAGCSSAYEKADAILSYLKYGDFIYSRDFVKSAADNVETFLFQ
FT                   NKTGVCYDYASAMTELCRAEGLIVRYVEGYSMSKPDERSFSGWDYAISTDEGHAFVDVF
FT                   IPGYGWMTMDPTSGNIVEEERADRANVISALQYSGLILFLAAIVLIGVFAYLVPFVRER
FT                   LFRRKFRRQMDAESVRAAYARLCRQWKANPAETPREVCAEQSKFLQLDLSELLAGVEET
FT                   VYAARCPKETAERVYRVYCAAHDAFAAAVKRRRRERRAERKQQFRKERLKT"
FT                   /product="hypothetical protein"
FT   CDS             complement(25256..26551)
FT                   /translation="MIGYLCAVGIALIFLFMMDGSIGVMMIAFLLLMPLISLIMTLLVR
FT                   KRVHLSLSLKPDITGKRGTASAEILLEKDSVLPLPFLRVNLQADAHFAPLNPEAEEMPD
FT                   KPLQGPGLAAAVRFRRQNQNYRRLRKIFLSPDTLPLCLSMGTAREKTYEIELSPEICGS
FT                   GAVKLSDPVLSDFLGIFKFRLKNEAAEAALLIMPEIPDMKTDSALFRTVATAIAAADEE
FT                   TEAEPNPLASSVPGYEHRDYIPGDSLKRINWKLSVKRRMLMVRKDEPVSLARMSVVLDF
```

```
FT                   RRAGDSRRLREKLAEEEQLIETALGFLMLCAQYGFPCSLSYADENGQWNSLSIDSGDQL
FT                   AAEAVTLLRGGFRDQEKLAGTDLMPAALLQDAGVLVMYFTTCGQEASAALEAVPTESYL
FT                   IVPSAQEAKCRAPKNGSLWLAMPDRSLEPGGH"
FT                   /product="FIG002343: hypothetical protein"
FT   CDS             complement(26548..27489)
FT                   /translation="MHSDILKLTESIGKVIVGKDRVIRCVTAGLLCGGHILIEDVPGVG
FT                   KTQLAAALSRSVGGTFNRIQLTPDVMPSDITGFTLVNPNDFSLEYRPGAAMCNFLLADE
FT                   INRASPKVQSALLEVMEEHQISIDGVTHKLPQPFMVLATQNPVETYGTYHLPEAQMDRF
FT                   FMRISMGYPTEEEEMQILTRTERRNPLENISEAALHPEDILNLQKEVENVFVSDEVRRY
FT                   ILRIVAATRTHDSLTLGASPRGSIALYKAAKANAFLSGRDFVLPDDVKAMAESVLAHRL
FT                   ILSPKGKSVHGSAEKVVADVLQSVPVPAGSGV"
FT                   /product="FIG022979: MoxR-like ATPases"
FT   CDS             complement(27520..27975)
FT                   /translation="MSITQSTFSGVPILNGIWYRDNSYVPYALTDGRLTAGARIPSLPE
FT                   EIHRRKGRPYYTNNQCFERSESLQCGFQCGVLEPMKCAYLAAIPANGEPILWIASFSYF
FT                   EQFEQIRFHGKRITVRTTEGQMVMFDAEDPMNVRVSLCAEPGMIVKN"
FT                   /product="hypothetical protein"
FT   CDS             complement(27972..28325)
FT                   /translation="MLKFYTNPETGEIVPESPETQGWEEVIANTVDASREKHVPVISRE
FT                   GQTVTVEVGSAAHPMIDAHYIGWILLETENGQERKELKPGEEPAAQFFAEEDDKIIAAY
FT                   AWCNLHGLWAAQA"
FT                   /product="Superoxide reductase (EC 1.15.1.2)"
FT                   /EC_number="1.15.1.2"
FT   CDS             complement(28388..29137)
FT                   /translation="MSGAYREIRRDAPDSGTAVLLLHGILSAPQFFDFLLNAVPDGTAV
FT                   SAPLLAGHGGTAGALSRAAMAQWKKQVRKTFLDLSRNRRRTVMIAHSMGTLFALEAARE
FT                   FPQSIRRMILLDVPLYAKMTLPGAFSFLKTAFGAPQDASRALAAAQAYSIAPDANLLHY
FT                   AGWIPRYLELFAEMRAVRGMISEIRVPAVIFQSEKDELVAARSARLLRGNPAFQVRLMR
FT                   NASHFDYPLREQERIRRAVRQAILSDS"
FT                   /product="hypothetical protein"
FT   CDS             29125..29307
FT                   /translation="MLRSLVEFDFSAVLAVHKAKSCRIGNPLQLLQKQIGRAPCIDAYA
FT                   DKRPAAPQTAQLSVC"
FT                   /product="hypothetical protein"
FT   CDS             complement(29343..30986)
FT                   /db_xref="GO:0004818"
FT                   /translation="MDYQKLAQLLFPDIDKRPEDFEQRYPKRNLPEGAVVTRIAPSPTG
FT                   FVHLGNLYNAIGERLARQTGGVFFLRIEDTDQKREVEGAVEAVIDAMSFYGVHFDEGAT
FT                   VDGEIGDYGPYRQRQRRELYQTIAKSLVERGMAYPCFCTEEELAAMHEAQEAAKDNYGY
FT                   YGKYAVWRDKSYDEIAAKIAAGEEWVLRFRSEGNEANTVTVQDGIRGELKLQENFTDFV
FT                   LLKSDGIPTYHFAHVCDDHFMRTTHVIRDESWLATLPIHVQLFDALGYERPVYCHTATL
FT                   MKMDGSSKRKLSKRKDPELALSYYRQAGFPQEAVWLYLLTVLNSNFEEWHLANPDADSR
FT                   EFTFSLEKMSISGSLFDLEKLDFIAKDYLSRQSVDFVYDGLLAWAKEYDPALAARMTER
FT                   AELYRGAIGVGRGGDKPRKDYGTWKQAADFLAFYDAETFTRTDACPECISDEDRKAILT
FT                   RYLETLDFSDAPEAWFAKVKALTEELGYAVQVKKYKKNPELYRGSIVDVTNLMRIALTG
FT                   RQNAPDINEISHVLGEAETRRRLQAYCDGE"
FT                   /product="Glutamyl-tRNA synthetase (EC 6.1.1.17)"
FT                   /EC_number="6.1.1.17"
FT   CDS             complement(31349..32278)
FT                   /translation="MKHPLKIAGALLGAAFLLGASVLPVSAAREFPELNSYCTVYDIED
FT                   AISADEEKMAAQRIQETCEKIDMYVAVYIVGEETPFSSDQAVERFADDAYDKLFNPVDG
FT                   KDTDGVLLVINDSTKYDYLTTSGTAQMYFENSDGGSRTWTILETITPNLKNKDYVGAIS
FT                   TFCAELEAFYKQGIPNGAFSYDSSTGEYYYYHKGQLQHAKKLPWWFGKNLGLLGGIAAI
FT                   CGAVAAFIASCFVKSGYHLVKSISPTNYVSEQESKYSVHEDHFLREHVARVHIDSGGGG
```

```
FT                   RTGGGGGGGGHSHMSSGGHSHGGGGHHR"
FT                   /product="hypothetical protein"
FT   CDS             complement(32275..33135)
FT                   /translation="MYSCSSCGAAIMADDNTAATFCYYCHNPVVLQGRVSGEYRPSAVL
FT                   PFQLDRAQAIAEFKKWCGEHRFLPSDFLSASQQEKIVGLYVPFWVTDCDIDVQMEALGK
FT                   KIRTYTSGREQITETKEFAVERNARVVIKGLPADGASHIDDDLMEAVEPFDYTKVRPFA
FT                   MQYLSGFLAEKYDMDFSKMFPRIQKRATNASDQLVRGSLKGYDSVSVTRSDIKVVKTDW
FT                   KYMLLPVWFMTYKYKDKIYEFALNGQSGKFVGDPPVSTGKLALFSGAIGLIIALIAFGL
FT                   GVLLS"
FT                   /product="hypothetical protein"
XX
SQ   Sequence 33187 BP; 6660 A; 10054 C; 9525 G; 6948 T; 0 other;
     gcatcgaaat agagcggaga ctggttgccg aagacggtgt cctccgaaaa ctgcagggtc        60
     tccgagagat tttcaatgta atcgatcaca gagcggtggc aggcgagaat gcccttcggc       120
     acgcctgtcg aaccggacgt aaagacaatg tagatcgggt caatgtcgag ctgctttgcg       180
     cggatggcct gcagcgcggc ttcgtcgggc gccgtctgaa tgagctcgtc aaacagtgcg       240
     gattcgccgg tatagccgaa gcctgcaagc tgcggctgcg tcacgctgtc gcagatgatg       300
     aggcgcggtg agagctgctg caggatcatt tcgatccgga atgcaggcat ttccgcgtcg       360
     agcggaacgt aatagcagcc ggcatagacc gtcccgaaga aggcgttgac cgccgccggg       420
     gacttttcca tgaagacggc aatcggttcg ccgtacagac cgcgctgcga aagcgccgtg       480
     ccgattgcct ttgcattccg ctgcagcgcc gaaaacgtca gctgcatatt gtcatcggca       540
     aaggcgattt tgtccggcac gcggggcgcc gtctgttcga gatactccaa aacatttgtt       600
     tgcatgatga tacccttcta tcaggctgtc agatttcaga agccggcatt tcccgattac       660
     gacggcgtcg gcggatagat gctcttgtcg cgttcgaagc agttgttgtt catcgcggag       720
     tagttgtcca tctgctcgtc cgtccagagg aagaattccg agcgattgac tctcggcgtg       780
     gtgtcgaagt gataccagcc gccgtcgacc ttgacgagat tccagtagtg cgcgttcgac       840
     cactcgttcg gcttctcgtg ctcgaccatc atgttttcaa agcctgcgca ggtcagcagg       900
     agacgcgcaa acgaatagta gttgcggcag tcgccgtagc caagcgtaat cgccacgaca       960
     gcagcgtgct cccagtcgag aatgtcgtga ttgtccttgt agaacatatg gttctgcacg      1020
     tagtagtaga tacggcgcag gtaggcgcgc ttgtcggtct tttccgtgcc gtccttgagc      1080
     gttccgacga tgtagtcgcc gatgcgggcg acgtcttcgg aggtcagctt ttcaacttcg      1140
     tccagactgg ccagcacgcg cagggttccg gtcgcttcgg cgcggttgcc ggccttgtcc      1200
     atgctggtgt agaggatttc gtagtcgccg ggggtgcgga tgctgaagtt cgcaggctct      1260
     gtgacctgaa cggcggtttc gccgtcattc gcgtcaatcg cctgcacgcc gcgcaggtag      1320
     tcgaccggca cgccggcaat cgtctgcatg acgccgacgc ccgtgaactc aggcggcaga      1380
     ccgtctccgc cgatgtagta gaccgcatcg gtggtcacgc tgttgcccgc ggcgtccgtc      1440
     gcgatgatct gaatcgagaa cgcgccctcc gctgccgtca ccggcacctc ggaaaaatgg      1500
     aactccactt cggagatatc ctgacaggat tcaatgaaat cgtcaaccgt gagcttctgg      1560
     ttgacataga aggcgaggtt ttccttcagc tgcacaaccg gcggcgtcgt gtcctgcgcg      1620
     acgagcgtga agtgcatggt cttgccgtcg atgtccacgt caatcgggta cgtgccgatt      1680
     tcggtgaacg agctcagcgg ctgagagaac tttgcgttct cgtagccctt gccgagcacc      1740
     tctgcggccg tttccgccga accgatttca tagcggatga ccggctcgcg gattgtcaga      1800
     acggcattct ctgtccgcgt cgtgccgtcg ccgagctgca tcagaatcgc gatgtcctga      1860
     tcgccgatgc tctcctccgg cgcaagcagg tagcgcgccg tgctgacaaa gctctttgta      1920
     ttgtcgttga ggaacgcact cggatccggt acgccctcgc catcccagac gctcagctga      1980
     ctgaccgcgt agtccggacc ggcaggaaga cgatccgctg ccggatctgc accgaagccc      2040
     gcaatcagcg agatgagcac cagtatcaat gccgcaagaa caatcaatcc gcccagcgca      2100
     taggccgttc ggaatatcga accgctcgaa agcctgcttg accgtttcat caaatgcaat      2160
     ccctttccag atagtttttc ggagccgtac attatattat aacgcttttt gcctgtgatg      2220
     tcaagtcgct tcaaagaaat gaaattgttt gcttgctttg caaaaagtgt gaaaaatgcc      2280
     cgtccttaaa gcggaatgac cggcagataa atcaggccgt catcctcttc cgtctgcacg      2340
     ggttccgtct tctgcgtctc cggcgcggcg gtcgttttcg tctgcgcggg agaagccgtg      2400
     ctctctttct gcttcggagc ggcggtttcc gttttctgcg gctctgtctc cgtcttggcg      2460
     gttttggcct ccgtgctccg cgcggttgtt tcggttttgg cagtctcaga ggagatttcc      2520
```

gtctccttct ttttggcctt ggcagtcgtt tccgcagcgg cgggagcggc ttcggtttcc 2580
gtcaaagcgg ttgtcacggt actgtccggc gcagcggtaa tctgaatcgg cgcttccgaa 2640
tccggattct gtccgccgca ggaagtcagc agtgcagcac agacaaataa agttccgatg 2700
atcttcacat caattcctcc tgttgaaaat gctttccctc tatttaacca catctgcggc 2760
agaatgtcaa gcgttttccg aaaaaagtcg gcagggccga cagccgtttc cgcgcttgtt 2820
tctgcgcggc tgttcattcc ggggccgcgg gacatacgct ccgacgaagc tgaatcagaa 2880
gtatcgctgc gcgatgattc gaaatggggc tccgccccaa accccgcctg ggacagtgtc 2940
cccggacccc gtttttgaga aatgctgatt tgattgctgt tcataaaaac aggattctta 3000
agagatgctg tcccctaaga atcctgatga caaaaacata tgaagattcg gttgtttatg 3060
catgcgctca atcgcctgcc gcgcggccga gaacgtcggg cacgtgcaga agcaagcgcg 3120
gcatcggcgg tcggcactgc cgaccgggtt tgggcggagc ccattatgaa tcatcgcgca 3180
gcgatacctc tttacggctt tgtgattgcc ccatagcact ccgggcggcg gtcgcggaag 3240
aggccccact ccagccgcgc ctgcgcgagc gcgtcgaggt catagctgtg cagaagcacg 3300
gtctcctcgg cagtgtccgc gctttccgca atctgaccga tgccgttcgt catgaacgac 3360
atgccgtaaa agctgagcgc cgattcctgt ccgccgtttt ccggacaggg cgtcactttc 3420
tccaggccga tgcggtttgc ggcaatcacc ggaatcatgt tcgccgccgc gtgtccctgc 3480
atggtcagct gccagtgcag gccgctctcc gtgccgagaa tcggctctgc gccgatggcc 3540
gtcgggaaga acatcagttc tgcgcccagc acggccagcg aacgcgccgt ctccggaaac 3600
cactgatccc agcagatgcc cgcgccgatg accgcatact tcgttttcca gaccttaaag 3660
cccgtattgc ccggcgagaa gtagaatttc tcctgataat agtgatcgtc ggggatgtgc 3720
gtcttgcggt acacgccgag caggctgccg tccgcgtcaa tcaccgcgag cgagttgaac 3780
agctgattgc cgtcccgctc ataaaagctc accggcagca caacgcccag ctctgccgca 3840
agctccgcgc agcgccggac agcggggttc tcctccgttc tctgtgcgaa ggcgtagtat 3900
tcgtatctgc gctcctggca gaagtactgc cgctcaaaca gctccggcag cagaatgatc 3960
tgcgcaccct tctccgcggc aaggcggatc agtctctccg cctttgccat gctctccttc 4020
gggtctctgc tgcacgccat ctgcacggcg gcgcaggtga tttttcggct catgctgatt 4080
ttcctccgtt agatccattg aatgaatgcc gtcttgtcgg tgcggttata gcccgcctgc 4140
tcgtaaaaat gcagcgtgcc ctcgtccttt gcgccggtca tgagcatcag cttatagcag 4200
ttttcgcgca gcgcaagctc cttcgcaaag ttcagacagg cactcgcaag cccgcgcctg 4260
cggtatgccg catccgtgac gacattctca accagcgcat acgggcgctg ttcgtgcgtg 4320
agattcggga cgatgacgca gacgcaggac gcggcgagcc tgccgtcctc gaatgcgccg 4380
atgatgtgat ggttcgggtc ctgcagaatg ctttgccaga gcgccgacag cgcagcgctt 4440
ttttcgggaa tcggattgcc gtgcagatgc gtgtacagct gaagcagggc atccagatcc 4500
tgcggttcca gttctctgac cattttcgcg ccctcaaatc attcctgcgc ggcggatttc 4560
gcggagccag ccgggaaatt cattcagaag gcggcggtag agctcttcgt tcgagaccgt 4620
gcggaaatcg tcgagcgcga agaagtttgc gttgtcgata aagcggccat ccatggtatc 4680
cagccgttcg agcacgccgt atccgctgcc gccgacgccg acaaactgcc agaaaatcgg 4740
gtaccgcgag gattcaacca gcagcttctt gatggcgttt tcgcttccga cgccgccgtc 4800
cgtcacgaaa atgacgtaag cgggaagccg gctgttcttg tattcgtcaa tgacctcctt 4860
catgacagca ggttcgtggt tgccgtaacc gagggagctc atcagctttt tccagttgtc 4920
cggaacagcg cccgtgtagt tcgccatggt gacttccggc atccgcctgc agcgcgaacc 4980
gtagtaccag aaatcgagcg cgccgtcgtc gtcgaactgc accgcgagcg gaagcacctt 5040
gttcacgatt tcctgcacgg tgccgttcgc gaagcgtccg gacatcgagc cggagatatc 5100
gagcacgagc gcaactcttg cgatttcatt ctccagaccg cagcgcttca gctcgttcat 5160
aatcggcttt gcgagcgaga caagcgccgg tgcgccgcgt tcgagcttct tttcgagcga 5220
aaccttgccc gaagagaccg gcgcctgcgc gggctgctgc gcggggaaag gctgctgcgc 5280
ggggaagggc gtctgcggct gtgcaggcgg agcgccgaac acgggcgtct gacgcgcggg 5340
atttgctgca ttgaatgccg gagagccgaa caccgggtgc gaattctgct gctgcggcga 5400
gggcatcggc gcagggcgcg gcggcgtcat cggctgcggc gaaggcatcg gtgcggggcg 5460
cggcggcgtc atcggctgcg gcgagggcat cggtgcgggg cgcggcggcg tcatcggctg 5520
cggcgagggc atcggtgcgg ggcgcggcgg cgctgccgga cgcggagcag acgccgaaac 5580
cggtgcgggc tgaggagcag cctccgcttc ttcgccgccg aagaatctca gcagatcgcc 5640
gaggccgccg ttaaagccgc gtgccgttgc ggcgagccgc cagacatcct tgcggtagat 5700
ttccacggaa ataatcgcgc gttcgttttt gaagtccgcg ccggaaagag aaagcgtgag 5760

gcgcggctga ccgccctgca gcagaacggc gctgtgcgag gcaatctgcc ccatcgtacc 5820
cgcgccgtca atgctgacgg taaacaccag cttgtcaatc gtctgcggca gccggtccag 5880
gctgacctgc cagcgcgacc cgccggggat ttctgtccat ttcagcgccc cttccggcga 5940
ctgcggctgg ttgtagaaga ccatgtaccg gtcatcggag agctttcctt ccttgtccac 6000
gccgaagcag ctgcagtcgt aaaccgcgcc gcccgccgtt ttcagctcca ctgtgaattc 6060
cgccgccgtg ctgagatgct ccggcagctt gactcttgtt ccgcgtgtca gttccatgcg 6120
tcacactcct ttttagtttg attgtgatat atgccttcag cgccggcgga aagacccgac 6180
cggcgctgag agatcattca gttttcggct ctgcctgcag tggcctcgag cagcgcttcc 6240
atcaccttcg ggaactgcac ggtgttggag tttctgttca tcaggccgaa gctctcgccg 6300
ctgttgtcat cgacgggcgc ctggaagtag ttgttgtccc accacacgca gacgatgccg 6360
tacttgtccg cattgctgac atagtacttt gcccacgccg cgcggtcatc gtcgttgtcc 6420
ttgttcatgg cgccgaattc gcccatgatg accggaatgc ccttgtcgat gtatctcgcc 6480
ttgaggtttg cgaatgcggt caggattgca ctggtgtcgc tggaattgct ctcgctccac 6540
ttgttcgtgc ccgtaacctg aagcgcgaag ggctgcgggt tgtaggcgtg tgcagaggcg 6600
atgatgtgat cgtcctccgg cagcacgacg ccgtcgaggt tggcctggtc ggcagatgcc 6660
gcgtaggagg gcagcatgac gaagcgcttt gcgttgttgc cgccggaatt gcggattgcc 6720
ttgaggcccg ctgcgttcag cttgttgacg cagtccattg cctccttgca gtcggcgctg 6780
ttcggatcaa tccaccactg gttccggtgg ccgatcatgc gcggttcgtt gagcgtttcg 6840
aaaatcagat ggtcgccgta gctcttgaag cgctcggaaa cctgcgtcca gacagcggtc 6900
acaaacttct cggaatgctc ccagtgcgcg ttggtcggat agaagtacgg gcaggagggc 6960
tccacgaact gctgattgat ttcgtagccg ttgtcgtggt gaatgttgag gatgcagtac 7020
atgccgttgt cgatgacata gtcgacaatc tcctgcacgc ggtccatcca cgcgccgtca 7080
atctcgtaat tgctgttcat cttctcgccc caggagaccg ggaggcggac ggtgttgaag 7140
cctgcggcct tgacgccgtc gatgagttcc cgcgtggtct ccgggttgcc ccatgcggtt 7200
tccgcctgct tcgggttctg ataggcgccc cggatctgcg cgtcgagcgt gttgccgagg 7260
ttccagccga gccgcaggtg cttgacgaat tccatcgccg tttcgccgct gtcgatggtc 7320
tgattgaagg agccgtccag aatgccgcgc ttgagcagcg tcaggtcgac agccgtcagc 7380
ttgttgtcgc ggtcgaggtc gccgttggcg ggcgctgcgg gctccgtgcc ctccgtgagc 7440
cacttcgcca gcgcttccgc gtccgcgagg ttcacggtct tgtcgtcgtt gaggtcgccc 7500
gtgcgggtcc ggatgccgat gtccgcaatg ccgctcgtca cgacgccgat gcccttgagc 7560
tgatacttgt caatcgtttt gtctttctcg ccccacatct gcacgctgaa ttcgttctgc 7620
agcgggtcct cctccggaat gtcgatcacg atcgttcctt cagaggcggg cggctgcagc 7680
caaaggccgt cgccgtacca gtcgttctgc tcgcccgtgg tcgggtcgat atgccccggc 7740
gacgtgtagc cgattgccgg catggcgggc aggccggatt caatgtccat ttcgaacttg 7800
atccagatct gcgtcacctt ttcggagagg ttgacgccca gtgccgcgag cgacgtgtcg 7860
tacggttctc cccgcttgac ttcaacgtcg agctccttgt aataattgca gacagacagt 7920
tccttcgacg tcatgctgcc tgctgcgttc accggaaacg ccgaaagcgg tgcgaatgca 7980
agaagtgccg ccgccgtcgc gccggcaaga cggatcagtt tcgcattctt catatcaaat 8040
ccccctaaaa gtttcaaatc ggacgcacgg tccatgttta ttataccaaa ttttcccagt 8100
tttgtaaagc ggcaaaactg ccaaaaagaa cgcaatccca aagtttatat tatacttttg 8160
aaacggcgca tcatcgcagt tttgacatt cacaatttgt tcatattaag ttcagagaaa 8220
ctgcacaact ctttgcgttg attctgctat actgaaagaa taaaatgggt atattgtgca 8280
gaaaggaggg agcagcttgt ctgaaccgca tgtgagaaga ttccggcgga aaaagccggt 8340
gccgaagcgc atcatcgacg ttgtcgccga tttcgtgtgc agctccgtca atccgacata 8400
caatcaggcg tatcggcagt ggcgcgcaaa cagcaaggct ccctgtatgt accgtttccg 8460
ggagaacaca gaagaacgtg tgttcgtttc cggaaaggcc gtcgagccgc cctctgccgc 8520
gcgcgcggaa acgcaggctc tgacccgttc cagcgaacgg atcggcgcgg cgctgctgct 8580
ctttctcgca gtcggcatcg ggagcgagct gctgatgaca gcgatttcc gcgcgttcgg 8640
ctccgatgcc cgcttcgacc tgttctcgct ctcgccgagc ggcagccagc cggtcgtcac 8700
gctgattctc tgcatcgcct cgctgctgaa atacggcgtt ccgattctct tcctgtgccg 8760
cacctgcaag ctcccgccgc gcgtgctgtt tccggtctcc gtcggcggtt tcccggagct 8820
catcgccgcg ttcggcttcg gcatgatcac ggcgtccgtc tatgcgctca cctctgcaga 8880
cgaaagcgtc cgcacggcgc agcgcctctt tgagtacagc gattccgcgg acgggctcgc 8940
cttcgcgctg tacggcgcgt tcgaaatgct cgccgtttcg ctcctcgcag agctctttct 9000

gcgcggcgcg gtcctgcccg tgctccggca gttcggcgac agcttcgccg ttgcctgcac 9060
agccgccgca gcattcttct tcccgaaccc cctgcacggc cgcatctgcg agctgctgct 9120
cggcctgttc tgcggctacc tgcttctgca gagcggctcg ctgctgaaat gcgtcctcgt 9180
gcgcatggtc tatgtcgcac tcgaattcgc aatgctgatt ctcgtctatg cggcgggccg 9240
gatgccgctt tggcagtacg cgctgctgct gctcaccgtc ggcttggtcc tgctctcctt 9300
ctacgcccgc atccgccgcg accgcctgct tctgcagaac cggcacacgc agctcagcga 9360
cgcgcgcaag accgcctgcg tcctgcaggc gccgaccgtg ctgccctggg ccgcaatctc 9420
cctgctgttc acggtgctgc agctgttccg ctgaggtgcg cccatgtttc tcgatgattt 9480
tgcatatatg cgcgaggcca tcgccctcgc ggaggaggcc gggcgcgcgg gggaagtccc 9540
cgtcggcgcg gttatcgtgg accggaatac cggcgccgtc atcggcagag gccgcaaccg 9600
cgtcgaagcc gaccgttctc cgactgcaca cgctgaaatg ctcgcaatcg cggaggctgc 9660
aaagctccgc ggttcgcgtc tcaccgactg cagcctctac gtgacgctcg aaccctgccc 9720
gatgtgcagc ggcgccattg tcatggcgcg cattacgcgg gtcattttcg gcgcgtacga 9780
cgagcggcag ggcggcgtct gctccgccgc aaagctgctt tcgctcccgc agagccaccg 9840
cccccgcgtc acggcaggcc tgctcgaaga gccctgcgcg gcgctcctgc agcagttttt 9900
ccgaaccatc cgcggcgaag aaagcgacaa accgcaggaa acggaacagt tctctgagtg 9960
aaagtttgtg cagtctgcac attccgcttt tgcagtgtgc gatgcacgat ggaaatattt 10020
atcaaatatt gtcaaagttt tatttgccgc ttgcatttcc ggaattgctg tgatataatg 10080
ataacaggat aatgggacaa gaaggacttt tgacgttttc cagacgactt gtgcattcca 10140
tgcccgacag attgatcaaa taaaagggcg ttaagaaagg tcgtgacaat atggcaacaa 10200
ttatggtaac cggcggctgc ggcatgatcg gcgatcatgt ctgctccggc ctgctcaaaa 10260
aaggaaacac cgtcattgcg gttgaccgga aggagagcga ctacaacaag ggaaaagaga 10320
atttctattt ccgccaggca ggccccgcag atcaggccaa gtacggtgaa atctttgatg 10380
aatttgaaat cgagtttgtt gtccatatcg cctgcacggc tgacaacgat atcggtccgg 10440
tcatcgaaaa gccgcagatg gacgagtgcg cggccatgga ccgtttcatt tacaaattcg 10500
ccctcggcaa gaaaatccgg aagttcatcc tcgtcagcac gacgcaggtc tacaagacgc 10560
cggaatcgcg cgaacccgtc cgcgaggatg acaccgtcaa gcctgtctcc aactacggca 10620
aaatgaaggc cgcttccgaa gccgcctttg ccgcggatac ccgcccgagc aaggaaatgc 10680
tccgctgcgt gatccgcgtg ccgcaggtct acacgctcaa cttcaccgcg aacctcatgt 10740
cgcgcatcac cgacccgaag gaccacacga tcttcgtctt ccgcaccggc gaatacggat 10800
tccatatgtg ctgcgtccac aacatctccg actttgtgat cggctatctc cggcaggcgg 10860
acgccgtctc ctattccggc gtctacaatg tcgcggattc ctcgctcatc atggcctccg 10920
aaatcatcac gttcatgaag gaacaccacc gcctcggccc ggtgctccag aaggcgaatc 10980
tcagcaggtc gctgaaggac cgcctcttcg gcagcgcgga ggagaagaac aactaccgct 11040
acctcgacac cgcgacgatt gacaagaact tcatgttcga cacgacgaag gcctcccgca 11100
tctgcccgtt ccgctggaac atcaccaaca ccaaataaac gcgctcacaa gcggcggttc 11160
tgttgttccg ccgctttgtt ttgctgtctg agtgctttca gcactcttca tttcagaaaa 11220
atcagaaagg tattttccgc ttatgattca gcctattctc gatcacgatt ttgtgcctgc 11280
ggtgtatttc aaccgcgatt ttctcgcaaa ggcaaaccgc ccgttctcca tcgccgtgga 11340
acgcgaggac ggtctggtct accgccgcga cacccgcctc ggcgatgact tcgcagagaa 11400
ctgccgctac gcagaacggc tcatcaagac cattctctgg tgcgcaggcg gctggaaggt 11460
catgctcgcg ggcgaccatg ccgtgtttga atacatccgc gatgcctaca cgaaggacgg 11520
tctccgcgcc ttcgacgtcg acttctggag cacgacctac gagcgtccct tcgaagtcga 11580
ggagcgcacg ttcgcgaacc tgccgcagag ctgcgcgcgg ccgctctcca tcgggcggaa 11640
cctcgcaggc tgcagagtcg gctttgacgc gggcggctcg gacatcaagg tggccgctgt 11700
cgtggacggc gaaaccgtct gggcagagga aatcatctgg ctcccgaagc tcgcggagga 11760
catctcctac cacaaagagc acatcaaggc cgccattctc aaggcggcgg agcatatgcc 11820
gcgcttcgat gccgtcggcg tcagcacggc gggcgtgctc gttgcgaacc gcgcaatggt 11880
ctcgcacctc ttcctgaagg tgccgaagga tacgcagggc gaagcctgca agaacgtcta 11940
cgtcgacgtt ctgaacgaac tcggcgtgcc gtacgcagtc gcaaatgacg gcgacgtcgc 12000
ggcgctcgcg gcctccatgg cgctcgacgt caacggcgtc ctcggcattg ccatgggcac 12060
ctcgcaggcg ggcggctacg tggacgcgaa cggcagcgtc accggctggc acaacgaact 12120
cgccttcgtg ccggtcgact acaaccccga ctcccccgtc gacgaatggt cgggcgatta 12180
cggctgcggc gtcaagtact tctcgcagga tgccgtcatc cgcctcgcac cgaaggccgg 12240

catcgaactc gatgagagcc tcaccctcgc cgaaaagctg aaggctgtgc agaatgtcct 12300
cgcggagggc cgcgagggcg cgaaggatat cttccgcacc atcggcacct acttcggcta 12360
cgcaatcgcg aactacgcgg acttctacga catctcctat cttcagattt ccggccgcgt 12420
cacctccggc ctcggcggcg atctgatcat tgaaaaggcg aaggaagtcc tgaacaagga 12480
attcccggaa ctcgccgcga agattcagtt tatcctgccg gacgaaatgg accggcgcgt 12540
cggacaggca gtcgcggcag cttccctgcc ggcaatcggc tgacaacctg cagaaaggcg 12600
gacggacatg attctgaaaa acctgcttct caacggcgaa ctgacggaca ttaccgtgga 12660
aaacggcaaa atcgctgcga tcgcacccgc ttctccgaaa acgggcggca tcgactgcag 12720
cggcctctcc gtcattcccg gcctggtcga tatgcacacg cacggctgcg tcggctatga 12780
cacgatggat gcgcacttcg aggagatgag cgtgttcctc gcgcagaacg gcacgacttc 12840
ctggctcccg acgaccatga ccatggattt caactccatt cggcgcgtca ccgaagcccc 12900
gcgcaatgtc accggcgcac agattctcgg cttccacatg gaaggaccgt acatcaacga 12960
aaagcgcaag ggcgcgcaga acggcaagtt catcaaggcg cccgacctcg cggaattccg 13020
gaagctgccg gatatgcgcg tcgtcaccat cgcgccggaa ctgcccggca gcgaggagtt 13080
cattcgggaa tgcggcgcac ttgtctgcat cggccacacc gacgcggact atgaaaccac 13140
gctgaaggcc atcgcggcgg gcgcaaactg cctgacccac accttcaacg ccatgcccgg 13200
catccaccac agagagcccg gccccatcgg cgcggccctc aaggaaaaca tctacgtgca 13260
ggccattacg gacggtctgc atctgcaccc ggccatcgtg ctgatgctct acaagctgtt 13320
cggcccggag cgcatggtca tcatcagcga tgcaatgcgc gcaaccggcc tcggcgacgg 13380
cgactatgaa ttcggcggac agaccatcac cgtcaaggac ggcgttgccc gcacgcagga 13440
cggcgccatt gccggctcga cctcgacgct gtggcagtgc gtgaagagag ctgcggaatt 13500
cggcgttccg tttgaagatg ccgtccgcat ggcgacccgc acccccgccg acatgatcgg 13560
cgcgccggaa aagggccgca ttgaagtcgg cgcggatgcc gatctcgttc tcctcgaccg 13620
gaacctggaa atcgagcgcg tcatgatcgc ggggcagttc atcgaagaac gcgatgagga 13680
acgccgccgc tatctcgaag aaaaagagca cgtgaagaaa atctgctcgc tcttccacac 13740
ggagggctcc ttcctcgcct atgaaacgct ccatcagggc aacatcaacg cgacctaccg 13800
cgtcgactac cgcctcgaaa gcggtcagct caagtcctat gtgctccaga aggtcaacac 13860
ctatgtgttc cgcaacccga aggaaatcat ggagaacatc cgcaaggtca cggagcatct 13920
ccgcggcaag gtccgccgcc tgaacaagcc ggcgctccac ttccacagaa ccgactccgg 13980
cgagaactac tatatcgaag acgatgcaac gttctggcgg ctcgaaaact ggattgaggg 14040
catcggctac gacacctgcg attccctcga cattctccgg aatgccggcg aggcgttcgg 14100
cgagttccag aatcagctct ccgatttcga cgcttcctcg ctcttcacga ccattcccga 14160
ttttcacaat acggccaagc gcctcgaaac cttcttcgca gcggttgagg ccgatccctg 14220
cggacgcgct gcacaggcac aggacgaaat cgcattcctg cgcgaaaacc gtgcgctcgc 14280
ctgcgaactg acggagatgc tggagcgcgg cgaagtcccg cttcgcgtca cccacaacga 14340
cacgaaaatc aacaacgtcc tgttcagccg cgaaaccggc gcaccgctga ccgtcatcga 14400
cctcgacacg gtcatgccgg gcctggccat gcacgacttc ggcgatgccg tccgctttgc 14460
ggcaagcacc gcgcgcgaag atgaaccgga tgtcagcaaa attgcatttg acctgcagaa 14520
ataccgcgct ttcgcggagg gcttcatcgg caagaccgcc gatgcgctca ccgaaaatga 14580
aatccgcacg atggcactcg gcgccattac cattacgatc gagcttgcat cgcgcttcct 14640
cgcagactac ctcaccggcg accagtactt caagaccgcg tatccggagc acaacctcgt 14700
gcgcgcccgc gcgcagattg cgctggcaaa ggatatgctt gcgaagcgga cccggatgga 14760
gagcattgtg cgggaaatcg tcgcaggcgt ataagcaaac cgcgattccc ggagaaatat 14820
gctccgggaa tcgaatcatg tcatatgcta aggaggcatt ctatcatgca aactgtactg 14880
acgctgttcg gcggcgtctc gccggaacac gacgtttcgc tcgtttccgt gaccgctgtg 14940
ctgaagaata tgcgcaagga gaaccggcgc gtccttccgg tcggcatcac gaaggagggc 15000
aagtggctgc tcttcaccgg cgagaactat gacgacatcg ccgcaaaccg ctggcaggaa 15060
cagagcaccc cggcattcct ctcgccggag cgcggcacgg gtctctgcat tctccgggac 15120
ggcgcgcttg aaaccgttcc ggtggacgtg gtctttcccg tgctgcacgg cgaaaacggc 15180
gaggacggct ccctgcaggg tctgctccag attgcgggaa tcccgtatgt cggctgcggc 15240
atctgcgcgt ccgcttcgtc catggacaag tccgtcacca agctgattgc cgcgacaacc 15300
ggcgtccggc aggcaaaatg gctcacgctc acccgccgcg cctttgcgga ggatgcacag 15360
gccatgtgca gaacagtcct cgaaacgctc ggcctgccgc tgttcgtgaa gccctgcagc 15420
accggctctt ccgtcggcgt ctcgaaggtc aagaccgaag cggaactgct gcccgcactc 15480

gaaaccgcct tccgcttcga taagaaaatt ctcgcggaag agttcatcaa cggcagagaa 15540
atcgaaaccg ccgtgctcgg caacgatgcc ccgaaggctg cggctgccgg cgaaatcgtc 15600
gcaggcgcgg agttttatga ctacgagacg aagtacatca cctcgacctc cgaagccatc 15660
atccccgcac cgattccggc ggacaaaatg caggaggtcc gcgaacgcgc cgtgcagatt 15720
tacagcgcgc tcggctgcgc gggtctctcc cgcgtcgact tcttcctcac gcgggacacg 15780
aacgagctcg ttttcaatga aatcaacacc atgcccggat tcacgccgat cagcatgtac 15840
ccgaagatgt gggcggcgga cggcctgccc ttcgacgaac tgattgaccg cgtgattgcg 15900
ctcgcgctgg aggactgaac atggctcagc tgcttgctcc ggcggacgcg gaaatgactg 15960
cggcggaact cgcttcggtg ctcggcacga aactgcggaa tgcggaaaat ccggatattg 16020
ccgtcaaagg cttcagcacc gacacgcgca ccgtcgccgc cggcaactgc tttctcgcac 16080
tgaagggcga gaatttcaac ggcaatctct atgcagaaca ggcagtcgaa aagggcgctg 16140
ttctgtgcat tctctcggaa gaacccaagg agcccatccc gttcttcctt gtctccgaca 16200
ccgtgcaggc gtatgcagac atcgcgcggc atttcatgca gaagcgcaaa gcggcgggac 16260
tccgcgttgt cggcgtgacc ggcagcagcg gcaaaacgac ggtcaaggac atgaccgcgc 16320
acgtcctttc gcatcacttc cgcacgatct gcacaaccgg aaacttcaac aaccacatcg 16380
gcgtcccgcg caccgtcctc ggcattcccg ctgatgcgga tgccgccgtc gtcgaaatgg 16440
gcatgaacca caccggcgaa atcgcgcacc tgacgcgcat cgctcagcct gatgccgcaa 16500
tcatcacgaa catcggcacc gcgcacatcg gcaacctcgg ctcgcaggaa aacattttcc 16560
gcgcgaaaat ggaaatcacg gagggcatgg gcgaaccgaa aaccatgatt ttcccggcgg 16620
aggacgcatt tctcagcacc gccgcagaac ggttcggcgc gcaccttgat ctgcacttca 16680
gttcccgcac cggcctgcag aatgcagaac tgaaggcaga gatcataacg gaaaatgctg 16740
acgatacgtg ctttaccgtt cgcttcggtg aagaggccgc agaggttcgt ctgccgatga 16800
ccggcctgca caatgtgacc gacgcgctgc ttgccgttca cgcgggactt ctgctcggca 16860
tgccgctcgc tgaatgcgcg gaagccctcg cagacttcac gcccggcgcc atgcggagcg 16920
aacgcctcag tgccgggaaa ttcacggtca tccgcgactg ctacaacgca aaccccgaag 16980
ccatggaggc cgcgctgaaa tcgctcggca cggcggcggc gggcgcgctc cgcatcgcgg 17040
tgctcggcaa catgaatgag ctcggcgcat tcgccggcga acgccaccgc gagctcggcg 17100
ccttctgcag aacgcacacg gatgcagcgt ttttctgcgg tgagaattac cgcgattttg 17160
cggagggtta cggcagttcc ggcacggcct tcgctgcgca ggaggagatg attcccgctg 17220
tgcttgcttt tctccgtcag cagacgcagc cggctttcgt gctcatcaaa gcctcgcgcg 17280
gcctgcacct cgaaaagagc tgtgaggctg ttctggcaga attcggcagt accgtgctgt 17340
aatctctctc cgaactcctc gctgctccct ccccaaagtg tctccgacgg atttataatg 17400
ggattccaaa ggcgactgcg tcggccgagg ggacagtatc cctttggcgg gtttggcgcg 17460
agcatgtttg ccttcggcaa actgcttagt ttcgctttgc gaaacataag cccattatta 17520
atcatcgcga agcgatacct ttttcgacaa actggattgc gccccgatca gatgaccggg 17580
gcgcgtcttt ttttgtatta caggaccttg gtgccgtcgc cgtcgatgtt cagcgaggtt 17640
gtgatcacat cctccgccgc aaacacagcg atttcaagct ccggagcttc gtaggtcttt 17700
ttcatcgtgc atcgtccttt ctatcagccg attgtctgat tattccataa actggttctc 17760
gccttccacg aatgcctcgc tcgaaattgc cgagctcaga tcaatgtcaa cggactcttc 17820
aatcacttcg ccgttgatgt ccgtgatgcg gaactcgtac gggccgctgt cgtcaagctt 17880
cggcagaccg tggtcgctgt ccgctgcaag gaacatattg tagtcgccct gccgttccat 17940
ctccacgtat tcgccgttct tcttgatttc cgccttcacg atcggatagc ggtggttctg 18000
aatctgaacc tcagtccacc aaatcgacga gccggatttg aagcggaaaa tcaccgggtc 18060
cgtcgtatcc agcggaatca cgcgccattt aatcgccgtt ttctgtccgg tcggcgtgcc 18120
gatggcgctg tagtctttct ccaggaaatc cagatcgccc agctgcatat tattgtttcc 18180
gtctttgttc gtgacaagca cattcacaac cgtgtcatca ttctcattgt ggacttccag 18240
ataagcaccg gccatcttgc cgcccgcata catttcatca ttcacggcgc aggtgatata 18300
gccgcggtct tcggcaaact gatcgagatt tgcagagccg ccctctgcac ttgctgtgta 18360
gaccgttgca tagccggtgt accatgtctc ttcttcgggc tcttcttcat ggaacgcatt 18420
gaccgcgttt ccgtaggcgt agagcgcttt gcagacattg gcaagcggct cttcttcctg 18480
cagcgccgcc gtaatgtact gcgtgaggcg gaaattctcc gtctgaccgt cgtagatgag 18540
ggaaaggtcc ttcgacagat ccgatgcggg aacgggaatg gaaacttcat agaagccctc 18600
tgccccgttc ggtgctgacg cacccatgct gagcgacaca tctttcatat acgcatctgc 18660
ttccgcatcc gtcgcggagt ccttcttctg gaagaacaga cggaatattg tctgtccgac 18720

gagcgaaagg ttcatgccgt aatagtccac cggcgcatct gcgaatttct cctgaaacgc 18780
tgccttgtca aagggatcgt ccggcactgc ggccaaatcg ctgtccgtca gctcgatgcc 18840
gctgtccgcg cagtccgctt tggcgatatc cggattaaaa taccgctgcg ccatgccgcc 18900
gtaaatgagc atggctcttg ccagtgcgcc gtactttttc gaatactgcg catcgttcat 18960
cagaacgttc aggtattccg caacgctgaa gctgtctgat ttcagcgtct cgccgtcctt 19020
ggtcagctta tacgagtatt gattcgtgaa ttcctttgca ggcacaagga gcggtgccgt 19080
ctgcgggctt tcggtgatct cgatttcttc cccgtccagc gtcagcacgg aaccgatacc 19140
gtttgtgaca accagctgaa tgcccactga tttgccgccc gcaatgacgc caaggcccag 19200
agaaggcgca tcgtcccccg ccgcaccggc aggaagcgca agcacccccg aagccgtcag 19260
gctgagcgct gccgctgcag ccgcggaaac aacccgtttg attttctgca ttccgtacac 19320
atcctttctg tttttcttca gtcttgaccg cagaacctgc ggatactgtc attataccac 19380
attcatacaa aaaatgcaag gagttcgcag aaaattcaca aataatgccg acagtaaatc 19440
gaaacattct ctacttgacg gaaccggaaa agtgtgttat actgaaaata cgggcaaacg 19500
caaccgcccg aatctgacag ggggaagaga aatatgaaac gaaaatggac gatatcggcg 19560
gctggagcgc tggcggcgtg ctgcgtctgc ccggcggttt ctgccgcaga aacagaagtc 19620
atgcactgcg gcttcgagga cggcattgcc ggctggcagg cgcgcggaag cggcacttcg 19680
gttgatgtct cagacgctgc atcggcggaa ggggcttctt ctgccgcagt cacagggcgc 19740
ggtgccgcat ggcacggaat tcagtatgcc gcggagctcg atgccggcgc gaaaatccgc 19800
gtttcggcgc aggtcatgca gaaatccggc gaatccgtgc atttcaaaat gaccatgcag 19860
tacggcagcg gcgattccgc cgtctatgac acgttcgcgg agggcgatgc cgcctccggg 19920
gaatggctga cgctctctgc ggaaaactac acagtcggaa gcggaaccag tcccgtgctg 19980
tactttgaaa cggaaagcgg ttcctgcgat ttctggctgg atgaggtcat tctcacgcag 20040
accggcgttt cgatcaagcc ttcgacgaag tacaggcgcg gcgatgccga ccacagcgga 20100
aaggtcgaac aagcggacgc cgaagccctc tgcagattcc tgcaggacgg cgaaaccgag 20160
gtctttgcgg acacggtcga cctcgacggc gatgaagccc tgactgccgc cgacctgaca 20220
attctcaagc ggatgctcct ctatcccgaa ccggagcccg aagccgatcc ggagtatatg 20280
tcgaaaatcc gcgatcaagt cacggcgaat gtgcccgatt ccgtcctgcg gggcgcgggc 20340
ggaaccacgg aacacattac ctatttctcg tccacggcga accgcgaaaa gggcgcgaat 20400
gtctggctgc cgccggggta cagtcaggcg gaaacctacc ccgtgttcta tgtcaaccac 20460
ggctacggcg gcgacgagta cgccatgatg gacggaaacg gcattcgcga aatcgcgacg 20520
aatctgattc aatccggcga tgccgttccg atgattatcg tgtttacgga ccagtattcc 20580
gacccggcgc atccgaagca gaccggaaac ggacaggcgg acgttccggg ctatgatttg 20640
tttgtcgagg acctgcccaa cagcctgatg ccctatatcg aatcgcacta cccggtcaag 20700
acgggacgcg aaaacaccgc tgttgcaggc ttttccatgg gcggcagaga atcgctctac 20760
atcggcatga agtgctgcgg caaagtcggc tatgtcggcg gtgctgcgcc ggcgccgggc 20820
atcttcccga cgaaggacca gttcatggac catccgggcg ttatgagcaa ggaagatatg 20880
cgcattgacc cgccgtatca gccctatctg ctgctgatcg cgggcggcac gaacgacggc 20940
atggtcaacg acttcccgct gcagtattcc aatctcttca cggagcacgg cacggaaaat 21000
atctacctct ccgtacccgg cggcaatcac gacaaatcca ccgtcatccc gctgatgtac 21060
aatttcatta agctcatttt caaggcgtga aaggggtatc gctctgcggt cggcagtgcc 21120
gacagccgtt tccgcgcttg tttctgcacg gatttgacgt tcttggccgc gcggcaggca 21180
tcatacaaaa caaggattct taaggggtac ttcctcttaa gaatcctttg ctttatgaaa 21240
agcatcaaca cataaaatcg ccgcatttcc ttaattttca aaaaagcggg attccaaagg 21300
ggcactgtct ctttggcggg cttggagcgg gcatgtttgc cttcggcaaa ctgcttagtt 21360
tcgctttgcg aaacataagc ccattattaa tcatcgcgaa gcgatacctt ttctacaaac 21420
gaaaagcacc cgcttttccg gcgggtgctt ttggctgctg tcagtgacgg ggggcttttg 21480
cggcgggctt tgcgcccttg cagccgtccc agtccttgtt cttgaagagc ttgctgcaga 21540
tccacagcgc gaggaatgcg taaacaatgc cgcagaacat accgaagagc acgtcggtcg 21600
gatagtgcac atacaggtac atgcgggaga acgcaaccag acacgccaga atcagcgcgg 21660
gaatgccggc tttccggttc cacatgcaga tggacgtcgc tgcggcaaag gaagcctgcg 21720
tatgccccga agggaagcag tagtccgaag gtgccttcac gagcagcgca atcagcgtgt 21780
gcccctccgg cgcagtcgtc gtcatggtct tgtgtccggc ggcgtactgc gtgatgccgt 21840
cctgaatcgt gtacgcgatg cgggcaagcg ggtggccgtt gtcaatccag aaggggcgcg 21900
aacgtgcaat cacattcttc agaatgaggt tgccgccgac cacgcagaaa atcagcgcga 21960

tggcagaaac atgcgcgtac ttgcgcgtcc gcttcggaat gaacagcagc agcgtcacga 22020
tgatccagaa gatgccgccg tctccgaaca tcgtaatgaa cggcatgaac cagtccataa 22080
acgcgttccg gatgtgatcc tgcaggtagt tgagaaattg gaattcccag tcgaaataca 22140
tagtggctta aaatcctttc atgttgattg gcacattatt ataccatatt caggggagtt 22200
tgtcaagccc gacgctcacg tcgcccgacg gttccttcga gcgcgtgatg gcccagccga 22260
acagcgtgcc gcagagaccg ccgatcagat ggcccatctg cgagacattg tcgctcttga 22320
acaggccgtc gacgatttcg ccgagcagga agatgatgac gacgagaatc agcgtgatcg 22380
gaatgcggcc gcgctggaaa ttcgcaaagg agctgagcag aatcagcatg aaagcgatgc 22440
cggacgcgcc gagcagcgcc gtgcccggga agaaaatcac ctgaacaatt cccgtgacaa 22500
ccgtcgtcac gagcatcatc ttcaggagca ccttcgagcc gtacttttcc tccagcatcg 22560
gcccgaccag cagaataatc gtgaaattgc cgatgaagtg ctcgtaattc gcatggccga 22620
acacgtacag aatcatccgg acgtacatca gcgggtcaag cagtctcgtc cggtacacgc 22680
agaagaggaa gtccgtcgac acgcctttcg tcagccaccc gaggagcagc gaaagcgccg 22740
acagcagcgc aaaggtcaga atgaccgggg aattgtactg gaattttgac aggaagtttt 22800
tcatgttttt aatctctcct ttctgaactg ctgcttccgt tccgccctgc gttcgcggcg 22860
tctccgcttc acggcggctg caaatgcgtc atgcgccgca cagtagacgc ggtaaacgcg 22920
ctctgcggtt tctttcggac agcgcgcggc gtagacggtt tcttccacgc ccgcaagcag 22980
ttcactcaaa tcaagctgca ggaatttgct ctgctctgca cagacctccc gcggcgtttc 23040
ggcgggattc gccttccact gtctgcagag ccgcgcatat gccgcacgga cactctccgc 23100
atccatttgt ctgcggaatt tccggcggaa cagccgctcc cgcacaaacg gcaccagata 23160
ggcaaacacg ccgatcagga cgatggccgc gagaaacaga atcagtccgg agtactgcag 23220
cgccgaaatc acattggcgc ggtccgcacg ctcctcttcc acgatgttgc cgcttgtcgg 23280
gtccatggtc atccagccgt agccgggaat gaacacatct acaaaagcgt ggccttcatc 23340
ggtcgaaatc gcgtagtccc agccggagaa ggagcgctca tccggcttgc tcatggaata 23400
tccttccacg taccggacaa tcagtccctc tgccctgcag agctccgtca tggcgctcgc 23460
atagtcgtag cacacacccg ttttgttttg gaacaaaaat gtctccacat tgtctgccgc 23520
ggatttcaca aaatcgcgcg aataaataaa atcgccgtac ttcaggtacg aaagaatcgc 23580
gtccgctttc tcgtatgccg agctgcagcc cgctgtcagc gaagccgcaa gttccgcgac 23640
gctgtccggc gtttcgctcg tcacggtcgt gctgtagagc agcgcttcga cgcagtcatc 23700
gtacgcagcc tgcgcaatct cctgctcttc cccttccgtc atgtctatca attcgaggag 23760
catgtagaag tattcatctt catcgagcga attgaccagc gtctgcacgg ctttcgtgta 23820
cgcaaactgc ggactgacgt agaccgtgct gtagttttcc tggaacggcc ggggaccgtt 23880
gaagcggaac ataatgcccg cacttgagaa gaacgggtca agctcgtccg ttgttgtcgc 23940
aagtcttgcg gtcagcgtcg cctgcttgac gtaattcggc accggataca cgaagtttcc 24000
ggagatttcc gcttcgacct ccaggtcgaa tgtatagtcc gtttccagtt ccgtctgcgc 24060
aaatgccgtc agattccacc gctctgcaaa cgctgcgtcg cgctgtgcgg ctgactgaat 24120
cacggccgca agctcccgcg ggtccgggga aacctgcagc gtccgcaggc cgcctttcat 24180
gatccacttt tcgttgactt ccggctttcg gtcgtattcc cctgcctgcc attcgtcatt 24240
ttcatagctg tagttcgtga aggtccgcat ccgcagattc agcggctctt tgcagcgcgc 24300
gtagtagagc gtccgccgga agttcatcgg cgaataattg ccgccgtcgg agctgtccgt 24360
gaagccgctg atcgcgttca tcaggtaatc cgaaagcgtg ctcatatcga gcgtcgtgtc 24420
gaggacggaa cggtccgcct gcacggtcgg cttcggaatg acgagaatca gaatgctcgc 24480
cgccgcaatg taaatgcccg cagaaccgag gaacgcgccg tcgagcatct ccggctttgt 24540
ccgggagaac tgcgatttct tcttcggcat cgaaagcttc gactgcgttt tcggcgcata 24600
gcgctgcacg acggtcttgt cctccgcgtg tgcctgccgg cagtagatca tgactgcgaa 24660
ccagcccgcg agcagaatga taatcatcgg cacgggcgcg ttctcgtcct ccttcgcgta 24720
gatcatcagc ggaaagacca tgaccatgaa ggacatcagc acgcggtact gcacaaatgt 24780
gaaatagtag gcgatgaatg cgataaagaa cacaaacagc agatacagcg cgagggtata 24840
gccgagatag agcgtttcca gaacggactg cggcgtcatg aaccagatga agaagtcaat 24900
ctgcttgctg ccgtctgcgg gtgagaatat cggttcggaa aagccgattt tcatgaaata 24960
cgtcgatgcc gcgagaaaca gaaagctgac aagggcgaag gcaagtccgc cgaggaagtg 25020
gtgctcccgc acgaagtcga agagccggaa cagccccgct gtcagcagca gcgtcagcgg 25080
aatccagagc cacagcagcc ccggcgcata gtggtaaaag atgccggcga caaccgtgac 25140
ggcatagtac atggggattc ggcgcggttc cgtcagatat ttcacgacgg gatgttcccg 25200

cagtgccgaa aacatggcat tccctccttt tccagagcgt tcagggcagg cgcgatcagt 25260
gcccgccggg ttcgagcgaa cggtcaggca ttgccagcca gagcgagccg tttttcggcg 25320
cacggcattt tgcctcctgc gccgacggaa caatcagata cgattctgtc ggaaccgcct 25380
cgagcgctgc agacgcctcc tgcccgcagg tcgtgaagta catcaccagc acgcccgcat 25440
cctgcagaag cgctgccggc atcagatccg ttccggcaag tttctcctgg tctcggaaac 25500
cgccgcgcag cagcgtcacg gcctccgccg cgagctggtc gccgctgtca atcgacaggc 25560
tgttccactg tccgttttcg tccgcatagg agagcgagca ggggaagccg tactgcgcgc 25620
agagcatcag aaagccgagc gccgtttcaa tcagctgttc ctcttcggcg agcttctccc 25680
gcagcctgcg cgaatcgccg gcgcgccgga aatccagcac caccgacatt cttgcaagcg 25740
aaaccggttc gtccttccgc accatcagca tgcgccgctt cacggagagc ttccagttga 25800
tgcgcttgag ggaatcgccg ggaatgtagt cccggtgttc gtagccgggc acggaggaag 25860
cgagcggatt cggctccgcc tccgtctctt cgtccgccgc ggcaatggcc gtcgcaactg 25920
tgcggaaaag cgcactgtcc gtcttcatgt ccggaatctc cggcataatc agcagcgcgg 25980
cctccgccgc ttcgttcttc agccggaact tgaaaatgcc gaggaagtcc gacagcaccg 26040
gatcagagag cttcactgcg ccgcttccgc agatttccgg ggaaagctcg atttcatacg 26100
tcttttcccg cgccgtcccc atcgagagac agagcggcaa agtgtccgga gagagaaaga 26160
ttttccgcag ccgccggtag ttctgattct gtctccgaaa gcggactgcg gcggcaagcc 26220
ccggcccctg caggggtttg tccggcattt cctccgcctc cggattcagc ggcgcgaaat 26280
gcgcgtcagc ctgcagattg acccgcagaa acggcagcgg cagaacgctg tccttttcca 26340
gcagaatctc cgccgacgcg gtcccgcgct ttccggtgat gtccggcttc agcgacaggg 26400
agagatgcac ccgcttccgg accaaaagcg tcataatcag ggaaatcagc ggcatcagca 26460
gcaggaacgc aatcatcatc acgccgatgc tgccgtccat catgaacagg aaaatcagcg 26520
cgatgccgac tgcacagaga tacccgatca tacgccgctg ccggcaggca cgggaaccga 26580
ctgcaggaca tcggcgacaa ccttttccgc agagccgtgc acggatttgc ccttcggcga 26640
gagaatcagc cggtgtgcaa ggacgctctc cgccattgcc ttcacgtcat ccggcaggac 26700
gaagtcgcgg ccggagagga aggcgtttgc cttcgcggcc ttgtagagcg cgatcgagcc 26760
gcgcgggctg gcgccgagcg tcagcgaatc gtgcgtgcgc gtcgccgcga caatgcgcag 26820
gatgtaccgc cgcacctcgt cgctgacaaa gacattttcg acttccttct gcagattcag 26880
gatgtcctcc ggatgcaggg ccgcttcgct gatgttttcc agcggattgc ggcgctccgt 26940
gcgcgtcaga atctgcatct cctcctcttc ggtcggatag cccatgctga tgcgcatgaa 27000
gaagcggtcc atctgcgctt cgggaaggtg gtaggttccg taggtttcca ccgggttctg 27060
cgttgcgagc accatgaacg gctgcggcag cttgtgcgtc acgccgtcaa tcgaaatctg 27120
atgctcctcc atgacttcga gcagtgccga ctgcaccttc ggagaagcgc ggttgatttc 27180
gtccgcgagc aggaaattgc acattgccgc gccgggccgg tattccagag aaaagtcgtt 27240
cgggttgacg agcgtgaagc cggtgatgtc cgaaggcatg acgtccggcg tcaactgaat 27300
gcggttgaag gtgccgccga ccgaacgcga gagcgccgcc gcgagctgcg tttttccgac 27360
gcccggaacg tcctcgatga gaatgtggcc gccgcagagc agacccgccg tgacgcagcg 27420
gatcacgcgg tccttgccga cgatgacttt tccgatgctt tcggtcagtt tcagaatatc 27480
agaatgcata gaatttgcgt tcctttcaaa tttgcaggat cagtttttca cgatcattcc 27540
aggctccgcg cagagggaga cgcggacatt catcggatcc tccgcatcga acatgaccat 27600
ctgcccctcc gtcgtccgga cggtgatgcg ctttccgtgg aagcggatct gctcaaactg 27660
ctcgaagtac gagaaggagg caatccagag aatcggctcg ccgttcgccg gaatcgcggc 27720
aagatacgcg catttcatcg gctcaagcac gccgcactga aagccgcact gcagactctc 27780
gctccgttca aagcactggt tgttcgtgta gtaggggcgc cccttccgcc ggtgaatctc 27840
ctccggcaga ctcggaatcc gcgcccctgc agtcagcctg ccgtccgtca gcgcgtacgg 27900
cacatagctg ttgtcccggt accagattcc gttcagaatc gggacgccgg aaaaggtgct 27960
ctgcgtgatg ctcatgcctg cgctgcccag aggccgtgca gattgcacca tgcgtatgct 28020
gcaatgatct tgtcatcctc ttccgcaaag aactgtgcgg cgggctcctc gcccggcttc 28080
agttccttgc gctcctggcc gttctcggtt tcgaggagaa tccagccgat gtagtgcgca 28140
tcaatcatcg ggtgcgcggc ggaaccgact tcgacggtca cggtctggcc ttcgcggctg 28200
atgaccggaa catgcttttc tctcgacgcg tcaaccgtgt tcgcgatgac ttcttcccag 28260
ccctgcgttt cggggctctc cggcacgatt tcaccggttt cgggatttgt gtagaacttc 28320
agcatcacaa tgctcctttc tgaaagccgc cccgtcattc tgcggaggcg gcgcaaatct 28380
tcggttttca ggaatcggac aggatcgcct gccggacagc gcgccggatg cgctcctgtt 28440

cgcgcagagg gtaatcgaaa tgggaggcgt tccgcatcag gcgcacctga aatgcgggat 28500
tgccgcgcag aagccgcgcc gaacgcgcag caacaagctc gtccttttcg gactggaaaa 28560
tcacagcagg gacacggatt tcagaaatca tgccccgcac cgcgcgcatc tccgcaaaga 28620
gctccagata gcgcggaatc cagcccgcat agtgcaggag attcgcatcg ggcgcaatgc 28680
tgtacgcctg tgccgcagca agcgcgcgcg aagcatcctg cggtgcgccg aacgctgttt 28740
tgaggaaaga gaatgcgccg gggagcgtca ttttcgcata gagcggcaca tcgagcagaa 28800
tcattctgcg gatgctctgc gggaattccc gcgccgcttc cagcgcaaag agcgttccca 28860
tggaatgcgc gatcatgacg gttctgcggc ggttccgcga aaggtcgaga aatgtcttcc 28920
gtacctgctt tttccactgc gccattgcgg cgcgtgagag cgctccggcg gtcccgccgt 28980
gtcccgcgag cagcggcgcg gaaactgcgg tgccgtccgg gactgcattc agcaggaaat 29040
caaaaaactg cggcgcgctg agaatgccgt ggagcagcag aaccgctgtc ccggaatccg 29100
gcgcatctct gcggatttcg cggtatgctc cgctcacttg tagaattcga tttttccgca 29160
gttctggcag tgcataaagc gaagtcctgc cgcatcggaa atccgctgca gctgcttcag 29220
aagcagattg gtcgggctcc atgtatcgac gcgtacgcgg acaagcggcc cgccgcaccg 29280
cagacagcgc agctgtccgt ctgctgattt tccgcgtacc ttctgtgatt ctgtcacgcc 29340
gttcactccc cgtcacagta cgcctgcagg cggcggcggg tttccgcttc gccgaggacg 29400
tgactgattt cattgatgtc cggtgcattc tgccggcccg tcagcgcaat gcgcatcagg 29460
ttcgtcacat cgacgatgct gccgcggtag agctccggat tttcttgta cttcttgacc 29520
tgcacggcat agccgagctc ttcggtcagc gctttgacct tcgcaaacca cgcttccggc 29580
gcgtcggaga agtccagcgt ttcgagatag cgcgtaagga tggcctttct gtcttcgtcg 29640
gaaatgcatt cagggcaggc gtccgtgcgg gtgaaggttt ccgcgtcgta gaatgcgagg 29700
aaatctgccg cctgcttcca tgtgccgtag tccttgcggg gcttgtcgcc gccgcgtccg 29760
acgccgatcg cgccgcggta cagctctgcg cgctccgtca tgcgggcagc aagcgccggg 29820
tcgtactcct tcgcccatgc gagcaggccg tcatagacga aatcaaccga ctgccgcgag 29880
aggtagtcct tggcgatgaa gtcgagcttt tcgaggtcga agagcgagcc ggagatgctc 29940
atcttctcaa gggagaatgt gaattcccgc gaatctgcgt ccggattggc aagatgccat 30000
tcctcgaaat tgctgttcag aaccgtcagc aggtagagcc agaccgcctc ctgcgggaag 30060
cccgcctgcc ggtagtagct gagcgcgagc tccgggtcct tgcgcttcga gagcttgcgc 30120
ttggagctgc cgtccatttt catcagcgtt gcggtgtggc agtagacggg acgctcatag 30180
ccgagcgcgt caaacagctg cacgtgaatc ggcagcgtcg cgagccagga ttcatcgcgg 30240
atgacatggg tcgtccgcat gaagtggtcg tcgcagacat gggcgaaatg gtaggtcgga 30300
atgccgtcgg acttcagcag aacaaagtcc gtgaagttct cctgcagctt cagttcgccg 30360
cggatgccgt cctgcaccgt gacggtgttc gcttcattgc cctcggagcg gaagcggagc 30420
acccactcct cgccggcggc aatcttcgcc gcgatttcgt cataggactt gtcgcgccag 30480
acggcgtatt tgccgtaata gccgtaattg tcctttgcgg cctcctgcgc ttcgtgcatg 30540
gcggcgagct cctcctccgt gcagaagcag gggtacgcca tgccgcgctc cacgagagat 30600
tttgcgatgg tctggtagag ctcccggcgc tgtctctgcc ggtaggggcc gtaatcgccg 30660
atttcaccgt cgaccgtcgc accctcgtcg aagtgtacgc cgtagaagga catcgcgtcg 30720
atgactgctt ccaccgcgcc ctccacctcg cgcttctggt cggtgtcttc gatgcgcagg 30780
aagaagacgc cgccggtctg ccgcgcaagg cgttcgccga ttgcgttgta gaggttgccg 30840
aggtgcacaa agcccgtcgg actcggcgca atgcgcgtca caactgcccc ttcgggcaga 30900
ttgcgcttgg gatagcgctg ttcaaagtcc tccggcctct tgtcgatgtc cgggaaaaga 30960
agctgtgcca gcttttggta atccatatca gaaagctcct gtctgtgaga taattcgtat 31020
aagaaaacgc tgaatataag tgtctccggc ggatgaaagt gtctccgacg gattttgaat 31080
gagggctccg ccctcaaact cccgccgggg gcagtgccca cagaccccat aataattgtt 31140
atgctgcaga caatgttcct gcaaaagcaa tgcggtttca tcaacgtttc cataaagtga 31200
agtacgcgca gtcaaaaggt cacgcgcacg cgagggggct tgggggattc ttcccccatt 31260
atcaatccgg gggcttgggg gattcttccc ccattatcaa tccgggggct tgggggattc 31320
ttcccccatt attaatcttc ttcccccatt accgatgatg cccgccgccg ccgtgagagt 31380
ggccgccgct cgacatgtgg gagtgaccgc cgccgcctcc gccgcctccg gtccgtccgc 31440
cgccgccgga atcaatgtgt acccgcgcga cgtgttcgcg aaggaaatga tcctcatgga 31500
cggaatactt gctctcctgt tcactgacgt agttcgtcgg ggaaatagac ttcacaagat 31560
gatagccgct cttgacgaag cacgatgcga tgaacgcagc aaccgcgccg cagatcgcag 31620
cgattccgcc cagcagtccg agatttttgc cgaaccacca cggcagcttt ttcgcgtgct 31680

```
     gcagctggcc cttgtgataa taatagtatt cgccggtcga actgtcatag gagaaggcgc     31740
     cgttcggaat gccctgcttg tagaaagctt ccagctctgc gcagaaggtc gaaatcgcgc     31800
     cgacataatc cttgttcttc aggttcggcg tgatcgtttc cagaatcgtc caagtgcggc     31860
     ttcctccgtc cgaattctca aagtacatct gcgccgtgcc ggaggtagtc aggtaatcgt     31920
     atttcgtcga atcgttgatg acgagcagca cgccgtccgt gtccttgccg tctaccggat     31980
     tgaacagctt gtcgtaggca tcgtccgcaa agcgctcaac ggcctgatcg ctgctgaacg     32040
     gcgtttcttc accgacaatg tacacagcaa cgtacatatc gattttttcg caggtttcct     32100
     gaatgcgctg cgcagccatc ttctcttcgt ccgcgctgat ggcatcctca atgtcgtaga     32160
     cggtgcaata gctgttgagt tcggggaact ctcttgcggc ggaaaccggg agaacggaag     32220
     cgccaagcag gaatgctgcg ccgagcagtg cgccggcaat tttcagcggg tgtttcatga     32280
     cagcagcacc cccaatccga atgcaatcaa ggcaataatc aggccgattg cgccggagaa     32340
     aagcgcgagc ttgcccgtgc tgaccggcgg atcgccgacg aattttccgg actgtccgtt     32400
     cagcgcaaat tcgtagattt tgtccttgta cttgtaggtc ataaaccaga ccggcagaag     32460
     catgtacttc cagtccgtct tcacgacctt gatgtccgaa cgcgtgacgg acacgctgtc     32520
     gtagcccttc agagagccgc ggaccagctg atcgctcgcg ttcgtcgcgc gcttctgaat     32580
     gcgcgggaac atcttgctga agtccatgtc gtacttctct gcaaggaagc cgctgagata     32640
     ctgcatcgca aacggacgca ccttcgtgta atcgaagggt tcaaccgctt ccatcaggtc     32700
     gtcgtcgatg tggctcgcgc cgtctgcggg caggcccttg atgacaacgc gcgcgttccg     32760
     ttcgacggcg aattcctttg tctcggtaat ctgttctctg cccgaagtgt aggtgcggat     32820
     tttcttgccg agtgcttcca tctgcacgtc aatgtcgcag tccgtcaccc agaacgggac     32880
     atacagaccg acgatctttt cctgctgcga agcggaaaga aaatcgctcg gcaggaagcg     32940
     gtgttcgccg caccatttct tgaattcggc aatcgcctgc gcgcggtcaa gctggaacgg     33000
     cagaaccgca ctcggacggt attcgccgga gacgcgcccc tgcagaacca cgggattgtg     33060
     gcagtagtag cagaaggtcg ccgcagtgtt gtcgtcagcc ataatcgccg cgccgcagct     33120
     gctgcaggaa tacacagagg tgtgtgcgtc ccattccgca tccgtctgga cttcttcctc     33180
     cggacgg                                                               33187
//
ID   XXX; XXX; linear; XXX; XXX; XXX;  XXX.
XX
AC   PRJEB67889
XX
DE   Uncultured bacterium DNA fragment from in vivo enriched cow rumen, clone
rum-51d10, contig 1
XX
FH   Key             Location/Qualifiers
FH
FT   source          1..9817
FT                   /organism="uncultured bacterium"
FT                   /environmental_sample
FT                   /mol_type="genomic DNA"
FT                   /isolation_source="Cow rumen"
FT                   /clone="rum-51d10"
FT                   /note="AZCL-xylan, AZCL-arabinoxylan active clone"
FT   CDS             709..1638
FT                   /translation="MSTFSVTFSNVLLTLMYLSAGFLLSKAKKVIPDHMSSISSILLYI
FT                   CGPCMFLDTIISMDYSASMTANIFLFLVITLAVMGLALLLIFLLLGKRRKDFAYRMMAI
FT                   ASVMGNAGFFGLPIVKAVFPAAPEAATFSCIYCVSMNIIAWTLCVFILTDDRKYMSVRA
FT                   SFLNPTVLSVLLAMILYALGAKNWMLRLVLRGIQSVGGISTPLCMFILGIRLSTMNLKS
FT                   FFTSPVPWLASFGKLLVFPLLCYGIVLLLPLSDVFRASILLLSATPCASIILSLSEIHH
FT                   HGQDIAARCALLSTLLSVLTIPLLSLLL"
FT                   /product="hypothetical protein"
FT   CDS             complement(1641..2786)
FT                   /translation="MNHTIKRVLSALVIILSITAVIIIAFSNPEMKNAWKALASMDLKW
FT                   VLGLFLCWAAYAGFEGFGTWSYLRSKGYKINLGRVIGVALIGFYYSNITPSAAGGQPMQ
```

```
FT                   VNSLHKAGIPVAYGTTAVTIRFVSNQLTICLMTLGLMIANKTFAYNQLGGLIWAARIGW
FT                   AINFIAVPLVLLAAFRRSLIQAVIEKLIALLAKLHLVRDKEATITKISEILDSYHAAMI
FT                   ALIHSPGQILLQILFSTLCMLGMTGSIVFTYYAFGLHGTPWYQILTMSCLLFVSASYTP
FT                   LPGASGAQEGGFMAYFAGIFTDGTIGVGLLIWRFFTYYMFLIVGVGTVLNDKRLIRLEK
FT                   KRRQAEERLSATEPAANGQPGNETGEKPEEPAEEPETPADA"
FT                   /product="Integral membrane protein"
FT   CDS             complement(2849..4924)
FT                   /db_xref="GO:0031216"
FT                   /translation="MVISQNVLSAVFSDETPDYRFPAEPDPGDSVTIRIRVAKNSAKRV
FT                   IVLFDSMSVGTQMENTRSDDYFDYYETSLVCNDQEVMYRFLIECHDDGRIAYDKSGARV
FT                   EEGTALDFNPVYAFRFIPGFHVPEWAKGAVQYQIFTDRFCNGDPSNDVTDNEYYYTIGH
FT                   AKHISDWDALPTDTDIRSFYGGDLQGIMNKLDYLQDIGIEVLYLNPIFVSPSSHKYDTQ
FT                   DYEHVDPHFGVITDDIEHEMQLWEKHNGFAPRYIRRVTSTENLEKSDELFARLCQEMHR
FT                   RGMRIILDGVFNHCGSFNKWMDHEGIYLGKAGFLPGAFQSVHSPYRSYFHFNDSGNGRS
FT                   PAYEGWWGYSTLPKLNYEESPELREQILKIAEKWMSPPYSIDGWRLDVAADLGHSKECN
FT                   HSFWQDFRKRVKAANPEALIIAEHYGDPTDWLNGREWDTVMNYDAFMEPVSWFLTGMEK
FT                   HSDSYRDDLYQNGSAFFGIMREKMAHLKYPSLITAMNELSNHDHSRFLTRTNRMVGRMT
FT                   TVGSGAAAAGINKGVFREAVTIQMTWPGAPTVYYADEAGQVGWTDPDNRRTYPWGHEDQ
FT                   SLISLHRDLIALRKKLPVLARGSLKPLLADYGRIAYARFDGNSRCVIAVNNTGGWTDFR
FT                   LFVRDAGAQEGEAFFRRILTMQESHEAGSVPLGSVTDGYLTFDIPPYSAAVISNAE"
FT                   /product="GH13"
FT                   /EC_number="3.2.1.135"
FT   CDS             complement(4961..5782)
FT                   /translation="MEGQPLVSIVIPAYNAEKTLKRMLDSVTGQTWQNIQIILVDDGSR
FT                   DNTLQIAHEAAGRDARIQVHTQENKGVSQARNTGMRFCTGKYIRFLDADDTLPPESTEI
FT                   LVNKAEEEASDLVIGGYKECWGTREFCFNLENRDDTLTTTEILPLLCRKANTYFYGVLW
FT                   NKLFRRDIIEDMGLAFQDGLTYGEDFAFHVTYLKEAERISFVKEYLYDYHRSMGSMTFR
FT                   QAFSCVTHPLRNIETKVRLYHSMRDMYIFRNEYGKYKNKLWLYLFKVGLDK"
FT                   /product="GT2"
FT   CDS             complement(5933..7459)
FT                   /db_xref="GO:0031176"
FT                   /translation="MMTGMTVAHAGQETMVYTSTFGAGTDLWHARGTEKVYHTTEATLR
FT                   TEGRTSDWNAPARDFELVAGTRYDMSVEVFQNDVDTATLMVSVQVTKEDGSPDYKNLFK
FT                   GEAKKGEWTTLTGSYTPDPSEDYIVYVETVGAPELSYEIRNFRLTAPDGVPDVQPTAAP
FT                   VTESEAVDEMPSLKDLYAGKFDFGTALPRNAFNDMQLLMLVKEQFSIITPENEMKPDAI
FT                   LDVYSSKKLAEEDETAVAVKFDACKTLLRFAQSNGIKVHGHTLLWHNQTPEALFHEGYD
FT                   TSKPLASREVMLGRMENYIRQVLTWTEENYPGVIVSWDVVNEAIDDGTNKLRTGANWYK
FT                   TVGPDYLARAFEYARKYAAEGVLLYYNDYNTAYSGKLYGIFDLLKSLKEEGNIDGYGFQ
FT                   MHHSLGEPSMDMITRAVEKIASLGLRLRVSELDINAGKPTEQNLQSQKNKYKQVMKLML
FT                   RFKDQTEAVQVWGVIDTMSWRRDGYPLLFDKNMNPKPAFFGVAEAAQEDE"
FT                   /product="CBM22-GH10"
FT                   /EC_number="3.2.1.8"
FT   CDS             complement(7650..9800)
FT                   /translation="MTTDISFTIDETGKVKTSGTTTTDEKGNKVILVEDAKTHVEVSKT
FT                   DIANGEELEGAHILVLREATDADKADENTKYADEEKKLVIVEEWTSTKDAHVIEGLLTD
FT                   TEYILRETVAPEGYTVTTDIGFTIDETGKVTTTGSTTTDEKGNTVILIEDAKTHIEVSK
FT                   VDVADGEELPGATIQILREATEADKADEKTKYADEEKTLVIVEEWISTEEAHVIEGLET
FT                   GVEYTLRETVAPHGYDIATDTKFTIDELGKVTTTGSVTENGILLVEDGVLPVTAAVKKI
FT                   WNDENNKDGVRPLTVRVNLLANGKKIDSVVLSIANNWMAAIGNLPMVDKNLKDIEYTWE
FT                   EEAPGNGYTLTSQEKKGTLTTITNSRPPENTGFAVQKIWEDDGNAYQTRPDEIRVQLYA
FT                   DGKACGEAVTLNAANSWKAEWTGLDKFANADGQSGHAAEVIYTVEELNVPEGYQATVTG
FT                   SAKAGFVIRNVLERGKLIIEKDFQFGEIEPEEPDDTPMDIPVTKTWNDNGNKDGNRPAS
FT                   ITVRLLADGTEVASAELTAAGGWKTVFTGLPRYNGEEKITYTITEDPVEWYTAEINGFN
FT                   IRNNYEPEVTSATVRKVWDDKNNAEKMRPTSIHVTLSNGTTVLLSEENNWTATINNLPT
```

```
FT                   KVNGKPVTYTWTEQTVIAYKLDKVEQQGTVTVFTNTFTQVPKIPEGPKQPTTPGGNWQK
FT                   FEEYETPLGVQSIINHVGDCFD"
FT                   /product="hypothetical protein"
XX
SQ   Sequence 9817 BP; 1819 A; 2994 C; 2452 G; 2552 T; 0 other;
     ctgtcctgcc ctcaaacctg tccccgtatg ttccgaatgc gatattgttc atcaggcggc        60
     tcctgtgctt tttcccgcag gctgtgccga tcacggatga cagcactttc ccggggcctg       120
     tcgccggatt ggggccgccg gttcccatga agatcgcgcc ggcgaatccg tccttcggat       180
     agcgggatgc atacatccgc acaatgaacg atcccatgga aaagcccatc agaatcagcc       240
     ggcatgccgg gtgctccttt ttggcataca gcaccatttc ccggtaatcc tcgatcaggg       300
     cattccatcc gccccgcctg ccgaaccatc ccttcagttt cggatcctgg ttggattctc       360
     cgtgattggc catatcatgc atatataccg caatcccgtg atcgcacagg aatcggatat       420
     attcctcgta tctcttctgg tgttccgcca ttccgtggtg aatcaccatc accgtctcga       480
     accagccgtc ttccggcgta tagatatggg cgctgatctc gcacacgcct gttttcgacg       540
     gaaaatgaaa cggtatgttg tttttcatgg ctcgtccctc caccggtttt gtttttggac       600
     tgtctcttgt tccatttgta ccatattttc cggcagcacg caaccgtttc tctgttgctt       660
     ttccgccggg atacagtgta tactgatgcc gtttcgggag gcgggaatgt gagtactttt       720
     tctgtaacct tcagcaatgt tctgctgaca ctgatgtatc tgtcagccgg ctttttgctg       780
     tccaaagcga aaaaggtcat accggatcat atgagcagca tcagttcgat cctgctctat       840
     atctgcggcc cctgcatgtt cctggacacg atcatcagca tggactattc cgcatccatg       900
     acggccaata tcttcctctt cctggtgatc acactggcag tcatgggact ggctctcctg       960
     ctgattttcc tgctgcttgg caaacggcgg aaggattttg cataccgcat gatggccatc      1020
     gcgtccgtca tgggaaacgc cggcttcttc ggcctgccga ttgtcaaggc cgtatttccc      1080
     gccgcgccgg aggctgccac tttttcctgc atctactgcg tttccatgaa tattatcgca      1140
     tggacgctgt gcgttttcat cctcacggat gaccggaaat acatgtccgt cagggcttct      1200
     ttcctgaatc cgaccgttct ttctgttctg ctcgcaatga ttctgtatgc cctgggtgca      1260
     aaaaactgga tgctccggct tgtactcagg ggcatccagt ccgtcggcgg tatttccacg      1320
     ccgctgtgta tgtttattct cgggatccgg ctttccacga tgaacctgaa gtcgttcttc      1380
     acttcgccgg ttccgtggct cgcttccttc ggcaaactcc ttgtcttccc tctgctatgc      1440
     tacggaatcg tgctgctgct tccgctgtcg gatgtgttcc gtgccagcat cctgctgctg      1500
     tccgccacac cgtgcgccag cattatcctc agtctttcgg agattcacca tcacggccag      1560
     gacatcgccg caaggtgcgc actcctgtcc acgctgctca gcgtactgac cattccgctc      1620
     ctgagcctgc tgctctgatc tcaggcgtct gcaggcgttt ccggttcttc cgcaggttcc      1680
     tccggtttct ccccggtttc atttcccggc tgtccattcg ctgccggttc ggttgctgac      1740
     aggcgttcct ccgcctgtct tcttttttc tccagccgga tcaggcgctt gtcgttcagc      1800
     acagttccca cgcccacgat caggaacata tagtaggtga agaaccgcca gatcagcagg      1860
     ccgacaccga tcgtaccgtc cgtaaatatg ccggcgaaat aggccatgaa tccgccttcc      1920
     tgtgcgccgc tcgcgccggg aaggggcgta tagctggctg acacaaacag caggcagctc      1980
     atcgtcagga tctggtacca cggggttccg tgcaggccga aagcataata ggtaaacacg      2040
     atggaaccgg tcattcccag catgcacagc gtgctgaaca gaatctgcag caggatctgc      2100
     ccgggagaat ggatcagggc gatcattgcg gcatgatagg aatccaggat ttcagaaatc      2160
     ttggtaatgg tggcttcctt atccctcacc aggtgcagtt tcgccagcag cgcgatcagc      2220
     ttttcaatca cagcctggat caggctccgc ctgaacgcgg caagcagcac cagcggaacc      2280
     gcgataaaat tgatcgccca gccgatgcgc gcagcccaga tcaggccgcc cagctgattg      2340
     taggcaaacg ttttgttggc aatcatgagt cccagcgtca tcaggcagat tgtcagctgg      2400
     ttggaaacga agcggatcgt gacagcggta gtaccgtatg ccacagggat ccccgcctta      2460
     tgcaggctgt tcacctgcat cggctgtccg ccggcagcgc tgggagtgat attgctgtaa      2520
     tagaaaccga tcagcgccac gccaatgacc cggcccaggt tgattttata acctttggac      2580
     cgcagatagc tccaagtgcc aaagccttcg aaaccggcat acgccgccca gcacaggaac      2640
     agaccgagca cccacttgag atccattgaa gcaagcgcct tccaggcgtt cttcatctcc      2700
     gggttgctga aagcgataat aatcactgca gtgatggata aaataatcac cagtgcactg      2760
     agtacccgct tgatcgtgtg attcatgtcc gactccttgc tgtcatgatc aatccgtgtc      2820
     tggaattagt tccttgtttt gatctgcctt attccgcgtt tgagatgact gccgcggaat      2880
     acggtggaat gtcaaatgtc agatatccgt ctgttacgct gcccagcggc acgctgccgg      2940
```

cttcatggga ttcctgcatc gtcaggatcc ggcggaagaa tgcttcccct tcctgcgcgc 3000
ccgcgtcccg gacaaacagg cggaaatccg tccatccgcc cgtattgttc accgcgataa 3060
cgcaccggct gtttccgtca aaccgggcat aggcaatccg gccgtaatcc gccagcagcg 3120
gcttcaggct cccgcgcgcc agcacaggca gcttcttccg cagcgcaatc aggtcccggt 3180
gcaggctgat caggctctgg tcctcatggc cccacggata tgtccgccgg ttgtccggat 3240
ccgtccagcc tacctgtcct gcttcatccg cgtaatacac ggtcggggcg cccggccagg 3300
tcatctggat ggttacggct tcccggaaaa cgcctttgtt gattccagca gccgcggcac 3360
cggaaccgac cgtcgtcatc cggcccacca tccggttggt ccgtgtcagg aaccgggaat 3420
ggtcatgatt gctcagctcg ttcatcgcgg taatcaggct cggatatttc aggtgtgcca 3480
tcttttcccg catgatgccg aagaatgcgc tgccgttctg gtacaggtca tcccggtagg 3540
aatcggaatg tttttccatt ccggtcagga accaggatac cggttccata aacgcgtcat 3600
agttcatcac ggtatcccat tcccggccgt tcagccagtc ggtgggatcc ccgtaatgct 3660
cggcaatgat cagagcttcc ggattggcgg ctttcacccg tttccggaaa tcctgccaga 3720
aactgtggtt gcactccttg ctgtgtccca gatccgccgc gacatccagc cgccatccgt 3780
caatgctgta aggcggcgac atccatttct cggcaatctt caggatctgt tcccgcagtt 3840
ccggcgattc ctcgtagttc agcttcggca gtgtggagta accccaccag ccttcatacg 3900
cgggagaccg gccgttgccg gaatcgttga agtggaaata gctccggtac gggctgtgca 3960
cgctttggaa agcacccggc aggaatccgg cctttcccag gtatattccc tcgtgatcca 4020
tccatttgtt gaacgatccg cagtggttga acacgccgtc caggataatc cgcatgccgc 4080
gccggtgcat ttcctggcac agcctggcga acagttcgtc actcttctcc aggttttccg 4140
tggacgtcac ccggcggatg taccggggcg caaaaccgtt atgcttctcc cacagctgca 4200
tttcatgctc gatgtcatcg gtaatcacgc cgaagtgggg atccacatgc tcgtaatcct 4260
gtgtatcata tttatggctc gacggggaaa cgaaaatcgg gttcaggtaa aggacttcaa 4320
ttccgatatc ctgcaggtaa tccaacttgt tcataattcc ctgcaggtcc ccgccgtaga 4380
aagaccggat atccgtgtcg gtcggcaaag catcccagtc gctgatatgc ttcgcgtgcc 4440
cgattgtata gtaatactcg ttgtccgtta cgtcgttcga cggatcaccg ttgcagaaac 4500
ggtccgtaaa aatctggtac tgtacggcac cctttgccca ttccggcaca tggaatccgg 4560
gaataaaccg gaacgcatac accggattaa agtccagcgc agttccttct tccacgcgtg 4620
ccccgctctt gtcataggca atccggccgt catcgtggca ctcaatcagg aatcggtaca 4680
tcacctcctg gtcgttgcag accaggctgg tctcataata atcaaaataa tcgtcgctgc 4740
gggtattctc catctgcgtg cctacgctca tggagtcaaa aagaacaatc acccgctttg 4800
cgctgttttt cgccacccgg atccggatgg taacggaatc gccgggatcc ggctcggccg 4860
gaaaacggta atccggtgtt tcatcgctga atacagcgga taaaacattc tgacttatca 4920
ccatgtatgc ctgctcctct tatccggatc tgtccggccg tcacttgtcg agcccaacct 4980
tgaacagata cagccacagt ttgtttttat atttcccgta ctcattccgg aagatataca 5040
tatcccgcat ggaatggtac aggcggactt ttgtttcaat gttccgcagc ggatgcgtca 5100
cacaactgaa cgcctgccgg aacgtcatgc tccccatgga ccggtggtaa tcgtacaggt 5160
attcctttac aaaggatatc cgctccgctt ccttcaggta ggtaacatgg aaggcgaagt 5220
cttcaccata ggtcagaccg tcctggaacg ccaatcccat atcctcaatg atgtcccgtc 5280
ggaacagttt gttccacagg acgccgtaaa aatatgtatt ggctttcctg cacagcagcg 5340
gcaggatctc tgttgtcgtc agcgtgtcat cccggttttc caggttgaag cagaattcac 5400
gcgttcccca gcattccttg tatccgccga tcaccaggtc ggatgcttct tcctccgcct 5460
tgttcaccag aatttcagtg gattccggcg gcagcgtgtc atccgcgtcc aggaaccgga 5520
tatatttccc cgtgcagaac cgcatcccgg tatttctcgc ctgggatacg cctttgttct 5580
cctgggtatg cacctgaatc cgggcatccc gtccggcagc ctcatgcgcg atctgcaggg 5640
tattgtccct gctgccgtcg tctaccagga taatctggat attctgccag gtctgtcctg 5700
tcacgctgtc cagcatgcgt ttcagggtct tctcggcgtt gtacgccgga atcacgatgc 5760
tgacaagggg ctgtccttcc aacttcgcat acctctctta aaagggatca tatgccaaat 5820
acatacagta tattgtacca tggttttccc ggaaaaggaa gtgcaaataa agagcgggga 5880
ctgtttgctc agtccccgcc gatacgcagc atccggtccg ggaagccccg tgttattcgt 5940
cttcctgcgc agcttcagcc actccgaaga acgcaggctt cggattcatg ttcttgtcaa 6000
acagcagcgg atagccgtcc ctgcgccagc tcatggtgtc aatcacaccc cagacctgta 6060
ccgcttcggt ctggtccttg aaccggagca tcagcttcat cacctgcttg tacttgttct 6120
tctgggactg gagattctgt tccgtgggct tgccggcatt gatatccagc tcgctcaccc 6180

gcagccggag ccccagggat gcgatctttt ccacggcacg ggtgatcata tccatggagg 6240
gttctcccag gctgtgatgc atctggaacc catacccgtc gatgtttccc tcttccttca 6300
ggctcttcag caggtcgaag attccgtaca gctttccgct gtatgcggtg ttatagtcgt 6360
tatagtacag cagtacgcct tccgccgcgt acttccgggc gtattcaaac gcccgtgcca 6420
ggtagtccgg tccgactgtc ttgtaccagt tggctccggt ccgcagcttg ttcgttccat 6480
catcaatcgc ttcattgacc acatcccagg atacgataac gcccggatag ttctcctcag 6540
tccaggtcag cacctggcgg atgtagttct ccatccggcc cagcatcact tcacggctgg 6600
caagcggttt ggaagtatca tatccctcat ggaagagcgc ttctggcgtc tggttatgcc 6660
acagcagcgt atggccgtgc accttgatgc cgttcgactg tgcaaaccgc aacagtgtct 6720
tgcaggcatc aaacttcaca gccaccgcgg tttcatcctc ttccgccagc tttttgctgc 6780
tgtacacgtc caggatggcg tccggcttca tctcattttc cggtgtaatg atgctgaact 6840
gttccttcac cagcatcagc agctgcatgt cgttgaatgc gttcctgggc agtgccgtac 6900
cgaagtcaaa ttttccggca tacagatcct tcagtgaagg catctcatcc accgcttcag 6960
actccgttac cggtgcggcg gtcggctgga catccggcac tccgtccggc gcggtcagcc 7020
ggaaattccg gatctcatag ctcagttccg gcgctccgac ggtctccaca tacacaatat 7080
aatcctcgga aggatccggc gtatagctgc cggtcaaggt ggtccattcg cctttcttcg 7140
cctcaccctt gaacaggttt ttgtagtccg gggacccgtc ttccttcgtc acctggacag 7200
agaccatcag cgtggcagta tccacgtcat tctggaacac ttcaacactc atgtcatacc 7260
gggtgccggc caccagttca aaatcccgtg ccggggcatt ccagtcgctg gtcctgcctt 7320
ctgtccgaag ggttgcctcc gtcgtatgat aaaccttctc ggttcccctg gcgtgccaca 7380
ggtccgttcc tgccccaaaa gtggacgtat ataccattgt ttcctgcccg gcatgggcaa 7440
ctgtcattcc tgtcatcatc aacatcatcg ctgtcagtat gatcccgatt cttttcatcg 7500
gtctttcctt tccccggtac ttccggcttc tgttgtgcct atatgataca cggaatttcc 7560
ctttgaatca atatttgttc gtacaaattc tgcgaaaaaa agcaaaaaag aggagccctt 7620
tcctcatcgg aaagggctcc gggattctgt tagtcgaagc agtcaccgac atggttgata 7680
atactctgaa cacccagcgg cgtctcgtat tcttcaaact tctgccagtt tccgccgggg 7740
gtggtcggct gcttcgggcc ttccggaatc ttcggcacct gtgtaaaggt gttggtgaag 7800
accgtcacag taccctgctg ttccaccttg tccagcttgt acgcgatcac cgtctgctct 7860
gtccatgtat aggttaccgg cttgccgttc accttcgtgg gcaggttgtt gatcgtcgca 7920
gtccagttgt tctcttcgct caggagcact gtcgttccgt tgctcagcgt cacatggatg 7980
ctggtcggcc gcatcttctc cgcgttgttc ttgtcatccc ataccttgcg gaccgttgcg 8040
ctggtcactt ccggctcata gttgttccgg atgttgaatc cgttgatttc cgccgtgtac 8100
cattccaccg gatcctcggt gatggtgtag gtgatctttt cctcgccgtt gtaacgcggc 8160
agtccggtga ataccgtctt ccagccgccg gccgctgtca gttcggcact ggcgacctct 8220
gttccgtctg ccagcaggcg gaccgtaatg ctcgccggcc ggttgccgtc cttgttcccg 8280
ttgtcattcc aggtcttggt caccggaata tccatcggag tgtcgtccgg ctcttcgggt 8340
tcaatttcgc caaactggaa gtccttttcg atgatcagct tcccgcgctc cagcacgttc 8400
cggatcacaa atccggcctt ggcgctgccg gtaactgtcg cctggtagcc ttccggcacg 8460
ttcagctctt caaccgtata gatcacttcc gccgcgtggc cgctctggcc gtctgcattg 8520
gcgaacttgt ccagtccggt ccactcagcc ttccagctgt tcgccgcatt cagcgtcacc 8580
gcttcgccgc aggccttgcc gtctgcgtac agctgcacac gaatttcatc aggtctggtc 8640
tgatacgcgt ttccgtcgtc ctcccagatc ttctggaccg cgaatccggt attctccggc 8700
gggcggctgt tggtaatggt cgtcagggta cccttcttct cctggctggt cagcgtatat 8760
ccgttgccgg gcgcttcctc ttcccaggta tattcgatat ccttcaggtt cttgtccacc 8820
atcggcaggt tgccgatcgc tgccatccag ttgttcgcga tactcagcac cacactgtcg 8880
atcttcttgc cgttcgccag caggttcacc cgcacggtca gcggccgtac gccgtccttg 8940
ttgttctcgt cgttccagat cttcttcacc gcggcggtta ccggcagcac gccgtcttcc 9000
accagcagga taccgttctc ggttaccgat ccggtcgtgg tcaccttgcc cagttcatcg 9060
atggtgaact tcgtatcggt cgcgatgtca tatccgtgcg gagccaccgt ctcacgcagt 9120
gtgtactcga cgcccgtctc caggccttcg atcacgtgcg cttcctcggt ggagatccac 9180
tcttcaacga taaccagcgt cttctcttcg tctgcgtact tcgtcttttc atctgccttg 9240
tcggcttcag ttgcttcacg caggatctgg atcgttgcgc ccgggagttc ctcaccgtcc 9300
gcgacatcca ccttgctcac ttcgatgtgg gtcttggcat cttcgatcag gatgaccgtg 9360
ttgcccttct catccgtggt cgtcgagccg gtcgttgtca ccttaccggt ttcatcgatc 9420

```
     gtaaagccga tatccgtggt cacggtgtat ccttccggag caaccgtctc acgcaggata      9480
     tactcggtat ccgtcagcag gccttcgatc acgtgcgcat ccttcgtcga ggtccactct      9540
     tcaacaatta cgagcttctt ctcttcgtca gcatacttcg tgttctcgtc tgccttatcc      9600
     gcgtcagtcg cttcacgcag caccaggata tgcgcgcctt ccagctcttc gccgttcgca      9660
     atgtccgtct tgcttacttc gacatgcgtc ttggcgtcct caaccaggat gaccttgttg      9720
     cccttttcat ccgtggtggt cgtgccgctc gtctttacct tgccggtctc gtcgatcgtg      9780
     aagctgatat ccgtcgtcac ggtgtatccg tccggag                               9817
//
ID   XXX; XXX; linear; XXX; XXX; XXX;  XXX.
XX
AC   PRJEB67889
XX
DE   Uncultured bacterium DNA fragment from in vivo enriched cow rumen, clone
rum-51d10, contig 2
XX
FH   Key             Location/Qualifiers
FH
FT   source          1..6846
FT                   /organism="uncultured bacterium"
FT                   /environmental_sample
FT                   /mol_type="genomic DNA"
FT                   /isolation_source="Cow rumen"
FT                   /clone="rum-51d10"
FT                   /note="AZCL-xylan, AZCL-arabinoxylan active clone"
FT   CDS             270..1331
FT                   /translation="MEARPILWMIIPCYNEEKVLPLTAPVFTQKLTQLADSGRISRNSR
FT                   VLFVNDGSTDRTWEIIRELAEQDERCIGISQSRNRGHQNAVLAGLMEAKDRCDITISID
FT                   CDGQDDINAMDEMVDQYLAGYEVVYGVRSKRETDTWFKRVTAEGFYKLMNTMGAEVVFN
FT                   HADYRLISARVLAEFADFEEVNIFLRGMVPLVGFPSTKVYYERSARLAGESHYPLRKML
FT                   ALAFNGITSLSVKPITLITTAGMIFSVIGLILMIWAVIQAIAGNTVAGWASTICIICLL
FT                   GGIQLISLGVLGQYIGKTYLETKHRPRYIISERTWESSQRRWIEQQGATAGQGSGSRRG
FT                   RKAVNPEETYKEQ"
FT                   /product="Glycosyltransferase"
FT   CDS             1522..3423
FT                   /translation="MKRKLQRIVSVVCALALMIGCGVFASLAEGADTVTRYITVEWVDE
FT                   DNYDNLRKDVSVSLGGQAPVVLSEANGWAGESSAPAGSEWEIPDVAGYVKSSSGNDITV
FT                   VTYTHAVKKTVAAATVKWNDSDNAAAVRPASVQVRLLADGKPFGAEKTANAGNSWTVTW
FT                   ENLPVTKKGTADAIAYSVEEVVPDGYTAAVSGLTITNTLQTGTLSLQASVSGAPAGADV
FT                   SGLKLTVTGPDPSMPRTLTYGQIAGGTYSFGSVLPGAYVVQEENADSLVEGYEMDPANS
FT                   RVGDAVLVKNGEAATLSFRYAYREPVAAEPNEDPMASVGALTFEIDGPDSRMPMTITYS
FT                   QFTGGQYTLENLAPGTYYVVERNAENLVEAYTLTSDSVTGMSITVGEGTATAVLFNRYT
FT                   PAVTPEPETELISIPVTKTWNDDNNKDGNRPQSVTVNLYADGALVDTAIITEAGGWVAT
FT                   FADKPLCYEDGTEIKYTVNEEPVEWYTAAVNGYNITNTYQPVTTSATVTKVWDDGNNYQ
FT                   RIRPKSIAVILQPVGTVYVLSAENGWTVTADKLPIRINGETVQYSWKEQESVGYVLSDV
FT                   LTNGNGTIFTNRVTRIPDTTGGEKKPTTPGGDFAYFEEYKTALGVETIINHVGDCFD"
FT                   /product="hypothetical protein"
FT   CDS             3459..5852
FT                   /translation="MRKGLINVNKLQRILSVLLAVMMIIACMPLAAAEEPADAAQEEIE
FT                   FPDELIVSHPTITKGDFFTEMFGNDTADIDVRALIHGYNLVNWDQNQGTYVFDPSVVRE
FT                   VLVGTDSEGYRQYTIRIADNLKYSDGTPITAWDYAFSFLLMMSPEIEEIGGKIYRAEHL
FT                   VDYDEYIAARRKAMAGETLDPDKDLDYLTAVTVLSDYQLRIWLDPDFLPYFFEIGLLMT
FT                   VPYPIDVIAPGCKVVYGFKDMEDNPLGIKIVNAEDSVEEPVLNAQGETVGYRIAEKSAP
FT                   IYTADLLRKTILDPETGYNTFPSKVSGPYVIKSWDGVTGHFEINPEFKGAWMVNNLPGG
FT                   ALATAWEKARAQGRVVKDTDSFGNPVQDADGNDIELVLPSIEKIAFTVADNDTVIEEIV
```

```
FT                   AGEKHLVNKVTYKETVEKGLGAYGEDGVEYQSYPRIGLAFLTFTYDWKTVNEKEVRQAI
FT                   AWCMDRDLLTEEYCGSNGLRVDGYFGLEQWEYQLLTRRIGFPVNFLPEVDAIGTPIEYE
FT                   EEELERMLKLKNRWVRSEADYEAAIKAWDALADEWDDGENSLLTKYTVDLDQANKLLDR
FT                   AGWKLDRNGEAYDPGVDDVRCKMIDDELVALDLKMMYPAGNHMAEFMSEEGNFVDNLAQ
FT                   VGIKLTLVPAAMEDLLRSYYREAERTTDMIYLATNFHVIVDPAITYSTDKTVGHIQWNN
FT                   TYSDDEYLFDLAVDLRKTEPGDVFQYVSKWMSFQRRYNEVLPTIPVYSNIYYDFYTPYL
FT                   QNYVITAQVTWSQAILLAYFGLPEGETEPVAVEETVGDDEMLFD"
FT                   /product="hypothetical protein"
FT   CDS             5948..6772
FT                   /translation="MQILCSTGALIGRPNGRDYRLLVAQAPMLHCDGFEFMMYDTWYPQ
FT                   ADEILQYLQGLRLNIPVLHCEKSLAEHISVGGQEELEEAFRLFHINCRMAAGLGARKMV
FT                   LHLWNGMISDTHFENNLAAYGDLRKIAEEYGIDLLVENVVCLEDPMSHWVELYREYTDI
FT                   HFIYDTKMADFHRQMDLLYMPEYDWLWKGNHIRHYHINDYDGGYMDWSNLKVLAPGKGH
FT                   IDFDRFFAFIDRIGYRDTFTFEATGFDKQGTIHTDMLNEQFDWARAHLNRMA"
FT                   /product="hypothetical protein"
XX
SQ   Sequence 6846 BP; 1664 A; 1785 C; 2042 G; 1355 T; 0 other;
     cccaggaagg caaacgccat gatccggctg atgctgacag cacggcgatg tttccactgc        60
     ctccgcgggc gacgggtctg ttttcctgtg gccatgattt tcccttcctt ccgggaaaac       120
     agtatacact ataacgcccc gccgaatcaa gcggggttga aacgccctgc gggatggaat       180
     ataatgaacg ggacgttata taaccggaaa gaaacagggc gcggaagccc cggcgggacc       240
     tctgcagccc ggaaggatgg accggaagca tggaagcaag accgattctg tggatgatta       300
     tcccctgtta taatgaagaa aaggtgctgc cgctgacagc acccgtgttt acccagaagc       360
     tgacacagct ggcggacagc ggccggatca gccggaacag ccgggtgctg tttgtaaatg       420
     acggctccac tgaccgtaca tgggagatta tccgggaact ggcggaacag gatgaacggt       480
     gcatcggcat cagccagagc cggaaccggg gccaccagaa cgcagtactg gccgggctga       540
     tggaggcgaa agaccgctgc gatatcacga tctccattga ctgcgacgga caggacgaca       600
     tcaacgcgat ggatgagatg gtagaccagt atctggccgg atatgaggtg gtgtacggtg       660
     tccgctccaa gcgggagacc gatacgtggt tcaagcgggt aaccgctgaa ggattctaca       720
     agctgatgaa caccatggga gcggaggtgg tgttcaacca cgcggactac cggctgattt       780
     ctgcccgggt tcttgcggaa ttcgcggact ttgaggaagt gaacatcttc ctgcgcggta       840
     tggtgccgct ggtcggattc ccgtccacga aggtctacta tgaacggtcg gcgcggctgg       900
     ccggtgaaag ccactatccc cttcggaaaa tgctcgcgct ggcgttcaac ggtattacca       960
     gcctgtccgt caagccgatt accctgatta ctacagccgg catgatcttc agcgtgatcg      1020
     gcctgatcct gatgatctgg gcggtgatcc aggccatcgc cggcaatacg gtagccggct      1080
     gggcttccac gatctgcatt atctgcctgc tgggcggcat ccagctgatc agcctgggcg      1140
     tgctgggaca gtatatcggg aagacctacc tggagacgaa gcaccggccc cggtacatca      1200
     tttccgagcg gacctgggaa tcatcacagc ggcgctggat tgaacagcag ggagcgaccg      1260
     ccggacaggg aagcggaagc cggcggggac ggaaagcggt gaatccggaa gaaacatata      1320
     aagaacagta aacgtacaat ttgcatgata caaatccggc agaatccggt cagaaagctt      1380
     accggatccc gccggatttg tatttttgc acttgctttt tgagtctcgt tctcatataa      1440
     taaactttga gtttatatgc cccaaaaagg gcggaaattt cattaaagct tttcggaata      1500
     tcaccgaagg gagataaaaa aatgaagaga aagctgcaga ggattgtgtc tgtcgtgtgc      1560
     gcgctcgcgc tgatgatcgg ctgcggagtc ttcgcttccc tggcggaagg cgctgatacc      1620
     gtgacccggt atatcaccgt ggagtgggtg gatgaggaca attacgataa cctgcggaaa      1680
     gacgtatcgg tttcgctcgg cgggcaggct cctgttgtcc tgagcgaggc gaacggctgg      1740
     gcgggagagt cttccgcgcc ggccggcagc gaatgggaaa tccccgatgt tgccggatat      1800
     gtcaagtctt cctccgggaa tgatattacc gttgtgacgt atacccacgc agtgaagaag      1860
     actgtcgcgg ctgcgacggt caaatggaac gacagcgaca atgccgccgc ggtacggcct      1920
     gcctccgtgc aggtgcgcct gctggcggac ggaaagccct tcggcgccga aaagaccgca      1980
     aacgccggca acagctggac cgtgacctgg gagaacctcc cggtgacgaa gaagggaaca      2040
     gcggacgcga ttgcctattc cgtggaggaa gtggttcccg acggatacac cgcggctgtt      2100
     tccggactga cgatcacgaa caccctgcag accggaaccc tgtctctcca ggcttctgtt      2160
     tccggcgccc cggcaggtgc ggatgtcagc ggactgaaac tgacggtcac cggtcccgat      2220
```

ccgtcgatgc ccagaaccct gacctacggc cagattgccg gaggcacata cagcttcggc 2280
agcgtcctgc ccggtgccta tgtggtgcag gaagaaaacg ccgattccct ggtggaagga 2340
tatgagatgg atcccgcaaa cagccgtgtc ggcgatgcgg tgctggtgaa gaacggggaa 2400
gccgcgacgc tgtccttccg ctatgcctac cgggagccgg tggccgcgga gccgaacgaa 2460
gatccgatgg cttccgtcgg tgcactgaca tttgagattg acggtccgga ttcccggatg 2520
ccgatgacga ttacctacag ccagttcacg ggcggacagt acaccctgga aaacctggca 2580
cccggcacct actatgtggt ggaacggaat gccgagaacc tggtggaagc ctataccctg 2640
acgagcgaca gcgtgaccgg catgagcatt acagtgggcg aaggcacggc gaccgcagtg 2700
ctgttcaacc gctatacacc ggcagtgaca ccggaaccgg agactgaact gatcagcatc 2760
ccggtgacca agacatggaa tgatgacaac aacaaggacg gcaaccgtcc tcagtccgtg 2820
acggtcaacc tgtatgccga cggtgcactg gtggataccg caattatcac cgaagccggc 2880
ggatgggtgg cgacctttgc ggataagccg ctctgctatg aggacggaac tgaaatcaaa 2940
tatacggtga atgaagagcc ggttgaatgg tacaccgccg cggtaaacgg atacaacatc 3000
acgaatactt accagccggt aacgacatcc gcgactgtga ccaaggtgtg ggatgacggc 3060
aacaactacc agaggatccg gccgaaatcc attgccgtga tcctgcagcc tgtcggcacg 3120
gtatatgtgc tgagcgcaga aaacggctgg accgtcacgg ctgacaaact gccgatccgc 3180
atcaacggtg agacggtgca gtattcctgg aaagaacagg aatcggtcgg atatgttctc 3240
tcggatgtgc tgaccaacgg aaacggaaca atctttacca accgggtgac gcgcattccc 3300
gatacgacgg gcggagagaa aaagccgacc actcccggcg gagactttgc ttactttgaa 3360
gaatacaaaa ccgcgctggg tgttgagaca atcatcaacc acgtgggtga ctgcttcgac 3420
tgattaccgg acagatattc atattcctga caaaacatat gagaaagggt ctgatcaacg 3480
tgaacaagct gcaaagaatc ctgagtgtgc tgctggctgt tatgatgata atcgcctgca 3540
tgccgctggc tgctgcggaa gagccggcag acgccgcaca ggaggaaatc gaatttccgg 3600
atgaactgat cgtctcccat ccgacgatta ccaagggcga tttcttcacc gaaatgttcg 3660
gcaacgacac cgcggatatc gacgtacgcg ctctgattca cggatacaac ctggtcaact 3720
gggaccagaa ccagggcacc tatgtttttg atccgagcgt tgtccgggaa gtgctggtcg 3780
gaacagacag cgagggttac cggcagtata ccatccgcat tgcggacaac ctgaagtact 3840
cggacggaac cccgatcact gcatgggatt acgctttctc cttcctgctc atgatgagcc 3900
cggagattga ggagatcggc ggaaaaatct accgtgccga gcacctggtg gactacgatg 3960
agtatattgc agcccgccgc aaggcgatgg ccggcgagac gctggatcct gacaaggacc 4020
tggattacct gaccgccgtg accgtgctga gcgattacca actgcggatc tggctggatc 4080
cggacttcct tccttatttc tttgagatcg gcctgctgat gaccgtcccg tatccgatcg 4140
atgtgatcgc tcccggatgc aaggtggtct atggtttcaa ggacatggag gacaatccgc 4200
tgggaatcaa gatcgtcaac gcggaagact ccgtggaaga gcctgtgctg aacgctcagg 4260
gcgaaaccgt gggataccgg atcgcggaga agtccgcgcc catctacacg gcagacctgc 4320
tccgcaagac catcctggac cccgaaaccg gatacaatac attcccctcc aaggtttccg 4380
gcccctatgt gatcaagtcc tgggacggag tgaccggcca ttttgagatc aaccctgaat 4440
tcaagggtgc ctggatggta aacaacctgc ccggcggcgc acttgcgacg gcatgggaaa 4500
aagcccgggc acagggccgc gtggtgaagg acacggattc cttcggaaat cccgtgcagg 4560
acgctgacgg caatgatatt gaactggtgc ttccctccat tgagaagatc gccttcaccg 4620
tggcggacaa cgataccgtg attgaggaaa tcgtcgcggg cgagaaacac ctggtcaaca 4680
aggtcaccta taaggaaacc gtggaaaagg gtctgggcgc ctatggcgaa gacggggttg 4740
aataccagag ctatccccgg atcggcctgg cctttctgac cttcacatat gattggaaga 4800
ccgtgaacga aaaagaggtc cgccaggcga ttgcgtggtg catggaccgg gatctgctga 4860
ccgaggaata ctgcggatcg aacggcctgc gtgtggatgg ctattttgga ctggaacagt 4920
gggaatacca gctgctgacc cgccggatcg gcttcccggt gaactttctg ccggaagtgg 4980
acgcgatcgg aacgcccatc gaatatgaag aggaagagct tgagcggatg ctgaagctga 5040
agaaccgctg ggtccgctcc gaggcggatt atgaagcggc catcaaggca tgggatgcgc 5100
tggcggatga gtgggacgac ggtgagaata gcctgctgac caagtatacc gttgacctgg 5160
atcaggcgaa caaactgctg gaccgtgccg gctggaagct ggaccggaac ggcgaagcct 5220
atgatcccgg tgtggacgat gtccgctgca agatgatcga tgatgagctg gttgccctgg 5280
acctgaagat gatgtatccg gccggcaacc atatggcgga gttcatgtct gaagaaggca 5340
acttcgtgga caacctggca caggttggca tcaagctgac gctggtgccc gctgccatgg 5400
aagacctgct caggtcctat taccgcgagg cggaacgcac cacggacatg atctacctgg 5460

```
        ccacgaactt ccacgtgatc gtggacccgg ctatcaccta cagcaccgac aagaccgtcg      5520
        gccatatcca gtggaacaat acttattcgg atgatgagta cctgtttgac ctggctgtgg      5580
        acctgcggaa gacagaaccc ggcgatgtat tccagtacgt cagcaagtgg atgtccttcc      5640
        agcggcgcta taacgaagtg cttccgacca ttccggttta ttcgaacatt tactacgatt      5700
        tctatacgcc ctatctgcag aactacgtga tcaccgccca ggtgacgtgg tcccaggcca      5760
        ttctgctggc ctacttcgga ctgccggagg gagaaacgga accggtagcc gtggaagaaa      5820
        cagtcggaga tgacgaaatg ctctttgact gattggtaaa ttgcagaaag tgttttcaga      5880
        ttcaccgggc gaatcatgcc cggtgaatct ttatacccgg caaaaacgtc ccgaaaggaa      5940
        taacaatatg cagattcttt gttccaccgg cgcactgatc ggccgcccga acgggcggga      6000
        ttaccgtctg ctggtagcgc aggcgcctat gctgcactgt gacggctttg aattcatgat      6060
        gtatgacaca tggtatcccc aggcggatga aatactgcag tacctgcagg gcctgcggct      6120
        gaatattccg gtgctgcact gcgagaaatc actggcggag catatttccg tcggcgggca      6180
        ggaagagctg gaggaggcgt tcaggctgtt ccatatcaac tgccgcatgg cagcgggcct      6240
        cggtgcccgg aagatggtcc tgcatctgtg gaacgggatg atctccgaca ctcatttcga      6300
        aaacaacctg gctgcctatg gagacctccg gaagattgca gaagagtatg gaatcgatct      6360
        gctggtggag aatgtggtgt gcctggaaga tcccatgagc cattgggtgg agctgtaccg      6420
        ggaatacacg gatatccact ttatttatga tacgaagatg gcggactttc accgccagat      6480
        ggacctgctc tatatgcctg aatatgactg gctctggaaa ggaaaccata tcaggcacta      6540
        ccatatcaac gactatgacg gcggatatat ggactggagc aacctgaagg tgcttgcgcc      6600
        ggggaaggga catatcgatt ttgaccgatt ctttgcattt atcgacagga tcggctaccg      6660
        ggacacattt acatttgagg caaccggttt tgacaagcag gggacgatcc atacagacat      6720
        gctgaatgag cagtttgact gggcacgtgc ccatctgaat cggatggcat aaacaaataa      6780
        tgctgcatag aataaattga gtcacagaag ctttgattca atatgaagta aacatagaag      6840
        gacaga                                                                6846
//
ID   XXX; XXX; linear; XXX; XXX; XXX;  XXX.
XX
AC   PRJEB67889
XX
DE   Uncultured bacterium DNA fragment from in vivo enriched cow rumen, clone
rum-52e22
XX
FH   Key             Location/Qualifiers
FH
FT   source          1..35389
FT                   /organism="uncultured bacterium"
FT                   /environmental_sample
FT                   /mol_type="genomic DNA"
FT                   /isolation_source="Cow rumen"
FT                   /clone="rum-52e22"
FT                   /note="AZO-galacto-mannan active clone"
FT   CDS             complement(417..1334)
FT                   /db_xref="SEED:fig|6666666.127280.peg.1"
FT                   /translation="MGTIQIGRTLLECKDYECSQDEVYVPRERLRLWLYINLTDACPAS
FT                   CPFCVHTVPGERGGHADPDQLSKVLSKTAPFISGVSLTGGEPMTDIPLMEEIIAVIREK
FT                   IPADIELDMVTNGLHIEKLPGLRGLERVTTVHISRHAAEDEQNAALMRWAGAPSKQTLK
FT                   QVFSALPDPGMTVLNCVLQKNGVHDAASVREYLETAAWIGAANVSLIGMFLANDYCTEQ
FT                   YVSPASIPFTEDPRFRVWNHYHDHDYCQCSTGSYNARSGPVRYYFRCPGNVQGPALCRQ
FT                   LVYGSDNRLRAGFGEAEVIDGIDD"
FT                   /product="hypothetical protein"
FT   CDS             complement(1336..1623)
FT                   /db_xref="SEED:fig|6666666.127280.peg.2"
FT                   /translation="MSFSRFIELLQQLITMTDPEDEDQVHYAAGVLQNLHELARRSDKI
FT                   DRRTLGIIYIAIVEFLDLLRMKEQFAGKPGDVAGNLAKRKRLAMALGPRC"
```

```
FT                   /product="hypothetical protein"
FT   CDS             complement(1620..2312)
FT                   /db_xref="SEED:fig|6666666.127280.peg.3"
FT                   /translation="MTEKIKKYMTPFQWAEVLAIIGFTVYFAITNKTDAWWYILIDSLA
FT                   AVCGVFCVVLCAGGKKSQYYWGFVNILAYIVIALINKYYGEVMLNGLYYLPTQFIGLWV
FT                   WGKHYRKEDDQVDGLRMDGRHIAVWSVCSVAGIFVYKLILDALGGNATLLDSMSTVFSI
FT                   VANILMVLRYREQWALWIIVDVVTVIMWIIAGDFLMTIMWAIYLVNAVYGWVMWTKMSS
FT                   EKREERTA"
FT                   /product="Ribosyl nicotinamide transporter, PnuC-like"
FT   CDS             complement(2305..3324)
FT                   /db_xref="SEED:fig|6666666.127280.peg.4"
FT                   /db_xref="GO:0000309"
FT                   /db_xref="GO:0050262"
FT                   /translation="MARYSCGMYGGSFNPLHLGHVRCMISAANQCDRLIVVISHGPNRG
FT                   EIDIRQRYRWVYQTVKHFPHVQIFVLEDDAPTKAAYSEETWYADAEKVKQFAGEPITAV
FT                   FFGDDYSRDSMWSKCYPEADVVVLPRDGISSTAIRRSPLACWDMMPAYVRSYYVKKVLL
FT                   IGTESTGKSTLTVSLARHYNTVFLEEVGRELSARSGTDLWMLPEDFTDILLEHKVKQLE
FT                   MISRANRVFFEDTNCLTTLFYVNFLEGEEKEKNRKLAEAISDLNTYDLVLLLGPDVAFV
FT                   QDGDRSEEIAADRAGFTALLKGMCEAHGLQVEELNGDYEQRYRKAVELVDRLLEEEHD"
FT                   /product="Nicotinamide-nucleotide adenylyltransferase,
FT                   NadR family (EC 2.7.7.1) / Ribosylnicotinamide kinase (EC
FT                   2.7.1.22)"
FT                   /EC_number="2.7.7.1"
FT                   /EC_number="2.7.1.22"
FT   CDS             complement(3533..4558)
FT                   /db_xref="SEED:fig|6666666.127280.peg.5"
FT                   /translation="MSKTRILSSAIPNLPWQPRPENDPHDAPLWRYSENPIIGRNPVPG
FT                   VARIFNSAAAPWQDGFIAVFRGEQVNGIPCVYMGTSTDGIHWDINPEKVPFTDEEGRPF
FT                   LPHYAYDPRLVKVDDTYYIMWCQDFYGASIGMAKTKDFKTFTRLENPFIPFNRNAVLFP
FT                   RKVGGNYLLLSRPSDSGHTPFGDIWLSRSPDMKFWGMHRHVMSPGGKWWESVKIGSGAA
FT                   PIETSEGWLLFYHGVSSTCNGFVYSFGAAVLDIDEPSIVKYRCENFLLTPEEWYEERGF
FT                   VPNVCFPCAALADADTGRIAIYYGAADTYLGLAFTEINPLMDYIKAHSCVSGNDTEPGI
FT                   R"
FT                   /product="GH130"
FT   CDS             complement(4732..6102)
FT                   /db_xref="SEED:fig|6666666.127280.peg.6"
FT                   /translation="MKRFTALFLALVMVLTMAASVSLADDARHLTFALWWDLYYDSDDE
FT                   SWEASPSATGKDSEIMRFENVKNIEETYNVTYEFVNATWAGVQDSINNSILAGEPDFDM
FT                   YMVDLMAFGVPAVMNGLATDLRNVLPADDPFLAHEDPVLDYIDLPDGSATLIYPKGAED
FT                   AVASTNPLAFNLQMIQEANLEDPRDLYERGEWTWDKFREYCQILTKDTNGDDVIDVYGY
FT                   GAWIGDALPAWYMSNGTYMAATPTENLSSSEIGETLKFLQDLWLTDKVAYPIPAENGWD
FT                   VCRYLYRDKKVAFTTTAVWIMANYDDYNWDGKAESTLDFDMVFVPYPIGPSGNAETNNS
FT                   KIASGNYWMLPAGVKDPKLVFDVFRAYQNWYHDDVTLRDDPEELEWWYTATSDKLDIQN
FT                   WNFSVLEKLGSKKMVDFYNSFSDSLPILEFLNGEYTPAQFQETYRQTVQDALDQIFD"
FT                   /product="ABC transporter substrate-binding protein -
FT                   sugar transport"
FT   CDS             complement(6189..7922)
FT                   /db_xref="SEED:fig|6666666.127280.peg.7"
FT                   /translation="MRKVLSWLLIVTMAVSLCAFASAETAAVDKIDFADGAFGFLGLST
FT                   AKGNADAAAELSVVDYHGAKALKIDSKSKAPYVAFNIEGLLGDQLSKVRAISATYGIDE
FT                   AGPDGKFYAASGVVYTYTGEENVETKHDWSVYLAKKNPRTIVTKLDESEVFVPGMNNSL
FT                   VISKETDNCQTATGTSLAVYLTDIQFLDAEGNALPLDLTAVYQSADTGRDTSNLFAISN
FT                   PVTFAGFETTGGGWGQNGFDWDPAVLEALVPGTVMEIEYESEDGSMWVVFPDAAAGWSR
FT                   IAQDQAYINGSKNIAQITYEQIAAVVGEDKMAWGGRLQCEAKTNWTVYSVKVGTCAPQV
```

```
FT                   TLANAVEFAGFETTGGGWGQNGFDWDPAVLEALVPGSAIEVTFESEDGSMWVVFPDAAA
FT                   GWSRIAQDTAAIQGGKAYITYDQIAAVVGEDKTAWGGRLQCEAKTNWTVYSVRVGQLVE
FT                   IPAFNKVVEFPGFETTGGGWGQNGFDWDPAVLEALVPGSALIVTFESEDNSMWVVFPDA
FT                   AAGWSRIAQDAAVINGGKAYITYDQIAAVVGEDKTAWGGRLQCEAKTNWTVYGVAVGQV
FT                   Q"
FT                   /product="hypothetical protein"
FT   CDS             complement(7985..8956)
FT                   /db_xref="SEED:fig|6666666.127280.peg.8"
FT                   /db_xref="GO:0005215"
FT                   /db_xref="GO:0006810"
FT                   /db_xref="GO:0016020"
FT                   /translation="MAKAQKQRISWQSADASSRSLDRQSRAVNGIKNTLLGIFRFVIIV
FT                   GISYVILAPVIGIIANSFFSRQDSINPMVFTIPINPTLERYTMAVKYLDYLPILGKTLL
FT                   YVLGVTLIQLLICSMVGYGFARYQFPLKKLLFGCVVVMIVVPLNTIQFPLYMTFRYFGI
FT                   GPNTVNLLKTPWPTVILSLFGCGLRSGLYIYIFNQFFRGLPKEIEEAALVDGAGSWYTY
FT                   FHIMLPNAAPALITVAVFSIVWQYNDSFFANTFNISDSIILTRKLDSLINVIANAEKIL
FT                   TISEQQLYFDAGVVLVLLPVVIIYLVLQRRFIEGVERSGIVG"
FT                   /product="N-Acetyl-D-glucosamine ABC transport system,
FT                   permease protein 2"
FT   CDS             complement(8971..9891)
FT                   /db_xref="SEED:fig|6666666.127280.peg.9"
FT                   /translation="MTAQKKKRFRLSYEQRNAMRGYAFILPWLIGFLVFYAGCLFQLVR
FT                   FSVMDIKIDAATGTRLDTFVGFDNFITAFTAHASFKQVLTSSLIDMVIDLPLIIFFSLF
FT                   VALLLNRKFRGRALVRAIFFLPIILSADAVGSAITRATELMNNGVSAQAAEMAGGGAVS
FT                   VSYYIALFGDLAIPPTILEYLVGAVNRIASIIQGSGVQIILFIAALQSIPGALYEVARI
FT                   EGATGYETLWKVTVPMVMPHIITNTIYTLVDRFTTSEVMRLANETYQKYNYGLSSVFSL
FT                   VATMATILILGVIVYLLNKRAFYYN"
FT                   /product="binding-protein-dependent transport systems
FT                   inner membrane component"
FT   CDS             complement(9888..12485)
FT                   /db_xref="SEED:fig|6666666.127280.peg.10"
FT                   /translation="MENAENTARLPSSVKRVLWILLAVLILALGAVIGWYVIHFRNYDA
FT                   YKAIPLRPAAYTEGTEFKPLEDSLKAVPGFALAAENETMALYFKEKTAEVAVYDKATGN
FT                   TMYSNPPAADKDPVAKATNLENLKSQFILSYLDGNAKEGTPWSSYAKSVSPGQVSCESI
FT                   ENGVRVIYLMSNEKMMLVPDQLDQAWYDILSTTGKKQAAKSYVFNEASGLYEIKTQGVT
FT                   ARNRQQIDADARAAGFTMEDYEAMQALRPAEEDDAEKAENLSFTVTLDWILKPDGVQVT
FT                   LPFEGISETGNGQIRAIQLLPFFGAAGSDEQGDLVIPEGSGGLIHFNNGKSTAAQYSKN
FT                   IYDLDLVDSDLTATQNLQTARLALFGICREDSSILATCERGASLAGITADVSGRNNSYN
FT                   YAYFTFALRRTDTLVISGENVIVAEKDLYPVDCSVRYTLLSPEKQGYSGIAQAYRERLI
FT                   REGLLAQETGEAQDIPFFYDVIGGVKETAHFLGIQYLRVLPMTTFSQAEEMIAEFRHQG
FT                   ITNQRMNLQGWMNGGYYHDPVTRLKVLGQLGGENGLKSLLGVMKRSGGTVYPDIALQQI
FT                   SDIARGFTASEEASRYYAQGYVVNLARISPVSLRRTDTMGYRELGFKLLSPKFLPRYAQ
FT                   AARTQADRLGLEALSLRDLANELHADKRRTNVISREAALDLVRSAFGTISEGGRELLVS
FT                   GGNDYSLAHAGYVINAPVQATPYPIMDEQIPLWEMIVHGSIGYAGSAMNMTQSESRQAD
FT                   LLHLIDYGASIHYTFTWRDAAEMKYTGLNSQYATTFSAWKDQAVEDYNYVNGALKHVSG
FT                   AAMLSHEKLSDTLSRSRYSNGVSIYVNTGSAQASADGHTVEALDYLVVGGEAQ"
FT                   /product="GH129"
FT   CDS             complement(12502..13137)
FT                   /db_xref="SEED:fig|6666666.127280.peg.11"
FT                   /translation="MKKGIFSGEKWKYLFYTVSHPNDGFYWIRHQDRGSVSIAVLLVIL
FT                   YGVCFSMNRIYASFVVNDIEPRTVNSIAELAGVLILYLVLCVGNWSVTCLMDGEGRFKD
FT                   IVIAIGYSMLPVIAATAAATLISQVVAENEEAFYTILIGIGTAYTVIMMLIGIMQIHNY
FT                   TLGKTLITLFLTVIAVLIILFLALLVINFITQVYSFLRSIYIELIFRI"
FT                   /product="FIG01031734: hypothetical protein"
```

```
FT   CDS             complement(13134..14609)
FT                   /db_xref="SEED:fig|6666666.127280.peg.12"
FT                   /translation="MKKIICCLLLLCLLPAAALANDSLPYDCYNYDYWNNILYTPAPYV
FT                   PDGSLGGSSLMWEGNPLGAFRSPQDLCVSPDGKIYLADTGNNRIVVIDGGLKTVLQVID
FT                   HFEADGGTQTFASPTGVAVSQNRQLYVADSQNHRIVVLELDGTFVKYVQNPQSEVLDEG
FT                   YIFTPLKVSVDYADRVYCVAQNMFQGIMVFTAEGDFTGFFGTISVKISLWEKFWRKFST
FT                   KEERSKQQLFIPTEFTGIDVDDEGFVYASNKDTDGIQAVRRLNPKGQDVIRKGLKENLG
FT                   GDLFFGTVGNYSGPSQIVDVVYRDHGIYSLLDARRGRVFTYDHEGNLLYIFGGIGTQEG
FT                   TFRTPAAIEQLGSRLLVLDAQQGNVNIFSETEYGRLINQAVSLRYNGDEALAVPLWSEV
FT                   LRLDENNELANAGIGKAYLSAGDNEKAMTYLRRGMNRAYYSVAFKRWRNEALRGNISWF
FT                   LTTLIVLIALTVVIRRVVLPRRAAKRKKGGNAA"
FT                   /product="NHL repeat containing protein"
FT   CDS             complement(14622..15551)
FT                   /db_xref="SEED:fig|6666666.127280.peg.13"
FT                   /translation="MSTASQHKPVDTRPAKIIKLEKKMRRKEGGKKLNRSMAGNTLLFI
FT                   IMGISGLFMVLPLVMIVNNALKPLDEIYQFPPRIFVRNPTLTNFSDLFVLMNESWVPFS
FT                   RYIFNTIIISLGGMLGHVIVASMAAYPLAKHKFPGKALLFSMVVLSMMFSWTVTQIPQY
FT                   LIISWLGINNTYAALVLPAWAFGMGLYLMKQFMEQLPDSLMESARLQGANEWQIFWQIV
FT                   MPNVRPAWLTLAIFQFQQMWGNTGGMFLRNEELKPLQYALQQITSGGVSRAGAGAAVTF
FT                   IIAAIPIIFFLICQSSILETMTTSGMKD"
FT                   /product="binding-protein-dependent transport systems
FT                   inner membrane component"
FT   CDS             complement(15553..16485)
FT                   /db_xref="SEED:fig|6666666.127280.peg.14"
FT                   /translation="MSRTEANRLPRVQKFTLRHEIWKNRASYLLMAPYFILFFLFTVIP
FT                   VLMSIYLSFTYFNMLEAPRFVGWSNYIKLLLEDDIFIISLRNTLIFAVVTGPVSYMLCL
FT                   LFAWIINEFRPKLRAVLTLVFYAPSICGNAYMVWNLILTGDRYGYLNGLLLRLDFINEP
FT                   VLWMQTEKFVLPVLIVVQLWLSLGTGFLSFIAGLQTVDKGMYEAAAIDGVKNRWQELWY
FT                   VTLPAMKPQLLFGAVMQITQSFAVADISINLAGNPSVNYAGATVVTHLLDYGSTRLDMG
FT                   YASAIATILFLLMVGTNKLVQKVLRRVGE"
FT                   /product="binding-protein-dependent transport systems
FT                   inner membrane component"
FT   CDS             complement(16487..19489)
FT                   /db_xref="SEED:fig|6666666.127280.peg.15"
FT                   /db_xref="GO:0005215"
FT                   /db_xref="GO:0006810"
FT                   /translation="MDRSEAIMKGWLSLAAVLIMLLAAVPSLAATDRSIAEYENLVSTY
FT                   SIDAGIPSYDDYRKARDEVRPGCRIVLDGADVVRYEEEGAAEPALLTDYAGMPGVSLLT
FT                   GEEALAEWEFTVEEEGWYDPCLEYYPYAGKNSEIQRAFFIDGQLPYDELSLIALTRVWR
FT                   NDLHDQVTRETDENGISVIRWRKDNQGNELKPSPEEAPEWVETALYDSNGYITDNLGVY
FT                   LTKGQHTLSVLSLREPMLIRRITLGQVEKPQPYAGRSDLPQAKGISVRIEAETAEKTSS
FT                   QMLYPVQDQGSSAVYPASARYLLNNTIGGNAWKNAGQWIEWKFSVPEEGDYCLTFYDKQ
FT                   NFVRGIDVYRKIYVDGQVPFQEFGALPFGYGQTWRMETAKTGSGEPYRIHLSAGEHKLR
FT                   MEVVLGSMAGIISQVQQCVLQLNSIYRQVLYITGVSPDVNRDYQIERNLPGLKQELEDV
FT                   RADLRVAIDTLEATAGHSSDKLTVLKTMDDQLIELIKDQERFTEVLSSYKTNVRACGNW
FT                   ITQVLGQPLQIDRIYVHTPDIDPKAENTSFGDTLSHEAQRLYNSFVIDYNQVGNIAEAG
FT                   GDSRVITLWIGTGRDQANVIKALIDENFTPETGINVNVQLVDMNTLLRATLSGQGPDVA
FT                   IQVANTTGIAGAVMNTGNDTPVNYGLRHAVLDLTGFSDFSEIAARFAPSAMVPFSFNGA
FT                   TYALPDTQTFPMMFYRKDILAEIGLEVPKTWDEVKVAMSVLSKNQMEFGMLPGEQTFAM
FT                   LLYQAGGEYYSENGDRSALDSDIAINVFRRYCEFYTDYKLDAATSVEERFRTGECPLII
FT                   SDYTTYNNLQVSAPDIKGLWDFTHVPGTVREDGTLNTATGCSGLADIIMSATKEPEACW
FT                   EFLKWWTSAATQTLYGREMESLMGSSARVATANQEALANLSWPIRDYKGLAAQFANVRG
FT                   IPQVPGGYYTWRNVNNAFYAVTTDNASKGRNESGNTPREELMDKIYYINAEITYKRTEF
FT                   GLPTAEQGEGE"
```

```
FT                   /product="N-Acetyl-D-glucosamine ABC transport system,
FT                   sugar-binding protein"
FT   CDS             complement(19515..20996)
FT                   /db_xref="SEED:fig|6666666.127280.peg.16"
FT                   /translation="MKRYERTSRRPLFTVAALAMCCLMMPAAPAAAQEDVVRLEAETGT
FT                   VCGHVKVVSSGEQTWVEGFQEAGDRVELTVHADQSGMYDISVILASADGGHKENPLLLD
FT                   GERVAGSIVEGKDFQTSVLSYIYLTEGDHTLALDTSWGWIRVDAVEIRPSQPLPADLYE
FT                   VSPVLVTKDPSPEAQRMFAWMCDIYGKKIITGQNCDGGMFGMENQAVWRATEGSYPALL
FT                   GLDMMDYSPSRVERGSQGKSVDHAIQYWEKGGIVTFCWHWNAPSPYLKADRWYSGFYTE
FT                   YSTINLKKIMDGEDPQGYDLLMKDLDAVAQQLGRLRDAGVPVLWRPLHEASGGWFWWGA
FT                   SGPNAYKWLWNLMFEKFTEEYHLNNLIWVWNGQHADWYPGDDTVDIIGEDIYPGERVYN
FT                   SQSSKFMDCLKYTSARKMIMLSENGCVPDPELVFRDGTVWSSYCTWGGDFVLKSSKFNK
FT                   PSEKYTEVWLLKKMYADERAVTRDDIPDLKSGPQE"
FT                   /product="CBM35-GH26"
FT                   /EC_number="3.2.1.78"
FT   CDS             21391..21594
FT                   /db_xref="SEED:fig|6666666.127280.peg.17"
FT                   /translation="MNSILVNKAGALLFTLTLLILSFATAVTGSPVFGGWTSTAVLAVF
FT                   SLLEAIDLRFTIRKIQRIREQA"
FT                   /product="hypothetical protein"
FT   CDS             complement(21877..22686)
FT                   /db_xref="SEED:fig|6666666.127280.peg.18"
FT                   /translation="MDSRKLIEAFQDTQNLIRQSPELMQETRRSQAESRLILPGFEDLL
FT                   VQEKSREEAVEILDDTTFHCAQQLMHPGEKCAVLNFANAYSPGGGVLAGNMAQEECLCR
FT                   SSNLYEVLTWPYLVRNYYKFNEKNVGQMGTDTLIYSPGIVVLKSDDEIPVLLDSPFRVD
FT                   VITCAAPRISPERKKKVQPEELERALRARIRNILEAASACGADLLVLGAFGCGAFANPP
FT                   ETVAEIFHDLLIGKHYADRFRKVVFAIKPDGETHNADIFRSVFFRWP"
FT                   /product="hypothetical protein"
FT   CDS             complement(22727..23929)
FT                   /db_xref="SEED:fig|6666666.127280.peg.19"
FT                   /translation="MKKKLQRPGLTAAVCLALCICLLCASAGADSFHGYTPTAITVGSS
FT                   VVFGRYEQDANWQNGQEPIVWDVLDIDGDHALLVSRMLLDCIIYHSFDTAVAWGDSYIS
FT                   WWLNNSFYSTAFNQTEQSCIIQSLVSGTMNYVFLPNRNQVLNYGLCDTGCDATVYAKSR
FT                   GVMIGKSSGKGCWWVRMDMTHANGNTQFVGTNGNVYDSNRVTVGDNGVRPCIVASVWAL
FT                   QNGGYGPGGSTPKTQTPQWTPVPEHNTEYYVTGTTNQKLATRCGPSTDYDETHTYNLGV
FT                   GTQVRVLRYQVTRGTAWVEVEFKYGGEWVRAWTGKKRIDCSDAVYLSGDYYVQDTGYVN
FT                   RETPGWFGPGSEYRILYTTIASGTGVDILASRGSWYLIEFMTYDSQLIHRAWVPKDRIS
FT                   K"
FT                   /product="hypothetical protein"
FT   CDS             23993..24151
FT                   /db_xref="SEED:fig|6666666.127280.peg.20"
FT                   /translation="MQFSFVHIKRLRARYSCLFRIFSRTLRRPAVFSLQALSHILSRCA
FT                   ISRFPHR"
FT                   /product="hypothetical protein"
FT   CDS             complement(24205..25533)
FT                   /db_xref="SEED:fig|6666666.127280.peg.21"
FT                   /translation="MIDRFVCAYAEDPDLAMRTLFYARDIRGGLGERRVFRVLLRYLAK
FT                   HEPQSVRKNLAYVAEMGRYDDLLVLLGTPCEVDVLALIDAQLKEDIAGMERGESISLLA
FT                   KWLPSVNTSCAESVRRGRYLARRLGMREAEYRRTLSRLRTHLKLLENYLRERNYTFSYE
FT                   EQPSSAMLKYHKAFERNDAKRYGNFLEKAARGEATLHTSSLNPCDLVQRAMAPSVMQDA
FT                   WERNALDVTWNALEDFTDDRNALCVVDTSGSMFYHGNPTPFAVAMSLGLYFAERNRGPF
FT                   RGRFITFSHTPRLIEILGRDLAEKVAYCRTLTEVSDTNIQAVFELILNAAVRWHLPQSQ
FT                   MPETIYIISDMEFNFCTQNASLTNFEYAKQLYASHHYRLPNVVFWNVQSRNTQQPVRMN
FT                   ERGVTLVSGYTPRLFRMVVSGESNPYDIMMGVLGSERYRAICA"
```

```
FT                   /product="Phage protein"
FT   CDS             25785..25931
FT                   /db_xref="SEED:fig|6666666.127280.peg.22"
FT                   /translation="MVAEGIFHAGSVPAAIRYRISRPDARKKSCELHLFPFFSKMPMYP
FT                   APS"
FT                   /product="hypothetical protein"
FT   CDS             complement(26230..26760)
FT                   /db_xref="SEED:fig|6666666.127280.peg.23"
FT                   /translation="MIAIFELVLGILLLVNPVGLTGVVMILVGAVMIALGGYHLYRYIR
FT                   LPREKAAETWNLATGAGLICVGITVISNQHWLVQMLGTLAAVYGIMTTVCSFMKLQMAV
FT                   DALRWKRPFWYLMAVSFVVTAVLAALLFIQPFAESAVWIFTGIVLLLLAVLDGTYFVLG
FT                   RKNRPAAGEAKQA"
FT                   /product="hypothetical protein"
FT   CDS             complement(26870..28345)
FT                   /db_xref="SEED:fig|6666666.127280.peg.24"
FT                   /db_xref="GO:0009011"
FT                   /translation="MSENHAGNHPVRVAFIGSECYPFIKTGGLGDVMYALPRALLREKC
FT                   EVRVLMPYYACIPEKFKNQMTFRGSFMMDLTQDGRSFFVGIMETEMDGVIYDFVENRDF
FT                   FDWGNPYTGLWEDIPKYCFFSKAALSVLNYLEWIPDVIHCHDWQAGLVPVYLRTEFAGT
FT                   PVGSAKSILTIHNLRFQGICSVDHLKYWTGLPDYLFDYSVFRIANDQANMFKGGLAYCD
FT                   RITTVSATYAREIQTDFYGENLNGHLWYHRAKLSGIVNGIDVDLWNSRTDKLLKAPYGV
FT                   EDVLEKKQENKAALQARLGLHTDTSRFVLGLISRLTDQKGLDLINTVFSKMMNANTQVV
FT                   ILGTGDARYENAFRYYESVYPGQVCAYIKYDESLAHQIYAGADALLVPSLFEPCGLTQL
FT                   IAMRYGTVPVVRETGGLKDTVEPYNQYTGEGNGFTFDYYNADLLLDAVERCRKVYREDR
FT                   DSWNAIVKRDMEKDVSWEHSASQYMDLYRELAQ"
FT                   /product="GT5"
FT                   /EC_number="2.4.1.21"
FT   CDS             complement(28356..28538)
FT                   /db_xref="SEED:fig|6666666.127280.peg.25"
FT                   /translation="MIRRGMRRGCAKWTTAEKPMEKSGKTSADAGKGGKRKYLLRNCTE
FT                   YAIFSIMALANEQET"
FT                   /product="hypothetical protein"
FT   CDS             28628..29410
FT                   /db_xref="SEED:fig|6666666.127280.peg.26"
FT                   /translation="MQILQRPKVKCMEYSVRVNGIEVTAHYEERAVHDIFLPLLRHLTG
FT                   LEREKGRRILVMLAAPPGAGKTTLLSFLETLSHQDPALCPVQVIGMDGFHRRQEYLLSH
FT                   TVFRNGEELPMVRIKGAPVTFDLERLTHFVRRAASGEVCTWPVYDRLLHNPIEDAVTVD
FT                   GRIVLLEGNYLLLDADGWRDLRSFADYTVSVRAPEDLLRRRLIDRRIRTGVEEDAARAF
FT                   VDFSDMPNVRVCLEKTMPADLELFLDDEGAYHITGSSS"
FT                   /product="Uridine kinase family protein"
FT   CDS             29559..30704
FT                   /db_xref="SEED:fig|6666666.127280.peg.27"
FT                   /translation="MKNSQKKASSGQWPGVLVWMLTGAACGIVSMQSMTSPEHADSLLP
FT                   LLLMIPFLLAAIVIQTIIHEGGHLLFGLMTGYRFISFRVFQFMWVMEGGKIRLRRMHLA
FT                   GTAGQCLMSPPEMENGTFPFRLYNYGGAILNLISALLFLGLSFLCAPSSPLRVFLLCLT
FT                   VLGTAFALLNGLPLPSGDVANDAQNVLDMSRHPEAVRAFWISMKVNEALSAGTRTKDMP
FT                   AEWFAVPSADAMKSSPLVAFLGDMACSRLIDEQRFQEADALMERILSEASGLTGLHRGL
FT                   MICDRIYLEAVGENRPEQLKAMRTKQQEKIMKSMASSPSVLRTEYACALLSDRNPEKAE
FT                   QLRRQFEKAAAVYPFPQEIESERGLIAVADEKAGRAAPDTA"
FT                   /product="hypothetical protein"
FT   CDS             30717..31187
FT                   /db_xref="SEED:fig|6666666.127280.peg.28"
FT                   /translation="MELKELTLRDRESMTKLYADVFTGEPWNDDWSCPGQLDAYITDLA
FT                   GQSNSLTFGYFDGDRLIGLAMGHIRHWHEGTEYCIDEFCIARDMQGRGHGGRFMDCVES
```

```
FT                   QLLERGIPWIFLLTDRHMPAYDFYRKHGFRELHDNAALVRKAGAPQPGRNTP"
FT                   /product="hypothetical protein"
FT   CDS             31184..31819
FT                   /db_xref="SEED:fig|6666666.127280.peg.29"
FT                   /translation="MIFPNAYFIIGTAYAGKSTMTRMLAEKHGGILLEENYHSRFFPTL
FT                   DPSAFPCLTYTRELKDWHDFIRRTPEEYEQWIDGAARECERLELEILPQFALGAQPVFV
FT                   DTNISAETLRQVALPGHVLVMLAPPEISVSRFFERQDEEKQFLHRLILEEPDPEKAMDN
FT                   YCRGLARINSPEKYERFLHSGFPVILRDENRTPAETLALVEEVFCLNG"
FT                   /product="hypothetical protein"
FT   CDS             31861..32052
FT                   /db_xref="SEED:fig|6666666.127280.peg.30"
FT                   /translation="MSKENNQLLWNISLAVLAICSLILGLTALFDGDLPDGLERLLGML
FT                   DLIALPVFVYASVQKARG"
FT                   /product="hypothetical protein"
FT   CDS             32077..32844
FT                   /db_xref="SEED:fig|6666666.127280.peg.31"
FT                   /translation="MTNQEILKIALQQSAEDIGCRAEDFLSDAPVLVPFRLGARARKYY
FT                   KEPIICQFVSYRANVVAASVPETAALAEEYIRKFPFYHCFETPNMDWLSTRLLEKGYKV
FT                   CFMAEYYLPDVHRLPDLPCPVETRVLGPEDFRGLYLPEWSNALCEERRELDVLGVGAYD
FT                   QGQLVGLAACSADCENMWQIGIDVLPRARGQGIAPALTSRLAKEILDRGKVPFYCSAWS
FT                   NIRSARNACRSGFLPAWAEMTVKPANLVDDMNR"
FT                   /product="hypothetical protein"
FT   CDS             32875..33657
FT                   /db_xref="SEED:fig|6666666.127280.peg.32"
FT                   /translation="MEKELFIREVRPGIFLMDEAHEATGYLVIGKEKACVIDTMNGYND
FT                   LYRAVRKLTDLPLVVVNTHGHPDHIYGNLYFEEAFLPPADLPLARSFAEDPEFLAFARQ
FT                   KGRSMPPFRDLLPGTRIDLGGRTLEAHALPGHTPGSTLLLLKEDRILFTGDSINHHLWL
FT                   QLDGCLPVHECAMQLDRVLFLMDKADIILHGHAGSEDDISLMRCLRAGMQEICDGKTEE
FT                   DAPYAWFGGTGLQHPFRCEEGKVYSSAQHVICYRPDNV"
FT                   /product="hypothetical protein"
FT   CDS             33694..33984
FT                   /db_xref="SEED:fig|6666666.127280.peg.33"
FT                   /translation="MIVLNVTYLCHPGMRDAFLQSIRAEGIDAASRAEDGNAQYEYFLP
FT                   ADGGDRLLLLEKWRDAEALSAHAHQAHFARLGELKAKYVRETVIEKYVSEA"
FT                   /product="hypothetical protein"
FT   CDS             34001..34447
FT                   /db_xref="SEED:fig|6666666.127280.peg.34"
FT                   /db_xref="GO:0008080"
FT                   /translation="MPGTDLRVRPMTMQDLEPLHALLSSPDVMRYLEPPYTRTQTTAFL
FT                   RRAISQHPPVLAAEADGTFIGYVIFHAWDETSMEIGWVLLPAYQGRGFATALTILLIRM
FT                   AREKGKQPVIECSPAQKATAHIARKFGFVRTETRDGLDVYRLSV"
FT                   /product="GCN5-related N-acetyltransferase"
FT   CDS             34515..34706
FT                   /db_xref="SEED:fig|6666666.127280.peg.35"
FT                   /translation="MDRTTPLYDTRLSDRERAEWLLSQLTLEEKFGFFTLNIRLERLGL
FT                   TLSTCGGEGAHGVQARAG"
FT                   /product="GH3"
FT   CDS             34810..35031
FT                   /db_xref="SEED:fig|6666666.127280.peg.36"
FT                   /translation="MDVMEKENRYTTQYPLAESPLENLRAIRETGMDAFLEKERAAWSC
FT                   GICGRLISVHTGICRGCGKQYGKPSRES"
FT                   /product="hypothetical protein"
XX
SQ   Sequence 35389 BP; 7448 A; 10440 C; 9845 G; 7656 T; 0 other;
```

agccctttg caggcccttc aagaggtacc ggggcagccc ggtacgggcg cagaatgatt 60
ccttctccgc tgttcgtttc cagaagctca tccatctctt tgtctgtgag ccctacaaag 120
gcaaaatcaa catgtttgag gcggaaggtc agcgggaaaa gcgaattcct ttctttgtcc 180
tcatccggcc agacactgct gaggatctct tcatagggaa caagcgtgtc gtgacgttcc 240
atcaggcact gcatgagctt ccattttttg cctttcagtt tcgttttctg tttcccgttg 300
accagctcct ggctgtttac atcaatgtag catgtctctg tcatatagta tcggttcttc 360
tgctgcatat ccgccgccat gttcttcctc ctcccgcctg ccggagaacc gggcactcaa 420
tcatctattc catcaataac ctctgcttcg ccaaaccctg ctctcagcct gttgtcggac 480
ccgtatacca gctgcctgca cagggccggt ccctgcacgt tgccggggca ccggaagtaa 540
taccgcacag gtccggagcg cgcattgtaa ctgcccgttg agcactgaca gtaatcatga 600
tcatgatagt ggttccagac cctgaaccgc ggatcctccg tgaatggaat cgaagcaggc 660
gaaacatact gttctgtgca gtaatcgttg gcgagaaaca tgccgatcag actgacattg 720
gccgccccga tccatgcggc cgtctccaga tattcccgga cagaggctgc atcatgcacg 780
ccgtttttct gaagcacaca gttgagcacg gtcatgcccg ggtccggaag ggccgagaaa 840
acctgcttca gggtctgttt cgatggggcc cctgcccatc gcatcagcgc ggcattctgc 900
tcatcctctg ccgcatgcct gctgatatgc acagtggtca cccgttctag gccgcgaagg 960
cccggcagtt tttcaatatg cagcccgttt gtgaccatat ccagttcaat gtccgccggg 1020
attttttcac gtatgacggc aatgatttcc tccatcagcg ggatatccgt catcggctcg 1080
ccgccggtca gcgaaacacc ggagatgaaa ggcgctgttt tggaaaggac cttcgacagc 1140
tgatcgggat ctgcatgccc tcccctttct ccgggtacgg tgtgcacgca aaaaggacag 1200
gatgcagggc atgcatccgt cagattgata tacagccaga gccgaagccg ttccctgggg 1260
acatagacct catcctggct gcattcatag tctttgcact ccagcagggt cctgccgatc 1320
tgaatggttc ccatctcagc atctgggacc gagcgccatg gccagccgct ttctctttgc 1380
caggttgccc gcaacatccc cgggtttccc tgcaaattgt tccttcatac gaagaaggtc 1440
caggaattca acgattgcaa tgtatatgat tccgagcgta cgccggtcga ttttatcgga 1500
tctgcgtgcc agttcatgca gattctgcag aaccccccgcg gcataatgca cctggtcctc 1560
atcctccggg tccgtcatcg taatcaactg ctggagcaac tcaataaacc tgctgaaact 1620
catgctgtcc tctcctctct tttttcactg ctcatttttg tccacattac ccagccatac 1680
actgcattga ccagatagat cgcccacatg atggtcatca ggaaatcgcc tgcgatgatc 1740
cacatgatca ccgtcacgac atccacaata atccacagtg cccactgttc cctgtagcga 1800
agcaccatca ggatgttcgc cacaatggaa aaaacggtac tcatgctgtc caggagcgtt 1860
gcgtttcctc caagcgcatc cagaatcagc ttatagacaa agatgcctgc aaccgaacac 1920
acagaccaga ccgcaatgtg tctcccgtcc atgcgcaggc cgtccacctg atcgtcttct 1980
ttcctgtagt gctttcccca tacccacagc ccgatgaact gtgtgggcag gtagtacagc 2040
ccgttgagca tcacctcgcc gtagtatttg ttgatcagcg caatgacgat ataggcgagg 2100
atattcacaa acccccagta atactggctc ttcttccctc cggcacagag cacaacgcag 2160
aagacaccgc acacggcggc cagcgagtca atgaggatat accaccaggc gtccgttttg 2220
ttcgtgatgg caaaatagac tgtaaagccg ataatggcaa ggacttcagc ccactgaaag 2280
ggcgtcatgt atttttttgat cttctcagtc atgctcttcc tccagcagcc ggtccaccag 2340
ctccaccgct ttccggtacc gctgttcata gtctccgttc agttcctcaa cctgcaggcc 2400
atgcgcctcg cacatgccct tcagcagagc cgtaaagccc gccctgtccg cagcgatctc 2460
ctcgctcctg tctccgtcct ggacaaaggc gacatccggg cccagcagaa gaaccaggtc 2520
atacgtattc agatcggata tcgcttctgc cagcttcctg ttcttttcct tctcttcccc 2580
ttcaaggaaa ttgacataga acagcgtggt cagacagttc gtatcctcaa agaacactct 2640
gttcgcgcgg gaaatcatct caagctgctt cactttgtgc tcgagcagga tatccgtaaa 2700
atcctccggc agcatccaca ggtccgtccc cgacctcgcc gaaagttccc tgccgacttc 2760
ttcgagaaag accgtattgt aatgtctggc cagactgacc gtcagcgtgg atttcccggt 2820
actctccgtg ccgatcagca gcaccttttt cacataatat gacctgacat acgccggcat 2880
catgtcccag caggcaagcg gagatctgcg gatggccgtc gaagaaatgc cgtccctcgg 2940
gagaaccacc acgtccgcct ccgggtaaca tttcgaccac atactgtctc tggaataatc 3000
atcgccgaaa aagacggctg tgatgggttc ccggcgaac tgtttcacct tttccgcgtc 3060
cgcataccag gtttcttcag aatacgctgc cttggtcggc gcatcatcct ccagcacaaa 3120
aatctggaca tgcgggaaat gttttacggt ctgatacacc cagcggtacc gctgcctgat 3180
atcaatctct cccctgttcg ggccgtgaga gatcacccaca atcagcctgt cgcactggtt 3240

cgccgcagag atcatgcagc ggacatgccc cagatgcagc ggattgaaac tgcctccgta 3300
catgccgcag gaataacgtg ccaaagccat tccccccgta caaaaggatc ctcagcatgc 3360
cggttatgcg catgcccctg tcgcagttat gttatgatct tattctatga aaatcccgcg 3420
gcctgtcaag gcgcgctgta tctcacacaa atccgaaggc ccaaagcaaa agagagagga 3480
agcaccagga atcacttccc tgctttctct ctcttttgat caggacacat tttcagcgga 3540
tgccgggttc ggtatcattc ccgcttacgc aggagtgcgc cttgatgtaa tccatcagcg 3600
ggttaatttc tgtgaaggcc aggcccaggt acgtatcggc tgcgccgtaa tagatcgcga 3660
tgcgcccggt gtcggcatct gccagcgcgg cacacgggaa gcagacattc gggacaaacc 3720
cgcgttcctc ataccactct tccggcgtga gcaggaagtt ctcgcagcgg tacttcacga 3780
tgctcggctc gtcgatatcc aggacggctg caccgaagga atatacaaag ccgttgcagg 3840
tgctggaaac accgtggtag aacagcagcc agccctcact ggtctcaatt ggggcggcgc 3900
cgcttccaat cttcacggat tcccaccatt ttcccccggg gctcatgaca tgccggtgca 3960
tgccccagaa cttcatgtcc gggctgcggc tgagccagat atccccgaac ggtgtatgcc 4020
ccgaatcgct cggcctggac agaagcaggt aattgccccc gacctttctc gggaacagaa 4080
cggcattgcg gttaaaggga atgaagggat tttccaggcg ggtaaacgtt ttaaaatcct 4140
ttgtctttgc catgccgatg gatgcaccgt aaaaatcctg gcaccacatg atgtaatagg 4200
tatcgtccac cttaaccagt ctcggatcat aggcgtaatg ggcagaaaa ggcctgccct 4260
cttcatccgt gaacgggact ttctcgggat tgatgtccca gtggatgccg tctgtgctcg 4320
tgcccatgta tacgcacgga attccgttca cctgttcccc tctgaacacg gcaatgaagc 4380
cgtcctgcca cggggcggca gcactgttga aaatacgcgc aacaccgggc accgggttcc 4440
gcccgatgat ggggttttcg ctgtagcgcc agagggggtgc atcatgggga tcgttttccg 4500
gacgcggctg ccagggcagg tttggaatgg cggacgaaag aattctggtc ttgctcatgc 4560
aatggtacct ccgtaatgct ttccattcgg gaattgcctt ttcgatgggt aaagaggccc 4620
gccccggtgc cctttggcgg catgcccggg gcaggccctg ctgtttcgga tgaccggtca 4680
tgcgttttcc ctgcggcagc tgcctcgggg gaaagcatgc gctctgtcag attagtcgaa 4740
gatctgatcc agggcatcct gcaccgtctg acgataggtc tcctggaact gtgcgggcgt 4800
atactcgccg ttgaggaact caaggatcgg cagagaatcg gagaaggaat tgtagaagtc 4860
aaccatcttc ttggagccga gcttttccag gacgctgaag ttccagttct ggatatccag 4920
cttgtcgctg gtggcggtgt accaccattc gagttcctcg ggatcgtcgc gaagagtgac 4980
gtcatcatga taccagttct gataggcgcg gaacacgtcg aagacgagct tgggatcctt 5040
cacgcctgcg ggcagcatcc agtagttgcc gctggcgatc ttggagttgt tggtttcagc 5100
attgccgctg gggccgatcg ggtaaggcac gaacaccatg tcgaagtcca gggtgctctc 5160
agcttttccg tcccagttat agtcgtcata gttggccatg atccacacag cggtcgtggt 5220
gaaggcaacc ttcttgtccc tgtagagata gcggcagaca tcccagccgt tttcggcagg 5280
aatcggatag gccaccttat cggtcagcca caggtcctgg aggaacttga gggtttcgcc 5340
gatctcactg gaggaaaggt tttcggtggg ggtagctgcc atataggtgc cgttggacat 5400
ataccaggca ggcagagcgt cgccgatcca tgcgccgtag ccgtacacgt cgatgacgtc 5460
gtcgccgttg gtgtccttcg tcaggatctg gcagtactca cggaacttgt cccaggtcca 5520
ttcgccgcgc tcatacaggt cgcggggatc ttccaggttg gcttcctgaa tcatctgcag 5580
gttgaatgcc agggggttgg tggaggcgac agcatcctct gcgcccttcg gatagatcag 5640
ggtcgcgctg ccgtcgggca ggtcaatgta gtccaggacg ggtcttcat gtgccaggaa 5700
gggatcatcc gcaggcagca cattgcgcag gtcggttgcc aggccgttca tgacggcagg 5760
cacaccgaag gccatgaggt cgaccatgta catgtcgaag tcaggctcac cggccaggat 5820
ggagttgttg atggaatcct gtacgccggc ccaggtcgca ttcacaaatt cataggtgac 5880
attataggtt tcttcaatgt tcttgacgtt ttcaaaacgc atgatctcag agtctttgcc 5940
ggttgcggaa ggactggctt cccaggattc atcatccgag tcatagtaca ggtcccacca 6000
cagggcgaag gtcagatgtc ttgcatcatc tgccagggaa acactggccg ccattgtcag 6060
aaccatgacc aaagccagga ataatgctgt gaaacgcttc attcgagttt cctcctttga 6120
gtttgtgtct ttcggtttct cagtgcctgc tgcaggatgc agcaggatta tccgatttca 6180
gcgcgggatt actggacctg gccgacagcc actccgtaga ccgtccagtt ggtcttggct 6240
tcgcactgca ggcgtccgcc ccaggccgtc ttgtcctcgc cgacaaccgc cgcgatctgg 6300
tcataggtga tgtaggcttt tccgccgttg atgacagcgg cgtcctgagc aatgcggctc 6360
cagccggctg ctgcgtccgg gaacacaacc cacatggagt tgtcttcgga ttcaaaggtc 6420
acgatcaggg cagaaccggg aacgagtgct tcgagcacag cgggatccca gtcaaagccg 6480

ttctggcccc agccgccgcc ggtggtctca aagccgggga attccacaac cttgttgaag 6540
gccgggattt caacgagctg accgacccgc acgctgtaca cggtccagtt ggtcttggct 6600
tcgcactgca ggcgtccgcc ccaggccgtc ttgtcttcgc cgacaacagc tgcgatctgg 6660
tcataggtga tgtaggcctt gccgccctgg atggctgcgg tatcctgagc gatacggctc 6720
cagccggctg ctgcatccgg gaacacgacc cacatgctgc cgtcttcgga ttcgaaggtc 6780
acttcgatgg cacttccggg aaccagtgct tcgagcacgg cgggatccca gtcaaagccg 6840
ttctggcccc agccgccgcc ggtggtctca aagcctgcga actcaaccgc gttggccagg 6900
gtcacctgag gtgcacaggt gccgaccttg acggagtaca cggtccagtt ggtcttggct 6960
tcgcactgca ggcgtccgcc ccaggccatc ttgtcttcgc cgacaacagc tgcgatctgt 7020
tcatacgtga tctgggcaat attcttgctg ccgttgatgt atgcctgatc ctgagcaata 7080
cggctccagc cggctgctgc gtccgggaac acaacccaca tgctgccgtc ttcggattca 7140
tactcgattt ccatcacggt gccgggaacc agtgcttcga gcacagcggg atcccagtca 7200
aagccgttct gtccccagcc gccgccggtg gtctcaaagc ctgcgaaggt cacaggattg 7260
gagatggcaa agaggttgga ggtgtcgcgg ccggtatcgg cactctgata gactgccgtc 7320
aggtccagag gcagtgcgtt gccctcggca tccaggaact gaatgtctgt caggtacacg 7380
gccagggagg ttccggtggc cgtctggcag ttgtcggttt ccttggagat gaccagggaa 7440
ttgttcatgc cgggaacaaa cacttcgctt tcatcgagct tggtcacgat ggtgcgcggg 7500
ttcttcttgg caagataaac gctccagtca tgcttggttt ccacgttttc ctcgccggtg 7560
taggtgtaga cgacaccgga agctgcgtag aacttgccgt cggggcccgc ttcatcaatg 7620
ccgtaggtcg cgctgatggc gcgcaccttg gacagctggt cgcccagcag gccttcaata 7680
ttgaaagcca catacggggc cttgctcttg gagtcgatct tcagtgcttt tgcgccatga 7740
taatcgacca cggaaagctc agccgcggca tctgcgttgc ccttggcagt ggaaaggccc 7800
aggaaaccaa acgcaccgtc ggcaaaatca atcttgtcga ccgctgccgt ttccgctgaa 7860
gcaaacgcgc acagactgac tgccatggtg acaatcagca accaggaaag gacttttctc 7920
atggtaatag cctccttgta atgatttaaa cccaccctgt gggcagagtc acgaattcag 7980
gtcgtcatcc cacaatgccg gagcgctcca cgccttcgat aaagcggcgc tgcagcacca 8040
ggtaaatgat gaccacgggg agaagcacca gcacgacgcc ggcgtcgaag tacagctgct 8100
gctcgctgat ggtgaggatt ttctcggcat tggcaatcac gttgatcagc gaatccagct 8160
tacgggtcag aatgatggag tcggaaatat taaaggtatt ggcaaagaac gaatcgttgt 8220
actgccagac aatggagaac acggcaacgg tgatcagggc cggggctgca ttgggcagca 8280
tgatatggaa gtaggtatac cacgatccgg cgccgtcgac cagggccgcc tcctcaatct 8340
ctttgggcag gccgcggaag aactgattga agatatagat gtacagaccg gagcgcaggc 8400
cgcagccaaa cagcgacagg atcaccgtgg gccacggggt tttcagaagg ttgaccgtgt 8460
tcgggccaat cccgaaatag cggaaggtca tgtacagagg gaactgaatg gtgttcagcg 8520
ggacaacgat catgaccacc acgcacccga agagcagttt cttgagcggg aactggtagc 8580
gtgcaaaccc gtacccgacc atggagcaga tgaggagctg gatcagggtg acacccagga 8640
catagagcag ggttttgccc aggatcggga ggtaatccag atacttgacc gccatcgtgt 8700
agcgctccag cgtcggattg atcgggatgg tgaacaccat ggggttgatg gaatcctgac 8760
gcgagaaaaa cgaattggca ataatgccta tcacgggggc caggatgaca tagctgatgc 8820
cgacaatgat cacgaaccgg aagatgccca gcagcgtgtt cttgatgccg ttcactgcgc 8880
ggctctgccg gtcgagggaa cgggaagaag catccgctga ctgccaggaa atacgctgtt 8940
tctgtgcttt cgccatgttt cttccccctc tcagttgtaa tagaaggcgc gtttgttgag 9000
cagatagaca atcacgccaa ggatcaggat cgtcgccatg gtcgcaacca gagaaaacac 9060
ggaggaaaga ccgtaattgt atttctggta ggtttcattg gcaaggcgca tcacttcgct 9120
ggttgtgaaa cggtcgacca gggtgtagat cgtgttggtg atgatatgcg gcatcaccat 9180
gggcaccgtc actttccata gggtttcata accggtggca ccctcgattc tcgccacctc 9240
atacagggcg ccgggaatcg actgcagggc ggcaatgaac aggatgatct gcacgccgga 9300
accctggata atgctggcga tgcggttgac cgcgccgacg aggtactcga gaatggtggg 9360
cgggatcgca aggtccccga agagcgcgat atagtagctc accgagaccg cgccgccccc 9420
ggccatttcc gccgcctgcg cggacacgcc gttgttcatg agctctgtgg cccttgtgat 9480
ggccgagccc acagcgtccg cgctcaggat aatggggagg aagaagatcg cgcgcaccag 9540
cgcacggccg cggaacttgc ggttcagaag cagtgcgaca aagaggctga agaagatgat 9600
caggggcagg tcaatgacca tgtcgatgag ggaggaggtc agcacctgct tgaaactggc 9660
gtgggccgta aacgccgtga tgaagttgtc aaaccctaca aacgtgtcca ggcgggtacc 9720

ggttgccgcg tcaattttga tgtccatcac gctgaaccgg accagctgga acagacagcc 9780
cgcgtagaag acgagaaagc cgatgagcca cggcaggatg aaggcatagc cgcgcatcgc 9840
attgcgctgt tcatagctca ggcggaatct tttctttttt tgtgcggtca ctgtgcctca 9900
cctcccacaa ccaggtaatc aagcgcctcc acggtatgac cgtcggcact cgcctgcgcg 9960
cttccggtat tcacatagat gctgacaccg ttggaatacc tgctgcgcga cagcgtatcg 10020
gacagctttt catgggacag catcgccgcg ccggacacat gcttcagtgc accgttcaca 10080
tagttgtaat cctcgaccgc ctggtccttc caggcagaga acgtggttgc atactggctg 10140
ttgagcccgg tgtatttcat ttcggcggca tcgcgccagg tgaatgtgta gtggatggaa 10200
gcaccgtaat cgatcagatg cagcaggtcg gcctgccggc tctcgctctg ggtcatgttc 10260
atggcactgc cggcatagcc gatggagccg tggacgatca tttcccacag gggatctgc 10320
tcatccatga tcggataggg cgttgcctgg acgggggcat tgatgacata ccccgcatgc 10380
gccaggctgt aatcgttccc gcccgatacc agcagttccc ttccgccttc cgagatggtc 10440
ccgaacgcgc tgcgcaccag gtccagtgcc gcctcgcggc tgatcacatt ggtgcgccgc 10500
ttgtctgcat gcagttcatt ggccaaatcg cgcagggaaa gggcttccag tcccagccgg 10560
tcggcctgtg tccgggccgc ctgcgcatag cgcggaagga acttgggaga caggagcttg 10620
aatcccagtt cccggtatcc catggtgtcg gtgcgccgca gggagacagg gctgatcctg 10680
gccaggttca ccacatagcc ctgggcgtaa taccttgagg cttcctcgct ggcggtgaag 10740
ccccgggcga tatcggaaat ctgctgcagc gcgatgtccg gataaacggt tcctccgctg 10800
cgcttcatga cgcccagcag gcttttcagc ccgttttcgc cgcccagctg gccgagcacc 10860
ttcaagcggg tcaccggatc gtggtaatag ccgccgttca tccagccctg caggttcatc 10920
cgctggttgg tgatgccctg gtgcctgaat tcggcgatca tttcctccgc ctgggagaag 10980
gtggtcatcg ggagcacgcg caggtactga atgcccagga agtgcgccgt ttccttgacg 11040
ccgccgatga catcatagaa gaagggaata tcctgcgcct ctccggtttc ctgcgccaga 11100
aggccttccc ggatcaggcg ctcacggtat gcctgtgcaa tcccggagta tccctgtttt 11160
tccggggaca ggagcgtata gcgcacgctg cagtccacgg gatacaggtc tttttccgcg 11220
acaatcacgt tctcgccgga aatgaccagc gtatccgttc tgcgcagggc aaaggtaaag 11280
taggcgtaat tgtagctgtt gttgcggccg gatacatcgg cggtaatgcc tgcaagcgac 11340
gcgccgcgtt cacaggtggc caggatggag ctgtcctccc ggcagatgcc gaacagggcc 11400
aggcgtgcgg tctgcaggtt ctgggtggcg gtcaggtcgc tgtccaccaa gtccaggtca 11460
tagatgtttt tgctgtactg ggccgcagtg ctttttccgt tgttgaaatg gatcagtccg 11520
ccgctgccct cgggaatcac caggtcgccc tgttcatcgc ttcccgcagc cccgaagaag 11580
ggcagaagct gaatggcgcg gatctggccg ttgccggtct cactgatccc ttcaaaggga 11640
agggtcacct gcacgccgtc gggcttgaga atccagtcca gagtgacggt aaaggacagg 11700
ttttctgcct tttccgcatc atcctcttcg gccgggcgga gcgcctgcat ggcctcataa 11760
tcctccatcg tgaacccggc agcacgcgca tcggcatcga tctgctgacg gttgcgggcc 11820
gtcacgccct gcgtcttgat ttcatacagc ccggatgcct cattgaatac atagcttttg 11880
gctgcctgtt tctttccggt ggtggagagg atgtcatacc atgcctggtc cagctggtcc 11940
gggacgagca tcatttctc attggacatg agatagatca cccggacgcc gttttcaatg 12000
gattcgcagg agacctggcc gggagagacg gacttggcat aggagctcca cggcgtgcct 12060
tccttggcgt tgccgtccag ataactgaga ataaactgac ttttcaggtt ttccagattg 12120
gtcgctttcg ccacgggatc cttgtcggcc gcgggcggat tggaatacat ggtgttcccg 12180
gtcgccttgt cgtagactgc cacttccgct gtttttttcct tgaaatacag cgccatggtt 12240
tcattctcgg cagccagtgc aaagcccggg accgctttca ggctgtcctc gagcggcttg 12300
aactccgtgc cctcggtgta cgccgcaggc ctgagaggga tggctttgta ggcgtcataa 12360
tttctgaaat gaatgacata ccagccgatc acggcaccca gggccaggat cagcacggcc 12420
agcaggatcc acaggacccg cttgacagag gaaggtagac gtgctgtgtt ttctgcattt 12480
tccatactgc cttccctcct ctcaaatccg gaaaatcagt tcgatgtaga tgcttctcag 12540
gaagctgtac acctgagtga taaaattgat gaccagcagg gcaaggaaaa ggatgatcag 12600
gacagcaatc acggtcagga acagtgtgat cagcgttttg cccagggtgt agttgtgaat 12660
ctgcatgatc ccgatgagca tcatgatcac ggtatacgcc gtcccgatcc cgatcaggat 12720
tgtgtaaaag gcttcctcgt tttctgcgac gacctgggag atcagggtgg ccgctgcagt 12780
ggctgcgatg acgggcagca tggagtagcc gatggcaatc acgatatcct tgaagcgccc 12840
ctcgccgtcc atcaggcagg tcacggacca gttgcccacg cacaggacaa gatagaggat 12900
cagaacgcct gccagttcgg caatggaatt cacggtcctg ggctcaatgt cgttgaccac 12960

aaagctggca tagatcctgt tcatggaaaa gcagacgccg tagaggatca ccaggaggac 13020
ggcaatgctg acgctgcccc ggtcctgatg gcggatccag taaaacccgt cattggggtg 13080
actgacggta tagaaaaggt acttccattt ttcaccgctg aaaattcctt ttttcatgcc 13140
gcatttcccc ctttcttccg tttggccgcg cgtcttggca gcaccacgcg ccggatcacg 13200
acagtcagcg cgatgagcac gatcagcgtg gtgaggaacc agctgatatt gccccgcagg 13260
gcctcattgc gccagcgctt gaatgccacc gaataatagg cacggttcat gccgcggcgc 13320
aggtaggtca tggccttttc gttgtccccg gcggagaggt aggctttccc gatgccggcg 13380
ttggccagct cattgttttc gtccaggcgg aggacttcgc tccacagcgg aaccgccagg 13440
gcttcatcgc cgttgtatcg caggctcacc gcctggttga tcagccttcc gtactcggtc 13500
tcggaaaaga tgttgacatt gccctgctgg gcatcgagca ccagcaggcg actgcccagc 13560
tgctcaatgg cggccggtgt gcggaaggtg ccttcctgcg tgccgatccc gccgaagata 13620
tagagcaggt tgccttcatg atcgtaggtg aacacgcgcc cgcgccgggc gtcgagcagg 13680
gaatagatgc cgtggtcccg gtacacgacg tccacaatct ggctggggcc ggagtaattg 13740
ccgaccgtgc cgaagaaaag gtcgccgccg aggttttcct tgagcccctt gcggatgacg 13800
tcctgtccct tgggattgag cctgcgcacg gcctgaatgc cgtcggtatc cttgttggag 13860
gcatacacaa agccttcatc gtccacatca atgccggtga attccgtggg aatgaacagc 13920
tgctgcttgg agcgctcttc cttggtggaa aacttcctcc agaatttttc ccacaggctg 13980
atcttgacgc tgatggtgcc gaagaagccg gtaaagtcgc cttcagcagt gaacaccatg 14040
atgccctgga acatgttctg ggcaacacag tacactctgt ccgcatagtc caccgacacc 14100
ttgagcgggg taaagatata gccctcatcg agcacctcgc tctgcgggtt ctgaacatac 14160
ttgacaaacg tcccgtcgag ttcgagcacc acaatgcggt ggttctggga gtccgccaca 14220
tacagctgcc ggttctggga gacggcgacg ccggtgggcg aggcaaaggt ttgcgtcccg 14280
ccgtccgcct caaagtggtc aatcacctga agcaccgtct tcaggccgcc gtcgatgaca 14340
acaatccggt tgtttcccgt gtccgcaaga tagatcttcc cgtccgggga aacgcacagg 14400
tcctgcggag agcggaaggc gccgagcggg tttccttccc acatcaggct gcttcccccc 14460
aggcttccgt ccgggacata cggcgcaggc gtatagagga tattgttcca gtagtcatag 14520
ttgtagcagt cgtagggcag agaatcgttt gccagggcag ctgcgggcag aaggcacagg 14580
agaagcagac agcagatgat ttttttcatt cttatgcccc cttaatcctt catgccggat 14640
gtcgtcatgg tttccagaat cgagctctga cagatcagga agaaaataat gggaatggcg 14700
gcaatgatga aggtgacggc cgcgcctgcg ccggcacggc tgacgccgcc ggaggtgatc 14760
tgctgcagcg catactgcag gggtttcagt tcctcattcc tcaggaacat gccgccggta 14820
ttgccccaca tctgctggaa ctggaagatg gccagtgtga gccaggcggg ccgcacattg 14880
ggcatgacaa tctgccagaa aatctgccat tcatttgcgc cctgcagccg cgcgctctcc 14940
atcagggagt ccggcagctg ctccataaac tgtttcatca ggtacagacc catgccgaag 15000
gcccaggcgg gcagcacgag cgcggcatag gtgttgttga tgcccagcca ggagataatc 15060
aggtactgcg ggatctgagt gacggtccag gagaacatca tcgacaggac caccatggag 15120
aacagcaggg ccttcccggg gaacttatgc ttggccagcg ggtaggccgc catggatgcg 15180
acaatgacat ggcccagcat gccgcccagg gaaatgatga tggtgttgaa aatgtagcgg 15240
gagaacggga cccagctttc gttcatcaga acgaacaggt cgctgaaatt ggtcagggtt 15300
ggattgcgga caaagatgcg cggcgggaac tgatagattt cgtcgagcgg tttgagcgca 15360
ttgttgacga tcatgaccag cggcaggacc atgaacagtc cgctgatgcc catgatgatg 15420
aaaagcaggg tgttgccggc catggaccgg ttcagctttt tcccgccttc cttcctgcgc 15480
attttctttt ccagcttaat gattttcgcc gggcgggtat ccacaggctt atgctgagac 15540
gctgtactca tcttattcac ccacccttct gagcaccttc tggaccagtt tgttggtgcc 15600
caccatcagc aggaagagaa tcgtggcgat ggccgaggca taacccatat ccagtcgggt 15660
ggacccgtag tccagcagat gggtcaccac ggtggcgccg gcatagttga cggatggatt 15720
gccggcgaga ttgatggaaa tatcggccac cgcgaaggac tgcgtgatct gcatcaccgc 15780
gccgaagagc agctgcggct tcatggcggg cagggtgaca taccacagtt cctgccagcg 15840
gtttttcacg ccgtcaatgg ccgcggcctc atacatgccc ttgtccacgg tctgcaggcc 15900
ggcaataaag gaaaggaagc cggtgcccag ggacagccac agctgaacca caatcagcac 15960
aggcagcaca aacttttcag tctgcatcca cagaaccggt tcattgatga aatccaggcg 16020
gagcagaaga ccgttcaggt atccgtagcg gtcaccggtc aggatcaggt tccagaccat 16080
gtaggcattg ccgcagatgg acggcgcata gaacaccagc gtcagcaccg cgcgcagttt 16140
gggcctgaac tcgttgataa tccacgcaaa caggaggcag agcatatagc tgaccgggcc 16200

tgtgacaacg gcaaagatca gggtgttgcg cagggagatg ataaaaatgt catcctccag 16260
aagcagcttg atgtaattgc tccagccgac aaagcggggt gcttccagca tattgaaata 16320
ggtaaaagac aggtaaatgg acatcagcac agggatgacc gtaaagagga aaaacaggat 16380
aaagtagggt gccatgagca gataggaggc cctgttcttc catatttcat gccgcagggt 16440
gaatttctga accctgggca ggcgatttgc ttcagtccgg ctcatgttac tccccctctc 16500
cctgttctgc ggtcggcagg ccgaattccg tgcgtttgta ggtgatctcc gcgttgatgt 16560
aataaatctt gtccatcagc tcctcgcggg gcgtattccc ggattcgttc cggcccttgc 16620
tggcgttgtc cgtggtcacc gcatagaacg cgttgttcac attgcgccag gtgtagtacc 16680
cgcccggcac ctgcggaatg ccccggacat tggcgaactg tgcggccagg cccttgtaat 16740
cccggatcgg ccaggacaga ttggccagcg cttcctggtt cgcggtcgcc acacgtgcgg 16800
aagaccccat cagggattcc atctcgcgtc cgtacagggt ctgggtggca gcgctggtcc 16860
accatttgag gaactcccag caggcttccg gttccttggt ggcgctcatg atgatgtcgg 16920
caagaccgga gcagccggtg gcggtattga gcgtgccgtc ctcgcgcacc gtgccgggaa 16980
catgcgtgaa gtcccacagt cccttgatgt ccggtgcaga gacctggagg ttgttgtagg 17040
tcgtgtaatc cgaaatgatg agcgggcact cgcctgtgcg gaagcgctct tccacggagg 17100
tggccgcgtc gagcttgtag tctgtgtaga attcgcagta tctgcggaat acattgattg 17160
cgatatcgga atcgagtgct gagcggtcgc cgttttcgct gtaatactcc ccgcccgcct 17220
ggtagagcag catggcgaag gtctgttcac cgggaagcat gccgaactcc atctgattct 17280
tggacagcac gctcatggcg accttcactt cgtcccatgt tttggggact tccaggccga 17340
tctccgcgag gatatccttt cggtagaaca tcatcgggaa ggtctgggta tcgggcagtg 17400
cgtacgtggc tccgttaaag gagaacggga ccatggcgct tggcgcgaaa cgggcggcga 17460
tctcggagaa atccgagaac ccggtcaggt ccagcaccgc atggcgcagt ccatagttga 17520
ccggcgtatc gttgccggta ttcatgaccg cgccggcaat accggtcgta ttggccacct 17580
gaatggccac atccggaccc tggcccgaca gcgtcgcacg caggagggta ttcatatcca 17640
ccagctgcac gttgacattg atgccggttt ccggtgtgaa gttttcatcg atcagggcct 17700
tgatcacatt ggcctggtcg cggccggtac cgatccacag cgtgatcacc cggctgtccc 17760
cgccggcttc agcaatattg ccgacctggt tgtaatcgat cacaaacgag ttgtacagtc 17820
tctgggcttc atggctcagg gtgtcgccga aggaggtgtt ctccgctttc ggatcgatat 17880
ccggcgtgtg cacatagatc ctgtcaatct gcagggggctg gccgaggacc tgagtgatcc 17940
agttgccgca ggcgcggaca ttggtcttat aggaggaaag gacctcggta aaccgctcct 18000
ggtctttgat cagttcgatc agctggtcat ccatggtttt gagcacggtc agcttatcgc 18060
tggagtgccc tgccgtcgcc tcgagggtat caatggcgac ccgaaggtca gcgcgcacat 18120
cttccagttc ctgcttcagg cccggaagat tccgctcgat ctgatagtcc cggttgacgt 18180
ccggggagac gccggtaatg tacagcacct gacggtaaat gctgttgagc tgcagcacgc 18240
actgctgcac ctggctgata atcccggcca tgctgcccag gaccacttcc atgcgcagct 18300
tgtgctcgcc cgccgagaga tggatgcggt acggttcccc gctcccggtt tttgcggtct 18360
ccatccgcca ggtctggcca tacccgaaag gaagcgcacc gaactcctgg aagggcacct 18420
gcccgtccac atagattttg cggtatacat caatgccccg gacgaaattc tgcttgtcat 18480
aaaaggtcag gcagtaatcg ccctcctcgg ggacggagaa tttccactcg atccactggc 18540
cggcattctt ccaggcattg ccgccaatgg tattgttcag aagatatctg gcactggcgg 18600
ggtacacggc ggaagacccc tgatcctgca ccggatagag catctgggat gaggtttttt 18660
ccgcagtctc cgcttcaatc cgtacggaaa tccccttcgc ctgcggcagg tcactgcgcc 18720
ctgcgtacgg ctggggcttt tccacctgtc cgagggtaat ccggcggatg agcatgggtt 18780
cgcgaaggga caggacggac agcgtgtgct gtccttttgt caggtagaca cccagattgt 18840
ccgtgatata tccgttgctg tcatagagcg cggtttccac ccattccggt gcttcctcag 18900
gagaaggctt cagttcgttg ccctggttgt ccttccgcca gcggatgacg ctgatcccgt 18960
tctcgtccgt ttcccgcgtg acctggtcat gcagatcgtt tctccatacc cgggtcagcg 19020
caatcaggga cagttcatca tagggcagct gcccgtcgat aaagaaagcg cgctggatct 19080
ctgaattttt tccggcgtag ggataatact ccagacaggg gtcataccag ccctcttcct 19140
ccaccgtaaa ttcccattct gcaagcgctt cctccccggt cagcagcgaa acgcccggca 19200
taccggcata atcggtaagc agcgcaggtt ctgctgcgcc ttcttcctca taccggacaa 19260
catctgcgcc atcgagcacg atccggcagc cgggccggac ctcatcccgg gccttccggt 19320
agtcatcata cgaaggaata cccgcgtcga tggagtatgt gctcaccagg ttctcatact 19380
ccgcaatgga cctgtctgtg gccgccaggg aaggcactgc cgccagaagc atgatcagca 19440

ctgcggccag actcaaccat cctttcatga ttgcttcgct cctgtccatg atttctttgt 19500
tgatgcatgt ccggttattc ctgcggaccg ctcttcaggt cgggaatgtc atcccgcgtc 19560
acagcccgtt catccgcata cattttttc agcagccaga cctcggtgta cttttcactg 19620
ggcttgttga atttgctgct cttgagcacg aaatcgccgc cccaagtgca gtaactcgac 19680
cagacggttc catcgcggaa caccagttcc gggtcaggca cgcatccatt ttccgagagc 19740
atgatcatct ttcttgcgct ggtatatttc aggcagtcca taaatttgga actctgcgag 19800
ttgtatacgc gctcgcccgg ataaatatcc tcgccgataa tatccaccgt atcatccccg 19860
gggtaccagt cggcatgctg tccgttccat acccagatca gattgttgag atgatattcc 19920
tccgtgaatt tctcaaacat cagattccac agccatttat aggcattcgg acccgaagcg 19980
ccccaccaga accagccccc gctcgcctca tgcagaggac gccagaggac cgggacgccg 20040
gcatcgcgca gccggcccag ctgctgagcg acagcatcca gatccttcat cagcaggtcg 20100
tatccctgcg ggtcttcccc gtccatgatt tttttcagat tgatggtgga atactccgta 20160
tagaacccgg agtaccaccg gtcggccttc agatagggg aaggcgcgtt ccagtgccag 20220
cagaaggtca caatgccgcc tttttcccag tactggatcg catggtccac tgacttgccc 20280
tgtgatccgc gctccacgcg cgaaggactg tagtccatca tatccaggcc cagcagcgca 20340
ggatacgatc cctccgtggc acgccagacg gcctggtttt ccatgccgaa catgcctccg 20400
tcacagttct gcccggtaat gatcttttg ccgtagatat cgcacatcca cgcaaacatc 20460
cgctgggctt ccggggatgg atcctttgtg acgagcaccg gtgaaacctc atagagatcg 20520
gccggcagcg gctgggaggg ccggatttcc acagcatcca ccctgatcca gccccagctg 20580
gtatccaggg ccagggtatg atccccctcg gtcaggtaga tataggaaag aacgcttgtc 20640
tgaaagtcct ttccctcgac gatgctgccc gccacccgct ctccgtccag aagcagcgga 20700
ttttccttgt gcccgccgtc tgcgcttgcc agaatgacag aaatatcata catgccgctc 20760
tgatccgcat gaaccgtcag ttccacacga tcacctgctt cctgaaatcc ttccacccag 20820
gtttgttccc cggaggatac gactttcaca tgaccgcaga cggtgcccgt ctcggcctcc 20880
aggcgcacaa catcctcctg cgcagctgcc ggggccgccg gcatcatcag gcagcacatt 20940
gccagtgccg caacggtgaa aagtggtctg cgggatgttc tttcatatct tttcatggcg 21000
ctgcctcccg attgcttctt ccggtgattc tgtgaaaaga tgtctgcctt ccgaccgggt 21060
ctgaaataaa aaacaatcct gtgttgaatg cagggataga aatgacaaaa aaacgggatc 21120
tgatctatga actgtccaac agacagcata atgaattcgg tgacatcatt ttacataata 21180
tatatttcgt cgtaaaggga gaaaagcgta tattttattg caaaaattga aaaaaacaaa 21240
aaaaacgaaa aaaaacaacg gcagctgccg cgcgcaaaaa aactgcgtat caattgatgc 21300
cggccgataa gaccggacac ggattcacat gcaccctgcc ggaaggcaga aaaaaacacc 21360
ggctgaccac cggcatttgg gaggtatcgt atgaattcca ttcttgtcaa taaagccgga 21420
gccctttgt tcacactgac cctgctcatt ctgagctttg ccaccgccgt caccggaagc 21480
cccgtgttcg ggggctggac gtccacggcc gtactggccg tcttcagtct ccttgaagcc 21540
attgaccttc gcttcaccat ccggaaaata cagcgcatca gggaacaggc ataaacggat 21600
tacaggatca tcttctgaca acggaccctt ccagggaagg attggccgcc ggcgagaaac 21660
gcacggaagc accgccgggg ctgacaaccg gaagcagtcc actgagcttt tccgcatccg 21720
ttcttctccg gattttccgg gcttcgcgcc ggggaaatca ggcgaggaac ggaaaaaaca 21780
gaagacaatc ctttcattca tcggctgacc gcttcagcca aaaaaaggct gcgaagcccg 21840
ggcacaacgc ccatcttcgc agctgttttt ttctcatcat ggccaccgga aaaagacgct 21900
tcggaaaata tccgcgttat gcgtttcccc gtccggcttg atggcaaaca ccactttcct 21960
gaagcggtct gcgtaatgct tcccgatcag caggtcatgg aaaatctctg ccacggtctc 22020
cggcgggttc gcaaacgcgc cgcatccaaa ggcacccagc accagcaggt ccgccccgca 22080
ggccgatgcc gcttccagga tattccggat gcgggcgcgc agcgcgcgct cgagttcctc 22140
cggctggacc ttctttttcc gctccggact gatgcgcggc gctgcacagg tgatgacgtc 22200
cacgcgaaag ggcgaatcca gcagcaccgg gatctcatca tctgacttga gcaccacaat 22260
gccgggcgaa tagatcagcg tgtccgttcc catctgcccg acgtttttt cgttgaattt 22320
gtagtagttc cggaccaggt acggccaggt cagcacttca tagagattgc tgctgcggca 22380
caggcattcc tcctgcgcca tgttgcccgc aagaacgccg cccccgggag aataggcatt 22440
ggcgaagttc agcaccgcac atttttcgcc cgggtgcatc agctgctgtg cgcagtggaa 22500
ggtggtatca tccaggatct ccaccgcttc ctccctgctc ttttcctgca caagcaggtc 22560
ttcaaacccg ggcaggatca gccgggactc cgcctgcgac cgcctggttt cctgcatcag 22620
ttcagggctc tggcggatca ggttctgcgt atcctgaaac gcctcaatca gcttccggct 22680

gtccatatga tctcccttcg tttccgtctc tgcacgcgca gggactttac ttcgaaatcc 22740
ggtccttcgg gacccaggcg cggtggatca gctggctgtc gtaggtcatg aattcaatca 22800
gataccagct gcctctggac gcgagaatat ccactccggt gcctgaagca atcgtcgtgt 22860
acaggatgcg gtattccgat cccggcccga accatcccgg cgtttcccgg ttgacatatc 22920
ccgtatcctg gacatagtaa tccccgctga gatacacagc gtcggaacag tcaatgcgtt 22980
tcttgcccgt ccaggcgcgc acccattctc cgccgtactt gaattccacc tcaacccagg 23040
ccgtgccgcg ggtcacctga tagcggagca cgcggacctg tgtgccgacg ccgagattgt 23100
aggtgtgcgt ctcatcataa tccgtggacg ggccgcagcg ggtcgccagt ttctgattgg 23160
tggttccggt gacataatat tccgtgttgt gctcgggcac cggggtccac tggggtgtct 23220
gggtcttcgg ggtgctgcct cccggtccgt atcccccgtt ctgcaaggcc cagacggagg 23280
ccacaatgca gggacgtacg ccgttgtcgc caacggtgac gcggttggag tcatagacat 23340
tgccgttcgt tccgacaaat tgggtgttcc cgttggcatg cgtcatatcc atgcgcaccc 23400
accagcagcc ttttccgctg ctcttaccga tcatcacgcc gcggcttttc gcatacacgg 23460
tcgcatcgca gccggtgtcg cacagtccgt aattgagcac ctgattgcgg ttgggcagga 23520
agacatagtt cattgtgccg gagaccaggg actggataat gcagctctgc tccgtctggt 23580
taaacgcggt cgaatagaac gaattgttca gccaccagct gatataggaa tccccccacg 23640
caaccgccgt gtcaaaggaa tgatagataa tgcagtccag cagcattctg gaaacgagca 23700
gggcatggtc cccgtcaata tccagcacat cccacacaat gggctcctgt ccgttctgcc 23760
agttggcatc ctgctcatac cggccgaaaa ccacagaact gcccaccgta atggctgtgg 23820
gcgtgtagcc gtggaaactg tccgctccgg cggatgcgca cagaagacag atacacaacg 23880
cgagacacac ggctgcggtc aaacccggcc gctgcagctt ctttttcata aacccttccc 23940
cctgttcctt ttttcctcac cggccgcctg tccatgcagg ccccttatcg gaatgcaatt 24000
ttctttcgta catataaaac gcctgagggc acggtattcc tgccttttcc ggattttttc 24060
ccgcacgctc cgccgccctg ccgttttttc tctgcaggca ttatctcaca tcctcagccg 24120
ctgtgcaatc tctcgttttc cgcacaggta aaaagcagtg aaagcaatcg ccttcactgc 24180
ccatgggttt ttcagaagaa tgccttatgc acagatcgcc cggtaccgtt cggagccgag 24240
cacgcccatc atgatgtcat acgggttgga ctctccggag acaaccatgc ggaacaggcg 24300
cggtgtatac ccgctcacca gggtaacccc ccgttcattc atgcgcaccg gctgctgggt 24360
attgcgggac tgcacattcc agaagaccac attcggcagg cggtagtggt gggaggcata 24420
cagctgcttt gcatattcaa agttggtgag ggatgcattc tgtgtgcaga agttgaactc 24480
catatccgaa atgatataga tggtttccgg catctgagac tgcgggagat gccaccggac 24540
ggccgcattc agaatcagtt caaagactgc ctggatgttg gtatccgaca cttccgtcag 24600
ggtgcggcag tatgccacct ttttccgccag gtcccgtccg aggatctcaa tgagccgcgg 24660
cgtgtgtgaa aaggtgatga accggccccg gaacggtccg cggttgcgct ctgcaaagta 24720
cagtccgagg gacatcgcca cagcaaaggg cgtcgggttc ccgtggtaaa acatcgagcc 24780
ggacgtatcc accacacaga gggcattgcg gtcatccgtg aaatcctcca gtgcattcca 24840
ggtgacatcc agcgcattcc gctcccacgc gtcctgcatc acagacggtg ccatggcgcg 24900
ctgcacaagg tcgcaggggt tcagggaaga cgtatggaga gtggcctcgc cgcgcgctgc 24960
cttttcgaga aagtttccgt agcgcttcgc gtcattgcgc tcaaaggcct tatggtactt 25020
gagcatggca gaggagggct gctcttcata ggaaaaggtg taattccgct cgcgcagata 25080
gttttccagc agcttcagat gcgtgcgcag acgggaaagt gtgcggcggt attccgcttc 25140
ccgcatgccc agccggcgcg ccagataccg gccccggcgg actgattccg cacaggatgt 25200
attcacagaa ggaagccact tggccagaag cgagatcgat tcaccgcgct ccatgccggc 25260
gatgtcttcc ttcagctgcg catcaatcag ggcaaggaca tcgacttcac agggtgtgcc 25320
gagaagcaca agcaggtcgt cataccggcc catctcggca acatacgcga gattcttccg 25380
gacagactgc ggttcatgct tcgcaaggta gcgcagcagg acacggaaga cacggcgctc 25440
gcccagtccg ccgcggatat cccgggcata gaacagcgtg cgcatggcaa ggtccggatc 25500
ctccgcatag gcgcagacaa aacggtcgat catcgtcttc tcgggcgctt cacgcagggc 25560
cccgatggtg ccgaagagat ccaggacgtg cgagcgggtc gacagaaggg ttgcagcacc 25620
gttttccgtc cgggtgcggc ggtgagctcc tgaagagccc gcagcatagg gtgaataaaa 25680
gacacaggac ttcctcctct cctgataaaa tcatgtccgg agccgctccg ggctccggac 25740
cgggaaaaca agatgccgtt ttcttcatta acagtgaagt acatttggtt gctgaaggca 25800
tctttcatgc aggttccgtc cctgcagcca tacgataccg catttcccgc cctgatgccc 25860
ggaaaaaaag ttgcgaactg catcttttcc cgttttttttc gaaaatgccc atgtatcccg 25920

ccccttcgtg acaaatgaca cgcccagcgc gtcctgtgca tgcacgattc caccttctgc 25980
ttccagaacc cgcatccgca cccctgcttc acgcccgact gaaagcatgt gctgtacgcg 26040
tccgatatcg ggggatacat caatctgtat ctggccgata tcccgtctta cgattccctg 26100
ccctgtctgg acgtcagagc ctgataggca cagaaaaaaa gaggatggtc ttctccctct 26160
tatgcagagc agaacgggaa agccatcctc tttttatatc tgtttttccg cgcggggtac 26220
tccccgtgct tatgcctgtt ttgcctctcc ggcggcaggg cggttctttc tgcccagcac 26280
gaagtaggtc ccgtcaagga cagccaggag gagcagtaca atgccggtaa agatccatac 26340
ggcactctcg gcaaagggct ggataaacag aagcgctgcc agaactgccg taacgacaaa 26400
gctgaccgcc atcaggtacc agaaaggccg cttccatctc aacgcgtcca cggccatctg 26460
aagcttcata aacgagcaga cggtggtcat aatcccgtag actgcagcca gcgtgcccag 26520
catctgcacc agccagtgct ggttgctgat aactgtaatc cccacgcaga tcagccctgc 26580
cccggtcgcc aggttccagg tctccgctgc cttttctctg ggcaggcgga tataccggta 26640
caggtgatag ccgcccaggg caatcatcac ggcgccgaca agaatcatga caacgccggt 26700
cagtcccacc gggttcacca gcagcagaat gccgagcacg agctcaaaga ttgcaatcat 26760
cagcgatacg ctgatttttc tgattcgatc cttcatgctg tgttcctcgc tttccttgtt 26820
ctttcagggc cgcagtcgct gaaaaacgaa atcagcatcc gaacgctgct cactgagcga 26880
gctccctgta cagatccatg tactggcttg cggaatgctc ccaggacaca tccttttcca 26940
tatcgcgttt gacgatcgca ttccagctgt ccctgtcttc tctgtacact tttctgcacc 27000
gctccacagc atccagaaga agatccgcat tgtaatagtc aaaggtaaaa ccgttgccct 27060
cgcctgtata ctgattatag ggttccacgg tatccttcag gccgcccgtc tcacgcacga 27120
ccggaaccgt tccgtagcgc atggcgatga gctgggtcag gccgcagggc tcaaagagac 27180
tgggcacgag gagtgcgtct gcgccggcat agatctgatg agcgagactc tcgtcatact 27240
tgatgtaggc gcagacctgt cccggataga cgctctcata atagcggaag gcatttttcat 27300
agcgcgcatc gccggtgccg agaatcacaa cctgcgtgtt cgcgttcatc atctttgaaa 27360
aaacagtatt gatcaggtcc aggcccttct gatcggtcag acgggagatc aggcccagga 27420
caaaccggct ggtgtccgtg tgcagtccga gccttgcctg cagggctgcc ttgttctcct 27480
gctttttctc cagcacgtcc tcaacgccgt agggtgcctt cagcagtttg tccgtccggc 27540
tgttccagag atccacgtca atgccgttga caatgccact gagcttggcc cggtgatacc 27600
acagatggcc gttgagattc tcgccgtaga agtctgtctg gatttcccgg gcataggtgg 27660
cgctcaccgt ggtgatgcgg tcgcagtagg caaggccgcc cttgaacatg ttcgcctgat 27720
cattggcgat gcggaagaca gaatagtcaa acagataatc cggaagcccg gtccagtatt 27780
tgaggtgatc cacactgcag atgccctgga agcggaggtt gtggatggtc aggatgctct 27840
tcgcactgcc caccggggtc cctgcaaact cggtacggag atacaccggt accaggcccg 27900
cctgccagtc atggcagtgg atgacatccg gaatccactc gagatagttg aggaccgaca 27960
gagctgcctt ggagaagaag cagtacttgg gaatgtcttc ccaaagcccg gtgtagggat 28020
tgccccagtc gaagaaatcg cggttttcaa cgaagtcata gatcacgccg tccatctcgg 28080
tctccatgat gcccacaaag aacgagcggc cgtcctgcgt gaggtccatc atgaatgagc 28140
cgcggaacgt catctggttt ttgaatttct caggaatgca ggcataatag ggcatgagga 28200
ctctgacctc acacttttcc cggagcagtg ccctgggcag ggcatacatg acatcgccca 28260
gaccaccggt cttgataaag ggatagcatt cagagccgat aaaggcgaca cggacaggat 28320
ggtttccagc atggttttcc gacatattcg atccctcacg tttcttgttc attggccaga 28380
gccattatac tgaatatagc gtattcggtg caattcctca agaggtattt tctttttccg 28440
ccctttcctg catccgctga agttttcccg ctttttttcca tcggtttctc agctgttgtc 28500
catttggcac atccccggcg cattcccctt cggatcattt acatcccccc tgcaatctgc 28560
tatgcttgcc atgcatccgc cgcacaatga aacagcatga aacagcctgt tgttttggca 28620
tgccgcggtg cagatcctgc agcgtccgaa ggtgaaatgc atggaataca gtgtccgcgt 28680
caacggcatc gaagtcacgg cgcactatga agagcgcgcg gtgcacgata tttttctccc 28740
cctgctcagg catctgacag gtcttgagcg cgaaaaggga cggcgcatcc tggtcatgct 28800
cgctgccccg cccggcgcgg gcaagaccac gctcctgagc tttctggaaa cgctctcaca 28860
ccaggacccc gcgctgtgtc cggtgcaggt catcggcatg gacggcttcc acaggcggca 28920
ggaatacctt ttgtcccata ccgtattccg taacggggag gagcttccca tggtccggat 28980
caaaggggcc ccggtcacct ttgacctcga gcgcctcaca cactttgtcc gccgcgccgc 29040
ctcaggcgag gtgtgcacct ggccggtcta tgaccgcctg ctccacaatc ccatcgagga 29100
tgctgtcacc gtcgacggcc gcattgtgct gctcgagggc aattatctgc tcctggatgc 29160

ggacggatgg cgggacctgc ggtcctttgc ggattacacc gtatccgtgc gcgcaccgga 29220
agacctgctc aggcgccggc tcattgaccg ccggatccgg accggcgtgg aagaggacgc 29280
cgcccgggcc ttcgtggact tcagtgatat gcccaatgtc agggtctgtc ttgaaaagac 29340
gatgcccgcc gacctcgagc tcttcctgga tgacgaaggc gcttaccata tcacaggctc 29400
ctcatcctga agatacagag aaaaccccga cagacggggc catttccctg cttttccgta 29460
tctgtgccct taggaacatg atatactcca tcttactgaa cagaaaggag agaagtctct 29520
cctcccacga cggaagtcat gggtgcccga gctttaccat gaaaaacagc cagaaaaaag 29580
cttcctccgg gcagtggccc ggtgtcctgg tctggatgct gaccggagcc gcctgcggca 29640
tcgtatccat gcagtccatg acctcccccg agcatgcaga ctcgctcctg ccgctcctgc 29700
tgatgatccc cttcctgctc gccgccatcg tcatccagac catcatccac gaaggtgggc 29760
acctcctttt cggcctgatg accggctacc gcttcatttc cttccgcgtc tttcagttta 29820
tgtgggtgat ggaaggcgga aagatccgcc tgcgcaggat gcacctggcc ggcacagccg 29880
gtcagtgcct gatgagcccg ccggagatgg agaacgggac gtttcccttc cgtctgtaca 29940
actatggcgg agccatcctg aacctgatct ccgcgctcct gttcctcggg ctttccttcc 30000
tctgcgcgcc gtcttccccg ctccgggtct tcctcctctg cctcaccgtc ctcggcacgg 30060
ccttcgccct gctcaacggg ctgcccctgc cctccggcga tgtcgccaat gacgcgcaga 30120
acgtgctgga tatgtcccgg catccggaag ccgtccgcgc cttctggatc tccatgaaag 30180
tcaatgaggc gctctctgcg ggcacgcgga cgaaggatat gccggcggaa tggttcgctg 30240
tgccctccgc ggatgccatg aagagcagcc cgctcgtcgc cttcctcggg gacatggcct 30300
gcagccgcct gatcgatgag cagcgctttc aggaagcaga tgccctcatg gagcgcatcc 30360
tctccgaagc ctccggactg accgggctgc accgcggcct catgatctgc gaccgcatct 30420
atctggaagc ggtcggggaa aaccggccgg agcagctgaa ggccatgcgc acaaagcagc 30480
aggagaagat catgaagtcc atggcctcct ccccttccgt cctgcgcaca gagtatgcct 30540
gtgccctgct ctcggacagg aatccggaaa aagcggagca gctccgcagg cagtttgaaa 30600
aggcagccgc cgtctatccc tttccccagg aaatcgagtc tgaacgcggg ctgatcgccg 30660
tcgcagacga aaaggcaggc cgtgcggccc cggatacagc atgagggggc tatggaatgg 30720
aactgaaaga actgacgctc cgggacaggg agagcatgac gaaactgtac gcggatgtat 30780
tcaccgggga gccctggaac gatgactgga gctgtcccgg acagctggat gcctatatca 30840
ccgaccttgc cggtcagagc aactccctca ccttcggata ctttgacggt gaccgcctca 30900
tcggcctcgc catgggccat atccgccact ggcacgaggg aaccgagtac tgcatcgatg 30960
aattctgcat tgcccgcgac atgcagggcc ggggccatgg cggccgcttt atggactgtg 31020
tggaatctca gcttctcgag cgcggaatcc cctggatctt cctgctcact gaccggcata 31080
tgcctgcata cgatttttac cggaaacacg gctttcgcga gctgcacgac aacgcggccc 31140
tggtcaggaa agccggtgcg ccgcagcctg ggaggaacac accatgatct ttcccaacgc 31200
ctatttcatc atcgggacag cctatgccgg gaaatcgacc atgacccgca tgctcgccga 31260
aaagcacggc ggcatcctgc tggaagaaaa ctatcacagc cgtttttttcc cgaccctcga 31320
tccctccgcc ttcccctgcc tgacctatac ccgcgagctg aaggactggc atgatttcat 31380
ccgcaggacg ccggaagaat atgaacaatg gatcgacggc gcggcgaggg aatgcgagcg 31440
gctggagctg gagattctcc ctcagttcgc ccttggcgca cagccggtct ttgtcgatac 31500
gaatatctcc gccgaaaccc tcaggcaggt cgccctcccg ggcatgtcc tggtcatgct 31560
cgcccctccg gagatctccg tcagccgctt ctttgagcgg caggatgagg aaaagcagtt 31620
cctccaccga ctgattctgg aggagccgga cccggaaaag gccatggaca actactgccg 31680
cgggctcgcg cggatcaatt ccccggaaaa atatgagcgc ttcctgcaca gcggtttccc 31740
ggtgatcctg cgggacgaaa acaggacacc cgcggaaacg cttgccctcg tggaggaggt 31800
attctgcctg aatggctgac cccactccca tataacatac agaaaggcac ctgaatcaac 31860
atgagtaagg aaaacaatca gcttctatgg aacatcagcc tggccgttct ggccatctgc 31920
tcgctcatcc tcggccttac cgccctgttc gacggcgacc tccccgacgg cctggagcgg 31980
ctgctcggca tgcttgatct gatcgcgctg ccggtctttg tgtatgccag cgtacagaag 32040
gcccgcggct gaaaaaaaat caatgaggtc acagagatga caaaccagga aattctgaag 32100
atcgccctgc agcagtccgc cgaggatatc ggctgccggg cggaggattt tctctcggac 32160
gcgccggtcc ttgtcccctt ccgtctcggg gcccgtgcgc ggaaatacta caaggaaccc 32220
atcatctgcc agtttgtctc ctaccgcgcc aacgtggtgg cagcctccgt cccggagacc 32280
gccgccctgg cggaggaata catccggaag tttcccttct accactgctt tgaaaccccg 32340
aacatggact ggctgagcac ccgcctgctc gaaaagggat ataaggtctg ttttatggcc 32400

gagtactatc tcccggacgt ccatcgcctc cctgacctcc cctgcccggt ggaaacgcgc 32460
gtgctgggac ctgaggactt cagagggctg tacctgcccg agtggagcaa tgccctgtgc 32520
gaagagcgcc gggaactgga tgtgctcggc gtcggtgcgt atgaccaggg acagctggtg 32580
ggcctcgcgg cctgctcggc tgactgtgaa aacatgtggc agatcgggat tgacgtgctc 32640
ccccgggcgc gcggacaggg cattgcccct gcgctgacca gccgtctggc aaaggaaatc 32700
ctggaccggg gcaaggtgcc gttctactgt tcggcctggt cgaatatccg ctcggcgagg 32760
aacgcctgcc gcagcgggtt cctcccggcc tgggccgaga tgacggtcaa gccggcaaat 32820
cttgttgatg acatgaaccg ctgaaaaaaa aacattccag ggaggcagaa tcatatggaa 32880
aaggaactct tcatccgcga agtccggccg ggcattttcc tcatggatga agcccatgag 32940
gccaccggct atctggtcat cggaaaggaa aaagcctgtg tcattgatac catgaacggg 33000
tataacgacc tgtaccgtgc tgtccggaag ctcacagacc ttccgcttgt cgtggtcaac 33060
acacacgggc atccggacca tatctacggc aatctctatt ttgaggaggc cttcctgccc 33120
cctgcggacc tgccgctcgc ccgcagtttt gcagaggacc cggaattcct ggcgtttgcc 33180
cggcagaagg gccgcagcat gccgcctttc cgggatctgc ttcccggcac ccggattgat 33240
ctgggcggaa ggacactgga agcgcatgcg ctccccgggc ataccccggg cagcacgctg 33300
ctcctgctca aagaagaccg catcctcttt accggggaca gcatcaacca tcacctgtgg 33360
ctgcagctgg acggctgcct gccggtgcac gaatgcgcca tgcagctcga ccgcgtcctg 33420
ttcctgatgg ataaggccga tatcatcctg cacgggcacg cgggctcgga ggatgatatc 33480
tccctcatgc gctgcctgag ggcgggcatg caggaaatct gcgacgggaa aacggaagag 33540
gacgcaccgt atgcatggtt cggcggcacc ggtctccagc acccgttccg ctgcgaagaa 33600
ggaaaagtct attccagcgc gcagcatgtc atctgttacc gtcccgacaa tgtctgaaac 33660
aaaagaccgc acctgcagat cggaggaaac aacatgattg tcctgaatgt gacctatctc 33720
tgccatcccg gcatgcggga tgcgtttctc cagagcatcc gggcagaggg catcgacgcc 33780
gccagccgcg cagaagacgg gaatgcgcag tatgagtatt tcctgcccgc agacggcggg 33840
gacaggcttc tgctgcttga aaaatggcgg gatgcggaag cgctttcagc gcacgcccat 33900
caggctcact ttgcccgcct cggggaactg aaggcaaagt atgtccggga gaccgtgatt 33960
gaaaagtatg tctccgaagc ctgagtgtct gccgcctgcc gtgccgggca cggacctgcg 34020
cgtccggccc atgacgatgc aggatcttga gcccctgcat gcgctcctct ccagccccga 34080
cgtcatgcgg tatcttgagc caccgtatac ccggacgcag accacagcct ttctccgccg 34140
cgccatctca cagcatccgc cggtcctggc cgcggaagca gacgggacct ttatcggcta 34200
tgtcattttt cacgcctggg atgagacctc catggaaatc ggctgggtgc tcctcccggc 34260
ttaccagggc agaggctttg ccacagcact gaccatcctc ctcatccgga tggcccggga 34320
aaagggaaaa cagccggtca ttgaatgctc tcccgcccag aaggcaaccg cacatatcgc 34380
ccggaagttc ggttttgtcc gtacagagac aagggacgga ctggatgtat accgcctcag 34440
cgtttaagct tccctccacg cacagcactt ccgccccggc cgctgccggt atcaaaaaaa 34500
cagggagaat ctgtatggac agaacgacac cgctctatga cacccgtctc tccgaccggg 34560
aacgcgctga atggctgctc tcccagctga cacttgagga gaaattcggt tttttcaccc 34620
tgaatatccg cctggagcgc ctgggactca ccctgtccac ctgcggcggg gaaggcgcgc 34680
acggcgtgca ggcccgggcc ggatgagcac aagccggaat tctgcgcgaa gcggtgcggc 34740
atcatcctgt gcacgaaaag gaagcagaac ggatggcggt tctgcgatgt ctgccctgac 34800
ttcccctgcg tggatgtcat ggaaaaagaa aaccgttata ccacgcagta cccgctcgca 34860
gaatcgccgc ttgaaaacct ccgcgccatc cgtgaaaccg gaatggatgc ctttctggaa 34920
aaagagcgag ccgcgtggag ctgcggcata tgcggccgtc tgatttcggt ccataccggc 34980
atctgcaggg gttgcgggaa acagtacgga aaaccgtccc gggagtcctg atcccccttt 35040
cgtccatgag caggccggag cctaacagca tatatatgag gaggagaatg cgttatgaaa 35100
aaacaggtat tcaatcccta tctgcccagc tatgagtaca tccccgacgg cgaaccccgt 35160
gtgtttgacg ggcggctcta catctatggc agccatgacc gctttaacgg cgatgtctac 35220
tgcgccaatg attatgtctg ctggtccgca ccggaagatg acctctcgga ctggcgctat 35280
gaaggcgtga tcttccgcaa gacgcaggac ccgcgcaacc cggacggcgc gcacgccctg 35340
tgggcaccgg atgtctgccg cggaaatgat ggaaagtact acctctatt 35389

//

ID XXX; XXX; linear; XXX; XXX; XXX;  XXX.

XX

AC PRJEB67889

```
XX
DE   Uncultured bacterium DNA fragment from in-reactor enriched cow rumen, clone
ferm-3f21
XX
FH   Key             Location/Qualifiers
FH
FT   source          1..30213
FT                   /organism="uncultured bacterium"
FT                   /environmental_sample
FT                   /mol_type="genomic DNA"
FT                   /isolation_source="Cow rumen"
FT                   /clone="ferm-3f21"
FT                   /note="AZCL-Barley beta glucan active clone"
FT   CDS             3083..4321
FT                   /protein_id=
FT                   /translation="MQSLGGDESSLENSIFADEVLLDKFLRMSTIPSAAFADTKVHYAL
FT                   LDGLRGVAALLVVWYHVFEGYAFAGGQAIIDGFNHGYLAVDFFFILSGFVISYAYDDRW
FT                   SKGFTVKDFFKRRLIRLHPMVVMGALLGVITFCIQRAVQWDGTQVAIPMIVLSLLGALF
FT                   LIPALPGSGQEVRGNGEMFPLNGPSWSLFFEYIGNILYAWFIRRLSNRALAVLVALLGV
FT                   ALTLFAVFDISGYGSIGVGWTMDGLNVAGGSLRMLFPFSMGMLLARHFKPIRVKGAFWI
FT                   CATALVALFAVPYIEGLEPICANGVYEAFCIVVVFPVLVWIGASGTTTDKQSTAICKFL
FT                   GDLSYPLYMVHYPFMYLFYAWLIQSQLFSFGETWPVALGVYGWNILLAWICLKYYDEPV
FT                   RRYLANRFLKRRR"
FT                   /product="Acyltransferase"
FT   CDS             4468..4971
FT                   /protein_id=
FT                   /translation="MLQNDIRPTVFYFNPNIFPKEEYEIRKAESKRYVRALGLDFVDAD
FT                   YDHADWREQVKGLENEPERGRRCVKCFMVRMLETARYASEHGFTVFTTTLASSRWKNLH
FT                   DIFAAGHAAEAEFQGVKFWDQNWRKGGLSERRLQIIAEYDFYNQQYCGCEFSFRKEKSP
FT                   TSSK"
FT                   /product="FIG053235: Diacylglucosamine hydrolase like"
FT   CDS             complement(5109..7262)
FT                   /db_xref="GO:0008422"
FT                   /protein_id=
FT                   /translation="MKVLKIFPACILLASVVSCKPSAEATGEAAVEQRVETLLKEMTLE
FT                   EKIGQMNQISPPEDVSQIADLIKKGEVGSILNLVDPEAINALQQIAVEETRMKIPLIVG
FT                   RDVIHGFKTIFPIPLGQAASFNPELVKAGARVAAVESASQGVHWTFAPMIDIARDPRWG
FT                   RIAESLGEDTYLTSVLGAAMIQGFQGDELSDPTAIAACPKHFVGYGAAESGRDYNSTFI
FT                   PERRLRNVYLPPFQAAVEAGAATFMTSFNDNDGVPSTGNRFILDEVLRKEWNFRGFVVS
FT                   DWASVSEMISHGFAKDGKEAAMKAINAGVDMEMVSYTYVTYVKELLAEKKISMESIDNA
FT                   VRNILRIKIQLGLFENPYVDTSAPSVLYAEEHLQKAKEAAIESAILLKNENGILPLNES
FT                   VKSIAVVGPMADAPHDQLGTWVFDGDKNYTQTPLKALQAAYGDQVRILYAPGTAFSRDT
FT                   RTEGIAAAVNAARNADVVLVFVGEEAILSGEAHSLADLNLQGAQSELIAAVAKTGKPVV
FT                   TIVMAGRPLTIGRDAALSQAVLYNFHPGTMGGPAIADLLFGKAVPSGKLPVTFPKMVGQ
FT                   IPIYYGHNSTGRPYHGNETLLADIPLEAGQTSLGNTSFHLDAGFAPLYSFGYGLSYTTF
FT                   DYSNLKISADRLKATDVMTVTADITNTGKYDATEIVQLYVQDCFGSVTRPVKRVERLCP
FT                   CLPEVRRNEDGRILITYRKARIF"
FT                   /product="GH3"
FT                   /EC_number="3.2.1.21"
FT   CDS             complement(7275..8324)
FT                   /protein_id=
FT                   /translation="MGCGGSEDTPVVDEKITCSPAAISAGYAGETATIEVVSTREWTAY
FT                   SNDEWLTCTPSSSIETTGTVTIQVDANPKTAIREGLVIVKAGSSRDTVRVTQAAKPETP
FT                   VDPNITVPEGYQLVWNDEFESGTAPGAEWWYETGDHGWGNNELQNYVAGSYGSEQLASV
```

```
FT                   KDGILTITAKKIGDTVYSIRMNTGQGWKYGYFEARLKLPTGKGTWPAFWMMPKNYTAWP
FT                   DDGEIDIMEEVGYNPNYVSSAIHCKAYNHSIGTQKTKDVLIPTAQTDFHVYALEWTEDY
FT                   IRTYVDGKELFYFANDKTGNKDTWPFNAAFYLKLNLAWGGNWGGAQGVDESVLPATYEI
FT                   DYVRVFQKK"
FT                   /product="GH16_3"
FT                   /EC_number="3.2.1.73"
FT   CDS             complement(8308..8442)
FT                   /protein_id=
FT                   /translation="MNKDDLEKKLVLEVIRIRKRENNHENKEYDSLINSNTCLYGLWR"
FT                   /product="hypothetical protein"
FT   CDS             complement(8446..9810)
FT                   /protein_id=
FT                   /translation="MKKRYNYAKSILMMLALALGITACSPDDFTSPNEAGIPLAANYED
FT                   AITILVNQETNWVTFSFNATGVMPLWIIDGTTYTSTFNMTKYYRKAGDYSVDVKIVNAN
FT                   GISDGVITKSFTINKTIMNGFGGFVYDSDFNLWKKATIATPTFWYAPGWSQIADPTFKS
FT                   DGASYSISLPEATTDTWQAQMLMGTDIATNANTHYDFSAILTATQDHPHVMVKLVDSTD
FT                   DNVFYFAETVKLTANEPVCFWKSDMEGIDIGSLKMVFDFGGNAAGTDITIENIVLKDHA
FT                   NDDGTIVPEEELVPEPNWSAVDSDDNLWKNVTFTNTFYYAPGWSQLADPDLTIEGTQYS
FT                   VDFPSATTDQWQNQVTFVTDNLSTTSAENYDFKVTLKASNDIQGVTVKLAQNDDDNVFL
FT                   VMERIDLLAGEDVVAKVINAEGVDITQGKLVFDFGGNPANTHVVIKDIILQKHKD"
FT                   /product="FIG00412289: hypothetical protein"
FT   CDS             complement(9831..11300)
FT                   /protein_id=
FT                   /translation="MLLTGCSESFLEVESMTQETIDTYFTTDEHIQEAVVAAYDPLHWS
FT                   DWGIGQYNPLCIMSDIMADDIWVGGADKNDNQYWHLMMNYEALPTNCMTGLWTVAYSGV
FT                   KRSNDVLTYIGWAGDNLTAENATYYEAQARVLRVLYYNWLWKFWGNIPYYEENLTAPYI
FT                   AEQLSADAVYANIIADLEGAIALDALPMKASSSDYGRVTKAMAYMLYAEIVLYQNDESR
FT                   YAKALAYMQEIINSGQYELVDDYTTIFKESGEWNSESIFEVNYKDDNAARSWGSPLVAG
FT                   GTVLPRLISPSGWADGTDGHDNGWGFCPVRLETYERYSNNDARREATCWNAAAIGSYNT
FT                   RYQDTGFFLEKYAAKTGDNADQIADADLNYNNNLRIYRFSETLLNAAELIVRGGGSGDA
FT                   KEYLNRVHKRAGLTIEVAATIDNIIEERHLEFVGEGKRYWDLIRTGKAASVLVPDSYGY
FT                   RTNAWTENKKYLPIPQSEIDAAQGTLTQNNY"
FT                   /product="RagB/SusD domain protein"
FT   CDS             11254..11373
FT                   /protein_id=
FT                   /translation="MDSTSKNDSEHPVSNKAEAMNPATILFLYLFITIYLRFE"
FT                   /product="hypothetical protein"
FT   CDS             complement(11374..14385)
FT                   /protein_id=
FT                   /translation="MVKGTTNGTMTDIDGQFILSAPDKSILTISYIGFKPIEVTADAST
FT                   TMNIVLREDTELLDEVVVVGYGVQKKSVVTASIAKVSSDDLAATAPVRMDNALKGLAAG
FT                   VTVTSSSGQPGAAAQIRIRGIGTINNSDPLYIVDGMPIEGGLDYLNPNDIASIEVLKDA
FT                   ASGAVYGARAANGVVLVTTKTGKEGKTKVTYDFSYGWQRPWKERDVLNATEYAILMNEG
FT                   AINSGQAPKYNDPYSYGVGTNWQKETFNYDAPVMSHQVSVSGAGEKINYLLSMGYYSQD
FT                   GIVGGNYGRSNYDRLTLRSNTGYTLFDESKNRNWLNSLKITSNVSYARIKSTGIETNSS
FT                   FGSPLGSALSLSPILGVYVEGQAAQDQLDYYSQYADFTPMYDPNNGKLYTMAGADYNEM
FT                   VNPLASLSLPGDKNWSHKFVANFSAELNIWDNLKFRTSYGADLSFWGYDGYTPLYYLTA
FT                   NNKSTRTNAYSNKQDGLVWQLENVLMYDKTLDKHSFSVLLGQSAKKSTGSQLGGNRYGI
FT                   INTDRPYIDASNGLQENGDMGVYGGPSDQATLASLFARASYNYDERYMLQVTVRRDGSS
FT                   RFGANNHYAVFPSFSLGWNLTNEAFMEKRPDWLTATKLRLSWGKNGNENIGNFRYTVLT
FT                   STGNNAIFGSGENLTNGVKASGLANPDLKWEESEQLDFGVDFGFFDNALTFTADYYKKT
FT                   TNGMLMEMNIPSYVGESKPIGNVGKMENSGFEFESAYRFHVNDWNFRVGGNLTYLKNKL
FT                   IEYGNDSGWANLDSFQGIGSITRAQNGKPFPYFYGYKTDGLFQNMEEVNAYVNNQGTLI
FT                   QPNAVPGDVRFVDMNGDGKITDDDRTDIGKGMPDFTYGMNFSVSWKNFDLNMMWQGTIG
```

```
FT                   NDVFDATRRTDISSSNLPSWMLGRWTGEGTSDKYPRFVLGDNTNWQSSDLYVYDGSYMR
FT                   LKNIQLGYTLPRQLTSKFMVSSLRFYVAAENLLTFTKYHGFDPEISSGGTSLGIDRGVY
FT                   PQAKTWTVGANLTF"
FT                   /product="TonB family protein / TonB-dependent receptor"
FT   CDS             complement(14689..17538)
FT                   /protein_id=
FT                   /translation="MFLGLMVLSSSAHAQWNHFITNFSKDLFGNGAQTWQINSYDDRFF
FT                   YFANKNGLLEYNGSDWKLFPIHNQTDLRSLHISRSRNRIYVGGESEFGYFDVNENGQLV
FT                   YKQLAENLNERYGLSGGYWGIYELDHIIYYVSDRYIVKQIADEFTVIPSENKIDCSSIV
FT                   NNVLYLGTFDGIKMLVGNTLLDMPKGELLKGKTIRNIIPYETGFLVATARSGLFLCQDN
FT                   QIVPLITGHEEFMRKNELFTLAVSEKYLAIGTIHKGLLLLDRKDFSEQYYNEDNGLQNN
FT                   TVLSIYFDRNQNIWLGLDNGIDYITLDSPLTNLYTYPNSRGTGYVSLLQNGKLFLGTNR
FT                   GLYFTDWPVRFTEKGVNLQFIPQLSGQVWELAAYGDDLFCLHDKGLFIVGDQGVELIPG
FT                   LRGAIHFVPHETDQNRCWVSTYDHFFLLEKTAGKWRIIHNYPEISNWPKNVVFESPEIL
FT                   WVRRLNDGMERILLDTTRNEIRESRIYTQKDGFDSITNLYAYKLNGEVVFTSDSGFYAY
FT                   DRITDKIVPALDYEIFRKPTQVLREWGNTVYALSGKNIQMARRNESATFLPPVTYSFSN
FT                   SKIDFIRYYESMDVIQDSLVIIPNEHGFALLNSHYPVQTNTHSLFIKNVHLTYPKDSLI
FT                   YEDNFLNYTVFPEIAYKHNGLRIEYDVRSMGQDKATKFRYRLSPDPIWSTLTASKVKEY
FT                   NHLREGDYTFEVEAYHPNGEVSAASFSFTILPPWYRSLYAWIAYSVLFLLFVRLLYLLD
FT                   ERRIARRKKKELAEKEKEILLMEQEFIKENLRKEQKIVELKNENLEQELNHKSQEMANL
FT                   MINLSRKNEILMAIKEELYKVMGEMRGEACVKPKRMIIALNNQIDTNIQSDDVLKKFEE
FT                   QFDLVHNKFITKLRAKHPDLSVGELKMCAYLKMNLTTKEIAPLLNLSVRGVETLRYRLR
FT                   KKIGLEREDNLLVYLNALS"
FT                   /product="FIG00417103: hypothetical protein"
FT   CDS             complement(17691..18398)
FT                   /protein_id=
FT                   /translation="MNLLPPDGKFCSFDCIYCECGLNEERKTTTRLPTREEARKQLEET
FT                   LLRMQADGMAPDVITFAGNGEPTMHPQFAGIIDDTVALRDQYFPKAKIAVLSNATMIKR
FT                   KSVFDALMKIDDRILKLDSVLDSRIQQLNAPYSPSFTYKWLVENLKPFAGNLIIQTMFL
FT                   NGTYKGESVTNMTEEEISGWINALKEINPRQVMIYTVERETPVKGLRKATKEELDAIAD
FT                   RAREAGFEVSVSY"
FT                   /product="Fe-S OXIDOREDUCTASE (1.8.-.-)"
FT   CDS             complement(18528..20591)
FT                   /protein_id=
FT                   /translation="MKALLITLELTLITYLITCFQSHAQSSHSKNDKWKEYIDNLSVSG
FT                   ASEEELETLYEDLSYLVENPFDLNQIKKEELERLPFLTDKQIAHILAYAGRYGKMYSVY
FT                   ELKNIDGLDYSTLELLIPFVHIVDTQVDKLPITVNNLLTYARNELAVRYDQGFQQKRGY
FT                   EALSDSVLQDNPNKKYLGEAFYQSIRYAYSFDERIQLGFVAEKDAGEPFLNKYHQGYDY
FT                   YSFNLLIKDWKSLKTFVAGDYKAAFGQGLVVSQDYSLSRSTILTQAERRNNGFRRHYST
FT                   NEFDFFRGVGVTLRMKGWDWSFFYSRRYLSGSTDSLTVTSLKMDGLHRLQRDWDKRNRV
FT                   GMQSFGTNLRYTRTNFYVGLTALHYDFGKRTYEPELKPYNLYYFRGKANTNMSVDYLFK
FT                   KGGIKFFGETAVSANKAVATLNGLQIQPASYASFLLLYRNYSVRYQAFFANAFSQGSSA
FT                   QNEEGVYLGMRWTPFPYWNCSAYLDVFRFPWLKYEVDAPSSGKEYSLQLEYNPGRLSSF
FT                   SLRYRRKVSEKNVTAENTTYIGQVDQHRLRFQFYQTVKDWQFRTALDGVGYGKESSGWK
FT                   KGWMIGQSAAYQPEKGKWRGDVFLAYFDTDHSSVTISSFERTPLYVYYRPTFYGQGLRW
FT                   ACSLRYDMSAAFDVVVKLASTRYFDREVIGTGLEEIDGSRKTDLNLVVRYKF"
FT                   /product="FIG00898434: hypothetical protein"
FT   CDS             complement(20602..21285)
FT                   /protein_id=
FT                   /translation="MSNPACLSLSTTQSIGFHYFNKYQLKELSAVSVIYENPVCALPFA
FT                   LHLSTFGYDKYRETLFRLALSKTLSSRWVIGVSFQYALLQTELYDEEAKKLSTDLGMVF
FT                   IPTENLLIGLSITDLPAVRLDEKNVHIEDFNYYSIQSGIQWTFINNLLIELTSKYVDRL
FT                   GWDFGFGMEYRFYEQVFFRTGFSTYPLIPGLGGGLQFAAFRLDVSANYHPQLGICPGVG
FT                   MRYSF"
```

```
FT                   /product="hypothetical protein"
FT   CDS             21518..22123
FT                   /protein_id=
FT                   /translation="MKGIRSYYLILTFLLGCTSLFGQSVKMNVLPKDHYKAYVEGSDTV
FT                   AMVNLREVYVYPKIKFKNKKEQQKYNKLVRDVKKTLPYAKMVYETLIETYEYMETLPTE
FT                   KERQAHLKRMEKELFKEYKPELKKLTYSQGKLLIKLIDRECNQSSYNLLKAYLGSFRAG
FT                   FWNLFAGMFGASLKTEYNPNGSDALTERVVVLVENGLI"
FT                   /product="FIG00896812: hypothetical protein"
FT   CDS             complement(22115..22645)
FT                   /db_xref="GO:0003723"
FT                   /db_xref="GO:0006396"
FT                   /db_xref="GO:0008173"
FT                   /protein_id=
FT                   /translation="MQKLKITELNRLSTEEFKKTDKIPLIVVLDHVRSLNNVGSVFRTS
FT                   DAFLLEAVYLCGITACPPHPEIHKTALGAEDSVDWLYFKDTLEAVDRLKQNGYTVCSIE
FT                   QVKGSIMLNKLELDKNKKYAVILGNEVKGVQQVVVDKSDCCIEIPQFGTKHSLNVSVTA
FT                   GIVIWDFFCQLSD"
FT                   /product="23S rRNA methyltransferase (EC 2.1.1.-)"
FT   CDS             23167..24165
FT                   /protein_id=
FT                   /translation="MKRIMTQNLSAGYVNVPIPKNIAIKEEIDRLRKEKNAVILAHYYQ
FT                   TGDLQDIADFVGDSLALAQWAAKTEADIIVLCGVHFMGETAKIICPDKKVLVPDLNAGC
FT                   SLADSCPAGDFSEFVRQHPGYTVISYVNTTAAVKAVTDVVVTSTNAKQIVDSFPKEQKI
FT                   IFGPDRNLGNYINSITGRNMLLWDGACHVHEQFSLEKIISLKKEYPEAEVITHPECKQP
FT                   IIQVSDFVGSTAALLKHTIVSPKKQFIVATESGVIHEMQKKSPYKEFIPAPPNDSTCAC
FT                   NECSFMRLNTMEKLYNCLKYELPEIFIDESIQKEAIKPIRRMLELSEKLGL"
FT                   /product="Quinolinate synthetase (EC 2.5.1.72)"
FT                   /EC_number="2.5.1.72"
FT   CDS             complement(24169..26457)
FT                   /protein_id=
FT                   /translation="MAVHAQDYESLAKHFKTYLSYYSSEKIAETNNLVYAVTNGSLLSY
FT                   NKTDNSIFTYSKQTGLSDNKITQIAYNPTVNSLLIVYSNGNIDLMDEAGNSYNISHLMT
FT                   NTNFNFKTVNQIDIYNQMAYLSMQFGVLALDMEKKEIKDTYRIVNDTVYSVSVLNNAIY
FT                   AATSKGVYKASLNDNLVDTNNWHAYSITLPASDDKIIKTVVFDNKLCFLVKNEKVKGVY
FT                   YLDNDNTTKNLLTNNTLVNIKMENGQLIAHSNNNAYIYSSWTQVDKLSGTINDISSYKD
FT                   QNTYWIAAGTGLIKGIKKNNSGGFDVLLSEININSPRNNNFYAMVTQGNKLLAVAGGKD
FT                   HDRYWRQGFFSTFENEEWSLFDQSKIEYYPGKFYPQDFTAVTYSQTDPTDYFISSYGEG
FT                   LFEIKDNEVVAAYNIHNSPLEGIDGKNQYVRISDIAFDSDNNLWMTNCQVESPIKVLKS
FT                   DGTWAGLSYKNYINKPQIMDKILITSKGHKWLNFPYTDQYNNKAGILIFQDANFEEDEE
FT                   NVLFLSSLQNSSGEDLGTSTYRCMVEDKKGYVWVGTNKGVVYTPVAANAIENPSRFYFS
FT                   RITREGDSGLYYFLDGDQVNTIAVDGGNRKWIGTQNSGVFLVSEDAQETIYNFTTENSP
FT                   LLSDCIVSLAINDMTGEVFIGTDKGLVSFMGEATEGKSSYSDVSVYPNPVRPEYEDRVT
FT                   ITGLMSDSNVKITNVSGQLIYQAKSIGGQLVWNCRNYKGNRVATGIYLVIAANADGNES
FT                   VVTKIMVVK"
FT                   /product="Immunoreactive 84kD antigen PG93"
FT   CDS             complement(26514..27089)
FT                   /protein_id=
FT                   /translation="MKLVFATNNKHKLDEVRKIVADKFEIVSLAEIGCHDDIPETAETL
FT                   EGNALQKANYIKDKFGFDCFADDTGLEVEALNNAPGVYSARYAGEAHNSEANMQKLLFE
FT                   LQGIANRKARFRTVIALVIGNETHTFEGIVNGHIIEEKKGNNGFGYDPVFVPEGYTETF
FT                   AELGDTIKNQISHRAEAVRQLNTYLSSI"
FT                   /product="Nucleoside 5-triphosphatase RdgB (dHAPTP, dITP,
FT                   XTP-specific) (EC 3.6.1.15)"
FT                   /EC_number="3.6.1.15"
```

```
FT   CDS             complement(27168..28043)
FT                   /protein_id=
FT                   /translation="METNNIGKLYNHAQDYLIIILGTLLYGFSFNAFILSNEIVTGGVS
FT                   GICAIIFFATSIPVSVSYFLINSVLLVIAFKILGIKFLIKTIVGVISLSLSLSFFEWVL
FT                   DGKPIIDGEPFMAILIGGALCGSGLGLIFSVSGSTGGTDIIAAIINKYKNISIGTALFF
FT                   CDFFIISSSYFLFQDVEKIVFGFVEMGVSNYVLDMIINRNKQSVQFLIFSQKYEEIAER
FT                   IIRDTQRGCTFLEGQGGYSKNPMKVIVLIARKSESLTIFRLVKDVDPNAFISQSFVRGV
FT                   YGEGFDQLKT"
FT                   /product="putative transporter"
FT   CDS             complement(28043..30211)
FT                   /db_xref="GO:0004823"
FT                   /protein_id=
FT                   /translation="MVNGLSERGGYPVEQRVMRQWCLRVSAYAQRLLDGLDQIDWTDSL
FT                   KETQRNWIGRSEGAEMVFRVKDSDVQFTIFTTRADTIFGVTFMVLAPESELVQALTTPE
FT                   QRADVDAYIESTKKRTERERIADRRVTGVFSGSYAINPLTNEAIPVWISDYVLAGYGTG
FT                   AIMAVPAHDSRDYAFAKHFNLPIIPLIEGCDVSEESFDAKEGIMMNSPVKGKEIVNGLV
FT                   LNGLSVPEAIAKTKAYIEANKLGKVKVNFRLRDAIFSRQRYWGEPFPVYYDANGLPRML
FT                   PEDKLPLELPEVDKFLPTETGEPPLGRATNWAWDTVNGKVVSTSLIDNKTIFQLELNTM
FT                   PGFAGSSAYYLRYMDPSNEQALVSRKIDKYWRNVDLYIGGTEHATGHLIYSRFWNKFLY
FT                   DIAISCEEEPFKKLINQGMIQGRSNFVYRINGTNKFVSYNLKDQYEVTPLHVDVNIVSN
FT                   DVLDIEAFRNWNPEYKTAEFDEELLENGKYICGWAVEKMSKSMYNVVNPDMIVEKYGAD
FT                   TLRLYEMFLGPLEQSKPWDTNGIDGVHRFLKKLWSIFFGNTDTLQLSVEAPTAQELKSL
FT                   HKLIKKVTFDIEHFSFNTSISAFMICLNELSSLKCNKQEIMEQFLILLAPFAPHISEEL
FT                   WHVIGHTTTICDAQWPEYQEEYLKEDVVTYAISFNGKARFSIELPADMPKEEVEKAALT
FT                   HEYSAKWLADKTPKKIIVVPQKIVNIVI"
FT                   /product="Leucyl-tRNA synthetase (EC 6.1.1.4)"
FT                   /EC_number="6.1.1.4"
XX
SQ   Sequence 30213 BP; 8344 A; 7110 C; 6290 G; 8469 T; 0 other;
     gtagagcacc ctatatataa atgatgtttt tactgcccct aagatacaga attgaaagct        60
     tatcacaact gaaacagagg cggccgaata taccggtgtg atgtttttac tgcccctaag       120
     atacagaatt gaaagcttat cacaacaatg tcgctgatgg actttgccga cgcgctgatg       180
     tttttactgc ccctaagata cagaattgaa agcttatcac aaccaaggcg tttaattaaa       240
     aaaaatagat tatgatgttt ttactgcccc taagatacag aattgaaagc ttatcacaac       300
     ctatcggaaa ctttaccagt caatacctgg atgtttttac tgcccctaag atacagaatt       360
     gaaagcttat cacaactaat ctgcgtagta cggggatccg tacggagatg tttttactgc       420
     ccctaagata cagaattgaa agcttatcac aacatcagac gaagagctaa acgcattcgt       480
     cgagatgttt ttactgcccc taagatacag aattgaaagc ttatcacaac ccgttgcagc       540
     attgccacgg aacgcaacat gatgttttta ctgcccctaa gatacagaat tgaaagctta       600
     tcacaaccga tacagataag gattttcgca ttacggagat gtttttactg cccctaagat       660
     acagaattga aagcttatca caacataacg attcacaaca attgatcgta ttcagatgtt       720
     tttactgccc ctaagataca gaattgaaag cttatcacaa cagttgaaat gccctcttcg       780
     tttacttcaa agatgttttt actgccccta agatacagaa ttgaaagctt atcacaactt       840
     accgcagcat tatagtagga ttctttgagg atgtttttac tgcccctaag atacagaatt       900
     gaaagcttat cacaactgac gctatcagcg tgtcatgtaa ggcactgatg tttttactgc       960
     ccctaagata cagaattgaa agcttatcac aactttattt tgcaggctat acattaacag      1020
     ccggatgttt ttactgcccc taagatacag aattgaaagc ttatcacaac taaaacagaa      1080
     agacactttg atattgattt gatgttttta ctgcccctaa gatacagaat tgaaagctta      1140
     tcacaacata tttgggcctg tgcaacccgt gttccttgat gtttttactg cccctaagat      1200
     acagaattga aagcttatca caacatattt gggcctgtgc aacccgtgtt ccttgatgtt      1260
     tttactgccc ctaagataca gaattgaaag cttatcacaa cccgggcttg gctctttcaa      1320
     agtcaaggtg agatgttttt actgccccta agatacagaa ttgaaagctt atcacaacga      1380
     agtggttgac atcgtacaag aaagaaggat gtttttactg cccctaagat acagaattga      1440
     aagcttatca caacggaata cggcagggac gttagatcac ttcgatgttt ttactgcccc      1500
```

taagatacag aattgaaagc ttatcacaac ttgttcgtta taaggtaagc gacaatatag 1560
atgtttttac tgcccctaag atacagaatt gaaagcttat cacaaccgtt cttaggcatt 1620
gagccggacg caagtagatg tttttactgc ccctaagata cagaattgaa agcttatcac 1680
aacatgaagc ggccgtttat ggtggaactt ccggatgttt ttactgcccc taagatacag 1740
aattgaaagc ttatcacaac caaaattgaa attattattc tatgattttt gatgttttta 1800
ctgcccctaa gatacagaat tgaaagctta tcacaacccg tcttgccgtg gttccagttg 1860
ctgaatggat gtttttactg cccctaagat acagaattga aagcttatca caacttaaac 1920
cattttattt gagtaatacc gtcggatgtt tttactgccc ctaagataca gaattgaaag 1980
cttatcacaa ctagctaaat caaccagctg tcgggaatat ggatgttttt actgccccta 2040
agatacagaa ttgaaagctt atcacaacac ttgatggttc gttaggagtg tactttggat 2100
gtttttactg cccctaagat acagaattga aagcttatca caacacttga tggttcgtta 2160
ggagtgtact ttggatgttt ttactgcccc taagatacag aattgaaagc ttatcacaac 2220
gcagccaggg cagccttacg agccagagcc gatgttttta ctgcccctaa gatacagaat 2280
tgaaagctta tcacaacaac aaggcgcaaa gatctgggtg atgctatgat gtttttactg 2340
cccctaagat acagaattga aagcttatca caacatttgt ctgatttttc cgacaaagat 2400
gaatgatgtt tttactgccc ctaagataca gaattgaaag cttatcacaa cgaaagatgc 2460
acagggtaat agcgatacca gatgttttta ctgcccctaa gatacagaat tgaaagctta 2520
tcacaacgaa agatgcacag ggtaatagcg ataccagatg tttttactgc ccctaagata 2580
cagaattgaa agcttatcac aacaatctta gggcttgcgt ggtagtcgtt agtgatgttt 2640
ttactgcccc taagatacag aattgaaagc ttatcacaac tgttattgcc ttgcttcggg 2700
aagcattaac gatgttttta ctgcccctaa gatacagaat tgaaagctta tcacaacgga 2760
atggtcaggt tgagatagca cttaaaggat gttattactg cccctaagat acagaattga 2820
aagcttatca caacaatatc tgccctaatg ccacaggaga ggccgatgtt tttactgccc 2880
ctaagataca gaattgaaag cttatcacaa ccggctcgcc ttgaaaaatc gtgaactggc 2940
ggatgttttt actgccccta agatacagaa ttgaaagctt atcacaacgc cagagacaag 3000
agctactata tagtatgtga tgtttttact gcccctaaga tacagaatta ttattgcagg 3060
gcggtaaata ttgagtgagg aattgcagtc attgggggggg gatgaaagtt cgttagaaaa 3120
cagtatattt gccgacgagg ttttactaga taaattttta cgaatgtcaa ctattccttc 3180
ggcggctttt gcggatacga aagtgcatta tgcacttctg gatggactgc gtggcgtggc 3240
agcgctgctc gtggtctggt atcatgtgtt cgagggctac gcttttgctg gtggccaggc 3300
gattattgat ggattcaacc atggttatct ggcggtggat ttctttttta tcctttcggg 3360
attcgttatc agttatgctt acgacgaccg atggagcaag ggtttacag taaaggattt 3420
ttttaagcgc cggttgatcc gtctgcatcc gatggtagtc atgggggctc ttctcggagt 3480
catcaccttt tgtatccaaa gggcagtgca atgggatgga acacaggtcg ccataccgat 3540
gatcgttttg tcgctgcttg gtgcattgtt tctgattcct gcacttcccg gcagcggaca 3600
ggaggttcgc ggaaatggtg aaatgttccc actgaacggg ccaagctggt cgctcttctt 3660
tgaatatatt ggcaatatct tgtacgcttg gtttatccga cgcttgtcaa accgggcact 3720
ggcggtgctg gtcgctttgc tgggggtggc cttgacgttg ttcgccgtct tcgatatctc 3780
cggctacgga agtatcgggg tgggctggac catggatggc ctcaacgtgg ccggcggttc 3840
gctaagaatg cttttcccgt tttccatggg tatgctgctc gcccgccatt tcaagccgat 3900
acgtgtgaaa ggggctttct ggatctgtgc gacggcttta gtcgcactgt ttgccgtgcc 3960
atatattgaa gggctggaac ccatctgcgc gaacggtgtc tacgaggcgt tttgcattgt 4020
cgtagttttt ccggtgcttg tctggattgg cgcttccgga acaacgacgg ataaacagtc 4080
gacagccatc tgcaagtttt tgggagacct ttcatatccg ctttatatgg tgcattatcc 4140
gtttatgtat ctgttctacg cttggctgat ccaatcgcag ctttttagtt ttggagagac 4200
ctggccggtc gctttgggcg tctacggatg gaatatactc ttggcctgga tatgcctgaa 4260
atattatgat gagcctgtgc gccgctactt ggcgaaccgc ttcttgaaaa ggagacgata 4320
aagccggtag cacaagttga cgcgatatac cttatctttg taccgaaaag ggattcggaa 4380
atgcttaaat tagagacacc caacggagag aagaacgtgt tgctgcatgc ttgctgcgca 4440
ccttgctcgt cggctatcat tgaatgtatg ttgcagaatg atatccggcc gacggtgttc 4500
tattttaatc cgaatatttt cccgaaggag gaatacgaga tccggaaggc ggaaagcaaa 4560
cgttatgtcc gggcactggg actcgatttc gtggatgcgg actatgacca tgcggattgg 4620
cgtgaacagg tgaaaggtct tgaaaacgag ccggaacgcg gacgacgctg cgtgaagtgt 4680
tttatggtgc gcatgcttga gacggctcgc tacgcttcgg aacacgggtt tacggtgttc 4740

accacgaccc ttgcctcatc gcgctggaaa aacctacatg atatttttgc ggccggtcat 4800
gcggcggaag cagaattcca gggggtgaaa ttctgggatc agaactggcg gaaaggggga 4860
ttaagcgaac gacgcttaca gataatcgca gaatacgatt tctataatca acaatactgc 4920
ggttgcgagt tcagcttccg caaagaaaaa tccccgacat cttcaaaata aaacatcggg 4980
gattttctac ctaggaacag tcggtcagtg tttattccac cacaaaggat gtttttaaac 5040
cctcctgact gttggggcct acccacagat taaaatcacc ggcttcaacg ccataattca 5100
tgtctaaatt aaaaaatgcg agcttttcta taggtaatga gaatacgacc gtcttcgttt 5160
cgccggactt caggaagaca cgggcaaagt ctttcaactc tttttacggg acgggtcacg 5220
gatccaaaac aatcttgtac gtacagctga acaatttctg ttgcgtcgta cttgcctgta 5280
tttgtgatat cggcggtaac ggtcatcacg tctgtcgctt tgagcctgtc tgccgagatt 5340
tttaaattac tgtaatcgaa cgtcgtatag gacaatccat atccaaagga atacagcgga 5400
gcgaatccag catccaaatg gaaagaggtg ttccccaaag aggtttgacc ggcttccaag 5460
gggatatccg ccaacaaggt ttcattgcca tgataggggc ggccggtgct attatgtccg 5520
taataaattg gaatctgtcc gaccatcttc gggaatgtga caggcaactt cccactgggt 5580
accgcttttc cgaaaaggag atcggctatc gccggaccgc ccatcgtacc gggatggaaa 5640
ttgtataaca ccgcctgcga aagcgccgca tcgcgtccga tcgttaaggg acgtcccgcc 5700
atcacaatgg tcactaccgg cttgccagtc ttggccaccg ccgcaatcaa ttcgctctgt 5760
gcaccctgca ggttcaagtc tgccagtgaa tgagcttctc cggaaaggat cgcttcttca 5820
ccgacaaaga ctaaaactac atccgcgttg cgggctgcgt ttacggctgc tgcaatgccc 5880
tcggtacggg tatcccgact aaacgccgtt cccggcgcgt agaggatacg tacctgatcg 5940
ccataagccg cctgcaaggc tttcaatgga gtctgcgtgt aattcttgtc tccatcaaaa 6000
acccatgtgc ctagttggtc gtgtggcgca tccgccatcg gaccgactac cgcgattgat 6060
tttacagatt cattaagtgg taaaatgccg ttttcgtttt ttaagaggat ggccgactct 6120
atcgccgctt ccttggcttt ctgcagatgc tcttctgcat acaaaacgga aggcgccgag 6180
gtgtcgacat aaggattctc gaacaagcct aattggattt ttatacgtag gatgttgcgt 6240
acggcattgt caatcgactc catgctgatt ttcttttcag ctagcaactc cttcacgtag 6300
gtgacatacg tgtaactcac catttccata tcaacaccgg cattgatcgc cttcatggcc 6360
gcttcttttc cgtctttcgc aaatccgtgc gagatcatct cgctgacaga agcccagtcg 6420
gaaacaacaa agccgcggaa gttccattct ttgcggagaa cttcatccag aataaagcga 6480
ttacccgtgg acggtacgcc gtcgttatcg ttgaacgagg tcatgaacgt cgccgcaccg 6540
gcttcaaccg cagcttggaa tggcggcaaa tagacgttgc gcaatcggcg ctcagggata 6600
aaggtcgaat tatagtcacg accactctcg gccgcaccgt aacctacgaa atgcttcgga 6660
caagcggcaa tggcagtcgg atcgctcagc tcgtcaccct gaaatccctg aatcatcgct 6720
gctccgagaa cagatgtcag gtaggtatct tcgcccaaac tttcggcgat tcgtccccaa 6780
cgcggatcgc gggcgatgtc aatcatcggt gcgaaagtcc aatgcacacc ttgcgaagcc 6840
gattcgaccg cagcgacacg tgcacccgct ttcaccagct caggattaaa cgaggctgcc 6900
tgacctaacg gaatggggaa gatggttttg aaaccgtgta tcacatcgcg accgacaatc 6960
aaagggatct tcatgcgggt ctcttcaacc gcaatctgct gaagcgcgtt gatggcttcc 7020
ggatcgacca gattcaaaat agaaccgact tcgcctttct tgatcaggtc ggctatttgg 7080
gatacatctt cgggaggact gatctgattc atctgaccga tcttttcctc caacgtcatt 7140
tctttcaata gggtttccac tcgttgctcg acggcagctt ctccggtcgc ctcggcagat 7200
ggtttacagg aaaccacgga tgccaacagg atacaagcgg gaaaaatctt taacactttc 7260
attttgtttc tctattattt tttctggaat acgcgtacat agtctatttc gtaagtggcc 7320
gggagaacgg attcatcgac tccctgcgcg ccgccccagt tgccgcccca agcgagattg 7380
agcttcaagt aaaacgctgc attaaagggc caggtatctt tatttcccgt tttgtcgttt 7440
gcaaaataga acaactcttt cccatccaca taggtacgta tataatcttc ggtccattcc 7500
aaggcgtaca catgaaaatc ggtttgggcg gttggaataa gcacatcttt cgttttctgc 7560
gtgccaatcg aatggttata ggccttgcaa tgaatggccg atgagacata gttcggattg 7620
tatccgactt cttccatgat atcgatctcg ccatcatcgg gccaagcggt atagttctta 7680
ggcatcatcc aaaatgccgg ccaggtgccc ttcccggtcg ggagttttaa gcgggcctca 7740
aaataaccgt atttccatcc ttggcctgtg ttcatacgga tggaatagac cgtgtcgcca 7800
atctttttgg cggtgatagt cagtatgccg tctttcacgg acgccaactg ctcggaacca 7860
tatgatccgg ctacataatt ctgcagttcg ttattccccc atccatggtc gccggtctcg 7920
taccaccact ccgcaccggg agctgtgccg gactcaaatt catcgttcca gaccagctga 7980

taaccttcgg gcacggtgat attggggtcg accggcgttt caggtttcgc tgcctgcgtc 8040
acccgaacgg tatcacgact actgccggct ttcacaataa ccagtccttc gcgtatcgct 8100
gtcttgggat tggcatcgac ctggatcgtt accgtacccg tcgtttcaat cgaactcgaa 8160
ggcgtacaag tcagccactc atcattactg taggccgtcc attcacgagt agaaacaacc 8220
tcaatcgttg ctgtttctcc ggcgtaaccg gcgctgatgg cagcgggtga acacgtgatt 8280
ttctcgtcaa cgacaggtgt gtcttcacta ccgccacagc ccatagaggc aagtattaga 8340
attaataagg gaatcatatt ctttattttc atgattattc tctcgtttgc gaatccggat 8400
gacctcaaga acgagttttt tctcaaggtc atccttattc atccattagt ctttatgctt 8460
ttgcaaaata atgtctttaa tgactacatg ggtattggcc ggattccctc cgaagtcgaa 8520
tacaagtttg ccttgcgtaa tatcaacccc ttcggcattg atcaccttgg ctacgacatc 8580
ttcacctgcg agtaaatcga tccgctccat caccagaaag acgttgtcgt catcgttctg 8640
ggccaatttt acggtcactc cctggatatc gttagaggct ttcaacgtca ctttaaaatc 8700
gtaattctca gccgaagtcg tggaaaggtt gtctgttaca aaggtaactt gattttgcca 8760
ttgatcggtc gttgcactcg ggaaatcgac cgaatattgt gtgccttcaa tcgttaagtc 8820
cggatccgcc aactgtgacc agcccggagc gtaatagaac gtatttgtga aggtgacgtt 8880
cttccacaga ttgtcgtcgc tgtcgacagc gctccaattg ggttcgggca ccaattcctc 8940
ttccggcacg atggtgccgt cgtcgttggc atgatctttc aggacaatat tttcgatggt 9000
gatatcggtt ccggcggcat tgcctccaaa atcgaatacc attttaagtg aaccgatatc 9060
gatgccttcc atgtcgctct tccagaaaca aaccggttca ttagctgtca acttcaccgt 9120
ttctgcaaaa tagaacacat tatcgtctgt gctgtctacg agtttcacca tgacatgggg 9180
atgatcctgt gtagcagtca gaatggcgga gaagtcataa tgcgtattgg cattggtagc 9240
gatatccgta cccatcagca tctgtgcctg ccaggtatcg gttgttgctt ccggcagact 9300
gattgaatag gaagcgccat ccgatttgaa ggtcgggtct gcaatctgtg accagcccgg 9360
agcgtaccag aatgtcggcg tggcaatcgt tgcctttttc cataagttga agtccgagtc 9420
gtaaacaaat ccgccgaacc cgttcatgat cgtcttattg atcgtaaatg atttggtgat 9480
cacgccgtca ctgattccgt tggcattgac aatcttcaca tccacactat agtcgcctgc 9540
cttacggtaa tacttcgtca tgttgaacgt agaggtatag gttgtcccgt caatgatcca 9600
gagcggcata acgcccgtcg cgttgaatga gaacgtgacc cagttggttt cttggttgac 9660
caggatcgtg attgcatctt cgtagttggc agccaatgga atgccggctt cattcggaga 9720
agtaaagtcg tccggagaac aagccgtaat ccccaaagcc aaggcaagca tcatcaaaat 9780
gcttttggca taattatatc gttttttcat gattcttcta ctttaataga ttaataattg 9840
ttttgagtca atgtcccctg agccgcgtcg atttcgctct gaggaatggg caggtatttt 9900
ttattctccg tccaagcgtt cgtacggtat ccgtagctgt cgggcaccaa tacactggcc 9960
gcctttccgg tacggatcaa atcccagtag cgcttacctt cacctacaaa ctccaggtga 10020
cgctcttcga tgatattatc gattgttgca gcgacctcga tcgtcagacc ggcccgttta 10080
tgaacccgat tcaggtattc cttcgcatca cctgaaccgc caccacgaac aatcagttcc 10140
gccgcattca aaagcgtttc cgaaaaacga tagatacgca ggttattatt gtagttcaag 10200
tcggcatccg caatctgatc ggcattatcc cccgtcttgg ctgcatattt ctccaagaag 10260
aatccggtat cctggtagcg ggtattgtaa cttccgatag ctgctgcgtt ccagcaggta 10320
gcctcacgac gcgcgtcgtt attggaatac cgttcgtagg tttccaaacg tacaggacag 10380
aatccccagc cgttatcgtg accgtctgtt ccgtcggccc agccactcgg actgatcaga 10440
cgcggtagaa cggttccccc ggccaccagc ggacttcccc atgagcgtgc cgcattatca 10500
tctttatagt tcacttcaaa gatggattcg ctgttccatt cgcccgactc cttgaagatc 10560
gtcgtatagt catcgactaa ctcatattga ccgctattga tgatttcctg catgtatgcc 10620
aaagctttgg catagcggct ttcatcattt tggtagagga cgatctcggc gtaaagcatg 10680
taagccatgg ctttcgtcac ccgaccgtaa tccgagctgc tcgctttcat aggaagcgca 10740
tccaaagcga tagccccttc caaatcggcg atgatattcg cataaacagc atccgcactc 10800
aattgttccg cgatataggg agcggtcagg ttttcttcgt aataaggaat attgccccag 10860
aatttccaaa gccagttata ataaagcacg cgaagaaccc ttgcctgcgc ttcataatag 10920
gtggcatttt ccgccgtcag gttatcaccg gcccaaccga tataggtcag cacgtcgttg 10980
cttcgtttca caccggaata agcgaccgtc caaaggcccg tcatgcaatt cgtcggcagt 11040
gcctcgtaat tcatcatcag gtgccagtac tgattgtcgt tcttatcggc accacccacc 11100
cagatatcat cggccatgat atcactcata atacataacg ggttatattg tccgataccc 11160
caatccgacc aatgcaaagg gtcgtaagcg gcgactaccg cttcttgaat atgttcatcg 11220

gtcgtaaaat acgtatcgat tgtttcttgc gtcatggact caacctccaa gaatgattcg 11280
gaacatcccg tcagcaacaa ggcggaagcc atgaatcctg ctactatatt gtttttatat 11340
cttttcataa cgatttattt gagatttgaa tgattaaaaa gtcaagtttg ctccgacagt 11400
ccaggtcttt gcctgaggat acaccccgcg gtcaatacct aatgaggtac cacccgatga 11460
gatctccgga tcaaaaccgt gatatttcgt gaaggttagc aggttctctg ccgccacata 11520
gaaacgcaaa gaagagacca tgaatttact tgtcagctga cgaggcaagg tataacccag 11580
ctgaatattc ttcagacgca tatagctgcc atcgtacaca tagaggtcgg acgactgcca 11640
gttcgtattg tcgcccaaca cgaaacgcgg gtatttgtcg gaagtacctt cgcccgtcca 11700
gcgacccagc atccaagaag gtaggttgct ggaagagata tccgtacggc gggtcgcgtc 11760
gaacacatcg tttccaatcg ttccttgcca catcatgttc aaatcaaagt tcttccagga 11820
tacgctgaaa ttcataccgt aagtaaagtc cggcatccct tttccgatat ccgtacgatc 11880
gtcgtccgtg atctttccat cgccattcat atcaacaaat cggacatcac cgggaacggc 11940
attgggctga atcaacgtgc cttggttatt cacataggca ttcacttcct ccatgttctg 12000
gaataggcca tccgttttat agccatagaa ataagggaaa ggtttgccgt tttgcgcacg 12060
cgtgatcgag ccgatacctt ggaacgagtc caggttggcc catccggaat catttccata 12120
ctcgatcaat ttattcttca ggtaggtcag gttacccccc acacggaagt tccaatcatt 12180
cacatggaac cggtaggctg attcaaactc aaaaccgctg ttttccatct tgccgacatt 12240
accgatcggt ttagattcac cgacatatga aggaatgttc atttccatca acataccatt 12300
cgttgttttc ttataataat cagccgtaaa ggtcaatgca ttgtcaaaga aaccgaagtc 12360
caccccgaag tcgagctgct cagactcttc ccatttcaaa tccgggttgg ccaatccgct 12420
tgctttcacc ccgttggtca ggttttcgcc cgaaccgaag atcgcgttat ttccggtcga 12480
tgtcaagacg gtatatctga aattaccgat gttctcgtta ccgttcttac cccaagacaa 12540
gcgaagctta gtagcagtca accagtccgg acgtttttcc atgaaagctt cgttagtcag 12600
gttccatccc aaagagaaag aagggaagac cgcataatgg ttattggccc caaaacggga 12660
agaaccatcc cgacgtacag ttacctgtaa catataacgt tcatcgtagt tatagcttgc 12720
gcgagcaaag agagacgcca atgttgcttg gtccgacgga cctccataaa cgcccatgtc 12780
gccgttttcc tgcaaaccgt tgctggcatc gatatagggc cggtcggtat tgatgattcc 12840
gtagcgatta ccacccaact gggaaccggt acttttcttg gccgactgac ccaaaagtac 12900
ggagaacgaa tgtttatcca gcgtcttatc gtacatcaat acattttcca gctgccacac 12960
cagaccgtct tgcttgtttg aatacgcatt cgtacgggtc gatttgttat tggctgtcag 13020
gtagtataat ggtgtataac cgtcatatcc ccaaaaggat aagtccgcgc catacgacgt 13080
tctaaatttc aaattgtccc agatattcag ttcggccgag aagttggcga cgaacttgtg 13140
actccagttt ttatctcccg gcaaggataa gctggccaac gggttcacca tttcattgta 13200
atcggcaccc gccatcgtat aaagcttgcc gttattcgga tcatacatcg gtgtgaagtc 13260
ggcatactgt gaataataat ccagctgatc ttgcgctgct tgcccttcta cataaacccc 13320
taagatcggt gacaacgaca acgcactgcc taacggcgaa ccgaaagaag agttggtttc 13380
tataccggtg ctcttgatac gggcataaga tacattcgac gtgatcttca aactgttcag 13440
ccagttcctg ttcttcgact cgtcgaacaa cgtataaccg gtattactgc gcagggtcag 13500
gcgatcgtaa ttggaacgcc cataattacc tccgacaata ccgtcctgcg aatagtatcc 13560
catggacagc aagtaattga tcttttcacc cgcaccgctg acactgacct ggtgactcat 13620
caccggtgcg tcataattga atgtttcttt ctgccagttc gttccaacgc cataggaata 13680
cggatcgttg tatttcggcg cctgacccga attgatcgca ccctcgttca tcaggatcgc 13740
gtattcagtc gcattcagta catcacgctc cttccaagga cgttgccatc cgtacgaaaa 13800
gtcgtacgtc actttcgtct tgccttcttt accggttttt gtcgtcacca aaactacccc 13860
gttcgctgca cgggctccat aaaccgcacc cgatgccgca tctttcaata cctcaataga 13920
agcgatgtcg ttcggattca aataatccag tccgccttcg atcggcatcc cgtctacaat 13980
atataacggg tcggaattat tgatcgtgcc gataccacga atacggatct gggcggctgc 14040
acccggctga cccgaagaag aggtaaccgt cacaccggca gccaatcctt tcaatgcatt 14100
atccatacgg accggcgccg tggcagccaa atcatcggaa gaaactttcg cgatggaggc 14160
cgtcaccaca cttttctttt gaacgccgta accgacaacc accacttcat ccaacaactc 14220
cgtatcctcc cgaaggacga tattcatcgt ggtcgaagcg tcggcggtga cttcgatcgg 14280
tttaaaacct atataagaaa tggttaagat acttttatcc ggagcggata aaataaactg 14340
accgtcgata tcggtcatcg ttccgttggt cgttcctttg accaccacat taactccagg 14400
caacggataa ttatcctcgc cagagataac aactccctta atctgtacat cttgttgcgc 14460

ttgtgcagag aaagacagac aaagcaataa agttattgct gcgaatatga gatttctcat 14520
aactaaatta aattaagatt tacatgttga aactgcgcaa acatagaccg attagtggca 14580
gtgaaacgaa aatctttaca ttctaaaata cgttatgagg cgaatataaa tacgtcaaca 14640
attcgtcatg aaatattcac agtaatgaaa cacagtgggt tacaaatatc aggacaaggc 14700
attcagatac accaacagat tatcttcccg ttcaagtccg atctttttac ggaggcgata 14760
gcggagcgtt tcaaccccac gaacggagag attcagcaag ggtgcaattt ctttggtagt 14820
cagattcatc ttcaagtagg cacacatttt taattcgccg accgacaaat cgggatgttt 14880
agcgcgcagt ttggtgatga atttgttgtg tacgaggtcg aactgttcct caaacttttt 14940
cagtacgtcg tcggattgga tattggtatc gatctgatta ttgagcgcga taatcatgcg 15000
tttaggtttg acacaagcct cgccccgcat ttcgcccatc accttatata actcctcctt 15060
gatcgccatc aggatctcgt ttttgcggga gaggttaatc atcaggttgg ccatctcctg 15120
actcttatga ttcagttcct gctccaggtt ttcgtttttc aattcgacaa tcttctgttc 15180
tttccgaaga ttctctttga taaactcctg ctccatcaac aaaatctcct tttccttttc 15240
cgccagttct tttttctttc gacgggcaat ccgacgttca tccaagagat acagcaatcg 15300
cacaaagagc aggaagagta ccgaataagc aatccaagca tacagcgaac ggtaccatgg 15360
cggaagaatc gtaaaagaaa acgaagcagc agacacttcc ccgttcgggt gatacgcctc 15420
cacttcgaag gtataatccc cttcgcgcag gtgattatat tctttgactt ttgaggcagt 15480
aagcgtcgac catatgggat ccggactaag gcggtaacgg aatttcgtgg ccttatcctg 15540
ccccatggaa cgcacgtcgt attcaatcct tagcccattg tgtttatagg caatctcagg 15600
gaaaaccgta taattcaaaa agttatcttc atagatcaac gaatctttcg gataggtaag 15660
atgcacattt ttaataaata aagaatgggt attggtctga accgggtaat ggctattcag 15720
taaggcaaaa ccgtgttcat tcggtataat caccaaggaa tcttggatga catccatcga 15780
ctcataataa cggataaaat caatcttcga gttggaaaat gaataggtaa cgggcggtaa 15840
gaaggtagcc gactcatttc tccgagccat ctggatattt ttaccggata aggcatagac 15900
cgtatttccc cattcccgta acacctgcgt aggttttctg aaaatctcat agtccaatgc 15960
aggcacgatc ttatcggtta tccggtcata cgcataaaaa cccgaatcgg aagtaaagac 16020
cacctcgcca ttcaatttat aagcatataa attcgtgatg gagtcaaacc cgtccttttg 16080
cgtatagata cgcgattctc gaatctcgtt acgggtcgta tcgagtagaa tccgttccat 16140
gccgtcattt aatcggcgca cccataaaat ctccggagat tcgaacacca cattcttcgg 16200
ccaattcgat atttcaggat aattgtggat gattctccat ttaccggccg tttttttccaa 16260
aaggaaaaaa tggtcatagg tactgaccca gcatcggttc tggtcggttt catgcggaac 16320
aaaatgaata gctccccgca aacccggaat cagttcgact ccttgatcgc caactataaa 16380
caagccttta tcgtgcaaac aaaacaaatc gtcgccatac gcagccaatt cccatacctg 16440
cccactgagt tgcgggatga actgcaggtt cacccctttc tccgtaaagc gaaccggcca 16500
atccgtgaaa tagagtcccc ggttcgttcc caaaaacaat ttaccatttt ggagtaagga 16560
aacataaccc gttcccctcg aattcggata cgtgtacaga ttcgtcaacg gcgaatccag 16620
ggtgatataa tcaatgccat tatctaaacc gagccagata ttctggttcc tatcaaaata 16680
gatggacagc accgtattat tttgtaagcc attatcctca ttataatatt gttccgaaaa 16740
atctttccga tccaatagaa gcaatccctt atgaatggtc ccgatggcca aatacttttc 16800
cgaaacagct agggtaaaca actcgttctt ccgcataaac tcctcgtgac cggtaatcaa 16860
cggaacgatt tggttatcct ggcataagaa taggccgctt cgggcggtag ccaccagaaa 16920
gcccgtttca tacggaataa tgttccggat cgtctttccc ttcagcagtt cccctttggg 16980
catatccagc agcgtgttgc ccaccagcat cttgatgccg tcgaaagtac ccagataaag 17040
cacattattg acaatcgacg aacaatcaat cttattctca gaaggaatga ctgtaaactc 17100
atccgctatc tgtttaacga tgtagcggtc cgacacataa taaataatat gatccaattc 17160
gtatatgccc caataccctc cggacaagcc ataccgctca ttcaagtttt cagccaactg 17220
tttataaacc agctgtccgt tttcattcac atcgaaataa ccgaactcac tctctccgcc 17280
cacataaatc cgattccgac tccgcgaaat atgtaaggac cgtaaatccg tttgattatg 17340
aatcgggaat aatttccaat cactcccatt gtattcaagg agtccgttct tattggcaaa 17400
gtagaaaaaa cggtcgtcgt aagaattaat ctgccaggtc tgggcaccat tgccgaacag 17460
gtctttcgag aagtttgtga taaaatgatt ccattgcgca tgagcagacg aggataagac 17520
cattaacccc aaaaacaacc tcaagaacct cataagaaaa cccattcctt tcgacctttt 17580
aaaaaagcaa atatacaggt tttgattctg aattcatacg ttatggcctg ataataacgc 17640
aatttatccg atcaggcagg catggctgag aaaggtgctt ctcttaggaa tcaataggaa 17700

accgacacct caaaacccgc ttctctggcc cgatccgcta tcgcatccaa ctcctctttg 17760
gtcgcttttc gcaaaccttt gaccggagtt tccctttcga ccgtatagat catgacctga 17820
cgcggattga tttctttcaa ggcgttgatc catccggaaa tttcttcttc cgtcatgttg 17880
gtgaccgact ccccttata cgtcccgttc agaaacatgg tttgtataat cagattccct 17940
gcaaacggtt tcaagttttc aaccagccat ttataggtaa aagaaggaga ataaggagcg 18000
ttcaattgtt gtatacggct gtccaatacc gaatccaact tcaatatccg gtcatcgatt 18060
ttcatcagtg cgtcgaacac ggatttacgt ttaatcatcg tcgcgttcga tagaacagca 18120
atcttcgctt tcgggaaata ctgatcacgt aaagcgaccg tatcgtcgat aatacccgca 18180
aactgcgggt gcatggttgg ctcaccattt ccggcaaagg taatcacatc cggcgccata 18240
ccatcggcct gcatcctcaa caacgtttcc tccaactgtt tcctggcctc ttctcgagtc 18300
ggcagtcgag tcgtcgtctt ccgctcctcg ttcaacccgc attcacaata aatacagtcg 18360
aacgaacaga acttcccatc cggaggaagc agattcattc ccaaagaaac tcctaagcgc 18420
ctgctcttca caggaccgaa aactatttta tcaaaaagga ttgtcgacat aaggcatgca 18480
ctattaataa ccgcttgcaa atgtaacaat aactccgccc gatcggttca aaatttgtag 18540
cgaaccacca aattcaagtc ggttttccgg ctgccgtcta tttcttccaa ccccgttccg 18600
atcacctcac gatcaaaata acgcgttgaa gccagtttca cgacaacatc gaaagccgct 18660
gacatgtcat aacggagcga acacgcccaa cgcagtccct gaccataaaa agtaggccga 18720
taatacacat agagcggcgt ccgttcaaaa gagctaatcg tcaccgacga atgatccgtg 18780
tcgaaatacg ctaaaaacac atctcctctc catttccctt tttccggttg ataagccgcg 18840
ctttgaccga tcatccaccc tttcttccat cccgacgatt ctttcccata gccgacgcca 18900
tccaatgccg tacggaattg ccagtctttc actgtttggt aaaactgaaa acgaagacga 18960
tgctgatcca cctgaccaat ataggtcgta ttctccgccg ttacattttt ttcggatact 19020
tttcgccggt accgaagcga gaacgacgat agccttcccg gattatattc gagctgcaga 19080
ctatattctt ttcccgacga aggcgcatcc acctcatact tcaaccaggg aaaacggaat 19140
acatccaggt aggccgaaca attccaatac gggaacggtg tccatcgcat acccagatac 19200
accccctctt cattctgtgc agaactgccc tgcgaaaaag catttgcgaa aaaggcctgg 19260
tatcggacac tataattgcg ataaagcaaa aggaacgaag catacgatgc cggctgtatc 19320
tgaagtccgt tcaaggtcgc caccgcctta ttggccgata ccgccgtttc gccgaagaat 19380
ttgattcccc cctttttaaa caagtaatcc acactcatgt tcgtattcgc cttaccccgg 19440
aaataataaa gattataagg ttttaactcg ggctcgtaag tccgtttgcc gaaatcataa 19500
tgcagggccg tcaatcccac gtaaaaattc gttctcgtat aacgcaggtt cgtaccgaaa 19560
ctctgcatac ccacccggtt ccgtttatcc caatcccgtt gcagccgatg caagccatcc 19620
attttcaaag aagtgaccgt caagctgtcc gtactgcccg atagatatcg gcgagaataa 19680
aaaaaactcc agtcccaccc tttcattcgc aaggtcaccc ctaccccacg gaaaaaatcg 19740
aattcattgg tcgaataatg cctccgaaaa ccgttattcc tcctttccgc ctgggtgagg 19800
atggtgctgc gtgaaagcga ataatcctga ctcaccacca acccttgtcc gaaagcagct 19860
ttataatcac cagccacgaa tgtcttcaga cttttccaat cctttatcaa caagttaaac 19920
gagtaataat catacccttg atgatactta ttcaggaatg gttcaccggc atccttttca 19980
gcaacaaaac ccagctgaat acgctcatcg aaactatagg catagcgtat agactgataa 20040
aatgcctcac ctaagtactt tttattggga ttatcctgta aaacactgtc agaaagcgcc 20100
tcataccctc gtttttgttg aaaaccctga tcatacctca cggccaactc attacgagca 20160
tatgttaaca agttattcac cgttattggc aacttatcca cttgtgtatc gacgatatga 20220
acaaatggaa tcagcaattc caaagtagaa taatccaatc cgtcgatatt tttcaattca 20280
tagaccgaat acattttccc gtaccgaccc gcgtatgcca aaatatgcgc tatttgttta 20340
tctgtcagaa aaggcaaacg ttccaattcc tccttcttaa tttgattcag atcaaacggg 20400
ttttccacca aatacgaaag atcttcgtat aaagtctcca gttcttcttc acttgctccg 20460
ctgaccgaca gattatcaat gtattccttc catttatcat tcttactgtg cgaggattgc 20520
gcatgcgatt ggaaacaagt tattaaataa gttatcaacg ttaattctaa ggttatcaac 20580
aaagctttca taagcagaaa attaaaaaga atacctcatt cccacacccg gacaaatccc 20640
caattgcgga tgataattgg ccgacacatc caaccgaaaa gccgcgaatt ggagtccccc 20700
gcccaatccc ggaatcaacg gatatgtaga aaaacccgtt cgaaagaata cctgctcata 20760
aaaccgatac tccataccaa aaccgaagtc ccatcctaac cggtccacat acttacttgt 20820
cagttctatt aacaggttgt tgataaacgt ccattggata ccggattgta tagaataata 20880
attaaaatcc tctatgtgaa cattttctc atccagtcta acagcaggca aatccgttat 20940

cgacaaccct atcaacaagt tttcagtagg tataaacacc atacccaaat ccgtagaaag 21000
cttcttcgct tcctcatcat agagttccgt ttgtaacaaa gcgtactgaa aactcacacc 21060
aataacccac ctcgaagaaa gagttttcga taaagccagg cgaaacaagg tttcccggta 21120
tttatcataa ccgaaagtcg aaagatgcaa agcaaaaggg agtgcgcaaa ccggattctc 21180
ataaatgaca gaaaccgcag acaactcttt gagctgatac ttgttaaaat aatgaaaccc 21240
aatagactgg gtggttgata aactcaaaca ggccgggttg gacaaaacgg aatgaactac 21300
gccattccct cccataccga gcgaacggat atcgttgaat cttaaattcg tttgtccgaa 21360
agtagtggta gaatgaacta ttcctaaaag gaacaggcag tgtattaaag ttctcatatg 21420
ttaagatcta tcttggttaa accaatacac aaaaataaac aaattataga ttattcagta 21480
ttacaagtgc ttaaaacatt tatttttgct acagtatatg aaaggtattc gatcctatta 21540
tttgattctg acattccttc ttggttgtac ttccctattt gggcaatcgg taaagatgaa 21600
tgtacttccg aaagaccatt acaaagcgta tgttgagggt tcggataccg ttgctatggt 21660
caatttaaga gaagtatacg tttatccaaa aatcaaattt aagaacaaga aagaacagca 21720
gaaatacaat aaactcgtac gcgatgtgaa gaaaacgctt ccctatgcca aatggtcta 21780
tgaaacgttg attgaaacct atgagtatat ggaaactctg ccgactgaaa aagaaagaca 21840
agcccatttg aaaagaatgg aaaaggaact gttcaaagaa tacaaaccgg aactaaaaaa 21900
actgacctat tcacaaggaa aactattgat aaaactgatt gatagggaat gtaaccaatc 21960
cagttataac ctattaaaag cctatcttgg aagtttccgt gccggatttt ggaatctatt 22020
cgccggcatg ttcggcgcca gcctgaaaac ggaatataat cccaatggaa gtgatgcctt 22080
gactgaacgg gtggtggttt tggttgaaaa cggattaatc tgaaagctga cagaaaaaat 22140
cccagattac aatacctgca gtaaccgaga cattcaaaga atgtttcgta ccgaattgcg 22200
gtatttcaat acaacagtca gatttatcca cgaccacctg ctgaacccct tttacctcat 22260
ttcctaaaat caccgcgtat ttcttatttt tatctaattc caatttattc agcatgatac 22320
tccccttcac ttgttcgatg ctgcagacgg tatacccatt ttgtttcaat cggtcgacgg 22380
cttccaacgt atccttaaaa tagagccaat caaccgaatc ttctgctccc aatgctgttt 22440
tatgaatttc cggatgcgga ggacaagccg taattccaca taaataaacc gcttcaagaa 22500
gaaaagcatc cgaagtacga aagacagaac ctacattatt taagctcctt acatgatcca 22560
aaaccacaat taatggtatt ttatccgttt ttttaaattc ttcagttgaa agtctattta 22620
attctgttat cttcagtttt tgcatgattg taactattat taccaacaaa agtagttctt 22680
tatttctttt tttgtacagt tgaaaaccta ttgataagaa gtgaaaagaa aatgaaaact 22740
agttcttgta attctgtctg aaaatgctat ataacttctt catattaaat gtcaatagga 22800
tgacaaagca aattactaga agatatccac ccgcatttta cacaggtttt ggataaaaaa 22860
atgactattt ttattgacta gacaagttat aaacagataa ctaataagta tacttaacta 22920
gctgaagata aggaacaaaa taagtatatt atctgttaat aagttgatga atatatagga 22980
tagactagta acaacgaaac aagcaacaaa aatgaaatat atttatcaat agtattaacc 23040
gtatatcata tacatcattt aatttttaaa gttttaaata aataataata ataatatgaa 23100
ttgttgataa ggtagaaaac taaggagtaa tgactaactt tgcatcggtt attcatctta 23160
tcataaatga agagaattat gacacagaat ttatccgccg gttatgtaaa cgttccgatc 23220
cccaaaaata tagctattaa agaagagata gacagactgc gtaaagaaaa aaacgcggtt 23280
atcttggccc attattacca gaccggtgat ttacaagata ttgccgattt tgttggagat 23340
agtttagcat tagcccaatg ggctgctaaa acggaagccg atatcatcgt tttatgcggg 23400
gtacatttca tgggtgaaac agccaaaatt atttgtccgg ataaaaaagt actcgtgccc 23460
gatctgaatg ccggctgttc attggccgat agttgtccgg ccggtgattt cagtgaattc 23520
gtcagacaac atccgggtta cacggtgatt tcttatgtga ataccaccgc cgcagttaaa 23580
gcagtcaccg atgttgtggt tacatcaacc aacgccaaac aaattgtaga cagttttccg 23640
aaagaacaaa aaatcatttt cggtccggat cgtaatctgg gaaattatat caatagtatt 23700
accggaagaa atatgttgtt atgggatggt gcttgccatg tacatgaaca attctccttg 23760
gagaaaataa tttcgctaaa aaaagaatat cccgaagcgg aagtgatcac gcatccggaa 23820
tgtaaacaac ccattatcca ggtatccgac ttcgtaggat cgacagctgc attgttaaag 23880
catacgatag tatcccctaa aaaacaattc attgtagcga ccgaaagtgg ggttattcac 23940
gaaatgcaga aaaagagtcc gtataaagaa tttataccgg ctccaccgaa tgacagtact 24000
tgtgcttgca atgaatgcag tttcatgcgc ttgaatacaa tggaaaaact gtataattgc 24060
cttaaatatg aattaccgga aattttcatt gacgaatcga ttcaaaaaga agccattaaa 24120
ccgatacgaa gaatgttgga attgtccgaa aaattaggtt tataatcctt atttaaccac 24180

catgattttg gtaactacac tttcgttgcc atctgcattg gctgcaatga ccagataaat 24240
gccggtagct acccggtttc ctttgtagtt acggcagttc catacgagtt ggccgcctat 24300
tgatttagcc tgataaatca gctgaccgct cacattggtt atcttcacat tcgagtcgct 24360
catcaatccg gtaatcgtta cccgatcttc gtattccgga cgaaccggat tcggatatac 24420
cgaaacatcc gaataagaag atttgccttc agtggcttct cccataaaag aaactaatcc 24480
tttatctgtt cctataaaaa cttcccctgt catatcattt atcgctaaac tgacgatgca 24540
atcggataag agaggagagt tttcggtcgt aaaattataa attgtttctt gggcatcttc 24600
gctgactaga aagacaccgg agttttgcgt acctatccat ttgcgattac cgccatcgac 24660
agcgattgta ttaacctgat caccatctag aaaataatac agtccgctgt ctccttcacg 24720
cgtgattctg gaaaaataaa agcgactcgg attctcgatt gcattggcag ctaccggcgt 24780
atagacgact cctttattgg ttcctaccca tacgtaacct tttttatctt ccaccataca 24840
acgataggtg ctggttccta gatcctcacc ggaactgttt tgcagggaac ttaaaaataa 24900
aacattctcc tcatcttctt caaaattcgc atcctgaaat ataagaatcc ccgctttatt 24960
gttgtactgg tccgtatacg gaaaattcaa ccacttatgg cctttagagg tgatgagaat 25020
cttatccatg atctgcggtt tattgatata atttttatac gataaacctg cccaagttcc 25080
gtcggatttg agcactttga tgggtgattc tacctgacaa ttggtcatcc aaagattatt 25140
atcactatca aaagcaatat cggatatacg gacatattgg ttttttccat cgattccttc 25200
taaggggcta ttatgtatat tatatgccgc aaccacttca ttatctttaa tttcaaataa 25260
accttctcca taagaagaga taaaatagtc agtaggatca gtttggctgt aggtaacggc 25320
tgtaaaatcc tgcggataga atttacctgg atagtattct attttggatt ggtcaaataa 25380
cgaccattct tcattttcga aagtagaaaa gaatccttgt ctccagtatc ggtcgtgatc 25440
ttttcctccg gctaccgcca acaatttatt tccttgcgta accatggcat aaaaattatt 25500
gtttcgggga ctgttgatgt tgatttcgga aagtaggacg tcgaatcctc cggaattgtt 25560
tttctttatt cctttaatta aaccggtgcc ggcagctatc caatacgtat tttggtcttt 25620
atacgaggaa atatcgttaa ttgttccgga taacttatcg acttgtgtcc aggaagaata 25680
aatatacgca ttgttgttgg aatgtgcgat tagttgaccg ttttccattt ttatattaac 25740
cagtgtattg ttagttaata agtttttcgt tgtgttgtcg ttatccaaat aatagactcc 25800
ttttactttc tcattttta ctagaaagca aagcttattg tcaaatacga ctgttttgat 25860
gattttatca tctgatgcag gcaaagtaat cgagtaagca tgccaattgt tcgtatccac 25920
taagttgtca ttgagggaag ctttgtatac tcctttggaa gtggctgcat atatcgcatt 25980
atttaatacc gaaacagaat agacggtatc attcacaatt cgataggtat cctttatttc 26040
ttttttttcc atatccagcg ccaatacacc gaattgcatg gataaatagg ccatctgatt 26100
atatatatca atctggttga ccgttttgaa gttaaaattg gtattggtca tcaggtgact 26160
gatattatag ctatttcccg cttcatccat caggtcgata tttccatttg agtacacaat 26220
caataaggag ttgaccgtcg gattataggc tatttgggtg attttgttgt cactaagacc 26280
agtttgtttg gaataggtaa atatactgtt atcggtttta ttataactga gtaatgagcc 26340
gtttgtgact gcgtaaacca agttgttagt ttccgctatt ttttcactgc tgtagtagga 26400
aagataggtt ttaaaatgtt tcgccagact ctcataatcc tgtgcatgaa cagccatttg 26460
gaaagaaaaa aggaatgaaa gtatgaaaag taggggtttc atctttgacc ggattaaatg 26520
gatgataaat acgtgtttaa ttgacggacg gcttctgccc gatgactgat ttggtttttg 26580
atggtatctc ccagttcggc gaaagtttct gtataccctt ccggaacgaa tacgggatca 26640
taaccaaaac cgttatttcc tttttctct tcgataatat ggccgtttac gattccttcg 26700
aaggtatggg tttcgtttcc tattactagg gcaatcacgg tacggaaacg ggcttttcta 26760
ttggctatgc cttggagttc gaaaagtaat ttttgcatat tggcttccga gttatgggct 26820
tcgccggcat accgggcgga atagactccc ggagcgttat ttaaagcctc cacttccaat 26880
ccggtatcat cggcaaagca atcgaatccg aacttatcct tgatatagtt ggctttctgc 26940
aaggcatttc cttccaaagt ttccgctgtt tccggaatat catcgtggca acctatttct 27000
gccagactga caatttcgaa tttatcggct acaatctttc tgacttcgtc gagtttatgt 27060
ttgttgttgg ttgcaaaaac tagtttcata cgtcctatag aataaaaaaa gcacttcgtt 27120
tacgtaataa taacgtgcag aagtgcttta aattgtttac tgtttgttta tgtttttaat 27180
tgatcgaaac cttctccgta gactcctcga acgaaactct gcgagatgaa tgcgttcgga 27240
tcgacatctt tcactaatcg gaaaatcgtc aaggattccg atttacgggc aatcaaaaca 27300
atcactttca tcgggttctt ggaatacccg ccttgacctt ccaggaaggt gcatcctctt 27360
tgggtatccc gtatgatccg ttcggctatt tcttcatatt tttgagagaa tatcaggaat 27420

```
        tgaaccgatt gtttgttccg gtttatgatc atgtccaaca catagttgct gacgcccatt     27480
        tccacaaaac cgaagacgat tttttccaca tcctggaaaa ggaaatatga agaagaaatg     27540
        ataaagaaat cacagaagaa caaagcggta ccgatagata tattttttgta tttattgatg     27600
        atggccgcaa tgatatctgt tcctcccgta cttccgctta ctgagaaaat caatcctaaa     27660
        ccggatccgc atagggcacc gccaattagg atagccataa aaggttcccc atcgataatg     27720
        ggttttccgt ccagaaccca ttcaaagaat gagagggaca acgacaagga tattacaccg     27780
        acaatcgttt tgataaggaa tttgatcccc aatattttaa aggcgatgac caataatacc     27840
        gagttgatta aaaaatagga aacggaaacg ggaatcgacg tggcaaagaa aatgatggcg     27900
        caaataccac tgactccccc cgtaacgatt tcattggaaa ggataaaagc gttgaagctg     27960
        aatccgtaca acaaggtacc caatataata atcaaataat cctgcgcgtg attatataat     28020
        tttccgatat tattggtttc catcaaatca caatatttac aattttttga ggcacaacga     28080
        ttatcttttt aggtgttttg tctgctagcc atttcgcgga atattcgtgt gtcagcgccg     28140
        ctttttcaac ttcttctttc ggcatgtcgg ccggcaattc gatgctgaac cgggcttttc     28200
        cattgaatga aatggcgtag gtgaccacgt cttctttcaa gtattcttcc tgatattccg     28260
        gccattgggc atcgcaaatg gtggttgtat gaccaatcac gtgccataat tcttccgaaa     28320
        tatggggtgc aaaaggagca agcagaatca ggaactgttc catgatttct tgcttgttac     28380
        atttcaaact acttaattcg ttcagacaga tcatgaacgc actgattgat gtattgaatg     28440
        agaaatgttc gatatcgaag gttactttct tgatcagttt atgcaatgat tttagttcct     28500
        gagcagtggg tgcttcaaca gaaagttgga gtgtatcggt attaccgaag aatatgctcc     28560
        ataatttttt caaaaaacgg tgtacaccgt ctatgccgtt cgtatcccag ggtttggatt     28620
        gttccaacgg accgaggaac atttcgtata gacgcaaggt gtcggctccg tatttctcga     28680
        caatcatatc gggattgact acgttgtaca tggatttgga cattttctcg actgcccaac     28740
        cgcagatata ctttccattt tccaagagtt cttcatcgaa ttcagcagtt ttatattccg     28800
        gattccaatt acggaacgcc tcaatgtcca acacatcatt gctcacaata ttgacatcca     28860
        cgtgaagcgg agtcacttcg tattggtctt tcaggttgta cgaaacgaac ttgttggtcc     28920
        cgttgatacg gtatacaaag ttggatcttc cttggatcat tccctggttg atcagttttt     28980
        tgaacggttc ttcctcgcag gatatggcga tgtcgtagag gaacttattc cagaaacggc     29040
        tatatatcaa gtgtcccgta gcatgttccg taccgccgat atataaatcc acattgcgcc     29100
        agtacttgtc aatctttcgg gaaaccagcg cctgctcatt gcttggatcc atataacgca     29160
        ggtaataggc ggaggatccc gcgaatcccg gcatcgtatt cagctctaac tggaaaatgg     29220
        ttttattgtc gatgagggag gtgctaacca ccttaccgtt gaccgtatcc caggcccaat     29280
        tcgttgcccg tccaagaggc ggttccccgg tttctgtcgg caagaactta tccacttccg     29340
        gcagttcgag cggtagcttg tcttccggca acattcgcgg cagtccgttg gcatcgtaat     29400
        aaacagggaa aggctctccc caataacgtt ggcgactgaa gatggcatcc cgtaagcgga     29460
        agttcacttt tacttttcct aatttgttcg cttcaatata agctttcgtc ttggcaatgg     29520
        cttccggaac agaaagtccg ttcaaaacca aaccgtttac tatttctttt cctttgaccg     29580
        gtgaattcat catgatgccc tctttggcat cgaaactttc ttcagacaca tcacatccct     29640
        cgataagcgg gatgatgggc aggttgaaat gtttcgcgaa ggcgtagtca cggctgtcat     29700
        gtgccggtac ggccatgatg gcgcctgtgc cgtatccggc caatacataa tcgctgatcc     29760
        ataccggtat cgcttcgttg gtgagcgggt tgatcgcgta ggatccggag aagacccccg     29820
        tcacccgacg gtcggcgata cgttcccttt cggttctctt tttcgtagat tcgatatagg     29880
        cgtccacatc ggcccgttgt tccggcgtgg tcaacgcttg cactaattca ctttccggag     29940
        ccaataccat gaaggtaact ccgaagattg tatcggcacg ggttgtgaat atggtaaact     30000
        gtacatcgga atctttcacc cggaagacca tctcggcccc ttcacttcga ccgatccagt     30060
        tccgttgcgt ttctttcaag gagtccgtcc agtcaatctg atccaatccg tccaataatc     30120
        gttgggcata ggccgaaaca cggaggcacc attgacgcat gacgcgctgt tccaccggat     30180
        acccaccccg ctcggaaagt ccgttgacca ctt                                  30213
//
ID   XXX; XXX; linear; XXX; XXX; XXX;  XXX.
XX
AC   PRJEB67889
XX
DE   Uncultured bacterium DNA fragment from in-reactor enriched cow rumen, clone
ferm-6f24
```

```
XX
FH   Key             Location/Qualifiers
FH
FT   source          1..21494
FT                   /organism="uncultured bacterium"
FT                   /environmental_sample
FT                   /mol_type="genomic DNA"
FT                   /isolation_source="Cow rumen"
FT                   /clone="ferm-6f24"
FT                   /note="AZO-CM cellulose, AZCL-Barley beta glucan active
clone"
FT   CDS             complement(1027..2538)
FT                   /protein_id=
FT                   /translation="MIEQLRQLMTLDANMNTSDIEERFEQIAKVLFECFAIQKGETMYL
FT                   FKEIEFYFYNKNHRDIITHPRVSKPLCWYVNDFGGIDLNFVSNIKRESKANSKGKNVEK
FT                   YVLDNGAYFGGILIRQLTSEDGCEVLNGPWACAELFRCYDATVIDKDLPILVELDNGMV
FT                   GYIREPRINLLTSKQTVEGKVDYILEEYHEVFERKCLYSDFSRFVDRRYRYVRCDTLMH
FT                   DKDTNLVYFSPWLKDNKEGHPEFYKLLKNVLNEMGIESKELKSTNDYWARDYMPIQLGE
FT                   NEFLKYRYYPDYLVKSKNKKDIETITDATKVLRGMGISCRSTNLIIDGGNMVPCGPYIV
FT                   MTDKVFMENGREKGDTDFKVLLESEIGHPVIIIPWTPHEDDVYGHSDGFIKWCGDNRIL
FT                   MGNHGDCYPEEAASIRRILESYGFEVTEMRFKGKVDKPCYDLNWAYINFLQVGKNIIMP
FT                   KFNIEEDPIAQQYIHEAFPDWNIRQIEMAEIAKEGGALHCISWNV"
FT                   /product="hypothetical protein"
FT   CDS             complement(2568..3572)
FT                   /protein_id=
FT                   /translation="MSYKDFQQKVYSYLIKYKKESLNINDKGISTRGVEHDCLLSKPYC
FT                   DAKIPVMLYNGIKETVKDIQASKYAYKAHLAASKHVASSQTACINLFVPILESEYADQI
FT                   LIGAGVVPKDFDHIDRNQLRKGYCFEYWESTLEGAKGLLGDHSPHAGTDSDVAIAYRNK
FT                   NDKLCLWLIEHKLTEQEFTTCGGYRSKAISHSEKEYCTSCGIEEILKNHDKCYYHKHCG
FT                   YFYWNIMDAQSSFFCGKFEGNGCPFRGGMNQLWRNQMLALELENRGVFEDVYFSVVSHP
FT                   DNSFLDKTMNEYRALTNNSPKFYDFKSNALVDVAVDYLPDWSNWYKKVYLGID"
FT                   /product="hypothetical protein"
FT   CDS             complement(3586..3792)
FT                   /protein_id=
FT                   /translation="MTEADKKFLISFESGQPEWEGYEFEYFKDYPSVQWKLLNLKKLAK
FT                   QNPLKLQMEVDKLRNLYNFSLNN"
FT                   /product="hypothetical protein"
FT   CDS             complement(3841..4011)
FT                   /protein_id=
FT                   /translation="MIEKIPPKGVSFSYFICTFAMPCYDFRLFSNCNTKISEKSDTAKS
FT                   WATFCCSGTYK"
FT                   /product="hypothetical protein"
FT   CDS             4169..5239
FT                   /protein_id=
FT                   /translation="MRHLSYHPTLRTCSSDTILRAIKELTQENISYTSDQGKTYDFNTA
FT                   DKLNTLLINALVSTGELKEIEEYDVDFDHQFLETEKYDAKPTYKKFLGYRPGVYVIGDK
FT                   IVYIENSDGNTNVRFHQADTHKRFFALLESQNIRVNRFRADCGSCSKEIVSEIEKHCKH
FT                   FYIRANRCSSLYNDIFALRGWKTEEINGVQFELNSILVEKWEGKCYRLVIQRQRRNSGD
FT                   LDLWEGEYTYRCILTNDYKSSTRDIVEFYNLRGGKERIFDDMNNGFGWSRLPKSFMAEN
FT                   TVFLLLTALIHNFYKTIMSRLDTKAFGLKKTSRIKAFVFRFISVPAKWIMTARQYVLNI
FT                   YTENRAYAKPFKTEFG"
FT                   /product="transposase"
FT   CDS             complement(5405..6115)
FT                   /protein_id=
```

```
FT                   /translation="MNDIKTSVYAQKVELLLRLIPVVMDEQVFAIHGGTAINLFLKDLP
FT                   RYSVDIDLTYIPLADRQTSINDINSHLKSIADKAKKAFKGMHIVPNFNTCKQLCEYSGK
FT                   QVKIEVNQTKRGIIGGEVQTISLSEKAQNEFSLFCEANVVPITQLYGGKIAAALSRQHP
FT                   RDLFDVKYMDIPLSECREGLIFCLLGSDRPIHESFAPSLIDQREAMENQFAGMTDIPFT
FT                   YEEFEETRANLINA"
FT                   /product="FIG00937925: hypothetical protein"
FT   CDS             complement(6108..6869)
FT                   /protein_id=
FT                   /translation="MATKIQQILEATPRDSVLFGSWLSSQGLDARGQYAYMKSGWLDRI
FT                   SKGVYKIHGVEPGLMAAVSSYNTQLGKSCIVGAYTALELQGYSHYLSMGKPMAYLFTDK
FT                   KNKLPSWLLKGEWDMTIKYMTTSFLGDELLGVETMTINQRDLLMSSPERAILECLNLPD
FT                   ASSSLLDIYYIMESMTTLRPKLVQALLEACTSHKVKRLFLYMSEKAGHSWFKALKLENI
FT                   NWGTSRLMITPTGKYINKYNMTISKELAEYE"
FT                   /product="Ynd"
FT   CDS             complement(7408..7665)
FT                   /protein_id=
FT                   /translation="MCLQCKKGEKILADFAQSDSCKNVNDAISTSNNPHIFFKKNDLYI
FT                   PYSTMLVRSKTHAWFMTHRTIVHWPSHHCSKALEALFIEA"
FT                   /product="hypothetical protein"
FT   CDS             complement(7640..7756)
FT                   /protein_id=
FT                   /translation="MKQQASLIDSSFLPHFFSLVFTKIIDDSKEYVPTVQEG"
FT                   /product="hypothetical protein"
FT   CDS             7910..8665
FT                   /protein_id=
FT                   /translation="MFGGLYFLGYLCGVSFIEEKGMDEKMYCYKYPHPSVTTDCVIFGF
FT                   DGDRLKVLLIERRNEPYKGKWAFPGGFLNPDETAEDGALRELQEETGLTGTYIEQFHTY
FT                   TVPDRDPRERVITIAYYALVRIQEVHGGDDATKAKWFPIDEVPQLAFDHERILRDAMRK
FT                   LRERIHFEPIGFELLPEKFTMKNLQMLYEAILDVKFDRRNFSKKMMHHKLLNQLDEKVW
FT                   PTAKREALLYSFNNEKYNEFKKKGFQIKF"
FT                   /product="Hypothetical Nudix-like regulator"
FT   CDS             8668..9465
FT                   /protein_id=
FT                   /translation="MKNLLLSAVIGDISGMPYEFEGRTKNYEAVNLLHSANTYTDDTVC
FT                   TFACAEALLHELDMAKNLWKRCRADFHRGFGGRFACWLMAEDIQPSYHSYGNGSGMRTS
FT                   AAGFMSKTKEECIALATRTALPTHDHPEGIKGAVATSLAIFHAMEGKGKDYIRKNVLDV
FT                   YYPDWSRKTYMEIKPEYYFDETCQLTIPAALICFLESKDYTDCLKLAISLGGDADTLAA
FT                   ISGPMAYAYYKMIPEELINHAKAKLPEWMLKINDEFDDYCNKL"
FT                   /product="hydrolase, putative"
FT   CDS             complement(9875..11110)
FT                   /protein_id=
FT                   /translation="MSKSTHFSGQPVVYSQVIKLLDKAEIVKISHQTKGSEAYVKRFDGY
FT                   QHLVVMLFGILKHFDSLRELEIGMKAEAHKLSHLGMDYLVRRSTLAEANIRRPQEFLAS
FT                   VYAYLLEKYAKFLADSRPAKSYKGQTHEPKDWEKLLYMMDSTTITLFDNILKGVGRHPK
FT                   SGKKKGGLKVHTVMKYHVGVPMVVQLTSAAKHDHYLLKEVHLPKDSTLAMDRGYMDVAQ
FT                   FQRLTEEGVCYVTKMKKNLKYKVLDSVTYVNAKGLVTHIDQRVHFTRGELTHEARRVEI
FT                   FHENKKPVALLSNNFEFSVEDISEIYRLRWAIESLYKQLKQNFPLHFFYGDSVNAIQIQ
FT                   TWVVLIANLLITVLSRSIKRRCAFSQVVSMVRLTLMYYIDFIAFMENPDKSWNNIRAKE
FT                   LQKAPPNPTLFD"
FT                   /product="Mobile element protein"
FT   CDS             complement(11571..13883)
FT                   /db_xref="GO:0008422"
FT                   /protein_id=
FT                   /translation="MKSFKAIFSILTMLPLAVYAQVSNPVIPRDDRIEAQVQQTLSRLT
```

```
FT                   LEEKVGQMMELVTDFFGSNDEKGVFHIDRQKTDSILARYKIGSILNAPNTCAPTAKEWE
FT                   VIIQDIQKSSMKQIGIPCVFGLDQNHGSTYTADGTLFPQNINVAATFNREIARRCAEAT
FT                   AYETRAISVPWTYSPTVDLGRDPRWPRIWENFGEDAYVNAEMAKAMVKGFQGENPNRID
FT                   QQHIAVSLKHYLGYGVPYSGKDRTPAYISLSELREKHFAPFLAGLEAGALTVMVNSASV
FT                   NGMPVHANKTLLTTWLKEETGWDGMLITDWADVNNLYTRERIAKDKKDALRIAINAGID
FT                   MVMEPYATDACDYLVELVKEGKIPMSRIDDAVSRILRLKYRLDLFKNPTQKLKNYPKFG
FT                   GEEFAKLSLEGAQESMILLKNQNNILPLKAGKRILVTGPNANQMRPLNGGWSYTWQGHR
FT                   TDEFTEKYNTIYEAVCQKYGKDNVVLSQGVTYNEKGKYYEENTPDIASAVAAAQNVDVI
FT                   FACIGENSYTETPGNLSDLNLSSNQKDLVKALAATGKPVILILNQGRPRLINDIEPLAQ
FT                   AVVNILIPGNYGADALVNLLAGEVNFSGKLPYTYPSEINSLATYDYKKSESVSTMEGAY
FT                   DYNAQITQQWGFGFGLSYTTYTYSNLKVDKTNFKHNDTIRVSVDVTNTGSMAGKESVLL
FT                   YSSDEIASIVPDNRRLRDFQKIELNPGETKTVSFLLPARNLAFVDYDGKWVIEEGTFAL
FT                   MAGNQLVKVNCSDTYYY"
FT                   /product="GH3"
FT                   /EC_number="3.2.1.21"
FT   CDS             complement(13883..14623)
FT                   /protein_id=
FT                   /translation="MNTFNKTIENKIFEGFSILAIIMVSLLCFSCSEDDPFFTATKDDA
FT                   PRILNTDIPEWRDGQPQTLQTVDFGDSLRFKVIATPRDYTTISWYIDDELVFEGDSIVY
FT                   PISEGKHLCKIVATTTQGLSTSRTCYVIGNPEPGKEKILWEGSFDVTWGTPFDVLKTSL
FT                   AENVQEGTVLRVYVEGDGQGTATSAWWNNILTGEGEDKRGDIAISGEQMLEFTLNSTSM
FT                   TLLSTQDGFLVVGNGYKITKITIM"
FT                   /product="hypothetical protein"
FT   CDS             complement(14762..16504)
FT                   /protein_id=
FT                   /translation="MKSFQKYIKSGLGASFALTLTVGAMSSCSDSYMSDLNTDDSKAMS
FT                   IDPNAQLTTSLLQTYGDFSLMDTYRSYITGFTQHFAGGWNVTNYAGSVHAFDDQTRLIW
FT                   DRYYTVSIKNMEDAISKTSEKPNINALLRIHKVYLMSVLTDTYGDVPCFNVGQGYMKGE
FT                   NNPAYDKQEDIYQWFFDELSLCESQLNAVGDRITGDVTSMNGDVAKWKKYCNSLRMRFA
FT                   MRISDVKPELAREQFEKALASENGYIAEASDDAYIKYLDQPFTLYKGAEKYDFRANALS
FT                   EILYGQDPTSPTFVSSTFYNIMKDSNDPRLYRICRHYINTTRSEIRADEAGNFDLTDEV
FT                   VQYLKDNGLSESPCDPGAAWYNNWINAPATKNIPTLQLLVDQNPAAGYDKNNFPARMMR
FT                   LFLSIDFEKGQNPGILFTSAEAEFLLAEAKIKGWNVEGSADEHYEAGVRASMKLLNTHY
FT                   LKEKDLINDAEIAAFIAANQLGENPKETINTQAWILHMTNPAEAWSNLRRSDYPVLMDR
FT                   TKLPTFPDFTYDDPDRSTPTRLKYPILEARYNSENYKEALERLGGTDDWHQRNWWDTAD
FT                   INVK"
FT                   /product="FIG00937602: hypothetical protein"
FT   CDS             complement(16616..19795)
FT                   /protein_id=
FT                   /translation="MMNKTILLSLLCTGMIGTPATTYAVSSSVKATLTAQQSDQEIKGS
FT                   VVDASGEALIGATVKIVGGTGGALTDMNGHFSLKCEPGTTLQISFMGYNPVTVKASNGM
FT                   KVTLQENVKVLGDVVVTALGIKRDKKALGYALEEVKGEELTKAKETNVINSLAGKVAGL
FT                   VVQNTAGGSSGSTRVLLRGNTEMGGDNQPLYVIDGVPLDNTNFGSVTTAGGHDLGDGIS
FT                   AINPDDIETMSVLKGPAASALYGSRASHGVILITTKKAEKDKVSVEYNGSITFDSQLAK
FT                   WNDIQQVYGMGNGGQYLETATSGTNMSWGPKADNYMVRYFDGVERPFLMYPNNTSNFFR
FT                   TGVTAQNTGILSFSSGKTGIRFSMTDMRNKDILPNTHMSRDNFNLRANTSSGAFDFDFT
FT                   ANYTREDVKNRPALGDSQSNVGKNLMTLAGTYNQDWLRYYQNPDGTYANWNGNDMYNKN
FT                   PYWDLYKNDNTSSKDVFRFTGKAVWNINKDWKVQGTIGTDINNMNFEEFIARTTPGSAS
FT                   GEMTNQVFQNRTLNAEILGLYGHTWDNFDVHATVGGNIYKVNNKTYNMLGHDQQISDMK
FT                   DIMNYSEHNVNRDTYVKQINSLYGSVSFGYNRTYYLDVTLRGDKSSTLPTSNNTYYVYPS
FT                   LSGSWIFSEYLKNKNILSYGKLRASWAKVGSDTDPFQLAINYSSGKYSYTGYMLGMINN
FT                   NNMPNGSLKPTMTNSFELGTELKFFNGRLGMDVTWYSQSSRNQIINLASSAPSGYPYRL
FT                   VNAGHIINRGVELAVNGRAVQYKDFAWDLGFNFSKNINKVKELVPGMDYFELEKATWAG
FT                   VSVGAMVGQNYGSIMGKDVARTDDGQVIIDRSTGMPVSAEGTHVIGNASWDWTGGFYST
```

```
FT                   FTYKNFHLSAGFDVKYGADLFSMSMRSAYQTGRATATLKGREAWYASEEARFAAGQTSE
FT                   DWRASGNMKGYVVPGVYLNDEGQYVKNDIPVNPEAYWKYATENCPAMFVYDNSYVKCRE
FT                   ITFGYTFPEKMLGKTVKGLTLSFVARNPFILWKNIPNIDPDSGYNTSGLGLEYGSLPSR
FT                   RSYGFNINLKF"
FT                   /product="FIG01289191: hypothetical protein"
FT   CDS             complement(19849..21096)
FT                   /protein_id=
FT                   /translation="MKTNRQIVTSLTLLLGMLFTTACEAHQPEAELPQKNEKEIKAPTA
FT                   QEWNREVTIGWNLGNSFECSAPGMNGESLLIGNPAGADNAETSWGNPVVSKKLIDAVKA
FT                   AGFDAIRIPVRWQCHITNAQAMTISRTWMNRIKEIVDYCLKCDMKVIINTHHEKWLESR
FT                   PTYKYKEENCQKLALLWMNIASEFKDYDYRLAFAGTNEVHERDNWGQPTAENLEVQNAY
FT                   NQTFVDIVRATGGNNAQRHLMVQTYVCNPEFGLNNGDFIIPTDLEENGNRYMSVEFHYY
FT                   QPWDYCGLGKLEYWGERYRAHGTAAPENEQTIIDTFTKVQQTWAFKGLGVVIGEWGVTD
FT                   HYKSSSADLMRENMAYYCQIYVAEAKKRGFATFIWDNNHFGNGAECFGIIDRTHGLQIK
FT                   APWVMEGIKAGLKAVE"
FT                   /product="GH5_4"
FT   CDS             complement(21306..21458)
FT                   /protein_id=
FT                   /translation="MRTIRLEQAAIYLKQGYPVLDVSIKTGFANVKYFSTVFKRHFGVQ
FT                   PSKYQ"
FT                   /product="FIG00938042: hypothetical protein"
XX
SQ   Sequence 21494 BP; 6072 A; 4704 C; 4400 G; 6318 T; 0 other;
     tgtccagcga cgtctgcgta catggaggac tgtcttcttg tcgcgtgtcg ggaaatccag        60
     aagacatgac tcgtcaccaa aaccatcgct cacagagcca ctcatgtctg gagtgcgatt       120
     gtcgcgttcg tcaagataga tatgaagctc agttgtgtac agaacttcgg tgtcctgtgt       180
     cggaacctca cagaagtcgg tcacttcaaa gtactcaagg acacctgtag ggaggaaata       240
     tgaggcaagg ccttcataca tgtttatatt gctgttacga ttcatactgc aaaggtacta       300
     actttttttc aatcgtgtat cgtttagcga cagtttttgg gttagcatcc gtacatcttc       360
     ccacagggtt tgtcgggtgc gcccattttt atcgaatttg gaagcaaaat acacgcatat       420
     aactctaaat atgagatatg atagccgtcc aaaagaaaga gaagccaatt attcagtttg       480
     atagcacaac cctttggttt agtttattcc catattatat atcaataacc taaactaaac       540
     cttaatgcaa gaatgtgcaa aaatctgcgt taaaaagtga aaacgtgcaa agaaatggaa       600
     aatcgatttt gcagatgttt gcattttctt tttttgaagg cagtcatatt tgcagatttt       660
     tgcctatttt acgtcccccc accgtccctc ggagtcactt tagttaccct aaaacaaagg       720
     catttgcgtt gattatcagc cacttatatt tgcagatagt aacttctgcg taggaaagta       780
     gattctgcgt gggaaagtga ttttttgcaa agaaaagcat actatctgca aatatggatc       840
     gtaaatcgct taatttcaag cattaagaat tacgcccatt tttgcagatt tttgattttt       900
     aacttcgatt tttgcagatt ttcgcccgtt ttgtccccca gctgtccccc gagatagtcg       960
     aaaatagccg agggacaaat agatctcata ggtggtgatt tatgttttta catgtgattt      1020
     tggaagttaa acattccaac ttatacaatg taaagcgcca ccttctttgg caatttctgc      1080
     catttctatt tgacgaatat tccaatcggg gaatgcttca tgaatgtatt gttgagcgat      1140
     tggatcttct tcaatgttga atttaggcat tatgatgttc ttgccaactt gcaggaagtt      1200
     gatatatgcc cagtttagat cataacaagg tttatcaact ttgcccttga aacgcatttc      1260
     tgtcacttca aagccataac tttcaagaat ccgacggatg gatgctgctt cttcaggata      1320
     gcaatcgcca tgattaccca tcagtattcg attatcgcca caccatttga tgaatccatc      1380
     cgaatgtcca tacacatcat cttcatgcgg agtccaagga ataatgataa caggatgccc      1440
     aatttcagat tccagcaaaa ccttgaaatc tgtatctcct ttctcacgtc cattttccat      1500
     aaatacttta tcagtcatca caatatatgg accgcatggc accatattac caccatcaat      1560
     gatgagattg gttgaacggc aactgatgcc cataccacgc aacaccttgg ttgcatcagt      1620
     aattgtctca atatcttttt tattcttgct cttcacaaga taatcaggat aatagcgata      1680
     cttcaagaac tcattttcgc ccaattgaat cggcatataa tcgcgagccc aataatcatt      1740
     tgtagatttc agttctttgg actcaattcc catttcatta agcacattct tcagaagctt      1800
     ataaaactct ggatgtcctt ctttattgtc ttttaaccaa ggagagaaat acacgaggtt      1860
```

tgtatcttta tcgtgcatta atgtatcaca acgcacatat ctataacgtc tatccacaaa 1920
tctcgaaaaa tcactatata gacatttcct ttcaaaaact tcatgatatt cttcaagaat 1980
ataatcaacc ttaccctcta ctgtttgttt agatgtaagt agattgatgc gaggttcacg 2040
aatatagccc accataccat tatcaagctc cacaagaata ggcaaatcct tgtcaatgac 2100
agtagcatca tagcacctaa acaattcagc acaagcccat ggtccattta gaacttcaca 2160
cccatcttca cttgttagtt gacgaatgag gatgccacca aaatatgcac cattatcgag 2220
aacatacttt tcaacattct ttcctttact attagcttta gactcacgct taatattact 2280
aacaaaattt aaatcaatcc ctccaaagtc atttacatac caacacaatg gttttgacac 2340
acgtggatgg gtgataatat cacgatgatt cttgttgtag aagtaaaact caatctcctt 2400
aaacagatac atcgtctcac ccttttgaat agcaaaacat tcaaatagca cctttgctat 2460
ttgttcaaaa cgctcttcga tgtctgatgt attcatgtta gcatcaagag tcattaattg 2520
tcggagttgt tctatcattt gtatattgtc ataatcacgg tgatacatta atcaattccc 2580
aaatacacct tcttgtacca atttgaccag tcaggcaaat aatcaactgc cacatccaca 2640
agtgcatttg atttgaagtc atagaacttt ggagagttat ttgttagtgc ccgatattca 2700
ttcatagtct tgtcaagaaa agaattatcc ggatgagaga caaccgagaa atagacatct 2760
tcaaatacgc ctctattctc aagttccaaa gccagcattt gatttctcca aagttgattc 2820
ataccacctc taaacggaca gccattgcct tcaaatttac cacagaagaa tgaagattgg 2880
gcatccataa tattccaata gaagtaaccg caatgcttgt ggtaataaca cttatcgtgg 2940
tttttcaaga tttcttcaat accacaagaa gtgcagtatt ccttctctga gtgagaaatt 3000
gcctttgaac ggtagcctcc acaagtcgtg aattcctgct ctgttaactt gtgctctatt 3060
agccaaaggc acagtttatc gtttttgttg cgatatgcaa tggcaacatc tgagtctgta 3120
ccagcatgag gactatggtc tccaagaagc ccttttgctc cttctagtgt tgactcccaa 3180
tactcaaagc aatatccttt gcgaagctgg tttctgtcaa tatggtcaaa atctttaggt 3240
acaactcctg cccctataag tatttggtct gcgtactcac tttccaagat tggcacaaat 3300
aggttgatac atgcagtttg agaacttgcc acatgtttag aagctgcaag gtgagcctta 3360
tatgcatact tgcttgcttg aatgtctttc actgtttctt ttataccatt ataaagcata 3420
acaggtattt tggcatcaca atatggcttg gataataaac aatcgtgttc aactccacgt 3480
gtagaaatgc ctttatcatt tatattaagg ctttccttct tgtatttaat caggtatgaa 3540
taaaccttct gctgaaaatc tttgtatgac ataagtagtg aactgttaat tattgagact 3600
aaaattatag agatttctta gtttatctac ttccatctgc aacttcagag gattctgctt 3660
ggcgagtttc ttcagattaa gcagtttcca ttgtactgac ggataatctt tgaaatattc 3720
aaattcatat ccttcccatt cgggctgacc agattcaaaa ctaatcagga acttcttatc 3780
cgcttcggtc atcaacaatt ttacatcatt accttaattc cgcaacacta taatttaact 3840
ttatttataa gttcctgagc aacaaaaagt tgcccaggat tttgccgtgt cagatttttc 3900
acttatctta gtgttgcaat tagaaaacaa gcgaaaatcg taacaaggca tggcaaaggt 3960
acaaataaaa tatgagaaac tcacaccttt tggaggaatt ttttcaatca tggagaaatt 4020
tgactccatg ctttcacccg ttatagactc aacactgggt cagagatgca gcagtatcat 4080
cggatatcag ttcagcgaga tagtccgttc gctgatgagc gtttatttct gtggcggctc 4140
atgcgtggaa gatgtaacgt cacacctgat gcgccatctc tcgtatcatc ctacccttcg 4200
tacatgcagc tctgatacca tcctcagagc catcaaggaa ctgacacagg aaaacatctc 4260
ctatacttcc gaccaaggca agacctatga tttcaatacc gcagacaaac tcaacacatt 4320
gcttataaac gctttggttt ctacaggcga gttgaaggaa attgaggaat acgatgttga 4380
ctttgaccat cagttccttg aaacggagaa gtatgatgca aaaccgacct acaaaaagtt 4440
cctcggctac aggcctggcg tatatgttat cggtgacaag atagtctata tcgagaacag 4500
cgatggcaac acgaatgtgc gttttcatca ggcagacacc cataagagat tcttcgctct 4560
tctggaatcc cagaacatcc gtgtaaatcg cttcagggca gactgcggtt cctgctcgaa 4620
ggaaatcgtc agtgagatag agaagcattg caaacatttc tacatccgtg ccaaccgatg 4680
cagttcgctc tacaatgaca tctttgctct gagaggatgg aagacggagg agattaacgg 4740
agtccagttc gaactcaatt ccattctcgt tgagaaatgg gaaggcaagt gctatcgtct 4800
tgtcatccag agacaaagac gcaacagtgg cgaccttgac ctgtgggaag gcgaatacac 4860
ttaccgttgt attctgacca acgattacaa gtcatcgaca agggacattg ttgaattcta 4920
caatctgcgt ggcggcaagg aacgtatctt tgacgacatg aacaacggat tcggttggag 4980
caggctcccc aagtcattca tggcggagaa tactgtcttt cttctgctta ctgcattgat 5040
acacaatttc tacaagacca tcatgagcag gcttgacacc aaggcttttg ggctcaagaa 5100

aacgagtcgc ataaaggctt ttgtcttcag attcatctcc gtacctgcca agtggatcat 5160
gactgcaagg caatacgtgc tgaatatcta cacagagaac cgcgcttatg caaaaccctt 5220
caaaacagaa ttcggataag aatcctttct ttccggttag aatatgcgta ttacctcaag 5280
tcgcatcgtg gggtaagggg atggtggcta catattgatg ttgtgctgtt gctttttact 5340
gaaagctgct actaacgact cataaatcta cccttaatca cgtattggat gatagttgcg 5400
gattttaggc attaataaga tttgcacggg tttcttcaaa ctcttcgtaa gtaaaaggaa 5460
tgtcagtcat tcccgcaaac tgattttcca tcgcttctcg ctggtctatc aaacttggag 5520
cgaatgattc atgaatgggt ctgtcactac caagtagaca gaaaataagc ccttcacgac 5580
attcactcaa tggaatatcc atgtatttta cgtcaaacaa atcacgagga tgctgacgag 5640
agagtgctgc tgcaatcttg ccaccgtata attgagttat cggaacaaca tttgcctcac 5700
aaaagagaga aaattcgttc tgtgccttct cactcaatga tatggtttgt acctcgccac 5760
ctattatacc tctctttgtc tgattaacct ctatcttcac ctgtttacca ctatactcac 5820
ataactgttt gcaagtattg aaattgggta cgatgtgcat ccctttgaat gctttctttg 5880
ctttatctgc aatggacttc agatgtgaat tgatatcatt tatacttgtc tgacggtctg 5940
ccagtgggat gtaggtcaga tcaatatcta cagaatagcg tggcaaatct ttcaagaaaa 6000
gattgatagc agtaccacca tgtatggcaa acacctgctc atccataaca acaggaatga 6060
gtctgagcaa cagttctacc ttctgagcgt aaacacttgt ttttatgtca ttcatattct 6120
gcaagttctt tggagattgt catattatat ttgttgatgt actttcctgt aggcgtaatc 6180
attaaccttg atgtacccca attgatattc tccagcttca aagccttaaa ccaggaatgc 6240
cctgcttttt ctgacatata taggaacagg cgcttcactt tatgggaagt gcatgcctcc 6300
agcaacgcct gtaccaactt gggacgtaat gttgtcatac tctccatgat gtaatagata 6360
tcaagcaatg aagatgatgc atccggcaaa ttcaagcact ccaatatggc tcgctctgga 6420
gatgacataa gcaaatcacg ctgattgata gtcatggttt ccacacctaa aagttcatca 6480
cccaggaaag aagtagtcat gtattttatc gtcatatccc actccccttt tagcaaccat 6540
gagggcaact tgtttttttt gtctgtaaac agataagcca taggtttccc catagatagg 6600
taatgagagt atccttgaag ttccaaagct gtatatgcac caactatgca actcttacca 6660
agttgtgtgt tataagacga tactgctgcc atcagacctg gttctacccc atgtatctta 6720
taaactcctt ttgatatacg gtcaagccat ccgcttttca tataagcata ttgaccacgt 6780
gcatccaatc cttgacttga aagccatgaa ccgaacaaaa cggaatctcg tggagtagcc 6840
tctaatattt gctgtatttt tgtagccatc ttcgttcttt ttatgaatac caggcaaaat 6900
ttgttgcaaa catactcttt ttattttgtc cagccaagca ttttcctcaa acattggaga 6960
acttttatag aaaaatctct ccaatttcat atttttagcg aattcaagca caaaaagtat 7020
gcatataacc atatacaacg agacgtttgc cgtctcaaaa aagaagaagc cagtttgaca 7080
agaggtttaa tttatcccac atataatata tccataaact aaactttaaa attcaacaat 7140
ttatgtcatc acgagatcag agggacagaa ttacaaccca gtagtgtacc cgatagacaa 7200
aagcataaaa cggcacttca ggacaataac aacactgtat ttcacattat tatgacgtga 7260
gttcgactaa ttcacgttat aattagcata acaaacgaat atccatctat aatttgttta 7320
tattactgtc cgctaaactt tgactttcca cctcaaaatt agggctgaca ccttgccttt 7380
ttgcatctcg catgtccgaa gtgtccttca tgcctcaatg aacaatgctt caagtgcctt 7440
tgaacaatgg tgcgatggcc aatgaacaat tgttcggtgg gtcatgaacc atgcgtgcgt 7500
ttttgacctc acaagcatgg tgctatatgg gatatataaa tcgttctttt tgaagaagat 7560
atgtggatta ttggatgtgg agatggcatc attcacattc ttgcaagagt cgctctgcgc 7620
aaagtctgcc agaatctttt caccccttctt gcactgtagg cacatactcc tttgaatcat 7680
ctatgatttt ggtaaaaaca agagaaaaga agtgtggcag aaaagaggaa tcaatcagtg 7740
aagcctgctg tttcaccttc tgaatttgtc gaactcccgt tgatttgcac ttgtttcaaa 7800
ttcatctaac tcaggttaaa aggttgcaca gggacttcat gtgttctttt tgactattac 7860
tctacatttt ttctataaac acaatgcttg ttgtgtactt ttcacacaaa tgtttggtgg 7920
tttatatttt ttaggttatc tttgcggtgt atccttcata gaagaaaaag gcatggatga 7980
aaagatgtat tgttataagt atccacatcc atctgtcaca acagattgtg tgatctttgg 8040
atttgacggt gacagactga aagttctact tatcgaacgt cgaaatgaac cttacaaagg 8100
gaaatgggct ttccctggcg gtttcttgaa tcctgatgaa acagccgaag atggtgcgct 8160
tcgcgagttg caggaggaga cagggctcac tggcacttat attgaacaat tccataccta 8220
cacagttcct gaccgtgacc ctcgtgaacg agttattacc attgcctatt atgcacttgt 8280
gcgcatacaa gaggttcatg gaggtgatga cgcaacgaaa gcgaaatggt ttcctattga 8340

tgaggttcca caactggctt ttgaccacga acgcatcctg agagacgcca tgcgtaaact 8400
tcgtgaaaga atccactttg agccaatagg atttgagctc ttaccagaga agttcaccat 8460
gaagaatctc cagatgcttt atgaagccat attagatgtg aagtttgaca gaagaaactt 8520
ctccaagaag atgatgcacc acaaacttct gaaccagttg gacgaaaaag tatggcctac 8580
cgcaaagaga gaagccctgc tctactcttt caacaacgag aaatacaatg aattcaagaa 8640
gaaaggattt caaattaagt tttaattatg aagaatctat tattaagtgc tgttattggc 8700
gatataagtg gcatgcccta tgagtttgaa ggcagaacaa agaactatga agcagtaaat 8760
cttctccatt cggcaaatac atacacagac gatacagtat gcaccttcgc ctgtgctgaa 8820
gctctgcttc atgagttaga catggcaaaa aatttatgga aaagatgccg tgccgacttc 8880
cacagaggat ttggaggccg ttttgcctgt tggctgatgg cagaagacat acaaccatct 8940
taccattcat acgggaatgg aagtggcatg agaacctcgg ctgccggttt tatgtccaag 9000
acaaaagagg aatgtattgc gcttgccacc agaacagcct tacccaccca tgaccatcca 9060
gaaggaatca aaggggcagt agccacctcg cttgccatct tccatgctat ggaagggaaa 9120
gggaaagatt acatacgcaa aaatgttctt gacgtgtatt atccagactg gagtagaaag 9180
acctatatgg aaatcaaacc agaatactat tttgatgaaa catgccaact aacaattcct 9240
gcagccctca tctgtttctt agagagcaag gattatacag attgcttgaa acttgccata 9300
tcccttggag gtgacgccga tactctcgcc gctatctctg gtcctatggc ctatgcttac 9360
tataaaatga ttccagaaga actcatcaat catgcaaaag cgaaactccc tgaatggatg 9420
ctgaagataa atgatgaatt tgatgattac tgcaacaaac tctaaaagaa gcatcatcac 9480
aatgagatga acaaacaaag catttcgttg caatcacctt ttaggcttta gaacaaaagc 9540
atcctcagta agaagagcct ccaaaagtta gacagaatac ttttggaggc tcttcaacgt 9600
tttactcaac aacgttaatt attcaagact cctatcgaag ccttatcgaa ggcatatcga 9660
aggcttgcag aagtatgggg agggtgtata gacaacatct tgttgtctca attgtctcaa 9720
tcacaatcac ccagatgcgc actgtggcac aagaggcaaa aaagttagaa tccgaattta 9780
ttgctgtccg gtaaaactca gaagtgcata gaaaccttgc ccgaagccca gtaaaatcag 9840
gcttcgggct ttctttttga aaaagtaagc cccccttaatc gaacaaagta ggatttggcg 9900
gtgccttttg cagctcttta gcacggatgt tgttccatga tttatctgga ttttccataa 9960
atgcgatgaa atcgatgtag tacataagtg ttagtcgcac catgctgact acctgagaga 10020
atgcacacct cctttttgatg cttcttgaga gcactgtgat gagtaaattg gcaatcaaaa 10080
ctacccaagt ctgtatttgt atggcattca cgctatcgcc atagaagaaa tgaaggggaa 10140
agttctgttt gagctgtttg taaagcgact caatggccca tcggaggcga taaatctcgg 10200
aaatatcttc cactgaaaac tcaaagttgt tgctcaacaa agccactggc ttcttgtttt 10260
cgtggaaaat ctctaccctt cgtgcctcat gcgtaagctc tcctcgtgta aagtggactc 10320
tttgatcaat gtgcgttacc aatcccttcg cattaacgta tgtaacggaa tccaggacct 10380
tatatttaag gttcttcttc attttggtaa catagcacac gccctcttcc gtaagccttt 10440
gaaattgtgc gacatccatg tatcctctat ccatggccaa ggtggaatcc tttggaagat 10500
gtacctcctt gagcaggtaa tggtcatgct ttgcggcaga agtgagctgt acgaccatag 10560
gaacgcccac atgatacttc atgaccgtat gcaccttcaa cccacctttc tttttacctg 10620
attttggatg ccgtccaacc cctttgagga tgttatcaaa gagcgttatc gtggtggaat 10680
ccatcatata gagcagtttc tcccagtcct tgggctcgtg tgtctgtcct ttataactct 10740
tagcaggacg gctgtccgct aaaaacttgg catacttctc caaaagatat gcgtatacgc 10800
tggctaagaa ctcctgagga cgccgtatat tggcttcagc cagcgtacta cgacgaacca 10860
agtagtccat gcccaagtgt gagagcttgt gcgcctcggc cttcatacca atttcaagtt 10920
ctcgtagaga atcaaagtgt ttgaggatgc cgaatagcat tacaaccaaa tgctgatagc 10980
catcgaaacg ctttacgtag gcctcgctac ccttggtttg atggctaatt ttgacaattt 11040
ctgccttatc gagtaatttt atgacctgac tatagaccgg ctgtccgcta aaatgtgtac 11100
ttttgctcat tgctagctgt tttttttgttc aaactcaaag ttagcaaaaa gttttaccgg 11160
acagcaatag ttgtttatat atacttccct ctcgtcgggg aaagtggcag cacagttggc 11220
ttctatgatt cctttataga aaaagaatcc ttcatgcttt atttactcat atcgtccgaa 11280
aacttacaca aggcaaagat ggataaggtt gtaaggggtg ttcacattcc atgttttctc 11340
cattgtcttt ctattgtgtc tctcctaccc ttctctactt ctttgtctct cctcatgaga 11400
tgaagtagta cagataccca gattgatgcg gaacaaaaaa ttattggtgt ctgcgaattt 11460
attttacaca aataaaatgc aaacaccaat aatcacaatt aattattgtc tgtgtgtttt 11520
tattaaattc ttttcaaatt taattttaat ggtataagat agtgtgtgag ttaataatag 11580

tatgtgtctg aacagttcac cttgaccaac tgattgcctg ccatgagtgc aaatgtgcct 11640
tcctcaatga cccacttgcc atcataatcc acaaaagcga ggttgcgagc aggaagaagg 11700
aaggagacag tcttggtctc acctggattg agctcaatct tctgaaagtc gcgcaagcga 11760
cgattgtcag gaacgatgct ggctatttca tcactactat agagcaacac gctttctttt 11820
cctgccatgc tacctgtgtt ggttacatcg acagagacac ggattgtgtc gttgtgctta 11880
aagttggtct tatcaacctt aaggttgctg taggtgtagg tagtgtagct gagaccaaag 11940
ccaaatcccc actgttgagt gatttgtgca ttgtagtcat aagcaccttc cattgtgctg 12000
acagactcac tcttcttgta gtcataggta gcaagagagt taatctcaga tggataagtg 12060
tagggaagtt taccgctgaa attaacttcg ccagcaagaa gattgacaag agcgtctgca 12120
ccataattac ctggaatgag gatgtttaca acagcctgtg caagtggttc tatatcgttg 12180
atgagacgag gacgcccttg attgagaatc aagatgacag gtttgccagt agcagcaagt 12240
gccttgacca aatccttctg attggaagat agattgaggt cggagagatt gccaggtgtt 12300
tctgtgtagg aattttcacc gatacaagca aaaatgacat ctacattctg tgcggctgct 12360
accgcagatg ctatgtcagg tgtattctct tcatagtact tgcccttctc gttgtaggtg 12420
actccttgag agagcacaac attgtcctta ccatacttct ggcagactgc ctcgtagatt 12480
gtgttatact tttcagtgaa ctcgtctgta cgatgacctt gccatgtgta gctccatcca 12540
ccattaagag ggcgcatctg attggcatta ggtccagtca caaggattct ttttcctgct 12600
ttgagaggaa ggatattgtt ctgattcttc aggaggatca tgctctcctg agcaccttcg 12660
agggaaagtt tggcaaactc ctcaccacca aacttaggat agttcttgag tttttgagta 12720
gggttcttga agaggtcaag acgatattta agacgaagaa tacggctcac agcgtcatca 12780
atacgtgaca tgggaatctt gccctcttta accagttcca caagataatc acatgcgtcg 12840
gtagcataag gttccatgac catatctata ccagcattga tggcgatacg aagggcgtcc 12900
ttcttgtcct tggcaatgcg ttcacgagtg tagaggttat tgacatctgc ccagtcagtg 12960
atgagcatgc cgtcccaacc tgtctcttct ttgagccaag tggtgagcaa ggtcttgttg 13020
gcatggacag gcataccatt gactgaagca gaattgacca tgacagtaag cgcacctgcc 13080
tcaagacctg cgaggaatgg agcaaaatgt ttttcacgca gttcagaaag cgaaatgtag 13140
gcgggagtgc ggtctttacc agaataaggt acgccataac ccagatagtg tttcaatgaa 13200
acagcaatat gttgttggtc aatgcggtta ggattttcgc cttgaaaacc tttgaccata 13260
gcctttgcca tttctgcatt aacataggca tcctcgccaa agttctccca gatgcgtggc 13320
caacgtgggt cgcgacctaa atctactgtg ggactataag tccaaggcac agagatggca 13380
cgtgtttcgt aggctgttgc ttctgcacaa cggcgagcga tctcacggtt gaatgtagcg 13440
gccacattga tgttctgagg gaaaagtgta ccatctgccg tgtaggtaga accgtggttt 13500
tgatcaagac caaacacaca tggaatgcca atctgcttca tggaactctt ctgaatatcc 13560
tgaatgatga cctcccactc ctttgctgta ggagcacaag tgttgggcgc attgaggata 13620
gagcctatct tgtagcgagc aaggatagag tctgtctttt gacgatcgat gtggaagaca 13680
cctttctcat cattagatcc gaagaagtct gtgaccagtt ccatcatttg cccaaccttc 13740
tcttcaagcg tgagacgaga gagggtttgt tgaacttgcg cctctatcct gtcatccctc 13800
gggatgacag gattagagac ttgagcataa actgccaaag gcagcattgt tagaatagag 13860
aaaattgctt tgaatgattt cattacatta ttgtgatctt agtgatttta tagccattgc 13920
caacaacgag gaatccgtct tgggtgctaa gaagagtcat ggaagtgctg ttgagtgtga 13980
attcgagcat ctgttcaccg ctgatagcta tatcgccacg cttgtcttca ccttcgcctg 14040
taagaatatt attccaccat gctgatgtag cagtgccctg tccatcacct tcaacataga 14100
ctctgaggac tgttccctct tgaacattct ctgccaatga ggtcttgagc acatcaaatg 14160
gagtgcccca tgtgacatcg aaagaacctt cccaaagaat cttctctttg ccaggctctg 14220
ggtttcctat cacatagcaa gtgcgagaag tggagagtcc ttgtgtggta gtagccacaa 14280
tcttgcagag atgcttgccc tcagatatag gatagacgat agagtcaccc tcgaaaacga 14340
gttcgtcgtc gatataccaa gagatggttg tatagtcacg aggagtggca atgaccttga 14400
agcgaaggct atcgccaaaa tccactgtct gaagtgtttg aggctgacca tccctccact 14460
caggaatgtc ggtgttgaga atgcgtggag catcgtcctt tgttgctgtg aaaaatgggt 14520
cgtcttcgga gcaagagaag cagagcaaag ataccatgat tatggcaaga atgctgaacc 14580
cttcgaagat tttgttttct atggtcttat taaatgtatt catgattttg atatttaata 14640
gggagtaaag gagtttaaag atggttgatg atacgagtgt ctaaggaaag aaacggtcat 14700
gagtccacct ttaactcctt ttctcatgtt tttaatgtat ttcaatcact tattcagcct 14760
attacttcac gttaatatcc gctgtatccc accagttgcg ctgatgccag tcgtctgttc 14820

cacccagacg ttcaagagcc tctttgtagt tttcgctatt gtagcgcgcc tcaaggattg 14880
ggtatttgag acgagtaggt gtagatctat cagggtcgtc ataagtgaag tctgggaatg 14940
tgggcaactt ggtgcgatcc atgagcacag gatagtccga gcgacgaagg ttagaccatg 15000
cctctgcagg attagtcatg tgaagaatcc aagcctgagt attgattgtc tctttaggat 15060
tctcgcccaa ctgattggca gcaatgaagg cagcaatttc tgcatcattg atcagatcct 15120
tctccttcaa ataatgggtg ttgagtagtt tcatagacgc tctcacacct gcttcatagt 15180
gctcatctgc actaccttcc acattccatc ccttaatctt tgcttcagca aggagaaact 15240
ctgcctcggc agaagtgaag agaatgccag gattctgccc cttctcaaag tcaatgctaa 15300
ggaacagtcg catcatgcgt gcagggaagt tgttcttgtc atagcccgcc gcggggttct 15360
ggtccactaa gagttggagg gttgggatgt tctttgtggc aggagcattg atccagttgt 15420
tgtaccaagc agcacctggg tcacaaggag actctgaaag tccattgtct ttaagatact 15480
gaaccacctc gtctgtcaag tcgaagttgc cagcctcgtc tgcacgaatc tcactacgag 15540
tagtatttat ataatgacgg caaatgcggt agaggcgtgg gtcattgctg tccttcataa 15600
tattgtagaa ggtactactc acgaaagttg gagaagtagg atcttgacca tagagaatct 15660
cactgagggc attagcacgg aagtcatatt tctctgcacc tttatagaga gtgaaaggct 15720
ggtcaagata cttgatgtaa gcatcgtcgg acgcttctgc aatatagcca ttctcacttg 15780
caagagcctt ttcgaattgt tcacgagcca gttctggttt tacatcagaa atacgcatgg 15840
caaaacgcat gcggagactg ttgcagtatt tcttccactt tgccacatca ccattcatac 15900
tggtcacatc acctgtgatt ctgtctccta ccgcattaag ttggctttcg cagagagaga 15960
gttcgtcaaa gaaccactgg tagatgtctt cctgtttgtc ataggcagga ttgttctctc 16020
ctttcatgta gccctgacca acattgaagc aaggaacatc tccataggtg tctgtgagga 16080
ctgacatcag ataaaccttg tgtatgcgta gaagtgcatt gatattaggc ttttcggacg 16140
tcttagagat agcatcttcc atattcttga tagacaccgt gtagtagcga tcccatataa 16200
gacgagtctg gtcatcgaag gcatgcacag agccagcata gttagttacg ttccaaccac 16260
ctgcaaaatg ctgggtgaat ccagtgatgt aactgcgata ggtgtccatc aggctgaagt 16320
cgccataggt ctggagaagg cttgttgtga gctgagcatt aggatcaatg ctcatagcct 16380
tactgtcatc agtgttaagg tcggacatat aggagtcaga acaactactc attgccccca 16440
cggtaagtgt cagagcaaat gaagcaccta aaccgctctt gatatatttt tgaaaacttt 16500
tcattgttgt ctatatattt aattccaaac acgatacatt gtaaaaatca aatggatgat 16560
tacatatcta tatttagaaa attaatgatc tgatgattct ttatcaaaaa atgatttaga 16620
acttaagatt gatattgaaa ccgtaactgc gacgagatgg aagtgaacca tactcaagtc 16680
cgagacctga agtgttgtag ccagagtctg ggtctatgtt aggaatgttc ttccacaaga 16740
tgaatggatt gcgagccaca aatgagagag taagtccctt tacagtcttg ccaagcatct 16800
tctctgggaa tgtgtagcca aaagtaatct cacggcactt cacataagag ttgtcataga 16860
cgaacatcgc tgggcagttt tctgtggcat acttccagta ggcttctgga ttgacaggaa 16920
tatcattctt cacatactga ccttcatcat taagatatac gccgggcaca acataaccct 16980
tcatgttgcc tgatgctctc caatcctctg atgtctgacc agcagcgaaa cgagcttctt 17040
ctgatgcgta ccatgcctca cgacctttga gagtggctgt ggcacgacct gtctgatatg 17100
ctgaacgcat agacattgag aagaggtctg caccatattt tacgtcaaaa cctgcagaga 17160
gatggaagtt cttgtaagtg aatgtgctgt agaaaccacc tgtccaatcc caagaagcat 17220
tacctattac gtgtgtgcct tcagcactta ctggcatgcc tgtgctacgg tcgatgatga 17280
cttgtccatc gtctgtgcga gcaacatcct tacccataat gctaccatag ttctgaccta 17340
ccatagcacc tacagaaaca cctgcccaag tagccttttc aagttcaaag taatccatgc 17400
cgggaacgag ttccttcact ttattgatgt tctttgagaa gttaaagccc aaatcccatg 17460
caaagtcctt atactgaacg gcacgaccat tcaccgccaa ctcaacaccg cgattgatga 17520
tgtgacctgc attgacaaga cgataaggat aaccagaagg agcagaggat gcaagattga 17580
taatctgatt gcgtgaactc tggctgtacc atgtaacatc catgccaagt cttccattga 17640
agaacttgag ctctgttcct aactcgaacg agttggtcat cgtaggcttg aggcttccgt 17700
tcggcatgtt gttgttattg atcataccaa gcatatagcc agtgtaacta tattttccgg 17760
aagaatagtt gatggcaagt tggaaaggat cagtgtcgct acctaccttt gcccaagatg 17820
cacgtaattt accataagaa aggatattct tgtttttgag atactctgaa aatatccagc 17880
ttcctgagag agaaggatat acataagtat tattgctggt tgggagagtt gatgacttgt 17940
caccacggag agtcacatcg aggtagtaag tgcggttgta gccgaaactc acactaccat 18000
agagtgagtt gatttgcttc acataagtgt ctcgattcac gttgtgttca ctgtagttca 18060

tgatgtcttt catgtcggaa atctgctgat catggccgag catgttatag gtcttgttat 18120
tgaccttata gatgttacca ccaacagtag catgaacatc aaagttgtcc caagtgtgac 18180
cgtagagacc gagaatctca gcattgagtg ttctattctg gaacacctga ttggtcattt 18240
caccagaagc acttccagga gtggtacgtg caatgaactc ctcaaagttc atgttgttga 18300
tgtcagtacc gattgtaccc tgaaccttcc agtctttgtt gatgttccat acagcctttc 18360
ctgtgaaacg gaaaacatcc ttagatgaag tgttgtcatt cttatagaga tcccagtatg 18420
ggttcttatt atacatgtca ttaccgttcc agttggcata agtgccatca ggattctggt 18480
aatagcgaag ccagtcctga ttgtaagtac cagccagtgt catgaggttc ttacctacat 18540
tactctgaga gtcaccaagg gcaggacgat tctttacgtc ctcacgagta tagttggcag 18600
tgaagtcaaa atcgaatgca ccagatgatg tgttggcacg cagattgaag ttatcacggc 18660
tcatgtgagt attaggtagg atgtccttat tacgcatgtc ggtcatagaa aaacggatac 18720
ctgtcttgcc agaactgaaa gaaaggatac ctgtgttttg agcagtgaca cccgtacgga 18780
agaagtttga tgtgttgtta ggatacataa ggaatggacg ctccacacca tcgaaataac 18840
gcaccatata gttgtccgcc ttaggtcccc aactcatgtt cgtacctgat gtggcagttt 18900
caagatactg tccgccatta cccataccat agacctgctg gatgtcattc cactttgcca 18960
actgagagtc gaaagtgata gaaccattgt attccacact caccttatct ttctcagcct 19020
tctttgtagt gataagaatc acaccatgag aggcacgact accatagagt gcagaagcag 19080
caggaccttt gagcacactc atagtctcta tgtcatcagg attgatagca gagataccat 19140
caccaagatc gtggccacct gctgtggtca cactaccgaa attagtattg tcaagcggaa 19200
caccgtcaat cacatagaga ggctgattat caccgcccat ctctgtgtta ccacgcaaaa 19260
gtacacgagt agagcctgat gaaccaccag ccgtgttctg cacaaccaaa cctgccactt 19320
tacctgcaag agagttgatc acatttgttt cctttgcctt agtgagttct tcgcccttca 19380
cctcctcaag tgcatagcca agagctttct tgtctcgctt aatgccgagg gctgttacaa 19440
ccacgtcacc aagcaccttg acgttttctt gaagtgtcac cttcatgcca ttagatgctt 19500
tcaccgtaac aggattataa cccataaagc tgatttgcag ggttgtacca ggctcgcatt 19560
tgagggagaa atgaccattc atgtcggtca aagcaccgcc tgtacctccc acaattttaa 19620
ctgtggctcc aatcaatgct tcacctgaag catcaaccac tgaacctttg atttcttgat 19680
cactctgctg agcagtaagt gttgctttca cagaagagga cacagcatag gttgttgcag 19740
gtgtcccgat catgcctgtg caaagcagac ttaaaagaat cgttttattc atcattattt 19800
atattggtaa tataggttga tatgataatt tatcaatttt tctttcaatt attctacggc 19860
tttaaggcca gccttgatac cttccatcac ccaaggcgct ttaatctgca agccatgcgt 19920
acggtcgata atgccaaagc attctgcacc attgccaaag tggttattgt cccaaatgaa 19980
agtggcaaac cctctcttct tggcctctgc cacataaatc tggcaatagt aagccatgtt 20040
ttcacgcatc aagtctgccg aagaagactt gtagtggtct gtcacccccc actctcctat 20100
cacgacacca agacccttaa aagcccaagt ctgctgaact ttggtgaagg tatcaatgat 20160
tgtctgctca ttctctggag cggcagtgcc atgcgcacga tagcgttctc cccaatactc 20220
cagttttcca agaccgcaat aatcccaagg ctgatagtaa tgaaactcca cactcatata 20280
acgattaccg ttctcttcca aatcagtagg aatgatgaag tctccattgt tgagtccaaa 20340
ttcagggtta catacatagg tctgcaccat aagatggcgc tgggcattat tgcctcctgt 20400
agcgcgcaca atatctacaa atgtctgatt ataggcattc tgcacttcaa ggttctctgc 20460
tgtaggctgt ccccagttgt ctctctcatg cacttcgttc gttccagcaa aagccaaacg 20520
ataatcatag tccttaaatt cagaagcgat attcatccag agcaaagcaa gcttttggca 20580
gttttcctct ttatatttat aagtaggacg actctcaagc catttctcat ggtgagtatt 20640
gatgatgacc ttcatgtcac acttcaagca gtaatcaact atttccttga tgcgattcat 20700
ccaagtccta ctgatggtca tcgcttgtgc atttgtgatg tgacattgcc aacggacagg 20760
aatacgaata gcatcaaaac ctgcggcctt cacagcatct atcagtttct tgcttaccac 20820
aggattgccc cacgacgtct ccgcattgtc tgctccagca ggattgccta taagcaaaga 20880
ttctccattc attccaggag cagaacattc aaatgaattg cccaagttcc aaccaattgt 20940
aacttctcta ttccattcct gagcagtggg agcctttatc tccttctcgt tcttctgggg 21000
taattctgcc tctggctgat gagcttcaca tgctgtagtg aatagcatgc caagcaaaag 21060
tgtgagtgat gttacgattt gtctgttagt tttcatcatt gaggtaatat gaattttatt 21120
gttaattgaa atcattgggg tattcaatta taggtaaccc tgttttagat gacgtgcatt 21180
ttagatttcg gtagcaaatt tattggtttt ttcaacgacc aaggttcgtt ttttatcgaa 21240
aaaggtttca aaactaacat ctgtcaattt tgaaaccttt tcctgtaatc tatcaacaca 21300

```
     ttgatttatt gatacttgct gggctgaaca ccaaagtgtc gcttaaaaac agtgctgaaa     21360
     tacttcacat ttgcaaatcc tgttttgatt gaaacatcaa gcacaggata gccctgcttg     21420
     aggtagatag ccgcctgctc aaggcggata gtacgcacaa aatcctgtgg tgtttgtccc     21480
     gtaaggcttt tcaa                                                       21494
//
ID   XXX; XXX; linear; XXX; XXX; XXX;  XXX.
XX
AC   PRJEB67889
XX
DE   Uncultured bacterium DNA fragment from in-reactor enriched cow rumen, clone
ferm-10d6
XX
FH   Key             Location/Qualifiers
FH
FT   source          1..21977
FT                   /organism="uncultured bacterium"
FT                   /environmental_sample
FT                   /mol_type="genomic DNA"
FT                   /isolation_source="Cow rumen"
FT                   /clone="ferm-10d6"
FT                   /note="AZO-CM cellulose, AZO Galactomannan active clone"
FT   CDS             104..265
FT                   /db_xref="SEED:fig|6666666.127227.peg.1"
FT                   /translation="MAEGCSTVVPSCAICQVRASPASTVTDSFPLGDASWYCCARTVPK
FT                   QRNNKQKV"
FT                   /product="hypothetical protein"
FT   CDS             complement(289..1293)
FT                   /db_xref="SEED:fig|6666666.127227.peg.2"
FT                   /translation="MINPIYSPAPERYDNGMKFRRCGQSGILLPEISLGLWHNFGDMNT
FT                   FANSQAMAHYAFDRGITHFDLANNYGPSYGSAEETFGMIMKKSFMPYRDELFISTKAGH
FT                   DMWPGPYGEWGSRKYLMSSLNQSLKRMNLEYVDIFYSHRYDPDTPLEETLQALVDIVRQ
FT                   GKALYVGISKYPKEAAEFAYKYLKERDVHCLLYQGRYNLFNREPEEEGILQQAKGNGTG
FT                   FIAFSPLAQGLLTNRYLNGIPEDSRMAKGAFLKKEALTDETLQKIKALNEVAASRGQTL
FT                   AEMSLAWLLKDDLVTSVIIGASSVAQLADNLKATENTDFSAEELEKIKKIIQK"
FT                   /product="Putative ion-channel protein"
FT   CDS             complement(1312..3798)
FT                   /db_xref="SEED:fig|6666666.127227.peg.3"
FT                   /translation="MQDKRFGYFDDDNREYVITDPKTPWPWINYLGNEDFFSLISNTAG
FT                   GYSFYKDAKFRRITRYRYNNVPMDNGGRYFYINEGGNVWSPTWKPCKTALDSYECRHGM
FT                   SYTRITGSKDGIEASLLFFVPLKTWGEVQKLTLRNTSAKVRKLKLFSFAEWCLWNAATD
FT                   MENFQRNFSTGEVEVDGSVIYHKTEYKERRNHYAFYSVNTPVDGFDTDRETFVGLYNEF
FT                   ADPERVVEGRPGNSIAHGWSPIASHYLEVELQPGESRDFIFLLGYVENKQEAKFEEREE
FT                   SLHEAFKQSVPSSPIINKVKAKAMISAFDTTAKVDAAFAELKAYWDRLLDIYVVKTDEE
FT                   KLDRMVNIWNQYQCMITFNMSRSASFFESGIGRGMGFRDSNQDLVGFVHQIPERARERI
FT                   IDIASTQFPDGGCYHQYQPLTKRGNNDIGGGFNDDPMWLIFGTVAYIKESGDFSILDEP
FT                   VPFDNKEGSEVSLFEHLRVSFNHVIENLGPHMLPLIGRADWNDCLNLNCFSWDPNESFQ
FT                   TTENQTEGSQAESLMIAGLFVVCGRDYVELCRRLGKDDEANRAQTYIDNMVEAVKKHGW
FT                   DGDWYLRAYDYFGHKVGSKENEEGQIFIESQGWCTMAGIGLEEGMVKKALDSVKERLDC
FT                   EHGIVLNNPPFTRYVVEYGEISTYPAGYKENAGIFCHNNPWVIIGETVLGRGDRSWEYF
FT                   RKICPSYTEEHSALHKVEPYVCCQMVAGKDAARPGEGKNSWLTGTAAWMWYAITQFILG
FT                   IKPSYEGLEINPCIPAGWKGFNVKRRFRGAEYHITVKNPDGVCKGIKSVTVDGQPIEGN
FT                   VVNHLPGTHTVEVIMG"
FT                   /product="GH94"
FT   CDS             4022..6466
```

```
FT                   /db_xref="SEED:fig|6666666.127227.peg.4"
FT                   /db_xref="GO:0004565"
FT                   /translation="MKHLFAFLFLLGITFSLSAQSVRERILLDDGWQFAFGNASSPEKD
FT                   FGCGTEYFNYLTKAASIHNEGPYSPKFNPEKWGVDWKTVNLPHDWVVDLPYAKEASHSH
FT                   GYKAVGYKFPENSVGWYRKSITVPQEDLGKHLYLQFDGIFRDARIWINGFYLGHEPSGY
FT                   ATQVYDITEYLNYGEENLICVRADATLEEGWFYEGAGIYRHVWLNKTAPLHVAPFGVFV
FT                   SSTLEAPFDKAVLVTEVQMKNSGKDPVSGKVWCRLLDADGQVQMEDEAEEITLLPQETR
FT                   SVKVRTKVLNPQLWSTEHPYLYKMETSVWQNGRQVDVVATPIGIRDIRFDADKGFLLNG
FT                   QPLKLKGVNMHQDHPGVGAGIPDALQVYRLKELKKFGCNACRSSHNPMTPEMLDACDSL
FT                   GILVIEENRLTGVNEEHTRLLKRMIDRDRNHPCIILWSVGNEEWGIEWKESGTRIAATM
FT                   REYCHRFDPTRLMTVASSSGPAILIPADVAGYNYILQNPVEQHRKDYPGRRALGSEETT
FT                   GCGTRGIYFDAHGKGHMVAHNRKPNGPDSLLNCIERGWKFYDERPYLAGLFYWTGFDYR
FT                   GEPNPMKFPATGSQFGILDYCGFPKDEAWYLKSWWTNEPVLHILPHWNLQGHEGDSIDI
FT                   WVYSNCDEVELTVNGKKLDRKPMPRNGHLSWKAVFQPGAVKAVGYKNGKKILARTVETT
FT                   GAPARISLTADRPVIQADNRDVAVCNIELQDKKKRFVPTACNDLTITVSGPVRILGVGN
FT                   GDPAYQATERPADADARTYQVKAFNGLAQVLLQSTGEAGEATLTVVSENLPETSCVLKL
FT                   QK"
FT                   /product="GH2"
FT                   /EC_number="3.2.1.23"
FT   CDS             6792..7658
FT                   /db_xref="SEED:fig|6666666.127227.peg.5"
FT                   /translation="MREITPLSERDCFYIADRRKTEFTYPIHCHAEYELNFTEHASGVR
FT                   RVVGDSAEIIGDYDLVLITGKELEHVWEQHECTSGDIREITIQFSPDLFFGNVVNKNQF
FT                   DSIRRMLERAQCGLSFPMQAIMKVYNWLDKLASEEQGFYAVMNFLRILYELSLYDNARV
FT                   LSSSSFAKIETFSDSRRVQKVQKYISTHYQEDIRLASLADMVGMTPVSFSRFFRLRTGK
FT                   TLSDYIIDIRLGFATRMLVDSTRTIAEVCYDCGFNNLSNFNRMFKRKKGCSPKEFRENY
FT                   RKKKLVI"
FT                   /product="Transcriptional regulator of various polyols
FT                   utilization, AraC family"
FT   CDS             7779..9014
FT                   /db_xref="SEED:fig|6666666.127227.peg.6"
FT                   /translation="MTAQIRGNEIRVVVSPDHSDWVYRLNEKCTFTVRVLKAQNLLSDV
FT                   KIDYELGPEMYPTEVKKDVVLKDGTLKLQGTMKTAGFLRCKVKAHVDGRTYEGLATSAY
FT                   APEQLQPVTKLPADFRDYWAKTLEEARKTPLNPLMTLLPERCTETDNVYQVSFQTKAWG
FT                   GRFYGILSIPKKEGKYPALLRVPGAGVRPYAGDTYTAPGKVITLEVGIHGIPVTMQQSV
FT                   YDALAGGALSNYWTFGRDSRDASYYNRVVVGALRAVDFICSLPQYNGKALGVTGSSQGG
FT                   ALSVITAALDSRVTFLAAVHPALCDHEAFFKKRACGWPHYFYYYGAPDEKQLETVRYYD
FT                   TANFARLLNVPGWFSWGYNDEVCPPTSMYAAYNVISSPKELHPYLETGHYWYQEQWDEW
FT                   LDWIRRQLNVPM"
FT                   /product="CE7"
FT   CDS             9106..10398
FT                   /db_xref="SEED:fig|6666666.127227.peg.7"
FT                   /translation="MKKQFLLLTVLLFLLGACAPKPAEHSFIKVNADGQFVRDGKPYYF
FT                   VGTNFWYGAILGSEGEGGNRERLHKELDFLKSIGINNLRVLVGADGENGIKTRVEPSLQ
FT                   VAPGVYNDTILAGLDYFMNELRERDMTAVLYLNNSWEWSGGYSVYLQWSGHGDAVVPAV
FT                   DGWPAYMEYVKQFPQSDSAKALFANHVNYIVSRTNRYNQIKYVDDPTIMSWQIGNEPRA
FT                   FSDENKEPFARWMADVAAQIKSLDPNHMVSSGSEGSWGCEMDMNLFEKIHADSNINYLN
FT                   IHIWPYNWSWVKADSLKELLPRAKENTKKYIDDHMVIARKYSKPIVLEEFGFPRDGFSF
FT                   SKEAPTTARDEYYRYVFDLIRQDRESGGLFAGCNFWAWGGFAGQNPDHVFWEKGDDYTG
FT                   DPAQEQQGLNSVFATDSTIEIIKAENRKLQN"
FT                   /product="GH5_7"
FT   CDS             10556..11482
FT                   /db_xref="SEED:fig|6666666.127227.peg.8"
FT                   /translation="MTIGCGTNPSKNENVKETLPALVVNGTQLMNTEGDTVVLHGVSYG
```

```
FT                   WHQFWPRFYNASSVAYLVNDWGAQVLRASMGVDLDSACYVNKPEFGIECVTKVVDAAIE
FT                   NGVYVIIDWHSHNLRQEEAKEFFTQMATRYKGVPNVIYEVFNEPVEDSWEQVKSYSVEV
FT                   IKTIRAIEPDAVILVGCPHWDQDIHLAADDPITGYSNIMYTLHFYANTHGQWLRDRADY
FT                   ALGKGLPIFVSECAGMEASGDGPISKEEWGNWLDWMQRHSISWVAWSLSDKDETCSMFY
FT                   PSASSEGNWKDADMKEWGRMVRDELKK"
FT                   /product="GH5_2"
FT                   /EC_number="3.2.1.4"
FT   CDS             11546..11659
FT                   /db_xref="SEED:fig|6666666.127227.peg.9"
FT                   /translation="MGKKEKRNYEDMDCLEVNLNVKAVIVYQGFLTQSHFC"
FT                   /product="hypothetical protein"
FT   CDS             11717..14983
FT                   /db_xref="SEED:fig|6666666.127227.peg.10"
FT                   /translation="MNLNFRKTALLAGICSVFSLGYAPQLFAVSSTDTVEAVQQTKKIT
FT                   GVVSDAMGPIIGANVLEKGTTNGVITDIDGNFALNVKPGATIVVSFIGYVSQEIKITNQ
FT                   TTVNITLKEDSEMLDEVVVVGYGTMKKSDLSGASVSMSEDKIKGSIITNLDQSLQGRAA
FT                   GVTAVSTSGAPGSSSSIRVRGQATINANAEPLYVIDGVIVQSQGQSGSSYGLGDALGNG
FT                   SVSTISPLSTINPADIVSMEILKDASATAIYGAQGSNGVVLITTKRGKAGEAKFTYDGM
FT                   FAVQRQTKRLDMMNLREYATYYNDLANMGEISDPSNYYADPSLLGVGTNWQDAIFQTAL
FT                   QHQHQISAQGGTEKIQYYVSASYMDQDGTIIGSNFNRYSFRANLDAQLKKWLKLGLTAT
FT                   YTATGDDLKLADSDEGLINYSLTTLPDIPIYNLDGSYATVVREGYTNPNPIALAMMDDI
FT                   LLDRQKLTGNIFFEVTPIKNLVWHAELGYDISSSKGERYKPMVDLGGWVRSSNESNIQK
FT                   NSSTFWQLKNYVTYSGNIGDKHHYTAMVGQECWESSYDFTSVFNSGLPSDEVHNPALGT
FT                   GTPTINAGFGSSSMASFFTRLTYNYADRYLGTYTYRYDGSSNFGPENRWAGFHSLATSW
FT                   RFSNETFFEPLKKVISNGKLRLGWGQTGNANIGGYAWGTSMSRMPSGLGMGYRPSNIPN
FT                   TSIKWESQEQWNVGVDLGFLHDRINLTVEWYKKTSSDMLMPLQLPSYMGTSGNASSALA
FT                   APKGNYGDIENTGFEISLNTHPITGKFEWDSDFQISFNKNKLKGLAGTANAQIVGYGQW
FT                   TDVVSVSSVGESLYNFYGFVCDGVYKDLEDLQTSPKPAKYPANGVFDRTNTVWVGDVKY
FT                   KDISGPDGVPDGVIDDYDKTNIGSPMPKFTFGWTNTFRYKNFDLSVFINGSYGNKVYNY
FT                   LSMKLTHMNSPWTNQLNAVNGRAQLVPINPNKVYEDGTMWYNDVSNVMVANPGASIPRA
FT                   TINDPNDNDRISDRYIEDGSYIRLKNISLGYTFPKKTIDKLGLSNLRIYANIQNLLTIT
FT                   GYDGYDPEIGASTSSVNVYGLDNGRYPSPTVYSFGLNVSF"
FT                   /product="TonB family protein / TonB-dependent receptor"
FT   CDS             15004..16725
FT                   /db_xref="SEED:fig|6666666.127227.peg.11"
FT                   /translation="MKIRYIKYWAFAAVLGMGTVSCEDFLNRPTEDNYNESNFYQNDEQ
FT                   CVQGVNYLYNSPWYDFQRGFIKVGEILSGNVYEGNSPYLNFTVNGTDENLVNMSYSLWA
FT                   VIGHANTVYTRLKTANASEAVKNQCMGECLAWKAMAYFYLVRSFGAVPIIHDNTDELNS
FT                   GNYNSKFKVEAADVYEYIIMTLEKAMELLPARGETGRIDRYCAEGLLAKVYLTKSGLSG
FT                   TRNADDLAKAAEYSKDVINNSGRNLLANYSDVFRLANNKNEECLFSWHWSAGRDPWTQQ
FT                   NTLQSDLAMVGFDEFGDCWGGYAGPSVDLQDAFGISALESPETRSDTDTRRKATMMMAG
FT                   DVYDYFWQDKGGFDYLRFIYDAEYGKGGPNGDYQSPTGANHVKHLYGNNNDHVLGLGVS
FT                   AGNMYSGLATHILRLSDIYLVYAEAKMGLATSTTDQSALDAFNAVRGRAIPGVTPKTSI
FT                   TWEDVWKERRLELACEGDRWYDYVRLAYYDSQRAINELKAQRRDVYYSLGTTYKAYYEN
FT                   GSWTVNPDETRYNPDAKAPNVTVSSFTLPFPTEDVVFNPNLMKDPVHVDVRSEFSY"
FT                   /product="putative outer membrane protein, probably
FT                   involved in nutrient binding"
FT   CDS             16730..18043
FT                   /db_xref="SEED:fig|6666666.127227.peg.12"
FT                   /translation="MLEKMKNNRYTNWFFAAAVVVAGFSMAACEDEPDKYEVAGGKPTL
FT                   KYVRVPDPAAADSLLTGAYMANTICLVGENLRSVYELYFNDQKAILNTSYITDHTLMVD
FT                   VPKEIPSVVTDKIYMVTKAKDTIDYDFKVLVPAPTVNSISCEFAKPGSEVTLIGDYFID
FT                   DPNVPLTITMAGNVEVTNITNITKTAVSFILPDNAPAGYINVKSIYGTGRSKFRYYDTR
FT                   NILFDWDGSHGGMAIAHGWRDGSKVLREADENSIDGAYICLTGALDGAIGATWAEDEFS
```

```
FT                   FNYWPEPSAGYPELSARPEFAELLEEYGFNGLQLKFEVNIPSSNPWQSCALQVMFTGND
FT                   VVTYATGTNAYFSDTNVPRGLWLPWKNTGSYDTGGKWTTVSMNLSEFNKTHEGNKCDRT
FT                   FDKTMLTGLTFFVWAGGVEGKDCNPVIMIDNIRVVPIE"
FT                   /product="FIG00417257: hypothetical protein"
FT   CDS             18065..20050
FT                   /db_xref="SEED:fig|6666666.127227.peg.13"
FT                   /translation="MNKTYKFPALWMMCLMLFSCLTFTACDNGDDEDTNQYKGGISLNV
FT                   FGPSPVSRGGVLRFLGSGMDKVTAVAIPGCDDITDIEVVSDTEIRVTVPQTAQPGLVVL
FT                   KTPKGDITTKTELTFTEPIALEAFAPAEVKPGSELTITGEYLNLIKEVIFADEVTVPAD
FT                   EFVSQSRQEIKVIVPDSAQTGKFILSDGAEIPNWIYSEGELEVTLPSVEAPLDLVDKKP
FT                   GDVIRVSGENFDLVKKVQMPNGDEVEFTMTASSEGDELTFTLPDNVSDGEVTVLPASDV
FT                   KVVVATVVVATPSNVVAVPAVNLRGGDMITLKGTNMDLVTDVTFPGVEEAVGLESQNST
FT                   EIKVLMPAAAISGDLQLNTNSGKATAVSIATAKPENISYSAATVPAGEALTVKGVNMDV
FT                   VSAVVFSGNVEVTVSDATATAISLTVPTTAETGALLLKMANGESVEAPSLTIEKPVCAY
FT                   LPALPDKLVRGRIVELEIVNADKLTNVLLNEAAVQYINDAAKGVLMLNVPAELDGTYSL
FT                   KLISSNGEIAYDVLVVANEETVWAGPLDISWGDGGRVLVPAVSFAKVTAGTVMKVYFDQ
FT                   KDQTWAQAQFNYGDWSGIAFSLFDTTMVPTDIYGWSFESRVMELTLAQEILDNIQAKQG
FT                   DCEDQTNVGIIIQGSDLTFTKITIVN"
FT                   /product="FIG00410329: hypothetical protein"
FT   CDS             20118..21815
FT                   /db_xref="SEED:fig|6666666.127227.peg.14"
FT                   /translation="MKKLNIKWMLSLIAAFTFASCDTDVDHDIPAVDAPVLVSTTPESG
FT                   AAKVKTGEITIEVKYDKNIFFATDNLSEIKFTGGELISADVLGASNILTVKVNVPGRET
FT                   ACSLSIPEGIVTGPNQMPAPAVSVQFSTVALDKALVAASSAKAVKLYNYLLDNFETKTL
FT                   SAMMANVAWNTEMSEKVYGWTGKYPAINCFDYVHLPASVAGADWINYGDITPVKDWSDK
FT                   GGIVAAMWHWNVPKKAVGEASSTRIWEGETVMPGDWSGNVQMTDDAAKTVFADAQVGQV
FT                   IRVAVKDVAEGAQGSFKNSGWSEIASGTDYFDISGDYTLVITEDILKSLQEGGLIIGGH
FT                   DYTAVAVYLESNGTALDPNKDYAFYKADTEFDAANATVEGTWENKVFTEDLKNAAAYLK
FT                   LLRDADIPVLWRPFHEAAGGWFWWGKDAASFKSLWIAMFNYFKTEGLDNLIWVWTTEGN
FT                   DSDWYPGDQYVDIVGRDVYNKETADCVSEYTSIAGNYGNKIVSLSECGTVGLISEQWAS
FT                   GARWSWFMPWYDGTNEDGSPAVHADEAWWKDAMSQEFVVSREELPSME"
FT                   /product="GH26"
FT                   /EC_number="3.2.1.78"
XX
SQ   Sequence 21977 BP; 5386 A; 4823 C; 5844 G; 5924 T; 0 other;
     gaaggaggac tccacggact tgcgttctgc acggattact tttacatttt taccgaaagt        60
     tattgctttc ttggctgatt tttcatctct gccttgcagg gcgatggcag agggttgcag       120
     caccgtggtt ccgtcgtgtg ctatctgcca ggtgagggct tctccggcat ctaccgtaac       180
     ggatagtttt ccgttgggtg atgcaagttg gtattgctgt gcacggaccg tgcccaagca       240
     aaggaataat aaacaaaagg tttgaataag gaggcttttc atgttgattt atttttgaat       300
     gattttcttt atcttctcca gttcctcggc agagaagtct gtattttcag tcgctttcaa       360
     gttatctgcc agttgggcaa cggagctcgc tccgatgatg acgcttgtca ccaggtcgtc       420
     cttcaacagc caggcaagtg acatctcggc aagggtttgc ccgcgtgagg cagcaacttc       480
     gttgagtgcc ttgatttttt gcagcgtctc gtcggtcagt gcttctttct tcaggaaagc       540
     ccccttttgcc atgcgcgaat cttcgggaat accattcagg tagcggttgg tcagcaggcc       600
     ctgcgccagc ggggagaagg cgatgaagcc ggttccgttc cctttggctt gctgcaggat       660
     accttcttct tccggttcgc ggttgaagag attgtatctg ccttgataga gcaggcagtg       720
     tacatcccgt tctttcaggt atttgtaggc aaactcggcc gcctctttgg gatatttgga       780
     gattcctaca tagagtgctt tgccttgccg tacgatatca accagtgctt gcagggtctc       840
     ttcaagcgga gtatccgggt cgtagcggtg ggagtagaag atgtccacat actccaggtt       900
     catccgtttc agactttggt tgaggctgct catcaggtac ttgcgtgagc cccactcccc       960
     gtaggggccg ggccacatgt cgtgtccggc cttggtggag atgaacagtt cgtcccgata      1020
     gggcatgaag gatttcttca taatcattcc gaaagtctct tcggcggaac cgtaggaagg      1080
     accgtaattg ttggcgaggt cgaaatgcgt gattccacgg tcgaaggcat agtgcgccat      1140
```

ggcctgggaa ttggcaaatg tgttcatgtc gccgaagttg tgccacaatc cgagtgatat 1200
ttccggcagc aagatgccgc tttgcccgca acggcggaat ttcatgccgt tgtcatagcg 1260
ttcgggcgct ggagaataaa tagggtttat catgatattg tattttgtgg gttaccccat 1320
tattacttct actgtatgcg tgcccggcag atggttcacc acattgcctt ctatcggctg 1380
gccgtctact gtgacggact tgatgccttt gcacacacca tcgggattct tcacggtaat 1440
atgatattct gccccgcgga acctacgctt cacgttgaag cctttccagc ctgccgggat 1500
acaaggattg atttccagac cttcatacga aggcttgatg ccgagtatga attgcgtgat 1560
ggcataccac atccaggcag ccgttcccgt cagccagctg ttcttgcctt caccggggcg 1620
ggcggcgtcc ttgccggcca ccatctggca gcatacgtaa ggttctactt tatgcagggc 1680
actgtgctct tcggtgtacg acgggcatat cttgcggaaa tactcccagg aacggtcacc 1740
ccgtcccagt acggtttctc cgatgatgac ccacgggttg ttgtggcaga agatacctgc 1800
gttctccttg tagccggcgg gataggtgga gatttctcca tactccacaa catagcgggt 1860
gaagggtgga ttgttgagta caataccgtg ctcgcagtcc aggcgttcct ttacggagtc 1920
cagcgctttt tttaccatgc cttcttccag tccgatgcct gccatggtgc accagccttg 1980
cgattcgatg aagatttgtc cttcttcgtt ctctttggag cctaccttgt ggccgaagta 2040
atcgtaggcg cgcagatacc agtctccgtc ccagccgtgt tttttcacgg cttccaccat 2100
attgtctatg taggtttgcg ctctgtttgc ttcgtcgtct ttacccaggc ggcggcacag 2160
ctccacgtag tcgcgtccgc ataccacgaa caagccggca atcatcagcg attctgcctg 2220
gctgccttcc gtctggtttt ccgtggtctg gaaggattca ttcggatccc aggagaagca 2280
gttcaggttc aggcagtcgt tccagtcggc acggccgatg aggggcaaca tgtgggggcc 2340
gaggttctcg atgacgtggt tgaaggagac gcgcaggtgc tcgaacagag acacttccga 2400
accttctttg ttgtcgaagg ggacgggctc atccaggata ctgaagtctc ccgattcctt 2460
gatgtatgcc accgtaccga agataagcca catggggtca tcgttgaagc cgccgccgat 2520
gtcgttgttg ccgcgtttgg tcagcggttg gtattggtgg tagcatcctc cgtcggggaa 2580
ttgggtagaa gctatgtcga tgattcgttc gcgggctcgt tccggtatct gatgcacgaa 2640
gcccacgagg tcctggttgg agtcgcggaa acccattcca cgaccgatgc cgctctcgaa 2700
gaaagaggcg gaacgggaca tgttgaaggt aatcatgcac tgatactggt tccagatgtt 2760
caccatgcgg tccaactttt cttcgtcggt cttcacgacg tagatgtcca gcaggcggtc 2820
ccagtaggct ttcagctcgg caaaggcagc atctactttg gcggtagtgt cgaaggccga 2880
aatcatcgcc ttagccttta ccttatttat aataggtgaa gagggtacac tctgcttgaa 2940
cgcttcatgc agagattctt cacgttcctc gaactttgcc tcctgcttgt tttccacata 3000
gcccagcagg aagatgaagt cgcggctttc gccgggttgc agttctactt ccagatagtg 3060
cgaggcgatg ggactccaac cgtgggcaat gctgttgccc ggtctgcctt ctactacccg 3120
ttccgggtcg gcaaactcgt tgtagagtcc cacgaatgtt tcgcggtcgg tatcgaaacc 3180
gtctacgggt gtattgacac tgtagaaggc gtagtgattg cggcgttcct tgtattccgt 3240
cttgtggtag atgacagacc cgtccacttc cacctctccc gtggagaagt tacgctggaa 3300
gttctccatg tccgtagcgg cattccacaa gcaccattcg gcaaaggaga atagtttgag 3360
ttttctcact ttggcagaag tattccgcaa ggtcagcttc tgcacctctc cccaggtctt 3420
cagtgggacg aagaagagca ggctggcttc gatgccgtct ttggatccgg tgatgcgggt 3480
gtagctcatg ccgtggcgac actcgtaact gtcgagtgca gtcttgcagg gcttccaggt 3540
gggagaccat acgttgccgc cttcattgat gtagaagtag cgtcccccgt tgtccatcgg 3600
cacgttgttg tagcggtagc gggtgatgcg gcggaacttg gcgtccttgt agaaggaata 3660
gcctccggca gtattggaga tgagtgagaa gaaatcttca ttgcccaggt agttaatcca 3720
tggccaaggt gtcttggggt cggtaatgac gtattcgcgg ttgtcgtcat cgaaatatcc 3780
gaatcttttg tcttgcatat tatttcttgt atttagtttg atacattctt cattagttgt 3840
gccatccggc cattttcccg gaagcacgct tcaaatttaa aaacttttct cctatcaccc 3900
ttgtttgccc ggcgtgtttt tttcccctcg gaaacatgaa gtaccggaaa atgatacaaa 3960
agtatcaccg cacgaatctt tttcgctcta tctttgtaaa cacattttat gattaaccga 4020
tatgaaacac ttattcgcat ttctgttcct tcttggaatt accttttctc tttctgccca 4080
atccgtccgc gaacgtattt tgctggacga cggctggcag tttgcttttg gcaatgcttc 4140
ctcaccggag aaagacttcg gctgcggaac ggagtacttc aactatctga ccaaggcagc 4200
ctctatccat aacgagggtc cttactcccc caagttcaat ccggagaagt ggggcgtgga 4260
ctggaagacg gtcaaccttc cgcatgactg ggtggttgat ttgccttatg ccaaagaggc 4320
gagccacagt catggttata aagccgtcgg ctataaattc cccgaaaact ctgtgggatg 4380

gtatcggaaa agtatcactg ttccgcaaga ggacctcggc aagcatcttt atcttcagtt 4440
cgacggtatc ttccgtgatg cccgtatctg gataaacggt ttctacctgg gacacgaacc 4500
cagcggctat gccacgcagg tatacgacat tacggaatac ctgaattacg gagaagagaa 4560
tcttatctgc gtccgtgcgg atgccacgct cgaggagggc tggttttacg agggtgcggg 4620
catctaccgc catgtatggc tgaacaagac ggcgccgttg catgtagctc ctttcggcgt 4680
atttgtatct tccacgctgg aagctccttt cgacaaggct gtgctggtga ccgaagtgca 4740
gatgaagaat tcgggaaaag accccgtctc cggcaaagta tggtgccgtc tgcttgatgc 4800
cgacggacaa gtgcagatgg aggacgaggc ggaagagata accctgctgc cgcaagaaac 4860
gcgtagcgta aaggtgcgta ccaaagtgct gaatccccag ctatggagta cagaacatcc 4920
ttacctttat aaaatggaaa ccagcgtctg gcaaaacggt aggcaggtgg acgtggtggc 4980
caccccgatt ggtattcgcg acatccgctt cgatgccgac aagggcttcc tcctgaacgg 5040
acagcctctg aaactgaaag gtgtgaacat gcaccaagac caccccggag taggtgccgg 5100
cattcccgat gccctgcaag tgtaccgact gaaggaactg aagaagttcg gctgcaatgc 5160
ctgccgttct tcccataacc cgatgactcc cgaaatgctc gatgcctgcg acagcctcgg 5220
cattcttgtg atagaggaaa accgtctgac gggcgtcaac gaggaacata cccgcctgct 5280
gaaacgtatg atagaccgcg accgcaacca tccgtgcatc atcctttgga gcgtaggcaa 5340
cgaggagtgg ggcatcgagt ggaaagaaag cggtacacgc attgctgcca cgatgcgcga 5400
atattgccac cgtttcgacc ctacgcgcct catgaccgtt gcttcgagca gtggccctgc 5460
catcttgatt cccgcagacg tggccggtta taactacatc cttcagaacc ctgtggagca 5520
gcaccgcaag gattatcccg gtcgccgggc gctcggttcg gaagagacca cgggctgtgg 5580
tacgcgcggc atctactttg atgcccacgg caaaggccac atggtggcgc acaaccgtaa 5640
accgaacggg cctgacagcc tgctgaactg catcgagcgt ggctggaaat tctatgacga 5700
acgcccctat ctggcaggtc ttttctattg gacgggtttc gactaccgcg gcgagcccaa 5760
ccccatgaag ttcccggcta cgggctcaca attcggcatt ctggactact gtggcttccc 5820
caaggacgaa gcctggtatc tgaagtcctg gtggacgaat gagccggtgc tgcacatcct 5880
tcctcactgg aatctgcaag gtcacgaagg tgatagcatc gacatttggg tatacagcaa 5940
ctgcgatgaa gtggaactga ctgtgaacgg caagaaactg gatcgcaaac cgatgccccg 6000
aaacggccat ctttcctgga aagccgtttt ccagcccggt gcagtgaagg ctgtaggcta 6060
taagaacggc aaaaaaatat tggcccgtac tgtggagact accggtgctc ctgcccgtat 6120
ctccctgacg gcagaccgtc ccgtcattca ggcagacaat cgggatgtgg cagtgtgtaa 6180
tattgagttg caggacaaaa agaaacgttt tgtccccact gcctgcaacg acttgacgat 6240
aaccgtttcc ggtccggtac gcatccttgg ggtggggaat ggagaccccg cttatcaggc 6300
tacggaacgt ccggccgacg ccgatgcccg cacctatcag gtaaaggcat tcaacggact 6360
tgcacaggtg ctgttgcaaa gtaccggcga agcgggagaa gccacactga cggtcgtttc 6420
ggaaaacctg ccggaaactt cctgtgtctt gaagttgcag aagtaagtgg cttatcattt 6480
atatcatccg gcgaagcgga gacctttctc ttgtctatgg ggcaagagag ggatctccgc 6540
ttctttcttc ctcaaactat agtgtgtaag tgtacgagat gcgcatctca tacatcctta 6600
cattagagcc tgagggggag aagaacggaa gttttgcttt tccgggtggg actttcccgg 6660
gtatctgctc atatataaca attattacct ctaaatgtaa caaatggtac tgttgtggga 6720
ttaaaaagaa tgagtatatt tgtggcatga tattttttta accaatgcct atggaaaacc 6780
agaactatat catgagggag ataacccctc tgtcggaacg tgactgtttc tatattgccg 6840
atcgtcgcaa aacagaattt acctacccca tccattgtca cgcagagtat gaactgaact 6900
tcacggaaca tgcttccggc gtacgccgtg tggtgggcga ttctgccgag attatcggtg 6960
attatgacct ggtgctgata accggtaagg agttggagca tgtatgggag caacatgaat 7020
gtacctccgg ggatattcgg gagattacca ttcagttctc tcccgacttg ttcttcggga 7080
acgttgtcaa caagaaccag ttcgacagta tccgtcggat gctggagcgt gcacagtgcg 7140
gacttagttt ccccatgcag gccatcatga aggtgtacaa ctggctggat aagctggctt 7200
ccgaggagca gggcttttat gccgtgatga actttctccg tatcctttac gaactttccc 7260
tctatgacaa tgcccgggtg ttgtccagtt cttcgtttgc caagatagag actttctctg 7320
acagccgccg ggtacagaag gtgcagaaat acatttccac ccattatcag gaggatatcc 7380
ggttggcttc attggccgat atggtgggca tgaccccggt ctctttcagc cgtttctttc 7440
gcctgcgcac cggcaagacg ctttccgact acattattga tatccgtctg ggctttgcca 7500
cgcgtatgtt ggtagattcc acgcgtacca ttgcagaggt gtgctatgac tgcggcttca 7560
acaatctgtc caatttcaat cgtatgttca agcggaagaa gggatgctcg ccgaaggaat 7620

tccgtgagaa ttatcggaaa aagaagctgg tgatatgata aaaaagtatt ggtattcttc 7680
attgaaagtg cctatttttg caccaatgat taatactaat atctgatatc atgaaaagag 7740
ttttttcctt atttttatgt ctgctgtgtg tatgtgctgt gacggcgcag atacgcggca 7800
atgaaatccg tgtagtcgtt tcacctgacc atagcgattg ggtgtatcgt ctgaatgaga 7860
agtgtacgtt taccgtccgg gtgctgaagg cacagaacct cttgtccgat gtcaagattg 7920
actatgaact gggccccgag atgtatccta ctgaagtaaa gaaagacgtg gtgctgaaag 7980
atggtacctt gaagttgcaa ggcacgatga agactgcagg cttcttgcgt tgcaaggtca 8040
aggcgcatgt ggacggccgg acttatgaag ggctggctac atcggcctat gctcccgagc 8100
agctgcaacc tgtgacgaaa cttcctgccg attttcggga ttattgggcc aagaccttgg 8160
aagaggcgcg caagactccg ttgaatcccc ttatgacgct gcttcccgaa cgctgtacgg 8220
aaacggataa tgtatatcag gtcagcttcc agacaaaagc ttggggagga cgtttttacg 8280
gtattctcag tattcccaag aaagaaggga agtatccggc tttgctgcgt gtgcccggtg 8340
cgggtgtccg cccttatgcg ggagatacgt atacggctcc tggaaaggtg attacattgg 8400
aggtgggtat ccacggcatt cccgtcacga tgcagcagtc tgtatatgat gcgctggcag 8460
gtggggcgct gagcaattac tggacgttcg gccgcgacag ccgggatgct tcttactata 8520
accgtgtggt ggtaggcgct ttgcgtgctg tggattttat ctgttcgctg ccgcagtaca 8580
acgggaaggc gctgggcgtt acgggctcca gccagggcgg tgccttgtct gtgattacgg 8640
cggctcttga ttcgcgcgtt actttcctgg cggctgttca tcctgcctta tgcgaccacg 8700
aagcgttctt caagaagcgc gcttgcggct ggccccatta tttctattat tacggtgctc 8760
ccgatgaaaa gcagcttgaa acggtgcgct attatgatac ggcaaacttt gcccgcctgc 8820
tgaatgtgcc cggttggttc agttgggggt ataacgatga ggtctgtcct cccacttcca 8880
tgtatgcagc ctacaatgtg atttcttctc cgaaggaact gcatccttat ctggaaaccg 8940
ggcattactg gtatcaggaa cagtgggatg aatggctgga ctggatacgg agacaactta 9000
atgtgccaat gtgattaatc acattggcac attaatctga taatatagta tcattttta 9060
tgatatgagc cgcgtatctt tgtgtatcgc ttaaaaactg ataatatgaa gaaacaattc 9120
cttttgctga cagtactttt attccttttg gggcatgtg cccctaagcc tgcggagcat 9180
tcgtttatta aggtcaatgc cgatggacag ttcgttcgcg acggaaaacc ctattatttt 9240
gtaggtacca atttctggta cggtgccatt cttggttccg aaggtgaagg cggcaatcgt 9300
gaacggttgc ataaggagct ggactttctg aaaagtattg gcatcaacaa cctgcgcgta 9360
ttggtgggtg ctgacggtga aaatggcata aagacgcggg tagagccctc tttacaagtg 9420
gcaccgggcg tctataatga tactatttta gccggattgg attattttat gaatgaactg 9480
cgtgaacgtg atatgacagc tgtcctctat ttgaacaatt cctgggaatg gagcggtggc 9540
tattccgttt acctgcaatg gtccggacac ggtgatgccg tggtacctgc cgtagacgga 9600
tggcccgcct atatggagta tgtcaagcag ttcccccaat cggacagcgc gaaggccttg 9660
tttgccaatc atgtcaatta tattgtttca cgcaccaata ggtataatca gataaaatat 9720
gtggatgacc ccactattat gtcttggcag attggtaacg agcctcgcgc tttctctgac 9780
gagaacaagg aaccgtttgc ccgctggatg gccgatgtgg ctgcacaaat caagtcactt 9840
gacccgaacc acatggtgtc ttctggcagt gagggctcgt ggggatgcga aatggatatg 9900
aatcttttcg agaagattca tgccgactcc aacatcaatt atctgaatat ccacatctgg 9960
ccttacaact ggagttgggt gaaggcggac agcctgaagg aacttcttcc gcgtgccaag 10020
gaaaatacta agaagtatat agatgaccac atggtcattg ctcggaagta cagcaagccg 10080
attgttttgg aggagttcgg ttttccccgt gacggtttca gcttttccaa agaggcaccg 10140
accactgccc gcgatgaata ttaccgttat gtattcgacc tcatccgcca ggaccgtgag 10200
agcggcggac tctttgccgg atgcaacttc tgggcatggg gtggttttgc cggacaaaat 10260
cccgaccatg tattctggga aaagggagat gactataccg gtgacccggc tcaagaacaa 10320
caaggactga actctgtctt tgccacagac agcactattg aaataattaa agcggaaaac 10380
cggaagctgc aaaattgaac ctatccgtgc ttttccgtgt aatccctgcc tgaaaattag 10440
aatgtaagcg tgtacaaagc gtttgtaccg ttatataaga aagattatta ctctaaatac 10500
taagtatgat atgaaaagca tatggaaagt tatgttggcg gtttgttgcc taggtatgac 10560
cattggttgt ggaacaaacc cgtcgaagaa tgaaaatgta aaggaaacgc taccggcatt 10620
ggtggtcaat ggtacgcagt tgatgaatac cgaaggagac acggtggtat tgcacggtgt 10680
cagctatggc tggcatcagt tctggccccg tttctacaat gcctcttcgg ttgcttatct 10740
ggtcaatgac tggggcgcac aagtgcttcg ggcctctatg ggagtggatt tggatagcgc 10800
ctgctatgtg aacaagcccg agttcggtat tgagtgtgtt acgaaagtgg tggatgctgc 10860

tatcgagaac ggcgtttacg tcatcatcga ctggcatagc cacaatctgc ggcaggaaga 10920
agcaaaggag ttcttcaccc aaatggctac ccgttacaaa ggggttccta atgtcatcta 10980
tgaggttttc aacgaaccgg tggaagacag ttgggagcag gtgaagtcat actctgtgga 11040
ggttatcaag acgattcgtg ccatagaacc ggatgccgtt atcctcgttg gttgtccgca 11100
ttgggaccag gacatccatt tggcagcgga cgatcctatt accggatata gcaatataat 11160
gtatacactc cacttttatg ccaacaccca cggacagtgg ctgcgcgacc gcgccgacta 11220
tgctctgggc aagggcttgc cgattttcgt ttccgaatgc gccggtatgg aggcaagtgg 11280
ggacggcccc atcagtaaag aagagtgggg gaattggctg gactggatgc agcggcattc 11340
catcagctgg gtggcctggt ctttgtctga caaggatgag acttgctcca tgttctatcc 11400
gtcggcttct tccgagggaa actggaaaga tgcagatatg aaggaatggg gaagaatggt 11460
acgggacgaa ctgaagaaat gaccctttcg ggaaaaagca atatgattta accattaata 11520
atttaagaaa aaaagaattg tgcctatggg aaaaaaggaa aaacggaact atgaagacat 11580
ggactgtttg gaagtaaacc taaatgtcaa ggctgtgata gtatatcagg gctttttgac 11640
tcagtctcat ttttgttaac taaaaaaatc ttcttttggg aggttttgtg tttgttttaa 11700
atctaagaaa tacagtatga atttgaattt tagaaagaca gcgctgcttg cggggatatg 11760
ctcggtgttc agtctgggtt atgctcctca gcttttcgca gtatcatcga cggataccgt 11820
tgaggcagtg cagcaaacaa agaaaattac gggtgtcgtc agtgatgcta tgggccctat 11880
tatcggtgcc aatgtattgg aaaaaggtac aacgaatggt gtaattaccg acattgacgg 11940
taactttgcc ttgaatgtga aaccgggagc taccattgta gtctccttta taggctatgt 12000
cagccaggaa ataaagataa cgaaccagac aacagtgaac atcacgctga aggaagactc 12060
tgagatgttg gacgaagtgg tggttgtggg gtacggtacg atgaagaaaa gcgacctttc 12120
cggtgcttcc gtttccatga gtgaagacaa gattaagggt tctattatca ccaatctcga 12180
ccagtccctt caagggcgtg ctgccggtgt gacggcagtc agcacttccg gtgctccggg 12240
ttcttcctcc tctatccgcg tacgtggtca ggccactatc aatgctaatg ccgagccgtt 12300
gtatgtaatt gacggtgtga ttgtacagag tcagggacaa agcggttcat cttatggttt 12360
gggcgatgca ctgggtaatg gctcggtttc caccatttct ccgctttcca ccatcaaccc 12420
tgctgatatc gtcagtatgg aaattctgaa agatgcttcg gcaaccgcca tttacggtgc 12480
gcaaggttcc aatggtgtgg ttttgattac gaccaaacgt ggtaaggcgg gtgaggctaa 12540
gtttacttat gacggtatgt ttgccgttca acgccagaca aagcgtctgg atatgatgaa 12600
tctgcgtgag tatgccactt attataatga tttggcaaac atgggtgaaa tctctgaccc 12660
tagtaattat tatgccgatc cttctctgct gggagtaggt acaaactggc aggacgccat 12720
cttccagact gctttgcagc atcagcacca gatttcggca caaggtggta cagagaagat 12780
acaatactac gtatcagctt cctatatgga tcaggacggt accatcatcg gttcaaactt 12840
taaccgttac agcttccgtg ccaacttgga tgcccaactg aagaagtggt tgaaactcgg 12900
tttgaccgcc acctatacag caacgggtga tgacctgaaa ctggcagaca gtgatgaagg 12960
tttgattaac tactcgttga ccactcttcc tgatatacct atctataatc tggatggcag 13020
ctatgccacc gttgttcgcg aaggctacac gaatcctaac ccgattgccc ttgccatgat 13080
ggatgacatt ctgctggacc gtcagaaact gacgggtaat attttctttg aggttactcc 13140
gattaaaaat ctggtatggc atgccgaatt gggttatgac atcagttcca gcaaaggtga 13200
gcgctacaag ccgatggttg atttgggtgg atgggtacgt tccagtaatg aaagcaatat 13260
ccagaaaaac agcagtacat tctggcaatt gaagaactat gtgacctatt ccggaaacat 13320
aggcgacaag catcattata cagccatggt gggccaggag tgctgggagt ccagctatga 13380
cttcacgagt gtgttcaata gcggtctgcc ttcggatgaa gtgcacaatc ctgccttggg 13440
tacgggaacc ccgactatca atgcaggttt tggaagctcg tctatggcat ctttctttac 13500
acgtctgact tataactatg ccgaccgtta tctgggtact tatacctatc gttatgacgg 13560
ctcttcaaac ttcggtcccg agaaccgttg ggcaggcttc cattcattgg ctacttcctg 13620
gcgcttttcc aacgaaacgt tctttgaacc gttgaagaag gtcatcagta acggtaagtt 13680
gcgtctggga tggggacaaa ctggtaatgc caacatcggt ggttatgctt ggggtacatc 13740
catgagtcgt atgccttccg gtctgggtat ggggtatcgc ccctccaata taccgaatac 13800
ttccatcaaa tgggagtccc aggagcaatg gaacgtcggt gtggatttgg gattcctgca 13860
cgaccgtatc aacctgacgg tagagtggta caagaaaact tccagtgata tgttgatgcc 13920
gttgcagttg ccttcttata tgggtacttc cggtaatgcc tcttctgcat tggctgctcc 13980
gaaaggtaac tatggtgaca tagagaatac cggttttgaa atctctctga acacacatcc 14040
tattacgggt aagtttgaat gggattctga cttccagatt tctttcaaca agaacaaact 14100

gaaaggattg gctggtacag ccaatgcgca gattgtgggt tacggacagt ggaccgatgt 14160
agtcagtgta tccagtgtag gcgaatcact ttataacttc tatggatttg tttgtgatgg 14220
tgtttacaag gacctggagg accttcagac ttctccgaaa cctgccaagt acccggcaaa 14280
cggtgtattc gaccgtacca ataccgtatg ggtgggtgat gtgaaatata aggatatcag 14340
tggtcctgac ggtgtgcccg atggcgttat tgacgactat gacaagacca atatcggttc 14400
tccgatgcct aagtttactt ttggctggac caatactttc cgttacaaga acttcgactt 14460
gtccgttttc atcaacggtt cctatggcaa caaggtgtac aactaccttt ccatgaagct 14520
gacacatatg aactctccct ggactaacca gctgaatgct gtaaacggac gtgcacagct 14580
tgttcctatc aatccgaaca aggtctacga agatggaacg atgtggtata atgatgtgtc 14640
taacgtcatg gtggcaaacc cgggagcttc cattccacgc gccactatca atgacccgaa 14700
tgacaacgac cgtatcagtg accgttatat tgaagatggt tcttatatcc gtctgaagaa 14760
catctctttg ggatatactt tcccgaaaaa gacgattgat aagctgggac tgagcaactt 14820
gcgtatctat gccaatatcc agaacctgct gaccattacc ggttatgatg gctatgaccc 14880
tgaaattggt gccagcactt caagcgtcaa tgtctatggt ttggacaacg gacgttatcc 14940
gtctcctaca gtttattcat tcggcttgaa cgtttcattc taacacttta aattgaagaa 15000
aacatgaaaa tcagatatat taaatattgg gcgtttgcag ctgtactcgg tatgggaaca 15060
gtttcatgcg aggatttctt gaaccgtcct accgaagaca attataacga atcgaacttc 15120
tatcagaatg acgaacaatg tgtgcagggg gttaattatc tttataactc accttggtat 15180
gactttcagc gcggttttat taaagtcggt gaaatcttgt ccggaaacgt atacgaggga 15240
aacagccctt acttgaattt tacggtgaat ggtacggatg aaaaccttgt gaacatgtcc 15300
tattctctat gggcagtgat tggccatgcc aatactgtat atactcgttt gaagacggca 15360
aatgcgtcag aagcagtgaa aaatcagtgt atgggcgagt gtctggcatg gaaagccatg 15420
gcttacttct atttggtacg tagttttggc gcagtaccca ttatccatga taatacggac 15480
gaattgaatt ccggtaatta taactccaaa ttcaaagtgg aggcagcaga tgtgtacgaa 15540
tatatcatta tgactctgga gaaagctatg gagttgcttc cggccagagg tgaaacggga 15600
cgtattgacc gctattgtgc cgaaggtctt ttggctaagg tgtatctgac caagagtggc 15660
ctttccggta cccgcaatgc ggatgatttg gcgaaagcgg cagaatattc caaggatgta 15720
attaataatt ccggccgtaa cttgctggct aactattccg atgttttccg tttggcaaac 15780
aacaagaatg aagagtgtct cttctcctgg cactggtcgg caggacgtga cccctggaca 15840
cagcagaata cgttgcagtc ggacttggca atggtgggat tcgatgaatt tggtgattgc 15900
tggggcggct atgcaggtcc gtcggttgat ttgcaggatg catttggaat ttccgctttg 15960
gaaagtcctg agacgcgttc ggatacggat acccgtcgta aggctactat gatgatggca 16020
ggtgatgtat acgattattt ctggcaggac aagggcggat ttgattatct gcggtttatt 16080
tatgatgccg aatatgggaa gggaggacct aatggcgatt accagtctcc taccggtgcc 16140
aatcacgtga agcatctcta tggtaacaat aatgaccatg tattgggctt gggcgtatcg 16200
gctggtaata tgtacagcgg cctggcaacc catatcctgc gcctgtccga tatctatctg 16260
gtatatgccg aggcgaaaat gggactggct acctctacta cagaccagag tgcactggat 16320
gctttcaatg cagtacgtgg tcgtgccatt ccgggagtta ctccgaaaac ctccattact 16380
tgggaggacg tctggaagga acgccggctg gaactggctt gcgaaggcga ccgttggtat 16440
gattatgtac gtttggcata ttatgattcc cagcgtgcta tcaatgagct gaaggcacag 16500
cgccgtgatg tttattatag tctgggtact acttataagg cttattatga aaacggcagt 16560
tggacggtga atccggacga gacccgttac aatccggatg caaaggcgcc gaatgtgaca 16620
gtctccagct ttacattgcc gttcccgaca gaagacgtgg tgttcaatcc taatttgatg 16680
aaagacccgg tacacgtaga tgtacgttct gaattcagtt attaatttaa tgcttgaaaa 16740
gatgaaaaat aatagatata cgaattggtt cttcgcggca gctgttgttg tagcaggctt 16800
ttccatggcg gcttgtgagg atgaaccgga taaatatgaa gtggcagggg gaaagcctac 16860
cttgaaatat gttcgtgtcc ctgatccggc agcggcagac tctttgctga cgggggctta 16920
tatggcgaat acgatttgtc tggtcggtga aaatctgcgg agtgtttacg agctctattt 16980
caatgaccag aaggcgatac ttaataccag ttacatcact gaccatacac ttatggtaga 17040
tgttccgaaa gagattcctt cggtagtgac tgataagatt tatatggtga ccaaagcaaa 17100
ggataccata gattatgatt tcaaggtgtt ggttccggct cctaccgtga actccatttc 17160
ttgtgagttt gcaaagccgg gtagtgaggt gacactgatt ggtgactatt tcattgatga 17220
cccgaatgtg ccgctgacca ttacgatggc cggtaatgtg gaagtgacta atattacgaa 17280
cataaccaag actgcggtca gctttattct tccggacaat gctcctgcag gctatatcaa 17340

cgtgaagtcc atttatggta ccggacgttc caagttccgc tattatgata cacgtaacat 17400
tctctttgac tgggacggtt ctcatggcgg tatggctatt gcccacggtt ggcgcgatgg 17460
tagcaaggtg ttgagagaag ccgatgaaaa cagtattgac ggtgcttata tctgtctcac 17520
cggagcactt gacggtgcta tcggagctac ttgggcagaa gatgagttct cgttcaacta 17580
ttggccggaa ccgtctgccg gttatcccga actttctgca cgtcctgagt ttgcagaact 17640
gcttgaggag tatggcttca atggcttgca actgaaattc gaggtgaata ttccttcttc 17700
caatccgtgg cagtcttgcg ctctacaagt gatgtttacg ggtaatgatg tggtgactta 17760
tgcgacggga actaatgctt attttagtga tacgaatgtg ccgcgcggtt tgtggttacc 17820
ttggaagaat acgggttcgt atgatacggg tggtaaatgg acaaccgttt ccatgaatct 17880
gtccgagttc aacaagacgc atgaaggtaa taagtgtgat agaacttttg acaagactat 17940
gctgaccgga ttgacattct ttgtctgggc aggtggtgtg gaaggtaagg attgcaatcc 18000
ggtgattatg attgacaaca tccgtgtggt acctattgaa taacctttta aatagaatac 18060
tgatatgaat aaaacatata aatttccggc actgtggatg atgtgcctga tgctcttcag 18120
ttgcttgacg tttacggcct gcgacaatgg ggatgatgaa gataccaacc aatataaagg 18180
gggaatcagt ctgaatgtat tcggaccgag tcccgtatcc cgcggcggtg tactccgttt 18240
cctaggcagt ggcatggata aagtgactgc ggtcgctatt cccggttgcg acgatataac 18300
agacatcgaa gtagtcagtg atacggaaat ccgtgtcact gtacctcaga cagcccagcc 18360
ggggttggtt gtcttgaaga ctccgaaagg ggatattaca acaaaaaccg agttgacatt 18420
tacggagccg attgctttgg aggcttttgc accggcagaa gtaaagcctg gtagtgaatt 18480
gaccattact ggtgaatatt tgaatttgat taaagaagtg atatttgctg acgaggtaac 18540
agtgcctgct gatgagtttg tcagtcaaag ccgtcaggag attaaagtta ttgttcctga 18600
ttctgctcag accggtaagt tcatcttgtc tgatggagct gaaatcccca attggattta 18660
ttctgaaggg gaactggagg ttactttgcc atcggtagag gctccactcg acctggtaga 18720
taagaagccg ggggatgtca ttcgagtttc tggtgaaaac tttgatttgg tgaagaaagt 18780
gcagatgccg aatggggatg aagttgaatt tacgatgact gcctcttccg aaggagatga 18840
acttactttt acattgccgg ataatgtctc ggacggagaa gttacggtac ttcctgcttc 18900
ggatgtaaaa gttgtggttg ctactgttgt agtagctacg ccttcgaacg tagtggctgt 18960
tcctgctgtg aacttgcgtg gtggagatat gattaccttg aaaggtacca atatggattt 19020
ggtgactgat gtaacgttcc ctggtgtaga agaagcagta ggattggaat cacaaaactc 19080
gactgaaata aaggtcttga tgccggctgc ggctattagt ggtgatttgc aattgaatac 19140
aaatagtgga aaagcaactg ccgtatctat tgcaacggca aagccggaaa acatttctta 19200
tagtgcggcg acggtcccgg caggagaggc tttgactgtc aaaggagtaa atatggatgt 19260
agtttcggct gttgtattca gtggtaatgt ggaggttacc gtatccgatg ccaccgctac 19320
tgccatttcc ttgactgtac ccacaactgc tgaaaccggg gccttactct tgaagatggc 19380
caatggtgaa tctgtggaag caccgtcttt gacaatagag aaaccggtat gtgcctattt 19440
gccggcattg cctgacaaat tggttcgggg acgaattgtg gaacttgaga ttgttaatgc 19500
tgataagctg acaaatgtat tgctgaatga ggctgctgtt cagtatatca atgatgcagc 19560
taaaggggta ttgatgttga atgtaccggc tgagctggat ggaacttata gcctaaaact 19620
gatttcttca aatggtgaaa ttgcttatga tgttcttgta gttgccaatg aagaaacagt 19680
ttgggctggt ccgctcgata ttagttgggg tgatggagga cgtgtgttgg taccggctgt 19740
atcttttgcc aaggtaacgg caggcactgt catgaaagtc tattttgatc agaaagacca 19800
gacttgggca caggctcagt ttaactatgg cgattggagt ggcatagctt tctctttgtt 19860
tgatacgact atggtaccga cagacattta cggctggagt ttcgagagcc gtgtgatgga 19920
gttgacgttg gctcaggaga tacttgataa tattcaagct aaacaaggag attgtgaaga 19980
ccagacaaat gtaggtatca ttattcaggg cagtgatttg acgtttacaa aaataacgat 20040
tgtaaattga tgaattgtgg cttgaagaat gtgactatgg atgccctgat taagaatgaa 20100
ataaaatcat aattaagatg aagaaactga atataaaatg gatgctttcg ttgatagcag 20160
cttttacttt tgcctcctgc gatactgatg tcgatcatga tatcccagca gtggatgcac 20220
cggtattggt gtctactact cctgaaagtg gagcagccaa agtgaaaacg ggagaaatca 20280
ctattgaggt gaaatatgac aagaatatat tctttgcaac agacaatctg agcgaaatca 20340
agtttaccgg tggtgaactg ataagtgccg atgtacttgg cgcaagcaat atactgactg 20400
taaaagtgaa tgttcccggg agagagacgg cttgttctct ctctatcccc gagggaatag 20460
taaccggacc gaaccagatg cctgctccgg ctgtatcggt tcagttctct actgtcgctt 20520
tggacaaggc actggttgcc gctagttcgg ccaaagctgt gaaactgtac aattatctat 20580

```
        tggataactt cgagacgaag acgctttcag ctatgatggc caatgttgcc tggaatacgg     20640
        aaatgtccga aaaggtgtac ggatggaccg gaaaatatcc ggctatcaac tgtttcgact     20700
        atgttcatct gccggcatct gtagctggag cagactggat taattacggt gacattactc     20760
        ccgttaagga ttggtctgac aaaggtggca tcgtagctgc catgtggcat tggaatgtac     20820
        ccaagaaagc tgtaggggaa gcttcttcta cccggatatg ggaaggagag acggtaatgc     20880
        cgggtgattg gagtggaaat gtacagatga cggatgatgc tgcaaaaact gtctttgccg     20940
        atgcacaagt ggggcaggtg atacgcgtgg ctgtgaaaga tgtggcagag ggggcgcaag     21000
        gttcgttcaa aaacagtggt tggagcgaga ttgcttcagg aactgattat tttgacatat     21060
        ccggagacta tacattggtg attaccgagg atattttgaa atccctgcaa gaaggtggcc     21120
        tcatcattgg cggacatgat tatacggcag ttgctgttta tcttgaaagt aacggtacag     21180
        ccctcgaccc gaataaggat tatgccttct ataaggcaga tacggagttt gacgctgcca     21240
        atgccacggt ggaaggcact tgggaaaaca aggtgtttac cgaagatttg aaaaatgctg     21300
        ccgcttactt gaagctgttg agagatgcgg atattccggt attgtggcgc ccattccatg     21360
        aggcagccgg tggctggttc tggtggggta aggatgctgc cagctttaaa tctctgtgga     21420
        ttgctatgtt caactacttc aagactgaag gattggataa tcttatctgg gtttggacta     21480
        ccgagggcaa tgattccgat tggtatccgg gagaccaata tgtagacatc gtgggacgtg     21540
        atgtttataa taaggaaact gcagattgtg tttcagaata tacttccatt gccggaaatt     21600
        atggaaacaa gattgtttcg ctgagtgaat gtggcacagt cggacttatt tccgaacaat     21660
        gggcatctgg tgcacgttgg agttggttca tgccgtggta tgatggcaca aatgaggatg     21720
        gtagtccggc agtccatgct gatgaggcgt ggtggaagga tgctatgagc caggaatttg     21780
        ttgtttcccg tgaagagctt ccttctatgg aatagcaatt tttaataata aaatagaaag     21840
        ggttgtctct ggttagtatc ggggcaaccc tttctgtctt ttatagcttt tagttgttgg     21900
        ttatagaatt catgctttat actttttagg aaaaatagta atggatgtta aatgagtgtc     21960
        gttatttgat aaaatag                                                    21977
//
ID   XXX; XXX; linear; XXX; XXX; XXX;  XXX.
XX
AC   PRJEB67889
XX
DE   Uncultured bacterium DNA fragment from in-reactor enriched cow rumen, clone
ferm-10n13
XX
FH   Key             Location/Qualifiers
FH
FT   source          1..46551
FT                   /organism="uncultured bacterium"
FT                   /environmental_sample
FT                   /mol_type="genomic DNA"
FT                   /isolation_source="Cow rumen"
FT                   /clone="ferm-10n13"
FT                   /note="ABTS active clone"
FT   CDS             38..664
FT                   /db_xref="SEED:fig|6666666.127229.peg.1"
FT                   /translation="MTALGIYPIGGKYDDGNGGKITSKDGSGCAPDKMLQAYSVPFMLA
FT                   ELVLAGEINGDAKKYLEDGIKASIAHVNAVTKLCDASAPLISSDDAQKLVDEVNEKFDA
FT                   ASAEGKLEIVMTEKWIANFYNPIEAYNDIRRTGYPKLFKGDENNMGWTPYAQTVEATPG
FT                   LSSFELVTILDYPRILWYPQNEIDTNPNVSNSSRNVAVKTVFWDK"
FT                   /product="hypothetical protein"
FT   CDS             690..1625
FT                   /db_xref="SEED:fig|6666666.127229.peg.2"
FT                   /translation="MKINKIFALIALGALFCSCDQKLPYDLEGTKHGVLISISKVQGTS
FT                   GILSTDMNAGNYQVKLDLPIYQGDYSMLEDAQIMAVYTGVDKTKKSAIVAEGIKAFPDT
FT                   VKLNIKDVASKLGISEINVGDKIEYTPCYTLKDGLQVNGWSELTGFNNKNFSSWKLEDG
FT                   SSFAYRVSYTAFAPLQLDKFQGDDIPFSMDGGDEGSMKVSQVDNKPEAKWIPAGANVED
```

```
FT                   LVGLKFEGDFWFGGDSFIIWINPADYSLIIPDQVMCPGFTYGSYGTFDAYMVSCDGEID
FT                   TLNNTIVFTFGTSWGPYSFGGDTVIFEFPR"
FT                   /product="hypothetical protein"
FT   CDS             2265..5618
FT                   /db_xref="SEED:fig|6666666.127229.peg.3"
FT                   /translation="MEKIKFFLAALLVFVSVGLMSAQNRTITVKGNVTTAVDGQPAPGA
FT                   YVSVEGTSVGTVTDLDGNFQIDNVPASAKWIVVSYLGYKDARVAITANIQVALEDDSEM
FT                   LEGAIATGMYNVDRRLNTGSAAKIDAAEAKLDGIADISRSLEGRVAGVSVQNVSGTFGT
FT                   APKIRVRGATSIYGSSKPLWVVDGVIMEDVVEVSADDLSSGDANTLISSAIAGLNSDDI
FT                   ESFDILKDGSATSIYGARAMAGVIVVTTKKGKAGQNRLSYTGEYTMRLKPSYSTFNIMN
FT                   SQDQMAVYQEIQQKGFLAYAGTANARNSGVYGKMYQLISQYDATLGKFGLQNTDEAKIA
FT                   YLRAAEYRNTDWFDRLFTYNIQHNHSVSLSGGTEKLNFYASLSLLDDAGWSLQSDVNRY
FT                   TLNLNASYKISDKLSLNVIGNGSHRTQRAPGTLNAEVDAVSGEVKRDFDINPYSYALNT
FT                   SRTLDPNVFYTRNYSPFNIIHELDNNYLDIDVDELRIQGELKFKPTKGLELSVLAATKS
FT                   TSTSQEHYIKDTANQALAYRELSTPDIRDNNPFLYRDPDHPNAYKITVLPVGGVFERKT
FT                   NAFKGWDFRTTASYSNTFNNVHILNLFAGAEANNVNRHATWFRGWGMMYDKGEKAMYDY
FT                   RIFKKGVEDNSDYYTLENTIARTAAFFANATYSYKGKYTINGTARYEGTNSLGKARSAR
FT                   WLPTWNVSGAWNAHEESFMRDLYPTVSHLSLKTSYSLTGDRGPSYVTNAEENINYRNPW
FT                   RPSASTKESALYIVDLANEELTYEKKHEFNIGIETGFLDNRINFAADWYTRNNYDLIGP
FT                   VITQGVGGQIEKYGNVASMKSDGYELSLTTTNIKNKDFSWTTNLVFSHSHNKITELENK
FT                   SRMMDLISGTGFAREGYPVRSLFSIKFMGLNSEGLPTFINEDGDVTISDIYFQTRDSKS
FT                   LENLVYSGSVDPTDVGSLGNIFTYKNFKLNVFVTFSFGNVIRLDPVFSDEYTDLVAMPK
FT                   EFADRWAVPGDENITNIPVISSRWMNHYYEANSNSRLSYAYNAYNYSTERIAKGDFIRM
FT                   KEISLSYDFPKKVTETLNVSNLGLKLQATNLFLIYADKKLNGQDPEFFNTGGVAVPLPK
FT                   QFTLTLRIGF"
FT                   /product="TonB family protein / TonB-dependent receptor"
FT   CDS             5627..7282
FT                   /db_xref="SEED:fig|6666666.127229.peg.4"
FT                   /translation="MKKMKNRIINIFASVIAVAAITSCDSFLDVTPDKRAVVDTQEEVQ
FT                   DLLVSAYPDVTYLAITEYMSDNVENLGDNISGSTRQIEQLYYWNEIVDENNDSPKDFWD
FT                   NCYKCIATANSAIAKINDLGGPEAGLQAEMAEALLCRAYAHFMLVNIFCQNYSSKTSGS
FT                   DLGIPYMTKSETTLNPKYERGTVKDVYEHIEEDILAALPYVTDAYYDVPKYHFTPSAAY
FT                   AFASRFYLFYEKWDLAIDYATKCLGDYPASVLRDWQYRSTLADSRQVKTLDFINESKKS
FT                   NLLLLTGVSQYGICFAYPAAYPKYSHTNYLGATETCFAKNVWGNTSVGYGGAARYYYDP
FT                   AASYVGGNFSQVALFKIPYLFEYTDPVNDIGYPHVVYPAFTTDEILLNRAEAYILTGDY
FT                   ERACEDLNIWIHNILNPSYFNKDLTPDMIKSFYDATPYARWDNFDPSTRKLQNNIKKQL
FT                   HPKFEIPSDPYAEPMLQCVLQFKRIENLTQGLRWFDIKRYGIEIWRRTLKSDSASPVGY
FT                   SPARLDDVLTVDDPRRAVQLPQEVILAGLDKNPR"
FT                   /product="FIG00937915: hypothetical protein"
FT   CDS             7301..8206
FT                   /db_xref="SEED:fig|6666666.127229.peg.5"
FT                   /translation="MKKKNILIAFVAAVSLVTVSCTEKKLSPDSVISAPSVEKNAFDLW
FT                   LDYNFLKPYNIEFRYRYDLNESDMYRMTIPADYDASIMYAHLVQYLCIDTYNEVAGVDF
FT                   TKAYFPKMFFLIGEWEYDNNGNIILGTAEGGKKIMLSGVNLLPGKFASFSGSGLKKELS
FT                   EYYIKTIHHEFTHILNQIKGYSESFKEITASTYQADSWSESDSKYRQRGYITKYAQKED
FT                   REDFAELLSEYITHDEDWWQKELTKAGDEGKSLILAKFIIVRDYMSSSWGIDVDELRDV
FT                   VNRRFDDVVEGRVNLTDLSI"
FT                   /product="FIG00938925: hypothetical protein"
FT   CDS             8218..9012
FT                   /db_xref="SEED:fig|6666666.127229.peg.6"
FT                   /translation="MNNRFFISLAAVAVSLVSLPSCQKDQDSVFDGSLTERQTQYLDGF
FT                   RSLITGAENGWAMYYFSGGAYATSRVFTVSFTDNEVTASSEDAPSVTAKSFYKVAVTSA
FT                   PSLIFDTYNKVLHTYAAPSHSYYQGRGGDFEFTIENYSEDKITLIGRRAYEQQILIPLK
FT                   ESPASYLNKIKSFAQSFNLTQVAGKVGNGDVVIYFDLRNRFLYIGRLGAEDTELEEIPY
```

```
FT                   IYTENSLLLQHPFWYNGTTFSEFSYNAEEHSITSEGITFKQF"
FT                   /product="hypothetical protein"
FT   CDS             9019..10353
FT                   /db_xref="SEED:fig|6666666.127229.peg.7"
FT                   /translation="MKKLFGILMVALSVFSLASCNGENEIDEKGAPTIVINSKDIQLPA
FT                   KGGESSFLVDSDVDFIVSVSHDWVSVSKDGNRVKVAAEKNSSLQGRYAELSVSNGKDGA
FT                   SIVINQLGYISSDVEFDDISVSYLGQFVELEYDYDDFISASTDVDWIILDNSRGTEGQR
FT                   ILRVNVGENKTGSQRSATIEWTLGLESGSFVVKQRKEMKEYSLNNSWNISYLGVDNSIY
FT                   KVGVSGVSGEYVVTAVSASKLAQGGFSNVAEYLTESDSDIAKAIADGEIAVYSGDKTVE
FT                   LSNLSGGENYAVVLGYKSSKSELTGEFNFVTFKVEGPDLRTPYEKWLGTWEIERESEID
FT                   VWTISKNVENVSYTILGVDGTSKENTPDVPVIAEFDSETGNLIIKAQHCGTFTSSSYGE
FT                   ADLSLYGLGPEYYYTYSSEPYTIATLSLSGKNSANLVRRRTSTHL"
FT                   /product="hypothetical protein"
FT   CDS             10710..12101
FT                   /db_xref="SEED:fig|6666666.127229.peg.8"
FT                   /translation="MLSENLEAAMKDYDKAISLDTSSTLAYYNRALIKYELQNYMGAME
FT                   DLNRVLRDDPGNSLTLYNRSLIYAKFGEYGSALDDMDRVIFNNPRNVLAYFNRASFFID
FT                   MGRWQDALEDYDKSIELYPDFAKAYKNRAYVELQLGRKAAAKKDYDTANRLVDEYRIKS
FT                   ADGTVSFADTTKKYNSLISFDTDFAKNNFDDEILLHRDIDIRLKPLYRFVITKDKDDVN
FT                   YALRNRYENPLIDKFIKESPVPVSIVNDIAVEKSSAEVEEELYGVRESSESVNSFLRGV
FT                   YEVSGKQFNLALNYFGDAISQASGSGDKTQDMYKSLYYMNRGVLRADMIDFISSIENNV
FT                   QTLTMDDQGTTRARVRDRVNRSYDYGDAISDMEAAASIVSDIPYVYFNLGNLYCLSSRL
FT                   VDAIGSYNKALELYPMMGDAYYNRGLVLIYLKDKEKGCIDLSRAGELGVGEAYSVIKRY
FT                   CEEAK"
FT                   /product="hypothetical protein"
FT   CDS             12083..12892
FT                   /db_xref="SEED:fig|6666666.127229.peg.9"
FT                   /translation="MRGGEMIKFMSFSSGSCGNCYWLGNDEKGILIDTGVSLRHLKKLM
FT                   LLHGLDENSFSSILVTHDHMDHIRSLGSFCKRLQKPVYATTLLHEALIRHSYAMPSIMG
FT                   CRRDLREGEWNDVDGFMVHYFVVPHDATQTVGYAIEFEGLRFVIMTDIGRMTDEAVEFA
FT                   SKADVVVVESNYDVDMLMGGSYPHDLKMRICQGNGHLSNDECAAAIKRFWHEGLRFIFL
FT                   CHLSENNNTPTLAYETSKRALDELGVSEKVQLAALPRQYPSRLYVIG"
FT                   /product="metallo-beta-lactamase family protein"
FT   CDS             complement(12889..13116)
FT                   /db_xref="SEED:fig|6666666.127229.peg.10"
FT                   /translation="MKLEGTYLPWIDCRSLGLKSEEIENSLLQNEKVWVNAGAMYGQDG
FT                   FIRINLATPASICREGLERIAAGLKRLSKA"
FT                   /product="Cystathionine beta-lyase, Bsu PatB (EC 4.4.1.8)"
FT                   /EC_number="4.4.1.8"
FT   CDS             complement(13321..13722)
FT                   /db_xref="SEED:fig|6666666.127229.peg.11"
FT                   /translation="MTIFSKIAQGEIPSYCCAQTDEFYAFLDINPLVKGHTLCIPRREV
FT                   DYIFDMEDDEIARFQQFAKHVALAIKKVCPCIKVGEAVLGLEVPHAHIHLVPMQSEHDM
FT                   QFGGPKKSFPAEEMQQVADAIRNEFEKLK"
FT                   /product="HIT family protein"
FT   CDS             complement(13728..14192)
FT                   /db_xref="SEED:fig|6666666.127229.peg.12"
FT                   /db_xref="GO:0003677"
FT                   /db_xref="GO:0003711"
FT                   /translation="MSYMSQEGYDKLRAQIKQLEEVERPEVIRQIQEAREKGDLSENAE
FT                   YDAAKEAQGKLETKIAELKAILMDAKIIDTSKVNTDTVQILTKVEMKNVKTGAVMTYTL
FT                   VSESEANLREGKISVKTPIAQGLLGKKVGDVATIKIPQGIIELEITNISI"
FT                   /product="Transcription elongation factor GreA"
FT   CDS             complement(14409..15716)
```

```
FT                   /db_xref="SEED:fig|6666666.127229.peg.13"
FT                   /db_xref="GO:0004069"
FT                   /translation="MKTPINKELIDNAISDFGIKDFAKATIREVVAIASNAEKASGVEF
FT                   IKMEMGVPGLPASKYGVEAQIQSLQNGIARLYPNINGLPELKEEASRFIKAFIGVDVNA
FT                   EGCVPVCGSMQGTFASFLTASQSDPKKNKILFIDPGFPVQKQQCVVMGAEYETFDVYDF
FT                   RGENLGPKLESYLKNGDIAAIVYSNPNNPSWICMTESELKTIGDLATKYDCIVFEDLAY
FT                   FAMDFRKDLSTPFQPPYQPTVARYTDNYVLLISGSKAFSYAGERIGVSCISNKLYHRSY
FT                   PALEKKYGNGTFGSVFIHRVLYALSSGTSHSAQFAMAAMLKAASDGIYDFRADISEYGR
FT                   RAHLLKEIFTRHGFIIVYDKDMDESIADGFYFTIGYPGMTGSQLARELMYYGVSAIALD
FT                   TTGSKQQGLRVCTSFIRDNQYDLLEQRMQIWEENHK"
FT                   /product="Aspartate aminotransferase (EC 2.6.1.1)"
FT                   /EC_number="2.6.1.1"
FT   CDS             complement(15697..16275)
FT                   /db_xref="SEED:fig|6666666.127229.peg.14"
FT                   /translation="MKTDIILAGVGGQGILSIATIIGEAATKAGLNLKQAEVHGMSQRG
FT                   GDVQSNLRLSNETIYSDLIPRGKADLIIAMEPMEALRYVEFLNPDGVIVTSSKPFINIP
FT                   NYPDQTELFKELDSQNKVVRLDIEEMAKEIKAPKSANVILLGMAAPFLDILSADELRSA
FT                   VGRVFSSKGEAIVEANYKAFDFGYENANK"
FT                   /product="Indolepyruvate oxidoreductase subunit IorB (EC
FT                   1.2.7.8)"
FT                   /EC_number="1.2.7.8"
FT   CDS             complement(16279..17931)
FT                   /db_xref="SEED:fig|6666666.127229.peg.15"
FT                   /translation="MEKRLLLGDEAIALGALHAGLSGVYAYPGTPSTEITEFIQNATGY
FT                   GKETYIIGGHELKANGKIHSRWCSNEKTAMEAALGMSYAGKRALVCMKHVGMNVCADAF
FT                   VNSAITGVNGGLVVLAADDPSMHSSQNEQDSRFYAKFAMIPIFEPSNQQEAYNMMEDAF
FT                   KLSESLKLPVLMRVVTRLAHSRANVEIKDKANEENQLNPDNGRHWVLLPAIARKNYLSL
FT                   IEKQKDMINASLSSKYNKLDTTTQNSKLGIIACGIGYNYVMESNIDGIEVMKISQYPLP
FT                   EQKIKDFAAKCDKILIVEDGQPVVEEAVHGILGADYKILGRLNGNLPRTGELNPDSVSL
FT                   AINSAITGEDPKEYPTATSPINVVGRPPALCQGCGHRDVYTALKEIVAEYKDARVFGDI
FT                   GCYTLGALPPFQAINSCVDMGASITMAKGAADAGVWPAISVIGDSTFTHSGMTGLLDAV
FT                   NDKSNITVIISDNLTTAMTGGQDSQGTYRFEQICLGLGVEPEHLHVVVPLPKNMDEIKQ
FT                   ILRDEINYKGVSVIIPRRECIQTHARKLRQAKK"
FT                   /product="Indolepyruvate oxidoreductase subunit IorA (EC
FT                   1.2.7.8)"
FT                   /EC_number="1.2.7.8"
FT   CDS             complement(17910..18692)
FT                   /db_xref="SEED:fig|6666666.127229.peg.16"
FT                   /translation="MKITLIQFEIAWANPTENIVRAEAAINRSPGSDLYIMPEMFTTGF
FT                   IMKPKGLAESKDSDSLKWMIKKSKEMNAAICGSIAVEDNGHFYNRMYFVKPDGEVTTYD
FT                   KSHLFIYGGEGEQYTSGNKRVVVNFRGVNILLEICYDLRFPMWCRNNMDENNNVLYDLI
FT                   VYSAEWPLARKLAWDTLLSARAIENQCYVAACSCVGADEWGEYFGGSKIIHPYGHVLAE
FT                   DELNKPSEITGEIDMEQLVKYRTKFPTLKDIIWKKDYC"
FT                   /product="Aliphatic amidase AmiE (EC 3.5.1.4)"
FT                   /EC_number="3.5.1.4"
FT   CDS             complement(18733..19743)
FT                   /db_xref="SEED:fig|6666666.127229.peg.17"
FT                   /db_xref="GO:0004073"
FT                   /translation="MKVAIVGASGAVGQEFLKVLDERQFPIDELVLFGSTRSAGTKYTF
FT                   RGKEYEVQLLKHGDDFKGVDIAFTSAGAGTSKEFAEDITRYGAVMIDNSSAFRMDDDVP
FT                   LVVPECNAEDALNRPRGIIANPNCTTIMMVVVLQPIEKLSHIKRIHIASYQSASGAGAA
FT                   AMAELQQQYKELTEGKPVTVNKFAYQLAYNVIPQIDVFQDNGYTKEEMKMFNETKKIMH
FT                   SDVRCSAMCVRISSLRAHSEAVWVETEKPLTVEEVQNALRNGEGVTLLDDPSTQTYPMP
FT                   LFTAGKDDVYVGRVRKDLANDNGITFWLSGDQIKKGAALNAVQIAEYLVKVGNIK"
```

```
FT                   /product="Aspartate-semialdehyde dehydrogenase (EC
FT                   1.2.1.11)"
FT                   /EC_number="1.2.1.11"
FT   CDS             20198..25573
FT                   /db_xref="SEED:fig|6666666.127229.peg.18"
FT                   /translation="MNTNKNYNDDHIDVLYKRIQNWLPNGSSQLKFTELKKRVLNKTII
FT                   NPEETLKSKDLKEIANYLSEYNKFKAERGDSKVKGMLNFSDIMTNVSEKQINNLINELD
FT                   SQNKPSTSLIPIKGISKVMQARLRDLKIVDVPTLLVKCKDANARKSLANKLGVDIRLVT
FT                   SWMKQAILWRVEGMTTDLAYLLVMVGIRNVEDLAQVDKAKILPILKGLVATHIDFEFEE
FT                   SELDTVLKNAQEMVVYNTPIKVNDLTSRLSDTMAGLTESLEEEFLNSIKEQLSSSLQEI
FT                   TTQSLRSARQPLSRQELHVTKDDEEDINDEEEDEEIDDEIYEDDEIEGDINEDDEDIDE
FT                   DEEDDEENEDDVNDEEENEGDVKRLELGVAHTYDLPNNFIANNYVHEDLREKIISEIIE
FT                   KEAQKLKQASAERIQEILKKEITGYQKENTVDISYVPNDINDVEPVHLFDEKLRKPLEI
FT                   LNGKNLKKGLDFLDDIQPSLPLPRKISGYIVIRKKDELIDISRNDSVYLQGCRRLSGAL
FT                   VEIQGIVSPSDDLKEAVTSPSCVTDGTGKFVIVLPERYSMKETITIVVSVGSKRQEFIK
FT                   TASEILESVEQHEILNRFYELDAIGDEIDYIESNIKTLTGQVAYLEDDVKLLISELDKL
FT                   NAELSNEETSDYQKILLKRDIDNKEKDRALKIKEKEQYTNQLIAYLQNIDILNEKYSTG
FT                   KAKLLSESFCAPAVDLKQAFDNFINNITSLEAKLSGDTLKPGETPKPFVVVEEIFNNER
FT                   MDLEKALPKVKLMGNDDKVIRLSTDTAPSKIYTYSMLQRLVEPEISTGDRQTLSHAIDV
FT                   MDFKESMITNIDKCPHACTLGMGYVLNMHQAWVPDGFALGDLLYSLVLAPGEEQRLIVR
FT                   ENKQTYEISDEANAIDADDESYALTQDDDTNAAFNYAMDQLSQGKSSYSYSASSWSIGG
FT                   SASGGAGGAMIGLSGGYSKSSGKGSASASQSNSHSETSSAAQNFQHNIKSASSKISQAK
FT                   RLSMEMATSDVTDSVATKIIANHNHSHAMTIQYWEVMRRYKLETCIDSVDLVLFVPLKL
FT                   INFLPTGDFYLGDTQNFDRTAFNNRYDTLLHYYDTLYSRLPSKYRSGLNLIKKYSELTN
FT                   WVMQKQEQSTKVLTFEIKVKLLTYDNLSATLVLKNGKGSIAGSCSDINNKRKKLTTSIY
FT                   QQYETSEDLRQGIIDFRNKVPDATDTLTFIFNIPDDTTDDDVSYIRLDYSCNECSYSLK
FT                   RDDNKLITIDYDNDDKHGNTTAGKLYQRYIDNIYDYSKNDKDTANEHKLEAYYERILPE
FT                   AYLHPIVKISSYEMRRLGTPTIMDAVLKIGSGKSVGSTSKDVETTYNIMLPSSKLSSSM
FT                   RLSISTSCYTLHNSELLKMESTLHHIASNPMKYSQIVWGSLSDDERVMMLEQFTINMNF
FT                   DSLDDDSSRYEKSNDLKIPLLNCVNVKNLLGFYGNCMILPFTFPQRLAKQLGKTAADIQ
FT                   NSLYRYHTNNFRVPTTTISLPTKGMIGEAVLGETNVSEEIDITRFWNWQDSPIDKMEID
FT                   SSYLNGNDYLQGKSTKDVTTLNMQGATAATPVTVPDLVSALVSKQTPTFDNITGLDQLK
FT                   EVLNAGTNSAAAGRDNAIKSSADVAKAALASLTPSSSSTTSSTDSSSSSSTPSTSSSST
FT                   PSTSSSSTPSTSSSSTPSTSSSSTPSTSSSSTPSTSSSSTPSTSSSSTPDTSDTGTPDP
FT                   SDTGTTDPTDTGTTDPETSDPETSETTSDIDMDLLSSIIKEAVEYIKTNGDDSVGFFKS
FT                   KVSDQYPDETVDDLANKFAEKHGLSLSDFIEKIFK"
FT                   /product="Probable exoglucanase a (1,4-beta-
FT                   cellobiosidase) protein (EC 3.2.1.91)"
FT                   /EC_number="3.2.1.91"
FT   CDS             25580..26773
FT                   /db_xref="SEED:fig|6666666.127229.peg.19"
FT                   /translation="MIIQIAILIQTFYTIKSLLDSRDKIEEWVKVVMAVYEKQMQIMDS
FT                   KKDPTILVNAFYKYLLEYADIFDADPSNNKPSLDSILKSSKKILESNSDRYVLSTTYAK
FT                   VSPFFPTPFRYLDNPKRNINPITAFTATEDDLKELLKKSNKDKNTDRFKELSKDIKSYR
FT                   ELLSKRSTLAQSTISALNPDNTMMTEYLFNTNSNTEYYWGGDKCNVYVGEAIYYALKKV
FT                   FLTYTNAKYDNLDGLLTDFLLDANHNLLTSSYYRYLQPSGIFEGYTEPIIGDDGKPKKG
FT                   DDGEDLKRLVNTTQIHSKIIFEKCPLNSVIEPGAFVCFITDGYKFGHIEMILSKIDVLQ
FT                   NKKTFVSIGAHPYGVYRKTQTFDSSKMQIVKVKESVIKSVVDIKPVAKEDDKKEAKN"
FT                   /product="hypothetical protein"
FT   CDS             26791..28212
FT                   /db_xref="SEED:fig|6666666.127229.peg.20"
FT                   /translation="MNEFELNRLKEIFNDFNLNDCLEQASSDSERGVCEDIAKIKEDWI
FT                   ESILATYTAKDNHCLFSIRTKEALDTGIYTKDNTYNNLEGLIYGKSYTFYILNKENNTT
FT                   VSDKSVEWSLYLSEDSNVKLIKKGTGNEIQFSVDNPEFLGKFLIVEVNYNGFTFPLNMK
```

```
FT                   GLFWIHYRFRHINFVCLAKDIVKRCLFPDGVSQRGTSLCGPAAVAYLFAKYQWDQYGRF
FT                   IIDIHHKGVATIKSTGYIVDLNKDRSLDYLVEDYKPETTFSQEYILNKSSEIPVVDFLF
FT                   LVVLKAYTNIAINVKPEDSFINNNAAGVTITSTEVEYMKNILGCKTITDDDYHEDCAYD
FT                   FCNQIQKWINRTTDKIASVAVFLVNHNAFTDTVSFSKRDSGYLTNWPTHWIVPESIEVD
FT                   LFKKEIGEIDMYAFTWGRIANIRLSSWLFAGGLHNYICAEIERTKMTPSDNIEHGIQYS
FT                   KTVCDELGLSTVVVP"
FT                   /product="hypothetical protein"
FT   CDS             28260..28559
FT                   /db_xref="SEED:fig|6666666.127229.peg.21"
FT                   /translation="MENELLIQAISQILQNEASEDTLKIKNKILERIALEADVKASRIP
FT                   APLNITQIGGYINLLRKLDKEEQERQTKQKTAYQQMLEQTLTSILGLPVQPPTE"
FT                   /product="hypothetical protein"
FT   CDS             29313..31316
FT                   /db_xref="SEED:fig|6666666.127229.peg.22"
FT                   /translation="MILAGVILGENGFNILERDGSFKIFGQVGIYYIMFLAGIEMDLVG
FT                   LKRNINKGLLFGVITSLIPFACGLAAGYYLLDYSFSASLLLACIFASHTIVAYPIVARY
FT                   GLNKHSAVTISISATMIALLFALLTLAVIASRYRGGGGLWFWVLFVVKCIAYFSGLFWL
FT                   FPIIIRWFFRRYTDRVLQYIFVIAMMFLAAGTANLCGIEGLLGAFLSGLIFNRFIPHMS
FT                   SLMNRIEFVGNALFIPYFLVGVGMLVNLKPIFTDMGAIVVVVIMVVVSTLSKLMASFIT
FT                   RKILHLPHSHGLVMFGLSEAHAAGAIAMVMVGTSLMVAPGEPLMNTAVLDGVVMMILLS
FT                   CIISSIATDKGARKLKLESDRETSQLTEKGDDEKIMVLVKESAKIRNLTQTAIMMRNPD
FT                   LNRGLICLNVVNDADPTEKSRRMSREILEQAEEICSANDVPVQVQSRLAVNLVNGVVHA
FT                   LRENDASEMILGLHEPRPHDTSYYGQFTEALVQSMNRQVMVVHYHQPVNTIGKIFVAVP
FT                   DRAEYETGFYRWVERVLRLAGQIGCKIVFYSSETTGRLIYQYSQTHHKSVRTDFKLMES
FT                   LTEVYSLRQVIHHDHMLVIVAARPSSLSYQRAFARLPKLLHDHFNDCSLMLIYPDQFGE
FT                   NNSETTFTEPHGNTTFEMSKTSAWLSKWLSKI"
FT                   /product="Na+/H+ antiporter"
FT   CDS             31345..32043
FT                   /db_xref="SEED:fig|6666666.127229.peg.23"
FT                   /translation="MIELRNINKSFGDLQVLKDISLTINDGEVVSIVGPSGAGKTTLLQ
FT                   IMGTLDNANSGSVIIDGIDINSLGSNQLAKFRNQHIGFVFQFHQLLPEFTAVENVMIPA
FT                   MIAGKSQRQAKQRALELLDMLGLKERADHKPAELSGGEKQRVAVARALINNPSVIMADE
FT                   PSGSLDTKNKTELHQLFFDLRDRLHCTFLIVTHDEQLASITDRTIHLKDGEIVANEIKN
FT                   VVDMPCETNE"
FT                   /product="Lipoprotein releasing system ATP-binding protein
FT                   LolD"
FT   CDS             32070..32936
FT                   /db_xref="SEED:fig|6666666.127229.peg.24"
FT                   /translation="MIQLGKQNRLQILRQVDFGMYLDGGEIGDILLPQRYIPKDAKVGD
FT                   FIDVFLYLDMEERLVATTEKPLVQVNQFAFLEVKWINQFGAFLDWGLMKDLFCPFREQK
FT                   MRMQQGRSYIVYCYIDTLTSRIVASAKIEKFLSKEMPPYNIGDKVELLIQQKTDLGFKA
FT                   IVDGKFSGLIYRNELFQEIHTGDRMSGYVKSVRNDGKLDISLQVTGKQHIEDFSSRLFR
FT                   YISNSEDGFCPFHDKSAAEDIYSEFQVSKKTFKKAVGDLYKRHLITIEDDGLRLTMEGR
FT                   ISGIDEE"
FT                   /product="FIG00938655: hypothetical protein"
FT   CDS             33028..33924
FT                   /db_xref="SEED:fig|6666666.127229.peg.25"
FT                   /db_xref="GO:0004311"
FT                   /translation="MSHSNPLLDKALKMIGARNGKMMRPILTLLIARLYGEIDEKALHS
FT                   ACAYEAFHTASLVHDDVVDESEQRRGTQSLNYSSGNKVAVLVGDYILSLALKHISMTGI
FT                   PRLVSIMSEAANRLSDGELLQLYNVNSRSIKEETYYKIIRNKTAALFSACAQSGAISKN
FT                   VSEDEISKAGLLGEYIGICFQIRDDIFDYTNDNIGKPTGNDMKEGKLTLPVIYALNNAS
FT                   NDEISLIAQRVKARVASQEEIAKLVDFTKENKGIEYAISSMKSYADKAKDIIMEYPDND
FT                   VRESLLAYVDYVVGRSY"
```

```
FT                   /product="Octaprenyl diphosphate synthase (EC 2.5.1.90) /
FT                   Dimethylallyltransferase (EC 2.5.1.1) / (2E,6E)-farnesyl
FT                   diphosphate synthase (EC 2.5.1.10) / Geranylgeranyl
FT                   pyrophosphate synthetase (EC 2.5.1.29)"
FT                   /EC_number="2.5.1.90"
FT                   /EC_number="2.5.1.29"
FT                   /EC_number="2.5.1.1"
FT                   /EC_number="2.5.1.10"
FT   CDS             complement(33986..34927)
FT                   /db_xref="SEED:fig|6666666.127229.peg.26"
FT                   /db_xref="GO:0004139"
FT                   /translation="MEHNCNCGCDHHHHHHAESSESKVQQAFEKYNFNISDEQVKKEVT
FT                   NILSKYLKENDTPEVKKFLFNSIELTSLKTTDSEDSIMKFVEKVNKFDDEYPELGHVAT
FT                   VCVYPCYAQICHDTLENEDVEIACVSACFPSSQSFIEVKIAETAMAIHDGATEIDIVQS
FT                   VGKFLAGDYETVADEISEIKDVCGDRKLKVILETGCLGNAENIKKASILAMYAGADYIK
FT                   TSTGKLEPAATPEAAYVMCQAIKEYYENTGIQIGFKPAGGMKTIQDALTYYTIVKEVLG
FT                   EKWLNNNLFRLGTSSLANLLLSDIIGKEVKFF"
FT                   /product="Deoxyribose-phosphate aldolase (EC 4.1.2.4)"
FT                   /EC_number="4.1.2.4"
FT   CDS             complement(34945..35397)
FT                   /db_xref="SEED:fig|6666666.127229.peg.27"
FT                   /translation="MRAVIQRVSKASVTINNEVKSSIGDGYLILLGCENADTQEDIEWL
FT                   SKKIVNLRVMNDENGVMNRSILDTGGDILVVSQFTLWASYKKGNRPSYLRAGSHEITIP
FT                   LYEKTCKQLEKDLGKPIFTGEFGADMKVELINDGPVTICMDTKNKE"
FT                   /product="D-tyrosyl-tRNA(Tyr) deacylase (EC 3.6.1.n1)"
FT   CDS             complement(35394..37256)
FT                   /db_xref="SEED:fig|6666666.127229.peg.28"
FT                   /db_xref="GO:0003677"
FT                   /db_xref="GO:0004518"
FT                   /db_xref="GO:0005622"
FT                   /db_xref="GO:0006281"
FT                   /db_xref="GO:0006289"
FT                   /translation="MFNKKQEQLKNIVLNLPEKPGCYQYFDKEGTIIYVGKAKNLKRRV
FT                   SSYFLKNQVSRKTQVLVSKIHDIQYIIVNSDEDALLLENNLIKKYKPRYNILLKDDKTY
FT                   PSVCVTNEEFPRIFKTRKIIPGAGTYYGPFSHAATLNNMLELINRLYPLRRCRQAITKD
FT                   RINEGKLKSCLEYHIKNCKAPCIGLQSHEEYMHDVAEVHEILKGNTRSICQKLLDKMQV
FT                   LASEMKFEEAETVKKKYLLALEFCEKSEVIANYSHDIDVFSIADDEKSAYINYLHVTNG
FT                   CINQAFTFEYSKRTEETPEDLLSLGIVEMRQRFKSHSKEIVIPFPINTELSDVTFTIPQ
FT                   RGDKKHLLDLSIANVRQYKFDKAKQAERLNPEQRTTNLLKELQQKLKLPKLPMHIECFD
FT                   NSNIQGADAVAACVVFKQAKPSKKDYRKYNIKTVEGPDDYASMAEVVFRRYSPLLPPEK
FT                   QAELGISHDTTDHSEHITPNLIIADGGKGQMEIIRKIVHEKLNLDIPIAGLAKDGKHKT
FT                   SELLAYNNGQIVTIGIESNSPLFLFLTRIQDEVHRFAIQFHREKRSKRQISSELDNIKG
FT                   IGETTKNTLLKHFKSVNRIKQASVEELSDVIGQTKAKLIKENLK"
FT                   /product="Excinuclease ABC subunit C"
FT   CDS             complement(37299..39194)
FT                   /db_xref="SEED:fig|6666666.127229.peg.29"
FT                   /translation="MIQEYDVIVVGAGHAGCEAASATARMGAKTCLITMDMNKIAQMSC
FT                   NPAVGGIAKGQIVREIDALGGMMGIVTDRCALQFRMLNISKGPAVWSPRAQCDRQKFIL
FT                   TWREILENTPNLNIWQDQVNELITSEDNSDSSHKHTIVGVRTIWGAEFHAKCVIITAGT
FT                   FLNGLMHIGRKQVTGGRIAEPAELELTKSINKLGIKSDRMKTGTPVRIDKRSVDFSLTE
FT                   RQDGDETFHRFSFMSEPRKLKQLPCWIFKTNKEVHDILFSGLKDSPLYNGQIKSIGPRY
FT                   CPSIETKLHTFPDKDEHQLFLEPEGETTNEMYLNGFSSSLPMDIQINALRKIPALRNVV
FT                   VYRPGYAIEYDYFDPTQLTHSLESKIVSGLFFAGQVNGTTGYEEAAGQGLIAGINAALK
FT                   CAGKPAFTLKRNEAYIGVLIDDLVTKGVDEPYRMFTSRAEYRVLLRQDDADMRLTEKSY
```

```
FT                   NIGLASKERYDFFVKKRDEIEHLIEFTKNFSIKPALINPALEKIGTTPLERGCKLFDLI
FT                   GRPQITIKGILPYIAAFNSEINKITERQDEIIEATEIRIKYKGYIEREQQVAEKMLRLE
FT                   NIRIKGKFDYSKLNQITIEARQKLAAIDPETLAQASRIPGVSPCDINVMLVLMGR"
FT                   /product="tRNA uridine 5-carboxymethylaminomethyl
FT                   modification enzyme GidA"
FT   CDS             complement(39303..39917)
FT                   /db_xref="SEED:fig|6666666.127229.peg.30"
FT                   /translation="MKTKNLIVLLFAMTMGTSCWAQNECKQTENKVIETIMQRRSIRKY
FT                   KDMPVEKEKLQLIADCGIAAPNAMNRQAWAVRIVDSKDFINGSTEAMKKVNPNITGNDK
FT                   NMYRNAPAIICVAVPSGDFSYIDAGLMGENMMLAAQSLGLGTCCLGGVAMQLNTRPELK
FT                   DYLLRLKLPNDYKIAYVLAVGYPDEQPQAKQRDKSKIMFVE"
FT                   /product="nitroreductase"
FT   CDS             40220..41872
FT                   /db_xref="SEED:fig|6666666.127229.peg.31"
FT                   /db_xref="GO:0004333"
FT                   /translation="MAQVPDFKYAPMFQLGKDDTEYYLLTKEGVSVGEFEGKPILKVTP
FT                   EALTTMANHAFKDVSFLLRRSHNLQVAKVLRDPEASENDKYVALTFLRNAEVASKGMLP
FT                   LCQDTGTAIIHGEKGQQVWTGFCDEEALSKGVYKTYTEENLRYSQNAPLSMYEEVNTKC
FT                   NLPAQIDLEATEGMEYRFLCVTKGGGSANKTYLYQETKARIQNPGTLIPFLVEKMKSLG
FT                   TAACPPYHIAIVIGGTSAEKNLLTVKLASTHYYDNLPTSGDETGRAFRDVELEKQLLQE
FT                   AYKIGLGAQFGGKYMAHDVRVVRLPRHGASCPIGLGVSCSADRNIKCKINKDGIWIEKL
FT                   DSNPGELIPEEYRKAGEGEVVKVDLNRPMSEVCKQLSQYPVSTRLSLSGTIIVGRDIAH
FT                   AKIKARLDAGEGMPQYLKDHPIYYAGPAKTPAGMACGSMGPTTAGRMDPYVDEFQDHGG
FT                   SLIMLAKGNRSIDVTNACKKHGGFYLGSIGGPAAILAQNNIKSIECVEYPELGMEAIWK
FT                   IEVEDFPAFILVDDKGNDFFKQLKPCEGCSILN"
FT                   /product="Fumarate hydratase class I, anaerobic (EC
FT                   4.2.1.2)"
FT                   /EC_number="4.2.1.2"
FT   CDS             41902..42465
FT                   /db_xref="SEED:fig|6666666.127229.peg.32"
FT                   /db_xref="GO:0006412"
FT                   /translation="MEEVKQYLNDAEESMEMATMALEEKLGRIRAGRANVHILDGIKVD
FT                   SYGAMMPLSNVANLSTPDARSIVIKPWDKNMLKVIEKAIIDSPIGIMPENNGEQLRLGI
FT                   PVLTEERRKELVKQCSKEAEQSKVSIRNARRDAIELLKKSVKGLGVPEDVEKDAEADCQ
FT                   KLHDKYIKKVDEMLEEKNKEIMTV"
FT                   /product="Ribosome recycling factor"
FT   CDS             42473..43402
FT                   /db_xref="SEED:fig|6666666.127229.peg.33"
FT                   /translation="MHGLVVKNTGSWYSVLTDEGTLEQCAVKGNFRIKGIKSTSPVVIG
FT                   DRVEISDGMITSVDDRKNYIVRKSINLSKQSHILAANLDQCFLIVTVNHPITSTTFIDR
FT                   FLATAEAYNIPVVLVFNKLDIYTDEDKEYMQLMKDLYENIGYKCVVISAITGEGMDEIR
FT                   MMLDSKVSLVSGNSGVGKSTFINSLLPDLNLKTADISDVHDTGMHTTTFSEMYPFKRNP
FT                   DSNEVPNSFIIDTPGIKGFGSFDMQREEISHYFREIFNYSHDCRFNNCTHTHEPGCAVL
FT                   AALERHDISQSRYNSYLSMLEEGDGKYR"
FT                   /product="Ribosome small subunit-stimulated GTPase EngC"
FT   CDS             complement(43788..44300)
FT                   /db_xref="SEED:fig|6666666.127229.peg.34"
FT                   /translation="MEPQEEKKVLLKLRLWIAFFMVALVLSGVTAFPVETELKTLCSVA
FT                   GLDSATVNDYPAIFGFLYSIKESIVELNQTTPQLSYGYDWLAFAHLVIALMFIGPFRDP
FT                   VRNKWVIQWGIIACVAVVPLALICGPIREIPLYWRFIDCSFGVFGVIPLIICLRLIKRL
FT                   ESDKHQD"
FT                   /product="hypothetical protein"
FT   CDS             complement(44544..46541)
FT                   /db_xref="SEED:fig|6666666.127229.peg.35"
```

```
FT                   /translation="MKQWASENGVTHYTHWFQPLTEGTAEKHDSFIEHDGKGGMIEEFS
FT                   GKLLVQQEPDASSFPSGGIRSTFEARGYSAWDPTSPVFIIDDTLCIPTVFISYSGEALD
FT                   YKAPLLRALHAVNVAATEVCHYFDPAVKKVTSNLGWEQEYFLVDEGLYAARPDLLLTGR
FT                   TLMGHDSAKNQQMDDHYFGSIPERVAAFMRELEIEALELGIPVKTRHNEVAPNQFEVAP
FT                   IFEETNLAVDHNMLLMSLMKRVARKHGFRALLHEKPFAGINGSGKHNNWSLSTDTGVLL
FT                   HAPGKTPEQNLRFAVFIVETLMGVWKHNGLLKASIMSATNAHRLGANEAPPAIISSFLG
FT                   KQLTELLDNIEKADKEDLFRIDGKQGMKLDITEIPELLIDNTDRNRTSPFAFTGNRFEF
FT                   RAVGSEANCASAMITLNAAVAEALVNFKARVDKRITEIADQYKENSGHTAKFHAIIDVL
FT                   REDIRACKPIRFDGNGYSDEWKEEAAKRGLDVETSCPVIFERYLDASSIEMFSSLGVMT
FT                   KKELEARNEVKWETYTKKIQIEARVLGDLSMNHIIPVATRYQSQLLNNVENMRTTFAPE
FT                   KANKLSARNLKIIEEIATRTETIESLVEELVSARKVANKIEDEHLKAIAYHDTVAPYFD
FT                   KIRRQIDKLELIVDDACWPLPKYRELLFIR"
FT                   /product="Glutamine synthetase type III, GlnN (EC
FT                   6.3.1.2)"
FT                   /EC_number="6.3.1.2"
XX
SQ   Sequence 46551 BP; 13534 A; 8876 C; 9758 G; 14383 T; 0 other;
     taactcagga tacaccagca gtactcagga atcctccatg actgctcttg gcatttatcc        60
     aatcggtggt aaatatgacg atggaaatgg cggcaagatc acttctaagg acggtagcgg       120
     ctgcgctccg gacaagatgc ttcaggctta ctctgttcct ttcatgcttg cggaacttgt       180
     tctcgcagga gaaatcaatg gagatgccaa aaaatacctc gaagatggta tcaaggcttc       240
     gattgcccat gtcaacgcag ttactaagct ttgtgacgct tccgccccgc ttatctcttc       300
     ggatgatgct cagaagcttg tggacgaagt caatgaaaaa tttgacgctg cttccgctga       360
     aggaaagctt gagatcgtca tgacagagaa gtggattgcc aacttctata accctatcga       420
     ggcttacaac gatatccgtc gtaccggtta tcctaagctt ttcaagggag acgagaataa       480
     tatgggatgg acaccttacg cccagactgt tgaggcaacc ccgggtcttt cctcttttga       540
     gcttgtcact atcctggatt atcctagaat actctggtat cctcagaatg aaatagacac       600
     aaaccctaac gtttccaact cttcaaggaa cgttgccgtc aagactgtgt tctgggacaa       660
     gtagttcaac caaagaaatt taaaagatta tgaaaatcaa caagatattt gcattgattg       720
     cgctcggagc cctgttctgc agctgcgatc agaagcttcc ttacgatctt gaagggacaa       780
     agcacggtgt cttaatatcc atcagcaagg ttcagggtac atccggaatc ctttcgacag       840
     atatgaacgc cgggaactac caggtcaagc ttgaccttcc aatttaccag ggagactact       900
     caatgcttga ggatgctcag ataatggctg tctatacagg tgtagacaaa accaagaaaa       960
     gcgcaatcgt tgcggaaggc atcaaggctt tccctgacac tgtcaaactt aacatcaaag      1020
     atgtggccag caagctcgga atcagcgaga taaacgttgg tgacaagatt gaatatactc      1080
     catgctacac tctcaaggac ggcctgcagg taaacggctg gtcagagctt actggtttca      1140
     acaacaagaa tttctcttcg tggaaacttg aagacggatc cagctttgca tatagagtat      1200
     cctatactgc tttcgctcca ttgcagctgg ataagttcca aggtgatgac attccgttct      1260
     cgatggatgg aggagacgaa ggttccatga aagtctcaca ggttgacaat aaaccagagg      1320
     caaaatggat tccagctgga gcaaatgtag aagatctggt tggactgaaa ttcgaaggcg      1380
     atttctggtt cggaggcgat tcatttatca tctggattaa tccggctgat tactccctta      1440
     tcatccctga tcaggtaatg tgtcctggtt tcacctatgg aagctatggt acattcgatg      1500
     cttatatggt atcatgcgac ggagaaattg acactctgaa caataccatc gtgtttacct      1560
     tcggtacaag ctggggtcca tactccttcg gaggtgacac agtaatattc gaatttcctc      1620
     gttaatagaa ttcaatgatc taaatatgaa gaaagagttt ggtgcatcac caaactcttt      1680
     cttttattat ccaaattatt gaaatcaaaa actaatgcag gtcaaagacg gcgtgaatga      1740
     agttggttcc gaagcagccc ttgaagatat actattacta ttcagttcta aatagagtcg      1800
     gaaaaaaggt tatctaatca accttctttt aaagaggtga caaaaagtca agttttccta      1860
     tttccggagc cattgatact caagatcagt tccttgaagt tatccattag gattactgac      1920
     tatgccaagc gggtcaaaaa aagacccgct tttttattt gcttttgttt tttatggatt      1980
     tggggtcttt tggttacaga ttgaaagtgc aattggtata aaagtcgatt ttttatatac      2040
     acaaaaacgt tttagagcct cgtggagcat agtttggcgc tgatttgcgc gtgttgcaaa      2100
     tatcataatt aataacttta tttgcagaaa tattgagaag atggcttata gatgtaggcg      2160
     tcggattaca gtttttaact tttcctcgct tacttttatt ggccattgct tttcactttt      2220
```

```
gggggaataa taatactcat ttaatattgt tttaatcatt gcttatggaa aaaatcaagt      2280
tttttctcgc ggcattgctg gtttttgtct ctgtaggact catgtcagcc cagaaccgta      2340
cgattacggt aaaaggtaac gtgactacag ctgtagacgg ccagccggct ccaggtgcct      2400
atgtatcggt agaaggcact tcagttggta cggttactga tctggacggt aatttccaga      2460
tcgacaatgt accagcttcc gctaagtgga tagttgtaag ctatcttggt tacaaggatg      2520
cccgtgtcgc cattaccgcc aacattcagg tagctctcga agatgattcc gagatgctcg      2580
aaggtgcaat cgctactggt atgtacaatg tggaccgtcg actcaacaca ggatctgccg      2640
ctaagatcga tgctgctgag gccaaactcg acggtatcgc cgatatcagc cgttccctcg      2700
agggacgtgt ggccggtgtg tctgtgcaga acgtatccgg tacattcggt actgccccta      2760
agatccgcgt ccgcggtgct acttctattt atggttcatc caagcctctt tgggttgtgg      2820
atggtgtaat catggaggat gtcgttgaag tcagcgctga cgatctctcg tcaggtgacg      2880
ccaacaccct tatctcttct gcaatcgctg gtctcaactc cgacgatatc gagagcttcg      2940
atatcctcaa ggatggttcc gctacttcca tttatggtgc gcgcgccatg gccggcgtga      3000
ttgtcgttac gaccaagaag ggtaaggccg gtcagaaccg cctcagctat acaggtgagt      3060
acaccatgcg tctgaagcca agctacagca cattcaacat aatgaactct caggaccaga      3120
tggctgttta tcaggaaatc cagcagaagg gcttccttgc ttatgccggt actgccaacg      3180
ccaggaactc cggagtatat ggaaagatgt accagcttat cagccagtat gatgctactt      3240
taggcaagtt cggtctccag aatactgacg aggccaagat tgcttatctc cgtgctgctg      3300
agtacaggaa taccgactgg ttcgacaggc tgtttacata taatatccag cacaatcatt      3360
ctgtcagcct ttccggcggt actgaaaagc tgaatttcta cgcttctctc tccctgctcg      3420
atgatgcagg atggtctttg cagagcgatg tcaacagata taccctcaac ctcaatgcgt      3480
cctataagat ttccgacaag ttgtctttga atgtaatcgg taacggttcg catcgtacac      3540
agagggctcc tggaactttg aatgcagagg tcgatgccgt ttcgggcgag gtaaagcgtg      3600
atttcgatat caacccttat tcttacgctc tgaatacatc caggactctc gatccgaatg      3660
tgttctatac ccgtaattat tctcctttca atattatcca cgagttggat aacaactatc      3720
tcgacattga cgtcgatgaa ctccgtatcc agggcgaact taagttcaag ccgacaaaag      3780
gtcttgaact cagcgttctc gccgctacca agtctacctc gacttcccag gagcattaca      3840
tcaaggatac cgccaaccag gctcttgctt atagagagct gagtactccg gacataaggg      3900
ataacaaccc tttcctttac agggatcctg atcatccgaa cgcgtataag atcaccgtcc      3960
ttcctgtcgg cggtgtgttc gagcgcaaaa cgaacgcttt caaaggctgg gacttccgta      4020
ctactgcaag ctacagcaat acattcaata atgtccatat actgaacctt ttcgccggcg      4080
ccgaggccaa taacgtgaac cgtcatgcta catggttccg tggctggggc atgatgtatg      4140
ataagggaga gaaggctatg tatgactata ggatattcaa gaagggtgtc gaggataaca      4200
gcgactacta tactctcgag aatactatag ccagaacagc agctttcttc gccaatgcga      4260
catattcata caagggaaaa tatacaatca acggtactgc ccgttatgag ggtaccaaca      4320
gcttgggtaa ggcccgtagc gccagatggc ttcctacatg gaacgtgtcc ggagcctgga      4380
atgctcatga ggaaagtttc atgagagacc tctatcctac tgtttctcac ctttccctca      4440
agacttcata ttccctcacc ggagaccgcg gtccttcata cgttacgaac gcagaggaga      4500
atatcaacta tcgtaatcct tggagaccgt ccgcctctac caaagagagc gctctctata      4560
ttgtcgatct tgccaacgaa gagcttactt atgagaagaa gcacgagttc aatatcggta      4620
tcgagactgg tttcctcgac aatagaatca actttgccgc agactggtat acccgtaaca      4680
actatgacct tatcggacct gtcatcactc agggtgtcgg tggtcagatc gaaaagtacg      4740
gtaacgtcgc ttcaatgaag tctgatggtt atgagttgtc tctcaccact accaacataa      4800
agaataaaga tttcagttgg acgaccaacc tggtcttctc ccattctcat aacaagatca      4860
ccgaacttga gaacaagtcc cgtatgatgg atcttatttc aggtactggt ttcgcaagag      4920
agggctaccc ggtacgaagc ctgttctcga tcaagttcat gggacttaat tccgagggac      4980
ttcctacttt catcaatgag gacggtgatg tcactatctc tgatatatat ttccagaccc      5040
gtgactccaa atctcttgag aacctcgttt attcaggcag tgtagaccct actgatgtgg      5100
gctccctggg aaatatcttc acttataaga atttcaaact caacgtattc gttacgttct      5160
cgttcggtaa tgtcatccgt ctcgatcctg tattctctga cgagtatact gatcttgtcg      5220
ccatgccgaa ggaattcgca gaccgttggg ctgtcccggg tgatgagaac ataactaaca      5280
tccctgtcat ttccagcaga tggatgaacc attattacga ggctaattct aattcaaggc      5340
tcagctatgc atacaatgca tacaactatt ctaccgagcg tatagccaag ggtgacttca      5400
tccgtatgaa ggagatatct ctttcttatg acttccctaa gaaggttaca gaaactctta      5460
```

acgtcagcaa ccttggtctg aagcttcagg cgaccaatct tttccttatc tatgctgata 5520
agaagctgaa tggccaggac cctgaattct tcaataccgg tggtgtggct gtcccacttc 5580
ctaagcagtt taccctcacc ttaagaatcg gtttttaatc tggaggatga aaaagatgaa 5640
aaatagaatt ataaatatat ttgcttcagt tattgctgta gcggcaatta cttcttgcga 5700
ctcttttctt gatgttactc ctgataagag agccgttgta gatactcagg aagaggtcca 5760
ggaccttctt gtttcggcat accctgacgt tacttatctg gctatcacag agtatatgtc 5820
tgacaatgta gagaatctcg gagacaatat ctccggctct acaagacaga tagagcagct 5880
ttattattgg aacgagattg tagatgagaa taacgattct cctaaggatt tctgggacaa 5940
ctgctacaaa tgcattgcca ccgccaattc cgctatcgcc aagatcaatg atctcggtgg 6000
tccggaggcc ggtctgcagg ctgagatggc cgaggctctg ctttgcaggg catacgctca 6060
tttcatgctt gtcaatatct tctgccagaa ctacagctcg aagacaagcg gttcagatct 6120
tggcatacct tatatgacaa aatccgagac cacgcttaac ccgaaatatg agcgtgggac 6180
agtcaaggac gtgtatgagc atatcgaaga ggatattctt gccgcccttc cttatgtgac 6240
cgatgcatac tatgacgtgc ctaagtacca ttttacccca agcgcggctt atgcattcgc 6300
ttcccgtttc tacctgttct acgaaaagtg ggaccttgcc atcgactatg ccactaaatg 6360
tcttggcgat tatccggcct ctgtcctcag agactggcag tacagatcca cccttgcgga 6420
tagcaggcag gtcaagactc ttgatttcat aaacgaatcc aagaagagca accttcttct 6480
tttgaccggt gtctctcaat atggtatttg tttcgcttat cctgcagcct accctaagta 6540
ctctcatacc aattatctcg gagcaactga gacttgtttt gccaagaacg tatggggcaa 6600
tacttctgta gggtacggcg gagccgcaag gtactattat gaccctgccg cttcttatgt 6660
cggcggtaat ttctcccagg tcgctctgtt caagatacct tacttattcg agtacactga 6720
cccggtcaac gatataggtt atcctcacgt cgtttatccg gcgttcacca ccgatgagat 6780
tctccttaac agggccgagg cttatatcct tacaggagat tatgagcgtg cttgcgagga 6840
cctcaatatc tggatccata atattctgaa tccaagttac ttcaataagg atcttactcc 6900
ggatatgatc aagtcattct atgatgctac accttatgca agatgggata acttcgaccc 6960
atcgacccgt aagcttcaga acaatatcaa gaagcagctg catcctaagt tcgagatccc 7020
ttccgatcct tatgccgagc ctatgcttca gtgcgttctc cagttcaaga gaatcgagaa 7080
ccttacccag ggactcagat ggtttgacat caagcgttat ggtatcgaaa tctggagaag 7140
gactcttaag tcagattcgg ccagccctgt cggttattct cctgcaaggc ttgacgatgt 7200
ccttacggta gacgatccgc gtagagctgt acagctccct caggaagtga ttcttgcagg 7260
tctggataag aacccacgtt aattcaaatc cggaattgtt atgaaaaaga aaaatatatt 7320
aattgcattt gtcgcagcag tctcacttgt gaccgtatcc tgcaccgaga aaaaactcag 7380
tccggacagc gtcatttctg cgccaagtgt cgaaaagaat gctttcgacc tctggcttga 7440
ttacaatttc ctcaagccat ataatatcga attccgttat cggtatgatc tcaatgagag 7500
tgatatgtac agaatgacga tccctgcgga ttatgatgct tcaatcatgt atgctcatct 7560
tgtacaatat ctttgtatcg atacttacaa cgaagttgcc ggcgtggatt tcaccaaggc 7620
ttatttccct aaaatgttct tccttatcgg agagtgggag tacgacaaca atggaaatat 7680
catcctcggt actgcggagg gcggaaaaaa gattatgctt tcaggcgtta atctccttcc 7740
ggggaaattc gcaagtttct cgggcagtgg tttgaagaag gagctcagtg agtattacat 7800
caagactatc catcatgagt tcacccatat cctcaaccag atcaagggtt attctgagtc 7860
tttcaaggag ataactgcat ctacgtatca ggctgattca tggtcggagt ctgattcaaa 7920
ataccgtcag agaggttata ttacaaaata cgcccagaaa gaggatcgcg aggatttcgc 7980
cgagcttctc tccgagtata tcactcatga tgaagactgg tggcagaaag agctgactaa 8040
agccggggat gaaggcaaaa gtcttatcct tgccaaattc attatcgtaa gagattatat 8100
gtcctcgtca tggggtatcg atgtcgatga gcttagagat gtggtcaacc gcaggttcga 8160
cgatgtcgtc gaaggacgtg tcaatcttac ggatctttca atctaacagg aggaaatatg 8220
aacaacagat tttcatttc attagccgca gtagcggtat cgctggtttc tcttccttcc 8280
tgccagaagg atcaggactc cgtttttgac gggtctctca cagagcgcca gacccagtat 8340
ctggatggct tccgtagtct tataacaggt gcggaaaatg gctgggcaat gtattatttc 8400
agtggcggag cctacgctac atcacgcgta ttcacagtat cttttactga caacgaagta 8460
accgcatcga gcgaagacgc tccgagcgtc accgccaaat cattttacaa ggttgcggta 8520
acttcagctc catccctgat tttcgataca tataacaaag tccttcatac ttatgcggca 8580
ccttctcatt cttattatca gggaagaggc ggtgactttg aattcactat agagaactat 8640
tccgaagata agatcactct tatcggaaga agagcttacg aacagcagat tctgatcccg 8700

ctgaaggagt ctcctgccag ctatctcaat aagatcaagt ctttcgccca aagcttcaat 8760
cttactcagg tcgccggcaa ggtaggtaat ggcgatgtcg tcatctattt cgatctcagg 8820
aacagattcc tttatatagg cagacttggc gctgaagata cagagcttga agaaattccg 8880
tatatctata ctgagaacag cttgttgctt cagcatcctt tctggtataa cggaacgacg 8940
ttctccgagt tctcctacaa tgctgaagag catagtataa cgtctgaggg tatcacattt 9000
aaacagtttt agacgcttat gaaaaagtta ttcggaatat tgatggtcgc cctgtctgtt 9060
ttctcgcttg cttcctgcaa cggcgaaaac gagatagacg agaaaggggc tcctacaatt 9120
gtaataaaca gcaaggatat ccagcttccg gcgaagggcg gtgagtcgag tttccttgta 9180
gactctgatg tggattttat cgtttcggtc agccatgact gggtttctgt cagcaaagat 9240
ggaaatcgcg taaaggttgc agccgagaag aactcatccc ttcagggcag atatgccgaa 9300
ctttctgtat ccaacggtaa ggacggtgct tccatagtga tcaaccagct tggctacatc 9360
tcttctgacg tcgaattcga cgatatatct gtcagctatc tcggacagtt tgtcgagctg 9420
gaatatgatt acgatgattt catttctgca tctaccgatg tggactggat tattcttgat 9480
aattccagag gtacggaagg tcaaaggatt ctccgtgtca atgtcggaga aaacaagacc 9540
ggaagccagc gtagtgcaac tatcgaatgg accctcggtc ttgagtccgg ctctttcgtc 9600
gtaaagcaga gaaaagaaat gaaagaatac tccttgaata attcatggaa tatctcatac 9660
ctcggcgtag acaattctat ctacaaagtt ggagtttccg gagtcagcgg agagtatgtt 9720
gtaacagctg tttctgcttc taaactggct cagggtggtt tctccaatgt ggcagaatat 9780
cttaccgagt ctgattctga tattgccaag gctatagctg acggcgagat cgctgtctat 9840
tctggcgaca agactgtcga actgagcaat ctgtctggcg gagagaatta tgctgtcgtt 9900
ctcggctaca aatcttctaa atccgagctt accggggaat ttaatttcgt tacctttaaa 9960
gtagaaggtc cggatctccg tactccttat gagaagtggc ttggtacctg ggagatagag 10020
agagagtctg aaattgacgt gtggacgata tccaagaatg tcgaaaatgt atcttatact 10080
atccttggcg tggatggaac tagcaaggaa aatactcctg acgttccggt cattgctgaa 10140
tttgattcag aaacaggcaa ccttattatc aaggcacagc attgcggtac atttactagc 10200
agctcttatg gcgaggcgga tttgtctctc tatggacttg gaccagagta ctactataca 10260
tattcatcag agccatatac aatcgctacc ctttctttga gtggcaagaa ttcagctaat 10320
cttgtgcgca gaaggacttc aactcatctg taaggctgaa tcccgtattc acgtccggct 10380
accattacag gggcatcacc aagagccgtt tcggagacta cgattcggcc ctcgaggatt 10440
ttgaacatgc aatcgcgctc cgcccgggat acaccggcct ttatttcagc cgtggagtaa 10500
catatttcct cgcccagcag ttcgacaagg ccatcgccga ctttgacaca tatatccgca 10560
aggagcctaa ggatccgtca gcttttctta acaggggagc gtcatacctt tatcaggggg 10620
acacgacgaa agctctgcag gattatgaca aggccattaa gatcgaccgt tacgaacctg 10680
aaggatatat aaggaggga aacctttaca tgctttcaga gaacctggaa gcggcgatga 10740
aggattacga caaggcgatt tctctcgata ccagcagcac tcttgcatat tacaaccgcg 10800
ccctgatcaa atatgaactc cagaactata tgggggccat ggaggatctc aaccgcgtcc 10860
ttcgcgacga tccgggcaac tccctgactt tgtacaacag gagtcttata tatgccaaat 10920
tcggggaata cgggtcggcc ctcgacgata tggacagagt gatcttcaat aatccccgca 10980
atgtgctggc ttatttcaac agggcttcct tcttcataga catggggcgc tggcaggatg 11040
ccctggagga ctatgacaag tcgatcgaat tatatcctga cttcgccaag gcatataaga 11100
acagggccta cgtggagctt cagctcggac gcaaggccgc ggccaagaaa gactacgata 11160
cagcgaacag acttgtcgat gagtacagga taaagtctgc cgacggtacg gtatcttttg 11220
ctgacaccac aaagaaatat aattctctca tatcgttcga taccgatttc gcgaagaaca 11280
atttcgatga cgagattctt cttcacaggg atatcgacat caggctcaag ccactttacc 11340
gcttcgtcat aacaaaggat aaggacgatg tcaactatgc gctccggaac cgttatgaga 11400
accctcttat cgacaagttc ataaaggagt ccccggttcc tgtctcaatc gtgaacgaca 11460
tcgccgttga gaagtcttcg gcagaggtgg aagaagaact ttatggagtc cgggaatctt 11520
cggaatccgt caactccttc ctcagaggcg tatatgaagt ctccggaaag cagttcaacc 11580
ttgctctgaa ttatttcggg gacgctatct cgcaggcctc gggctcggga gacaagaccc 11640
aggatatgta caaatccctt tactacatga acaggggcgt tctcagggca gatatgatcg 11700
atttcatttc ttcgatagag aataacgtac agactcttac gatggatgac cagggtacga 11760
ccagggccag agtaagggac agggtcaacc gctcatacga ttacggggat gccatttctg 11820
atatggaagc tgccgcttca atagtatcgg atatccctta cgtttatttc aatctgggta 11880
atctgtactg tctgtcatcc cgtcttgtag atgccatcgg cagctacaat aaggctctgg 11940

agctttatcc gatgatggga gacgcctact ataacagggg actcgtgctt atctatctga 12000
aggataagga gaaaggctgt atagaccttt cccgtgccgg agagcttgga gtaggcgagg 12060
cttatagtgt aatcaagagg tattgcgagg aggcgaaatg ataaagttca tgagtttttc 12120
cagcggaagc tgcggtaatt gctactggct cggaaatgac gagaaaggaa tactgataga 12180
tacaggcgtt agtctgagac atctcaagaa actgatgctg ctgcacggac tggacgagaa 12240
ttcgttcagt tcgattcttg tgacccatga ccacatggac catatccgca gtctcggctc 12300
attctgcaaa agactccaga agcccgtcta tgcgactacg ctgctccatg aggccctgat 12360
ccgccactca tatgctatgc cttcgatcat gggatgccgc cgcgatctcc gggaaggcga 12420
atggaacgat gtggacggtt tcatggtcca ttatttcgtc gtccctcatg atgctaccca 12480
gaccgtagga tatgccatcg aattcgaggg gctgcggttc gttataatga ccgatatcgg 12540
ccgtatgaca gacgaggccg tcgaattcgc gtcgaaggcg gacgtcgttg tcgtggagtc 12600
gaattatgat gtcgacatgc tgatgggggg ctcatatccc catgatctca aaatgcgtat 12660
ctgccagggg aacggtcatc tcagcaatga cgagtgcgcc gccgccatca agcgtttctg 12720
gcatgagggg ctgcggttca tattcctctg ccatctcagc gaaaacaaca atactcccac 12780
tcttgcctac gagacctcga aaagggctct ggacgagttg ggagtatcgg agaaggtcca 12840
gctggcggct ctgccgaggc agtatccgtc gaggctgtat gtgataggct aggccttgga 12900
aagccgtttc agaccggcgg ctatcctttc cagaccttcg cggcagattg atgccggcgt 12960
ggcgagattg attctgatga agccgtcctg tccgtacatc gctcctgcgt ttacccatac 13020
tttttcattc tggagcaggc tgttctcgat ttcttcggat ttgaggccca ggctgcgaca 13080
gtctatccac ggcaggtatg tgccttccag tttcactatc ctggcgaacg gaatctcttt 13140
ggcgaatatt tccttcagag tacggtaatt accgtagata taggcgttga gttcgtcgag 13200
ccatgcatct ccttcattgc tgttgtgata aaatacaaat aggaagttac tatcatcgta 13260
atatctccct atttgtatat attgtatgtt ctgaaatttt attgaagtat ataagtttta 13320
ttatttcagt ttttcaaatt catttcttat agcatcggca acctgttgca tttcttctgc 13380
agggaatgat tttttaggtc ctccaaactg catatcgtgt tctgattgca ttggaacgag 13440
gtgtatgtgt gcgtgtggca cttcaagtcc aagaacggct tcacctactt ttatgcaagg 13500
gcacactttt ttgattgcaa gtgctacatg ttttgcaaat tgctggaaac gcgcaatttc 13560
gtcgtcttcc atgtcgaaga tatagtcaac ctcgcgtctt ggaatacata aagtatgtcc 13620
ttttaccaat gggttgatat ctaagaaagc atagaactca tctgtttgtg cgcagcaata 13680
acttggaatt tctccctgtg ctatttttga gaatatagtc ataatgatta gatgctaata 13740
ttagtgattt caagttcaat aataccttgt ggtattttga ttgtagccac atcacctact 13800
ttcttgccaa gtaagccttg tgcgatagga gttttgacag agatttttcc ttcgcgcaga 13860
tttgcttcgc tttcgcttac aagagtgtat gtcattacag cacctgtttt aacattcttc 13920
atttctactt tagtaagaat ctgaacagtg tctgtattta ctttagatgt atcaataatc 13980
tttgcatcca tcaagatggc cttgagttct gcgatttttg tttcgagttt gccttgtgct 14040
tcttttgcag catcgtattc ggcattctcg ctcaaatcgc ctttttctct ggcttcttgt 14100
atttgacgga taacttcagg gcgttcaacc tcttctaatt gcttaatttg ggctctgagt 14160
ttatcatagc cctcttgtga catatatgac attgtataaa agtttttttag taattaatat 14220
gtattaagtt tctcttataa ttattcgctt atattaaaaa gaataagaat tccaatacta 14280
actgtattgg aaccctttttt atcttaaatg cgttgcaaag ataatacttt tattaatacg 14340
gacaaaattt ttattgattt tttagattta gcttatcttg ggtaaatcga taatgattat 14400
taatcaggtt atttgtgatt ttcttcccag atttgcatgc gttgttctaa taaatcatat 14460
tgattgtcac gtatgaacga agtacaaact ctaagtcctt gctgtttgct tcctgttgta 14520
tccagtgcaa tagcactaac tccataatac atcagttcgc gtgcaagttg gcttcctgtc 14580
atgcctggat atcctattgt gaaatagaaa ccatcagcga ttgattcatc catgtctttg 14640
tcataaacaa tgatgaagcc atgacgtgtg aaaatctcct ttaacaagtg tgctcttctg 14700
ccgtattcac tgatatctgc gcggaaatca taaatgccgt cgcttgcagc tttaagcatt 14760
gctgccatgg caaattgtgc actatgactt gttccacttg agagtgcgta taagactctg 14820
tgaataaata cactaccgaa tgttccgttt ccatatttct tctccaaagc aggataagat 14880
ctgtggtaga gtttgtttga aatacagcta acacctattc tttcgccggc atatgagaat 14940
gctttacttc cacttatgag caatacatag ttatctgtgt atcgtgctac tgttggttgg 15000
tatggaggtt ggaatggtgt acttaagtct ttacggaagt ccattgcgaa gtatgccaaa 15060
tcctcaaaaa cgatacaatc gtattttgtt gctaagtcac cgattgtttt gagttctgat 15120
tcagtcatac atatccaact tgggttattt gggttggaat atacgatggc tgcgatatct 15180

ccgtttttga gatatgattc aagttttgga ccgagatttt ctccgcggaa gtcatataca 15240
tcgaatgtct cgtattcagc tcccattaca acacattgtt gtttttgtac agggaatcct 15300
gggtcgatga ataatatctt attcttttta ggatcgcttt ggcttgctgt gaggaatgaa 15360
gcaaatgttc cttgcatgct tccgcaaaca ggtacacagc cttctgcgtt tacatctact 15420
ccgataaatg cttttatgaa tcggcttgct tcttctttta gttctggaag tccgtttata 15480
ttaggataca agcgtgctat tccgttttgg agagattgta tttgtgcttc aactccatat 15540
ttagatgcag gcagtcctgg aactcccatt tccatcttta tgaattctac tcctgaagct 15600
ttttctgcat tagaggcaat ggcaaccact tcgcgaatag ttgcttttgc aaagtctttt 15660
attccaaagt cagaaatagc gttgtcgatt agttctttat ttattggcgt tttcatatcc 15720
gaaatcaaat gctttgtaat tagcttcaac gatggcttct ccttttgatg agaatactct 15780
gcctactgca ctgcgaagtt catcagcaga aagaatatcc aggaatggtg ctgccattcc 15840
aagaaggatt acgtttgcgc tttttggtgc ttttatttct tttgccattt cctcaatatc 15900
aagacgtaca accttgtttt ggctgtcgag ttccttgaat aactcagttt ggtcaggata 15960
gttaggaata tttatgaaag gcttagaact tgttactatt actccatcag gatttaggaa 16020
ttctacataa cgtagtgctt ccattggttc cattgcgatg ataagatctg cttttccgcg 16080
aggaataaga tcgctatata ttgtttcgtt agagaggcga agattacttt gtacatcacc 16140
gccacgctga ctcattccgt gaacctcagc ttgtttaaga ttaagtcctg cttttgtggc 16200
agcttcacct atgattgttg cgatggagag gattccttgt cctcctacac cagctaatat 16260
gatatctgtt ttcataactt atttcttagc ttgacggagt ttacgtgcat gagtttgtat 16320
gcattcacgt cgtggaatga taacgctcac gcccttataa ttgatttcat cacgaaggat 16380
ttgtttgatt tcatccatgt ttttaggaag tggaacaact acatgcaagt gttctggttc 16440
aactcctaaa ccgagacata tttgttcgaa tcggtatgta ccttgagagt cttgtccacc 16500
agtcattgca gttgtaaggt tgtcgcttat gataacagtt atgttagatt tatcattaac 16560
tgcatcaagt aatcctgtca ttcctgaatg agtgaatgta gagtctccaa ttactgatat 16620
tgcaggccat acaccagcat cagcggcacc ttttgccatg gtaattgatg cacccatgtc 16680
aacacagcta ttgattgctt ggaatggagg aagtgctcct aatgtgtagc aaccgatatc 16740
tccaaatact cttgcgtctt tgtattcagc tacaatttct tttaaggcag tgtaaacatc 16800
tctgtgaccg cagccttggc atagtgcagg aggacgtcct acaacattta ttggacttgt 16860
tgctgttgga tattctttag ggtcttcgcc tgtgatagca gaattgatag caagtgaaac 16920
agagtcaggg ttaagttcgc cagtacgtgg aagatttccg ttgagtctgc ctaagatctt 16980
ataatcagca ccgagtattc cgtgtacagc ttcttcaact acaggttgac catcttcaac 17040
aataaggatt ttgtcgcatt tagcagcgaa gtccttaatt ttctgttcag gaagcggata 17100
ttgtgaaatc ttcattactt cgattccatc aatattgctt tccatcacat agttgtatcc 17160
aataccgcat gcgatgatac cgagttttga gttctgtgta gttgtatcta atttattata 17220
ttttgaagaa agagatgcgt taatcatatc tttctgtttt tcaattaggc taagatagtt 17280
ctttcgtgca atagcaggaa gaagtaccca atgacgtccg ttgtctggat taagttgatt 17340
ttcttcatta gctttatctt tgatttccac gtttgcgcgt gaatgagcca aacgagttac 17400
aacacgcatg agaacaggta attttaaaga ttcagacaat ttgaatgcat cttccatcat 17460
gttgtatgct tcttgttggt ttgatggttc aaaaatagga atcatcgcaa atttagcata 17520
aaaacgtgag tcttgttcat tttgtgaaga gtgcattgat gggtcatcgg cagcgagaac 17580
cacaagtcct ccgtttacac cagttatagc gctgtttaca aaggcatcgg cacatacgtt 17640
cattcctacg tgtttcatac ataccagtgc acgtttacca gcatatgaca ttcctaacgc 17700
agcttccatt gctgtttttt cgttagagca ccagcggcta tgtatttttc cgttggcttt 17760
cagttcatgt ccgccgatta tatatgtttc ttttccgtat ccggtagcgt tctgtataaa 17820
ttctgttatt tcagttgatg gtgtaccagg ataggcgtac acaccggaga gtccggcatg 17880
aagtgctccg agtgcaatag cctcatcgcc taacaatagt ctttttttcca tattatatct 17940
tttagtgtag ggaatttagt gcgatatttc accagttgtt ccatatctat ttcccctgtt 18000
atttctgatg gtttattgag ttcatcttca gccaatacgt gtccgtaagg gtgaattatt 18060
tttgagcctc cgaagtattc tccccactcg tcagcaccga cacaactaca tgcagctaca 18120
taacattggt tttctatggc tcttgctgat aggagagtgt cccatgcaag ttttcttgct 18180
aaaggccatt ctgcagaata tacaataagg tcgtaaagta cgttgttgtt ttcgtccatg 18240
ttattacggc accacatagg gaagcgtaag tcatagcata tttcaagcaa gatatttaca 18300
ccgcggaaat ttacgacaac acgtttgttt cctgatgtat attgttctcc ttctccacca 18360
taaatgaaaa gatgactctt atcgtatgtt gttacttctc cgtcgggttt tacgaaatac 18420

atacgattat agaaatgtcc gttgtcttca actgcgatac ttccacagat agcagcgttc 18480
atttctttag attttttat catccattta agtgaatcgg agtctttcga ttctgccagt 18540
ccttttggtt tcataatgaa tccagtggta aacatttctg gcataatata taggtcagag 18600
ccgggactgc ggtttattgc ggcttcggct ctgactatat tttctgtggg attagcccac 18660
gctatttcaa attgtatgag agtaattttc atcttctctg tcttgttctg tattgcaggg 18720
tctgagtgga tattatttta tgttacccac tttgacgagg tattctgcaa tctgaacggc 18780
attaagtgct gcacctttct tgatttggtc gccgctgagc cagaaagtga taccattgtc 18840
gtttgcgaga tctttacgaa cgcgtcctac gtatacatcg tctttacctg cagtgaacag 18900
aggcattgga tatgtttgtg tagaaggatc atcgagaaga gtaacccctt cgccatttct 18960
caatgcgttt tgtacttctt caactgttaa tggtttttca gtttctaccc atacagcttc 19020
actatgagcg cgaagagaag agatacgtac acacatagca ctgcatcgaa cgtcagagtg 19080
catgattttc tttgtttcgt taaacatctt catttcctct tttgtgtatc cattgtcttg 19140
gaatacatca atttgtggaa tgacgttgta tgcaagttgg tatgcaaact tatttacagt 19200
tacaggtttt ccttctgtta gttctttgta ttgctgttga agttcagcca tagcagcagc 19260
tccagctcct gatgcagact gataacttgc gatatgaata cgtttgatat gacttaactt 19320
ctcgattggt tgaagaacga caaccatcat gattgttgta cagtttggat ttgcgatgat 19380
accacgtggg cggttgagtg catcttcagc attgcattca ggtacaacca atggcacatc 19440
atcatccata cggaatgcac ttgagttgtc aatcatgact gcaccatatc gtgtgatgtc 19500
ttccgcaaat tcctttgatg taccagctcc agcagatgta aatgctatat ctacaccctt 19560
gaaatcgtca ccatgtttaa gaagttgtac ttcatattcc tttccacgga aagtgtattt 19620
tgttccagca ctacgtgtag aaccgaataa tactaattca tcaattggga attgtctctc 19680
gtctagaact ttaaggaact cttgtcctac ggcgccactg gcaccaacga tagctacttt 19740
catatttatt ttatatacgt tttatttgtg tgcgcaaaga taatgattat ttatgactta 19800
agatttatcc tttgcatttt ttttcacttt attcatgttt taatataata ataaggtgta 19860
tttgctcatt gaacttaata ataaataaag caatttttat tatacgatta ttgtttttct 19920
cgtgcgtatt acataataaa ataagcattg tcggtgtttt tctattctat gtttattttt 19980
tttgttaaaa tttacggaaa aatttgataa atttagaaat tagtactaaa tttgtgcatc 20040
aaaataaaca ttagatattc gccgattaag aattacagat tcagtagtac gagtcaagca 20100
gacttaaaag tttacataaa tctacatttt ctttgtcata atccttttgt ttaccctttt 20160
tttgaataca tacttaagaa aaaatatact ataaaaaatg aatactaaca agaattacaa 20220
tgatgatcat atagacgtat tgtataaaag aatacagaat tggttaccaa atggatcatc 20280
acaattaaaa tttactgaac ttaaaaaacg tgttttaaat aaaacaatca tcaatccgga 20340
ggagacgtta aagtcgaaag atttgaaaga gattgcgaat tatctttcag aatataataa 20400
gtttaaggct gaacgtggtg attcaaaagt gaaaggtatg ttgaattttt cggatataat 20460
gactaatgta tctgaaaaac aaataaataa tctcataaat gaattggata gtcagaacaa 20520
accaagtact tcgttaattc ctataaaagg aatcagcaag gttatgcaag cacgattacg 20580
tgatcttaag attgttgatg ttcctacttt attggtaaaa tgtaaggatg caaatgccag 20640
aaaatcactt gccaacaagc ttggtgtaga cattcgttta gttacttcat ggatgaagca 20700
agctatatta tggcgagtgg aaggtatgac cacagacttg gcttatctgc ttgtgatggt 20760
aggtatcaga aatgttgagg accttgctca agttgataaa gcaaagattc ttccaatatt 20820
gaaagggtta gttgcaactc atatagattt tgaatttgag gaatctgaat tagacactgt 20880
tctgaagaat gcccaagaga tggtagtgta taacactcct atcaaggtta atgatttgac 20940
ttctcgtttg tcggacacaa tggctggttt gacagagtca ttagaggagg aattcttgaa 21000
ttcaataaaa gagcaattat ccagttcttt acaagaaatc acaactcaat ctttacgttc 21060
agcaagacaa cctttgtctc gacaagagtt acatgtaact aaagatgatg aagaggatat 21120
caatgatgag gaggaagatg aagaaataga tgatgaaata tatgaggatg atgagattga 21180
aggtgatata aatgaagatg atgaggacat cgatgaggat gaagaagatg atgaagagaa 21240
tgaggatgat gtaaatgacg aggaagaaaa tgagggtgat gtaaaacgat tagaattagg 21300
agttgctcat acatatgatc ttccaaataa ttttattgct aataattatg ttcacgaaga 21360
tcttcgtgaa aaaattattt ctgagattat tgagaaagag gctcagaaac tgaaacaggc 21420
atctgcagag agaattcaag agattcttaa aaaagaaata actggttatc aaaaagaaaa 21480
tacagtagat atctcatatg tccctaatga tataaacgat gttgaacctg tacatctttt 21540
tgatgaaaaa ctcagaaagc cacttgaaat tcttaatggt aagaatttga agaagggatt 21600
agatttcttg gatgatatcc aaccatcatt acctcttcca cgaaagataa gtggttatat 21660

cgttattaga aaaaaagacg aactaataga tatttccaga aacgatagtg tctatctcca 21720
aggttgcaga cgtttgtcgg gggctcttgt tgaaattcaa ggaattgtaa gtccttcaga 21780
tgatttaaaa gaggctgtaa cttctccatc atgcgttaca gatggtacag gtaaatttgt 21840
tattgttctg cctgaacgtt atagtatgaa ggagacaata acaatcgttg tttctgtagg 21900
ttcaaaaaga caagaattta tcaagactgc ttctgaaata ttagaatcag ttgagcaaca 21960
tgagatattg aacagattct atgaacttga tgcaattggc gacgagattg actatataga 22020
atctaatata aaaacactta ctggtcaggt tgcatatctt gaggacgatg ttaaactctt 22080
gatttcggaa ttagataagt taaatgctga attatctaat gaggaaacct ctgattatca 22140
gaaaattctc ttaaagcgtg atattgataa taaggagaaa gaccgtgccc ttaagattaa 22200
ggaaaaagaa caatatacaa atcagttaat agcatatctg cagaatattg atattctgaa 22260
tgaaaaatat agtacaggta aggcaaaact gttgtcagag agtttttgtg cgccagcggt 22320
tgatcttaaa caggcatttg ataatttcat aaataatatt accagtttag aagcaaaact 22380
gagtggagat acattaaaac caggtgaaac tcctaagcct tttgttgttg ttgaggaaat 22440
attcaataac gaacgtatgg acttggagaa agcattacca aaggttaaat tgatgggtaa 22500
tgatgataag gtgataagat tatcaaccga tacagctcca tcaaagattt atacatatag 22560
tatgcttcaa cgtttggtag aaccagagat tagtacaggt gaccgacaaa cgttatccca 22620
tgcaattgat gtcatggatt tcaaggaatc tatgattacc aatatagata aatgtcctca 22680
tgcatgtact ttaggcatgg gatatgtact gaatatgcat caggcatggg ttccagatgg 22740
ttttgcatta ggcgacttat tgtattcatt agtattggct ccaggtgagg aacaacgttt 22800
gattgtccgt gagaacaagc agacatacga gatttctgat gaagcgaatg ctattgatgc 22860
agatgatgaa agctatgctt tgactcaaga tgatgataca aatgcagcat ttaattatgc 22920
aatggatcaa ttgtctcaag gtaaatcaag ttatagttat agtgcatcat catggagtat 22980
cggaggaagt gccagtggtg gtgctggtgg cgcaatgatc ggactctcag gaggttattc 23040
taaatcaagc ggaaaaggct cggcttctgc aagtcaaagt aactcccata gtgagacctc 23100
ttcggcagca cagaatttcc aacataatat caagagtgca tcaagtaaga tttctcaagc 23160
taaacgtctg agcatggaga tggcgacaag tgatgtcact gattctgtag ctaccaagat 23220
aattgccaac cataaccata gtcatgctat gactattcag tattgggagg tgatgcgcag 23280
atataagctc gagacatgta tcgacagtgt agatttagtt ttgtttgttc cattaaaatt 23340
aataaatttc cttcctacag gtgatttcta tttaggtgat acacagaact ttgatcgtac 23400
tgcatttaat aatcgttatg atactttgtt gcattactat gatacacttt actcaagact 23460
tccaagtaag tatcgtagtg gtttaaatct gattaagaag tattcagaat tgaccaattg 23520
ggtaatgcag aagcaagagc agtcaactaa ggttcttact ttcgaaataa aggttaaact 23580
gcttacttat gataatctgt ctgctacatt agtattgaag aatggtaaag gttctattgc 23640
tggatcttgt agtgacatca ataataagag aaagaaatta accacttcaa tatatcagca 23700
atatgaaaca tcagaagatt taagacaagg tataatagat ttcagaaata aagtcccaga 23760
tgcaacagat accttgacat ttatattcaa tatacctgat gatacaaccg atgatgatgt 23820
ttcatacatc cgattagact attcatgcaa tgaatgtagt tatagtttga aacgcgatga 23880
taataaactc atcactatcg actatgataa cgacgataaa catggaaata cgacagccgg 23940
taagctttat cagcgttata tagacaatat ctatgattat tcaaagaatg ataaagatac 24000
tgcaaatgaa cataaactgg aggcttacta tgaacgtata ttgccagagg catatcttca 24060
ccctatcgta aagatctctt cttatgagat gagacgtttg ggaacccta ctataatgga 24120
tgcggttttg aagatcggtt caggaaagtc tgttggttca acaagtaagg acgttgagac 24180
tacatataat ataatgttgc catctagtaa gttgagttca tctatgcgac ttagtatctc 24240
aacatcttgc tatacattgc acaattcaga attgcttaag atggagagca ctcttcatca 24300
cattgcttct aacccaatga aatattctca gatagtttgg ggttcattaa gtgatgacga 24360
acgtgtgatg atgttggaac aattcactat aaatatgaat tttgattcat tagatgatga 24420
ttcttcaaga tatgagaaat ctaatgactt aaagattcct ttacttaatt gtgtaaacgt 24480
aaagaatttg ttgggattct atggtaactg tatgatttta ccattcacat tcccacaacg 24540
tttagcaaaa caattgggta agactgcggc agacattcag aattctcttt atcgttatca 24600
cactaataac ttccgtgtac ctacaacaac aatctctctc ccaacaaaag gtatgattgg 24660
tgaggctgta ctcggtgaga caaatgttag tgaggaaatc gatatcacac gcttctggaa 24720
ctggcaggat tctccaattg acaagatgga aatagatagt agttatctta atggaaatga 24780
ctatctgcaa ggaaagagta caaaagatgt tactactttg aatatgcaag gtgcgacagc 24840
tgctactcca gtaactgttc cagatcttgt ttcggcattg gtaagcaaac aaactccaac 24900

atttgacaat ataacaggtc ttgatcagtt gaaggaagta cttaatgctg gtacaaatag 24960
tgctgcagcg ggtcgtgaca atgctatcaa gtcatctgct gatgttgcaa aagcagcttt 25020
agcttctttg acaccaagta gtagttctac tacatcaagt acagatagtt caagcagttc 25080
aagtactcca agtacaagca gttcaagcac tccaagcaca agtagctcaa gcactccaag 25140
tacaagcagc tcaagtactc caagcacaag tagctcaagc actccaagta caagcagttc 25200
aagcactcca agcacaagta gctcaagcac tccaagtaca agtagctcaa gtactcctga 25260
tacaagtgat acaggtactc ctgatccaag tgatacagga actacagatc caacagatac 25320
aggaactaca gacccagaaa catctgatcc agagacaagc gaaacaacct ctgatatcga 25380
tatggatctc ttgtcaagta ttattaaaga ggcagtagag tatatcaaga ctaatggcga 25440
tgattctgtt ggattcttta agtcaaaagt atcagatcaa tatccagatg aaactgtaga 25500
tgacttggct aataaatttg ctgagaaaca tggattatca ttaagcgatt ttattgaaaa 25560
aatttttaaa tagttgatta tgattataca aatagctata ttaatacaga cattttatac 25620
aataaagtcg ttgctcgatt ctcgtgataa gatcgaggaa tgggttaagg ttgtcatggc 25680
tgtttatgag aagcaaatgc agataatgga ttctaaaaag gaccctacta tacttgtcaa 25740
tgctttctat aaatatttat tggaatacgc agatattttt gatgcagacc cttcaaataa 25800
taagccgtca ttagatagta tattaaagag tagcaagaaa atacttgaat ccaattcaga 25860
tcgttatgtg ttgagtacaa cttacgcaaa ggtttctcct ttcttcccta caccatttcg 25920
ttatttggat aatcctaaaa gaaacattaa tccaatcaca gcatttactg ctacagaaga 25980
tgatcttaaa gaactattga agaaatctaa taaagataag aatactgata gatttaagga 26040
gctatctaaa gatattaagt catatcgtga gttgttatca aagagaagta cgcttgcaca 26100
aagtacaata agtgcgttaa atcctgataa tacaatgatg actgaatacc tttttaatac 26160
taattcaaat actgaatact attggggagg tgataagtgt aatgtgtatg taggtgaagc 26220
aatttactat gcactcaaga aagtttttct tacatataca aatgcgaaat atgataactt 26280
agatggtctt ttaacagatt tcttgttaga tgcaaatcac aatcttttaa cttcaagtta 26340
ctatcgatat ttgcaaccaa gtggtatatt cgagggttat acagaaccga taattggcga 26400
tgatggcaaa ccgaaaaaag gcgatgatgg tgaagactta aaacgtttag taaatactac 26460
ccaaatacat agtaagatta ttttcgagaa atgtcctctt aactcggtaa tagagccagg 26520
tgcttttgtt tgctttataa ctgatggata caaatttggt cacattgaga tgatattatc 26580
taagattgat gttttgcaaa ataaaaaaac atttgtttct ataggtgctc atccatatgg 26640
agtatataga aagacacaga cttttgattc ttctaaaatg caaatcgtca aagttaagga 26700
gagtgttatt aaaagtgtcg ttgacataaa gccagtggct aaagaagatg ataagaaaga 26760
agccaaaaat tgattaaaaa gagtttacag atgaatgaat ttgaattaaa cagattaaaa 26820
gaaatcttta acgattttaa tcttaatgat tgtcttgagc aagccagttc tgattctgaa 26880
cgtggcgtct gtgaggatat cgctaagatt aaagaagatt ggattgaatc aatattggca 26940
acctatacgg caaaggataa tcattgttta ttctccataa ggactaaaga ggcgttagat 27000
acaggtatat ataccaaaga taatacatat aataatcttg agggattgat atatggtaag 27060
tcttacactt tttatatcct aaataaggaa aataatacta cagtgtcgga taagagcgta 27120
gaatggtctc tgtatttatc agaggactca aacgtgaaac ttataaagaa aggaacaggt 27180
aatgaaattc agttctcagt tgataatcct gagttcttgg gtaagttttt gatcgttgag 27240
gtaaattata atggctttac tttcccgtta aacatgaaag gattgttctg gattcattat 27300
cgattcagac atattaattt cgtttgcctt gccaaagata ttgtcaaaag atgtctcttc 27360
cctgatggtg tcagtcaacg cggtacatct ttatgtggcc ctgctgcggt agcatatcta 27420
tttgctaaat accaatggga ccaatatggc cgttttatta tagatataca tcataagggt 27480
gttgcaacta tcaagagtac aggatacatt gttgacttaa acaaagatag gagtttagat 27540
tatctggtag aggattataa accagaaact acttttagcc aagaatatat acttaataag 27600
tcttcagaaa tccctgttgt tgatttcttg ttcctggttg tgcttaaggc atatactaat 27660
atagccataa atgtaaaacc tgaggatagt tttattaata acaatgctgc aggagtgacc 27720
atcacaagta cagaagttga atatatgaaa aatattcttg gttgtaagac aatcactgac 27780
gatgattacc atgaggattg cgcttatgac ttctgtaatc agatccaaaa atggattaat 27840
agaactacag acaaaattgc atcggttgca gtctttttgg ttaatcataa tgcattcaca 27900
gatactgttt ctttttcgaa aagagattca ggatatttga ctaattggcc aacccattgg 27960
attgtaccgg aaagtatcga ggttgatttg tttaaaaagg aaatcggtga gattgatatg 28020
tatgcattta catggggaag aatcgctaat attcgtctta gttcgtggtt attcgctggt 28080
ggattgcata attacatatg tgctgagatt gaaagaacca agatgacgcc aagcgataat 28140

atagaacatg gtatacaata ttcaaagact gtttgtgatg aattaggatt aagcactgtg 28200
gtggtaccat aatcacaatc cgtaatttaa cgaagaataa atattaaata gaaataaaaa 28260
tggaaaacga attattaata caggctattt ctcagattct tcagaatgag gcatctgaag 28320
atactttaaa gataaagaat aagattcttg aaagaatagc attggaagca gatgttaagg 28380
catcacgtat tccagctcca ttgaatataa ctcagatagg cggatatatc aacctgctga 28440
gaaaattaga taaagaagaa caagaaagac agacaaagca gaaaactgct tatcaacaga 28500
tgttagaaca aacattaact tctatactcg gattacctgt gcaaccgcct acagagtaaa 28560
gcccgtgggg ttagtgcgat tagtctgatt cccgaagttt attaacgttc tacatatgta 28620
taatgtacga taatgcctcg tgttttgtat ggattcggtt gtgacggtgt tatgacagaa 28680
gaagcttatc ctagacagaa tagaaatgaa ggcgatgtca taggataatg gagcttaggc 28740
ttccgcttac ctgctgagag tatcagtgtg tgagttttac aaagagcatt attagtgtca 28800
ggaatgacgc aaggacgaca tttgtagtta agataccctg caacgggtat ggaggaaggt 28860
tgtatgaact tctatgtatt gaagagagta agtacagaat ttgggacaac tccaatcgca 28920
taggtgaacg agtcgcttat ccttatggat ggcttcagtg agggtgtgtc ataacaataa 28980
ttgacacacc ctctttttag taatatccag aatacctgaa gattgataaa tataggagag 29040
tagaagtggt gaacgaatga agaattattc tcgaggtgta aacatcgaca atgtaaagtc 29100
ttttgtttgt ttttgtagat tgtgaataat tttgtacctt tgctgccgaa tttgaatgaa 29160
tgtcactcaa aacccgaaac agcatggact caatcaccca atatctgaat tttccgatcc 29220
aagatcctac atgggttttc tttatcgtcc ttcttataat attatttgcg cctttgttgt 29280
ttcataaact gcacattcca cacataatag gaatgatatt ggcaggagta attcttgggg 29340
aaaatggttt caatatctta gagcgtgacg gtagttttaa gatttttggt caagtaggaa 29400
tatattatat catgtttctt gctggaatag agatggatct tgttggcttg aaacgcaaca 29460
tcaataaagg tctgcttttt ggtgtgataa cttcacttat cccatttgca tgcggattag 29520
ccgcagggta ttatctgctt gattattcat tttcagcatc tttgttgtta gcatgtatat 29580
ttgcttcaca tacaattgtg gcatatccta ttgtcgcaag atatggatta aacaaacatt 29640
cagctgtaac tatcagtatc tcagccacaa tgatagcatt gttgtttgct ttgttgacat 29700
tggcagtgat agccagtaga tacagaggcg gtggaggctt gtggttctgg gtattgtttg 29760
tagtaaaatg tattgcttat ttttctggtt tattttggct tttcccaata ataatccgct 29820
ggtttttcag aagatataca gatagagtgt tacaatatat atttgtaatc gcaatgatgt 29880
ttctcgcagc aggaacagct aatctgtgtg gaatagaggg acttttaggc gcattcttgt 29940
caggattgat attcaatagg tttatacctc atatgtcgtc cttgatgaac cgtattgagt 30000
ttgttggaaa cgcattgttt atcccttatt tccttgtggg agttggtatg ttggttaatt 30060
taaaaccgat atttaccgat atgggagcaa tcgtcgtggt tgtaatcatg gttgtggtga 30120
gtacgctgtc aaaacttatg gcttcattta taactagaaa gattcttcat ctgcccccatt 30180
cccatggact cgtaatgttt ggtttgtctg aagctcatgc agcgggtgct atagcgatgg 30240
ttatggttgg aaccagtttg atggttgccc caggagagcc attgatgaat acggcagtgt 30300
tagacggagt ggtaatgatg attctgcttt catgtatcat cagttctata gcgacagata 30360
agggcgctcg taaactgaaa ttagaatcag atagggaaac gagccagttg accgaaaagg 30420
gtgatgatga aaagataatg gttttggtaa aagaatcagc taaaatacgt aatctgaccc 30480
aaaccgccat aatgatgcgt aatccggatt tgaacagagg attaatttgt ttgaatgtag 30540
taaatgatgc ggatccaacg gaaaaatcac gtcgtatgag tcgtgaaata ttggaacaag 30600
ccgaagagat ctgttcagca aatgatgtcc ctgtacaggt tcaaagccga ttggctgtta 30660
atcttgtgaa tggtgttgtt catgcattaa gagagaacga tgcgtcagag atgattcttg 30720
gactacatga accacgtccg cacgacacgt cctattatgg acaatttaca gaagctttgg 30780
tacagagtat gaatcgtcag gtaatggttg tacattatca tcagcctgtt aataccattg 30840
gtaagatttt tgttgcggtt cctgatagag ctgaatatga aacaggtttt tatcgttggg 30900
ttgaacgagt attacgactt gcaggtcaaa taggttgtaa gatcgttttt tattcatcgg 30960
aaactaccgg aagactgatt tatcagtatt cccaaactca tcataagagt gtacgtaccg 31020
actttaaact aatggaatcc ttgactgagg tttattcttt gcgtcaagtg attcatcacg 31080
atcatatgtt ggtaatagtt gcagctcgtc ctagttcgtt aagttatcag agggcatttg 31140
cacgattgcc aaaactatta cacgatcatt ttaatgattg cagtcttatg ttgatttatc 31200
ccgatcaatt cggcgaaaat aactccgaga caacatttac agaaccacac ggaaatacaa 31260
catttgagat gtcaaagact tcagcgtggc tttcaaaatg gttaagtaag atataaatgt 31320
caatttttga aagaagcgac agaaatgata gaattaagaa atataaataa gagtttcggt 31380

gatttgcaag tactaaaaga tattagtctt acaatcaacg atggtgaggt ggttagtatt 31440
gttggtccaa gtggagcagg caagactaca ttacttcaga ttatgggcac cttggataat 31500
gctaattcag gctctgtgat tattgatgga attgatatta acagtttggg ttctaatcaa 31560
ttggcaaagt ttcgtaatca gcatataggt tttgtttttc agtttcatca acttctccct 31620
gaatttactg cagttgagaa tgttatgatt ccggctatga ttgcaggaaa atcacaacgt 31680
caggcaaagc aacgtgcatt ggagttattg gatatgttag gtcttaaaga acgagcagat 31740
cataaacctg cagagttgtc gggtggagaa aaacaacgtg tagcagttgc cagagctctt 31800
attaataatc catcggttat aatggcagat gagccttcag gaagtcttga tacgaagaat 31860
aagacagaac tacatcaatt attctttgat cttcgtgata gattacattg tacatttctt 31920
attgtaaccc atgatgaaca attggcgtca atcacagaca gaaccattca tttaaaagat 31980
ggtgagattg tggcaaatga gataaagaat gtagtggata tgccatgtga aacaaacgaa 32040
tgatatttgt aaagaaataa gaatataaga tgatacagtt aggtaaacaa aatagacttc 32100
agatattacg tcaggttgac tttggaatgt acttagatgg aggtgagatt ggagacattc 32160
tgcttcctca gcgatatatc ccaaaagatg ctaaagttgg ggatttcatc gatgttttcc 32220
tttatcttga tatggaggag agattggtag ctacaactga aaaaccattg gttcaggtca 32280
accagtttgc atttctcgaa gtgaaatgga ttaaccagtt tggtgcattt cttgattggg 32340
gactgatgaa agacttgttt tgtcctttcc gtgagcagaa gatgcgtatg caacaaggta 32400
gaagttatat agtgtattgt tatatagata cacttacttc acgcattgtg gcatctgcaa 32460
agatagagaa gtttctttct aaggagatgc ctccatataa tatcggcgat aaggtggaat 32520
tgttaatcca gcaaaagact gatttaggtt ttaaagccat tgtggatgga aaattcagtg 32580
gtttgatata cagaaacgag ttgtttcagg aaatacacac aggagataga atgtctggtt 32640
atgtgaaatc tgtaagaaat gacggaaaat tagatatctc tcttcaagta acaggaaagc 32700
aacatataga agatttctca agtagattgt tcagatatat atccaactct gaggacggat 32760
tttgtccatt tcatgataag agtgctgccg aggacatata ttcagagttt caagtgagca 32820
agaaaacttt caaaaaggct gtaggagatc tatacaaacg tcatctcata acaattgaag 32880
atgatggatt gagattaact atggaaggac gaatttctgg tattgacgaa gaataaaagc 32940
caagttatta atatggatat atttcaacag atacaacaac caatatcaaa ggagttggag 33000
ttgtgtagaa acatgttatc agactcaatg tctcactcta atcctttatt agataaagcc 33060
ttgaagatga ttggagcccg aaatggtaag atgatgcgtc cgatattaac gcttttgatt 33120
gcgagattat acggtgaaat tgatgaaaag gcacttcatt cagcatgtgc gtatgaggct 33180
tttcacactg caagtcttgt acatgatgac gtagttgacg agagcgagca gcgacgtgga 33240
acacagtcgc ttaattattc atctggtaat aaggtagcag tgttagtagg agattatatt 33300
ctatctttag cgttgaaaca tatctcaatg acaggcattc cacgattggt ttctataatg 33360
tcagaagccg caaatcgttt atctgatggt gaacttcttc aattatacaa tgtaaatagt 33420
cgttctatca aagaggaaac atattataag ataattcgta ataaaacagc agcacttttt 33480
tctgcatgtg ctcaatcggg agcaatatca aaaaatgtat cagaagacga gataagcaag 33540
gcaggtttat taggagaata tatcggtatc tgttttcaaa ttcgtgacga tatatttgat 33600
tatacaaatg ataatatagg taagcctaca ggcaacgaca tgaaagaggg aaaactcact 33660
ttgcccgtta tctatgctct aaacaatgcc tctaatgatg aaatatcgtt gattgctcaa 33720
agggtaaaag ctagagttgc ttctcaagag gaaatagcca aactggttga tttcacaaaa 33780
gaaaataaag gtatagaata tgcgatttca tcgatgaaat catatgcaga taaagcaaag 33840
gacatcatta tggaatatcc agataatgat gtccgtgagt cactgcttgc ctatgtcgat 33900
tatgttgtag gccgcagtta ttagtttgtt tcatatcata tcctgattat ggatatagat 33960
attacttaga ttttacgatt tataatcaga agaatttaac ttcttttccg attatgtcgc 34020
ttaaaagaag attggcaaga ctgcttgtgc ctaagcggaa taagttgtta ttaagccatt 34080
tttcacccaa aacttctttt acgatagtgt agtatgtcag agcatcttgt attgtcttca 34140
ttcctccagc aggtttgaat ccaatctgta ttcctgtatt ttcgtaatat tccttaattg 34200
cttgacacat tacatatgca gcttcaggtg ttgcagcagg ttcaagtttg cctgtagatg 34260
tcttaatata gtcagcacca gcatacatag ccagtataga tgctttcttt atgttttcag 34320
catttccaag gcatcctgtt tcaagaatga ctttaagttt gcggtcgcca catacatcct 34380
taatttcgct tatctcatca gcaacagtct cgtagtcgcc agcaagaaat ttgcctacag 34440
actgaacgat atctatttct gtggctccgt cgtgaatagc cattgcagtt tcagctattt 34500
ttacttcaat aaatgattgt gatgaaggga aacatgcaga aacacaagca atttcaacgt 34560
cctcattttc taatgtatca tgacagattt gagcatagca tgggtaaaca catactgttg 34620

caacatgtcc taattcaggg tattcatcat caaatttatt tactttctct acaaatttca 34680
tgatactatc ttctgaatcg gttgttttca acgatgtcag ttcaatgctg ttgaataaga 34740
atttctttac ttcaggagta tcgttttcct tgagatattt tgaaaggatg tttgtcactt 34800
cctttttac ttgttcatcg ctgatgttga aattatattt ctcaaaagct tgttgaactt 34860
tactttctga tgattctgcg tgatgatgat ggtggtgatc acatccacaa ttgcaattat 34920
gttccatatt ctatattttg ttttttattc tttattcttt gtatccatac atattgtaac 34980
gggcccgtcg ttgattaatt ccactttcat gtcagctccg aattcaccag tgaaaatcgg 35040
ttttcctaaa tccttttcga gttgtttaca tgtcttctca tatagtggaa ttgttatttc 35100
atgacttcca gcccttaaat agcttggacg attccctttc ttataggatg cccagagtgt 35160
gaattggcta acaacaagta tgtcgcctcc tgtgtcgagt attgaacggt tcattactcc 35220
attttcatcg ttcatcactc ttaggttcac aatctttttg cttagccatt ctatgtcttc 35280
ttgtgtgtca gcattttcac atcccaatag gatgaggtat ccatctccaa tcgaagattt 35340
tacttcgtta ttgatagtta ctgatgcctt actaactctt tggattactg ctctcatttt 35400
aagttttctt taatcaattt ggcttttgtc tgccctataa catctgataa ttcctcaaca 35460
gatgcttgtt ttatcctgtt tacactctta aagtgtttca gtaatgtatt ctttgttgtc 35520
tctcctatgc cttttatgtt atccagttcg ctggatattt gtcgtttgct tcttttttca 35580
cgatggaatt gaattgcaaa cctatgaacc tcatcttgta ttcttgtgag gaacaagaat 35640
aaaggactgt tgctttctat tcctattgtt actatttgtc cgttgttata cgcaagtaat 35700
tcagaagtct tatgtttacc gtcctttgct aaaccagcaa tcgggatatc caagtttagc 35760
ttttcatgta caatcttacg tattatctcc atctgccctt ttcctccgtc tgcgataatc 35820
agatttgggg tgatatgctc actatggtct gttgtgtcgt gtgatatgcc caattcagct 35880
tgcttttcag gaggaagcag aggtgaatat ctacggaata ccacttcagc catagatgca 35940
taatcatcgg ggccttcaac ggtcttaatg ttgtatttgc gatagtcttt ttttgacggt 36000
tttgcttgtt tgaaaactac gcatgcagca acagcatcgg ctccttgtat gtttgagttg 36060
tcaaagcatt cgatatgcat tggcagtttc ggaagtttta gcttttgttg cagttctttc 36120
aacaggtttg tggttctttg ttctgggtta agacgttcag cctgtttagc tttgtcgaat 36180
ttatattgtc ttacatttgc tatagataga tcaagcaagt gtttttgtc gcctctttgt 36240
ggaatagtaa atgtaacatc tgacaattct gtattgattg gaaatggtat cacgatttct 36300
ttagagtgac ttttgaatcg ttgtcgcatt tccacaatgc ccaaggaaag caagtcttct 36360
ggcgtttcct ctgtacgttt tgagtattca aaggtgaatg cctgatttat acatccgttg 36420
gtgacatgca gatagttaat atatgcagat ttctcatcat cagcgatgga gaaaacatcg 36480
atatcatgac tataatttgc gatgacttcg ctttttcgc agaattccaa tgcaagaaga 36540
tacttttttt tcacggtctc agcttcttca aatttcattt cagaagcaag aacttgcatc 36600
ttatcaagta atttttggca tatactgcgt gtattaccct ttagtatctc atgaacctcg 36660
gcaacatcgt gcatgtattc ttcatgactt tgaagtccga tgcatggtgc tttacagttt 36720
tttatgtgat attcaagaca tgatttcagt tttccttcat ttatacgatc ttttgtgatt 36780
gcttgtcggc atctacgaag agggtataat cgatttatta gttccagcat gttgttgagt 36840
gtagcagcat gactgaatgg tccgtaatat gtgccggctc caggtattat tttccttgtt 36900
ttaaatattc ttgggaattc ttcatttgtg acacataccg aagggtatgt cttatcgtct 36960
tttagaagga tattgtatct tggcttgtat ttctttataa ggttattttc aaggagaaga 37020
gcatcctcgt cagagttcac gatgatatat tggatgtcgt gaattttgct aacaaggact 37080
tgtgtctttc ttgacacttg attctttagg aaataggaac taactcggcg tttcaggttt 37140
ttagccttgc caacatatat tattgtgcct tctttgtcaa aatattgata acatcctggt 37200
ttttcaggaa ggtttagcac aatattcttg agttgttcct gtttcttgtt gaacattgat 37260
gaataaaaat agtgttctga taaatcatct tcaggatatt atctacccat aagtacaagc 37320
atgacgttga tatcgcatgg tgagactcct ggaatgcggc tggcttgtgc caatgtttca 37380
gggtcgatag cagccaattt ctggcgagct tcaatggtta tttgattcag ttttgagtaa 37440
tcaaacttgc ccttgattct tatattttcc agacgaagca ttttctcagc cacttgttgt 37500
tcacgttcga tatatccttt atattttatg cgtatttctg tagcttcgat gatttcgtct 37560
tgacgttctg ttatcttgtt gatttcagag ttgaatgcgg caatataagg aagtattcct 37620
tttatagtga tttgtggtct gccgattaaa tcgaacaatt tacatccacg ttccagaggt 37680
gtagttccta ttttctcaag tgcaggattg attaatgcag gtttaattga gaaattcttg 37740
gtgaattcga tgagatgttc gatttcatcg cgttttttta caaagaaatc ataacgttcc 37800
ttagatgcca atccgatatt atatgatttc tcggttaagc gcatgtcggc gtcgtcttgg 37860

cgtaagagaa ctcgatattc agctctactt gtaaacattc gataaggctc gtccacaccc 37920
tttgttacaa gatcgtcgat aagcactccg atataagctt cgtttctttt aagtgtgaat 37980
gctggttttc cagcgcattt caatgccgca ttgattcctg caatcagtcc ttgtcctgct 38040
gcttcttcat atccagttgt tccgttaact tgtccggcga agaatagtcc gctaacgatt 38100
tttgactcaa gagagtgagt tagttgagtt gggtcgaaat agtcatattc tatggcatat 38160
ccaggtcggt acacaacaac atttcttagt gcaggaattt ttcttaatgc gttaatctgt 38220
atatccattg gtaaagacga agagaatcca ttgaggtaca tctcgttggt tgtttctcct 38280
tcaggttcaa ggaatagctg atgttcgtct ttatcaggga aagtgtgcaa ttttgtttct 38340
atacttgggc agtatcgtgg tccgatagat ttaatctgtc cgttgtagag aggtgagtcc 38400
tttaatccac taaaaagaat atcgtgaact tccttgttag tcttgaatat ccaacaagga 38460
agttgtttta gtttgcgagg ttcgctcatg aaagagaatc gatggaaggt ctcatctccg 38520
tcttgtcttt cagttaaaga gaagtcaaca gaacgcttgt cgattctgac aggtgttccg 38580
gtcttcattc tatcgctttt gattccaagt ttattgatag acttggtcag ttcaagttct 38640
gcaggttcgg caattcttcc tcctgtaact tgttttctac cgatgtgcat aagtccattg 38700
aggaatgttc cagctgtaat gatcacacat tttgcatgaa attcagcgcc ccagatggta 38760
cgtactccta cgatagtgtg tttatgagag gaatcggaat tatcttccga ggttataagt 38820
tcgtttactt gatcctgcca gatattcagg ttgggagtat tttccaatat ctccctccat 38880
gtcaatatga atttctgtcg gtcgcactga gctctcggac tccacacagc aggtcctttt 38940
gagatattca gcatgcgaaa ttgtaaggca catctatctg tgacaatgcc catcattcct 39000
cctaaggcat caatttcgcg tacaatttgt ccttttgcta ttccaccaac agccggattg 39060
cacgacattt gcgcaatctt attcatatcc atagttatca gacaagtctt agctcccatt 39120
cgtgctgtag cagatgcagc ttcgcatcca gcatgacctg ctccgacaac tattacgtca 39180
tattcctgta tcatgaacgt tatttcttag cgaataattt ttagaaattc tccttaaaat 39240
ccttaaaacc atgcaaagtt atgaaatgtt attgagatac agaaataatc tgtatcaaat 39300
tgctattcca cgaacatgat tttgctttta tcgcgttgtt ttgcttgtgg ttgttcatca 39360
gggtaaccaa cagccaaaac atatgcaatc ttgtaatcgt ttggtaattt gagtcgcaag 39420
agatagtctt tcagttcagg tcgggtgttc aattgcatcg ccacgcctcc aagacaacaa 39480
gttcccagtc cgagtgattg tgcagcaagc atcatatttt ctcccatcaa tccagcatcg 39540
atataactga aatctccact tggtacagca acacaaatga tagcaggagc attgcggtac 39600
atgtttttat cgttgccagt tatgttagga ttaacctttt tcatggcttc agttgaacca 39660
ttgataaagt ccttagaatc aacaattcta acggcccatg cttgcctgtt cattgcgtta 39720
ggagcagcaa tgccacagtc ggcaatcaat tgcagttttt ctttctctac aggcatgtct 39780
ttgtacttac gtattgatct gcgctgcatg atagtttcaa taaccttgtt ttctgtttgt 39840
ttgcattcat tttgtgccca acagcttgtt cccattgtca tggcgaaaag caaaactata 39900
agattttttg ttttcatgac tttataattt ataattatta attgtattta tattcatgat 39960
atcttttaca aagatactca aaaaaaatga ataatgcaaa taataagaaa aatgtacatg 40020
cttttgggta ttgcaaaaga gtgtgtaaag tgttgttttt gctgtaagca acttgtaaaa 40080
tgggttaaaa atgcgattaa aagtttgtga tgtgattttt ttgaattatc tttgtaaact 40140
gaatgtattc ctctgtaaat atcggagaag gatgcgtttg tgaacgtgaa attataaaac 40200
aaacaaaata agatataaaa tggcacaagt tccagatttt aagtatgctc ctatgtttca 40260
gttaggcaag gatgatactg aatattattt gcttactaaa gaaggagtga gcgttggtga 40320
gtttgaagga aaacctattt taaaggtgac accagaggca ttgacaacaa tggctaatca 40380
tgcattcaag gatgttagtt ttcttttacg cagaagtcat aatttacagg ttgcaaaggt 40440
gttacgcgat ccagaagcaa gtgaaaacga taaatatgta gcacttacgt ttcttcgtaa 40500
cgcggaggta gcatctaaag gtatgcttcc attatgtcag gatacaggaa ctgccattat 40560
tcatggtgag aaaggtcagc aagtgtggac aggtttctgt gacgaagagg ctctttcaaa 40620
aggtgtttat aaaacataca ctgaagagaa tctgcgttat agtcagaatg ctccattgtc 40680
aatgtatgag gaagtaaaca caaaatgtaa cttacctgct cagattgacc ttgaagcaac 40740
agaaggtatg gagtatcgtt tcctttgtgt tactaaaggt ggaggaagtg caaataagac 40800
ttatctttat caggaaacaa aggctcgtat tcagaaccca ggaacattaa ttcctttctt 40860
agtggagaag atgaagagct taggtacagc tgcttgtcct ccatatcaca ttgcaattgt 40920
tattggtggt acaagtgcag aaaagaacct tcttacagtt aaacttgcat ctacacacta 40980
ttatgataat ttacctacaa gtggagacga gacaggtcgt gcattccgtg atgtagaatt 41040
ggaaaagcaa ttgcttcagg aagcatataa gattggtctt ggtgcccaat tcggtggtaa 41100

gtacatggcg catgatgtac gtgtggtacg tttacctcgc catggtgcaa gttgcccaat 41160
aggacttgga gtaagttgtt cagcagatag aaatatcaag tgtaagataa ataaggacgg 41220
tatttggatt gagaaattgg atagtaatcc aggcgaattg attccagaag aatatcgtaa 41280
agcaggtgag ggtgaagttg tcaaagttga tttgaatcgt cctatgagtg aagtgtgcaa 41340
gcaattgagt caatatcctg taagtacacg tttgtcactc tcaggaacaa taatcgtagg 41400
tcgtgatatt gctcatgcaa aaatcaaggc tcgtttggat gcaggcgaag gtatgccaca 41460
atacttgaaa gatcatccaa tttattatgc aggtcctgca aagacaccag caggtatggc 41520
atgtggttca atgggaccaa caacagcagg tcgtatggac ccatatgtag atgagttcca 41580
agatcatgga ggcagcctta taatgcttgc aaagggaaac cgcagtatcg atgtgacaaa 41640
tgcatgtaag aagcacggag gtttctatct cggaagcatc ggtggtccgg ctgcaatctt 41700
agctcagaac aacataaaat caatagagtg tgtagaatat ccagaattag gaatggaagc 41760
aatctggaag attgaggttg aagatttccc agcatttatt cttgtagatg ataaaggcaa 41820
cgacttcttc aagcaactca aaccatgtga gggatgttca atacttaact aataataaat 41880
aatcattaat aaagaattat tatggaagaa gttaagcaat atcttaacga tgcagaagaa 41940
agcatggaga tggctacaat ggctttggaa gaaaaacttg gccgtattcg tgccggacgt 42000
gcaaatgtac atatcttgga cggaataaag gttgattcat atggagcaat gatgccattg 42060
tcaaatgtag caaacctaag tacaccagat gctcgcagta tagtaatcaa gccatgggat 42120
aagaatatgc ttaaggtaat agagaaagca attatagatt ctcctatcgg tattatgcca 42180
gagaataatg gtgaacaact tcgtcttggt attcctgttc ttacagaaga gcgtcgtaag 42240
gaactcgtaa aacaatgttc aaaagaagcc gagcaatcaa aggttagtat ccgtaatgct 42300
cgtcgcgatg ctattgaact tcttaagaag agcgtgaagg gtcttggagt ccctgaggac 42360
gtagagaagg acgccgaagc agattgtcag aaacttcatg acaaatacat caagaaggtt 42420
gacgagatgc ttgaagagaa gaacaaagaa ataatgacag tataaaagag ctatgcatgg 42480
acttgtagtt aaaaatacag gtagttggta tagtgttctc actgatgaag gtacccttga 42540
gcaatgtgct gtcaagggta actttcgtat taaaggcatc aaatctacca gtcctgtcgt 42600
tataggcgat cgtgttgaga tttctgacgg gatgataacc tctgttgacg atcgaaagaa 42660
ctacatagta cgcaaatcca ttaatctaag taagcagtcg catatacttg cagcaaactt 42720
agatcagtgt tttcttattg tcacagtaaa tcatccgata acttcaacca cattcattga 42780
taggtttctt gccacggcag aagcatataa tataccagtt gtactggtgt ttaataagtt 42840
ggatatctat accgacgaag ataaggaata tatgcaatta atgaaggacc tgtacgagaa 42900
tatcggatat aagtgtgttg ttatctcggc aataacaggt gagggaatgg atgagatacg 42960
catgatgctc gattcaaaag tatcattggt ctcaggtaat agcggagtgg gaaaatccac 43020
ttttatcaat tccttactcc ccgacttgaa tctaaaaacg gcagatataa gtgatgttca 43080
tgacacagga atgcacacca caacattctc tgaaatgtat ccgtttaaaa gaaatcccga 43140
tagtaatgaa gtccctaata gttttataat agatactcca ggaatcaagg gatttggctc 43200
gtttgatatg caacgtgagg aaataagtca ttattttcgc gagatattca actatagtca 43260
tgattgtaga ttcaataact gtacgcatac tcacgagcca ggatgtgccg ttcttgctgc 43320
attagaacga catgacataa gccaaagtcg atataactca tacctctcaa tgttggaaga 43380
aggcgacggc aaatacaggt gacatatccg ttgtttttga tttatatgct aaaaataaat 43440
aatgtgcagt cgtaataggc tgcacattat tgttatatag acaatatctt attcctatat 43500
tgtcaggatt aaaacaacgt gtttgtgttt ttggacgttc taatcgtgtc atttcaattc 43560
cactatggta cgattaaaat atgtacactt agaatcccat ataaaagata aacttttttta 43620
catatcctga caactgctaa ctcagccaaa tcttattatc aaaaaataat gaaccacata 43680
gaaaatcacg aagtcccgat aatgctgttg tattcctaag ccttaattaa ataatttgtt 43740
acaatacata aataataata atattttgaa taagagagaa aaaactttca atcttgatgt 43800
ttgtcactct ccaacctctt aattaatcgc aaacaaataa tcagcggaat aacgccaaac 43860
actccgaacg aacagtcgat aaacctccaa tagagcggaa tctctctgat aggtccgcag 43920
ataagggcaa gcggaacaac tgccacacag gcgattattc ccactgaat cacccatttg 43980
ttgcgtactg gatcacgaaa cggaccaata aacatcaatg caataaccaa atgcgcgaat 44040
gccaaccaat catatccgta cgacaactga ggagtggttt gattgagttc tacaatactt 44100
tccttaatag aatacagaaa accaaaaatt gcaggataat cgtttaccgt agccgaatcc 44160
aaccctgcaa cagagcagag agttttcaac tcagtttcca cagggaaagc cgttactccc 44220
gacagcacta acgccaccat aaagaacgca atccataagc gtaacttcaa caatactttt 44280
ttctcttctt gtggttccat aactttactc tttctaaaac taatactgca aatttacaat 44340

```
     taataattat aaataataat aacctaatat tagaatattt catgaagaca tctttttctt     44400
     ttattgctgc aaatatacga aaatagtttt atgttgcgta tattaattgt tactttataa     44460
     taatccccca gaagtttcac tttaacttct gggggataac gtctcgttat ggataacgag     44520
     atgaatagat gataaagatg ggtttatcgg ataaataata actctctgta tttaggtaga     44580
     ggccagcaag cgtcgtctac tataagttcg agcttatcta tttgtctacg aatcttatcg     44640
     aagtatggtg ctactgtgtc atgatatgca atagccttaa gatgttcgtc ttcaatcttg     44700
     tttgcaactt tacgtgctga aacgagttct tcaacgagag attctattgt ttctgtgcgt     44760
     gtagcaattt cttcgattat cttcaagtta cgtgcgctga gtttgtttgc tttttctggt     44820
     gcaaatgtag tgcgcatgtt ttcaacgttg ttgagcaact gactttggta gcgtgtagca     44880
     acagggatga tgtggttcat tgacaagtcg ccgagaacgc gtgcttcaat ctgaatcttc     44940
     ttagtgtatg tttcccactt aacttcgtta cgtgcttcaa gttcttttt  agtcataact     45000
     ccgagtgatg agaacatctc aatgctgctt gcatcaaggt aacgttcgaa gataactggg     45060
     cagcttgttt ctacatcaag accacgtttt gcagcttctt ccttccattc gtcgctgtat     45120
     ccgtttccat cgaatctgat tggcttacaa gcgcgaatat cttcacgaag aacgtcgatg     45180
     atagcgtgga acttagcagt atgtccagag ttttctttat actgatctgc aatttctgtg     45240
     attctcttgt caacacgtgc tttaaagtta acgagtgctt cagcaacagc tgcgttgagg     45300
     gtaatcattg cagatgcgca gtttgcctca ctaccgactg cacggaattc gaaacgatta     45360
     ccagtgaaag caaatggtga agttctgtta cggtcagtat tgtcaatgag caattctgga     45420
     atctctgtta tatcgagttt catacсttgt ttaccatcaa tacggaagag atcttctttg     45480
     tctgccttct cgatgttatc aagtaattca gtaagctgtt tgcccaagaa acttgagatg     45540
     attgctggtg gagcttcgtt tgcgccaagt cggtgagcat ttgttgcact catgatactt     45600
     gctttaagca aaccgttgtg tttccataca cccatcagtg tttctacgat gaacactgcg     45660
     aaacgtaagt tctgttctgg tgttttgcct ggtgcatgta agagaacacc tgtgtcggtt     45720
     gacaaactcc agttgttatg cttaccgcta ccatttatac ctgcaaatgg tttttcgtgt     45780
     aataatgcgc ggaaaccatg ctttctggca acacgtttca taagtgacat taataacatg     45840
     ttgtggtcaa cggcaaggtt ggtttcttca aagattggag ctacttcgaa ctggttagga     45900
     gcaacctcat tatgacgtgt cttaacagga atgccgagct caagagcttc gatttccaat     45960
     tcacgcatga atgcagcaac tcgttcaggg atgcttccga aatagtggtc atccatctgc     46020
     tggttctttg cactgtcgtg tcccatcaat gtacgacctg taagaagtaa gtctggacgt     46080
     gcagcatata atccttcatc tacaaggaaa tattcttgtt cccatcctaa attacttgta     46140
     accttcttga ctgctgggtc aaagtaatga catacttctg tagctgctac attaacagca     46200
     tgaagagcac gtaacaatgg tgctttatag tctaatgctt caccactata actgatgaat     46260
     acagttggga tacaaagtgt atcgtctata atgaatactg gtgatgtagg atcccatgca     46320
     gaataaccac gagcttcaaa tgtagagcgt atacctccac ttgggaaaga tgaagcgtca     46380
     ggttcctgct gaactaagag ttttccacta aactcttcta tcattcctcc tttaccatca     46440
     tgttcaataa atgaatcgtg tttctctgct gtaccttcag tcaatggctg gaaccagtgt     46500
     gtatagtgtg ttacaccatt ctcgcttgcc cattgtttca taccagcagc a               46551
//
ID   XXX; XXX; linear; XXX; XXX; XXX;  XXX.
XX
AC   PRJEB67889
XX
DE   Uncultured bacterium DNA fragment from in-reactor enriched cow rumen, clone
ferm-11m20, contig-1
XX
FH   Key             Location/Qualifiers
FH
FT   source          1..17653
FT                   /organism="uncultured bacterium"
FT                   /environmental_sample
FT                   /mol_type="genomic DNA"
FT                   /isolation_source="Cow rumen"
FT                   /clone="ferm-11m20, contig 1"
FT                   /note="Tween 20, pNPA, pNPB, pNPP active clone"
```

```
FT   CDS             complement(3..1190)
FT                   /db_xref="SEED:fig|6666666.127230.peg.1"
FT                   /translation="MHLWSQNGPLPALASREMRYVLTCQAERLKALGLTMEESIEHTEG
FT                   EITGGKARSGETIDAATMYREGVRLVTFRQNGRILRKIRKPVTFYATMLNKRGYQDFSC
FT                   TCPNCGHTDMASRMREKCAYCGTVFEIEDAWPVFSSYYFVPGVVERSNLGSNLKRVLIR
FT                   IFLAAAVILAVVSLILNRDQELAERILGAVFGGLAAGAAITFFCYMFYSFFLLGKMFKE
FT                   AGRSMPLLKGLRTEKKLEERMRQYEPDFSYPYFEGRLISLLRAVAYSGRRETLSIYEGS
FT                   QDLSFLDTLVDMQYRGVLQLLNFQVKDGVIRVRLRAFMDDLRYGRRIRRKDENFIVTME
FT                   RDIEAGDDPGFQLHQVKCRSCAGSFDAYHRKTCPYCGSAYRLSQDDWMITSIERG"
FT                   /product="hypothetical protein"
FT   CDS             complement(1231..2178)
FT                   /db_xref="SEED:fig|6666666.127230.peg.2"
FT                   /db_xref="GO:0004617"
FT                   /translation="MKISLIEPLNVKQEVIDRLAAPLKDAGHTFVSYDDLAADEADQAE
FT                   RMKDSDIVMIANHPVAAEAIRGAKDLKMIDVAFTGIDHVPADVCRECGVMICNAAGYST
FT                   ETVAELVIAMAVDALRNVAKANEVVRQGGTAAGIGGREIAGRTVGIIGLGAIGMRCAEL
FT                   FQAFGAKVIAFNRSRKEEAVARGIEYKTLDEVLAESDIISLNLPNTPETRGLISREKIA
FT                   LMKKDAVFINCARGPIVDNAALADALNADELGFACIDVYDMEPPLPADYPLLSAKNTLL
FT                   TPHQGFISEEAMIRRAEIVFRNVYAFLEGKPENV"
FT                   /product="D-3-phosphoglycerate dehydrogenase (EC
FT                   1.1.1.95)"
FT                   /EC_number="1.1.1.95"
FT   CDS             complement(2231..2665)
FT                   /db_xref="SEED:fig|6666666.127230.peg.3"
FT                   /translation="MTASVGAIAGVKYVGGELRLEVLGAAKPKGICGSGFIDILGILYE
FT                   EDILDEIGHILDPEDVDSELARYIGEADDRPVFYLTDDRSICISQADISKFQMAKAAVS
FT                   AALSEEARAELTRIRERVRVIDLPTRPSFSDLFTEEIFFE"
FT                   /product="Methyltransferase corrinoid activation protein"
FT   CDS             2846..4186
FT                   /db_xref="SEED:fig|6666666.127230.peg.4"
FT                   /translation="MVFNDFYFLFLFFPVFYVVYLMLRKTCAKWLIVIGSVVFYTIGVW
FT                   GHPFQLVLLLAVTAVGTLGCMLFQQPRFCRKWLLAVFIVLVAAPLGYVKLAGLVLPSSP
FT                   ALPLGLSFYTFQMIAFLVYAWREGQVDPLSVAAGILFFPKLLSGPIAEPRPLLRDMWKP
FT                   KLDKKLIDASLQEFIIGLAYKVILADHLGGVLGKIRMRTVQGVSVQLAWLGVICYGLQL
FT                   YFDFCGYSRMAMGIAGMLGIRLPENFRHPYCSTSIREFWRRWHITLGTWFRNHVYIPLG
FT                   GSRNGAARMVLATLVVWLLTGIWHGAGWNFVFWGLTMFVLLMGEYFIWGKWLEKVPVIG
FT                   HIYVPFIIMLSWIFFITTGPAEAGHYFIRLFGVTNMPFDAGDWKIVLSQCWMYLVPAVI
FT                   FATPLPQMLFKKVSKKTVGWVILAAVFVASIFFMSTAASEPFLYFSF"
FT                   /product="Probable poly(beta-D-mannuronate) O-acetylase
FT                   (EC 2.3.1.-)"
FT   CDS             4196..5122
FT                   /db_xref="SEED:fig|6666666.127230.peg.5"
FT                   /db_xref="GO:0008781"
FT                   /translation="MAERKKQKKGKFYAFKFLILAEVFALALLVSGMVLRRTLLAGMPQ
FT                   YDSVQTIALPMMVLKDHRQIMAQRAHMAPAGPGGASGTVIDGSETEAETAAVQTPAETE
FT                   PQTETAAPEPVTVDEHYFDNTLFIGDSRTEELSLWGRLGTARYYCGVGFSVFNIFGETV
FT                   TNDDFSGQTLEGALAEHQYDQIYIQLGYNESGYPLESLMSQYTVLLDTVREAQPDARIV
FT                   LHAVMHASRGAVERWDNYAPENLDAINARIRELAESRDRVYYVDVNDLFIDENGYMREG
FT                   LSNDGEHLDIEWTEVWAQALIDRAVVE"
FT                   /product="N-Acetylneuraminate cytidylyltransferase (EC
FT                   2.7.7.43)"
FT                   /EC_number="2.7.7.43"
FT   CDS             complement(5190..5765)
FT                   /db_xref="SEED:fig|6666666.127230.peg.6"
```

```
FT                   /translation="MKKDIGAFPAVYPMPVLMVAAYDANEKVNVMNVAWGQICDTDKII
FT                   LFIGEGKRTWLNIRESRAFTVALADEAHMDVADFFGIASGNKMDDKFERTGYHAVKSDK
FT                   VHAPIIEEFPLVMECELLEFLESEHVQGIVGKIVNVKAEEAVLNDAGKVDPEKLHALIF
FT                   DQFLHGYYTTGAKAGTAWNAGAALMKKS"
FT                   /product="Flavoredoxin"
FT   CDS             complement(5838..6158)
FT                   /db_xref="SEED:fig|6666666.127230.peg.7"
FT                   /translation="MKEETIARIRKFTEDRDWQQYHTPANLAKSIAIEAGELLECFQWS
FT                   ETEYDTEHVKEELADVMVYCRNLLDELGLDEDEIVNAKMDKNEAKYPVEKARGRNTKYD
FT                   RL"
FT                   /product="hypothetical protein"
FT   CDS             complement(6169..7125)
FT                   /db_xref="SEED:fig|6666666.127230.peg.8"
FT                   /db_xref="GO:0004034"
FT                   /translation="MITKTPFASSSHPSAAKFRITNAAGDFAEIIEYGARVSRICVSGR
FT                   DLCLRYETPEEYERDPFYLGAAVGRAANRIRGAAFPLNGETVRLQPNEGRNLLHGGAVN
FT                   FSTALWTGEPAGEDAVRFRLTMEDDGFPGRLDAAVTYSWTDDGVLGILFEGRSDADTVF
FT                   SPTNHMYFNLSGDFSTPVSDHIVEIRSEAYLPVDEEHIPSGAAAPVGGTPFDFRVPREI
FT                   GPGGYDHHFVCSGEPGTLRTLAEAYSPASGIRLICRSDLPGLQFYTGDYLTGFTPRSGF
FT                   CFEAQFAPDALNREDEPKPLLRAGGTVRYRTEYIIST"
FT                   /product="Aldose 1-epimerase (EC 5.1.3.3)"
FT                   /EC_number="5.1.3.3"
FT   CDS             7277..9229
FT                   /db_xref="SEED:fig|6666666.127230.peg.9"
FT                   /translation="MNRYYPHLGSSLKVGGVMLKNRIVNAPMAPPEMGPIGDMTPENIG
FT                   FYGLRAKGGAAVVTVSEGIVHTATGRSHTKHLSLDNPMMLPSLAEVAREIHYHNALASI
FT                   ELSHGGKYAGNRVQGQDPNTVTRYGPIDEVLSDGREVHAMPEELILEIADCFGKAAAMC
FT                   KQAGFDMVLVHAAHGWLLSQFMSPGTNKRTDKWGGSRENRMRFPLLVVQKIREAVGPNF
FT                   PIEYRLSGSEYVEGGYDIDECVEMCKMLDGKVDILYISAGVHENRDSFVLTHPSMFLPH
FT                   GSNVHLAAAVKKHMKHTPVGCVGGINDPAQAEEILATGQADLVCMARQLLVDPDFPKKA
FT                   LSGHAEDITKCCRCFTCFASTLTNRITSCALNPLIGRELDQRYGRPATTPKRLVIVGGG
FT                   PGGMQAALTAREQGHEVILIEKSDRLGGQLNSEEFVPFKADLYAYARQQARRVEQSGTE
FT                   VRLNTEATPELIESLHPDALVLAVGAKPVVPPIPGIDSPKVKGLSALHQADPDLGQRVV
FT                   ILGGGLVGSETAVWLNSLGRDVTVVEMKDDFAAEANEMHKIALEHEFKNSRIRLHLNTS
FT                   AVEVTDEGLVCRGPEGTFTVPCDSILLAAGMRADWDSVERLRDAAPWYWAIGDCVRAGQ
FT                   VVNATEQGHNVVLDL"
FT                   /product="2,4-dienoyl-CoA reductase [NADPH] (EC 1.3.1.34)"
FT                   /EC_number="1.3.1.34"
FT   CDS             complement(9312..10736)
FT                   /db_xref="SEED:fig|6666666.127230.peg.10"
FT                   /translation="MNRKAVDLLHGPVLGKLLAFSIPLVLTQWLQTLFNSADSMIVGRW
FT                   GGEGALASVGATFSLNLLIICFFGGLAAGVMVCAGTDWGAGDRESFDRTAHTAILLAFL
FT                   VGLCATVIIELLAAPVLTAMHVPEDIRPGAILYFRLYLLVIPGQMVYNAGASIMRARGD
FT                   SQRPLIFLTVAGVCNVGLNLLFVIAFRWGVFGVGIATVITQYLSAVLALIILFRPGSEY
FT                   GMQFSRLRLHKDKVFRIIRVGLPAGLQASMLAASDIPLQTCVNSLGSLAVAGNAAALNL
FT                   DGFTFSAMMSLGQGCMVFTSQCVGARDFARARKVLWNGMLMIVGIGAAIGWLCVLFRMQ
FT                   LLGLFQPGNAAAIAFGEQRVMAVCTLAFIYGFLESMNSSMRGYAISVPQAIINGLGLFG
FT                   FRTLWSIFYFPHHQDLFHLYLSYPVAWSLCIAAMAAIYVPMVRRRVASITAEDAARDAV
FT                   PADSGSGGALPENSGS"
FT                   /product="hypothetical protein"
FT   CDS             complement(10813..11448)
FT                   /db_xref="SEED:fig|6666666.127230.peg.11"
FT                   /translation="MSKKITWIGHSCFKIEADGFAIVTDPYEDGSVPGLGPVREQANLV
FT                   LASHEHFDHNARQLVKVEEKADCPFRITEIAAWHDEVQGAKRGPNTIYIMETEGFKIAH
```

```
FT                   LGDLGCDLEEEQIEQLKGLDVCLIPVGGFFTIDGLQAAELIKKIQPRAAVPMHFRDDAA
FT                   GFGFDAISEVDVFAGQFDSVCRLPGSVLDPDELPDAQVVILRPQNAEK"
FT                   /product="Zn-dependent hydrolases of the metallo-beta-
FT                   lactamase superfamily"
FT   CDS             complement(11497..11970)
FT                   /db_xref="SEED:fig|6666666.127230.peg.12"
FT                   /translation="MTNEKAAWMFRKLEENIREDRSTLNGMIDIRFVSCSVEDRTAEFL
FT                   FTAEDWERNVFGGIHGGMIATVLDSCMGITSGVFSEKPASTVSMNVQYLQPGTGAEYRV
FT                   LVSLPHKGRRIIHAAAEMKDAAGGASCAMAEAVFTLRESMDMADRLGGSAPEW"
FT                   /product="hypothetical protein"
FT   CDS             complement(12023..12280)
FT                   /db_xref="SEED:fig|6666666.127230.peg.13"
FT                   /translation="MSELNLNDLEHVSGGASAEGARRTYIVQPGDTLSEIAAKYGTTTK
FT                   KLSLLNADILIAGAEERGIKINDSSEYADHIYAGQILRLP"
FT                   /product="hypothetical protein"
FT   CDS             complement(12309..12821)
FT                   /db_xref="SEED:fig|6666666.127230.peg.14"
FT                   /translation="MADYVMNREGKYGYLFVQDMSGAPESRRSPEALAGYRQFGARQLW
FT                   IDGNCVPGAFQMNTCFWTKANKELMLQHPDAPVVTGHSHPYPEILGFVGTDPDDPLDLG
FT                   GEVEFWVEDEPHIITRSTMVFLPPNMKHCPMFILNVKDEKTPILHFSVVMNELYGYEQG
FT                   GEKVDHV"
FT                   /product="hypothetical protein"
FT   CDS             complement(13048..14448)
FT                   /db_xref="SEED:fig|6666666.127230.peg.15"
FT                   /translation="MQTLSNIVNTIVGGVNNVLWNYLLLILLVGTGVFFTIRTRFVQIR
FT                   RFGAAMKRVFGGIKLHGDKAGKEGMSSFQALATAIAAQVGTGNVAGCATALVSGGPGAI
FT                   FWMWVAAFFGMATIYGEAFLAQITKTRDKEGHVIGGPVFYIRKAFGEKGFGKFLAVFFS
FT                   IAIIIALGFMGNMVQSNSIGDAFFTAFGVPKIAVGIAVAAVAAVIFIGGIGRIASVTEK
FT                   IVPIMAALYIVGSLVVIIINVRAVPGAFASIFVGAFSPRAVGGAVAGIAVRQAIRYGVA
FT                   RGLFSNEAGMGSTPHAHAMAKVDTPEHQGETAMVGVFIDTFVVLTLTALVILTSGVLNG
FT                   MIEGGISGIPVAQAAFSTGFGSRIGPAFVAICLLFFAFSTIIGWYFFAKTNITILFGKK
FT                   GVLPFTVIALFFILLGSVLKVNLVWDLSDMFNGLMVIPNLIALIALSGLVAKAAIDGHR
FT                   KREGEKIR"
FT                   /product="Na(+)-linked D-alanine glycine permease"
FT   CDS             complement(14859..15605)
FT                   /db_xref="SEED:fig|6666666.127230.peg.16"
FT                   /translation="MSLYAIGDLHLHFQSELKAPGQLHDRVWKKHEQKFQAYCGKHLTD
FT                   EDTLVLVGDHSWGKSLEQAETDLQYICDLPGRKILTRGNHDMFWDANKTASLNARYAGK
FT                   LAFLQGNYYAYRDYALVGTKGFTFEGPFRVDRRGRIVDWDDAAEEHAKKIVERELKRLR
FT                   DSFEAASRDGYRKFIMFLHYPPTSVIEEESGFTRMAEEYGAEQVIYAHCHGEARFHDSI
FT                   EGLFHDIRYSLVSGDFLKWRPYKVLD"
FT                   /product="Phosphohydrolase"
FT   CDS             complement(15700..16689)
FT                   /db_xref="SEED:fig|6666666.127230.peg.17"
FT                   /translation="MRYRHFDRMDRDVAEINVGTWGLGGDGYGETNDEDAVQAIRAMLD
FT                   RGVNAIDTAPAYGDGHAETLVGRAIAGYDREKIIVATKCGIMNHTLKAKRGAAGEVRDA
FT                   SFGNILYECEKSLRRLNTDYIDILYVHWPDYDTPLEETAEAFNVLKAEGKIRHIGLSNF
FT                   SREQIEEISKYCRVDALQLPYSMIVREKEELMRWASAQGIDTFTYGSLGSGILSGKYRT
FT                   LPEFGPRDPRAMGFYPYFREPGFSKVMELLKVLDEVAEETGRPVIEVSLNWTASKPFVS
FT                   TVLCGVRNAAQAEGLVRAFEWELTEAQTAKIDAAIGRYIDFDGGNGMK"
FT                   /product="L-fuco-beta-pyranose dehydrogenase (EC
FT                   1.1.1.122)"
FT                   /EC_number="1.1.1.122"
FT   CDS             complement(16705..17280)
```

```
FT                   /db_xref="SEED:fig|6666666.127230.peg.18"
FT                   /translation="MTPKVFLDILSQAGKLKTTLRHCWLEADRRESVSDHCWRLALMAM
FT                   LLAPEPEFAETDMDRVMRMCLIHDLGESFTGDIPVFRKQDSQEHAEEDVLLKWIAGFPE
FT                   PQRAEWTALLAEMTAMETREARTYKALDALEALISHDESDIDTWLPLECDLQLTYRQEN
FT                   MKVSPFFEELRACVDAWTRAKIAGSGKY"
FT                   /product="Hydrolase (HAD superfamily)"
XX
SQ   Sequence 17653 BP; 3466 A; 5630 C; 4917 G; 3640 T; 0 other;
     gctcatcccc gctcgatcga tgtgatcatc cagtcatcct gcgacagccg atacgcggat        60
     ccgcagtacg ggcaggtctt ccgatgatac gcgtcgaagc tgccggcaca ggaacgacac       120
     tttacctgat gaagctgaaa tccgggatca tctcccgcct cgatatccct ctccatcgtg       180
     acgataaagt tctcatcttt tctccggatc cgcctcccat accgcagatc atccatgaat       240
     gctctgagcc ggacgcgtat cacaccgtct ttcacctgaa aattcagaag ctgcagcacg       300
     cccctgtact gcatatccac cagcgtatcc agaaaagaca gatcctgtga cccttcatag       360
     atgctgaggg tctctctccg gccgctgtaa gcgactgccc gcagcagcga gatgagccgc       420
     ccttcgaaat aggggtagga aaaatccggt tcgtactgcc gcatccgctc ttccagcttt       480
     ttctccgtcc gcaagccttt tagcagcggc atggaccgac ccgcttcttt gaacatcttc       540
     cccagaagga agaaactgta aaacatgtag cagaaaaagg tgatcgcagc gccggcggcc       600
     aggccgccga acacagcgcc caggatccgc tccgccaatt cctgatcgcg gttcaggatc       660
     agagacacca ccgcgaggat caccgcggcg gccagaaaga tccgaatcag gacacgtttc       720
     agattcgaac cgaggttcga ccgttccacg acgccaggca cgaaataata agaagaaaac       780
     acgggccatg catcttcgat ctcgaacacg gtcccgcagt aagcacactt ctccctcatc       840
     cgggacgcca tgtccgtatg accgcagttc gggcaggtgc aggaaaaatc ctgatagccg       900
     cgcttattaa gcatcgtcgc ataaaaagtg accggttttc ggatcttgcg caggatcctt       960
     ccgttctgcc ggaatgtgac cagacgcacg ccttcgcgat acatggtcgc agcgtcgatg      1020
     gtctctccgc ttctggcttt cccgccggtg atctcacctt ccgtgtgttc gatgctctct      1080
     tccatggtca ggccgagggc ttttaaacgt tccgcctgac aggtcagcac ataacgcatc      1140
     tcccgcgatg ccaatgccgg aagcggtccg ttctgcgacc agagatgcat ttcgtcggta      1200
     aaagtattga gtggattcct gctgtctcgg tcaaacattc tcgggctttc cttccagaaa      1260
     cgcgtacaca ttccggaaca cgatctccgc gcgccggatc atggcctctt ccgagataaa      1320
     gccctggtgc ggcgtcagaa gcgtattctt cgcggaaagg agcggataat ccgcaggcag      1380
     cggcggctcc atgtcataga cgtcgatgca ggcgaagccc agctcgtccg cgttcagcgc      1440
     gtccgcgagc gcggcattgt ccacgatggg gccgcgcgca cagttgatga acaccgcatc      1500
     cttcttcatg agcgcgatct tctcgcggct tatgagccct ctggtctcgg gcgtgttcgg      1560
     gaggttcagc gaaatgatat cgctctccgc gaggacctca tccagcgtct tatactcgat      1620
     gccgcgcgcg accgcctctt ccttccggct gcggttgaac gcgatcacct tcgcgccgaa      1680
     cgcctgaaac agctccgcgc agcgcatgcc gatcgcaccg aggccgatga tgccgaccgt      1740
     acggcccgcg atctcgcgtc cgccgatgcc tgcagcggtg cctccctgcc ggacgacttc      1800
     gttcgccttc gcgacgttcc gcagagcgtc caccgccatg gcgatcacga gctccgcgac      1860
     cgtctcggtg gaatacccgg cggcgttgca gatcatgacg ccgcactcac ggcagacgtc      1920
     cgcggggacg tggtcgatcc cggtgaacgc cacgtcgatc atcttcagat ccttcgctcc      1980
     gcggatcgcc tccgcggcga ccgggtggtt cgcgatcatc acgatgtcgc tgtctttcat      2040
     ccgctccgcc tgatccgctt cgtcagcggc caggtcatca tacgacacga acgtgtgacc      2100
     cgcgtccttc agcgggggcgg ccaggcggtc aatgacctcc tgcttcacgt tcaggggttc      2160
     gatcagactg atcttcatat tctggtacct ctcttaaaaa cattgtcctg ttccgccgta      2220
     ttccggcgta tcattcgaaa aagatctcct cggtaaacag gtcgctgaag gacggacgcg      2280
     tcggcaggtc gatcacacgg acccgttccc ggatcctggt cagctcagcc cgcgcctctt      2340
     ccgacagcgc ggccgagacc gcggccttcg ccatctggaa tttactgatg tccgcctggg      2400
     aaatgcagat gctccggtcg tccgtcagat aaaacacggg gcggtcgtcc gcctcgccga      2460
     tatagcgggc cagctccgag tccacgtcct ccggatccag gatatggccg atctcgtcca      2520
     gaatgtcttc ctcatacagg attccgagaa tatcgataaa accggatccg cagatccctt      2580
     tcggcttcgc cgcgcccagc acttcgaggc gcagttcgcc gccaacatat ttgacgccgg      2640
     cgatggcgcc gaccgacgct gtcatgccgt gctcgagcag ggagccgtaa tgctctgaat      2700
     accggcatgc tgcagaacac tcagaatggt ctcgccctct tcgaaaggca gttcgcgctg      2760
```

```
tatctgcgac tatcatacta tattgtgtgt ttttttgctc tatccctcgc attttcccgc      2820
cgaatatcgt ataataacag ctgttatggt atttaacgac ttctacttcc tgtttctttt      2880
cttcccggtc ttctacgtag tctatctcat gctgcggaag acctgtgcga agtggctgat      2940
cgtgatcggg agcgtcgtct tctacacgat cggcgtgtgg gggcatccct tccagctggt      3000
gctccttctc gcggtgacgg cggtcggcac gctcggctgc atgctgttcc agcagccgcg      3060
gttctgcagg aagtggctgc tcgcagtttt catcgtcctc gtggctgcgc cgctcggcta      3120
cgtgaagctg gccgggctgg tcctgccgag ttctcccgcc cttcccctgg gtctgagttt      3180
ctatacgttc cagatgatcg cgttcctggt gtatgcctgg cgcgagggtc aggtggaccc      3240
gctgagcgtg gccgcgggca tcctgttctt cccgaagctg ctgtccggcc cgatcgcgga      3300
gccgcggccg ctcctcaggg acatgtggaa gccgaagctc gataagaagc tgatcgacgc      3360
gagccttcag gagttcatca ttggtctcgc ttataaagtg atcctggccg accacctggg      3420
cggcgtgctc ggaaagatcc gaatgcgtac ggtccagggc gtctccgtgc agctcgcatg      3480
gctgggcgtg atctgctacg gtctgcagct gtatttcgat ttctgcggct attcgaggat      3540
ggccatgggc atcgccggta tgctcggcat ccgcctgccg gagaacttcc ggcatcccta      3600
ctgctcgacc agcatccgcg agttctggcg cagatggcat atcacgctgg gaacctggtt      3660
tcggaatcac gtctacattc cgctgggcgg cagccgtaac ggcgcggccc gcatggtcct      3720
cgccacgctg gtggtctggc ttctcactgg catctggcac ggcgccggct ggaacttcgt      3780
gttctgggga ctgacgatgt tcgtgctcct gatgggcgaa tatttcatct gggggaaatg      3840
gctggagaaa gtccccgtga tcggacacat ctacgtgccg ttcatcatca tgctctcgtg      3900
gatcttcttc atcacgacgg gtcccgcgga ggccgggcac tatttcattc ggctgttcgg      3960
cgtaacgaac atgccgttcg acgcaggcga ctggaagatc gtgctctcgc agtgctggat      4020
gtacctcgtg cccgcggtca tcttcgccac tccgctgccg cagatgctct ttaaaaaagt      4080
gtcgaagaag acggtcggct gggtcatcct cgcggcggtc ttcgtggcca gcatcttctt      4140
catgtcgacc gcggcttccg agccgttcct gtatttcagc ttttaagggg gcgtcatggc      4200
ggaaaggaaa aaacagaaaa aaggcaagtt ctatgctttt aagttcctga tcctcgcgga      4260
ggtcttcgcg ctggcgcttc tggtgtcggg aatggtcctg cggcgtacgc tcctcgcagg      4320
catgccgcag tatgactcgg tccagacgat cgcgctgccg atgatggtac tgaaagacca      4380
caggcagatc atggcacagc gggcgcatat ggcgccggcc gggcccgggg gcgcgtccgg      4440
gaccgtcatt gacgggtctg agaccgaggc cgagacagcg gcggtccaga ctcccgccga      4500
gaccgaaccg cagaccgaga cggccgcgcc ggaaccggtc acggtcgacg agcattattt      4560
cgacaatacg ctgttcatcg gagactcgag gaccgaggag ctctcgctgt ggggcaggct      4620
cggaacggca cgctattact gcggcgtggg cttctccgtg ttcaacatct tcggtgagac      4680
cgtgacgaac gacgatttct cggggcagac cctcgaaggc gcgctcgcgg agcatcagta      4740
cgaccagatc tacatccagc tcggctataa cgagtcgggc tacccgctcg agagcctgat      4800
gtcgcagtac acggtccttc tggacaccgt gcgcgaggca cagcccgacg cacgcatcgt      4860
tctgcatgcg gtcatgcacg cgagccgcgg cgcggtggag cgctgggata attacgcgcc      4920
ggaaaacctg gatgcgatca atgcccggat ccgtgagctc gcggaaagcc gcgaccgcgt      4980
ctactacgtg gatgtgaacg acctgttcat cgatgagaac ggctacatgc gcgagggcct      5040
gagcaatgac ggcgagcacc tcgacatcga gtggaccgag gtctgggcgc aggcattaat      5100
agaccgggcc gtcgtggagt aaggcccggt cccggtattc ttcatttcat tcaactgtta      5160
agctgagcga cccggctccg gccggctgct cagcttttct tcatcagagc agcgccggca      5220
ttccatgccg tgccggcttt cgcgccggtg gtgtaatagc cgtgcaggaa ctggtcgaag      5280
atcagcgcgt gcagcttctc cggatcgacc tttcccgcat cgttaagaac ggcctcctcg      5340
gccttcacat tcacgatctt tccgacgatc ccctgaacat gctcgctctc caggaactcc      5400
agcagctcgc attccatgac cagcgggaat tcctcgatga tcggcgcgtg gaccttatcg      5460
ctctttaccg cgtgataacc ggtccgctca aatttgtcgt ccatcttatt tccggaggcg      5520
atgccgaaga aatccgcgac gtccatgtgc gcttcatctg ccagtgccac cgtgaacgca      5580
cggctctccc ggatgttgag ccaggtcctc ttcccctcgc cgatgaacag gatgatctta      5640
tccgtgtcgc agatctgtcc ccaggccacg ttcatgacat tgaccttctc attcgcgtcg      5700
taggccgcca ccatcagcac cggcatgggg tacacggcgg gaaatgctcc gatatctttt      5760
ttcataattc tgctctcctt tcaggctgtt ttgatcattt atcttaaacg gcagatgctc      5820
tttttgcagc gcgtccctca cagacggtca tacttcgtgt tccgaccccg ggccttttcg      5880
acgggatact tcgcctcgtt cttatccatc ttggcattga cgatctcatc ctcgtccagg      5940
ccgagctcgt ccagcaggtt ccggcagtag accatcacat cggccagctc ttccttcaca      6000
```

tgctccgtgt catattccgt ctcgctccac tggaagcact ccagcagctc tcccgcctcg 6060
atggcaatgg acttggccag gttcgcgggc gtgtgatact gctgccagtc tctgtcctct 6120
gtgaatttgc gaatgcgggc aatggtttcc tctttcatat catcatcctc acgtgcttat 6180
aatgtattcc gtcctgtacc ggacggtccc gccggccctc agcagaggtt tcggctcatc 6240
ttcccgattc aatgcatccg gcgcgaactg tgcctcgaag cagaagcccg accgcggcgt 6300
gaagccggtc aggtagtctc cggtataaaa ctgcagtccg ggcaggtccg accggcagat 6360
cagacgtatc ccgctcgcgg gcgaatacgc ctccgccagc gtgcgcaggg tcccgggctc 6420
cccggaacag acaaaatggt ggtcgtaacc gcccggcccg atctccctcg gaacgcggaa 6480
gtcgaagggc gtgcccccga ccggcgccgc tgctccgctg ggaatatgtt cctcgtccac 6540
cggcagatac gcctccgacc gtatctccac gatgtgatcc gagaccggcg tcgagaagtc 6600
ccccgacaga ttaaaataca tgtggttcgt cgggctgaag accgtgtcgg cgtcgcttct 6660
cccctcgaag aggatcccga gtacgccgtc atccgtccag ctgtatgtga ccgccgcgtc 6720
tagccgcccc ggaaaaccgt cgtcttccat cgtcagccga aaccggaccg catcttctcc 6780
cgccggttct cccgtccaga gagccgtact gaaattcacc gcgccgccgt gaagcaggtt 6840
ccgcccttca ttcggctgca ggcgcacggt ctctccgttc aggggaaacg ccgccccgcg 6900
gatgcggttc gcggcgcggc cgaccgccgc gcccagataa aacgggtcgc gctcgtactc 6960
ttctggggtc tcataccgca ggcacaggtc ccggccgctc acgcagatgc gcgagacccg 7020
ggcgccgtat tcgatgatct ccgcaaaatc gcccgcggcg ttcgtgatgc ggaacttcgc 7080
ggccgaaggg tgcgatgatg aggcaaatgg cgttttcgtg atcataatat ccccgatact 7140
tctgtccgtt cggaaaccgt ttctctgcag tgcgccgcat ttattatagc acagaaccgg 7200
ctccgctttc ttttcatgct gtttatggta aaatgagcca tatcgaaccc gaattctttt 7260
actacggagg atccctatga accgttatta tccgcatctg ggaagcagcc tgaaagtggg 7320
cggcgtcatg ctgaagaacc gtatcgtgaa tgcaccgatg gctccgcccg agatgggccc 7380
gatcggcgat atgacgccgg agaatatcgg tttttacggc ctgcgcgcga aaggcggcgc 7440
cgccgtcgtc acggtcagcg aaggcatcgt gcacacggcc accggccgca gccacacgaa 7500
gcacctgtcg ctcgataacc cgatgatgct cccctcgctg gccgaggtcg ccagggagat 7560
ccattatcac aatgcgctgg ccagcatcga gctctcgcac ggcggaaagt acgccggaaa 7620
ccgggtgcag ggacaggatc cgaacacggt cacccgctac ggcccgatcg acgaggtgct 7680
ctccgacgga cgcgaggtcc acgccatgcc cgaggagctg atcctggaga tcgcggactg 7740
cttcggaaag gccgccgcga tgtgcaagca ggcgggattt gacatggtgc tcgtccacgc 7800
cgcgcacggc tggcttctga gccagttcat gtctcccggc acgaataagc gcacggacaa 7860
atggggcggc agccgtgaga accggatgcg cttccctctt ctcgtcgtgc agaagatccg 7920
cgaggccgtg ggcccgaact tcccgatcga gtaccgcctc agcggctcgg agtatgtcga 7980
aggcggctac gatatcgacg agtgcgtcga aatgtgcaag atgctcgacg gaaaggtcga 8040
catcctctac atttccgccg gcgtccacga gaaccgcgac agcttcgtgc tgactcatcc 8100
ctcgatgttc cttccgcacg ggagcaacgt ccatctggcc gctgccgtga agaagcacat 8160
gaagcacacc ccggtcggat gcgtcggcgg aatcaatgac ccggcgcagg ccgaggagat 8220
cctcgccacc ggtcaggccg acctggtctg catggcgcgc cagctgctgg tggatccgga 8280
tttcccgaag aaagccctct ccggccatgc cgaagacatc acaaagtgct gccgctgctt 8340
cacctgcttc gcctcgaccc tgacgaaccg catcacctcc tgcgccttga acccgctgat 8400
cgggcgggag ctggaccagc gttacggaag gcccgcgacc acgccgaagc gcctggtcat 8460
cgtgggcggc ggccccggcg gcatgcaggc ggccctcacc gcccgtgagc agggtcatga 8520
agtcatcctg atcgagaagt ccgaccgtct gggcgggcag ttaaacagcg aggagttcgt 8580
cccgttcaag gccgacctgt acgcctatgc caggcagcag gccagacgcg tggaacagtc 8640
cggcacggag gtgcgcctga acacggaggc cacgccggag ctgatcgagt cgctgcaccc 8700
ggacgctctg gtgctcgcgg tgggcgcgaa gcctgtggtc ccgccgatcc cggggatcga 8760
ttccccgaag gtgaaggggc tttccgccct gcatcaggca gatcccgatc tgggtcagcg 8820
cgtcgtgatc ctcggcgggg gcctggtggg cagcgagacc gcggtctggc tgaacagtct 8880
gggccgggac gtgaccgtgg tggaaatgaa ggacgatttc gccgcggaag cgaatgagat 8940
gcacaagatc gcgctggaac acgagtttaa gaacagccgg atccggctgc atctgaatac 9000
gtccgccgtg gaagtcacgg acgaagggct cgtctgccgc gggccggagg ggacctttac 9060
cgttccctgc gattccatcc tgctggccgc gggcatgcgt gcagactggg acagcgtcga 9120
gcggctgcgc gacgcggcgc cctggtactg ggcgatcggc gactgcgtcc gggccggtca 9180
ggtcgttaac gcgaccgagc agggacacaa cgtcgtgctc gatctttgat acgcgcaccc 9240

gttcaaccaa taagagaccc gggccgccag gcccgggtct ctttgtctca ggtcgtctga 9300
gattctctct ttcatgatcc gctgttttcc gggagtgcac cgccggaccc gctgtctgcc 9360
ggaacagcat cccgggcggc atcttccgct gtgatactcg cgacccttcg ccggaccatc 9420
ggcacataga tggcagccat ggcggcgatg cagagcgacc acgccacagg ataagacaga 9480
tagagatgga acagatcctg atggtgcgga aaataaaaga tcgaccacag cgtgcggaag 9540
ccgaacagcc ccagtccgtt gatgatcgcc tgcggcactg agatcgcgta accgcgcatg 9600
gatgaattca tgctctccag gaagccatag atgaaagcca gagtacagac cgccatgacg 9660
cgctgctctc cgaaagcgat cgcggccgcg ttcccgggtt ggaaaagtcc gagaagctgc 9720
atgcggaaca ggacgcagag ccatccgatc gccgcgccta ttcccacgat catgagcatg 9780
ccgttccaca acactttcct cgcgcgtgcg aaatctctgg ctccgacgca ctgactggta 9840
aagaccatac acccctggcc gagggacatc atcgcggaga aggtaaagcc gtcgaggttc 9900
agggccgcgg cattgcccgc gaccgcgaga gaccccagac tgtttacgca ggtctgcagc 9960
ggaatatccg aagccgccag catggaggcc tgaaggcccg cgggcagtcc gacccgtatg 10020
atgcggaaca ctttatcctt atgaaggcgc agacggctga actgcatgcc gtattccgag 10080
cccggtcgga acaggatgat caatgccagc accgctgaca gatattgcgt aatgaccgta 10140
gcgatgccca ctccgaagac gccccagcgg aatgcgatga cgaagagaag gttcaggccg 10200
acgttgcaga cgccggcgac cgtaaggaag atcagcggac gctgcgagtc gccccgcgcg 10260
cgcataatcg aagcgccggc attatagacc atctggcccg gaatgacgag caaatacagc 10320
cgaaaataaa ggatggcgcc cggccggatg tcctccggca catgcatggc cgtcagcaca 10380
ggcgcggcca gcagctcgat gatcaccgta gcgcacaggc cgaccagaaa cgcgagcagg 10440
atggctgtgt gggcggtccg gtcgaaactt tcccggtctc ccgctcccca gtctgtgccc 10500
gcgcagacca tgacacccgc ggcaagtccg ccgaaaaagc agatgatcag caggttcagg 10560
gagaacgtcg cccccacgga tgcgagcgcg ccttcgccgc cccaacggcc gacgatcatg 10620
ctgtctgcgg agttaaacag cgtctgcagc cactgcgtca ggaccaatgg gatcgaaaat 10680
gccagaagct tccccaggac aggcccgtgc agcagatcga cagcttttct gttcatagga 10740
atcctccgtt cgaaaaactt cgtggtttcg actcagatgc cttaactgaa atgatctatg 10800
aaaatgatgc ttttactttt ccgcgttctg cgggcgcagg atcacgacct gcgcatccgg 10860
cagctcatcc ggatccagca cacttccggg cagcctgcag acagaatcga actgtccggc 10920
gaacacgtcg acctcgctga tcgcgtcaaa accgaacccg gccgcgtcgt cccggaaatg 10980
catcgggacc gcggccctcg gctgaatctt tttgatcagc tccgccgcct gcagcccgtc 11040
gatcgtaaag aagccgccga ccgggatcag gcagacgtcg aggcctttca gctgctcgat 11100
ctgctcttct tccaggtcgc agcccagatc gccgagatgt gcgatcttaa agccttccgt 11160
ctccatgata tagatcgtgt tcggcccgcg cttcgcgccc tgaacctcgt catgccaggc 11220
cgcgatctcc gtgatccgga acggacagtc cgctttctct tcgaccttca ccagctgccg 11280
ggcgttatga tcgaaatgtt catgactggc cagcacgagg ttcgcctgct cgcgcaccgg 11340
cccgagcccg ggaaccgagc cgtcctcgta aggatccgtc acgatcgcaa acccgtctgc 11400
ttcgatctta aaacaagaat gaccgatcca tgtgatcttt ttactcatga tccacctgtc 11460
ctttctctaa tgactccgat tctgcccggc gcgccttcac cattccggag cgctgcctcc 11520
cagacggtct gccatatcca tgctctcccg cagggtaaac acggcttccg ccatggcgca 11580
gctcgctccg ccggcagcat ccttcatctc ggctgccgcg tgtatgatac ggcgcccttt 11640
gtgcggcagc gagaccagaa cacggtactc cgctcccgtt ccgggctgca ggtactgcac 11700
attcatcgat accgtgctcg caggcttttc gctgaagaca cccgaggtga tgcccatgca 11760
ggagtccagc acggtcgcga tcatgccgcc gtggatgccg ccgaacacgt ttctctccca 11820
gtcctccgcg gtaaacagaa attccgccgt ccgatcttcc acagaacatg aaacgaaacg 11880
gatatcgatc atgccgttca gggtcgatct gtcttcccgg atgttctcct ccagtttccg 11940
gaacatccat gccgcttttt cattcgtcat aatcccaacc ttcggtcttt ttctgcccgg 12000
atccttcggc cgttctcccg cctcagggca gccgcaggat ctgccccgca taaatgtggt 12060
cggcgtattc cgaggaatcg tttatcttga tgccccgctc ctctgcaccc gcgataagga 12120
tatccgcgtt aagaagcgac agttttttcg tggtggtccc gtatttcgcc gcgatctcgg 12180
aaagcgtatc tcccggctgc acgatatagg tcctccgggc gccttcggcg gaagctccgc 12240
cggaaacatg ttccagatca ttcaggttta attccgacat aactgttcct ccttttatct 12300
ctgaccggtc agacgtgatc gaccttttct ccgccctgct cgtatccgta gagctcgttc 12360
atcacgaccg agaagtgaag gatcggcgtc ttctcgtcct tgacgttcag tatgaacatc 12420
gggcaatgct tcatgttcgg gggaaggaag accatcgtgg atctggtaat gatatgcggt 12480

tcgtcctcta cccagaactc cacctcgccg ccgagatcca gagggtcatc gggatctgtg 12540
cccacaaatc cgaggatctc cggatagggg tgcgaatggc ccgtgaccac aggcgcgtcg 12600
gggtgctgaa gcatgagctc cttgttcgcc ttcgtccaga agcaggtgtt catctggaac 12660
gcgcccggca cgcagttgcc gtcgatccag agctgtctgg ctccgaactg cctgtatccc 12720
gcgagagctt ccggggatct tctgctctcc ggggcgccgc tcatgtcctg cacgaaaagg 12780
tagccgtatt tgccttccct gttcattaca tagtcagcca ttttttgatc tccttatgct 12840
ttaataacag ataacctctc gatctgtatg tataccattt tactccatta ccccggttcg 12900
catatacatt tttgagcatc gtttttttatc cagtttcagg ccgcataaaa agcagaccgc 12960
atacctctgg atccggaaaa gaaaaattgc agatgatccc atggatcacc tgcaatctgt 13020
tatctctttt ttgtaccggc gaaccgatca gcggatcttc tcaccctcac gtttcctgtg 13080
cccgtcgatc gcagcctttg cgaccagacc ggacagcgcg atcagcgcga tcaggttcgg 13140
aatgaccatg aggccattga acatatcgga cagatcccat accaggttca ccttcaggac 13200
cgaacccagc aggatgaaga acagcgcgat gaccgtgaac ggcagaacgc ccttcttccc 13260
gaaaaggatg gtaatgttcg tcttcgcgaa gaagtaccac ccgatgatag tcgaaaacgc 13320
gaagaacagc aggcagatcg ccacaaacgc gggaccgatt cgcgagccga agcccgtgga 13380
aaacgcggcc tgtgccacgg ggatgccgct gatgccgccc tcgatcatgc cgttcaggac 13440
gcccgaagtc aggatgacca gcgcggtcag tgtcagcacg acgaacgtat cgatgaacac 13500
gccgaccatg gcggtctcgc cctgatgctc cggagtgtcg accttcgcca tcgcatgcgc 13560
gtgcggggtc gagcccatgc cggcctcatt cgagaacagg ccgcgcgcca caccgtaacg 13620
gatggcctga cgcaccgcga tgcccgcgac cgcaccgccg accgcgcggg ggctgaacgc 13680
gcccacgaag atggaggcga aagcgccggg gaccgcacgc acattgatga tgatcacgac 13740
caggctgccc acgatgtaaa gcgccgccat gattgggacg atcttctcgg tgaccgaggc 13800
gatacgcccg atgccgccga tgaaaatgac cgccgccaca gccgcgaccg cgatgcccac 13860
cgcgatcttc ggcacaccga acgccgtgaa gaaagcgtcg ccgatggagt tcgactggac 13920
catattgccc atgaaaccga gggcaatgat gatcgcgatc gagaaaaaca ccgcaaggaa 13980
ctttccgaag cccttctcgc cgaaagcctt gcggatatag aagaccgggc cgccgatgac 14040
gtggccttcc ttatcacggg tcttcgtgat ctgtgcgagg aatgcctcgc cgtagatggt 14100
cgccatgccg aaaaaggccg cgacccacat ccagaagatg gcgccggggc cgccggaaac 14160
gagcgcggtg gcgcagccgg cgacattgcc cgtgccgacc tgggccgcga tggcggtcgc 14220
cagagcctgg aatgagctca tgccttcctt gcccgcttta tcgccgtgca gcttgatgcc 14280
gccgaagaca cgcttcatgg cggcgccgaa cctgcgaatc tggacgaagc gggtgcggat 14340
cgtgaagaac acgcccgtgc cgaccagcag aatcagcagc aggtagttcc agaggacgtt 14400
gttcacgccc cccacaatgg tgttgacaat gttcgaaagt gtctgcatac aagttcccct 14460
tagccggtag taccttcgga cgaagtgatg ccgaggccgt gatgtcgcgc gaaaacgccg 14520
tacgaaaaaa ccggatcctc atgagagaat ccggttcgac gcctcgtgtt cagcccgctg 14580
cgcgatacgc ctgtccgcgc ctgcggcagc tgatcaaaag ctctgtcctt ttggcctgag 14640
agattcacgc ggcctgcgcc gcgcttgccc cttcggcacc cgactgaacg ggattctcca 14700
gaggtgctcc gctgccggtc gggaaggatc cgttctggca gcatccgcat attgggtttt 14760
cgcaagcgtt atcttaacag cccgggcacg atatgacaat cggttttttt acccgggccc 14820
gatcgtaccc ggcgcgttcc gccttttcgt gctgtccgtc agtccagcac cttatacggc 14880
cgccatttca ggaagtcccc cgagaccagg ctgtatcgga tgtcatgaaa cagcccctcg 14940
atgctgtcat ggaacctggc ctcaccatgg cagtgcgcat agatcacctg ctctgcgccg 15000
tactcctcgg ccatccgcgt aaagccgctc tcctcttcga tcacgctggt gggcggatag 15060
tgcaggaaca tgatgaactt ccggtatccg tcccgggacg ccgcctcaaa agaatcccgc 15120
aggcgtttca gttcgcgctc cacgatcttc ttcgcgtgct cttccgccgc gtcgtcccaa 15180
tcaacgatgc ggccgcgccg gtcgacccgg aacggtccct cgaacgtaaa gcccttcgtc 15240
ccgacgagcg cataatcccg atacgcatag taatttccct gcagaaatgc cagtttcccg 15300
gcataccgcg cattcagcga ggcggtctta ttcgcgtccc agaacatgtc gtgattgccc 15360
cgggtcagga tcttccgtcc gggcaggtca cagatatact gcaggtccgt ttccgcctgc 15420
tccaggctct ttccccagct gtggtctccg accaggacca gcgtgtcctc atccgtcaga 15480
tgtttcccgc aatatgcctg aaacttctgt tcatgcttct tccatacgcg gtcatgcagc 15540
tgtcccggcg ccttcagttc cgactgaaaa tgcagatgca ggtcgccgat cgcatataaa 15600
ctcataaata cgtcactccg atgcggcctt tcggctcatg cggctcgatg ggataatccg 15660
cgaaacatgc acagtctgtc actcgaattc agaaatccgt tatttcatcc cgttcccgcc 15720

```
     gtcaaaatcg atatagcggc cgatcgccgc gtcgatcttc gcggtctgcg cttcggtcag     15780
     ctcccattcg aacgcgcgca cgagcccctc ggcctgcgcg gcattccgca caccgcagag     15840
     caccgtcgac acgaacggtt tcgacgcggt ccagttcagg ctgacctcga tgaccggacg     15900
     gcccgtctcc tccgcgactt cgtccaggac tttcagcagc tccatcactt tcgagaaccc     15960
     gggctcccgg aaatacggat aaaagcccat ggcgcgcgga tcacgcgggc cgaattctgg     16020
     cagcgtccgg tactttcccg acaggatgcc ggagcccagc gaaccgtagg taaaagtgtc     16080
     gataccctgc gcggacgccc agcgcatcag ctcctctttt tcacgcacga tcatcgaata     16140
     cggaagctgc aatgcatcga cccggcaata cttcgagatc tcctcgatct gctcccgcga     16200
     gaagttcgac agaccgatgt ggcggatctt tccttccgcc tttaagacat tgaacgcctc     16260
     cgccgtttcc tcgagcggcg tatcgtaatc cggccagtgc acatacagaa tatcgatata     16320
     atccgtgttc agccggcgca gtgatttctc acattcatag aggatgttcc caaacgacgc     16380
     gtcgcggacc tcgcccgccg caccgcgctt cgccttcagc gtgtggttca tgatcccgca     16440
     cttcgtcgcc acgatgatct tctcccggtc atagcccgcg atggcgcgcc ccacgagcgt     16500
     ctcggcatgg ccgtccccgt aagcgggcgc ggtgtcgatc gcgttaacgc cgcggtccag     16560
     catggcgcgg atggcctgca ctgcatcctc gtcattcgtc tctccgtatc cgtctccgcc     16620
     gaggccccag gtccccacat tgatctccgc tacgtcgcgg tccatacggt cgaaatgtct     16680
     gtatcgcatc ttggatctcc tttctcaata cttccctgat cccgcgatct tcgcccgcgt     16740
     ccaggcgtcc acacaagccc gcagttcttc gaaaaacggc gagaccttca tattctcctg     16800
     ccggtacgtc agctgcagat cgcattccag aggaagccag gtgtcgatgt ccgactcatc     16860
     gtgcgagatc agggcctcga gtgcatccag tgccttatac gtccgggctt cccgggtctc     16920
     catcgcagtc atctcggcga gcaatgccgt ccactcggcc cgctgcggct ccggaaaacc     16980
     cgcgatccac tttagcagaa cgtcctcctc cgcatgttcc tgcgaatcct gctttcgaaa     17040
     caccggaata tccccggtga aagactcacc cagatcatgg atcaaacaca tgcgcatcac     17100
     gcggtccatg tccgtctcag cgaattccgg ctccggcgcc agcagcatcg ccatcagagc     17160
     cagccgccag cagtggtccg acacgctctc gcggcggtcc gcctccagcc agcagtgccg     17220
     aagcgtggtc ttcagttttc cggcctgcga caggatatcc aaaaacactt tcggtgtcat     17280
     acgccctcca tccattctct catgtttcgg atcttgtata tcgttaggcc atgtcgcaca     17340
     tggcagcatt tggggagaca ttacttgtct gctgtaccgg atgaaagccc ggcgtccccc     17400
     ggcagcacaa gtttgcaaga taattttacc aaagatacag ctgcattgaa aagcgtgaaa     17460
     ggccgacctc atttgggggа ctgtcgttca ggaagaagac gttcccaaaa actcccccgc     17520
     agaatgatcc gcaggggagt ttcccgtaca tattcaccgt gcggaaggtt tccgcggtcc     17580
     ggccgcctca tcctctctgc cgcaccattt tcaggatggc cttgatcacg ccgaagccca     17640
     ccgccgcgat cgg                                                         17653
//
ID   XXX; XXX; linear; XXX; XXX; XXX;  XXX.
XX
AC   PRJEB67889
XX
DE   Uncultured bacterium DNA fragment from in-reactor enriched cow rumen, clone
ferm-11m20, contig-2
XX
FH   Key             Location/Qualifiers
FH
FT   source          1..15380
FT                   /organism="uncultured bacterium"
FT                   /environmental_sample
FT                   /mol_type="genomic DNA"
FT                   /isolation_source="Cow rumen"
FT                   /clone="ferm-11m20, contig 2"
FT                   /note="Tween 20, pNPA, pNPB, pNPP active clone"
FT   CDS             25..777
FT                   /db_xref="SEED:fig|6666666.127232.peg.1"
FT                   /db_xref="GO:0005524"
FT                   /db_xref="GO:0006810"
```

```
FT                   /db_xref="GO:0016021"
FT                   /db_xref="GO:0016887"
FT                   /translation="MKEVVPFIEGVKEITLIDAADNRVDLKIDGMLYAKDLPENAFKSL
FT                   AGDPVRVIVPEGDARYVTWFCDPADDEKFTVFARSVLDEKYPLLTEDTYYDLGYNVQIS
FT                   RANQMIKVMNIAIILGEIILTGLIILILLMGFASVISTLSSNIRVRRREFAVLKSVGMT
FT                   RGALEKMLYGESLCCILKAVGKGTVFGILIPWAINLFIRRVFAVRYELPVAYLAVGIVV
FT                   VALIVVFITRVEIGKLGKQNIIEDIRME"
FT                   /product="ABC transporter ATP-binding protein"
FT   CDS             complement(1058..2212)
FT                   /db_xref="SEED:fig|6666666.127232.peg.2"
FT                   /translation="MKKRYLPIVLIAALTLTACGKGAEGSSDVVDELTSEETVATDFTA
FT                   TDSEEPVASFVSDPAKEAETTDDAAYAIGENVVDGKYVFNAHVFSDNDTAAWGTNGRDA
FT                   LFNLCDALRKGEDTFECANEQAYMNAIGGRLINYYFPVASNCVMEVEDGGFSDGVGKIE
FT                   YLIPKEEFLEREKAFEAQIVTILNENVKPDYSDFEKALALYVYISQNYTYDYDKADRLG
FT                   PDTDDVLTYSPLALGKGICQEFSGLYSYLLMQCGVQADVVCGDMYDATTPGHQWDFVRI
FT                   NGTDYYVDTTFALHDPEVCDDHTPLEYFLMTDDMRSFYGNFDPGTYQMCGQGETSRSYF
FT                   DIYAKDDRYSELWNGYFLEMDTDNNVVKYVEYGSEEIKEFHYED"
FT                   /product="putative cell wall-binding protease"
FT   CDS             complement(2228..3262)
FT                   /db_xref="SEED:fig|6666666.127232.peg.3"
FT                   /translation="MALTACGKEAEGALDTTEVVEATTPEPTPEITPEPTEEPEPEPAK
FT                   FEFNPHVYSPILAERFGPEYWESFYNLCDALRAGEDTFECASQEVYDWCTEEVILCALF
FT                   PAACCKVCGKSDDGTVPFENGVGRIYYKMPKEEFLQREKDFEDMITEILNDNVRTDYSD
FT                   FEKTLAIHDYFAREYTYDEDIFDHEDDGASYSTFVYKSGVCSELAIAYAYLLMQCGVDA
FT                   ISAGSFDIFHSFDYIRLNGKGYFIDPTWSLWSEYNTDHTPLHYFLITEKDREIDGCTIE
FT                   NNKVELLRYFNAVLKIDDYVVSATDDTYAAFRRTHFLGMDTEKNIVKLLDLETGKTREF
FT                   HYED"
FT                   /product="S-layer protein"
FT   CDS             3549..4289
FT                   /db_xref="SEED:fig|6666666.127232.peg.4"
FT                   /translation="MYIYDDKTRLNAFLDMPKDGADKCPLCIIIHGFTGYATETHIIGV
FT                   QEAMNEIGIATLRVEMYGHGESDGEFRNHTLHKWISNAMAAFDYVKTLDFVTEVYLCGH
FT                   SQGGLLSMLVAPMEKEVVKLLIPLSPAICIPKGARAGNLLGIPFDPDHVPDELAWDDRT
FT                   LDGKYIRVAQTIDTDRAIAGYKGKVLIVHADTDEVVPYEDSVDAAAKYADCELVTIKND
FT                   DHCYNAHLDQVKEAVKEFLLRQRG"
FT                   /product="Alpha/beta superfamily hydrolase"
FT   CDS             4321..5163
FT                   /db_xref="SEED:fig|6666666.127232.peg.5"
FT                   /translation="METIKISDLNTEELSIFGETSEIQLYRYYEPEPGIFIAESPNVVM
FT                   RAIEAGHVPFRMLVEDKLYDKEIPPIVNALESLHGPEATKNLTVYVGDHETLKDLTGYA
FT                   LVKGLWCAFRRSPVMGLRGFCEGKRRLAVLFDVVNPTNVGAIVRSAAAMGVDGVVLTHG
FT                   SVNPLYRRASRVSMGTVFQIPWTMATKDESEGVNLIGLLHEFGFETAAMALTDKSVGIN
FT                   DTRVKNAEKLAIVLGTEGDGLPTEVIEACTHTIKIPMYHGVDSLNVAAASAVAFWELV"
FT                   /product="putative rRNA methylase"
FT   CDS             5294..5746
FT                   /db_xref="SEED:fig|6666666.127232.peg.6"
FT                   /translation="MRPSRNEKENAMREKLIAELYENQFNKFVGFEVLELGPTYSKAKV
FT                   KCSEKIQNPYGSIHGGALLALVDTMAGATGCMNGYYVTTVSANLNFLLPAVNTEYIYCE
FT                   CMKLKTGKHILVFDIRITDDEGNLLDSGEFSFFASKTPLLDNPKIY"
FT                   /product="Phenylacetic acid degradation protein PaaD,
FT                   thioesterase"
FT   CDS             5773..6462
FT                   /db_xref="SEED:fig|6666666.127232.peg.7"
FT                   /translation="MAILGFYVLSIICFLGVVFQAIFIAVEHKEKYVPAVIFKGLASCC
```

```
FT                   FVIMGFLGYKNAQNGDFAKLVLLGLIFGALGDILLNLRFVFAKKGQLIFLIGIAAFLAG
FT                   HILYLAALIPLSSSLLVSVIAGVVLAAILLAIIFKKFEVKIAFKIFGIFYIGAIAIMTV
FT                   VSIGNLIALQDTNRILYAVGAVAFLISDVVLIFNTFGSETKFSMRITNLAFYYVGQLLI
FT                   AFSLFFI"
FT                   /product="hypothetical protein"
FT   CDS             6592..6882
FT                   /db_xref="SEED:fig|6666666.127232.peg.8"
FT                   /translation="MHIMRRNAMFYVPDGTERAGFYECEDCGTRFLDIRIAPRITCPYC
FT                   GEEPDMEIGPDDEMPSASEAAKLIQMVEGEDVEKYDVLLSLAITGGDYEWI"
FT                   /product="hypothetical protein"
FT   CDS             6896..8857
FT                   /db_xref="SEED:fig|6666666.127232.peg.9"
FT                   /translation="MKKKLLKFTGLFITFIMVMWFAGCGNKTAETSETNSGDNQTAETS
FT                   AGTETETSTEEPAKEPEHIETYALTQGYNGDLIYIIDSEGNKLKTIDRADIAKNVPDNE
FT                   KEQYRFASEETYMRCTLAAEGDGFLFFSDNVPGEGSDQYEHSVFAVKEDDLKIYPIAKI
FT                   PNEHYVISTDYYEGALFVDFSFGYGENNEYLGVGEKKFVYDENTDVFTEKDSEYNGVFE
FT                   SAKVMGVRLNGTTRVMGYSQDSFTRTMHECGFILGSTDNGYAMVDDRGAITPLVNFEGG
FT                   YINMYSKGHMFFINSSYDTGVAEAFIYDVDGKALNQISDDWSLIQSIGRNGNDYYYYTQ
FT                   DEESYGIKINHVFKYNAESGLTTALYEAKSIPGVSIQPGVEDFRLCGDKMYAICYDDGA
FT                   TKWMRIDSSNGVATYTDIDCPVSTVDIFNYGKISSTSIVDTCPNCGTPLVKIYAERFEL
FT                   DSKYSKNADKINETLKDVQDRELSYSTDGDDYYGSDCTEHQEHPTWYTITDDYTVSAVK
FT                   IINDKYLAVDMGGYWYGGGAHGMPMRDQHLFDLETGEEKTLEDFYTMTNEDFKALVAYK
FT                   TKEDFLKYENGESPYFAADADTVFNDAYEYAELDRGTVDFAEDGIYYYYPPYEMGPYAA
FT                   GYIDVFISYEELLGRKDL"
FT                   /product="hypothetical protein"
FT   CDS             complement(9327..9485)
FT                   /db_xref="SEED:fig|6666666.127232.peg.10"
FT                   /translation="MLPLEPGLEISALSELAPQPEIIRADIPRIAIACFLTNIGTHQVL
FT                   LGFYISC"
FT                   /product="hypothetical protein"
FT   CDS             complement(9570..9866)
FT                   /db_xref="SEED:fig|6666666.127232.peg.11"
FT                   /translation="MLGLEFPEGVGLEVGVGVAVPEGVGVGVTVPVGVGVAVPADVGDG
FT                   DGVDVDPPAVEDVPEGVEFDADGVLKLPPVSLPSNSPPLYMAAASLAILSSSL"
FT                   /product="hypothetical protein"
FT   CDS             9880..11007
FT                   /db_xref="SEED:fig|6666666.127232.peg.12"
FT                   /translation="MEVIFYDIGQGDCTLVKCDGHSLLIDCGETSKGSEVRLKLLQKQN
FT                   VENLDFLMLTHPDSDHIGGAASVIDNVKIDAIWMSDFVKTNDVYTRLMDTIRKKDVYYY
FT                   TPKPGNEYYLGDAVIKVIAPNNTYDNPNDSSIGIIIEHGENRFLFAGDATTNAEADIIK
FT                   NGMDISADVYKVNHHGSASSTSQAFLDAVNPTYAVISCGKNNDYGHPTKEVLDKLKAKN
FT                   VSVFRTDEQGTVTVTSDGSNLTWSTEPSTTWAAGTGTGKTYGEQAHAGATATPTPTPTP
FT                   APSPTPTPASTPFIVNNVGDTGNMDRSLPNTSGSSTVKRDLPEPEPGPGAVWIPTDGGK
FT                   KYHKDPTCSGMTNPIQVSVHTAQEYGFTACKKCYD"
FT                   /product="Late competence protein ComEC, DNA transport"
FT   CDS             11033..11581
FT                   /db_xref="SEED:fig|6666666.127232.peg.13"
FT                   /translation="MQQKTITKIKSFFAFALIIALLASFAGCDSSELRTSVVQKNTSNI
FT                   TREVVETPTPKPTVVETPTPKPTTIETPTPEPTPIETPTPEPTPIETPTPEPTAVETPT
FT                   PEPTAIETPTPEPTIVEAAPVQEESPAQEYILNKNTKKFHYPNCKSVGQMKEKNKIYSN
FT                   ESRDTIIGKGYVPCKNCNP"
FT                   /product="hypothetical protein"
FT   CDS             11782..12519
FT                   /db_xref="SEED:fig|6666666.127232.peg.14"
```

```
FT                   /translation="MRSVRKRVFEIIEVSRDNDRASLAYDIAMLVCIVVSLTPLAFKNP
FT                   PKFFKTTDYVTTGVFIVDYVLRLATADYKLNKKGVISFIKYPFTLWAIVDLLSILPGLD
FT                   VLSSGFKLFRVFRVIRTFRIFRVFKAFRYSKNIEKIARVIKNSKDALIAVCTLAIGYIL
FT                   VSALIIFNVEGDSFSSFFDAIYWATVSLTTVGYGDIYPVTTAGRIITMISSIFGIAIVA
FT                   LPAGIITAGYMDVINEEKNNKEE"
FT                   /product="Potassium voltage-gated channel subfamily KQT;
FT                   possible potassium channel, VIC family"
FT   CDS             12628..12843
FT                   /db_xref="SEED:fig|6666666.127232.peg.15"
FT                   /translation="MDVAVRIHWDDEAEVWVAINDELGIALESESYDALVERLKNAIPE
FT                   MAELNGLSNIKSIIITTDTRQMTYGR"
FT                   /product="hypothetical protein"
FT   CDS             12951..13853
FT                   /db_xref="SEED:fig|6666666.127232.peg.16"
FT                   /db_xref="GO:0008677"
FT                   /translation="MRKYLVIGAGGTGGPTGAYLAKAGKDVTFIVRGENLRVMKEKGLK
FT                   VLRPEDEFTVTNFKACEFKDYNDTPDVIFVCVKAYSLDAILPDIKRVVGTNTIIIPILN
FT                   GFGIGEKIQKSIPGSLVTDGCTYVAAELKEPGVIQMNGGIFRVIFGVRDKADYRDELKE
FT                   IEKELNEADIIGIYSENIVRDVLVKFSYVSPAGACGLYLNVAAGDMQKPGESRDLFAGL
FT                   IAEVSEIGNAMGIEFEEDLVERNLKIMDSLIPTATTSMQRDVMAGKSSEMDSLVYEVVR
FT                   LADRYGVEAPLYRKVAAAL"
FT                   /product="2-dehydropantoate 2-reductase (EC 1.1.1.169)"
FT                   /EC_number="1.1.1.169"
FT   CDS             13862..14749
FT                   /db_xref="SEED:fig|6666666.127232.peg.17"
FT                   /db_xref="GO:0016020"
FT                   /translation="MRNLKVTNNHIKGILCILAAALGFSFMTFFVRISGDLPTMQKAFF
FT                   RNAVAAVVASVALLRSPEKFKIKKTSYGDIAFRCIFGTSGLICNFYAIDRLGIADANML
FT                   NKLSPFFAILLSIPLLKEVPKVRDIVATIIAFIGALFIIRPGGDMSVLPALIGLYGGFG
FT                   AGTAYVFVRRLGGKGERTPIIVFCFSMFSCLVTAPFLIFNFVPMTGWQWFCLVMAGVSA
FT                   SLGQFSITTAYKFAPAKEISVFDYTQVIFAAILGMIFLSETPSYLSVIGYVIIIGTAVV
FT                   RWYLTLKREREVQA"
FT                   /product="Permease of the drug/metabolite transporter
FT                   (DMT) superfamily"
FT   CDS             14868..15080
FT                   /db_xref="SEED:fig|6666666.127232.peg.18"
FT                   /translation="MKVFDANTFGEVIKRQRKKMGYTQKYICEVSGISASYISDLENGK
FT                   ATIELGKAIQLANLLGIDVELTERG"
FT                   /product="hypothetical protein"
XX
SQ   Sequence 15380 BP; 4376 A; 3325 C; 3716 G; 3963 T; 0 other;
     tactcttata atgacgatgg ccgaatgaaa gaggttgtgc cctttattga gggtgtcaaa        60
     gaaattacat tgattgatgc ggcggataac agggtggatt taaagattga cgggatgctc       120
     tatgcaaaag atcttcccga aaatgccttc aaaagcctgg ccggggatcc ggtgagggtt       180
     attgtgcctg agggtgatgc cagatatgtt acctggtttt gtgaccccgc agatgatgag       240
     aaatttacgg tttttgccag aagtgtgtta gatgagaagt atccgctttt gacagaggat       300
     acatattatg acctggggta caatgtgcaa atcagccgcg ccaaccagat gattaaggtt       360
     atgaatatcg caattatcct tggggagatt attctgacgg ggcttattat tctcattctt       420
     cttatgggat ttgccagcgt tatcagcact cttagctcca atatcagagt caggaggcgg       480
     gagtttgcgg tgcttaagag cgtcggaatg accagaggtg ccctggagaa gatgctttac       540
     ggcgagagtc tttgctgcat tctcaaggct gtgggaaagg gaacggtttt tgggatcctg       600
     attccctggg caattaacct ctttatcagg agggtcttcg cagtccgata tgaactgcct       660
     gtggcttacc tcgccgtggg aattgttgtc gtagctttga ttgtggtgtt catcacccgg       720
     gtggaaatcg gcaagcttgg gaagcagaac atcattgaag acatcaggat ggagtaggtg       780
```

caggatgggg acaggcccaa attgcacatt ttccgcagaa tgtcatcagc gggatgatat 840
tttacacggt agacacgttc gcacttagat ctcgcacgta gcggtacata aggtgctaaa 900
ggacagcacc taagtaccga cggctcgata aagtctacct tagtaaaaca tcatcccgct 960
gattctgttc tcgaatattt ggtgcaatta gggcctgtcc ccatcctgca cggtgcaaaa 1020
tagggcctgt ccccaacctg cacatcaggc tcaaaattta gtcttcatag tggaattctt 1080
tgatttcctc tgagccgtac tcaacatact tcacgacatt gttgtcggtg tccatctcaa 1140
gaaaatatcc gttccacagc tcactgtagc gatcatcttt ggcgtaaata tcaaagtagg 1200
agcggcttgt ttcgccctgg ccgcacatct gataggtgcc ggggtcaaag tttccataga 1260
atgaacgcat gtcatcagtc ataaggaaat actcaagggg tgtgtgatcg tcacacactt 1320
ccgggtcatg aagtgcaaag gtggtgtcaa cataataatc agtgccatta attctgacga 1380
aatcccactg gtggccggga gtggtagcat catacatgtc tccgcacact acatcggcct 1440
ggacgccgca ttgcatcaga agatatgagt ataatcccga aaactcctgg cagattccct 1500
tgccgagggc gagggggctg taggtcagaa catcgtctgt atcgggaccg agacggtctg 1560
ctttatcata gtcatatgtg taattttgtg agatgtatac atataaagcc aaggcttttt 1620
caaaatcgct gtaatcgggc ttaacatttt catttaaaat ggttacaatc tgcgcctcaa 1680
aagctttttc cctttcgaga aattcttcct tgggaatgag atattcgatt tttcccacac 1740
catcagaaaa accaccatcc tctacttcca tgacacagtt cgaggcaacc gggaagtagt 1800
agttgataag tcgtccacca atggcattca tgtaggcctg ctcattggca cattcgaagg 1860
tatcctcgcc ctttcttaag gcatcacaga gattgaaaag agcgtctctg ccgttggtgc 1920
cccaggctgc tgtatcatta tcggaaaaaa catgtgcgtt aaaaacatac ttgccgtcaa 1980
caacattttc accgatggca taggctgcat cgtcagtggt ttcggcttct tttgccgggt 2040
cgcttacgaa ggatgccaca ggctcctcgg aatcggtggc ggtaaaatcg gtggctacag 2100
tttcttcaga tgtaagctcg tccaccacgt cggaagaccc ttccgctccc ttgccgcaag 2160
cagtaagtgt aagtgccgct ataagcacta tgggtaaata tctttttttc atttttttct 2220
ctcctcatta atcttcatag tgaaactcac gtgtcttacc tgtttcaaga tcaagcagtt 2280
tcacaatatt cttttccgtg tccatgccaa gaaaatgggt tcttctgaag gccgcatacg 2340
tatcatccgt ggcggaaacc acgtaatcat caatctttaa taccgcattg aaatatctca 2400
aaagctccac cttgttattc tcgatggtgc agccgtcgat ttccctgtct tttcagtaa 2460
ttaaaaagta atgaagcggg gtgtgatccg tgttgtattc cgaccataaa gaccaggtgg 2520
gatcgataaa atagcccttt ccgttgagcc ttatgtagtc aaaagaatgg aagatatcga 2580
aacttccggc tgaaatggca tcaacgccgc actgcatcaa aagataggca taggctatgg 2640
ccagttcgga gcacacaccg cttttataca caaaggtgct gtaactggct ccgtcatcct 2700
cgtggtcgaa gatgtcctca tcgtatgtat attctctggc aaaataatca tgaatcgcca 2760
gagtcttctc aaaatcactg taatccgtac ggacattatc attaagaatc tcagtgatca 2820
tgtcctcaaa atccttttcc ctctgaagga attcctcttt tggcatcttg tagtaaatcc 2880
tgccgacacc attctcaaaa ggtacggttc cgtcgtcgct tttgccgcat actttacagc 2940
aggcagcggg gaaaagtgca cacaaaatta cttcctcggt acaccagtcg taaacctcct 3000
ggcttgcaca ttcgaaggta tcctcgcctg cacgaagtgc atcgcaaaga ttataaaagc 3060
tctcccagta ctcaggcccg aatctttcgg ccaaaatcgg cgagtacacg tgggggttaa 3120
actcaaactt cgcaggctcc ggttccggct cctcagtagg ttccggggta atctcgggag 3180
tcggctcagg agtagttgcc tccactacct cagtggtatc caaagcccct tcagcttctt 3240
tgccacaggc cgtcagcgcc aacgataatg ccagcaatac aggtatcaat cgattcctca 3300
tatttattct cctctcgaag atgtgcaatt tggggcctgt ccccaacctg cacggtgcag 3360
attgggcctg tccccaaatt gcacggtgca aaacggggac aggccccaaa ttgcacatag 3420
gctcaaaact gcaccttcgc attgactata tcaagtggat tgaaagaaaa aggggcaaat 3480
ttgattaaga tttgtttaat tcatgctata attcactagg aattattttt cgaggaaggt 3540
atagcacaat gtatatttac gacgacaaaa caagactcaa cgcatttctt gacatgccaa 3600
aggacggggc tgataagtgc ccgctttgca ttataattca cggcttcacg ggttacgcta 3660
cagagactca cattattggg gttcaggaag caatgaacga aatcggcatc gctaccctta 3720
gggttgaaat gtacgggcac ggcgaaagtg acggggagtt tagaaaccac acccttcaca 3780
agtggatttc aaacgccatg gcggcttttg attacgtaaa gaccttggat ttcgttacgg 3840
aggtttatct ttgcggccac tcccagggcg gactcctttc catgcttgtt gcccctatgg 3900
aaaaagaggt ggtaaagctt ctcattccct tgtctcctgc catctgtatc cccaagggcg 3960
cccgagcagg aaatcttctt gggattccct ttgacccgga tcatgtgccc gacgaactgg 4020

cctgggatga cagaactttg gacggaaagt acataagagt tgcacagacc atcgatacag 4080
acagggctat tgccggatat aagggcaagg ttttgattgt tcatgcggac acagacgagg 4140
ttgttcccta tgaggattca gtggacgcag cggcaaagta tgccgattgc gagctggtga 4200
caatcaaaaa tgatgatcac tgctacaatg ctcaccttga tcaggtgaaa gaagcggtta 4260
aagaattttt attaagacag cgcgggtaag aaaacagaca gagaataaga agcagtattt 4320
atggaaacaa taaaaatcag tgatttaaat acagaagaac tttcgatttt tggcgaaaca 4380
tcggagattc agctctacag atactatgag ccggagcccg gaattttcat tgccgagagc 4440
cccaatgtgg tgatgcgcgc cattgaggcg gggcatgtgc cctttcgaat gttagtggaa 4500
gataagcttt atgacaagga gattccgccc atagtaaacg cacttgaaag tcttcacggg 4560
cccgaggcca cgaagaatct tactgtttac gtgggagacc acgaaaccct aaaggacctt 4620
acgggctatg ctttggtaaa aggcctctgg tgcgccttta gacgctcacc tgttatggga 4680
ttacgcgggt tttgcgaagg aaaaaggcgt ctggccgtat tattcgatgt ggtcaacccc 4740
acaaatgtag gcgccatagt ccgttccgca gccgcaatgg gtgtggacgg agtggttctt 4800
acccacggga gtgtgaatcc tttatacaga agagcttcaa gagtgagtat gggtacggtt 4860
tttcagattc cctggaccat ggccacaaag gatgaaagtg aaggggtgaa tttgattgga 4920
cttcttcatg aatttggctt tgaaacagcc gccatggcat tgactgacaa gtcagtcggc 4980
ataaatgaca cacgtgtcaa gaatgccgaa aagctggcaa tagtgctggg tacagagggc 5040
gacggcttgc ccacagaggt aatcgaagcc tgcacccaca ccataaagat tccaatgtac 5100
cacggagtgg attccttaaa tgttgccgcc gccagcgcag tggcattctg ggaacttgta 5160
tagaaactgt tgggccgatg tgcagaatgg ggatgtgcag aatggggaca ggcccaaatt 5220
gcaccgtgca aaacggggac aggccccaaa ttgcacatcg gcccaaaccg caaatacatg 5280
gtataataga cgcatgagac catcaagaaa tgaaaaagaa aacgcaatga gggaaaagct 5340
tattgcggaa ctttacgaaa atcaatttaa taaattcgtc ggatttgagg ttttggagct 5400
tggccccacg tacagtaagg ccaaggtgaa atgctcggag aaaatccaaa atccctacgg 5460
aagtatacac gggggagctt tgcttgccct ggttgatacc atggcgggag caacgggttg 5520
catgaacggc tactatgtta ccaccgtttc tgccaacctt aatttccttt tgcccgcggt 5580
caataccgaa tacatctact gcgagtgcat gaagctcaag accggcaagc atattctggt 5640
attcgatatc agaataacag acgatgaagg aaatcttctc gacagcggag agttctcgtt 5700
ttttgccagc aaaactccgc ttcttgacaa tccaaagata tactagcaaa aaatgctata 5760
tgggagggta aaatggctat actcggattt tatgtgttat caatcatttg ttttctggga 5820
gttgtctttc aggctatatt cattgcggtg gagcacaaag aaaaatatgt tccggcagtg 5880
attttcaagg gcctggcatc ctgttgcttc gtgattatgg gctttctggg gtataaaaat 5940
gcccagaatg gagactttgc caaactggta ctccttggcc tgatattcgg agctttggga 6000
gatatacttc tgaatctgag atttgtattc gctaagaagg ggcagctgat tttctaata 6060
ggaatcgcag cattcctggc aggacatatc ctttatcttg cagcgctgat tcccctgtca 6120
tcgagcctgc ttgtttcggt tatcgcaggc gtggtgctgg cagcgattct tctcgccatc 6180
atcttcaaga agtttgaggt taagattgct tttaaaatct tcggtatctt ctatatcggt 6240
gccattgcca taatgacggt ggtttccatc ggaaacctca tcgcccttca ggataccaac 6300
agaatcctct atgcggtggg cgcagtggca ttcctgattt cagacgtggt gctgattttc 6360
aataccttcg gatccgaaac caagttttcc atgagaatta caaacctggc gttctactat 6420
gtgggccagc tgctgattgc attttcgtta tttttcatat aaagcaaaat aatgtgcagt 6480
ttgggggcctg tccccaaatt gcaccgtgca gaatggggac aggcccaaat tgcaccgtgc 6540
agaatgggga caggcccaaa ttgcaccgtg cagaatgggg acagacccaa attgcacata 6600
atgaggagaa atgccatgtt ttacgtaccg gatggcactg agcgtgccgg attttatgag 6660
tgtgaggatt gcggaacccg ctttctggat ataagaattg ccccgaggat tacctgcccc 6720
tattgcgggg aggagcctga tatggagatt gggcctgatg atgagatgcc atctgcatcc 6780
gaggcggcta aactcatcca gatggttgaa ggagaagatg tggaaaaata cgatgttctt 6840
ctaagtctcg ccatcaccgg cggagactac gagtggatat agggggaaag gcactatgaa 6900
gaagaaatta ttaaagttca ccggattatt tatcacgttt atcatggtga tgtggtttgc 6960
gggatgcggc aataaaaccg ccgagacctc agaaaccaac tccggggaca accaaactgc 7020
ggaaacttcc gccgggactg agaccgaaac gtccacagag gaaccggcaa aagagcccga 7080
acacattgaa acctacgccc ttacccaggg ctataacggg gacctcatat acatcattga 7140
ctccgaggga aacaagctaa aaaccatcga cagagcggat atagcaaaga atgttccgga 7200
taatgagaag gagcagtatc gttttgcctc agaagaaacc tacatgagat gcactctggc 7260

tgccgaaggc gatggattcc ttttcttcag cgacaacgtt cccggtgagg gcagcgacca 7320
atatgaacat tctgtttttg cggtaaaaga agatgactta aaaatctacc ctatcgcaaa 7380
gattcccaat gagcactatg ttatttcaac agattactac gaaggtgccc tgtttgtgga 7440
cttcagtttt ggctacggag agaacaatga gtatcttgga gtaggcgaga agaagtttgt 7500
ttatgatgag aacactgacg tttttaccga aaaggattcc gagtataacg gagttttcga 7560
gtcggcgaaa gttatgggtg tcagattaaa cggaaccacc agagttatgg gatacagcca 7620
ggactccttt acccgtacaa tgcatgagtg tggcttcatc ctcggaagca ccgacaacgg 7680
ctatgcgatg gtggatgaca ggggcgcgat taccccttg gtgaatttcg aaggcggcta 7740
catcaacatg tacagcaagg gccacatgtt tttcataaat tccagttacg acacaggagt 7800
tgccgaggct tttatctatg atgtggacgg aaaggccctg aaccagattt ccgacgactg 7860
gagcctgatt cagtccatcg gccgaaacgg aaacgattat tattactaca ctcaggacga 7920
ggaaagttac ggaataaaga taaaccacgt gtttaagtac aatgcggagt cagggctcac 7980
cacggccctt tacgaggcaa agagcatccc tggggtcagc attcagccgg gcgttgagga 8040
cttccggctt tgcggcgata agatgtacgc catttgctac gacgacggcg ccaccaagtg 8100
gatgcgaatc gacagcagca acggtgtggc gacctacacc gacatcgact gtcccgtttc 8160
caccgtcgat atctttaatt atggtaaaat cagctcaacc tccatagtgg acacctgccc 8220
caattgcggc accccttgg taaaaatata tgcggagcgc tttgaacttg acagcaagta 8280
ctcaaagaac gcagacaaga taaatgagac ccttaaagat gttcaggaca gggagctttc 8340
ttatagcacc gacggcgatg actactatgg atccgattgt acggaacatc aggagcatcc 8400
cacctggtac accattaccg acgactacac cgtgtcagcc gtaaaaatca tcaatgataa 8460
gtaccttgcc gtggacatgg gcggttactg gtatggaggc ggcgctcacg gtatgcccat 8520
gcgtgatcag cacctgtttg accttgaaac cggtgaggaa aaaacgctcg aagatttcta 8580
caccatgact aacgaggact ttaaagccct tgtggcatac aagaccaaag aggacttcct 8640
caaatatgaa aatggcgaat cgccttattt tgcagctgat gcagatactg tattcaatga 8700
cgcatatgaa tatgcagagc ttgacagagg taccgtagat tttgccgagg atggcatata 8760
ttattactat cctccctacg agatgggccc ctatgccgca ggatatatcg atgttttcat 8820
ttcgtatgag gagctgctgg gaaggaaaga tttgtaaatt tgtaaatgat ttgaggattc 8880
ttgcgggatt tgtgggattc ttgcgggggt ctcgagcatt cccagcttcg gtggccgcga 8940
aatactactt aaatgctgaa acggcggatt tgttagtacg gtttttgctg atgggtacta 9000
ctgaaatttc aaaaatgctg aattcattag tacggtttct gcggcgggta ctactgaaat 9060
atcggaaacg gcagattcat tagtacagtt tcgccgtgct gtactactga aatgtagaaa 9120
tgttgaattc attagtacaa ccctgacggc gcgtactact gaaattataa ggatgttgaa 9180
ttcattagta taattttcaa cgtgctggac tattgaaatg tagaattatt gaattcatta 9240
gtacaacctc gatggcgcgt actactgaaa tgcagaagtg taaaattcta tagtatgact 9300
tcgctgaaaa ctgcaggacg tagtgattag caagaaatgt aaaaccccaa aaggacttgg 9360
tgagttccta tgtttgtgag aaaacaggca atagcaattc tgggaatatc tgctctaatt 9420
atctcgggct gcggtgcgag ctctgaaagt gcacttattt ccaatcccgg ttctaacggc 9480
agcattggcg gacagtatat cagtgagcaa aacagagaag agaatcttgc tgtaagcaat 9540
gagcctcagg atgtaattcc tgaggcttat tatagtgatg atgaaagaat tgccaagctg 9600
gcagctgcca tgtataacgg cggagaattt gagggaagcg agaccggagg aagcttcagg 9660
acaccatccg catcaaactc cacgccttcg ggaacatcct ccactgcagg aggatcgaca 9720
tccactccat ctccatcccc cacatcggct ggcactgcca cacccacccc gacaggtacg 9780
gttacaccaa caccgactcc ctctggtacg gctaccccga caccaacttc caggccaaca 9840
ccctcaggga actctaaacc caacacagac attgccccca tggaagtgat attttacgac 9900
atcggacagg gagattgtac tttggtaaaa tgcgacggtc actccttact tattgactgt 9960
ggcgaaacca gcaaaggttc agaggttcga ctgaaattat tacagaagca aaatgttgag 10020
aacctggatt tcctgatgct tacccaccct gattcggatc atatcggcgg tgcagcttct 10080
gttatcgata acgtgaaaat agatgccatc tggatgagtg acttcgttaa gactaacgat 10140
gtttacacca gattgatgga cacaatcagg aaaaaagacg tgtactacta tactcctaag 10200
cccgggaatg agtattatct tggggatgcc gtgattaagg taattgctcc aaataacaca 10260
tacgataatc ccaatgattc ttctatagga atcataattg aacacggaga aaaccgcttc 10320
ctcttcgcag gggatgcgac caccaacgcc gaagcagaca taataaaaaa tggaatggat 10380
atctctgcag atgtctacaa agttaatcat cacggaagcg cttcatccac ttcccaggct 10440
ttcctggacg cggttaatcc tacatatgca gtaatcagct gtggaaagaa caatgattac 10500

ggtcatccta ccaaggaagt gcttgataag cttaaggcaa agaacgtaag tgttttcagg 10560
acagatgagc aaggtacagt tacggtcact tccgatggca gtaaccttac ctggagcact 10620
gaaccttcca ccacctgggc agcaggtacg ggaaccggaa aaacatatgg tgagcaggca 10680
catgcagggg caactgcaac gcctacaccc acgcctacac ctgcgccttc tcccacacct 10740
actcctgctt ctacaccttt catagtgaat aatgtaggag ataccggcaa tatggacagg 10800
tcattgccaa atacaagcgg ctcgagcaca gtaaaacggg acttgcctga accagagcca 10860
ggacctggag cagtatggat acccaccgat ggaggaaaga aatatcataa ggacccaacc 10920
tgcagtggaa tgacgaatcc aatacaagtt tcggtacaca ctgcgcaaga atatggcttc 10980
actgcctgca agaaatgcta cgactgattg ctattaaaag ggaggatggg aaatgcaaca 11040
aaagactatt acaaaaatca aatctttctt cgcatttgca ttgataattg ctttattagc 11100
aagctttgcc ggatgtgatt caagtgaact tagaacttcg gtagttcaaa aaaatacaag 11160
caacattacc agagaagttg ttgagacacc aactccaaag ccaacagtag ttgagacacc 11220
gactccgaag cctacaacca tagagacgcc caccccggag ccaacccta tagaaacgcc 11280
aactccggag ccaacaccca tagagacgcc cacgccagaa ccgacagcgg tagagacacc 11340
tactccggaa ccaacggcta ttgagacacc gaccccggaa ccaacaatag tagaggcggc 11400
gccagttcag gaagaatccc ctgctcaaga atatattttg aataaaaaca ctaagaaatt 11460
tcattaccca aattgcaaaa gcgtggggca gatgaaggaa aagaacaaga tttattccaa 11520
tgagtcaaga gatacaataa ttggtaaggg atatgttcca tgtaagaact gcaatccata 11580
gatgatgcgg aacaagcgga ggaatacgta tggcaaaaaa gaaattcaag attccgggac 11640
tttcatttag ttggaaaaga gcaacgggac ttactaaaga aaaaagaaag gtctcaaaag 11700
ccacaggtgt gcccacctca aagcagggaa gacgaaacaa aatagcgaag tggctgggaa 11760
ttaaatagtt agggagatag tatgaggtcc gtgagaaaaa gagtttttga gattattgag 11820
gtatccagag ataacgaccg agcaagcttg gcttatgata ttgctatgct cgtttgtata 11880
gttgtgagcc tgactccatt agcattcaaa aacccaccta agtttttaa aactacagat 11940
tatgttacaa caggggtatt tattgttgat tatgttttac ggcttgcaac tgctgattat 12000
aagttgaaca aaaagggagt tatatcattt atcaagtatc cctttacatt atgggcaatt 12060
gtggacttgc tttcaatttt gccgggactg gatgttctta gcagtggatt caaactgttt 12120
agggtattta gagtaataag gacatttaga atttttagag tctttaaagc ttttagatat 12180
tccaaaaaca ttgagaagat tgcaagggta atcaaaaact cgaaagatgc gctgattgcc 12240
gtatgtacct tggctatagg ttatattctt gtttctgcat taataatatt taacgttgaa 12300
ggagattcgt tttcttcatt ctttgatgcc atttattggg cgaccgtttc gctcacaaca 12360
gttggatatg gtgatattta cccagttact acagcaggaa gaataatcac tatgatttcc 12420
tcgatttttg gtatagctat agtagcatta cctgcgggca ttattactgc gggatacatg 12480
gatgtaataa atgaagaaaa aaacaataaa gaagaataga aaagtatgat ataattactt 12540
tacttattct tcgttatttc atgctaagta ccgtggctat tgtagtacgc aatgttgcgg 12600
gaaaggagaa agtctatgag gtgttttatg gacgtagcag ttagaatcca ttgggatgat 12660
gaagctgaag tttgggtggc tataaatgat gaattgggta tagcacttga atctgaatct 12720
tatgacgctc ttgtagaaag actgaaaaat gccattcctg aaatggctga acttaatgga 12780
ttatcgaaca ttaagtcaat aattattaca accgatacta gacagatgac atacggcaga 12840
taaagcaacc cgctttacct gccgttcctt ttttcatata attgatatca gattgaatta 12900
acttgaacaa atacacacac aacacgacat taagtgaagg gagtaaacag atgaggaaat 12960
acctggtaat cggagcgggc ggaacaggag gccccacagg agcatatctg gcgaaggccg 13020
gcaaggacgt gacttttatc gtgcgaggcg agaatttaag ggttatgaag gaaaaaggac 13080
ttaaagttct ccgccccgag gacgaattta cagtaacgaa ttttaaggcc tgcgagttta 13140
aggactacaa cgacacgccg gacgtgattt ttgtatgcgt aaaggcctat tcattggacg 13200
caattcttcc ggatattaaa agggttgtgg gcacgaatac aattatcatc cccattctca 13260
acggctttgg tatcggagaa aagattcaaa agagcatccc cggttccctg gttaccgacg 13320
gctgcaccta cgtggcagca gagcttaagg agcccggcgt tatccagatg aacggcggta 13380
tcttcagggt tatctttgga gtgagagata aggccgacta tcgcgatgaa ctcaaggaaa 13440
tcgagaagga gcttaacgaa gccgatatca tcggaattta ctccgagaac atagttcgag 13500
atgttttggt aaaattctcg tatgtctctc ctgcgggtgc ctgcggcttg taccttaatg 13560
tcgctgcagg agatatgcaa aagcccgggg aatcaagaga ccttttgcg ggactgatag 13620
ctgaagtaag tgaaatcgga aatgccatgg gaatcgagtt tgaagaggat cttgtagaga 13680
gaaatttaaa gattatggac agtcttattc ccactgccac aacttccatg cagcgtgacg 13740

```
     ttatggccgg aaagagctct gagatggata gcctggttta cgaggttgtg agacttgcag     13800
     acaggtatgg ggttgaagca cccttataca gaaaggtcgc tgcggcgtta tagaatcgga     13860
     aatgcgaaat ttaaaggtaa caaacaacca cattaaggga atcctgtgca tcctcgcggc     13920
     cgcactggga ttctctttta tgactttttt tgtaaggatt tcaggagatc tgcccacaat     13980
     gcaaaaggcg tttttcagaa atgccgtggc agcagtggtg gcatcggtcg ctcttctcag     14040
     gtcccccgaa aagtttaaga ttaagaagac gagttacgga gatattgcat tcaggtgcat     14100
     ctttggcacc agcggtttga tctgtaactt ctacgccata gacaggctgg gaatagctga     14160
     tgcgaacatg cttaataagc tgtccccctt ctttgcaatt ttgctttcga ttcctcttct     14220
     taaggaggtc cccaaagtaa gggatattgt ggcaacaatc atagctttta taggcgcgct     14280
     ctttattatc cgtcccggcg gagatatgtc ggttttgcct gccctcattg gcctttacgg     14340
     aggttttgga gcaggaacag cctatgtatt tgtgagacgc cttggaggca aaggagagag     14400
     aacgcctata atagttttct gcttctccat gttttcctgc ctcgtcaccg cgcctttctt     14460
     gatatttaac tttgtgccaa tgaccggctg gcagtggttc tgccttgtaa tggcgggagt     14520
     atccgcatcc cttggccagt tctcaattac cacggcatat aaatttgccc ctgccaagga     14580
     gatttccgtt ttcgattaca cccaggttat ttttgctgcc atattgggaa tgattttttt     14640
     aagcgagact ccatcttatt taagtgtcat tggatatgtc attatcatcg gaaccgccgt     14700
     tgtaaggtgg tatttgacgt taaagaggga gcgggaagta caagcttaga atagtattga     14760
     catagccgta cgactatatt tataatagaa acataggaat atagtcgaac ggctatattt     14820
     atgccctcaa accattcata tagccgtacg gctatgaaag gagaaccatg aaggtttttg     14880
     acgcaaatac atttggtgaa gtcataaaaa ggcagcgaaa gaaaatgggc tatacccaga     14940
     aatacatatg tgaggtcagc ggaatcagcg caagctacat ttctgatttg gagaatggca     15000
     aggcaaccat tgagttaggc aaagccattc aactggccaa ccttttgggc atcgatgttg     15060
     aactgacgga gagagggtag tctatgagaa atcttggagt atatctggag ataaacggca     15120
     gcttttgtca tgtgggaaat atcgtaggca attcatctgc tgatgcctgc tttcagtatt     15180
     caaaggcata tttggaatca tcggacagcc atcccatatc aatatcactt cctttgtcag     15240
     gtgagatatt tacgcctgac agaaccaaaa ccttttttga gggccttctt ccggaaggct     15300
     attcaagaag ggcagtgtcg gattggatca agacagatga gaatgattat ctctccatac     15360
     tggcgatgct tggaagagag                                                 15380
//
ID   XXX; XXX; linear; XXX; XXX; XXX;  XXX.
XX
AC   PRJEB67889
XX
DE   Uncultured bacterium DNA fragment from in-reactor enriched cow rumen, clone
ferm-12e13, contig-1
XX
FH   Key             Location/Qualifiers
FH
FT   source          1..13214
FT                   /organism="uncultured bacterium"
FT                   /environmental_sample
FT                   /mol_type="genomic DNA"
FT                   /isolation_source="Cow rumen"
FT                   /clone="ferm-12e13, contig 1"
FT                   /note="AZO-galacto-mannan, AZCL-debranched arabinan active
clone"
FT   CDS             15..929
FT                   /db_xref="SEED:fig|6666666.127233.peg.1"
FT                   /db_xref="GO:0046556"
FT                   /translation="MYRRYSTYCRNYDGNSLYKIASGASDYDYNWTEVLMKNVGHRMDG
FT                   LSLHYYTVTGWSGSKGAATKFNKDDYYWTMGKCLEIEDVVKKHIAIMDQYDPKKHIGLL
FT                   VDEWGTWWDEEPGTTRGHLYQQNTLRDAFVASLTLDIFHKYTDRIKMANIAQIVNVLQS
FT                   MILTKGKEMILTPTYHVFNMYKVHQGATYLPLNLISDKINVRDNRTVPFISATASKDKN
FT                   GIIHISLSNVDADEYQEVSIELPGVKVNNISGQILTSNNLTDYNSFENPNVIKPQDFKD
```

```
FT                   AKITKSLLKVKLPAKSIVTLTLK"
FT                   /product="GH51"
FT   CDS             999..3011
FT                   /db_xref="SEED:fig|6666666.127233.peg.2"
FT                   /db_xref="GO:0004802"
FT                   /translation="MNDNKLMNRAADNIRILAASMVEKANSGHPGGAMGGADFVNVLFS
FT                   EFLVFDPKNPKWEGRDRFFLDPGHMSPMLYSVLALTGKYNMNELSQFRQWGSPTPGHPE
FT                   VDVMRGVENTSGPLGQGHTFAVGAAIAAKFLKARFGEVMNQTIYAYISDGGVQEEISQG
FT                   SGRIAGHLGLDNLVMFYDSNDIQLSTETDAVTSEDVAKKYEAWNWKVITINGNDPDAIR
FT                   AALKEAKATEKQPTLIIGKTIMGKGARKADNSSFESNCATHGAPLGGDAYKNTIINLGG
FT                   NPENPFVIFPEVAELYAKRAKELESIVAEKYAAKANWAKANPELALKMETFFQGNAPAV
FT                   DWNGIAQKAGSATRAASATVLAALATQVENMIVASADLSNSDKTDGFLKKTHAFTKDDF
FT                   SGAFLQAGVSELTMACISIGMSLHGGVIPACATFFVFSDYMKPAVRMAALMEQPVKFIW
FT                   THDAFRVGEDGPTHEPVEQEAQIRLMEKLKNHKGHNSMLVLRPADAEETTQAWKLAMEN
FT                   TSTPTALILSRQNIANLPAGTDYAQTAKGAYVIAGSDSNPDVILVASGSEVSTLVAGAE
FT                   LLRKDGIKLSIVSAPSEGLFRAQDKEYQETVIPANAKVFGLTAGLPVNLQGLVGANGKV
FT                   FGLESFGFSAPYEVLDEKLGFTAENVYNQVKALLA"
FT                   /product="Transketolase (EC 2.2.1.1)"
FT                   /EC_number="2.2.1.1"
FT   CDS             3113..3544
FT                   /db_xref="SEED:fig|6666666.127233.peg.3"
FT                   /db_xref="GO:0004751"
FT                   /translation="MKTIGICSDHAGYELKQFVKKWLEEQGWPYKDFGTYTAESCDYPD
FT                   YAHALASSVEAGECYPGIAICGSGNGINMTLNKHQGIRAALCWIPEISSLARKHNNANV
FT                   LVMPGRFVSEEEAAKIMTEFFHTEFEGGRHQKRIDKIPL"
FT                   /product="Ribose 5-phosphate isomerase B (EC 5.3.1.6)"
FT                   /EC_number="5.3.1.6"
FT   CDS             complement(3663..5552)
FT                   /db_xref="SEED:fig|6666666.127233.peg.4"
FT                   /db_xref="GO:0009044"
FT                   /translation="MFCTLILACSNNDDPPLSQKPSLENPVVTNLTVSSVDLSAVISGS
FT                   QTDVIKRGFCYSITKLTPTTTDNVKEAVGDKLELTLSALQSATTYYVRAYLLTSEGTIY
FT                   SAVTSFKTLAEYEDPLLAQYQPPVYSDNYTSIAAWSQRDQWNLANVHDPTVFKADDGYY
FT                   YMYQTDASYGNAHSGYGHFHGRRSKDLVNWEYLGATMPENPPSWAKDKLNEYRSEMGLP
FT                   LIDNPRYGYWAPVARKVRTGLYRMYYCLVPGHYIKTGKYDDATNFDNSWTERALIGVME
FT                   CSDPASNQWVDKGYVICSASNKGINEWTRSSTNDWEAYFMWNAIDPSYVITPEGEHWLV
FT                   YGSWHSGFTAVQLNAETGKPLNELGKPWSINGDHSSYGDLIARRGYGRWQASEAPEIIY
FT                   NSKTGYYYLFMAYDELSVAYNTRVARSKNIKGPYLDMVGKNITENGGGDAYPVVTHPYK
FT                   FNNSYGWVGISHCAIFNDGQDNWFYASQGRLPANITGINASNAVMMGHVRSIRWTDEGW
FT                   PLVMPERYGLVPSVAIKEEELVGEWEHINMGYNYQKQFTSVSLKLNSNHISEGALTGSW
FT                   NFDATKQTVTIGGVKLYLQREVDWEISPRKLTIVYAGYSSDGKTTYWGKKIVN"
FT                   /product="GH43_4"
FT                   /EC_number="3.2.1.37"
FT   CDS             complement(5622..7421)
FT                   /db_xref="SEED:fig|6666666.127233.peg.5"
FT                   /translation="MKKILIYITVGLLATVSSSCSDYLDKEVDLTLSEEQIFSKYENTR
FT                   GFLANVYTYLPDAFAGYSDGQFLAASRDCMTDNSLSFWNVHYYHSVLTDAYSATSHPFA
FT                   ERFWPNDLKGIRAANQFMKNARESVIGNAEKTGDDNHLYDRNMAEARLLRAIFHFDLAG
FT                   WFGDIPVIGNDENGVPIVFEAGNASAMNMSRTACSEALKWIADECDAVKDILPFRYSNE
FT                   NENWGRVNGAAAYALKSRALLYRASALNNPEGKTDWWQEAAQAAKDFIAKNNQQAHPYK
FT                   LFTTEDNDPNKNYYQCFVSTPHLNNEYILSRSEWNTREIELFLAPCGFSGNVNSTGRTN
FT                   PTQNLVDAYEMRNGKPIDDAASGYDEQNPYANRDPRLEQTILHHGSIWGDKVQEEERAV
FT                   DVSYPDGKDYQDLHGGTTTGYYTKKFLNNMSFKNPTTYVHACPIFRYAEILLNAAEAIN
FT                   ESEGPTNAYQYVNQVRGRVGMPAYSGMTKEQLRERIRNERRIELCFEDHRFFDERRWKL
```

```
FT                   FEGKTAASEKSLPRYQQVYNIYSVVVTPGASSVYSYEADGTHPSRAFNSPKNYRFPIPD
FT                   SEVKKAPLLKQNPGWELTTSAAE"
FT                   /product="Putative outer membrane protein, probably
FT                   involved in nutrient binding"
FT   CDS             complement(7446..10580)
FT                   /db_xref="SEED:fig|6666666.127233.peg.6"
FT                   /translation="MQTIALAQNEQVLKGRIVDSKGAPIAGAVVNVAEQSRIALSDKDG
FT                   YFILKNVKSQDEICVSSVGYLNATANADFNEGFQIVLAEDLDEYLHTMPVPFGRKAKKL
FT                   MTEATSVVTGQELQKHPVTILQNAFTATVNGMETYEWSSEPGWSETAMYIRGIRTMNQN
FT                   ARAPLIIVDNVERDLSFLDAFPIENITVLKDAASTAIYGMRGANGAILVTTKRGHAGKT
FT                   NIDFTQEIGYQMLSNKMELQNSYNMALTRNQVRYLSGSDPMYTDEQIEKYRRVSNGETL
FT                   EGVDRYRYFDTNWFEQLYRDAAPMYKTNMQISGGNNRARYYVSFSYLRQEGMWNETGTG
FT                   WNKNFDTQHMLNRWNLRSNIDIDVTKFLNVSLDLGGRIDNINQPTQSVFDLTTFGAVEA
FT                   NPMEPVYCPNGELYASSTANNPLRYLAASGQEKNRRRNLYSTLSVNGDLSSLLKGLKVN
FT                   ATVSFDSYETFESTQTNAINTYNYDYTNVDVTDPSEFTYTKYTSYSALTNPSANQREYY
FT                   YHLNFNGGLNYSNKFGKHAVDARAFIRTYQNVVSGSASSERTLSWNGQAMYAYDNKYIF
FT                   SGNISRMASDNFAPEERWGTFYGTSLGWVASEESWLKNKNIDLLKLRASYGRAGQSSTG
FT                   AGRYPYQGTYSSGTGYSFGYSQSNIGGYYESKAGNSNNKWEISDMLNIGVDFDFWGKKL
FT                   YGAVDVFKEWRSGILVTRSTIPSIIGVSVSSDSYGKAESKGLEVTLGHRGKIGKVNYYL
FT                   EGLLTYNTNKITEMDETEPNVEWQRKTGRRIYDGTEVAALYESSFNNTVGGWNIYKFMQ
FT                   WANDPDKVATSQQDAIDHPEKYPYHSASNGKQTLGTAVFKDLNGDRQIDSNDMLPDSYT
FT                   IIPEIIPALNIGLEWNGFDVRAVFNAYLNRSVFLSPAISFSGWSNMGTHEVTKAWGYYT
FT                   DNPADPRNINATYPRPLYGGFNAIDSDRATGTYKNDIWVRNGNYLSLRNVEIGYSLPKE
FT                   LIAKISMTKCRFYFSGYNLYNWSDLPDDVDPEKPMSYCWWYPKTRSFTFGVNIGF"
FT                   /product="TonB-dependent receptor"
FT   CDS             complement(10674..12440)
FT                   /db_xref="SEED:fig|6666666.127233.peg.7"
FT                   /translation="MKKTILYSLLALTISTSVLTSCEDAFGSFLDKQPSNELTEEQVFG
FT                   DWSLMEQFHFDTYNFLRHGACRINNSWLDAATDLAETSYATGGVRSTFNIGNYYGSAGA
FT                   AELTDTWEHYYRGIRKCNMIINRIESVPKSADLSNDKYLQDKTNYTSEARFLRAWFYWE
FT                   LFLRYGPVPIVTEVLDPDGDLLSGYTDRPTVKEYVVDFLLKELSECESGLLTYEDAWNA
FT                   TYAGRIGQPMARALASRIKLYMASPRYSAESGITWQEAADAAKSFMDTYGSKFALFTEE
FT                   TTDGILPYNNALLRTTYLDSNKEIIFFRNDVAVGWSAIQNDTPVGEGGKGGLCPSQNLV
FT                   DMYDMIDGSSPFTTYDATGAPVYNNGQPTINTGSGYSDATMWTGRDKRLEATVLYHGTE
FT                   WGNGTINVIKGQRDNPVGNANATPTGYYVRKYIPEAILGSVHTGTSRRLWTFIRYAEIM
FT                   LNYAEALNEVQGPSTEVYNMLDQIRRRSGITGSVANRSDLTSSKEAMRNFIHKERTVEL
FT                   AFEEHRAWDVRRWSVAVEALSRPIYGIDVAANGAISRKTAQGRVFENKMYLYPIPEGEV
FT                   WKTNIENNPDWK"
FT                   /product="putative outer membrane protein, probably
FT                   involved in nutrient binding"
FT   CDS             complement(12446..13165)
FT                   /db_xref="SEED:fig|6666666.127233.peg.8"
FT                   /translation="MDEPELKTAYRKREGNPISQYFGLIAEGFVTQADLDNPDFPVSTF
FT                   GAVKVGDLKYKDANGDGFIDDRDESRIGYSDIPENTYALSLGVNYKGWGFSVMFQGVDH
FT                   VSRYYDAEAMYAFVSGGKVKEHHLERWNPAQSEAYNLQHANYPLLHYDNYGDHNQRTNS
FT                   FFLKNGSFVRLKNIELSYTLPENWSKVAGMSNCRLYVNANNLITWDHLNNLTDPESNGS
FT                   NRYPIMKTVNFGVNIKF"
FT                   /product="TonB-dependent receptor"
XX
SQ   Sequence 13214 BP; 3951 A; 2990 C; 2506 G; 3767 T; 0 other;
     aatactatgc tgatttgtac cgtcgttatt ctacctactg tcgcaattat gatggcaata        60
     gcctctataa gatagctagt ggggccagcg attacgatta caactggaca gaagtgttaa       120
     tgaaaaacgt aggtcaccgc atggacggtc tctctttaca ctattacact gttaccggat       180
     ggagtggaag caaaggagca gcaaccaagt ttaacaagga tgattactat tggacaatgg       240
```

gtaagtgctt ggaaattgag gatgtagtaa agaagcacat tgctatcatg gatcaatatg 300
accctaaaaa acatatcggc ttactagttg atgaatgggg aacttggtgg gatgaagaac 360
ctggaacaac gcgtgggcat ttatatcagc aaaacacatt gcgagatgca tttgtcgctt 420
cgttaaccct tgatattttt cataaatata cggatcgtat aaagatggct aatatagcgc 480
aaatagtaaa cgtcctacaa tctatgattt tgaccaaagg caaagaaatg atactcactc 540
cgacttacca tgttttcaac atgtataaag tacatcaagg tgcaacttat cttccgctga 600
atctcatttc cgacaaaatc aatgttcgag acaacagaac cgttcctttt atcagcgcta 660
ccgcttcaaa agacaaaaac ggtatcatac acatctcact atctaatgtt gatgcagatg 720
aatatcaaga ggtaagcatt gaattaccag gggttaaggt aaacaatatc tcagggcaga 780
tactaacatc aaataatctt acagactata actcattcga aaatccaaat gttataaaac 840
cgcaagattt taaagatgca aaaataacaa aaagcctatt aaaggttaag ttgcctgcta 900
aatcaattgt tacactgaca ctaaaataaa tagttttttc agaaagataa cttacaataa 960
agtattatct ttgcagacaa acttaatgat tatatattat gaatgataat aagctaatga 1020
atcgcgcggc tgataatatt cgtatcttgg ctgcatcaat ggttgagaaa gcaaactcag 1080
gtcacccggg tggtgctatg ggaggagctg attttgtaaa cgtactcttc tcagagtttc 1140
ttgtattcga tcccaaaaac cctaaatggg aaggtcgtga ccgcttcttt cttgacccag 1200
gccacatgtc tccaatgtta tattctgtgt tagcattgac cggtaaatac aacatgaatg 1260
aactttctca atttcgtcaa tggggaagtc ctactcccgg ccatccggaa gtagatgtaa 1320
tgcgtggtgt tgagaacaca tcaggaccat taggtcaagg acatactttt gcagtaggag 1380
ctgccattgc tgctaaattc ttaaaagctc gttttggtga agttatgaac caaaccattt 1440
acgcttatat atcagacggt ggtgttcaag aagaaatatc gcaaggttca ggacgtattg 1500
caggacactt aggacttgac aatttagtta tgttctacga ttcaaacgat attcagctgt 1560
ctaccgaaac ggatgctgtt accagcgagg atgttgccaa gaagtatgaa gcgtggaatt 1620
ggaaagtcat cacaatcaac ggtaatgacc ccgatgcaat acgcgccgct ttaaaagagg 1680
ccaaagcaac tgaaaaacaa cctacactta tcatcggtaa aaccatcatg ggtaagggtg 1740
ctcgcaaggc agacaattca agctttgaaa gtaactgtgc tactcatgga gctcctctgg 1800
gcggtgacgc atataaaaat acaattatca acttgggcgg aaatcctgaa aatccatttg 1860
tgatattccc tgaagttgcc gagttatatg caaaacgcgc aaaagaatta gaatctattg 1920
tagctgaaaa atacgcagca aaagctaatt gggcaaaagc caatccggaa ctagcattga 1980
aaatggaaac attcttccaa ggcaatgctc ctgctgttga ttggaatggc attgcacaga 2040
aagctggttc ggctacacgt gctgcttctg caactgtttt agctgctctt gctacacaag 2100
ttgagaacat gattgttgcc tctgctgacc tttctaattc agacaaaact gatggcttcc 2160
tcaaaaaaac tcatgcattt actaaggatg atttcagtgg tgcgttttta caagcgggtg 2220
tgtctgaatt aaccatggca tgtatttcaa tcggtatgtc gcttcatggc ggtgtgattc 2280
ctgcatgtgc tactttcttt gtattctcag actacatgaa accggcagta cgtatggccg 2340
cactaatgga gcaaccggtt aaattcattt ggacacatga tgcgttccgt gtaggtgaag 2400
atggtcctac tcacgaacct gttgaacaag aggcacaaat tcgcttaatg gagaaactta 2460
agaatcacaa aggacacaac tccatgttag ttttacgtcc ggctgatgca gaagaaacca 2520
ctcaggcatg gaaattagca atggaaaata cgagcactcc aacagctctt attctttcaa 2580
gacaaaacat tgccaatctt ccggctggaa cagattatgc acaaactgca aaaggcgcat 2640
atgtaattgc cggttctgat tctaaccccg atgttatatt agtagcttcc ggttcagagg 2700
tttcaacttt agttgcaggt gctgaattac ttcgtaaaga cggaatcaaa ttaagcattg 2760
tatcggctcc ttcagaaggc ttattccgtg ctcaggataa agagtatcaa gagactgtta 2820
taccagccaa tgcaaaggtc ttcggattaa ccgctggttt gcctgttaat ctgcaaggat 2880
tagtaggagc aaatgggaaa gtttttggtc tggaatcatt tggtttctca gctccttacg 2940
aagtgttgga tgagaaactt ggcttcactg ctgaaaacgt ttacaaccaa gtaaaagcat 3000
tattagctta aaagtatgat tcatgcttta ggtgatttgc tatcagtaaa atcacctaaa 3060
gtaatgagtc agcaaaacat aaatagtaca aaccgcttct ttatatttta gaatgaaaac 3120
aataggaatc tgttccgatc acgcaggcta cgaattgaaa caattcgtaa agaaatggct 3180
ggaagaacaa ggatggccat acaaagactt tggcacatac actgcagaga gttgtgatta 3240
tcctgattat gcacatgctc ttgcttcatc tgtagaggca ggagaatgtt atccgggtat 3300
tgctatttgc ggaagtggca acggcatcaa catgacttta aacaagcatc agggaattcg 3360
ggcagcactt tgctggattc ctgagatttc gtctctcgcc cgcaaacaca acaacgcaaa 3420
tgtattggtc atgcccggac gatttgtgtc ggaagaagag gctgctaaaa taatgactga 3480

gttctttcac accgaatttg aaggaggacg tcatcaaaag cgaatagata aaattccttt 3540
gtagaaagcg attggaaaaa attgagtata taaaaagggc tgtaacttca ggttgcagcc 3600
ctttttatat atactctcac taaataaaat tgttccttta tctagcaaat agcttgattc 3660
ctttagttca caatcttctt tccccaataa gtagttttcc catcggaact atatccagca 3720
taaacgatgg tcaacttacg tggagaaatt tcccagtcaa cctctcgttg taaatacaat 3780
ttcacccccc caatcgtcac agtttgcttg gtagcatcaa agttccagga accagttaag 3840
gccccttcgg aaatatgatt ggaattaagt ttcaatgaaa cagacgtaaa ttgcttctga 3900
taattatatc ccatatttat atgttcccac tcaccgacta attcttcctc cttaatagca 3960
acagatggaa ccaatccata gcgttcaggc atcaccaaag gccaaccttc atccgtccat 4020
ctgatactac gcacatgacc catcataaca gcattactag cattaattcc ggtaatgtta 4080
gcaggtaagc gcccctgtga agcataaaac caattatctt gtccgtcatt aaaaatagca 4140
caatgagaaa ttcccaccca accatagcta ttattaaatt tatacggatg cgtcacaaca 4200
ggataagcat ctcctccccc attttctgtt atattcttgc caaccatatc aagataagga 4260
cctttatat tctttgagcg agccacacgc gtattatatg ctacggacaa ttcatcatat 4320
gccataaaca aataatagta acctgtttta gaattataaa taatttcggg agcttcagaa 4380
gcctgccacc ttccatatcc cctacgagct atcaagtcgc cgtaagaact atgatcgcca 4440
ttaatactcc aaggtttacc caattcattc aatggtttac cagtttctgc attaagctga 4500
acagcagtaa acccactgtg ccaggaacca taaacaagcc aatgctctcc ctccggagta 4560
atgacatacg aaggatctat cgcattccac ataaaatagg cttcccaatc atttgtactc 4620
gagcgagtcc attcatttat acctttattg gaagcagaac aaattacata acctttatca 4680
acccattggt tgctagccgg atccgaacac tccatcacac cgatgagtgc acgttcggtc 4740
catgaattat caaagttagt tgcatcgtca tacttacccg tttttatata atgtcccggc 4800
actaaacaat aatacattct atacaatccg gttcttacct ttctggccac gggagcccaa 4860
tatccataac gcggattatc aataagaggt aaccccatct ccgaacgata ttcatttaat 4920
ttatccttgg cccaactagg aggattctca ggcattgttg ctcccaaata ttcccaatta 4980
accaaatctt tagatctacg tccatgaaaa tggccatagc cggaatgcgc attaccatat 5040
gaagcatcag tttgatacat gtaataatat ccatcatcag ctttaaaaac agttggatca 5100
tgtacattgg ccaagttcca ttgatcccgc tgactccagg cagctataga ggtataatta 5160
tctgagtaaa caggtggttg atactgagcc agtaaaggat cttcatattc tgctaatgtt 5220
ttaaaactag ttacagcaga gtaaatagtt ccttcagaag tcagaagata agcacgtaca 5280
taataggttg tagcagattg caaggccgat aaggtcagct caagtttatc acccacagct 5340
tcctttacat tatcagtagt tgtaggtgtt aatttggtta tgctataaca aaaaccacgt 5400
ttaatgacat ctgtttgtga accagaaatg actgcgctta aatcgacaga agaaacagtt 5460
aagttcgtaa ctacaggatt ctccaaggac ggtttttgtg ataaaggagg gtcatcattg 5520
ttgctgcacg ctaaaattag cgtgcagaac attgagaaga aatataggtt aaaatacctc 5580
atatctattt attcagttaa taacatatta gatttaaatt atcattcagc tgcagaagtc 5640
gttaactccc accctggatt ttgcttcaac aaaggtgctt tcttaacttc tgaatccggt 5700
ataggaaaac gataattctt cggactatta aacgctctgg aaggatgcgt tccgtccgct 5760
tcataagaat agacagaaga tgccccagga gtaactacta cactataaat gttgtagacc 5820
tgctgataac gaggtagaga tttctcactg gcagcagttt tcccttcaaa caatttccag 5880
cgacgttcgt caaagaagcg atgatcttcg aaacaaagtt caatgcgacg ttcgttacga 5940
atgcgttcgc gaagctgttc tttcgtcatc ccactataag caggcattcc cacgcgtcca 6000
cgaacctgat tgacatattg atatgcatta gtcggtccct cagattcatt gatggcttca 6060
gctgcgttca acaatatctc ggcataacgg aaaatgggac aagcatgaac atacgtggta 6120
ggattcttga aagacatgtt attcaagaat tttttggtat aataacctgt tgttgttcca 6180
ccatgcaagt cttgataatc ctttccatcg ggatatgata catcaacagc tcgttcctcc 6240
tcttgtactt tatcacccca aatagagcca tgatgcagaa ttgtttgttc caaacgcgga 6300
tcacgattag cataaggatt ttgctcatcg taaccggaag cagcatcgtc aataggctta 6360
ccattcctca tttcatacgc atcaaccagg ttttgtgttg gattggtacg tcctgtagaa 6420
tttacatttc ctgaaaagcc acaaggtgcc aagaaaagct caatttcgcg agtattccac 6480
tcggaacggc tcagaatgta ttcattgttt aagtgaggcg tagacacaaa gcactgataa 6540
tagtttttat tgggatcgtt atcctcagta gtaaacagtt tataaggatg cgcctgttga 6600
ttattcttcg caataaaatc tttggcagct tgtgcggcct cctgccacca gtcagtcttg 6660
ccctccggat tgttcaatgc agacgctctg taaagcagag cccttgattt cagagcataa 6720

gcagcagcgc catttacacg tccccagttc tcattctcat tactataacg aaatggaagt 6780
atatctttaa cagcatcaca ttcatcagca atccatttca gagcttcact acaagcagtg 6840
cgcgacatat tcatggccga agcattcccc gcttcaaaaa cgatagggac accgttctca 6900
tcattaccga taaccggtat atcgccaaac cagccggcaa gatcaaagtg aaagatagcc 6960
ctgagtaatc tggcttcagc catattacgg tcatacaaat gattatcatc acctgttttt 7020
tcagcattac caataacact ttctctggcg tttttcataa attggttggc ggctctaatc 7080
cccttcaaat cgttgggcca aaatcgttcc gcaaacgggt gactcgtagc agaataagca 7140
tctgtcagca cactgtgata ataatgtacg ttccagaatg agagtgaatt atcagtcatg 7200
cagtcacgtg aagcagcaag aaactgtcca tcgctatatc cggcaaaagc atcaggaagg 7260
tatgtgtata cattcgctaa aaagccacgt gtattttcgt acttgctaaa tatctgttcc 7320
tcagacaagg tcaaatctac ttctttatcc aaataatcgc tacaagaact ggaaacagtt 7380
gccagtaatc ctacagtgat atatatcaat atctttttca tcatttcata atttaaagta 7440
caacattaaa atccgatgtt tacaccaaat gtaaacgaac gggtcttggg ataccaccaa 7500
caataactca tcggtttttc aggatctacg tcatccggaa gatcactcca gttatataaa 7560
ttatatccgc taaagtaaaa tcggcacttc gtcatgctaa ttttggcaat aagttctttg 7620
ggcaacgaat aaccaatctc cacattacgt aaagacagat aattgccatt tctaacccaa 7680
atatcattct tgtatgtacc tgtagcacga tctgagtcta tggcattaaa accaccgtat 7740
agcggacgtg gataagttgc gtttatgtta cgcggatcag cgggattatc agtataatat 7800
ccccatgcct tcgttacttc gtgtgttccc atgttactcc agccactaaa gctgatagca 7860
ggtgaaagaa aaacactacg attcaggtaa gcattaaata cagcacgtac atcaaaacca 7920
ttccattcca aaccaatatt caatgccggt ataatttcgg gaataatggt atagctatca 7980
ggtaacatgt cgttagaatc aatttgtcgg tcaccgttca aatccttgaa aacagccgta 8040
cctaatgtct gcttaccgtt agatgctgaa tgatacggat atttctcggg atgatcaatt 8100
gcatcttgct gactggtagc tactttatca ggatcgttcg cccattgcat gaacttatag 8160
atattccatc cgcccactgt gttattaaat gagctctcat ataaagctgc tacttccgta 8220
ccatcataaa ttcgacgacc cgttttacgt tgccattcca cattgggctc cgtttcatcc 8280
atttcagtta tcttattcgt attgtaagtt agtaaacctt ctaagtaata atttacctta 8340
cctatcttac cacgatggcc taaagtcacc tcaagtccct tgctctctgc ttttccataa 8400
gaatcagaac ttactgaaac tccaataatc gaaggaatag tggagcgtgt taccaagatg 8460
cccgaacgcc actctttaaa cacatctacc gcaccgtaaa gtttcttacc ccagaaatca 8520
aagtccaccc caatattcaa catatcggat atttcccatt tattgttact attgcctgct 8580
ttgctctcgt agtaacctcc gatattgctt tgcgaataac caaagctata accagttcca 8640
ctagagtaag taccctgata aggatatctt ccggcacctg tgctcgactg cccggcacgg 8700
ccataagaag cacgtagctt caacaaatct atatttttgt tttcaacca tgactcctca 8760
gaagcaaccc aaccgagtga tgtaccgtaa aaagtacccc aacgttcttc cggagcaaag 8820
ttatcactag ccattctact tatgtttccg gagaagatat acttattatc gtaagcatac 8880
attgcttgtc cgttccatga cagcgtacgt tccgacgatg cacttccgct taccacattc 8940
tggtaagtgc gaataaaagc acgtgcatcg accgcatgct taccgaactt attgctataa 9000
ttcaacccac cgttaaagtt caaatgatag taatactcac gctgattagc cgatggatta 9060
gtcaatgccg aatatgaagt atacttcgta tacgtaaatt cggacggatc agtaacatcc 9120
acattagtat agtcgtaatt atacgtattg atagcatttg tttgtgtaga ttcaaatgtt 9180
tcgtaagagt cgaaacttac ggtagcattc accttcaatc ctttcaacaa actactgagg 9240
tctccgttca cactcaatgt actatacaaa ttacgtcgac ggtttttctc ttgtccacta 9300
gcggccaaat agcgtaaggg gttattggct gtactactgg catacagttc accgttggga 9360
caataaaccg gttccatcgg gttagcctcc actgctccga acgttgttaa atcgaacaca 9420
ctctgtgtag gctgattgat gttatcaata cgtccaccta aatctaatga tacatttaaa 9480
aacttcgtca catcaatatc aatgttagaa cgcaaattcc atcgattgag catgtgctgc 9540
gtatcaaagt tcttgttcca accggttcct gtctcgttcc acataccttc ttgtcgaaga 9600
taagagaaag acacataata acgagcccgg ttattacccc cgcttatctg catgttcgtt 9660
ttgtacatgg gagctgcatc ccgatacaac tgttcaaacc aattcgtatc gaagtatcta 9720
tagcgatcaa cgccctctaa tgtttcacca ttacttaccc gacgatactt ttcaatctgt 9780
tcatccgtat acatcggatc acttccactc aggtaacgca cttggttacg agtcaatgcc 9840
atattatagg aattctgcaa ttccattttg ttgctcagca tttgataacc aatttcctga 9900
gtaaagtcaa tattggtttt accagcatgc ccacgcttag tggttacaag tatagcacca 9960

ttggctcctc tcataccgta aatagcagta gacgcagcat ctttcaatac agtaatgttc 10020
tcaatcggaa aagcatcgag aaaagagaga tctctctcta cattgtccac aatgataagc 10080
ggagcacgcg cattctgatt catggtacgt ataccacgaa tgtacatggc agtttcactc 10140
caccccggct cactactcca ttcgtaagtt tccataccgt taacggtagc tgtaaatgca 10200
ttttgtaaaa tggtaacagg gtgcttttgt aattcttgcc cggttactac agaagttgcc 10260
tcagtcatca gctttttcgc cttgcgtcca aaggaaccg gcatggtatg caagtattca 10320
tccaagtcct ccgccaatac aatctgaaag ccttcattga aatcggcatt tgccgtagca 10380
tttaaatagc caacactact cacgcatatc tcatcctgag acttcacatt cttgagaata 10440
aaataaccat ctttgtctga cagggcaata cgactttgtt cggctacatt gactacagca 10500
ccggcaatag gagcaccttt gctgtctaca atccgccctt tcaatacttg ctcgttctgt 10560
gcaagggcaa ttgtttgcaa ccagagaaag cctagcagca acagaaacct gcctgatgct 10620
gtgcattttc ttatactata ttttatgtta ctcataatta ttcaatttaa gaattatttc 10680
caatccgggt tattctcaat gttggtcttc catacctcac cttcgggtat tggatagagg 10740
tacatcttat tttcaaagac acgtccttgt gccgttttac ggctaatagc cccattggcc 10800
gctacatcaa taccataaat gggacgagaa agagcctcaa cagctacact ccagcgacgc 10860
acatcccaag cacgatgttc ctcgaaagca agctcaaccg tacgttcttt gtgaatgaag 10920
tttcgcatgg cttcttttga actggtcaag tcactacgat tggccaccga accggtaatt 10980
cccgaacgcc gacgaatctg gtcgagcata ttataaacct ctgtagaagg tccttgcact 11040
tcattcagcg cctctgcata gttcagcata atttcagcgt aacgaatgaa agtccacaaa 11100
cgacgtgaag taccggtatg taccgaacct aaaatagctt ccggaatgta tttgcgaaca 11160
taataacccg taggcgtagc atttgcatta cccaccggat tgtcgcgttg tcctttgatc 11220
acattgattg ttccgttgcc ccattccgta ccgtgataaa gaacagtagc ctccaatcgt 11280
ttatcgcgac ctgtccacat agtagcatca ctatatccac tgcctgtgtt aatggtaggt 11340
tgaccgttgt tgtaaaccgg tgcacccgtg gcgtcataag tagtgaaagg cgaagaaccg 11400
tcaatcatgt cgtacatgtc gactaagttc tgcgagggac aaagacctcc tttaccacct 11460
tctcccaccg gcgtgtcatt ttgtatggcg ctccatccta ctgctacatc atttctgaag 11520
aatataattt ctttgttgct gtccaaatag gttgtacgaa gcaacgcatt attatatggc 11580
aaaataccat ctgttgtttc ctctgtaaaa agagcaaact tagaaccata agtatccata 11640
aaactcttcg cagcatctgc cgcttcctgc caggttattc cactctcagc actataccgg 11700
ggactggcca tgtacaactt aatgcgtgaa gcaagtgccc gtgccattgg ctgcccaatg 11760
cgacctgcat aagttgcatt ccacgcatct tcataagtaa gcaagccact ttcacattca 11820
ctcagttcct tcaataagaa atcgactaca tattccttca ccgttggccg gtcggtatat 11880
ccgctcagca aatcaccatc cggatcaagt acttcggtaa caatgggtac cgggccgtaa 11940
cgcaaaaaca actcccagta aaaccatgcg cgcaagaagc gagcttcaga agtatagttt 12000
gttttgtcct gcaagtactt atcattactt aagtcggcac ttttgggcac actttctatg 12060
cgattaatga tcatgttgca tttgcggata ccgcgatagt aatgttccca agtatcagtg 12120
agttcggcag caccggccga tccgtaataa ttaccgatat taaaagtact acgcacacca 12180
ccagttgcat aactggtttc ggccaaatca gtagccgcat ccagccaaga attgtttatg 12240
cgacatgctc cgtgacgcag gaagttatac gtgtcaaagt ggaattgttc catcaagctc 12300
caatcaccga agacctgttc ctcggttaat tcgttactgg gctgcttgtc caagaaacta 12360
ccaaatgcat cttcgcatga agtaagaaca cttgtagaaa tggttagtgc aagcaagctg 12420
tataatattg ttttcttcat gtgtgtcaga atttaatgtt tacaccaaag ttcactgttt 12480
tcataatagg gtaacgattt gaaccattac tttcaggatc ggttaagttg tttaaatgat 12540
cccaagtaat taagttgttc gcattcacgt aaagacgaca attgctcatg ccggccacct 12600
tagaccaatt ctcgggcaga gtatagctta actctatgtt tttcagacga acaaatgatc 12660
cgttcttcaa gaagaaagag tttgtacgct gattgtgatc accgtaattg tcgtagtgca 12720
acaacggata gttggcatgt tgcagattgt aagcttcgct ctgcgcagga ttccagcgtt 12780
ccaaatgatg ttctttcacc ttaccgccac taacgaaagc atacatcgct tcggcatcgt 12840
agtaacgact tacgtgatcc acaccttgga acatcacgct gaatccccat cctttatagt 12900
taacgcccaa agacaatgcg tacgtatttt cgggaatatc actatatcct atacgacttt 12960
catcgcgatc gtcaatgaaa ccatcaccgt ttgcatcctt atatttaaga tcgcccacct 13020
ttacggcacc aaacgtagat acggggaaat caggattatc taaatcagcc tgcgtaacaa 13080
agccctcggc aatcagcccg aaatactgac tgataggatt accctcgcgc ttgcgataag 13140
cggtcttcag ttcgggttca tccatgctga taatttcatt gcgtgcgtgc gaatacgtta 13200

```
     atccaacgtt gtag                                                       13214
//
ID   XXX; XXX; linear; XXX; XXX; XXX;  XXX.
XX
AC   PRJEB67889
XX
DE   Uncultured bacterium DNA fragment from in-reactor enriched cow rumen, clone
ferm-12e13, contig-2
XX
FH   Key             Location/Qualifiers
FH
FT   source          1..10713
FT                   /organism="uncultured bacterium"
FT                   /environmental_sample
FT                   /mol_type="genomic DNA"
FT                   /isolation_source="Cow rumen"
FT                   /clone="ferm-12e13, contig 2"
FT                   /note="AZO-galacto-mannan, AZCL-debranched arabinan active
clone"
FT   CDS             318..1505
FT                   /db_xref="SEED:fig|6666666.127234.peg.1"
FT                   /translation="MSPLSAVTALAAFALRRRCSSRSFQFRSVVSFCTCFSLSEKAISC
FT                   PASFKSRCCCKVISFFFINIFIKMIIGRFRGRESSFLSSRLFAKVTTCRGRFANHSPLI
FT                   LQKSFLPTNILIYAAEPFIFSTFAWFFTSERPGSSNRLTRVVQSNGPSCSFGRPESFTY
FT                   RKTLPREETNTLNALKPIDMAILYDWYENPGTNEEAPEERGLHPRPLLNGKVTMRQLYN
FT                   RVHARSSLTVGDVMNAIDCLAQICGEELRDGHEVHIEGLGYFAPTLEATQKVTRSTPNK
FT                   HLKLRLKSISFRPDVRLKQAMTGVSAIRSKYTRHSLKLSEVEIDMKLKEYFADHDVLIR
FT                   SDFQALCGMARTTANVHLKRLQQEGKLKNVGKPTQPIYRAMPGYYGVSRDAKAGR"
FT                   /product="putative DNA-binding protein"
FT   CDS             1548..1661
FT                   /db_xref="SEED:fig|6666666.127234.peg.2"
FT                   /translation="MGEGGEGDAGIKRNILRPEIRGNTKRNVNYIRKGNRE"
FT                   /product="hypothetical protein"
FT   CDS             complement(1828..2472)
FT                   /db_xref="SEED:fig|6666666.127234.peg.3"
FT                   /translation="MDSNVPPTPTGYARVVLPIGGADASVAVSITVCDESGQQYVLRTT
FT                   TPNAAPYCYFLAYGVYRVVALEDCTAQSNWGALTVGTIFEVTGGGYISLNYIGTISTPS
FT                   IVQASGTDDNVPPSKVGYNIMKVYGIETNGGGVLVDSDKKEYGIFNYTGHIGGAHYFYI
FT                   KPGTYTIKSLSATNGNYIYLEYGNVRQPLSAGRTFTVPASFFSIVFSKKAL"
FT                   /product="hypothetical protein"
FT   CDS             complement(2621..3490)
FT                   /db_xref="SEED:fig|6666666.127234.peg.4"
FT                   /translation="MKKDMKKILFAIPLFAMLLVACDPIVDEISPDPNVTPENLTFELV
FT                   AKNQGNNNIQIVPTPSRYVKVYDATSDIKLAEGTAPSVQVAPPNKELQVYVTTINSDGS
FT                   ITKSASKSIAVTEYTDLPAIYADVFGTTPDGGYGTTTWVWNTEAADGCWGNGGYLGNTG
FT                   PGWWICDAVGDVGNGRGQIDEQAVGKGLPDDGLNGWFSLSLTGVQTSRGESGSVKVSED
FT                   VVQSGWDVGTMEFSGTIPLLGIQPNQGNARQYEYHILKITEDELRLCAPEPGAGAWGTA
FT                   WFWNFKKK"
FT                   /product="FIG00416167: hypothetical protein"
FT   CDS             complement(3505..5274)
FT                   /db_xref="SEED:fig|6666666.127234.peg.5"
FT                   /translation="MKKIAYLIGAALMIAGMFASCSLDSMSLTEKDTTNFPKTAEDANQ
FT                   ALAAIYQNLNNVNAFPQQSFLYNSLLACDDMLGGGGPNDKLMQSLDLICNTGTNATQDY
FT                   WVARYQGINRANTLLEALAGIEMDPEVRAQAEGEAKFLRAFYYYELASMYGRVPLVITA
```

```
FT                   LLPEDTTPPTAAVIWGQILQDLRDAAANMPAVRKSDGHVDKYTAEAMLGRAWLYYTGMY
FT                   GNGETLNDLVSTNYSPLTSVDLPDGTTLTKQDVVGYIDDCVNNSGYSLVPEFRNLWAYT
FT                   NKCTVEDFPFTAGQGLKWVEDDAAVNPESMFAIKYNKLASWSTTIGYGNGYALHFGVRG
FT                   NDQANSFPFGQGWGAGPVAPNLYNDWSVAEQDDARRPASVFKTEDMPSYNKGGGDGFIQ
FT                   ETDYYQMKIGSIMAYSTDAAGNKTIEPVFEKIMYGADGWINDNLMQTGSIHDLVLIRFA
FT                   DVLLMQSELKEDVSGINRVRSRAGLEPIGTYSLAALQNERRWELCFEGTRWNDIRRWHI
FT                   AADALERQTGAKVYTNGAESRNVAQNGGYAARYKATAGFQKIPENQVLLNEGKVEQNEG
FT                   YTDASAEYNGWQE"
FT                   /product="putative outer membrane protein, probably
FT                   involved in nutrient binding"
FT   CDS             complement(5287..8535)
FT                   /db_xref="SEED:fig|6666666.127234.peg.6"
FT                   /translation="MNLNFRRTALLVGICSAVSLTYTPQLFAASVDAIDAVQQAKKITG
FT                   TVTDAMGPVIGANVLEKGTTNGVITDIDGNFTLNVQPGATIVVSFIGYQPQEIVVGNQT
FT                   SFKIQLKEDTELLDEVVVVGYGVQKKKLVTGATVEVKGEDIAKLNTTQALGALQSQSPG
FT                   VNIQAVSGQPGDGFKINIRGAGTNGNTAPVYVIDGVAGGDINALNPADIERIDVLKDAA
FT                   SCAIYGSSGANGVILITTKQGKVGKVSVSYDGNIGWANIYKMPDMLNAKEYMAVMDQVA
FT                   YNNGGQPYDWSKFVDADLLTAYQNGTNPGTDWVKEFRNKNAVVTNHALNVTGGSEFSKF
FT                   STGIGYQYQDGAFGGPVKTDYRRFTFRINSEHVIYRKGDMDVIKFGENIYYQHKQNQGV
FT                   QLGNQYSNDLSNALRAIPLIPVYNENGDFFMYDDLKNFGTSANGILDYTAYASNPMAHM
FT                   VYNQAGNNKNKNFNLNTAAYLEVQPLKNLIYKGQVSYKQWSSSWRSYLPVYQINNQGDS
FT                   RDKDQTINNVSLGWNWSLTNTLSYRFDITDLHHFDVLAGTEYSKSRPTYGESVEATGYN
FT                   SAFGDFTHAYLHNTERKATATVNGYPSDYGSKMSYFGRLNYDFKETYMFSAIIRADGSS
FT                   KFAKGNQWGYFPSFSAGWVISNEAFMKNTASWLGFLKVRAGWGQNGNDNIPNSNWRAGY
FT                   EFGDYGLYTFGSDKNGGTTGAYPNRLANPDLTWETSEQTNIGIDARFLDNRLSFTMDWY
FT                   SKQTKDLLVEVGTNAASGFATQYQNAGTVKNTGLELSMGWRDQIGKEFKYGVNVNMAYN
FT                   KNEVTEVNNANHFIEGGNDLLAQSTGRFVRMEEGHPIGYFYGYKTDGVIQNQNDLQAYL
FT                   DQNCKGNAANSKQGASIKPGDLKFVDVDGNGVINDDDKTDLGNPHPDVTMGLTLSAEYK
FT                   GFDFSVTTYGAFGAQVARSWRKFSDGQYENYTTEVYDYWHGEGTSNRYPLLAPGNSGQN
FT                   FQAISDIYIDDADYVRIQNLTIGYDFKRIWKNCPFQQLRVYAAAQNLFTFTGYKGMDPE
FT                   NGRALNDKEPWVTGVDVGNYPQPRTYMVGVNVKF"
FT                   /product="SusC, outer membrane protein involved in starch
FT                   binding"
FT   CDS             complement(8941..10638)
FT                   /db_xref="SEED:fig|6666666.127234.peg.7"
FT                   /translation="MKKLNIKWMLSLIAAFTFASCDTDVDHDIPAVDAPVLVSTTPESG
FT                   AAKVKTGEITIEVKYDKNIFFATDNLSEIKFTGGELISADVLGASNILTVKVNVPGRET
FT                   ACSLSIPEGIVTGPNQMPAPAVSVQFSTVALDKALVAASSAKAVKLYNYLLDNFETKTL
FT                   SAMMANVAWNTEMSEKVYGWTGKYPAINCFDYVHLPASVAGADWINYGDITPVKDWSDK
FT                   GGIVAAMWHWNVPKKAVGEASSTRIWEGETVMPGDWSGNVQMTDDAAKTVFADAQVGQV
FT                   IRVAVKDVAEGAQGSFKNSGWSEIASGTDYFDISGDYTLVITEDILKSLQEGGLIIGGH
FT                   DYTAVAVYLESNGTALDPNKDYAFYKADTEFDAANATVEGTWENKVFTEDLKNAAAYLK
FT                   LLRDADIPVLWRPFHEAAGGWFWWGKDAASFKSLWIAMFNYFKTEGLDNLIWVWTTEGN
FT                   DSDWYPGDQYVDIVGRDVYNKETADCVSEYTSIAGNYGNKIVSLSECGTVGLISEQWAS
FT                   GARWSWFMPWYDGTNEDGSPAVHADEAWWKDAMSQEFVVSREELPSME"
FT                   /product="GH26"
FT                   /EC_number="3.2.1.78"
XX
SQ   Sequence 10713 BP; 3072 A; 2703 C; 2030 G; 2908 T; 0 other;
     atccccgcac ccagtggagc cgggaagaga cgcggcaggc aagcgcagcc agccggtcct        60
     tgccgtccca atcgggcagt tcgtccatgt agagcaggaa gggatggtag gcagggatgt       120
     aagtggaata gacgtatcgg ttcaagtcct tgtcccagca aggaatacct tcggcatggg       180
     cttcgatgca gaaagtgttc aactcgcgct tgcccacggg ggtgaaggtt gcggtgcgtt       240
     gcccggcggg gcggaattcg gtttcgtccg tcagcaggtt gtagcgaaag tcgtagcgcg       300
```

aggtgaggaa ctcctcgatg tcgccgctca gtgcagtcac ggcgcttgcg gctttcgcgc 360
tccggcgacg ctgctcttcc cgctccttcc agtttcggtc ggtcgtttct ttttgtacgt 420
gtttcagcct ttcggaaaag gctatcagtt gtcccgcttc cttcaagagt cggtgttgct 480
gtaaagtcat cagtttttc ttcataaata tttttattaa aatgattatt gggaggttcc 540
ggggaaggga atcatccttt ctttcctctc gcctcttcgc caaagttact acctgccgag 600
ggcgttttgc aaaccattct ccgcttattc ttcaaaaatc attcctgccg acaaacatac 660
ttatttatgc tgcggaaccc tttattttca gtacctttgc atggttcttt acaagtgaac 720
gacctgggtc gtccaatcgg ctgacccggg tcgtccaatc gaacggcccg agttgttcgt 780
tcggccgacc ggagtcgttc acttatagaa agacacttcc gagggaagaa acaaatacgt 840
tgaacgcact aaaacctatt gatatggcaa tactatatga ctggtacgag aatccaggca 900
ccaatgagga agccccggaa gagagaggcc tccatccccg cccgttgctc aacgggaaag 960
taaccatgcg gcagctctac aaccgcgtgc atgcccgcag ctcgctcacg gtgggcgacg 1020
tgatgaacgc gatagactgc ctggcgcaaa tctgcggcga agaactgcgc gacggccacg 1080
aggtgcacat cgaagggctg ggctacttcg cccccaccct ggaagccaca cagaaagtga 1140
cccgcagcac cccgaacaag cacctgaaac tccggctaaa aagcatcagt ttccgtcccg 1200
acgtcaggct gaaacaagcg atgacgggag tcagtgctat ccgcagcaaa tacacccgcc 1260
actccctgaa gctttcggaa gtggagatag acatgaagct gaaagaatac tttgccgacc 1320
acgacgtgct gattcgcagt gacttccaag cgctctgcgg catggcgcgc accaccgcga 1380
acgtccacct gaaacggttg cagcaagagg gcaagctgaa gaatgtgggg aagcccacgc 1440
aacccatcta ccgggccatg ccgggatact acggcgtgtc gcgcgatgca aaggcggggc 1500
gatgagggtg gaaagcggtt tcaccgggaa agccgatgaa caggaggatg ggagaaggag 1560
gtgaaggaga tgcagggata aaaagaaaca ttctgcgccc tgaaatcagg ggtaatacaa 1620
aaaggaatgt gaattacatc cggaaaggca atagagaatg aatgcgaaaa ggcgaatctc 1680
acgactcgcc ttttcctgat ttaaccatta aacaattgta ctattcggtt aatgcattta 1740
caattatccg atgttacggg cgactaataa agtcgggggg ggtaacgaca actgctttga 1800
gttgaataat tcatttatag attattctta aagtgccttt ttagaaaaca caatggagaa 1860
gaatgacgca ggaactgtga aagtcctacc agcagaaagc ggttgtctaa catttccata 1920
ctccaaataa atataatttc cattggtagc actcaaagat tttatagtgt atgtgcctgg 1980
cttaatgtaa aagtaatgag caccgcctat atgtccggta taattaaaga taccatattc 2040
ttttttgtca gaatcaacca acaccccacc gccgtttgtc tctattccat acactttcat 2100
tatattgtaa cctactttgg aaggaggaac attatcatca gtacccgatg cttgaacaat 2160
acttggagta gagattgttc ctatataatt taaagaaata tagccacctc ctgtcacttc 2220
gaaaattgta ccaaccgtaa gtgcgcccca attagattgt gctgtacaat cctctaaagc 2280
gaccacgcgg taaacgccat aagccaagaa ataacaatat ggagcagcat ttggggtagt 2340
agttctcaaa acatactgtt gaccactttc gtcacaaaca gttatgctta cggctaccga 2400
agcgtcagca cctcctatcg gaagtacaac ccgagcatat ccggttggtg tgggaggtac 2460
attactatcc attgtatacg gtcccaaaac atccgcttta acaaaaagag cacagaagaa 2520
cagtaatatc aaaaagaaag ttcgtttcat tgttgcattt ttttaattag taataaaagt 2580
atacgagtgg tcgttaccca cccgtacctt tcaatattaa ctattttttc ttgaagttcc 2640
agaaccatgc cgtaccccaa gcgccagcac cgggttcagg agcacaaaga cgaagttcat 2700
cttctgtaat cttcaaaata tgatactcat actgacgtgc atttccttga ttcggctgta 2760
tacctaacaa tggaatagta ccagagaact ccatagtacc tacatcccaa ccggattgca 2820
ccacatcttc tgacaccttg acagaaccac tttcaccacg acttgtctga acaccagtca 2880
aactaagtga gaaccaaccg tttaatccat catcaggcaa tccttttcct acagcttgtt 2940
catcaatttg ccctctaccg tttccgacat ctcccacagc atcacaaatc caccaaccgg 3000
ggccggtatt gccgagataa cctccattac cccaacaacc gtctgcagct tctgtattcc 3060
agacccaagt ggttgttcca taaccaccgt ccggagttgt gccgaataca tcagcataaa 3120
tagcgggaag atctgtatac tccgttactg ctattgattt ggaagccgat tttgttatgg 3180
agccgtccga attaatggtt gttacatata cttgcaattc tttattagga ggagctactt 3240
gaaccgacgg agccgtacct tctgccaact tgatatcaga cgtagcatca taaaccttca 3300
cataacgtga cggagtagga acaatctgaa tgttgttgtt tccctggttc ttggccacaa 3360
gttcaaaagt aagattttcc ggtgtgacat tgggatccgg actgatttca tccactatcg 3420
gatcgcaagc caccagcagc attgcaaaca aaggtattgc gaataatatc tttttcatat 3480
ctttttttcat tttaaattca tattctattc ttgccaacca ttatactcag cacttgcatc 3540

cgtataacct tcattttgtt ctacctttcc ttcattcaag aggacttggt tttcgggaat 3600
cttctggaaa cctgccgtag ccttataacg tgcggcataa ccaccgttct gcgccacatt 3660
acggctttca gccccatttg tataaacctt tgctcccgtc tgtctttcaa gggcatcggc 3720
agcaatatgc caacggcgaa tatcgttcca gcgagtacct tcaaaacata gttcccaacg 3780
gcgttcattc tgcaatgcag ccaacgaata tgtaccaata ggttcaagtc cggcacgact 3840
acgcactctg tttataccgg aaacatcttc cttcaattca gactgcatca acaatacatc 3900
cgcaaaacga atcaaaacca aatcatgaat agaaccggtt tgcatcagat tatcattaat 3960
ccagccgtcg gcaccataca ttatcttctc aaataccggt tcaatcgttt tatttccagc 4020
cgcatccgta ctataagcca tgatagaacc tattttcatc tgataataat ccgtttcttg 4080
gataaagccg tcaccaccac ctttattata actcggcata tcctccgtct tgaatacgga 4140
ggccggacga cgggcatcat cttgttctgc cacactccaa tcattataaa gattgggagc 4200
tacaggaccg gcaccccatc cttgtccgaa ggggaaagaa ttggcctgat cattgccacg 4260
tacaccaaaa tgcagagcat aaccgttacc ataaccgata gtagtgctcc aagatgcaag 4320
tttattatat ttaatggcaa acatggattc gggatttacc gctgcatcat cttctaccca 4380
cttcaatccc tgaccggcag taaacggaaa atcttctacc gtacatttat tggtatatgc 4440
ccacaaatta cgaaactcag gaacaagact ataaccagaa ttattcacac aatcatcaat 4500
atacccaacc acatcctgct tggtcaaagt tgtcccatca ggcaaatcaa ctgaagtcaa 4560
cggactatag ttcgtactaa ccaaatcgtt cagagtttca ccattgccat acataccagt 4620
ataatacaac caagcacgtc ctaacatagc ttcagccgta tatttatcaa catgcccatc 4680
cgacttgcga acagccggca tatttgcagc cgcatcgcga agatcctgca gaatctgccc 4740
ccagataaca gcggcagtcg gaggggtagt atcttcgggc aacaaagcag taatcaccaa 4800
aggtacgcgc ccatacatgg aagccaactc gtagtaataa aaagcacgaa ggaacttggc 4860
ttcgccctca gcctgcgctc ttacttccgg atccatttca ataccagcaa gagcctccaa 4920
caacgtattg gcacggttga ttccttgata gcgagccacc cagtaatcct gcgtagcatt 4980
tgtacccgta ttacaaataa ggtccaaaga ttgcatcaac ttatcattgg gaccaccacc 5040
acccaacata tcatcgcaag ccagcagaga attatacaag aaagactgct gtggaaaagc 5100
attgacatta ttcaggttct gatatatggc agctaaagcc tgattggcat cttcagccgt 5160
tttaggaaaa ttggtcgtat ctttttccgt cagactcatc gagtccaagc tacaagaagc 5220
aaacattcct gctatcatca aagctgcacc tattaaatat gctatttttt tcatcgtgta 5280
tcaattttag aatttaacat taacacctac catataagta cgaggttgag gatagttacc 5340
tacatcaaca ccggttaccc acggttcttt gtcattgagg gctctaccat tctccggatc 5400
catcccttta taaccggtaa aagtaaatag attctgtgca gcagcataca cacgaagttg 5460
ctggaacgga cagttcttcc aaattctctt gaaatcgtat cctatagtca gattttgaat 5520
gcgaacatag tcggcatcat caatataaat atctgaaatg gcctggaaat tctgtccact 5580
atttcccggc gcaagcagcg gatatctgtt tgaggttcct tcgccatgcc agtaatcgta 5640
aacttccgtc gtatagtttt catactgtcc gtctgagaac tttcgccaag agcgagctac 5700
ctgtgcacca aaagcgccat aggttgttac agagaaatca aatcctttat attccgcact 5760
tagcgtaaga cccattgtca cgtcaggatg aggattaccc aaatcagtct tatcgtcatc 5820
attaatcacc ccatttccat caacatccac aaacttcaaa tcgcccggtt taatacttgc 5880
accttgctta gagtttgccg catttccttt acaattctgg tcaagatatg cctgcaagtc 5940
attctggttc tgaatgacac catccgtctt ataaccataa aaatagccaa taggatgtcc 6000
ttcttccata cggacaaacc gacccgtact ctgagccaac aaatcattac cgccttcaat 6060
aaaatgattg gcgttattaa cctccgttac ttcattttta ttataggcca tattgacatt 6120
caccccatac ttgaactcct taccaatctg atctctccag cccattgaaa gttcaaggcc 6180
agtattcttt accgtaccag cattttgata ttgagtagcg aaaccggaag ccgcattggt 6240
acccacttcc actaacaagt ctttcgtctg cttgctatac cagtccatgg taaaactcag 6300
acggttatcc aagaaacggg catcaatacc gatattggtc tgttctgaag tttcccatgt 6360
caaatcgggg ttagccaaac gattcgggta agctccggtt gtacctccat tcttatccga 6420
gccaaatgta taaagaccat aatcgccaaa ctcgtaaccg gcacgccaat ttgaattagg 6480
gatattgtca ttaccattct gtccccaacc agcacggact ttcaagaaac caagccaaga 6540
agcagtattc ttcataaaag cctcgtttga aataacccaa ccggcagaga atgatgggaa 6600
atatccccat tgattgcctt tcgcaaactt agacgagcca tcggcacgaa tgatagcaga 6660
aaacatataa gtttccttaa agtcataatt caaacgtcca aaataagaca tcttagaacc 6720
ataatccgag ggataaccat ttactgtagc agtagcttta cgttctgtat tatgcaaata 6780

agcatgtgta aagtcaccaa aagcactatt ataaccggtt gcctctactg attcgccata 6840
agttggtctt gatttagaat actctgtacc agccaaaaca tcaaaatgat gtaagtctgt 6900
aatatcaaac cgataactca acgtattagt caaactccag ttccaaccca aagacacgtt 6960
atttatcgtt tggtctttat cacggctatc tccctgatta ttaatttggt aaacgggaag 7020
atatgaacgc catgaacttg accattgttt gtaactaacc tgacctttat aaataagatt 7080
cttaagtggt tgtacttcca gatatgccgc tgtgttcaga ttaaaattct tattcttatt 7140
attcccagcc tgattatata ccatgtgagc catcggatta gaagcatatg cagtataatc 7200
aagtatacca ttagcactcg ttccaaaatt cttcaaatca tcatacatga agaaatcacc 7260
attttcatta tatacaggaa tcagaggaat agcacgcaaa gcattagaaa ggtcattcga 7320
atattgatta cctaactgca caccttgatt ctgtttgtgc tgataatata tattctcacc 7380
aaactttatc acatccatat ctcccttacg atatatcaca tgctcagaat taatgcggaa 7440
agtaaaacga cgatagtcag tcttcaccgg accaccgaat gcaccatcct gatactgata 7500
gcctatacct gtcgaaaatt tggagaattc agaaccacct gttacattca acgcatgatt 7560
agtaaccaca gcatttttat tccggaattc ttttacccaa tctgtaccag gattggtacc 7620
attctgataa gcagtcagca aatcagcatc tacgaatttt gaccaatcat acggttggcc 7680
tccattatta taggctactt gatccattac cgccatatat tctttggcat tcagcatatc 7740
aggcatctta tatatatttg cccaacctat gtttccatca taagatacag aaacctttcc 7800
taccttaccc tgctttgtag taataaggat aacgccgttg gcaccgctcg aaccgtaaat 7860
tgcacaagat gcagcatctt tcaacacgtc aatacgttca atatcggcag gattcaaggc 7920
gttaatatcg ccaccggcta caccatcaat aacgtaaact ggagctgtat ttccgttagt 7980
accagcacca cgaatattaa ttttaaaccc atcacccggc tgaccggaaa cggcctgaat 8040
atttacaccc ggagactggc tttgtaaagc gcccaatgcc tgcgtagtat ttaattttgc 8100
gatgtcttcg ccttttacct caaccgtagc acctgtcacc agtttcttct tctgtacacc 8160
gtaacccact acgactactt catccaacaa ttccgtatct tcctttagct gaatcttgaa 8220
actagtctgg ttacctacca ctatttcttg cggctgataa cctataaaag acacaacaat 8280
cgttgctccc ggctgaacat tcaatgtgaa attaccgtca atgtcggtaa tgacaccatt 8340
tgtggttcct ttctccaata cattggcacc gatgaccggt cccattgcat ctgttacagt 8400
acctgtgatt ttttttgctt gctgcacagc atctattgca tcgaccgatg cggcgaaaag 8460
ttgcggagta taagtcaggc tgacagccga acagataccc acgagcagcg cggttcttct 8520
gaaatttaaa ttcatacttg tatttttctt agattaatga attggtattg cctggcggca 8580
tctaccgttt aatataaggt taaacttctc tttttgagga aaaattaaat ataccttatt 8640
ctataaagtg gttggtatac atatcactcc tttctcatag gcatttttca tttcattagc 8700
ggttaaacat tatatgatta ttatcaaatt ttaattaggc ttgacaaaag tagactttta 8760
gcagcaaacc ttataatact attttatcaa ataacgacac tcatttaaca tccattacta 8820
tttttcctaa aaagtataaa gcatgaattc tataaccaac aactaaaagc tataaaagac 8880
agaaagggtt gccccgatac taaccagaga caaccctttc tattttatta ttaaaaattg 8940
ctattccata gaaggaagct cttcacggga aacaacaaat tcctggctca tagcatcctt 9000
ccaccacgcc tcatcagcat ggactgccgg actaccatcc tcatttgtgc catcatacca 9060
cggcatgaac caactccaac gtgcaccaga tgcccattgt tcggaaataa gtccgactgt 9120
gccacattca ctcagcgaaa caatcttgtt tccataattt ccggcaatgg aagtatattc 9180
tgaaacacaa tctgcagttt ccttattata aacatcacgt cccacgatgt ctacatattg 9240
gtctcccgga taccaatcgg aatcattgcc ctcggtagtc caaacccaga taagattatc 9300
caatccttca gtcttgaagt agttgaacat agcaatccac agagatttaa agctggcagc 9360
atccttaccc caccagaacc agccaccggc tgcctcatgg aatgggcgcc acaataccgg 9420
aatatccgca tctctcaaca gcttcaagta agcggcagca tttttcaaat cttcggtaaa 9480
caccttgttt tcccaagtgc cttccaccgt ggcattggca gcgtcaaact ccgtatctgc 9540
cttatagaag gcataatcct tattcgggtc gagggctgta ccgttacttt caagataaac 9600
agcaactgcc gtataatcat gtccgccaat gatgaggcca ccttcttgca gggatttcaa 9660
aatatcctcg gtaatcacca atgtatagtc tccggatatg tcaaaataat cagttcctga 9720
agcaatctcg ctccaaccac tgtttttgaa cgaaccttgc gccccctctg ccacatcttt 9780
cacagccacg cgtatcacct gccccacttg tgcatcggca aagacagttt ttgcagcatc 9840
atccgtcatc tgtacatttc cactccaatc acccggcatt accgtctctc cttcccatat 9900
ccgggtagaa gaagcttccc ctacagcttt cttgggtaca ttccaatgcc acatggcagc 9960
tacgatgcca cctttgtcag accaatcctt aacgggagta atgtcaccgt aattaatcca 10020

```
     gtctgctcca gctacagatg ccggcagatg aacatagtcg aaacagttga tagccggata     10080
     ttttccggtc catccgtaca ccttttcgga catttccgta ttccaggcaa cattggccat     10140
     catagctgaa agcgtcttcg tctcgaagtt atccaataga taattgtaca gtttcacagc     10200
     tttggccgaa ctagcggcaa ccagtgcctt gtccaaagcg acagtagaga actgaaccga     10260
     tacagccgga gcaggcatct ggttcggtcc ggttactatt ccctcgggga tagagagaga     10320
     acaagccgtc tctctcccgg gaacattcac ttttacagtc agtatattgc ttgcgccaag     10380
     tacatcggca cttatcagtt caccaccggt aaacttgatt tcgctcagat tgtctgttgc     10440
     aaagaatata ttcttgtcat atttcacctc aatagtgatt tctcccgttt tcactttggc     10500
     tgctccactt tcaggagtag tagacaccaa taccggtgca tccactgctg ggatatcatg     10560
     atcgacatca gtatcgcagg aggcaaaagt aaaagctgct atcaacgaaa gcatccattt     10620
     tatattcagt ttcttcatct taattatgat tttatttcat tcttaatcag ggcatccata     10680
     gtcacattct tcaagccaca attcatcaat tta                                 10713
//
ID   XXX; XXX; linear; XXX; XXX; XXX;  XXX.
XX
AC   PRJEB67889
XX
DE   Uncultured bacterium DNA fragment from in-reactor enriched cow rumen, clone
ferm-13f22, contig-1
XX
FH   Key             Location/Qualifiers
FH
FT   source          1..24657
FT                   /organism="uncultured bacterium"
FT                   /environmental_sample
FT                   /mol_type="genomic DNA"
FT                   /isolation_source="Cow rumen"
FT                   /clone="ferm-13f22, contig 1"
FT                   /note="AZCL-arabinoxylan active clone"
FT   CDS             4..1998
FT                   /db_xref="SEED:fig|6666666.127235.peg.1"
FT                   /translation="MHKLITAQADANVTFYDSYAASVDWWEQAATLRPHLVSPFIPLSY
FT                   IEETDQNSLNAVLNSSYIKDGKYFFGGTQQNPTNPVADAYAAGDSKLVSRQFQFNTRFD
FT                   INLSPLLKGLYFRAKYGIDYASTYNQGYTSKYATFSPVWANYNGKDVIASITQYGKDEK
FT                   SGSENIDNSAYRYTYNVSGQFDYQRTINADHNVFAMILANAWQTQRSGHYHRTTNANLG
FT                   FQASYNYQHRYYADFSVAMPYSTKLPEGKRIAYSPTVTLGWNLAKESFMENSIFDDLML
FT                   TASAGIINQDLDITASDNEDGYYLYKAVIQSGGWYSWGDNGGLAATEFQRGNNPDMTYV
FT                   KRKELTAGLRGSMWDKLISFDVNFFTSKMDGGLVRTNSLYPNYFTQTGYPSSSIIPYVN
FT                   FNVDQRTGFDFSVYANKKVGEVELTLGVSGMYYKSTAKKRDENVEFNYLSAVGRALNGH
FT                   WGLQSEGFFQNQSEIDAAPDQTFGDVKPGDIRYKDQNNDNKIDNNDRVFLGRWDSPFMS
FT                   GVNLTAKWRNFTLYAMGNLYMGGYGMKDNSYYWVKGDGKYSEVVRNRWTPETAATATYP
FT                   RLTTTDGANNFRTSDFWMYKNDRFNLTQVQLTYQMPEKVLEGTFIKGLSFFANANNLLT
FT                   IAKERKALELNVGSAPQTRFYQLGFKGTF"
FT                   /product="putative outer membrane protein, probably
FT                   involved in nutrient binding"
FT   CDS             2041..3837
FT                   /db_xref="SEED:fig|6666666.127235.peg.2"
FT                   /translation="MKLKNIILSAALLVAFSACDDLFEPTIENHKDAADLENMPSWAVG
FT                   LLGHAYISNPLGENANSWKFTDVATDDAVSNDIGNSYLKIATGSWRADNNPMDNWQYLR
FT                   ASWQYLNQFLEISENVEWARDQVASDLFKKRFQGDAYGMRALYMYHLLMSCAGWSADGQ
FT                   LLGIPILTESENIDSDFNKPRNTFRECLDQLNLDVDKAIEMLPEEYGDVDGANNVPAKY
FT                   TQMGADPSQYNRVFGDHAKNRMSARVARAIRAQAALLAASPAYSEGSGVTWEQAANSMA
FT                   EVLANLGSDPIANIDPTGNKWYEDANSIKNLVAGYNRPEFLWRSNKNESADLEKDQYPP
FT                   SLFGKGRVNPSQNLVDAFPSLTGYPISDSRSGYDAQNPYANRDPRLALYIIYNGSKAGT
```

```
FT                   DKSVITTAFDGSNNDAMNKFDGASTRTGYYLRKFLDMNVNANPSNTTNGYHIKPWIRYT
FT                   EIFLGYAEAANEAWGPQGKGSNSYSAYDVIKAIRKRAGIGENGGDPYLESVKGDKDKMR
FT                   ELIRNERRLELCFEGFRFWDLRRWKVDISKLNETVKGMKITGTNHEVVDVEKRDYKDYM
FT                   YYGPVPYSEMLKFDALIQNKGW"
FT                   /product="FIG00938343: hypothetical protein"
FT   CDS             3868..4902
FT                   /db_xref="SEED:fig|6666666.127235.peg.3"
FT                   /translation="MKIAMKKSFLIIAAMMGALFTACENGDWEFPDYKYSAVYFAYQSP
FT                   IRTITLGEDKTVDNSLDNEHKCQIMATIGGVYENKGDVEIGIRVDNSLCDGYTFESTGG
FT                   DVEPLPASHYTLSADKIVIKKGQVLGGVTVNLTDAFFNDPKSTSLNYVIPVVMTNVKNA
FT                   DSILVGKPSVDTPHRIKKTDWDVQPKDYVLYAVKYISKYDSHYLRRGIDTYSGNKTGTE
FT                   VRHAAYVEKDETIDGFTTMAMNKVQWARPTRNTAGKIIESKLLLTFDESGNCIITSNSQ
FT                   GVVASGSGKFVVKGEKKSWGNEDRDALYLDYTMDYDGIHCATKDTLVVHYRGVKAEWFT
FT                   PLKK"
FT                   /product="FIG00939126: hypothetical protein"
FT   CDS             4965..7139
FT                   /db_xref="SEED:fig|6666666.127235.peg.4"
FT                   /db_xref="GO:0031176"
FT                   /translation="MLLTAAAVAVTSCVDDSKLLFAFEKPASIAGMEYLNEYDALKTYV
FT                   NSSANPNFKLGIALTADDYIAGGLVTRLANSNFQEMTAGNAMKYASCVADDGTMNFGTV
FT                   ENFVAAAKAGGLTIYGHTLAWHAQQNNKYLNSLIKDKEIEIDPNQKLDVVDKIVDFSAM
FT                   TKFPYYVMGYEPDIVNGILVSKNPGSWYQYFVADGVPTVSGKEYKVTAMIKSSKEGEFI
FT                   VKMGDWDASTDKNIKVTTEWKEQTIDFGTLAAKSSFVIFQPGGFDGEIQIKWIKVSHPE
FT                   APSVSFYVNQLDNSEMKAGGTMKNFVVRENGKGDIPGPIFEGQGPDGINCTKITSIANP
FT                   ANEWDTQFFIYTPDKEWQGGQKYKISFWYKASAEADCSTQCHGDPGAYKHGAMLPQNPH
FT                   FTTKWQHYEATGTIPSEGSGMKSIAFNLNVSKDAVTYYFANIVWESEEKGNTIPLTPAE
FT                   KKDTLTWAMNNWINGMMKATGGYVTAWDVVNEAIAGGGDDGEGFYPLQSATNVSADDAK
FT                   NNFYWQDYLGSEDYVRIAVAAARKYYAENGGTNPLRLFINDYNLESDWDDNKKVKSLVH
FT                   WIEKWEADGVTKIDGIGTQMHVSCHANAETQKSKEDHVVKMFEILAESGKLVKITELDM
FT                   GYVDEEGNSVKTADMTQAQHKAMSEYYKFIVKKYFEIIPVAQQYGITQWCITDSPTGSG
FT                   WRGGEPVGLWDANYNRKHTYAGFADGLAGK"
FT                   /product="GH10-CBM4"
FT   CDS             complement(7146..7274)
FT                   /db_xref="SEED:fig|6666666.127235.peg.5"
FT                   /translation="MHIFLLQKLALLIVRMTSNKLYYPTVSDTDGNTDYGVMRTII"
FT                   /product="hypothetical protein"
FT   CDS             7228..9450
FT                   /db_xref="SEED:fig|6666666.127235.peg.6"
FT                   /db_xref="GO:0031176"
FT                   /translation="MRTINKASFCKRKICNKKPFKNYKMKHLFKFSVCALTFAMSTATG
FT                   TAQNSGTGLKDAYKDYFSIGVAVNMRNITTPEQIAIIKNDFNSITAENDMKPQPTEPAY
FT                   GQFNWENADKIADFCRSNGIKLRGHCLIWHSQIGQWMYQDEKGNPVSKEKLFANMKHHI
FT                   TAIMERYKDVVYAWDVVNEAISDGGWQGSRPGMGQQSGPYRNSPLYQIAGDEFIKKAFI
FT                   YAREVDPDVLLFYNDYNAADPGKRDRIYNMVKAMKEEGVPIDGIGMQGHYNIYGPSMED
FT                   VDAALTKYSTIVKHIHITELDIRANEEMGGQLNFSREAGNISPVVKTLQEDQYARLFRV
FT                   LRKHKDVVDNVTFWNLSDRDSWLGARNYPLPYDENYKPKPVYNIIKDFDSALDVAEVKE
FT                   DFRPSTLNQPGQQYPMVNSQGYARFRVVAPDAKSVIVSLGLGGRGGTVLRKDKEGVWVG
FT                   TTDGPMDEGFHYYHLTIDGGVFNDPGTLNYYGSCRWESGIEIPAHDSDFYAMKQVPHGN
FT                   VQQVYFYSKSTNAHRRAFVYTPPTYGKDKKKYPVLYLQHGWGEDETAWSNQGYANLIMD
FT                   NLIAEGKIAPFIIVMTYGMTNDIKFGRLNEFTAKEFETVLVDELIPYIDSNFRTLADKK
FT                   HRAMAGLSMGGFETKLITLRRPEIFNYYGLLSGGTYAPSDIKDQNQVKSIFISCGSKEN
FT                   PDGVTKAVNDLKAAGLKATSYVSPGTAHEFLTWRRSLYHMAQELFK"
FT                   /product="GH10-CBM48"
FT   CDS             complement(9595..13485)
```

```
FT                   /db_xref="SEED:fig|6666666.127235.peg.7"
FT                   /translation="MIWIATEDGLNKYDGVKFTVYKHKKNDEHSLCHNYVRTLFEDSKG
FT                   NLIIGTYTGIQMYDPATDRFTPQARWENGDLFDSNIIAILERQNGEIWVSGNILCKLII
FT                   KKEKLVVKMLNLPIATTSTDYMIEDKKRNIWITKGENGIYKISDKKKVTHYLKDEKGIT
FT                   IVDFCEDLNGNIYVGTMGKGLFVYDQLSDSFKPIPYKGKQNLPIKSLYPDNQNELYIGT
FT                   DGKGMKVYNIKQQKITDAPFDNSYFDSNKSKVHSILKDNTGNFWLAIYQKGVMMIPAQP
FT                   NSFKYIGNKAIDKNIIGSNSVTSLCRDHEGTLWVGTDNDGIYGITQELKQKKHFYPADT
FT                   DSEENVPSTVFGLYEDSERNMWIGSFTKGMCRLDRETGRCIYLKELLDQEGNRVQRVYD
FT                   FVEDKNKRLWIATMGAGLYYYELKTNKLKYPEKANKNNKWINCLLYSKDNKLYTGTYDG
FT                   VYCINLNSDEFITKKVLTRRIVFSIYEDKKGIIWIGTSDGLSSWNPKNQQLVTSTTENG
FT                   LPSDAVYAIVGDEQDNLWISTNAGISQYNRSSHKFINYFVGDGLQGNEFSKNASFKDAN
FT                   GIIWFGGTDGITYFNPQEITNPAKKWNIRITDFYLHNQPVRQGVLSAGREIINEPVFEA
FT                   TRFDLSHSDNSFTIEFSTLEMNNPERITYLYAMNDNNWVSLPIGNNRVSFSNLSPGTYH
FT                   FKVKVQDYMVDSDVKEITIYIHPAWWASWWAKLIYALLTGGTIWFIILQAKHRYRTKQE
FT                   MLQHIHAEQINEAKLQFFMNISHEIRTPMSLIISPLQKLISTDGDPSRQKTYHTIYRNA
FT                   ERILRLINQLMDIRKIDKGQMSLMFREIDMVAFIQDLCNTFAGQTVNKHISLTFHREDF
FT                   DTFGLWIDPTNFDKVILNILSNAFKFTPEGGRVDIYLKSGHSEDTETTLSNYAEIIIAD
FT                   TGIGIAPEEIEHIFERFYQIRNHLNNSNIGTGIGLHLTRSLVELHHGHIHAENNPEGIG
FT                   SRFVIRLPMGCSHLKQEELDNSTDNELAYTPAETIPAIEESGDEEELKVRKKTKYRVLI
FT                   VEDDEEIRRYIKQELFADYHILESCNGKEALDMLFKRKPDVVISDIMMPEMDGLTLCKK
FT                   IKQNVNLNHIPIILLTAKTREEDNIEGLEVGADAYLTKPFHIEILRKTVQNLINSRERL
FT                   RNTFSGNQDHEDKMEEIKLQSPDEKLMERVMNAINKNLSNPNLTVELIASEIGISRVHL
FT                   HRKLKELTNQTTRDFIRNTRLKQAAILLTQKRYTMAEISDLTGFANPTIFSTAFKELYG
FT                   VPPTKYMEEHLK"
FT                   /product="DNA-binding response regulator, AraC family"
FT   CDS             13889..16414
FT                   /db_xref="SEED:fig|6666666.127235.peg.8"
FT                   /translation="MKINRCQISFFTIGFIGYIVLSGFQKETVYMNENHLIVNDSDYFE
FT                   TRGLNYFVFSNIYDATFDDSKISAVEIIHHGIRTATNGDVRLNPTPGQWDKLPRFIDRK
FT                   VDQKNKRIEVSLSYPEYAFDYKLVAEAKDGGLYLSVYTEKALPDKLCGIAGLNLEFVPP
FT                   VFWGHSYILDDIHGLFPTSPADFMTTIQGTVEPEPIATGSKIEIAPDDPEKHVSIRTTD
FT                   GNSLMLFDGRNKQQNGNFVVRTLLPGKKTGKIAEWFIQAETDTQWIRKPLVAYSQVGYH
FT                   PAQKKMAVIELDKNDEVSHEVTLLKVGSDGRLTKVLSAMPADWGSYTRYKYKQFDFTAI
FT                   NEPGIYVLEYDKLRTAPFPIGKNVFQKAWFPTLDVFFPVQMDHMCVREAYRIWHGAPHL
FT                   DDALQAPLNRNHWDGWHQGPTTDNKYKPLEHIEGLNVGGWFDAGDFDIQTPSQQSVVQT
FT                   FADLWSDFRVSRDQTSINQQTRYTEIHVPDGKPDLLQQLEHGVLQLIGQVNAVGYAIPG
FT                   ITESHLYQYRHLGDAVNKTDNKVYNANLDSLQTDGPTSGTFDDRWAFTNRNPYLNYGTA
FT                   ISLAAAARCLKEYNPQLSMEASRVAQFIWKDEHSRPTPEKIEFGGRSFNSADFMNSVEC
FT                   RAAFELWRTVGDTVYKNRMQKLFPILLTQFDRNVSVIARILPFMDNQIKKQVEPMVQAY
FT                   VKQVTEQNKENPFGVQISRFSWAGNHGIVQEAINAYQLHLLYPDIVKPDLIYRSLDYLF
FT                   GCHPFHNLSFVSGVGAQPKKVAYGNNRADYSFIPGGVVPGIRIVKPDFPENRDDYPFLW
FT                   SENEYVIDLAASYIYLVNAADHLLNNSGK"
FT                   /product="GH9"
FT   CDS             16459..17583
FT                   /db_xref="SEED:fig|6666666.127235.peg.9"
FT                   /db_xref="GO:0031176"
FT                   /translation="MEKRLILLVTFLSVVMIAMAQPVSDSDTSGSVRSVYCVRGAQYPR
FT                   ITPDHRAQFKVVAPEAKLVQIDLGKKYDLVRNEKGEWLGTTEPLGPGFHYYFLIVDGVR
FT                   VADPSSESFYGCSMHTSGIEIPYAKDVLQYELADVPHGEVRMKRYFSKTANAWRRMFVY
FT                   TPPCYEKSEKEYPVLYLQHGGGEDERGWALQGRTDIILDNLIAQQKAVPMVIVMCDGNS
FT                   QDFTAELLNDCIPFVEENYRVKKDRSHRALAGLSMGGIHTLNTAIPHPELFAYVGVFSS
FT                   GWWAKAPMPGLMKMDTETYYSLLQEKKDEYNASFKEFWISMGGKEDIAYENCRIMRERF
FT                   DKIGIKYTYYETNGGHTWPVWRESLYQFAPLLFK"
FT                   /product="CBM48-CE1"
```

```
FT   CDS             17638..20157
FT                   /db_xref="SEED:fig|6666666.127235.peg.10"
FT                   /db_xref="GO:0031176"
FT                   /translation="MVHLSGAQNPIIRHQFSADPTARVFDGKVYVYPSHDIPSPVERLK
FT                   EWFCMADYHVFSSNNLVDWEDHGVILSQENVPWVNAESYSMWAPDCVFANGKYYFYFPA
FT                   SPKGENQRGFKVGVAVADKPTGPFTPLAEPIKGINGIDPCVLIDKNGEAYIYWSGRGMY
FT                   VARLKSNMTELASEPVQIKNLPEGFMEGPFAFERNGKYYFTFPWVQDKTETLAYAMGDS
FT                   PMGPFEFKGLIMDQSATGCWTNHHSLVEYNKQWYLFYHHNDYSPAFDKNRSVRVDSLSF
FT                   NADGTIKKVVPTLRGVGLTTASSKIQLDRYSQISNQGAAIAFVDENNKFEGWKAVFTKP
FT                   GAWLRYNRVDFGDGGYRKMQMRVNSSTGGVVEIRTADKAAKLLASLVVPKSDGWIEKEY
FT                   DLKLALRNVHDLSVSLKGEGQVEIDWMRFGQNAGEFAVQSRASIKPWEQGAFETRKYRN
FT                   LFAEAGYKQADIDAKLKSVFNDIFYGPNKVYFETNDSMAYVSDIKNHDVRTEGMSYGLM
FT                   IAVQFNRKDIFDRLWRWCKKYMQHQEGPLEGYFAWSCKTDGTRNAQGPASDGELYYVTS
FT                   LIFASNSWGNDSDINYLGEAQHILNCSMKKDGTNRVMPLINMEHKLITFVPDTFGGRFT
FT                   DPSYHVPAFYEVWARWANDGRADFWRECAARSREYLHKSIHPVTGLNPDYNNYDGSLLN
FT                   MKRGIIGDAFRFDSWRVPMNIALDYSWACADKEWQQGYGNKIQNFLYSQGIDTFVDQYN
FT                   VDGTTVAEILDAGGYKKLRHSLGLVATAAAASLVTTHTKSYEFVDKLWNAKHEPYEDGY
FT                   FDAYYDGLLRLFAFMHLSGNYRIIFPQ"
FT                   /product="GH43_2-CBM6-GH8"
FT   CDS             complement(20221..21183)
FT                   /db_xref="SEED:fig|6666666.127235.peg.11"
FT                   /translation="MSCSAQNSLPVFKNTGNPLIKNKFTADPAPMVHNGRLYLYVGHDE
FT                   YRDGQDTASGGQEFNITEWLCYSTDDMQNWTDHGSVLKPTDFKWAVGEAWASQVVERNG
FT                   KFYYYTTVQAGAPYNSKTVGVAVSDSPTGPFVDAIGKPLVTDDMTSNGARGWWNDIDPT
FT                   VLIDDDGQAWLCWGNGTCFLAKLKSNMIELDGDIEVINVPRYVEGPWLHKRGDMYYLTY
FT                   ASMGKGKETIAYATASSMKGPWTPQGELTGMAENSFTIHPGIIEFNKKWYLFYHNATLT
FT                   INGRAGAIGRRSVCVDELQYAPDGKMIYVNQTKEGVAKK"
FT                   /product="GH43_29"
FT   CDS             21674..22165
FT                   /db_xref="SEED:fig|6666666.127235.peg.12"
FT                   /db_xref="GO:0004733"
FT                   /translation="MFREMRRKKQLLSAADTIAVLNRMTSGVLALLGDNDYPYAVPISY
FT                   VYHENTLYFHSALAGHKVDAIAQNSKASFCVIEQDQIVPEEYTTYFRSAIAFGKISMVN
FT                   DDVEKRTAIELLAEKYSPAQEEGRLKEIEKGFHHMCVIKLDIEHLTGKEAVELVRAGSK
FT                   "
FT                   /product="Pyridoxamine 5'-phosphate oxidase (EC 1.4.3.5)"
FT                   /EC_number="1.4.3.5"
FT   CDS             complement(22187..22603)
FT                   /db_xref="SEED:fig|6666666.127235.peg.13"
FT                   /translation="MLLRDPDIFPSEEVLRNALGETNYNLFQGFVQNITSPTGGLAMEW
FT                   RYYNDGKAWLCKVVNKKKTVFWLSIWEHYFKTSFYFTEKHLESIDALSIADSLKQQLAQ
FT                   AKSIGKLIPLIIDVHNDEAMTDLLTLAHFKKSLK"
FT                   /product="hypothetical protein"
FT   CDS             22852..23058
FT                   /db_xref="SEED:fig|6666666.127235.peg.14"
FT                   /translation="MIQSFAQMSKSCIFTVDVFKGRYDFASENFSQLFGYDPQCISNIR
FT                   QEGDLLEERIHPDDRAQLLDYQV"
FT                   /product="transcriptional regulator"
XX
SQ   Sequence 24657 BP; 7121 A; 4703 C; 5478 G; 7355 T; 0 other;
     cagttgcata aacttatcac cgcacaggct gatgccaatg taactttcta tgactcttat        60
     gcggcaagtg tggattggtg ggagcaagct gctacgttgc gcccgcactt ggtgtcgccg       120
     tttatccctt tgagttacat cgaagaaact gaccagaatt cattgaatgc tgtgcttaat       180
     agtagttata tcaaagatgg taaatacttt tttggcggta cgcagcagaa ccctactaat       240
```

ccggttgccg atgcttatgc ggctggagac agtaagttag tgagtcgcca gtttcagttt 300
aatacacgct ttgatattaa tctttctcca ttgcttaaag gactttattt tcgtgctaaa 360
tatggtatag actatgcttc tacctacaac caagggtata ccagcaaata tgctactttt 420
tcacctgtgt gggccaacta caatgggaag gatgtcattg ctagtattac gcagtatggt 480
aaggatgaga agagtggcag tgaaaatatt gataattcag cttatagata tacctataat 540
gtatccggtc agtttgatta ccagcgtact attaacgccg accataatgt ttttgccatg 600
attttagcta atgcttggca gacacaacga agcgggcatt atcatcgcac taccaatgcc 660
aatttaggtt ttcaggcatc gtataactat caacatagat actatgcaga ctttagtgtt 720
gcaatgcctt attcgaccaa gttgccagaa ggcaagcgca tcgcttattc tcccactgtt 780
actttgggtt ggaatttagc taaagagtca tttatggaaa actccatctt tgacgacttg 840
atgcttactg cttcggcagg tatcattaat caggatcttg acatcactgc ttccgataac 900
gaagatggat attaccttta taaggcagtt atacaatcgg gtggttggta tagttggggt 960
gataacggag gtttggctgc tactgagttt caacgtggta acaatcctga tatgacctat 1020
gtgaaacgta aagagttgac tgcaggtttg cgtggttcta tgtgggataa gttgataagt 1080
tttgatgtta actttttcac aagcaagatg gatggtggat tggttcgtac caattcgttg 1140
tatcctaatt actttacgca aaccggttat ccgtcatcat ctatcattcc ttacgtaaac 1200
tttaatgtag atcagcgcac gggatttgat ttcagtgtgt atgctaataa aaaggtaggt 1260
gaagttgaac taactcttgg agtaagcggt atgtactata agagcacggc taaaaaacgc 1320
gatgaaaatg tcgagttcaa ttatttatca gctgtaggtc gtgccttgaa tggacattgg 1380
gggttgcaga gcgaaggttt tttccagaat caatctgaaa tcgacgctgc tcctgatcag 1440
acatttggag atgtgaagcc aggcgatatt cgttataagg atcaaaataa tgataataag 1500
attgacaaca atgaccgcgt gtttttaggt cgttgggatt ctccttttat gagtggcgtc 1560
aaccttaccg ctaagtggcg taattttact ctctatgcga tgggtaacct ctacatgggt 1620
ggttacggta tgaaagacaa ttcatattat tgggtcaagg gagacggcaa atattcggaa 1680
gtggtgcgca atcgttggac accagagaca gcagctactg ctacctatcc acggttgact 1740
acaactgatg gtgccaataa tttccgtact tctgacttct ggatgtacaa gaatgaccgc 1800
tttaacctca ctcaagtaca actaacatac cagatgcctg aaaaagtgtt ggaaggaact 1860
ttcatcaaag gtttgagctt ttttgccaat gcaaacaatc tgcttaccat tgcgaaagaa 1920
cgaaaagctt tggaactgaa tgtgggcagt gcacctcaaa cacgcttcta ccaacttggt 1980
tttaagggaa ctttctaaaa aatgataaat ttaataatca agattgaaaa ataagaagtt 2040
atgaaactaa aaaatatcat cttatcagca gccttattgg tggcattctc tgcttgcgac 2100
gatttgtttg agccgacaat agagaaccat aaagatgctg cagatttgga aaacatgcct 2160
tcgtgggctg ttggcttgct gggtcatgcc tatatcagta acccgctggg tgaaaatgcc 2220
aactcttgga agtttactga tgtagctacc gatgacgccg tatccaatga tataggaaat 2280
tcttatctga agatagcaac aggttcttgg cgtgctgata ataatcccat ggacaattgg 2340
caatatttgc gtgcatcatg gcaatatctt aatcagtttt tggaaatctc cgaaaatgta 2400
gaatgggcta gagatcaagt cgcttccgac ctcttcaaaa aacgtttcca aggtgatgcg 2460
tatggtatgc gtgctctcta tatgtatcac ttgttgatga gttgtgccgg atggagtgct 2520
gatggacaac tattgggtat tcccatcttg acagagtcag aaaacataga ttcggacttt 2580
aataaacctc gtaacacgtt tcgtgaatgt cttgaccaac tgaatctaga tgtagacaaa 2640
gccattgaaa tgcttcccga agagtatggt gatgtagatg gtgcaaacaa tgtgcccgca 2700
aaatacactc agatgggtgc tgatcctagt cagtacaatc gcgtgtttgg tgaccatgcc 2760
aagaatcgta tgagtgctcg tgtggctcgt gctattcgtg cccaagcagc tctgctcgca 2820
gcaagtcctg cctacagcga agggtctgga gttacttggg aacaggctgc taatagcatg 2880
gctgaagtgc ttgccaatct tggtagtgac cctatagcga atattgatcc cacaggtaac 2940
aaatggtacg aggatgctaa cagtatcaag aatctggtag caggctacaa tcgtcccgaa 3000
ttcctgtggc gcagcaacaa gaatgaaagt gctgaccttg aaaaagatca atatcctcct 3060
tcactctttg gcaaggggcg tgttaatcct tcgcagaact tggttgacgc atttccatca 3120
ctgaccggtt acccgattag cgactcgcgg agtggatacg atgcacagaa tccgtatgct 3180
aatcgtgacc ctcgcttagc gctatatata atatataatg gtagcaaggc aggtacggat 3240
aagagtgtta ttacgactgc tttcgatggt agcaacaacg atgcaatgaa caagttcgac 3300
ggtgcttcta cacgcactgg ttactatttg cgcaagttct tggatatgaa cgtgaatgct 3360
aatcctagca acacaaccaa tggttaccat attaaacctt ggattcgtta caccgagatt 3420
tttcttggtt atgctgaagc agccaatgag gcttggggac cacaaggcaa aggaagtaat 3480

agctattcgg cttacgatgt gattaaagct attcgcaaac gtgccggtat cggtgaaaat 3540
ggtggtgatc cttatctgga atcagtgaaa ggcgataaag acaagatgcg tgaacttatt 3600
cgcaatgaac gtcgcctaga actttgcttc gaaggtttcc gcttttggga tttgcgtcgc 3660
tggaaggtgg atatcagcaa gttgaacgaa acggtaaagg gcatgaagat tacgggaact 3720
aaccatgaag tggtagatgt agaaaaacgc gactacaagg actatatgta ttacggtcct 3780
gtaccttata gcgaaatgtt gaagtttgat gctctcatcc agaataaagg atggtaacgt 3840
gtaaacgaat ttatataaac tgaataaatg aaaatagcaa tgaaaaaatc attcctaata 3900
atagccgcta tgatgggagc cctattcact gcatgtgaaa acggtgactg ggagttccct 3960
gattataagt attctgcggt ttactttgcc tatcaaagtc ctatacgcac gattacattg 4020
ggcgaggata agactgtgga caactcgttg gacaatgaac acaaatgtca gattatggca 4080
acaattggcg gtgtatacga gaacaagggc gatgtagaaa ttggtattcg cgtagataat 4140
tcgttgtgcg atggttacac ttttgaaagc acaggtggtg atgtagagcc tttgccggct 4200
tctcactata cattgagtgc tgacaagatt gtcatcaaga agggacaagt actaggcggt 4260
gtcactgtaa atctaaccga tgcattcttt aatgacccca aatcgacgag tttgaattat 4320
gtcatccctg tagtgatgac caatgttaag aacgcagact ctattttggt gggaaaacct 4380
tcagttgata ctccccatcg cattaaaaaa accgattggg atgtgcagcc taaagattac 4440
gtcctttatg ctgtgaagta cattagcaag tatgattcac actacctgcg acggggtatt 4500
gatacatata gtggtaataa aaccggtacg gaggttcgtc atgccgctta tgtagagaaa 4560
gatgagacta ttgacggatt cactactatg gccatgaata aagtgcagtg ggctcgtccc 4620
actcgcaata cagccggaaa gattattgaa agcaagctgt tgctgacttt tgatgaaagt 4680
ggcaactgta tcattacttc caattctcaa ggtgtagtag ctagcggtag tggtaagttc 4740
gttgttaaag gtgagaaaaa aagctggggt aacgaagacc gtgatgcact ttatctggat 4800
tataccatgg attatgacgg tattcattgt gctactaaag acaccttagt cgtacattat 4860
cgtggtgtga aagctgaatg gttcactcct ttgaagaaat aatgatcagg tttcttatta 4920
aaaaaagtat ttatatgaaa catacaaaaa atattttagg aacaatgctt ttaacagctg 4980
ctgctgttgc tgtcacttct tgtgtggacg acagcaagtt gctgtttgct tttgagaagc 5040
ctgcctctat tgcaggtatg gaatatctca acgagtatga cgcattgaaa acttatgtaa 5100
acagcagtgc taatcccaac tttaagttgg gtattgcact gactgctgat gactatattg 5160
caggtggttt ggtaacacgt ctagctaatt ctaacttcca ggaaatgacg gctggaaacg 5220
ctatgaaata tgcttcgtgc gtggcagatg acggcacgat gaattttgga accgtagaga 5280
attttgtagc tgctgctaaa gcaggaggac ttaccattta tggacacacc ttagcatggc 5340
atgcgcagca gaataataag tatttaaatt cgctgattaa agataaagag atagagattg 5400
atcccaatca aaagcttgat gtggtagaca agattgttga cttttctgcg atgacaaaat 5460
ttccatatta tgtaatgggc tatgagccgg atattgtaaa cggtatactc gtcagtaaga 5520
atcctggtag ttggtatcag tatttcgtgg cagatggtgt tcctaccgtt tctggtaagg 5580
aatacaaagt tactgccatg attaaatctt ctaaagaagg tgaatttatt gtcaaaatgg 5640
gtgattggga tgcatctact gataagaata tcaaggttac tactgaatgg aaagaacaaa 5700
ccattgattt tggtaccctc gctgctaaaa gctcgtttgt aatattccaa ccgggcggct 5760
ttgatggtga aattcaaatt aagtggatta aagtgagtca tccggaagct ccttctgtct 5820
ctttctatgt taatcagctt gataatagcg aaatgaaagc aggtggtaca atgaaaaatt 5880
tcgttgttcg cgaaaatggc aagggggata ttcccggtcc tatttttgaa ggacaaggac 5940
ctgatggtat aaactgtacg aagatcacgt ctattgctaa tcctgctaat gagtgggata 6000
ctcagttctt tatctataca ccggataaag aatggcaagg tggtcagaag tataaaatat 6060
ctttttggta caaagcttcc gctgaagctg attgttctac tcagtgccac ggagatcctg 6120
gtgcttacaa acatggggct atgcttccgc aaaatcctca cttcactact aaatggcaac 6180
attatgaagc cactggaaca attcctagtg agggttcagg tatgaaatcc atagctttca 6240
atcttaatgt aagcaaagat gcggtaacct actactttgc aaatatagtt tgggaatcgg 6300
aagaaaaagg taacaccatt cctcttactc ctgcagagaa aaaagacacg cttacatggg 6360
caatgaacaa ctggataaac ggaatgatga aagctactgg tggttatgtc accgcatggg 6420
atgtcgtgaa tgaagcgatt gctggtggag gtgatgacgg cgaaggtttt tatcctttgc 6480
agtcagcgac taatgttagt gcagatgatg ctaagaacaa tttctactgg caagactatt 6540
taggtagcga agactacgtg cgtattgctg tggctgctgc tcgcaaatat tatgccgaga 6600
atggtggcac caatcctttg agactcttta tcaatgatta caatttggaa tcagattggg 6660
atgacaacaa aaaggtcaag agtttggtac attggattga gaagtgggaa gctgacggtg 6720

tgacgaaaat tgatggtatc ggaacacaga tgcatgtgag ttgtcatgcc aatgccgaga 6780
cgcagaagag caaggaggat catgtggtga agatgttcga aattctggca gagagtggaa 6840
aactcgtgaa gattactgaa cttgatatgg gctatgtaga tgaggaaggc aatagtgtga 6900
agactgcgga tatgactcaa gcacaacaca aggcgatgtc tgaatactac aagttcattg 6960
tgaaaaagta cttcgagatt attcctgttg ctcaacaata tggtattact cagtggtgta 7020
tcaccgatag tcctaccgga tcgggttggc gcggtggtga accggtaggt ctctgggatg 7080
caaactacaa ccgtaaacat acctatgctg gatttgcaga cggattagca ggtaagtaag 7140
cctctctaaa tgattgtccg catcacgccg taatctgtgt ttccatctgt gtcggaaaca 7200
gtaggataat acagtttatt ggatgtcatg cggacaatta ataaagcgag cttctgtaaa 7260
aggaagatat gcaataaaaa accttttaaa aactataaaa tgaaacactt gtttaagttt 7320
tccgtctgtg cattgacatt tgccatgagc acagctaccg gaactgcaca aaatagcgga 7380
acgggattga aggatgctta taaagattac ttctctatcg gtgtggctgt taatatgcgc 7440
aacattacga cacctgaaca aatagcgatc attaaaaatg actttaatag tattacggcg 7500
gaaaacgata tgaagcccca acccactgaa cctgcttatg ggcagtttaa ctgggaaaat 7560
gctgataaaa ttgcagactt ctgccgcagt aacggaatca aactccgtgg acactgtttg 7620
atatggcact ctcaaatagg acaatggatg tatcaggacg aaaaaggaaa tcctgtgtcg 7680
aaagaaaaat tgttcgccaa catgaagcat catattacag ctatcatgga acgttacaaa 7740
gatgttgtat atgcgtggga tgtggtgaac gaggctattt ctgacggtgg atggcagggt 7800
agtagaccgg gtatgggaca gcagtctggt ccatatcgca attcgcctct ttatcagata 7860
gcgggtgacg agtttattaa gaaagccttt atatatgctc gtgaagttga tcctgatgtg 7920
ttactctttt acaacgatta taatgcggcg gacccaggaa agagagaccg tatttacaac 7980
atggttaaag ccatgaaaga agagggcgtg cccatcgatg gtattggtat gcagggacat 8040
tataatatct atggtccgag tatggaagat gtggatgccg cattgacaaa gtattctacg 8100
atagtgaagc atatccatat taccgagttg gacattcgtg ccaacgagga gatgggaggt 8160
caacttaact ttagccggga agctggtaat atcagtccgg tagtgaaaac acttcaggaa 8220
gatcaatatg cgcgtctgtt ccgtgttctt cgtaagcata aagatgtggt ggataatgtc 8280
actttctgga atctttcaga tcgcgattca tggcttggtg cacgcaacta tcccctccct 8340
tatgacgaga attacaagcc aaaacccgtt tataatataa tcaaggattt tgattcggca 8400
cttgacgttg ctgaagtaaa agaggatttc cgtccttcca cgcttaacca gcccggacaa 8460
cagtacccga tggttaattc gcaaggatat gcacgtttcc gtgtagttgc tcctgatgcc 8520
aaatctgtta tcgtcagtct cggattaggc ggccgtggcg gcacagtgct tcgtaaagat 8580
aaagaaggag tatgggtcgg aactacagat ggtccaatgg atgaaggatt tcattattat 8640
cacctgacta ttgacggcgg cgtgtttaac gatccgggta ctctgaatta ttacggctct 8700
tgtcggtggg agagcggtat cgaaattccg gctcatgata gtgatttcta tgcaatgaag 8760
caggtacctc atggaaatgt tcagcaagtt tatttctatt ctaaaagtac aaacgctcat 8820
cgtcgtgcgt ttgtctatac acctcctact tacggtaaag acaagaagaa atatcctgtt 8880
ctctaccttc agcacggatg gggagaagat gaaacagcat ggtccaatca gggatacgca 8940
aatttaataa tggataatct gattgctgaa ggcaagattg ctccgttcat catcgtaatg 9000
acgtatggca tgacgaatga tattaaattt ggacgtctaa atgaatttac cgccaaagaa 9060
tttgaaacag tattggtgga cgagctgatt ccttatattg atagtaattt tcgtacgctg 9120
gccgacaaga aacaccgtgc catggcaggg ctttctatgg gaggcttcga aacgaaattg 9180
ataaccttga gacgaccgga gatatttaac tattacggac tgttgagcgg aggtacatat 9240
gcaccaagcg acatcaaaga ccagaatcag gtgaaatcta tctttatcag ttgtgggagt 9300
aaagaaaatc cggatggtgt gactaaggcg gtgaacgatc ttaaggctgc cggattaaag 9360
gcgacatctt atgtttctcc tggtacagct catgaattcc tgacttggcg tagaagccta 9420
tatcatatgg cgcaagagct tttcaaataa gatatgaacc caatgaaatt tggattattg 9480
tatttcattg ggctcacaat aaagaaagta tttaatttaa aggttaaaac ataatatatt 9540
atttgtgctt tttatacttg gcctgcactg tctgtgaaga tcgtgcaggt ttttctattt 9600
caggtgttct tccatatatt ttgttggagg aacaccgtaa agctctttga aagcagttga 9660
gaaaatagtg ggattcgcaa aaccagttaa gtcagatatt tctgccatcg tataacgctt 9720
ttgagtaagt aaaatggcag cttgctttaa gcgggtattg cgaatgaaat ctctagtagt 9780
ctgattggta agctctttga gtttacgatg taaatgtact ctacttatgc caatctcact 9840
tgcaattaat tctactgtta gattaggatt acttaagttc ttattaatcg cattcataac 9900
acgctccatc aatttttcgt cgggagactg taacttaatc tcctccatct tatcttcgtg 9960

atcttgatta ccactgaaag tattgcgcaa tctttctcgg ctattgatca ggttttgtac 10020
tgtttttctg agaatttcaa tatggaaagg tttagtcaga taagcatctg caccaacttc 10080
caagccttca atattatctt cttctcttgt ttttgcagtc agcaagataa tgggaatatg 10140
gtttaaattt acattctgtt ttattttctt gcatagtgtc aaaccgtcca tttcaggcat 10200
cattatatca ctgatgacta catccggttt gcgtttgaaa agcatatcca aagcttcttt 10260
tccgttacag ctttccaata tgtgataatc ggcaaaaagt tcttgcttaa tgtatctgcg 10320
aatttcttca tcatcttcta ctatcaatac gcgatatttt gtttttttgc gaactttgag 10380
ttcttcttca tctccactct cttctatagc tggtattgtc tctgcaggtg tatatgcgag 10440
ctcattatca gtagaattat ctagctcttc ctgttttaaa tggctgcaac ccataggcag 10500
gcgaatgaca aaacggctac caataccttc cggattgttt tcggcgtgaa tatgtccgtg 10560
atgtaattca actaatgagc gagtaaggtg cagaccaata cctgtgccta tgtttgagtt 10620
atttaaatga ttgcgtattt gatagaatcg ttcaaatatg tgctcaattt cttctggtgc 10680
aattccgata cctgtgtcgg caattataat ttcggcataa ttgcttaaag tagtttctgt 10740
atcttcgcta tggccagatt tgaggtatat atcaacacgt ccaccttccg gagtaaattt 10800
aaaggcattt gaaagaatgt ttaagatcac tttatcaaag tttgtaggat caatccataa 10860
accaaatgta tcgaaatctt cacggtgaaa ggtcaatgaa atatgtttgt tgactgtttg 10920
cccagcaaaa gtattgcata gatcctgaat gaatgccacc atatctattt ctcgaaacat 10980
aagagacatt tgccctttat caattttacg gatatccatt aactgattga tgagtcgaag 11040
aatgcgttcc gcatttctat aaatagtatg atatgttttc tgtcttgaag gatcgccgtc 11100
agtagagatt agtttctgaa gaggactaat aataagggac attggcgtgc gaatttcatg 11160
tgaaatattc atgaaaaatt gcaatttggc ttcgttgatt tgctctgcat ggatgtgttg 11220
taacatttct tgtttggtgc ggtatcgatg tttggcttgc aatataataa accagattgt 11280
tccgcccgtc aacaatgcat aaatgagttt ggcccaccaa gaagcccacc aggccggatg 11340
aatatatatt gtgatttctt tcacgtctga atcgaccatg taatcttgta ctttcacctt 11400
gaagtgatag gttcctggcg acagattact gaatgagact ctattattac caattggtaa 11460
actaacccaa ttgttatcat tcattgcata taaataagtt atacgctcgg gattattcat 11520
ttctagtgtg gagaattcta tagtaaatga attatcactg tgcgaaagat caaatcttgt 11580
ggcttcaaat accggttcat taataatttc tcttcctgcg gataatacac cttgtcttac 11640
cggttgatta tgtaaataga agtcagtgat gcgtatattc cactttttgg cggggttagt 11700
aatctcttga gggttgaaat aggtaatgcc atcagttcct ccaaaccaga ttattccatt 11760
agcgtctttg aatgaggcgt tcttactaaa ttcattcccc tgcaatccat cacctacaaa 11820
ataattgata aatttatggc tgcttcggtt atactgtgaa atgccggcat ttgtacttat 11880
ccataaatta tcttgttcat cgccaacaat agcataaaca gcatcgctgg gtaacccatt 11940
ctctgtggtg gatgttacta attgttgatt ctttggattc caactagata atccatcgga 12000
tgttcctatc caaataatgc ctttttttatc ttcataaata gaaaatacga tgcgacgagt 12060
aagcaccttt ttagttataa actcatctga gtttaagttg atgcaatata caccatcgta 12120
agtgcctgtg tagagtttat tgtctttaga ataaagcaaa caattaatcc atttgttgtt 12180
cttgttggcc ttttcgggat actttagttt attcgttttg agctcataat agtacagacc 12240
agctcccata gtagctatcc aaagcctttt atttttgtct tcaacaaaat catatacacg 12300
ttgaactcta ttcccttctt gatccagtag ctcttttaaa taaatgcaac ggcctgtttc 12360
acgatcaagt ctgcacattc ctttagtaaa agaaccgatc cacatatttc gttcggaatc 12420
ttcatagagt ccgaatacag tagacggtac attctcttca ctatctgtat ctgccggata 12480
gaaatgtttt ttttgtttta gttcttgtgt aataccatat atgccatcat tgtctgtccc 12540
tacccacaag gtaccttcat ggtcacggca aagagacgtg acactatttg aaccaatgat 12600
atttttatct atggccttat ttccaatata cttgaaacta ttaggttgtg cgggaatcat 12660
cattacccct ttttgataga tggcgagcca gaaattaccg gtattgtctt ttagaataga 12720
gtgtactttt gatttgttgg aatcaaaata actattgtca aatggtgcat ccgttatctt 12780
ctgctgttta atgttataca ctttcattcc tttcccatct gttccaatgt ataactcatt 12840
ttggttgtcc gggtatagcg attttatagg taagttctgt tttcctttgt atggaatggg 12900
tttaaatgaa tcacttaatt gatcatatac aaaaagccct ttacccattg tacctacata 12960
tatattgcca ttcagatctt cgcagaaatc aacgatagtt attccctttt catcttttaa 13020
ataatgagtc actttcttct tatccgatat tttgtatatg ccattttctc ctttagttat 13080
ccatatattc cgctttttat cttcaatcat ataatctgtt gaggtggtag ctatgggcaa 13140
attgagcatt ttcacaacca gtttttcttt ctttataatc agtttacata aaatatttcc 13200

tgatacccat atttctccat tttgtcgttc taaaatggca ataatgttgc tgtcaaacag 13260
atccccgttt tcccatctgg cttgcggtgt aaaacgatca gttgccggat cgtacatttg 13320
aataccagta taagtaccga taatcaggtt tcctttgcta tcttcaaaca gcgtgcgaac 13380
ataattgtga cagagtgaat gttcgtcatt cttcttgtgt ttgtaaacgg taaatttgac 13440
tccatcgtat ttattcagtc catcttcggt ggcaatccat atcataccat ttctatcttg 13500
atatattttg ttgatcaagc tgcttgacaa tttactattc gtggtgaaca gtttgcatga 13560
ctgactaaaa gccccgattg aaaagaatag aataaagaag atacttaatg tgtttctcat 13620
attgataata tctgtttgtt actgcaaaat taagcatatt ttcaaataaa aaaaggattg 13680
tctccgctaa tgtaacagag aaaccgtatg ttgtaacatc ttcaaaaggt tatgtaacat 13740
gatgtggctg ataagtcgaa acagtacact aatattgttg ctgtaaagaa tggaagaaga 13800
acacatttat ttcctgccca ttcatttact caatttgcca taaccatagg tcagtaaact 13860
aattgtgtta tataatactt aatctaatat gaagatcaac agatgtcaaa taagtttttt 13920
tactatcgga tttattggtt acatagtatt aagtggattt cagaaagaga cagtctacat 13980
gaatgaaaat cacttgatag tgaatgattc tgactatttt gagacgcgtg gattaaacta 14040
ttttgttttc agtaacatat atgatgctac ttttgatgat tcgaaaatca gtgccgtaga 14100
gatcatacat catggtattc gtaccgcaac gaatggtgac gtgcgattga accccactcc 14160
cggacaatgg gataagctgc ctcgattcat tgaccgtaag gttgatcaaa aaaataagag 14220
aattgaggtt tcattaagct acccggagta tgcatttgat tataaattgg ttgcagaagc 14280
taaggatgga ggcctatatc ttagcgtata tacggagaaa gcattacctg ataaattgtg 14340
tggcatagcc ggacttaatc ttgaatttgt gcctccagtg ttttggggac actcatacat 14400
acttgatgac atacacggtt tgtttccaac ttctccggca gatttcatga ccactattca 14460
aggaactgtg gagccggaac cgatagcaac cggtagcaaa atagagatag cgcccgatga 14520
tcctgaaaaa cacgtgtcta tacgtacaac agatggtaat agtctcatgt tgtttgatgg 14580
acggaacaaa caacaaaatg gaaattttgt ggttcgaaca ttacttccgg gaaagaagac 14640
gggaaaaatt gccgaatggt ttattcaggc tgaaacggat acccagtgga tacgcaaacc 14700
gcttgttgct tattctcagg tgggatatca tccggctcaa aagaagatgg ctgtaattga 14760
attggataaa aatgatgaag tatcacatga agtcacattg ttgaaagtag gcagcgacgg 14820
aaggctgacg aaggtattat cagcaatgcc tgcagattgg ggaagctata cccgttataa 14880
gtataaacaa tttgacttta ctgcaataaa tgaaccaggt atctatgttt tggaatatga 14940
caaactgcgt accgctcctt ttcccattgg aaaaaatgta tttcaaaaag catggttccc 15000
cacgttagat gtattttttc ctgtccaaat ggatcacatg tgtgttcgag aagcctatcg 15060
aatctggcat ggagcgccac atctcgacga tgccttgcaa gcacccctta accgtaatca 15120
ttgggatgga tggcaccaag gacctactac cgacaataaa tataagccgc tggaacacat 15180
tgaaggttta aatgtaggag ggtggtttga tgcgggcgac tttgacatcc aaacaccttc 15240
tcagcaatct gttgttcaga cttttgcaga tctttggtca gactttagag tttctcgtga 15300
tcaaacaagc atcaatcaac aaacccgtta cacagaaata catgttcccg atggcaaacc 15360
tgatttgttg cagcaattgg aacacggagt tttgcaacta ataggacagg ttaatgctgt 15420
aggatatgcg atacccggca ttaccgagtc acatttgtat cagtacagac atttgggtga 15480
tgcagtgaat aaaacggata ataaggtata taatgctaat ttagatagtt tgcaaaccga 15540
cggaccaacc agcggtacct ttgatgatcg ttgggcattt actaatagaa atccgtatct 15600
gaattacggt acagctattt ctttggcagc tgcagcccgt tgtttgaaag agtataatcc 15660
acaattaagc atggaagcct ccagagtcgc tcagtttatt tggaaagacg aacattcacg 15720
tcccactccc gaaaagattg aatttggtgg gcggagtttt aatagtgctg atttcatgaa 15780
tagcgtggaa tgcagagccg cttttgagct ttggcgtaca gtaggtgata ctgtctataa 15840
aaacagaatg caaaaactgt ttccgattct tttgacgcaa tttgatagaa acgtttctgt 15900
cattgcaaga atattacctt ttatggataa tcagataaag aaacaggtgg aacctatggt 15960
gcaagcgtat gtcaaacaag tgacagaaca aaataaagaa aacccgtttg gagttcaaat 16020
cagtcgtttc tcttgggcgg gaaatcacgg aattgttcaa gaagccatta acgcttacca 16080
acttcatctg ctatatcctg atatcgttaa accagatttg atttaccgaa gccttgatta 16140
tcttttcgga tgtcatcctt ttcataatct ttcgtttgtg tcgggtgttg gagcgcagcc 16200
caagaaagtg gcgtacggca ataaccgggc cgattattcc tttattcccg gtggagttgt 16260
gcccggaata cgtattgtaa agcccgactt tcctgaaaat agagatgatt atcccttctt 16320
gtggagcgaa aacgaatacg ttatcgattt ggcggccagt tatatttatt tggtgaatgc 16380
agcagatcat ctattaaaca atagcggcaa atagttatgc agtctatttt ataaattata 16440

ccaaactcaa aattaagtat ggaaaaaaga ttaattcttt tagtaacgtt tctctcggtg 16500
gtcatgattg ctatggctca acctgtttct gacagtgata cttcagggag tgtacgttca 16560
gtgtattgtg tcagaggtgc acaataccct cggattacac cggatcatcg tgctcagttt 16620
aaagtagtag cccctgaagc taaattggtg caaattgatt taggcaaaaa atacgatttg 16680
gttcgtaacg aaaaaggtga gtggttgggt accactgaac cgttaggccc gggatttcat 16740
tattactttc tcattgttga tggcgtacgt gtggccgatc cttccagtga gtctttctat 16800
ggttgcagta tgcacaccag tggcatagag ataccttatg ccaaagatgt gcttcaatat 16860
gaactggccg atgtacctca cggcgaagtt cgtatgaaga gatatttttc taaaacggcc 16920
aatgcctgga gacgcatgtt tgtttatact cctccctgct acgaaaagag cgaaaaggaa 16980
tatccggtgc tgtatcttca acacggtggc ggtgaagatg aaagaggatg ggcgttgcaa 17040
ggccgtaccg atatcattct cgacaacctt attgctcaac agaaagcagt tcctatggtc 17100
attgtaatgt gcgatggtaa ttctcaagac ttcacggcgg aacttttaaa tgattgtatt 17160
cctttttgtag aagaaaacta tcgcgtgaag aaagatcgct ctcatcgtgc tttagccgga 17220
ctttccatgg gaggtattca cacattaaat accgctattc ctcaccctga actgtttgct 17280
tatgttggtg tctttagctc cggatggtgg gcgaaggctc ctatgcccgg cttaatgaag 17340
atggatacgg aaacatatta ttcacttctt caggaaaaaa aggacgagta taatgctagt 17400
tttaaagaat tctggatatc gatgggaggg aaagaagaca ttgcttacga aaactgtcgg 17460
atcatgagag aacgttttga taagataggc atcaaatata cgtattatga aaccaatgga 17520
ggacatacat ggcctgtgtg gcgtgaaagc ctttatcagt ttgctccgct tttatttaaa 17580
taaaaaacct tctatatgaa tactagaaaa atttggttga ccttttttatt atgcctgatg 17640
gttcatttgt cgggagctca gaatcccatt atccggcatc aattctcggc cgaccctacg 17700
gcaagagtgt ttgatggaaa agtatatgtt tacccgtcgc acgacattcc cagtcccgtt 17760
gaacggttaa aggaatggtt ttgcatggca gattatcacg tgttttcttc caacaatctt 17820
gttgattggg aagatcacgg agttattctt agtcaggaaa atgttccttg ggtgaatgcg 17880
gagtcatatt ccatgtgggc tcccgactgt gtctttgcca acggaaaata ttatttctat 17940
tttccagcgt ctcctaaagg agaaaatcag cgagggttca aagttggcgt agccgttgcc 18000
gataaaccta ccggtccttt tactccattg gctgaaccta ttaaaggcat aaatggcata 18060
gacccttgcg tgctgattga taaaaatggt gaggcttata tttattggtc aggaagagga 18120
atgtatgttg cgcgcttgaa gagcaacatg acagaactgg cttcggagcc tgtacagatt 18180
aaaaatttgc ccgaaggttt tatggaaggc ccttttgcct ttgaaagaaa cggaaagtat 18240
tatttcactt tcccgtgggt gcaagataaa accgagacgt tggcctatgc catgggcgac 18300
agtccgatgg gaccgtttga attcaaaggg ctgatcatgg atcagtcggc cacgggctgc 18360
tggaccaatc atcattcttt agtagaatac aacaaacaat ggtatttgtt ttaccatcac 18420
aacgattatt ctcctgcttt tgataagaat agatccgtac gcgttgattc tctttctttc 18480
aacgcagacg gaacgatcaa gaaggtagtg cctactttac gcggagtggg actcacgact 18540
gcaagcagta aaattcaatt agacaggtat agtcaaatca gcaaccaggg tgcagcgatt 18600
gctttcgttg atgaaaacaa taaatttgaa ggatggaaag ccgtctttac taagcccgga 18660
gcttggttgc gatataacag agtcgatttt ggcgacggtg gctatcgaaa aatgcaaatg 18720
agagtaaact cctctaccgg gggtgtggtt gaaataagaa cggcagataa ggctgccaaa 18780
ttgcttgcca gtctggttgt tccgaagtcc gacggatgga ttgaaaagga gtatgatcta 18840
aaacttgctt tacgaaacgt gcacgactta tctgtcagtt tgaaaggaga aggacaagtt 18900
gaaatagact ggatgcgctt tggtcaaaac gccggtgagt ttgccgtaca aagcagggct 18960
tccataaaac catgggagca gggagccttc gaaacccgta aatatcgaaa tttatttgct 19020
gaggccggat acaaacaagc cgatattgat gccaaattaa agtcggtttt caatgacatt 19080
ttttacgggc ccaataaagt ttattttgag acaaatgatt caatggccta cgtcagtgac 19140
ataaaaaacc acgatgtgcg caccgaaggt atgtcttacg gattgatgat tgccgtacag 19200
tttaaccgaa aagatatttt tgatcgtttg tggcgttggt gcaaaaaata tatgcaacac 19260
caagaaggtc cgctcgaagg ctatttcgcc tggagttgca agaccgacgg tacgcgtaat 19320
gctcagggac cggcttccga tggcgagtta tattacgtca cttcgttgat tttcgcttcc 19380
aacagttggg gcaatgactc ggacattaat tacttaggag aggcacaaca catcctcaac 19440
tgttccatga agaaagacgg aaccaatcgg gttatgcctc ttattaatat ggagcacaag 19500
ttaataacct ttgttcccga tacttttgga ggacgattta ccgacccttc ttatcacgtt 19560
cctgcctttt atgaagtgtg ggcccgctgg gcaaacgacg gaagagccga cttctggaga 19620
gaatgtgccg cgcgtagccg agaatatttg cataaaagta ttcatccggt aacgggactc 19680

aatcccgatt acaataacta cgatggcagc cttcttaaca tgaaacgcgg aattataggc 19740
gatgctttcc gctttgactc gtggcgtgtt ccgatgaaca tcgccttgga ttattcgtgg 19800
gcttgtgcag ataaagaatg gcagcagggt tacggcaata aaatacagaa cttcttgtac 19860
agtcaaggta ttgatacatt tgtagatcaa tacaatgtag atgggacaac tgttgcagaa 19920
atactagacg caggagggta taaaaagtta agacactctc tgggactggt tgccactgcc 19980
gcagcagctt cgctggttac tacgcacaca aagagttatg agtttgttga taagttatgg 20040
aatgctaaac atgaacctta cgaagacggg tattttgatg cctactatga tggtttgtta 20100
cgtttgtttg ctttcatgca tttgagtgga aactaccgca ttattttccc ccaatgagtt 20160
tatgcattaa gaggtgagca tttgcccacc tcttaatcta tatataataa atgttgatcg 20220
ttacttcttg gctacaccct ctttggtttg atttacataa atcatcttac catcgggtgc 20280
atactgcaat tcatccacac acacactgcg ccttcctatt gctccggcac gtccgttaat 20340
ggtcaacgta gcattgtgat aaaacaaata ccatttctta ttaaattcaa tgattcccgg 20400
gtgaattgta aagctgtttt cagccattcc ggtcagctca ccttgtggag tccacgggcc 20460
tttcatggaa gaagccgtag cataagcaat ggtttccttg ccttttccca ttgaagcgta 20520
agtgaggtaa tacatgtcgc cacgtttgtg caaccacggt ccctcaacat atctgggcac 20580
attaatcacc tcaatatcac catccagctc aatcatgttg ctttttaatt tggcaagaaa 20640
acaagttccg ttgccccagc ataaccacgc ttgcccgtca tcatcaatca gtacggtagg 20700
atcgatgtca ttccaccagc cgcgtgcacc gtttgaagtc atatcatcgg tcaccaaagg 20760
cttgccaatg gcatcaacaa aaggaccggt aggactatcg ctcactgcaa ctccaaccgt 20820
tttactgtta taaggagcac ctgcctgaac ggtcgtataa taatagaact taccgtttct 20880
ctccactacc tgcgacgccc aagcttcgcc cactgcccac ttaaaatctg tcggcttcaa 20940
aactgatccg tggtccgtcc agttttgcat atcgtctgtt gaataacaaa gccattcggt 21000
aatgttaaac tcttgcccgc ccgaggccgt atcttgtccg tccctatact catcgtgccc 21060
tacgtataaa tagagtcgcc cgttgtgtac catcggggca ggatctgccg taaatttgtt 21120
tttaattaaa ggattgcctg tgtttttaaa tacaggaaga ctgttttgtg cactgcaact 21180
catcaatatg agtatggcat tgactagaaa taagttgatt cgtttcataa aatattatgt 21240
ttatggttca ctggaacaaa tataggttct ttgtgattca taaacccctt ttgaatgata 21300
cggatatgtt tgtacatgta acatcctgcc tttttttatc ccatcagaca ctttatctct 21360
tcgttgacgc tgtacttatt cattttctca tgcaaactaa gtgatttctc ctttattgac 21420
agttcgtccc cgctacccgt gacgaatggc accgctagcg gggacgtaag gcaccgcatg 21480
ctgtgacgaa tggtttccat acctgtgacg aactaattat aaggagcaat caacttagtt 21540
tgtgctaaaa gagagagtag ggaatgcttt atttttaaat aggaagtgct gggtaaaaga 21600
agagattgta tctttgccgg gaatttaaat cagatgattg gtctaaagat tgtttcattt 21660
taaaatagaa tatatgttca gagagatgcg ccgtaagaaa cagttgttgt cggctgccga 21720
cacaatagcc gtactaaacc gcatgacatc cggagtgctg gcactcttgg gtgataacga 21780
ttacccgtat gcagtgccca ttagctatgt gtatcacgaa aataccctat actttcacag 21840
tgcgctggcc ggtcacaagg tcgatgccat tgctcaaaac agcaaagcct ctttttgtgt 21900
cattgagcaa gaccagattg tgccagaaga gtacaccacg tatttcagaa gcgccattgc 21960
ctttggaaaa ataagcatgg tgaatgacga tgttgaaaaa cgaacagcca ttgaactact 22020
ggccgaaaaa tattctccgg cgcaggaaga agggcgtttg aaggagatag aaaaaggatt 22080
tcatcacatg tgcgtgatca aactcgacat agaacaccta accggtaaag aagccgtgga 22140
actggttcgt gccggtagta aataaaaata agaaaggatt atcgatttat tttaagctct 22200
tcttgaaatg agccaaggtc agtaaatcgg tcattgcttc atcgttatgc acatcaataa 22260
tcagaggaat taatttccct atgcttttgg cttgtgcaag ttgctgtttg agagagtcgg 22320
caatacttag cgcatcgatg ctttccaagt gtttttcggt gaaataaaag ctggttttga 22380
aatagtgttc ccaaatggaa agccaaaaaa cggtcttctt cttgttaacc accttgcaaa 22440
gccatgcttt tccgtcgtta tagtatcgcc attccatggc taagccgcca gttggggagg 22500
tgatgttttg aacgaatccc tgaaataaat tgtaatttgt ttcgcctaaa gcgtttcgca 22560
atacttcctc ggaaggaaat atatcgggat ctcttaataa catttgtgtt tccataaaga 22620
tgattctttg tttttgacgt ccaaagatag tttttttcaa tcaaatgatg tagttttgca 22680
ataaatgttt aatagtgatt tgacactgtt gccaagaata acagtcagaa atataaaaag 22740
aacatgaatc cgacaatgag tgtaaagaac acaactcctc aagaattatg gtcgaggcag 22800
cgcattagca gtctcgatgt ggattatgac ttttagaacc ccaagagaga gatgattcaa 22860
tcatttgcac aaatgagtaa aagttgcatt tttacagttg atgtttttaa aggaagatat 22920

```
     gactttgcgt ccgaaaactt ttcgcagctc tttggatacg atccgcaatg catcagcaac     22980
     atccggcagg aaggggactt gctggaggaa agaattcacc ctgacgaccg ggcgcaattg     23040
     cttgactatc aggtgtaacg cgggctatgc atcaattggt gctgacttat tgcggctctc     23100
     ctagtttatt tattggttct gtgtaacaca ggctaaggta tatgtaacag aactctttta     23160
     tgatgtggat gataaatccc tatatttgtg ccactttata atcttgttat cattaaaatc     23220
     atgagaaaaa tacatgtaat cgtctttttg ttgtttttgg cttttgccgc tcaggcacag     23280
     aatttaaaat tatggtacag caaaccggca caacattgga ctgaagctct tcctttaggg     23340
     aattctcgtt taggggtgat ggttttcggg ggtacctcgc gtgaggaact tcagttgaac     23400
     gaggagacgt tttggtcggg cggaccttac aataacaata atccgaaagg acgtgaggcg     23460
     cttcctaaag tgcgcgaact cgttttccaa aataaatttg ttgaggcgca aaaggtgatt     23520
     gatgagaatt tcttcactcc acaacacgga atgcgttacc tgactttggg cagtctcttt     23580
     attcatcaac ccggacacga gcaggctcgt aactattacc gggacttaaa tctggagaat     23640
     gccacagcca ctgttcgcta tgaagtgaat ggggttaatt atactcgtac ggtgttttct     23700
     tcattctcgg ataatgtgat catggttcgt ctggaagcag ataaatcaaa ggcgttgagt     23760
     ttttctcttt cttacgattc tcctttgaaa tcggccgtaa cggtgaaggg gcgcaaactg     23820
     ttggttcgat gtgagggagc agagcatgag gggattcctg ccgcaatgca tgccgaatgc     23880
     cgggttgagg ttaaaacaga cggtgtgcag aacacggccg ataagcagtt ggtcattacg     23940
     gatgctacga ctgctactat atatatatcg gccgctacta attttgtgaa ttaccgtgat     24000
     gtgagcgccg atgcttccaa aagagctgag aattatttga aagctgctgt tcgtaagcct     24060
     tacgagacat cattggcagc gcatgtttcg ctttataaaa agcaatttga cagggtttct     24120
     ttttctttgc ctgcaacgga acagtcgctg gcggaaacgg atacgcgcgt gttgaatttc     24180
     cacaagggga atgatttgaa tctggcggct cttctgtttc agtacggaag gtatttgttg     24240
     atttcttctt ctcaaccggg tggacagcct gctaatttac aaggaatttg gaacaatagt     24300
     ccgtatgcac cgtgggatag taaatatacg gtgaacatca acgcggaaat gaattattgg     24360
     ccggcggagg taacaaacct aagcgaaacg catgatcctc tttttaaaat gattgccgaa     24420
     ctggccgtcg caggtagtga aaccgcccga gttctttatg gtgcaaaagg ttgggttaca     24480
     catcataata cggacttgtg gcgggcttgc ggcccggtcg acggggcttt ctttggcatg     24540
     tggcctaacg gtggtgcatg gcttacgcaa catctgtggc agcattatct gtttacgggc     24600
     gacaaagatt ttttgaagca atattatccg gtatttaaag gtagcgctga tttcttt        24657
//
ID   XXX; XXX; linear; XXX; XXX; XXX;  XXX.
XX
AC   PRJEB67889
XX
DE   Uncultured bacterium DNA fragment from in-reactor enriched cow rumen, clone
ferm-13f22, contig-2
XX
FH   Key             Location/Qualifiers
FH
FT   source          1..4084
FT                   /organism="uncultured bacterium"
FT                   /environmental_sample
FT                   /mol_type="genomic DNA"
FT                   /isolation_source="Cow rumen"
FT                   /clone="ferm-13f22, contig 2"
FT                   /note="AZCL-arabinoxylan active clone"
FT   CDS             complement(1282..3024)
FT                   /db_xref="SEED:fig|6666666.127236.peg.1"
FT                   /translation="MKKIIIPVLLAVATLFSACSESLLDIERKGVVDVDSFYKTDADAE
FT                   AALVAAYQGFLWNICTKEGGNIYAPIHAVFNLCGDDMYAAGESFGDNDFMGAMNEFRYD
FT                   TGSEVVRNAFQNIYYAMYYANLVIDHVKDADSKTKKRCVAEARVLRAYEHLLLAIGWNC
FT                   PPRIDHVLAATDLPYNCDKDPENPMTHNELLEWIAKECDDAEADLDERESLTDKKGAVK
FT                   VTKGFANAVAGKARLFAGDYAGAKASLEKVIKSEKYKLVPPERYAENFHASGDANEEKI
FT                   FEANLQTADGISVWWDIVNRSTWMEANIWNWRGDHFKVSPSVAYSSIDGWGGCGVPQEF
```

```
FT                   GDEFYANDGDSPRFKASLISIEDVVYNLNYPAVVDSKTGKAWADMTLEEKKVCPEIGID
FT                   NRGVYGQSKWLPLKPICTPEDLVGPGQNIRVNNFVIMRYAEVLLMYAEACLQTGDTQGA
FT                   KDAINLVQKHAGSATISDEVNMEVLEREKKYELWLEGCRWADMVRWGHFDGVKTAGQRV
FT                   PTLYDAMSKDGESAHRFYTVYSYPAMEIRKTETGFKAGKHEYFPYPEAETSLNPNLTQN
FT                   PGWE"
FT                   /product="SusD, outer membrane protein"
FT   CDS             complement(3043..3864)
FT                   /db_xref="SEED:fig|6666666.127236.peg.2"
FT                   /translation="MLNRGIEIELGWKDHIGDLNYSINANMSTLHNEVTYLHSSLARQE
FT                   GQVGGVSGLNNKVRTAFEVGHKVWYFRGYKFAGVDSEGHATYYDKNGKVVNSVGDEDLQ
FT                   DIGAALPDVLYGFTINLDYKGFDLSIYGSGTIGNDIFSLLYNADRPKANCLSHYWKDSW
FT                   TPKNTNAKYPAASIVANDWMFWSSDASIFNGSYLKIKQIQLGYTLPAKISRKALIENLR
FT                   CFVSFDDFFTFSSYPGCDPETATTASAQNMGYDCGNYPTTKKIVLGLNITF"
FT                   /product="putative outer membrane protein, probably
FT                   involved in nutrient binding"
XX
SQ   Sequence 4084 BP; 994 A; 963 C; 1082 G; 1045 T; 0 other;
     aggtgtatag aaaccggaat cgacatcgtt gaaggtggac catgagcctt ccttcgcgag        60
     ggtaagatac cagccggcga agtagttggt agtatcatgg tactcgacag aagcctcgaa       120
     ggcatcgctg taacggtcaa cctggatcat ggactcggcc tttacaaggc ccacgccacg       180
     gagagtaggg gtgacaggat tgatggttcc gtcttcgttg aaggtgactc tgtcgattct       240
     taccgaacgg ttcttgtcga aattcggaga atagtcattg tgatggtaga aaagatacca       300
     ctctcctttg tattcgacaa aggaatggtg gttggtccag cagccggtag gagactcgac       360
     catgaataca cctttgtatt cgaacgggcc catcggattg tcggacatag cgtatgcgag       420
     agtctcagtg ccgttcttct cgcgtaccca tggataagta agataatagt ggccgttacg       480
     ctcgaaagcg aacgggcctt ctttctggcc tgcagggaag tccgcgtcga gacgggtcgg       540
     acggccttcg atagccttga ggtcgtcagt catctttgcc acgcagatgc ccatgccctg       600
     ccagtatagg taggcgctgc cgtccttgtc gacgagtatg caagggtcga tgccgaacac       660
     gcctttgatt gcttcaggtt caggggtgaa cggtccttcc gggctgtcgg cagtagcgat       720
     gccgattccg aatccgcggc cgttcttcgg ggcgttaggg aaagcgaaat aatacttccc       780
     gtccttatat acgcagtcag gcgcccacat cgaatatccg gccgggttgc cccaagggac       840
     attctcctgg ctgaggatta cgccgtggtc cctccagttg acaaggtctt cggaggagaa       900
     tacatggtaa tcctccatac agaaccactt tctctctgga gcgaccggag agatgatgtc       960
     atgggacggg tagagatata cctttccgtt gaacactctg gctgtaggat cagcagtgta      1020
     ctgattacgg atcacaggat tctgtgctag gcacgcgagc gaaagcagaa gtgccgggat      1080
     aattattagt ttctttatca ttgtatagag tgttgattat ttacaggtgt cccgagtcgt      1140
     cccggtctat cattagacaa ttatcagcgt gagaggttta gtcttttcaa gaaaaaaatg      1200
     ggcggatccc ggattggaga tccgcccaaa cacaactaac acaaacaaac taatatactg      1260
     tcttattcaa aagctgaagg actattccca acccgggttc tgggtaagat tagggttaag      1320
     agaggtttca gcctcaggat aagggaagta ttcatgcttt ccggccttga atccggtttc      1380
     agtcttacgg atctccattg caggatatga gtagactgtg tagaatctat gtgcagactc      1440
     accatcctta gacattgcgt cataaagggt aggaacgcgc tgacctgcag tcttgacacc      1500
     gtcgaagtga ccccagcgga ccatgtcagc ccagcggcag ccttcgagcc agagctcata      1560
     cttcttctct ctctcgagaa cttccatatt tacttcgtcg ctgatggttg cagatccggc      1620
     atgtttctga actaagttga tggcatcctt agcaccctgg gtgtcaccgg tctggagaca      1680
     agcctcagca tacatgagga ggacctcagc ataacgcatt atgacgaagt tgttgacacg      1740
     gatattctgt ccaggaccaa cgaggtcttc aggagtacag ataggcttga gaggaagcca      1800
     ttttgactga ccgtaaacac ctctgttatc gattccaatc tctgggcata ccttcttttc      1860
     ttcgagagtc atgtcagccc aggccttacc ggtcttggag tcgactactg cagggtagtt      1920
     gaggttatag actacatctt caatgctgat gagagaagct ttgaaacgag gtgagtcgcc      1980
     atcgttcgca tagaactcgt caccgaactc ctgaggcacg ccgcagccgc cccatccgtc      2040
     gatagaagaa taagctacag aaggagatac cttgaagtgg tcgccacgcc agttccagat      2100
     gttagcttcc atccaggttg aacggttgac gatatcccac catacgctga taccatcagc      2160
     tgtctggagg ttggcctcga agatcttctc ctcgttggcg tcaccggaag cgtggaagtt      2220
```

```
     ctcggcatat ctctcaggtg gaacaagctt atatttttct gatttgatga ccttctcgag      2280
     agaagccttg gcacctgcat agtcgcctgc aaaaagacgt gcctttccgg cgacagcgtt      2340
     tgcaaagccc ttggtgacct tgacggcgcc cttcttgtcg gtcagtgatt cacgctcatc      2400
     gaggtctgct tcagcgtcat cgcactcttt tgcaatccac tcgaggagct cgttatgggt      2460
     cataggattc tcaggatcct tgtcgcagtt gtaagggagg tcggtagctg caagaacatg      2520
     gtctattctt ggagggcagt tccagccgat agcgagaagg agatgctcgt atgcacggag      2580
     aacgcgggcc tcggcgacgc aacgcttctt ggtcttgctg tcagcgtcct tgacgtggtc      2640
     gattacgagg ttggcgtaat acattgcata gtagatattc tggaaagcgt ttctgacgac      2700
     ttcgctaccg gtatcgtaac ggaactcgtt catggcgccc atgaagtcat tatcaccgaa      2760
     gctttcacct gcagcataca tatcgtcgcc gcagaggttg aagactgcgt ggataggagc      2820
     gtagatatta ccaccttcct tggtacagat attccagagg aagccctggt aagcagcaac      2880
     aagagcggct tctgcatcgg catcggtctt atagaaagaa tcgacatcga ctacaccttt      2940
     cctctcgata tcgagcaggc tttcactgca ggcagagaag agagtcgcaa cggccagaag      3000
     tactggtatg attatttttt tcataatata ttcctattga aattagaatg ttatgttaag      3060
     accgaggacg atcttcttgg ttgtagggta gttaccgcag tcataaccca tgttctgagc      3120
     acttgctgta gtagcggtct cagggtcgca gccagggtag ctggagaatg tgaagaagtc      3180
     gtcaaaggaa acgaagcatc tgaggttttc aatgagagcc tttctggaga tcttggcagg      3240
     aagggtatat ccgagctgaa tctgtttgat cttaagataa gagccattga agatagaggc      3300
     gtcagaactc cagaacatcc agtcattagc tacgattgaa gctgccggat acttggcatt      3360
     ggtattctta ggagtccaag agtccttcca gtaatggcta aggcagtttg ccttaggacg      3420
     gtcggcattg tagaggaggg agaagatatc gtttccgata gttcctgatc cataaatgct      3480
     gaggtcgaat cccttgtagt cgagattgat cgtgaaacca taaagtacgt caggaagagc      3540
     tgctccgata tcctggagat cctcgtcacc cacactgttc acgactttac cattcttgtc      3600
     gtaataagta gcatgaccct ctgagtctac acctgcaaac ttatagccgc ggaagtacca      3660
     taccttgtgg ccaacctcaa atgccgtacg gaccttgtta ttaagaccgg atacaccacc      3720
     gacctgacct tcctggcgag ccagagaaga gtggaggtaa gtaacctcgt tgtgcagggt      3780
     tgacatgttg gcgttgatgc tgtagttgag atctccgata tggtctttcc atccgagttc      3840
     aatctcaata cctctgttga gcacgctacc agcatttatg gtagtactct tttcaccgat      3900
     gaagcaaggt ggtttgatag tagtaagaag gtctttggtc ttcttgtcat agtaaactac      3960
     ggatgcggaa agtctgtctc tgaagaagcg gccttcgata ccgaggtcga tctgctcgga      4020
     ggtctcccag gtaaccttag ggttagaaag gcctgaagga gctgaaccga gggtctgggc      4080
     gcct                                                                   4084
//
ID   XXX; XXX; linear; XXX; XXX; XXX;  XXX.
XX
AC   PRJEB67889
XX
DE   Uncultured bacterium DNA fragment from in-reactor enriched cow rumen, clone
ferm-14j10
XX
FH   Key             Location/Qualifiers
FH
FT   source          1..27156
FT                   /organism="uncultured bacterium"
FT                   /environmental_sample
FT                   /mol_type="genomic DNA"
FT                   /isolation_source="Cow rumen"
FT                   /clone="ferm-14j10"
FT                   /note="AZCL-Barley beta glucan, AZO-CM cellulose,Tween 20
active clone"
FT   CDS             32..625
FT                   /db_xref="SEED:fig|6666666.127237.peg.1"
FT                   /translation="MDGEEYNLAKISGKDLRKYRSRVGFVFQNFNLYGNKTALGNVTLG
FT                   LTVARKMDKKEATEIGKKLLDRVGLSDRVDFYPSKLSGGQQQRVAIARALATNPDVIFF
```

```
FT                   DEPTSALDPELTGEVLSVIKQLADDGITMIVVTHELEFARKVANRVVFMENGEIVEENT
FT                   AESFFSNPKTERTKEFICKSSVWTSKNQYEGSGI"
FT                   /product="Amino acid ABC transporter, ATP-binding protein"
FT   CDS             644..1156
FT                   /db_xref="SEED:fig|6666666.127237.peg.2"
FT                   /translation="MSVISLIDIDNASDEVKEAVKKHLDDGHTITNEKRTLLHNVPAFW
FT                   ALEGKSYEMSAELKKFITPRAAALFEYAISVENDCLVCTTYYRKYYESLGIDIDHVEYT
FT                   PEEELVIEYARAIVKDRKHVPDELFDRLKDRLGEEGLIVLTTMGMFMIANNYFNDILHV
FT                   QSEFLQK"
FT                   /product="putative transmembrane protein"
FT   CDS             1179..1565
FT                   /db_xref="SEED:fig|6666666.127237.peg.3"
FT                   /translation="MYLKNVGIFVKDLEGAKKFFESYFGAKVLKTFDNPDKGYYSYILE
FT                   FEGQAWLELMTKPEIVDSPKDPNRTGIAHVCFKIDTRDKLNEIIAGFKKDGYKIQYEPE
FT                   LPEGPGEVRAVTFEDIVIEANYGY"
FT                   /product="Possible glyoxylase family protein
FT                   (Lactoylglutathione lyase) (EC 4.4.1.5)"
FT                   /EC_number="4.4.1.5"
FT   CDS             1754..3049
FT                   /db_xref="SEED:fig|6666666.127237.peg.4"
FT                   /db_xref="GO:0003961"
FT                   /translation="MSNHEFKFETKQLHVGQEQADPVTDSRAVPIYQTTSYVFHSAQHA
FT                   ADRFNLRDAGNIYGRLTNSTQGVLEERVAALEGGVAALALASGASAVTYSILAVAGAGD
FT                   NIVAQNSIYGGSYNLLDNTLPTYGIDTKFVDIHDHAAVAAAVDEKTKAIFIETLGNPGS
FT                   DIPDIEKIAEIAHSHKIPLIIDNTFGTPYLIRPLEHGADIVVHSATKFIGGHGTTLGGI
FT                   IVDGGKFDWAASGKFPRFTEPNASYHGVSFVGAVGPAAYVTYIRAILLRDTGAAISPFN
FT                   AWALIQGLETLSLRLERHVENTLKVVEFLANHPLVEKVSHPSLKDHPQHDLYERYFPNG
FT                   GGSIFTFDIKGGQKEAYAFIDGLKIFSLLANVADVKSLVIHPYDTTHAEMTPEQLAEAG
FT                   ISQATIRLSIGTEHIDDIIWDLEQGFEAVKAL"
FT                   /product="O-acetylhomoserine sulfhydrylase (EC 2.5.1.49) /
FT                   O-succinylhomoserine sulfhydrylase (EC 2.5.1.48)"
FT                   /EC_number="2.5.1.48"
FT                   /EC_number="2.5.1.49"
FT   CDS             3190..4122
FT                   /db_xref="SEED:fig|6666666.127237.peg.5"
FT                   /db_xref="GO:0004124"
FT                   /translation="MAKIYKSAIDLIGNTPLLELTHIEKKYNLKARLLAKVEYFNPAGS
FT                   VKDRIAKEMIEAAEADGSLKPGATIIEPTSGNTGIGLAAIAAAKGYKLIIVLPETMSVE
FT                   RRNIIKAYGAELVLTEGSKGMKGAIAKANELHEQIENSFIPAQFENPANPAAHYKTTGP
FT                   EIWNDTDGEVDAFIAGVGTGGTITGVGKFLKEKKSDVKIIAVEPASSPVLSTGTAGAHK
FT                   IQGIGAGFVPNTLDTQIYDEIITIENEVAFEGGKEVSREEGFQVGISSGAALKAGIQVA
FT                   QRPEFAGKTVVVLLPDSGDRYYSTDLFTN"
FT                   /product="Cysteine synthase (EC 2.5.1.47)"
FT                   /EC_number="2.5.1.47"
FT   CDS             complement(4174..5907)
FT                   /db_xref="SEED:fig|6666666.127237.peg.6"
FT                   /translation="MSFSERLDSYLKELGVSAKELSLESGVSQVVISRYRTGQRIPSPD
FT                   NRGVLEELAYGIEALSHKNDREPISAVDVLEAFDSDLSLLNVPFNYENFNFLIRELSLN
FT                   ASEMSRVMNYDPSYISRIRTGKRRPPDPEEFVSTLCKFIYPALGKNVSVTALKDLLSVP
FT                   ASEKFESYDSFCTLLKGYILNNDNISEDKNVSVFLKYVDEFDLNDYIRSVHFDEIKIPN
FT                   VPFTIPKSKVYYGPDGHKAAELDILKLIISSKSNEPVFFFSDFELSDLINDSEFTKKFM
FT                   TGLAMILKKGVKLQYVLNLDSTPKEILFALEALLPLYMTGQVEVYYLPERHNDIFKHLI
FT                   RVAGGVAMIGTGVSGDQKGYCFRTTTSKAFVAGVRQTADAIIANASPLMEMYMDYNQPE
FT                   FIEFLKINTYQKRPRYNKLASLPMCTIHEKTLKNVLKRNNVPDETISKEIEYLRGLKTM
```

```
FT                   MEVWIRDMEVVDEVPLVSKEDFEKCPLFMQLPGGPLPFDLPYTYEEYVEQLEASREFAK
FT                   KNKNYRLVENPNSTFRNINIIIYRGEFIWISKMRAPSVQLIVRNAELREAFEKMYFSVE
FT                   E"
FT                   /product="hypothetical protein"
FT   CDS             6107..7135
FT                   /db_xref="SEED:fig|6666666.127237.peg.7"
FT                   /translation="MREFKGYVHGVDLGGWFSQCDHSQDRYDNFIKEEDFATIASWGLD
FT                   HVRLPIDYELVEDENGGYRQDGFDRIRKALEWAGKYNLNTVLDLHKTYGYSFDVGYGES
FT                   GFFENEKYQERFYKLWEKIASEFGGYGDALAFELLNEVTKKEYCDTWNRIAKECIGRIR
FT                   AILPDVKILIGGYYNNSIVALKDLEAPYDENIVYNFHCYDPLVFTHQGAGWVDRMSLDF
FT                   RIPVDVTYGELAKASDEQTNGIGGDLKLLDPNEKLSSKYFDKCFAEAVAVAEERNVPLY
FT                   CGEYGVIECASAEDTLKWYEMIGESFNRFNIGRAAWSYKEMNFGLTAPELDSVRDKLIK
FT                   LL"
FT                   /product="GH5_37"
FT                   /EC_number="3.2.1.4"
FT   CDS             7463..8911
FT                   /db_xref="SEED:fig|6666666.127237.peg.8"
FT                   /db_xref="GO:0004049"
FT                   /translation="MTIEELKAFKDQGKYDIAPVSCEMLSDMRTPIQVLRILKNVSDHC
FT                   YLLESAKDNDNWGRYTFLGFDPKLEITCLNGKMKIGNESFDTDCPDKHIRELLSRYRSP
FT                   KVASLPPFTGGLVGYFAYDYIKYAEKSLHLTEEDEEQFNDIDLMLFDKVIAFDHLKQKI
FT                   IIIVNIHLADIDAEYARAQAELSKLKDLIENGKMCADTAGRATGEFTPVFDKEAYCAMV
FT                   EKAKHHIVEGDIFQIVLSNKLSAPFEGSLLNTYRMLRTINPSPYMFYFSSNDLELAGAS
FT                   PETLVKLQDGVLHTFPLAGTRKRGKDEEEDKVLVEDLLADPKELAEHNMLVDLGRNDLG
FT                   RISKFGTVEVEKLREVQFFSHVMHIGSTVRGEIRDDEDAMSAIGAVLPAGTLSGAPKIR
FT                   ACELIDELEGNKRGVYGGAIGYIDFTGNLDTCISIRLAYKKKGKVFVRSGAGIVADSVP
FT                   ESEFIECINKASAVIKAITQASEV"
FT                   /product="Anthranilate synthase, aminase component (EC
FT                   4.1.3.27)"
FT                   /EC_number="4.1.3.27"
FT   CDS             8913..9482
FT                   /db_xref="SEED:fig|6666666.127237.peg.9"
FT                   /db_xref="GO:0004049"
FT                   /translation="MILLIDNYDSFSYNLFQLVGEINPDIRVIRNDAHTVEEIRQMKPS
FT                   HIILSPGPGRPEDAGVCIDVVKELSGEIPILGVCLGHQSICAAFGAVVTYAKTLMHGKS
FT                   SIVTLDTDSALFKGFGKTTEVARYHSLAADENTMPECLKVTARTDDGEVMAVEHKEFKV
FT                   YGVQFHPESILTPEGRKMLRNYLDVK"
FT                   /product="Anthranilate synthase, amidotransferase
FT                   component (EC 4.1.3.27)"
FT                   /EC_number="4.1.3.27"
FT   CDS             9707..14236
FT                   /db_xref="SEED:fig|6666666.127237.peg.10"
FT                   /db_xref="GO:0004355"
FT                   /translation="MDNNLSYDREHDACGIGAVVKIDGVPEYSVLDDALSIVEKLEHRA
FT                   GKDASGKVGDGVGIMLQISHSFFASALKKAGIKVGHAGDYGIGMFFLPQDELKRTFAKR
FT                   LFEVIADKEGVEVKGWREVPTNPKILGERAFDCMPSIWQCVVGRPENIQKGIDFDRKLY
FT                   VIRREFEQSSDDTYICSLSSRTIVYKGMFLVSQLRRFYDDLQSKEYTTAVALVHSRFST
FT                   NTLPSWERAHPYRFIAHNGEINTIRGNSDRMLAREETMHSALLESEKSKIYPVISASGS
FT                   DSAMLDNTLEFLYMNGMDLPLAMMALIPEPWKHNRFMSESKKDFYHYYATMMEPWDGPA
FT                   AILFTDGELFGATLDRNGLRPSRYYVTDDNRLILSSEVGVLDIAPEHIVEKSRLTPGRM
FT                   LLVDMRKKRIISDEECKEFYASSNPYGEWLDAHLIRLKDLTIPNHQVPVHTQPERDRLY
FT                   KVFGYSYEEVKTYILEAAKNGAEPTVSMGHDVPLASLSDNYPLMYRFFQQQFAQVTNPP
FT                   IDSLREKIVTDTTVYIGSDGDLLCEKSDNCTVLEVNNPILTGVDVMKIRSMKRKGFKTE
FT                   TISLLYYKHTSLSRAVEQLYLSVDRAVSQGANIIILSDRGVDENHVPIPALLAVSAVEQ
```

```
FT                   HLVHTKKRTAVSLILETAEVRDVHQTACILGFGARAVNPYLAHECIGELINLGLLDKDL
FT                   HTAIDDFDKALLSGVVKIAAKMGISTIQSYQSARIFEAIGLSDEVIRDYFAGVASRVGG
FT                   IGVKELEEGISYRHDRAFDPMGLPVDPVLDSMGYHNLRSGNGKEDHLYSPKVIVLLQQA
FT                   VRTGDYEKFKEYTSLVDDAKHPHTLRGLLEFVPAGKSIDIDKVEPVSEIVKRFQVGAMS
FT                   YGSLSREAHETIATAMNRLGGRSNTGEGGEASDRFGTERNSAVKQVASGRFGVTEEYLL
FT                   SANEIQIKMAQGAKPGEGGNLPAAKVYPWVAKTRCSTPGVQLISPPPHHDIYSIEDLAQ
FT                   LIYDLKNANRNARISVKLVSEAGVGTIASGVAKAGAQVVLISGYDGGTGAAPYSSVHGA
FT                   GLPWELGVAETHQALIESGLRSRVRIETDGKLMTGRDVAIAALLGAEEFGFATAPLVSL
FT                   GCMMMRVCSKDTCPVGIATQNEILRKRFAGKPEHVINFMTFAAMQLREIMASLGFKTVD
FT                   EMIGRTDCLKKRASFDQPHAHTISFDAILKPEYAHIEENRFDPKQAYDFELEKTLDVNK
FT                   LLPWFEKASKTKSASSKVSVKVSSTDRSLGTILGSEIVKKYGEGCRGLAEERYVVECEG
FT                   GGGQSFGAFIPHGLTIKLKGDANDGFGKGLSGGKLVVTPPEGSTFEASKNIIVGNVALY
FT                   GATSGKAYITGVAGERFCVRNSGATAVAEGCGDHGLEYMTGGRVVILGSTGKNFAAGMS
FT                   GGIAYVLDKDHSLYRRINKDMIELEEVSEKVDREELRTILEDYVKCTGSKLGKEILDNF
FT                   DEELLCFKKVIARDYQRMIRTIAKYEEEGASRDRAELEAFKEVAGTAV"
FT                   /product="Glutamate synthase [NADPH] large chain (EC
FT                   1.4.1.13)"
FT                   /EC_number="1.4.1.13"
FT   CDS             14253..15716
FT                   /db_xref="SEED:fig|6666666.127237.peg.11"
FT                   /db_xref="GO:0004355"
FT                   /translation="MGKQTGFLEYERTENEEIPAKERIKGYKEFHTFQTEETRRKQGAR
FT                   CMNCGVPFCQSAMSLSGMVVGCPLHNLIPEWNDAIYKGEDAHALARLLKTNRFPEFTGR
FT                   VCPALCEKACICGQYDDPVTVRDNELFLIESGYKNGWMKAQPPVKESGKSVAVIGSGPS
FT                   GLSAADALRKRGHKVTVFEREDRIGGLLMYGIPNMKLDKSVIARRQKLMEEEGIEFKTG
FT                   VKVGVDIKASELRKKYDAIVLACGAKKARALTAEGIEGTDGVYFAVDFLTAATKNVLGI
FT                   KKLPAHFDAKGKNVIIVGGGDTGNDCIGTVIRMGCKSVTAIEMMPKPPVERAANNPWPE
FT                   WPKTLRTDYGHAEAIEVFGEDPRVFKTTVTKVFSKKGKISEVELSEVYFDKGKLCTKKG
FT                   TERKVKCDLLLVAAGFVGAESEVPEDFGLDMTARGVVATPEGKYSTNVPGVFSAGDMRR
FT                   GQSLVVWALSEGKECAAEVDEYLMGYRNG"
FT                   /product="Glutamate synthase [NADPH] small chain (EC
FT                   1.4.1.13)"
FT                   /EC_number="1.4.1.13"
FT   CDS             15803..17020
FT                   /db_xref="SEED:fig|6666666.127237.peg.12"
FT                   /db_xref="GO:0004356"
FT                   /translation="MNYTSQEVKQFVEEEDVKFIRLAFSDVFGKLKNVSIMPTELDRAF
FT                   TEGIAIDAWSITGFDCFDRSDLFLHPDPDTLAILPWRPEHGRVIRMYCDITWPDGTPFE
FT                   CDSRTFLKGAIEAAENKGYTFRFGTEQEFYLFRNDLEGEPTDNPYDNAGYMDVGPADRC
FT                   ANIRREICITLEQMGIMPESSHHEEGPGQNEIDFRYCDALKAADDAMTFRSVVGAIANR
FT                   NGATAVFTPKPFKGKPGNGMHINFSVKKGNEDVNLEPVIAGILEHIAEMTLILNPIRES
FT                   YDRIGALKAPGYITWAKENRSALIRIPAAFGEYVRGELRSPDACANPYLAFGLLIYAGL
FT                   EGLEKGLKAPEPVNTNLSTAGEEVTKNLKKLPMSLEEAVSTASKGNFVVEKLPSEILAS
FT                   YLDLNR"
FT                   /product="Glutamine synthetase type I (EC 6.3.1.2)"
FT                   /EC_number="6.3.1.2"
FT   CDS             17041..17622
FT                   /db_xref="SEED:fig|6666666.127237.peg.13"
FT                   /translation="MQAERVYSVLLVSAAEKMNTRLATFFSGGGFEPVTVVDSVAKAQR
FT                   RLLNREYDLVIVNSPLPDDFGKKLAVDVVAESGGVSLLLVKSDMYDEISSSMIPRGVMV
FT                   AKRPLDDSMIGQLLNIMCSVCERLKGIRKKTMSLEDKMEEIRLVNRAKWALIKSCHMTE
FT                   EDAHRFIQKQAMDLCLSKKETAENILKTYG"
FT                   /product="Response regulator receiver and ANTAR domain
FT                   protein"
```

```
FT   CDS             complement(17661..18488)
FT                   /db_xref="SEED:fig|6666666.127237.peg.14"
FT                   /db_xref="GO:0008800"
FT                   /db_xref="GO:0030655"
FT                   /db_xref="GO:0046677"
FT                   /translation="MALLQVSYLSKALFRTVPLYVILPSDKIDYRTFGYANDRGKKFKT
FT                   LYLLHGLLGDCSDWVNGTRIRRWAEEKNLAVVMPSGDNAFYIDYPGTKNNYSTFIGKEL
FT                   VEVTRNMFPLSDKRDDTYIAGLSMGGYGALRNGIVYSDTFSHVAGLSSATHLFDGKEHD
FT                   AMLMDEKAIFGDMEKAAKTDLNPTVAFDNLLKENRPVPKFYISCGTEDGLFDANVEFKN
FT                   YLVGKGADVTWDEEKRGHDWDFWDSQIKKVLDWLPLEGENLGLGSGNVRVGED"
FT                   /product="putative esterase"
FT   CDS             18687..20168
FT                   /db_xref="SEED:fig|6666666.127237.peg.15"
FT                   /translation="MAKVIIVGGGIAGLSAGIYALKAGYEAEIYEKNPLVGGECMGWNR
FT                   KGYHIDNCIHWLTGTDPKTGLWEVWKTVGAIDEKTEYAKTSKFYSSRAEGREVTLWNDL
FT                   DKTEKELIAESPEDEEEIKKFIQHVRYAEDCVISSIKPLDMMGIRDYIEMGKGMANMPK
FT                   VMKEYGKINCFELGKRFKSPVIRKMMSDYLPGEYTAYSLIVSYATMTSGNGNIPMKGSL
FT                   AMSLRIAERFKEMGGVIHTATPVSEIVIKGKKTEGIKLEDGSFIPADFVISAVDTDFLF
FT                   GRLIDRKYMPKDLVKAYEKRDAYPVTTGFQVAYAIDDTFSGEDTLFFDCEPMQIGTRTF
FT                   SRLSVKNYAYDKSFAPEGKAVLQANLIQTDADYEYWASLSKEEYRAKKEELTSMITDRI
FT                   VKEFPELVGKIELLDSWSPMTYNRYCNAYHGSYMGFVTTVGNKQMRFKGVVKGVDNLYV
FT                   AGQWVMSPGGLPIAVVSGKFAVQRILKKAGKSIII"
FT                   /product="Carotenoid cis-trans isomerase (EC 5.2.-.-)"
FT   CDS             20178..20741
FT                   /db_xref="SEED:fig|6666666.127237.peg.16"
FT                   /translation="MTRKEQKEERRNAILMTALNLFVERGYYDTKIADIAAAVPMSTGL
FT                   LFHYFESKEELLLLELVKMGLQGPGAIGDVPGVPPDVYLTGFLKQIFAFAKEQPWVFNMF
FT                   VFMGQVRRAGMPEEARRLAQSVDAVTPTVKLIKKGQKDGVFHKGDADTMARCFWASVQG
FT                   IMEQMSADKNMKTPDPEWIVSILK"
FT                   /product="hypothetical protein"
FT   CDS             20871..22973
FT                   /db_xref="SEED:fig|6666666.127237.peg.17"
FT                   /translation="METHSVPDYFGELVFDDRQMRARLSSKIYASLRKTIDEGKKLDPK
FT                   VAEYVAEAMKEWALEKGATHYTHWFQPLTGVTAEKHDSFISPSPDGSVIMEFSGKELIQ
FT                   GEPDASSFPTGGIRATFEARGYTAWDPTSYAFIKDKTLCIPTAFCSYSGEALDKKTPLL
FT                   RSMEALNKQALRILALFGNTTAKNVRTSVGPEQEYFLITKEMYDQRPDLKFTGRTLFGA
FT                   KSPKGQEMDDHYFGVIKPRVQAFMNDLNKELWKLGILAKTEHNEVAPAQHELAPIYSTT
FT                   NIATDQNQLTMEIMQKVASKHGLVCLLHEKPFAGVNGSGKHNNWSMATDTGINLLSPGD
FT                   TPAENAQFLLFLCAVIKAVDDYQDLLRLSVATAGNDHRLGANEAPPAIISVFLGDELSA
FT                   ILDAIESDTPYKGTEQKIMKLGVDVLPAFPRDNTDRNRTSPFAFTGNKFEFRSLGSSNS
FT                   IACANIMLNTVVAESLKIYADRLEGAEDFEKTLHEMIRKTIKDHKRILFNGNGYDDAWV
FT                   KEATEVRGLCNYRTTPDCMPHILDEKNVKLLTSHGVFTEKEIESRCEIMLDNYCKQVRI
FT                   EANTMIDMALKQILPAITCYADKLSTGVIAKKSACEGIACEYEKSVISRLSVLADQIFS
FT                   RTGELEKAVLKLADAKDIIEESAMIRDTVLPKMGELRACADEAETLVAKECWPLPSYGD
FT                   LLFGVR"
FT                   /product="Glutamine synthetase type III, GlnN (EC
FT                   6.3.1.2)"
FT                   /EC_number="6.3.1.2"
FT   CDS             23031..24617
FT                   /db_xref="SEED:fig|6666666.127237.peg.18"
FT                   /db_xref="GO:0004066"
FT                   /translation="MCSIMAWCKKAADEKKFTAALDEMTSRGPDDGRVIKTEGGLLGFR
FT                   RLSIMGLTPEGMQPFELKGCSVVCNGEIYGFTPIRDELKSKGYTFKSDSDCEILLPMYF
FT                   EYGTEMFKKLDAEFACVIYDARTDEWIAARDPIGIRPLYYTFDKGGSIIFASEPKSLMP
```

```
FT                   LTEEHIYPFPPGHFYKDGKFTCFNNIAKPDKVIKDDLETVCANIKEKLIAGVEKRLIAD
FT                   AKVGFLLSGGLDSSLVCSIAARKSDKPIRTFAIGMNEDAIDLKYAKQTADFLGADHTEV
FT                   IITKEDVLEALPKVIHLLGTYDITTIRASMGMYLLCKAIHEQTDIRVILTGEVSDEIFG
FT                   YKYTDFAPSAEAFQAEAEKRVSELHMYDVLRADRCISVNSLEARVPFGDLDFVKYCMSI
FT                   DPEMKRNHYNKGKYLLRHAFEGGDYLPDEILMREKAAFSDAVGHSMVDHLKEYAEEQYT
FT                   EEEFVEGCKKYSHARPFTKESLLYREIFEKFYPGQSRMVKDFWMPNKEWEGCNVNDPSA
FT                   RVLSNYGASGV"
FT                   /product="Asparagine synthetase [glutamine-hydrolyzing]
FT                   (EC 6.3.5.4)"
FT                   /EC_number="6.3.5.4"
FT   CDS             24829..25860
FT                   /db_xref="SEED:fig|6666666.127237.peg.19"
FT                   /db_xref="GO:0003883"
FT                   /translation="MTKYIFVTGGVVSGLGKGITAASLGRLLKARGLKVAAQKLDPYIN
FT                   VDPGTMSPYQHGEVFVTEDGAETDLDLGHYERFIDEDLNKYSNLTSGRVYWNVLNRERR
FT                   GEYLGSTVQIIPHITDEIKDFVYSVGKKTEADVVITEIGGTVGDIESQPFIEAVRQISL
FT                   EVGRENSLFVHVTLVPFLHGSDEHKSKPTQHSVKELQGMGISPNILILRSDEPLEDEIF
FT                   KKIAHFCNVKEDCVIENITLPNLYEAPLMLEDSGFSQIVCRELGIEAGEPDLDHWREMV
FT                   DRIKNLKKTAVIGLVGKYVQLHDAYLSVAEALRHAGYENEGKGSDQVDRFREDHRRKCG
FT                   RDF"
FT                   /product="CTP synthase (EC 6.3.4.2)"
FT                   /EC_number="6.3.4.2"
FT   CDS             25844..26437
FT                   /db_xref="SEED:fig|6666666.127237.peg.20"
FT                   /db_xref="GO:0003883"
FT                   /translation="MEETFKGIDGIIVPGGFGDRGIEGMITSAKYARTHDIPYFGICLG
FT                   MQIACIEYARNVLGYEDANSGEFDELCAHKVIDFMPGQSDSINKGGTLRLGAYPCKISE
FT                   GSKMKEFYGKEEISERHRHRYEFNNDFKEAFEKAGATFAGMSPDGKLVEEMEITDKDFY
FT                   LGVQYHPEFKSRPDKAHPLFKGFVREACKRNGNS"
FT                   /product="CTP synthase (EC 6.3.4.2)"
FT                   /EC_number="6.3.4.2"
XX
SQ   Sequence 27156 BP; 8012 A; 5898 C; 6456 G; 6790 T; 0 other;
     tcttacaaaa gccgacagcg gaaccttgat catggacgga gaggaataca atctggccaa        60
     aatctccggt aaggatctga gaaaatacag aagcagagtc ggatttgttt ttcagaattt       120
     taatttatac ggcaacaaga ctgcactagg aaatgtaaca ctcggactta ccgttgccag       180
     aaaaatggat aagaaagaag ccacagagat aggcaaaaaa ctgcttgata gagtaggcct       240
     gtcagacaga gtggattttt atccttcgaa attgtcgggt ggtcagcaac aaagagttgc       300
     aatagccaga gcgttggcta caaaccctga tgtaatattt tttgatgaac ctacatcagc       360
     acttgatccg gaacttaccg gagaggttct gagtgttata aagcaactgg ctgatgacgg       420
     aataacgatg atagttgtta ctcatgaact tgagtttgcg agaaaagtgg caaacagagt       480
     tgtgttcatg gaaaacggag agatagtcga agaaaatact gccgaaagtt ttttcagtaa       540
     tccaaaaacc gaaagaacaa aggaattcat atgtaaatcc tctgtttgga cttccaagaa       600
     tcagtatgaa ggaagcggaa tctgataagt aagggagata tttatgtcag ttataagctt       660
     aattgatatt gacaatgcat ccgatgaagt aaaagaagct gtaaaaaaac atttggatga       720
     cggtcatacc ataaccaatg aaaaaagaac tctacttcac aatgtgcctg cgttttgggc       780
     gcttgaagga aaatcatatg agatgagtgc ggaacttaag aaattcatca ctcccagggc       840
     ggcggctctt ttcgagtatg caatatctgt ggagaatgat tgtcttgtat gtactacata       900
     ttacaggaag tattatgaat cactgggcat agatatcgat catgttgaat atactccgga       960
     agaagagctt gtgatcgaat atgccagggc aatcgtaaaa gacagaaagc atgtaccgga      1020
     tgaactgttt gaccgattaa aagatcgctt gggtgaggaa ggactgatag tccttaccac      1080
     aatgggaatg tttatgattg cgaataatta ctttaatgat attcttcatg tgcaatcaga      1140
     gtttcttcaa aaataacgtt ctgtaaagga gtatagatat gtatctcaag aatgtaggaa      1200
     tattcgtaaa ggaccttgaa ggtgccaaga agtttttgga aagctatttc ggtgccaagg      1260
```

ttttgaaaac ctttgataat cccgataagg gttattattc atatatactt gagtttgaag 1320
gccaggcatg gctggaacta atgaccaagc cagagattgt tgactctcca aaggatccca 1380
accgcaccgg tatagcccat gtatgtttca agatagatac cagagataaa ttaaatgaga 1440
tcatagcggg cttcaagaaa gacggatata agatccagta tgagcctgaa ttacctgaag 1500
gccccggaga agtacgtgca gtcacttttg aagatatcgt aatagaggct aattacggtt 1560
attgatttct atatatatga gtattcagga tttactgata aagcgggcta taagaaaaac 1620
ttatagcccg ctcttaattt tatgaattat acatcatttt gaaaaaggtg tatcttaaca 1680
tcatcaaaca aaacaacaac ttcttacaca gttaagaaaa gcatttgaaa gaaagtttca 1740
ggaggtacac attatgagca accacgaatt caaatttgaa accaagcagc ttcatgtagg 1800
acaggaacag gcagatcccg taaccgattc cagagcggtt cccatttacc agactacctc 1860
ctatgtattc cattcagctc agcatgcagc tgacagattc aatttaagag atgcaggtaa 1920
catctacgga agactcacca actcaactca gggcgttctc gaagaaagag tagcagctct 1980
tgaaggcgga gtcgcagcac tcgctcttgc atccggtgca tccgcagtaa cttattcaat 2040
ccttgcagtc gcaggcgcag gcgataatat cgtagcacag aacagcatat acggcggcag 2100
ctacaacctc cttgataata cgcttcctac ctacggcatt gataccaagt tcgttgatat 2160
ccatgatcat gcagcagttg ccgccgctgt tgatgagaag accaaggcaa tcttcatcga 2220
aacactcgga aatcccggat cggatattcc cgatatcgag aagattgcag agatcgctca 2280
ctcacataag attcctctta ttatagataa tactttcgga actccttatc tcatccgtcc 2340
ccttgagcac ggcgcagaca ttgtagttca ttccgctacc aagttcatcg gcggacacgg 2400
aaccacccttt ggcggcatca tcgtagacgg cggtaagttt gactgggcag caagcggaaa 2460
gttccccaga tttaccgaac ccaatgcaag ctatcacgga gtatcctttg taggcgctgt 2520
aggacctgca gcatatgtta cttacatcag agcaatcctt ttaagagata ccggagcagc 2580
aatttctcct ttcaacgcat gggcactcat tcagggactt gagacacttt cactcagact 2640
tgagcgtcat gtagagaata ctcttaaggt tgttgagttc cttgcaaatc atcctcttgt 2700
agagaaggta agccatcctt cacttaagga tcatcctcag catgatcttt acgagagata 2760
tttccctaac ggcggaggtt ctattttcac cttcgatatc aagggcggac agaaagaggc 2820
ttatgcattc atagacggtc tcaagatctt ctcacttctt gcaaatgttg cggatgttaa 2880
gagccttgtc atccatcctt atgacaccac tcacgcagag atgactcccg agcagcttgc 2940
agaggcaggc atcagccagg ctaccatccg tctttccatc ggtacagagc atatagatga 3000
catcatctgg gaccttgaac agggctttga ggcagtaaag gctctctaaa ttttgggagg 3060
agaaagagta tgtttacaag tggatttcga tgtagtacat atatttaata gataaagaaa 3120
aagcttcgac ctcgcaaaat actacattat gaaatttgaa ggatcacatg atccggtaag 3180
gagataatta tggctaagat ttataaaagt gcaattgacc tgattggaaa cactcctctt 3240
ctggagctta ctcatattga aaagaaatat aacctcaagg caagacttct tgcaaaggtt 3300
gagtatttta accccgcagg atccgttaag gatcgtatcg caaaggagat gatcgaggca 3360
gcagaggctg acggaagcct taagcccggc gcaaccatca tcgagcccac ttccggaaat 3420
accggtatcg gacttgcagc tatcgcagca gccaagggat acaagcttat cattgttctt 3480
cccgagacca tgagcgttga gcgcagaaac attatcaagg catacggcgc agaacttgtt 3540
cttactgaag gctccaaggg tatgaagggt gcaattgcca aggctaacga gcttcacgag 3600
cagatcgaga acagctttat tcctgcacag ttcgaaaatc ccgcaaaccc tgcagctcat 3660
tacaagacca caggtcccga gatctggaac gataccgacg gagaagttga tgcatttatt 3720
gcaggcgtag gaaccggcgg aaccatcacc ggcgttggta aattccttaa ggaaaagaaa 3780
tccgacgtta agatcatcgc tgttgagccc gcatcttcac ctgttctttc tacaggaaca 3840
gcaggagcac ataagattca gggtatcggt gccggatttg ttcccaacac tcttgatact 3900
cagatctatg atgagatcat taccattgaa aatgaagttg catttgaagg cggtaaggaa 3960
gtatccagag aagaaggctt ccaggttggt atctcttcag gcgctgcact taaggccggc 4020
attcaggttg cacagcgtcc cgagtttgca ggaaagaccg tagttgtact tcttcccgac 4080
agcggagaca gatattactc aaccgacctt ttcaccaact aattaaatga gaatctaaaa 4140
cgcacgcctc attcggcgtg cgtttttttgt tctttattct tctacagaaa aatacatttt 4200
ctcaaaagcc tccctcaatt cggcattcct tactatcagt tggactgagg gagctctcat 4260
tttagagatc catatgaatt cgcctctgta aatgataatg ttgatgttcc tgaaagtact 4320
gttgggattt tcgacaagtc ggtagttctt gttcttcttg gcaaattctc ttgatgcttc 4380
aagctgctct acatattctt catatgtata gggaaggtcg aaaggaagcg gacctccggg 4440
aagctgcatg aagaggggac acttctcaaa gtcttctttt gataccaaag gaacttcgtc 4500

aactacttcc atatccctga tccagacttc catcatagtt ttgagacctc ttaagtattc 4560
gatctctttt gaaatggttt catcgggaac gttatttctc ttaagaacat tcttaagggt 4620
cttctcatgg atggtacaca tgggaaggct ggccagctta ttatatctgg gtctcttctg 4680
gtaagtgttt attttcagga actcgataaa ttcaggctgg ttataatcca tatacatctc 4740
cataagagga gaagcattgg cgattatagc atcggcagtt tgtcttacac ctgctacaaa 4800
agccttggat gtggttgtgc ggaagcagta tcccttctgg tctccggata caccggtacc 4860
gatcattgca actccgcccg caactctgat gagatgcttg aatatatcat tatgtctttc 4920
gggcagatag taaacttcaa cctgacctgt catatataaa ggaagaagtg cttcaagcgc 4980
aaagaggatc tctttgggag ttgagtccag attgagaaca tactggagct ttactccttt 5040
tttgaggatc attgcaaggc cggtcatgaa cttcttggtg aattcggaat cattgataag 5100
atccgacaat tcgaagtccg agaagaagaa tacaggctcg tttgacttgg atgaaataat 5160
gagcttgaga atatccagct ccgcagcttt atgcccgtca ggtccgtagt aaactttgga 5220
cttgggaatt gtaaaaggaa cattgggaat cttaatctcg tcgaaatgaa ccgatctgat 5280
atagtcgttg agatcaaact catccacata tttaaggaag acggagacgt tcttgtcttc 5340
ggaaatatta tcgttattaa ggatatatcc cttcaaaaga gtacagaaag aatcatagga 5400
ttcgaacttc tctgatgcgg gcacgcttaa aagatctttt aaagccgtaa ccgatacgtt 5460
ctttcctaga gcgggatata tgaacttgca aagtgtgctt acgaattcct ccggatcggg 5520
agggcgacgc tttcctgtcc tgattcttga aatataagaa gggtcatagt tcatgactct 5580
ggacatttcg gaagcattga gagagagttc tcttataagg aagttaaaat tttcgtaatt 5640
gaaaggaacg ttcagaagag aaagatctga atcgaatgcc tcaagtacat cgacagcact 5700
tatcggttcg cggtcatttt tgtgcgacag tgcttcaatg ccgtatgcga gttcctccaa 5760
taccccctctg ttgtcggggg atggaattct ctgaccggta cggtacctac tgatgacgac 5820
ctgtgataca ccgctttcaa gggacaattc cttagcagaa acaccgagtt ctttcaaata 5880
tgagtccaaa cgttctgaaa atgacatagg cctatcctta gaaaaaaagt caatgacaat 5940
attggtaatt ccaatattat taagcatata ttacattata acgcacgtta aataaagagt 6000
tttttgaagc attttttttat ctatttttta ttggctgtcg ggaacaggat tttgatataa 6060
taatcgggta gttaaaaaac gtttttttcat aatatatggg agtactatga gagaatttaa 6120
agggtatgtt cacggggtcg atcttggcgg ttggttttca cagtgtgacc attctcagga 6180
tcgttatgac aacttcataa aggaagaaga ttttgccact attgcatcat ggggactcga 6240
tcacgtaaga cttcccatcg actatgaatt ggttgaggat gaaaacggcg gatacagaca 6300
ggatggtttc gaccgcatca gaaaagctct tgagtgggcc ggaaaatata accttaacac 6360
cgttctcgat cttcacaaga cctatgggta ttcttttgac gtaggatacg gagagagcgg 6420
ctttttttgag aatgagaagt atcaggagag attctacaaa ctatgggaga agattgcatc 6480
cgaattcgga ggatatggcg acgctcttgc ttttgagctt ctcaatgaag ttaccaagaa 6540
ggaatactgc gatacctgga acaggattgc aaaagaatgc atcggaagaa tccgtgcaat 6600
acttcccgat gttaaaattc ttatcggcgg atattacaac aacagcatcg tagcacttaa 6660
ggatcttgaa gctccttacg atgagaatat agtttacaac ttccattgct atgatcctct 6720
tgtcttcacc catcagggtg cgggctgggt tgacaggatg agccttgatt tccggattcc 6780
tgtggatgtt acatacggag aactggcaaa agcatcggat gagcagacaa acggaatagg 6840
aggggatctt aagcttcttg atcctaatga gaaactcagt tccaagtatt tcgacaagtg 6900
ttttgctgag gctgttgcag tcgcagaaga gagaaatgtt cctctttatt gcggagaata 6960
cggagttatc gagtgtgcat ccgcagaaga tactctcaag tggtatgaaa tgatcggtga 7020
gtctttcaac agatttaaca tcggaagagc agcatggtca tataaggaga tgaacttcgg 7080
acttacagct cccgaacttg attcggttcg tgacaagctg atcaaactct tataattttt 7140
ccgaaaaaga ttgcaaagag tttacacatg tattattctg aataaaatca tatataaacc 7200
gatgacgaag aggagtagca aaggatgctt gctcagagag agcccggggt ggtgtgatcc 7260
ggacagtaaa ggcttttgtg aatggacttc ggagggcatg gaggaaatac catgacgtaa 7320
ggcatacgtt agttgccggg gatatgtagg tatcccgcga gagcatgggt caaaccatga 7380
agcagggtgg caccgcgaat tgtagattcg tccctgacag tatagttgta ctgtcaggga 7440
tttttttttat ggaggaaagc acatgactat cgaagaatta aaagctttta aagaccaggg 7500
aaaatatgac atcgcaccgg ttagctgtga aatgctttcc gatatgagaa caccgataca 7560
ggttctcagg atactaaaga atgtatccga tcactgctat cttcttgagt ctgccaagga 7620
caatgacaat tggggaagat atactttcct tggctttgat cccaagcttg agatcacttg 7680
ccttaacgga aagatgaaga tcggaaatga aagctttgat accgattgtc ccgacaagca 7740

tataagggaa ctactttccc gttacagaag ccccaaggta gcctcacttc ctccttttac 7800
cggaggcctt gtaggatact ttgcatatga ctacattaag tatgcggaga aatcccttca 7860
ccttacagag gaagatgaag aacagttcaa tgatatagac cttatgctct ttgataaggt 7920
aatcgcattc gatcatctca agcagaagat cattatcata gtaaacattc atttagcgga 7980
tatcgatgca gagtatgcaa gagctcaggc tgaactgtct aagttaaaag atctcatcga 8040
aaacggcaag atgtgcgcag ataccgcagg aagggctaca ggagagttta ctccggtatt 8100
cgataaagaa gcatactgtg caatggttga aaaggctaag catcatatcg ttgaaggaga 8160
tatcttccag atagttctct ccaataaact cagcgctcct tttgaaggat ctctacttaa 8220
tacatacaga atgcttagaa ccatcaatcc ttcaccttat atgttttatt tcagttcaaa 8280
cgatctggaa cttgcgggtg cttctcccga gactcttgta aagcttcagg acggcgtgct 8340
tcatacattc cctcttgcgg gaacaagaaa gcgcggtaag gacgaagagg aagataaggt 8400
acttgtagaa gatctcctgg cagatcccaa ggaacttgca gagcacaaca tgctggtgga 8460
cctgggaaga aacgatctgg gaagaatctc gaagttcgga accgttgaag tagaaaagct 8520
tcgtgaagta cagttcttct cacatgtaat gcatataggc tctaccgtac gaggtgagat 8580
aagagatgat gaggatgcaa tgtccgcaat aggtgcggtt cttcctgcag gaactctctc 8640
cggtgcaccc aagatcaggg catgtgaact tatcgatgaa cttgagggta ataagagagg 8700
ggtatacgga ggcgcaatcg gatatatcga ctttaccgga aacctcgata catgtatctc 8760
cataagactt gcatataaga aaaagggcaa ggtatttgta agaagcggcg caggtattgt 8820
tgccgattcc gtacccgagt cagagtttat cgaatgcatc aacaaggcat ccgcagttat 8880
taaagccatc acacaggcat cggaggtgta agatgatcct tcttatagat aattacgaca 8940
gcttttcata taacctgttc cagcttgtag gggagatcaa tcccgacatt cgtgtcataa 9000
gaaatgatgc acataccgtc gaggagataa ggcagatgaa gccttctcac ataatcttaa 9060
gtcccggccc cggaagaccc gaggatgcgg gagtatgcat cgatgttgtt aaagagcttt 9120
ccggcgagat tcccattctt ggggtttgtc tcgggcatca gtcgatctgc gcggcattcg 9180
gtgcagtggt aacttacgca aagaccctta tgcacggaaa gagttcgatt gttacccttg 9240
ataccgactc agcacttttt aaaggattcg gtaagacaac ggaggttgca agatatcact 9300
ctcttgctgc ggatgaaaat acaatgcctg aatgccttaa ggttaccgca cgtaccgatg 9360
acggcgaagt aatggctgtt gagcataaag agtttaaggt atacggagta cagttccacc 9420
ctgaatccat ccttactccc gagggcagaa agatgctcag aaactacctt gatgtaaagt 9480
aatgtcaatc cggctttgca ctttgcacag tgtatcgcat taaagtaata aagtttttg 9540
ttgacaaatc ttttataaaa gtctatttta ttgagaaaga ggcaaaggcg tctctaatgt 9600
tttaacgaca ttagaggcgt ctttttcttt ttttctgcgg attgaaatgt atgcctgact 9660
gagatgggca ttgatcatta aatgctttgg atgaaaggaa gaagacatgg acaataatct 9720
ttcatatgac agagaacatg atgcgtgtgg tatcggtgcg gtagttaaga ttgatggcgt 9780
acccgaatat tccgttctcg atgatgctct ttccatcgtt gaaaagcttg agcacagagc 9840
aggtaaagat gcgagcggta aggtcggaga cggcgtaggt atcatgcttc agatctctca 9900
ttcttttttc gcgtcagcgc ttaagaaggc tggaataaag gttggtcatg cgggagatta 9960
cggtatcggt atgttcttcc ttcctcagga tgaattaaag cgcactttcg caaagagact 10020
ctttgaagtt atcgcagaca aagagggcgt agaggtaaag ggctggagag aggttcccac 10080
caatcccaag atcctcggcg agagagcctt tgattgcatg ccttcaatct ggcagtgcgt 10140
tgtcggaaga cctgagaata ttcagaaggg aatcgacttc gacagaaagc tttatgtcat 10200
cagaagagag tttgagcagt cttccgatga tacttatatc tgttcacttt cttcaagaac 10260
aatagtctat aagggcatgt tccttgtttc acagctcaga cgtttctatg acgatcttca 10320
gagcaaggaa tatacaaccg cagttgctct tgttcattca agatttcta ccaatacact 10380
tcccagctgg gagagagcac atccctacag attcatcgct cacaacggtg agatcaatac 10440
cataagaggt aactctgaca gaatgcttgc ccgtgaggag accatgcact ctgcacttct 10500
cgagtccgaa aagagcaaga tctatcccgt tatctccgca agcggatccg actctgcaat 10560
gctggacaat actcttgaat tcctctacat gaacggaatg gatcttcctc ttgcaatgat 10620
ggcactgatc cccgagcctt ggaagcataa cagattcatg agcgaatcca agaaggattt 10680
ctatcactac tatgcaacca tgatggagcc ctgggacggc cctgctgcaa ttcttttcac 10740
cgacggtgaa ctcttcggtg caactctcga ccgtaacgga cttcgtcctt caagatacta 10800
tgttaccgat gacaacagac tgatcctctc atcagaggtc ggtgttcttg atattgctcc 10860
cgagcacatc gtagaaaaat caagacttac tcccggaaga atgcttcttg tggatatgcg 10920
caagaagaga ataatttccg atgaagagtg caaggaattc tatgcatcat ccaatcctta 10980

cggtgagtgg cttgatgcac atctgatcag acttaaggac ctgaccattc ccaatcatca 11040
ggttcccgtt cacactcagc ccgaaagaga cagactctat aaggtattcg gatacagcta 11100
tgaagaggtt aagacctata tcctcgaagc tgccaaaaac ggcgcagaac ccaccgtatc 11160
aatgggtcac gacgtgcctc ttgcatctct ttccgacaat taccctctga tgtacagatt 11220
cttccagcag cagtttgctc aggttacgaa tcctcccatc gactcactca gagaaaagat 11280
tgtcacagac acaaccgtat acatcggatc cgacggagac cttctctgtg aaaagagtga 11340
taactgcacc gtgcttgagg ttaacaaccc catccttacc ggcgtagatg tcatgaagat 11400
ccgttcaatg aagagaaaag gtttcaagac cgagaccatt tcccttctgt attacaagca 11460
cacctccctg tcgcgcgcgg tcgagcagct ctatctctcc gtagaccgcg cggtatcgca 11520
gggtgcaaat atcatcatcc tgtcagaccg cggagttgac gaaaaccacg tacccatccc 11580
cgcacttctt gcagtatccg cagttgagca gcatctcgtt cacacgaaaa agagaacggc 11640
agtatcgctc atcctcgaga ccgctgaagt aagagatgtc caccagaccg catgtatact 11700
gggctttggc gcacgtgccg ttaaccctta tcttgcacat gagtgtatcg gagaacttat 11760
taaccttgga cttctcgata aggatcttca caccgcaatc gatgatttcg acaaggcact 11820
tttatcaggc gttgttaaga ttgcagctaa gatgggtatt tccactatcc agtcatatca 11880
gtccgcaaga atcttcgaag ctatcggact ttccgatgaa gttatcaggg attattttgc 11940
aggcgttgca tcaagagtcg gcggaatcgg tgttaaggag cttgaagagg gtatctctta 12000
cagacacgat agagcattcg atcccatggg tcttcccgtt gatccggtac tggacagcat 12060
gggctatcac aacttaagaa gcggaaacgg caaggaagat catctctatt ctcccaaggt 12120
tattgttctt ctccagcagg ctgtaagaac aggggattac gagaagttca aagaatacac 12180
ttcgcttgta gatgatgcaa agcatcctca tacactcaga ggtcttcttg aattcgttcc 12240
cgcaggtaag agcattgata tcgacaaagt agaacctgta agcgagatcg ttaaacgttt 12300
ccaggtcggt gcaatgagct acggttcact ttcaagagaa gcacacgaga ccatcgcaac 12360
cgcaatgaac agactgggtg gaagatccaa taccggtgaa ggcggtgagg catccgacag 12420
attcggtacc gagagaaact ctgcagttaa gcaggttgca tccggaagat tcggtgttac 12480
cgaagagtat cttctcagcg caaatgagat tcagatcaag atggcacagg gtgctaagcc 12540
cggtgaaggc ggaaaccttc ccgcagctaa ggtttaccct tgggttgcaa agaccagatg 12600
ttcaacaccc ggcgtacagc tgatctcccc tccacctcat cacgatatct attccattga 12660
ggatctggct cagcttattt acgaccttaa gaacgcaaac agaaacgcca gaatatccgt 12720
aaagcttgtt tcggaagcag gcgtaggaac catagcatca ggtgttgcca aggcaggtgc 12780
acaggttgta cttatctccg gttatgacgg aggtacaggt gcggcccctt acagctccgt 12840
acacggtgca ggacttccct gggagcttgg tgttgccgag acacatcagg cacttatcga 12900
aagcggactt cgttcaagag taagaatcga gacagacgga aaactcatga caggacgtga 12960
cgttgcaatc gcagcacttc tcggtgcgga ggaattcgga tttgcaacag ctccccttgt 13020
atccctggga tgcatgatga tgagagtctg ctccaaggat acatgtcccg taggtatcgc 13080
tacccagaac gagatcttaa gaaagagatt tgcaggaaag cccgagcatg taattaactt 13140
catgaccttt gcggcaatgc agcttcgtga gatcatggca agcttaggat tcaagaccgt 13200
tgatgagatg atcggtcgta ccgactgcct gaaaaagaga gcaagcttcg atcagcccca 13260
tgctcatacg atttcttttg atgcaatact taagcctgaa tatgctcaca ttgaagagaa 13320
cagattcgat cccaagcagg catatgattt tgaactcgag aagactctcg atgtaaataa 13380
gcttcttccc tggtttgaaa aggcatccaa gacaaagtct gcttcatcaa aagtatccgt 13440
caaggtaagc tcgacagata gaagcctcgg aaccatcctt ggatcagaga tcgttaaaaa 13500
gtacggcgaa ggatgcagag ggcttgctga ggaaagatac gttgttgaat gcgaaggcgg 13560
cggtggccag tcattcggtg cattcattcc tcacggtctt accattaagc ttaagggtga 13620
tgcgaacgac ggattcggta agggactctc aggcggaaag cttgtagtta ctcctcccga 13680
aggaagcacc ttcgaagcat ccaagaacat catagttggt aacgtagctc tttatggcgc 13740
aacttcaggt aaggcatata taaccggtgt tgcaggcgaa agattctgtg taagaaattc 13800
cggagctacc gcagttgctg aaggttgcgg tgaccacggc cttgagtata tgaccggcgg 13860
aagagtagta attctcggaa gcacaggtaa gaacttcgca gccggtatga gcggcggtat 13920
cgcatatgtt cttgataagg atcacagtct ctatcgcagg atcaacaagg atatgattga 13980
actcgaagaa gtatccgaga aggtggacag agaagaactt agaacaatcc ttgaagatta 14040
cgtaaagtgc accggatcaa agctcggtaa agagatactc gataacttcg atgaagaact 14100
tctttgcttc aagaaagtaa tcgcaagaga ttatcagaga atgatcagaa ccattgctaa 14160
gtacgaagag gaaggcgcat caagagatag agccgaactc gaagcgttca aagaagtagc 14220

aggcacagcg gtttaaaagg cagggagata aaatgggaaa acagacaggc tttcttgaat 14280
acgaaagaac agaaaacgaa gagatcccgg ctaaggaaag gatcaaaggc tacaaagaat 14340
tccatacctt tcagaccgaa gagactagga gaaaacaggg cgcaagatgc atgaactgtg 14400
gcgttccctt ttgccagtcg gcaatgagcc tttcgggaat ggtagtagga tgccctcttc 14460
acaatctaat ccccgagtgg aatgatgcca tatacaaggg agaggacgct catgcacttg 14520
caagactcct taagacaaac agattccccg agttcacagg aagagtatgc cccgcacttt 14580
gcgagaaggc atgtatctgc ggacagtacg acgaccccgt caccgtaaga gacaatgagc 14640
tcttccttat cgagtcgggt tacaagaacg gctggatgaa ggctcaacct ccggtaaaag 14700
aaagcggaaa gagcgtagct gttatcggta gtggcccttc cggactttcc gcagcggatg 14760
cacttcgtaa aagaggacat aaggtaaccg tatttgaaag agaggaccgt atcggcggac 14820
ttcttatgta cggaatcccc aacatgaaac ttgacaagag tgtcattgca agaagacaaa 14880
agctcatgga agaggaaggt atcgagttta agaccggggt caaagtcggc gtagatatca 14940
aggcttccga acttcgtaag aaatatgatg ccatcgttct tgcgtgcggt gccaagaaag 15000
caagagcact taccgcagaa ggaatcgaag gaaccgacgg agtatacttt gcggttgatt 15060
tcttaaccgc agctacaaag aacgtactcg gtattaagaa acttcccgcg catttcgatg 15120
caaagggaaa gaacgtaata atcgtaggcg gcggtgatac cggtaatgac tgtatcggaa 15180
ctgttatcag aatgggatgc aaatccgtaa cagcaatcga gatgatgccc aagcctccgg 15240
tagaaagagc tgcgaataac cctggcccg agtggcccaa gactctccgc acagactacg 15300
gtcatgcaga ggctatcgaa gtattcggag aagatcccag agtatttaag accacggtta 15360
ccaaagtatt ttccaagaag ggtaagataa gcgaagtaga actttcagag gtttattttg 15420
acaagggaaa gctttgcacc aagaagggaa ccgagagaaa ggttaagtgc gatctgcttc 15480
ttgttgccgc aggatttgta ggcgcggagt ctgaagttcc cgaggatttc ggtcttgata 15540
tgactgcaag gggagttgtg gcaactcccg aaggaaagta ctctacaaat gttcccggtg 15600
tcttttccgc aggagacatg agaagaggac agagccttgt tgtatgggcg ctcagcgaag 15660
gaaaagagtg cgctgccgag gttgacgagt atctgatggg atacagaaac ggctgatgat 15720
ttgccggcaa gcgttaattg acaggtagcc ttgcatatag aataattaat taaaacctat 15780
cagtaaccat cttggaggat atatgaatta cacatctcag gaagtgaaac agttcgtcga 15840
agaagaagac gtcaaattta tccgtcttgc tttttccgat gtattcggaa aactgaagaa 15900
tgtatctatt atgcccacgg agcttgatcg tgcatttacc gaaggcatag caatcgatgc 15960
atggagcata accggatttg actgcttcga cagatcggat ctcttccttc atcccgatcc 16020
cgacaccctg gcaattcttc cctggagacc cgagcacgga agagttatca gaatgtattg 16080
cgatataaca tggcccgacg gtactccttt tgaatgtgat tccagaacat tccttaaggg 16140
cgccatcgaa gctgccgaga ataagggata tactttcaga ttcggaaccg agcaggaatt 16200
ctatctgttc agaaacgatc ttgagggaga acccaccgat aatccttatg acaatgcagg 16260
atatatggat gtaggtcccg cagacagatg tgcgaacatc agaagagaaa tctgcattac 16320
tctcgagcag atgggaatca tgcccgaaag ctctcatcat gaagaaggtc ccggacaaaa 16380
cgagatcgac ttcagatatt gtgatgccct taaagctgcc gatgatgcaa tgacattcag 16440
aagtgtggtt ggtgcgatcg caaacagaaa cggagctacc gcagtattta cacccaagcc 16500
tttcaagggt aagcccggta acggaatgca cattaacttc tccgttaaaa agggaaatga 16560
agatgtaaat cttgagcctg tcatcgcagg aatactggag catattgcgg agatgactct 16620
tatcctgaat cccataaggg aaagctatga cagaataggt gcgctcaaag cacccggata 16680
tatcacctgg gccaaggaga accgttccgc tcttatcaga attcccgcag cattcggcga 16740
gtacgtaaga ggagaactca gatctcccga tgcttgtgca aatccttatc ttgcattcgg 16800
acttttgatc tatgcgggac ttgaaggtct cgaaaaagga cttaaagcac ccgaaccagt 16860
aaacaccaat ctctccacag ccggagaaga agtaacaaag aatcttaaga agttacccat 16920
gtctcttgag gaggcggtat caactgcttc aaagggtaat tttgtggtcg aaaaacttcc 16980
ttccgagatt cttgcttcat atcttgatct gaacagataa atagaaaagg gttgctttaa 17040
atgcaggctg aaagagtata cagcgtgctg cttgtatctg cagcggaaaa aatgaatacc 17100
aggctggcta cctttttttc cggaggggga tttgaaccgg taacggttgt agacagtgtt 17160
gcaaaggcgc agcgtagatt attgaacaga gagtacgatc tggtcatagt caactcgcct 17220
cttcccgacg acttcggaaa aaagcttgcg gttgatgtgg tggccgaaag cggaggagta 17280
tcacttctcc tggtaaagag cgatatgtac gatgagattt catcaagcat gattccaagg 17340
ggcgtaatgg tggcaaaaag gcctcttgat gattcgatga taggtcagct gcttaatatc 17400
atgtgcagcg tttgtgagcg tctcaagggc atcagaaaga agactatgag ccttgaggat 17460

aagatggaag agatcagact cgtcaacagg gctaaatggg cacttatcaa gtcctgccac 17520
atgaccgagg aagatgctca cagatttatt cagaagcagg ccatggatct gtgcctgagt 17580
aaaaaagaaa ccgcagaaaa tattcttaaa acttacggat aaaaaatgtg acacggggtt 17640
taccccgtgt cgcatttatt ttagtcttcg cctactctta cattgccgct tcccagtccg 17700
aggttttcac cttcgagagg aagccagtcc agaaccttct tgatctggga atcccagaag 17760
tcccagtcgt gacctctctt ttcttcatcc caggtaacat ccgcaccttt accgacaaga 17820
tagtttttaa attctacatt tgcatcaaag aggccgtctt cggttccgca ggagatatag 17880
aacttcggaa cgggacggtt ttcttttaac aaattatcga atgcaacggt gggattcagg 17940
tccgtcttgg cggctttttc catatcgccg aatattgctt tctcatccat cagcatggcg 18000
tcatgttctt tgccgtcgaa taaatgcgta gcggatgaaa gccccgcaac atggctgaag 18060
gtatccgaat aaactatgcc gtttctgaga gcaccgtatc cgcccatgga aaggcctgcg 18120
atgtatgtat catccctttt atccgaaaga gggaacatgt ttctcgtaac ctcaacaagt 18180
tccttgccga taaaggtgct gtaattattc ttggttccgg gatagtcgat gtagaaagcg 18240
ttgtcgcccg agggcatgac aaccgcaaga tttttctctt cggcccatct tctgatacgg 18300
gttccgttta cccagtcact gcagtctccg agaagaccgt ggagaagata gagagttttg 18360
aacttcttgc cacgatcgtt agcgtatccg aaagttctgt agtcaatctt gtccgaagga 18420
aggatcacat agagcgggac agttctgaaa agagcttttg ataagtaact tacctgaagc 18480
agtgccatat ctgtacctcc gtttgcttta ttttccccat atttaaattt tatcttagca 18540
gggatgaaaa gccaataaat gatttcgctg aattaaaatt caataaaagt attgacaaat 18600
gatttgcgtc gatttatatt gaatacagat tcagtgaata tgaattcaac aaacgacaaa 18660
acatttgaat aagtgaggat aaagatatgg caaaggtaat aatcgtaggt ggcggaattg 18720
caggactttc agcaggtata tatgcgctta aggcaggata tgaggctgag atttatgaga 18780
agaacccccct ggtgggcgga gagtgcatgg gatggaaccg caaaggttat cacatcgata 18840
actgcatcca ctggctcacc ggcaccgacc ctaagaccgg actttgggaa gtatggaaga 18900
cggtgggtgc catcgatgaa aaaacagagt atgccaagac ttcaaagttc tattcaagca 18960
gagctgaggg aagggaagtt actctctgga acgatcttga taaaacagag aaagagctta 19020
ttgcagagtc acccgaggac gaagaagaga tcaagaaatt cattcagcac gtaagatatg 19080
cggaggattg cgtaatttcc tccatcaaac cccttgatat gatgggcata cgtgattata 19140
tcgaaatggg taagggaatg gcgaatatgc ccaaggttat gaaagagtac ggcaagatca 19200
actgtttcga actgggcaag agatttaaat cccccgtcat caggaagatg atgagcgatt 19260
atctccccgg agaatacact gcttactccc ttatcgtttc ttacgcaacc atgacaagcg 19320
gaaacggcaa tatccccatg aaaggatccc ttgcaatgtc acttcgcatc gcagagagat 19380
tcaaggaaat gggtggcgtg atccataccg ctactcccgt aagtgaaatt gtgatcaagg 19440
gaaaaaagac cgagggcatt aagcttgagg acggctcatt cattcctgcg gatttcgtaa 19500
taagcgcagt agatacggat ttcctcttcg gaagacttat cgacagaaaa tacatgccta 19560
aggatcttgt taaggcttac gaaaagagag atgcatatcc cgttaccaca ggcttccagg 19620
ttgcatatgc aatcgatgat acattctcgg gagaagatac gctctttttt gactgcgagc 19680
ccatgcagat tggaacaaga actttctcaa gactttccgt taagaactat gcatatgaca 19740
agagttttgc tcccgaggga aaagcagttc ttcaggcaaa ccttatccag acagatgcgg 19800
actatgaata ctgggcatca ctcagtaagg aagagtacag agcaaagaag gaagaactca 19860
cttccatgat cacagacagg atcgtaaaag agttccccga gcttgtcgga aaaatagaac 19920
tccttgattc atggtctcct atgacttata acagatattg taatgcttat cacggctcct 19980
atatgggatt tgttacgacc gtgggaaata agcagatgag atttaaaggt gtagttaagg 20040
gcgtggataa tctttacgta gccggccagt gggtaatgag ccccggcgga ttacccatag 20100
ctgttgtatc cggaaaattc gcagtacaga ggatccttaa gaaagccgga aagtccatca 20160
ttatctgagg aaataatatg accagaaaag aacagaagga agaaagaaga aacgcaatac 20220
tgatgacggc gcttaatctt ttcgtcgaaa gaggttatta cgacaccaag atcgcagata 20280
ttgcggcagc tgtgcccatg agcacgggtc ttttattcca ctattttgaa tcgaaggaag 20340
aacttcttct tgagcttgtg aaaatgggct tgcagggacc gggtgccatc ggggatgtgc 20400
ccggagttcc gcccgatgtg tatcttacgg gattccttaa acagatcttt gcattcgcca 20460
aagagcagcc ctgggtattt aacatgttcg ttttcatggg acaggtaaga agagcgggaa 20520
tgcccgagga agcaaggcgt ctggcgcagt ccgtggatgc cgttaccccc acggtaaaat 20580
tgatcaaaaa aggtcagaaa gacggggtat ttcataaagg cgatgcggat accatggcca 20640
ggtgtttctg ggcttccgta cagggtatca tggagcagat gtctgcggat aagaacatga 20700

aaactcccga ccccgaatgg attgtttcta ttttgaaata ataccttgac aacttaaatt 20760
tacagtagta atttatcact tgaaaaggca aaggcgtctt ttctgtaaac gtacagaaaa 20820
ggcacctttg tctttttctg tttccaaaaa catttaggag gaacaatact atggaaacac 20880
attcagtacc tgattacttc ggagaactcg tatttgacga tcgtcagatg agagcaagac 20940
tttcatccaa gatctatgca tccctcagaa agaccatcga cgagggtaag aaacttgatc 21000
ccaaggttgc ggaatatgta gcagaggcca tgaaagaatg ggcactcgaa aaaggcgcaa 21060
ctcactatac acattggttc cagcctctta ccggcgtaac cgcagaaaag cacgattcat 21120
tcataagccc ttcacctgac ggaagtgtca taatggaatt ctcgggcaag gaactcatcc 21180
agggcgagcc cgacgcatct tcattcccta ccggcggtat ccgcgcaaca ttcgaagcaa 21240
gaggatatac ggcatgggat cccacctctt acgcattcat caaagataag accctttgca 21300
ttcccaccgc attctgttca tacagcggtg aggctctcga taagaagact cctctccttc 21360
gttccatgga agctctcaac aagcaggctc ttcgtatcct tgcacttttc ggaaacacca 21420
ctgccaagaa cgtaagaacc tccgttggtc ccgagcagga atacttcctt atcaccaagg 21480
aaatgtatga tcagcgtccc gaccttaagt tcaccggaag aactcttttt ggtgccaagt 21540
ctcccaaggg tcaggagatg gacgatcatt acttcggagt tatcaagccc cgtgtacagg 21600
cattcatgaa cgacctcaac aaggaactct ggaaactcgg aatccttgca aagacagagc 21660
acaacgaagt tgcacccgct cagcacgagc ttgctcccat ctattcaacc actaacatcg 21720
caaccgacca gaaccagctc accatggaga tcatgcagaa ggttgcttcc aagcacggac 21780
ttgtatgcct gcttcatgag aagcctttcg caggcgtaaa cggatcaggt aagcacaaca 21840
actggtcaat ggcaaccgat accggcatca accttctctc cccggagat acccccgcag 21900
agaatgcaca gttccttctt ttcctgtgtg cggtaatcaa ggctgttgat gattatcagg 21960
atctccttag actctccgta gcaaccgcag gaaacgatca cagactcgga gctaacgagg 22020
cacctcccgc aatcatctcc gtattcctcg gcgatgagct ttccgcaatc cttgatgcca 22080
tcgagagcga cactccttac aagggaacag agcagaagat catgaagctc ggcgttgatg 22140
tccttcccgc attccccaga gacaacaccg accgtaacag aacttcacca ttcgcattca 22200
ccggtaacaa gttcgagttc agaagccttg gttcatccaa ctcgattgca tgtgcaaaca 22260
tcatgctcaa caccgtagtt gcagagtcac ttaagatcta cgctgacaga cttgaaggcg 22320
ctgaagactt tgagaagacc cttcacgaga tgatccgtaa gaccattaag gatcacaaga 22380
gaatcctctt caacggcaac ggatacgacg atgcatgggt taaggaagct accgaagtca 22440
gaggactttg caactacaga acaactcccg actgcatgcc tcatatcctc gatgagaaga 22500
acgttaagct ccttacttct cacggagtat ttaccgaaaa agaaatcgag tcacgttgcg 22560
agatcatgct tgataactac tgcaagcagg ttcgcatcga agctaacacg atgatcgaca 22620
tggctcttaa gcagatcctt cccgcaatca cctgctacgc tgacaagctt tccaccggcg 22680
tcatcgctaa gaagtcagct tgcgaaggaa ttgcttgtga atatgaaaag agcgtaatct 22740
caagactttc cgttcttgca gatcagatct tctcaagaac cggagaactt gaaaaggcag 22800
ttcttaaact cgctgatgct aaagacataa tcgaagaaag tgcaatgatc cgtgacaccg 22860
tgcttcccaa gatgggcgag ctcagagcat gtgcggatga agcagagaca ctcgtagcta 22920
aggagtgctg gcctcttcct tcgtacggcg atctcctctt cggagtaaga taataaaaat 22980
tcttcacctg catcgaaaga tgcaggtgaa ctaattcgga ggatgaaaaa atgtgttcga 23040
taatggcgtg gtgtaaaaaa gccgctgatg agaaaaagtt caccgccgct ctggatgaga 23100
tgacatcaag aggaccggac gacggaagag taattaaaac cgaaggaggt cttctcggat 23160
tcagaagact ctcgatcatg ggtcttaccc ctgaaggaat gcagcccttc gaattgaaag 23220
gctgctctgt tgtttgtaac ggggaaatat acggattcac tcctataagg gatgaactga 23280
agagcaaagg ctacaccttt aagagtgaca gtgactgcga gatcctgctt cccatgtatt 23340
ttgaatacgg aacagagatg ttcaaaaagc ttgatgcgga atttgcatgt gtaatctatg 23400
acgcaaggac agatgagtgg attgctgcaa gagatcccat cggaatcaga cctttatatt 23460
acacattcga taagggcgga agcataatct ttgcaagtga gcccaagagt ctcatgcctc 23520
ttaccgaaga acatatctat cctttccctc ccggacactt ctataaggat gggaagttta 23580
cctgcttcaa taatattgca aagcctgaca aggttatcaa agacgacctg gagacggttt 23640
gtgcaaacat caaagaaaag cttattgcag gcgttgagaa aaggcttatc gcggatgcta 23700
aagtcggttt cctcctttcc ggcggacttg attcatccct tgtatgcagc attgctgcaa 23760
gaaagtccga taagcccatc cgtacatttg cgatcggcat gaacgaagat gccatcgacc 23820
ttaaatatgc taagcagacc gcagatttcc tcggtgcaga ccacaccgaa gttatcatta 23880
caaaagagga tgtactcgaa gcacttccca aggtaattca cctgcttgga acttatgaca 23940

taacaaccat cagagcttcg atgggaatgt atcttctctg taaagcaatt cacgagcaga 24000
ccgatatcag agtaatcctt accggtgagg tttccgatga aatcttcgga tacaagtaca 24060
ccgactttgc tcccagtgca gaggctttcc aggctgaagc cgaaaagaga gtgagcgagc 24120
tccacatgta tgacgtactc cgtgcggaca gatgtatctc agtaaactca cttgaagcaa 24180
gagttccttt cggtgatctg gacttcgtaa aatattgcat gtccatcgat cccgagatga 24240
agcgcaatca ctacaacaag ggtaagtacc tccttcgtca tgcattcgaa ggcggagatt 24300
atcttcccga tgagattctc atgcgtgaga aggcagcatt ctccgatgca gtcggacatt 24360
caatggtaga tcatctgaaa gagtacgcag aggagcagta caccgaagag gaattcgtag 24420
agggatgcaa gaagtactca cacgctcgtc ccttcaccaa agaatcgctt ctctacagag 24480
agatcttcga gaagttctat cccggacagt cccgtatggt taaggacttc tggatgccga 24540
ataaagagtg ggagggatgt aacgtaaacg atccttcagc tcgagttctt tcaaactacg 24600
gtgcaagcgg agtataaata tatagtcacg gcaccgtcta cctttacatg tttttccgcg 24660
ggatgcgctt gtcgccccgt cctcgcgtag aggtactaag ctcgaacttc gcagagctcg 24720
ttctcactaa gtacctgcgc tccggaggtc ccgcagaaac acatgtaaag ctcccggtgc 24780
ccgatctaag aaataatcag catcttttac gtatacaggt aggaatttat gacgaaatat 24840
atctttgtaa caggcggtgt tgtatccggt ctcggaaaag gaataactgc agcttcactc 24900
ggaagacttc tcaaagcaag aggtcttaag gttgctgccc agaaactcga tccctacatc 24960
aatgtagatc ccggaaccat gagcccttat cagcatggtg aagttttcgt aaccgaagac 25020
ggtgcggaga cagaccttga tctcggacat tatgagagat tcatcgatga ggacctcaac 25080
aaatattcga atcttacttc cggaagagtt tactggaatg ttcttaaccg tgagagaaga 25140
ggcgagtatc tgggatccac cgttcagatc attcctcata tcaccgatga gatcaaagat 25200
ttcgtctaca gcgtaggtaa aaagaccgaa gctgatgttg ttatcaccga aatcggagga 25260
accgtaggtg atatcgaatc acagcctttc atagaagctg ttcgtcagat ttcacttgaa 25320
gtcggaagag aaaacagcct ctttgttcat gttactttag ttcccttcct tcacggatcg 25380
gatgagcaca aatcaaagcc cacccagcac tctgttaagg aacttcaggg aatgggaata 25440
tcccctaaca tcctgatctt aagaagcgat gagcctcttg aggatgagat cttcaagaag 25500
atcgcacatt tctgcaacgt taaggaagat tgcgtaatcg aaaacattac ccttcctaac 25560
ctgtatgaag cacctcttat gcttgaagat tccggtttct cacagattgt ttgcagagaa 25620
ctcggtatcg aagcaggtga acccgatctt gatcactgga gagaaatggt cgatcgcatc 25680
aagaatctta agaagactgc ggttatcggc cttgtaggaa aatatgtcca gcttcacgat 25740
gcatatctgt cagttgccga ggcccttcgt cacgcaggat atgagaacga aggcaaaggt 25800
tcagatcaag tggatcgatt ccgagaagat cacagaagaa aatgtggaag agacttttaa 25860
gggtatcgac ggaatcattg ttcccggagg attcggtgac agaggaatcg agggaatgat 25920
cacatcagcc aagtatgcca gaacccacga tattccttat ttcggaatat gtctcggaat 25980
gcagatcgca tgtatcgagt atgcaagaaa tgttctcgga tatgaggatg caaactcagg 26040
tgaattcgat gagctctgtg cgcataaggt catcgatttc atgcccggac aaagcgacag 26100
cattaataaa ggcggaacgc tcagactcgg agcttatccc tgcaagataa gcgaaggctc 26160
caagatgaaa gagttctacg gcaaggaaga aatctcagag cgtcatcgcc acagatatga 26220
attcaataac gatttcaagg aagcttttga gaaagcggga gctacattcg ccggaatgtc 26280
acccgacgga aaactcgtag aggagatgga aatcacggat aaagatttct acctcggagt 26340
tcagtatcat cccgaattca agtcaagacc cgacaaagct catcctcttt tcaagggatt 26400
cgtaagggag gcatgtaaac gaaatggcaa ttcatgaaga atgcggtgta ttcggagtag 26460
tcggcagtaa aaataatgat attgcacgtg acatttacca cgggctgtat gctcttcagc 26520
acagagggca ggaaagctgc ggaatcgtgg taaatgatga cggtgtattt gcaacacata 26580
aagacctcgg cctagtcggc gaggtctttt ccgagactga attaaataat cttccgggcg 26640
gatgcatggc gatcggccat gtcagatacg gcacaaccgg cggaaatatc gcaagtaaca 26700
tccagcccat ggtcgtcagc caccagaagg gtaatcttgc tcttgcacat aacggaaatc 26760
tggcaaattc cctggatctt cgcgattctc tcgagctttc cggtgctatt ttccactcaa 26820
ccagcgatac cgagaccatc tgctacatga tcaccaggga aaggctcaaa acagatagca 26880
tagaaaaggc ggttcttgcg gctatgaagc accttgaggg cgcatacagc cttgttatga 26940
tgtcttccac caagctcctt gcagtcaggg atccccacgg tttcagaccc ctttgctacg 27000
gtaagaggga ggacggaagc tttgttgttg cttcagagag ctgtgccctg agcgctgttg 27060
gcgcaacctt cgtaagagac atcgctcccg gcgaaatgtt ggtattccag tccgataaac 27120
cgttttcgga gcccgtttcc tatacagaaa actgcg 27156

```
//
ID   XXX; XXX; linear; XXX; XXX; XXX;  XXX.
XX
AC   PRJEB67889
XX
DE   Uncultured bacterium DNA fragment from in-reactor enriched cow rumen, clone
ferm-15f15 contig1
XX
FH   Key             Location/Qualifiers
FH
FT   source          1..11647
FT                   /organism="uncultured bacterium"
FT                   /environmental_sample
FT                   /mol_type="genomic DNA"
FT                   /isolation_source="Cow rumen"
FT                   /clone="ferm-15f15"
FT                   /note="AZCL-Barley beta glucan, AZO-CM cellulose, AZCL
Tamarind xyloglucan, AZCL-HE cellulose active clone"
FT   CDS             1140..3074
FT                   /db_xref="SEED:fig|6666666.127239.peg.1"
FT                   /translation="MLSYYMLSKNVLRERYCAIMLITIKRTVKQMSESTHRYSSIIKPA
FT                   ISVIFSNFSVAAFGIIYNFITPKGYYGDDIFSFTPVSLMVFSLTIILVLRFLNASVLKN
FT                   IARVVISCILGISMSATALLSTLMLYGTNTIYQNGLSIRTQILTILGMSIFFIPLVSEL
FT                   TGVVDAISGKRNYTSDTSHIFFYEKNNTIYFFFVWLLTFLCFVPLFLYTWPGNFIADAG
FT                   FQVFNYLDKSINTHHPILHTLLMGFLYKTGFEHNNISFWFQFYTILQMLLLSASFAFFM
FT                   SHIKKMQVSRRFRAIIFLLFAFNPVVSYYAVSTVKETLCVAFFIFALTFLLRLLEGKST
FT                   ACDFVFFVLSAVMSCHFRNNMLYALMTTALVIVIFKRGIKAKLIALILAVVIITMYRLS
FT                   LSLIIKLSGAYDKDSAVESMSVPLMCFAHVMIDEKNSLSPDLYDEMCLYIDEGSLQNYS
FT                   FIIADPIKGKANESLIKTNKINFFKLFAKVGLRYPGKYLESFVGLTAGYYTPITSPYFI
FT                   TGTTKLYTVPVEGWYPSPAVDNKLPFGSAFFDYMYGERDGRLRLPILGWLWRSAPYFWS
FT                   FAFAFLYLIYKKKYNNLIVLLFPFIYMLTCLLGPTAFLRYMFINIATMPIALYLVIYRD
FT                   TTSAKEVLG"
FT                   /product="hypothetical protein"
FT   CDS             complement(3046..4215)
FT                   /db_xref="SEED:fig|6666666.127239.peg.2"
FT                   /db_xref="GO:0008784"
FT                   /translation="MESTLKRTWSEIDLDALVHNYKTLRARIGDKVKFLGVVKADAYGH
FT                   GSVMVAQTLEKEGADYLAVSSIDEAMELRVNDIKMPILILGHTPKEQVDRLIEYNITQA
FT                   VTCKAKAVEYSEEATRLNKTLKIHIKVDTGMSRLGYLCDGDFFDTGVQGIVDAVTLPGL
FT                   DAEGIFTHFAVADEEGPECLEYTRHQFKLFTDVCSKVEEKIGKPFLIKHCANTGAVARF
FT                   PETYLDMVRPGLLLYGYGEYAKELGLKPVMTLKTVVSTVKVYPAGTAISYGGIYKTDKK
FT                   ERIGVLPYGYADGFFRSLSNKCSLMTDEGPAPQRGKICMDMCMIDITDKYGVDVGSEIE
FT                   IFGKRNPIETLSDIAGTIPYELTCAVSKRVPRLYINDGEVINRELLLRM"
FT                   /product="Alanine racemase (EC 5.1.1.1)"
FT                   /EC_number="5.1.1.1"
FT   CDS             complement(4248..4871)
FT                   /db_xref="SEED:fig|6666666.127239.peg.3"
FT                   /translation="MNDNRVIIEAIGYVGTALVLISFLMTSVYKLRIINTAGSLISVIY
FT                   GVIMQVYPTVVLNGALALINIFFLWRHLSYKNQKIYTAGKTYAEDPIVKLFLDKYREDI
FT                   LKFFPSFGVAHLEPNIAYLIMCEDTMAGVMLGNLKENGELDIYLDFVTGTYRDFSVGKY
FT                   LYRELTRDGVKKCRFMTDIPKSYVYLRKVGYKRVGDVMELTLNE"
FT                   /product="hypothetical protein"
FT   CDS             complement(4881..6245)
FT                   /db_xref="SEED:fig|6666666.127239.peg.4"
```

```
FT                   /translation="MTTIKKRIIATMLIAALALTGCESTKASVSSADTVSTETVSASST
FT                   EEVKEEVKDETQETVSASASTEEKKEEKKDENVRPDNLINVLVSKHKAPESEALSFTAD
FT                   MGAGINLGNTFDASDAKNIFDDDLKFETYWQKDVTTQDMIKAIKNEGFKTIRIPVAWHN
FT                   HVDDDFNINEKWMNRVYEVVNWALEEDMYVILNIHHDNLKKYMYPSYECRENSLKYVTK
FT                   IWAQVAEKFGDCDEHLVFESLNEPRLAGTDIEWNLPDLNAKEAKETLDCINQYNQAIVD
FT                   TIRQTPGAYNKTRYITVPGYAGSPDYILCDQFVVPTDTYGEKENRILLTVHAYRPYLFA
FT                   LADPGKGDTKTFDTEGAVSELNALFMNLFVKYVKNGTGVIIDEFGARDRGGNTEERAKF
FT                   TAYFVAYARSYRMPVCWWDNNNFSGSGEVFGIFKRSACKFEYQDIVSQLVVYYGNR"
FT                   /product="GH5_4"
FT   CDS             complement(6299..7171)
FT                   /db_xref="SEED:fig|6666666.127239.peg.5"
FT                   /translation="MQGKIIKGIAGFYYVNIDGKGTFECKAKGSFRNEGTKPLVGDNVE
FT                   IDIIDEEKSLGNVTRVYKSKSRMIRPACANVDQALIVFAIKSPDPNFNLLDRFLIYMES
FT                   ESVPCIIVFNKDDLASPEDIETIREAYVGAGYPIFFVSARENAGVEALRDLLKGKTTVM
FT                   AGPSGVGKSSLINLLCNCDIMETGDISVKTQRGKHTTRHAELLPLGEGTYVMDTPGFTS
FT                   LDIFGADRQNLKDYYNEFSSVEPCRFADCVHVNEPDCAVKAAVEAGKISKLRYDNYLEL
FT                   FSQIGTKTR"
FT                   /product="Ribosome small subunit-stimulated GTPase EngC"
FT   CDS             complement(7183..9264)
FT                   /db_xref="SEED:fig|6666666.127239.peg.6"
FT                   /translation="MVKIGMMIGERYEVLEKIGTGGMSDVYKAKDHKLNRFVAVKVLKQ
FT                   EFAENANFVSKFKIEAQAAAGLMHPNIVNVYDVGEENGTNYIVMELVEGITLKKYIEKK
FT                   ARLSVKEAVTIAIQIGMGIEAAHNNHIIHRDIKPQNVIISKEGKVKVTDFGIAKAATSN
FT                   TITSNVMGSVHYTSPEQARGGYSDEKSDIYSLGITLFEMLTGRVPFNGETTVAIAIKQI
FT                   QEEMPSPREYVDDIPVSVEQIVLKCTQKSPDRRYQSMGALIEDLKKTFITPDEDFVKIK
FT                   EDDEFGATRPVTAGELEVLKKSTKGTEELTGELPKVNPNASQDDSEYDAKMDKLTTILA
FT                   IFMGAAILVIAIFLISKGIGVLKLGKENTEKVPTTVSMEKPEQIVMPKVVGKSQEDAQK
FT                   ILEKNNLKYFVKYTESDSVKEGYVISASVQDGLKVDAGSEIVLNVSSGSTSVIVPDVVG
FT                   KALEAANKSIMDAGLKVEIKKDYSDTVPEGNIISQFPEGGEKTPEESTVLIIVSQGVAE
FT                   NLTKVPNVLGKDNDDAIIILTEAGLTVNPVAPVYDNIFPEGQICYQSLEADKEVPKGTA
FT                   IDLHASMGPEPPDTVMYRFNGSINAPSTAEDPEFEMGTKVEITVSDANGKVLQSTWTTS
FT                   FPVSVNISNIAGVNSGTITLSYEVKIPAEKDVSTGEEITPAKTEKKVIKRDINFEVDE"
FT                   /product="Serine/threonine protein kinase PrkC, regulator
FT                   of stationary phase"
FT   CDS             complement(9292..10020)
FT                   /db_xref="SEED:fig|6666666.127239.peg.7"
FT                   /translation="MLKTFSLSDVGKKRTMNQDYIFTSEEPVGNLPNLFALADGMGGYN
FT                   GGEYASLRTVETVTSFIDKSKSTTPRKLFTEAIKEANRIIRDEASQNPEYDKMGTTIVL
FT                   ASVVGDCLQVANVGDSRLYVIGDDIKQVTVDHSYVEEMIKMGSLDRESARTHPQKNIIT
FT                   RAVGADDDIDTDFFTVRLHKGDMVLMCSDGLTNMLEDSEILMILKKERDIASKAEALVA
FT                   AANNSGGKDNISVILIEPFA"
FT                   /product="Protein serine/threonine phosphatase PrpC,
FT                   regulation of stationary phase"
FT   CDS             complement(10066..11103)
FT                   /db_xref="SEED:fig|6666666.127239.peg.8"
FT                   /translation="MDKIDIKSLDFEELENSIKEMGLPKFRASQIYDWLHKKLVTSFGE
FT                   MTNLSADLRNKLSETYYITACEVARMQESREDGTRKYLFKLRDGNYVESVFMRYHHGNS
FT                   VCISSQVGCRMGCKFCASTLDGLTRCLEPSEMLEQIYRIQALTGERVSNVVVMGTGEPM
FT                   DNYDNLLKFIKLLTDGHGLNISQRNLTVSTCGIVENMKRLADEKLSITLALSLHSVTDE
FT                   KRRELMPIANKYSINELMDACEYYFDKTGRRITFEYALVRDVNDRDEDAKGLIKLAGRV
FT                   KAHVNLIPVNPIKERDYRETARDDVLAFKNKLEKSRINVTIRREMGRDIDGACGQLRKK
FT                   TMDSE"
FT                   /product="Ribosomal RNA large subunit methyltransferase N
FT                   (EC 2.1.1.-)"
```

```
FT   CDS             complement(11103..11627)
FT                   /db_xref="SEED:fig|6666666.127239.peg.9"
FT                   /translation="MYPEGEVVSCDVSDTKVSMIADNAERLHLSNLKPTLQDATEFNEE
FT                   YEASFDIVIADVPCSGLGVMSRKSDIKYNITNEAMKDICDLQKAIVTNVSRYVKPGGVL
FT                   LYSTCTIHKAENEKMVKFITENLPFKEDSLKPFVPFLFDKPRKSECHIQLRPDTDGTDG
FT                   FFVARFVKSED"
FT                   /product="Ribosomal RNA small subunit methyltransferase B
FT                   (EC 2.1.1.-)"
XX
SQ   Sequence 11647 BP; 2929 A; 2733 C; 2428 G; 3557 T; 0 other;
     tcgagagaca aatcaacacc cgcataggcg cccatggcgg tttcaagttc cgtaaaggta        60
     tcggaaatct gaagtaccgc attgccgtct gcatctatat tgaggtgacc cagtattctg       120
     tcatcgctct taacgtctat tacatatcct tccgggcgcg cacatacctt gggaatggag       180
     ggaaccttgt aatgctgtcc cgccaccaca tggaatcccg gaatcttgtc attaatcttg       240
     gcgcccgcgg gattaatgtg cagaggctca ttgaaagtat aagttacctt tattgtgtca       300
     tccaaagaaa taacggtatc gcccgattca tcaagaggct taacattacc ctccgaatcc       360
     tggaacataa ccgtcatgga ctttatcaaa gcattctcat cgggaatgtt tgaatcgtcc       420
     tcatcctcca aagaaaccac aactcccagt gttcccgccg aagctgtctt ttgtgaagac       480
     aacaaaagcc ccgcgaagtc aagcttctca gagtttaaat cttcttcatt caattcgttt       540
     aatctgtcta tggaaaacaa agccaaagaa ttttgacttt ctaaagtatc atccgctaaa       600
     gatacctcat tgcccgatat gtcgccgcct gatacatcat tgcctgaagt atcattgtcg       660
     gaaatatcat tgtccgatac ggaaggctcg ataacaacag gcgcatcacc cgaaacatct       720
     tccgttaccg aaggcatctc atcgtctgca gaaacatcgc gagactccag tgatgaaatt       780
     acgggctcaa cagtttccga aatctcttct tcctcttcct caatcgcttc atcggttgaa       840
     gtcagcgtcg gttctgcgtt ttcatcctcg cttatcagga tttcttctgt gctttcttgc       900
     tcctcataac ctaagtcggg atcggccgca tatacgctta tgacgtcatt tctcagcacg       960
     cagaaaacaa cgaccatact taaaagcacg gccaaaatac gtcgccccat tatacccatt      1020
     cctttcactt gaattcagtt taaaattccg atgagtaccc ctactctaca aaagtgaaat      1080
     attatcgtaa cagattataa cacttttccc cagaaataaa acattttttt acaaatttga      1140
     tgctaagtta ttatatgctg agtaaaaatg ttttacgaga acgatactgt gctataatgc      1200
     ttataacaat aaaaaggacg gttaaacaaa tgtcagaatc aacacatcgt tattcttcca      1260
     taattaaacc tgctatttca gttatttttt caaattttag cgttgcggct ttcggtatca      1320
     tatacaattt tattacccca aaaggatact atggagatga tattttttct tttacgcctg      1380
     taagtctcat ggttttctca ttgacaatta ttttagtttt aagattttta aacgcttcag      1440
     tgcttaagaa catcgccaga gtagtgatat cttgtattct cggaatatcc atgtcggcca      1500
     ctgcactttt atccacgctt atgttatatg gtaccaatac tatatatcaa aatggattaa      1560
     gcatccgaac tcagattctg acaatactgg gaatgtctat tttcttcatt cctttggtat      1620
     cagaacttac aggagtggtc gatgccatct ccggcaaaag aaactatact tccgatactt      1680
     cacatatatt tttttacgag aaaaacaata cgatttattt ctttttgtt tggctcttaa       1740
     ccttcttatg cttcgttcct ctctttcttt acacttggcc cggcaatttt atcgcagatg      1800
     ccggatttca agtattcaat tatcttgaca aaagtataaa cactcatcat cccatactac      1860
     atactctgct tatgggattc ttatacaaaa caggcttcga gcataataac atatccttct      1920
     ggttccagtt ttacacaatt ttgcagatgc ttcttctttc tgcttctttt gcattcttta      1980
     tgagccacat caagaaaatg caggtttcac ggagatttag ggctattatt ttcttattgt      2040
     ttgcttttaa ccccgttgtt tcttattatg ccgtttcaac agttaaagag actctttgtg      2100
     tggcattttt tatatttgcg ctgacttttc ttcttcgact gcttgaaggg aaatctacgg      2160
     catgtgattt tgttttcttt gtccttagcg cagttatgtc ttgtcatttt aggaataata      2220
     tgttgtatgc tctcatgact accgctctgg tcattgttat ttttaagcgc ggaattaagg      2280
     ccaaacttat tgcccttata ctggctgtgg taatcattac catgtacaga ttaagtcttt      2340
     ctctcataat caaattgtcc ggagcatatg ataaagacag cgctgttgaa agtatgtcag      2400
     taccacttat gtgctttgct catgtaatga tcgatgaaaa aaatagcctg agtcctgacc      2460
     tttacgacga aatgtgtttg tatatcgacg aagggtctct tcaaaactat tcattcataa      2520
     tagcggatcc gattaaaggc aaagccaatg aaagtcttat taaaaccaac aaaattaact      2580
     tttttaagct tttgcaaaa gtcggactga gatatccggg taaatatttg gaatcatttg       2640
```

taggtcttac agcaggttac tatacaccca taacttctcc gtattttatt accggtacca 2700
ccaagcttta caccgtgccc gtagaaggtt ggtatccttc gcctgccgtt gataataaac 2760
ttcctttcgg aagcgctttt tttgattaca tgtatggtga acgtgatggg agactccgcc 2820
ttccgatact gggctggctg tggcgctcgg ctccctactt ttggagcttt gctttcgcat 2880
ttttatattt aatatataaa aagaagtaca acaatctgat tgtacttctc tttccattta 2940
tttatatgct aacctgtctt ctcggaccca cggcatttct ccgctacatg tttatcaata 3000
tagcgaccat gccaattgct ctatatcttg tcatatatcg agatactaca tccgcaaaag 3060
aagttctcgg ttaataacct caccatcatt aatgtataag cggggaactc tcttgctgac 3120
cgcacatgtg agctcgtaag gaatggtgcc cgctatatcg gaaagtgttt ctatgggatt 3180
tcttttgccg aaaatttcta tttcggaacc tacgtcaaca ccgtatttgt cggtgatgtc 3240
tatcatgcac atgtccatgc atatcttgcc gcgctgaggt gcgggacctt cgtcggtcat 3300
gagactgcat ttattggaga ggcttctgaa gaagccgtcg gcatatccgt aagggagtac 3360
gcctatgcgc tcttttttgt ctgttttgta gataccgccg tagcttatgg ctgttccggc 3420
aggatatacc ttaacggtgc ttaccacggt ctttaaggtc ataacgggct taaggcccag 3480
ttctttggca tactcgccgt atccgtaaag aaggagtccc ggacgaacca tatccagata 3540
agtctcaggg aatcttgcca cggcaccggt attggcgcaa tgcttgatta aaaaaggctt 3600
accaatcttc tcttcaacct tagaacatac atcggtaaac agtttaaact gatgtctcgt 3660
atattcaagg cactcgggac cttcttcatc cgcaacagcg aaatgagtaa agataccttc 3720
cgcatcaagc cccggcaatg taaccgcatc cactattccc tgaacgcctg tgtcaaagaa 3780
atctccgtcg cacagataac cgaggcggga catgcccgta tccaccttta tatgaatctt 3840
cagggtctta ttaagtcttg tggcttcctc ggagtactca accgctttag ccttgcaggt 3900
tactgcctga gtgatattgt attcgataag tctgtccacc tgctccttgg gcgtatgtcc 3960
caggattaaa atgggcatct taatatcgtt aaccctaagc tccatggctt cgtcaatact 4020
gctgacagcc aggtaatcag cgccttcttt ctcaagggtc tgtgccacca taacggaacc 4080
gtgtccgtag gcatcagcct taaccacacc taagaattta accttatcac cgattcttgc 4140
tcttaaagtc ttgtaattgt ggaccaaagc atcaaggtcg atttccgacc atgttctttt 4200
taaagtggat tccatgataa ataaacccat cctcccaaaa tcatctctta ttcgttaagc 4260
gtaagctcca taacgtcgcc cacacgcttg taaccgactt ttcttaagta aacgtaactc 4320
ttgggaatgt ctgtcataaa ccggcatttc ttgacaccgt ctcttgtaag ctccctgtag 4380
aggtatttgc ccacggaaaa gtccctgtag gttcccgtta caaaatcaag gtagatgtcg 4440
agttcaccgt tttcttttaa attacctaac attacacccg ccatggtgtc ttcgcacata 4500
ataagatatg cgatgttggg ttcaaggtgc gccacaccga agcttgggaa gaatttaagt 4560
atatcctccc tgtatttgtc aagaaacagt ttgactatgg gatcctcggc gtaggtcttg 4620
cctgccgtat aaatcttttg attcttgtaa gataagtggc gccataaaaa gaaaatattt 4680
ataagcgcca gagctccgtt taaaacaacc gtagggtaca cctgcataat aacgccgtaa 4740
ataaccgaaa taagggaacc tgcggtattt atgatgcgaa gcttgtatac cgaagtcata 4800
aggaaggata taagcaccaa agcggtgcca acataaccta tagcttcaat aatcactctg 4860
ttatcgttca taagagaatc ctatctgttt ccgtaataaa ccaactgact tacgatgtcc 4920
tgatactcga atttgcatgc ggaacgctta aatataccga atacttcgcc gcttcccgag 4980
aagttgttgt tgtcccacca gcatacgggc atacggtagc ttctggcata tgcaacgaaa 5040
taagccgtaa atttggcacg ttcctcggta ttgccgcctc tgtctcttgc accgaattcg 5100
tcgatgataa cgccggtacc gttcttaacg tacttaacga acaggttcat gaataaagcg 5160
ttaagttcgg atacagcgcc ctccgtatca aaggtcttgg tatctccctt gccgggatcg 5220
gcaagtgcga aaagataagg tctgtatgca tgaaccgtca aaagaatgcg gttttctttt 5280
tcaccgtatg tatctgtggg aaccacaaac tggtcacaga ggatataatc gggtgaaccc 5340
gcataaccgg gaacggtgat atatctggtc ttattgtaag caccgggagt ttgtctgatg 5400
gtatctacaa ttgcctgatt gtactgattg atgcagtcaa gggtttcctt ggcttctttg 5460
gcatttaagt cgggaaggtt ccactcaatg tcggtaccgg cgagtctggg ctcattcagt 5520
gattcaaata ccagatgctc gtcgcagtcg ccgaatttct ccgcaacctg cgcccagatc 5580
ttggtaacgt acttaaggga gttctcacgg cactcatagg aaggatacat atacttcttt 5640
aagttgtcgt gatgaatgtt aaggataacg tacatatcct cttcaagggc ccagttaaca 5700
acttcgtaga ctctgttcat ccacttttcg ttgatgttaa agtcatcgtc aacatgattg 5760
tgccatgcta cgggaattct gatggtctta aagccttcgt tcttaatggc cttaatcatg 5820
tcctgagtgg taacgtcctt ctgccagtat gtctcaaatt ttaagtcatc gtcaaaaatg 5880

ttcttggcat cgcttgcgtc gaaggtgttg ccaaggttaa tgcccgcgcc catgtcagcg 5940
gtaaaagaaa gtgcctcgct ctcgggtgct ttgtgtttgg ataccaatac attgataaga 6000
ttgtcgggtc tcacgttttc gtccttttc tcttcttttt tctcttctgt ggatgcggaa 6060
gcggaaacgg tttcttgtgt ctcgtccttt acttcctctt taacttcttc tgttgaagaa 6120
gctgatacgg tttcggtgct tactgtgtct gcggatgata cgcttgcttt ggtggattcg 6180
cagccggtca gagcaagggc cgcaataagc attgttgcaa taattctctt tttgatggta 6240
gtcataataa tcctctacat tataaagttt gtgtccggcg tggccggtat atgaaccgtt 6300
acctggtctt cgtccctatc tgggaaaaaa gttcaaggta attgtcataa cgaagtttag 6360
aaatcttacc cgcttccact gccgccttaa cagcacagtc gggttcgttg acatgtacgc 6420
agtctgcgaa tctgcagggc tctacggaag aaaattcatt gtagtaatcc ttaaggttct 6480
gtctgtctgc gccaaaaatg tcaagggacg taaaaccggg ggtgtccata acataagtgc 6540
cctcaccgag gggtaacaat tccgcatgcc tggtggtatg tttgcccctt tgggtcttca 6600
cacttatatc accggtttcc attatatcgc aattgcataa cagatttatt agagatgatt 6660
tgcccacacc gctggggccg gccataaccg ttgttttgcc ttttaacagg tctcttaagg 6720
cctcaactcc cgcattctct ctcgcactga caaagaaaat ggggtagccc gctcccacat 6780
atgcctctct tatggtctca atgtcctccg gggatgccag atcgtccttg ttaaagacta 6840
tgatgcaggg aaccgattcg ctctccatgt agattaaaaa cctgtcgaga aggttgaaat 6900
tggggtcggg gcttttaatt gcaaaaacaa ttaaagcctg gtcaacgttg gcacaggcag 6960
gccgtatcat acggcttttg ctcttgtaaa cccgtgtcac gtttcctaag cttttctctt 7020
cgtcgattat gtctatttca acattgtcgc ctaccaatgg cttggtgcct tcattacgaa 7080
aagaaccttt ggccttacac tcaaaggttc ctttaccgtc gatattaacg taataaaatc 7140
cggcaatgcc tttaataatc tttccttgca ttaatctccc gattattcgt caacttcaaa 7200
atttatgtct cttttaatga ccttttctc ggtcttggca ggggtaatct cctcgcccgt 7260
ggatacgtcc ttctccgcag gaatcttaac ttcatacgaa agggtaatgg taccgctgtt 7320
aactcccgca atgttggaaa tgttaacgga aacggggaaa gaagtggtcc aggtagactg 7380
aagcaccttg ccgtttgcgt cagataccgt tatctcaacc ttggtaccca tttcaaattc 7440
gggatcttct gccgtggagg gtgcatttat gcttccgtta aatctgtaca taacggtatc 7500
cggaggttcg ggacccatgc tggcatgaag gtcaatagcg gttcccttgg gaacttcttt 7560
gtccgcttca aggctctggt agcaaatctg tccttcgggg aagatattgt cgtatacggg 7620
agctacgggg ttaaccgtaa gacccgcttc ggtcagaata ataatagcat cgtcattgtc 7680
cttaccaaga acattgggaa ccttggtgag gttctcggca acgccctggc ttacgataat 7740
gagcacggta gattcttcgg gagtcttctc tccgccttcg ggaaactgtg aaatgatgtt 7800
gccttcggga acggtatcgg aatagtcctt tttaatctct acctttaagc cggcatccat 7860
aatcgattta ttggctgcct ccaaagcttt gcccactaca tcgggcacaa taacactggt 7920
ggaaccggag cttacgttaa gcacgatttc cgagcccgca tctaccttta agccgtcctg 7980
aacgcttgct gaaataacat agccttcttt gacggaatcg gattccgtat attttacaaa 8040
gtatttaaga ttgttcttct caagtatttt ctgagcgtct tcctggctct tgcccacaac 8100
cttgggcatt actatctgct caggcttctc catggaaact gtggtgggaa ccttctccgt 8160
gttttctttg ccgagcttca gtactccgat gcccttgctg ataaggaaaa tggctataac 8220
aagaatggcc gcacccataa agatggcaag aatggtggtg agtttgtcca tcttggcatc 8280
gtattcgctg tcatcctgag aggcgttcgg attaaccttg ggaagttcgc ccgtaagctc 8340
ttccgtgcct tttgtactct tcttaaggac ctcaagttca cctgcggtga cgggtcttgt 8400
ggcgccgaat tcgtcgtctt ctttaatctt tacaaaatct tcgtcgggag tgataaaggt 8460
cttcttaagg tcctcgataa gggctcccat gctctgatag cgtctgtcgg ggctcttctg 8520
ggtacactta agcacaatct gctcaacact tacgggaata tcgtcaacgt attccctggg 8580
ggagggcatt tcttcctgaa tctgtttgat agctatagct acggtggtct ctccgttaaa 8640
aggaactctt cctgtaagca tttcaaagag agtgatacca agtgaataga tatcgctctt 8700
ctcatcggag tatccgccgc gagcctgctc gggagatgta tagtgaacag accccataac 8760
gttactggta atggtgttgc tggtagcggc cttggcgatg ccaaagtctg ttaccttaac 8820
cttgccttct ttggaaataa taacgttctg aggtttaatg tctctgtgaa taatatgatt 8880
gttgtgggcg gcttcgatac ccatgcctat ctggatggct atggtaacgg cttctttgac 8940
cgaaaggcgg gccttcttct caatatattt cttgagagtg ataccctcaa caagttccat 9000
tacaatgtaa ttggtgccgt ttcttcgcc tacgtcatag acgttgacga tattggggtg 9060
cataagacct gcggcggcct gagcctcgat tttaaacttc gagacgaagt tggcattctc 9120

```
     cgcaaactcc tgcttaagca ccttaacggc aacaaaacga ttcaatttat gatctttagc      9180
     tttatagacg tcggacatac cgccggttcc gattttttca agaacctcgt atctctcgcc      9240
     tatcatcata ccgattttaa ccatgtgtat cctctgtcag acgattcttt atcaagcaaa      9300
     gggctctatt aagataaccg aaatgttatc cttgccgccg gagttattgg cagctgccac      9360
     aagtgcttct gctttgcttg ctatatcccg ctctttcttg agtatcataa ggatttcact      9420
     gtcctcaagc atgttggtga ggccgtccga gcacataagc accatatcgc ccttatgaag      9480
     ccttacggta aagaaatccg tgtcgatgtc atcgtccgca ccgacagctc ttgttatgat      9540
     attcttctgc gggtgggtcc tggcactttc tctgtcaaga cttcccattt tgatcatctc      9600
     ctcaacatag gagtgatcga cagtaacctg tttgatatcg tccccgatta catataacct      9660
     tgagtcgcca acattggcaa cctgaaggca atccccccaca acagatgcaa gcactatggt      9720
     agtacccatc ttgtcatatt cgggattctg agaggcttcg tctctgatga tacggttggc      9780
     ctctttaata gcttccgtga aaagtttcct cggcgtggtg ctcttggact tgtcaataaa      9840
     agaagtgaca gtctccacgg tacgaagaga tgcgtactct ccgccgttat aacctcccat      9900
     gccgtcggca agggcaaaaa ggttgggcag atttcccaca ggctcctcgc ttgtgaaaat      9960
     gtagtcctgg ttcattgttc tcttttttcc cacatccgat aaggaaaaag ttttaagcac     10020
     ccctatgcct cctgattaat accatcgctt caaaagaaaa caagtctatt cactgtccat     10080
     tgtcttcttg cgaagctgtc cgcaggcccc gtctatatcg cgacccattt cccttctaat     10140
     agtaacattt attcggcttt tttcaagttt atttttgaat gccagcacgt cgtctctggc     10200
     cgtttccctg taatcacgct cttttatggg attgaccgga ataaggttta catgagcctt     10260
     taccettccc gcaagcttaa taaggccttt ggcatcctcg tccctgtcat tgacatctcg     10320
     caccagagca tactcgaagg tgatacgtct gccggtcttg tcaaaataat attcacaggc     10380
     atccataagc tcgtttatag agtatttatt ggctattggc atgagctcgc gacgtttttc     10440
     gtccgtaacc gaatgaaggg acaaagcaag tgttatggag agtttctcat cagccagacg     10500
     cttcatgttc tcaacgattc cgcaggttga tacggtaagg tttctttggc ttatattgag     10560
     tccgtgcccg tcggttaaaa gctttatgaa tttaagcaga ttgtcgtaat tgtccatggg     10620
     ctcgccggta cccattacca ctacgttgga tactctctcg ccggtgaggg cctgaatacg     10680
     gtaaatctgc tcaagcatct cagagggctc aaggcatctg gtaagtccgt ccaaagtgga     10740
     ggcgcagaac ttacagccca tgcggcagcc tacctgggaa gaaatgcaca cggaattgcc     10800
     gtgatgatag cgcatgaaaa cgctctccac gtaattgccg tcgcgaagct taaagaggta     10860
     ctttctggta ccgtcttccc ttgactcctg cattctcgcc acttcgcagg cggttatgta     10920
     gtacgtttcc gaaagtttat tacgaaggtc tgccgagaga ttggtcattt cgccaaaaga     10980
     agtaaccagt ttcttgtgga gccagtcata aatctgtgag gctctgaatt tgggaagacc     11040
     catttctttt atggaatttt caagttcttc aaagtccaat gacttaatgt cgattttatc     11100
     cattaatctt cactcttaac aaatcttgcc acaaagaagc cgtcggtgcc gtccgtatcg     11160
     ggacgaagct gaatgtggca ttcgctctta cgaggcttgt caaacaggaa cggaacaaaa     11220
     ggtttaaggg aatcttcttt gaagggaagg ttctccgtta taaacttaac catttttca      11280
     ttttcagctt tatgtatggt gcaggtggaa taaagaagca ctccgcccgg cttaacgtag     11340
     cgtgaaacat tggtaacgat ggcctttga aggtcacata tgtccttcat ggcttcgttg      11400
     gtaatattgt atttaatatc gctcttacgt gacattacgc ccagtcccga acatggaacg     11460
     tcggctatta ctatgtcaaa agaagcctca tattcttcat tgaattccgt ggcatcctgc     11520
     agtgtaggct taagattgct taagtgaagc ctctctgcat tgtcggctat catggatact     11580
     ttggtatccg aaacatcgca gcttacgacc tcgccctcag gatacatgcg cgaagccacg     11640
     tacgagg                                                                11647
//
ID   XXX; XXX; linear; XXX; XXX; XXX;  XXX.
XX
AC   PRJEB67889
XX
DE   Uncultured bacterium DNA fragment from in-reactor enriched cow rumen, clone
ferm-15f15-contig2
XX
FH   Key             Location/Qualifiers
FH
FT   source          1..16634
```

```
FT                   /organism="uncultured bacterium"
FT                   /environmental_sample
FT                   /mol_type="genomic DNA"
FT                   /isolation_source="Cow rumen"
FT                   /clone="ferm-15f15"
FT                   /note="AZCL-Barley beta glucan, AZO-CM cellulose, AZCL
Tamarind xyloglucan, AZCL-HE cellulose active clone"
FT   CDS             30..266
FT                   /db_xref="SEED:fig|6666666.127240.peg.1"
FT                   /db_xref="GO:0003715"
FT                   /db_xref="GO:0003723"
FT                   /db_xref="GO:0005524"
FT                   /db_xref="GO:0006353"
FT                   /db_xref="GO:0015986"
FT                   /db_xref="GO:0016469"
FT                   /translation="MELQLDRNLSNKRIFPAVNIVASSTRRDDLLQDKQTLDRMWILRK
FT                   YLADMNPIEAMDFVKDRLEKTKDNDEFLMSMNS"
FT                   /product="Transcription termination factor Rho"
FT   CDS             421..1866
FT                   /db_xref="SEED:fig|6666666.127240.peg.2"
FT                   /db_xref="GO:0047753"
FT                   /translation="MGMALCSFPVMAQQSPNFLIIQCDHLTQRVVGAYGADPSCTPPID
FT                   RIASQGVTFANAYVGCPLSQPSRAALWSGLMPHQTNVRSNSAEHINPPLPDSIPTLGSL
FT                   FTANGYEAVHFGKTNDMGSLRGFRHKEPVAKPFTDPEFPVNNDSFLDVGTCEDAVAYLS
FT                   NPPQEPFICIADFQNPHNICGYVGENKGEHIDSPISTPLPLLPENFEVEDWEKLPLPIQ
FT                   YICCSHRRMMQAAHWNEGNYRHYVAAFQHYTRIVARQIESVLEALYSTPAGKNTIVIVL
FT                   ADHGDGMGSHRMVTKQVSFYEEMTNVPFIIAGPGIHPRQKPVEDLLTQPTIDLLPTLCE
FT                   LAGIPVPSSKPGISLAPFLKGEKQKRQHPYAASEWHSEYEVTVSPGRMIRSPRYKYTHY
FT                   LEGNGEELYDMLKDKGERRNLAHDPKYSKILAEHRALLDDYIRRTQDDYRSLKVEADKR
FT                   WRSHAPGYPNHEGPCARDIINRK"
FT                   /product="Choline-sulfatase (EC 3.1.6.6)"
FT                   /EC_number="3.1.6.6"
FT   CDS             2060..3382
FT                   /db_xref="SEED:fig|6666666.127240.peg.3"
FT                   /db_xref="GO:0003723"
FT                   /db_xref="GO:0005525"
FT                   /db_xref="GO:0006605"
FT                   /db_xref="GO:0006614"
FT                   /db_xref="GO:0016020"
FT                   /translation="MFDNLSERLERSFKILKGEGKITEINVAETLKDVRKALLDADVNY
FT                   KVAKNFTDTVKEKALGQNVLTAVKPSQLMVKIVHDELTQLMGGETSEINLDSKPAIILM
FT                   SGLQGSGKTTFSGKLARMLKSKKNKKPLLVACDVYRPAAIEQLRVLAEQIDVPMYSEAD
FT                   SKNPVAIAQNAIQEAKAKGYDLVIVDTAGRLAVDEEMMNEIAAIKKAINPDEILFVVDS
FT                   MTGQDAVNTAKEFNERLDFNGVVLTKLDGDTRGGAALSIRSVVNKPIKFVGTGEKLDAI
FT                   DQFHPSRMADRILGMGDIVSLVERAQEQYDEEEAKRLQKKIAKNQFDFNDFLSQIAQIK
FT                   KMGNLKELASMIPGVGKAIKDIDIDDNAFKSIEAIIYSMTPEERSNPAILNGSRRQRIA
FT                   KGSGTNIQEVNRLLKQFDQTRKMMKMVTGSKMGKMMPKLKR"
FT                   /product="Signal recognition particle, subunit Ffh SRP54
FT                   (TC 3.A.5.1.1)"
FT   CDS             3394..4275
FT                   /db_xref="SEED:fig|6666666.127240.peg.4"
FT                   /db_xref="GO:0004477"
FT                   /db_xref="GO:0004488"
FT                   /translation="MTLIDGKAISEQVKQEIAAEVAEIVARGGKRPHLAAILVGHDGGS
```

```
FT                   ETYVAAKVKACEVCGFKSSLIRYEADVTEEELLAKVRELNEDADVDGFIVQLPLPKHIS
FT                   EQKVIETIDYRKDVDGFHPINVGRMSIGLPCYVSATPNGILELLKRYQIETSGKKCVVL
FT                   GRSNIVGKPMAALMMQKAYPGDATVTVCHSRSKDLVKECQEADIIIAALGQPNFVKAEM
FT                   VKEGAVVIDVGTTRVPDASKKSGFKLTGDVKFDEVAPKCSFITPVPGGVGPMTIVSLMK
FT                   NTLLAGKKAIYQ"
FT                   /product="Methylenetetrahydrofolate dehydrogenase (NADP+)
FT                   (EC 1.5.1.5) / Methenyltetrahydrofolate cyclohydrolase (EC
FT                   3.5.4.9)"
FT                   /EC_number="1.5.1.5"
FT                   /EC_number="3.5.4.9"
FT   CDS             4283..5428
FT                   /db_xref="SEED:fig|6666666.127240.peg.5"
FT                   /translation="MKTLAVFIAVLLYSVILSSQERITLLFVGDLMQHQAQIDAAHVSE
FT                   GKYDYSSCFSLIKEQISQADLAIGNLEVTLGGRPYRGYPMFSAPDEYLQAIKDAGFDIL
FT                   LTANNHCLDRGKKGMERTIQLLDSSGIRYAGTYKNLSERRQRYPLFINRNGFRIALLNY
FT                   TYGTNGIKATSPNIVNYIDKNTILQDIQSAKARQPDAIIACMHWGEEYQSLPNREQREL
FT                   ANWLLQQGVTHIIGSHPHVIQPMELRTNGTEQHIIVYSLGNFISNMSKANTDGGLIFTL
FT                   QLEKHPLPQPIRPSYTGQSQTPLPDSNPLPFPYCRVSYCGYNLVWTGRPELTKEKNYIL
FT                   YPASFPTEHLTPEARKHLEIFVKSSRKLLQQHNIGIYEKKK"
FT                   /product="Capsule biosynthesis protein capA"
FT   CDS             6053..7747
FT                   /db_xref="SEED:fig|6666666.127240.peg.6"
FT                   /translation="MKILWIDLNSSYAHSSLALPALHAQIAKNNSVKWEIVSATINENV
FT                   GMIVDEIYRHQPDIIAATTWLFNHEQLLHITARLKALLPNSCFILGGPEFLGNNEDFLH
FT                   KNPFIDCVFRGEGEEAFPQWLSCWNQPEKWNSIPGLCYMNPQKQYIDNGIARVLNFSRL
FT                   VPPENSCFFNWSKPFVQLETTRGCFNTCAFCVSGEEKPVRTLSIESIRERLQIIHSHGI
FT                   KNVRVLDRTFNYNPRRAKELLQLFLEFNPDICFHLEIHPALLSEELKKELKKLPQGLLH
FT                   LEAGIQSLREPVLQKSRRMGSLEDSLQGLRFLCSLPNMETHADLIAGLPLYRLSEIFED
FT                   VRTLAGYNAGEIQLESLKLLPGTEMRRRAEELGIRYSPLPPYEVLQTNEISVNELQTAR
FT                   QLSRLLDGFYNTTAWQAITRKLILDDNDFLRRFLEFLIDKNLIDQPMSLEKRGLVLYEF
FT                   CSMHYPAYKIMVTIAWIEAGMSLKKKPAEKVKTKRQMPPEYWEVIYGNYKESLRLCFLP
FT                   IDDNTQNGYWFGFESEIQKAEPVFKAKGIMERYQNTQSPQINTDKSS"
FT                   /product="Fe-S oxidoreductase"
FT   CDS             complement(7809..9311)
FT                   /db_xref="SEED:fig|6666666.127240.peg.7"
FT                   /db_xref="GO:0004356"
FT                   /translation="MNQELSMSPNLLVAALQKPASEFTKADIVSYIKENDIRMVNFMYP
FT                   AADGRLKTLNFVINNEAYLDAILTCGERVDGSSLFPFIEAGSSDLYVVPRFRTAFIDPF
FT                   SELPTLSMLCSYFNKDGEPLESSPEYTLRKACQAFTTVTGMEFQAMGELEYYVISENDG
FT                   LFPATDQRGYHESAPYAKFNDFRTECMSYIAQAGGQIKYGHSEVGNFTLDDKIYEQNEI
FT                   EFLPVRAEEAADQLVIAKWVIRNLASQYGYNITFAPKITAGKAGSGLHIHMRIMKDGQN
FT                   QMLKDGALSETARKAIAGMMKLASSITAFGNTNPTSYFRLVPHQEAPTNICWGDRNRSV
FT                   LVRVPLGWSAKTDMCMLANPLETPSHYDTTQKQTVEMRSPDGSADLYQLIAGLAVACRY
FT                   GFEIDDALGIAEKTYVNVNIHKKENEDKLKQLEQLPDSCAASADCLERQRAVFEQYHVF
FT                   SPAMVDGVISKLRSYEDRTLRAEIQDSPEEMLKLVEKYFHCG"
FT                   /product="Glutamine synthetase type I (EC 6.3.1.2)"
FT                   /EC_number="6.3.1.2"
FT   CDS             9340..9486
FT                   /db_xref="SEED:fig|6666666.127240.peg.8"
FT                   /translation="MVNKTHAIFCHNKKLLFRKDTKSCIRKKTNNLNLIREPQPFPSYP
FT                   HFT"
FT                   /product="hypothetical protein"
FT   CDS             complement(9527..10453)
FT                   /db_xref="SEED:fig|6666666.127240.peg.9"
```

```
FT                   /translation="MENLNAVITGVGGYVPDYVLTNDEISKMVDTNDEWIMGRIGIKER
FT                   RILHEEGLGTSYMARKAVKQLMQRTKTSPNDIDLVIVATTTPDYHFPSTASILCDKLGL
FT                   NNAFAFDMQAVCSGFLYAMEAGANFIRSGKYKKVVIVGADKMSSVIDYTDRATCPIFGD
FT                   GAAAFMLEPTREDMGVMDAVLRTDGKGLPFLHIKAGGSVCAPSYYTLDNRMHYIYQEGR
FT                   TVFKYAVANMADACEAVMERNHLAKEDIDWVIPHQANMRIITAVSQRLGVSPEKVMVNI
FT                   ERYGNTSAGTLPLCVWDFEDKLKRGIN"
FT                   /product="3-oxoacyl-[acyl-carrier-protein] synthase,
FT                   KASIII (EC 2.3.1.180)"
FT                   /EC_number="2.3.1.180"
FT   CDS             complement(10622..10792)
FT                   /db_xref="SEED:fig|6666666.127240.peg.10"
FT                   /translation="MQAADFLFFIALFIVYASKNKERRIHSSCLQKGYTEVLLQGSDSS
FT                   ESRCVLKCYYL"
FT                   /product="hypothetical protein"
FT   CDS             11135..11305
FT                   /db_xref="SEED:fig|6666666.127240.peg.11"
FT                   /translation="MVILELTVDVTDLQNASSICAGCMMHTEQTDDALGANARCVFADK
FT                   ALMTIKGCYII"
FT                   /product="hypothetical protein"
FT   CDS             complement(11558..11692)
FT                   /db_xref="SEED:fig|6666666.127240.peg.12"
FT                   /translation="MDKAGQITLQKSLLFRYLFEILENKKLIYRRLDVWTQYPPFPLN"
FT                   /product="hypothetical protein"
FT   CDS             11912..12238
FT                   /db_xref="SEED:fig|6666666.127240.peg.13"
FT                   /db_xref="GO:0003700"
FT                   /db_xref="GO:0005622"
FT                   /translation="MKEKQYTTEQEQIARFAKAMGHPARMAILAFLAKQDSCFFGDIHE
FT                   ELPIAKATVSQHLKELKEAGLIQGEIETPKVRYCINRENWELARKLFAAFLGDCKCKGT
FT                   SCCK"
FT                   /product="Arsenical resistance operon repressor"
FT   CDS             12309..12548
FT                   /db_xref="SEED:fig|6666666.127240.peg.14"
FT                   /db_xref="GO:0015446"
FT                   /translation="MTMEIKVLGTGCSSCKALYETTKQAISELGCDATLIKEEDLLKIM
FT                   EYNILGLPALVIDEKVVSAGKKLSLAEVKELITK"
FT                   /product="Arsenical pump-driving ATPase (EC 3.6.3.16)"
FT                   /EC_number="3.6.3.16"
FT   CDS             12558..13001
FT                   /db_xref="SEED:fig|6666666.127240.peg.15"
FT                   /translation="MRKLFYLLIATLVLVSCGNGSKKKTGENQAEEIQSNRIEVLYFHG
FT                   AQRCITCRAIETNTVALLDSLYSKEKADGKIIYKVIDISKKENEAIADRYEVTWSSLFV
FT                   NGWKDGKENVNNMTEFSFSNAKNAPDKFKEGIKSKIDELLKQL"
FT                   /product="hypothetical protein"
FT   CDS             13003..13710
FT                   /db_xref="SEED:fig|6666666.127240.peg.16"
FT                   /translation="MDFLQSLLDNSSIPAITAFILGILTAVSPCPLATNITAIGFISKD
FT                   IENHHRIFINGLLYTFGRIVSYTVLGFILIPILREGASMFMVQKAVSKYGEMLIAPALI
FT                   IIGIFMLDIIKFNLPKINIGGESLKKRTKGGWGAMLLGVLFALAFCPTSGVFYFGMLMP
FT                   LSAAETGGYLLPVIYAIATGLPVILVAWILAYSVAGLGKFYNRMQIFEKWFRKIVAILF
FT                   IAVGIYYAVVFYL"
FT                   /product="Cytochrome c biogenesis protein"
FT   CDS             13745..14071
FT                   /db_xref="SEED:fig|6666666.127240.peg.17"
```

```
FT                   /translation="MKKIEIFDPAMCCPTGLCGTNIDPELMRIAVVIETLKKQGVIVTR
FT                   HNLRDEPQVYVSNKTVNEHLQKHGADALPITLVDGEIAVSQTYPTTKQMSEWTGINLDF
FT                   MPVK"
FT                   /product="Arsenical resistance operon trans-acting
FT                   repressor ArsD"
FT   CDS             14097..15809
FT                   /db_xref="SEED:fig|6666666.127240.peg.18"
FT                   /db_xref="GO:0015446"
FT                   /translation="MKTFNLSDIDLTKYLFFTGKGGVGKTSIACATAVGLADNGKKILL
FT                   ISTDPASNLQDVFNQTLNGHGTDIQEVPGLTVVNLDPEQAAAEYRESVIAPFRGQLPES
FT                   VIQNMEEQLSGSCTVEIAAFNQFSDFITDADKAKEYDHIIFDTAPTGHTLRMLQLPSAW
FT                   STFISESTHGASCLGQLSGLEERKGIYKQAVETLSDANATRLVLVSRPEIAPLKEAARS
FT                   SHELQLLGIKNQLLVINGLLLQLDEADNVSKQIYDRQQNALKQTPAELLEYPSYYVPLR
FT                   SYNLSNIANIRRMLYNDNLTNDVNYQRITDAKGMDELVNDLYQSGKRVVFTMGKGGVGK
FT                   TTLATEIALKLTKLGAKVHLTTTDPANHLNYNLAVQAGITVSRIDEAEVLEAYKNEVRS
FT                   KAAETMTAEDMEYIEEDLRSPCTQEIAVFRAFAEIVDKAENEVVVIDTAPTGHTLLLLD
FT                   ATESYHKEVQRTHGDTPASVRKLLPRLRNQQETEVVIVTLPEATPVFEAERLQMDLQRA
FT                   GINNKWWVVNACLSLTDTENSFLRAKAQNELVWIKKVEELSKGNAALIAWKNN"
FT                   /product="Arsenical pump-driving ATPase (EC 3.6.3.16)"
FT                   /EC_number="3.6.3.16"
FT   CDS             15814..16242
FT                   /db_xref="SEED:fig|6666666.127240.peg.19"
FT                   /db_xref="GO:0008794"
FT                   /translation="MKILILCTGNSCRSQMAHGFLQSFDNKLDVFSAGTKPAEKVNPMA
FT                   VKVMDEMGIDLTHHTPKSVNLYIGQEWDYVITVCGGANESCPMFTGEVRNRLHIGFDDP
FT                   SEATGTPEFINSEFHRVRDEIKARFYDFYLKELRPQLS"
FT                   /product="Arsenate reductase (EC 1.20.4.1)"
FT                   /EC_number="1.20.4.1"
XX
SQ   Sequence 16634 BP; 5029 A; 3810 C; 3565 G; 4230 T; 0 other;
     tcttcgaaga attcaagggt acgggtaata tggagttgca gctcgaccgc aacttgtcca        60
     acaagcgtat cttcccggct gtcaacatcg ttgcctccag cactcgccgc gatgacctgc       120
     tgcaagacaa gcagacactg gaccgcatgt ggattctccg caagtatctt gccgacatga       180
     atccgataga agccatggac ttcgtaaaag accgtctgga gaagaccaaa gacaacgacg       240
     aattcctgat gagcatgaac agttaacaac aactgcatac agatattgta caaaagcttc       300
     catttacgtt cataaacgta ggtggaagct ttttttatat ccgtttttc atacatttgc       360
     caccatacaa taaccttgaa caaataaaat ccaaatctga tgaaaacgat tcactctttt       420
     atgggaatgg cattgtgcag ctttccggta atggcacagc aatcgcccaa ttttcttatt       480
     atccaatgcg accatctgac ccagcgggta gtaggcgcat acggtgccga ccccagctgc       540
     actccaccta tagaccgaat tgcctcccaa ggtgtaacat ttgccaacgc atacgtggga       600
     tgccccctca gccaaccgtc acgcgctgcc ttatggagcg gactgatgcc gcatcaaacc       660
     aatgtacgtt ccaactcggc cgaacacatc aatcctccgc tgcccgacag cattccgacc       720
     ctgggaagcc tcttcaccgc caacggttat gaggccgtcc actttggcaa aaccaatgac       780
     atgggttccc taaggggatt ccgccacaag gaaccggttg ccaagccctt caccgacccc       840
     gaatttcccg tcaacaatga cagctttctg gatgtaggaa cctgcgagga tgccgtggcc       900
     tatctcagca atccgccgca agaacctttc atctgcatag ccgatttcca aaacccgcac       960
     aatatctgcg gatatgtagg cgaaaataaa ggggagcaca tagatagccc catcagtacc      1020
     ccactcccgc tactacccga aaactttgaa gtggaagact gggaaaaact tccgttgccc      1080
     attcagtaca tctgctgcag ccaccggagg atgatgcaag ccgcccactg gaacgagggc      1140
     aattaccggc actacgtcgc tgccttccag cattacacgc ggatagtggc cagacagata      1200
     gagagtgtac tggaagccct ctactccact cccgccggca aaaatacaat tgtcattgta      1260
     ctggcagacc acggagacgg aatggggtca caccgcatgg taaccaagca ggtcagcttt      1320
     tacgaagaaa tgacaaatgt gcccttcatc atagccggtc ccggtattca tccccgtcaa      1380
     aaaccggtgg aagatttact gacccaaccc actatcgacc tgctgcccac tctctgtgaa      1440
```

ctggcaggca ttccggtgcc ttcctccaag cccggcatca gccttgcccc tttcctgaaa 1500
ggcgagaagc aaaagaggca gcacccctat gcagcctccg aatggcacag tgaatacgaa 1560
gtgaccgtca gcccgggacg gatgatacgc agtcccagat acaaatacac acactatctg 1620
gaaggcaacg gggaagaact gtatgatatg ctgaaggata aaggagaaag acgaaacctt 1680
gcccatgacc ccaaatactc aaagattctg gccgagcacc gggcactgct ggacgactac 1740
atccgccgca cacaggacga ctaccgaagc ctgaaagttg aagccgacaa gagatggcgc 1800
agccacgcac cgggatatcc caaccacgaa ggaccctgcg ccagagacat tatcaataga 1860
aagtaagcaa aaagaactaa aaaaaacaga ataatataga tgattatcag tgccttgacc 1920
ggcacttcca ttcttttcgc ccatcccgct gcaagtcttc aataatattc tgcttcaaaa 1980
gaaggcagtt aaagggattt tttgtacttt tgcgcctcaa taagagtaat ttgtaattta 2040
gtaattgtaa ttagatacta tgttcgataa tttaagcgag agacttgaaa ggtcgtttaa 2100
aatttgaaa ggtgagggca aaatcaccga aatcaatgtt gcggaaaccc tgaaagacgt 2160
acgtaaggca cttttggatg ccgacgtgaa ctataaggta gccaaaaact tcaccgatac 2220
cgttaaggaa aaagccttgg gacaaaacgt gctgacagcc gttaagccca gccagctgat 2280
ggtaaaaatc gtgcacgatg aactgaccca gctcatgggc ggcgagactt ctgaaataaa 2340
cctcgactcc aagcctgcca ttatcctaat gtccggtttg caaggctccg gtaaaacaac 2400
tttctccgga aagctggccc gcatgctgaa gagcaagaag aacaaaaagc cgttattggt 2460
ggcttgtgac gtgtaccgtc ctgccgctat cgagcagttg cgtgtactgg ctgaacagat 2520
tgacgtaccg atgtactcgg aagcggacag caaaaatccg gtagccattg cccaaaatgc 2580
catccaagaa gccaaagcca aaggttacga ccttgtaatt gtcgatacag ccggtcgtct 2640
ggcagtggat gaggagatga tgaacgagat tgccgccatc aagaaggcca tcaatccgga 2700
cgagattctg ttcgtagtgg attcaatgac cggtcaggat gcagtgaata ccgctaagga 2760
attcaacgaa cggctcgact tcaatggtgt agtattgacc aaactggatg gtgatacccg 2820
cggtggtgcc gccctttcta tccgttcggt ggtaaacaag cccatcaagt ttgtaggtac 2880
cggtgaaaag ctggatgcca tcgaccagtt ccatccctca cgtatggccg accgtattct 2940
cggtatgggt gacatcgtat ctttggtgga acgtgcacag gaacaatatg acgaggaaga 3000
agccaaacgc ctgcaaaaga agattgccaa gaaccagttc gacttcaacg acttcctgag 3060
ccagattgca caaatcaaga aaatgggtaa cttgaaggaa cttgcctcca tgataccggg 3120
cgtaggcaaa gccatcaagg atatagacat tgacgacaat gccttcaaga gcattgaagc 3180
catcatctac tccatgactc cggaagaacg cagcaatccg gccattctga acggttcacg 3240
ccgtcagcgc attgccaaag gtagcggtac gaatattcag gaagtaaacc gcctgctgaa 3300
acagtttgac cagacccgca agatgatgaa gatggtcacc ggcagcaaga tgggtaagat 3360
gatgccgaaa ttaaaaagat aagataatag cctatgacac tgattgatgg aaaagcaatt 3420
tcagagcaag taaagcaaga gattgcagca gaagtagccg agatagtggc tcgtgggggt 3480
aaacgtccgc atctggccgc cattcttgta ggacacgatg gaggtagcga aacctacgtt 3540
gccgccaaag taaaagcatg cgaagtttgc ggattcaagt cctcacttat ccgttacgag 3600
gcagacgtta ccgaagagga gctgcttgcc aaggtacgcg aattgaacga ggatgcggat 3660
gtggacggtt tcatcgtaca gcttcccttg cccaagcata tctccgagca gaaagtgatt 3720
gaaaccatcg actaccgtaa ggatgtggac ggtttccatc ccatcaatgt gggacggatg 3780
tccattggtc ttccctgcta tgtatccgct acccccaacg gtatcctcga actgctgaag 3840
cgctatcaga tagagacttc aggcaagaaa tgcgtggtat tgggacgcag caacattgta 3900
ggcaaaccca tggctgccct gatgatgcag aaagcctatc ccggtgatgc cacagtaacc 3960
gtatgccaca gccgcagcaa agacctggta aaggaatgcc aggaagcgga catcatcatt 4020
gctgcgttgg gacaaccgaa cttcgttaag gcggaaatgg tcaaggaagg tgctgtcgtt 4080
attgatgtag gtactacccg tgtgcccgat gcctccaaga aatccgggtt caagttgacc 4140
ggtgatgtga aattcgatga agtagctcct aaatgttcat tcattacccc cgtaccggga 4200
ggcgtaggtc cgatgaccat cgtttccctt atgaagaata cattgcttgc cggcaagaaa 4260
gcaatctacc aataacaacg taatgaaaac gctcgccgtc ttcattgctg tcttactgta 4320
tagcgtaatc ctttcctccc aggaaaggat tacgctatta tttgtcggtg acctgatgca 4380
gcaccaggca caaatagacg ctgcacatgt atctgaagga aaatatgact actcctcctg 4440
cttctccctc ataaaagagc aaatatccca ggcagacctc gccataggaa atctggaagt 4500
cacattgggg ggcaggccct atcggggcta tcctatgttc agcgcccccg atgaatatct 4560
tcaggccata aaggacgccg gattcgatat tttactgaca gccaacaacc actgtctgga 4620
ccgtggaaaa aagggaatgg agcgtaccat acaattgctt gactcctccg gtattcgtta 4680

cgcaggcacc tacaagaatc tgtcggagcg caggcaacgc tatcccctgt tcatcaacag 4740
aaacggcttt cgcattgccc tactcaacta tacttacggc accaatggta tcaaggctac 4800
ctctcccaat atcgtaaact atattgataa gaataccatc ctgcaagata tccaaagtgc 4860
caaggcacgc caaccggatg ccattattgc ctgcatgcat tggggagagg aatatcagtc 4920
ccttcccaac cgtgagcaac gcgaactggc caactggttg ctccaacaag gcgtaacaca 4980
tattatcggt tcacaccctc acgtcattca accgatggaa ctacgtacca acggtaccga 5040
acagcacatc atagtctact ccttgggaaa cttcatttcc aatatgagca aggccaacac 5100
tgacggcggc ctcatcttca ctttgcagtt ggagaaacat cctctccccc aacccatccg 5160
tccctcttac accggtcaga gtcagacgcc cctgccagac tccaatcctc tgccttttcc 5220
atactgtcgt gtttcctatt gtggatataa tctggtctgg accggccggc cggaacttac 5280
taaagaaaaa aattacattc tttatcctgc cagttttcct acagaacatt taacgccgga 5340
agctcgtaaa cacctggaaa tctttgtcaa aagctctcga aagctgctcc agcagcacaa 5400
tataggtatc tacgagaaga aaaaataatg cagaaatctc aaaaattttc gtctcattct 5460
ttgcaatatt aaagattatc cctatctttg cactgcattt aagaaatgca ctaacatggt 5520
ggctgtagtt cagttggtta gagcgtcaga ttgtggttct gaatgtcgtg ggttcgagtc 5580
ccatctgcca cccgcaaaga agaggaaatt cgaaagaata ttcctctttt tttgttattg 5640
tccaaacaat cccccaaagg gaggcaacgt ggaatcaatg taaaaaattg tagggatact 5700
cctcacagag tatccctaca tcataggaag gaatcaatta cctaaaccag tccttgtaca 5760
aaagagttta ctttctcaaa atgttcaaaa tagctggaaa cttattccga tattttgcat 5820
tctgtatcaa tcacacttaa tagtacttcc ttccttactt tatcaagcca gtgataagac 5880
aacccgttat caccttatag aaataagaat acaaaaagaa tatttcatct ttcaaaaaac 5940
attctatacc aataactcta ctattttcga tttccgacat gcaaatataa cccttttttgt 6000
ttatacgaaa agcaaaatca gttatctttg caaagatgaa attcaatcaa atatgaaaat 6060
tctctggata gacctcaata gctcgtatgc acactcttcg ttggcactcc ctgcactgca 6120
cgcacaaata gcaaagaaca attccgtaaa atgggaaatc gtatctgcca ccatcaatga 6180
aaacgtagga atgattgtcg acgaaatata ccggcaccaa cccgacatca ttgccgcaac 6240
gacatggttg ttcaaccacg aacaactgtt gcacatcaca gcgagattaa aagcattgct 6300
tcccaatagc tgcttcatct tgggtggacc tgaattctta ggaaacaatg aagactttct 6360
gcataaaaac cctttcatcg actgtgtatt caggggagag ggagaagaag cattccccca 6420
atggctgtct tgctggaacc agccggaaaa atggaacagt attccgggac tctgctatat 6480
gaatcctcaa aagcaatata tagacaacgg catagcccgt gtacttaact tttcacggct 6540
cgtaccaccg gaaaacagct gtttttttcaa ctggagcaaa ccattcgtac aactggagac 6600
tacccgaggc tgtttcaaca catgcgcatt ctgcgtcagc ggtgaagaaa aaccggtgcg 6660
cacactgtcc attgaaagta tccgtgaacg ccttcaaatt atccacagcc atggaataaa 6720
aaacgtacgc gtcctcgacc ggacctttaa ttataatccg cgccgtgcta aagaattact 6780
gcaacttttc ctggaattca acccggacat atgctttcac ttagaaatac accctgcact 6840
attatccgaa gagctgaaaa aggaattgaa aaagctacct caaggattgc tccatctcga 6900
ggccggtatt caaagcctgc gcgaaccggt cctccagaaa agccgtcgga tgggaagtct 6960
ggaggattca ctgcaaggac ttcgtttcct atgttcccta cccaatatgg aaacacacgc 7020
cgacctcatt gccggcttgc ctctataccg cctatccgaa atatttgaag atgtgcgtac 7080
acttgccgga tacaatgccg gcgagataca gcttgaatcc ctgaagttac ttcccggcac 7140
agaaatgaga agaagagcgg aagaactggg catccgatat tctcctctcc cccttatga 7200
agtgcttcaa acaaatgaaa tcagtgtcaa cgagcttcaa acagcccggc aactgtcacg 7260
tcttctggat ggattctata acacaactgc ttggcaagcc atcacccgca agctgattct 7320
ggatgacaac gacttcctac gtcgctttct ggaattttta atcgataaaa atcttataga 7380
ccagcccatg agcctggaaa agcgaggcct cgtactctat gaattctgca gtatgcatta 7440
tccggcctat aaaatcatgg tcaccattgc ctggatagag gcaggcatgt cattaaagaa 7500
aaaaccagca gagaaagtca agaccaagcg ccagatgcca ccggaatatt gggaagtcat 7560
ctatggaaac tacaaagaaa gcctccgctt atgttttcta cccattgacg acaataccca 7620
aaatggttat tggttcggct tcgaatcgga aatacaaaaa gcagaaccag tattcaaagc 7680
caaaggaata atggagaggt atcaaaatac acaatcaccg cagattaaca cagataaaag 7740
ttcttgatta tcagaaattc tatagcaatc tgtgtaacct gcggcgagtt atgccctcat 7800
ttcttctcct atccgcagtg gaaatacttc tcgaccagtt tcagcatctc ctccggactg 7860
tcttgtatct ctgcacgcaa ggttctatcc tcgtaagaac gcagtttgga aataacccca 7920

tcaaccattg caggactgaa cacatgatat tgctcaaaga cagcacgctg tctttccaaa 7980
caatcggcag aagcggcaca actgtcagga agctgttcca gttgtttcaa cttatcttca 8040
ttttctttct tgtggatatt tacattgaca tatgtttttt ctgcaattcc taaagcatcg 8100
tctatctcaa atccataacg gcaagctaca gcaagtccgg caatcaattg atacaaatcg 8160
gccgaaccgt caggcgaacg catttccacc gtctgtttct gtgtagtatc ataatgactt 8220
ggggtctcca acggatttgc caacatgcac atatccgttt ttgccgacca gcccaacgga 8280
acgcgtacca ataccgaacg gttgcggtcg ccccagcaaa tgttggtcgg tgcttcctga 8340
tgaggaacca gacgaaaata ggaagtagga tttgtattac cgaaagccgt aatggaagaa 8400
gccagcttca tcatgcccgc aatggcctta cgggctgttt cagaaagtgc cccatctttc 8460
agcatctggt tctgcccgtc cttcatgata cgcatatgta tatgtaatcc ggaaccggct 8520
tttcctgccg tgattttcgg tgcaaaagta atgttatatc catactggct agccagatta 8580
cgaataaccc attttgctat caccagttgg tcagcagcct cctcggcacg caccggcagg 8640
aactctattt cattctgctc ataaatctta tcgtccagcg taaaattacc gacttcagaa 8700
tgcccatact ttatctgtcc gcctgcctgc gcaatatatg acatacattc cgtacggaaa 8760
tcattgaact tggcataagg ggccgattcg tgataaccac gctggtcggt agccgggaac 8820
aatccgtcat tttcagaaat gacataatat tccaattctc ccatagcctg aaattccatc 8880
cccgtaaccg tagtgaatgc ctgacatgcc ttacgcaggg tatattccgg agaactttcc 8940
agcggttcac catccttatt gaaataagaa cagagcattg acagtgtggg caattcagaa 9000
aaagggtcta taaaagcggt ccggaaacgc ggcactacat acaaatcact gctaccggcc 9060
tctataaacg gaaacaagct ggaaccatct acgcgttcac cacatgtaag tatggcatca 9120
aggtatgctt cattattgat gacaaagttc agtgtcttca gtctcccgtc tgccgcagga 9180
tacatgaagt tcaccatacg aatgtcgttc tcctttatat aggaaacaat gtctgctttc 9240
gtaaattcgg aagccggttt ttgaagggca gccaccaata gattggggct catggataat 9300
tcctggttca taatacgctg attgttttta gttttctata tggttaacaa aacacatgca 9360
attttctgtc acaataaaaa actgctgttt cgcaaagata ctaaaagttg tatcagaaaa 9420
aaaacaaaca acctaaatct tatcagagag cctcaacctt ttccgtcgta tccccacttc 9480
acgtaaatag ccccccaagc aaatcctgct ccgaatgctg taaaaatcag tttatccccc 9540
ttttcagctt atcctcgaaa tcccatacac acaatggaag cgtgccggca cttgtatttc 9600
cataccgttc tatattgacc attacctttt caggagatac tcccaagcgt tgcgaaacag 9660
cagtaattat acgcatgttt gcctgatggg gaatcaccca gtcaatgtcc tccttcgcca 9720
agtgattcct ctccatgacc gcctcacaag catccgccat attggccaca gcatatttaa 9780
agactgtacg tccttcctga taaatataat gcatccgatt gtccagtgta tagtaagatg 9840
gagcacatac ggagcctccc gctttgatat gtaaaaaggg cagacctttc ccgtctgttc 9900
tcaatacggc atccattact cccatatcct cccgggtggg ttccagcatg aaagcggcag 9960
caccgtcacc aaaaatagga caggtagcgc ggtctgtata atcaatcacc gatgacatct 10020
tatcagctcc tactatgaca accttcttat attttccgga acgtatgaaa ttcgcccctg 10080
cctccattgc atacagaaat ccactgcaaa cagcctgcat atcaaaagca aaagcattgt 10140
tcaggcccaa tttgtcacat aagatggatg cggtagaagg aaagtgataa tcgggagtag 10200
tagtagctac gataaccaag tcaatgtcat tcgggctggt ctttgtgcgc tgcatcagtt 10260
gtttcacagc cttacgagcc atatatgaag ttccaagccc ctcctcatgc aaaatgcgtc 10320
tttcctttat accaatgcgc cccataatcc attcatcatt ggtatccacc attttagata 10380
tttcatcatt agtcaagaca tagtccggaa catatcctcc aactcccgta atcactgcat 10440
tcaaattctc cattaatcct atccagttta gtgtaaaatc aggaataacc ctatttaaat 10500
atgttatcag aatatcacac ttttttcaga acagtcgtgt gacaatcttt aattaatggg 10560
cgcaaatata cgaaaagaga atagaaatca atgaatctga caacaaaaag acattaactg 10620
tctacaaata ataacacttt aaaacacaac gggattcact actatccgaa ccctgcagga 10680
gtacttccgt ataacctttc tgcaaacaag aagaatgaat acgtctttcc ttattctttg 10740
atgcatatac aataaacaaa gctataaaga acaaaaaatc cgctgcctgc aacgatggcg 10800
cacgattggc cgcacaaccg gccacatttc cttttccagc gaatggagca tcgcaggatg 10860
cggtttatgc attatatacg atttcttttt catctgaact aacaaactca cccccctaccc 10920
caattcctcc ctctccctct tgaagtagcc cagcaccgtg gctagcgtga gacggcagcg 10980
ttccacgctt ttcaggtggc cctaaaaggt gtagcgagag cacagcacga ggtagaacct 11040
atagttcaag ctacggtgca agcggtgctg aaggcaggct ctgacatctt cggcggagag 11100
tttcattcct ttgtttgggg ttactttcat tgtaatggtt attttagaat tgactgttga 11160

tgtgacggat ttgcaaaacg catcatccat ctgtgccgga tgcatgatgc acaccgagca 11220
aacggatgat gcgttgggag caaatgcacg atgcgttttc gctgataaag ccttaatgac 11280
cattaagggt tgttatataa tatgatatta atcgctgccg gtttggaggt tagcggaaaa 11340
catgtacatt tgcaatgtgt ttttcatagt attagattta aggttaacaa aagattggct 11400
gtctgggata gatagccttt ttttatgttg ttttctgacc tgcttcgtgg aattaagttt 11460
ttaatagaaa atactttaag acagacacaa ggggataaaa aaaacataaa tcattgaatt 11520
tgtgggactt gtctgcttac tgttcggttt tgtccggtta gttcagcgga aagggagggt 11580
attgagtcca gacatctaac cgcctataaa tcaacttttt attttccaaa atttcaaata 11640
ggtaacgaaa cagtaacgat ttttgcaggg ttatttgtcc ggctttgtcc actgtttgtc 11700
tgcctatctc acgggttcaa agatacaaca catttttttta attacgcaaa tctctgtata 11760
taaatgtttc agcttaggtg caaggtgcaa gctatatata tagatatata cttgcacctt 11820
gcacctatcc gaaaaaaata tgccataaaa tttggctact aaaaataatg ttcgtagttt 11880
tgcgaacaac aaataagaaa ggagaattga agtgaaagag aaacagtaca caacagagca 11940
ggaacagatt gctcgttttg ccaaagcaat gggacatccg gcacggatgg ctattcttgc 12000
tttcttggca aaacaagata gttgtttctt cggtgatatt cacgaagaac tgcctatcgc 12060
taaagcaacc gtttcgcagc atttgaaaga actgaaagaa gcagggttaa ttcaggggga 12120
aatagaaacg ccaaaagtcc ggtattgtat aaaccgtgag aattgggaac ttgcccgaaa 12180
attgtttgca gctttttttag gggactgtaa atgtaaaggt acatcgtgct gtaaatagca 12240
gtaccttttt ttggaataat tgttcgtagt tctacgaact acgattatta tattaacttt 12300
aaagtagaat gacaatggaa attaaagtat taggaactgg gtgttcaagc tgtaaagcct 12360
tgtacgaaac taccaaacaa gctatttcag aattaggttg cgatgcaacc cttatcaaag 12420
aggaagattt attgaaaatc atggagtata acattttagg gcttccggct ttggtgattg 12480
acgagaaagt cgtatcagcc gggaaaaagt tatcccttgc agaagtgaaa gagctgataa 12540
ccaaataaaa atgaactatg agaaaattat tttatctatt gattgcaacg ctggtcttag 12600
tttcctgtgg taacggttca aagaaaaaga ccggagaaaa ccaagcggaa gaaatacagt 12660
ctaaccgcat agaggttctt tattttcatg gagcgcaacg gtgtatcact tgccgggcga 12720
tagaaacaaa tacagtagcc cttttggata gcctttattc aaaggaaaaa gcagacggta 12780
aaattatcta caaagtgata gatatttcaa agaaagaaaa tgaagcaatt gcagacaggt 12840
acgaagttac atggtcgtct ttgtttgtga atggttggaa agacggcaaa gagaatgtaa 12900
acaacatgac agagtttagt ttttccaatg cgaaaaatgc accggacaag tttaaagagg 12960
gaattaaaag caagattgac gaattgttaa aacagttgta agatggattt tcttcaatcg 13020
ctattagata acagttctat tcctgctatt acagctttta tcttgggcat tttaacagcg 13080
gtcagcccgt gtccgttagc gaccaacata acggcaatag gatttattag taaggatata 13140
gaaaatcatc accggatatt tataaacggc ttgctttata ctttcggcag aatagtaagc 13200
tatacggttc ttggttttat ccttatccct attctccgtg agggggcaag tatgtttatg 13260
gttcaaaaag ccgtaagcaa gtacggggaa atgctgattg ctccggcgtt aatcatcatt 13320
ggaatattta tgctcgatat aataaaattc aatttgccga aaatcaacat cggcggtgaa 13380
agtttgaaaa agaggacgaa aggcggttgg ggggctatgc tattgggtgt tttattcgcc 13440
cttgcttttt gtcctaccag tggagtattt tatttcggta tgcttatgcc tttgtcagca 13500
gcagaaacgg gcgggtatct cttacctgta atttacgcta ttgctacggg attacccgta 13560
atccttgttg catggatatt agcgtacagt gttgccggac tgggtaagtt ttataaccgg 13620
atgcaaatat tcgagaaatg gtttcgtaaa atagttgcca tcctctttat tgcggtagga 13680
atttattatg cagtagtctt ttatctataa ataaacatag agtatcattt aaaataaaaa 13740
cagtatgaaa aagatagaaa ttttcgaccc ggcaatgtgt tgccctacgg gtctttgtgg 13800
aacaaatatc gaccctgaat taatgcgtat tgcggttgtt attgaaacat tgaagaaaca 13860
gggtgtcatt gttacccgcc acaatttacg tgacgaaccg caagtgtatg taagtaataa 13920
aacagtaaac gagcatctgc aaaaacatgg ggcggatgca ctgcctatta ctttagtgga 13980
cggcgaaatc gctgtttcac aaacctatcc taccaccaaa caaatgagtg aatggacggg 14040
tatcaatttg gattttatgc cagtaaaata agtattcacc agctaattat ataaaaatga 14100
aaacatttaa tttatcagat atagatttga cgaaatacct tttcttcacc ggaaagggcg 14160
gtgtaggaaa aacttcgatt gcctgtgcta ctgcggtagg cttggcagat aatggaaaga 14220
aaatacttct tatcagtaca gacccggctt caaacctgca agatgtgttt aatcaaacac 14280
tgaacggaca cggcacggat attcaagaag tacccggctt gacggtcgtt aatctcgacc 14340
cggaacaagc cgcagccgaa tacagggaaa gtgttatcgc tcctttccgg ggacaactgc 14400

```
     ccgaaagtgt tattcagaat atggaagaac aactttcagg ttcttgtacg gtagaaattg     14460
     cagctttcaa tcagttttcc gattttatca cggatgccga taaagcaaaa gaatacgacc     14520
     atattatctt tgatacagca cccacgggac acacattgcg aatgctacaa ctgccatcag     14580
     cgtggagtac atttattagt gagagtacgc acggtgcatc ttgcttaggt caattatccg     14640
     gtttggaaga acggaaaggc atctataagc aagcagttga aacactttcg gatgcaaacg     14700
     caacccgctt agtattggtt agccgtcctg aaatcgcacc attaaaagaa gccgcccgtt     14760
     cttcacatga attgcaatta ctgggaatta aaaaccagct attggtaatc aacgggcttt     14820
     tgctgcaatt agatgaagcc gataacgtat cgaaacagat atatgacagg cagcaaaatg     14880
     ccctgaaaca gacccctgcg gaactgctgg aatatccgtc ttattatgtt cctttacggt     14940
     catacaattt atctaatatt gcaaatatcc gccggatgct ttacaatgat aacttaacaa     15000
     atgatgtgaa ctatcagcgg attaccgatg ccaaagggat ggatgaactg gtaaacgacc     15060
     tctatcaatc aggaaaaagg gttgttttta ctatgggaaa aggcggcgtg ggaaaaacca     15120
     cgttagccac tgaaatagcc ctgaaattaa caaaattggg tgcaaaggtg catcttacca     15180
     ccactgaccc ggcaaaccat ctgaattata accttgctgt tcaggctggc attacggtaa     15240
     gccggattga cgaagcggaa gtgttggaag cctataaaaa cgaggttcgc agtaaagctg     15300
     cggaaacaat gacagccgaa gatatggaat atatagagga agatttacgt tcaccgtgta     15360
     cgcaggaaat tgcagtattc cgggctttcg ctgaaattgt cgataaagca gaaaatgaag     15420
     tcgtagtaat tgacaccgca cctaccggac acactttgtt attgctggat gcaacggaaa     15480
     gttaccataa agaagtacaa cgtacacatg gcgacactcc cgcttccgta agaaagctgt     15540
     taccacgttt gagaaaccag caggaaacag aagtcgttat tgtgaccctg cccgaagcaa     15600
     cacccgtatt tgaagccgaa cgtttgcaaa tggatttaca acgtgccggg attaataata     15660
     aatggtgggt cgtgaatgct tgcttgtcat taactgacac cgaaaattct ttcttgcggg     15720
     caaaggcgca aaatgaattg gtttggatta agaaagtaga agaattgtca aagggaaatg     15780
     ctgccctgat agcttggaaa aataactaaa aacatgaaga ttttaattct ttgtacaggg     15840
     aatagctgcc gcagccaaat ggcacatggc tttctacaat catttgacaa taaactggat     15900
     gtcttttcag caggaacaaa acctgctgaa aaggttaatc cgatggcggt aaaggttatg     15960
     gatgaaatgg gtatagattt aacccaccat acccctaaaa gtgtaaacct atatatagga     16020
     caggaatggg actatgttat tacggtctgc ggtggggcaa acgaaagctg cccgatgttt     16080
     acgggtgaag taaggaacag gctgcatata ggatttgacg acccctcgga agcaaccgga     16140
     acgccggagt ttataaacag tgaatttcat cgggtacggg atgagataaa agcccgtttc     16200
     tatgacttct acctgaaaga attaagacca caattaagct aaatagttgt tgttatggaa     16260
     aagaaacaag gaattggatt ttttgaaaaa tacctgacta tctgggttgc cttgtgcatc     16320
     attattggaa ttgccgtggg acaatggctt ccggcaattc cgcaaacatt aagcaaattt     16380
     gaatatgcca acgtgtctat acccgtggca atcctgattt ggttaatgat ttatccgatg     16440
     atgctaaaag tggattttca aagtgttaag aatgtcggca agcgtccgaa aggaataata     16500
     gtcacttgcg ttacaaattg gctgataaag ccatttacca tgttcggaat agcttatttg     16560
     ttcttctatg tggttttcaa agcctttata cctgctggat tagccgaaga atatttagcc     16620
     ggggcggtat tatt                                                       16634
//
ID   XXX; XXX; linear; XXX; XXX; XXX;  XXX.
XX
AC   PRJEB67889
XX
DE   Uncultured bacterium DNA fragment from in-reactor enriched cow rumen, clone
ferm-15k21
XX
FH   Key             Location/Qualifiers
FH
FT   source          1..39574
FT                   /organism="uncultured bacterium"
FT                   /environmental_sample
FT                   /mol_type="genomic DNA"
FT                   /isolation_source="Cow rumen"
FT                   /clone="ferm-15k21"
```

```
FT                   /note="AZO-CM cellulose, AZCL-HE cellulose, AZCL-Barley
beta glucan, AZCL-Tamarind xyloglucan, AZCL-arabinoxylan active clone"
FT   CDS             complement(490..684)
FT                   /db_xref="SEED:fig|6666666.127241.peg.1"
FT                   /db_xref="GO:0003735"
FT                   /db_xref="GO:0005622"
FT                   /db_xref="GO:0005840"
FT                   /db_xref="GO:0006412"
FT                   /translation="MIIVPVKEGENIERALKRFKRKFEKTGAIRELRARKNFEKPSVKK
FT                   RIQMQKAIYVQHLQLMDEQ"
FT                   /product="SSU ribosomal protein S21p"
FT   CDS             992..2002
FT                   /db_xref="SEED:fig|6666666.127241.peg.2"
FT                   /translation="MSLIQQTLLNIRVTGMQFVVGQFNLLNVRTIASFLQRYGSTAIRT
FT                   VASTPEEYVSIVAEYLKDTHFVFDQDRYSADDEILSLNWDIDDNDIAAILRLPFLLSSD
FT                   RNVKIIMIIDEFQNVEYTEDGERVLKAMEKLFSEMRSAGKTDCVFLLCGSMVNAMKHIF
FT                   EEKHYFYRQVEHLSLRKIDEREIIEYIHRGFMLSGKEIDRSLLHGTCQLFRCNMWYINH
FT                   FISICDSLSKGYIVESTLIEALNIILSIHEPRFMAMMAGLTSFQVNMLRAVVDGNVRFS
FT                   AADVIRNYGLNSSANVKRVREALVKKEIITFNEKEEPLIMDPLFEYWVTKYYFEK"
FT                   /product="hypothetical protein"
FT   CDS             1999..2706
FT                   /db_xref="SEED:fig|6666666.127241.peg.3"
FT                   /translation="MKKKIAVLTGAGVSAESGISTFRDSGGLWKNYDPQVVASIDGWYR
FT                   DKKLMLDFYNMLRKDLAAAKPNAAHYAIADLEKDYDVTVVTQNVDNLHERAGSTRIIHL
FT                   HGELTKVRPEHGVYDELFSEKEVFDIGYEAVNLGDRAPNGAQLRPHIVWFGEAVPKIDA
FT                   AIDVFEAADVVLIVGTSLQVYPAAGLYRYAKHDVPIYIIDPQDVPVSDPRITHIKDVAT
FT                   RGMDRFMQILREK"
FT                   /product="NAD-dependent protein deacetylase of SIR2
FT                   family"
FT   CDS             2830..3141
FT                   /db_xref="SEED:fig|6666666.127241.peg.4"
FT                   /db_xref="GO:0003723"
FT                   /db_xref="GO:0003735"
FT                   /db_xref="GO:0005622"
FT                   /db_xref="GO:0005840"
FT                   /db_xref="GO:0006412"
FT                   /translation="MYAIVDIAGQQFKVEAGSVLYVNRLAAKKDEAVEFDKVLLLDNEG
FT                   AVKVGAPYVEGAVVKATVLKDDVKGNKVIVFKKIRRKGFQKLNGHRQKLSLIKVDAIA"
FT                   /product="LSU ribosomal protein L21p"
FT   CDS             3161..3421
FT                   /db_xref="SEED:fig|6666666.127241.peg.5"
FT                   /db_xref="GO:0003735"
FT                   /db_xref="GO:0005622"
FT                   /db_xref="GO:0005840"
FT                   /db_xref="GO:0006412"
FT                   /translation="MAHKKGQSSSKNGRESQSKRLGLKKFGGEVVKAGNIIVRQRGTVH
FT                   NPDKNVGMGKDHTLYALIDGTVKFTKKRNDKSYVSVLPLEN"
FT                   /product="LSU ribosomal protein L27p"
FT   CDS             3505..4776
FT                   /db_xref="SEED:fig|6666666.127241.peg.6"
FT                   /db_xref="GO:0004828"
FT                   /translation="MLTLKLLRDDPEFVIKKLAVKNFDAKEIVAKILELDTNRRSLQTE
FT                   SDALLAQQKQKAAVIGGLMKEGKKAEAEEAKKEVAALKERSNELLAKAEENNKELESQL
FT                   VLLPNIPCDLVPEGKSAEDNLVVKMGGNDPVLPENALPHWELAKKYDIIDFDLGVKITG
```

FT                   AGFPVYKGKGARLQRALINFFLDYNTKAGYKEVEPPVVVNAASGFGTGQLPDKEAQMYY
FT                   CGLDDFYLVPTAEVPVTNIFRDEILSAGEIPQKLTAYTPCFRREAGSYGKDVRGLNRLH
FT                   QFDKVEIVRVEKPEDSYKALDEMVAHVESIVNALGLKYRILRLCGGDMSFTSAITYDFE
FT                   LWSAAQQRWLEISSVSNFESYQANRLHLRYRDENGKTQLAHTLNGSSLALPRVVAALLE
FT                   DNQKEDRIVIPEVLRPYTGFDYID"
FT                   /product="Seryl-tRNA synthetase (EC 6.1.1.11)"
FT                   /EC_number="6.1.1.11"
FT   CDS             4783..5346
FT                   /db_xref="SEED:fig|6666666.127241.peg.7"
FT                   /translation="MAENLQERIIVGIDPGTNILGFGVIRAVGRSVSYVDMGILDLRKE
FT                   KDAYSKLRQIYDTVSEVCRTYHATEMALESPFYGKNAQVILKLGRAQGAAMIAAMEQGI
FT                   DVVEYAPRKAKEAITGNGAASKEQVSTMIQKTLGIPLDDKFLDATDALAIALCHFYQTT
FT                   NPLASVKGSSSWEKFVAANPDRVK"
FT                   /product="Crossover junction endodeoxyribonuclease RuvC
FT                   (EC 3.1.22.4)"
FT                   /EC_number="3.1.22.4"
FT   CDS             complement(5339..6085)
FT                   /db_xref="SEED:fig|6666666.127241.peg.8"
FT                   /translation="MSIAGNSIFNRTGLLFGEKGLERLSAAKVIIFGVGGVGSWCAEGL
FT                   IRSGICNLTIVDSDKVCVTNINRQLMATTRTIGQMKVDALKERLLEINPEASITAIPEI
FT                   YGEETADKFNLSEYDYIIDAIDSLSNKMLLMLRACETSAVFFSSMGAALKVDPTRIRVA
FT                   EFWEVRGCPLGAALRKKFKRAKLCPARKFLCVYDDEVLPNVGQAEDDGSCTFDKVVANG
FT                   TTASVTAIFGFTLSGLVVKDIYSKLI"
FT                   /product="ThiF domain protein"
FT   CDS             complement(6082..7587)
FT                   /db_xref="SEED:fig|6666666.127241.peg.9"
FT                   /translation="MERIAEVGKIKRFWIWHYSTFVLWNAATTFWVCISTVGGGLFAIF
FT                   ANALQMSLIFGLFRWSKKTFRGSLPYIFLALMWIAWERFYFSAEISWPWLTLGNSFART
FT                   ISLVQWYEYTGTLGGSLWIWACNLAVFGLLCLIADDDWRRLGWKARLAAITGTVVVFVA
FT                   PVAVSEVMYKNYEETGEPLDVLIAQPNIDPYHKFVALSQDSQDAILDGLIQDAMKGRDT
FT                   ADTSPLLILGPETFTSGIDTDNIGNSKTFRRLSRIAREYPGTNVLFGASTWTYIFSDER
FT                   PSETARKVMDGIWVECHNSAIITDGSDDYDIFHKSRLVVGVELMPYPKIFRPLDEALGG
FT                   VIGRDIGQDEISLLNVRTQDGGKLPLGCAICYESVYGEYCTGYVRKGAKALTVITNDAW
FT                   WADTPGYRQHLSYASLRAIETRRDIARCANTGISAIIDQRGDIVSRTTWWEPEVLSGTI
FT                   NLTDGETFFVRNGDVTGKFCTFIFLLLLLALVTRSFIPKSLRK"
FT                   /product="Apolipoprotein N-acyltransferase (EC 2.3.1.-)"
FT   CDS             7855..8421
FT                   /db_xref="SEED:fig|6666666.127241.peg.10"
FT                   /translation="MAILVLQKSLSGGHRAGFFTGLGATFTDTLYSVIAIFALAITQQF
FT                   ITEHEVFIKITGGIAIILLGANMLFKDPFRKMKDQSGPSYSIKDFLQAVATGLSNPFAI
FT                   LIIMSLFGFFGLAPESRDLTVAPVILAVSGGSMTYWFFYSLIFSRIRKNFKLSTLLWIG
FT                   RISGAIIMIIGIALASDGLMKLIFE"
FT                   /product="Putative threonine efflux protein"
FT   CDS             8423..9205
FT                   /db_xref="SEED:fig|6666666.127241.peg.11"
FT                   /translation="MPEILKNWIVKNLLLAIGLLLALVLVSALFLNLLTRHGKELVVPD
FT                   FTNMKISEATYTASVDGMRIEVVDSVYIRRMGRGLVYSQNPKPGSKVKKGRRILLTINS
FT                   VTPKKVQMPNLVGYSMRQAKAELSSRGLALGKLMYVSDIATNNVLRQLVGNMEITPGRM
FT                   IESGTAVNLVVGLNSSDSQTYIPDVKGLKYLRAVDVVHDNSLNIKRLVFDESVKNYSDS
FT                   LDAIVVRQGPEASMAPIVMGSDVTLYLSVDPAKTIMKK"
FT                   /product="hypothetical protein"
FT   CDS             9207..10238
FT                   /db_xref="SEED:fig|6666666.127241.peg.12"
FT                   /translation="MEFDDLQDDGMFEHFRLNVDPGQTPMRVDKYMACHLEDTSRHRVQ

```
FT                   NAIKSGYVRVNDKVAKANVIVRPGDVIRFVMPYERHGFEVLPENIPLNIVYEDDDFLIV
FT                   NKPAGMVVHPGHGHFDGTLINALAYHLGISQGPDAEDDRMGILVHRIDMDTSGLLVVAK
FT                   NDEAQLNLAQQFFVHSIERKYVAVVWGNIKEDEGTIVGNIGRDPNDRLKFRVYQEEDKG
FT                   KHAVTHYKVLERFGFVTVVECRLETGRTHQIRVHMSYIGHPLFNDARYGGAEIRKGTIY
FT                   AKYKQFIQNCFEICPRQALHAKTLGFVHPRTGKPVSFDSPLPEDMTALLEKWRKYAGDM
FT                   PKE"
FT                   /product="Ribosomal large subunit pseudouridine synthase D
FT                   (EC 4.2.1.70)"
FT                   /EC_number="4.2.1.70"
FT   CDS             complement(10256..11827)
FT                   /db_xref="SEED:fig|6666666.127241.peg.13"
FT                   /db_xref="GO:0003723"
FT                   /translation="MDSEKELLKDLIVDVQATEVQIALQENHRLIELNKESSSGRSFAV
FT                   GDAYLGRVKKVMPALNAAFVDIGDEKEAFIHYLDLGLYFGAFDEFVKKTNPNTNPSRLF
FT                   GSIKLGPILEKEGRIENVLTQGQMIIVQIVKEPISTKGSRLTAEISLAGRNIVLLPFAE
FT                   KVSISQKISSKEEKKRLETLVRSILPKNYGAIIRTAAEGKNAAILVAELKSLIGKWEDS
FT                   WENLSKNKSIQPLFTEYSKTTTILRDLLNDSFSNIYVNNEKEYEEIRRYISLISPEQEK
FT                   IVKLYQGKEPIFDHFEITRQIKSSFGKVVPLKQGAYLVIETTEALNVVDVNSGIRAKTK
FT                   EQEENTFEVNKFAAEEIARQLRLRDMGGIVIVDFIDMESVEHRNNLFKYMQELMSKDRA
FT                   KHNVLPLTKFGLMQITRQRVRPVTEINTTEVCPVCHGTGKIASSLLIDEQIERRLSYYV
FT                   NDKGIRNFTLKVSPILGAYITRGLFNSYVGKWKKKYKCKIKVVEITDFAVLQNELFNEK
FT                   GDKLDS"
FT                   /product="Cytoplasmic axial filament protein CafA and
FT                   Ribonuclease G (EC 3.1.4.-)"
FT   CDS             complement(12056..12328)
FT                   /db_xref="SEED:fig|6666666.127241.peg.14"
FT                   /translation="MTKAEIVNEVAKATGIEKLAVQSVVESFMESVEASLAKGEPVYLR
FT                   GFGSFIIKHRAEKAARNITKNTTLTIPAHNIPAFKPAKVFVSKVK"
FT                   /product="DNA-binding protein HU"
FT   CDS             12572..14956
FT                   /db_xref="SEED:fig|6666666.127241.peg.15"
FT                   /translation="MFPASSHEVNIIPVMSGEPALNFDASELPGSLPILALRNAVIFPG
FT                   VVFPITIGREKSMLLVDKAQGNGMFIGAVPQKNVAEENPLSLADFSEFGTVVKVLKTFE
FT                   MPDGTVTALLQGYKRLRMLELTDTDPYHMARVSYLDDILPQNDSKVPVIADILKENASS
FT                   IIRSSSFVPKEALEALKNIDDFNFLVNFIATTIDMDDFMEKVDLLKYDDVKERAMALLG
FT                   VLSRQLEIQRLKQEISSKVKTELDQQQREYFLNNQLRTIQEELGMDEMDDIARFREAAE
FT                   KKEWPASVGETFEKELAKLERFNPSTPEYSIQYNFLQFMIDLPWYHTSDDNLDLKHAQK
FT                   VLDEDHFGLENVKDRIIEYLAVLKLKGNMKSPILCLYGPPGVGKTSLGKSIARALGRKY
FT                   VRISLGGMHDEAEIRGHRKTYVGALPGRILNGINKAGTSNPVMVLDEIDKLGSDFKGDP
FT                   SSALLEVLDPEQNTAFHDNYLDIDYDLSNVMFITTANTINGIQPALLDRMELINVTGYL
FT                   AEEKVQIAKKFLVPKQLEEHGIPKKELKIDKAALEEIIDKYTHESGVRGLEKQIARIAR
FT                   VSAKKMALGEDYPKTIKKEHLKEYLGLQTAFHDSQKGNEAPGVVTGLAWTSMGGEILFI
FT                   ETSVSDGKGVLTMTGNLGDVMKESATIGYQYIKAHPQLAGLTSEEFAKKDVHVHVPEGA
FT                   VPKDGPSAGITMVSSMVSALRGKKVRSGIAMTGEMTLRGRVLPVGGIKEKILAAKRAGV
FT                   STIIISEENRKDIEDIKEIYVKGLTFVYVKTIDDVIGYIFE"
FT                   /product="ATP-dependent protease La (EC 3.4.21.53) Type I"
FT                   /EC_number="3.4.21.53"
FT   CDS             14956..15624
FT                   /db_xref="SEED:fig|6666666.127241.peg.16"
FT                   /translation="MDVKIEQTWKDALRDEFEKPYFASLVRFLHNEKEAGKVIYPPGGL
FT                   IFRAFELTPLPEVKVVILGQDPYHGPRQAMGLSFSVPSDVPAPPSLKNIFKEIQTDLGI
FT                   TMSGSPNLEGWARQGVLLLNAMLTVQAGVAGSHKAIGWQEFTDAVIKCISDRCNGVVFL
FT                   LWGNFARGKKELIDTSRHYVLEAAHPSPLAGGAFFGCRHFSKTNELLIKEGKTPIDWKL
FT                   "
```

```
FT                   /product="Uracil-DNA glycosylase, family 1"
FT   CDS             15621..16094
FT                   /db_xref="SEED:fig|6666666.127241.peg.17"
FT                   /db_xref="GO:0004595"
FT                   /translation="MKRKALFPGSFDPFTAGHLNILRRALTMFDEIVIAVGVNQDKRGF
FT                   FSMDQRMDIIRQAVRGMDGIEIIKYDNLTIDTCRELGIAHIIRGVRNMLDFENERAIAD
FT                   ANRRLAPEIETIIIPTAQEFAHISSSAVRDILKHKGDTSLFLPEGVVLPDHDA"
FT                   /product="Phosphopantetheine adenylyltransferase (EC
FT                   2.7.7.3)"
FT                   /EC_number="2.7.7.3"
FT   CDS             complement(16096..16290)
FT                   /db_xref="SEED:fig|6666666.127241.peg.18"
FT                   /translation="MVSYIVSRARRNEEIALGPLYIQRLGLQRRKILLKLCIQGGHLDG
FT                   IRILGAQGKRKQRKKDYED"
FT                   /product="hypothetical protein"
FT   CDS             16333..17565
FT                   /db_xref="SEED:fig|6666666.127241.peg.19"
FT                   /translation="MGDEAVAVHLADNWFIPGKVSFPDEIDLMNAKIFAEFNRQSLLGC
FT                   KAPGLVAEDSLGRKVEVPLGLSDRYRILYIYSPDCATCSLQTMLMDSLFSRKSYPVDLI
FT                   AFNSDDNTEAWVQYRRNRLSFDGVRVMHFRDPSLESDFQRKYGVLQTPRMFLIDKNGII
FT                   VGRGLDAAALEKLLDATPDLGIGYGSKSSADYFDAVLESFGDELRGSDIRYFIGRLEEA
FT                   TLESADTAGFRQMTGDMLYYLAGKRGEAFKDAEAFVVDSMVLSRPDVWRFADDSLTVVD
FT                   MAMSLKHMLSLAPVGARVPDVRVHATVLRAGRKPATVHKKLRSLLKGKRNYVLFHTAGC
FT                   TFCEAELAAADSVASEKGVRVILVDMDKLTSLYPEEADRLLDAFDLSVLPFIMETDNKG
FT                   CVVRKYLSLNK"
FT                   /product="hypothetical protein"
FT   CDS             complement(17554..18126)
FT                   /db_xref="SEED:fig|6666666.127241.peg.20"
FT                   /translation="MSIMGKAYIFLADGFEETEALATLDILRRGKIEAYTVSITEEVYI
FT                   TSSHGVTVVADMGRSDFSEIDIDEKDAMIFPGGMPGSKKLAADTPLIGLMNVHYGNGGI
FT                   VAAICAAPGLVATQIKGIEGKKATCFDGFEENLAASGMEYVKAPAVTDGQLITGRGAAF
FT                   AIDFGLAILEKMGGKEAVAKVKAALLI"
FT                   /product="DJ-1/YajL/PfpI superfamily, includes chaperone
FT                   protein YajL (former ThiJ), parkinsonism-associated
FT                   protein DJ-1, peptidases PfpI, Hsp31"
FT   CDS             complement(18236..19348)
FT                   /db_xref="SEED:fig|6666666.127241.peg.21"
FT                   /db_xref="GO:0008784"
FT                   /translation="MNRVDDFSRLELNLTGCMENYRYFRSKLEQTTKLLVLVKANAYGH
FT                   GAVEFAAMMQRAGANYLAVAYPVEGMELRKAGISLPILVLTAGTDFFNEIIDNRLEPGI
FT                   PNLYALKILCGILKERGIKDYPVHIKLDTGMHRLGFMTSELQDLTDFLKTCEEVKVKSV
FT                   YSHLAVSEDPEQDEFTLGQIGMFKTNADRITSEIGYKPMYHILNSAGIERFPQYQFDMV
FT                   RLGIGIYGISALPDVRLAHVASLKCKILQIKTLKPEDGTIGYGRYGHIAPEGTVIATIP
FT                   VGYADGIDRHLGRGHASFSVNGHRAPTIGNICMDMCMLDVTGIDAKVGDTVTIFGEDPT
FT                   VNELADILGTIPYEILTSVPRRIERIVVRK"
FT                   /product="UDP-N-acetylmuramoylalanyl-D-glutamyl-2,6-
FT                   diaminopimelate--D-alanyl-D-alanine ligase (EC 6.3.2.10) /
FT                   Alanine racemase (EC 5.1.1.1)"
FT                   /EC_number="6.3.2.10"
FT                   /EC_number="5.1.1.1"
FT   CDS             19707..20381
FT                   /db_xref="SEED:fig|6666666.127241.peg.22"
FT                   /translation="MDMIWFIIIAVMVLSFVIQQVLNSKFNKYSKVGLHSGLTGAQVAA
FT                   KMLEDHGITDVSITCIKGKLTDHYNPVNKTVNLSEEVYYGNSVAAAAVAAHECGHVVQH
```

```
FT                   ATGYAPLKLRSALVPVVSFSSNAVQWVLLIGVMIIKITPTLLYLGIAMFALTTLFSVVT
FT                   LPVEINASARAVQWLDKAGITGPATKAKAVDALKWAAYTYVIAAIGSIATLIYYIAMAR
FT                   RR"
FT                   /product="probable metal-dependent peptidase"
FT   CDS             complement(20378..20671)
FT                   /db_xref="SEED:fig|6666666.127241.peg.23"
FT                   /translation="MTGDDYKIISDTTENGVRHITAIPSSLVCSQQIDFDIIDGKVRNL
FT                   AYVRGCNGNLQAIGRLVEGMDIKKIVDTLSGVNCNFRGTSCTDQLARILSSL"
FT                   /product="hypothetical protein"
FT   CDS             complement(20688..21122)
FT                   /db_xref="SEED:fig|6666666.127241.peg.24"
FT                   /db_xref="GO:0008270"
FT                   /db_xref="GO:0016787"
FT                   /translation="MEKDELYMKEALREAQAALAEDEIPIGAVIVCKGRIVGKGHNMTE
FT                   RLHDPTAHAEMIAITAATEALGGKYLNECTLYVTVEPCPMCAGALNWAQIGRIVYGAID
FT                   PKRGHSLFTPSLLHPKTEVEAGVLAEECGSMVTEFFKKKR"
FT                   /product="tRNA-specific adenosine-34 deaminase (EC 3.5.4.-
FT                   )"
FT   CDS             complement(21243..21473)
FT                   /db_xref="SEED:fig|6666666.127241.peg.25"
FT                   /translation="MFKIPAGNQRKLIFAYLFDTVAYDSPDALAMLDEVQLVFPVPVEG
FT                   EDEFGLMSLDYVEAVPVRKRCYFRYYRGHIH"
FT                   /product="hypothetical protein"
FT   CDS             21558..22130
FT                   /db_xref="SEED:fig|6666666.127241.peg.26"
FT                   /translation="MMRNQFSSIQHMFEQARNGLTFSLAAIMKVYSLLDALPSEIDKFD
FT                   QFQHFMRLLYDLSLAGDSRILASTSFAHNDADKESRRVQKIKQYVGDHFAENITLEELA
FT                   EMVGMSPSAFSRFFKSRTGKTVIDYIIDIRLGNAARLLVDTTQNISEICYSCGFNNLSN
FT                   FNRLFKSKRGYTPREFRTLFKKNKVFV"
FT                   /product="Transcriptional regulator, AraC family"
FT   CDS             22237..24699
FT                   /db_xref="SEED:fig|6666666.127241.peg.27"
FT                   /translation="MIKRLFPALFIALFAILSCQPMKMDIVKVDDGSFVVDGHFYYYIG
FT                   TNFWYGGLLGSEYGDRSRLAAELDSLHALGLDNLRVLVGSDGPDGVATRVYPTLQKEPG
FT                   VYDEKVFEGLDYLMAELGKRNMRAVLYINNSWEWSGGYGMYLEWAGAGKAIIPSVGGYG
FT                   PFMEQMAQYATNARAQELFENHLRTVVSRVNSITGKPYSEDPAIFSWQIGNEPRCFSSD
FT                   PAVQDGFVSWIWRAASIIKSIDPNHMVSTGNEGIWGCEENPELFRRLNECPDIDYITAH
FT                   IWPYNWSWVGKESPQLDIDRAEENTGRYIDAHLEVAESLGKPLVIEEFGFPRDDMQFAK
FT                   GTPVSGRDRYYRYVFSKVMENQKNGGRLAGANFWSWGGLAVPAHRYWQEGDDYAGDPAQ
FT                   EEQGLNSVFADDYSTLAIIKHTTARLSAVANASAVLENDWMFTTGDQKPLRVAVQSQNG
FT                   FKGSVPVYVRVSTDKGVSVSVSGKKARLGKGADTLDFDLGLAPGFYKVCVSLNENDPLD
FT                   ENFCVGCDPEKIVSPQDKQPDFDEFWEDNLRQLAMVPMNARMTPLKDHSNADRTAYRVD
FT                   MKSFGGKEISGLLYLPNKEGKFAARISYMGYGSDVWFADPSAEPDMIEFTLSVRDQGFN
FT                   KTPGQGWVTRGLADKDSYYYRGAYLDCVRAIDFVSSLPQTDPDHIYAEGGSQGGAFTLI
FT                   AASLDNRLRAIAPFVPFLSDFPDYFAIADWPGGEVLSAAKAQGISEDELYKTLSYFDVK
FT                   NFTDRIGCPVLMGFGLQDPVCPPHTNFSGYNHITADKQWLCVPVSGHHLEQFPEWWAAR
FT                   TEFFTKNK"
FT                   /product="GH5_7-CE7"
FT   CDS             24787..27936
FT                   /db_xref="SEED:fig|6666666.127241.peg.28"
FT                   /translation="MAQNKVTGRITDNSGQGVINASVMVKGTNNGVVADLDGYYEIVVG
FT                   QGSTLVYSSIGYKDQEVVVGTQSVINVLLVENSELLDEIVVVGYGTMKRSDLAGSSVSM
FT                   KEDDLKGSIITNLDQSLQGRAAGVTAVQTSGAPGSSSSIRVRGQATINANAEPLYVIDG
FT                   VIMQGGGNSGADFGLGDALGNGKVSTISPLSTINPADIVSMEILKDASATAIYGAQGAN
```

```
FT                   GVVLITTKHGSRGDAKFSYDGMFAVSRQTTRIDMMNLREYAEFYNSMVETGEISEANPY
FT                   YATPGLLGKGTNWQDEIFRTAIQHQHQISAQGGTDKVQYYVSGSFMDQEGTIRGSEFGR
FT                   YTFRTNLDAQLKEWFKLGLNLTFANTNDDLKLADGSEGLITYSLTTIPDIPVYDVYGNF
FT                   SSTVREGYTSPNPVALAMMDEILLDRQKLNGNIYAELTPVKHFTWRTELGFDVSGSTAT
FT                   RFRPTVNLGGWTRDTNSMSYQKNSNRFWQIKNYLTYANSFGKHSVTAMLGQEAWESSWS
FT                   YVQGANTKLPSNDVHNVALGGGTPSLSSGFGSSAMASFFTRETYNFDDRYLLTYTYRYD
FT                   GSSNFGPEKRWAGFHSLAVAWKFTNEKFMDFLKPVVNNGKLRLGWGQTGNSSIGGYAWG
FT                   ASISRMPSALGMGYRPANIPNPAIHWESQTQQNIGLDLNMFDNRLNLTVDAYLKKSDNM
FT                   LMSMQLASYMGTQGNSSSALAAPKGNFGSIENKGLEITLDAHPVQIKDFAWDSNFQISF
FT                   NRNKLVSLDGTTNAALIGYGQWSDVVCVSEIGKPLYNFYGYKVAGVYTSLEDIENSPKP
FT                   EKYPSNGEYARGNTVWVGDIKYEDISKDGVINEKDRTDIGNPLPKFTFGWSNTFTYKDF
FT                   DLNIFINGSYGNKVLNYNAISLTHMNSAWTNQINDAVSGRAMLAPIDASVVYADGSKWY
FT                   DHIDNVRVLNPGTKTPRASINDPNDNDRISDRYVEDGSYIRIKNITLGYNVPKSFLNKF
FT                   KVDNVRVYVNIQNLYTFTKYTGYDPEVGASTQDSTGLTYGVDNGRYPSPTVYSCGLNIT
FT                   F"
FT                   /product="TonB family protein / TonB-dependent receptor"
FT   CDS             27949..29697
FT                   /db_xref="SEED:fig|6666666.127241.peg.29"
FT                   /translation="MKISNMMKGFAVATAVFALTSCEGFLNRPNEDNYNVDNFYKTDEQ
FT                   CEQGVNYLYNSPWYDFQRGFIKIGEVMSGNMYWGSSPYLDFSTNGTDQDLVNMSYSLWA
FT                   VNGHANTVIYNLLNSQGPSKAAIDKCIGEALTWKAFAYFFMVRTFGDVPIVHSNSDLLS
FT                   SGEYNSISKVERADVYEYIITTLEKAMEMLPKKTSGFNNYERIDYYAAEALLAKVYLTK
FT                   AGLSGSLNQDDLKQAAKYAKDVIDNSGRTLLPVYSDVFRLTGFNATGECLISWEWSVGR
FT                   NPWTQQNTLQSDLAMVGFSDQGDCWGGWGGPSVDLADAFGVSATQNPAERVGEKDSRRK
FT                   ATLMLPGDKYDYFWTDKGGFDYLRFIYDTEYGKGGPGGSMQCPTGSNNVKHLYGDNFDH
FT                   ENAGLGTPENMAYQLPTHLLRLSDVYLIYAEASLLTGSNGPALEYVNKVRERAGADALA
FT                   SVTFEDIWKERRLELAGEGDRWYDYVRRSYYDMDACIAELKAQRRSYWDGLDDVWKSYI
FT                   GDAFDYSGTWDASEVQYNNAEDNPNVTAASFKLPFPTEDVVFNGNMASDAVPVHVDVRE
FT                   EYSYNF"
FT                   /product="putative outer membrane protein, probably
FT                   involved in nutrient binding"
FT   CDS             29710..31014
FT                   /db_xref="SEED:fig|6666666.127241.peg.30"
FT                   /translation="MKSINRIITGSLAILAMASFASCDKVDYPDRYEVADGKPTVYSVR
FT                   YADKDVNITQAYMDEVVCLLGKNLKSIVELWFNDQQATLNTSFITDNTLLVSVPKNMPV
FT                   VQTDKIYMITSGKDTLAYDFAVLPPTPKVTSMSNEWAAEGETVTIYGDFLIDKADKPLQ
FT                   IAFAGAEVDHASMVFNGSSSVSFPVPAGAQPGYVTVTSLSGTGKSKFMYKDTRNILFDW
FT                   DGSHGGFATGHGWRNGVIHAPGADAWDAIDGSYLYFGGADLGPGSADAWAEDQYSFNYW
FT                   PEAGTDYDALNLRPEFAEYIDKYGVSGLVLKFEVLVPSSNPWSASGMQLMFSSDAVVTY
FT                   GTATNSYFSDETFPRAIWQPWTTNGGSYDTADKWVTVSFPLSSFNKNQENKECATAFDK
FT                   SYLTGLTFFVWNGGIDGTACSPVFAIDNIRVVPNE"
FT                   /product="FIG00417257: hypothetical protein"
FT   CDS             31021..33447
FT                   /db_xref="SEED:fig|6666666.127241.peg.31"
FT                   /translation="MKLIMKKIFSISAILAACVLALGLSGCSSKELDTDQYSDKDVKLA
FT                   SFGPNPVMRGATLRFFGSNLDRIVEVSVPGLDPITDIEVVTSGNPSEIRIQLPAEGAQV
FT                   GKISLKSSDGKVFTTQVDVEYTEPIVFDSFSPAEAMPGDVITVKGDYMNLVQSVTFEGG
FT                   AIVTEINHNGRHEASFKVPSKAITGKIILSDEGAIANLLYSEEDLVIGDPTVKSLTVAV
FT                   AKPGENAVVKGAYLDMIDKVVFEGGAEVSDLALNEDATEFNLVIPAGAQSGEVKAVSHA
FT                   AKEFVAGEIELVIPTDVAAAPSPVKAGADLVISGNDLDVVTSVDFPGASGVEFVFAEDA
FT                   ITVTVPATATEGDIALNMASGEAVNVAFTLVHPVVSAIAPVELMAGETITVTGTDLDLV
FT                   TGVTLGGKPVEFTAEAEKIVVQTANTSVSGKLVLSLANGEKIEPEDAITLSYDSFIIVN
FT                   DMPAAEHIGAIVTLKGENFMMIENIYIGEEKVSKYNLRSDNEIQFVMPWNKVGTYSMYF
FT                   HLLSGDVETCPQTIEVLLEQDIMTVWEGSLQITWNDGGRVTVPAASFTGIPAGSKLRFY
```

```
FT                   YTQVDQQWDQAQLNYGNWTGINFDDPAGTVINGTLVPTDVYGWFTDGILDRCTEVVLTK
FT                   EILENIQANKGDCEDKTACGIIIQGSGLTFTKIEIVCDIPQEVTIYEGPTTLTWGDDGR
FT                   FGLAMTYFEAAKPGSQLIFYLEQTDAWGQVQLNDGWWANADMNFPEIGGAYITTDNLGG
FT                   KDVTRIALTLTADILDHILATSGDYFGLNSQFQGDGRVAMVIQGSDMIINKVCIL"
FT                   /product="hypothetical protein"
FT   CDS             33542..35068
FT                   /db_xref="SEED:fig|6666666.127241.peg.32"
FT                   /translation="MIFLAACSCSKDDAEKKEPSVPAPKAVTVDVASVSAPVEGTSVVV
FT                   AISSPSKPAVSVSGDWIKAELGPYNPDTYKVKLTVTVGENPTYESRSGSVTLTCQGFSP
FT                   VGISVSQDAAAKPVGPSANPAYDLLGLGWNMGNQFDAFYNGSWAGDKYNYPDETVWGNA
FT                   KATQATFDGVKKAGFTSVRIPVTWLNMIGPGPEYNIDETWMNRIYEVVGYAEKAGLKVI
FT                   INTHHDENNNDDHWLDILSASKDARTNEAIKAEIAAVWTHIANKFKDKGDFLIMESFNE
FT                   IQDGGWGWSAAFQADPSKQCDVLNEWNQVFVDAVRATGGNNATRWLGVPTYAAGTSWTK
FT                   YFRLPEDPAGKVMMAVHCYDPYEYTIGNIQYPEWGHTGAAGKKTPEGEKEITDLFSRLN
FT                   TEFISKGIPVYLGEFGCSMRSKGDARAWMFYKYYMEYVSKAAHTYSLPGILWDNGSTKT
FT                   GKECHGYIDHGTGKYIGNSKEIVDLLVKGMLSTDESYTLQSVYDRAPKTN"
FT                   /product="GH5_4"
FT   CDS             35185..36213
FT                   /db_xref="SEED:fig|6666666.127241.peg.33"
FT                   /translation="MDRMAASVKAGKILYGHQDDLVYGHSWCVDPDSAATDALNRSDVK
FT                   AVAGDYPAVVGFELGGLELGWDWSIDKVPFLLMRRAAVEHAGRGGIVTFSWHPHNPYTG
FT                   GDAWDISSDKAVASVLEGGEKAEMFGVWLERLGDFMDSLRDDKGNLIPVIFRPWHENIG
FT                   SWFWWGGALCTADEYIALFRRTVEYLRNDRGLDNIIMCYSPNSGITAEQYMSRYPGDDV
FT                   VDILGYDHYETVMEGESLEDAGKRYAGQLDLGLSIVKSLASAHGKLMTLSETGMEGLPD
FT                   RSWWTSVLYPAVKDFPIVYVLTWRNAWDRETHFYAPYPGNEGDFREFASLPEISLLSSF
FT                   RP"
FT                   /product="GH26"
FT   CDS             36291..37253
FT                   /db_xref="SEED:fig|6666666.127241.peg.34"
FT                   /translation="MENMSFEERLAWLEARHQELIAKKNSPIYGNGVFERYENPVLTGD
FT                   HAPLFWRYDLDKNSNPFLEERFGIHAAFNAGAIKWKDKYVVCARVEGADRKSFFAIAES
FT                   PNGIDNFRFWDHPVTMPDYGEPATNVYDMRLTAHEDGWIYGIFCVERKDPDAPEGDLSS
FT                   AVAAAGIARTKDLVNWERLPDLVSDSQQRNVVLHPEFVNGKYALYTRPQDGFIDAGNGG
FT                   GIGWALVDSMENAVVTSEKIINCRRYHTIKETKNGEGPAPIKTDRGWLHLAHGVRGCAS
FT                   GLRYVLYMYMTSLEDPSKVIAELPVSSWSRKGRSMWVMS"
FT                   /product="GH130"
FT   CDS             37214..37465
FT                   /db_xref="SEED:fig|6666666.127241.peg.35"
FT                   /translation="MVPEGEEYVGDVMNVLFSNGWIADEDGKVFIYYASCDTRLHVATS
FT                   TVDRLVDYCMNTEPDLFTTSASVRRLNALIDKNLSLSM"
FT                   /product="GH130"
FT   CDS             37486..38913
FT                   /db_xref="SEED:fig|6666666.127241.peg.36"
FT                   /translation="MDIITAPAKPTLKEKIGYGFGDMSSSMFWKIFSYYLPIFYSDVFG
FT                   LKPQHAAFLLLITKLYDAISDPVMGIIADRTRSRWGKYRPYLLWVALPFSLIGLLSFYT
FT                   PDASYGLKHVYAYVMYILMMTVYTAINVPYGAMLGVVTPDSREKSVFSSYRMFFAYIGS
FT                   FIAMGIFGVFEKLISGTVRVMPDGSERVVRGVGDAAPGQWTLVVGIVSLCCFVLFLLCF
FT                   LMTRERVGTENKAVSEGGSSIAEDLKALGRNRPWWLLLGAGIGILLFNSIRGGAAAYYF
FT                   SNILGTSVFLTCAVYLFIGEVAQMVGVLFTVPLSERIGKKATFISVLLTVSVLSVAIFF
FT                   LPETPAGFWGLLVLQVLICIAFGVQSPLLWSMFADVADYSEEKNGSASTGLIFSSSSMA
FT                   QKFGGALGAFLLMQILAIFAYDKDAAVQAPQTLAAIRGLMSYIPAIGALLGALCLLFYP
FT                   LTESRMASIREKLSARR"
FT                   /product="Xyloside transporter XynT"
XX
```

SQ   Sequence 39574 BP; 9470 A; 10001 C; 10231 G; 9872 T; 0 other;

```
     gagggtcatg taattgacgg agacggtcag gttgccgaac gtggcgagag tatcctcggg        60
     ctcgctccgg ccttcttcgc gaagatgtcc ctggaccccg acggtatcgt aggccgcgaa       120
     tttttcatgc gggtaggtta tgctggcttc atcgaccgaa atattgaggt aggggaaact       180
     cttgaacacg gaggcgctta tgcttccgaa acggaggttt ccgtcgatga attcggcggc       240
     atacttgttg gccgtcttag tgagaaatct ggtatcaagt attatctgca gggctattat       300
     gattatggcc catagtccca caaaaattcc gagggctatc ttgagtcccc tgaatctgcg       360
     ttttttttcg gattcagcca ttaatgattg ttatatattt tacaaatata actatttagg       420
     cggtatatag tacaaaaaaa agaagccaat gtgattgact tcttctttgt atctatagca       480
     tcggggagat tactgctcgt ccatcagctg aagatgctgc acatagatag ccttctgcat       540
     ctgaattctc ttctttacag atggcttctc aaagttcttc cttgcgcgaa gctcacggat       600
     tgcgccggtc ttctcgaatt tccttttgaa tctcttgaga gctctctcta tgttctcacc       660
     ttcttttaca ggtactatga tcatttgcta tatcaccccc tttttgcttg tttttctttt       720
     ttagcgggtg caaaggtact aaaacttttt tcatttcaaa aaaaaccgct aaaaaaatta       780
     cagatagata taaaatatta tgaaataatt tattattttt gctaacagaa atgttttaca       840
     accattaaca cccatggaaa gtcccttcgt ttattacagg tatgtgaccg ggaaaaactt       900
     cctcggacgc aaggacgatt gccggattct tggcaatctt ttactacagg gagaacatgt       960
     ggcgatatat gagagcccga aatcgggtaa aatgtcattg attcagcaga ccctacttaa      1020
     cattcgtgtt accggaatgc agttcgtcgt cggacagttc aatctgctca atgtcaggac      1080
     tatcgcttct tttctccaac gctacggatc gaccgccatc aggacagttg cttcaactcc      1140
     ggaggaatac gtttctatcg ttgccgaata tcttaaagat acccatttcg tcttcgatca      1200
     ggatcgttac tccgctgatg atgagatcct ttcactcaac tgggatatag atgacaatga      1260
     cattgccgct atcctccgtc ttccgttcct tctctcgtcc gacaggaatg tcaagatcat      1320
     aatgatcata gacgaattcc agaatgtcga atataccgag gacggcgaac gtgtcctcaa      1380
     ggccatggag aagctcttct ccgagatgcg ctcggccggg aagactgact gcgtcttcct      1440
     tctctgcggc tctatggtca atgccatgaa gcatatcttc gaggagaagc attatttcta      1500
     ccgtcaggtg gagcatcttt ccctccggaa gatcgacgag cgtgagatca tcgagtacat      1560
     ccaccgcggt ttcatgctca gcggaaagga gatcgaccgg tccctgcttc acggcacttg      1620
     ccagctcttc cgctgcaaca tgtggtatat caaccatttc atatcgatct gcgactccct      1680
     ctccaagggc tatatcgtcg aatctactct tatcgaggcc ctgaacataa tactgtccat      1740
     ccatgaaccg agattcatgg cgatgatggc aggactgact tcgttccagg tcaatatgct      1800
     ccgtgccgtc gtggatggca atgtgcgctt cagcgccgcc gatgtcatcc ggaattacgg      1860
     cctcaactct tcggccaatg tgaagagagt cagggaggcc cttgtcaaga aagagatcat      1920
     tactttcaat gagaaggaag agccgttgat catggatcct ctcttcgaat actgggtaac      1980
     caaatattat ttcgagaaat gaaaaagaag atagcagtcc ttaccggtgc cggagtaagc      2040
     gccgagagcg ggataagcac tttccgcgac agcgggggat tgtggaagaa ctatgatccc      2100
     caggtagttg cgtcgataga tggctggtat cgggacaaga aactgatgct ggatttctat      2160
     aacatgttgc gtaaggacct tgctgcggcc aagcctaatg cggcccatta tgccatcgct      2220
     gatctcgaga aggattacga tgtcacagtg gtgacgcaga atgtcgacaa tctccatgag      2280
     agggccggca gcaccaggat catccacctc cacggcgaac ttaccaaagt ccgcccggag      2340
     catggagtct atgacgagct tttctccgag aaggaggtct tcgacatagg atacgaggcg      2400
     gtcaacctgg gggacagggc accgaacggc gcccagctcc gccctcatat cgtatggttc      2460
     ggggaagccg tgcctaagat tgacgccgct atagacgtct tcgaggctgc cgacgtggtc      2520
     cttatcgtag gtacttcttt gcaggtttat cctgccgccg gactgtaccg ctatgcgaag      2580
     catgacgtcc ctatctatat aatagatcca caggatgttc cggttagcga cccccgtatt      2640
     acccatataa aggacgtcgc cacccgggga atggaccgtt ttatgcagat tttgagggag      2700
     aaatagttgt aaaaccagaa agttttacta cctttgtggt cccaaaaatg taaatggccg      2760
     tcgagcaagt aaagatctga catttagggc aggcgcttac ggctctaaac tttaagagtt      2820
     tattaaacaa tgtacgcaat tgtagatatc gcaggacagc agttcaaagt tgaggctggt      2880
     tccgtgcttt atgtaaatcg tctcgcagca aagaaagatg aggctgtaga gttcgacaag      2940
     gtgctccttc tcgataacga gggtgcagta aaggttggcg ctccatatgt agagggtgct      3000
     gtagtaaagg caacagtcct taaggacgac gtcaagggta acaaggtgat cgtgttcaag      3060
     aagatccgtc gtaaaggctt ccagaagctc aacggccatc gtcagaaact ttcccttatc      3120
     aaggtagacg ctatcgcata aattgaggag gaattaaatt atggcacata agaaaggaca      3180
```

atcaagttca aagaacggtc gtgagtcaca aagcaaacgt ctcggactca agaaattcgg 3240
tggcgaagtc gtaaaggccg gtaacatcat cgtccgccag agaggtacag tccacaatcc 3300
tgacaagaac gtcggaatgg gcaaggacca cacactttac gctctcatcg acggtaccgt 3360
taaattcacc aagaagagaa acgacaaatc ttacgtttcg gttcttcctt tagagaacta 3420
agactcggag gtttatgaat agaacgaaag gacagcactt cgagccgatg tgcagtcctt 3480
ttttcattat aaaacaggat acagatgctt acactgaaat tactccgtga cgatccggag 3540
tttgtgatca agaaactcgc cgtcaagaat tttgacgcaa aggagattgt agcaaagatt 3600
ctcgagctcg ataccaaccg tcgctccctc cagacagaga gtgacgccct tctcgcacag 3660
cagaagcaga aagctgccgt aatcggcggt ctcatgaagg aaggcaagaa agccgaggct 3720
gaagaggcta agaaagaagt ggccgctctc aaggagcgct ccaacgaact tctcgcaaag 3780
gcagaagaga ataacaagga actcgagtct cagctcgtgc ttcttcctaa catcccgtgc 3840
gacctcgtac cggaaggcaa gagcgctgag gataatcttg tcgtcaagat gggcggcaat 3900
gatcctgtac tccctgagaa tgctcttcct cactgggagc ttgcaaagaa atatgatatc 3960
atcgacttcg acctcggcgt gaagatcacc ggagccggat tccctgtata taaaggcaaa 4020
ggcgcccgtc tccagagggc tctcatcaac ttcttcctgg actacaatac aaaggcaggt 4080
tacaaggaag tggagccgcc tgtagtggta aacgccgctt caggtttcgg tacaggccag 4140
cttcctgaca aggaggccca gatgtactac tgcggactcg acgatttcta cctcgttcct 4200
accgcagagg ttcctgtgac caacatcttc agggatgaga tcctctcagc aggcgagatc 4260
cctcagaagc ttacagccta tactccttgc ttccgccgcg aggccggttc atacggcaag 4320
gacgtacgcg gactcaaccg tctgcaccag ttcgacaaag tcgagatcgt gcgcgtagag 4380
aagcctgaag attcatacaa ggctctcgac gagatggtag cgcatgtgga gtcaatcgtg 4440
aatgctctcg gtcttaagta ccgtattctc cgtctctgcg gcggcgacat gagcttcacc 4500
tctgccatca cttatgactt cgaactctgg tcagcagccc agcagcgctg gctcgagatt 4560
tcttcagtat ccaacttcga gagctaccag gccaaccgtc tccatctccg ttacagggat 4620
gagaacggca agacccagct cgcccatact ctcaacggaa gctcgctcgc ccttccacgc 4680
gtcgttgcag ctctcctcga ggacaaccag aaggaagaca gaattgtcat ccctgaggtg 4740
ctcagaccat acaccggctt cgattacatc gactagtctg ccatggcgga gaaccttcag 4800
gaaaggatta tagtcggcat agaccccgga accaacatac ttggtttcgg ggttatccgt 4860
gccgtcggcc gcagtgtcag ctatgttgac atgggcatcc ttgacctccg caaagaaaag 4920
gacgcatatt ccaagctgcg tcagatttac gataccgtat cagaagtctg ccgtacctac 4980
catgccacgg aaatggctct ggaatctcct ttctacggga agaacgccca ggttatcctt 5040
aagctcggac gtgcccaggg agcggctatg attgccgcga tggagcaggg gattgatgtt 5100
gtggaatatg ccccgagaaa ggctaaagag gctattacgg gcaatggtgc agcttccaag 5160
gagcaggtca gcaccatgat ccagaagact cttggtattc cgctggacga caagttcctg 5220
gacgcgacgg atgcgcttgc aatcgcgctc tgccacttct accagacgac aaacccgctg 5280
gcttcggtca agggatcttc aagttgggag aaattcgtgg ccgccaatcc ggaccgcgtc 5340
aaataagctt ggagtagata tctttcacta ccagccctga aagggtaaat ccgaaaatag 5400
cggtaacgga cgccgttgtg ccgtttgcca cgactttatc gaaagtgcat gagccgtcat 5460
cttcggcctg accgacatta ggcagcactt cgtcatcata gacacatagg aacttgcgcg 5520
caggacagag ttttgcgcgc ttgaattttt ttctgagagc ggctccgagc gggcagccgc 5580
gcacttccca gaattcggct acgcgtatcc tggtagggtc taccttgagc gccgccccca 5640
tggaggagaa gaatacagcg gaagtctcgc aggcgcgcag catcagcagc atcttgttgc 5700
tgagactgtc gatagcgtcg attatatagt catattccga cagattgaac ttgtcagccg 5760
tttcttcgcc gtagatttcc gggatggcgg taatcgaagc ctccggattg atctccagaa 5820
gtctctcctt gagggcgtcc accttcatct ggccgatggt cctggtggtg gccatcagct 5880
ggcggttgat gttggtgacg catactttgt ccgaatccac tatcgtcagg ttgcatattc 5940
cggagcgtat cagcccttct gcacaccagc ttccgacacc tccgacgccg aagattatga 6000
ctttggcggc agagagcctc tccaggccct tctctccgaa gagaagcccg gtcctgttga 6060
atatggaatt gccggcaatg ctcatttctt gagagactta ggaatgaaac tgcgggttac 6120
gagggccagc agaagaagca ggaaaatgaa agtgcagaat tttccggtca catctccgtt 6180
cctgacgaaa aacgtctctc cgtcggtaag attaatggta ccggaaagga cttccggttc 6240
ccaccatgtc gtgcggctga cgatgtctcc gcgctggtcg attatcgccg aaattccggt 6300
gttggcgcag cgggcgatgt ccctgcgggt ttcgatcgcc ctgagggaag catagctcag 6360
atgctgtctg tatccgggag tgtcggccca ccaggcgtca ttcgttatca cagtcagagc 6420

tttcgctccc tttctgacgt aaccggtgca gtactcgccg tagactgatt cgtagcatat 6480
tgcgcagccc agaggaagct ttcctccgtc ctgcgtcctg acattgagca gagaaatctc 6540
gtcctgccct atatcccggc cgataacccc tccgagggcc tcgtccagcg gcctgaatat 6600
cttcgggtac ggcatcagtt cgactccgac tacaagccgg cttttatgga agatgtcgta 6660
gtcgtcgctt ccgtcggtaa tgatggcgga gttgtgacac tcgacccata tcccgtccat 6720
gactttccgc gcagtctcgg aaggcctttc gtcactaaag atatatgtcc atgtagaagc 6780
cccgaaaagg acgttcgttc ccgggtattc cctcgcaatc cttgacaggc ggcggaaggt 6840
cttgctgttg ccgatgttat ctgtatcgat gccggaagtg aacgtttccg gccccaggat 6900
caggagaggg gaggtgtctg cggtgtcgcg gcctttcatc gcgtcctgga tcagtccgtc 6960
gaggatggca tcctggctgt cctggctgag ggcgacgaat ttatggtaag gatcaatgtt 7020
cggctgggcg ataaggacat ccagcggctc tccggtctct tcatagttct tgtacatgac 7080
ttcggacact gcgaccggag cgacgaaaac gacgacagtc ccggtaatcg cggcgaggcg 7140
ggctttccat cccaggcggc gccagtcgtc gtccgcaatc aggcagagca gcccgaatac 7200
ggcgaggttg catgcccata tccatagcga cccgccgagc gtgcctgtat attcgtacca 7260
ttggacaagg cttatcgtgc gggcgaagga gttgcccaga gtcagccatg gccatgaaat 7320
ctcggcgctg aagtagaatc tttcccatgc gatccacatt aatgcaagga agatataagg 7380
cagcgagccc cggaaagttt tcttgctcca gcggaacagc ccgaagatca gggacatctg 7440
gagggcgttg gcgaagattg cgaaaaggcc gcctccgaca gtcgagatgc agacccagaa 7500
tgtggtcgcg gcattccaca gtacgaatgt ggaataatgc cagatccaga atctttttat 7560
ctttccgacc tcggcgatcc gttccatgca caatagcggg acgagtccta tgagggccac 7620
gatgccgcag tgcggtacca ggaaaggtat cgacataagt gctgccgata gcagcatcag 7680
aataatgagc gagacagtct ctttacgcat atccaccaac ttaggtttta caaatttaca 7740
tagaattatg cgaaaaaata gctaattttg taaattgctt gattacgaaa tatgttattg 7800
gatatactaa aggcttttct tgtaggacta tgcgcctccg ccccgctcgg acctatggcg 7860
atacttgtcc ttcagaaatc tctgagcgga ggacacaggg ccggtttctt taccggtctc 7920
ggagcgactt tcacggatac gctttattcc gtgatcgcta tcttcgccct tgccattacc 7980
cagcagttca taacagaaca cgaggtattc atcaagataa ccggaggtat agcgataata 8040
ctgcttggag caaacatgct tttcaaggac cctttccgca agatgaagga ccagagcggt 8100
ccgtcatatt cgatcaagga tttcctgcag gcggtagcga ccggactgag caatccgttc 8160
gccatactta tcataatgtc ccttttcggc tttttcggcc ttgctcctga aagccgtgac 8220
ctgacagtcg ctccggtgat acttgcagtg agcggcggct cgatgactta ctggttcttc 8280
tattcgctga ttttcagcag gataagaaag aacttcaaac ttagtaccct gctttggatc 8340
ggaaggattt ccggcgccat cataatgata atcggtatcg cgctggcttc tgacggtctt 8400
atgaagttaa tattcgaatg atatgcctga aatactgaaa aactggattg taaagaacct 8460
gttgctggcg ataggtctgt tactcgcgct tgtgctggta tcggcacttt tcctcaatct 8520
gcttacccgt cacggtaagg aactggtcgt gccggacttc accaatatga agatatccga 8580
agctacttat actgcctccg tcgacggaat gcgtatagaa gtcgtggact ctgtatatat 8640
ccgcaggatg ggcaggggac tggtatacag ccagaatcct aagccgggca gcaaagtcaa 8700
gaaaggacgc cgtatcctcc ttacgatcaa ttccgtgacc cctaagaagg tccagatgcc 8760
taacctcgtg ggctattcta tgaggcaggc caaggcggaa ctttcgtcaa ggggacttgc 8820
cctcggaaaa ctcatgtatg tcagcgatat cgctaccaac aatgtgctcc gtcagctcgt 8880
cggcaatatg gagattactc cgggccgtat gatcgagagc ggaactgccg tcaatctggt 8940
agtcggactc aactccagcg acagtcagac ttatatccct gatgtcaagg gactaaagta 9000
tctgcgtgct gtcgatgtag tgcatgacaa ctcgctgaat atcaagaggc ttgtcttcga 9060
cgaaagcgtc aagaactatt ccgattcgct tgatgccata gtagtccgtc agggaccgga 9120
ggcctcgatg gccccgatag tgatgggctc cgatgtcact ctttatctct cggtcgatcc 9180
ggccaagaca ataatgaaga aataatatgg aattcgacga tcttcaggac gacggaatgt 9240
tcgagcactt caggctcaac gtcgatcccg gacagacccc gatgagggtg gacaaatata 9300
tggcctgcca tctcgaggat acatcgagac atagggtgca gaatgccatc aagagcggat 9360
atgtccgcgt caatgacaag gtcgctaagg ccaatgtcat cgtacgcccg ggcgatgtca 9420
tccgtttcgt gatgccatat gagcgccatg gcttcgaggt cctgccggag aatattcccc 9480
tgaatatcgt atatgaagat gatgatttcc tgatcgtcaa caagccggcc ggaatggttg 9540
tccatccggg ccatggccat ttcgacggca ccctcatcaa tgctctggcc taccatctcg 9600
ggatcagcca gggaccggat gccgaggatg acaggatggg cattcttgtc catagaatcg 9660

atatggacac ttcaggcctg ctggtcgtgg ccaagaatga cgaggctcag cttaatctgg 9720
cgcagcagtt ctttgtccat tcgatagagc gaaagtatgt cgccgtcgtc tgggggaaca 9780
taaaggagga cgaggggacg attgtcggta atatcggcag ggaccctaac gaccgcctca 9840
agttccgcgt atatcaggag gaggacaagg gaaagcatgc cgtaactcat tacaaggtgc 9900
ttgagcgttt cggattcgtt accgtggtgg aatgccgcct ggagaccgga cgtacccacc 9960
agataagggt acacatgtcc tatataggtc atccgctttt caatgatgcc agatacggcg 10020
gggcggagat ccgcaagggg actatctatg cgaaatataa gcagttcatc cagaactgtt 10080
tcgagatttg cccgcgtcag gcgctgcatg caaagactct gggtttcgtg caccctcgca 10140
cgggaaagcc ggtcagcttc gattctccgc ttccggagga catgaccgcg ctgctggaga 10200
agtggcgcaa gtatgccggc gacatgccga aagaataaaa aaaggggggac acccttttatg 10260
aatcaagttt gtccctttt tcgttgaata gttcattctg cagaactgca aaatcggtaa 10320
tttctaccac tttgatcttg catttgtact ttttcttcca cttgcctaca taagagttga 10380
acaggcctcg ggtaatatat gcccctaaaa tagggcttac cttcagagtg aagttcctga 10440
ttcccttgtc atttacgtag taggaaagcc ttcgttcaat ctgctcgtcg atgagcaggc 10500
ttgatgctat ttttcctgta ccgtggcata ccgggcagac ttccgtcgtg ttgatctcgg 10560
ttaccggccg tacgcgctga cgggtgatct gcataagacc gaactttgtc agcggcagca 10620
cattgtgctt agccctgtct ttgctcataa gctcctgcat gtacttgaag aggttgttcc 10680
tgtgctctac cgattccata tcgatgaagt ccacaatcac tattcctccc atatcgcgga 10740
gtctcagctg tcttgcgatc tcttccgccg caaacttgtt gacttcaaag gtgttctctt 10800
cctgttcctt tgtcttggca cggatgccac tgttgacgtc aacaacgttt aaggcttctg 10860
tcgtctcgat tacaagatag gctccctgtt tgagcggtac cacttttccg aaagaactct 10920
ttatctgtct tgtgatctca aagtggtcga agataggttc ctttccctga tagagcttta 10980
ctatcttctc ctgttccgga gaaataaggg agatatacct tctgatctct tcgtattctt 11040
tctcgttgtt cacatagatg ttcgagaaag agtcgttaag cagatctctc agaattgtag 11100
tggtcttgct gtactctgta aacagaggct gtatgctctt gttcttcgag agattttccc 11160
aggaatcttc ccatttaccg ataagcgact tcagctctgc aaccagaata gctgcgttct 11220
tgccttcagc cgcggtgcgg atgattgcac cgtaattctt gggaagaata ctcctgacaa 11280
gtgtctcaag tctcttcttc tcctccttgg aggagatttt ctgggatatg cttaccttct 11340
cggcgaaggg aagcagtact atgtttcgtc ctgccaatga aatctctgcg gtcagtcgtg 11400
atccctttgt ggaaataggc tccttgacta tctggactat tatcatctgg ccctgggtga 11460
gcacattctc gatccggcct tccttttcga ggataggacc gagctttatg cttccgaata 11520
gcctcgacgg gttggtatta gggtttgtct tcttgacgaa ctcgtcgaaa gcgccgaaat 11580
aaagtccgag gtccagataa tggatgaaag cctctttctc gtcaccgata tccacaaagg 11640
cagcgttgag tgcaggcatc actttcttta ctcttccgag gtaggcatcg cctacagcga 11700
agctgcgacc cgagctggac tctttgttca gttcgataag cctgtgattt tcctgtaagg 11760
ctatctgaac ttcagtggcc tgaacgtcaa cgatcaggtc tttgagtagt tccttttcag 11820
aatccattgg ataatttgtg aaacaataac aaaagggccg tccagatctc actgtcggcc 11880
cttctctttt agcagactgc taaaatggca gacacaggag aattatttct tcttgtgtct 11940
attcttacgc aagcgttttt tgcgcttgtg tgtcgccatt ttatgtctct ttcttttctt 12000
accgcttggc ataattgaaa aaaagttaaa aaattattaa atattatacc ggtaattact 12060
taaccttgct tacgaatacc tttgctggct tgaaagccgg gatattgtga gctggaatgg 12120
taagggtagt attcttggtg atgttccttg ctgccttctc agcacggtgc ttgataatga 12180
agctgccgaa gccacgaaga taaaccggtt caccccttcgc aagtgaagcc tctacacttt 12240
ccataaaaga ttcaacgact gactgtactg ccagtttctc aataccggta gcttttgcta 12300
cttcattaac gatctctgcc ttagtcataa tagagtaatt tatataaagt tatgtgtgtg 12360
caaattttaa gactgcaaat atagactttt ttttgcatag ggcaaattat ttcgttaaaa 12420
atctttgtct tacatatttt gatattacct ttgtgcttgg cataaaggtt gacattatag 12480
tatctgcctt aaaaacgcca tccttctctg acaaaatgtc agcagggagg attgtataca 12540
ctgataaatt atgatagatt ttaataaaga aatgtttcct gcctcgtccc acgaggtcaa 12600
tataataccg gtaatgtctg gagagcccgc tctcaacttc gacgcctcgg aactcccggg 12660
atcccttccg atcctggcac tccgcaacgc cgtgatcttt ccgggtgtcg tcttccctat 12720
aactatcggc cgcgagaagt ctatgcttct cgtcgacaag gcccagggaa acggcatgtt 12780
cataggcgca gtccctcaga agaatgtcgc cgaggagaac ccgctgtctc tcgccgactt 12840
ctcggaattc ggtactgtcg taaaggtcct gaagactttc gagatgcctg acggcactgt 12900

cacggcgctt ctccagggct acaagcgtct ccgcatgctg gaactgacgg acaccgatcc 12960
ttaccacatg gcaagagtca gctatctcga cgatatcctg ccgcagaacg actcgaaagt 13020
tccggtcata gccgatatcc tcaaggagaa tgcgtcaagc atcatccgtt cgtcttcgtt 13080
cgttcctaag gaggccctcg aggccctcaa gaacatcgac gacttcaatt tcctcgtcaa 13140
tttcatcgct acgacgatcg acatggacga tttcatggaa aaagtcgacc tgctgaaata 13200
tgacgacgta aaggaaaggg ccatggctct tctcggcgtg ctcagccgtc agctggagat 13260
ccagaggctc aagcaggaga tcagctccaa ggtcaagacc gagctcgacc agcagcagcg 13320
cgagtatttc cttaataatc agctccgtac gatccaggag gagctcggca tggatgagat 13380
ggacgacatc gcacgcttcc gcgaggctgc cgagaagaaa gagtggcctg catccgtagg 13440
ggagaccttc gaaaaggaac tcgcaaagct cgagagattc aatccttcta ctccggaata 13500
cagcatccag tataacttcc tgcagttcat gatcgacctg ccgtggtatc atacttcaga 13560
cgacaatctc gacctgaagc atgcccagaa agttctcgat gaggaccatt tcggactcga 13620
gaacgtcaag gacaggatca tcgagtatct cgccgtcctc aagctgaagg gtaacatgaa 13680
atccccgatc ctctgcctct atggccctcc gggagtcggc aagacaagcc tcggaaagtc 13740
gatcgccaga gccctcggac gcaagtatgt ccgtatctcc ctcggcggca tgcatgacga 13800
ggccgagatc cgtggccaca ggaaaacata cgtaggcgcc cttccgggac gtatccttaa 13860
cggaatcaac aaggccggca cttccaaccc ggtaatggtt ctcgacgaga tcgacaaact 13920
cggaagcgac ttcaaaggcg atccgtcttc cgctcttctc gaggtcctcg acccggagca 13980
gaacacggct ttccacgaca actacctcga tatcgattac gacctttcga acgtaatgtt 14040
cataacaacg gccaatacta taaacggtat ccagccggct ctgctcgaca ggatggagct 14100
tatcaatgtg accggttatc tggcagagga gaaggtccag atcgccaaga agttcctcgt 14160
gcctaagcag ctcgaggagc atggaatccc taagaaggaa ctgaagatcg acaaggctgc 14220
ccttgaagag atcatcgaca aatataccca tgagtcaggc gtccgcggcc tggagaagca 14280
gatcgcccgt atcgccagag tctctgccaa gaagatggcc ctcggcgagg actatccgaa 14340
gaccataaag aaagagcatc tcaaagagta tctcggcctg cagacggcct tccacgacag 14400
ccagaagggc aacgaggctc caggagtcgt gaccggtctc gcatggacct ccatgggcgg 14460
cgagatactg ttcatagaga cttccgtcag cgatggcaag ggagtcctga caatgacggg 14520
aaatctcggc gatgtcatga aagagtccgc tactatcggc tatcagtata tcaaggctca 14580
tccgcagctc gccggcctta cttcggaaga gttcgcgaag aaggatgtcc atgtccatgt 14640
ccctgaagga gccgttccta aggacggtcc ttcggccgga atcaccatgg tcagctcgat 14700
ggtgtcagcc ctccgcggaa agaaagtccg cagcggaatc gccatgaccg gagaaatgac 14760
cctgaggggc cgtgtgctgc ctgtaggtgg catcaaggag aagattctag cggcaaagag 14820
ggccggagtc tctaccataa tcatttctga agaaaacagg aaagatattg aagatatcaa 14880
ggaaatttac gtaaaaggtc ttacctttgt gtatgtcaag acaattgacg acgtaatcgg 14940
atatatcttc gaataatgga cgttaagata gaacagactt ggaaagacgc actgcgcgac 15000
gagttcgaaa aaccgtactt cgcctcgctg gtgcgttttc tgcataatga aaaagaggcc 15060
gggaaggtca tctatcctcc cggaggcctg atattcaggg cattcgaact cactccgctg 15120
ccggaggtga aagtcgttat cctaggccag gatccgtatc atgggccccg ccaggccatg 15180
ggactttcgt tctccgtgcc gtcggacgtg ccggcccctc cgtccctgaa gaatatattc 15240
aaggagatcc agacggatct gggcataacc atgagcggaa gcccgaatct tgaaggctgg 15300
gcccgtcagg gcgtgcttct gctgaacgcc atgctgacag tccaggccgg agtcgccggc 15360
tcgcacaagg ctatcggctg gcaggagttc acggatgccg tgatcaagtg catctctgac 15420
cgctgcaacg gggtcgtgtt cctgctctgg ggtaatttcg cccgcggaaa gaaggaactc 15480
atagatacgt cccgtcatta tgtgctcgaa gccgcccatc cttctccgct tgccgggggga 15540
gccttcttcg ggtgcagaca tttttctaag actaacgaac ttctcataaa agagggcaaa 15600
acccctatcg actggaaatt atgaaacgta aagcactttt cccgggatct ttcgatcctt 15660
ttaccgccgg ccatctgaat attctccgca gagcgctgac gatgttcgat gaaatcgtca 15720
tcgcagtcgg tgtcaaccag gacaaacgcg gattctttag catggatcag cgtatggata 15780
tcatacgtca ggctgtccgc ggtatggacg ggatcgagat aatcaaatac gacaacctta 15840
ctatcgatac ctgccgcgag ctgggtatcg cccatataat aaggggcgtc aggaacatgc 15900
ttgatttcga gaacgagcgt gccattgcgg atgccaaccg ccgtcttgct ccggaaatcg 15960
agactatcat catccctacc gcacaggaat tcgcgcatat ctcgtcttca gccgtcaggg 16020
atatccttaa acataagggg gatacttctc tcttccttcc ggagggagtc gtccttccgg 16080
atcacgatgc gtaggttaat cctcataatc tttctttctc tgctttcttt tgccttgggc 16140

gcccagaatg cggattccgt caagatggcc gccctggatg caaagcttga ggagtatttt 16200
gcggcgctgg agaccgagtc tctggatgta aagaggaccg agtgcgattt cctcatttct 16260
tcgtgcacga ctgacgatgt acgaaaccat gtcgccgtac gtctttactc gcatttcatg 16320
cagtcgaagc ttatgggggga tgaggctgtc gccgtccatc ttgccgacaa ctggttcatt 16380
cccggcaagg tttctttccc ggatgagata gatctgatga atgcgaagat tttcgctgag 16440
ttcaacagac agtcgctgct cggatgcaag gctcccggtc tggtagccga ggatagtctc 16500
ggcaggaagg tcgaggtccc tctgggattg tctgacagat accggattct ttatatctat 16560
tctccggact gtgccacatg cagtcttcag actatgctga tggactctct tttcagcagg 16620
aagagctatc cggtcgacct tatcgcattc aattcagacg ataatactga agcctgggtg 16680
cagtaccgca ggaaccgcct ttcttttgac ggagtccgtg tcatgcattt cagggatcct 16740
tcgctggagt cggatttcca gcggaaatac ggagtgcttc agactccccg catgttcctt 16800
atcgacaaga acgggattat agtaggccgc ggactggacg ctgctgcgct ggagaaattg 16860
ctggatgcaa ctcctgacct tggcataggg tatggttcga agtcgtctgc cgactatttc 16920
gacgctgtgc tggagagctt cggggatgag ctgcgcggct ctgatatccg ctattttatc 16980
ggaagacttg aagaggctac tctcgaaagc gccgataccg caggattccg tcagatgacc 17040
ggtgacatgc tgtattatct tgccggaaag aggggagagg cgttcaagga tgccgaggcc 17100
ttcgtcgtcg attcgatggt cctgtcccgt ccggatgtgt ggcgctttgc cgatgattcg 17160
ttgacggtcg ttgacatggc catgagcctg aaacatatgc tctcgctagc tccagtaggt 17220
gccagggttc cggatgtacg tgtccatgcc actgtccttc gcgccggacg aaagccggcc 17280
accgtccata agaaattgcg gtcgttgctt aagggaaagc ggaattatgt gcttttccat 17340
actgccggct gcactttctg cgaggctgaa cttgccgccg ccgattctgt cgcttcggag 17400
aagggtgtcc gtgtcatcct ggtcgatatg gacaagctga caagcttata cccggaggag 17460
gctgatcggc ttctggacgc attcgacctg tccgttctcc cgttcataat ggagacggat 17520
aataaaggtt gtgtcgtacg taagtatctg agtttaaata agtagggctg cctttacctt 17580
agctacggcc tctttgcctc ccatcttttc gaggatggcg agtccgaagt ctatcgcgaa 17640
ggctgcgcct ctgccggtga tgagctggcc gtcggtaact gccggggctt ttacgtattc 17700
cattccgctg gcggcgaggt tctcttcgaa gccgtcaaag caggtggctt tcttgccttc 17760
gattcccttg atctgggtgg cgacgaggcc cggggcggcg cagattgcgg ccacaatgcc 17820
tccgttgccg tagtggacgt tcatcagacc gataagcggg gtgtcggctg cgagtttctt 17880
cgagcctggc attcctcccg ggaagatcat ggcgtctttt tcgtcgatat ctatttcgct 17940
gaagtcggag cggcccatgt ctgccaccac cgttactccg tgggagctgg tgatatagac 18000
ttcttcggtg attgatactg tgtatgcttc gattttacct ctgcggagaa tgtcgagggt 18060
tgcgagggct tcggtctctt caaaaccgtc agccaggaag atgtatgcct ttcccatgat 18120
gctcatgtgg tttagattgg ttgaggtaag agggttcgaa cctctactga gggaaccaaa 18180
atccctagtg ctaccattac accatacctc agtggggtgc aaatatacga aaaaattatt 18240
tgcggacaac aattctctct atacgtctag ggacgctagt gagaatttcg taagggattg 18300
tcccgagaat gtcggcgagt tcgttgactg tagggtcttc tccgaagata gtcacggtgt 18360
ctccgacttt agcgtcaatt cctgtcacat caagcatgca catgtccatg cagatgttcc 18420
cgatcgtcgg cgctctgtgc ccgttgacgg agaatgaggc gtgaccgcgt ccgagatggc 18480
ggtcgatgcc gtcggcgtat cctaccggga tcgtggcgat tacggtgcct tcaggggcga 18540
tatgaccgta tcttccgtag cctatcgtgc cgtcttccgg cttgagggtc tttatctgca 18600
atattttgca tttgagggac gcgacgtggg cgaggcggac atcaggcagg gcgctgattc 18660
cgtagatacc gattccgagg cggaccatgt cgaactggta ttgcgggaat ctttcgattc 18720
cggcggagtt gaggatatgg tacatcggct tgtagcctat ctcggaggtg atcctgtcgg 18780
cgtttgtctt gaacatgccg atctggccga gggtgaattc gtcctgctca gggtcttcgg 18840
atactgcaag gtgcgaatat acggacttga ctttgacttc ttcgcaggtc ttgaggaagt 18900
cggtgagatc ctggagctcg ctcgtcatga acccgaggcg gtgcattccg gtgtcgagtt 18960
tgatatggac cggataatcc ttgatgccgc gttctttgag tattccgcag agtatcttga 19020
gggcgtacag gttcgggatt cccggctcga ggcggttgtc gatgatttcg ttgaagaagt 19080
ccgtgcctgc ggtcaggacg aggatcggga ggctgatacc ggctttgcgg agttccatgc 19140
cttcgacagg gtaggcgact gcgaggtagt tcgctcccgc acgctgcatc atggctgcga 19200
attcgacggc tccgtgcccg taagcatttg cctttacgag tacaagtaat ttagtcgtct 19260
gctccagttt ggacctgaaa tatctgtagt tttccataca tcctgtcagg ttcagctcca 19320
gccttgaaaa atcatcaacc ctgttcattc cacatcaatt tagaatacaa atttatcaaa 19380

tttctccgac ataggttaca gtttttcgct tttctttccc cgtaaagacc cttacagtac 19440
gtccgctgtc gccgccgaca gcgtctgctt tggcgggaat cctcgtggcc tcggctctgt 19500
tccgccgact ctgtcggctc catccctccc aatgtctact tcgtcctgct tcgcagaact 19560
cgtatccatc gggcccctcc ctctaacgtg gccgagggtg gccacgctcg ccgagaccag 19620
aggacagccg ccatcaatta agacgctgac ggcaagtgag aggttggcat ggatttggaa 19680
gcaaccagaa ggattgttaa aattttatgg atatgatttg gtttattatc attgctgtga 19740
tggtgctgag ctttgtgatt cagcaggttc tgaacagtaa attcaataaa tattcgaagg 19800
tgggattgca ttccgggctg accggggcgc aagtcgctgc caaaatgttg gaagaccatg 19860
gcataaccga tgtctctatt acctgcatca aaggaaagct gacagaccat tacaaccctg 19920
tcaacaagac agtcaacctc agcgaagaag tctattacgg caattcggta gccgctgcgg 19980
ccgttgcagc ccatgaatgt ggccatgtag tgcagcatgc cacgggatat gctccgctga 20040
aactgagatc ggctctcgtt ccggtagtct cattctccag caatgccgtg cagtgggtac 20100
ttcttattgg cgtaatgatc atcaagatta ctccgaccct cctgtatctc ggaatcgcca 20160
tgtttgcctt gacgaccttg ttcagcgtcg tgacacttcc tgtggaaatc aacgccagcg 20220
ccagagccgt gcaatggctc gacaaggccg ggataacagg tcccgctaca aaagcgaaag 20280
ctgtcgatgc cctcaaatgg gcagcatata catacgtcat tgccgccatc ggttcgatag 20340
cgacattgat ctactacatt gccatggcca ggcgacgcta gagactcgaa agaatccttg 20400
caagctggtc cgtgcaggat gtaccacgga aattgcagtt gacgcccgaa agagtgtcaa 20460
ctattttttt tatgtccatg ccctcgacga gacggccgat cgcctgaaga ttaccgttgc 20520
agccccgcac gtatgcgaga ttgcggacct tgccgtctat tatgtcgaaa tcgatctgct 20580
ggctgcatac caggctgctc gggatggcag ttatgtgtct gacaccgttc tcggtggtgt 20640
cgctgatgat cttgtagtcg tctcctgtca ttgttgtagt ccgtttttta tctttttttc 20700
ttgaaaaact ccgtcaccat ggaaccgcat tcctccgcga gcactcctgc ctcgacctcg 20760
gtcttcggat gcagcagcga aggagtgaaa agtgagtgcc cccgcttcgg atcgatcgct 20820
ccatagacga tccggccgat ctgggcccag ttcagggctc cggcgcacat cggacacggc 20880
tcgaccgtga cgtagagcgt gcattcgttg aggtatttgc cccgagcgc ctccgtagcg 20940
gcggtgatgg ctatcatctc ggcatgggcc gtaggatcat gtaaacgttc ggtcatattg 21000
tgaccctttc ccacgatccg gcctttgcat acgataactg ccccgatcgg gatttcatcc 21060
tcggccaggg ccgcctgcgc ctcgcgcaga gcttccttca tatatagctc gtctttttcc 21120
ataattacag caaagatagc aattcttgga caatcccaat agttgataaa acaattaaga 21180
aaatttgcga aatgatactt ttttgacatt tttatcatat atttgccacg tgataatata 21240
agttaatgga tatgacctcg gtaataacgg aaataacacc tctttcggac agggactgct 21300
tctacatagt cgagagacat aagaccgaat tcgtcttccc cctccacagg cacagggaat 21360
acgagctgaa cttcatcgag catggcaagg gcgtccggcg agtcgtaggc gacagtgtcg 21420
aagagatagg cgaatatgag cttacgctga ttgccggcgg ggatcttgaa catgcctggg 21480
agcagggcaa ttgtacctcc gaggacgtac gcgagatcac tatccagttc gatccggaga 21540
tattctccag cggcctcatg atgagaaacc agttctcatc gatacagcat atgttcgagc 21600
aggcgaggaa cgggctgacc ttttcgctgg cggccataat gaaggtgtat tcgcttcttg 21660
acgccctgcc gtcggaaata gataaattcg atcagttcca gcactttatg aggctgcttt 21720
atgacctttc cctggcgggt gactccagga tactggccag cacgtcattc gcccataacg 21780
atgccgacaa ggaaagccgt cgggtgcaga agatcaagca atatgtcgga gaccatttcg 21840
cggaaaacat tacccttgaa gaattggcgg aaatggtcgg catgagcccg tcggcgttca 21900
gccgcttttt caagagcagg accgggaaga cggtgataga ctatataatc gacatccgac 21960
tcgggaatgc ggcaaggctg ctggtcgaca ccacccagaa tatctctgaa atctgctata 22020
gctgcgggtt caacaacctg tcgaatttca atcgtctttt caagtcaaaa agaggctaca 22080
cgccgcgcga attcaggact ctgttcaaga agaacaaggt ttttgtataa cgtgcaaaat 22140
agtatcggtt attgacaaat ttgtatttgt tgggtaaaac gttgcatgct aaatttgcaa 22200
cgtactcaga caagtatgga tttatcgata aaccgcatga ttaaaagatt attcccggca 22260
ctgtttatcg cccttttcgc gattctttcg tgtcagccta tgaaaatgga catcgtcaaa 22320
gttgacgatg gctcttttgt cgttgatggt cacttctatt attatatcgg tacaaatttc 22380
tggtacggag gacttctcgg ctccgaatat ggtgacagaa gcaggcttgc cgccgaactt 22440
gattccctgc atgccctcgg actggacaac ctccgcgtac tggtaggcag cgacggcccc 22500
gacggggtgg cgaccagagt ctatccgacc ctgcagaagg aaccgggcgt ctatgacgaa 22560
aaggtcttcg aaggtctgga ctaccttatg gccgaacttg ggaaaaggaa catgagggca 22620

gtcctctata tcaataattc ctgggaatgg tccggcggct atggcatgta cctcgaatgg 22680
gccggagccg gcaaggcgat aattccttcg gtcggaggat atggtccgtt catggaacag 22740
atggcccagt atgcgacgaa cgccagggcg caggaactct tcgagaatca tctccgtact 22800
gtcgtttcaa gagtcaacag catcaccgga aagccttata gcgaggatcc ggcgatattc 22860
tcatggcaga tcggcaacga gccgcgctgc ttctcctccg atcctgccgt ccaggacgga 22920
ttcgtatcat ggatatggcg cgctgcctcg ataatcaaat cgatagaccc taatcatatg 22980
gtttctaccg gcaatgaggg catatggggc tgcgaagaga atccggagct attccgccgg 23040
ctgaatgaat gtcctgacat tgattatatt accgcccata tatggccata caactggagc 23100
tgggtaggaa aggaatcccc tcagctggat atagaccgcg ccgaggagaa taccgggaga 23160
tacatcgacg cccatctgga ggtggccgag agtctgggca agcctctcgt gatcgaggaa 23220
ttcggcttcc cgagagatga tatgcagttt gccaaaggga ctcccgtaag cggacgcgac 23280
cgctattaca gatacgtttt ctccaaggtc atggagaatc agaaaaatgg cggcagactc 23340
gccggcgcca atttctggag ctggggaggt cttgcggtgc ctgcccacag atattggcag 23400
gaaggagacg attacgccgg agatccggcc caggaagaac aggggctcaa ttctgtattc 23460
gccgacgatt acagtaccct tgctattata aaacatacaa cggcgagact ttctgccgtg 23520
gctaatgcaa gcgcagtgct tgaaaatgat tggatgttca ctaccggaga ccagaaacct 23580
ctcagagttg ctgtacagtc tcagaatggt tttaagggaa gtgtcccggt ttatgtgcgg 23640
gtgtcaacag acaaaggggt ttcagtgtcg gtctcaggaa agaaagcccg tctcggcaaa 23700
ggcgccgata ctctggattt cgatcttgga ctagccccag gcttttataa ggtgtgtgtc 23760
agcctgaacg agaatgaccc gcttgatgaa aacttctgtg tcggctgtga tcctgagaag 23820
atagtatctc ctcaggataa acagcctgat ttcgatgaat tctgggaaga caacctcagg 23880
cagcttgcca tggtcccgat gaatgccagg atgactccgt tgaaggatca ttcaaacgct 23940
gacaggactg cctacagggt tgatatgaag tctttcggag gcaaagagat ctccggtctg 24000
ctttatcttc cgaacaaaga gggaaaattc gctgccagga tctcatatat gggctacggt 24060
tcggacgtct ggttcgctga tccgtcggcg gaaccggaca tgatcgagtt cactctcagc 24120
gtccgtgacc agggtttcaa caagactccg ggacaaggct gggtgacaag gggacttgcc 24180
gataaggact cgtattatta cagaggcgca tatctggact gtgtgcgggc cattgatttc 24240
gtctcttctc ttcctcagac tgatccggac catatctacg ccgaaggtgg aagtcagggc 24300
ggagcattta ccctgatagc tgcatctctg gacaaccgtc tccgcgcaat cgctccgttc 24360
gtgcctttcc tttccgattt cccggactat ttcgccatcg cagactggcc gggcggggaa 24420
gtgctctcgg cggcaaaggc tcagggcatt tctgaagacg aactttataa gaccctgtct 24480
tatttcgacg tgaagaactt tacggacagg ataggttgtc ctgtgcttat gggtttcggc 24540
ctgcaggatc ccgtctgtcc tcctcatacc aatttttcgg ggtataacca tattactgca 24600
gataaacaat ggctttgcgt tcctgtctca ggccaccatc tggagcaatt tcccgagtgg 24660
tgggcagcaa ggacggaatt cttcacaaaa aataaataac acaactaatt attcacattc 24720
aaaatcagta gcatgaaaag acagctatta acatgtatgc tgatgctctt tggagcattg 24780
tcacttatgg cgcagaacaa agtgaccgga agaattacag ataattccgg acagggcgtc 24840
ataaatgcca gcgtaatggt gaaaggcacc aacaatggtg tcgttgcaga tcttgatgga 24900
tactacgaga ttgttgtcgg ccaaggctcc actcttgtat attcttccat cggttacaaa 24960
gatcaggaag ttgttgtagg cacccagagt gtcatcaacg ttcttcttgt cgaaaactcc 25020
gagctgctcg acgaaatagt cgtagtcggt tacggtacta tgaagagaag cgacctcgcc 25080
ggatcatccg tttccatgaa ggaggatgac ctcaagggtt cgatcatcac caacctcgac 25140
cagtctctcc agggacgtgc tgccggtgtc acggcagtcc agacttccgg agctcctggt 25200
tcgtcctcaa gtatccgtgt ccgcggtcag gcgacaatca atgccaacgc agaacctctc 25260
tacgtcatcg atggcgtaat catgcaggga ggcggtaact ccggtgcaga cttcggtctc 25320
ggagatgctc tcggcaatgg taaagtctct acgatttctc cactttccac tatcaaccct 25380
gccgatatcg taagcatgga aatcctcaag gatgcttccg caaccgctat ctacggtgcc 25440
cagggtgcga atggtgttgt cctcattact accaagcatg gctccagagg tgatgcgaaa 25500
ttctcatacg acggtatgtt cgccgtttca aggcagacaa cccgtatcga tatgatgaat 25560
ctccgtgaat acgcagagtt ctacaactct atggttgaaa ccggagaaat cagtgaggcc 25620
aatccttact atgctactcc gggcctgctc ggaaagggta ccaactggca ggacgaaatc 25680
ttccgcacag ccatccagca ccagcatcag atcagcgccc agggcggtac ggacaaagtc 25740
cagtactatg tttccggatc attcatggat caggagggta ctatcagggg ttccgaattc 25800
ggccgttata ccttcaggac caaccttgac gcccagttga aggaatggtt caagctcggt 25860

ctcaacctta cctttgccaa caccaatgat gaccttaagc ttgccgacgg tagcgaaggt 25920
cttatcacat actcccttac cacaatccct gacatccctg tttatgatgt ctatggcaat 25980
ttctcatcta cagtacgcga aggctatact tctccgaacc ctgtagctct tgcaatgatg 26040
gatgagattc tgctggaccg tcagaaactc aacggtaata tctacgcgga acttactcct 26100
gtaaagcatt tcacctggag aacagaactc ggtttcgatg tcagcggctc tacggctacc 26160
cgtttcagac caacagtcaa tctcggtggc tggacaagag ataccaactc tatgagctat 26220
cagaagaaca gcaacagatt ctggcagatc aagaactatc tcacttatgc caactccttc 26280
ggaaagcaca gcgttaccgc catgcttggt caggaggcct gggagtcttc gtggagttac 26340
gtacagggcg caaataccaa gctcccttca aacgatgtcc ataacgtcgc cctcggtgga 26400
ggaactccgt cgctttcttc cggtttcgga agctctgcga tggcgtcatt cttcactcga 26460
gaaacttata atttcgacga tcgttacctc cttacctata cctaccgtta tgatggatca 26520
tccaacttcg gtcctgagaa acgttgggcc ggtttccatt cattggcagt ggcttggaaa 26580
ttcaccaatg agaaattcat ggatttcctc aagcctgtgg tcaacaacgg aaaactccgt 26640
ctcggatggg ggcagacagg taactcaagt atcggtggtt atgcatgggg agcatccata 26700
agccgtatgc cttcggctct cggcatgggt taccgtcctg ctaacatccc taatcctgct 26760
atccattggg agagccagac gcagcagaac atcggtctcg acctcaatat gttcgacaac 26820
aggctgaatc ttacagtaga tgcatacctg aagaagtctg acaacatgct tatgtcaatg 26880
cagcttgctt cctatatggg aacccagggt aacagttctt ctgctcttgc agctcctaaa 26940
ggtaacttcg gttctatcga gaacaagggt cttgaaatta ctctcgatgc ccatcctgtc 27000
cagatcaagg atttcgcctg ggacagcaac ttccagattt ccttcaacag aaacaagctt 27060
gtcagccttg acggaacgac taatgccgct ctcataggtt acggacagtg gtctgacgtt 27120
gtgtgtgtgt ctgaaatcgg caagcctctc tataacttct atggttacaa ggtcgctggc 27180
gtatatactt ctctcgagga tatcgagaat tctccgaaac ctgagaaata tccttctaat 27240
ggagagtatg cccgtggaaa taccgtgtgg gtaggtgata tcaagtacga agatatcagc 27300
aaggacggag taatcaatga aaaggaccgt accgatatcg gcaatcctct tcctaagttc 27360
acctttggct ggagcaatac ctttacctac aaggacttcg acctcaatat cttcatcaac 27420
ggaagctatg gtaacaaagt cctcaactac aatgccattt ctcttaccca tatgaacagc 27480
gcatggacca accagatcaa tgacgctgtc tccggcaggg caatgcttgc tcctatagac 27540
gcaagcgtag tctatgctga cggttccaag tggtatgacc atatcgacaa tgtcagggtc 27600
ctcaaccctg gcacgaagac tcctcgtgca agcatcaacg accctaacga caacgaccgt 27660
atcagcgacc gttatgtgga ggatggttca tatatccgta tcaagaacat cactctcgga 27720
tataatgtcc ctaagtcttt cctgaacaag ttcaaggtgg ataatgtacg tgtttacgtc 27780
aacatccaga acctttatac tttcacgaag tacaccggat atgatcctga agtaggtgcc 27840
agcacccagg attccaccgg tcttacctat ggcgttgata acggacgtta tccttctcct 27900
acagtatatt cttgcggttt gaacattaca ttctaaaatt gaaggaatat gaaaatctct 27960
aatatgatga aaggttttgc tgtggcgacc gcagtctttg ctctgacttc ctgcgaaggg 28020
ttcctcaaca gacctaacga agataattat aacgtagata atttctacaa gaccgacgag 28080
cagtgcgaac agggcgtgaa ttacctgtac aactctccat ggtatgactt ccagagaggc 28140
ttcatcaaga tcggtgaggt catgtccgga aacatgtact ggggctcgtc tccatacctt 28200
gatttctcca ccaacggtac cgaccaggac ctcgtcaaca tgtcctattc tctctgggct 28260
gtgaacggcc atgccaatac cgtgatctac aaccttctta attcgcaggg cccgtccaag 28320
gcagcgatcg acaagtgtat cggagaagct cttacctgga aggcattcgc ctatttcttc 28380
atggtacgaa ccttcggtga cgttcctatc gtccacagca attccgatct cctctccagc 28440
ggcgagtaca actctatctc caaggtggag cgcgccgatg tctatgagta catcatcacg 28500
accctcgaaa aggcaatgga aatgcttcct aagaagactt ccggcttcaa taattacgag 28560
cgtatcgatt actatgcagc agaggccctt ctcgcaaaag tatatcttac caaggccgga 28620
ctttccggat ccttgaacca ggatgacctc aagcaggctg ccaagtatgc caaggatgtg 28680
atcgacaatt ccggacgtac ccttcttcct gtatattcgg acgtcttccg tcttaccgga 28740
ttcaacgcca caggcgagtg ccttatctcc tgggaatgga gcgtaggccg taacccttgg 28800
actcagcaga ataccctcca gagcgacctc gcaatggtcg gattcagcga ccagggagac 28860
tgctggggag gctggggcgg tccttcagtg gatcttgcag acgcattcgg cgtaagtgcg 28920
acccaaaatc ctgcagagcg tgtaggagag aaggacagcc gccgcaaggc aaccttgatg 28980
ctcccgggtg acaagtatga ttacttctgg actgacaagg gaggtttcga ttatctccgc 29040
ttcatctatg atacggaata tggcaaaggc ggtcctggcg gttccatgca gtgccctaca 29100

ggttccaaca acgtaaagca tctctatggc gataactttg atcatgagaa cgccggtctg 29160
ggaacccctg agaacatggc ttatcagctt cctactcatc ttctccgtct ttccgacgtg 29220
tatcttatct acgcagaggc aagccttctt accggaagca acggccctgc tctcgagtat 29280
gtcaacaagg ttcgcgaacg tgctggcgcc gatgctctcg catccgtcac tttcgaggat 29340
atctggaaag agcgtcgtct tgagcttgca ggcgagggcg accgctggta tgactatgta 29400
cgccgttcat actacgatat ggacgcatgc atcgctgaac tcaaagctca gagaaggtca 29460
tactgggatg gactcgacga tgtttggaaa tcatatatcg gagatgcttt cgattattcc 29520
ggaacctggg atgcttcaga agtacagtac aacaacgcag aggacaatcc taacgttacc 29580
gctgcaagct tcaagcttcc gttccctaca gaggacgtag tgttcaatgg caacatggca 29640
tctgacgccg tgcctgtcca tgtcgatgtc cgcgaggaat acagctataa cttttaatct 29700
gacgaaaaga tgaaatctat aaatagaata atcaccggtt cactggcaat cctcgcaatg 29760
gcgtcattcg cttcctgcga caaggttgac tatccggacc gttatgaagt agctgacggc 29820
aagccgacgg tttattccgt cagatacgca gacaaggacg tcaacatcac ccaggcatat 29880
atggatgagg tcgtatgtct cctcggcaag aatctcaaga gtattgtgga actctggttc 29940
aacgaccagc aggcaaccct caataccagc ttcattacag acaatactct tctggtatcc 30000
gtgcctaaga atatgcctgt ggtacagaca gacaagatat atatgattac ttcgggcaag 30060
gatacgctcg cgtatgattt cgccgtgctt cctccgactc cgaaggttac ttccatgtcc 30120
aatgagtggg ctgcggaagg ggagaccgtc acaatctacg gcgacttcct tattgacaag 30180
gctgacaagc ctctccagat agcctttgca ggagccgagg tcgatcatgc ttcaatggtg 30240
ttcaacggtt cgtcttcagt cagcttccct gttcctgcag gagcccagcc tggatatgtt 30300
accgtgactt cactctccgg taccggaaag agcaagttca tgtacaagga taccagaaac 30360
atcctcttcg actgggatgg cagccacggc ggctttgcta ccggtcacgg atggcgtaac 30420
ggcgtgatcc acgctcctgg cgctgatgca tgggatgcga tcgacggttc atacctctac 30480
ttcggaggcg ccgatcttgg tcctggaagt gcagatgcat gggcagagga ccagtacagc 30540
ttcaactact ggccagaggc cggcaccgat tacgatgccc tcaacctccg tccggagttc 30600
gcagagtata tcgacaagta tggagtaagc ggcctcgtgc tcaagttcga ggttctcgtt 30660
ccttcatcca acccttggtc tgcttcaggt atgcagctca tgttctcgag cgacgctgtc 30720
gttacctacg gtaccgctac gaacagctac ttcagtgacg aaaccttccc tcgtgctatc 30780
tggcagccat ggactaccaa cggaggttct tatgataccg ctgacaagtg ggttaccgta 30840
agtttccctc tcagctcatt caacaagaac caggaaaaca aggagtgtgc aaccgcattt 30900
gacaagtctt accttaccgg tctcacattc ttcgtatgga acggcggtat cgacggcaca 30960
gcctgctctc cggtcttcgc gatcgacaat atccgtgtgg ttcctaatga ataatattaa 31020
atgaagctca ttatgaaaaa gatattcagt atttcagcca ttcttgccgc ctgtgtcctg 31080
gcactcgggc tctccggatg ttcatccaaa gagcttgata ccgaccagta ttccgacaag 31140
gacgtgaagc tggcatcatt cggccctaac ccggtgatgc gtggagctac actccggttc 31200
ttcggaagta atctcgacag gattgtcgaa gtttcagttc cgggactcga tcctatcacc 31260
gacatcgaag tcgttacctc cggtaatcct tcagaaatcc gcatccagct tcctgcagag 31320
ggcgcacagg tcggaaagat ctctctcaag tcatccgacg gaaaggtctt tactacccag 31380
gtggatgtag aatatacaga acctatcgta tttgacagtt tctctcctgc agaggctatg 31440
cctggcgacg tgataacggt caagggcgat tatatgaacc tcgtgcagag cgttaccttc 31500
gaaggcggcg ccatcgtgac cgaaatcaac cataacggcc gtcatgaggc ttccttcaag 31560
gtcccttcaa aggctattac cggaaagatc atcctttctg acgaaggagc catcgccaat 31620
ctcctctact cggaggaaga tcttgtgatc ggcgatccga ccgtcaagtc tcttaccgta 31680
gctgttgcta aacctggcga gaacgcagtt gtgaagggag catatctcga tatgatcgac 31740
aaggtcgtat tcgaaggtgg tgcagaggtt tccgaccttg ctctcaatga ggatgcaacc 31800
gaattcaatc tcgtaatccc tgccggagca cagtccggtg aagtaaaggc tgtcagccac 31860
gctgccaagg aattcgtcgc cggcgaaatc gaacttgtca tccctacgga tgtcgcagct 31920
gcaccttcgc ctgtaaaggc tggtgccgac ctcgtgattt ccggtaacga tcttgatgtg 31980
gtaacttctg ttgatttccc tggcgccagc ggagtagaat tcgtatttgc cgaggacgcc 32040
attacagtga ccgttcctgc tactgctact gaaggtgata tcgccctcaa tatggccagc 32100
ggagaagctg tgaatgtggc attcaccctt gtgcacccgg tagtttctgc catagctcct 32160
gtcgagctca tggccggcga gaccatcacc gtgacaggta ccgacctcga ccttgtcact 32220
ggagtaactc tcggcggaaa gccggtcgag ttcactgcag aggctgagaa gattgtcgta 32280
cagaccgcca atacttccgt ttcaggaaag ctcgtgcttt ctcttgcgaa tggtgagaag 32340

attgaaccgg aggatgccat caccctcagc tatgattcat tcatcatcgt gaacgacatg 32400
cctgccgcag agcatatcgg ggctatcgta accctcaagg gcgagaactt catgatgatc 32460
gagaacatct atatcggcga agagaaagta tcgaagtaca accttcgttc tgataatgaa 32520
atccagttcg taatgccttg gaacaaggtc ggaacataca gcatgtactt ccatcttctc 32580
tccggcgatg tcgagacatg tcctcagaca atcgaagttc tcctcgagca ggacatcatg 32640
actgtatggg aaggctctct gcagatcacc tggaatgacg gcggccgtgt aaccgtgccg 32700
gcagcatcat tcaccggtat ccctgcaggc agcaagctca gattctacta cactcaggtg 32760
gaccagcagt gggatcaggc ccagctcaac tatggtaact ggaccggaat caacttcgac 32820
gatcctgcag gtaccgtcat taacggaacc ctcgttccga ctgacgtcta tggatggttc 32880
actgacggaa tccttgaccg ctgcacagag gtcgtgctta caaaggaaat cctcgaaaac 32940
atccaggcta acaagggtga ctgcgaagac aagactgcct gcggtatcat catccagggc 33000
tctggcctta ccttcacgaa gatcgagatc gtatgcgata ttccgcagga ggttacgatc 33060
tacgaaggtc ctacaaccct tacatggggt gacgacggcc gtttcggtct cgccatgact 33120
tacttcgagg cagcaaagcc gggctctcag cttatcttct atctcgagca gacggacgca 33180
tggggccagg ttcagctcaa cgacggctgg tgggccaacg ccgacatgaa cttccctgag 33240
atcggcggtg catacattac caccgacaac ctcggaggaa aggacgtcac acgtatcgcc 33300
cttaccctga ctgctgatat ccttgaccat atacttgcta catccggtga ctatttcggt 33360
ctcaatagcc agttccaggg tgacggacgc gtcgccatgg ttatccaggg atctgatatg 33420
attatcaata aagtttgtat attgtaacac ttttattgcc ggaggcgggt gatgccgcct 33480
ccggtagcca gtaaacaaat tttatataat gaaagcacac gtcagtttct ttgtattatc 33540
tatgattttt cttgcggcat gttcctgctc aaaggatgat gccgaaaaga aggaaccgtc 33600
cgtcccggcc cctaaggcgg taacggtaga tgtcgcgtct gtcagcgctc ccgtagaggg 33660
aacgtcggtc gtcgtcgcca tctcctcccc gtccaagcct gcggtctctg tctccggaga 33720
ctggatcaaa gccgaactgg gtccgtacaa tccggataca tataaagtga aacttacggt 33780
gaccgtcggc gagaatccga catacgagtc acgcagcggt agtgtgaccc ttacttgcca 33840
gggcttttct ccggtcggga taagcgtatc ccaggatgcc gcggcgaagc cggtcggacc 33900
gtcggctaat cctgcatacg atcttctcgg tctcggctgg aatatgggta accagttcga 33960
tgctttctac aacggctcat gggccggaga taaatacaat tatccggacg agaccgtctg 34020
gggtaatgcc aaggcgaccc aggctacttt cgacggggtg aagaaagccg gattcacatc 34080
agtgcgaatc cctgtcacgt ggctcaatat gataggtcca ggacctgagt acaatattga 34140
cgagacttgg atgaaccgta tatatgaggt ggtaggatat gccgagaaag ccggactcaa 34200
ggtcatcatc aatacccatc acgacgaaaa caacaatgac gaccattggc tggatatcct 34260
ttctgcatct aaggatgcca ggaccaatga agcgataaag gcggaaatcg cggctgtatg 34320
gacccatatc gccaataagt tcaaggataa gggagatttc cttatcatgg agtccttcaa 34380
tgaaattcag gacggaggat ggggctggag cgctgctttc caggctgacc cttccaagca 34440
gtgtgatgtg cttaacgaat ggaatcaggt tttcgtggat gccgtacgtg ctacaggagg 34500
caacaacgct acccgctggc tgggtgttcc tacatacgcc gcaggtactt catggacgaa 34560
gtatttcaga ctgccggagg atcctgccgg aaaggtcatg atggcagtgc attgctatga 34620
tccgtatgaa tatactatag ggaacataca gtatcctgag tggggccata ccggcgccgc 34680
aggaaagaag accccggaag gggagaaaga gataacggat ctcttctcca gacttaatac 34740
ggaattcatc agcaagggga tacctgtata tcttggcgaa ttcggctgtt cgatgagaag 34800
caaaggggat gcccgggcat ggatgttcta taaatactat atggagtatg tctccaaagc 34860
tgcgcatacc tattccttac cgggaatcct gtgggataac ggaagtacta agaccggaaa 34920
ggagtgccac ggctacattg accatggtac cggcaagtat atagggaatt ctaaggagat 34980
cgtggatctt ctcgtaaagg gaatgctcag cacggacgaa agctatactc tccagagtgt 35040
atatgacagg gctccgaaaa ccaattgatt atgaatctaa tccttaaact tagcatcatt 35100
ttatcattag tagttgctgt atccggctgc ggacgcaata aagaagcggc cggaacagct 35160
tttaacgaag aaacccggat catgatggat cgtatggcgg cttcagtaaa ggccggcaag 35220
attctttatg gccatcagga tgacctcgtt tatggccatt catggtgtgt ggatcctgat 35280
tccgcagcga cagatgcgtt gaaccgttcg gatgtcaagg ctgtcgccgg agattatccg 35340
gctgtcgtgg gattcgaact cggcggtctg gaactgggat gggactggag cattgacaaa 35400
gtcccgttcc tgctgatgcg ccgggctgct gtagagcatg ccggaagggg aggaatcgtg 35460
acgttctcat ggcacccgca caatccttat acaggcggtg atgcctggga tatcagttcc 35520
gacaaggcag tggcttctgt cctcgaagga ggggagaagg cggagatgtt cggagtctgg 35580

ctggagaggc tgggcgattt catggactcg ctccgtgacg ataagggtaa tctgataccg 35640
gtcattttca ggccgtggca tgagaatatc ggcagctggt tctggtgggg cggagccctt 35700
tgtactgcgg atgagtatat cgcccttttc aggaggacgg tcgagtatct gagaaatgac 35760
aggggactgg acaatatcat aatgtgttat tctccgaatt ccgggattac ggcggagcag 35820
tatatgtccc gttatccggg cgatgatgtc gttgatatcc tggggtatga ccattatgag 35880
actgtcatgg agggcgagag tctggaagat gccggaaagc gttatgccgg gcagttggat 35940
cttggccttt ccatcgtcaa gtctcttgct tcggcccacg gaaagctgat gactctttcc 36000
gaaaccggaa tggagggcct cccggacagg agctggtgga cttcggtgct ttatcctgcg 36060
gtaaaggatt ttccgattgt ctatgtgctg acgtggagaa atgcatggga cagggagacg 36120
catttttatg cgccgtatcc tggtaatgag ggcgacttca gggaattcgc ttcgctgccg 36180
gagatctctc tcctgtcgtc gttccggcct tagtccgtca ttcccggcga agtgtcgtcc 36240
tttccggtcg gaccgggaat ctgtaacctt aaaacaataa ataacaatat atggagaata 36300
tgagttttga agaacgtctg gcgtggctgg aagcacgcca ccaggaactt attgctaaga 36360
aaaactcgcc tatatacggc aatggtgtgt ttgagagata tgagaaccct gttcttaccg 36420
gtgaccatgc cccttttgttc tggagatatg atctggataa gaactccaat ccttttctgg 36480
aagagagatt cggcatccat gccgcattca atgctggagc tataaagtgg aaggacaaat 36540
atgtcgtctg cgcccgtgtc gaaggtgctg accgcaaatc attcttcgct atcgccgaga 36600
gcccgaacgg aatcgacaat ttccggttct gggaccaccc cgtgactatg cctgattatg 36660
gcgaacctgc cacgaatgta tatgacatga gacttactgc ccatgaagac gggtggatct 36720
atggtatatt ctgcgtcgag cgtaaggatc cggatgctcc ggaaggcgat ctttcgtctg 36780
ccgtcgcagc tgcgggaatc gcccgtacca aggatctggt caattgggaa agactgccgg 36840
atctggtttc tgattctcag cagcgcaatg tcgtacttca tcctgagttc gtgaatggga 36900
aatatgctct ttacacccgt cctcaggatg gattcataga tgccggaaac ggcggcggca 36960
tcggctgggc cctggtggac agcatggaga atgccgttgt cacttccgag aagatcatca 37020
actgcaggag ataccatact atcaaggaga ccaagaatgg cgaaggccct gctcctatca 37080
agactgacag gggctggctg catctggccc atggcgtcag ggctgcgct tccggcctgc 37140
gttatgtgct ttatatgtac atgacttcgc tggaggatcc ttcgaaagtg attgccgaac 37200
tgccggtttc ttcatggtcc cggaagggga ggagtatgtg ggtgatgtca tgaacgtcct 37260
gttctctaac ggttggattg ccgatgagga cggaaaggtc tttatatatt atgcttcctg 37320
cgatacgcgt ctccatgtgg cgacttctac cgtcgacaga ctcgtggatt actgtatgaa 37380
tactgagccg gatctgttta ccacttctgc ttcagtacga cgtctgaacg cgcttataga 37440
caagaatctt tcgctatcta tgtaggacta ataatcaggt ttcagatgga tatcattact 37500
gcaccggcta agccgacatt gaaggaaaag atcggatatg gcttcggaga catgtcgtct 37560
tcgatgttct ggaagatttt ttcgtattat ctgccgatct tctattcgga tgtgttcggg 37620
cttaagcctc agcatgcggc tttcctgctt cttatcacaa aactgtatga tgcaatatcg 37680
gaccctgtca tgggaatcat cgccgacagg acccggtcca gatggggaaa gtatcgtccg 37740
tatcttctct gggtggctct gccgttctcg cttataggcc ttctgtcttt ctatactcct 37800
gatgccagct atggcctgaa gcatgtatat gcgtatgtga tgtatatcct catgatgact 37860
gtctatacgg ctataaacgt gccttacggg gccatgctgg gagtcgtcac tccggacagc 37920
cgtgagaaat cggttttctc gtcttacagg atgttctttg cgtatatcgg cagtttcatc 37980
gccatgggta tattcggcgt gttcgagaag ctgatttccg gtaccgtgcg tgttatgccg 38040
gacggtagcg agagggtggt gcgcggagtc ggcgatgctg ctcccggaca gtggactctt 38100
gtcgtcggca tcgtttctct gtgctgtttc gtgctgtttc tgctttgctt ccttatgacc 38160
agggagcggg tggggactga aaataaggcg gtttccgagg gaggctcttc tatcgcagag 38220
gatctcaagg ctctaggccg taacaggcct tggtggctgc ttcttggcgc cggtatcggt 38280
atattgctgt tcaattcgat ccgaggcgga gccgctgcat attatttctc gaatatcctc 38340
ggtacgagcg tgttcctgac ttgcgctgtc tatcttttca ttggcgaggt ggctcagatg 38400
gttggggtgc tgtttactgt gccgctttcg gagcgtatcg gcaagaaggc gacttttatc 38460
tctgtgcttc tgacggtttc tgtgctgagt gtggcgatat tcttcctgcc tgagactcct 38520
gccggtttct ggggacttct ggtgctgcag gtgcttatat gcattgcttt tggtgtccag 38580
tctcctctgc tgtggtcgat gtttgccgat gtcgccgatt attcggagga gaagaacggc 38640
tcggcttcta ccggacttat tttctcgtct tcttctatgg ctcagaagtt cggcggggcc 38700
ctcggcgctt tcttgctgat gcagattctt gcaatcttcg catatgacaa ggatgctgca 38760
gtccaggcgc ctcagactct tgctgccatc cgtggcctga tgagctatat cccggcaatc 38820

```
     ggcgccctcc tcggagccct ctgcctgctg ttctatcctc ttacggaatc gcgtatggct     38880
     tcaattagag agaaactttc tgctcgcaga taacatcttc attcatgtag gtatttgtta     38940
     ttccaaagga tggccctctt cggccatcct ttttcgtctg gctgattcga gtcgggcctt     39000
     tctgctggcg ggagaaggac gggacggctc cgtcaagttc ggaaaagctg cttggtcgct     39060
     gtcgccgcct aaacggctaa cttctccctt ttacgatggg cgagccatcg taacggtcgt     39120
     caaacacacg ccgttcttca cggcgccgac ggccgctccc gcgctctccg cttttcctca     39180
     attcctccgc cgcctccgcc ctcctcccgc cgcaggactg ctgaccaggg ttccggggac     39240
     gtggcccccc tcggccacgg gaagagggac ggggcccgca agatgcaagt tatcgcgaag     39300
     cgatgacgcg gcagattgtg ggagggtacg gagtgagcga tagcgaacga agttccccgg     39360
     gaccctgcgt cagcaggact gtggcaaagg aagcagaatg caggactgcg gcaaagatga     39420
     gggactggag ggaattgaaa aaaatgttat atttgtattt cgttccttaa tccggagaac     39480
     tatggcagac aaactatacc tgattgacgg tcacaaccta atcttcaaga tgtactacgc     39540
     attcctgcgt cgtcctatga tcaattccaa ggga                                39574
//
ID   XXX; XXX; linear; XXX; XXX; XXX;  XXX.
XX
AC   PRJEB67889
XX
DE   Uncultured bacterium DNA fragment from in-reactor enriched cow rumen, clone
ferm-20g16
XX
FH   Key             Location/Qualifiers
FH
FT   source          1..35118
FT                   /organism="uncultured bacterium"
FT                   /environmental_sample
FT                   /mol_type="genomic DNA"
FT                   /isolation_source="Cow rumen"
FT                   /clone="ferm-20g16"
FT                   /note="AZO-CM cellulose, AZCL-HE cellulose, AZCL-Barley
beta glucan, AZCL-Tamarind xyloglucan, AZCL-arabinoxylan active clone"
FT   CDS             complement(10..870)
FT                   /db_xref="SEED:fig|6666666.127242.peg.1"
FT                   /translation="MRNAYLILAHNEWSLLNILLSCLDDERNDIYVHIDKKVRTLPKLH
FT                   TDKAGLRILENRADVRWGDLSMVEAEYALFEEALKKGGYDYYHLLSGVDLPLKSQDYIN
FT                   DFLEKNKGKEFIGYTLTQITPEVVRKVQRWHLFPKDFRKRSGAKRIIRAGFLRIQEVLG
FT                   IRRNKGIDFKKGSQWCSISEDMARYFVSRKDWVMKVFRHTFCSDEIVMQTLCWESPYRK
FT                   NIYCTTDDGRGCMRAIGWRNGELVDWCAKDYSMLCESDAIFARKFNSSDMSFIRRITEL
FT                   QQTHD"
FT                   /product="GT14"
FT   CDS             complement(871..1950)
FT                   /db_xref="SEED:fig|6666666.127242.peg.2"
FT                   /translation="MARMDNHSGRLVDDQEIIIFIYDIQGNILRQDLEAQSAPFTGGEP
FT                   LGWQLYHYISELISHNIGAVAIPMFFMISGYLFFAGCADRNLQFSDFITKWKKRVRTLL
FT                   VPYIIWNVLYIVAILAKKVIFTRLGLENDDKDQLLARGFGYLFWTGPLDFPLWYVRSLM
FT                   VMSLLAPVYYWLIRKALPLFIVFLGVMCLLPWDIECPDASSLIFFGTGACLSLKDVDVL
FT                   SFCRKFRIPSFILSVILLAVTLFSENYEYRYIFLRISCFIWFIAVFNIFDSVCDKKKIS
FT                   EILTWLAGASFFVYALHAVYLTDWVRGFLQRVLSTGLFGTSLSYVLITPTVCLICLAFY
FT                   TFLKKFMPHTLSVLCGTRI"
FT                   /product="Succinoglycan biosynthesis protein"
FT   CDS             1930..2634
FT                   /db_xref="SEED:fig|6666666.127242.peg.3"
FT                   /translation="MVVHPGHGHFDGTLINALAYHLGISQGPDAEDDRMGILVHRIDMD
FT                   TSGLLVVAKNDEAQLNLAQQFFVHSIERKYVAVVWGNIKEDEGTIVGNIGRDPNDRLKF
```

```
FT                   RVYQEEDKGKHAVTHYKVLERFGFVTVVECRLETGRTHQIRVHMSYIGHPLFNDARYGG
FT                   AEIRKGTIYAKYKQFIQNCFEICPRQALHAKTLGFVHPRTGKPVSFDSPLPEDMTALLE
FT                   KWRKYAGDMPKE"
FT                   /product="Ribosomal large subunit pseudouridine synthase D
FT                   (EC 4.2.1.70)"
FT                   /EC_number="4.2.1.70"
FT   CDS             complement(2652..4223)
FT                   /db_xref="SEED:fig|6666666.127242.peg.4"
FT                   /db_xref="GO:0003723"
FT                   /translation="MDSEKELLKDLIVDVQATEVQIALQENHRLIELNKESSSGRSFAV
FT                   GDAYLGRVKKVMPALNAAFVDIGDEKEAFIHYLDLGLYFGAFDEFVKKTNPNTNPSRLF
FT                   GSIKLGPILEKEGRIENVLTQGQMIIVQIVKEPISTKGSRLTAEISLAGRNIVLLPFAE
FT                   KVSISQKISSKEEKKRLETLVRSILPKNYGAIIRTAAEGKNAAILVAELKSLIGKWEDS
FT                   WENLSKNKSIQPLFTEYSKTTTILRDLLNDSFSNIYVNNEKEYEEIRRYISLISPEQEK
FT                   IVKLYQGKEPIFDHFEITRQIKSSFGKVVPLKQGAYLVIETTEALNVVDVNSGIRAKTK
FT                   EQEENTFEVNKFAAEEIARQLRLRDMGGIVIVDFIDMESVEHRNNLFKYMQELMSKDRA
FT                   KHNVLPLTKFGLMQITRQRVRPVTEINTTEVCPVCHGTGKIASSLLIDEQIERRLSYYV
FT                   NDKGIRNFTLKVSPILGAYITRGLFNSYVGKWKKKYKCKIKVVEITDFAVLQNELFNEK
FT                   GDKLDS"
FT                   /product="Cytoplasmic axial filament protein CafA and
FT                   Ribonuclease G (EC 3.1.4.-)"
FT   CDS             complement(4452..4724)
FT                   /db_xref="SEED:fig|6666666.127242.peg.5"
FT                   /translation="MTKAEIVNEVAKATGIEKLAVQSVVESFMESVEASLAKGEPVYLR
FT                   GFGSFIIKHRAEKAARNITKNTTLTIPAHNIPAFKPAKVFVSKVK"
FT                   /product="DNA-binding protein HU"
FT   CDS             5010..7352
FT                   /db_xref="SEED:fig|6666666.127242.peg.6"
FT                   /translation="MSGEPALNFDASELPGSLPILALRNAVIFPGVVFPITIGREKSML
FT                   LVDKAQGNGMFIGAVPQKNVAEENPLSLADFSEFGTVVKVLKTFEMPDGTVTALLQGYK
FT                   RLRMLELTDTDPYHMARVSYLDDILPQNDSKVPVIADILKENASSIIRSSSFVPKEALE
FT                   ALKNIDDFNFLVNFIATTIDMDDFMEKVDLLKYDDVKERAMALLGVLSRQLEIQRLKQE
FT                   ISSKVKTELDQQQREYFLNNQLRTIQEELGMDEMDDIARFREAAEKKEWPASVGETFEK
FT                   ELAKLERFNPSTPEYSIQYNFLQFMIDLPWYHTSDDNLDLKHAQKVLDEDHFGLENVKD
FT                   RIIEYLAVLKLKGNMKSPILCLYGPPGVGKTSLGKSIARALGRKYVRISLGGMHDEAEI
FT                   RGHRKTYVGALPGRILNGINKAGTSNPVMVLDEIDKLGSDFKGDPSSALLEVLDPEQNT
FT                   AFHDNYLDIDYDLSNVMFITTANTINGIQPALLDRMELINVTGYLAEEKVQIAKKFLVP
FT                   KQLEEHGIPKKELKIDKAALEEIIDKYTHESGVRGLEKQIARIARVSAKKMALGEDYPK
FT                   TIKKEHLKEYLGLQTAFHDSQKGNEAPGVVTGLAWTSMGGEILFIETSVSDGKGVLTMT
FT                   GNLGDVMKESATIGYQYIKAHPQLAGLTSEEFAKKDVHVHVPEGAVPKDGPSAGITMVS
FT                   SMVSALRGKKVRSGIAMTGEMTLRGRVLPVGGIKEKILAAKRAGVSTIIISEENRKDIE
FT                   DIKEIYVKGLTFVYVKTIDDVIGYIFE"
FT                   /product="ATP-dependent protease La (EC 3.4.21.53) Type I"
FT                   /EC_number="3.4.21.53"
FT   CDS             7352..8020
FT                   /db_xref="SEED:fig|6666666.127242.peg.7"
FT                   /translation="MDVKIEQTWKDALRDEFEKPYFASLVRFLHNEKEAGKVIYPPGGL
FT                   IFRAFELTPLPEVKVVILGQDPYHGPRQAMGLSFSVPSDVPAPPSLKNIFKEIQTDLGI
FT                   TMSGSPNLEGWARQGVLLLNAMLTVQAGVAGSHKAIGWQEFTDAVIKCISDRCNGVVFL
FT                   LWGNFARGKKELIDTSRHYVLEAAHPSPLAGGAFFGCRHFSKTNELLIKEGKTPIDWKL
FT                   "
FT                   /product="Uracil-DNA glycosylase, family 1"
FT   CDS             8017..8490
FT                   /db_xref="SEED:fig|6666666.127242.peg.8"
```

```
FT                   /db_xref="GO:0004595"
FT                   /translation="MKRKALFPGSFDPFTAGHLNILRRALTMFDEIVIAVGVNQDKRGF
FT                   FSMDQRMDIIRQAVRGMDGIEIIKYDNLTIDTCRELGIAHIIRGVRNMLDFENERAIAD
FT                   ANRRLAPEIETIIIPTAQEFAHISSSAVRDILKHKGDTSLFLPEGVVLPDHDA"
FT                   /product="Phosphopantetheine adenylyltransferase (EC
FT                   2.7.7.3)"
FT                   /EC_number="2.7.7.3"
FT   CDS             complement(8492..8710)
FT                   /db_xref="SEED:fig|6666666.127242.peg.9"
FT                   /translation="MRVKTYGDMVSYIVSRARRNEEIALGPLYIQRLGLQRRKILLKLC
FT                   IQGGHLDGIRILGAQGKRKQRKKDYED"
FT                   /product="hypothetical protein"
FT   CDS             8714..9961
FT                   /db_xref="SEED:fig|6666666.127242.peg.10"
FT                   /translation="MQSKLMGDEAVAVHLADNWFIPGKVSFPDEIDLMNAKIFAEFNRQ
FT                   SLLGCKAPGLVAEDSLGRKVEVPLGLSDRYRILYIYSPDCATCSLQTMLMDSLFSRKSY
FT                   PVDLIAFNSDDNTEAWVQYRRNRLSFDGVRVMHFRDPSLESDFQRKYGVLQTPRMFLID
FT                   KNGIIVGRGLDAAALEKLLDATPDLGIGYGSKSSADYFDAVLESFGDELRGSDIRYFIG
FT                   RLEEATLESADTAGFRQMTGDMLYYLAGKRGEAFKDAEAFVVDSMVLSRPDVWRFADDS
FT                   LTVVDMAMSLKHMLSLAPVGARVPDVRVHATVLRAGRKPATVHKKLRSLLKGKRNYVLF
FT                   HTAGCTFCEAELAAADSVASEKGVRVILVDMDKLTSLYPEEADRLLDAFDLSVLPFIME
FT                   TDNKGCVVRKYLSLNK"
FT                   /product="Thiol:disulfide interchange protein tlpA"
FT   CDS             complement(9950..10522)
FT                   /db_xref="SEED:fig|6666666.127242.peg.11"
FT                   /translation="MSIMGKAYIFLADGFEETEALATLDILRRGKIEAYTVSITEEVYI
FT                   TSSHGVTVVADMGRSDFSEIDIDEKDAMIFPGGMPGSKKLAADTPLIGLMNVHYGNGGI
FT                   VAAICAAPGLVATQIKGIEGKKATCFDGFEENLAASGMEYVKAPAVTDGQLITGRGAAF
FT                   AIDFGLAILEKMGGKEAVAKVKAALLI"
FT                   /product="DJ-1/YajL/PfpI superfamily, includes chaperone
FT                   protein YajL (former ThiJ), parkinsonism-associated
FT                   protein DJ-1, peptidases PfpI, Hsp31"
FT   CDS             complement(10632..11744)
FT                   /db_xref="SEED:fig|6666666.127242.peg.12"
FT                   /db_xref="GO:0008784"
FT                   /translation="MNRVDDFSRLELNLTGCMENYRYFRSKLEQTTKLLVLVKANAYGH
FT                   GAVEFAAMMQRAGANYLAVAYPVEGMELRKAGISLPILVLTAGTDFFNEIIDNRLEPGI
FT                   PNLYALKILCGILKERGIKDYPVHIKLDTGMHRLGFMTSELQDLTDFLKTCEEVKVKSV
FT                   YSHLAVSEDPEQDEFTLGQIGMFKTNADRITSEIGYKPMYHILNSAGIERFPQYQFDMV
FT                   RLGIGIYGISALPDVRLAHVASLKCKILQIKTLKPEDGTIGYGRYGHIAPEGTVIATIP
FT                   VGYADGIDRHLGRGHASFSVNGHRAPTIGNICMDMCMLDVTGIDAKVGDTVTIFGEDPT
FT                   VNELADILGTIPYEILTSVPRRIERIVVRK"
FT                   /product="UDP-N-acetylmuramoylalanyl-D-glutamyl-2,6-
FT                   diaminopimelate--D-alanyl-D-alanine ligase (EC 6.3.2.10) /
FT                   Alanine racemase (EC 5.1.1.1)"
FT                   /EC_number="6.3.2.10"
FT                   /EC_number="5.1.1.1"
FT   CDS             12103..12777
FT                   /db_xref="SEED:fig|6666666.127242.peg.13"
FT                   /translation="MDMIWFIIIAVMVLSFVIQQVLNSKFNKYSKVGLHSGLTGAQVAA
FT                   KMLEDHGITDVSITCIKGKLTDHYNPVNKTVNLSEEVYYGNSVAAAAVAAHECGHVVQH
FT                   ATGYAPLKLRSALVPVVSFSSNAVQWVLLIGVMIIKITPTLLYLGIAMFALTTLFSVVT
FT                   LPVEINASARAVQWLDKAGITGPATKAKAVDALKWAAYTYVIAAIGSIATLIYYIAMAR
FT                   RR"
```

```
FT                   /product="probable metal-dependent peptidase"
FT   CDS             complement(12774..13067)
FT                   /db_xref="SEED:fig|6666666.127242.peg.14"
FT                   /translation="MTGDDYKIISDTTENGVRHITAIPSSLVCSQQIDFDIIDGKVRNL
FT                   AYVRGCNGNLQAIGRLVEGMDIKKIVDTLSGVNCNFRGTSCTDQLARILSSL"
FT                   /product="hypothetical protein"
FT   CDS             complement(13084..13518)
FT                   /db_xref="SEED:fig|6666666.127242.peg.15"
FT                   /db_xref="GO:0008270"
FT                   /db_xref="GO:0016787"
FT                   /translation="MEKDELYMKEALREAQAALAEDEIPIGAVIVCKGRIVGKGHNMTE
FT                   RLHDPTAHAEMIAITAATEALGGKYLNECTLYVTVEPCPMCAGALNWAQIGRIVYGAID
FT                   PKRGHSLFTPSLLHPKTEVEAGVLAEECGSMVTEFFKKKR"
FT                   /product="tRNA-specific adenosine-34 deaminase (EC 3.5.4.-
FT                   )"
FT   CDS             complement(13639..13869)
FT                   /db_xref="SEED:fig|6666666.127242.peg.16"
FT                   /translation="MFKIPAGNQRKLIFAYLFDTVAYDSPDALAMLDEVQLVFPVPVEG
FT                   EDEFGLMSLDYVEAVPVRKRCYFRYYRGHIH"
FT                   /product="hypothetical protein"
FT   CDS             13954..14526
FT                   /db_xref="SEED:fig|6666666.127242.peg.17"
FT                   /translation="MMRNQFSSIQHMFEQARNGLTFSLAAIMKVYSLLDALPSEIDKFD
FT                   QFQHFMRLLYDLSLAGDSRILASTSFAHNDADKESRRVQKIKQYVGDHFAENITLEELA
FT                   EMVGMSPSAFSRFFKSRTGKTVIDYIIDIRLGNAARLLVDTTQNISEICYSCGFNNLSN
FT                   FNRLFKSKRGYTPREFRTLFKKNKVFV"
FT                   /product="Transcriptional regulator, AraC family"
FT   CDS             14696..17095
FT                   /db_xref="SEED:fig|6666666.127242.peg.18"
FT                   /translation="MKMDIVKVDDGSFVVDGHFYYYIGTNFWYGGLLGSEYGDRSRLAA
FT                   ELDSLHALGLDNLRVLVGSDGPDGVATRVYPTLQKEPGVYDEKVFEGLDYLMAELGKRN
FT                   MRAVLYINNSWEWSGGYGMYLEWAGAGKAIIPSVGGYGPFMEQMAQYATNARAQELFEN
FT                   HLRTVVSRVNSITGKPYSEDPAIFSWQIGNEPRCFSSDPAVQDGFVSWIWRAASIIKSI
FT                   DPNHMVSTGNEGIWGCEENPELFRRLNECPDIDYITAHIWPYNWSWVGKESPQLDIDRA
FT                   EENTGRYIDAHLEVAESLGKPLVIEEFGFPRDDMQFAKGTPVSGRDRYYRYVFSKVMEN
FT                   QKNGGRLAGANFWSWGGLAVPAHRYWQEGDDYAGDPAQEEQGLNSVFADDYSTLAIIKH
FT                   TTARLSAVANASAVLENDWMFTTGDQKPLRVAVQSQNGFKGSVPVYVRVSTDKGVSVSV
FT                   SGKKARLGKGADTLDFDLGLAPGFYKVCVSLNENDPLDENFCVGCDPEKIVSPQDKQPD
FT                   FDEFWEDNLRQLAMVPMNARMTPLKDHSNADRTAYRVDMKSFGGKEISGLLYLPNKEGK
FT                   FAARISYMGYGSDVWFADPSAEPDMIEFTLSVRDQGFNKTPGQGWVTRGLADKDSYYYR
FT                   GAYLDCVRAIDFVSSLPQTDPDHIYAEGGSQGGAFTLIAASLDNRLRAIAPFVPFLSDF
FT                   PDYFAIADWPGGEVLSAAKAQGISEDELYKTLSYFDVKNFTDRIGCPVLMGFGLQDPVC
FT                   PPHTNFSGYNHITADKQWLCVPVSGHHLEQFPEWWAARTEFFTKNK"
FT                   /product="GH5_7-CE7"
FT   CDS             17153..20332
FT                   /db_xref="SEED:fig|6666666.127242.peg.19"
FT                   /translation="MLMLFGALSLMAQNKVTGRITDNSGQGVINASVMVKGTNNGVVAD
FT                   LDGYYEIVVGQGSTLVYSSIGYKDQEVVVGTQSVINVLLVENSELLDEIVVVGYGTMKR
FT                   SDLAGSSVSMKEDDLKGSIITNLDQSLQGRAAGVTAVQTSGAPGSSSSIRVRGQATINA
FT                   NAEPLYVIDGVIMQGGGNSGADFGLGDALGNGKVSTISPLSTINPADIVSMEILKDASA
FT                   TAIYGAQGANGVVLITTKHGSRGDAKFSYDGMFAVSRQTTRIDMMNLREYAEFYNSMVE
FT                   TGEISEANPYYATPGLLGKGTNWQDEIFRTAIQHQHQISAQGGTDKVQYYVSGSFMDQE
FT                   GTIRGSEFGRYTFRTNLDAQLKEWFKLGLNLTFANTNDDLKLADGSEGLITYSLTTIPD
FT                   IPVYDVYGNFSSTVREGYTSPNPVALAMMDEILLDRQKLNGNIYAELTPVKHFTWRTEL
```

```
FT                   GFDVSGSTATRFRPTVNLGGWTRDTNSMSYQKNSNRFWQIKNYLTYANSFGKHSVTAML
FT                   GQEAWESSWSYVQGANTKLPSNDVHNVALGGGTPSLSSGFGSSAMASFFTRETYNFDDR
FT                   YLLTYTYRYDGSSNFGPEKRWAGFHSLAVAWKFTNEKFMDFLKPVVNNGKLRLGWGQTG
FT                   NSSIGGYAWGASISRMPSALGMGYRPANIPNPAIHWESQTQQNIGLDLNMFDNRLNLTV
FT                   DAYLKKSDNMLMSMQLASYMGTQGNSSSALAAPKGNFGSIENKGLEITLDAHPVQIKDF
FT                   AWDSNFQISFNRNKLVSLDGTTNAALIGYGQWSDVVCVSEIGKPLYNFYGYKVAGVYTS
FT                   LEDIENSPKPEKYPSNGEYARGNTVWVGDIKYEDISKDGVINEKDRTDIGNPLPKFTFG
FT                   WSNTFTYKDFDLNIFINGSYGNKVLNYNAISLTHMNSAWTNQINDAVSGRAMLAPIDAS
FT                   VVYADGSKWYDHIDNVRVLNPGTKTPRASINDPNDNDRISDRYVEDGSYIRIKNITLGY
FT                   NVPKSFLNKFKVDNVRVYVNIQNLYTFTKYTGYDPEVGASTQDSTGLTYGVDNGRYPSP
FT                   TVYSCGLNITF"
FT                   /product="TonB family protein / TonB-dependent receptor"
FT   CDS             20345..22093
FT                   /db_xref="SEED:fig|6666666.127242.peg.20"
FT                   /translation="MKISNMMKGFAVATAVFALTSCEGFLNRPNEDNYNVDNFYKTDEQ
FT                   CEQGVNYLYNSPWYDFQRGFIKIGEVMSGNMYWGSSPYLDFSTNGTDQDLVNMSYSLWA
FT                   VNGHANTVIYNLLNSQGPSKAAIDKCIGEALTWKAFAYFFMVRTFGDVPIVHSNSDLLS
FT                   SGEYNSISKVERADVYEYIITTLEKAMEMLPKKTSGFNNYERIDYYAAEALLAKVYLTK
FT                   AGLSGSLNQDDLKQAAKYAKDVIDNSGRTLLPVYSDVFRLTGFNATGECLISWEWSVGR
FT                   NPWTQQNTLQSDLAMVGFSDQGDCWGGWGGPSVDLADAFGVSATQNPAERVGEKDSRRK
FT                   ATLMLPGDKYDYFWTDKGGFDYLRFIYDTEYGKGGPGGSMQCPTGSNNVKHLYGDNFDH
FT                   ENAGLGTPENMAYQLPTHLLRLSDVYLIYAEASLLTGSNGPALEYVNKVRERAGADALA
FT                   SVTFEDIWKERRLELAGEGDRWYDYVRRSYYDMDACIAELKAQRRSYWDGLDDVWKSYI
FT                   GDAFDYSGTWDASEVQYNNAEDNPNVTAASFKLPFPTEDVVFNGNMASDAVPVHVDVRE
FT                   EYSYNF"
FT                   /product="putative outer membrane protein, probably
FT                   involved in nutrient binding"
FT   CDS             22106..23410
FT                   /db_xref="SEED:fig|6666666.127242.peg.21"
FT                   /translation="MKSINRIITGSLAILAMASFASCDKVDYPDRYEVADGKPTVYSVR
FT                   YADKDVNITQAYMDEVVCLLGKNLKSIVELWFNDQQATLNTSFITDNTLLVSVPKNMPV
FT                   VQTDKIYMITSGKDTLAYDFAVLPPTPKVTSMSNEWAAEGETVTIYGDFLIDKADKPLQ
FT                   IAFAGAEVDHASMVFNGSSSVSFPVPAGAQPGYVTVTSLSGTGKSKFMYKDTRNILFDW
FT                   DGSHGGFATGHGWRNGVIHAPGADAWDAIDGSYLYFGGADLGPGSADAWAEDQYSFNYW
FT                   PEAGTDYDALNLRPEFAEYIDKYGVSGLVLKFEVLVPSSNPWSASGMQLMFSSDAVVTY
FT                   GTATNSYFSDETFPRAIWQPWTTNGGSYDTADKWVTVSFPLSSFNKNQENKECATAFDK
FT                   SYLTGLTFFVWNGGIDGTACSPVFAIDNIRVVPNE"
FT                   /product="FIG00417257: hypothetical protein"
FT   CDS             23417..25843
FT                   /db_xref="SEED:fig|6666666.127242.peg.22"
FT                   /translation="MKLIMKKIFSISAILAACVLALGLSGCSSKELDTDQYSDKDVKLA
FT                   SFGPNPVMRGATLRFFGSNLDRIVEVSVPGLDPITDIEVVTSGNPSEIRIQLPAEGAQV
FT                   GKISLKSSDGKVFTTQVDVEYTEPIVFDSFSPAEAMPGDVITVKGDYMNLVQSVTFEGG
FT                   AIVTEINHNGRHEASFKVPSKAITGKIILSDEGAIANLLYSEEDLVIGDPTVKSLTVAV
FT                   AKPGENAVVKGAYLDMIDKVVFEGGAEVSDLALNEDATEFNLVIPAGAQSGEVKAVSHA
FT                   AKEFVAGEIELVIPTDVAAAPSPVKAGADLVISGNDLDVVTSVDFPGASGVEFVFAEDA
FT                   ITVTVPATATEGDIALNMASGEAVNVAFTLVHPVVSAIAPVELMAGETITVTGTDLDLV
FT                   TGVTLGGKPVEFTAEAEKIVVQTANTSVSGKLVLSLANGEKIEPEDAITLSYDSFIIVN
FT                   DMPAAEHIGAIVTLKGENFMMIENIYIGEEKVSKYNLRSDNEIQFVMPWNKVGTYSMYF
FT                   HLLSGDVETCPQTIEVLLEQDIMTVWEGSLQITWNDGGRVTVPAASFTGIPAGSKLRFY
FT                   YTQVDQQWDQAQLNYGNWTGINFDDPAGTVINGTLVPTDVYGWFTDGILDRCTEVVLTK
FT                   EILENIQANKGDCEDKTACGIIIQGSGLTFTKIEIVCDIPQEVTIYEGPTTLTWGDDGR
FT                   FGLAMTYFEAAKPGSQLIFYLEQTDAWGQVQLNDGWWANADMNFPEIGGAYITTDNLGG
FT                   KDVTRIALTLTADILDHILATSGDYFGLNSQFQGDGRVAMVIQGSDMIINKVCIL"
```

```
FT                   /product="hypothetical protein"
FT   CDS             25938..27464
FT                   /db_xref="SEED:fig|6666666.127242.peg.23"
FT                   /translation="MIFLAACSCSKDDAEKKEPSVPAPKAVTVDVASVSAPVEGTSVVV
FT                   AISSPSKPAVSVSGDWIKAELGPYNPDTYKVKLTVTVGENPTYESRSGSVTLTCQGFSP
FT                   VGISVSQDAAAKPVGPSANPAYDLLGLGWNMGNQFDAFYNGSWAGDKYNYPDETVWGNA
FT                   KATQATFDGVKKAGFTSVRIPVTWLNMIGPGPEYNIDETWMNRIYEVVGYAEKAGLKVI
FT                   INTHHDENNNDDHWLDILSASKDARTNEAIKAEIAAVWTHIANKFKDKGDFLIMESFNE
FT                   IQDGGWGWSAAFQADPSKQCDVLNEWNQVFVDAVRATGGNNATRWLGVPTYAAGTSWTK
FT                   YFRLPEDPAGKVMMAVHCYDPYEYTIGNIQYPEWGHTGAAGKKTPEGEKEITDLFSRLN
FT                   TEFISKGIPVYLGEFGCSMRSKGDARAWMFYKYYMEYVSKAAHTYSLPGILWDNGSTKT
FT                   GKECHGYIDHGTGKYIGNSKEIVDLLVKGMLSTDESYTLQSVYDRAPKTN"
FT                   /product="GH5_4"
FT   CDS             27578..28609
FT                   /db_xref="SEED:fig|6666666.127242.peg.24"
FT                   /translation="MMDRMAASVKAGKILYGHQDDLVYGHSWCVDPDSAATDALNRSDV
FT                   KAVAGDYPAVVGFELGGLELGWDWSIDKVPFLLMRRAAVEHAGRGGIVTFSWHPHNPYT
FT                   GGDAWDISSDKAVASVLEGGEKAEMFGVWLERLGDFMDSLRDDKGNLIPVIFRPWHENI
FT                   GSWFWWGGALCTADEYIALFRRTVEYLRNDRGLDNIIMCYSPNSGITAEQYMSRYPGDD
FT                   VVDILGYDHYETVMEGESLEDAGKRYAGQLDLGLSIVKSLASAHGKLMTLSETGMEGLP
FT                   DKSWWTSVLYPAVKDFPIVYVLTWRNAWDRETHFYAPYPGNEGDFREFASLPEISLLSS
FT                   FRP"
FT                   /product="GH26"
FT   CDS             28667..29842
FT                   /db_xref="SEED:fig|6666666.127242.peg.25"
FT                   /translation="MENMSFEERLAWLEARHQELIAKKNSPIYGNGVFERYENPVLTGD
FT                   HAPLFWRYDLDKNSNPFLEERFGIHAAFNAGAIKWKDKYVVCARVEGADRKSFFAIAES
FT                   PNGIDNFRFWDHPVTMPDYGEPATNVYDMRLTAHEDGWIYGIFCVERKDPDAPEGDLSS
FT                   AVAAAGIARTKDLVNWERLPDLVSDSQQRNVVLHPEFVNGKYALYTRPQDGFIDAGNGG
FT                   GIGWALVDSMENAVVTSEKIINCRRYHTIKETKNGEGPAPIKTDRGWLHLAHGVRGCAS
FT                   GLRYVLYMYMTSLEDPSKVIAEPAGFFMVPEGEEYVGDVMNVLFSNGWIADEDGKVFIY
FT                   YASCDTRLHVATSTVDRLVDYCMNTEPDLFTTSASVRRLNALIDKNLSLSM"
FT                   /product="COG2152 predicted glycoside hydrolase"
FT   CDS             29863..31290
FT                   /db_xref="SEED:fig|6666666.127242.peg.26"
FT                   /translation="MDIITAPAKPTLKEKIGYGFGDMSSSMFWKIFSYYLPIFYSDVFG
FT                   LKPQHAAFLLLITKLYDAISDPVMGIIADRTRSRWGKYRPYLLWVALPFSLIGLLSFYT
FT                   PDASYGLKHVYAYVMYILMMTVYTAINVPYGAMLGVVTPDSREKSVFSSYRMFFAYIGS
FT                   FIAMGIFGVFEKLISGTVRVMSDGSERVVRGVGDAAPGQWTLVVGIVSLCCFVLFLLCF
FT                   LMTRERVGTEHKAVSEGGSSIAEDLKALGRNRPWWLLLGAGIGILLFNSIRGGAAAYYF
FT                   SNILGTSVFLTCAVYLFIGEVAQMVGVLFTVPLSERIGKKATFISVLLTVSVLSVAIFF
FT                   LPETPAGFWGLLVLQVLICIAFGVQSPLLWSMFADVADYSEEKNGSASTGLIFSSSSMA
FT                   QKFGGALGAFLLMQILAIFAYDKDAAVQAPQTLAAIRGLMSYIPAIGALLGALCLLFYP
FT                   LTESRMASIREKLSARR"
FT                   /product="Xyloside transporter XynT"
FT   CDS             31846..34494
FT                   /db_xref="SEED:fig|6666666.127242.peg.27"
FT                   /db_xref="GO:0003887"
FT                   /translation="MADKLYLIDGHNLIFKMYYAFLRRPMINSKGMDMSILYGFTKYVL
FT                   ELIEREKPTHMAIAFDPPGGTFRNEMYPEYKANRSETPQLVIDALEPLLEICKALNIKT
FT                   LMIPGYEADDVIGTVAKSYGKPGTDVYMVTPDKDYGQLVAEHVYQVKPGKAGGADETVD
FT                   VKAICEKYGIKTPAQVIEMLTLCGDTADNVPGVHGIGEVGAGKLIAKYGTVENIYAHLD
FT                   ELTDRQKTQFEEARGHIGLSHDLVTIKTDVPVETSLDEMRLDLDYAPEIADLFEKYEFN
FT                   SLRKYIAHSSRAAKAVAEVKRLDYSETGAEELVKMAPEKISLIVEPEGPGIFADIRRVI
```

```
FT                   VAAEKKDGGIVVASGAAKDFRKILEGEAVKYGYDLKLQSNLLENAGIKLGGRLYDIELM
FT                   HYLINPEKSHKVEILAKSYLGVDIEEAAPVEEAPVSLFDVAEESPANGPDRNLEAAAVL
FT                   KLGEVVRAGMDEAGVGALYEDMEAPLAAVLARMERAGVKIDLGQLRTFADELKVEMDAL
FT                   EAQVRKLADEPGLNVSSPKQIGIVLFEKLKLDPKMKKSASGSYSTDETTLTAIADRHPI
FT                   IDKILEFRAVKKLLSTYIEPFPTYVSPVTGRVHTTFNQALTATGRLSSSNPNLQNIPIR
FT                   TERGKEIRKAFIPATPDGIIMSADYSQIELRIMAHLCGDEHLISSFKAGLDVHKATAAK
FT                   IFGIPVEEVTSDQRRIAKTANFGIMYGISSFGLAQRLHVSRTEAKKIIDDYFAAFPSIR
FT                   TFIDKTIADAREKGYVETVFGRRRYLPDITAKNATIRALAERNAVNAPIQGTSADIIKL
FT                   AMISVDKRIREAGLKSKMVLQIHDELLFDAVPEEIEELGRIVVAEMENVITLSIPLTVE
FT                   CNYGKNWLEAH"
FT                   /product="DNA polymerase I (EC 2.7.7.7)"
FT                   /EC_number="2.7.7.7"
FT   CDS             34466..35071
FT                   /db_xref="SEED:fig|6666666.127242.peg.28"
FT                   /db_xref="GO:0003677"
FT                   /db_xref="GO:0003700"
FT                   /db_xref="GO:0006352"
FT                   /db_xref="GO:0016987"
FT                   /translation="MAKTGLKLTDHDDGSLVEMALGGGPEAEAAFFTIYARYHKGVFAH
FT                   ISRFIQEQEEVEDICMESFEKAFKQLESYKPSNKFSTWIFTIARNTAFDHLDKDKVRGR
FT                   NKDKMPIDYSDLEHIDVPEDNVSPEDEIISSQDHENFLSCVQGLPDLYREVASMCFVDN
FT                   LGYKEISEKAGLPINTVKTRISRAKGLIVQMMLDMEEE"
FT                   /product="RNA polymerase sigma factor RpoE"
XX
SQ   Sequence 35118 BP; 8432 A; 8876 C; 9159 G; 8651 T; 0 other;
     atgattcttt taatcatgtg tctgttgtaa ttcggttatt cttctgataa acgacatgtc        60
     cgaactattg aatttccttg caaaaatagc gtcggattca catagcatgg aatagtcttt       120
     cgcacaccag tcgaccagct ctccatttcg ccatccaatg gccctcatgc accctcttcc       180
     gtcgtcggta gtgcagtaaa tgttcttgcg atatggagat tcccagcaaa gggtctgcat       240
     tacgatctcg tcagagcaga aggtgtgtcg gaatactttc atcacccagt ccttgcgtga       300
     gacgaaatat ctggccatat cttcagaaat gctgcaccac tgggagcctt tcttgaaatc       360
     tatgcccttg ttccgtctga ttccgagaac ttcctgaatt ctcaggaatc cggctcgtat       420
     tatccgtttt gccccggatc tcttcctgaa atccttaggg aacagatgcc atctctggac       480
     tttacgtact acttccgggg ttatttgtgt caaggtatat cctatgaatt ctttcccttt       540
     attcttttcc agaaagtcgt ttatatagtc ttgggatttc aatgggagat cgacgccgga       600
     caggagatgg tagtaatcgt atccgccttt cttcagggcc tcttcgaaaa gtgcatactc       660
     cgcttctacc atgctcagat ccccccagcg cacgtcggct ctgttttcca gaatcctcag       720
     tcctgctttg tcggtatgga gtttcggcag agtgcggact ttcttgtcga tatggacata       780
     gatgtcgttc ctctcgtcgt ccaggcaaga aagcaggatg ttcagcaggc tccattcgtt       840
     atgggcaagt atgaggtatg cgtttctcat ctatattctg gtgccgcaaa gcactgaaag       900
     ggtgtgcggc atgaacttct tcaagaatgt atagaatgca aggcagatca ggcagactgt       960
     cggagtaatc agcacatagc tcaagctggt tccgaacagg ccggtgctga gaactctttg      1020
     caggaagccc cttacccagt cggtgaggta tacggcgtgt agggcatata cgaagaatga      1080
     tgctcctgcc agccaggtaa gtatttccga tatctttttc ttgtcgcaga cactgtcgaa      1140
     gatgttgaag acagcgatga accatataaa gcatgagatc cgcaggaaaa tgtatctgta      1200
     ctcgtagttt tcggaaaaca gcgtcaccgc cagtaaaatg accgagagga tgaaagatgg      1260
     tatcctgaat ttcctgcaga acgatagcac gtcaacgtcc ttcaggctga ggcaggcacc      1320
     cgtcccgaag aatatcagtg atgaggcgtc cggacactca atgtcccatg gcagcaggca      1380
     cattacgcca aggaagacaa tgaatagcgg cagggctttg cggatgagcc aataatagac      1440
     cggagccagc agagacatta ccatcaagct cctgacatac cacagtggga aatcgagcgg      1500
     tcccgtccag aacagatatc cgaagcctct ggcgagcagc tggtccttgt catcgttttc      1560
     cagacccagc cgggtgaaaa tgactttctt cgccaggatc gccacgatgt atagtacatt      1620
     ccatatgatg tacggaacca gaagcgtccg gactctcttc ttccattttg tgatgaaatc      1680
     ggagaactgg aggttcctgt ctgcacagcc ggcaaagaac aggtagcctg atatcatgaa      1740
```

gaacatcggg atggcgaccg ctccgatgtt gtgcgagatc agttcggaga tgtaatgata 1800
aagctgccat ccgagaggtt ctcctcccgt gaacggagca gactgggctt cgaggtcctg 1860
ccggagaata ttcccctgaa tatcgtatat gaagatgatg atttcctgat cgtcaacaag 1920
ccggccggaa tggttgtcca tccgggccat ggccatttcg acggcaccct catcaatgct 1980
ctggcctacc atctcgggat cagccaggga ccggatgccg aggatgacag gatgggcatt 2040
cttgtccata gaatcgatat ggacacttca ggcctgctgg tcgtggccaa gaatgacgag 2100
gctcagctta atctggcgca gcagttcttt gtccattcga tagagcgaaa gtatgtcgcc 2160
gtcgtctggg ggaacataaa ggaggacgag gggacgattg tcggtaatat cggcagggac 2220
cctaacgacc gcctcaagtt ccgcgtatat caggaggagg acaagggaaa gcatgccgta 2280
actcattaca aggtgcttga gcgtttcgga ttcgttaccg tggtggaatg ccgcctggag 2340
accggacgta cccaccagat aagggtacac atgtcctata taggtcatcc gcttttcaat 2400
gatgccagat acggcggggc ggagatccgc aaggggacta tctatgcgaa atataagcag 2460
ttcatccaga actgtttcga gatttgcccg cgtcaggcgc tgcatgcaaa gactctgggt 2520
ttcgtgcacc ctcgcacggg aaagccggtc agcttcgatt ctccgcttcc ggaggacatg 2580
accgcgctgc tggagaagtg gcgcaagtat gccggcgaca tgccgaaaga ataaaaaaag 2640
ggggacaccc tttatgaatc aagtttgtcc ccttttcgt tgaatagttc attctgcaga 2700
actgcaaaat cggtaatttc taccactttg atcttgcatt tgtacttttt cttccacttg 2760
cctacataag agttgaacag gcctcgggta atatatgccc ctaaaatagg gcttaccttc 2820
agagtgaagt tcctgattcc cttgtcattt acgtagtagg aaagccttcg ttcaatctgc 2880
tcgtcgatga gcaggcttga tgctattttt cctgtaccgt ggcataccgg gcagacttcc 2940
gtcgtgttga tctcggttac cggccgtacg cgctgacggg tgatctgcat aagaccgaac 3000
tttgtcagcg gcagcacatt gtgcttagcc ctgtctttgc tcataagctc ctgcatgtac 3060
ttgaagaggt tgttcctgtg ctctaccgat tccatatcga tgaagtccac aatcactatt 3120
cctcccatat cgcggagtct cagctgtctt gcgatctctt ccgccgcaaa cttgttgact 3180
tcaaaggtgt tctcttcctg ttcctttgtc ttggcacgga tgccactgtt gacgtcaaca 3240
acgtttaagg cttctgtcgt ctcgattaca agataggctc cctgtttgag cggtaccact 3300
tttccgaaag aactctttat ctgtcttgtg atctcaaagt ggtcgaagat aggttccttt 3360
ccctgataga gctttactat cttctcctgt tccggagaaa taagggagat ataccttctg 3420
atctcttcgt attctttctc gttgttcaca tagatgttcg agaaagagtc gttaagcaga 3480
tctctcagaa ttgtagtggt cttgctgtac tctgtaaaca gaggctgtat gctcttgttc 3540
ttcgagagat tttcccagga atcttcccat ttaccgataa gcgacttcag ctctgcaacc 3600
agaatagctg cgttcttgcc ttcagccgcg gtgcggatga ttgcaccgta attcttggga 3660
agaatactcc tgacaagtgt ctcaagtctc ttcttctcct ccttggagga gattttctgg 3720
gatatgctta ccttctcggc gaagggaagc agtactatgt ttcgtcctgc caatgaaatc 3780
tctgcggtca gtcgtgatcc ctttgtggaa ataggctcct tgactatctg gactattatc 3840
atctggccct gggtgagcac attctcgatc cggccttcct tttcgaggat aggaccgagc 3900
tttatgcttc cgaatagcct cgacgggttg gtattagggt ttgtcttctt gacgaactcg 3960
tcgaaagcgc cgaaataaag tccgaggtcc agataatgga tgaaagcctc tttctcgtca 4020
ccgatatcca caaaggcagc gttgagtgca ggcatcactt tctttactct tccgaggtag 4080
gcatcgccta cagcgaagct gcgacccgag ctggactctt tgttcagttc gataagcctg 4140
tgattttcct gtaaggctat ctgaacttca gtggcctgaa cgtcaacgat caggtctttg 4200
agtagttcct tttcagaatc cattggataa tttgtgaaac aataacaaaa gggccgtcca 4260
gatctcactg tcggcccttc tcttttagca gactgctaaa atggcagaca caggagaatt 4320
atttcttctt gtgtctattc ttacgcaagc gttttttgcg cttgtgtgtc gccattttat 4380
gtctctttct tttcttaccg cttggcataa ttgaaaaaaa gttaaaaaat tattaaatat 4440
tataccggta attacttaac cttgcttacg aataccttttg ctggcttgaa agccgggata 4500
ttgtgagctg gaatggtaag ggtagtattc ttggtgatgt tccttgctgc cttctcagca 4560
cggtgcttga taatgaagct gccgaagcca cgaagataaa ccggttcacc cttcgcaagt 4620
gaagcctcta cactttccat aaaagattca acgactgact gtactgccag tttctcaata 4680
ccggtagctt ttgctacttc attaacgatc tctgccttag tcataataga gtaatttata 4740
taaagttatg tgtgtgcaaa ttttaagact gcaaatatag acttttttttt gcatagggca 4800
aattatttcg ttaaaaatct ttgtcttaca tattttgata ttacctttgt gcttggcata 4860
aaggttgaca ttatagtatc tgccttaaaa acgccatcct tctctgacaa aatgtcagca 4920
gggaggattg tatacactga taaattatga tagattttaa taaagaaatg tttcctgcct 4980

cgtcccacga ggtcaatata ataccggtaa tgtctggaga gcccgctctc aacttcgacg 5040
cctcggaact cccgggatcc cttccgatcc tggcactccg caacgccgtg atctttccgg 5100
gtgtcgtctt ccctataact atcggccgcg agaagtctat gcttctcgtc gacaaggccc 5160
agggaaacgg catgttcata ggcgcagtcc ctcagaagaa tgtcgccgag gagaacccgc 5220
tgtctctcgc cgacttctcg gaattcggta ctgtcgtaaa ggtcctgaag actttcgaga 5280
tgcctgacgg cactgtcacg gcgcttctcc agggctacaa gcgtctccgc atgctggaac 5340
tgacggacac cgatccttac cacatggcaa gagtcagcta tctcgacgat atcctgccgc 5400
agaacgactc gaaagttccg gtcatagccg atatcctcaa ggagaatgcg tcaagcatca 5460
tccgttcgtc ttcgttcgtt cctaaggagg ccctcgaggc cctcaagaac atcgacgact 5520
tcaatttcct cgtcaatttc atcgctacga cgatcgacat ggacgatttc atggaaaaag 5580
tcgacctgct gaaatatgac gacgtaaagg aaagggccat ggctcttctc ggcgtgctca 5640
gccgtcagct ggagatccag aggctcaagc aggagatcag ctccaaggtc aagaccgagc 5700
tcgaccagca gcagcgcgag tatttcctta ataatcagct ccgtacgatc caggaggagc 5760
tcggcatgga tgagatggac gacatcgcac gcttccgcga ggctgccgag aagaaagagt 5820
ggcctgcatc cgtaggggag accttcgaaa aggaactcgc aaagctcgag agattcaatc 5880
cttctactcc ggaatacagc atccagtata acttcctgca gttcatgatc gacctgccgt 5940
ggtatcatac ttcagacgac aatctcgacc tgaagcatgc ccagaaagtt ctcgatgagg 6000
accatttcgg actcgagaac gtcaaggaca ggatcatcga gtatctcgcc gtcctcaagc 6060
tgaagggtaa catgaaatcc ccgatcctct gcctctatgg ccctccggga gtcggcaaga 6120
caagcctcgg aaagtcgatc gccagagccc tcggacgcaa gtatgtccgt atctccctcg 6180
gcggcatgca tgacgaggcc gagatccgtg gccacaggaa aacatacgta ggcgcccttc 6240
cgggacgtat ccttaacgga atcaacaagg ccggcacttc caacccggta atggttctcg 6300
acgagatcga caaactcgga agcgacttca aaggcgatcc gtcttccgct cttctcgagg 6360
tcctcgaccc ggagcagaac acggctttcc acgacaacta cctcgatatc gattacgacc 6420
tttcgaacgt aatgttcata acaacggcca atactataaa cggtatccag ccggctctgc 6480
tcgacaggat ggagcttatc aatgtgaccg gttatctggc agaggagaag gtccagatcg 6540
ccaagaagtt cctcgtgcct aagcagctcg aggagcatgg aatccctaag aaggaactga 6600
agatcgacaa ggctgccctt gaagagatca tcgacaaata tacccatgag tcaggcgtcc 6660
gcggcctgga gaagcagatc gcccgtatcg ccagagtctc tgccaagaag atggccctcg 6720
gcgaggacta tccgaagacc ataaagaaag agcatctcaa agagtatctc ggcctgcaga 6780
cggccttcca cgacagccag aagggcaacg aggctccagg agtcgtgacc ggtctcgcat 6840
ggacctccat gggcggcgag atactgttca tagagacttc cgtcagcgat ggcaagggag 6900
tcctgacaat gacgggaaat ctcggcgatg tcatgaaaga gtccgctact atcggctatc 6960
agtatatcaa ggctcatccg cagctcgccg gccttacttc ggaagagttc gcgaagaagg 7020
atgtccatgt ccatgtccct gaaggagccg ttcctaagga cggtccttcg gccggaatca 7080
ccatggtcag ctcgatggtg tcagccctcc gcggaaagaa agtccgcagc ggaatcgcca 7140
tgaccggaga aatgaccctg aggggccgtg tgctgcctgt aggtggcatc aaggagaaga 7200
ttctagcggc aaagagggcc ggagtctcta ccataatcat ttctgaagaa aacaggaaag 7260
atattgaaga tatcaaggaa atttacgtaa aaggtcttac ctttgtgtat gtcaagacaa 7320
ttgacgacgt aatcggatat atcttcgaat aatggacgtt aagatagaac agacttggaa 7380
agacgcactg cgcgacgagt tcgaaaaacc gtacttcgcc tcgctggtgc gttttctgca 7440
taatgaaaaa gaggccggga aggtcatcta tcctcccgga ggcctgatat tcagggcatt 7500
cgaactcact ccgctgccgg aggtgaaagt cgttatccta ggccaggatc cgtatcatgg 7560
gccccgccag gccatgggac tttcgttctc cgtgccgtcg gacgtgccgg cccctccgtc 7620
cctgaagaat atattcaagg agatccagac ggatctgggc ataaccatga gcggaagccc 7680
gaatcttgaa ggctgggccc gtcagggcgt gcttctgctg aacgccatgc tgacagtcca 7740
ggccggagtc gccggctcgc acaaggctat cggctggcag gagttcacgg atgccgtgat 7800
caagtgcatc tctgaccgct gcaacggggt cgtgttcctg ctctggggta atttcgcccg 7860
cggaaagaag gaactcatag atacgtcccg tcattatgtg ctcgaagccg cccatccttc 7920
tccgcttgcc gggggagcct tcttcgggtg cagacatttt tctaagacta acgaacttct 7980
cataaaagag ggcaaaaccc ctatcgactg gaaattatga aacgtaaagc acttttcccg 8040
ggatctttcg atccttttac cgccggccat ctgaatattc tccgcagagc gctgacgatg 8100
ttcgatgaaa tcgtcattgc agtcggtgtc aaccaggaca aacgcggatt ctttagcatg 8160
gatcagcgta tggatatcat acgtcaggct gtccgcggta tggacgggat cgagataatc 8220

aaatacgaca accttactat cgatacctgc cgcgagctgg gtatcgccca tataataagg 8280
ggcgtcagga acatgcttga tttcgagaac gagcgtgcca ttgcggatgc caaccgccgt 8340
cttgctccgg aaatcgagac tatcatcatc cctaccgcac aggaattcgc gcatatctcg 8400
tcttcagccg tcagggatat ccttaaacat aagggggata cttctctctt ccttccggag 8460
ggagtcgtcc ttccggatca cgatgcgtag gttaatcctc ataatctttc tttctctgct 8520
ttcttttgcc ttgggcgccc agaatgcgga ttccgtcaag atggccgccc tggatgcaaa 8580
gcttgaggag tattttgcgg cgctggagac cgagtctctg gatgtaaaga ggaccgagtg 8640
cgatttcctc atttcttcgt gcacgactga cgatgtacga aaccatgtcg ccgtacgtct 8700
ttactcgcat ttcatgcagt cgaagcttat gggggatgag gctgtcgccg tccatcttgc 8760
cgacaactgg ttcattcccg gcaaggtttc tttcccggat gagatagatc tgatgaatgc 8820
gaagattttc gctgagttca acagacagtc gctgctcgga tgcaaggctc ccggtctggt 8880
agccgaggat agtctcggca ggaaggtcga ggtccctctg ggattgtctg acagataccg 8940
gattctttat atctattctc cggactgtgc cacatgcagt cttcagacta tgctgatgga 9000
ctctcttttc agcaggaaga gctatccggt cgaccttatc gcattcaatt cagacgataa 9060
tactgaagcc tgggtgcagt accgcaggaa ccgcctttct tttgacggag tccgtgtcat 9120
gcatttcagg gatccttcgc tggagtcgga tttccagcgg aaatacggag tgcttcagac 9180
tccccgcatg ttccttatcg acaagaacgg gattatagta ggccgcggac tggacgctgc 9240
tgcgctggag aaattgctgg atgcaactcc tgaccttggc atagggtatg gttcgaagtc 9300
gtctgccgac tatttcgacg ctgtgctgga gagcttcggg gatgagctgc gcggctctga 9360
tatccgctat tttatcggaa gacttgaaga ggctactctc gaaagcgccg ataccgcagg 9420
attccgtcag atgaccggtg acatgctgta ttatcttgcc ggaaagaggg gagaggcgtt 9480
caaggatgcc gaggccttcg tcgtcgattc gatggtcctg tcccgtccgg atgtgtggcg 9540
ctttgccgat gattcgttga cggtcgttga catggccatg agcctgaaac atatgctctc 9600
gctagctcca gtaggtgcca gggttccgga tgtacgtgtc catgccactg tccttcgcgc 9660
cggacgaaag ccggccaccg tccataagaa attgcggtcg ttgcttaagg gaaagcggaa 9720
ttatgtgctt ttccatactg ccggctgcac tttctgcgag gctgaacttg ccgccgccga 9780
ttctgtcgct tcggagaagg gtgtccgtgt catcctggtc gatatggaca agctgacaag 9840
cttataccca gaggaggctg atcggcttct ggacgcattc gacctgtccg ttctcccgtt 9900
cataatggag acggataata aaggttgtgt cgtacgtaag tatctgagtt taaataagta 9960
gggctgcctt taccttagct acggcctctt tgcctcccat cttttcgagg atggcgagtc 10020
cgaagtctat cgcgaaggct gcgcctctgc cggtgatgag ctggccgtcg gtaactgccg 10080
gggcttttac gtattccatt ccgctggcgg cgaggttctc ttcgaagccg tcaaagcagg 10140
tggctttctt gccttcgatt cccttgatct gggtggcgac gaggcccggg gcggcgcaga 10200
ttgcggccac aatgcctccg ttgccgtagt ggacgttcat cagaccgata agcggggtgt 10260
cggctgcgag tttcttcgag cctggcattc ctcccgggaa gatcatggcg tctttttcgt 10320
cgatatctat ttcgctgaag tcggagcggc ccatgtctgc caccaccgtt actccgtggg 10380
agctggtgat atagacttct tcggtgattg atactgtgta tgcttcgatt ttacctctgc 10440
ggagaatgtc gagggttgcg agggcttcgg tctcttcaaa accgtcagcc aggaagatgt 10500
atgcctttcc catgatgctc atgtggttta gattggttga ggtaagaggg ttcgaacctc 10560
tactgaggga accaaaatcc ctagtgctac cattacacca tacctcagtg gggtgcaaat 10620
atacgaaaaa attatttgcg gacaacaatt ctctctatac gtctagggac gctagtgaga 10680
atttcgtaag ggattgtccc gagaatgtcg gcgagttcgt tgactgtagg gtcttctccg 10740
aagatagtca cggtgtctcc gactttagcg tcaattcctg tcacatcaag catgcacatg 10800
tccatgcaga tgttcccgat cgtcggcgct ctgtgcccgt tgacggagaa tgaggcgtga 10860
ccgcgtccga gatggcggtc gatgccgtcg gcgtatccta ccgggatcgt ggcgattacg 10920
gtgccttcag ggcgatatg accgtatctt ccgtagccta tcgtgccgtc ttccggcttg 10980
agggtcttta tctgcaatat tttgcatttg agggacgcga cgtgggcgag gcggacatca 11040
ggcagggcgc tgattccgta gataccgatt ccgaggcgga ccatgtcgaa ctggtattgc 11100
gggaatcttt cgattccggc ggagttgagg atatggtaca tcggcttgta gcctatctcg 11160
gaggtgatcc tgtcggcgtt tgtcttgaac atgccgatct ggccgagggt gaattcgtcc 11220
tgctcagggt cttcggatac tgcaaggtgc gaatatacgg acttgacttt gacttcttcg 11280
caggtcttga ggaagtcggt gagatcctgg agctcgctcg tcatgaaccc gaggcggtgc 11340
attccggtgt cgagtttgat atggaccgga taatccttga tgccgcgttc tttgagtatt 11400
ccgcagagta tcttgagggc gtacaggttc gggattcccg gctcgaggcg gttgtcgatg 11460

atttcgttga agaagtccgt gcctgcggtc aggacgagga tcgggaggct gataccggct 11520
ttgcggagtt ccatgccttc gacagggtag gcgactgcga ggtagttcgc tcccgcacgc 11580
tgcatcatgg ctgcgaattc gacggctccg tgcccgtaag catttgcctt tacgagtaca 11640
agtaatttag tcgtctgctc cagtttggac ctgaaatatc tgtagttttc catacatcct 11700
gtcaggttca gctccagcct tgaaaaatca tcaaccctgt tcattccaca tcaatttaga 11760
atacaaattt atcaaatttc tccgacatag gttacagttt ttcgcttttc tttccccgta 11820
aagaccctta cagtacgtcc gctgtcgccg ccgacagcgt ctgctttggc gggaatcctc 11880
gtggcctcgg ctctgttccg ccgactctgt cggctccatc cctcccaatg tctacttcgt 11940
cctgcttcgc agaactcgta tccatcgggc ccctccctct aacgtggccg agggtggcca 12000
cgctcgccga gaccagagga cagccgccat caattaagac gctgacggca agtgagaggt 12060
tggcatggat ttggaagcaa ccagaaggat tgttaaaatt ttatggatat gatttggttt 12120
attatcattg ctgtgatggt gctgagcttt gtgattcagc aggttctgaa cagtaaattc 12180
aataaatatt cgaaggtggg attgcattcc gggctgaccg gggcgcaagt cgctgccaaa 12240
atgttggaag accatggcat aaccgatgtc tctattacct gcatcaaagg aaagctgaca 12300
gaccattaca accctgtcaa caagacagtc aacctcagcg aagaagtcta ttacggcaat 12360
tcggtagccg ctgcggccgt tgcagcccat gaatgtggcc atgtagtgca gcatgccacg 12420
ggatatgctc cgctgaaact gagatcggct ctcgttccgg tagtctcatt ctccagcaat 12480
gccgtgcagt gggtacttct tattggcgta atgatcatca agattactcc gaccctcctg 12540
tatctcggaa tcgccatgtt tgccttgacg accttgttca gcgtcgtgac acttcctgtg 12600
gaaatcaacg ccagcgccag agccgtgcaa tggctcgaca aggccgggat aacaggtccc 12660
gctacaaaag cgaaagctgt cgatgccctc aaatgggcag catatacata cgtcattgcc 12720
gccatcggtt cgatagcgac attgatctac tacattgcca tggccaggcg acgctagaga 12780
ctcgaaagaa tccttgcaag ctggtccgtg caggatgtac cacggaaatt gcagttgacg 12840
cccgaaagag tgtcaactat ttttttatg tccatgccct cgacgagacg gccgatcgcc 12900
tgaagattac cgttgcagcc ccgcacgtat gcgagattgc ggaccttgcc gtctattatg 12960
tcgaaatcga tctgctggct gcataccagg ctgctcggga tggcagttat gtgtctgaca 13020
ccgttctcgg tggtgtcgct gatgatcttg tagtcgtctc ctgtcattgt tgtagtccgt 13080
tttttatctt tttttcttga aaaactccgt caccatggaa ccgcattcct ccgcgagcac 13140
tcctgcctcg acctcggtct tcggatgcag cagcgaagga gtgaaaagtg agtgcccccg 13200
cttcggatcg atcgctccat agacgatccg gccgatctgg gcccagttca gggctccggc 13260
gcacatcgga cacggctcga ccgtgacgta gagcgtgcat tcgttgaggt atttgccccc 13320
gagcgcctcc gtagcggcgg tgatggctat catctcggca tgggccgtag gatcatgtaa 13380
acgttcggtc atattgtgac cctttcccac gatccggcct ttgcatacga taactgcccc 13440
gatcgggatt tcatcctcgg ccagggccgc ctgcgcctcg cgcagagctt ccttcatata 13500
tagctcgtct ttttccataa ttacagcaaa gatagcaatt cttggacaat cccaatagtt 13560
gataaaacaa ttaagaaaat ttgcgaaatg atactttttt gacattttta tcatatattt 13620
gccacgtgat aatataagtt aatggatatg acctcggtaa taacggaaat aacacctctt 13680
tcggacaggg actgcttcta catagtcgag agacataaga ccgaattcgt cttcccctc 13740
cacaggcaca gggaatacga gctgaacttc atcgagcatg gcaagggcgt ccggcgagtc 13800
gtaggcgaca gtgtcgaaga gataggcgaa tatgagctta cgctgattgc cggcggggat 13860
cttgaacatg cctgggagca gggcaattgt acctccgagg acgtacgcga gatcactatc 13920
cagttcgatc cggagatatt ctccagcggc ctcatgatga gaaaccagtt ctcatcgata 13980
cagcatatgt tcgagcaggc gaggaacggg ctgacctttt cgctggcggc cataatgaag 14040
gtgtattcgc ttcttgacgc cctgccgtcg gaaatagata aattcgatca gttccagcac 14100
tttatgaggc tgctttatga cctttccctg gcgggtgact ccaggatact ggccagcacg 14160
tcattcgccc ataacgatgc cgacaaggaa agccgtcggg tgcagaagat caagcaatat 14220
gtcggagacc atttcgcgga aaacattacc cttgaagaat tggcggaaat ggtcggcatg 14280
agcccgtcgg cgttcagccg ctttttcaag agcaggaccg ggaagacggt gatagactat 14340
ataatcgaca tccgactcgg gaatgcggca aggctgctgg tcgacaccac ccagaatatc 14400
tctgaaatct gctatagctg cgggttcaac aacctgtcga atttcaatcg tcttttcaag 14460
tcaaaaagag gctacacgcc gcgcgaattc aggactctgt tcaagaagaa caaggttttt 14520
gtataacgtg caaaatagta tcggttattg acaaatttgt atttgttggg taaaacgttg 14580
catgctaaat ttgcaacgta ctcagacaag tatggattta tcgataaacc gcatgattaa 14640
aagattattc ccggcactgt ttatcgccct tttcgcgatt ctttcgtgtc agcctatgaa 14700

aatggacatc gtcaaagttg acgatggctc ttttgtcgtt gatggtcact tctattatta 14760
tattggtaca aatttctggt acggaggact tctcggctcc gaatatggtg acagaagcag 14820
gcttgccgcc gaacttgatt ccctgcatgc cctcggactg gacaacctcc gcgtactggt 14880
aggcagcgac ggccccgacg gggtggcgac cagagtctat ccgaccctgc agaaggaacc 14940
gggcgtctat gacgaaaagg tcttcgaagg tctggactac cttatggccg aacttgggaa 15000
aaggaacatg agggcagtcc tctatatcaa taattcctgg gaatggtccg gcggctatgg 15060
catgtacctc gaatgggccg gagccggcaa ggcgataatt ccttcggtcg gaggatatgg 15120
tccgttcatg gaacagatgg cccagtatgc gacgaacgcc agggcgcagg aactcttcga 15180
gaatcatctc cgtactgtcg tttcaagagt caacagcatc accggaaagc cttatagcga 15240
ggatccggcg atattctcat ggcagatcgg caacgagccg cgctgcttct cctccgatcc 15300
tgccgtccag gacggattcg tatcatggat atggcgcgct gcctcgataa tcaaatcgat 15360
agaccctaat catatggttt ctaccggcaa tgagggcata tggggctgcg aagagaatcc 15420
ggagctattc cgccggctga atgaatgtcc tgacattgat tatattaccg cccatatatg 15480
gccatacaac tggagctggg taggaaagga atcccctcag ctggatatag accgcgccga 15540
ggagaatacc gggagataca tcgacgccca tctggaggtg gccgagagtc tgggcaagcc 15600
tctcgtgatc gaggaattcg gcttcccgag agatgatatg cagtttgcca aagggactcc 15660
cgtaagcgga cgcgaccgct attacagata cgttttctcc aaggtcatgg agaatcagaa 15720
aaatggcggc agactcgccg gcgccaattt ctggagctgg ggaggtcttg cggtgcctgc 15780
ccacagatat tggcaggaag gagacgatta cgccggagat ccggcccagg aagaacaggg 15840
gctcaattct gtattcgccg acgattacag taccottgct attataaaac atacaacggc 15900
gagactttct gccgtggcta atgcaagcgc agtgcttgaa aatgattgga tgttcactac 15960
cggagaccag aaacctctca gagttgctgt acagtctcag aatggtttta agggaagtgt 16020
cccggtttat gtgcgggtgt caacagacaa aggggtttca gtgtcggtct caggaaagaa 16080
agcccgtctc ggcaaaggcg ccgatactct ggatttcgat cttggactag ccccaggctt 16140
ttataaggtg tgtgtcagcc tgaacgagaa tgacccgctt gatgaaaact tctgtgtcgg 16200
ctgtgatcct gagaagatag tatctcctca ggataaacag cctgatttcg atgaattctg 16260
ggaagacaac ctcaggcagc ttgccatggt cccgatgaat gccaggatga ctccgttgaa 16320
ggatcattca aacgctgaca ggactgccta cagggttgat atgaagtctt tcggaggcaa 16380
agagatctcc ggtctgcttt atcttccgaa caaagaggga aaattcgctg ccaggatctc 16440
atatatgggc tacggttcgg acgtctggtt cgctgatccg tcggcggaac cggacatgat 16500
cgagttcact ctcagcgtcc gtgaccaggg tttcaacaag actccgggac aaggctgggt 16560
gacaagggga cttgccgata aggactcgta ttattacaga ggcgcatatc tggactgtgt 16620
gcgggccatt gatttcgtct cttctcttcc tcagactgat ccggaccata tctacgccga 16680
aggtggaagt cagggcggag catttaccct gatagctgca tctctggaca accgtctccg 16740
cgcaatcgct ccgttcgtgc ctttcctttc cgatttcccg gactatttcg ccatcgcaga 16800
ctggccgggc ggggaagtgc tctcggcggc aaaggctcag ggcatttctg aagacgaact 16860
ttataagacc ctgtcttatt tcgacgtgaa gaactttacg gacaggatag gttgtcctgt 16920
gcttatgggt ttcggcctgc aggatcccgt ctgtcctcct cataccaatt tttcggggta 16980
taaccatatt actgcagata aacaatggct ttgcgttcct gtctcaggcc accatctgga 17040
gcaatttccc gagtggtggg cagcaaggac ggaattcttc acaaaaaata aataacacaa 17100
ctaattattc acattcaaaa tcagtagcat gaaaagacag ctattaacat gtatgctgat 17160
gctctttgga gcattgtcac ttatggcgca gaacaaagtg accggaagaa ttacagataa 17220
ttccggacag ggcgtcataa atgccagcgt aatggtgaaa ggcaccaaca atggtgtcgt 17280
tgcagatctt gatggatact acgagattgt tgtcggccaa ggctccactc ttgtatattc 17340
ttccatcggt tacaaagatc aggaagttgt tgtaggcacc cagagtgtca tcaacgttct 17400
tcttgtcgaa aactccgagc tgctcgacga aatagtcgta gtcggttacg gtactatgaa 17460
gagaagcgac ctcgccggat catccgtttc catgaaggag gatgacctca agggttcgat 17520
catcaccaac ctcgaccagt ctctccaggg acgtgctgcc ggtgtcacgg cagtccagac 17580
ttccggagct cctggttcgt cctcaagtat ccgtgtccgc ggtcaggcga caatcaatgc 17640
caacgcagaa cctctctacg tcatcgatgg cgtaatcatg cagggaggcg gtaactccgg 17700
tgcagacttc ggtctcggag atgctctcgg caatggtaaa gtctctacga tttctccact 17760
ttccactatc aaccctgccg atatcgtaag catggaaatc ctcaaggatg cttccgcaac 17820
cgctatctac ggtgcccagg gtgcgaatgg tgttgtcctc attactacca agcatggctc 17880
cagaggtgat gcgaaattct catacgacgg tatgttcgcc gtttcaaggc agacaacccg 17940

tatcgatatg atgaatctcc gtgaatacgc agagttctac aactctatgg ttgaaaccgg 18000
agaaatcagt gaggccaatc cttactatgc tactccgggc ctgctcggaa agggtaccaa 18060
ctggcaggac gaaatcttcc gcacagccat ccagcaccag catcagatca gcgcccaggg 18120
cggtacggac aaagtccagt actatgtttc cggatcattc atggatcagg agggtactat 18180
caggggttcc gaattcggcc gttatacctt caggaccaac cttgacgccc agttgaagga 18240
atggttcaag ctcggtctca accttacctt tgccaacacc aatgatgacc ttaagcttgc 18300
cgacggtagc gaaggtctta tcacatactc ccttaccaca atccctgaca tccctgttta 18360
tgatgtctat ggcaatttct catctacagt acgcgaaggc tatacttctc cgaaccctgt 18420
agctcttgca atgatggatg agattctgct ggaccgtcag aaactcaacg gtaatatcta 18480
cgcggaactt actcctgtaa agcatttcac ctggagaaca gaactcggtt tcgatgtcag 18540
cggctctacg gctacccgtt tcagaccaac agtcaatctc ggtggctgga caagagatac 18600
caactctatg agctatcaga agaacagcaa cagattctgg cagatcaaga actatctcac 18660
ttatgccaac tccttcggaa agcacagcgt taccgccatg cttggtcagg aggcctggga 18720
gtcttcgtgg agttacgtac agggcgcaaa taccaagctc ccttcaaacg atgtccataa 18780
cgtcgccctc ggtggaggaa ctccgtcgct ttcttccggt ttcggaagct ctgcgatggc 18840
gtcattcttc actcgagaaa cttataattt cgacgatcgt tacctcctta cctataccta 18900
ccgttatgat ggatcatcca acttcggtcc tgagaaacgt tgggccggtt tccattcatt 18960
ggcagtggct tggaaattca ccaatgagaa attcatggat ttcctcaagc ctgtggtcaa 19020
caacggaaaa ctccgtctcg gatgggggca gacaggtaac tcaagtatcg gtggttatgc 19080
atggggagca tccataagcc gtatgccttc ggctctcggc atgggttacc gtcctgctaa 19140
catccctaat cctgctatcc attgggagag ccagacgcag cagaacatcg gtctcgacct 19200
caatatgttc gacaacaggc tgaatcttac agtagatgca tacctgaaga agtctgacaa 19260
catgcttatg tcaatgcagc ttgcttccta tatgggaacc cagggtaaca gttcttctgc 19320
tcttgcagct cctaaaggta acttcggttc tatcgagaac aagggtcttg aaattactct 19380
cgatgcccat cctgtccaga tcaaggattt cgcctgggac agcaacttcc agatttcctt 19440
caacagaaac aagcttgtca gccttgacgg aacgactaat gccgctctca taggttacgg 19500
acagtggtct gacgttgtgt gtgtgtctga aatcggcaag cctctctata acttctatgg 19560
ttacaaggtc gctggcgtat atacttctct cgaggatatc gagaattctc cgaaacctga 19620
gaaatatcct tctaatggag agtatgcccg tggaaatacc gtgtgggtag gtgatatcaa 19680
gtacgaagat atcagcaagg acggagtaat caatgaaaag gaccgtaccg atatcggcaa 19740
tcctcttcct aagttcacct ttggctggag caataccttt acctacaagg acttcgacct 19800
caatatcttc atcaacggaa gctatggtaa caaagtcctc aactacaatg ccatttctct 19860
tacccatatg aacagcgcat ggaccaacca gatcaatgac gctgtctccg gcagggcaat 19920
gcttgctcct atagacgcaa gcgtagtcta tgctgacggt tccaagtggt atgaccatat 19980
cgacaatgtc agggtcctca accctggcac gaagactcct cgtgcaagca tcaacgaccc 20040
taacgacaac gaccgtatca gcgaccgtta tgtggaggat ggttcatata tccgtatcaa 20100
gaacatcact ctcggatata atgtccctaa gtctttcctg aacaagttca aggtggataa 20160
tgtacgtgtt tacgtcaaca tccagaacct ttatactttc acgaagtaca ccggatatga 20220
tcctgaagta ggtgccagca cccaggattc caccggtctt acctatggcg ttgataacgg 20280
acgttatcct tctcctacag tatattcttg cggtttgaac attacattct aaaattgaag 20340
gaatatgaaa atctctaata tgatgaaagg ttttgctgtg gcgaccgcag tctttgctct 20400
gacttcctgc gaagggttcc tcaacagacc taacgaagat aattataacg tagataattt 20460
ctacaagacc gacgagcagt gcgaacaggg cgtgaattac ctgtacaact ctccatggta 20520
tgacttccag agaggcttca tcaagatcgg tgaggtcatg tccggaaaca tgtactgggg 20580
ctcgtctcca taccttgatt tctccaccaa cggtaccgac caggacctcg tcaacatgtc 20640
ctattctctc tgggctgtga acggccatgc caataccgtg atctacaacc ttcttaattc 20700
gcagggcccg tccaaggcag cgatcgacaa gtgtatcgga gaagctctta cctggaaggc 20760
attcgcctat ttcttcatgg tacgaacctt cggtgacgtt cctatcgtcc acagcaattc 20820
cgatctcctc tccagcggcg agtacaactc tatctccaag gtggagcgcg ccgatgtcta 20880
tgagtacatc atcacgaccc tcgaaaaggc aatggaaatg cttcctaaga agacttccgg 20940
cttcaataat tacgagcgta tcgattacta tgcagcagag gcccttctcg caaaagtata 21000
tcttaccaag gccggacttt ccggatcctt gaaccaggat gacctcaagc aggctgccaa 21060
gtatgccaag gatgtgatcg acaattccgg acgtaccctt cttcctgtat attcggacgt 21120
cttccgtctt accggattca acgccacagg cgagtgcctt atctcctggg aatggagcgt 21180

aggccgtaac ccttggactc agcagaatac cctccagagc gacctcgcaa tggtcggatt 21240
cagcgaccag ggagactgct ggggaggctg gggcggtcct tcagtggatc ttgcagacgc 21300
attcggcgta agtgcgaccc aaaatcctgc agagcgtgta ggagagaagg acagccgccg 21360
caaggcaacc ttgatgctcc cgggtgacaa gtatgattac ttctggactg acaagggagg 21420
tttcgattat ctccgcttca tctatgatac ggaatatggc aaaggcggtc ctggcggttc 21480
catgcagtgc cctacaggtt ccaacaacgt aaagcatctc tatggcgata actttgatca 21540
tgagaacgcc ggtctgggaa cccctgagaa catggcttat cagcttccta ctcatcttct 21600
ccgtctttcc gacgtgtatc ttatctacgc agaggcaagc cttcttaccg gaagcaacgg 21660
ccctgctctc gagtatgtca acaaggttcg cgaacgtgct ggcgccgatg ctctcgcatc 21720
cgtcactttc gaggatatct ggaaagagcg tcgtcttgag cttgcaggcg agggcgaccg 21780
ctggtatgac tatgtacgcc gttcatacta cgatatggac gcatgcatcg ctgaactcaa 21840
agctcagaga aggtcatact gggatggact cgacgatgtt tggaaatcat atatcggaga 21900
tgctttcgat tattccggaa cctgggatgc ttcagaagta cagtacaaca acgcagagga 21960
caatcctaac gttaccgctg caagcttcaa gcttccgttc cctacagagg acgtagtgtt 22020
caatggcaac atggcatctg acgccgtgcc tgtccatgtc gatgtccgcg aggaatacag 22080
ctataacttt taatctgacg aaaagatgaa atctataaat agaataatca ccggttcact 22140
ggcaatcctc gcaatggcgt cattcgcttc ctgcgacaag gttgactatc cggaccgtta 22200
tgaagtagct gacggcaagc cgacggttta ttccgtcaga tacgcagaca aggacgtcaa 22260
catcacccag gcatatatgg atgaggtcgt atgtctcctc ggcaagaatc tcaagagtat 22320
tgtggaactc tggttcaacg accagcaggc aaccctcaat accagcttca ttacagacaa 22380
tactcttctg gtatccgtgc ctaagaatat gcctgtggta cagacagaca agatatatat 22440
gattacttcg ggcaaggata cgctcgcgta tgatttcgcc gtgcttcctc cgactccgaa 22500
ggttacttcc atgtccaatg agtgggctgc ggaaggggag accgtcacaa tctacggcga 22560
cttccttatt gacaaggctg acaagcctct ccagatagcc tttgcaggag ccgaggtcga 22620
tcatgcttca atggtgttca acggttcgtc ttcagtcagc ttccctgttc ctgcaggagc 22680
ccagcctgga tatgttaccg tgacttcact ctccggtacc ggaaagagca agttcatgta 22740
caaggatacc agaaacatcc tcttcgactg ggatggcagc cacggcggct ttgctaccgg 22800
tcacggatgg cgtaacggcg tgatccacgc tcctggcgct gatgcatggg atgcgatcga 22860
cggttcatac ctctacttcg gaggcgccga tcttggtcct ggaagtgcag atgcatgggc 22920
agaggaccag tacagcttca actactggcc agaggccggc accgattacg atgccctcaa 22980
cctccgtccg gagttcgcag agtatatcga caagtatgga gtaagcggcc tcgtgctcaa 23040
gttcgaggtt ctcgttcctt catccaaccc ttggtctgct tcaggtatgc agctcatgtt 23100
ctcgagcgac gctgtcgtta cctacggtac cgctacgaac agctacttca gtgacgaaac 23160
cttccctcgt gctatctggc agccatggac taccaacgga ggttcttatg ataccgctga 23220
caagtgggtt accgtaagtt tccctctcag ctcattcaac aagaaccagg aaaacaagga 23280
gtgtgcaacc gcatttgaca agtcttacct taccggtctc acattcttcg tatggaacgg 23340
cggtatcgac ggcacagcct gctctccggt cttcgcgatc gacaatatcc gtgtggttcc 23400
taatgaataa tattaaatga agctcattat gaaaaagata ttcagtattt cagccattct 23460
tgccgcctgt gtcctggcac tcgggctctc cggatgttca tccaaagagc ttgataccga 23520
ccagtattcc gacaaggacg tgaagctggc atcattcggc cctaacccgg tgatgcgtgg 23580
agctacactc cggttcttcg gaagtaatct cgacaggatt gtcgaagttt cagttccggg 23640
actcgatcct atcaccgaca tcgaagtcgt tacctccggt aatccttcag aaatccgcat 23700
ccagcttcct gcagagggcg cacaggtcgg aaagatctct ctcaagtcat ccgacggaaa 23760
ggtctttact acccaggtgg atgtagaata tacagaacct atcgtatttg acagtttctc 23820
tcctgcagag gctatgcctg gcgacgtgat aacggtcaag ggcgattata tgaacctcgt 23880
gcagagcgtt accttcgaag gcggcgccat cgtgaccgaa atcaaccata acggccgtca 23940
tgaggcttcc ttcaaggtcc cttcaaaggc tattaccgga aagatcatcc tttctgacga 24000
aggagccatc gccaatctcc tctactcgga ggaagatctt gtgatcggcg atccgaccgt 24060
caagtctctt accgtagctg ttgctaaacc tggcgagaac gcagttgtga agggagcata 24120
tctcgatatg atcgacaagg tcgtattcga aggtggtgca gaggtttccg accttgctct 24180
caatgaggat gcaaccgaat tcaatctcgt aatccctgcc ggagcacagt ccggtgaagt 24240
aaaggctgtc agccacgctg ccaaggaatt cgtcgccggc gaaatcgaac ttgtcatccc 24300
tacggatgtc gcagctgcac cttcgcctgt aaaggctggt gccgacctcg tgatttccgg 24360
taacgatctt gatgtggtaa cttctgttga tttccctggc gccagcggag tagaattcgt 24420

atttgccgag gacgccatta cagtgaccgt tcctgctact gctactgaag gtgatatcgc 24480
cctcaatatg gccagcggag aagctgtgaa tgtggcattc acccttgtgc acccggtagt 24540
ttctgccata gctcctgtcg agctcatggc cggcgagacc atcaccgtga caggtaccga 24600
cctcgacctt gtcactggag taactctcgg cggaaagccg gtcgagttca ctgcagaggc 24660
tgagaagatt gtcgtacaga ccgccaatac ttccgtttca ggaaagctcg tgctttctct 24720
tgcgaatggt gagaagattg aaccggagga tgccatcacc ctcagctatg attcattcat 24780
catcgtgaac gacatgcctg ccgcagagca tatcggggct atcgtaaccc tcaagggcga 24840
gaacttcatg atgatcgaga acatctatat cggcgaagag aaagtatcga agtacaacct 24900
tcgttctgat aatgaaatcc agttcgtaat gccttggaac aaggtcggaa catacagcat 24960
gtacttccat cttctctccg gcgatgtcga gacatgtcct cagacaatcg aagttctcct 25020
cgagcaggac atcatgactg tatgggaagg ctctctgcag atcacctgga atgacggcgg 25080
ccgtgtaacc gtgccggcag catcattcac cggtatccct gcaggcagca agctcagatt 25140
ctactacact caggtggacc agcagtggga tcaggcccag ctcaactatg gtaactggac 25200
cggaatcaac ttcgacgatc ctgcaggtac cgtcattaac ggaaccctcg ttccgactga 25260
cgtctatgga tggttcactg acggaatcct tgaccgctgc acagaggtcg tgcttacaaa 25320
ggaaatcctc gaaaacatcc aggctaacaa gggtgactgc gaagacaaga ctgcctgcgg 25380
tatcatcatc cagggctctg gccttacctt cacgaagatc gagatcgtat gcgatattcc 25440
gcaggaggtt acgatctacg aaggtcctac aacccttaca tggggtgacg acggccgttt 25500
cggtctcgcc atgacttact tcgaggcagc aaagccgggc tctcagctta tcttctatct 25560
cgagcagacg gacgcatggg gccaggttca gctcaacgac ggctggtggg ccaacgccga 25620
catgaacttc cctgagatcg gcggtgcata cattaccacc gacaacctcg gaggaaagga 25680
cgtcacacgt atcgccctta ccctgactgc tgatatcctt gaccatatac ttgctacatc 25740
cggtgactat ttcggtctca atagccagtt ccagggtgac ggacgcgtcg ccatggttat 25800
ccagggatct gatatgatta tcaataaagt ttgtatattg taacactttt attgccggag 25860
gcgggtgatg ccgcctccgg tagccagtaa acaaatttta tataatgaaa gcacacgtca 25920
gtttctttgt attatctatg atttttcttg cggcatgttc ctgctcaaag gatgatgccg 25980
aaaagaagga accgtccgtc ccggccccta aggcggtaac ggtagatgtc gcgtctgtca 26040
gcgctcccgt agagggaacg tcggtcgtcg tcgccatctc ctccccgtcc aagcctgcgg 26100
tctctgtctc cggagactgg atcaaagccg aactgggtcc gtacaatccg gatacatata 26160
aagtgaaact tacggtgacc gtcggcgaga atccgacata cgagtcacgc agcggtagtg 26220
tgacccttac ttgccagggc ttttctccgg tcgggataag cgtatcccag gatgccgcgg 26280
cgaagccggt cggaccgtcg gctaatcctg catacgatct tctcggtctc ggctggaata 26340
tgggtaacca gttcgatgct ttctacaacg gctcatgggc cggagataaa tacaattatc 26400
cggacgagac cgtctggggt aatgccaagg cgacccaggc tactttcgac ggggtgaaga 26460
aagccggatt cacatcagtg cgaatccctg tcacgtggct caatatgata ggtccaggac 26520
ctgagtacaa tattgacgag acttggatga accgtatata tgaggtggta ggatatgccg 26580
agaaagccgg actcaaggtc atcatcaata cccatcacga cgaaaacaac aatgacgacc 26640
attggctgga tatcctttct gcatctaagg atgccaggac caatgaagcg ataaaggcgg 26700
aaatcgcggc tgtatggacc catatcgcca ataagttcaa ggataaggga gatttcctta 26760
tcatggagtc cttcaatgaa attcaggacg gaggatgggg ctggagcgct gctttccagg 26820
ctgacccttc caagcagtgt gatgtgctta acgaatggaa tcaggttttc gtggatgccg 26880
tacgtgctac aggaggcaac aacgctaccc gctggctggg tgttcctaca tacgccgcag 26940
gtacttcatg gacgaagtat ttcagactgc cggaggatcc tgccggaaag gtcatgatgg 27000
cagtgcattg ctatgatccg tatgaatata ctatagggaa catacagtat cctgagtggg 27060
gccataccgg cgccgcagga aagaagaccc cggaaggggga gaaagagata acggatctct 27120
tctccagact taatacggaa ttcatcagca aggggatacc tgtatatctt ggcgaattcg 27180
gctgttcgat gagaagcaaa ggggatgccc gggcatggat gttctataaa tactatatgg 27240
agtatgtctc caaagctgcg catacctatt ccttaccggg aatcctgtgg gataacggaa 27300
gtactaagac cggaaaggag tgccacggct acattgacca tggtaccggc aagtatatag 27360
ggaattctaa ggagatcgtg gatcttctcg taaagggaat gctcagcacg gacgaaagct 27420
atactctcca gagtgtatat gacagggctc cgaaaaccaa ttgattatga atctaatcct 27480
taaacttagc atcattttat cattagtagt tgctgtatcc ggctgcggac gcaataaaga 27540
agcggccgga acagctttta acgaagaaac ccggatcatg atggatcgta tggcggcttc 27600
agtaaaggcc ggcaagattc tttatggcca tcaggatgac ctcgtttatg gccattcatg 27660

gtgtgtggat cctgattccg cagcgacaga tgcgttgaac cgttcggatg tcaaggctgt 27720
cgccggagat tatccggctg tcgtgggatt cgaactcggc ggtctggaac tgggatggga 27780
ctggagcatt gacaaagtcc cgttcctgct gatgcgccgg gctgctgtag agcatgccgg 27840
aaggggagga atcgtgacgt tctcatggca cccgcacaat ccttatacag gcggtgatgc 27900
ctgggatatc agttccgaca aggcagtggc ttctgtcctc gaaggagggg agaaggcgga 27960
gatgttcgga gtctggctgg agaggctggg cgatttcatg gactcgctcc gtgacgataa 28020
gggtaatctg ataccggtca ttttcaggcc gtggcatgag aatatcggca gctggttctg 28080
gtggggcgga gccctttgta ctgcggatga gtatatcgcc cttttcagga ggacggtcga 28140
gtatctgaga aatgacaggg gactggacaa tatcataatg tgttattctc cgaattccgg 28200
gattacggcg gagcagtata tgtcccgtta tccgggcgat gatgtcgtcg atatcctggg 28260
gtatgaccat tatgagactg tcatggaggg cgagagtctg gaagatgccg gaaagcgtta 28320
tgccgggcag ctggatcttg gcctttccat agtcaagtct cttgcttcgg cccacggaaa 28380
gctgatgact ctttccgaaa ccggaatgga gggcctcccg gacaagagct ggtggacttc 28440
ggtgctttat cctgcggtaa aggattttcc gattgtctat gtgctgacgt ggagaaatgc 28500
atgggacagg gagacgcatt tttatgcgcc gtatcctggg aatgagggcg acttcaggga 28560
attcgcttcg ctgccggaga tctctctcct gtcgtcgttc cggccttagt ccgtcattcc 28620
cgacctgacc ggggatatgt aagcttaaaa caataaataa caatatatgg agaatatgag 28680
ttttgaagaa cgtctggcgt ggctggaagc acgccaccag gaacttattg ctaagaaaaa 28740
ctcgcctata tacggcaatg gtgtgtttga gagatatgag aaccctgttc ttaccggtga 28800
ccatgcccct ttgttctgga gatatgatct ggataagaac tccaatcctt ttctggaaga 28860
gagattcggc atccatgccg cattcaatgc tggagctata aagtggaagg acaaatatgt 28920
cgtctgcgcc cgtgtcgaag gtgctgaccg caaatcattc ttcgctatcg ccgagagccc 28980
gaacggaatc gacaatttcc ggttctggga ccaccccgtg actatgcctg attatggcga 29040
acctgccacg aatgtatatg acatgagact tactgcccat gaagacgggt ggatctatgg 29100
tatattctgc gtcgagcgta aggatccgga tgctccggaa ggcgatcttt cgtctgccgt 29160
cgcagctgcg ggaatcgccc gtaccaagga tctggtcaat tgggaaagac tgccggatct 29220
ggtttctgat tctcagcagc gcaatgtcgt acttcatcct gagttcgtga atgggaaata 29280
tgctctttac acccgtcctc aggatggatt catagatgcc ggaaacggcg gcggcatcgg 29340
ctgggccttg gtggacagca tggagaatgc cgtggtcact tccgagaaga taatcaactg 29400
caggagatac catactatca aggagaccaa gaatggcgaa ggccctgctc ctataaagac 29460
tgacaggggc tggctgcatc tggcccatgg cgtcaggggc tgcgcttccg gcctgcgtta 29520
tgtgctttat atgtacatga cttcgctgga ggatccttcg aaagtgattg ccgaacctgc 29580
cggtttcttc atggtcccgg aaggggagga gtatgtgggt gatgtcatga acgtcctgtt 29640
ctctaacggt tggattgccg atgaggacgg aaaggtcttt atatattatg cttcctgcga 29700
tacgcgtctc catgtggcga cttctaccgt cgacagactc gtggattact gtatgaatac 29760
tgagccggat ctgtttacca cttctgcttc ggtacgacgt ctgaacgcgc ttatagacaa 29820
gaatctttcg ctatctatgt aggactaata atcaggtttc agatggatat cattactgca 29880
ccggctaagc cgacattgaa ggaaaagatc ggatatggct tcggagacat gtcgtcttca 29940
atgttctgga agattttttc gtattatctg ccgatcttct attcggatgt gttcgggctt 30000
aagcctcagc atgcggcttt cctgcttctt atcacaaaac tgtatgatgc aatatcggac 30060
cctgtcatgg gaatcatcgc cgacaggacc cggtccagat ggggaaagta tcgtccgtat 30120
cttctctggg tggctctgcc gttctcgctt ataggccttc tgtctttcta tactcctgat 30180
gccagctatg gcctgaagca tgtatatgcg tatgtgatgt atatcctcat gatgactgtc 30240
tatacggcta tcaacgtgcc ttacggggcc atgctgggag tcgtcactcc ggacagccgt 30300
gagaaatcgg ttttctcgtc ttacaggatg ttctttgcgt atatcggcag tttcatcgcc 30360
atggggatat tcggcgtgtt cgagaagctg atttccggta ccgtgcgtgt tatgtcggac 30420
ggtagcgaga gggtggtgcg cggagtcggc gatgcagctc ccggacagtg gactcttgtc 30480
gtcggtatcg tttctctgtg ctgtttcgtg ctgtttctgc tttgcttcct tatgaccagg 30540
gagcgggtgg ggactgaaca taaggcggtt tccgagggcg gctcttctat cgcagaggat 30600
ctcaaggctc taggccgtaa caggccttgg tggctgcttc ttggcgccgg tatcggtata 30660
ttgctgttca attcgattag aggcggagcc gctgcatatt atttctcgaa tatcctcggt 30720
acgagcgtgt tcctgacttg cgctgtctat cttttcattg gcgaggtggc gcagatggtc 30780
ggggtgctgt ttactgtgcc gctttcggag cgtatcggca agaaggcgac ttttatctct 30840
gtgcttctga cggtttctgt gctgagtgtg gcgatattct tcctgcctga gactcctgcc 30900

ggtttctggg gacttctggt gctgcaggtg cttatatgca ttgcttttgg tgtccagtct 30960
cctctgctgt ggtcgatgtt tgccgatgtc gccgattatt cggaggagaa gaacggctcg 31020
gcttctaccg gacttatttt ctcgtcttct tctatggctc agaagttcgg cggggccctc 31080
ggcgctttct tgctgatgca gattcttgca atcttcgcat atgacaagga tgctgcagtc 31140
caggcgcctc agactcttgc tgccatccgt ggcctgatga gctatatccc ggcaatcggc 31200
gcccttctcg gtgccctctg cctgctgttc tatcctctta cggaatcgcg tatggcttca 31260
attagagaga aactttctgc tcgcagatag catcttcatt catgtaggta tttgttattc 31320
caaaggatgg ccttttcggc catccttttt cgtctggctg attcgagtcg ggcctttctg 31380
ctggcgggag aagggcggag gcggcggagg aattgaggaa aagcggagag cgcgggagcg 31440
gccgtcgacg ccgtgaagaa cggcgtgtgt ttgacgaccg ttacgatggc tcgcccatcg 31500
taaaagggag aagttagccg tttaggcggc gacagcgacc aagcagcttt tccgaacttg 31560
acggagccgt cccgtccttc tccagccagc agacatctcg ctacagtcct gctgacgcag 31620
ggtcccgggg aacttctttc gctgacgctc actccgtacc ctcccacaat ctgccgcgtc 31680
atcgcttcgc gataacttgc atcttgcggg ccccgtccct cttcccgtgg ccgagggggg 31740
ccacgtcccc gagaccctgg tcagcaggac tgcggcaaag atgagggact ggagggaatt 31800
gaaaaaaatg ttatatttgt atttcgttcc ttaatccgga gaactatggc agacaaacta 31860
tacctgattg acggtcacaa cctaatcttc aagatgtact acgcattcct gcgccgtcct 31920
atgatcaatt ccaagggaat ggacatgtca atcctgtatg gcttcacgaa atatgtgctc 31980
gaactcatcg agagagagaa gcctacccat atggccatag cgttcgatcc tcccggcggc 32040
accttcagga acgaaatgta ccccgaatac aaggcaaaca gaagcgaaac tccgcagctt 32100
gtgatcgatg ccctcgaacc gctgctggaa atatgcaaag ccttgaacat aaagaccttg 32160
atgattccgg gatatgaggc cgacgacgtg atcggaaccg ttgcgaagag ttacgggaaa 32220
cccgggacgg acgtctatat ggtcaccccg gacaaagatt acggccagct ggtcgccgaa 32280
catgtgtacc aggtgaagcc tggaaaggcc ggcggcgcag acgagaccgt cgatgtcaag 32340
gcaatctgcg aaaaatacgg aatcaagact ccggcgcagg tgatcgaaat gctgaccctc 32400
tgcggagata cggccgacaa tgtgcccgga gtgcatggca tcggggaagt cggggccggc 32460
aagctcatcg caaaatatgg caccgtcgag aatatctacg cccatctgga cgagctgact 32520
gaccgccaga agacgcagtt cgaggaggcc aggggacata tcggcctgag ccatgacctg 32580
gtgacaataa agaccgacgt gccggtagag acaagcctcg acgagatgag gctggacctg 32640
gattatgctc ccgaaatcgc cgaccttttc gaaaaatacg aattcaactc gctgagaaaa 32700
tatatagccc atagctcccg tgctgccaag gcagtggcgg aggtcaagcg cctggattat 32760
tccgagacgg gggcggaaga gctcgtgaag atggctccgg aaaagatttc gctgatagtg 32820
gaaccggagg gaccgggcat attcgccgat atccgcaggg tgatcgtggc cgccgagaag 32880
aaagacggcg gaatcgtcgt cgcttcaggc gctgcgaagg attttcgcaa gatactggag 32940
ggcgaagccg tgaaatacgg atatgatctc aagctccagt cgaatcttct ggagaatgcc 33000
ggaataaagc tcggaggccg gttatatgac atcgaactga tgcattatct catcaatccg 33060
gagaaaagcc ataaggtcga aatcctcgcc aagagctatc tcggcgtcga tattgaagag 33120
gccgctcccg tagaagaagc tcctgtttcc ctgtttgacg tcgcggaaga gagcccggca 33180
aacggaccgg acaggaatct cgaagccgcg gcagtgctga aactcggaga agtggtccgc 33240
gcaggcatgg acgaagccgg cgtaggagcc ctttatgagg atatggaagc ccctctcgca 33300
gccgtgctcg cccgcatgga aagggccggc gtgaagatag acctcggaca gctcaggacg 33360
tttgccgatg aactcaaagt cgaaatggat gccctcgaag cccaggtcag gaaactggcc 33420
gacgagccgg ggctgaacgt ttcttccccg aagcagatag gcatcgtgct cttcgaaaag 33480
ctgaaactcg acccgaagat gaagaaatcg gccagcggta gctatagcac cgacgaaacg 33540
acgcttactg ccatcgccga caggcaccct ataattgaca agatacttga attcagagcc 33600
gtcaagaagc tgctctccac atacatagag ccgttcccga cctacgtctc ccccggttacg 33660
ggaagggtcc atacgacctt caaccaggcg ctgacggcga cgggaagact cagctcctcc 33720
aaccctaacc tgcagaacat tccgatccgc actgagaggg gaaaggagat caggaaagcc 33780
ttcataccgg cgactccgga cggaatcata atgtccgccg actactccca gatcgagctc 33840
aggatcatgg cccatctctg cggcgacgag catctgataa gctcattcaa ggcagggctg 33900
gatgtccata aggccaccgc agccaagatt ttcggcatac cggtagagga ggtcacatcc 33960
gaccagaggc gcatcgcgaa gaccgccaac ttcggaatca tgtacggcat ctcatctttc 34020
gggctcgccc agaggctgca tgtctccagg acggaagcca agaagatcat cgacgattac 34080
ttcgccgcct tcccttcgat ccggaccttc atcgacaaga ctattgcgga cgcccgcgaa 34140

```
     aagggatatg tcgagacagt cttcggacgc aggaggtatc ttccggacat aacggcgaag     34200
     aacgctacca taagggctct ggccgagagg aatgccgtaa atgctcctat ccagggcaca     34260
     tcagccgata taatcaagct tgcgatgata agcgtcgaca agcgcatccg cgaggcgggc     34320
     ctgaagagca agatggtact tcagatccat gatgagcttc ttttcgatgc cgtgccggaa     34380
     gaaatcgagg aactcggcag gatagttgta gcggaaatgg agaacgtaat taccttgtca     34440
     atacctttaa ctgtagaatg taactatggc aaaaactggc ttgaagctca ctgatcatga     34500
     cgacggatct cttgtggaga tggcgttagg tggcggcccc gaggccgagg ccgctttctt     34560
     cacgatttat gcacgttacc ataagggcgt attcgcccac atctcacgct tcatccagga     34620
     acaggaggaa gttgaggata tctgtatgga gagcttcgag aaggccttca agcagctgga     34680
     atcatacaag ccgagcaaca agttctcgac atggatattc actatagccc gcaacacggc     34740
     cttcgaccac ctcgacaagg acaaagtacg cggccgcaac aaagacaaga tgccgatcga     34800
     ctactccgac ctggagcata tagacgtccc tgaggacaat gtcagcccgg aggacgagat     34860
     cataagttcg caggaccatg agaacttcct gagttgtgtc cagggtcttc cggacctgta     34920
     tcgcgaggtc gcatccatgt gctttgtgga caatctcgga tacaaggaga tttcggagaa     34980
     ggcggggctg cctatcaata cagtaaaaac ccgcatcagc agggctaagg gcctgatagt     35040
     ccagatgatg cttgatatgg aagaagaatg aatttcaaag aattcgaaga tatagtcgtc     35100
     gctgaatttc cggagctg                                                   35118
//
ID   XXX; XXX; linear; XXX; XXX; XXX;  XXX.
XX
AC   PRJEB67889
XX
DE   Uncultured bacterium DNA fragment from in-reactor enriched cow rumen, clone
ferm-21e20
XX
FH   Key             Location/Qualifiers
FH
FT   source          1..37465
FT                   /organism="uncultured bacterium"
FT                   /environmental_sample
FT                   /mol_type="genomic DNA"
FT                   /isolation_source="Cow rumen"
FT                   /clone="ferm-21e20"
FT                   /note="lignin alkali active clone"
FT   CDS             complement(2606..3292)
FT                   /db_xref="SEED:fig|6666666.127243.peg.1"
FT                   /translation="MKKLIPIALIALVLVPGCGNMSNLAKDSLIGSGAGAAVGAAVGAI
FT                   IGKDAKGALIGAAIGTAVGGGAGAVIGRKMDKKAEELAALQDAQIEKIYDANGLEALKV
FT                   TFASGILFQTNKADLSAASKKELAEFASKMNDLKDTDITIYGHTDNTGSAAVNEKLSAE
FT                   RAASVAAYLKSCGIAGDRMKTQGLSYTDPVADNSTVEGRAKNRRVEIYITADENMIAAA
FT                   QNGTLQ"
FT                   /product="OmpA family outer membrane protein"
FT   CDS             complement(3374..5296)
FT                   /db_xref="SEED:fig|6666666.127243.peg.2"
FT                   /db_xref="GO:0004342"
FT                   /translation="MKHFVLPYEGGLITDKLPQDILHSYEKVNAEIYSESREASSKVAD
FT                   IIVNGIKATKGRLFRLGLTTGVTPVSLYNELARRCAAGEVSFRNVEIVSIDEYYPFRAD
FT                   QHQSRNSRLHEALIDKVDILKENVHIPDGTVPAEEISAYCASFDKLARGLDLLVIGVGE
FT                   QGQLGFNEAGSTEKSRTRTVPLPYKSRKRQAHNFNGDISATPAAAITIGISTMLSASRI
FT                   IVMAWGEDKAEAVKAIAEGEVTPLCPASYLQLHDNASIFVDETAGSLLTRVSAPWLVGS
FT                   CDWTPKFIRKAVVWLCQQVGKPILKLTERDYLENSLGELIEKYGPFDKININVFNDLQH
FT                   TITGWPGGKPNADDSTRPVSAEPFPKTVLIFSPHPDDDVISMGGTFIRLVAQGHNVHVA
FT                   YETSGDVAVYDDVVLQHMDAAYQLGFADKFDEVKALIDSKRPGEAEPRALLDIKAAIRR
FT                   SEARSAVRSFGLNDKTNAHFLNLPFYESGGVKKMPRTQVDVDIIKKLMNELKPDMIFMA
```

```
FT                   GDLADPHGTHRVCTEAAMDAFEQLREEGAKWTENAHIWLYRGAWMEWELGRVDMAVPLS
FT                   PGEVVSKRHAIFRHLSQKDIVPFPGDDPREFWQRAEERTQNTARLYDELGMAEYQAMEV
FT                   FLKLF"
FT                   /product="Glucosamine-6-phosphate deaminase (EC 3.5.99.6)"
FT                   /EC_number="3.5.99.6"
FT   CDS             complement(5535..6131)
FT                   /db_xref="SEED:fig|6666666.127243.peg.3"
FT                   /translation="MKKAALIIFLSLAIASCGKENPDQESGTTGLREDYVVTKIEYHLD
FT                   DSAVIEQLPDYVASDQLHNNTPELILASRTFTFEVKESSSFLSSGDVSVPEGFYAPVPY
FT                   LMPETNSIFLTEPRYNTWGEDSTTSDVRNVEVSLNTPPYSSVNVTVSIKRYRMTVRYTA
FT                   YLKGVMTGMETSVDGIWEGVSTSGTETIWTQQDLD"
FT                   /product="hypothetical protein"
FT   CDS             complement(6214..7995)
FT                   /db_xref="SEED:fig|6666666.127243.peg.4"
FT                   /translation="MKNYLKLILTVSCAAVAFSGCEKFLDTESYTTRNTSNFPSSETDA
FT                   IQLVTGVYAALNGSVYEPSASYWMAAMIASDECYGGGGIDDYDCQADDHLLYYEPDVHS
FT                   DYWKEFFAGITRANMAIANLDKVENEELRNQLLGEAYFLRSYFLFELTQCFGEIPLTSS
FT                   VPETVEEAKKYPAQASIEEIYGTIAAGLKKAIEIMPDKPWNQCVSGLRHATKWDAEGIL
FT                   ARVYLFYTGFYGDKNGTTLTTLPLVDLETGELTSETVSKDYVVSKLKDCIDNSGHSLVS
FT                   DFRLQWSYTNSATKADYPYAQTVEGTWIIDSENPEEMFSICCTNLASSSRFCNKYDQYL
FT                   GVRKRSNWEVNAFPFGRGYGWATVNPYVWKQWDIEEPGDLRKVASIYNYEDECVPASAY
FT                   TWGNDQQYEETGLWQKKYQTWGAMVDGKFFYEFSSVPEYGGTGTDLGRGHSPQNIQLLR
FT                   FADVLLMHSELTRTVEGINKVRARVNLAPIAAYTDEALRMERKHELAFEAVRWGDMRRY
FT                   GKAYAIAALETQLGQEIRNAGGETVMKDQGAGYKARYEATYGFRPIPRSEISLSNGVLK
FT                   QNEGWTDTASAQYAGWK"
FT                   /product="putative outer membrane protein, probably
FT                   involved in nutrient binding"
FT   CDS             complement(8002..11127)
FT                   /db_xref="SEED:fig|6666666.127243.peg.5"
FT                   /translation="MFPAGVFAQVATVKGTVTDAGTGDPVPYATVVVKGTSSWTTTDKD
FT                   GLYSIESPANGVLSVSLLGYSNVEEKVNGRTLVDITLVPEFDQLMETVVVGYGVQQKKL
FT                   VTGSTIQVKGDDLMKLSTTSALGALQSQTPGVNITQNSGQPGQGYKVNIRGIGTTGDSE
FT                   PLYVIDGVAGGSLSSLNPSDIESIDVLKDAASAAIYGARAANGVVLVTTKQGKEGRVIT
FT                   SYDAYLGAQYIAKMPDLCNAQEYIQLQDLAKTNSGSPLYDWQTLLPQALYQSVLNGTWK
FT                   GTDWVNEMYNKGARTENHSVNIVGGNADHKFSMGFSCTAQDGILGYNKIQPVNAEYRRY
FT                   TFRINTDNVVKRINGREVFKVGETLNFSYGTNNGIAEGDIYWNSLHNAITANPLLPAYT
FT                   YDEQGNITGFYDEAARDAAGWDFDGTAKHPLGMDFYTSRGRNISKSYNLQASVYAEISP
FT                   IKNFKFRSQFGYRMSGSSSRSYTMLYRLSSGQYSDNDEVSQSLNLNQRYTWENYASYHF
FT                   DINKDHVFDVVLGQSIEKWGFGEKVGASAFNSIFLNDFDRAYISNTKPEALNQLELSGS
FT                   PSSRGAIASFFGRVNYSLKDTYLLSAIVRTDGSSNFARGHRWGIFPSVSAGWIISNEAF
FT                   MANTKDWLSFLKLRASWGQNGNCDIDNFQYLTTIKLSSSAGYYFANKAVPTTGAIPGVL
FT                   ANPDVSWETSEQTNIGLDARFFNSALGVTLDGYVKTTRDWLVQAPIASVYGFEAPYVNG
FT                   GDVRNSGVEIAIDYGKYTGDFTYGVKFNGSYNKNKVTRIANSEGIIHGEDDVLSQGTSE
FT                   MYRAQVGYPIGYFYGYKTAGIFQNDAQVAATAVKYEDAKPGDVIFVDTDGDGAITEEDR
FT                   TMIGNPHPDFTTGLNLWFAYKNFDFSVTGYGAFGQQIAKSYRSFFDSPRENYTRDFFDC
FT                   WHGEGTSNRYPTLTLANDRNWSNISDIYIENGDYFKISNVTIGYDFSRLVKKSILSKAR
FT                   IYFSAQNLYTFTGYSGMDPEIGYGFDEDWVSGIDLGYYPGARTYMIGVNLQF"
FT                   /product="SusC, outer membrane protein involved in starch
FT                   binding"
FT   CDS             complement(11212..12045)
FT                   /db_xref="SEED:fig|6666666.127243.peg.6"
FT                   /translation="MWLGIDLGGTNLRVGVVDGSAVVESVTVPSIPKEYSLESSIGYLV
FT                   EVIRPLCSSDIRGIGIGAPSVVDAEQGIVYDVTNIPQWVEVPLKAILEKAFPSKKVSVN
FT                   NDSNCFTLGEHRFGKGRGTSSMVGITLGTGVGSGIIIDGKLYGGVNTGAGEIGCIPYLG
```

```
FT                   KTLEDYCASKFFTERGLTAKDAAEAANGGDSAMIRLWSEFGRHMAFLVGVAMYAYDPEM
FT                   IVFGGGLTKAEPLFRDSMNESLKDFAFQKSLEKIRIEYSVLKDVAVLGAASLVMD"
FT                   /product="Hypothetical sugar kinase, ROK family"
FT   CDS             complement(12133..13290)
FT                   /db_xref="SEED:fig|6666666.127243.peg.7"
FT                   /translation="MTDFTEAFAHYLLQGIYKYASTPGDWIVCRMPPAYKERFGLKAVV
FT                   TWAKKWKADAIIGQFSDTDDVSLFRKNGIIALAQDYKSRFSVIPNITSDYFATGAKAAD
FT                   FFIKRRYRHFAFFGYRNVVWSDERYMGYRRRLEDLGMEVSPSYQEQDISELWHYDAEPI
FT                   RRWLDSLPKPVAIFACDDNQGTKLIEVCNVAGYKVPEDVAVLGVDNDETVCQMSFPPQS
FT                   SIVTDIVQGGYDAAKLIDDILAGRVEPSGETNVVIRPLYIVSRTSTNVIATEDADVQKV
FT                   LRFIYDNVHRKINVDDVLATVPMSRKLLEIRFRKAMGDSIYHFIGKVKLERFRSYLINM
FT                   DRPVGEIAGLVGEEDPNNIFRRFRKVYGCTPMEYRRKHNSQNASD"
FT                   /product="Xylose operon regulatory protein"
FT   CDS             complement(13394..15634)
FT                   /db_xref="SEED:fig|6666666.127243.peg.8"
FT                   /db_xref="GO:0004563"
FT                   /translation="MNITPAPVEAVRTDGEFEITKHTVISSADAPEVAEFFAEKIGKST
FT                   GYKIRTSTRGGQIELSISAGAGLPEEGYYLEVTPGKVTAVASDKAGLFYAMQTFLQLLP
FT                   PQIESLSKVKGVKWTAQCARVTDYPRFKWRGFHLDVCRHFMSMDAIKRQIDIMASYKIN
FT                   TMHWHLTEDQGWRIEIKKYPRLTEVGAWRTEFDGSVYGGFYTQEQAKEIVAYAAERFVT
FT                   VIPEIELPGHTIAAVVSYPELSCDGEQINPSYSWGVMDRVLCPGNDKVFEFIEGVFEEL
FT                   IPIFPSVYYHVGGDECPKTRWKVCPKCQARIKAEGLFADENHTAEEKLQSYCIQRVEKI
FT                   LDKFGKKLIGWDEILEGGLSGEAAVMSWRGEQGGIAAAQMHHYVVMTPSHEGCYFDSYQ
FT                   GDPKAEPLGFGRYIPLKKIYDYNPVPAELQKDGLEKYVMGVQACNWSEYMYDDDYRDYM
FT                   LFPRGFALSEVAWSQLSSKNWEDFSVRVNDACERLDMRGVTYHIPLPEQPDGSCNNLVI
FT                   TGESEIAFTTTRPEKMYYTLDGSEPDGNSTEYTAPIKVDGNSLIKICTVLPSGKKSMVR
FT                   EIKVEKQTLAPAVKLENPSNGLKFRKVYGRFLSASELDKVKEGWIESTVTELKQIERQE
FT                   PRQNNMQDVNFYGAVAEGYINISEDGVWTVSSDYDQVWIDGKKLIDNGTEVKRYSRHDA
FT                   QVALEKGLHEIKVVFISNIIGGHATARNNTTIQMRRPGGKGFHAVNADQLYR"
FT                   /product="Beta-hexosaminidase (EC 3.2.1.52)"
FT                   /EC_number="3.2.1.52"
FT   CDS             complement(15667..18771)
FT                   /db_xref="SEED:fig|6666666.127243.peg.9"
FT                   /db_xref="GO:0004035"
FT                   /translation="MGIVRDVTLFETGSVAVTDLFVKPELNLDTDESADLEVRVTLRNM
FT                   SRKPVNGEVRGRWAGGEFVFPVSLAASETKEVVLTPKECKNLHTAKPSIWWTRDLGKPY
FT                   LYTMQASFVAGKETTDSREVKFGIRKIESRIDSHGHRVFLLNGKKILIKGAGWTDNVNL
FT                   RDTHESLDAQMEHVLNMGLNTIRFENIWGKDQYIYDLCDQNGVMALVGWSCQWEWENYC
FT                   GLPETDKYGCINDETTMKLAARYFRDQVKWLRNHPSVIGWMTGSDRIPNPELETEYLRI
FT                   YSELDYRPYINSAKAMTSKISGKSGTKMEGPYEYVGPDYWYYDTEAGGAFGFNTETGVG
FT                   ANWPQIETIRTMIPESELWPLGPAYSKYCTTSSSAMNSTKMLESVVNAQYGEADGIEDF
FT                   TRKAHAADYDATRAMFEAFRVNLPNTTGIVQWMLNSACPSLYWQLYDNNLTPTASYYGV
FT                   KKACAPIQLIFNYKDYKVYAVNESSKDKDIKAYIKVYDAQSNLLVDTSDHVGTERRHPK
FT                   VAFDLDSLRGRDIFVALRIRDGQDNFYCIPAKMTTYKFNKTNWYVTPAEEYADMHFVSD
FT                   MPRAAVSYKVVESVDNLGKLWTVTLTNDSDYISYQNVLKLVRPDGSLLEPAIWSDNFVA
FT                   VLPHSEKTVTCRCDRKENARVAMTGWNTDITLIPDDNTRDKSKYAEHVYGDGDKYVATA
FT                   RYDVPVIKEPKGKKVKNVIMMIGDGMGLEQISCAWVANGGHLYLDNFPVVGLSRTYATD
FT                   KLITDSSAGGSALATGVKTRYGFIAVDEDGLPVKSILADAHERGLRTGVSVVCRIGDAT
FT                   PAAFCNHSISRYDEEGLAAQYLDCGADFLIGGGIKFFKDRSDGRDIVEEMKAKGYNFVD
FT                   TREELAASEKLPILGLFADTEMAPALDRGPILEESAVKALELLDNDKGFFLMIEGSSID
FT                   DHCHHNQVGYAMEELFDFDKTIGKVLEWAAKDGETLVIVTADHATGGLTLLGGSLEKGE
FT                   IKVNFSTSGHNGIMVPVYAFGPHSEDFSGTMENAEVANRIKAIFK"
FT                   /product="Alkaline phosphatase (EC 3.1.3.1)"
FT                   /EC_number="3.1.3.1"
```

```
FT   CDS             18870..19316
FT                   /db_xref="SEED:fig|6666666.127243.peg.10"
FT                   /translation="MCQDFTQVMIAALRGIGIPAGYVSGFLRTFPPPGQPRLEGADAMH
FT                   AWVRAWVGRDAGWIEFDPTNNQFAGEDYIAVAFGRDYSDVAPVRGVLRSSGGQNSHQAV
FT                   DVIPLDPDTAQPLVRSVPARRAGTRAENDMETQAEPDAAVPPEI"
FT                   /product="Small protein containing transglutaminase-like
FT                   domain"
FT   CDS             complement(19343..20497)
FT                   /db_xref="SEED:fig|6666666.127243.peg.11"
FT                   /db_xref="GO:0008915"
FT                   /translation="MKVFLIAGEASGDKLGAALMAGLKSLVPGVGFAGVGGPLMQAEGL
FT                   TSLFPMEDLSVMGLAEILPKYPLLRRRVQEAADAAIATGPDVLITIDSPDFCLRVARKV
FT                   KDARPDLFCTHYVAPSVWAWRAGRAKRMAQVIDHVLALLPFEPPYMRAAGMSCDFVGHP
FT                   VVSEPRATGAEAAAWRTRAGIAADEKLLLVLPGSRRGEVSRLAPKFGATLGLLKDRVPG
FT                   LRVVLPTVRGVSDMVRAAVADWPVPALIVEDRADRLAAFAAADAALAASGTVSLELAAN
FT                   RVPMVIGYDANRLTWAIVWRMMKIDTVTLVNLVSETRAVPEFLGPACQPEAMADAVVAL
FT                   FGDAGARAAQLQAMEITMTRLGEGGEAPGLRAARSVLGALSGGA"
FT                   /product="Lipid-A-disaccharide synthase (EC 2.4.1.182)"
FT                   /EC_number="2.4.1.182"
FT   CDS             complement(20494..21312)
FT                   /db_xref="SEED:fig|6666666.127243.peg.12"
FT                   /translation="MSQGDIAAPDLALIAGTGALPALLMRARPDMLLCELEGFASGIPG
FT                   AEPLRFRVERLVPFMTELTRRGIRKVCFAGAIRRPKIEPAKFDLRTALLVPRIMGAIQQ
FT                   GDDATLRAVIALFEERGLEVIGADHIAPDLLGGEGVLCGKPTDTDRKDMARAAEIVAAL
FT                   GAVDVGQGAVVARGLCLAVEALPGTDAMLDFVATTRPEAHRGGVLMKAPKPGQDLRIDM
FT                   PAIGPRTVEGAARARLSCLCWRAGGVLLLDRAAVIAAAEKHGIALWGVQA"
FT                   /product="Protein of unknown function DUF1009 clustered
FT                   with KDO2-Lipid A biosynthesis genes"
FT   CDS             complement(21309..22103)
FT                   /db_xref="SEED:fig|6666666.127243.peg.13"
FT                   /db_xref="GO:0008780"
FT                   /translation="MSIDPGAEIHPSAVVEDGATVAAGVRIGPFCHVGPEVTLHAGVEL
FT                   KSHVVIGGWTEVGEDTVIFPFASIGQIPQDLKFKGEHTRLIIGKRNRIREYVTMNPGTE
FT                   GGGGVTRVGDDGLFMSSCHVAHDVIIGDRVILVNSVAIAGHCHIEDDVIVGGLSGVHQF
FT                   VRIGKGAIIGALSMVAADVIPHGLVAAPRAGLEGLNLVGLKRHGVSRAEIGALREAYMA
FT                   LSVGNFREQARKLADEGTDSEMVREVLDFILGPSDRSFLAPK"
FT                   /product="Acyl-[acyl-carrier-protein]--UDP-N-
FT                   acetylglucosamine O-acyltransferase (EC 2.3.1.129)"
FT                   /EC_number="2.3.1.129"
FT   CDS             complement(22135..22602)
FT                   /db_xref="SEED:fig|6666666.127243.peg.14"
FT                   /translation="MIEPAPYSSVDLSLIKQILPHRYPFLMIDKVRDIVPFEGGVGIKM
FT                   VTSNEPHFQGHFPEEPVMPGVLIVEAMAQTAAIVVGVSTNIIGGDMLTYFMGIDSCKFR
FT                   RKVVPGDILELNVKCLRPGGKVWKFRGEATVDGELAAEAEFMAMMAPKPKA"
FT                   /product="3-hydroxyacyl-[acyl-carrier-protein]
FT                   dehydratase, FabZ form (EC 4.2.1.59)"
FT                   /EC_number="4.2.1.59"
FT   CDS             complement(22745..23458)
FT                   /db_xref="SEED:fig|6666666.127243.peg.15"
FT                   /translation="MAQRGGTRIWRAGMAVLLCLAAPAVAQQATAQPTGGADDYPTGVA
FT                   PGSAVVAASRDPGDALIRLPSPVLTIDWEQLFDGSRWGQRILADMQRDTGAQKAENDRI
FT                   ADQLIAEERALTEARGKIDAATFQARAEEFDARATKIRAEQSAKGKALEQRYNDARNAF
FT                   FGAMVPLLDEVLHRRGAVVVLDQRAVIRALPEADVTPDMIALVDARMGEGPADTASGSH
FT                   DSDPAGDPVPAPASP"
```

```
FT                   /product="Outer membrane protein H precursor"
FT   CDS             complement(23467..25848)
FT                   /db_xref="SEED:fig|6666666.127243.peg.16"
FT                   /db_xref="GO:0019867"
FT                   /translation="MRRLPAVALALTTASPLPGALPLTVAAGIAAGALVPVAAMAEDFR
FT                   FSAVKIEGNKRIEPATILAYAKVAKGQTVSGDQLNDAYQRLVGSGLFETVDLVPQGGTL
FT                   VIRVKEYPMVGVVAFEGNRVLKDDAIAKSVKTKPNHVFSPAQVEADANLIAEGYTQAGR
FT                   INVRVDPKVIRRGDDRVDVAFEIKEGYTSEVHRISFTGNRSYSDFTLRQALSTKQAGIL
FT                   HRLIQSDNFVPDRLEQDKQNLIDFYQSRGYPDVKVTGISSDMARARDGYFVTFNIEEGQ
FT                   QYKFGNITTVSEVPDLNVDEFAPLAKIKTGSVYTPTAIDLAITRMEQLALKKGANFVRV
FT                   DPRVTRDPATQTLNVQFALVRGPRIFVERIDIKGNTTTLDSVIRREFRTDEGDPFNPRE
FT                   IRNSADRIRALGFFSKADVNTKPGTNPDQVVVDVNVEEQPTGSLTLGASYGAASGFGVL
FT                   VGVQEDNFLGRGQSLGVQIGTTADNQQSSIDFVEPYFLGRDVKARMHLWYDTTSNDNAD
FT                   YNTRIQGLQPSVEFPVAENSRLEARYAYGKKKLFGVTDYTAASGTTSASGSSPVLSYEQ
FT                   SLGERTASSVGYTYTWDTMTTNLDPRYGYTFKFGQDYYGLGGDVDAISTTALLAAQRKV
FT                   MHESVRLRAELEGGVITPRSGSDLTILDRFSGYGKVRGFESHGWGPRDVGAPNGDALGG
FT                   QYFWALRTETEFPIGLPEEYGITGGLFWDTGALWGVSSSERALAMGLDDSDKMHIRSAV
FT                   GFSFFWRSALGPLRIDFSKAIKKETYDKEQNFDLTISTKF"
FT                   /product="Outer membrane protein assembly factor YaeT
FT                   precursor"
FT   CDS             complement(26009..27379)
FT                   /db_xref="SEED:fig|6666666.127243.peg.17"
FT                   /translation="MDLLPDFGNVLYTAAAFVVALSIIVFIHEFGHYIVGRWSGIGAEV
FT                   FSLGFGPVLVSRVDRRGTRWQIAAIPLGGYVRFNGDSNAASAGAADEAELDAMSPEERR
FT                   RTLAGAPLWARFATVAAGPVFNFILSIAIFAGAVAWQGVATEAPTVGRLATLPGGEGAL
FT                   QPGDRILALDGHETPDYRAFYAALADLPPTPTVRWSIERDGTARDIDGPQPMPPLLSGV
FT                   MPSSAAAGAGLKAGDVITKVDGAPIWRFEDLRSRVGASAGAPLDLTVWRPLPAGAAGGT
FT                   AATGAQLLTLSLTPKRMDLPLPNGDFETRWMIGATGDMAFAPETRAAGPLEALERGAGQ
FT                   VWTIVTSSVSAFEHMILGKISSCNLHGAVGIAQGSAAAAQAGAMDFIWFIGVLSTAVGF
FT                   LNLFPVPVLDGGHLMFYLWEAVTRRPPSPAVLNVLTAAGMVAVLSLMLFGLWNDVTC"
FT                   /product="Membrane-associated zinc metalloprotease"
FT   CDS             complement(27383..28606)
FT                   /db_xref="SEED:fig|6666666.127243.peg.18"
FT                   /db_xref="GO:0030604"
FT                   /translation="MRRVSVFGSTGSIGVNTLDLLERQGGAGAFQVVALTGGRNVDLLA
FT                   SQARALRADIAVTAYEECLPALRAALHGSEIEAAAGPAALIEAAERPADWIMSAIVGAA
FT                   ALAPGYAALRHGTTLALANKESMVAAGPLFLEAARRHGARLLPVDSEHSAVFQALHGED
FT                   MASVERVIITASGGAFRDWPMERIARATVAEASTHPNWAMGQRITIDSASMFNKGMELI
FT                   EAKELFGLRPDQIEAVIHPESVIHALVGFRDGALMAHMGVPDMRHAIGYALNWPERAEL
FT                   PVKRLDLAALGSLSFRPADPVRYPALRLAREVMATGGLAGAVYNAAKEVALDGFIAGAI
FT                   GFLDMAAVVEQVLSDMSARQGLGNPATSLDEVLATDAEARALGRARIAGRHGAGRHGAS
FT                   GDTPRGRI"
FT                   /product="1-deoxy-D-xylulose 5-phosphate reductoisomerase
FT                   (EC 1.1.1.267)"
FT                   /EC_number="1.1.1.267"
FT   CDS             complement(28608..29420)
FT                   /db_xref="SEED:fig|6666666.127243.peg.19"
FT                   /db_xref="GO:0004605"
FT                   /translation="MTQDGSGRWGDLRTRLVSGLVMLVLGGAAVWRGGFAMEALAIALC
FT                   AAMIWELTTLCAPGRGRLGYEIGVGAAAVSFAVLELRSLFPPSLGLLVLYPAAAAFLVT
FT                   RDRLLFAVYGFAILLTGYGFVALREGYGAGEGLWIIVWILAVVGVSDVAGYFAGRTLGG
FT                   PKFWPALSPKKTWSGTVAGWIGAVIVGLCLVAAGSGWGIVLLSPLVAFAGQMGDIAESW
FT                   IKRRAGVKDSSGLIPGHGGALDRFDALIGAILLVLLVVQFGALPQIGG"
FT                   /product="Phosphatidate cytidylyltransferase (EC
```

```
FT                   2.7.7.41)"
FT                   /EC_number="2.7.7.41"
FT   CDS             complement(29417..30169)
FT                   /db_xref="SEED:fig|6666666.127243.peg.20"
FT                   /translation="MTQMAATEISPAPADVDYGVGHVAIIMDGNGRWAKNRGWPRLVGH
FT                   RKGAERVKQIVRACPDMGVHWLTVYAFSTENWKRSTEEVLGLMGIFARYIEKEADALSA
FT                   EGVRLRFIGARGRLDRKLQDLMAGIEARTAGNCRLNLTVAINYGGRDEIRRAVQAIAAE
FT                   VAEGRLNPDQIDDATISAHLDTADIPDPDLVIRTSGETRTSNFLPWQAAYAEYEFTPAL
FT                   WPDFTPELLAGILSRFGQRERRFGGVKG"
FT                   /product="Undecaprenyl diphosphate synthase (EC 2.5.1.31)"
FT                   /EC_number="2.5.1.31"
FT   CDS             complement(30185..30748)
FT                   /db_xref="SEED:fig|6666666.127243.peg.21"
FT                   /db_xref="GO:0006412"
FT                   /translation="MADEFEIDTDDLQRRMDGAMASLKTEFASLRTGRASASILEPVMV
FT                   DAYGSMTPINQVGTVNVPEPRMVTINVWDKGLVNKVEKAIRESGLGINPQTNGTIIMLP
FT                   IPELNEERRRELTKVAAQYAESTRVAIRNVRRDGMDQVKKAKAAGMSEDDVKFWEGEVD
FT                   DLTKKNIALVDKALEHKQAEIMQV"
FT                   /product="Ribosome recycling factor"
FT   CDS             complement(30868..31599)
FT                   /db_xref="SEED:fig|6666666.127243.peg.22"
FT                   /translation="MTAVAEPTVKYKRVLLKISGEALMGDQGYGLHPPTVARIAREVKS
FT                   VHELGVEICMVIGGGNIFRGLAGAAQGMERTTADYMGMLATVMNALGMQAQLEALGLHT
FT                   RVISAIPMDQVCEPYIRRRAVRHLEKKRICIFAAGTGNPYFTTDTAATLRANEMNCEAI
FT                   FKGTKVDGVYDKDPKKFSDAKRYGEVSYDECLQKNLGVMDASAIALARDNNLPIIVFSL
FT                   DEPGGFRGILAGQGTYTTVHA"
FT                   /product="Uridine monophosphate kinase (EC 2.7.4.22)"
FT                   /EC_number="2.7.4.22"
FT   CDS             31681..32580
FT                   /db_xref="SEED:fig|6666666.127243.peg.23"
FT                   /translation="MGGMDNFARISPDRPVLIAGATASGKSALALEIARAQGGVVVNAD
FT                   ALQVWSCWRLLSARPTPEDEAAAPHLLYGHRAPGEEYSVGHWLREAGEILSAWKAGTGE
FT                   RPIFVGGTGLYFTALTAGLAEIPPTPAHVRAEADARMQAEGPAALLAEIDAETRARIDI
FT                   RNPVRIQRAWEVMRATGRDLASWQDATPAPLLALSEAEALVLDADRDWLASRIDRRFRI
FT                   MMEQGALDEVRAVAPFWQPAAPWAKAIGAPELMAYLQGDIGLECAIARAQAASRQYAKR
FT                   QRTWFRARMRDWNHLPRP"
FT                   /product="tRNA dimethylallyltransferase (EC 2.5.1.75)"
FT                   /EC_number="2.5.1.75"
FT   CDS             32774..33535
FT                   /db_xref="SEED:fig|6666666.127243.peg.24"
FT                   /translation="MPIARLAQGGRWRVEALRALPEACLLWFTRGQGRVTLDGVTRGYG
FT                   PHNAIFVPAGVIHGFEIAPQTIGFAVFFGFNLPDPLPGKALHLRVRDAAPQVELSALIE
FT                   AASRELDGGRPGAARAARFQLGLIGVWLERQAAVQPNEVEANDAARRLVARYSALLERD
FT                   FRTGQGVGDYAAKLGVTATHLTRSCRATSGRAALEMLHDRRLYEARRLLSETATPVQEI
FT                   AAGLGFSSAAYFTRAFHAATGKTPSDFRRAG"
FT                   /product="Transcriptional regulator, AraC family"
FT   CDS             complement(33545..33913)
FT                   /db_xref="SEED:fig|6666666.127243.peg.25"
FT                   /translation="MSESDTTAVPEAVPALEATDAARALLAAIVADHGPVLLILSGGCC
FT                   DGTAPMCYPEGEFLLAPDDLRIGAVDGAAVYLPAGLAPLWQGGLTLDAVPGRGGGVFSL
FT                   DNGREAHFVARSAVCLKP"
FT                   /product="hypothetical protein"
FT   CDS             complement(34122..35651)
FT                   /db_xref="SEED:fig|6666666.127243.peg.26"
```

```
FT                   /db_xref="GO:0004029"
FT                   /translation="MATDMSAQFQGVMVSPFRARYDNFINGTFKAPNAGRYFDNITPLT
FT                   GAKICECARSDESDVNDALDAAHAAKELWGHKMSPAERAGALLRIADRMEANAELLATC
FT                   ETWDNGKPIRETRAADVPLAIDHFRYFAGVLRAQEGTMSEIDADTVAYHFHEPLGVVGQ
FT                   IIPWNFPLLMAAWKLAPALAAGNCVVLKPAEQTPATILVFAELIADLLPPGVLNIVNGF
FT                   GLEAGKPLASSKRIAKIAFTGETTTGRLIMQYASQNLIPVTLELGGKSPNIFFSDVARE
FT                   DDAFFDKAIEGFVMFALNQGEVCTCPSRALIQEDIYDKFMEKAIARTRAIVQGDPRDAA
FT                   TMIGAQASEEQKNKILSYFDIGRAEGAEVLTGGAAAALSGDLAGGYYIQPTILRGHNKM
FT                   RVFQEEIFGPVVSVTTFKTQEDALEIANDTLFGLGAGVWSRDANTCYRFGRGIQAGRVW
FT                   VNNYHAYPAHAAFGGYKQSGIGRETHKMMLDHYQQTKNMLVSYNPNKLGFF"
FT                   /product="Aldehyde dehydrogenase (EC 1.2.1.3)"
FT                   /EC_number="1.2.1.3"
FT   CDS             35706..35867
FT                   /db_xref="SEED:fig|6666666.127243.peg.27"
FT                   /translation="MPASDPSGNASAARRPGTETVPREKTHIRYATMEVQRGRAIEGQA
FT                   NFPQVIVY"
FT                   /product="hypothetical protein"
FT   CDS             complement(35988..37379)
FT                   /db_xref="SEED:fig|6666666.127243.peg.28"
FT                   /db_xref="GO:0004777"
FT                   /translation="MSYATINPYTGETVATFPEATDAEVANALDRADAAFHEWRKTSFA
FT                   HRAAVLQKAADLLRRDADEHARILTLEMGKLFAEAKAEVELSAKIFEYYVRNAEKLLAP
FT                   EKLPVLDPEEGEAVLVHEPLGILLAIEPWNFPYYQIARILAPQLSAGNVLILKHASNVP
FT                   QAAARFEKLMAEAGLLAGGFTNLYATRDQINTIINDPRVHGVALTGSEAAGAVVAAEAG
FT                   KALKKSTMELGGADAFVVLKDADLDKTIDWAVFGRHWNGGQVCVSSKRMIVVDEVYDKF
FT                   VEGYKKGVAKLVAGDPMDPATTLAPLSSQKAADDLKVQIAAAIAHGAKAEEVGPAVPNK
FT                   GAFVQPTILTGMDGDNPARLWEFFGPVSMIFRARDEADAIRIANDTPFGLGGSVFTKDE
FT                   KRGFEVASQIATGMVFVNHPTKVEADLPFGGIKRSGYGRELFGLGLKEFVNHKLIDVVK
FT                   IDAHF"
FT                   /product="Succinate-semialdehyde dehydrogenase [NAD] (EC
FT                   1.2.1.24); Succinate-semialdehyde dehydrogenase [NAD(P)+]
FT                   (EC 1.2.1.16)"
FT                   /EC_number="1.2.1.16"
FT                   /EC_number="1.2.1.24"
XX
SQ   Sequence 37465 BP; 7134 A; 11244 C; 11444 G; 7643 T; 0 other;
     cctcgacctg cgggcccgga taaacgtaat cagctatagg atagaccttt gtcgaatcaa        60
     aatcgaatgg cttgtacatt acgccgtaaa ggtctgtgac gccgtcggcc gccttgacct       120
     tgaatcgctc cgggaacttg tatccggctg cgaagagcag ggagaggtcg gccttttcga       180
     ggtccattat cttcttaccg gcagcattgt agagggctac cgcaggagca tagtccacgc       240
     gggaataatt ggccacgaag tacttggcat cgtcgctggc ggcggacatc acgtccatgt       300
     ccctgatatc gagaggctgg agggctccgc cggaaaggga tactttgtag gtatggttct       360
     ggtaaggatc ttcatccttg actactccgc aggcgcttac gagcatatag ccggcggctt       420
     cgttgactgc aagtacgtca gagatgtgga aggctccttc cgtaaggttc ttgacaaggg       480
     ttccgtcaga gttgtagagg tagatgttgg cccatccgtt gcgctcggac caccagagga       540
     gctgttttcc gcctttgagg agacgtactc cgcgggtttc cacgtatgtg ttgaggcgtt       600
     ccttgacgat cgccttggta gagtcggcgt ccacgcctat ccagcagagg tcgatacgct       660
     tgaggtcgcg gctgagacgg tcgatataga aaccggtatt gtcgccgagc catttcgggc       720
     tatagcactc gagataactg tcttcggccg tttctgcagg agtctcgaag gcaaggtcct       780
     ggtctttgaa agcatggttc tttatctccc tgtacttgcc gtccggcatc gagaagacat       840
     agatatagct gacaggtccc ggctctcccg gcatctggta tttgtatgtc tcgagtttcg       900
     ggcgggggct gctgactgag ttgattaccc agagatcctt taccgggctc atgtcgaagc       960
     ggaaggctgc gaagtgcctg gagtccgggg accagcggat catcgggatg tttcttgcgg      1020
     tcgagtctgt ctcggtatct ccggaataat tgtctccgtt ccagctgaag tcggcttttc      1080
```

cgtcaaaggt taggcggcgc tcgacgaggg tgctgtcttt gtcatcctcg gcagcctttg 1140
ccagattctc tgcatccata cagtagagat catggttctt tacatagacg ccggtctttc 1200
cgtccgggga gacggctgcc cagtccgggt atttatcttt ctcgccttct tcgagctcgg 1260
tgagcttccc ggaggctatg tcatatttga agtggaagat tttcttgacg gtcttcgtgc 1320
tgtctttctt gtccttgacg tcgatgctgg agtggatatc gaagaggaaa tagctgtcgt 1380
ctttgagctt gaggttgtcg atcgggagat ggttggcttc gaaggggtcg cgtacgattt 1440
cggatatctg catcgcgagg cgttctttat cgaagatttc agtcttggtt ccggctgcgg 1500
gatcgacaat gtaatagcgg gtgccggaag gggtcttcca tgagtaccag aagcggtccg 1560
agttgcggaa ccaggcgggt ctcatctggg ttgagaagac catggaattg attttctttg 1620
cagagaatct ttctgcaagt ccgtagttcg gcttctgcgc ggagagcgtc agcgctgccg 1680
cgagagccag tgcaatagcg agtattcttt tcataaaaac ctgaatattc tgcaagataa 1740
gaatttttcg cagaaaacga tacatatatc ggccggaatg gcgacgtcgt aaagaacgtc 1800
gtagaccatt taaatggcgc ggaccttttt atgttgaggg ggcgccatcc cttaatacag 1860
gccactcagc ggcacattgc tgcaccctca atgccggaaa atcaaggttg aggggacgcc 1920
atccttcagt acagggcact cagtggcaca gtgctgcacc ctcaatgctt tgtccgacgg 1980
attagaggac tcgtaaagcg agggcgagac aattgtccgg aagtcggacg ccgctggacg 2040
ggtgacggag acgccggagg cattgaggaa aagcgatgag cgagggagcg gccgtcgccg 2100
ccgtgaagaa cggcgtgtgt ttgaaggacg ttacggcgac ttcgagcgga gcgacgaagg 2160
aggcgtataa ggactgagtt agccgtttag gcggcgacag cgactgagca gcttttccga 2220
acctgacgga ggcggcccgg cagctggacg cagcggagac agcaggacag cggagaacaa 2280
gcaggcggac aggtaacgga gacgccgtag gaagtcagag ggttggtagc ggcgatagat 2340
tgacgagggc agcagtggcg aaaaaacggc gggggtcctc tggtctcggc gagcgtggcc 2400
accctcggcc acgttagagg gaggggcccg atggatacga gttctgcgaa gcaggacgaa 2460
gtagacattg ggagggatgg agccgacgga gtcggcggaa cagagccgag accacgaggc 2520
ttcccggaga gggaggacaa aaaaaagccg gcccgggcgg ggtcggctta taagtatcag 2580
gatggccggt cggccatgac gcagattact ggagagttcc gttctgggcg gcagcgatca 2640
tgttctcgtc ggcagtgatg tagatctcca cacgacggtt ctttgcgcgg ccctcaacag 2700
tactgttgtc agcaacagga tcggtatatg aaagaccctg ggtcttcata cgatcgcctg 2760
cgataccaca actcttaaga taagcggcaa cggaagcggc tctctctgca gaaagctttt 2820
cgttgacagc ggcagaaccg gtattgtcgg tatggccata aatcgtgata tccgtatcct 2880
tgagatcgtt catcttgctg gcgaactctg cgagttcctt ctttgaagca gcagaaaggt 2940
cagccttgtt ggtctggaag agaattccgc tagcgaaagt aactttcaga gcctcgagac 3000
cgttggcatc gtagatcttc tcgatctggg catcctggag agctgcaagt tcttcagcct 3060
tcttgtccat cttgcggccg ataacagcgc cggcacctcc acctacagcg gtaccgattg 3120
cagctccgat aagagctccc tttgcatcct taccgatgat agcgcctacg gctgcaccga 3180
cagctgcgcc ggcacctgaa ccgataagac tatctttagc gagattggac atgttaccgc 3240
aacccggtac gagaaccagt gcgatcaatg cgataggaat taactttttc ataatatagt 3300
ttatttaaca gtataggtct taaaacaacg taaatgtatg tattttttttc taaaactgca 3360
aataatcaat tatttaaaaa agcttgagga agacctccat cgcctggtat tccgccattc 3420
cgagctcatc gtagagacgc gcagtattct gggtcctctc ctcggcacgc tgccagaact 3480
cacgaggatc gtctccaggg aatggaacga tgtccttctg gctgaggtga cggaagatag 3540
catgacgctt gcttacgacc tctcccgggc tgagcggaac ggccatgtcc acgcggccga 3600
gctcccactc catccatgcg cccctgtaga gccagatatg ggcgttctcg gtccacttgg 3660
caccttcctc gcggagctgc tcgaaagcgt ccatggctgc ttcagtgcag accctgtggg 3720
ttccgtgagg gtcggcgagg tcgccggcca tgaagatcat gtcaggcttg agctcattca 3780
tgagcttctt gattatgtcg acgtccacct gggtcctcgg catcttcttg acgccgccgg 3840
actcatagaa cggaaggttc aggaagtggg cgttggtctt gtcgttgaga ccgaatgagc 3900
gaacggcgct tctggcttcg ctgcggcgga tggctgcctt gatgtcgaga agggcgcgag 3960
gttcggcctc tccaggacgc ttggagtcga tgagggcctt gacttcgtcg aacttgtcgg 4020
cgaatccgag ctggtatgcg gcatccatgt gctggagtac cacgtcgtca tagactgcta 4080
cgtctccgga ggtctcgtat gctacatgga cattgtggcc ctgggctacg agacggatga 4140
atgtaccgcc catggagatc acgtcgtcat ccggatgcgg agagaagatc aggacggtct 4200
tagggaacgg ttcggcgctg acaggacggg ttgaatcgtc tgcattcggc tttccgcccg 4260
gccatcctgt gatggtgtgc tggaggtcgt tgaagacgtt tatgttgatc ttgtcgaacg 4320

gtccgtattt ttcgatgagc tcgccgaggg agttttcgag ataatcgcgc tcggtgagtt 4380
tgaggatagg ctttccaacc tgctggcaga gccatacgac ggccttgcgg atgaacttag 4440
gagtccagtc gcagctgcct acgagccatg gggcgctcac gcgggtgagc agggatccgg 4500
cagtctcgtc tacgaagatg gatgcgttgt catggagctg gagatatgaa gccgggcaga 4560
gaggggtaac ctcgccttct gcgatggctt tgacggcttc agccttatcc tcgccccagg 4620
ccatcacgat aatcctggat gcgctgagca ttgtgctgat gccgatggtg attgctgcgg 4680
caggggtggc ggagatgtct ccgttgaagt tatgggcctg tctcttgcgg gacttgtatg 4740
ggagcggaac ggtgcgggtg cggctctttt cggtcgaacc tgcctcgttg aagccgagct 4800
ggccctgctc tcctactccg attacgagga ggtcgaggcc acgggcgagt ttgtcgaagg 4860
atgcgcagta ggcggagatc tcttcggcag ggacagtccc gtcagggata tgtacatttt 4920
ccttgaggat atcgaccttg tcgatcaggg cctcgtggag tcggctgttg cggctctggt 4980
gctggtctgc gcggaatgga taatactcgt cgattgagac gatctctacg ttgcggaagg 5040
agacttctcc ggcggcgcaa cggcgcgcga gttcattgta aagggaaaca ggggtaacgc 5100
ctgtagtcag tccgagacgg aagaggcggc ccttagtagc cttgattcca ttaacgataa 5160
tgtctgcgac tttcgagctt gcttcgcgag attcgctgta gatttcagcg tttactttct 5220
cgtagctgtg caggatgtct tggggcagct tgtctgttat aagaccgccc tcataaggta 5280
aaacaaagtg tttcattctt ctgtgtcaat gttagctgaa agtgaacttc aaaggtagta 5340
attttttttg aaatagaaac aaaatgggag ctccaggcct tttttgctac ggtgcgccgc 5400
gaaatgcatc ctgctgatat tcaggcatat acaattttag agatgcaccg cggtacatct 5460
ctaaaatcat aaagcgctaa cttctaagtg tttacagttt acaccagata gataaatctg 5520
ttcatattac tttgctaatc aagatcctgc tgagtccata tggtttccgt gccggaggtg 5580
ctgacaccct cccagatacc gtccacactg gtttccatgc cggtcatcac tcctttcaga 5640
tatgcagtat accggacagt catacgatat ctttttatac tgactgtgac attcacactg 5700
ctgtaaggag gggtgttcag ggagacttca acgttcctga catcggatgt cgtcgagtct 5760
tctccccacg tattgtagcg cggttccgtc aaaaatatgg aatttgtctc cggcatcaga 5820
taaggaacag gcgcataaaa cccttccgga acggacacgt ccccggagct taagaaggat 5880
gagctttcct taacttcaaa ggtgaaagtc ctacttgcca aaatgagctc gggagtgtta 5940
ttatgtagct gatccgaggc aacgtagtcg ggcagctgtt caatgacggc ggaatcgtca 6000
agatgatact ctatcttggt tacgacataa tcttccctca gcccggtcgt gccggactcc 6060
tggtcgggat tttcttttcc gcaagatgca atagcgagcg acaaaaaaat aatcaatgca 6120
gctttcttca taaggaatca gtagttttat atgcgaaaat ataaaaaacc gggcaatctg 6180
tcaaaggttg cccggccttt ttcttatgtc agactatttc cagccagcgt actgtgcaga 6240
ggcggtatct gtccagcctt cattctgctt gagaacaccg ttggagaggc tgatctcgga 6300
acgagggata ggcctgaagc cgtaggttgc ctcgtagcgg gccttgtagc ctgcaccctg 6360
atctttcatg acagtctcgc cacctgcgtt gcgaatctcc tgaccgagct gggtctcgag 6420
ggcggcgata gcataagcct tgccataacg tctcatgtca ccccatctta cggcctcgaa 6480
tgcgagctca tgctttctct ccatgcggag ggcttcatcg gtgtaagcgg ctatagggggc 6540
gaggttgacc cttgcgcgca ccttgttgat gccttccacg gtcctggtaa gctcggagtg 6600
catcaggagc acgtcagcga agcggaggag ctggatattc tgaggagaat ggccgcggcc 6660
gaggtcggta ccggtaccgc cgtactcagg aacagatgag aattcataga agaatttgcc 6720
atcgaccatg gcgccccatg tctggtattt cttctgccag agtccggtct cttcatactg 6780
ctggtcgtta ccccaggtat atgctgatgc aggaacgcac tcgtcttcgt agttgtagat 6840
tgaagccacc ttgcggaggt cgcccggctc ctcgatatcc cactgtttcc atacatatgg 6900
attgacagtt gcccatccgt atccacggcc gaacgggaag gcgttgacct cccagttgga 6960
gcgcttgcgg actccgagat actggtcgta cttgttgcag aaacgggatg aagaagcgag 7020
gttggtgcag cagatgctga acatctcctc agggttctct gagtcaatga tccatgttcc 7080
ctcgactgtc tgggcataag gatagtcagc cttggtggcg gagttggtgt atgaccactg 7140
gaggcggaag tcgctcacga ggctatggcc agagttgtcg atacagtctt tgagcttgga 7200
gacaacatag tccttgctga cagtttctga ggtgagctct ccggtctcga ggtctacgag 7260
aggaagggtt gtaagagtag tgccgttctt gtcaccatag aagccggtgt agaagagata 7320
tactcttgcg aggatacctt cggcgtccca cttggttgcg tgacggagac ctgaaacaca 7380
ctggttccaa ggcttgtcag gcatgatctc gatagccttc ttgaggccgg ctgcgatagt 7440
accgtatatc tcttcgattg aagcctgtgc aggatacttc tttgcctctt caacagtctc 7500
aggtacgctg ctggtcagag gaatctcgcc gaagcactgg gtgagttcga agaggaagta 7560

tgagcggagg aagtaagcct cgccgagaag ctggttgcgc agctcttcat tctcgacctt 7620
gtcaagattg gcaatggcca tgttggcgcg ggtgattccg gcgaagaatt ctttccagta 7680
gtcactatgg acatccggct catagtagag gaggtgatcg tcggcctggc agtcgtagtc 7740
gtcgataccg ccgccgccgt agcattcgtc ggaagcgatc atggcagcca tccagtagga 7800
tgcgctcggc tcatatactg aaccgttgag agcggcatag acaccggtca cgagctggat 7860
cgcgtctgtc tcggacgaag ggaagttact tgtattcctt gtcgtatagc tttccgtatc 7920
gaggaacttc tcgcagccgg aaaaagcgac tgcggcgcat gatacggtga gtataagttt 7980
cagataattt ttcatgacaa tctagaattg aaggttaaca ccaatcatgt atgttctcgc 8040
acccggatag taaccgaggt cgataccgga tacccagtcc tcgtcgaaac cgtaaccgat 8100
ctcaggatcc attccggagt agcctgtgaa ggtgtagagg ttctgggcag agaaatagat 8160
acgagccttg ctgaggattg actttttgac aagacggctg aagtcgtatc cgatagtgac 8220
gttggaaatc ttgaagtagt ctccgttctc gatgtagatg tcggagatgt tggaccagtt 8280
cctgtcgttt gcgagggtaa gggtaggata cctgttagag gtaccttcgc cgtgccagca 8340
gtcgaagaag tccctggtgt agttctcgcg agggctgtcg aagaatgaac ggtaagactt 8400
ggcgatctgc tggccgaatg cgccgtaacc ggtcacgctg aagtcgaagt tcttgtatgc 8460
gaaccagagg ttgagacctg tggtgaagtc aggatgaggg ttaccgatca tggtgcggtc 8520
ctcttctgtg atagcgccgt caccgtcggt atcgacgaag atcacgtcgc ccggcttggc 8580
atcctcatac ttgactgcgg ttgctgcaac ctgggcatcg ttctggaaga taccggcggt 8640
cttgtaaccg tagaagtaac cgataggata gcctacctgg gcgcggtaca tctcggaggt 8700
accctggctg agcacgtcgt cctcgccatg gatgataccc tcggagttgg cgatacgggt 8760
caccttgttc ttgttgtaag aaccgttgaa cttgacgccg taggtgaagt ctccggtgta 8820
tttgccgtag tcgatggcaa tttcgacacc tgagttgcgc acgtcaccac cgttcacgta 8880
tggagcttcg aagccatata cggaagcgat aggagcctgt acgagccagt cgcgggtagt 8940
cttgacataa ccgtcgaggg ttactccgag agctgagttg aagaatcttg cgtcgagacc 9000
gatattggtc tgctctgagg tctcccagct tacgtcaggg ttggcaagga cgcccggaat 9060
ggcaccggta gtaggcactg ccttgttggc gaagtagtag ccggcgcttg aggagagctt 9120
gatggtggta aggtactgga agttgtcgat gtcgcagtta ccgttctgac cccatgatgc 9180
acggagcttg aggaagctga gccagtcctt cgtgtttgcc atgaaggctt cgttgctgat 9240
gatccagccg gcggatactg aagggaagat accccagcgg tgaccgcggg cgaagttgga 9300
agaaccatcg gtacgtacga tagccgagag aaggtatgta tctttaaggc tatagttgac 9360
acggccgaag aatgaagcga tggcgcccct tgatgatggc gaacctgaga gttcgagctg 9420
gttgagggcc tccggcttgg tgttggagat atatgccctg tcaaagtcgt tgaggaagat 9480
tgagttgaaa gcgctggctc caactttctc tccgaagccc cacttctcga ttgactgacc 9540
gaggacgacg tcgaagacat ggtccttatt gatatcgaag tggtatgaag cgtagttctc 9600
ccaggtatat ctctggttga ggttcaggct ctggctgact tcgtcgttgt cagaatactg 9660
accgctggaa agtctgtaga gcatggtgta tgagcgcgag ctcgaaccgg acatgcggta 9720
accgaactgg ctcctgaact tgaagttctt gataggagat atttctgcgt atacgctagc 9780
ctgaaggttg tatgacttgc tgatgttacg tcctcttgag gtataaaagt ccatgccgag 9840
agggtgcttt gcagtaccgt caaaatccca gcctgcagca tcgcgggcag cctcgtcata 9900
gaaacctgtg atattaccct gctcgtcata tgtatatgca ggaaggagag ggttggcagt 9960
gatggcgttg tggagggagt tccagtagat gtcaccttct gcaataccgt tattggtgcc 10020
gtagctgaag ttgagagtct cgccgacctt gaagacttcg cgaccgttga ttctctttac 10080
tacgttgtcc gtgttgatac ggaaggtata ccttctgtat tcagcgttta ccggctggat 10140
cttgttgtat ccgaggatac cgtcctgggc agtacaagag aaacccatcg aaaacttgtg 10200
gtcagcatta ccgccgacga tgttgacgga gtggttctcg gtccttgctc ccttgttgta 10260
catctcgttg acccagtcgg ttcctttcca tgtgccgttg agcactgact ggtagagagc 10320
ctgaggaagg agtgtctgcc agtcatagag aggagaaccg gaattcgtct ttgcaaggtc 10380
ctggagctgg atatattcct gagcattgca gagatccggc atctttgcga tatactgggc 10440
gccgagatat gcgtcataag atgtgatcac gcggccttcc ttgccctgct ttgtggttac 10500
aagcacgaca ccgttggctg cacgggcacc gtagatagca gcagatgctg catccttgag 10560
cacgtcgatg gactcgatat cagaagggtt gagagagctc aggcttcctc ctgcgactcc 10620
gtcgatcaca tagagaggct cggaatcgcc cgtggtaccg ataccacgga tattgacttt 10680
gtagccctgg ccaggctgac ctgaattctg ggtaatattg acaccagggg tctggctctg 10740
gagagcaccg agagcggaag tagtgctcag tttcatgagg tcgtcaccct ttacctggat 10800

ggtagaacca gtgacgagtt ttttctgctg gacgccgtat cccacgacaa ctgtttccat 10860
gagctggtca aattcaggga cgagagtgat atcgaccaac gtccttccat tgactttttc 10920
ctcgacattg ctgtatccga gaagggatac agagaggact ccgtttgcag gggactcaat 10980
agaatacaga ccatccttat cagtggtggt ccagctggag gtacctttca cgacaacggt 11040
tgcgtaaggg acaggatctc cagtccccgc gtccgtcacc gtacctttaa cagttgcaac 11100
ctgtgcgaaa acgcctgcag ggaacaacgc taaaagcaat aatagactta atcttttcat 11160
acgagaaaat ttttaatggt ttattagttt tcggttctgc ccacttatcg tttagtccat 11220
gacaagggaa gctgcgccga gcacagcgac atccttgagc acggaatact cgattctgat 11280
tttttcgaga gacttctgga acgcaaaatc tttcagcgac tcgttcatgg aatcgcggaa 11340
gagaggttcg gccttcgtca gaccgccgcc gaatacgatc atttcaggat cataggcata 11400
catggcaacc ccgaccagga atgccatatg ccgaccgaat tcggaccaga gtcggatcat 11460
agcagaatca ccgccattgg cggcctccgc tgcatctttt gctgtcaagc cacgttctgt 11520
aaagaacttg ctggcgcaat agtcttcgag agtcttcccg agatacggaa tgcatccgat 11580
ctccccggcg ccggtattga cgccgccgta gagtttgccg tcgattatga ttcccgagcc 11640
cacgccggta ccgagagtga ttccgaccat gctggatgtg ccgcgaccct tgccgaaacg 11700
gtgttcgcca agagtaaagc agttcgaatc attgttgacg ctgactttct ttgaagggaa 11760
ggctttttcg agtatcgctt tgagcggaac ctcaacccac tgcgggatat tcgtcacatc 11820
ataaacaatc ccctgttccg catccaccac cgaaggcgca ccaattccta tacctctgat 11880
atcggatgaa cataaaggtc gaatcacttc gacaagataa ccaatggaag actctagcga 11940
gtattccttc gggatagagg gaactgtaac gctttccacg acagcggatc catcaactac 12000
acctactctc agattggttc cgcccaaatc gatacctaac cacataaaaa actattgact 12060
tgattactat tagtgttaaa tgtcttcgca aagttatata atgaaaaaag tgattaatct 12120
gcaatttgcg gattaatcac ttgcattttg agaattatgc ttccgtctgt attccattgg 12180
tgtacaacca taaacttttc ggaaccgtcg gaatatattg ttcgggtcct cttccccgac 12240
aagcccggct atttccccga ccgggcggtc catgtttatg agataggacc tgaacctttc 12300
cagcttgacc ttgccgatga aatggtaaat cgaatccccc atcgctttcc ggaaccttat 12360
ctccagcagt ttcctcgaca tcggcaccgt ggcgagcacg tcgtcgacat ttatcttccg 12420
gtgcacattg tcataaatga atcgcaggac cttctgcaca tcggcgtctt cggtagcgat 12480
gacgttggta gaggtgcggc tgacgatata gagaggccgg atgacgacat ttgtctctcc 12540
ggaaggttct acccggccgg caagtatgtc atcgataagc ttcgctgcat catatccccc 12600
ctgcacaata tcagtcacaa tgctcgactg cggcgggaat gacatctggc agaccgtctc 12660
gtcgttatcg accccgagca cggcgacatc ctccgggacc ttgtagccgg cgacgttgca 12720
gacttcgatg agcttcgtgc cctgattgtc gtcgcaggcg aagatggcaa ccggtttcgg 12780
caggctgtcc agccatctgc ggataggttc ggcatcataa tgccagagct cggatatgtc 12840
ctgctcctga taggacggag agacttccat acccagatcc tccagccggc ggcggtagcc 12900
catgtacctt tcatcggacc agacgacgtt gcggtagccg aagaaggcga aatgccggta 12960
gcggcgcttg atgaagaagt cggccgcctt ggctcctgtc gcgaaatagt ccgacgtaat 13020
gttcgggatt acagaaaacc gtgatttgta gtcctgcgcc agggcgatga ttccgttctt 13080
gcggaagaga gagacgtcgt cggtgtcact gaactggccg atgatagcgt cggctttcca 13140
ttttttcgcc catgtcacga ctgccttcag accgaacctt tccttgtagg ccggcggcat 13200
gcggcagact atccagtctc cgggagtgga ggcatattta tagatgccct gcagcagata 13260
gtgcgcaaag gcctctgtaa aatcggtcat cagtattagt cttaccatag caaagtttaa 13320
aaaattgccg cacggagacc ggttccggca tccgtgcggc ctgatcgtag aaaaaatatg 13380
gattaatgca gttttacctg tataactggt cagcgttcac tgcgtggaat ccctttccac 13440
caggacggcg catctggatg gtagtgttgt tgcgcgctgt agcatgacct ccgataatgt 13500
tgcttatgaa cactaccttg atctcgtgga gacccttctc gagagcgacc tgggcgtcgt 13560
gacgggagta tctcttgacc tcggtaccgt tgtcgatgag cttcttgccg tcgatccata 13620
cctggtcgta gtcagaagaa acagtccata cgccatcctc ggagatattg atgtagcctt 13680
cagctacggc tccgtagaaa ttgacgtcct gcatgttgtt ctggcgaggc tcctggcgct 13740
cgatctgctt gagctcggta acggttgact cgatccagcc ttccttgact ttgtcaagct 13800
cgctggctga aaggaagcgg ccatagacct tgcggaactt cagtccgttt gaaggattct 13860
caagcttgac agctggagcg agggtctgct tctcgacctt gatctcgcgg accatgctct 13920
tctttccgga aggaaggaca gtgcagatct tgatgagaga gtttccgtcc accttgatag 13980
gagcggtgta ctcggtcgaa tttccgtcag gctccgatcc gtcgagggtg tagtacatct 14040

tctcaggacg agtggtagtg aatgcaattt cagactcgcc ggtgatgacg aggttgttgc 14100
atgaaccgtc cggctgctca gggagcggaa tatgataggt aactccgcgc atgtcaagac 14160
gctcgcaggc atcgttgacc cttacgctga agtcttccca gttcttggaa gagagctggc 14220
tccatgcgac ctcgctgaga gcgaatccgc gagggaagag catgtaatcc ctgtagtcgt 14280
catcatacat gtactctgac cagttgcatg cctgtactcc catgacgtac ttctcgagac 14340
cgtctttctg gagctctgcc ggcaccggat tgtaatcata gatcttcttc agaggaatgt 14400
agcggccgaa accgagaggc tctgccttag ggtcgccctg atagctgtcg aagtagcagc 14460
cttcgtggct cggggtcatg accacatagt ggtgcatctg tgcagcggca ataccgcctt 14520
gctcgcctct ccaggacatc acggctgctt cgccgctgag tccgccttcg aggatttcgt 14580
cccagccgat gagcttcttg ccgaacttgt caaggatttt ctccacgcgc tggatgcagt 14640
agctctggag cttctcctca gcagtgtggt tctcgtctgc gaacaggcct tcggccttga 14700
tcctggcctg gcacttaggg cagactttcc atcttgtctt agggcattcg tcgccgccga 14760
catggtagta gacagaaggg aagataggaa tgagttcttc gaacactccc tcgatgaatt 14820
caaatacttt atcatttcca gggcagagga cacggtccat tacgccccat gaatagcttg 14880
gattgatttg ctctccgtcg caggacagtt ccggataaga aacgacggct gcgatggtat 14940
ggccagggag ctcgatctca gggatcacgg tcacgaatct ctcggctgca taagcgacga 15000
tttccttggc ctgttcctgg gtatagaagc cgccatatac ggaaccgtcg aactcagtcc 15060
tccatgctcc tacttcggta agacgaggat acttctttat ctctatcctc catccctggt 15120
cctcggtgag gtgccagtgc atggtgttga tcttgtatga ggccatgata tcgatctggc 15180
gcttgattgc atccatgctc atgaagtgac ggcatacatc gagatggaat cctctccatt 15240
tgaatctcgg ataatcggtc actctggcgc actgggcagt ccatttcacg ccctttacct 15300
tagagaggct ttcgatctga ggaggaagga gctggaggaa ggtctgcatc gcatagaaca 15360
gaccggcctt gtcgctggct accgctgtga cttttcctgg agtcacttcg aggtagtagc 15420
cttcttccgg aagtccggcg ccggccgaaa tcgaaagttc gatctggcct ccgcgggttg 15480
atgtgcggat cttgtagccg gtagatttgc cgattttttc agcaaagaat tcagcaacct 15540
caggtgcgtc ggcgctgctg attaccgtgt gcttggtaat ttcgaattcc ccgtcggtac 15600
ggaccgcttc gacaggggca ggggttatgt tcaaaagccc tgcgacgagt agtgtcagta 15660
aataattcat ttaaaaatag cttttatacg gttagcgact tcggcatttt ccattgtgcc 15720
ggagaaatct tctgaatgtg gaccgaaggc atataccggg accatgattc cattatggcc 15780
cgaggtcgag aagttaacct tgatttcgcc cttctccagg cttccgccaa ggagggtaag 15840
acctccggta gcatggtccg cagtgacgat cacgagggtt tcgccatcct tggcggccca 15900
ctcgagaacc ttacctattg tcttgtcgaa atcaaagagc tcttccatgg catagccgac 15960
ctggttgtgg tggcaatgat catcgatgct ggatccctcg atcataagga aaaagccctt 16020
atcgttgtcg agaagctcga gagccttcac ggcgctctct tcaagaatcg ggccgcgatc 16080
cagggccgga gccatctcgg tatcggcaaa aagaccgagt ataggaagtt tctccgaggc 16140
tgcgagctct tcacgtgtat ccacgaaatt gtagcccttt gctttcattt cctcgactat 16200
gtcacggccg tcactgcggt ctttgaagaa cttgatgcct ccgccgataa ggaagtcggc 16260
gccacagtcg agatactgcg cagcaagtcc ctcttcgtca tagcggctga tcgaatggtt 16320
gcagaatgct gcaggagtag cgtcgcctat acggcagacc acgctgacac cggttctgag 16380
gcctctttca tgggcatcgg caaggatgct cttgacagga agtccgtctt cgtcgactgc 16440
aatgaagcca tagcgggtct tgacgcctgt agcgagagca gagccgccgg cagaagagtc 16500
tgtgataagc ttatcggtag cgtaggtgcg ggaaaggccg actacaggga agttgtcgag 16560
atagagatga ccgccgttgg cgacccaggc gcagctgatc tgctcgaggc ccatgccgtc 16620
gccgatcatc ataatcacgt tctttacctt cttgccctta ggctccttga ttacagggac 16680
atcgtagcgg gcagttgcga catatttgtc gccgtcgccg tagacatgct cagcatattt 16740
ggacttgtcg cgggtgttgt cgtcaggaat gagggtgatg tcggtattcc agccggtcat 16800
ggcgacacgg gcgttctcct tgcggtcgca acggcaggtg acggtcttct cggagtgcgg 16860
gaggactgcc acgaagttgt cgctccagat tgcaggctcg aggagactgc cgtcaggacg 16920
gacaagcttg aggacgttct ggtaagagat atagtcggag tcgttggtga gggtgacagt 16980
ccagagtttt ccgagattgt cgactgactc tacgactttg tatgacactg cagcccttgg 17040
catgtcagat acgaagtgca tgtcggcgta ttcctctgca ggggtgacat accagttggt 17100
cttgttgaac ttgtaggtgg tcatcttcgc agggatgcag tagaagttgt cctggccgtc 17160
cctgatgcgg agagctacga agatgtcgcg gccgcggaga gagtcgaggt cgaaggcaac 17220
cttaggatgg cgtctttcgg tacctacatg gtcgctggta tccaccagga ggtttgactg 17280

ggcgtcgtaa actttgatat aagccttgat atccttgtcc ttggagcttt cgttgactgc 17340
atagacctta tagtctttgt agttgaatat aagctggatc ggggcgcagg ctttcttgac 17400
tccgtagtag gatgcagtcg gggtcagatt gttgtcataa agctgccagt agagcgacgg 17460
gcaggcggag ttgagcatcc actggacgat gccggtggtg ttagggagat tgacgcggaa 17520
tgcctcgaac atggcgcggg tagcgtcata gtcggctgca tgggccttac gggtgaagtc 17580
ctcgatgcca tcagcctcgc cgtactgggc gttgacgacg ctctcgagca ttttggtgct 17640
gttcatcgcg gatgaagagg tcgtgcagta tttgctgtag gccgggccga gaggccagag 17700
ctctgattca gggatcatgg tccttatggt ctcaatctga ggccagttgg cgccgacgcc 17760
tgtctccgtg ttgaagccga atgcgccgcc agcttcggta tcgtagtacc agtagtcagg 17820
accgacgtat tcgtatggac cctccatctt ggtgccggac ttgccggaaa tcttgctggt 17880
catcgcctta gcggagttga tgtatgggcg atagtcgagc tccgaataga tgcggagata 17940
ttctgtctcg agttccggat tagggatgcg gtcgctgccg gtcatccagc cgatgactga 18000
aggatggttg cggagccatt tcacctggtc gcggaaatag cgggcggcga gtttcatggt 18060
cgtctcgtcg ttgatgcatc cgtatttgtc ggtctcagga aggccgcagt agttttccca 18120
ttcccactgg cagctccagc ctacgagggc catgactccg ttctggtcgc agaggtcata 18180
gatatactgg tccttgcccc agatattctc gaaacggatt gtgttcaggc ccatgttcag 18240
gacgtgctcc atctgggcgt cgagggattc gtgggtgtcg cggaggttga cgttgtcggt 18300
ccagcctgcg ccctttatga ggatcttctt tccgttgagg aggaagactc tgtggccgtg 18360
gctgtcgatg cggctctcga ttttgcggat accgaatttg acttcacggg agtcggtcgt 18420
ctcttttccg gcgacaaagg aagcctgcat ggtatagagg taaggctttc cgaggtctct 18480
tgtccaccag atcgaaggct tagcggtatg caggttcttg cactccttcg gagtaaggac 18540
tacctctttt gtctcggaag cggcgagaga gaccgggaac acgaattcgc cgccggccca 18600
acggccacga acttcgccgt tgacgggctt tctggacatg ttcctaaggg tcacgcggac 18660
ttccaggtct gcgctctcgt ctgtgtcgag gttgagctca ggcttgacga agaggtcggt 18720
tacggctaca ctgccggtct cgaagagggt aacgtctctt acgataccca tgctttcgtc 18780
ggcaggacgc ggattccagt cgacgaagcg atcagaaagc caccaccgtc gacaccgccc 18840
cggccgaggc cttcgcaaag cggcgcggcg tgtgtcagga cttcacccag gtgatgattg 18900
cggcgctgcg tggcatcggc attccggcgg gctatgtctc ggggtttctg cgcaccttcc 18960
cgccgcccgg ccagccgcgc cttgaagggg cggatgcgat gcacgcctgg gtccgcgcct 19020
gggtcggccg cgatgccggc tggatcgagt tcgacccgac gaataaccag ttcgccggcg 19080
aggattacat cgccgtagcc ttcggacgcg actattccga cgtcgccccg gtgcgcggcg 19140
tgctgcgctc ttcgggcggg cagaacagcc atcaggcggt cgatgtgatc ccgctcgacc 19200
cggacacggc gcagcctctg gtgcgcagcg tgcccgcgcg gcgggccggg acccgcgccg 19260
aaaacgacat ggagacgcag gcagagccgg acgccgcggt gcctcctgaa atctgaaagc 19320
ccccgtgctt cggcccggtc gctcaggccc cgcccgacag cgccccgagc accgagcgcg 19380
cggcgcgcag ccccggtgcc tcgccgccct cgccaaggcg ggtcatggtg atctccatcg 19440
cctgcagctg cgccgcgcgc gccccggcat cgccaaacag cgcgaccacc gcatcggcca 19500
tcgcctccgg ctgacaggcc gggccgagga attccggcac cgcacgggtt tccgacacca 19560
gattgaccag cgtcaccgtg tcgatcttca tcatgcgcca cacaatggcc caggtcagcc 19620
ggttggcgtc atagccgatg accatcggca cgcgattggc ggcaagttcc agactgaccg 19680
tgcccgaagc ggcaagtgcg gcatcggcgg ccgcgaaagc cgccagacgg tcggcgcgat 19740
cctcgacgat cagcgcgggc acgggccagt ccgccaccgc cgcgcgcacc atgtcagaca 19800
cgccgcgcac cgtcggcaac accacccgca gccccggcac ccggtccttc agcagcccga 19860
gcgtggcacc gaacttcggc gcaaggcgcg aaacctcgcc gcgccgcgag ccgggcagca 19920
ccagaagcag tttctcatcg gcggcgatac cggcgcgggt gcgccatgcg gccgcctcgg 19980
ccccggtcgc gcggggttcg gacaccaccg gatgaccgac gaaatcgcag ctcatgcccg 20040
ccgcgcgcat atagggcggc tcgaacggca acagcgccag aacgtgatcg atcacctgtg 20100
ccatgcgctt cgcccgcccc gcccgccagg cccagaccga gggcgcgacg taatgggtgc 20160
agaacagatc cggacgcgca tccttgacct tgcgtgcgac gcgcaggcag aaatccgggc 20220
tgtcgatggt gatcagcaca tccggcccgg tggcgatcgc ggcatcggcc gcttcctgca 20280
cccggcggcg cagcagcgga tatttcggca agatctcggc cagacccatg accgacagat 20340
cttccatcgg gaacaggctg gtcagcccct cggcctgcat cagcggcccg ccgacgccgg 20400
cgaagccgac accgggcacc agcgacttca gccccgccat cagcgccgcc ccgagcttgt 20460
cgccagacgc ctcgcccgcg atcaggaaga ccttcatgcc tgcactcccc acaaggcgat 20520

gccgtgtttt tcggccgccg cgatcaccgc cgcgcggtcg agcagcagca ccccgcccgc 20580
gcgccagcac aggcacgaca gacgcgcacg cgcagccccc tcgacggtgc gcggaccgat 20640
cgccggcatg tcgatgcgca ggtcctgacc gggcttcggc gccttcatca gcacgccgcc 20700
gcgatgcgcc tcggggcggg tggtggcgac aaaatcgagc atcgcatcgg tgccgggcaa 20760
tgcctcgacc gcaaggcaca gcccgcgcgc gacgaccgcg ccctgcccga catcgacagc 20820
gccgagcgcc gcgacgatct cggccgcgcg ggccatgtcc ttgcggtcgg tgtcggtggg 20880
tttgccgcac agcacaccct cgccgcccag cagatccggc gcgatatggt ccgcgccgat 20940
gacctccagc ccgcgttcct cgaacagcgc gatcaccgcg cgcagcgtcg catcgtcgcc 21000
ctgctggatc gcccccatga tccgcggcac cagaagcgcg gtgcgcagat cgaatttcgc 21060
cggttcgatc ttcgggcggc ggatcgcacc ggcgaagcac accttgcgga tgccgcgccg 21120
ggtcagctcc gtcatgaacg gaaccagacg ttccaccctg aagcgcaagg gctcggcccc 21180
cgggatcccg gaggcgaagc cctccagctc gcacagcagc atgtcagggc gcgcgcgcat 21240
cagcaacgcc ggcaacgccc ccgtccccgc gatcagcgca aggtccggcg cggcgatgtc 21300
gccctggctc atttcggcgc caggaaactg cgatccgacg gcccgaggat gaaatcgagc 21360
acctcgcgca ccatttcgct gtcggtccct tcgtcggcga gcttgcgggc ctgctcgcgg 21420
aagttgccga ccgaaagcgc catgtaggcc tcgcgcagcg cgccgatctc ggcgcggctg 21480
acgccgtggc gcttcagccc gacgaggttc agcccttcga gcccggcgcg cggtgccgcg 21540
acgagaccgt gcgggatcac atcggccgcc accatcgaca gcgcaccgat gatcgccccc 21600
ttgccgatgc gcacgaactg atgcacgccc gacagaccgc cgacgatcac gtcgtcctcg 21660
atatggcaat gcccggcgat ggcgacggaa ttcaccagga tcacccggtc gccgatgatc 21720
acgtcatggg cgacgtggca actcgacatg aacaggccgt cgtcgccgac gcgggtcacg 21780
ccgccgccgc cttcggttcc ggggttcatc gtcacgtatt cgcggatgcg gttgcgtttg 21840
ccgatgatca gacgggtatg ctcgcccttg aacttcaggt cctgcgggat ctgcccgatc 21900
gaggcgaagg ggaagatcac cgtgtcctcg ccgacctcgg tccagccgcc gatcacgaca 21960
tgcgacttca gttcgacgcc cgcgtgaagc gtgacctcgg ggccgacatg gcaaaacggt 22020
ccgatgcgca ccccggcggc aacggtggcg ccgtcctcga cgacggcaga ggggtggatc 22080
tcggcacccg gatcaatcga catgcgtctt ctcctcgatg cggtggggtg cggctcaggc 22140
cttgggcttc ggggccatca tggccatgaa ctcggcctcg gcggcaagct cgccatcgac 22200
cgtagcctcg ccacggaatt tccacacctt gccgcccggg cgcaggcatt tcacgttcag 22260
ttccagaatg tcgcccggca ccaccttgcg gcggaacttg cacgagtcga tgcccatgaa 22320
ataggtcagc atgtcgccgc cgatgatatt ggtcgacacc cccaccacga tcgccgcggt 22380
ctgcgccatc gcctcgacga tcagcacgcc cggcatcacc ggctcttccg ggaaatggcc 22440
ctggaaatgc ggctcgttcg aggtgaccat cttgataccg acgccgcctt cgaagggaac 22500
gatgtcgcgc accttgtcga tcatcaggaa cgggtaacgg tgcggcagga tctgcttgat 22560
cagcgacaga tccacggagc tgtaaggagc gggttcgatc atctcatgtc cttgaactgc 22620
aaggcgcccg gacaggggcg cgggggatgt gcaaagcgcc cgggccgggg cgcgcgggct 22680
ttccgggtct tgctagcaac cggggggcgc cttggcaagc cgggccgagg ggatggcgcc 22740
gcagtcaggg cgaggccgga gccggaaccg gatcgcccgc cggatcgctg tcgtgcgacc 22800
cggatgccgt gtcggccggc ccctcgccca tgcgcgcatc gacgagcgcg atcatgtccg 22860
gggtcacgtc ggcttcgggc agcgcccgga tcaccgcgcg ctgatcgaga acgaccaccg 22920
caccgcgccg gtgcaaaacc tcatcgagca gcggcaccat cgccccgaag aaggcattgc 22980
gcgcatcgtt gtagcgctgt tcgagcgcct tgcccttggc gctctgttcg gcccggatct 23040
tcgtcgcgcg ggcgtcgaac tcctcggcgc gggcctgaaa ggtggccgcg tcgatcttgc 23100
cccgggcctc ggtcagggcg cgttcctcgg caatcagctg atcggcgatg cgatcgttct 23160
cggccttctg ggcgccggta tcgcgctgca tatcggccag gatccgctgt ccccagcgtg 23220
atccgtcgaa cagctgctcc cagtcgatgg tgagcaccgg gctcggcagg cggatcagcg 23280
catcgccggg atcgcgggac gcagcgacga ccgcagagcc cggcgcgacg ccggtcggat 23340
agtcgtcagc cccgccggtc ggctgagccg tggcctgctg cgccacggcc ggcgcagcaa 23400
ggcacagcag caccgccatg cccgcccgcc agatccgcgt tccgccccgc tgtgccatcg 23460
ccgtcgtcag aacttcgtcg agatcgtcag gtcgaagttc tgctccttgt cataggtttc 23520
cttcttgatc gccttcgaga agtcgatccg cagcggaccg agcgccgaac gccagaagaa 23580
cgagaagccg acggccgaac ggatgtgcat cttgtcgctg tcgtcgagcc ccatcgccag 23640
cgcgcgctcg ctgctcgata cgccccacag cgcgccggta tcccagaaca gaccgccggt 23700
gatgccgtat tcctcgggca gaccgatcgg gaattcggtc tcggtgcgca gcgcccagaa 23760

atactgaccg ccgagcgcat cgccattggg tgcgcccacg tcgcgcgggc cccagccatg 23820
cgattcgaaa ccgcgcacct tgccgtaacc gctgaaccgg tcgaggatgg tcaggtcgct 23880
gccgctgcgc ggcgtgatca cgcccccctc cagctcggcg cgcaggcgga cggattcgtg 23940
catcaccttg cgctgcgccg ccagaagcgc agtggtgctg atcgcgtcga catcgccgcc 24000
gaggccgtaa taatcctgac cgaacttgaa cgtataccca tagcgcgggt cgaggttcgt 24060
ggtcatcgtg tcccaggtgt aggtataccc caccgaagag gccgtgcgct cgcccaggct 24120
ctgctcatag ctcagaaccg gagaagagcc cgacgccgac gtcgtgccgc tggccgcggt 24180
ataatcggtg acgccaaaca gcttcttctt gccataggca tagcgcgcct cgaggcggga 24240
attctcggcc accgggaatt cgacagaagg ctgcagcccc tggatacggg tattgtaatc 24300
ggcattgtcg ttgcttgtgg tgtcatacca caggtgcatc cgcgccttga catcgcgacc 24360
gaggaaatac ggctccacga agtcgatgga gctttgctga ttgtccgcgg tggtgccgat 24420
ctgcacgccg agcgactggc cgcggccgag gaagttgtcc tcctgcacgc cgaccagaac 24480
gccgaagccc gaggccgcgc catagctcgc accgagcgtc agcgaaccgg tgggctgttc 24540
ctcgacattg acgtcgacga cgacctgatc ggggttggtg cccggcttgg tgttcacgtc 24600
cgccttgctg aagaacccga gcgcacggat ccggtcggcg gagttgcgga tctcgcgcgg 24660
gttgaagggg tcgccttcgt cggtgcggaa ttcgcgccgg atcaccgaat cgagcgtggt 24720
ggtattgccc ttgatgtcga tacgttcgac gaagatacgc ggtccgcgca ccagggcgaa 24780
ctgcacgttc agggtctgcg tcgccggatc gcgggtcacc cgcggatcga cgcggacgaa 24840
attcgcgcct ttcttcagcg ccagctgctc catccgggtg atcgcgaggt cgatcgccgt 24900
cggcgtgtag acagacccgg tcttgatctt cgccagcggt gcgaactcat cgacgttcag 24960
atccggcact tcggaaaccg tcgtgatatt gccgaactta tactgctggc cctcttcgat 25020
gttgaaggtg acgaaatagc cgtcgcgcgc ccgcgccatg tcggacgaaa tccccgtcac 25080
cttgacgtcg gggtagccgc gcgactgata gaagtcgatc aggttctgct tgtcctgctc 25140
cagccggtcg gggacgaaat tgtcgctctg gatcagccgg tgaagaatgc cggcctgctt 25200
ggtcgacaac gcctgacgca gggtgaaatc ggaatagctg cggttaccgg tgaagctgat 25260
gcggtgaact tccgaggtat agccttcctt gatctcgaag gcgacgtcga cccggtcgtc 25320
gccgcgccgg atcaccttcg gatcgacgcg cacgttgatg cgcccggcct gcgtgtagcc 25380
ctcggcgatc aggttggcat cggcctcgac ctgagcgggc gaaaacacgt ggttgggctt 25440
ggtcttgacg ctcttggcga tggcatcgtc cttgagcacg cggttgccct cgaaggcgac 25500
gacgccgacc atcgggtatt ccttgacgcg gatcaccagc gtgccgccct gcggcaccag 25560
gtcgaccgtc tcgaacaggc ccgagcccac cagccgctga taggcatcgt tgagctgatc 25620
gcccgagacc gtctgcccct tcgcgacctt ggcataggcg agaatcgtcg ccggctcgat 25680
gcgcttgttg ccctcgatct tcaccgcgga gaaccggaaa tcctcggcca tcgcggcgac 25740
gggaaccagt gccccggcag cgatcccggc cgccaccgtc aggggcagcg ccccgggcag 25800
agggctcgcc gtcgtcagtg ccagcgccac cgccggcaga cgacgcacgg ccgaccgccg 25860
tgcaggtgct gcatgttgcc gccgcttcgt cattccctca ccccgtctat tgggacagtt 25920
ctttgtcccg tgagtagcgg caacgccgcc gattgtcaaa agcgccgagc caggcagagg 25980
ctgccggacc ggcgaaatgc cgcgaagctc agcaggtcac gtcgttccac agcccgaaca 26040
gcatcagcga cagaaccgcc accatccccg ccgcggtcag cacgttcagc accgccggcg 26100
acggcgggcg gcgcgtcacc gcctcccaaa gataaaacat cagatgcccg ccatcgagca 26160
ccggcaccgg aaacaggttc agaaacccca ccgccgtcga caacacgccg atgaaccaga 26220
tgaaatccat cgcccccgcc tgcgccgccg cggcagagcc ctgcgcgatg cccaccgcgc 26280
catgcagatt gcacgacgaa atcttgccga ggatcatatg ctcgaacgcc gagaccgagg 26340
aggtcacgat ggtccacacc tgacccgcgc cgcgctccag cgcctcaagc gggccggccg 26400
cgcgggtttc gggggcgaaa gccatgtcgc cagtcgcccc gatcatccag cgggtctcga 26460
aatcgccgtt gggcaagggc aggtccatcc gcttcggcgt caggctcagc gtcaggagct 26520
gcgcccccgt cgccgccgtg ccgcccgccg cccccgcggg aagcggacgc cacaccgtca 26580
gatcgagggg ggcgcccgcg cttgcaccga cgcggctgcg cagatcctcg aaccgccaga 26640
tcggcgcgcc gtcgactttg gtgatcacgt cgcccgcctt cagcccggcc cccgccgcgg 26700
ccgaggacgg catgaccccg ctcagcagcg gcggcatcgg ctgcggcccg tcgatgtcgc 26760
gggcggtgcc gtcgcgctcg atgctccagc gcaccgtggg cgtcggcggc agatcggcga 26820
gcgcggcata gaaggcgcga taatccgggg tctcatgccc gtcgagcgca aggatccggt 26880
cgcccggctg caacgcgccc tcgcccccgg gcagcgtcgc gagacgcccg acggtcggcg 26940
cttcggtcgc gacaccctgc caggcgacgg cgccggcgaa gatcgcgatc gacaggatga 27000

agttgaacac cggccccgcc gccacggtgg cgaaacgcgc ccagagaggc gcgcccgcaa 27060
gcgtgcgccg gcgctcctcg gggctcatcg cgtcaagctc cgcctcgtcg gcggccccgg 27120
ccgaggcggc gttgctgtcg ccgttgaacc gcacatagcc gccaagcggg atcgccgcga 27180
tctgccagcg cgtgccgcgc ctgtcgacgc gcgacaccag caccggcccg aagccgagcg 27240
agaacacctc ggccccgatc ccggaccagc gcccgacgat gtaatgaccg aactcatgga 27300
tgaagacgat gatcgacagc gccacgacga aagccgcggc ggtgtagagc acattgccga 27360
aatcgggaag caggtccaag gctcagatcc ttccgcgggg ggtatcgccg ctggcgccgt 27420
gccttccggc accgtgtctg ccggcgatgc gggcccggcc gagcgcgcgc gcctcggcat 27480
cggtggcgag aacctcatcg aggctggtgg ccggattgcc cagcccctga cgggcggaca 27540
tgtcggacag cacctgttcg accactgccg ccatatcgag aaaaccgatg gcgccggcga 27600
tgaacccgtc gagcgcgacc tctttcgcgg cgttgtagac cgcgccggcg agaccgcccg 27660
tcgccatcac ctcgcgtgcg aggcgcagcg ccggataacg cacgggatcg gccgggcgga 27720
agctgaggct gccgagcgcg gcaagatcga gccgcttcac cggcagctcg gcgcgctcgg 27780
gccagttcag cgcatagccg atggcgtggc gcatatcggg cacacccata tgcgccatca 27840
gcgcaccgtc gcggaagccg accagcgcat ggatcacgct ttcggggtgg atcaccgcct 27900
cgatctggtc gggacgcagg ccgaacagct ccttcgcctc gatcagttcc atccccttgt 27960
tgaacatcga ggcggaatcg atggtgatac gctgccccat cgcccagttc ggatgggtcg 28020
aggcctcggc cacggtcgcg cgggcgatgc gctccatcgg ccagtcacga aaggcgccgc 28080
ccgaggcggt gatgatgacg cgctcgacgc tggccatgtc ctcgccgtga agcgcctgga 28140
acaccgcgga atgttcgctg tcgaccggca gcagacgcgc cccgtgccgc cgcgcggcct 28200
cgaggaacag cggacccgcc gcgaccatgc tttccttgtt ggcaagcgcg agcgtcgtgc 28260
catggcgcag cgcggcataa ccgggcgcca gcgccgccgc cccgacgatg gcactcatga 28320
tccagtcggc cggacgttcc gccgcctcta tcagcgccgc ggggccggcc gcggcctcga 28380
tctcggagcc atgcagcgcc gcgcgcagcg ccggcaggca ctcttcataa gcggtcaccg 28440
cgatatcggc gcgcagcgcg cgtgcctgcg aggccagcag atccacgttt cgcccgccgg 28500
tcagcgcgac gacctgaaac gcccccgctc cgccctgacg ctccagcaag tcaagcgtgt 28560
tgaccccgat ggagccggtc gatccgaaca ccgataccct gcgcatctca gcctccgatc 28620
tgcggcagag ccccgaactg aacgaccagc agcaccagca agatcgcacc gatcagcgcg 28680
tcgaaccggt cgagcgctcc gccatggccc gggatcaggc ccgagctgtc cttgacgcct 28740
gcgcgccgct tgatccagct ttccgcgata tcgcccatct ggccggcaaa ggcgaccagc 28800
ggggacagca ggacaattcc ccagcccgag cccgccgcga cgaggcacag ccccacgatc 28860
accgcgccga tccagcccgc gacagtgccc gaccaggttt tcttcgggct gagcgcgggc 28920
cagaacttcg gcccgccaag cgtgcgcccg gcgaaatagc ctgccacatc cgatacgccg 28980
accacggcca ggatccagac gatgatccac agcccttcgc ccgcgccata gccttcgcgc 29040
agcgcgacga atccatagcc cgtcagcagg attgcaaagc cgtagaccgc gaaaagcagc 29100
cggtcgcgcg tcaccagaaa cgccgctgcc gccggataaa gcaccaacag cccgagcgag 29160
ggcggaaaca gagagcgcag ctccagcacc gcgaaggaga ccgccgccgc ccccacgccg 29220
atctcgtagc cgagacgccc gcgcccgggt gcgcacagcg tcgtcagctc ccagatcatg 29280
gcagcgcaca gcgcgatggc cagcgcctcc atcgcgaaac cgccgcgcca gaccgccgcc 29340
ccgcccagca ccagcatcac cagccccgag acaagacggg tccgcaggtc gccccagcga 29400
ccggaaccgt cctgcgtcat cccttgaccc cgccaaagcg ccgctcgcgc tgcccgaagc 29460
gcgacaggat gcccgccagc aactcgggcg tgaaatcggg ccagagcgcg ggggtgaact 29520
catattccgc ataagcggcc tgccagggca ggaaattcga ggtccgcgtc tcgccagagg 29580
tgcggatcac cagatccgga tcggggatgt ccgcggtgtc gagatgcgcc gaaatcgtcg 29640
catcgtcgat ctgatcgggg ttcagccggc cctcggccac ctcggcggca atcgcctgca 29700
cggcgcggcg gatctcgtcg cggccgccgt agttgatcgc cacggtcagg ttcagccggc 29760
agttgcccgc cgtgcgcgcc tcgatccccg ccatcagatc ctgcagcttc cggtcgagcc 29820
gcccgcgcgc gccgatgaag cgcaaccgca ccccctcggc cgagagcgcg tcggcttcct 29880
tctcgatgta gcgcgcgaaa atgcccatca gaccgaggac ttcctcggtc gagcgtttcc 29940
agttctcggt ggagaacgca tagacggtca gccagtgcac gcccatgtcg ggacaggcgc 30000
gcacgatctg cttgacgcgc tcggcgccct tgcggtgacc gacgagacgc ggccagccgc 30060
ggttcttcgc ccaccggccg tttccgtcca tgatgatcgc gacatgtccg acgccataat 30120
cgacgtcggc aggcgcggga gaaatctcgg tggcggccat ctgggtcact ttctgccggt 30180
cctctcagac ctgcatgatc tccgcctgct tgtgctcgag cgccttgtcg acgagcgcga 30240

tgttcttctt ggtgaggtca tcgacctcgc cttcccagaa cttcacgtca tcctcggaca 30300
tgccggcggc tttcgccttc ttcacctggt ccatgccgtc gcggcgcacg ttgcggatcg 30360
cgacgcgggt gctctcggca tattgcgcgg cgaccttggt cagctcgcgg cggcgctcct 30420
cgttcagctc ggggatcggc agcatgatga tggtgccgtt ggtctgcgga ttgatgccga 30480
ggcccgactc gcggatcgcc ttctcgacct tgttcaccag ccccttgtcc cagacgttga 30540
tggtgaccat ccgcggctcg ggacgttga cggtgccgac ctggttgatc ggggtcatcg 30600
aaccgtaggc gtcgaccatc accggctcca gaatggaggc cgaggcgcgc ccggtgcgca 30660
gcgaggcgaa ctccgtcttg agcgaggcca tcgcgccatc catgcggcgc tgcagatcat 30720
cggtatcgat ttcgaattcg tcggccatcg gtcagggctc cttggtgtcg gtgttctttt 30780
agccgctctt gcatgggaag catagggcaa agcaggggcg gcggggtgcg cacaggcact 30840
ccgccgccga ttgaggggga aaccggctca ggcgtggacg gtggtgtaag tgccctgccc 30900
ggccagaatg ccgcggaacc cgcccggctc gtcgagcgag aacacgatga tcggcaggtt 30960
gttgtcgcgt gcgagcgcga tcgccgaggc atccatcacg ccaaggttct tctgcaggca 31020
ctcgtcgtag ctgacttcac cgtagcgctt cgcatccgaa aacttcttcg gatccttgtc 31080
gtagacgcca tcgaccttgg tgcccttgaa gatcgcctcg cagttcatct cgttggcgcg 31140
cagcgtcgcg gcggtatcgg tggtgaaata cgggttgccg gtgccggcgg cgaagatgca 31200
gatccgcttc ttctccagat ggcgcacggc gcggcggcgg atgtagggct cgcagacctg 31260
atccatcgga atggccgaga tcacccgcgt atgcagcccg agcgcctcga gctgcgcctg 31320
catcccgagg gcgttcatca ccgtcgccag cattcccatg taatcggcgg tggtgcgctc 31380
catcccctga gcggcaccgg cgaggccgcg gaagatgttg ccgccgccga tcaccatgca 31440
aatctcgacg ccgagttcat ggaccgactt cacctcgcgg gcgatgcgcg cgacggtcgg 31500
cggatgcagc ccgtagccct gatcgcccat cagtgcctcg cccgagatct tcagaagcac 31560
acgtttgtat ttcacagtcg gctcggccac tgcagtcatc ccgggattcc catttgcggt 31620
tgtcatttcg gcgacaaatg tcgcaaatgg ccgccggaat caacctcggc cttgcaccgg 31680
atgggcggca tggacaactt tgcgcggata tcccccgacc gcccggtgct gatcgcgggc 31740
gcgaccgcct cgggcaagtc ggcgctggcg ctggaaatcg cgcgcgcgca gggcggcgtg 31800
gtggtgaatg ccgatgcgct gcaggtctgg tcgtgctggc ggctgctgtc ggcccgcccc 31860
acccccgagg acgaggccgc cgcgccgcat ctcctctacg gtcaccgcgc accgggcgag 31920
gaatactccg tcgggcactg gctgcgcgag gcgggcgaga tcctgagcgc gtggaaggcc 31980
ggcaccggtg agcggccgat cttcgtcggg ggcacggggc tttacttcac cgcgctgacc 32040
gcgggactgg ccgagatccc gccgaccccg gcccatgtgc gcgccgaggc ggacgcccgg 32100
atgcaggccg aaggcccggc ggcgctgctc gccgaaatcg acgccgaaac gcgcgcccgc 32160
atcgacatcc ggaacccggt ccggatccag cgcgcctggg aggtgatgcg cgccaccggg 32220
cgcgacctcg caagctggca ggacgccacg cccgcgccgc ttctggccct gagcgaggcc 32280
gaggcgctgg tcctcgacgc cgaccgcgac tggctggcca gccgcatcga ccgccggttc 32340
cggatcatga tggagcaagg cgcgctcgac gaggtgcgcg ccgtggcgcc tttctggcag 32400
cccgccgccc cctgggcgaa agccatcggc gcgcccgagc tgatggcgta tctgcaaggc 32460
gacatcggtc ttgaatgcgc catcgcccgc gcgcaggccg cctcgcgcca atatgccaaa 32520
cgccagcgca cctggtttcg cgcccggatg cgcgactgga accacctgcc ccggccctga 32580
ggcccggca atcccgactc ctttgcgcgc attccgccca cgcaaattcc gattggaagc 32640
cgcgcccggc ccttctatct tcgggcgagc attcttttcg ccccttcctg ttcatggccc 32700
cgtcgtgccc ccagcctgcg ccccctctgc cccgatcctg cccgacgctc cggcgcctcg 32760
ggaaccgcgc gtcgtgccaa tcgcccggct ggcccagggc gggcgctggc gggtcgaggc 32820
gctgcgcgcc ctgcctgaag cctgcctgtt gtggttcacc cgcggacagg gccgggtgac 32880
gctggacggg gtaacccgcg gctacgggcc gcacaacgcg atcttcgtgc ccgccggcgt 32940
catccacggc ttcgagatcg ccccgcaaac aattggcttt gcggtatttt tcggattcaa 33000
cttgcccgac cccctgcccg gaaaggcgct gcacctgcgg gtgcgtgacg ccgccccccca 33060
ggtcgagctg tcggccctga tcgaggccgc ctcgcgcgag ctggacggcg gacggcccgg 33120
tgccgcacgc gcggcgcggt ttcagctcgg actgatcggt gtgtggctgg aacgtcaggc 33180
cgcggtgcag ccgaacgagg tcgaggcgaa cgatgccgca cggcggctgg tggcgcgcta 33240
ctcggcgctt ctcgaacgcg atttccgcac cggtcagggg gtcggcgact acgcggcaaa 33300
actcggggtg acggcgacgc atctgacgcg ttcgtgccgg gccacctcgg ggcgcgccgc 33360
gctggagatg ctgcacgacc gccgcctcta cgaggcgcgc cggctgctga gcgaaaccgc 33420
caccccggtg caggagatcg cggccgggct cgggttcagc tctgccgcct atttcacccg 33480

cgccttccat gccgcaaccg ggaaaacccc gtcggatttc cgccgcgccg gctgaaaccg 33540
gagatcaggg tttcagacac accgcggagc gggcgacgaa atgcgcctcc cgcccgttgt 33600
cgaggctgaa caccccgccg ccccgcccgg gcaccgcatc gagggtcagc ccgccctgcc 33660
acagcggcgc aagccccgcc ggaagataga ccgccgcacc gtcgaccgcc ccgatgcgca 33720
gatcgtccgg cgccaacaga aactcgcctt ccgggtagca catcggcgcc gtcccgtcgc 33780
agcatccgcc cgacaggatc agcagcaccg gcccgtgatc cgccacgatc gcggccagca 33840
aggcgcgcgc ggcatccgtc gcctccagcg ccgggacggc ttcggggacc gctgtcgtgt 33900
cactctcgct catgatccgc ctttccggcg ccggggacga aaagcccatg cgccctacgc 33960
cgaaaacggg cggcgcgggg ggcgcgccgc ccgtccgggc cggacctcgg gatcggggga 34020
ggatcagcgt tcgcgccgcg aacgatgcgc ccgtcgatcc tgcccgtccg atccggcacc 34080
cgggcgaagc cccgccccgg tgccgtccgt tcccgcccgc ctcagaagaa gccgagcttg 34140
ttcgggttgt agctgaccag catgttcttg gtctgctgat agtggtcgag catcatcttg 34200
tgggtctcgc gcccgatgcc cgactgcttg tagccgccga aggccgcatg ggccggatag 34260
gcgtggtagt tgttcaccca aacacgcccg gcctggatgc cgcgcccgaa gcggtagcag 34320
gtgttcgcat cgcgcgacca cacgccggca ccgaggccga acagcgtgtc attggcgatc 34380
tcgagcgcgt cttcctgggt cttgaaggtg gtcaccgaca cgaccggccc gaagatttcc 34440
tcctggaaca cccgcatctt gttgtgaccg cgcaggatcg tcggctgaat gtagtagccg 34500
cccgccagat cgcccgagag cgcggccgcc gcaccgccgg tgagcacctc ggcgccctcc 34560
gcccggccga tgtcgaaata gctgaggatc ttgttcttct gctcttccga ggcctgcgca 34620
ccgatcatcg tcgccgcatc gcgcgggtcg ccctgcacga tggcccgggt gcgcgcgatc 34680
gccttctcca tgaacttgtc atagatgtct tcctggatca gggcgcgcga cgggcaggtg 34740
cagacctcgc cctggttcag cgcaaacatc acgaagccct cgatcgcctt gtcgaagaag 34800
gcgtcatcct cgcgcgcgac atccgagaag aagatgttgg gcgacttgcc gccgagttcg 34860
agcgtcaccg ggatcaggtt ctgcgaggcg tattgcatga tcaggcggcc cgtcgtcgtc 34920
tcgccggtga aggcgatctt ggcgatccgc ttcgagctgg cgagcggctt gcccgcctcg 34980
aggccgaaac cgttgacgat gttgagcaca ccgggcggca gcagatcggc gatcagctcg 35040
gcgaagacga ggatggtggc cggggtctgc tcggccggtt tgagcaccac gcagttgccc 35100
gccgcaagcg ccggagcgag cttccatgcc gccatcagca gcgggaagtt ccacgggatg 35160
atctggccga cgacgccgag ggctcgtgg aagtgatagg cgacggtatc ggcgtcgatt 35220
tccgacatcg tgccttcctg cgcccgcagc acgcccgcga agtaacggaa gtggtcgatg 35280
gcgagcggca cgtcggcggc gcgggtctcg cggatcggct tgccgttgtc ccaggtctcg 35340
caggtggcga gaagctcggc attcgcctcc atccggtcgg cgatgcgcag cagcgcgccc 35400
gcgcgctcgg ccggggacat cttgtggccc cacagctctt tcgcggcatg ggcggcgtcg 35460
agcgcgtcat tcacgtcgct ctcgtccgag cgggcgcatt cgcagatctt cgcgccggtg 35520
agcggcgtga tgttgtcgaa atagcgccct gcgttcggcg ccttgaaggt gccgttgatg 35580
aagttgtcgt agcgcgccct gaaaggcgat accatcacgc cctgaaactg ggcagacata 35640
tccgtggcca tgggtctttc tcctcctctg gggcgggccc tccccggccc gcgactgagg 35700
aaagaatgcc tgcttcggat ccctccggca acgcttctgc ggcccgccgt cccggaactg 35760
agacagtccc tcgcgaaaaa acgcatatcc ggtatgcaac gatggaagtg cagcgcgggc 35820
gggccataga aggtcaggcg aattttccac aagtgattgt ttattaatga aattgagaaa 35880
gagcctgcat cccgcgccgc actctggccg ggcacgggac atgaaacggc cggccccgtt 35940
tcgggggccg gccatcgctt gcgtgctgtt tctgtgcctg gaagggctca gaaatgggcg 36000
tcgatcttca ccacgtcgat cagcttgtgg ttgacgaatt ccttcagccc gaggccgaag 36060
agctcgcgcc cgtagcccga gcgcttgatc ccgccgaagg gcagatccgc ctcgaccttg 36120
gtcgggtggt tgacgaagac catgcccgta gcgatctggc tggcgacttc gaaaccgcgc 36180
ttctcgtcct tggtgaagac cgagccgccc agcccgaagg gcgtgtcatt ggcgatgcgg 36240
atcgcgtcgg cttcgtcccg ggcacggaag atcatcgaca ccgggccgaa gaattcccaa 36300
agccgggcgg ggttgtcgcc gtccatgccg gtcaggatcg tcggctgcac gaaggcgccc 36360
ttgttcggca cggcgggacc gacttcctcg gccttcgcac cgtgcgcgat ggcggcggcg 36420
atctgcacct tcagatcgtc ggcggccttt tgcgacgaca gcggcgcaag cgtggtggcc 36480
gggtccatcg gatcgcccgc gacgagcttc gcgacgccct tcttgtagcc ctcgacgaac 36540
ttgtcataga cctcgtcgac cacgatcatg cgcttcgacg agacgcaaac ctgaccgccg 36600
ttccagtgcc ggccgaacac cgcccagtcg atggtcttgt cgagatcggc atccttcagc 36660
acgacgaaag catcggcacc gccgagttcc atcgtcgact tcttcagcgc cttgcctgcc 36720

```
     tcggccgcga cgacggcacc cgccgcctcc gacccggtca gcgccacgcc atgcacgcgc     36780
     gggtcgttga tgatcgtgtt gatctgatcg cgcgtcgcgt agaggttggt gaagcccccc     36840
     gccagcagac cggcttcggc catcagcttc tcgaagcgcg cggccgcctg cggcacgttc     36900
     gaggcatgtt tcaggatcag cacgttgccg gccgagagct gcggcgcaag gatccgcgcg     36960
     atctgataat aggggaagtt ccacggctcg atcgccagca ggatgccgag cggctcgtgc     37020
     accagcaccg cctcgccttc ctcgggatcg aggaccggca gcttctcggg ggcgagcagc     37080
     ttctcggcgt tgcggacgta atattcgaag atcttcgccg aaagctcgac ctcggccttg     37140
     gcctcggcga acagcttgcc catttcaagg gtcaggatcc gcgcatgttc atcggcatcg     37200
     cgacgcagca ggtcagcggc cttttgcagc accgccgcac gatgcgcgaa cgaggtcttg     37260
     cgccattcat ggaacgccgc atccgcccgg tcgagcgcat tcgccacttc ggcatcggtg     37320
     gcctcgggga aagtcgcgac ggtttcgccg gtgtagggat tgatcgttgc gtaggacatg     37380
     ttgttcctcc tttgcgtcgg ccggagcgag tgcgccggtc accctgagcg aagcatcctc     37440
     ggatcgcgat cctctcgggc ttcgt                                           37465
//
ID   XXX; XXX; linear; XXX; XXX; XXX;  XXX.
XX
AC   PRJEB67889
XX
DE   Uncultured bacterium DNA fragment from in-reactor enriched cow rumen, clone
ferm-23o13
XX
FH   Key             Location/Qualifiers
FH
FT   source          1..35397
FT                   /organism="uncultured bacterium"
FT                   /environmental_sample
FT                   /mol_type="genomic DNA"
FT                   /isolation_source="Cow rumen"
FT                   /clone="ferm-23o13"
FT                   /note="AZCL-xylan, AZCL-arabinoxylan active clone"
FT   CDS             347..652
FT                   /db_xref="SEED:fig|6666666.127244.peg.1"
FT                   /translation="MKKYAANTFGDMIKCVADVEKGLLAIDADLHADLERMMLENGSEQ
FT                   TSLWGFNLYPDEIGDDFIEYDSLINIRSWQGNPSRDVLDKDVRERIASIVSKFITE"
FT                   /product="hypothetical protein"
FT   CDS             652..981
FT                   /db_xref="SEED:fig|6666666.127244.peg.2"
FT                   /translation="MQHQVLENGRWAQLTFPQQMANIGSETSRIYRALEAGKDSRAESA
FT                   FARLQELVDLTIKYGRSDASPMLRSALLEELCRFRELYCKAYLERDLSELAAFNRYLDK
FT                   FAKLS"
FT                   /product="hypothetical protein"
FT   CDS             complement(1210..1545)
FT                   /db_xref="SEED:fig|6666666.127244.peg.3"
FT                   /translation="MSAKDHGIDHQHLIGNGGEHPHYDQDIQHLIYAEDNAGLGKRIPL
FT                   AFKIYLEGFRKSPRQREQDRRTAECDKIFRFYAERSQMRDKDNPPYDNGSHDKVQALFL
FT                   TSRTILL"
FT                   /product="hypothetical protein"
FT   CDS             1528..1725
FT                   /db_xref="SEED:fig|6666666.127244.peg.4"
FT                   /translation="MIFGAHLMPFSWLYKSRTYLVVSIVVPILSLGIGLLFPPAVLAAV
FT                   MVLIELLFSFLLYMECKAIQ"
FT                   /product="FIG00523589: hypothetical protein"
FT   CDS             2236..2790
FT                   /db_xref="SEED:fig|6666666.127244.peg.5"
```

```
FT                   /translation="MGIGSYFIRQVSPQDHWIWPFGIIPMPLEPAHYLQYVMMFVIGVL
FT                   AGRCGWLENMSNRVGIASLVLGALFAIGNYVRGDGSWNLFVWRWFGIYESLMCIFISFG
FT                   LLWLFREYANVSGRFLRWCSTQSYGAYIFHMILMLALQNATDGVWMGALGKFFFIGIVS
FT                   TVISFAFTGLVRMIPGVKKVI"
FT                   /product="hypothetical protein"
FT   CDS             2899..3219
FT                   /db_xref="SEED:fig|6666666.127244.peg.6"
FT                   /translation="MSNLNSALTEAIFYILLALQEPLHGYGIMQRTSTLSKGRLILSAG
FT                   TLYGAISNLLDKGWIMPAGESAENDGRRKMYLITDAGKAALVEELERLEELVKNGRNYI
FT                   KR"
FT                   /product="Transcriptional regulator, PadR family"
FT   CDS             3225..3794
FT                   /db_xref="SEED:fig|6666666.127244.peg.7"
FT                   /translation="MKTKKFALRWFVDFDKEEKWLNDMSKAGWCFWHTNGVIYRFKACA
FT                   PGEEIFQIDFDEKKGLSGEDYVAFRNTCGDTFVHQWKSKIYWKRAASSGPFEGEGNVAA
FT                   KLRLTNKAYNYHIKSLLGLTLIAAIAFLICVPVGKYMLPSGGFSKWLVDVGVGLTAGIL
FT                   FTEIVILLPIVKKLKKKMNMLISQLF"
FT                   /product="hypothetical protein"
FT   CDS             complement(3914..4777)
FT                   /db_xref="SEED:fig|6666666.127244.peg.8"
FT                   /translation="MKQEISFWLVTTEHLEDRIWFRDEDDFKVGMNYVAVQVAIGQVLL
FT                   LAFILMSNHVHFVLHGYYKDALAFVNGFKMRFAKYLRNKYGAKEFLRDNGVDIRALPSG
FT                   EEAVERAIAYVHMNSVAANICANASQYPWGSGDVLFNRTIQNGTRVDKMSNRERERILH
FT                   CSLDIPSHWILENGYILPSSYLSVKKAESIFRTPKRMNFFLSSSSKARKIKSVEDGIPS
FT                   FRDQIIAAAIPDLCMTLYGRSSLTGLSTEEMVEVLRQIRYRFSSNVNQIARVTGLSYAE
FT                   AARLLD"
FT                   /product="hypothetical protein"
FT   CDS             4886..5515
FT                   /db_xref="SEED:fig|6666666.127244.peg.9"
FT                   /translation="MKVEFGKMRRGEVRQAAELAARAFDDYEYFTNWFPEAESRHRVLF
FT                   SIIWHEYKTNFNVADYLVARQDGRLVAVAQLDAPTYRKPSDLRYLLHGWLKVYGTGDRK
FT                   TIDDWLAMDAAAGKPCHDYQKTGAGIWYASSLTVDPGAQGTGIGTRFLEYWEEYIRERG
FT                   GDQIVFFTNSRANLDFYTKRGYQVFDEREFGYNGKTIGSWSLRKKL"
FT                   /product="hypothetical protein"
FT   CDS             5584..7191
FT                   /db_xref="SEED:fig|6666666.127244.peg.10"
FT                   /db_xref="GO:0031176"
FT                   /translation="MLIPRIRVKILYFTIIVVTAVVLASGCDKKNSTPEKDTPFIDNHE
FT                   KASDAVRNMKLGWNIVNTLDATASYGMDSETSWGQPRVTRELIRAMADAGFGAIRVPVT
FT                   WYNHMDEVGNVDIQWMDRVEEVVGYVLDANLYCILNVHHDTGEGSQWLHADLENMDSIL
FT                   PKFDTLWRQISVRFASYGDRLIFEGWNEILDSKNSWNFARDKKAYDAIDTLASTFVTAV
FT                   RSTGGNNVGRNLIISTYAAASGGVSWAPTPDDVLEGFSLPEDSVEDHLIVEVHSYDPWL
FT                   WGADHGKWTASCEQTNLAMFSRLEKWFTSKGIPVIIGEFGPSGGDSHSIEDRAEQAKYM
FT                   RFFVHNARERGIGLFAWVNFMDGLDRMTLNWTFPDLAEAALKAWYGDGYSLPESGFTPS
FT                   AINVRFDNLWSEYCLYLGDGLDMREYKGFRLELAEAPAPGDFQIKMQTQAQFHGETTSP
FT                   DGQYPRIRNAVSEILFDKTCFDGDFLWNINLQACRSDVSASVKRMALIRNDGTEEEIPT
FT                   GLDGAHSGWGCTIVSVSR"
FT                   /product="GH5_4-CBM72"
FT   CDS             complement(7262..8719)
FT                   /db_xref="SEED:fig|6666666.127244.peg.11"
FT                   /translation="MKNVFKYLGIFLTVALAAGFTACDPTEENESAEAGLSIKVFFPTK
FT                   VIAGQEVTINGTGLSDVTEIIFPDNVSVKEFTHAGDGMIRVIAPAGIAAEGGNIIVRTE
FT                   DDEAVSKLPLTLGKVNVTGYSNRAGDTIVGGSPFTIFGKDLEFLKKVELLDEDGEPLLI
FT                   SDLAFDRKSTNSVSFTIPQNIYTGDIEGKVYACDGSVITMPLLSFKPAPTGHWETVKTL
```

```
FT                   IWKNDGTHGNINWSSEYRFAPESNSTGEEIYTVPQDIWDKMKETTFYMSFAEPGAQIRV
FT                   TTGWWSTTWTKDDIFVGNELITDNGDGTYSLAVTLAGSPILDVLDAQHLLFTGGGYTPL
FT                   EVYFSEQVWVENGSGPKEVVIWENDGSHGNINWSSEYRFAPESNSTGEEIYTVPQDIWD
FT                   NVIKKGTFYMQFAEPGAQVRVTTGWWSTTWTKDDIFVGNELITENEDGTYSLAVNLAGS
FT                   PILDVLDVQHLLFTGGGYTVLKLYYLE"
FT                   /product="Potassium uptake protein TrkH"
FT   CDS             complement(8750..10426)
FT                   /db_xref="SEED:fig|6666666.127244.peg.12"
FT                   /translation="MKLDKISLFILACVIPAACVDLDYTEENTRDEEWTYEYYENGIKN
FT                   MVFDVYAQVYNTEFDDNSAYFLAGATDEAQYALETGAVNNYVNGGWSAANPYSRTWTKS
FT                   YTAIADICMYLEKIEQTDISDWKYNSSYEIWVQQLELFPYELRFLRAYFYFELFKTYGD
FT                   VPLVTTTLTNAQANNISRNKSEEIVDFIVNEIDAVAPHLPLTYANIPGAEVGRATRIAA
FT                   LALKARTLLYAASPLYNKNNDTAKWAKAAEACKFILDNANAWGLKLSDYGALWGHDAFF
FT                   NPEIIFGLGRGESNSFEMANYPVGVENGSSGNCPTQSFVDQYEYQNNGETFAKRNKVEP
FT                   IDLSKNPYDGLDPRFALTVVKNGDQWPSNGSQKKVIETFTGGFNAAPKYGATPTGYYLR
FT                   KYVDGSCVTTADNQTSRRHTWIIFRLAEIYLDYAEAVFNATGSANDDTYGMTANDAINV
FT                   LRERPSIQMPEFTEDGEAWVERYERERLVELAFENHRFWDVRRWKKADKYFKTIQVASI
FT                   SSDLKLTRKAVTRKWDDKYYFYPIPQTEIKKNPNLTQNEGW"
FT                   /product="Putative outer membrane protein, probably
FT                   involved in nutrient binding"
FT   CDS             complement(10440..13223)
FT                   /db_xref="SEED:fig|6666666.127244.peg.13"
FT                   /translation="MKLRNILIYSLLSFCAAPFAANAQIQLSDKEAQKVDLGYGVESSE
FT                   FLTTAAAYTITAEELGRTSAISLADALYGKLLGLTAVQNTGFKGEEGKGASFSIRGAQT
FT                   TGEKDILILVDGVERPIDRLTVEEVESVTVLRDAAAVALYGHEGINGVLLVKTKRGIAE
FT                   SGFKIKAGASHKIQFDPQTADMVSAYDYAKALNIARRNDGGAVAYNDAELQKFADGSDP
FT                   FVYPDVNWKDVVFKNIAHETNGFVSMYGGSENVQYYSILDYTDARGLYKNPDLSEFNSQ
FT                   LKYSKMNIRTNVDFQVSSTTRVKTNILAVFMETNGVPGINSSDAWWHLYKVPALAFPIR
FT                   TSNGVWGGSQAYGDFNLLAKESAGGAKTHQRQIWADVTLEQDLDFLLKGLKFYVGGSYD
FT                   NASITDESHTRGYQYGWNYYDSANAMQEVVMGTVKKDATFNYELSSHWRTGTVNGGFRY
FT                   ATRLDNGDNISANANYNMKSEIRKGRSNTWYRENWNLAAHYDHANKFMADLVLAVNGSN
FT                   RSYPAKWAFSPTLGLGWIYANAPEAGFLNYGKLRLSSGIQHTDYVPENGLWLAAWDNSH
FT                   GQFWYGQNFASSTGAFLTQFPTTDFNQEAAYKANLGTDLRIANALDFTVDVFYQQRRNI
FT                   LVSAASLNSDVVGIQSSYDDKGRVASYGIETGLRYAKTFSNGLNLNFGGNFSFVKSEIL
FT                   EWIEAPAFPNLSRIGEPADASRGLVAIGFFQDQADIDNSPLQEFGQVKVGDIKYKDVNN
FT                   DNVINENDFVAMDHGNAFPALNYAFSLGIEYKGFGINATFQGAGCQTKNLLDVDGVWGA
FT                   LSDNRNLSQEYFDNCFDIAGANAKYPRLSTEKTANNTQQSTNWYRNVSWFKLRDCEVYY
FT                   KLPKSFLNSVKVADAKVFVQGQNLLSFDNIPAMDAENLNTGYPVMKAINVGLSVVF"
FT                   /product="TonB-dependent receptor"
FT   CDS             complement(13245..15311)
FT                   /db_xref="SEED:fig|6666666.127244.peg.14"
FT                   /translation="MNMKKILNYILGLGICVTALCMSSCVDRFAVGDDFLEKAPGVDVN
FT                   IDTVFSSAEFTRNFLWSAYGQLYCTYTSGNMMNGAPIDVLSDSYHCYVSWGGPIQSYYP
FT                   GLLTENAQDTDDGNLQGKFSYSTKNPASTADGRVSIYETVRKCWLLIENIDRVPDMSED
FT                   EKSRLRGEAYTIMASRYFDAFRNFGGLCLVKKAYAVGEPFTSGRSTAWETVEFIDSLIN
FT                   CAIIEPGYIWNIPNSDIGQWSGRLTRASARALRAKVWMFAASPLFNSNKPYMEYSPDKV
FT                   TEFTNIEHVWFGKEDRSLWDRCLKACNDFFEDNAANGNYYQLIQPESDDEGGYRMAFRR
FT                   AYRYRNDETHHEKIFDVNPTQLMSSSVVDGNINNGWGWGWAGFALDCYRQGGAVPTNEL
FT                   VECFGMADGTNFDVKYPDIYGPGKNPGGADIFADRDPRLYETVLVPRRNIPTDKVGDYN
FT                   GYNYADTWVNGAMYFNKDNFGDITSDDAHSGYRKFKWMLDYSSSKMDDEYIGVAYIRLA
FT                   EMYLIRAEAKAETGDLQGALADVNIVRARVGLGKIEDKNTDLHLGSNKENLIDEILRER
FT                   NCEIGSECGDRIYDMIRRKRQDIFTKPLHEIKIYRLGNDGKRLSLFNPDGTPADLSWDP
FT                   QTDWPKFEYEKPEITIGHRRWWDAGYWTNKWYLDPVSRDEIQKGYGLTQNPGW"
FT                   /product="FIG00405972: hypothetical protein"
```

```
FT   CDS             complement(15319..15717)
FT                   /db_xref="SEED:fig|6666666.127244.peg.15"
FT                   /translation="MAALRYFRDQFGSTNTSALTQWVADNSWTEDNPDAILPRISFTNK
FT                   GNNNLNSRAWMINSKYVRLKNVEIGYTINKPKSMPFFNYIRVFVSGQNLLTFADFKGND
FT                   PEAPGQGLDYGIRYPMTRVINIGAQVNF"
FT                   /product="TonB-dependent receptor"
FT   CDS             15733..16059
FT                   /db_xref="SEED:fig|6666666.127244.peg.16"
FT                   /db_xref="GO:0003700"
FT                   /translation="MAAYFQKQNISIGIATIYRQIEKFVAMGVVQKYFLGEQNAACFQY
FT                   MGEECHKEVSHFHLKCEKCGTLIHLECHDLEQLSSHLMAEHGFALDPFRTVFYGLCENC
FT                   RLAS"
FT                   /product="Ferric uptake regulation protein FUR"
FT   CDS             16096..17253
FT                   /db_xref="SEED:fig|6666666.127244.peg.17"
FT                   /translation="MKKNKKPVVLITGYLGSGKTTLLNNLLKQEKRKVALIVNDMGSIN
FT                   VDAEILKKNGSNVTECPMFELQNGCICCTLRDEFIQQIEKISNLDSIEVVFVEASGISD
FT                   PGAVSASFLAYEEDNPNTNVYLTSIVTVVDGDRIYREFLSELKQKKEERDSLAEKYDLS
FT                   DEEISTLIVDQIEFCNFIILNKCDLLSADQLKEVESIVRDFQPRAPIIHSVNGDIDIEK
FT                   IMTTKPFNYDQIDSSSAIQKATATLTQPGRKNNGCVDEYGITSFVFEARRPFNRTRFME
FT                   FVNNRYPSQLIRSKGYIWFSDASKDVQLFEQAGRNSSILPVSYWIDALREDVKQSYIAE
FT                   NPEIKENWDTQFGDRENQVVFIGKGYNKDDIVVELEKCLDERAYA"
FT                   /product="Putative metal chaperone, involved in Zn
FT                   homeostasis, GTPase of COG0523 family"
FT   CDS             17347..18252
FT                   /db_xref="SEED:fig|6666666.127244.peg.18"
FT                   /translation="MNYKQINLNDWELQGEGAFGESYFSKTDSSIMLKLMKPEGRKEDI
FT                   IAEFENSRKIASVGFKTPAAIELVKESQSGRLGIIYEKVQEKISFTRMIHDNPSDLPRI
FT                   AKIHAEEAKKFHGINCDPIQFLCYKETIRKAIPKLFTFKKYKDILKAAIELVPDSYGCL
FT                   QYDFQPGNIVHSNKTGENYWIDMGDFGYGHYLFDIAFLYMFTNILCTKKSVQQIFHMTE
FT                   QQLRDYWVEFTAAYFGRSIQPDEMFGKKIRLHLAMAVTIKYHMIPAPGPIKYLILNVML
FT                   RNALKGITLEDVADIMAAEV"
FT                   /product="aminoglycoside phosphotransferase"
FT   CDS             18567..21662
FT                   /db_xref="SEED:fig|6666666.127244.peg.19"
FT                   /translation="MEKKRYRLGIDLGTASIGTALVELDGPAEHPYPTNILHMGVRIFD
FT                   DGRDPKTKVPLGVARRMAKQNRKMLDRYLRRRKRLTDALIKFKIWPADETQRKSLCLLN
FT                   PYDIRTRALDEKISLEELGRAIFHLNQRRGFKSNRKTVGGEDGKIRQAVAKLEETFQKE
FT                   GCRTLGEFYQKRLEKGQSVRARLEGAGAQASYPFYPSRQMIECEFDKIWDRQVEFYPDI
FT                   LTEIAKKEIRATLLYQRPLKPAKPGKCFLEPSEDRAPIASPYFQRFRILQTINELRIIS
FT                   RYGDQRPLTLKERNILLEDLSQKKELKFSAAKKLLSLSVYDRFSIQDERRDKFQGDQIA
FT                   VLLGNKKYFGKNWKLLPLENQNAIVAELINQDDTEKLIKKLCAFGLSNESAAQLSEETL
FT                   PTGYGKLSLVALKKIVPFLEEAVITYSEAAAKAGYKHSDNYTGEIFHYLPYYGKVLENR
FT                   VMRGSGNEADNDEDRYGKVGNPTVHIALNQLRRVVNCVIKHFGSPEQIVIEVARDLKMT
FT                   QEEIKDYNKRQAENEKRNERIDNELKKLGQSINRENRERYKLWEELADNPMNRCCPYTG
FT                   KAIGIEKLFSEEVEIEHILPFAQTYEDSLANKTLSMRYANRLKQNRSPYEAFAEHSYKG
FT                   ILSYQEMLDKLPKDKAWENKRWRFEKGAMDRYNANGDFQARQLTDTAYASRLALEYLKF
FT                   VCPKDSVWAIPGKLTSMLRRQWGLNSILGRELPDGSFEKSRDHHGHHAIDGLVVAMTDR
FT                   AMLKKMSDATKINWEHQERILSHIDPPIVDIYDQAKERLSHLVVSHRKDHGLGGALHND
FT                   TAYGIIGEYKEGVVSTVRHKIPITSITGSNFNQIQNESIRYGIQELMANSSSDAALKDA
FT                   INEYASKTGIRRVTLVEKLSVVPIKNIKGQIYKAFKPDGNYCYDIWANGDKWEGKIVSR
FT                   FEANTKGAKKPQNLLMRIFVSDTLCCEINGKKTIYHVIGITDKKITMVEHFEAGNLRER
FT                   DRDKNDYLKYIVKSPNTLKPLKARIVHVDECGFVYDPGFKEK"
FT                   /product="CRISPR-associated protein, Csn1 family"
```

```
FT   CDS             21659..22579
FT                   /db_xref="SEED:fig|6666666.127244.peg.20"
FT                   /translation="MIGRILEIGTPGKFLKLFRGFCIIEEKGSELGRIALDDLESVILH
FT                   SPDGIISYQLLAEFVNRNIPVVFCNEKHMPIGIYWGLNGHYQGSKRINQQAQLSLPKRK
FT                   QIWQQIVSEKIKNQATILKAAKLDYLDLEHFANNVKSGDSDNREGIAAQAYWSRLFGSE
FT                   FRRNPDLDGLNAFLNYGYAILRASVTRGIVAVGLHPAFSLFHHNSEDPTPLANDLMEPW
FT                   RPIIDIAVYKLYLAERTELTPEEKKFLTQVLEWDLELNNQHSPLRICIQRLCNSFVDIC
FT                   EGTSSNLQFAKIPSIEEIQEEWSEV"
FT                   /product="CRISPR-associated protein Cas1"
FT   CDS             22612..22968
FT                   /db_xref="SEED:fig|6666666.127244.peg.21"
FT                   /translation="MVLFDLPVVEELDRKAATAFRQFLLDEGFEMAQFSVYMRHCSGKE
FT                   NAEKYYRHIEMNLPPGGKVHIITLTDKQFENIVRFQNNIQEPRKKIQTSYACSRFSARF
FT                   SGCLVVLKYVLSIS"
FT                   /product="CRISPR-associated protein Cas2"
FT   CDS             complement(23531..24682)
FT                   /db_xref="SEED:fig|6666666.127244.peg.22"
FT                   /translation="MQSSSSSEAQSSVAESSSSQSSSSSSEAPSSSSAQPIKIDFYNGA
FT                   DISEVQEYERKNTKFYDVDGKESDIFTILKNHGFNSIRLRTFVSPKAKYGYAASGCGHD
FT                   SEAYGDKDHVVAFAKKVKAAGMGLLVDIHYSDVWADPGKQIIPERWRNVNNANAMADSV
FT                   YAYTKDLMIALKNAGATPDMVQVGNETTPGILIHKPNSKTDCWGNGVDKAATSVNGDMG
FT                   TSAGKANAAKYFNAGIKAVKEVSPTTKTVLHIESIRKEETVKWWMGVIFDDYKIPADVM
FT                   GFSAYTAYGDGTPDKWKNLFNTITTKYSKLEFIVAEYNGGDSDNHYNFDKSRQKTRESV
FT                   REMNRWIGSFFWEPTIGGAWGAGLFDWRGNDVYANAKAFEEFF"
FT                   /product="GH53"
FT   CDS             25119..26456
FT                   /db_xref="SEED:fig|6666666.127244.peg.23"
FT                   /translation="MFNADLLTSAKVQDFIKLATKKKLDALQVSTQLAKEFSNEERAAI
FT                   MDYMALVPKFREKFGIENTAFLLCDKLALEQSTAQDIGRWKASLWPSGPEAVGKTVHDL
FT                   CCGMGGDSFFLPKELTAIGVDLDENRLAMYRYNSSVMRGVSPESSNAHTILGDVREVAT
FT                   HSNAASDAQHADYFTIDPARRAIEGENQRDLRNLTPTFEEVIEISKHYKGGMAKIPPGY
FT                   PICEIPRGTEILYIGSRTDCRECLVLFGELAQNPDHVRAVMVDKTGNEIANWTFARDEK
FT                   REARNESEQSQYDHNYELEGRDRVYRTSSSESDLPLGGISKFFAEPAPVLLRSHLFGVV
FT                   ALAHDETTHLISPGIAYVTSEKPLPAPAFANYEILESSEISTGTVRAMLKSHDIGKLTL
FT                   KLRGVKVDPDQEIKRLKPKGKNSATLFYTRLDGEKIAILANPVKNL"
FT                   /product="hypothetical protein"
FT   CDS             26473..27150
FT                   /db_xref="SEED:fig|6666666.127244.peg.24"
FT                   /translation="MSVKLEESWLKLLADQFEQPYFKQIKAKLLQEHAEHHVVYPPGPQ
FT                   IFAALDYCPVDKVKAVIIGQDPYHNPGQAHGLCFSVPFGIEPPPSLINIFQELHDDLGI
FT                   TPPPHGNLEGWAHQGILLLNASLTVRAHMAASHAGIGWQQFTDTIIQRLSQVRENLVFL
FT                   LWGSFAIKKQVLIAPNRGHLILTAPHPSPLSAYRGFFGCKHFSKANNYLISKGLEPIDW
FT                   SIK"
FT                   /product="Uracil-DNA glycosylase, family 1"
FT   CDS             27271..28407
FT                   /db_xref="SEED:fig|6666666.127244.peg.25"
FT                   /db_xref="GO:0006812"
FT                   /db_xref="GO:0008324"
FT                   /db_xref="GO:0016020"
FT                   /translation="MAKNCNEEKHNEENRNKVIVRTSVVGIVTNVILASVKAVIGLMAN
FT                   SIAVVLDAVNNLSDALSSIITIVGNKLSRKLPNEKHPLGYGRIEYLTAMVIASIVIYAG
FT                   ITSCVESIKKIIHPEEADYSTISLIIIAIAVVVKVILGRYVKSQGERVNSGSLIASGSD
FT                   ALFDAILSLSVLASALIFIFTGLSLEAYVGVLISIFIIKAGFEMLKDTVSDLLGKKNDT
FT                   ELTKQIKSLIRSEEGVHGAYDLVINDYGPEKKLASVHVELPDTMTVADLDFLTRKLEKK
```

```
FT                   VYRETGVILTAIGVYSYNTQNDESAKIRDTILDKVKSHDWALQMHGFYVDIEAKEMRFD
FT                   VVIKFGVNAQEALETIKSEVAQLYPDYKIQVSPDIDLG"
FT                   /product="Cobalt-zinc-cadmium resistance protein"
FT   CDS             28458..28784
FT                   /db_xref="SEED:fig|6666666.127244.peg.26"
FT                   /translation="MTHKNSLGRLLLFIVVGLILGGILSEALGALFGELGVLLNAGGYD
FT                   NIVHHFFTAGPDFSIGFSGDTPQPVVIDLYFVKFALGLGFKGNALGLVGMVLAIYIMKW
FT                   SGER"
FT                   /product="hypothetical protein"
FT   CDS             28784..30202
FT                   /db_xref="SEED:fig|6666666.127244.peg.27"
FT                   /translation="MQTIQELQAQLASGSTTAEKLAQASLEKIESTKNLNAYISVLNDR
FT                   ALERAKESDKRRTEGKSLGALDGIPVAVKDNMCLTGTRTTAASKILDNFVAPYTATALE
FT                   KLEAAGAIIVGKTNMDEFAMGSSNETSYYGPVKNPLDESRVPGGSSGGSAVAVASGTVP
FT                   CALGSDTGGSIRQPAACTGVVGLKPTYGRVSRYGLLAYASSLDQIGPFGATVRDCATLL
FT                   GAICGIDPHDNTTSTRPTEDFTAKLGTGVKGKVIGVPKEYFGEGLDAECKAAIENMLKK
FT                   MEAEGATIKEVSLPHISYAVSSYYIIATAEASSNLSRYDGVRYGYRSPEARKLFDLYAK
FT                   SRSEGFGKEVQRRILLGSYVLSAGFYDAYYVQAQKVRRLITDDFTEAFKTCDVIASPTM
FT                   PGLPLKCGMNESDPMAVYLSDIYTVSLNLAGLPGVSVPCGMAGGLPVGLQWIGKPFQEA
FT                   DLLAVAEATERLNK"
FT                   /product="Aspartyl-tRNA(Asn) amidotransferase subunit A
FT                   (EC 6.3.5.6) @ Glutamyl-tRNA(Gln) amidotransferase subunit
FT                   A (EC 6.3.5.7)"
FT                   /EC_number="6.3.5.6"
FT                   /EC_number="6.3.5.7"
FT   CDS             complement(30268..30957)
FT                   /db_xref="SEED:fig|6666666.127244.peg.28"
FT                   /translation="MLVENGGKGSVTTYGSITAKETSFGGACNYGQTNIQYYAAIHVNV
FT                   SPGDDKGPWDGGAACGGCVHVKAKTPDGWKEVTVRITDRCPDANCGVDLGGAPASDIMG
FT                   IQVGRYYGEWEFVSCEGVEGVWGDSTSIWVKEGASEFWSIIQVRNPKDAVRKIVFNELD
FT                   GDKSYTLEFVVGTENFWTVPQEILQSNKKYSVVVTYRSGTDDEWILESSELTIPEANLY
FT                   LYEHRKE"
FT                   /product="hypothetical protein"
FT   CDS             30989..31192
FT                   /db_xref="SEED:fig|6666666.127244.peg.29"
FT                   /translation="MQTPLSDDSAGFSEEEDCTGFSELELDDCATFSEEDSASFSELEL
FT                   SDFTELELAEIFAELLNGFADS"
FT                   /product="hypothetical protein"
FT   CDS             complement(31532..32989)
FT                   /db_xref="SEED:fig|6666666.127244.peg.30"
FT                   /db_xref="GO:0003938"
FT                   /translation="MKLLPEALTFDDVLLVPAESSVLPAQTDVSTQLAPNIKLNIPIIS
FT                   AAMDTVTTAPLAISLALQGGLGIIHKNMSIEDQAEEVRKVKRWQSGIVTNPVTLDADEP
FT                   VSAAFEQRARNKVSGFPILSKGKLVGMLTSRDLRTVSDMNVKISTVMTKNPVTASPKVS
FT                   LAKAKEILAEKRIEKLPLVDASGALKGLITMTDILKRENNPNASLDKNGQLLVGAAVST
FT                   SANTLERVAALVDAGVDLLIIDTAHGHHIGVRNMVKTVRKKYPKLTICAGNVCTPEAVE
FT                   ELAKCGANIVKVGIGPGSICTTRIIAGVGYPQFSAVVECGKMARKVGVKIIADGGLKFS
FT                   GDIVKALAAGGHAVMVGSLFAGTEEAPGEVILADGRSYKSYRGMGSLGAMKAGSADRYF
FT                   QGGVQEPRKFVPEGIEGRVPYKGPLRDTVYQLIGGIHSAMGYAGAANLEELYKKATFVR
FT                   ITGAGLRESHPHDVTITKEAPNYRTGD"
FT                   /product="Inosine-5'-monophosphate dehydrogenase (EC
FT                   1.1.1.205)"
FT                   /EC_number="1.1.1.205"
FT   CDS             complement(33261..34256)
```

```
FT                   /db_xref="SEED:fig|6666666.127244.peg.31"
FT                   /translation="MGLTAGVLAQDFCNAAKHTGASKKVNGNEVSSFNGVGYELWYDRA
FT                   TSGSLTIYEDGSMACSFQNAGDYLCRAGLSFDSDKKYNELGGDMLAEFKLVKQNINGAD
FT                   YSYVGVYGWMEEVSKSPSKLVEYYVVDNWLTGWRPGDWVASHKMGDYTIDGAKYSVYWG
FT                   EHTGPAIKTQGNTTFLQYFSIRENARDCGVINISAHMRQWEKLGMEMGKLYEAKVLGEA
FT                   GSMSNGVSGTADFPHIKVYVKGEEVPSSSSVALSSSSVASSSSKMVPSSSSEKVAKSSS
FT                   SVGGSSSSGTDAIPFTVQRFDKAGIYQVFDMQGNFLGKVELKSGLSLKQL"
FT                   /product="GH11"
FT                   /EC_number="3.2.1.8"
FT   CDS             complement(34327..35394)
FT                   /db_xref="SEED:fig|6666666.127244.peg.32"
FT                   /translation="MTVVFDRYKDSTVSAFIDDFAKNGKSVDDSTIYRIAKIAFNSGVV
FT                   TVSGDVYLPTELNRFRNSFDRWKSVYGTGFDMTSIAYHGMIPEFEEVVENFWRSEFGLE
FT                   KCDKEHLGTVGKTRFSEYYTCAYDNMVGMMRSSDYFWRDANSLEANSYGFKCDSADYFI
FT                   TPGGVNSSLIVCDFGSWRYALDVEKQVGLCTNENEGSLEDTDKYGYVCKSNMWEVFTDT
FT                   LEDERDGKNYAYSKAAGLYWMIDDLGDTTFAWAEVLCPEGWRLPTVDEWNNLQKKLGGW
FT                   RTPTKDGHQMNLLDGMPSFKMWWTSNESTENDSLAIAFAASYSPMDYQQAQYFSMMTRE
FT                   EDKNSKLGVRCVKEP"
FT                   /product="hypothetical protein"
XX
SQ   Sequence 35397 BP; 9483 A; 8758 C; 8343 G; 8813 T; 0 other;
     gcgtgatcaa tgtcaagaat cttgcttcgg gagagcagaa gcagttctct aaggacgata        60
     ttgcgggtat tctggcgttt gtcaggtagc ctcggacgct atccggtggc agattttcaa       120
     aaaaaatttg cagttaaaaa aataatgctt acctttgcag tcccaaaaca tcgcggggta       180
     gagcagttgg tagctcgtcg ggctcataac ccggaggccg gaggttcgag tccttccccc       240
     gctactatat atagaggatt tctgaaaggg aatcctcttt ttttgttata tttgcaaaaa       300
     caggtttacg atgattatct tggagcatcc tatatctaga gccgaattga aaaagtatgc       360
     cgccaacacg tttggagaca tgattaagtg tgtcgcagat gtggagaaag gcctgctggc       420
     cattgatgcg gatttgcacg ccgatttgga aagaatgatg cttgaaaacg gctcggagca       480
     aacctcattg tggggattta acctgtatcc agatgagatt ggggatgatt ttatcgagta       540
     cgattccctc attaacatcc gttcctggca gggcaatcct tccagagatg tgctggataa       600
     ggatgtgaga gagaggattg caagcatcgt tagcaagttc attactgaat aatgcagcat       660
     caagttcttg aaaatggccg ttgggcccaa ttgacgttcc cgcagcagat ggcaaacatc       720
     ggcagcgaga cttcccgtat ctatcgggcg ctggaggctg ggaaagattc cagggccgaa       780
     agtgctttcg cccgtcttca ggagctcgta gaccttacca taaaatatgg tcgttcagat       840
     gcttccccga tgcttcgttc agccttgcta gaggaactct gccgtttcag ggaactttac       900
     tgcaaagcgt accttgaaag agatctttcc gaactggccg ctttcaacag gtatctagac       960
     aagtttgcta aattatcata agcccttcat tgaagcggag gtcctataca ggagtgtatg      1020
     gtctactata gaggatttct gaaagggaat cctctatttt tgttataatt gctttatgga      1080
     aaacgataga atactgttgc ggccttggtt ggaaagcgac gccgaggctc tttataaatt      1140
     ttttgaaatc aacaggaaaa aattctatat ttgactgcta attaacagat aattatggaa      1200
     ctgagcgaat tacagaagga ttgtgcgaga cgtcagaaaa agggcctgca ctttatcatg      1260
     gcttccgttg tcatatgggg gattatcctt gtcgcgcatc tgactccttt cagcatagaa      1320
     acgaaaaatc ttatcacatt ctgcagttcg gcggtcttgt tccctttggc gtggggactt      1380
     gcgaaagccc tcaagataga ttttgaaggc aaggggaatc cgcttaccaa ggccggcatt      1440
     atcttcagca taaatcagat gctgtatatc ctgatcgtaa tgtgggtgtt cgccgccgtt      1500
     cccaataaga tgctgatggt ctatgccatg atctttggcg ctcatcttat gcctttctca      1560
     tggctctata aatccagaac atatctggta gtctccattg ttgtgcctat tctttctctc      1620
     ggcatcgggc tgcttttccc gccggctgtg cttgccgcgg tgatggtttt aatcgaactg      1680
     ctttttttcat tcctcctata tatggaatgt aaggcaatcc aataataatt tactaattat      1740
     aattaacgta ttatggaaac aaacgcaaat gcaaaccgac tgtcttatct tgacaatgta      1800
     aaggttttcc tgacggtcct ggtaatcttt caccacgcgg gtcaggctta tgggcaaggc      1860
     gggggctggg cgtatatgcc gagcaatcct gctgaaacgc taccttggat ctggcacttc      1920
     ttttcgacca atgccagctt tttcatggga ctatatttct tgatctctgg ctactttgtc      1980
```

ccgaggagct acgacagaca tggcggaagg actttcgtcc ggaagaaact cgtcaggcta 2040
ggtattcctc tgatcgtgat gggcggactt atgattgtat tgtcaggcaa attggagatt 2100
ggacatatgt ggttcgtgga aagcttgttg gtattctgcc tcttctatgc gctgatccgc 2160
tgtttcgttc cgccgatggg caatagccat gtacgacgga tcccgttttg gggactgttg 2220
gcgactggac tcgtgatggg tatcggcagc tatttcattc gtcaggtcag cccgcaggat 2280
cattggatat ggccgttcgg gatcattccg atgccgctcg agccggctca ttatcttcag 2340
tatgtaatga tgtttgtcat tggtgtgctg gcgggccgtt gcggatggct cgagaacatg 2400
agcaatcgtg tgggcatcgc gtcgcttgtc ctgggcgctc ttttcgccat cggcaactat 2460
gtccgtggcg acggttcatg gaaccttttc gtatggcgtt ggttcggcat ctatgagtcg 2520
ctgatgtgca tattcatttc gttcggcctg ctgtggctgt tccgcgaata tgccaatgtc 2580
agcggacgct tcttgcgctg gtgctccaca cagtcatacg gcgcctatat attccatatg 2640
atactgatgc tcgcgctgca gaatgctacg gacggtgttt ggatgggcgc gcttggaaag 2700
ttcttcttta tcggtatagt aagcacggtg atttcgtttg cattcacagg gcttgtacgt 2760
atgatcccgg gagtcaagaa ggtcatatag ctctttcaga cttagcacct tcaaattttt 2820
ttgaaaacag tttgtttttt caaatatatc gtctaccttt gtatcggcat acgatataac 2880
gttaaacaat atattatcat gagtaacttg aattcagctc tgacagaagc gatcttctat 2940
atcctcctgg ctctccagga gccgcttcat ggctatggaa tcatgcaaag gacctctaca 3000
ttaagtaagg gtcggcttat tctttcagca ggaacccttt acggcgcaat atccaatctt 3060
ctggacaagg gatggatcat gccggccggg gagtctgcag aaaatgatgg acgtcgtaag 3120
atgtacttga taacagatgc ggggaaagcc gctctggtcg aggaacttga gcgtcttgaa 3180
gaactggtta aaaatgggag aaactatatt aaacgataac ggttatgaaa acaaagaaat 3240
ttgcgctaag gtggtttgtc gattttgaca aggaggaaaa gtggctcaac gacatgagta 3300
aggccggatg gtgcttctgg catactaatg gggtgattta tagattcaag gcgtgtgctc 3360
ctggggaaga gatcttccag attgatttcg atgagaagaa gggcctgtct ggcgaagatt 3420
atgtcgcttt caggaatacc tgcggagata cattcgtcca ccagtggaag agcaagatct 3480
attggaagag ggcggcatcc tcgggtcctt tcgaaggaga aggaaacgtc gcagcaaaac 3540
tgcgccttac taacaaggcc tacaactatc acatcaagag tcttttggga ctcacgttga 3600
ttgccgcaat cgctttcctg atatgcgtcc cggtaggaaa atatatgttg ccgtcaggag 3660
ggttttccaa atggctcgta gatgtcgggg taggccttac cgcaggaata ctgtttacgg 3720
aaatcgtaat actccttcct attgttaaga aactcaagaa gaagatgaac atgctgattt 3780
ctcaactgtt ttgacatttt gtttttttctc agtttacatt agtgtgcggg tatcgtccat 3840
gttttccgga cgatgcccgc gtttgagccc gtttttgcgg ttatcgtccg tagtttcagg 3900
acgatgaccg cacctagtca agcaaccttg cagcttcagc gtaggacagc ccggtaaccc 3960
gggcaatctg attgacgttg gaagagaacc tgtaacgtat ctgtctgagc acttcgacca 4020
tttcttcagt agatagtccg gtcagagaag accgtccata tagtgtcata cacagatcag 4080
gtattgcagc ggctatgatc tgatccctaa aggaggggat accatcttcc acggatttga 4140
ttttgcgtgc ctttgaggaa gatgaaagaa agaaattcat ccttttaggc gtcctgaata 4200
tgctttccgc tttttttaacc gacagataag aagacgggag gatatatccg ttttcgagaa 4260
tccaatgtga cggaatatca agagaacagt gcaaaatccg ttcacgctcc ctattcgaca 4320
tcttgtccac acgagtacca ttctggattg tccggttgaa gagaacgtca ccgcttcccc 4380
aagggtattg cgaggcattg gcgcagatat tcgctgcgac actattcata tggacatatg 4440
cgatagctct ttcgacagct tcttctcctg aagggagagc ccggatatcg acgccgttat 4500
cccgcagaaa ttccttcgcg ccgtatttgt tacgtaaata cttggcaaaa cgcatcttga 4560
agccgttgac aaaagccaag gcatctttgt agtaaccgtg cagtacaaaa tgcacgtgat 4620
tagacataag aatgaaggcc agcaacaaca cctgacctat agccacctga acggccacat 4680
aattcattcc aaccttaaaa tcatcttcat ctcggaacca gatgcggtcc tctaagtgct 4740
cggtagtaac gagccagaaa gaaatctcct gcttcatatc tttttctcat cctgaaatga 4800
ctggacttga gacaaatata caaaaatagt tagtatattt acaaccattc gaagtgatta 4860
aaatgtcagg aaaaaagaaa gtgcgatgaa ggtggaattc ggaaagatga gacgtggtga 4920
ggtgcgtcag gcggcggagc tcgcggcaag ggcgttcgat gactacgagt atttcacgaa 4980
ctggttcccg gaggcggaga gccggcatcg tgtacttttc tccataatct ggcatgaata 5040
caagaccaac ttcaacgtcg ccgactacct cgtagccaga caggacgggc gtcttgtggc 5100
cgtagctcag ctggacgccc cgacctaccg gaaaccatcg gatctgcgat accttcttca 5160
cgggtggctc aaagtctatg gaaccggcga ccggaaaacc attgacgact ggctggccat 5220

ggatgctgca gccgggaagc cttgccatga ctatcagaaa acgggagccg gaatatggta 5280
tgccagctcc ctgaccgtgg acccgggggc gcaaggcacg ggaatcggaa cccggtttct 5340
ggaatactgg gaagaatata tccgtgaacg cggaggagac cagatagtct tcttcaccaa 5400
ctccagagcc aacctggatt tctacacaaa acgcggctac caggtcttcg acgagcgcga 5460
attcggatac aacggaaaga caatcggaag ctggagcctc agaaagaaac tgtaatatcg 5520
ctggtaagaa tttttttact atctttaccc cgtacctgtc aatctaactt agcttaacga 5580
gttatgctta taccacgtat cagagtaaag attctttact ttaccatcat cgtagtcaca 5640
gccgtcgtac ttgcttccgg atgtgataag aagaattcta ctcccgaaaa agataccccg 5700
tttatagata atcatgaaaa agcctccgat gcagtccgga atatgaagct cggctggaat 5760
atcgtcaaca ctctcgacgc cacggcctct tacggaatgg acagcgagac ttcatgggga 5820
cagccgcgtg tgacccgcga gctgataaga gccatggccg atgccggatt cggcgcaatc 5880
cgagtcccgg tgacgtggta caaccatatg gatgaagtcg gcaacgtcga catacagtgg 5940
atggaccggg tcgaagaagt cgtcggatac gtcctcgacg cgaatctcta ctgcatactt 6000
aacgtccatc atgataccgg agagggctct cagtggctgc atgccgacct cgaaaacatg 6060
gattcgatcc tcccgaaatt cgatactctc tggcgccaga tctcggtccg tttcgcctca 6120
tacggcgacc gcctgatctt cgaaggctgg aatgaaatcc tggactcgaa aaacagctgg 6180
aacttcgccc gcgacaagaa agcctatgat gctatcgaca ctctcgcatc gacctttgtc 6240
acggccgtac gctcgaccgg cggcaacaat gtcggcagga acctgatcat aagcacatat 6300
gccgccgctt ccggcggggt atcatgggca ccaacgcctg acgacgtact cgagggcttt 6360
tctctcccgg aggacagtgt cgaggaccat ctgatcgtcg aggtccattc ctacgaccca 6420
tggctatggg gcgcggacca cggcaaatgg acggcatcat gcgaacagac gaatctggcc 6480
atgttcagcc ggctcgaaaa atggttcacc tccaaaggaa tccccgtaat catagggggag 6540
ttcggtccta gcggcggcga cagtcatagc atcgaggacc gggcggagca ggcgaaatac 6600
atgcgattct ttgtccataa cgcccgcgag cgcggaatcg gcctgttcgc ctgggtcaat 6660
ttcatggacg gactggaccg gatgaccctt aactggacct ttccggatct tgccgaggcg 6720
gctctgaagg cgtggtatgg cgacggttat tctcttcctg agagcgggtt cacccсttct 6780
gccatcaatg tacgtttcga caatctatgg tcagaatact gcctgtatct cggagatggc 6840
ctcgacatga gagaatacaa gggattccgt cttgaacttg cagaagcgcc ggctccaggc 6900
gatttccaga taaagatgca gactcaagcc cagttccatg gcgagacgac gtccccggac 6960
ggccagtatc cacgaatccg caatgctgtc agcgaaatcc ttttcgacaa gacatgtttc 7020
gacggagact tcctctggaa cataaacctg caggcctgcc gcagcgatgt ctctgcttcc 7080
gtgaaacgca tggctcttat aaggaatgac gggacggagg aggaaatccc taccggactt 7140
gacggagccc acagcggctg gggctgcacg atagtttccg taagccgttg atatagcgga 7200
aaagataaaa aaatcaggct gcccgacaag gcagcctgat tcagtatagt agagtataag 7260
actactcgag gtaatagagt ttcagcacag tatatccgcc accggtaaag agcagatgct 7320
gaacatccag gacatccaga ataggactgc ctgcaagatt tacggcaagc gagtaggtcc 7380
cgtcctcatt ctcggtaatc agctcgtttc ctacgaagat gtcgtctttt gtccaggttg 7440
tgctccacca tccggtagtt acgcggacct gagcgcccgg ctctgcaaac tgcatgtaga 7500
aggtaccttt cttgataacg ttgtcccaga tatcctgagg aacggtatag atctcctcgc 7560
cggtagaatt gctctcagga gcaaagcggt attcgctgct ccagttgatg ttgccgtgag 7620
agccatcgtt ctcccagata actacctctt taggtcctga gccgttctca acccaaacct 7680
gctcagagaa ataaacctcc aaaggagtgt atccgccacc ggtaaagagc agatgctgag 7740
catccaggac atccagaata ggactgccag caagcgttac ggcaagcgag taggttccgt 7800
caccgttgtc tgtgataagc tcgtttccta cgaagatgtc gtcttttgtc caggtcgtgc 7860
tccaccatcc ggtagttacg cggatctgag cgcccggttc ggcgaatgac atgtagaacg 7920
tcgtctcttt catcttgtcc cagatatcct gaggaacggt atagatttcc tcgccggtag 7980
aattgctctc aggagcaaag cggtactcgc tgctccagtt gatgttgccg tgggtgccat 8040
cgttcttcca gatcaaggtc ttgacagtct cccagtgacc tgtaggggct ggcttaaaac 8100
tcaacagagg cattgtgatt acagatccgt cgcaggcata tactttaccc tcgatatcgc 8160
cggtataaat attctgcggg atcgtgaaag atacgctgtt cgtactctta cggtcgaaag 8220
ccaaatcgga aatcagaagg ggctctccgt cttcgtcgag aagctcgacc tttttgagga 8280
actcaaggtc ctttccaaag atagtgaacg ggctgccgcc gacgatggta tcgccggcac 8340
ggttagagta accggtaacg ttgacttttc cgagagtcag aggcagtttc gatacagcct 8400
cgtcatcttc ggtacggaca attatattac ctccttcggc agcgatgccg gcaggggcga 8460

ttacccttat cattccgtcg ccggcatggg tgaactcctt gacggaaaca ttgtcaggga 8520
atatgatttc cgtgacatcg ctgagaccgg ttccattgat agtcacctcc tggccggcaa 8580
tcactttggt cgggaagaac accttgatgc tcagtccggc ctcagctgat tcgttttctt 8640
cggtagggtc gcatgccgtg aaaccggctg caagggctac tgtcagaaat attcctagat 8700
atttaaatac attcttcata attgaaatcc ttttaaaata agtagaatac taccagcctt 8760
cattctgggt gaggttaggg tttttcttaa tttcagtctg agggatcgga tagaagtagt 8820
acttgtcatc ccatttacgg gtaacagcct ttctggtaag cttgagatca ctgctgatag 8880
aagcaacctg gatggtcttg aagtacttgt cagccttctt ccagcggcgt acatcccaga 8940
aacgatggtt ctcgaatgcg agttccacaa gacgctcgcg ctcgtagcgc tcaacccacg 9000
cctctccgtc ctctgtgaat tctggcatct ggatgctcgg acgttcgcga agtacgttga 9060
tagcatcgtt ggctgtcatt ccgtaggtat cgtcatttgc gcttccggta gcgttgaata 9120
cagcctcggc atagtcgaga tatatctcgg caagacggaa gatgatccat gtgtgacggc 9180
gtgaagtctg gttgtccgcg gttgtcacgc aggaaccgtc cacgtacttc ctgaggtaat 9240
aacctgtagg agtagcaccg tacttaggag ccgcattgaa gccgccggtg aaggtttcga 9300
tgaccttctt ctgtgaaccg ttgctcggcc actggtcgcc gttctttacg actgtaagag 9360
cgaatcgtgg atcgagaccg tcataaggat tcttgctgag gtctataggt tctactttgt 9420
ttctttttgc gaaggtttcg ccgttattct gatactcata ctggtctacg aagctctggg 9480
tagggcagtt gcctgaggaa ccgttctcga cgccgacagg atagttggcc atctcgaagg 9540
agttgctctc accacgtccg agtccgaata tgatttcagg gttgaagaat gcatcgtggc 9600
cccagagagc gccataatcg ctgagcttca agccccatgc gttggcgttg tccaggatga 9660
acttgcaggc ttcagccgct tttgcccact tggccgtgtc attattcttg ttataaagag 9720
gtgatgcggc atagaggagc gtgcgagcct tgagagcgag agcagcgata cgggtagcgc 9780
ggccgacttc tgcgccaggg atatttgcat aagtcagagg aagatgaggc gcgacagcat 9840
cgatttcgtt tacgatgaag tcgacgattt cttcagactt gttgcgggat atgttgttgg 9900
cctgagcgtt cgtaagagtg gtcgtcacca gaggaacgtc gccataagtc ttgaacagct 9960
cgaaatagaa gtatgccctg aggaaacgga gttcataagg gaacagttcc aactgctgaa 10020
cccagatttc atagctggag ttgtacttcc agtccgagat atcggtctgc tcgatcttct 10080
ccagatacat acatatgtcc gcaatagcag tataggactt cgtccatgta cgtgaatatg 10140
ggtttgctgc gctccagcct ccgtttacgt agttgtttac agccccggtc tcgagagcgt 10200
actgcgcctc gtcggttgca cctgcaagga agtaagcgct gttatcatcg aactcggtgt 10260
tgtagacctg ggcgtaaaca tcgaatacca tgttctttat accattctcg tagtattcgt 10320
aggtccattc ctcgtcacgt gtgttttcct cggtatagtc gaggtctacg cacgcagccg 10380
ggatgacgca ggcgagaatg aataaactta ttttatctag tttcataatt caatctttat 10440
tagaatacta ctgaaagacc cacgttgata gccttcatta ctggataacc ggtattgagg 10500
ttttctgcat ccattgcagg gatattgtcg aaagacagaa ggttctgtcc ctggacaaat 10560
accttggcat cggcaacctt gacgctgttg aggaatgact tcggaagctt gtagtagacc 10620
tcgcagtcac gaagcttgaa ccagcttacg ttgcggtacc agttggtaga ttgctgggtg 10680
ttgttggcgg tcttctcggt cgagagacga ggatacttgg cgttggctcc ggcaatgtcg 10740
aagcagttgt caaaatactc ctgggaaagg ttcctgttgt cagaaagggc accccatacg 10800
ccatcgacgt caaggagatt cttggtctgg cagccagcgc cctggaacgt agcgttgata 10860
ccgaatccct tatactctat accaagactg aaagcatagt tgagagcagg gaaggcattg 10920
ccgtgatcca tagcgacaaa gtcgttctcg ttgatgacat tgtcattgtt gacatccttg 10980
tacttgatgt caccgacctt gacctgtccg aattcctgca gagggctgtt gtcgatgtca 11040
gcctgatcct ggaagaatcc gatggcaaca agtccgcggg aagcgtcggc aggctctccg 11100
atcctggaaa ggttagggaa tgcaggggcc tcgatccact cgagaatctc gctcttgacg 11160
aatgagaagt ttccgccgaa gttgaggttg agcccgttgc tgaaggtctt cgcatagcgg 11220
agtccggtct cgataccgta gcttgccaca cgtcccttgt catcgtaaga agactggata 11280
cctacaacgt cagagttgag ggatgcggca cttacgagaa tattgcgtct ctgctggtag 11340
aatacatcga cggtgaagtc gagggcgttg gctattctca ggtcggtacc gaggttggcc 11400
ttgtaggcag cctcctgatt gaaatcagtg gtagggaact gggtaaggaa tgctccggtt 11460
gagcttgcga aattctggcc ataccagaac tgtccgtggg agttgtccca ggctgcgagc 11520
cacagtccgt tttcagggac atagtcggta tgctggatac cactggagag acggagctta 11580
ccgtaattca ggaagcctgc ctcaggggca ttggcataga tccagccgag tccgagagtt 11640
ggggagaacg cccattttgc aggataagaa cggttagaac cgttgactgc gagcacgagg 11700

tcagccatga atttgttggc atggtcgtag tgcgcagcca gattccagtt ctcgcggtac 11760
caagtattgc ttcgtccttt acggatttcg ctcttcatgt tgtagttggc gttggcagag 11820
atattgtcgc cattatcgag tctggtagca tatctgaatc cgccgttgac agttccggta 11880
cgccaatgtg aagaaagttc gtaattgaag gtcgcatcct tcttgacggt acccataaca 11940
acttcctgca ttgcgtttgc agagtcgtag tagttccatc catactgata tccgcgggtg 12000
tgagattcgt cggtgatgga agcgttgtca taagaaccgc ctacgtagaa cttcagtcct 12060
ttgagcagga agtcaagatc ttgctccaga gtcacatcag cccagatctg acgctggtga 12120
gtcttggcgc cgccggcact ttctttagcc agaaggttga aatctccgta tgcctggctg 12180
ccgccccaga ctccattgct ggtcctgata gggaaggcaa gtgcaggaac cttatagaga 12240
tgccaccatg cgtcgctgga gttgatgcca ggtacgccgt tggtttccat gaatactgcc 12300
agaatattag tctttactct ggttgtgctg gaaacctgga aatccacatt tgtgcggata 12360
ttcatcttgg aatatttcag ctgggagttg aactcgctca gatcaggatt cttgtaaaga 12420
ccgcgtgcgt ccgtatagtc aagtatgcta tagtactgca cgttttctga tccgccatac 12480
atggaaacga atccgttggt ttcatgggcg atattcttga agacaacgtc cttccagttt 12540
acgtcaggat aaacgaacgg gtcgctgccg tcagcgaatt tctggagttc tgcatcattg 12600
tatgcaactg caccgccgtc gttcctgcgg gcgatattga gggctttggc ataatcatat 12660
gcgcttacca tgtcggcagt ctgaggatcg aactgtatct tgtgggaagc tcccgccttg 12720
atcttgaaac cgctttcggc aattcctctc tttgtcttga caaggagtac tccgttgata 12780
ccttcatgac cataaagagc cacggcggca gcatctctga ggacagtgac gctttccacc 12840
tcttcgacag taagacggtc aatagggcgt tctacaccgt cgacaagaat aagaatatcc 12900
ttctcgcctg tagtctgggc tccacggatg ctgaatgacg cgcctttgcc ttcttcaccc 12960
ttgaaacctg tattctggac tgcagtaagt ccgagcaatt tgccatagag tgcatcggcc 13020
aggcttatgg cagaggtacg gccaagttcc tcggcggtaa tagtgtaagc tgccgcggta 13080
gttagaaatt ccgaactctc gacaccatag cctaaatcca ctttctgtgc ctccttatcg 13140
ctcaactgaa tctgagcgtt tgctgcaaaa ggagcggcac agaagctcaa aaggctgtat 13200
ataagtatat tacgtaattt cataatcgtg aaaatttctg tttattacca tccaggattc 13260
tgagtaagac catatccctt ctgaatttca tcacgggata ccgggtcgag ataccatttg 13320
ttcgtccagt aacctgcatc ccaccagcgg cgatgaccta tggttatctc aggcttttcg 13380
tattcgaatt taggccaatc agtctgagga tcccagctaa ggtcagcagg agtcccatcc 13440
ggattgaaca aagaaagacg cttgccgtcg ttgccaagac gatagatctt gatctcgtga 13500
agaggctttg tgaagatgtc ctgacgtttg cggcgtatca tatcatagat acggtcgcca 13560
cactcgctgc caatctcgca gttgcgttcg cggagaattt cgtcgataag gttctccttg 13620
ttggatccaa gatgaagatc cgtgttttta tcttctattt taccgagacc tacacgtgca 13680
cggacgatat ttacgtcggc aagagctccc tggagatctc cggtttctgc cttggcttcc 13740
gcacggatga gatacatctc ggcaagtcgt atataggcaa ctcctatata ctcgtcatcc 13800
atcttggacg aagaatagtc gagcatccat ttgaatttcc tgtagccgga atgagcatcg 13860
tcggaggtga tatcaccaaa gttgtccttg ttgaagtaca tagctccgtt gacccaggta 13920
tcggcataat tgtatccatt atagtctccg accttgtcag tagggatatt tctgcgtgga 13980
acaagcacgg tctcgtaaag acgagggtcg cggtcggcga agatatcggc tcctccagga 14040
ttcttcccag gaccatagat gtcagggtat ttgacatcga agttggtacc gtcggccatt 14100
ccgaagcatt ctacgagctc attggtagga acagcgccac cctgacgata gcagtcgagg 14160
gcgaagccgg cccatcccca gccccagccg ttgttgatgt ttccgtcaac tacggaactt 14220
gacataagct gagtcggatt gacatcgaat atcttctcgt ggtgagtctc gtcgttacgg 14280
taacggtatg cgcggcggaa agccatacgg tagccaccct catcatcgga ttcaggttgg 14340
ataagctggt aatagttgcc gttggcagcg ttgtcttcga agaaatcgtt gcatgctttc 14400
agacagcggt cccagaggct gcgatcttcc ttgccgaacc atacatgctc gatgttggtg 14460
aactcggtaa ccttgtcagg agaatattcc atataaggtt tattgctgtt gaacagaggt 14520
gatgccgcga acatccatac ttttgcacgg agtgcacgcg cagacgctct tgtaagtcga 14580
cctgaccact ggcctatgtc ggaattagga atattccaga tatagccagg ttcgatgatt 14640
gcgcagttga taagtgaatc gatgaattcg acagtctccc atgcggtaga acggccactt 14700
gtgaatggct ctcctactgc ataagccttc ttgacaaggc aaaggcctcc gaagttacgg 14760
aaagcatcga aataacggct ggccatgatg gtataggcct cgccgcgcag acggctcttt 14820
tcatcctcgg acatatccgg aactctgtcg atattctcta tcagcagcca gcatttacgc 14880
acggtttcgt agattgagac acgtccgtct gcagtggaag ccgggttctt tgtagaatag 14940

gagaatttgc cctgaaggtt accgtcatcg gtatcctgcg cattctcggt aagaagtcca 15000
gggtagtagc tctggatagg accaccccat gaaacatagc agtgataaga gtctgaaagc 15060
acgtcgatag gcgcgccatt catcatatta cctgaggtat aggtgcagta gagctgacca 15120
taggcactcc acaggaagtt cctggtaaat tcagcgcttg agaatactgt gtcgatattt 15180
acgtcgacac ccggggcctt ctcgaggaaa tcgtcaccga cggcaaagcg gtcgacgcag 15240
gacgacatgc agagagcggt gacacatatt cccagaccca gtatgtaatt caatatcttt 15300
ttcatattca caattctttt agaagttaac ctgggctcct atgttgatga cacgggtcat 15360
agggtaacgt ataccatagt cgagtccctg gccaggagct tcagggtcat ttcccttgaa 15420
gtcggcgaag gtcaacaggt tctgtccgga aacaaataca cggatgtaat tgaagaacgg 15480
catgctctta ggtttgttta tggtatagcc gatctccacg ttcttcagac ggacatactt 15540
cgaattgatc atccatgctc tggagttaag gttgttgtta cctttattcg tgaaggaaat 15600
tcttggaaga attgcatcag gattgtcttc cgtccaggag ttgtcagcga cccactgggt 15660
gagggctgaa gtgttggtgg aaccgaactg gtcgcggaaa taccgcaggg ccgccatttc 15720
acagcagaag acgtggcggc ttatttccag aagcagaaca tttctatcgg cattgcaact 15780
atctatcgcc agattgaaaa gtttgtggcg atgggcgttg ttcaaaagta tttccttggc 15840
gaacagaatg ccgcctgctt ccaatacatg ggcgaagaat gccacaaaga agtttcgcat 15900
tttcacctga agtgtgaaaa atgcggcacg ctcatccatc tcgaatgtca cgatcttgaa 15960
cagctgagtt cgcacctgat ggcagaacac ggttttgcac ttgatccatt ccgcactgtt 16020
ttttacggac tctgtgagaa ctgccgccta gcgagctaac aacataatta cataattttt 16080
taacaaagga ttcttatgaa aaagaataag aaaccggtag tactcatcac cgggtatttg 16140
gggtcgggta aaaccacgct tttgaacaat cttctcaagc aagaaaagag aaaagtcgcc 16200
ctcatcgtaa acgatatggg tagcatcaac gtcgatgccg aaatcttgaa aaagaacggt 16260
tcgaacgtca ccgaatgccc gatgttcgaa ttgcaaaacg gctgcatctg ctgcaccttg 16320
cgcgatgaat tcatccagca gattgaaaaa atttccaacc tcgattccat tgaagtcgtg 16380
tttgtcgaag catccggcat cagcgacccg ggcgcagtca gcgccagttt ccttgcttac 16440
gaagaagaca atccgaatac gaatgtctac ttgacttcca tcgtcacagt cgtcgatggc 16500
gatagaattt accgtgaatt ccttagcgaa ttaaagcaga aaaaagaaga aagagattcg 16560
ctcgccgaaa aatacgattt aagcgacgaa gaaatttcga cactgattgt agaccaaatt 16620
gaattctgca atttcatcat tttgaacaag tgcgacctac tcagcgcaga ccagctcaaa 16680
gaagtggaat ctatcgttcg cgatttccag cctcgcgctc cgattatcca ttccgtcaac 16740
ggcgatatcg acatcgaaaa aatcatgacg acaaagccgt tcaactatga tcaaattgat 16800
tcctcttcgg ccatccagaa ggctaccgca acattaaccc agccaggtcg caagaataat 16860
ggttgcgttg acgaatatgg gattacgtca ttcgttttcg aagccaggcg cccgttcaac 16920
agaacccgct ttatggaatt tgtcaacaac agatacccgt cacagctcat ccgttccaag 16980
ggctacattt ggttctcgga tgccagcaag gacgtgcaac ttttcgaaca ggcaggtcgt 17040
aattcctcga ttttgcccgt ttcgtactgg attgatgctc tgcgcgaaga cgttaagcaa 17100
tcatacatcg ccgaaaatcc agaaatcaaa gaaaactggg acactcaatt tggcgaccgc 17160
gagaaccagg tcgtgttcat cggcaaaggc tacaacaagg atgacatcgt tgtcgagctt 17220
gaaaaatgcc ttgacgaaag agcgtatgct taacgggtat cacaaaatat ttgtttaacc 17280
gcaaaaaaat cgaaaaaagg gcgaccagac aatcactgag tcgccttttt gtttatattc 17340
gtaattatga attacaaaca gatcaattta aatgactggg aattgcaagg ggaaggggct 17400
ttcggagaaa gctacttttc taaaacagat tcgagcatta tgctcaagct catgaaacct 17460
gaggggcgca aggaagacat tatcgcagag tttgaaaact ctagaaagat cgcttccgtc 17520
ggctttaaaa ctcctgcagc aatcgaacta gttaaagaaa gtcaatcggg acgccttggc 17580
ataatctatg agaaagtgca agaaaaaatt tcgttcacaa ggatgattca cgacaatcca 17640
agtgacctgc cccgcatcgc aaaaatccac gccgaagaag cgaaaaaatt tcatggcata 17700
aactgcgacc caatccaatt cttgtgctat aaagaaacca tacgcaaggc cattccaaag 17760
ctattcacat tcaagaagta caaagatatc ttgaaggccg ctatagagct agttcctgac 17820
agctatggat gcctacagta cgacttccaa cccggcaata tcgtacattc aaacaaaacc 17880
ggtgaaaatt actggataga catgggcgat ttcggctatg ggcattacct ttttgatata 17940
gcctttctgt acatgtttac caacattctt tgtaccaaaa agagcgtgca acaaatattt 18000
cacatgacag aacaacagtt gcgcgattac tgggtggaat ttacagccgc atattttggc 18060
agatcaattc aaccagacga aatgttcggg aaaaagattc ggcttcactt agccatggct 18120
gtgactataa aataccacat gatccctgca ccaggcccca taaagtatct cattttaaat 18180

gtcatgctta gaaacgccct caagggaatc actctggaag acgttgccga cataatggcg 18240
gcggaagttt aagaaaatct aaaaaaactg tcctagagtc aaaaattttg actctaggat 18300
ttcgcaataa tgttccaatt gaaatataaa attttcatca aatagaacaa taattaaatt 18360
ggtctattaa aaatgacatt atttgtcgtc taaaaatttt atataaaaaa aaagggaatt 18420
tgcataaaaa aacgagggtt cccttgcgag aaccctattt agtgctagtt tttcagctta 18480
ttactggcga aaaacctcat ttcctaagtc actaatgacc atattatagt atttttattt 18540
gagaaagtca atagaaggaa taagaaatgg agaaaaaacg atatcgttta ggaatcgatc 18600
ttggcacagc aagcattgga acagcccttg ttgaattaga tggaccagcc gagcacccat 18660
atccaaccaa tatccttcat atgggagtcc gcatctttga tgatggtcgt gaccccaaga 18720
caaaagttcc actcggtgtc gcccgtagaa tggccaagca gaaccgaaaa atgcttgacc 18780
gttatcttag gcgacgcaag cgccttaccg acgcacttat caaattcaaa atttggccag 18840
cagatgaaac acaacgaaag tctctctgtc ttttaaatcc gtacgatatt cggactcgtg 18900
ccctagatga aaagatttct ctcgaggaac tcggaagagc tattttccat ttaaaccaga 18960
gacgtggttt taaaagcaac agaaaaactg ttggcggtga agacggcaag attagacaag 19020
ccgtagccaa actcgaagaa accttccaaa aagaaggctg tcgaacatta ggcgaatttt 19080
atcagaaaag acttgaaaag ggacaatctg tccgagcaag actagaaggc gctggagcac 19140
aagcatctta tccattttat ccttcccgtc aaatgattga atgtgaattt gataaaatat 19200
gggatagaca agtagaattt tatccagaca ttttgacgga aatcgccaaa aaagagattc 19260
gagcaacact cctctatcag cgtccgctta aaccagccaa accgggggaaa tgctttttag 19320
aacccagcga agatcgcgct cccattgcaa gcccttattt tcaacgtttc aggattctgc 19380
aaacaatcaa tgaattgaga attatcagcc gatacggaga ccaaaggcca ctaaccctaa 19440
aagaacgcaa tattctttta gaagatcttt cgcagaaaaa agaacttaaa ttctccgctg 19500
ccaagaagct cctttctttg tccgtttacg acagattcag catccaagat gagcgccgag 19560
acaaattcca aggagaccaa attgcagttc tacttggcaa caagaaatat ttcggtaaaa 19620
attggaaatt attgccatta gaaaaccaaa atgcaattgt tgcagaactc atcaatcaag 19680
atgatactga aaagctaata aaaaaattgt gcgcctttgg actatccaat gaatcagcgg 19740
cgcagttgtc cgaggaaaca ctccccactg gatatggcaa actttctctc gttgcattaa 19800
agaaaatagt tcccttcctt gaagaagctg tcattacata ttccgaagct gcagcaaaag 19860
ctggctataa gcacagcgac aactatacag gcgaaatatt ccactaccta ccctactatg 19920
gtaaagttct agaaaataga gttatgcgcg gaagcggcaa cgaagccgac aacgacgagg 19980
accgttatgg gaaagtcggt aatccaacag ttcatatagc attaaaccaa ctccgtcgcg 20040
tagtaaattg cgtcataaag cattttggca gtccagaaca aatcgtcatt gaagtcgctc 20100
gcgatttgaa gatgacacaa gaagaaatta aggattacaa taagcgccaa gcagaaaatg 20160
aaaaacgaaa tgagcgcatc gataacgaac taaaaaaact aggtcaatcc atcaatcgag 20220
aaaacagaga acgttataag ctttgggaag aattagccga taatccaatg aatcgctgct 20280
gtccctatac aggcaaagcc atcggtatag aaaagctctt ttctgaagaa gtcgaaatag 20340
aacatattct cccctttgcg caaacatatg aggactcttt ggcaaataaa acgctttcca 20400
tgcgatatgc caacagactt aagcaaaacc gttctcctta cgaagcattt gcggagcatt 20460
cctataaagg gattttgtct taccaagaaa tgcttgataa acttccaaaa gacaaagctt 20520
gggaaaacaa acgttggcgt tttgaaaaag gagcgatgga cagatataac gcaaatggag 20580
acttccaagc aagacaatta acagatacag cttacgcttc ccgtttagcc cttgaatact 20640
taaaattcgt atgtccaaag gattccgttt gggctattcc tggaaaatta acatccatgc 20700
tcaggcgcca atggggttta aacagcattt taggccgaga acttcccgat ggttcatttg 20760
aaaaaagccg cgaccaccat gggcatcacg ccattgatgg acttgtcgtc gctatgaccg 20820
atcgtgctat gctgaaaaaa atgtctgacg caacgaaaat taattgggaa catcaggaaa 20880
ggattctttc tcatattgat cctcctattg ttgatattta cgatcaagca aaagaacgac 20940
tttctcattt ggtagtttct catcgcaaag accatggttt gggaggagcc ctccataatg 21000
atacagccta tggaataatc ggtgaataca aagaaggagt cgtttcaaca gttcgacaca 21060
aaataccaat cacatctata acaggctcaa atttcaacca aattcaaaat gaatctattc 21120
gttatggcat tcaagagcta atggcaaatt catcttcaga tgcggctctg aaagatgcta 21180
tcaacgaata cgcaagcaag acaggaatca gaagagttac ccttgttgaa aagctgtcgg 21240
ttgtgccaat taaaaacata aagggccaaa tatataaggc attcaagccc gatggtaatt 21300
actgctatga catttgggcg aatggagata aatgggaagg taagattgtc agcagatttg 21360
aagcaaatac aaaaggagca aagaagccac aaaatcttct aatgagaatt ttcgtaagtg 21420

acactctctg ttgtgagatt aatggtaaaa agaccattta tcatgttatt ggaataactg 21480
ataaaaagat tactatggtt gagcactttg aagcaggaaa tttaagggaa cgtgatcgcg 21540
ataaaaacga ctacctcaaa tacattgtaa aatcacccaa cactctaaaa ccactcaaag 21600
cccgaatcgt ccatgttgat gagtgtggct ttgtctatga ccctggattt aaggagaaat 21660
gattggtagg attctagaaa taggaactcc aggaaaattt ctaaagcttt tccgaggatt 21720
ctgtattatc gaagaaaaag gttctgagtt gggcagaata gcccttgatg acttagaatc 21780
tgttattcta cattctcctg atggaataat ttcataccaa ttactagcgg aatttgtaaa 21840
tcgaaatata cccgtcgttt tctgcaacga aaagcacatg ccaataggaa tctattgggg 21900
gcttaatggt cactatcaag gatctaaacg aatcaatcaa caggctcaat taagtttacc 21960
caaaagaaaa caaatatggc aacaaatcgt tagtgaaaaa ataaaaaacc aagcaacgat 22020
attaaaagcg gcaaaacttg actatctaga cctagagcat ttcgcaaaca acgtaaaatc 22080
tggagattca gataaccgag aaggaattgc agcacaagcc tactggtctc gactttttgg 22140
atctgaattt cgaaggaatc ctgatttaga tggtctaaac gcattcttaa attacggcta 22200
cgccatttta cgcgctagcg taactagagg tatcgttgca gtcggcctac atccagcatt 22260
ttccttattt catcataatt cagaagaccc gacaccttta gccaatgatc taatggaacc 22320
atggcgtccc attatagata ttgccgttta taaattatat ctagccgaaa gaacagaact 22380
aactccagaa gaaaaaaagt ttctgaccca agtcttagaa tgggatctag aattgaataa 22440
tcaacattcc cccctacgca tttgtattca acgtttgtgc aatagttttg ttgatatttg 22500
tgaagggact tcttcaaatc tccaatttgc caagataccc tccattgaag aaatccaaga 22560
agaatggtct gaagtttgaa acggctgagt ggatatcgaa ttttgtggtt gatggttctt 22620
tttgatttgc ctgttgtaga agaactcgac cgaaaagcgg ctacagcttt tcgccaattt 22680
ctgttagacg aaggttttga gatggcgcaa ttttcagtct acatgcgcca ctgttccggc 22740
aaagagaatg ccgaaaaata ttatcgccat atagaaatga acctacctcc aggaggaaag 22800
gttcatatta tcactttaac cgacaaacaa ttcgaaaaca ttgttcgttt tcagaacaat 22860
atccaagaac cacgcaaaaa aatccagacc agttatgcct gttctaggtt ttcggcccgt 22920
ttttctggct gtttagtcgt gctaaaatac gttttatcaa tatcttagcg cgaaactaat 22980
atagaaactt aggaaatgag gtatcgccag aacattatat gtgtccggac ggcacgggtc 23040
ttgaatatag aaacttagga aatgaggtat cgccagaact atctacaacg gcaacgtccc 23100
ggcgtctaga atatagaaac ttaggaaatg aggtatcgcc agaaccactt gcatgtcgtg 23160
tcaacgttct tccagaatat agaaacttag gaaatgaggt atcgccagaa cttcttccga 23220
gtcgcgcacc atgtagccgt taatatagaa acttaggaaa tgaggtatcg ccagaacgga 23280
tacgaaagcg tgggcgtggt cggtgcaaat atagaaactt aggaaatgag gtttcgccag 23340
aactgagggt gatggacgga gaaaacggga tgtaatatag aaacttagga aatgaagttt 23400
cgccagaact agtggttctg cgggttcttc tcgccgttca atatagaaac ttaggaaatg 23460
aggtttcgcc agaacgggtt ctggttacca ccgaaaaggc cgccaaatat agaaacttag 23520
gaattcattt ctagaaaaac tcttcaaaag ctttggcatt agcgtagacg tcgttgccac 23580
gccagtcaaa aagccctgcg ccccaggcac cgccaattgt cggttcccag aagaaggagc 23640
caatccagcg gttcatttcg cggacggatt cacgagtctt ttggcgggat ttgtcaaaat 23700
tgtagtggtt atcggaatca ccgccgttgt attccgcaac gataaattcc aattttgaat 23760
actttgtcgt aatcgtattg aacaaattct tccacttatc aggtgtgccg tcgccgtatg 23820
ccgtgtaagc ggagaagccc atcacgtcag cagggatttt atagtcatca aagatgacgc 23880
ccatccacca ctttacggtt tcttctttgc ggatgctttc gatgtggagc accgttttcg 23940
ttgtcggcga aacttccttg accgccttga tgcccgcatt gaaatacttg gcggcattcg 24000
ccttgcccgc cgaagtcccc atatcgccat tgacggaagt tgcagcctta tcgacgccat 24060
taccccagca atctgttttg ctgtttggct tgtggatcaa gatgcccgga gtcgtttcat 24120
tgcccacttg caccatgtcc ggcgttgccc ccgcattctt aagtgcaatc atcaaatcct 24180
tggtgtatgc gtacacggaa tccgccatcg cattggcatt attcacgtta cgccagcgtt 24240
cggggataat ctgcttgcca ggatccgccc acacatcgct gtaatggata tcaacgagaa 24300
gccccatgcc cgcggccttg actttcttgg caaaagcaac aacgtggtcc ttgtcgccat 24360
acgcttcgga atcgtggccg caaccgcttg ccgcatagcc gtactttgcc ttgggcgaaa 24420
caaacgtgcg caagcgaatg gagttaaatc cgtgattttt caggattgtg aagatgtcgc 24480
tttctttgcc atcgacatcg taaaactttg tgtttttttcg ttcgtattcc tggacttcgg 24540
aaatatcagc gccattgtaa aagtcaattt ttatgggttg agcggaagat gaacttggcg 24600
cttccgaaga agaactggat gattgcgatg acgaagattc tgccaccgat gactgagctt 24660

cggacgaaga ggaactctgc aacgtttcag atgaagtcgg ttctaccgca gacgatgtca 24720
ccaaagacga agacgattca gtccctgccg acaaagaagt ttccgaggac gaggaaccac 24780
ccacatcttg actagaggac gataaatttt caacagacga cgaggattct tgagccgagg 24840
actccggcac gctagaagat gattccggat caagtccgga atgacatgac gaagattctt 24900
gcacagacga agaactaaaa gcataaacag gtgacaatgc agtatcgtca ccacaagatt 24960
gcattagcaa agcaacagaa gcgcaagaca cgcacgccgc aagatttttc aaaaacttca 25020
taccaaccca gtccttttct tccttaaaaa tacatcttaa tcggcaatcc tcaaaatttt 25080
aaattgtaaa attcccgcat aaaaattaaa tttctagcat gttcaacgcg gatttactca 25140
cttcggctaa agttcaggat ttcatcaagc ttgcaaccaa gaaaaaattg gatgcattgc 25200
aagtttcgac gcaacttgcc aaggaattca gcaacgaaga acgagccgcc attatggact 25260
atatggctct cgtgccaaaa tttcgcgaaa aattcggaat cgagaataca gcgttcctcc 25320
tttgcgacaa attagcgttg gaacagagta cggcgcaaga tatcgggcgt tggaaagcaa 25380
gcttgtggcc aagtggaccc gaagccgtag gcaaaaccgt tcacgacctt tgctgtggca 25440
tgggtggcga ttccttcttt ttgccgaagg aactgactgc aattggcgtt gacctcgacg 25500
aaaaccgcct tgcaatgtac cgctacaaca gcagcgtgat gcgtggagtc tcgccagaat 25560
ccagcaacgc acacaccatt ctcggagatg tccgcgaagt cgcgacgcat tccaacgcag 25620
catcagacgc ccaacacgcc gactatttca cgattgaccc tgcccgccga gccatcgaag 25680
gcgaaaacca gcgcgatctg cgcaatctta cgcccacatt cgaagaagtc atcgagattt 25740
caaagcatta caaaggcggc atggctaaaa ttccgcctgg ctaccccatt tgcgaaattc 25800
cgcgaggcac agaaattctg tacatcggct cccgcacgga ctgccgcgaa tgcttagtgc 25860
tgttcggcga actcgcccaa aatccagacc acgtgcgcgc tgtgatggtc gataaaacag 25920
ggaatgaaat cgccaactgg acattcgccc gcgacgaaaa gcgcgaagcc cgcaacgaga 25980
gcgaacaatc gcaatacgac cacaactatg aactcgaagg ccgcgaccgc gtttaccgca 26040
catcgagcag cgaatccgac ttgcctcttg gcgggatttc aaagttcttc gcagaacccg 26100
caccggtgtt gctccgcagc cacctctttg gagtggtcgc cctcgctcac gacgaaacga 26160
cgcatctgat ttcgccgggc atcgcttacg tgacaagcga aaagccgctc cctgcccccg 26220
ctttcgccaa ctacgagatt ttggaatcgt ccgagatttc gacaggcacc gtccgcgcga 26280
tgctcaaatc gcacgacatc ggaaaactca cgctcaagct gcgaggcgtc aaagtcgacc 26340
ccgaccaaga aatcaagcgc ctcaagccca aaggcaaaaa cagcgcaaca ctattctaca 26400
cacgcctcga cggcgagaaa atcgcaatac tcgcaaatcc tgtaaagaac ctataaccca 26460
ataacagcta ttatgtccgt caaactcgaa gaatcttggc taaaactgct cgccgaccag 26520
ttcgaacagc cgtattttaa gcaaatcaag gcaaaacttt tgcaggaaca cgcggaacat 26580
cacgtggtgt accctcccgg accgcagatt tttgcagccc tcgactattg ccccgtcgat 26640
aaagtcaagg ccgtcatcat cggacaagac ccgtaccaca atccgggcca ggctcacggt 26700
ctttgcttct cggtgccgtt tggcatcgag ccgccgccgt ccctcatcaa cattttccag 26760
gaactccacg acgatctcgg tatcacgccg ccgccacacg ggaatttgga aggctgggct 26820
caccagggaa ttttgctgtt aaacgcatcg ctcacggtgc gcgcccacat ggcggcaagt 26880
cacgcaggca ttggctggca gcagttcacg gacacgatta tccagcgcct ttcgcaagtc 26940
cgcgaaaatc tcgtcttttt gctctggggc agtttcgcca tcaaaaagca agtgctcatc 27000
gcacccaatc gcggccacct gattctcacc gccccgcacc cgagcccgct ctcagcatac 27060
cgcggattct tcggttgcaa gcactttagc aaggctaata actatcttat cagcaagggg 27120
ctcgaaccca ttgactggag tattaaatag acgagagacg acaaatgcga aagccgagag 27180
tcgcgacaaa acttgttttg gcatgaccga ggttagcatt tgggattaga gcaaagcgat 27240
aatccaaaga aaaattagta ggagaatcat atggcaaaga actgcaatga agaaaagcac 27300
aacgaagaaa accgcaataa ggtcatcgtc agaactagcg ttgtcgggat tgtcaccaac 27360
gtcattctcg cgagcgtcaa ggcggttatc ggccttatgg caaactccat tgccgtcgtt 27420
ctcgatgccg tgaacaacct ctcggacgca ctctcgtcca tcatcacgat tgtcggcaac 27480
aagctctcgc gaaaattgcc caacgaaaag cacccgctcg gttacggccg tattgaatac 27540
ctgaccgcca tggtcatcgc ctccatcgtg atttacgcgg gcatcacctc ttgcgtggaa 27600
tccatcaaga aaatcatcca tcccgaagaa gccgattact ccaccatttc acttatcatc 27660
atcgccatcg ccgtcgtcgt gaaagtcatc ctcggacgtt atgtgaaatc gcaaggagaa 27720
cgtgtaaatt ccggttcgct catcgcctcc ggttccgacg cgctcttcga cgccatcctc 27780
tccctttccg ttctcgcctc ggcactcatc ttcattttca caggtctctc gctcgaagcc 27840
tacgttggcg tcctcatttc catattcatt atcaaagccg gctttgaaat gttgaaggac 27900

acagtgagcg acttgctcgg caagaaaaat gacaccgaac tcaccaagca aatcaagagc 27960
ctcatccgca gcgaagaagg cgtccacggt gcatacgacc tcgtcatcaa cgattacggt 28020
cctgaaaaga aactcgcctc cgtgcacgtg gaactccccg acaccatgac tgtcgccgac 28080
ctggatttcc tcacccgcaa actcgaaaag aaagtctacc gcgaaacagg cgtcatcctc 28140
accgctatcg gtgtttactc ttacaacacc caaaacgacg aatccgccaa aatccgcgac 28200
acgattttag acaaagtcaa gtcacacgat tgggcgctcc agatgcacgg tttttacgtg 28260
gacatcgaag ccaaggaaat gcgctttgac gtggtcatca aattcggcgt gaacgcccag 28320
gaagccctcg aaaccatcaa gagcgaagtt gcgcaactct accccgacta caagattcaa 28380
gtttcacccg atatcgattt gggttaatga gattatagac gagagacgag agacgagaga 28440
aatactatat ttgcggcatg actcacaaga actcactagg acgtctctta ctattcatcg 28500
ttgtcggcct tattttaggt ggcatcttga gcgaagctct cggagctttg tttggcgaac 28560
ttggagtcct cttgaacgca ggtggctacg ataacattgt acaccacttc tttacagcgg 28620
gtcccgactt cagtatcggt ttttctggcg atacaccgca gcccgtcgtc attgatttgt 28680
actttgtcaa atttgcactt ggtttaggct tcaaagggaa cgccttaggc cttgttggca 28740
tggttttagc tatttacatc atgaaatggt ccggagaaag ataatgcaga ctattcagga 28800
acttcaggct caacttgcta gcggcagcac cactgccgaa aagctcgctc aggcctccct 28860
cgaaaaaatt gaatctacaa agaatttgaa tgcctatatc tcggtgctga acgaccgcgc 28920
tctcgaacgc gctaaggaaa gcgacaagcg ccgcaccgaa ggcaagtctc tgggcgccct 28980
cgatggcatc ccggtcgccg tgaaggacaa catgtgcttg accggcactc gcaccacggc 29040
tgcctccaag attctcgata actttgttgc cccctacact gcaacagcac tcgaaaagct 29100
cgaagccgca ggagccatca tcgtcggcaa gacgaacatg gacgaattcg ccatgggttc 29160
cagcaacgaa acctcttact acggcccggt caagaatccg ctggacgaat cccgcgttcc 29220
gggaggttcc tcgggtggtt ccgcagtcgc cgtcgcctca ggcacggttc cctgcgctct 29280
cggttctgat acgggtggat ctatccgcca gccggctgcc tgcacaggcg ttgtcggttt 29340
gaagccgacc tacggccgcg tttcccgtta cggcctcctc gcctacgcaa gctctcttga 29400
ccagattggt ccgttcggtg caaccgttcg tgactgcgca acgctcctcg gtgctatttg 29460
cggcattgac ccgcacgaca acaccacgag cactcgcccg accgaagact ttaccgcaaa 29520
gctcggcacc ggcgttaagg gcaaggtcat cggcgttccg aaggaatact ttggcgaagg 29580
cctcgatgca gaatgcaagg ccgctatcga aaacatgctt aagaagatgg aagccgaagg 29640
cgccaccatc aaggaagtga gcctcccgca catcagctac gcagtttcga gctactacat 29700
cattgcaacg gcagaagcaa gctccaacct cagccgctac gacggcgtgc gttacggcta 29760
ccgcagtccc gaagctcgta agctcttcga cctttacgcc aagtcccgta gcgaaggttt 29820
cggcaaggaa gtgcagcgcc gcatcctcct cggaagctac gttctctccg ccggtttcta 29880
tgacgcttac tacgtgcagg cccagaaggt tcgccgcctc atcacggacg acttcaccga 29940
agcattcaag acttgcgacg tgattgcaag cccgacgatg ccgggtctcc cgctcaagtg 30000
cggcatgaac gaatccgatc cgatggccgt ttacctctcc gacatctaca ccgttagctt 30060
gaacctcgcc ggcctcccgg gcgtgagcgt tccgtgcggt atggcaggcg gactcccggt 30120
tggcctccag tggattggca agccgttcca ggaagcagac ttgctcgcag ttgccgaagc 30180
aaccgagcgc ttgaacaagt aaaaaaagcg cgactttaaa aagccgcgac tttagcaaaa 30240
aacgacgccc accaaggcgc cgttttttta ttccttcctg tgttcgtaca gataaaggtt 30300
cgcctcagga attgttaatt cggagctttc taaaatccat tcatcgtcag taccgctgcg 30360
atacgtaacg actacgctat attttttatt ggactgcaaa atttcttgcg gaacggtcca 30420
gaagttttcc gtccccacaa caaactcaag cgtataagac ttgtcaccat ccaactcatt 30480
gaatacaatt ttgcggacag catctttcgg attgcgcacc tggataatgc tccaaaattc 30540
gcttgctcct tccttcaccc agatagatgt agaatcgccc cacacgcctt cgacaccttc 30600
gcagctcacg aattcccatt caccatagta gcgaccgacc tggattccca tgatatcgct 30660
agcgggcgca ccgcccaaat ccacgccgca gtttgcatcc gggcaacggt ccgtaatgcg 30720
taccgtcact tctttccagc catccggcgt ttttgccttc acatgaacgc agccaccgca 30780
agcagcgccg ccatcccacg ggcctttatc atcccccggc gagacattca catggatagc 30840
cgcataatat tgaatattcg tctgtccgta attgcaggca ccaccgaaac tcgtttcttt 30900
cgcagtgatg ctcccgtaag tcgtcacgga acccttgccg ccattttcca caagcatcgg 30960
gtctgccgcg gtatagctat cgcagttcgt gcaaacaccg ctatccgacg attccgcagg 31020
cttctcggaa gaagaggatt gcacaggttt ctcggaactg gaacttgatg attgcgcgac 31080
cttttcagaa gaagattccg caagcttttc cgaacttgaa ctttcggact tcaccgagct 31140

ggaacttgca gaaattttcg cagagctgct caatggcttc gcggattctt gagaagacga 31200
agattccgcc aacatctcgg aagaagataa atcctcgccc atagcaaccg gcgcaccatc 31260
agtcgaatcg ccacatgcca caaaagcaaa cggcaacgca atcgaacctg cgagcaccgc 31320
cgccaacaac caactcaaac gcgacctttt attcataaaa aacgctacct tgctgtcatg 31380
cccgtcgatc ccgtcacttc gttccaggat gacagagcat ctccttttca ccaatatata 31440
tcaaaagacc cgcaacatca agttacgggt ctcttaaata atcatttcca aaaaggcgcg 31500
ctttcccgat taagtctcgg gcatttagcg attagtcgcc tgtgcggtag ttcggagctt 31560
ccttcgtgat ggtcacatcg tgcggatgag attcacggag gcctgcgccc gtgatacgga 31620
caaacgttgc cttcttgtag agttcttcaa ggttagcagc accggcataa cccatagcag 31680
agtggatacc gccaatgagc tggtaaacgg tgtcgcggag cggtcccttg tacggaacac 31740
gaccttcaat gccttccgga acgaacttgc gcggttcctg aacgccaccc tggaagtaac 31800
ggtcggcaga accggccttc atcgcaccga gggagcccat gccgcggtag ctcttgtagc 31860
tacggccatc agcgaggatg acttcgcccg gagcttcttc agtaccggca aacaaggagc 31920
ccaccatcac agcgtgaccg ccagcagcca aagccttgac gatatcgcca gagaacttga 31980
ggccaccgtc agcgataatc ttcacgccga ccttgcgagc catcttgccg cattccacga 32040
cagcggagaa ctgcgggtaa ccgacaccag caatgatacg ggtcgtgcag atggaacccg 32100
gaccgatacc caccttgacg atgttggcgc cgcacttggc gagttcttcg acagcttccg 32160
gcgtgcaaac gttaccggcg caaatcgtga gcttcggata tttcttgcga acggtcttca 32220
ccatgttacg cacaccgatg tggtggccat gagccgtatc gataatcaac aagtcaacgc 32280
ctgcgtccac gagggctgca acgcgttcaa gagtatttgc agaagtgcta acagcagcac 32340
caacgagcaa ctgaccattc ttgtcgaggc ttgcgttcgg gttgttttcg cgcttcaaga 32400
tgtccgtcat cgtgatgagg cccttcaaag caccagaagc gtccacgagc gggagctttt 32460
caatgcgctt ttcggcgagg atttccttcg ccttggcgag gctcaccttc gggcttgccg 32520
tcaccggatt cttggtcatg accgtgctga ttttcacgtt catgtcgcta acggtgcgga 32580
ggtcgcggct cgtgagcata ccgacgagct tacccttgga gagaatcggg aaaccgctca 32640
ccttgttgcg agcacgctgt tcaaaagcag cagacaccgg ttcatcggca tcaagcgtca 32700
ccggattggt cacaataccg gactgccagc gcttgacctt gcgaacttct tcggcctggt 32760
cttcaatgga catgttcttg tggataatgc cgaggccacc ctgcaaagca agggaaatag 32820
ccaacggagc cgtcgtaacg gtatccattg cagcactgat tataggaatg ttcagcttga 32880
tattcggggc gagctgggtg ctcacatccg tctgagccgg aagaacagaa gattcagcgg 32940
gaacgagtaa gacgtcatca aacgtcaaag cttctggcaa aagtttcata tagtacctct 33000
tttgcgcatt gaatatagta attttcggtt ttaaatccta gctctaaacc gctagaatta 33060
tgtaaaaagt ggttattcgt tagcgtttta aaatggggga agcctccccc tggttcgcac 33120
tacgtcgctc tccacccctt cgaacgggcg ctcagtcgcc gccccgtaac gccccggctt 33180
gtattagcgc tcgacagcaa tcgtcttggc aaaggcccca cgcttgacca gataaacgga 33240
cgaacgagca aacttgtttg ctacagctgc ttaagcgaca aaccactctt cagttcaacc 33300
ttgccgagga aatttccctg catatcgaac acttgataaa tccccgcctt gtcaaagcgc 33360
tgtaccgtaa acggaatcgc atcggtcccc gaactcgaag aaccgccaac actgctcgaa 33420
gatttcgcta ccttctccga tgacgaactc ggcaccattt ttgacgaact cgacgcaaca 33480
gaacttgaac ttaaagcaac ggaactggac gacggaactt cttcaccttt gacgtaaacc 33540
tttatatgcg gaaaatcggc cgtaccggaa acgccgttac tcatgcttcc ggcttcgcca 33600
aggaccttgg cttcgtagag ttttcccatc tccatgccca gcttttccca ctgtctcatg 33660
tgagcgctaa tattgatgac tccgcaatca cgggcatttt cacgaatgct gaaatactgc 33720
aaaaaagtcg tattgccctg tgtcttgatg gccggccccg tatgttcacc ccaataaacg 33780
gagtatttgg caccatcgat tgtgtaatcg cccatcttat ggctagcgac ccaatcgcca 33840
ggcctccacc ccgtcagcca gttatcgaca acataatatt cgaccagttt gctcggggat 33900
ttcgaaactt cttccatcca accgtagaca ccgacatagg agtaatcggc accattgata 33960
ttctgcttca cgagcttgaa ttccgcaagc atatcgccgc caagttcgtt atacttcttg 34020
tcgctatcga acgaaagccc cgcacggcaa aggtagtctc cagcattctg gaaagaacat 34080
gccatggatc cgtcttcata aattgtaagc gaacccgaag tagccctatc gtaccaaagc 34140
tcgtaaccga caccgttgaa cgagctcact tcgtttccat tcactttttt gcttgcacct 34200
gtatgtttag ctgcattgca aaaatcctgt gccaagaccc ccgccgtgag tcccatactc 34260
aaggcgataa taattttatt caccatacat acctccacca aaagtttaat ttcaatataa 34320
caacttttag ggttctttca cacagcggac gccaagcttg ctgtttttat cctcctcacg 34380

```
     ggtcatcatt gaaaaatact gggcctgctg atagtccatt ggactatatg aagcagcaaa     34440
     tgcaatggca agcgaatcgt tttccgtaga ttcattggaa gtccaccaca ttttaaagga     34500
     aggcatccca tcaaggagat tcatttgatg cccatctttt gtaggagtcc tccaaccgcc     34560
     caatttttc tgcaaattat tccattcatc cacagtcggg agcctccaac cctcagggca     34620
     tagcacctcc gcccacgcaa aggtggtatc gcccaaatcg tcaatcatcc aataaaggcc     34680
     agccgccttg ctataggcat agtttttgcc atcacgttca tcttcaagcg tatcggtaaa     34740
     cacttcccac atattcgact tgcagacata accatattta tcagtatctt caagcgagcc     34800
     ttcattttca tttgtacaca acccaacctg tttttcgaca tccagagcat aacgccatga     34860
     accaaaatcg cacacaatca aagaagaatt cacaccaccc ggcgttataa aataatccgc     34920
     gctatcgcat ttgaatccat aagaattagc ttcaagacta tttgcatcac gccaaaaata     34980
     atctgaactg cgcatcatac ccaccatatt atcgtacgcg cacgtataat attcgctaaa     35040
     tcgcgttttt ccaacagtcc ctagatgttc cttatcgcac ttttcaagac caaactcact     35100
     acgccaaaaa ttttcaacaa cctcttcaaa ttcgggaatc atgccgtggt aagcaatgga     35160
     tgtcatatcg aatccagtac cataaacgct tttccaacga tcaaaactat tgcggaagcg     35220
     attcagctca gtcggcaaat aaacatctcc cgacaccgtg acgacgccac tattaaaagc     35280
     aatttttgca atccgataaa ttgtagaatc atccaccgat ttaccgttct tggcaaaatc     35340
     atctataaaa gcgctgaccg tcgaatcctt atacctgtca aaaacaacgg tcaaccc        35397
//
ID   XXX; XXX; linear; XXX; XXX; XXX;  XXX.
XX
AC   PRJEB67889
XX
DE   Uncultured bacterium DNA fragment from in-reactor enriched cow rumen, clone
ferm-26i17
XX
FH   Key             Location/Qualifiers
FH
FT   source          1..41253
FT                   /organism="uncultured bacterium"
FT                   /environmental_sample
FT                   /mol_type="genomic DNA"
FT                   /isolation_source="Cow rumen"
FT                   /clone="ferm-26i17"
FT                   /note="ABTS active clone"
FT   CDS             33..506
FT                   /db_xref="SEED:fig|6666666.127245.peg.1"
FT                   /translation="MLPILQKEVVEKRGWVSEADVMDYYAISQCMPGIIAVNTSLFIGH
FT                   KIKGARGSVMAALGVVFPSLVIIMIIAAFISSFAHLPVVQNAFAGIRVCVFVLIVNAIM
FT                   KLGKKSVTDWVSGAICLAVLLLSFLTEVSPVLLVVASGLAGVLYMGRKGTEKA"
FT                   /product="Chromate transporter"
FT   CDS             503..1078
FT                   /db_xref="SEED:fig|6666666.127245.peg.2"
FT                   /translation="MIYLRLFIEFFQIGLFAIGGGLATLPFLEKLSQSTGWFTLADLTD
FT                   MIAISESTPGPIGINMATYAGFKAAGVPGSIIATIGEVLPCVILVLIVARFLKRFSENR
FT                   IVQAVFYGLRAASVGLIVAAGLSVLQISVLNVPLWRQSGQILDLIQWKGVVLAAALFFV
FT                   IKKWKPHPVVTIAASAVVGIVFRFAGVG"
FT                   /product="probable chromate transport protein, putative"
FT   CDS             1307..2515
FT                   /db_xref="SEED:fig|6666666.127245.peg.3"
FT                   /translation="MPCVWREHMKKTDSTLIVFTLMNFVLSMTAFVFSAILDKAAVDIG
FT                   VSVAEAGLLNTMFSLGAAIGVPVTLIVFRKIDRTALLKVMLLFAALLNLLLTASANFQT
FT                   MLLIRLLSGIAVNGYGVLAVAAVVALVSKERMGRSLAFLIAGNSLSLVVGIPLARALSD
FT                   MLNWRIIFLALSLLMAGCLVYFQLHLKPQRQDQPRLDFRSELELLKNPRVLLILLFTLI
FT                   MFLGYHGAYNYVAPYLMEMFPSAPQGMSVFLMLFGLGSFFGNYLGGLVSDRIGYARSML
```

```
FT                   LGAAFQTVVTLLMLLLRSSLAANVILVTLWVMSAWFMGLQLNAGIAGETNNRSSFLLSL
FT                   NGSAIQLGSAMGTSLAALLIANGLMRQLFCMSLLTALAVTLLQVYSLKTYHHKREDAAG
FT                   QAE"
FT                   /product="putative integral membrane transport protein"
FT   CDS             2560..3453
FT                   /db_xref="SEED:fig|6666666.127245.peg.4"
FT                   /translation="MEERSMNIGIGTYSFGGLASYLGLGPTLREQFGQIQALGFDSVEL
FT                   LPVDLEEDPKDILKWLEETGLKVVSVHAEPTEEMVKKLAAVGGKAVIWAGTPFNSKAEA
FT                   IEVARLLDSMAEMAAPYGIKVGYHNHSQEFYVDEGKALLEHLLDNGSKFYLQLDCGWAM
FT                   NAGMYPPYFIRKYRNRILSIHVKENDKVNGPGERPASRHAEPPAFAKAFANARSLPLEE
FT                   RKKMLEQFTSGASDDHSMDNQCRMGAPESNLDWKEIKKALDEQDFEAFWVVEREWFYDE
FT                   HDKCIREDCEWLRKNL"
FT                   /product="Sugar phosphate isomerases/epimerases"
FT   CDS             3802..3933
FT                   /db_xref="SEED:fig|6666666.127245.peg.5"
FT                   /translation="MGLHFTYSATGGVQFRPIRRSESEAKRRMAGSPPVDKLGLSTA"
FT                   /product="hypothetical protein"
FT   CDS             complement(4003..4836)
FT                   /db_xref="SEED:fig|6666666.127245.peg.6"
FT                   /translation="MKKQFWLGTFALVMLISACNNKPKDGFTDTYTSGVIAIAADESFR
FT                   PIIQEEIDVFEGTFALAGIVPKFTTEVNAINLLLKDSVRLAIATRTLSAEEMSSFHSRK
FT                   FFPREIKLATDGLALIINRANKDSLISLRDIKRMLTGEVKTWKEIYPSSSLKNIQVVFD
FT                   NKNSSTVRFAVDSICKGKKLSKDLKALNNNQEVIDFVAQNSHAIGVIGVNWLGNRSDTT
FT                   NLSFRNEIRVMSVSEDDIATKDNSYKPYQAYLFYGDYPLTRSIYILLNDPRNGWA"
FT                   /product="Putative phosphate ABC transporter, phosphate-
FT                   binding component"
FT   CDS             complement(4839..5654)
FT                   /db_xref="SEED:fig|6666666.127245.peg.7"
FT                   /db_xref="GO:0005381"
FT                   /db_xref="GO:0006826"
FT                   /translation="MAKIDLTSSEWCELIFNGKNKAYGAYKMRGDSSKRHNMAMIIVVV
FT                   IALIGFSIPTLIRLATPKQKEVMTEVTALSQLEEPEVKQDEMKKVEPVAPPPPALKSSI
FT                   KFTAPVIKKDSEVADEDEIKSQEELTQTKVSISIADVKGNDEANGKDIADLKQVVTQAP
FT                   VEEEKVFDMVEQMPTFPGGQAELLSFIGKNLKYPTIAQENGTQGRVICQFIVGKDGTVR
FT                   DVQVVKSLDPYCDKEAIRVIRSMPRWIPGKQNGKPVTVKYTVPITFRLQ"
FT                   /product="Ferric siderophore transport system, periplasmic
FT                   binding protein TonB"
FT   CDS             complement(5676..6332)
FT                   /db_xref="SEED:fig|6666666.127245.peg.8"
FT                   /db_xref="GO:0005215"
FT                   /db_xref="GO:0006810"
FT                   /db_xref="GO:0016020"
FT                   /translation="MSAEVQESGNKKKGGKGKQKKMTVRVDFTPMVDMNMLLITFFMLC
FT                   TSLSKPQTMEISMPSNDKDITEEQQTKVKASQAITLLLGSDDKLYYYEGEPDYKDYSSL
FT                   KETSYKPDGIRSLLLKRNKEAVRKVNELKQKKLNMEISEEEYTQQLSEIKNGKNTPVVI
FT                   IKATEKSKYVNLIDALDEMQICSIGKYVIVDITSGDEFLLKNFESRGGLSQNVAD"
FT                   /product="Biopolymer transport protein ExbD/TolR"
FT   CDS             complement(6347..6952)
FT                   /db_xref="SEED:fig|6666666.127245.peg.9"
FT                   /db_xref="GO:0005215"
FT                   /db_xref="GO:0006810"
FT                   /db_xref="GO:0016020"
FT                   /translation="MGRAKIKKKSTFIDMTAMSDVTVLLLTFFMLTSTFVKKEPVQVTT
FT                   PASVSEIKIPEKDILQILVNPDGKIFMSLDKQADLKATLEGMGQEYGITFSPEQIKKFM
```

```
FT                   VSTTFGVPIKSMKTYLDLPAEKQDAVLKNEGIPCDSVDNQFKSWVRNARLVNSELRIAI
FT                   KADESTPYSVIKKVMNSLQDLRENRYNLITSLKTTSEN"
FT                   /product="Biopolymer transport protein ExbD/TolR"
FT   CDS             complement(6978..7790)
FT                   /db_xref="SEED:fig|6666666.127245.peg.10"
FT                   /db_xref="GO:0006810"
FT                   /db_xref="GO:0008565"
FT                   /db_xref="GO:0016020"
FT                   /translation="METNQKKAYSFTGIRSAGLVIIFCFIIAVCIYQFLLGNPANFMNN
FT                   DPNNHPLPGNFLGTIYKGGFIVPIIQTLLLTVLALSIERYFALRSAFGKGSLVKFVKNI
FT                   KEALTAGDLKKAQAICDKQRGSVANVVTSTLQKYDEMERDATLSKDQKLLAIQKELEEA
FT                   TALELPMMEQNLPIIGTITTLGTLMGLLGTVIGMIRSFAALAAGGAADSMALSQGISEA
FT                   LINTAFGILTGALAVIAYNYYTNKIDKLTYSLDEVGFSIVQTFAATHK"
FT                   /product="MotA/TolQ/ExbB proton channel family protein"
FT   CDS             complement(7910..8311)
FT                   /db_xref="SEED:fig|6666666.127245.peg.11"
FT                   /translation="MLIALLVLPLSAGANTVCAVHACNEAAESFSSEDESASLYSSLNM
FT                   DMLYKTLTVSYADLSIKGVSESLLHANHYRARRAAEYLSAIRDLLLHLSIRENQLVTDR
FT                   SKLFYSNTSHHCLSGVCDYYVYLLRRILI"
FT                   /product="hypothetical protein"
FT   CDS             11678..11797
FT                   /db_xref="SEED:fig|6666666.127245.peg.12"
FT                   /translation="MLLYIMSKIVVGSKEPEVENNGFEPLTLCVQGRCSSQLS"
FT                   /product="hypothetical protein"
FT   CDS             14289..14420
FT                   /db_xref="SEED:fig|6666666.127245.peg.13"
FT                   /translation="MAYLILYKGKCKQKKKRVIAIVRRKARMEEKDSKQSGGIKKTK"
FT                   /product="hypothetical protein"
FT   CDS             complement(14832..16247)
FT                   /db_xref="SEED:fig|6666666.127245.peg.14"
FT                   /translation="MIFSSGFFLWLFTAFISVYVLLTQKNTARILFVTAFSYFFYYKSS
FT                   GVYFLLLALVTVSDFFLARRMNITASAVQRKLLVFFSLLINLGLLCYFKYSNFFGTILA
FT                   DITGGSFSPIDVFLPVGISFFTFQSLSYTIDVYRKQIEPLNHLMDYAFYVSFFPQLVAG
FT                   PIVRARDFIPQIRKPLMVTSEMFGRGVFLILCGLFKKSVISDYISVNFVERVFDAPSLY
FT                   SGVENLVAVYGYALQIYCDFSGYSDMAIGIALLLGFHFNLNFDSPYKSLSITEFWRRWH
FT                   ISLSSWLKDYLYISLGGNRKGKLRQYVNLIVTMFLGGLWHGASWNFVLWGVLHGLALAV
FT                   HKIWLELLVVSRLSIKIPNFVGAFFTFHFVCFCWIFFRNTSFENSLNMIRQIFVNFHVD
FT                   VLPQMIIGYWPVFVLMVIGFLLHFAPDNWEKACCSFITRIPLWAKALIIILVVFVIIQV
FT                   KSADIQPFIYFQF"
FT                   /product="Probable poly(beta-D-mannuronate) O-acetylase
FT                   (EC 2.3.1.-)"
FT   CDS             complement(16288..17133)
FT                   /db_xref="SEED:fig|6666666.127245.peg.15"
FT                   /translation="MSSQDKLPQKAARNLISRIPKPALKVKWPADTIPYTQLLPEIFRN
FT                   VKSNEIVDSTDVLEPVMKQLQEMRLDMEVDTFRILHIGDSHVRGHIYPQSAGLVLTQEF
FT                   HSLLYYDMGVNGATCLTFTHPARISAIVEAKPSLLILSFGTNESHGKGYKTNVHYNQMH
FT                   ELIELLKDSLPGVPMLLTSPPGSFERRRLSRRRSTYSPNPRTIMAAKTIREYASRNGLA
FT                   FWDLYTVVGGEQWACKNWDGAKLMRPDHVHYLPEGYTLQGKLLAQAIIKAYNNYVVNRH
FT                   "
FT                   /product="putative periplasmic protein"
FT   CDS             complement(17180..18439)
FT                   /db_xref="SEED:fig|6666666.127245.peg.16"
FT                   /translation="MLSDIRSSKTDLSVPDTLPLVAPKSKILVDTCKSGITCVEDFTDS
FT                   IGQGMALFYNALSKLDAGEKRIVRIAVFGDSFIEGDIFTADLREMLQRKFGGCGVGFIP
```

```
FT                   ITSMTSGYRPTVKHSFGGWNSHAATDTTYFNKKKQGISGHYFIADSAAFVELRGQAQYA
FT                   SLLDSCTCAYLLFANKAPVKVGIRLNREIEKLVDVPSSPSNGSLRQITLKGHIGSVRWN
FT                   IYKADSTVFYGAALEGSKGIVLDNFSLRGSSGVSLRYIPEQTLKAFRSKRPYDLIILQY
FT                   GLNVATEKGRNYDYYRIGMNKSIAHIKKCFPEAAILMLGVGDRSYKTEEGELKTMPGVK
FT                   NLIQYQKVIASESSVAFWNMFQAMGGEGAMVKMVHAKPSLANYDYTHINFRGGKYLAGL
FT                   LYESLMYGYEQHKRREDYAK"
FT                   /product="putative periplasmic protein"
FT   CDS             complement(18618..18911)
FT                   /db_xref="SEED:fig|6666666.127245.peg.17"
FT                   /translation="MKSVFITFDQAYYEQIIGTLDRLNCRGFSYFEQVKGRGSKTGEPH
FT                   FGSHAWPSMCSAIITMVDDNKVDPLLDALHKLDTRTEKLGLRAFVWNIEKTI"
FT                   /product="FIG00936253: hypothetical protein"
FT   CDS             complement(18908..22066)
FT                   /db_xref="SEED:fig|6666666.127245.peg.18"
FT                   /translation="MSLYEGAVKKPIMTSLCFLAVVIFGLFSLSKLPVDLYPDIDTNTI
FT                   MVMTTYQGASASDIENNVTRPLENVLNSVSNLKHITSKSSENISVITLEFEYGYDIDVL
FT                   TNDVRDKLDMVNSALPENVETPIIFKFSTDMIPIVLLSVQAEESQSALYKILDDNVVNP
FT                   LARIPGVGTVSISGAPKREIQVYCDPNKLEAYNLSIETISSIIGAENKNIPGGNFDIGN
FT                   ETYALRVEGEFNDTKQLENIVVGTRNEANIFLRDVAKVVDTVEERAQETYNNGQKGAMI
FT                   IVQKQSGANSVNISKEVLKALPKLQKSLPSDVKLGIIVNTSDNILNTIDSLTETVAYAL
FT                   LFVVVVVFLFLGRWRATLIISITIPLSLIASFIYLAITNNTINIISLSSLSIAIGMVVD
FT                   DAIVVLENVTTHIERGSAPKQAAVHGTNEVAISVIASTLTMIAVFFPLTMITGMTGVMF
FT                   RQLGWMMCTIMFISTVAALSLTPMLCSQLLKLEKKQTKSFQKFYAPIERILNGFDNWYA
FT                   HVLNKAVRHRMLVLISCIGLFLLSLLCANGISTEFFPTQDNARISVALELPIGSRKELA
FT                   QNLADRLTKQWRSKYKDEILVCNYTVGQADTDNAWAVMQNNGSHIISFNIRLSDPGDRK
FT                   KTLEQVCDEMRQDLKSYPELSKAQVILGGSNTGLTGQSTADFEIYGYDMSVTDSVAANL
FT                   KRQLYQIKGVSEVNISRSDYQPEYQVDFDREKLAMHGLNLSTAGTYLRNRVNGAIASQY
FT                   REDGDEYDIKVRYAPEFRTSLESLENIIIYNSKGEAVRVKELGKVVERFTPPTIERKDR
FT                   ERIITVSAVISGEALGDVVAAGNDVIDNMDIPAGVTVQVAGSFEDQQESFQDLGVLGVL
FT                   IVILVFIVMAAQFESLTYPFIIMFSLPFAFSGVLIALFTSQNTLNVMSMLGGIMLIGIV
FT                   VKNGIVLIDYISLCRERGQAVINSVVTAGRSRLRPVIMTTATTVLGMVPMAISTGQGSE
FT                   MWRPMAIAVIGGLSISTILTLILIPVLYCVFAGTGIKRQRKVMKRRRALSDYFQEHKSE
FT                   IVKK"
FT                   /product="Cobalt-zinc-cadmium resistance protein CzcA;
FT                   Cation efflux system protein CusA"
FT   CDS             complement(22079..23095)
FT                   /db_xref="SEED:fig|6666666.127245.peg.19"
FT                   /translation="MRKYIQLVSFMFILLMSACTESKDKKGVDVISEKPRVKLATVTAR
FT                   QVDQILEYTATVEAEVKNNIAPASPVRIDHIYVEVGDKVSKGQKLVQMDAASLKQLKLQ
FT                   LDNQEIEFRRLDELYKVGGVSKSEWDASKMSLDVKKTSYRNLLENTSLLSPISGIITAR
FT                   NYDNGDMYNGNTPVLVVEQIVPVKLLINISENYFSKIKKGSPVKVKFDVFEGEVFNGKI
FT                   SLIYPTINAATRTFPVELVLDNEEMKVRPGMFARVEVNFGSENHVVVPDLAIVKQAGSG
FT                   DRYVFVYEKGKVRYQKVELGRRMGTEYELISGVSNNSQVVVAGHTRLVNGTEVVIEK"
FT                   /product="Membrane fusion protein of RND family multidrug
FT                   efflux pump"
FT   CDS             complement(23130..24467)
FT                   /db_xref="SEED:fig|6666666.127245.peg.20"
FT                   /translation="MFLDKILLLIVLLALFTNSVEAQDSSLSQDKDTLVLTIESAMKIA
FT                   LSESPIVKISEKEVVLKKEINKEAYAALFPDINLSGTYSRTIKKQTMAMQFGEETTTIQ
FT                   VGRDNTYNGGMIISLPIFAPALYKTINLTKTDIDLATEKARFSKLDLINQIKKAFYQLL
FT                   LAQDSYSVLTRSYRQAEANFNIVNAKFIHGTVSEFDKIRAEVQMRNLKPSVVSAHNGVN
FT                   LAKLQLKVLMGIDANYELVINGGLKEYEMEMFISKENNVYYNLSNNSDLKQLDLSNKLL
FT                   QNTLKLQQTNFMPTLSASFNYLYMSMNDNFRISSYDWYPYSTFGLSLTIPLYKASNFNK
FT                   LKQTKIQIQQLALNRLNLERQLNMKISSYLDSMNASSEQVMSNKESVAQAEKGRQIASK
```

```
FT                   RYDVGKGTILELNDSELALTQAQLVYNQSIYDFLVAKADLELVLGE"
FT                   /product="Outer membrane efflux protein precursor"
FT   CDS             complement(24502..25134)
FT                   /db_xref="SEED:fig|6666666.127245.peg.21"
FT                   /translation="MYEKENNRTELKHKTVTLAMNAFATKGIKGVTMDDIAMGLGISKR
FT                   TLYEMFEDKEELLIACMLEHQREKNVFVNEVISNSVNVLEVILKCYKKSVEDYHKTNVR
FT                   FLEEIKKYPKVCELMNDHKKKNSNYVISFFRKGVEQGLFRADIKFDIVHDLLHEQIDML
FT                   IHSETFKKYSFLEVYESIVLIFVRGISTDKGIGILENFLIEYNAKSV"
FT                   /product="Transcriptional regulator, TetR family"
FT   CDS             complement(25263..25895)
FT                   /db_xref="SEED:fig|6666666.127245.peg.22"
FT                   /translation="MSMKRVLILLGFALFSVLIFAQNKFYLSVKDVYIHRGDSLSKVVL
FT                   EKRTKNQLMLTPGGDYKISASSPSTCKRIKGRYFAIEIDGNLYLNCRKLRFNRFRFGTC
FT                   YAPAMWVKDKIYFSALPIGPAAVSITSATNYNMGDVGRAIASSSLLTERVYYFIDPITY
FT                   KVEFVSRDVMSELLIDQPDLRAEYLSSHSELANVTGKYLKILAGVDQ"
FT                   /product="FIG00405371: hypothetical protein"
FT   CDS             complement(26423..26914)
FT                   /db_xref="SEED:fig|6666666.127245.peg.23"
FT                   /db_xref="GO:0003735"
FT                   /db_xref="GO:0005622"
FT                   /db_xref="GO:0005840"
FT                   /db_xref="GO:0006412"
FT                   /translation="MRHNKKFNHLGRTASHRAAMLSNMASSLIKHKRITTTVAKAKALK
FT                   KFVEPLITKSKNDTTNSRRVVFSNLQDKIAVTELFKEIATKIADRPGGYTRIIKTGNRL
FT                   GDNAEMCFIELVDYNDNMSKDIVAKKSPRTRRSKKVVNADSTDLAQDAQSTEENNASVE
FT                   "
FT                   /product="LSU ribosomal protein L17p"
FT   CDS             complement(26918..27910)
FT                   /db_xref="SEED:fig|6666666.127245.peg.24"
FT                   /db_xref="GO:0003899"
FT                   /translation="MAILAFQKPDKVLMLESDPKFGKFEFRPLEPGFGITVGNALRRIL
FT                   LSSLEGFAITTIRIEGVEHEFSSVPGVKEDVTNIILNLKQVRFKQVVEEFETEKVTITV
FT                   ENSSEFKAGDIGKYLTGFEVLNPELVICHLDSKATMQIDITINKGRGYVPADENREFCT
FT                   DVNVIPIDSIYTPIRNVKYLVENFRVEQKTDYEKLILEITTDGSINPKDALKEAAKILI
FT                   YHFMLFSDEKITLENTDIDGNEEFDEEVLHMRQLLKTKLVDMDLSVRALNCLKAADVET
FT                   LGDLVQFNKTDLLKFRNFGKKSLTELDDLLESLNLTFGTDISKYKLEKE"
FT                   /product="DNA-directed RNA polymerase alpha subunit (EC
FT                   2.7.7.6)"
FT                   /EC_number="2.7.7.6"
FT   CDS             complement(27922..28527)
FT                   /db_xref="SEED:fig|6666666.127245.peg.25"
FT                   /db_xref="GO:0003723"
FT                   /db_xref="GO:0003735"
FT                   /db_xref="GO:0005622"
FT                   /db_xref="GO:0005840"
FT                   /db_xref="GO:0006412"
FT                   /translation="MARYTGPKSRIARKFGEGIFGADKVLSKKNYPPGQHGNSRKRKTS
FT                   EYGIQLREKQKAKYTYGVLEKQFRNLFEKAETAKGITGEILLQLLEGRLDNVVFRLGIA
FT                   PSRAAARQLVSHKHITVDGSVVNIPSYAVKPGQVIGVRERSKSLEVIGNSLAGFNHSKY
FT                   PWLEWDDNAKVGKLLHIPERTDIPENIKEHLIVELYSK"
FT                   /product="SSU ribosomal protein S4p (S9e)"
FT   CDS             complement(28634..29023)
FT                   /db_xref="SEED:fig|6666666.127245.peg.26"
FT                   /db_xref="GO:0003735"
```

```
FT                   /db_xref="GO:0005622"
FT                   /db_xref="GO:0005840"
FT                   /db_xref="GO:0006412"
FT                   /translation="MAKKTVAAKKRNVKVDANGQLHVHSSFNNIIVSLANSEGQIISWS
FT                   SAGKMGFRGSKKNTPYAAQMAAQDCAKIAFDLGLRKVKAYVKGPGNGRESAIRTIHGAG
FT                   IEVTEIVDVTPLPHNGCRPPKRRRV"
FT                   /product="SSU ribosomal protein S11p (S14e)"
FT   CDS             complement(29035..29415)
FT                   /db_xref="SEED:fig|6666666.127245.peg.27"
FT                   /db_xref="GO:0003723"
FT                   /db_xref="GO:0003735"
FT                   /db_xref="GO:0005622"
FT                   /db_xref="GO:0005840"
FT                   /db_xref="GO:0006412"
FT                   /translation="MAIRIVGVDLPQNKRGEIALTYVYGIGRSSSAKILEKAGVDKNLK
FT                   VKDWSDDQAASIREIIGAEFKVEGDLRSEVQLNIKRLMDIGCYRGVRHRIGLPVRGQST
FT                   KNNARTRKGRKKTVANKKKATK"
FT                   /product="SSU ribosomal protein S13p (S18e)"
FT   CDS             complement(29449..29565)
FT                   /db_xref="SEED:fig|6666666.127245.peg.28"
FT                   /db_xref="GO:0003735"
FT                   /db_xref="GO:0005622"
FT                   /db_xref="GO:0005840"
FT                   /db_xref="GO:0006412"
FT                   /translation="MKVRASLKKRTSECKIVRRNGRLYVINKKNPKFKQRQG"
FT                   /product="LSU ribosomal protein L36p"
FT   CDS             complement(29574..29792)
FT                   /db_xref="SEED:fig|6666666.127245.peg.29"
FT                   /db_xref="GO:0003723"
FT                   /db_xref="GO:0003743"
FT                   /db_xref="GO:0006413"
FT                   /translation="MAKQSAIEQDGIIVEALSNAMFRVELDNGHEITAHISGKMRMHYI
FT                   KILPGDKVRVEMSPYDLSKGRIVFRYK"
FT                   /product="Translation initiation factor 1"
FT   CDS             complement(29807..30598)
FT                   /db_xref="SEED:fig|6666666.127245.peg.30"
FT                   /db_xref="GO:0004239"
FT                   /translation="MIFLKTEEEIEFLRESNLLVSKTLGYVASLIKPGVTTLELDKSAE
FT                   TYIRDHGGVPTFKGYPNNFGTPFPATLCTSINDQIVHGIPSGAVTLKEGDIISVDCGVR
FT                   LNGFCGDSAYTFGVGEINSNVKHLLKVTKQALLEGIDNAVVGKRIGDIGFAIQNLCENS
FT                   GYGVVREFVGHGIGRNMHEDPQVPNYGRKGYGPLLKTGMCIAIEPMITLGSRLIIMERD
FT                   GWTVRTKDCSYSAHFEHTVAITNKEADVLSTFDYIDEVLRI"
FT                   /product="Methionine aminopeptidase (EC 3.4.11.18)"
FT                   /EC_number="3.4.11.18"
FT   CDS             complement(30607..31956)
FT                   /db_xref="SEED:fig|6666666.127245.peg.31"
FT                   /db_xref="GO:0009306"
FT                   /db_xref="GO:0015450"
FT                   /db_xref="GO:0016020"
FT                   /translation="MRKAIETLKNIWKIEDLRQRILITILFVAIYRFGSYVVLPGINPG
FT                   MLTQLHQQTSEGLLALLNMFSGGAFSNASIFALGIMPYISASIVIQLLGIAVPYFQKMQ
FT                   REGESGRRKMNQYTRYLTILILLVQAPSYLLNLKMQAGPSLNASLDWTLFMFTSTIILA
FT                   AGSMFILWLGERITDKGIGNGISFIILVGIIARLPQSLFQELISRMSDKTGGLIMFLIE
FT                   IVFLLLVIAAAILLVQGTRKIPVQYAKKIVGNKQYGGARQYIPLKVNAAGVMPIIFAQA
```

```
FT                   IMFIPITFIGFSNANTSNSIIQAFTDHTSLWYNLVFAVMIILFTYFYTAITINPTQMAE
FT                   DMKRNNGFIPGIKPGKSTAEYIDDIMSRITFPGSFFLAIVAIMPAFAGLLGVKAEFAQF
FT                   FGGTSLLILVGVVLDTLQQIESHLLMRHYDGLLKSGRIKGRSASSTVNAY"
FT                   /product="Preprotein translocase secY subunit (TC
FT                   3.A.5.1.1)"
FT   CDS             complement(31961..32407)
FT                   /db_xref="SEED:fig|6666666.127245.peg.32"
FT                   /db_xref="GO:0003735"
FT                   /db_xref="GO:0005622"
FT                   /db_xref="GO:0005840"
FT                   /db_xref="GO:0006412"
FT                   /translation="MDLSNLKPAFGSTKTRKRVGRGSGSGLGGTSTRGHKGAKSRSGYS
FT                   KKIGFEGGQMPLQRRVPKFGFKNISRVEYKAINIDVVQKLAEKKNLTVVGFSDLVDAGF
FT                   VSANQLVKILGNGTISIKLEVQAHAFSKSAISAIEALDGNVVKL"
FT                   /product="LSU ribosomal protein L15p (L27Ae)"
FT   CDS             complement(32435..32611)
FT                   /db_xref="SEED:fig|6666666.127245.peg.33"
FT                   /db_xref="GO:0003735"
FT                   /db_xref="GO:0005622"
FT                   /db_xref="GO:0005840"
FT                   /db_xref="GO:0006412"
FT                   /translation="MSTIKIKQIKSRIGAPIDQKRTLDALGLRKLNRVVEHESTPSILG
FT                   MVSKVKHLVAIVK"
FT                   /product="LSU ribosomal protein L30p (L7e)"
FT   CDS             complement(32623..33144)
FT                   /db_xref="SEED:fig|6666666.127245.peg.34"
FT                   /db_xref="GO:0003735"
FT                   /db_xref="GO:0005622"
FT                   /db_xref="GO:0005840"
FT                   /db_xref="GO:0006412"
FT                   /translation="MIMAGINKVKVTNDLELKDRLVAINRVTKVTKGGRTFSFSAIVVV
FT                   GNEEGVIGWGLGKAGEVTAAIAKGVEAAKKNLTKVPLLKGTVPHEQVAKFGGAEVFIKP
FT                   ASHGTGVVAGGAMRAVLESVGITDVLAKSKGSSNPHNLVKATILALSEMRDARMVAQNR
FT                   GISVEKVFRG"
FT                   /product="SSU ribosomal protein S5p (S2e)"
FT   CDS             complement(33144..33488)
FT                   /db_xref="SEED:fig|6666666.127245.peg.35"
FT                   /db_xref="GO:0003735"
FT                   /db_xref="GO:0005622"
FT                   /db_xref="GO:0005840"
FT                   /db_xref="GO:0006412"
FT                   /translation="MTTKVERRTKIKYRVRSKISGTSDCPRMSVFRSNKQIYVQIIDDL
FT                   SGKTLASASSLSFDEKVTKKEQAAKVGEVIAKRAQEAGIMTVVFDRNGYLYHGRIKEVA
FT                   DAARNGGLNF"
FT                   /product="LSU ribosomal protein L18p (L5e)"
FT   CDS             complement(33511..34065)
FT                   /db_xref="SEED:fig|6666666.127245.peg.36"
FT                   /db_xref="GO:0003735"
FT                   /db_xref="GO:0005622"
FT                   /db_xref="GO:0005840"
FT                   /db_xref="GO:0006412"
FT                   /translation="MSRIGKLPITVPAGVTLEFKDNVITVKGPKGELRQYVNPVISVNL
FT                   DNNEITLTRSTDDKQERAYHGLYRSLINNMIIGVSVGYKKELELVGVGYRASNNGNIID
FT                   LSLGYTHNIFIQLPPEIKVETKSERNKNPLIILESCDKQLLGQVCSKIRSFRKPEPYKG
```

```
FT                   KGIKFVGEIIRRKSGKSAGAK"
FT                   /product="LSU ribosomal protein L6p (L9e)"
FT   CDS             complement(34083..34478)
FT                   /db_xref="SEED:fig|6666666.127245.peg.37"
FT                   /db_xref="GO:0003735"
FT                   /db_xref="GO:0005622"
FT                   /db_xref="GO:0005840"
FT                   /db_xref="GO:0006412"
FT                   /translation="MTDPIADYLTRLRNAISANHRTVEVPASNLKKEITKILFEKGYIL
FT                   NYKFVSDGPQGTIKIALKYDSVNKVNAIKKLIRVSSPGMRKYTGYKDMPRVINGLGIAI
FT                   ISTSRGVMTNKEASDLKVGGEVLCYVY"
FT                   /product="SSU ribosomal protein S8p (S15Ae)"
FT   CDS             complement(34531..34800)
FT                   /db_xref="SEED:fig|6666666.127245.peg.38"
FT                   /db_xref="GO:0003735"
FT                   /db_xref="GO:0005622"
FT                   /db_xref="GO:0005840"
FT                   /db_xref="GO:0006412"
FT                   /translation="MAKESMKAREVKRAKLVEKYAEKRALLKKEGNFEALQALPKNSSP
FT                   VRLHNRCKLTGRPKGYIRQFGVSRIQFREMASNGLIPGVKKASW"
FT                   /product="SSU ribosomal protein S14p (S29e) @ SSU
FT                   ribosomal protein S14p (S29e), zinc-independent"
FT   CDS             complement(34805..35362)
FT                   /db_xref="SEED:fig|6666666.127245.peg.39"
FT                   /db_xref="GO:0003735"
FT                   /db_xref="GO:0005622"
FT                   /db_xref="GO:0005840"
FT                   /db_xref="GO:0006412"
FT                   /translation="MSNTASLKKEYVERIAPELKSQFQYSSTMQIPMLKKIVINQGLGM
FT                   AVADKKIIEVAINEMTTITGQKAVATISRKDIANFKLRKKMPIGVMVTLRRERMYEFLE
FT                   RLVKVALPRIRDFKGIESKFDGKGNYTLGIQEQIIFPEINIDSITRILGMNITFVTTAK
FT                   TDEEGYALLKEFGLPFKNAKKD"
FT                   /product="LSU ribosomal protein L5p (L11e)"
FT   CDS             complement(35365..35682)
FT                   /db_xref="SEED:fig|6666666.127245.peg.40"
FT                   /translation="MNKLHIKKGDTVYVNSGDDKGKTGKVLKVLVKEGRAIVDGINLVS
FT                   KSAKPSAKYPQGGIIKQEASVHISNLNVVDAKTNKPTRIGRRKSSEGTLVRYSKKSGEE
FT                   I"
FT                   /product="LSU ribosomal protein L24p (L26e)"
FT   CDS             complement(35702..36067)
FT                   /db_xref="SEED:fig|6666666.127245.peg.41"
FT                   /db_xref="GO:0003735"
FT                   /db_xref="GO:0005622"
FT                   /db_xref="GO:0005840"
FT                   /db_xref="GO:0006412"
FT                   /translation="MIQAESRLVVCDNSGAREALCIRVLGGTGRRYASVGDIIVVSVKN
FT                   VIPSSDVKKGAVSKALIVRTKKEIRRADGSYIRFDDNACVLLNNAGEIRGSRIFGPVAR
FT                   ELRSTNMKVVSLAPEVL"
FT                   /product="LSU ribosomal protein L14p (L23e)"
FT   CDS             complement(36070..36327)
FT                   /db_xref="SEED:fig|6666666.127245.peg.42"
FT                   /db_xref="GO:0003735"
FT                   /db_xref="GO:0005622"
FT                   /db_xref="GO:0005840"
```

```
FT                   /db_xref="GO:0006412"
FT                   /translation="MEARNLRKERTGIVLGNKMDKTITVAAKFKEKHPIYGKFVSKTKK
FT                   YHAHDEKNECNVGDTVCIMETRPLSKTKRWRLVEIIERAK"
FT                   /product="SSU ribosomal protein S17p (S11e)"
FT   CDS             complement(36341..36535)
FT                   /db_xref="SEED:fig|6666666.127245.peg.43"
FT                   /db_xref="GO:0003735"
FT                   /db_xref="GO:0005622"
FT                   /db_xref="GO:0005840"
FT                   /db_xref="GO:0006412"
FT                   /translation="MKNSEIKELSTVDLVERIEAEKISYSQLVLNHSISPLDNPAKIKV
FT                   LRRTIARMKTELRDRKLNK"
FT                   /product="LSU ribosomal protein L29p (L35e)"
FT   CDS             complement(36541..36975)
FT                   /db_xref="SEED:fig|6666666.127245.peg.44"
FT                   /db_xref="GO:0003735"
FT                   /db_xref="GO:0005622"
FT                   /db_xref="GO:0005840"
FT                   /db_xref="GO:0006412"
FT                   /translation="MLQPKKTKFRRQQKGRMKGNAQRGNQLAFGSFGIKALQNKWITGR
FT                   QIEAARIAVTRYMQRQGQIWIRIFPDKPITKKPAEVRMGKGKGSPEGFVAPITPGRIII
FT                   EVEGVSYEIAKEALRLAAQKLPVTTKFIVRRDYDVQNQNA"
FT                   /product="LSU ribosomal protein L16p (L10e)"
FT   CDS             complement(36998..37723)
FT                   /db_xref="SEED:fig|6666666.127245.peg.45"
FT                   /db_xref="GO:0003676"
FT                   /db_xref="GO:0003735"
FT                   /db_xref="GO:0005622"
FT                   /db_xref="GO:0005840"
FT                   /db_xref="GO:0006412"
FT                   /translation="MGQKVNPISNRLGIIRGWDSNWYGGNDYGDSLLEDSKIRKYLNAR
FT                   LAKASISRIIIERTLKLVTITVCTARPGIIIGKGGQEVDKLKEELKKITDKDVQINIFE
FT                   VKRPELDAVIVANNIARQVEGKIAYRRAIKMAIANTMRMGAEGIKVQISGRLNGAEMAR
FT                   SEMYKEGRTPLHTFRADIDYCHAEALTKVGLLGIKVWICRGEVFGKRDLAPNFTQTKES
FT                   ARSTNNSGGKNFKRKKNNR"
FT                   /product="SSU ribosomal protein S3p (S3e)"
FT   CDS             complement(37735..38139)
FT                   /db_xref="SEED:fig|6666666.127245.peg.46"
FT                   /db_xref="GO:0003735"
FT                   /db_xref="GO:0005622"
FT                   /db_xref="GO:0005840"
FT                   /db_xref="GO:0006412"
FT                   /translation="MGTRKKISADKRKEALKATYFAKLQNVPTSPRKMRLVVDMIRGME
FT                   VNKALGVLRFSSKEASSRVEKLLRSAIANWEQKNERKAEGGELFVTKVFVDCGSTLKRM
FT                   RPAPQGRGYRIRKRSNHVTLFVDAKSNNED"
FT                   /product="LSU ribosomal protein L22p (L17e)"
FT   CDS             complement(38166..38435)
FT                   /db_xref="SEED:fig|6666666.127245.peg.47"
FT                   /db_xref="GO:0003735"
FT                   /db_xref="GO:0005622"
FT                   /db_xref="GO:0005840"
FT                   /db_xref="GO:0006412"
FT                   /translation="MSRSLKKGPYINVKLEGKVLAMNESGKKAVVKTWARASMISPDFV
FT                   GHTVAVHNGNKFIPVYVTENMVGHKLGEFAPTRTFRGHAGNKKK"
```

```
FT                   /product="SSU ribosomal protein S19p (S15e)"
FT   CDS             complement(38455..39279)
FT                   /db_xref="SEED:fig|6666666.127245.peg.48"
FT                   /db_xref="GO:0003735"
FT                   /db_xref="GO:0005622"
FT                   /db_xref="GO:0005840"
FT                   /db_xref="GO:0006412"
FT                   /translation="MAVRKFKPTTPGQRHKIIGTFDDITAQVPEKSLVYGKKSSGGRNN
FT                   VGKMTMRYIGGGHKKVIRVVDFKRNKDGVPAVVKTIEYDPNRSARIALLYYADGEKRYI
FT                   IAPNGLQVGVTLLSGADAAPEIGNTLPLQNIPVGTVIHNIELRPGQGAAFVRSAGNFAQ
FT                   LTSREGKYCVIKLPSGEVRQILSTCKATIGSVGNSDHALESSGKAGRSRWMGRRPRNRG
FT                   VVMNPVDHPMGGGEGRASGGHPRSRKGLYAKGLKTRAPKKQSSKYIIERRKK"
FT                   /product="LSU ribosomal protein L2p (L8e)"
FT   CDS             complement(39281..39571)
FT                   /db_xref="SEED:fig|6666666.127245.peg.49"
FT                   /translation="MGIIIKPLVTEKMTAITDKLNRFGFIVRPEANKLEIKREVEMHYN
FT                   VTVVDVNTTRYAGKTKSRYTRAGIIKGRTNAFKKAIVTLKDGDSIDFYSNI"
FT                   /product="LSU ribosomal protein L23p (L23Ae)"
FT   CDS             complement(39584..40210)
FT                   /db_xref="SEED:fig|6666666.127245.peg.50"
FT                   /db_xref="GO:0003735"
FT                   /db_xref="GO:0005622"
FT                   /db_xref="GO:0005840"
FT                   /db_xref="GO:0006412"
FT                   /translation="MEVNVYNINGEATGRKVTLNESIFGIEPNDHAIYLDVKQFMANQR
FT                   QGTHKSKERSEISGSTRKIGRQKGGGGARRGDMNSPVLVGGGRVFGPRPRDYFFKLNKK
FT                   VKALARKSALSYKAKYDAIIVLEDFTFEVPKTREFVSLTKKLNISNKKLLMILLETNKN
FT                   VYLSARNIAGASVQTTSGLNTYRVLNSDVVVFTESSLSYFDKVSI"
FT                   /product="LSU ribosomal protein L4p (L1e)"
FT   CDS             complement(40210..40827)
FT                   /db_xref="SEED:fig|6666666.127245.peg.51"
FT                   /db_xref="GO:0003735"
FT                   /db_xref="GO:0005622"
FT                   /db_xref="GO:0005840"
FT                   /db_xref="GO:0006412"
FT                   /translation="MPGLLGKKIGMTSVFSADGKNVPCTVIEAGPCVVTQVKTSKVDGY
FT                   EALQLGFQEKKEKHTTKPLMGHFKRAGVTPKRHLAEFKDFDQVFNLGDTINVSLFSDED
FT                   FVDVIGTSKGKGFQGVVKRHGFGGVGQSTHGQHNRARKPGSIGACSYPAKVFKGMRMGG
FT                   QLGGDRVTVQNLKVLKVIEEHNLLLIKGSVPGCKGSIVIIEK"
FT                   /product="LSU ribosomal protein L3p (L3e)"
FT   CDS             complement(40848..41153)
FT                   /db_xref="SEED:fig|6666666.127245.peg.52"
FT                   /db_xref="GO:0003735"
FT                   /db_xref="GO:0005622"
FT                   /db_xref="GO:0005840"
FT                   /db_xref="GO:0006412"
FT                   /translation="MSQKIRIKLKSYDHNLVDKSAEKIVKTVKSTGAIVSGPIPLPTHK
FT                   RIFTVNRSTFVNKKSREQFELSSYKRLIDIYSSTAKTVDALMKLELPSGVEVEIKV"
FT                   /product="SSU ribosomal protein S10p (S20e)"
XX
SQ   Sequence 41253 BP; 12303 A; 9297 C; 7001 G; 12652 T; 0 other;
     tgggcgggct gaccttcggc gggggctatg ccatgctgcc catcctgcaa aaagaagtgg        60
     tggagaagcg gggctgggtc agcgaggccg acgtgatgga ttattacgcc atcagccagt       120
     gcatgcccgg catcatcgcc gtcaacacct cgctgttcat cgggcacaag atcaagggcg       180
```

cacgcggcag cgtcatggcg gccctcggcg tcgtgttccc ttcgctggtc atcatcatga 240
tcatcgcggc cttcatcagc agcttcgccc atctgcccgt cgtgcaaaac gccttcgcgg 300
gtatccgcgt gtgtgtattc gtattgattg tcaatgccat tatgaagctg ggtaaaaagt 360
cggtgaccga ctgggtcagc ggcgccatct gcctggcggt gctgctgctc tctttcctga 420
cggaggtcag ccccgtgctg ctggtcgtgg cctccggcct tgccggcgtg ctgtatatgg 480
gccgaaaggg gactgagaag gcatgattta tctgcgcctg tttattgagt tctttcagat 540
cggccttttc gccatcggcg gcggccttgc caccctgccc ttccttgaga aactgtccca 600
aagcacgggt tggttcacgt tggccgattt gaccgacatg atcgccattt ccgagtcgac 660
cccggcccg atcggcatca atatggccac ctacgcgggc ttcaaggcgg ccggcgtgcc 720
gggcagcatc atcgccacca tcggcgaggt gcttccctgc gtcattctgg tgctgatcgt 780
ggcgcgcttc ctgaaacgct tcagcgagaa ccgcatcgtg caggctgtct tctacggcct 840
gcgcgcggct tcggtaggcc tgatcgtggc cgcaggcctc agtgtgctgc agatttccgt 900
gctcaacgtg cccctgtggc gccagtccgg tcagattctc gacctgatcc agtggaaggg 960
cgttgtgctg gccgccgcgc tgtttttcgt catcaagaag tggaaacccc accccgtggt 1020
gaccatcgcc gcctccgccg tggtgggcat cgtgttccgg tttgccggcg tcggctgagc 1080
cgcgaatccg tttgccttct ttaaaatcct gtctctgctg tcgaaacaga gacaggattt 1140
ttgatgttct gttaaagaaa atgcgtagag catatcttct atcaatctat atgcctgaaa 1200
aattgctttt cctgttttct tttcatgaag gactgtccct ttttcgtctg gatagggtat 1260
actggggaca gataagggat cgtattttga cgaatgattt gatggcatgc catgtgtctg 1320
gagagagcat atgaagaaaa ccgactccac gctgattgtg tttactctga tgaattttgt 1380
cctgtctatg accgcctttg ttttcagcgc catactggat aaggcggccg tggatatagg 1440
cgtcagcgtg gcggaggcag gtttgctcaa taccatgttt tcacttggcg cggcgatcgg 1500
tgtgcccgtc acgctgattg ttttccgcaa aatcgacagg accgcgctcc tcaaggtcat 1560
gctgttgttc gctgccctgc tgaatctcct gctgacggcg agcgcgaatt ttcagacaat 1620
gctgctgatt cgcctgctgt ccggcatcgc ggtcaacggc tacggggtgc tggcggtcgc 1680
tgcggtggtg gcgctcgttt cgaaggaacg tatggggcgc tctctggcct ttttgattgc 1740
cggcaattct ctgtcccttg tcgtgggcat accgcttgca cgggcgcttt ccgatatgct 1800
gaactggcgc atcattttc ttgcgctcag cctgctgatg gcaggctgcc ttgtgtattt 1860
tcagctgcat ctgaagccgc agcggcagga tcagccgagg ctggattttc gcagcgagct 1920
ggagctgctc aaaaacccgc gcgtgctcct gattcttctg tttaccctca tcatgtttct 1980
gggctatcac ggagcctaca attatgtagc cccttatctg atggaaatgt tcccctcggc 2040
tccgcagggg atgagcgtct ttctgatgct ctttggtctt ggcagcttct ttggcaatta 2100
tctgggcggg ctggtgtcgg accgcatcgg ctacgcccga tccatgctgc tgggcgccgc 2160
ttttcaaacg gttgttaccc tgctgatgct tttgctgcgt tcgagccttg ctgcaaacgt 2220
cattctcgtg acgctgtggg taatgagcgc gtggtttatg ggctgcagc tgaacgcggg 2280
cattgccggg gagaccaata accgctccag ctttctctta agcctcaacg gttcggccat 2340
ccagctgggc tccgctatgg gcacaagcct ggccgcgttg ctgatcgcaa acggcttgat 2400
gcggcagctg ttctgcatgt ccctgctgac tgcgctggcg gtgacgctct tgcaggttta 2460
ttcattaaaa acatatcatc acaaacggga agacgctgcc gggcaagctg aatgatttat 2520
cccgtcggtt tcttccgctg cgcgtgattg cataaagtaa tggaggaaag aagcatgaac 2580
atcggtatcg gtacctattc ttttggtggg ctggccagct atctcggtct cggtccgacg 2640
ctgcgcgagc agttcgggca gattcaggca ctcggctttg acagcgtgga gctgctgccc 2700
gtcgaccttg aggaagaccc gaaggatatt ctcaaatggc tggaggagac aggcctgaag 2760
gtcgtttccg tgcacgcgga accgaccgag gagatggtca aaaagctggc cgccgtcggc 2820
ggcaaggcgg tcatctgggc cggcacgccc tttaacagca aggcggaagc catcgaagtg 2880
gcgcgcctgc ttgacagcat ggccgaaatg gccgcgccct atggcatcaa ggtcggctac 2940
cataaccaca gccaggagtt ttatgtggac gagggcaagg cgctgctcga gcacctgctg 3000
gataacggct ccaagttcta cctgcagctc gactgcggct gggccatgaa tgccggcatg 3060
tatccgccct actttatccg caagtataga aaccgtatcc tctccatcca cgttaaggaa 3120
aatgacaagg tcaacggccc gggggaaagg ccggcctccc gccatgccga gccgcccgcc 3180
tttgccaagg cctttgccaa tgcccgctcc ctgccgcttg aagagcgcaa aaaaatgctg 3240
gagcagttta cctctggggc ctcggatgac cactcgatgg acaaccagtg caggatgggc 3300
gcccccgaat ccaaccttga ctggaaagaa atcaaaaagg cgctcgacga gcaggatttt 3360
gaagccttct gggttgtgga gcgcgagtgg ttttacgatg agcacgacaa gtgcatccgg 3420

gaagactgcg agtggctgag gaagaacctg taatctcata tcagacgcat cgtaacaaaa 3480
agggactgta gcatttttgt tttgctacag ccccttcaaa ctattgacaa accctgtggg 3540
tggtacagga gggccgcagc cctcctgttg ggggtgggcc tcggcggtct caatttgcaa 3600
tggtcaaggg cttcgcggcg caggcaccgc tctcgctttc ccttgcttat tgggctgacg 3660
gttcgcttca tcccgcacag cgggcgctca cctctgccct tgcgcggcga ggccgtaagc 3720
cttgcgtagc aagggtttcc ttgtatcaat tcgcaatagt cgccttcttc gcgcctacgc 3780
gcttgaaggt tcctttgctc attggggctt cacttcactt actccgccac aggcggcgtt 3840
cagttccgcc ccatccgccg ctcggagagc gaagcgaaaa ggcggatggc cggttcccct 3900
cccgttgaca aacttggttt gtcaacggct tgagcccccct gaagcagttt cgcttcaggg 3960
ggctttgctt attcgccggg gacaacggtc atgctcagag cctcaggccc agccattgcg 4020
aggatcgttg agtaaaatgt aaatgctacg agttaacgga taatcaccat aaaacaaata 4080
agcttgataa ggtttatagc tattatcctt cgttgcaata tcgtcttcgc tcacagacat 4140
taccctgatt tcgtttctga atgataaatt cgttgtgtca ctccgattac ctaaccaatt 4200
aactccaatc accctattg catgcgagtt ttgagcaaca aaatcaataa cctcttgatt 4260
attattcaat gctttgagat cttttgaaag ctttttgcct ttacagatag agtctactgc 4320
aaaacgaacc gtacttgagt ttttattatc aaatacaact tgaatgtttt ttaaagatga 4380
agaagggtaa atttctttcc aggttttcac ctcaccagtc agcatacgtt ttatatcacg 4440
cagactaatc aatgaatctt tattggctcg gttaataatc agagccaaac catccgttgc 4500
caattttatt tctcgaggaa aaaacttacg actatggaac gaactcattt cttccgccga 4560
tagtgtacgt gtggcaatag ccaaacgtac actatctttt agcagaagat taatagcatt 4620
tacttctgtt gtaaatttgg gaacaattcc agccaaggca aaagtacctt cgaaaacatc 4680
aatttcctct tgtataattg gccggaaact ttcatcagca gcaatagcaa ttacccctga 4740
agtataagta tccgtaaaac cgtctttcgg tttattatta cacgctgaaa ttaacataac 4800
aagcgcaaaa gtgccaagcc aaaactgttt cttcatattt actgcaatct aaatgtaata 4860
ggtactgtat attttactgt aacgggttta ccgttttgct taccaggaat ccatcttggc 4920
atagaacgaa taactcgaat tgcttcttta tcacaatatg gatcaagaga tttaacgact 4980
tgaacatctc gcaccgtacc atctttccct acgataaact gacagataac tctaccctga 5040
gttccgtttt cctgtgcaat agtaggatat ttaaggttct tccctataaa agagagcaac 5100
tcagcttgtc ctccggggaa agttggcatc tgctcaacca tatcaaaaac cttctcttct 5160
tcaacaggag cttgtgtaac tacctgtttc aagtctgcaa tatctttacc attagcttca 5220
tcattaccct taacatcggc aatagaaata cttaccttag tctgtgttaa ctcttcctga 5280
ctcttaattt catcttcatc agctacctca gaatcttttt ttataactgg ggcagtgaac 5340
ttgatagaac ttttcagagc aggaggtggc ggagccacag gttctacttt tttcatttca 5400
tcttgcttaa cttcgggttc ttccaattgc gacaaagcag ttacttctgt catcacctct 5460
ttctgtttag gagtagccaa ccgtatcaaa gttggtatac taaaaccaat caaagcaatg 5520
acgacaacaa taatcatagc catattatga cgtttagaag aatctccacg cattttatat 5580
gcaccatagg ctttattctt gccattaaat atcagctcac accactcaga agatgtcaaa 5640
tcaatttttg ccattttttta ctttaatttt aggtgttaat cagctacatt ttgtgaaagt 5700
ccacctcttg attcaaaatt cttcagcaag aattcgtcac cactagtaat atcaactatc 5760
acatatttac ctatactgca tatttgcatt tcatcaagcg catcaataag atttacatac 5820
tttgatttct cagtagcttt aatgataaca accggagtat ttttaccatt ttttatttca 5880
ctgagctgtt gggtatattc ttcttctgat atttccatat tcaacttttt ctgtttcagc 5940
tcattaacct tacgtacagc ttctttatta cgtttaagca ataaagaacg aattccgtcg 6000
ggtttgtaag atgtttcttt caaagaagaa taatctttgt agtctggttc gccttcgtaa 6060
taatataact tatcatcact tcctaacagt aaagttatag cctgtgaggc ttttacttta 6120
gtttgttgtt cttcagttat atctttatca ttactgggca tacttatctc catcgtttga 6180
ggtttgctca acgaggtaca taacataaag aaagtaataa gaagcatatt catatcaacc 6240
ataggagtaa aatcgacacg tacggtcatt tttttctgct taccctttcc gcctttcttt 6300
ttattgccac tctcttgtac ttcagcactc atagtttatt cgatttttaa ttttcagaag 6360
ttgttttcaa cgaagtaatc aagttgtatc gattctctct gagatcctga agtgaattca 6420
taaccttctt aattaccgaa tagggagttg actcatctgc tttgatagca atacgcaatt 6480
cggaattaac cagacgtgca ttcctaaccc atgatttaaa ttgattatct acactatcac 6540
aaggaattcc ttcattcttt agcacggcat cttgtttctc tgccggtaaa tcaagatatg 6600
ttttcatact tttgattggt acaccaaatg tagttgaaac cataaacttc tttatttgtt 6660

ccggagaaaa tgtaatacca tactcttgcc ccatcccttc taaagttgct tttaaatcag 6720
cttgtttatc caagctcata aagatcttgc catcgggatt taccaatatt tgcaagatgt 6780
ccttttcagg gatcttaatt tcagagacag acgctggagt tgttacttgt accggttcct 6840
tctttacaaa tgttgaagtc aacatgaaga aggtaagcaa tagcacagta acatcactca 6900
tagctgtcat atctatgaaa gtactttttt tcttaatttt ggctctaccc ataattttaa 6960
gcgtttaaaa tgaactctta cttatgagtt gctgcaaatg tttgtacaat tgaaaagcct 7020
acttcatcaa ggctataagt cagtttatca attttattag tataatagtt ataggctatt 7080
actgctaaag cacctgttaa gataccaaac gctgtattta ttaacgcttc agaaatacct 7140
tgcgataatg ccattgaatc agcagcccca ccagcggcca aagcagcaaa tgaacgaatc 7200
ataccgataa ccgtaccaag caaccccatc aaagtaccca aagtcgttat agtaccgata 7260
ataggtaaat tttgttccat cattggcaat tccaaagcgg tagcttcttc cagttctttt 7320
tgaatagcca atagtttttg gtctttagaa agagtcgcat cacgttccat ttcatcatac 7380
ttctgaagag tagacgtaac tacatttgca acagaacctc tttgtttgtc acaaatagct 7440
tgagcctttt taagatcacc agctgtcaaa gcttctttga tattcttaac gaactttaca 7500
agagaacctt tgccaaaagc agaacgaaga gcaaagtaac gttcaatact tagcgcaagt 7560
actgtaagta acaaggtctg aataatcgga acaataaaac cacctttata aatagtacct 7620
aaaaaatttc ctggcagcgg atgattgtta ggatcattgt tcataaagtt agccgggtta 7680
cctaataaga actgataaat acatactgca attataaagc agaagataat taccaaacct 7740
gcagatctaa tacctgtaaa tgaataagct tttttttgat tagtttccat aatgtgtaag 7800
tataaaaaga tatgtttaaa aataaaataa ttctgataag cataatatct cctagcagac 7860
aagataatac aagcctgcta aacatattag agataaaaaa gtcttggaac taaataagaa 7920
ttcgtcttaa gaggtagacg taataatcgc acacacctga aagacaatgg tgtgaggtat 7980
ttgagtagaa taatttgctt ctgtcagtaa ctaactgatt ctctctgata ctcaaatgca 8040
ataataaatc tcttatcgca ctaagatact cagcagcacg acgtgcacga taatgatttg 8100
catgtaagag agattccgaa actcctttta tacttaaatc agcatatgat acggtcagtg 8160
ttttatagag catatccata ttcagcgatg aatataagct agcactttca tcctctgatg 8220
aaaacgattc agctgcttca ttgcatgcat gtactgcaca tacagtattt gcaccagcac 8280
tcaaaggaag taccagcaat gcaattaaca aaagacacct aagcataaac ttcatcactc 8340
tctgctattc gaattttcat tggcaaaatt acgaagtata ttaataccaa attcattatt 8400
cacatctttt aagtcagata taatcgtata aaaaaacata aataggaagt ctttattcta 8460
ataattataa cctccttaaa cgcaaaaaag tcccacagct ttgagctgca ggacttcttc 8520
aaaaaacggc gactacctac tctcccactg ttacgcagta ccatcggcgt gacaaggctt 8580
aacttctctg ttcggaatgg gaagaggtgg aacccttgtg ctatagtcac cttaataagg 8640
tagacataat gtacacaaaa aataaacaat cagtcagcta aacgtatata aaaaccatac 8700
tgacaagaag aaagtgaacg ggcaattagt attgctcggc tgtgatattt ctatctatac 8760
acctgcaacc tatcaacgtc atcgtctttg acgaccctaa gaaatctaat cttgtggctg 8820
gcttcgtact tagatgcttt cagcacttat ccaatcccga cttagatacc cggcgatgca 8880
cctggcggca caaccggtaa accagaggtc agtccaacac ggtcctctcg tactagtgtc 8940
agagccacgc aaatttcata cgcccacgat agatagagac cgaactgtct cacgacgttc 9000
tgaacccagc tcgcgtgcca ctttaatggg cgaacagccc aacccttggg accttctcca 9060
gccccaggat gtgacgagcc gacatcgagg tgccaaaccc ctccgtcgat atgagctctt 9120
gggagggatc agcctgttat ccccggagta ccttttatcc tttgagcgat gtcctttcca 9180
tacagaaaca ccggatcact atgctctagt ttcctacctg atcgacttgt cggtctccca 9240
gtcaagcacc cttatgccat tacactctac ggacggttac caatcgtcct gagggtacct 9300
ttagaagcct ccgttacact tttggaggcg accaccccag tcaaactacc caccaaacag 9360
tgtccccatt aaaaacgggt tagaattcaa acaatcaaag ggtcgtattt caacagcgac 9420
tccacgaata ctagcgtacc cgcttcatag tctccgacct atcctacaca tcaatggccc 9480
aaattcaatg ttaagctata gtaaaggttc acggggtctt ttcgtcccat cgcgggtaat 9540
cggcatcttc accgatacta caatttcact gagctcacgg ttgagacagt gtccagatca 9600
ttacaccatt cgtgcaggtc ggaacttacc cgacaaggaa tttcgctacc ttaggaccgt 9660
tatagttacg gccgccgttt actggggctt caattcaatg cgtctcttgc gataacatct 9720
cctcttaacc ttccagcacc gggcaggtgt caggctgtat acgtgatctt tcgattttgc 9780
acagccctgt gtttttgtta aacagttgcc tggacctatt ctctgcgccc tctcttgcga 9840
ggagggaccc tttatcccga agttacaggg tcaatttgcc tagttcctta accgtgattc 9900

actcaagcgc cttagtatat tcaacccgac tacgtgtgtc cgtttacggt acgggtacct 9960
gtaagattaa gtttagcgga ttttcttgga agtctgttta catgcactat tggattgtcc 10020
gaagacgctc catactatca ggttcgaatc ggaatgtgga tttgcctgca ttcctcaact 10080
tctacaccct ttaacgaact attccgtcag ttcgcggcac tatcactact ccgtccccac 10140
atcactctta aaggtagtaa tggaatatta accatttcag ccatcggcat agcctttcgg 10200
cggagcctta ggacccgact aaccctgatc cgattagcgt tgatcaggaa accttagtct 10260
ttcggcgagg gggtttctca ccccctttat cgttacttat acctacattt gcgtttccac 10320
acgctccaga gaagctgacg cttcgccttc aacgcagagt ggaatgctcc cctaccaacc 10380
ataaatggtt ccatagcttc ggtaaaccgc ttatgcccga ttattatcca cgccaaattc 10440
ctcgactagt gagctgttac gcactcttta aatgaatggc tgcttccaag ccaacatcct 10500
agctgtctta gcaatctgac ttcgttagtt caactcagcg gttatttggg gaccttagct 10560
gatggtctgg attcttctcc tctcggacgc ggaccttagc acccgcgccc tcactcctgt 10620
tctcaaacta atgcgcattc ggagtttatc aagacttgat aggcggtgaa gccctcgcat 10680
cttatcagtc gctctacctc acattagtaa taaacaaggc tgcacctaaa tgcatttcgg 10740
ggagtacgag ctatctccaa gtttgattag cctttcaccc ctaccctcaa ctcatccgga 10800
agcttttcaa cgcttatcgg ttcggtcctc cagttagtgt tacctaacct tcaacctggc 10860
caagggtaga tcacttggtt tcgcgtctac tccttccgac taaatcgccc tgttaagact 10920
cgctttcgct tcggctgcag gtctcaagac ccttaacctt gccggaaaaa gtaactcgta 10980
ggttcattat gcaaaaggca cgccgtcaca gcacaaggct gctccgaccg cttgtaggca 11040
catggtttca ggaactattt cactcttcta ttcgaagtgc ttttcacctt tccttcacag 11100
tactggttca ctatcggtct ctcaggagta tttagcctta ccggatggtc ccggcagatt 11160
cacacagaat ttcacgtgtt ccgcgctact caggatacca ctagggtcaa acagtattcg 11220
aataaggggc tatcacccgc tatggcgtaa ctttccagtt acttcttctc tactgttgag 11280
atcccacatc gtggtcctac aaccccatat atgccgtaac atatatggtt tgggctattc 11340
cgcgttcgct cgccactact agcggaatca ttattatttt cttttcctgc aggtactaag 11400
atgtttcagt tccctgcgtt agcctactgc atagcagtat aatatctctt caagatattg 11460
ggttgtccca ttcggaaatc ttcggatcaa aggtcatttg cacctacccg aagcttatcg 11520
cagcttatca cgtccttcat cgcctctgag agccaaggca tccaccatgt gcccttgctt 11580
actttcttct tatcaaacac ttcttaatcc aatataaata tcgaatcaaa acgtatggtt 11640
gatatatact tttagctctt actaaaaaat ttactttttg cttttgtaca ttatgtcaaa 11700
gatcgttgtc ggttctaaag aacctgaagt ggagaataac ggattcgaac cgttgaccct 11760
ctgcgtgcaa ggcagatgct ctagccagct gagctaatcc cccgaatcaa gagattcgta 11820
gtcccaggca gagttgaact gccgacctct acattatcag tgtagcgctc taaccaactg 11880
agctatagga ctgtgttcaa acccttgcct ttcggctcgg cttctttttt ttctcttgtg 11940
ttatctaaca actatatatt aaataaacaa gtaccagtag tacaacagat aaatctaacc 12000
ttaatcagac ttaaagtaat aatcataaaa ccttttacga aatcactcca gaaaggaggt 12060
gttccagccg caccttccgg tacggctacc ttgttacgac ttagccccag tcaccagttt 12120
taccctagga cgctccttgc ggttacgtac ttcaggtacc cccggctccc atggcttgac 12180
gggcggtgtg tacaaggccc gggaacgtat tcaccgcgcc gtggctgatg cgcgattact 12240
agcgaatcca gcttcacgaa gtcgggttgc agacttcgat ccgaactgag agaggttttt 12300
gggattggca tccggtcgcc cggtagctgc cctctgtacc ccccattgta acacgtgtgt 12360
agccccggac gtaagggccg tgctgatttg acgtcatccc caccttcctc acatcttacg 12420
acggcagtct cactagagtc ctcagcatga cctgttagta actaatgata agggttgcgc 12480
tcgttatggc acttaagccg acacctcacg gcacgagctg acgacaacca tgcagcacct 12540
tcacagagac cttgcggcca taatctttca actatgttcc tctgcaattc aagcccgggt 12600
aaggttcctc gcgtatcatc gaattaaacc acatgttcct ccgcttgtgc gggcccccgt 12660
caattccttt gagtttcacc gttgccggcg tactccccag gtggaatact taatgctttc 12720
gcttggccgc ttactgtata tcgcaaacag cgagtattca tcgtttactg tgtggactac 12780
cagggtatct aatcctgttt gatacccaca ctttcgagcc tcagtgtcag ttgtacccca 12840
gtgagctgcc ttcgcaattg gagttcttcg tgatatctaa gcatttcacc gctacaccac 12900
gaattccgcc cacctcttgt acactcaaga catacagtat caactgcaat ttcacggttg 12960
agccgtaaac tttcacagct gacttatatg tcaacctacg ctccctttaa acccaataaa 13020
tccggataac gcttggatcc tccgtattac cgcggctgct ggcacggagt tagccgatcc 13080
ttattcataa ggtacataca aaaagggaca cgtccctcac tttattccct tataaaagaa 13140

gtttacaacc catagggcag tcatccttca cgctacttgg ctggttcagg ctctcgccca 13200
ttgaccaata ttcctcactg ctgcctcccg taggagtttg gaccgtgtct cagttccaat 13260
gtgggggacc ttcctctcag aacccctatc catcgtagac ttggtgggcc gttaccccgc 13320
caacaatcta atggaacgca tccccatcgc ttaccgaaat tctttaataa agagaacatg 13380
cgaactcttt ataccatcgg gtattaatct ttctttcgaa aggctatccc cgagtaagcg 13440
gcaggttgga tacgtgttac tcacccgtgc gccggtcgcc atctgaagta ttgctacccc 13500
aatgctgccc ctcgacttgc atgtgttaag cctgtagcta gcgttcatcc tgagccagga 13560
tcaaactctt cattgtaaaa gtattgtgta cctatgaaaa atacccgaaa gtatcaatca 13620
taagcagttt ttagctctgt tcaggatttc gtaaatttat ttaagttcac accctgctga 13680
aagcaggata ttgacggttc taaattttac ccaaaattac ttctcttgca atcatcagtt 13740
tccgaagaaa ccaaacaaga agaaaagcat tttgctattg tactacttgt attgtttatg 13800
taattctatc aaagaacgct tcttttattt ttgctttgtt ttctaaagcg ggtgcaaagg 13860
taaggagttt aaaagtaaca accaaatgtt ttcggaagtt tttttttaaa agttttcttt 13920
atcgctttct ccttctaaaa ctcttttagc aagagcgaag gcctaaacaa aaaaaagatc 13980
ctttttttgcg aaccggactg caaagataaa gactttttag cttcacttcc aaactttcag 14040
agaagttttt ttttaaaagt aaaaccgcat aagtcaaacc ccacaggcgc tttatcaaca 14100
tttcaagaaa acactttgtc tctcttgcaa agcgggtgca aaagtagaag cttttgccat 14160
acaaagcaag aagatcatac tctttttttta gagaagaaag gagcgaaaag gcttaaagcg 14220
ctagacatca agggataaca gcaggaaaat aaatgcaaag aaagaaggta tgagggaaat 14280
accaagaaat ggcatacctt atattatata aggggaaatg taagcagaag aaaaaaaggg 14340
ttattgcaat agtaagaagg aaggcaagga tggaagaaaa agactcgaaa caaagtgggg 14400
gtatcaagaa aacaaagtag gaaggtggta gttgaaaggc gaaaggcaga cattagcgaa 14460
gaagaaagaa gaaggaagga caggatagac tattcaatag ctcaagccaa aagagcccga 14520
agaaaaaaga gcggctaacc ttggttggac gactggctaa ccatcgttgg gcgactggct 14580
aaccatcgtt gggcgactgg ctaaccttgg ttattcaact agctaacctt ggtcagccga 14640
taaatgcaaa aggaagatac tcgaaagaga agaaaagaga gaagacaaag cggatgaagc 14700
taaaaaagag ataagagaat acgaaagatt ctagattttg gagaaccatt gtttaataac 14760
aaagcacata aacaactagg cgaatctgtt gattcgccta aagtaaaaag atcatctttg 14820
atgtttaagc gtcagaattg aaaataaata aagggctgaa tatctgcact cttaacctga 14880
ataataacga atacgactag aataataatg agggccttag cccataaggg tattcgtgta 14940
ataaatgaac aacaagcttt ttcccaatta tcaggtgcaa aatgcaataa gaacccaatc 15000
accattaaca caaaaacggg ccaatagcca attatcattt gaggtaacac atctacatga 15060
aagttcacaa atatctgtcg aatcatgtta agcgaatttt caaagctggt atttctaaaa 15120
aatatccagc agaagcacac aaagtggaaa gtgaaaaagg ctccaacaaa atttggaatc 15180
ttaatactaa gtcgagaaac gaccaatagc tccaaccata ttttgtgaac cgctaaggcc 15240
aggccatgta gcactcccca caaaacaaag ttccaagaag caccatgcca taagccgcca 15300
agaaacatgg taacaatcaa gttcacatat tgacgtaatt tgcccttgcg gttacctccc 15360
aacgatatgt ataaataatc tttcaaccaa cttgaaagcg atatgtgcca gcggcgccag 15420
aactctgtaa tagacaatga tttatagggt gaatcaaaat tgagattaaa atgaaacccc 15480
aacaaaagag ctatacctat agccatgtcg ctataacccg agaaatcaca ataaatttga 15540
agtgcatatc catacacagc aactaaattt tcaacaccgg aataaagcga aggagcatca 15600
aacacacgtt ctacaaaatt cacacttata taatcagaga tgactgactt cttaaatagt 15660
ccgcacagaa taaggaacac gcctctgcca aacatttcag aagtcaccat caaaggcttt 15720
cttatctgcg gaataaagtc gcgcgcgcgt acaatagggc cagctactaa ctgcgggaaa 15780
aaagacacgt agaaagcata atccatcaaa tgatttaaag gttctatctg tttgcgataa 15840
acatctatgg tatagcttaa ggattgaaac gtaaaaaaag aaatcccaac cggtagaaac 15900
acgtcaatcg gactaaaaga tcctcccgta atatcagcca aaatggttcc aaagaagtta 15960
ctatatttaa aatagcaaag aagtcctagg tttattaaca agctaaaaaa caccaaaagc 16020
tttctttgaa cagcagatgc agtaatattc attcttcgag ccaaaaagaa atcactaacg 16080
gttaccaatg ccaaaagcag aaaataaacg ccactacttt tataatagaa gaaataagaa 16140
aatgccgtta cgaacaaaat tctggctgta ttcttctgtg ttagtaaaac gtatacagag 16200
ataaatgcgg taaataacca gagaaaaaaa ccgctactaa aaatcatcgg tgcctgcgaa 16260
tcatagctaa acacccctat tagtcgacta atgtctattg acaacataat tattataagc 16320
ttttataatg gcttgagcca aaagttttcc ttgcaaagta tatccttcag gcaaataatg 16380

cacatggtcg ggacgcataa gttttgcccc atcccagttt ttacaagccc attgctctcc 16440
tcctaccaca gtgtatagat cccaaaacgc aagtccatta cgagatgcat attctctgat 16500
tgtctttgct gccataatag tacgcggatt aggagaataa gtgctgcgcc tgcgacttag 16560
ccgacgtctt tcaaatgacc ccggaggaga agtaagcaac atgggaacac ccggtaaact 16620
atccttcaac aactctatta actcatgcat ctgattatag tgcacatttg ttttgtaacc 16680
ttttccatgg ctctcattgg taccaaacga taagattaac aacgacggct tagcttcaac 16740
aatagccgat attcgcgccg gatgcgtaaa agtaagacag gtagctccgt tgactcccat 16800
gtcataataa agaagagaat gaaattcttg agtaagtact aaacctgccg attgaggata 16860
aatatggcca cgaacgtgac tatcaccgat atgaagaatt cggaatgtat cgacttccat 16920
gtccagcctc atctcttgca gctgtttcat gacaggctcc aatacatctg tagaatctac 16980
aatttcattc gacttaacgt tcctaaaaat ttcaggcagt aattgagtat acggaatcgt 17040
atcagccggc cattttactt tcagtgcagg tttgggtatt ctacttatta aattccgtgc 17100
cgctttctgc ggcagtttat cttgagacga catgcccgat gcatagagca gcaggcaaag 17160
gaggcacttc attctaaagc tacttagcat aatcttccct ccgtttatgt tgctcgtatc 17220
catacattaa ggattcatat agtaaacctg ccaaatattt gccgccccga aaattgatgt 17280
gcgtataatc ataattggcc aaagaaggtt tagcatgtac cattttgacc atagcacctt 17340
ccccacccat ggcctgaaac atattccaaa aagcaaccga agactctgag gctatcacct 17400
tttgatattg aatgaggttc tttacacccg gcatagtttt caactcccct tcctctgtct 17460
tatagcttct atcacccacg cccaacatta gaatagccgc ttcgggaaag cattttttta 17520
tatgagcaat gcttttgttc atacctatgc ggtagtagtc atagttacgt cctttttcgg 17580
tagcaacatt tagtccgtat tgcagaatta ttaaatcata aggtctctta gatctaaaag 17640
ctttcaatgt ttgctccgga atatatcgca gtgatacccc cgaactgccc ctcaaagaaa 17700
aattatcaag gacaattccc tttgaccctt ctaacgccgc tccataaaac acggtagaat 17760
cggccttata aatgttccac ctcacagagc cgatgtgtcc ctttagagta atctgccgca 17820
atgatccatt tgagggagaa gatgggacat cgacaagttt ttctatttcc cgattaagtc 17880
tgatccccac tttgacggga gccttattgg caaacaaaag gtaagcgcat gtacaagaat 17940
ctagtaaaga ggcatattga gcctgccccc tgagttctac aaatgctgct gaatcggcaa 18000
taaaataatg gccggatatc ccctgtttct ttttattaaa ataagtagta tcagttgcag 18060
catgactatt ccagcctcca aacgaatgtt tcaccgtagg ccgatagcca ctggtcatag 18120
aggttatggg aataaagccc actccgcatc ctccaaattt tcgttgaagc atttctctta 18180
aatcggcagt gaatatatct ccctctataa aagaatctcc gaacacagca attctcacta 18240
tacgtttttc tccggcatca agcttcgaca acgcattata aaacagcgcc attccctgtc 18300
ctatagaatc tgtgaaatct tcaacacaag taatgccgct cttacatgta tctactaaaa 18360
tctttgattt gggagcaaca agaggtagcg tgtcaggtac agacagatcc gttttgctac 18420
tacgtatatc actcaacata tcaaccttcc gtaaagtgtg accacagatt attgtagaag 18480
gcaaataatg tagcaaaaga agtgccgaca gaacaattac tgtcagccaa aacgtatctt 18540
taatatgatt catacagcca tgttacgatt ttaaggctgc aaagatatat aaaaggaaga 18600
gaaagatcga tttacaatca aatcgttttt tcaatattcc aaacgaacgc acgcagacct 18660
aatttttcag ttctcgtatc caacttatgc aacgcatcaa gcaatggatc aactttatta 18720
tcatctacca ttgttataat ggccgaacac atagaaggcc atgcatgact cccgaaatgc 18780
ggttcgccgg ttttgctccc acgccctttt acttgttcaa aatagctaaa cccacgacaa 18840
ttcaaacgat ctagggttcc tattatctgt tcataatagg cttggtcaaa ggttataaat 18900
acagacttca ttttttaaca atttcacttt tatgttcttg aaaataatca gatagagcac 18960
gcctgcgctt catcacttta cgttgcctct tgattcctgt acctgcaaac acacagtaga 19020
gaactgggat taatatcaaa gtaagaatag tagagatact taatccaccg ataacagcaa 19080
tagccattgg tcgccacatt tctgaacctt gtcccgttga gattgccata ggtaccatac 19140
ctaacacggt tgtagccgtt gtcattataa ccgggcgaag acgactacgg cctgccgtaa 19200
caacagaatt tattaccgcc tgcccccttt ctcgacataa actaatataa tcaatcaata 19260
caataccatt cttaacaacg ataccaataa gcataatgcc tcccaacata ctcatcacat 19320
tcagcgtgtt ttggctggta aacaaagcaa tcaatacacc actaaatgca aatggcaatg 19380
aaaacatgat gataaacgga taggtaagcg attcaaattg ggcagccatc acaataaaaa 19440
caagaataac aataagcacc ccaagcactc ctaaatcttg gaacgactct tgttgatctt 19500
caaaagaacc agcaacttgt actgttactc ccgcaggaat atccatattg tcaataacgt 19560
cattgccagc tgcaacaaca tctcccaaag cctctcctga aataacggca gagactgtta 19620

ttatacgctc cctgtctttt cgctcgattg ttggcggcgt aaatcgctca acgactttac 19680
caagttcttt aactcgcact gcctcacctt tactattata tatgataatg ttctcaaggc 19740
tttccaaact agtcctgaat tcaggggcat aacgcacctt aatatcatat tcatcaccat 19800
cttcacgata ttgcgatgca attgcgccat taactctatt ccttaaataa gttcctgcgg 19860
tagatagatt tagcccatgc atggctagct tttctcggtc aaaatcaacc tgatattctg 19920
gttgataatc actcctacta atattaacct ctgaaacacc ttttatctgg taaagctgac 19980
gtttcagatt agcggcaaca ctatctgtga cagacatatc ataaccgtaa atttcaaagt 20040
cagcagtaga ttggccggtt aagcctgtat tactaccacc taatatcact tgagctttac 20100
ttaattcggg ataagacttt aagtcttgac gcatttcatc gcaaacctgt tccaacgtct 20160
tttttctatc tcccggatca gacaatcgga tattaaatga aataatatgc gaaccattat 20220
tctgcattac tgcccacgca ttatctgtat cggcttggcc aacggtataa ttacacacca 20280
atatttcatc cttatatttt gatcgccact gcttggttaa tcgatctgcc aaattttgcg 20340
ctaactcttt tcgagatccg ataggcaact ctagtgcaac ggatatgcgt gcattgtctt 20400
gtgtcggaaa gaattccgta cttataccat tggcgcaaag aaggctcaag agaaataagc 20460
cgatacaaga aatgaggacc aacattctat ggcgcactgc tttgtttaat acatgcgcat 20520
accaattgtc aaaaccatta agaatgcgtt caatcggagc atagaatttc tgaaaagatt 20580
tggtctgctt cttctctaat ttcagtaact gcgaacaaag cattggcgtc aatgaaagag 20640
ctgcaacagt tgatataaac atgattgtac acatcatcca accaagttgt ctaaacataa 20700
ctcctgtcat accagtaatc attgtaagcg gaaaaaacac agcaatcatg gttagtgtag 20760
aagcaattac tgatatggca acttcattag taccatgaac agctgcttgc ttaggggcag 20820
aacctctttc gatgtgggta gtgacatttt caaggacaac aatcgcatca tcaacaacca 20880
taccaattgc aatggataaa gaagataacg atataatgtt aattgtatta ttagtaatag 20940
caagatatat aaatgatgca atcagtgaaa ggggaatcgt aattgatata ataagtgttg 21000
ccctccatct tcctaaaaag agaaatacaa caacgactac aaacaataac gcatatgcta 21060
ccgtttctgt taaactgtct atcgtattta atatgttatc ggacgtatta acaatgatac 21120
caagtttcac gtcactcggc agtgatttct gtagtttggg taaagcttta agtacttctt 21180
ttgatatatt gactgaattt gctccagatt gcttttgtac aataatcata gcacctttct 21240
gcccattatt atacgtctct tgagctcgct cttcaactgt gtcaaccact tttgctacat 21300
cacgaagaaa aatattcgct tcattacgcg tgccaactac aatattctct aactgtttgg 21360
tatcattaaa ctctccttca acacgtaaag cataggtttc atttccaata tcaaaattgc 21420
ctccggggat atttttgttt tctgccccta ttatagagct aatagtttca atactgagat 21480
tataggcttc cagtttattg ggatcacaat aaacctgtat ttctctctta ggtgcaccac 21540
taattgatac agttccaaca cctgggattc gggctaaagg attaacaaca ttatcatcta 21600
gtattttata cagagcagat tggctttctt cagcttgcac agagagcagt actataggaa 21660
tcatatctgt gctgaacttg aaaataattg gagtctcaac attttcgggc aaagctgagt 21720
taaccatatc taacttatca cgcacgtcat tagttaagac atcaatatca taaccatatt 21780
cgaactctag cgtgatgact gaaatgttct ctgatgattt tgacgttatg tgctttaagt 21840
tactaactga attaagaaca ttctcaagag gacgtgtgac attattttca atatcagaag 21900
cactagcccc ttgatacgtt gtcattacca ttatagtgtt cgtatcaatg tcaggataca 21960
aatctacagg caacttagac aaagagaaaa gaccaaaaat cactactgcc agaaagcaaa 22020
gcgaagtcat aattggcttt tttacagctc cttcgtataa actcatagcc tagctttatt 22080
atttttcaat aactacttca gttccattaa ccaaacgtgt atgtccagct acaacaacct 22140
gcgagttgtt gcttactcct gaaattaact catattcagt tcccattctg cgtcccaact 22200
ctaccttttg gtaacgaacc tttcccttct catagacaaa tacataacgg tcgccagatc 22260
cagcctgttt aacaatggca aggtcaggaa ctactacatg attttccgaa ccgaaattta 22320
cttcaacacg cgcaaacata cctggacgta ctttcatttc ttcattatct agcacaagct 22380
caacaggaaa agtgcgtgtt gctgcgttaa tggttggata gattaaacta attttgccat 22440
taaagacttc tccttcaaaa acatcaaact taactttaac tggagatcct ttctttattt 22500
tagaaaaata gttttctgat atattaatta gcagctttac aggaacaatt tgttctacta 22560
ctaaaacagg agtattgcca ttatacatat ctccattatc ataattacga gccgttatta 22620
ttccgcttat gggactaagt aaagacgtat tttccaatag attccgatag gaagtctttt 22680
ttacgtctaa agacattttt gaagcatccc actcagactt cgaaactcct ccgactttat 22740
acaattcatc aagacgtcta aattctattt cttggttatc caactgaagt tttaattgtt 22800
ttaaacttgc agcatccatt tgaacaagtt tttgcccttt acttacctta tcgcctactt 22860

caacatagat atgatcaatt ctaacaggag aagcaggagc aatattattt tttacttctg 22920
cctctacagt tgctgtatat tccaaaattt gatcaacttg tcgagctgta acagttgcaa 22980
gttttaccct cggcttttca ctaattacat caactccttt cttatctttg ctttctgtgc 23040
atgcactcat tagcaaaata aacatgaagg atactaattg aatatacttt ctcatataat 23100
acttttaatt gaaattactc tatttgtcgt tattcgccta gcactagctc taaatcggct 23160
ttggcaacta aaaaatcata tatagattgg ttatatacca attgagcttg cgttaaagcc 23220
aattcagaat cattcaattc taaaatagtt cctttaccta catcatatcg ctttgatgca 23280
atttgacgac ctttttccgc ctgagccaca ctttctttgt tactcattac ttgctcggaa 23340
ctagcattca tactatctaa ataactacta attttcatat tcagctgacg ttccaaattg 23400
agtctattca gcgccagttg ctggatttga atttttgttt gcttcagttt attaaaattg 23460
ctagctttat ataacggtat agtcagactt aatccaaacg tagagtatgg ataccaatcg 23520
taactagata ttctgaaatt atcattcatc gacatatata aataattgaa agaagccgat 23580
aaggtaggca tgaaattagt ttgttgtagt ttaagagtat tctgaagcaa cttattacta 23640
aggtctaatt gctttaggtc tgaattatta ctaaggttat agtagacatt attctcttta 23700
gatatgaaca tctccatttc atattctttt agacctccat tgataactag ttcataattt 23760
gcatcaatcc ccattaaaac tttgagttgc agttttgcta aattaacgcc attatgggct 23820
gacacaacag aaggttttaa attgcgcatc tggacttctg cacgaatttt atcaaactca 23880
ctaacagtgc catgaataaa ctttgcatta acaatattaa aattagcctc agcctgacgg 23940
taacttctag tcaaaacgct ataactgtcc tgagctagaa gcaactgata aaatgctttt 24000
ttaatttgat ttataagatc caattttgag aatcgagctt tctcagtagc aagatcaata 24060
tctgttttcg taagattaat agttttgtat aatgcgggag caaaaattgg taaactaatt 24120
atcataccac cattataagt gttgtcacgt ccaacttgta tagttgtagt ttcctcaccg 24180
aattgcattg ccattgtttg tttttgattt gttcttgaat aagtaccact caagtttata 24240
tcagggaata aggcagcgta tgcttcttta ttaatttctt tcttcaaaac aacttctttc 24300
tcggatatct ttacgatagg actttcagaa agtgcaattt tcatagcaga ttcgatagtt 24360
agaaccaatg tatccttatc ttgtgaaaga ctcgaatctt gcgcttccac tgaattagta 24420
aataaagcta gcaagacaat caataataaa attttatcca aaaacatttt cttaatcatc 24480
atatactcac aggatattgc attatacaga tttagcatta tattctatca agaagttttc 24540
cagtattcca atccccttat cagttgatat gcctcgtacg aatatcaaaa ctattgattc 24600
ataaacctct agaaaagaat attttttaaa agtctcactg tgaattaaca tatcgatctg 24660
ctcatgcaac aaatcatgaa caatatcaaa cttaatatct gctctaaaaa gcccttgctc 24720
aactcctttc ctaaaaaaag aaataacgta gttactattt ttctttttat gatcattcat 24780
tagctcacaa actttcggat atttctttat ttcctctaaa aaacgtacgt tcgttttatg 24840
ataatcctca acactcttct tataacattt gagaatgacc tctagaacat tcactgaatt 24900
agatataact tcattaacaa atacattttt ttctctttga tgctccagca tacatgctat 24960
caagagttcc tctttatctt cgaacatttc atatagagtt ctttttgata tacccaaccc 25020
catagcaatg tcatccattg tcactccttt tataccttta gtcgcaaaag cgttcatcgc 25080
taaagtgacg gttttatgct tcagctctgt tctgttgttt tctttttcat acataattat 25140
atactgttta ttattatata gttatatctg atttctaaat taaaaaatcg catgagtaac 25200
acactaaaag atgtagtata aagatattat taaaaaatag gaaagtatta attgaagaat 25260
gattattgat caacacctgc taatatttta agatacttac ctgtcacatt agctaattct 25320
gaatgggaag ataagtattc tgctcgcaaa tcaggttgat caatcaataa ttcgctcata 25380
acatcacgac taacaaattc aaccttataa gttataggat caataaaata gtagacacgt 25440
tcagttagta gtgaggaaga agcaattgcc cgtccaacat ctcccatatt ataatttgtc 25500
gctgatgtta tactaactgc tgcaggtcct atcggcaaag ccgagaaata aattttatct 25560
ttcacccaca tagcaggggc ataacaagta ccaaacctaa aacggttaaa gcgcaatttt 25620
ctgcaattca aatataaatt tccatctatc tcaatcgcaa aataccgtcc tttgattctc 25680
ttgcaagttg aaggagaaga ggcactaatt ttataatccc cgccaggtgt aagcataagc 25740
tgattctttg ttcttttctc taaaacaact ttggagagag aatcaccacg atgtatgtag 25800
acatctttaa cactcaaata aaacttattt tgagcaaata taagcactga aaaaagtgca 25860
aatcccaata atattaaaac ccttttcata gacataaaca atcagcatat taaaaacttt 25920
ggaaagatac gaattaaagg tgaaatgaca ataaatgaat ataataaatc aaggtgtata 25980
ttaaatatga tgttcaggca taaattagag ctaacataca cttaaaggaa aaaacaactt 26040
tagattctga aataggaagg tgagtattat aaatccattc aagaaaagca ttatctagtt 26100

gatttttaaa cagagaaatc tattttgatg aagcaatttt tgctataata aaataataat 26160
catctaattt tggctggcaa tcaagatcaa tataagtcta acaactagtg aatcaaaaaa 26220
atacattaac cgccgcatct aggatatcga atcaatcgat ataaaacatt tcatctaaaa 26280
gatttgtagg ctaatactaa aattacaagc atagtcaacc ctaaacatta gtcattacta 26340
aaaaaataac tcgagtttgt gactcgagtt atttttagt atggtcaaca tatctcatgt 26400
gacccttata agaaaatctc tactattcaa cggaagcatt attttcctct gtagattgag 26460
cgtcttgggc taaatctgta gaatcagcat taactacttt ctttgaacgg cgtgttcgag 26520
gtgatttttt agcaacaata tctttcgaca tgttatcgtt ataatcaacc aattctataa 26580
aacacatttc agcattatca cccaatcgat tccctgtctt aataattctc gtatagccac 26640
ctggtcgatc tgcaatttta gtcgcaattt ctttaaataa ttcagtaact gcaattttat 26700
cttgtaaatt gctaaaaact acacgacgag aatttgttgt atcatttttt gatttagtta 26760
tcaatggttc aacaaatttc tttaatgctt tagccttagc aacggtcgta gtgattcttt 26820
tgtgctttat caaagaagat gccatattag ataacatagc agctctatga gaagcagtac 26880
ggcctaaatg attgaatttt ttattatgtc tcattttta ttctttttct aatttgtatt 26940
tcgagatgtc agtcccaaat gttagattca gactttccag caaatcatca agctcggtaa 27000
gcgatttctt tccaaaattt ctaaacttca gtaaatcagt cttattaaat tgtacaagat 27060
ctcctaaagt ttcaacatca gctgctttta agcagttaag ggcacgaaca gacaaatcca 27120
tatcaactaa cttagtctta agcaattgtc tcatgtgcaa aacttcctca tcaaattcct 27180
cattaccatc tatatctgta ttttccaacg taatcttttc atcagaaaaa agcataaaat 27240
gataaattag aattttggca gcttctttta aagcatcttt tggattaatg gaaccatctg 27300
tagtaatctc aagtattaat ttctcatagt cagtttttg ctcaacacgg aaattttcta 27360
ctaaatactt gacattacga ataggagtat aaattgaatc aattggtata acattaacat 27420
ctgtacagaa ctccctgttt tcatcagcag ggacataacc tctaccttta ttaattgtta 27480
tatcaatttg catagttgcc ttagaatcta aatgacaaat aaccaattca ggatttaaca 27540
cttcaaatcc agtcaaatac ttacctatgt caccagcctt aaattcacta gagttttcaa 27600
ctgtaatagt taccttttct gtttcgaact cttcaactac ttgcttaaat cttacttgtt 27660
taagatttaa gattatgttg gtaacatctt ccttgacacc aggaacacta gaaaattcat 27720
gctctacacc ttcaatccta attgttgtta tagcaaaacc ttccaacgaa gaaagaagaa 27780
ttcggcgcaa tgcattacct actgtaatac caaaacccgg ctctaaggga cgaaattcaa 27840
attttccaaa tttcggatcc gactccaaca ttaatacttt atcaggtttt tgaaatgcta 27900
atatcgccat ggaactaatt attatttaga atacaactca acaatcaaat gctctttaat 27960
attctcagga atatctgtac gctcaggaat atgaagcaat ttacctactt tagcattgtc 28020
atcccattct aaccatggat atttactatg attaaaacct gcaagagaat taccaattac 28080
ttctaaagac tttgaacgct ctcttacacc aataacttgc ccaggtttca ccgcatatga 28140
aggaatatta acaacactac cgtcaacagt tatgtgttta tgacttacta actgacgagc 28200
tgcggcacgc gatggagcaa ttccaagacg gaacactaca ttatccaaac gcccttctag 28260
taattgaagc aagatctcac ctgtaatacc tttagcggtc tcagcctttt caaacaaatt 28320
acgaaattgt tttccaaaaa cgccataagt atattttgcc ttttgcttct cacgaagttg 28380
gattccgtat tcagaagttt ttctttttct agaattacca tgttgtcctg gaggatagtt 28440
cttctttgac aacaccttat cggccccaaa aatgccttca ccaaatttac gcgcaattct 28500
agattttggt cctgtatatc tagccattat tatttttatt aaattatagt aatcatcaac 28560
ttaatttgtt gcagccgcga ttacagagaa gaaagagtgc aatccataac aaaattaagt 28620
ctcattttat taattaaact ctacgtcttt taggaggacg acatccatta tgtggcaatg 28680
gagttacatc gacaatctct gtaacctcaa taccagcacc atgaatagta cgaattgctg 28740
actctctacc attaccgggt cctttaacat aagccttaac ttttctaagt ccaagatcaa 28800
aagcaatttt tgcacaatct tgtgcagcca tttgagctgc gtaaggggta ttctttttag 28860
agcctctaaa gcccattttt ccagcagaag accacgaaat tatctgccct tcactattag 28920
caagtgaaac aattatgtta ttgaatgagg aatgaacatg taattgtcca ttagcatcaa 28980
ctttaacatt cctttttttt gctgcaacag ttttttttgc catatcaaga attattattt 29040
agtagctttt ttcttattag caacggtttt ctttctccct ttacgagttc gtgcattatt 29100
cttagtactt tgaccacgaa caggtaagcc aatacgatga cggacaccac gatagcaacc 29160
aatgtccatt aatcttttaa tatttaattg aacttcagaa cgaagatcac cttcaacctt 29220
aaattctgca ccaataatct cacgaattga tgccgcttga tcatctgacc aatctttgac 29280
tttaaggttt ttatctacac cagctttctc tagaatttta gctgaactac tgcgtcctat 29340

accatataca taggttaacg ctatttctcc tctcttattt tgaggtaaat ctacaccaac 29400
tattcttata gccataaaat aaaatattta tcttgcaaaa ataagaaatt atccttgacg 29460
ttgtttaaac ttaggatttt tcttgttaat aacatacaaa cgaccgttac gtctaacaat 29520
tttacattct gacgtgcgtt tttttaatga tgctcttact ttcatatcta aatttattta 29580
tatctaaaca ctattcttcc tttggataaa tcatagggag acatttcaac tctaacttta 29640
tctccaggta aaatctttat ataatgcatt ctcattttac ctgaaatatg agccgtaatc 29700
tcatgtccat tatctaactc aactcgaaac attgcattag acaacgcctc aactattatt 29760
ccatcttgtt ctattgcaga ttgctttgcc ataaatatta aaatactcaa attcttaaaa 29820
cctcatcaat gtaatcaaaa gtagataaaa catcagcttc tttgttcgtt atagctacag 29880
tatgttcaaa atgtgcagaa taggaacaat ctttggttct cacagtccaa ccatcacgct 29940
ccataataat aagtctactg cctaaagtta tcatgggttc aattgcgatg cacattcctg 30000
tttttagtaa aggtccataa ccttttctcc cataatttgg aacctgagga tcttcgtgca 30060
tatttcgacc aataccatga cctacaaact cacgaacaac tccgtaccca ctattttcac 30120
acaaattctg tattgcgaaa ccaatatcac caatgcgttt acctacgaca gcattatcta 30180
taccttctag caacgcttgt tttgtaactt tcagcaaatg tttcacattg ctgttaattt 30240
cacccacacc aaacgtatat gcagaatcgc cacagaaacc attgaggcga actccacaat 30300
caacagaaat aatatcacct tcttttaaag ttacagcacc acttggaata ccgtgaacta 30360
tttggtcatt aatagatgta caaagtgttg ctggaaaagg agtaccaaaa ttatttggat 30420
aacctttaaa agttggtact cctccgtgat ctcttatata cgtttctgct gatttatcta 30480
actctaaagt agttacacca ggtttaataa gtgaagcaac ataacctaaa gttttactaa 30540
ccaatagatt actctcccta agaaactcaa tttcttcttc agttttaaga aaaatcattt 30600
taacaatcaa taagcattaa cggttgagct agcactccga cctttaatac gaccagactt 30660
caacaatcca tcataatgcc tcattaacaa atggctttca atctgctgta aagtatctaa 30720
aacgacacca actaagatta aaagcgatgt accaccaaaa aattgagcaa attcagcttt 30780
aacccccaat aagccagcaa aagctggcat gatagccaca attgccagaa agaaagagcc 30840
tggaaaggta atgcgtgaca taatatcatc tatatactcc gcagtactct tacccggttt 30900
aatacctgga atgaaaccat tattcctttt catatcttca gccatttgag tcggattaat 30960
agttattgct gtataaaaat aagtaaacaa aataatcatc acagcaaaaa ctaaattata 31020
ccaaagactc gtatgatctg taaaagcctg aataatacta ttagaagtgt tagcatttga 31080
aaaaccgatg aaagttatag gtatgaacat aattgcctga gcaaatatga ttggcataac 31140
accggcagca ttaactttta atggaatata ttgacgtgca cctccatatt gcttattacc 31200
aacaattttc ttagcatact gaacaggaat ttttcttgtt ccttgaacca ataaaatagc 31260
agcagcaata accaacagta ggaaaacaat ttctattaaa aacataatca agccaccagt 31320
cttatcactc atacgagaaa ttaactcctg aaataaagat tgcggcagcc ttgcgataat 31380
tcccactaat ataataaaag aaataccatt gccaatacct ttatcagtaa ttctctcacc 31440
taaccaaagt ataaacatgc ttcctgcagc caatataatc gtagaggtaa acatgaacaa 31500
ggtccaatct aatgaagcat ttaaagaagg acctgcttgc attttcaaat taagtaaata 31560
ggaaggtgct tgtactaaaa gtattagaat cgtcaaataa cgagtatatt gattcatttt 31620
ccttctacca ctctcacctt cacgctgcat tttctgaaaa taggggacag caatacccaa 31680
taattgaatt acaatcgatg ctgaaatata tggcataatt cccaaagcaa aaattgatgc 31740
atttgaaaac gcaccgccag aaaacatatt taacagcgca agaagaccct cactagtttg 31800
ctgatgcagt tgagttaaca taccaggatt aattcctggt aaaacaacat atgatccaaa 31860
gcgatatatc gcaacaaaca atatggttat gaggatcctt tgtctaagat cctcaatttt 31920
ccatatattc tttaatgttt caatagcttt tctcattgat ctagagtttt acaacatttc 31980
cgtccaacgc ttcaatggca ctaattgcag atttggaaaa agcatgagct tgtacctcca 32040
atttaataga gatggtaccg ttacccaaaa tcttaactaa ttgattagcc gaaacaaatc 32100
ccgcatcaac taaatcagaa aaaccgacaa ccgttaaatt tttcttttca gcaagttttt 32160
gtacgacatc aatattaata gctttgtact ctactcggct aatattttta aatccaaatt 32220
taggcactct acgttgaaga ggcatctgac ctccttcaaa acctattttt tttgaatatc 32280
cagatcttga cttagcacct ttatgtcctc tagtcgaagt accacctaaa ccagaaccag 32340
aaccacgtcc tactcttttc cttgttttgg ttgagccaaa agctggtttt aaattactta 32400
aatccatatg gtaaaatttt aaatttcaca atgtttattt aacaatagca accaagtgct 32460
ttaccttact caccattcct aagatagaag gtgtagactc atgttctacc actctattta 32520
gttttcggag accaagagca tctaaggtac gcttttggtc aattggagca ccaattctac 32580

tttttatttg tttaatcttt atagttgaca tattatatcc tcctaacctc tgaaaacttt 32640
ttcaacacta atacctctgt tctgtgcaac catacgcgca tctctcatct ccgataacgc 32700
taagattgta gccttaacca aattatgagg attcgaagat cccttagatt ttgctaaaac 32760
gtcggtaatt ccaacacttt ctaaaacagc acgcatagca ccaccagcaa ctactccggt 32820
accatgtgac gcaggtttaa taaaaacttc ggcaccacca aattttgcaa cttgctcgtg 32880
aggaactgtt ccttttaaaa gaggcacctt agtaagattc ttttttgcag cttctacgcc 32940
tttagcaatt gctgcagtaa cttcacctgc tttaccaagt ccccagccaa ttacaccttc 33000
ttcattacca acaactacta tcgctgaaaa actaaaagtt ctaccacctt tagtcacctt 33060
tgttacacga ttaatggcaa ctaatctatc ttttagttct aaatcgttag taactttaac 33120
tttattaatt cctgccataa tcattaaaaa ttaagtccac cattacgtgc tgcatcagca 33180
acctctttaa tgcgtccgtg atacaaataa ccattgcgat caaaaacaac agtcataatt 33240
cctgcttctt gagctctttt agcaataacc tcaccaactt tcgcagcctg ttctttctta 33300
gttacttttt catcaaaaga taaagaagaa gcagaagcta aagtctttcc cgatagatca 33360
tcaattatct gtacatagat ttgtttatta cttctaaaca cactcatacg agggcaatcc 33420
gatgttccag atattttact acggactcta tatttgattt tggtacgtct ttctactttt 33480
gttgtcataa tattatctat tatttaaaac ttatttagca ccagctgatt ttcctgattt 33540
tctacgaata atttctccca caaatttaat acctttacct ttgtaaggct caggcttacg 33600
gaaagaacga attttagaac aaacctgccc taaaagttgt ttatcgcaag attccaaaat 33660
aataagcgga tttttattcc tttcagattt agtttcaact tttatctcag gtggcaactg 33720
aataaatata ttatgcgtat aacctaaaga taaatcaatt atattaccat tatttgaagc 33780
tctataaccg acacccacaa gttcaagttc ctttttgtag ccaacagaaa ctccaataat 33840
catattgttt atcaaggaac gatacaatcc atgatatgca cgttcttgtt tgtcatcagt 33900
cgaacgagta agagtaatct cattattatc taagttaaca gatataactg gatttacata 33960
ctgacgtaac tctcctttag gacccttaac agtaataaca ttatctttaa actctaatgt 34020
aacacctgct ggcacagtaa taggtaattt tccaattcta gacattgctt aatcctccct 34080
tattaatata cataacacaa tacctcacca ccaactttca ggtcagaagc ttctttgttt 34140
gtcatcactc ccctagaagt agatattata gcaataccta aaccattaat tactctaggc 34200
atatctttat aacctgtgta cttcctcata ccagggggaag atacccctaat aagtttcttg 34260
attgcgttaa ctttattaac agaatcatac ttcaaggcaa tcttaattgt accttgaggt 34320
ccatcactta caaacttata attaagaatg taaccttttt caaagagtat tttagtaatc 34380
tctttttttca aattagaggc tggaacttct actgttctat gatttgcact aatcgcattt 34440
cttagcctcg tcaaataatc agctatcgga tctgtcatat gaaataaatt aatcaggtct 34500
atcctgacaa tatttaaaat ttaacaataa ttaccagctt gctttcttta cgccagggat 34560
aagcccatta gaagccattt ctctaaactg gattcttgat acaccaaact gacgtatata 34620
acctttggga cgccctgtaa gtttgcaacg attatgcaaa cgaacaggtg aagaattttt 34680
aggcaaagct tgtaaagctt caaaattacc ttctttcttc aataatgcac gtttttctgc 34740
atacttttcc actaacttag cacgttttac ctcacgagct ttcattgatt cttttgccat 34800
aatatcaatc ctttttagcg tttttaaaag gtaatccaaa ttccttcagg agtgcataac 34860
cttcttcatc agtcttagcc gttgttacaa aagttatatt catccctaaa attcttgtta 34920
tactatcaat atttatttca ggaaaaataa tttgttcttg aatgcctaaa gtatagtttc 34980
ccttaccatc aaatttactt tcaatccctt taaagtctct aatacgaggt aaagcaactt 35040
tcactaagcg ctcaagaaat tcgtacatcc tctctctgcg taaagtaacc ataacaccaa 35100
ttggcatttt ctttctcaat ttgaaattgg caatatcttt acgagaaatt gttgcaacag 35160
ccttttgacc tgtaatagta gtcatttcat taattgcaac ctcaataatt tttttatcag 35220
ccacagccat acctaatcct tgattaatga caatcttttt taacataggt atttgcatgg 35280
tagaagaata ttgaaattga gacttcaatt caggagctat acgctcaaca tattcttttt 35340
ttaaactagc agtattactc attattaaat ctcctctcca ctttttttag aataacgtac 35400
taaagtccct tcagaactct ttcttctacc aatacgagta ggcttgtttg tcttagcatc 35460
aacaacattt aagttagaga tgtgtacaga cgcctcttgc ttaattatac caccttgagg 35520
atactttgca ctaggcttag cacttttgga aactaagtta atgccatcaa cgattgctcg 35580
cccctcttta actaaaactt ttaaaacctt acctgttttg cctttatcat cacctgaatt 35640
gacataaaca gtatcacctt tttttatatg taacttattc atgactaaag tcttttctaa 35700
attataaaac ttcaggtgca agcgaaacaa ctttcatatt tgtagaacgt agttctctag 35760
caactggacc aaaaatacgg cttccacgta tctccccagc attattcaac agcacacatg 35820

cattgtcatc aaagcgtata taagaaccat cagcacgacg aatttctttc tttgtcctaa 35880
caattaaagc cttagaaact gcaccttttt ttacatcact agaggggata acattttta 35940
ctgaaacaac aataatatcc ccaaccgaag cataacgacg acctgtaccg ccaagaacgc 36000
ggatacacaa agcctcccta gctccactat tatcacacac tactaatcta gattctgcct 36060
gtatcataac tacttagctc tttcgattat ttctactaat ctccatctct tagttttact 36120
taagggacga gtctccataa tgcaaactgt atcaccaaca ttgcattcat tcttttcatc 36180
atgtgcatgg tattttttg tcttactaac aaacttacca tatatgggat gtttctcttt 36240
aaacttagcg gccacagtaa tggtcttatc catcttgtta cctagcacaa tcccagttct 36300
ttcttttctt aaatttcttg cttccattta aggcaatcat ttatttattt aattttctgt 36360
cacgaagctc agttttcatc ctagcaatcg ttctacgcaa aactttaatt tttgctggat 36420
tatctaaagg agagattgaa tgattcaaaa ccaattggct ataactaatt ttctcagctt 36480
cgattctttc tactaggtca actgtagaca actcttttat ttcagaattc ttcatatgtc 36540
ttacgcattt tgattttgaa catcataatc acgcctaaca ataaattttg tagtaacagg 36600
aagtttctga gcagctaacc ttaaagcttc ttttgcaatc tcatatgaaa ctccttcaac 36660
ttcaattata attctaccag gagtaatagg cgcaacaaaa ccttctggag aacctttttcc 36720
tttacccata cgtacttctg caggtttctt agtaataggc ttatcaggga aaattcttat 36780
ccagatttgt ccttgacgtt gcatatatct agtcacagca atacgcgcag cttctatttg 36840
cctaccagta atccatttat tctggagtgc tttaattcca aaagatccaa aagcaagctg 36900
attgccgcgc tgggcattgc ccttcatgcg tcccttttgt tgtcttctga attttgtctt 36960
tttcggttgt aacataattc ctaaaattca aattcgttta gcgattattt ttctttcttt 37020
tgaagttttt tccgccactg ttatttgtag aacgtgcact ttctttagtt tgagtaaagt 37080
ttggagccaa atctcttttt ccaaatactt ctcctctaca aatccagacc ttaattccaa 37140
gcagcccaac ttttgtcaaa gcctcagcat gacaataatc tatatcagct ctaaaagtgt 37200
gtaacggagt ccttccttct ttatacatct cggaacgagc catttcagca ccattaagac 37260
gtcctgaaat ttgcactttt attccttcag ctcccattcg catcgtattt gctattgcca 37320
tcttaatagc acgcctatag gctattttac cttcaacctg tcttgcgata ttattcgcaa 37380
caataacagc atctaactca ggccttttaa cctcaaaaat attgatttga acatctttgt 37440
cggttatttt cttaagctct tccttaagtt tatcaacctc ttgtcctccc ttaccaataa 37500
taataccagg acgagctgtg catactgtaa tagtaaccaa ttttagcgtc ctttcaatta 37560
taatacgaga tatgctcgct tttgctaacc tagcatttaa atatttacga atcttgctat 37620
cttcaagcaa agaatcacca taatcatttc caccatacca gttagaatcc caacctctaa 37680
tgattcccaa acgattacta attggattaa ctttttgtcc catgtagcat ttatttaatc 37740
ttcattatta ctcttagcat caacgaacaa agttacgtga tttgaacgtt tgcgaattct 37800
gtaacctctc ccttgaggag ctggtctcat tctctttaat gttgaaccac aatcaacaaa 37860
aacttttgta acaaacagtt ctccaccttc agctttacgt tcgtttttct gttcccaatt 37920
ggcaatcgct gagcgtaata gtttctctac tctcgaagat gcttccttag atgaaaatcg 37980
taacacacca agtgctttat ttacttccat tccacgaatc atgtcaacaa caagacgcat 38040
tttccgagga gaagtaggaa cattttgcag ctttgcaaaa tatgtagctt taagggcttc 38100
ttttctttta tctgccgata ttttttttct tgttcccatt ataacttaat ttttccttat 38160
aatcattact ttttcttatt acctgcatgt cctctgaagg tacgtgttgg ggcaaattct 38220
ccaagtttat gacctaccat attctcggta acatatacag gtataaattt attcccatta 38280
tgtactgcaa cagtgtggcc aacaaagtct ggcgatatca ttgaagccct agcccatgtt 38340
ttaacaacag ccttttttccc agattcattc attgcaagaa cttttccttc aagcttaaca 38400
ttaatatagg gacctttttt taatgaacga ctcataactt actaattaat cagattactt 38460
tttccttctc tcaataatat acttagaaga ttgcttttta ggagctctcg ttttaagtcc 38520
ttttgcatat agacctttac gagatcttgg gtgacctcca gaagcacgtc cttcaccacc 38580
acccattggg tgatcaacag ggttcattac aactccacga tttcgaggac ggcgtcccat 38640
ccatcttgat cggcctgctt taccagaact ctcaagtgca tgatctgaat taccaacact 38700
acctatagtc gctttacaag tactcaaaat ctgcctaact tccccagaag gcaatttaat 38760
aacgcaatac ttgccttctc ttgatgttaa ttgggcaaaa ttaccagcag aacgaacaaa 38820
agctgctcct tgaccggggc gtaactctat attatgaatc acagtaccaa caggaatatt 38880
ttgaagaggt aaagtattcc ctatttcagg tgcagcatcc gcacctgaca ataatgtaac 38940
acctacttgc aaaccattag gcgcaatgat ataacgtttc tcaccatcag catagtacaa 39000
caacgcaatg cgagctgaac gatttgggtc atactcaatt gtcttaacta ccgcaggaac 39060

```
     accatcctta tttctcttga agtcaacaac acgaataacc ttcttatgac cgcctccaat     39120
     gtaacgcatt gtcattttac cgacattatt acggcctcca gaagacttct taccatatac     39180
     aagagatttt tctggtactt gagcagttat atcatcaaaa gtaccaataa tcttatgtct     39240
     ttgccctggt gttgtgggct taaatttacg tactgccatt ttaaatattg ctataaaaat     39300
     ctatagaatc accatctttt aaagttacta ttgctttctt aaaagcatta gtacgacctt     39360
     tgataatacc agcgcgcgtg taacgacttt tagttttgcc agcatatctt gttgtattaa     39420
     catccacaac agtaacattg taatgcattt caacttctct ctttatttct aatttattag     39480
     cttcaggacg tacaataaaa ccgaaacgat taagtttatc agttattgca gtcattttct     39540
     ccgtcaccaa tggtttaatt ataattccca ttatactact cctttaaatt gaaactttat     39600
     caaaatatga tagcgaactt tcagtaaaca caacaacatc agaattcaac actctataag     39660
     tattgagccc agatgtagtt tgcacagaag caccagcaat gttacgagcc gacaaatata     39720
     cgtttttatt cgtttccaat agaatcatta acagcttttt attggaaata ttaagttttt     39780
     ttgtcaaact tacaaattct ctagtcttag gaacctcaaa agtgaaatct tcaagcacta     39840
     tgatcgcatc atacttagcc ttataagaaa gggctgattt acgagccaac gctttcactt     39900
     ttttattgag cttaaagaaa tagtcgcgag gtcttggacc gaaaactcta ccaccaccaa     39960
     ctaaaacagg agaattcata tcgcctcgtc gagctccccc accgcccttt tgacgaccaa     40020
     ttttgcgggt agaaccgcta atttcacttc tttccttaga tttatgtgta ccttgacgct     40080
     gatttgccat aaattgcttt acgtccagat aaatagcatg atcattgggt tcaataccga     40140
     agatagattc gtttaacgta acctttcttc cagtggcttc accatttata ttatatacgt     40200
     taacttccat tacttttcaa ttattacgat tgaaccttta caaccaggga cagaaccttt     40260
     aatcaacaaa agattgtgtt cctcaatcac ttttaaaact ttaagatttt gcactgtaac     40320
     tctgtctcct ccaagttggc ctcccattcg cattccttta aaaacctttg caggatacga     40380
     gcaagcgcca atagaacccg gtttacgagc acgattatgc tgtccatgtg tcgattgccc     40440
     aacacctcca aaaccatggc gctttacaac accttgaaaa cctttacctt tagatgttcc     40500
     aattacatct acaaaatcct catcactaaa taaggaaaca ttaatcgtgt ctcctaaatt     40560
     aaagacctga tcgaaatcct taaactcagc caagtgtctc ttgggcgtaa ctccagctct     40620
     cttaaaatgg cccatcaagg gcttcgtagt atgtttctct tttttttctt gaaaacctaa     40680
     ctgcaaagct tcataaccat ctactttaga cgttttcacc tgagtaacta cacaaggacc     40740
     tgcttcgata acagtgcatg gtacattttt accatcggca ctgaaaacgg atgtcattcc     40800
     gattttcttt cctaataatc ctggcatttc actaacttat taaattatca aactttaatt     40860
     tcaacctcaa ctccacttgg caactcaagc ttcattaatg catcaactgt tttagcagtc     40920
     gaactataaa tatctatcaa ccttttgtac gaagaaagtt caaattgctc cctagacttc     40980
     ttgtttacaa aagtagatct atttaccgta aaaatacgtt tatgagtagg caacggtata     41040
     ggcccactaa caatagcacc agtacttttt acagttttta caatcttttc agccgactta     41100
     tcaaccaaat tatgatcgta tgattttaat ttaattctaa ttttttgact catcttcagt     41160
     tttaattaaa tattaataaa tagtagtgac ttaagagtca ttcactaagt cactacatta     41220
     attgattata ataaatctgt ccttcctttc act                                 41253
//
ID   XXX; XXX; linear; XXX; XXX; XXX;  XXX.
XX
AC   PRJEB67889
XX
DE   Uncultured bacterium DNA fragment from in-reactor enriched cow rumen, clone
ferm-28d5, contig-1
XX
FH   Key             Location/Qualifiers
FH
FT   source          1..17825
FT                   /organism="uncultured bacterium"
FT                   /environmental_sample
FT                   /mol_type="genomic DNA"
FT                   /isolation_source="Cow rumen"
FT                   /clone="ferm-28d5, contig 1"
FT                   /note="AZCL-Barley beta glucan,  AZCL-arabinoxylan active
```

```
clone"
FT   CDS             167..2233
FT                   /db_xref="SEED:fig|6666666.127246.peg.1"
FT                   /translation="MDVEYTEPIVFDSFSPAEAMPGDVITVKGDYMNLVQSVTFEGGAI
FT                   VTEINHNGRHEASFKVPSKAITGKIILSDEGAIANLLYSEEDLVIGDPTVKSLTVAVAK
FT                   PGENAVVKGAYLDMIDKVVFEGGAEVSDLALNEDATEFNLVIPAGAQSGEVKAVSHAAK
FT                   EFVAGEIELVIPTDVAAAPSPVKAGADLVISGNDLDVVTSVDFPGASGVEFVFAEDAIT
FT                   VTVPATATEGDIALNMASGEAVNVAFTLVHPVVSAIAPVELMAGETITVTGTDLDLVTG
FT                   VTLGGKPVEFTAEAEKIVVQTANTSVSGKLVLSLANGEKIEPEDAITLSYDSFIIVNDM
FT                   PAAEHIGAIVTLKGENFMMIENIYIGEEKVSKYNLRSDNEIQFVMPWNKVGTYSMYFHL
FT                   LSGDVETCPQTIEVLLEQDIMTVWEGSLQITWNDGGRVTVPAASFTGIPAGSKLRFYYT
FT                   QVDQQWDQAQLNYGNWTGINFDDPAGTVINGTLVPTDVYGWFTDGILDRCTEVVLTKEI
FT                   LENIQANKGDCEDKTACGIIIQGSGLTFTKIEIVCDIPQEVTIYEGPTTLTWGDDGRFG
FT                   LAMTYFEAAKPGSQLIFYLEQTDAWGQVQLNDGWWANADMNFPEIGGAYITTDNLGGKD
FT                   VTRIALTLTADILDHILATSGDYFGLNSQFQGDGRVAMVIQGSDMIINKVCIL"
FT                   /product="hypothetical protein"
FT   CDS             2328..3854
FT                   /db_xref="SEED:fig|6666666.127246.peg.2"
FT                   /translation="MIFLAACSCSKDDAEKKEPSVPAPKAVTVDVASVSAPVEGTSVVV
FT                   AISSPSKPAVSVSGDWIKAELGPYNPDTYKVKLTVTVGENPTYESRSGSVTLTCQGFSP
FT                   VGISVSQDAAAKPVGPSANPAYDLLGLGWNMGNQFDAFYNGSWAGDKYNYPDETVWGNA
FT                   KATQATFDGVKKAGFTSVRIPVTWLNMIGPGPEYNIDETWMNRIYEVVGYAEKAGLKVI
FT                   INTHHDENNNDDHWLDILSASKDARTNEAIKAEIAAVWTHIANKFKDKGDFLIMESFNE
FT                   IQDGGWGWSAAFQADPSKQCDVLNEWNQVFVDAVRATGGNNATRWLGVPTYAAGTSWTK
FT                   YFRLPEDPAGKVMMAVHCYDPYEYTIGNIQYPEWGHTGAAGKKTPEGEKEITDLFSRLN
FT                   TEFISKGIPVYLGEFGCSMRSKGDARAWMFYKYYMEYVSKAAHTYSLPGILWDNGSTKT
FT                   GKECHGYIDHGTGKYIGNSKEIVDLLVKGMLSTDESYTLQSVYDRAPKTN"
FT                   /product="GH5_4"
FT   CDS             3968..4999
FT                   /db_xref="SEED:fig|6666666.127246.peg.3"
FT                   /translation="MMDRMAASVKAGKILYGHQDDLVYGHSWCVDPDSAATDALNRSDV
FT                   KAVAGDYPAVVGFELGGLELGWDWSIDKVPFLLMRRAAVEHAGRGGIVTFSWHPHNPYT
FT                   GGDAWDISSDKAVASVLEGGEKAEMFGVWLERLGDFMDSLRDDKGNLIPVIFRPWHENI
FT                   GSWFWWGGALCTADEYIALFRRTVEYLRNDRGLDNIIMCYSPNSGITAEQYMSRYPGDD
FT                   VVDILGYDHYETVMEGESLEDAGKRYAGQLDLGLSIVKSLASAHGKLMTLSETGMEGLP
FT                   DRSWWTSVLYPAVKDFPIVYVLTWRNAWDRETHFYAPYPGNEGDFREFASLPEISLLSS
FT                   FRP"
FT                   /product="GH26"
FT   CDS             5077..6252
FT                   /db_xref="SEED:fig|6666666.127246.peg.4"
FT                   /translation="MENMSFEERLAWLEARHQELIAKKNSPIYGNGVFERYENPVLTGD
FT                   HAPLFWRYDLDKNSNPFLEERFGIHAAFNAGAIKWKDKYVVCARVEGADRKSFFAIAES
FT                   PNGIDNFRFWDHPVTMPDYGEPATNVYDMRLTAHEDGWIYGIFCVERKDPDAPEGDLSS
FT                   AVAAAGIARTKDLVNWERLPDLVSDSQQRNVVLHPEFVNGKYALYTRPQDGFIDAGNGG
FT                   GIGWALVDSMENAVVTSEKIINCRRYHTIKETKNGEGPAPIKTDRGWLHLAHGVRGCAS
FT                   GLRYVLYMYMTSLEDPSKVIAEPAGFFMVPEGEEYVGDVMNVLFSNGWIADEDGKVFIY
FT                   YASCDTRLHVATSTVDRLVDYCMNTEPDLFTTSASVRRLNALIDKNLSLSM"
FT                   /product="GH130"
FT   CDS             6273..7700
FT                   /db_xref="SEED:fig|6666666.127246.peg.5"
FT                   /translation="MDIITAPAKPTLKEKIGYGFGDMSSSMFWKIFSYYLPIFYSDVFG
FT                   LKPQHAAFLLLITKLYDAISDPVMGIIADRTRSRWGKYRPYLLWVALPFSLIGLLSFYT
FT                   PDASYGLKHVYAYVMYILMMTVYTAINVPYGAMLGVVTPDSREKSVFSSYRMFFAYIGS
FT                   FIAMGIFGVFEKLISGTVRVMPDGSERVVRGVGDAAPGQWTLVVGIVSLCCFVLFLLCF
```

```
FT                   LMTRERVGTENKAVSEGGSSIAEDLKALGRNRPWWLLLGAGIGILLFNSIRGGAAAYYF
FT                   SNILGTSVFLTCAVYLFIGEVAQMVGVLFTVPLSERIGKKATFISVLLTVSVLSVAIFF
FT                   LPETPAGFWGLLVLQVLICIAFGVQSPLLWSMFADVADYSEEKNGSASTGLIFSSSSMA
FT                   QKFGGALGAFLLMQILAIFAYDKDAAVQAPQTLAAIRGLMSYIPAIGALLGALCLLFYP
FT                   LTESRMASIREKLSARR"
FT                   /product="Xyloside transporter XynT"
FT   CDS             8185..9375
FT                   /db_xref="SEED:fig|6666666.127246.peg.6"
FT                   /db_xref="GO:0003887"
FT                   /translation="MQDCGKDEGLEGIEKNVIFVFRSLIRRTMADKLYLIDGHNLIFKM
FT                   YYAFLRRPMINSKGMDMSILYGFTKYVLELIEREKPTHMAIAFDPPGGTFRNEMYPEYK
FT                   ANRSETPQLVIDALEPLLEICKALNIKTLMIPGYEADDVIGTVAKSYGKPGTDVYMVTP
FT                   DKDYGQLVAEHVYQVKPGKAGGADETVDVKAVCEKYGISSPEQVIEMLTLCGDTADNVP
FT                   GVHGIGEVGAGKLIAKYGTVENIYAHLDELTDRQKTQFEEARGHIGLSHDLVTIKTDVP
FT                   VETSLDEMRLDLDYAPEIADLFEKYEFNSLRKYIAHSSRAAKAVAEVKRLDYTETGAEE
FT                   LVKMAPEKISLIVEPEGPGIFADIRRVIVAAEKKDGGIVVLQALRRIFARYWRAKP"
FT                   /product="DNA polymerase I (EC 2.7.7.7)"
FT                   /EC_number="2.7.7.7"
FT   CDS             9372..10916
FT                   /db_xref="SEED:fig|6666666.127246.peg.7"
FT                   /db_xref="GO:0003887"
FT                   /translation="MKYGYDLKLQSNLLENAGIKLGGRLYDIELMHYLINPEKSHKVEI
FT                   LAKSYLGVDIEEAAAVEEAPVSLFDVAEESPANGPDRNLEAAAVLKLGEVVRAGMDEAG
FT                   VGALYEDMEAPLAAVLARMERAGVKIDLGQLRTFADELKVEMDALEAQVRKLADEPGLN
FT                   VSSPKQIGIVLFEKLKLDPKMKKSASGSYSTDETTLTAIADRHPIIDKILEFRAVKKLL
FT                   STYIEPFPTYVSPVTGRVHTTFNQALTATGRLSSSNPNLQNIPIRTERGKEIRKAFIPA
FT                   TPDGIIMSADYSQIELRIMAHLCGDEHLISSFKAGLDVHKATAAKIFGIPVEEVTSDQR
FT                   RIAKTANFGIMYGISSFGLAQRLHVSRTEAKKIIDDYFAAFPSIRTFIDKTIADAREKG
FT                   YVETVFGRRRYLPDITAKNATIRALAERNAVNAPIQGTSADIIKLAMISVDKRIREAGL
FT                   KSKMVLQIHDELLFDAVPEEIEELGRIVVAEMENVITLSIPLTVECNYGKNWLEAH"
FT                   /product="DNA polymerase I (EC 2.7.7.7)"
FT                   /EC_number="2.7.7.7"
FT   CDS             10888..11493
FT                   /db_xref="SEED:fig|6666666.127246.peg.8"
FT                   /db_xref="GO:0003677"
FT                   /db_xref="GO:0003700"
FT                   /db_xref="GO:0006352"
FT                   /db_xref="GO:0016987"
FT                   /translation="MAKTGLKLTDHDDGSLVEMALGGGPEAEAAFFTIYARYHKGVFAH
FT                   ISRFIQEQEEVEDICMESFEKAFKQLESYRPSNKFSTWIFTIARNTAFDHLDKDKVRGR
FT                   NKDKMPIDYSDLEHIDVPEDNVSPEDEIISSQDHENFLSCVQGLPDLYREVASMCFVDN
FT                   LGYKEISEKAGLPINTVKTRISRAKSLIVQMMLDMEEE"
FT                   /product="RNA polymerase sigma factor RpoE"
FT   CDS             11490..12152
FT                   /db_xref="SEED:fig|6666666.127246.peg.9"
FT                   /translation="MNFKEFEDIVVAEFPELTPEMLGKYEAMEPLYEEWNAKINVISRK
FT                   DIGGLYLHHVLHSLAIASYLKRRRPEVYSAWTEGASVLDLGTGGGFPGIPLAVMFPKVR
FT                   FTLCDSVGKKIIVASEVSKALSLGNVTTVNARAESLPQVFDHVVSRAVTSLDNFYPWVK
FT                   GKFSGSILYLKGGDIVPEIAELMKKNRLRTGAVSTWPIETWLKDEYFKEKFVIEIEK"
FT                   /product="rRNA small subunit 7-methylguanosine (m7G)
FT                   methyltransferase GidB"
FT   CDS             12631..15408
FT                   /db_xref="SEED:fig|6666666.127246.peg.10"
FT                   /translation="MQVWLNEDHSQPVVHGAVVVKAGGRDCAGTGMAHYLEHMLFKGTE
```

```
FT                   DIGTIDYASEKVYLDSIETCYDLIAATSSSVKRDSLQKEINRLSIKALDYAIPNEFDRL
FT                   TANIGGTKLNAFTSYDCTCYHNSFSPQFIAQWAELNSHRLLNPVFRLFQSELEAVYEEK
FT                   NAAADSQFDMILEKMLEKLGDENPYSKSIIGTTDNLKNPSLSEMKKFFNKYYVGCNMGL
FT                   ILAGDIDASSILPLLESTFGRIPRGDKPERIPVPVPDFSSREDYPIKVNFPMIKASLNV
FT                   FNGPAAKDADYPAMELALSLLNNSFSTGSLDSLINNGKVLAGEVSVLDFFDEMGLIAMI
FT                   TIPGIPFGSLNKAEARCFKEVERLKAGDFAEKDFESLKYDLVKSYKRCLETSESRFSIM
FT                   IDVMSQDRSWDSYLSQVESISSLTSEDIVRVSNKYLGDNRIRFHKVRGTYPKDNLQAPS
FT                   YETLSPKNGNAVSDYATRLMEMPYDTLPPRLVDMNNDIRKVAVNDHVTLYYNGNPVNDV
FT                   FELTVRINKGIFEEPMICPLTQLLGLSGTDSLSARDFSMSLAELGSDLSFKDKNHTLEM
FT                   VLTGFDDRIGPSLDLLRHFMKDFKPEKSALNTVISDTKMSRNAFRSSGNSAIFNGMFNY
FT                   SLLGNQSPLLRQPTEKDLKSLGLKGMKELFNSLVEYDFDVVYSGGLSPEEVVNEISGHL
FT                   DVECPSGKPESSFFAYRKYEQPVVYIYNVPGARQVMSITYQNVATPENDHDSALLGLMQ
FT                   ECYGGSMASIMFQEVREFRSLAYTSSSTLQKPLPNVSSPSAFITYYGTQADKFPQALEL
FT                   VDSLIRTMPMTDLCFDSAVRSIISRANNGYPSFRALPEYVANLELFGYGEDDDASVVRG
FT                   LPGITKQDVLDYHEKSIGNAPCSYFIIGDKKNLPMDALRERGRIVELKLSDIYR"
FT                   /product="ZINC PROTEASE (EC 3.4.99.-)"
FT   CDS             complement(15422..16081)
FT                   /db_xref="SEED:fig|6666666.127246.peg.11"
FT                   /translation="MRIKDLNIGHRGKTLVSGISFETEPGEFILLCGPNGSGKSTLLRA
FT                   LAGKDAIMVPTGVPKVKGFTLREFIATTLYSESDWRGRISPAMASDISKSLECLGISSL
FT                   ADRDISTLSDGEFQKGCIASALSRHAKTLLLDEPTSFLDVDNKSLILQALRDIARSTGT
FT                   TVIMASHDLAPALSCADRVFGITADHRFLASSSRLSDSGLSASEVISACFSTYSSR"
FT                   /product="iron (III) ABC transporter ATP-binding"
FT   CDS             complement(16078..17079)
FT                   /db_xref="SEED:fig|6666666.127246.peg.12"
FT                   /translation="MAIKRRNRLITAAGATALLLLSVCDLILGGSGIGAAEALGGLFGG
FT                   EGIASSIVWSIRLPRVLTAIAAGACLALSGAQMQSVFRNPLADPHIMGVSGGAGLGAAI
FT                   ATLLISGTGLLSGFSSMAGLTVAGAAFVGAILASALIVLVSSRFRSGTTLLVFGVMLGF
FT                   IFSAITSILEYSASEESLKLFYSWSAGSFSGTRTFETLTIFAALLVGTGLYTSGAKGLD
FT                   ILLFGEEYATLAGASVRRIRGVAMVSSCLMTAAATAFCGPLGFVGIVSPHIARRLLRSS
FT                   GHMSILPATLITGSAVSVLADIISQAPGLPLPVGSTMALIGIPIILLILMKK"
FT                   /product="Vitamin B12 ABC transporter, permease component
FT                   BtuC"
FT   CDS             complement(17088..17759)
FT                   /db_xref="SEED:fig|6666666.127246.peg.13"
FT                   /translation="MAGVVGHHGLSLEVHSFGQAGVLQDVGEEHFAEISLDLGIAGKRV
FT                   GEIAGLVPDGFAAGVEILNALFEAGVPLYGVFLAFLDGVAELFEFLLDRFHQFRDCLGA
FT                   LCLEGLACLGRKVLEDFLHLLELFGMPFRGLLRGFAHLILIFLLSLQATLQYGFFALDI
FT                   GQSVLESVFDLFGTRLETENQYRSYDNDCDKKNYQNCFGAHPQCILIFQIYENPLKFLS
FT                   L"
FT                   /product="hypothetical protein"
XX
SQ   Sequence 17825 BP; 4338 A; 4459 C; 4853 G; 4175 T; 0 other;
     cgacaggatt gtcgaagttt cagttccggg actcgatcct atcaccgaca tcgaagtcgt        60
     tacctccggt aatccttcag aaatccgcat ccagcttcct gcagagggcg cacaggtcgg       120
     aaagatctct ctcaagtcat ccgacggaaa ggtctttact acccaggtgg atgtagaata       180
     tacagaacct atcgtatttg acagtttctc tcctgcagag gctatgcctg gcgacgtgat       240
     aacggtcaag ggcgattata tgaacctcgt gcagagcgtt accttcgaag gcggcgccat       300
     cgtgaccgaa atcaaccata acggccgtca tgaggcttcc ttcaaggtcc cttcaaaggc       360
     tattaccgga aagatcatcc tttctgacga aggagccatc gccaatctcc tctactcgga       420
     ggaagatctt gtgatcggcg atccgaccgt caagtctctt accgtagctg ttgctaaacc       480
     tggcgagaac gcagttgtga agggagcata tctcgatatg atcgacaagg tcgtattcga       540
     aggtggtgca gaggtttccg accttgctct caatgaggat gcaaccgaat tcaatctcgt       600
     aatccctgcc ggagcacagt ccggtgaagt aaaggctgtc agccacgctg ccaaggaatt       660
```

cgtcgccggc gaaatcgaac ttgtcatccc tacggatgtc gcagctgcac cttcgcctgt 720
aaaggctggt gccgacctcg tgatttccgg taacgatctt gatgtggtaa cttctgttga 780
tttccctggc gccagcggag tagaattcgt atttgccgag gacgccatta cagtgaccgt 840
tcctgctact gctactgaag gtgatatcgc cctcaatatg gccagcggag aagctgtgaa 900
tgtggcattc acccttgtgc acccggtagt ttctgccata gctcctgtcg agctcatggc 960
cggcgagacc atcaccgtga caggtaccga cctcgacctt gtcactggag taactctcgg 1020
cggaaagccg gtcgagttca ctgcagaggc tgagaagatt gtcgtacaga ccgccaatac 1080
ttccgtttca ggaaagctcg tgctttctct tgcgaatggt gagaagattg aaccggagga 1140
tgccatcacc ctcagctatg attcattcat catcgtgaac gacatgcctg ccgcagagca 1200
tatcggggct atcgtaaccc tcaagggcga gaacttcatg atgatcgaga acatctatat 1260
cggcgaagag aaagtatcga agtacaacct tcgttctgat aatgaaatcc agttcgtaat 1320
gccttggaac aaggtcggaa catacagcat gtacttccat cttctctccg gcgatgtcga 1380
gacatgtcct cagacaatcg aagttctcct cgagcaggac atcatgactg tatgggaagg 1440
ctctctgcag atcacctgga atgacggcgg ccgtgtaacc gtgccggcag catcattcac 1500
cggtatccct gcaggcagca agctcagatt ctactacact caggtggacc agcagtggga 1560
tcaggcccag ctcaactatg gtaactggac cggaatcaac ttcgacgatc ctgcaggtac 1620
cgtcattaac ggaaccctcg ttccgactga cgtctatgga tggttcactg acggaatcct 1680
tgaccgctgc acagaggtcg tgcttacaaa ggaaatcctc gaaaacatcc aggctaacaa 1740
gggtgactgc gaagacaaga ctgcctgcgg tatcatcatc cagggctctg gccttacctt 1800
cacgaagatc gagatcgtat gcgatattcc gcaggaggtt acgatctacg aaggtcctac 1860
aacccttaca tggggtgacg acggccgttt cggtctcgcc atgacttact tcgaggcagc 1920
aaagccgggc tctcagctta tcttctatct cgagcagacg gacgcatggg gccaggttca 1980
gctcaacgac ggctggtggg ccaacgccga catgaacttc cctgagatcg gcggtgcata 2040
cattaccacc gacaacctcg gaggaaagga cgtcacacgt atcgccctta ccctgactgc 2100
tgatatcctt gaccatatac ttgctacatc cggtgactat ttcggtctca atagccagtt 2160
ccagggtgac ggacgcgtcg ccatggttat ccagggatct gatatgatta tcaataaagt 2220
ttgtatattg taacactttt attgccggag gcgggtgatg ccgcctccgg tagccagtaa 2280
acaaatttta tataatgaaa gcacacgtca gtttctttgt attatctatg atttttcttg 2340
cggcatgttc ctgctcaaag gatgatgccg aaaagaagga accgtccgtc ccggccccta 2400
aggcggtaac ggtagatgtc gcgtctgtca gcgctcccgt agagggaacg tcggtcgtcg 2460
tcgccatctc ctccccgtcc aagcctgcgg tctctgtctc cggagactgg atcaaagccg 2520
aactgggtcc gtacaatccg gatacatata aagtgaaact tacggtgacc gtcggcgaga 2580
atccgacata cgagtcacgc agcggtagtg tgacccttac ttgccagggc ttttctccgg 2640
tcgggataag cgtatcccag gatgccgcgg cgaagccggt cggaccgtcg gctaatcctg 2700
catacgatct tctcggtctc ggctggaata tgggtaacca gttcgatgct ttctacaacg 2760
gctcatgggc cggagataaa tacaattatc cggacgagac cgtctggggt aatgccaagg 2820
cgacccaggc tactttcgac ggggtgaaga aagccggatt cacatcagtg cgaatccctg 2880
tcacgtggct caatatgata ggtccaggac ctgagtacaa tattgacgag acttggatga 2940
accgtatata tgaggtggta ggatatgccg agaaagccgg actcaaggtc atcatcaata 3000
cccatcacga cgaaaacaac aatgacgacc attggctgga tatcctttct gcatctaagg 3060
atgccaggac caatgaagcg ataaaggcgg aaatcgcggc tgtatggacc catatcgcca 3120
ataagttcaa ggataaggga gatttcctta tcatggagtc cttcaatgaa attcaggacg 3180
gaggatgggg ctggagcgct gctttccagg ctgacccttc caagcagtgt gatgtgctta 3240
acgaatggaa tcaggttttc gtggatgccg tacgtgctac aggaggcaac aacgctaccc 3300
gctggctggg tgttcctaca tacgccgcag gtacttcatg gacgaagtat ttcagactgc 3360
cggaggatcc tgccggaaag gtcatgatgg cagtgcattg ctatgatccg tatgaatata 3420
ctatagggaa catacagtat cctgagtggg gccataccgg cgccgcagga aagaagaccc 3480
cggaaggga gaaagagata acggatctct tctccagact taatacggaa ttcatcagca 3540
aggggatacc tgtatatctt ggcgaattcg gctgttcgat gagaagcaaa ggggatgccc 3600
gggcatggat gttctataaa tactatatgg agtatgtctc caaagctgcg catacctatt 3660
ccttaccggg aatcctgtgg gataacggaa gtactaagac cggaaaggag tgccacggct 3720
acattgacca tggtaccggc aagtatatag ggaattctaa ggagatcgtg gatcttctcg 3780
taaagggaat gctcagcacg gacgaaagct atactctcca gagtgtatat gacagggctc 3840
cgaaaaccaa ttgattatga atctaatcct taaacttagc atcattttat cattagtagt 3900

tgctgtatcc ggctgcggac gcaataaaga agcggccgga acagctttta acgaagaaac 3960
ccggatcatg atggatcgta tggcggcttc agtaaaggcc ggcaagattc tttatggcca 4020
tcaggatgac ctcgtttatg gccattcatg gtgtgtggat cctgattccg cagcgacaga 4080
tgcgttgaac cgttcggatg tcaaggctgt cgccggagat tatccggctg tcgtgggatt 4140
cgaactcggc ggtctggaac tgggatggga ctggagcatt gacaaagtcc cgttcctgct 4200
gatgcgccgg gctgctgtag agcatgccgg aaggggagga atcgtgacgt tctcatggca 4260
cccgcacaat ccttatacag gcggtgatgc ctgggatatc agttccgaca aggcagtggc 4320
ttctgtcctc gaaggagggg agaaggcgga gatgttcgga gtctggctgg agaggctggg 4380
cgatttcatg gactcgctcc gtgacgataa gggtaatctg ataccggtca ttttcaggcc 4440
gtggcatgag aatatcggca gctggttctg gtggggcgga gccctttgta ctgcggatga 4500
gtatatcgcc cttttcagga ggacggtcga gtatctgaga aatgacaggg gactggacaa 4560
tatcataatg tgttattctc cgaattccgg gattacggcg gagcagtata tgtcccgtta 4620
tccgggcgat gatgtcgttg atatcctggg gtatgaccat tatgagactg tcatggaggg 4680
cgagagtctg gaagatgccg gaaagcgtta tgccgggcag ttggatcttg gcctttccat 4740
cgtcaagtct cttgcttcgg cccacggaaa gctgatgact ctttccgaaa ccggaatgga 4800
gggcctcccg gacaggagct ggtggacttc ggtgctttat cctgcggtaa aggattttcc 4860
gattgtctat gtgctgacgt ggagaaatgc atgggacagg gagacgcatt tttatgcgcc 4920
gtatcctggt aatgagggcg acttcaggga attcgcttcg ctgccggaga tctctctcct 4980
gtcgtcgttc cggccttagt ccgtcattcc cggcgaagtg tcgtcctttc cggtcggacc 5040
gggaatctgt aaccttaaaa caataaataa caatatatgg agaatatgag ttttgaagaa 5100
cgtctggcgt ggctggaagc acgccaccag gaacttattg ctaagaaaaa ctcgcctata 5160
tacggcaatg gtgtgtttga gagatatgag aaccctgttc ttaccggtga ccatgcccct 5220
ttgttctgga gatatgatct ggataagaac tccaatcctt ttctggaaga gagattcggc 5280
atccatgccg cattcaatgc tggagctata aagtggaagg acaaatatgt cgtctgcgcc 5340
cgtgtcgaag gtgctgaccg caaatcattc ttcgctatcg ccgagagccc gaacggaatc 5400
gacaatttcc ggttctggga ccaccccgtg actatgcctg attatggcga acctgccacg 5460
aatgtatatg acatgagact tactgcccat gaagacgggt ggatctatgg tatattctgc 5520
gtcgagcgta aggatccgga tgctccggaa ggcgatcttt cgtctgccgt cgcagctgcg 5580
ggaatcgccc gtaccaagga tctggtcaat tgggaaagac tgccggatct ggtttctgat 5640
tctcagcagc gcaatgtcgt acttcatcct gagttcgtga atgggaaata tgctctttac 5700
acccgtcctc aggatggatt catagatgcc ggaaacggcg gcggcatcgg ctgggccctg 5760
gtggacagca tggagaatgc cgttgtcact tccgagaaga tcatcaactg caggagatac 5820
catactatca aggagaccaa gaatggcgaa ggccctgctc ctatcaagac tgacaggggc 5880
tggctgcatc tggcccatgg cgtcaggggc tgcgcttccg gcctgcgtta tgtgctttat 5940
atgtacatga cttcgctgga ggatccttcg aaagtgattg ccgaacctgc cggtttcttc 6000
atggtcccgg aaggggagga gtatgtgggt gatgtcatga acgtcctgtt ctctaacggt 6060
tggattgccg atgaggacgg aaaggtcttt atatattatg cttcctgcga tacgcgtctc 6120
catgtggcga cttctaccgt cgacagactc gtggattact gtatgaatac tgagccggat 6180
ctgtttacca cttctgcttc agtacgacgt ctgaacgcgc ttatagacaa gaatctttcg 6240
ctatctatgt aggactaata atcaggtttc agatggatat cattactgca ccggctaagc 6300
cgacattgaa ggaaaagatc ggatatggct tcggagacat gtcgtcttcg atgttctgga 6360
agatttttttc gtattatctg ccgatcttct attcggatgt gttcgggctt aagcctcagc 6420
atgcggcttt cctgcttctt atcacaaaac tgtatgatgc aatatcggac cctgtcatgg 6480
gaatcatcgc cgacaggacc cggtccagat ggggaaagta tcgtccgtat cttctctggg 6540
tggctctgcc gttctcgctt ataggccttc tgtctttcta tactcctgat gccagctatg 6600
gcctgaagca tgtatatgcg tatgtgatgt atatcctcat gatgactgtc tatacggcta 6660
taaacgtgcc ttacggggcc atgctgggag tcgtcactcc ggacagccgt gagaaatcgg 6720
ttttctcgtc ttacaggatg ttctttgcgt atatcggcag tttcatcgcc atgggtatat 6780
tcggcgtgtt cgagaagctg atttccggta ccgtgcgtgt tatgccggac ggtagcgaga 6840
gggtggtgcg cggagtcggc gatgctgctc ccggacagtg gactcttgtc gtcggcatcg 6900
tttctctgtg ctgtttcgtg ctgtttctgc tttgcttcct tatgaccagg gagcgggtgg 6960
ggactgaaaa taaggcggtt tccgagggag gctcttctat cgcagaggat ctcaaggctc 7020
taggccgtaa caggccttgg tggctgcttc ttggcgccgg tatcggtata ttgctgttca 7080
attcgatccg aggcggagcc gctgcatatt atttctcgaa tatcctcggt acgagcgtgt 7140

tcctgacttg cgctgtctat cttttcattg gcgaggtggc tcagatggtt ggggtgctgt 7200
ttactgtgcc gctttcggag cgtatcggca agaaggcgac ttttatctct gtgcttctga 7260
cggtttctgt gctgagtgtg gcgatattct tcctgcctga gactcctgcc ggtttctggg 7320
gacttctggt gctgcaggtg cttatatgca ttgcttttgg tgtccagtct cctctgctgt 7380
ggtcgatgtt tgccgatgtc gccgattatt cggaggagaa gaacggctcg gcttctaccg 7440
gacttatttt ctcgtcttct tctatggctc agaagttcgg cggggccctc ggcgctttct 7500
tgctgatgca gattcttgca atcttcgcat atgacaagga tgctgcagtc caggcgcctc 7560
agactcttgc tgccatccgt ggcctgatga gctatatccc ggcaatcggc gccctcctcg 7620
gagccctctg cctgctgttc tatcctctta cggaatcgcg tatggcttca attagagaga 7680
aactttctgc tcgcagataa catcttcatt catgtaggta tttgttattc caaaggatgg 7740
ccctcttcgg ccatcctttt tcgtctggct gattcgagtc gggcctttct gctggcggga 7800
gaaggacggg acggctccgt caagttcgga aaagctgctt ggtcgctgtc gccgcctaaa 7860
cggctaactt ctccctttta cgatgggcga gccatcgtaa cggtcgtcaa acacacgccg 7920
ttcttcacgg cgccgacggc cgctcccgcg ctctccgctt ttcctcaatt cctccgccgc 7980
ctccgccctc ctcccgccgc aggactgctg accagggttc cggggacgtg gcccccctcg 8040
gccacgggaa gagggacggg gcccgcaaga tgcaagttat cgcgaagcga tgacgcggca 8100
gattgtggga gggtacggag tgagcgatag cgaacgaagt tccccgggac cctgcgtcag 8160
caggactgtg gcaaaggaag cagaatgcag gactgcggca aagatgaggg actggaggga 8220
attgaaaaaa atgttatatt tgtatttcgt tccttaatcc ggagaactat ggcagacaaa 8280
ctatacctga ttgacggtca caacctaatc ttcaagatgt actacgcatt cctgcgtcgt 8340
cctatgatca attccaaggg aatggacatg tcaatcctgt atggcttcac gaaatatgtg 8400
ctcgaactca tcgagagaga gaagcctacc catatggcca tagcgttcga tcctcccggc 8460
ggcaccttca ggaacgaaat gtaccccgaa tacaaggcaa acagaagcga aactccgcag 8520
ctcgtgatcg atgccctcga accgctgctg gaaatatgca aagccttgaa cataaagacc 8580
ttgatgattc cgggatatga ggccgacgac gtgatcggaa ccgttgcgaa gagttacggg 8640
aaacccggga cggacgtcta tatggtcacc ccggacaaag attacggcca gctggtcgcc 8700
gaacatgtgt accaggtgaa gcctggaaag gccggcgggg ctgacgagac cgtcgatgtc 8760
aaggcagtct gcgaaaaata cggaatctcc agtccggaac aggtcatcga aatgctgacc 8820
ctctgcggag atacggccga caatgtgccc ggagtgcatg gcatcgggga agtcggggca 8880
ggcaagctca tcgcaaaata tggcaccgtc gagaatatct atgcccatct ggacgagctg 8940
actgaccgcc agaagacgca gttcgaggag gccaggggac atatcggcct gagccatgac 9000
ctggtgacaa taaagaccga cgtgccggta gagacaagcc tcgacgagat gaggctggac 9060
ctggattatg ctcccgaaat cgccgacctt ttcgaaaaat acgaattcaa ctcgctgaga 9120
aaatatatag cccatagctc ccgcgctgcc aaggcagtgg cggaggtcaa gcgcctggat 9180
tataccgaga caggggcgga agagctcgtg aagatggctc cggaaaagat ttcgctgata 9240
gtcgaaccgg agggaccggg catattcgcc gatatccgca gggtgatcgt ggccgcggag 9300
aagaaagacg gcggaatcgt cgtccttcag gcgctgcgaa ggattttcgc aagatactgg 9360
agggcgaagc cgtgaaatac ggatatgacc tcaagctcca gtcgaatctt ctggagaatg 9420
ccggaataaa gctcggaggc cggctatatg acatcgagct gatgcattat ctcatcaatc 9480
cggagaagag ccacaaggtc gaaatccttg ccaagagcta tctcggcgtc gatatcgaag 9540
aggccgctgc tgtggaagaa gctcctgttt ccctgtttga cgtcgcggaa gaaagcccgg 9600
caaacggacc ggacaggaat ctcgaagccg cggcagtgct gaaactcgga gaagtggtcc 9660
gcgcaggcat ggacgaggcc ggcgtaggag ccctgtatga ggatatggaa gcccctctcg 9720
cagccgtgct cgcccgcatg gaaagggcag gcgtgaagat agacctcgga cagctcagga 9780
cgtttgccga tgaactcaaa gtcgaaatgg atgccctcga agcccaggtc aggaaactgg 9840
ccgacgagcc ggggctgaac gtttcttccc cgaagcagat tggcatcgtg ctcttcgaaa 9900
agctgaaact cgacccgaag atgaagaaat cggccagcgg tagctatagc accgacgaaa 9960
cgacgcttac tgccatcgcc gacaggcacc ctataattga caagatactt gaattcagag 10020
ccgtaaagaa gctgctctcc acatacatag agccatttcc gacctacgtc tccccggtta 10080
cgggaagggt ccatacgacc ttcaaccagg cgctgacggc gacgggaaga ctcagttcct 10140
ccaaccctaa cctgcagaac attccgatcc gcacggagag gggaaaggag atcaggaaag 10200
ccttcatacc ggcgactccg gacggaatca taatgtccgc cgactactcc cagatcgagc 10260
tcaggatcat ggcccatctc tgcggcgacg agcatctgat aagctcattc aaggcagggc 10320
tggatgtcca taaggccacc gcagccaaga ttttcggcat accggtagag gaggtcacat 10380

ccgaccagag gcgcatcgcg aagaccgcaa acttcggaat catgtacggc atctcatcct 10440
tcgggctcgc ccagaggctg catgtctcca ggacggaagc caagaagata atagacgatt 10500
acttcgccgc cttcccttcg atccggactt tcatcgacaa gactattgcg gacgcccgcg 10560
aaaaagggta tgtcgagaca gtcttcggac gcaggaggta tcttccggac ataacggcga 10620
agaacgctac cataagggct ctggccgaga ggaatgccgt aaatgctcct atccagggca 10680
catcagccga tataatcaag cttgcgatga taagcgtcga caagcgcatc cgcgaggcgg 10740
gactgaagag caagatggtg ctccagatcc atgatgagct tcttttcgat gccgtgccgg 10800
aagaaatcga ggaactcggc aggatagttg tagcggaaat ggagaacgta attaccttgt 10860
caataccttt aactgtagaa tgtaactatg gcaaaaactg gcttgaagct cactgatcat 10920
gacgacggat ctcttgtgga gatggcgtta ggtggcggcc ccgaagccga ggccgctttc 10980
ttcacgattt atgcacgtta ccataagggc gtattcgccc acatctcacg cttcatccag 11040
gaacaggagg aagttgagga tatctgtatg gagagcttcg agaaggcctt caagcagctg 11100
gaatcataca ggccgagcaa caagttctcg acttggatat tcactatagc ccgcaacacg 11160
gccttcgacc acctcgacaa ggacaaagtg cgcggccgca acaaagacaa gatgccgatc 11220
gactactccg acctggagca tatagacgtc cctgaggaca atgtcagccc ggaggacgag 11280
atcataagtt cgcaggacca tgagaacttc ctgagttgtg tccaggggct tccggacctg 11340
tatcgcgaag tcgcatccat gtgctttgtg gacaatctcg gatacaagga gatttcggag 11400
aaggcagggc tgcctatcaa tacagtaaaa acccgcatca gcagggctaa gagcctgata 11460
gtccagatga tgcttgatat ggaagaagaa tgaatttcaa agaattcgaa gatatagttg 11520
ttgctgagtt tccggagctg actccggaga tgcttggaaa gtacgaggcc atggagcccc 11580
tgtatgagga gtggaacgcc aagatcaacg taatttcccg aaaagacata ggcggccttt 11640
atctgcacca tgtgctgcat tctcttgcga tcgcctcgta tctgaagaga cgcaggccgg 11700
aggtgtattc cgcatggacc gaaggcgcgt ccgtgctcga cctgggcacc ggcggcggct 11760
ttccggggat tcctctcgcg gtaatgttcc cgaaagttcg ctttaccctc tgcgattccg 11820
tgggcaagaa gataatcgtg gcatcggaag tgtccaaggc tctttccctg ggcaacgtga 11880
ccactgtcaa tgcccgtgca gaatcgctgc cgcaggtttt cgatcatgtc gtttcccgcg 11940
ccgtgacctc gctggataat ttctatccat gggtgaaggg caagttctcc ggaagcattc 12000
tgtatctcaa gggaggggac atagtcccgg aaattgcaga actgatgaag aagaacagac 12060
tccgtaccgg agcagtttcg acctggccga ttgaaacttg gctgaaggac gaatatttta 12120
aggaaaagtt tgtgattgaa atagaaaaat aatttgcaaa tccacataaa ataactacct 12180
ttgcagtcct atttccggaa aactggtttt ttcggcctga taagaataaa taaaaaaaca 12240
gataacgatg aaaagaacat tccaaccatc aaagcgcaag cgtataaaca agcacggttt 12300
ccgcgagcgt atgtcaaccc ctaacggccg tcgtgtactt gctgcacgcc gtcgtcatgg 12360
ccgcaaaaaa ctcagcgttt cttcagagga tagattctag tcatctctct ccggacgttg 12420
tttgtttgga tcgaaattca ggaggcttcc gaaaggatgc ctcctttttt cattatggca 12480
gattatttga tagacgaggg acgtaatcat taagtctcaa tggaaaagtt tttttatca 12540
atcttgtctt gcgcgctttc agcgacaata gctgtttctg cgcagagtag cggaccatta 12600
gttgtccacc agtatgaatt gtcaaacgga atgcaagtct ggcttaacga agaccattcg 12660
cagccggtag tccacggagc cgtggtcgtc aaggccggag gccgggattg cgccgggacc 12720
ggaatggccc attatctgga gcatatgctc ttcaaaggta cggaggacat cggtaccatc 12780
gactatgctt cagagaaggt ctatcttgat tctatcgaga cttgttacga cctcatcgca 12840
gcaacatcct ccagcgtaaa aagggattct ctccagaagg agatcaaccg tcttagcatc 12900
aaggcgctgg actatgcgat tccgaatgaa ttcgaccgtc ttacggccaa tatcggaggc 12960
acaaagctca atgctttcac gtcttatgac tgtacatgct accataattc tttttctccg 13020
caatttatcg cgcagtgggc cgaactcaat agccataggc tgctgaaccc tgttttcagg 13080
cttttccaga gcgaacttga agcagtatat gaggagaaga acgcggcagc cgattcccag 13140
tttgacatga ttctcgagaa gatgctggag aaacttgggg atgaaaatcc atacagcaaa 13200
tctataatcg gaactacaga taacctgaag aatccaagtc tttccgagat gaagaagttc 13260
ttcaacaagt attatgtggg ctgcaacatg ggacttatcc ttgccggaga catagatgct 13320
tcgtcgatat tgcctctgct ggaatctacc ttcggacgga ttccgcgtgg cgataagccg 13380
gaaaggattc ctgtacctgt gccggatttc tcgtcccggg aagattatcc tataaaagtt 13440
aatttcccta tgatcaaggc gtcgctgaac gtgttcaacg gtcctgcagc caaggatgcc 13500
gattatcctg caatggaact cgctctcagc ctcctgaaca acagttttag taccggaagt 13560
ctggattccc tcataaataa cggcaaggtt ctggctggag aggtctcagt cttggacttt 13620

tttgacgaaa tggggcttat tgccatgata acgatacctg gaatcccttt cggatctctc 13680
aacaaggcgg aggcgcggtg cttcaaggaa gtggaaagac tgaaggccgg cgattttgcg 13740
gagaaggatt tcgagtcgtt gaaatatgat cttgtcaaat catataaaag gtgcctggaa 13800
acgtccgagt caaggttctc tatcatgata gacgtgatgt cccaggacag gtcgtgggat 13860
tcttatcttt cccaggtgga gtcgatatct tcgcttactt cggaggatat agtcagggta 13920
agcaataaat atctgggcga caatcgtata cgtttccata aagtcagagg aacctatccg 13980
aaggataatc tgcaagctcc ttcatacgag actctttcgc cgaagaatgg caatgcggta 14040
tcggattatg ccaccagact gatggagatg ccttatgaca ctcttcctcc aagactggtg 14100
gatatgaata atgatatccg gaaagtcgca gttaatgacc atgtgacgct ttactataac 14160
gggaacccgg tcaatgatgt ctttgaactg accgtccgga tcaacaaggg tattttcgag 14220
gaacctatga tctgtccttt gacacagctt cttggcctct cggggaccga cagcctgtcc 14280
gcccgtgatt tttcaatgag tcttgccgag ctgggttcgg acttgtcttt taaagataag 14340
aaccatacct tggagatggt tctgactggc tttgatgacc gtataggacc ttccctcgac 14400
ctgctccgcc actttatgaa ggatttcaag ccggaaaaaa gtgcgctcaa tacggtgatt 14460
tccgatacga agatgtccag gaatgcattc aggagcagtg gcaattcagc catcttcaat 14520
ggtatgttta attattctct attagggaac cagtcacctc ttctgaggca gccgacagaa 14580
aaagatctca aatctctagg tctgaaaggg atgaaagagc ttttcaacag cctggttgaa 14640
tatgatttcg atgtagtcta ttccggaggc ctttctccag aggaagtcgt caatgagatt 14700
tccggccatc ttgatgtgga atgtccttcc ggaaaacctg agagttcatt tttcgcttac 14760
agaaaatatg aacagcctgt ggtctatatt tacaatgtcc cgggagcgcg ccaggtgatg 14820
tcgataactt accagaatgt cgctactccg gagaatgacc atgacagcgc actgctcggc 14880
cttatgcagg aatgctatgg cggttccatg gcttcgatca tgttccagga agtccgtgaa 14940
ttccgttctc ttgcatatac ttcttccagt acgcttcaga agccacttcc gaatgtatcc 15000
tctccttctg cattcataac ctactatggt actcaggctg ataaattccc tcaggctctt 15060
gaactggtgg attccctcat ccgtacgatg ccgatgactg atctgtgctt cgattcagca 15120
gtccgctcta tcatttcccg cgccaataac ggctatcctt cgttcagggc tttgcctgaa 15180
tatgtcgcga atttggaatt gttcggatat ggagaggatg atgacgcttc cgtagttaga 15240
ggccttccgg gaataaccaa gcaagacgtc ctggattacc atgaaaagag tatcggcaat 15300
gctccttgct cttatttcat aatcggagac aagaagaatc tgcctatgga cgcgcttcgg 15360
gaacgcggcc gcatagtcga gctgaaactc tctgacatct accgttagtt cgctgaatcc 15420
cttatcggga cgaataggtg gagaagcagg cggagattac ttcggaagcc gaaagtcccg 15480
agtcggagag gcggctggag gaggcaagga aacggtggtc ggcggtgatg ccgaagacgc 15540
ggtccgcgca gctgagtgcg ggtgccaggt cgtgagaagc catgatcacg gtggtgcccg 15600
tacttcttgc gatgtccctg agagcctgta gtatcagcga cttgttgtcg acgtccagga 15660
atgaagtcgg ctcgtccagc agcagcgtct tcgcatggcg gctgagcgcc gacgcgatgc 15720
agcctttctg gaattctccg tcgctcaggg tgctgatgtc cctgtctgcg agagatgaaa 15780
ttccgagaca ctccagcgac ttcgagatgt ccgaagccat ggccggagaa atgcggccgc 15840
gccagtcgct ttccgagtat agggtggtgg cgatgaactc cctgagagtg aatcccttga 15900
ctttcgggac ccctgtcggt accatgattg cgtccttgcc ggcgagagct ctcagcaggg 15960
tgctctttcc gctgccgttg ggaccgcaga gaagaatgaa ctctccgggc tcagtctcga 16020
aactgatgcc cgaaaccagg gtcttgcccc tgtgtccgat attcaagtcc tttattctca 16080
ttttttcatc agtatcagga gtattatcgg tatgcctatc agggccattg tgctgccgac 16140
cggaagcggg agtccgggcg cctgcgaaat gatgtccgcc agcaccgaga cggccgagcc 16200
tgtaatgaga gtggccggca gtatggacat atgtccggaa ctccggagca gcctgcgggc 16260
gatatgaggg gagacgattc cgacgaagcc cagaggcccg cagaatgcag tcgccgctgc 16320
cgtcatcaga cagctgctga ccatggcgac gccgcgtatc cggcgcaccg aggcgccagc 16380
gagagtggcg tattcctccc cgaaaagaag tatgtccaga cccttggctc cgcttgtgta 16440
aagccccgtt cctaccagca acgcggcaaa tatcgtcaat gtctcgaaag ttcttgtgcc 16500
ggagaaactt ccggcggacc agctatagaa gagcttgagg ctttcttcgc tggctgaata 16560
ctcaagtatg gaggtgatgg cgctgaatat gaagccgagc atcacgccga aaaccagcag 16620
ggtggtaccg ctgcggaaac ggcttgagac cagcactatc agggctgatg ccaggatcgc 16680
accgacgaat gcggctccgg ccacggtgag cccggccatg gacgagaagc ccgaaagcaa 16740
cccggtgcct gagatcagca gggtagctat cgcggctccg aggcctgcgc ctccgcttac 16800
gcccatgata tgagggtcgg cgagagggtt gcggaaaact gactgcatct gggctccgga 16860

```
     gagggccagg catgcgcctg ctgcaatggc cgtcagtacc ctcgggagac ggatgctcca     16920
     gactatcgag gaagctatcc cttctcctcc gaacaggcct ccaagcgcct cggcagcccc     16980
     gattccggaa cctccgagta tcaggtcgca tacggagagc agcagaagcg ccgttgctcc     17040
     tgccgccgtg ataagtctgt ttctgcgttt tatggccata ttaactctta caaagataga     17100
     aacttcaagg gattttcgta tatttggaag attaaaatac attgcggatg agcaccgaaa     17160
     caattctgat aattcttctt gtcgcagtca ttgtcatagc tgcgatactg attttccgtc     17220
     tccaggcgcg tgccgaaaag atcaaagacg gattccagga cagactggcc gatgtccagc     17280
     gcgaaaaagc cgtactggag agtcgcctgc aggctaagga ggaatatgag aatcagatgc     17340
     gcgaagccgc gaaggagacc tctgaaaggc atgccaaaga gctcgagcag atgaaggaag     17400
     tcttcaagaa cctttcgacc gagacaagcg aggccttcaa ggcaaagagc gcccagacaa     17460
     tctctgaact gatgaaacct gtccaggaga aattcaaaga gttcagcgac gccgtcaagg     17520
     aaagccagaa agacaccata gagaggcact cccgcctcga acagagcatt aaggatctcg     17580
     acacccgcag caaatccgtc ggggacgagg cccgcaatct cgccaacgcg cttaccggct     17640
     attccaaggt ccagggagat ttcggcgaaa tgctcctcac cgacgtcctg aagaacgccg     17700
     gcctgaccga aggagtgcac ttcgagactc agtccgtgat gaccgacaac tccggccatg     17760
     aaatcaaatc cgagagcggc aggaccatga ttccggacgt caaagtttat tacccggacg     17820
     acact                                                                 17825
//
ID   XXX; XXX; linear; XXX; XXX; XXX;  XXX.
XX
AC   PRJEB67889
XX
DE   Uncultured bacterium DNA fragment from in-reactor enriched cow rumen, clone
ferm-28d5, contig-2
XX
FH   Key             Location/Qualifiers
FH
FT   source          1..14524
FT                   /organism="uncultured bacterium"
FT                   /environmental_sample
FT                   /mol_type="genomic DNA"
FT                   /isolation_source="Cow rumen"
FT                   /clone="ferm-28d5, contig 2"
FT                   /note="AZCL-Barley beta glucan,  AZCL-arabinoxylan active
clone"
FT   CDS             422..1354
FT                   /db_xref="SEED:fig|6666666.127247.peg.1"
FT                   /translation="MMSKSFRCLSYTEVIISIFKSLGDRLGFCIKRHIAEVAVFLKLVI
FT                   ITVHRRGLHGIVCLFPVESLDVLDNHVSKYRNRMITDHAPGFVGIERPYRKHLLDTLVI
FT                   MRKHGSRDIRVDVLMHQVHPWMEGPVGIPKREYSIVRKPVRRGDILVESPVSAVRILEN
FT                   EWMERRMVCRSIEGLHCLAVSLELISGELIFPEVHAFLVKAVEGLSFSLCEILKGTFQA
FT                   YAGNRHAYCNLLAFRGLQTDPGGQFPAPGFTGAAEAHRIIDYGLSVLIEQHSLQAAGEC
FT                   GREIYMPFRSPSVNQLISGYRRVVLHAKT"
FT                   /product="hypothetical protein"
FT   CDS             complement(1869..2651)
FT                   /db_xref="SEED:fig|6666666.127247.peg.2"
FT                   /translation="MKRYIIIPLCIIAAATAACGQKNSGKTSGDWSDYNIGNVIFEDKA
FT                   KGTEGSAIYAGIIKDPVAYISEQARDVLATLYYSPEDSIPPVENIYYTLEDVEGISAKG
FT                   DEGNDAFVFYSTRHIEHSFADADTAKVLFETRGVLLHELTHVYQLSPQGIGNYDDNRVF
FT                   WCFTEGMADAVRVANGGFTSDDRPRGGSYMDGYRITGFFLVWLRDNKDPDFLRKFNAST
FT                   LEVIPWSFNGAIKHILGENADIDDLWHEYQVAVGDIKL"
FT                   /product="putative secretory protein"
FT   CDS             complement(2661..4937)
FT                   /db_xref="SEED:fig|6666666.127247.peg.3"
```

```
FT                   /translation="MAGFLVSAQPSRFVDPFIGTDGMGHTFPGACVPFGGIQLSPDTDI
FT                   VPHNIAGKYQQRTYDYCAGYRHSDNTIVGFSHRHLSGTGHSDLGDILIMPTTGPVQLTP
FT                   GTADHPEDGYRSRFSHDTEVARPGLYEVTLADYGIRARLSATKRVGIHEYTYPSYQKQN
FT                   IILDLTHSIYGYDGKVCRATLRVENDTLVTGFRLTDGWARTNYTYFAISFSKPIRDYGY
FT                   EDKKPSAYSGFWRRFDIGHNFPDMAGRDVAAWFSFDEGEPLTVKVAISAVDAAGAIRNL
FT                   EAEAAGSFDEIASMAAQEWDKALGRFEVEGTESEKTMFFTSVYHTLINPSVYMDVDGKY
FT                   RGVDGNIHQAEGFTNYTVFSLWDTYRAEHPFLGLLWPEENSDMIKSMVEHWKQNAVGML
FT                   PVWSLMGNENWCMSGYHAVPVLADAIVKGTYKGSAEEALAAMTATSKVRIYEGLGFYMD
FT                   HGFVPLDESGTAASTTLEYSYDDWTIHAAALKAGSSEIADEYGRRAMNYANVYDPSIGY
FT                   ARPRYSDGSFKKDFDIDQTYGEGFIEGNSRNFSFHVPQDVKGVMDLMGGEKAFLGKLEN
FT                   LFAMDLDPKYYEDNEDIEKECLVGGYVHGNEPSHHVPYLFAWTSKPWLTQYWVREIMNK
FT                   MYREGIDGLGGNDDCGQMSAWYLFSAMGFYPVCPGTDQYVLGAPYLPYIKITLPDGNTL
FT                   EIKAKGVSDTNRYVRKIRINGKTLDRRYVTHKEILEGGVWEFEMGSRPSRNNRLEKPYS
FT                   MSDNN"
FT                   /product="GH92"
FT   CDS             complement(4990..6924)
FT                   /db_xref="SEED:fig|6666666.127247.peg.4"
FT                   /translation="MIMASVSSCNDFFPTIPGTQYNLEDTFSDRNKTEEFLNNVYSYVP
FT                   DETCERWPTYYAGGIWTGGSIEGDITWSGDGEEATSWSLGSVYASTNWIGYWYNEYYKG
FT                   IAKASTFISNVDKCVEASASDRKYWKAQARALRAYYYFNIFRCYGPVTIIGEEAMALDT
FT                   PLDDLLKERNTVDENIDFIVSELDKAAEDLPAKYGDANLGRIDRGTCKALKAKALLYAA
FT                   SPLFNCNPDYAEIRNLDGTQIFPQDKSQEQAKWERARDAYKEFLDEFDGKVYSLQIVRT
FT                   PAGNIDHYESYRQAASISDYTRNTEQIFIRLSDHDLIQYQLTPYHRDCQDESIRGGLGF
FT                   GTTQEMVDMYFTDKGLRICEDPDYDEYTGVPDPSHYGWDEDYNDPFNPERNLFKANTDK
FT                   TLKQWAHREPRFYVCVTFNGSTWLNTNTNDGEVTTELTVNGNSGFNAANWDAPYTGYGE
FT                   RRMAPMNGDEQGAHCAVLLRLGDMYLGYAECLSACGQYDEAMKYVNLIRARAGVPGYGN
FT                   GGGTDANGMNYISYPVSRDEVDKRIRRERLVELSFEWNHFFDVRRWKVADMAVGDDWIY
FT                   PSYHRGGEGGPIHAMDFMADAPDFFKKVEFETRVCSKQHFLMPIPDAEMRRNPKMVQNY
FT                   GWTSEATER"
FT                   /product="putative outer membrane protein, probably
FT                   involved in nutrient binding"
FT   CDS             complement(6972..10127)
FT                   /db_xref="SEED:fig|6666666.127247.peg.5"
FT                   /translation="MELSKIKLLAGKIWLAVVIFITTGIMMSAQTRNVSGRVSDDKGEM
FT                   LAGIAVYEKGTTNGTLTDDNGQYSLTLKSTNPILVFSALGFEEQEIEVGKQTLIDVALV
FT                   TVVSELDESVVVGYGTQRKISVVGSQSSLKSDQIKMPSASLSSVVAGRLAGVVSVQRTG
FT                   EPGHDETDIWIRGISTFTNQPSRPLVLVDGVERSFNNIDPEDIESFTVLKDASATAVYG
FT                   VRGANGVIIIKTKPGRVGKPKFTVDYYESITRLTRIPKLANAYEYMDAANEAYLNSRGA
FT                   LLFTPQYIEATKKADGIIPNDDPKAYNSYLYPNVNWIDELFRPLGHNRRAQISVRGGVP
FT                   NATYYVSLTYYNETGLTRNAKMENYNANMRYDRYNYTANLNLRPTETTTIDLGFSGYLS
FT                   EGNYPQQSTSSLFGACFQINPVYLPKEMPDGSIPGISSMGDLRNPLADLTKRGYRNEIN
FT                   NQLNSNIRIAQDLGFFDWSKGLSANALLAFDVNNWRQNKYEKRDDTYYFSGSKNTETGL
FT                   WNNDVFTDDEWAITRTYTGSRELGYGRDQTMDRNVYFEASLNYDRTFGKHGVTGLLLYN
FT                   QNVYAAGHPDSLIESLNYKQQGLAARATYAYDDRYFAEVNAGYNGSENFDPKRRFGFFP
FT                   AAGLGWAVSNEPFWKPLKDVWSFFKIRYTIGQVGSDSAAGRRFMYQDEMGSADGTRFGT
FT                   SNSPKDGWGYKKYGSNVGWSTSLKQNLGFDFKFFKDELSLTVELFKEHRTGIFLQRATI
FT                   PDYCGFIEMPYANLGIVDNKGIDMNIEYNKQLATKTFLTLRGNLTWNEDRIIEDDQPEP
FT                   AYPWLETRGTNVLARWGWIAEGLFTSDDEIIDHARQFGEEYPGQISRPGDIKYKDLNGD
FT                   GVVDDYDKCVIGQGDVPKLYYGFGADLQVGQFSFGVIFSGNAFADRCLSGDGIHPFSDQ
FT                   AGLSNLFSNITDRWSAENPENTNVFYPRLHYGKTDNLNNTKESTWWQKDVSFLRLKQAT
FT                   ITYDLPGRIVDKTFLDAASVYITGTNLLTFSKFKIWDPELNTNNGCSYPNAMTISVGLN
FT                   VKF"
FT                   /product="TonB-dependent receptor"
FT   CDS             complement(10312..14082)
```

```
FT                   /db_xref="SEED:fig|6666666.127247.peg.6"
FT                   /translation="MNKTLYPLLLAVLNIFISVFSADAADNGGEYVFRRVGYSDGLSHS
FT                   AVLSIFQDNTDLVWMGTYDGLDCYDGSSMHVYRSDFSSGGSLTNNIIYSICQADGDNLW
FT                   VNTFQSLNRLSKGSGEITKAFSFKGEAYISSNSKGNTWLVCRDSLYYFNTVKDDFILVD
FT                   DSFIHGEDPEHRFFVNEDGEMMYFGRDDGKVRIYSVSSWKSAPSSVTIKYKEENFHPKP
FT                   ISDIFYQTGMMCFVDQDKDLYMYDTARNTKVYIRNISALIEEYGSIGGIVPFYEDFLIG
FT                   FNANGLFRLKAFSKYEAEPVDRNLRIFCLYHDLRQGVIWIGSDGQGALMYEKKYSIADN
FT                   IILSSVSENLSRPVRAILTDSRGGMWVGTKGDGIVYFPDYKKGTSGGEVITPSGRRSVV
FT                   GYSRESTEFKVYAIKESRFRDGVWIGAGNEGLMFCPYGGIPVKIAFPAGVYAPDEVHDI
FT                   YEKDKNTLYISCAGMGIHKLILSGNKVINDLNYSFYFEGREIGLFYAMKAVGDSLLYAG
FT                   SRGDGIVRLDMRTNEYSVISMRYLYNEAIDDILSLCPASDGSLYVGTVSGMLRLYNSGD
FT                   DVRAEFIGKKNGLINDMIHGILEDNDGMLWLGTDRGLVKYNPSNGLAQTYLFTAGLEFG
FT                   EFSDDAYYKDPRTGKLYLGGNRGLISLAPHTPEMVQYYSNIQLRSLEVAHEEVRFSDYT
FT                   DGDGNLVLKGSPVSFAVTFAVPDYRASGLVEYSAFLEGYDKEWSVYTLDNKVSYTDVPA
FT                   GTYTLRLRYKKDVYDTLPEEFSIKVRIRRGFFRSSLFMILVGLAFLGLLGYSVFSYRNN
FT                   RRKIQAARDSDKAMADFRKRLSIAYADTFDNASVERSTIKELSDSVTSLLAARGAQTEG
FT                   ISFIGPEVPFPVYTNYMKRLLLVVYDHILSSGGSADISCEIEDKSLHMVVSAEKKLLEP
FT                   LYRAVSGGSAHNLSSRFCEDSTTQLDFLCIAAQQGIPERIDFDGKNLVFVFNPAIQRNS
FT                   GATERDRLLLLESRPEMYWLLSDILSSDYEVVYASEPAAALKSLKHSDANLIVVDAHLG
FT                   IKDENAFRDTLSKNRTTISDIPFVVILGNDNPGHNLGSELVSWSDSYVFDPEGMHLVKD
FT                   VISRAVDGKKDFRRIKMEELGSLADDIVCTNEEDMAFVRKTIQVLSENIAREDLGTALI
FT                   ADRMAMTPRSFYRRFRSISHITPGTLIKIYRLEMASRLLLDDSKQIQDVIAEVGISSRS
FT                   YFYKEFTARFGMTPGNWKSSHGLSAEPMSSK"
FT                   /product="DNA-binding response regulator, AraC family"
FT   CDS             14192..14503
FT                   /db_xref="SEED:fig|6666666.127247.peg.7"
FT                   /translation="MSKKKQDITLENVVIEAVAAEGKALAHVDGAVLFVQFAVPGDVVN
FT                   VKVFKKKKNYMEGFITEIVKPSPDRLEPFCSHFGICGGCKWQPLPLQDAARSKAAAGL"
FT                   /product="RNA methyltransferase, TrmA family"
XX
SQ   Sequence 14524 BP; 3525 A; 3671 C; 3621 G; 3707 T; 0 other;
     tctcgcggat cctcttctgg cacttccacg gctgcctgaa ataattgtag atataaggga        60
     catggtgaca tggctcattg ccgtgccagt atgcaccgat cctgccctga atatcatgag       120
     cgcccggaat gtcctcagga agctcgtaag taaacaggct gtcaagtcgg gaaagaagct       180
     tcttctcacc tgcaatatcc atataaccct gcatatcgtg aggaacatac caggtatact       240
     gtacggtaaa cccttccgta atgtcgcccc agccgttgac tgagcccggt ccctgataat       300
     cgagaggtga gaacggagtt ctccactccc ctttcgaatc ccttcctctc atgaatccgg       360
     tctccggatc gataaggttc ctatatgaga gggacctttt cagatagtac tcgtaatcct       420
     catgatgtcc aagagctttc gctgcctgag ctatacagaa gtcatcataa gcatattcaa       480
     gagtcttgga gaccgactcg gattctgtat caaaaggcac atagccgagg ttgcggtatt       540
     cctcaagctt gtcataatta cggttcatcg ccgtggtctt catggcatcg tatgcctttt       600
     ccctgtcgaa tcccttgacg tccttgacaa tcatgtcagc aagtaccgaa accgcatgat       660
     aaccgatcat gcaccaggtt tcgttggcat agaaagacca tatcggaagc atcttctcga       720
     cactcttgtc ataatgcgca agcatggaag tcgcgatatc cgcgttgatg tcctgatgca       780
     ccaggttcat ccatggatgg aaggccctgt aggtatccca aagcgagaat acagtatagt       840
     tcgtaaaccc gtcagacgtg gcgatattct tgtcgagtcc cctgtatctg ccgtccgcat       900
     cctcgaaaac gaatggatgg agcgccgcat ggtatgcaga agtatagaag gtctccattg       960
     tcttgcggtc agccttgagc tcatatctgg agagctcatt ttcccagagg tccacgcctt      1020
     tctcgtgaag gctgtcgaag gtctctcctt cagtctctgc gagattctta aaggcacctt      1080
     ccaggcttac gccggaaacc gccacgcgta ctgtaacctg ctcgccttcc gtggtcttca      1140
     gaccgatcct ggcggacagt ttcctgcccc aggcttcacc ggtgctgcgg aagcgcatcg      1200
     tatcatagat tacgggcttt ccgtcctcat agaacagcat tcccttcagg ccgcaggaga      1260
     atgtggccgc gaaatataca tgcctttccg gtccccatcc gtcaaccagc ttatatccgg      1320
     ttatcgtcga gtcgttctcc acgcgaagac ttgaccagac acatttctgg taaagcgtat      1380
```

ggtccatatc gacggcgatg aaggccgagt cggactccgg aaatgtaaaa cggaatattc 1440
cgcttcgcag gcccgaggtc atctctgcct ttacaccata atcatccaac acgacgctgt 1500
aataatccgg tgtagcaact tccttgtcat ggctgtaggc agaggctggt ctcagctccc 1560
cagtaccggg gatgaacagg aaatctccga gatcagggat gcctgtgccg ctaagatggg 1620
taaggctgaa tccctggatc gtattgtggg aatacttgta tccggaagcc gcatcataga 1680
aattgtcatc ggtatcaggg ctgatctgga tacctccgaa cggaatctgg gcacctggat 1740
agacattccc gtatccatct gttccaacaa gtggctttac ataatcagta agcttctgag 1800
ccgtcatcgg agcgatgccg gccagaagcc ctaatactgc cattgtcagt ctgtatgttc 1860
tcattgcatt agagtttgat gtccccgaca gcgacctgat actcatgcca gagatcatcg 1920
atatcggcat tttctccaag gatatgcttg atggctccat tgaatgacca tggaatcacc 1980
tcaagcgtgc tggcattgaa tttcctgaga aaatcaggat ccttgttgtc cctgagccat 2040
acaaggaaga atccggtaat acggtatccg tccatatagc tgccgcctct aggcctgtcg 2100
tcagaggtaa aacctccgtt tgcgacacgt acggcatcag ccattccctc ggtgaagcac 2160
cagaatacac ggttgtcatc atagtttccg attccctgag gcgaaagctg atatacatgg 2220
gtgagttcgt ggagcaggac gccgcgggtc tcgaagagga ccttggcggt atcggcatca 2280
gcaaaactat gctcgatatg ccttgtgctg tagaaaacga aggcatcatt accctcgtct 2340
cccttggcag agattccttc tacgtcctca agggtataat agatgttctc gacaggaggg 2400
atgctgtctt caggagagta ataaagggtt gcaagcacgt ccctggcctg ttccgaaata 2460
taggctacag ggtcctttat gattccggca tagatcgcgg aaccctcggt gcccttagcc 2520
ttgtcttcga atattacgtt tccgatatta tagtctgacc agtctccgga tgtctttccg 2580
gaatttttct gtccgcaggc cgcagtggcc gcagctatga tacaaagcgg gataatgata 2640
tatctcttca tattatgtta ttagttgttg tctgacattg aatatggctt ttcaagccta 2700
ttgtttctgg aaggtctgga gcccatttcg aactcccata cgccgccttc aaggatctcc 2760
ttatgggtga cgtatctcct gtccagagtt tttccgttaa ttcttatttt tcttacatac 2820
cggttggtat cgctgacgcc tttcgccttg atctcaagtg tattgccatc aggaagggtt 2880
atctttatgt aaggaagata aggggcgccg agcacatact ggtccgttcc tgggcataca 2940
ggatagaatc ccatagccga gaacagatac caggcgctca tctggccgca atcatcattg 3000
ccgccgagac cgtcaatccc ctctctatac attttgttca tgatttcgcg aacccagtac 3060
tgggtaagcc acggcttgga tgtccaggcg aaaagatatg gcacatggtg gctcggttcg 3120
ttgccatgta catatccgcc gaccagacat tctttttcga tatcctcgtt atcctcgtaa 3180
tatttcgggt caaggtccat cgcaaacaga ttctcaagtt ttccaaggaa agccttttcg 3240
ccgcccatca ggtccatgac acctttgacg tcctggggaa catggaaaga gaaattacgc 3300
gaattccctt cgatgaaacc ttccccgtaa gtctggtcga tatcgaaatc tttcttgaaa 3360
gaaccgtcgg aatatctcgg acgcgcataa ccgatcgaag gatcatatac attggcatag 3420
ttcatggccc tgcgaccgta ttcatcggct atctccgaag atccggcttt cagcgcggcg 3480
gcatggatag tccagtcgtc atagctgtat tccagagtcg tcgaagccgc agtcccgctt 3540
tcatcgagcg gcacgaatcc atggtccata taaaagccga gaccttcgta gatcctgacc 3600
ttcgaggtcg cggtcatggc cgcgagagcc tcttcggccg agcctttata tgttcctttg 3660
acgatggcgt cggcgagtac cgggacggca tgataaccgc tcatgcacca gttttcgttg 3720
cccatgaggc tccagaccgg gagcatgccg acagcattct gcttccagtg ctcgaccatt 3780
gactttatca tatccgagtt ctcctcgggc cagagaaggc ccaggaaagg atgctcggcc 3840
ctgtaggtat cccagagtga aaatactgtg tagttggtga atccttcggc ctggtggatg 3900
tttccgtcca ctcccctgta cttgccgtcc acgtccatat agactgacgg attgatcagg 3960
gtatgataca cggaggtaaa gaacatcgtc ttttcgcttt cggtgccttc cacctcgaat 4020
ctgccgagag ccttgtccca ttcctgagcc gccatggagg cgatctcgtc gaatgagcct 4080
gcagcttcag cctcgagatt ccggatcgcg cctgcagcgt cgactgccga aatcgctact 4140
tttaccgtga gcggttctcc ttcgtcgaaa gagaaccatg ccgctacgtc ccggccggcc 4200
atatcgggga aattatggcc aatgtcgaaa cgtctccaga atccgctgta ggcagaaggt 4260
ttcttgtcct cgtaaccata atcgcggatc ggcttggaga atgaaattgc aaaataagtg 4320
tagttggtac gggcccatcc gtctgtgaga cggaagccgg taaccagagt gtcattctcg 4380
actctcaagg ttgcacggca aactttgccg tcgtatccgt aaatgctgtg tgtcaggtcc 4440
agaataatat tctgtttttg gtaggatgga taagtgtatt cgtgtatgcc gaccctttttt 4500
gttgctgaaa ggcgggccct gatcccgtaa tccgcgagag taacctcata gagacctggt 4560
cttgcgactt cggtatcatg cgaaaaacgg gatctgtatc cgtcttcagg gtgatctgcc 4620

gttcccggag tcagctgtac gggaccggtg gtaggcatga tcagtatatc cccgaggtct 4680
gaatgacctg ttccactgag atgcctgtgg gagaagccta cgatcgtgtt gtcggaatgt 4740
ctgtatccgg cgcaatagtc ataggttctc tgctggtatt tgccagcgat gttatgggga 4800
acgatatcgg tatcaggact caactggata cctccgaaag gcacgcatgc cccagggaat 4860
gtatgcccca ttccgtcggt tccgatgaac ggatcgacga aacgggatgg ttgtgcggaa 4920
accaggaatc ctgccattgc cagagaagct gataccaata ttttcctcat gatttgtgtt 4980
agctttaaat tatctttcag ttgcttcaga agtccagccg tagttctgca ccatcttagg 5040
attccttctc atttctgcat ctggaatcgg cataaggaag tgctgttttg agcagactct 5100
ggtctcgaac tcgactttct tgaagaaatc aggcgcatcg gccatgaaat ccatggcgtg 5160
gataggaccg ccttcgccac ctctgtggta agaaggatag atccagtcat caccgacagc 5220
catgtcagcc actttccagc gacgtacatc gaagaagtga ttccactcga atgaaagctc 5280
tacgagtctc tcgcgacgga ttcgcttgtc gacttcatca cggctgacag gataagagat 5340
atagttcatg ccgttggcat ccgtaccacc gccattgcca tatccaggca cacctgctct 5400
cgcacggatg agattgacat acttcatggc ctcgtcatac tgaccgcagg cgctgaggca 5460
ttctgcatat ccgagataca tgtcgccgag gcggagaagg actgcgcagt gagcgccctg 5520
ctcgtcacca ttcataggag ccatcctacg ttcgccgtag ccggtataag gagcatccca 5580
gttggctgcg ttgaagccgg agttaccgtt tactgtaagc tcggtggtca cctcgccgtc 5640
gttggtattg gtgttgagcc atgtggatcc gttgaaggtc acgcatacat agaagcgagg 5700
ctcacggtgt gcccactgct tgagggtctt gtcggtatta gccttgaaga ggtttctctc 5760
agggttgaac gggtcattgt aatcttcgtc ccagccatag tgggaaggat ccggaacgcc 5820
tgtgtactca tcatagtcag gatcttcgca gatacggaga cctttgtctg taaagtacat 5880
gtcgaccatc tcctgggttg ttccgaaacc gagacctccg cggatacttt cgtcctggca 5940
atccctgtgg taaggggtga gctggtactg gataagatcg tggtcggaaa gcctgatgaa 6000
tatctgctct gtgttgcggg tatagtcaga gatagacgct gcctgacgat atgactcata 6060
atggtcgata tttcctgcag gagtcctcac gatctggagt gagtagacct taccgtcgaa 6120
ctcgtcgagg aattccttgt atgcatctct tgccctttcc cactttgcct gctcctggct 6180
cttgtcctga gggaagatct gggtaccgtc gagattcctg atctccgcat agtcagggtt 6240
gcagttgaaa agagggctgg cggcataaag aagggcttta gccttgagag ccttgcaggt 6300
acccctgtct atacggccga ggttggcatc gccgtacttt gcagggagat cctcggcagc 6360
cttgtcaagc tcggacacga tgaaatcgat attctcgtcg accgtattcc tttccttgag 6420
aaggtcatca agaggagtat cgagagccat ggcttcttca ccgatgatag tcacaggacc 6480
gtagcaacgg aagatgttga agtaatagta ggcacggaga gctcttgcct gggccttcca 6540
gtacttacgg tctgaagcgc tggcttcgac gcatttatct acattgctga tgaatgtgct 6600
ggcctttgca ataccettgt agtattcatt ataccagtat ccgatccagt tggtagaagc 6660
gtataccgag ccgagagacc agctggtagc ttcttcgcca tcaccgctcc aggtgatatc 6720
gccttcaatt gatccgccgg tccagatacc gccggcataa taggtaggcc agcgttcgca 6780
ggtctcgtca ggaacataac tatagacgtt gttgaggaac tcctctgtct tgttacggtc 6840
tgagaaggtg tcttccagat tgtactgcgt acccggaatc gtcgggaaaa agtcgttgca 6900
ggacgaaaca gacgccatga tcatgaccgc agctgcaatt ttatataatg ttttcataat 6960
gacatccggt tttagaattt gacattgaga cctactgaga tcgtcatcgc gttaggatat 7020
gaacatccgt tgttggtgtt gagctcagga tcccagatct tgaatttaga gaaggtcaga 7080
aggttggttc ctgtgatata tacacttgcg gcatcgagga aggtcttgtc gacaattctt 7140
ccaggaaggt cgtatgtgat ggtagcctgc ttcaggcgga ggaaacttac gtccttctgc 7200
caccaggtgc tttccttggt gttattgagg ttgtctgtct ttccgtagtg gagacgtggg 7260
tagaatacgt tggtgttctc agggttttct gcagaccatc tgtcggtaat gttgctgaac 7320
aggttggaaa gaccggcctg gtcagagaac ggatggatac cgtcaccgga gagacagcgg 7380
tcagcgaagg catttccgga gaaaatcacg ccgaatgaga actggcctac ctggaggtct 7440
gcgccaaagc cgtaataaag cttcggaacg tcaccctggc cgattacgca cttgtcataa 7500
tcatcgacta cgccgtcacc gttgagatcc ttatacttga tatcaccagg acgtgagatc 7560
tgtccaggat attcctcgcc gaactgtctt gcatggtcga tgatctcgtc atcgcttgtg 7620
aagaggcctt cagcgatcca tccccagcgt gcaaggacgt tggttccgcg ggtctcgagc 7680
catggatatg caggttccgg ctgatcatcc tcgatgatgc ggtcctcgtt ccaggtgagg 7740
ttgccccctga gggtaaggaa ggtcttggta gcaagctgct tgttgtattc gatgttcatg 7800
tcgataccct tgttgtccac gataccgagg ttggcataag gcatctcgat gaaaccgcag 7860

tagtcaggaa tagtagcacg ctggaggaag attccggtac ggtgctcctt gaagagctct 7920
actgtgagcg agagttcgtc cttgaagaac ttgaagtcga aaccgaggtt ctgcttgagg 7980
gatgttgacc agcccacgtt ggaaccatat ttcttgtatc cccaaccgtc tttagggctg 8040
tttgaagtac cgaaacgggt accgtctgcg ctgcccatct cgtcctgata catgaaacgg 8100
cgtcctgctg cggagtcgga tcctacctga ccgatggtat agcggatctt gaagaatgac 8160
catacgtcct tgagaggttt ccagaacggt tcatttgaaa cagcccatcc gagaccggct 8220
gccgggaaga aaccgaacct gcgcttaggg tcgaagttct cggaaccgtt gtaaccggca 8280
ttgacttctg cgaaataacg gtcgtcgtaa gcatatgtcg ctcttgcggc aaggccctgc 8340
tgcttgtagt tgagcgactc gataaggctg tcagggtgac ctgcggcata gacattctgg 8400
ttgtagagaa gaagtccggt gactccgtgc ttgccgaagg tcctgtcata attgagggag 8460
gcctcgaaat agacgttgcg gtccatggtc tggtccctgc catagccaag ctcgcggcta 8520
ccggtatatg tacgggtgat agcccactcg tcatcggtga atacatcatt gttccagaga 8580
cctgtctcag tgttcttcga tcctgagaag tagtaggtat cgtcacgttt ctcgtatttg 8640
ttctgacgcc agttgtttac gtcgaaagcg agaagggcgt tggcggaaag acctttgctc 8700
cagtcgaaga atccgagatc ctgggcgatt ctgatattgg aattgagctg gttgttgatc 8760
tcgttcctgt agcctctctt tgtaaggtct gcgagagggt tgcggaggtc acccatggat 8820
gagattccag ggatggagcc gtcaggcatc tccttaggaa gatatacagg gttgatctgg 8880
aagcatgccc cgaagaggct ggaggtactc tgctgaggat agttgccctc ggagagatat 8940
ccgctgaatc caagatcgat ggtagtagtc tctgtaggac ggagattcag gttggcagta 9000
tagttatacc tgtcataacg catgttggcg ttgtagttct ccatcttggc gttacgtgta 9060
agacctgtct cattataata tgtgagagac acataatatg tagcgttagg cacgccacct 9120
cttacgctga tctgcgcacg gcggttatgt cccaatggtc tgaaaagctc gtcgatccag 9180
ttgacattag gataaagata gctgttatag gccttaggat catcgttagg gatgattccg 9240
tcagccttct tggtagcctc gatatactga ggagtgaaga gaagcgcacc acggctgttc 9300
aggtaagcct cgtttgctgc gtccatatac tcatatgcgt tcgcaagctt agggattctg 9360
gtgagtctgg tgatactctc gtaataatcg acagtgaact taggcttgcc taccctaccc 9420
ggcttagtct tgatgatgat gacaccgttg gcgccacgta caccgtagac agcagttgca 9480
gatgcatctt tgaggacagt gaaactctcg atatcctcag ggtcgatatt gttgaagcta 9540
cgctccacac cgtcgacaag cacgagaggc cttgaaggct gattcgtaaa ggtcgaaata 9600
ccacggatcc agatgtcggt ctcatcatga ccaggctctc ctgtcctctg gacagagaca 9660
acgcctgcca gtcgtccggc gacaacagat gaaagacttg ccgaaggcat ctttatctgg 9720
tctgacttga gggatgactg agatccgaca acggagatct ttctctgggt accataacct 9780
acgacaacgc tctcgtcaag ttcacttact actgtaacga gggcaacgtc gataagggtc 9840
tgtttgccga cttcgatctc ctgctcttcg aaaccaaggg ctgagaatac aagaataggg 9900
ttagtgctct taagagtaag ggagtactga ccgttgtcat cggtaagggt accgtttgtg 9960
gtacctttct cgtacaccgc aattcctgca agcatttcac ccttgtcatc agatacacga 10020
ccggagacat tccttgtctg tgcagacatc attatcccgg tggtgatgaa aatcacaact 10080
gccagccaga tcttaccggc aagaagttta attttagata attccataaa tgggtgtttt 10140
ggttacatta gctttatgtt gcactttcat gcataacaaa gaaaagaaga tataattaca 10200
atcatttcct catgcgcgac aaaatagaaa gggtgtccgt aaagaggaca ccctatcaaa 10260
attgagcaca aggcaattat ttctatgtca aaagtttact ctatccgctg gtcatttaga 10320
agacatcggt tcagcactca gcccatggga acttttccag tttcccggag tcattccgaa 10380
gcgggcggtg aactccttgt aaaaatatga acggctcgag attcccacct ccgcgatcac 10440
atcctggatc tgtttcgagt catccagaag cagacgcgac gccatctcca gacgataaat 10500
cttgatcaat gttcccggag ttatatgcga gatgctgcgg aaacgtctgt agaacgagcg 10560
cggagtcatg gccatcctgt cggcgataag ggccgttccg agatcctcgc gggcgatatt 10620
ctcggaaagc acctggattg tcttgcggac aaaggccata tcctcttcat tcgtgcatac 10680
tatgtcatcg gccagactgc cgagctcttc catcttgatg cgccggaaat ccttctttcc 10740
gtcgacagcg cgcgatatca cgtccttgac aagatgcatt ccttcagggt cgaatacata 10800
gctgtcggac catgagacaa gctccgaacc aagattatgg cccgggttgt cattcccaag 10860
gatcacgacg aacgggatat ccgaaatagt cgtcctgttc ttggacaatg tatccctgaa 10920
cgcattttcg tcttttattc ccaaatgggc gtcgacgact atcaggttgg catccgaatg 10980
tttcagcgac ttgagtgcag ctgccggctc cgacgcatat acgacttcat aatctgaaga 11040
aagaatgtcc gaaagaagcc agtacatctc aggccggctc tccagcagga gaagcctgtc 11100

cctttcagtc gccccgctgt tacgctggat ggccggattg aatacgaaaa caagattctt 11160
tccgtcgaag tctattctct ccggaatccc ctgctgagca gctatgcaga gaaaatccag 11220
ctgagtcgta ctgtcctcgc agaatcttga agaaagatta tgtgcagaac cacctgaaac 11280
ggccctgtac agaggctcca gcagtttctt ttcagcagaa acgaccatgt gcagagactt 11340
gtcttctatt tcgcaggaaa tgtcggctga gcctcctgac gaaagaatat ggtcatagac 11400
tacaagcagc agccgcttca tgtaattcgt atagaccggg aacggaactt ccgggccaat 11460
gaaactgatt ccctcggtct gcgctcccct tgcagcaagc agactggtca cggaatcgga 11520
aagttccttg atagtgcttc tttcaacgga tgcattgtcg aaagtatcgg catatgcaat 11580
cgaaagcctc ttcctgaaat cggccattgc tttgtcagaa tcccttgccg cctgaatctt 11640
gcgcctgtta ttgcggtatg agaatacact gtatccgaga agtcccagga aagccagccc 11700
gacgaggatc atgaagagcg aagacctgaa gaagcccctg cgtatacgga cctttatact 11760
gaactcttcc ggaagagtat catatacatc cttcttatag cggagacgca aggtataagt 11820
gccggcagga acgtctgtat aggacacttt gttgtcgagg gtatatacag accattcctt 11880
gtcatatcct tcgaggaaag ccgaatactc cacaagtccg gatgccctgt aatcaggtac 11940
ggcgaacgtg accgcgaaag ataccggact gcctttcagg acaaggtttc cgtcgccgtc 12000
ggtatagtcc gaaaaccgca cctcctcatg ggccacctcg agacttctca gctggatatt 12060
gctgtaatac tgcaccatct ccggagtatg aggggccaga cttatgagcc ctctgttacc 12120
gccaagatag agttttccgg tcctcggatc cttgtaatac gcgtcgtcac tgaattcccc 12180
gaattccagg ccggcggtaa agagataagt ctgtgcaagg ccgttggaag ggttgtactt 12240
gaccagtccc ctgtccgtcc cgagccagag catgccgtca ttatcttcga ggattccatg 12300
gatcatatca ttgataagac cgttcttctt tcctatgaat tcagcgcgca catcatctcc 12360
ggagttatac agacgcaaca ttcccgagac tgtaccgaca tataggcttc cgtcggatgc 12420
aggacagaga ctcagaatgt cgtctatcgc ctcattatag agatagcgca tagagatgac 12480
gctatattca ttcgtcctca tatctagcct aactattccg tctccacggc tgccggcata 12540
cagcagcgaa tccccgacag ccttcatggc atagaaaagt cctatctctc gtccctcgaa 12600
atagaaagaa taattcaagt cattgatcac cttgttcccg gaaagaatca gcttgtggat 12660
acccattccg gcgcaggaga tataaagagt gttcttatcc ttctcataaa tatcgtggac 12720
ctcatcaggc gcatagactc ccgcaggaaa tgcgattttg acagggatgc cgccatatgg 12780
acagaacatc agaccttcgt taccggcccc gatccagact ccgtcgcgga acctgctctc 12840
ctttatagca tagaccttga attccgtaga ttccctcgaa taaccgacaa cgctcctgcg 12900
tcccgaagga gtaatcacct cccctccgga cgttcccttc ttgtagtcag ggaaatagac 12960
tattccgtcc ccctttgtgc ctacccacat tccgcctctg gaatcagtaa gtatagccct 13020
gacaggccgg gaaagatttt cggagacaga agacaggatg atattgtcag cgatagaata 13080
cttcttctcg tacatcagag cgccctgacc gtcggacccg atccaaatga caccctgacg 13140
aagatcatga tatagacaga aaatcctcag gttcctgtca actggttctg cctcatactt 13200
gctgaacgcc ttcagccgga aaaggccgtt ggcattgaat cctataagga agtcctcata 13260
aaaaggcacg atacctccga tagacccata ctcttcgata agggcggata tgttccttat 13320
atatacttta gtgttcctgg cggtatcata catataaagg tccttgtcct gatccacgaa 13380
acacatcatt ccggtctggt agaagatatc ggagataggt ttcgggtgga aattctcctc 13440
cttatacttg attgtcacgg aagaaggggc ggacttccag gatgaaacgg aatagatcct 13500
cacttttccg tcgtcccggc cgaaatacat catctcgccg tcttcgttca cgaaaaacct 13560
gtgctccgga tcttcgccat ggataaaaga gtcgtcgacc aggatgaaat catccttgac 13620
agtattgaaa taatacaggg aatccctgca tacaagccag gtattgccct tgctgttcga 13680
agagatatat gcctcaccct tgaaagagaa agccttggta atctctcccg accctttcga 13740
gagacggttg agagactgga aagtatttac ccagagattg tccccgtccg cctgacagat 13800
ggaatagatt atattgttgg tcagcgaacc tccggaagag aaatcggacc tgtagacatg 13860
catcgaggaa ccgtcatagc agtccagtcc atcgtacgtt cccatccaca caagatccgt 13920
attgtcctgg aaaatagaaa gaacggcact gtgggacaac ccgtcggagt atccgaccct 13980
tctgaagaca tactccccgc cattgtcggc agcatctgca gagaagaccg agataaatat 14040
attcagaacc gccagcagaa gcggatacag agttttattc ataaaatcaa ggcaattagg 14100
tatgcaaaag gtgcaaatac aaacttacac aaaaatttgt tttgacgcaa gagttaccta 14160
actttgtatt taattacagc ataaaaggaa tatgtcgaaa aagaaacagg atataactct 14220
cgaaaacgtt gtcattgaag cagttgcagc agaaggaaaa gcattggccc atgtagatgg 14280
cgcagtactc ttcgtacagt tcgcagtgcc cggcgacgta gtcaacgtaa aagtattcaa 14340

```
     gaaaaagaaa aactatatgg aaggcttcat cacagagatt gtgaagccgt ccccggaccg     14400
     cctcgaaccg ttctgctctc acttcggaat atgcggagga tgcaaatggc agccgctccc     14460
     cctacaagat gcagctcgaa gcaaagcagc agcaggtcta tgaccagctt gtgcgcatag     14520
     gcca                                                                  14524
//
ID   XXX; XXX; linear; XXX; XXX; XXX;  XXX.
XX
AC   PRJEB67889
XX
DE   Uncultured bacterium DNA fragment from in-reactor enriched cow rumen, clone
ferm-30d14
XX
FH   Key             Location/Qualifiers
FH
FT   source          1..23779
FT                   /organism="uncultured bacterium"
FT                   /environmental_sample
FT                   /mol_type="genomic DNA"
FT                   /isolation_source="Cow rumen"
FT                   /clone="ferm-30d14"
FT                   /note="AZO-CM cellulose, AZCL-Barley beta glucan,
AZCL-arabinoxylan active clone"
FT   CDS             13..1584
FT                   /db_xref="SEED:fig|6666666.127248.peg.1"
FT                   /translation="MALLDNLPEKAKTYVAMAQRNGEKLFRMVSELMDFQKKEEGIHCV
FT                   KWQAVDLPSFLRVQIDKFADSAASKNITLLLGDVPDVSIMTDVSIADLVFDNLISNAVK
FT                   YTPEYGKVDISAGLLHNNVEIYVRDNGIGIPETETKDVFQNFFRASNAVNSQEMGSGLG
FT                   LSFTRNAVKRLKGSLTFESVEHRGTVFTLTLPTTDKEPVALEKIGTTHSFLDADGDWSI
FT                   TEDVMEKDTILFVDDNEDLRKYVRLVFSNNYNVVDVPNAQAALDYLSKDVCDIVVSDVM
FT                   MPGMQGDELCQCIKENPNTSWLPVILLTAKVGRDFMISGLKQGADDYINKPFDSTVLGS
FT                   KIDSMLKNRRRMAAYYLKRSRAIVRGEAISAEIPEEMMMDPTQPSSQVPENILQKRVEE
FT                   ALATDDNSTPLDKVSTGEELQAQNEFIDQATRIVMQNLEDTEFNIDNLCREMAMSRTLF
FT                   YGKLKSLTGQTPQDFVRTIRLEQAAIYLKQGYPVLDVSIKTGFANVKYFSTVFKRHFGV
FT                   QPSKYQ"
FT                   /product="FIG00938042: hypothetical protein"
FT   CDS             1791..3041
FT                   /db_xref="SEED:fig|6666666.127248.peg.2"
FT                   /translation="MMKTNRQIVTSLTLLLGMLFTTACEAHQPEAELPQKNEKEIKAPT
FT                   AQEWNREVTIGWNLGNSFECSAPGMNGESLLIGNPAGADNAETSWGNPVVSKKLIDAVK
FT                   AAGFDAIRIPVRWQCHITNAQAMTISRTWMNRIKEIVDYCLKCDMKVIINTHHEKWLES
FT                   RPTYKYKEENCQKLALLWMNIASEFKDYDYRLAFAGTNEVHERDNWGQPTAENLEVQNA
FT                   YNQTFVDIVRATGGNNAQRHLMVQTYVCNPEFGLNNGDFIIPTDLEENGNRYMSVEFHY
FT                   YQPWDYCGLGKLEYWGERYRAHGTAAPENEQTIIDTFTKVQQTWAFKGLGVVIGEWGVT
FT                   DHYKSSSADLMRENMAYYCQIYVAEAKKRGFATFIWDNNHFGNGAECFGIIDRTHGLQI
FT                   KAPWVMEGIKAGLKAVE"
FT                   /product="GH5_4"
FT                   /EC_number="3.2.1.4"
FT   CDS             3095..6274
FT                   /db_xref="SEED:fig|6666666.127248.peg.3"
FT                   /translation="MMNKTILLSLLCTGMIGTPATTYAVSSSVKATLTAQQSDQEIKGS
FT                   VVDASGEALIGATVKIVGGTGGALTDMNGHFSLKCEPGTTLQISFMGYNPVTVKASNGM
FT                   KVTLQENVKVLGDVVVTALGIKRDKKALGYALEEVKGEELTKAKETNVINSLAGKVAGL
FT                   VVQNTAGGSSGSTRVLLRGNTEMGGDNQPLYVIDGVPLDNTNFGSVTTAGGHDLGDGIS
FT                   AINPDDIETMSVLKGPAASALYGSRASHGVILITTKKAEKDKVSVEYNGSITFDSQLAK
```

```
FT                   WNDIQQVYGMGNGGQYLETATSGTNMSWGPKADNYMVRYFDGVERPFLMYPNNTSNFFR
FT                   TGVTAQNTGILSFSSGKTGIRFSMTDMRNKDILPNTHMSRDNFNLRANTSSGAFDFDFT
FT                   ANYTREDVKNRPALGDSQSNVGKNLMTLAGTYNQDWLRYYQNPDGTYANWNGNDMYNKN
FT                   PYWDLYKNDNTSSKDVFRFTGKAVWNINKDWKVQGTIGTDINNMNFEEFIARTTPGSAS
FT                   GEMTNQVFQNRTLNAEILGLYGHTWDNFDVHATVGGNIYKVNNKTYNMLGHDQQISDMK
FT                   DIMNYSEHNVNRDTYVKQINSLYGSVSFGYNRTYYLDVTLRGDKSSTLPTSNNTYVYPS
FT                   LSGSWIFSEYLKNKNILSYGKLRASWAKVGSDTDPFQLAINYSSGKYSYTGYMLGMINN
FT                   NNMPNGSLKPTMTNSFELGTELKFFNGRLGMDVTWYSQSSRNQIINLASSAPSGYPYRL
FT                   VNAGHIINRGVELAVNGRAVQYKDFAWDLGFNFSKNINKVKELVPGMDYFELEKATWAG
FT                   VSVGAMVGQNYGSIMGKDVARTDDGQVIIDRSTGMPVSAEGTHVIGNASWDWTGGFYST
FT                   FTYKNFHLSAGFDVKYGADLFSMSMRSAYQTGRATATLKGREAWYASEEARFAAGQTSE
FT                   DWRASGNMKGYVVPGVYLNDEGQYVKNDIPVNPEAYWKYATENCPAMFVYDNSYVKCRE
FT                   ITFGYTFPEKMLGKTVKGLTLSFVARNPFILWKNIPNIDPDSGYNTSGLGLEYGSLPSR
FT                   RSYGFNINLKF"
FT                   /product="FIG01289191: hypothetical protein"
FT   CDS             6386..8128
FT                   /db_xref="SEED:fig|6666666.127248.peg.4"
FT                   /translation="MKSFQKYIKSGLGASFALTLTVGAMSSCSDSYMSDLNTDDSKAMS
FT                   IDPNAQLTTSLLQTYGDFSLMDTYRSYITGFTQHFAGGWNVTNYAGSVHAFDDQTRLIW
FT                   DRYYTVSIKNMEDAISKTSEKPNINALLRIHKVYLMSVLTDTYGDVPCFNVGQGYMKGE
FT                   NNPAYDKQEDIYQWFFDELSLCESQLNAVGDRITGDVTSMNGDVAKWKKYCNSLRMRFA
FT                   MRISDVKPELAREQFEKALASENGYIAEASDDAYIKYLDQPFTLYKGAEKYDFRANALS
FT                   EILYGQDPTSPTFVSSTFYNIMKDSNDPRLYRICRHYINTTRSEIRADEAGNFDLTDEV
FT                   VQYLKDNGLSESPCDPGAAWYNNWINAPATKNIPTLQLLVDQNPAAGYDKNNFPARMMR
FT                   LFLSIDFEKGQNPGILFTSAEAEFLLAEAKIKGWNVEGSADEHYEAGVRASMKLLNTHY
FT                   LKEKDLINDAEIAAFIAANQLGENPKETINTQAWILHMTNPAEAWSNLRRSDYPVLMDR
FT                   TKLPTFPDFTYDDPDRSTPTRLKYPILEARYNSENYKEALERLGGTDDWHQRNWWDTAD
FT                   INVK"
FT                   /product="FIG00937602: hypothetical protein"
FT   CDS             8267..9007
FT                   /db_xref="SEED:fig|6666666.127248.peg.5"
FT                   /translation="MNTFNKTIENKIFEGFSILAIIMVSLLCFSCSEDDPFFTATKDDA
FT                   PRILNTDIPEWRDGQPQTLQTVDFGDSLRFKVIATPRDYTTISWYIDDELVFEGDSIVY
FT                   PISEGKHLCKIVATTTQGLSTSRTCYVIGNPEPGKEKILWEGSFDVTWGTPFDVLKTSL
FT                   AENVQEGTVLRVYVEGDGQGTATSAWWNNILTGEGEDKRGDIAISGEQMLEFTLNSTSM
FT                   TLLSTQDGFLVVGNGYKITKITIM"
FT                   /product="hypothetical protein"
FT   CDS             9007..11319
FT                   /db_xref="SEED:fig|6666666.127248.peg.6"
FT                   /db_xref="GO:0008422"
FT                   /translation="MKSFKAIFSILTMLPLAVYAQVSNPVIPRDDRIEAQVQQTLSRLT
FT                   LEEKVGQMMELVTDFFGSNDEKGVFHIDRQKTDSILARYKIGSILNAPNTCAPTAKEWE
FT                   VIIQDIQKSSMKQIGIPCVFGLDQNHGSTYTADGTLFPQNINVAATFNREIARRCAEAT
FT                   AYETRAISVPWTYSPTVDLGRDPRWPRIWENFGEDAYVNAEMAKAMVKGFQGENPNRID
FT                   QQHIAVSLKHYLGYGVPYSGKDRTPAYISLSELREKHFAPFLAGLEAGALTVMVNSASV
FT                   NGMPVHANKTLLTTWLKEETGWDGMLITDWADVNNLYTRERIAKDKKDALRIAINAGID
FT                   MVMEPYATDACDYLVELVKEGKIPMSRIDDAVSRILRLKYRLDLFKNPTQKLKNYPKFG
FT                   GEEFAKLSLEGAQESMILLKNQNNILPLKAGKRILVTGPNANQMRPLNGGWSYTWQGHR
FT                   TDEFTEKYNTIYEAVCQKYGKDNVVLSQGVTYNEKGKYYEENTPDIASAVAAAQNVDVI
FT                   FACIGENSYTETPGNLSDLNLSSNQKDLVKALAATGKPVILILNQGRPRLINDIEPLAQ
FT                   AVVNILIPGNYGADALVNLLAGEVNFSGKLPYTYPSEINSLATYDYKKSESVSTMEGAY
FT                   DYNAQITQQWGFGFGLSYTTYTYSNLKVDKTNFKHNDTIRVSVDVTNTGSMAGKESVLL
FT                   YSSDEIASIVPDNRRLRDFQKIELNPGETKTVSFLLPARNLAFVDYDGKWVIEEGTFAL
FT                   MAGNQLVKVNCSDTYYY"
```

```
FT                   /product="Beta-glucosidase (EC 3.2.1.21)"
FT                   /EC_number="3.2.1.21"
FT   CDS             11780..13015
FT                   /db_xref="SEED:fig|6666666.127248.peg.7"
FT                   /translation="MSKSTHFSGQPVYSQVIKLLDKAEIVKISHQTKGSEAYVKRFDGY
FT                   QHLVVMLFGILKHFDSLRELEIGMKAEAHKLSHLGMDYLVRRSTLAEANIRRPQEFLAS
FT                   VYAYLLEKYAKFLADSRPAKSYKGQTHEPKDWEKLLYMMDSTTITLFDNILKGVGRHPK
FT                   SGKKKGGLKVHTVMKYHVGVPMVVQLTSAAKHDHYLLKEVHLPKDSTLAMDRGYMDVAQ
FT                   FQRLTEEGVCYVTKMKKNLKYKVLDSVTYVNAKGLVTHIDQRVHFTRGELTHEARRVEI
FT                   FHENKKPVALLSNNFEFSVEDISEIYRLRWAIESLYKQLKQNFPLHFFYGDSVNAIQIQ
FT                   TWVVLIANLLITVLSRSIKRRCAFSQVVSMVRLTLMYYIDFIAFMENPDKSWNNIRAKE
FT                   LQKAPPNPTLFD"
FT                   /product="Mobile element protein"
FT   CDS             complement(13425..14222)
FT                   /db_xref="SEED:fig|6666666.127248.peg.8"
FT                   /translation="MKNLLLSAVIGDISGMPYEFEGRTKNYEAVNLLHSANTYTDDTVC
FT                   TFACAEALLHELDMAKNLWKRCRADFHRGFGGRFACWLMAEDIQPSYHSYGNGSGMRTS
FT                   AAGFMSKTKEECIALATRTALPTHDHPEGIKGAVATSLAIFHAMEGKGKDYIRKNVLDV
FT                   YYPDWSRKTYMEIKPEYYFDETCQLTIPAALICFLESKDYTDCLKLAISLGGDADTLAA
FT                   ISGPMAYAYYKMIPEELINHAKAKLPEWMLKINDEFDDYCNKL"
FT                   /product="hydrolase, putative"
FT   CDS             complement(14225..14980)
FT                   /db_xref="SEED:fig|6666666.127248.peg.9"
FT                   /translation="MFGGLYFLGYLCGVSFIEEKGMDEKMYCYKYPHPSVTTDCVIFGF
FT                   DGDRLKVLLIERRNEPYKGKWAFPGGFLNPDETAEDGALRELQEETGLTGTYIEQFHTY
FT                   TVPDRDPRERVITIAYYALVRIQEVHGGDDATKAKWFPIDEVPQLAFDHERILRDAMRK
FT                   LRERIHFEPIGFELLPEKFTMKNLQMLYEAILDVKFDRRNFSKKMMHHKLLNQLDEKVW
FT                   PTAKREALLYSFNNEKYNEFKKKGFQIKF"
FT                   /product="Hypothetical Nudix-like regulator"
FT   CDS             15134..15250
FT                   /db_xref="SEED:fig|6666666.127248.peg.10"
FT                   /translation="MKQQASLIDSSFLPHFFSLVFTKIIDDSKEYVPTVQEG"
FT                   /product="hypothetical protein"
FT   CDS             15210..15482
FT                   /db_xref="SEED:fig|6666666.127248.peg.11"
FT                   /translation="MIQRSMCLQCKKGEKILADFAQSDSCKNVNDAISTSNNPHIFFKK
FT                   NDLYIPYSTMLVRSKTHAWFMTHRTIVHWPSHHCSKALEALFIEA"
FT                   /product="hypothetical protein"
FT   CDS             16102..16782
FT                   /db_xref="SEED:fig|6666666.127248.peg.12"
FT                   /translation="MDARGQYAYMKSGWLDRISKGVYKIHGVEPGLMAAVSSYNTQLGK
FT                   SCIVGAYTALELQGYSHYLSMGKPMAYLFTDKKNKLPSWLLKGEWDMTIKYMTTSFLGD
FT                   ELLGVETMTINQRDLLMSSPERAILECLNLPDASSSLLDIYYIMESMTTLRPKLVQALL
FT                   EACTSHKVKRLFLYMSEKAGHSWFKALKLENINWGTSRLMITPTGKYINKYNMTISKEL
FT                   AEYE"
FT                   /product="Ynd"
FT   CDS             16775..17485
FT                   /db_xref="SEED:fig|6666666.127248.peg.13"
FT                   /translation="MNDIKTSVYAQKVELLLRLIPVVMDEQVFAIHGGTAINLFLKDLP
FT                   RYSVDIDLTYIPLADRQTSINDINSHLKSIADKAKKAFKGMHIVPNFNTCKQLCEYSGK
FT                   QVKIEVNQTKRGIIGGEVQTISLSEKAQNEFSLFCEANVVPITQLYGGKIAAALSRQHP
FT                   RDLFDVKYMDIPLSECREGLIFCLLGSDRPIHESFAPSLIDQREAMENQFAGMTDIPFT
FT                   YEEFEETRANLINA"
FT                   /product="FIG00937925: hypothetical protein"
```

```
FT   CDS             complement(17651..18721)
FT                   /db_xref="SEED:fig|6666666.127248.peg.14"
FT                   /translation="MRHLSYHPTLRTCSSDTILRAIKELTQENISYTSDQGKTYDFNTA
FT                   DKLNTLLINALVSTGELKEIEEYDVDFDHQFLETEKYDAKPTYKKFLGYRPGVYVIGDK
FT                   IVYIENSDGNTNVRFHQADTHKRFFALLESQNIRVNRFRADCGSCSKEIVSEIEKHCKH
FT                   FYIRANRCSSLYNDIFALRGWKTEEINGVQFELNSILVEKWEGKCYRLVIQRQRRNSGD
FT                   LDLWEGEYTYRCILTNDYKSSTRDIVEFYNLRGGKERIFDDMNNGFGWSRLPKSFMAEN
FT                   TVFLLLTALIHNFYKTIMSRLDTKAFGLKKTSRIKAFVFRFISVPAKWIMTARQYVLNI
FT                   YTENRAYAKPFKTEFG"
FT                   /product="transposase"
FT   CDS             19098..19304
FT                   /db_xref="SEED:fig|6666666.127248.peg.15"
FT                   /translation="MTEADKKFLISFESGQPEWEGYEFEYFKDYPSVQWKLLNLKKLAK
FT                   QNPLKLQMEVDKLRNLYNFSLNN"
FT                   /product="hypothetical protein"
FT   CDS             19318..20322
FT                   /db_xref="SEED:fig|6666666.127248.peg.16"
FT                   /translation="MSYKDFQQKVYSYLIKYKKESLNINDKGISTRGVEHDCLLSKPYC
FT                   DAKIPVMLYNGIKETVKDIQASKYAYKAHLAASKHVASSQTACINLFVPILESEYADQI
FT                   LIGAGVVPKDFDHIDRNQLRKGYCFEYWESTLEGAKGLLGDHSPHAGTDSDVAIAYRNK
FT                   NDKLCLWLIEHKLTEQEFTTCGGYRSKAISHSEKEYCTSCGIEEILKNHDKCYYHKHCG
FT                   YFYWNIMDAQSSFFCGKFEGNGCPFRGGMNQLWRNQMLALELENRGVFEDVYFSVVSHP
FT                   DNSFLDKTMNEYRALTNNSPKFYDFKSNALVDVAVDYLPDWSNWYKKVYLGID"
FT                   /product="hypothetical protein"
FT   CDS             20322..21863
FT                   /db_xref="SEED:fig|6666666.127248.peg.17"
FT                   /translation="MYHRDYDNIQMIEQLRQLMTLDANMNTSDIEERFEQIAKVLFECF
FT                   AIQKGETMYLFKEIEFYFYNKNHRDIITHPRVSKPLCWYVNDFGGIDLNFVSNIKRESK
FT                   ANSKGKNVEKYVLDNGAYFGGILIRQLTSEDGCEVLNGPWACAELFRCYDATVIDKDLP
FT                   ILVELDNGMVGYIREPRINLLTSKQTVEGKVDYILEEYHEVFERKCLYSDFSRFVDRRY
FT                   RYVRCDTLMHDKDTNLVYFSPWLKDNKEGHPEFYKLLKNVLNEMGIESKELKSTNDYWA
FT                   RDYMPIQLGENEFLKYRYYPDYLVKSKNKKDIETITDATKVLRGMGISCRSTNLIIDGG
FT                   NMVPCGPYIVMTDKVFMENGREKGDTDFKVLLESEIGHPVIIIPWTPHEDDVYGHSDGF
FT                   IKWCGDNRILMGNHGDCYPEEAASIRRILESYGFEVTEMRFKGKVDKPCYDLNWAYINF
FT                   LQVGKNIIMPKFNIEEDPIAQQYIHEAFPDWNIRQIEMAEIAKEGGALHCISWNV"
FT                   /product="hypothetical protein"
FT   CDS             22629..23006
FT                   /db_xref="SEED:fig|6666666.127248.peg.18"
FT                   /translation="MYEGLASYFLPTGVLEYFEVTDFCEVPTQDTEVLYTTELHIYLDE
FT                   RDNRTPDMSGSVSDGFGDESCLLDFPTRDKKTVLHVRRRRWTMPDGTTKSVDLDKKIQL
FT                   KHPGTRYSKDFALFLKKADGQ"
FT                   /product="hypothetical protein"
XX
SQ   Sequence 23779 BP; 6993 A; 4875 C; 5305 G; 6606 T; 0 other;
     cctatcaacg atttggctct tctcgacaat ctgccagaaa aggctaagac ctatgtggct        60
     atggctcaac gcaatggcga gaaactcttc aggatggtgt ctgagctcat ggatttccag       120
     aagaaagaag agggcattca ttgtgtgaaa tggcaggcag tggatcttcc atccttcttg       180
     cgtgtgcaga tagacaagtt tgctgattct gcagcaagca agaatatcac tctcttgttg       240
     ggtgatgtgc cagatgtgtc gattatgact gacgtctcta ttgcagatct cgtgtttgac       300
     aatcttatct ctaatgccgt gaagtacact ccagaatatg gaaaggtgga catctctgcg       360
     ggattgttac acaataatgt ggaaatctat gtgagagaca atggtatagg tattccagag       420
     acagagacaa aagatgtctt ccagaatttc ttccgtgcct ctaatgccgt aaattctcag       480
     gagatgggta gcggattggg actttccttt actcgcaatg cagtgaagag actcaagggt       540
     tctcttacct ttgagagtgt ggaacataga gggactgtct tcactcttac cctgccaaca       600
```

acagataaag agcctgttgc acttgaaaag attggcacga cccattcctt tttagatgcg 660
gacggcgatt ggtctataac agaggatgtg atggagaaag acaccatact ttttgtggat 720
gacaatgagg atcttcgcaa gtatgttcgt ttggtgttca gcaacaacta caatgttgtg 780
gatgttccaa atgcacaggc ggctctggat tatctctcca aggatgtgtg tgacattgtg 840
gtgagtgatg tgatgatgcc tggtatgcaa ggagatgaac tctgccagtg cattaaggag 900
aatcctaaca cctcatggct gcctgtcatt ctcctcactg ccaaggtggg acgagacttc 960
atgatttcag gtttgaagca aggtgctgac gattatatca acaaaccatt cgactctact 1020
gtgttgggca gcaagattga cagcatgctc aagaaccgtc gtcgcatggc cgcttactat 1080
ctcaaaagga gtagagcaat tgtgagggga gaagcgatta gtgctgaaat accagaagag 1140
atgatgatgg atcctaccca gccatccagc caagtgcctg agaacatatt gcagaaaagg 1200
gtggaagagg ctcttgctac tgatgataat agcactccac ttgacaaggt ttctacaggt 1260
gaagagttgc aggcacaaaa tgagtttatt gatcaagcga ctcgcattgt gatgcaaaac 1320
ctggaggata cggagtttaa tattgacaat ctttgtcgtg agatggcaat gagtcgcaca 1380
ctcttctatg gaaaattgaa aagccttacg ggacaaacac cacaggattt tgtgcgtact 1440
atccgccttg agcaggcggc tatctacctc aagcagggct atcctgtgct tgatgtttca 1500
atcaaaacag gatttgcaaa tgtgaagtat ttcagcactg tttttaagcg acactttggt 1560
gttcagccca gcaagtatca ataaatcaat gtgttgatag attacaggaa aaggtttcaa 1620
aattgacaga tgttagtttt gaaacctttt tcgataaaaa acgaaccttg gtcgttgaaa 1680
aaaccaataa atttgctacc gaaatctaaa atgcacgtca tctaaaacag ggttacctat 1740
aattgaatac cccaatgatt tcaattaaca ataaaattca tattacctca atgatgaaaa 1800
ctaacagaca aatcgtaaca tcactcacac ttttgcttgg catgctattc actacagcat 1860
gtgaagctca tcagccagag gcagaattac cccagaagaa cgagaaggag ataaaggctc 1920
ccactgctca ggaatggaat agagaagtta caattggttg gaacttgggc aattcatttg 1980
aatgttctgc tcctggaatg aatggagaat ctttgcttat aggcaatcct gctggagcag 2040
acaatgcgga gacgtcgtgg ggcaatcctg tggtaagcaa gaaactgata gatgctgtga 2100
aggccgcagg ttttgatgct attcgtattc ctgtccgttg gcaatgtcac atcacaaatg 2160
cacaagcgat gaccatcagt aggacttgga tgaatcgcat caaggaaata gttgattact 2220
gcttgaagtg tgacatgaag gtcatcatca atactcacca tgagaaatgg cttgagagtc 2280
gtcctactta taaatataaa gaggaaaact gccaaaagct tgctttgctc tggatgaata 2340
tcgcttctga atttaaggac tatgattatc gtttggcttt tgctggaacg aacgaagtgc 2400
atgagagaga caactgggga cagcctacag cagagaacct tgaagtgcag aatgcctata 2460
atcagacatt tgtagatatt gtgcgcgcta caggaggcaa taatgcccag cgccatctta 2520
tggtgcagac ctatgtatgt aaccctgaat ttggactcaa caatggagac ttcatcattc 2580
ctactgattt ggaagagaac ggtaatcgtt atatgagtgt ggagtttcat tactatcagc 2640
cttgggatta ttgcggtctt ggaaaactgg agtattgggg agaacgctat cgtgcgcatg 2700
gcactgccgc tccagagaat gagcagacaa tcattgatac cttcaccaaa gttcagcaga 2760
cttgggcttt taagggtctt ggtgtcgtga taggagagtg gggggtgaca gaccactaca 2820
agtcttcttc ggcagacttg atgcgtgaaa acatggctta ctattgccag atttatgtgg 2880
cagaggccaa gaagagaggg tttgccactt tcatttggga caataaccac tttggcaatg 2940
gtgcagaatg ctttggcatt atcgaccgta cgcatggctt gcagattaaa gcgccttggg 3000
tgatggaagg tatcaaggct ggccttaaag ccgtagaata attgaaagaa aaattgataa 3060
attatcatat caacctatat taccaatata aataatgatg aataaaacga ttcttttaag 3120
tctgctttgc acaggcatga tcgggacacc tgcaacaacc tatgctgtgt cctcttctgt 3180
gaaagcaaca cttactgctc agcagagtga tcaagaaatc aaaggttcag tggttgatgc 3240
ttcaggtgaa gcattgattg gagccacagt taaaattgtg ggaggtacag gcggtgcttt 3300
gaccgacatg aatggtcatt tctccctcaa atgcgagcct ggtacaaccc tgcaaatcag 3360
ctttatgggt tataatcctg ttacggtgaa agcatctaat ggcatgaagg tgacacttca 3420
agaaaacgtc aaggtgcttg gtgacgtggt tgtaacagcc ctcggcatta agcgagacaa 3480
gaaagctctt ggctatgcac ttgaggaggt gaagggcgaa gaactcacta aggcaaagga 3540
aacaaatgtg atcaactctc ttgcaggtaa agtggcaggt ttggttgtgc agaacacggc 3600
tggtggttca tcaggctcta ctcgtgtact tttgcgtggt aacacagaga tgggcggtga 3660
taatcagcct ctctatgtga ttgacggtgt tccgcttgac aatactaatt tcggtagtgt 3720
gaccacagca ggtggccacg atcttggtga tggtatctct gctatcaatc ctgatgacat 3780
agagactatg agtgtgctca aaggtcctgc tgcttctgca ctctatggta gtcgtgcctc 3840

tcatggtgtg attcttatca ctacaaagaa ggctgagaaa gataaggtga gtgtggaata 3900
caatggttct atcactttcg actctcagtt ggcaaagtgg aatgacatcc agcaggtcta 3960
tggtatgggt aatggcggac agtatcttga aactgccaca tcaggtacga acatgagttg 4020
gggacctaag gcggacaact atatggtgcg ttatttcgat ggtgtggagc gtccattcct 4080
tatgtatcct aacaacacat caaacttctt ccgtacgggt gtcactgctc aaaacacagg 4140
tatcctttct ttcagttctg gcaagacagg tatccgtttt tctatgaccg acatgcgtaa 4200
taaggacatc ctacctaata ctcacatgag ccgtgataac ttcaatctgc gtgccaacac 4260
atcatctggt gcattcgatt ttgacttcac tgccaactat actcgtgagg acgtaaagaa 4320
tcgtcctgcc cttggtgact ctcagagtaa tgtaggtaag aacctcatga cactggctgg 4380
tacttacaat caggactggc ttcgctatta ccagaatcct gatggcactt atgccaactg 4440
gaacggtaat gacatgtata ataagaaccc atactgggat ctctataaga atgacaacac 4500
ttcatctaag gatgttttcc gtttcacagg aaaggctgta tggaacatca acaaagactg 4560
gaaggttcag ggtacaatcg gtactgacat caacaacatg aactttgagg agttcattgc 4620
acgtaccact cctggaagtg cttctggtga aatgaccaat caggtgttcc agaatagaac 4680
actcaatgct gagattctcg gtctctacgg tcacacttgg gacaactttg atgttcatgc 4740
tactgttggt ggtaacatct ataaggtcaa taacaagacc tataacatgc tcggccatga 4800
tcagcagatt tccgacatga aagacatcat gaactacagt gaacacaacg tgaatcgaga 4860
cacttatgtg aagcaaatca actcactcta tggtagtgtg agtttcggct acaaccgcac 4920
ttactacctc gatgtgactc tccgtggtga caagtcatca actctcccaa ccagcaataa 4980
tacttatgta tatccttctc tctcaggaag ctggatattt tcagagtatc tcaaaaacaa 5040
gaatatcctt tcttatggta aattacgtgc atcttgggca aaggtaggta gcgacactga 5100
tcctttccaa cttgccatca actattcttc cggaaaatat agttacactg gctatatgct 5160
tggtatgatc aataacaaca acatgccgaa cggaagcctc aagcctacga tgaccaactc 5220
gttcgagtta ggaacagagc tcaagttctt caatggaaga cttggcatgg atgttacatg 5280
gtacagccag agttcacgca atcagattat caatcttgca tcctctgctc cttctggtta 5340
tccttatcgt cttgtcaatg caggtcacat catcaatcgc ggtgttgagt tggcggtgaa 5400
tggtcgtgcc gttcagtata aggactttgc atgggatttg ggctttaact tctcaaagaa 5460
catcaataaa gtgaaggaac tcgttcccgg catggattac tttgaacttg aaaaggctac 5520
ttgggcaggt gtttctgtag gtgctatggt aggtcagaac tatggtagca ttatgggtaa 5580
ggatgttgct cgcacagacg atggacaagt catcatcgac cgtagcacag gcatgccagt 5640
aagtgctgaa ggcacacacg taataggtaa tgcttcttgg gattggacag gtggtttcta 5700
cagcacattc acttacaaga acttccatct ctctgcaggt tttgacgtaa aatatggtgc 5760
agacctcttc tcaatgtcta tgcgttcagc atatcagaca ggtcgtgcca cagccactct 5820
caaaggtcgt gaggcatggt acgcatcaga agaagctcgt ttcgctgctg gtcagacatc 5880
agaggattgg agagcatcag gcaacatgaa gggttatgtt gtgcccggcg tatatcttaa 5940
tgatgaaggt cagtatgtga agaatgatat tcctgtcaat ccagaagcct actggaagta 6000
tgccacagaa aactgcccag cgatgttcgt ctatgacaac tcttatgtga agtgccgtga 6060
gattactttt ggctacacat tcccagagaa gatgcttggc aagactgtaa agggacttac 6120
tctctcattt gtggctcgca atccattcat cttgtggaag aacattccta acatagaccc 6180
agactctggc tacaacactt caggtctcgg acttgagtat ggttcacttc catctcgtcg 6240
cagttacggt ttcaatatca atcttaagtt ctaaatcatt ttttgataaa gaatcatcag 6300
atcattaatt ttctaaatat agatatgtaa tcatccattt gatttttaca atgtatcgtg 6360
tttggaatta aatatataga caacaatgaa aagttttcaa aaatatatca agagcggttt 6420
aggtgcttca tttgctctga cacttaccgt gggggcaatg agtagttgtt ctgactccta 6480
tatgtccgac cttaacactg atgacagtaa ggctatgagc attgatccta atgctcagct 6540
cacaacaagc cttctccaga cctatggcga cttcagcctg atggacacct atcgcagtta 6600
catcactgga ttcacccagc attttgcagg tggttggaac gtaactaact atgctggctc 6660
tgtgcatgcc ttcgatgacc agactcgtct tatatgggat cgctactaca cggtgtctat 6720
caagaatatg gaagatgcta tctctaagac gtccgaaaag cctaatatca atgcacttct 6780
acgcatacac aaggtttatc tgatgtcagt cctcacagac acctatggag atgttccttg 6840
cttcaatgtt ggtcagggct acatgaaagg agagaacaat cctgcctatg acaaacagga 6900
agacatctac cagtggttct ttgacgaact ctctctctgc gaaagccaac ttaatgcggt 6960
aggagacaga atcacaggtg atgtgaccag tatgaatggt gatgtggcaa agtggaagaa 7020
atactgcaac agtctccgca tgcgttttgc catgcgtatt tctgatgtaa aaccagaact 7080

ggctcgtgaa caattcgaaa aggctcttgc aagtgagaat ggctatattg cagaagcgtc 7140
cgacgatgct tacatcaagt atcttgacca gcctttcact ctctataaag gtgcagagaa 7200
atatgacttc cgtgctaatg ccctcagtga gattctctat ggtcaagatc ctacttctcc 7260
aactttcgtg agtagtacct tctacaatat tatgaaggac agcaatgacc cacgcctcta 7320
ccgcatttgc cgtcattata taaatactac tcgtagtgag attcgtgcag acgaggctgg 7380
caacttcgac ttgacagacg aggtggttca gtatcttaaa gacaatggac tttcagagtc 7440
tccttgtgac ccaggtgctg cttggtacaa caactggatc aatgctcctg ccacaaagaa 7500
catcccaacc ctccaactct tagtggacca gaaccccgcg gcgggctatg acaagaacaa 7560
cttccctgca cgcatgatgc gactgttcct tagcattgac tttgagaagg ggcagaatcc 7620
tggcattctc ttcacttctg ccgaggcaga gtttctcctt gctgaagcaa agattaaggg 7680
atggaatgtg gaaggtagtg cagatgagca ctatgaagca ggtgtgagag cgtctatgaa 7740
actactcaac acccattatt tgaaggagaa ggatctgatc aatgatgcag aaattgctgc 7800
cttcattgct gccaatcagt tgggcgagaa tcctaaagag acaatcaata ctcaggcttg 7860
gattcttcac atgactaatc ctgcagaggc atggtctaac cttcgtcgct cggactatcc 7920
tgtgctcatg gatcgcacca agttgcccac attcccagac ttcacttatg acgaccctga 7980
tagatctaca cctactcgtc tcaaataccc aatccttgag gcgcgctaca atagcgaaaa 8040
ctacaaagag gctcttgaac gtctgggtgg aacagacgac tggcatcagc gcaactggtg 8100
ggatacagcg gatattaacg tgaagtaata ggctgaataa gtgattgaaa tacattaaaa 8160
acatgagaaa aggagttaaa ggtggactca tgaccgtttc tttccttaga cactcgtatc 8220
atcaaccatc tttaaactcc tttactccct attaaatatc aaaatcatga atacatttaa 8280
taagaccata gaaaacaaaa tcttcgaagg gttcagcatt cttgccataa tcatggtatc 8340
tttgctctgc ttctcttgct ccgaagacga cccatttttc acagcaacaa aggacgatgc 8400
tccacgcatt ctcaacaccg acattcctga gtggagggat ggtcagcctc aaacacttca 8460
gacagtggat tttggcgata gccttcgctt caaggtcatt gccactcctc gtgactatac 8520
aaccatctct tggtatatcg acgacgaact cgttttcgag ggtgactcta tcgtctatcc 8580
tatatctgag ggcaagcatc tctgcaagat tgtggctact accacacaag gactctccac 8640
ttctcgcact tgctatgtga taggaaaccc agagcctggc aaagagaaga ttctttggga 8700
aggttctttc gatgtcacat ggggcactcc atttgatgtg ctcaagacct cattggcaga 8760
gaatgttcaa gagggaacag tcctcagagt ctatgttgaa ggtgatggac agggcactgc 8820
tacatcagca tggtggaata atattcttac aggcgaaggt gaagacaagc gtggcgatat 8880
agctatcagc ggtgaacaga tgctcgaatt cacactcaac agcacttcca tgactcttct 8940
tagcacccaa gacggattcc tcgttgttgg caatggctat aaaatcacta agatcacaat 9000
aatgtaatga aatcattcaa agcaattttc tctattctaa caatgctgcc tttggcagtt 9060
tatgctcaag tctctaatcc tgtcatcccg agggatgaca ggatagaggc gcaagttcaa 9120
caaaccctct ctcgtctcac gcttgaagag aaggttgggc aaatgatgga actggtcaca 9180
gacttcttcg gatctaatga tgagaaaggt gtcttccaca tcgatcgtca aaagacagac 9240
tctatccttg ctcgctacaa gataggctct atcctcaatg cgcccaacac ttgtgctcct 9300
acagcaaagg agtgggaggt catcattcag gatattcaga agagttccat gaagcagatt 9360
ggcattccat gtgtgtttgg tcttgatcaa aaccacggtt ctacctacac ggcagatggt 9420
acactttcc ctcagaacat caatgtggcc gctacattca accgtgagat cgctcgccgt 9480
tgtgcagaag caacagccta cgaaacacgt gccatctctg tgccttggac ttatagtccc 9540
acagtagatt taggtcgcga cccacgttgg ccacgcatct gggagaactt tggcgaggat 9600
gcctatgtta atgcagaaat ggcaaaggct atggtcaaag gttttcaagg cgaaaatcct 9660
aaccgcattg accaacaaca tattgctgtt tcattgaaac actatctggg ttatggcgta 9720
ccttattctg gtaaagaccg cactcccgcc tacatttcgc tttctgaact gcgtgaaaaa 9780
cattttgctc cattcctcgc aggtcttgag gcaggtgcgc ttactgtcat ggtcaattct 9840
gcttcagtca atggtatgcc tgtccatgcc aacaagacct tgctcaccac ttggctcaaa 9900
gaagagacag gttgggacgg catgctcatc actgactggg cagatgtcaa taacctctac 9960
actcgtgaac gcattgccaa ggacaagaag gacgcccttc gtatcgccat caatgctggt 10020
atagatatgg tcatggaacc ttatgctacc gacgcatgtg attatcttgt ggaactggtt 10080
aaagagggca agattcccat gtcacgtatt gatgacgctg tgagccgtat tcttcgtctt 10140
aaatatcgtc ttgacctctt caagaaccct actcaaaaac tcaagaacta tcctaagttt 10200
ggtggtgagg agtttgccaa actttccctc gaaggtgctc aggagagcat gatcctcctg 10260
aagaatcaga acaatatcct tcctctcaaa gcaggaaaaa gaatccttgt gactggacct 10320

aatgccaatc agatgcgccc tcttaatggt ggatggagct acacatggca aggtcatcgt 10380
acagacgagt tcactgaaaa gtataacaca atctacgagg cagtctgcca gaagtatggt 10440
aaggacaatg ttgtgctctc tcaaggagtc acctacaacg agaagggcaa gtactatgaa 10500
gagaatacac ctgacatagc atctgcggta gcagccgcac agaatgtaga tgtcattttt 10560
gcttgtatcg gtgaaaattc ctacacagaa acacctggca atctctccga cctcaatcta 10620
tcttccaatc agaaggattt ggtcaaggca cttgctgcta ctggcaaacc tgtcatcttg 10680
attctcaatc aagggcgtcc tcgtctcatc aacgatatag aaccacttgc acaggctgtt 10740
gtaaacatcc tcattccagg taattatggt gcagacgctc ttgtcaatct tcttgctggc 10800
gaagttaatt tcagcggtaa acttccctac acttatccat ctgagattaa ctctcttgct 10860
acctatgact acaagaagag tgagtctgtc agcacaatgg aaggtgctta tgactacaat 10920
gcacaaatca ctcaacagtg gggatttggc tttggtctca gctacactac ctacacctac 10980
agcaacctta aggttgataa gaccaacttt aagcacaacg acacaatccg tgtctctgtc 11040
gatgtaacca acacaggtag catggcagga aaagaaagcg tgttgctcta tagtagtgat 11100
gaaatagcca gcatcgttcc tgacaatcgt cgcttgcgcg actttcagaa gattgagctc 11160
aatccaggtg agaccaagac tgtctccttc cttcttcctg ctcgcaacct cgcttttgtg 11220
gattatgatg gcaagtgggt cattgaggaa ggcacatttg cactcatggc aggcaatcag 11280
ttggtcaagg tgaactgttc agacacatac tattattaac tcacacacta tcttatacca 11340
ttaaaattaa atttgaaaag aatttaataa aaacacacag acaataatta attgtgatta 11400
ttggtgtttg cattttattt gtgtaaaata aattcgcaga caccaataat tttttgttcc 11460
gcatcaatct gggtatctgt actacttcat ctcatgagga gagacaaaga agtagagaag 11520
ggtaggagag acacaataga aagacaatgg agaaaacatg gaatgtgaac accccttaca 11580
accttatcca tctttgcctt gtgtaagttt tcggacgata tgagtaaata aagcatgaag 11640
gattcttttt ctataaagga atcatagaag ccaactgtgc tgccactttc cccgacgaga 11700
gggaagtata tataaacaac tattgctgtc cggtaaaact ttttgctaac tttgagtttg 11760
aacaaaaaaa cagctagcaa tgagcaaaag tacacatttt agcggacagc cggtctatag 11820
tcaggtcata aaattactcg ataaggcaga aattgtcaaa attagccatc aaaccaaggg 11880
tagcgaggcc tacgtaaagc gtttcgatgg ctatcagcat ttggttgtaa tgctattcgg 11940
catcctcaaa cactttgatt ctctacgaga acttgaaatt ggtatgaagg ccgaggcgca 12000
caagctctca cacttgggca tggactactt ggttcgtcgt agtacgctgg ctgaagccaa 12060
tatacggcgt cctcaggagt tcttagccag cgtatacgca tatcttttgg agaagtatgc 12120
caagtttttta gcggacagcc gtcctgctaa gagttataaa ggacagacac acgagcccaa 12180
ggactgggag aaactgctct atatgatgga ttccaccacg ataacgctct ttgataacat 12240
cctcaaaggg gttggacggc atccaaaatc aggtaaaaag aaaggtgggt tgaaggtgca 12300
tacggtcatg aagtatcatg tgggcgttcc tatggtcgta cagctcactt ctgccgcaaa 12360
gcatgaccat tacctgctca aggaggtaca tcttccaaag gattccacct tggccatgga 12420
tagaggatac atggatgtcg cacaatttca aaggcttacg gaagagggcg tgtgctatgt 12480
taccaaaatg aagaagaacc ttaaatataa ggtcctggat tccgttacat acgttaatgc 12540
gaagggattg gtaacgcaca ttgatcaaag agtccacttt acacgaggag agcttacgca 12600
tgaggcacga agggtagaga ttttccacga aaacaagaag ccagtggctt tgttgagcaa 12660
caactttgag ttttcagtgg aagatatttc cgagatttat cgcctccgat gggccattga 12720
gtcgctttac aaacagctca aacagaactt tccccttcat ttcttctatg gcgatagcgt 12780
gaatgccata caaatacaga cttgggtagt tttgattgcc aatttactca tcacagtgct 12840
ctcaagaagc atcaaaagga ggtgtgcatt ctctcaggta gtcagcatgg tgcgactaac 12900
acttatgtac tacatcgatt tcatcgcatt tatggaaaat ccagataaat catggaacaa 12960
catccgtgct aaagagctgc aaaaggcacc gccaaatcct actttgttcg attaaggggg 13020
cttacttttt caaaaagaaa gcccgaagcc tgattttact gggcttcggg caaggtttct 13080
atgcacttct gagttttacc ggacagcaat aaattcggat tctaactttt ttgcctcttg 13140
tgccacagtg cgcatctggg tgattgtgat tgagacaatt gagacaacaa gatgttgtct 13200
atacaccctc cccatacttc tgcaagcctt cgatatgcct tcgataaggc ttcgatagga 13260
gtcttgaata attaacgttg ttgagtaaaa cgttgaagag cctccaaaag tattctgtct 13320
aacttttgga ggctcttctt actgaggatg cttttgttct aaagcctaaa aggtgattgc 13380
aacgaaatgc tttgtttgtt catctcattg tgatgatgct tcttttagag tttgttgcag 13440
taatcatcaa attcatcatt tatcttcagc atccattcag ggagtttcgc ttttgcatga 13500
ttgatgagtt cttctggaat cattttatag taagcatagg ccataggacc agagatagcg 13560

gcgagagtat cggcgtcacc tccaagggat atggcaagtt tcaagcaatc tgtataatcc 13620
ttgctctcta agaaacagat gagggctgca ggaattgtta gttggcatgt ttcatcaaaa 13680
tagtattctg gtttgatttc catataggtc tttctactcc agtctggata atacacgtca 13740
agaacatttt tgcgtatgta atctttccct ttcccttcca tagcatggaa gatggcaagc 13800
gaggtggcta ctgccccttt gattccttct ggatggtcat gggtgggtaa ggctgttctg 13860
gtggcaagcg caatacattc ctcttttgtc ttggacataa aaccggcagc cgaggttctc 13920
atgccacttc cattcccgta tgaatggtaa gatggttgta tgtcttctgc catcagccaa 13980
caggcaaaac ggcctccaaa tcctctgtgg aagtcggcac ggcatctttt ccataaattt 14040
tttgccatgt ctaactcatg aagcagagct tcagcacagg cgaaggtgca tactgtatcg 14100
tctgtgtatg tatttgccga atggagaaga tttactgctt catagttctt tgttctgcct 14160
tcaaactcat agggcatgcc acttatatcg ccaataacag cacttaataa tagattcttc 14220
ataattaaaa cttaatttga aatcctttct tcttgaattc attgtatttc tcgttgttga 14280
aagagtagag cagggcttct ctctttgcgg taggccatac tttttcgtcc aactggttca 14340
gaagtttgtg gtgcatcatc ttcttggaga agtttcttct gtcaaacttc acatctaata 14400
tggcttcata aagcatctgg agattcttca tggtgaactt ctctggtaag agctcaaatc 14460
ctattggctc aaagtggatt ctttcacgaa gtttacgcat ggcgtctctc aggatgcgtt 14520
cgtggtcaaa agccagttgt ggaacctcat caataggaaa ccatttcgct ttcgttgcgt 14580
catcacctcc atgaacctct tgtatgcgca caagtgcata ataggcaatg gtaataactc 14640
gttcacgagg gtcacggtca ggaactgtgt aggtatggaa ttgttcaata taagtgccag 14700
tgagccctgt ctcctcctgc aactcgcgaa gcgcaccatc ttcggctgtt tcatcaggat 14760
tcaagaaacc gccagggaaa gcccatttcc ctttgtaagg ttcatttcga cgttcgataa 14820
gtagaacttt cagtctgtca ccgtcaaatc caaagatcac acaatctgtt gtgacagatg 14880
gatgtggata cttataacaa tacatctttt catccatgcc tttttcttct atgaaggata 14940
caccgcaaag ataacctaaa aaatataaac caccaaacat ttgtgtgaaa agtacacaac 15000
aagcattgtg tttatagaaa aaatgtagag taatagtcaa aaagaacaca tgaagtccct 15060
gtgcaacctt ttaacctgag ttagatgaat ttgaaacaag tgcaaatcaa cgggagttcg 15120
acaaattcag aaggtgaaac agcaggcttc actgattgat tcctcttttc tgccacactt 15180
cttttctctt gtttttacca aaatcataga tgattcaaag gagtatgtgc ctacagtgca 15240
agaagggtga aaagattctg gcagactttg cgcagagcga ctcttgcaag aatgtgaatg 15300
atgccatctc cacatccaat aatccacata tcttcttcaa aaagaacgat ttatatatcc 15360
catatagcac catgcttgtg aggtcaaaaa cgcacgcatg gttcatgacc caccgaacaa 15420
ttgttcattg gccatcgcac cattgttcaa aggcacttga agcattgttc attgaggcat 15480
gaaggacact tcggacatgc gagatgcaaa aaggcaaggt gtcagcccta attttgaggt 15540
ggaaagtcaa agtttagcgg acagtaatat aaacaaatta tagatggata ttcgtttgtt 15600
atgctaatta taacgtgaat tagtcgaact cacgtcataa taatgtgaaa tacagtgttg 15660
ttattgtcct gaagtgccgt tttatgcttt tgtctatcgg gtacactact gggttgtaat 15720
tctgtccctc tgatctcgtg atgacataaa ttgttgaatt ttaaagttta gtttatggat 15780
atattatatg tgggataaat taaacctctt gtcaaactgg cttcttcttt tttgagacgg 15840
caaacgtctc gttgtatatg gttatatgca tactttttgt gcttgaattc gctaaaaata 15900
tgaaattgga gagatttttc tataaaagtt ctccaatgtt tgaggaaaat gcttggctgg 15960
acaaaataaa aagagtatgt ttgcaacaaa ttttgcctgg tattcataaa aagaacgaag 16020
atggctacaa aaatacagca aatattagag gctactccac gagattccgt tttgttcggt 16080
tcatggcttt caagtcaagg attggatgca cgtggtcaat atgcttatat gaaaagcgga 16140
tggcttgacc gtatatcaaa aggagtttat aagatacatg ggtagaacc aggtctgatg 16200
gcagcagtat cgtcttataa cacacaactt ggtaagagtt gcatagttgg tgcatataca 16260
gctttggaac ttcaaggata ctctcattac ctatctatgg ggaaacctat ggcttatctg 16320
tttacagaca aaaaaaacaa gttgccctca tggttgctaa aaggggagtg ggatatgacg 16380
ataaaataca tgactacttc tttcctgggt gatgaacttt taggtgtgga aaccatgact 16440
atcaatcagc gtgatttgct tatgtcatct ccagagcgag ccatattgga gtgcttgaat 16500
ttgccggatg catcatcttc attgcttgat atctattaca tcatggagag tatgacaaca 16560
ttacgtccca agttggtaca ggcgttgctg gaggcatgca cttcccataa agtgaagcgc 16620
ctgttcctat atatgtcaga aaaagcaggg cattcctggt ttaaggcttt gaagctggag 16680
aatatcaatt ggggtacatc aaggttaatg attacgccta caggaaagta catcaacaaa 16740
tataatatga caatctccaa agaacttgca gaatatgaat gacataaaaa caagtgttta 16800

cgctcagaag gtagaactgt tgctcagact cattcctgtt gttatggatg agcaggtgtt 16860
tgccatacat ggtggtactg ctatcaatct tttcttgaaa gatttgccac gctattctgt 16920
agatattgat ctgacctaca tcccactggc agaccgtcag acaagtataa atgatatcaa 16980
ttcacatctg aagtccattg cagataaagc aaagaaagca ttcaaaggga tgcacatcgt 17040
acccaatttc aatacttgca aacagttatg tgagtatagt ggtaaacagg tgaagataga 17100
ggttaatcag acaaagagag gtataatagg tggcgaggta caaaccatat cattgagtga 17160
gaaggcacag aacgaatttt ctctcttttg tgaggcaaat gttgttccga taactcaatt 17220
atacggtggc aagattgcag cagcactctc tcgtcagcat cctcgtgatt tgtttgacgt 17280
aaaatacatg gatattccat tgagtgaatg tcgtgaaggg cttattttct gtctacttgg 17340
tagtgacaga cccattcatg aatcattcgc tccaagtttg atagaccagc gagaagcgat 17400
ggaaaatcag tttgcgggaa tgactgacat tccttttact tacgaagagt ttgaagaaac 17460
ccgtgcaaat cttattaatg cctaaaatcc gcaactatca tccaatacgt gattaagggt 17520
agatttatga gtcgttagta gcagctttca gtaaaaagca acagcacaac atcaatatgt 17580
agccaccatc cccttacccc acgatgcgac ttgaggtaat acgcatattc taaccggaaa 17640
gaaaggattc ttatccgaat tctgttttga agggttttgc ataagcgcgg ttctctgtgt 17700
agatattcag cacgtattgc cttgcagtca tgatccactt ggcaggtacg gagatgaatc 17760
tgaagacaaa agcctttatg cgactcgttt tcttgagccc aaaagccttg gtgtcaagcc 17820
tgctcatgat ggtcttgtag aaattgtgta tcaatgcagt aagcagaaga aagacagtat 17880
tctccgccat gaatgacttg gggagcctgc tccaaccgaa tccgttgttc atgtcgtcaa 17940
agatacgttc cttgccgcca cgcagattgt agaattcaac aatgtccctt gtcgatgact 18000
tgtaatcgtt ggtcagaata caacggtaag tgtattcgcc ttcccacagg tcaaggtcgc 18060
cactgttgcg tctttgtctc tggatgacaa gacgatagca cttgccttcc catttctcaa 18120
cgagaatgga attgagttcg aactggactc cgttaatctc ctccgtcttc catcctctca 18180
gagcaaagat gtcattgtag agcgaactgc atcggttggc acggatgtag aaatgtttgc 18240
aatgcttctc tatctcactg acgatttcct tcgagcagga accgcagtct gccctgaagc 18300
gatttacacg gatgttctgg gattccagaa gagcgaagaa tctcttatgg gtgtctgcct 18360
gatgaaaacg cacattcgtg ttgccatcgc tgttctcgat atagactatc ttgtcaccga 18420
taacatatac gccaggcctg tagccgagga actttttgta ggtcggtttt gcatcatact 18480
tctccgtttc aaggaactga tggtcaaagt caacatcgta ttcctcaatt tccttcaact 18540
cgcctgtaga aaccaaagcg tttataagca atgtgttgag tttgtctgcg gtattgaaat 18600
cataggtctt gccttggtcg gaagtatagg agatgttttc ctgtgtcagt tccttgatgg 18660
ctctgaggat ggtatcagag ctgcatgtac gaagggtagg atgatacgag agatggcgca 18720
tcaggtgtga cgttacatct tccacgcatg agccgccaca gaaataaacg ctcatcagcg 18780
aacggactat ctcgctgaac tgatatccga tgatactgct gcatctctga cccagtgttg 18840
agtctataac gggtgaaagc atggagtcaa atttctccat gattgaaaaa attcctccaa 18900
aaggtgtgag tttctcatat tttatttgta cctttgccat gccttgttac gattttcgct 18960
tgttttctaa ttgcaacact aagataagtg aaaaatctga cacggcaaaa tcctgggcaa 19020
ctttttgttg ctcaggaact tataaataaa gttaaattat agtgttgcgg aattaaggta 19080
atgatgtaaa attgttgatg accgaagcgg ataagaagtt cctgattagt tttgaatctg 19140
gtcagcccga atgggaagga tatgaatttg aatatttcaa agattatccg tcagtacaat 19200
ggaaactgct taatctgaag aaactcgcca agcagaatcc tctgaagttg cagatggaag 19260
tagataaact aagaaatctc tataatttta gtctcaataa ttaacagttc actacttatg 19320
tcatacaaag attttcagca gaaggtttat tcatacctga ttaaatacaa gaaggaaagc 19380
cttaatataa atgataaagg catttctaca cgtggagttg aacacgattg tttattatcc 19440
aagccatatt gtgatgccaa aatacctgtt atgctttata atggtataaa agaaacagtg 19500
aaagacattc aagcaagcaa gtatgcatat aaggctcacc ttgcagcttc taaacatgtg 19560
gcaagttctc aaactgcatg tatcaaccta tttgtgccaa tcttggaaag tgagtacgca 19620
gaccaaatac ttataggggc aggagttgta cctaaagatt ttgaccatat tgacagaaac 19680
cagcttcgca aaggatattg ctttgagtat tgggagtcaa cactagaagg agcaaaaggg 19740
cttcttggag accatagtcc tcatgctggt acagactcag atgttgccat tgcatatcgc 19800
aacaaaaacg ataaactgtg cctttggcta atagagcaca agttaacaga gcaggaattc 19860
acgacttgtg gaggctaccg ttcaaaggca atttctcact cagagaagga atactgcact 19920
tcttgtggta ttgaagaaat cttgaaaaac cacgataagt gttattacca caagcattgc 19980
ggttacttct attggaatat tatggatgcc caatcttcat tcttctgtgg taaatttgaa 20040

ggcaatggct gtccgtttag aggtggtatg aatcaacttt ggagaaatca aatgctggct 20100
ttggaacttg agaatagagg cgtatttgaa gatgtctatt tctcggttgt ctctcatccg 20160
gataattctt ttcttgacaa gactatgaat gaatatcggg cactaacaaa taactctcca 20220
aagttctatg acttcaaatc aaatgcactt gtggatgtgg cagttgatta tttgcctgac 20280
tggtcaaatt ggtacaagaa ggtgtatttg ggaattgatt aatgtatcac cgtgattatg 20340
acaatataca aatgatagaa caactccgac aattaatgac tcttgatgct aacatgaata 20400
catcagacat cgaagagcgt tttgaacaaa tagcaaaggt gctatttgaa tgttttgcta 20460
ttcaaaaggg tgagacgatg tatctgttta aggagattga gttttacttc tacaacaaga 20520
atcatcgtga tattatcacc catccacgtg tgtcaaaacc attgtgttgg tatgtaaatg 20580
actttggagg gattgattta aattttgtta gtaatattaa gcgtgagtct aaagctaata 20640
gtaaaggaaa gaatgttgaa aagtatgttc tcgataatgg tgcatatttt ggtggcatcc 20700
tcattcgtca actaacaagt gaagatgggt gtgaagttct aaatggacca tgggcttgtg 20760
ctgaattgtt taggtgctat gatgctactg tcattgacaa ggatttgcct attcttgtgg 20820
agcttgataa tggtatggtg ggctatattc gtgaacctcg catcaatcta cttacatcta 20880
aacaaacagt agagggtaag gttgattata ttcttgaaga atatcatgaa gtttttgaaa 20940
ggaaatgtct atatagtgat ttttcgagat ttgtggatag acgttataga tatgtgcgtt 21000
gtgatacatt aatgcacgat aaagatacaa acctcgtgta tttctctcct tggttaaaag 21060
acaataaaga aggacatcca gagttttata agcttctgaa gaatgtgctt aatgaaatgg 21120
gaattgagtc caaagaactg aaatctacaa atgattattg ggctcgcgat tatatgccga 21180
ttcaattggg cgaaaatgag ttcttgaagt atcgctatta tcctgattat cttgtgaaga 21240
gcaagaataa aaaagatatt gagacaatta ctgatgcaac caaggtgttg cgtggtatgg 21300
gcatcagttg ccgttcaacc aatctcatca ttgatggtgg taatatggtg ccatgcggtc 21360
catatattgt gatgactgat aaagtattta tggaaaatgg acgtgagaaa ggagatacag 21420
atttcaaggt tttgctggaa tctgaaattg ggcatcctgt tatcattatt ccttggactc 21480
cgcatgaaga tgatgtgtat ggacattcgg atggattcat caaatggtgt ggcgataatc 21540
gaatactgat gggtaatcat ggcgattgct atcctgaaga agcagcatcc atccgtcgga 21600
ttcttgaaag ttatggcttt gaagtgacag aaatgcgttt caagggcaaa gttgataaac 21660
cttgttatga tctaaactgg gcatatatca acttcctgca agttggcaag aacatcataa 21720
tgcctaaatt caacattgaa gaagatccaa tcgctcaaca atacattcat gaagcattcc 21780
ccgattggaa tattcgtcaa atagaaatgg cagaaattgc caaagaaggt ggcgctttac 21840
attgtataag ttggaatgtt taacttccaa aatcacatgt aaaaacataa atcaccacct 21900
atgagatcta tttgtccctc ggctattttc gactatctcg ggggacagct gggggacaaa 21960
acgggcgaaa atctgcaaaa atcgaagtta aaaatcaaaa atctgcaaaa atgggcgtaa 22020
ttcttaatgc ttgaaattaa gcgatttacg atccatattt gcagatagta tgcttttctt 22080
tgcaaaaaat cactttccca cgcagaatct actttcctac gcagaagtta ctatctgcaa 22140
atataagtgg ctgataatca acgcaaatgc ctttgtttta gggtaactaa agtgactccg 22200
agggacggtg gggggacgta aaataggcaa aaatctgcaa atatgactgc cttcaaaaaa 22260
agaaaatgca aacatctgca aaatcgattt tccatttctt tgcacgtttt cactttttaa 22320
cgcagatttt tgcacattct tgcattaagg tttagtttag gttattgata tataatatgg 22380
gaataaacta aaccaaaggg ttgtgctatc aaactgaata attggcttct ctttcttttg 22440
gacggctatc atatctcata tttagagtta tatgcgtgta ttttgcttcc aaattcgata 22500
aaaatgggcg cacccgacaa accctgtggg aagatgtacg gatgctaacc caaaaactgt 22560
cgctaaacga tacacgattg aaaaaaagtt agtacctttg cagtatgaat cgtaacagca 22620
atataaacat gtatgaaggc cttgcctcat atttcctccc tacaggtgtc cttgagtact 22680
ttgaagtgac cgacttctgt gaggttccga cacaggacac cgaagttctg tacacaactg 22740
agcttcatat ctatcttgac gaacgcgaca atcgcactcc agacatgagt ggctctgtga 22800
gcgatggttt tggtgacgag tcatgtcttc tggatttccc gacacgcgac aagaagacag 22860
tcctccatgt acgcagacgt cgctggacaa tgccagacgg cacaaccaaa tcagtggatc 22920
tcgacaagaa gatacagttg aagcatcctg gcacgcgata ctcgaaagac ttcgcgcttt 22980
ttttaaagaa ggcggatgga caatgagacg ataaccgctc gtcaaatagg ctacacatac 23040
atgattaacg acgccacgtt tgcgaagaga tacaaagata ctctgagtga cttcgagacg 23100
tgggagcaga aagagcatgc ctccgaatgg atacttcttc cgcaaaacgt aggcaaggag 23160
caaggcatag acgagacatc cctccaaggc gagctgtata ccattgtcca caacaaggaa 23220
gcccatggtc gcaagggcgc aatcgttgcg gtagtcaagg gaacgaatcc tgctgatgtc 23280

```
     ctgagagttc tcatgcaact gcctgcagat aagcgtgaaa tggtcgagag tattacgatg     23340
     gacctttctg attgcatgcg tgccatagca cgggaagcat tcccacaggc cactgttata     23400
     cgtgactgct tccatgttgt cagacgaggc ggtgagggat gcgaggagat acgcctgcgt     23460
     ctgaaacgag aagcaataaa agaactgaac aggcagaagg cagagttccg caaatacctc     23520
     gaaggactgg cagcgcagag aaaggcttac cgggagatga tgagggcaaa acacggaaag     23580
     aagtggaaga aaagcaaacg tggcaagaag ccaaagaggc tgaacactcg atttgaaccg     23640
     ccaaggctta ctaacggaga aacactcgtg gaagcgttaa cccggagtcg caagcaactg     23700
     gccatgagcc gagataaatg gagtgcaacg cagggaaaac gtgcgacgat tctctttgaa     23760
     ctatatccaa aattagagg                                                 23779
//
ID   XXX; XXX; linear; XXX; XXX; XXX;  XXX.
XX
AC   PRJEB67889
XX
DE   Uncultured bacterium DNA fragment from in-reactor enriched cow rumen, clone
ferm-38a17
XX
FH   Key             Location/Qualifiers
FH
FT   source          1..37749
FT                   /organism="uncultured bacterium"
FT                   /environmental_sample
FT                   /mol_type="genomic DNA"
FT                   /isolation_source="Cow rumen"
FT                   /clone="ferm-38a17"
FT                   /note="AZO-CM cellulose, AZCL Tamarind xyloglucan, AZCL-HE
cellulose, AZCL-Barley beta glucan, AZCL-arabinoxylan active clone"
FT   CDS             267..833
FT                   /db_xref="SEED:fig|6666666.127251.peg.1"
FT                   /translation="MAILVLQKSLSGGHRAGFFTGLGATFTDTLYSVIAIFALAITQQF
FT                   ITEHEVFIKITGGIAIILLGANMLFKDPFRKMKDQSGPSYSIKDFLQAVATGLSNPFAI
FT                   LIIMSLFGFFGLAPESRDLTVAPVILAVSGGSMTYWFFYSLIFSRIRKNFKLSTLLWIG
FT                   RISGAIIMIIGIALASDGLMKLIFE"
FT                   /product="Putative threonine efflux protein"
FT   CDS             835..1617
FT                   /db_xref="SEED:fig|6666666.127251.peg.2"
FT                   /translation="MPEILKNWIVKNLLLAIGLLLALVLVSALFLNLLTRHGKELVVPD
FT                   FTNMKISEATYTASVDGMRIEVVDSVYIRRMGRGLVYSQNPKPGSKVKKGRRILLTINS
FT                   VTPKKVQMPNLVGYSMRQAKAELSSRGLALGKLMYVSDIATNNVLRQLVGNMEITPGRM
FT                   IESGTAVNLVVGLNSSDSQTYIPDVKGLKYLRAVDVVHDNSLNIKRLVFDESVKNYSDS
FT                   LDAIVVRQGPEASMAPIVMGSDVTLYLSVDPAKTIMKK"
FT                   /product="hypothetical protein"
FT   CDS             1619..2650
FT                   /db_xref="SEED:fig|6666666.127251.peg.3"
FT                   /translation="MEFDDLQDDGMFEHFRLNVDPGQTPMRVDKYMACHLEDTSRHRVQ
FT                   NAIKSGYVRVNDKVAKANVIVRPGDVIRFVMPYERHGFEVLPENIPLNIVYEDDDFLIV
FT                   NKPAGMVVHPGHGHFDGTLINALAYHLGISQGPDAEDDRMGILVHRIDMDTSGLLVVAK
FT                   NDEAQLNLAQQFFVHSIERKYVAVVWGNIKEDEGTIVGNIGRDPNDRLKFRVYQEEDKG
FT                   KHAVTHYKVLERFGFVTVVECRLETGRTHQIRVHMSYIGHPLFNDARYGGAEIRKGTIY
FT                   AKYKQFIQNCFEICPRQALHAKTLGFVHPRTGKPVSFDSPLPEDMTALLEKWRKYAGDM
FT                   PKE"
FT                   /product="Ribosomal large subunit pseudouridine synthase D
FT                   (EC 4.2.1.70)"
FT                   /EC_number="4.2.1.70"
```

```
FT   CDS             complement(2668..4239)
FT                   /db_xref="SEED:fig|6666666.127251.peg.4"
FT                   /db_xref="GO:0003723"
FT                   /translation="MDSEKELLKDLIVDVQATEVQIALQENHRLIELNKESSSGRSFAV
FT                   GDAYLGRVKKVMPALNAAFVDIGDEKEAFIHYLDLGLYFGAFDEFVKKTNPNTNPSRLF
FT                   GSIKLGPILEKEGRIENVLTQGQMIIVQIVKEPISTKGSRLTAEISLAGRNIVLLPFAE
FT                   KVSISQKISSKEEKKRLETLVRSILPKNYGAIIRTAAEGKNAAILVAELKSLIGKWEDS
FT                   WENLSKNKSIQPLFTEYSKTTTILRDLLNDSFSNIYVNNEKEYEEIRRYISLISPEQEK
FT                   IVKLYQGKEPIFDHFEITRQIKSSFGKVVPLKQGAYLVIETTEALNVVDVNSGIRAKTK
FT                   EQEENTFEVNKFAAEEIARQLRLRDMGGIVIVDFIDMESVEHRNNLFKYMQELMSKDRA
FT                   KHNVLPLTKFGLMQITRQRVRPVTEINTTEVCPVCHGTGKIASSLLIDEQIERRLSYYV
FT                   NDKGIRNFTLKVSPILGAYITRGLFNSYVGKWKKKYKCKIKVVEITDFAVLQNELFNEK
FT                   GDKLDS"
FT                   /product="Cytoplasmic axial filament protein CafA and
FT                   Ribonuclease G (EC 3.1.4.-)"
FT   CDS             complement(4468..4740)
FT                   /db_xref="SEED:fig|6666666.127251.peg.5"
FT                   /translation="MTKAEIVNEVAKATGIEKLAVQSVVESFMESVEASLAKGEPVYLR
FT                   GFGSFIIKHRAEKAARNITKNTTLTIPAHNIPAFKPAKVFVSKVK"
FT                   /product="DNA-binding protein HU"
FT   CDS             4963..7368
FT                   /db_xref="SEED:fig|6666666.127251.peg.6"
FT                   /translation="MIDFNKEMFPASSHEVNIIPVMSGEPALNFDASELPGSLPILALR
FT                   NAVIFPGVVFPITIGREKSMLLVDKAQGNGMFIGAVPQKNVAEENPLSLADFSEFGTVV
FT                   KVLKTFEMPDGTVTALLQGYKRLRMLELTDTDPYHMARVSYLDDILPQNDSKVPVIADI
FT                   LKENASSIIRSSSFVPKEALEALKNIDDFNFLVNFIATTIDMDDFMEKVDLLKYDDVKE
FT                   RAMALLGVLSRQLEIQRLKQEISSKVKTELDQQQREYFLNNQLRTIQEELGMDEMDDIA
FT                   RFREAAEKKEWPASVGETFEKELAKLERFNPSTPEYSIQYNFLQFMIDLPWYHTSDDNL
FT                   DLKHAQKVLDEDHFGLENVKDRIIEYLAVLKLKGNMKSPILCLYGPPGVGKTSLGKSIA
FT                   RALGRKYVRISLGGMHDEAEIRGHRKTYVGALPGRILNGINKAGTSNPVMVLDEIDKLG
FT                   SDFKGDPSSALLEVLDPEQNTAFHDNYLDIDYDLSNVMFITTANTINGIQPALLDRMEL
FT                   INVTGYLAEEKVQIAKKFLVPKQLEEHGIPKKELKIDKAALEEIIDKYTHESGVRGLEK
FT                   QIARIARVSAKKMALGEDYPKTIKKEHLKEYLGLQTAFHDSQKGNEAPGVVTGLAWTSM
FT                   GGEILFIETSVSDGKGVLTMTGNLGDVMKESATIGYQYIKAHPQLAGLTSEEFAKKDVH
FT                   VHVPEGAVPKDGPSAGITMVSSMVSALRGKKVRSGIAMTGEMTLRGRVLPVGGIKEKIL
FT                   AAKRAGVSTIIISEENRKDIEDIKEIYVKGLTFVYVKTIDDVIGYIFE"
FT                   /product="ATP-dependent protease La (EC 3.4.21.53) Type I"
FT                   /EC_number="3.4.21.53"
FT   CDS             7368..8036
FT                   /db_xref="SEED:fig|6666666.127251.peg.7"
FT                   /translation="MDVKIEQTWKDALRDEFEKPYFASLVRFLHNEKEAGKVIYPPGGL
FT                   IFRAFELTPLPEVKVVILGQDPYHGPRQAMGLSFSVPSDVPAPPSLKNIFKEIQTDLGI
FT                   TMSGSPNLEGWARQGVLLLNAMLTVQAGVAGSHKAIGWQEFTDAVIKCISDRCNGVVFL
FT                   LWGNFARGKKELIDTSRHYVLEAAHPSPLAGGAFFGCRHFSKTNELLIKEGKTPIDWKL
FT                   "
FT                   /product="Uracil-DNA glycosylase, family 1"
FT   CDS             8033..8506
FT                   /db_xref="SEED:fig|6666666.127251.peg.8"
FT                   /db_xref="GO:0004595"
FT                   /translation="MKRKALFPGSFDPFTAGHLNILRRALTMFDEIVIAVGVNQDKRGF
FT                   FSMDQRMDIIRQAVRGMDGIEIIKYDNLTIDTCRELGIAHIIRGVRNMLDFENERAIAD
FT                   ANRRLAPEIETIIIPTAQEFAHISSSAVRDILKHKGDTSLFLPEGVVLPDHDA"
FT                   /product="Phosphopantetheine adenylyltransferase (EC
FT                   2.7.7.3)"
```

```
FT                   /EC_number="2.7.7.3"
FT   CDS             complement(8508..8726)
FT                   /db_xref="SEED:fig|6666666.127251.peg.9"
FT                   /translation="MRVKTYGDMVSYIVSRARRNEEIALGPLYIQRLGLQRRKILLKLC
FT                   IQGGHLDGIRILGAQGKRKQRKKDYED"
FT                   /product="hypothetical protein"
FT   CDS             8730..9977
FT                   /db_xref="SEED:fig|6666666.127251.peg.10"
FT                   /translation="MQSKLMGDEAVAVHLADNWFIPGKVSFPDEIDLMNAKIFAEFNRQ
FT                   SLLGCKAPGLVAEDSLGRKVEVPLGLSDRYRILYIYSPDCATCSLQTMLMDSLFSRKSY
FT                   PVDLIAFNSDDNTEAWVQYRRNRLSFDGVRVMHFRDPSLESDFQRKYGVLQTPRMFLID
FT                   KNGIIVGRGLDAAALEKLLDATPDLGIGYGSKSSADYFDAVLESFGDELRGSDIRYFIG
FT                   RLEEATLESADTAGFRQMTGDMLYYLAGKRGEAFKDAEAFVVDSMVLSRPDVWRFADDS
FT                   LTVVDMAMSLKHMLSLAPVGARVPDVRVHATVLRAGRKPATVHKKLRSLLKGKRNYVLF
FT                   HTAGCTFCEAELAAADSVASEKGVRVILVDMDKLTSLYPEEADRLLDAFDLSVLPFIME
FT                   TDNKGCVVRKYLSLNK"
FT                   /product="Thiol:disulfide interchange protein tlpA"
FT   CDS             complement(9966..10538)
FT                   /db_xref="SEED:fig|6666666.127251.peg.11"
FT                   /translation="MSIMGKAYIFLADGFEETEALATLDILRRGKIEAYTVSITEEVYI
FT                   TSSHGVTVVADMGRSDFSEIDIDEKDAMIFPGGMPGSKKLAADTPLIGLMNVHYGNGGI
FT                   VAAICAAPGLVATQIKGIEGKKATCFDGFEENLAASGMEYVKAPAVTDGQLITGRGAAF
FT                   AIDFGLAILEKMGGKEAVAKVKAALLI"
FT                   /product="DJ-1/YajL/PfpI superfamily, includes chaperone
FT                   protein YajL (former ThiJ), parkinsonism-associated
FT                   protein DJ-1, peptidases PfpI, Hsp31"
FT   CDS             complement(10648..11760)
FT                   /db_xref="SEED:fig|6666666.127251.peg.12"
FT                   /db_xref="GO:0008784"
FT                   /translation="MNRVDDFSRLELNLTGCMENYRYFRSKLEQTTKLLVLVKANAYGH
FT                   GAVEFAAMMQRAGANYLAVAYPVEGMELRKAGISLPILVLTAGTDFFNEIIDNRLEPGI
FT                   PNLYALKILCGILKERGIKDYPVHIKLDTGMHRLGFMTSELQDLTDFLKTCEEVKVKSV
FT                   YSHLAVSEDPEQDEFTLGQIGMFKTNADRITSEIGYKPMYHILNSAGIERFPQYQFDMV
FT                   RLGIGIYGISALPDVRLAHVASLKCKILQIKTLKPEDGTIGYGRYGHIAPEGTVIATIP
FT                   VGYADGIDRHLGRGHASFSVNGHRAPTIGNICMDMCMLDVTGIDAKVGDTVTIFGEDPT
FT                   VNELADILGTIPYEILTSVPRRIERIVVRK"
FT                   /product="UDP-N-acetylmuramoylalanyl-D-glutamyl-2,6-
FT                   diaminopimelate--D-alanyl-D-alanine ligase (EC 6.3.2.10) /
FT                   Alanine racemase (EC 5.1.1.1)"
FT                   /EC_number="6.3.2.10"
FT                   /EC_number="5.1.1.1"
FT   CDS             12119..12793
FT                   /db_xref="SEED:fig|6666666.127251.peg.13"
FT                   /translation="MDMIWFIIIAVMVLSFVIQQVLNSKFNKYSKVGLHSGLTGAQVAA
FT                   KMLEDHGITDVSITCIKGKLTDHYNPVNKTVNLSEEVYYGNSVAAAAVAAHECGHVVQH
FT                   ATGYAPLKLRSALVPVVSFSSNAVQWVLLIGVMIIKITPTLLYLGIAMFALTTLFSVVT
FT                   LPVEINASARAVQWLDKAGITGPATKAKAVDALKWAAYTYVIAAIGSIATLIYYIAMAR
FT                   RR"
FT                   /product="probable metal-dependent peptidase"
FT   CDS             complement(12790..13083)
FT                   /db_xref="SEED:fig|6666666.127251.peg.14"
FT                   /translation="MTGDDYKIISDTTENGVRHITAIPSSLVCSQQIDFDIIDGKVRNL
FT                   AYVRGCNGNLQAIGRLVEGMDIKKIVDTLSGVNCNFRGTSCTDQLARILSSL"
FT                   /product="hypothetical protein"
```

```
FT   CDS             complement(13100..13534)
FT                   /db_xref="SEED:fig|6666666.127251.peg.15"
FT                   /db_xref="GO:0008270"
FT                   /db_xref="GO:0016787"
FT                   /translation="MEKDELYMKEALREAQAALAEDEIPIGAVIVCKGRIVGKGHNMTE
FT                   RLHDPTAHAEMIAITAATEALGGKYLNECTLYVTVEPCPMCAGALNWAQIGRIVYGAID
FT                   PKRGHSLFTPSLLHPKTEVEAGVLAEECGSMVTEFFKKKR"
FT                   /product="tRNA-specific adenosine-34 deaminase (EC 3.5.4.-
FT                   )"
FT   CDS             complement(13655..13885)
FT                   /db_xref="SEED:fig|6666666.127251.peg.16"
FT                   /translation="MFKIPAGNQRKLIFAYLFDTVAYDSPDALAMLDEVQLVFPVPVEG
FT                   EDEFGLMSLDYVEAVPVRKRCYFRYYRGHIH"
FT                   /product="hypothetical protein"
FT   CDS             14003..14542
FT                   /db_xref="SEED:fig|6666666.127251.peg.17"
FT                   /translation="MFEQARNGLTFSLAAIMKVYSLLDALPSEIDKFDQFQHFMRLLYD
FT                   LSLAGDSRILASTSFAHNDADKESRRVQKIKQYVGDHFAENITLEELAEMVGMSPSAFS
FT                   RFFKSRTGKTVIDYIIDIRLGNAARLLVDTTQNISEICYSCGFNNLSNFNRLFKSKRGY
FT                   TPREFRTLFKKNKVFV"
FT                   /product="Transcriptional regulator, AraC family"
FT   CDS             14649..17111
FT                   /db_xref="SEED:fig|6666666.127251.peg.18"
FT                   /translation="MIKRLFPALFIALFAILSCQPMKMDIVKVDDGSFVVDGHFYYYIG
FT                   TNFWYGGLLGSEYGDRSRLAAELDSLHALGLDNLRVLVGSDGPDGVATRVYPTLQKEPG
FT                   VYDEKVFEGLDYLMAELGKRNMRAVLYINNSWEWSGGYGMYLEWAGAGKAIIPSVGGYG
FT                   PFMEQMAQYATNARAQELFENHLRTVVSRVNSITGKPYSEDPAIFSWQIGNEPRCFSSD
FT                   PAVQDGFVSWIWRAASIIKSIDPNHMVSTGNEGIWGCEENPELFRRLNECPDIDYITAH
FT                   IWPYNWSWVGKESPQLDIDRAEENTGRYIDAHLEVAESLGKPLVIEEFGFPRDDMQFAK
FT                   GTPVSGRDRYYRYVFSKVMENQKNGGRLAGANFWSWGGLAVPAHRYWQEGDDYAGDPAQ
FT                   EEQGLNSVFADDYSTLAIIKHTTARLSAVANASAVLENDWMFTTGDQKPLRVAVQSQNG
FT                   FKGSVPVYVRVSTDKGVSVSVSGKKARLGKGADTLDFDLGLAPGFYKVCVSLNENDPLD
FT                   ENFCVGCDPEKIVSPQDKQPDFDEFWEDNLRQLAMVPMNARMTPLKDHSNADRTAYRVD
FT                   MKSFGGKEISGLLYLPNKEGKFAARISYMGYGSDVWFADPSAEPDMIEFTLSVRDQGFN
FT                   KTPGQGWVTRGLADKDSYYYRGAYLDCVRAIDFVSSLPQTDPDHIYAEGGSQGGAFTLI
FT                   AASLDNRLRAIAPFVPFLSDFPDYFAIADWPGGEVLSAAKAQGISEDELYKTLSYFDVK
FT                   NFTDRIGCPVLMGFGLQDPVCPPHTNFSGYNHITADKQWLCVPVSGHHLEQFPEWWAAR
FT                   TEFFTKNK"
FT                   /product="GH5_7-CE7"
FT   CDS             17169..20348
FT                   /db_xref="SEED:fig|6666666.127251.peg.19"
FT                   /translation="MLMLFGALSLMAQNKVTGRITDNSGQGVINASVMVKGTNNGVVAD
FT                   LDGYYEIVVGQGSTLVYSSIGYKDQEVVVGTQSVINVLLVENSELLDEIVVVGYGTMKR
FT                   SDLAGSSVSMKEDDLKGSIITNLDQSLQGRAAGVTAVQTSGAPGSSSSIRVRGQATINA
FT                   NAEPLYVIDGVIMQGGGNSGADFGLGDALGNGKVSTISPLSTINPADIVSMEILKDASA
FT                   TAIYGAQGANGVVLITTKHGSRGDAKFSYDGMFAVSRQTTRIDMMNLREYAEFYNSMVE
FT                   TGEISEANPYYATPGLLGKGTNWQDEIFRTAIQHQHQISAQGGTDKVQYYVSGSFMDQE
FT                   GTIRGSEFGRYTFRTNLDAQLKEWFKLGLNLTFANTNDDLKLADGSEGLITYSLTTIPD
FT                   IPVYDVYGNFSSTVREGYTSPNPVALAMMDEILLDRQKLNGNIYAELTPVKHFTWRTEL
FT                   GFDVSGSTATRFRPTVNLGGWTRDTNSMSYQKNSNRFWQIKNYLTYANSFGKHSVTAML
FT                   GQEAWESSWSYVQGANTKLPSNDVHNVALGGGTPSLSSGFGSSAMASFFTRETYNFDDR
FT                   YLLTYTYRYDGSSNFGPEKRWAGFHSLAVAWKFTNEKFMDFLKPVVNNGKLRLGWGQTG
FT                   NSSIGGYAWGASISRMPSALGMGYRPANIPNPAIHWESQTQQNIGLDLNMFDNRLNLTV
FT                   DAYLKKSDNMLMSMQLASYMGTQGNSSSALAAPKGNFGSIENKGLEITLDAHPVQIKDF
```

```
FT                   AWDSNFQISFNRNKLVSLDGTTNAALIGYGQWSDVVCVSEIGKPLYNFYGYKVAGVYTS
FT                   LEDIENSPKPEKYPSNGEYARGNTVWVGDIKYEDISKDGVINEKDRTDIGNPLPKFTFG
FT                   WSNTFTYKDFDLNIFINGSYGNKVLNYNAISLTHMNSAWTNQINDAVSGRAMLAPIDAS
FT                   VVYADGSKWYDHIDNVRVLNPGTKTPRASINDPNDNDRISDRYVEDGSYIRIKNITLGY
FT                   NVPKSFLNKFKVDNVRVYVNIQNLYTFTKYTGYDPEVGASTQDSTGLTYGVDNGRYPSP
FT                   TVYSCGLNITF"
FT                   /product="TonB family protein / TonB-dependent receptor"
FT   CDS             20361..22109
FT                   /db_xref="SEED:fig|6666666.127251.peg.20"
FT                   /translation="MKISNMMKGFAVATAVFALTSCEGFLNRPNEDNYNVDNFYKTDEQ
FT                   CEQGVNYLYNSPWYDFQRGFIKIGEVMSGNMYWGSSPYLDFSTNGTDQDLVNMSYSLWA
FT                   VNGHANTVIYNLLNSQGPSKAAIDKCIGEALTWKAFAYFFMVRTFGDVPIVHSNSDLLS
FT                   SGEYNSISKVERADVYEYIITTLEKAMEMLPKKTSGFNNYERIDYYAAEALLAKVYLTK
FT                   AGLSGSLNQDDLKQAAKYAKDVIDNSGRTLLPVYSDVFRLTGFNATGECLISWEWSVGR
FT                   NPWTQQNTLQSDLAMVGFSDQGDCWGGWGGPSVDLADAFGVSATQNPAERVGEKDSRRK
FT                   ATLMLPGDKYDYFWTDKGGFDYLRFIYDTEYGKGGPGGSMQCPTGSNNVKHLYGDNFDH
FT                   ENAGLGTPENMAYQLPTHLLRLSDVYLIYAEASLLTGSNGPALEYVNKVRERAGADALA
FT                   SVTFEDIWKERRLELAGEGDRWYDYVRRSYYDMDACIAELKAQRRSYWDGLDDVWKSYI
FT                   GDAFDYSGTWDASEVQYNNAEDNPNVTAASFKLPFPTEDVVFNGNMASDAVPVHVDVRE
FT                   EYSYNF"
FT                   /product="putative outer membrane protein, probably
FT                   involved in nutrient binding"
FT   CDS             22122..23426
FT                   /db_xref="SEED:fig|6666666.127251.peg.21"
FT                   /translation="MKSINRIITGSLAILAMASFASCDKVDYPDRYEVADGKPTVYSVR
FT                   YADKDVNITQAYMDEVVCLLGKNLKSIVELWFNDQQATLNTSFITDNTLLVSVPKNMPV
FT                   VQTDKIYMITSGKDTLAYDFAVLPPTPKVTSMSNEWAAEGETVTIYGDFLIDKADKPLQ
FT                   IAFAGAEVDHASMVFNGSSSVSFPVPAGAQPGYVTVTSLSGTGKSKFMYKDTRNILFDW
FT                   DGSHGGFATGHGWRNGVIHAPGADAWDAIDGSYLYFGGADLGPGSADAWAEDQYSFNYW
FT                   PEAGTDYDALNLRPEFAEYIDKYGVSGLVLKFEVLVPSSNPWSASGMQLMFSSDAVVTY
FT                   GTATNSYFSDETFPRAIWQPWTTNGGSYDTADKWVTVSFPLSSFNKNQENKECATAFDK
FT                   SYLTGLTFFVWNGGIDGTACSPVFAIDNIRVVPNE"
FT                   /product="FIG00417257: hypothetical protein"
FT   CDS             23433..25859
FT                   /db_xref="SEED:fig|6666666.127251.peg.22"
FT                   /translation="MKLIMKKIFSISAILAACVLALGLSGCSSKELDTDQYSDKDVKLA
FT                   SFGPNPVMRGATLRFFGSNLDRIVEVSVPGLDPITDIEVVTSGNPSEIRIQLPAEGAQV
FT                   GKISLKSSDGKVFTTQVDVEYTEPIVFDSFSPAEAMPGDVITVKGDYMNLVQSVTFEGG
FT                   AIVTEINHNGRHEASFKVPSKAITGKIILSDEGAIANLLYSEEDLVIGDPTVKSLTVAV
FT                   AKPGENAVVKGAYLDMIDKVVFEGGAEVSDLALNEDATEFNLVIPAGAQSGEVKAVSHA
FT                   AKEFVAGEIELVIPTDVAAAPSPVKAGADLVISGNDLDVVTSVDFPGASGVEFVFAEDA
FT                   ITVTVPATATEGDIALNMASGEAVNVAFTLVHPVVSAIAPVELMAGETITVTGTDLDLV
FT                   TGVTLGGKPVEFTAEAEKIVVQTANTSVSGKLVLSLANGEKIEPEDAITLSYDSFIIVN
FT                   DMPAAEHIGAIVTLKGENFMMIENIYIGEEKVSKYNLRSDNEIQFVMPWNKVGTYSMYF
FT                   HLLSGDVETCPQTIEVLLEQDIMTVWEGSLQITWNDGGRVTVPAASFTGIPAGSKLRFY
FT                   YTQVDQQWDQAQLNYGNWTGINFDDPAGTVINGTLVPTDVYGWFTDGILDRCTEVVLTK
FT                   EILENIQANKGDCEDKTACGIIIQGSGLTFTKIEIVCDIPQEVTIYEGPTTLTWGDDGR
FT                   FGLAMTYFEAAKPGSQLIFYLEQTDAWGQVQLNDGWWANADMNFPEIGGAYITTDNLGG
FT                   KDVTRIALTLTADILDHILATSGDYFGLNSQFQGDGRVAMVIQGSDMIINKVCIL"
FT                   /product="hypothetical protein"
FT   CDS             25954..27480
FT                   /db_xref="SEED:fig|6666666.127251.peg.23"
FT                   /translation="MIFLAACSCSKDDAEKKEPSVPAPKAVTVDVASVSAPVEGTSVVV
FT                   AISSPSKPAVSVSGDWIKAELGPYNPDTYKVKLTVTVGENPTYESRSGSVTLTCQGFSP
```

```
FT                   VGISVSQDAAAKPVGPSANPAYDLLGLGWNMGNQFDAFYNGSWAGDKYNYPDETVWGNA
FT                   KATQATFDGVKKAGFTSVRIPVTWLNMIGPGPEYNIDETWMNRIYEVVGYAEKAGLKVI
FT                   INTHHDENNNDDHWLDILSASKDARTNEAIKAEIAAVWTHIANKFKDKGDFLIMESFNE
FT                   IQDGGWGWSAAFQADPSKQCDVLNEWNQVFVDAVRATGGNNATRWLGVPTYAAGTSWTK
FT                   YFRLPEDPAGKVMMAVHCYDPYEYTIGNIQYPEWGHTGAAGKKTPEGEKEITDLFSRLN
FT                   TEFISKGIPVYLGEFGCSMRSKGDARAWMFYKYYMEYVSKAAHTYSLPGILWDNGSTKT
FT                   GKECHGYIDHGTGKYIGNSKEIVDLLVKGMLSTDESYTLQSVYDRAPKTN"
FT                   /product="GH5_4"
FT   CDS             27594..28625
FT                   /db_xref="SEED:fig|6666666.127251.peg.24"
FT                   /translation="MMDRMAASVKAGKILYGHQDDLVYGHSWCVDPDSAATDALNRSDV
FT                   KAVAGDYPAVVGFELGGLELGWDWSIDKVPFLLMRRAAVEHAGRGGIVTFSWHPHNPYT
FT                   GGDAWDISSDKAVASVLEGGEKAEMFGVWLERLGDFMDSLRDDKGNLIPVIFRPWHENI
FT                   GSWFWWGGALCTADEYIALFRRTVEYLRNDRGLDNIIMCYSPNSGITAEQYMSRYPGDD
FT                   VVDILGYDHYETVMEGESLEDAGKRYAGQLDLGLSIVKSLASAHGKLMTLSETGMEGLP
FT                   DRSWWTSVLYPAVKDFPIVYVLTWRNAWDRETHFYAPYPGNEGDFREFASLPEISLLSS
FT                   FRP"
FT                   /product="GH26"
FT   CDS             28703..29878
FT                   /db_xref="SEED:fig|6666666.127251.peg.25"
FT                   /translation="MENMSFEERLAWLEARHQELIAKKNSPIYGNGVFERYENPVLTGD
FT                   HAPLFWRYDLDKNSNPFLEERFGIHAAFNAGAIKWKDKYVVCARVEGADRKSFFAIAES
FT                   PNGIDNFRFWDHPVTMPDYGEPATNVYDMRLTAHEDGWIYGIFCVERKDPDAPEGDLSS
FT                   AVAAAGIARTKDLVNWERLPDLVSDSQQRNVVLHPEFVNGKYALYTRPQDGFIDAGNGG
FT                   GIGWALVDSMENAVVTSEKIINCRRYHTIKETKNGEGPAPIKTDRGWLHLAHGVRGCAS
FT                   GLRYVLYMYMTSLEDPSKVIAEPAGFFMVPEGEEYVGDVMNVLFSNGWIADEDGKVFIY
FT                   YASCDTRLHVATSTVDRLVDYCMNTEPDLFTTSASVRRLNALIDKNLSLSM"
FT                   /product="GH130"
FT   CDS             29899..31326
FT                   /db_xref="SEED:fig|6666666.127251.peg.26"
FT                   /translation="MDIITAPAKPTLKEKIGYGFGDMSSSMFWKIFSYYLPIFYSDVFG
FT                   LKPQHAAFLLLITKLYDAISDPVMGIIADRTRSRWGKYRPYLLWVALPFSLIGLLSFYT
FT                   PDASYGLKHVYAYVMYILMMTVYTAINVPYGAMLGVVTPDSREKSVFSSYRMFFAYIGS
FT                   FIAMGIFGVFEKLISGTVRVMPDGSERVVRGVGDAAPGQWTLVVGIVSLCCFVLFLLCF
FT                   LMTRERVGTENKAVSEGGSSIAEDLKALGRNRPWWLLLGAGIGILLFNSIRGGAAAYYF
FT                   SNILGTSVFLTCAVYLFIGEVAQMVGVLFTVPLSERIGKKATFISVLLTVSVLSVAIFF
FT                   LPETPAGFWGLLVLQVLICIAFGVQSPLLWSMFADVADYSEEKNGSASTGLIFSSSSMA
FT                   QKFGGALGAFLLMQILAIFAYDKDAAVQAPQTLAAIRGLMSYIPAIGALLGALCLLFYP
FT                   LTESRMASIREKLSARR"
FT                   /product="Xyloside transporter XynT"
FT   CDS             31895..34543
FT                   /db_xref="SEED:fig|6666666.127251.peg.27"
FT                   /db_xref="GO:0003887"
FT                   /translation="MADKLYLIDGHNLIFKMYYAFLRRPMINSKGMDMSILYGFTKYVL
FT                   ELIEREKPTHMAIAFDPPGGTFRNEMYPEYKANRSETPQLVIDALEPLLEICKALNIKT
FT                   LMIPGYEADDVIGTVAKSYGKPGTDVYMVTPDKDYGQLVAEHVYQVKPGKAGGADETVD
FT                   VKAVCEKYGISSPEQVIEMLTLCGDTADNVPGVHGIGEVGAGKLIAKYGTVENIYAHLD
FT                   ELTDRQKTQFEEARGHIGLSHDLVTIKTDVPVETSLDEMRLDLDYAPEIADLFEKYEFN
FT                   SLRKYIAHSSRAAKAVAEVKRLDYTETGAEELVKMAPEKISLIVEPEGPGIFADIRRVI
FT                   VAAEKKDGGIVVASGAAKDFRKILEGEAVKYGYDLKLQSNLLENAGIKLGGRLYDIELM
FT                   HYLINPEKSHKVEILAKSYLGVDIEEAAAVEEAPVSLFDVAEESPANGPDRNLEAAAVL
FT                   KLGEVVRAGMDEAGVGALYEDMEAPLAAVLARMERAGVKIDLGQLRTFADELKVEMDAL
FT                   EAQVRKLADEPGLNVSSPKQIGIVLFEKLKLDPKMKKSASGSYSTDETTLTAIADRHPI
FT                   IDKILEFRAVKKLLSTYIEPFPTYVSPVTGRVHTTFNQALTATGRLSSSNPNLQNIPIR
```

```
FT                   TERGKEIRKAFIPATPDGIIMSADYSQIELRIMAHLCGDEHLISSFKAGLDVHKATAAK
FT                   IFGIPVEEVTSDQRRIAKTANFGIMYGISSFGLAQRLHVSRTEAKKIIDDYFAAFPSIR
FT                   TFIDKTIADAREKGYVETVFGRRRYLPDITAKNATIRALAERNAVNAPIQGTSADIIKL
FT                   AMISVDKRIREAGLKSKMVLQIHDELLFDAVPEEIEELGRIVVAEMENVITLSIPLTVE
FT                   CNYGKNWLEAH"
FT                   /product="DNA polymerase I (EC 2.7.7.7)"
FT                   /EC_number="2.7.7.7"
FT   CDS             34530..35120
FT                   /db_xref="SEED:fig|6666666.127251.peg.28"
FT                   /db_xref="GO:0003677"
FT                   /db_xref="GO:0003700"
FT                   /db_xref="GO:0006352"
FT                   /db_xref="GO:0016987"
FT                   /translation="MKLTDHDDGSLVEMALGGGPEAEAAFFTIYARYHKGVFAHISRFI
FT                   QEQEEVEDICMESFEKAFKQLESYRPSNKFSTWIFTIARNTAFDHLDKDKVRGRNKDKM
FT                   PIDYSDLEHIDVPEDNVSPEDEIISSQDHENFLSCVQGLPDLYREVASMCFVDNLGYKE
FT                   ISEKAGLPINTVKTRISRAKSLIVQMMLDMEEE"
FT                   /product="RNA polymerase sigma factor RpoE"
FT   CDS             35117..35779
FT                   /db_xref="SEED:fig|6666666.127251.peg.29"
FT                   /translation="MNFKEFEDIVVAEFPELTPEMLGKYEAMEPLYEEWNAKINVISRK
FT                   DIGGLYLHHVLHSLAIASYLKRRRPEVYSAWTEGASVLDLGTGGGFPGIPLAVMFPKVR
FT                   FTLCDSVGKKIIVASEVSKALSLGNVTTVNARAESLPQVFDHVVSRAVTSLDNFYPWVK
FT                   GKFSGSILYLKGGDIVPEIAELMKKNRLRTGAVSTWPIETWLKDEYFKEKFVIEIEK"
FT                   /product="rRNA small subunit 7-methylguanosine (m7G)
FT                   methyltransferase GidB"
FT   CDS             complement(36417..36683)
FT                   /db_xref="SEED:fig|6666666.127251.peg.30"
FT                   /translation="MAIEFGPLRDKLRRKRIMVACTVIRRESIELCASDIGRKTVEFIR
FT                   NRIVQRLDAKTVDLLLERIPFYAGGCCCDEVVTSLDRIKIDLL"
FT                   /product="hypothetical protein"
XX
SQ   Sequence 37749 BP; 9136 A; 9489 C; 9828 G; 9296 T; 0 other;
     cacaatagcg ggacgagtcc tatgagggcc acgatgccgc agtgcggtac caggaaaggt        60
     atcgacataa gtgctgccga tagcagcatc agaataatga gcgagacagt ctctttacgc       120
     atatccacca acttaggttt tacaaattta catagaatta tgcgaaaaaa tagctaattt       180
     tgtaaattgc ttgattacga aatatgttat tggatatact aaaggctttt cttgtaggac       240
     tatgcgcctc cgccccgctc ggacctatgg cgatacttgt ccttcagaaa tctctgagcg       300
     gaggacacag ggccggtttc tttaccggtc tcggagcgac tttcacggat acgctttatt       360
     ccgtgatcgc tatcttcgcc cttgccatta cccagcagtt cataacagaa cacgaggtat       420
     tcatcaagat aaccggaggt atagcgataa tactgcttgg agcaaacatg cttttcaagg       480
     accctttccg caagatgaag gaccagagcg gtccgtcata ttcgatcaag gatttcctgc       540
     aggcggtagc gaccggactg agcaatccgt tcgccatact tatcataatg tcccttttcg       600
     gctttttcgg ccttgctcct gaaagccgtg acctgacagt cgctccggtg atacttgcag       660
     tgagcggcgg ctcgatgact tactggttct tctattcgct gattttcagc aggataagaa       720
     agaacttcaa acttagtacc ctgctttgga tcggaaggat ttccggcgcc atcataatga       780
     taatcggtat cgcgctggct tctgacggtc ttatgaagtt aatattcgaa tgatatgcct       840
     gaaatactga aaaactggat tgtaaagaac ctgttgctgg cgataggtct gttactcgcg       900
     cttgtgctgg tatcggcact ttcctcaat ctgcttaccc gtcacggtaa ggaactggtc        960
     gtgccggact tcaccaatat gaagatatcc gaagctactt atactgcctc cgtcgacgga      1020
     atgcgtatag aagtcgtgga ctctgtatat atccgcagga tgggcagggg actggtatac      1080
     agccagaatc ctaagccggg cagcaaagtc aagaaaggac gccgtatcct ccttacgatc      1140
     aattccgtga cccctaagaa ggtccagatg cctaacctcg tgggctattc tatgaggcag      1200
     gccaaggcgg aactttcgtc aaggggactt gccctcggaa aactcatgta tgtcagcgat      1260
```

atcgctacca acaatgtgct ccgtcagctc gtcggcaata tggagattac tccgggccgt 1320
atgatcgaga gcggaactgc cgtcaatctg gtagtcggac tcaactccag cgacagtcag 1380
acttatatcc ctgatgtcaa gggactaaag tatctgcgtg ctgtcgatgt agtgcatgac 1440
aactcgctga atatcaagag gcttgtcttc gacgaaagcg tcaagaacta ttccgattcg 1500
cttgatgcca tagtagtccg tcagggaccg gaggcctcga tggccccgat agtgatgggc 1560
tccgatgtca ctctttatct ctcggtcgat ccggccaaga caataatgaa gaaataatat 1620
ggaattcgac gatcttcagg acgacggaat gttcgagcac ttcaggctca acgtcgatcc 1680
cggacagacc ccgatgaggg tggacaaata tatggcctgc catctcgagg atacatcgag 1740
acatagggtg cagaatgcca tcaagagcgg atatgtccgc gtcaatgaca aggtcgctaa 1800
ggccaatgtc atcgtacgcc cgggcgatgt catccgtttc gtgatgccat atgagcgcca 1860
tggcttcgag gtcctgccgg agaatattcc cctgaatatc gtatatgaag atgatgattt 1920
cctgatcgtc aacaagccgg ccggaatggt tgtccatccg ggccatggcc atttcgacgg 1980
caccctcatc aatgctctgg cctaccatct cgggatcagc cagggaccgg atgccgagga 2040
tgacaggatg ggcattcttg tccatagaat cgatatggac acttcaggcc tgctggtcgt 2100
ggccaagaat gacgaggctc agcttaatct ggcgcagcag ttctttgtcc attcgataga 2160
gcgaaagtat gtcgccgtcg tctgggggaa cataaaggag gacgagggga cgattgtcgg 2220
taatatcggc agggacccta acgaccgcct caagttccgc gtatatcagg aggaggacaa 2280
gggaaagcat gccgtaactc attacaaggt gcttgagcgt ttcggattcg ttaccgtggt 2340
ggaatgccgc ctggagaccg gacgtaccca ccagataagg gtacacatgt cctatatagg 2400
tcatccgctt ttcaatgatg ccagatacgg cggggcggag atccgcaagg ggactatcta 2460
tgcgaaatat aagcagttca tccagaactg tttcgagatt tgcccgcgtc aggcgctgca 2520
tgcaaagact ctgggtttcg tgcaccctcg cacgggaaag ccggtcagct tcgattctcc 2580
gcttccggag gacatgaccg cgctgctgga gaagtggcgc aagtatgccg gcgacatgcc 2640
gaaagaataa aaaaaggggg acacccttta tgaatcaagt ttgtcccctt tttcgttgaa 2700
tagttcattc tgcagaactg caaaatcggt aatttctacc actttgatct tgcatttgta 2760
ctttttcttc cacttgccta cataagagtt gaacaggcct cgggtaatat atgcccctaa 2820
aatagggctt accttcagag tgaagttcct gattcccttg tcatttacgt agtaggaaag 2880
ccttcgttca atctgctcgt cgatgagcag gcttgatgct atttttcctg taccgtggca 2940
taccgggcag acttccgtcg tgttgatctc ggttaccggc cgtacgcgct gacgggtgat 3000
ctgcataaga ccgaactttg tcagcggcag cacattgtgc ttagccctgt ctttgctcat 3060
aagctcctgc atgtacttga agaggttgtt cctgtgctct accgattcca tatcgatgaa 3120
gtccacaatc actattcctc ccatatcgcg gagtctcagc tgtcttgcga tctcttccgc 3180
cgcaaacttg ttgacttcaa aggtgttctc ttcctgttcc tttgtcttgg cacggatgcc 3240
actgttgacg tcaacaacgt ttaaggcttc tgtcgtctcg attacaagat aggctccctg 3300
tttgagcggt accacttttc cgaaagaact ctttatctgt cttgtgatct caaagtggtc 3360
gaagataggt tcctttccct gatagagctt tactatcttc tcctgttccg gagaaataag 3420
ggagatatac cttctgatct cttcgtattc tttctcgttg ttcacataga tgttcgagaa 3480
agagtcgtta agcagatctc tcagaattgt agtggtcttg ctgtactctg taaacagagg 3540
ctgtatgctc ttgttcttcg agagattttc ccaggaatct tcccatttac cgataagcga 3600
cttcagctct gcaaccagaa tagctgcgtt cttgccttca gccgcggtgc ggatgattgc 3660
accgtaattc ttgggaagaa tactcctgac aagtgtctca agtctcttct tctcctcctt 3720
ggaggagatt ttctgggata tgcttacctt ctcggcgaag ggaagcagta ctatgtttcg 3780
tcctgccaat gaaatctctg cggtcagtcg tgatcccttt gtggaaatag gctccttgac 3840
tatctggact attatcatct ggccctgggt gagcacattc tcgatccggc cttcctttc 3900
gaggatagga ccgagcttta tgcttccgaa tagcctcgac gggttggtat tagggtttgt 3960
cttcttgacg aactcgtcga aagcgccgaa ataaagtccg aggtccagat aatggatgaa 4020
agcctctttc tcgtcaccga tatccacaaa ggcagcgttg agtgcaggca tcactttctt 4080
tactcttccg aggtaggcat cgcctacagc gaagctgcga cccgagctgg actctttgtt 4140
cagttcgata agcctgtgat tttcctgtaa ggctatctga acttcagtgg cctgaacgtc 4200
aacgatcagg tctttgagta gttccttttc agaatccatt ggataatttg tgaaacaata 4260
acaaaagggc cgtccagatc tcactgtcgg cccttctctt ttagcagact gctaaaatgg 4320
cagacacagg agaattattt cttcttgtgt ctattcttac gcaagcgttt tttgcgcttg 4380
tgtgtcgcca ttttatgtct ctttcttttc ttaccgcttg gcataattga aaaaaagtta 4440
aaaaattatt aaatattata ccggtaatta cttaaccttg cttacgaata cctttgctgg 4500

cttgaaagcc gggatattgt gagctggaat ggtaagggta gtattcttgg tgatgttcct 4560
tgctgccttc tcagcacggt gcttgataat gaagctgccg aagccacgaa gataaaccgg 4620
ttcacccttc gcaagtgaag cctctacact ttccataaaa gattcaacga ctgactgtac 4680
tgccagtttc tcaataccgg tagcttttgc tacttcatta acgatctctg ccttagtcat 4740
aatagagtaa tttatataaa gttatgtgtg tgcaaatttt aagactgcaa atatagactt 4800
ttttttgcat agggcaaatt atttcgttaa aaatctttgt cttacatatt ttgatattac 4860
ctttgtgctt ggcataaagg ttgacattat agtatctgcc ttaaaaacgc catccttctc 4920
tgacaaaatg tcagcaggga ggattgtata cactgataaa ttatgataga ttttaataaa 4980
gaaatgtttc ctgcctcgtc ccacgaggtc aatataatac cggtaatgtc tggagagccc 5040
gctctcaact tcgacgcctc ggaactcccg ggatcccttc cgatcctggc actccgcaac 5100
gccgtgatct ttccgggtgt cgtcttccct ataactatcg gccgcgagaa gtctatgctt 5160
ctcgtcgaca aggcccaggg aaacggcatg ttcataggcg cagtccctca gaagaatgtc 5220
gccgaggaga acccgctgtc tctcgccgac ttctcggaat tcggtactgt cgtaaaggtc 5280
ctgaagactt tcgagatgcc tgacggcact gtcacggcgc ttctccaggg ctacaagcgt 5340
ctccgcatgc tggaactgac ggacaccgat ccttaccaca tggcaagagt cagctatctc 5400
gacgatatcc tgccgcagaa cgactcgaaa gttccggtca tagccgatat cctcaaggag 5460
aatgcgtcaa gcatcatccg ttcgtcttcg ttcgttccta aggaggccct cgaggccctc 5520
aagaacatcg acgacttcaa tttcctcgtc aatttcatcg ctacgacgat cgacatggac 5580
gatttcatgg aaaaagtcga cctgctgaaa tatgacgacg taaaggaaag ggccatggct 5640
cttctcggcg tgctcagccg tcagctggag atccagaggc tcaagcagga gatcagctcc 5700
aaggtcaaga ccgagctcga ccagcagcag cgcgagtatt tccttaataa tcagctccgt 5760
acgatccagg aggagctcgg catggatgag atggacgaca tcgcacgctt ccgcgaggct 5820
gccgagaaga aagagtggcc tgcatccgta ggggagacct tcgaaaagga actcgcaaag 5880
ctcgagagat tcaatccttc tactccggaa tacagcatcc agtataactt cctgcagttc 5940
atgatcgacc tgccgtggta tcatacttca gacgacaatc tcgacctgaa gcatgcccag 6000
aaagttctcg atgaggacca tttcggactc gagaacgtca aggacaggat catcgagtat 6060
ctcgccgtcc tcaagctgaa gggtaacatg aaatccccga tcctctgcct ctatggccct 6120
ccgggagtcg gcaagacaag cctcggaaag tcgatcgcca gagccctcgg acgcaagtat 6180
gtccgtatct ccctcggcgg catgcatgac gaggccgaga tccgtggcca caggaaaaca 6240
tacgtaggcg cccttccggg acgtatcctt aacggaatca acaaggccgg cacttccaac 6300
ccggtaatgg ttctcgacga gatcgacaaa ctcggaagcg acttcaaagg cgatccgtct 6360
tccgctcttc tcgaggtcct cgacccggag cagaacacgg ctttccacga caactacctc 6420
gatatcgatt acgacctttc gaacgtaatg ttcataacaa cggccaatac tataaacggt 6480
atccagccgg ctctgctcga caggatggag cttatcaatg tgaccggtta tctggcagag 6540
gagaaggtcc agatcgccaa gaagttcctc gtgcctaagc agctcgagga gcatggaatc 6600
cctaagaagg aactgaagat cgacaaggct gcccttgaag agatcatcga caaatatacc 6660
catgagtcag gcgtccgcgg cctggagaag cagatcgccc gtatcgccag agtctctgcc 6720
aagaagatgg ccctcggcga ggactatccg aagaccataa agaaagagca tctcaaagag 6780
tatctcggcc tgcagacggc cttccacgac agccagaagg gcaacgaggc tccaggagtc 6840
gtgaccggtc tcgcatggac ctccatgggc ggcgagatac tgttcataga gacttccgtc 6900
agcgatggca agggagtcct gacaatgacg ggaaatctcg gcgatgtcat gaaagagtcc 6960
gctactatcg gctatcagta tatcaaggct catccgcagc tcgccggcct tacttcggaa 7020
gagttcgcga agaaggatgt ccatgtccat gtccctgaag gagccgttcc taaggacggt 7080
ccttcggccg gaatcaccat ggtcagctcg atggtgtcag ccctccgcgg aaagaaagtc 7140
cgcagcggaa tcgccatgac cggagaaatg accctgaggg gccgtgtgct gcctgtaggt 7200
ggcatcaagg agaagattct agcggcaaag agggccggag tctctaccat aatcatttct 7260
gaagaaaaca ggaaagatat tgaagatatc aaggaaattt acgtaaaagg tcttaccttt 7320
gtgtatgtca agacaattga cgacgtaatc ggatatatct tcgaataatg gacgttaaga 7380
tagaacagac ttggaaagac gcactgcgcg acgagttcga aaaaccgtac ttcgcctcgc 7440
tggtgcgttt tctgcataat gaaaaagagg ccgggaaggt catctatcct cccggaggcc 7500
tgatattcag ggcattcgaa ctcactccgc tgccggaggt gaaagtcgtt atcctaggcc 7560
aggatccgta tcatgggccc cgccaggcca tgggactttc gttctccgtg ccgtcggacg 7620
tgccggcccc tccgtccctg aagaatatat tcaaggagat ccagacggat ctgggcataa 7680
ccatgagcgg aagcccgaat cttgaaggct gggcccgtca gggcgtgctt ctgctgaacg 7740

ccatgctgac agtccaggcc ggagtcgccg gctcgcacaa ggctatcggc tggcaggagt 7800
tcacggatgc cgtgatcaag tgcatctctg accgctgcaa cggggtcgtg ttcctgctct 7860
ggggtaattt cgcccgcgga aagaaggaac tcatagatac gtcccgtcat tatgtgctcg 7920
aagccgccca tccttctccg cttgccgggg gagccttctt cgggtgcaga catttttcta 7980
agactaacga acttctcata aaagagggca aaacccctat cgactggaaa ttatgaaacg 8040
taaagcactt ttcccgggat ctttcgatcc ttttaccgcc ggccatctga atattctccg 8100
cagagcgctg acgatgttcg atgaaatcgt catcgcagtc ggtgtcaacc aggacaaacg 8160
cggattcttt agcatggatc agcgtatgga tatcatacgt caggctgtcc gcggtatgga 8220
cgggatcgag ataatcaaat acgacaacct tactatcgat acctgccgcg agctgggtat 8280
cgcccatata ataaggggcg tcaggaacat gcttgatttc gagaacgagc gtgccattgc 8340
ggatgccaac cgccgtcttg ctccggaaat cgagactatc atcatcccta ccgcacagga 8400
attcgcgcat atctcgtctt cagccgtcag ggatatcctt aaacataagg gggatacttc 8460
tctcttcctt ccggagggag tcgtccttcc ggatcacgat gcgtaggtta atcctcataa 8520
tctttctttc tctgctttct tttgccttgg gcgcccagaa tgcggattcc gtcaagatgg 8580
ccgccctgga tgcaaagctt gaggagtatt ttgcggcgct ggagaccgag tctctggatg 8640
taaagaggac cgagtgcgat ttcctcattt cttcgtgcac gactgacgat gtacgaaacc 8700
atgtcgccgt acgtctttac tcgcatttca tgcagtcgaa gcttatgggg gatgaggctg 8760
tcgccgtcca tcttgccgac aactggttca ttcccggcaa ggtttctttc ccggatgaga 8820
tagatctgat gaatgcgaag attttcgctg agttcaacag acagtcgctg ctcggatgca 8880
aggctcccgg tctggtagcc gaggatagtc tcggcaggaa ggtcgaggtc cctctgggat 8940
tgtctgacag ataccggatt ctttatatct attctccgga ctgtgccaca tgcagtcttc 9000
agactatgct gatggactct cttttcagca ggaagagcta tccggtcgac cttatcgcat 9060
tcaattcaga cgataatact gaagcctggg tgcagtaccg caggaaccgc ctttcttttg 9120
acggagtccg tgtcatgcat ttcagggatc cttcgctgga gtcggatttc cagcggaaat 9180
acggagtgct tcagactccc cgcatgttcc ttatcgacaa gaacgggatt atagtaggcc 9240
gcggactgga cgctgctgcg ctggagaaat tgctggatgc aactcctgac cttggcatag 9300
ggtatggttc gaagtcgtct gccgactatt tcgacgctgt gctggagagc ttcggggatg 9360
agctgcgcgg ctctgatatc cgctatttta tcggaagact tgaagaggct actctcgaaa 9420
gcgccgatac cgcaggattc cgtcagatga ccggtgacat gctgtattat cttgccggaa 9480
agaggggaga ggcgttcaag gatgccgagg ccttcgtcgt cgattcgatg gtcctgtccc 9540
gtccggatgt gtggcgcttt gccgatgatt cgttgacggt cgttgacatg gccatgagcc 9600
tgaaacatat gctctcgcta gctccagtag gtgccagggt tccggatgta cgtgtccatg 9660
ccactgtcct tcgcgccgga cgaaagccgg ccaccgtcca taagaaattg cggtcgttgc 9720
ttaagggaaa gcggaattat gtgcttttcc atactgccgg ctgcactttc tgcgaggctg 9780
aacttgccgc cgccgattct gtcgcttcgg agaagggtgt ccgtgtcatc ctggtcgata 9840
tggacaagct gacaagctta tacccggagg aggctgatcg gcttctggac gcattcgacc 9900
tgtccgttct cccgttcata atggagacgg ataataaagg ttgtgtcgta cgtaagtatc 9960
tgagtttaaa taagtagggc tgcctttacc ttagctacgg cctctttgcc tcccatcttt 10020
tcgaggatgg cgagtccgaa gtctatcgcg aaggctgcgc ctctgccggt gatgagctgg 10080
ccgtcggtaa ctgccggggc ttttacgtat tccattccgc tggcggcgag gttctcttcg 10140
aagccgtcaa agcaggtggc tttcttgcct tcgattccct tgatctgggt ggcgacgagg 10200
cccggggcgg cgcagattgc ggccacaatg cctccgttgc cgtagtggac gttcatcaga 10260
ccgataagcg gggtgtcggc tgcgagtttc ttcgagcctg gcattcctcc cgggaagatc 10320
atggcgtctt tttcgtcgat atctatttcg ctgaagtcgg agcggcccat gtctgccacc 10380
accgttactc cgtgggagct ggtgatatag acttcttcgg tgattgatac tgtgtatgct 10440
tcgattttac ctctgcggag aatgtcgagg gttgcgaggg cttcggtctc ttcaaaaccg 10500
tcagccagga agatgtatgc ctttcccatg atgctcatgt ggtttagatt ggttgaggta 10560
agagggttcg aacctctact gagggaacca aaatccctag tgctaccatt acaccatacc 10620
tcagtggggt gcaaatatac gaaaaaatta tttgcggaca acaattctct ctatacgtct 10680
agggacgcta gtgagaattt cgtaagggat tgtcccgaga atgtcggcga gttcgttgac 10740
tgtagggtct tctccgaaga tagtcacggt gtctccgact ttagcgtcaa ttcctgtcac 10800
atcaagcatg cacatgtcca tgcagatgtt cccgatcgtc ggcgctctgt gcccgttgac 10860
ggagaatgag gcgtgaccgc gtccgagatg gcggtcgatg ccgtcggcgt atcctaccgg 10920
gatcgtggcg attacggtgc cttcaggggc gatatgaccg tatcttccgt agcctatcgt 10980

gccgtcttcc ggcttgaggg tctttatctg caatattttg catttgaggg acgcgacgtg 11040
ggcgaggcgg acatcaggca gggcgctgat tccgtagata ccgattccga ggcggaccat 11100
gtcgaactgg tattgcggga atctttcgat tccggcggag ttgaggatat ggtacatcgg 11160
cttgtagcct atctcggagg tgatcctgtc ggcgtttgtc ttgaacatgc cgatctggcc 11220
gagggtgaat tcgtcctgct cagggtcttc ggatactgca aggtgcgaat atacggactt 11280
gactttgact tcttcgcagg tcttgaggaa gtcggtgaga tcctggagct cgctcgtcat 11340
gaacccgagg cggtgcattc cggtgtcgag tttgatatgg accggataat ccttgatgcc 11400
gcgttctttg agtattccgc agagtatctt gagggcgtac aggttcggga ttcccggctc 11460
gaggcggttg tcgatgattt cgttgaagaa gtccgtgcct gcggtcagga cgaggatcgg 11520
gaggctgata ccggctttgc ggagttccat gccttcgaca gggtaggcga ctgcgaggta 11580
gttcgctccc gcacgctgca tcatggctgc gaattcgacg gctccgtgcc cgtaagcatt 11640
tgcctttacg agtacaagta atttagtcgt ctgctccagt ttggacctga aatatctgta 11700
gttttccata catcctgtca ggttcagctc cagccttgaa aaatcatcaa ccctgttcat 11760
tccacatcaa tttagaatac aaatttatca aatttctccg acataggtta cagtttttcg 11820
cttttctttc cccgtaaaga cccttacagt acgtccgctg tcgccgccga cagcgtctgc 11880
tttggcggga atcctcgtgg cctcggctct gttccgccga ctctgtcggc tccatccctc 11940
ccaatgtcta cttcgtcctg cttcgcagaa ctcgtatcca tcgggcccct ccctctaacg 12000
tggccgaggg tggccacgct cgccgagacc agaggacagc cgccatcaat taagacgctg 12060
acggcaagtg agaggttggc atggatttgg aagcaaccag aaggattgtt aaaattttat 12120
ggatatgatt tggtttatta tcattgctgt gatggtgctg agctttgtga ttcagcaggt 12180
tctgaacagt aaattcaata aatattcgaa ggtgggattg cattccgggc tgaccggggc 12240
gcaagtcgct gccaaaatgt tggaagacca tggcataacc gatgtctcta ttacctgcat 12300
caaaggaaag ctgacagacc attacaaccc tgtcaacaag acagtcaacc tcagcgaaga 12360
agtctattac ggcaattcgg tagccgctgc ggccgttgca gcccatgaat gtggccatgt 12420
agtgcagcat gccacgggat atgctccgct gaaactgaga tcggctctcg ttccggtagt 12480
ctcattctcc agcaatgccg tgcagtgggt acttcttatt ggcgtaatga tcatcaagat 12540
tactccgacc ctcctgtatc tcggaatcgc catgtttgcc ttgacgacct tgttcagcgt 12600
cgtgacactt cctgtggaaa tcaacgccag cgccagagcc gtgcaatggc tcgacaaggc 12660
cgggataaca ggtcccgcta caaaagcgaa agctgtcgat gccctcaaat gggcagcata 12720
tacatacgtc attgccgcca tcggttcgat agcgacattg atctactaca ttgccatggc 12780
caggcgacgc tagagactcg aaagaatcct tgcaagctgg tccgtgcagg atgtaccacg 12840
gaaattgcag ttgacgcccg aaagagtgtc aactattttt tttatgtcca tgccctcgac 12900
gagacggccg atcgcctgaa gattaccgtt gcagccccgc acgtatgcga gattgcggac 12960
cttgccgtct attatgtcga aatcgatctg ctggctgcat accaggctgc tcgggatggc 13020
agttatgtgt ctgacaccgt tctcggtggt gtcgctgatg atcttgtagt cgtctcctgt 13080
cattgttgta gtccgttttt tatcttttttt tcttgaaaaa ctccgtcacc atggaaccgc 13140
attcctccgc gagcactcct gcctcgacct cggtcttcgg atgcagcagc gaaggagtga 13200
aaagtgagtg cccccgcttc ggatcgatcg ctccatagac gatccggccg atctgggccc 13260
agttcagggc tccggcgcac atcggacacg gctcgaccgt gacgtagagc gtgcattcgt 13320
tgaggtattt gcccccgagc gcctccgtag cggcggtgat ggctatcatc tcggcatggg 13380
ccgtaggatc atgtaaacgt tcggtcatat tgtgaccctt tcccacgatc cggcctttgc 13440
atacgataac tgccccgatc gggatttcat cctcggccag ggccgcctgc gcctcgcgca 13500
gagcttcctt catatatagc tcgtcttttt ccataattac agcaaagata gcaattcttg 13560
gacaatccca atagttgata aaacaattaa gaaaatttgc gaaatgatac tttttgaca 13620
tttttatcat atatttgcca cgtgataata taagttaatg gatatgacct cggtaataac 13680
ggaaataaca cctctttcgg acagggactg cttctacata gtcgagagac ataagaccga 13740
attcgtcttc cccctccaca ggcacaggga atacgagctg aacttcatcg agcatggcaa 13800
gggcgtccgg cgagtcgtag gcgacagtgt cgaagagata ggcgaatatg agcttacgct 13860
gattgccggc ggggatcttg aacatgcctg ggagcagggc aattgtacct ccgaggacgt 13920
acgcgagatc actatccagt tcgatccgga gatattctcc agcggcctca tgatgagaaa 13980
ccagttctca tcgatacagc atatgttcga gcaggcgagg aacgggctga ccttttcgct 14040
ggcggccata atgaaggtgt attcgcttct tgacgccctg ccgtcggaaa tagataaatt 14100
cgatcagttc cagcacttta tgaggctgct ttatgacctt tccctggcgg gtgactccag 14160
gatactggcc agcacgtcat tcgcccataa cgatgccgac aaggaaagcc gtcgggtgca 14220

gaagatcaag caatatgtcg gagaccattt cgcggaaaac attacccttg aagaattggc 14280
ggaaatggtc ggcatgagcc cgtcggcgtt cagccgcttt ttcaagagca ggaccgggaa 14340
gacggtgata gactatataa tcgacatccg actcgggaat gcggcaaggc tgctggtcga 14400
caccacccag aatatctctg aaatctgcta tagctgcggg ttcaacaacc tgtcgaattt 14460
caatcgtctt ttcaagtcaa aaagaggcta cacgccgcgc gaattcagga ctctgttcaa 14520
gaagaacaag gtttttgtat aacgtgcaaa atagtatcgg ttattgacaa atttgtattt 14580
gttgggtaaa acgttgcatg ctaaatttgc aacgtactca gacaagtatg gatttatcga 14640
taaaccgcat gattaaaaga ttattcccgg cactgtttat cgcccttttc gcgattcttt 14700
cgtgtcagcc tatgaaaatg gacatcgtca aagttgacga tggctctttt gtcgttgatg 14760
gtcacttcta ttattatatc ggtacaaatt tctggtacgg aggacttctc ggctccgaat 14820
atggtgacag aagcaggctt gccgccgaac ttgattccct gcatgccctc ggactggaca 14880
acctccgcgt actggtaggc agcgacggcc ccgacggggt ggcgaccaga gtctatccga 14940
ccctgcagaa ggaaccgggc gtctatgacg aaaaggtctt cgaaggtctg gactacctta 15000
tggccgaact tgggaaaagg aacatgaggg cagtcctcta tatcaataat tcctgggaat 15060
ggtccggcgg ctatggcatg tacctcgaat gggccggagc cggcaaggcg ataattcctt 15120
cggtcggagg atatggtccg ttcatggaac agatggccca gtatgcgacg aacgccaggg 15180
cgcaggaact cttcgagaat catctccgta ctgtcgtttc aagagtcaac agcatcaccg 15240
gaaagcctta tagcgaggat ccggcgatat tctcatggca gatcggcaac gagccgcgct 15300
gcttctcctc cgatcctgcc gtccaggacg gattcgtatc atggatatgg cgcgctgcct 15360
cgataatcaa atcgatagac cctaatcata tggtttctac cggcaatgag ggcatatggg 15420
gctgcgaaga gaatccggag ctattccgcc ggctgaatga atgtcctgac attgattata 15480
ttaccgccca tatatggcca tacaactgga gctgggtagg aaaggaatcc cctcagctgg 15540
atatagaccg cgccgaggag aataccggga gatacatcga cgcccatctg gaggtggccg 15600
agagtctggg caagcctctc gtgatcgagg aattcggctt cccgagagat gatatgcagt 15660
ttgccaaagg gactcccgta agcggacgcg accgctatta cagatacgtt ttctccaagg 15720
tcatggagaa tcagaaaaat ggcggcagac tcgccggcgc caatttctgg agctggggag 15780
gtcttgcggt gcctgcccac agatattggc aggaaggaga cgattacgcc ggagatccgg 15840
cccaggaaga acaggggctc aattctgtat tcgccgacga ttacagtacc cttgctatta 15900
taaaacatac aacggcgaga ctttctgccg tggctaatgc aagcgcagtg cttgaaaatg 15960
attggatgtt cactaccgga gaccagaaac ctctcagagt tgctgtacag tctcagaatg 16020
gttttaaggg aagtgtcccg gtttatgtgc gggtgtcaac agacaaaggg gtttcagtgt 16080
cggtctcagg aaagaaagcc cgtctcggca aaggcgccga tactctggat ttcgatcttg 16140
gactagcccc aggcttttat aaggtgtgtg tcagcctgaa cgagaatgac ccgcttgatg 16200
aaaacttctg tgtcggctgt gatcctgaga agatagtatc tcctcaggat aaacagcctg 16260
atttcgatga attctgggaa gacaacctca ggcagcttgc catggtcccg atgaatgcca 16320
ggatgactcc gttgaaggat cattcaaacg ctgacaggac tgcctacagg gttgatatga 16380
agtctttcgg aggcaaagag atctccggtc tgctttatct tccgaacaaa gagggaaaat 16440
tcgctgccag gatctcatat atgggctacg gttcggacgt ctggttcgct gatccgtcgg 16500
cggaaccgga catgatcgag ttcactctca gcgtccgtga ccagggtttc aacaagactc 16560
cgggacaagg ctgggtgaca aggggacttg ccgataagga ctcgtattat tacagaggcg 16620
catatctgga ctgtgtgcgg gccattgatt tcgtctcttc tcttcctcag actgatccgg 16680
accatatcta cgccgaaggt ggaagtcagg gcggagcatt taccctgata gctgcatctc 16740
tggacaaccg tctccgcgca atcgctccgt tcgtgccttt cctttccgat ttcccggact 16800
atttcgccat cgcagactgg ccgggcgggg aagtgctctc ggcggcaaag gctcagggca 16860
tttctgaaga cgaactttat aagaccctgt cttatttcga cgtgaagaac tttacggaca 16920
ggataggttg tcctgtgctt atgggtttcg gcctgcagga tcccgtctgt cctcctcata 16980
ccaatttttc ggggtataac catattactg cagataaaca atggctttgc gttcctgtct 17040
caggccacca tctggagcaa tttcccgagt ggtgggcagc aaggacggaa ttcttcacaa 17100
aaaataaata acacaactaa ttattcacat tcaaaatcag tagcatgaaa agacagctat 17160
taacatgtat gctgatgctc tttggagcat tgtcacttat ggcgcagaac aaagtgaccg 17220
gaagaattac agataattcc ggacagggcg tcataaatgc cagcgtaatg gtgaaaggca 17280
ccaacaatgg tgtcgttgca gatcttgatg gatactacga gattgttgtc ggccaaggct 17340
ccactcttgt atattcttcc atcggttaca aagatcagga agttgttgta ggcacccaga 17400
gtgtcatcaa cgttcttctt gtcgaaaact ccgagctgct cgacgaaata gtcgtagtcg 17460

gttacggtac tatgaagaga agcgacctcg ccggatcatc cgtttccatg aaggaggatg 17520
acctcaaggg ttcgatcatc accaacctcg accagtctct ccagggacgt gctgccggtg 17580
tcacggcagt ccagacttcc ggagctcctg gttcgtcctc aagtatccgt gtccgcggtc 17640
aggcgacaat caatgccaac gcagaacctc tctacgtcat cgatggcgta atcatgcagg 17700
gaggcggtaa ctccggtgca gacttcggtc tcggagatgc tctcggcaat ggtaaagtct 17760
ctacgatttc tccactttcc actatcaacc ctgccgatat cgtaagcatg gaaatcctca 17820
aggatgcttc cgcaaccgct atctacggtg cccagggtgc gaatggtgtt gtcctcatta 17880
ctaccaagca tggctccaga ggtgatgcga aattctcata cgacggtatg ttcgccgttt 17940
caaggcagac aacccgtatc gatatgatga atctccgtga atacgcagag ttctacaact 18000
ctatggttga aaccggagaa atcagtgagg ccaatcctta ctatgctact ccgggcctgc 18060
tcggaaaggg taccaactgg caggacgaaa tcttccgcac agccatccag caccagcatc 18120
agatcagcgc ccagggcggt acggacaaag tccagtacta tgtttccgga tcattcatgg 18180
atcaggaggg tactatcagg ggttccgaat tcggccgtta taccttcagg accaaccttg 18240
acgcccagtt gaaggaatgg ttcaagctcg gtctcaacct taccttttgcc aacaccaatg 18300
atgaccttaa gcttgccgac ggtagcgaag gtcttatcac atactccctt accacaatcc 18360
ctgacatccc tgtttatgat gtctatggca atttctcatc tacagtacgc gaaggctata 18420
cttctccgaa ccctgtagct cttgcaatga tggatgagat tctgctggac cgtcagaaac 18480
tcaacggtaa tatctacgcg gaacttactc ctgtaaagca tttcacctgg agaacagaac 18540
tcggtttcga tgtcagcggc tctacggcta cccgtttcag accaacagtc aatctcggtg 18600
gctggacaag agataccaac tctatgagct atcagaagaa cagcaacaga ttctggcaga 18660
tcaagaacta tctcacttat gccaactcct tcggaaagca cagcgttacc gccatgcttg 18720
gtcaggaggc ctgggagtct tcgtggagtt acgtacaggg cgcaaatacc aagctccctt 18780
caaacgatgt ccataacgtc gccctcggtg gaggaactcc gtcgctttct tccggtttcg 18840
gaagctctgc gatggcgtca ttcttcactc gagaaactta taatttcgac gatcgttacc 18900
tccttaccta tacctaccgt tatgatggat catccaactt cggtcctgag aaacgttggg 18960
ccggtttcca ttcattggca gtggcttgga aattcaccaa tgagaaattc atggatttcc 19020
tcaagcctgt ggtcaacaac ggaaaactcc gtctcggatg gggcagaca ggtaactcaa 19080
gtatcggtgg ttatgcatgg ggagcatcca taagccgtat gccttcggct ctcggcatgg 19140
gttaccgtcc tgctaacatc cctaatcctg ctatccattg ggagagccag acgcagcaga 19200
acatcggtct cgacctcaat atgttcgaca acaggctgaa tcttacagta gatgcatacc 19260
tgaagaagtc tgacaacatg cttatgtcaa tgcagcttgc ttcctatatg ggaacccagg 19320
gtaacagttc ttctgctctt gcagctccta aaggtaactt cggttctatc gagaacaagg 19380
gtcttgaaat tactctcgat gcccatcctg tccagatcaa ggatttcgcc tgggacagca 19440
acttccagat ttccttcaac agaaacaagc ttgtcagcct tgacggaacg actaatgccg 19500
ctctcatagg ttacggacag tggtctgacg ttgtgtgtgt gtctgaaatc ggcaagcctc 19560
tctataactt ctatggttac aaggtcgctg gcgtatatac ttctctcgag gatatcgaga 19620
attctccgaa acctgagaaa tatccttcta atggagagta tgcccgtgga aataccgtgt 19680
gggtaggtga tatcaagtac gaagatatca gcaaggacgg agtaatcaat gaaaaggacc 19740
gtaccgatat cggcaatcct cttcctaagt tcacctttgg ctggagcaat acctttacct 19800
acaaggactt cgacctcaat atcttcatca acggaagcta tggtaacaaa gtcctcaact 19860
acaatgccat ttctcttacc catatgaaca gcgcatggac caaccagatc aatgacgctg 19920
tctccggcag ggcaatgctt gctcctatag acgcaagcgt agtctatgct gacggttcca 19980
agtggtatga ccatatcgac aatgtcaggg tcctcaaccc tggcacgaag actcctcgtg 20040
caagcatcaa cgaccctaac gacaacgacc gtatcagcga ccgttatgtg gaggatggtt 20100
catatatccg tatcaagaac atcactctcg gatataatgt ccctaagtct ttcctgaaca 20160
agttcaaggt ggataatgta cgtgtttacg tcaacatcca gaacctttat actttcacga 20220
agtacaccgg atatgatcct gaagtaggtg ccagcaccca ggattccacc ggtcttacct 20280
atggcgttga taacggacgt tatccttctc ctacagtata ttcttgcggt ttgaacatta 20340
cattctaaaa ttgaaggaat atgaaaatct ctaatatgat gaaaggtttt gctgtggcga 20400
ccgcagtctt tgctctgact tcctgcgaag ggttcctcaa cagacctaac gaagataatt 20460
ataacgtaga taatttctac aagaccgacg agcagtgcga acagggcgtg aattacctgt 20520
acaactctcc atggtatgac ttccagagag gcttcatcaa gatcggtgag gtcatgtccg 20580
gaaacatgta ctggggctcg tctccatacc ttgatttctc caccaacggt accgaccagg 20640
acctcgtcaa catgtcctat tctctctggg ctgtgaacgg ccatgccaat accgtgatct 20700

acaaccttct taattcgcag ggcccgtcca aggcagcgat cgacaagtgt atcggagaag 20760
ctcttacctg gaaggcattc gcctatttct tcatggtacg aaccttcggt gacgttccta 20820
tcgtccacag caattccgat ctcctctcca gcggcgagta caactctatc tccaaggtgg 20880
agcgcgccga tgtctatgag tacatcatca cgaccctcga aaaggcaatg gaaatgcttc 20940
ctaagaagac ttccggcttc aataattacg agcgtatcga ttactatgca gcagaggccc 21000
ttctcgcaaa agtatatctt accaaggccg gactttccgg atccttgaac caggatgacc 21060
tcaagcaggc tgccaagtat gccaaggatg tgatcgacaa ttccggacgt acccttcttc 21120
ctgtatattc ggacgtcttc cgtcttaccg gattcaacgc cacaggcgag tgccttatct 21180
cctgggaatg gagcgtaggc cgtaaccctt ggactcagca gaataccctc cagagcgacc 21240
tcgcaatggt cggattcagc gaccagggag actgctgggg aggctggggc ggtccttcag 21300
tggatcttgc agacgcattc ggcgtaagtg cgacccaaaa tcctgcagag cgtgtaggag 21360
agaaggacag ccgccgcaag gcaaccttga tgctcccggg tgacaagtat gattacttct 21420
ggactgacaa gggaggtttc gattatctcc gcttcatcta tgatacggaa tatggcaaag 21480
gcggtcctgg cggttccatg cagtgcccta caggttccaa caacgtaaag catctctatg 21540
gcgataactt tgatcatgag aacgccggtc tgggaacccc tgagaacatg gcttatcagc 21600
ttcctactca tcttctccgt ctttccgacg tgtatcttat ctacgcagag gcaagccttc 21660
ttaccggaag caacggccct gctctcgagt atgtcaacaa ggttcgcgaa cgtgctggcg 21720
ccgatgctct cgcatccgtc actttcgagg atatctggaa agagcgtcgt cttgagcttg 21780
caggcgaggg cgaccgctgg tatgactatg tacgccgttc atactacgat atggacgcat 21840
gcatcgctga actcaaagct cagagaaggt catactggga tggactcgac gatgtttgga 21900
aatcatatat cggagatgct ttcgattatt ccggaacctg ggatgcttca gaagtacagt 21960
acaacaacgc agaggacaat cctaacgtta ccgctgcaag cttcaagctt ccgttcccta 22020
cagaggacgt agtgttcaat ggcaacatgg catctgacgc cgtgcctgtc catgtcgatg 22080
tccgcgagga atacagctat aacttttaat ctgacgaaaa gatgaaatct ataaatagaa 22140
taatcaccgg ttcactggca atcctcgcaa tggcgtcatt cgcttcctgc gacaaggttg 22200
actatccgga ccgttatgaa gtagctgacg gcaagccgac ggtttattcc gtcagatacg 22260
cagacaagga cgtcaacatc acccaggcat atatggatga ggtcgtatgt ctcctcggca 22320
agaatctcaa gagtattgtg gaactctggt tcaacgacca gcaggcaacc ctcaatacca 22380
gcttcattac agacaatact cttctggtat ccgtgcctaa gaatatgcct gtggtacaga 22440
cagacaagat atatatgatt acttcgggca aggatacgct cgcgtatgat ttcgccgtgc 22500
ttcctccgac tccgaaggtt acttccatgt ccaatgagtg ggctgcggaa ggggagaccg 22560
tcacaatcta cggcgacttc cttattgaca aggctgacaa gcctctccag atagcctttg 22620
caggagccga ggtcgatcat gcttcaatgg tgttcaacgg ttcgtcttca gtcagcttcc 22680
ctgttcctgc aggagcccag cctggatatg ttaccgtgac ttcactctcc ggtaccggaa 22740
agagcaagtt catgtacaag gataccagaa acatcctctt cgactgggat ggcagccacg 22800
gcggctttgc taccggtcac ggatggcgta acggcgtgat ccacgctcct ggcgctgatg 22860
catgggatgc gatcgacggt tcatacctct acttcggagg cgccgatctt ggtcctggaa 22920
gtgcagatgc atgggcagag gaccagtaca gcttcaacta ctggccagag gccggcaccg 22980
attacgatgc cctcaacctc cgtccggagt tcgcagagta tatcgacaag tatggagtaa 23040
gcggcctcgt gctcaagttc gaggttctcg ttccttcatc caacccttgg tctgcttcag 23100
gtatgcagct catgttctcg agcgacgctg tcgttaccta cggtaccgct acgaacagct 23160
acttcagtga cgaaaccttc cctcgtgcta tctggcagcc atggactacc aacggaggtt 23220
cttatgatac cgctgacaag tgggttaccg taagtttccc tctcagctca ttcaacaaga 23280
accaggaaaa caaggagtgt gcaaccgcat ttgacaagtc ttaccttacc ggtctcacat 23340
tcttcgtatg gaacggcggt atcgacggca cagcctgctc tccggtcttc gcgatcgaca 23400
atatccgtgt ggttcctaat gaataatatt aaatgaagct cattatgaaa aagatattca 23460
gtatttcagc cattcttgcc gcctgtgtcc tggcactcgg gctctccgga tgttcatcca 23520
aagagcttga taccgaccag tattccgaca aggacgtgaa gctggcatca ttcggcccta 23580
acccggtgat gcgtggagct acactccggt tcttcggaag taatctcgac aggattgtcg 23640
aagtttcagt tccgggactc gatcctatca ccgacatcga agtcgttacc tccggtaatc 23700
cttcagaaat ccgcatccag cttcctgcag agggcgcaca ggtcggaaag atctctctca 23760
agtcatccga cggaaaggtc tttactaccc aggtggatgt agaatataca gaacctatcg 23820
tatttgacag tttctctcct gcagaggcta tgcctggcga cgtgataacg gtcaagggcg 23880
attatatgaa cctcgtgcag agcgttacct tcgaaggcgg cgccatcgtg accgaaatca 23940

accataacgg ccgtcatgag gcttccttca aggtcccttc aaaggctatt accggaaaga 24000
tcatcctttc tgacgaagga gccatcgcca atctcctcta ctcggaggaa gatcttgtga 24060
tcggcgatcc gaccgtcaag tctcttaccg tagctgttgc taaacctggc gagaacgcag 24120
ttgtgaaggg agcatatctc gatatgatcg acaaggtcgt attcgaaggt ggtgcagagg 24180
tttccgacct tgctctcaat gaggatgcaa ccgaattcaa tctcgtaatc cctgccggag 24240
cacagtccgg tgaagtaaag gctgtcagcc acgctgccaa ggaattcgtc gccggcgaaa 24300
tcgaacttgt catccctacg gatgtcgcag ctgcaccttc gcctgtaaag gctggtgccg 24360
acctcgtgat ttccggtaac gatcttgatg tggtaacttc tgttgatttc cctggcgcca 24420
gcggagtaga attcgtattt gccgaggacg ccattacagt gaccgttcct gctactgcta 24480
ctgaaggtga tatcgccctc aatatggcca gcggagaagc tgtgaatgtg gcattcaccc 24540
ttgtgcaccc ggtagtttct gccatagctc ctgtcgagct catggccggc gagaccatca 24600
ccgtgacagg taccgacctc gaccttgtca ctggagtaac tctcggcgga aagccggtcg 24660
agttcactgc agaggctgag aagattgtcg tacagaccgc caatacttcc gtttcaggaa 24720
agctcgtgct ttctcttgcg aatggtgaga agattgaacc ggaggatgcc atcaccctca 24780
gctatgattc attcatcatc gtgaacgaca tgcctgccgc agagcatatc ggggctatcg 24840
taaccctcaa gggcgagaac ttcatgatga tcgagaacat ctatatcggc gaagagaaag 24900
tatcgaagta caaccttcgt tctgataatg aaatccagtt cgtaatgcct tggaacaagg 24960
tcggaacata cagcatgtac ttccatcttc tctccggcga tgtcgagaca tgtcctcaga 25020
caatcgaagt tctcctcgag caggacatca tgactgtatg ggaaggctct ctgcagatca 25080
cctggaatga cggcggccgt gtaaccgtgc cggcagcatc attcaccggt atccctgcag 25140
gcagcaagct cagattctac tacactcagg tggaccagca gtgggatcag gcccagctca 25200
actatggtaa ctggaccgga atcaacttcg acgatcctgc aggtaccgtc attaacggaa 25260
ccctcgttcc gactgacgtc tatggatggt tcactgacgg aatccttgac cgctgcacag 25320
aggtcgtgct tacaaaggaa atcctcgaaa acatccaggc taacaagggt gactgcgaag 25380
acaagactgc ctgcggtatc atcatccagg gctctggcct taccttcacg aagatcgaga 25440
tcgtatgcga tattccgcag gaggttacga tctacgaagg tcctacaacc cttacatggg 25500
gtgacgacgg ccgtttcggt ctcgccatga cttacttcga ggcagcaaag ccgggctctc 25560
agcttatctt ctatctcgag cagacggacg catggggcca ggttcagctc aacgacggct 25620
ggtgggccaa cgccgacatg aacttccctg agatcggcgg tgcatacatt accaccgaca 25680
acctcggagg aaaggacgtc acacgtatcg cccttaccct gactgctgat atccttgacc 25740
atatacttgc tacatccggt gactatttcg gtctcaatag ccagttccag ggtgacggac 25800
gcgtcgccat ggttatccag ggatctgata tgattatcaa taaagtttgt atattgtaac 25860
acttttattg ccggaggcgg gtgatgccgc ctccggtagc cagtaaacaa attttatata 25920
atgaaagcac acgtcagttt ctttgtatta tctatgattt ttcttgcggc atgttcctgc 25980
tcaaaggatg atgccgaaaa gaaggaaccg tccgtcccgg cccctaaggc ggtaacggta 26040
gatgtcgcgt ctgtcagcgc tcccgtagag ggaacgtcgg tcgtcgtcgc catctcctcc 26100
ccgtccaagc ctgcggtctc tgtctccgga gactggatca aagccgaact gggtccgtac 26160
aatccggata catataaagt gaaacttacg gtgaccgtcg gcgagaatcc gacatacgag 26220
tcacgcagcg gtagtgtgac ccttacttgc cagggctttt ctccggtcgg gataagcgta 26280
tcccaggatg ccgcggcgaa gccggtcgga ccgtcggcta atcctgcata cgatcttctc 26340
ggtctcggct ggaatatggg taaccagttc gatgctttct acaacggctc atgggccgga 26400
gataaataca attatccgga cgagaccgtc tggggtaatg ccaaggcgac ccaggctact 26460
ttcgacgggg tgaagaaagc cggattcaca tcagtgcgaa tccctgtcac gtggctcaat 26520
atgataggtc caggacctga gtacaatatt gacgagactt ggatgaaccg tatatatgag 26580
gtggtaggat atgccgagaa agccggactc aaggtcatca tcaataccca tcacgacgaa 26640
aacaacaatg acgaccattg gctggatatc ctttctgcat ctaaggatgc caggaccaat 26700
gaagcgataa aggcggaaat cgcggctgta tggacccata tcgccaataa gttcaaggat 26760
aagggagatt tccttatcat ggagtccttc aatgaaattc aggacggagg atggggctgg 26820
agcgctgctt tccaggctga cccttccaag cagtgtgatg tgcttaacga atggaatcag 26880
gttttcgtgg atgccgtacg tgctacagga ggcaacaacg ctacccgctg gctgggtgtt 26940
cctacatacg ccgcaggtac ttcatggacg aagtatttca gactgccgga ggatcctgcc 27000
ggaaaggtca tgatggcagt gcattgctat gatccgtatg aatatactat agggaacata 27060
cagtatcctg agtggggcca taccggcgcc gcaggaaaga agaccccgga aggggagaaa 27120
gagataacgg atctcttctc cagacttaat acggaattca tcagcaaggg gatacctgta 27180

tatcttggcg aattcggctg ttcgatgaga agcaaagggg atgcccgggc atggatgttc 27240
tataaatact atatggagta tgtctccaaa gctgcgcata cctattcctt accgggaatc 27300
ctgtgggata acggaagtac taagaccgga aaggagtgcc acggctacat tgaccatggt 27360
accggcaagt atatagggaa ttctaaggag atcgtggatc ttctcgtaaa gggaatgctc 27420
agcacggacg aaagctatac tctccagagt gtatatgaca gggctccgaa aaccaattga 27480
ttatgaatct aatccttaaa cttagcatca ttttatcatt agtagttgct gtatccggct 27540
gcggacgcaa taaagaagcg gccggaacag cttttaacga agaaacccgg atcatgatgg 27600
atcgtatggc ggcttcagta aaggccggca agattcttta tggccatcag gatgacctcg 27660
tttatggcca ttcatggtgt gtggatcctg attccgcagc gacagatgcg ttgaaccgtt 27720
cggatgtcaa ggctgtcgcc ggagattatc cggctgtcgt gggattcgaa ctcggcggtc 27780
tggaactggg atgggactgg agcattgaca aagtcccgtt cctgctgatg cgccgggctg 27840
ctgtagagca tgccggaagg ggaggaatcg tgacgttctc atggcacccg cacaatcctt 27900
atacaggcgg tgatgcctgg gatatcagtt ccgacaaggc agtggcttct gtcctcgaag 27960
gaggggagaa ggcggagatg ttcggagtct ggctggagag gctgggcgat ttcatggact 28020
cgctccgtga cgataagggt aatctgatac cggtcatttt caggccgtgg catgagaata 28080
tcggcagctg gttctggtgg ggcggagccc tttgtactgc ggatgagtat atcgcccttt 28140
tcaggaggac ggtcgagtat ctgagaaatg acaggggact ggacaatatc ataatgtgtt 28200
attctccgaa ttccgggatt acggcggagc agtatatgtc ccgttatccg ggcgatgatg 28260
tcgttgatat cctggggtat gaccattatg agactgtcat ggagggcgag agtctggaag 28320
atgccggaaa gcgttatgcc gggcagttgg atcttggcct ttccatcgtc aagtctcttg 28380
cttcggccca cggaaagctg atgactcttt ccgaaaccgg aatggagggc ctcccggaca 28440
ggagctggtg gacttcggtg ctttatcctg cggtaaagga ttttccgatt gtctatgtgc 28500
tgacgtggag aaatgcatgg gacagggaga cgcatttttta tgcgccgtat cctggtaatg 28560
agggcgactt cagggaattc gcttcgctgc cggagatctc tctcctgtcg tcgttccggc 28620
cttagtccgt cattcccggc gaagtgtcgt cctttccggt cggaccggga atctgtaacc 28680
ttaaaacaat aaataacaat atatggagaa tatgagtttt gaagaacgtc tggcgtggct 28740
ggaagcacgc caccaggaac ttattgctaa gaaaaactcg cctatatacg gcaatggtgt 28800
gtttgagaga tatgagaacc ctgttcttac cggtgaccat gcccctttgt tctggagata 28860
tgatctggat aagaactcca atccttttct ggaagagaga ttcggcatcc atgccgcatt 28920
caatgctgga gctataaagt ggaaggacaa atatgtcgtc tgcgcccgtg tcgaaggtgc 28980
tgaccgcaaa tcattcttcg ctatcgccga gagcccgaac ggaatcgaca atttccggtt 29040
ctgggaccac cccgtgacta tgcctgatta tggcgaacct gccacgaatg tatatgacat 29100
gagacttact gcccatgaag acgggtggat ctatggtata ttctgcgtcg agcgtaagga 29160
tccggatgct ccggaaggcg atctttcgtc tgccgtcgca gctgcgggaa tcgcccgtac 29220
caaggatctg gtcaattggg aaagactgcc ggatctggtt tctgattctc agcagcgcaa 29280
tgtcgtactt catcctgagt tcgtgaatgg gaaatatgct ctttacaccc gtcctcagga 29340
tggattcata gatgccggaa acggcggcgg catcggctgg gccctggtgg acagcatgga 29400
gaatgccgtt gtcacttccg agaagatcat caactgcagg agataccata ctatcaagga 29460
gaccaagaat ggcgaaggcc ctgctcctat caagactgac aggggctggc tgcatctggc 29520
ccatggcgtc aggggctgcg cttccggcct gcgttatgtg ctttatatgt acatgacttc 29580
gctggaggat ccttcgaaag tgattgccga acctgccggt ttcttcatgg tcccggaagg 29640
ggaggagtat gtgggtgatg tcatgaacgt cctgttctct aacggttgga ttgccgatga 29700
ggacggaaag gtctttatat attatgcttc ctgcgatacg cgtctccatg tggcgacttc 29760
taccgtcgac agactcgtgg attactgtat gaatactgag ccggatctgt ttaccacttc 29820
tgcttcagta cgacgtctga acgcgcttat agacaagaat ctttcgctat ctatgtagga 29880
ctaataatca ggtttcagat ggatatcatt actgcaccgg ctaagccgac attgaaggaa 29940
aagatcggat atggcttcgg agacatgtcg tcttcgatgt tctggaagat tttttcgtat 30000
tatctgccga tcttctattc ggatgtgttc gggcttaagc ctcagcatgc ggctttcctg 30060
cttcttatca caaaactgta tgatgcaata tcggaccctg tcatgggaat catcgccgac 30120
aggacccggt ccagatgggg aaagtatcgt ccgtatcttc tctgggtggc tctgccgttc 30180
tcgcttatag gccttctgtc tttctatact cctgatgcca gctatggcct gaagcatgta 30240
tatgcgtatg tgatgtatat cctcatgatg actgtctata cggctataaa cgtgccttac 30300
ggggccatgc tgggagtcgt cactccggac agccgtgaga aatcggtttt ctcgtcttac 30360
aggatgttct ttgcgtatat cggcagtttc atcgccatgg gtatattcgg cgtgttcgag 30420

aagctgattt ccggtaccgt gcgtgttatg ccggacggta gcgagagggt ggtgcgcgga 30480
gtcggcgatg ctgctcccgg acagtggact cttgtcgtcg gcatcgtttc tctgtgctgt 30540
ttcgtgctgt ttctgctttg cttccttatg accagggagc gggtggggac tgaaaataag 30600
gcggtttccg agggaggctc ttctatcgca gaggatctca aggctctagg ccgtaacagg 30660
ccttggtggc tgcttcttgg cgccggtatc ggtatattgc tgttcaattc gatccgaggc 30720
ggagccgctg catattattt ctcgaatatc ctcggtacga gcgtgttcct gacttgcgct 30780
gtctatcttt tcattggcga ggtggctcag atggttgggg tgctgtttac tgtgccgctt 30840
tcggagcgta tcggcaagaa ggcgactttt atctctgtgc ttctgacggt ttctgtgctg 30900
agtgtggcga tattcttcct gcctgagact cctgccggtt tctggggact tctggtgctg 30960
caggtgctta tatgcattgc ttttggtgtc cagtctcctc tgctgtggtc gatgtttgcc 31020
gatgtcgccg attattcgga ggagaagaac ggctcggctt ctaccggact tattttctcg 31080
tcttcttcta tggctcagaa gttcggcggg gccctcggcg ctttcttgct gatgcagatt 31140
cttgcaatct tcgcatatga caaggatgct gcagtccagg cgcctcagac tcttgctgcc 31200
atccgtggcc tgatgagcta tatcccggca atcggcgccc tcctcggagc cctctgcctg 31260
ctgttctatc ctcttacgga atcgcgtatg gcttcaatta gagagaaact ttctgctcgc 31320
agataacatc ttcattcatg taggtatttg ttattccaaa ggatggccct cttcggccat 31380
cctttttcgt ctggctgatt cgagtcgggc ctttctgctg gcgggagaag gacgggacgg 31440
ctccgtcaag ttcggaaaag ctgcttggtc gctgtcgccg cctaaacggc taacttctcc 31500
cttttacgat gggcgagcca tcgtaacggt cgtcaaacac acgccgttct tcacggcgcc 31560
gacggccgct cccgcgctct ccgcttttcc tcaattcctc cgccgcctcc gccctcctcc 31620
cgccgcagga ctgctgacca gggttccggg gacgtggccc ccctcggcca cgggaagagg 31680
gacgggggccc gcaagatgca agttatcgcg aagcgatgac gcggcagatt gtgggagggt 31740
acggagtgag cgatagcgaa cgaagttccc cgggaccctg cgtcagcagg actgtggcaa 31800
aggaagcaga atgcaggact gcggcaaaga tgagggactg gagggaattg aaaaaaatgt 31860
tatatttgta tttcgttcct taatccggag aactatggca gacaaactat acctgattga 31920
cggtcacaac ctaatcttca agatgtacta cgcattcctg cgtcgtccta tgatcaattc 31980
caagggaatg gacatgtcaa tcctgtatgg cttcacgaaa tatgtgctcg aactcatcga 32040
gagagagaag cctacccata tggccatagc gttcgatcct cccggcggca ccttcaggaa 32100
cgaaatgtac cccgaataca aggcaaacag aagcgaaact ccgcagctcg tgatcgatgc 32160
cctcgaaccg ctgctggaaa tatgcaaagc cttgaacata aagaccttga tgattccggg 32220
atatgaggcc gacgacgtga tcggaaccgt tgcgaagagt tacgggaaac ccgggacgga 32280
cgtctatatg gtcaccccgg acaaagatta cggccagctg gtcgccgaac atgtgtacca 32340
ggtgaagcct ggaaaggccg gcggggctga cgagaccgtc gatgtcaagg cagtctgcga 32400
aaaatacgga atctccagtc cggaacaggt catcgaaatg ctgaccctct gcggagatac 32460
ggccgacaat gtgcccggag tgcatggcat cggggaagtc ggggcaggca agctcatcgc 32520
aaaatatggc accgtcgaga atatctatgc ccatctggac gagctgactg accgccagaa 32580
gacgcagttc gaggaggcca ggggacatat cggcctgagc catgacctgg tgacaataaa 32640
gaccgacgtg ccggtagaga caagcctcga cgagatgagg ctggacctgg attatgctcc 32700
cgaaatcgcc gaccttttcg aaaaatacga attcaactcg ctgagaaaat atatagccca 32760
tagctcccgc gctgccaagg cagtggcgga ggtcaagcgc ctggattata ccgagacagg 32820
ggcggaagag ctcgtgaaga tggctccgga aaagatttcg ctgatagtcg aaccggaggg 32880
accgggcata ttcgccgata tccgcagggt gatcgtggcc gcggagaaga aagacggcgg 32940
aatcgtcgtc gcttcaggcg ctgcgaagga ttttcgcaag atactggagg gcgaagccgt 33000
gaaatacgga tatgacctca agctccagtc gaatcttctg gagaatgccg gaataaagct 33060
cggaggccgg ctatatgaca tcgagctgat gcattatctc atcaatccgg agaagagcca 33120
caaggtcgaa atccttgcca agagctatct cggcgtcgat atcgaagagg ccgctgctgt 33180
ggaagaagct cctgtttccc tgtttgacgt cgcggaagaa agcccggcaa acggaccgga 33240
caggaatctc gaagccgcgg cagtgctgaa actcggagaa gtggtccgcg caggcatgga 33300
cgaggccggc gtaggagccc tgtatgagga tatggaagcc cctctcgcag ccgtgctcgc 33360
ccgcatggaa agggcaggcg tgaagataga cctcggacag ctcaggacgt ttgccgatga 33420
actcaaagtc gaaatggatg ccctcgaagc ccaggtcagg aaactggccg acgagccggg 33480
gctgaacgtt tcttccccga agcagattgg catcgtgctc ttcgaaaagc tgaaactcga 33540
cccgaagatg aagaaatcgg ccagcggtag ctatagcacc gacgaaacga cgcttactgc 33600
catcgccgac aggcacccta taattgacaa gatacttgaa ttcagagccg taaagaagct 33660

gctctccaca tacatagagc catttccgac ctacgtctcc ccggttacgg gaagggtcca 33720
tacgaccttc aaccaggcgc tgacggcgac gggaagactc agttcctcca accctaacct 33780
gcagaacatt ccgatccgca cggagagggg aaaggagatc aggaaagcct tcataccggc 33840
gactccggac ggaatcataa tgtccgccga ctactcccag atcgagctca ggatcatggc 33900
ccatctctgc ggcgacgagc atctgataag ctcattcaag gcagggctgg atgtccataa 33960
ggccaccgca gccaagattt tcggcatacc ggtagaggag gtcacatccg accagaggcg 34020
catcgcgaag accgcaaact tcggaatcat gtacggcatc tcatccttcg ggctcgccca 34080
gaggctgcat gtctccagga cggaagccaa gaagataata gacgattact tcgccgcctt 34140
cccttcgatc cggactttca tcgacaagac tattgcggac gcccgcgaaa aagggtatgt 34200
cgagacagtc ttcggacgca ggaggtatct tccggacata acggcgaaga acgctaccat 34260
aagggctctg gccgagagga atgccgtaaa tgctcctatc cagggcacat cagccgatat 34320
aatcaagctt gcgatgataa gcgtcgacaa gcgcatccgc gaggcgggac tgaagagcaa 34380
gatggtgctc cagatccatg atgagcttct tttcgatgcc gtgccggaag aaatcgagga 34440
actcggcagg atagttgtag cggaaatgga gaacgtaatt accttgtcaa tacctttaac 34500
tgtagaatgt aactatggca aaaactggct tgaagctcac tgatcatgac gacggatctc 34560
ttgtggagat ggcgttaggt ggcggccccg aagccgaggc cgctttcttc acgatttatg 34620
cacgttacca taagggcgta ttcgcccaca tctcacgctt catccaggaa caggaggaag 34680
ttgaggatat ctgtatggag agcttcgaga aggccttcaa gcagctggaa tcatacaggc 34740
cgagcaacaa gttctcgact tggatattca ctatagcccg caacacggcc ttcgaccacc 34800
tcgacaagga caaagtgcgc ggccgcaaca aagacaagat gccgatcgac tactccgacc 34860
tggagcatat agacgtccct gaggacaatg tcagcccgga ggacgagatc ataagttcgc 34920
aggaccatga gaacttcctg agttgtgtcc aggggcttcc ggacctgtat cgcgaagtcg 34980
catccatgtg ctttgtggac aatctcggat acaaggagat ttcggagaag gcagggctgc 35040
ctatcaatac agtaaaaacc cgcatcagca gggctaagag cctgatagtc cagatgatgc 35100
ttgatatgga agaagaatga atttcaaaga attcgaagat atagttgttg ctgagtttcc 35160
ggagctgact ccggagatgc ttggaaagta cgaggccatg gagcccctgt atgaggagtg 35220
gaacgccaag atcaacgtaa tttcccgaaa agacataggc ggcctttatc tgcaccatgt 35280
gctgcattct cttgcgatcg cctcgtatct gaagagacgc aggccggagg tgtattccgc 35340
atggaccgaa ggcgcgtccg tgctcgacct gggcaccggc ggcggctttc cggggattcc 35400
tctcgcggta atgttcccga aagttcgctt taccctctgc gattccgtgg gcaagaagat 35460
aatcgtggca tcggaagtgt ccaaggctct ttccctgggc aacgtgacca ctgtcaatgc 35520
ccgtgcagaa tcgctgccgc aggttttcga tcatgtcgtt tcccgcgccg tgacctcgct 35580
ggataatttc tatccatggg tgaagggcaa gttctccgga agcattctgt atctcaaggg 35640
aggggacata gtcccggaaa ttgcagaact gatgaagaag aacagactcc gtaccggagc 35700
agtttcgacc tggccgattg aaacttggct gaaggacgaa tattttaagg aaaagtttgt 35760
gattgaaata gaaaaataat ttgcaaatcc acataaaata actacctttg cagtcctatt 35820
tccggaaaac tggttttttc ggcctgataa gaataaataa aaaaacagat aacgatgaaa 35880
agaacattcc aaccatcaaa gcgcaagcgt ataaacaagc acggtttccg cgagcgtatg 35940
tcaaccccta acggccgtcg tgtacttgct gcacgccgtc gtcatggccg caaaaaactc 36000
agcgtttctt cagaggatag attctagtca tctctctccg gacgttgttt gtttggatcg 36060
aaattcagga ggcttccgaa aggatgcctc ctttttttcat tatggcagat tatttgatag 36120
acgagggacg taatcattaa gtctcaatgg aaaagttttt tttatcaatc ttgtcttgcg 36180
cgctttcagc gacaatagct gtttctgcgc agagtagcgg accattagtt gtccaccagt 36240
atgaattgtc aaacggaatg caagtctggc ttaacgaaga ccattcgcag ccggtagtcc 36300
acggagccgt ggtcgtcaag gccggaggcc gggattgcgc cgggaccgga atggcccatt 36360
atctggagca tatgctcttc aaaggtacgg aggacatcgg taccatcgac tatgcttcag 36420
agaaggtcta tcttgattct atcgagactt gttacgacct catcgcagca acatcctcca 36480
gcgtaaaaag ggattctctc cagaaggaga tcaaccgtct tagcatcaag gcgctggact 36540
atgcgattcc gaatgaattc gaccgtctta cggccaatat cggaggcaca aagctcaatg 36600
ctttcacgtc ttatgactgt acatgctacc ataattcttt ttctccgcaa tttatcgcgc 36660
agtgggccga actcaatagc cataggctgc tgaaccctgt tttcaggctt ttccagagcg 36720
aacttgaagc agtatatgag gagaagaacg cggcagccga ttcccagttt gacatgattc 36780
tcgagaagat gctggagaaa cttggggatg aaaatccata cagcaaatct ataatcggaa 36840
ctacagataa cctgaagaat ccaagtcttt ccgagatgaa gaagttcttc aacaagtatt 36900

```
     atgtgggctg caacatggga cttatccttg ccggagacat agatgcttcg tcgatattgc     36960
     ctctgctgga atctaccttc ggacggattc cgcgtggcga taagccggaa aggattcctg     37020
     tacctgtgcc ggatttctcg tcccgggaag attatcctat aaaagttaat ttccctatga     37080
     tcaaggcgtc gctgaacgtg ttcaacggtc ctgcagccaa ggatgccgat tatcctgcaa     37140
     tggaactcgc tctcagcctc ctgaacaaca gttttagtac cggaagtctg gattccctca     37200
     taaataacgg caaggttctg gctggagagg tctcagtctt ggactttttt gacgaaatgg     37260
     ggcttattgc catgataacg atacctggaa tccctttcgg atctctcaac aaggcggagg     37320
     cgcggtgctt caaggaagtg gaaagactga aggccggcga ttttgcggag aaggatttcg     37380
     agtcgttgaa atatgatctt gtcaaatcat ataaaaggtg cctggaatcg tccgagtcaa     37440
     ggttctctat catgatagac gtgatgtccc aggacaggtc gtgggattct tatctttccc     37500
     aggtggagtc gatatcttcg cttacttcgg aggatatagt cagggtaagc aataaatatc     37560
     tgggcgacaa tcgtatacgt ttccataaag tcagaggaac ctatccgaag gataatctgc     37620
     aagctccttc atacgagact ctttcgccga agaatggcaa tgcggtatcg gattatgcca     37680
     ccagactgat ggagatgcct tatgacactc ttcctccaag actggtggat atgaataatg     37740
     atatccgga                                                             37749
//
ID   XXX; XXX; linear; XXX; XXX; XXX;  XXX.
XX
AC   PRJEB67889
XX
DE   Uncultured bacterium DNA fragment from in-reactor enriched cow rumen, clone
ferm-40d8
XX
FT   source          1..38624
FT                   /organism="uncultured bacterium"
FT                   /environmental_sample
FT                   /mol_type="genomic DNA"
FT                   /isolation_source="Cow rumen"
FT                   /clone="ferm-40d8"
FT                   /note="AZCL-Barley beta glucan active clone"
FT   CDS             103..846
FT                   /db_xref="SEED:fig|6666666.127252.peg.1"
FT                   /translation="MNWIIVQRQDLFWQFGVNGAYNHQKITKMTTNDGDGYKGVDVGGI
FT                   AGAVGNTIQKHMTGYAPNTFYVFQQIYDVDGRPIMGAYVDRNEDGVVDEEDKYYYKSPH
FT                   PKVTLGFNTSLNWKQWNFAASAHANLGNYVYDNNSSQLSLLTDLWTNNFIANRIPQGVK
FT                   DGFIKAQYFSDYYIKDASFFKLDNVTVGYTFALKNDMALNLFGTVQNVLIISPYDGLDP
FT                   EIFGGIDSNIWPRPRTFVIGAKFNF"
FT                   /product="SusC, outer membrane protein involved in starch
FT                   binding"
FT   CDS             865..2451
FT                   /db_xref="SEED:fig|6666666.127252.peg.2"
FT                   /translation="MKLYRIAISIFVALGLTACVGDLDVTPIDPSVTLPEEVLNSEDAY
FT                   AQLLAKCYAGLTVSGSDGPDGAPDIDGIDGGFGQYMRALFYMQEFTTDEALIVWNDQTV
FT                   KDLHALKWTTTDVFVTAMFSRVFYQIGLCNEFIRRAQASEFASSANMKTMIAEARALRA
FT                   LSYYHAIDIFGNVPFSTETSTVGAVGPDQISRADLFKWIDDECADLIAGGELKGAKQNV
FT                   YGRLDVNFVKMVRAHLNLNAAVYLGISASEAKPYYDVVGTLCKEIKNDYPALHAKYTDL
FT                   FGADNDKMTDEMIFCIPQDGTNIQNYGGTNFIIKSSCKGGDKATAAALGIDDGWGGTLV
FT                   TPEFIAKFDDADARKLFWGVGGDMSASQAIDVLEFDYGWSAHKFSNVNSDGTTPSVINF
FT                   MSTDVPLFRAADAYLMLAEAQLRGASTVSETEGKEAWNAVRERAGLGECTYSLDELLDE
FT                   RGRELYWESWRRSDLVRFDKLTSGDYLWTWKGGVQEGTSVDSHYNLMPIPTSELNSNSK
FT                   LSQNAGYVANK"
FT                   /product="SusD, outer membrane protein"
FT   CDS             2461..4842
FT                   /db_xref="SEED:fig|6666666.127252.peg.3"
```

```
FT                   /translation="MKRIKSIIFAGLAAIAAISCDREELATFNPVNADAPVLLEYSATE
FT                   DAISASFSPAVFNMGFNDKVAANHFLVIVSVNDAPVEKALTASAKDGVITVKNSALSKT
FT                   LMGLGYEEGSKVSFDMVVRASLQTNGDNGKNGYIDSEGRISVKDFEVTIPVAQGNPWEN
FT                   FTEVSTWSVIGDIASAGLKWNGDIVMYTDGSRHVARNVKLTTADQFKFRNNGGWDENFG
FT                   GTDESDPFVASTGVEYTAVQKGKNISVAEDGSYDLVLDVDNQTFTVYAAFQTYPGFDQE
FT                   SAWSVIGSIASFEMVWNKDIAMTTDGEWHVAEGVELTPDDQFKFRKDQSWDVNIGAVGD
FT                   TEPYIAELDTEIEGLEKGKNLAVAEAGTYDLLVNPDAKLFKVVKSLGGKSGLVGEGGGE
FT                   TPPGPTVKGWNIIGLNGDWDNDVLATASGNVWTAFVTAKEATSFKWRKDGGWDDNYGGT
FT                   MVALGEAFDAVAGGDNIELSAAGFYKVVLDLDKLKITVSDGNVWSLIGDFNSWSGDVDM
FT                   TLTDGKWVSPVTKISAGGFKIRRNHDWTENVGGVMDALGIPFAAIAGGDDIAIEEENDY
FT                   VVTYDPEDQTILVEVIGWGVVGTINSWGGSADLPMKEENGMFVRKNVTLTADDEIKVRY
FT                   RNSWDVNYGGRTDVGFAVEAISGGDNLKPGAGTYDIWLDADKKALFVMTAGAAPEYWGV
FT                   VGTINGWGGTPDAVMYHDAEGYFVAKGITLAADDEIKIRKNSDWTDNRGASDAAPDTAL
FT                   AAVAGGDNIKVAAGTYDIYYQSASEKIYVMTAGKTPADAQ"
FT                   /product="hypothetical protein"
FT   CDS             5108..7348
FT                   /db_xref="SEED:fig|6666666.127252.peg.4"
FT                   /translation="MKYIMAVAATIAVLSSCGGDNKETPDVKVTLSATPTTVTFAPKAT
FT                   GTKIIAVSTTGDWTATPSATWFHLDKTSGKGKGSIFVTADENTGSDERSGSIRISVSGA
FT                   PDVTVSVSQSADNMVPLTATPAAFDGTKRASTTYQLLVYSFADSNGDGIGDFNGITSKL
FT                   DYLSDVMGVTALWLSPVHPAMSYHGYDVLDYFSLNPEYGTEDDFRNLLAKAKAKGIDIY
FT                   LDYVLNHAGKDHPWFKDACASEESQYRDYFLFSKTPSADIQAGKLPMFPKNGFDAGQWF
FT                   AVGGEKVGAKGRFRFDLDWKSDSAPVVTVTETEAAVETGSSEKYIYYGNDDKCRAFVED
FT                   GDGKYHIVVDFDSDWGFLIRTSSVTWENGTKYGGKAGDVIAFGEPFTLNNTEAADIVFK
FT                   EVTYYHSHLWTDWFADFNYGKASEAETSPAFVDLANSADKWIDMGVAGLRLDAVKHIYW
FT                   SETSSENPTFLDKWYDRCNATYKAAGGEGDIFMVGEVFSDAPATYSYYDGLPSVFEFSF
FT                   WWGLRDVLNNRRAGKFVADIIGYHDAYKARRSDAIPALKLSNHDEDRAASDLGKNLAKE
FT                   KQAAAFLLTAGGKPFIYQGEELGYWGTKSKGDEYVRTPMMWTKSNTGLASKRLGSKIDK
FT                   SMLTASISVESQLADEESLLNVYKTFSALRNTYPALAEGQMSIMSIDDPAIASWYMTSG
FT                   SSKLLVIHNVANSDKTIEVDDSMAHCVALLGTAYVDGTSLKLGANSSVVFEL"
FT                   /product="GH13_36-CBM58"
FT                   /EC_number="3.2.1.1"
FT   CDS             7362..10502
FT                   /db_xref="SEED:fig|6666666.127252.peg.5"
FT                   /db_xref="GO:0031216"
FT                   /translation="MKRLLLAAVAAIVALAGCRSNVEKTIATHPEWAYNSVVYEINIRQ
FT                   FSEEGTFAAVEAQLPRLKELGVDVLWLMPMYKIGEEGRKGTLGSYYAISDYCSTNPEFG
FT                   TMEDFQSLLDAAHSMGFKVILDWVANQTAPDHIWMTEKPADFYERDAEGNAIYEYDWTD
FT                   TRSLNYENKEVWAAQDSCMRFWLEKGVDGFRCDAAGEVPADFWYGILPGIRRDYPEIYL
FT                   LAEAETPKLSSANSEFEATYAWKLHHLLNDIAQGKAKASAIRKYVVEDDKEMGKEGFRL
FT                   MFTSNHDENSWSGSEFERMGDAANVMEVLCFTLPKGHPLIYTGQEVGVTRRIEFFEKDP
FT                   VADWSENEYTTFFKTLVKMRHDNPALQAGSRGGDMRFFDNVPEDVLAFSRSLGGNEVVV
FT                   IANLSAQKQTVSLSLEGEYTSVFSGVGMDSPKSVDLAPWGYMVLAKNSESPVGRVEPLS
FT                   WWTGMKMPLQLMVKGDRIASCNVRIEGGKGVSVKSVTPGDSQDYLFVDVAISASAEPGT
FT                   YDLVFSDDEREFSYPYRIEARAEGSAERKSFTTADMIYLVFPDRFANGDPANDSVKDMA
FT                   EPGDRNLTGGRHGGDIQGIIDHLDYIAGLGATAIWSTPLLVDDEPEGSYHGYACGDYYH
FT                   IDPRFGSNGLYREMVSKAHEKGLKVIMDIVTNHCGTAHWWMDDLPFKDWIHVFPEFTGS
FT                   NVAFSTNMDPNASKYDLNLQESGWFVPSMPDMNLDNPFMLKYFQQWAIWWIEYSGLDGF
FT                   RVDTYPYNEKVPMSKWCAAVLEEYPDFNIVGECWTSSHSQLAYWQAGNGNKDGFNSNLP
FT                   TIMDFPLQEAIVAAMCERGDNPGWGQGMARVYDCLSHDFVYHDLSHMMIFLANHDHARL
FT                   GDIFGSDPSKMKVALAMLATMRGIPQLYNGDEMMFKAFNGEQSDPARRVDFPGGWADDK
FT                   TDLFSADGRKGVSAELHDYTARLFNWRKEKSVLHDGKTLHFLGRDNTYAYFRYNETEAV
FT                   FVFVNNTGHDVIVPWNHYAEFVSGPVSATDVVSGMPVTLGPGLTVPAGTALIAEFDR"
FT                   /product="GH13-GH13"
```

```
FT                   /EC_number="3.2.1.135"
FT   CDS             10559..12838
FT                   /db_xref="SEED:fig|6666666.127252.peg.6"
FT                   /translation="MMFAAAVMFSVAAYSGETLKSPDGNLELDFRLDERGAPVYNLSYK
FT                   GSPVVLDSRMGFLLVFNNDPSMHRMPYAGRFGGPWSLYDGFSMTDSKTSSFDETWTPVW
FT                   GEEAEIRNHYNELAVTLKQESGRIMIVRFRLYDDGLGFRYEFPAENALTYFLISEEVTE
FT                   FALAGDHLAWWLPGDYDTQEYNYTESRLSEVRKLVDRARTANASQAAVPGVSVQTSLQM
FT                   KTDDGLYVNIHEAEVVNFPTANLELDDENMKLRVWLTPDAQGVKGRMQAPCQSPWRTVM
FT                   VVDDARKVLASRLILNLNEPCALEDVSWIHPVKYMGVWWEMITNKSLWSYTNDFPSVQL
FT                   GKTDYSAAKPHGRHGANNENVRRYIDFAAENHFDALLIEGWNEGWEDWFGHQKDYVFDF
FT                   VTPYPDFDLPGLNAYAHSKGIRLIMHHESSSSTVNYERHIDKAYDLMNEYGYDAVKSGY
FT                   VGDIIPYGEHHYSQPIINHYALCIKKAAEKHIMVNAHEAVRPTGLCRTYPNMIGNESAM
FT                   GTEFRSDINPGHTTILPFTRLQGGPMDYTPGIFEMDMSKLNPRERGHMRMTICNQLGLY
FT                   VTFYSPLQMAADLPEHYAPFMDAFQFIKDVAVDWDKSLYLAAEPGAYVVTARHPKLSTL
FT                   NKSQEGVASLSDGKKDALSGAARFVYALGEDATSADLEGAVARDVWYVGGITDESARDI
FT                   TVALDFLQPKVKYEAVIYADGPDADFETNPQSYTITRKTVTSKSKLNLHMARSGGFAIS
FT                   LREIRK"
FT                   /product="GH97"
FT                   /EC_number="3.2.1.20"
FT   CDS             12851..13171
FT                   /db_xref="SEED:fig|6666666.127252.peg.7"
FT                   /translation="MTLIQKIRLSERYGVPFYVISALFSIAAAIVCFFYPVAIPVCMTM
FT                   KLLCIPVILYLCESFQNKHAIYFYLNLGISKIEYYLIPVVTEFLFFIVLISVSCFISNV
FT                   VR"
FT                   /product="hypothetical protein"
FT   CDS             13182..13823
FT                   /db_xref="SEED:fig|6666666.127252.peg.8"
FT                   /db_xref="GO:0005524"
FT                   /db_xref="GO:0006810"
FT                   /db_xref="GO:0016021"
FT                   /db_xref="GO:0016887"
FT                   /translation="MLVDSVHLEFGQQIVLRSAFITSETGKVTGLFGRNGVGKSCLFRC
FT                   IMGGIKAQNLFVRFNDEMDTDYNSIGKRVKYLPQGLFMPARMTLKEAFDLYHVDYDGLV
FT                   SFDVRFLNKQGKRFHELSGGEARLAEIYLVLRSTSEFCLLDEPFSYIAPKDVEKIQNLI
FT                   LAEKKRKGIIVSDHMYEALLPVSDELYLIKDGYTFPIQTKADLIKHGYII"
FT                   /product="ABC transporter ATP-binding protein"
FT   CDS             complement(13816..14001)
FT                   /db_xref="SEED:fig|6666666.127252.peg.9"
FT                   /translation="MSTLIYLAGIVLAIVAVLDIVKKPIKTSGKIIASVILLLTSWVGL
FT                   IIYYLYAKDHLTDWFK"
FT                   /product="hypothetical protein"
FT   CDS             complement(14036..15754)
FT                   /db_xref="SEED:fig|6666666.127252.peg.10"
FT                   /db_xref="GO:0004849"
FT                   /translation="MTSIRILCKNTGKYISVEAGSSLYEISEQFCKVVKDPKTGVEYRV
FT                   ISALVDNKLKELDFTVFHPHEVEFLGYNHADGRRTYIRSLAFMLQNAVRKIYPDKILVV
FT                   DHSLPSGLYCEILSYDENDTTDIHAHSKPVVPDLELLKQEMKNLSSRDLRFTKKKIPAE
FT                   EGIKIFENNKQFHKARLEESLGRFFLSVYYLDGQADTFHGPLAYSTGMIKVFNLVHFGK
FT                   GFCLQYPSIADPQKVIPYKEQSKVASSLEEYSQWCKTIGILGVGTLNKAIRNGRAIELI
FT                   NISEARMERHYADIADKIFAEKSRIRIVFIAGPSSSGKTSSSLRIAQQCKVLGMKPKVI
FT                   ELDNYFVSRKFTPRDEDGNLDFECLEAMDLKLLNKHLNALLAGEKVELPRFDFLRGEPV
FT                   FEGNFMQLEDNDILIMEGIHALDPAMVPEVDNNRIFRVYASALTSLNLDENNNISTSDN
FT                   RILRRMVRDYNYRGHTPENTINGWQSVRRGEEKYIFPFQENADAQINTALMYELPLLKY
FT                   YAEPLLRRITPNSPAYTEAVRLLKFLDYIVALQPNEIAAIPPTSIMREFIGGQTL"
```

```
FT                   /product="Uridine kinase (EC 2.7.1.48)"
FT                   /EC_number="2.7.1.48"
FT   CDS             complement(15751..18540)
FT                   /db_xref="SEED:fig|6666666.127252.peg.11"
FT                   /db_xref="GO:0005524"
FT                   /db_xref="GO:0016887"
FT                   /translation="MGKEYIEIRGAKAHNLKGASLSIPRNEFVVVTGLSGSGKSSLAFD
FT                   TIYAEGQRRYMDTLSTYAKHFMGILEKPEVEEITGLSPIIAIEQKTTGNNPRSTVGTIT
FT                   EIYDFLRLLYARASRAYSPSTGKEMVRYTDEQITDLILTNYGGRKCYLLAPLVRGRKGH
FT                   YKELFDQMRKRGYTQVRIDGELKDLSEATQLDRYKAHFIELVIDKLKPDASDAKRIRES
FT                   VLTALTRGKGSLAVMDMDTGALQHYSRHLVDPDSGLSLQEPAPHSFSFNSPEGYCPHCK
FT                   GLGMIVDVNIDAIIPDRNVSIADGGIVPLGKIRETRKFDIIRSIARKYNFSLYDPIAAI
FT                   PDEAVSLIVFGSDELFRVGDGALSEMVAFPGIVDDIDDKIVCPVCKGTRLNEQTNCFRI
FT                   DGKTIAEVSSMEISALKEWLDSLPEKMTGRENRIGRDILKELRERVRFMLDVGLDYLSL
FT                   DRATASLSGGESQRIRLATQIGSKLVNVLYILDEPSIGLHQRDNRKLINSLKELRDEGN
FT                   SIMVVEHDAETMYASDWLVDVGPGAGEEGGRVCLSAPADYLLKGKKPDPDTLKHCIVGP
FT                   SLTLEYLRGASEIAVPKHRRLGNGNSLGLRGASGNNLKNIDIDFPLGCFIGISGVSGSG
FT                   KSSLINETLMPILKNKFYNAKLKPLPYTALHGIRNIDKLIEIDQSPIGRTPRSNPATFT
FT                   GVFNDIRTLFAETVDAKVRGFGSGRFSFNVPGGRCEACKGAGIKVIEMNFLPSVHVPCD
FT                   ECRGRRYKDDTLAVRYKGKNINDVLEMPISEAYEFFKPVPKIAQKLKALLDVGLGYVHL
FT                   GQSAVTLSGGESQRMKLAAELFRKATGNTLYILDEPTTGLHFEDIKVLLGVLQQLVDQG
FT                   NTVIIIEHNLDVLKSVDYIFDLGPDGGRRGGYIVAEGTPEQLAKNPDSVTGPYLKEVL"
FT                   /product="Excinuclease ABC subunit A"
FT   CDS             complement(18540..19247)
FT                   /db_xref="SEED:fig|6666666.127252.peg.12"
FT                   /translation="MLNIRNFTFNLFQEVCTLLWDDTMEGAIVDPGCLGEAECSRLTDY
FT                   VRDNGIRIKMILLTHGHFDHIYGVPAMVAEYGVPVYMNPADKVLLENNHLLSGVFGMPD
FT                   APKDFATKDIKEGDRLSFGKTEFEVIETPGHTPGCVCFYDKADKLLISGDTLFAGSIGR
FT                   TDSAWGDYDKEIVSIMDKIMGLDGDVAVIPGHGPKTDIAYERTHNPFLQPFNEPDDDLN
FT                   WDGEGISIDGDFQ"
FT                   /product="Hypothetical metal-binding enzyme, YcbL homolog"
FT   CDS             19317..19964
FT                   /db_xref="SEED:fig|6666666.127252.peg.13"
FT                   /db_xref="GO:0004136"
FT                   /db_xref="GO:0004136"
FT                   /db_xref="GO:0004138"
FT                   /db_xref="GO:0004138"
FT                   /translation="MFIGVAGNIGAGKTTLTRMLVDYYGWTPKYESVTYNPYLEDYYKD
FT                   IQRWSFNLEIYFLEQRFQDLMEIAKSDNVIIQDRTIFEGVYVFVANNYEQGNLSKRDFE
FT                   TYMRLFNLMMSMVKAPDLMIYLKSSVPHLVAQIQKRGREYEQGMNLSYLQGLNEKYEKF
FT                   INEDYKGHVLTIDVDGLDFESRPEDFAMITERIDSELFSLFGQTNNIQGLCI"
FT                   /product="Deoxyadenosine kinase (EC 2.7.1.76) /
FT                   Deoxyguanosine kinase (EC 2.7.1.113)"
FT                   /EC_number="2.7.1.76"
FT                   /EC_number="2.7.1.113"
FT   CDS             19955..20575
FT                   /db_xref="SEED:fig|6666666.127252.peg.14"
FT                   /db_xref="GO:0004136"
FT                   /db_xref="GO:0004136"
FT                   /db_xref="GO:0004138"
FT                   /db_xref="GO:0004138"
FT                   /translation="MHIAIAGNIGSGKTTLTKMLAAHYGWTPKFESVDFNPYLSDFYED
FT                   MERWSFNLQIYFLNKRFKDVVDISKCDDVIIQDRTIYEDARIFAPNLHAMGLMSTRDFE
FT                   NYSDLFDLMMSLVNPPDLLIYLKSSIPNLVSQIQKRGREYEKSIRIDYLTGLNERYENW
```

```
FT                   IKDYKGNLLVIDADRIKFGNRPEDFQTVTDMIDGQMFGLFPQK"
FT                   /product="Deoxyadenosine kinase (EC 2.7.1.76) /
FT                   Deoxyguanosine kinase (EC 2.7.1.113)"
FT                   /EC_number="2.7.1.76"
FT                   /EC_number="2.7.1.113"
FT   CDS             20582..21364
FT                   /db_xref="SEED:fig|6666666.127252.peg.15"
FT                   /translation="MDGGIGGIALGLLIPFAGTVLGSAFVFFMKKEMSEKLQKALLGFA
FT                   SGVMVAASVWSLLIPAMEMGDGKMSVVPATVGLLAGFAFLLLIDYITPHLHLDGDQEGP
FT                   ESHLSKTTMLSLAVTIHNLPEGMAVGVAIAGALSSDVGMSVAGALALSLGIAIQNIPEG
FT                   AIISMPLRAVGNTRLRSFGIGSLSGIVEPIGGLLVLLLASIVTPMMPYLLAFAAGAMLY
FT                   VVIEELIPEASTGKHSNISTVGFALGFALMMVLDVVLG"
FT                   /product="Metal transporter, ZIP family"
FT   CDS             complement(21586..22800)
FT                   /db_xref="SEED:fig|6666666.127252.peg.16"
FT                   /translation="MKRENYIRTWLKDFRGIGWFELRQIFSDWGVMLIFFVAGLIYPLL
FT                   YNVIYLNGVLNDTPIAVVDDANCDVSRRYIREMDATREVAVEYRCANMAEAEKLMQERK
FT                   VKGIVMFPSDFNEKLARMETATLSVYCDMSSFLYYKNALMAANHVMLDEIGHIQIERYA
FT                   AAGFTGEEAAQLVQAIPYEENNPYNSAFSYSFFLISAILFLIIQQTMFYGMSLLVGTFR
FT                   EQGRSFATLPDRLDGIGLGRVVLGRGAAYWLVYMGISMYIAFIVPAIFGLPQRGEFWDI
FT                   VVLLLFFVTDCVFFSMTWSTVVTRRETVFVLFLFMSPICLFLTGTSWPTAAFPAFWKYF
FT                   SYIFPSTFGCQAFINLNAAGGDLAAIRSQMFAMTIQSVAYYFLASAAVFLENRILKYKI
FT                   KTLSQ"
FT                   /product="ABC-type multidrug transport system, permease
FT                   component"
FT   CDS             complement(22802..23971)
FT                   /db_xref="SEED:fig|6666666.127252.peg.17"
FT                   /translation="MGMWQQIFAVTGRERRIWRHRPIYLIGSAGVIAFCAIFFLTFFRD
FT                   GLPHDLPIGIVDNDNSSLSRNFIQQLDATQLGKTVQFDDFEQARLAMQGGKITSICVIP
FT                   ENMYADVSANRRPVFTYYLNSMYFVGGALAYKDILTMINLTNGAVQRQVLRAKGVNERE
FT                   IMGRIRPIDIDTHQIGNVYTNYGYYLTNIILPAVLELVIIIVLIYSLGAEMKYEKSPEL
FT                   MKIAGGSITNALTGKLILYTALFSGLGFLIVTLLYHWMHFPIAGSIWNMFIAIFLMVTA
FT                   SEAIAIFIIGCLPVPRLALSIGALYSVLGFSLSGFTLPIEAMPAWIQGWASAFPIRHYY
FT                   LFYVQEVIFGAGFAGWWQEIVHLLFFVLLPLPVLWRLKGAYVNLNFPKN"
FT                   /product="ABC-type multidrug transport system, permease
FT                   component"
FT   CDS             complement(23976..24965)
FT                   /db_xref="SEED:fig|6666666.127252.peg.18"
FT                   /translation="MTKEQKTYNITIGVAALLIIILIISIIGYFVSRPKPLVIQGQVEA
FT                   SEYRVSGKVPGRVEELYVKEGQEVHKGDTVVYIDSPEVRAKLAQAKGAVAAARAQKTKA
FT                   ANGARQEQITGAYELWQQARVQEEVMRKSFDRVQKLYDQSVISAQKYDETKAKYDASVA
FT                   QAKAAKSQYDMAVNGARIEDKEAAEALVERAMGALQEVEGYLGELYLTSPADGIISATF
FT                   PKVGELVGTGSPIMTVTDTHDIWFTFNVREDYLHGLKQGDKITLTVPALDNKEITAVVS
FT                   YRAVRESYATWKATKETGMYDAKTFEVRAVPEEYTEGLYPGMTVILKQ"
FT                   /product="FIG00937622: hypothetical protein"
FT   CDS             complement(24979..26457)
FT                   /db_xref="SEED:fig|6666666.127252.peg.19"
FT                   /translation="MAIESNKDLDQARTKVEMAGYDRKIARANYFPEISATGAYLYNNR
FT                   NLSLVTDSQSAMLTNLGTSAQGALDQAVGAISGKVNGAVTQLSTAIMSDPRLAAEYMNS
FT                   PMWQTVLNMLKESGSGSATAPDIATPINAIGQDIDKSMHLDIENVFIGAVTLKQPVFVG
FT                   GKIVLSNQIAKMAEELARAEYDMKYADVLVDIDQAYWQIVSIANKKKLAESYSDLLHQL
FT                   EKDVAISVQEGVATKSDELQIKVKANEADMMLTKATNGLELAKMLLCKQIGLPLDSQIT
FT                   LSDEALDAIPVPVPGEEKSMEDIWEDRAETRSLELATKIYDKKIGIARADMMPQIALVA
FT                   GYSVTNPNAFHGYENSWNGGLFSAGVMVNVPIFHGTEAYQRTRKAKAEATLYRDKLEDA
```

```
FT                   RNMINLQVEQSRQHLEEARERLLMSENNLSSAEENMRMATVGFEAGVIETNTVLAAQTA
FT                   WLQAHSEYIDAGIDLQMANANLRKAEGYYNIDNE"
FT                   /product="Type I secretion system, outer membrane
FT                   component LapE"
FT   CDS             26745..27662
FT                   /db_xref="SEED:fig|6666666.127252.peg.20"
FT                   /translation="MEETLQTIDIKGFKSLFPIEYDAELGDDFFIADITYDRALRMLQY
FT                   PTRFDGFLGIFCVHGHLSMDINLSTYKVTENSLLFSVPGYIIRVSDVGTDDLKNLRFIV
FT                   IAVSRGFMSSMRMDFTRLFNESMSVLSNSCIRLEPDELALCSKYMTLAIEIVKNGVTSY
FT                   KKETIGSLISSVCYFLGGMWADKLSKARRESNPDISARAKLVFDHFLQLVTDNHTSQRN
FT                   MAFYADKLCLTPKYLSKLVKNVSGRSAPEWIDSFVILEAKNMLKYSEISIKEIVYKLNF
FT                   PNQSVFYKFFKAHTGLTPSEYRNS"
FT                   /product="Transcriptional regulator, AraC family"
FT   CDS             27665..28435
FT                   /db_xref="SEED:fig|6666666.127252.peg.21"
FT                   /translation="MQNLTASFAIHALNTLGYKVSPPIGERTFGQWSFCYLISGEMFME
FT                   SAPDSYLIRGHEFVLVPPGVPFRISWYRDSIGYMGGFDETILKDRSYTCLQARRPLRVS
FT                   VSADDEGFVDELMYKLLRKKDDMPVVLGCLDVMLSQIDADTPESRMPDKPLVNLFLDRI
FT                   FDREKEFLGVAGYAEELNVTPNYLNRLVRKSTGRSANEWIEISRLTYAKLLLRRKELAI
FT                   IDVAAMVGLPDQSYFSRFFKKMEGVTPSEFRKDL"
FT                   /product="Transcriptional regulator, AraC family"
FT   CDS             28541..30790
FT                   /db_xref="SEED:fig|6666666.127252.peg.22"
FT                   /translation="MHYLAIIITILGGAFAAEPESEVFDTLAVSVVSSSAKLVQPLEQL
FT                   SSPSTSVLYKDFESHGLNSPKLLSSIVPNLHIPDYGTSMTSSIYMRGFGSRIDNPVLGL
FT                   YIDDIPVLDKNNYEFELMDIRRADLLRGPQGTLFGRNSLTGVLSISTLDPSVISGGKAV
FT                   LEYGSANTMLAKGSWYGAGLGVSAAYRHSDGFYVNDYDGSDIDKYDSFSLRLTHGKDLN
FT                   PRLHMDNTLSASWLVQGGYAYRRLVESGEGDPAYGGVLLPVNYNGDSGYRRVSVLGGTK
FT                   FRYEHDRYQLRSVTSIQLLFDRMNMDQDFTPQSVFTLQQKQKQAAVTQEFIFRPKEDVE
FT                   WWDSQTGVFAFYRYNYMDAPVNFLYDGVNTLILGNANKGIPDPDYLDFLEDEFLIATDF
FT                   KIHTYNTAFYHESYFTFGNWHFTAGLRLDHEGNFMRYCSGTELTFIYSPIMKAYEPFET
FT                   VYKGLERNFYFELLPKLSALYDFGSLNVFATVSKGYKSGGFNTQIFSDILQNKLMEGMM
FT                   NHIGVHSVDSGEAPEASATCYKPESCFNYEAGLRYSHVSESGHRISASASAFYIDCHNQ
FT                   QITIFPPGKNTGRRMANVGKSRSYGAEAEFSYSYKNFSVNGSYGYVNAKFTEYSDGKND
FT                   YSGNRIPYSPEQTAFMRASYTLPLSGGILDSISFSADANCTGRIWWNEDNSYVEPVHVL
FT                   FGGDITLDFGKFDLYVRGRNLSNRDYNTFYFKSMDNSFFQKAKPVSYTAGIRFIF"
FT                   /product="TonB-dependent receptor"
FT   CDS             30808..31260
FT                   /db_xref="SEED:fig|6666666.127252.peg.23"
FT                   /translation="MKKLVFGLMIALTVFAACSKDDDNKEVTVEDGSYVGTVTVVEPHF
FT                   SASDIKVSVTPIEGTTTLDIKMFEVRFSERMPKTIDFTIPKASYTIDGDTINIFGENII
FT                   PTMGAGYEKPEYTVTGLKGTIKDGNIEFSLNFGSFPTKYSGTLAKK"
FT                   /product="hypothetical protein"
FT   CDS             complement(31580..32608)
FT                   /db_xref="SEED:fig|6666666.127252.peg.24"
FT                   /translation="MTACNNTPKVTTIKGTINPADSLVAVYYYDWNESELQRQSKAVRV
FT                   NNGSFETTIPVCALRMASVRDLSWAKQVDFVSDGSTIMIDVDSCTAISSDPEGIHSRYV
FT                   KYTNYYINHVYLDEALEIWKRDTESVHRFKEICDSLNLDMLREAYTENTDNVICDEIIY
FT                   WLENLVSLEELQTYLDMASEDMKKDTRIGLKLKNMPAKLNTAPGKMYSDIEGIKDPDNP
FT                   ESKARLSDYVGKGKYVLADFCGAGCGPCRQMTKDDLVPNYERLKDKVEFLTIMVWDETL
FT                   ETTQSFVETYGITWPVIHRCLEARDIYAIEGVPTLIIFAPDGTIVERGHSDLFLRLGDY
FT                   IK"
FT                   /product="Thiol:disulfide interchange protein tlpA"
FT   CDS             complement(32785..33744)
```

```
FT                   /db_xref="SEED:fig|6666666.127252.peg.25"
FT                   /db_xref="GO:0004340"
FT                   /translation="MAQELVIGIDVGGQTTKCGVIDARGTVLAQTVIRTDTYTEIEPYI
FT                   AELAEALQHIIAESKAEGLIRGIGVGAPNGNYYTGEIENAVNLTWGGTKKIAFSKLLSD
FT                   AMNGIPVSLTNDANAAAMGEMTYGAARGMKNFIMITLGTGVGSGIVIDGKIVYGHDGFA
FT                   GELGHTCAVRNNGRLCNCGKTGCLETYCSATGVARTAREWLDMTDEPSTLRSLDKVSSK
FT                   DVYEAAKDGDKLALKIFEYTGNILGQKFADFIEFSAPEAIVLFGGLARAKDFLREPIER
FT                   SMNANVLPLWRNKVKLVFSQLKESDAAILGASALAWEL"
FT                   /product="Glucokinase (EC 2.7.1.2)"
FT                   /EC_number="2.7.1.2"
FT   CDS             complement(33771..35051)
FT                   /db_xref="SEED:fig|6666666.127252.peg.26"
FT                   /translation="MAKKNGNIIAIVTMIFLFGMISFVTNLAAPIGNIWKMQPQIAGSN
FT                   ALGMMGNMMNFLAYLFMGVPAGKMLTKYGYKKTALIAIAVGFIGVFIQFLSGVVASFPA
FT                   NFACYLLGAFISGFSVCMLNTVVNPMLTLIGGEKHGNQYVQIGGTFNSLMGTLTPMLVG
FT                   ALIGTLTAQTVIKDVNPVLYIAMGVFLAAFIALSFIPIEDPEEVADAANAESPWNYKHF
FT                   IYGVIAIFVYVGVEVGIPGTLNFYLADPENPAQMGANAATVAGFVAGTYWFLMLIGRFL
FT                   SSLISGKVSAKAQITAVSAVALCLVIGAMVVSPEVRASMPAFTGSNFEMVNVPLSALLL
FT                   VLCGLCTSVMWGAIFALSVDGLGSATNKASGLFMMMVVGGGIIPLLQNFIADHSTYMFS
FT                   YILPAIGLAYILFFGLIGSKPAKRVKA"
FT                   /product="Predicted glucose transporter in maltodextrin
FT                   utilization gene cluster"
FT   CDS             complement(35158..37410)
FT                   /db_xref="SEED:fig|6666666.127252.peg.27"
FT                   /db_xref="GO:0008422"
FT                   /translation="MRHNINLLLPFVAIAITISCSGPSASTVGSTPEIEKRIDKLLSQM
FT                   TLQEKIGQMNQISAGGSVEPFSDAVRNGQVGSILNETDPGRVNELQRIAVEESRLGIPL
FT                   LVARDVIHGFNTVFPIPLGIAATFDPQLAEEGARVAAIEATSVGVRWTFSPMLDIARDP
FT                   RWGRIAEGSGEDPYLDARMGEALVRGYQGEPGDSTSMAACIKHFVGYGAAEGGRDYNST
FT                   YLTERQLRNVYLPPFEAGVKAGAMTLMTSFNDNDGVPSTGNRFIVNDILRDEWGYDGFV
FT                   VTDWNSMGEMMVHGFAADHKNSAALAIGAGVDMDMMTYGFIANLETLVKEGKVSEKQID
FT                   AAVRNILRVKFLLGLFENPYVDPARAGKVLYSEENLDAARRTAEEAAILLKNDGVLPLD
FT                   ASRIRTILVTGPMADAPHDQLGTWAFDGDKTHTVTPLAALRERFGGKARIVYVPGLEYS
FT                   RQEKDSFADVVSAARNADVVLAFLGEEAILSGEAHSLADLDLKGSQSALLAALKSSCKP
FT                   IVATIMAGRPLTIEKDLVNCDAMLYEFHPGTMGGPATVNLLFGDAVPSGKTPVTFLRTV
FT                   GQAPMYYSHNNTGRPYNGELLLDQIPLEAGQTSLGNTSYYLDYGPAPLFPFGYGLSYTT
FT                   FSYSGISLDKKVYSAEDTVRATFTLSNTGKYNGVEVAQLYIRDLVGSVTRPVKELKYFE
FT                   RIALEVGESREITFELPVRELAFWNMDMEKVVEPGAFRLWIAGDSASGEAAEFSVE"
FT                   /product="GH3"
FT                   /EC_number="3.2.1.21"
FT   CDS             complement(37422..38405)
FT                   /db_xref="SEED:fig|6666666.127252.peg.28"
FT                   /translation="MELSGEAGSASVTTEANTTWTAETDGSFWLKISARAIRSGSQEFT
FT                   VTVLKNYSEEPRTAKVIFTSGSSKVELEVIQAKGEPFVDGAFVPEGYQLAWHDEFDGEK
FT                   VDPDNWRFEDWAPGRVNHELQRYVAGGERDGKKTAFVEDGALNIRAMKHNGEVISARMN
FT                   SKAAWKYGYMEARIWLPKGLGTWPAFWMMPTDQSLGWPKCGEIDILEEVGANPEYTSSS
FT                   IHTESYNHVKNTQKTHEQLTEGAENEYHIYSCRWTAEALEFYVDGKLHLTFRNDGAGNV
FT                   STWPFDKEFFIILNLAWGGDWGGYKGIDESALPTTMKIDYVRVYQK"
FT                   /product="GH16_3"
FT                   /EC_number="3.2.1.73"
XX
SQ   Sequence 38624 BP; 8954 A; 9942 C; 10113 G; 9615 T; 0 other;
     tcggatctgc tcaattatac tcctgtcgct tccggcgcaa acctcaagaa ctacctcaac        60
     gccaatatcg gaaccctcgt caacaagggt atcgaggtcg acatgaactg gatcatagtc       120
```

cagaggcagg acctcttctg gcagttcggc gtgaacgggg cgtacaacca ccagaagatc 180
accaagatga ccaccaacga cggcgacggc tataagggtg tcgatgtcgg aggtattgct 240
ggtgctgtcg gcaacactat ccagaagcat atgacgggct acgctcctaa taccttctat 300
gtgttccagc agatatacga cgttgacggc cgtccgatca tgggcgctta tgtagaccgc 360
aacgaagacg gcgtagtaga tgaggaggac aagtactatt acaagagccc tcatccgaag 420
gttactctcg gtttcaacac ttctctcaac tggaagcagt ggaacttcgc cgcatcggcc 480
cacgccaatc tcggcaacta tgtctatgac aacaattcat ctcagctgag tcttcttacc 540
gacctctgga ccaacaactt catagccaac cgtattccgc agggcgtgaa ggacggcttc 600
atcaaggcac agtatttctc agactactat atcaaggacg cttcgttctt caagctcgac 660
aatgttaccg taggctatac cttcgctctc aagaacgata tggctctcaa cctcttcggt 720
actgtccaga acgtattgat catatctcct tacgacggtc tcgatcctga aatcttcggc 780
ggtatagact ccaacatctg gccgcgtccg agaacattcg tcatcggagc caagtttaac 840
ttttaatcaa ggaggaaaag aatcatgaaa ctttatagaa tcgcaatatc catctttgtc 900
gcccttgggc ttactgcctg cgtaggcgat cttgatgtca ctcctatcga tccttcggtt 960
accctccctg aggaagtgct caactccgaa gacgcatatg cccagctcct tgcgaagtgc 1020
tatgccggcc ttacggtctc aggctccgac ggccctgacg gtgctccgga tatcgacggt 1080
atcgacggcg gtttcggcca gtacatgcgt gcgctcttct acatgcagga gttcactacc 1140
gacgaggccc ttatcgtatg gaatgaccag acagtaaagg atctccacgc tctcaagtgg 1200
actaccaccg acgtattcgt aaccgcaatg ttctccaggg tattctacca gatcggactt 1260
tgcaacgaat ttatccgccg cgcccaggct tcggagttcg cttcatccgc aaacatgaag 1320
accatgattg cagaagcccg cgctctccgc gccctcagct actaccacgc tatcgatatc 1380
ttcggtaacg ttccgttctc taccgagacc agcacagtag gtgctgtcgg tcctgaccag 1440
atcagccgcg ccgatctctt caagtggatc gatgacgagt gcgccgacct tatcgccggc 1500
ggagaactca agggtgccaa gcagaatgtc tacggacgtc tcgacgtaaa cttcgtcaag 1560
atggtacgtg cgcacctgaa cctcaacgct gccgtatatc tcggaatctc cgcttctgag 1620
gccaagcctt actacgatgt agtcggcacc ctctgcaagg agatcaagaa cgattatccg 1680
gctctccatg ctaaatatac cgacctcttc ggcgccgaca acgacaagat gaccgacgag 1740
atgatcttct gcatacctca ggatggtacc aacatccaga actatggcgg aaccaacttc 1800
atcatcaagt cttcatgcaa gggtggtgac aaggccactg ccgccgctct cggtatcgat 1860
gacggctggg gaggtactct cgtaactcct gagttcatcg ccaagttcga cgatgccgac 1920
gcccgcaaac ttttctgggg cgtaggcgga gacatgagtg cttctcaggc aatcgacgtc 1980
ctcgagttcg actatggctg gtcagcccac aagttctcca acgtcaacag cgacggaaca 2040
actccgtcag taatcaactt catgtcaacc gacgttcctc ttttcagggc agccgatgca 2100
tacctcatgc ttgctgaggc tcagctccgc ggcgcttcta cagtaagcga gaccgagggt 2160
aaggaggcgt ggaacgcagt acgagaaaga gccggcctgg gcgagtgcac atactctctc 2220
gacgaacttc tcgacgagcg tggccgcgag ctttactggg agtcatggag aagaagcgac 2280
ctagtccgtt tcgacaagct caccagcggt gactatctct ggacatggaa aggtggcgtc 2340
caggaaggta cttccgtaga cagccactac aacctcatgc ctatcccgac ttccgagctc 2400
aactccaaca gcaagctcag ccagaatgca ggttatgtcg ccaataaata gactatcgtt 2460
atgaagagaa ttaaatcaat catatttgcg ggacttgcag ccattgccgc aatctcctgc 2520
gaccgggaag aactggcgac attcaatcct gtgaatgccg acgcccctgt ccttcttgag 2580
tattcagcca cagaggatgc catctcggct tcgttctctc cggctgtgtt caacatgggc 2640
ttcaatgata aagttgcagc caaccacttc cttgtcatcg tgtcagtaaa cgatgcgccg 2700
gtagaaaaag ccctcaccgc ttccgcaaag gacggcgtga tcactgtcaa gaattccgcc 2760
ctcagcaaga ctctcatggg cctcggatat gaggaaggtt cgaaagtctc tttcgacatg 2820
gtagtcaggg caagcctgca gaccaacggt gacaacggca agaacggata tatcgattcc 2880
gaaggcagaa tatctgtcaa ggacttcgaa gtgacgattc ctgtcgccca gggcaatcct 2940
tgggagaact tcacggaggt cagcacatgg agcgtgatcg gcgatatcgc cagcgccggc 3000
ctgaaatgga atggtgacat cgtcatgtac actgacggtt ctcgccacgt cgcccgcaac 3060
gtcaaactta ccaccgcaga ccagttcaag ttccgtaaca acggaggctg ggacgagaac 3120
ttcggcggta ctgacgagtc cgatccattc gtggcatcga ctggtgtaga atacactgcc 3180
gtacaaaaag gaaagaacat ctccgttgca gaagacggtt cgtatgacct cgtgctcgac 3240
gtagacaacc agaccttcac ggtttacgca gccttccaga cctatccggg ctttgaccag 3300
gagagcgcat ggagcgtcat aggctcgatt gcaagctttg agatggtttg gaacaaggat 3360

atcgctatga ctaccgacgg agagtggcat gtcgcagaag gtgtagaact taccccctgac 3420
gatcagttta agttccgtaa ggatcagagc tgggatgtca atatcggcgc tgtcggtgat 3480
accgagcctt atatcgcaga actcgataca gagatcgagg gcttggagaa aggcaagaat 3540
ctggcagtcg cagaggctgg tacatatgac cttctcgtca accctgatgc caagctcttc 3600
aaggttgtca agtctctcgg aggcaagagc ggtcttgtgg gcgaaggcgg cggcgaaacc 3660
cctccgggac ctacagtaaa gggctggaac atcatcggcc tcaacggtga ctgggacaac 3720
gacgtgctcg ccactgcttc cggcaacgta tggactgcat ttgtaactgc taaagaagct 3780
acatcattca agtggcgcaa ggatggcggc tgggacgata actatggcgg aaccatggtt 3840
gctctcggag aagccttcga tgcagttgcc ggtggcgaca acatcgagct ctcagcagcc 3900
ggtttctaca aggtagtcct cgacctcgac aagcttaaga ttactgtatc agacggtaat 3960
gtatggtccc tcattggaga cttcaacagc tggagcggcg acgtggacat gacccttacc 4020
gacggcaagt gggtaagccc tgtcaccaag atttctgcag gcggcttcaa gatccgtcgc 4080
aaccatgact ggaccgagaa tgtcggcggt gtcatggatg ctctcggaat cccgttcgca 4140
gctatcgccg gtggtgacga tatcgccatc gaagaggaaa atgactatgt ggtgacatac 4200
gaccctgagg atcagaccat ccttgtcgag gtaataggct ggggcgtcgt aggtaccatc 4260
aacagctggg gcggctctgc cgaccttcct atgaaagagg agaacggcat gttcgtaagg 4320
aagaacgtca ctcttaccgc tgacgatgag atcaaagtcc gttaccgcaa ctcatgggat 4380
gtcaactatg gcggccgcac cgatgtgggc ttcgcagtag aggctatctc cggcggtgac 4440
aacctcaagc cgggcgctgg cacctatgac atctggctgg atgccgataa gaaggccctg 4500
ttcgtgatga cggcaggcgc tgctcctgag tactggggcg tggtcggcac cattaacggc 4560
tggggtggaa ctcctgatgc cgtgatgtac catgacgctg aaggatattt cgttgcgaaa 4620
ggcatcactc ttgcagcaga cgacgagata aagatcagga agaacagcga ttggacagac 4680
aacagaggtg cttccgacgc agctccggat accgctctcg ctgcagttgc gggcggcgac 4740
aatatcaagg tggctgccgg tacgtacgac atctattacc agagcgcatc tgaaaagata 4800
tacgtaatga cagcgggcaa gacccctgct gacgctcagt agcatatttt tcggactgga 4860
gcccctgacc ggactccagt ctgccaatct atagacagtg gtctatatat ttttaaaatt 4920
ctgttttgct tcgtaatatt gattacggca aaggtactat agggttcgga gaccgtggcc 4980
gcccgcggcc atcgtgagcg ggaggggccc agccgtcagg ctgggaggga tagggccgaa 5040
ggcccgtagt ctcagggccc tatagtacct ttgccgggaa caaataaaac tatatatgaa 5100
aaaaatattg aagtatataa tggctgtggc cgcaactatt gcggtattga gctcctgtgg 5160
cggcgacaac aaagagactc cggacgtaaa ggtcaccctc tcggcaaccc cgacaaccgt 5220
cacctttgct ccgaaagcca ccggcacaaa aataatcgcc gtaagtacca ccggagactg 5280
gacggcaacc ccgtccgcca cctggttcca tctcgacaag acttccggca aaggcaaggg 5340
ttccattttc gtcacagctg acgagaatac gggctcggac gaacgcagcg gttcaatccg 5400
catctccgtt tcaggagctc ctgatgtcac tgtgtccgtg agccagagcg ccgataacat 5460
ggtccctctg accgccactc ctgcagcctt cgacggtacc aagagagctt cgacgaccta 5520
tcagcttctc gtgtacagct tcgcggacag caacggcgac ggcatcggtg atttcaacgg 5580
aatcacctca aagctggatt atctttcaga cgtgatggga gtgacagccc tctggctctc 5640
gccggtccat ccggccatgt cttatcatgg atacgatgtc ctggactatt tctccctcaa 5700
tccggaatat ggtaccgagg acgatttccg caaccttctg gcgaaagcca aggcaaaggg 5760
catagacatc tacctcgact atgtgctcaa ccatgccgga aaggaccatc catggttcaa 5820
ggacgcctgc gcaagcgaag agagccagta cagggattac ttcctcttca gcaagactcc 5880
ttccgcagat atccaggccg gaaagttacc gatgttcccg aagaacggat tcgatgccgg 5940
ccaatggttc gcggtaggtg gagagaaggt cggggcgaag ggacgtttcc gtttcgacct 6000
cgactggaag agcgattccg ctccggtagt gactgttacc gaaacggagg ctgcagtcga 6060
aacgggctcg tcggagaaat atatctatta tggcaatgac gataaatgta gggctttcgt 6120
agaagacggg gacggaaaat accatatcgt cgtcgatttc gattctgact ggggcttcct 6180
cataaggaca tcgagtgtaa cctgggaaaa cggtaccaag tatggcggaa aggccggaga 6240
tgtaatcgcc ttcggagagc ctttcaccct caacaatacc gaagccgcag atatagtctt 6300
caaggaagtg acatactatc attcccatct ctggaccgac tggttcgccg atttcaatta 6360
cggcaaggct tcggaggcgg agacttctcc tgcatttgtc gatcttgcga attcagctga 6420
caaatggatc gacatgggtg tggcgggact ccgtctcgac gccgtcaagc atatctactg 6480
gagcgagaca tcctccgaga accctacctt cctcgacaaa tggtatgaca ggtgcaacgc 6540
tacctataag gccgccggcg gagaagggga tatctttatg gtcggcgaag tcttcagtga 6600

cgctcccgca acctacagct actacgacgg ccttccgtca gtcttcgaat tcagcttctg 6660
gtgggggctc agggacgtgc tcaacaacag gcgcgcagga aagtttgtcg ctgacataat 6720
tgggtaccat gatgcctata aggcccgcag gagcgatgcc attcctgctc ttaagctctc 6780
caaccatgac gaggacaggg ctgccagcga tctcggcaag aatctcgcaa aggagaaaca 6840
ggctgctgct ttcctactca ccgcgggcgg caagccgttc atttatcagg gcgaggaact 6900
cggatactgg ggtaccaagt ccaagggaga cgagtatgtg cgtaccccta tgatgtggac 6960
caagagcaac accggccttg cttctaagcg ccttggatcg aagattgaca agagcatgct 7020
gaccgcttcc atttccgtgg agagccagct tgccgacgaa gagtccctgc tgaacgttta 7080
caagactttc tcggcacttc gtaataccta tccggctctt gcagaaggtc agatgagcat 7140
catgtcaatt gacgacccgg ctattgcctc atggtacatg acctccggca gctccaaact 7200
tctggtcatc cacaatgtgg ccaactccga caagacgata gaggtggacg acagcatggc 7260
acattgcgtg gccctgctcg gtaccgcata cgtcgatgga acttcgctca aactcggggc 7320
gaactcttca gttgtatttg aactatagat aacgaaaata tatgaaaaga cttttactcg 7380
ccgcagttgc cgcgatcgtt gcgctggcag gctgccggag caatgttgaa aagaccattg 7440
cgacccatcc tgaatgggcc tataattctg tggtctatga aatcaatata cgccagttct 7500
ccgaagaggg cacgttcgct gccgtcgagg cgcagctccc gagactcaag gaactcggag 7560
tcgacgtact ttggcttatg cctatgtata agatcggcga agagggccgt aagggtacgc 7620
tcggctcgta ttatgccatc tccgactact gcagcaccaa tcctgagttc ggtacgatgg 7680
aggatttcca gtcgcttttg gatgccgccc attccatggg attcaaggtg attctcgact 7740
gggtcgccaa ccagactgct ccggaccata tatggatgac ggagaaaccg gcagatttct 7800
acgagaggga tgccgaaggc aatgccatct atgagtatga ctggaccgat acccgcagtc 7860
tcaactacga gaacaaggaa gtctgggcag cccaggacag ttgcatgagg ttctggctgg 7920
agaagggcgt ggacggtttc cgctgcgatg ccgccggaga agttcctgcc gatttctggt 7980
acggaatcct tcctggtatc cgcagggatt atccagaaat ctacctgctt gcagaggccg 8040
agacgccgaa gctcagcagc gccaattctg aatttgaagc tacttatgca tggaaactgc 8100
accatcttct caacgatatt gcgcagggta aggccaaggc ctctgcgatc cgcaagtacg 8160
tcgttgaaga tgataaggag atgggcaaag agggtttccg tctcatgttt acgtccaacc 8220
atgacgagaa ctcatggagc ggaagcgagt ttgagcgtat gggcgatgcc gccaatgtga 8280
tggaggtgct ctgcttcacc cttcctaagg gccatccgct tatctatacc ggacaggaag 8340
taggagttac gcgccgtatc gagttcttcg aaaaagaccc tgtcgctgac tggtcggaga 8400
atgagtatac taccttcttc aagactctgg tgaagatgcg ccacgacaat ccggcacttc 8460
aggccggatc ccgcggcggc gatatgcgtt tctttgacaa tgtcccggaa gacgtgcttg 8520
cattcagccg ttctctcggt ggaaacgaag tcgtggtgat cgccaatctt tcggcgcaga 8580
agcagacagt cagcctttct cttgaaggag aatatacttc tgtcttctcc ggtgtcggca 8640
tggattctcc gaagtcagtc gatctcgctc cgtggggcta tatggtcctt gcgaaaaact 8700
ctgagtcacc ggtcggaaga gtcgagcctc tcagctggtg gacaggcatg aagatgcctc 8760
tccagcttat ggtcaagggc gacagaatcg catcctgcaa tgtccgtata gagggcggaa 8820
agggtgtctc tgtcaaatcc gtaactcctg gcgacagcca ggattatctt tttgtggacg 8880
tcgctatctc tgcttcggca gagcccggga cttatgatct cgtcttctcc gacgacgagc 8940
gcgagttcag ctatccctac aggatagagg caagagctga aggttcggct gagcgtaaga 9000
gttttactac ggcggacatg atatatctgg tattcccaga ccgcttcgcc aacggcgatc 9060
ctgccaacga ttctgtcaag gacatggctg aaccgggcga ccgcaatctc accggaggcc 9120
gtcacggcgg cgatatccag ggtatcatcg accatctgga ctatatcgcc ggactcggcg 9180
cgaccgccat ctggagcact ccgctgcttg tcgacgacga gcctgagggt tcatatcatg 9240
gctacgcctg cggcgactat taccatatcg acccgcgttt cggcagcaac ggactttatc 9300
gcgagatggt atcgaaggcc catgaaaaag gcctgaaagt catcatggat atcgtcacca 9360
accattgcgg aacggcgcac tggtggatgg atgacctgcc tttcaaggac tggatccacg 9420
tcttcccgga gtttaccggc tctaatgtgg ctttctccac caacatggat ccgaacgctt 9480
cgaaatatga ccttaatctt caggagagcg gctggttcgt gccttccatg cctgacatga 9540
atctggataa tccgtttatg ctgaagtatt tccagcagtg ggcgatatgg tggatcgagt 9600
attccgggct ggacggtttc agagttgaca cttatccata taatgagaag gttcctatga 9660
gcaagtggtg tgcagcagtg cttgaggagt accctgattt caatatcgta ggggagtgct 9720
ggacttcttc tcactcgcag ctggcctatt ggcaggcagg gaacgggaac aaggacggct 9780
tcaattccaa cttgcctacg attatggatt ttcctctcca ggaggctatc gtagcggcta 9840

```
tgtgcgagcg cggtgacaat ccggggtggg gacagggaat ggcccgtgtc tatgattgtc      9900
tgagccatga tttcgtatat catgatctct cgcatatgat gatctttctg gcaaatcatg      9960
accatgcccg tcttggggat atcttcggca gcgatccttc gaagatgaaa gtggcgctgg     10020
ccatgctcgc taccatgcgc ggaatcccgc agctgtacaa cggcgacgag atgatgttca     10080
aagcattcaa tggcgaacag agcgatcctg cccgcagggt tgatttcccg ggcggatggg     10140
ccgatgacaa gaccgatctt ttctctgcag acggccgcaa gggggtgtct gccgagttgc     10200
atgactatac cgccaggctt ttcaactggc gcaaggaaaa gtctgtgctg catgatggca     10260
agactctcca tttcctcggc agggacaata cttatgctta tttccgttac aatgagacgg     10320
aggctgtatt tgtcttcgtc aacaataccg gccatgatgt tattgtccct tggaaccatt     10380
atgccgaatt cgtgtccggt ccggtttctg ccacggatgt ggtttcaggt atgcctgtga     10440
ctctcggtcc tggcctgaca gtgcctgccg ggactgcgct tattgctgaa tttgacagat     10500
aattttatta tattggacaa ttgtttgaac agttagaaat aatatgaaga aagttttgat     10560
gatgtttgcc gctgccgtca tgttttcggt agctgcctat tctggagaga ctctgaagag     10620
tccggatggt aatcttgaac tggatttccg tctggacgag cgtggagctc ctgtctataa     10680
tctctcttat aagggcagtc cggtggtact tgacagccgc atgggtttcc ttcttgtatt     10740
caataatgat ccttccatgc ataggatgcc gtatgctggc cgattcggcg ggccatggag     10800
cctgtatgac ggtttcagca tgactgattc caagacttct tcttttgacg agacgtggac     10860
tcctgtctgg ggcgaggagg ctgagataag gaatcattat aatgagcttg cagtcactct     10920
gaagcaggaa tccggcagga tcatgattgt ccgtttccgc ctttatgacg acggtctggg     10980
attccggtat gaattccctg ccgagaatgc tctgacttat ttccttatca gcgaggaagt     11040
taccgagttt gctcttgccg gagaccatct cgcctggtgg ctccccggag attacgatac     11100
ccaggagtac aattataccg agagccgtct ttcggaggta cgcaaacttg tagatcgcgc     11160
gcgtactgcc aatgcttcgc aggctgccgt gcctggtgtt tctgtccaga cttctcttca     11220
gatgaagaca gacgacggcc tgtatgtcaa tatccatgaa gccgaggtcg tgaatttccc     11280
taccgccaat ctggaactgg atgacgagaa catgaagctt cgtgtgtggc ttacccctga     11340
tgcccagggt gtcaaaggac gtatgcaggc tccatgccag agtccgtgga ggacggtcat     11400
ggtcgtggat gatgcgcgta aggtgctcgc ttcgagactt atcctgaatc tcaatgagcc     11460
ttgtgccctg gaggatgtct cctggatcca tccggtaaag tatatgggtg tctggtggga     11520
gatgatcacc aacaagagcc tctggtcata tacaaatgac ttcccgtcag tccagctcgg     11580
aaagaccgat tattccgctg ccaagcctca tggccgtcat ggcgccaaca atgagaatgt     11640
gcgccgttac atcgacttcg ccgccgagaa ccatttcgat gctcttctta tcgaaggctg     11700
gaatgaaggc tgggaggact ggttcggcca ccagaaggat tatgtcttcg atttcgtcac     11760
tccttatccg gatttcgatc ttccggggct gaatgcatat gcccattcca agggtatcag     11820
gcttataatg caccatgagt cttcgtcttc gacggtcaat tatgagcgtc atatcgacaa     11880
ggcttacgat ctgatgaacg agtatggcta tgatgccgtg aagagcggat acgtgggcga     11940
tataatccct tacggcgagc accattacag ccagcctata atcaaccatt atgctctctg     12000
catcaagaaa gccgccgaga aacatatcat ggtaaacgcc catgaggctg tacgtcctac     12060
cggtctctgc aggacttatc ctaacatgat cggcaacgag tctgccatgg gtacggagtt     12120
ccgttcggat atcaatccgg gccataccac catcctgccg ttcacgcgtc tgcagggcgg     12180
tcctatggac tatactccgg gtatcttcga gatggacatg agcaagctca atcctaggga     12240
gcgcggacat atgaggatga ctatctgcaa tcagcttggt ctttatgtga cattctattc     12300
tcctttgcag atggctgcgg accttccgga gcattatgct cctttcatgg acgcattcca     12360
gttcatcaag gatgtggctg tcgactggga caagagcctt taccttgctg ccgagccggg     12420
ggcttatgtg gtgactgcgc gtcatcctaa gctttctacg cttaacaagt cccaggaggg     12480
cgtggcttcc ctttctgacg gaaagaagga tgccctcagc ggtgccgcaa ggtttgtgta     12540
tgctctcgga gaggatgcca cttcggctga cctggaggga gccgtggcac gtgacgtatg     12600
gtatgtgggc ggcatcacgg atgagtctgc ccgtgacatt acggtggccc tggatttcct     12660
ccagcctaag gtgaagtatg aggctgtaat ctatgcggat ggtccggatg ctgatttcga     12720
gaccaatcct cagtcttaca ccataacccg caagactgtc acctccaagt ccaagctcaa     12780
tctccacatg gcccgctccg gcggcttcgc catctccctc cgcgaaatcc gcaaataagg     12840
agctgtcctg atgactctga tacagaaaat acgtctttcg gaaagatacg gagtgccttt     12900
ctatgtgata tcggcattgt tttccattgc tgccgccata gtgtgcttct tctatcccgt     12960
ggcgataccg gtatgcatga cgatgaaact gttatgcata ccagtaatcc tgtatctgtg     13020
cgagtcgttc cagaacaagc atgctattta tttctatctg aatctgggca tcagcaagat     13080
```

cgagtactat ctgatcccgg tcgtgacgga gtttttattc ttcatagtcc tgatttctgt 13140
ttcatgcttc atcagtaatg tcgtcagata ggataaataa gatgctggtt gacagcgtgc 13200
atctcgaatt cggtcagcag atcgtcctcc ggagtgcgtt cataacttcc gagaccggca 13260
aggttaccgg cctcttcgga aggaacgggg tcggaaagtc ctgtctgttc aggtgcataa 13320
tgggcgggat caaggcccag aacctgttcg tgcggttcaa cgatgagatg gataccgact 13380
acaacagtat cggcaaaagg gtaaagtatc ttccccaggg cttgttcatg ccggcgagga 13440
tgacgctgaa ggaagcgttt gacttgtacc atgtcgacta cgacggtctg gtgtcgtttg 13500
acgtacgttt cctgaataaa caggggaagc gcttccatga gctctccgga ggagaggcca 13560
ggctcgcgga aatatacctt gtgctcaggt ctacctccga attctgcctt ctggacgagc 13620
cattctccta catcgccccg aaggacgtgg agaagataca gaatctcatc ctcgcggaaa 13680
agaagcgcaa gggaatcata gtctccgacc atatgtatga ggcactgcta cccgtgtcgg 13740
acgagctcta cctgatcaag gacggctaca ccttccccat ccagacaaaa gccgacctca 13800
tcaagcacgg ctacattatt tgaaccagtc ggtgaggtgg tccttggcgt agaggtagta 13860
gatgataaga ccgacccagc tggtcagcag aaggatgact gaagcgatta tctttccgct 13920
tgtcttgata ggcttcttga cgatgtcaag aactgcgacg attgccagga cgatgcctgc 13980
aaggtaaatt aatgtactca tagttttatg ctttagtgtt tagtaaatgg ggttcctaga 14040
gggtctggcc gccaatgaat tcgcgcatta tcgaagtcgg aggaatggcg gcgatctcgt 14100
taggctgaag agcgacgata tagtcaagga acttcagcag gcgtacggcc tcggtgtagg 14160
ccgggctgtt aggagtgata cggcggagca gaggctcggc atagtacttc agaagaggaa 14220
gctcgtacat caaagccgtg tttatctgcg catcggcatt ctcctggaac gggaagatat 14280
acttctcctc gccacggcgc acgctctgcc acccgttaat agtattctcc ggagtatgcc 14340
cgcggtaatt gtaatcccgg accatgcggc ggaggatacg gttgtcggaa gtagagatat 14400
tgttgttctc gtcaaggttc agggacgtca gggctgaagc gtagaccctg aagatacggt 14460
tgttgtccac ctcagggacc atcgccggat caagagcatg gataccctcc atgataagga 14520
tgtcattatc ttccagctgc atgaaattgc cctcgaagac aggctcgcca cggaggaaat 14580
caaaacgcgg aagctcgacc ttctctccgg cgagcaaagc attcagatgc ttgttcagaa 14640
gcttaaggtc catggcctcc agacactcga aatcaagatt cccgtcctca tcccgcggag 14700
tgaacttacg cgaaacgaaa taattgtcca gctcgattac tttaggcttc atgcccagta 14760
ccttgcactg ctgggcgata cgcagcgaag aagaagtctt gccgcttgaa gaaggacctg 14820
cgatgaacac aatccttatc cgggacttct cggcgaagat cttgtcggcg atatcggcat 14880
aatgacgctc catccgcgcc tcggaaatat tgataagctc gattgcgcgg ccgttacgga 14940
tagccttgtt gagagtgccc actccgagta tgccgatagt cttgcaccac tgggaatatt 15000
cctcgagaga agacgcaacc ttggactgct ccttgtaagg gatgaccttc tgtggatcgg 15060
cgatggaagg gtactgaagg cagaatcctt tgccgaaatg caccagattg aacaccttga 15120
tcattccggt agaatatgcg agaggcccat ggaaagtatc ggcctggcca tcaagataat 15180
acacgctgag gaagaatcgt ccgagagact cctcaagacg tgccttgtgg aactgtttgt 15240
tgttctcgaa tatcttaatc ccttcctctg ccggaatctt cttcttggtg aaacggagat 15300
cccttgaaga aaggttcttc atctcctgct tcagcagctc cagatccgga acgacaggct 15360
tcgaatgggc atggatatct gtcgtatcat tctcatcata ggagagaatc tcgcagtaga 15420
gaccgctcgg cagcgaatgg tccaccacca gaatcttgtc cggatagatc ttgcgcactg 15480
cgttctggag catgaaagcc agagaacgga tatatgtcct ccggccgtcg gcatggttat 15540
agccgaggaa ctccacctca tgcggatgga agacggtaaa gtcaagctcc ttgagcttgt 15600
tgtccacaag ggccgaaatg accctgtatt cgactcccgt cttcgggtcc tttacaacct 15660
tgcagaactg ctcggagatc tcgtaaagag aggaaccggc ctctacactg atgtacttgc 15720
ccgtattctt gcataagatt ctgatagatg tcatagtact tcctttaaat atggtccggt 15780
aaccgaatcc ggattcttgg caagctgctc cggagtaccc tcggcaacta tatatccgcc 15840
cctgcgtccg ccgtccggac cgagatcgaa aatatagtcc accgacttga gcacgtccag 15900
attatgctcg attataatta cagtattacc ctggtccaca agctgctgca gaaccccgag 15960
caacactttt atatcttcaa aatgcaggcc tgtagtaggt tcgtcaagaa tataaagcgt 16020
atttcccgta gcctttcgga aaagttcggc tgcaagtttc atgcgctggc tttcacctcc 16080
ggaaagcgtc acggccgact ggcccagatg cacatatccg agcccgacat caagcagggc 16140
cttaagcttc tgcgcaatct tcggcacagg cttgaagaac tcgtaggcct cgctgatcgg 16200
catctccagt acatcattta tattttttcc cttgtaacgc actgcaagag tatcgtcctt 16260
gtacctgcgc ccgcggcact cgtcgcatgg aacatgtacg gacggcagga agttcatctc 16320

gatgaccttg atacccgctc ccttgcaggc ctcgcaccgt cctcccggga cattgaacga 16380
gaaccttccc gaaccgaaac cgcgcacctt ggcatcgacg gtttccgcaa acagagtgcg 16440
gatatcgttg aatactccag taaaggtggc cggattgctc cggggagtac gtccgatagg 16500
actctggtcg atctcgataa gcttgtcaat attcctgatg ccgtgcagag ccgtatacgg 16560
aagcggctta agcttggcat tgtagaactt gttcttcagt ataggcatca gcgtctcgtt 16620
gatgagagaa gacttgccgc tgccgctgac tccgcttatg ccgatgaagc atcccagcgg 16680
aaaatctatg tctatattct ttagattgtt gcccgacgct cctcgcagtc cgagagaatt 16740
cccgttcccc agtcgcctgt gcttagggac ggcgatctcg gaggccccgc gcagatactc 16800
cagcgtcagg gacgggccga cgatacaatg cttcagcgtg tccggatcag gcttctttcc 16860
cttcaggaga tagtccgccg gagcgctcag gcagacccgg cctccttcct ctccggcgcc 16920
cggaccgaca tctacgagcc agtcggacgc atacatggtc tcggcatcat gttcgacgac 16980
catgatcgag ttgccctcgt cgcgcagctc cttcagcgaa ttgatcagct tccggttgtc 17040
tctctgatgc aggccgatgc tcggctcatc gaggatgtag agcacattga ccagcttcga 17100
gcctatctgt gtggccagac ggatgcgctg gctttcacct ccggaaaggg atgcagtagc 17160
cctgtccaga gaaagatagt ccagaccgac gtccagcatg aaacggaccc tttcccggag 17220
ctccttgaga atgtcgcgtc cgatcctgtt ctccctgccg gtcatcttct ccggcagcga 17280
atccagccac tccttcaggg cgctgatctc catcgacgaa acttccgcga tcgtctttcc 17340
gtcgatccgg aaacagttgg tctgctcatt cagacgcgtg cctttgcaga ccggacatac 17400
gatcttgtcg tctatgtcat ccacaatccc ggggaaggcc accatctccg agagggcgcc 17460
gtccccgacc ctgaacagtt catccgaacc gaaaactatc agagacacgg cttcgtccgg 17520
aatcgcggct ataggatcat aaagggagaa attgtatttg cgcgctatag agcgtataat 17580
atcgaatttt ctggtttccc gtatctttcc gagagggacg atacctccgt cggcgatcga 17640
gacgttgcgg tcagggatta tcgcgtcgat attgacatcg acaatcattc cgaggccttt 17700
gcagtgcggg cagtagccct ccggagagtt gaacgagaac gaatgaggcg ccggctcctg 17760
tagcgaaagc ccgctgtcag gatccacgag atgtcttgaa taatgctgca gggcgccggt 17820
atccatgtcc attactgcca gggagccttt gcctctcgta agggcggtaa gtaccgactc 17880
gcggatacgc ttggcatccg aggcgtccgg cttgagcttg tcgatcacga gctcgatgaa 17940
atgggccttg tacctgtcca gctgggtcgc ctcggaaaga tccttcagct cgccgtcaat 18000
gcgtacctgc gtatatccgc gcttgcgcat ctggtcgaag agctccttgt aatggccttt 18060
acgccctctg acgagcggag cgagaaggta acatttacgg cctccgtagt tcgtaagtat 18120
gaggtctgtg atctgctcgt cggtatagcg gaccatctcc ttcccggtgg aaggggaata 18180
ggccctcgaa gccctggcgt atagcagtcg caggaaatcg tatatctccg ttatcgtgcc 18240
tacggtcgaa cgcgggttgt tgccggtagt cttctgctcg attgcgatta tcgggctgag 18300
accggtgatc tcctcgacct caggcttctc cagaatgccc atgaaatgct tcgcataggt 18360
gctcagcgta tccatgtacc ttctctggcc ttctgcgtat atcgtgtcga atgcaagcga 18420
agacttgccg ctgccgctga gtccggtgac gacgacaaat tcgttgcgcg ggatgctcag 18480
gctcgctcct ttgaggttgt gggccttcgc tcctcgtatt tcgatatatt ctttacccat 18540
tattgaaaat ctccgtcgat gctgatgccc tcgccatccc agttgagatc gtcgtcgggc 18600
tcgttgaaag gctgtagaaa cggattgtgc gtgcgctcgt aagcgatatc ggtcttgggc 18660
ccgtggcccg gaatcacggc gacatccccg tccagcccca tgatcttgtc catgatactt 18720
acgatctcct tgtcgtagtc gccccaggcg ctgtcggtgc ggccgatgct gccggcgaac 18780
agagtatctc cgctgatgag cagtttgtca gccttgtcgt agaagcagac gcagccggga 18840
gtatgcccgg gcgtctctat gacctcgaac tcagttttgc cgaaactgag cctgtcgcct 18900
tcctttatgt ccttcgtggc gaagtctttc ggagcgtcgg gcatcccgaa gactccggag 18960
agtaaatggt tgttctccag caggactttg tcggccgggt tcatatagac gggaaccccg 19020
tattcggcga ccatggccgg gactccgtag atatggtcga agtgcccgtg ggtcagaagt 19080
atcatcttga ttcttatgcc gttgtccctg acgtagtccg tcagccggct gcattcggcc 19140
tctccgagac agcccggatc cacgatcgct ccttccatag tgtcatccca gaggagggtg 19200
cagacttcct ggaaaagatt gaatgtgaaa tttcggatat tcagcataag tgatggaaca 19260
tatttttaca aaattacaaa aaatcctcta aatttgtaga atcataaata catactatgt 19320
tcattggtgt tgccggaaat atcggtgcag ggaagaccac tctgacgaga atgcttgtgg 19380
attattatgg atggaccccg aaatacgagt ctgtcacata taacccatac ctggaggatt 19440
actacaagga tatccagaga tggtcgttca accttgagat atatttcctc gagcagcgtt 19500
tccaggacct catggagata gccaagtcgg ataatgtgat aatccaggac aggacaatct 19560

tcgagggcgt ctatgtattc gtcgccaaca actatgaaca ggggaatctt tcgaagaggg 19620
atttcgagac ttacatgaga cttttcaacc tgatgatgtc gatggtgaag gctcctgacc 19680
tgatgatcta cctgaagagt tccgtcccgc atcttgtcgc ccagatccag aagagagggc 19740
gcgagtacga gcaggggatg aacctcagct atctgcaggg gctgaatgaa aagtacgaga 19800
aattcatcaa tgaggactat aaggggcatg tgctcacgat cgatgtggac gggctggatt 19860
tcgagagcag acctgaggac ttcgcgatga ttacggaaag gatcgattcc gaactcttca 19920
gcttatttgg tcagacaaac aatattcaag gattatgcat atagcgatag caggaaacat 19980
cggttcgggt aagacgactc ttaccaagat gctggcggcc cattatggct ggacgcctaa 20040
gttcgagtct gtggatttca atccatatct ctctgatttc tatgaggata tggagaggtg 20100
gagcttcaat ctccagattt acttcctcaa caagaggttc aaggatgtcg tcgacatctc 20160
caaatgcgac gacgtcataa tccaggacag gactatctac gaggacgcca ggatattcgc 20220
gccgaacctg catgctatgg gccttatgag cacaagggat ttcgagaact attcggatct 20280
cttcgacctg atgatgtccc tcgtcaatcc gcctgacctg cttatctatc tcaagagctc 20340
gatccctaac ctcgtgtccc agatccagaa gaggggccgc gagtacgaga agagcatccg 20400
tatagactat cttaccggtc tcaacgagag atacgagaac tggatcaagg attataaggg 20460
taatctgcta gtcatagatg ctgaccgcat caagttcgga aaccgtccgg aggacttcca 20520
gaccgtcacc gatatgatcg acggacagat gttcggtctt ttccctcaaa aataacagtc 20580
tatggacggt gggatcggcg gcatagcttt aggtctgctc ataccttttg ccggaactgt 20640
gctcggctct gccttcgtgt tcttcatgaa gaaggagatg tcggaaaagc ttcagaaagc 20700
tctgctcggc ttcgcgtccg gagtcatggt cgccgcgtct gtatggtccc ttctcatccc 20760
cgccatggag atgggggatg gaaaaatgtc ggtagtccct gcgactgtcg gccttcttgc 20820
cggtttcgcg ttcctgctcc ttatagacta tattaccccg cacctgcatc tcgatggtga 20880
tcaggaaggc cctgagagcc acctgtcgaa gacgacgatg ctgtctcttg ccgtgaccat 20940
ccataatctc ccggagggaa tggccgtcgg agtcgctatc gccggcgctc tcagctcgga 21000
tgtgggcatg tctgtagccg gtgctctggc cctgtccctc ggtatagcta tccagaatat 21060
ccccgagggg gcgatcatat cgatgcctct tcgcgcagtc ggcaacacac gtctccgttc 21120
cttcggaatc ggcagcctca gcggtatcgt cgagcctatc ggcggattgc ttgtcctgct 21180
tctcgcttcg atcgttaccc cgatgatgcc ttaccttctg gcattcgctg ccggagcgat 21240
gctttatgtc gtaatagagg agctgattcc ggaagcttca acgggtaaac actccaatat 21300
cagtactgtc ggctttgctc tcggcttcgc cctgatgatg gtcctggatg tggttttggg 21360
ctagtctcat aataatttaa cgtcccgaat gtcacttttt tacgatttgg gacgggaatt 21420
tcaaatctat ttttctatat ttgttcccag ataggaaata attaaactgg caaactatga 21480
aaaatagaat ggctttcgtg ggagtgcttt tgttgttttt tatggtaagc aactgctctc 21540
atgaaccggg attcgagatc tcgtcaaatg aagaagactc tgtcgttatt gtgacagagt 21600
ctttatttta tatttaagta tcctgttctc caggaataca gcagcactcg caaggaaata 21660
atatgctacg ctctggatgg tcatcgcgaa catctggctg cgtatcgcag caagatctcc 21720
gccggcagcg ttgaggttga tgaaagcctg acagccgaac gtggacggga agatatatga 21780
gaagtacttc cagaatgccg ggaaggcagc ggtaggccaa gaggttccgg taaggaagag 21840
acaaatcggg gacatgaaca ggaacaatac gaagactgtc tccctgcggg tcaccaccgt 21900
gctccaggtc atgctgaaga agacgcagtc cgtcacgaag aacagcagca ggacgacgat 21960
atcccagaac tcgcctctct gaggcagccc gaagattgcc ggcactataa aggctatgta 22020
catggaaatg cccatatata cgagccagta ggcggctccc cggcccagta cgacgcgccc 22080
gagaccgatt ccgtccagtc tgtcaggaag ggttgcgaaa cttctgccct gctctctgaa 22140
tgttccgaca agcagggaca ttccgtagaa cattgtctgc tggattatga ggaacagaat 22200
cgccgagatc aggaagaaac tgtatgagaa agcgctgttg tacgggttgt tctcctcgta 22260
tggaatcgcc tgcacaagct gggccgcctc ttctccggta aagccggctg ccgcatatct 22320
ttctatctgg atatggccga tttcgtcaag catgacatga ttcgcggcca tcagagcatt 22380
cttatagtat aggaaactgc tcatgtcgca atagacagac aaagtagccg tctccattct 22440
tgcgagcttc tcattgaagt ctgacgggaa catcactatt cccttgacct tcctctcctg 22500
cataagcttc tctgcctccg ccatgtttgc gcagcggtat tcgacagcga cttctctggt 22560
cgcatccatc tcccggatgt atctccggct gacatcgcag ttggcgtcgt cgactacggc 22620
tataggggtg tcattcagga ctccgttgag gtagatcaca ttgtagagca gaggatatat 22680
cagacctgcc acgaagaaga tcagcattac tccccagtca gagaaaatct gcctgagttc 22740
gaaccatccg atgccccgga agtccttcag ccatgttctt atataatttt ctcttttcat 22800

tctaattctt agggaagtta agattgacat atgctccttt cagcctccat agtaccggaa 22860
gaggaagcag gacaaagaac agcaggtgca ctatttcctg ccaccatcct gcgaatccgg 22920
ccccgaagat cacttcctgc acatagaaca ggtagtaatg ccttatcgga aaagctgaag 22980
cccagccctg gatccacgcc ggcatcgcct ctatcgggag agtgaagccc gagagggaga 23040
agcccaggac gctgtagagg gctccgatac tcagcgccag cctcggtacc gggaggcagc 23100
cgattatgaa tatggctatc gcctcgctgg ctgtcaccat caggaagatt gctatgaaca 23160
tgttccagat cgacccggct atagggaaat gcatccagtg gtacagcagg gtcactatca 23220
ggaatcccag accggagaac agggctgtat agaggatgag ttttcctgtc aaggcgttcg 23280
ttatcgaacc gccggctatc ttcatcagct ccggcgactt ttcgtatttc atctctgctc 23340
cgagagagta gatcagcact attatgatga ccagctccag tactgcgggc aggattatgt 23400
tcgtaagata gtagccgtag ttggtataga cgttgccgat ctggtgggtg tctatgtcga 23460
tcgggcgtat gcggcccatg atctcccttt cgttgacgcc tttagctctg agcacctgac 23520
gctggacggc tccgttggtc aggttgatca tcgtcagtat gtccttatat gcaagggcgc 23580
cgccgacgaa gtacatgctg ttgagatagt atgtgaagac cggacggcgg tttgcgctga 23640
cgtccgcata catgttctcc ggaatcacgc agatagaagt gatcttgccg ccctgcatgg 23700
ccaggcgggc ctgctcgaag tcgtcgaact gtacggtctt tccgagctgt gtcgcgtcga 23760
gctgctggat gaagttcctc gaaagggaag agttgtcgtt gtcgactatc cctatcggca 23820
ggtcatgcgg cagaccgtcc cggaagaagg tcaggaagaa gatggcgcag aatgcgatca 23880
ccccggccga gccgatcaga tatatcggcc tgtgcctcca tatcctccgc tcacggcctg 23940
taaccgcgaa tatttgttgc cacattccca tttcgctact gctttagaat aactgtcatt 24000
ccaggataaa gtccttcggt atattcctcc ggaacggcgc ggacttcaaa ggtcttggca 24060
tcatacattc ccgtctcctt ggtagccttc catgttgcgt aggactcgcg tactgcgcgg 24120
tagctgacaa ctgcggtgat ttccttgttg tcgagagcag gcacagtaag ggttatcttg 24180
tctccttgtt tcagaccatg gagataatct tcacgtacgt tgaaagtaaa ccatatgtca 24240
tgggtatccg taacggtcat gatagggggag ccggtgccga caagttcgcc gaccttaggg 24300
aaggtggcgg aaataatacc gtcggccgga gaagtcagat agagctcacc gagatagcct 24360
tcgacctcct ggagagcgcc catggctctt tctacaaggg cctcggctgc ttccttgtct 24420
tcgatacgcg cgccgttgac agccatgtca tactggctct ttgcggcttt tgcctgtgct 24480
accgaagcgt catacttggc ttttgtctcg tcatacttct gggcggagat cacgctctgg 24540
tcgtagagtt tctgcactct gtcgaaggat ttgcgcatga cctcttcctg aaccctggcc 24600
tgctgccaga gttcgtatgc gcccgtgatc tgctcctggc gtgctccgtt ggctgcctta 24660
gtcttctggg cgcgggctgc tgcgacggcg cctttagcct gtgcgagttt ggcgcggacc 24720
tcaggggaat cgatatatac tacggtatcg cccttgtgga cttcctgtcc ctcttttaca 24780
tagagttcct cgactctgcc cggcaccttg ccggatacgc ggtactcgct ggcctctacc 24840
tggccctgga ttacaagcgg cttcggtctg gatacgaaat aaccgattat cgagatgatc 24900
aggattatga tcagcaaagc cgcaacacct atcgtaatat tgtaggtttt ctgttctttt 24960
gtcatagttt atgtattttt attcattatc aatattgtag taaccttcag ccttgcggag 25020
attggcattg gccatctgca ggtcgattcc cgcatcgatg tactccgaat gggcctggag 25080
ccatgcggtc tgcgccgcga ggacggtatt ggtctcgatg accccggcct cgaagccgac 25140
tgtggccatc ctcatgttct cctcggcgct gctgagattg ttctcgctca tgagaagtct 25200
ctcgcgggcc tcctccagat gctggcggct ctgttcgacc tggagattga tcatgttcct 25260
ggcatcctcg agcttgtccc tgtagagggt cgcctctgcc ttggctttgc gggtccgctg 25320
gtaggcttca gtcccgtgga atatagggac attgaccatg actccggcgc tgaagagtcc 25380
tccgttccaa ctgttctcgt atccgtggaa ggcgttcgga ttggtcacgc tgtatccggc 25440
aacgagggcg atctgcggca tcatgtcagc acgcgcgatt ccgattttct tgtcgtagat 25500
cttggtggcg agttcgagac tgcgggtttc ggcgcggtct tcccagatat cctccatgct 25560
cttctcttct cccggaaccg ggaccggaat cgcgtcgagg gcctcgtcgg agagggttat 25620
ctgagagtcg agaggcaggc cgatctgctt gcacagaagc atctttgcga gttcgaggcc 25680
gttggtggct ttggtaagca tcatgtcggc ttcgttggcc tttaccttta tctggagttc 25740
gtcagacttc gtggcgactc cttcctggac agaaattgcc acgtccttct cgagctgatg 25800
cagcaggtcg gaatatgatt cggcgagttt ctttttgttg gctatcgata ctatctgcca 25860
gtaggcctgg tcgatatcga cgagaacgtc ggcatatttc atgtcatatt cggctctggc 25920
gagttcctca gccatctttg cgatctggtt ggaaaggact atctttcctc cgacgaatac 25980
cggctgcttg agggtgacgg ccccgatgaa gacgttctcg atgtcgagat gcatggattt 26040

gtcgatatcc tggccgatag cgttgatagg agtggcgatg tccggagcgg tggcgctgcc 26100
ggagccggat tccttaagca tgttcagcac cgtctgccac atcgggctgt tcatgtactc 26160
tgcggcaagt ctcgggtctg acattatggc cgtagagagt tgtgtgactg ctccgttgac 26220
tttgccggag attgcgccga cggcctggtc gagggctccc tgggcgctcg tgcccaagtt 26280
tgtcaacatg gcagactggc tgtcagtgac cagagacaag ttcctgttat tgtaaaggta 26340
cgctcccgtg gcgcttattt ccgggaagta atttgcacgg gcgatctttc tgtcgtaccc 26400
ggccatctcg accttagtcc ttgcctgatc gaggtctttg ttgctttcta tcgccatctg 26460
acggcattcc tccagcgtga gtacttgctg ggcattcatc tgaagaggta gaatgcatgt 26520
cgctgcgata atggggatga agattcttct catatttagt ttattctgtt tttccggatt 26580
gtttttttcct gcaaaaaagt tgccgcttta tgccctaaaa actgcctttt ctggctaaat 26640
attgttcaaa aggtcgaaaa ttgcgaagaa ttgatacttt tgcactttat ttgcaggaca 26700
gagggaggat aaacttggta aactatatag cctgattaca taaaatggaa gagactttgc 26760
agacaattga catcaaaggc tttaagtcgt tgtttccgat agagtatgac gctgaactcg 26820
gagacgattt cttcattgcc gacatcactt acgaccgggc attgagaatg ctccagtatc 26880
ctacccgctt cgacgggttc ctcgggatct tctgtgtgca cgggcatctc agcatggata 26940
taaacctcag cacctataag gtgaccgaga attccctgct gttcagcgtg ccgggctata 27000
taataagggt ttcggacgtc gggaccgatg acctgaagaa tctcaggttc atcgtgatcg 27060
ccgtctcccg cggttttatg tccagcatga gaatggactt caccaggctt ttcaacgaga 27120
gtatgagcgt cctgtctaat tcatgcatca ggcttgagcc ggatgagctt gccctttgca 27180
gcaagtacat gactctggcc atagaaatag tcaagaatgg agtgacctct tacaagaaag 27240
aaactatcgg atcgctcatt tcttcggtat gctacttcct cggcggaatg tgggctgaca 27300
agctgtccaa ggcgcgccgg gagagcaatc ctgacatctc cgccagggcc aagcttgtct 27360
tcgaccattt cctgcagctg gtgacagaca atcatacttc ccagaggaac atggcttttt 27420
atgctgacaa actgtgtctt actccgaagt atctctccaa gctcgtaaag aacgtaagcg 27480
gccgttcggc tccggaatgg atcgactcct ttgtcatcct tgaggccaag aacatgctga 27540
aatactccga gatttccatc aaagagatag tatacaagct caatttcccg aatcagagcg 27600
tcttttataa gttcttcaag gcccatacag gcctgactcc gagcgaatac cgtaacagtt 27660
agccatgcag aatcttaccg catccttcgc catccatgcc ctgaatacct tgggctacaa 27720
agtgtctcct cctatcggag aaaggacttt cggccagtgg tcgttctgct atctgatttc 27780
cggcgaaatg ttcatggaat cggcgccgga ttcttatctg atccggggcc atgagttcgt 27840
gctcgtccct ccgggagttc cgttccgcat cagctggtac cgggacagta tcggttatat 27900
gggaggattc gacgagacaa tcctaaagga tcgcagctac acctgtcttc aggccaggag 27960
gccgctccgg gtgtctgtaa gcgccgatga tgaaggcttc gtcgacgagc tcatgtacaa 28020
actgctccgg aagaaagacg atatgcccgt cgttctgggt tgcctggatg taatgctgtc 28080
gcagatcgat gctgataccc cggagagccg catgccggac aagccgcttg tgaatttgtt 28140
cctcgacagg atcttcgaca gggagaagga gttccttggg gttgccggat acgccgaaga 28200
actgaatgtc acgcctaatt atctcaatcg tcttgtccgc aaaagtaccg gccgcagtgc 28260
gaatgaatgg atagagattt cgcgcctgac ttatgcgaag ttgttgctcc ggcgaaagga 28320
actggccatc atcgatgtcg cggccatggt gggtcttcct gaccagtcct atttctcccg 28380
cttcttcaag aaaatggagg gcgtcactcc ttcagagttc cggaaagact tatagatctc 28440
tattgaaacg tttaggattg atttgtccta atctgcccgg ttcttgtcct aaaatagcta 28500
tacggattat cataattttg caggcgtaaa acaatatgcc atgcactatc ttgctataat 28560
aattacaatt cttggcggag cgtttgctgc tgaacctgag agtgaagtgt tcgatactct 28620
cgctgtttcg gtagtatctt cttccgcaaa actggtacaa cctctcgaac agttgtcttc 28680
accttcgaca tctgtcctct ataaagactt tgaaagtcat gggctgaata gtcctaaact 28740
tctgtcttct atcgttccga atcttcatat tcctgattac ggaacttcta tgacttcctc 28800
tatatatatg cgtggattcg gctccagaat cgacaatccg gtgctaggtc tttatataga 28860
tgacatcccg gtgcttgaca agaacaacta cgagttcgag ctgatggaca tccgtcgtgc 28920
cgatctcctc agaggtcctc agggcactct tttcggccgt aattccctta ccggagttct 28980
ttcaatcagt actctcgacc cttcagtcat cagtggaggc aaggctgtac tggaatatgg 29040
ctccgccaac accatgcttg caaaagggtc atggtatggt gccggtctcg gcgtctctgc 29100
cgcctacagg cacagcgatg gcttttatgt caatgactac gatggcagcg atatagataa 29160
atatgattct ttctcgctga ggctgactca tgggaaagat ctgaatccgc gactccatat 29220
ggacaatact ctttcggcat cttggctggt gcagggcggc tatgcgtacc gccgtctggt 29280

ggagtcggga gagggtgatc cggcttatgg cggagtgctg cttccggtca attacaacgg 29340
cgacagcggt taccggaggg tttctgtcct cggcggaacg aagtttcgtt atgagcatga 29400
ccgctaccag ctgcgaagcg tgaccagtat ccagctgctc ttcgaccgta tgaatatgga 29460
ccaggatttc actccgcagt ctgtgtttac tctgcagcag aagcagaagc aggcagcagt 29520
cactcaggag tttatcttcc gtccaaagga ggatgtcgaa tggtgggaca gccagaccgg 29580
agtttttgct ttctatcgct acaattatat ggatgcacct gtcaatttcc tgtatgacgg 29640
agtcaatact cttattctgg gaaatgcgaa taaaggtata ccggatccag attacctgga 29700
tttcctggag gatgaatttt tgatagctac tgattttaaa atacacactt acaacacagc 29760
gttctatcac gagtcatatt ttactttcgg aaactggcat ttcactgcag gattacgtct 29820
ggatcacgaa ggtaatttta tgagatattg ctccgggacc gagttgactt ttatctactc 29880
cccgattatg aaggcttacg aacctttcga gacagtgtat aaggggttgg aaaggaattt 29940
ctattttgaa ctgcttccta aattatcggc cttatatgat tttggcagtc tgaatgtgtt 30000
tgcaactgtc tcgaagggtt ataagtctgg cggattcaac acccagattt tttccgatat 30060
cctgcagaac aagttgatgg aagggatgat gaatcatatc ggagtgcatt cagtggacag 30120
cggtgaggct cctgaggcgt ctgcgacttg ctataagcct gagtcatgtt tcaactatga 30180
agccggtctg cgttattctc atgtttcgga atccggccac cgtatctctg catcggcatc 30240
ggctttctat atcgattgcc ataaccagca gatcaccatc tttcctccgg gaaagaatac 30300
tggacgccgt atggcaaatg tgggcaaatc ccgcagttac ggagctgaag cggagttctc 30360
ttattcatac aagaatttca gtgtcaatgg ttcgtatggc tatgtgaatg caaagtttac 30420
cgagtattcc gacggaaaaa atgattattc cggaaaccgt atcccttatt ctccggagca 30480
gacagctttc atgcgtgctt cctatactct tcctttaagc ggaggtatcc tggatagcat 30540
cagtttctcg gccgatgcca attgcacggg cagaatctgg tggaacgaag ataactctta 30600
tgtcgagccg gtccatgtcc ttttcggtgg tgatatcacc ctggatttcg gaaagtttga 30660
cttatatgtc cgcggacgta acctttcaaa ccgcgattat aatactttttt atttcaagtc 30720
gatggacaac agctttttcc agaaggcgaa acctgtcagc tatacggctg gaatcagatt 30780
tatttttttaa tatatataat agctattatg aagaaattag ttttcgggct tatgattgcc 30840
ctcacagtgt tcgctgcatg cagcaaagat gatgacaaca aggaagtaac tgtagaggac 30900
ggttcttacg ttggtaccgt caccgttgtc gagcctcatt tttccgcttc tgatatcaag 30960
gttagcgtta ctccaataga gggaacaacg actctggata tcaagatgtt tgaagttaga 31020
ttcagcgaga gaatgcctaa aaccatagat tttaccatac ctaaggcttc ttataccatc 31080
gatggtgata ccatcaacat cttcggtgaa aacattattc ctacaatggg cgccggatat 31140
gagaaacctg agtatactgt taccggactc aaaggaacaa tcaaggatgg aaatatagaa 31200
ttctccctca acttcggctc tttccctacc aaatattcag gaacgctcgc caagaagtaa 31260
cgatttaata acgaacttat cggcatttaa agcgcattta gttgctattt cgaaaaattt 31320
tagcaaaaaa aattaaaaat tgttttgtta aaatgattta aagaattata tttgcaccag 31380
acattcgaaa agattttttc atgtctattt gttaagttcg ttttatatac gggatcaggc 31440
tttggggaaa gcctgatccc ttcttttttt tcacctctgc agctttgaat atgcaaaata 31500
acctggtcga tattgaccag gttactttgc gtaaatcgtt gatgacaagc agaatccctt 31560
tcacgcttga aaccggcggc tatttgatat aatccccaag cctgagaaac agatcagagt 31620
gaccgcgctc aacgattgtg ccatcaggag cgaagataat cagggttggg acccccttcaa 31680
tagcatagat gtcacgggct tccagacacc tgtgtattac gggccatgtg atgccataag 31740
tctcgacaaa cgactgggtc gtttctaaag tctcgtccca taccattatg gtaagaaatt 31800
cgactttatc ttttaacctc tcgtagtttg gtactaaatc gtccttggtc atttgccggc 31860
aagggccgca tccggcgccg cagaaatccg cgagcacata ttttcctttt cctacatagt 31920
cagacagcct tgcctttgat tccgggttgt caggatcctt gatgccttct atatcggaat 31980
acatctttcc tggagcggtg ttgagttttg ccggcatgtt tttaagtttc aatccgatcc 32040
tcgtgtcttt tttcatatct tcggaagcca tatccaggta tgtctgaagc tcctcaagag 32100
agacaaggtt ctcaagccag tatattattt cgtcgcatat tacgttatct gtgttctcgg 32160
tataggcctc tcgtaacata tccagattca agctgtcaca tatttccttg aaccgatgca 32220
cagattcggt atctcttttc cagatttcca gggcctcatc aagataaaca tgatttatat 32280
aataattcgt gtacttgacg tagcggctgt gtattccttc cggatcggag gaaatggccg 32340
tgcaactgtc gacatctatc atgatggtcg atccgtccga gacaaaatca acttgtttcg 32400
cccaagatag atccctgacg gatgccattc tgagcgcgca taccggaatc gtggtctcaa 32460
agcttccatt atttacacga accgccttcg actggcgttg tagttcagac tcattccaat 32520

cgtaatagta tactgcgacc agggagtctg cagggttgat tgtgcccttt atagtggtga 32580
cttttggagt gttgttgcac gctgtcataa gcaaaagcaa cataagaata gaacgataca 32640
tggttttagt gtttattatt gtatggcaat gataggataa aaaattatgt cccggatggc 32700
aattcacgta tttccgtgct actttttacg aaaaaaaagc cgactcgatt tcgagccggc 32760
cttaggtgtg ctatatagtc ccggttagag ctcccaggcg agggcggagg cgccgaggat 32820
ggcggcgtcg gactccttga gctgggagaa cacgagcttt accttgtttc tccaaagagg 32880
aagcacattg gcgttcatgc tcctttcgat aggctcgcgg aggaaatcct ttgcccttgc 32940
gagaccgccg aaaagcacaa ttgcctcagg cgcagagaac tcgatgaaat cggcaaactt 33000
ctgtcccagg atatttccgg tatattcgaa aatcttgagg gcaagcttgt caccatcctt 33060
cgcagcctca tatacatcct tcgaagaaac cttgtcaaga ctacgcagcg tagaaggctc 33120
atcagtcatg tcaagccact cgcgtgccgt acgagcgaca ccagtggcag aacagtaagt 33180
ctccagacag ccggtctttc cgcagttgca gagacggccg ttgttgcgga ctgcgcaagt 33240
atgtccaagc tcgccggcga aaccgtcgtg gccatagaca atctttccgt caatgacaat 33300
gcctgaacct acgcccgtgc cgagagtgat catgatgaaa ttcttcatgc ccctcgcggc 33360
gccgtatgtc atctctccca ttgctgccgc gttggcatca ttggtaagag aaacaggaat 33420
cccgttcatc gcgtcggaaa gcaacttcga gaaagcgatc ttcttggtgc caccccatgt 33480
aaggttgaca gcattctcga tctctcccgt ataatagttg ccgttaggag ctccgacgcc 33540
gataccgcgg ataagtccct cggccttgga ttcagcgatg atatgctgga gagcttcggc 33600
gagctcggcg atataaggtt cgatttcagt atatgtgtcg gtacggatga cggtttgagc 33660
cagcactgta ccacgggcat caatgacgcc gcatttggtg gtctgtccgc cgacatcaat 33720
tccgattaca agttcttgtg ccataatgat aatagagttt tttgtgtagt ttatgcttta 33780
actctcttgg ccggctttga gccgataaga ccgaagaaga ggatatatgc aagtccgatg 33840
gcaggaagga tatagctgaa catataggtg ctgtggtcag ctatgaagtt ctgcaggagc 33900
ggaatgatac caccgccgac aaccatcatc atgaacaggc cggaagcctt gttggtcgca 33960
gagccgagac cgtcgacaga aagtgcgaag attgctcccc acatgactga agtacagaga 34020
ccacaaagta ccagaagaag ggctgagagc ggaacgttca ccatctcgaa gtttgaacct 34080
gtgaatgcag gcatagaagc ccttacctca ggagaaacta ccatagcgcc gattacgagg 34140
cagagagcta ctgcagaaac tgcggtgatc tgggccttgg cagaaacctt gccgctgata 34200
aggctggaaa ggaaacggcc gataagcatc aggaaccagt aggtacctgc aacgaaacct 34260
gcgacagtag cggcgttggc acccatctgg gcaggattct caggatcagc aaggtagaag 34320
ttgagggttc cagggatgcc gacctcgaca cccacgtata cgaagatggc gattacgcca 34380
tagatgaaat gcttgtaatt ccaagggctc tcggcattgg ctgcatcagc gacctcctca 34440
ggatcttcga tagggatgaa ggagagggcg atgaaggctg caaggaacac acccatggcg 34500
atgtacagaa ccggattgac atccttgatc actgtctggg ctgtaagggt tccgataaga 34560
gcgccgacga gcataggggt aagggtaccc atgagagagt tgaaagtacc tccgatctgg 34620
acatactggt tgccgtgctt ctcaccaccg ataagagtaa gcatagggtt tactacggtg 34680
ttgagcatac ataccgagaa tccggaaatg aatgcgccga gaagatagca agcaaagttg 34740
gcagggaagg atgcgacgac gccggaaagg aactggatga agactccgat gaaaccgacg 34800
gcgatagcga taagcgctgt cttcttgtag ccgtatttgg taagcatctt tcctgcaggg 34860
acacccatga agaggtatgc gaggaagttc atcatgttac ccatcattcc gagggcgttg 34920
gatccagcaa tctgaggctg catcttccag atgttgccga ttggagctgc gagattagtt 34980
acgaaggaga tcattccgaa gaggaagatc atagtaacga ttgcgataat gttaccgttt 35040
ttctttgcca tttgtctatc tgtattagta attagtttca gcatgtacgc agatttaccc 35100
tgcggagtca caaatttaag aaaaataaat ggaaagccac tattatacgg taaattttta 35160
ctcgactgaa aactcggctg cttcgcctga tgcgctgtcc ccggcaatcc atagccggaa 35220
tgccccgggc tccactactt tttccatatc catattccag aaggcaagct ccctgaccgg 35280
aagttcgaag gtgatttccc tgctctctcc cacctccaga gcgattctct cgaaatattt 35340
cagctccttg accggacggg ttacagaccc gaccagatct ctgatataaa gctgggcgac 35400
ctcgacgccg ttatatttcc cggtattgga aagggtaaac gtagcgcgga cagtatcttc 35460
ggccgaatat accttcttgt caagagatat tccggaataa gagaatgtag tatagctcag 35520
tccatatccg aacgggaaca gcggagccgg accatagtcc agataatatg aagtattgcc 35580
aagagaagtc tggccggcct ccagaggaat ctggtccagc aggagctctc cgttatacgg 35640
acgtccggta ttgttatggc tataatacat cggcgcctgt cctactgtac gcaggaatgt 35700
gaccggagtc tttccgctcg gcacggcatc cccgaaaaga agattgacgg ttgcgggacc 35760

```
     gcccatggtt cctggatgga actcataaag catggcatcg caattgacga gatccttctc     35820
     gatcgtaaga ggccttccgg ccatgatcgt ggcaactatc ggcttgcagg aactcttcag     35880
     tgcggcgaga agagcgctct gcgagccctt caggtcaagg tctgccagag aatgggcctc     35940
     tccggaaagg atggcctctt ctcccaggaa cgccagcacg acatccgcat tgcgggcggc     36000
     gctgacgaca tcggcgaaac tgtctttctc ctgacggctg tattccaggc cgggaacata     36060
     tacaatcctg gccttccctc cgaatctctc cctgagagcc gcgagcggag tgacggtatg     36120
     agtcttgtcg ccgtcgaagg cccatgtgcc cagctggtca tgaggggcat cggccatagg     36180
     gcctgtcacc agaatcgtcc tgatgcggga agcgtcgagc ggcagcacgc catcgttctt     36240
     cagcagaatc gcagcctcct cggccgtccg tctggcggca tcgaggttct cctcgctata     36300
     gaggactttc ccggccctag cgggatcgac atatggattc tcgaacagtc cgagcagaaa     36360
     tttcactctc agaatgttgc gcacggcggc gtcgatctgc ttctcgctga cctttccctc     36420
     cttgaccaga gtctccaggt tggcgatgaa tccgtaggtc atcatgtcca tgtcaacgcc     36480
     ggcgccgatg gccagcgcgg cagaattctt atggtcggcc gcgaatccat ggaccatcat     36540
     ctctcccatc gaattccagt ccgtgactac gaatccgtcg tagccccatt cgtcccgaag     36600
     gatgtcgttg acgatgaaac ggttgccggt cgaagggacg ccgtcattgt cgttgaatga     36660
     tgtcatcaga gtcatagcac cggctttgac tccggcctcg aacggaggaa gatatacgtt     36720
     gcggagctgg cgttcggtaa gataggtgct gttgtagtcg cgtccgcctt ccgccgcgcc     36780
     atagccgacg aaatgcttga tgcaggcggc cattgacgtg gaatctcccg gttcgccctg     36840
     atagcctctg accagcgctt cacccatgcg cgcgtcgaga tatggatcct cgccggaacc     36900
     ttcggcgatg cggccccatc ttgggtcgcg ggcgatgtcg agcatcgggg agaatgtcca     36960
     tctgactccg accgaagtgg cctcgatggc ggcgacgcgg gccccttcct cagcgagctg     37020
     cggatcgaag gtcgccgcga tgcccagtgg aatagggaag actgtattga atccatggat     37080
     cacgtctctg gcgaccagaa gcggaatccc gaggcgggat tcttcgaccg cgatgcgctg     37140
     gagttcattg actctgcccg gatcggtctc gttgagtatg gatccgacct gaccgttgcg     37200
     tacggcatcg gagaaaggtt cgaccgagcc tccggctgag atctggttca tctggccgat     37260
     cttctcctgg agagtcatct gcgacaggag cttatctatg cgcttttcga tttccggggt     37320
     cgacccgact gtcgaggcgg aaggcccgga acaggatatt gttatcgcta tcgccacgaa     37380
     aggcaacagc aggtttatgt tatgtctcat ctctatatgt tttatttctg atataccctg     37440
     acgtagtcga ttttcatggt ggtcgggaga gcactctcgt ctattccttt gtagccaccc     37500
     cagtctccgc cccaagccag gttcaggata atgaaaaatt ctttgtcgaa cggccatgtc     37560
     gatacgttgc cggcgccgtc attcctgaag gtcaggtgca gcttcccgtc aacatagaac     37620
     tccagagctt cggccgtcca tctgcatgaa tagatatggt attcattttc tgcgccttcg     37680
     gtaagctgct cgtgggtctt ctgggtattc ttgacatggt tgtagctttc tgtatggatc     37740
     gatgaggaag tatattccgg atttgcgccg acctcttcca ggatatcgat ctcgccgcat     37800
     ttcggccatc cgagactctg gtctgtcggc atcatccaga atgccggcca tgtgcccagg     37860
     cctttcggga gccagatcct ggcttccata taaccgtatt tccatgcggc cttgctgttc     37920
     attcttgcgg aaatcacttc tccgttgtgt ttcatcgcgc gtatgttcag cgcaccgtct     37980
     tctacgaagg cggtcttctt tccgtcgcgt tctcctcctg cgacatatct ctggagctca     38040
     tggtttactc ttcccggggc ccagtcttcg aacctccagt tgtccggatc gactttttca     38100
     ccatcgaatt cgtcgtgcca tgccagctgg tagccttcag gaacgaatgc accgtccacg     38160
     aacggttcgc ccttggcctg gattacctcc agctcgactt tgctgctgcc tgaggtgaat     38220
     atcacctttg ccgtcctcgg ctcttccgag tagttcttga ggaccgtaac ggtaaactcc     38280
     tggctgccgg atctgatcgc tcttgcggag atcttgagcc agaaggatcc gtccgtctct     38340
     gccgtccagg tcgtattggc ttcagtggtc actgatgcgc tgccggcttc tccgctgagt     38400
     tccaatctgg tggaagaaag ggtcagttcc ggtacgaggc atggctgtat aacctgcagt     38460
     tccttctttt cctttccgga ggtgaagctg actgtggcct cgcgggttgt tcccgtaagg     38520
     ttttcgccgg cggagaggtt cagtatgctt tcgcctttgt aaagttcttt tgcggatagg     38580
     gtgagccagg actggtctgt ctttatcgac caggccgtgt ttac                      38624
//
ID   XXX; XXX; linear; XXX; XXX; XXX;  XXX.
XX
AC   PRJEB67889
XX
DE   Uncultured bacterium DNA fragment from in-reactor enriched cow rumen, clone
```

```
ferm-41j20,contig1
XX
FT   source          1..23867
FT                   /organism="uncultured bacterium"
FT                   /environmental_sample
FT                   /mol_type="genomic DNA"
FT                   /isolation_source="Cow rumen"
FT                   /clone="ferm-41j20"
FT                   /note="AZO-galactomannan active clone"
FT   CDS             25..2022
FT                   /db_xref="SEED:fig|6666666.127253.peg.1"
FT                   /translation="MSFTACDNGDEEDTNQYVGGIQLNVFGPSPVARGGELRFLGSGMN
FT                   KVTGVVIPGCDEITDIQVISDNEIRVTVPQIAEPGLVVLHIPNGEIVTKTKLTYTEPVG
FT                   LDAITPKSIKAGQELTLTGEYLNLVKEVIFADNVVVKSAAFTKHERKEIKLIVPNEAQT
FT                   GKIIISDGAEIPNWIYSAESLTVVLPSVSKTLDLTSKKPGDVITIEGEDLDLVKQILMP
FT                   NEKPVDFTIVKMDIAATKSTAGAKYKLNFVLPEDASDGAVVMVAHSGVKVAIANIGMAV
FT                   PTSLVATPAADIKGGDVISIKGVNMELVTSVTFPGVTDKVTPNSKTATEIKVTMPDAAI
FT                   SGNLVLHTASGYSASVAISTLKPEVMAYNPSPVAAGNDVVLQGKHLDLVSAVTFAGDKK
FT                   VSVSPTTSTELKVTVPVDAESGEVILTMKNGETVKCATLEVSKPVFCYIPVLPDASQEI
FT                   HAGKILAISVQNGDKLTNVQVNGATTQYIFQGGTLNVLIPSNAGGETALKLISSNGEVT
FT                   YTIKVIASGVTETVLFTGPLLITWSDGGRVILPATAFTGLTAGAILKIYFTQNDNWGQI
FT                   QINNGNWSTISFAELGNGGYITTNTYNDKTITSQELVLTTDVLSIINSNASNGNAIIMQ
FT                   GSDFVIDKVSVITKGGVSAKARLYKIKNLK"
FT                   /product="FIG00410329: hypothetical protein"
FT   CDS             2244..3524
FT                   /db_xref="SEED:fig|6666666.127253.peg.2"
FT                   /translation="MGSTPTNGATDVSASNISIVLTFNQNVTAPTTGHDKITLSGATIT
FT                   SVVAYLTKVTIEVTGLEKGKTYTLRIPKGVILGPSKVEAPEISISFTTTSPQQITNKLC
FT                   TANPLTQAQNVYDFLIANYGTKIISGSMASDSWNVNEAEWVFKHTGKYPALNGFDYGHL
FT                   YASPANWIDYGNTTIIENWWNNHGLVAATWHWNVPKSSGSTEYGFYYTGKNSGKGETSF
FT                   DISKAVQDGTSENQIVKADLAKIADYLLLLKAKNIPVIWRPLHEAAGGWFWWGAKGATS
FT                   FKALWKLMFETFQSKGVNNLIWVWTSETGDEDWYPGDAYVDIVGRDVYNKILASQSVEE
FT                   FRKLSTAYSHKLITLSECGNVSKISLQWSASANWSWFMPWYDYSRTNKMDDPAFNETSH
FT                   QFANIEWWNDALNTEAVLSRDEMPDLR"
FT                   /product="GH26"
FT                   /EC_number="3.2.1.78"
FT   CDS             3635..5917
FT                   /db_xref="SEED:fig|6666666.127253.peg.3"
FT                   /db_xref="GO:0008422"
FT                   /translation="MTMAIAATVGSGSAEKKEMYKDPSVPVKDRVEDLLSRMTLEEKVG
FT                   QMNQFVGIEHIKANSAVMTADELKNNTANAFYPGVTDKDVERWTAQGLIGSFLHVLTIE
FT                   EANHLQSLAMKSRLQIPIIFGIDAIHGNANAPDNTVYPTNINLACSFDTAMAYKIARQT
FT                   AKEMRAMNMHWTFNPNVEVARDARWGRVGETYGEDPYLVTLMGVESVKGYQGSLNGKED
FT                   VLACIKHFVGGSEPINGTNGSPTDLSERTLREVFFPPFKAGVEAGAMSLMTAHNELNGI
FT                   PCHSNEWLMQDVLRGEWKFPGFVVSDWMDIEHIHDLHATAENIKEAFYQSIMAGMDMHM
FT                   HGIHWNEMVVELVKEGRIPESRIDESVHRILDIKFRLGLFEQPFADTSKSMKVRLSKEH
FT                   RETALEAARNGIVLLKNEGVLPLDASKYKKVLVTGINANDMNVLGDWSAVQKEENVTTI
FT                   LEGLRQVAPGTAFDFVDQGWDPRNMDPAKVAEAEKRAKESDLNIVVAGEYMMRFRWTDR
FT                   TDGEDTDRSDLDLVGLQNELIRKVAASGKPTILILVNGRPLGVEWAAQHLPAIVEAWAP
FT                   GMYAGQAVAEILYGKVNPSAKLAITIPRSVGQLQMIYNHKPSQYFHPYAVKPSTPLWPF
FT                   GYGLSYTTYKYDDLKLSAKEVSQNQNVEVSVKVTNTGQRDGVEIVQLYIRDKYSSVTRP
FT                   VKELKDFARVALKAGESKVVTFTITPDKLAFYDKKMKWIVEPGEFVVMAGASSADESLL
FT                   KETFILK"
FT                   /product="GH3"
```

```
FT   CDS             6010..7044
FT                   /db_xref="SEED:fig|6666666.127253.peg.4"
FT                   /translation="MKMQSSERTVETKNLLINLGTIHQKGFMFGHHDDPVYGIGWEGDA
FT                   DRSDVKSVCGDYPAVMSFDLGRIELGGDKNLDKVPFDKIRREILAQYARGGMVSLSWHV
FT                   DNPLTGKDSWDVSDTTVVSSVLSGGANHQKFLGWLDKVADFMNSLTTDKGVKVPVLFRP
FT                   WHEHSGSWFWWGQNLCTAAQYKALWKMTYDRMQEKGVNNLLYAYSPGTEPQTVEEYLER
FT                   YPGNDIIDLVGFDTYQFDRQKYIDQLNKSLTILTEVGKKQGKPIAITETGFETIPDATW
FT                   WTETLLPVISQYPISYVLVWRNAREKDNHYYAPYPGQVSAPDFVKFYQNSKTLFARDIK
FT                   KLYD"
FT                   /product="GH26"
FT   CDS             7077..8267
FT                   /db_xref="SEED:fig|6666666.127253.peg.5"
FT                   /translation="MSLFKDKVAKLLTDHEAFLSQKNEPLAVDNGVVTRYKNPVLTAEH
FT                   TPVFWRYDLNETTNPYLMERIGMNATMNSGAIKWNGKYVLMVRVEGSDRKSFFAVAESP
FT                   NGIDNFRFWDYPVQMPEDLIPATNIYDMRLTAHEDGWVYGVFCAERHDDNAPAGDLSSA
FT                   TATAAIARTKDLIHWERLPDLKTKSQQRNVVLHPEFVDGKYALYTRPQDGFIDAGSGGG
FT                   IGWALIDDITHAVVKEEKIIDPRYYHTIKEVKNGEGPHPIKTPKGWLHLAHGVRGCAAG
FT                   LRYVLYMYMTALEDPARLIASPAGYFLAPQGEERIGDVSNVLFTNGWIADEDGTVFIYY
FT                   ASSDTRMHVATSTVDKLVDYCLHTPEDGLSSSASVATLSKLIEKNLALLNQFSLKK"
FT                   /product="GH130"
FT   CDS             8287..9720
FT                   /db_xref="SEED:fig|6666666.127253.peg.6"
FT                   /translation="MNKMGNTASKIVADKERISLKEKIGYGFGDMASSMFWKLFGSYLM
FT                   IFYTDVFGLPAAVVGTMFLITRVWDSAFDPIVGVVADRTHSRWGKFRPYLLYLAVPFAL
FT                   IGILTFTTPALSDTGKLIYAYVTYSLMMMVYSAINVPYASLLGVMSPDPKERNTLSTYR
FT                   MTFAYIGSFIALLLFMPMANLFAGSEADAVHEQRGWMLAVVVIAVMCAVLFYFCFALTR
FT                   ERVKPIKEKQSPLKEDLKNLLGNGPWWILLGAGVAALVFNSIRDGATVYYFKYFVVEAD
FT                   YATVSLFGVSFVLSGLYLAVGQAANIVGVILAAPLSNRIGKKMTYMGAMVIATVLSIIF
FT                   YWFDKDDIALIFVFQILISVCAGSIFPLLWSMYADCADYSELRTGNRATGLIFSSSSMS
FT                   QKFGWAIGSALTGWLLAYFGFRANEIQSEEAIHGIKMFLSFLPAVGTVLSVIFISMYPL
FT                   SEKKMKVITEELEAKRKQQ"
FT                   /product="Cation symporter"
FT   CDS             9717..10907
FT                   /db_xref="SEED:fig|6666666.127253.peg.7"
FT                   /translation="MNTKITHLKGEVTEVLTTNILPYWINKMVDTTYGGFYGRIDGTEV
FT                   CRPEAEKGAILHARMLWTFSAAYRLLKKEEYLDMATRAKRYMIDSFYDKTYGGIYWSLT
FT                   HQGDPLDTKKQIYAIGFAIYGLSEYYRATADAEALTYAIHLFESIETHSFDKIKNGYCE
FT                   ALTREWGEIADMRLSEKDANERKTMNTHLHILEPYTNLYRVWKDERLKNQLRNLIFLFT
FT                   DKILNKETHHLELFFDDDWNSKYSIVSYGHDIEASWLIHEAALELKDPAVLQYVEPLVQ
FT                   RIAEAATEGFTPEGGMIYERNLTNGHVDADRHWWVQAETVVGYINLYQHFGDEASLDKA
FT                   LACWEFTKSHLIDRVNGEWFWSLRADGTINHTDDKAGFWKCPYHNGRMCMEIMERF"
FT                   /product="N-acylglucosamine 2-epimerase (EC 5.1.3.8)"
FT                   /EC_number="5.1.3.8"
FT   CDS             complement(10914..11399)
FT                   /db_xref="SEED:fig|6666666.127253.peg.8"
FT                   /translation="MAAYIFEDVIKEIEQIIEKEELFLNGLDEETISTRLNSQNRSIKQ
FT                   IVGHMIDSASNNLHRVVHLHYGANPLVFPNYATNGNNDRWIGIQHFQEEDWQTLVQLWK
FT                   YSNLHFVHVIKHVDAHKLNNKWIAGEGEEVSLQDMIRDYPRHLELHMGEVHALHELR"
FT                   /product="hypothetical protein"
FT   CDS             complement(11507..12526)
FT                   /db_xref="SEED:fig|6666666.127253.peg.9"
FT                   /db_xref="GO:0005506"
FT                   /db_xref="GO:0006118"
FT                   /db_xref="GO:0009055"
```

```
FT                   /translation="MEKSLCIYYFSGTGNARNVAQIIEREMWRHNVAVQINDISKTDGL
FT                   HIKSPQSHELLVLVSPVHGFNYPPIMLHYLLRFPKGNNEVMLMNNTRAGMRIGNWVTPGI
FT                   SGISLYLSALILRWKGYRIQSLYPVDFPSNWISLHPALNKKTVDFIYHQVTPKIEKAAR
FT                   GFLNGKREFTCVRECIIDLLVAPIAILYYFIGRFVIAKTFYASQDCTKCGLCERNCAIH
FT                   AIKTVNGKPFWTFRCESCMNCMSNCPQKAIETAHGFIIVIAIAFSILVSSVFDRYLPVI
FT                   TDIKWLKFLLENMLFIAYIAVFYRLLHFMLRFRWFERMVVFTSLTKLSFWGRYKAPKQ"
FT                   /product="Ferredoxin"
FT   CDS             complement(12538..13347)
FT                   /db_xref="SEED:fig|6666666.127253.peg.10"
FT                   /translation="MNVLPSLLLAPYVKYYWVVKADETTAIQTVPSGCIHLVFHRGGSM
FT                   YFSDGEQQPQSFIRGQLSGPGVLQSKGGIDMVAVIFHPLGFNVFFSLPLQLIYNQYVDV
FT                   DSMEDAGLKELKGNILHEEDSLACIQLIEHFLLSRITDWHDYNFKRMFHSVEVISRQAE
FT                   IDVASLASQTCLGVRHFNREFSKYIGTSPKEYLRIVRFQRLLHLLQNNPGTELTELAYG
FT                   CGFCDSSHLVKEFRSLTGCTPTEYLQSRKPHSTFFSKGSRLNIIKKI"
FT                   /product="Transcriptional regulator, AraC family"
FT   CDS             complement(13351..13908)
FT                   /db_xref="SEED:fig|6666666.127253.peg.11"
FT                   /db_xref="GO:0008908"
FT                   /translation="MVSMNKEALLIIDMQNDYFPGGANELVGAEKASQNAKAILERFRG
FT                   CNKPVVHIQHVSTRPEATFFIPNTHGVEIHPNVSPGESETVVTKGFPNSFINTPLLEKL
FT                   KEKGVDSLVIVGMMTHMCVDATTRAAKDLGFTCSVIGDACATKDLFIHDQSVKAKDVQN
FT                   AFLAALSYFYAAVVTTEEYLNL"
FT                   /product="Isochorismatase (EC 3.3.2.1)"
FT                   /EC_number="3.3.2.1"
FT   CDS             complement(14921..16189)
FT                   /db_xref="SEED:fig|6666666.127253.peg.12"
FT                   /translation="MVIKKITYSILYKEVDYNNLIIIMEIQNKILDSLFDDIDRELKLA
FT                   DFLGSANFKRIMNQMKAYEAYCHIYNEVESNSKVIMFGMDQLYRVDAFKILLDRLYSSS
FT                   KNDFYGVLASILSVYEVWSKTKININEIMLDLELLPEFNEEKKKWFVGIGTMTENEGDI
FT                   EDVENNRELEINIENVGYLETEYRELRDRKGDYTNSQDTINAYHDWFNLSITLFSRYFD
FT                   DTNSDYQKFKSVDNSGNGHKLSCNFNQIQGVYSILCDKIKNKISENSKINKIENKKIFI
FT                   VHGHDNTMKLEVDSFLKQLEFETIILNNQANNGNTIIEKLEENSNVGFAIVLLSPCDEG
FT                   RRKGTNELRSRARQNVILELGYFIGLLGRTRVCTLKKTDVEEPSDFTGIVYTDYDNAGG
FT                   WKFLLVRELRSAGYNLDMNKIGV"
FT                   /product="Predicted nucleotide-binding protein"
FT   CDS             complement(16203..16391)
FT                   /db_xref="SEED:fig|6666666.127253.peg.13"
FT                   /translation="MQFNGLIFYLFGWQQVGWKQFVCSYQTNSFVLGNSESHPHETGET
FT                   LNGPEIVTFNIFILNNL"
FT                   /product="hypothetical protein"
FT   CDS             complement(16741..17868)
FT                   /db_xref="SEED:fig|6666666.127253.peg.14"
FT                   /translation="MKKWTYAILALTLTAGLAACNGSNDETDNNIVFGDNQIGNGEQEF
FT                   IITHNQTIKKGVYLLKGWCYVADGATLVIEAGTVIKGDSETRAALIVEPGGQIIARGTA
FT                   ELPIVFTSEKAPGQRKPGDWGGLILCGNARNNQGIMQVEGGPRTMHGGIKDTDNSGILS
FT                   YVRIEFAGSPLRTNQEINGLTLGSVGSNTQIDHIQVSYSNDDAFEWFGGTVSPNYLVAY
FT                   HTWDDDFDADNGYRGTATNLLGIRHPRIADASTSHGFECSSSSNAEPQTAPTFSNVNLY
FT                   GPSSTDAEFENTTTYINGAGLLPDNQSLLGLYGAAIALSSNTAVNFKNANINGYSLNWE
FT                   GTAATQENVTFGTATVSFPTWTNGWCNFDPQNTVY"
FT                   /product="FIG00896557: hypothetical protein"
FT   CDS             complement(17980..20352)
FT                   /db_xref="SEED:fig|6666666.127253.peg.15"
FT                   /translation="MKMTNWKLATLAMAATLFAFTSCSDNDEPNNGGGVDAGSTLTGEL
FT                   TKDLTLKSGNSYYLSGGYHVKSGTTLTIQPGVNIIAKDDDIVDYILIEQGAKIDAQGTA
```

```
FT                   SSPIIMTSEKKEAGAWGGLHICGYAHTNAGTGAKSEIGDAAYGGSVENDNSGTLRYIRL
FT                   EYTGYAFSEEKEANGVSFYGVGNGTTVEYLQAYKGSDDGFEFFGGSVNVKYLVATDCSD
FT                   DSFDWTEGWNGYGQFLVAYQGDKATIGYDCDCLIEADNNGKNAAATPVSAPVLANMTLV
FT                   GNSSTANKRGVRLRAGTQAKIYNTLVKGKTSCLTTETVETETALVNGTSVLDYVYLESG
FT                   VGTSGNGYTEALFTGNTNNGISKIIPLTSNYIGTIDGGKDMSTVNSFFTKATYKGAVSS
FT                   SNDWTAGWTLASNAVQQSETLEGSLTSGTKTLTANKVYYLTGGYTVKSGATLKIEAGVT
FT                   VIAKNDNIVDYILIEQGGKIDAQGTASSPIVMTSEKKEPGAWGGIHICGYAHTNLGATS
FT                   KSEIGDATYGGTNDADNSGTLRYIRLEYTGYAFSEEKEANGVSFYGVGNGTTVEYLQAY
FT                   KGSDDGFEFFGGSVNVKYLVATSCSDDSFDWTEGWNGKGQFLVAYQQPSTTLGYECDCL
FT                   IEADNLGSNAAATPISQPTLANLTLIGNGSTANTRGIRLRAGTYAKIYNALVTGKTNNL
FT                   TTETKETEEALVGGTKSILNYIHVANDIKVSGLGGYTSALFAAATGNAINQTLTFTDTF
FT                   FGTVDGGADLSADSFFTKATYKGAISTANNWTAGWIR"
FT                   /product="FIG00403577: hypothetical protein"
FT   CDS             complement(20404..23046)
FT                   /db_xref="SEED:fig|6666666.127253.peg.16"
FT                   /translation="MAENIKGIIRDKSSKEPLTGATVQIVGTNIGVIADLDGNYNLAGL
FT                   KNGTYNLEIKYVSYKTIQIRNIKVSGSTPTVLNFEMEPESQVLGDVTVVARKNLEGEKA
FT                   LLQERQKATLAIENIGAKEMSLKGVSSVADGVKKITGVSLASAGQLIVRGLGDRYSTTT
FT                   LNGLPIASPNPDNKLIPLDIFPTSTVKNITVSKVYEAGAFADYSGAHVDIGTRENTGED
FT                   FFSISINAGGKINTLFKDFYSSDRKGSLFSTGNLNKSLLTMNKSEFEDSYVKNNDPFGT
FT                   SFSISKNTSLPVIGGSFGVGKSWKLNNGQKLSLLASVGISNDSQILKDAYVTTLTAQGT
FT                   NLNEFSYDKYTSELKMAGLASVGYTFRKADRINYTLFYARNAVDNYMRREGYDSEGVNL
FT                   IGSNSVFHAYTLLNNQLLGHHEFGKQWDLNWSASYGETGSDEPDRRQVMFRKDDAGALS
FT                   LFKLNKQETMRYYGELDEKEVVGDLRLNYKFGESNKIKVGGTYKDKDRDFTSARFYYNL
FT                   NAINPTITDIYDTDGYLNQSNLANGTISITRDFQPKYTYYAGSEIAAAFAEAEYYPIQS
FT                   LLVSLGVRYEHSTQWVRYWDDASNEKRSELNTDDFFPALNMKYNFNPANSLRLSASRTV
FT                   TRPSFIEMAPFLYKESYGSSEIRGNENLKNGYNYNIDLRYEFFPQNSSDMFSITGYYKK
FT                   LESPIERVQESSGGSAVHSFRNADNGMAAGVEIEIRKEIIKNLRFGINGSYMYTDVKLP
FT                   EGEGIYTDSQRQLQGASPYLINADLSYTPSFGEERRMSLAMMYNVQGPRIHSVGIYNLG
FT                   NVEQQALHTFDFVGNYEFNKHFSAKVQLKDLLNSTVRFKQDVPSTGKVIEVESFKPGTS
FT                   AEIGFTYKF"
FT                   /product="TonB-dependent receptor"
FT   CDS             complement(23311..23688)
FT                   /db_xref="SEED:fig|6666666.127253.peg.17"
FT                   /db_xref="GO:0008137"
FT                   /translation="MEMDDYNGLYQSNPKLAFLMTLSLFSLAGIPPFAGFFSKFFIFMS
FT                   AFRSGFHLLVFIALVNTVISLYYYLLIVKAMYINKNETPVAAFRSDAYSKAGMIICLIG
FT                   IIGLGIASIVYETINQFSFGI"
FT                   /product="NADH-ubiquinone oxidoreductase chain N (EC
FT                   1.6.5.3)"
FT                   /EC_number="1.6.5.3"
XX
SQ   Sequence 23867 BP; 6485 A; 4888 C; 5412 G; 7082 T; 0 other;
     ctatgcctga taattcttgg tagtttgtcg tttaccgctt gtgacaatgg agatgaggaa        60
     gacaccaatc aatatgtcgg tggcattcag ctaaacgtct ttggtccgag tccggtagca       120
     cgtggaggag aacttcgttt cttgggaagc ggaatgaata aagtgacagg agtcgttatt       180
     ccgggatgcg atgaaataac agatattcaa gtaattagtg ataatgaaat ccgtgttact       240
     gtgcctcaaa tagcagaacc gggtttggtt gtactacata ttcctaacgg agaaattgta       300
     acgaaaacca agctgactta tacagaacct gtcggtttgg atgctattac tcctaaatca       360
     ataaaagcag gtcaggaatt aactcttacg ggtgagtacc ttaatttagt taaagaagtg       420
     attttcgctg ataatgtggt ggtgaaaagc gcagctttta ccaaacatga gcgtaaggaa       480
     atcaaactga tagttcctaa tgaggctcaa acgggtaaga taatcatttc tgatggcgca       540
     gaaataccga actggattta ttctgctgaa tccttaacgg ttgtattgcc ttcggtatct       600
     aaaactttag atctgacgtc aaagaaacca ggcgatgtga ttacgattga aggcgaagac       660
```

ttggatttgg ttaaacaaat attaatgcct aatgaaaaac cggtagactt taccatcgta 720
aagatggata ttgctgcaac taagagtact gcaggtgcta aatataaact aaactttgta 780
ttaccggaag atgcctctga tggtgctgtt gttatggttg ctcattcagg cgtaaaagtt 840
gctatagcca acattggtat ggctgttcca acgtctttgg ttgcaacacc ggctgctgat 900
attaaagggg gtgatgtgat tagtataaaa ggcgttaata tggaattagt aacatccgta 960
accttccctg gagttacaga caaggttacg cctaattcaa aaacagctac tgagattaaa 1020
gttactatgc ccgatgctgc gattagtggt aatctcgtct tgcatacagc tagtggctat 1080
tcagcgtcag ttgctatctc gactttgaag cctgaagtta tggcatataa tccttcacca 1140
gttgctgccg gaaacgatgt tgttttgcaa ggtaaacact tagatctagt atctgctgtg 1200
acctttgctg gcgataaaaa agtatctgtc agtccaacta cttctactga gttgaaggtc 1260
accgttcctg ttgatgcaga aagtggagag gtcattctga ctatgaagaa tggtgaaact 1320
gtaaaatgtg ccacattaga agtttcaaaa cctgtattct gttatattcc tgtgttaccg 1380
gacgccagtc aagaaatcca tgctggaaaa attctggcta tttcagtaca gaatggcgat 1440
aaactaacga atgttcaggt gaatggggcc actacgcagt atatcttcca aggaggtaca 1500
ttgaatgtgc ttattccgtc taatgcggga ggagagacag ccttgaaact gatttcttca 1560
aatggtgaag ttacttatac gattaaagtt attgctagtg gcgttacaga aactgttcta 1620
tttacaggac cattgctcat cacatggagt gatggcggac gtgttatttt gccggcaaca 1680
gcctttaccg gtttaacagc aggagctatt ctcaaaatct attttactca aaatgataac 1740
tggggacaga ttcagattaa taatggcaac tggtctacta tttcttttgc agagctagga 1800
aatggtggct atattactac taatacatac aatgataaga cgattacttc tcaggaattg 1860
gttcttacca ctgacgtatt gagcattatc aatagtaatg caagcaatgg caatgctatt 1920
attatgcaag gttccgattt cgtaatagac aaagtttctg ttataacgaa aggtggagtc 1980
tcggcaaagg cacggttgta taagattaaa aatttgaaat aatatcgttt taataatcta 2040
agaagacttg ctcaggacaa taacttgagc aagtcctttt ttgcctctta catatactga 2100
agggaacata taatacaaat cactcaatca tgaaaagtct tttttatcgt tttttgctct 2160
tgtttttagt aatctcattt tttctagcgt gtagtggagg tagtgacaca cccgatacac 2220
ctttgcttga agctcctcaa ctgatgggaa gtactccgac aaatggtgct acagatgtat 2280
cggcaagcaa cataagtatt gtactgactt ttaatcagaa tgtaacagct cctactaccg 2340
gacatgataa aataacgttg agtggagcaa ccataaccag tgtcgttgct tatctgacta 2400
aggttacgat tgaggttacc ggattggaaa agggtaaaac ttatactctt agaattccca 2460
aaggagttat attgggtccc tcaaaagtcg aagctcctga aatctctatt agtttcacaa 2520
caacttctcc ccaacaaata accaataaac tctgtacggc caacccgttg acacaagcac 2580
agaatgtata tgactttctc attgctaatt atggtacaaa aataatttca ggtagcatgg 2640
ccagcgacag ctggaacgtt aatgaagctg agtgggtatt taaacataca ggtaaatatc 2700
cggctttgaa tggttttgat tatggacacc tgtatgcttc gcctgcaaat tggattgatt 2760
acggcaatac aactattatt gaaaactggt ggaataatca tggccttgta gctgctacat 2820
ggcattggaa tgtccctaaa agttcaggta gtacggaata tggattttac tatacgggta 2880
agaattcggg caaaggtgaa acttcttttg atataagtaa agccgttcaa gatggaactt 2940
cagaaaatca aatagtgaaa gctgatttgg ctaaaatagc tgattatctt ttattgttaa 3000
aggctaaaaa tatccctgta atttggcgtc cgctgcatga ggcagccggt ggctggttct 3060
ggtggggagc aaaaggggca acttctttta aagcgctttg gaagttgatg tttgaaacct 3120
ttcaatctaa aggagttaat aatcttatct gggtatggac tagtgaaacg ggagatgaag 3180
attggtatcc cggagatgcg tatgttgaca ttgttgggcg tgatgtatac aataaaatat 3240
tagcgtctca atctgtagaa gaatttcgga aattatcgac cgcatacagt cataaactta 3300
tcacattgag cgaatgcggt aatgtatcga aaatatcact tcaatggagc gcgtctgcaa 3360
actggtcatg gtttatgcct tggtatgatt atagtcgtac aaataagatg gatgatccgg 3420
cttttaatga aacgagtcat cagtttgcta atattgaatg gtggaacgat gctctcaaca 3480
ccgaagccgt gctttcacga gacgagatgc cagatttgag ataatattat ctttggataa 3540
aattgtattt gtttgattcc tcgtacactc gtacttttgt cgaatagaat gatgcgttac 3600
attgaaactg atataaacca tgaaaaactt tttaatgaca atggcaattg cggcaacggt 3660
gggaagtgga tccgccgaaa aaaaggagat gtataaagat ccttcagttc cggtgaaaga 3720
tcgtgtggaa gacttgctga gtcggatgac tttggaagaa aaagtggggc agatgaatca 3780
atttgtcggt attgaacaca tcaaggcaaa tagcgctgtg atgactgcgg atgagttgaa 3840
aaacaatacc gcaaatgcat tttatccggg agtgaccgat aaggatgtag aacgatggac 3900

agcacaagga ctcatcggtt ctttcctgca tgtgcttacc attgaagaag ccaaccattt 3960
gcagtcactg gcaatgaaaa gccgcctgca aattcctatt atttttggta tcgatgccat 4020
tcacggcaat gcaaatgctc ccgataacac ggtttatccg accaatataa atctggcttg 4080
ttcctttgat acagccatgg cttacaaaat agcccggcaa actgcaaagg aaatgcgggc 4140
catgaatatg cactggacgt ttaatcccaa tgtagaggta gcacgcgatg cccgctgggg 4200
aagagttggt gaaacctatg gtgaagaccc ttatctggta actttgatgg gcgttgaatc 4260
tgtaaaagga tatcaaggaa gtttaaacgg caaagaagat gttttggcat gcattaaaca 4320
tttcgttggc ggtagcgaac ctattaacgg aacaaacgga tcaccgacag atctgtcgga 4380
gcgcactttg cgcgaagtgt tttttcccacc attcaaagca ggagtagaag ccggtgccat 4440
gtcattgatg actgcgcaca atgagctcaa cggcattccg tgtcacagca atgaatggtt 4500
gatgcaggat gtgcttcgtg gtgaatggaa gttccctggt tttgtggtga gcgactggat 4560
ggatattgaa cacattcatg acttacatgc cacagcggaa aacatcaaag aggcttttta 4620
tcaatccatt atggctggca tggatatgca catgcatggc attcactgga atgaaatggt 4680
cgttgaacta gtcaaagaag gacgtattcc cgaatcacgt atcgacgagt cggtgcatag 4740
gatattggac atcaaattcc gtttaggact gtttgaacaa ccttttgccg atacttctaa 4800
aagcatgaaa gtgcgtttat ccaaagaaca tcgcgaaaca gctttggaag cagcccgaaa 4860
cggtatcgta ttacttaaaa atgaaggtgt acttccttta gacgcttcaa aatataaaaa 4920
agtattggtt acaggcatca acgccaatga catgaatgtg ttgggcgatt ggagtgcggt 4980
gcagaaagaa gaaaatgtta ccaccatact ggaaggtctt cgtcaggtgg caccgggtac 5040
ggcatttgat tttgtagatc aaggttggga tcctcgtaat atggatcctg ccaaggtggc 5100
tgaggctgag aaacgtgcca aggagtccga cctaaatatt gttgttgccg gagaatacat 5160
gatgcgtttt cgctggacgg atcgtaccga cggtgaagat accgatcgtt cggatcttga 5220
tcttgtgggg cttcaaaatg aattgatccg taaagttgct gcttcgggaa agcctacaat 5280
tcttattttg gtcaacggtc gcccgttggg tgtggaatgg gcagctcaac atcttccggc 5340
catcgtagaa gcttgggcac cgggcatgta tgccggtcag gcagtggccg aaatactata 5400
tggaaaagta aatccatcgg ctaagttggc aatcactatc ccacggagcg ttggccaatt 5460
gcagatgatt tacaatcata aaccgtcgca atactttcac ccgtatgccg ttaaacccag 5520
cactcctttg tggccgtttg gttatgggct ttcatatacc acttacaaat atgatgattt 5580
aaaactctct gccaaagagg tgtcgcaaaa ccaaaacgtt gaagtgagtg tgaaagtgac 5640
caataccggt cagcgtgatg gagttgaaat agtacaactg tatatacgag acaaatacag 5700
ttctgtaact cgtccggtca aagaactgaa agattttgcg cgggtagcac tgaaagccgg 5760
agagagtaaa gtcgtcacct ttaccatcac gcccgataaa ctggcatttt atgataaaaa 5820
gatgaaatgg atagttgaac cgggagaatt tgtggtgatg gcgggtgctt cctctgccga 5880
cgaatcatta ttaaaagaaa cattcatcct taaataagtc gccttatgag acatattaag 5940
tttataaccg caagcatgct aatagctgcg gggctatctt cttgcaacct gtttggacag 6000
aaaggtacaa tgaaaatgca atcatcagaa cgaacagttg agaccaaaaa cttacttata 6060
aacctaggga caatccatca aaaaggtttt atgtttggtc accacgatga tccggtttac 6120
ggcattggct gggaaggtga tgcagacaga agcgacgtga aaagcgtatg tggagattat 6180
ccggccgtga tgtcttttga tttgggaaga atagaattgg gaggcgataa aaacctggat 6240
aaagtacctt ttgataagat caggcgagaa attctggccc aatacgcacg gggtggtatg 6300
gtttcattga gttggcatgt tgataaccct cttacaggta aggactcttg ggatgtgagt 6360
gatacaactg ttgtgtcctc tgtattgtcc ggtggcgcta atcatcaaaa gtttttgggt 6420
tggttggata aggtggccga ctttatgaat tcgctcacta ccgataaagg tgtgaaagtt 6480
cctgtacttt tccgcccctg gcatgagcac agcggcagtt ggttttggtg gggacagaat 6540
ctttgcacgg ctgcccagta caaagccttg tggaagatga cttatgaccg tatgcaggaa 6600
aagggagtaa acaacctgct ttatgcatac tctcccggaa cggaaccaca aacagtagaa 6660
gaatatttgg aacggtatcc gggtaatgac atcattgact tggtgggttt cgatacgtat 6720
caattcgacc gtcagaaata catcgatcaa ttaaacaaat cactgaccat cctgactgaa 6780
gtcggtaaga aacaggggaa gcctattgcc attaccgaaa cagggtttga gactattccc 6840
gatgcaactt ggtggacaga aacattgttg cccgttatca gtcaataccc tatctcatac 6900
gtgttggtgt ggcgcaatgc ccgtgagaaa gataaccatt attacgctcc gtaccccgga 6960
caggtttcgg cacccgattt tgtgaagttc taccagaact ccaaaacctt attcgcccga 7020
gacataaaga agctatatga ctaacgaaat tcaataacga aatcaataac ctaattatga 7080
gcttgtttaa agataaagtg gcgaaattac tgactgacca cgaagctttt ttatcacaga 7140

agaacgaacc tcttgcggta gataacggtg tggtaacccg ttacaagaac cctgtgctca 7200
cagcagagca cacccccgta ttctggcgat atgacttaaa cgaaacaacc aatccgtact 7260
taatggaacg catcgggatg aatgcaacga tgaattcggg agccatcaag tggaatggaa 7320
aatatgtact gatggtacgt gtggaaggat ccgacaggaa atcatttttc gccgtggccg 7380
agagccctaa cggaattgat aacttccgat tctgggacta tccggtacaa atgcccgagg 7440
acctgattcc cgctaccaat atatacgata tgcggcttac agcccatgaa gacggctggg 7500
tatacggagt gttttgtgcc gaacgacacg atgataatgc cccggcgggc gacttgtcgt 7560
cggcaacagc aacagcagct attgcacgca ccaaagatct tattcactgg gagcgcttac 7620
ccgatttaaa gaccaagagt cagcaacgca acgtggtgtt acaccccgaa tttgttgacg 7680
gtaaatacgc tctttacacc cgtccccaag acggatttat tgatgcgggc agtggcggag 7740
gtattggctg ggcattgatt gacgatataa cccatgcggt ggtaaaggaa gaaaaaataa 7800
tcgatccgcg ttactatcac accattaagg aagtgaagaa tggagaaggt ccgcacccca 7860
taaaaacacc taaaggatgg ttgcatttgg cgcacggcgt acgcggttgt gcagccggtt 7920
tacgttatgt gctctacatg tatatgaccg ctttggaaga cccggcccga ttgatcgctt 7980
cacctgccgg ttactttttg gcacctcagg gagaagaacg tatcggcgat gtgtctaacg 8040
ttcttttcac caatggttgg attgccgatg aagacggaac cgtgtttatt tactatgcat 8100
cctcagacac acgcatgcac gtagctacct caactgtcga caaattagtt gattattgtc 8160
tgcatacgcc ggaagacggt ttgagctctt cggcatctgt ggctacatta agtaaattaa 8220
ttgaaaaaaa tctggcctta cttaatcagt tttcactgaa aaaatagcag atttgtatta 8280
taatgaatga ataagatggg aaacacagca agtaagatag tagcagataa ggaacgaata 8340
tcactcaaag aaaaaattgg ttacggtttt ggtgatatgg cctcgtctat gttttggaaa 8400
cttttcgggt catacttaat gatttttat acagacgtgt tcggtttacc ggctgcggta 8460
gtaggtacta tgtttttaat tacccgcgtc tgggactcag ctttcgaccc cattgtagga 8520
gtagtggctg acagaaccca tagtcgctgg gggaaatttc gtccttatct gctctatctg 8580
gctgtgcctt ttgcacttat cggcattctg acctttacta caccggcact aagtgatacc 8640
ggaaaactga tttatgccta cgtcacctac tcattgatga tgatggttta ctcggctatc 8700
aatgttccct atgcatcttt gctgggcgtg atgagtcccg acccgaaaga gcgcaacaca 8760
ctttcaacct atcgaatgac ttttgcctat atcggtagct tcatagccct gttgcttttt 8820
atgccgatgg ctaatctctt tgctggcagc gaagcagatg cggtgcacga acagcgcgga 8880
tggatgctgg ctgtagtcgt gattgcggtg atgtgcgccg tgctgtttta cttctgcttt 8940
gccttgactc gcgaacgggt aaagcccatc aaagaaaaac aaagtccact gaaagaagac 9000
cttaaaaacc tgcttggtaa tgggccgtgg tggattctgc tgggtgccgg cgtagcagca 9060
ctggtgttca actccattcg tgacggagcc actgtgtatt actttaaata ttttgttgtt 9120
gaggccgact atgcaaccgt atcccttttt ggtgtgtcgt ttgtgttgag cggattgtat 9180
ttggcagtgg ggcaggcggc caacattgtt ggagtcatac tggccgctcc gttaagcaat 9240
cgcattggaa agaagatgac ctacatgggc gccatggtta ttgccaccgt tctcagcatt 9300
attttctact ggtttgataa agatgatatt gcgctgatct ttgtgttcca gatactgatc 9360
agtgtgtgtg ccggcagcat cttcccgtta ttgtggtcta tgtatgccga ttgtgccgat 9420
tattcagaac tgagaacagg caatagagcc accggcctta tcttttcctc ctcctccatg 9480
agtcagaagt tcggttgggc cataggcagc gcacttaccg gctggctatt agcttatttc 9540
ggattccggg ccaacgaaat acaaagcgaa gaagccattc acggcattaa aatgttcttg 9600
agtttcttgc cggctgtggg taccgtgtta tctgttatat ttattagcat gtatcccctc 9660
agtgaaaaga aaatgaaagt cataaccgaa gagctcgaag caaagagaaa acaacaatga 9720
atacaaagat aactcactta aagggagaag taaccgaagt ccttaccaca aatatacttc 9780
cgtattggat taataaaatg gtcgacacca cttacggtgg cttttacggc cgtattgacg 9840
gaacagaagt gtgccgcccc gaagctgaga agggcgcaat actccatgcc cgtatgttgt 9900
ggaccttctc agccgcttac cgtttgctga agaaagaaga atatttagac atggccacac 9960
gtgccaaacg gtatatgata gactcttttt atgataaaac gtacggagga atctattggt 10020
cgcttaccca ccagggcgat ccgttggata ccaaaaagca aatctacgcc attggctttg 10080
ccatttacgg gttgagcgaa tattaccgcg ccacggcaga tgccgaggca cttacttatg 10140
ccattcatct ctttgaaagc atagaaactc atagctttga taagataaag aacggttatt 10200
gtgaagccct gactcgtgag tggggcgaga ttgccgatat gcgtttgagt gaaaaagatg 10260
ccaacgaaag aaagaccatg aatacacacc tgcacatatt ggaaccctat accaatttgt 10320
accgtgtgtg gaaagacgaa cgcttgaaga atcagttgcg taacctgatt tttcttttta 10380

cagataagat actcaataaa gaaacccatc acctggaact attttttgat gacgactgga 10440
acagcaaata ctccatcgta tcttacgggc atgacattga agcatcgtgg ttgattcacg 10500
aggctgcact ggagctgaaa gaccctgccg tattgcaata cgtcgagccg ttggtacaga 10560
gaattgctga ggctgctacc gaaggattta cacccgaagg aggaatgatt tacgaacgca 10620
atctcactaa cgggcatgtt gatgccgacc gccattggtg ggtgcaagcc gaaaccgtag 10680
taggatatat taatttgtac cagcactttg gtgatgaggc ttcacttgat aaagcactcg 10740
cctgttggga atttaccaaa tcgcatctga ttgaccgtgt caacggagaa tggttttgga 10800
gccttcgtgc agatggcacc ataaaccaca ccgatgacaa agcggggttc tggaaatgcc 10860
cctatcacaa cggtcgcatg tgcatggaaa tcatggaacg attctgatac tcgttagcgg 10920
agttcatgta atgcgtgaac ttcgcccatg tgaagttcaa gatgtcgtgg gtagtcccta 10980
atcatatctt gcagagacac ttcttctccc tcaccggcaa tccacttgtt attcaactta 11040
tgtgcatcaa catgctttat gacgtgcaca aaatgcaggt tactgtattt ccatagctgc 11100
accaatgttt gccaatcttc ctcctgaaaa tgctgaatgc caatccatct gtcattgttc 11160
ccgttggtgg cataattagg aaacacgaga ggatttgcgc cgtaatgcaa gtgaactacg 11220
cggtgcaagt tattggaagc cgagtcaatc atgtgcccca caatctgttt gatggaacga 11280
ttctgactat tcaaacgggt agagattgtt tcttcatcta acccattcag gaatagttcc 11340
tccttctcaa taatttgctc aatttccttg ataacatctt caaaaatata cgctgccata 11400
gtcttctttt ttatgattca acaactttaa ctaattcata cttctgcaaa agtaaagcga 11460
ttaatgccat ctgagttgta aaaatcggac tttgtattgt tgtccgttat tgtttaggag 11520
ctttatacct tccccagaaa cttaactttg ttaatgaagt aaagaccacc atacgttcaa 11580
accatcgaaa acgtagcatg aaatgaagta acctataaaa tacagcaatg tatgcaatga 11640
aaagcatgtt ttccagaaga aacttaagcc attttatatc tgtaattaca ggcaaataac 11700
gatcaaagac agatgataca agaatgctga atgctatggc tatgactatg ataaaaccat 11760
gtgccgtttc aatagccttt tgaggacaat tgctcatgca attcatgcaa ctctcacacc 11820
gaaaagtcca aaacggtttg ccattaacgg ttttgatagc atgtattgca cagtttcgtt 11880
cacataaacc gcacttagta caatcttgtg acgcataaaa agttttagct atcacaaagc 11940
gacctatgaa ataatataag atggcaatag gagctaccag taaatcgata atgcattctc 12000
ttacgcaggt aaattctctt tttccgttaa gaaaacctcg agcagctttt tctattttcg 12060
gagttacctg atgatagatg aaatctactg tctttttatt taatgccgga tgcagtgata 12120
tccagttcga cgggaaatca acagggtata acgattgtat tctataccct ttccatctta 12180
gtatcaatgc cgataaatag agactgatgc ccgatatgcc gggcgtaacc cagttcccaa 12240
ttctcatacc tgcccgcgta ttcatcaaca ttacttcatt gttacctttg gggaagcgca 12300
gcagataatg caacattatt ggtggatagt taaacccatg aacgggtgaa accaaaacca 12360
acagttcatg gctttgtggc gatttaatat gtaggccatc tgtcttagaa atatcgttta 12420
tttgtaccgc cacattgtga cgccacattt ctcgttctat tatttgtgcc acattgcggg 12480
catttcctgt gccagagaaa tagtatatgc aaaggctctt ttccatctgt tatatatcta 12540
tatctttttg ataatattga gcctgcttcc tttagagaag aaagtggagt gaggctttct 12600
tgattgcaga tattcagtag gcgtgcatcc ggtgaggctt ctaaactcct tcaccagatg 12660
agaagagtcg cagaacccgc aaccataggc tagttctgtt aattctgtgc cgggattgtt 12720
ttgcaataga tgcaatagcc gttgaaaacg aacaatgcgc aaatattctt taggacttgt 12780
gcctatgtat ttagaaaatt ccctgttgaa gtgtctcaca cccaagcaag tttgggaggc 12840
taaagaagcc acatcaattt ctgcctgccg actgattact tcaacggagt gaaacattct 12900
cttgaagttg taatcatgcc aatcagttat gcgagacagc agaaaatgct ctatcagttg 12960
gatacaggca agagaatctt cttcatgcaa aatgttgcct ttcagttcct ttagtccggc 13020
atcctccatg gaatcaacat ccacatattg attgtaaata agctgcaaag gtaatgaaaa 13080
gaatacattg aaacccagcg gatggaaaat cacggcaacc atgtctattc cacctttgaa 13140
ttgaagaact cctggacccg aaagctgtcc tctgataaaa gactgcggct gttgctctcc 13200
atcagaaaag tacatacttc ctcctctgtg aaacaccaaa tgaatacatc ccgacggtac 13260
ggtctgaatg gctgttgttt catccgcttt cactacccag taatacttta cgtaaggtgc 13320
cagtaacaga gaaggaagta cgttcatgga ttataaattt aaatactctt cagtagtaac 13380
aacggctgca tagaaataac ttaaagcagc aagaaaagca ttttgaacat ccttagcttt 13440
aaccgattga tcatgaatga acaaatcttt cgttgcacat gcatcgccaa tgacggaaca 13500
agtaaaacct aaatcttttg cggctcgggt agtggcatca acacacatgt gcgtcatcat 13560
tcccacaata acaagactgt caactccctt ctccttcagc ttttctaata aaggagtatt 13620

tataaaactg tttgggaagc ctttcgttac aactgtctcg ctctcgcccg gagatacgtt 13680
ggggtgtatt tccactccat gcgtatttgg aatgaaaaat gtggcttccg gtcgggttga 13740
aacatgttga atgtgaacaa ccggtttgtt gcaacctcta aatcgttcaa gaattgcttt 13800
tgcattttgg cttgctttct cagcacccac cagctcgttg gcaccacccg gaaagtaatc 13860
attctgcatg tcgatgatta ataatgcttc tttattcata gataccatgc ttattatctt 13920
tttgatgtat tatttgtatc ctatacacat actttttcat gtactttga atttccaaat 13980
tcatttgcgt atgtataata gcctgactaa taaacagtgc aagttataga aatcttttta 14040
gatgacaaag tgtgtaaacg tatataatct tgcctcctta tgaaaatcgc actggttttt 14100
cctcataaaa tgatagaaaa aaagtatcac caagtatcac caactttgca atgctttgat 14160
tatcagtttt ttatgtgtga cacttggcta aaaaaagcct caccaagtat caccaagtat 14220
caccaacttt tttggtgatt attccttata tgcatctatc tgtttattgt ttgtatatag 14280
ggtttttcgt tggttccttt atgtatttat tttgtggatt ggcggttttc gttacattcg 14340
aaacaccacg gtttgtgccc taataataac ctgataaaag agccacaatg atttggaatt 14400
gacagtataa tgtattatct ttgtacttat aatcttatag tatattcaaa ttattatagg 14460
cgcaaaggca ggagctcata ctcttgcctt tttttatgtc ttaacggttt atgattgcat 14520
ttacacagta tctcgcgccg gtttgcacgg tagcaaatgg tcaataagaa accgtctgat 14580
ggcgaagaga cggtttctcc cgaaccaaaa gacggtttat gatttaaaat tgctcgttat 14640
ctttttgaaa aagtcggtta tgtttttcaa aaagttggtt atgttttacc aaagtcttgc 14700
tgttccttgt tgcttccctg tttacccgtt aataatatat aaaactgctc ttgacaggat 14760
ttgtctgatt catgatgtta aggggtaaat ttgtaaaata agttctacta cgtggccaac 14820
atagtagaaa taaatgttac tctagactcg atggaatcca attatcaagg catgagaaat 14880
ttgatattta aagtgagagc tctaagcata gatagtttag ttacactcct attttattca 14940
tatcaagatt gtaacccgct gatcttaatt ctcttactag aagaaatttc cagccaccag 15000
cattgtcata atcagtataa actatacctg taaaatcaga aggttcttca acatctgttt 15060
tttttagtgt acaaactcgt gttctaccta gaagtccaat aaaatagcct aactcaagga 15120
ttacgttctg tcgcgctcta ctacgaagct cattagtgcc ctttcttctt ccttcatcac 15180
agggagatag aagcacaata gcaaagccta catttgaatt ttcttcaagt ttttctatta 15240
tggtattgcc attattagcc tgattattta atattatggt ttcaaattca agctgtttta 15300
aaaaactatc tacttccaat ttcatggtat tatcatgccc gtgaactata aatatctttt 15360
tattttctat tttatttatt ttgctatttt cagatatttt atttttaatt ttatcgcata 15420
aaattgaata tactccttga atttgattaa aattgcagct taatttatgc ccattcccac 15480
tattgtcaac cgatttaaat ttctgatagt ccgaatttgt gtcatcaaaa tacctactaa 15540
atagagtaat ggataaatta aaccagtcgt gataagcatt aatagtatct tgagaatttg 15600
tataatcacc ttttcgatct ctcaattccc tatactctgt ttctaaatat cccacattct 15660
ctatattgat ctcaagttcc ctattatttt ctacgtcttc gatatcacct tcattctcag 15720
tcattgttcc aatccctaca aaccattttt tcttttcttc attaaattcg ggtaataatt 15780
caaggtctaa catgatttca tttatattaa tctttgtctt gctccatacc tcataaacgc 15840
ttaaaattga agctaacacg ccataaaagt catttttaga tgaagaatag aggcgatcta 15900
ataagatttt aaatgcatca acgcggtata attgatccat tccaaacatt ataacttttg 15960
aatttgattc aacttcatta taaatatggc aatacgcctc gtatgctttc atttggttca 16020
taattctttt aaaattagct gatcctaaaa aatcggctaa tttaagttct ctatctatat 16080
catcaaatag agagtctagt attttatttt gaatttccat aattattatt aaattgttat 16140
agtctacttc tttatataag attgaatatg ttattttttt aattaccaaa gatactaatt 16200
atctataaat tatttaatat aaatatgttg aatgtaacta tttcaggacc attcagagtt 16260
tctccagttt catggggatg agactccgag tttcctagta caaaactgtt ggtttgatag 16320
gagcaaacaa actgcttcca acccacttgc tgccaaccaa ataaatagaa tattaaacca 16380
ttaaattgca tattattacg ctttacccta tcttttctaa tagcgaatca accccaaaaa 16440
gcgtttagtc aacccaatat gctctcaaaa tgaagagtta tgaagcgaaa aaagagtgtt 16500
tttcatcgaa aacgggctta aatgagcaaa acgtcgacta atttatagtt aaatggtagt 16560
taacgaatct gtatacatgg ttaaaagtga gagatagtga tagtaaaaag cctgttttag 16620
tggtagacat attttactat cactcctctg taaccgctta cagtaagccg tttctttcca 16680
gttggtgagg gtaagtgagg gtaaaccaca cttttttttc tttaggagga aagtggtgta 16740
ttaatataca gtattctgtg ggtcaaagtt acaccaaccg ttggtccaag tagggaaaga 16800
aactgttgcc gtgccaaagg ttacgttttc ctgagtagcg gcagtacctt cccaattcag 16860

cgagtatccg ttgatgtttg cgttcttgaa attaaccgca gtgttactgc tcaaagcaat 16920
ggctgccccg taaagaccta gcagactctg attatcgggc agcaatccgg ctccgttgat 16980
gtaggtcgtg gtgttttcaa attcggcatc ggtggaggag gggccgtaca ggtttacgtt 17040
gctgaaggtt ggggcggtct gtggttcggc gttgctgcta ctgctgcact caaaaccgtg 17100
tgaggtggag gcatcggcta tccgtgggtg acgaatgccc agcaggttgg ttgcggtacc 17160
gcggtatccg ttgtcggcat cgaagtcatc gtcccacgta tggtaggcta ccaggtagtt 17220
gggacttact gttccgccaa accattcaaa ggcgtcatcg ttggagtaag atacttgtat 17280
gtggtctatc tgcgtgttac tgcctactga tcctagtgtt aatccgttaa tttcttggtt 17340
agtgcgcaga ggtgatccgg caaactctat gcgtacgtag cttagtatgc ccgagttgtc 17400
agtgtctttg atgccgccgt gcatggttcg cggaccgcct tctacttgca taatgccttg 17460
gttgttgcgg gcgttgccac agagaataag gccaccccag tcgcccggct tgcgttgtcc 17520
gggagctttc tcggaagtaa atacaatagg aagttcggct gttccgcggg ctatgatttg 17580
tccgcccggc tccacaataa gtgctgcacg tgtttcgctg tcacctttga taacggtgcc 17640
cgcctcaatg actagtgtag caccgtctgc tacgtagcac cagcctttga gcaggtaaac 17700
tcctttttta attgtctggt tgtgggtaat gataaactct tgttcaccgt tacctatctg 17760
gttatcacca aaaacaatgt tgttgtcggt ttcgtcgtta ctgccgttgc aggcggcaag 17820
gccggcagtc agtgtaagtg ccaggatggc ataagtccat tttttcatct cgtttttatt 17880
taataacgaa agccgatggg aaacagttca ccatcggctt aagcaagtta gttgaaaaga 17940
aacgctatta taatgatatc aacaaactaa tttgcatgat taacgaatcc aacctgctgt 18000
ccagttatta gctgtactga tggctccttt gtaagtagct ttggtaaaga agctgtcggc 18060
gctcaggtct gcacctccgt caacggttcc gaagaacgta tctgtaaatg tcagcgtctg 18120
attaatggca ttgccggtag ctgctgcgaa taaagccgat gtgtatccgc ctaaaccgct 18180
taccttgatg tcattggcta catgtatgta gttcaaaatt gattttgttc cgcctaccaa 18240
tgcttcttct gtttctttgg tttctgtggt gaggttattg gtcttaccgg ttaccaaagc 18300
gttatatatc ttagcatagg tacctgcacg caaacgaatg ccgcgtgtgt tggctgtcga 18360
tccgttacca atcagtgtga ggttggccaa tgtgggttgt gagataggtg ttgctgctgc 18420
attgcttccc aagttgtcgg cttcaatcag acagtcgcac tcataaccca gtgtggtgct 18480
cggttgctgg taagcaacga ggaactgacc tttaccgttc cagccttcag tccagtcgaa 18540
cgagtcgtcg ctacagctgg ttgccacaag gtatttcacg tttaccgatc cgccaaagaa 18600
ctcaaatccg tcatcagatc ctttgtaggc ttgcaggtat tcaacggttg ttccgttacc 18660
cacgccgtag aatgatacgc cgttagcttc tttctcttcg ctgaaagcat aaccggtgta 18720
ctccaaacgg atgtagcgaa gtgttcctga gttgtcggca tcattggttc cgccataagt 18780
tgcgtcaccg atttctgatt ttgaggtagc tcccaggttg gtgtgtgcat atccgcaaat 18840
gtgtatgcca ccccatgctc cgggttcttt cttttcagaa gtcatcacga tgggtgaact 18900
ggctgttccc tgtgcatcaa ttttaccgcc ttgttcaata aggatgtaat ctactatatt 18960
atcgttcttg gcaataacag ttacgccggc ttcgatttta agtgtagcac cgctcttcac 19020
ggtatatccg cctgtgaggt agtacacttt gttggctgtc agtgttttgg taccgctggt 19080
tagtgaacct tctagtgttt ccgactgttg cacagcgttg ctggctagtg tccaaccggc 19140
tgtccagtcg ttgctgctac ttacggctcc tttataagtg gctttggtga agaagctgtt 19200
tacggtgctc atgtctttac caccgtcaat ggtaccgatg tagttgctgg tcaacggaat 19260
gattttgctg atgccgttgt tggtattacc ggtgaaaaga gcttctgtat aaccgttgcc 19320
gctagtgcct acaccgcttt ccagatacac atagtcgagc acggaagtac cgttaaccaa 19380
tgccgtttct gtttcaacgg tttcagtggt caagcagctg gtttttcctt ttaccaacgt 19440
gttgtagatt ttagcctgag taccggcacg cagacgaaca ccgcgtttgt tggctgtgga 19500
tgagttaccc accaatgtca tgttagccaa tacgggtgct gaaacaggag ttgcagcagc 19560
gttcttgccg ttgttgtcag cttcaatgag acagtcgcag tcgtaaccga tggtagcttt 19620
atcgccctga taagcaacca agaattgtcc gtaaccgttc cagccttctg tccagtcgaa 19680
tgagtcatcg ctacagtcag ttgccacgag gtatttcacg tttaccgatc cgccgaagaa 19740
ttcaaatccg tcgtcagaac ctttgtaagc ttgcaggtat tcaacagttg taccgttacc 19800
tacaccgtag aatgatacgc cgtttgcttc tttttcttcg ctgaaagcat aaccggtgta 19860
ctccaaacgg atgtagcgaa gtgttcctga gttgtcgttc tcaacgcttc cgccgtaagc 19920
agcgtcacca atttctgatt tggctccggt tccggcattg gtatgtgcat atccgcagat 19980
gtgcaaaccg ccccatgctc cggcttcttt cttttcggaa gtcatgataa taggtgagct 20040
ggcagtacct tgcgcatcaa ttttagctcc ttgttcaatg aggatgtaat caacgatgtc 20100

atcgtctttg gctatgatgt ttacgcccgg ctgaatggtc agtgtagtac cgctcttcac 20160
gtggtaaccg cccgacaggt agtaggagtt accgcttttc agcgtgaggt ctttggtgag 20220
ttcgccggtt aaggtcgatc ctgcatccac tccgcccccg ttattgggtt catcgttatc 20280
gctgcacgaa gtgaaagcga acagggtggc tgccatggcc agtgttgcta atttccaatt 20340
tgtcattttc atacgaataa aagttagtgt tactttttaa ttttttactt taataatcta 20400
tgcttagaat ttgtaagtga atccaatctc agcacttgtt ccgggtttga aagattctac 20460
ctcaattacc ttgcctgtac tgggcacgtc ttgcttgaag cgtacggtac tgttcagcag 20520
gtctttgagt tgcactttgg cactgaaatg tttgttgaat tcgtagttgc ctacaaagtc 20580
gaatgtatgc aaggcttgtt gttccacatt tcccaggttg tagataccca ctgaatgtat 20640
gcggggacct tgtacgttgt acatcatagc gagtgacatg cgtctttctt cgccaaagct 20700
gggtgtgtaa ctcaggtcgg cattgatcag gtaaggcgat gctccctgta gctgacgctg 20760
actgtcggtg tagattcctt ctccttcggg cagtttcacg tcggtgtaca tgtatgatcc 20820
gttgatgccg aagcgaaggt tcttgataat ttctttgcgg atttctattt ctactccggc 20880
tgccataccg ttatcggcat tgcggaagga gtgtacggcc gatccgcccg atgattcttg 20940
tacgcgttca atagggcttt ccaatttctt gtagtaaccg gtgatagaga acatgtcgct 21000
gctgttttgc gggaagaact cgtaacggag gtcgatgttg tagttgtatc cgttcttcag 21060
gttttcgtta ccgcgtattt ctgaacttcc gtatgactct ttgtagagga acggtgccat 21120
ctcaatgaat gaggggcgtg ttacggttct ggaagccgat aagcgaaggc tgttggccgg 21180
gttgaagttg tacttcatgt tcagtgcggg gaagaagtcg tccgtgttca actcggaacg 21240
tttctcgttg gaagcatcgt cccagtagcg tacccactgg gtagagtgtt cgtaacgaac 21300
gcccagacta accaagagcg actgaatggg atagtattcg gcttcagcaa aggctgctgc 21360
tatttcactg cctgcgtagt aggtgtactt cggctgaaaa tctcgtgtga tggagatggt 21420
accgttggcc aggttcgact ggttgaggta gccgtctgta tcataaatat cggtgatggt 21480
agggttgatg gcgttgaggt tgtaatagaa gcgagcagag gtgaagtcgc ggtctttgtc 21540
tttataggtt ccgcccacct tgattttgtt gctttcacca aacttgtagt tcaggcgaag 21600
atcgcctact acttcttttt cgtctaactc accatagtat ctcatggttt cttgcttgtt 21660
gagtttgaag aggctcaatg caccggcatc gtctttgcgg aacatcacct gacggcggtc 21720
gggttcgtca cttccggttt cgccgtatga ggcgctccag ttcaaatccc attgcttgcc 21780
gaattcgtgg tgacccagca actgattgtt gagcagtgtg taggcgtgaa acacgctgtt 21840
gctaccgatg aggtttacac cttctgaatc atagccttca cgacgcatgt agttgtcaac 21900
agcgttgcgg gcatagaaca gggtgtagtt gatgcggtcg gccttgcgga aggtgtaacc 21960
cacgctggcc agacctgcca ttttcaactc ggatgtgtat ttgtcataac tgaattcgtt 22020
gaggttggtg ccttgtgcgg tgagggtggt tacgtaggcg tctttcaaaa tctgactgtc 22080
gttgctgata cctacggatg ccaggagact aagtttctgt ccgttattca gcttccagct 22140
tttacctact ccgaatgatc cgccgataac gggtagtgag gtgttctttg aaatggagaa 22200
gctggtaccg aacgggtcgt tgttttttac gtagctgtct tcgaactcgc ttttattcat 22260
ggtaagcagg cttttgttga ggttgcctgt gctgaataaa gagcctttgc gatcactgct 22320
gtagaagtct ttgaaaagtg tgtttatctt tcctccggca ttgatggaaa tagagaagaa 22380
gtcttctccg gtattttcac gcgtaccgat gtcaacgtgt gctcccgagt aatcggcaaa 22440
tgcgccggct tcgtacacct tactcacggt gatgtttttt acggttgagg tagggaagat 22500
gtcgaggggt atgagcttgt tgtcggggtt gggtgaagca ataggtagcc cgttgagagt 22560
ggttgtgctg tagcggtcgc ccagtccgcg aacaataagc tgtcctgcgc tggctagtga 22620
tacgccggtg attttcttca ctccgtcggc cacactcgat acgcctttca aactcatttc 22680
tttcgctcct atgttctcaa tggccaacgt cgctttctgg cgctcctgca agagggcttt 22740
ctcgccttcg aggtttttac gtgccacaac ggttacgtcg cccaagactt gtgactcggg 22800
ttccatctcg aagttcagaa cggttggggt gctgcccgat actttaatgt ttctaatctg 22860
gatggttttg taagatacgt acttgatttc cagattgtac gttccgtttt tgagtccggc 22920
taagttgtaa ttaccatcta agtcagctat tacaccgatg ttggtaccca cgatttgtac 22980
ggtggctccg gtgaggggtt ctttggagct tttatcccga ataattcctt tgatgttttc 23040
tgccatcagt tgagtggata gcagactaag taatgtgaca atgaataaaa cttttttttct 23100
aactgttttc atacttaact ataatgcgtt tctttttcga tgcaaagatg cgggtatgag 23160
attacaatag taagacgggt tttatacatt taattgacag atgtgttaca gtttggttaa 23220
gaacgaggag ggtgcgaata caaaaaaggc ccggccgaca ggggtactgt ctgttccgga 23280
ccttctgact ttaggtttgg ttgtatgtgg ttagatgccg aagctaaact ggtttatcgt 23340

```
     ttcgtaaaca atgctggcta tccctaaacc gatgatgccg atgaggcaga taatcatgcc     23400
     tgctttgctg tacgcatcac tgcggaatgc cgcaacaggt gtttcatttt tgttaatgta     23460
     catggccttt acaatgagca ggtagtagta caaggatata acggtgttga ccaacgcaat     23520
     gaataccaac aggtggaagc cgctgcggaa tgccgacatg aaaatgaaga acttactaaa     23580
     aaaacctgca aacggtggaa ttccggcgag tgagaacaaa ctcagtgtca tgaggaatgc     23640
     cagtttgggg ttgctttggt acagaccgtt atagtcgtcc atctccaact ttccgctgcg     23700
     tcactcggag acgggcattg acattcaccc tggagccacg attggcgagt atttcacaat     23760
     cgaccacggc acaggtgtgg tgattggcga gacaagcgtg attggcaaca atgtgaagct     23820
     ctatcagggt gtgacattgg gcgcaaagag ttttccgctt gacgaag                   23867
//
ID   XXX; XXX; linear; XXX; XXX; XXX;  XXX.
XX
AC   PRJEB67889
XX
DE   Uncultured bacterium DNA fragment from in-reactor enriched cow rumen, clone
ferm-41j20,contig2
XX
FT   source          1..4322
FT                   /organism="uncultured bacterium"
FT                   /environmental_sample
FT                   /mol_type="genomic DNA"
FT                   /isolation_source="Cow rumen"
FT                   /clone="ferm-41j20"
FT                   /note="AZO-galactomannan active clone"
FT   CDS             86..424
FT                   /db_xref="SEED:fig|6666666.127254.peg.1"
FT                   /translation="MVEISSDVIRGYNDTMILCILMKGPSYGYEISKQIKTISEEKYVI
FT                   KETTLYSAFTRMEKNEFIESFVADPGPDGNGKKRTYYRITEKGKEYYREKCEEWALTKD
FT                   VIEKFIEH"
FT                   /product="Transcriptional regulator, PadR family"
FT   CDS             435..1415
FT                   /db_xref="SEED:fig|6666666.127254.peg.2"
FT                   /translation="METIKNYLESMFASMPNTPEVRKAKAELLQMMEDKYNELISEGTS
FT                   ENAAVGTVISEFGNLDELAESLGLTKEVEVARGMETEMPRRFISLDEAKAYLSNRNKKA
FT                   LFRALGILLCIISVSFVIILGPKGVEVLSGVGLMILSIAIGIGFLVYSGFTDHEWTFIG
FT                   HEYCSIDMNTAAYVKEKQRSFEPIRALCVSLGIVLCIVCWTPNILIPFRYQPLGAGIMF
FT                   IVVGFGVFLIVYSNSIARGFKRLLNVNEKSTISGQYANGSGYNKPVKYKSKTAQTIVEV
FT                   YWPTVTCIYLIISFLTFSWGSTWIIWVIAGVFHRVVMINCVDDED"
FT                   /product="hypothetical protein"
FT   CDS             1425..2180
FT                   /db_xref="SEED:fig|6666666.127254.peg.3"
FT                   /translation="MTKLYITVISIVTVVAIMFGCFIHIFRRGNFSWNVNNLEAVSDTV
FT                   ELSGDLDTIEVNMSYAGITIEYGDDFHVEYDLPRGYVPEIDLNNGTLSIENKNTLTIGN
FT                   HGWNDFTMVIVIPEGTELEKLDVKLNAGNLEIEDIPVSNFKVNCDAGNIELENIECAEM
FT                   DVEVDAGNIEIRDCNITKFTVSADAGNIEMHNSTIDTIKADADAGNIESHNCTINSGSV
FT                   ETDLGNINLRGDIGDVTTKTSLGIVNTDN"
FT                   /product="hypothetical protein"
FT   CDS             complement(2233..4257)
FT                   /db_xref="SEED:fig|6666666.127254.peg.4"
FT                   /translation="MCACLSAVLLIINLFTGCVYDIDEANAYHRGLFFIVNYLLVFAGP
FT                   VLDLTIVLEFRKKISKKLVKSFLLFIIVPIAAGVIQAAKYGVSLINISLAIVSIFLYTF
FT                   AYEDINDKISDEKDTEVEHLKSDQIKIQNLFEQTIKAFVNAIDARVDYAEGHSVRVADY
FT                   SYMLAQMNGKNDDECKEIYQAALLHDVGKIGISDEILKKEGNLTKEEIDSMRQEPLIGE
FT                   RILSGIAEYPYLSLGAKYHMERYDGSGFPKGLKGTDIPEAARIIAIAECYDSMTTRNSY
```

```
FT                   REPLPQSVVREELIKESGYKYDPLYTGIMLQLIDSDVDYQMKEETENVDTIFKNEFTCE
FT                   EYRSAISYGIRIRPEKKIIAFCATPVDEENAYSSPAIIIFDSLDERVHDTDRAIKDTRY
FT                   TEFGEIWFDGHFVCSEARNIKLISIEPIEQDEDETYFIEMARNRDHVLLKLEGGGYRSE
FT                   VVVALPDTSKYAYISMTGENCHISSIEVEKTDKRVPEDEIPRIAEEINYINRLESDIPN
FT                   VQIDSNRSDYSEGVKIKDGLRLIFHTMSLPSANLVWHCPFLLLYSSEDAKVMGRGYKEY
FT                   AMIRLDGEVQGGGPDSENKTEAVKGKDFTNWDAWKKLNRKGFECTVSFTRFKNKIITET
FT                   ENGGISITNITTLNDKKADVYISITGDQVAVTDIRIKNR"
FT                   /product="two-component response regulator"
XX
SQ   Sequence 4322 BP; 1265 A; 981 C; 859 G; 1217 T; 0 other;
     ctaaaacaaa cctgacgagg tgcgtgagcg gtccggccgg ccgcagtacc atgtaaaaca        60
     atcacctatt aacaggagga agaaaatggt agagatcagc agtgatgtga tccgcggtta       120
     caacgataca atgattctct gcattctcat gaagggacct tcctacggct acgaaatttc       180
     caagcagatc aaaacgatct cagaagaaaa atacgtcatc aaggaaacaa ccctctactc       240
     ggccttcacg agaatggaaa aaaatgaatt catcgagtca ttcgtggcag accccggtcc       300
     cgacggtaac ggtaaaaaga gaacgtatta ccggataact gaaaagggta aggagtacta       360
     cagggaaaaa tgtgaagaat gggccctgac caaagatgtt atcgagaagt tcatcgaaca       420
     ttaaggagag aaaaatggaa actataaaaa attatctgga atcgatgttt gcctccatgc       480
     ccaacactcc cgaagtgcgt aaggcaaagg ctgaactcct tcagatgatg gaagataaat       540
     acaacgaact catctccgaa ggaacaagcg aaaatgctgc agtcggaaca gtcatctccg       600
     aattcggaaa tcttgacgaa cttgccgaga gcttaggcct caccaaagaa gtcgaagttg       660
     caagaggaat ggaaaccgaa atgccccgca gatttatctc tctcgacgag gcaaaagcat       720
     acctctcaaa cagaaataaa aaagctcttt tcagggctct cggaatactc ttatgcatca       780
     tatcggtatc ctttgtcata atccttggcc ccaaaggagt ggaagttctg agcggcgtag       840
     gtttgatgat actcagtatc gcgatcggta tcggttttct ggtttattcc ggttttacag       900
     accacgaatg gaccttcatc ggtcatgaat actgcagcat cgacatgaac accgcagctt       960
     acgttaaaga aaagcaacgc agtttcgaac ccataagagc actttgcgta tccctcggta      1020
     ttgtcctgtg catagtctgc tggacaccta atatcctgat tcccttcaga taccagcctc      1080
     ttggcgccgg gatcatgttc attgtagtcg gattcggagt attcctcatc gtatactcca      1140
     attccatagc tagaggtttc aaaagacttc tcaatgtcaa cgaaaaaagc accatcagcg      1200
     gacaatatgc taacggaagc ggttacaaca aacctgttaa gtataagagc aagaccgcac      1260
     agactatcgt tgaagtatac tggccgaccg ttacctgcat ttatctgata atcagcttcc      1320
     ttactttcag ctggggatca acctggatca tctgggttat agccggagta ttccacagag      1380
     tggtaatgat caattgtgtt gacgatgaag actaggagga aatcatgact aagttatata      1440
     taacagtaat ttcaattgtc acagttgtgg ctatcatgtt cggatgtttt atccatatat      1500
     tccgcagagg aaacttctca tggaacgtta ataacctcga agccgtaagc gatacggttg      1560
     agctttcagg agatcttgat accatcgaag taaatatgag ctatgcaggt atcaccatcg      1620
     aatacggtga tgactttcat gtagaatatg atcttcccag aggttatgtt cccgagattg      1680
     acttaaacaa cggaactctt tccatcgaga acaaaaacac attaaccatc ggaaatcacg      1740
     gatggaatga ttttacaatg gttatagtca tccctgaagg cacagagctg gaaaaattag      1800
     atgtaaaact caatgcagga aatcttgaga tcgaggatat acccgtaagc aacttcaagg      1860
     ttaactgtga tgcaggtaac atcgagcttg aaaatatcga atgtgccgaa atggatgttg      1920
     aagttgatgc aggtaacatc gagatcagag actgcaatat caccaagttc acagtaagtg      1980
     cggatgcagg taatatcgag atgcacaatt caaccatcga taccattaag gccgatgcgg      2040
     atgcaggaaa tatcgagtct cacaactgca ccatcaatag cggatccgtt gagacagatt      2100
     taggcaacat taaccttcgc ggagatatcg gagatgtaac taccaagact tccctcggaa      2160
     tcgtaaatac cgacaactga tatatacaaa aagtgacacg gggttcactt cagatcccgt      2220
     gtcacttttt tattatctgt tctttattct gatatccgta acagctacct gatcacccgt      2280
     gatcgatata taaacatccg ccttcttatc attaagagtt gtgatatttg tgatcgatat      2340
     tccgccgttt tccgtttcgg tgatgatctt attcttaaac cttgtaaagc tgacagtaca      2400
     ttcaaaacct tttctgttca gcttcttcca ggcatcccag tttgtgaaat ccttgccttt      2460
     aaccgcttcc gtcttatttt ccgaatcggg tcctccgccc tgcacttcac cgtccaggcg      2520
     gatcatggca tattccttat agccccttcc cattaccttc gcatcttccg aagagtagag      2580
     caagaggaaa ggacaatgcc agacaagatt tgcggaagga agactcatgg tgtggaaaat      2640
```

```
     aagtctcaag ccgtctttta tcttcacacc ttcggagtaa tcacttctgt tactgtcgat      2700
     ctgaacattg ggaatatcgc tttcaagcct gtttatatag ttaatctctt cggctattct      2760
     gggtatttca tcttcaggaa ctctcttatc agtcttctca acctctatgg aagagatgtg      2820
     acagttttct cccgtcatac tgatatacgc atacttggaa gtatcgggga gtgcaaccac      2880
     aacttcgctt ctgtatccgc cgccttcaag cttaagcaga acatgatccc tgtttcttgc      2940
     catttctata aaataggtct catcctcatc ctgctctatc ggttcaattg agatgagctt      3000
     aatgtttctg gcttcagagc atacaaaatg tccgtcaaac cagatctctc cgaattcagt      3060
     atatctggta tctttgattg ctctgtcggt gtcatgaact ctttcatcaa gcgaatcaaa      3120
     tatgatgata gcaggagaag aatatgcatt ctcctcatca accggagttg cgcaaaacgc      3180
     aataatcttc ttctcgggtc tgatccttat tccgtatgat atcgcgcttc tgtactcctc      3240
     gcacgtaaat tcgttcttaa atatggtatc tacattttcc gtttcttctt tcatctgata      3300
     atcaacatca ctgtctatca gctgaagcat aattccggta tataaaggat cgtacttgta      3360
     tcctgattct ttgatcagtt cttctcttac aaccgactga ggaagaggtt ccctgtaact      3420
     gtttctggtc gtcatcgaat catagcattc cgcaattgca atgatccttg cagcttcggg      3480
     aatatcagtc ccctttaatc ctttgggaaa tccgcttccg tcatatctct ccatgtgata      3540
     tttggcaccg agtgaaagat acggatactc ggcaatgccc gacagaattc tttctccgat      3600
     caggggttcc tgcctcatcg aatcgatttc ttctttggta agatttcctt ctttttttaag      3660
     aatttcatcg gagattccta tcttacctac atcatgcaaa agcgcagcct gatatatctc      3720
     cttacattca tcgtcatttt ttccgttcat ctgggcaagc atgtaagaat aatccgcaac      3780
     tctgacagaa tgtccttccg catagtcgac tcttgcatct atagcattaa caaaagcctt      3840
     gatggtctgc tcaaacagat tctggatctt gatctggtcg cttttgagat gctcaacttc      3900
     ggtgtctttc tcgtcactta tcttgtcgtt gatatcttca tacgcaaatg tgtacaggaa      3960
     tatcgaaact atggcaagtg aaatattaat aagcgaaaca ccgtattttg ccgcctgaat      4020
     tacacctgcc gcaatcggaa cgataataaa gagcaggaat gatttaacaa gttttttact      4080
     aatctttttt ctaaactcaa gaacaatagt aagatcaaga acaggtccgg caaataccag      4140
     aagataattt acaatgaaga ataatcctct gtgatatgca tttgcttcat cgatatcata      4200
     aacgcatccc gtaaagagat tgatgatcaa taaaactgca ctaagacacg cacacatagc      4260
     agccgccctt aatctcttcg gtgcgttctc tcccgtatcc tctaatacta ccgttcctac      4320
     at                                                                4322
//
ID   XXX; XXX; linear; XXX; XXX; XXX;  XXX.
XX
AC   PRJEB67889
XX
DE   Uncultured bacterium DNA fragment from in-reactor enriched cow rumen, clone
ferm-41o1, contig-1
XX
FH   Key             Location/Qualifiers
FH
FT   source          1..22594
FT                   /organism="uncultured bacterium"
FT                   /environmental_sample
FT                   /mol_type="genomic DNA"
FT                   /isolation_source="Cow rumen"
FT                   /clone="ferm-41o1, contig 1"
FT                   /note="AZCL-Barley beta glucan active clone"
FT   CDS             347..1810
FT                   /db_xref="SEED:fig|6666666.127255.peg.1"
FT                   /translation="MKLMKKSFYYIITGLLLALLIVFGWQVFWLHGLYMTLEEELKRDL
FT                   IKQIEVADQFDLLRRLKMVENRYKQKNFEKVISISGKTDDDGQYEFEKEERYVDRRKDD
FT                   KDTTVIISQVNPKGPISISEHLIKSINESIHDAVDIEEPIDLQRIDSVYRQNLLNSQIH
FT                   ITHYYTELYDQKNDSLLSTTFDASQMVKKRETPTFTYSFGAAGRYCYRVYMEPLTQTVL
FT                   IRMGGILLTTVLIMLILGFTFWYLIRTLFKQKTLEEMKDDFTNNMTHELKTPIAVAYAA
FT                   TDSLLNFDLSKDEEKRRKYLQICRDQLMRLSHLVEQILSMSMEQRKSFQLHPEEIPVKE
```

```
FT                   LINGLVEQHKLKSDKPVTFTVSCATDSEIVYADRTHLSNMLSNLIDNAIKYSGEQVNIR
FT                   IDARRKNNRCEFRVSDNGPGIPHDKQAHIFDKYYRIPNGNLHTIKGYGLGLFYVKVMAE
FT                   KHGGQVAVKSAPGKGSTFIIQLPINPEIS"
FT                   /product="Osmosensitive K+ channel histidine kinase KdpD
FT                   (EC 2.7.3.-)"
FT   CDS             1807..2499
FT                   /db_xref="SEED:fig|6666666.127255.peg.2"
FT                   /db_xref="GO:0000156"
FT                   /db_xref="GO:0003677"
FT                   /translation="MKNKHIQVLLVEDEATLAMIIKDTLEESNLFKIQIAEDGEKGLRL
FT                   FFETKSDIVVTDVMMPKVDGFEMVRRLRQKDPTTPILFLTARSAIKDVVAGFELGANDY
FT                   LRKPFGMQELIVRIKALVQKTQRNTETEDAFVIGDYRLDPVAQKLYHLGEIDEELSYRE
FT                   TEILKRLCRNKDQIVNSQDILLDLWGDDSFFNSRSLHVFITKLRHKLSKDPRIRILNIR
FT                   GIGYKLTI"
FT                   /product="Two-component system response regulator"
FT   CDS             2724..7178
FT                   /db_xref="SEED:fig|6666666.127255.peg.3"
FT                   /translation="MKNILGLDLGTNSIGWALIDKDAHKIVGMGSRIIPMGQDILGDFE
FT                   KGNSISQTAERTRYRGIRRLRERHLLRRERLHRILHILGFLPEHYENCIDFEKRLGQFI
FT                   AETEPKLANNNQNFIFKKSFEEMVEDFEKRNPTLFRVKTNGKKTKIPYDWTIYYLRKKA
FT                   LTQKIEKEELAWIILNFNQKRGYYQLRGEDIEENPNKLVEYHSLRIVEVVADEKPNNKG
FT                   ETWYSLHLENGWIYRRSSKNPLFDWKDKVRDFIVTTDLDNDGHIKLDKEGNEKRSFRAP
FT                   NEDDWTLLKKKTEAEIENAHKTVGTYIYDCLIDNPTQKIRGKLVRTIERKYYKNELIAI
FT                   LKKQIELQPELFSEDVYNDCIRELYRNNETHLFNLSKRDFIHLFIDDIIFYQRPLKSKK
FT                   SSISNCSLEYRLYQDKNKNWTKEYLKAIPKSHPIYQEFRLWQWLYNIKIFRKSDDQDIT
FT                   SEYLANAQDKEDCFDFLNSRKEIDQKALLKYFKLTVKTHRWNYVEDKIYPCNETGTQIK
FT                   TRLAKVKNYAPDFLTPEIEHNLWHIIYSVTDKIAYKNALKTFARKHNLDEDTFVENFEK
FT                   YPPFKNEYGTYSEKAIKKLLPLMRLGKYWDWDAIDITTQNRINKILTGEFDPSIKDRVR
FT                   EKTITLTNEHDFQGLPLWLAQYVVYDRHAEAGDIKCWRTPEEIDNYLKEFKQHSLRNPI
FT                   VEQVITETLRVVRDIWKKYGDISEIHVELGREMKKTANERKQLTSQITENENTNIRIKN
FT                   LLMELKQNSDGKLQVDNVRPYSPIQQELLKIYEEGVIKSNIEIPDDILRISKTAQPSTS
FT                   ELQRYKLWLEQKYRSPYTGEFIPLGKLFTTEYEIEHIIPQSRYFDDSFSNKVICESAVN
FT                   KLKDRQLGLEFIKNHHGEKVQLGLKKIATVFSEEKYVAFVKENYANSRSKMKKLLLEEI
FT                   PESFVDRQMNDTRYISKIVKGLLSNIVREEDEQEATSKNVLTSNGSITSTLKRDWGLDA
FT                   VWNSLIISRFERLNILTNSTAFTSWNEQYQKFIPTVPLELSKGFQKKRIDHRHHAMDAL
FT                   VIACTTIDHVNLLNNQSANSDLKRYDLQRKLRKFEQTSYFDKTTNKQVVRNIPKEFIKP
FT                   WDSFTTDAKKELDKIIISFKQNLRIINKSTNKYQAYENEKKVFKSQTKGDSWSIRKPLH
FT                   KDTVYGKVSLQKTKRVRLSEALKNPESIVDKALKKEIQRIAALYGKFDTDWINKYFKDR
FT                   KYLFEKTDISKVDIYFFESENAAVRKPLDTSFTEKTIKESITDSGNQKILLNYLATKDH
FT                   KPELAFSPEGIEEMNENIAAFNDNKPHQPIRKVRVYEPIGYKFQVGYTGNKKVKYVEAA
FT                   KGTNLFFAIYVDESGNRSYESIPLNIIIERMKQGLYAVPLKNEKGYELLFHLSPNDLVY
FT                   VPTEEELASGQNISMEDLDISRIYKMVSCTGNRSFFIPNNIACPIEQTTELGPNNKAEK
FT                   SWDGIMIKQVCIKIITDRLGYIM"
FT                   /product="CRISPR-associated protein, Csn1 family"
FT   CDS             7252..8193
FT                   /db_xref="SEED:fig|6666666.127255.peg.4"
FT                   /translation="MMIKKTLYFGNPTYLSLRNNQLVVKLPEIENNDSLPDKFKKESEV
FT                   SFPIEDIGVIILDHKQITVTHGLIEQLLDNKSALITCNSSRMPVGLMLPLSGNKIQTEK
FT                   FNAQIQATDPLKKNLWQQTVKAKIKNQTEVLRKYRRCNIRNMQYWASEVKSGDPDNYEA
FT                   RAAAYYWKEAFPQIENFVRGREELPPNNLLNYGYAILRAVVARGLVSSGLLPTLGIHHH
FT                   NRYNAYCLADDIMEPYRPFVDAIVLSIVDRGDDFNELSKSIKTELLRIPTIDVIINGKK
FT                   SPLMVAVSQTTSSLTQCFDGKLRKIKYPEFII"
FT                   /product="CRISPR-associated protein Cas1"
FT   CDS             8235..8540
```

```
FT                   /db_xref="SEED:fig|6666666.127255.peg.5"
FT                   /translation="MWIFVLFDLPTETKKDRKIYADFRKNLQKDGFTMFQFSIYLRHCM
FT                   SMENANVHINRVKKFLPEKGHVGIICITDKQFGMMEVFYGKTAEAKKTEGQQLELF"
FT                   /product="CRISPR-associated protein Cas2"
FT   CDS             12475..12966
FT                   /db_xref="SEED:fig|6666666.127255.peg.6"
FT                   /translation="MQSLGGDESSLENSIFADEVLLDKFLRMSTIPSAAFADTKVHYAL
FT                   LDGLRGVAALLVVWYHVFEGYAFAGGQAIIDGFNHGYLAVDFFFILSGFVISYAYDDRW
FT                   SKGFTVKDFFKRRLIRLHPMVVMGALLGVITFCIQRAVQWDGTQVAIPMIVLSCLVHCF
FT                   "
FT                   /product="Acyltransferase"
FT   CDS             12957..13712
FT                   /db_xref="SEED:fig|6666666.127255.peg.7"
FT                   /translation="MFLIPALPGSGQEVRGNGEMFPLNGPSWSLFFEYIGNILYAWFIR
FT                   RLSNRALAVLVALLGVALTLFAVFDISGYGSIGVGWTMDGLNVAGGSLRMLFPFSMGML
FT                   LARHFKPIRVKGAFWICATALVALFAVPYIEGLEPICANGVYEAFCIVVVFPVLVWIGA
FT                   SGTTTDKQSTAICKFLGDLSYPLYMVHYPFMYLFYAWLIQSQLFSFGETWPVALGVYGW
FT                   NILLAWICLKYYDEPVRRYLANRFLKRRR"
FT                   /product="Acyltransferase"
FT   CDS             13859..14362
FT                   /db_xref="SEED:fig|6666666.127255.peg.8"
FT                   /translation="MLQNDIRPTVFYFNPNIFPKEEYEIRKAESKRYVRALGLDFVDAD
FT                   YDHADWREQVKGLENEPERGRRCVKCFMVRMLETARYASEHGFTVFTTTLASSRWKNLH
FT                   DIFAAGHAAEAEFQGVKFWDQNWRKGGLSERRLQIIAEYDFYNQQYCGCEFSFRKEKSP
FT                   TSSK"
FT                   /product="FIG053235: Diacylglucosamine hydrolase like"
FT   CDS             complement(14403..16652)
FT                   /db_xref="SEED:fig|6666666.127255.peg.9"
FT                   /db_xref="GO:0008422"
FT                   /translation="MKVLKIFPACILLASVVSCKPSAEATGEAAVEQRVETLLKEMTLE
FT                   EKIGQMNQISPPEDVSQIADLIKKGEVGSILNLVDPEAINALQQIAVEETRMKIPLIVG
FT                   RDVIHGFKTIFPIPLGQAASFNPELVKAGARVAAVESASQGVHWTFAPMIDIARDPRWG
FT                   RIAESLGEDTYLTSVLGAAMIQGFQGDELSDPTAIAACPKHFVGYGAAESGRDYNSTFI
FT                   PERRLRNVYLPPFQAAVEAGAATFMTSFNDNDGVPSTGNRFILDEVLRKEWNFRGFVVS
FT                   DWASVSEMISHGFAKDGKEAAMKAINAGVDMEMVSYTYVTYVKELLAEKKISMESIDNA
FT                   VRNILRIKIQLGLFENPYVDTSAPSVLYAEEHLQKAKEAAIESAILLKNENGILPLNES
FT                   VKSIAVVGPMADAPHDQLGTWVFDGDKNYTQTPLKALQAAYGDQVRILYAPGTAFSRDT
FT                   RTEGIAAAVNAARNADVVLVFVGEEAILSGEAHSLADLNLQGAQSELIAAVAKTGKPVV
FT                   TIVMAGRPLTIGRDAALSQAVLYNFHPGTMGGPAIADLLFGKAVPSGKLPVTFPKMVGQ
FT                   IPIYYGHNSTGRPYHGNETLLADIPLEAGQTSLGNTSFHLDAGFAPLYSFGYGLSYTTF
FT                   DYSNLKISADRLKATDVMTVTADITNTGKYDATEIVQLYVQDCFGSVTRPVKELKDFAR
FT                   VFLKSGETKTVVFSLPIEKLAFFNLDMNYGVEAGDFNLWVGPNSQEGLKTSFVVE"
FT                   /product="GH3"
FT                   /EC_number="3.2.1.21"
FT   CDS             complement(16665..17762)
FT                   /db_xref="SEED:fig|6666666.127255.peg.10"
FT                   /translation="MKIKNMIPLLILILASMGCGGSEDTPVVDEKITCSPAAISAGYAG
FT                   ETATIEVVSTREWTAYSNDEWLTCTPSSSIETTGTVTIQVDANPKTAIREGLVIVKAGS
FT                   SRDTVRVTQAAKPETPVDPNITVPEGYQLVWNDEFESGTAPGAEWWYETGDHGWGNNEL
FT                   QNYVAGSYGSEQLASVKDGILTITAKKIGDTVYSIRMNTGQGWKYGYFEARLKLPTGKG
FT                   TWPAFWMMPKNYTAWPDDGEIDIMEEVGYNPNYVSSAIHCKAYNHSIGTQKTKDVLIPT
FT                   AQTDFHVYALEWTEDYIRTYVDGKELFYFANDKTGNKDTWPFNAAFYLKLNLAWGGNWG
FT                   GAQGVDESVLPATYEIDYVRVFQKK"
FT                   /product="GH16_3"
```

```
FT                   /EC_number="3.2.1.73"
FT   CDS             complement(17836..19200)
FT                   /db_xref="SEED:fig|6666666.127255.peg.11"
FT                   /translation="MKKRYNYAKSILMMLALALGITACSPDDFTSPNEAGIPLAANYED
FT                   AITILVNQETNWVTFSFNATGVMPLWIIDGTTYTSTFNMTKYYRKAGDYSVDVKIVNAN
FT                   GISDGVITKSFTINKTIMNGFGGFVYDSDFNLWKKATIATPTFWYAPGWSQIADPTFKS
FT                   DGASYSISLPEATTDTWQAQMLMGTDIATNANTHYDFSAILTATQDHPHVMVKLVDSTD
FT                   DNVFYFAETVKLTANEPVCFWKSDMEGIDIGSLKMVFDFGGNAAGTDITIENIVLKDHA
FT                   NDDGTIVPEEELVPEPNWSAVDSDDNLWKNVTFTNTFYYAPGWSQLADPDLTIEGTQYS
FT                   VDFPSATTDQWQNQVTFVTDNLSTTSAENYDFKVTLKASNDIQGVTVKLAQNDDDNVFL
FT                   VMERIDLLAGEDVVAKVINAEGVDITQGKLVFDFGGNPANTHVVIKDIILQKHKD"
FT                   /product="FIG00412289: hypothetical protein"
FT   CDS             complement(19221..20645)
FT                   /db_xref="SEED:fig|6666666.127255.peg.12"
FT                   /translation="MTQETIDTYFTTDEHIQEAVVAAYDPLHWSDWGIGQYNPLCIMSD
FT                   IMADDIWVGGADKNDNQYWHLMMNYEALPTNCMTGLWTVAYSGVKRSNDVLTYIGWAGD
FT                   NLTAENATYYEAQARVLRVLYYNWLWKFWGNIPYYEENLTAPYIAEQLSADAVYANIIA
FT                   DLEGAIALDALPMKASSSDYGRVTKAMAYMLYAEIVLYQNDESRYAKALAYMQEIINSG
FT                   QYELVDDYTTIFKESGEWNSESIFEVNYKDDNAARSWGSPLVAGGTVLPRLISPSGWAD
FT                   GTDGHDNGWGFCPVRLETYERYSNNDARREATCWNAAAIGSYNTRYQDTGFFLEKYAAK
FT                   TGDNADQIADADLNYNNNLRIYRFSETLLNAAELIVRGGGSGDAKEYLNRVHKRAGLTI
FT                   EVAATIDNIIEERHLEFVGEGKRYWDLIRTGKAASVLVPDSYGYRTNAWTENKKYLPIP
FT                   QSEIDAAQGTLTQNNY"
FT                   /product="RagB/SusD domain protein"
FT   CDS             20644..20763
FT                   /db_xref="SEED:fig|6666666.127255.peg.13"
FT                   /translation="MDSTSKNDSEHPVSNKAEAMNPATILFLYLFITIYLRFE"
FT                   /product="hypothetical protein"
FT   CDS             complement(20764..22581)
FT                   /db_xref="SEED:fig|6666666.127255.peg.14"
FT                   /translation="MVNPLASLSLPGDKNWSHKFVANFSAELNIWDNLKFRTSYGADLS
FT                   FWGYDGYTPLYYLTANNKSTRTNAYSNKQDGLVWQLENVLMYDKTLDKHSFSVLLGQSA
FT                   KKSTGSQLGGNRYGIINTDRPYIDASNGLQENGDMGVYGGPSDQATLASLFARASYNYD
FT                   ERYMLQVTVRRDGSSRFGANNHYAVFPSFSLGWNLTNEAFMEKRPDWLTATKLRLSWGK
FT                   NGNENIGNFRYTVLTSTGNNAIFGSGENLTNGVKASGLANPDLKWEESEQLDFGVDFGF
FT                   FDNALTFTADYYKKTTNGMLMEMNIPSYVGESKPIGNVGKMENSGFEFESAYRFHVNDW
FT                   NFRVGGNLTYLKNKLIEYGNDSGWANLDSFQGIGSITRAQNGKPFPYFYGYKTDGLFQN
FT                   MEEVNAYVNNQGTLIQPNAVPGDVRFVDMNGDGKITDDDRTDIGKGMPDFTYGMNFSVS
FT                   WKNFDLNMMWQGTIGNDVFDATRRTDISSSNLPSWMLGRWTGEGTSDKYPRFVLGDNTN
FT                   WQSSDLYVVYDGSYMRLKNIQLGYTLPRQLTSKFMVSSLRFYVAAENLLTFTKYHGFDPE
FT                   ISSGGTSLGIDRGVYPQAKTWTVGANLTF"
FT                   /product="TonB family protein / TonB-dependent receptor"
XX
SQ   Sequence 22594 BP; 6761 A; 5014 C; 4755 G; 6064 T; 0 other;
     ccaacaaacg cggaacatca tacgttaact taccatcttt cactttcaca accggtcgtt        60
     caccactaac catgatatca ggtaagttca gcatagtaat caacgtatcc cgatcccatg       120
     cgacagtctg ttccacccct tcctgtgctt ttaccgatgt caaaatcatc accccaacca       180
     gcataaaacc caaaatccgt ttcatattca tttcgtttac tattttctgc aaaaaaaggt       240
     attcgagaaa caaccctatg ttatcattct cttacgaagt gttatttagt cttatcgatt       300
     tccggatgaa caggatttac cccgtatatt tgttatcaga caaaagatga agcttatgaa       360
     aaagtctttt tactacatca tcaccggcct cttgctcgcc ttactgattg tcttcggttg       420
     gcaggtcttt tggctccacg ggctgtatat gacactcgag gaggagctca aacgcgattt       480
     gatcaaacag atagaagtgg ccgatcagtt tgaccttctg cgaaggttga aaatggttga       540
     aaaccgttat aaacagaaaa acttcgaaaa ggtaatcagc atttccggaa agacagatga       600
```

tgacggacag tacgagtttg aaaaagaaga acgctacgta gaccgacgaa aagacgataa 660
agacactacc gtgataatca gtcaggttaa cccgaaaggg ccaatttcga tttcagaaca 720
cctgatcaaa agcataaacg aatctattca cgacgcggta gatatagaag agccgatcga 780
cttacagcgc attgacagcg tttatcgtca aaatctgctc aacagccaga ttcatattac 840
gcattattac acggagttat atgaccagaa aaacgacagt ttgctaagca ccactttcga 900
tgcttcccaa atggtcaaaa agcgggaaac gcccactttc acctattcct ttggagcagc 960
aggacgctat tgctaccggg tctatatgga gcccctcacc caaacggtac tgatccgtat 1020
gggaggcatc ttgttaacga ccgtgctgat catgctgatt ttgggcttta ccttttggta 1080
tctcatccgt actcttttta aacaaaagac actggaagag atgaaagacg acttcaccaa 1140
caacatgaca cacgaattga aaacgcctat tgctgtcgcc tatgctgcaa cggactcgct 1200
actcaatttc gatctgtcga aagacgaaga gaaaaggcga aaataccttc aaatatgccg 1260
ggaccagttg atgcgtctaa gtcacctggt cgaacaaatc ctgtccatga gtatggagca 1320
aaggaaatcc ttccagttgc atccggaaga gattccggta aaggaattga taaacgggct 1380
agtcgaacaa cacaagctta agtccgacaa acccgtgacg tttacagtaa gctgtgccac 1440
cgattcggaa atcgtctacg cagatcgcac ccatttgagc aacatgttga gcaacctgat 1500
cgacaacgcc ataaaataca gtggcgagca agtaaacatc cggatcgacg caaggagaaa 1560
aaacaatcga tgcgaattcc gggtttctga caacgggccc ggtattcccc acgataaaca 1620
agcacatatt ttcgacaagt attaccgcat tccgaacgga aatctgcata ccatcaaagg 1680
atatggcctg ggactctttt atgtgaaagt catggcggaa aagcatggcg gacaggttgc 1740
cgtcaagagt gcgccgggca aaggctctac ttttattatt cagctaccta tcaatccgga 1800
aatctcatga agaacaagca catacaggtt ttattggtcg aagacgaagc gactctagca 1860
atgatcatca aagatacgct ggaggaaagc aatctgttta aaatccagat tgcggaagac 1920
ggagaaaaag ggctacgact cttcttcgag acgaaatccg acattgttgt caccgatgtg 1980
atgatgccca aggtagatgg tttcgaaatg gtacgtcgac tcagacaaaa agatccgacc 2040
acccccatcc tgtttctgac tgcccgatca gccatcaaag atgtcgtagc cggattcgag 2100
ttgggagcca acgactacct ccggaaacct ttcggtatgc aagaactaat cgtccgcatc 2160
aaggcgctcg tccaaaaaac ccaacgaaac accgaaacgg aagatgcgtt tgtgattggc 2220
gactaccgac tcgatcccgt cgcacaaaaa ctttaccacc taggagaaat tgatgaagaa 2280
ctctcctacc gggaaaccga aatattaaaa cgactctgcc ggaacaaaga ccaaattgtc 2340
aactcacaag acatcctgct cgacctctgg ggcgatgaca gtttcttcaa ctcgcgcagc 2400
ctccatgtct ttatcaccaa gcttcgccac aagctgtcca aagacccacg catacgcatc 2460
ctcaacatcc gcggaatagg ctataaattg accatctaaa acctccaatc cccaaaatca 2520
cttcgagaac agacagccca attcgtcaaa accagaaaga agacaaaaca aaaaaggcga 2580
gaaataatcc ccgccttaca tgttttcaca acggaataat ccttattgtg ataagctttc 2640
aattactaca aaaatacatt tttatttaga aagcacacca tattcttaaa aataattatc 2700
tttaagtgtt aattatttac attatgaaaa atattttagg gctagattta ggcacaaaca 2760
gtatcggatg ggcgttaata gacaaagatg ctcataaaat agttggcatg ggaagccgaa 2820
taattccaat gggacaggat attctcggtg acttcgaaaa agggaattca atatcacaga 2880
cagcagagag aacgagatat agaggaatca gacgattacg ggaaaggcac ctacttcgcc 2940
gggaaagatt gcatcgcatt ttacacattc ttggttttct ccctgaacat tatgaaaact 3000
gcattgattt tgaaaaacgc ttaggacagt ttattgccga aacagaacct aaattagcta 3060
ataataatca aaactttatt ttcaaaaaat cttttgagga aatggtcgaa gattttgaaa 3120
agagaaatcc gaccttgttt cgtgttaaaa cgaatggtaa aaagacaaag attccatacg 3180
attggactat ttattattta cgcaaaaaag ccctaacaca aaagattgaa aaagaagaac 3240
ttgcttggat aatattaaat tttaatcaaa aacgaggata ttaccaatta agaggagaag 3300
atatagaaga aaatcccaat aaattagtcg aatatcattc cttaaggatt gttgaagtcg 3360
tagcggatga aaagccaaac aacaagggag aaacctggta ttcattacat ttggaaaatg 3420
gctggattta tagaaggtca agcaagaatc cattattcga ctggaaagat aaagtccgag 3480
attttattgt taccaccgac cttgacaacg atggccatat aaaattagat aaagaaggaa 3540
acgagaaacg ctcatttcgt gctccaaatg aagatgattg gactttgtta aaaaagaaaa 3600
cagaagcaga aatagagaat gcccataaaa cggttggtac atatatttat gactgtttga 3660
ttgataatcc aacccaaaag attagaggga aattagtacg cactatcgaa cgaaagtatt 3720
ataagaatga actaatagcc atattgaaaa aacaaattga gctacagcca gagttatttt 3780
ccgaagatgt gtataacgat tgcattcggg aattgtatcg caacaatgaa actcatctgt 3840

tcaacctcag taaaagagat tttatccacc tatttattga tgatattatt ttctatcaac 3900
gtcccctaaa aagtaaaaaa tcgtctattt caaattgctc tctagaatat aggctatatc 3960
aagataaaaa caaaaactgg acgaaagaat atctgaaagc cataccaaag tcacatccta 4020
tataccaaga gttccgccta tggcaatggt tgtataatat taaaatattt agaaaatcgg 4080
atgatcaaga tatcacttca gaatatttag caaatgctca agataaagaa gactgctttg 4140
atttcctgaa tagccgtaaa gagattgatc aaaaagcatt actgaaatat tttaagctaa 4200
cagtcaaaac gcatcgttgg aactatgtag aagacaagat ctatccttgt aatgaaaccg 4260
gaacccagat aaagacccgt ttggcaaaag taaaaaacta tgcacctgat ttcttaacac 4320
cagaaatcga acataactta tggcatatca tctattccgt aactgataag atagcatata 4380
aaaatgcatt gaaaactttt gcacggaaac ataatctgga tgaagatact tttgttgaga 4440
attttgaaaa atatccacct ttcaagaatg aatacggaac atattctgaa aaagcgataa 4500
aaaaactgtt gccgttaatg cgtttaggga aatattggga ttgggacgcc attgatataa 4560
caacccaaaa tagaatcaac aaaatcctaa caggagaatt tgatccaagt atcaaagatc 4620
gcgttcgtga aaaaacaata accctaacta atgaacatga ttttcaaggt cttcctttat 4680
ggctagctca atacgtcgta tatgacagac atgccgaagc tggcgatatt aagtgttgga 4740
gaactccaga ggagattgat aattatctta aggagtttaa gcagcattcg cttcgcaatc 4800
cgattgtaga acaagtgata acagaaaccc tgagagtcgt tcgtgatatc tggaagaaat 4860
acggtgatat ttctgaaata catgttgagc tgggacgtga aatgaaaaaa acagcgaatg 4920
aacgaaagca gctcacttct caaattacag aaaatgaaaa cacgaacatt cgcatcaaga 4980
acctattgat ggagttgaaa caaaattcag acggaaaact gcaagttgat aatgttcgac 5040
cctattctcc tattcaacag gaactcttga aatttatgga agaaggcgtc atcaaatcaa 5100
atattgaaat tccggatgat atcctgagaa taagcaaaac ggcacaacct tcgacctcgg 5160
aacttcagcg ttataaatta tggttagagc aaaaatatcg ctctccatat accggtgagt 5220
ttattccgtt gggtaaatta tttaccacag aatacgaaat agaacatatc attccacaaa 5280
gtcggtactt cgatgatagt ttcagcaata aagtcatctg cgaatcagcg gtaaacaaac 5340
tcaaagacag acaactagga ttagagttta tcaaaaatca ccatggcgaa aaagtccaat 5400
taggattaaa gaaaattgcg acggtttttt cagaagagaa atacgttgcc tttgtaaaag 5460
agaattatgc caatagtcgc tccaagatga aaaaactact attggaagaa attccggaat 5520
catttgttga ccgccaaatg aatgacacac gctatatcag caagattgta aaaggacttt 5580
tatccaacat cgtccgtgaa gaagatgagc aagaggcgac ttcaaaaaat gtattgactt 5640
ccaatggtag cattacctct acattaaaac gcgattgggg tttggacgct gtgtggaaca 5700
gtctcattat ttcacgcttc gagcggctaa atattttaac taatagtacg gcttttactt 5760
cttggaacga acaatatcag aaatttattc caactgttcc attggaatta tcaaaagggt 5820
ttcaaaaaaa acgaatcgac catcgccacc atgcaatgga tgcattagtt attgcctgca 5880
caacgataga ccatgtaaat ctgctaaaca accaatcagc caattccgat ttaaaacgat 5940
atgaccttca aagaaaatta aggaaatttg aacagacttc gtatttcgac aagacaacaa 6000
ataagcaagt cgttcggaat attcccaaag agtttattaa gccatgggat agctttacaa 6060
ccgatgccaa aaaggagctg gataaaataa tcatcagttt caaacaaaac ctgcgtataa 6120
taaataaatc aaccaataaa taccaggcat atgaaaacga aaaaaaagtt tttaaatcgc 6180
aaacgaaagg agacagttgg tccatacgaa aaccactaca taaagacacc gtttatggga 6240
aagtttcctt acagaaaacg aagagggtta gactaagtga agctttgaag aacccggaaa 6300
gcatcgttga caaagcatta aaaaaagaaa ttcaaagaat tgctgcccta tacggcaagt 6360
tcgatacaga ctggattaat aaatacttta aagacagaaa atatttattt gaaaaaacgg 6420
atatttcaaa ggtggatatc tatttttttg aatccgagaa tgccgctgtt cgaaaacctt 6480
tggacacttc attcacagaa aagacaatca aagagtctat aactgatagc gggaaccaaa 6540
aaatccttct gaattattta gcgaccaaag atcataagcc agaattagca ttttctccag 6600
aaggaatcga agagatgaat gaaaatattg ccgcttttaa tgataacaaa ccccatcaac 6660
ctatacgcaa agttagggta tacgaaccta ttggttataa atttcaagtc ggatatacag 6720
ggaataaaaa agttaaatat gtagaagcag cgaaaggaac aaacctattc tttgctattt 6780
atgttgatga aagcggaaac cgctcctacg agagcattcc actaaacatt atcattgaac 6840
ggatgaaaca aggactctat gcagtcccgc ttaaaaacga gaaggggtat gaattactat 6900
ttcacttatc gccaaatgat ttagtctatg ttccaacaga ggaggaatta gccagcggtc 6960
aaaacatctc aatggaagat ttggatatca gtcggattta taaaatggtc agttgcactg 7020
gtaacagaag tttctttatt cctaataata tcgcttgccc gatcgaacaa acaacagaat 7080

taggaccaaa caataaggca gaaaaatctt gggatggaat aatgattaaa caagtctgca 7140
tcaaaatcat aacagacagg ttaggatata ttatgtaatt atatcaatcg catataagcg 7200
aggctattat tggccccatt tttaaacaaa gctatttatc aaacgtgaat aatgatgatt 7260
aaaaaaacgc tttatttcgg aaacccaacg tatctgagtc tgcggaacaa tcagttagtt 7320
gtaaaacttc ctgagattga gaataatgat tctttacctg ataaattcaa aaaggaaagt 7380
gaagtcagtt tccctattga agatataggt gttatcattt tggatcataa acaaatcacc 7440
gtgactcatg gattgattga acagttgttg gataataaat cggcacttat tacctgtaat 7500
agcagccgaa tgccagttgg tctcatgcta cctttatctg gaaataaaat acaaacagaa 7560
aaatttaatg cgcagattca ggcaacagac ccattgaaaa aaaatctttg gcaacaaaca 7620
gtaaaagcta aaataaaaaa tcaaactgag gttttacgaa agtaccgtag atgtaacatt 7680
cgaaatatgc aatactgggc gagtgaagta aaaagcggtg atccggataa ttatgaagct 7740
cgtgctgcag cttactactg gaaagaagct tttcctcaaa ttgaaaattt tgttagaggc 7800
cgagaagaac ttccaccaaa caacctatta aattatggat atgcaatttt aagagctgtt 7860
gttgcgagag gtttagtgtc aagcggatta ttgccaacat tgggtattca tcatcataac 7920
cgttacaatg cttattgctt agcagacgat ataatggaac cttatcgccc gtttgtcgac 7980
gcaatagtac tttccattgt tgatcgtgga gatgatttta acgaattatc caaatccata 8040
aaaacagaac tgttacgaat tccaacaatc gatgtaataa ttaatgggaa gaagagtcct 8100
ttaatggttg cagtaagcca aaccacatcg tcattaacac agtgttttga tgggaaatta 8160
agaaaaataa aatatcctga atttataatc taactattgt tatgaaacat gacaggttaa 8220
atcaatatcg aattatgtgg atttttgttt tatttgatct ccctactgaa acgaaaaaag 8280
acaggaaaat ttatgcggat ttccgaaaaa atcttcaaaa agatggattt acaatgtttc 8340
agttttcgat ttaccttcgc cattgcatga gtatggaaaa tgcaaatgtt catataaatc 8400
gggtaaagaa atttctaccg gaaaagggac atgtcgggat catttgtatt actgacaaac 8460
aatttggaat gatggaggta ttttatggca aaacggctga agccaagaaa actgaaggac 8520
aacaattgga actcttttga aaataaaaat cccgacaaaa atcgggattt ttattgaaaa 8580
cacatcattt ttctaaatct taacggtcta cattagatac tgtatctcag caaattaaac 8640
acgcctatga tgtttttact gcccctaaga tacagaattg aaagcttatc acaacaaaat 8700
cggcaaaaca gacatgggaa ttgctgatgt ttttactgcc cctaagatac agaattgaaa 8760
gcttatcaca acagaagcta tggagtggct tgaggaggct ggatgttttt actgccccta 8820
agatacagaa ttgaaagctt atcacaaccc agggctggta tgtatctccc atccagtaga 8880
tgtttttact gcccctaaga tacagaattg aaagcttatc acaacagaaa gatcagcaac 8940
ctgttcacta atgcgatgtt tttactgccc ctaagataca gaattgaaag cttatcacaa 9000
ccgccgtaaa agccgcgccg aaattgttta cgatgttttt actgccccta agatacagaa 9060
ttgaaagctt atcacaaccg gaacttatta cgggcgttag caactacgat gtttttactg 9120
cccctaagat acagaattga aagcttatca caacatgggg atcatcatcc aagcgaacac 9180
ataggatgtt tttactgccc ctaagataca gaattgaaag cttatcacaa cggaagtggt 9240
agagcaccct atatataaat gatgttttta ctgcccctaa gatacagaat tgaaagctta 9300
tcacaactga aacagaggcg gccgaatata ccggtgtgat gtttttactg cccctaagat 9360
acagaattga aagcttatca caacaatgtc gctgatggac tttgccgacg cgctgatgtt 9420
tttactgccc ctaagataca gaattgaaag cttatcacaa ccaaggcgtt taattaaaaa 9480
aaatagatta tgatgttttt actgccccta agatacagaa ttgaaagctt atcacaacct 9540
atcggaaact ttaccagtca atacctggat gtttttactg cccctaagat acagaattga 9600
aagcttatca caactaatct gcgtagtacg gggatccgta cggagatgtt tttactgccc 9660
ctaagataca gaattgaaag cttatcacaa catcagacga agagctaaac gcattcgtcg 9720
agatgttttt actgccccta agatacagaa ttgaaagctt atcacaaccc gttgcagcat 9780
tgccacggaa cgcaacatga tgtttttact gcccctaaga tacagaattg aaagcttatc 9840
acaaccgata cagataagga ttttcgcatt acggagatgt ttttactgcc cctaagatac 9900
agaattgaaa gcttatcaca acataacgat tcacaacaat tgatcgtatt cagatgtttt 9960
tactgcccct aagatacaga attgaaagct tatcacaaca gttgaaatgc cctcttcgtt 10020
tacttcaaag atgtttttac tgcccctaag atacagaatt gaaagcttat cacaacttac 10080
cgcagcatta tagtaggatt ctttgaggat gtttttactg cccctaagat acagaattga 10140
aagcttatca caactgacgc tatcagcgtg tcatgtaagg cactgatgtt tttactgccc 10200
ctaagataca gaattgaaag cttatcacaa ctttattttg caggctatac attaacagcc 10260
ggatgttttt actgccccta agatacagaa ttgaaagctt atcacaacta aaacagaaag 10320

acactttgat attgatttga tgtttttact gcccctaaga tacagaattg aaagcttatc 10380
acaacatatt tgggcctgtg caacccgtgt tccttgatgt tttactgcc cctaagatac 10440
agaattgaaa gcttatcaca acatatttgg gcctgtgcaa cccgtgttcc ttgatgtttt 10500
tactgcccct aagatacaga attgaaagct tatcacaacc cgggcttggc tctttcaaag 10560
tcaaggtgag atgtttttac tgcccctaag atacagaatt gaaagcttat cacaacgaag 10620
tggttgacat cgtacaagaa agaaggatgt ttttactgcc cctaagatac agaattgaaa 10680
gcttatcaca acggaatacg gcagggacgt tagatcactt cgatgttttt actgccccta 10740
agatacagaa ttgaaagctt atcacaactt gttcgttata aggtaagcga caatatagat 10800
gtttttactg cccctaagat acagaattga aagcttatca caaccgttct taggcattga 10860
gccggacgca agtagatgtt tttactgccc ctaagataca gaattgaaag cttatcacaa 10920
catgaagcgg ccgtttatgg tggaacttcc ggatgttttt actgccccta agatacagaa 10980
ttgaaagctt atcacaacca aaattgaaat tattattcta tgatttttga tgtttttact 11040
gcccctaaga tacagaattg aaagcttatc acaacccgtc ttgccgtggt tccagttgct 11100
gaatggatgt ttttactgcc cctaagatac agaattgaaa gcttatcaca actctcctta 11160
gaatatgagc atttcggttt aagatgtttt tactgcccct aagatacaga attgaaagct 11220
tatcacaacc cgtcttgccg tggttccagt tgctgaatgg atgtttttac tgcccctaag 11280
atacagaatt gaaagcttat cacaacttaa accattttat ttgagtaata ccgtcggatg 11340
tttttactgc ccctaagata cagaattgaa agcttatcac aactagctaa atcaaccagc 11400
tgtcgggaat atggatgttt ttactgcccc taagatacag aattgaaagc ttatcacaac 11460
acttgatggt tcgttaggag tgtactttgg atgtttttac tgcccctaag atacagaatt 11520
gaaagcttat cacaacactt gatggttcgt taggagtgta ctttggatgt ttttactgcc 11580
cctaagatac agaattgaaa gcttatcaca acgcagccag ggcagcctta cgagccagag 11640
ccgatgtttt tactgcccct aagatacaga attgaaagct tatcacaaca acaaggcgca 11700
aagatctggg tgatgctatg atgtttttac tgcccctaag atacagaatt gaaagcttat 11760
cacaacattt gtctgatttt tccgacaaag atgaatgatg tttttactgc ccctaagata 11820
cagaattgaa agcttatcac aacgaaagat gcacagggta atagcgatac cagatgtttt 11880
tactgcccct aagatacaga attgaaagct tatcacaacg aaagatgcac agggtaatag 11940
cgataccaga tgtttttact gcccctaaga tacagaattg aaagcttatc acaacaatct 12000
tagggcttgc gtggtagtcg ttagtgatgt ttttactgcc cctaagatac agaattgaaa 12060
gcttatcaca actgttattg ccttgcttcg ggaagcatta acgatgtttt tactgcccct 12120
aagatacaga attgaaagct tatcacaacg gaatggtcag gttgagatag cacttaaagg 12180
atgttattac tgcccctaag atacagaatt gaaagcttat cacaacaata tctgccctaa 12240
tgccacagga gaggccgatg tttttactgc ccctaagata cagaattgaa agcttatcac 12300
aaccggctcg ccttgaaaaa tcgtgaactg gcggatgttt ttactgcccc taagatacag 12360
aattgaaagc ttatcacaac gccagagaca agagctacta tatagtatgt gatgttttta 12420
ctgcccctaa gatacagaat tattattgca gggcggtaaa tattgagtga ggaattgcag 12480
tcattggggg gggatgaaag ttcgttagaa aacagtatat ttgccgacga ggttttacta 12540
gataaatttt tacgaatgtc aactattcct tcggcggctt ttgcggatac gaaagtgcat 12600
tatgcacttc tggatggact gcgtggcgtg gcagcgctgc tcgtggtctg gtatcatgtg 12660
ttcgagggct acgcttttgc tggtggccag gcgattattg atggattcaa ccatggttat 12720
ctggcggtgg atttcttttt tatcctttcg ggattcgtta tcagttatgc ttacgacgac 12780
cgatggagca aggggtttac agtaaaggat ttttttaagc gccggttgat ccgtctgcat 12840
ccgatggtag tcatgggggc tcttctcgga gtcatcacct tttgtatcca aagggcagtg 12900
caatgggatg gaacacaggt cgccataccg atgatcgttt tgtcctgctt ggtgcattgt 12960
ttctgattcc tgcacttccc ggcagcggac aggaggttcg cggaaatggt gaaatgttcc 13020
cactgaacgg gccaagctgg tcgctcttct ttgaatatat tggcaatatc ttgtacgctt 13080
ggtttatccg acgcttgtca aaccgggcac tggcggtgct ggtcgctttg ctgggggtgg 13140
ccttgacgtt gttcgccgtc ttcgatatct ccggctacgg aagtatcggg gtgggctgga 13200
ccatggatgg cctcaacgtg gccggcggtt cgctaagaat gcttttcccg ttttccatgg 13260
gtatgctgct cgcccgccat ttcaagccga tacgtgtgaa aggggctttc tggatctgtg 13320
cgacggcttt agtcgcactg tttgccgtgc catatattga agggctggaa cccatctgcg 13380
cgaacggtgt ctacgaggcg ttttgcattg tcgtagtttt tccggtgctt gtctggattg 13440
gcgcttccgg aacaacgacg gataaacagt cgacagccat ctgcaagttt ttgggagacc 13500
tttcatatcc gctttatatg gtgcattatc cgtttatgta tctgttctac gcttggctga 13560

tccaatcgca gctttttagt tttggagaga cctggccggt cgctttgggc gtctacggat 13620
ggaatatact cttggcctgg atatgcctga aatattatga tgagcctgtg cgccgctact 13680
tggcgaaccg cttcttgaaa aggagacgat aaagccggta gcacaagttg acgcgatata 13740
ccttatcttt gtaccgaaaa gggattcgga aatgcttaaa ttagagacac ccaacggaga 13800
gaagaacgtg ttgctgcatg cttgctgcgc accttgctcg tcggctatca ttgaatgtat 13860
gttgcagaat gatatccggc cgacggtgtt ctattttaat ccgaatattt tcccgaagga 13920
ggaatacgag atccggaagg cggaaagcaa acgttatgtc cgggcactgg gactcgattt 13980
cgtggatgcg gactatgacc atgcggattg gcgtgaacag gtgaaaggtc ttgaaaacga 14040
gccggaacgc ggacgacgct gcgtgaagtg ttttatggtg cgcatgcttg agacggctcg 14100
ctacgcttcg gaacacgggt ttacggtgtt caccacgacc cttgcctcat cgcgctggaa 14160
aaacctacat gatatttttg cggccggtca tgcggcggaa gcagaattcc agggggtgaa 14220
attctgggat cagaactggc ggaaaggggg attaagcgaa cgacgcttac agataatcgc 14280
agaatacgat ttctataatc aacaatactg cggttgcgag ttcagcttcc gcaaagaaaa 14340
atccccgaca tcttcaaaat aaaacatcgg ggatttttcta cctaggaaca gtcggtcagt 14400
gtttattcca ccacaaagga tgtttttaaa ccctcctgac tgttggggcc tacccacaga 14460
ttaaaatcac cggcttcaac gccataattc atgtctaaat taaaaaatgc gagcttttct 14520
ataggtaatg agaatacgac cgtcttcgtt tcgccggact tcaggaagac acgggcaaag 14580
tctttcaact cttttacggg acgggtcacg gatccaaaac aatcttgtac gtacagctga 14640
acaatttctg ttgcgtcgta cttgcctgta tttgtgatat cggcggtaac ggtcatcacg 14700
tctgtcgctt tgagcctgtc tgccgagatt tttaaattac tgtaatcgaa cgtcgtatag 14760
gacaatccat atccaaagga atacagcgga gcgaatccag catccaaatg gaaagaggtg 14820
ttccccaaag aggtttgacc ggcttccaag gggatatccg ccaacaaggt ttcattgcca 14880
tgatagggc ggccggtgct attatgtccg taataaattg gaatctgtcc gaccatcttc 14940
gggaatgtga caggcaactt cccactgggt accgcttttc cgaaaaggag atcggctatc 15000
gccggaccgc ccatcgtacc gggatggaaa ttgtataaca ccgcctgcga aagcgccgca 15060
tcgcgtccga tcgttaaggg acgtcccgcc atcacaatgg tcactaccgg cttgccagtc 15120
ttggccaccg ccgcaatcaa ttcgctctgt gcaccctgca ggttcaagtc tgccagtgaa 15180
tgagcttctc cggaaaggat cgcttcttca ccgacaaaga ctaaaactac atccgcgttg 15240
cgggctgcgt ttacggctgc tgcaatgccc tcggtacggg tatcccgact aaacgccgtt 15300
cccggcgcgt agaggatacg tacctgatcg ccataagccg cctgcaaggc tttcaatgga 15360
gtctgcgtgt aattcttgtc tccatcaaaa acccatgtgc ctagttggtc gtgtggcgca 15420
tccgccatcg gaccgactac cgcgattgat tttacagatt cattaagtgg taaaatgccg 15480
ttttcgtttt ttaagaggat ggccgactct atcgccgctt ccttggcttt ctgcagatgc 15540
tcttctgcat acaaaacgga aggcgccgag gtgtcgacat aaggattctc gaacaagcct 15600
aattggattt ttatacgtag gatgttgcgt acggcattgt caatcgactc catgctgatt 15660
ttcttttcag ctagcaactc cttcacgtag gtgacatacg tgtaactcac catttccata 15720
tcaacaccgg cattgatcgc cttcatggcc gcttcttttc cgtctttcgc aaatccgtgc 15780
gagatcatct cgctgacaga agcccagtcg gaaacaacaa agccgcggaa gttccattct 15840
ttgcggagaa cttcatccag aataaagcga ttacccgtgg acggtacgcc gtcgttatcg 15900
ttgaacgagg tcatgaacgt cgccgcaccg gcttcaaccg cagcttggaa tggcggcaaa 15960
tagacgttgc gcaatcggcg ctcagggata aaggtcgaat tatagtcacg accactctcg 16020
gccgcaccgt aacctacgaa atgcttcgga caagcggcaa tggcagtcgg atcgctcagc 16080
tcgtcaccct gaaatccctg aatcatcgct gctccgagaa cagatgtcag gtaggtatct 16140
tcgcccaaac tttcggcgat tcgtccccaa cgcggatcgc gggcgatgtc aatcatcggt 16200
gcgaaagtcc aatgcacacc ttgcgaagcc gattcgaccg cagcgacacg tgcacccgct 16260
ttcaccagct caggattaaa cgaggctgcc tgacctaacg gaatggggaa gatggttttg 16320
aaaccgtgta tcacatcgcg accgacaatc aaagggatct tcatgcgggt ctcttcaacc 16380
gcaatctgct gaagcgcgtt gatggcttcc ggatcgacca gattcaaaat agaaccgact 16440
tcgcctttct tgatcaggtc ggctatttgg gatacatctt cgggaggact gatctgattc 16500
atctgaccga tcttttcctc caacgtcatt tctttcaata gggtttccac tcgttgctcg 16560
acggcagctt ctccggtcgc ctcggcagat ggtttacagg aaaccacgga tgccaacagg 16620
atacaagcgg gaaaaatctt taacactttc attttgtttc tctattattt tttctggaat 16680
acgcgtacat agtctatttc gtaagtggcc gggagaacgg attcatcgac tccctgcgcg 16740
ccgccccagt tgccgcccca agcgagattg agcttcaagt aaaacgctgc attaaagggc 16800

caggtatctt tatttcccgt tttgtcgttt gcaaaataga acaactcttt cccatccaca 16860
taggtacgta tataatcttc ggtccattcc aaggcgtaca catgaaaatc ggtttgggcg 16920
gttggaataa gcacatcttt cgttttctgc gtgccaatcg aatggttata ggccttgcaa 16980
tgaatggccg atgagacata gttcggattg tatccgactt cttccatgat atcgatctcg 17040
ccatcatcgg gccaagcggt atagttctta ggcatcatcc aaaatgccgg ccaggtgccc 17100
ttcccggtcg ggagttttaa gcgggcctca aaataaccgt atttccatcc ttggcctgtg 17160
ttcatacgga tggaatagac cgtgtcgcca atctttttgg cggtgatagt cagtatgccg 17220
tctttcacgg acgccaactg ctcggaacca tatgatccgg ctacataatt ctgcagttcg 17280
ttattccccc atccatggtc gccggtctcg taccaccact ccgcaccggg agctgtgccg 17340
gactcaaatt catcgttcca gaccagctga taaccttcgg gcacggtgat attggggtcg 17400
accggcgttt caggtttcgc tgcctgcgtc acccgaacgg tatcacgact actgccggct 17460
ttcacaataa ccagtccttc gcgtatcgct gtcttgggat tggcatcgac ctggatcgtt 17520
accgtacccg tcgtttcaat cgaactcgaa ggcgtacaag tcagccactc atcattactg 17580
taggccgtcc attcacgagt agaaacaacc tcaatcgttg ctgtttctcc ggcgtaaccg 17640
gcgctgatgg cagcgggtga acacgtgatt ttctcgtcaa cgacaggtgt gtcttcacta 17700
ccgccacagc ccatagaggc aagtattaga attaataagg gaatcatatt ctttattttc 17760
atgattattc tctcgtttgc gaatccggat gacctcaaga acgagttttt tctcaaggtc 17820
atccttattc atccattagt ctttatgctt ttgcaaaata atgtctttaa tgactacatg 17880
ggtattggcc ggattccctc cgaagtcgaa tacaagtttg ccttgcgtaa tatcaacccc 17940
ttcggcattg atcaccttgg ctacgacatc ttcacctgcg agtaaatcga tccgctccat 18000
caccagaaag acgttgtcgt catcgttctg ggccaatttt acggtcactc cctggatatc 18060
gttagaggct ttcaacgtca ctttaaaatc gtaattctca gccgaagtcg tggaaaggtt 18120
gtctgttaca aaggtaactt gattttgcca ttgatcggtc gttgcactcg ggaaatcgac 18180
cgaatattgt gtgccttcaa tcgttaagtc cggatccgcc aactgtgacc agcccggagc 18240
gtaatagaac gtatttgtga aggtgacgtt cttccacaga ttgtcgtcgc tgtcgacagc 18300
gctccaattg ggttcgggca ccaattcctc ttccggcacg atggtgccgt cgtcgttggc 18360
atgatctttc aggacaatat tttcgatggt gatatcggtt ccggcggcat tgcctccaaa 18420
atcgaatacc attttaagtg aaccgatatc gatgccttcc atgtcgctct tccagaaaca 18480
aaccggttca ttagctgtca acttcaccgt ttctgcaaaa tagaacacat tatcgtctgt 18540
gctgtctacg agtttcacca tgacatgggg atgatcctgt gtagcagtca gaatggcgga 18600
gaagtcataa tgcgtattgg cattggtagc gatatccgta cccatcagca tctgtgcctg 18660
ccaggtatcg gttgttgctt ccggcagact gattgaatag gaagcgccat ccgatttgaa 18720
ggtcgggtct gcaatctgtg accagcccgg agcgtaccag aatgtcggcg tggcaatcgt 18780
tgcctttttc cataagttga agtccgagtc gtaaacaaat ccgccgaacc cgttcatgat 18840
cgtcttattg atcgtaaatg atttggtgat cacgccgtca ctgattccgt tggcattgac 18900
aatcttcaca tccacactat agtcgcctgc cttacggtaa tacttcgtca tgttgaacgt 18960
agaggtatag gttgtcccgt caatgatcca gagcggcata acgcccgtcg cgttgaatga 19020
gaacgtgacc cagttggttt cttggttgac caggatcgtg attgcatctt cgtagttggc 19080
agccaatgga atgccggctt cattcggaga agtaaagtcg tccggagaac aagccgtaat 19140
ccccaaagcc aaggcaagca tcatcaaaat gcttttggca taattatatc gtttttttcat 19200
gattcttcta ctttaataga ttaataattg ttttgagtca atgtcccctg agccgcgtcg 19260
atttcgctct gaggaatggg caggtatttt ttattctccg tccaagcgtt cgtacggtat 19320
ccgtagctgt cgggcaccaa tacactggcc gcctttccgg tacggatcaa atcccagtag 19380
cgcttacctt cacctacaaa ctccaggtga cgctcttcga tgatattatc gattgttgca 19440
gcgacctcga tcgtcagacc ggcccgttta tgaacccgat tcaggtattc cttcgcatca 19500
cctgaaccgc caccacgaac aatcagttcc gccgcattca aaagcgtttc cgaaaaacga 19560
tagatacgca ggttattatt gtagttcaag tcggcatccg caatctgatc ggcattatcc 19620
cccgtcttgg ctgcatattt ctccaagaag aatccggtat cctggtagcg ggtattgtaa 19680
cttccgatag ctgctgcgtt ccagcaggta gcctcacgac gcgcgtcgtt attggaatac 19740
cgttcgtagg tttccaaacg tacaggacag aatccccagc cgttatcgtg accgtctgtt 19800
ccgtcggccc agccactcgg actgatcaga cgcggtagaa cggttccccc ggccaccagc 19860
ggacttcccc atgagcgtgc cgcattatca tctttatagt tcacttcaaa gatggattcg 19920
ctgttccatt cgcccgactc cttgaagatc gtcgtatagt catcgactaa ctcatattga 19980
ccgctattga tgatttcctg catgtatgcc aaagctttgg catagcggct ttcatcattt 20040

```
        tggtagagga cgatctcggc gtaaagcatg taagccatgg ctttcgtcac ccgaccgtaa     20100
        tccgagctgc tcgctttcat aggaagcgca tccaaagcga tagccccttc caaatcggcg     20160
        atgatattcg cataaacagc atccgcactc aattgttccg cgatataggg agcggtcagg     20220
        ttttcttcgt aataaggaat attgccccag aatttccaaa gccagttata ataaagcacg     20280
        cgaagaaccc ttgcctgcgc ttcataatag gtggcatttt ccgccgtcag gttatcaccg     20340
        gcccaaccga tataggtcag cacgtcgttg cttcgtttca caccggaata agcgaccgtc     20400
        caaaggcccg tcatgcaatt cgtcggcagt gcctcgtaat tcatcatcag gtgccagtac     20460
        tgattgtcgt tcttatcggc accacccacc cagatatcat cggccatgat atcactcata     20520
        atacataacg ggttatattg tccgataccc caatccgacc aatgcaaagg gtcgtaagcg     20580
        gcgactaccg cttcttgaat atgttcatcg gtcgtaaaat acgtatcgat tgtttcttgc     20640
        gtcatggact caacctccaa gaatgattcg gaacatcccg tcagcaacaa ggcggaagcc     20700
        atgaatcctg ctactatatt gtttttatat cttttcataa cgatttattt gagatttgaa     20760
        tgattaaaaa gtcaagtttg ctccgacagt ccaggtcttt gcctgaggat acaccccgcg     20820
        gtcaatacct aatgaggtac cacccgatga gatctccgga tcaaaaccgt gatatttcgt     20880
        gaaggttagc aggttctctg ccgccacata gaaacgcaaa gaagagacca tgaatttact     20940
        tgtcagctga cgaggcaagg tataacccag ctgaatattc ttcagacgca tatagctgcc     21000
        atcgtacaca tagaggtcgg acgactgcca gttcgtattg tcgcccaaca cgaaacgcgg     21060
        gtatttgtcg gaagtacctt cgcccgtcca gcgacccagc atccaagaag gtaggttgct     21120
        ggaagagata tccgtacggc gggtcgcgtc gaacacatcg tttccaatcg ttccttgcca     21180
        catcatgttc aaatcaaagt tcttccagga tacgctgaaa ttcataccgt aagtaaagtc     21240
        cggcatccct tttccgatat ccgtacgatc gtcgtccgtg atctttccat cgccattcat     21300
        atcaacaaat cggacatcac cgggaacggc attgggctga atcaacgtgc cttggttatt     21360
        cacataggca ttcacttcct ccatgttctg gaataggcca tccgttttat agccatagaa     21420
        ataagggaaa ggtttgccgt tttgcgcacg cgtgatcgag ccgatacctt ggaacgagtc     21480
        caggttggcc catccggaat catttccata ctcgatcaat ttattcttca ggtaggtcag     21540
        gttacccccc acacggaagt tccaatcatt cacatggaac cggtaggctg attcaaactc     21600
        aaaaccgctg ttttccatct tgccgacatt accgatcggt ttagattcac cgacatatga     21660
        aggaatgttc atttccatca acataccatt cgttgttttc ttataataat cagccgtaaa     21720
        ggtcaatgca ttgtcaaaga aaccgaagtc caccccgaag tcgagctgct cagactcttc     21780
        ccatttcaaa tccgggttgg ccaatccgct tgctttcacc ccgttggtca ggttttcgcc     21840
        cgaaccgaag atcgcgttat ttccggtcga tgtcaagacg gtatatctga aattaccgat     21900
        gttctcgtta ccgttcttac cccaagacaa gcgaagctta gtagcagtca accagtccgg     21960
        acgtttttcc atgaaagctt cgttagtcag gttccatccc aaagagaaag aagggaagac     22020
        cgcataatgg ttattggccc caaaacggga agaaccatcc cgacgtacag ttacctgtaa     22080
        catataacgt tcatcgtagt tatagcttgc gcgagcaaag agagacgcca atgttgcttg     22140
        gtccgacgga cctccataaa cgcccatgtc gccgttttcc tgcaaaccgt tgctggcatc     22200
        gatatagggc cggtcggtat tgatgattcc gtagcgatta ccacccaact gggaaccggt     22260
        acttttcttg gccgactgac ccaaaagtac ggagaacgaa tgtttatcca gcgtcttatc     22320
        gtacatcaat acattttcca gctgccacac cagaccgtct tgcttgtttg aatacgcatt     22380
        cgtacgggtc gatttgttat tggctgtcag gtagtataat ggtgtataac cgtcatatcc     22440
        ccaaaaggat aagtccgcgc catacgacgt tctaaatttc aaattgtccc agatattcag     22500
        ttcggccgag aagttggcga cgaacttgtg actccagttt ttatctcccg gcaaggataa     22560
        gctggccaac gggttcacca tttcattgta atcg                                 22594
//
ID   XXX; XXX; linear; XXX; XXX; XXX;  XXX.
XX
AC   PRJEB67889
XX
DE   Uncultured bacterium DNA fragment from in-reactor enriched cow rumen, clone
ferm-41o1, contig-2
XX
FH   Key             Location/Qualifiers
FH
FT   source          1..10243
```

```
FT                   /organism="uncultured bacterium"
FT                   /environmental_sample
FT                   /mol_type="genomic DNA"
FT                   /isolation_source="Cow rumen"
FT                   /clone="ferm-41o1, contig 2"
FT                   /note="AZCL-Barley beta glucan active clone"
FT   CDS             17..808
FT                   /db_xref="SEED:fig|6666666.127257.peg.1"
FT                   /translation="MLWGVGGLTFGLSMRYLGIALGQSIALGTCAAFGTLFPALLGGMD
FT                   LLHGDGLILLIGVSITLAGIAVIGYAGSLRARNMTEEEKKAAVKDFALTKGLLVALLAG
FT                   VMSACFNLGLEAGSGIVERVKAAGANELFALNPVILLVTMGGFVTNAVYCIYQNIKNKS
FT                   GKEYLSVSGSVLVNNLLFCALAGVLWYSQFFGLGMGKSYFADSPVMLAFSWSILMSLNV
FT                   LFSNLWGILLKEWKGAGEKTITVLVIGMGILIFSLLFPNLV"
FT                   /product="L-rhamnose-proton symporter"
FT   CDS             845..1639
FT                   /db_xref="SEED:fig|6666666.127257.peg.2"
FT                   /db_xref="GO:0008994"
FT                   /translation="MIRDNKELMAEVGRVAEVAGYLWTKGWAERNGGNISVNVTHLMSD
FT                   EEKALPAISAPVALPAKMEALCGHVFFVTGTGKRMRYVAQAPFDNGSLIRISGDGLSFE
FT                   IIAEKPIAPTSELPSHLSMHNYLRLQGRDNRVVLHTHPTDLIGLTHGPKFLNSEYLTRT
FT                   LWAMIPECRIIVPKGIGIVRYEIPGTMDLARATIKQLEKHDVVFWEKHGILAVGEDLIE
FT                   CFDAIDTLSKSAQIYFSARMAGFEPEGMTDEQLDGLVDAFNL"
FT                   /product="Rhamnulose-1-phosphate aldolase (EC 4.1.2.19)"
FT                   /EC_number="4.1.2.19"
FT   CDS             1650..4871
FT                   /db_xref="SEED:fig|6666666.127257.peg.3"
FT                   /translation="MEREMYNKLLSGLFILACCLISCGKQQQAASSMDEIEANFKRVPD
FT                   VQTSVYWYWINGNISKEGVVKDLHAMKEVGINRAFIGNIGDQAGIPHGAVKLFSDEWWE
FT                   IMHTALKTAGELDIEIGIFNCPGWSQSGGPWVKPEQAMRYLASSETRLKGPQSLQIKLD
FT                   KPTSEFEDVKLLAYPIPASYRDNLLEKQGTVITCSSASMRNGTRFLLPEGESSMSITLA
FT                   TDEPVRSLVFYPENIVYTDVDFQLFEAGAYKTVRSFTVNRTNANLNVGFDPFAPVAIAL
FT                   PETQAQTYRLLFRNTNKGATFRRIQLSATPVVERYAEKSLAKMFQTPLPYWHDYLWEVQ
FT                   PEVTDASLLIDPASVLDLSDKLDAEGNLTWEVPAGEWLVMRTGMTPTGVVNAPADPEGR
FT                   GLEVDKMSKSHVAAHFDAFLGELLRRIPEADRKTWRVVVEDSYETGGQNFTDGFLTEFR
FT                   EQYGYDPVAFLPVLKGYTIGSPELSDRFLWDLRRLVADKVAYDYVGGLKEVSNKHGLTT
FT                   WLENYGHWGFPGEFLQYGGQSDEIGGEFWSEGELGNIENRAASSAAHIYGKTKVSAESF
FT                   TSGGPHFARYPKTMKQRGDRFFTEGINNTLLHVYIQQPYEEKNPGMNASFGNEFNRKNT
FT                   WFSHLDLFTDYLKRCNYLLQQGTYVADAAYFIGEDVPKMTGVCDPALPKGYSFDYINRE
FT                   VLMTRASVVDGKLTLPDGLQYRVLVLPKQETIRPDLLARIAAFVRDGAVVLGPRPSRSP
FT                   SMQNYPEADKEVQAIAAELWGAVDGVQVKQARVGKGLVCSGMTMEEVFDLIGVKPDMET
FT                   NAADPLLFIHRRIGDADMYFVSNQSEERVNVSPRFRISGRQPELWNPIEASVRDLPAFQ
FT                   MDGQTTAVPLQLKPLESAFIVFRRSAGAGNEKVLSVNFPNPNLLVTLDGPWEVSFDPAK
FT                   GGPKEPVRMETLSDWTLSENADVKYYSGEAVYRNQFDLSNIPEGEVYLDLGEAAVMAKV
FT                   SLNGEYVGGAWTAPWRVRITDMLKTGNNAVEIRVVNNWVNRLIGDSRLPESKRTTWAPV
FT                   NPYTPESPLQRSGLLGPVTVVAYNR"
FT                   /product="GH106"
FT   CDS             4916..6715
FT                   /db_xref="SEED:fig|6666666.127257.peg.4"
FT                   /translation="MMRQLKILFSVSLVLLPLLVNAQLPAVVENANKHIRKTELTRSYL
FT                   TPTRIVWKSDASGKYVQNESVLLKPGIGQSDLNTKTGFKLISDKEQKGGIVLDFGCEIQ
FT                   GGLEIVTTIVNQNPAARVRVRFGESVSEVMSDVKGEKGATNDHAIRDQVVTLPWLGCIQ
FT                   LGQTGFRFVRLDLVDADTALDIKEISAIFTYRDIPYRGAFRCNDERLNKIWMTGAYTVH
FT                   VNMQEFLWDGIKRDRLVWIGDMHPEVMTVNTVFGYNEVVPKSLDLIRDITPSSDWMNGI
FT                   SSYSIWWLLIHRDWYYYQGDLEYLKEQREYLKGLLTHLISKIDKDGREQLDGNRFLDWP
```

```
FT                   SSENQAGVDAGLQALMLWAMKSGNELCRVLGENDLAESCASMADKLEKAGRKIAKNYPN
FT                   TQSVPDAPGVKQGAALMALAGLMKSEEANRNFLAVNGARGFSTFYGYYMLKAMAEAGNY
FT                   QGAIDVIREFWGAMIDLGATTFWEDFNIDWLPNASRIDELVPEGKKDIHGDYGAYCYVG
FT                   LRHSLCHGWASGPTSWMSEYVLGVKVLEPGCRVVQIRPNLGDLEWVEGRFPTPYGDIQI
FT                   RHEKVNGQIESRIDAPAEVRVVR"
FT                   /product="GH78"
FT                   /EC_number="3.2.1.40"
FT   CDS             6785..9319
FT                   /db_xref="SEED:fig|6666666.127257.peg.5"
FT                   /translation="MGVWLLTGCMNNTVPVNLRTEYLTEPLGLDVRNPRFTWEYAGEEE
FT                   TFSPSRYEIKIGTDPKNLQPYSDDFQLRPHTRYYWQVAVWNEEGERSRWSDVTWFETAK
FT                   LSPSDWNGRWITDGYDQEFEPAPLLRKVFTAEKKVKSARLYIASAGYHEVFLNGRRIGE
FT                   NYLDPGYTHFDKRVLYVTHDVTEEVKQGENALTAVLGNGWYNEQSVAVWNFHEARWRNR
FT                   PRMLCELRITYTDDSVETIVSDRSWKTHIGAYTYNNIYSGDQYDARQEEAGWQTAAFDD
FT                   AQWEEAVETESPAPLLVAQQMPGIRIVEEIRPVSVKAFSDTLYVYSFPKNMSGLCRLKV
FT                   TGEAGTRVTLKHGELLKADGRLEQGNIDVYYKPVKPGEVFQMDAYTLKGEGEEVFMPSF
FT                   TYHGFQYVEVESTRPIQLDEDNLTALFMHTDLRPVGNYSCSDSLLNKVWKATMQAYRSN
FT                   LHSIPTDCPQREKNGWTADAHVSVDLGLLGYDGITLYEKWMNDFIDNQTEAGDISGIIP
FT                   SSGWGYGKWIGPVWDAAMFIIPDALYKYYNDTQAIEKLYPTMRNYLDYLKTQEEDGLLT
FT                   YGLGDWVYWKSTTPNAYTSTAYYYLDHTLMAKFASLLGEDPTPYLEKAEALKKRINAQF
FT                   FQEETGTYANGTQTAQALALYLHLVPEGKRQRVADKLHEVVAANDYFLDFGLIGSKTVP
FT                   AMLTQYGYVEDVMKMLKKTEAPSWGYWVETMGYTTLPETWTLSPEFKDASLNHVFMGDI
FT                   SAWMMNHLAGIQYDEEKPGFAHIRIEPHFVKDLDWVHGEYHSPKGLIRSEWKREGERYG
FT                   LSVTIPVGCTATVSVDEKEFQLPSGTHQLLFR"
FT                   /product="GH78"
FT                   /EC_number="3.2.1.40"
XX
SQ   Sequence 10243 BP; 2474 A; 2236 C; 3036 G; 2497 T; 0 other;
     gtcggttatc tacgggatgc tttggggagt cggcggcttg acctttggct tgagcatgcg        60
     ctacctgggg attgccctgg ggcaaagtat tgccttgggg acttgtgccg cgttcggcac       120
     tttgttcccg gctttattgg ggggcatgga tttgttgcat ggcgacggac tgatcctgct       180
     gatcggcgtt tccattactt tagcgggtat cgcggtgatc ggctatgccg gcagtttgcg       240
     tgcccggaac atgacggagg aggaaaagaa agcagcggtc aaggatttcg cgctgacgaa       300
     aggactgctg gtggctttgc tggccggtgt gatgagcgcc tgtttcaatc tcggtctgga       360
     agcgggtagt ggtatcgtgg aaagggttaa agcggccggc gctaatgaac tattcgcctt       420
     gaatccggtg attttgctgg tgacgatggg cgggtttgtg acgaatgccg tctattgtat       480
     ttatcagaat ataaagaata agtcgggtaa agaatatctt tctgtttccg gctccgtgtt       540
     ggtgaataat ctcttatttt gtgccttggc cggagtcttg tggtattctc aattctttgg       600
     attgggcatg ggtaagagtt attttgccga ttcgccggtg atgctggctt tctcgtggag       660
     catactgatg tcgctgaatg tgttgttcag caatctgtgg ggcatcttgc tgaaggaatg       720
     gaaaggggcc ggtgaaaaga ccatcacggt tttggtgatt ggcatgggca ttctgatttt       780
     ctccctgctt ttcccgaatt tggtgtaaat cttggaagtg aacgagtatt aattcaacat       840
     aacaatgata agagataata aagaattgat ggcggaagta ggtcgtgtgg ccgaagtagc       900
     cggttatttg tggacaaaag gatgggcaga acggaacggt ggaaacattt cggtgaatgt       960
     gacccatttg atgagtgatg aggagaaagc gttacctgcg atatctgcgc cggttgcctt      1020
     gcctgctaag atggaagcgt tgtgcggaca tgtctttttc gtcaccggaa cgggtaagcg      1080
     catgcgctat gtggctcagg cacctttgga taacggatcg ctgatacgga tcagtggcga      1140
     cggactttca ttcgagatca tcgcggagaa gccgattgcg ccgacatccg aattgccgtc      1200
     gcatctgtcc atgcataatt acctgcgcct gcagggacgg gataaccgtg tggtacttca      1260
     tacgcacccg acggatttga tcggtctgac gcatggaccg aaattcttaa attcggaata      1320
     cctgacgcgc acgctttggg cgatgattcc ggagtgccgc atcatcgttc ccaagggaat      1380
     cggtattgtg cgctacgaaa tacccggaac gatggacttg gcacgggcga cgatcaagca      1440
     attggagaaa cacgatgtcg tattttggga gaaacacggt attttggctg tcggcgaaga      1500
     cttgatcgag tgttttgatg ctattgatac gttgagcaaa tccgcccaga tatatttcag      1560
```

tgcccggatg gcgggatttg agccggaagg aatgactgat gaacaattag acggactcgt 1620
cgacgcattt aatttgtaaa atagaatgga tggagagaga gatgtataat aaactattat 1680
ccggtttatt tattctggcc tgctgtctga tttcctgcgg taaacagcag caggccgctt 1740
catctatgga tgagattgaa gcgaatttta agcgggtgcc ggatgtgcag accagtgtgt 1800
attggtattg gattaatggg aatatctcga aggagggggt cgtcaaagac ctccacgcga 1860
tgaaggaggt cgggattaac cgggcattca tcggtaatat cggagaccag gcgggaattc 1920
ctcatggtgc ggtcaaactt ttttccgacg aatggtggga gattatgcat acggccctga 1980
aaacggcggg tgaactggat atcgaaatcg gtatctttaa ctgtccgggc tggagtcagt 2040
cgggcggccc ctgggtgaaa ccggagcagg ccatgcgcta tctggcttct tctgagacgc 2100
gcctgaaagg tccacagtcc ttacaaatca aactggataa accgacatcc gaatttgagg 2160
atgtcaagct gctggcttat ccgattccag cttcttaccg cgataacctg ctggaaaagc 2220
aagggaccgt catcacgtgc tcgtctgctt ccatgcggaa tggaacacgc tttttgctgc 2280
ccgagggtga atcatcgatg tcgattacgc ttgcgaccga tgaacctgtt cgcagtttgg 2340
tattctaccc cgaaaatatc gtttatacgg atgtcgattt tcagctattc gaagccggcg 2400
cgtataagac ggttcgctcg tttaccgtga atcggacgaa tgccaacttg aatgtcgggt 2460
ttgatccgtt tgcgcctgtt gcgattgcct tgccggaaac ccaagcacaa acataccgtc 2520
tcttgttcag gaatacgaat aagggggcga cctttcgtcg gattcagctg agtgccacac 2580
ccgtagtgga acgatacgcg gagaaatcgc tggcgaaaat gttccaaacc ccgctgcctt 2640
attggcatga ttatctttgg gaggttcagc cggaagtgac cgacgcctcg ttgctgatcg 2700
atccggcctc ggtcttggat ctctcggaca agttggatgc cgaggggaat ctgacttggg 2760
aagtgcctgc cggcgaatgg ctggtgatgc gtacaggtat gacacctacg gggtcgtca 2820
acgccccggc cgatccggaa ggaaggggcc tggaagtgga caagatgagc aagtcgcatg 2880
tcgctgcaca tttcgatgcc tttctgggag aattgctgcg ccgtattccg gaggccgaca 2940
ggaagacttg gcgggtggtg gtggaagata gctacgagac aggaggacag aactttacgg 3000
atggattttt aacggaattc cgagagcaat acggctatga tcctgtggcg tttctcccgg 3060
tattgaaagg atatacgatt ggaagtccgg aattatccga ccgctttctg tgggatctac 3120
gtcgtctggt ggcggataag gtggcttatg attatgtcgg cggactcaaa gaggtaagca 3180
acaaacacgg tctgactact tggttggaga actacggaca ttgggggttc ccgggagaat 3240
tcctgcagta cggcggacag tcggatgaaa tcggcggtga gttctggagc gaaggggagt 3300
tgggcaatat agaaaaccga gccgcctctt ctgcagcgca tatttacggg aagaccaagg 3360
tctcagcgga atcgttcacc agcggtggcc cacattttgc ccgttatccg aagacaatga 3420
aacaacgggg ggatcgtttc tttacggaag ggatcaacaa tacgctgctc catgtctata 3480
ttcagcagcc ctatgaagag aaaaatccgg ggatgaatgc ctccttcgga aatgaattca 3540
atcgcaaaaa tacttggttc agtcacctgg acttgtttac cgactacctg aaaaggtgta 3600
actacctgct gcagcaagga acttatgtag ccgatgcggc ttattttatc ggagaggatg 3660
tgccgaagat gaccggcgta tgtgatccgg ccttgccgaa agggtattca tttgattata 3720
tcaatcggga ggtgctgatg actcgcgcgt ctgtggtcga cggcaagctg accttgcccg 3780
acgggcttca gtaccgtgtg cttgtcttgc ctaaacagga gacgattcgt ccggatttat 3840
tggcccggat cgccgcattc gtgcgtgatg gagctgttgt ccttggtccg cgtccgtctc 3900
gctcgccgag tatgcaaaac tatccggagg cggataagga agtacaggcg attgcagccg 3960
agctctgggg cgccgttgat ggagtgcagg tgaaacaggc acgggtaggc aaagggctgg 4020
tttgctcggg tatgactatg gaggaggtgt ttgacctgat tggcgtgaag ccggatatgg 4080
agacgaatgc agcagatccc ttgctcttta tccatcgtcg gataggtgac gcggatatgt 4140
attttgtgtc taaccaatct gaagaaaggg tgaatgtgtc tccgcgtttc cgcatttccg 4200
gcagacaacc cgaattatgg aacccgattg aggcgagtgt tcgcgacctt cctgcttttc 4260
aaatggatgg gcagacgact gcggttccgt tgcaactgaa acctttggaa agtgccttca 4320
tcgtctttcg ccggtcggcc ggcgcaggaa acgagaaagt cctttctgtc aatttcccga 4380
atccgaattt attggttacc ttagatgggc cttgggaggt gagcttcgac ccggccaaag 4440
gaggaccgaa agagcctgtt cggatggaaa cgttgtcaga ctggacgctt tcggagaatg 4500
cggacgtgaa gtattattcc ggagaggctg tttaccggaa tcaatttgac ttgtctaaca 4560
ttccggaagg agaggtctac ttggatttgg gtgaagcggc cgtcatggcg aaggtcagcc 4620
taaacggtga atatgtcgga ggtgcttgga cggcgccttg gcgggtgcga ataacggaca 4680
tgctgaaaac cgggaacaac gcggtggaga ttcgcgtggt gaataactgg gtgaatcgcc 4740
tgataggaga cagccgtttg ccggaatcga agcggacgac ttgggcaccg gtgaatccct 4800

atacaccgga gagtcccctg caacgctcgg gattattggg cccagttacg gttgtggcgt 4860
ataaccgata aaataaaata taaacgaatt cagccttggg cacttgttat agattatgat 4920
gagacaattg aaaatccttt ttagcgtcag tttggtgttg ctcccgcttt tggtcaatgc 4980
gcagcttccg gctgtcgtgg aaaatgcgaa taagcatatc cgaaaaacgg aattaacccg 5040
atcgtatctc acgccgacaa ggattgtgtg gaagtcggac gcctccggta aatacgtgca 5100
gaacgagtcg gtcttgttaa agccgggaat cggtcagtcc gacctcaata cgaaaacggg 5160
gtttaaactg attagcgata aggaacaaaa ggggggaatc gtgctggatt tcggttgcga 5220
gatccaaggc ggactcgaga ttgtgacgac gatcgtgaat caaaatccgg cggcccgggt 5280
gcgggtgcgt ttcggcgagt cggtctctga ggtgatgagt gacgtgaaag gcgaaaaggg 5340
ggctaccaac gaccatgcga tacgggatca ggtggtcacc ctcccttggc tgggatgtat 5400
tcagctggga cagaccggct ttcgctttgt gcgtctggat ctggtcgatg cggatacggc 5460
tttggacatc aaagaaattt ccgcaatctt tacctatcgg gacattcctt accgggggc 5520
tttccgatgc aatgacgagc gactcaataa aatctggatg accggtgcgt atacggtgca 5580
tgtcaatatg caggagttcc tttgggacgg tatcaaacga gaccgcctag tctggatcgg 5640
tgatatgcac cccgaggtga tgactgtaaa tacggtgttc gggtataacg aggtggttcc 5700
taaaagtctg gatctgatcc gggacattac gccttcttcc gattggatga acgggatcag 5760
ctcgtattcg atctggtggc tcttgattca tcgcgactgg tattattatc agggcgatct 5820
ggagtatctg aaggaacagc gggagtactt gaaaggcttg ttgactcatc tgatcagcaa 5880
gatcgataaa gacggtcgtg aacaattgga cggcaatcgc ttcttggatt ggccttccag 5940
cgaaaatcaa gcgggcgtgg atgccggttt gcaggctttg atgctgtggg caatgaagtc 6000
gggcaacgag ctatgccgtg tcctcggcga gaatgacctg gccgaatcct gcgcttcgat 6060
ggcggataag ttggaaaaag cgggcagaaa aatcgctaaa aactacccaa atacacagtc 6120
ggtgccggat gcgcccggag tcaaacaagg agctgctctg atggccctgg ccggattgat 6180
gaagagcgaa gaggcgaacc ggaatttcct ggccgtgaat ggggcgcggg gtttctcgac 6240
cttttacgga tattatatgt tgaaagcgat ggcagaggcc ggtaattatc agggagctat 6300
cgatgtgatc cgtgaattct ggggggctat gattgatttg ggagcaacca cctttttggga 6360
agattttaac atcgactggc taccgaacgc gagtcggata gacgaacttg ttccggaagg 6420
gaaaaaggat attcatggag attatggcgc ttattgctac gtcggactcc gtcacagtct 6480
ttgtcacggt tgggcttcgg gacctacttc gtggatgagc gaatacgtac tgggagtcaa 6540
ggtgcttgaa ccgggttgcc gggtcgtgca gattcgtccg aatctcggcg atctggaatg 6600
ggtggaagga cgtttcccta ctccttatgg cgatattcaa atacgccatg agaaggtgaa 6660
cggacagata gaaagtcgga ttgatgcacc cgctgaggta cgtgtggtcc ggtaatccga 6720
caacatttcg aaaagcgtaa acgattagcg tatgatatca attaataaag ttcttgttgc 6780
cgtaatgggc gtatggttac tgaccggctg tatgaataat acggttccgg tcaatctgcg 6840
tacggaatac ctgactgaac ccttgggtct ggatgtgcgg aatccgcgtt ttacttggga 6900
atatgccggc gaggaggaga ccttctctcc aagccgctac gagatcaaga ttggaacaga 6960
tccgaagaat ttacagcctt atagcgacga cttccaactc cggccgcata cccgttacta 7020
ttggcaggtg gccgtttgga acgaggaggg agagcggtcg cgctggtcgg acgtaacctg 7080
gtttgagacg gccaagttgt ccccgtcgga ttggaacggt cgctggataa cggacgggta 7140
tgatcaggag tttgagccgg caccgctttt gcggaaggta ttcaccgccg aaaagaaggt 7200
gaaaagcgcc cgactttaca ttgcttcggc aggatatcat gaggtttttc tgaatggccg 7260
tcggataggg gaaaattacc tcgatccggg ttatacgcat tttgataagc gggtgttgta 7320
tgtgacgcac gatgtgaccg aagaggtcaa gcaaggagag aatgccctga cggccgtatt 7380
aggcaatggc tggtataatg agcagtctgt cgccgtctgg aatttccacg aagcccggtg 7440
gcggaaccgt ccccgtatgc tctgtgaatt gcggattacc tatacggacg actcggtcga 7500
gacgattgtg tcggatcgtt cgtggaaaac ccatatcggt gcgtatactt acaataatat 7560
ctatagtggt gaccaatacg atgcccgtca ggaggaggcg ggttggcaaa cagccgcatt 7620
tgatgacgcc cagtgggaag aagccgtgga aaccgagtcg cccgccccat tactcgttgc 7680
gcagcaaatg ccgggcataa ggattgttga agagattcgc cctgtctcgg taaaggcctt 7740
tagtgacacc ctgtatgtct attcgttccc caagaacatg tccggacttt gccgtttgaa 7800
agtgacggga gaagccggaa cgcgggttac gctaaaacac ggggaactgc tgaaagcgga 7860
cggccgcttg gagcaaggga atattgatgt ctactacaaa cccgtgaaac cgggtgaggt 7920
gttccaaatg gatgcctata cactcaaggg agagggagag gaagtcttca tgccttcctt 7980
cacctatcac ggttttcagt atgtcgaagt ggaaagcact cgcccgatcc aactggacga 8040

```
     ggataatctt accgccttgt tcatgcacac cgacctgcgt ccggtgggga attactcctg      8100
     ttccgattcg ttactgaata aagtatggaa ggcaacgatg caggcgtatc gcagtaacct      8160
     gcacagcatt ccgaccgact gtccgcagcg tgagaagaac ggttggacgg ccgatgcaca      8220
     tgtctcggtc gacttaggac tgctgggtta tgacggcatc acgctctatg agaaatggat      8280
     gaacgacttt atcgataatc aaaccgaggc cggcgatatc tccggaatca tcccttcgag      8340
     cgggtgggga tacggaaaat ggatcggacc ggtttgggat gcggccatgt ttattattcc      8400
     cgatgccttg tataaatact ataacgatac gcaggccatc gaaaaactgt atcctacgat      8460
     gcggaactat ttggactatt tgaaaacgca ggaggaggac ggactgctga cttacggctt      8520
     gggtgactgg gtgtattgga aatcaacgac gccgaatgcg tatacttcga cggcctatta      8580
     ttatctggat catacgttga tggccaagtt tgcttcgctg ctgggcgaag atccaacgcc      8640
     ttatttggaa aaagccgaag cattgaaaaa acggattaac gcgcaattct ttcaggagga      8700
     gacgggtacg tatgcgaatg gcacgcagac cgcccaagcg ttggcgctct atctgcacct      8760
     cgttcccgaa ggtaaacggc aacgggtggc ggataagttg cacgaagtgg tcgcggccaa      8820
     cgattatttc ttggacttcg gattgatcgg cagcaagacg gtgccggcga tgctcaccca      8880
     atacgggtat gtagaggatg tgatgaagat gctgaagaaa acggaagcgc cttcgtgggg      8940
     atattgggtc gagaccatgg gttatacgac gttgccggaa acgtggacgc tcagtcccga      9000
     attcaaggat gcttcgttga accatgtatt catgggcgat atatctgcct ggatgatgaa      9060
     tcatttagcc ggaattcagt atgacgaaga gaaaccgggc ttcgcgcata tccgcatcga      9120
     accgcatttt gtgaaggatc tggattgggt gcatggtgag taccattctc ccaaaggact      9180
     catacgtagt gaatggaaac gcgagggtga acgctatgga ttgtcagtca cgattcccgt      9240
     cggttgtacg gcgacggtct cggtcgatga aaaagaattc caactgccat caggcacaca      9300
     tcaattactt ttccgttaaa ctaaatctat agaacgatga aaaacacaat tgctattatt      9360
     cttttattgg gaagtctttg gatatacccg ctccgtgcgg agaattggaa agggcattgg      9420
     atcaatacgg aacaatgtca aagtgcgacg aacacgtggc tggcttatcg gaaaacagtt      9480
     tcgctcgagc gagttcccca acatctcacg gccaagatag ccgccgacac caaatattgg      9540
     ctgtggatca acgaccaatt agtcgtcttc gagggcggat tgaaacgggg gccggctccg      9600
     ggagcgattt attatgaccg ggtggatatc gctccatttc ttaaaatagg cgataatcaa      9660
     atcgccgtcc tgctttggca ttttgggaaa gacggattct gccatatcaa cagcggtaaa      9720
     gccgcgttgt tgtttgatgc ccggggggag ggtgttgaaa tcgtgtcgga cgcgtcgtgg      9780
     caatgtgccc tgtatgaagc ctaccaaaac accggggcgc ctttcccgaa ttaccgtttg      9840
     ccggaaagca atatccgttt cgatgcacgt aaggaaatgc gcggatggac ttcgcccgat      9900
     tataaaagac atatgccggc cgccgtatgt gtggccgatg ccaacgacct gcctttcggc      9960
     agattggtag aacggccgat tccgttgtgg aaagactacg gtttgaaggc gtatgctgaa     10020
     gtgaggcggt cggcctcgaa tgatacgctc ttttgccgtt taccctataa ttgccaggtc     10080
     accccttatc tgaaagtaaa cgcggaagcc ggaaaaacaa tccatatgtt gacggacaac     10140
     taccgcgggg gaggttcgcc gaacgtcaga ggggagtaca ttacccggaa aggggttcag     10200
     gaatatgaaa gctacggctg gatgaacgga catgaggtat ggt                       10243
//
ID   XXX; XXX; linear; XXX; XXX; XXX;  XXX.
XX
AC   PRJEB67889
XX
DE   Uncultured bacterium DNA fragment from in-reactor enriched cow rumen, clone
ferm-42a14, contig1
XX
FH   Key             Location/Qualifiers
FH
FT   source          1..22822
FT                   /organism="uncultured bacterium"
FT                   /environmental_sample
FT                   /mol_type="genomic DNA"
FT                   /isolation_source="Cow rumen"
FT                   /clone="ferm-42a14"
FT                   /note="ABTS active clone"
```

```
FT   CDS             complement(122..448)
FT                   /db_xref="SEED:fig|6666666.127258.peg.1"
FT                   /translation="MAKKTNLENLRDAVSKQAKNLDPAQKEFVLAEFDHYEWNAKRIAE
FT                   LETSLENGIIDSEGLHNYDAEGKLFRQRHQLVAEQSSLFSHIMRWLKGTATEESPIDDF
FT                   LNGN"
FT                   /product="hypothetical protein"
FT   CDS             complement(448..645)
FT                   /db_xref="SEED:fig|6666666.127258.peg.2"
FT                   /translation="MKKLKKCPWCKKHAHYMEWIDKGDGTKLYSVACKTLRCFGSSSRM
FT                   FFTSKKDARKAWNKRNGKVQ"
FT                   /product="hypothetical protein"
FT   CDS             complement(642..1004)
FT                   /db_xref="SEED:fig|6666666.127258.peg.3"
FT                   /translation="MAKTATWRNTPFAHKFYHSRAWKNARAFVWTRAHGLCERCAEQGE
FT                   LKPADVVHHKIPLNPVNINDPYISLNADNLIALCNECHTEVHKELGIGAFENNKEVEAR
FT                   VGFDEDGNIIRLESEQ"
FT                   /product="Prophage LambdaBa04, Gp54"
FT   CDS             complement(1410..1589)
FT                   /db_xref="SEED:fig|6666666.127258.peg.4"
FT                   /translation="MTNRADIIITVLVIISIVQLLLAGALSLLVSMASGTFVGILGCYT
FT                   ILASIFLEIYSQRG"
FT                   /product="hypothetical protein"
FT   CDS             complement(1586..1915)
FT                   /db_xref="SEED:fig|6666666.127258.peg.5"
FT                   /translation="MHSNCPNCGAPITDDYCHYCGTVFNPASKMMIGKTIKIEFEHEGL
FT                   VHKFDMVLDQFNLDYNSDATTFYADGAAYYHATINTFCEALLRGRIKPKEVNGKQVFMT
FT                   TEVAR"
FT                   /product="hypothetical protein"
FT   CDS             complement(1905..2117)
FT                   /db_xref="SEED:fig|6666666.127258.peg.6"
FT                   /translation="MDKPIFIVKLKGMHRAAEIEAWKSKLKEQLDGTVIVIDEQIEYIE
FT                   QLNPADKSRLLSLLEAELKGADDAQ"
FT                   /product="hypothetical protein"
FT   CDS             complement(2110..3081)
FT                   /db_xref="SEED:fig|6666666.127258.peg.7"
FT                   /db_xref="GO:0016020"
FT                   /db_xref="GO:0019350"
FT                   /db_xref="GO:0047355"
FT                   /translation="MASKVLFVSQNKLGRGENITALYNSYDGDKVFKHGLVNMLDAELK
FT                   GYDVVVCDALPQYIAFKDKCKSIVICHGMTGNKVYGADEDRDVDIEAFKQTDIATAASK
FT                   ASIPIVAQQLRIPENRVKAIGFPRTDTYIGKKKGEGFTKLGNKRAYLFAPTFRDADKGG
FT                   WIPWIDWEIVDNLLEEDEIVAVKRHYFTDRPLLGANFKRVIEIEPDKPLTPYLIDCDVL
FT                   ITDYSSCMTDGYLLGKPCILTVDDMDEYLPDRPMYYPYPDTYSSRWIVGEGHEEELVKM
FT                   MREAASNGLTDVEKHYMELTASECDGHSIERTVALIKEMANG"
FT                   /product="CDP-glycerol: N-acetyl-beta-D-mannosaminyl-1,4-N-
FT                   acetyl-D-glucosaminyldiphosphoundecaprenyl
FT                   glycerophosphotransferase"
FT   CDS             complement(3081..3674)
FT                   /db_xref="SEED:fig|6666666.127258.peg.8"
FT                   /translation="MAKVLLAIPTGKGFIWADASDALYGLNRGQNDVVQATKSFYAVDE
FT                   ARNALVRFALKNDFDYVFFVDDDTIVPPDALLNLMENNVDVCLGYYIKSDSDNLTPMVE
FT                   PRGNNYRTYYSVEEIEEMRKDGQTLVEIGGGGFGCALIKVSALKILTDPWFKFNKHLSE
FT                   DYYFCHQWKSIRGKIYLDTRVACDHIKLERKKAM"
FT                   /product="hypothetical protein"
```

```
FT   CDS             complement(3667..4386)
FT                   /db_xref="SEED:fig|6666666.127258.peg.9"
FT                   /translation="MKYALYSGSRNLYGDMQTATKSLIANSDVDKVYLLVEDDDYDFWL
FT                   PDMVEVIDVSSQTYFPEDSPSMKTPFSYLSLMRAALALMPELSHVDKVLSLDVDTICVR
FT                   DVSNAWNMPIEGKYFSASIEPCSCYKNMLYCNTGVALYNLEMLRNGKAQEVVDALNERL
FT                   YPNIEQDVFSFMCCGKIHDMPSEYNCTNFTKPTNIVRIMHYAGYKTEQWRKEALVGKYR
FT                   DMSWDEALKMHERLKNG"
FT                   /product="hypothetical protein"
FT   CDS             complement(4383..4781)
FT                   /db_xref="SEED:fig|6666666.127258.peg.10"
FT                   /translation="MAQLKHTPKFKRERRSIHRRHCPYCDGQRVLYEVVPDTRINVEGH
FT                   DLPFRTIMHVFSNYLALDIKAPDRPFKFYKREYQVGNPERLTPDSLLKFEEIDIKPKTQ
FT                   RLLTPIKYCPMCGRDFKGTREKNWRKFK"
FT                   /product="hypothetical protein"
FT   CDS             complement(4782..4979)
FT                   /db_xref="SEED:fig|6666666.127258.peg.11"
FT                   /translation="MDANTFWEIVKPCPFCKRHGAQRLRERHGKHYVACSRNKCGCGAR
FT                   GPMARTQEVAVRIWNSATEE"
FT                   /product="hypothetical protein"
FT   CDS             complement(4988..5194)
FT                   /db_xref="SEED:fig|6666666.127258.peg.12"
FT                   /translation="MDNFDDDDWKTLMPDSGDYGLSIQETLKEVGLFLARIVGVLLLIA
FT                   LIIGLFAYSILREYLFINFLLGN"
FT                   /product="hypothetical protein"
FT   CDS             complement(5203..6918)
FT                   /db_xref="SEED:fig|6666666.127258.peg.13"
FT                   /translation="MTASKRTISEAEQYLLDVQAGKITVCKRIKQLAEMMLPRFKDGYK
FT                   KWHFDIDKARRPVEFIEKFCVLPESGEPMVLEPYELVVIETAFGFVDDEGYRQFHEVLV
FT                   MWARKQGKSSLGAAIELYLLVADGEGAPQVYNVATSKAQASLAYGIVVKMLKRSKELSK
FT                   VLRKGTVVDRDQDGVIFDRNDGYITVLTNQTRHLDGLNTHGCLFDEMHANTNRDQFDLI
FT                   WQSMSTRRQPLLIAITTNGFERGNLFDDQYEYACGILDGKIEDDRMLPFLYELDDRSEW
FT                   LREECWIKANPGLGTVKSWDYMRDAVNKAMQDPSFLPTVMTKDFNMPESRASAWLSFDE
FT                   AVNTEPFPELPPSGKLSDLGFRYGVAGIDASDTTDLSAAKILMMRQDDPKIYELSMYWL
FT                   PEDALKKDDGYRRERDDVPYRIWEQKGYIRTVPGNIVPKSVFIDWFDEVKRELDVWLFA
FT                   IGYDRWGFSDEDIARFEQYVGKDRSEIVRQGPMTFSSPMKELKAMYGSNQIVDGHNPTN
FT                   EWNRMNVQIKQDSNANIRPIKAEGKAVHRIDGFMAELNGFIAMQRHLEEYKGII"
FT                   /product="Phage terminase, large subunit"
FT   CDS             complement(6905..7063)
FT                   /db_xref="SEED:fig|6666666.127258.peg.14"
FT                   /translation="MVEKNCDTCAYKQMESKDENGNRIIDCEANELQIFAPFAEDCKHW
FT                   EKSVDCK"
FT                   /product="hypothetical protein"
FT   CDS             complement(7072..7341)
FT                   /db_xref="SEED:fig|6666666.127258.peg.15"
FT                   /translation="MATKSKLEQAVEQQMAHLNPAQKEIVKSQLSRHKKNKARIFEIDE
FT                   TLKSPMSTQLHPAERMALTSERSQLSTSNTEIERNLFNQLKDTD"
FT                   /product="hypothetical protein"
FT   CDS             7554..8876
FT                   /db_xref="SEED:fig|6666666.127258.peg.16"
FT                   /translation="MSKILDKILGKETAPSATAVQTVGNKPYTSSWNGSMYQQVLVRSV
FT                   IERIALACSKLKPVIKGSARPRIRRAIETCPNEFQTWPQFLHRCATLYMNNTTVCVVPE
FT                   FKPGSQVQTGFYPVPLANAEVVDYGGEYWLRYTTRDGDSRAIELKYVAIVTRFQYESDW
FT                   FGDGNILANTLSMLKAQEEAQKNAINDSAQLKFLGQLTGQVREEDMKKKRDRFYQDNLT
FT                   SDNNTPLMLYDATFASFTQLKAENWTIPNEEMERIENNVFDYFGINRRILQNAYDENAW
```

```
FT                   DAFYEGIIEPFGIMLGEALTKATFTMRERPANQILFSANRLEYAAASSKRNINKDMTDR
FT                   GIMTLNEAREILQLPPIEGGDVFILRGEYKVGHTFDEIFQAQQAKAEAKASGRVASPDE
FT                   DRDGADGDNIRGDSDGYGSPGDTDTGDVTSTSQDRWSENAS"
FT                   /product="Phage portal protein"
FT   CDS             8892..9581
FT                   /db_xref="SEED:fig|6666666.127258.peg.17"
FT                   /translation="MPAKPEERQYRMMSTPLFIPDAAPETITDENGEEQTSYQNRFNSP
FT                   CYVEGFATTFDDPYTLFEDSWDGWKYVEIIDRHALDDADMSDVIFQYDHEGRVYARNTN
FT                   NTLFFEANDHGLYIAADLSKTSLARQMYEDIAVGNVTRMSWAFIPTEEEYVSDMENKVF
FT                   TTRIMRVKKVFDVSAVSYPADPNTEISARHLVNGEIESRRQQELLQRELDRKRKEIAMR
FT                   AKSLSIR"
FT                   /product="Phage head maturation protease"
FT   CDS             9602..10906
FT                   /db_xref="SEED:fig|6666666.127258.peg.18"
FT                   /translation="MDFTAMDARAYRSLNADQYAERRSLVLSLAEDMPEDFTEEQMRSL
FT                   DEELGFIKSEDARRDQLAEVRNHKAVAVASGSGKVVGETEAPEVRTKQRKDNSLGARAW
FT                   DAIQERGYKRDERFQVSDIAFRSASDVQTVAELDGENSPNYYDYALTQVDLEIKEGYRR
FT                   PMSIWDLFNHETTEKDSVTWFVEGTDEGDAGMTAENGAFSQYHVNDPVAVTTPLKKVTA
FT                   LWRQTDEILSDAPRFVSHVNSRAGYKLDCKVEDQLLAGTGSSNQITGLVNVSGIHTASA
FT                   SAFDMAFIESLLDERTKIRKATPNFNVDTLLLADEDYDTLMKLKNNADQYVLGGPTGFV
FT                   YGNGVTIGKNLWNTITMVPTPALSSGTAILGAFKDGATVYEHVTGRRFDIGYDGEDFSH
FT                   GRVSFRAYQRLALAVEYPAAFCKYTVAGGASGLNA"
FT                   /product="Phage major capsid protein #Fam0024"
FT   CDS             10918..11148
FT                   /db_xref="SEED:fig|6666666.127258.peg.19"
FT                   /translation="MQVIALVDFHDNLKNVDRKRGEQFVVTKERFEEINAVGMRAINGP
FT                   IVEEKRIETPKETPESRAKKAPAKRKTAVAK"
FT                   /product="hypothetical protein"
FT   CDS             11217..11549
FT                   /db_xref="SEED:fig|6666666.127258.peg.20"
FT                   /translation="MIDAARFDMERAGVDPTLLALNPVTCDLDNEYVKQAVRLYAKANF
FT                   GFDNSEAERLQESYEQHMRNLLNSSHNIAAIALEPASQSTIGSYPTSEPTQEPEVQEPE
FT                   TQEVEG"
FT                   /product="hypothetical protein"
FT   CDS             11550..11918
FT                   /db_xref="SEED:fig|6666666.127258.peg.21"
FT                   /translation="MRWNETVTLIAKAYEPDDEGVPQVTDTKTEVFCNPRTMGANTWSS
FT                   MYEIGISPVAEIQVRTCDYDGQRDVFYRGKYLSVEVVKEQGDFTVLVLRHQQSDSDDSA
FT                   EESSSEIPNETPEEGEGE"
FT                   /product="hypothetical protein"
FT   CDS             11915..12319
FT                   /db_xref="SEED:fig|6666666.127258.peg.22"
FT                   /translation="MTHDITCGIDAFAKSIEELVGKIPVACQEECGKAVAKSTRQGVKK
FT                   VKEKAAKGGKHEWSDEYVNGFSSRVTKGAETTGEIGNKAKPGLVHLLEKGHATLTGRRT
FT                   TAYPHMAPAFEEIQDDFIKEVEKAVERAIS"
FT                   /product="hypothetical protein"
FT   CDS             12319..12651
FT                   /db_xref="SEED:fig|6666666.127258.peg.23"
FT                   /translation="MAHKDVYETVSAIVPTCHMEWPNDSAPELPWALYHGEETPIVADD
FT                   VQIAVRRRWTVELYEKRRNKTIEKKLGDALRARFGSVRRSESYVENDNMLEVIYTFYEI
FT                   EGEFDG"
FT                   /product="hypothetical protein"
FT   CDS             12644..13249
FT                   /db_xref="SEED:fig|6666666.127258.peg.24"
```

```
FT                   /translation="MANKVRFGLRNTKYAIYNAQTGEYGALKDMPGAVSLSLSREGGDS
FT                   SDFYADDGIYYTFAGTNGGYSGDLTLARITDEVRKDLLNEIVDDATGVQFETTTAEPAE
FT                   FALITEMQGDQGPIGFAFYNCKASRPEISANTKNDSPSVDQDTLSLRIASRSFTYDGVA
FT                   TPFVQGHIEKTQENATKFAAFFQSVVTPGASGGSGLSA"
FT                   /product="Phage major tail protein"
FT   CDS             13262..13624
FT                   /db_xref="SEED:fig|6666666.127258.peg.25"
FT                   /translation="MFEVEINGKKVKAEVTFYTAQLYEAEFRADLIKELFGEQSLEPQI
FT                   ETDGENIIKIDFTKINWLATAKVLWAAVKTANEATPSYVSWMKETSGVNMWLVSEQLSA
FT                   EVADCFFRTEAPRTQE"
FT                   /product="hypothetical protein"
FT   CDS             13668..13835
FT                   /db_xref="SEED:fig|6666666.127258.peg.26"
FT                   /translation="MALNGLEAGLTWRDMRHMKYTHLMQFLYEWEDMHGADVDEVVDAT
FT                   PADVMALTRL"
FT                   /product="hypothetical protein"
FT   CDS             13847..18889
FT                   /db_xref="SEED:fig|6666666.127258.peg.27"
FT                   /translation="MGDAFRGLTIRLGADARPLNSAINSITRAASGAQKQLNAMNKALK
FT                   FDNTNVAAMQSRLELVEDKALLTARAANKIEVAMRQASQETLQFSARSGLGVKNLKEMA
FT                   AETKAVHAANEKLHADFNHVNTDLAHIYEAAARVRAEAKGISFEKALKQVKSLASHMNS
FT                   TTAAGKAAKDAVYSLVRSAAMRTDIGEKFGLQKTIGHASKLLGVFLELRRASRDFSSDI
FT                   ELMNKVEGFRAAQTRLISFKADLMQAASEAARFKSELYALGTNGGLVSALNEIKLIDAA
FT                   VAKSTAHAREMAEAYALAPRHLGAAKERLNSYVAAQMALNNQLTSIKSALRRIESDPAF
FT                   DKQAANSAHAYTNAVKVESAYTKVKSKIALATAALQRLEEEQRELSLVQTKTDKQSARY
FT                   EQLTSHINKVNSAINRLKSELASLDQKHAAASLVIEFKELTAAQAKCVAETERLKQALS
FT                   SANRVFGNLRTMGYGLYSTITPAILMAGRYAVQSADEVDAAYRDMRKTVNGTEEEFEHL
FT                   RDAALAFGRTHVTSADTILEIEAIGGQLSIQAKNLEAFSRVVSNLDIATNLDTETISQN
FT                   IGQLSNIMRDMNQDLASGPGSMEAFSDSLVRLGNNSAAQEDKIMSVMMRIASMGTISNM
FT                   TTPDLLALATATAATGQGAEAAGTALSRTFSNIEAAVGKGGAKLEKFAEISGMSASEFA
FT                   DAWNTRPMEAFTAFINGLKRVDEEGGSVANTLSELSINSVRQRQTLMGLTTTTDVLTKS
FT                   LIMSNDAWNGVGDTWGDAGDAAREAQRKAEGFSGQLQIMKNAAQELGVQLLDGAVPMLK
FT                   ELGGALQFVAGLLAAAPSGVTQLITRFAVLAAAAGPVLVAVGAIGDGITKLSEAVKAAK
FT                   AFWLLKDAARSSLAEINTAIAKNNAAIEQNIIAQERKVAAIENVNKAELENYNVTTEST
FT                   AVKERYIAGLNQETEALKAQQIQLEINRDKLLAQKEATVAAANNAKKLGTAVKAIGSVA
FT                   IFMVMTELITKSVEGIGETVERIKQLENATHGLKVAALDFTEAANSQATAIKTVGLATE
FT                   DLQSITEDAIKGNNSLIDSISSNKESASANSALVQEYANTILELSGQCKGSAESLALLR
FT                   AAIKSYNSLTGESVAITDEFTGAINVNEDSIRKHAEAFKKQAWADAVKKNAESATQELA
FT                   ALDTKIDALYKRRDTMKAQNYDLDEINATIKNLESMRDSAQAVYDQSWSDYKKLVTDGA
FT                   NEIKKAGEGVSSVSDTISDVTDELDEANDELEELSDTIDSLSEKYPEFGEKFAQWGLGV
FT                   EAFSNLLADLGLSMEDVVQGVEEMADKVSDGFNRIEENSEISLEKYMENLQHNRELTAA
FT                   WGDNLKKLYDKSSDETYRAWVNEMAEAGPEYAALIAEMAEQDAATLEEEARAWQEAKST
FT                   GVDAYLASIGLTSEEANTKVKLMVSGIADALGGLEAVEAKVVVTDDGTVQVLLDKLTEV
FT                   DRKRLADKRLLVTDDGTIIDMQDKCKSLNVYEINGKHFYVTDDGTIYDEKGKADQLNDV
FT                   VNRLSKTVTTYFKGDTSNLDDAVTRANDLISRVSGGITAIFRRSATGNIFPNQAIKAIP
FT                   RHADGGINGIVTHPTLTNIGWVGENGTEALLHMKNAGGAVIPLSNRTYVRPFARAVAME
FT                   MGGTQAPNITYNVTVNGAPTDSPEQFARRFTRAVAMQQRTKA"
FT                   /product="Phage tail length tape-measure protein"
FT   CDS             18901..22011
FT                   /db_xref="SEED:fig|6666666.127258.peg.28"
FT                   /translation="MATPKNHKIERSSSAPTNMSASWDLSLQETDTFSVRNFHVFGKNV
FT                   DTGEEKRFYTVPPNDTDWVDTNQSNNDVEVIFPAQFYPTTRWRLSFVQHHCYGRGWWLS
FT                   NIDVTAFNFEEPLPPKVTLTRSDLKLSWKVEAGDMPNTSKYYRTETQYWLQRIEHGQSW
FT                   PSYSKYRMPTPTGTFDESDDSYDYTSQSTYTPDKPVRIRLLAINRGPSGASEQAFEEHV
```

```
FT                   FAQVNTPTVSVKSVRSSSSNNVTAFSVNPNADDWHPIDHVILQRQIASSMNPSGSWDDI
FT                   VDIADVAANVNLMTDDTAAQVPDEDEATFWRVMTWHDDKQANAAYGYPANQPVRAGKPK
FT                   APSSVTTTINSDTSDDEYGYITASWTINSACSPNHIAELWGKTQGGVMERIESKTVAAG
FT                   ATSVTFDHDYSSLAAYSVRVQAYKPWSQSQKLSSDWAESTSMASVSIDTVSLLDDGVSL
FT                   YVEESHDNDDANYTEYSWSDRGDGWSCTEEPKTYNRLSTGSASHVSIAGLEEGKEYYVQ
FT                   ARRYDSETGAYGPYSAQVSAHSGATPSMPTLTASGRVAAGSPIEYEWDFPGGNQTAAML
FT                   FLEHDIARDETFIGDGETDTFELDLVPDSTPTVLVDGEVATFTFSGNTVTLNEAPADGA
FT                   WVEVSYIAPSGITVIGIDNDNMAYAYDTDESWDGEIAAYVKVTTGGNWSVASPIVKTEI
FT                   VQPPTCEVSATGTLVSDSSSYGGYKLTALPLTLDLGGTAGIFDVQVTCVNGTSIPSPRG
FT                   ISKVAAGSNVAWVPSCAPGETDIVPERGVLIGGGQYDVTVTGRDPSTGAKSEPVTISFA
FT                   VDWTNKALAPTAVAVVSDGEGKITVTKNPLSPITDRLDIWRKTADGAVLCAADAAFGME
FT                   YTDQVPPYGWEGNAYIAQVTTIDGDTNWVEAPYTLKSSGATVNWGDSSVTLPWNLDVSS
FT                   KYASTFERRAHMDGSRVGYWQAGVDRDSTMSGELIKGRDTEIINLFRQLGKHDGISFVR
FT                   SGGGLAFACNVDVDMQEDYMSQAVPVTLTITEVDDDGTWRARTELVE"
FT                   /product="hypothetical protein"
XX
SQ   Sequence 22822 BP; 5969 A; 5334 C; 5673 G; 5846 T; 0 other;
     acaaataaag gggcgcgaaa atttttcgtt ttttggtcga gtgcgcaaga gcgcactctc        60
     ccttctcaca acttagttgc gagttaagga gagttaacta ttagattatt ttcgtgtggg       120
     gttaattgcc gttcaggaag tcatcaatcg gcgattcctc agtggcggtg cctttcaacc       180
     aacgcatgat gtgactgaat aggcttgact gctctgcgac taattggtgc cgctgtctaa       240
     atagcttacc ttctgcatcg taattgtgca acccctcaga atcaataatt ccgttttcca       300
     aggatgtttc cagttcagcg atgcgttttg cgttccattc atagtggtca aactcagcca       360
     atacaaattc tttttgtgct gggtcgaggt tctttgcttg cttcgataca gcgtctcgca       420
     ggttttcaag attggttttc tttgccatta ctgtaccttt ccgtttcgtt tgttccaagc       480
     cttgcgtgca tccttcttac tcgtgaaaaa catcctactg ctggaaccaa aacagcgcag       540
     agtcttacac gcaaccgagt agagttttgt tccgtctcct ttatcaatcc attccatgta       600
     gtgagcgtgc ttcttacacc acggacattt ttttagtttt ttcattgttc gctctccaat       660
     ctgattatgt ttccgtcctc gtcaaatcca actcgcgctt caacttcctt gttgttttca       720
     aatgcgccga tacctaattc cttgtgtact tctgtgtgac actcattaca tagtgcaatc       780
     aagttgtcag cgttaagcga gatgtaaggg tcgttgatgt ttacagggtt gagcgggatt       840
     ttatggtgaa ctacatctgc tggtttaagt tctccttgtt cagcgcatct ttcacaaagc       900
     ccgtgtgctc gcgtccagac aaatgctcgt gcgtttttcc aggctcgtga gtggtagaac       960
     ttgtgcgcga atggcgtgtt tcgccaggtt gctgtttttg ccattggctt tctcccatcg      1020
     tggctagtaa accttttggt tggttagaag tatctccata tagatatatg tcagccatca      1080
     tctttgcgcg gcgttcgttt agtatttctc ccgccttaat actgatgtag ttgagaatta      1140
     cctttcgttc ttctggtggc aggtcaacaa gccattgtgg tgcttcgatt ctgatttcgt      1200
     acttactctc aaattcattc atgcctaatt accccttcag cttctttgat tagcttgtgt      1260
     cgtgctattg ggtatccaca aaagggacag tagaacggga ttactcggct tgcgcatata      1320
     tgcacgccat caggtttttc tactagagtc aggtgtattg ctgaagctaa ttccttacac      1380
     ctggggcatt tctcggtatg gtgcataagc taacctcgtt gagagtagat ttctaagaag      1440
     atggatgcca ggatggtgta gcagccgaga atacctacga aggttccaga tgccatacta      1500
     accaagagag acaaagcacc tgcaagtagt agctgtacga tagagataat gacgagtacc      1560
     gtaataataa tgtcagctct gtttgtcatc gcgctacctc cgtggtcata aacacttgtt      1620
     tgccgtttac ctctttaggc ttaatccttc ctctgagtag agcttcgcag aaggtgttta      1680
     tggttgcatg gtaataagct gctccatcag cgtagaacgt agtcgcgtcg ctgttatagt      1740
     cgaggttgaa ttggtctaac accatatcga acttatgaac aagaccttcg tgctcaaact      1800
     caatcttgat ggtcttacca atcatcatct ttgacgcagg attaaagaca gtcccgcagt      1860
     aatggcaata gtcatcagtg ataggagcgc cgcagttagg gcaattactg tgcatcgtca      1920
     gctcctttca gttctgcttc aagcaaggat aggagccgtg acttgtcagc agggtttagt      1980
     tgctcgatgt attcaatctg ctcatctatt acgattaccg taccgtcaag ctgctctttt      2040
     agcttagact tccatgcttc aatttcagcg gcacggtgca tccctttcaa cttgacaatg      2100
     aagataggct tatccatttg ccatctcctt aatcaaagca actgtacgct cgatgctatg      2160
     accatcacac tcactagctg tgagttccat gtagtgcttc tcaacgtctg taagaccatt      2220
```

agacgctgct tctctcatca tcttgacgag ttcttcttcg tgtccttctc caacaatcca 2280
acgagagctg taagtgtcgg ggtaggggta atacatcggg cggtcaggta ggtattcatc 2340
catatcgtca acggtaagga tgcaaggctt acctaagagg tatccgtctg tcatgcagct 2400
agagtagtca gtgattaaaa cgtcgcagtc aatcaggtag ggtgttagtg gtttgtcagg 2460
ctcaatctcg attactcgct tgaagttagc accgagtaat ggtctgtcgg taaagtagtg 2520
acgcttgact gctacgattt cgtcttcttc aagcaggttg tcaacaatct cccagtcaat 2580
ccagggaatc cagccgccct tgtcagcgtc tctaaatgta ggagcaaata ggtaagcacg 2640
cttattgccg agcttagtga atccttctcc tttcttcttt ccgatgtagg tgtcggttct 2700
ggggaatccg attgccttta ccctgttttc agggatacgc aactgctgtg cgacgatagg 2760
gatactagct tttgacgctg cggtagcaat atctgtttgc ttgaacgctt caatgtcaac 2820
gtctctatct tcgtctgctc catagacctt attgcccgtc atgccgtggc aaatcacgat 2880
ggacttacac ttatccttga acgcgatgta ttgagggagt gcgtcgcata ctacaacgtc 2940
atatcctttg agttcagcgt cgagcatgtt tactagcccg tgcttaaaca ccttgtcccc 3000
gtcgtaggag ttgtataagg ctgtgatgtt ttccccccta ccaagtttgt tctgagaaac 3060
aaacagtacc ttagaagcca ttacatcgcc ttctttcttt ctaacttgat gtggtcacaa 3120
gctactcgcg tgtcgagata tatctttcct cggatgcttt tccattgatg acagaagtaa 3180
tagtcttcgc tcaggtgctt attgaatttg aaccacgggt ctgtaaggat tttgagagca 3240
ctaaccttaa tcagagcgca cccaaagcca ccgccgccga tttccacaag ggtttgtccg 3300
tccttgcgca tctcttcaat ttcttctact gagtagtagg ttcgataatt gttgccgcga 3360
ggttcgacca tcggagtcaa attgtcacta tcactcttga tgtagtagcc aagacataca 3420
tcaacattgt tctccatgag atttagtagt gcatctgggg gaacaatggt gtcatcatcg 3480
acaaaaaaga cgtaatcgaa gtcgtttttt agggcaaatc gcaccagagc gtttctcgct 3540
tcgtctacgg cgtagaagct cttagttgct tgtactacat cattttgacc tctgttcagg 3600
ccgtagagag cgtctgatgc gtccgcccaa atgaagccct ttccagtcgg aattgcaagg 3660
aggactttag ccatttttca gcctttcgtg catctttaag gcttcatccc aactcatatc 3720
acggtactta ccgacgagag cttcctttct ccattgttct gtcttatatc ctgcgtagtg 3780
cataatgcga acaatgttgg ttggcttagt aaagttggtg cagttgtact cagatggcat 3840
gtcgtgaatc ttgccgcaac acatgaacga gaatacgtct tgttcgatgt ttggatacag 3900
acgttcgttt agagcatcaa ctacttcttg tgcctttccg ttgcgcaaca tttcgagatt 3960
gtagagagcc acacctgtat tgcagtagag catgttcttg tagcaactac atggctcaat 4020
agatgccgag aaatactttc cttcgattgg catgttccag gcattagata catctcgtac 4080
acagatagta tccacgtcca aggataatac cttatcaaca tgagacaact caggcattaa 4140
ggctagtgca gcacgcatga gcgaaagata cgagaacgga gtcttcatgc ttggtgaatc 4200
ttcagggaag taggtttgac tagacacgtc gattacctca accatatcag gcaaccagaa 4260
gtcatagtca tcgtcttcta ccagaagata aaccttgtct acgtctgagt tagcaatcaa 4320
agacttggtt gctgtttgca tatctccgta gagattgcga gaacctgagt agagcgcgta 4380
cttcatttga atttcctcca attcttttcc cttgtcccct taaagtccct gccgcacatc 4440
gggcagtatt taatgggtgt taataaccgc tgagtcttag gcttaatgtc tatttcctca 4500
aacttcaaaa ggctgtcagg cgtaagtctt tcgggattac caacttgata ttcgcgcttg 4560
tagaatttga acggtctgtc aggcgcttta atatcaagtg ctaggtaatt tgagaacacg 4620
tgcatgatgg ttcggaatgg caagtcgtgt ccttcaacgt tgatacgtgt atcggggaca 4680
acttcgtaca gcactctctg accgtcgcag tagggacaat gccttctatg aatcgaccta 4740
cgctctcgct tgaacttagg tgtgtgcttc agttgtgcca tctattcctc cgttgcgcta 4800
ttccagatac gaactgctac ttcttgcgtg cgtgccattg gccctcttgc tccacatccg 4860
cacttgtttc tactgcaagc tacatagtgc ttcccgtgac gttctctgag acgctgtgcg 4920
ccgtgacgct tacagaacgg acaaggctta acgatttccc agaaggtgtt agcatccatg 4980
tgttctccta gttgccaagt aagaagttga taaacaggta ttcacgtaag attgagtatg 5040
cgaataaacc gatgattaga gcgataagca gaagtacgcc tacgatacga gcaagaaaca 5100
agccgacttc tttcagtgtt tcttgaattg acagcccgta gtcacctgaa tctggcatca 5160
gtgttttcca gtcatcgtca tcgaagttat ccataagtcc tcctagatga ttcccttgta 5220
ctcttcaagg tgcctttgca tagcgataaa gccgttcagt tcagccatga aaccgtcgat 5280
acggtgaaca gctttgcctt cagctttaat gggtctgatg ttggcgttgg aatcctgctt 5340
gatttggacg ttcatgcggt tccattcgtt tgtggggtta tgtccgtcaa caatctggtt 5400
agacccgtac atagctttca gttccttcat tggggatgag aaagtcattg ggccttgtct 5460

```
tacaatctct gaacggtctt tcccaacgta ctgctcgaat cgagcaatat cctcgtcgct      5520
gaatccccaa cggtcatagc cgattgcaaa tagccatacg tccaactcgc gctttacttc      5580
atcgaaccag tcaatgaata cgctcttagg cacgatgttt cctggtactg ttcttatgta      5640
gcccttctgt tcccaaatac ggtatggaac atcgtcacgc tcgcgcctat acccgtcatc      5700
cttctttagc gcgtcttcgg ggagccaata catagacagc tcgtaaatct tcgggtcatc      5760
ctgtctcatc atgagaatct ttgcagcact aaggtcagtg gtatcactag cgtcgattcc      5820
agccacgcca tagcgaaagc ccaggtcaga gagctttcct gatggtggaa gctctgggaa      5880
tggctcagtg tttacagctt cgtcaaacga gagccatgca gacgcacggg actcaggcat      5940
gttgaagtct tttgtcatga ccgttgggag gaagctaggg tcttgcatag ccttgttgac      6000
agcatccctc atgtaatccc acgacttaac ggttccaagt ccaggattag cttttatcca      6060
acactcttca cggagccatt cactcctgtc atctaactcg taaaggaagg ggagcatacg      6120
gtcatcttca atcttgccgt cgaggatacc gcacgcatat tcatactggt catcgaaaag      6180
gttgccgcgc tcaaagccat ttgtggtaat cgctataagg aggggctggc gtctggtaga      6240
cattgactgc caaattaaat cgaactggtc acgattcgta tttgcgtgca tctcatcgaa      6300
caggcacccg tgagtgttaa gaccgtcgag gtgtcgtgtc tgattggtga gcaccgtgat      6360
gtagccgtcg ttgcggtcga atatcacacc gtcttggtcg cggtctacaa cggttccttt      6420
tctcaggact ttagatagtt ccttagaacg tttgagcatc tttactacga tgccataggc      6480
tagagacgct tgtgcttttg aggttgccac attatagacc tggggcgctc cttcaccatc      6540
agcaacgagc agatataact cgattgcagc accaaggcta gacttgcctt gcttacgcgc      6600
ccacataacg aggacttcat ggaattgcct ataaccttca tcatcgacaa agccaaaggc      6660
cgtttcaata acaacaagct catacggctc caagaccata ggctcaccag attcagggag      6720
cacacagaac ttctcgataa actctactgg ccttcttgct ttatcaatat caaaatgcca      6780
tttcttgtat ccatctttga agcgcggaag catcatctct gcaagctgtt taatacgctt      6840
acagaccgta atctttcccg cttgtacgtc gagtaggtat tgctcggctt ctgatatggt      6900
tcgcttactt gcagtcaacg gacttctccc agtgtttgca gtcctcagcg aacggggcga      6960
agatttgtaa ctcattggct tcgcagtcaa taatgcggtt gccgttctcg tctttagatt      7020
ccatctgctt ataggcgcag gtgtcgcagt tcttctcaac catgattact cctaatctgt      7080
gtctttgagc tggttaaaca gatttctttc aatctcagta ttagaggtgg agagttgaga      7140
acgctctgat gtaagagcca tgcgttctgc tggatggagc tgggttgaca taggggactt      7200
taaggtttcg tcaatctcga aaatccgtgc cttattcttc ttgtgcctag ataactgaga      7260
tttgacaatt tctttttgag cagggttgag atgtgccatt tgttgttcaa cggcttgctc      7320
tagtttgctc ttcgttgcca ttcggctcct ttcccgaatc gctaaatcct gcggtgccag      7380
tcaacttgtt cgccaggaaa aacgtgaaag tcaggaaaga ctaattcgcc agaaagcccc      7440
aaatcgccct aaacgacgtt aatacgcccc gttttgatat gagcgcaaat gtcagagttt      7500
agggacataa actcttcaag taagtgtacc acaaacagga cactaaggaa cacatgagca      7560
aaatacttga taagatactc ggcaaagaga cagctccgag cgcaactgct gtccaaactg      7620
tcggtaataa accctataca tcgtcatgga atggctcgat gtatcagcag gtgcttgttc      7680
gtagtgttat tgagcgtatc gcacttgcat gttccaagct caaaccagta attaagggga      7740
gcgcacgccc acgtattcga cgtgctattg agacttgtcc taacgagttt cagacgtggc      7800
cccagttcct tcatcgttgc gcaactcttt atatgaataa cactacggtt tgtgtggttc      7860
ctgagtttaa gcctggttca caagtccaaa cggggttcta tcccgtacct ctagccaacg      7920
ccgaagtcgt tgactacggt ggtgagtatt ggcttcggta cacaactcgt gacggtgact      7980
cacgcgctat tgaattgaag tacgtggcta tcgttacacg tttccaatat gaaagtgact      8040
ggtttggcga tggaaatatc ctcgccaaca cgctttctat gctcaaagca caagaagaag      8100
cccagaagaa cgctatcaac gactctgctc agttgaagtt cttgggtcag ctaacgggtc      8160
aggtacgcga agaggatatg aagaagaagc gcgaccgctt ctaccaggac aatcttacga      8220
gcgacaacaa cactccgctc atgctctatg atgccacgtt cgcttccttt actcagttga      8280
aggctgagaa ctggacaatc cctaacgagg agatggagcg cattgagaac aatgtgttcg      8340
actactttgg aatcaaccgc cgtattcttc agaacgcata cgacgagaac gcctgggacg      8400
cattttacga gggaattatt gagcctttcg gaatcatgct aggtgaagca ctaaccaagg      8460
ctacattcac catgagagag cgtcctgcga accagattct ctttagtgct aacagacttg      8520
aatacgcagc agcttcttca aaacgcaaca tcaacaaaga catgaccgac cgaggaatca      8580
tgacgcttaa cgaagcgcgt gaaatcctac aactcccgcc gattgaaggg ggagacgtat      8640
tcattcttcg cggtgaatac aaggttggtc acaccttcga tgaaatcttc caagcccagc      8700
```

aagccaaggc tgaagccaag gctagtgggc gtgttgcaag tcctgatgag gacagagatg 8760
gagccgacgg agacaacatc cgtggtgact cagatggtta cggctcacct ggcgataccg 8820
atacgggtga cgtgacctca accagccaag accgatggag cgagaacgca tcctagttaa 8880
aggagactat catgccagcg aaaccagaag agcgtcaata cagaatgatg agcacaccgc 8940
tctttattcc agatgccgca cccgaaacaa ttacagatga aaacggcgaa gagcaaactt 9000
cgtatcaaaa ccgattcaat agcccgtgct acgttgaggg ttttgctacg acgtttgatg 9060
acccttatac cttgttcgag gattcctggg acggctggaa gtacgttgag attatcgacc 9120
gtcacgcatt ggatgacgct gatatgtctg acgttatctt ccagtatgac catgaaggtc 9180
gcgtctatgc tcgaaacacc aacaacacgc ttttctttga agcaaatgac cacggcttgt 9240
atatcgcagc cgacctttct aaaacctcac tcgcacgtca gatgtatgag gatattgcag 9300
ttggcaacgt tacccgcatg agctgggctt ttattccaac tgaggaagag tacgtcagcg 9360
acatggagaa caaagttttc accacgcgaa ttatgcgtgt gaaaaaggta ttcgacgtaa 9420
gtgcggttag ctaccccgct gacccgaata ccgagataag cgcacgtcac ctcgtcaacg 9480
gagagattga gagcagacgg cagcaggagt tgttgcagcg cgaacttgac cgcaagcgca 9540
aggagattgc gatgcgtgcc aagagtttgt caatccgtta gtaaagagga aaggagaagt 9600
catggacttc accgctatgg acgcgagggc ttatcgtagc ctgaacgccg accagtacgc 9660
agagcgtcgt tctctggtac tgtctctggc agaggatatg ccagaagact tcaccgaaga 9720
gcagatgcgc tcgcttgacg aggaacttgg ctttatcaag tccgaagatg ctcgtcgtga 9780
ccagctcgcc gaagtacgca accacaaggc tgtcgctgtt gcaagtggct ctggcaaggt 9840
tgttggcgaa acggaagctc ccgaagttcg cacgaagcag cgcaaggaca actcccttgg 9900
cgctcgcgct tgggacgcaa ttcaggaacg tggctacaag cgtgatgaac gcttccaggt 9960
ttccgatatt gctttccgtt cggcatccga tgttcagacg gtagcagaac ttgacggcga 10020
gaatagtccg aactattacg actatgctct cacgcaggtt gaccttgaaa tcaaggaagg 10080
ctatcgccgc ccaatgtcca tttgggactt gttcaaccac gaaaccaccg agaaggattc 10140
ggtcacgtgg ttcgtggaag gcaccgacga aggtgacgct ggcatgaccg ctgagaacgg 10200
cgcatttagc cagtatcacg tcaatgaccc tgttgctgtt acgaccccgc tgaagaaggt 10260
cacggctctg tggcgacaga ccgacgagat tctgtccgat gctccgcgct ttgtgagcca 10320
cgtgaactct cgcgctggct acaagctgga ttgcaaggtt gaagaccagc tccttgctgg 10380
tactggttct tctaaccaga ttacgggtct tgtcaatgta tctggtatcc acaccgcttc 10440
tgcttctgcc ttcgacatgg ctttcatcga gtcgctgctt gatgaacgca cgaagattcg 10500
caaggctacg ccgaacttca acgttgacac cctgcttctg gctgacgagg actatgacac 10560
cctcatgaag ctgaagaaca acgctgacca gtacgtgctc ggtggcccta cgggcttcgt 10620
gtacggcaac ggtgtgacca ttggtaagaa cctgtggaac accatcacga tggttccgac 10680
ccctgctctg tccagcggta ctgccattct tggtgctttc aaggatggtg caacggttta 10740
cgagcatgtc acaggccgtc gcttcgacat tggctatgac ggcgaagact tcagccacgg 10800
tcgcgtttcg ttccgtgctt atcagcgtct cgctctggct gtcgaatatc ctgcggcctt 10860
ctgcaagtac acggtcgctg gcggcgcttc tggtctgaac gcttaaagga ggttcacatg 10920
caagtcatcg ccttagtaga ttttcatgac aacttgaaga acgttgaccg caagcgcggt 10980
gagcagttcg tcgtgaccaa ggaacgcttc gaggaaatca acgctgttgg catgagggca 11040
atcaatgggc ctatcgttga agagaagcgc attgaaaccc cgaaggaaac gcctgaatcc 11100
cgtgcgaaga aggcacctgc caagcgcaag acggcggtgg caaagtgacc ctcttagccg 11160
acattaaggc ttcgctccga gttacctccg atgcttttga ctcggaggta cgaggattga 11220
ttgacgcagc acgcttcgat atggagcgtg ctggcgttga cccgacgctt cttgcattga 11280
atccagtgac gtgcgacttg gataacgagt atgtcaagca agcagttcgt ttgtatgcaa 11340
aggctaactt tggatttgat aactccgaag cagagcgttt gcaggaaagc tacgaacagc 11400
acatgcgcaa cttgctcaac tcgtcgcaca acatcgcagc aatcgcgctt gaaccagcaa 11460
gccaaagcac gattggtagc tatccaacct ctgaaccaac tcaggagcct gaagtccagg 11520
agcctgaaac tcaggaggtc gaaggctaga tgcgctggaa tgagacagtc actcttatcg 11580
ctaaggctta cgagcctgat gatgagggtg tgccgcaagt gactgacaca aaaactgagg 11640
tattttgcaa cccgcgcact atgggcgcta acacatggtc ttccatgtat gagattggca 11700
tttctcctgt cgctgaaatt caggtacgca cctgtgatta cgacggtcag cgagacgtgt 11760
tctatcgcgg caagtatctc tcggtcgaag tggtgaaaga gcagggtgac ttcactgttc 11820
tagtcttgcg acaccaacag tctgattccg atgattctgc cgaagaatct tccagtgaga 11880
ttcctaatga aactcccgaa gaaggtgaag gcgaatgaca cacgatatta cgtgcgggat 11940

tgatgcgttt gcaaaaagca tcgaggaact tgtaggtaag attcctgtgg cttgccaaga 12000
agagtgcggt aaggccgtag ccaagtctac acgtcaaggc gtgaagaagg ttaaggaaaa 12060
ggcagcaaag ggcggtaagc acgaatggtc tgatgagtac gttaatgggt tctctagtcg 12120
agttactaag ggcgcagaga cgacgggtga aattggaaac aaggcaaagc ctggtctagt 12180
tcacctgctt gaaaaaggac acgcgacact aacaggccga cgcactactg cttatccaca 12240
catggcaccc gcatttgaag agatacaaga tgactttatt aaggaagtcg agaaggcagt 12300
agagagggct atttcgtaat ggcacacaaa gacgtatatg aaacagtctc agcgattgtt 12360
cctacgtgcc acatggagtg gcctaacgac tctgctcctg aacttccttg ggctttatac 12420
cacggggaag aaacacctat tgtcgctgac gatgtgcaga ttgcagtacg tcggcgttgg 12480
actgtcgagc tgtatgaaaa gcgtcgcaat aagaccattg aaaagaaact cggagacgcg 12540
ctaagagcac gttttgggtc tgtaaggcga agtgaatcat acgtcgaaaa cgacaacatg 12600
ctcgaagtaa tttatacatt ctacgagatt gaaggagagt tcgatggcta acaaagttcg 12660
tttcggtttg cgaaacacga aatacgctat ctacaacgct cagactggcg agtatggcgc 12720
tttgaaggat atgcctggtg ctgtttctct gagcctgtcc cgcgaaggtg gcgatagctc 12780
cgacttttat gctgacgatg gtatctacta taccttcgct ggcaccaacg gcggttactc 12840
tggcgatttg acgctggctc gtatcaccga cgaggttcgt aaggacttgc ttaacgaaat 12900
cgttgatgat gctactggcg ttcagtttga aaccacgacc gctgaacctg ctgagtttgc 12960
tctcatcact gagatgcagg gtgaccaggg gccgattggc tttgcgttct ataactgcaa 13020
ggcttcgcgt cctgaaatca gcgcaaacac gaagaacgat tcgccgagcg ttgaccagga 13080
taccctgagt ctgcgtatcg cttcccgttc gttcacctac gacggcgtgg caactccgtt 13140
cgtccagggt cacattgaaa agacccagga aaacgcaacg aagttcgcgg cgttcttcca 13200
gtctgttgtt accccctggcg cttctggcgg ttctggtctg tctgcttaaa ggagatacta 13260
aatgtttgaa gtcgaaatca atggcaagaa agtcaaagct gaagtcacgt tctacacggc 13320
tcaactctat gaagccgagt tccgcgccga cctcattaaa gaactgttcg gcgagcagag 13380
tttagagccg cagattgaga ctgatggtga gaatatcatc aagattgact ttaccaagat 13440
taactggctt gccacagcaa aagtgctttg ggcggcagtt aagacagcca acgaagcgac 13500
tccatcctac gtctcctgga tgaaagaaac gtcgggagtc aatatgtggc tcgtctctga 13560
gcaactgtcg gctgaggttg ccgactgctt ctttcgtacc gaagctccca gaacgcaaga 13620
gtaacagcga tagcaaacga gagcctactt ccaggccgta cacggctatg gcactaaacg 13680
gcctggaagc gggtcttaca tggcgcgaca tgcgccacat gaagtacacg cacctgatgc 13740
agttccttta tgaatgggaa gacatgcacg gtgctgatgt tgatgaagtt gtagatgcca 13800
cacccgcaga tgtaatggca ttaaccagat tgtaaggggt tgagatatgg gtgacgcttt 13860
ccgtggcttg actattcggc ttggtgccga cgctaggccg ctcaactccg caatcaactc 13920
aatcacacgc gctgcatctg gcgcacaaaa gcaacttaac gccatgaaca aggcgttgaa 13980
gtttgataat accaatgttg ccgctatgca gtctcgcctt gaacttgttg aggataaggc 14040
gcttctaaca gcaagggcag caaataaaat cgaagttgct atgcgtcagg cttcgcagga 14100
gacacttcag tttagcgccc gaagtggtct aggtgtaaaa aatctcaaag agatggcagc 14160
ggaaacaaag gctgtccatg ctgctaatga gaaattacac gcagatttta accatgtaaa 14220
tactgacctt gctcatatct atgaagctgc tgccagagtt agagctgaag ctaaaggcat 14280
ttcgtttgaa aaagcactaa agcaggttaa gtcgcttgcg tctcacatga atagcacgac 14340
tgctgctggc aaagctgcta aagatgctgt ctattctttg gttcgctctg ctgctatgag 14400
aacagacatt ggcgaaaagt ttgggcttca aaaaacaata ggccatgcga gtaagttact 14460
cggtgttttc ctagagctaa gaagagcatc aagggacttt agcagcgata tagagttgat 14520
gaataaggtc gaaggattta gagctgcgca aacaaggctc atttcgttca aagctgattt 14580
aatgcaagct gcttcagaag cagctcgctt taagtcagag ttatatgcgc ttggaacgaa 14640
tggtggtctt gttagtgctc tgaatgaaat taagctgata gatgctgctg ttgcaaaatc 14700
gactgcacat gctagagaaa tggctgaagc atacgcttta gcccctcgtc accttggagc 14760
tgcgaaagag cgtcttaact catacgttgc agcacagatg gcgctcaata atcagcttac 14820
ttcaatcaaa tctgcactta gaagaattga gagcgaccca gcatttgata agcaagcggc 14880
aaactcagct catgcctaca caaatgcagt taaggttgaa agcgcatata caaaggttaa 14940
atcaaagatt gctcttgcaa ccgcagccct tcaaaggctt gaagaagaac agcgtgagct 15000
tagtcttgtt cagacaaaaa ccgataagca gtctgctcgc tacgagcaac ttacttcgca 15060
cataaacaag gtaaacagcg caatcaatag gcttaaatct gagcttgcat cgcttgacca 15120
aaaacatgcc gcagcttctt tagtcattga gttcaaagaa ttgacagcag cacaggcaaa 15180

atgcgttgct gaaacagagc gactaaaaca ggcgcttagt tctgcaaata gggtttttgg 15240
caaccttaga actatgggct atggtcttta ttcaaccatt accccagcta tcttaatggc 15300
tggtagatac gctgttcaat cggctgatga agttgacgct gcatatcgtg acatgcgtaa 15360
gaccgtcaat ggaactgaag aagagttcga gcacttacgc gatgcagctc tagcctttgg 15420
tcgtacccac gtcacttccg ctgatacgat tttagagatt gaagcaatag gcggtcagct 15480
ttcgattcaa gcaaagaatc tcgaagcatt tagtcgtgtt gtttcaaatt tggatattgc 15540
tacgaatctc gatacggaga ctatttctca gaacatcggt cagctttcta atatcatgag 15600
ggacatgaac caagacctag cttctggccc tggttcaatg gaagcatttt ctgactcgct 15660
cgttcgtctt ggtaataact cagcagccca agaagacaaa atcatgtcag ttatgatgcg 15720
tattgcatca atgggtacta tttccaacat gacaactcct gatttgctcg cattggcaac 15780
tgctaccgca gcaacaggcc aaggtgctga agccgctggt actgcattat cccgtacttt 15840
ttctaacatt gaagcagcgg ttggaaaagg tggcgcaaag ttagagaagt ttgctgaaat 15900
atcaggcatg agtgcaagtg agtttgcaga tgcttggaat actcgcccga tggaagcgtt 15960
tacagccttc atcaacggct tgaagcgtgt tgacgaagaa ggcggctctg ttgctaatac 16020
tctgtcagaa cttagtatca atagcgttcg tcagcgtcaa acgttgatgg gcttaactac 16080
taccaccgac gttctaacca agtcgctaat tatgagtaac gatgcctgga atggcgttgg 16140
agacacctgg ggcgatgcag gagacgcagc gagagaagcc cagagaaaag ctgaaggctt 16200
ctctggtcag ttgcaaatca tgaaaaacgc agctcaggaa ctaggagtac aacttcttga 16260
tggcgctgtg ccaatgctaa aagagctggg tggcgcttta caatttgtag cagggttgct 16320
tgctgctgct cctagcgggg ttactcagtt gattacaaga tttgcggtgc ttgctgcggc 16380
agctggccca gttttagttg ctgttggtgc tattggcgat ggcatcacaa agctaagtga 16440
agccgtgaaa gctgcaaagg ctttctggct gttaaaggat gcagcaaggt ctagcctagc 16500
tgagataaat actgcaatag caaagaacaa tgctgctatt gaacagaaca taattgctca 16560
ggaaagaaaa gtagccgcaa tcgaaaatgt caacaaggct gaattagaga attacaatgt 16620
gactaccgaa tctactgctg tgaaagagag gtatatagct gggcttaacc aggaaacaga 16680
agctctaaag gcgcagcaaa ttcaacttga aataaacaga gataaattac ttgcgcaaaa 16740
ggaagccacc gtagcggctg ctaataacgc gaagaaactg ggaactgctg taaaggcaat 16800
cggttcggta gctatcttta tggttatgac agaactgatt acgaagagcg tagaaggtat 16860
aggcgaaaca gttgagagaa tcaaacagct tgaaaacgca acgcacgggc tgaaagttgc 16920
tgctcttgat ttcactgaag ccgctaactc tcaggctact gcaatcaaga cggttggcct 16980
agcaactgaa gatttacaat ctattacaga agacgcaatc aagggcaata acagcctgat 17040
tgattctata tcatctaata aagaatctgc ttccgctaat tctgcgcttg ttcaggagta 17100
cgctaacaca atcctagagt tgtctggtca gtgtaaggga agtgctgaaa gccttgcgct 17160
gttgagggct gctattaagt cctataacag ccttacaggt gaatcggttg ctatcacaga 17220
tgagttcact ggcgcaatca atgtaaatga agattcaatt aggaagcacg ctgaagcgtt 17280
caaaaagcaa gcatgggctg atgccgtaaa gaagaacgct gaatccgcaa cacaggaact 17340
cgctgctctt gatactaaga tagatgcttt atataagcgc cgagatacaa tgaaggctca 17400
aaattatgac cttgacgaga taaatgcaac aatcaagaat ctcgaatcta tgcgcgattc 17460
tgctcaggcg gtttatgacc agtcgtggag cgattacaag aaactcgtaa cagatggtgc 17520
taatgaaatt aaaaaggctg gcgaaggtgt atctagtgta agcgacacca tatctgatgt 17580
tactgatgag ctagatgaag cgaatgacga actggaagaa cttagcgata cgattgattc 17640
tctaagcgag aagtatcctg agtttggcga gaagtttgct cagtggggtc ttggagtaga 17700
agcgtttagt aatttgttgg ctgaccttgg attaagcatg gaagatgtag tccaaggcgt 17760
tgaagagatg gctgacaagg tatctgatgg cttcaaccgc atagaagaga actccgaaat 17820
ctccctcgaa aagtacatgg agaaccttca gcacaacagg gagcttacgg ctgcttgggg 17880
cgataacctg aaaaagctat acgataagtc ttctgatgaa acatatagag catgggtcaa 17940
tgagatggct gaagctggcc ctgagtatgc agccctaatt gctgaaatgg ctgaacaaga 18000
tgctgcgaca ctagaagagg aagctagggc gtggcaagaa gcaaagagca caggtgttga 18060
tgcgtactta gcttcaattg gcctgacaag cgaagaagcc aataccaagg ttaagctaat 18120
ggtgagcgga atcgctgatg cgcttggcgg tctggaagcc gttgaagcaa aagttgtagt 18180
aactgatgat ggtacggttc aggttctact tgataagctg acagaggttg accgaaagcg 18240
tcttgcagac aagagacttc ttgtaactga tgatggcact atcatcgaca tgcaagataa 18300
gtgcaagtcg ctcaatgttt atgaaatcaa cggcaagcac ttctacgtta ccgatgatgg 18360
aacaatttac gacgagaagg gtaaagcaga ccagttaaat gacgttgtaa atcgactcag 18420

taagacggta actacctatt tcaagggaga cacaagtaac cttgacgatg ctgttactcg 18480
tgcaaatgac cttatttcaa gagtatctgg cggtattacg gcgatattta gaaggtctgc 18540
gactggaaac attttcccga accaagctat caaagcaatc cctcgtcacg cagatggcgg 18600
catcaacggc atagttactc acccgactct taccaatatc ggctgggtag gagagaacgg 18660
cacagaagca ttactccaca tgaagaacgc aggtggcgct gttattcctc tgtctaaccg 18720
cacttatgtc agaccatttg ctagggcagt agctatggaa atgggcggta ctcaggcacc 18780
caacatcaca tacaacgtaa ctgtcaacgg cgcaccaaca gactctcctg aacagtttgc 18840
gagaaggttc actcgtgcgg tagcgatgca gcagagaacg aaggcataag gagatattaa 18900
atggcaacac ctaagaacca taagatagaa cgaagctcat cggctcccac gaacatgagt 18960
gcatcgtggg acttatcgtt gcaggagaca gacacgtttt cggtgcgcaa ctttcacgtc 19020
tttggaaaga acgttgacac tggcgaagaa aagcgattct acactgttcc cccaaacgat 19080
acagactggg ttgacaccaa tcagtcaaac aacgatgtag aggttatatt ccctgctcag 19140
ttctatccga caactaggtg gcggttaagt ttcgtgcagc accattgtta cgggcgtggt 19200
tggtggctca gtaacattga cgttacagcg ttcaactttg aagaaccact cccaccaaag 19260
gtcacgctta caaggagtga cctaaagctg tcatggaagg tcgaagctgg cgatatgcct 19320
aatacttcca agtattacag aaccgagact caatactggt tacagcgcat tgaacacggt 19380
cagtcgtggc cttcttactc caagtatcgt atgccgacac ctactggcac gttcgatgag 19440
tctgatgatt cttatgatta cacgagtcag agtacataca cgcctgataa gccagtcaga 19500
atacgccttc tagcaatcaa tagggggccg tctggcgcgt ctgagcaagc gtttgaagaa 19560
cacgtatttg cgcaggtcaa cacgcctacg gtatcggtta agtctgttcg ctcgtctagt 19620
tccaataacg tcacagcttt ctctgtcaat cctaatgcgg atgactggca tcctattgac 19680
cacgtaattc tgcaaaggca gatagcgtcc tcaatgaatc cgtctggctc gtgggatgac 19740
attgttgaca tagcggacgt tgccgcgaac gtgaacctta tgactgatga cacggcggcg 19800
caagtcccag atgaggacga agctacattc tggcgtgtca tgacgtggca cgatgataaa 19860
caagctaatg ccgcctacgg ataccctgcg aatcagcctg tgagggcagg taagccaaag 19920
gcaccatcat ctgtgacaac aacaatcaac tcagatacca gcgatgatga gtacggttac 19980
atcacggcga gttggactat caactctgcg tgttcaccga atcatattgc cgagttgtgg 20040
ggtaagacac aaggcggcgt gatggagcgc attgagtcaa agacagtggc ggcgggagca 20100
acgagtgtta cgtttgacca cgactattca tctttagcag cgtactctgt gcgtgtgcaa 20160
gcgtataagc cctggtcaca gtcgcaaaaa ctttcgtctg actgggctga gtcaacttct 20220
atggcatcgg tcagcataga tacggtgtcg cttttagatg atggcgtgtc gctctacgtc 20280
gaagaatcac atgataacga cgatgcaaac tatacggagt atagctggtc tgacagaggg 20340
gatggctggt cttgcacaga ggaacccaag acatacaaca ggctttccac gggtagcgca 20400
tctcatgtat ctatcgctgg gttagaagaa ggcaaggagt attacgtcca agcaaggcgt 20460
tacgatagcg aaacgggagc ttacggccca tacagcgcac aggtatctgc tcattcaggc 20520
gcaacaccaa gtatgccaac tcttaccgct tctggcaggg ttgcagcagg tagtccaatt 20580
gagtatgagt gggacttccc aggtggcaac cagacggcag ctatgctgtt tttggaacat 20640
gacatagcga gggatgaaac attcattgga gacggcgaaa ccgatacgtt tgagcttgac 20700
cttgttcctg acagtacccc aacggtactt gttgatggag aggttgctac gtttacgttc 20760
tctggtaaca cggttacgct taatgaagca cccgcagatg gagcgtgggt cgaggtttcg 20820
tatatagcac catctggaat cacagttata ggcatagaca acgacaacat ggcatacgct 20880
tatgacactg atgaaagctg ggacggcgaa attgcagcct atgtgaaggt gactacgggc 20940
ggcaactggt cagtcgcatc gcctatcgta aagacagaga ttgttcagcc acctacttgt 21000
gaggtatcgg caacaggaac actcgtctct gactcatcta gctatggcgg ttataaactg 21060
acagccttac cactcacctt agacctcggc ggtactgcgg ggatatttga tgttcaggtt 21120
acgtgtgtga atggcacatc cattccttcc cctcgtggca tctcgaaggt tgcggctggg 21180
tcgaatgtcg cttgggtgcc gtcgtgcgcc ccaggagaga cagatattgt ccctgagagg 21240
ggcgtgttaa taggtggcgg tcagtacgac gttaccgtta caggccgtga cccatcaaca 21300
ggtgctaagt ctgagcctgt aactatcagc ttcgctgttg actggacgaa caaggcgctt 21360
gcaccaaccg cagtagccgt tgtgtctgac ggtgaaggca aaataaccgt aactaagaac 21420
ccgctatctc caatcacaga ccgcctagat atatggcgca aaactgctga cggagctgtg 21480
ctatgtgccg ctgacgcagc ttttggtatg gagtacacag accaagttcc cccgtatggc 21540
tgggaaggaa acgcttatat cgcacaggtc acaacgattg atggtgacac gaactgggtt 21600
gaagcacctt acacgctgaa atcaagcggc gcaactgtca actgggggga tagttcagta 21660

```
     acgctccctt ggaacctaga cgtgtcgagc aagtatgcat ccacattcga gcgcagagcg     21720
     cacatggacg gttcgcgtgt tggatattgg caagctggtg tagaccgcga ctctaccatg     21780
     tctggtgaac tcataaaggg aagagatacc gagattatca atctgttcag gcagcttggc     21840
     aagcatgatg gcatctcctt tgtccgctct ggcggtggac tcgcgttcgc gtgcaacgtc     21900
     gatgtagaca tgcaggaaga ttatatgtcg caagcagtac cagtgaccct gacaattact     21960
     gaagtagacg atgacggcac ctggcgtgct aggacggagt tagtggaata acatggactt     22020
     ctccaaagga tacacagcat cgtttagggc ggttttgctt gacccatcgt ggagtgaggg     22080
     tgagtctctt acgggcttaa cttcgatttc gattacgaaa gacgtatcgt ctgatgttcc     22140
     gctcattgac tcggctgaca ttgacctcga ctctccaccc gcagacgagt cttatgtgag     22200
     gatatatctt gatgccgtac aggacggcgg cgtaacgtca tacccactag ggacgttcct     22260
     tgtcgaaagc tcagatgaag aaacgggcgc agggtcagac tggtcaacag ccgaactgtg     22320
     gtctgtcctt aaaccagcag atgagcttat gctaccaact ggctggtatg cacagaaggg     22380
     agctgcatca gcacttaaag cagcgtctct actagcagag tgcatctcag cacccgttga     22440
     agcacagggc ggttctccgt cacttgcgga acactacgtt tctgatggaa cacagagtca     22500
     tcttgcagcg gcttgggcgc ttctaccgtc tggctggggc atagcagttg atggcatggg     22560
     taacgtaacc gttcgtgaga atccaagtga ccccgtacca atacttacgt ctgagattat     22620
     cggcacagtc aaagagtcat ggaaccttgc aggtgtccct aacagggtaa cggtatcaga     22680
     tgacgctggg aacgaagcta cggctgtgaa cacagacccg tcatctccta catcgaccgt     22740
     tgctagaggt cgaatcatcg acggagacgg cggtggagta cgtgagtctg gcgagtcttt     22800
     gagggcgttt gcaaagagac tc                                              22822
//
ID   XXX; XXX; linear; XXX; XXX; XXX;  XXX.
XX
AC   PRJEB67889
XX
DE   Uncultured bacterium DNA fragment from in-reactor enriched cow rumen, clone
ferm-42a14, contig2
XX
FT   source          1..6910
FT                   /organism="uncultured bacterium"
FT                   /environmental_sample
FT                   /mol_type="genomic DNA"
FT                   /isolation_source="Cow rumen"
FT                   /clone="ferm-42a14"
FT                   /note="ABTS active clone"
FT   CDS             complement(477..1052)
FT                   /db_xref="SEED:fig|6666666.127259.peg.1"
FT                   /translation="MINRIWLGMMFIGLVFGLCNGQGAALSEAVIDGSTQAVNLSIAMV
FT                   GGMMFWMGLMEIVEKSNLTQILARGLGRILRPLFREVQHDEKAMSQISLNIAANMLGLG
FT                   NAATPFGLCAMQTLQAHNLHPKTATCAQIMLICIDCSAVQFLPTSIMTLLAAAGSAAPT
FT                   ALVLPILAATSATTLTAVVACLMAQRNK"
FT                   /product="Spore maturation protein A"
FT   CDS             1184..2125
FT                   /db_xref="SEED:fig|6666666.127259.peg.2"
FT                   /translation="MAWKDRFVHAAEPADSADQAAQEQAYARRRAAFYQQIAQARALPE
FT                   RTLYYHEPDTRTLSVLGLGVNRKEGMFTHMQAPMTAHEPDGTKLYAFEWHAAPIADLMS
FT                   LRVSDGSTPVFSITAERDAANKDIRPGFFMTTSLLTRKPVFRETSAVTLDLTFRDPAKM
FT                   PPIHIMASFDRLEERSIEKDPLLAPDDQSHLLPISRLAPAFEDMINHTENLNRIKPLIE
FT                   KWVLDLAVLMQEQAQKRLSGESAAPTVNTDIPSAPLAASVHPAVDGAAHSGQAEGVDAV
FT                   LATIEKLAALHSAGILTDDEFAEKKKELLAKI"
FT                   /product="hypothetical protein"
FT   CDS             complement(2199..2675)
FT                   /db_xref="SEED:fig|6666666.127259.peg.3"
FT                   /translation="MQIQVLSVGKVKEPFFRQACAEYEKRLSRYAKLNCIEVADESIPE
```

```
FT                   KAGPAECRRVVQKEGERLRAHCKPGLTIALCIQGRTMDSPAFAGLLRQAEQSGISCVQF
FT                   LIGGSLGLDASVVQQADIKISFSPMTFPHNLAKVMLLEQIYRAMKINRGETYHK"
FT                   /product="LSU m3Psi1915 methyltransferase RlmH"
FT   CDS             complement(2687..3505)
FT                   /db_xref="SEED:fig|6666666.127259.peg.4"
FT                   /translation="MERDEGLKLAFCPLFSGSSGNAVFIGSQQARVLVDAGVSATAICQ
FT                   ALVEIDVRPDQLDGILVTHEHTDHTRALQTLARKYGIPIYANRATFDALGKKTEAIPTG
FT                   LARVFETGQEFYIGNLAVRSFSIPHDAADPVGFAVHANGKKACVMTDLGHVPTRLLDQA
FT                   AQADVMLLESNHDVHLLEQGKYPAYLKRRILSTKGHLSNETAGHTAVRLAQRGVKRFFL
FT                   GHLSQENNREDVALSAVCGALTAQQIQPGRDVALAMAHRNCRSTLVVLDR"
FT                   /product="Zn-dependent hydrolase (beta-lactamase
FT                   superfamily)"
FT   CDS             3634..4569
FT                   /db_xref="SEED:fig|6666666.127259.peg.5"
FT                   /translation="MASAQIVVQQVLVMFLLLGVGILIRRLGWVTEEAARQFSNLLLMI
FT                   VFPALLIEVYQRPFIQEQFIDQLISMALAVIFHLLAIAVSHLCIRPREHHAYRIERLAA
FT                   VYSNCGFMGFPLLSALLGEEGLFYGVAFSGVFNLVFWIHGARTCSLDDRLEKKRIVLNP
FT                   GCIGFLTGFILYITQLPLPSFLCETITHVANLNTPLAMIITGVFLADIRWKKLWDDASI
FT                   WRACLLRNMLLPLGMFVLLRCIGFASWFSGAPEIALAHLIACACPSAASTILLAAKRNM
FT                   DSAHGAKILALSTLLSIVTLPAIVLLAQSV"
FT                   /product="Malate permease"
FT   CDS             complement(4615..5880)
FT                   /db_xref="SEED:fig|6666666.127259.peg.6"
FT                   /db_xref="GO:0008760"
FT                   /translation="MEDTLIIQGGRPLRGVVSPSGAKNAAVALLPAALLCDEPCTVENL
FT                   PDIRDVRVLASILEQMGASVDYRSGVMTLDPRCVTSREATYDLIRHMRASYYLLGAMLG
FT                   KFGEAKIALPGGCEIGQRPIDQHLKGMEALGAEVEVSHGFVSAKAPQGRLVGGEVYLDV
FT                   VSVGATANVMMAAVRAQGRTTIVNAAKEPHIVDLANFLNAMGANVKGAGTDVIRVRGVE
FT                   RLHGCTYTVIPDQIETGTLMIAAAATRGDVVIENMIPAHMESISAKLEEMGVIVEEGES
FT                   SIRVADSGKPFHQTNIKTMPYPGFPTDLQQPISALLCVTQGSSMVEENIFEDRFKHLGQ
FT                   LQRMGAQYRVSGRVAMIDGVKSLSGSRVVATDLRAGACLVIAGLMAQGTTEITGLRHID
FT                   RGYDHLEEKLRSLGADIRRVQK"
FT                   /product="UDP-N-acetylglucosamine 1-
FT                   carboxyvinyltransferase (EC 2.5.1.7)"
FT                   /EC_number="2.5.1.7"
FT   CDS             6396..6755
FT                   /db_xref="SEED:fig|6666666.127259.peg.7"
FT                   /db_xref="GO:0004526"
FT                   /translation="MTRNAEFQYVYRRGRSQADGRMVLLVARSRTGTRAGFTVGKKVGN
FT                   SVVRSRVKRLMRESYRLMAGQVQPRTLLVFIARPSMAGSSFAQVQASMKKLMVRSGCLR
FT                   EQTDENASDCDDSRL"
FT                   /product="Ribonuclease P protein component (EC 3.1.26.5)"
FT                   /EC_number="3.1.26.5"
XX
SQ   Sequence 6910 BP; 1529 A; 1938 C; 1836 G; 1607 T; 0 other;
     aatgaatgaa gtgggtacag tataagataa gcagaagtac agatgtgata atgataagtc        60
     acagctgaca ctggaataca atgctgggag caatgagctt gggaaccctg gtaacctaaa       120
     gagagatgtt ctttgtaaag ggcagtcatc atgtactttt tgctgactgt tgagtacagc       180
     caatgccgcg ctgcccgaaa aggggcgcac caccgcgacg ggtaacgctt caggcggcaa       240
     cccaatcgct tgtgccaggg gcgtacatat atgcaccaac cagtccaata cgccgctggc       300
     acgcaatacc gatacggcta ccagcatggc aatcagcgtg ggcaacacat gcagacaggt       360
     ctgcaggccc tttggcgccc cttgcgaaaa cgctgcaaaa ctatccacct ttttaacact       420
     ggcgtaaaaa agcgtgccgc cgatcaaaat cgcgaccgcg gcggacgaaa ggccgctcat       480
     ttgttccgtt gcgccatcag gcaggcgacc acagcggtca gcgtcgtagc ggacgtagcc       540
```

gcaagaatcg gcaacaccag cgcggtgggc gcagcgctgc ccgccgctgc cagcagcgtc 600
atgatggacg tgggtaaaaa ctggacggct gaacagtcga tacagatgag catgatttgc 660
gcacacgtgg cagttttagg atgcagatta tgcgcctgca gcgtttgcat cgcgcacaaa 720
ccgaacggcg ttgccgcatt ccccaggccg agcatattag cggcgatatt caaactgatt 780
tggctcatcg ccttttcatc atgttggacc tccctgaaca gtggccgcag tatacgccct 840
aaaccacgtg ccagaatctg cgtcaggttg ctttttttcca cgatctccat caggcccatc 900
cagaacatca ttcctccaac catggcgatg gatagattga ccgcctgcgt ggaaccgtca 960
atgaccgcct cactcagagc cgcgccctgg ccgttgcaca agccgaacac caggccgata 1020
aacatcattc ccaaccagat tcggttgatc atctccatcc ctcctccttg ggttcactgt 1080
atgcggctgg caaacaaaaa aggacggaat cgtcctaggg gattttcatt gtgtacaatg 1140
gcgtagtttg atatactgaa taaaaagaaa gcgaggggtt tgcatggctt ggaaggatcg 1200
tttcgttcac gcagccgaac cggcagacag cgcggaccag gctgcgcagg aacaagcata 1260
tgcgcgccgg cgtgccgcct tttatcagca gatcgcacag gcgcgagcct tgccggagcg 1320
cacgttgtat tatcatgaac ctgatacccg cacgttgagc gttctggggc ttggcgtcaa 1380
ccgcaaggag ggcatgttca cgcatatgca ggccccgatg accgcgcacg agccggacgg 1440
taccaaactg tatgcatttg aatggcacgc tgcgcccatc gcggatttaa tgtccttgcg 1500
ggtcagcgac ggcagcacgc ccgtgttttc cattacagcg gaacgggacg ccgccaacaa 1560
ggatattcgt cccggctttt tcatgaccac ttccctgttg acccgcaaac ccgttttccg 1620
ggaaacgtcg gccgtaacgc tggacttgac ctttcgcgat cccgccaaga tgccgccaat 1680
tcatatcatg gcttcgttcg accggctgga agaacgcagc atcgaaaaag atccgttgtt 1740
ggcgccggat gaccaatcgc atcttctgcc tatcagccgg ttggcgccgg cctttgaaga 1800
tatgattaat cacacggaaa atctgaaccg cattaagccg ttgattgaaa aatgggtatt 1860
ggacctggca gtgttgatgc aggaacaggc gcaaaagcgg ctcagcggcg aatcggccgc 1920
ccccacagtt aacaccgata taccctcggc accccctggcc gcgtccgtgc atcctgccgt 1980
ggacggcgcc gcgcatagtg gacaggcgga aggagtcgat gctgtgctgg ccaccatcga 2040
aaagctggct gcgctgcata gcgccggcat tctgaccgac gatgagtttg cggaaaagaa 2100
aaaagagttg ttggcaaaga tttaaaaaac ggatgcaaag agccggctga cgtgatgtca 2160
gccggctctt ttgtctttac ggcagatctt taccatgcct atttatgata ggtttccccg 2220
cgattgatct tcatcgcccg atagatctgt tccagcagca tcaccttggc caggttgtgt 2280
ggaaatgtca tcggagaaaa ggaaatcttt atatcggctt gctgcacgac cgatgcatcc 2340
aaccccagcg agccgccgat aagaaactgc acacagctaa tgccgctttg ttcagcttga 2400
cgcagcaaac cggcaaaagc aggcgagtcc attgtccgcc cctgtatgca aagggcgatg 2460
gtcagcccgg gtttgcaatg cgcacgcagc cgttcccctt ctttttgcac gacgcgccgg 2520
cattccgccg gacccgcttt ttctgggatg ctttcgtcgg ctacctcgat acaattcagc 2580
ttcgcataac ggctgagcct cttttcatat tccgcacagg cctgtctgaa aaatggttcc 2640
ttgacctttc ccacgctcaa tacttggatt tgcaaaatag gcctcctcag cggtcgagta 2700
cgacgagcgt actgcggcag tttcgatgcg ccatggcaag cgccacatcc cggcctggtt 2760
gtatctgttg ggcggttaac gctccgcaga cggcggagag cgccacatct tcccggttgt 2820
tttcctgcga aagatggcca agaaaaaaac gcttgacgcc gcgctgtgcc aaccgaacgg 2880
ccgtatggcc ggcggtttcg ttgctcaggt ggcctttggt ggaaagaatg cgccgtttta 2940
aatacgccgg atatttgccc tgttcaagca gatgtacgtc atggttgctt tccagcagca 3000
tcacatccgc ctgcgccgcc tgatccagca ggcgcgtggg cacatggccc agatccgtca 3060
tcacgcacgc cttcttgcca ttagcatgga cggcaaaccc gactggatcc gccgcatcgt 3120
gcggaatgga aaacgaccga acggccagat taccgatata aaattcctgt ccggtttcaa 3180
agacgcgcgc caggcccgtg ggaatggctt cggtcttttt gcccagcgca tcgaacgtgg 3240
cacggttggc gtagatggga atcccgtatt tacgcgccaa ggtctgcaga gcgcgcgtat 3300
gatcggtatg ttcatgggtg actagaatgc cgtccagttg atccgggcgg acatcgatct 3360
ccaccagcgc ttgacagatg gcggtcgcag acaccccggc gtccactaaa acacgcgctt 3420
gttgcgaacc aatgaaaacc gcgtttccgc ttgaaccgga aaaaagcgga caaaaagcca 3480
gtttcaatcc ttcatccctc tccattcatt cgacacgatt gtaccataaa ccgtggcctt 3540
cgtcataacg ggattttttc ttgtcagcgg gacgcctgtg cggtacaatg aacgcgtttt 3600
gaactcgttt tataataaag aggtgttcgt cccatggctt cggcacaaat tgtggtgcaa 3660
caggtgttgg tcatgttcct tttgctgggc gtcgggatat tgatccgccg gctgggatgg 3720
gtgacggaag aggctgcgcg ccagttcagc aatctacttt tgatgatcgt cttccccgcc 3780

ttgttgattg aagtatatca gcgcccattt atacaagaac aatttattga ccagttgatt 3840
tccatggcgc tggcggtgat ctttcatcta ctggcgattg ccgttagcca tctatgcatc 3900
cgaccgcgtg agcatcatgc ctaccggatc gaacgcctgg ccgccgttta ttccaactgt 3960
ggattcatgg ggttcccgct gctgtccgcg ctgctgggcg aagaggggct gttttatggg 4020
gttgcattct ccggcgtatt caatctggta ttctggattc atggcgcaag aacgtgttcg 4080
ttggatgacc ggctcgaaaa aaagcggatc gtactcaatc ccggctgcat tggcttttta 4140
accggcttta ttctttacat cacgcagctg ccgcttccgt cctttctctg tgaaaccatc 4200
acccatgtgg ccaacctgaa tacgccgctg gctatgatca tcacaggcgt atttctcgct 4260
gatatccgct ggaagaaact atgggacgac gcatctattt ggcgggcctg cctattgcgc 4320
aacatgctct tgccgttggg aatgtttgtg ctgcttcgct gcatcggatt tgcttcttgg 4380
ttttccggcg cgccggagat tgcgttggcg catctgatcg cctgcgcctg tccgagcgcg 4440
gcttccacca ttttgctggc ggcgaaacgc aacatggata gcgcgcatgg cgcgaaaatc 4500
ctcgcccttt ccacgctgtt atcgattgtg actttgcccg ccatcgtatt gctggcccag 4560
tccgtataaa cgtatagaaa aaggaaggcc aacgccttcc tttttttcatc atggctattt 4620
ctgtacgcgc cggatgtcgg cgcccagact gcgcagcttt tcctccaggt ggtcatagcc 4680
ccggtcaata tgccgcaggc cagtgatttc cgtcgttcct tgtgccataa gccccgcgat 4740
caccaaacag gcgccggcgc gtaaatccgt cgcaaccacg cggcttccgc ttaggctctt 4800
tacgccgtcg atcatcgcaa cgcgcccgct gacgcgatat tgcgcgccca tgcgctgcaa 4860
ctgaccgagg tgtttaaagc ggtcctcaaa gatattctct tctaccatgc tgctcccctg 4920
cgttacgcac agcagtgcgg aaatcggctg ctgcaaatcc gtgggaaagc ccggatatgg 4980
catggtctta atattggtct gatggaacgg tttgccgctg tccgccacgc ggatgctcga 5040
ctcgccctct tcaacgatga cgcccatctc ttccaatttg gcggaaatgc tttccatatg 5100
ggccggaatc atgttttcaa tcacgacgtc tccgcgggtt gccgcggcgg cgatcattag 5160
cgttcccgtt tcaatctgat ccggaatgac ggtatacgta cagccatgca gccgttcgac 5220
tccgcgaacg cgaatcacat ccgtaccggc gcctttaaca ttggcgccca tggcgttgag 5280
aaaattcgcc aaatccacaa tgtgcggctc tttggctgcg ttgacgatgg ttgtacgtcc 5340
ctgcgcacga accgccgcca tcatcacatt ggcggttgcg ccgacactga cgacatccag 5400
atatacttcg ccgccgacca aacgcccctg cggagctttg gcgctgacaa agccatggct 5460
gacctctact tccgccccca gcgcttccat gcccttgaga tgctggtcaa tcggccgctg 5520
gccaatttca cagccgccgg gaagagcgat tttcgcctcg ccaaattttc ccagcatcgc 5580
gcccagcaga taataggatg cgcgcatgtg acgaatcaaa tcataggtag cttcgcgcga 5640
cgtcacacag cggggatcca acgtcataac gccgctgcga taatcgaccg aagcgcccat 5700
ctgttccaga atgctggcaa gcacccgcac atcccgaata tccggcagat tctccaccgt 5760
acagggttca tcgcacaaca gcgccgccgg cagaagcgct accgccgcat ttttcgctcc 5820
gctgggactg acgacgccgc gcaagggccg tcctccctgg atgatcaatg tatcttccac 5880
cctgcatcct ccttgaaata tttcattgat gcaaaccttt tgttgcaata ttgttacata 5940
cgtttcttag tataatcgtt atgctttttt tgtgcaatgg atgtttctct atataaaatt 6000
tgtatttttc gtcatacgaa aagagctgct gctgtttgaa gaaagggatt tacgggatgt 6060
ttcttgattg gatggattga cagcctgtcg tccatccttt ataattacaa agaattgcca 6120
tactctgctc tatgtgtgga ggagggcgca aaattttgtc atctacggcg ggtatccgct 6180
taattagaag gagtgcttaa catgttacag acgtatcaac cgaaaaaacg gcatcgcaaa 6240
cgcgtacacg gtttccgttc ccgtatgaaa agtgcaaccg gtgcaaaagt catttcgcgc 6300
aggcgccgta aaggccgcaa ggtgctctct gcttaaagtc gcgcaacgcg gcttttttca 6360
tcatggcacg atattctttt cagcgacccc aacgtatgac gcgcaacgcc gaatttcaat 6420
atgtgtaccg gcgcggccgt agccaggcgg atggacgcat ggtattgctc gtggcacgca 6480
gccgaaccgg cacgcgcgcc ggttttaccg tgggaaaaaa ggtaggcaac agcgtggtac 6540
gcagccgtgt gaaacgactg atgcgggaaa gttatcgttt gatggccgga caggtgcagc 6600
cgcgtacgct cctggtattc attgcacggc cgagcatggc cggcagctcg tttgcccagg 6660
tgcaggccag catgaaaaaa ttgatggtgc gcagcggatg cctgcgggag caaacggatg 6720
aaaacgcttc tgattgcgat gattcgcggc tataaaaagt atatctcccc tgcattgccc 6780
aacgcttgcc ggttctcgcc gacgtgttcg gaatatgcca tgcaggcgct ggaagaatgg 6840
ggagttatcc gggggttggg cctggccata tggcgcatat tgcgttgtaa ccctttgtgc 6900
aagggcggct 6910

//

```
ID   XXX; XXX; linear; XXX; XXX; XXX;  XXX.
XX
AC   PRJEB67889
XX
DE   Uncultured bacterium DNA fragment from in-reactor enriched cow rumen, clone
ferm-44C18, contig1
XX
FH   Key             Location/Qualifiers
FH
FT   source          1..17351
FT                   /organism="uncultured bacterium"
FT                   /environmental_sample
FT                   /mol_type="genomic DNA"
FT                   /isolation_source="Cow rumen"
FT                   /clone="ferm-44c18"
FT                   /note="AZCL-Barley beta glucan, AZO-CM cellulose, AZCL
Tamarind xyloglucan, AZCL-HE cellulose active clone"
FT   CDS             18..665
FT                   /db_xref="SEED:fig|6666666.127261.peg.1"
FT                   /translation="MIDEKNSLSPDLYDEMCLYIDEGSLQNYSFIIADPIKGKANESLI
FT                   KTNKINFFKLFAKVGLRYPGKYLESFVGLTAGYYTPITSPYFITGTTKLYTVPVEGWYP
FT                   SPAVDNKLPFGSAFFDYMYGERDGRLRLPILGWLWRSAPYFWSFAFAFLYLIYKKKYNN
FT                   LIVLLFPFIYMLTCLLGPTAFLRYMFINIATMPIALYLVIYRDTTSAKEVLG"
FT                   /product="hypothetical protein"
FT   CDS             complement(637..1821)
FT                   /db_xref="SEED:fig|6666666.127261.peg.2"
FT                   /db_xref="GO:0008784"
FT                   /translation="MGLFIMESTLKRTWSEIDLDALVHNYKTLRARIGDKVKFLGVVKA
FT                   DAYGHGSVMVAQTLEKEGADYLAVSSIDEAMELRVNDIKMPILILGHTPKEQVDRLIEY
FT                   NITQAVTCKAKAVEYSEEATRLNKTLKIHIKVDTGMSRLGYLCDGDFFDTGVQGIVDAV
FT                   TLPGLDAEGIFTHFAVADEEGPECLEYTRHQFKLFTDVCSKVEEKIGKPFLIKHCANTG
FT                   AVARFPETYLDMVRPGLLLYGYGEYAKELGLKPVMTLKTVVSTVKVYPAGTAISYGGIY
FT                   KTDKKERIGVLPYGYADGFFRSLSNKCSLMTDEGPAPQRGKICMDMCMIDITDKYGVDV
FT                   GSEIEIFGKRNPIETLSDIAGTIPYELTCAVSKRVPRLYINDGEVINRELLLRM"
FT                   /product="Alanine racemase (EC 5.1.1.1)"
FT                   /EC_number="5.1.1.1"
FT   CDS             complement(1839..2462)
FT                   /db_xref="SEED:fig|6666666.127261.peg.3"
FT                   /translation="MNDNRVIIEAIGYVGTALVLISFLMTSVYKLRIINTAGSLISVIY
FT                   GVIMQVYPTVVLNGALALINIFFLWRHLSYKNQKIYTAGKTYAEDPIVKLFLDKYREDI
FT                   LKFFPSFGVAHLEPNIAYLIMCEDTMAGVMLGNLKENGELDIYLDFVTGTYRDFSVGKY
FT                   LYRELTRDGVKKCRFMTDIPKSYVYLRKVGYKRVGDVMELTLNE"
FT                   /product="hypothetical protein"
FT   CDS             complement(2472..3836)
FT                   /db_xref="SEED:fig|6666666.127261.peg.4"
FT                   /db_xref="GO:0031176"
FT                   /translation="MTTIKKRIIATMLIAALALTGCESTKASVSSADTVSTETVSASST
FT                   EEVKEEVKDETQETVSASASTEEKKEEKKDENVRPDNLINVLVSKHKAPESEALSFTAD
FT                   MGAGINLGNTFDASDAKNIFDDDLKFETYWQKDVTTQDMIKAIKNEGFKTIRIPVAWHN
FT                   HVDDDFNINEKWMNRVYEVVNWALEEDMYVILNIHHDNLKKYMYPSYECRENSLKYVTK
FT                   IWAQVAEKFGDCDEHLVFESLNEPRLAGTDIEWNLPDLNAKEAKETLDCINQYNQAIVD
FT                   TIRQTPGAYNKTRYITVPGYAGSPDYILCDQFVVPTDTYGEKENRILLTVHAYRPYLFA
FT                   LADPGKGDTKTFDTEGAVSELNALFMNLFVKYVKNGTGVIIDEFGARDRGGNTEERAKF
FT                   TAYFVAYARSYRMPVCWWDNNNFSGSGEVFGIFKRSACKFEYQDIVSQLVYYGNR"
```

```
FT                   /product="GH5_4"
FT                   /EC_number="3.2.1.8"
FT   CDS             complement(3890..4762)
FT                   /db_xref="SEED:fig|6666666.127261.peg.5"
FT                   /translation="MQGKIIKGIAGFYYVNIDGKGTFECKAKGSFRNEGTKPLVGDNVE
FT                   IDIIDEEKSLGNVTRVYKSKSRMIRPACANVDQALIVFAIKSPDPNFNLLDRFLIYMES
FT                   ESVPCIIVFNKDDLASPEDIETIREAYVGAGYPIFFVSARENAGVEALRDLLKGKTTVM
FT                   AGPSGVGKSSLINLLCNCDIMETGDISVKTQRGKHTTRHAELLPLGEGTYVMDTPGFTS
FT                   LDIFGADRQNLKDYYNEFSSVEPCRFADCVHVNEPDCAVKAAVEAGKISKLRYDNYLEL
FT                   FSQIGTKTR"
FT                   /product="Ribosome small subunit-stimulated GTPase EngC"
FT   CDS             complement(4774..6855)
FT                   /db_xref="SEED:fig|6666666.127261.peg.6"
FT                   /translation="MVKIGMMIGERYEVLEKIGTGGMSDVYKAKDHKLNRFVAVKVLKQ
FT                   EFAENANFVSKFKIEAQAAAGLMHPNIVNVYDVGEENGTNYIVMELVEGITLKKYIEKK
FT                   ARLSVKEAVTIAIQIGMGIEAAHNNHIIHRDIKPQNVIISKEGKVKVTDFGIAKAATSN
FT                   TITSNVMGSVHYTSPEQARGGYSDEKSDIYSLGITLFEMLTGRVPFNGETTVAIAIKQI
FT                   QEEMPSPREYVDDIPVSVEQIVLKCTQKSPDRRYQSMGALIEDLKKTFITPDEDFVKIK
FT                   EDDEFGATRPVTAGELEVLKKSTKGTEELTGELPKVNPNASQDDSEYDAKMDKLTTILA
FT                   IFMGAAILVIAIFLISKGIGVLKLGKENTEKVPTTVSMEKPEQIVMPKVVGKSQEDAQK
FT                   ILEKNNLKYFVKYTESDSVKEGYVISASVQDGLKVDAGSEIVLNVSSGSTSVIVPDVVG
FT                   KALEAANKSIMDAGLKVEIKKDYSDTVPEGNIISQFPEGGEKTPEESTVLIIVSQGVAE
FT                   NLTKVPNVLGKDNDDAIIILTEAGLTVNPVAPVYDNIFPEGQICYQSLEADKEVPKGTA
FT                   IDLHASMGPEPPDTVMYRFNGSINAPSTAEDPEFEMGTKVEITVSDANGKVLQSTWTTS
FT                   FPVSVNISNIAGVNSGTITLSYEVKIPAEKDVSTGEEITPAKTEKKVIKRDINFEVDE"
FT                   /product="Serine/threonine protein kinase PrkC, regulator
FT                   of stationary phase"
FT   CDS             complement(6883..7611)
FT                   /db_xref="SEED:fig|6666666.127261.peg.7"
FT                   /translation="MLKTFSLSDVGKKRTMNQDYIFTSEEPVGNLPNLFALADGMGGYN
FT                   GGEYASLRTVETVTSFIDKSKSTTPRKLFTEAIKEANRIIRDEASQNPEYDKMGTTIVL
FT                   ASVVGDCLQVANVGDSRLYVIGDDIKQVTVDHSYVEEMIKMGSLDRESARTHPQKNIIT
FT                   RAVGADDDIDTDFFTVRLHKGDMVLMCSDGLTNMLEDSEILMILKKERDIASKAEALVA
FT                   AANNSGGKDNISVILIEPFA"
FT                   /product="Protein serine/threonine phosphatase PrpC,
FT                   regulation of stationary phase"
FT   CDS             complement(7657..8694)
FT                   /db_xref="SEED:fig|6666666.127261.peg.8"
FT                   /translation="MDKIDIKSLDFEELENSIKEMGLPKFRASQIYDWLHKKLVTSFGE
FT                   MTNLSADLRNKLSETYYITACEVARMQESREDGTRKYLFKLRDGNYVESVFMRYHHGNS
FT                   VCISSQVGCRMGCKFCASTLDGLTRCLEPSEMLEQIYRIQALTGERVSNVVVMGTGEPM
FT                   DNYDNLLKFIKLLTDGHGLNISQRNLTVSTCGIVENMKRLADEKLSITLALSLHSVTDE
FT                   KRRELMPIANKYSINELMDACEYYFDKTGRRITFEYALVRDVNDRDEDAKGLIKLAGRV
FT                   KAHVNLIPVNPIKERDYRETARDDVLAFKNKLEKSRINVTIRREMGRDIDGACGQLRKK
FT                   TMDSE"
FT                   /product="Ribosomal RNA large subunit methyltransferase N
FT                   (EC 2.1.1.-)"
FT   CDS             complement(8694..9026)
FT                   /db_xref="SEED:fig|6666666.127261.peg.9"
FT                   /translation="MSRKSDIKYNITNEAMKDICDLQKAIVTNVSRYVKPGGVLLYSTC
FT                   TIHKAENEKMVKFITENLPFKEDSLKPFVPFLFDKPRKSECHIQLRPDTDGTDGFFVAR
FT                   FVKSED"
FT                   /product="16S rRNA (cytosine(967)-C(5))-methyltransferase
FT                   (EC 2.1.1.176)"
```

```
FT                   /EC_number="2.1.1.176"
FT   CDS             complement(9026..10042)
FT                   /db_xref="SEED:fig|6666666.127261.peg.10"
FT                   /translation="MSVNTREIITDILLELDKEGSKSHLLIREVLEKYDYLDSRDKGLI
FT                   KRISEGTVFHRITLDFVLDKYSQKPMNKCKPVVAAILRMSLYQILFMDKIPDNAICDEA
FT                   VKLCRKKSFEAFCPFVNGVLRNILRDKENALNFDGIDDKVQRISVMYSCPEWIVKMLLK
FT                   EQEDGESLVKALSEIRPAFVKVVNEADTEALTKAWGDAGISFEPARYIKNAYKIEDFDG
FT                   MESVPGFSDGKLIVQDESSMICAECAGVQKGSNLRILDVCAAPGGKSSYVASRMYPEGE
FT                   VVSCDVSDTKVSMIADNAERLHLSNLKPTLQDATEFNEEYEASFDIVIADVPCSGWA"
FT                   /product="Ribosomal RNA small subunit methyltransferase B
FT                   (EC 2.1.1.-)"
FT   CDS             complement(10039..10983)
FT                   /db_xref="SEED:fig|6666666.127261.peg.11"
FT                   /db_xref="GO:0004479"
FT                   /translation="MKTVFMGTPEFAASILKALKEAGHEVTAVYTQPDKPQGRSGALIA
FT                   PPVKVYAEENGIPVYQPAKIKAPESVEVLKTIPADVYVVAAYGQILSQEILDIPKFGCI
FT                   NVHGSLLPKLRGAAPIQRAIANGDRETGVSIMKMDKGMDTGAVYSSVSFPIEPEDNEDS
FT                   VYLKMADYGAKELLKVLKDVEAGTAVATPQDPDLATYAPPLKKEEGLIDFGRSAESIDC
FT                   LVRGFLKWPTAYTTVNGKLLKIFGTKVIDSIPGVDTADYEPGALIVTKKNVYVSTGDGI
FT                   LELLEVQLEGKKRMSACDFVRGMHLGGGDFLGQ"
FT                   /product="Methionyl-tRNA formyltransferase (EC 2.1.2.9)"
FT                   /EC_number="2.1.2.9"
FT   CDS             complement(10983..11468)
FT                   /db_xref="SEED:fig|6666666.127261.peg.12"
FT                   /db_xref="GO:0042586"
FT                   /translation="MALRKIREWGDEVLTKKCKEVTEITPRTKLLIDDMIDTMRAADGV
FT                   GLAACQVGVLKRIVVIDTTGEDLHVLINPVILETSGEQEGYEGCLSVPGKSGIVKRPYY
FT                   VKVKAFDEHLEPFELEGEELLARAICHELEHLDGHLYTERVIGELVDNSMLHEEEAE"
FT                   /product="Peptide deformylase (EC 3.5.1.88)"
FT                   /EC_number="3.5.1.88"
FT   CDS             complement(11487..13709)
FT                   /db_xref="SEED:fig|6666666.127261.peg.13"
FT                   /translation="MENFSYAKVIINISHESVDRPFTYIIPEDLKSRVDIGTAVTVPFG
FT                   AGNTLKKGYVVELTDTTDVAADKLKSIAEVSDRDISLSDKRIALAAWMKRNYGSTMIAA
FT                   LKTVIPVKEKVKESVKKTIVRLKSPEEIREYIALNKRNASQIRLLTALLDEERIPKSIL
FT                   DDKLHIAPGAYKKLSEAGFIDICENRVLRGTVNLDVAHDFHLDLSPEQQLVVDSVRKDF
FT                   DAEVNNTYLIHGITGSGKTEVYMQIIDDVIRRGRQAIVLIPEISLTYQTLMRFYKRFGA
FT                   RVGVMNSTLSKGEKYDLFEQAVNGDIDIVIGPRSALFTPFNNLGIIIIDEEHENSYKSE
FT                   TMPKYHARETAIELARMSKASVILGSATPSIDSYYRAKTGEYKLFTLTRRLTGHQLPNV
FT                   TVADLRQELREGNRSILSRDLQALIEDRLNKNEQIMLFLNRRGVAGFVSCRACGFVVKC
FT                   PHCDVSLSLHRNGKMVCHYCGYTTDSYKTCPQCGSKYILGFKAGTEQVEEIVNKMYPKA
FT                   RTLRMDADTTRKKESYEEILSRFLDHEADILIGTQMIVKGHDFKNVTLMGVLAADMSLF
FT                   AADYRAPEKTFDLLTQAAGRAGRGDKPGDVVIQTYQPDHYAISFSVTQDYEGFYEEEIG
FT                   YRDLMDYPPAAHMLSVLVTGADEKACEALATNLVQGIREHFKILVIGPAPATVSKINDI
FT                   YRQVFYLKHKSYDALVQIKDFLEEALSSRKQFKENVFFDFDPQNGV"
FT                   /product="Helicase PriA essential for oriC/DnaA-
FT                   independent DNA replication"
FT   CDS             complement(13755..15530)
FT                   /db_xref="SEED:fig|6666666.127261.peg.14"
FT                   /translation="MENWVEIRKSADFMKMASDYAISPITARILRNRDLTDPSEIRNFL
FT                   SPGPEAFHDPITLTGMDRVILVLKEKISAGLKIRIIGDYDVDGITSTYIFYKGLKYFGA
FT                   DVDYVIPHRIEDGYGINIRLIDDAYKAGVDTILTCDNGIAAYDQTEHATSLGMTMVITD
FT                   HHEVPFTEDEKGEKIYKIPRAAAVTDPKLPGSEYAFEGICGAFVAFKTIMAYAREMGED
FT                   STDSFAGLRDELTEFAALATVCDVMELKDENRSLVCLGLKLMQHSRNLGMRALLRVTGM
```

```
FT                   EGKVLSPYHMGFVIGPCLNASGRLDTSLKAVDLFLETDEARATDKATELKMLNEERKQM
FT                   TLDETAKAFEIVDGRPSLDNVLVVYLPDCHESLAGIIAGKVRERYVRPAFVITKTDKGL
FT                   KGSGRSIDAYNMYEELTKVKDCLSKFGGHKLAAGISLADDTPDAFREALNANCTLTDKD
FT                   FTETVRIDMELPLEYADLNLARDLMKLEPYGVGNEKPLFAVRHVTIISGRLMGKNGNVA
FT                   KYKVRNDAGRVYEMIYFGDVSAFEQFVADIYDEETARRLHSSVAVNVVLDICYRVEINT
FT                   YMGRDSVSLQMVHYR"
FT                   /product="Single-stranded-DNA-specific exonuclease RecJ
FT                   (EC 3.1.-.-)"
FT   CDS             complement(15534..17303)
FT                   /db_xref="SEED:fig|6666666.127261.peg.15"
FT                   /translation="MADAQAGTTRNDMGNTQIMVNLTLTKEGTDSFAKATEAAFNRGKD
FT                   VIGIYYDGQFISVPAVQGVISGGTAQITGQKDAEEAQKLASTIRIGGLSLELEELRSNV
FT                   VGASLGQDAIQSSLKAGLVGFVIVVIFMLVVYYFLGLTSAIALTLYAALEIMLINGFAI
FT                   TLTLPGIAGIILSIGMAVDANVIIFARLREELATGKTMDSAIKVAYSKAMSAILDGNIT
FT                   TLIAAAVLWIMGSGSIVGFAQTLTLGICLSMFTAIVITKALLKAFVAAGIKDPKFFGVA
FT                   KQRKTIDFLSKRGLFFGISGLVIAAGLVFMLAVNGGNANRGHSLNYSLEFLGGTQTVVP
FT                   FEQNYELEEINDLMVADIEKITGSANVVTTKVQGSNEVIFKTVTLNADQRAELNNYFNE
FT                   KFGIDKETIKTESISSTISKEAQRETIIAVVVACICMLIYIWFRFKDIRFGASSVLALT
FT                   HDVLVTLAFYAIAYVSVGSSFVACMLTIVGYSINATIVIFDRIRENLKEAAPRTALKDI
FT                   VNDSITTTLSRSIFTSLTTFIMVVALYVFGVSSIKEFALPLMVGIVCGTYSSICLAGSM
FT                   WFVFKNKFSKKEK"
FT                   /product="Protein-export membrane protein SecD (TC
FT                   3.A.5.1.1) / Protein-export membrane protein SecF (TC
FT                   3.A.5.1.1)"
XX
SQ   Sequence 17351 BP; 4301 A; 4173 C; 3874 G; 5003 T; 0 other;
     tgtgctttgc tcatgtaatg atcgatgaaa aaaatagcct gagtcctgac ctttacgacg        60
     aaatgtgttt gtatatcgac gaagggtctc ttcaaaacta ttcattcata atagcggatc       120
     cgattaaagg caaagccaat gaaagtctta ttaaaaccaa caaaattaac tttttaagc        180
     tttttgcaaa agtcggactg agatatccgg gtaaatattt ggaatcattt gtaggtctta       240
     cagcaggtta ctatacaccc ataacttctc cgtattttat taccggtacc accaagcttt       300
     acaccgtgcc cgtagaaggt tggtatcctt cgcctgccgt tgataataaa cttcctttcg       360
     gaagcgcttt ttttgattac atgtatggtg aacgtgatgg gagactccgc cttccgatac       420
     tgggctggct gtggcgctcg gctccctact tttggagctt tgctttcgca tttttatatt       480
     taatatataa aaagaagtac aacaatctga ttgtacttct ctttccattt atttatatgc       540
     taacctgtct tctcggaccc acggcatttc tccgctacat gtttatcaat atagcgacca       600
     tgccaattgc tctatatctt gtcatatatc gagatactac atccgcaaaa gaagttctcg       660
     gttaataacc tcaccatcat taatgtataa gcggggaact ctcttgctga ccgcacatgt       720
     gagctcgtaa ggaatggtgc ccgctatatc ggaaagtgtt tctatgggat ttcttttgcc       780
     gaaaatttct atttcggaac ctacgtcaac accgtatttg tcggtgatgt ctatcatgca       840
     catgtccatg catatcttgc cgcgctgagg tgcgggacct tcgtcggtca tgagactgca       900
     tttattggag aggcttctga agaagccgtc ggcatatccg taagggagta cgcctatgcg       960
     ctcttttttg tctgttttgt agataccgcc gtagcttatg gctgttccgg caggatatac      1020
     cttaacggtg cttaccacgg tctttaaggt cataacgggc ttaaggccca gttctttggc      1080
     atactcgccg tatccgtaaa gaaggagtcc cggacgaacc atatccagat aagtctcagg      1140
     gaatcttgcc acggcaccgg tattggcgca atgcttgatt aaaaaaggct taccaatctt      1200
     ctcttcaacc ttagaacata catcggtaaa cagtttaaac tgatgtctcg tatattcaag      1260
     gcactcggga ccttcttcat ccgcaacagc gaaatgagta aagatacctt ccgcatcaag      1320
     ccccggcaat gtaaccgcat ccactattcc ctgaacgcct gtgtcaaaga aatctccgtc      1380
     gcacagataa ccgaggcggg acatgcccgt atccaccttt atatgaatct tcagggtctt      1440
     attaagtctt gtggcttcct cggagtactc aaccgcttta gccttgcagg ttactgcctg      1500
     agtgatattg tattcgataa gtctgtccac ctgctccttg ggcgtatgtc ccaggattaa      1560
     aatgggcatc ttaatatcgt taaccctaag ctccatggct tcgtcaatac tgctgacagc      1620
     caggtaatca gcgccttctt tctcaagggt ctgtgccacc ataacggaac cgtgtccgta      1680
```

ggcatcagcc ttaaccacac ctaagaattt aaccttatca ccgattcttg ctcttaaagt 1740
cttgtaattg tggaccaaag catcaaggtc gatttccgac catgttcttt ttaaagtgga 1800
ttccatgata aataaaccca tcctcccaaa atcatctctt attcgttaag cgtaagctcc 1860
ataacgtcgc ccacacgctt gtaaccgact tttcttaagt aaacgtaact cttgggaatg 1920
tctgtcataa accggcattt cttgacaccg tctcttgtaa gctccctgta gaggtatttg 1980
cccacggaaa agtccctgta ggttcccgtt acaaaatcaa ggtagatgtc gagttcaccg 2040
ttttctttta aattacctaa cattacaccc gccatggtgt cttcgcacat aataagatat 2100
gcgatgttgg gttcaaggtg cgccacaccg aagcttggga agaatttaag tatatcctcc 2160
ctgtatttgt caagaaacag tttgactatg ggatcctcgg cgtaggtctt gcctgccgta 2220
taaatctttt gattcttgta agataagtgg cgccataaaa agaaaatatt tataagcgcc 2280
agagctccgt ttaaaacaac cgtagggtac acctgcataa taacgccgta aataaccgaa 2340
ataagggaac ctgcggtatt tatgatgcga agcttgtata ccgaagtcat aaggaaggat 2400
ataagcacca aagcggtgcc aacataacct atagcttcaa taatcactct gttatcgttc 2460
ataagagaat cctatctgtt tccgtaataa accaactgac ttacgatgtc ctgatactcg 2520
aatttgcatg cggaacgctt aaatataccg aatacttcgc cgcttcccga gaagttgttg 2580
ttgtcccacc agcatacggg catacggtag cttctggcat atgcaacgaa ataagccgta 2640
aatttggcac gttcctcggt attgccgcct ctgtctcttg caccgaattc gtcgatgata 2700
acgccggtac cgttcttaac gtacttaacg aacaggttca tgaataaagc gttaagttcg 2760
gatacagcgc cctccgtatc aaaggtcttg gtatctccct tgccgggatc ggcaagtgcg 2820
aaaagataag gtctgtatgc atgaaccgtc aaaagaatgc ggttttcttt ttcaccgtat 2880
gtatctgtgg gaaccacaaa ctggtcacag aggatataat cgggtgaacc cgcataaccg 2940
ggaacggtga tatatctggt cttattgtaa gcaccgggag tttgtctgat ggtatctaca 3000
attgcctgat tgtactgatt gatgcagtca agggtttcct tggcttcttt ggcatttaag 3060
tcgggaaggt tccactcaat gtcggtaccg gcgagtctgg gctcattcag tgattcaaat 3120
accagatgct cgtcgcagtc gccgaatttc tccgcaacct gcgcccagat cttggtaacg 3180
tacttaaggg agttctcacg gcactcatag gaaggataca tatacttctt taagttgtcg 3240
tgatgaatgt taaggataac gtacatatcc tcttcaaggg cccagttaac aacttcgtag 3300
actctgttca tccacttttc gttgatgtta aagtcatcgt caacatgatt gtgccatgct 3360
acgggaattc tgatggtctt aaagccttcg ttcttaatgg ccttaatcat gtcctgagtg 3420
gtaacgtcct tctgccagta tgtctcaaat tttaagtcat cgtcaaaaat gttcttggca 3480
tcgcttgcgt cgaaggtgtt gccaaggtta atgcccgcgc ccatgtcagc ggtaaaagaa 3540
agtgcctcgc tctcgggtgc tttgtgtttg gataccaata cattgataag attgtcgggt 3600
ctcacgtttt cgtccttttt ctcttctttt ttctcttctg tggatgcgga agcggaaacg 3660
gtttcttgtg tctcgtcctt tacttcctct ttaacttctt ctgttgaaga agctgatacg 3720
gtttcggtgc ttactgtgtc tgcggatgat acgcttgctt tggtggattc gcagccggtc 3780
agagcaaggg ccgcaataag cattgttgca ataattctct ttttgatggt agtcataata 3840
atcctctaca ttataaagtt tgtgtccggc gtggccggta tatgaaccgt tacctggtct 3900
tcgtccctat ctgggaaaaa agttcaaggt aattgtcata acgaagttta gaaatcttac 3960
ccgcttccac tgccgcctta acagcacagt cgggttcgtt gacatgtacg cagtctgcga 4020
atctgcaggg ctctacggaa gaaaattcat tgtagtaatc cttaaggttc tgtctgtctg 4080
cgccaaaaat gtcaagggac gtaaaaccgg gggtgtccat aacataagtg ccctcaccga 4140
ggggtaacaa ttccgcatgc ctggtggtat gtttgcccct ttgggtcttc acacttatat 4200
caccggtttc cattatatcg caattgcata acagatttat tagagatgat ttgcccacac 4260
cgctggggcc ggccataacc gttgttttgc cttttaacag gtctcttaag gcctcaactc 4320
ccgcattctc tctcgcactg acaaagaaaa tggggtagcc cgctcccaca tatgcctctc 4380
ttatggtctc aatgtcctcc ggggatgcca gatcgtcctt gttaaagact atgatgcagg 4440
gaaccgattc gctctccatg tagattaaaa acctgtcgag aaggttgaaa ttggggtcgg 4500
ggcttttaat tgcaaaaaca attaaagcct ggtcaacgtt ggcacaggca ggccgtatca 4560
tacggctttt gctcttgtaa acccgtgtca cgtttcctaa gcttttctct tcgtcgatta 4620
tgtctatttc aacattgtcg cctaccaatg gcttggtgcc ttcattacga aaagaacctt 4680
tggccttaca ctcaaaggtt cctttaccgt cgatattaac gtaataaaat ccggcaatgc 4740
ctttaataat ctttccttgc attaatctcc cgattattcg tcaacttcaa aatttatgtc 4800
tcttttaatg accttttttct cggtcttggc aggggtaatc tcctcgcccg tggatacgtc 4860
cttctccgca ggaatcttaa cttcatacga aagggtaatg gtaccgctgt taactcccgc 4920

aatgttggaa atgttaacgg aaacggggaa agaagtggtc caggtagact gaagcacctt 4980
gccgtttgcg tcagataccg ttatctcaac cttggtaccc atttcaaatt cgggatcttc 5040
tgccgtggag ggtgcattta tgcttccgtt aaatctgtac ataacggtat ccggaggttc 5100
gggacccatg ctggcatgaa ggtcaatagc ggttcccttg ggaacttctt tgtccgcttc 5160
aaggctctgg tagcaaatct gtccttcggg gaagatattg tcgtatacgg gagctacggg 5220
gttaaccgta agacccgctt cggtcagaat aataatagca tcgtcattgt ccttaccaag 5280
aacattggga accttggtga ggttctcggc aacgccctgg cttacgataa tgagcacggt 5340
agattcttcg ggagtcttct ctccgccttc gggaaactgt gaaatgatgt tgccttcggg 5400
aacggtatcg gaatagtcct ttttaatctc tacctttaag ccggcatcca taatcgattt 5460
attggctgcc tccaaagctt tgcccactac atcgggcaca ataacactgg tggaaccgga 5520
gcttacgtta agcacgattt ccgagcccgc atctaccttt aagccgtcct gaacgcttgc 5580
tgaaataaca tagccttctt tgacggaatc ggattccgta tattttacaa agtatttaag 5640
attgttcttc tcaagtattt tctgagcgtc ttcctggctc ttgcccacaa ccttgggcat 5700
tactatctgc tcaggcttct ccatggaaac tgtggtggga accttctccg tgttttcttt 5760
gccgagcttc agtactccga tgcccttgct gataaggaaa atggctataa caagaatggc 5820
cgcacccata aagatggcaa gaatggtggt gagtttgtcc atcttggcat cgtattcgct 5880
gtcatcctga gaggcgttcg gattaacctt gggaagttcg cccgtaagct cttccgtgcc 5940
ttttgtactc ttcttaagga cctcaagttc acctgcggtg acgggtcttg tggcgccgaa 6000
ttcgtcgtct tctttaatct ttacaaaatc ttcgtcggga gtgataaagg tcttcttaag 6060
gtcctcgata agggctccca tgctctgata gcgtctgtcg ggctcttcct gggtacactt 6120
aagcacaatc tgctcaacac ttacgggaat atcgtcaacg tattccctgg gggagggcat 6180
ttcttcctga atctgtttga tagctatagc tacggtggtc tctccgttaa aaggaactct 6240
tcctgtaagc atttcaaaga gagtgatacc aagtgaatag atatcgctct tctcatcgga 6300
gtatccgccg cgagcctgct cgggagatgt atagtgaaca gaccccataa cgttactggt 6360
aatggtgttg ctggtagcgg ccttggcgat gccaaagtct gttaccttaa ccttgccttc 6420
tttggaaata ataacgttct gaggtttaat gtctctgtga ataatatgat tgttgtgggc 6480
ggcttcgata cccatgccta tctggatggc tatggtaacg gcttctttga ccgaaaggcg 6540
ggccttcttc tcaatatatt tcttgagagt gataccctca acaagttcca ttacaatgta 6600
attggtgccg ttttcttcgc ctacgtcata gacgttgacg atattggggt gcataagacc 6660
tgcggcggcc tgagcctcga ttttaaactt cgagacgaag ttggcattct ccgcaaactc 6720
ctgcttaagc accttaacgg caacaaaacg attcaattta tgatctttag ctttatagac 6780
gtcggacata ccgccggttc cgattttttc aagaacctcg tatctctcgc ctatcatcat 6840
accgatttta accatgtgta tcctctgtca gacgattctt tatcaagcaa agggctctat 6900
taagataacc gaaatgttat ccttgccgcc ggagttattg gcagctgcca caagtgcttc 6960
tgctttgctt gctatatccc gctctttctt gagtatcata aggatttcac tgtcctcaag 7020
catgttggtg aggccgtccg agcacataag caccatatcg cccttatgaa gccttacggt 7080
aaagaaatcc gtgtcgatgt catcgtccgc accgacagct cttgttatga tattcttctg 7140
cgggtgggtc ctggcacttt ctctgtcaag acttcccatt ttgatcatct cctcaacata 7200
ggagtgatcg acagtaacct gtttgatatc gtccccgatt acatataacc ttgagtcgcc 7260
aacattggca acctgaaggc aatcccccac aacagatgca agcactatgg tagtacccat 7320
cttgtcatat tcgggattct gagaggcttc gtctctgatg atacggttgg cctctttaat 7380
agcttccgtg aaaagtttcc tcggcgtggt gctcttggac ttgtcaataa aagaagtgac 7440
agtctccacg gtacgaagag atgcgtactc tccgccgtta taacctccca tgccgtcggc 7500
aagggcaaaa aggttgggca gatttcccac aggctcctcg cttgtgaaaa tgtagtcctg 7560
gttcattgtt ctcttttttc ccacatccga taaggaaaaa gttttaagca cccctatgcc 7620
tcctgattaa taccatcgct tcaaaagaaa acaagtctat tcactgtcca ttgtcttctt 7680
gcgaagctgt ccgcaggccc cgtctatatc gcgacccatt tcccttctaa tagtaacatt 7740
tattcggctt ttttcaagtt tatttttgaa tgccagcacg tcgtctctgg ccgtttccct 7800
gtaatcacgc tcttttatgg gattgaccgg aataaggttt acatgagcct ttacccttcc 7860
cgcaagctta ataaggcctt tggcatcctc gtccctgtca ttgacatctc gcaccagagc 7920
atactcgaag gtgatacgtc tgccggtctt gtcaaaataa tattcacagg catccataag 7980
ctcgtttata gagtatttat tggctattgg catgagctcg cgacgttttt cgtccgtaac 8040
cgaatgaagg gacaaagcaa gtgttatgga gagtttctca tcagccagac gcttcatgtt 8100
ctcaacgatt ccgcaggttg atacggtaag gtttctttgg cttatattga gtccgtgccc 8160

gtcggttaaa agctttatga atttaagcag attgtcgtaa ttgtccatgg gctcgccggt 8220
acccattacc actacgttgg atactctctc gccggtgagg gcctgaatac ggtaaatctg 8280
ctcaagcatc tcagagggct caaggcatct ggtaagtccg tccaaagtgg aggcgcagaa 8340
cttacagccc atgcggcagc ctacctggga agaaatgcac acggaattgc cgtgatgata 8400
gcgcatgaaa acgctctcca cgtaattgcc gtcgcgaagc ttaaagaggt actttctggt 8460
accgtcttcc cttgactcct gcattctcgc cacttcgcag gcggttatgt agtacgtttc 8520
cgaaagttta ttacgaaggt ctgccgagag attggtcatt tcgccaaaag aagtaaccag 8580
tttcttgtgg agccagtcat aaatctgtga ggctctgaat ttgggaagac ccatttcttt 8640
tatggaattt tcaagttctt caaagtccaa tgacttaatg tcgattttat ccattaatct 8700
tcactcttaa caaatcttgc cacaaagaag ccgtcggtgc cgtccgtatc gggacgaagc 8760
tgaatgtggc attcgctctt acgaggcttg tcaaacagga acggaacaaa aggtttaagg 8820
gaatcttctt tgaagggaag gttctccgtt ataaacttaa ccattttttc attttcagct 8880
ttatgtatgg tgcaggtgga ataaagaagc actccgcccg gcttaacgta gcgtgaaaca 8940
ttggtaacga tggccttttg aaggtcacat atgtccttca tggcttcgtt ggtaatattg 9000
tatttaatat cgctcttacg tgacattacg cccatcccga acatggaacg tcggctatta 9060
ctatgtcaaa agaagcctca tattcttcat tgaattccgt ggcatcctgc agtgtaggct 9120
taagattgct taagtgaagc ctctctgcat tgtcggctat catggatact ttggtatccg 9180
aaacatcgca gcttacgacc tcgccctcag gatacatgcg cgaagccacg tacgaggact 9240
tgcctccggg ggccgcacac acatcaagga tgcgtaaatt gctgcctttt tgaactccag 9300
cgcactcggc acatatcatg gagctttcgt cctgaacgat aagtttgcca tctgagaatc 9360
ccggcacgga ttccatgccg tcaaaatcct cgattttgta ggcattctta atatatcgtg 9420
cgggttcaaa agaaatccct gcatcacccc aggccttagt aagtgcctct gtgtctgctt 9480
cattgacaac cttcacgaaa gcaggtctta tctcgctcaa tgcctttaca agactctctc 9540
cgtcctcctg ctcttttaaa agcatcttca ctatccattc ggggcaggaa tacatgacac 9600
tgattctttg gactttgtcg tctatgccgt caaagttaag ggcgttttct ttgtccctta 9660
atatgttgcg aaggacaccg tttacaaagg ggcagaaagc ctcaaagctc ttcttgcggc 9720
agagcttgac cgcttcgtca catatggcgt tgtcggggat tttgtccata aaaagaatct 9780
ggtaaagact catgcgaaga atggcagcca caacaggctt acacttgttc atgggctttt 9840
gagagtattt gtccagcaca aaatcgaggg taatcctgtg aaagacggta ccctcgctga 9900
tacgctttat aagtcctttg tcccttgaat ccaggtaatc gtatttttca agcacttctc 9960
ttatgaggag atggctcttc gatccttctt tgtcaagttc caaaagaata tccgtaatta 10020
tttctcttgt attaacactc attgacccaa aaagtctccg ccgcccaagt gcatgcctct 10080
tacaaagtca catgcgctca tgcgtttttt gccttcaagc tgaacttcaa gcagttccaa 10140
aattccgtct ccggtggaaa cgtaaacatt tttcttggta acgataaggg caccgggttc 10200
ataatccgca gtatccacgc ccggaatgct gtcaatcact ttggtgccga atatctttaa 10260
gagtttgccg ttaacggtgg tataggctgt gggccacttc aaaaagcctc ttacaagaca 10320
gtctatgctc tctgcggatc ggccgaaatc tatgagacct tcctccttct taagaggagg 10380
cgcataagtg gcaaggtcag ggtcctgagg tgtggcaaca gctgtgccgg cctcaacgtc 10440
ctttaaaacc tttaagagtt cttttgcacc gtaatccgcc atttttaagt aaaccgagtc 10500
ttcattgtct tcaggctcaa tgggaaaaga aaccgagctg taaactgcac cggtatccat 10560
tcctttgtcc attttcatga tggaaacgcc ggtttcacga tcaccgttgg cgatggcgcg 10620
ctgaatgggt gcggcaccgc gaagcttggg aagaagtgaa ccgtgaacat taatgcagcc 10680
aaacttggga atatccagta tctcctggga aagaatctgt ccgtaagccg ccacaacata 10740
aacgtctgcg ggaatggtct ttaaaacctc cacagattcg ggagccttaa tcttagcggg 10800
ctgatatacg gggataccgt tctcctcggc atagacttta accgggggag caataagcgc 10860
tccgcttctg ccctgaggct tgtcgggctg tgtgtaaacc gccgttactt cgtgacctgc 10920
ttcttttaac gctttaagta tagacgccgc gaattcgggc gtacccataa atactgtctt 10980
cattattcgg cttcctcttc atgcagcatt gaattgtcta ccagttcgcc gataaccctt 11040
tcggtgtaga ggtgaccgtc gaggtgctca agttcgtggc agatggcacg ggccaaaagc 11100
tcctcaccct caagttcgaa tggctcaaga tgctcgtcga aagccttaac tttaacataa 11160
taggggcgct taacgatacc gctcttgccc ggaacgctta agcatccttc atatccctcc 11220
tgttcgccgg aggtctcgag tattacgggg ttaataagaa catgcaggtc ttcacccgta 11280
gtgtcgatta caacgattct tttaagcact cccacctggc atgccgcaag gcctacgccg 11340
tcggcagcgc gcatggtatc aatcatatcg tcgattaaaa gctttgtgcg gggagtaatc 11400

tccgttactt ccttgcattt cttggtaaga acttcgtcgc cccattctct aatttttctg 11460
agtgccatag tatcttcctt tctaaatcag accccgtttt gagggtcaaa gtcaaaaaat 11520
acgttttctt taaactgttt cctcgaagaa agggcttcct cgagaaaatc ctttatttga 11580
accaaagcat cataggattt gtgctttagg taaaatacct gtctgtatat gtcgttaatc 11640
ttggaaacgg tggcgggagc aggtccgatt acaagaatct taaagtgctc tctgatgccc 11700
tgcacgagat tggtcgcaag ggcctcacag gctttctcat ccgcgcccgt aaccagtacc 11760
gacagcatat gtgccgccgg agggtagtcc attaagtctc tgtagcctat ctcctcttca 11820
taaaagcctt cgtagtcctg agttacggaa aaagaaattg catagtgatc cggctggtag 11880
gtctgaataa ccacatcacc gggcttgtcg cctcttccgg ctcttcccgc agcctgggta 11940
agaaggtcga aggttttctc gggcgccctg taatctgccg caaataaaga catatcggcg 12000
gccagaaccc ccattaatgt aacgttctta aagtcatgtc ccttgactat catctgtgtg 12060
ccgataagaa tgtcggcttc atggtcaagg aaacggctga gaatttcttc gtagctttct 12120
tttttcctgg tggtatccgc gtccatgcga agggtacggg ctttgggata cattttgttg 12180
actatctctt ctacctgttc ggtgccggct ttgaagccaa ggatatattt ggacccgcac 12240
tgtgggcagg tcttgtagga atcggtggta tagccgcaat agtggcatac catcttaccg 12300
tttctgtgaa gggacaaaga aacgtcgcag tgcggacatt ttactacaaa gccgcaggct 12360
ctgcaagata cgaagcccgc cacgcctctt ctgttcaaaa agagcattat ctgctcgttc 12420
ttattgagac ggtcctcaat aagtgcctgc aggtctcttg aaagaatgct tctgttgcct 12480
tcccgcaatt cctgcctcaa atcggccaca gtcacattgg gaagctgatg gccggttaaa 12540
cgtctggtaa gagtgaaaag cttatactct cccgtcttcg ccctgtaata ggaatctatt 12600
gaaggggttg cggagccgag tatcacactt gccttactca tgcgcgcaag ttcaatggcg 12660
gtttcacgag cgtgatattt gggcattgtt tcgctcttat aggagttttc gtgctcctcg 12720
tctattatta taatcccaag gttattaaaa ggcgtaaaaa gcgctgacct ggggcctatt 12780
actatatcaa tgtctccatt aaccgcctgc tcgaaaaggt cgtatttctc gccctttgaa 12840
agagttgagt tcatgacgcc cactcttgcg ccgaatcgtt tgtaaaagcg catgagggtc 12900
tggtatgtca aagaaatctc cggtatcaaa acgatggcct gtctgccccg tctgataaca 12960
tcgtcgatta tctgcatgta aacttctgtt ttgccgcttc ccgtgatacc gtgaatcaga 13020
taagtgttat tgacttccgc atcaaaatcc ttgcgaaccg aatccaccac cagctgctgt 13080
tcgggactta agtcaagatg aaagtcatga gccacatcaa gattgactgt gccgcgaaga 13140
acacgattct cgcagatatc tataaatcct gcttccgaaa gctttttgta agcgccgggt 13200
gctatgtgga gcttatcgtc aaggatgctc ttgggaatcc tctcctcgtc gagtaatgcg 13260
gtcagaagac gaatctgtga tgcgtttctt ttgttaagtg cgatatactc acgaatttct 13320
tcgggggatt taaggcgcac tatggtcttt ttgacgcttt ctttgacctt ttctttgaca 13380
ggaatgacgg tttttaaagc ggctatcatg gtggagccgt aattacgctt catccaggca 13440
gccagagcta ttcttttgtc ggacaaagaa atgtccctgt cagagacttc ggctatggat 13500
ttgagtttgt ctgcggctac atcggtggtg tctgtgagtt ccaccacata accctttttg 13560
agggtattcc ccgctccgaa gggtacggtc acggcggtac cgatgtccac ccggctcttt 13620
aagtcctccg gaataatgta ggtaaaagga cggtccacgc tttcatgaga aatatttata 13680
atcacttttg cgtatgaaaa attctccacg tgtcctccct aatccataat ctgattcata 13740
ttattttatc atacttatcg gtaatgcacc atctgaaggg atacgctgtc cctgcccata 13800
taagtattga tttccacacg ataacagata tccagcacaa cgttaaccgc aaccgaagaa 13860
tggagcctac gggcagtttc ttcgtcgtag atatccgcca caaattgttc aaaagcagac 13920
acgtctccga aatatatcat ctcgtatacg cggcctgcat cgtttctgac cttgtatttg 13980
gcaacgtttc cgtttttgcc cattaagcgt cccgaaatta tcgtcacatg tcttacggca 14040
aacagaggct tttcattgcc cacaccgtaa ggctcaagct tcattaaatc cctggcaaga 14100
tttaaatccg catactcaag ggaagctcc atgtcgattc gaaccgtttc cgtaaaatcc 14160
ttgtccgtaa gggtacagtt ggcgttaagc gcctccctga aggcatccgg agtatcatca 14220
gcaagggaaa tgcctgcggc aagcttatgc cctccgaact tggaaagaca gtctttgacc 14280
ttcgtaagct cttcgtacat gttataggca tctatacttc ttccgctgcc ctttaaccct 14340
ttgtcggttt tcgtaatgac aaaagcagga cgcacgtaac gctcacgcac cttgcccgct 14400
ataatgcccg caaggctttc gtgacagtcc ggaagataca ccaccagaac attgtcaaga 14460
gaaggccttc cgtccactat ttcaaaggct ttggcggtct cgtcaagagt catctgcttt 14520
ctttcttcat taagcatttt aagttctgta gccttatctg tagcacgggc ttcatcggtc 14580
tcaagaaata agtcaacagc cttcagcgaa gtgtcaagac gtccgctggc attcaggcag 14640

```
     ggaccgatta caaagcccat atgataagga ctcagcactt ttccttccat gccggttacc     14700
     cgcaacagcg cccgcattcc gaggtttctg ctgtgctgca taagtttgag gcccagacat     14760
     accaaagaac ggttctcgtc cttaagctcc attacatcgc aaacagtggc aagtgcggca     14820
     aattccgtaa gttcgtccct taagcctgca aaagaatccg tactgtcctc tcccatttcg     14880
     cgggcgtaag ccattatggt cttaaaagcc acaaaagctc cgcagatacc ttcaaaagca     14940
     tattcagagc ccggaagctt ggggtccgtg acagctgccg cacggggaat cttataaatc     15000
     ttttctccct tttcatcctc ggtaaaaggc acttcatggt ggtcggtaat aaccattgtc     15060
     atgccaaggg atgttgcgtg ttcggtctgg tcatatgccg caataccgtt atcgcaggta     15120
     agaatggtat ccactccagc tttatatgca tcatcgataa gtcgaatatt gataccgtat     15180
     ccgtcttcta ttctgtgcgg aattacataa tccacatccg cgccgaaata cttaagaccc     15240
     ttatagaaaa tataagtgga tgtgataccg tccacatcgt agtcgcctat tatgcggatt     15300
     ttaaggcccg ccgagatttt ttcttttaac actaaaataa cccgatccat tccggtgagt     15360
     gtaatcgggt catgaaaagc ctcgggcccc ggagataaaa agtttcttat ctccgagggg     15420
     tcggtgaggt ctctgtttct taatattctt gcggtaatgg gagaaattgc gtagtccgag     15480
     gccatcttca taaagtcggc gctctttctt atttcaaccc aattttccat agtttatttt     15540
     tccttcttag aaaacttatt tttaaataca aaccacatgg aacctgcaag gcagatggat     15600
     gagtatgtac cgcacacaat acctaccata agaggaagtg cgaactcctt aatggagctt     15660
     acgccgaaga cgtaaagagc cacaaccatg atgaaggtgg tgagggaagt aaatatactt     15720
     cttgaaagtg tggtggtaat actgtcgttg acaatgtcct taagagccgt tctgggagct     15780
     gcctccttta agttctcacg gattctgtca aatataacga tggttgcgtt gatggagtaa     15840
     cctacgattg taagcataca tgctacaaaa gaagaaccaa cgcttacata tgctatggcg     15900
     tagaaagcaa gagttacaag cacgtcatgg gtaagtgcaa gaaccgagga agcgccaaag     15960
     cggatgtcct tgaatctgaa ccagatgtaa ataagcatgc agatgcaggc tacaaccaca     16020
     gcgatgatgg tctcgcgctg agcttcttta ctgatggtgg agctgatgga ttcggtcttg     16080
     atggtttctt tgtcaatgcc gaacttttcg ttaaagtaat tgttaagctc cgctctctgg     16140
     tctgcgttaa gggttactgt cttaaatatt acttcattgg agccctgaac cttggtggta     16200
     acaacgtttg cgctgccggt aatcttctcg atatcggcta ccataaggtc gttaatctct     16260
     tcaagttcgt aattctgttc gaaaggaact accgtctgag taccgccgag gaactcaagt     16320
     gagtagttga gggaatgtcc tctgtttgcg ttgccgccgt taacagcaag catgaataca     16380
     agacctgcgg caatcacaag gccggaaatg ccaaagaaca agcctctctt ggaaaggaag     16440
     tcgatggtct ttctctgctt ggcaacacca aagaacttgg gatccttgat gccggctgca     16500
     acaaaggcct taagaagtgc cttggtaata acgatggctg tgaacatgga caagcagata     16560
     ccgagggtaa gtgtctgagc aaaacctacg atggaaccgg agcccatgat ccaaagaacc     16620
     gctgctgcaa taagggtggt gatgttaccg tccagaattg cgctcatagc cttggagtat     16680
     gcaaccttaa tggctgaatc catggtcttg cctgtagcaa gttcttctct gagtcttgcg     16740
     aaaataataa cgtttgcgtc aaccgccata ccgattgaaa ggatgatacc tgcaataccg     16800
     ggaagggtaa gggtaatggc gaagccgttt ataagcatta tttcaagtgc tgcgtaaaga     16860
     gtaagtgcaa tggcactggt aagtccgagg aagtagtaaa ccacaagcat gaagattact     16920
     acgattacaa aacctacgag acctgccttt aagctgctct gaatagcgtc ctgtccgagg     16980
     gatgcgccta caacgttgga acgaagttct tcaagttcga gggaaagacc accgatacgg     17040
     atggtgctgg caagcttctg tgcttcttcg gcatccttct gacccgtaat ctgagcggtt     17100
     ccgccggaga taacgccctg aactgcggga acgcttatga actgaccgtc atagtaaata     17160
     ccgataacgt ccttgcctct gttgaaggct gcttccgtag ccttggcaaa agaatccgtt     17220
     ccttcttttg tcagtgtcag gttaaccatg atctgggtgt tgcccatatc gttacgggta     17280
     gtacctgcct gagcgtctgc cacgtcggta ccttccagaa cgatgccgcc gtcagcctta     17340
     atctcgtcaa t                                                          17351
//
ID   XXX; XXX; linear; XXX; XXX; XXX;  XXX.
XX
AC   PRJEB67889
XX
DE   Uncultured bacterium DNA fragment from in-reactor enriched cow rumen, clone
ferm-44C18, contig2
XX
```

```
FT   source          1..6528
FT                   /organism="uncultured bacterium"
FT                   /environmental_sample
FT                   /mol_type="genomic DNA"
FT                   /isolation_source="Cow rumen"
FT                   /clone="ferm-44c18"
FT                   /note="AZCL-Barley beta glucan, AZO-CM cellulose, AZCL
Tamarind xyloglucan, AZCL-HE cellulose active clone"
FT   CDS             25..2934
FT                   /db_xref="SEED:fig|6666666.127262.peg.1"
FT                   /translation="MTGAISSVNGSDILAAPVSNISNALIGNTPGVAGLQTSGEPGLNA
FT                   ANIFVRGVSTYGDATPLIIIDGVEQASEQAFAEFNAMDANEIQGISVLKDAASTAVYGV
FT                   RGANGVIIVTTKHGSTGKPVINFSANYGLTSATQLQEGVTAYEYALFRNEAIRNEQKGY
FT                   PGNDGLSSYIYDDYDLWKFKNNRDFTPQEVEAMTWLTPEQKKALNESPALYYGDHNLYK
FT                   EQFNKVAPQYQANLNVSGGTERVKYFVSFGYFRQEGITHNFHYFGADTRSLFNRYNFRS
FT                   NFDIKITDRTSVTIHTAGQFGETTGPGNGADVYDLSARYKIIMQYIYDCNPFICPGIVE
FT                   GHLISGFAGIAGSAQNPIASKTNFSIGNQNAVYNLLTSGTATIYNSLLDNTLRLNHKMD
FT                   YLLPELKAHVTVSYQDNYNRYVKYTPSLPSYLIQRNAENPLKYDFFGGSRGSGSFESWG
FT                   YSNWNKIYFDAGLDWARSFGKHNVSALLLGKASKYTMPSDSYHVPSGIMGFVGRVTYNY
FT                   ADRYMAEVNMGYNGTEQFAKGRRFGLFPAVSAGWVPTLEPWFPKNDWVSFIKIRGSYGE
FT                   VGNDQLGASRRYYYLPNTWNLKQSGYWLGNSDGSAPNPYFEGAREGSLGNAAITWERSR
FT                   KYDVGLETKFFKSRLSADFDYFYEKRDNILTTLGIIPAIYGVSQSDVPPANVGITTNRG
FT                   YEVVLNWTDKVGSLMYTLGADLSYSRNKIIYMAEAENPYEWMNKTGHMIGQRYGLTSDG
FT                   LFNTKEELDARPYNTYTGNQATLGDIRYVSLDGDDKIDQNDVGPIGFPNYALYHFNFKV
FT                   QLAYKGFDLKLLFTGSKDGSYYLDDGYTIPFFKNGGNAWKWQYDGRWTPEKYAAGEKIT
FT                   FPRAAYSATSSHNNYLVSDYWMVSTDHVKLKNIELGYTFDLSGRESILQAVRLYANANN
FT                   VYTFRNALTPYGIDPETTDGSTYVYPLTRIFTLGLSFRF"
FT                   /product="putative outer membrane protein, probably
FT                   involved in nutrient binding"
FT   CDS             2971..4782
FT                   /db_xref="SEED:fig|6666666.127262.peg.2"
FT                   /translation="MTAVLILGSATSCSYFLEKPDTTGTVDQEAVFSTAKNAKSALMTC
FT                   YRDALRHGWPGGIGIGHSTLGALCGELGRGYSWHGSYTITQQGLSVNGIDGSDAGAENY
FT                   GNNWRIIRECYIVYENVDKVTDLTEAEKQNMKGEALALIAYRYMGMFYRYGGVPIVRKS
FT                   FEASDDLTAGRATLQEMVDFIMELCDEAIGLLPEKWNAANTGRMSKGAALAIKARTLQF
FT                   AARPLFNSDRPYLDNGADNDLICFGNASVERWKAAIDANEAVLSWAAANGVVLINTGDA
FT                   GVGEPNPHAFDDYGRATSEPSNEELILAYKFNDNGGTDALVNFINCSNYQTYNHFDTAE
FT                   SGILTNHLEKYWDKDGNDIDWPKVGDAAPKSGADWRANVAKTEPRAHADIKFGGFDSDN
FT                   NPGNYNWTNAGWGRGGYSADKNKKDVFPNAISGGRGIGERTKFYYHAGSRLWFEPPLFR
FT                   LAETYLYLAEAYNEYGNTAKALENLNMVHNRAGLPKITETDQAKLRAIIQRECAVELFQ
FT                   EGGHRYYDAKHWKLADIADGVCGGPMRMLQFNIKNDNSVVWPYPADAVEDYWDSVVYES
FT                   FWSDAMYLEPFPQTEVNKKTITQNPGY"
FT                   /product="putative outer membrane protein, probably
FT                   involved in nutrient binding"
FT   CDS             4779..4913
FT                   /db_xref="SEED:fig|6666666.127262.peg.3"
FT                   /translation="MIVNEHYYDTFCYKISYSGRGTASRQSEPSGPGLWDGSNFDGFG"
FT                   /product="hypothetical protein"
XX
SQ   Sequence 6528 BP; 1594 A; 1930 C; 1699 G; 1305 T; 0 other;
     ttcgccaccc agaaaaaggt caatgtgacc ggagcgatat cttccgtgaa cggaagtgac        60
     atcctcgcag ctccggtatc caacatctcc aatgccctta tcggcaatac cccgggcgtc       120
     gccggtctcc agacatcagg tgagccggga ctcaatgccg ccaacatctt cgtccgcggc       180
     gtatcaacct atggagatgc taccсctctg atcattatcg acggcgtcga gcaggcttcc       240
```

gagcaggcct tcgccgagtt caacgcgatg gacgcgaatg agatacaggg catcagcgtg 300
ctcaaggacg ccgcctctac ggccgtctat ggagtccgcg gcgccaatgg agtaatcatc 360
gtgaccacga aacatggctc caccggaaag ccggtgatca atttctccgc aaactatggt 420
cttacgtcgg ccactcagct gcaggagggc gtgaccgcct atgaatatgc cctcttccgc 480
aacgaggcca tccgcaacga gcagaaaggc tatccgggca acgacggact gtcatcatat 540
atctatgacg actatgacct ctggaaattc aagaacaacc gcgacttcac tccgcaggag 600
gtcgaggcca tgacatggct cacccccgag cagaagaagg cgctcaacga atcgccggct 660
ctctactacg gcgaccataa cctctacaag gagcagttca acaaagtcgc tccgcagtac 720
caggccaacc tgaacgtctc cggaggtacc gagagggtga agtatttcgt gtcattcgga 780
tacttccgcc aggagggtat cacccacaac ttccactact tcggggcaga caccaggtcc 840
ctcttcaacc gctacaactt ccgaagcaac ttcgacatca agatcaccga caggacgtcc 900
gtgactatcc atactgccgg acagttcgga gagaccacgg gcccgggcaa cggcgccgac 960
gtatatgacc tctccgcccg ttacaagatc ataatgcagt acatctatga ctgtaacccg 1020
ttcatctgtc cgggtatcgt cgaggggcat cttatcagcg gtttcgcagg tatcgccggt 1080
tccgcccaga acccgatcgc ctcgaagacc aacttctcca tcggcaacca gaacgctgtg 1140
tacaacttgc tgaccagcgg taccgcgacc atctacaaca gcctcctgga caacaccctc 1200
aggctgaacc acaagatgga ctacctgctt cctgagctga aggcccatgt cacggtctca 1260
tatcaggaca actacaaccg ctatgtgaaa tataccccgt ccctgccttc atacctgatc 1320
cagcgcaatg ccgagaatcc tctcaagtat gatttcttcg gcggatcgag gggcagcggc 1380
tcattcgagt catggggcta ttccaactgg aacaagatct atttcgacgc cggtctcgac 1440
tgggcgaggt cattcgggaa gcacaatgtt tccgccctcc tgctcggaaa ggcgtccaaa 1500
tatacaatgc cttcggattc atatcatgtg ccttcgggca tcatgggttt cgtaggccgt 1560
gtcacataca attacgctga ccgctacatg gctgaggtga acatgggcta caacggcacc 1620
gagcagtttg caaagggccg ccgtttcggc ctgttcccgg cagtctctgc aggttgggtc 1680
cctaccctgg agccatggtt ccctaagaac gactgggtaa gcttcatcaa gatcagaggc 1740
tcatacggag aggtcggaaa cgaccagctc ggagcaagcc gccgctacta ctatcttcca 1800
aacacatgga atctcaagca gagcggctac tggctcggaa actctgacgg ctctgccccg 1860
aacccttatt tcgagggtgc aagggaaggc agcctgggca atgccgccat cacatgggag 1920
cgatcccgca aatatgatgt cggcctggag accaaattct tcaagagccg cctctcggca 1980
gacttcgact atttctatga gaaacgagac aacatcctta cgacgctggg catcattccg 2040
gccatctatg gcgtgagcca gagtgatgtc cctccggcaa acgtcggcat caccaccaac 2100
cgtggttatg aagtcgtgct caactggact gacaaggtcg ggtctctcat gtacacattg 2160
ggcgcagacc tcagctactc gcgcaacaag atcatctaca tggccgaggc cgagaatcca 2220
tatgaatgga tgaacaagac cgggcacatg atcggccagc gctacggtct gacaagcgac 2280
ggactgttca ataccaagga agagctcgac gcccgtccgt acaacacata caccggaaac 2340
caggccaccc tgggcgacat ccgctatgtt tccctggacg gggatgacaa gatcgaccag 2400
aacgacgtcg gcccgatcgg attcccgaac tatgccctgt accacttcaa cttcaaggtg 2460
cagctcgcat acaagggctt cgacctcaag ctcctcttca ccggctccaa ggacggctca 2520
tactacctgg acgacggcta taccatcccg ttcttcaaga atggaggaaa tgcctggaaa 2580
tggcagtatg acggccgctg gacccctgag aaatatgccg ccggcgagaa gatcaccttc 2640
ccgagggctg cgtacagcgc gacttcttcg cacaacaact atctggtcag cgactactgg 2700
atggtctcta ctgaccatgt caagctcaag aacatagaac ttggctacac tttcgacctc 2760
agcggccgcg aatcgatcct ccaggctgtc aggctatatg caaatgccaa caacgtgtat 2820
actttccgca atgccctcac tccttacggc atcgacccgg agaccaccga cggcagtact 2880
tacgtctatc cgctcacccg tatctttacg cttggcctga gcttccgttt ttaatccacg 2940
agcattatga aaaagattac aagcatattg atgacagcgg tccttatact gggatccgcc 3000
acctcgtgca gttacttcct ggagaaaccg gatacgaccg gaactgtcga ccaggaagca 3060
gtgttctcga cggccaagaa tgccaagtct gccttgatga cctgctaccg agacgccctg 3120
cgtcacggct ggccgggggg catcggcatc ggccacagca cgcttggagc tctctgcggc 3180
gaactcggcc gcggctacag ctggcacggc agttacacca tcacccagca gggcctttcc 3240
gtcaacggaa tcgacggctc cgatgcgggc gcagagaact atggcaacaa ctggcgcatc 3300
atccgcgagt gctacatagt atatgagaat gtagataagg tgactgacct caccgaggcc 3360
gagaagcaga acatgaaggg agaggccctg gcccttatcg cctatcgcta tatgggcatg 3420
ttctaccgtt acggcggagt ccctatagtc cgcaagtcct tcgaggcgag cgatgacctg 3480

```
actgcaggcc gcgctaccct ccaggagatg gtagacttca taatggaact ctgcgacgag      3540
gccatcggac tgcttcctga gaaatggaac gcagccaata ccggacgaat gagcaaggga      3600
gctgccctgg caatcaaggc ccgcaccctg cagttcgccg cccgccctct gttcaacagc      3660
gacaggccgt atctcgacaa cggcgccgac aatgacctca tatgtttcgg caacgcgagc      3720
gtcgagaggt ggaaggccgc gatcgatgcc aacgaggccg tgctctcatg ggccgcagcc      3780
aacggagtcg tgctcatcaa taccggcgat gccggagtcg gagagccgaa tccacacgca      3840
ttcgacgact atggacgcgc tacttcagag ccttcgaacg aggagctgat acttgcgtac      3900
aagttcaacg acaatggtgg gactgacgct ctggtcaact ttatcaactg ttccaactat      3960
cagacataca atcatttcga tacggccgag agcggaatcc ttaccaacca tctcgagaag      4020
tattgggata aggacggcaa tgatatagac tggcctaagg tcggggatgc tgctccaaaa      4080
tccggcgcgg actggcgggc caatgtcgcc aagacggagc cgagagccca tgccgacatc      4140
aagttcggcg gctttgattc tgataataat ccgggcaact ataactggac gaatgcagga      4200
tggggccgcg gaggttattc ggccgacaag aacaagaagg atgtattccc taacgccata      4260
agtggcggac gaggaatcgg ggaacgtacc aagttctatt accatgccgg cagccgtctc      4320
tggttcgaac cgcctctgtt ccgccttgcc gagacttatc tctatctcgc agaggcatac      4380
aacgagtacg gcaatactgc aaaagccctc gagaacctga acatggttca caacagggca      4440
ggacttccta agatcaccga gaccgaccag gccaagctcc gcgcgatcat tcagcgcgag      4500
tgcgcagtgg aacttttcca ggagggcggc caccgttact atgacgccaa gcactggaag      4560
cttgccgaca tagcagacgg agtctgcgga ggccctatga ggatgctcca gttcaacatc      4620
aagaatgaca attcggtcgt atggccatat cctgcagatg ctgtggagga ctattgggac      4680
tccgtagtat atgagtcatt ctggtcagat gccatgtatc tggaaccatt ccctcagacc      4740
gaggtgaaca agaagactat cacccagaac ccggggtatt gattgtaaat gaacactact      4800
atgatacatt ctgttacaag atatcttatt ctggcagggg cactgcttct cgccagtctg      4860
agccttccgg cccaggatta tgggacggga gcaatttcga cggcttcggg tgacgatctg      4920
tacaagaccg tgaatccgaa cctgacgaac accttcgtcg gcaacttctc cggtatgaat      4980
gtttatcagg gtaccggcga gctcggaggc aacaacggca aatggatgat ccggggaatc      5040
ggcagctatg gaatgggctc atggtccatg gccaagttct tcgtggacgg cttcgaggtg      5100
aacgccgagt atgtatgcgc catctctcct tccgaaatcg agaaggttga ggtattgaag      5160
gatgccgccg cccttgccct ctacggagag aagggagcca acggaatcat ccgcataact      5220
acccgcaggg gcgtcgaagg caagacttcg gtccaggccc gcgtgcggta cggaatccag      5280
gccccacaga gcatgaacaa gcctcttggc tcattcgaat acgccagtct ctacaaccag      5340
gccgtctcca acgacaacgg catgagatgg actcctgtct attcggagga gcagctcgcc      5400
gcctacatgg acggcaccgg catcgatgtc gactggtatg acaaggcgat gcgggattat      5460
ggtacattct ccgacgtgga cgtgatcctc aacggcggta ccaagggcgc caggtacaat      5520
atcaatctgg actatgtcgg caaccagggc cttctttccg caaagaattc ggactcgacc      5580
aagaatctcg gatacaaccg cttcaacgta agggccaacc tggatttcag cgtgctgaag      5640
atcttcgagg tccgcttcga catgggcggc cgtatcgaga tgctgcaccg tccgaactac      5700
gccatctcca gcctgttctc gaacctccag aaatatccgt caaacatata cgatgtctat      5760
gacgacgagg cctgcgagaa ttacagcggt accgctgtct atccgaacaa cccttatgct      5820
tcagtccacg gactcggctg gtactcatac aagggccgca gcctccagac aaatctctcc      5880
gtcagggaga atctggacat ggtaaccccg ggcctctacc tggaagaagc cgtctccctg      5940
tacagctata cgctgagcac atattccaag actgcgaatt atgcccgctg gcacaacggc      6000
tcgaccacca ctaccgacca gaccaccacg ctcaccgcga gcggctatgg ctccgccggg      6060
atgcaggacc gcaagcaggg caggatcacg gccggttatg accatgcctt cgggaagcac      6120
cacctgacga cggccctgaa cttctatctc tcggccttca agggagacgg atacttctca      6180
tacaaataca acgtcgccaa tctcaacggc ttcatcaact atgtatatga cggccgctac      6240
gtcgtcgagg cagcgttctc cgaattcggc aacgacgcat atgctcccgg acaccgctgg      6300
gcattctatc cgactgtctc tgcggcctgg atagtctcca atgagggatt catgaaagga      6360
agcgccttcg actggctcaa ggtcagggca tctgccggac tctccggtta ctctgattca      6420
gctgccacgt cggtactcgc ggactactcc tccaagggtc gttacctgtt caaggattac      6480
tatacataca gctacatcgg cagcttctac actggagcga cttcaggt                   6528
//
ID   XXX; XXX; linear; XXX; XXX; XXX;  XXX.
XX
```

```
AC   PRJEB67889
XX
DE   Uncultured bacterium DNA fragment from in-reactor enriched cow rumen, clone
ferm-44d8
XX
FH   Key             Location/Qualifiers
FH
FT   source          1..43948
FT                   /organism="uncultured bacterium"
FT                   /environmental_sample
FT                   /mol_type="genomic DNA"
FT                   /isolation_source="Cow rumen"
FT                   /clone="ferm-44d8"
FT                   /note="AZO-CM cellulose, AZCL-Barley beta glucan, AZCL
Tamarind xyloglucan, AZCL-arabinoxylan, AZCL-HE cellulose active clone"
FT   CDS             complement(373..1944)
FT                   /db_xref="SEED:fig|6666666.127265.peg.1"
FT                   /db_xref="GO:0003723"
FT                   /translation="MDSEKELLKDLIVDVQATEVQIALQENHRLIELNKESSSGRSFAV
FT                   GDAYLGRVKKVMPALNAAFVDIGDEKEAFIHYLDLGLYFGAFDEFVKKTNPNTNPSRLF
FT                   GSIKLGPILEKEGRIENVLTQGQMIIVQIVKEPISTKGSRLTAEISLAGRNIVLLPFAE
FT                   KVSISQKISSKEEKKRLETLVRSILPKNYGAIIRTAAEGKNAAILVAELKSLIGKWEDS
FT                   WENLSKNKSIQPLFTEYSKTTTILRDLLNDSFSNIYVNNEKEYEEIRRYISLISPEQEK
FT                   IVKLYQGKEPIFDHFEITRQIKSSFGKVVPLKQGAYLVIETTEALNVVDVNSGIRAKTK
FT                   EQEENTFEVNKFAAEEIARQLRLRDMGGIVIVDFIDMESVEHRNNLFKYMQELMSKDRA
FT                   KHNVLPLTKFGLMQITRQRVRPVTEINTTEVCPVCHGTGKIASSLLIDEQIERRLSYYV
FT                   NDKGIRNFTLKVSPILGAYITRGLFNSYVGKWKKKYKCKIKVVEITDFAVLQNELFNEK
FT                   GDKLDS"
FT                   /product="Cytoplasmic axial filament protein CafA and
FT                   Ribonuclease G (EC 3.1.4.-)"
FT   CDS             complement(2173..2445)
FT                   /db_xref="SEED:fig|6666666.127265.peg.2"
FT                   /translation="MTKAEIVNEVAKATGIEKLAVQSVVESFMESVEASLAKGEPVYLR
FT                   GFGSFIIKHRAEKAARNITKNTTLTIPAHNIPAFKPAKVFVSKVK"
FT                   /product="DNA-binding protein HU"
FT   CDS             2731..5073
FT                   /db_xref="SEED:fig|6666666.127265.peg.3"
FT                   /translation="MSGEPALNFDASELPGSLPILALRNAVIFPGVVFPITIGREKSML
FT                   LVDKAQGNGMFIGAVPQKNVAEENPLSLADFSEFGTVVKVLKTFEMPDGTVTALLQGYK
FT                   RLRMLELTDTDPYHMARVSYLDDILPQNDSKVPVIADILKENASSIIRSSSFVPKEALE
FT                   ALKNIDDFNFLVNFIATTIDMDDFMEKVDLLKYDDVKERAMALLGVLSRQLEIQRLKQE
FT                   ISSKVKTELDQQQREYFLNNQLRTIQEELGMDEMDDIARFREAAEKKEWPASVGETFEK
FT                   ELAKLERFNPSTPEYSIQYNFLQFMIDLPWYHTSDDNLDLKHAQKVLDEDHFGLENVKD
FT                   RIIEYLAVLKLKGNMKSPILCLYGPPGVGKTSLGKSIARALGRKYVRISLGGMHDEAEI
FT                   RGHRKTYVGALPGRILNGINKAGTSNPVMVLDEIDKLGSDFKGDPSSALLEVLDPEQNT
FT                   AFHDNYLDIDYDLSNVMFITTANTINGIQPALLDRMELINVTGYLAEEKVQIAKKFLVP
FT                   KQLEEHGIPKKELKIDKAALEEIIDKYTHESGVRGLEKQIARIARVSAKKMALGEDYPK
FT                   TIKKEHLKEYLGLQTAFHDSQKGNEAPGVVTGLAWTSMGGEILFIETSVSDGKGVLTMT
FT                   GNLGDVMKESATIGYQYIKAHPQLAGLTSEEFAKKDVHVHVPEGAVPKDGPSAGITMVS
FT                   SMVSALRGKKVRSGIAMTGEMTLRGRVLPVGGIKEKILAAKRAGVSTIIISEENRKDIE
FT                   DIKEIYVKGLTFVYVKTIDDVIGYIFE"
FT                   /product="ATP-dependent protease La (EC 3.4.21.53) Type I"
FT                   /EC_number="3.4.21.53"
FT   CDS             5073..5741
```

```
FT                   /db_xref="SEED:fig|6666666.127265.peg.4"
FT                   /translation="MDVKIEQTWKDALRDEFEKPYFASLVRFLHNEKEAGKVIYPPGGL
FT                   IFRAFELTPLPEVKVVILGQDPYHGPRQAMGLSFSVPSDVPAPPSLKNIFKEIQTDLGI
FT                   TMSGSPNLEGWARQGVLLLNAMLTVQAGVAGSHKAIGWQEFTDAVIKCISDRCNGVVFL
FT                   LWGNFARGKKELIDTSRHYVLEAAHPSPLAGGAFFGCRHFSKTNELLIKEGKTPIDWKL
FT                   "
FT                   /product="Uracil-DNA glycosylase, family 1"
FT   CDS             5738..6211
FT                   /db_xref="SEED:fig|6666666.127265.peg.5"
FT                   /db_xref="GO:0004595"
FT                   /translation="MKRKALFPGSFDPFTAGHLNILRRALTMFDEIVIAVGVNQDKRGF
FT                   FSMDQRMDIIRQAVRGMDGIEIIKYDNLTIDTCRELGIAHIIRGVRNMLDFENERAIAD
FT                   ANRRLAPEIETIIIPTAQEFAHISSSAVRDILKHKGDTSLFLPEGVVLPDHDA"
FT                   /product="Phosphopantetheine adenylyltransferase (EC
FT                   2.7.7.3)"
FT                   /EC_number="2.7.7.3"
FT   CDS             6204..7682
FT                   /db_xref="SEED:fig|6666666.127265.peg.6"
FT                   /translation="MRRLILIIFLSLLSFALGAQNADSVKMAALDAKLEEYFAALETES
FT                   LDVKRTECDFLISSCTTDDVRNHVAVRLYSHFMQSKLMGDEAVAVHLADNWFIPGKVSF
FT                   PDEIDLMNAKIFAEFNRQSLLGCKAPGLVAEDSLGRKVEVPLGLSDRYRILYIYSPDCA
FT                   TCSLQTMLMDSLFSRKSYPVDLIAFNSDDNTEAWVQYRRNRLSFDGVRVMHFRDPSLES
FT                   DFQRKYGVLQTPRMFLIDKNGIIVGRGLDAAALEKLLDATPDLGIGYGSKSSADYFDAV
FT                   LESFGDELRGSDIRYFIGRLEEATLESADTAGFRQMTGDMLYYLAGKRGEAFKDAEAFV
FT                   VDSMVLSRPDVWRFADDSLTVVDMAMSLKHMLSLAPVGARVPDVRVHATVLRAGRKPAT
FT                   VHKKLRSLLKGKRNYVLFHTAGCTFCEAELAAADSVASEKGVRVILVDMDKLTSLYPEE
FT                   ADRLLDAFDLSVLPFIMETDNKGCVVRKYLSLNK"
FT                   /product="Thiol:disulfide interchange protein tlpA"
FT   CDS             complement(7671..8243)
FT                   /db_xref="SEED:fig|6666666.127265.peg.7"
FT                   /translation="MSIMGKAYIFLADGFEETEALATLDILRRGKIEAYTVSITEEVYI
FT                   TSSHGVTVVADMGRSDFSEIDIDEKDAMIFPGGMPGSKKLAADTPLIGLMNVHYGNGGI
FT                   VAAICAAPGLVATQIKGIEGKKATCFDGFEENLAASGMEYVKAPAVTDGQLITGRGAAF
FT                   AIDFGLAILEKMGGKEAVAKVKAALLI"
FT                   /product="DJ-1/YajL/PfpI superfamily, includes chaperone
FT                   protein YajL (former ThiJ), parkinsonism-associated
FT                   protein DJ-1, peptidases PfpI, Hsp31"
FT   CDS             complement(8353..9465)
FT                   /db_xref="SEED:fig|6666666.127265.peg.8"
FT                   /db_xref="GO:0008784"
FT                   /translation="MNRVDDFSRLELNLTGCMENYRYFRSKLEQTTKLLVLVKANAYGH
FT                   GAVEFAAMMQRAGANYLAVAYPVEGMELRKAGISLPILVLTAGTDFFNEIIDNRLEPGI
FT                   PNLYALKILCGILKERGIKDYPVHIKLDTGMHRLGFMTSELQDLTDFLKTCEEVKVKSV
FT                   YSHLAVSEDPEQDEFTLGQIGMFKTNADRITSEIGYKPMYHILNSAGIERFPQYQFDMV
FT                   RLGIGIYGISALPDVRLAHVASLKCKILQIKTLKPEDGTIGYGRYGHIAPEGTVIATIP
FT                   VGYADGIDRHLGRGHASFSVNGHRAPTIGNICMDMCMLDVTGIDAKVGDTVTIFGEDPT
FT                   VNELADILGTIPYEILTSVPRRIERIVVRK"
FT                   /product="UDP-N-acetylmuramoylalanyl-D-glutamyl-2,6-
FT                   diaminopimelate--D-alanyl-D-alanine ligase (EC 6.3.2.10) /
FT                   Alanine racemase (EC 5.1.1.1)"
FT                   /EC_number="6.3.2.10"
FT                   /EC_number="5.1.1.1"
FT   CDS             9824..10498
FT                   /db_xref="SEED:fig|6666666.127265.peg.9"
```

```
FT                   /translation="MDMIWFIIIAVMVLSFVIQQVLNSKFNKYSKVGLHSGLTGAQVAA
FT                   KMLEDHGITDVSITCIKGKLTDHYNPVNKTVNLSEEVYYGNSVAAAAVAAHECGHVVQH
FT                   ATGYAPLKLRSALVPVVSFSSNAVQWVLLIGVMIIKITPTLLYLGIAMFALTTLFSVVT
FT                   LPVEINASARAVQWLDKAGITGPATKAKAVDALKWAAYTYVIAAIGSIATLIYYIAMAR
FT                   RR"
FT                   /product="probable metal-dependent peptidase"
FT   CDS             complement(10495..10788)
FT                   /db_xref="SEED:fig|6666666.127265.peg.10"
FT                   /translation="MTGDDYKIISDTTENGVRHITAIPSSLVCSQQIDFDIIDGKVRNL
FT                   AYVRGCNGNLQAIGRLVEGMDIKKIVDTLSGVNCNFRGTSCTDQLARILSSL"
FT                   /product="hypothetical protein"
FT   CDS             complement(10805..11239)
FT                   /db_xref="SEED:fig|6666666.127265.peg.11"
FT                   /db_xref="GO:0008270"
FT                   /db_xref="GO:0016787"
FT                   /translation="MEKDELYMKEALREAQAALAEDEIPIGAVIVCKGRIVGKGHNMTE
FT                   RLHDPTAHAEMIAITAATEALGGKYLNECTLYVTVEPCPMCAGALNWAQIGRIVYGAID
FT                   PKRGHSLFTPSLLHPKTEVEAGVLAEECGSMVTEFFKKKR"
FT                   /product="tRNA-specific adenosine-34 deaminase (EC 3.5.4.-
FT                   )"
FT   CDS             11369..12247
FT                   /db_xref="SEED:fig|6666666.127265.peg.12"
FT                   /translation="MTSVITEITPLSDRDCFYIVERHKTEFVFPLHRHREYELNFIEHG
FT                   KGVRRVVGDSVEEIGEYELTLIAGGDLEHAWEQGNCTSEDVREITIQFDPEIFSSGLMM
FT                   RNQFSSIQHMFEQARNGLTFSLAAIMKVYSLLDALPSEIDKFDQFQHFMRLLYDLSLAG
FT                   DSRILASTSFAHNDADKESRRVQKIKQYVGDHFAENITLEELAEMVGMSPSAFSRFFKS
FT                   RTGKTVIDYIIDIRLGNAARLLVDTTQNISEICYSCGFNNLSNFNRLFKSKRGYTPREF
FT                   RTLFKKNKVFV"
FT                   /product="Transcriptional regulator of various polyols
FT                   utilization, AraC family"
FT   CDS             12417..14816
FT                   /db_xref="SEED:fig|6666666.127265.peg.13"
FT                   /translation="MKMDIVKVDDGSFVVDGHFYYYIGTNFWYGGLLGSEYGDRSRLAA
FT                   ELDSLHALGLDNLRVLVGSDGPDGVATRVYPTLQKEPGVYDEKVFEGLDYLMAELGKRN
FT                   MRAVLYINNSWEWSGGYGMYLEWAGAGKAIIPSVGGYGPFMEQMAQYATNARAQELFEN
FT                   HLRTVVSRVNSITGKPYSEDPAIFSWQIGNEPRCFSSDPAVQDGFVSWIWRAASIIKSI
FT                   DPNHMVSTGNEGIWGCEENPELFRRLNECPDIDYITAHIWPYNWSWVGKESPQLDIDRA
FT                   EENTGRYIDAHLEVAESLGKPLVIEEFGFPRDDMQFAKGTPVSGRDRYYRYVFSKVMEN
FT                   QKNGGRLAGANFWSWGGLAVPAHRYWQEGDDYAGDPAQEEQGLNSVFADDYSTLAIIKH
FT                   TTARLSAVANASAVLENDWMFTTGDQKPLRVAVQSQNGFKGSVPVYVRVSTDKGVSVSV
FT                   SGKKARLGKGADTLDFDLGLAPGFYKVCVSLNENDPLDENFCVGCDPEKIVSPQDKQPD
FT                   FDEFWEDNLRQLAMVPMNARMTPLKDHSNADRTAYRVDMKSFGGKEISGLLYLPNKEGK
FT                   FAARISYMGYGSDVWFADPSAEPDMIEFTLSVRDQGFNKTPGQGWVTRGLADKDSYYYR
FT                   GAYLDCVRAIDFVSSLPQTDPDHIYAEGGSQGGAFTLIAASLDNRLRAIAPFVPFLSDF
FT                   PDYFAIADWPGGEVLSAAKAQGISEDELYKTLSYFDVKNFTDRIGCPVLMGFGLQDPVC
FT                   PPHTNFSGYNHITADKQWLCVPVSGHHLEQFPEWWAARTEFFTKNK"
FT                   /product="GH5_7-CE7"
FT   CDS             14904..18053
FT                   /db_xref="SEED:fig|6666666.127265.peg.14"
FT                   /translation="MAQNKVTGRITDNSGQGVINASVMVKGTNNGVVADLDGYYEIVVG
FT                   QGSTLVYSSIGYKDQEVVVGTQSVINVLLVENSELLDEIVVVGYGTMKRSDLAGSSVSM
FT                   KEDDLKGSIITNLDQSLQGRAAGVTAVQTSGAPGSSSSIRVRGQATINANAEPLYVIDG
FT                   VIMQGGGNSGADFGLGDALGNGKVSTISPLSTINPADIVSMEILKDASATAIYGAQGAN
FT                   GVVLITTKHGSRGDAKFSYDGMFAVSRQTTRIDMMNLREYAEFYNSMVETGEISEANPY
```

```
FT                   YATPGLLGKGTNWQDEIFRTAIQHQHQISAQGGTDKVQYYVSGSFMDQEGTIRGSEFGR
FT                   YTFRTNLDAQLKEWFKLGLNLTFANTNDDLKLADGSEGLITYSLTTIPDIPVYDVYGNF
FT                   SSTVREGYTSPNPVALAMMDEILLDRQKLNGNIYAELTPVKHFTWRTELGFDVSGSTAT
FT                   RFRPTVNLGGWTRDTNSMSYQKNSNRFWQIKNYLTYANSFGKHSVTAMLGQEAWESSWS
FT                   YVQGANTKLPSNDVHNVALGGGTPSLSSGFGSSAMASFFTRETYNFDDRYLLTYTYRYD
FT                   GSSNFGPEKRWAGFHSLAVAWKFTNEKFMDFLKPVVNNGKLRLGWGQTGNSSIGGYAWG
FT                   ASISRMPSALGMGYRPANIPNPAIHWESQTQQNIGLDLNMFDNRLNLTVDAYLKKSDNM
FT                   LMSMQLASYMGTQGNSSSALAAPKGNFGSIENKGLEITLDAHPVQIKDFAWDSNFQISF
FT                   NRNKLVSLDGTTNAALIGYGQWSDVVCVSEIGKPLYNFYGYKVAGVYTSLEDIENSPKP
FT                   EKYPSNGEYARGNTVWVGDIKYEDISKDGVINEKDRTDIGNPLPKFTFGWSNTFTYKDF
FT                   DLNIFINGSYGNKVLNYNAISLTHMNSAWTNQINDAVSGRAMLAPIDASVVYADGSKWY
FT                   DHIDNVRVLNPGTKTPRASINDPNDNDRISDRYVEDGSYIRIKNITLGYNVPKSFLNKF
FT                   KVDNVRVYVNIQNLYTFTKYTGYDPEVGASTQDSTGLTYGVDNGRYPSPTVYSCGLNIT
FT                   F"
FT                   /product="TonB family protein / TonB-dependent receptor"
FT   CDS             18081..19814
FT                   /db_xref="SEED:fig|6666666.127265.peg.15"
FT                   /translation="MMKGFAVATAVFALTSCEGFLNRPNEDNYNVDNFYKTDEQCEQGV
FT                   NYLYNSPWYDFQRGFIKIGEVMSGNMYWGSSPYLDFSTNGTDQDLVNMSYSLWAVNGHA
FT                   NTVIYNLLNSQGPSKAAIDKCIGEALTWKAFAYFFMVRTFGDVPIVHSNSDLLSSGEYN
FT                   SISKVERADVYEYIITTLEKAMEMLPKKTSGFNNYERIDYYAAEALLAKVYLTKAGLSG
FT                   SLNQDDLKQAAKYAKDVIDNSGRTLLPVYSDVFRLTGFNATGECLISWEWSVGRNPWTQ
FT                   QNTLQSDLAMVGFSDQGDCWGGWGGPSVDLADAFGVSATQNPAERVGEKDSRRKATLML
FT                   PGDKYDYFWTDKGGFDYLRFIYDTEYGKGGPGGSMQCPTGSNNVKHLYGDNFDHENAGL
FT                   GTPENMAYQLPTHLLRLSDVYLIYAEASLLTGSNGPALEYVNKVRERAGADALASVTFE
FT                   DIWKERRLELAGEGDRWYDYVRRSYYDMDACIAELKAQRRSYWDGLDDVWKSYIGDAFD
FT                   YSGTWDASEVQYNNAEDNPNVTAASFKLPFPTEDVVFNGNMASDAVPVHVDVREEYSYN
FT                   F"
FT                   /product="putative outer membrane protein, probably
FT                   involved in nutrient binding"
FT   CDS             19875..21131
FT                   /db_xref="SEED:fig|6666666.127265.peg.16"
FT                   /translation="MASFASCDKVDYPDRYEVADGKPTVYSVRYADKDVNITQAYMDEV
FT                   VCLLGKNLKSIVELWFNDQQATLNTSFITDNTLLVSVPKNMPVVQTDKIYMITSGKDTL
FT                   AYDFAVLPPTPKVTSMSNEWAAEGETVTIYGDFLIDKADKPLQIAFAGAEVDHASMVFN
FT                   GSSSVSFPVPAGAQPGYVTVTSLSGTGKSKFMYKDTRNILFDWDGSHGGFATGHGWRNG
FT                   VIHAPGADAWDAIDGSYLYFGGADLGPGSADAWAEDQYSFNYWPEAGTDYDALNLRPEF
FT                   AEYIDKYGVSGLVLKFEVLVPSSNPWSASGMQLMFSSDAVVTYGTATNSYFSDETFPRA
FT                   IWQPWTTNGGSYDTADKWVTVSFPLSSFNKNQENKECATAFDKSYLTGLTFFVWNGGID
FT                   GTACSPVFAIDNIRVVPNE"
FT                   /product="FIG00417257: hypothetical protein"
FT   CDS             21138..23564
FT                   /db_xref="SEED:fig|6666666.127265.peg.17"
FT                   /translation="MKLIMKKIFSISAILAACVLALGLSGCSSKELDTDQYSDKDVKLA
FT                   SFGPNPVMRGATLRFFGSNLDRIVEVSVPGLDPITDIEVVTSGNPSEIRIQLPAEGAQV
FT                   GKISLKSSDGKVFTTQVDVEYTEPIVFDSFSPAEAMPGDVITVKGDYMNLVQSVTFEGG
FT                   AIVTEINHNGRHEASFKVPSKAITGKIILSDEGAIANLLYSEEDLVIGDPTVKSLTVAV
FT                   AKPGENAVVKGAYLDMIDKVVFEGGAEVSDLALNEDATEFNLVIPAGAQSGEVKAVSHA
FT                   AKEFVAGEIELVIPTDVAAAPSPVKAGADLVISGNDLDVVTSVDFPGASGVEFVFAEDA
FT                   ITVTVPATATEGDIALNMASGEAVNVAFTLVHPVVSAIAPVELMAGETITVTGTDLDLV
FT                   TGVTLGGKPVEFTAEAEKIVVQTANTSVSGKLVLSLANGEKIEPEDAITLSYDSFIIVN
FT                   DMPAAEHIGAIVTLKGENFMMIENIYIGEEKVSKYNLRSDNEIQFVMPWNKVGTYSMYF
FT                   HLLSGDVETCPQTIEVLLEQDIMTVWEGSLQITWNDGGRVTVPAASFTGIPAGSKLRFY
FT                   YTQVDQQWDQAQLNYGNWTGINFDDPAGTVINGTLVPTDVYGWFTDGILDRCTEVVLTK
```

```
FT                   EILENIQANKGDCEDKTACGIIIQGSGLTFTKIEIVCDIPQEVTIYEGPTTLTWGDDGR
FT                   FGLAMTYFEAAKPGSQLIFYLEQTDAWGQVQLNDGWWANADMNFPEIGGAYITTDNLGG
FT                   KDVTRIALTLTADILDHILATSGDYFGLNSQFQGDGRVAMVIQGSDMIINKVCIL"
FT                   /product="hypothetical protein"
FT   CDS             23626..25185
FT                   /db_xref="SEED:fig|6666666.127265.peg.18"
FT                   /translation="MKAHVSFFVLSMIFLAACSCSKDDAEKKEPSVPAPKAVTVDVASV
FT                   SAPVEGTSVVVAISSPSKPAVSVSGDWIKAELGPYNPDTYKVKLTVTVGENPTYESRSG
FT                   SVTLTCQGFSPVGISVSQDAAAKPVGPSANPAYDLLGLGWNMGNQFDAFYNGSWAGDKY
FT                   NYPDETVWGNAKATQATFDGVKKAGFTSVRIPVTWLNMIGPGPEYNIDETWMNRIYEVV
FT                   GYAEKAGLKVIINTHHDENNNDDHWLDILSASKDARTNEAIKAEIAAVWTHIANKFKDK
FT                   GDFLIMESFNEIQDGGWGWSAAFQADPSKQCDVLNEWNQVFVDAVRATGGNNATRWLGV
FT                   PTYAAGTSWTKYFRLPEDPAGKVMMAVHCYDPYEYTIGNIQYPEWGHTGAAGKKTPEGE
FT                   KEITDLFSRLNTEFISKGIPVYLGEFGCSMRSKGDARAWMFYKYYMEYVSKAAHTYSLP
FT                   GILWDNGSTKTGKECHGYIDHGTGKYIGNSKEIVDLLVKGMLSTDESYTLQSVYDRAPK
FT                   TN"
FT                   /product="GH5_4"
FT   CDS             25188..26330
FT                   /db_xref="SEED:fig|6666666.127265.peg.19"
FT                   /translation="MNLILKLSIILSLVVAVSGCGRNKEAAGTAFNEETRIMMDRMAAS
FT                   VKAGKILYGHQDDLVYGHSWCVDPDSAATDALNRSDVKAVAGDYPAVVGFELGGLELGW
FT                   DWSIDKVPFLLMRRAAVEHAGRGGIVTFSWHPHNPYTGGDAWDISSDKAVASVLEGGEK
FT                   AEMFGVWLERLGDFMDSLRDDKGNLIPVIFRPWHENIGSWFWWGGALCTADEYIALFRR
FT                   TVEYLRNDRGLDNIIMCYSPNSGITAEQYMSRYPGDDVVDILGYDHYETVMEGESLEDA
FT                   GKRYAGQLDLGLSIVKSLASAHGKLMTLSETGMEGLPDRSWWTSVLYPAVKDFPIVYVL
FT                   TWRNAWDRETHFYAPYPGNEGDFREFASLPEISLLSSFRP"
FT                   /product="GH26"
FT   CDS             26408..27583
FT                   /db_xref="SEED:fig|6666666.127265.peg.20"
FT                   /translation="MENMSFEERLAWLEARHQELIAKKNSPIYGNGVFERYENPVLTGD
FT                   HAPLFWRYDLDKNSNPFLEERFGIHAAFNAGAIKWKDKYVVCARVEGADRKSFFAIAES
FT                   PNGIDNFRFWDHPVTMPDYGEPATNVYDMRLTAHEDGWIYGIFCVERKDPDAPEGDLSS
FT                   AVAAAGIARTKDLVNWERLPDLVSDSQQRNVVLHPEFVNGKYALYTRPQDGFIDAGNGG
FT                   GIGWALVDSMENAVVTSEKIINCRRYHTIKETKNGEGPAPIKTDRGWLHLAHGVRGCAS
FT                   GLRYVLYMYMTSLEDPSKVIAEPAGFFMVPEGEEYVGDVMNVLFSNGWIADEDGKVFIY
FT                   YASCDTRLHVATSTVDRLVDYCMNTEPDLFTTSASVRRLNALIDKNLSLSM"
FT                   /product="GH130"
FT   CDS             27604..29031
FT                   /db_xref="SEED:fig|6666666.127265.peg.21"
FT                   /translation="MDIITAPAKPTLKEKIGYGFGDMSSSMFWKIFSYYLPIFYSDVFG
FT                   LKPQHAAFLLLITKLYDAISDPVMGIIADRTRSRWGKYRPYLLWVALPFSLIGLLSFYT
FT                   PDASYGLKHVYAYVMYILMMTVYTAINVPYGAMLGVVTPDSREKSVFSSYRMFFAYIGS
FT                   FIAMGIFGVFEKLISGTVRVMPDGSERVVRGVGDAAPGQWTLVVGIVSLCCFVLFLLCF
FT                   LMTRERVGTENKAVSEGGSSIAEDLKALGRNRPWWLLLGAGIGILLFNSIRGGAAAYYF
FT                   SNILGTSVFLTCAVYLFIGEVAQMVGVLFTVPLSERIGKKATFISVLLTVSVLSVAIFF
FT                   LPETPAGFWGLLVLQVLICIAFGVQSPLLWSMFADVADYSEEKNGSASTGLIFSSSSMA
FT                   QKFGGALGAFLLMQILAIFAYDKDAAVQAPQTLAAIRGLMSYIPAIGALLGALCLLFYP
FT                   LTESRMASIREKLSARR"
FT                   /product="Xyloside transporter XynT"
FT   CDS             29516..32248
FT                   /db_xref="SEED:fig|6666666.127265.peg.22"
FT                   /db_xref="GO:0003887"
FT                   /translation="MQDCGKDEGLEGIEKNVIFVFRSLIRRTMADKLYLIDGHNLIFKM
FT                   YYAFLRRPMINSKGMDMSILYGFTKYVLELIEREKPTHMAIAFDPPGGTFRNEMYPEYK
```

```
FT                   ANRSETPQLVIDALEPLLEICKALNIKTLMIPGYEADDVIGTVAKSYGKPGTDVYMVTP
FT                   DKDYGQLVAEHVYQVKPGKAGGADETVDVKAVCEKYGISSPEQVIEMLTLCGDTADNVP
FT                   GVHGIGEVGAGKLIAKYGTVENIYAHLDELTDRQKTQFEEARGHIGLSHDLVTIKTDVP
FT                   VETSLDEMRLDLDYAPEIADLFEKYEFNSLRKYIAHSSRAAKAVAEVKRLDYTETGAEE
FT                   LVKMAPEKISLIVEPEGPGIFADIRRVIVAAEKKDGGIVVASGAAKDFRKILEGEAVKY
FT                   GYDLKLQSNLLENAGIKLGGRLYDIELMHYLINPEKSHKVEILAKSYLGVDIEEAAAVE
FT                   EAPVSLFDVAEESPANGPDRNLEAAAVLKLGEVVRAGMDEAGVGALYEDMEAPLAAVLA
FT                   RMERAGVKIDLGQLRTFADELKVEMDALEAQVRKLADEPGLNVSSPKQIGIVLFEKLKL
FT                   DPKMKKSASGSYSTDETTLTAIADRHPIIDKILEFRAVKKLLSTYIEPFPTYVSPVTGR
FT                   VHTTFNQALTATGRLSSSNPNLQNIPIRTERGKEIRKAFIPATPDGIIMSADYSQIELR
FT                   IMAHLCGDEHLISSFKAGLDVHKATAAKIFGIPVEEVTSDQRRIAKTANFGIMYGISSF
FT                   GLAQRLHVSRTEAKKIIDDYFAAFPSIRTFIDKTIADAREKGYVETVFGRRRYLPDITA
FT                   KNATIRALAERNAVNAPIQGTSADIIKLAMISVDKRIREAGLKSKMVLQIHDELLFDAV
FT                   PEEIEELGRIVVAEMENVITLSIPLTVECNYGKNWLEAH"
FT                   /product="DNA polymerase I (EC 2.7.7.7)"
FT                   /EC_number="2.7.7.7"
FT   CDS             32235..32825
FT                   /db_xref="SEED:fig|6666666.127265.peg.23"
FT                   /db_xref="GO:0003677"
FT                   /db_xref="GO:0003700"
FT                   /db_xref="GO:0006352"
FT                   /db_xref="GO:0016987"
FT                   /translation="MKLTDHDDGSLVEMALGGGPEAEAAFFTIYARYHKGVFAHISRFI
FT                   QEQEEVEDICMESFEKAFKQLESYRPSNKFSTWIFTIARNTAFDHLDKDKVRGRNKDKM
FT                   PIDYSDLEHIDVPEDNVSPEDEIISSQDHENFLSCVQGLPDLYREVASMCFVDNLGYKE
FT                   ISEKAGLPINTVKTRISRAKSLIVQMMLDMEEE"
FT                   /product="RNA polymerase sigma factor RpoE"
FT   CDS             32822..33484
FT                   /db_xref="SEED:fig|6666666.127265.peg.24"
FT                   /translation="MNFKEFEDIVVAEFPELTPEMLGKYEAMEPLYEEWNAKINVISRK
FT                   DIGGLYLHHVLHSLAIASYLKRRRPEVYSAWTEGASVLDLGTGGGFPGIPLAVMFPKVR
FT                   FTLCDSVGKKIIVASEVSKALSLGNVTTVNARAESLPQVFDHVVSRAVTSLDNFYPWVK
FT                   GKFSGSILYLKGGDIVPEIAELMKKNRLRTGAVSTWPIETWLKDEYFKEKFVIEIEK"
FT                   /product="rRNA small subunit 7-methylguanosine (m7G)
FT                   methyltransferase GidB"
FT   CDS             33963..35471
FT                   /db_xref="SEED:fig|6666666.127265.peg.25"
FT                   /translation="MQVWLNEDHSQPVVHGAVVVKAGGRDCAGTGMAHYLEHMLFKGTE
FT                   DIGTIDYASEKVYLDSIETCYDLIAATSSSVKRDSLQKEINRLSIKALDYAIPNEFDRL
FT                   TANIGGTKLNAFTSYDCTCYHNSFSPQFIAQWAELNSHRLLNPVFRLFQSELEAVYEEK
FT                   NAAADSQFDMILEKMLEKLGDENPYSKSIIGTTDNLKNPSLSEMKKFFNKYYVGCNMGL
FT                   ILAGDIDASSILPLLESTFGRIPRGDKPERIPVPVPDFSSREDYPIKVNFPMIKASLNV
FT                   FNGPAAKDADYPAMELALSLLNNSFSTGSLDSLINNGKVLAGEVSVLDFFDEMGLIAMI
FT                   TIPGIPFGSLNKAEARCFKEVERLKAGDFAEKDFESLKYDLVKSYKRCLESSESRFSIM
FT                   IDVMSQDRSWDSYLSQVESISSLTSEDIVRVSNKYLGDNRIRFHKVRGTYPKDNLQAPS
FT                   YETLSPKNGNAVSDYATRLMEMPYDTLPPRLVDMNNDIRKVAVK"
FT                   /product="ZINC PROTEASE (EC 3.4.99.-)"
FT   CDS             complement(35530..37128)
FT                   /db_xref="SEED:fig|6666666.127265.peg.26"
FT                   /translation="MELWDKAERLWTRVKNWKTVRGWHIWKLKLVDARLYFVSMFVGLL
FT                   TGLVAVPYHYLLYYLFHLRSGFFASHPAWYWHIPLFLFSWGILVFVMWLVGKMPLIGGG
FT                   GIPQTRGVINGRITYRHPFIEMVSKFVGGILSFSAGLSLGREGPSVQIGSYVGSLVSRW
FT                   THILKGEQKQLLAAGAGAGLAAAFAAPLASSLIVIESIERFDAPKTAITTLLAGVVAGA
FT                   VASMVFPVNPYHLIEVTEPVFAFFVQMKYFLILAVIVSVCGKFYSSTMNAFRHVYAKVN
```

```
FT                   SPAYVKMFYLVLVAYIISLMQVNLTGGGEQFLLAQAQNGSTQIMWVTAVMLLHFVFSVF
FT                   SVSSGLPGGSFIPTLVTGGLLGQIVGLVGVRYGVIGQENVSYIMLVSMSAFLVAVVRTP
FT                   LTAIVLITEITGHFEVFYPSVVVGGLTYYFTELLQMKPSNVSLYEDMIHAPDFKEEKRY
FT                   TLSVEIMTDSYLDGKLVDELSLPERCAIINVHRDGKNLVPAGTRLIPGDQVQIEMDSQD
FT                   IEKLYEPLVSMANIY"
FT                   /product="Voltage-gated chloride channel family protein"
FT   CDS             complement(37201..37353)
FT                   /db_xref="SEED:fig|6666666.127265.peg.27"
FT                   /translation="MLYSFINMKLRDFYPSLDALCEDMNVEKEDIVRKLKLVGFEYNPA
FT                   KNRFW"
FT                   /product="FIG00405231: hypothetical protein"
FT   CDS             complement(37497..39041)
FT                   /db_xref="SEED:fig|6666666.127265.peg.28"
FT                   /translation="MKCFILHFSLFIFFLAGSARADGGDVLDRIVRLPGMKGTVYALLG
FT                   KVSEQSGYLFIYDSKVIHNDSVVRVRKQECSVRQAIHRITGSRNLELKVLGNHILITRA
FT                   AERKSMEETVLLPPRPAYSTIAGILLDKETGEPVVSASVIAQGTSLGNITNQNGEFRLN
FT                   LPDSLRRCMLSFSHLGYVGQTVEASALEGRSNVLSLEPKVISLQEVLIRLVEPKKLLRE
FT                   MIEHRDRNCSTSPVYLTTFYREGVQLKNKFQSLTEAVFKVYKSPTMEPGLKDQVKLLKM
FT                   SKIDNREQTDSVLAKISSGVEACLQLDIMKNLPDFLLLESGEELYTYTSGDIVSVDDRT
FT                   ANVVYFEQKRGVKEPLFCGELYIDSENSALLRARFEIHPRYVKAATRLFVIRQAPKIRL
FT                   TTEKLVYTVSYKPWNGTYYVHHIRGDLYFKMKRKRMLFSNPTLYTWFEMVTCRIDGENV
FT                   TRFPRAERLPVHTVFSETDFKYDADFWGDFNVIPLEEELGKIIEKVSLKIEQTEAP"
FT                   /product="putative outer membrane protein, probably
FT                   involved in nutrient binding"
FT   CDS             complement(39047..39907)
FT                   /db_xref="SEED:fig|6666666.127265.peg.29"
FT                   /translation="MMEVDKKKTRTDEAWVRLYARFETDCLLPADAEHAPAMHRRLYLK
FT                   WGSVAAAVIAGIVCFAAWWAVPEQGGDSRSLLTEENREKPTLVKTLEDGSVVYLAQEST
FT                   LKYPEHFAEDKREVNLQGEAFFDVAKKPEQAFTIETARVRVEVLGTAFNVRSNEGAPFS
FT                   LSVKRGKVKVLLKQEEQTVFVEAGETVILQGGSLLVSETENPELFDRYTSNIRFKDECL
FT                   EDVLRAINKESAGWQIEAASPTLGKRRLTVEFSNNSPESVAELICWTFDLKCTRQGNRL
FT                   ILSEQ"
FT                   /product="putative anti-sigma factor"
FT   CDS             complement(39904..40455)
FT                   /db_xref="SEED:fig|6666666.127265.peg.30"
FT                   /translation="MLDELMILARIKEGDIKAFEGIFRRYYSPLCWYAAGITGDTASAE
FT                   EIVEELFYVLWKNRERLQVFQSVKSYLYRSVRNEAVQFCEHQEVKERYRASVRTESGPG
FT                   YSSDPHAHLEYEELRTLIRHTLDKQPRRRQQIFRMHRMEGKKYAEIALSLSLSVKTVEA
FT                   EMTKVLRTLRNEIDNYIRTQ"
FT                   /product="RNA polymerase ECF-type sigma factor"
FT   CDS             complement(40507..41700)
FT                   /db_xref="SEED:fig|6666666.127265.peg.31"
FT                   /db_xref="GO:0004455"
FT                   /translation="MAQDAAVNELVITGTVKNKDSRKKLENVNVSVVGNNIGTVTNADG
FT                   TFSLKVAEAEAFRGLEVSHIGYLTTHLSLEELEKTGGLTIWMIPAPNLLSEIVVYGNNP
FT                   RVIVEEAIKKIPVNYSGNDNMLTAFYRETVQKRRRYISVSEAVMDVYKTDYNSRDVDRD
FT                   KVQLLKGRRLLSQKQSDTLAVKVVGGPNLSLYLDIVKNGDALLSTDNLDYYEFRMEDPV
FT                   NLDNRMQYVVSFRPRVSLMYALFIGKLYIDYERLSFTRAEFGLDMANRVKAVEAILHKK
FT                   PVGLRFRPQEVTYLVTYKQQGAKTYLNYIRNVIRFKCDWKKRLFSSSYTAFSEMVVTDR
FT                   TESPLSGISNKNTFKRKQVFYDLVDEYWNEDFWRNYNIIEPTESLENAVSKLRKQSN"
FT                   /product="Ketol-acid reductoisomerase (EC 1.1.1.86)"
FT                   /EC_number="1.1.1.86"
FT   CDS             complement(42003..43031)
FT                   /db_xref="SEED:fig|6666666.127265.peg.32"
```

```
FT                   /translation="MSKGWKYGIGLGVVILLLFAGNLLVGSVSIPPADVFRILLGGEGE
FT                   KASWSFILWESRLPQALTALLCGGALAVCGLMLQTAFKNPLAGPSILGINAGASLGVAF
FT                   VMLLFGGSITAGAFSLSGFFSVLLGAFIGAMLIMALILFFSTLIKSNIMLLITGIMIGY
FT                   IASSAIALLNFFATAEGVQSYMIWGLGNFGGVSLQQMPAFALVTIVGLFGSLLLIKPLN
FT                   ALLLGERYAENLGVNIRRVRNWLLIITGLLTAVTTAFCGPVAFIGLAVPHVARMILGTT
FT                   NHNSLLPVTILSGGAVALLCNLICVLPGEAGIIPLNAVTPIIGAPVIIYVILSQRKPQQ
FT                   FN"
FT                   /product="Vitamin B12 ABC transporter, permease component
FT                   BtuC"
FT   CDS             complement(43113..43931)
FT                   /db_xref="SEED:fig|6666666.127265.peg.33"
FT                   /db_xref="GO:0005381"
FT                   /db_xref="GO:0006827"
FT                   /translation="MDNLGVFGSIGGVCDLKYIKLPTVHEACRRGTIADCGDGMNPDME
FT                   RIIDLHPDAILLSPFENSGGYGRIEKLDIPIIECADYMETSALGRAEWMRFYGLLFGVA
FT                   PQADNLFAEVDSCYQRLKMRAQLSSTSFSVVSELKSGSAWYVPGGRSTIGRLFQDACGR
FT                   YAFADDTHSGSVPLAFETVFDKAGDADVWLVKYNRDRDMSYADLEADYIGYTGFKAFKT
FT                   RNVYGCNTAKVPFYEETPFRPDYLLSDLIQILHPEIGDLGGLRYFCRLKE"
FT                   /product="Vitamin B12 ABC transporter, B12-binding
FT                   component BtuF"
XX
SQ   Sequence 43948 BP; 10702 A; 11304 C; 11272 G; 10670 T; 0 other;
     cggcctgaag agcgtttcga cccgcctccc ccgaccactg cctatttgat attgattcgc        60
     attacggctc gcactaggtc gtcgtccgga agggggaccg ataggtcatc cgcttttcaa       120
     tgatgccaga tacggcgggg cggagatccg caaggggact atctatgcga aatataagca       180
     gttcatccag aactgtttcg agatttgccc gcgtcaggcg ctgcatgcaa agactctggg       240
     tttcgtgcac cctcgcacgg gaaagccggt cagcttcgat tctccgcttc cggaggacat       300
     gaccgcgctg ctggagaagt ggcgcaagta tgccggcgac atgccgaaag aataaaaaaa       360
     gggggacacc ctttatgaat caagtttgtc cccttttttcg ttgaatagtt cattctgcag     420
     aactgcaaaa tcggtaattt ctaccacttt gatcttgcat ttgtactttt tcttccactt       480
     gcctacataa gagttgaaca ggcctcgggt aatatatgcc cctaaaatag ggcttacctt       540
     cagagtgaag ttcctgattc ccttgtcatt tacgtagtag gaaagccttc gttcaatctg       600
     ctcgtcgatg agcaggcttg atgctatttt tcctgtaccg tggcataccg ggcagacttc       660
     cgtcgtgttg atctcggtta ccggccgtac gcgctgacgg gtgatctgca taagaccgaa       720
     ctttgtcagc ggcagcacat tgtgcttagc cctgtctttg ctcataagct cctgcatgta       780
     cttgaagagg ttgttcctgt gctctaccga ttccatatcg atgaagtcca caatcactat       840
     tcctcccata tcgcggagtc tcagctgtct tgcgatctct tccgccgcaa acttgttgac       900
     ttcaaaggtg ttctcttcct gttcctttgt cttggcacgg atgccactgt tgacgtcaac       960
     aacgtttaag gcttctgtcg tctcgattac aagataggct ccctgtttga gcggtaccac      1020
     ttttccgaaa gaactcttta tctgtcttgt gatctcaaag tggtcgaaga taggttcctt      1080
     tccctgatag agctttacta tcttctcctg ttccggagaa ataagggaga tataccttct      1140
     gatctcttcg tattctttct cgttgttcac atagatgttc gagaaagagt cgttaagcag      1200
     atctctcaga attgtagtgg tcttgctgta ctctgtaaac agaggctgta tgctcttgtt      1260
     cttcgagaga ttttcccagg aatcttccca tttaccgata agcgacttca gctctgcaac      1320
     cagaatagct gcgttcttgc cttcagccgc ggtgcggatg attgcaccgt aattcttggg      1380
     aagaatactc ctgacaagtg tctcaagtct cttcttctcc tccttggagg agattttctg      1440
     ggatatgctt accttctcgg cgaagggaag cagtactatg tttcgtcctg ccaatgaaat      1500
     ctctgcggtc agtcgtgatc cctttgtgga aataggctcc ttgactatct ggactattat      1560
     catctggccc tgggtgagca cattctcgat ccggccttcc ttttcgagga taggaccgag      1620
     ctttatgctt ccgaatagcc tcgacgggtt ggtattaggg tttgtcttct tgacgaactc      1680
     gtcgaaagcg ccgaaataaa gtccgaggtc cagataatgg atgaaagcct ctttctcgtc      1740
     accgatatcc acaaaggcag cgttgagtgc aggcatcact ttctttactc ttccgaggta      1800
     ggcatcgcct acagcgaagc tgcgacccga gctggactct ttgttcagtt cgataagcct      1860
     gtgattttcc tgtaaggcta tctgaacttc agtggcctga acgtcaacga tcaggtcttt      1920
```

gagtagttcc ttttcagaat ccattggata atttgtgaaa caataacaaa agggccgtcc 1980
agatctcact gtcggccctt ctcttttagc agactgctaa aatggcagac acaggagaat 2040
tatttcttct tgtgtctatt cttacgcaag cgtttttttgc gcttgtgtgt cgccatttta 2100
tgtctctttc ttttcttacc gcttggcata attgaaaaaa agttaaaaaa ttattaaata 2160
ttataccggt aattacttaa ccttgcttac gaataccttt gctggcttga aagccgggat 2220
attgtgagct ggaatggtaa gggtagtatt cttggtgatg ttccttgctg ccttctcagc 2280
acggtgcttg ataatgaagc tgccgaagcc acgaagataa accggttcac ccttcgcaag 2340
tgaagcctct acactttcca taaaagattc aacgactgac tgtactgcca gtttctcaat 2400
accggtagct tttgctactt cattaacgat ctctgcctta gtcataatag agtaatttat 2460
ataaagttat gtgtgtgcaa attttaagac tgcaaatata gacttttttt tgcatagggc 2520
aaattatttc gttaaaaatc tttgtcttac atattttgat attacctttg tgcttggcat 2580
aaaggttgac attatagtat ctgccttaaa aacgccatcc ttctctgaca aaatgtcagc 2640
agggaggatt gtatacactg ataaattatg atagatttta ataaagaaat gtttcctgcc 2700
tcgtcccacg aggtcaatat aataccggta atgtctggag agcccgctct caacttcgac 2760
gcctcggaac tcccgggatc ccttccgatc ctggcactcc gcaacgccgt gatctttccg 2820
ggtgtcgtct tccctataac tatcggccgc gagaagtcta tgcttctcgt cgacaaggcc 2880
cagggaaacg gcatgttcat aggcgcagtc cctcagaaga atgtcgccga ggagaacccg 2940
ctgtctctcg ccgacttctc ggaattcggt actgtcgtaa aggtcctgaa gactttcgag 3000
atgcctgacg gcactgtcac ggcgcttctc cagggctaca agcgtctccg catgctggaa 3060
ctgacggaca ccgatcctta ccacatggca agagtcagct atctcgacga tatcctgccg 3120
cagaacgact cgaaagttcc ggtcatagcc gatatcctca aggagaatgc gtcaagcatc 3180
atccgttcgt cttcgttcgt tcctaaggag gccctcgagg ccctcaagaa catcgacgac 3240
ttcaatttcc tcgtcaattt catcgctacg acgatcgaca tggacgattt catggaaaaa 3300
gtcgacctgc tgaaatatga cgacgtaaag gaaagggcca tggctcttct cggcgtgctc 3360
agccgtcagc tggagatcca gaggctcaag caggagatca gctccaaggt caagaccgag 3420
ctcgaccagc agcagcgcga gtatttcctt aataatcagc tccgtacgat ccaggaggag 3480
ctcggcatgg atgagatgga cgacatcgca cgcttccgcg aggctgccga gaagaaagag 3540
tggcctgcat ccgtagggga gaccttcgaa aaggaactcg caaagctcga gagattcaat 3600
ccttctactc cggaatacag catccagtat aacttcctgc agttcatgat cgacctgccg 3660
tggtatcata cttcagacga caatctcgac ctgaagcatg cccagaaagt tctcgatgag 3720
gaccatttcg gactcgagaa cgtcaaggac aggatcatcg agtatctcgc cgtcctcaag 3780
ctgaagggta acatgaaatc cccgatcctc tgcctctatg gccctccggg agtcggcaag 3840
acaagcctcg gaaagtcgat cgccagagcc ctcggacgca agtatgtccg tatctccctc 3900
ggcggcatgc atgacgaggc cgagatccgt ggccacagga aaacatacgt aggcgccctt 3960
ccgggacgta tccttaacgg aatcaacaag gccggcactt ccaacccggt aatggttctc 4020
gacgagatcg acaaactcgg aagcgacttc aaaggcgatc cgtcttccgc tcttctcgag 4080
gtcctcgacc cggagcagaa cacggctttc cacgacaact acctcgatat cgattacgac 4140
ctttcgaacg taatgttcat aacaacggcc aatactataa acggtatcca gccggctctg 4200
ctcgacagga tggagcttat caatgtgacc ggttatctgg cagaggagaa ggtccagatc 4260
gccaagaagt tcctcgtgcc taagcagctc gaggagcatg gaatccctaa gaaggaactg 4320
aagatcgaca aggctgccct tgaagagatc atcgacaaat atacccatga gtcaggcgtc 4380
cgcggcctgg agaagcagat cgcccgtatc gccagagtct ctgccaagaa gatggccctc 4440
ggcgaggact atccgaagac cataaagaaa gagcatctca aagagtatct cggcctgcag 4500
acggccttcc acgacagcca gaagggcaac gaggctccag gagtcgtgac cggtctcgca 4560
tggacctcca tgggcggcga gatactgttc atagagactt ccgtcagcga tggcaaggga 4620
gtcctgacaa tgacgggaaa tctcggcgat gtcatgaaag agtccgctac tatcggctat 4680
cagtatatca aggctcatcc gcagctcgcc ggccttactt cggaagagtt cgcgaagaag 4740
gatgtccatg tccatgtccc tgaaggagcc gttcctaagg acggtccttc ggccggaatc 4800
accatggtca gctcgatggt gtcagccctc cgcggaaaga aagtccgcag cggaatcgcc 4860
atgaccggag aaatgaccct gaggggccgt gtgctgcctg taggtggcat caaggagaag 4920
attctagcgg caaagagggc cggagtctct accataatca tttctgaaga aaacaggaaa 4980
gatattgaag atatcaagga aatttacgta aaaggtctta cctttgtgta tgtcaagaca 5040
attgacgacg taatcggata tatcttcgaa taatggacgt taagatagaa cagacttgga 5100
aagacgcact gcgcgacgag ttcgaaaaac cgtacttcgc ctcgctggtg cgttttctgc 5160

ataatgaaaa agaggccggg aaggtcatct atcctcccgg aggcctgata ttcagggcat 5220
tcgaactcac tccgctgccg gaggtgaaag tcgttatcct aggccaggat ccgtatcatg 5280
ggccccgcca ggccatggga ctttcgttct ccgtgccgtc ggacgtgccg gcccctccgt 5340
ccctgaagaa tatattcaag gagatccaga cggatctggg cataaccatg agcggaagcc 5400
cgaatcttga aggctgggcc cgtcagggcg tgcttctgct gaacgccatg ctgacagtcc 5460
aggccggagt cgccggctcg cacaaggcta tcggctggca ggagttcacg gatgccgtga 5520
tcaagtgcat ctctgaccgc tgcaacgggg tcgtgttcct gctctggggt aatttcgccc 5580
gcggaaagaa ggaactcata gatacgtccc gtcattatgt gctcgaagcc gcccatcctt 5640
ctccgcttgc cgggggagcc ttcttcgggt gcagacattt ttctaagact aacgaacttc 5700
tcataaaaga gggcaaaacc cctatcgact ggaaattatg aaacgtaaag cacttttccc 5760
gggatctttc gatcctttta ccgccggcca tctgaatatt ctccgcagag cgctgacgat 5820
gttcgatgaa atcgtcatcg cagtcggtgt caaccaggac aaacgcggat tctttagcat 5880
ggatcagcgt atggatatca tacgtcaggc tgtccgcggt atggacggga tcgagataat 5940
caaatacgac aaccttacta tcgatacctg ccgcgagctg ggtatcgccc atataataag 6000
gggcgtcagg aacatgcttg atttcgagaa cgagcgtgcc attgcggatg ccaaccgccg 6060
tcttgctccg gaaatcgaga ctatcatcat ccctaccgca caggaattcg cgcatatctc 6120
gtcttcagcc gtcagggata tccttaaaca taagggggat acttctctct tccttccgga 6180
gggagtcgtc cttccggatc acgatgcgta ggttaatcct cataatcttt ctttctctgc 6240
tttcttttgc cttgggcgcc cagaatgcgg attccgtcaa gatggccgcc ctggatgcaa 6300
agcttgagga gtattttgcg gcgctggaga ccgagtctct ggatgtaaag aggaccgagt 6360
gcgatttcct catttcttcg tgcacgactg acgatgtacg aaaccatgtc gccgtacgtc 6420
tttactcgca tttcatgcag tcgaagctta tgggggatga ggctgtcgcc gtccatcttg 6480
ccgacaactg gttcattccc ggcaaggttt ctttcccgga tgagatagat ctgatgaatg 6540
cgaagatttt cgctgagttc aacagacagt cgctgctcgg atgcaaggct cccggtctgg 6600
tagccgagga tagtctcggc aggaaggtcg aggtccctct gggattgtct gacagatacc 6660
ggattcttta tatctattct ccggactgtg ccacatgcag tcttcagact atgctgatgg 6720
actctctttt cagcaggaag agctatccgg tcgaccttat cgcattcaat tcagacgata 6780
atactgaagc ctgggtgcag taccgcagga accgcctttc ttttgacgga gtccgtgtca 6840
tgcatttcag ggatccttcg ctggagtcgg atttccagcg gaaatacgga gtgcttcaga 6900
ctccccgcat gttccttatc gacaagaacg ggattatagt aggccgcgga ctggacgctg 6960
ctgcgctgga gaaattgctg gatgcaactc ctgaccttgg catagggtat ggttcgaagt 7020
cgtctgccga ctatttcgac gctgtgctgg agagcttcgg ggatgagctg cgcggctctg 7080
atatccgcta ttttatcgga agacttgaag aggctactct cgaaagcgcc gataccgcag 7140
gattccgtca gatgaccggt gacatgctgt attatcttgc cggaaagagg ggagaggcgt 7200
tcaaggatgc cgaggccttc gtcgtcgatt cgatggtcct gtcccgtccg gatgtgtggc 7260
gctttgccga tgattcgttg acggtcgttg acatggccat gagcctgaaa catatgctct 7320
cgctagctcc agtaggtgcc agggttccgg atgtacgtgt ccatgccact gtccttcgcg 7380
ccggacgaaa gccggccacc gtccataaga aattgcggtc gttgcttaag ggaaagcgga 7440
attatgtgct tttccatact gccggctgca ctttctgcga ggctgaactt gccgccgccg 7500
attctgtcgc ttcggagaag ggtgtccgtg tcatcctggt cgatatggac aagctgacaa 7560
gcttataccc ggaggaggct gatcggcttc tggacgcatt cgacctgtcc gttctcccgt 7620
tcataatgga gacggataat aaaggttgtg tcgtacgtaa gtatctgagt ttaaataagt 7680
agggctgcct ttaccttagc tacggcctct ttgcctccca tcttttcgag gatggcgagt 7740
ccgaagtcta tcgcgaaggc tgcgcctctg ccggtgatga gctggccgtc ggtaactgcc 7800
ggggctttta cgtattccat tccgctggcg gcgaggttct cttcgaagcc gtcaaagcag 7860
gtggctttct tgccttcgat tcccttgatc tgggtggcga cgaggcccgg ggcggcgcag 7920
attgcggcca caatgcctcc gttgccgtag tggacgttca tcagaccgat aagcggggtg 7980
tcggctgcga gtttcttcga gcctggcatt cctcccggga agatcatggc gtctttttcg 8040
tcgatatcta tttcgctgaa gtcggagcgg cccatgtctg ccaccaccgt tactccgtgg 8100
gagctggtga tatagacttc ttcggtgatt gatactgtgt atgcttcgat tttacctctg 8160
cggagaatgt cgagggttgc gagggcttcg gtctcttcaa aaccgtcagc caggaagatg 8220
tatgcctttc ccatgatgct catgtggttt agattggttg aggtaagagg gttcgaacct 8280
ctactgaggg aaccaaaatc cctagtgcta ccattacacc atacctcagt ggggtgcaaa 8340
tatacgaaaa aattatttgc ggacaacaat tctctctata cgtctaggga cgctagtgag 8400

aatttcgtaa gggattgtcc cgagaatgtc ggcgagttcg ttgactgtag ggtcttctcc 8460
gaagatagtc acggtgtctc cgactttagc gtcaattcct gtcacatcaa gcatgcacat 8520
gtccatgcag atgttcccga tcgtcggcgc tctgtgcccg ttgacggaga atgaggcgtg 8580
accgcgtccg agatggcggt cgatgccgtc ggcgtatcct accgggatcg tggcgattac 8640
ggtgccttca ggggcgatat gaccgtatct tccgtagcct atcgtgccgt cttccggctt 8700
gagggtcttt atctgcaata ttttgcattt gagggacgcg acgtgggcga ggcggacatc 8760
aggcagggcg ctgattccgt agataccgat tccgaggcgg accatgtcga actggtattg 8820
cgggaatctt tcgattccgg cggagttgag gatatggtac atcggcttgt agcctatctc 8880
ggaggtgatc ctgtcggcgt ttgtcttgaa catgccgatc tggccgaggg tgaattcgtc 8940
ctgctcaggg tcttcggata ctgcaaggtg cgaatatacg gacttgactt tgacttcttc 9000
gcaggtcttg aggaagtcgg tgagatcctg gagctcgctc gtcatgaacc cgaggcggtg 9060
cattccggtg tcgagtttga tatggaccgg ataatccttg atgccgcgtt ctttgagtat 9120
tccgcagagt atcttgaggg cgtacaggtt cgggattccc ggctcgaggc ggttgtcgat 9180
gatttcgttg aagaagtccg tgcctgcggt caggacgagg atcgggaggc tgataccggc 9240
tttgcggagt tccatgcctt cgacagggta ggcgactgcg aggtagttcg ctcccgcacg 9300
ctgcatcatg gctgcgaatt cgacggctcc gtgcccgtaa gcatttgcct ttacgagtac 9360
aagtaattta gtcgtctgct ccagtttgga cctgaaatat ctgtagtttt ccatacatcc 9420
tgtcaggttc agctccagcc ttgaaaaatc atcaaccctg ttcattccac atcaatttag 9480
aatacaaatt tatcaaattt ctccgacata ggttacagtt tttcgctttt ctttccccgt 9540
aaagaccctt acagtacgtc cgctgtcgcc gccgacagcg tctgctttgg cgggaatcct 9600
cgtggcctcg gctctgttcc gccgactctg tcggctccat ccctcccaat gtctacttcg 9660
tcctgcttcg cagaactcgt atccatcggg cccctccctc taacgtggcc gagggtggcc 9720
acgctcgccg agaccagagg acagccgcca tcaattaaga cgctgacggc aagtgagagg 9780
ttggcatgga tttggaagca accagaagga ttgttaaaat tttatggata tgatttggtt 9840
tattatcatt gctgtgatgg tgctgagctt tgtgattcag caggttctga acagtaaatt 9900
caataaatat tcgaaggtgg gattgcattc cgggctgacc ggggcgcaag tcgctgccaa 9960
aatgttggaa gaccatggca taaccgatgt ctctattacc tgcatcaaag gaaagctgac 10020
agaccattac aaccctgtca acaagacagt caacctcagc gaagaagtct attacggcaa 10080
ttcggtagcc gctgcggccg ttgcagccca tgaatgtggc catgtagtgc agcatgccac 10140
gggatatgct ccgctgaaac tgagatcggc tctcgttccg gtagtctcat tctccagcaa 10200
tgccgtgcag tgggtacttc ttattggcgt aatgatcatc aagattactc cgaccctcct 10260
gtatctcgga atcgccatgt ttgccttgac gaccttgttc agcgtcgtga cacttcctgt 10320
ggaaatcaac gccagcgcca gagccgtgca atggctcgac aaggccggga taacaggtcc 10380
cgctacaaaa gcgaaagctg tcgatgccct caaatgggca gcatatacat acgtcattgc 10440
cgccatcggt tcgatagcga cattgatcta ctacattgcc atggccaggc gacgctagag 10500
actcgaaaga atccttgcaa gctggtccgt gcaggatgta ccacggaaat tgcagttgac 10560
gcccgaaaga gtgtcaacta ttttttttat gtccatgccc tcgacgagac ggccgatcgc 10620
ctgaagatta ccgttgcagc cccgcacgta tgcgagattg cggaccttgc cgtctattat 10680
gtcgaaatcg atctgctggc tgcataccag gctgctcggg atggcagtta tgtgtctgac 10740
accgttctcg gtggtgtcgc tgatgatctt gtagtcgtct cctgtcattg ttgtagtccg 10800
tttttttatct ttttttcttg aaaaactccg tcaccatgga accgcattcc tccgcgagca 10860
ctcctgcctc gacctcggtc ttcggatgca gcagcgaagg agtgaaaagt gagtgccccc 10920
gcttcggatc gatcgctcca tagacgatcc ggccgatctg ggcccagttc agggctccgg 10980
cgcacatcgg acacggctcg accgtgacgt agagcgtgca ttcgttgagg tatttgcccc 11040
cgagcgcctc cgtagcggcg gtgatggcta tcatctcggc atgggccgta ggatcatgta 11100
aacgttcggt catattgtga ccctttccca cgatccggcc tttgcatacg ataactgccc 11160
cgatcgggat ttcatcctcg gccagggccg cctgcgcctc gcgcagagct tccttcatat 11220
atagctcgtc tttttccata attacagcaa agatagcaat tcttggacaa tcccaatagt 11280
tgataaaaca attaagaaaa tttgcgaaat gatacttttt tgacattttt atcatatatt 11340
tgccacgtga taatataagt taatggatat gacctcggta ataacggaaa taacacctct 11400
ttcggacagg gactgcttct acatagtcga gagacataag accgaattcg tcttcccccct 11460
ccacaggcac agggaatacg agctgaactt catcgagcat ggcaagggcg tccggcgagt 11520
cgtaggcgac agtgtcgaag agataggcga atatgagctt acgctgattg ccggcgggga 11580
tcttgaacat gcctgggagc agggcaattg tacctccgag gacgtacgcg agatcactat 11640

ccagttcgat ccggagatat tctccagcgg cctcatgatg agaaaccagt tctcatcgat 11700
acagcatatg ttcgagcagg cgaggaacgg gctgaccttt tcgctggcgg ccataatgaa 11760
ggtgtattcg cttcttgacg ccctgccgtc ggaaatagat aaattcgatc agttccagca 11820
ctttatgagg ctgctttatg acctttccct ggcgggtgac tccaggatac tggccagcac 11880
gtcattcgcc cataacgatg ccgacaagga aagccgtcgg gtgcagaaga tcaagcaata 11940
tgtcggagac catttcgcgg aaaacattac ccttgaagaa ttggcggaaa tggtcggcat 12000
gagcccgtcg gcgttcagcc gctttttcaa gagcaggacc gggaagacgg tgatagacta 12060
tataatcgac atccgactcg ggaatgcggc aaggctgctg gtcgacacca cccagaatat 12120
ctctgaaatc tgctatagct gcgggttcaa caacctgtcg aatttcaatc gtcttttcaa 12180
gtcaaaaaga ggctacacgc cgcgcgaatt caggactctg ttcaagaaga acaaggtttt 12240
tgtataacgt gcaaaatagt atcggttatt gacaaatttg tatttgttgg gtaaaacgtt 12300
gcatgctaaa tttgcaacgt actcagacaa gtatggattt atcgataaac cgcatgatta 12360
aaagattatt cccggcactg tttatcgccc ttttcgcgat tctttcgtgt cagcctatga 12420
aaatggacat cgtcaaagtt gacgatggct cttttgtcgt tgatggtcac ttctattatt 12480
atatcggtac aaatttctgg tacggaggac ttctcggctc cgaatatggt gacagaagca 12540
ggcttgccgc cgaacttgat tccctgcatg ccctcggact ggacaacctc cgcgtactgg 12600
taggcagcga cggcccgac ggggtggcga ccagagtcta tccgaccctg cagaaggaac 12660
cgggcgtcta tgacgaaaag gtcttcgaag gtctggacta ccttatggcc gaacttggga 12720
aaaggaacat gagggcagtc ctctatatca ataattcctg ggaatggtcc ggcggctatg 12780
gcatgtacct cgaatgggcc ggagccggca aggcgataat tccttcggtc ggaggatatg 12840
gtccgttcat ggaacagatg gcccagtatg cgacgaacgc cagggcgcag gaactcttcg 12900
agaatcatct ccgtactgtc gtttcaagag tcaacagcat caccggaaag ccttatagcg 12960
aggatccggc gatattctca tggcagatcg gcaacgagcc gcgctgcttc tcctccgatc 13020
ctgccgtcca ggacggattc gtatcatgga tatggcgcgc tgcctcgata atcaaatcga 13080
tagaccctaa tcatatggtt tctaccggca atgagggcat atggggctgc gaagagaatc 13140
cggagctatt ccgccggctg aatgaatgtc ctgacattga ttatattacc gcccatatat 13200
ggccatacaa ctggagctgg gtaggaaagg aatcccctca gctggatata gaccgcgccg 13260
aggagaatac cgggagatac atcgacgccc atctggaggt ggccgagagt ctgggcaagc 13320
ctctcgtgat cgaggaattc ggcttcccga gagatgatat gcagtttgcc aaagggactc 13380
ccgtaagcgg acgcgaccgc tattacagat acgttttctc caaggtcatg gagaatcaga 13440
aaaatggcgg cagactcgcc ggcgccaatt tctggagctg gggaggtctt gcggtgcctg 13500
cccacagata ttggcaggaa ggagacgatt acgccggaga tccggcccag gaagaacagg 13560
ggctcaattc tgtattcgcc gacgattaca gtacccttgc tattataaaa catacaacgg 13620
cgagactttc tgccgtggct aatgcaagcg cagtgcttga aaatgattgg atgttcacta 13680
ccggagacca gaaacctctc agagttgctg tacagtctca gaatggtttt aagggaagtg 13740
tcccggttta tgtgcgggtg tcaacagaca aaggggtttc agtgtcggtc tcaggaaaga 13800
aagcccgtct cggcaaaggc gccgatactc tggatttcga tcttggacta gccccaggct 13860
tttataaggt gtgtgtcagc ctgaacgaga atgacccgct tgatgaaaac ttctgtgtcg 13920
gctgtgatcc tgagaagata gtatctcctc aggataaaca gcctgatttc gatgaattct 13980
gggaagacaa cctcaggcag cttgccatgg tcccgatgaa tgccaggatg actccgttga 14040
aggatcattc aaacgctgac aggactgcct acagggttga tatgaagtct ttcggaggca 14100
aagagatctc cggtctgctt tatcttccga acaaagaggg aaaattcgct gccaggatct 14160
catatatggg ctacggttcg gacgtctggt tcgctgatcc gtcggcggaa ccggacatga 14220
tcgagttcac tctcagcgtc cgtgaccagg gtttcaacaa gactccggga caaggctggg 14280
tgacaagggg acttgccgat aaggactcgt attattacag aggcgcatat ctggactgtg 14340
tgcgggccat tgatttcgtc tcttctcttc ctcagactga tccggaccat atctacgccg 14400
aaggtggaag tcagggcgga gcatttaccc tgatagctgc atctctggac aaccgtctcc 14460
gcgcaatcgc tccgttcgtg cctttccttt ccgatttccc ggactatttc gccatcgcag 14520
actggccggg cggggaagtg ctctcggcgg caaaggctca gggcatttct gaagacgaac 14580
tttataagac cctgtcttat ttcgacgtga agaactttac ggacaggata ggttgtcctg 14640
tgcttatggg tttcggcctg caggatcccg tctgtcctcc tcataccaat ttttcggggt 14700
ataaccatat tactgcagat aaacaatggc tttgcgttcc tgtctcaggc caccatctgg 14760
agcaatttcc cgagtggtgg gcagcaagga cggaattctt cacaaaaaat aaataacaca 14820
actaattatt cacattcaaa atcagtagca tgaaaagaca gctattaaca tgtatgctga 14880

tgctctttgg agcattgtca cttatggcgc agaacaaagt gaccggaaga attacagata 14940
attccggaca gggcgtcata aatgccagcg taatggtgaa aggcaccaac aatggtgtcg 15000
ttgcagatct tgatggatac tacgagattg ttgtcggcca aggctccact cttgtatatt 15060
cttccatcgg ttacaaagat caggaagttg ttgtaggcac ccagagtgtc atcaacgttc 15120
ttcttgtcga aaactccgag ctgctcgacg aaatagtcgt agtcggttac ggtactatga 15180
agagaagcga cctcgccgga tcatccgttt ccatgaagga ggatgacctc aagggttcga 15240
tcatcaccaa cctcgaccag tctctccagg gacgtgctgc cggtgtcacg gcagtccaga 15300
cttccggagc tcctggttcg tcctcaagta tccgtgtccg cggtcaggcg acaatcaatg 15360
ccaacgcaga acctctctac gtcatcgatg gcgtaatcat gcagggaggc ggtaactccg 15420
gtgcagactt cggtctcgga gatgctctcg gcaatggtaa agtctctacg atttctccac 15480
tttccactat caaccctgcc gatatcgtaa gcatggaaat cctcaaggat gcttccgcaa 15540
ccgctatcta cggtgcccag ggtgcgaatg gtgttgtcct cattactacc aagcatggct 15600
ccagaggtga tgcgaaattc tcatacgacg gtatgttcgc cgtttcaagg cagacaaccc 15660
gtatcgatat gatgaatctc cgtgaatacg cagagttcta caactctatg gttgaaaccg 15720
gagaaatcag tgaggccaat ccttactatg ctactccggg cctgctcgga aagggtacca 15780
actggcagga cgaaatcttc cgcacagcca tccagcacca gcatcagatc agcgcccagg 15840
gcggtacgga caaagtccag tactatgttt ccggatcatt catggatcag gagggtacta 15900
tcaggggttc cgaattcggc cgttatacct tcaggaccaa ccttgacgcc cagttgaagg 15960
aatggttcaa gctcggtctc aaccttacct ttgccaacac caatgatgac cttaagcttg 16020
ccgacggtag cgaaggtctt atcacatact cccttaccac aatccctgac atccctgttt 16080
atgatgtcta tggcaatttc tcatctacag tacgcgaagg ctatacttct ccgaaccctg 16140
tagctcttgc aatgatggat gagattctgc tggaccgtca gaaactcaac ggtaatatct 16200
acgcggaact tactcctgta aagcatttca cctggagaac agaactcggt ttcgatgtca 16260
gcggctctac ggctacccgt ttcagaccaa cagtcaatct cggtggctgg acaagagata 16320
ccaactctat gagctatcag aagaacagca acagattctg gcagatcaag aactatctca 16380
cttatgccaa ctccttcgga aagcacagcg ttaccgccat gcttggtcag gaggcctggg 16440
agtcttcgtg gagttacgta cagggcgcaa ataccaagct cccttcaaac gatgtccata 16500
acgtcgccct cggtggagga actccgtcgc tttcttccgg tttcggaagc tctgcgatgg 16560
cgtcattctt cactcgagaa acttataatt tcgacgatcg ttacctcctt acctatacct 16620
accgttatga tggatcatcc aacttcggtc ctgagaaacg ttgggccggt ttccattcat 16680
tggcagtggc ttggaaattc accaatgaga aattcatgga tttcctcaag cctgtggtca 16740
acaacggaaa actccgtctc ggatgggggc agacaggtaa ctcaagtatc ggtggttatg 16800
catggggagc atccataagc cgtatgcctt cggctctcgg catgggttac cgtcctgcta 16860
acatccctaa tcctgctatc cattgggaga gccagacgca gcagaacatc ggtctcgacc 16920
tcaatatgtt cgacaacagg ctgaatctta cagtagatgc atacctgaag aagtctgaca 16980
acatgcttat gtcaatgcag cttgcttcct atatgggaac ccagggtaac agttcttctg 17040
ctcttgcagc tcctaaaggt aacttcggtt ctatcgagaa caagggtctt gaaattactc 17100
tcgatgccca tcctgtccag atcaaggatt tcgcctggga cagcaacttc cagatttcct 17160
tcaacagaaa caagcttgtc agccttgacg gaacgactaa tgccgctctc ataggttacg 17220
gacagtggtc tgacgttgtg tgtgtgtctg aaatcggcaa gcctctctat aacttctatg 17280
gttacaaggt cgctggcgta tatacttctc tcgaggatat cgagaattct ccgaaacctg 17340
agaaatatcc ttctaatgga gagtatgccc gtggaaatac cgtgtgggta ggtgatatca 17400
agtacgaaga tatcagcaag gacggagtaa tcaatgaaaa ggaccgtacc gatatcggca 17460
atcctcttcc taagttcacc tttggctgga gcaatacctt tacctacaag gacttcgacc 17520
tcaatatctt catcaacgga agctatggta acaaagtcct caactacaat gccatttctc 17580
ttacccatat gaacagcgca tggaccaacc agatcaatga cgctgtctcc ggcagggcaa 17640
tgcttgctcc tatagacgca agcgtagtct atgctgacgg ttccaagtgg tatgaccata 17700
tcgacaatgt cagggtcctc aaccctggca cgaagactcc tcgtgcaagc atcaacgacc 17760
ctaacgacaa cgaccgtatc agcgaccgtt atgtggagga tggttcatat atccgtatca 17820
agaacatcac tctcggatat aatgtcccta agtctttcct gaacaagttc aaggtggata 17880
atgtacgtgt ttacgtcaac atccagaacc tttatacttt cacgaagtac accggatatg 17940
atcctgaagt aggtgccagc acccaggatt ccaccggtct tacctatggc gttgataacg 18000
gacgttatcc ttctcctaca gtatattctt gcggtttgaa cattacattc taaaattgaa 18060
ggaatatgaa aatctctaat atgatgaaag gttttgctgt ggcgaccgca gtctttgctc 18120

tgacttcctg cgaagggttc ctcaacagac ctaacgaaga taattataac gtagataatt 18180
tctacaagac cgacgagcag tgcgaacagg gcgtgaatta cctgtacaac tctccatggt 18240
atgacttcca gagaggcttc atcaagatcg gtgaggtcat gtccggaaac atgtactggg 18300
gctcgtctcc ataccttgat ttctccacca acggtaccga ccaggacctc gtcaacatgt 18360
cctattctct ctgggctgtg aacggccatg ccaataccgt gatctacaac cttcttaatt 18420
cgcagggccc gtccaaggca gcgatcgaca agtgtatcgg agaagctctt acctggaagg 18480
cattcgccta tttcttcatg gtacgaacct tcggtgacgt tcctatcgtc cacagcaatt 18540
ccgatctcct ctccagcggc gagtacaact ctatctccaa ggtggagcgc gccgatgtct 18600
atgagtacat catcacgacc ctcgaaaagg caatggaaat gcttcctaag aagacttccg 18660
gcttcaataa ttacgagcgt atcgattact atgcagcaga ggcccttctc gcaaaagtat 18720
atcttaccaa ggccggactt tccggatcct tgaaccagga tgacctcaag caggctgcca 18780
agtatgccaa ggatgtgatc gacaattccg gacgtaccct tcttcctgta tattcggacg 18840
tcttccgtct taccggattc aacgccacag gcgagtgcct tatctcctgg gaatggagcg 18900
taggccgtaa cccttggact cagcagaata ccctccagag cgacctcgca atggtcggat 18960
tcagcgacca gggagactgc tggggaggct ggggcggtcc ttcagtggat cttgcagacg 19020
cattcggcgt aagtgcgacc caaaatcctg cagagcgtgt aggagagaag gacagccgcc 19080
gcaaggcaac cttgatgctc ccgggtgaca agtatgatta cttctggact gacaagggag 19140
gtttcgatta tctccgcttc atctatgata cggaatatgg caaaggcggt cctggcggtt 19200
ccatgcagtg ccctacaggt tccaacaacg taaagcatct ctatggcgat aactttgatc 19260
atgagaacgc cggtctggga acccctgaga acatggctta tcagcttcct actcatcttc 19320
tccgtctttc cgacgtgtat cttatctacg cagaggcaag ccttcttacc ggaagcaacg 19380
gccctgctct cgagtatgtc aacaaggttc gcgaacgtgc tggcgccgat gctctcgcat 19440
ccgtcacttt cgaggatatc tggaaagagc gtcgtcttga gcttgcaggc gagggcgacc 19500
gctggtatga ctatgtacgc cgttcatact acgatatgga cgcatgcatc gctgaactca 19560
aagctcagag aaggtcatac tgggatggac tcgacgatgt ttggaaatca tatatcggag 19620
atgctttcga ttattccgga acctgggatg cttcagaagt acagtacaac aacgcagagg 19680
acaatcctaa cgttaccgct gcaagcttca agcttccgtt ccctacagag gacgtagtgt 19740
tcaatggcaa catggcatct gacgccgtgc ctgtccatgt cgatgtccgc gaggaataca 19800
gctataactt ttaatctgac gaaaagatga aatctataaa tagaataatc accggttcac 19860
tggcaatcct cgcaatggcg tcattcgctt cctgcgacaa ggttgactat ccggaccgtt 19920
atgaagtagc tgacggcaag ccgacggttt attccgtcag atacgcagac aaggacgtca 19980
acatcaccca ggcatatatg gatgaggtcg tatgtctcct cggcaagaat ctcaagagta 20040
ttgtggaact ctggttcaac gaccagcagg caaccctcaa taccagcttc attacagaca 20100
atactcttct ggtatccgtg cctaagaata tgcctgtggt acagacagac aagatatata 20160
tgattacttc gggcaaggat acgctcgcgt atgatttcgc cgtgcttcct ccgactccga 20220
aggttacttc catgtccaat gagtgggctg cggaaggggа gaccgtcaca atctacggcg 20280
acttccttat tgacaaggct gacaagcctc tccagatagc ctttgcagga gccgaggtcg 20340
atcatgcttc aatggtgttc aacggttcgt cttcagtcag cttccctgtt cctgcaggag 20400
cccagcctgg atatgttacc gtgacttcac tctccggtac cggaaagagc aagttcatgt 20460
acaaggatac cagaaacatc ctcttcgact gggatggcag ccacggcggc tttgctaccg 20520
gtcacggatg gcgtaacggc gtgatccacg ctcctggcgc tgatgcatgg gatgcgatcg 20580
acggttcata cctctacttc ggaggcgccg atcttggtcc tggaagtgca gatgcatggg 20640
cagaggacca gtacagcttc aactactggc cagaggccgg caccgattac gatgccctca 20700
acctccgtcc ggagttcgca gagtatatcg acaagtatgg agtaagcggc ctcgtgctca 20760
agttcgaggt tctcgttcct tcatccaacc cttggtctgc ttcaggtatg cagctcatgt 20820
tctcgagcga cgctgtcgtt acctacggta ccgctacgaa cagctacttc agtgacgaaa 20880
ccttccctcg tgctatctgg cagccatgga ctaccaacgg aggttcttat gataccgctg 20940
acaagtgggt taccgtaagt ttccctctca gctcattcaa caagaaccag gaaaacaagg 21000
agtgtgcaac cgcatttgac aagtcttacc ttaccggtct cacattcttc gtatggaacg 21060
gcggtatcga cggcacagcc tgctctccgg tcttcgcgat cgacaatatc cgtgtggttc 21120
ctaatgaata atattaaatg aagctcatta tgaaaaagat attcagtatt tcagccattc 21180
ttgccgcctg tgtcctggca ctcgggctct ccggatgttc atccaaagag cttgataccg 21240
accagtattc cgacaaggac gtgaagctgg catcattcgg ccctaacccg gtgatgcgtg 21300
gagctacact ccggttcttc ggaagtaatc tcgacaggat tgtcgaagtt tcagttccgg 21360

gactcgatcc tatcaccgac atcgaagtcg ttacctccgg taatccttca gaaatccgca 21420
tccagcttcc tgcagagggc gcacaggtcg gaaagatctc tctcaagtca tccgacggaa 21480
aggtctttac tacccaggtg gatgtagaat atacagaacc tatcgtattt gacagtttct 21540
ctcctgcaga ggctatgcct ggcgacgtga taacggtcaa gggcgattat atgaacctcg 21600
tgcagagcgt taccttcgaa ggcggcgcca tcgtgaccga aatcaaccat aacggccgtc 21660
atgaggcttc cttcaaggtc ccttcaaagg ctattaccgg aaagatcatc ctttctgacg 21720
aaggagccat cgccaatctc ctctactcgg aggaagatct tgtgatcggc gatccgaccg 21780
tcaagtctct taccgtagct gttgctaaac ctggcgagaa cgcagttgtg aagggagcat 21840
atctcgatat gatcgacaag gtcgtattcg aaggtggtgc agaggtttcc gaccttgctc 21900
tcaatgagga tgcaaccgaa ttcaatctcg taatccctgc cggagcacag tccggtgaag 21960
taaaggctgt cagccacgct gccaaggaat tcgtcgccgg cgaaatcgaa cttgtcatcc 22020
ctacggatgt cgcagctgca ccttcgcctg taaaggctgg tgccgacctc gtgatttccg 22080
gtaacgatct tgatgtggta acttctgttg atttccctgg cgccagcgga gtagaattcg 22140
tatttgccga ggacgccatt acagtgaccg ttcctgctac tgctactgaa ggtgatatcg 22200
ccctcaatat ggccagcgga gaagctgtga atgtggcatt caccсttgtg cacccggtag 22260
tttctgccat agctcctgtc gagctcatgg ccggcgagac catcaccgtg acaggtaccg 22320
acctcgacct tgtcactgga gtaactctcg gcggaaagcc ggtcgagttc actgcagagg 22380
ctgagaagat tgtcgtacag accgccaata cttccgtttc aggaaagctc gtgctttctc 22440
ttgcgaatgg tgagaagatt gaaccggagg atgccatcac cctcagctat gattcattca 22500
tcatcgtgaa cgacatgcct gccgcagagc atatcggggc tatcgtaacc ctcaagggcg 22560
agaacttcat gatgatcgag aacatctata tcggcgaaga gaaagtatcg aagtacaacc 22620
ttcgttctga taatgaaatc cagttcgtaa tgccttggaa caaggtcgga acatacagca 22680
tgtacttcca tcttctctcc ggcgatgtcg agacatgtcc tcagacaatc gaagttctcc 22740
tcgagcagga catcatgact gtatgggaag gctctctgca gatcacctgg aatgacggcg 22800
gccgtgtaac cgtgccggca gcatcattca ccggtatccc tgcaggcagc aagctcagat 22860
tctactacac tcaggtggac cagcagtggg atcaggccca gctcaactat ggtaactgga 22920
ccggaatcaa cttcgacgat cctgcaggta ccgtcattaa cggaaccctc gttccgactg 22980
acgtctatgg atggttcact gacggaatcc ttgaccgctg cacagaggtc gtgcttacaa 23040
aggaaatcct cgaaaacatc caggctaaca agggtgactg cgaagacaag actgcctgcg 23100
gtatcatcat ccagggctct ggccttacct tcacgaagat cgagatcgta tgcgatattc 23160
cgcaggaggt tacgatctac gaaggtccta caacccttac atggggtgac gacggccgtt 23220
tcggtctcgc catgacttac ttcgaggcag caaagccggg ctctcagctt atcttctatc 23280
tcgagcagac ggacgcatgg ggccaggttc agctcaacga cggctggtgg gccaacgccg 23340
acatgaactt ccctgagatc ggcggtgcat acattaccac cgacaacctc ggaggaaagg 23400
acgtcacacg tatcgccctt accctgactg ctgatatcct tgaccatata cttgctacat 23460
ccggtgacta tttcggtctc aatagccagt tccagggtga cggacgcgtc gccatggtta 23520
tccagggatc tgatatgatt atcaataaag tttgtatatt gtaacacttt tattgccgga 23580
ggcgggtgat gccgcctccg gtagccagta aacaaatttt atataatgaa agcacacgtc 23640
agtttctttg tattatctat gatttttctt gcggcatgtt cctgctcaaa ggatgatgcc 23700
gaaaagaagg aaccgtccgt cccggcccct aaggcggtaa cggtagatgt cgcgtctgtc 23760
agcgctcccg tagagggaac gtcggtcgtc gtcgccatct cctccccgtc caagcctgcg 23820
gtctctgtct ccggagactg gatcaaagcc gaactgggtc cgtacaatcc ggatacatat 23880
aaagtgaaac ttacggtgac cgtcggcgag aatccgacat acgagtcacg cagcggtagt 23940
gtgacccttа cttgccaggg cttttctccg gtcgggataa gcgtatccca ggatgccgcg 24000
gcgaagccgg tcggaccgtc ggctaatcct gcatacgatc ttctcggtct cggctggaat 24060
atgggtaacc agttcgatgc tttctacaac ggctcatggg ccggagataa atacaattat 24120
ccggacgaga ccgtctgggg taatgccaag gcgacccagg ctactttcga cggggtgaag 24180
aaagccggat tcacatcagt gcgaatccct gtcacgtggc tcaatatgat aggtccagga 24240
cctgagtaca atattgacga gacttggatg aaccgtatat atgaggtggt aggatatgcc 24300
gagaaagccg gactcaaggt catcatcaat acccatcacg acgaaaacaa caatgacgac 24360
cattggctgg atatcctttc tgcatctaag gatgccagga ccaatgaagc gataaaggcg 24420
gaaatcgcgg ctgtatggac ccatatcgcc aataagttca aggataaggg agatttcctt 24480
atcatggagt ccttcaatga aattcaggac ggaggatggg gctggagcgc tgctttccag 24540
gctgaccctt ccaagcagtg tgatgtgctt aacgaatgga atcaggtttt cgtggatgcc 24600

gtacgtgcta caggaggcaa caacgctacc cgctggctgg gtgttcctac atacgccgca 24660
ggtacttcat ggacgaagta tttcagactg ccggaggatc ctgccggaaa ggtcatgatg 24720
gcagtgcatt gctatgatcc gtatgaatat actataggga acatacagta tcctgagtgg 24780
ggccataccg gcgccgcagg aaagaagacc ccggaagggg agaaagagat aacggatctc 24840
ttctccagac ttaatacgga attcatcagc aaggggatac ctgtatatct tggcgaattc 24900
ggctgttcga tgagaagcaa aggggatgcc cgggcatgga tgttctataa atactatatg 24960
gagtatgtct ccaaagctgc gcatacctat tccttaccgg gaatcctgtg ggataacgga 25020
agtactaaga ccggaaagga gtgccacggc tacattgacc atggtaccgg caagtatata 25080
gggaattcta aggagatcgt ggatcttctc gtaaagggaa tgctcagcac ggacgaaagc 25140
tatactctcc agagtgtata tgacagggct ccgaaaacca attgattatg aatctaatcc 25200
ttaaacttag catcatttta tcattagtag ttgctgtatc cggctgcgga cgcaataaag 25260
aagcggccgg aacagctttt aacgaagaaa cccggatcat gatggatcgt atggcggctt 25320
cagtaaaggc cggcaagatt ctttatggcc atcaggatga cctcgtttat ggccattcat 25380
ggtgtgtgga tcctgattcc gcagcgacag atgcgttgaa ccgttcggat gtcaaggctg 25440
tcgccggaga ttatccggct gtcgtgggat tcgaactcgg cggtctggaa ctgggatggg 25500
actggagcat tgacaaagtc ccgttcctgc tgatgcgccg ggctgctgta gagcatgccg 25560
gaaggggagg aatcgtgacg ttctcatggc acccgcacaa tccttataca ggcggtgatg 25620
cctgggatat cagttccgac aaggcagtgg cttctgtcct cgaaggaggg gagaaggcgg 25680
agatgttcgg agtctggctg gagaggctgg gcgatttcat ggactcgctc cgtgacgata 25740
agggtaatct gataccggtc attttcaggc cgtggcatga gaatatcggc agctggttct 25800
ggtggggcgg agccctttgt actgcggatg agtatatcgc ccttttcagg aggacggtcg 25860
agtatctgag aaatgacagg ggactggaca atatcataat gtgttattct ccgaattccg 25920
ggattacggc ggagcagtat atgtcccgtt atccgggcga tgatgtcgtt gatatcctgg 25980
ggtatgacca ttatgagact gtcatggagg gcgagagtct ggaagatgcc ggaaagcgtt 26040
atgccgggca gttggatctt ggcctttcca tcgtcaagtc tcttgcttcg gcccacggaa 26100
agctgatgac tctttccgaa accggaatgg agggcctccc ggacaggagc tggtggactt 26160
cggtgcttta tcctgcggta aaggattttc cgattgtcta tgtgctgacg tggagaaatg 26220
catgggacag ggagacgcat ttttatgcgc cgtatcctgg taatgagggc gacttcaggg 26280
aattcgcttc gctgccggag atctctctcc tgtcgtcgtt ccggccttag tccgtcattc 26340
ccggcgaagt gtcgtccttt ccggtcggac cgggaatctg taaccttaaa acaataaata 26400
acaatatatg gagaatatga gttttgaaga acgtctggcg tggctggaag cacgccacca 26460
ggaacttatt gctaagaaaa actcgcctat atacggcaat ggtgtgtttg agagatatga 26520
gaaccctgtt cttaccggtg accatgcccc tttgttctgg agatatgatc tggataagaa 26580
ctccaatcct tttctggaag agagattcgg catccatgcc gcattcaatg ctggagctat 26640
aaagtggaag gacaaatatg tcgtctgcgc ccgtgtcgaa ggtgctgacc gcaaatcatt 26700
cttcgctatc gccgagagcc cgaacggaat cgacaatttc cggttctggg accaccccgt 26760
gactatgcct gattatggcg aacctgccac gaatgtatat gacatgagac ttactgccca 26820
tgaagacggg tggatctatg gtatattctg cgtcgagcgt aaggatccgg atgctccgga 26880
aggcgatctt tcgtctgccg tcgcagctgc gggaatcgcc cgtaccaagg atctggtcaa 26940
ttgggaaaga ctgccggatc tggtttctga ttctcagcag cgcaatgtcg tacttcatcc 27000
tgagttcgtg aatgggaaat atgctcttta cacccgtcct caggatggat tcatagatgc 27060
cggaaacggc ggcggcatcg gctgggccct ggtggacagc atggagaatg ccgttgtcac 27120
ttccgagaag atcatcaact gcaggagata ccatactatc aaggagacca agaatggcga 27180
aggccctgct cctatcaaga ctgacagggg ctggctgcat ctggcccatg gcgtcagggg 27240
ctgcgcttcc ggcctgcgtt atgtgcttta tatgtacatg acttcgctgg aggatccttc 27300
gaaagtgatt gccgaacctg ccggtttctt catggtcccg gaaggggagg agtatgtggg 27360
tgatgtcatg aacgtcctgt tctctaacgg ttggattgcc gatgaggacg gaaaggtctt 27420
tatatattat gcttcctgcg atacgcgtct ccatgtggcg acttctaccg tcgacagact 27480
cgtggattac tgtatgaata ctgagccgga tctgtttacc acttctgctt cagtacgacg 27540
tctgaacgcg cttatagaca agaatctttc gctatctatg taggactaat aatcaggttt 27600
cagatggata tcattactgc accggctaag ccgacattga aggaaaagat cggatatggc 27660
ttcggagaca tgtcgtcttc gatgttctgg aagatttttt cgtattatct gccgatcttc 27720
tattcggatg tgttcggget taagcctcag catgcggctt tcctgcttct tatcacaaaa 27780
ctgtatgatg caatatcgga ccctgtcatg ggaatcatcg ccgacaggac ccggtccaga 27840

tggggaaagt atcgtccgta tcttctctgg gtggctctgc cgttctcgct tataggcctt 27900
ctgtctttct atactcctga tgccagctat ggcctgaagc atgtatatgc gtatgtgatg 27960
tatatcctca tgatgactgt ctatacggct ataaacgtgc cttacggggc catgctggga 28020
gtcgtcactc cggacagccg tgagaaatcg gttttctcgt cttacaggat gttctttgcg 28080
tatatcggca gtttcatcgc catgggtata ttcggcgtgt tcgagaagct gatttccggt 28140
accgtgcgtg ttatgccgga cggtagcgag agggtggtgc gcggagtcgg cgatgctgct 28200
cccggacagt ggactcttgt cgtcggcatc gtttctctgt gctgtttcgt gctgtttctg 28260
ctttgcttcc ttatgaccag ggagcgggtg gggactgaaa ataaggcggt ttccgaggga 28320
ggctcttcta tcgcagagga tctcaaggct ctaggccgta acaggccttg gtggctgctt 28380
cttggcgccg gtatcggtat attgctgttc aattcgatcc gaggcggagc cgctgcatat 28440
tatttctcga atatcctcgg tacgagcgtg ttcctgactt gcgctgtcta tcttttcatt 28500
ggcgaggtgg ctcagatggt tggggtgctg tttactgtgc cgctttcgga gcgtatcggc 28560
aagaaggcga cttttatctc tgtgcttctg acggtttctg tgctgagtgt ggcgatattc 28620
ttcctgcctg agactcctgc cggtttctgg ggacttctgg tgctgcaggt gcttatatgc 28680
attgcttttg gtgtccagtc tcctctgctg tggtcgatgt ttgccgatgt cgccgattat 28740
tcggaggaga agaacggctc ggcttctacc ggacttattt tctcgtcttc ttctatggct 28800
cagaagttcg gcggggccct cggcgctttc ttgctgatgc agattcttgc aatcttcgca 28860
tatgacaagg atgctgcagt ccaggcgcct cagactcttg ctgccatccg tggcctgatg 28920
agctatatcc cggcaatcgg cgccctcctc ggagccctct gcctgctgtt ctatcctctt 28980
acggaatcgc gtatggcttc aattagagag aaactttctg ctcgcagata acatcttcat 29040
tcatgtaggt atttgttatt ccaaaggatg gccctcttcg gccatccttt ttcgtctggc 29100
tgattcgagt cgggcctttc tgctggcggg agaaggacgg gacggctccg tcaagttcgg 29160
aaaagctgct tggtcgctgt cgccgcctaa acggctaact tctccctttt acgatgggcg 29220
agccatcgta acggtcgtca aacacacgcc gttcttcacg gcgccgacgg ccgctcccgc 29280
gctctccgct tttcctcaat tcctccgccg cctccgccct cctcccgccg caggactgct 29340
gaccagggtt ccggggacgt ggccccctc ggccacggga agagggacgg ggcccgcaag 29400
atgcaagtta tcgcgaagcg atgacgcggc agattgtggg agggtacgga gtgagcgata 29460
gcgaacgaag ttccccggga ccctgcgtca gcaggactgt ggcaaaggaa gcagaatgca 29520
ggactgcggc aaagatgagg gactggaggg aattgaaaaa aatgttatat ttgtatttcg 29580
ttccttaatc cggagaacta tggcagacaa actatacctg attgacggtc acaacctaat 29640
cttcaagatg tactacgcat tcctgcgtcg tcctatgatc aattccaagg gaatggacat 29700
gtcaatcctg tatggcttca cgaaatatgt gctcgaactc atcgagagag agaagcctac 29760
ccatatggcc atagcgttcg atcctcccgg cggcaccttc aggaacgaaa tgtaccccga 29820
atacaaggca aacagaagcg aaactccgca gctcgtgatc gatgccctcg aaccgctgct 29880
ggaaatatgc aaagccttga acataaagac cttgatgatt ccgggatatg aggccgacga 29940
cgtgatcgga accgttgcga agagttacgg gaaacccggg acggacgtct atatggtcac 30000
cccggacaaa gattacggcc agctggtcgc cgaacatgtg taccaggtga agcctggaaa 30060
ggccggcggg gctgacgaga ccgtcgatgt caaggcagtc tgcgaaaaat acggaatctc 30120
cagtccggaa caggtcatcg aaatgctgac cctctgcgga gatacggccg acaatgtgcc 30180
cggagtgcat ggcatcgggg aagtcggggc aggcaagctc atcgcaaaat atggcaccgt 30240
cgagaatatc tatgcccatc tggacgagct gactgaccgc cagaagacgc agttcgagga 30300
ggccagggga catatcggcc tgagccatga cctggtgaca ataaagaccg acgtgccggt 30360
agagacaagc ctcgacgaga tgaggctgga cctggattat gctcccgaaa tcgccgacct 30420
tttcgaaaaa tacgaattca actcgctgag aaaatatata gcccatagct cccgcgctgc 30480
caaggcagtg gcggaggtca agcgcctgga ttataccgag acaggggcgg aagagctcgt 30540
gaagatggct ccggaaaaga tttcgctgat agtcgaaccg gagggaccgg gcatattcgc 30600
cgatatccgc agggtgatcg tggccgcgga gaagaaagac ggcggaatcg tcgtcgcttc 30660
aggcgctgcg aaggattttc gcaagatact ggagggcgaa gccgtgaaat acggatatga 30720
cctcaagctc cagtcgaatc ttctggagaa tgccggaata aagctcggag gccggctata 30780
tgacatcgag ctgatgcatt atctcatcaa tccggagaag agccacaagg tcgaaatcct 30840
tgccaagagc tatctcggcg tcgatatcga agaggccgct gctgtggaag aagctcctgt 30900
ttccctgttt gacgtcgcgg aagaaagccc ggcaaacgga ccggacagga atctcgaagc 30960
cgcggcagtg ctgaaactcg gagaagtggt ccgcgcaggc atggacgagg ccggcgtagg 31020
agccctgtat gaggatatgg aagcccctct cgcagccgtg ctcgcccgca tggaaagggc 31080

aggcgtgaag atagacctcg gacagctcag gacgtttgcc gatgaactca aagtcgaaat 31140
ggatgccctc gaagcccagg tcaggaaact ggccgacgag ccggggctga acgtttcttc 31200
cccgaagcag attggcatcg tgctcttcga aaagctgaaa ctcgacccga agatgaagaa 31260
atcggccagc ggtagctata gcaccgacga aacgacgctt actgccatcg ccgacaggca 31320
ccctataatt gacaagatac ttgaattcag agccgtaaag aagctgctct ccacatacat 31380
agagccattt ccgacctacg tctccccggt tacgggaagg gtccatacga ccttcaacca 31440
ggcgctgacg gcgacgggaa gactcagttc ctccaaccct aacctgcaga acattccgat 31500
ccgcacggag aggggaaagg agatcaggaa agccttcata ccggcgactc cggacggaat 31560
cataatgtcc gccgactact cccagatcga gctcaggatc atggcccatc tctgcggcga 31620
cgagcatctg ataagctcat tcaaggcagg gctggatgtc cataaggcca ccgcagccaa 31680
gattttcggc ataccggtag aggaggtcac atccgaccag aggcgcatcg cgaagaccgc 31740
aaacttcgga atcatgtacg gcatctcatc cttcgggctc gcccagaggc tgcatgtctc 31800
caggacggaa gccaagaaga taatagacga ttacttcgcc gccttccctt cgatccggac 31860
tttcatcgac aagactattg cggacgcccg cgaaaaaggg tatgtcgaga cagtcttcgg 31920
acgcaggagg tatcttccgg acataacggc gaagaacgct accataaggg ctctggccga 31980
gaggaatgcc gtaaatgctc ctatccaggg cacatcagcc gatataatca agcttgcgat 32040
gataagcgtc gacaagcgca tccgcgaggc gggactgaag agcaagatgg tgctccagat 32100
ccatgatgag cttcttttcg atgccgtgcc ggaagaaatc gaggaactcg gcaggatagt 32160
tgtagcggaa atggagaacg taattacctt gtcaatacct ttaactgtag aatgtaacta 32220
tggcaaaaac tggcttgaag ctcactgatc atgacgacgg atctcttgtg gagatggcgt 32280
taggtggcgg ccccgaagcc gaggccgctt tcttcacgat ttatgcacgt taccataagg 32340
gcgtattcgc ccacatctca cgcttcatcc aggaacagga ggaagttgag gatatctgta 32400
tggagagctt cgagaaggcc ttcaagcagc tggaatcata caggccgagc aacaagttct 32460
cgacttggat attcactata gcccgcaaca cggccttcga ccacctcgac aaggacaaag 32520
tgcgcggccg caacaaagac aagatgccga tcgactactc cgacctggag catatagacg 32580
tccctgagga caatgtcagc ccggaggacg agatcataag ttcgcaggac catgagaact 32640
tcctgagttg tgtccagggg cttccggacc tgtatcgcga agtcgcatcc atgtgctttg 32700
tggacaatct cggatacaag gagatttcgg agaaggcagg gctgcctatc aatacagtaa 32760
aaacccgcat cagcagggct aagagcctga tagtccagat gatgcttgat atggaagaag 32820
aatgaatttc aaagaattcg aagatatagt tgttgctgag tttccggagc tgactccgga 32880
gatgcttgga aagtacgagg ccatggagcc cctgtatgag gagtggaacg ccaagatcaa 32940
cgtaatttcc cgaaaagaca taggcggcct ttatctgcac catgtgctgc attctcttgc 33000
gatcgcctcg tatctgaaga gacgcaggcc ggaggtgtat tccgcatgga ccgaaggcgc 33060
gtccgtgctc gacctgggca ccggcggcgg ctttccgggg attcctctcg cggtaatgtt 33120
cccgaaagtt cgctttaccc tctgcgattc cgtgggcaag aagataatcg tggcatcgga 33180
agtgtccaag gctctttccc tgggcaacgt gaccactgtc aatgcccgtg cagaatcgct 33240
gccgcaggtt ttcgatcatg tcgtttcccg cgccgtgacc tcgctggata atttctatcc 33300
atgggtgaag ggcaagttct ccggaagcat tctgtatctc aagggagggg acatagtccc 33360
ggaaattgca gaactgatga agaagaacag actccgtacc ggagcagttt cgacctggcc 33420
gattgaaact tggctgaagg acgaatattt taaggaaaag tttgtgattg aaatagaaaa 33480
ataatttgca aatccacata aaataactac ctttgcagtc ctatttccgg aaaactggtt 33540
ttttcggcct gataagaata aataaaaaaa cagataacga tgaaaagaac attccaacca 33600
tcaaagcgca agcgtataaa caagcacggt ttccgcgagc gtatgtcaac ccctaacggc 33660
cgtcgtgtac ttgctgcacg ccgtcgtcat ggccgcaaaa aactcagcgt ttcttcagag 33720
gatagattct agtcatctct ctccggacgt tgtttgtttg gatcgaaatt caggaggctt 33780
ccgaaaggat gcctcctttt ttcattatgg cagattattt gatagacgag ggacgtaatc 33840
attaagtctc aatggaaaag ttttttttat caatcttgtc ttgcgcgctt tcagcgacaa 33900
tagctgtttc tgcgcagagt agcggaccat tagttgtcca ccagtatgaa ttgtcaaacg 33960
gaatgcaagt ctggcttaac gaagaccatt cgcagccggt agtccacgga gccgtggtcg 34020
tcaaggccgg aggccgggat tgcgccggga ccggaatggc ccattatctg gagcatatgc 34080
tcttcaaagg tacggaggac atcggtacca tcgactatgc ttcagagaag gtctatcttg 34140
attctatcga gacttgttac gacctcatcg cagcaacatc ctccagcgta aaaagggatt 34200
ctctccagaa ggagatcaac cgtcttagca tcaaggcgct ggactatgcg attccgaatg 34260
aattcgaccg tcttacggcc aatatcggag gcacaaagct caatgctttc acgtcttatg 34320

actgtacatg ctaccataat tctttttctc cgcaatttat cgcgcagtgg gccgaactca 34380
atagccatag gctgctgaac cctgttttca ggcttttcca gagcgaactt gaagcagtat 34440
atgaggagaa gaacgcggca gccgattccc agtttgacat gattctcgag aagatgctgg 34500
agaaacttgg ggatgaaaat ccatacagca aatctataat cggaactaca gataacctga 34560
agaatccaag tctttccgag atgaagaagt tcttcaacaa gtattatgtg ggctgcaaca 34620
tgggacttat ccttgccgga gacatagatg cttcgtcgat attgcctctg ctggaatcta 34680
ccttcggacg gattccgcgt ggcgataagc cggaaaggat tcctgtacct gtgccggatt 34740
tctcgtcccg ggaagattat cctataaaag ttaatttccc tatgatcaag gcgtcgctga 34800
acgtgttcaa cggtcctgca gccaaggatg ccgattatcc tgcaatggaa ctcgctctca 34860
gcctcctgaa caacagtttt agtaccggaa gtctggattc cctcataaat aacggcaagg 34920
ttctggctgg agaggtctca gtcttggact tttttgacga aatggggctt attgccatga 34980
taacgatacc tggaatccct ttcggatctc tcaacaaggc ggaggcgcgg tgcttcaagg 35040
aagtggaaag actgaaggcc ggcgattttg cggagaagga tttcgagtcg ttgaaatatg 35100
atcttgtcaa atcatataaa aggtgcctgg aatcgtccga gtcaaggttc tctatcatga 35160
tagacgtgat gtcccaggac aggtcgtggg attcttatct ttcccaggtg gagtcgatat 35220
cttcgcttac ttcggaggat atagtcaggg taagcaataa atatctgggc gacaatcgta 35280
tacgtttcca taaagtcaga ggaacctatc cgaaggataa tctgcaagct ccttcatacg 35340
agactctttc gccgaagaat ggcaatgcgg tatcggatta tgccaccaga ctgatggaga 35400
tgccttatga cactcttcct ccaagactgg tggatatgaa taatgatatc cggaaagtcg 35460
cagttaagtg aattacgagc tgacggatgc cgacatccag aagaatctgg acgagatagg 35520
caatctggct taatagatat tcgccatact gacaagcggt tcatagagtt tctctatgtc 35580
ctgcgagtcc atctcaatct gtacctggtc ccccggaatc aaacgagttc ctgcggggac 35640
cagatttttt ccatcccgat gtacattgat gatggcgcaa cgttccggca gggacagttc 35700
gtccaccagt ttcccgtcca gataagaatc cgtcataatc tccactgaaa gcgtatagcg 35760
tttctcttct ttaaaatcgg gagcatgaat catatcttca tagagggaga cgttggaagg 35820
cttcatttgc agaagttccg taaaatagta ggtcaaccca ccaacaacca ccgaagggta 35880
aaagacctca aaatgccccg taatctccgt aatcagcaca atggccgtga gcggtgtacg 35940
gaccacggcc accagaaaag ccgacatgga aacaagcatg atataactga cattctcttg 36000
cccgatgaca ccgtacctca ctcctaccag accgacaatc tgtcccaaca accctcccgt 36060
aaccagcgtc ggaatgaaac tgccgcccgg caatccggaa gagacagaga aaacggaaaa 36120
gacgaaatgc agcaacatga cagccgtcac ccacataatc tgcgtactgc cgttctgtgc 36180
ctgggccagc agaaactgct ccccgcctcc agtcagattg acctgcatca gcgaaatgat 36240
gtaggccacc agcaccagat agaacatctt gacatacgcc ggagaattga ctttggcata 36300
cacatgcctg aaagcattca tggtggagga gtagaacttc ccgcatacgg agacaatcac 36360
cgccaatatc aagaaatact tcatctggac aaagaaggcg aaaaccggtt ccgttacttc 36420
tatcagatgg taaggattca caggaaagac catgcttgcc acggctcctg ccacaacacc 36480
cgccagcaag gtagtgatgg ccgtcttagg ggcatcgaag cgttcaatgg attctattac 36540
aatgagcgag gaagccagtg gcgctgcaaa agccgcagcg agccccgctc ccgccccggc 36600
tgccagcaac tgtttctgtt cgcctttcag gatatgagtc cagcgagata ccaggctacc 36660
cacataagag cctatctgca cggaaggccc ttcgcgcccc aaagacaacc ccgccgagaa 36720
ggacagaatg ccccctacaa acttggatac catctcgatg aaaggatgcc ggtaagtgat 36780
gcgcccgtta atcacgccgc gtgtctgcgg aatacctccc ccgccgataa gcggcatttt 36840
ccccacgagc cacatgacga agacgagaat gccccaagaa aacagaaaga gtggaatgtg 36900
ccagtaccag gccggatgcg aagcgaagaa accactgcgg aggtggaaca gataatagag 36960
aagatagtga tagggcacag cgaccagtcc cgtcagcaaa ccgacgaaca tgctgacgaa 37020
atacaaccga gcatccacca gcttcaactt ccagatgtgc cagccgcgta ctgttttcca 37080
gttcttgacc ctcgtccaaa gacgttctgc cttatcccat agttccataa caaaacaact 37140
cccgttaact tgcgctttac aacaacaagc aacgggagtt gttttataca gaaacattat 37200
ttaccaaaat ctgttctttg caggattata ctcaaatccc acgagcttca attttcgtac 37260
aatgtcctct ttctccacgt tcatatcctc gcagagcgcg tccagcgaag gatagaaatc 37320
acgcagtttc atgttgatga agctgtacag catcaccggg tcctcaggta atctcatagt 37380
tacaatcttt aaatttaaac taataccggg gacaaaatta caataaaaaa acaaaataga 37440
gataatttta gtgtagaata tacaacctac ccaccgcaaa cgattgaggg ccatgtttac 37500
ggagcttccg tctgctctat cttcagggag accttctcga taatctttcc aagctcctct 37560

tccagcggaa tcacattgaa gtcgccccag aaatcggcgt catacttgaa gtcggtttca 37620
gagaagacgg tatgcaccgg caaccgttcc gcacgcggga agcgcgtcac gttctccccg 37680
tcaatccggc aggtcaccat ctcgaaccag gtatacagag tgggattact gaagagcatg 37740
cgttttctct tcatcttgaa gtaaaggtcg ccacggatgt ggtggacata ataggtaccg 37800
ttccagggtt tgtatgagac ggtatagaca agtttctccg ttgtcagcct gattttcgga 37860
gcctggcgta tcacgaaaag ccgtgtggcg gccttcacat accggggatg aatctcgaag 37920
cgggcacgca gcaaggcgct gttctccgag tctatataga gttctccgca gaacaaaggc 37980
tcctttaccc cacgtttctg ctcaaaatag acgacgttgg cagtacggtc gtccaccgat 38040
acgatgtcgc ccgaagtata ggtataaagt tcctcgccgg attccagcaa gaggaagtcg 38100
ggaagattct tcatgatgtc caactgcaga cacgcctcca ccccggagct gattttggca 38160
agcacactgt ccgtctgctc ccggttgtct atcttgctca tcttgagcaa cttcacctgg 38220
tccttcagtc cgggctccat cgtgggcgat ttgtagacct tgaaaaccgc ctccgtcagg 38280
ctttggaact tgttcttcag ttgtacgcct tcccggtaga aggtggtcag atagaccgga 38340
gaagtggagc agttcctatc ccggtgctct atcatctcgc gaagcagctt tttaggctcc 38400
accaggcgga taagcacttc ttgtaaggaa atgactttcg gctcaaggct cagaacattg 38460
ctgcgtcctt ccaaagcaga ggcttccacc gtttgcccca cataacccag atgggagaag 38520
gagagcatgc agcgcctcaa cgagtcgggc aggttgagcc ggaattcacc gttctggttg 38580
gtaatgttac ccagtgaggt tccttgtgcg atgaccgacg cactgacgac gggctctccc 38640
gtctccttgt ccaacaggat tccggcaatg gtggaatacg ccggtcgggg cggcaggagt 38700
accgtttctt ccatactctt cctttctgcc gcccgtgtta tcaatatatg gttgcctaat 38760
accttcaatt cgaggttccg actgccggtt atcctatgaa tggcctgtct cacggaacac 38820
tcctgcttcc taaccctcac tactgagtcg ttatggatga ccttgctgtc atagataaac 38880
agatagccgg actgctccga gactttcccc aacagcgcat aaaccgttcc cttcattccg 38940
ggcagccgga cgatgcggtc cagcacatcg cctccgtccg ctcttgcact gcccgccagg 39000
aagaaaatga agagtgaaaa atggagaatg aagcatttca ttgaattcat tgttccgaca 39060
atatcagtct attgccctga cgtgtacact tcaagtcgaa cgtccagcat atcagctcgg 39120
ccacagactc cggggagtta ttcgaaaatt cgaccgtcag ccgccttttg cccagggttg 39180
gcgacgcggc ttctatctgc cagccggcag actccttatt gatagcacgc agcacatcct 39240
ccaggcactc gtccttgaaa cggatgttgc tggtgtaacg gtcaaagagc tccggattct 39300
ccgtctcgct gacgagcaga ctgcccccct gcaagataac cgtttctccc gcctctacaa 39360
agaccgtctg ttcctcttgt ttcagcagca ccttcacctt cccgcgcttc acggacaagc 39420
tgaagggggc cccttcattg ctcctgacat tgaaagccgt gcccagcacc tccacacgca 39480
ccctcgccgt ctcaatcgta aaagcctgct cgggtttctt tgccacatcg aagaaggctt 39540
caccctgcag gttcacttcc cgcttgtctt ccgcaaaatg ttcgggatac ttcagcgtgc 39600
tttcctgcgc cagatatacc accgaaccgt cttcaagggt cttgaccagt gtgggcttct 39660
cccggttctc ctccgtcagc aagctacgac tgtccccacc ttgttccggt acggcccacc 39720
atgcggcaaa acatacgatt ccggcaatca cggctgccgc aacgctcccc cacttgaggt 39780
aaagcctgcg gtgcattgcc ggagcatgtt ccgcatctgc cggcagcagg cagtccgttt 39840
cgaagcgggc ataaagccgt acccatgcct cgtcggtcct tgtcttcttt ttatctactt 39900
ccatcattgt gtccggatat aattatcaat ttcatttcgt aaggttcgga gtaccttggt 39960
catttccgct tccaccgtct tgacggacaa cgaaagcgac aaggcaatct ctgcatattt 40020
cttcccctcc atgcggtgca tcctgaaaat ctgctgccgg cgccggggct gcttgtcgag 40080
cgtatgccgg atgagagtcc gtagttcctc gtactccaaa tgcgcgtggg ggtcggacga 40140
atagcccggc ccggactccg tacggacaga tgcccgatag cgctccttga cctcctgatg 40200
ctcacaaaac tgcacggctt cattcctcac agaacggtag agatagcttt tcacggactg 40260
gaatacttgt aaccgctccc ggttcttcca cagcacgtag aacagctcct ccacaatttc 40320
ctcggcagac gccgtatcac ccgtgatgcc cgcagcatac cagcaaagag gggaataata 40380
ccggcggaag attccttcga acgcctttat atcgccctcc tttattctgg ccagtatcat 40440
cagttcgtcc agcatacacg gtgttttttc tcggttgctc caaatataag gaaaactatc 40500
agataatcaa ttggattgtt tccgcaattt gctgactgca ttctccaggg attccgtagg 40560
ttctatgata ttgtagttcc tccagaagtc ttcgttccag tactcatcca ccaggtcgta 40620
gaatacctgc ttccgtttga aggtgttctt gttggagatg ccggagagag ggctttccgt 40680
cctgtcggtc accaccattt cagagaaagc ggtgtagctg gaggagaaca gccttttttt 40740
ccagtcgcat ttaaagcgga tgacattccg tatatagttc aggtaagtct tggccccctg 40800

ctgcttgtag gtcaccagat aggtgacttc ctgcgggcgg aaccgtaagc cgacgggctt 40860
cttatgcaga atggcctcca ctgccttcac ccggtttgcc atgtccaggc caaactcggc 40920
acgtgtaaag gaaagccgtt cgtagtctat atacagtttc ccgatgaaca aggcatacat 40980
caggctgacc cgcggacgga agctgaccac atactgcatg cggttatcca aattgacggg 41040
gtcctccata cggaattcat aatagtccag attgtccgtg ctcaacagcg catcgccatt 41100
cttcacaatg tccagataaa gggacaaatt ggggcctccc accactttca cagccagcgt 41160
gtcactctgc ttttggctca gcaggcggcg gcctttcagc agctgcacct tgtcgcggtc 41220
cacatcgcgg gagttataat ccgtcttata gacatccatc actgcctcgg acacgctgat 41280
atagcggcgg cgcttctgca cggtttcccg atagaaggcg gtcagcatgt tgtcattgcc 41340
agaatagttt actggaattt tcttgatggc ctcttccact atgacgcggg gattattgcc 41400
atagaccaca atctcgctca aaagattggg agccggtatc atccagatgg tcagcccgcc 41460
cgtttttttca agttcttcca gagagaggtg tgtagtgaga tagccgatat gggagacttc 41520
cagtccacgg aaagcttccg cttccgctac tttcaaggaa aatgtgccgt cggcattggt 41580
caccgttccg atattgttgc cgacaacaga gacgttcacg ttttcgagtt tcttcctgct 41640
gtccttattc ttgaccgtac cggtgatgac gagctcgttc actgccgcgt cctgcgccat 41700
caacggggtg atacccgaag ataagaatgc aaacaaacac ccaagtgtaa atcctacaaa 41760
gtttttcata aggcatgttt ttaaggggtt tctattaata agagcggtga ggaagataaa 41820
accctatgga atggagagaa aaatacaaaa aaagtagcac gttgataatc agagagaaat 41880
tttgccaaga cattgccata ggtttggcaa aacgttgcca tgcctatggc aaaagtttgc 41940
caacgcggtg gcaaagtctt gccaatagta aggcaaaaac ttgtcaagct gtcggcaata 42000
gcttagttga attgttgggg cttccgctgg ctcagaataa cgtagatgat gaccggcgca 42060
ccgataatgg gcgtgacggc attcaaggga ataatacccg cttcgccggg caagacgcaa 42120
atcagattgc aaagcagtgc cactgcaccg ccactcagga tggtgaccgg caatagggaa 42180
ttgtgattgg ttgtccccaa tatcatgcgg gccacatgcg gcacggccag gccaataaag 42240
gcgaccggac cacagaaggc ggtggtgaca gccgtcaaga gcccggtgat aatcaaaagc 42300
cagttgcgca cacggcggat attgacacct agatttcgg catagcgctc tcccagcaac 42360
aaggcgttca ggggctttat caacaataat gagccgaaca gtcccacaat ggtgaccagc 42420
gcaaaggccg gcatctgctg tagggaaact ccaccgaagt tgcccagccc ccatatcata 42480
taggactgca ccccctctgc cgtagcaaag aagttcagca gggcaatggc cgaagaagcg 42540
atgtagccta tcatgatacc cgttatcagc agcatgatat tgctctttat cagcgtggaa 42600
aagaaaagga taagtgccat aatcagcatg gctccgataa atgcgccaag cagtacggag 42660
aagaagccgg acagactgaa cgcccctgcc gtgatgctac cgccaaacag cagcatgacg 42720
aaggccactc ccagacttgc cccggcgttg atgcccagaa tggatggtcc tgccagcgga 42780
ttcttgaaag ccgtctgcaa catcagtccg cagacagcca atgcgccacc gcaaagcagc 42840
gccgtcagcg cttggggcaa acgggattcc caaaggatga agctccagct tgccttctcc 42900
ccctctccac cgagcaggat acggaaaaca tcggcaggcg gaatcgaaac cgaccctacc 42960
agcagattgc ccgcaaaaag caataagatt actactccca acccgatacc atatttccag 43020
cctttactca tagttacatt atttagcaca cagagacacg gaggcacaga gttttctatt 43080
actttcacaa aagttccgta tctctgtgtt catcactcct tcagtctgca aaaatagcgc 43140
aagcctccca aatctcctat ttcgggatgc agaatctgta tcaggtcgga taacaaataa 43200
tcgggacgga acggagtctc ctcgtagaaa ggaactttcg ccgtattaca accatagaca 43260
ttccgggtct tgaaagcctt gaaaccggtg tagccgatat aatccgcctc caaatcagca 43320
tagctcatgt cgcggtcacg gttgtacttc accagccaga cgtcggcatc gcccgccttg 43380
tcgaaaacgg tctcgaaagc cagaggaaca gagccgctgt gggtatcgtc ggcaaaagcg 43440
taacgcccgc aagcatcctg aaaaagcctt cctatggtgc tacgccctcc cggcacgtac 43500
catgccgagc cgcttttcaa ctcactgaca acggagaacg aggtggatga aagttgcgca 43560
cgcatcttga ggcgttggta acaactgtcc acttcagcaa aaaggttgtc ggcttgtggc 43620
gcaaccccga agagcagtcc gtagaaacgc atccactcgg cacgtcccaa agcggaggtt 43680
tccatgtagt cggcacattc gatgatggga atatccagtt tctcgatacg gccataaccg 43740
ccactgttct cgaacggaga cagaaggatg gcatcgggat gcaggtcgat gatgcgctcc 43800
atatcgggat tcataccgtc gccacaatcg gcgatggtgc cccggcggca ggcttcgtga 43860
acggtgggca gtttgatgta cttcaagtcg cagacaccgc cgatgctacc aaacacaccc 43920
agattatcca tcagtccgca gtggacgg 43948

//

```
ID   XXX; XXX; linear; XXX; XXX; XXX;  XXX.
XX
AC   PRJEB67889
XX
DE   Uncultured bacterium DNA fragment from in-reactor enriched cow rumen, clone
ferm-44f17
XX
FH   Key             Location/Qualifiers
FH
FT   source          1..35340
FT                   /organism="uncultured bacterium"
FT                   /environmental_sample
FT                   /mol_type="genomic DNA"
FT                   /isolation_source="Cow rumen"
FT                   /clone="ferm-44f17"
FT                   /note="AZO-CM cellulose, AZCL-Barley beta glucan, AZCL
Tamarind xyloglucan, AZCL-arabinoxylan, AZCL-HE cellulose active clone"
FT   CDS             complement(171..1694)
FT                   /db_xref="SEED:fig|6666666.127266.peg.1"
FT                   /db_xref="GO:0008815"
FT                   /translation="MTNEQLIREAAAKLGKKVYQGPGQKIPTFRRDLQKESCKVVSLED
FT                   AIRKSGLKDGMTISFHHHFRGGDKIVNMVVDKIAEMGIKGLTLAASSLSDVHAPLVGHI
FT                   KNGVISKITTSGVRGELAEQISRGLMDEPVIFRSHGTRGAAVASGELHIDVAFLGAPSC
FT                   DPLGNACGYSRSENAKSICGSLGYALPDARYADKVVILTDDLVPYPNTPNSISEHDVDY
FT                   VVVVDSVGDSSKISSGAIRDSKNPRDILLARLAAKAIINSGYFKDGFSIQTGTGGAALA
FT                   AVKFIRESMIEKGIKASYALGGITAHMVRLHEEGLIERLIDVQSFDKVAAESVKNDPAH
FT                   KEVASYEYASGMEPGSAVHYLDIVILSALEVDVNFNANVLVGSDGIIRGAIGGHPDTAA
FT                   DSALSIVVCPLLRGRIPCIVDHVTTLITPGASVDVVITEYGTVVNPRRPEIAERLKAAG
FT                   MKLVDMQYLRDCARKIIGEPAPIPFGDKVVGVVLGRDGSVQDVIRNVLK"
FT                   /product="Citrate lyase alpha chain (EC 4.1.3.6)"
FT                   /EC_number="4.1.3.6"
FT   CDS             complement(1691..2581)
FT                   /db_xref="SEED:fig|6666666.127266.peg.2"
FT                   /db_xref="GO:0008815"
FT                   /translation="MERLRRTMMFIPGNNPGMMQDAYIYGPDSIMLDLEDSVTMAEKDA
FT                   ARLLVYNALKTVDYGNIERVVRINPLSTPYGKKDVEAVVKAGVDVIRMPKTETADEVRE
FT                   LEAEILKVEEEIGCVGRTQIMAAIESALGIVNAYQIASASSRMMGIALGAEDYTANLKT
FT                   MRSNEGIELQLARETIVVAARAAGISCFDTVFSNLEDMEIFRKEVEFIKTLGFDGKSII
FT                   NPRQIDVVNEVFTPTEKEIAKAQTIIAAIKEARQRGSGVISVNGKMVDRPVVIRAERII
FT                   ALAVAAGVIPEEDAL"
FT                   /product="Citrate lyase beta chain (EC 4.1.3.6)"
FT                   /EC_number="4.1.3.6"
FT   CDS             complement(2588..2866)
FT                   /db_xref="SEED:fig|6666666.127266.peg.3"
FT                   /db_xref="GO:0008815"
FT                   /translation="MELKNASAGTFESGDILIRIDPAEKKGLEIELESSVADQFGVQIK
FT                   KVISETLEGLGISDAKVKAIDKGALDCTIRARVTAAAVRASGKDVWK"
FT                   /product="Citrate lyase gamma chain, acyl carrier protein
FT                   (EC 4.1.3.6)"
FT                   /EC_number="4.1.3.6"
FT   CDS             complement(2979..4298)
FT                   /db_xref="SEED:fig|6666666.127266.peg.4"
FT                   /translation="MSTETILIILLVAVIVIAAILIFRLQARAEKIKDGFQDRLADVQR
FT                   EKAVLESRLQAKEEYENQMREAAKETSERHAKELEQMKEVFKNLSTETSEAFKAKSAQT
```

```
FT                   ISELMKPVQEKFKEFSDAVKESQKDTIERHSRLEQSIKDLDTRSKSVGDEARNLANALT
FT                   GYSKVQGDFGEMLLTDVLKNAGLTEGVHFETQSVMTDNSGHEIKSESGRTMIPDVKVYY
FT                   PDDTIVIIDSKVSLTAYNEYMAATTVEDRTRLAKAHVRSIEDHVNELKVKDYAGYIPEG
FT                   RRKVDYNIMFIPVEGAFRLMLEEAPRLWQVAKDNNILIVSQMTLIIVLNMIQMAWKQHE
FT                   QEKNISDVYKTAEELMSQLKGWMESYVKIGDYLGKASSAYEDSKKKLSDSNQSVIKKIG
FT                   KLEKLGIAPKRSNAKIKTGSRLSGPESVIPKELSDDEITD"
FT                   /product="DNA recombination protein RmuC"
FT   CDS             4597..5367
FT                   /db_xref="SEED:fig|6666666.127266.peg.5"
FT                   /translation="MQSVFRNPLADPHIMGVSGGAGLGAAIATLLISGTGLLSGFSSMA
FT                   GLTVAGAAFVGAILASALIVLVSSRFRSGTTLLVFGVMLGFIFSAITSILEYSASEESL
FT                   KLFYSWSAGSFSGTRTFETLTIFAALLVGTGLYTSGAKGLDILLFGEEYATLAGASVRR
FT                   IRGVAMVSSCLMTAAATAFCGPLGFVGIVSPHIARRLLRSSGHMSILPATLITGSAVSV
FT                   LADIISQAPGLPLPVGSTMALIGIPIILLILMKK"
FT                   /product="Vitamin B12 ABC transporter, permease component
FT                   BtuC"
FT   CDS             5364..6023
FT                   /db_xref="SEED:fig|6666666.127266.peg.6"
FT                   /translation="MRIKDLNIGHRGKTLVSGISFETEPGEFILLCGPNGSGKSTLLRA
FT                   LAGKDAIMVPTGVPKVKGFTLREFIATTLYSESDWRGRISPAMASDISKSLECLGISSL
FT                   ADRDISTLSDGEFQKGCIASALSRHAKTLLLDEPTSFLDVDNKSLILQALRDIARSTGT
FT                   TVIMASHDLAPALSCADRVFGITADHRFLASSSRLSDSGLSASEVISACFSTYSSR"
FT                   /product="iron (III) ABC transporter ATP-binding"
FT   CDS             complement(6037..8814)
FT                   /db_xref="SEED:fig|6666666.127266.peg.7"
FT                   /translation="MQVWLNEDHSQPVVHGAVVVKAGGRDCAGTGMAHYLEHMLFKGTE
FT                   DIGTIDYASEKVYLDSIETCYDLIAATSSSVKRDSLQKEINRLSIKALDYAIPNEFDRL
FT                   TANIGGTKLNAFTSYDCTCYHNSFSPQFIAQWAELNSHRLLNPVFRLFQSELEAVYEEK
FT                   NAAADSQFDMILEKMLEKLGDENPYSKSIIGTTDNLKNPSLSEMKKFFNKYYVGCNMGL
FT                   ILAGDIDASSILPLLESTFGRIPRGDKPERIPVPVPDFSSREDYPIKVNFPMIKASLNV
FT                   FNGPAAKDADYPAMELALSLLNNSFSTGSLDSLINNGKVLAGEVSVLDFFDEMGLIAMI
FT                   TIPGIPFGSLNKAEARCFKEVERLKAGDFAEKDFESLKYDLVKSYKRCLETSESRFSIM
FT                   IDVMSQDRSWDSYLSQVESISSLTSEDIVRVSNKYLGDNRIRFHKVRGTYPKDNLQAPS
FT                   YETLSPKNGNAVSDYATRLMEMPYDTLPPRLVDMNNDIRKVAVNDHVTLYYNGNPVNDV
FT                   FELTVRINKGIFEEPMICPLTQLLGLSGTDSLSARDFSMSLAELGSDLSFKDKNHTLEM
FT                   VLTGFDDRIGPSLDLLRHFMKDFKPEKSALNTVISDTKMSRNAFRSSGNSAIFNGMFNY
FT                   SLLGNQSPLLRQPTEKDLKSLGLKGMKELFNSLVEYDFDVVYSGGLSPEEVVNEISGHL
FT                   DVECPSGKPESSFFAYRKYEQPVVYIYNVPGARQVMSITYQNVATPENDHDSALLGLMQ
FT                   ECYGGSMASIMFQEVREFRSLAYTSSSTLQKPLPNVSSPSAFITYYGTQADKFPQALEL
FT                   VDSLIRTMPMTDLCFDSAVRSIISRANNGYPSFRALPEYVANLELFGYGEDDDASVVRG
FT                   LPGITKQDVLDYHEKSIGNAPCSYFIIGDKKNLPMDALRERGRIVELKLSDIYR"
FT                   /product="ZINC PROTEASE (EC 3.4.99.-)"
FT   CDS             complement(9293..9955)
FT                   /db_xref="SEED:fig|6666666.127266.peg.8"
FT                   /translation="MNFKEFEDIVVAEFPELTPEMLGKYEAMEPLYEEWNAKINVISRK
FT                   DIGGLYLHHVLHSLAIASYLKRRRPEVYSAWTEGASVLDLGTGGGFPGIPLAVMFPKVR
FT                   FTLCDSVGKKIIVASEVSKALSLGNVTTVNARAESLPQVFDHVVSRAVTSLDNFYPWVK
FT                   GKFSGSILYLKGGDIVPEIAELMKKNRLRTGAVSTWPIETWLKDEYFKEKFVIEIEK"
FT                   /product="rRNA small subunit 7-methylguanosine (m7G)
FT                   methyltransferase GidB"
FT   CDS             complement(9952..10542)
FT                   /db_xref="SEED:fig|6666666.127266.peg.9"
FT                   /db_xref="GO:0003677"
FT                   /db_xref="GO:0003700"
```

```
FT                   /db_xref="GO:0006352"
FT                   /db_xref="GO:0016987"
FT                   /translation="MKLTDHDDGSLVEMALGGGPEAEAAFFTIYARYHKGVFAHISRFI
FT                   QEQEEVEDICMESFEKAFKQLESYRPSNKFSTWIFTIARNTAFDHLDKDKVRGRNKDKM
FT                   PIDYSDLEHIDVPEDNVSPEDEIISSQDHENFLSCVQGLPDLYREVASMCFVDNLGYKE
FT                   ISEKAGLPINTVKTRISRAKSLIVQMMLDMEEE"
FT                   /product="RNA polymerase sigma factor RpoE"
FT   CDS             complement(10529..13261)
FT                   /db_xref="SEED:fig|6666666.127266.peg.10"
FT                   /db_xref="GO:0003887"
FT                   /translation="MQDCGKDEGLEGIEKNVIFVFRSLIRRTMADKLYLIDGHNLIFKM
FT                   YYAFLRRPMINSKGMDMSILYGFTKYVLELIEREKPTHMAIAFDPPGGTFRNEMYPEYK
FT                   ANRSETPQLVIDALEPLLEICKALNIKTLMIPGYEADDVIGTVAKSYGKPGTDVYMVTP
FT                   DKDYGQLVAEHVYQVKPGKAGGADETVDVKAVCEKYGISSPEQVIEMLTLCGDTADNVP
FT                   GVHGIGEVGAGKLIAKYGTVENIYAHLDELTDRQKTQFEEARGHIGLSHDLVTIKTDVP
FT                   VETSLDEMRLDLDYAPEIADLFEKYEFNSLRKYIAHSSRAAKAVAEVKRLDYTETGAEE
FT                   LVKMAPEKISLIVEPEGPGIFADIRRVIVAAEKKDGGIVVASGAAKDFRKILEGEAVKY
FT                   GYDLKLQSNLLENAGIKLGGRLYDIELMHYLINPEKSHKVEILAKSYLGVDIEEAAAVE
FT                   EAPVSLFDVAEESPANGPDRNLEAAAVLKLGEVVRAGMDEAGVGALYEDMEAPLAAVLA
FT                   RMERAGVKIDLGQLRTFADELKVEMDALEAQVRKLADEPGLNVSSPKQIGIVLFEKLKL
FT                   DPKMKKSASGSYSTDETTLTAIADRHPIIDKILEFRAVKKLLSTYIEPFPTYVSPVTGR
FT                   VHTTFNQALTATGRLSSSNPNLQNIPIRTERGKEIRKAFIPATPDGIIMSADYSQIELR
FT                   IMAHLCGDEHLISSFKAGLDVHKATAAKIFGIPVEEVTSDQRRIAKTANFGIMYGISSF
FT                   GLAQRLHVSRTEAKKIIDDYFAAFPSIRTFIDKTIADAREKGYVETVFGRRRYLPDITA
FT                   KNATIRALAERNAVNAPIQGTSADIIKLAMISVDKRIREAGLKSKMVLQIHDELLFDAV
FT                   PEEIEELGRIVVAEMENVITLSIPLTVECNYGKNWLEAH"
FT                   /product="DNA polymerase I (EC 2.7.7.7)"
FT                   /EC_number="2.7.7.7"
FT   CDS             complement(13746..15173)
FT                   /db_xref="SEED:fig|6666666.127266.peg.11"
FT                   /translation="MDIITAPAKPTLKEKIGYGFGDMSSSMFWKIFSYYLPIFYSDVFG
FT                   LKPQHAAFLLLITKLYDAISDPVMGIIADRTRSRWGKYRPYLLWVALPFSLIGLLSFYT
FT                   PDASYGLKHVYAYVMYILMMTVYTAINVPYGAMLGVVTPDSREKSVFSSYRMFFAYIGS
FT                   FIAMGIFGVFEKLISGTVRVMPDGSERVVRGVGDAAPGQWTLVVGIVSLCCFVLFLLCF
FT                   LMTRERVGTENKAVSEGGSSIAEDLKALGRNRPWWLLLGAGIGILLFNSIRGGAAAYYF
FT                   SNILGTSVFLTCAVYLFIGEVAQMVGVLFTVPLSERIGKKATFISVLLTVSVLSVAIFF
FT                   LPETPAGFWGLLVLQVLICIAFGVQSPLLWSMFADVADYSEEKNGSASTGLIFSSSSMA
FT                   QKFGGALGAFLLMQILAIFAYDKDAAVQAPQTLAAIRGLMSYIPAIGALLGALCLLFYP
FT                   LTESRMASIREKLSARR"
FT                   /product="Xyloside transporter XynT"
FT   CDS             complement(15194..16360)
FT                   /db_xref="SEED:fig|6666666.127266.peg.12"
FT                   /translation="MSFEERLAWLEARHQELIAKKNSPIYGNGVFERYENPVLTGDHAP
FT                   LFWRYDLDKNSNPFLEERFGIHAAFNAGAIKWKDKYVVCARVEGADRKSFFAIAESPNG
FT                   IDNFRFWDHPVTMPDYGEPATNVYDMRLTAHEDGWIYGIFCVERKDPDAPEGDLSSAVA
FT                   AAGIARTKDLVNWERLPDLVSDSQQRNVVLHPEFVNGKYALYTRPQDGFIDAGNGGGIG
FT                   WALVDSMENAVVTSEKIINCRRYHTIKETKNGEGPAPIKTDRGWLHLAHGVRGCASGLR
FT                   YVLYMYMTSLEDPSKVIAEPAGFFMVPEGEEYVGDVMNVLFSNGWIADEDGKVFIYYAS
FT                   CDTRLHVATSTVDRLVDYCMNTEPDLFTTSASVRRLNALIDKNLSLSM"
FT                   /product="GH130"
FT   CDS             complement(16447..17466)
FT                   /db_xref="SEED:fig|6666666.127266.peg.13"
FT                   /translation="MAASVKAGKILYGHQDDLVYGHSWCVDPDSAATDALNRSDVKAVA
FT                   GDYPAVVGFELGGLELGWDWSIDKVPFLLMRRAAVEHAGRGGIVTFSWHPHNPYTGGDA
```

```
FT                   WDISSDKAVASVLEGGEKAEMFGVWLERLGDFMDSLRDDKGNLIPVIFRPWHENIGSWF
FT                   WWGGALCTADEYIALFRRTVEYLRNDRGLDNIIMCYSPNSGITAEQYMSRYPGDDVVDI
FT                   LGYDHYETVMEGESLEDAGKRYAGQLDLGLSIVKSLASAHGKLMTLSETGMEGLPDRSW
FT                   WTSVLYPAVKDFPIVYVLTWRNAWDRETHFYAPYPGNEGDFREFASLPEISLLSSFRP"
FT                   /product="GH26"
FT   CDS             complement(17592..19118)
FT                   /db_xref="SEED:fig|6666666.127266.peg.14"
FT                   /translation="MIFLAACSCSKDDAEKKEPSVPAPKAVTVDVASVSAPVEGTSVVV
FT                   AISSPSKPAVSVSGDWIKAELGPYNPDTYKVKLTVTVGENPTYESRSGSVTLTCQGFSP
FT                   VGISVSQDAAAKPVGPSANPAYDLLGLGWNMGNQFDAFYNGSWAGDKYNYPDETVWGNA
FT                   KATQATFDGVKKAGFTSVRIPVTWLNMIGPGPEYNIDETWMNRIYEVVGYAEKAGLKVI
FT                   INTHHDENNNDDHWLDILSASKDARTNEAIKAEIAAVWTHIANKFKDKGDFLIMESFNE
FT                   IQDGGWGWSAAFQADPSKQCDVLNEWNQVFVDAVRATGGNNATRWLGVPTYAAGTSWTK
FT                   YFRLPEDPAGKVMMAVHCYDPYEYTIGNIQYPEWGHTGAAGKKTPEGEKEITDLFSRLN
FT                   TEFISKGIPVYLGEFGCSMRSKGDARAWMFYKYYMEYVSKAAHTYSLPGILWDNGSTKT
FT                   GKECHGYIDHGTGKYIGNSKEIVDLLVKGMLSTDESYTLQSVYDRAPKTN"
FT                   /product="GH5_4"
FT   CDS             complement(19213..21639)
FT                   /db_xref="SEED:fig|6666666.127266.peg.15"
FT                   /translation="MKLIMKKIFSISAILAACVLALGLSGCSSKELDTDQYSDKDVKLA
FT                   SFGPNPVMRGATLRFFGSNLDRIVEVSVPGLDPITDIEVVTSGNPSEIRIQLPAEGAQV
FT                   GKISLKSSDGKVFTTQVDVEYTEPIVFDSFSPAEAMPGDVITVKGDYMNLVQSVTFEGG
FT                   AIVTEINHNGRHEASFKVPSKAITGKIILSDEGAIANLLYSEEDLVIGDPTVKSLTVAV
FT                   AKPGENAVVKGAYLDMIDKVVFEGGAEVSDLALNEDATEFNLVIPAGAQSGEVKAVSHA
FT                   AKEFVAGEIELVIPTDVAAAPSPVKAGADLVISGNDLDVVTSVDFPGASGVEFVFAEDA
FT                   ITVTVPATATEGDIALNMASGEAVNVAFTLVHPVVSAIAPVELMAGETITVTGTDLDLV
FT                   TGVTLGGKPVEFTAEAEKIVVQTANTSVSGKLVLSLANGEKIEPEDAITLSYDSFIIVN
FT                   DMPAAEHIGAIVTLKGENFMMIENIYIGEEKVSKYNLRSDNEIQFVMPWNKVGTYSMYF
FT                   HLLSGDVETCPQTIEVLLEQDIMTVWEGSLQITWNDGGRVTVPAASFTGIPAGSKLRFY
FT                   YTQVDQQWDQAQLNYGNWTGINFDDPAGTVINGTLVPTDVYGWFTDGILDRCTEVVLTK
FT                   EILENIQANKGDCEDKTACGIIIQGSGLTFTKIEIVCDIPQEVTIYEGPTTLTWGDDGR
FT                   FGLAMTYFEAAKPGSQLIFYLEQTDAWGQVQLNDGWWANADMNFPEIGGAYITTDNLGG
FT                   KDVTRIALTLTADILDHILATSGDYFGLNSQFQGDGRVAMVIQGSDMIINKVCIL"
FT                   /product="hypothetical protein"
FT   CDS             complement(21646..22902)
FT                   /db_xref="SEED:fig|6666666.127266.peg.16"
FT                   /translation="MASFASCDKVDYPDRYEVADGKPTVYSVRYADKDVNITQAYMDEV
FT                   VCLLGKNLKSIVELWFNDQQATLNTSFITDNTLLVSVPKNMPVVQTDKIYMITSGKDTL
FT                   AYDFAVLPPTPKVTSMSNEWAAEGETVTIYGDFLIDKADKPLQIAFAGAEVDHASMVFN
FT                   GSSSVSFPVPAGAQPGYVTVTSLSGTGKSKFMYKDTRNILFDWDGSHGGFATGHGWRNG
FT                   VIHAPGADAWDAIDGSYLYFGGADLGPGSADAWAEDQYSFNYWPEAGTDYDALNLRPEF
FT                   AEYIDKYGVSGLVLKFEVLVPSSNPWSASGMQLMFSSDAVVTYGTATNSYFSDETFPRA
FT                   IWQPWTTNGGSYDTADKWVTVSFPLSSFNKNQENKECATAFDKSYLTGLTFFVWNGGID
FT                   GTACSPVFAIDNIRVVPNE"
FT                   /product="FIG00417257: hypothetical protein"
FT   CDS             complement(22963..24711)
FT                   /db_xref="SEED:fig|6666666.127266.peg.17"
FT                   /translation="MKISNMMKGFAVATAVFALTSCEGFLNRPNEDNYNVDNFYKTDEQ
FT                   CEQGVNYLYNSPWYDFQRGFIKIGEVMSGNMYWGSSPYLDFSTNGTDQDLVNMSYSLWA
FT                   VNGHANTVIYNLLNSQGPSKAAIDKCIGEALTWKAFAYFFMVRTFGDVPIVHSNSDLLS
FT                   SGEYNSISKVERADVYEYIITTLEKAMEMLPKKTSGFNNYERIDYYAAEALLAKVYLTK
FT                   AGLSGSLNQDDLKQAAKYAKDVIDNSGRTLLPVYSDVFRLTGFNATGECLISWEWSVGR
FT                   NPWTQQNTLQSDLAMVGFSDQGDCWGGWGGPSVDLADAFGVSATQNPAERVGEKDSRRK
FT                   ATLMLPGDKYDYFWTDKGGFDYLRFIYDTEYGKGGPGGSMQCPTGSNNVKHLYGDNFDH
```

```
FT                   ENAGLGTPENMAYQLPTHLLRLSDVYLIYAEASLLTGSNGPALEYVNKVRERAGADALA
FT                   SVTFEDIWKERRLELAGEGDRWYDYVRRSYYDMDACIAELKAQRRSYWDGLDDVWKSYI
FT                   GDAFDYSGTWDASEVQYNNAEDNPNVTAASFKLPFPTEDVVFNGNMASDAVPVHVDVRE
FT                   EYSYNF"
FT                   /product="putative outer membrane protein, probably
FT                   involved in nutrient binding"
FT   CDS             complement(24724..27873)
FT                   /db_xref="SEED:fig|6666666.127266.peg.18"
FT                   /translation="MAQNKVTGRITDNSGQGVINASVMVKGTNNGVVADLDGYYEIVVG
FT                   QGSTLVYSSIGYKDQEVVVGTQSVINVLLVENSELLDEIVVVGYGTMKRSDLAGSSVSM
FT                   KEDDLKGSIITNLDQSLQGRAAGVTAVQTSGAPGSSSSIRVRGQATINANAEPLYVIDG
FT                   VIMQGGGNSGADFGLGDALGNGKVSTISPLSTINPADIVSMEILKDASATAIYGAQGAN
FT                   GVVLITTKHGSRGDAKFSYDGMFAVSRQTTRIDMMNLREYAEFYNSMVETGEISEANPY
FT                   YATPGLLGKGTNWQDEIFRTAIQHQHQISAQGGTDKVQYYVSGSFMDQEGTIRGSEFGR
FT                   YTFRTNLDAQLKEWFKLGLNLTFANTNDDLKLADGSEGLITYSLTTIPDIPVYDVYGNF
FT                   SSTVREGYTSPNPVALAMMDEILLDRQKLNGNIYAELTPVKHFTWRTELGFDVSGSTAT
FT                   RFRPTVNLGGWTRDTNSMSYQKNSNRFWQIKNYLTYANSFGKHSVTAMLGQEAWESSWS
FT                   YVQGANTKLPSNDVHNVALGGGTPSLSSGFGSSAMASFFTRETYNFDDRYLLTYTYRYD
FT                   GSSNFGPEKRWAGFHSLAVAWKFTNEKFMDFLKPVVNNGKLRLGWGQTGNSSIGGYAWG
FT                   ASISRMPSALGMGYRPANIPNPAIHWESQTQQNIGLDLNMFDNRLNLTVDAYLKKSDNM
FT                   LMSMQLASYMGTQGNSSSALAAPKGNFGSIENKGLEITLDAHPVQIKDFAWDSNFQISF
FT                   NRNKLVSLDGTTNAALIGYGQWSDVVCVSEIGKPLYNFYGYKVAGVYTSLEDIENSPKP
FT                   EKYPSNGEYARGNTVWVGDIKYEDISKDGVINEKDRTDIGNPLPKFTFGWSNTFTYKDF
FT                   DLNIFINGSYGNKVLNYNAISLTHMNSAWTNQINDAVSGRAMLAPIDASVVYADGSKWY
FT                   DHIDNVRVLNPGTKTPRASINDPNDNDRISDRYVEDGSYIRIKNITLGYNVPKSFLNKF
FT                   KVDNVRVYVNIQNLYTFTKYTGYDPEVGASTQDSTGLTYGVDNGRYPSPTVYSCGLNIT
FT                   F"
FT                   /product="TonB family protein / TonB-dependent receptor"
FT   CDS             complement(27961..30354)
FT                   /db_xref="SEED:fig|6666666.127266.peg.19"
FT                   /translation="MDIVKVDDGSFVVDGHFYYYIGTNFWYGGLLGSEYGDRSRLAAEL
FT                   DSLHALGLDNLRVLVGSDGPDGVATRVYPTLQKEPGVYDEKVFEGLDYLMAELGKRNMR
FT                   AVLYINNSWEWSGGYGMYLEWAGAGKAIIPSVGGYGPFMEQMAQYATNARAQELFENHL
FT                   RTVVSRVNSITGKPYSEDPAIFSWQIGNEPRCFSSDPAVQDGFVSWIWRAASIIKSIDP
FT                   NHMVSTGNEGIWGCEENPELFRRLNECPDIDYITAHIWPYNWSWVGKESPQLDIDRAEE
FT                   NTGRYIDAHLEVAESLGKPLVIEEFGFPRDDMQFAKGTPVSGRDRYYRYVFSKVMENQK
FT                   NGGRLAGANFWSWGGLAVPAHRYWQEGDDYAGDPAQEEQGLNSVFADDYSTLAIIKHTT
FT                   ARLSAVANASAVLENDWMFTTGDQKPLRVAVQSQNGFKGSVPVYVRVSTDKGVSVSVSG
FT                   KKARLGKGADTLDFDLGLAPGFYKVCVSLNENDPLDENFCVGCDPEKIVSPQDKQPDFD
FT                   EFWEDNLRQLAMVPMNARMTPLKDHSNADRTAYRVDMKSFGGKEISGLLYLPNKEGKFA
FT                   ARISYMGYGSDVWFADPSAEPDMIEFTLSVRDQGFNKTPGQGWVTRGLADKDSYYYRGA
FT                   YLDCVRAIDFVSSLPQTDPDHIYAEGGSQGGAFTLIAASLDNRLRAIAPFVPFLSDFPD
FT                   YFAIADWPGGEVLSAAKAQGISEDELYKTLSYFDVKNFTDRIGCPVLMGFGLQDPVCPP
FT                   HTNFSGYNHITADKQWLCVPVSGHHLEQFPEWWAARTEFFTKNK"
FT                   /product="GH5_7-CE7"
FT   CDS             complement(30530..31414)
FT                   /db_xref="SEED:fig|6666666.127266.peg.20"
FT                   /translation="MDMTSVITEITPLSDRDCFYIVERHKTEFVFPLHRHREYELNFIE
FT                   HGKGVRRVVGDSVEEIGEYELTLIAGGDLEHAWEQGNCTSEDVREITIQFDPEIFSSGL
FT                   MMRNQFSSIQHMFEQARNGLTFSLAAIMKVYSLLDALPSEIDKFDQFQHFMRLLYDLSL
FT                   AGDSRILASTSFAHNDADKESRRVQKIKQYVGDHFAENITLEELAEMVGMSPSAFSRFF
FT                   KSRTGKTVIDYIIDIRLGNAARLLVDTTQNISEICYSCGFNNLSNFNRLFKSKRGYTPR
FT                   EFRTLFKKNKVFV"
FT                   /product="Transcriptional regulator of various polyols
```

```
FT                   utilization, AraC family"
FT   CDS             31538..31972
FT                   /db_xref="SEED:fig|6666666.127266.peg.21"
FT                   /db_xref="GO:0008270"
FT                   /db_xref="GO:0016787"
FT                   /translation="MEKDELYMKEALREAQAALAEDEIPIGAVIVCKGRIVGKGHNMTE
FT                   RLHDPTAHAEMIAITAATEALGGKYLNECTLYVTVEPCPMCAGALNWAQIGRIVYGAID
FT                   PKRGHSLFTPSLLHPKTEVEAGVLAEECGSMVTEFFKKKR"
FT                   /product="tRNA-specific adenosine-34 deaminase (EC 3.5.4.-
FT                   )"
FT   CDS             31989..32282
FT                   /db_xref="SEED:fig|6666666.127266.peg.22"
FT                   /translation="MTGDDYKIISDTTENGVRHITAIPSSLVCSQQIDFDIIDGKVRNL
FT                   AYVRGCNGNLQAIGRLVEGMDIKKIVDTLSGVNCNFRGTSCTDQLARILSSL"
FT                   /product="hypothetical protein"
FT   CDS             complement(32279..32947)
FT                   /db_xref="SEED:fig|6666666.127266.peg.23"
FT                   /translation="MIWFIIIAVMVLSFVIQQVLNSKFNKYSKVGLHSGLTGAQVAAKM
FT                   LEDHGITDVSITCIKGKLTDHYNPVNKTVNLSEEVYYGNSVAAAAVAAHECGHVVQHAT
FT                   GYAPLKLRSALVPVVSFSSNAVQWVLLIGVMIIKITPTLLYLGIAMFALTTLFSVVTLP
FT                   VEINASARAVQWLDKAGITGPATKAKAVDALKWAAYTYVIAAIGSIATLIYYIAMARRR
FT                   "
FT                   /product="probable metal-dependent peptidase"
FT   CDS             33312..34424
FT                   /db_xref="SEED:fig|6666666.127266.peg.24"
FT                   /db_xref="GO:0008784"
FT                   /translation="MNRVDDFSRLELNLTGCMENYRYFRSKLEQTTKLLVLVKANAYGH
FT                   GAVEFAAMMQRAGANYLAVAYPVEGMELRKAGISLPILVLTAGTDFFNEIIDNRLEPGI
FT                   PNLYALKILCGILKERGIKDYPVHIKLDTGMHRLGFMTSELQDLTDFLKTCEEVKVKSV
FT                   YSHLAVSEDPEQDEFTLGQIGMFKTNADRITSEIGYKPMYHILNSAGIERFPQYQFDMV
FT                   RLGIGIYGISALPDVRLAHVASLKCKILQIKTLKPEDGTIGYGRYGHIAPEGTVIATIP
FT                   VGYADGIDRHLGRGHASFSVNGHRAPTIGNICMDMCMLDVTGIDAKVGDTVTIFGEDPT
FT                   VNELADILGTIPYEILTSVPRRIERIVVRK"
FT                   /product="UDP-N-acetylmuramoylalanyl-D-glutamyl-2,6-
FT                   diaminopimelate--D-alanyl-D-alanine ligase (EC 6.3.2.10) /
FT                   Alanine racemase (EC 5.1.1.1)"
FT                   /EC_number="6.3.2.10"
FT                   /EC_number="5.1.1.1"
FT   CDS             34534..35106
FT                   /db_xref="SEED:fig|6666666.127266.peg.25"
FT                   /translation="MSIMGKAYIFLADGFEETEALATLDILRRGKIEAYTVSITEEVYI
FT                   TSSHGVTVVADMGRSDFSEIDIDEKDAMIFPGGMPGSKKLAADTPLIGLMNVHYGNGGI
FT                   VAAICAAPGLVATQIKGIEGKKATCFDGFEENLAASGMEYVKAPAVTDGQLITGRGAAF
FT                   AIDFGLAILEKMGGKEAVAKVKAALLI"
FT                   /product="DJ-1/YajL/PfpI superfamily, includes chaperone
FT                   protein YajL (former ThiJ), parkinsonism-associated
FT                   protein DJ-1, peptidases PfpI, Hsp31"
FT   CDS             complement(35095..35331)
FT                   /db_xref="SEED:fig|6666666.127266.peg.26"
FT                   /translation="MLFHTAGCTFCEAELAAADSVASEKGVRVILVDMDKLTSLYPEEA
FT                   DRLLDAFDLSVLPFIMETDNKGCVVRKYLSLNK"
FT                   /product="hypothetical protein"
XX
SQ   Sequence 35340 BP; 8392 A; 9394 C; 8922 G; 8632 T; 0 other;
```

atcaatcccg gcctccgcta tctttatcgt atattcatca cgcttgacat tccctggcat 60
caccacggtc atgcttatga gcgtagagcc atagctatgc ttgagcagat attcctgctc 120
tttctgcctg gcctcgcggg aggctaatat catatcaagc gtaacttcca ttatttcagc 180
acattcctta tcacatcctg tacggatccg tctcttccga ggacgacgcc cacgaccttg 240
tctccgaacg gaatcggagc aggctcccct attatcttgc gggcgcagtc acgaaggtac 300
tgcatatcca ccagcttcat tccggccgcc ttcagccttt cggcaatctc cggcctgcga 360
ggattgacca cggtgccata ctcggtgatc accacgtcca cggacgctcc cggagtgatc 420
agggtggtga catggtccac gatacatgga atcctgcccc tgagcagagg gcagaccaca 480
atcgacaggg ccgagtcagc cgccgtatcc ggatggcctc cgatggcgcc tctgattatg 540
ccgtcgctgc ctaccagaac gttggcgttg aaattgacat ccacctcgag cgccgacagg 600
attacaatat ccagataatg cacggccgaa ccgggttcca tgccgcttgc atattcatac 660
gaggccactt ccttgtgggc cggatcattc tttacggact cggcagccac tttgtcgaaa 720
gactgcacgt cgatgagcct ttcgatgagt ccctcttcat ggagcctgac catgtgggcg 780
gtgatgccgc cgagggcata gcttgccttg atgcctttct ctatcatgct ctcgcggatg 840
aacttgacgg cagcaagggc tgcgcctccg gttccggtct ggatggagaa tccgtccttg 900
aagtagccgc tgtttatgat tgcctttgcg gcaagccttg cgagaagtat gtcacgaggg 960
ttcttggagt cacggatggc tccggaagat atctttgagg aatcgcctac cgaatccacc 1020
actaccacat aatccacgtc atgctccgag atagagttcg gagtattcgg ataaggtacc 1080
aggtcatcgg taaggatcac gaccttgtca gcatagcgcg catccggcag ggcatatccc 1140
agggagccgc agattgactt ggcgttttcg gaacggctgt atccgcaggc gtttccgaga 1200
gggtcgcatg aaggtgcgcc gaggaaggcg acgtctatgt gaagctctcc gctggccacg 1260
gcagcgcctc tggtaccatg ggaccggaat atcaccggtt catccatgag gccgcgggag 1320
atctgttccg cgagttctcc gcgcacgccg gaggtcgtga tcttcgagat cacgccgttc 1380
ttgatatggc cgacaagcgg agcatgtacg tcagagagac tggatgcagc caatgtcagg 1440
cctttgatgc ccatttccgc gatcttgtcg actaccatgt tgacgatctt gtcgccgccg 1500
cggaaatgat gatggaaaga aatcgtcatt ccgtccttga gaccgctctt gcggatcgca 1560
tcctcaaggg agacgacctt gcagctttcc ttctgcaggt ccctgcggaa tgtaggtatc 1620
ttctggccgg ggccctggta aaccttcttg ccaagctttg cagcggcctc tcttataagc 1680
tgttcatttg tcatagcgca tcctcctctg gtattacccc ggccgcaacg gcaagggcga 1740
ttattctttc agctcttatc actaccggac ggtcgaccat cttgccattg accgaaatca 1800
ctccgctacc cctctgacgg gcttccttga ttgcggcgat tatggtctgg gccttggcga 1860
tctccttttc ggtcggagtg aagacttcgt tgaccacgtc gatctggcga gggttgatga 1920
tggattttcc gtcgaagccg agagtcttga tgaactctac ctccttacgg aagatttcca 1980
tgtcctcgag attggagaat actgtatcga agcaagagat tccggcggcg cgggcggcga 2040
ctactatcgt ctctcgggcg agctggagct cgatgccttc gttggagcgc atagtcttca 2100
ggttggcggt gtagtcctcg gcaccgaggg cgatacccat catcctggac gatgcggagg 2160
ctatctggta agcgttgact ataccgaggg cgctttcgat tgcggccatt atctgggtcc 2220
tgccgacgca gcctatttct tcctcgacct tgagtatctc ggcttcgagt tcgcggactt 2280
cgtcagccgt ttcggtcttt ggcatacgaa taacgtcgac tccggctttg acaacggctt 2340
cgacgtcttt ctttccatac ggagtgctca gaggattgat cctgacgact ctttcgatat 2400
ttccgtagtc tacggtcttg agggcattgt agacaagaag acgggcggca tccttttccg 2460
ccatcgtcac agaatcctca agatcgagca ttatggaatc agggccatag atataggcgt 2520
cctgcatcat tccgggatta ttgcccggaa tgaacatcat tgttcttctt agtctttcca 2580
tgattcctta tttccaaacg tccttgcctg aggcgcgcac ggcggctgcc gtgactctcg 2640
cccttatagt acagtcgagg gcgcccttgt cgattgcctt gactttggcg tcactaattc 2700
cgagtccttc gagcgtctca gaaataactt tctttatctg tacgccgaac tggtccgcta 2760
ccgaactctc aagctctatc tcgagcccct tcttctcggc cggatcaatc ctgataagga 2820
tatctccaga ttcgaacgta cctgctgatg catttttttag ttccatgttc tagcaatagt 2880
gttattaaac actgctaaat tagccaattt tattaataaa aagaaatctc aagcacggat 2940
tttaaagcag aatataagca atacggctta tttccctatt aatccgtaat ctcatcatcg 3000
cttaattcct taggaatcac cgattccggt ccgctcaacc tgctgccggt cttgatcttg 3060
gcgttggagc gcttcggagc gatgccaagc ttctccagct ttcctatctt cttgatcacc 3120
gactggttag agtcgctgag tttcttcttt gagtcttcat aggccgaaga agccttgccc 3180
agatagtccc ctatcttgac ataagattcc atccatccct tgagctggga catcagttcc 3240

tccgcggtct tgtatacgtc ggagatgttc ttctcctgct catgctgctt ccaggccatc 3300
tggatcatgt tcagcacaat tataagagtc atctggctga cgataaggat gttgttgtcc 3360
ttcgccacct gccagagcct cggggcttct tcaagcataa ggcggaaagc tccctcgaca 3420
gggatgaaca tgatgttata atcaactttc ctgcggcctt cagggatgta gcccgcgtaa 3480
tcctttacct tcagttcgtt gacatgatcc tcgatgctgc ggacatgagc cttggccagc 3540
ctggtcctgt cctcgaccgt cgtcgccgcc atgtactcgt tataggcggt caggctgacc 3600
ttagagtcga tgatgactat agtgtcgtcc gggtaataaa ctttgacgtc cggaatcatg 3660
gtcctgccgc tctcggattt gatttcatgg ccggagttgt cggtcatcac ggactgagtc 3720
tcgaagtgca ctccttcggt caggccggcg ttcttcagga cgtcggtgag gagcatttcg 3780
ccgaaatctc cctggacctt ggaatagccg gtaagcgcgt tggcgagatt gcgggcctcg 3840
tccccgacgg atttgctgcg ggtgtcgaga tccttaatgc tctgttcgag gcgggagtgc 3900
ctctctatgg tgtctttctg gctttccttg acggcgtcgc tgaactcttt gaatttctcc 3960
tggacaggtt tcatcagttc agagattgtc tgggcgctct ttgccttgaa ggcctcgctt 4020
gtctcggtcg aaaggttctt gaagacttcc ttcatctgct cgagctcttt ggcatgcctt 4080
tcagaggtct ccttcgcggc ttcgcgcatc tgattctcat attcctcctt agcctgcagg 4140
cgactctcca gtacggcttt ttcgcgctgg acatcggcca gtctgtcctg gaatccgtct 4200
ttgatctttt cggcacgcgc ctggagacgg aaaatcagta tcgcagctat gacaatgact 4260
gcgacaagaa gaattatcag aattgtttcg gtgctcatcc gcaatgtatt ttaatcttcc 4320
aaatatacga aaatcccttg aagtttctat ctttgtaaga gttaatatgg ccataaaacg 4380
cagaaacaga cttatcacgg cggcaggagc aacgggcttc tgctgctctc cgtatgcgac 4440
ctgatactcg gaggttccgg aatcggggct gccgaggcgc ttggaggcct gttcggagga 4500
gaagggatag cttcctcgat agtctggagc atccgtctcc cgagggtact gacggccatt 4560
gcagcaggcg catgcctggc cctctccgga gcccagatgc agtcagtttt ccgcaaccct 4620
ctcgccgacc ctcatatcat gggcgtaagc ggaggcgcag gcctcggagc cgcgatagct 4680
accctgctga tctcaggcac cgggttgctt tcgggcttct cgtccatggc cgggctcacc 4740
gtggccggag ccgcattcgt cggtgcgatc ctggcatcag ccctgatagt gctggtctca 4800
agccgtttcc gcagcggtac caccctgctg gttttcggcg tgatgctcgg cttcatattc 4860
agcgccatca cctccatact tgagtattca gccagcgaag aaagcctcaa gctcttctat 4920
agctggtccg ccggaagttt ctccggcaca agaactttcg agacattgac gatatttgcc 4980
gcgttgctgg taggaacggg gctttacaca agcggagcca agggtctgga catacttctt 5040
ttcggggagg aatacgccac tctcgctggc gcctcggtgc gccggatacg cggcgtcgcc 5100
atggtcagca gctgtctgat gacggcagcg gcgactgcat tctgcgggcc tctgggcttc 5160
gtcggaatcg tctcccctca tatcgcccgc aggctgctcc ggagttccgg acatatgtcc 5220
atactgccgg ccactctcat tacaggctcg gccgtctcgg tgctggcgga catcatttcg 5280
caggcgcccg gactcccgct tccggtcggc agcacaatgg ccctgatagg cataccgata 5340
atactcctga tactgatgaa aaaatgagaa taaaggactt gaatatcgga cacaggggca 5400
agaccctggt ttcgggcatc agtttcgaga ctgagcccgg agagttcatt cttctctgcg 5460
gtcccaacgg cagcggaaag agcaccctgc tgagagctct cgccggcaag gacgcaatca 5520
tggtaccgac aggggtcccg aaagtcaagg gattcactct cagggagttc atcgccacca 5580
ccctatactc ggaaagcgac tggcgcggcc gcatttctcc ggccatggct tcggacatct 5640
cgaagtcgct ggagtgtctc ggaatttcat ctctcgcaga cagggacatc agcaccctga 5700
gcgacggaga attccagaaa ggctgcatcg cgtcggcgct cagccgccat gcgaagacgc 5760
tgctgctgga cgagccgact tcattcctgg acgtcgacaa caagtcgctg atactacagg 5820
ctctcaggga catcgcaaga agtacgggca ccaccgtgat catggcttct cacgacctgg 5880
cacccgcact cagctgcgcg gaccgcgtct tcggcatcac cgccgaccac cgtttccttg 5940
cctcctccag ccgcctctcc gactcgggac tttcggcttc cgaagtaatc tccgcctgct 6000
tctccaccta ttcgtcccga taagggattc agcgaactaa cggtagatgt cagagagttt 6060
cagctcgact atgcggccgc gttcccgaag cgcgtccata ggcagattct tcttgtctcc 6120
gattatgaaa taagagcaag gagcattgcc gatactcttt tcatggtaat ccaggacgtc 6180
ttgcttggtt attcccggaa ggcctctaac tacggaagcg tcatcatcct ctccatatcc 6240
gaacaattcc aaattcgcga catattcagg caaagccctg aacgaaggat agccgttatt 6300
ggcgcgggaa atgatagagc ggactgctga atcgaagcac agatcagtca tcggcatcgt 6360
acggatgagg gaatccacca gttcaagagc ctgagggaat ttatcagcct gagtaccata 6420
gtaggttatg aatgcagaag gagaggatac attcggaagt ggcttctgaa gcgtactgga 6480

agaagtatat gcaagagaac ggaattcacg gacttcctgg aacatgatcg aagccatgga 6540
accgccatag cattcctgca taaggccgag cagtgcgctg tcatggtcat tctccggagt 6600
agcgacattc tggtaagtta tcgacatcac ctggcgcgct cccgggacat tgtaaatata 6660
gaccacaggc tgttcatatt ttctgtaagc gaaaaatgaa ctctcaggtt ttccggaagg 6720
acattccaca tcaagatggc cggaaatctc attgacgact tcctctggag aaaggcctcc 6780
ggaatagact acatcgaaat catattcaac caggctgttg aaaagctctt tcatcccttt 6840
cagacctaga gatttgagat ctttttctgt cggctgcctc agaagaggtg actggttccc 6900
taatagagaa taattaaaca taccattgaa gatggctgaa ttgccactgc tcctgaatgc 6960
attcctggac atcttcgtat cggaaatcac cgtattgagc gcactttttt ccggcttgaa 7020
atccttcata aagtggcgga gcaggtcgag ggaaggtcct atacggtcat caaagccagt 7080
cagaaccatc tccaaggtat ggttcttatc tttaaaagac aagtccgaac ccagctcggc 7140
aagactcatt gaaaaatcac gggcggacag gctgtcggtc cccgagaggc caagaagctg 7200
tgtcaaagga cagatcatag gttcctcgaa aatacccttg ttgatccgga cggtcagttc 7260
aaagacatca ttgaccgggt tcccgttata gtaaagcgtc acatggtcat taactgcgac 7320
tttccggata tcattattca tatccaccag tcttggagga agagtgtcat aaggcatctc 7380
catcagtctg gtggcataat ccgataccgc attgccattc ttcggcgaaa gagtctcgta 7440
tgaaggagct tgcagattat ccttcggata ggttcctctg actttatgga aacgtatacg 7500
attgtcgccc agatatttat tgcttaccct gactatatcc tccgaagtaa gcgaagatat 7560
cgactccacc tgggaaagat aagaatccca cgacctgtcc tgggacatca cgtctatcat 7620
gatagagaac cttgactcgg acgtttccag gcacctttta tatgatttga caagatcata 7680
tttcaacgac tcgaaatcct tctccgcaaa atcgccggcc ttcagtcttt ccacttcctt 7740
gaagcaccgc gcctccgcct tgttgagaga tccgaaaggg attccaggta tcgttatcat 7800
ggcaataagc cccatttcgt caaaaaagtc caagactgag acctctccag ccagaacctt 7860
gccgttattt atgagggaat ccagacttcc ggtactaaaa ctgttgttca ggaggctgag 7920
agcgagttcc attgcaggat aatcggcatc cttggctgca ggaccgttga acacgttcag 7980
cgacgccttg atcataggga aattaacttt tataggataa tcttcccggg acgagaaatc 8040
cggcacaggt acaggaatcc tttccggctt atcgccacgc ggaatccgtc cgaaggtaga 8100
ttccagcaga ggcaatatcg acgaagcatc tatgtctccg gcaaggataa gtcccatgtt 8160
gcagcccaca taatacttgt tgaagaactt cttcatctcg gaaagacttg gattcttcag 8220
gttatctgta gttccgatta tagatttgct gtatggattt tcatccccaa gtttctccag 8280
catcttctcg agaatcatgt caaactggga atcggctgcc gcgttcttct cctcatatac 8340
tgcttcaagt tcgctctgga aaagcctgaa aacagggttc agcagcctat ggctattgag 8400
ttcggcccac tgcgcgataa attgcggaga aaaagaatta tggtagcatg tacagtcata 8460
agacgtgaaa gcattgagct ttgtgcctcc gatattggcc gtaagacggt cgaattcatt 8520
cggaatcgca tagtccagcg ccttgatgct aagacggttg atctccttct ggagagaatc 8580
ccttttacg ctggaggatg ttgctgcgat gaggtcgtaa caagtctcga tagaatcaag 8640
atagaccttc tctgaagcat agtcgatggt accgatgtcc tccgtacctt tgaagagcat 8700
atgctccaga taatgggcca ttccggtccc ggcgcaatcc cggcctccgg ccttgacgac 8760
cacggctccg tggactaccg gctgcgaatg gtcttcgtta agccagactt gcattccgtt 8820
tgacaattca tactggtgga caactaatgg tccgctactc tgcgcagaaa cagctattgt 8880
cgctgaaagc gcgcaagaca agattgataa aaaaaacttt tccattgaga cttaatgatt 8940
acgtccctcg tctatcaaat aatctgccat aatgaaaaaa ggaggcatcc tttcggaagc 9000
ctcctgaatt tcgatccaaa caaacaacgt ccggagagag atgactagaa tctatcctct 9060
gaagaaacgc tgagtttttt gcggccatga cgacggcgtg cagcaagtac acgacggccg 9120
ttaggggttg acatacgctc gcggaaaccg tgcttgttta tacgcttgcg ctttgatggt 9180
tggaatgttc ttttcatcgt tatctgtttt tttatttatt cttatcaggc cgaaaaaacc 9240
agttttccgg aaataggact gcaaaggtag ttattttatg tggatttgca aattattttt 9300
ctatttcaat cacaaacttt tccttaaaat attcgtcctt cagccaagtt tcaatcggcc 9360
aggtcgaaac tgctccggta cggagtctgt tcttcttcat cagttctgca atttccggga 9420
ctatgtcccc tcccttgaga tacagaatgc ttccggagaa cttgcccttc acccatggat 9480
agaaattatc cagcgaggtc acggcgcggg aaacgacatg atcgaaaacc tgcggcagcg 9540
attctgcacg ggcattgaca gtggtcacgt tgcccaggga aagagccttg gacacttccg 9600
atgccacgat tatcttcttg cccacggaat cgcagagggt aaagcgaact ttcgggaaca 9660
ttaccgcgag aggaatcccc ggaaagccgc cgccggtgcc caggtcgagc acggacgcgc 9720

cttcggtcca tgcggaatac acctccggcc tgcgtctctt cagatacgag gcgatcgcaa 9780
gagaatgcag cacatggtgc agataaaggc cgcctatgtc ttttcgggaa attacgttga 9840
tcttggcgtt ccactcctca tacaggggct ccatggcctc gtactttcca agcatctccg 9900
gagtcagctc cggaaactca gcaacaacta tatcttcgaa ttctttgaaa ttcattcttc 9960
ttccatatca agcatcatct ggactatcag gctcttagcc ctgctgatgc gggtttttac 10020
tgtattgata ggcagccctg ccttctccga aatctccttg tatccgagat tgtccacaaa 10080
gcacatggat gcgacttcgc gatacaggtc cggaagcccc tggacacaac tcaggaagtt 10140
ctcatggtcc tgcgaactta tgatctcgtc ctccgggctg acattgtcct cagggacgtc 10200
tatatgctcc aggtcggagt agtcgatcgg catcttgtct ttgttgcggc cgcgcacttt 10260
gtccttgtcg aggtggtcga aggccgtgtt gcgggctata gtgaatatcc aagtcgagaa 10320
cttgttgctc ggcctgtatg attccagctg cttgaaggcc ttctcgaagc tctccataca 10380
gatatcctca acttcctcct gttcctggat gaagcgtgag atgtgggcga atacgccctt 10440
atggtaacgt gcataaatcg tgaagaaagc ggcctcggct tcggggccgc cacctaacgc 10500
catctccaca agagatccgt cgtcatgatc agtgagcttc aagccagttt ttgccatagt 10560
tacattctac agttaaaggt attgacaagg taattacgtt ctccatttcc gctacaacta 10620
tcctgccgag ttcctcgatt tcttccggca cggcatcgaa aagaagctca tcatggatct 10680
ggagcaccat cttgctcttc agtcccgcct cgcggatgcg cttgtcgacg cttatcatcg 10740
caagcttgat tatatcggct gatgtgccct ggataggagc atttacggca ttcctctcgg 10800
ccagagccct tatggtagcg ttcttcgccg ttatgtccgg aagatacctc ctgcgtccga 10860
agactgtctc gacataccct ttttcgcggg cgtccgcaat agtcttgtcg atgaaagtcc 10920
ggatcgaagg gaaggcggcg aagtaatcgt ctattatctt cttggcttcc gtcctggaga 10980
catgcagcct ctgggcgagc ccgaaggatg agatgccgta catgattccg aagtttgcgg 11040
tcttcgcgat gcgcctctgg tcggatgtga cctcctctac cggtatgccg aaaatcttgg 11100
ctgcggtggc cttatggaca tccagccctg ccttgaatga gcttatcaga tgctcgtcgc 11160
cgcagagatg ggccatgatc ctgagctcga tctgggagta gtcggcggac attatgattc 11220
cgtccggagt cgccggtatg aaggctttcc tgatctcctt tcccctctcc gtgcggatcg 11280
gaatgttctg caggttaggg ttggaggaac tgagtcttcc cgtcgccgtc agcgcctggt 11340
tgaaggtcgt atggaccctt cccgtaaccg gggagacgta ggtcggaaat ggctctatgt 11400
atgtggagag cagcttcttt acggctctga attcaagtat cttgtcaatt atagggtgcc 11460
tgtcggcgat ggcagtaagc gtcgtttcgt cggtgctata gctaccgctg gccgatttct 11520
tcatcttcgg gtcgagtttc agcttttcga agagcacgat gccaatctgc ttcggggaag 11580
aaacgttcag ccccggctcg tcggccagtt tcctgacctg ggcttcgagg gcatccattt 11640
cgactttgag ttcatcggca aacgtcctga gctgtccgag gtctatcttc acgcctgccc 11700
tttccatgcg ggcgagcacg gctgcgagag gggcttccat atcctcatac agggctccta 11760
cgccggcctc gtccatgcct gcgcggacca cttctccgag tttcagcact gccgcggctt 11820
cgagattcct gtccggtccg tttgccgggc tttcttccgc gacgtcaaac agggaaacag 11880
gagcttcttc cacagcagcg gcctcttcga tatcgacgcc gagatagctc ttggcaagga 11940
tttcgacctt gtggctcttc tccggattga tgagataatg catcagctcg atgtcatata 12000
gccggcctcc gagctttatt ccggcattct ccagaagatt cgactggagc ttgaggtcat 12060
atccgtattt cacggcttcg ccctccagta tcttgcgaaa atccttcgca gcgcctgaag 12120
cgacgacgat tccgccgtct ttcttctccg cggccacgat caccctgcgg atatcggcga 12180
atatgcccgg tccctccggt tcgactatca gcgaaatctt ttccggagcc atcttcacga 12240
gctcttccgc ccctgtctcg gtataatcca ggcgcttgac ctccgccact gccttggcag 12300
cgcgggagct atgggctata tattttctca gcgagttgaa ttcgtatttt tcgaaaaggt 12360
cggcgatttc gggagcataa tccaggtcca gcctcatctc gtcgaggctt gtctctaccg 12420
gcacgtcggt ctttattgtc accaggtcat ggctcaggcc gatatgtccc ctggcctcct 12480
cgaactgcgt cttctggcgg tcagtcagct cgtccagatg ggcatagata ttctcgacgg 12540
tgccatattt tgcgatgagc ttgcctgccc cgacttcccc gatgccatgc actccgggca 12600
cattgtcggc cgtatctccg cagagggtca gcatttcgat gacctgttcc ggactggaga 12660
ttccgtattt ttcgcagact gccttgacat cgacggtctc gtcagccccg ccggcctttc 12720
caggcttcac ctggtacaca tgttcggcga ccagctggcc gtaatctttg tccggggtga 12780
ccatatagac gtccgtcccg ggtttcccgt aactcttcgc aacggttccg atcacgtcgt 12840
cggcctcata tcccggaatc atcaaggtct ttatgttcaa ggctttgcat atttccagca 12900
gcggttcgag ggcatcgatc acgagctgcg gagtttcgct tctgtttgcc ttgtattcgg 12960

ggtacatttc gttcctgaag gtgccgccgg gaggatcgaa cgctatggcc atatgggtag 13020
gcttctctct ctcgatgagt tcgagcacat atttcgtgaa gccatacagg attgacatgt 13080
ccattccctt ggaattgatc ataggacgac gcaggaatgc gtagtacatc ttgaagatta 13140
ggttgtgacc gtcaatcagg tatagtttgt ctgccatagt tctccggatt aaggaacgaa 13200
atacaaatat aacatttttt tcaattccct ccagtccctc atctttgccg cagtcctgca 13260
ttctgcttcc tttgccacag tcctgctgac gcagggtccc ggggaacttc gttcgctatc 13320
gctcactccg taccctccca caatctgccg cgtcatcgct tcgcgataac ttgcatcttg 13380
cgggccccgt ccctcttccc gtggccgagg ggggccacgt ccccggaacc ctggtcagca 13440
gtcctgcggc gggaggaggg cggaggcggc ggaggaattg aggaaaagcg gagagcgcgg 13500
gagcggccgt cggcgccgtg aagaacggcg tgtgtttgac gaccgttacg atggctcgcc 13560
catcgtaaaa gggagaagtt agccgtttag gcggcgacag cgaccaagca gcttttccga 13620
acttgacgga gccgtcccgt ccttctcccg ccagcagaaa ggcccgactc gaatcagcca 13680
gacgaaaaag gatggccgaa gagggccatc ctttggaata acaaatacct acatgaatga 13740
agatgttatc tgcgagcaga aagtttctct ctaattgaag ccatacgcga ttccgtaaga 13800
ggatagaaca gcaggcagag ggctccgagg agggcgccga ttgccgggat atagctcatc 13860
aggccacgga tggcagcaag agtctgaggc gcctggactg cagcatcctt gtcatatgcg 13920
aagattgcaa gaatctgcat cagcaagaaa gcgccgaggg ccccgccgaa cttctgagcc 13980
atagaagaag acgagaaaat aagtccggta gaagccgagc cgttcttctc ctccgaataa 14040
tcggcgacat cggcaaacat cgaccacagc agaggagact ggacaccaaa agcaatgcat 14100
ataagcacct gcagcaccag aagtccccag aaaccggcag gagtctcagg caggaagaat 14160
atcgccacac tcagcacaga aaccgtcaga agcacagaga taaaagtcgc cttcttgccg 14220
atacgctccg aaagcggcac agtaaacagc accccaacca tctgagccac ctcgccaatg 14280
aaaagataga cagcgcaagt caggaacacg ctcgtaccga ggatattcga gaaataatat 14340
gcagcggctc cgcctcggat cgaattgaac agcaatatac cgataccggc gccaagaagc 14400
agccaccaag gcctgttacg gcctagagcc ttgagatcct ctgcgataga agagcctccc 14460
tcggaaaccg ccttattttc agtccccacc cgctccctgg tcataaggaa gcaaagcaga 14520
aacagcacga aacagcacag agaaacgatg ccgacgacaa gagtccactg tccgggagca 14580
gcatcgccga ctccgcgcac caccctctcg ctaccgtccg gcataacacg cacggtaccg 14640
gaaatcagct tctcgaacac gccgaatata cccatggcga tgaaactgcc gatatacgca 14700
aagaacatcc tgtaagacga gaaaaccgat ttctcacggc tgtccggagt gacgactccc 14760
agcatggccc cgtaaggcac gtttatagcc gtatagacag tcatcatgag gatatacatc 14820
acatacgcat atacatgctt caggccatag ctggcatcag gagtatagaa agacagaagg 14880
cctataagcg agaacggcag agccacccag agaagatacg gacgatactt tccccatctg 14940
gaccgggtcc tgtcggcgat gattcccatg acagggtccg atattgcatc atacagtttt 15000
gtgataagaa gcaggaaagc cgcatgctga ggcttaagcc cgaacacatc cgaatagaag 15060
atcggcagat aatacgaaaa aatcttccag aacatcgaag acgacatgtc tccgaagcca 15120
tatccgatct tttccttcaa tgtcggctta gccggtgcag taatgatatc catctgaaac 15180
ctgattatta gtcctacata gatagcgaaa gattcttgtc tataagcgcg ttcagacgtc 15240
gtactgaagc agaagtggta aacagatccg gctcagtatt catacagtaa tccacgagtc 15300
tgtcgacggt agaagtcgcc acatggagac gcgtatcgca ggaagcataa tatataaaga 15360
cctttccgtc ctcatcggca atccaaccgt tagagaacag gacgttcatg acatcaccca 15420
catactcctc cccttccggg accatgaaga aaccggcagg ttcggcaatc actttcgaag 15480
gatcctccag cgaagtcatg tacatataaa gcacataacg caggccggaa gcgcagcccc 15540
tgacgccatg ggccagatgc agccagcccc tgtcagtctt gataggagca gggccttcgc 15600
cattcttggt ctccttgata gtatggtatc tcctgcagtt gatgatcttc tcggaagtga 15660
caacggcatt ctccatgctg tccaccaggg cccagccgat gccgccgccg tttccggcat 15720
ctatgaatcc atcctgagga cgggtgtaaa gagcatattt cccattcacg aactcaggat 15780
gaagtacgac attgcgctgc tgagaatcag aaaccagatc cggcagtctt tcccaattga 15840
ccagatcctt ggtacgggcg attcccgcag ctgcgacggc agacgaaaga tcgccttccg 15900
gagcatccgg atccttacgc tcgacgcaga atataccata gatccacccg tcttcatggg 15960
cagtaagtct catgtcatat acattcgtgg caggttcgcc ataatcaggc atagtcacgg 16020
ggtggtccca gaaccggaaa ttgtcgattc cgttcgggct ctcggcgata gcgaagaatg 16080
atttgcggtc agcaccttcg acacgggcgc agacgacata tttgtccttc cactttatag 16140
ctccagcatt gaatgcggca tggatgccga atctctcttc cagaaaagga ttggagttct 16200

tatccagatc atatctccag aacaaagggg catggtcacc ggtaagaaca gggttctcat 16260
atctctcaaa cacaccattg ccgtatatag gcgagttttt cttagcaata agttcctggt 16320
ggcgtgcttc cagccacgcc agacgttctt caaaactcat attctccata tattgttatt 16380
tattgtttta aggttacaga ttcccggtcc gaccggaaag gacgacactt cgccgggaat 16440
gacggactaa ggccggaacg acgacaggag agagatctcc ggcagcgaag cgaattccct 16500
gaagtcgccc tcattaccag gatacggcgc ataaaaatgc gtctccctgt cccatgcatt 16560
tctccacgtc agcacataga caatcggaaa atcctttacc gcaggataaa gcaccgaagt 16620
ccaccagctc ctgtccggga ggccctccat tccggtttcg gaaagagtca tcagctttcc 16680
gtgggccgaa gcaagagact tgacgatgga aaggccaaga tccaactgcc cggcataacg 16740
ctttccggca tcttccagac tctcgccctc catgacagtc tcataatggt catacccag 16800
gatatcaacg acatcatcgc ccggataacg ggacatatac tgctccgccg taatcccgga 16860
attcggagaa taacacatta tgatattgtc cagtcccctg tcatttctca gatactcgac 16920
cgtcctcctg aaaagggcga tatactcatc cgcagtacaa agggctccgc ccaccagaa 16980
ccagctgccg atattctcat gccacggcct gaaaatgacc ggtatcagat tacccttatc 17040
gtcacggagc gagtccatga aatcgcccag cctctccagc cagactccga acatctccgc 17100
cttctcccct ccttcgagga cagaagccac tgccttgtcg gaactgatat cccaggcatc 17160
accgcctgta taaggattgt gcgggtgcca tgagaacgtc acgattcctc cccttccggc 17220
atgctctaca gcagcccggc gcatcagcag gaacgggact ttgtcaatgc tccagtccca 17280
tcccagttcc agaccgccga gttcgaatcc cacgacagcc ggataatctc cggcgacagc 17340
cttgacatcc gaacggttca acgcatctgt cgctgcggaa tcaggatcca cacaccatga 17400
atggccataa acgaggtcat cctgatggcc ataaagaatc ttgccggcct ttactgaagc 17460
cgccatacga tccatcatga tccgggtttc ttcgttaaaa gctgttccgg ccgcttcttt 17520
attgcgtccg cagccggata cagcaactac taatgataaa atgatgctaa gtttaaggat 17580
tagattcata atcaattggt tttcggagcc ctgtcatata cactctggag agtatagctt 17640
tcgtccgtgc tgagcattcc ctttacgaga agatccacga tctccttaga attccctata 17700
tacttgccgg taccatggtc aatgtagccg tggcactcct ttccggtctt agtacttccg 17760
ttatcccaca ggattcccgg taaggaatag gtatgcgcag ctttggagac atactccata 17820
tagtatttat agaacatcca tgcccgggca tccccttttgc ttctcatcga acagccgaat 17880
tcgccaagat atacaggtat ccccttgctg atgaattccg tattaagtct ggagaagaga 17940
tccgttatct ctttctcccc ttccggggtc ttctttcctg cggcgccggt atggccccac 18000
tcaggatact gtatgttccc tatagtatat tcatacggat catagcaatg cactgccatc 18060
atgacctttc cggcaggatc ctccggcagt ctgaaatact tcgtccatga agtacctgcg 18120
gcgtatgtag gaacacccag ccagcgggta gcgttgttgc ctcctgtagc acgtacggca 18180
tccacgaaaa cctgattcca ttcgttaagc acatcacact gcttggaagg gtcagcctgg 18240
aaagcagcgc tccagcccca tcctccgtcc tgaatttcat tgaaggactc catgataagg 18300
aaatctccct tatccttgaa cttattggcg atatgggtcc atacagccgc gatttccgcc 18360
tttatcgctt cattggtcct ggcatcctta gatgcagaaa ggatatccag ccaatggtcg 18420
tcattgttgt tttcgtcgtg atgggtattg atgatgacct tgagtccggc tttctcggca 18480
tatcctacca cctcatatat acggttcatc caagtctcgt caatattgta ctcaggtcct 18540
ggacctatca tattgagcca cgtgacaggg attcgcactg atgtgaatcc ggctttcttc 18600
accccgtcga aagtagcctg ggtcgccttg gcattacccc agacggtctc gtccggataa 18660
ttgtatttat ctccggccca tgagccgttg tagaaagcat cgaactggtt acccatattc 18720
cagccgagac cgagaagatc gtatgcagga ttagccgacg gtccgaccgg cttcgccgcg 18780
gcatcctggg atacgcttat cccgaccgga gaaaagccct ggcaagtaag ggtcacacta 18840
ccgctgcgtg actcgtatgt cggattctcg ccgacggtca ccgtaagttt cactttatat 18900
gtatccggat tgtacggacc cagttcggct ttgatccagt ctccggagac agagaccgca 18960
ggcttggacg gggaggagat ggcgacgacg accgacgttc cctctacggg agcgctgaca 19020
gacgcgacat ctaccgttac cgccttaggg gccgggacgg acggttcctt cttttcggca 19080
tcatcctttg agcaggaaca tgccgcaaga aaaatcatag ataatacaaa gaaactgacg 19140
tgtgctttca ttatataaaa tttgtttact ggctaccgga ggcggcatca cccgcctccg 19200
gcaataaaag tgttacaata tacaaacttt attgataatc atatcagatc cctggataac 19260
catggcgacg cgtccgtcac cctggaactg gctattgaga ccgaaatagt caccggatgt 19320
agcaagtata tggtcaagga tatcagcagt cagggtaagg gcgatacgtg tgacgtcctt 19380
tcctccgagg ttgtcggtgg taatgtatgc accgccgatc tcagggaagt tcatgtcggc 19440

gttggcccac cagccgtcgt tgagctgaac ctggccccat gcgtccgtct gctcgagata 19500
gaagataagc tgagagcccg gctttgctgc ctcgaagtaa gtcatggcga gaccgaaacg 19560
gccgtcgtca ccccatgtaa gggttgtagg accttcgtag atcgtaacct cctgcggaat 19620
atcgcatacg atctcgatct tcgtgaaggt aaggccagag ccctggatga tgataccgca 19680
ggcagtcttg tcttcgcagt caccccttgtt agcctggatg ttttcgagga tttcctttgt 19740
aagcacgacc tctgtgcagc ggtcaaggat tccgtcagtg aaccatccat agacgtcagt 19800
cggaacgagg gttccgttaa tgacggtacc tgcaggatcg tcgaagttga ttccggtcca 19860
gttaccatag ttgagctggg cctgatccca ctgctggtcc acctgagtgt agtagaatct 19920
gagcttgctg cctgcaggga taccggtgaa tgatgctgcc ggcacggtta cacggccgcc 19980
gtcattccag gtgatctgca gagagccttc ccatacagtc atgatgtcct gctcgaggag 20040
aacttcgatt gtctgaggac atgtctcgac atcgccggag agaagatgga agtacatgct 20100
gtatgttccg accttgttcc aaggcattac gaactggatt tcattatcag aacgaaggtt 20160
gtacttcgat actttctctt cgccgatata gatgttctcg atcatcatga agttctcgcc 20220
cttgagggtt acgatagccc cgatatgctc tgcggcaggc atgtcgttca cgatgatgaa 20280
tgaatcatag ctgagggtga tggcatcctc cggttcaatc ttctcaccat tcgcaagaga 20340
aagcacgagc tttcctgaaa cggaagtatt ggcggtctgt acgacaatct tctcagcctc 20400
tgcagtgaac tcgaccggct ttccgccgag agttactcca gtgacaaggt cgaggtcggt 20460
acctgtcacg gtgatggtct cgccggccat gagctcgaca ggagctatgg cagaaactac 20520
cgggtgcaca agggtgaatg ccacattcac agcttctccg ctggccatat tgagggcgat 20580
atcaccttca gtagcagtag caggaacggt cactgtaatg gcgtcctcgg caaatacgaa 20640
ttctactccg ctggcgccag ggaaatcaac agaagttacc acatcaagat cgttaccgga 20700
aatcacgagg tcggcaccag cctttacagg cgaaggtgca gctgcgacat ccgtagggat 20760
gacaagttcg atttcgccgg cgacgaattc cttggcagcg tggctgacag cctttacttc 20820
accggactgt gctccggcag ggattacgag attgaattcg gttgcatcct cattgagagc 20880
aaggtcggaa acctctgcac caccttcgaa tacgaccttg tcgatcatat cgagatatgc 20940
tcccttcaca actgcgttct cgccaggttt agcaacagct acggtaagag acttgacggt 21000
cggatcgccg atcacaagat cttcctccga gtagaggaga ttggcgatgg ctccttcgtc 21060
agaaaggatg atctttccgg taatagcctt tgaagggacc ttgaaggaag cctcatgacg 21120
gccgttatgg ttgatttcgg tcacgatggc gccgccttcg aaggtaacgc tctgcacgag 21180
gttcatataa tcgcccttga ccgttatcac gtcgccaggc atagcctctg caggagagaa 21240
actgtcaaat acgataggtt ctgtatattc tacatccacc tgggtagtaa agacctttcc 21300
gtcggatgac ttgagagaga tctttccgac ctgtgcgccc tctgcaggaa gctggatgcg 21360
gatttctgaa ggattaccgg aggtaacgac ttcgatgtcg gtgataggat cgagtcccgg 21420
aactgaaact tcgacaatcc tgtcgagatt acttccgaag aaccggagtg tagctccacg 21480
catcaccggg ttagggccga atgatgccag cttcacgtcc ttgtcggaat actggtcggt 21540
atcaagctct ttggatgaac atccggagag cccgagtgcc aggacacagg cggcaagaat 21600
ggctgaaata ctgaatatct ttttcataat gagcttcatt taatattatt cattaggaac 21660
cacacggata ttgtcgatcg cgaagaccgg agagcaggct gtgccgtcga taccgccgtt 21720
ccatacgaag aatgtgagac cggtaaggta agacttgtca aatgcggttg cacactcctt 21780
gttttcctgg ttcttgttga atgagctgag agggaaactt acggtaaccc acttgtcagc 21840
ggtatcataa gaacctccgt tggtagtcca tggctgccag atagcacgag ggaaggtttc 21900
gtcactgaag tagctgttcg tagcggtacc gtaggtaacg acagcgtcgc tcgagaacat 21960
gagctgcata cctgaagcag accaagggtt ggatgaagga acgagaacct cgaacttgag 22020
cacgaggccg cttactccat acttgtcgat atactctgcg aactccggac ggaggttgag 22080
ggcatcgtaa tcggtgccgg cctctggcca gtagttgaag ctgtactggt cctctgccca 22140
tgcatctgca cttccaggac caagatcggc gcctccgaag tagaggtatg aaccgtcgat 22200
cgcatcccat gcatcagcgc caggagcgtg gatcacgccg ttacgccatc cgtgaccggt 22260
agcaaagccg ccgtggctgc catcccagtc gaagaggatg tttctggtat ccttgtacat 22320
gaacttgctc tttccggtac cggagagtga agtcacggta acatatccag gctgggctcc 22380
tgcaggaaca gggaagctga ctgaagacga accgttgaac accattgaag catgatcgac 22440
ctcggctcct gcaaaggcta tctggagagg cttgtcagcc ttgtcaataa ggaagtcgcc 22500
gtagattgtg acggtctccc cttccgcagc ccactcattg gacatggaag taaccttcgg 22560
agtcggagga agcacggcga aatcatacgc gagcgtatcc ttgcccgaag taatcatata 22620
tatcttgtct gtctgtacca caggcatatt cttaggcacg gataccagaa gagtattgtc 22680

tgtaatgaag ctggtattga gggttgcctg ctggtcgttg aaccagagtt ccacaatact 22740
cttgagattc ttgccgagga gacatacgac ctcatccata tatgcctggg tgatgttgac 22800
gtccttgtct gcgtatctga cggaataaac cgtcggcttg ccgtcagcta cttcataacg 22860
gtccggatag tcaaccttgt cgcaggaagc gaatgacgcc attgcgagga ttgccagtga 22920
accggtgatt attctattta tagatttcat cttttcgtca gattaaaagt tatagctgta 22980
ttcctcgcgg acatcgacat ggacaggcac ggcgtcagat gccatgttgc cattgaacac 23040
tacgtcctct gtagggaacg gaagcttgaa gcttgcagcg gtaacgttag gattgtcctc 23100
tgcgttgttg tactgtactt ctgaagcatc ccaggttccg gaataatcga aagcatctcc 23160
gatatatgat ttccaaacat cgtcgagtcc atcccagtat gaccttctct gagctttgag 23220
ttcagcgatg catgcgtcca tatcgtagta tgaacggcgt acatagtcat accagcggtc 23280
gccctcgcct gcaagctcaa gacgacgctc tttccagata tcctcgaaag tgacggatgc 23340
gagagcatcg gcgccagcac gttcgcgaac cttgttgaca tactcgagag cagggccgtt 23400
gcttccggta agaaggcttg cctctgcgta gataagatac acgtcggaaa gacggagaag 23460
atgagtagga agctgataag ccatgttctc aggggttccc agaccggcgt tctcatgatc 23520
aaagttatcg ccatagagat gctttacgtt gttggaacct gtagggcact gcatggaacc 23580
gccaggaccg cctttgccat attccgtatc atagatgaag cggagataat cgaaacctcc 23640
cttgtcagtc cagaagtaat catacttgtc acccgggagc atcaaggttg ccttgcggcg 23700
gctgtccttc tctcctacac gctctgcagg attttgggtc gcacttacgc cgaatgcgtc 23760
tgcaagatcc actgaaggac cgccccagcc tccccagcag tctccctggt cgctgaatcc 23820
gaccattgcg aggtcgctct ggagggtatt ctgctgagtc caagggttac ggcctacgct 23880
ccattcccag gagataaggc actcgcctgt ggcgttgaat ccggtaagac ggaagacgtc 23940
cgaatataca ggaagaaggg tacgtccgga attgtcgatc acatccttgg catacttggc 24000
agcctgcttg aggtcatcct ggttcaagga tccggaaagt ccggccttgg taagatatac 24060
ttttgcgaga agggcctctg ctgcatagta atcgatacgc tcgtaattat tgaagccgga 24120
agtcttctta ggaagcattt ccattgcctt ttcgagggtc gtgatgatgt actcatagac 24180
atcggcgcgc tccaccttgg agatagagtt gtactcgccg ctggagagga gatcggaatt 24240
gctgtggacg ataggaacgt caccgaaggt tcgtaccatg aagaaatagg cgaatgcctt 24300
ccaggtaaga gcttctccga tacacttgtc gatcgctgcc ttggacgggc cctgcgaatt 24360
aagaaggttg tagatcacgg tattggcatg gccgttcaca gcccagagag aataggacat 24420
gttgacgagg tcctggtcgg taccgttggt ggagaaatca aggtatggag acgagcccca 24480
gtacatgttt ccggacatga cctcaccgat cttgatgaag cctctctgga agtcatacca 24540
tggagagttg tacaggtaat tcacgccctg ttcgcactgc tcgtcggtct tgtagaaatt 24600
atctacgtta taattatctt cgttaggtct gttgaggaac ccttcgcagg aagtcagagc 24660
aaagactgcg gtcgccacag caaaaccttt catcatatta gagattttca tattccttca 24720
attttagaat gtaatgttca aaccgcaaga atatactgta ggagaaggat aacgtccgtt 24780
atcaacgcca taggtaagac cggtggaatc ctgggtgctg gcacctactt caggatcata 24840
tccggtgtac ttcgtgaaag tataaaggtt ctggatgttg acgtaaacac gtacattatc 24900
caccttgaac ttgttcagga aagacttagg gacattatat ccgagagtga tgttcttgat 24960
acggatatat gaaccatcct ccacataacg gtcgctgata cggtcgttgt cgttagggtc 25020
gttgatgctt gcacgaggag tcttcgtgcc agggttgagg accctgacat tgtcgatatg 25080
gtcataccac ttggaaccgt cagcatagac tacgcttgcg tctataggag caagcattgc 25140
cctgccggag acagcgtcat tgatctggtt ggtccatgcg ctgttcatat gggtaagaga 25200
aatggcattg tagttgagga ctttgttacc atagcttccg ttgatgaaga tattgaggtc 25260
gaagtccttg taggtaaagg tattgctcca gccaaaggtg aacttaggaa gaggattgcc 25320
gatatcggta cggtcctttt cattgattac tccgtccttg ctgatatctt cgtacttgat 25380
atcacctacc cacacggtat ttccacgggc atactctcca ttagaaggat atttctcagg 25440
tttcggagaa ttctcgatat cctcgagaga agtatatacg ccagcgacct tgtaaccata 25500
gaagttatag agaggcttgc cgatttcaga cacacacaca acgtcagacc actgtccgta 25560
acctatgaga gcggcattag tcgttccgtc aaggctgaca agcttgtttc tgttgaagga 25620
aatctggaag ttgctgtccc aggcgaaatc cttgatctgg acaggatggg catcgagagt 25680
aatttcaaga cccttgttct cgatagaacc gaagttacct ttaggagctg caagagcaga 25740
agaactgtta ccctgggttc ccatatagga agcaagctgc attgacataa gcatgttgtc 25800
agacttcttc aggtatgcat ctactgtaag attcagcctg ttgtcgaaca tattgaggtc 25860
gagaccgatg ttctgctgcg tctggctctc ccaatggata gcaggattag ggatgttagc 25920

aggacggtaa cccatgccga gagccgaagg catacggctt atggatgctc cccatgcata 25980
accaccgata cttgagttac ctgtctgccc ccatccgaga cggagttttc cgttgttgac 26040
cacaggcttg aggaaatcca tgaatttctc attggtgaat ttccaagcca ctgccaatga 26100
atggaaaccg gcccaacgtt tctcaggacc gaagttggat gatccatcat aacggtaggt 26160
ataggtaagg aggtaacgat cgtcgaaatt ataagtttct cgagtgaaga atgacgccat 26220
cgcagagctt ccgaaaccgg aagaaagcga cggagttcct ccaccgaggg cgacgttatg 26280
gacatcgttt gaagggagct tggtatttgc gccctgtacg taactccacg aagactccca 26340
ggcctcctga ccaagcatgg cggtaacgct gtgctttccg aaggagttgg cataagtgag 26400
atagttcttg atctgccaga atctgttgct gttcttctga tagctcatag agttggtatc 26460
tcttgtccag ccaccgagat tgactgttgg tctgaaacgg gtagccgtag agccgctgac 26520
atcgaaaccg agttctgttc tccaggtgaa atgctttaca ggagtaagtt ccgcgtagat 26580
attaccgttg agtttctgac ggtccagcag aatctcatcc atcattgcaa gagctacagg 26640
gttcggagaa gtatagcctt cgcgtactgt agatgagaaa ttgccataga catcataaac 26700
agggatgtca gggattgtgg taagggagta tgtgataaga ccttcgctac cgtcggcaag 26760
cttaaggtca tcattggtgt tggcaaaggt aaggttgaga ccgagcttga accattcctt 26820
caactgggcg tcaaggttgg tcctgaaggt ataacggccg aattcggaac ccctgatagt 26880
accctcctga tccatgaatg atccggaaac atagtactgg actttgtccg taccgccctg 26940
ggcgctgatc tgatgctggt gctggatggc tgtgcggaag atttcgtcct gccagttggt 27000
accctttccg agcaggcccg gagtagcata gtaaggattg gcctcactga tttctccggt 27060
ttcaaccata gagttgtaga actctgcgta ttcacggaga ttcatcatat cgatacgggt 27120
tgtctgcctt gaaacggcga acataccgtc gtatgagaat ttcgcatcac ctctggagcc 27180
atgcttggta gtaatgagga caacaccatt cgcaccctgg gcaccgtaga tagcggttgc 27240
ggaagcatcc ttgaggattt ccatgcttac gatatcggca gggttgatag tggaaagtgg 27300
agaaatcgta gagactttac cattgccgag agcatctccg agaccgaagt ctgcaccgga 27360
gttaccgcct ccctgcatga ttacgccatc gatgacgtag agaggttctg cgttggcatt 27420
gattgtcgcc tgaccgcgga cacggatact tgaggacgaa ccaggagctc cggaagtctg 27480
gactgccgtg acaccggcag cacgtccctg gagagactgg tcgaggttgg tgatgatcga 27540
acccttgagg tcatcctcct tcatggaaac ggatgatccg gcgaggtcgc ttctcttcat 27600
agtaccgtaa ccgactacga ctatttcgtc gagcagctcg gagttttcga caagaagaac 27660
gttgatgaca ctctgggtgc ctacaacaac ttcctgatct ttgtaaccga tggaagaata 27720
tacaagagtg gagccttggc cgacaacaat ctcgtagtat ccatcaagat ctgcaacgac 27780
accattgttg gtgcctttca ccattacgct ggcatttatg acgccctgtc cggaattatc 27840
tgtaattctt ccggtcactt tgttctgcgc cataagtgac aatgctccaa agagcatcag 27900
catacatgtt aatagctgtc ttttcatgct actgattttg aatgtgaata attagttgtg 27960
ttatttattt tttgtgaaga attccgtcct tgctgcccac cactcgggaa attgctccag 28020
atggtggcct gagacaggaa cgcaaagcca ttgtttatct gcagtaatat ggttataccc 28080
cgaaaaattg gtatgaggag gacagacggg atcctgcagg ccgaaaccca taagcacagg 28140
acaacctatc ctgtccgtaa agttcttcac gtcgaaataa gacagggtct tataaagttc 28200
gtcttcagaa atgccctgag cctttgccgc cgagagcact tccccgcccg gccagtctgc 28260
gatggcgaaa tagtccggga aatcggaaag gaaaggcacg aacggagcga ttgcgcggag 28320
acggttgtcc agagatgcag ctatcagggt aaatgctccg ccctgacttc caccttcggc 28380
gtagatatgg tccggatcag tctgaggaag agaagagacg aaatcaatgg cccgcacaca 28440
gtccagatat gcgcctctgt aataatacga gtccttatcg gcaagtcccc ttgtcaccca 28500
gccttgtccc ggagtcttgt tgaaaccctg gtcacggacg ctgagagtga actcgatcat 28560
gtccggttcc gccgacggat cagcgaacca gacgtccgaa ccgtagccca tatatgagat 28620
cctggcagcg aattttccct ctttgttcgg aagataaagc agaccggaga tctctttgcc 28680
tccgaaagac ttcatatcaa ccctgtaggc agtcctgtca gcgtttgaat gatccttcaa 28740
cggagtcatc ctggcattca tcgggaccat ggcaagctgc ctgaggttgt cttcccagaa 28800
ttcatcgaaa tcaggctgtt tatcctgagg agatactatc ttctcaggat cacagccgac 28860
acagaagttt tcatcaagcg ggtcattctc gttcaggctg acacacacct tataaaagcc 28920
tggggctagt ccaagatcga aatccagagt atcggcgcct ttgccgagac gggctttctt 28980
tcctgagacc gacactgaaa ccccttttgtc tgttgacacc cgcacataaa ccgggacact 29040
tcccttaaaa ccattctgag actgtacagc aactctgaga ggtttctggt ctccggtagt 29100
gaacatccaa tcattttcaa gcactgcgct tgcattagcc acggcagaaa gtctcgccgt 29160

tgtatgtttt ataatagcaa gggtactgta atcgtcggcg aatacagaat tgagcccctg 29220
ttcttcctgg gccggatctc cggcgtaatc gtctccttcc tgccaatatc tgtgggcagg 29280
caccgcaaga cctccccagc tccagaaatt ggcgccggcg agtctgccgc catttttctg 29340
attctccatg accttggaga aaacgtatct gtaatagcgg tcgcgtccgc ttacgggagt 29400
ccctttggca aactgcatat catctctcgg gaagccgaat tcctcgatca cgagaggctt 29460
gcccagactc tcggccacct ccagatgggc gtcgatgtat ctcccggtat tctcctcggc 29520
gcggtctata tccagctgag gggattcctt tcctacccag ctccagttgt atggccatat 29580
atgggcggta atataatcaa tgtcaggaca ttcattcagc cggcggaata gctccggatt 29640
ctcttcgcag ccccatatgc cctcattgcc ggtagaaacc atatgattag ggtctatcga 29700
tttgattatc gaggcagcgc gccatatcca tgatacgaat ccgtcctgga cggcaggatc 29760
ggaggagaag cagcgcggct cgttgccgat ctgccatgag aatatcgccg gatcctcgct 29820
ataaggcttt ccggtgatgc tgttgactct tgaaacgaca gtacggagat gattctcgaa 29880
gagttcctgc gccctggcgt tcgtcgcata ctgggccatc tgttccatga acggaccata 29940
tcctccgacc gaaggaatta tcgccttgcc ggctccggcc cattcgaggt acatgccata 30000
gccgccggac cattcccagg aattattgat atagaggact gccctcatgt tccttttccc 30060
aagttcggcc ataaggtagt ccagaccttc gaagaccttt tcgtcataga cgcccggttc 30120
cttctgcagg gtcggataga ctctggtcgc caccccgtcg ggccgtcgc tgcctaccag 30180
tacgcggagg ttgtccagtc cgagggcatg cagggaatca agttcggcgg caagcctgct 30240
tctgtcacca tattcggagc cgagaagtcc tccgtaccag aaatttgtac cgatataata 30300
atagaagtga ccatcaacga caaaagagcc atcgtcaact ttgacgatgt ccattttcat 30360
aggctgacac gaaagaatcg cgaaaagggc gataaacagt gccgggaata atcttttaat 30420
catgcggttt atcgataaat ccatacttgt ctgagtacgt tgcaaattta gcatgcaacg 30480
ttttacccaa caaatacaaa tttgtcaata accgatacta ttttgcacgt tatacaaaaa 30540
ccttgttctt cttgaacaga gtcctgaatt cgcgcggcgt gtagcctctt tttgacttga 30600
aaagacgatt gaaattcgac aggttgttga acccgcagct atagcagatt tcagagatat 30660
tctgggtggt gtcgaccagc agccttgccg cattcccgag tcggatgtcg attatatagt 30720
ctatcaccgt cttcccggtc ctgctcttga aaaagcggct gaacgccgac gggctcatgc 30780
cgaccatttc cgccaattct tcaagggtaa tgttttccgc gaaatggtct ccgacatatt 30840
gcttgatctt ctgcacccga cggctttcct tgtcggcatc gttatgggcg aatgacgtgc 30900
tggccagtat cctggagtca cccgccaggg aaaggtcata aagcagcctc ataaagtgct 30960
ggaactgatc gaatttatct atttccgacg gcagggcgtc aagaagcgaa tacaccttca 31020
ttatggccgc cagcgaaaag gtcagcccgt tcctcgcctg ctcgaacata tgctgtatcg 31080
atgagaactg gtttctcatc atgaggccgc tggagaatat ctccggatcg aactggatag 31140
tgatctcgcg tacgtcctcg gaggtacaat tgccctgctc ccaggcatgt tcaagatccc 31200
cgccggcaat cagcgtaagc tcatattcgc ctatctcttc gacactgtcg cctacgactc 31260
gccggacgcc cttgccatgc tcgatgaagt tcagctcgta ttccctgtgc ctgtggaggg 31320
ggaagacgaa ttcggtctta tgtctctcga ctatgtagaa gcagtccctg tccgaaagag 31380
gtgttatttc cgttattacc gaggtcatat ccattaactt atattatcac gtggcaaata 31440
tatgataaaa atgtcaaaaa agtatcattt cgcaaatttt cttaattgtt ttatcaacta 31500
ttgggattgt ccaagaattg ctatctttgc tgtaattatg gaaaaagacg agctatatat 31560
gaaggaagct ctgcgcgagg cgcaggcggc cctggccgag gatgaaatcc cgatcggggc 31620
agttatcgta tgcaaaggcc ggatcgtggg aaagggtcac aatatgaccg aacgtttaca 31680
tgatcctacg gcccatgccg agatgatagc catcaccgcc gctacggagg cgctcggggg 31740
caaatacctc aacgaatgca cgctctacgt cacggtcgag ccgtgtccga tgtgcgccgg 31800
agccctgaac tgggcccaga tcggccggat cgtctatgga gcgatcgatc cgaagcgggg 31860
gcactcactt ttcactcctt cgctgctgca tccgaagacc gaggtcgagg caggagtgct 31920
cgcggaggaa tgcggttcca tggtgacgga gtttttcaag aaaaaaagat aaaaaacgga 31980
ctacaacaat gacaggagac gactacaaga tcatcagcga caccaccgag aacggtgtca 32040
gacacataac tgccatcccg agcagcctgg tatgcagcca gcagatcgat ttcgacataa 32100
tagacggcaa ggtccgcaat ctcgcatacg tgcggggctg caacggtaat cttcaggcga 32160
tcggccgtct cgtcgagggc atggacataa aaaaaatagt tgacactctt tcgggcgtca 32220
actgcaattt ccgtggtaca tcctgcacgg accagcttgc aaggattctt tcgagtctct 32280
agcgtcgcct ggccatggca atgtagtaga tcaatgtcgc tatcgaaccg atggcggcaa 32340
tgacgtatgt atatgctgcc catttgaggg catcgacagc tttcgctttt gtagcgggac 32400

```
     ctgttatccc ggccttgtcg agccattgca cggctctggc gctggcgttg atttccacag     32460
     gaagtgtcac gacgctgaac aaggtcgtca aggcaaacat ggcgattccg agatacagga     32520
     gggtcggagt aatcttgatg atcattacgc caataagaag tacccactgc acggcattgc     32580
     tggagaatga gactaccgga acgagagccg atctcagttt cagcggagca tatcccgtgg     32640
     catgctgcac tacatggcca cattcatggg ctgcaacggc cgcagcggct accgaattgc     32700
     cgtaatagac ttcttcgctg aggttgactg tcttgttgac agggttgtaa tggtctgtca     32760
     gctttccttt gatgcaggta atagagacat cggttatgcc atggtcttcc aacattttgg     32820
     cagcgacttg cgccccggtc agcccggaat gcaatcccac cttcgaatat ttattgaatt     32880
     tactgttcag aacctgctga atcacaaagc tcagcaccat cacagcaatg ataataaacc     32940
     aaatcatatc cataaaattt taacaatcct tctggttgct tccaaatcca tgccaacctc     33000
     tcacttgccg tcagcgtctt aattgatggc ggctgtcctc tggtctcggc gagcgtggcc     33060
     accctcggcc acgttagagg gaggggcccg atggatacga gttctgcgaa gcaggacgaa     33120
     gtagacattg ggagggatgg agccgacaga gtcggcggaa cagagccgag gccacgagga     33180
     ttcccgccaa agcagacgct gtcggcggcg acagcggacg tactgtaagg gtctttacgg     33240
     ggaaagaaaa gcgaaaaact gtaacctatg tcggagaaat ttgataaatt tgtattctaa     33300
     attgatgtgg aatgaacagg gttgatgatt tttcaaggct ggagctgaac ctgacaggat     33360
     gtatggaaaa ctacagatat ttcaggtcca aactggagca gacgactaaa ttacttgtac     33420
     tcgtaaaggc aaatgcttac gggcacggag ccgtcgaatt cgcagccatg atgcagcgtg     33480
     cgggagcgaa ctacctcgca gtcgcctacc ctgtcgaagg catggaactc cgcaaagccg     33540
     gtatcagcct cccgatcctc gtcctgaccg caggcacgga cttcttcaac gaaatcatcg     33600
     acaaccgcct cgagccggga atcccgaacc tgtacgccct caagatactc tgcggaatac     33660
     tcaaagaacg cggcatcaag gattatccgg tccatatcaa actcgacacc ggaatgcacc     33720
     gcctcgggtt catgacgagc gagctccagg atctcaccga cttcctcaag acctgcgaag     33780
     aagtcaaagt caagtccgta tattcgcacc ttgcagtatc cgaagaccct gagcaggacg     33840
     aattcaccct cggccagatc ggcatgttca agacaaacgc cgacaggatc acctccgaga     33900
     taggctacaa gccgatgtac catatcctca actccgccgg aatcgaaaga ttcccgcaat     33960
     accagttcga catggtccgc ctcggaatcg gtatctacgg aatcagcgcc ctgcctgatg     34020
     tccgcctcgc ccacgtcgcg tccctcaaat gcaaaatatt gcagataaag accctcaagc     34080
     cggaagacgg cacgataggc tacggaagat acggtcatat cgcccctgaa ggcaccgtaa     34140
     tcgccacgat cccggtagga tacgccgacg gcatcgaccg ccatctcgga cgcggtcacg     34200
     cctcattctc cgtcaacggg cacagagcgc cgacgatcgg gaacatctgc atggacatgt     34260
     gcatgcttga tgtgacagga attgacgcta aagtcggaga caccgtgact atcttcggag     34320
     aagaccctac agtcaacgaa ctcgccgaca ttctcgggac aatcccttac gaaattctca     34380
     ctagcgtccc tagacgtata gagagaattg ttgtccgcaa ataatttttt cgtatatttg     34440
     caccccactg aggtatggtg taatggtagc actagggatt ttggttccct cagtagaggt     34500
     tcgaaccctc ttacctcaac caatctaaac cacatgagca tcatgggaaa ggcatacatc     34560
     ttcctggctg acggttttga agagaccgaa gccctcgcaa ccctcgacat tctccgcaga     34620
     ggtaaaatcg aagcatacac agtatcaatc accgaagaag tctatatcac cagctcccac     34680
     ggagtaacgg tggtggcaga catgggccgc tccgacttca gcgaaataga tatcgacgaa     34740
     aaagacgcca tgatcttccc gggaggaatg ccaggctcga agaaactcgc agccgacacc     34800
     ccgcttatcg gtctgatgaa cgtccactac ggcaacggag gcattgtggc cgcaatctgc     34860
     gccgccccgg gcctcgtcgc cacccagatc aagggaatcg aaggcaagaa agccacctgc     34920
     tttgacggct tcgaagagaa cctcgccgcc agcggaatgg aatacgtaaa agccccggca     34980
     gttaccgacg gccagctcat caccggcaga ggcgcagcct tcgcgataga cttcggactc     35040
     gccatcctcg aaaagatggg aggcaaagag gccgtagcta aggtaaaggc agccctactt     35100
     atttaaactc agatacttac gtacgacaca acctttatta tccgtctcca ttatgaacgg     35160
     gagaacggac aggtcgaatg cgtccagaag ccgatcagcc tcctccgggt ataagcttgt     35220
     cagcttgtcc atatcgacca ggatgacacg gacacccttc tccgaagcga cagaatcggc     35280
     ggcggcaagt tcagcctcgc agaaagtgca gccggcagta tggaaaagca cataattccg     35340
//
ID   XXX; XXX; linear; XXX; XXX; XXX;  XXX.
XX
AC   PRJEB67889
XX
```

```
DE   Uncultured bacterium DNA fragment from in-reactor enriched cow rumen, clone
ferm-45h9, contig1
XX
FH   Key             Location/Qualifiers
FH
FT   source          1..14385
FT                   /organism="uncultured bacterium"
FT                   /environmental_sample
FT                   /mol_type="genomic DNA"
FT                   /isolation_source="Cow rumen"
FT                   /clone="ferm-45h9"
FT                   /note="AZO-CM cellulose active clone"
FT   CDS             complement(286..1284)
FT                   /db_xref="SEED:fig|6666666.127267.peg.1"
FT                   /translation="MEKRENFTVISEEEIQKAIGIASAINKYYYEKMVGQKHLGTDLLA
FT                   TMMVNGHLLIESVPGLAKTTAAKTMTEAVAGKFSRIQCTPDLLPSDIIGTQILNYKTNE
FT                   FETKLGPVHANFVLLDEINRSSAKTQSAMLEVMQEHQTTIGGEIYPVPDVFMVIATQNP
FT                   IEQEGTYLLSEAQLDRFMMKTIITYPSQEEELTVLDRIEADVFSESRSVATIDDIRYLQ
FT                   ELTKRVYMDEAIKKYIISLVDATRYPEKFISKQLAEYVTIGSSIRGAICLMEVSKAIAL
FT                   LKGRDYVTPDDVKSHIRPVLRHRVTLNFAAAADGVNEDMIIDAITGAIPTP"
FT                   /product="widely conserved MoxR-like protein in magnesium
FT                   chelatase family"
FT   CDS             complement(1397..2362)
FT                   /db_xref="SEED:fig|6666666.127267.peg.2"
FT                   /translation="MITLKFLAAYFIIYAVAGWCLEVVYATTVTGKFVNRGFLNGPLCP
FT                   IYGLGAIVVYVSLNPFGENLLILYGCSVILTTILEFITGLVLEKIFHQKWWDYSKRPFN
FT                   LMGYVCLQFSLLWGVACVIVVKVLQPAVIAFVTFIDNLGNNALNILLCIAYGILIADIA
FT                   VTVIALLNISIRLRLIADMDKYIRSVSDSVGMKLSDETLKTMNETQKIKARYDRLLEKL
FT                   KTKYGENYAEYVGKLTKVQARIQNAFPNLDFSGIAKISEKMDKITSYMEKQKELFGKKM
FT                   GDTKDQLVKNVKNVINGIEEAISPQMDIEIADVDNNDDKG"
FT                   /product="FIG028593: membrane protein"
FT   CDS             complement(2377..2835)
FT                   /db_xref="SEED:fig|6666666.127267.peg.3"
FT                   /db_xref="GO:0008892"
FT                   /translation="MNKEYMNEAIKEAYDGIKNGHGGPFGSVVVKNGKIVGRGHNRVLL
FT                   KNDPTCHGEVEAIRDACSNLGTHSLEGCELYTTAEPCPMCLGAILWANIGKVYQGCNRV
FT                   DTDIIGFRDDRFYDHLSGKVKLLDIEETGREDCLKLFNDYNAMNAERY"
FT                   /product="Guanine deaminase (EC 3.5.4.3)"
FT                   /EC_number="3.5.4.3"
FT   CDS             complement(2850..3833)
FT                   /db_xref="SEED:fig|6666666.127267.peg.4"
FT                   /translation="MPRGKKTKLLYIIRLFFERSDENNMLTIDDIIDYLDENGIGAERK
FT                   SLYDDLKELNDFGIKVKKKKVGKECYYYLSERFLETAEIKLLVDSIQSSKFISENKSRT
FT                   LIQKLESFAGVSERKKLSRQVYVQGRVKTMNESVYKSIDTIHSAIADGNQLSFQYLQWN
FT                   MNKELEPKYDGKTYVVSPWALSCDEQNYYLIAYDPGHGEIRHYRVDKMKNVKETSDKRE
FT                   GKTQYRETNVAEYVKGNIGMMGGEDTPVTVSIPSYMIGVFIDRFGKEIRVFKTGEDKCE
FT                   VTFHVRLSSMFYGWVAGLGKDTVIISPAAAVEGAKQFAKENFYKYK"
FT                   /product="hypothetical protein"
FT   CDS             complement(3850..5085)
FT                   /db_xref="SEED:fig|6666666.127267.peg.5"
FT                   /db_xref="GO:0004813"
FT                   /translation="MDDRIGRVNTKLLYEEDAYIREFEATVTGCVKAEPGDKKKDLYII
FT                   ELDKTAFFPEGGGQSSDTGFIEDAAVSHVTLEGGRVLHTTDKAFEEGTVVKCRIDFAKR
FT                   YDRMQNHSAEHLLCGLIHNATGFDNVGFHLGDDETILDLNGPVSAELLSELELKANRII
```

```
FT                   YENVPITAAIYSGDELKSIDYRSKLDITENVRIVTIEGYDKCACCAPHVRTTGEIGSIK
FT                   VVDFAPHRGGTRIAIIAGERAFKDHVMLDESNRKIMALLSTKRPEGAKGTEQLLNRNLA
FT                   LNGELVSIKKELTSMVFNGVKNGLVGKTDEKPVLVFSENFDEIMMRNLINDTLKLTEAA
FT                   VAGFMGNDEKGYRYIIGAHKEKGILYGGEYEALPAFAKKFNESLSGRGGGSPLMIQGSV
FT                   TGRRKDIEEFFG"
FT                   /product="Alanyl-tRNA synthetase (EC 6.1.1.7)"
FT                   /EC_number="6.1.1.7"
FT   CDS             complement(5381..5557)
FT                   /db_xref="SEED:fig|6666666.127267.peg.6"
FT                   /translation="MTLEKILQDYGFEILGNKSIFTKEVSTEHFLEYKTMLERLRGTIA
FT                   DRGRKELSGRTMG"
FT                   /product="hypothetical protein"
FT   CDS             complement(5801..7261)
FT                   /db_xref="SEED:fig|6666666.127267.peg.7"
FT                   /db_xref="GO:0003938"
FT                   /translation="MGTIVKEGITFDDVLLIPGYSEVIPNQVDLSTNLTKNIKLNIPVM
FT                   SAGMDTVTEYRMAIAMARQGGIGIIHKNMSIDEQADEVDKVKRSENGVITDPFSLSQEH
FT                   TLADADALMAKYRISGVPITEGRKLIGIITNRDLKFETDFTKKIKESMTTAKDLVTAKE
FT                   GITLEEAKKILADSRKEKLPIVDDDFTLKGLITIKDIEKAIKYPLAAKDSKGRLLCGAA
FT                   VGITANVLDRVDALVKVGVDVVVLDSAHGHSLNVINCVKMIKEKYPELQLIAGNVATGD
FT                   ATKALIEAGVDAVKVGIGPGSICTTRVVAGIGVPQVTAIMDCYEVAKQYNVPIIADGGI
FT                   KYSGDITKAIAAGGSVCMMGSMFAGCEESPGSFELFQGRKYKVYRGMGSISAMENGSKD
FT                   RYFQENAKKLVPEGVEGRVAYKGTLEDTVFQLMGGLRSGMGYCGAKDIPTLRETGKFVK
FT                   ISAASLKESHPHDIHITKEAPNYSTDDK"
FT                   /product="Inosine-5'-monophosphate dehydrogenase (EC
FT                   1.1.1.205)"
FT                   /EC_number="1.1.1.205"
FT   CDS             complement(7312..7827)
FT                   /db_xref="SEED:fig|6666666.127267.peg.8"
FT                   /translation="MDSYMEYMVKRGKTKGDFIIKGVCVWAIIVLCMLSWMFGGVPLIK
FT                   YLLLALTIGMLFLSWYVIFPRTDVEYEYLYCDKTITVDVIYSKSKRKNLATYDLDRIEI
FT                   LAPVNSHRLDEYKNKNLTVKDYASGMESDAHKPYALIYGGNEKVILDLPSEFVKMVQNN
FT                   GPRKVFFD"
FT                   /product="hypothetical protein"
FT   CDS             complement(7860..9128)
FT                   /db_xref="SEED:fig|6666666.127267.peg.9"
FT                   /db_xref="GO:0031176"
FT                   /translation="MRFKNLKRILPIILTTAVFVLGCGGNIPAVSENTTDVMRTDPEKV
FT                   LSTDLYVAEVVCDSSIDVPVYDFEKSAVPDSDAFAFVSDMGLGWNLGNTFDAVTTGEVQ
FT                   GAGLSLEMSWCGINTTKEMIDSIKTAGFNTIRIPVSWHNHLKDNYEIAERWMDRIEEIV
FT                   KIAYEDDLYVIINIHHDIDEKYYYPDRDHFANSSEYMYKIWEQIAERFKDYDHHVIFEA
FT                   VNEPRLVGTDVEWWYMPGNGKATGAIECINDLNQIFVDTVRESGGYNSDRYLMIPSYDQ
FT                   SVQGTLDSAFVLPTDTVEGRMILSVHAYTPYDFALNISGTSVFDESGKGEIDSFLSDLY
FT                   GKYVCNGIPVVMTEFGALDKNNLQDRVNFTAYYTARAASYGIPVVWWDNNAFAGSGENF
FT                   GIFNRGMLTFTFGDILNAMLLNK"
FT                   /product="GH5_4"
FT                   /EC_number="3.2.1.8"
FT   CDS             complement(9137..10744)
FT                   /db_xref="SEED:fig|6666666.127267.peg.10"
FT                   /translation="MKLTFVGAAHEVTGSCHYLNAAGKHILVDYGMQQGIDVLESVPLP
FT                   IDESQVDYVLLTHAHIDHSGMLPLLVKNGFKGQIIATDATVDLCSIMLRDCAHIQMQEA
FT                   EWKSRKAKRSASQAPIEPIYNMEDAEGAIHRLTPFPYNEVIDLCEGIKIRFTDVGHLLG
FT                   SASIEVWATEGEDTRKIVFSGDIGNKKGTLIKDPEYVKDADYVVMESTYGDRYHEEHRP
FT                   DYVAELTEIISETLAKGGNVVIPSFAVGRTQEMLYYIREIKEKNLIPENPDFPVYVDSP
```

```
FT                   LAVEATGVFQKNIQNCFSDEAKAMVNAGINPLTFPNLHLSITSDESKLINFDETPKVII
FT                   SASGMCEAGRIRHHLKHNLWRQECVILFVGYQANGTLGRFIIEGAKEVKLFGETIDVRA
FT                   KIMTLPGMSGHADKNGLIEWITNVGSDIKKVFIVHGEDTVTKLFSETLRAEYGYDTMAP
FT                   YSGSSYDLIKNEPIYIAKPVYVQKKKPGRISDVFARLLAAGQRLISVIHKNEGCANKDL
FT                   AKFADQVNSLCDKWDREI"
FT                   /product="Metallo-beta-lactamase family protein, RNA-
FT                   specific"
FT   CDS             complement(10981..11877)
FT                   /db_xref="SEED:fig|6666666.127267.peg.11"
FT                   /db_xref="GO:0004143"
FT                   /db_xref="GO:0007205"
FT                   /translation="MSDFKRLLFIVNPNAGMKKETPFESICNVFEDYGYEVTVCFTRAS
FT                   GDGTKIVKEHGVGDFDLIVCMGGDGTLNETFAGAYETGWRKPIGYIPAGTTNDFATSIG
FT                   LPSDHVEAAKHIMTHEPRALDLGVFNKRMFVYTACCGLFTKTSYETPQKIKNVFGHLAY
FT                   VLEGVKDLTQIHSIHMKINTGKEKLEDNYIFVGLCNTFSLGGVMTLDENQVSLDDGEFE
FT                   LLTISMPRDILQLNTIITCLTNQKYDEAKDLVHFTKAKKAEILFPQFEDWSLDGERGEG
FT                   REKNRFEVVHDAVRLIY"
FT                   /product="Transcription regulator [contains diacylglycerol
FT                   kinase catalytic domain]"
FT   CDS             complement(11890..12042)
FT                   /db_xref="SEED:fig|6666666.127267.peg.12"
FT                   /translation="MLVSFLNLKLRDEYESFDSLCDSLDLDKAEIIDKLKKAGFVYRED
FT                   QKQFR"
FT                   /product="hypothetical protein"
FT   CDS             complement(12098..13036)
FT                   /db_xref="SEED:fig|6666666.127267.peg.13"
FT                   /db_xref="GO:0019164"
FT                   /translation="MENTFNFKEIQGRPDRLAPGHRMCAGCGGTITVRNVLKALHPGDK
FT                   AVVGNATGCLEVSTFMYPFTAYEDSYIHNAFENAGATLSGVETAYNVLKAKGKISENYK
FT                   FITFGGDGGTYDIGLQSLSGAMERGHDMVYVCYDNGAYMNTGTQRSSATPMYADTTTTP
FT                   AGKCSAGKPQYRKDLAAIMAAHNIPYVGQTTFIGNMKDLYVKSEKAIYTHGPAFLNVMS
FT                   PCPRGWRYDAPEIMNICKLAVETCYWPLFEVENGRWILNYQPKNKLPIEDFLKTQGRFK
FT                   HLFKPENEELIVLFQKELDRRWEELLTRCNA"
FT                   /product="Pyruvate:ferredoxin oxidoreductase, beta subunit
FT                   (EC 1.2.7.1)"
FT                   /EC_number="1.2.7.1"
FT   CDS             complement(13065..14246)
FT                   /db_xref="SEED:fig|6666666.127267.peg.14"
FT                   /db_xref="GO:0019164"
FT                   /translation="MSKKNRMSGNEALGEAIRQINPDVMPAFPITPSTEVPQFVSNMIA
FT                   NGQVDTEFIPVESEHSAMSAAMGSEAAGARTITATSSCGLALMWEVLYVAASSRLPLAM
FT                   CVVNRGLTGPLNINAEHSDSMGARDSGWLQIYAENNQEVYDNMLMAYRIAEHKDVRLPI
FT                   MICQDGFITSHAVENIVLEDDEPIKKFVGTYEPEHYLLNENEPMAMGPYANSPFCMEAK
FT                   RAQVQGMMNAKQVILDVAKEYAKISGREYGFYEGYKLDDADYVLVCIGSACGTIKDAVD
FT                   EMRDKGVKAGLLKVRVFRPFPGEEFAKALENAKAVAVMDRAESYTGHGGPLGAELMAAM
FT                   YRNKNKAEAINVIYGLSGRDLTVDMVCGLYDQLADIAVNGSEGKQYQYLGLRE"
FT                   /product="Pyruvate:ferredoxin oxidoreductase, alpha
FT                   subunit (EC 1.2.7.1)"
FT                   /EC_number="1.2.7.1"
XX
SQ   Sequence 14385 BP; 3695 A; 3413 C; 2675 G; 4602 T; 0 other;
     ctccgccgag agtctcaccc ttcatctttt ttcccgtatc ggcaacgatg aggatattgt        60
     gcttcttctc tgcgatataa cgcttaacaa gaagtgaacc gtttcttgaa gacgctttcc       120
     agtctatgtc ctttatgctg tctcccggta cgtactcgcg aaggtcctcg aaattgaggc       180
```

ttcgtcccat gaagacagat aaatacgaac cgtcaagaat actgttggac ttcttggtcg 240
catatattcc cgtattcgct tttacctttt tgatatagtc ttttatcatg gtgtaggtat 300
tgctccggta attgcatcaa taatcatgtc ttcatttacg ccgtctgctg ctgctgcaaa 360
gttaagagta acgcggtgac gaagaacagg tctaatgtga gacttaacat catcgggagt 420
aacataatct cttcccttga gaagagctat ggcctttgaa acttccataa ggcagattgc 480
gcctcggata cttgatccga ttgttacgta ctctgcaagc tgctttgaaa taaacttctc 540
aggatatctt gtagcatcaa caagagagat aatatatttc ttgatagctt catccatgta 600
aactctcttt gtgagttcct ggagataacg gatatcatcg attgttgcaa cgcttcttga 660
ctcagagaat acgtcagcct cgatacggtc aagaaccgta agttcctctt cctgtgaagg 720
atatgtgatg attgtcttca tcatgaatcg gtcaagttgc gcctctgaaa gaaggtatgt 780
accttcctgc tcgatagggt tctgtgttgc gattaccatg aacacgtcgg gaacgggata 840
aatctcgccg cctatggttg tctggtgctc ctgcatgacc tcaagcatgg ctgactgtgt 900
ctttgcactt gatctgttaa tctcatcaag aagaacaaag tttgcatgaa caggtccgag 960
ctttgtctcg aactcatttg tcttgtagtt aagaatctgg gttccgataa tatcgctggg 1020
aagaaggtcg ggcgtacact gaatacgcga gaactttcct gctacagcct ctgtcattgt 1080
ctttgcggca gttgtctttg caagaccggg aaccgactcg ataagaagat ggccgtttac 1140
catcattgtt gcaagaagat ctgttccgag gtgtttctga cctaccatct tctcgtagta 1200
atacttgttg atagctgaag ctatccctat ggctttctgt atctcttcct cagagattac 1260
cgtgaaattt tcccgttttt ccattttttt aaatttcctc tcttttttta tttcccttca 1320
taaatttttt ataatcgccc ctctgattat cagtaattat atcatcaggc aaatatttta 1380
ttcttaaaag aacattttac cctttatcat cattattgtc cacatcggca atctcgatgt 1440
ccatctgagg gcttattgct tcttctattc cgttaatcac atttttaacg tttttttacaa 1500
gctgatcttt tgtatctccc atcttctttc caaacagttc tttctgtttt tccatataag 1560
atgtgatttt atccatcttc tcggatatct ttgcaattcc cgagaaatca aggttaggaa 1620
atgcattctg aatacgtgcc tgaacctttg taagttttcc aacatactcg gcataattct 1680
cgccgtactt tgtcttgagt ttttcaagaa gtctgtcata tcttgccttt atcttctgag 1740
tctcattcat tgtcttaaga gtctcatccg aaagtttcat tccgacagaa tccgatacgc 1800
ttcttatata tttatccata tcggcgataa ggcgaagtct gatgctgata ttaagaagtg 1860
caattaccgt aacggctata tccgcgatga ggatgccgta ggctatgcag agaagaatat 1920
taagcgcatt attaccgaga ttatcgataa acgtcacaaa tgcaattacc gcaggctgca 1980
gcacctttac aacaatgacg caagccactc cccataaaag cgaaaactgc aggcatacgt 2040
atcccataag attgaaaggt ctctttgaat aatcccacca tttctgatgg aatatcttct 2100
caagtacaag acccgtaata aattcgagaa tagttgtgag gatcaccgaa catccgtaaa 2160
gaatcaaaag gttttcacca aacggattta atgagacata tacaacaatg gcacccagtc 2220
cgtatatggg acataagggc ccgtttaaaa atcctctgtt aacaaatttg ccggttaccg 2280
tcgttgcata caccacctca agacaccatc cggctacggc ataaataata aaatatgccg 2340
ccaaaaaattt taatgtaatc atttattgtt cccgtcctaa tacctttcgg cattcatagc 2400
gttatagtca ttaaaaagtt tgagacagtc ttctcttccc gtctcttcta tatcaagaag 2460
ttttaccttg cccgataaat gatcgtaaaa tctgtcatcg cggaaaccga ttatatctgt 2520
atcaaccctg ttgcatccct ggtatacttt tccgatattg gcccagagta tggcgccaag 2580
acacatgggg cagggttctg ccgttgtata aagttcgcat ccttcaaggg aatgggtacc 2640
caggttgctg catgcatctc ttattgcttc cacttccccg tgacatgttg gatcgttttt 2700
aagaagcact ctgttatgac ctcttcccac aatcttgccg ttctttacca caacagagcc 2760
gaaaggtccg ccgtgtccgt tttttattcc gtcatatgct tctttgattg cttcattcat 2820
gtattcttta ttcatatggc ttctccgatt tatttatatt tataaaagtt ttcttttgca 2880
aactgctttg cgccttcaac cgcagcagcg ggacttatga ttacggtatc cttgccaaga 2940
cctgctaccc atccgtaaaa catggaactt aatcttacat ggaaagtaac ttcacattta 3000
tcttctcccg ttttgaaaac tcttatttcc ttgccgaatc tgtcaatgaa gacgcctatc 3060
atatatgagg gaatgcttac cgttacggga gtatcctcgc ctcccatcat tccgatattt 3120
ccttttacat actcggcaac attggtctcc ctgtactgag tcttgccttc acgcttatca 3180
gaagtctctt ttacgttctt catcttatcg actctgtaat gacgtatctc gccgtgaccg 3240
ggatcgtaag ctataagata ataattctgc tcatcacatg aaagtgccca aggacttaca 3300
acatacgtct tgccgtcata cttcggttcc agttctttat tcatattcca ctgaagatac 3360
tggaaagaca gctgattgcc gtcagctatt gcagagtgaa tcgtatcgat tgatttatat 3420

acgctctcgt tcatggtctt aactcggccc tgaacgtata cctgtcttga aagcttttta 3480
cgttcactta ctcccgcaaa tgattcaagt ttctgtataa gcgttcttga tttgttctcc 3540
gaaataaatt tcgatgactg tattgagtct accagaagtt ttatctctgc tgtctcaaga 3600
aaacgctcgg acaggtagta ataacattcc tttccgactt tcttcttttt gactttaatg 3660
ccgaaatcgt taagttcttt taaatcatca taaagacttt ttctctcggc accgataccg 3720
ttttcatcaa gataatcgat aatgtcatcg atggtcagca tattgttttc atctgacctt 3780
tcaaagaaca gcctgattat ataaagcagt tttgttttct ttcctctcgg catgttaacc 3840
cctccgttat taaccgaaaa attcttctat gtcttttctt ctgcctgtta cggagccctg 3900
tatcatcaag ggacttccgc cgcctcttcc cgataatgat tcgttaaact tttttgcaaa 3960
cgcaggaaga gcttcatatt caccgccgta taagattccc ttttccttat gggcgcctat 4020
tatgtagcgg tatccctttt catcattacc cataaatccg gcaacggctg cctccgtaag 4080
tttaagagta tcgttaataa gatttctcat cataatctca tcaaaattct ccgaaaagac 4140
aagcaccggc ttttcatctg tcttgcccac aagaccattt ttgacaccgt taaataccat 4200
ggatgtaagt tctttcttta tacttacaag ttctccgtta agggccagat ttctgtttaa 4260
aagctgttct gttccctttg caccttcagg tctttttgtt gaaagaagcg ccatgatctt 4320
tctgtttgat tcatcaagca tcacatgatc tttaaatgct ctctcacctg ctattatggc 4380
tattcttgtt ccgcctctgt ggggagcaaa gtccacgact ttaatgctgc cgatttcccc 4440
tgttgtcctt acatggggag cacagcatgc acacttgtca tagccctcta ttgtgacgat 4500
tctcacattc tctgtaatat caagcttgct tctgtaatca atggatttaa gttcatcccc 4560
ggaatatatg gctgccgtta tgggaacgtt ttcatatatt atcctgtttg ccttaagttc 4620
aagttcggat aaaagctccg cagatacagg tccgtttaaa tcaagaattg tctcatcatc 4680
tccaagatga aatcctacat tatcaaaccc tgtggcatta tgaataagac cgcagagaag 4740
atgttccgca ctgtgattct gcatcctgtc atatctcttt gcaaaatcaa tccggcactt 4800
tacaacagta ccttcctcaa aagctttatc ggtcgtatga aggactcttc cgccttcaag 4860
ggttacgtgt gacacggcag catcttcaat gaagcctgta tcacttgact ggccgcctcc 4920
ctcagggaaa aaagctgttt tatcaagttc gataatgtat aagtctttct ttttatcacc 4980
gggctccgct tttacgcatc cggttactgt tgcctcaaat tcccttatat atgcatcttc 5040
ttcatagaga agttttgtat ttacacgtcc aatacggtca tccattgact gtctccttta 5100
ttacggatct taagattatc cgttccttat tattttcaaa gggtaatttt ttactctttg 5160
actctttact gattatactg caaatcggta ttttcttcga taaaaacagt aaaaacgaca 5220
aaaaaatatc ccgcagctta tggctgcggg atatcttata aattttacgg aagtatattc 5280
cgaatatttt cttcgattaa tgcttagggg ggattttgtg taactaaggt aactaatgtt 5340
tgtaaggagc agtttatcga agctttatcg aaatatgtta ttaacccatt gttctgcctg 5400
acagttcttt acgtcctctg tcagcgattg ttcctctgag gcgctcaagc attgtcttgt 5460
attcgaggaa atgctctgtt gaaacctctt tggtaaaaat actcttgtta ccgagaattt 5520
caaatccgta atcctgtaaa atcttttcca atgtcatttt cttccccctc ttcgcttcct 5580
gcaggtatat cttttattgt ttatcaacaa taacccgcca cttctattgt tcaaatttac 5640
tatgaccata gtctacatct tcactcttaa caattactta accaatatat tgtgtttttg 5700
atataccttg ttcatttatg tgataaaaaa aacaggaatc cgtaaattcc tgtttttata 5760
tatcaaatat ttaatatttt ttactgtcag acttaaaaat ttatttatcg tctgtgctgt 5820
agttgggagc ctcttttgtg atgtgaatat catgaggatg tgactcctta agtgaagcag 5880
cggaaatctt aacgaatttt cctgtctcac ggagtgtggg gatatccttt gccccgcaat 5940
atcccatacc tgaacgaaga ccgcccataa gctggaatac tgtatcttca agagttccct 6000
tatatgcaac tcgtccctca actccttcag gcacaagttt ctttgcattc tcctggaaat 6060
aacggtcttt tgatccgttc tccatagctg agatagaacc cattccacgg tataccttgt 6120
attttcttcc ctggaaaagt tcgaagcttc cgggtgattc ctcgcatccc gcaaacattg 6180
aacccatcat acatacgctg ccgcctgctg cgattgcctt tgtgatatct ccggaatact 6240
tgataccacc gtctgcgata ataggtacgt tatactgctt tgctacctcg tagcaatcca 6300
taatagctgt aacctgagga acaccgatac ctgcaacgac acgtgttgta cagatagatc 6360
cgggtccgat tcccacctta actgcatcaa cgccggcttc gataagagcc tttgtagcat 6420
ctcccgtagc aacgttacct gcgataagct gaagctcggg atatttttcc tttatcatct 6480
ttacgcagtt gattacgtta agtgaatggc catgtgcaga gtcaagtact actacgtcaa 6540
ctccaacctt tacaagtgcg tcaactctgt cgagaacgtt tgctgtaata cctacggcag 6600
ctccgcagag aagtcttccc tttgaatcct ttgctgcaag gggatacttg atagccttct 6660

caatatcctt aattgtaata agtcccttta atgtgaaatc atcatcaacg atgggaagtt 6720
tctcttttct tgaatctgca agaatcttct ttgcttcctc aagagtgatt ccttcctttg 6780
ctgtaacaag atcttttgct gtagtcattg actctttaat cttctttgtg aagtcagtct 6840
caaacttaag atctctgttt gtgatgatac ctataagctt tctgccctct gtgataggca 6900
caccggaaat acggtacttt gccataagag catctgcatc tgcaagagtg tgctcctggg 6960
aaagtgaaaa cggatcagtg ataacgccgt tctccgaacg ttttacctta tctacctcat 7020
ctgcctgctc atcaatgctc atattcttgt ggatgatacc tataccaccc tggcgcgcca 7080
tcgctatcgc catacggtac tccgtaaccg tatccatacc ggcactcata accggtatat 7140
taagtttgat gtttttggtg aggtttgtcg aaagatctac ctgattgggg ataacttcag 7200
aataacccgg aatcaaaagc acatcatcga aggtaatgcc ctctttcact attgttccca 7260
ttgctctcct cctaactata tttttttatat atttaaaggt cgcaaaaccc tttaatcaaa 7320
gaaaactttc ctcggtccgt tgttttgtac cattttaaca aactcactcg gtaaatcaag 7380
tataactttt tcgttaccgc cgtaaataag agcataaggc ttatgggcat cggactccat 7440
accgcttgca taatccttta cggtgaggtt tttgtttttg tattcatcga gtctgtgtga 7500
atttacaggc gccagaattt cgatcctgtc aagatcgtat gtggcaagat tctttctctt 7560
tgactttgaa tagattacat cgactgtgat tgtcttgtcg caataaagat attcgtactc 7620
aacatctgtt ctggggaaaa tcacatacca tgaaagaaaa agcattccta tggtaagtgc 7680
aagaagtaaa tatttaatga gaggaacacc tccaaacatc catgaaagca tgcagaggac 7740
aattattgcc caaacgcaca ctccttttat gataaagtcg cctttggtct tgcctctctt 7800
aaccatatat tccatgtatg aatccatttt tgttattcct cttctttcaa tttatttttt 7860
tatttattca aaagcatggc atttaagata tcaccgaatg taaatgtcag catgcctctg 7920
ttaaatattc cgaagttttc gccgcttccc gcaaacgcgt tgttgtccca ccatacaacg 7980
gggattccgt atgaagctgc tctggccgtg taataagccg tgaaattaac tctgtcctga 8040
agattgtttt tatcaagagc tccaaactca gtcatcacaa cgggaattcc gttgcagaca 8100
tatttgccgt aaaggtctga taaaaagctg tcaatctcgc cttttccgct ttcgtcaaat 8160
accgacgttc cgcttatatt aagagcaaaa tcatagggcg tataagcatg tacgcttaat 8220
atcattctgc cttccaccgt atcggtgggc agtacaaatg ccgaatccag tgtaccctgt 8280
acgctctgat cgtagcttgg aatcataaga tatctgtctg aattgtatcc gcctgattct 8340
cttaccgtat cgacaaaaat ctgattaagg tcgttaatgc attctatcgc ccggttgcc 8400
ttaccgttgc ccggcatata ccaccattcc acatcagtgc ctacaagtct cggctcgttt 8460
acggcctcaa atatcacatg atgatcgtaa tctttaaacc gctccgctat ctgctcccag 8520
attttataca tatactccga tgagtttgcg aaatgatctc tgtcgggata ataatacttt 8580
tcatctatgt cgtgatgaat gttgatgatt acatagaggt catcctcgta tgcaatcttt 8640
acaatctcct cgatcctgtc catccatctc tcggctattt cataattatc ctttaaatgg 8700
ttgtgccatg aaaccggaat tcttattgta ttaaaccctg cggtttttat ggaatcaatc 8760
atttccttgg tcgtattgat tccgcaccag ctcatttcaa gggaaagacc tgctccctga 8820
acctcacctg ttgtcaccgc atcaaacgta ttaccaaggt tccagccaag tcccatgtcg 8880
ctcacaaacg caaatgcatc gctgtcagga acagccgact tttcaaaatc atagacaggc 8940
acatctatac tgctgtcgca tacaacttca gccacataaa gatctgtgga aagcaccttt 9000
tcagggtcag tccgcataac atcagtcgta ttctcactga ctgcggggat gttacctccg 9060
cagccaagaa caaatacggc agttgtaagt ataatcggta aaatacgttt taagttttta 9120
aatctcatat aatactctaa atctctctat cccacttgtc gcaaagagaa ttgacctgat 9180
ccgcaaattt cgcaagatct ttgtttgcac atccctcgtt cttgtgaata acggagataa 9240
gcctctgtcc tgcggcaaga agtcttgcaa atacatcgga tatgcggccc ggctttttct 9300
tctgaacgta tacgggcttt gcaatgtata taggctcgtt cttgataaga tcataacttg 9360
atccgctgta gggagccatc gtatcgtatc cgtactcagc gcgaagtgtc tctgaaaaca 9420
gctttgttac cgtatcttcg ccgtgaacaa taaatacttt cttaatatcg gatccgacgt 9480
ttgtaatcca ctcgatgagt ccgttcttgt cagcatgtcc gctcattccg ggaagtgtca 9540
ttatcttggc tctgacatcg attgtctcac caaagagctt gacttctttt gcaccctcga 9600
taatgaatcg tccgagtgta ccgtttgcct ggtatcctac aaagaggatt acacactcct 9660
gtctccagag gttgtgcttt aaatggtgtc tgatacgacc tgcctcacac attccggatg 9720
cggagataat taccttcggt gtctcatcaa agtttatgag ttttgattcg tctgatgtaa 9780
ttgaaagatg aaggttagga aatgtaagag ggtttatgcc tgcattaacc attgcctttg 9840
cttcatcgga aaagcagttc tgaatgtttt tctggaatac acccgtagcc tcaacggcaa 9900

gaggtgaatc cacgtaaacg gggaaatcgg gattctcggg aataagattc ttttccttga 9960
tttcccttat gtaatagagc atctcctgag tacgtcctac ggcaaatgac gggataacaa 10020
cgttaccgcc ttttgcaaga gtctcggaaa taatctccgt aagctccgct acgtaatcgg 10080
gacgatgttc ctcgtgatat ctgtcaccgt atgttgattc cataacaaca taatcggcat 10140
ccttgacata ctcgggatct ttaataagtg tacccttctt gttaccgata tctccggaga 10200
atacgatttt tctcgtatcc tctccttcgg tagcccacac ttcaatagaa gccgatccga 10260
gaagatggcc cacatctgta aatcttattt ttataccttc gcaaaggtct ataacttcgt 10320
tgtagggaaa aggagtaagc ctgtgaattg cgccttctgc atcctccatg ttgtagatag 10380
gctctatcgg tgcctgactg gcactacgtt ttgcctttct tgatttccat tcagcttcct 10440
gcatctgtat atgagcacag tcacgaagca ttatggagca aagatccacc gtggcgtcgg 10500
tggcgataat ctgtccttta aaaccgttct ttacaagaag cggcagcata cccgaatgat 10560
cgatatgagc atgggtaaga agaacatagt caacctgaga ctcatcaatg ggaagcggga 10620
cggattcaag tacatctata ccctgttgca taccgtagtc aacaagaatg tgctttcccg 10680
cagcattaag ataatggcag gatcccgtaa cctcatgagc agccccgaca aaagtcagtt 10740
tcatgtataa tacctcctct tatgaaaaag taatccccaa aatactgttt ctattatatc 10800
ataattcctc ggcactcgtc atccgtgact cgttgtaccc tcggaattac aacataacat 10860
ccttgttata tcataattcc tcggcactcg tcatccgtga ctcgttgtac cctcggaatt 10920
acaacataac atccttgtta tatcataatt cctgactttt taatacttat gacaacctaa 10980
tcagtaaata agtctgacgg cgtcatgtac gacttcgaac ctgtttttct ctcttccttc 11040
ccctctctct ccgtcaagag accagtcctc aaactgaggg aagagaatct cggctttctt 11100
tgcctttgtg aaatgtacaa ggtcctttgc ctcgtcatat ttctgatttg tgaggcatgt 11160
gataatcgta tttaactgca gaatatccct gggcattgag atagtcagaa gttcaaactc 11220
tccgtcatca agacttacct gattttcatc aagagtcatc actccgccga gggagaatgt 11280
attgcaaaga cctacgaaga tataattatc ttcaagtttt tccttgcccg tatttatctt 11340
catgtgaatc gaatgaatct gtgtaaggtc cttcacacct tcaaggacat aggcaaggtg 11400
gccgaatacg tttttaatct tctgaggtgt ctcataagat gtcttcgtaa agaggccgca 11460
acatgccgta tagacaaaca ttcttttgtt gaatacacca agatcaagcg cccttggttc 11520
atgtgtcata atatgctttg ccgcttccac gtggtcagag ggaagtccga tgcttgtggc 11580
aaaatcgttt gtcgtaccgg caggaatata acctatgggt tttctccagc ctgtctcata 11640
tgcgccggca aaagtctcgt taagagtacc gtctccgccc atgcacacaa taagatcaaa 11700
atcgcccacg ccgtgctctt ttacgatttt cgtgccgtca ccggatgctc tcgtaaagca 11760
cacggttacc tcatatccgt aatcttcgaa tacattgcaa atcgattcaa acggagtctc 11820
ttttttcatt ccggcatttg gattaactat aaaaagcagt cttttaaaat cactcatacc 11880
cgcacacttt catctgaact gtttctgatc ttccctgtat acaaagccag ctttctttaa 11940
cttgtcgatt atctcagcct tatcaaggtc aagagaatcg caaagcgaat caaaactttc 12000
atattcgtct ctgagcttca agttaagaaa acttacaagc ataacagggt cctgtggtat 12060
catcggttat caaagctcct ttataacgtt tcctgcatta agcattacat cttgttaaga 12120
gttcttccca tcttctgtca agttccttct ggaagagaac aatgagttcc tcattctcag 12180
gcttgaagag atgcttgaat cttccctgag tctttaagaa gtcttcaatg ggaagtttat 12240
tcttgggctg gtagttgaga atccatctgc cgttctctac ttcaaacaaa ggccagtagc 12300
atgtctcaac agcaagcttg cagatattca taatctcggg agcgtcatat ctccatcctc 12360
tggggcaagg gctcattacg ttaaggaatg cgggaccatg tgtgtaaata gccttctctg 12420
actttacata gaggtccttc atgtttccaa taaatgttgt ctgtcccaca taaggaatat 12480
tatgtgctgc cataattgca gcaagatctt ttctgtactg aggcttacct gcagagcact 12540
ttcctgcggg agttgtcgtt gtatcggcgt acatgggagt tgctgaagat ctctgtgtac 12600
ccgtgttcat gtaggcgccg ttgtcatagc acacgtaaac catgtcatgg ccacgctcca 12660
ttgctccgga aagtgactga agaccgatat catatgtacc gccgtcaccg ccaaatgtta 12720
taaatttata attctcgctg atttttccct ttgccttgag tacgttgtat gcagtctcaa 12780
cacctgaaag agttgctccg gcattttcga atgcattatg gatatagctg tcctcatatg 12840
cagtaaaggg atacataaat gttgatactt caagacatcc tgttgcatta cccacaacag 12900
ccttatctcc cggatgaaga gccttaagta cgtttcttac cgtgatggta ccgccgcatc 12960
ccgcacacat tctgtgtccc ggtgcaagtc tgtcggggcg tccctgtatt tccttaaaat 13020
tgaatgtatt ttccatcgtc atttacctct ttatcctttg aagattactc gcgaagtcca 13080
agatactgat actgctttcc ttcagaaccg ttaacggcta tatcggcaag ctgatcgtaa 13140

```
     agtccgcata ccatatcaac cgtaaggtct cttcctgaaa gaccgtagat aacatttatt     13200
     gcttcagcct tgttcttgtt tctgtacatt gctgccataa gttctgcgcc aagaggtccg     13260
     ccgtgtcccg tgtagctctc tgctctgtcc attacggcta ccgcttttgc attctcaaga     13320
     gcctttgcga attcttcgcc ggggaagggt ctgaatactc ttactttaag aagtcctgcc     13380
     ttaacaccct tgtctctcat ctcgtcaacg gcatccttga ttgttccgca tgcagatcca     13440
     atacatacaa gtacgtaatc tgcatcatcg agtttatatc cctcgtagaa tccgtactca     13500
     cggcctgaaa tctttgcgta ttcctttgca acatcaagga taacctgctt tgcattcatc     13560
     atgccctgaa cctgcgcacg ctttgcttcc atacagaagg ggctgtttgc gtaaggtccc     13620
     attgccatcg gctcattttc gttaagaaga taatgctcgg gctcatatgt tccgacaaac     13680
     ttctttatgg gctcatcgtc ttcaagaaca atgttctcaa ccgcatgtga tgtgatgaaa     13740
     ccgtcctggc atatcataat gggaagtctg acatccttat gctctgcaat tctgtatgcc     13800
     ataagcatgt tgtcatatac ttcctgattg ttctctgcat aaatctgaag ccaacccgaa     13860
     tctcttgcac ccattgaatc cgaatgctcg gcgttgatgt taagaggtcc tgtaagtcct     13920
     ctgtttacaa cacacatagc aaggggaagt ctcgaagaag ccgcaacgta aagaacctcc     13980
     cacataagtg caagtccgca tgatgatgtg gctgtgattg ttcttgctcc tgctgcctct     14040
     gatcccatgg ctgccgacat ggcactgtgc tcactctcta cggggataaa ctccgtatca     14100
     acctgtccgt tggctatcat atttgaaaca aactgaggca cctctgttga aggtgttatg     14160
     ggaaatgcag gcattacatc agggtttatc tgtctgattg cctctcccag agcctcgttt     14220
     cccgacattc tgttcttttt actcatctta gtattctccc ttaatcctga ttatttaccc     14280
     ggtctcatgg taattgctcc gaaaggacat gccttctcgc atataccgca tcccttgcag     14340
     tgcataaggt caaagcttct gcgctcgccg ttcacaacag gaata                     14385
//
ID   XXX; XXX; linear; XXX; XXX; XXX;  XXX.
XX
AC   PRJEB67889
XX
DE   Uncultured bacterium DNA fragment from in-reactor enriched cow rumen, clone
ferm-45h9, contig2
XX
FH   Key             Location/Qualifiers
FH
FT   source          1..11613
FT                   /organism="uncultured bacterium"
FT                   /environmental_sample
FT                   /mol_type="genomic DNA"
FT                   /isolation_source="Cow rumen"
FT                   /clone="ferm-45h9"
FT                   /note="AZO-CM cellulose active clone"
FT   CDS             1552..2385
FT                   /db_xref="SEED:fig|6666666.127269.peg.1"
FT                   /translation="MRKEGPLLNKKLIAIDLDGTTLNQDSMITSKTKETLKKAINAGHH
FT                   VSIATGRPFRMSHQFYQQLELTTPMVNFNGALVHIPNQHWDGERETLINREIAFEILSQ
FT                   KKQLNLDFIAAENRDTFFIDSFDFFDEKIFASSRPGEKNLLSPKNLTTNPTSLLVRTDK
FT                   RFAEAVSAELTRQFGSYVDVRTWGGPTAILEIVSKGIQKAKGVQEIANYLSIDQQDVIA
FT                   FGDEHNDLELLDYAGWGVAMANGTDQLKGIANDVTPLSNQEDGLAVYLEKLLKLQ"
FT                   /product="Cof protein:HAD-superfamily hydrolase, subfamily
FT                   IIB"
FT   CDS             complement(2688..4841)
FT                   /db_xref="SEED:fig|6666666.127269.peg.2"
FT                   /translation="MMKDFMKKIVNTSWIEKIRKGAQEPKNKTFRRSHVPFRLNFLFFI
FT                   IFTLFVALIARLGYLQIVNGEEMEARVKSTSTITVQESTPRGMIYDSTGKALVTNKANQ
FT                   AITYTRGTQVTAKDLLEIATKLNSLIDVPADENLTDRDKKDFWLADPEHLKEVTKRLSA
FT                   KEKQLGNSEQYSAMVNKVTDDEINLNEEQLKIATIFKRMNSAYELNTVYIKNSGVTDQE
FT                   LAVVAERASELPGVSTGTDWTREYNAADSLKSILGSVTTEKQGLPADEAEKYLAKGYSR
```

```
FT                   NDRVGQSYLEKQYEDVLQGTKTQYEVSLDNEGNVSNQKEIFSGEKGSNLMLSMNAEFQS
FT                   KVEEILKRNYQTLINNGKAQYSPGAYAVAMNPQTGEVLAMTGFSHEQGSKEITENALGT
FT                   ITSAFAPGSVVKAGTLTAGWASNAISGNQVLIDEPIRLQGASEKSSVFNRNGQVALDAV
FT                   KALELSSNTYMIKVALKMLGLDYTPGMGLPSLDEEAKAYQQLRDSFKEFGLGTTTGIDL
FT                   PNESPGISRSVEYMKKFNADNGKEWYTPGNFTDLAFGQFDTYTPIQLAQYASTVANGGK
FT                   RVQPHLVKAIYGNDENGNLGEVKKEISTTVENTVNISAENMAILREGFHQVVHGTDPYT
FT                   TAKPLASAKMDLSAKTGTAETVAEGHPDITTVNSNIVAYGPTDNPEIAISVVLPNLLDE
FT                   QDHMNLTIAKEIMDAYYDMFMAK"
FT                   /product="Cell division protein FtsI [Peptidoglycan
FT                   synthetase] (EC 2.4.1.129)"
FT                   /EC_number="2.4.1.129"
FT   CDS             complement(4874..4990)
FT                   /db_xref="SEED:fig|6666666.127269.peg.3"
FT                   /translation="MQLQRFAVKQKVDFILKLGKEFLAPLIFSFLLRSFWVQ"
FT                   /product="hypothetical protein"
FT   CDS             complement(5147..6007)
FT                   /db_xref="SEED:fig|6666666.127269.peg.4"
FT                   /translation="MEELMSVISRIESMMEDYSSAEKKLANYIIGNVEKVPTMTANELA
FT                   EASGLSAPTVVRFSKKIGFQSLTDFKITISTELQTGIDEGFSDIEANESFYSIKNKLGN
FT                   NAQVAIKETVDILEEETIQKVVECLETAESVFLYGVGASSLVVEDILQKWSRVGKPIIF
FT                   EKDIHVLLPQLVSNEKKKVLWLVSNSGRSADVVALAELAKNMNIEIIALTQFGNNPLSK
FT                   IADVLVQTSRPKEITNRSAATNSLLAQFVTIDIIFYFYMAKNESLSEKVKKTREAIQDY
FT                   FMDSK"
FT                   /product="Transcriptional regulator, RpiR family"
FT   CDS             complement(6217..8160)
FT                   /db_xref="SEED:fig|6666666.127269.peg.5"
FT                   /db_xref="GO:0004829"
FT                   /translation="MIKITFPDGAVKEFEAGTTTLAIAESISKSLAKKALAGKVNGQLI
FT                   DLTRPIEEDASIEIITPDHEDALGLVRHSAAHLMAQAMRRLYPNIHFGVGPAIDSGFYY
FT                   DTDNGQNQVTAEDLPAIEAEMMKIVKENLPIERRVLSKQEALEIFASDPYKVELISELP
FT                   EEEVITAYQQGEFIDLCRGPHVPSTGRIQVFKLLSVAGAYWRGNSNNQMMQRVYGTAFF
FT                   DKKALKEYIRLREEAKERDHRKLGKELDLFMVSPEVGSGLPFWLPKGATIRRTIERYIV
FT                   DKEVSLGYQHVYTPIMGDVELYKTSGHWDHYQEDMFPPMDMGDGEMLVLRPMNCPHHMM
FT                   VYKNAIHSYRELPIRIAELGMMHRYEKSGALSGLQRVREMTLNDGHTFVRPDQIKDEFK
FT                   RTLELMVAVYADFNITDYRFRLSYRDPNNTDKYFDDDAMWEKAQAMLKAAMDELELDYF
FT                   EAEGEAAFYGPKLDVQVKTALGTEETLSTIQLDFLLPERFDLTYVGEDGENTHRPVVIH
FT                   RGIVSTMERFVAYLTEVYKGAFPTWLAPIQATIIPVSVDAHGDYAYEIKERLQMKGLRV
FT                   EVDDRNEKMGYKIRASQTQKIPYQLVVGDKELEDATVNVRRYGSKETAVEDLNIFIDAM
FT                   EAEVKNYSRENK"
FT                   /product="Threonyl-tRNA synthetase (EC 6.1.1.3)"
FT                   /EC_number="6.1.1.3"
FT   CDS             complement(8157..8270)
FT                   /db_xref="SEED:fig|6666666.127269.peg.6"
FT                   /translation="MEPRTFRPSISFQEMLGFFVFQLDDLINQNISRRKLL"
FT                   /product="hypothetical protein"
FT   CDS             complement(8496..8711)
FT                   /db_xref="SEED:fig|6666666.127269.peg.7"
FT                   /db_xref="GO:0006725"
FT                   /db_xref="GO:0016853"
FT                   /translation="MKKERSERGENIMPFVHVELVEGRSPEQLENMMKDITEAVHKNTQ
FT                   APKEHIHVIINEMKKGTYGVNGEWKK"
FT                   /product="4-oxalocrotonate tautomerase (EC 5.3.2.-);
FT                   Xylose transport system permease protein xylH"
FT   CDS             complement(8722..10545)
```

```
FT                   /db_xref="SEED:fig|6666666.127269.peg.8"
FT                   /translation="MTHLKKFLLTLFIGGVALLSEFVFHQPGLAFMIVAFTGGILAFVM
FT                   FLEMIKTLRSGRYGVDILAITAILATLAVKEYWASLMILIMLTGGDSLEDYANKRAGKE
FT                   LQSLLDNTPRIAHRFKEGRWQDIVIDEINVDDQLLVKPGELVPVDGRVIKGESSVDESS
FT                   LTGESRPETKKPGDQIMSGSLNGDGSIQMIAEKRAADSQYQMIIKLVEESKEKPARFVR
FT                   LADRYAVPFTLIAYLIGGISWWFSKDPLRFAEVLVVASPCPLILAAPIALVAGMSRSSK
FT                   NGIVVKTGTTIEKLARTKTIAFDKTGTLTKGILEVEAIQPVNGMKKEELLMYAASAEQE
FT                   SSHILARSLVNNVKNSQLYPVSQLKEVTGQGIQAVVNNHDVKIGRASFVSVEEIETSET
FT                   AIYVAIDQSFAGTILFSDQIRSEAAKTIKELRNLGVKNLMMITGDGSAIAQSIAEEVKL
FT                   DSVHAHCLPQDKLTILEHIPKDQRPVAMIGDGVNDAPALAAADIGIAMGAHGSTAASES
FT                   ADAVILKDDLLRVAKAVVISKDTMRIAKQSVLIGIFICIFLMLIASTGLIPALFGAVLQ
FT                   EVVDTVSILSALRAKKDPAIFPDSVPKTNIV"
FT                   /product="cadmium-translocating P-type ATPase"
XX
SQ   Sequence 11613 BP; 3349 A; 2464 C; 2057 G; 3743 T; 0 other;
     ataaagatcg atgcaaagaa ggctggtaaa gcttcttcca tatctagcca tttgatgtct        60
     gcgaaagaag ccatcatcat tactccgacc aaaatcagtg ctggcgcagt tgcttgattt       120
     ggtacaatgg cgatcagcgg ggagaacaga ctacttaatg cgaatagcac tgccactact       180
     actgaagtca aaccagtacg tccgccggcc ccgattccag ctgcagattc gacataagtt       240
     gttgtatttg atgtaccaaa gatagcacct acggaagtag cgattgcatc ggcaaatagc       300
     gctttatcca tttttgtttt gaacccgcgg ccatcttcca atgcgatctc gtcctctttt       360
     gagaaaatac ctgtgcggcg tcctgtaccg ataaatgttc cgatcgtatc aaatgtatca       420
     gataaactga atgcgatgat cgtcatcaat acttgaggga tcttggaaga atcactgaat       480
     aaagattgca ttccttcaga accaaatgct gcaccgaacg ttgttcctaa ttctttaaat       540
     gaattgccta acgaattcgt ttgccaatca atggaactta aatcaacaac acccataaaa       600
     atcccaagga ctgtggtcgc tacgataccg ataagaatag caccacggac gtttttttaca       660
     actaaaatgg ctgtcagtac caaaccaatc acagctaaca agataggtgc gttgtcaaag       720
     ttggctaacg cgggaacgat tccgccattt atcgtaacat ttgttgcttt tccaccttca       780
     acgacagaac tcgaaatagc agattggtct gctgaaaaag ataaaaatcc tgcatttttg       840
     atcccgacat acgcaacgaa aataccgatc ccgccaccaa tcgcatgttg aagactttca       900
     ggaatcgccc gaatgatcat ctttcggata tttgtaacag taataaaaat attgattaat       960
     ccgcaaataa aaaccattgc caaggcttgt tgccaagaat agccaagccc gaaaacgacc      1020
     gtaaaagtga aaaatgcgtt caatcccata ccaggtgcct gagcatatgg tacattcgcg      1080
     aacaatccca tgactagcgt accaatgatt gatgcaatga ttgtggctaa aaatactgct      1140
     tggaacggca tacctgatgc tgataaaatc gtcggattta cgaataagat atagctcatg      1200
     gcaaagaatg tcgttactcc cgccattatt tcagttgata catttgtatt gtttttcttt      1260
     aaaccgaaaa acttttccat aaaaagttcc actccctgaa tttttttctg atggaaaact      1320
     ataataataa aaaatatgca aaatatggca aatccatcga acaatttcaa attatataga      1380
     cgttttattc gtttttcaac gtttttttaa caaaaaaacc gaaaacaata cttaatgttc      1440
     gtgtttagct cgtatttatt tatttacata gccataactg tattttttgt taggaataac      1500
     tgattcctct tgttaattaa ctaaaatatg ctagactaaa acaaatgcta gttgaggaag      1560
     gaaggacctt tattgaataa aaaattaatt gccatcgatc tggacggaac caccctcaat      1620
     caagattcga tgattacctc taaaacaaaa gaaaccctga aaaaagcaat caatgctggt      1680
     catcatgtca gtatcgctac tggacgccca tttcgaatga gccatcaatt ttaccaacaa      1740
     ttagagttaa caacaccgat ggttaatttt aacggcgcac tggtgcatat ccctaatcaa      1800
     cattgggatg gcgaaagaga aacattgatt aatcgtgaaa ttgcttttga aatcttatcg      1860
     caaaaaaaac aattgaattt agactttatc gctgctgaaa atcgggatac ttttttttatc      1920
     gacagctttg acttctttga tgaaaaaatt tttgcttctt cacgtcctgg agaaaaaaac      1980
     ttgctttctc ctaaaaatct gacgacaaat cctacatctt tgcttgtccg aacagacaag      2040
     cgttttgcag aagctgtatc tgctgaatta acacgtcaat ttggcagtta tgtggatgtt      2100
     cgtacgtggg gcggtcctac ggcgatctta gagattgttt caaaaggaat acaaaaagca      2160
     aaaggtgtgc aagagattgc taactaccta tctattgatc aacaagacgt gattgctttt      2220
     ggtgatgagc ataacgattt ggagctgctg gattatgcag gctggggagt ggctatggct      2280
     aatggtactg atcagctcaa aggcatcgca aatgatgtca ctcctttgag caatcaagaa      2340
```

gacggtttag ctgtttatct ggaaaaatta cttaaactac aatagagaat tagattccac 2400
taaaaaaggt tgtgacaatt tttgtcacaa cctttttttgc cgagtaaacg gtgctcaatc 2460
agctgtcctt cggtattagc tcaaaaacag caaaatatga aaatctattt tcccattttg 2520
tttctctaat actcagatca gggtgctgat ttcacaccct cttgataaat agtgttcaaa 2580
agccagtttc tcggtatgaa gtcaaattta agtaaaaata tgagaagcta ttttcacaaa 2640
atttttcttc atactcgaaa ctaagtggct ttttcacgac ctctttttta ttttgccatg 2700
aacatatcat aatacgcatc cattatttct ttagcaatcg tcaagttcat gtggtcttgt 2760
tcatctaata agttcggcaa gaccacactg attgcgattt ccggattgtc tgttggtccg 2820
tatgccacga tattactatt gactgtcgta atatctgggt gaccttccgc aaccgtttct 2880
gcagtccctg tttttgcaga aaggtccatt ttcgcactgg ctaatggttt agctgttgtg 2940
taagggtcgg taccatgaac aacctgatgg aatccttcac ggaggattgc catattttct 3000
gcagatatat tgaccgtatt ttctactgtc gtgctaattt ctttttttcac ttcacctaaa 3060
ttcccatttt cgtcattgcc atagattgct ttgactaggt gaggctgtac tcgtttcccg 3120
ccattagcga ctgttgaagc atattgagct aattggatcg gtgtatatgt gtcaaactgt 3180
ccaaatgcca aatcggtaaa gttaccagga gtataccatt ctttgccatt atccgcattg 3240
aacttcttca tatactcaac agagcgtgaa atcccaggag attcattcgg caaatcaatc 3300
ccagtcgttg taccaagacc aaattctttg aaagagtctc tcaattgttg ataagcttta 3360
gcttcttcgt ctaaactagg tagtcccatt ccaggagtat agtctagtcc tagcatcttt 3420
aatgccactt tgatcatata cgtattggaa gacaattcca atgctttgac tgcatcaaga 3480
gcaacttgtc cgtttcggtt aaagacagag ctttttttctg aagcaccttg tagacgaatc 3540
ggttcatcaa tcaacacttg attcccgctg atcgcgttac ttgcccaacc tgctgtcaaa 3600
gtaccagcct tgactacaga ccctggagca aaggcgctag tgatcgtacc caaagcattc 3660
tcggtgattt ccttggaacc ttgctcatga ctgaaaccag tcatggctaa cacttctcct 3720
gtctgaggat tcatggctac tgcataagca ccaggcgagt actgtgcttt accattattg 3780
atcaatgttt gataatttct ttttaagatt tcttctactt ttgattggaa ttctgcattc 3840
atcgataaca ttaaattaga acctttttcc ccagagaaga tttctttttg atttgaaaca 3900
ttgccttcat tatcaagaga aacttcatat tgggttttag ttccttgtag gacatcttca 3960
tattgttttt ccaaataact ttgtcccaca cgatcatttc gagaataacc ttttgctaaa 4020
tacttctctg cttcatctgc cggaagtcct tgtttttcag tcgtcacact acctaaaata 4080
cttttcagtg aatctgctgc gttatattcg cgtgtccagt ctgtgcctgt cgaaactcct 4140
ggaagctcag aagcacgttc cgccacaaca gctaattctt ggtcagtaac accagagttc 4200
ttgatataga ccgtgtttaa ttcatatgca ctgttcatcc gtttaaagat ggtagcgatc 4260
tttaattgtt cctcgtttaa gttgatctca tcatctgtta ctttattgac catcgcgctg 4320
tattgttcgc tgtttcctaa ttgcttttct tttgccgaaa gtctttttgt gacttccttc 4380
aaatgttccg gatcagccag ccaaaaatct tttttatctc tatctgtcag attttcatct 4440
gccggcacat caatcagcga gttcagcttc gttgcgattt caagtaggtc tttagccgtc 4500
acttgtgtgc ccctcgtata ggtgattgcc tgattcgctt tatttgtgac taacgctttt 4560
cctgtggaat cgtagatcat tccacgaggg gtactttctt ggactgtaat cgtagaagta 4620
gatttcactc gcgcttccat ctcttcacca ttcacgatct gtaaatagcc caatcgagca 4680
atcaacgcaa caaataacgt aaatataatg aaaaaaagaa aatttaagcg aaaaggaaca 4740
tgcgaccggc gaaacgtttt gtttttaggc tcttgtgccc cttttcttat tttttctatc 4800
caagaagtat ttactatttt tttcataaaa tctttcatca tttcctatcc ttccgcctat 4860
ttacattcat gcttcattgt acccaaaaac ttcttaataa aaaagaaaaa atgagcggag 4920
caagaaattc tttacctaat ttcaagataa aatcaacttt ttgtttcaca gcaaacctct 4980
gaagttgcat atccaactat ttttttcatt aaaaactaaa cttatttttg aaaattatgc 5040
tttctaaagt cgtgaaacac tttaatagaa catttctttg tggaaaacta taagttgtta 5100
ttctaaaaaa acttatatca aaaatagtaa ggtacattta gttttttttat tttgaatcca 5160
taaaatagtc ttgaattgct tctcttgttt ttttcacttt ttcactaagt gattcgtttt 5220
tagccatata aaaataaaaa atgatatcaa tcgtaacgaa ttgggccaaa agtgaattgg 5280
ttgccgcact tcgattggtg atttcttttg gtcttgatgt ttgtacgagt acatcggcta 5340
tcttgctaag tgggttattt ccaaattgcg tcaacgcaat gatctcgata ttcatatttt 5400
ttgctagctc agctaatgca actacatctg cgcttctccc tgagttcgaa acaagccaaa 5460
gtactttctt cttctcattg ctgacaagtt gtgggagcaa gacatggata tctttttcaa 5520
aaatgatggg cttaccgaca cgagaccatt tttgcagaat gtcttctacc actaaagaag 5580

aggcacccac accatataaa aaaacagact cagccgtttc caaacattct acgacttttt 5640
ggatcgtttc ttcttccaga atatctactg tttcttttat cgctacttgc gcattgttcc 5700
caagtttgtt tttgatagag taaaaggact catttgcttc gatatcgcta aatccttcat 5760
cgattccagt ctgtaattct gttgaaatcg tgattttgaa gtccgtcaaa ctttgaaacc 5820
cgattttttt tgaaaaacgt acaactgtcg gcgcactcaa acctgaagct tccgctaatt 5880
cattagctgt cattgtgggg accttctcaa catttccaat aatgtagttg gccaactttt 5940
tttctgctga agaataatct tccatcatac tttcaatacg actaattacg ctcataagtt 6000
cctccatcct tttaataatc gtctatgtaa gtataccata tcagtcccct gttaacggtc 6060
gtgtactgga taaaaaaaca aaaaaacaat ccgaacaaca ggatagctca ttccatcaat 6120
cccacaaatt ctattgtcca tctgtgggat tgtggaatat ctgttctcta ttgtttgaat 6180
tgtttttat ttagtgcttt tttataggca gcttatctat ttgttttctc tgctgtaatt 6240
tttgacttct gcttccatag catcaataaa gatattcaaa tcttctacag ctgtttcttt 6300
gcttccgtaa cgacgtacat tcactgttgc atcttccaat tctttgtcac cgacaactaa 6360
ttggtaagga attttttgtg tttgagaggc acgaatctta tagcccattt tttcattgcg 6420
gtcatcgact tctacacgta accctttcat ttgcaagcgt tctttgattt cataagcata 6480
atcgccatgt gcatcaacag aaacagggat gattgtcgct tgtattggtg ctagccatgt 6540
tggaaaagca cctttgtata cttctgtcaa ataggcgaca aaacgttcca tcgtagaaac 6600
gattccgcgg tggataacta ctggacgatg cgtattttct ccatcttctc caacatatgt 6660
caaatcaaag cgttccggca ataagaagtc taattgtata gttgacaatg tttcttctgt 6720
cccaagtgct gttttgactt ggacatctaa ttttggtccg taaaaggctg cttccccttc 6780
tgcttcaaag taatctagct ctaattcatc catagcagct ttcagcattg cttgggcttt 6840
ttcccacatt gcatcatcat caaaatattt atctgtattg tttggatcgc ggtagcttaa 6900
gcggaaacga taatctgtaa tattgaagtc agcatatacc gcgaccatca attcaagcgt 6960
acgcttaaat tcatctttga tttggtcagg acgaacaaat gtatgaccat cgttcaacgt 7020
catttctcgt acacgttgta aacctgacaa agcacctgat ttttcataac gatgcatcat 7080
tcctaattca gcgatacgaa ttggtaattc acggtaagaa tggatggcgt ttttgtagac 7140
catcatgtga tgagggcagt tcattggacg aagaacaagc atttcgccat ctcccatatc 7200
cattggcggg aacatgtctt cttgataatg atcccagtgt ccagatgttt tgtacaactc 7260
aacatctccc atgatcggtg tatacacatg ttggtaacct agactaactt ctttgtctac 7320
gatataacgt tcgatcgtgc gtcggattgt tgccccttt ggaagccaga atggcaaacc 7380
agaaccaact tctggagaaa ccataaacaa gtcaagttct ttccctagtt tacgatgatc 7440
tctttctttt gcttcttcac gcagacggat atattctttc aaagcttttt tatcaaaaaa 7500
ggctgttccg taaactcgtt gcatcatttg attgtttgag ttcccacgcc agtaagcacc 7560
tgccactgat aataatttga atacttgtat acgaccagta gacggaacat gtggaccacg 7620
gcacaaatcg atgaattcac cttgttgata ggcagtaatc acttcttctt ctggcaattc 7680
actaatcagt tctactttgt atggatcact tgcaaaaatt tctaatgctt cttgtttcga 7740
aagaacacgg cgttcgattg gcagattttc tttgacgatt ttcatcatct ctgcttcgat 7800
tgcgggaaga tcttctgcag ttacttgatt ttgtccattg tctgtatcat agtaaaaccc 7860
tgaatcaata gctggcccaa caccaaagtg gatatttgga taaaggcggc gcattgcttg 7920
agccatcaaa tgagctgctg aatgtcggac taatccaagt gcatcttcat ggtctggtgt 7980
gatgatctca atacttgcat cttcctcgat tggacgagtc aaatcaataa gttggccgtt 8040
cactttacca gctaaagctt tttttgctaa gctttttgaa atactttcag cgatggccaa 8100
tgttgttgta ccagcctcaa attctttgac tgcaccatct ggaaaagtaa tttttatcat 8160
aatagtttcc tccttgaaat gttttgattg atcagatcat caagctgaaa aacaaaaaat 8220
cctagcatct cttgaaaaga gatactagga cgaaaagttc gtggttccac ctaatgttca 8280
aacgcacatg cgttcctcaa acgttgtaac ggtccgaccg cccttgcttc agcaagggtt 8340
tcgaaagtgg tcatcctttg gttttctcct ggaaaagttt cagcctagct tttcctctct 8400
tatcgaagac tctcaaattc agttggcttt ctcatcaatc aattatgtgt ttatttaacc 8460
actaataaat agaaaaatca agtgtgaatc taaacttatt ttttccattc cccatttaca 8520
ccgtatgttc cttttttcat ctcgttgatg atgacatgga tgtgttcttt tggtgcttgc 8580
gtgtttttat gaacagcttc cgtgatatct ttcatcatat tttctaattg ttccggactg 8640
cgaccttcta ctaattcgac atgtacgaat ggcataatat tctctcctct ttctgatcgt 8700
tctttttca tctcgattct atcatactat gtttgtcttg ggtactgaat ccggaaaaat 8760
ggctgggtct ttttttgctc tcaacgcact caggatagag actgtatcca cgacttcttg 8820

```
        caaaacagct ccgaataacg caggaatcag acccgtactt gctattaaca tcaaaaaaat       8880
        acaaataaag atgccaatca gtacagattg tttcgctatt ctcatcgtat cttttgagat       8940
        tactactgct tttgctacac gcaaaagatc atctttcaag atgaccgcat ctgcactttc       9000
        actagcagct gtagaaccat gagcgcccat tgcaatgcca atatctgcag cagcaagtgc       9060
        tggtgcgtca ttcacaccgt ctcctatcat cgccacaggt cgttgatctt tagggatatg       9120
        ttctaaaatg gtcaatttgt cttgaggcaa gcaatgagca tgtacagaat ctaatttgac       9180
        ttcctctgca atcgattgag caatggctga tccatccccg gtaatcatca tcaaattttt       9240
        cacaccaaga tttcttagct ctttgattgt ctttgctgct tctgagcgaa tctgatccga       9300
        aaacaggatg gttcctgcaa aggattgatc aatggctaca tagattgctg tttcagaagt       9360
        ttcgatttct tccactgaaa caaaagaagc gcgtccaatc tttacgtcat gattattgac       9420
        tactgcttgg attccttgac ctgtcacttc tttcagttga ctaacaggat acagttggct       9480
        gtttttcacg ttgttcacta acgaacgagc taagatatgg ctagattcct gttctgcact       9540
        tgctgcatac ataagtagct cttctttttt cattccattg actggttgaa tcgcctctac       9600
        ttctaatatt cctttggtaa gcgttcctgt tttatcaaaa gcaatcgttt ttgtacgtgc       9660
        gagcttttca atcgttgttc cagttttaac aacaatcccg tttttacttg aacggctcat       9720
        cccagcaact aaagcgatcg gcgcagctaa aatcaatgga caaggcgatg ctacgaccaa       9780
        tacttctgca aaacgcaaag gatcttttga gaaccaccaa gaaataccgc caatcaaata       9840
        tgcgatcaat gtaaagggaa ccgcataacg atcagctaaa cggacgaatc gggcaggttt       9900
        ttcttttgat tcttcgacta atttaatgat catttgatat tggctatctg cagcgcgttt       9960
        ttcagcaatc atttggattg atccgtctcc atttaaagac ccagacatga tctgatctcc      10020
        aggtttcttt gtttctggac gagactctcc tgtcaaagaa gattcatcga ctgatgattc      10080
        tcctttgatc actcgaccat caactggtac aagctcgcca ggcttgacta gcagctgatc      10140
        atccacattt atttcatcaa ttactatatc ctgccatctt ccttctttga atcgatgggc      10200
        gattcttggg gtgttgtcta ataaagattg caattcttta cctgctcgtt tattcgcgta      10260
        atcttctaga ctatctccgc ctgtaagcat gatcaatatc atcaaactag cccaatattc      10320
        tttgacggct aatgttgcaa ggattgcggt aatcgctaat atatctacac catatcttcc      10380
        tgaacgcaat gttttgatca tttccaaaaa catcacgaag gcgagaatcc ctccagtaaa      10440
        tgcaacaatc ataaaagcaa gtcccggctg atggaaaaca aattcactta acaaagctac      10500
        tccgccaata aataaagtaa gtaaaaattt ttttaaatgt gtcatagttt cttctcctct      10560
        tttttgtgac ttagaggtta gtcatcttca attgcctctc tttctcttaa gcatagtcct      10620
        tgcttaacac tttttaaaac aataggcctt aattgaaaac aactatcact atcagcgttt      10680
        ccaatagtcg ccagtccgac aaacgaatct atcgagcaaa gaggactttt ttcgattcat      10740
        ccgttaaaaa gtaccttttt tcgaaaaact gtgctatact tttaattcgt tatttacggc      10800
        ggaaggagca aacatgtgat gaatgagaaa caactagaac aaacgcacgt ggatgaaacg      10860
        atcgaactaa tccagatcga acaaaatatg ctggatacaa aacaacgaga attgactcaa      10920
        gaaatgaatc attcgacaaa agacacagct aatcacaaga ttcgagctgg ttccaatgaa      10980
        gctttttatg aatctgctgt tgaatatcgt caacatgagc aagaattgct tttaaaatat      11040
        catactcttg aggcgcagca aaaacgccta caaacgttgt ctgttatgaa agacagtcct      11100
        tactttgccc gtatcgattt taaagaagac cgtgaaacag aaacattgta tctaggtatc      11160
        gcttctttaa gagatctagc agaagaaaca attgttattg actggcgcgc accaattgcc      11220
        aacctttatt atgaaggtga gattggtccc gcattttatg aaactgaaac agatagaata      11280
        gatgtcgatc tgcagttgaa acgacaattt aaaatccaag aaggacgtat tttatctatg      11340
        gtagatacgt cggaagtaat caacgatgac tttttgctag agatattgga cgaagcatca      11400
        agcaaccata tgaaaaatat cgtctctacg atacaaaaag cacaaaatgc aatcattcgg      11460
        gacacaacaa gtaaagtgat gctgattgaa ggaatcgccg gaagcggaaa aacgtctgct      11520
        ctgcttcaac gtattgcttt tctgttgtat cataatcgca aatggctaga ggcagaaaac      11580
        gtgcttctat tttcaccaaa ccatttgttt tct                                   11613
//
ID   XXX; XXX; linear; XXX; XXX; XXX;  XXX.
XX
AC   PRJEB67889
XX
DE   Uncultured bacterium DNA fragment from in-reactor enriched cow rumen, clone
ferm-46a3
```

```
XX
FH   Key             Location/Qualifiers
FH
FT   source          1..35712
FT                   /organism="uncultured bacterium"
FT                   /environmental_sample
FT                   /mol_type="genomic DNA"
FT                   /isolation_source="Cow rumen"
FT                   /clone="ferm-46a3"
FT                   /note="lignin alkali, ABTS active clone"
FT   CDS             56..379
FT                   /db_xref="SEED:fig|6666666.127270.peg.1"
FT                   /translation="MQLMFSSDAVVTYGTATNSYFSDETFPRAIWQPWTTNGGSYDTAD
FT                   KWVTVSFPLSSFNKNQENKECATAFDKSYLTGLTFFVWNGGIDGTACSPVFAIDNIRVV
FT                   PNE"
FT                   /product="hypothetical protein"
FT   CDS             398..2812
FT                   /db_xref="SEED:fig|6666666.127270.peg.2"
FT                   /translation="MKKIFSISAILAACVLALGLSGCSSKELDTDQYSDKDVKLASFGP
FT                   NPVMRGATLRFFGSNLDRIVEVSVPGLDPITDIEVVTSGNPSEIRIQLPAEGAQVGKIS
FT                   LKSSDGKVFTTQVDVEYTEPIVFDSFSPAEAMPGDVITVKGDYMNLVQSVTFEGGAIVT
FT                   EINHNGRHEASFKVPSKAITGKIILSDEGAIANLLYSEEDLVIGDPTVKSLTVAVAKPG
FT                   ENAVVKGAYLDMIDKVVFEGGAEVSDLALNEDATEFNLVIPAGAQSGEVKAVSHAAKEF
FT                   VAGEIELVIPTDVAAAPSPVKAGADLVISGNDLDVVTSVDFPGASGVEFVFAEDAITVT
FT                   VPATATEGDIALNMASGEAVNVAFTLVHPVVSAIAPVELMAGETITVTGTDLDLVTGVT
FT                   LGGKPVEFTAEAEKIVVQTANTSVSGKLVLSLANGEKIEPEDAITLSYDSFIIVNDMPA
FT                   AEHIGAIVTLKGENFMMIENIYIGEEKVSKYNLRSDNEIQFVMPWNKVGTYSMYFHLLS
FT                   GDVETCPQTIEVLLEQDIMTVWEGSLQITWNDGGRVTVPAASFTGIPAGSKLRFYYTQV
FT                   DQQWDQAQLNYGNWTGINFDDPAGTVINGTLVPTDVYGWFTDGILDRCTEVVLTKEILE
FT                   NIQANKGDCEDKTACGIIIQGSGLTFTKIEIVCDIPQEVTIYEGPTTLTWGDDGRFGLA
FT                   MTYFEAAKPGSQLIFYLEQTDAWGQVQLNDGWWANADMNFPEIGGAYITTDNLGGKDVT
FT                   RIALTLTADILDHILATSGDYFGLNSQFQGDGRVAMVIQGSDMIINKVCIL"
FT                   /product="hypothetical protein"
FT   CDS             2907..4433
FT                   /db_xref="SEED:fig|6666666.127270.peg.3"
FT                   /translation="MIFLAACSCSKDDAEKKEPSVPAPKAVTVDVASVSAPVEGTSVVV
FT                   AISSPSKPAVSVSGDWIKAELGPYNPDTYKVKLTVTVGENPTYESRSGSVTLTCQGFSP
FT                   VGISVSQDAAAKPVGPSANPAYDLLGLGWNMGNQFDAFYNGSWAGDKYNYPDETVWGNA
FT                   KATQATFDGVKKAGFTSVRIPVTWLNMIGPGPEYNIDETWMNRIYEVVGYAEKAGLKVI
FT                   INTHHDENNNDDHWLDILSASKDARTNEAIKAEIAAVWTHIANKFKDKGDFLIMESFNE
FT                   IQDGGWGWSAAFQADPSKQCDVLNEWNQVFVDAVRATGGNNATRWLGVPTYAAGTSWTK
FT                   YFRLPEDPAGKVMMAVHCYDPYEYTIGNIQYPEWGHTGAAGKKTPEGEKEITDLFSRLN
FT                   TEFISKGIPVYLGEFGCSMRSKGDARAWMFYKYYMEYVSKAAHTYSLPGILWDNGSTKT
FT                   GKECHGYIDHGTGKYIGNSKEIVDLLVKGMLSTDESYTLQSVYDRAPKTN"
FT                   /product="GH5_4"
FT   CDS             4559..5578
FT                   /db_xref="SEED:fig|6666666.127270.peg.4"
FT                   /translation="MAASVKAGKILYGHQDDLVYGHSWCVDPDSAATDALNRSDVKAVA
FT                   GDYPAVVGFELGGLELGWDWSIDKVPFLLMRRAAVEHAGRGGIVTFSWHPHNPYTGGDA
FT                   WDISSDKAVASVLEGGEKAEMFGVWLERLGDFMDSLRDDKGNLIPVIFRPWHENIGSWF
FT                   WWGGALCTADEYIALFRRTVEYLRNDRGLDNIIMCYSPNSGITAEQYMSRYPGDDVVDI
FT                   LGYDHYETVMEGESLEDAGKRYAGQLDLGLSIVKSLASAHGKLMTLSETGMEGLPDRSW
FT                   WTSVLYPAVKDFPIVYVLTWRNAWDRETHFYAPYPGNEGDFREFASLPEISLLSSFRP"
FT                   /product="GH26"
```

```
FT   CDS             5665..6831
FT                   /db_xref="SEED:fig|6666666.127270.peg.5"
FT                   /translation="MSFEERLAWLEARHQELIAKKNSPIYGNGVFERYENPVLTGDHAP
FT                   LFWRYDLDKNSNPFLEERFGIHAAFNAGAIKWKDKYVVCARVEGADRKSFFAIAESPNG
FT                   IDNFRFWDHPVTMPDYGEPATNVYDMRLTAHEDGWIYGIFCVERKDPDAPEGDLSSAVA
FT                   AAGIARTKDLVNWERLPDLVSDSQQRNVVLHPEFVNGKYALYTRPQDGFIDAGNGGGIG
FT                   WALVDSMENAVVTSEKIINCRRYHTIKETKNGEGPAPIKTDRGWLHLAHGVRGCASGLR
FT                   YVLYMYMTSLEDPSKVIAEPAGFFMVPEGEEYVGDVMNVLFSNGWIADEDGKVFIYYAS
FT                   CDTRLHVATSTVDRLVDYCMNTEPDLFTTSASVRRLNALIDKNLSLSM"
FT                   /product="GH130"
FT   CDS             7057..8280
FT                   /db_xref="SEED:fig|6666666.127270.peg.6"
FT                   /translation="MGIIADRTRSRWGKYRPYLLWVALPFSLIGLLSFYTPDASYGLKH
FT                   VYAYVMYILMMTVYTAINVPYGAMLGVVTPDSREKSVFSSYRMFFAYIGSFIAMGIFGV
FT                   FEKLISGTVRVMPDGSERVVRGVGDAAPGQWTLVVGIVSLCCFVLFLLCFLMTRERVGT
FT                   ENKAVSEGGSSIAEDLKALGRNRPWWLLLGAGIGILLFNSIRGGAAAYYFSNILGTSVF
FT                   LTCAVYLFIGEVAQMVGVLFTVPLSERIGKKATFISVLLTVSVLSVAIFFLPETPAGFW
FT                   GLLVLQVLICIAFGVQSPLLWSMFADVADYSEEKNGSASTGLIFSSSSMAQKFGGALGA
FT                   FLLMQILAIFAYDKDAAVQAPQTLAAIRGLMSYIPAIGALLGALCLLFYPLTESRMASI
FT                   REKLSARR"
FT                   /product="Predicted beta-glucoside transporter, GPH
FT                   family"
FT   CDS             8765..11497
FT                   /db_xref="SEED:fig|6666666.127270.peg.7"
FT                   /db_xref="GO:0003887"
FT                   /translation="MQDCGKDEGLEGIEKNVIFVFRSLIRRTMADKLYLIDGHNLIFKM
FT                   YYAFLRRPMINSKGMDMSILYGFTKYVLELIEREKPTHMAIAFDPPGGTFRNEMYPEYK
FT                   ANRSETPQLVIDALEPLLEICKALNIKTLMIPGYEADDVIGTVAKSYGKPGTDVYMVTP
FT                   DKDYGQLVAEHVYQVKPGKAGGADETVDVKAVCEKYGISSPEQVIEMLTLCGDTADNVP
FT                   GVHGIGEVGAGKLIAKYGTVENIYAHLDELTDRQKTQFEEARGHIGLSHDLVTIKTDVP
FT                   VETSLDEMRLDLDYAPEIADLFEKYEFNSLRKYIAHSSRAAKAVAEVKRLDYTETGAEE
FT                   LVKMAPEKISLIVEPEGPGIFADIRRVIVAAEKKDGGIVVASGAAKDFRKILEGEAVKY
FT                   GYDLKLQSNLLENAGIKLGGRLYDIELMHYLINPEKSHKVEILAKSYLGVDIEEAAAVE
FT                   EAPVSLFDVAEESPANGPDRNLEAAAVLKLGEVVRAGMDEAGVGALYEDMEAPLAAVLA
FT                   RMERAGVKIDLGQLRTFADELKVEMDALEAQVRKLADEPGLNVSSPKQIGIVLFEKLKL
FT                   DPKMKKSASGSYSTDETTLTAIADRHPIIDKILEFRAVKKLLSTYIEPFPTYVSPVTGR
FT                   VHTTFNQALTATGRLSSSNPNLQNIPIRTERGKEIRKAFIPATPDGIIMSADYSQIELR
FT                   IMAHLCGDEHLISSFKAGLDVHKATAAKIFGIPVEEVTSDQRRIAKTANFGIMYGISSF
FT                   GLAQRLHVSRTEAKKIIDDYFAAFPSIRTFIDKTIADAREKGYVETVFGRRRYLPDITA
FT                   KNATIRALAERNAVNAPIQGTSADIIKLAMISVDKRIREAGLKSKMVLQIHDELLFDAV
FT                   PEEIEELGRIVVAEMENVITLSIPLTVECNYGKNWLEAH"
FT                   /product="DNA polymerase I (EC 2.7.7.7)"
FT                   /EC_number="2.7.7.7"
FT   CDS             11484..12074
FT                   /db_xref="SEED:fig|6666666.127270.peg.8"
FT                   /db_xref="GO:0003677"
FT                   /db_xref="GO:0003700"
FT                   /db_xref="GO:0006352"
FT                   /db_xref="GO:0016987"
FT                   /translation="MKLTDHDDGSLVEMALGGGPEAEAAFFTIYARYHKGVFAHISRFI
FT                   QEQEEVEDICMESFEKAFKQLESYRPSNKFSTWIFTIARNTAFDHLDKDKVRGRNKDKM
FT                   PIDYSDLEHIDVPEDNVSPEDEIISSQDHENFLSCVQGLPDLYREVASMCFVDNLGYKE
FT                   ISEKAGLPINTVKTRISRAKSLIVQMMLDMEEE"
FT                   /product="RNA polymerase sigma factor RpoE"
```

```
FT   CDS             12071..12733
FT                   /db_xref="SEED:fig|6666666.127270.peg.9"
FT                   /translation="MNFKEFEDIVVAEFPELTPEMLGKYEAMEPLYEEWNAKINVISRK
FT                   DIGGLYLHHVLHSLAIASYLKRRRPEVYSAWTEGASVLDLGTGGGFPGIPLAVMFPKVR
FT                   FTLCDSVGKKIIVASEVSKALSLGNVTTVNARAESLPQVFDHVVSRAVTSLDNFYPWVK
FT                   GKFSGSILYLKGGDIVPEIAELMKKNRLRTGAVSTWPIETWLKDEYFKEKFVIEIEK"
FT                   /product="rRNA small subunit 7-methylguanosine (m7G)
FT                   methyltransferase GidB"
FT   CDS             13212..15989
FT                   /db_xref="SEED:fig|6666666.127270.peg.10"
FT                   /translation="MQVWLNEDHSQPVVHGAVVVKAGGRDCAGTGMAHYLEHMLFKGTE
FT                   DIGTIDYASEKVYLDSIETCYDLIAATSSSVKRDSLQKEINRLSIKALDYAIPNEFDRL
FT                   TANIGGTKLNAFTSYDCTCYHNSFSPQFIAQWAELNSHRLLNPVFRLFQSELEAVYEEK
FT                   NAAADSQFDMILEKMLEKLGDENPYSKSIIGTTDNLKNPSLSEMKKFFNKYYVGCNMGL
FT                   ILAGDIDASSILPLLESTFGRIPRGDKPERIPVPVPDFSSREDYPIKVNFPMIKASLNV
FT                   FNGPAAKDADYPAMELALSLLNNSFSTGSLDSLINNGKVLAGEVSVLDFFDEMGLIAMI
FT                   TIPGIPFGSLNKAEARCFKEVERLKAGDFAEKDFESLKYDLVKSYKRCLESSESRFSIM
FT                   IDVMSQDRSWDSYLSQVESISSLTSEDIVRVSNKYLGDNRIRFHKVRGTYPKDNLQAPS
FT                   YETLSPKNGNAVSDYATRLMEMPYDTLPPRLVDMNNDIRKVAVNDHVTLYYNGNPVNDV
FT                   FELTVRINKGIFEEPMICPLTQLLGLSGTDSLSARDFSMSLAELGSDLSFKDKNHTLEM
FT                   VLTGFDDRIGPSLDLLRHFMKDFKPEKSALNTVISDTKMSRNAFRSSGNSAIFNGMFNY
FT                   SLLGNQSPLLRQPTEKDLKSLGLKGMKELFNSLVEYDFDVVYSGGLSPEEVVNEISGHL
FT                   DVECPSGKPESSFFAYRKYEQPVVYIYNVPGARQVMSITYQNVATPENDHDSALLGLMQ
FT                   ECYGGSMASIMFQEVREFRSLAYTSSSTLQKPLPNVSSPSAFITYYGTQADKFPQALEL
FT                   VDSLIRTMPMTDLCFDSAVRSIISRANNGYPSFRALPEYVANLELFGYGEDDDASVVRG
FT                   LPGITKQDVLDYHEKSIGNAPCSYFIIGDKKNLPMDALRERGRIVELKLSDIYR"
FT                   /product="ZINC PROTEASE (EC 3.4.99.-)"
FT   CDS             complement(16003..16662)
FT                   /db_xref="SEED:fig|6666666.127270.peg.11"
FT                   /translation="MRIKDLNIGHRGKTLVSGISFETEPGEFILLCGPNGSGKSTLLRA
FT                   LAGKDAIMVPTGVPKVKGFTLREFIATTLYSESDWRGRISPAMASDISKSLECLGISSL
FT                   ADRDISTLSDGEFQKGCIASALSRHAKTLLLDEPTSFLDVDNKSLILQALRDIARSTGT
FT                   TVIMASHDLAPALSCADRVFGITADHRFLASSSRLSDSGLSASEVISACFSTYSSR"
FT                   /product="iron (III) ABC transporter ATP-binding"
FT   CDS             complement(16659..17660)
FT                   /db_xref="SEED:fig|6666666.127270.peg.12"
FT                   /translation="MAIKRRNRLITAAGATALLLLSVCDLILGGSGIGAAEALGGLFGG
FT                   EGIASSIVWSIRLPRVLTAIAAGACLALSGAQMQSVFRNPLADPHIMGVSGGAGLGAAI
FT                   ATLLISGTGLLSGFSSMAGLTVAGAAFVGAILASALIVLVSSRFRSGTTLLVFGVMLGF
FT                   IFSAITSILEYSASEESLKLFYSWSAGSFSGTRTFETLTIFAALLVGTGLYTSGAKGLD
FT                   ILLFGEEYATLAGASVRRIRGVAMVSSCLMTAAATAFCGPLGFVGIVSPHIARRLLRSS
FT                   GHMSILPATLITGSAVSVLADIISQAPGLPLPVGSTMALIGIPIILLILMKK"
FT                   /product="Vitamin B12 ABC transporter, permease component
FT                   BtuC"
FT   CDS             17729..19048
FT                   /db_xref="SEED:fig|6666666.127270.peg.13"
FT                   /translation="MSTETILIIILLVAVIVIAAILIFRLQARAEKIKDGFQDRLADVQR
FT                   EKAVLESRLQAKEEYENQMREAAKETSERHAKELEQMKEVFKNLSTETSEAFKAKSAQT
FT                   ISELMKPVQEKFKEFSDAVKESQKDTIERHSRLEQSIKDLDTRSKSVGDEARNLANALT
FT                   GYSKVQGDFGEMLLTDVLKNAGLTEGVHFETQSVMTDNSGHEIKSESGRTMIPDVKVYY
FT                   PDDTIVIIDSKVSLTAYNEYMAATTVEDRTRLAKAHVRSIEDHVNELKVKDYAGYIPEG
FT                   RRKVDYNIMFIPVEGAFRLMLEEAPRLWQVAKDNNILIVSQMTLIIVLNMIQMAWKQHE
FT                   QEKNISDVYKTAEELMSQLKGWMESYVKIGDYLGKASSAYEDSKKKLSDSNQSVIKKIG
FT                   KLEKLGIAPKRSNAKIKTGSRLSGPESVIPKELSDDEITD"
```

```
FT                   /product="DNA recombination protein RmuC"
FT   CDS             19161..19439
FT                   /db_xref="SEED:fig|6666666.127270.peg.14"
FT                   /db_xref="GO:0008815"
FT                   /translation="MELKNASAGTFESGDILIRIDPAEKKGLEIELESSVADQFGVQIK
FT                   KVISETLEGLGISDAKVKAIDKGALDCTIRARVTAAAVRASGKDVWK"
FT                   /product="Citrate lyase gamma chain, acyl carrier protein
FT                   (EC 4.1.3.6)"
FT                   /EC_number="4.1.3.6"
FT   CDS             19467..20336
FT                   /db_xref="SEED:fig|6666666.127270.peg.15"
FT                   /db_xref="GO:0008815"
FT                   /translation="MMFIPGNNPGMMQDAYIYGPDSIMLDLEDSVTMAEKDAARLLVYN
FT                   ALKTVDYGNIERVVRINPLSTPYGKKDVEAVVKAGVDVIRMPKTETADEVRELEAEILK
FT                   VEEEIGCVGRTQIMAAIESALGIVNAYQIASASSRMMGIALGAEDYTANLKTMRSNEGI
FT                   ELQLARETIVVAARAAGISCFDTVFSNLEDMEIFRKEVEFIKTLGFDGKSIINPRQIDV
FT                   VNEVFTPTEKEIAKAQTIIAAIKEARQRGSGVISVNGKMVDRPVVIRAERIIALAVAAG
FT                   VIPEEDAL"
FT                   /product="Citrate lyase beta chain (EC 4.1.3.6)"
FT                   /EC_number="4.1.3.6"
FT   CDS             20333..21856
FT                   /db_xref="SEED:fig|6666666.127270.peg.16"
FT                   /db_xref="GO:0008815"
FT                   /translation="MTNEQLIREAAAKLGKKVYQGPGQKIPTFRRDLQKESCKVVSLED
FT                   AIRKSGLKDGMTISFHHHFRGGDKIVNMVVDKIAEMGIKGLTLAASSLSDVHAPLVGHI
FT                   KNGVISKITTSGVRGELAEQISRGLMDEPVIFRSHGTRGAAVASGELHIDVAFLGAPSC
FT                   DPLGNACGYSRSENAKSICGSLGYALPDARYADKVVILTDDLVPYPNTPNSISEHDVDY
FT                   VVVVDSVGDSSKISSGAIRDSKNPRDILLARLAAKAIINSGYFKDGFSIQTGTGGAALA
FT                   AVKFIRESMIEKGIKASYALGGITAHMVRLHEEGLIERLIDVQSFDKVAAESVKNDPAH
FT                   KEVASYEYASGMEPGSAVHYLDIVILSALEVDVNFNANVLVGSDGIIRGAIGGHPDTAA
FT                   DSALSIVVCPLLRGRIPCIVDHVTTLITPGASVDVVITEYGTVVNPRRPEIAERLKAAG
FT                   MKLVDMQYLRDCARKIIGEPAPIPFGDKVVGVVLGRDGSVQDVIRNVLK"
FT                   /product="Citrate lyase alpha chain (EC 4.1.3.6)"
FT                   /EC_number="4.1.3.6"
FT   CDS             21856..22359
FT                   /db_xref="SEED:fig|6666666.127270.peg.17"
FT                   /db_xref="GO:0050519"
FT                   /translation="MEVTLDMILASREARQKEQEYLLKHSYGSTLISMTVVMPGNVKRD
FT                   EYTIKIAEAGIDALLNAFDGGINRIVQRDLETGYEAFLLVKAPLEETKRVVSLIEETHP
FT                   LGRLFDIDVIGEDGVPVTRTGYGMTPRKCLLCDNEARYCMRNHTHTPEQLKARIREMVD
FT                   EYVR"
FT                   /product="Apo-citrate lyase phosphoribosyl-dephospho-CoA
FT                   transferase (EC 2.7.7.61)"
FT                   /EC_number="2.7.7.61"
FT   CDS             22349..24079
FT                   /db_xref="SEED:fig|6666666.127270.peg.18"
FT                   /db_xref="GO:0046917"
FT                   /translation="MYGEYQIIDLPLSAGVVRSKVETFLAENGLRLEAVDRYVGIEDSD
FT                   ERLIAGGGICKDVLKCIAVSEEARSGGLTGKLVARLMQLGEEAGQHDLKVFTKPEYEDI
FT                   FASLGFSVIAKAGKAILMENGDGLKRYCSDLEKHRRPGRCGAVVMNAAPLTWGHDYLVE
FT                   KACESVDALFVIPVAEDTPGFSYRDRYQALASAWLEELKVEVCSGSPYAVSKATFPTYF
FT                   LKDLSDAAETQMTLDLDIFARHIAPALGVTVRFVGSEPSDALTARYNELMKKILPEHGI
FT                   EVVEVPRLESEGRPVSASEVRKAVAEGSFRRADAIAHPESMPMLVGALASRALRMELDL
FT                   TPKPGLVDKNGSGAHKDMDYVTMSASIDAIVPFIHAMASINPDDDVPSDMVELGIKAEQ
```

```
FT                   KMLAATGGVNTHKGAIFSLGLMATAFAQCYWEGETVGEKALKSKIVALACAVPKSKDTH
FT                   GAKVCSKFKAKGALQNARDGYPALFKEWLPFYRKVKAEKNGLLKTLLLIVSQLEDTNVL
FT                   YRKGPAVAEKVRSMAREALKNFSEDKVRNICGRFTSEGISPGGSADMLALTVFADSLLS
FT                   "
FT                   /product="2-(5''-triphosphoribosyl)-3'-dephosphocoenzyme-A
FT                   synthase (EC 2.7.8.25)"
FT                   /EC_number="2.7.8.25"
FT   CDS             24113..26350
FT                   /db_xref="SEED:fig|6666666.127270.peg.19"
FT                   /translation="MVDLINHGVYLLDGVVSDAAPGHMSPDEARENTITYGILRSHDVD
FT                   GGKGKKMRIRFDAMVSHDITYVGIIQTARASGLEKFPLPYAMTNCHNSLCAVGGTINED
FT                   DHVFGLSAAKKYGGIYVPANQAVIHQYAREALTKCGNMILGSDSHTRYGSLGNMGVGEG
FT                   GGELVKQLLRNTWDINAPEVVLVWLEGTPRHGIGPHDVAISLVKAVFENGFVKNKVLEF
FT                   AGPGVASLPIDFRNGIDVMTTETTCLSSIWITDEKVREYYEMHGRAADYKKLEPGKVAY
FT                   YDSMIRIDLSTQESMIALPFHPSNAYTIHELQADPERILRAVEEDARKRFGDKISIDLL
FT                   SKIHDGKVVADQGIIAGCSGGTYDNLSEAATILKGKSIGNDYFTMSAYPQSTPVYLATT
FT                   RNHIAEELLEAGVVIKPAFCGPCFGAGDVPSNNGFSIRHTTRNFPNREGSKPGQGQISL
FT                   VSLMDARSIAATAANGGVITAATDIEYNDTHKPYAFDDRIYKSRIYNGFGHADLSQELR
FT                   YGPNIKDWPKIYPMEKNMLVELAAVIHDPVTTTDELIPSGETSSYRSNPLKLSEFALSR
FT                   RVPEYVCLSKRIQSEDLRRRAGDVPANVAAALAKVGASASDTSFGSCVFANRPGDGSAR
FT                   EQAASCQKVLGGDANICYEYATKRYRSNCINWGLVPFTIAPETAFDYKPGDFVFVPGIR
FT                   DAILSGKEEIDAKVITASGVSDIHLYFNNLTPVEREILADGCLMNYYMTHK"
FT                   /product="Putative enzyme"
FT   CDS             26414..26530
FT                   /db_xref="SEED:fig|6666666.127270.peg.20"
FT                   /translation="MSGRGPAVRLGGIAPKARSFQAIVSADKTIDKTIKENK"
FT                   /product="hypothetical protein"
FT   CDS             26544..27755
FT                   /db_xref="SEED:fig|6666666.127270.peg.21"
FT                   /db_xref="GO:0004450"
FT                   /translation="MRKIQMTTPLVEMDGDEMTRILWKMIKDELILPFVDLKSEYYDLG
FT                   LVNRDKTADQVTVDSALATKKYGVAVKCATITPNVQRMSEYNLHEMWKSPNGTIRAMLD
FT                   GTVFRCPITIPSIHPAVKFWKKPITIARHAYGDVYKGVEIKTDEPGTAKLVFEGESGKK
FT                   EEVVIQKFDGPGVIQGMHNTDKSIRSFAHSCFKYALDVKQTIWFATKDTISKKYDGRFK
FT                   AIFEEVFQEYKDRFEAAGIEYFYTLIDDAVARVMRSEGGFIWACKNYDGDVMSDMVSTA
FT                   FGSLAMMTSVLVSPDGKYEYEAAHGTVTRHYYKYLKGEETSTNPMATIFAWSGAFRKRA
FT                   ELDNLPELVNYADQLEASCFDTLNSGIATKDLVNLMEGVEAKAVNSLDFLKAIRANLEK
FT                   RLQN"
FT                   /product="Isocitrate dehydrogenase [NADP] (EC 1.1.1.42)"
FT                   /EC_number="1.1.1.42"
FT   CDS             27755..29431
FT                   /db_xref="SEED:fig|6666666.127270.peg.22"
FT                   /db_xref="GO:0005524"
FT                   /db_xref="GO:0006810"
FT                   /db_xref="GO:0016021"
FT                   /db_xref="GO:0016887"
FT                   /translation="MADEKIIFSMNGVGKVLPHNNKVILKDIYLSFFYGAKIGIIGLNG
FT                   SGKSTLMKIIAGVEKSYKGEVVWAPGYSVGYLEQEPQMDPDKTVIEVVQEGVQEVMDLL
FT                   NEYNEISAKFCEPLEDDEMNKLIERQGELTDKIEHCGGWEIDSKLERAMDALQCPPSEA
FT                   KIATLSGGEKRRVALCRLLLRQPDVLLLDEPTNHLDTESIQWLEAHLQQYKGTVIAVTH
FT                   DRYFLDNVAGWILELDRGEGIPWKGNYSSWLDQKTKRLAMEEKQESKRRKTLERELEWV
FT                   RMAPKARQAKQKARLGAYEKLASADTKEKESNLEIYIPNGPHLGNKVIEFHNVSKSYGD
FT                   KLLFENLDFVLPPAGIVGVIGPNGAGKTTLFRLIMGLEKPDTGTIEIGETVKISYVDQQ
FT                   HRSIDPDLTVYETIAGNSEWIRLGNKDINARSWVSRFNFSGADQEKLCGVLSGGERNRL
```

```
FT                   HLALTLKDEGNVLLLDEPTNDVDVNTIRALEEGLDGFAGCAVVVSHDRWFLDRIATHIL
FT                   AFEGDSKVVFFDGNYSEYEENKKMRLGDVEPKRFKYRKLME"
FT                   /product="ABC transporter ATP-binding protein"
FT   CDS             complement(29440..30465)
FT                   /db_xref="SEED:fig|6666666.127270.peg.23"
FT                   /translation="MISFVCDYLEGAHPEIMKRLAETNLEQTPGYGQDKYSASAISKIR
FT                   EACGCPEAQVYFLAGGTQTNSTVIGSLLNGNEGVISATSGHINVHEAGAVEYTGHKVLT
FT                   VPAHLGKMDATDLDSYLRGFFADPAWDHMVIPGMVYISHPTEYGTIYSLAELSEISAVA
FT                   HKYGLQLYLDGARLGYGLMATGTDVSLKDIARLCDVFYIGGTKVGALFGEAVVFPRGNA
FT                   PRHMFSMIKQHGALLAKGRIAGLQFDTLFTDDLYFRIGRHAIEMAEKMKAIFIEKGYEF
FT                   FIDSPTNQQFIILTSEQREMLSALVDFEFWEALPDGRCAVRFATSWATTPESIDYLRTI
FT                   L"
FT                   /product="Low-specificity L-threonine aldolase (EC
FT                   4.1.2.48)"
FT                   /EC_number="4.1.2.48"
FT   CDS             complement(30470..30598)
FT                   /db_xref="SEED:fig|6666666.127270.peg.24"
FT                   /translation="MKYDTIAHLLQYWFLEVTNLQNILHIPISFHMTEKAYICAMF"
FT                   /product="hypothetical protein"
FT   CDS             30575..31717
FT                   /db_xref="SEED:fig|6666666.127270.peg.25"
FT                   /db_xref="GO:0004017"
FT                   /translation="MSNSIVLHDKVFKPYLSYDRIISAIDGVAARINADFEGCTDIPVI
FT                   LCVLNGSIMFTGELLKRLKFNVELVSMKLTSYDGVRSTGKVRQVMGLSGSVKGKRVIVV
FT                   EDIVDTGNTIIDLMDIVRGEGASDVKICTMLLKPEVYKKDVKLDYVGMEIPNAFIVGFG
FT                   LDYNEIGRNYRDIYVLDPDATEQLNEKMKYYILFGPPGAGKGTQAAAMVEKYNLCHIST
FT                   GELLRKEISAGSELGLKAKALIDAGALVPDEVVEGMIENQFKTVKGVKGFLLDGFPRTI
FT                   AQAEALDAMLAKNSEEVTSVVSIMIPDEMITARIKHRASIEGRADDANDATIANRIKTY
FT                   HDKTEPLIGYYRKAGKYAEIDGTGSIDEVRGKIFATMDKF"
FT                   /product="Adenylate kinase (EC 2.7.4.3)"
FT                   /EC_number="2.7.4.3"
FT   CDS             31948..33498
FT                   /db_xref="SEED:fig|6666666.127270.peg.26"
FT                   /translation="MFSPQAPPETLLGVTDMEELNEEAAERLDFLERHPLKVNYATEAR
FT                   MAASGLMSRYQAASLADYRSLHGDVLSVAELAAVDGFGPAVAEAMRPYLSFESSRLPGQ
FT                   ADEPAGLSHEIELRAGTKFQRTTFAGRYRLEYGSRIEAAVSYRDRLNGYASVSGRRWKA
FT                   VAGCFNARFGQGLALWSGMSIGGVTTPEAAGRRASGIAPTWSFSNAGFRGLAGEYAVGK
FT                   LRMSAFLDARGRVSPGGNILWLMKNGELGLTATRSKASADFRLTLRGSDLFGEAAYDFR
FT                   DHVIAGIVGTAFPLWERGRAAVVFRHYPTGFDGKDSGAMRSSTKVTDETGVFAALRLGS
FT                   ILTSADAVVHPGTGRRQLKLLLLHEWKISPSVSLKTRATERLRDYDRRTRTDVRTDLGI
FT                   TRGSMVMNFRLNALKCREIGLAGYAEAGYKREKSALWLRWTAFKVDNWDDRIYVYERDA
FT                   PGNFSVPACYGSGFTFSLYGSWKIRRLRLYLKSGTTFKERSGTRSLSSDLHFQLSYSL"
FT                   /product="hypothetical protein"
FT   CDS             complement(33500..34918)
FT                   /db_xref="SEED:fig|6666666.127270.peg.27"
FT                   /translation="MVSSVAPLDSYAQSESIFDKITQPFRKNGPKKKNKDLKKENERLQ
FT                   AEINALRQQLNDYIEQHEAADSIAAEMIEIYEENEDRIGAGIAADDYTPEVTDSLLNLW
FT                   YSTRKSMNMENNSYNMDSVHFTSNVPDSVYIKRLRKMNSYITLPYNETVKNYIILYSEK
FT                   MPTRMGHMLGLAQYYFPIFEETLNKYNMPEELKYMAVIESALNPTAVSRANAKGMWQFM
FT                   YSTAKNYGLKINSYVDERLDPVKAADAAARYLRDAYSIFGDWSLAISAYNCGAGNVNKA
FT                   IRRAGGNKNFWAIYEYLPRETRGYVPAFVGAMYAFTYYKEHGIIPEPTAIPSHLDTFAI
FT                   RKNLHFAQINGVINVPVDMLKELNPQYMHEIVPGNEGVSILTLPSQYTTAFIDYEDSIY
FT                   NYQADKYLSPKTLEDIKKARTEESGRIVYKVRNGDYLGRIAARYHVTVAQIKRWNNLRS
FT                   NNIRVGQRLVIYRR"
```

```
FT                   /product="GH23-CBM50"
FT   CDS             complement(34966..35712)
FT                   /db_xref="SEED:fig|6666666.127270.peg.28"
FT                   /translation="MFSFRQYFRAMRHKEEGKIGTMFAGSTIVIGGQQIYNRDYWKLPV
FT                   IYGGLAATAGLGSYYIHKDGKKDLGKYLLAGAGAIYWGSLMDGVVSYDTKGEHSAGRAT
FT                   LYSLLVPGLGQIYNKEYWKLPLYYSGLMVSASLLIENSANYHRFKRIHNEITDENSTYS
FT                   GHYTESSALYMRNTYRRYRDYSVVALVGVYVLQIIDANVFSYMLDFDIDDEIAVDVSPA
FT                   IITPHSQFAMGMPVDNAYGISIGIKF"
FT                   /product="hypothetical protein"
XX
SQ   Sequence 35712 BP; 8621 A; 9081 C; 9801 G; 8209 T; 0 other;
     cgtgctcaag ttcgaggttc tcgttccttc atccaaccct tggtctgctt caggtatgca        60
     gctcatgttc tcgagcgacg ctgtcgttac ctacggtacc gctacgaaca gctacttcag       120
     tgacgaaacc ttccctcgtg ctatctggca gccatggact accaacggag gttcttatga       180
     taccgctgac aagtgggtta ccgtaagttt ccctctcagc tcattcaaca agaaccagga       240
     aaacaaggag tgtgcaaccg catttgacaa gtcttacctt accggtctca cattcttcgt       300
     atggaacggc ggtatcgacg gcacagcctg ctctccggtc ttcgcgatcg acaatatccg       360
     tgtggttcct aatgaataat attaaatgaa gctcattatg aaaaagatat tcagtatttc       420
     agccattctt gccgcctgtg tcctggcact cgggctctcc ggatgttcat ccaaagagct       480
     tgataccgac cagtattccg acaaggacgt gaagctggca tcattcggcc ctaacccggt       540
     gatgcgtgga gctacactcc ggttcttcgg aagtaatctc gacaggattg tcgaagtttc       600
     agttccggga ctcgatccta tcaccgacat cgaagtcgtt acctccggta atccttcaga       660
     aatccgcatc cagcttcctg cagagggcgc acaggtcgga aagatctctc tcaagtcatc       720
     cgacggaaag gtctttacta cccaggtgga tgtagaatat acagaaccta tcgtatttga       780
     cagtttctct cctgcagagg ctatgcctgg cgacgtgata acggtcaagg gcgattatat       840
     gaacctcgtg cagagcgtta ccttcgaagg cggcgccatc gtgaccgaaa tcaaccataa       900
     cggccgtcat gaggcttcct tcaaggtccc ttcaaaggct attaccggaa agatcatcct       960
     ttctgacgaa ggagccatcg ccaatctcct ctactcggag gaagatcttg tgatcggcga      1020
     tccgaccgtc aagtctctta ccgtagctgt tgctaaacct ggcgagaacg cagttgtgaa      1080
     gggagcatat ctcgatatga tcgacaaggt cgtattcgaa ggtggtgcag aggtttccga      1140
     ccttgctctc aatgaggatg caaccgaatt caatctcgta atccctgccg gagcacagtc      1200
     cggtgaagta aaggctgtca gccacgctgc caaggaattc gtcgccggcg aaatcgaact      1260
     tgtcatccct acggatgtcg cagctgcacc ttcgcctgta aaggctggtg ccgacctcgt      1320
     gatttccggt aacgatcttg atgtggtaac ttctgttgat ttccctggcg ccagcggagt      1380
     agaattcgta tttgccgagg acgccattac agtgaccgtt cctgctactg ctactgaagg      1440
     tgatatcgcc ctcaatatgg ccagcggaga agctgtgaat gtggcattca cccttgtgca      1500
     cccggtagtt tctgccatag ctcctgtcga gctcatggcc ggcgagacca tcaccgtgac      1560
     aggtaccgac ctcgaccttg tcactggagt aactctcggc ggaaagccgg tcgagttcac      1620
     tgcagaggct gagaagattg tcgtacagac cgccaatact tccgtttcag gaaagctcgt      1680
     gctttctctt gcgaatggtg agaagattga accggaggat gccatcaccc tcagctatga      1740
     ttcattcatc atcgtgaacg acatgcctgc cgcagagcat atcggggcta tcgtaaccct      1800
     caagggcgag aacttcatga tgatcgagaa catctatatc ggcgaagaga aagtatcgaa      1860
     gtacaacctt cgttctgata atgaaatcca gttcgtaatg ccttggaaca aggtcggaac      1920
     atacagcatg tacttccatc ttctctccgg cgatgtcgag acatgtcctc agacaatcga      1980
     agttctcctc gagcaggaca tcatgactgt atgggaaggc tctctgcaga tcacctggaa      2040
     tgacggcggc cgtgtaaccg tgccggcagc atcattcacc ggtatccctg caggcagcaa      2100
     gctcagattc tactacactc aggtggacca gcagtgggat caggcccagc tcaactatgg      2160
     taactggacc ggaatcaact tcgacgatcc tgcaggtacc gtcattaacg gaaccctcgt      2220
     tccgactgac gtctatggat ggttcactga cggaatcctt gaccgctgca cagaggtcgt      2280
     gcttacaaag gaaatcctcg aaaacatcca ggctaacaag ggtgactgcg aagacaagac      2340
     tgcctgcggt atcatcatcc agggctctgg ccttaccttc acgaagatcg agatcgtatg      2400
     cgatattccg caggaggtta cgatctacga aggtcctaca acccttacat ggggtgacga      2460
     cggccgtttc ggtctcgcca tgacttactt cgaggcagca aagccgggct ctcagcttat      2520
     cttctatctc gagcagacgg acgcatgggg ccaggttcag ctcaacgacg gctggtgggc      2580
```

caacgccgac atgaacttcc ctgagatcgg cggtgcatac attaccaccg acaacctcgg 2640
aggaaaggac gtcacacgta tcgcccttac cctgactgct gatatccttg accatatact 2700
tgctacatcc ggtgactatt tcggtctcaa tagccagttc cagggtgacg gacgcgtcgc 2760
catggttatc cagggatctg atatgattat caataaagtt tgtatattgt aacactttta 2820
ttgccggagg cgggtgatgc cgcctccggt agccagtaaa caaattttat ataatgaaag 2880
cacacgtcag tttctttgta ttatctatga tttttcttgc ggcatgttcc tgctcaaagg 2940
atgatgccga aaagaaggaa ccgtccgtcc cggcccctaa ggcggtaacg gtagatgtcg 3000
cgtctgtcag cgctcccgta gagggaacgt cggtcgtcgt cgccatctcc tccccgtcca 3060
agcctgcggt ctctgtctcc ggagactgga tcaaagccga actgggtccg tacaatccgg 3120
atacatataa agtgaaactt acggtgaccg tcggcgagaa tccgacatac gagtcacgca 3180
gcggtagtgt gacccttact tgccagggct tttctccggt cgggataagc gtatcccagg 3240
atgccgcggc gaagccggtc ggaccgtcgg ctaatcctgc atacgatctt ctcggtctcg 3300
gctggaatat gggtaaccag ttcgatgctt tctacaacgg ctcatgggcc ggagataaat 3360
acaattatcc ggacgagacc gtctggggta atgccaaggc gacccaggct actttcgacg 3420
gggtgaagaa agccggattc acatcagtgc gaatccctgt cacgtggctc aatatgatag 3480
gtccaggacc tgagtacaat attgacgaga cttggatgaa ccgtatatat gaggtggtag 3540
gatatgccga gaaagccgga ctcaaggtca tcatcaatac ccatcacgac gaaaacaaca 3600
atgacgacca ttggctggat atcctttctg catctaagga tgccaggacc aatgaagcga 3660
taaaggcgga aatcgcggct gtatggaccc atatcgccaa taagttcaag gataagggag 3720
atttccttat catggagtcc ttcaatgaaa ttcaggacgg aggatggggc tggagcgctg 3780
ctttccaggc tgacccttcc aagcagtgtg atgtgcttaa cgaatggaat caggttttcg 3840
tggatgccgt acgtgctaca ggaggcaaca acgctacccg ctggctgggt gttcctacat 3900
acgccgcagg tacttcatgg acgaagtatt tcagactgcc ggaggatcct gccggaaagg 3960
tcatgatggc agtgcattgc tatgatccgt atgaatatac tatagggaac atacagtatc 4020
ctgagtgggg ccataccggc gccgcaggaa agaagacccc ggaagggggag aaagagataa 4080
cggatctctt ctccagactt aatacggaat tcatcagcaa ggggataccт gtatatcttg 4140
gcgaattcgg ctgttcgatg agaagcaaag ggatgcccg ggcatggatg ttctataaat 4200
actatatgga gtatgtctcc aaagctgcgc atacctattc cttaccggga atcctgtggg 4260
ataacggaag tactaagacc ggaaaggagt gccacggcta cattgaccat ggtaccggca 4320
agtatatagg gaattctaag gagatcgtgg atcttctcgt aaagggaatg ctcagcacgg 4380
acgaaagcta tactctccag agtgtatatg acagggctcc gaaaaccaat tgattatgaa 4440
tctaatcctt aaacttagca tcattttatc attagtagtt gctgtatccg gctgcggacg 4500
caataaagaa gcggccggaa cagcttttaa cgaagaaacc cggatcatga tggatcgtat 4560
ggcggcttca gtaaaggccg gcaagattct ttatggccat caggatgacc tcgtttatgg 4620
ccattcatgg tgtgtggatc ctgattccgc agcgacagat gcgttgaacc gttcggatgt 4680
caaggctgtc gccggagatt atccggctgt cgtgggattc gaactcggcg gtctggaact 4740
gggatgggac tggagcattg acaaagtccc gttcctgctg atgcgccggg ctgctgtaga 4800
gcatgccgga aggggaggaa tcgtgacgtt ctcatggcac ccgcacaatc cttatacagg 4860
cggtgatgcc tgggatatca gttccgacaa ggcagtggct tctgtcctcg aaggagggga 4920
gaaggcggag atgttcggag tctggctgga gaggctgggc gatttcatgg actcgctccg 4980
tgacgataag ggtaatctga taccggtcat tttcaggccg tggcatgaga atatcggcag 5040
ctggttctgg tggggcggag ccctttgtac tgcggatgag tatatcgccc ttttcaggag 5100
gacggtcgag tatctgagaa atgacagggg actggacaat atcataatgt gttattctcc 5160
gaattccggg attacggcgg agcagtatat gtcccgttat ccgggcgatg atgtcgttga 5220
tatcctgggg tatgaccatt atgagactgt catggagggc gagagtctgg aagatgccgg 5280
aaagcgttat gccgggcagt tggatcttgg cctttccatc gtcaagtctc ttgcttcggc 5340
ccacggaaag ctgatgactc tttccgaaac cggaatggag ggcctcccgg acaggagctg 5400
gtggacttcg gtgctttatc ctgcggtaaa ggattttccg attgtctatg tgctgacgtg 5460
gagaaatgca tgggacaggg agacgcattt ttatgcgccg tatcctggta atgagggcga 5520
cttcagggaa ttcgcttcgc tgccggagat ctctctcctg tcgtcgttcc ggccttagtc 5580
cgtcattccc ggcgaagtgt cgtcctttcc ggtcggaccg ggaatctgta accttaaaac 5640
aataaataac aatatatgga gaatatgagt tttgaagaac gtctggcgtg gctggaagca 5700
cgccaccagg aacttattgc taagaaaaac tcgcctatat acggcaatgg tgtgtttgag 5760
agatatgaga accctgttct taccggtgac catgcccctt tgttctggag atatgatctg 5820

gataagaact ccaatccttt tctggaagag agattcggca tccatgccgc attcaatgct 5880
ggagctataa agtggaagga caaatatgtc gtctgcgccc gtgtcgaagg tgctgaccgc 5940
aaatcattct tcgctatcgc cgagagcccg aacggaatcg acaatttccg gttctgggac 6000
caccccgtga ctatgcctga ttatggcgaa cctgccacga atgtatatga catgagactt 6060
actgcccatg aagacgggtg gatctatggt atattctgcg tcgagcgtaa ggatccggat 6120
gctccggaag gcgatctttc gtctgccgtc gcagctgcgg gaatcgcccg taccaaggat 6180
ctggtcaatt gggaaagact gccggatctg gtttctgatt ctcagcagcg caatgtcgta 6240
cttcatcctg agttcgtgaa tgggaaatat gctctttaca cccgtcctca ggatggattc 6300
atagatgccg gaaacggcgg cggcatcggc tgggccctgg tggacagcat ggagaatgcc 6360
gttgtcactt ccgagaagat catcaactgc aggagatacc atactatcaa ggagaccaag 6420
aatggcgaag gccctgctcc tatcaagact gacaggggct ggctgcatct ggcccatggc 6480
gtcaggggct gcgcttccgg cctgcgttat gtgctttata tgtacatgac ttcgctggag 6540
gatccttcga aagtgattgc cgaacctgcc ggtttcttca tggtcccgga aggggaggag 6600
tatgtgggtg atgtcatgaa cgtcctgttc tctaacggtt ggattgccga tgaggacgga 6660
aaggtcttta tatattatgc ttcctgcgat acgcgtctcc atgtggcgac ttctaccgtc 6720
gacagactcg tggattactg tatgaatact gagccggatc tgtttaccac ttctgcttca 6780
gtacgacgtc tgaacgcgct tatagacaag aatctttcgc tatctatgta ggactaataa 6840
tcaggtttca gatggatatc attactgcac cggctaagcc gacattgaag gaaaagatcg 6900
gatatggctt cggagacatg tcgtcttcga tgttctggaa gattttttcg tattatctgc 6960
cgatcttcta ttcggatgtg ttcgggctta agcctcagca tgcggctttc ctgcttctta 7020
tcacaaaact gtatgatgca atatcggacc cctgtcatgg gaatcatcgc cgacaggacc 7080
cggtccagat ggggaaagta tcgtccgtat cttctctggg tggctctgcc gttctcgctt 7140
ataggccttc tgtctttcta tactcctgat gccagctatg gcctgaagca tgtatatgcg 7200
tatgtgatgt atatcctcat gatgactgtc tatacggcta taaacgtgcc ttacggggcc 7260
atgctgggag tcgtcactcc ggacagccgt gagaaatcgg ttttctcgtc ttacaggatg 7320
ttctttgcgt atatcggcag tttcatcgcc atgggtatat tcggcgtgtt cgagaagctg 7380
atttccggta ccgtgcgtgt tatgccggac ggtagcgaga gggtggtgcg cggagtcggc 7440
gatgctgctc ccggacagtg gactcttgtc gtcggcatcg tttctctgtg ctgtttcgtg 7500
ctgtttctgc tttgcttcct tatgaccagg gagcgggtgg ggactgaaaa taaggcggtt 7560
tccgagggag gctcttctat cgcagaggat ctcaaggctc taggccgtaa caggccttgg 7620
tggctgcttc ttggcgccgg tatcggtata ttgctgttca attcgatccg aggcggagcc 7680
gctgcatatt atttctcgaa tatcctcggt acgagcgtgt tcctgacttg cgctgtctat 7740
cttttcattg gcgaggtggc tcagatggtt ggggtgctgt ttactgtgcc gctttcggag 7800
cgtatcggca agaaggcgac ttttatctct gtgcttctga cggtttctgt gctgagtgtg 7860
gcgatattct tcctgcctga gactcctgcc ggtttctggg gacttctggt gctgcaggtg 7920
cttatatgca ttgcttttgg tgtccagtct cctctgctgt ggtcgatgtt tgccgatgtc 7980
gccgattatt cggaggagaa gaacggctcg gcttctaccg gacttatttt ctcgtcttct 8040
tctatggctc agaagttcgg cggggccctc ggcgctttct tgctgatgca gattcttgca 8100
atcttcgcat atgacaagga tgctgcagtc caggcgcctc agactcttgc tgccatccgt 8160
ggcctgatga gctatatccc ggcaatcggc gccctcctcg gagccctctg cctgctgttc 8220
tatcctctta cggaatcgcg tatggcttca attagagaga aactttctgc tcgcagataa 8280
catcttcatt catgtaggta tttgttattc caaaggatgg ccctcttcgg ccatcctttt 8340
tcgtctggct gattcgagtc gggcctttct gctggcggga gaaggacggg acggctccgt 8400
caagttcgga aaagctgctt ggtcgctgtc gccgcctaaa cggctaactt ctccctttta 8460
cgatgggcga gccatcgtaa cggtcgtcaa acacacgccg ttcttcacgg cgccgacggc 8520
cgctcccgcg ctctccgctt ttcctcaatt cctccgccgc ctccgccctc ctcccgccgc 8580
aggactgctg accagggttc cggggacgtg gcccccctcg gccacgggaa gagggacggg 8640
gcccgcaaga tgcaagttat cgcgaagcga tgacgcggca gattgtggga gggtacggag 8700
tgagcgatag cgaacgaagt tccccgggac cctgcgtcag caggactgtg gcaaaggaag 8760
cagaatgcag gactgcggca aagatgaggg actggaggga attgaaaaaa atgttatatt 8820
tgtatttcgt tccttaatcc ggagaactat ggcagacaaa ctatacctga ttgacggtca 8880
caacctaatc ttcaagatgt actacgcatt cctgcgtcgt cctatgatca attccaaggg 8940
aatggacatg tcaatcctgt atggcttcac gaaatatgtg ctcgaactca tcgagagaga 9000
gaagcctacc catatggcca tagcgttcga tcctcccggc ggcaccttca ggaacgaaat 9060

gtaccccgaa tacaaggcaa acagaagcga aactccgcag ctcgtgatcg atgccctcga 9120
accgctgctg gaaatatgca aagccttgaa cataaagacc ttgatgattc cgggatatga 9180
ggccgacgac gtgatcggaa ccgttgcgaa gagttacggg aaacccggga cggacgtcta 9240
tatggtcacc ccggacaaag attacggcca gctggtcgcc gaacatgtgt accaggtgaa 9300
gcctggaaag gccggcgggg ctgacgagac cgtcgatgtc aaggcagtct gcgaaaaata 9360
cggaatctcc agtccggaac aggtcatcga aatgctgacc ctctgcggag atacggccga 9420
caatgtgccc ggagtgcatg gcatcgggga agtcggggca ggcaagctca tcgcaaaata 9480
tggcaccgtc gagaatatct atgcccatct ggacgagctg actgaccgcc agaagacgca 9540
gttcgaggag gccaggggac atatcggcct gagccatgac ctggtgacaa taaagaccga 9600
cgtgccggta gagacaagcc tcgacgagat gaggctggac ctggattatg ctcccgaaat 9660
cgccgacctt ttcgaaaaat acgaattcaa ctcgctgaga aaatatatag cccatagctc 9720
ccgcgctgcc aaggcagtgg cggaggtcaa gcgcctggat tataccgaga caggggcgga 9780
agagctcgtg aagatggctc cggaaaagat ttcgctgata gtcgaaccgg agggaccggg 9840
catattcgcc gatatccgca gggtgatcgt ggccgcggag aagaaagacg gcggaatcgt 9900
cgtcgcttca ggcgctgcga aggattttcg caagatactg gagggcgaag ccgtgaaata 9960
cggatatgac ctcaagctcc agtcgaatct tctggagaat gccggaataa agctcggagg 10020
ccggctatat gacatcgagc tgatgcatta tctcatcaat ccggagaaga gccacaaggt 10080
cgaaatcctt gccaagagct atctcggcgt cgatatcgaa gaggccgctg ctgtggaaga 10140
agctcctgtt tccctgtttg acgtcgcgga agaaagcccg gcaaacggac cggacaggaa 10200
tctcgaagcc gcggcagtgc tgaaactcgg agaagtggtc cgcgcaggca tggacgaggc 10260
cggcgtagga gccctgtatg aggatatgga agcccctctc gcagccgtgc tcgcccgcat 10320
ggaaagggca ggcgtgaaga tagacctcgg acagctcagg acgtttgccg atgaactcaa 10380
agtcgaaatg gatgccctcg aagcccaggt caggaaactg gccgacgagc cggggctgaa 10440
cgtttcttcc ccgaagcaga ttggcatcgt gctcttcgaa aagctgaaac tcgacccgaa 10500
gatgaagaaa tcggccagcg gtagctatag caccgacgaa acgacgctta ctgccatcgc 10560
cgacaggcac cctataattg acaagatact tgaattcaga gccgtaaaga agctgctctc 10620
cacatacata gagccatttc cgacctacgt ctccccggtt acgggaaggg tccatacgac 10680
cttcaaccag gcgctgacgg cgacgggaag actcagttcc tccaacccta acctgcagaa 10740
cattccgatc cgcacggaga ggggaaagga gatcaggaaa gccttcatac cggcgactcc 10800
ggacggaatc ataatgtccg ccgactactc ccagatcgag ctcaggatca tggcccatct 10860
ctgcggcgac gagcatctga taagctcatt caaggcaggg ctggatgtcc ataaggccac 10920
cgcagccaag attttcggca taccggtaga ggaggtcaca tccgaccaga ggcgcatcgc 10980
gaagaccgca aacttcggaa tcatgtacgg catctcatcc ttcgggctcg cccagaggct 11040
gcatgtctcc aggacggaag ccaagaagat aatagacgat tacttcgccg ccttcccttc 11100
gatccggact ttcatcgaca agactattgc ggacgcccgc gaaaaagggt atgtcgagac 11160
agtcttcgga cgcaggaggt atcttccgga cataacggcg aagaacgcta ccataagggc 11220
tctggccgag aggaatgccg taaatgctcc tatccagggc acatcagccg atataatcaa 11280
gcttgcgatg ataagcgtcg acaagcgcat ccgcgaggcg ggactgaaga gcaagatggt 11340
gctccagatc catgatgagc ttcttttcga tgccgtgccg gaagaaatcg aggaactcgg 11400
caggatagtt gtagcggaaa tggagaacgt aattaccttg tcaatacctt taactgtaga 11460
atgtaactat ggcaaaaact ggcttgaagc tcactgatca tgacgacgga tctcttgtgg 11520
agatggcgtt aggtggcggc cccgaagccg aggccgcttt cttcacgatt tatgcacgtt 11580
accataaggg cgtattcgcc cacatctcac gcttcatcca ggaacaggag gaagttgagg 11640
atatctgtat ggagagcttc gagaaggcct tcaagcagct ggaatcatac aggccgagca 11700
acaagttctc gacttggata ttcactatag cccgcaacac ggccttcgac cacctcgaca 11760
aggacaaagt gcgcggccgc aacaaagaca agatgccgat cgactactcc gacctggagc 11820
atatagacgt ccctgaggac aatgtcagcc cggaggacga gatcataagt tcgcaggacc 11880
atgagaactt cctgagttgt gtccaggggc ttccggacct gtatcgcgaa gtcgcatcca 11940
tgtgctttgt ggacaatctc ggatacaagg agatttcgga gaaggcaggg ctgcctatca 12000
atacagtaaa aacccgcatc agcagggcta agagcctgat agtccagatg atgcttgata 12060
tggaagaaga atgaatttca aagaattcga agatatagtt gttgctgagt ttccggagct 12120
gactccggag atgcttggaa agtacgaggc catggagccc ctgtatgagg agtggaacgc 12180
caagatcaac gtaatttccc gaaaagacat aggcggcctt tatctgcacc atgtgctgca 12240
ttctcttgcg atcgcctcgt atctgaagag acgcaggccg gaggtgtatt ccgcatggac 12300

cgaaggcgcg tccgtgctcg acctgggcac cggcggcggc tttccgggga ttcctctcgc 12360
ggtaatgttc ccgaaagttc gctttaccct ctgcgattcc gtgggcaaga agataatcgt 12420
ggcatcggaa gtgtccaagg ctctttccct gggcaacgtg accactgtca atgcccgtgc 12480
agaatcgctg ccgcaggttt tcgatcatgt cgtttcccgc gccgtgacct cgctggataa 12540
tttctatcca tgggtgaagg gcaagttctc cggaagcatt ctgtatctca agggagggga 12600
catagtcccg gaaattgcag aactgatgaa gaagaacaga ctccgtaccg gagcagtttc 12660
gacctggccg attgaaactt ggctgaagga cgaatatttt aaggaaaagt ttgtgattga 12720
aatagaaaaa taatttgcaa atccacataa aataactacc tttgcagtcc tatttccgga 12780
aaactggttt tttcggcctg ataagaataa ataaaaaaac agataacgat gaaaagaaca 12840
ttccaaccat caaagcgcaa gcgtataaac aagcacggtt tccgcgagcg tatgtcaacc 12900
cctaacggcc gtcgtgtact tgctgcacgc cgtcgtcatg gccgcaaaaa actcagcgtt 12960
tcttcagagg atagattcta gtcatctctc tccggacgtt gtttgtttgg atcgaaattc 13020
aggaggcttc cgaaaggatg cctccttttt tcattatggc agattatttg atagacgagg 13080
gacgtaatca ttaagtctca atggaaaagt tttttttatc aatcttgtct tgcgcgcttt 13140
cagcgacaat agctgtttct gcgcagagta gcggaccatt agttgtccac cagtatgaat 13200
tgtcaaacgg aatgcaagtc tggcttaacg aagaccattc gcagccggta gtccacggag 13260
ccgtggtcgt caaggccgga ggccgggatt gcgccgggac cggaatggcc cattatctgg 13320
agcatatgct cttcaaaggt acggaggaca tcggtaccat cgactatgct tcagagaagg 13380
tctatcttga ttctatcgag acttgttacg acctcatcgc agcaacatcc tccagcgtaa 13440
aaagggattc tctccagaag gagatcaacc gtcttagcat caaggcgctg gactatgcga 13500
ttccgaatga attcgaccgt cttacggcca atatcggagg cacaaagctc aatgctttca 13560
cgtcttatga ctgtacatgc taccataatt ctttttctcc gcaatttatc gcgcagtggg 13620
ccgaactcaa tagccatagg ctgctgaacc ctgttttcag gcttttccag agcgaacttg 13680
aagcagtata tgaggagaag aacgcggcag ccgattccca gtttgacatg attctcgaga 13740
agatgctgga gaaacttggg gatgaaaatc catacagcaa atctataatc ggaactacag 13800
ataacctgaa gaatccaagt ctttccgaga tgaagaagtt cttcaacaag tattatgtgg 13860
gctgcaacat gggacttatc cttgccggag acatagatgc ttcgtcgata ttgcctctgc 13920
tggaatctac cttcggacgg attccgcgtg gcgataagcc ggaaaggatt cctgtacctg 13980
tgccggattt ctcgtcccgg gaagattatc ctataaaagt taatttccct atgatcaagg 14040
cgtcgctgaa cgtgttcaac ggtcctgcag ccaaggatgc cgattatcct gcaatggaac 14100
tcgctctcag cctcctgaac aacagtttta gtaccggaag tctggattcc ctcataaata 14160
acggcaaggt tctggctgga gaggtctcag tcttggactt ttttgacgaa atggggctta 14220
ttgccatgat aacgatacct ggaatccctt tcggatctct caacaaggcg gaggcgcggt 14280
gcttcaagga agtggaaaga ctgaaggccg gcgattttgc ggagaaggat ttcgagtcgt 14340
tgaaatatga tcttgtcaaa tcatataaaa ggtgcctgga atcgtccgag tcaaggttct 14400
ctatcatgat agacgtgatg tcccaggaca ggtcgtggga ttcttatctt tcccaggtgg 14460
agtcgatatc ttcgcttact tcggaggata tagtcagggt aagcaataaa tatctgggcg 14520
acaatcgtat acgtttccat aaagtcagag gaacctatcc gaaggataat ctgcaagctc 14580
cttcatacga gactctttcg ccgaagaatg gcaatgcggt atcggattat gccaccagac 14640
tgatggagat gccttatgac actcttcctc caagactggt ggatatgaat aatgatatcc 14700
ggaaagtcgc agttaatgac catgtgacgc tttactataa cgggaacccg gtcaatgatg 14760
tctttgaact gaccgtccgg atcaacaagg gtattttcga ggaacctatg atctgtcctt 14820
tgacacagct tcttggcctc tcggggaccg acagcctgtc cgcccgtgat ttttcaatga 14880
gtcttgccga gctgggttcg gacttgtctt ttaaagataa gaaccatacc ttggagatgg 14940
ttctgactgg ctttgatgac cgtataggac cttccctcga cctgctccgc cactttatga 15000
aggatttcaa gccggaaaaa agtgcgctca atacggtgat ttccgatacg aagatgtcca 15060
ggaatgcatt caggagcagt ggcaattcag ccatcttcaa tggtatgttt aattattctc 15120
tattagggaa ccagtcacct cttctgaggc agccgacaga aaaagatctc aaatctctag 15180
gtctgaaagg gatgaaagag cttttcaaca gcctggttga atatgatttc gatgtagtct 15240
attccggagg cctttctcca gaggaagtcg tcaatgagat ttccggccat cttgatgtgg 15300
aatgtccttc cggaaaacct gagagttcat ttttcgctta cagaaaatat gaacagcctg 15360
tggtctatat ttacaatgtc ccgggagcgc gccaggtgat gtcgataact taccagaatg 15420
tcgctactcc ggagaatgac catgacagcg cactgctcgg ccttatgcag gaatgctatg 15480
gcggttccat ggcttcgatc atgttccagg aagtccgtga attccgttct cttgcatata 15540

cttcttccag tacgcttcag aagccacttc cgaatgtatc ctctccttct gcattcataa 15600
cctactatgg tactcaggct gataaattcc ctcaggctct tgaactggtg gattccctca 15660
tccgtacgat gccgatgact gatctgtgct tcgattcagc agtccgctct atcatttccc 15720
gcgccaataa cggctatcct tcgttcaggg ctttgcctga atatgtcgcg aatttggaat 15780
tgttcggata tggagaggat gatgacgctt ccgtagttag aggccttccg ggaataacca 15840
agcaagacgt cctggattac catgaaaaga gtatcggcaa tgctccttgc tcttatttca 15900
taatcggaga caagaagaat ctgcctatgg acgcgcttcg ggaacgcggc cgcatagtcg 15960
agctgaaact ctctgacatc taccgttagt tcgctgaatc ccttatcggg acgaataggt 16020
ggagaagcag gcggagatta cttcggaagc cgaaagtccc gagtcggaga ggcggctgga 16080
ggaggcaagg aaacggtggt cggcggtgat gccgaagacg cggtccgcgc agctgagtgc 16140
gggtgccagg tcgtgagaag ccatgatcac ggtggtgccc gtacttcttg cgatgtccct 16200
gagagcctgt agtatcagcg acttgttgtc gacgtccagg aatgaagtcg gctcgtccag 16260
cagcagcgtc ttcgcatggc ggctgagcgc cgacgcgatg cagcctttct ggaattctcc 16320
gtcgctcagg gtgctgatgt ccctgtctgc gagagatgaa attccgagac actccagcga 16380
cttcgagatg tccgaagcca tggccggaga aatgcggccg cgccagtcgc tttccgagta 16440
tagggtggtg gcgatgaact ccctgagagt gaatcccttg actttcggga cccctgtcgg 16500
taccatgatt gcgtccttgc cggcgagagc tctcagcagg gtgctctttc cgctgccgtt 16560
gggaccgcag agaagaatga actctccggg ctcagtctcg aaactgatgc ccgaaaccag 16620
ggtcttgccc ctgtgtccga tattcaagtc ctttattctc atttttttcat cagtatcagg 16680
agtattatcg gtatgcctat cagggccatt gtgctgccga ccggaagcgg gagtccgggc 16740
gcctgcgaaa tgatgtccgc cagcaccgag acggccgagc ctgtaatgag agtggccggc 16800
agtatggaca tatgtccgga actccggagc agcctgcggg cgatatgagg ggagacgatt 16860
ccgacgaagc ccagaggccc gcagaatgca gtcgccgctg ccgtcatcag acagctgctg 16920
accatggcga cgccgcgtat ccggcgcacc gaggcgccag cgagagtggc gtattcctcc 16980
ccgaaaagaa gtatgtccag acccttggct ccgcttgtgt aaagccccgt tcctaccagc 17040
aacgcggcaa atatcgtcaa tgtctcgaaa gttcttgtgc cggagaaact tccggcggac 17100
cagctataga agagcttgag gctttcttcg ctggctgaat actcaagtat ggaggtgatg 17160
gcgctgaata tgaagccgag catcacgccg aaaaccagca gggtggtacc gctgcggaaa 17220
cggcttgaga ccagcactat cagggctgat gccaggatcg caccgacgaa tgcggctccg 17280
gccacggtga gcccggccat ggacgagaag cccgaaagca acccggtgcc tgagatcagc 17340
agggtagcta tcgcggctcc gaggcctgcg cctccgctta cgcccatgat atgagggtcg 17400
gcgagagggt tgcggaaaac tgactgcatc tgggctccgg agagggccag gcatgcgcct 17460
gctgcaatgg ccgtcagtac cctcgggaga cggatgctcc agactatcga ggaagctatc 17520
ccttctcctc cgaacaggcc tccaagcgcc tcggcagccc cgattccgga acctccgagt 17580
atcaggtcgc atacggagag cagcagaagc gccgttgctc ctgccgccgt gataagtctg 17640
tttctgcgtt ttatggccat attaactctt acaaagatag aaacttcaag ggattttcgt 17700
atatttggaa gattaaaata cattgcggat gagcaccgaa acaattctga taattcttct 17760
tgtcgcagtc attgtcatag ctgcgatact gattttccgt ctccaggcgc gtgccgaaaa 17820
gatcaaagac ggattccagg acagactggc cgatgtccag cgcgaaaaag ccgtactgga 17880
gagtcgcctg caggctaagg aggaatatga gaatcagatg cgcgaagccg cgaaggagac 17940
ctctgaaagg catgccaaag agctcgagca gatgaaggaa gtcttcaaga acctttcgac 18000
cgagacaagc gaggccttca aggcaaagag cgcccagaca atctctgaac tgatgaaacc 18060
tgtccaggag aaattcaaag agttcagcga cgccgtcaag gaaagccaga aagacaccat 18120
agagaggcac tcccgcctcg aacagagcat taaggatctc gacacccgca gcaaatccgt 18180
cggggacgag gcccgcaatc tcgccaacgc gcttaccggc tattccaagg tccagggaga 18240
tttcggcgaa atgctcctca ccgacgtcct gaagaacgcc ggcctgaccg aaggagtgca 18300
cttcgagact cagtccgtga tgaccgacaa ctccggccat gaaatcaaat ccgagagcgg 18360
caggaccatg attccggacg tcaaagttta ttacccggac gacactatag tcatcatcga 18420
ctctaaggtc agcctgaccg cctataacga gtacatggcg gcgacgacgg tcgaggacag 18480
gaccaggctg gccaaggctc atgtccgcag catcgaggat catgtcaacg aactgaaggt 18540
aaaggattac gcgggctaca tccctgaagg ccgcaggaaa gttgattata acatcatgtt 18600
catccctgtc gagggagctt tccgccttat gcttgaagaa gccccgaggc tctggcaggt 18660
ggcgaaggac aacaacatcc ttatcgtcag ccagatgact cttataattg tgctgaacat 18720
gatccagatg gcctggaagc agcatgagca ggagaagaac atctccgacg tatacaagac 18780

cgcggaggaa ctgatgtccc agctcaaggg atggatggaa tcttatgtca agatagggga 18840
ctatctgggc aaggcttctt cggcctatga agactcaaag aagaaactca gcgactctaa 18900
ccagtcggtg atcaagaaga taggaaagct ggagaagctt ggcatcgctc cgaagcgctc 18960
caacgccaag atcaagaccg gcagcaggtt gagcggaccg gaatcggtga ttcctaagga 19020
attaagcgat gatgagatta cggattaata gggaaataag ccgtattgct tatattctgc 19080
tttaaaatcc gtgcttgaga tttcttttta ttaataaaat tggctaattt agcagtgttt 19140
aataacacta ttgctagaac atggaactaa aaaatgcatc agcaggtacg ttcgaatctg 19200
gagatatcct tatcaggatt gatccggccg agaagaaggg gctcgagata gagcttgaga 19260
gttcggtagc ggaccagttc ggcgtacaga taaagaaagt tatttctgag acgctcgaag 19320
gactcggaat tagtgacgcc aaagtcaagg caatcgacaa gggcgccctc gactgtacta 19380
taagggcgag agtcacggca gccgccgtgc gcgcctcagg caaggacgtt tggaaataag 19440
gaatcatgga aagactaaga agaacaatga tgttcattcc gggcaataat cccggaatga 19500
tgcaggacgc ctatatctat ggccctgatt ccataatgct cgatcttgag gattctgtga 19560
cgatggcgga aaaggatgcc gcccgtcttc ttgtctacaa tgccctcaag accgtagact 19620
acggaaatat cgaaagagtc gtcaggatca atcctctgag cactccgtat ggaaagaaag 19680
acgtcgaagc cgttgtcaaa gccggagtcg acgttattcg tatgccaaag accgaaacgg 19740
ctgacgaagt ccgcgaactc gaagccgaga tactcaaggt cgaggaagaa ataggctgcg 19800
tcggcaggac ccagataatg gccgcaatcg aaagcgccct cggtatagtc aacgcttacc 19860
agatagcctc cgcatcgtcc aggatgatgg gtatcgccct cggtgccgag gactacaccg 19920
ccaacctgaa gactatgcgc tccaacgaag gcatcgagct ccagctcgcc cgagagacga 19980
tagtagtcgc cgcccgcgcc gccggaatct cttgcttcga tacagtattc tccaatctcg 20040
aggacatgga aatcttccgt aaggaggtag agttcatcaa gactctcggc ttcgacggaa 20100
aatccatcat caaccctcgc cagatcgacg tggtcaacga agtcttcact ccgaccgaaa 20160
aggagatcgc caaggcccag accataatcg ccgcaatcaa ggaagcccgt cagaggggta 20220
gcggagtgat ttcggtcaat ggcaagatgg tcgaccgtcc ggtagtgata agagctgaaa 20280
gaataatcgc ccttgccgtt gcggccgggg taataccaga ggaggatgcg ctatgacaaa 20340
tgaacagctt ataagagagg ccgctgcaaa gcttggcaag aaggtttacc agggccccgg 20400
ccagaagata cctacattcc gcagggacct gcagaaggaa agctgcaagg tcgtctccct 20460
tgaggatgcg atccgcaaga gcggtctcaa ggacggaatg acgatttctt tccatcatca 20520
tttccgcggc ggcgacaaga tcgtcaacat ggtagtcgac aagatcgcgg aaatgggcat 20580
caaaggcctg acattggctg catccagtct ctctgacgta catgctccgc ttgtcggcca 20640
tatcaagaac ggcgtgatct cgaagatcac gacctccggc gtgcgcggag aactcgcgga 20700
acagatctcc cgcggcctca tggatgaacc ggtgatattc cggtcccatg gtaccagagg 20760
cgctgccgtg gccagcggag agcttcacat agacgtcgcc ttcctcggcg caccttcatg 20820
cgaccctctc ggaaacgcct gcggatacag ccgttccgaa aacgccaagt caatctgcgg 20880
ctccctggga tatgccctgc cggatgcgcg ctatgctgac aaggtcgtga tccttaccga 20940
tgacctggta ccttatccga atactccgaa ctctatctcg gagcatgacg tggattatgt 21000
ggtagtggtg gattcggtag gcgattcctc aaagatatct tccggagcca tccgtgactc 21060
caagaaccct cgtgacatac ttctcgcaag gcttgccgca aaggcaatca taaacagcgg 21120
ctacttcaag gacggattct ccatccagac cggaaccgga ggcgcagccc ttgctgccgt 21180
caagttcatc cgcgagagca tgatagagaa aggcatcaag gcaagctatg ccctcggcgg 21240
catcaccgcc cacatggtca ggctccatga agagggactc atcgaaaggc tcatcgacgt 21300
gcagtctttc gacaaagtgg ctgccgagtc cgtaaagaat gatccggccc acaaggaagt 21360
ggcctcgtat gaatatgcaa gcggcatgga acccggttcg gccgtgcatt atctggatat 21420
tgtaatcctg tcggcgctcg aggtggatgt caatttcaac gccaacgttc tggtaggcag 21480
cgacggcata atcagaggcg ccatcggagg ccatccggat acggcggctg actcggccct 21540
gtcgattgtg gtctgccctc tgctcagggg caggattcca tgtatcgtgg accatgtcac 21600
caccctgatc actccgggag cgtccgtgga cgtggtgatc accgagtatg gcaccgtggt 21660
caatcctcgc aggccggaga ttgccgaaag gctgaaggcg gccggaatga agctggtgga 21720
tatgcagtac cttcgtgact gcgcccgcaa gataataggg gagcctgctc cgattccgtt 21780
cggagacaag gtcgtgggcg tcgtcctcgg aagagacgga tccgtacagg atgtgataag 21840
gaatgtgctg aaataatgga agttacgctt gatatgatat tagcctcccg cgaggccagg 21900
cagaaagagc aggaatatct gctcaagcat agctatggct ctacgctcat aagcatgacc 21960
gtggtgatgc cagggaatgt caagcgtgat gaatatacga taaagatagc ggaggccggg 22020

attgatgctc tcctgaacgc ctttgacggc ggaatcaaca ggatagtgca gcgcgacctc 22080
gaaacaggat acgaagcctt cttgcttgtg aaagctcctt tggaagaaac caagagggtg 22140
gtcagcctta tcgaggagac tcatcctctc ggaaggcttt tcgatattga cgtaataggc 22200
gaggacggtg tgccggtgac caggacaggt tacggcatga ctccccgcaa atgcctgctc 22260
tgcgacaatg aggcccggta ttgtatgagg aatcataccc atactccgga gcagctcaag 22320
gctcgtatca gggaaatggt agacgaatat gtacggtgag tatcagataa ttgatctgcc 22380
gctttcggca ggagtcgtcc gcagcaaggt tgagaccttt cttgcggaga acggcctgcg 22440
tctggaagct gtggaccgct atgtcggcat tgaagactcc gatgaaaggc ttattgccgg 22500
cggcggaatc tgcaaggacg tgctcaagtg catcgccgtc tccgaggaag cccgctcagg 22560
cggacttacc ggaaaactgg tcgccagact gatgcagctg ggcgaagagg cgggtcagca 22620
tgatctcaag gtcttcacta agcccgagta tgaggatatc ttcgcgagtc tgggattcag 22680
cgtcatcgcg aaagccggga aagccatact catggagaat ggcgacggac tgaagagata 22740
ctgctccgat ctggagaaac atcgccgtcc gggccgctgc ggcgccgttg tcatgaacgc 22800
ggctccgttg acatggggcc atgattatct cgtcgagaaa gcatgcgagt cggtggatgc 22860
ccttttcgtg attccggttg ccgaagatac tccgggattc agctacaggg accgttatca 22920
ggccctcgca agcgcctggc tggaggaact taaggtcgaa gtatgcagcg gcagtcctta 22980
tgccgtgtcg aaggccacgt ttccgaccta tttcctgaag gatctcagcg atgctgccga 23040
gactcagatg actctcgatc tcgatatttt cgcccgtcat atagctccgg cgctgggtgt 23100
gacggtccgt ttcgtcggca gcgagccgtc ggatgccctg accgcccgtt acaacgagct 23160
catgaaaaag atacttcctg agcatggaat cgaagtcgtc gaagtgccgc ggctggaatc 23220
ggagggacgg cctgtatcgg cctccgaagt ccgtaaagcc gtcgccgaag gctcgttccg 23280
ccgcgccgat gccatcgctc atcctgagtc gatgccgatg cttgtcggcg ctcttgcctc 23340
cagggctctg cgtatggagc tcgaccttac tcccaagccg ggactggttg acaagaacgg 23400
ctccggggcc cataaggaca tggattatgt caccatgagc gcgagcatag acgctatcgt 23460
gccgtttatc catgcgatgg ccagcatcaa tcctgacgac gacgttcctt ccgacatggt 23520
ggaactggga atcaaggccg agcagaagat gcttgcggcg accggaggag tgaacaccca 23580
caagggagct atattcagtc tcggattaat ggcgacagcc tttgcccagt gctactggga 23640
aggggagact gtcggggaaa aggctctcaa gtcgaagatc gtggctctgg catgtgcagt 23700
tccgaaatcg aaggataccc acggggccaa ggtctgctct aagtttaagg ccaagggagc 23760
gctgcagaat gcccgtgacg gctatcctgc cctttttaag gaatggctgc ctttctaccg 23820
gaaggtgaag gcagagaaga atggcttgct gaagactttg ctgctgattg tttctcagct 23880
tgaggatacg aatgtgcttt accgtaaggg acctgccgtt gcggagaagg tccgctcgat 23940
ggcgcgcgaa gctctgaaga acttcagtga ggataaggta agaaatattt gcggacggtt 24000
tacgtctgag ggcatctctc cgggcggaag cgcggacatg ctggctctga ccgtatttgc 24060
tgattcgcta ttgtcatgat acaacaatta ttaatataaa acggatttga ttatggtaga 24120
tctgattaac catggagttt atctacttga cggtgttgta agcgacgccg ctccgggaca 24180
tatgtctccg gacgaggccc gcgagaacac cattacctac ggaattctgc gctctcatga 24240
cgtggacgga ggcaaaggca agaaaatgag gattcgtttc gacgctatgg tgtcccacga 24300
catcacttac gtcggaatca tccagacggc ccgcgccagc ggactagaga agttcccgct 24360
tccttacgcc atgaccaact gccacaattc cctctgcgca gtgggcggaa cgatcaacga 24420
ggacgaccat gtcttcggtc tctccgcggc gaagaagtac ggcggcatat acgtgcctgc 24480
caaccaggct gtcatccatc agtacgcccg cgaggcgctc accaagtgcg gtaacatgat 24540
tctgggctct gacagccata cccgctacgg ctctctcggc aacatgggag tcggcgaggg 24600
cggcggcgag ctcgtgaagc agcttctccg caatacctgg gacatcaacg ctcccgaggt 24660
tgtgcttgta tggcttgaag gaactcctcg ccacggcatc ggtccccacg atgtcgcaat 24720
ctccctggtc aaggctgtgt tcgagaacgg tttcgtgaag aacaaggtgc tcgagttcgc 24780
aggacctggc gtggcttccc ttccgatcga tttccgcaac ggaatcgacg tgatgactac 24840
cgagacgacc tgtctcagct ccatctggat aactgacgag aaggtcaggg aatactacga 24900
gatgcacgga cgcgcggccg actacaagaa gcttgagccg ggcaaggtgg catattacga 24960
cagcatgatc cgcatcgacc tgagcactca ggagagcatg atcgccctgc cgttccaccc 25020
ttcgaatgca tatacgatac atgaactcca ggctgatccg gagaggattc tccgcgctgt 25080
cgaggaagac gcgcgcaagc gtttcggaga caagataagc atagatcttc tcagcaagat 25140
ccatgacgga aaggtcgtgg ctgaccaggg tatcatagcc ggctgctccg gaggtacata 25200
tgacaacctc agcgaggctg ccaccatact caagggcaag tctatcggca atgactattt 25260

caccatgtcc gcctatcctc agtccactcc ggtatatctc gcaactacca ggaaccatat 25320
cgcagaggaa ctgctcgagg ccggcgtggt catcaagcct gccttctgcg ggccttgctt 25380
cggagcagga gacgtgcctt ctaacaacgg cttctctatc cgccacacca ccaggaattt 25440
ccctaaccgc gagggctcca agcctggaca gggccagatt tctctggttt cgctcatgga 25500
tgcgcgctcg attgccgcaa ctgcagccaa cggcggagtc atcactgccg caaccgatat 25560
cgaatacaat gatactcaca agccatacgc cttcgacgac cgtatctaca agagccgtat 25620
ctacaatgga ttcggccatg ccgatctctc gcaggaactc cgctatggac cgaatatcaa 25680
ggactggccg aagatctatc ctatggagaa gaacatgctc gtagagcttg ccgccgtgat 25740
ccatgatcct gtgactacta ccgacgagct cattccttcg ggcgagactt caagctaccg 25800
ttcgaatcct ctgaagctca gcgagttcgc cctgtcacgc cgcgtgcctg agtatgtgtg 25860
ccttagcaag aggatccaga gtgaggatct ccgccgtcgt gcaggtgacg ttcctgccaa 25920
tgttgcggct gccctcgcga aagtgggcgc ttctgcctcc gacacctcct tcggttcatg 25980
cgtgttcgcc aacaggccgg gtgacggttc tgcccgtgag caggctgcct catgccagaa 26040
ggtgctcggc ggcgatgcca acatctgcta tgagtatgct accaagagat accgcagcaa 26100
ctgtatcaac tggggtctgg tgcctttcac catagctcct gagactgctt ttgactataa 26160
gccgggcgac ttcgtgtttg tgccgggtat cagggatgcc atactttcag gcaaggagga 26220
gattgatgcc aaggtgatca ccgcttccgg tgtctccgac atccacctct acttcaacaa 26280
cctcactcct gtcgagcggg aaattcttgc cgacggttgc ctcatgaact attatatgac 26340
tcataaatag ttatatttgt atgttatgtc ggcagagact atggtctgaa agccgtggcc 26400
gcccgcggcc atcgtgagcg ggaggggccc ggccgtcagg ctgggaggga tagcgccgaa 26460
ggcgcgtagc tttcaggcca tagtctctgc cgacaagaca atcgacaaga cgataaaaga 26520
aaataaataa ttgatataaa gatatgagaa aaatacagat gactacgccg cttgtcgaga 26580
tggacggcga cgagatgaca aggatcctct ggaaaatgat caaggatgag ctgatcctcc 26640
ctttcgtcga cctcaagtcc gaatattacg acctcggcct ggtcaacagg gacaagacag 26700
cagaccaggt gactgtggat tctgcccttg ccacaaagaa atatggcgtg gccgtgaaat 26760
gcgccaccat cactcctaac gtacagcgca tgagcgagta caaccttcat gagatgtgga 26820
agagccctaa cggcaccatc cgcgccatgc ttgacggtac cgttttccgc tgcccgatca 26880
ccatcccttc catccatcct gcagtgaagt tctggaagaa gcctatcacc atcgcccgtc 26940
atgcatacgg cgacgtctac aagggcgtcg agatcaagac tgacgagcct ggtacagcca 27000
agctggtgtt cgagggcgag agcggcaaga aggaggaggt cgtgatccag aagttcgacg 27060
gcccgggcgt gatccagggc atgcataaca ctgacaagtc catcaggtca ttcgcacatt 27120
cctgcttcaa atacgcgctg gacgtgaaac agaccatctg gtttgcaacc aaggatacca 27180
tttccaagaa atatgacggc cgcttcaagg ctatcttcga agaagtgttc caggagtata 27240
aggacaggtt cgaggccgct ggaatagagt atttctatac cctcatcgat gatgccgtag 27300
cccgcgtgat gcgcagcgag ggcggattca tatgggcctg caagaactat gacggcgacg 27360
tcatgtccga catggtttcg accgcattcg gctctctcgc catgatgact tcggtgcttg 27420
tcagcccgga cggcaaatat gaatatgagg ccgcccacgg aacggtgaca cgtcactatt 27480
acaagtatct caagggtgag gagacatcta cgaaccctat ggctacaata ttcgcctggt 27540
ccggcgcctt ccgcaagaga gcagagctgg acaacctccc ggaactggtc aactatgcag 27600
accagctcga ggcctcatgc ttcgataccc tcaattccgg catcgctaca aaagaccttg 27660
ttaaccttat ggaaggtgta gaagccaagg ctgtcaactc ccttgacttc ctgaaggcta 27720
tccgcgccaa cctcgaaaaa cgccttcaga actaatggca gacgaaaaga taatattttc 27780
catgaacggg gtgggaaaag tccttcccca caacaataaa gtaatcctca aggatattta 27840
cctgagcttt ttctatggag ctaagatcgg catcatcggt ctcaacggtt ccggtaagtc 27900
aaccctcatg aagatcatcg ccggagtgga gaaatcctac aagggcgagg tagtgtgggc 27960
tcccggctat tcggtcggtt acctggagca ggagcctcag atggatccgg acaagactgt 28020
gatcgaagtc gtgcaggagg gcgtccagga agtgatggac cttctgaacg agtataacga 28080
gatctcggcc aagttctgcg agcctctcga ggatgacgag atgaacaagc tgatcgagcg 28140
tcagggcgaa cttaccgaca agattgagca ttgcgggggc tgggagatcg attccaagct 28200
cgaaagggcc atggatgcgc tccagtgccc tccgtcagag gccaagatcg caactctctc 28260
cggaggagag aaacgccgcg tggctctctg ccgccttttg ttgaggcagc ctgacgtgtt 28320
gcttctggac gagcctacca accacctcga taccgagagt atccagtggc tcgaggcaca 28380
tctccagcag tacaaaggca cggtcatcgc cgtgacccac gaccgttact tcctcgacaa 28440
cgtggccggc tggatccttg agcttgaccg cggcgagggt attccatgga agggaaatta 28500

cagctcatgg ctcgaccaga agaccaagcg cctggccatg gaggagaagc aggagagcaa 28560
gcgccgcaag actctcgaga gggagctgga gtgggtgcgc atggctccta aggcccgtca 28620
ggccaagcag aaggcccgtc tcggagcata tgagaaactg gcttctgccg ataccaagga 28680
gaaagagtcc aatcttgaaa tttacatccc gaacggaccg catctcggca acaaggtcat 28740
cgagttccat aatgtcagca agtcatatgg agacaagctg ctgttcgaga atctcgattt 28800
cgtgctgcct ccggccggca ttgtcggtgt cataggtcct aacggagccg gcaagaccac 28860
gttgttccgt cttatcatgg gactggagaa accggatacc ggcacgatcg agatagggga 28920
gaccgtcaag atatcctatg tggaccagca gcatcgcagc atcgatccag accttacggt 28980
atatgagacc attgccggca actcggaatg gataagactc ggcaacaagg acatcaatgc 29040
ccgctcatgg gtgtccaggt tcaatttcag cggcgctgac caggagaaac tctgcggagt 29100
gctttccggt ggcgagagaa accgtctgca ccttgccctg acgctcaagg acgagggcaa 29160
cgtgctgctt ctcgacgagc ctaccaacga tgtggacgtg aatacgatcc gggccctcga 29220
ggaaggtctc gacgggtttg ccggctgcgc cgtggtcgtc agccatgacc gctggttcct 29280
cgaccgtatc gcaacccata tcctggcctt cgagggtgac tcaaaggttg tcttcttcga 29340
cggtaactac tcggagtatg aggagaacaa gaaaatgcgg ctcggtgatg tcgaacctaa 29400
gcgtttcaag tatcgtaaac tgatggaata gcctggaaat cagaggatag tccggaggta 29460
atctatggat tccggggtgg tggcccagct ggtggcgaag cgtacggcgc agcggccgtc 29520
agggagcgcc tcccagaatt caaagtcgac aagggcggac agcatttcgc gctgctcgga 29580
agtgagaata atgaattgct ggttggtagg ggagtcgatg aagaactcgt agcccttttc 29640
gatgaagatt gccttcatct tttctgccat ttcaatcgcg tgacggccga tcctgaaata 29700
aaggtcgtcc gtaaacaatg tatcgaactg caatcccgcg atacgaccct tagccagaag 29760
agctccatgc tgcttgatca tactgaacat gtggcgtgga gcatttccgc gcgggaatac 29820
tacggcctcg ccgaagagcg cgccgacttt cgtgcctccg atatagaaaa catcgcagag 29880
acgcgctata tccttcaagg agacatcggt tcccgtcgcc atgagcccgt aacccaggcg 29940
ggcaccgtcc agatacagct gaagcccata tttatgggcg acagccgaaa tctccgaaag 30000
ttcagcaaga gaatagatgg ttccgtactc agtcggatgc gagatataga ccattcccgg 30060
gatgaccata tggtcccagg ccgggtcggc aaagaagccg cgaaggtatg agtccagatc 30120
cgtcgcatcc atctttccga gatgggcagg aacggtcagc accttgtgtc cggtatattc 30180
gaccgcaccg gcttcatgga cgttgatatg gccgctggta gcggagatga ctccttcgtt 30240
gccgttcagc agggacccga tgactgtcga gttggtctgg gtgccgccgg cgagaaagta 30300
gacctgcgcc tcgggacaac cgcaggcttc gcgtatcttg gaaatagctg aagctgaata 30360
tttatcctga ccatagcccg gagtctgctc caggttggtc tcggcaagtc ttttcatgat 30420
ctcgggatgg gctccctcga gatagtcgca aacaaaggat atcataaagt caaaacattg 30480
cacaaatata agctttttct gtcatatgaa atgatattgg aatatgaagg atattttgta 30540
aatttgtcac ttctaaaaac caatattgta acagatgagc aatagtatcg tacttcacga 30600
caaggttttc aagccttatc tgtcttatga ccggatcatt tcggctatcg atggcgtcgc 30660
cgccaggatc aatgcagatt ttgaaggatg taccgatatc ccggttatcc tgtgtgtctt 30720
gaacggctcc ataatgttca ctggtgaact tctgaagaga ctcaagttca atgtggagct 30780
ggtttccatg aagctgactt cttatgacgg ggtccgttcg accggcaagg tgcgccaggt 30840
catgggcctg agcggaagtg tgaagggcaa gagagtgatt gtcgtcgagg atatcgtcga 30900
tactggcaat acgatcattg accttatgga cattgtaagg ggcgagggcg cttcggatgt 30960
gaagatctgc accatgcttc tcaagcctga agtctataag aaagacgtga agctcgatta 31020
tgtcggaatg gagattccta atgcttttat cgtaggcttc ggtctggatt acaatgagat 31080
cggaagaaac tacagggata tctatgtgct cgatcccgat gcaacggaac aacttaacga 31140
aaagatgaaa tattatatcc tctttggccc tccgggagcc ggtaaaggca cccaggccgc 31200
agctatggtt gagaaataca acctttgtca tatctctaca ggtgagcttc tccgcaagga 31260
gatctctgca ggaagcgaac tcggtctcaa ggccaaggct cttatcgatg ccggcgcact 31320
cgttcctgac gaggttgtcg aaggaatgat cgaaaaccag ttcaagactg tcaagggcgt 31380
caagggcttc cttctcgatg gtttccctcg tactatcgcg caggcagaag ctctcgatgc 31440
aatgctcgca aagaactctg aggaggtgac ttcggttgtc tctatcatga ttcctgacga 31500
gatgatcact gcccgcatca agcatcgcgc ttctatcgag ggccgcgctg acgatgccaa 31560
cgacgctacg atcgccaacc gtatcaagac ttatcatgac aagaccgaac ctcttatcgg 31620
ttactaccgc aaggccggca aatatgccga gatcgacggt accggttcta tagacgaagt 31680
ccgcggcaag attttcgcca caatggacaa gttctaatcc ttagcgaatg gatttccggc 31740

cccggccggt taaaaataag gctgaccctc ccgggccagc tttttttga tacaaatgta 31800
tccgttccga caatccatcg aaaaaatcat aaaaactatt tataaaatca taaaaacctt 31860
acttgtgcaa attgactgac cttatccttt ttcttttctt cgcggaaaac aaaggctatg 31920
ataaggttag ttttattgct ggctctgatg ttttcgccgc aggcgccgcc ggagacgttg 31980
cttggggtta cggatatgga ggagctgaac gaggaggcgg cggagaggct ggactttctg 32040
gaaaggcatc cgcttaaggt gaattacgcg acggaggcaa ggatggccgc gtccgggctt 32100
atgtccaggt atcaggcggc gtcgctggcg gattaccgca gcctgcatgg ggatgtgctt 32160
tcggtagcgg aactggcggc tgtcgacggg ttcgggcctg cagttgccga ggccatgaga 32220
ccttatcttt ctttcgagtc ttcgcgtctt cccggacagg cggacgaacc tgcgggactg 32280
agtcatgaga tcgaactccg ggccgggacg aagttccagc ggactacttt cgcggggcgg 32340
tatcgtctgg agtatgggag caggatcgag gctgcggtat cgtataggga tcgcctgaac 32400
gggtatgcat cagttagcgg aaggcgttgg aaggccgtag cgggctgttt caatgcccgg 32460
ttcgggcagg gactggcctt gtggagcggg atgtcgatag gcggagtcac gactccggag 32520
gcggccggcc gccgcgcttc cggcatcgct cctacatggt ccttctccaa tgccggattc 32580
aggggcctgg caggggagta tgccgtcgga aagctcagga tgtcggcgtt tctggatgcc 32640
cgcgggagag tctctcctgg cggaaatatt ctctggctca tgaaaaacgg cgaactggga 32700
ctgacggcaa cccgttcaaa ggcttccgcg gatttccggc tgactctccg gggctcggat 32760
cttttcggcg aggcggcgta tgacttccgg gaccatgtaa tcgcggggat cgtcgggact 32820
gctttcccgt tgtgggagag gggccgggcg gcggttgttt tccgacacta cccgacaggt 32880
ttcgacggaa aggattccgg ggccatgaga agttcgacga aggtgaccga cgagaccgga 32940
gtcttcgctg ctctcaggct tgggagcatt ctgaccagtg ctgatgctgt cgtccatccc 33000
gggaccgggc ggcggcaact gaaactgctg ctactgcatg aatggaaaat atcgccgtcg 33060
gtcagtctta agactcgtgc gacggagagg ctgcgggatt atgaccgacg gaccaggact 33120
gacgtccgga ccgatctcgg aatcactcgg gggagtatgg tcatgaattt ccggttgaat 33180
gccctgaagt gcagggaaat agggcttgcc ggctatgcgg aggctggcta taagagggag 33240
aaatcggcac tatggttgcg ctggactgcg tttaaagtcg ataactggga tgaccgtatc 33300
tacgtttatg aacgtgatgc tcccggcaat ttctctgttc ctgcctgcta cgggagtgga 33360
tttacatttt ctctttatgg ttcatggaag atccgtcgtc tccgtcttta tctgaaatcc 33420
ggaacgacgt tcaaggagcg ctccggcacc cggtcgctct cctctgacct gcacttccaa 33480
ctcagttatt ctctgtagtt tacctccggt agatcaccag tcgctgacca acacggatat 33540
tgttgctgcg gagattgttc cacctcttga tctgcgctac ggtgacatgg tatcttgcag 33600
cgatgcggcc cagataatcg ccgttgcgca ccttgtagac tatcctgccg gactcttccg 33660
tacgggcctt cttgatatcc tccagagtct tcggactcag atatttatcg gcctggtaat 33720
tatagattga atcctcgtaa tcgataaagg cagtagtgta ctgcgaagga agggtaagaa 33780
tgctgacacc ctcattgccc ggcactatct catgcatata ctgagggtta agttccttga 33840
gcatgtcgac tgggacgttg atcacgccat tgatctgggc aaaatgcagg ttcttcctga 33900
tggcaaaagt atccagatgg ctcggaatgg cagtcggctc aggaattatc ccgtgctcct 33960
tgtaatacgt aaaggcatac atcgctccta caaaggcagg cacataacct ctggtctcgc 34020
gcggaagata ctcgtagata gcccagaaat tcttgttgcc tcctgcacgg cggatcgcct 34080
tgttgacatt gccggctccg cagttgtatg cggaaatagc cagactccag tcaccgaaga 34140
tactgtaggc atccctgagg tatctggcag cagcgtcggc agccttgacc gggtcaagcc 34200
tctcgtcgac ataggagttt atcttcaggc cgtaattctt tgccgtagaa tacatgaact 34260
gccacattcc ctttgcattc gccctggata ccgccgtagg attgagggca gactcgatca 34320
cagccatgta cttcagctct tcaggcatgt tgtacttgtt aagggtctct tcgaagatag 34380
ggaaataata ctgggccaga ccgagcatgt gccccatgcg ggtcggcatt ttctcggaat 34440
agagtattat gtagttcttt accgtttcgt tgtaggggag ggtaatatag gagttcatct 34500
tccgcagcct cttgatatag acgctgtccg gaacgttcga ggtgaaatgg acagaatcca 34560
tgttatagga attgttttcc atgttcatgg atttgcgggt gctgtaccag agattcagaa 34620
gactgtcggt cacctccggc gtataatcgt cggcagctat gcccgctccg atgcggtcct 34680
cgttttcctc atatatctcg atcatctctg cggcgatgct gtccgcagcc tcgtgctgct 34740
caatataatc gttgagctgc tgcctgagcg catttatttc agcctgcagt ctctcattct 34800
ccttcttgag atccttattc tttttcttag gcccgttttt gcggaagggc tgggttatct 34860
tgtcaaatat actctccgac tgggcgtatg agtccagcgg agcaactgaa gaaaccatga 34920
caaccgttaa tgcggttgcc gctatcaatt tatttaatct catttctaga attttattcc 34980

```
gatgcttatt ccgtaggcat tgtcgaccgg catgcccatc gcgaactggc tgtgcggggt     35040
tatgatggcc ggactgacat cgacagctat ttcgtcatcg atgtcgaaat caagcatata     35100
cgagaatacg ttggcatcta ttatctgaag tacatatact ccgaccagag ctactaccga     35160
atagtccctg tatcttctgt atgtatttct catatacaaa gcggaagatt cggtgtagtg     35220
gcccgaatat gtgctgttct catctgttat ttcgttgtgg atcctcttga agcggtggta     35280
attggcgctg ttctctatca gaaggctcgc cgaaaccatc agcccggagt agtaaagagg     35340
caatttccaa tattccttgt tgtatatctg gccaagaccc gggacaagca gggaatagag     35400
cgtagccctt ccggccgaat gctcgccttt tgtgtcgtag cttaccactc cgtccatcag     35460
ggagccccaa tagatagccc ctgcgcctgc gagaagatac tttccgaggt ccttcttgcc     35520
gtccttatgg atataatatg atccgagacc ggctgtcgcg gcaaggcctc cgtatatcac     35580
cggcagtttc cagtaatccc tgttatatat ctgctggccg ccgatgacga tggtggagcc     35640
ggcgaacatg gtgccgatct tgccttcctc tttatgtctc atggccctga aatactgcct     35700
gaaggagaac at                                                         35712
//
```
